# A higher burden of multiple sclerosis genetic risk confers an earlier onset

**DOI:** 10.1101/2020.12.18.423477

**Authors:** Elina Misicka, Mary F. Davis, Woori Kim, Steven W. Brugger, Jeremy Beales, Stephanie Loomis, Paola G. Bronson, Farren B.S. Briggs

## Abstract

Multiple sclerosis is a neurodegenerative, autoimmune disease characterized by inrreversible neurological disability. The age at onset of multiple sclerosis is an objective and influential predictor of the evolution of multiple sclerosis independent of disease duration. Little is known about the mechanisms contributing to variation in onset age of multiple sclerosis, though *HLA-DRB1*15:01*, the predominant risk variant, confers an earlier onset. Here we present an age at onset genome-wide association analysis for 9.2 million variants, including gene-based and pathway enrichment analyses, for 3,495 cases who were non-Latinx white with onset ≥18 years. We investigated whether a higher burden of multiple sclerosis risk variants conferred an earlier age at onset for combinations of *HLA-DRB1*15:01* alleles and quintiles of a genetic risk score for 200 risk variants that reside outside the major histocompatibility complex. The study population had a mean age at onset of 32 years, 29% was male, and 46% were *HLA-DRB1*15:01* carriers. *HLA-DRB1*15:01* carriers were on average one year younger at onset than non-carriers (*p*<0.001); a similar effect was observed for a 10-risk-allele increase in the genetic risk score (*p*<1×10^-8^). Those in the highest genetic risk score quintile (n=717) were on average 2.5 years younger at onset than those in the lowest quintile (n=698; p=1.2×10^-7^). For those with the greatest genetic risk burden (highest genetic risk score quintile with two *HLA-DRB1*15:01* alleles) were on average five years younger at onset (*p*=0.002) than those with the lowest genetic risk burden (lowest genetic risk score quintile with no *HLA-DRB1*15:01* alleles). There was an apparent inverse relationship between the genetic multiple sclerosis risk burden and age at onset of multiple sclerosis (*p*<5×10^-8^). We did not observe any individual variants reaching genome-wide significance in the genome-wide association analysis of age at onset. The most significantly associated independent genic loci (*p*<5×10^-6^) were located within *HLA-DQB1, COL21A1, LINC01484, UBR3*, and *CSMD1*. At the gene-level, the most significant associations (*p*<5×10^-5^) were for *SSB, TRAFD1, HECTD2, MMP8, NAA25* and *UBR3*. There was an enrichment of genes involved in adaptive and innate immunity, specifically genes in the complement pathway, and genes involved in synapses and collagen biosynthesis. In summary, we demonstrated a significant gradient between elevated genetic risk burden and an earlier onset of multiple sclerosis.

## Introduction

Multiple sclerosis (MS) is a demyelinating autoimmune disease of the central nervous system. It is a leading cause of neurological disability in young and middle-aged adults, affecting near 900,000 individuals in the United States,^1^ who variably accrue neurological impairments across multiple functional domains ^2^. There is currently no cure for MS, and existing treatments primarily aim to modulate disease activity due to inflammation with modest effect on neurodegeneration ^3^. As such, it is important for new and evolving research to identify and characterize factors that influence disease presentation and severity, in order to create opportunities for the development of novel therapeutic targets.

MS can occur at any age, but for the majority of persons with MS (PwMS), age at onset (AAO) is between 20 to 40 years of age. MS AAO is an objective and influential predictor of the evolution of MS independent of disease duration. An earlier AAO is associated with a higher frequency of relapses prior the transition to progressive disease ^4-6^. In addition, PwMS with an older AAO reach major disability milestones in a shorter time span, such as Expanded Disability Status Scale scores of 3 and 6; however, those with an earlier AAO still reach these milestones at younger ages ^4-12^. Of particular importance is the influence of AAO on the transition from relapsing remitting (RR) to secondary progressive (SP) MS. PwMS with an older AAO transition to SPMS in a shorter time span, but those with an early AAO do transition to SPMS at younger ages, and this is important to note since those who experience SP conversion at earlier ages attain advanced disability milestones in less time ^9-12^.

Factors influencing AAO in MS are not well established, though it appears to be multifactorial ^13^. It is clear that females have a significantly younger AAO ^13, 14^, and that there is a strong genetic component demonstrated by positive AAO correlations within families.

Additionally, consistent associations exist between the primary MS risk allele (*HLA-DRB1*15:01*) and earlier AAO, and a genetic risk score (GRS) capturing a higher burden of MS risk variants residing outside the major histocompatibility complex (MHC) also confers an earlier onset ^13, 15-18^. In a recent modestly sized study, those with the highest genetic risk burden for MS (carriers of *HLA-DRB1*15:01* and in the highest quintile of the non-MHC GRS; n=102) were significantly younger (3.7 years) at MS onset compared to those with the lowest genetic risk burden (non-carriers of *HLA-DRB1*15:01* and in the lowest quintile of the non-MHC GRS; n=121), with gradated increases in AAO with decreasing genetic risk burden ^13^. These findings need to be confirmed in a large population. It is also unclear if there are prominent associations for individual non-MHC risk variants. In 2011, a genome-wide association (GWA) study of ~465,000 SNPs observed no significant associations (p<5.0×10^-5^) with AAO outside the MHC ^18^. Additionally, a modestly-sized candidate gene study which focused on only five SNPs in four vitamin D level-associated genes and did not observe any significant associations with AAO;^19^ thus, this absence of association should also be confirmed.

In the current study, we add significant resolution to our understanding of the genetic component underlying MS AAO through an expansive GWA study of >9.7 million SNPs, that incorporated gene-based and pathway enrichment analyses. We also add resolution to relationship between AAO and MS genetic risk burden by investigating specific combinations of copies of *HLA-DRB1*15:01* alleles and quintiles of the non-MHC GRS. Confirming a gradient between decreasing genetic risk burden and delayed AAO of MS would importantly suggest that a higher genetic risk burden accelerates onset of MS.

## Materials and Methods

### Study Population and Association Analysis

The study population consisted of 3,495 MS cases with an AAO of at least 18 years and of non-Latinx, European ancestry from five cohorts (Table 1). 1,268 MS cases were from the Accelerated Cure Project for MS (ACP), a biorepository of PwMS and other demyelinating diseases who were recruited from the communities of 10 MS specialty clinics across the United States. A neurologist confirmed all MS cases met standard diagnostic criteria at the time of enrollment ^20, 21^. Inclusion and exclusion criteria have been described (Supplementary Table 1A) ^22^. ACP participants comprised two cohorts (ACP1 and ACP2), which were defined by their genotyping method (Table 1). ACP1 included 1,061 PwMS genotyped on a customized Illumina iSelect platform, the Illumina Exome Chip, as part of a recent international effort to identify MS risk variants ^23^. ACP2 included 207 PwMS who were genotyped on the Illumina MEGAEx BeadChip. ACP1 and ACP2 were similarly imputed using the Michigan Imputation Server, and SNPs with an imputation quality score (r^2^) of <0.8 info score were removed; this has also been described ^24^. Multidimensional scaling (MDS) components were generated for a subset of independent SNPs to determine genetic outliers and cryptic relatives, who were removed from the data ^24^. The first five MDS dimensions were used to adjust for population substructure. For all ACP participants, AAO of MS was defined as the age of first neurological symptom suggestive of MS.

**Table 1.** Study population and study cohort characteristics

| Trait<br>N, Mean (SD) or % | Overall | ACP1 | ACP2 | ADVANCE | ASCEND | DECIDE |
| --- | --- | --- | --- | --- | --- | --- |
| N | 3,495 | 1,061 | 207 | 655 | 555 | 1,017 |
| Male | 29.0% | 21.9% | 24.2% | 27.4% | 39.1% | 32.7% |
| Onset Age (years) | 32.0 (8.9) | 33.3 (9.6) | 34.0 (9.9) | 31.4 (8.5) | 32.0 (8.2) | 30.7 (8.1) |
| Genotyping Platform | NA | Illumina<br>Exome<br>Chip | Illumina<br>MegaEX | Affymetrix UK Biobank Axiom Array |  |  |
| Imputation Approach | NA | University of Michigan Imputation Server |  |  |  |  |
| No. SNPs <sup>1</sup> | 9.2×10<sup>6</sup> | 1.2×10<sup>6</sup> | 7.3×10<sup>6</sup> | 9.6×10<sup>6</sup> | 9.6×10<sup>6</sup> | 9.6×10<sup>6</sup> |
| *HLA-DRB1\*15:01* carriers <sup>2</sup> | 45.7% | 46.2% | 46.9% | 43.2% | 45.9% | 46.3% |
| Genetic Risk Score <sup>3</sup> | 209.2 (8.7) | 208.4 (8.6) | 207.5 (9.0) | 209.7 (8.6) | 209.7 (8.6) | 209.8(8.7) |

1. The number of SNPs listed for each cohort are those with minor allele frequencies >1% and HWE p>0.0001. The number of SNPs listed for the overall study population, are those included in the final meta-analysis (occurring in at least two cohorts and I<sup>2</sup> p>0.05)
2. Determined by the tagging SNP rs3135388A
3. Sum of the risk alleles spanning 198 non-MHC MS variants (Supplementary Table 2).

The remaining 2,227 MS cases were participants in three MS clinical trials: ADVANCE (N=655) ^25^, ASCEND (N=555) ^26^, and DECIDE (N=1,017) ^27, 28^. Genotyping was performed on the Affymetrix UK Biobank Axiom Array in two batches; samples from ASCEND were genotyped separately from samples from ADVANCE and DECIDE. Imputation was performed using the University of Michigan imputation server for each batch, and Eigensoft v7.2.1 was used to conduct principal components analysis and identify genetic outliers (see Supplementary Methods^29^). Inclusion and exclusion criteria for each study is presented in Supplementary Table 1A. AAO was generated by subtracting years since onset of symptoms from age at the enrollment in ADVANCE and by subtracting the birth year from year of first study indication symptom in ASCEND and DECIDE.

### Statistical Analysis

#### GWA meta-analysis

A schematic of the analytical framework is presented in Figure 1. In each cohort, SNPs with a minor allele frequency <1% and Hardy-Weinberg Equilibrium p<0.0001 were excluded (Supplementary Table 1B), and GWA linear regression models with AAO (years; Figure 2) as the dependent variable and each of the retained SNPs as the independent variable of interest (additive genetic model), adjusting for population substructure and sex, were conducted. Meta-analysis of the GWA AAO results from the five cohorts was conducted using METAL, with a standard-error based approach ^30^. Genomic control correction was applied, automatically adjusting for unaccounted-for relatedness between subjects, and inflation of the test statistics was estimated through comparison of the median test statistic to the value expected from random chance. Heterogeneity of effect estimates for each SNP was calculated using the I^2^ statistic. METAL discarded variants with nonmatching allele pairs, invalid standard error values, and extra copies of duplicated variants across cohorts, resulting in 10.3 million SNPs (Supplementary Table 1B). SNPs observed in only one cohort were also excluded, resulting in 9.7 million SNPs. Finally, only the results for 9.2 million SNPs with heterogeneity I^2^ p>0.05 were retained for interpretation and subsequent analyses. SNPs were annotated using Seattle Seq Variation

**Figure 1.**
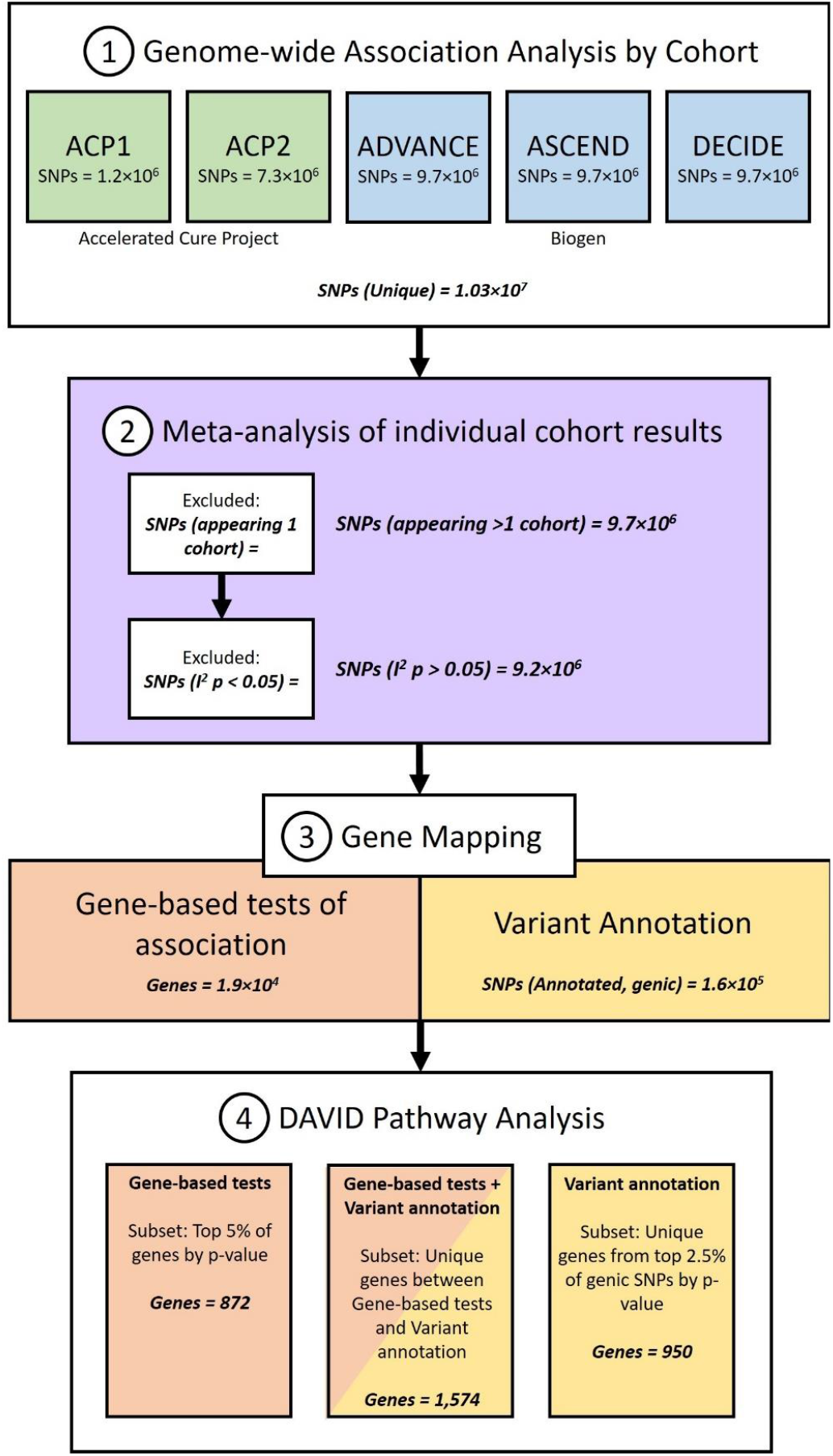
Flow Chart of Study Analyses

**Figure 2.**
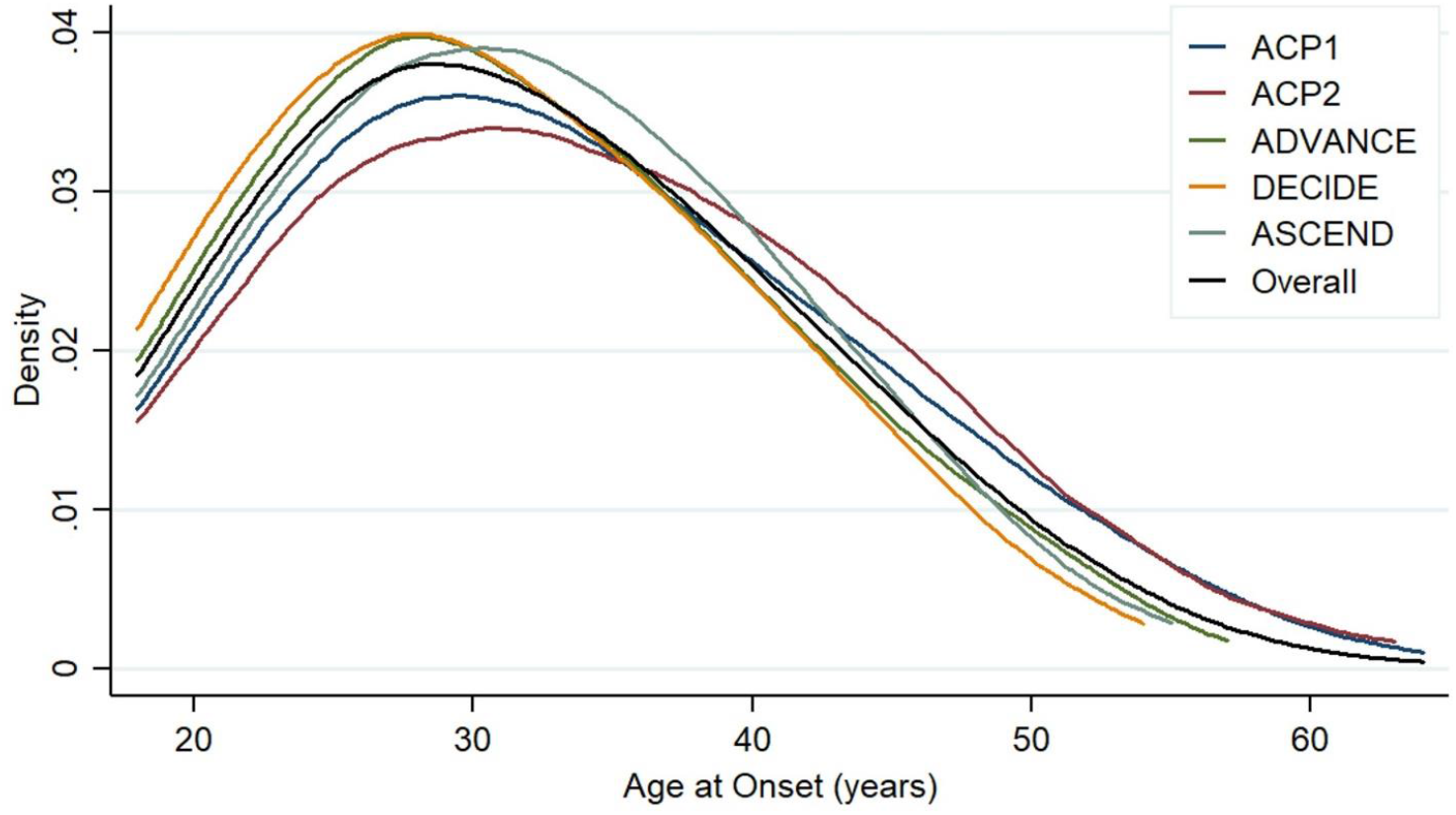
Distribution of Age at Onset of MS across study cohorts

Annotation 138 ^31^ and genome build hg19 and were determined genic if within 3’-5’ UTR gene boundaries.

Independent lead variants were identified by FUMA ^32^, using the linkage disequilibrium (LD) structure for the European (EUR) 1000 Genomes population. Independent significant SNPs were first identified by having a discovery significance threshold of p<5×10^-6^ and being independent at a level of r^2^<0.6. SNPs in LD (r^2^>0.6) with the independent variants were considered candidate SNPs. Finally, independent lead variants were selected from the independent significant SNPs if they remained independent from each other at r^2^<0.1. Lead SNPs were visualized through FUMA’s annotation plot software, where lead variants and their candidate variants and other nearby variants were plotted based on genomic location and -log p-value.

#### MS genetic risk burden analyses

Given prior associations between *HLA-DRB1:15:01* and AAO, in addition to evidence that a higher non-MHC GRS, particularly amongst carriers of *HLA-DRB1*15:01*, was associated with earlier AAO in ACP1 ^13^, we sought to confirm the relationship between MS genetic risk and AAO. For *HLA-DRB1*15:01*, we used the tagging SNP rs3135388A. For the 200 non-MHC MS risk variants ^23^, 198 variants were available across all cohorts. These 198 variants were either the original risk variant or a LD proxy identified by using the 1000 Genomes CEU population when the original variant was not available (Supplementary Table 2). LD proxies had a r^2^≥0.6 with the risk variant, favoring variants in higher LD and closer to the risk variants. In the case that the risk variant had no nearby variant in high LD, the discovery SNP from the original study was used. In each cohort, we constructed a non-MHC GRS by summing the risk alleles across these variants (Supplementary Figure 1). A binary variable capturing *HLA-DRB1*15:01* carrier status and a categorical variable capturing quintiles of the GRS were constructed. Using STATA v13.1 (StataCorp, College Station, TX), a fixed-effect meta-analysis with AAO as the dependent variable and *HLA-DRB1*15:01*, sex, and population substructure as predictors was conducted. This model was repeated for *HLA-DRB1*15:01* carrier status, the GRS, GRS quintiles, and combinations of number of *HLA-DRB1*15:01* alleles and GRS quintiles.

#### Gene-Based Tests

Gene-based tests of association were performed using MAGMA implemented in FUMA ^32, 33^, using three gene ranges: exact gene boundaries, gene boundaries +/-10kb, and gene boundaries +/-25kb. MAGMA creates a gene-based test statistic by converting p-values of SNP-level summary statistics to χ^2^ values, which are then grouped by SNPs mapped to a specific gene and gene range ^32, 33^. For the group of SNPs in each gene, the mean χ^2^ is determined, and converted into a p-value to indicate significance of the gene’s association with AAO. LocusZoom visualized the individual SNPs results for the top gene-based associations ^34^.

#### Pathway Enrichment Analysis

Pathway enrichment analyses was completed for three gene sets. The first gene set (GS1) consisted of the genes with a SNP-level association amongst the top 2.5% of unique genic associations based on p-value ranking. The second gene set (GS2) consisted of genes amongst the top 5% of associations from the gene-based test (exact gene boundary analysis) based on p-value ranking. The ranking thresholds for GS1 and GS2 were selected, as they resulted in similar number of genes. Lastly, the union of GS1 and GS2 comprised the third gene set (GS3).

Pathway enrichment analyses were performed using the DAVID Functional Annotation database version 6.8 and all human genes as the reference and the Reactome, KEGG, and BioCarta databases^35, 36^. Enrichment was determined using the EASE (Expression Analysis Systematic Explorer) score, which can be considered as a conservatively modified Fisher Exact Test ^37^ and is implemented in DAVID. In addition to pathway enrichment, functional annotation clusters of genes were identified, where genes in the gene set are clustered based on the degree of co-association between heterogeneous annotation terms to account for annotation redundancy across databases; all available annotation databases were used. An enrichment score is calculated; it is a geometric mean of EASE scores of terms involved in the annotation cluster.

### Data Availability Statement

Individual level data may be made available to qualified persons with the approval of a data use agreement between the respective institutions. Summary level statistics are available by contacting specific authors of the manuscript (ACP via FB, Biogen via PB).

## Results

Five cohorts of non-Latinx white PwMS were included in this study (Table 1). The total study population consisted of 3,495 individuals, of whom 29% were male. The overall mean AAO was 32.0 years (standard deviation= 8.9; Figure 2), which is representative of the general MS population. While inclusion and exclusion criteria for the ACP and Biogen cohorts differed, the mean AAO for each cohort were within 3.3 years with similar standard errors. The cohorts were also genetically similar, with ~45% of participants positive for *HLA-DRB1*15:01*, and all cohorts had a similar burden and distribution of the GRS (Table 1; Figure 3).

**Figure 3.**
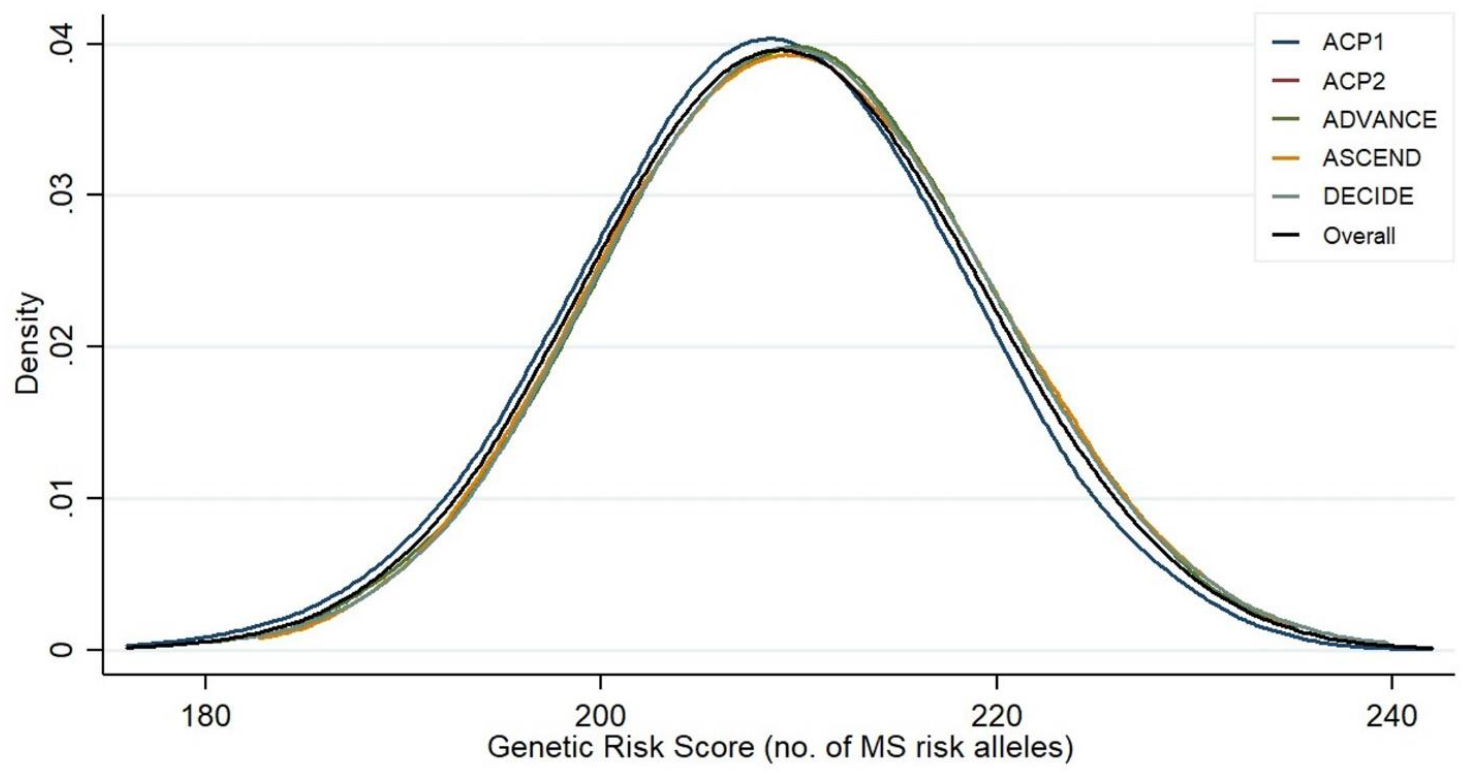
Distribution of genetic risk score across study cohorts

### MS genetic risk burden results

The MS genetic risk component significantly conferred an earlier AAO of MS (Table 2). Increasing copies of *HLA-DRB1*15:01* was associated with earlier AAO (β (95% CI) = −0.67 (−1.16, −0.19); p=0.0062), with carriers of *HLA-DRB1*15:01* experiencing MS onset a full year earlier than non-carriers (β (95% CI) = −1.00 (−1.58, −0.41), p=0.0009). The non-MHC GRS was also significantly associated with AAO (p=9.8×10^-9^), where an increase of 10 and 50 risk alleles would confer an earlier onset of MS by one and five years, respectively. This trend for increasing non-MHC genetic risk burden and earlier AAO is also demonstrated by the results for GRS quintiles (p_trend_=2.1×10^-7^); those in the highest quintile (n=698) were 2.5 years younger at onset than those in the lowest quintile (n=717; β (95%CI) = −2.49 (−3.41, −1.57), p=1.2×10^-7^). For combinations of *HLA-DRB1*15:01* alleles and GRS quintiles (Table 2), those in the highest GRS quintile with two *HLA-DRB1*15:01* alleles (those with highest genetic risk burden for MS) were on average five years younger at MS onset than those in the lowest quintile with zero *HLA-DRB1*15:01* alleles (n=31, β (95%CI) = −4.98 (−8.19, −1.78), p=0.0023; similar to unadjusted observed AAO differences in the data). There was an apparent gradient between increasing genetic risk burden and earlier AAO. Interestingly, of the non-MHC risk variants, nine (4.5%) were associated with AAO (p<0.05), including genic variants in *PHGDH, SH2B3*, and *ATXN1*, which were associated with earlier AAO (β= −0.5 to −0.6; Supplementary Table 2). Thus, the effect of the GRS on AAO appears to be driven by the cumulative effect of risk alleles and not by large effects for specific risk alleles.

**Table 2.** The association between genetic risk factors and age of onset of MS ^1^.

| Genetic Predictor | | | Mean<br>AAO<br>(Years) | AAO<br>Standard<br>Deviation | $\beta$ (95% CI) | P | p trend |
| --- | --- | --- | --- | --- | --- | --- | --- |
| *HLA-DRB1\*15:01* | | | - | - | -0.67 (-1.16, -0.19) | $6.2 \times 10^{-3}$ | - |
| Carrier of *HLA-DRB1\*15:01* | | | 31.60 | 8.64 | -1.00 (-1.58, -0.41) | $9.0 \times 10^{-4}$ | - |
| GRS | | | - | - | -0.10 (-0.13, -0.06) | $9.8 \times 10^{-9}$ | - |
| GRS Quintile | Q1 (n=717) | | 34.00 | 9.40 | Ref | - | $2.1 \times 10^{-7}$ |
| | Q2 (n=722) | | 32.04 | 8.98 | -1.84 (-2.74, -0.94) | $6.8 \times 10^{-5}$ | |
| | Q3 (n=658) | | 31.34 | 8.67 | -2.27 (-3.20, -1.34) | $1.7 \times 10^{-6}$ | |
| | Q4 (n=700) | | 31.39 | 8.64 | -2.32 (-3.24, -1.41) | $6.7 \times 10^{-7}$ | |
| | Q5 (n=698) | | 31.20 | 8.46 | -2.49 (-3.41, -1.57) | $1.2 \times 10^{-7}$ | |
| Combinations<br>of genetic risk<br>factors | 0 copies of<br>*HLA-*<br>*DRB1\*15:01* | Q1 (n=388) | 34.43 | 9.63 | Ref | - | $2.9 \times 10^{-8}$ |
| | | Q2 (n=384) | 32.42 | 8.88 | -1.93 (-3.17, -0.70) | $2.1 \times 10^{-3}$ | |
| | | Q3 (n=332) | 32.20 | 8.86 | -2.08 (-3.36, -0.80) | $1.5 \times 10^{-3}$ | |
| | | Q4 (n=399) | 31.49 | 8.94 | -2.64 (-3.87, -1.42) | $2.4 \times 10^{-5}$ | |
| | | Q5 (n=396) | 31.62 | 8.85 | -2.44 (-3.67, -1.21) | $1.0 \times 10^{-4}$ | |
|  | 1 copy of<br>*HLA-*<br>*DRB1\*15:01* | Q1 (n=273) | 33.23 | 9.12 | -1.32 (-2.68, 0.04) | 0.057 |  |
| | | Q2 (n=300) | 31.64 | 9.04 | -2.72 (-4.04, -1.40) | $5.5 \times 10^{-5}$ | |
| | | Q3 (n=289) | 30.83 | 8.46 | -3.47 (-4.80, -2.13) | $3.6 \times 10^{-7}$ | |
| | | Q4 (n=256) | 30.93 | 8.11 | -3.37 (-4.76, -1.99) | $1.8 \times 10^{-6}$ | |
| | | Q5 (n=271) | 30.81 | 8.00 | -3.55 (-4.91, -2.19) | $3.2 \times 10^{-7}$ | |
|  | 2 copies of<br>*HLA-*<br>*DRB1\*15:01* | Q1 (n=56) | 34.86 | 9.02 | 0.01 (-2.45, 2.46) | 0.996 |  |
|  |  | Q2 (n=38) | 31.26 | 9.60 | -3.53 (-6.44, -0.61) | 0.018 |  |
|  |  | Q3 (n=37) | 31.11 | 8.34 | -3.47 (-6.42, -0.52) | 0.021 |  |
|  |  | Q4 (n=45) | 33.13 | 8.80 | -1.22 (-3.92, 1.48) | 0.376 |  |
| | | Q5 (n=31) | 29.16 | 6.87 | -4.98 (-8.19, -1.78) | $2.3 \times 10^{-3}$ | |

1. Results from individual fixed effect meta-analyses with AAO as the dependent variable, and each genetic predictor (i.e. GRS Quintile) as the sole independent variable of interest, adjusted for sex and genetic ancestry.

### GWA meta-analysis results

In the meta-analysis of 9.24 million SNPs, no association met genome-wide significance (p<5.0×10^-8^; Figure 4). At a discovery significance level of p<5×10^-6^, 7 lead independent risk variants were identified from 12 independent significant SNPs (Table 3). 1,235 variants were considered candidate variants for the 7 lead risk loci (Supplementary Table 3). Three of the lead risk variants were located on chromosome 6: rs28672722 in *HLA-DQB1* (which is ~70 kb downstream of *HLA-DRB1*; however, it is not in LD with *HLA-DRB1*15:01* [r^2^<0.03]), rs149847639 in *COL21A1*, and an intergenic variant rs17066212 (Table 2). The other four variants were located on chromosomes 2 (rs145201293 in *UBR3*), 5 (rs17076315 in *LINC01484*), 8 (rs74402157 in *CSMD1*), and 19 (rs34132828, intergenic). The most significant association was for *LINC01484* rs17076315 (p=6.9×10^-7^), while three less common variants (MAF<5%) had large effect sizes for increasing copies of the minor allele, including AAO betas of −7.5 years for *COL21A1* rs149847639 and −4.5 years for *CSMD1* rs74402157. The complete list of associations with p<0.001 is shown in Supplementary Table 3.

**Figure 4.**
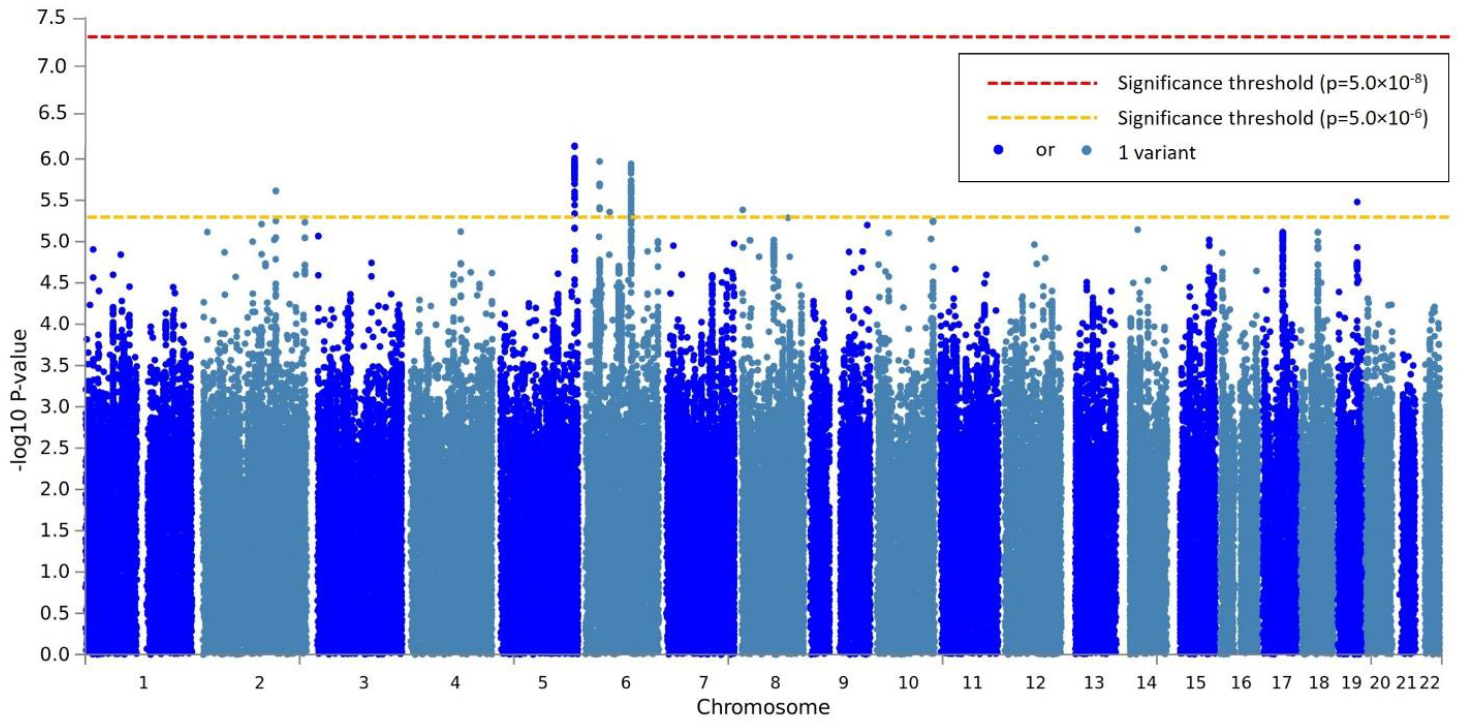
Manhattan Plot of Meta-Analyzed GWAS Results

**Table 3.** Genomic risk loci from the MS AAO meta-analysis at p<5.0×10^-6^.

| Chr | Base pair position | rsID | MAF <sup>1</sup> | A1 <sup>2</sup> | A2 | N <sup>3</sup> | Cohorts <sup>4</sup> | $\beta$ (SE) | p | Gene | SNP Function |
| --- | --- | --- | --- | --- | --- | --- | --- | --- | --- | --- | --- |
| 2 | 170812811 | rs145201293 | 0.30 | T | TTA | 2,434 | 3 | 1.3 (0.28) | $2.4 \times 10^{-6}$ | *UBR3* | Indel |
| 5 | 173141042 | rs17076315 | 0.35 | A | G | 2,434 | 4 | -1.3 (0.26) | $6.9 \times 10^{-7}$ | *LINC01484* | Intron variant |
| 6 | 32626537 | rs28672722 | 0.26 | T | G | 3,495 | 5 | 1.2 (0.25) | $1.1 \times 10^{-6}$ | *HLA-DQB1* | Upstream transcript variant |
| 6 | 56121561 | rs149847639 | 0.015 | A | C | 1,210 | 2 | -7.5 (1.6) | $4.4 \times 10^{-6}$ | *COL21A1* | Intron variant, genic upstream transcript variant |
| 6 | 106149587 | rs17066212 | 0.022 | A | G | 2,434 | 4 | -3.9 (0.80) | $1.1 \times 10^{-6}$ | | Intergenic |
| 8 | 3655967 | rs74402157 | 0.028 | T | G | 2,434 | 3 | -4.5 (0.98) | $4.1 \times 10^{-6}$ | *CSMD1* | Intron variant, genic upstream transcript variant |
| 19 | 46163870 | rs34132828 | 0.24 | A | G | 3,495 | 5 | 1.2 (0.25) | $3.3 \times 10^{-6}$ | | Intergenic (7.6 kb upstream of *GIPR*) |

1. Minor allele frequency in the 1000 Genomes European (EUR) reference. 2. A1 is the minor allele. 3. Due the differences in the genotyping platforms, not all variants were able across all studies even after imputation. 4. Number of cohorts in which the SNP was available

Prominent vitamin D level-associated variants in *GC* (rs7041, rs4588, rs2282679), *CYP27B1* (rs12368653), and *CYP24A1* (rs2248359) were not associated with AAO (p>0.05; data not shown). *CYP2R1* rs10741657 was marginally associated with AAO (p=0.04; Supplementary Table 3). We do note that ~20 intronic *GC* SNPs were associated with AAO (p<0.05), as were several SNPs within 25kb of *CYP27B1* (e.g. rs113625101), *CYP24A1* (e.g. rs2248817), and *CYP2R1* (e.g. rs11819875) (Supplementary Table 3). Similar modestly significant relationships were present for SNPs near the *DHCR7-NADSYN1* locus (e.g. rs61885923A associated with a 2.9 year earlier AAO) but not *CYP27A1*. Within *VDR*, five intronic variants had strong effects (p=0.05-0.005), including a less common variant rs142161130T (MAF=2%) which was associated with an earlier onset of MS by 2.9 years (p=0.037; Supplementary Table 3).

### Gene-Based test results

Gene-based tests of associations were conducted for >19,000 genes (Supplementary Table 4). Top gene-based results were for *SSB* on chromosome 2 (p=2.6×10^-5^), *MMP8* on chromosome 11 (p=2.9×10^-5^), and *TRAFD1* (p=2.7×10^-5^), *HECTD4* (p=2.8×10^-5^), and *NAA25* (p=4.5×10^-5^) located on chromosome 12. Of the genes in which the lead independent SNP associations were located, *UBR3* was significantly associated with AAO at the gene-level as well, across all gene boundary definitions (p_0kb_ = 4.6×10^-5^, p_10kb_=4.0×10^-5^, p_25kb_=1.3×10^-5^). *COL21A1* was trending in association (p<0.10), as were *HLA-DQB1* (p<0.15) and *HLA-DRB1* (p<0.10), while *HLA-DRA* (where rs3135388, which tags *HLA-DRB1*15:01*, resides) was associated with AAO (p=0.0023). The associations for the HLA genes were not unexpected, given the complex extended LD spanning the region. None of the genes associated with vitamin D levels demonstrated any association with AAO, except for *CYP2R1* when considering a 25kb window around the gene boundary (p=0.03; Supplementary Table 4).

### Pathway enrichment analyses

GS1, GS2, and GS3 consisted of 986 genes, 952 genes, and 1,679 genes, respectively (Supplementary Table 5); for these sets, DAVID recognized 950 (96.3%), 872 (91.6%), and 1,574 (93.7%) of the submitted gene symbols, respectively.

For GS1, there was an enrichment in cell membrane and adhesion proteins, as well as pathways involving numerous HLA genes, suggesting a prominent role for adaptive immune response (Table 4, Supplementary Table 6). The most significant pathway identified from enriched genes in this list was NCAM1 interactions with a fold enrichment (FE) of 4.5 (p=6.7×10^-4^; FDR p=0.0099). Other enriched pathways included the generation of second messenger molecules (FE=4.7; p=1.3×10^-3^; FDR p=0.019), production of cell adhesion molecules (FE=2.4; p=1.7×10^-3^; FDR=0.022), the translocation of the ZAP-70 kinase to immunological synapses (FE=5.8; p=3.1×10^-3^; FDR=0.045), and collagen biosynthesis (FE=3.1; p=4.0×10^-3^; FDR=0.059). When applying functional annotation clustering, the top cluster had an enrichment score of 4.99 and included genes involved presynaptic and postsynaptic cell membranes.

**Table 4.**
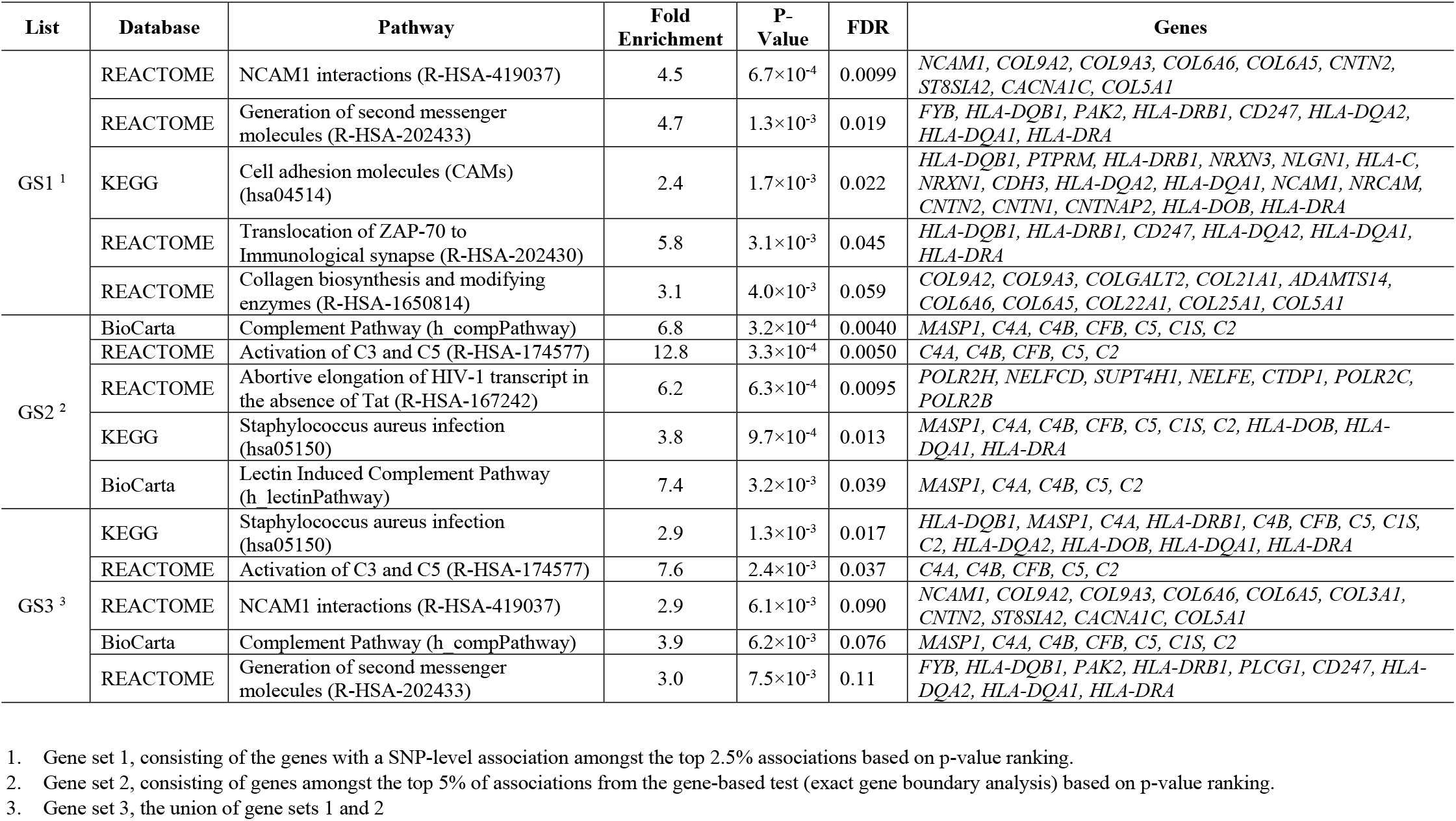
Top 5 Pathway Enrichment Results per Gene Set

For GS2, there was an enrichment of complement pathways and gene clusters that suggested a role for innate immune response (Table 4, Supplementary Table 7). The most significant enrichment was observed for complement response (FE=6.8; p=3.2×10^-4^; FDR p=0.004) and activation of C3 and C5 (FE=12.8; p=3.3×10^-4^; FDR p=0.005). The pathway for the abortive elongation of the HIV-1 transcript in the absence of Tat was also enriched (FE=6.2; p=6.3×10^-4^; FDR p=0.0095), as was immune response to Staphylococcus aureus infection (FE=3.8; p=9.7×10^-4^, FDR = 0.013), which included complement system and HLA genes. The top gene cluster by functional annotation was centered around the innate immune response, with an enrichment score of 3.55.

For GS3, significant pathway enrichment involved genes involved in both adaptive and innate immune response, as well as cell adhesion interactions (Table 4, Supplementary Table 8). Top enriched pathways included the NCAM1 interactions (FE=2.9; p=6.1×10^-3^, FDR =0.090) and generation of second messenger molecules (FE=3.0; p=7.5×10^-3^, FDR=0.11), both enriched in GS1 alone. Pathways from the Staphylococcus aureus infection (FE=3.8; p=9.7×10^-4^, FDR=0.013), C3 and C5 activation (FE=7.6; p=2.4×10^-3^, FDR = 0.037), and complement pathway (FE=3.9; p=6.2×10^-3^, FDR = 0.076) were also enriched. The top gene clusters by functional annotation entirely consisted of genes involved in the function of the cell/plasma membrane (enrichment score of 3.90) and synaptic cell member (enrichment score of 3.65) (Supplementary Table 9).

## Discussion

Here we present a comprehensive and updated genome-wide investigation of AAO of MS. AAO is an objective predictor of the evolution of MS independent of disease duration, influencing the accrual of neurological disability and MS progression, which both substantially impact quality of life and long-term care decisions. The study’s objective was to identify genetic factors and biological processes contributing to variation in the AAO of MS, as these associations may underline relationships contributing to disability accrual and MS progression, and present opportunities for the development of novel drug targets. While no non-MS risk SNP or gene-level association met genome-wide significance, a higher burden of MS risk variants significantly contributed to an earlier onset (p<5×10^-8^). Those with the highest genetic burden (highest non-MHC GRS quintile and homozygous for *HLA-DRB1*15:01*) were on average five years younger at MS onset than those with the lowest genetic burden. The results also suggest a prominent role for variation spanning the MHC and beyond *HLA-DRB1*15:01*, with strong evidence supporting a role for the complement system, which is a functional bridge between innate and adaptive immune response to pathogenic challenges ^38^ and the structure of cell membranes at synapses.

### MS Genetic Risk Component and AAO

To date, the identified MHC and 200 non-MHC genetic risk variants for MS additively explain 21% and 18%, respectively, of the 19% of the liability for MS explained by common genetic variation ^23^. Prior studies established that presence of *HLA-DRB1*15:01* conferred earlier onset, which we also demonstrated. There have been two studies that have investigated the relationship between a MS non-MHC GRS and AAO. The first study constructed a GRS based on 106 variants and the second was based on the complete 200 non-MHC risk variants (which is the ACP1 cohort in this study) – in both, a higher GRS was associated with earlier onset ^13, 15^. We add resolution to these prior observations, by reporting the individual associations for the most recent list of non-MHC risk variants. Of these variants, only nine (4.5%) had a significant effect on AAO at p<0.05, including *ATXN1* rs719316 (p=0.020). Ataxin-1, encoded by *ATXN1*, aggregates in the brain and spinal cord ^39^, and is associated with spinocerebellar ataxia, a fatal, progressive neurodegenerative disorder ^40^. Other genes in which significant variants were mapped include *SH2B3*, a lymphocyte adapter protein associated with various autoimmune and vascular disorders ^41^, and *PHGDH*, a catalyst involved in the biosynthesis of L-serine. Mutations in PHGDH can cause Neu-laxova syndrome, a congenital disorder that affects the abnormal development of the brainstem and spinal cord ^42^. While there was limited evidence for pronounced effects for individual non-MHC risk variants, collectively, a higher burden of these variants significantly conferred an earlier MS onset (p<1 ×10^-9^). Thus, we importantly confirm that there is a gradient between decreasing genetic risk burden and delayed AAO of MS.

Given that MS genetic risk is associated with AAO, and that AAO is a strong predictor of long term outcomes in PwMS, one would hypothesize that MS genetic risk may also have a direct effect on these outcomes; however, there has been no robust evidence to support this hypothesis. Previous work saw no evidence for association between the *HLA-DRB1*15:01*, the non-MHC GRS and time to transition to SPMS, even when stratifying by 10-year increments to account for time-varying effects ^9^. There is also no evidence that MS genetic risk contributes to accrual of disability captured by the Multiple Sclerosis Severity Score (MSSS). A GWA study found no variants associated with MSSS at a genome-wide significance threshold ^43^. A second MSSS study of 52 MS risk variants and their composite GRS, reported no significant associations upon correction for multiple testing ^44^. Furthermore, given the relationship between vitamin D insufficiency and MS ^45^ and that a prior study focused only on five related variants ^19^, our results do appear to support relationships between genetic variation in related genes (particularly *VDR* and the *DHCR7-NADSYN1* locus) and AAO and warrants closer inspection.

### Adaptive and Complement Immune Systems and MS AAO

Genetic variants associated with several aspects of the complement system were enriched amongst the AAO associations. The lead variant rs74402157 resides within *CSMD1*, which encodes a complement regulatory protein that contributes indirectly to the innate immune function ^46^. *CSMDI* is highly expressed in the brain ^47^, and mutations in *CSMD1* are associated with several neurological diseases including Parkinson’s Disease and Schizophrenia ^48-50^. Genes from the complement system significantly associated with AAO at both the SNP and gene-level, and they were strongly enriched in pathways derived from GS2 and GS3. Activation of the complement pathway enhances and expedites the innate and adaptive immune response, playing various roles in autoimmune responses ^51, 52^. Elevated levels of C3 and C4a have been reported in the cerebrospinal fluid (CSF) of relapsing-remitting patients compared to healthy controls ^53^, with elevated levels for C4a in those with active versus stable disease ^54, 55^. Both of these factors have also been positively correlated with the Expanded Disability Status Scale ^55, 56^. C3 was recently associated with atrophy of the ganglion cell layer of the inner retina in patient with MS, and *C1QA* and *CR1* variants were associated with increased loss of low-contrast letter acuity ^57^. Additionally, earlier AAO has been associated with higher levels of visual symptoms at onset ^58^. Considering MS AAO is an established predictor of reaching disability milestones, these findings suggest that dysregulation in the complement system may be a shared biological mechanism.

### Innate Immune System and MS AAO

In the gene-based tests, *TRAFD1* and *SSB* were identified as a gene significantly associated with AAO. *TRAFD1* is a negative feedback regulator of the innate immune system, suppressing over-reactive immune responses, and variation in this gene has been associated with body mass index and variation in systolic and diastolic blood pressure ^59-62^. *SSB* encodes a small RNA-binding exonuclease protection factor that is associated with Sjögren’s Syndrome and systemic lupus erythematosus ^63, 64^. Additionally, pathways involved in innate immune response were enriched in all gene sets, but most dominantly in GS1. These pathways, such as the cell adhesion molecule pathway (hsa04514), contain many MHC genes, including *HLA-DRB1.* Additionally, the lead variant rs28672722 is <1kb upstream of *HLA-DQB1*, and variation in DQB1 has been associated with MS risk ^23^.

### Cellular Matrix, Adhesion, and MS AAO

Many of the enriched pathways were involved in cell membrane construction and the creation and transportation of messenger molecules. For example, *COL21A1* and *MMP8* were associated with AAO at both the gene and variant level. COL21A1 is a fibril that bonds to other molecules in the cellular matrix ^65^, and genetic variation in *COL21A1* has been associated with brain volume ^66^, systolic and diastolic blood pressure ^67^, pulse pressure ^67^, and heel bone mineral density ^68^. *MMP8* is a matrix metalloproteinase, specifically named neutrophil collagenase for breaking down collagen in both neutrophils and leukocytes. The cell adhesion molecule pathway is involved in the formation of the nodes of Ranvier along the axons of neurons ^69^. Nodes of Ranvier lie between the myelin sheathes on neuronal axons, which, when damaged, result in a substantial portion of neurological disability seen in PwMS. Other pathways of note involved NCAM1 interactions, which contribute to neuronal development by inducing cell differentiation^70, 71^.

### Study Strengths and Limitations

A key strength of this study is the inclusion of multiple cohorts who were representative of the non-Latinx white MS population by sex, AAO, and genetic predisposition. A second strength is the opportunity to characterize the relationship between MS genetic risk burden and AAO, and the expansion genome-wide coverage of genetic variants. Another strength is the conducting of enrichment analysis for multiple gene sets (variant-based and gene-based tests), and the inclusion of multiple reference databases. A notable limitation is that while there was extensive genetic overlap across data sets, several of our lead findings were not available in all samples. We also acknowledge that our findings may not be generalizable to diverse and understudied populations nor to those with pediatric-onset MS.

## Conclusions

Given that AAO is an important predictor of long-term outcomes, it is important to characterize factors influencing heterogeneity in MS presentation. This study demonstrated that an increasing load of genetic risk variants contributes to earlier onset of MS, suggesting that multiple aberrant processes accelerates the initiation of etiologic events. Amongst the genic associations, there was an enrichment of genes involved in the adaptive and innate immunity, specifically in the complement systems, and genes involved in cell membrane adhesion and cell signaling.

## Acknowledgements

The authors appreciated Mr. Justin Yu’s contributions on imputing the ACP data.

## Supplementary Material

See additional files for Supplementary Tables (.xlsx), Supplementary Figures (.docx), and Supplementary Methods (.docx).

