## Supplementary Methods for "A higher burden of multiple sclerosis genetic risk confers an earlier onset"

*Biogen Sample Genotyping and Quality Control*

All samples were of white, non-Latinx ancestry. Serum samples were collected at baseline, and were genotyped in two batches on the Affymetrix UK Biobank Axiom array by Thermo Fisher Scientific (Santa Clara, CA). Batch 1 consisted of the ASCEND cohort, genotyped in 2017. Batch 2 consisted of the ADVANCE and DECIDE cohorts, genotyped in 2018. Quality control for each batch was performed in PLINK v1.9, including removing samples with >2% missingness, minor allele frequency<1%, Hardy-Weinberg equilibrium $p<1x10^{-4}$, and SNP missingness<1%. Sex checks were performed using LD-pruned data with default options. Identity by descent analyses were conducted and samples with excess heterozygosity (>6 standard deviations from the mean) compared to the LD-pruned dataset were excluded. Pairs of individuals were considered related with a pi-hat >0.4. One individual from each related pair was randomly selected for exclusion.

Two rounds of principal components analysis (PCA; SmartPCA, Eigensoft v7.2.1) were performed, using an LD-pruned dataset merged with ancestry information markers. Samples were excluded if they were 6 standard deviations from the top 10 principal components with a maximum of 10 outlier removal iterations. Tracy-Widom statistics were used to identify significant principle components at p<0.05. Imputation to the 1000G phase 3 v5 was performed after quality control exclusions using the University of Michigan imputation server. Phasing was performed with ShapeIT v2.r790. The pseudo-autosomal region of chromosome X was excluded, as well as any variants with MAF<0.01, and info score <0.8.
