## Supplementary Table 1 for "A higher burden of multiple sclerosis genetic risk confers an earlier onset"

**Supplementary Table 1A**: Inclusion and Exclusion Criteria for ACP, ADVANCE, ASCEND, and DECIDE

| | | ACP | ADVANCE |
|---|---|---|---|
| Inclusion | MS Diagnosis | Standard MS diagnostic criteria defined at the time of enrollment, confirmed by a neurologist | RRMS as defined by McDonald criteria, at least 2 relapses in the 3 years before study enrollment |
| | Age Range | ≥18 years | 18-65 years |
| | Disability Scores | *NA* | A score of 0-5 on the EDSS |
| Exclusion | Other forms of MS | *NA* | A diagnosis for a progressive form of MS |
| | Clinical characteristics | Clinical or radiological evidence of a stroke, or a history of meningitis, neoplastic, peripheral nervous system, primary muscle disease, or other non-demyelinating disease of the central nervous system, or history of bloodborne pathogens, or allogenic blood marrow transplant, or weight < 37lbs | Any pre-specified laboratory abnormalities |
| | Treatment Status | *NA* | Treatment with interferon for MS for 4+ weeks or discontinuation <6 months before ASCEND study baseline. |

**Supplementary Table 1B**. Sample size by number of variants at each stage of the MS AAO meta-analysis.

| Cohort | Number of Autosomal Variants after Quality Control (MAF>1%, HWE p>0.0001) | Number of Variants Retained by METAL (i.e. matching alleles, valid SE, removal of duplicates) | Number of Variants in final Meta-analysis (i.e. present in more than one cohort) |
|---|---|---|---|
| ACP1 | 1,240,687 | 1,239,788 | 1,229,423 |
| ACP2 | 7,347,748 | 7,329,355 | 7,108,052 |
| ADVANCE | 9,660,356 | 9,602,863 | 9,478,535 |
| ASCEND | 9,689,382 | 9,629,972 | 9,513,822 |
| DECIDE | 9,689,411 | 9,630,084 | 9,555,065 |
| All Cohorts | | 10,264,634 | 9,717,633 |

| ASCEND | DECIDE |
|---|---|
| SPMS for two years or more before study enrollment | RRMS as defined by the McDonald criteria, as well as at least 2 relapses in the 3 years before study enrollment OR |
| 18-58 years | 18-55 years |
| A score of 3.0-6.5 on the EDSS, a score of ≥4 on the MSSS | A score of 0-5 on the EDSS |
| Clinically confirmed relapses ≤3 months before trial randomization | Relapse ≤50 days before trial randomization |
| *NA* | No cranial MRI evidence of lesions associated with MS |
| Treatment with natalizumab | Treatment with mitoxantrone, cyclophosphamide, fingolimod, or natalizumab ≤1 year before trial randomization OR treatment with intravenous or oral corticosteroids or glatiramer acetate ≤30 days before trial randomization |

| Number of non-heterogeneous variants ($pI^2$>0.05) |
|---|
| 1,167,913 |
| 6,758,542 |
| 9,012,609 |
| 9,045,548 |
| 9,085,036 |
| 9,239,744 |
