## Supplementary Table 2 for "A higher burden of multiple sclerosis genetic risk confers an earlier onset"

**Supplementary Table 2. Meta-Analysis AAO results for 198 non-MHC variants  associated with MS risk**

| Column Header | Key |
|---|---|
| **rsID** | dbSNP identifier (rsID) or chr:bp location if no rsID |
| **chromosome** | Chrosomome location |
| **position** | Position location (hg19) |
| **Minor Allele** | Minor allele |
| **Minor Allele is Risk Allele** | Column denoting if tested allele corresponded with the risk allele from IMSGC 2019 (1 = risk allele match, NA=tested variant was in LD with effect variant, nonmatching alleles) |
| **IMSGC Risk Allele** | Second Allele |
| **LD Proxy** | Identifier of variants that served as proxies for IMSGC effect SNPs not available in the meta-analyzed dataset |
| **Proxy LD r2** | Linkage disequilibrium r2 of the proxy SNP with the original IMSGC effect SNP |
| **Proxy LD D'** | Linkage disequilibrium D' of the proxy SNP with the original IMSGC effect SNP |
| **Effect** | Effect estimate of the risk allele in β form |
| **StdErr** | Standard error of the effect estimate |
| **P-Value** | Significance level of the effect estimate |
| **HetISq** | I-squared heterogeneity measure |
| **HetPVal** | Significance level of the Heterogeneity I-squared measure |
| **functionGVS** | GVS class of function variation, using only sumitted alleles and hg19 genome build |
| **functionDBSNP** | dbSNP class of variation function |
| **geneList** | HUGO names, any for which the transcription region overlaps the variation |
| **EuropeanHapMapFreq** | MAF in HapMap EUR population |

| rsID | chromosome | position | Minor Allele | Minor Allele is Risk Allele | IMSGC Risk Allele | LD Proxy | Proxy LD r2 | Proxy LD D' | Effect | StdErr | P-Value | HetISq | HetPVal | functionGVS | functionDBSNP | geneList |
|---|---|---|---|---|---|---|---|---|---|---|---|---|---|---|---|---|
| rs4325907 | 3 | 101749022 | t | 1 | t | | | | 0.6761 | 0.2149 | 0.001653 | 0 | 0.9741 | intergenic | | |
| rs6032662 | 20 | 44734310 | t | 0 | c | | | | -0.6782 | 0.2305 | 0.003261 | 20.2 | 0.2862 | intergenic | | |
| rs483180 | 1 | 120267505 | c | 0 | g | | | | -0.5887 | 0.2371 | 0.01305 | 8.1 | 0.3605 | intron | intron-variant | PHGDH |
| rs438613 | 3 | 28072086 | t | 0 | c | | | | -0.4996 | 0.2082 | 0.01639 | 0 | 0.4738 | intergenic | | |
| rs3184504 | 12 | 111884608 | t | 1 | t | | | | -0.4935 | 0.2071 | 0.0172 | 41.5 | 0.1447 | missense | missense | SH2B3 |
| rs719316 | 6 | 16672760 | t | 1 | t | | | | -0.491 | 0.211 | 0.01994 | 69.9 | 0.009916 | intron | intron-variant | ATXN1 |
| rs6564681 | 16 | 79652720 | t | 1 | t | | | | 0.5066 | 0.2258 | 0.02488 | 23.8 | 0.2628 | intergenic | | |
| rs72922276 | 1 | 65429319 | a | 0 | g | | | | -0.8935 | 0.3986 | 0.02499 | 5.9 | 0.3734 | intron | intron-variant | JAK1 |
| rs9878602 | 3 | 71535338 | t | 1 | t | | | | 0.4631 | 0.2124 | 0.0292 | 0 | 0.667 | intron | intron-variant | FOXP1 |
| rs6498163 | 16 | 11213951 | t | 1 | t | | | | 0.4189 | 0.2194 | 0.05619 | 0 | 0.7223 | intron | intron-variant | CLEC16A |
| rs1323292 | 1 | 192541021 | a | 1 | a | | | | -0.5681 | 0.3001 | 0.05832 | 59.7 | 0.04158 | upstream-gene | | |
| rs10245867 | 7 | 28142186 | t | 0 | g | | | | -0.4097 | 0.2168 | 0.05876 | 0 | 0.6839 | intron | intron-variant | JAZF1 |
| rs32658 | 5 | 118703662 | t | 0 | g | | | | 0.4027 | 0.215 | 0.06112 | 0 | 0.8809 | intron | intron-variant | TNFAIP8 |
| rs9308424 | 1 | 212877776 | a | 1 | a | | | | 0.4381 | 0.2348 | 0.06212 | 42.4 | 0.1386 | upstream-gene | | |
| rs10063294 | 5 | 35877505 | a | 1 | a | | | | 0.3881 | 0.2086 | 0.06278 | 0 | 0.9892 | 3-prime-UTR | utr-variant-3-prime | IL7R |
| rs9568402 | 13 | 50961957 | a | 1 | a | | | | 0.6019 | 0.3267 | 0.06543 | 0 | 0.5246 | intergenic | | |
| rs62420820 | 6 | 137438057 | a | 1 | a | | | | -0.4196 | 0.2402 | 0.08073 | 0 | 0.5603 | intergenic | | |
| rs249677 | 5 | 141539339 | a | 0 | c | | | | -0.3878 | 0.2226 | 0.08148 | 25.9 | 0.2487 | intergenic | | |
| rs1250551 | 10 | 81059335 | t | 0 | g | | | | 0.3825 | 0.2216 | 0.08429 | 10 | 0.3494 | intron | intron-variant | ZMIZ1 |
| rs1059091 | 11 | 309127 | a | *NA* | a | *Discovery SNP for IMSGC effe* | 0.102 | 0.744 | 0.3844 | 0.2277 | 0.09138 | 0 | 0.4633 | missense | missense | IFITM2 |
| rs4808760 | 19 | 18301979 | c | 0 | g | | | | 0.3991 | 0.2367 | 0.09178 | 0 | 0.9697 | upstream-gene | | |
| rs1112718 | 10 | 94479107 | a | 1 | a | | | | 0.3598 | 0.2176 | 0.09817 | 0 | 0.4523 | intergenic | | |
| rs760517 | 22 | 37258986 | t | 0 | c | | | | -0.3786 | 0.2297 | 0.09929 | 0 | 0.955 | intron | intron-variant | NCF4 |
| rs17741873 | 10 | 75653800 | t | 0 | g | | | | 0.4314 | 0.2707 | 0.111 | 0 | 0.8599 | intergenic | | |
| rs2269434 | 11 | 47360412 | t | 1 | t | | | | 0.3464 | 0.2227 | 0.1199 | 0 | 0.4624 | intron | intron-variant | MYBPC3 |
| rs2027982 | 22 | 31593435 | t | *NA* | t | *Discovery SNP for IMSGC effe* | 0.223 | 0.617 | -0.3316 | 0.2147 | 0.1225 | 0 | 0.8841 | intron | intron-variant | RNF185 |
| rs12614091 | 2 | 204632861 | a | 1 | a | | | | -0.3801 | 0.2475 | 0.1245 | 68.5 | 0.01289 | intergenic | | |
| rs12147246 | 14 | 103265844 | a | 0 | g | | | | 0.3302 | 0.2164 | 0.1269 | 72.4 | 0.005877 | intron | intron-variant | TRAF3 |
| rs4939490 | 11 | 60793651 | c | 0 | g | | | | -0.3424 | 0.2248 | 0.1277 | 0 | 0.7499 | intergenic | | |
| rs1014486 | 3 | 159691112 | t | 0 | c | | | | 0.3328 | 0.2188 | 0.1283 | 0 | 0.4814 | intergenic | | |
| rs6990534 | 8 | 128814091 | a | 1 | a | | | | 0.36 | 0.24 | 0.1336 | 0 | 0.6424 | intergenic | | |
| rs138433213 | 3 | 112693983 | t | 1 | t | | | | -1.9458 | 1.3095 | 0.1373 | 0 | 0.4587 | upstream-gene | upstream-variant-2KB | |
| rs10271373 | 7 | 138729795 | a | 0 | c | | | | 0.3098 | 0.214 | 0.1478 | 0 | 0.6135 | 3-prime-UTR | utr-variant-3-prime | ZC3HAV1 |
| rs12133753 | 1 | 92222089 | t | 0 | c | | | | -0.4275 | 0.3008 | 0.1553 | 45.6 | 0.1185 | intron | intron-variant | TGFBR3 |
| rs7819665 | 8 | 129177769 | t | 1 | t | | | | 0.38 | 0.2692 | 0.1582 | 46.4 | 0.1133 | intergenic | | |
| rs4728142 | 7 | 128573967 | a | 0 | g | | | | 0.2952 | 0.2105 | 0.1608 | 0 | 0.4913 | upstream-gene | | |
| rs7260482 | 19 | 45143942 | a | 1 | a | | | | -0.3118 | 0.2255 | 0.1667 | 41.3 | 0.1459 | upstream-gene | | |
| rs11578655 | 1 | 101412902 | t | 1 | t | | | | 0.4875 | 0.3675 | 0.1846 | 0 | 0.7972 | intron | intron-variant | SLC30A7 |
| rs9808753 | 21 | 34787312 | a | 0 | g | | | | -0.4004 | 0.3067 | 0.1917 | 0 | 0.4746 | missense | missense | IFNGR2 |
| rs72989863 | 4 | 164493807 | a | 0 | g | | | | -0.2974 | 0.228 | 0.192 | 0 | 0.5174 | intron | intron-variant | MTARC1 |
| rs2255214 | 3 | 121770739 | t | *NA* | t | *Discovery SNP for IMSGC effe* | 0.784 | 0.958 | 0.2706 | 0.2093 | 0.1961 | 0 | 0.4063 | upstream-gene | | |
| rs146566517 | 16 | 11353879 | t | 0 | c | | | | 0.7848 | 0.6073 | 0.1963 | 72.7 | 0.005481 | upstream-gene | | |
| rs11125803 | 2 | 25052177 | t | 1 | t | | | | 0.3147 | 0.2454 | 0.1997 | 40.9 | 0.1488 | intron | intron-variant | ADCY3 |
| rs34695601 | 14 | 76014298 | t | 1 | t | | | | 0.3359 | 0.2646 | 0.2042 | 0 | 0.4127 | downstream-gene | | |
| rs2286974 | 16 | 11114512 | a | 0 | g | | | | 0.2772 | 0.2204 | 0.2085 | 34.3 | 0.1925 | intron | intron-variant | CLEC16A |
| rs2331964 | 3 | 121542898 | t | 0 | c | | | | -0.2725 | 0.2168 | 0.2089 | 14 | 0.3252 | intron | intron-variant | IQCB1 |

| | | | | | | | | | | | | | |
|---|---|---|---|---|---|---|---|---|---|---|---|---|---|
| rs11749040 | 5 | 40396425 | a | 0 | g | 0.3723 | 0.2989 | 0.2129 | 0 | 0.6221 | intergenic | | |
| rs5756405 | 22 | 37310954 | a | 1 | a | -0.2553 | 0.2089 | 0.2218 | 0 | 0.4733 | intron | intron-variant | CSF2RB |
| rs1415069 | 1 | 93426869 | c | 1 | c | -0.3708 | 0.3106 | 0.2325 | 0 | 0.4565 | intron | intron-variant | FAM69A |
| rs67111717 | 5 | 176790162 | a | 0 | g | -0.2593 | 0.2202 | 0.239 | 56.2 | 0.05785 | intron | intron-variant | RGS14 |
| rs35486093 | 1 | 85729820 | a | 1 | a | 0.3829 | 0.3272 | 0.2419 | 0 | 0.8145 | upstream-gene | | |
| rs4262739 | 11 | 128421175 | a | 1 | a | 0.2434 | 0.2116 | 0.25 | 0 | 0.6472 | intron | intron-variant | ETS1 |
| rs6742 | 20 | 62374441 | t | 0 | c | -0.3233 | 0.283 | 0.2532 | 0 | 0.5266 | 3-prime-UTR | utr-variant-3-prime | SLC2A4RG |
| rs6738544 | 2 | 191989356 | a | 0 | c | -0.2575 | 0.2253 | 0.2532 | 0 | 0.9876 | intron | intron-variant | STAT4 |
| rs17724508 | 16 | 79350204 | t | 1 | t | 0.4942 | 0.4334 | 0.2542 | 35.4 | 0.1851 | intergenic | | |
| rs10191360 | 2 | 136884679 | t | 1 | t | 0.2378 | 0.2107 | 0.2593 | 0 | 0.4947 | intergenic | | |
| rs61884005 | 11 | 14402930 | c | 1 | c | 0.3716 | 0.3363 | 0.2692 | 0 | 0.8553 | intergenic | | |
| rs75937181 | 3 | 121783015 | t | 1 | t | -0.4358 | 0.3946 | 0.2694 | 65.4 | 0.02081 | intron | intron-variant | CD86 |
| rs1087056 | 10 | 31395761 | a | 0 | g | -0.283 | 0.2568 | 0.2704 | 0 | 0.7209 | intergenic | | |
| rs10801908 | 1 | 117090493 | t | 0 | c | -0.3901 | 0.3552 | 0.2721 | 40.8 | 0.1493 | intron | intron-variant | CD58 |
| rs6672420 | 1 | 25291010 | a | 0 | t | -0.2327 | 0.2164 | 0.282 | 0 | 0.5162 | missense | missense | RUNX3 |
| rs11899404 | 2 | 12607893 | t | 1 | t | 0.2321 | 0.2176 | 0.2862 | 0 | 0.5683 | intergenic | | |
| rs11852059 | 14 | 52306091 | a | 1 | a | 0.2756 | 0.2595 | 0.2883 | 0 | 0.6369 | intergenic | | |
| rs11231749 | 11 | 64095178 | t | 0 | c | -0.2375 | 0.2239 | 0.2889 | 52.3 | 0.07853 | intergenic | | |
| rs35703946 | 16 | 86021505 | a | 0 | g | -0.3708 | 0.3522 | 0.2925 | 76.8 | 0.001755 | intergenic | | |
| rs11256593 | 10 | 6117322 | t | 1 | t | -0.2242 | 0.2166 | 0.3006 | 0 | 0.6034 | intergenic | | |
| rs4812772 | 20 | 42579051 | t | 0 | c | -0.2456 | 0.2376 | 0.3013 | 0.1 | 0.4052 | intron | intron-variant | TOX2 |
| rs71252597 | 2 | 112492986 | t | 1 | t | 0.3256 | 0.3151 | 0.3014 | 0 | 0.4751 | intergenic | | |
| rs6911131 | 6 | 143865221 | a | 1 | a | 0.4351 | 0.4224 | 0.3031 | 71.3 | 0.007479 | intergenic | | |
| rs10230723 | 7 | 50239880 | a | 1 | a | 0.2993 | 0.2924 | 0.3061 | 0 | 0.8238 | intergenic | | |
| rs3809627 | 16 | 30103160 | a | 1 | a | 0.22 | 0.2151 | 0.3065 | 0.9 | 0.4013 | intron | intron-variant | TBX6 |
| rs12609500 | 19 | 11173928 | t | 0 | c | 0.2527 | 0.2477 | 0.3076 | 53 | 0.07455 | downstream-gene | | |
| rs62013236 | 15 | 79247482 | t | 0 | c | -0.3149 | 0.3138 | 0.3155 | 0 | 0.7936 | downstream-gene | | |
| rs61863928 | 10 | 64449549 | t | 0 | g | 0.2372 | 0.2363 | 0.3156 | 0 | 0.5073 | intergenic | | |
| rs34681760 | 5 | 6712834 | t | 0 | c | -0.2294 | 0.2288 | 0.3162 | 0 | 0.561 | upstream-gene | upstream-variant-2KB | |
| rs2726479 | 4 | 106255589 | t | 1 | t | 0.2083 | 0.2121 | 0.3262 | 30 | 0.2218 | intergenic | | |
| rs2469434 | 18 | 67544046 | t | 1 | t | 0.208 | 0.212 | 0.3265 | 0 | 0.4506 | intron | intron-variant | CD226 |
| rs11079784 | 17 | 45702280 | t | 0 | c | -0.2054 | 0.2126 | 0.3339 | 0 | 0.6819 | intron | | LOC101929046 |
| rs354033 | 7 | 149289464 | a | 0 | g | -0.2374 | 0.2457 | 0.334 | 0 | 0.5248 | intron | intron-variant | ZNF767 |
| rs701006 | 12 | 58106836 | a | 0 | g | -0.2061 | 0.2142 | 0.3358 | 0 | 0.9053 | intron | intron-variant | OS9 |
| rs9591325 | 13 | 50811220 | t | 1 | t | -0.4 | 0.4232 | 0.3446 | 0 | 0.8627 | intergenic | | |
| rs74449127 | 1 | 101290432 | a | 1 | a | 0.2366 | 0.2535 | 0.3507 | 37 | 0.1747 | intergenic | | |
| rs12622670 | 2 | 68646536 | t | 1 | t | -0.1922 | 0.2101 | 0.3602 | 0 | 0.5441 | intergenic | | |
| rs17780048 | 6 | 138179146 | t | 0 | c | -0.5031 | 0.5532 | 0.3631 | 0 | 0.9741 | intron | intron-variant | LOC100130476 |
| rs6670198 | 1 | 2520527 | t | 0 | c | -0.1994 | 0.2263 | 0.3784 | 7.5 | 0.3639 | intron | intron-variant | FAM213B |
| rs9909593 | 17 | 37970149 | a | 1 | a | 0.182 | 0.2108 | 0.3879 | 0 | 0.4415 | intron | intron-variant | IKZF3 |
| rs1177228 | 2 | 61242410 | a | 0 | g | -0.2124 | 0.2464 | 0.3887 | 0 | 0.9537 | intron | intron-variant | PUS10 |
| rs58166386 | 19 | 16559421 | a | 0 | g | -0.1978 | 0.2339 | 0.3979 | 0 | 0.6802 | intron | intron-variant | EPS15L1 |
| rs735542 | 8 | 128175696 | a | 1 | a | 0.1811 | 0.2198 | 0.4098 | 0 | 0.5986 | intergenic | | |
| rs111635774 | 6 | 14691215 | t | 0 | c | -0.4235 | 0.5143 | 0.4102 | 0 | 0.6871 | intergenic | | |
| rs6837324 | 4 | 48127262 | a | 1 | a | 0.1775 | 0.2155 | 0.4102 | 0 | 0.6624 | intron | intron-variant | TXK |
| rs2150879 | 17 | 57859210 | a | 0 | g | -0.1674 | 0.2086 | 0.4221 | 22.5 | 0.2712 | intron | intron-variant | VMP1 |
| rs73414214 | 7 | 105706462 | a | 0 | c | 0.3055 | 0.3849 | 0.4273 | 0 | 0.7519 | intergenic | | |
| rs12365699 | 11 | 118743286 | a | 0 | g | -0.2324 | 0.2962 | 0.4328 | 30.9 | 0.2156 | intergenic | | |
| rs12588969 | 14 | 103230758 | c | 1 | c | 0.1774 | 0.2261 | 0.4328 | 0 | 0.4345 | intergenic | | |
| rs12925972 | 16 | 79111297 | t | 0 | c | -0.1681 | 0.2145 | 0.4333 | 24.8 | 0.2558 | intron | intron-variant | WWOX |
| rs10951042 | 7 | 3139417 | t | 0 | c | -0.1667 | 0.2132 | 0.4341 | 35.4 | 0.1855 | intergenic | | |
| rs56095240 | 11 | 95421830 | a | 0 | t | 0.2375 | 0.3086 | 0.4415 | 0 | 0.7931 | intergenic | | |
| rs962052 | 2 | 151644203 | t | 0 | c | -0.1801 | 0.2383 | 0.4499 | 0 | 0.5996 | intergenic | | |
| rs57116599 | 2 | 112770799 | a | 1 | a | 0.1955 | 0.2655 | 0.4615 | 55.2 | 0.06273 | intron | intron-variant | MERTK |
| rs2986736 | 1 | 6512547 | t | 1 | t | 0.1845 | 0.2524 | 0.4648 | 0 | 0.8566 | intron | intron-variant | ESPN |
| rs6589706 | 11 | 118747813 | a | 1 | a | -0.1513 | 0.2084 | 0.468 | 0 | 0.9556 | intergenic | | |
| rs11919880 | 3 | 32962051 | a | 1 | a | -0.1581 | 0.2212 | 0.4746 | 0 | 0.6796 | intergenic | | |
| rs2705616 | 4 | 87862396 | c | 0 | g | 0.1479 | 0.2099 | 0.4812 | 0 | 0.5667 | intron | intron-variant | AFF1 |
| rs4896153 | 6 | 135833463 | a | 1 | a | 0.1765 | 0.2585 | 0.4947 | 51.3 | 0.1284 | intron | intron-variant | LINC00271 |
| rs4940730 | 18 | 56269737 | a | 1 | a | 0.1471 | 0.2171 | 0.4979 | 38.8 | 0.1624 | intron | intron-variant | ALPK2 |
| rs9610458 | 22 | 22205353 | t | 1 | t | -0.1533 | 0.2264 | 0.4983 | 32.6 | 0.2039 | intron | intron-variant | MAPK1 |
| rs7731626 | 5 | 55444683 | a | 0 | g | -0.1496 | 0.2211 | 0.4986 | 0 | 0.6147 | intron | intron-variant | ANKRD55 |
| rs59655222 | 1 | 200875897 | t | 1 | t | -0.1623 | 0.2462 | 0.5098 | 8.5 | 0.3581 | intron | intron-variant | C1orf106 |
| rs140522 | 22 | 50971266 | t | 0 | c | 0.1422 | 0.2173 | 0.5129 | 0 | 0.9921 | upstream-gene | upstream-variant-2KB | |
| rs61708525 | 12 | 94661453 | a | 0 | g | -0.1493 | 0.2317 | 0.5193 | 0 | 0.6174 | intron | intron-variant | PLXNC1 |
| rs116899835 | 14 | 88523488 | t | 0 | c | -0.3323 | 0.5174 | 0.5207 | 33.6 | 0.197 | intron | intron-variant | LOC283587 |
| rs13327021 | 3 | 27783015 | t | 0 | c | 0.1337 | 0.216 | 0.5359 | 58 | 0.04922 | intergenic | | |
| rs17051321 | 4 | 122119449 | t | 0 | c | -0.1517 | 0.2481 | 0.5408 | 0 | 0.4581 | intron | intron-variant | TNIP3 |
| rs116877451 | 7 | 50328339 | a | 1 | a | -0.4241 | 0.694 | 0.5412 | 38.3 | 0.1659 | intergenic | | |

| | | | | | | | | | | | | | | | |
|---|---|---|---|---|---|---|---|---|---|---|---|---|---|---|---|
| rs12722559 | 10 | 6070273 | a | | 0 | c | | -0.1901 | 0.3127 | 0.5433 | 6.9 | 0.3671 | intron | intron-variant | IL2RA |
| rs4409785 | 11 | 95311422 | t | | 1 | t | | 0.1619 | 0.2673 | 0.5448 | 0 | 0.823 | intergenic | | |
| rs58394161 | 1 | 92939959 | t | | 1 | t | | 0.1634 | 0.2754 | 0.5531 | 0 | 0.5818 | downstream-gene | downstream-variant-500B | |
| rs1465697 | 19 | 49837246 | t | | 0 | c | | 0.1359 | 0.2297 | 0.5539 | 0 | 0.9428 | upstream-gene | upstream-variant-2KB | |
| rs1801133 | 1 | 11856378 | a | | 1 | a | | -0.1327 | 0.2292 | 0.5626 | 34.4 | 0.192 | missense | missense | MTHFR |
| rs3737798 | 1 | 160389984 | a | | 1 | a | | -0.1161 | 0.2107 | 0.5816 | 25.8 | 0.2493 | intron | intron-variant | VANGL2 |
| rs9955954 | 18 | 56348044 | a | | 0 | g | | -0.1657 | 0.3173 | 0.6015 | 14 | 0.3127 | intron | intron-variant | MALT1 |
| rs12434551 | 14 | 69253364 | a | | 1 | a | | 0.1083 | 0.2091 | 0.6044 | 49.3 | 0.09598 | downstream-gene | | |
| rs4796224 | 17 | 34842521 | a | | 1 | a | | -0.1069 | 0.2156 | 0.6199 | 0 | 0.4279 | 5-prime-UTR | | ZNHIT3 |
| rs6789653 | 3 | 141150990 | a | | 0 | g | | -0.2044 | 0.4184 | 0.6251 | 0 | 0.6505 | intron | intron-variant | ZBTB38 |
| rs1076928 | 6 | 36348689 | t | | 0 | c | | 0.1026 | 0.2108 | 0.6263 | 43.2 | 0.1334 | intron | intron-variant | ETV7 |
| rs2289746 | 3 | 105455955 | t | | 0 | c | | 0.1119 | 0.2308 | 0.6278 | 0 | 0.7738 | intron | intron-variant | CBLB |
| rs9992763 | 4 | 109058718 | t | | 1 | t | | 0.1022 | 0.2126 | 0.6306 | 0 | 0.9565 | intron | intron-variant | LEF1 |
| rs1026916 | 17 | 40529835 | a | | 0 | g | | -0.1006 | 0.2148 | 0.6398 | 54.5 | 0.06652 | intron | intron-variant | STAT3 |
| rs9843355 | 3 | 119228508 | a | | 0 | g | | 0.134 | 0.2871 | 0.6406 | 36.9 | 0.1755 | intron | intron-variant | TIMMDC1 |
| rs2546890 | 5 | 158759900 | a | | 1 | a | | -0.0961 | 0.2092 | 0.6461 | 0 | 0.7076 | non-coding-exon | nc-transcript-variant | LOC285626 |
| rs34536443 | 19 | 10463118 | c | | 0 | g | | -0.2626 | 0.5722 | 0.6463 | 0 | 0.4709 | missense | missense | TYK2 |
| rs6427540 | 1 | 160634588 | t | | 0 | c | | -0.133 | 0.3079 | 0.6659 | 67.9 | 0.01413 | intergenic | | |
| rs7222450 | 17 | 43407670 | a | | 1 | a | | 0.1058 | 0.2511 | 0.6736 | 0 | 0.383 | intergenic | | |
| rs12211604 | 6 | 7100029 | a | | 1 | a | | 0.1077 | 0.2582 | 0.6765 | 0 | 0.7817 | intergenic | | |
| rs983494 | 1 | 160703965 | a | | 0 | g | | -0.1118 | 0.268 | 0.6766 | 56.6 | 0.05593 | upstream-gene | | |
| rs531612 | 11 | 65705432 | t | | 0 | c | | -0.0883 | 0.2118 | 0.6768 | 0 | 0.572 | intergenic | | |
| rs6020055 | 20 | 48422095 | a | | 1 | a | | 0.2179 | 0.5355 | 0.684 | 0 | 0.9402 | intergenic | | |
| rs35540610 | 2 | 231121829 | t | | 0 | c | | 0.0941 | 0.2341 | 0.6876 | 43.8 | 0.1298 | intron | | SP140 |
| rs34947566 | 16 | 11412926 | a | | 0 | c | | -0.1181 | 0.298 | 0.6918 | 0 | 0.8338 | intergenic | | |
| rs9900529 | 17 | 73335776 | c | | 1 | c | | 0.094 | 0.2387 | 0.6936 | 0 | 0.4087 | intron | intron-variant | GRB2 |
| rs7977720 | 12 | 9866349 | t | | 0 | c | | -0.0835 | 0.2132 | 0.6953 | 0 | 0.477 | downstream-gene | | |
| rs55858457 | 7 | 2443302 | t | | 0 | g | | 0.1006 | 0.2603 | 0.6991 | 0 | 0.5291 | intron | intron-variant | CHST12 |
| rs13385171 | 2 | 65661843 | t | | 0 | c | | -0.0829 | 0.2153 | 0.7003 | 0 | 0.5106 | upstream-gene | | |
| rs1077667 | 19 | 6668972 | t | | 0 | c | | -0.099 | 0.2591 | 0.7024 | 0 | 0.6206 | intron | intron-variant | TNFSF14 |
| rs60600003 | 7 | 37382465 | t | | 1 | t | | 0.1327 | 0.3482 | 0.7032 | 24.1 | 0.2608 | intron | intron-variant | ELMO1 |
| rs10951154 | 7 | 27135314 | t | | 1 | t | | -0.1206 | 0.3201 | 0.7063 | 0 | 0.8137 | missense | | HOXA1 |
| rs72928038 | 6 | 90976768 | a | | 0 | g | | -0.0946 | 0.2675 | 0.7237 | 0 | 0.9411 | intron | intron-variant | BACH2 |
| rs2317231 | 1 | 157686337 | t | | 0 | g | | -0.0762 | 0.2193 | 0.7282 | 0 | 0.9416 | intergenic | | |
| rs11542663 | 6 | 119215402 | a | | 1 | a | | -0.0803 | 0.2321 | 0.7294 | 20.1 | 0.2866 | 5-prime-UTR | utr-variant-5-prime | ASF1A,MCM9 |
| rs2585447 | 20 | 52744437 | t | | 1 | t | | -0.1003 | 0.2906 | 0.7299 | 0 | 0.9354 | intergenic | | |
| rs9863496 | 3 | 18798848 | t | | 1 | t | | 0.082 | 0.2401 | 0.7328 | 0 | 0.4144 | intergenic | | |
| rs7855251 | 9 | 100868189 | t | | 1 | t | | -0.0811 | 0.2513 | 0.7468 | 13.9 | 0.3255 | intron | intron-variant | TRIM14 |
| rs3923387 | 8 | 144986793 | t | | 1 | t | | 0.0701 | 0.2173 | 0.7471 | 0 | 0.4845 | downstream-gene | | |
| rs2364485 | 12 | 6514963 | a | | 0 | c | | 0.1088 | 0.3409 | 0.7496 | 5 | 0.3785 | intergenic | | |
| rs244656 | 5 | 133449827 | a | | 1 | a | | -0.0991 | 0.3138 | 0.752 | 0 | 0.7046 | upstream-gene | upstream-variant-2KB | |
| rs2836438 | 21 | 39864727 | a | | 0 | g | | 0.0991 | 0.3177 | 0.755 | 0 | 0.9189 | intron | intron-variant | ERG |
| rs77654077 | 13 | 100026952 | a | | 1 | a | | 0.2231 | 0.7316 | 0.7604 | 45.2 | 0.1206 | intron | intron-variant | UBAC2 |
| rs28703878 | 8 | 79417222 | a | | 1 | a | | 0.0714 | 0.2442 | 0.7699 | 47.4 | 0.107 | intergenic | | |
| rs6589939 | 11 | 122518525 | a | | 1 | a | | 0.0619 | 0.2154 | 0.7737 | 60.5 | 0.03825 | intergenic | | |
| rs767455 | 12 | 6450935 | t | *NA* | | t | *Discovery SNP for IMSGC effect SNP 12:6441622* | 0.0598 | 0.2121 | 0.7779 | 54.5 | 0.06663 | synonymous | synonymous-codon | TNFRSF1A |
| rs2327586 | 6 | 135495226 | t | | 1 | t | | -0.0679 | 0.2439 | 0.7807 | 0 | 0.5003 | intergenic | | |
| rs13414105 | 2 | 30472442 | c | | 1 | c | | -0.0704 | 0.2687 | 0.7933 | 58.1 | 0.092 | intron | intron-variant | LBH |
| rs137955 | 22 | 40291807 | t | | 1 | t | | -0.0561 | 0.2182 | 0.797 | 59.4 | 0.04319 | upstream-gene | | |
| rs112741635 | 1 | 24207504 | a | | 1 | a | | 0.0708 | 0.2769 | 0.7981 | 0 | 0.924 | intron | intron-variant | CNR2 |
| rs149114341 | 11 | 118783424 | a | | 0 | g | | -0.1802 | 0.7201 | 0.8024 | 0 | 0.6448 | intron | | BCL9L |
| rs111430408 | 3 | 100848597 | t | | 0 | c | | -0.1182 | 0.4781 | 0.8048 | 0 | 0.4776 | intergenic | | |
| rs2590438 | 3 | 187565968 | t | | 1 | t | | -0.0526 | 0.2138 | 0.8058 | 0 | 0.6477 | intergenic | | |
| rs6533052 | 4 | 103911781 | a | | 1 | a | | -0.0513 | 0.209 | 0.806 | 0 | 0.9826 | intron | intron-variant | SLC9B1 |
| rs28834106 | 19 | 10592144 | t | | 1 | t | | 0.0581 | 0.2404 | 0.809 | 27.1 | 0.241 | downstream-gene | | |
| rs139504223 | 1 | 93152635 | t | | 0 | c | | -0.0538 | 0.2292 | 0.8145 | 0 | 0.6324 | intron | intron-variant | EVI5 |
| rs112344141 | 1 | 154983036 | t | | 1 | t | | -0.1196 | 0.5265 | 0.8203 | 0 | 0.4519 | intron | intron-variant | ZBTB7B |
| rs6880809 | 5 | 40429250 | a | | 1 | a | | -0.0449 | 0.2097 | 0.8303 | 0 | 0.6048 | intergenic | | |
| rs6496663 | 15 | 90887584 | a | | 1 | a | | -0.0474 | 0.2309 | 0.8372 | 0 | 0.4689 | downstream-gene | | |
| rs75191738 | 6 | 130348257 | t | | 1 | t | | 0.0567 | 0.2986 | 0.8494 | 63.6 | 0.02677 | intron | intron-variant | L3MBTL3 |
| rs2248137 | 20 | 52789743 | c | | 0 | g | | 0.0395 | 0.2228 | 0.8592 | 0 | 0.5448 | intron | intron-variant | CYP24A1 |
| rs405343 | 16 | 1067832 | t | | 0 | g | | 0.0463 | 0.2629 | 0.8601 | 0 | 0.856 | intergenic | | |
| rs79979643 | 1 | 32738415 | a | | 0 | g | | 0.0479 | 0.2799 | 0.864 | 0 | 0.5761 | intron | intron-variant | LCK |
| rs11083862 | 19 | 47638539 | a | | 1 | a | | -0.0325 | 0.21 | 0.8771 | 0 | 0.6771 | intron | intron-variant | SAE1 |
| rs1800693 | 12 | 6440009 | t | | 1 | t | | 0.0296 | 0.2104 | 0.888 | 36.3 | 0.1794 | intron | intron-variant | TNFRSF1A |
| rs34026809 | 11 | 118480695 | c | | 1 | c | | 0.069 | 0.5994 | 0.9084 | 0 | 0.7629 | intron | intron-variant | PHLDB1 |
| rs11161550 | 1 | 85682020 | a | | 1 | a | | -0.0224 | 0.2088 | 0.9147 | 0 | 0.6036 | intergenic | | |
| rs1738074 | 6 | 159465977 | t | | 1 | t | | 0.0219 | 0.2198 | 0.9207 | 0 | 0.8721 | 5-prime-UTR | utr-variant-5-prime | TAGAP |
| rs13136820 | 4 | 40307564 | t | | 0 | c | | -0.0211 | 0.2252 | 0.9252 | 0 | 0.9779 | intergenic | | |

| | | | | | | | | | | | | | |
|---|---|---|---|---|---|---|---|---|---|---|---|---|---|
| rs1365120 | 11 | 36438075 | t | 1 | t | -0.0278 | 0.3345 | 0.9338 | 0 | 0.7902 | intron | intron-variant | PRR5L |
| rs8062446 | 16 | 57077094 | t | 1 | t | -0.0177 | 0.2171 | 0.9351 | 49.9 | 0.0922 | intron | intron-variant | NLRC5 |
| rs10936602 | 3 | 169536637 | t | 1 | t | 0.0198 | 0.2444 | 0.9354 | 15.8 | 0.314 | intron | | LRRIQ4 |
| rs12971909 | 19 | 4466466 | a | 0 | g | -0.0239 | 0.3127 | 0.9391 | 0 | 0.7103 | intergenic | | |
| rs6072343 | 20 | 39968188 | a | 0 | g | -0.0211 | 0.2981 | 0.9436 | 0 | 0.6566 | upstream-gene | upstream-variant-2KB | |
| rs883871 | 17 | 38252660 | a | 0 | g | 0.0201 | 0.2915 | 0.9451 | 35.9 | 0.1822 | intron | intron-variant | NR1D1 |
| rs13066789 | 3 | 187987624 | t | 0 | c | -0.0112 | 0.2098 | 0.9575 | 0 | 0.8645 | intron | intron-variant | LPP |
| rs1399180 | 10 | 8098719 | t | 1 | t | -0.014 | 0.2923 | 0.9619 | 64.9 | 0.0224 | intron | intron-variant | GATA3 |
| rs631204 | 6 | 137959455 | a | 1 | a | 0.0079 | 0.2154 | 0.9707 | 0 | 0.4727 | intergenic | | |
| rs12478539 | 2 | 43355324 | c | 0 | g | -0.0086 | 0.244 | 0.9719 | 69 | 0.01181 | intergenic | | |
| rs2084007 | 5 | 133891282 | t | 0 | c | 0.0067 | 0.2095 | 0.9744 | 0 | 0.6468 | intron | intron-variant | PHF15 |
| rs802730 | 6 | 128280104 | t | 1 | t | 0.0055 | 0.2329 | 0.9813 | 0 | 0.9864 | intergenic | | |
| rs7975763 | 12 | 123604053 | t | 0 | c | 0.0051 | 0.2475 | 0.9835 | 0 | 0.8539 | intron | | PITPNM2 |
| rs78727559 | 8 | 95851818 | t | 1 | t | 0.0051 | 0.4039 | 0.9899 | 0 | 0.5522 | intron | intron-variant | INTS8 |
