## Supplementary Table 3 for "A higher burden of multiple sclerosis genetic risk confers an earlier onset"

**Supplementary Table 3. Meta-Analysis results for variants with p<0.05, I-squared p>0.05.**

| Column Header | Key |
|---|---|
| **rsID** | dbSNP identifier (rsID) or chr:bp location if no rsID |
| **chromosome** | Chrosomome location |
| **position** | Position location (hg19) |
| **Total Cohorts** | Number of cohorts in which the observed variant appeared/was retained |
| **IndSigSNP** | Variant flag for independent significant SNP (1=indepentent sigificant SNP, 0=no). Independent SNPs were first identified by having a p-value<5x10-6 and being independent from each other at a level of r2<0.6. SNPs at a level of r2>0.6 with the independent variants were considered candidate SNPs. Finally, independent lead variants were selected from the independent significant SNPs if they remained independent from each other at r2<0.1. |
| **CandidateSNP** | Variant flag for candidate for significant SNPs (1=candidate, 0=no) |
| **Allele1** | Tested Allele |
| **Allele2** | Second Allele |
| **Effect** | Effect estimate in β form |
| **StdErr** | Standard error of the effect estimate |
| **P-Value** | Significance level of the effect estimate |
| **Direction** | Direction of the effect estimate in each of the individual five cohorts, listed in order: ADVANCE, ASCEND, DECIDE, ACP1, ACP2. "?" Refers to an unknown effect estimate, meaning the variant was not present in the specific cohort |
| **HetISq** | I-squared heterogeneity measure |
| **HetPVal** | Significance level of the Heterogeneity I-squared measure |
| **functionGVS** | GVS class of function variation, using only sumitted alleles and hg19 genome build |
| **functionDBSNP** | dbSNP class of variation function |
| **geneList** | HUGO names, any for which the transcription region overlaps the variation |

| rsID | chromosome | position | Total Cohorts | IndSigSNP | CandidateSNP | Allele1 | Allele2 | Effect | StdErr | P-value | Direction | HetISq | HetPVal | functionGVS | geneList |
|---|---|---|---|---|---|---|---|---|---|---|---|---|---|---|---|
| rs17076315 | 5 | 173141042 | 4 | 1 | 1 | A | G | -1.3059 | 0.2631 | 6.91E-07 | ---?- | 48.5 | 0.1203 | intergenic | |
| rs17076316 | 5 | 173141093 | 4 | 0 | 1 | A | G | -1.3031 | 0.2627 | 7.07E-07 | ---?- | 48.9 | 0.1183 | intergenic | |
| rs12520579 | 5 | 173158992 | 4 | 1 | 1 | T | C | -1.3777 | 0.2814 | 9.77E-07 | ---?- | 32 | 0.2202 | intergenic | |
| rs9313646 | 5 | 173155866 | 4 | 0 | 1 | A | G | 1.3026 | 0.2667 | 1.04E-06 | +++?+ | 43.4 | 0.1511 | intergenic | |
| rs28672722 | 6 | 32626537 | 5 | 1 | 1 | T | G | 1.2363 | 0.2534 | 1.07E-06 | +++++ | 0 | 0.6686 | downstream-gene | |
| rs57487085 | 5 | 173152486 | 4 | 0 | 1 | C | G | -1.3009 | 0.267 | 1.10E-06 | ---?- | 32.7 | 0.2161 | intergenic | |
| rs34605315 | 5 | 173155060 | 4 | 0 | 1 | T | C | -1.3105 | 0.269 | 1.11E-06 | ---?- | 43.6 | 0.1501 | intergenic | |
| rs17066212 | 6 | 106149587 | 4 | 1 | 1 | A | G | -3.8943 | 0.8006 | 1.15E-06 | ---?- | 22.2 | 0.2774 | intergenic | |
| rs59436056 | 5 | 173152799 | 4 | 0 | 1 | T | G | -1.3003 | 0.2673 | 1.15E-06 | ---?- | 28.8 | 0.2394 | intergenic | |
| rs10223185 | 5 | 173156307 | 4 | 0 | 1 | C | G | 1.292 | 0.2659 | 1.18E-06 | +++?+ | 44.4 | 0.145 | intergenic | |
| rs6860531 | 5 | 173159426 | 4 | 0 | 1 | T | C | 1.3973 | 0.2877 | 1.19E-06 | +++?+ | 44.7 | 0.1434 | intergenic | |
| rs35805769 | 5 | 173153723 | 4 | 0 | 1 | A | C | -1.2975 | 0.2672 | 1.20E-06 | ---?- | 31.7 | 0.2224 | intergenic | |
| rs34131421 | 5 | 173153002 | 4 | 0 | 1 | T | C | 1.2963 | 0.2672 | 1.23E-06 | +++?+ | 30.4 | 0.2301 | intergenic | |
| rs35238759 | 5 | 173152905 | 4 | 0 | 1 | T | C | 1.2958 | 0.2672 | 1.24E-06 | +++?+ | 30.1 | 0.2317 | intergenic | |
| rs17066216 | 6 | 106150158 | 4 | 0 | 1 | T | C | 3.8765 | 0.8 | 1.26E-06 | +++?+ | 22.6 | 0.2753 | intergenic | |
| rs61359098 | 5 | 173152707 | 4 | 0 | 1 | A | T | 1.2942 | 0.2672 | 1.28E-06 | +++?+ | 29.2 | 0.2367 | intergenic | |
| rs10079152 | 5 | 173159091 | 4 | 0 | 1 | T | C | 1.3653 | 0.282 | 1.29E-06 | +++?+ | 31.5 | 0.2232 | intergenic | |
| rs10079151 | 5 | 173159090 | 4 | 0 | 1 | T | G | 1.3642 | 0.2818 | 1.29E-06 | +++?+ | 31.5 | 0.2235 | intergenic | |
| rs79534154 | 6 | 106154295 | 4 | 0 | 1 | A | G | 3.9171 | 0.8106 | 1.35E-06 | +++?+ | 33.2 | 0.2128 | intergenic | |
| rs2002546 | 5 | 173150340 | 4 | 0 | 1 | A | G | 1.2907 | 0.2672 | 1.36E-06 | +++?+ | 30.6 | 0.2289 | intergenic | |
| rs2002548 | 5 | 173150415 | 4 | 0 | 1 | A | T | 1.2894 | 0.2671 | 1.38E-06 | +++?+ | 30.7 | 0.2283 | intergenic | |
| rs1820084 | 5 | 173151538 | 4 | 0 | 1 | T | G | -1.2847 | 0.2665 | 1.43E-06 | ---?- | 31.4 | 0.2236 | intergenic | |
| rs73775828 | 6 | 106158312 | 4 | 0 | 1 | T | G | -3.8785 | 0.8058 | 1.49E-06 | ---?- | 35 | 0.2025 | intergenic | |
| rs35105788 | 5 | 173144163 | 4 | 0 | 1 | A | G | -1.271 | 0.2643 | 1.52E-06 | ---?- | 25.8 | 0.2566 | intergenic | |
| rs56794815 | 5 | 173152723 | 4 | 0 | 1 | T | C | -1.2832 | 0.2669 | 1.52E-06 | ---?- | 29.2 | 0.2368 | intergenic | |
| rs12054964 | 5 | 173149709 | 4 | 0 | 1 | T | C | -1.2966 | 0.2699 | 1.55E-06 | ---?- | 38.8 | 0.179 | intergenic | |
| rs34840467 | 5 | 173144222 | 4 | 0 | 1 | T | C | 1.264 | 0.2634 | 1.59E-06 | +++?+ | 26.6 | 0.2523 | intergenic | |
| rs59711172 | 5 | 173143857 | 4 | 0 | 1 | T | C | 1.2678 | 0.2644 | 1.62E-06 | +++?+ | 26.1 | 0.2554 | intergenic | |
| rs10067485 | 5 | 173144583 | 4 | 0 | 1 | A | C | 1.2595 | 0.2634 | 1.73E-06 | +++?+ | 24.2 | 0.2663 | intergenic | |
| rs1107406 | 5 | 173151427 | 4 | 0 | 1 | T | C | 1.2728 | 0.2665 | 1.78E-06 | +++?+ | 33.1 | 0.2135 | intergenic | |
| rs76541020 | 6 | 106160924 | 4 | 0 | 1 | T | C | -3.8714 | 0.8111 | 1.82E-06 | ---?- | 35.5 | 0.1993 | intergenic | |
| rs73775831 | 6 | 106173349 | 4 | 0 | 1 | A | G | -3.8402 | 0.8055 | 1.87E-06 | ---?- | 37.4 | 0.1878 | intergenic | |
| rs17076322 | 5 | 173142582 | 4 | 0 | 1 | A | T | -1.2585 | 0.2646 | 1.98E-06 | ---?- | 26.9 | 0.2507 | intergenic | |
| rs17076328 | 5 | 173143508 | 4 | 0 | 1 | A | G | -1.268 | 0.2667 | 2.00E-06 | ---?- | 35.5 | 0.199 | intergenic | |

| | | | | | | | | | | | | | | | |
|---|---|---|---|---|---|---|---|---|---|---|---|---|---|---|---|
| rs28718232 | 6 | 32626750 | 5 | 0 | 1 | A | G | -1.3213 | 0.278 | 2.00E-06 | ----- | 0 | 0.6942 | downstream-gene |  |
| rs28414666 | 6 | 32626451 | 5 | 0 | 1 | A | G | 1.32 | 0.2778 | 2.02E-06 | +++++ | 0 | 0.6927 | downstream-gene | |
| rs184408348 | 6 | 106164607 | 4 | 0 | 1 | A | G | 3.8426 | 0.8096 | 2.07E-06 | +++?+ | 36.2 | 0.1951 | intergenic | |
| rs28407950 | 6 | 32626348 | 5 | 0 | 1 | T | C | 1.178 | 0.2484 | 2.11E-06 | +++++ | 22.8 | 0.2692 | downstream-gene | |
| rs74544066 | 6 | 106143945 | 4 | 0 | 1 | T | C | 3.9164 | 0.8261 | 2.13E-06 | +++?+ | 41.6 | 0.1623 | intergenic | |
| rs72931572 | 6 | 106241543 | 4 | 1 | 1 | A | C | -3.3908 | 0.7169 | 2.25E-06 | ---?- | 59.7 | 0.05898 | intergenic | |
| rs75148708 | 6 | 106131257 | 4 | 0 | 1 | A | C | -3.9326 | 0.8324 | 2.31E-06 | ---?- | 50.5 | 0.1087 | intergenic | |
| rs74450414 | 6 | 106175652 | 4 | 0 | 1 | T | C | -3.8286 | 0.8107 | 2.33E-06 | ---?- | 37.8 | 0.1853 | intergenic | |
| rs57223568 | 6 | 106142953 | 4 | 0 | 1 | T | C | 3.8416 | 0.8146 | 2.40E-06 | +++?+ | 33.6 | 0.2106 | intergenic | |
| rs72931565 | 6 | 106238455 | 4 | 0 | 1 | A | C | -3.3775 | 0.7164 | 2.43E-06 | ---?- | 60.4 | 0.05576 | intergenic | |
| 2:170812811 | 2 | 170812810 | 3 | 1 | 0 | T | TTA | 1.3059 | 0.277 | 2.43E-06 | +++?? | 56.6 | 0.09974 | | |
| rs72933607 | 6 | 106259396 | 4 | 1 | 1 | T | C | -2.4828 | 0.5268 | 2.44E-06 | ---?- | 23.4 | 0.2708 | intergenic | |
| rs9284986 | 5 | 173158157 | 4 | 0 | 1 | T | C | 1.3109 | 0.2784 | 2.49E-06 | +++?+ | 30.9 | 0.2266 | intergenic | |
| 3:187898598 | 3 | 187898598 | 3 | 0 | 0 | <ins:me: | T | 3.3855 | 0.7199 | 2.57E-06 | +++?? | 0 | 0.4434 | | |
| rs79228492 | 6 | 106129623 | 4 | 0 | 1 | A | G | 3.9537 | 0.841 | 2.59E-06 | +++?+ | 43.5 | 0.1507 | intergenic | |
| rs76855436 | 6 | 106127703 | 4 | 0 | 1 | T | C | 3.947 | 0.8399 | 2.61E-06 | +++?+ | 43.4 | 0.1514 | intergenic | |
| rs74891492 | 6 | 106127508 | 4 | 0 | 1 | A | G | -3.9464 | 0.8398 | 2.61E-06 | ---?- | 43.4 | 0.1513 | intergenic | |
| rs35065704 | 5 | 173153149 | 4 | 0 | 1 | T | C | -1.2633 | 0.2693 | 2.71E-06 | ---?- | 16.5 | 0.3091 | intergenic | |
| rs75878272 | 6 | 106123582 | 4 | 0 | 1 | A | G | -3.9124 | 0.8367 | 2.93E-06 | ---?- | 50.3 | 0.1099 | intergenic | |
| rs9313647 | 5 | 173156099 | 4 | 0 | 1 | T | C | -1.2516 | 0.2677 | 2.93E-06 | ---?- | 28.7 | 0.2397 | intergenic | |
| rs10223202 | 5 | 173156504 | 4 | 0 | 1 | T | C | -1.2474 | 0.267 | 3.00E-06 | ---?- | 29.8 | 0.2336 | intergenic | |
| rs59969576 | 5 | 173152800 | 4 | 0 | 1 | A | G | -1.2567 | 0.269 | 3.00E-06 | ---?- | 13 | 0.3276 | intergenic | |
| rs77269080 | 6 | 106178623 | 4 | 0 | 1 | T | C | -3.7533 | 0.8058 | 3.20E-06 | ---?- | 25.3 | 0.2598 | intergenic | |
| rs34132828 | 19 | 46163870 | 5 | 1 | 1 | A | G | 1.1596 | 0.2494 | 3.32E-06 | ++++- | 0 | 0.4828 | intergenic | |
| rs17486144 | 6 | 106256999 | 4 | 0 | 1 | T | C | 2.3444 | 0.505 | 3.44E-06 | +++?+ | 48.2 | 0.1221 | intergenic | |
| rs10051062 | 5 | 173159651 | 4 | 0 | 1 | T | C | 1.3754 | 0.2969 | 3.61E-06 | +++?+ | 53.1 | 0.09368 | intergenic | |
| rs78899175 | 6 | 106126691 | 4 | 0 | 1 | A | G | 3.8922 | 0.8408 | 3.68E-06 | +++?+ | 44.2 | 0.1461 | intergenic | |
| rs62404084 | 6 | 32603321 | 5 | 1 | 1 | T | C | 1.3826 | 0.2994 | 3.88E-06 | +++++ | 29.5 | 0.2246 | upstream-gene | |
| rs17499411 | 6 | 32665002 | 5 | 0 | 1 | C | G | -1.6769 | 0.3638 | 4.04E-06 | ----- | 27.2 | 0.2405 | intergenic |  |
| rs56296353 | 6 | 106180611 | 4 | 0 | 1 | T | C | -3.6972 | 0.8023 | 4.06E-06 | ---?- | 28 | 0.2442 | intergenic | |
| rs74402157 | 8 | 3655967 | 3 | 1 | 1 | T | G | -4.5215 | 0.9819 | 4.13E-06 | ---?? | 0 | 0.6538 | intron | CSMD1 |
| rs80246006 | 6 | 106178480 | 4 | 0 | 1 | C | G | -3.73 | 0.8105 | 4.18E-06 | ---?- | 24.9 | 0.2618 | intergenic | |
| rs149847639 | 6 | 56121561 | 2 | 1 | 1 | A | C | -7.4704 | 1.6266 | 4.38E-06 | ?--?? | 0 | 0.6093 | intergenic | |
| rs17747385 | 5 | 173145040 | 4 | 0 | 1 | A | G | 1.2233 | 0.2669 | 4.58E-06 | +++?+ | 39.7 | 0.1734 | intergenic | |
| rs61057753 | 6 | 106180964 | 4 | 0 | 1 | T | C | -3.6678 | 0.8006 | 4.62E-06 | ---?- | 29 | 0.2381 | intergenic | |
| rs12660518 | 6 | 106255758 | 4 | 0 | 1 | A | G | 2.3049 | 0.5036 | 4.72E-06 | +++?+ | 41.4 | 0.163 | intergenic | |
| rs57136171 | 6 | 106180996 | 4 | 0 | 1 | A | T | -3.6616 | 0.8003 | 4.75E-06 | ---?- | 29.2 | 0.2367 | intergenic | |
| rs72931562 | 6 | 106237540 | 4 | 0 | 1 | T | G | -3.2247 | 0.7049 | 4.77E-06 | ---?- | 55.4 | 0.08122 | intergenic | |
| rs12660517 | 6 | 106255713 | 4 | 0 | 1 | A | G | 2.2976 | 0.5028 | 4.90E-06 | +++?+ | 41.5 | 0.1627 | intergenic | |
| 8:109012762 | 8 | 109012762 | 3 | 0 | 0 | G | GT | 2.3734 | 0.5206 | 5.14E-06 | +++?? | 26 | 0.2587 | | |
| rs72931594 | 6 | 106253897 | 4 | 0 | 1 | A | G | -2.2866 | 0.5019 | 5.22E-06 | ---?- | 38 | 0.1843 | intergenic | |
| rs12660507 | 6 | 106255631 | 4 | 0 | 1 | T | G | 2.2799 | 0.5011 | 5.38E-06 | +++?+ | 38.1 | 0.1835 | intergenic | |
| rs112666352 | 6 | 106088034 | 4 | 0 | 1 | A | C | -3.9779 | 0.8746 | 5.41E-06 | ---?- | 32.4 | 0.2181 | intergenic | |
| rs111420040 | 6 | 106210842 | 4 | 0 | 1 | A | G | -3.2281 | 0.71 | 5.45E-06 | ---?- | 60.4 | 0.05572 | intergenic | |
| rs72931548 | 6 | 106225773 | 4 | 0 | 1 | T | C | 3.2273 | 0.7101 | 5.49E-06 | +++?+ | 58.9 | 0.06314 | intergenic | |
| rs41282882 | 10 | 131499645 | 4 | 0 | 0 | T | C | 1.8956 | 0.4175 | 5.61E-06 | +++?+ | 0 | 0.9068 | intron | MGMT |
| rs10754972 | 2 | 170875105 | 4 | 0 | 1 | A | G | -1.1879 | 0.2616 | 5.62E-06 | ---?- | 60.3 | 0.05591 | intron | UBR3 |
| rs77159423 | 10 | 131498735 | 4 | 0 | 0 | T | C | 1.8956 | 0.4176 | 5.63E-06 | +++?+ | 0 | 0.9051 | intron | MGMT |
| rs78579101 | 10 | 131493784 | 4 | 0 | 0 | T | C | 1.8868 | 0.4161 | 5.78E-06 | +++?+ | 0 | 0.8865 | intron | MGMT |
| rs73999196 | 2 | 238549024 | 4 | 0 | 0 | A | C | 4.393 | 0.9693 | 5.84E-06 | +++?+ | 0 | 0.589 | intron | LRRFIP1 |
| rs73775833 | 6 | 106181597 | 4 | 0 | 1 | A | G | -3.6013 | 0.7964 | 6.12E-06 | ---?- | 31.1 | 0.2256 | intergenic | |
| rs116533843 | 2 | 137646848 | 4 | 0 | 0 | T | C | 4.5524 | 1.0068 | 6.14E-06 | +++?+ | 0 | 0.7617 | intron | THSD7B |
| rs41297215 | 9 | 133231049 | 4 | 0 | 0 | T | G | 5.3779 | 1.1909 | 6.31E-06 | +++?+ | 42.2 | 0.1586 | intron | HMCN2 |
| rs10076158 | 5 | 173141811 | 4 | 0 | 1 | T | C | -1.2093 | 0.2688 | 6.82E-06 | ---?- | 46.7 | 0.1314 | intergenic | |
| rs10076268 | 5 | 173141810 | 4 | 0 | 1 | A | G | -1.2083 | 0.2688 | 6.96E-06 | ---?- | 46.7 | 0.1314 | intergenic | |
| rs28469403 | 14 | 38488576 | 4 | 0 | 0 | T | C | 1.4205 | 0.3165 | 7.17E-06 | +++?+ | 0 | 0.7573 | intergenic | |
| rs72931584 | 6 | 106251139 | 4 | 0 | 1 | A | G | -2.3349 | 0.5206 | 7.28E-06 | ---?- | 47.1 | 0.1288 | intergenic | |
| rs12664161 | 6 | 106229573 | 4 | 0 | 1 | A | G | 3.1607 | 0.7051 | 7.36E-06 | +++?+ | 58 | 0.0674 | intergenic | |
| rs76607806 | 6 | 106181089 | 4 | 0 | 1 | T | C | 3.6099 | 0.8053 | 7.37E-06 | +++?+ | 29.2 | 0.2368 | intergenic | |

| | | | | | | | | | | | | | | | |
|---|---|---|---|---|---|---|---|---|---|---|---|---|---|---|---|
| rs12663858 | 6 | 106229562 | 4 | 0 | 1 | T | C | 3.1599 | 0.705 | 7.40E-06 | +++?+ | 58 | 0.06727 | intergenic | |
| rs78831300 | 6 | 106140677 | 4 | 0 | 1 | A | G | -3.7588 | 0.8391 | 7.48E-06 | ---?- | 42.7 | 0.1551 | intergenic | |
| rs79161041 | 6 | 106140679 | 4 | 0 | 1 | T | C | -3.7588 | 0.8391 | 7.49E-06 | ---?- | 42.7 | 0.1551 | intergenic | |
| rs75004031 | 6 | 106139783 | 4 | 0 | 1 | A | G | 3.7554 | 0.8387 | 7.54E-06 | +++?+ | 42.8 | 0.155 | intergenic | |
| rs9995547 | 4 | 117281523 | 4 | 0 | 0 | T | G | -3.898 | 0.8707 | 7.58E-06 | ---?- | 26.8 | 0.2511 | intergenic | |
| rs73775834 | 6 | 106182146 | 4 | 0 | 1 | A | C | -3.5488 | 0.7927 | 7.58E-06 | ---?- | 32.6 | 0.2166 | intergenic | |
| rs73775836 | 6 | 106182450 | 4 | 0 | 1 | A | T | 3.5366 | 0.7902 | 7.62E-06 | +++?+ | 32.7 | 0.2164 | intergenic | |
| rs78688982 | 2 | 11050238 | 3 | 0 | 0 | T | C | -3.3299 | 0.7442 | 7.66E-06 | ---?? | 0 | 0.5548 | upstream-gene | |
| rs917051 | 17 | 47383945 | 5 | 0 | 0 | C | G | -1.1395 | 0.2547 | 7.68E-06 | ----+ | 0.7 | 0.4021 | intron | ZNF652 |
| 18:39974520 | 18 | 39974520 | 3 | 0 | 0 | AT | ATT | 1.6169 | 0.3615 | 7.71E-06 | +++?? | 0 | 0.7949 | | |
| rs77671147 | 17 | 47353803 | 3 | 0 | 0 | A | G | 1.3385 | 0.2994 | 7.80E-06 | +++?? | 0 | 0.8685 | intergenic | |
| 10:28230252 | 10 | 28230252 | 3 | 0 | 0 | G | GA | -1.1134 | 0.2491 | 7.85E-06 | ---?? | 30.4 | 0.2375 | | |
| rs57637664 | 6 | 106180774 | 4 | 0 | 1 | T | C | -3.6202 | 0.8102 | 7.89E-06 | ---?- | 39.5 | 0.175 | intergenic | |
| rs62076405 | 17 | 47393605 | 5 | 0 | 0 | A | G | -1.1166 | 0.25 | 7.97E-06 | ----+ | 0 | 0.4872 | intron | ZNF652 |
| 17:47353791 | 17 | 47353791 | 3 | 0 | 0 | G | GA | -1.3349 | 0.2993 | 8.18E-06 | ---?? | 0 | 0.8624 | | |
| rs4794040 | 17 | 47392718 | 5 | 0 | 0 | A | G | 1.1149 | 0.2502 | 8.37E-06 | ++++- | 0 | 0.4882 | intron | ZNF652 |
| rs2269932 | 17 | 47386688 | 3 | 0 | 0 | T | C | 1.3263 | 0.2977 | 8.41E-06 | +++?? | 0 | 0.8876 | intron | ZNF652 |
| rs72931552 | 6 | 106226772 | 4 | 0 | 1 | T | C | 3.1386 | 0.7047 | 8.43E-06 | +++?+ | 58.8 | 0.06346 | intergenic | |
| 17:47351759 | 17 | 47351759 | 3 | 0 | 0 | <Cn0> | C | -1.3332 | 0.2994 | 8.45E-06 | ---?? | 0 | 0.8507 | | |
| rs7639143 | 3 | 2468938 | 3 | 0 | 0 | A | T | 1.2971 | 0.2915 | 8.58E-06 | +++?? | 0 | 0.5951 | intron | CNTN4 |
| rs7649185 | 3 | 2468939 | 3 | 0 | 0 | T | G | -1.2971 | 0.2915 | 8.58E-06 | ---?? | 0 | 0.5949 | intron | CNTN4 |
| rs75498623 | 6 | 106181603 | 4 | 0 | 1 | A | T | -3.5771 | 0.8044 | 8.71E-06 | ---?- | 31.3 | 0.2244 | intergenic | |
| rs1050118 | 6 | 31237776 | 3 | 0 | 0 | T | C | 1.2163 | 0.2736 | 8.74E-06 | +++?? | 0 | 0.563 | missense | HLA-C |
| rs17485214 | 6 | 106218143 | 4 | 0 | 1 | A | G | -3.131 | 0.7042 | 8.75E-06 | ---?- | 61.5 | 0.0503 | intergenic | |
| rs76156757 | 17 | 47391807 | 5 | 0 | 0 | A | C | -1.1128 | 0.2504 | 8.85E-06 | ----+ | 0 | 0.4894 | intron | ZNF652 |
| rs7221336 | 17 | 47387604 | 5 | 0 | 0 | T | C | -1.1154 | 0.251 | 8.87E-06 | ----+ | 0 | 0.5005 | intron | ZNF652 |
| rs76225746 | 17 | 47384750 | 5 | 0 | 0 | A | G | -1.1179 | 0.2517 | 8.93E-06 | ----+ | 0 | 0.4981 | intron | ZNF652 |
| rs6747881 | 2 | 170857653 | 4 | 0 | 1 | T | G | -1.1685 | 0.2631 | 8.93E-06 | ---?- | 61.3 | 0.05148 | intron | UBR3 |
| rs2189785 | 17 | 47385544 | 5 | 0 | 0 | T | C | 1.1157 | 0.2513 | 8.99E-06 | ++++- | 0 | 0.4997 | intron | ZNF652 |
| rs113035845 | 2 | 238538300 | 4 | 0 | 0 | A | T | 4.3532 | 0.9804 | 8.99E-06 | +++?+ | 0 | 0.573 | intron | LRRFIP1 |
| rs111549905 | 2 | 238538203 | 4 | 0 | 0 | T | C | -4.3527 | 0.9804 | 9.02E-06 | ---?- | 0 | 0.573 | intron | LRRFIP1 |
| 17:47347152 | 17 | 47347152 | 3 | 0 | 0 | CT | C | 1.3274 | 0.2991 | 9.08E-06 | +++?? | 0 | 0.8195 | | |
| 17:47408251 | 17 | 47408251 | 3 | 0 | 0 | A | AAAAC | -1.3428 | 0.3028 | 9.23E-06 | ---?? | 0 | 0.9444 | | |
| 10:126829709 | 10 | 126829709 | 3 | 0 | 0 | CA | C | -1.9209 | 0.4333 | 9.29E-06 | ---?? | 0 | 0.7767 | | |
| rs78410390 | 6 | 106182656 | 4 | 0 | 1 | T | C | 3.4924 | 0.7884 | 9.43E-06 | +++?+ | 34.1 | 0.2074 | intergenic | |
| 2:167270363 | 2 | 167270363 | 2 | 0 | 0 | T | C | 7.1506 | 1.6148 | 9.50E-06 | ???++ | 8.4 | 0.296 | intron | SCN7A |
| rs72754076 | 15 | 87480575 | 4 | 0 | 0 | T | C | 3.6174 | 0.8169 | 9.51E-06 | +++?+ | 0 | 0.975 | intron | AGBL1 |
| rs33917769 | 8 | 76139316 | 4 | 0 | 0 | T | G | 1.3409 | 0.3029 | 9.54E-06 | +++?+ | 0 | 0.5533 | intron | CASC9 |
| rs118105237 | 8 | 20167335 | 4 | 0 | 0 | C | G | -4.1805 | 0.9447 | 9.63E-06 | ---?- | 0 | 0.8214 | intergenic | |
| rs4794039 | 17 | 47378729 | 5 | 0 | 0 | A | G | 1.1159 | 0.2522 | 9.65E-06 | ++++- | 0 | 0.4965 | intron | ZNF652 |
| rs78763027 | 17 | 47361402 | 3 | 0 | 0 | T | C | -1.3274 | 0.3 | 9.67E-06 | ---?? | 0 | 0.8882 | intergenic | |
| rs35805352 | 8 | 21039132 | 4 | 0 | 0 | A | G | -1.9021 | 0.4299 | 9.68E-06 | ---?- | 0 | 0.6407 | intergenic | |
| rs6504604 | 17 | 47389935 | 5 | 0 | 0 | T | G | -1.1087 | 0.2507 | 9.77E-06 | ----+ | 0 | 0.4914 | intron | ZNF652 |
| rs4794038 | 17 | 47377876 | 5 | 0 | 0 | A | T | 1.1157 | 0.2523 | 9.79E-06 | ++++- | 0 | 0.4966 | intron | ZNF652 |
| rs7744123 | 6 | 168620659 | 4 | 0 | 0 | T | C | -1.5887 | 0.3595 | 9.92E-06 | ---?- | 0 | 0.9688 | intergenic | |
| 18:40019473 | 18 | 40019473 | 3 | 0 | 0 | A | AT | -1.5216 | 0.3444 | 9.95E-06 | ---?? | 0 | 0.8532 | | |
| rs116564918 | 2 | 116598778 | 3 | 0 | 0 | A | G | -5.1348 | 1.1625 | 1.00E-05 | ---?? | 59.3 | 0.08546 | intron | DPP10 |
| rs72931545 | 6 | 106224252 | 4 | 0 | 1 | A | G | 3.1078 | 0.7039 | 1.01E-05 | +++?+ | 59.8 | 0.05852 | intergenic | |
| rs7760810 | 6 | 168621020 | 4 | 0 | 0 | T | C | 1.5852 | 0.359 | 1.01E-05 | +++?+ | 0 | 0.9677 | intergenic | |
| rs9455996 | 6 | 168622133 | 4 | 0 | 0 | T | C | -1.582 | 0.3589 | 1.05E-05 | ---?- | 0 | 0.9645 | intergenic | |
| rs56277478 | 7 | 158624126 | 2 | 0 | 0 | A | G | -7.4107 | 1.6823 | 1.06E-05 | ??-?- | 0 | 0.5094 | upstream-gene | |
| rs1397086 | 8 | 76107814 | 4 | 0 | 0 | T | G | 1.3327 | 0.3026 | 1.06E-05 | +++?+ | 0 | 0.5232 | intergenic | |
| rs7823733 | 8 | 76107231 | 4 | 0 | 0 | A | T | 1.3324 | 0.3026 | 1.07E-05 | +++?+ | 0 | 0.5231 | intergenic | |
| rs11079869 | 17 | 47373417 | 5 | 0 | 0 | T | C | 1.1112 | 0.2525 | 1.08E-05 | ++++- | 0 | 0.4924 | 3-prime-UTR | ZNF652 |
| rs11178134 | 12 | 70597450 | 3 | 0 | 0 | T | C | -1.2652 | 0.2876 | 1.09E-05 | ---?? | 51.3 | 0.1285 | intergenic | |
| rs72760176 | 15 | 87614911 | 4 | 0 | 0 | A | G | 3.2839 | 0.7468 | 1.10E-05 | +++?+ | 0 | 0.6773 | intergenic | |
| rs16947943 | 17 | 47368946 | 5 | 0 | 0 | A | G | -1.1103 | 0.2525 | 1.10E-05 | ----+ | 0 | 0.496 | 3-prime-UTR | ZNF652 |
| rs864699 | 7 | 16836477 | 4 | 0 | 0 | A | G | -1.6876 | 0.3842 | 1.12E-05 | ---?- | 0 | 0.8361 | intron | AGR2 |
| rs140097284 | 15 | 87546523 | 4 | 0 | 0 | A | G | -3.6465 | 0.8302 | 1.12E-05 | ---?- | 0 | 0.7353 | intron | AGBL1 |

| | | | | | | | | | | | | | | | |
|---|---|---|---|---|---|---|---|---|---|---|---|---|---|---|---|
| rs8086170 | 18 | 40105420 | 4 | 0 | 0 | A | G | -1.4553 | 0.3318 | 1.15E-05 | ---?+ | 25.9 | 0.256 | intron | LINC00907 |
| rs7237867 | 18 | 40103034 | 4 | 0 | 0 | A | G | -1.454 | 0.3315 | 1.15E-05 | ---?+ | 25.9 | 0.2562 | intron | LINC00907 |
| rs10111361 | 8 | 76101212 | 4 | 0 | 0 | T | G | -1.3252 | 0.3022 | 1.16E-05 | ---?- | 0 | 0.5164 | intergenic | |
| rs4794037 | 17 | 47369302 | 5 | 0 | 0 | A | G | 1.1071 | 0.2525 | 1.16E-05 | ++++- | 0 | 0.4907 | 3-prime-UTR | ZNF652 |
| rs112880267 | 6 | 106124004 | 4 | 0 | 1 | T | C | -3.8641 | 0.8813 | 1.16E-05 | ---?- | 27.9 | 0.2446 | intergenic | |
| rs8107376 | 19 | 46163769 | 5 | 0 | 1 | T | C | 0.9936 | 0.2267 | 1.17E-05 | +++++ | 0 | 0.813 | intergenic | |
| rs8109244 | 19 | 46163796 | 5 | 0 | 1 | A | C | 0.9935 | 0.2267 | 1.17E-05 | +++++ | 0 | 0.8129 | intergenic | |
| rs72931544 | 6 | 106223827 | 4 | 0 | 1 | T | C | 3.0837 | 0.7037 | 1.18E-05 | +++?+ | 59.9 | 0.05817 | intergenic | |
| 8:3642167 | 8 | 3642167 | 3 | 0 | 0 | CT | C | 5.3238 | 1.215 | 1.18E-05 | +++?? | 0 | 0.617 | | |
| rs181025707 | 8 | 3642169 | 3 | 0 | 0 | T | G | 5.3237 | 1.2149 | 1.18E-05 | +++?? | 0 | 0.6167 | intron | CSMD1 |
| rs7231412 | 18 | 40105657 | 4 | 0 | 0 | T | C | 1.4529 | 0.3316 | 1.18E-05 | +++?- | 25.9 | 0.2565 | intron | LINC00907 |
| rs13255145 | 8 | 76120341 | 4 | 0 | 0 | A | G | -1.3389 | 0.3056 | 1.18E-05 | ---?- | 0 | 0.556 | intergenic | |
| rs77859392 | 18 | 40104002 | 4 | 0 | 0 | A | G | -1.4534 | 0.3318 | 1.18E-05 | ---?+ | 25.9 | 0.2564 | intron | LINC00907 |
| rs7237672 | 18 | 40103183 | 4 | 0 | 0 | A | G | 1.4534 | 0.3318 | 1.19E-05 | +++?- | 25.9 | 0.2564 | intron | LINC00907 |
| rs77546496 | 18 | 40103192 | 4 | 0 | 0 | A | T | 1.4534 | 0.3318 | 1.19E-05 | +++?- | 25.9 | 0.2564 | intron | LINC00907 |
| rs78235502 | 18 | 40103339 | 4 | 0 | 0 | A | G | 1.4534 | 0.3318 | 1.19E-05 | +++?- | 25.9 | 0.2564 | intron | LINC00907 |
| rs73775838 | 6 | 106183047 | 4 | 0 | 1 | T | C | -3.4314 | 0.7836 | 1.19E-05 | ---?- | 35.6 | 0.1987 | intergenic | |
| rs73775839 | 6 | 106183068 | 4 | 0 | 1 | A | G | 3.431 | 0.7837 | 1.20E-05 | +++?+ | 35.6 | 0.1987 | intergenic | |
| rs58882737 | 17 | 47413672 | 3 | 0 | 0 | A | G | -1.3325 | 0.3044 | 1.20E-05 | ---?? | 0 | 0.9134 | intron | ZNF652 |
| rs16976281 | 18 | 40104100 | 4 | 0 | 0 | C | G | -1.4511 | 0.3316 | 1.21E-05 | ---?+ | 25.8 | 0.2569 | intron | LINC00907 |
| rs79155636 | 18 | 40103345 | 4 | 0 | 0 | A | C | 1.4511 | 0.3316 | 1.21E-05 | +++?- | 25.8 | 0.2569 | intron | LINC00907 |
| rs10102195 | 8 | 76135582 | 4 | 0 | 0 | T | C | -1.321 | 0.302 | 1.22E-05 | ---?- | 0 | 0.5981 | non-coding-exon | CASC9 |
| rs141063723 | 17 | 47385017 | 5 | 0 | 0 | A | G | 1.0979 | 0.2511 | 1.23E-05 | ++++- | 0 | 0.5204 | intron | ZNF652 |
| rs7011621 | 8 | 76090988 | 4 | 0 | 0 | A | G | -1.3198 | 0.3019 | 1.23E-05 | ---?- | 0 | 0.5238 | intergenic | |
| rs56398835 | 6 | 168625719 | 4 | 0 | 0 | A | G | -1.5623 | 0.3574 | 1.24E-05 | ---?- | 0 | 0.9519 | intergenic | |
| rs17066223 | 6 | 106183476 | 4 | 0 | 1 | T | C | 3.4282 | 0.7845 | 1.24E-05 | +++?+ | 35.6 | 0.1987 | intergenic | |
| rs115296667 | 1 | 18339303 | 3 | 0 | 0 | A | G | 6.8854 | 1.5757 | 1.24E-05 | ++??+ | 18.2 | 0.2943 | intergenic | |
| rs17066225 | 6 | 106184035 | 4 | 0 | 1 | T | C | 3.4272 | 0.7849 | 1.26E-05 | +++?+ | 35.6 | 0.1987 | intergenic | |
| rs28729931 | 8 | 76093983 | 4 | 0 | 0 | T | C | 1.3178 | 0.3019 | 1.27E-05 | +++?+ | 0 | 0.5252 | intergenic | |
| rs9886511 | 8 | 76153209 | 4 | 0 | 0 | A | C | -1.3209 | 0.3027 | 1.28E-05 | ---?- | 0 | 0.4815 | intron | CASC9 |
| rs17066226 | 6 | 106184215 | 4 | 0 | 1 | T | C | -3.4258 | 0.7852 | 1.29E-05 | ---?- | 35.5 | 0.1992 | intergenic | |
| rs17701489 | 17 | 47397493 | 5 | 0 | 0 | T | C | -1.0936 | 0.2507 | 1.29E-05 | ----+ | 0 | 0.4775 | intron | ZNF652 |
| rs10073233 | 5 | 173159652 | 4 | 0 | 1 | A | G | -1.3084 | 0.3 | 1.30E-05 | ---?- | 45.3 | 0.1398 | intergenic | |
| rs73775843 | 6 | 106184425 | 4 | 0 | 1 | A | C | 3.4248 | 0.7855 | 1.30E-05 | +++?+ | 35.5 | 0.199 | intergenic | |
| rs73775841 | 6 | 106184073 | 4 | 0 | 1 | A | G | 3.4221 | 0.785 | 1.30E-05 | +++?+ | 35.5 | 0.1994 | intergenic | |
| 6:144621222 | 6 | 144621222 | 3 | 0 | 0 | A | AT | 1.9527 | 0.4481 | 1.32E-05 | +++?? | 65.6 | 0.0547 | | |
| rs1360690 | 9 | 122624757 | 3 | 0 | 0 | A | G | 1.3523 | 0.3104 | 1.32E-05 | +++?? | 0 | 0.5305 | intergenic | |
| rs35017111 | 8 | 76154837 | 4 | 0 | 0 | T | C | -1.3197 | 0.303 | 1.33E-05 | ---?- | 0 | 0.4821 | intron | CASC9 |
| rs111343227 | 9 | 90729704 | 2 | 0 | 0 | A | G | 4.601 | 1.0566 | 1.33E-05 | +?+?? | 0 | 0.7068 | intergenic | |
| rs72931531 | 6 | 106215511 | 4 | 0 | 1 | A | G | -3.0706 | 0.7052 | 1.34E-05 | ---?- | 61.3 | 0.05126 | intergenic | |
| rs115212420 | 6 | 106184758 | 4 | 0 | 0 | T | C | 3.4229 | 0.7862 | 1.34E-05 | +++?+ | 35.6 | 0.1989 | intergenic | |
| rs12677743 | 8 | 76148146 | 4 | 0 | 0 | A | G | 1.3186 | 0.303 | 1.35E-05 | +++?+ | 0 | 0.4747 | intron | CASC9 |
| 2:51173075 | 2 | 51173075 | 3 | 0 | 0 | G | GTATTA[illegible] | 4.4839 | 1.0303 | 1.35E-05 | +++?? | 0 | 0.8033 | | |
| rs141339729 | 6 | 106184821 | 4 | 0 | 0 | A | G | 3.4215 | 0.7864 | 1.36E-05 | +++?+ | 35.5 | 0.199 | intergenic | |
| rs151272052 | 6 | 106184916 | 4 | 0 | 0 | A | G | -3.4217 | 0.7865 | 1.36E-05 | ---?- | 35.5 | 0.1991 | intergenic | |
| rs10102416 | 8 | 76135649 | 4 | 0 | 0 | T | G | -1.3173 | 0.3029 | 1.37E-05 | ---?- | 0 | 0.5914 | non-coding-exon | CASC9 |
| rs34493352 | 16 | 5459204 | 4 | 0 | 0 | T | G | -1.0606 | 0.2439 | 1.37E-05 | ---?- | 0 | 0.7164 | intron | RBFOX1 |
| rs10282969 | 8 | 76134424 | 4 | 0 | 0 | T | C | -1.317 | 0.3029 | 1.37E-05 | ---?- | 0 | 0.586 | downstream-gene | |
| 6:106150219 | 6 | 106150219 | 3 | 0 | 0 | G | GGAT | 3.5906 | 0.8259 | 1.38E-05 | +++?? | 0 | 0.3752 | | |
| rs10094554 | 8 | 76133763 | 4 | 0 | 0 | T | C | -1.3175 | 0.3031 | 1.39E-05 | ---?- | 0 | 0.5764 | downstream-gene | |
| 6:106184712 | 6 | 106184712 | 3 | 0 | 0 | T | TC | 3.6332 | 0.8365 | 1.41E-05 | +++?? | 47.3 | 0.1499 | | |
| rs58410208 | 6 | 106185710 | 4 | 0 | 1 | A | C | 3.4201 | 0.7875 | 1.41E-05 | +++?+ | 35.3 | 0.2002 | intergenic | |
| 6:106184715 | 6 | 106184715 | 3 | 0 | 0 | A | T | 3.6333 | 0.8367 | 1.41E-05 | +++?? | 47.3 | 0.1499 | intergenic | |
| rs143738042 | 2 | 137669900 | 3 | 0 | 0 | A | T | -5.0611 | 1.166 | 1.42E-05 | ---?? | 0 | 0.5847 | intron | THSD7B |
| rs13253586 | 8 | 76089712 | 4 | 0 | 0 | A | G | -1.3086 | 0.3015 | 1.42E-05 | ---?- | 0 | 0.507 | intergenic | |
| rs13279464 | 8 | 76089121 | 4 | 0 | 0 | A | C | -1.3076 | 0.3014 | 1.44E-05 | ---?- | 0 | 0.506 | intergenic | |
| rs77198376 | 1 | 82663128 | 3 | 0 | 0 | A | G | 5.4778 | 1.2627 | 1.44E-05 | +++?? | 0 | 0.53 | intergenic | |
| 6:55142458 | 6 | 55142458 | 3 | 0 | 0 | CT | C | 1.3473 | 0.3106 | 1.44E-05 | +++?? | 48.8 | 0.1416 | | |
| rs2061867 | 8 | 76088980 | 4 | 0 | 0 | A | T | -1.3072 | 0.3014 | 1.44E-05 | ---?- | 0 | 0.5054 | intergenic | |

| | | | | | | | | | | | | | | | |
|---|---|---|---|---|---|---|---|---|---|---|---|---|---|---|---|
| rs17485319 | 6 | 106219328 | 4 | 0 | 1 | A | G | -3.0426 | 0.7015 | 1.44E-05 | ---?- | 61.6 | 0.05015 | intergenic | |
| rs28867768 | 8 | 76159205 | 4 | 0 | 0 | T | C | -1.3166 | 0.3036 | 1.45E-05 | ---?- | 0 | 0.4832 | intron | CASC9 |
| rs12335321 | 8 | 76088493 | 4 | 0 | 0 | T | C | -1.3059 | 0.3013 | 1.46E-05 | ---?- | 0 | 0.5049 | intergenic | |
| rs967844 | 8 | 76159829 | 4 | 0 | 0 | A | G | 1.3162 | 0.3037 | 1.47E-05 | +++?+ | 0 | 0.4833 | intron | CASC9 |
| rs12600968 | 17 | 47403726 | 5 | 0 | 0 | C | G | -1.0935 | 0.2524 | 1.48E-05 | ----+ | 0 | 0.4853 | intron | ZNF652 |
| rs62076440 | 17 | 47405067 | 5 | 0 | 0 | T | C | -1.0954 | 0.253 | 1.49E-05 | ----+ | 0 | 0.4611 | intron | ZNF652 |
| rs35243817 | 17 | 47359858 | 5 | 0 | 0 | A | G | 1.0903 | 0.2518 | 1.49E-05 | ++++- | 0 | 0.4179 | intergenic | |
| rs72807378 | 5 | 173152647 | 3 | 0 | 1 | C | G | 1.2045 | 0.2784 | 1.51E-05 | +++?? | 0 | 0.6089 | intergenic | |
| rs76043254 | 17 | 47359114 | 5 | 0 | 0 | A | G | -1.0894 | 0.2518 | 1.52E-05 | ----+ | 0 | 0.4138 | intergenic |  |
| 8:41753419 | 8 | 41753419 | 3 | 0 | 0 | G | GC | -4.391 | 1.0153 | 1.53E-05 | ---?? | 0 | 0.3687 | | |
| rs145768829 | 8 | 113211629 | 3 | 0 | 0 | T | C | -1.2016 | 0.2778 | 1.53E-05 | ---?? | 0 | 0.8923 | intergenic | |
| rs2898858 | 17 | 47405413 | 5 | 0 | 0 | A | C | 1.0947 | 0.2531 | 1.53E-05 | ++++- | 0 | 0.4614 | intron | ZNF652 |
| 6:106161654 | 6 | 106161654 | 3 | 0 | 0 | CA | C | 3.6862 | 0.8533 | 1.56E-05 | +++?? | 5.6 | 0.3468 | | |
| rs16939041 | 8 | 76079317 | 4 | 0 | 0 | T | C | -1.3084 | 0.303 | 1.57E-05 | ---?- | 0 | 0.5039 | intergenic | |
| rs28719789 | 8 | 76083388 | 4 | 0 | 0 | T | C | 1.2974 | 0.3006 | 1.58E-05 | +++?+ | 0 | 0.4997 | intergenic | |
| 12:95921614 | 12 | 95921614 | 3 | 0 | 0 | T | TC | 1.2622 | 0.2924 | 1.58E-05 | +++?? | 0 | 0.8323 | | |
| rs10105244 | 8 | 76078401 | 4 | 0 | 0 | A | T | -1.3109 | 0.3039 | 1.61E-05 | ---?- | 0 | 0.506 | intergenic | |
| rs62076399 | 17 | 47364991 | 5 | 0 | 0 | A | G | -1.0903 | 0.2528 | 1.61E-05 | ----+ | 0 | 0.4678 | downstream-gene |  |
| rs62522815 | 8 | 76163309 | 4 | 0 | 0 | A | G | -1.3117 | 0.3042 | 1.62E-05 | ---?- | 0 | 0.4779 | intron | CASC9 |
| rs13262359 | 8 | 76082094 | 4 | 0 | 0 | A | G | 1.295 | 0.3004 | 1.62E-05 | +++?+ | 0 | 0.498 | intergenic | |
| rs73971708 | 2 | 170580483 | 4 | 0 | 0 | A | G | 1.378 | 0.3198 | 1.64E-05 | +++?+ | 0 | 0.9928 | intron | PHOSPHO2-KLHL23 |
| rs55921605 | 8 | 76163672 | 4 | 0 | 0 | A | G | 1.3113 | 0.3043 | 1.64E-05 | +++?+ | 0 | 0.4778 | intron | CASC9 |
| rs9747090 | 17 | 47408645 | 5 | 0 | 0 | A | G | -1.0928 | 0.2538 | 1.66E-05 | ----+ | 0 | 0.4636 | intron | ZNF652 |
| rs17212553 | 6 | 32665687 | 5 | 0 | 1 | T | G | 1.6566 | 0.3847 | 1.67E-05 | +++++ | 0 | 0.4307 | intergenic | |
| rs4420575 | 17 | 47410127 | 5 | 0 | 0 | A | C | -1.0939 | 0.2541 | 1.67E-05 | ----+ | 0 | 0.4681 | intron | ZNF652 |
| rs9747404 | 17 | 47408729 | 5 | 0 | 0 | T | C | -1.0926 | 0.2538 | 1.67E-05 | ----+ | 0 | 0.4638 | intron | ZNF652 |
| 5:173141342 | 5 | 173141342 | 3 | 0 | 0 | A | ATTTTTT | 1.1684 | 0.2715 | 1.68E-05 | +++?? | 0 | 0.466 | | |
| rs13249617 | 8 | 76160368 | 4 | 0 | 0 | A | G | 1.3121 | 0.3051 | 1.70E-05 | +++?+ | 0 | 0.4777 | intron | CASC9 |
| rs2411984 | 17 | 47445751 | 3 | 0 | 0 | A | G | -1.1976 | 0.2785 | 1.71E-05 | ---?? | 0 | 0.8291 | intergenic | |
| rs62076442 | 17 | 47410679 | 5 | 0 | 0 | A | C | 1.0931 | 0.2543 | 1.71E-05 | ++++- | 0 | 0.469 | intron | ZNF652 |
| rs4794041 | 17 | 47410841 | 5 | 0 | 0 | A | G | -1.0928 | 0.2543 | 1.73E-05 | ----+ | 0 | 0.4694 | intron | ZNF652 |
| rs11751024 | 6 | 32586236 | 5 | 0 | 0 | A | C | 0.9669 | 0.2253 | 1.78E-05 | +++++ | 0 | 0.7815 | intergenic | |
| rs8104845 | 19 | 46161326 | 5 | 0 | 1 | A | G | 0.9721 | 0.2266 | 1.78E-05 | +++++ | 0 | 0.7738 | intergenic | |
| rs10409882 | 19 | 46160935 | 5 | 0 | 1 | T | C | -0.971 | 0.2265 | 1.81E-05 | ----- | 0 | 0.7734 | intergenic |  |
| rs112107334 | 3 | 126542229 | 4 | 0 | 0 | A | T | -3.7677 | 0.8788 | 1.81E-05 | ---?- | 0 | 0.966 | intron | CHCHD6 |
| rs10404527 | 19 | 46160703 | 5 | 0 | 1 | A | G | -0.9708 | 0.2264 | 1.81E-05 | ----- | 0 | 0.7732 | intergenic |  |
| rs10404142 | 19 | 46160698 | 5 | 0 | 1 | A | C | -0.9707 | 0.2264 | 1.81E-05 | ----- | 0 | 0.7731 | intergenic |  |
| rs55872740 | 19 | 46160323 | 5 | 0 | 1 | A | G | 0.9694 | 0.2262 | 1.83E-05 | +++++ | 0 | 0.7736 | intergenic | |
| rs35242582 | 6 | 32600057 | 5 | 0 | 1 | A | G | -1.1519 | 0.2688 | 1.83E-05 | ----- | 22.5 | 0.2712 | intergenic |  |
| rs1948195 | 4 | 117280580 | 3 | 0 | 0 | C | G | -3.8327 | 0.8947 | 1.84E-05 | ---?? | 52.9 | 0.1196 | intergenic | |
| rs72913363 | 4 | 117279296 | 4 | 0 | 0 | A | G | -3.8304 | 0.8942 | 1.84E-05 | ---?- | 32.3 | 0.2183 | intergenic | |
| rs16825716 | 2 | 146658515 | 4 | 0 | 0 | T | G | -3.8293 | 0.894 | 1.84E-05 | ---?- | 0 | 0.8382 | intergenic | |
| rs968155 | 6 | 32380715 | 5 | 0 | 0 | T | C | 0.9078 | 0.212 | 1.85E-05 | +++++ | 0 | 0.9141 | intergenic | |
| rs58069157 | 19 | 46160168 | 5 | 0 | 1 | A | G | 0.9684 | 0.2261 | 1.85E-05 | +++++ | 0 | 0.7734 | intergenic | |
| rs28881587 | 8 | 76165240 | 4 | 0 | 0 | T | C | -1.3086 | 0.3056 | 1.85E-05 | ---?- | 0 | 0.4807 | intron | CASC9 |
| rs55864746 | 19 | 46160246 | 5 | 0 | 1 | A | G | -0.9684 | 0.2262 | 1.85E-05 | ----- | 0 | 0.7733 | intergenic |  |
| rs55698417 | 19 | 46160138 | 5 | 0 | 1 | T | C | -0.9681 | 0.2261 | 1.86E-05 | ----- | 0 | 0.7731 | intergenic |  |
| rs79798982 | 12 | 76032156 | 3 | 0 | 0 | T | C | -4.7387 | 1.1068 | 1.86E-05 | ---?? | 55.9 | 0.1038 | intergenic | |
| rs55892531 | 6 | 106185884 | 4 | 0 | 1 | A | C | -3.3716 | 0.7876 | 1.86E-05 | ---?- | 37.3 | 0.1884 | intergenic | |
| rs59721711 | 6 | 106185873 | 4 | 0 | 1 | T | C | -3.3711 | 0.7876 | 1.87E-05 | ---?- | 37.2 | 0.1887 | intergenic | |
| rs59086866 | 6 | 106185961 | 4 | 0 | 1 | A | T | -3.3713 | 0.7877 | 1.87E-05 | ---?- | 37.3 | 0.1885 | intergenic | |
| rs8107236 | 19 | 46158436 | 5 | 0 | 1 | A | G | 0.9666 | 0.2259 | 1.87E-05 | ++++- | 0 | 0.7657 | intergenic | |
| rs8107527 | 19 | 46158417 | 5 | 0 | 1 | A | G | -0.9668 | 0.2259 | 1.87E-05 | ----+ | 0 | 0.767 | intergenic |  |
| rs72913359 | 4 | 117277155 | 4 | 0 | 0 | A | G | -3.8463 | 0.8988 | 1.87E-05 | ---?- | 32.1 | 0.2199 | intergenic | |
| rs11098340 | 4 | 117278263 | 4 | 0 | 0 | A | T | 3.8398 | 0.8973 | 1.87E-05 | +++?+ | 32.3 | 0.2185 | intergenic | |
| rs71352280 | 19 | 46159778 | 5 | 0 | 1 | T | C | 0.967 | 0.226 | 1.88E-05 | +++++ | 0 | 0.773 | intergenic | |
| rs4340404 | 18 | 40123296 | 4 | 0 | 0 | A | T | -1.4419 | 0.3371 | 1.90E-05 | ---?+ | 26.1 | 0.2554 | intron | LINC00907 |
| 10:4409289 | 10 | 4409289 | 3 | 0 | 0 | CTGTGT( | C | 1.1927 | 0.2789 | 1.90E-05 | +++?? | 57 | 0.09759 | | |
| rs10107017 | 8 | 76076922 | 4 | 0 | 0 | A | G | -1.3006 | 0.3042 | 1.90E-05 | ---?- | 0 | 0.4797 | intergenic | |

| | | | | | | | | | | | | | | | |
|---|---|---|---|---|---|---|---|---|---|---|---|---|---|---|---|
| rs77418080 | 2 | 238546723 | 3 | 0 | 0 | A | G | -4.5305 | 1.0598 | 1.91E-05 | -?-?- | 0 | 0.4087 | intron | LRRFIP1 |
| rs11872433 | 18 | 40057666 | 3 | 0 | 0 | A | T | 1.407 | 0.3292 | 1.92E-05 | +++?? | 0 | 0.9716 | intron | LINC00907 |
| rs190441245 | 6 | 106165643 | 3 | 0 | 1 | A | G | -3.5619 | 0.8335 | 1.93E-05 | ---?? | 27.8 | 0.2505 | intergenic | |
| rs114211941 | 2 | 238547863 | 3 | 0 | 0 | A | G | -4.5231 | 1.0586 | 1.93E-05 | -?-?- | 0 | 0.4067 | intron | LRRFIP1 |
| rs17205177 | 6 | 32602518 | 5 | 0 | 1 | A | G | -1.1573 | 0.2709 | 1.94E-05 | ----- | 14.4 | 0.3224 | upstream-gene |  |
| rs17211510 | 6 | 32602430 | 5 | 0 | 1 | A | C | 1.1541 | 0.2702 | 1.94E-05 | +++++ | 13 | 0.3313 | upstream-gene | |
| rs7258890 | 19 | 46159609 | 5 | 0 | 1 | T | C | 0.9648 | 0.226 | 1.96E-05 | ++++- | 0 | 0.7588 | intergenic | |
| rs8108269 | 19 | 46158513 | 5 | 0 | 1 | T | G | 0.9635 | 0.2257 | 1.96E-05 | ++++- | 0 | 0.7625 | intergenic | |
| rs10422601 | 19 | 46159414 | 5 | 0 | 1 | C | G | -0.9644 | 0.2259 | 1.96E-05 | ----+ | 0 | 0.7591 | intergenic |  |
| 17:47333988 | 17 | 47333988 | 3 | 0 | 0 | CA | C | 1.341 | 0.3141 | 1.96E-05 | +++?? | 0 | 0.5984 | | |
| rs1600283 | 8 | 76167155 | 4 | 0 | 0 | A | C | 1.3054 | 0.3058 | 1.97E-05 | +++?+ | 0 | 0.4829 | intron | CASC9 |
| rs9447925 | 6 | 77655297 | 4 | 0 | 0 | T | C | 1.1593 | 0.2716 | 1.97E-05 | +++?+ | 0 | 0.5023 | intergenic | |
| rs10403772 | 19 | 46160659 | 5 | 0 | 1 | A | C | 0.9672 | 0.2266 | 1.97E-05 | +++++ | 0 | 0.7774 | intergenic | |
| rs10403962 | 19 | 46160638 | 5 | 0 | 1 | A | C | -0.967 | 0.2266 | 1.98E-05 | ----- | 0 | 0.7775 | intergenic |  |
| rs16825718 | 2 | 146658569 | 4 | 0 | 0 | T | C | 4.0557 | 0.9505 | 1.98E-05 | +++?+ | 0 | 0.8396 | intergenic | |
| rs10421566 | 19 | 46159123 | 5 | 0 | 1 | T | C | -0.963 | 0.2258 | 1.99E-05 | ----+ | 0 | 0.7585 | intergenic |  |
| rs10415769 | 19 | 46158643 | 5 | 0 | 1 | T | G | -0.9617 | 0.2255 | 2.00E-05 | ----+ | 0 | 0.7598 | intergenic |  |
| rs17211580 | 6 | 32602754 | 5 | 0 | 1 | A | G | 1.1563 | 0.2712 | 2.00E-05 | +++++ | 14.8 | 0.3203 | upstream-gene | |
| rs28633132 | 6 | 32626053 | 5 | 0 | 1 | A | T | -1.6074 | 0.377 | 2.01E-05 | ----- | 18.2 | 0.2989 | downstream-gene |  |
| rs78596106 | 6 | 168605635 | 4 | 0 | 0 | A | G | -2.2531 | 0.5284 | 2.01E-05 | ---?- | 0 | 0.9151 | intergenic | |
| rs74684772 | 10 | 131470000 | 4 | 0 | 0 | T | C | 1.8473 | 0.4333 | 2.01E-05 | +++?+ | 0 | 0.8821 | intron | MGMT |
| rs72762272 | 15 | 87690493 | 3 | 0 | 0 | T | C | 3.5355 | 0.8301 | 2.05E-05 | +++?? | 0 | 0.9065 | intergenic | |
| rs76526967 | 6 | 106187225 | 4 | 0 | 1 | A | G | 3.3927 | 0.7967 | 2.06E-05 | +++?+ | 35.4 | 0.2 | intergenic | |
| rs9447903 | 6 | 77615816 | 4 | 0 | 0 | C | G | -1.1569 | 0.2717 | 2.07E-05 | ---?- | 0 | 0.4389 | intergenic | |
| rs28407322 | 6 | 32603355 | 5 | 0 | 1 | T | C | -1.1569 | 0.2718 | 2.07E-05 | ----- | 15 | 0.3191 | upstream-gene |  |
| rs10421259 | 19 | 46158860 | 5 | 0 | 1 | A | G | -0.9608 | 0.2257 | 2.07E-05 | ----+ | 0 | 0.7611 | intergenic |  |
| rs10420680 | 19 | 46158828 | 5 | 0 | 1 | A | G | 0.9607 | 0.2257 | 2.07E-05 | ++++- | 0 | 0.7612 | intergenic | |
| rs9447905 | 6 | 77616528 | 4 | 0 | 0 | C | G | 1.1566 | 0.2717 | 2.08E-05 | +++?+ | 0 | 0.4402 | intergenic | |
| rs10421206 | 19 | 46158786 | 5 | 0 | 1 | A | G | -0.9605 | 0.2257 | 2.08E-05 | ----+ | 0 | 0.7613 | intergenic |  |
| 9:119051495 | 9 | 119051495 | 3 | 0 | 0 | T | TGTAGC | 3.6367 | 0.8546 | 2.09E-05 | +++?? | 57.4 | 0.09549 | | |
| rs9447908 | 6 | 77619399 | 4 | 0 | 0 | A | G | 1.156 | 0.2717 | 2.10E-05 | +++?+ | 0 | 0.444 | intergenic | |
| 14:100095847 | 14 | 100095847 | 3 | 0 | 0 | A | AT | -1.1614 | 0.273 | 2.10E-05 | ---?? | 0 | 0.4865 | | |
| rs111766349 | 17 | 47350442 | 5 | 0 | 0 | A | T | 1.073 | 0.2523 | 2.11E-05 | ++++- | 4.8 | 0.3792 | intergenic | |
| rs1335435 | 6 | 77622102 | 4 | 0 | 0 | A | C | -1.1555 | 0.2717 | 2.11E-05 | ---?- | 0 | 0.4467 | intergenic | |
| 17:47434041 | 17 | 47434041 | 3 | 0 | 0 | T | TC | -1.3084 | 0.3077 | 2.11E-05 | ---?? | 0 | 0.9247 | | |
| rs1414189 | 6 | 77625266 | 4 | 0 | 0 | A | T | -1.1554 | 0.2717 | 2.12E-05 | ---?- | 0 | 0.4502 | intergenic | |
| rs9447911 | 6 | 77623867 | 4 | 0 | 0 | T | C | -1.1551 | 0.2717 | 2.12E-05 | ---?- | 0 | 0.4492 | intergenic | |
| rs9443323 | 6 | 77624386 | 4 | 0 | 0 | A | G | -1.155 | 0.2717 | 2.13E-05 | ---?- | 0 | 0.4495 | intergenic | |
| rs9443324 | 6 | 77624637 | 4 | 0 | 0 | T | G | -1.155 | 0.2717 | 2.13E-05 | ---?- | 0 | 0.4497 | intergenic | |
| rs4596448 | 6 | 77621749 | 4 | 0 | 0 | T | C | -1.155 | 0.2717 | 2.13E-05 | ---?- | 0 | 0.4472 | intergenic | |
| rs7772149 | 6 | 77623125 | 4 | 0 | 0 | A | G | -1.1552 | 0.2718 | 2.14E-05 | ---?- | 0 | 0.4467 | intergenic | |
| rs9443325 | 6 | 77627860 | 4 | 0 | 0 | T | C | -1.1545 | 0.2717 | 2.14E-05 | ---?- | 0 | 0.4526 | intergenic | |
| rs9443326 | 6 | 77628331 | 4 | 0 | 0 | A | C | -1.1544 | 0.2717 | 2.14E-05 | ---?- | 0 | 0.4534 | intergenic | |
| rs78072691 | 11 | 34365785 | 3 | 0 | 0 | T | G | 5.7581 | 1.3553 | 2.15E-05 | +++?? | 0 | 0.8399 | intron | ABTB2 |
| rs1335437 | 6 | 77629837 | 4 | 0 | 0 | T | G | 1.1539 | 0.2716 | 2.16E-05 | +++?+ | 0 | 0.4553 | intergenic | |
| rs7767954 | 6 | 77622777 | 4 | 0 | 0 | T | C | 1.1543 | 0.2717 | 2.16E-05 | +++?+ | 0 | 0.4449 | intergenic | |
| rs28558693 | 19 | 46161036 | 5 | 0 | 0 | A | T | -0.966 | 0.2276 | 2.19E-05 | ----- | 0 | 0.8483 | intergenic |  |
| rs145003984 | 2 | 238579272 | 3 | 0 | 0 | T | C | -5.358 | 1.2631 | 2.22E-05 | -?-?- | 0 | 0.4963 | intron | LRRFIP1 |
| rs114011447 | 2 | 238525915 | 3 | 0 | 0 | A | G | 4.4717 | 1.0545 | 2.23E-05 | +?+?+ | 0 | 0.5548 | intergenic | |
| rs9447919 | 6 | 77644459 | 4 | 0 | 0 | A | G | -1.1515 | 0.2716 | 2.24E-05 | ---?- | 0 | 0.4571 | intergenic | |
| rs1335438 | 6 | 77636017 | 4 | 0 | 0 | A | G | 1.1516 | 0.2717 | 2.24E-05 | +++?+ | 0 | 0.4588 | intergenic | |
| rs79239212 | 6 | 106187487 | 4 | 0 | 1 | A | T | -3.3681 | 0.7947 | 2.25E-05 | ---?- | 35.1 | 0.2018 | intergenic | |
| rs59255914 | 7 | 141157415 | 4 | 0 | 0 | T | C | 6.0894 | 1.437 | 2.26E-05 | +++?+ | 0 | 0.9765 | intron | TMEM178B |
| rs7748134 | 6 | 77638120 | 4 | 0 | 0 | A | T | -1.1511 | 0.2717 | 2.27E-05 | ---?- | 0 | 0.4596 | intergenic | |
| rs62048636 | 16 | 85215308 | 4 | 0 | 0 | T | C | 1.5122 | 0.3569 | 2.27E-05 | +++?+ | 0 | 0.7826 | intron | GSE1 |
| rs199681785 | 6 | 106167451 | 3 | 0 | 1 | A | G | 3.5432 | 0.8364 | 2.28E-05 | +++?? | 27.9 | 0.2498 | intergenic | |
| rs28675927 | 6 | 32603488 | 5 | 0 | 1 | A | G | -1.1525 | 0.2721 | 2.28E-05 | ----- | 16.1 | 0.3118 | upstream-gene |  |
| rs7074695 | 10 | 20509520 | 2 | 0 | 0 | T | C | 1.7989 | 0.4249 | 2.30E-05 | ???++ | 0 | 0.7701 | intron | PLXDC2 |
| rs2127222 | 8 | 76075619 | 4 | 0 | 0 | A | G | 1.2912 | 0.305 | 2.30E-05 | +++?+ | 0 | 0.487 | intergenic | |

| | | | | | | | | | | | | | | | |
|---|---|---|---|---|---|---|---|---|---|---|---|---|---|---|---|
| rs62031641 | 15 | 87693262 | 4 | 0 | 0 | A | T | -3.3166 | 0.7834 | 2.30E-05 | ---?- | 0 | 0.7106 | intergenic | |
| rs1999700 | 6 | 77658878 | 4 | 0 | 0 | T | C | -1.1512 | 0.2719 | 2.30E-05 | ---?- | 0 | 0.4688 | intergenic | |
| rs75557050 | 18 | 40106834 | 4 | 0 | 0 | T | C | -1.4154 | 0.3343 | 2.30E-05 | ---?+ | 22.8 | 0.2743 | intron | LINC00907 |
| rs2052004 | 16 | 5460330 | 4 | 0 | 0 | A | G | 1.2184 | 0.2879 | 2.31E-05 | +++?+ | 0 | 0.8178 | intron | RBFOX1 |
| rs114164410 | 2 | 238524877 | 3 | 0 | 0 | A | G | -4.6363 | 1.0958 | 2.33E-05 | -?-?- | 0 | 0.4449 | intergenic | |
| rs1932210 | 6 | 77643424 | 4 | 0 | 0 | A | T | 1.1498 | 0.2718 | 2.33E-05 | +++?+ | 0 | 0.4607 | intergenic | |
| rs1577429 | 6 | 77647177 | 4 | 0 | 0 | A | C | 1.1488 | 0.2715 | 2.33E-05 | +++?+ | 0 | 0.4596 | intergenic | |
| rs10943397 | 6 | 77658340 | 4 | 0 | 0 | A | G | -1.1498 | 0.2718 | 2.34E-05 | ---?- | 0 | 0.4679 | intergenic | |
| rs4708305 | 6 | 77647287 | 4 | 0 | 0 | T | C | -1.1486 | 0.2715 | 2.34E-05 | ---?- | 0 | 0.4597 | intergenic | |
| rs6925365 | 6 | 77658252 | 4 | 0 | 0 | T | G | 1.1496 | 0.2718 | 2.34E-05 | +++?+ | 0 | 0.4678 | intergenic | |
| rs2210774 | 6 | 77647451 | 4 | 0 | 0 | A | T | 1.1485 | 0.2715 | 2.34E-05 | +++?+ | 0 | 0.4599 | intergenic | |
| rs9443331 | 6 | 77657239 | 4 | 0 | 0 | T | C | 1.1491 | 0.2717 | 2.35E-05 | +++?+ | 0 | 0.4669 | intergenic | |
| rs72735355 | 9 | 101605241 | 4 | 0 | 0 | A | G | -3.3481 | 0.7918 | 2.35E-05 | ---?- | 0 | 0.4008 | intron | GALNT12 |
| rs7755762 | 6 | 77657092 | 4 | 0 | 0 | A | G | 1.149 | 0.2717 | 2.35E-05 | +++?+ | 0 | 0.4667 | intergenic | |
| rs9447929 | 6 | 77657360 | 4 | 0 | 0 | A | T | 1.1491 | 0.2718 | 2.36E-05 | +++?+ | 0 | 0.4671 | intergenic | |
| rs77816582 | 4 | 140544823 | 4 | 0 | 0 | A | C | 2.5076 | 0.5931 | 2.36E-05 | +++?+ | 0 | 0.7444 | intergenic | |
| rs9447930 | 6 | 77657681 | 4 | 0 | 0 | A | C | 1.1491 | 0.2718 | 2.36E-05 | +++?+ | 0 | 0.4675 | intergenic | |
| rs7755032 | 6 | 77656898 | 4 | 0 | 0 | A | C | -1.1489 | 0.2717 | 2.36E-05 | ---?- | 0 | 0.4664 | intergenic | |
| rs6453952 | 6 | 77648771 | 4 | 0 | 0 | C | G | 1.1478 | 0.2715 | 2.36E-05 | +++?+ | 0 | 0.4605 | intergenic | |
| rs7771149 | 6 | 77656619 | 4 | 0 | 0 | T | C | -1.1486 | 0.2717 | 2.36E-05 | ---?- | 0 | 0.4662 | intergenic | |
| rs1577424 | 6 | 77658024 | 4 | 0 | 0 | T | C | 1.1491 | 0.2718 | 2.37E-05 | +++?+ | 0 | 0.4683 | intergenic | |
| rs1591436 | 6 | 77649041 | 4 | 0 | 0 | A | G | -1.1476 | 0.2715 | 2.37E-05 | ---?- | 0 | 0.4607 | intergenic | |
| rs28431396 | 7 | 153662454 | 4 | 0 | 0 | T | C | -1.5457 | 0.3657 | 2.37E-05 | ---?- | 0 | 0.5509 | intron | DPP6 |
| rs9447927 | 6 | 77656082 | 4 | 0 | 0 | T | C | -1.1482 | 0.2716 | 2.37E-05 | ---?- | 0 | 0.4651 | intergenic | |
| rs9443327 | 6 | 77649924 | 4 | 0 | 0 | T | G | -1.1472 | 0.2714 | 2.38E-05 | ---?- | 0 | 0.4611 | intergenic | |
| rs9443328 | 6 | 77650689 | 4 | 0 | 0 | A | G | -1.1473 | 0.2715 | 2.38E-05 | ---?- | 0 | 0.4605 | intergenic | |
| rs9447924 | 6 | 77655105 | 4 | 0 | 0 | A | G | 1.1476 | 0.2716 | 2.38E-05 | +++?+ | 0 | 0.4644 | intergenic | |
| rs9447921 | 6 | 77650274 | 4 | 0 | 0 | A | T | -1.1469 | 0.2714 | 2.39E-05 | ---?- | 0 | 0.4609 | intergenic | |
| rs11735872 | 4 | 190010868 | 4 | 0 | 0 | A | G | 1.0129 | 0.2398 | 2.40E-05 | +++?+ | 0 | 0.5806 | intergenic | |
| rs13194552 | 6 | 77651025 | 4 | 0 | 0 | T | C | -1.1464 | 0.2714 | 2.40E-05 | ---?- | 0 | 0.4611 | intergenic | |
| rs9447922 | 6 | 77653724 | 4 | 0 | 0 | A | C | 1.1468 | 0.2715 | 2.40E-05 | +++?+ | 0 | 0.4631 | intergenic | |
| rs9447923 | 6 | 77654813 | 4 | 0 | 0 | A | G | 1.147 | 0.2716 | 2.41E-05 | +++?+ | 0 | 0.4646 | intergenic | |
| rs4257828 | 6 | 77651478 | 4 | 0 | 0 | T | C | -1.1462 | 0.2714 | 2.41E-05 | ---?- | 0 | 0.4611 | intergenic | |
| rs34004019 | 6 | 32626403 | 4 | 0 | 1 | A | G | -1.0922 | 0.2586 | 2.41E-05 | ----? | 5.4 | 0.366 | downstream-gene |  |
| rs4510626 | 6 | 77651906 | 4 | 0 | 0 | A | C | 1.1461 | 0.2714 | 2.41E-05 | +++?+ | 0 | 0.461 | intergenic | |
| rs76743140 | 2 | 238524492 | 3 | 0 | 0 | T | C | -4.6535 | 1.1021 | 2.42E-05 | -?-?- | 0 | 0.4429 | intergenic | |
| rs4472308 | 6 | 77652140 | 4 | 0 | 0 | A | G | -1.1459 | 0.2714 | 2.42E-05 | ---?- | 0 | 0.4614 | intergenic | |
| rs4426969 | 6 | 77652131 | 4 | 0 | 0 | T | C | -1.1456 | 0.2714 | 2.43E-05 | ---?- | 0 | 0.4611 | intergenic | |
| rs112792621 | 17 | 47349948 | 5 | 0 | 0 | A | G | -1.0645 | 0.2522 | 2.44E-05 | ----+ | 7.4 | 0.3645 | intergenic |  |
| rs299372 | 5 | 134340030 | 4 | 0 | 0 | A | G | -1.1397 | 0.2701 | 2.45E-05 | ---?- | 0 | 0.7742 | intron | CATSPER3 |
| rs13422023 | 2 | 238526178 | 4 | 0 | 0 | A | C | -3.9864 | 0.9448 | 2.45E-05 | ---?- | 0 | 0.5415 | intergenic | |
| rs13386886 | 2 | 238526071 | 4 | 0 | 0 | T | C | 3.986 | 0.9448 | 2.46E-05 | +++?+ | 0 | 0.5417 | intergenic | |
| 6:106150903 | 6 | 106150903 | 3 | 0 | 0 | CT | C | 3.4734 | 0.8235 | 2.47E-05 | +++?? | 19 | 0.2909 | | |
| rs114851365 | 2 | 238524269 | 3 | 0 | 0 | A | T | 4.6636 | 1.1061 | 2.48E-05 | +?+?+ | 0 | 0.4422 | intergenic | |
| 7:155151434 | 7 | 155151434 | 2 | 0 | 0 | T | C | 6.5381 | 1.5512 | 2.50E-05 | ++??? | 0 | 0.487 | non-coding-exon | BLACE |
| 7:35942450 | 7 | 35942450 | 3 | 0 | 0 | CT | C | 4.8653 | 1.1544 | 2.50E-05 | +++?? | 35.5 | 0.212 | | |
| rs7587286 | 2 | 218359846 | 4 | 0 | 0 | T | C | -1.0679 | 0.2535 | 2.52E-05 | ---?- | 0 | 0.6347 | intron | DIRC3 |
| rs74399905 | 4 | 100549105 | 4 | 0 | 0 | A | T | -1.7469 | 0.4146 | 2.52E-05 | ---?- | 36.9 | 0.1904 | downstream-gene | |
| rs17126437 | 1 | 64811160 | 4 | 0 | 0 | T | G | 1.5693 | 0.3725 | 2.52E-05 | +++?+ | 0 | 0.5379 | intergenic | |
| 11:106764191 | 11 | 106764191 | 3 | 0 | 0 | A | AAT | -1.2886 | 0.3059 | 2.52E-05 | ---?? | 0 | 0.5065 | | |
| rs16976160 | 18 | 40040474 | 4 | 0 | 0 | A | G | -1.413 | 0.3354 | 2.53E-05 | ---?+ | 0 | 0.4356 | intron | LINC00907 |
| rs756872 | 7 | 107384849 | 4 | 0 | 0 | T | C | -1.0164 | 0.2414 | 2.56E-05 | ---?- | 17.3 | 0.3048 | intron | CBLL1 |
| rs11129148 | 3 | 2486763 | 4 | 0 | 0 | T | C | -1.1596 | 0.2755 | 2.56E-05 | ---?+ | 18 | 0.3008 | intron | CNTN4 |
| 17:47324505 | 17 | 47324505 | 3 | 0 | 0 | CGCCAG | C | 1.2819 | 0.3046 | 2.57E-05 | +++?? | 0 | 0.6525 | | |
| rs951915 | 15 | 98044012 | 4 | 0 | 0 | T | C | 1.1469 | 0.2726 | 2.59E-05 | +++?+ | 0 | 0.8235 | intergenic | |
| rs12447061 | 16 | 5460977 | 4 | 0 | 0 | A | G | -1.2022 | 0.2859 | 2.61E-05 | ---?- | 0 | 0.8053 | intron | RBFOX1 |
| rs6509231 | 19 | 46158293 | 5 | 0 | 1 | A | G | 0.9552 | 0.2272 | 2.62E-05 | ++++- | 0 | 0.7712 | intergenic | |
| rs60670643 | 7 | 107391322 | 4 | 0 | 0 | A | C | -1.0334 | 0.2458 | 2.62E-05 | ---?- | 8.8 | 0.3489 | intron | CBLL1 |
| rs112011161 | 3 | 126358691 | 4 | 0 | 0 | T | C | 3.0994 | 0.7374 | 2.63E-05 | +++?+ | 0 | 0.8481 | intron | TXNRD3 |

| | | | | | | | | | | | | | | | |
|---|---|---|---|---|---|---|---|---|---|---|---|---|---|---|---|
| rs2395887 | 7 | 107385825 | 4 | 0 | 0 | T | G | -1.0135 | 0.2412 | 2.64E-05 | ---?- | 16.6 | 0.3086 | intron | CBLL1 |
| 10:28197685 | 10 | 28197685 | 3 | 0 | 0 | CAGGGC | C | -1.0371 | 0.2468 | 2.65E-05 | ---?? | 0 | 0.5181 | | |
| rs7573386 | 2 | 77090914 | 3 | 0 | 0 | A | G | -5.1055 | 1.2156 | 2.67E-05 | ---?? | 43.5 | 0.1702 | intron | LRRTM4 |
| rs10255946 | 7 | 107387595 | 4 | 0 | 0 | T | C | -1.0193 | 0.2428 | 2.68E-05 | ---?- | 14 | 0.3222 | intron | CBLL1 |
| rs10274201 | 7 | 107386810 | 4 | 0 | 0 | T | C | -1.0153 | 0.2418 | 2.68E-05 | ---?- | 14.9 | 0.3175 | intron | CBLL1 |
| rs114655864 | 1 | 18360590 | 2 | 0 | 0 | T | C | 8.9114 | 2.1242 | 2.73E-05 | +???+ | 0 | 0.4816 | intergenic | |
| rs2023711 | 7 | 107390207 | 4 | 0 | 0 | T | C | -1.0242 | 0.2443 | 2.77E-05 | ---?- | 10.9 | 0.3385 | intron | CBLL1 |
| rs10225292 | 7 | 156173238 | 4 | 0 | 0 | T | C | -1.0341 | 0.2467 | 2.77E-05 | ---?- | 22.7 | 0.2747 | intergenic | |
| rs11073559 | 15 | 98044496 | 4 | 0 | 0 | T | G | 1.1431 | 0.2728 | 2.79E-05 | +++?+ | 0 | 0.8211 | intergenic | |
| rs34677551 | 6 | 32663936 | 5 | 0 | 1 | A | C | -1.6379 | 0.3909 | 2.79E-05 | ----- | 3.1 | 0.3887 | intergenic |  |
| 8:76146017 | 8 | 76146017 | 3 | 0 | 0 | CAT | C | -1.3203 | 0.3152 | 2.80E-05 | ---?? | 20.1 | 0.286 | | |
| rs10274831 | 7 | 156175708 | 4 | 0 | 0 | A | C | 1.0376 | 0.2478 | 2.81E-05 | +++?+ | 22.5 | 0.2755 | intergenic | |
| rs1442655 | 18 | 40025574 | 4 | 0 | 0 | A | C | -1.4042 | 0.3353 | 2.81E-05 | ---?+ | 0 | 0.4483 | intron | LINC00907 |
| rs537576 | 16 | 5461604 | 4 | 0 | 0 | T | C | 1.2183 | 0.291 | 2.83E-05 | +++?+ | 0 | 0.7887 | intron | RBFOX1 |
| rs36186647 | 6 | 32666431 | 5 | 0 | 1 | T | C | 1.5963 | 0.3813 | 2.83E-05 | +++++ | 6.2 | 0.3712 | intergenic | |
| rs12606301 | 18 | 40023782 | 4 | 0 | 0 | A | G | -1.4024 | 0.3352 | 2.87E-05 | ---?+ | 0 | 0.4511 | intron | LINC00907 |
| rs4552541 | 4 | 117281499 | 4 | 0 | 0 | A | G | -3.5516 | 0.849 | 2.88E-05 | ---?- | 23.1 | 0.2724 | intergenic | |
| rs13178761 | 5 | 173144887 | 4 | 0 | 1 | T | C | 1.0592 | 0.2533 | 2.89E-05 | +++?+ | 20.9 | 0.2845 | intergenic | |
| rs59269598 | 15 | 85513033 | 4 | 0 | 0 | T | C | 1.034 | 0.2473 | 2.89E-05 | +++?- | 54.8 | 0.08464 | intergenic | |
| rs34593937 | 6 | 32666271 | 5 | 0 | 1 | T | C | -1.5917 | 0.3807 | 2.90E-05 | ----- | 7.3 | 0.3653 | intergenic |  |
| rs35836258 | 6 | 32666296 | 5 | 0 | 1 | A | G | 1.5917 | 0.3807 | 2.90E-05 | +++++ | 7.3 | 0.3653 | intergenic | |
| rs34389498 | 6 | 32666348 | 5 | 0 | 1 | T | C | -1.5917 | 0.3807 | 2.90E-05 | ----- | 7.3 | 0.3653 | intergenic |  |
| rs148539701 | 19 | 49115774 | 4 | 0 | 0 | T | G | 1.7924 | 0.4288 | 2.91E-05 | +++?+ | 0 | 0.8972 | intron | FAM83E |
| rs13194877 | 6 | 32655548 | 5 | 0 | 1 | A | G | 1.5892 | 0.3802 | 2.92E-05 | +++++ | 6 | 0.3723 | intergenic | |
| rs113476886 | 17 | 47348147 | 5 | 0 | 0 | T | G | 1.0552 | 0.2525 | 2.92E-05 | ++++- | 10.6 | 0.3458 | intergenic | |
| rs62076392 | 17 | 47347674 | 5 | 0 | 0 | T | G | -1.055 | 0.2524 | 2.92E-05 | ----+ | 10.7 | 0.3448 | intergenic |  |
| rs62076391 | 17 | 47347607 | 5 | 0 | 0 | A | G | -1.0549 | 0.2524 | 2.92E-05 | ----+ | 10.8 | 0.3447 | intergenic |  |
| rs34475663 | 6 | 32666563 | 5 | 0 | 1 | A | T | 1.591 | 0.3807 | 2.93E-05 | +++++ | 7.5 | 0.3638 | intergenic | |
| rs140713539 | 6 | 99747959 | 4 | 0 | 0 | C | G | -3.568 | 0.8539 | 2.93E-05 | ---?- | 0 | 0.977 | intron | FAXC |
| rs12528368 | 6 | 32653643 | 5 | 0 | 1 | T | C | -1.5899 | 0.3806 | 2.94E-05 | ----- | 7 | 0.3666 | intergenic |  |
| rs217667 | 14 | 62358847 | 4 | 0 | 0 | T | C | -1.2531 | 0.3 | 2.96E-05 | ---?- | 0 | 0.693 | intron | SYT16 |
| rs17212525 | 6 | 32665539 | 5 | 0 | 1 | T | C | -1.5959 | 0.3822 | 2.97E-05 | ----- | 7.9 | 0.3614 | intergenic |  |
| rs12526612 | 6 | 32653598 | 5 | 0 | 1 | A | G | 1.5882 | 0.3805 | 3.00E-05 | +++++ | 7.3 | 0.3652 | intergenic | |
| 15:85559615 | 15 | 85559615 | 3 | 0 | 0 | A | AT | 1.0357 | 0.2482 | 3.01E-05 | +++?? | 62.4 | 0.06994 | | |
| rs17212518 | 6 | 32665475 | 5 | 0 | 1 | A | C | -1.5916 | 0.3816 | 3.04E-05 | ----- | 6.3 | 0.3707 | intergenic |  |
| rs77267725 | 8 | 86248605 | 4 | 0 | 0 | A | C | -2.1544 | 0.5168 | 3.06E-05 | ---?- | 43.3 | 0.152 | intron | CA1 |
| 10:131469694 | 10 | 131469694 | 3 | 0 | 0 | CT | C | -1.7167 | 0.4118 | 3.07E-05 | ---?? | 39.6 | 0.1912 | | |
| 13:45630983 | 13 | 45630983 | 3 | 0 | 0 | CGGAGT | C | -2.2314 | 0.5356 | 3.09E-05 | ---?? | 0 | 0.6036 | | |
| rs6842339 | 4 | 100547579 | 4 | 0 | 0 | T | G | 1.7222 | 0.4135 | 3.12E-05 | +++?+ | 42.9 | 0.154 | downstream-gene | |
| rs112048442 | 4 | 100547715 | 4 | 0 | 0 | A | G | -1.7224 | 0.4136 | 3.12E-05 | ---?- | 42.9 | 0.1542 | downstream-gene | |
| 7:130135619 | 7 | 130135619 | 3 | 0 | 0 | G | GTTTTT | 1.253 | 0.3009 | 3.12E-05 | +++?? | 55.5 | 0.1059 | | |
| rs4515336 | 6 | 77612396 | 3 | 0 | 0 | C | G | 1.1689 | 0.2807 | 3.13E-05 | +++?? | 27.1 | 0.2536 | intergenic | |
| rs9447894 | 6 | 77605806 | 3 | 0 | 0 | A | G | 1.1679 | 0.2805 | 3.14E-05 | +++?? | 28.4 | 0.2472 | intergenic | |
| rs11663061 | 18 | 47752648 | 4 | 0 | 0 | T | C | 1.292 | 0.3104 | 3.15E-05 | +++?+ | 0.4 | 0.3898 | downstream-gene | |
| rs79293902 | 4 | 100548163 | 4 | 0 | 0 | T | C | -1.7225 | 0.4139 | 3.16E-05 | ---?- | 42.7 | 0.1552 | downstream-gene | |
| rs11670462 | 19 | 46148386 | 5 | 0 | 1 | T | C | -0.9555 | 0.2296 | 3.16E-05 | ----+ | 0 | 0.5074 | intron | EML2 |
| rs61684991 | 14 | 28206082 | 4 | 0 | 0 | T | C | 1.8596 | 0.4469 | 3.17E-05 | +++?+ | 0 | 0.7412 | intergenic | |
| rs17099462 | 11 | 102600706 | 5 | 0 | 0 | T | G | 0.9043 | 0.2174 | 3.19E-05 | +++++ | 20.2 | 0.2857 | upstream-gene | |
| rs4708297 | 6 | 77614293 | 3 | 0 | 0 | T | C | 1.1673 | 0.2807 | 3.21E-05 | +++?? | 26.4 | 0.2568 | intergenic | |
| 15:98044155 | 15 | 98044155 | 3 | 0 | 0 | G | GA | 1.1639 | 0.2799 | 3.21E-05 | +++?? | 0 | 0.6614 | | |
| rs72642413 | 18 | 40049909 | 4 | 0 | 0 | A | G | 1.3972 | 0.3361 | 3.22E-05 | +++?- | 0 | 0.4557 | intron | LINC00907 |
| rs56364844 | 15 | 91586112 | 4 | 0 | 0 | T | C | -2.2497 | 0.5412 | 3.22E-05 | ---?- | 0 | 0.5601 | intergenic | |
| rs115225609 | 6 | 32599272 | 5 | 0 | 0 | A | G | -1.1182 | 0.269 | 3.23E-05 | ----- | 9.4 | 0.3528 | intergenic |  |
| rs78117460 | 6 | 32599337 | 5 | 0 | 0 | T | C | 1.1174 | 0.2688 | 3.23E-05 | +++++ | 9.6 | 0.3518 | intergenic | |
| rs1335434 | 6 | 77616335 | 3 | 0 | 0 | T | C | 1.1669 | 0.2808 | 3.24E-05 | +++?? | 25.6 | 0.2608 | intergenic | |
| rs3731674 | 2 | 170668813 | 5 | 0 | 0 | T | C | 0.9448 | 0.2274 | 3.25E-05 | +++++ | 57.5 | 0.05177 | intron | METTL5 |
| 2:170513008 | 2 | 170513008 | 3 | 0 | 0 | GGGA | GGGAGC | -1.3823 | 0.3327 | 3.26E-05 | ---?? | 0 | 0.9345 | | |
| rs72691350 | 4 | 161047853 | 4 | 0 | 0 | A | G | -4.1617 | 1.0018 | 3.27E-05 | ---?- | 0 | 0.5354 | intergenic | |
| rs10282626 | 7 | 141168003 | 4 | 0 | 0 | A | G | -5.5592 | 1.3385 | 3.28E-05 | ---?- | 0 | 0.8326 | intron | TMEM178B |

| | | | | | | | | | | | | | | | |
|---|---|---|---|---|---|---|---|---|---|---|---|---|---|---|---|
| rs35282906 | 6 | 32657201 | 5 | 0 | 1 | A | T | 1.5753 | 0.3793 | 3.28E-05 | +++++ | 9 | 0.3553 | intergenic | |
| rs60601875 | 7 | 107391310 | 3 | 0 | 0 | A | T | 1.0556 | 0.2542 | 3.28E-05 | +++?? | 37.2 | 0.2033 | intron | CBLL1 |
| 7:107391311 | 7 | 107391311 | 3 | 0 | 0 | G | GACC | -1.0549 | 0.2541 | 3.30E-05 | ---?? | 36.9 | 0.2049 | | |
| rs10516447 | 4 | 100551372 | 4 | 0 | 0 | A | G | 1.7244 | 0.4154 | 3.30E-05 | +++?+ | 42.1 | 0.1587 | intergenic | |
| 6:106170180 | 6 | 106170180 | 3 | 0 | 0 | G | GA | 3.4489 | 0.8309 | 3.31E-05 | +++?? | 24.2 | 0.2673 | | |
| rs35740559 | 6 | 32666132 | 5 | 0 | 1 | T | C | 1.6186 | 0.39 | 3.32E-05 | +++++ | 8.8 | 0.3564 | intergenic | |
| 17:47379654 | 17 | 47379654 | 3 | 0 | 0 | A | AT | 1.3178 | 0.3175 | 3.32E-05 | +++?? | 0 | 0.7883 | | |
| rs7450344 | 6 | 77607688 | 4 | 0 | 0 | T | G | 1.134 | 0.2732 | 3.32E-05 | +++?+ | 13.2 | 0.3267 | intergenic | |
| 6:37308748 | 6 | 37308748 | 3 | 0 | 0 | A | AT | 1.1044 | 0.2661 | 3.32E-05 | +++?? | 0 | 0.4267 | | |
| 5:173154706 | 5 | 173154706 | 3 | 0 | 0 | G | GA | 1.1492 | 0.2769 | 3.33E-05 | +++?? | 3.2 | 0.3559 | | |
| rs6844547 | 4 | 100551756 | 4 | 0 | 0 | T | C | -1.7243 | 0.4155 | 3.33E-05 | ---?- | 42 | 0.1594 | intergenic | |
| 7:107391318 | 7 | 107391318 | 3 | 0 | 0 | T | TGG | -1.0547 | 0.2542 | 3.33E-05 | ---?? | 37.1 | 0.204 | | |
| 7:107391316 | 7 | 107391316 | 3 | 0 | 0 | T | TA | -1.0545 | 0.2541 | 3.33E-05 | ---?? | 37.2 | 0.2036 | | |
| rs6457614 | 6 | 32651900 | 5 | 0 | 1 | T | G | -1.5865 | 0.3824 | 3.34E-05 | ----- | 8.1 | 0.3605 | intergenic |  |
| rs6845301 | 4 | 100551902 | 4 | 0 | 0 | A | G | -1.7242 | 0.4156 | 3.34E-05 | ---?- | 42 | 0.1596 | intergenic | |
| 7:107358829 | 7 | 107358829 | 3 | 0 | 0 | G | GT | 1.0762 | 0.2594 | 3.35E-05 | +++?? | 26.2 | 0.2577 | | |
| rs3998180 | 6 | 32650897 | 5 | 0 | 1 | A | C | 1.5887 | 0.3831 | 3.37E-05 | +++++ | 8.1 | 0.3606 | intergenic | |
| 7:107387366 | 7 | 107387366 | 3 | 0 | 0 | T | TA | 1.0387 | 0.2505 | 3.38E-05 | +++?? | 41.3 | 0.1821 | | |
| 8:135156885 | 8 | 135156885 | 3 | 0 | 0 | T | C | 4.5515 | 1.0981 | 3.40E-05 | +++?? | 30.8 | 0.2359 | intergenic | |
| rs10829615 | 10 | 131471179 | 4 | 0 | 0 | T | C | 1.5496 | 0.3741 | 3.45E-05 | +++?+ | 0 | 0.469 | intron | MGMT |
| rs12907910 | 15 | 91588698 | 4 | 0 | 0 | T | G | -1.6829 | 0.4063 | 3.45E-05 | ---?- | 0 | 0.9935 | intergenic | |
| rs34388057 | 6 | 32675951 | 5 | 0 | 1 | A | T | -1.5801 | 0.3816 | 3.46E-05 | ----- | 7.3 | 0.3651 | intergenic |  |
| 8:85638889 | 8 | 85638889 | 3 | 0 | 0 | A | ATTACT( | -2.6717 | 0.6452 | 3.46E-05 | ---?? | 0 | 0.7733 | | |
| rs10947332 | 6 | 32677440 | 5 | 0 | 1 | A | G | 1.58 | 0.3816 | 3.46E-05 | +++++ | 7.3 | 0.3651 | intergenic | |
| rs17206378 | 6 | 32675161 | 5 | 0 | 1 | T | G | 1.5799 | 0.3816 | 3.47E-05 | +++++ | 7.3 | 0.365 | intergenic | |
| rs111944102 | 6 | 32677818 | 5 | 0 | 1 | A | T | 1.5799 | 0.3816 | 3.47E-05 | +++++ | 7.3 | 0.3652 | intergenic | |
| rs35472821 | 6 | 32674529 | 5 | 0 | 1 | C | G | 1.5799 | 0.3816 | 3.47E-05 | +++++ | 7.3 | 0.3649 | intergenic | |
| rs145281025 | 6 | 32674252 | 5 | 0 | 1 | A | G | 1.5798 | 0.3816 | 3.47E-05 | +++++ | 7.3 | 0.3649 | intergenic | |
| 13:45631455 | 13 | 45631455 | 3 | 0 | 0 | G | GT | 2.2177 | 0.5357 | 3.47E-05 | +++?? | 0 | 0.6055 | | |
| rs9442770 | 6 | 73048415 | 4 | 0 | 0 | A | G | 1.6856 | 0.4072 | 3.48E-05 | +++?+ | 52.8 | 0.09543 | intron | RIMS1 |
| rs77651188 | 12 | 91532271 | 3 | 0 | 0 | T | C | -5.0355 | 1.2165 | 3.49E-05 | ---?? | 0 | 0.9385 | intergenic | |
| 7:107386855 | 7 | 107386855 | 3 | 0 | 0 | A | AT | -1.0356 | 0.2502 | 3.50E-05 | ---?? | 42.2 | 0.1774 | | |
| rs79685911 | 6 | 32679053 | 5 | 0 | 0 | A | T | 1.5825 | 0.3824 | 3.50E-05 | +++++ | 9.5 | 0.3523 | intergenic | |
| 1:103097544 | 1 | 103097544 | 3 | 0 | 0 | A | T | -4.7265 | 1.1424 | 3.51E-05 | ---?? | 59.2 | 0.0864 | intergenic | |
| rs62433561 | 6 | 77721102 | 4 | 0 | 0 | C | G | 1.2346 | 0.2985 | 3.53E-05 | +++?+ | 21 | 0.2841 | intergenic | |
| rs12535876 | 7 | 156165232 | 4 | 0 | 0 | T | C | 1.0655 | 0.2576 | 3.53E-05 | +++?+ | 5.2 | 0.3669 | intergenic | |
| rs12530545 | 7 | 156165191 | 4 | 0 | 0 | C | G | 1.0654 | 0.2576 | 3.54E-05 | +++?+ | 5.3 | 0.3666 | intergenic | |
| 6:83854840 | 6 | 83854840 | 3 | 0 | 0 | T | TTG | -4.4807 | 1.0834 | 3.54E-05 | ---?? | 37.5 | 0.202 | | |
| rs17212483 | 6 | 32665250 | 5 | 0 | 1 | T | C | 1.5707 | 0.3798 | 3.54E-05 | +++++ | 12.6 | 0.3338 | intergenic | |
| rs17212504 | 6 | 32665369 | 5 | 0 | 1 | A | G | -1.5707 | 0.3798 | 3.54E-05 | ----- | 12.6 | 0.3338 | intergenic |  |
| rs372244887 | 15 | 93873158 | 3 | 0 | 0 | C | G | 1.4318 | 0.3463 | 3.55E-05 | +++?? | 32.1 | 0.2292 | intergenic | |
| rs11857301 | 15 | 42086382 | 5 | 0 | 0 | A | G | -1.2584 | 0.3044 | 3.56E-05 | ----- | 32.4 | 0.2056 | intron | MAPKBP1 |
| rs1572995 | 1 | 205024012 | 5 | 0 | 0 | T | C | 1.2438 | 0.3009 | 3.56E-05 | +++++ | 31.1 | 0.2141 | intron | CNTN2 |
| rs62480450 | 7 | 156167239 | 4 | 0 | 0 | A | G | -1.0637 | 0.2573 | 3.57E-05 | ---?- | 5.1 | 0.3675 | intergenic | |
| 8:76088879 | 8 | 76088879 | 3 | 0 | 0 | CA | C | -1.2437 | 0.3009 | 3.59E-05 | ---?? | 43.5 | 0.1705 | | |
| rs6922148 | 6 | 77615656 | 4 | 0 | 0 | T | C | 1.1313 | 0.2738 | 3.60E-05 | +++?+ | 9.2 | 0.3469 | intergenic | |
| rs115255544 | 6 | 32664628 | 5 | 0 | 1 | A | G | -1.567 | 0.3792 | 3.60E-05 | ----- | 10.9 | 0.3437 | intergenic |  |
| rs114361720 | 6 | 32664635 | 5 | 0 | 1 | C | G | 1.5669 | 0.3792 | 3.60E-05 | +++++ | 10.9 | 0.3437 | intergenic | |
| rs72844343 | 6 | 32660993 | 5 | 0 | 1 | A | T | -1.5669 | 0.3793 | 3.61E-05 | ----- | 11 | 0.3434 | intergenic |  |
| rs72844311 | 6 | 32659536 | 5 | 0 | 1 | T | C | 1.5667 | 0.3793 | 3.61E-05 | +++++ | 11 | 0.3433 | intergenic | |
| rs17206119 | 6 | 32665104 | 5 | 0 | 1 | A | T | 1.5667 | 0.3792 | 3.61E-05 | +++++ | 11 | 0.3433 | intergenic | |
| rs17212462 | 6 | 32665125 | 5 | 0 | 1 | T | C | 1.5667 | 0.3792 | 3.61E-05 | +++++ | 11 | 0.3433 | intergenic | |
| rs35472814 | 6 | 32659897 | 5 | 0 | 1 | T | C | -1.5667 | 0.3793 | 3.61E-05 | ----- | 11 | 0.3432 | intergenic |  |
| 6:77658933 | 6 | 77658933 | 3 | 0 | 0 | T | TAATA | -1.1605 | 0.2809 | 3.61E-05 | ---?? | 20.6 | 0.2839 | | |
| rs17212469 | 6 | 32665153 | 5 | 0 | 1 | T | G | 1.5665 | 0.3792 | 3.62E-05 | +++++ | 11 | 0.3431 | intergenic | |
| rs114169440 | 6 | 32662921 | 5 | 0 | 1 | A | C | 1.5665 | 0.3792 | 3.62E-05 | +++++ | 11 | 0.3431 | intergenic | |
| rs6923022 | 6 | 32664289 | 5 | 0 | 1 | T | G | 1.5665 | 0.3792 | 3.62E-05 | +++++ | 11 | 0.3431 | intergenic | |
| rs10484561 | 6 | 32665420 | 5 | 0 | 1 | T | G | -1.5665 | 0.3792 | 3.62E-05 | ----- | 11 | 0.3431 | intergenic |  |
| rs17427564 | 6 | 32667067 | 5 | 0 | 1 | T | C | -1.5665 | 0.3792 | 3.62E-05 | ----- | 11 | 0.3431 | intergenic |  |

| | | | | | | | | | | | | | | | |
|---|---|---|---|---|---|---|---|---|---|---|---|---|---|---|---|
| rs34492195 | 6 | 32670833 | 5 | 0 | 1 | A | G | 1.5665 | 0.3792 | 3.62E-05 | +++++ | 11 | 0.3431 | intergenic | |
| rs35900855 | 6 | 32664038 | 5 | 0 | 1 | A | G | -1.5665 | 0.3792 | 3.62E-05 | ----- | 11 | 0.3431 | intergenic |  |
| rs66490218 | 6 | 32668105 | 5 | 0 | 1 | A | G | 1.5665 | 0.3792 | 3.62E-05 | +++++ | 11 | 0.3431 | intergenic | |
| rs71202902 | 6 | 32667175 | 5 | 0 | 1 | T | C | 1.5665 | 0.3792 | 3.62E-05 | +++++ | 11 | 0.3431 | intergenic | |
| rs72847158 | 6 | 32664708 | 5 | 0 | 1 | T | C | -1.5664 | 0.3792 | 3.62E-05 | ----- | 11 | 0.3429 | intergenic |  |
| rs72847174 | 6 | 32668026 | 5 | 0 | 1 | T | C | 1.5665 | 0.3792 | 3.62E-05 | +++++ | 11 | 0.3431 | intergenic | |
| rs78825896 | 6 | 32667181 | 5 | 0 | 1 | T | G | -1.5665 | 0.3792 | 3.62E-05 | ----- | 11 | 0.3431 | intergenic |  |
| rs35169534 | 6 | 32663791 | 5 | 0 | 1 | A | G | -1.5665 | 0.3793 | 3.62E-05 | ----- | 11 | 0.343 | intergenic |  |
| rs35473146 | 6 | 32663713 | 5 | 0 | 1 | A | G | -1.5665 | 0.3792 | 3.62E-05 | ----- | 11 | 0.3431 | intergenic |  |
| rs35998847 | 6 | 32666997 | 5 | 0 | 1 | A | G | 1.5665 | 0.3792 | 3.62E-05 | +++++ | 11 | 0.343 | intergenic | |
| rs16888213 | 6 | 77600686 | 4 | 0 | 0 | A | C | 1.1297 | 0.2735 | 3.62E-05 | +++?+ | 11.7 | 0.3342 | intergenic | |
| rs17427445 | 6 | 32663764 | 5 | 0 | 1 | A | G | 1.5664 | 0.3793 | 3.63E-05 | +++++ | 10.9 | 0.3436 | intergenic | |
| rs10808033 | 7 | 145171914 | 4 | 0 | 0 | T | C | 1.3029 | 0.3155 | 3.64E-05 | +++?+ | 0 | 0.507 | intergenic | |
| rs72691338 | 4 | 161022208 | 4 | 0 | 0 | T | C | -3.8951 | 0.9435 | 3.66E-05 | ---?- | 0 | 0.408 | intergenic | |
| rs1346987 | 18 | 40102641 | 4 | 0 | 0 | A | C | -1.3786 | 0.3341 | 3.69E-05 | ---?+ | 21.9 | 0.2794 | intron | LINC00907 |
| rs1870894 | 6 | 77600134 | 4 | 0 | 0 | T | C | -1.1287 | 0.2736 | 3.69E-05 | ---?- | 11.7 | 0.3341 | intergenic | |
| rs7001087 | 8 | 76089347 | 4 | 0 | 0 | A | C | 1.1459 | 0.2777 | 3.69E-05 | +++?+ | 0 | 0.5261 | intergenic | |
| 6:77648613 | 6 | 77648613 | 3 | 0 | 0 | CAT | C | -1.157 | 0.2805 | 3.70E-05 | ---?? | 22.1 | 0.277 | | |
| rs35986284 | 6 | 77656347 | 3 | 0 | 0 | A | C | -1.1576 | 0.2807 | 3.72E-05 | ---?? | 21.1 | 0.2815 | intergenic | |
| rs12995559 | 2 | 170631368 | 4 | 0 | 0 | A | G | -1.0032 | 0.2433 | 3.73E-05 | ---?- | 0 | 0.7891 | intergenic | |
| rs4708306 | 6 | 77655637 | 3 | 0 | 0 | A | G | -1.1569 | 0.2806 | 3.73E-05 | ---?? | 21.4 | 0.2803 | intergenic | |
| 7:107378746 | 7 | 107378746 | 3 | 0 | 0 | CTTTA | C | -1.0374 | 0.2516 | 3.73E-05 | ---?? | 46.1 | 0.1565 | | |
| rs35332745 | 6 | 32674225 | 5 | 0 | 1 | T | C | -1.5739 | 0.3818 | 3.74E-05 | ----- | 8.3 | 0.3591 | intergenic |  |
| rs4706640 | 6 | 77655821 | 3 | 0 | 0 | A | G | 1.157 | 0.2806 | 3.74E-05 | +++?? | 20.9 | 0.2823 | intergenic | |
| rs7772268 | 6 | 77623152 | 4 | 0 | 0 | A | G | -1.129 | 0.2739 | 3.75E-05 | ---?- | 7.1 | 0.3575 | intergenic | |
| rs11980149 | 7 | 107396941 | 4 | 0 | 0 | A | G | 1.0214 | 0.2478 | 3.75E-05 | +++?+ | 30.3 | 0.2302 | intron | CBLL1 |
| rs113984677 | 6 | 32666636 | 5 | 0 | 1 | A | G | 1.5752 | 0.3822 | 3.76E-05 | +++++ | 4.4 | 0.3815 | intergenic | |
| rs111848638 | 6 | 32666630 | 5 | 0 | 1 | A | T | -1.5752 | 0.3822 | 3.76E-05 | ----- | 4.4 | 0.3815 | intergenic |  |
| rs13208645 | 6 | 77650945 | 3 | 0 | 0 | A | G | -1.1554 | 0.2804 | 3.77E-05 | ---?? | 22 | 0.2775 | intergenic | |
| rs75040946 | 10 | 131470783 | 4 | 0 | 0 | C | G | 1.771 | 0.4298 | 3.79E-05 | +++?+ | 0 | 0.9183 | intron | MGMT |
| 6:77648149 | 6 | 77648149 | 3 | 0 | 0 | T | TA | -1.1562 | 0.2808 | 3.82E-05 | ---?? | 22.1 | 0.2771 | | |
| rs10502789 | 18 | 40070199 | 4 | 0 | 0 | A | G | 1.3725 | 0.3334 | 3.83E-05 | +++?- | 17.1 | 0.3057 | intron | LINC00907 |
| rs10266547 | 7 | 107402435 | 4 | 0 | 0 | A | G | 1.0295 | 0.2501 | 3.85E-05 | +++?+ | 0 | 0.4295 | downstream-gene | |
| rs146616726 | 18 | 40126913 | 4 | 0 | 0 | A | G | 1.3999 | 0.3401 | 3.86E-05 | +++?- | 22.6 | 0.2751 | intron | LINC00907 |
| 13:45632397 | 13 | 45632397 | 3 | 0 | 0 | T | TTTTTG | 2.2055 | 0.536 | 3.87E-05 | +++?? | 0 | 0.6037 | | |
| rs323233 | 6 | 77669395 | 3 | 0 | 0 | A | C | 1.1606 | 0.282 | 3.87E-05 | +++?? | 19.8 | 0.2875 | intergenic | |
| rs113259939 | 17 | 8761017 | 4 | 0 | 0 | A | G | -3.3277 | 0.8087 | 3.87E-05 | ---?- | 0 | 0.8576 | intron | PIK3R6 |
| rs7768254 | 6 | 77637933 | 4 | 0 | 0 | T | C | -1.1287 | 0.2744 | 3.90E-05 | ---?- | 4.9 | 0.3686 | intergenic | |
| rs1932212 | 6 | 77631909 | 4 | 0 | 0 | A | G | 1.1268 | 0.2739 | 3.90E-05 | +++?+ | 5.2 | 0.3672 | intergenic | |
| rs565799 | 16 | 5461361 | 4 | 0 | 0 | C | G | -1.179 | 0.2867 | 3.90E-05 | ---?- | 0 | 0.784 | intron | RBFOX1 |
| rs16976051 | 18 | 39981024 | 4 | 0 | 0 | A | G | -1.43 | 0.3477 | 3.92E-05 | ---?+ | 0 | 0.4367 | intron | LINC00907 |
| rs35992469 | 6 | 77641958 | 4 | 0 | 0 | T | C | 1.1281 | 0.2744 | 3.94E-05 | +++?+ | 5 | 0.3681 | intergenic | |
| 10:28240358 | 10 | 28240358 | 3 | 0 | 0 | A | AG | 1.0053 | 0.2445 | 3.94E-05 | +++?? | 0 | 0.524 | | |
| rs11753149 | 6 | 77597224 | 4 | 0 | 0 | C | G | -1.1233 | 0.2733 | 3.95E-05 | ---?- | 10.2 | 0.3419 | intergenic | |
| rs1381677 | 6 | 77598255 | 4 | 0 | 0 | A | G | -1.1199 | 0.2725 | 3.96E-05 | ---?- | 10.8 | 0.3387 | intergenic | |
| rs114487668 | 1 | 32005997 | 3 | 0 | 0 | T | G | 4.5404 | 1.1048 | 3.96E-05 | +++?? | 0 | 0.8587 | intergenic | |
| rs10849939 | 12 | 111811604 | 5 | 0 | 0 | A | G | 1.1202 | 0.2726 | 3.97E-05 | +-+++ | 56.1 | 0.05834 | upstream-gene | |
| rs17206196 | 6 | 32665629 | 4 | 0 | 1 | T | G | 1.6451 | 0.4003 | 3.97E-05 | ++++? | 47 | 0.1293 | intergenic | |
| rs79423472 | 13 | 103355510 | 4 | 0 | 0 | T | C | 2.8729 | 0.6992 | 3.98E-05 | +++?+ | 0 | 0.7246 | intergenic | |
| rs9393525 | 6 | 24070942 | 5 | 0 | 0 | A | G | -1.5721 | 0.3828 | 4.00E-05 | ----+ | 0 | 0.61 | intergenic |  |
| rs13204857 | 6 | 77641181 | 4 | 0 | 0 | T | G | -1.1259 | 0.2741 | 4.01E-05 | ---?- | 4.6 | 0.3698 | intergenic | |
| rs34485365 | 6 | 77644585 | 4 | 0 | 0 | T | C | 1.127 | 0.2744 | 4.01E-05 | +++?+ | 4.6 | 0.3701 | intergenic | |
| rs35725597 | 6 | 32659112 | 5 | 0 | 1 | T | C | 1.554 | 0.3784 | 4.01E-05 | +++++ | 11.4 | 0.3409 | intergenic | |
| rs12913355 | 15 | 91589625 | 4 | 0 | 0 | A | T | 1.6926 | 0.4122 | 4.02E-05 | +++?+ | 0 | 0.9948 | intergenic | |
| rs7743922 | 6 | 77591293 | 4 | 0 | 0 | T | C | 1.1212 | 0.2731 | 4.03E-05 | +++?+ | 14.1 | 0.3217 | intergenic | |
| rs76039124 | 5 | 166153365 | 2 | 0 | 0 | A | G | -5.1385 | 1.2518 | 4.04E-05 | -???- | 0 | 0.6654 | intergenic | |
| rs182753584 | 6 | 168613527 | 3 | 0 | 0 | T | C | -2.2216 | 0.5412 | 4.05E-05 | ---?? | 0 | 0.9739 | intergenic | |
| 19:3699361 | 19 | 3699361 | 3 | 0 | 0 | A | ACT | -2.0022 | 0.4879 | 4.07E-05 | ---?? | 0 | 0.5325 | | |
| rs2411979 | 17 | 47427243 | 5 | 0 | 0 | A | G | -1.0522 | 0.2564 | 4.07E-05 | ----+ | 0 | 0.4074 | intron | ZNF652 |

| | | | | | | | | | | | | | | | |
|---|---|---|---|---|---|---|---|---|---|---|---|---|---|---|---|
| rs7236611 | 18 | 40125717 | 4 | 0 | 0 | T | C | 1.3932 | 0.3397 | 4.11E-05 | +++?- | 22.6 | 0.2754 | intron | LINC00907 |
| rs148342015 | 6 | 106172769 | 4 | 0 | 1 | A | C | 4.0101 | 0.978 | 4.13E-05 | +++?+ | 36 | 0.1964 | intergenic | |
| rs9379637 | 6 | 24078475 | 5 | 0 | 0 | A | G | -1.5854 | 0.3868 | 4.16E-05 | ----+ | 0 | 0.5504 | intergenic |  |
| 14:43264333 | 14 | 43264333 | 3 | 0 | 0 | G | GT | 1.607 | 0.3922 | 4.17E-05 | +++?? | 53.4 | 0.1169 | | |
| rs1356998 | 6 | 24055673 | 5 | 0 | 0 | T | G | -1.4598 | 0.3563 | 4.18E-05 | ----+ | 0 | 0.7044 | intergenic |  |
| rs72748343 | 1 | 209020811 | 3 | 0 | 0 | A | G | 4.5186 | 1.1031 | 4.20E-05 | ?++?+ | 10.7 | 0.3261 | intergenic | |
| 7:107372359 | 7 | 107372359 | 3 | 0 | 0 | T | TA | 1.0435 | 0.2547 | 4.20E-05 | +++?? | 44.5 | 0.1649 | | |
| rs62542 | 16 | 5461624 | 4 | 0 | 0 | A | G | 1.0835 | 0.2646 | 4.21E-05 | +++?+ | 0 | 0.8598 | intron | RBFOX1 |
| rs112498857 | 8 | 3094110 | 3 | 0 | 0 | T | C | 4.0844 | 0.9974 | 4.22E-05 | +++?? | 0 | 0.9703 | intron | CSMD1 |
| rs73489806 | 9 | 90836739 | 4 | 0 | 0 | A | G | 2.1207 | 0.5179 | 4.22E-05 | +++?+ | 0 | 0.7106 | intergenic | |
| rs117642721 | 7 | 158561813 | 4 | 0 | 0 | T | C | -4.5772 | 1.118 | 4.24E-05 | ---?- | 24 | 0.2671 | intron | ESYT2 |
| rs7785204 | 7 | 10146653 | 4 | 0 | 0 | T | G | -1.2065 | 0.2947 | 4.25E-05 | ---?- | 52.1 | 0.09958 | intergenic | |
| 6:144619637 | 6 | 144619637 | 3 | 0 | 0 | A | ATG | 1.9692 | 0.4811 | 4.25E-05 | +++?? | 34.1 | 0.2192 | | |
| rs149185459 | 18 | 40126915 | 4 | 0 | 0 | T | C | -1.3918 | 0.34 | 4.26E-05 | ---?+ | 22.5 | 0.2755 | intron | LINC00907 |
| rs74783222 | 10 | 131467811 | 4 | 0 | 0 | T | C | 1.8209 | 0.4449 | 4.26E-05 | +++?+ | 0 | 0.8826 | intron | MGMT |
| rs144961538 | 2 | 117045480 | 4 | 0 | 0 | A | G | -4.1529 | 1.0147 | 4.27E-05 | ---?- | 6 | 0.3629 | intergenic | |
| rs144243324 | 6 | 36927769 | 3 | 0 | 0 | T | G | 4.4959 | 1.0986 | 4.27E-05 | +++?? | 29.6 | 0.2415 | intron | PI16 |
| rs28708580 | 7 | 107370532 | 3 | 0 | 0 | A | G | -1.0476 | 0.2561 | 4.30E-05 | ---?? | 40.5 | 0.1863 | intergenic | |
| rs299371 | 5 | 134339715 | 4 | 0 | 0 | T | C | 1.0945 | 0.2676 | 4.32E-05 | +++?+ | 0 | 0.5879 | intron | CATSPER3 |
| rs73173307 | 3 | 172322918 | 4 | 0 | 0 | T | C | 2.3113 | 0.5652 | 4.32E-05 | +++?- | 21.1 | 0.2834 | intergenic | |
| rs4510635 | 6 | 77651472 | 4 | 0 | 0 | T | C | -1.1209 | 0.2741 | 4.32E-05 | ---?- | 3.1 | 0.3769 | intergenic | |
| rs6965530 | 7 | 141189752 | 4 | 0 | 0 | A | G | -5.516 | 1.3491 | 4.34E-05 | ---?- | 0 | 0.7922 | intergenic | |
| rs9874917 | 3 | 77395423 | 4 | 0 | 0 | A | C | 1.0257 | 0.2509 | 4.34E-05 | +++?+ | 0 | 0.532 | intron | ROBO2 |
| rs299370 | 5 | 134339065 | 4 | 0 | 0 | T | C | 1.0971 | 0.2683 | 4.35E-05 | +++?+ | 0 | 0.5942 | intron | CATSPER3 |
| rs3769769 | 2 | 170675765 | 5 | 0 | 0 | T | C | 0.92 | 0.225 | 4.35E-05 | +++++ | 56.5 | 0.05644 | intron | METTL5 |
| 6:144619248 | 6 | 144619248 | 3 | 0 | 0 | A | AT | -1.9676 | 0.4815 | 4.38E-05 | ---?? | 34 | 0.2196 | | |
| rs2162516 | 2 | 170677492 | 5 | 0 | 0 | A | G | -0.9207 | 0.2253 | 4.38E-05 | ----- | 56.3 | 0.05748 | intron | METTL5 |
| rs77282998 | 2 | 52008457 | 3 | 0 | 0 | T | C | -4.2999 | 1.0524 | 4.39E-05 | ---?? | 0 | 0.7452 | intergenic | |
| rs7748939 | 6 | 77638489 | 4 | 0 | 0 | T | C | 1.1195 | 0.274 | 4.40E-05 | +++?+ | 0.1 | 0.391 | intergenic | |
| rs2023709 | 7 | 107390163 | 4 | 0 | 0 | A | T | -0.9991 | 0.2446 | 4.41E-05 | ---?- | 13 | 0.3275 | intron | CBLL1 |
| rs60150543 | 18 | 39938729 | 4 | 0 | 0 | A | G | -1.5943 | 0.3904 | 4.42E-05 | ---?- | 0 | 0.7507 | intron | LINC00907 |
| rs117917800 | 6 | 83866365 | 4 | 0 | 0 | A | G | 3.5486 | 0.8689 | 4.43E-05 | +++?+ | 0 | 0.407 | intron | DOPEY1 |
| 18:40003866 | 18 | 40003866 | 3 | 0 | 0 | CT | CTT | -1.3753 | 0.3368 | 4.43E-05 | ---?? | 0 | 0.5138 | | |
| rs73518625 | 7 | 141181486 | 4 | 0 | 0 | A | G | -6.2715 | 1.5363 | 4.46E-05 | ---?- | 0 | 0.851 | downstream-gene | |
| rs2098982 | 2 | 170674920 | 5 | 0 | 0 | T | C | -0.9198 | 0.2253 | 4.47E-05 | ----- | 56.8 | 0.05496 | intron | METTL5 |
| rs10237263 | 7 | 107402466 | 4 | 0 | 0 | T | C | -1.0215 | 0.2503 | 4.47E-05 | ---?- | 0 | 0.4159 | downstream-gene | |
| rs72748854 | 15 | 85889204 | 3 | 0 | 0 | C | G | -6.4171 | 1.5722 | 4.47E-05 | --??- | 0 | 0.9035 | intergenic | |
| rs74399289 | 4 | 155657965 | 4 | 0 | 0 | T | G | 3.9113 | 0.9583 | 4.47E-05 | +++?+ | 0 | 0.8374 | intergenic | |
| rs4577935 | 8 | 140375578 | 4 | 0 | 0 | A | G | -1.489 | 0.3648 | 4.48E-05 | ---?- | 0 | 0.9889 | intergenic | |
| rs59840594 | 11 | 2549743 | 3 | 0 | 0 | T | G | -7.1755 | 1.7582 | 4.48E-05 | ---?? | 0 | 0.3862 | intron | KCNQ1 |
| rs16997350 | 4 | 117278898 | 4 | 0 | 0 | T | G | -3.8629 | 0.9467 | 4.50E-05 | ---?- | 12.9 | 0.3281 | intergenic | |
| rs62076449 | 17 | 47441678 | 5 | 0 | 0 | T | C | -1.0421 | 0.2554 | 4.51E-05 | ----+ | 29.3 | 0.2264 | upstream-gene |  |
| rs11644062 | 16 | 6541029 | 4 | 0 | 0 | A | G | -1.094 | 0.2682 | 4.51E-05 | ---?- | 0 | 0.6755 | intron | RBFOX1 |
| rs11756314 | 6 | 24074700 | 5 | 0 | 0 | A | C | 1.5679 | 0.3844 | 4.52E-05 | ++++- | 0 | 0.5734 | intergenic | |
| rs6457615 | 6 | 32651964 | 5 | 0 | 1 | T | C | -1.5567 | 0.3817 | 4.54E-05 | ----- | 23.3 | 0.2659 | intergenic |  |
| rs117404686 | 6 | 83870271 | 4 | 0 | 0 | A | G | 3.5346 | 0.8668 | 4.55E-05 | +++?+ | 1.6 | 0.3843 | intron | DOPEY1 |
| rs76667225 | 17 | 47442527 | 5 | 0 | 0 | T | C | -1.0434 | 0.2559 | 4.55E-05 | ----+ | 32.8 | 0.2029 | upstream-gene |  |
| rs75267052 | 7 | 158603864 | 2 | 0 | 0 | A | T | -6.6819 | 1.6387 | 4.55E-05 | ??-?- | 0 | 0.6256 | intron | ESYT2 |
| 10:126848527 | 10 | 126848527 | 3 | 0 | 0 | CT | C | 1.4534 | 0.3565 | 4.56E-05 | +++?? | 36.7 | 0.206 | | |
| rs78724452 | 17 | 47432212 | 5 | 0 | 0 | T | C | -1.0464 | 0.2567 | 4.57E-05 | ----+ | 0 | 0.4142 | intron | ZNF652 |
| rs75760547 | 12 | 91448747 | 3 | 0 | 0 | C | G | -5.2772 | 1.2945 | 4.57E-05 | ---?? | 0 | 0.9262 | intron | KERA |
| rs34984116 | 6 | 32636392 | 5 | 0 | 1 | A | T | 1.5553 | 0.3816 | 4.58E-05 | +++++ | 4.3 | 0.3825 | upstream-gene | |
| rs17053975 | 9 | 90836065 | 4 | 0 | 0 | A | C | -2.1196 | 0.5201 | 4.59E-05 | ---?- | 0 | 0.71 | intergenic | |
| rs72913376 | 4 | 117280946 | 4 | 0 | 0 | A | T | -3.8418 | 0.9427 | 4.59E-05 | ---?- | 12.7 | 0.3291 | intergenic | |
| rs1827266 | 15 | 98046060 | 3 | 0 | 0 | T | C | 1.1433 | 0.2805 | 4.60E-05 | +++?? | 0 | 0.6431 | intergenic | |
| rs6926512 | 6 | 83870537 | 4 | 0 | 0 | A | G | -3.5333 | 0.867 | 4.60E-05 | ---?- | 1.1 | 0.3864 | intron | DOPEY1 |
| rs12511242 | 4 | 117281692 | 4 | 0 | 0 | C | G | 3.8373 | 0.9418 | 4.61E-05 | +++?+ | 13 | 0.3277 | intergenic | |
| rs1197704 | 15 | 42083094 | 5 | 0 | 0 | T | C | -1.239 | 0.3041 | 4.61E-05 | -+--- | 34.3 | 0.1924 | intron | MAPKBP1 |
| rs55979242 | 9 | 90837462 | 4 | 0 | 0 | A | G | -2.1259 | 0.5218 | 4.61E-05 | ---?- | 0 | 0.7124 | intergenic | |

| | | | | | | | | | | | | | | | |
|---|---|---|---|---|---|---|---|---|---|---|---|---|---|---|---|
| rs55938995 | 2 | 170484235 | 4 | 0 | 0 | A | T | -1.3106 | 0.3217 | 4.61E-05 | ---?- | 0 | 0.9964 | intron | PPIG |
| rs79315064 | 8 | 86476247 | 4 | 0 | 0 | T | C | -2.4211 | 0.5943 | 4.62E-05 | ---?- | 47.8 | 0.1243 | intergenic | |
| rs192227308 | 5 | 166754182 | 2 | 0 | 0 | C | G | -6.4705 | 1.5883 | 4.62E-05 | -???- | 0 | 0.6652 | intron | TENM2 |
| rs4877420 | 9 | 90837001 | 4 | 0 | 0 | T | G | -2.1221 | 0.5209 | 4.63E-05 | ---?- | 0 | 0.7107 | intergenic | |
| rs10506216 | 12 | 43130885 | 4 | 0 | 0 | A | G | 1.7042 | 0.4184 | 4.64E-05 | +++?+ | 0 | 0.419 | intergenic | |
| 10:28229190 | 10 | 28229190 | 3 | 0 | 0 | A | ATATG | 1.0051 | 0.2468 | 4.65E-05 | +++?? | 0 | 0.5456 | | |
| rs12910149 | 15 | 91590259 | 4 | 0 | 0 | A | G | -1.6955 | 0.4163 | 4.65E-05 | ---?- | 0 | 0.9946 | intergenic | |
| rs2050427 | 6 | 77587889 | 4 | 0 | 0 | A | C | 1.1138 | 0.2735 | 4.66E-05 | +++?+ | 9.1 | 0.3476 | intergenic | |
| rs72913374 | 4 | 117280930 | 4 | 0 | 0 | A | G | 3.839 | 0.9428 | 4.67E-05 | +++?+ | 12.8 | 0.3287 | intergenic | |
| rs12924480 | 16 | 6539449 | 4 | 0 | 0 | A | G | -1.089 | 0.2675 | 4.68E-05 | ---?- | 0 | 0.6527 | intron | RBFOX1 |
| rs75982363 | 10 | 6409909 | 4 | 0 | 0 | A | G | 3.8055 | 0.935 | 4.70E-05 | +++?+ | 0 | 0.7378 | intergenic | |
| rs4238868 | 16 | 6540014 | 4 | 0 | 0 | T | C | 1.0894 | 0.2678 | 4.73E-05 | +++?+ | 0 | 0.6626 | intron | RBFOX1 |
| rs3898202 | 6 | 77608692 | 4 | 0 | 0 | T | C | -1.1073 | 0.2722 | 4.73E-05 | ---?- | 25.5 | 0.2587 | intergenic | |
| rs43142 | 16 | 6539775 | 4 | 0 | 0 | T | G | 1.0889 | 0.2677 | 4.73E-05 | +++?+ | 0 | 0.6585 | intron | RBFOX1 |
| rs142419323 | 6 | 83867590 | 4 | 0 | 0 | T | C | 3.5359 | 0.8692 | 4.74E-05 | +++?+ | 0 | 0.392 | intron | DOPEY1 |
| rs2074492 | 6 | 31239869 | 3 | 0 | 0 | T | C | -1.0331 | 0.254 | 4.76E-05 | ---?? | 56.6 | 0.09969 | 5-prime-UTR | HLA-C |
| rs28414211 | 7 | 141164149 | 4 | 0 | 0 | A | T | 5.7625 | 1.417 | 4.77E-05 | +++?+ | 0 | 0.7986 | intron | TMEM178B |
| rs4332649 | 13 | 60015474 | 4 | 0 | 0 | T | C | -0.9999 | 0.2459 | 4.79E-05 | ---?- | 0 | 0.7449 | intergenic | |
| rs299374 | 5 | 134340942 | 4 | 0 | 0 | T | C | -1.0949 | 0.2693 | 4.80E-05 | ---?- | 0 | 0.6111 | intron | CATSPER3 |
| rs34709043 | 6 | 32676486 | 5 | 0 | 1 | A | C | -1.553 | 0.382 | 4.80E-05 | ----- | 4.1 | 0.3831 | intergenic |  |
| rs917647 | 16 | 6538699 | 4 | 0 | 0 | T | C | 1.0855 | 0.2672 | 4.84E-05 | +++?+ | 0 | 0.6112 | intron | RBFOX1 |
| rs76491206 | 10 | 131476494 | 4 | 0 | 0 | A | G | 1.6918 | 0.4164 | 4.85E-05 | +++?+ | 0 | 0.8844 | intron | MGMT |
| 11:30670984 | 11 | 30670984 | 3 | 0 | 0 | G | GCTCA | 1.1538 | 0.2841 | 4.89E-05 | +++?? | 9.7 | 0.3304 | | |
| rs9470028 | 6 | 35418227 | 5 | 0 | 0 | T | C | 3.9256 | 0.9668 | 4.90E-05 | +++++ | 0 | 0.7955 | upstream-gene | |
| rs6116900 | 20 | 5681944 | 4 | 0 | 0 | A | T | -1.201 | 0.2958 | 4.91E-05 | ---?- | 0 | 0.5097 | intergenic | |
| rs9348631 | 6 | 24072212 | 5 | 0 | 0 | A | G | 1.554 | 0.383 | 4.95E-05 | ++++- | 0 | 0.5914 | intergenic | |
| rs9832713 | 3 | 77393019 | 4 | 0 | 0 | T | C | -1.0199 | 0.2513 | 4.96E-05 | ---?- | 0 | 0.4456 | intron | ROBO2 |
| rs10879452 | 12 | 41369271 | 4 | 0 | 0 | T | C | 1.2564 | 0.3097 | 4.97E-05 | +++?+ | 0 | 0.8549 | intron | CNTN1 |
| rs177252 | 5 | 134343799 | 4 | 0 | 0 | A | G | 1.0771 | 0.2655 | 4.99E-05 | +++?+ | 0 | 0.5626 | synonymous | CATSPER3 |
| rs11813363 | 10 | 131469926 | 4 | 0 | 0 | T | C | -1.5365 | 0.3788 | 4.99E-05 | ---?- | 4.6 | 0.3701 | intron | MGMT |
| rs12589648 | 14 | 43230092 | 3 | 0 | 0 | T | C | -1.6348 | 0.4031 | 5.00E-05 | ---?? | 51.9 | 0.1248 | intergenic | |
| rs323239 | 6 | 77698853 | 4 | 0 | 0 | A | C | -1.1167 | 0.2754 | 5.02E-05 | ---?- | 9.7 | 0.3444 | intergenic | |
| rs76971763 | 5 | 131413145 | 2 | 0 | 0 | A | G | -9.1586 | 2.259 | 5.03E-05 | ?-??- | 32.5 | 0.2235 | downstream-gene | |
| rs7177999 | 15 | 87687819 | 4 | 0 | 0 | C | G | 2.4079 | 0.5941 | 5.05E-05 | +++?+ | 0 | 0.6546 | intergenic | |
| rs9393523 | 6 | 24067664 | 5 | 0 | 0 | A | G | -1.5514 | 0.3828 | 5.07E-05 | ----+ | 0 | 0.6231 | intergenic |  |
| rs7235528 | 18 | 40058882 | 4 | 0 | 0 | T | C | 1.3003 | 0.3209 | 5.07E-05 | +++?- | 0 | 0.4704 | intron | LINC00907 |
| rs141257196 | 6 | 83859341 | 4 | 0 | 0 | T | C | 3.539 | 0.8736 | 5.10E-05 | +++?+ | 0 | 0.43 | intron | DOPEY1 |
| 4:21081261 | 4 | 21081261 | 3 | 0 | 0 | G | GT | -2.6687 | 0.6588 | 5.10E-05 | ---?? | 0 | 0.5652 | | |
| 6:83865373 | 6 | 83865373 | 3 | 0 | 0 | G | GC | 3.6216 | 0.894 | 5.10E-05 | +++?? | 27.7 | 0.2508 | | |
| rs28557291 | 18 | 40058675 | 4 | 0 | 0 | T | C | -1.2996 | 0.3208 | 5.11E-05 | ---?+ | 0 | 0.4705 | intron | LINC00907 |
| rs2197109 | 3 | 77392452 | 4 | 0 | 0 | C | G | 1.0188 | 0.2516 | 5.12E-05 | +++?+ | 0 | 0.4459 | intron | ROBO2 |
| rs112478066 | 18 | 39942532 | 4 | 0 | 0 | T | C | -1.5795 | 0.39 | 5.13E-05 | ---?- | 0 | 0.7435 | intron | LINC00907 |
| rs4749288 | 10 | 28255835 | 3 | 0 | 0 | T | C | -1.0145 | 0.2506 | 5.15E-05 | ---?? | 0 | 0.48 | intron | ARMC4 |
| rs4449296 | 3 | 77394540 | 4 | 0 | 0 | T | C | 1.0156 | 0.2509 | 5.17E-05 | +++?+ | 0 | 0.4515 | intron | ROBO2 |
| 6:83871941 | 6 | 83871941 | 3 | 0 | 0 | G | GTT | 3.6088 | 0.8917 | 5.18E-05 | +++?? | 32.3 | 0.2285 | | |
| rs7453887 | 6 | 77608156 | 4 | 0 | 0 | T | G | -1.1023 | 0.2724 | 5.20E-05 | ---?- | 25 | 0.2617 | intergenic | |
| rs1028547 | 6 | 83862242 | 4 | 0 | 0 | T | G | 3.5331 | 0.8732 | 5.20E-05 | +++?+ | 0 | 0.4084 | intron | DOPEY1 |
| rs9447896 | 6 | 77608251 | 4 | 0 | 0 | T | G | 1.1022 | 0.2724 | 5.20E-05 | +++?+ | 24.9 | 0.2619 | intergenic | |
| rs17069793 | 8 | 4126283 | 4 | 0 | 0 | A | C | -1.3142 | 0.3248 | 5.21E-05 | ---?- | 0 | 0.5433 | intron | CSMD1 |
| rs9443311 | 6 | 77608293 | 4 | 0 | 0 | C | G | -1.1022 | 0.2724 | 5.21E-05 | ---?- | 24.9 | 0.2621 | intergenic | |
| rs9447897 | 6 | 77608338 | 4 | 0 | 0 | T | C | 1.1022 | 0.2724 | 5.21E-05 | +++?+ | 24.9 | 0.2621 | intergenic | |
| rs7312913 | 12 | 111793552 | 5 | 0 | 0 | A | G | -1.0705 | 0.2646 | 5.21E-05 | -+--- | 51.2 | 0.08463 | downstream-gene | |
| rs1953343 | 6 | 77606795 | 4 | 0 | 0 | T | C | 1.1021 | 0.2724 | 5.22E-05 | +++?+ | 24.7 | 0.2631 | intergenic | |
| rs7356848 | 6 | 77607554 | 4 | 0 | 0 | A | C | -1.1017 | 0.2723 | 5.22E-05 | ---?- | 25.2 | 0.2603 | intergenic | |
| rs6453944 | 6 | 77607625 | 4 | 0 | 0 | A | C | 1.1016 | 0.2723 | 5.23E-05 | +++?+ | 25.1 | 0.2607 | intergenic | |
| rs3118466 | 1 | 85203104 | 5 | 0 | 0 | T | C | -0.9311 | 0.2302 | 5.23E-05 | ----- | 25.6 | 0.2509 | intergenic |  |
| rs5012254 | 6 | 77607806 | 4 | 0 | 0 | A | G | 1.1015 | 0.2723 | 5.23E-05 | +++?+ | 25.1 | 0.2611 | intergenic | |
| rs5012255 | 6 | 77607823 | 4 | 0 | 0 | T | C | 1.1015 | 0.2723 | 5.23E-05 | +++?+ | 25.1 | 0.2611 | intergenic | |
| rs5012256 | 6 | 77607854 | 4 | 0 | 0 | A | G | 1.1015 | 0.2723 | 5.24E-05 | +++?+ | 25 | 0.2613 | intergenic | |

| | | | | | | | | | | | | | | | |
|---|---|---|---|---|---|---|---|---|---|---|---|---|---|---|---|
| 5:179985620 | 5 | 179985620 | 3 | 0 | 0 | G | GCT | -1.3994 | 0.346 | 5.24E-05 | ---?? | 42.9 | 0.1736 | | |
| rs6832927 | 4 | 100544867 | 4 | 0 | 0 | A | C | -1.6848 | 0.4165 | 5.24E-05 | +--?- | 41.4 | 0.1632 | 3-prime-UTR | MTTP |
| rs9443310 | 6 | 77607111 | 4 | 0 | 0 | A | T | -1.1014 | 0.2723 | 5.25E-05 | ---?- | 25.1 | 0.2608 | intergenic | |
| rs150722044 | 6 | 83867998 | 4 | 0 | 0 | A | G | 3.5224 | 0.871 | 5.25E-05 | +++?+ | 0 | 0.4115 | intron | DOPEY1 |
| rs6832119 | 4 | 100544511 | 4 | 0 | 0 | A | G | 1.6847 | 0.4166 | 5.25E-05 | -++?+ | 41.3 | 0.1641 | 3-prime-UTR | MTTP |
| rs928460 | 9 | 7824232 | 4 | 0 | 0 | T | C | 1.8025 | 0.4457 | 5.26E-05 | +++?+ | 36.1 | 0.1953 | intergenic | |
| rs113653192 | 4 | 100545232 | 4 | 0 | 0 | T | C | -1.6846 | 0.4166 | 5.26E-05 | +--?- | 41.4 | 0.1635 | downstream-gene | |
| 10:28224631 | 10 | 28224631 | 3 | 0 | 0 | T | TAA | 0.9917 | 0.2453 | 5.27E-05 | +++?? | 0 | 0.575 | | |
| rs6817699 | 4 | 190021887 | 4 | 0 | 0 | T | G | 0.9763 | 0.2414 | 5.27E-05 | +++?+ | 0 | 0.437 | intergenic | |
| rs74542928 | 4 | 100544188 | 4 | 0 | 0 | T | C | -1.6846 | 0.4166 | 5.27E-05 | +--?- | 41.1 | 0.165 | 3-prime-UTR | MTTP |
| rs6974826 | 7 | 107404440 | 4 | 0 | 0 | T | C | 1.0179 | 0.2518 | 5.27E-05 | +++?+ | 0 | 0.4128 | downstream-gene | |
| 6:18340608 | 6 | 18340608 | 3 | 0 | 0 | T | TA | 2.1223 | 0.5249 | 5.28E-05 | +++?? | 0 | 0.6767 | | |
| rs7693865 | 4 | 100545820 | 4 | 0 | 0 | T | C | -1.6849 | 0.4168 | 5.28E-05 | +--?- | 41.2 | 0.1642 | downstream-gene | |
| rs12916539 | 15 | 91588040 | 4 | 0 | 0 | A | G | 1.6194 | 0.4006 | 5.28E-05 | +++?+ | 0 | 0.9562 | intergenic | |
| rs12722051 | 6 | 32609147 | 3 | 0 | 0 | A | T | -1.5359 | 0.3799 | 5.28E-05 | ---?? | 0 | 0.7923 | missense | HLA-DQA1 |
| rs56010116 | 17 | 47436514 | 5 | 0 | 0 | A | G | -1.0388 | 0.257 | 5.29E-05 | ----+ | 0 | 0.4232 | intron | ZNF652 |
| rs13096695 | 3 | 77403512 | 4 | 0 | 0 | A | T | -1.0031 | 0.2482 | 5.30E-05 | ---?- | 0 | 0.5635 | intron | ROBO2 |
| rs9443309 | 6 | 77606341 | 4 | 0 | 0 | T | C | 1.1009 | 0.2724 | 5.31E-05 | +++?+ | 24.9 | 0.2618 | intergenic | |
| rs11531337 | 18 | 39957838 | 4 | 0 | 0 | A | G | -1.4349 | 0.355 | 5.31E-05 | ---?+ | 0 | 0.4372 | intron | LINC00907 |
| rs112769434 | 4 | 100542921 | 4 | 0 | 0 | A | G | 1.6841 | 0.4167 | 5.31E-05 | -++?+ | 40.7 | 0.1673 | intron | MTTP |
| rs4794030 | 17 | 47338890 | 4 | 0 | 0 | C | G | -1.1797 | 0.2919 | 5.32E-05 | ---?+ | 16.7 | 0.3079 | downstream-gene | |
| rs9447895 | 6 | 77606115 | 4 | 0 | 0 | A | G | 1.1008 | 0.2724 | 5.32E-05 | +++?+ | 24.9 | 0.2621 | intergenic | |
| rs6919763 | 6 | 83859945 | 4 | 0 | 0 | T | C | -3.5275 | 0.8729 | 5.32E-05 | ---?- | 0 | 0.4411 | intron | DOPEY1 |
| rs16948042 | 17 | 47436040 | 5 | 0 | 0 | T | C | 1.0381 | 0.257 | 5.34E-05 | ++++- | 0 | 0.4201 | intron | ZNF652 |
| rs79837054 | 4 | 100546600 | 4 | 0 | 0 | T | C | 1.6848 | 0.417 | 5.34E-05 | -++?+ | 41.1 | 0.1651 | downstream-gene | |
| rs1814246 | 12 | 111801951 | 5 | 0 | 0 | A | T | 1.0183 | 0.2521 | 5.35E-05 | +-+++ | 48.4 | 0.1011 | intron | FAM109A |
| rs1491244 | 4 | 100542167 | 4 | 0 | 0 | A | T | 1.6835 | 0.4167 | 5.36E-05 | -++?+ | 40.4 | 0.1693 | intron | MTTP |
| rs877811 | 5 | 134346485 | 4 | 0 | 0 | T | G | 1.0796 | 0.2673 | 5.36E-05 | +++?+ | 0 | 0.5922 | intron | CATSPER3 |
| rs7399113 | 12 | 111792848 | 5 | 0 | 0 | A | G | 1.0665 | 0.2641 | 5.37E-05 | +-+++ | 50.7 | 0.08729 | downstream-gene | |
| rs9447893 | 6 | 77605453 | 4 | 0 | 0 | T | C | -1.1003 | 0.2725 | 5.38E-05 | ---?- | 24.7 | 0.2634 | intergenic | |
| rs1819167 | 6 | 77705465 | 4 | 0 | 0 | A | G | 1.1767 | 0.2914 | 5.38E-05 | +++?+ | 26.6 | 0.252 | intergenic | |
| rs113462175 | 17 | 47339826 | 4 | 0 | 0 | T | C | -1.1789 | 0.292 | 5.40E-05 | ---?+ | 16.5 | 0.3087 | downstream-gene | |
| rs7356913 | 6 | 77604953 | 4 | 0 | 0 | T | C | -1.1 | 0.2725 | 5.41E-05 | ---?- | 24.6 | 0.2638 | intergenic | |
| rs111702517 | 17 | 47339836 | 4 | 0 | 0 | C | G | -1.1787 | 0.292 | 5.42E-05 | ---?+ | 16.5 | 0.309 | downstream-gene | |
| rs35169156 | 2 | 2762533 | 4 | 0 | 0 | A | G | 1.7488 | 0.4333 | 5.43E-05 | +++?+ | 0 | 0.4132 | intergenic | |
| 6:23856928 | 6 | 23856928 | 3 | 0 | 0 | T | TAGAGA | 2.9878 | 0.7402 | 5.43E-05 | +++?? | 0 | 0.4121 | | |
| rs2447561 | 8 | 140465090 | 4 | 0 | 0 | A | T | -1.699 | 0.421 | 5.44E-05 | ---?- | 0 | 0.741 | intergenic | |
| rs1486950 | 13 | 60000182 | 4 | 0 | 0 | A | G | 1.0083 | 0.2499 | 5.46E-05 | +++?+ | 0 | 0.7367 | intergenic | |
| rs2011768 | 17 | 47340311 | 4 | 0 | 0 | T | G | -1.1785 | 0.2921 | 5.46E-05 | ---?+ | 16.4 | 0.3095 | downstream-gene | |
| rs331120 | 2 | 150851790 | 4 | 0 | 0 | A | G | -1.096 | 0.2717 | 5.48E-05 | ---?- | 0 | 0.6654 | intergenic | |
| rs7300400 | 12 | 111792179 | 5 | 0 | 0 | A | C | -1.0649 | 0.264 | 5.49E-05 | -+--- | 50.5 | 0.08845 | downstream-gene | |
| rs9447906 | 6 | 77616796 | 4 | 0 | 0 | T | C | 1.1003 | 0.2728 | 5.51E-05 | +++?+ | 21.5 | 0.2812 | intergenic | |
| rs9447907 | 6 | 77616915 | 4 | 0 | 0 | A | T | -1.1003 | 0.2728 | 5.51E-05 | ---?- | 21.5 | 0.2815 | intergenic | |
| rs933399 | 12 | 111807499 | 5 | 0 | 0 | A | G | 1.0223 | 0.2535 | 5.52E-05 | +-+++ | 48.4 | 0.1014 | upstream-gene | |
| 20:7956304 | 20 | 7956304 | 2 | 0 | 0 | CTT | C | 6.0579 | 1.5023 | 5.52E-05 | ?++?? | 0 | 0.6803 | | |
| rs112707021 | 4 | 100524148 | 4 | 0 | 0 | A | G | -1.6783 | 0.4162 | 5.53E-05 | +--?- | 38.8 | 0.179 | intron | MTTP |
| rs111303866 | 4 | 100524149 | 4 | 0 | 0 | A | C | -1.6782 | 0.4162 | 5.53E-05 | +--?- | 38.8 | 0.179 | intron | MTTP |
| rs11753230 | 6 | 77620561 | 4 | 0 | 0 | A | G | -1.0999 | 0.2729 | 5.56E-05 | ---?- | 20.8 | 0.2853 | intergenic | |
| rs7840428 | 8 | 76121004 | 4 | 0 | 0 | A | G | -1.1038 | 0.2738 | 5.56E-05 | ---?- | 23.2 | 0.2717 | intergenic | |
| rs7316698 | 12 | 111790511 | 5 | 0 | 0 | T | C | 1.0646 | 0.2641 | 5.56E-05 | +-+++ | 50.5 | 0.08885 | downstream-gene | |
| rs117770737 | 15 | 87650327 | 4 | 0 | 0 | T | C | -2.4144 | 0.599 | 5.57E-05 | ---?- | 0 | 0.7567 | intergenic | |
| rs299377 | 5 | 134344902 | 4 | 0 | 0 | T | C | 1.0721 | 0.266 | 5.58E-05 | +++?+ | 0 | 0.5634 | intron | CATSPER3 |
| rs12540351 | 7 | 107405738 | 4 | 0 | 0 | A | T | 1.0167 | 0.2523 | 5.58E-05 | +++?+ | 0 | 0.415 | downstream-gene | |
| 2:117195559 | 2 | 117195559 | 3 | 0 | 0 | CAG | C | 1.7345 | 0.4304 | 5.58E-05 | +++?? | 24.9 | 0.264 | | |
| rs17207599 | 5 | 66231022 | 4 | 0 | 0 | T | C | 2.4453 | 0.6068 | 5.59E-05 | +++?+ | 0 | 0.8265 | intron | MAST4 |
| rs7958312 | 12 | 111789977 | 5 | 0 | 0 | T | C | -1.0642 | 0.2641 | 5.60E-05 | -+--- | 50.4 | 0.08922 | downstream-gene | |
| rs12916522 | 15 | 91588006 | 4 | 0 | 0 | A | G | 1.6185 | 0.4017 | 5.61E-05 | +++?+ | 0 | 0.9699 | intergenic | |
| rs2249596 | 5 | 134347956 | 4 | 0 | 0 | T | C | 1.0774 | 0.2675 | 5.63E-05 | +++?+ | 0 | 0.5127 | downstream-gene | |
| rs6553277 | 4 | 190012421 | 4 | 0 | 0 | T | C | -0.9864 | 0.2449 | 5.65E-05 | ---?- | 0 | 0.4065 | intergenic | |

| | | | | | | | | | | | | | | | |
|---|---|---|---|---|---|---|---|---|---|---|---|---|---|---|---|
| 15:98048184 | 15 | 98048184 | 3 | 0 | 0 | TTAAG | TTATAA( | 1.1267 | 0.2798 | 5.66E-05 | +++?? | 0 | 0.5593 | | |
| rs75164113 | 12 | 43133486 | 4 | 0 | 0 | A | G | 1.6822 | 0.4178 | 5.67E-05 | +++?+ | 0 | 0.4078 | intergenic | |
| rs72761306 | 15 | 91584219 | 4 | 0 | 0 | A | G | 2.2392 | 0.5562 | 5.67E-05 | +++?+ | 0 | 0.5873 | intergenic | |
| rs323241 | 6 | 77700830 | 4 | 0 | 0 | T | C | 1.1107 | 0.2759 | 5.68E-05 | +++?+ | 3.9 | 0.3733 | intergenic | |
| rs9527981 | 13 | 60001040 | 4 | 0 | 0 | A | G | 1.0049 | 0.2496 | 5.68E-05 | +++?+ | 0 | 0.7389 | intergenic | |
| rs1601643 | 13 | 60001417 | 4 | 0 | 0 | A | G | -1.0041 | 0.2495 | 5.70E-05 | ---?- | 0 | 0.7399 | intergenic | |
| 2:202947572 | 2 | 202947572 | 3 | 0 | 0 | A | ATTT | -2.4554 | 0.6101 | 5.70E-05 | ---?? | 0 | 0.4521 | | |
| rs1843751 | 13 | 60001514 | 4 | 0 | 0 | T | C | -1.0038 | 0.2495 | 5.72E-05 | ---?- | 0 | 0.7398 | intergenic | |
| rs1224232 | 6 | 77695933 | 4 | 0 | 0 | A | G | 1.1092 | 0.2757 | 5.74E-05 | +++?+ | 4.9 | 0.3684 | intergenic | |
| rs3207561 | 6 | 31236685 | 3 | 0 | 0 | A | G | -1.025 | 0.2548 | 5.74E-05 | ---?? | 55.8 | 0.1039 | 3-prime-UTR | HLA-C |
| rs76704606 | 12 | 43137222 | 4 | 0 | 0 | C | G | 1.6659 | 0.4141 | 5.76E-05 | +++?+ | 0 | 0.4099 | intergenic | |
| 6:31236682 | 6 | 31236682 | 3 | 0 | 0 | A | AC | 1.0249 | 0.2548 | 5.76E-05 | +++?? | 55.8 | 0.104 | | |
| rs1916289 | 17 | 47333271 | 4 | 0 | 0 | T | C | 1.1714 | 0.2913 | 5.78E-05 | +++?- | 27.5 | 0.2469 | intron | FLJ40194 |
| rs142821198 | 12 | 104450172 | 4 | 0 | 0 | A | C | 4.607 | 1.1456 | 5.79E-05 | +++?+ | 26.7 | 0.2514 | intron | GLT8D2 |
| rs67294117 | 6 | 32504774 | 5 | 0 | 1 | A | G | 1.4409 | 0.3583 | 5.79E-05 | +++++ | 4.4 | 0.3817 | intergenic | |
| rs6955419 | 7 | 141167443 | 4 | 0 | 0 | A | G | 5.4783 | 1.3624 | 5.80E-05 | +++?+ | 0 | 0.8505 | intron | TMEM178B |
| rs12591344 | 15 | 98041445 | 4 | 0 | 0 | A | G | 1.0829 | 0.2693 | 5.80E-05 | +++?+ | 0 | 0.9498 | intergenic | |
| rs10124631 | 9 | 7821868 | 4 | 0 | 0 | A | G | -1.8575 | 0.462 | 5.80E-05 | ---?- | 15.5 | 0.3144 | intergenic | |
| rs111526997 | 6 | 23865881 | 3 | 0 | 0 | T | G | 2.9811 | 0.7414 | 5.80E-05 | +++?? | 0 | 0.4552 | intergenic | |
| 3:191150895 | 3 | 191150895 | 3 | 0 | 0 | A | AT | 1.5945 | 0.3966 | 5.80E-05 | +++?? | 49.2 | 0.1399 | | |
| rs11700220 | 20 | 60770931 | 2 | 0 | 0 | A | G | -6.9586 | 1.7309 | 5.81E-05 | ???-- | 0 | 0.8005 | missense | MTG2 |
| rs80107531 | 20 | 60792972 | 2 | 0 | 0 | T | C | 6.9586 | 1.7309 | 5.81E-05 | ???++ | 0 | 0.8005 | intron | HRH3 |
| rs9563724 | 13 | 60003049 | 4 | 0 | 0 | T | C | 1.0005 | 0.2489 | 5.83E-05 | +++?+ | 0 | 0.7636 | intergenic | |
| rs13206018 | 6 | 32663497 | 5 | 0 | 1 | T | C | 1.5004 | 0.3733 | 5.84E-05 | +++++ | 6 | 0.3724 | intergenic | |
| rs213506 | 5 | 134338223 | 3 | 0 | 0 | T | C | -1.1302 | 0.2812 | 5.84E-05 | ---?? | 0 | 0.541 | intron | CATSPER3 |
| rs68067315 | 6 | 32663340 | 5 | 0 | 1 | A | G | 1.5004 | 0.3733 | 5.84E-05 | +++++ | 6 | 0.3723 | intergenic | |
| rs7123658 | 11 | 102593979 | 5 | 0 | 0 | A | T | 0.8407 | 0.2092 | 5.84E-05 | +++++ | 0 | 0.8512 | intron | MMP8 |
| rs11121617 | 1 | 10842021 | 3 | 0 | 0 | A | G | -1.1533 | 0.287 | 5.85E-05 | ---?? | 33.8 | 0.2209 | intron | CASZ1 |
| rs112383833 | 6 | 83854803 | 4 | 0 | 0 | T | C | 3.5253 | 0.8774 | 5.87E-05 | +++?+ | 0 | 0.4563 | intron | DOPEY1 |
| 15:98048414 | 15 | 98048414 | 3 | 0 | 0 | CT | C | -1.1269 | 0.2805 | 5.88E-05 | ---?? | 0 | 0.5651 | | |
| rs141116581 | 6 | 83853720 | 4 | 0 | 0 | A | T | 3.5272 | 0.8781 | 5.90E-05 | +++?+ | 0 | 0.4568 | intron | DOPEY1 |
| rs141084113 | 20 | 54141479 | 3 | 0 | 0 | T | C | -5.1653 | 1.286 | 5.90E-05 | ---?? | 0 | 0.67 | intergenic | |
| rs67813730 | 6 | 77609247 | 4 | 0 | 0 | T | C | -1.0948 | 0.2726 | 5.90E-05 | ---?- | 25.1 | 0.2612 | intergenic | |
| rs596783 | 17 | 47329705 | 4 | 0 | 0 | T | C | 1.1608 | 0.289 | 5.91E-05 | +++?- | 29.7 | 0.234 | non-coding-exon | FLJ40194 |
| 17:52303160 | 17 | 52303160 | 3 | 0 | 0 | G | GT | -2.6232 | 0.6532 | 5.92E-05 | ---?? | 0 | 0.8434 | | |
| 12:66001646 | 12 | 66001646 | 3 | 0 | 0 | G | GCAGAT | -3.6832 | 0.9172 | 5.93E-05 | ---?? | 0 | 0.8676 | | |
| rs2315641 | 6 | 77642689 | 4 | 0 | 0 | T | C | -1.0961 | 0.273 | 5.94E-05 | ---?- | 18 | 0.3007 | intergenic | |
| rs1932741 | 13 | 60015433 | 4 | 0 | 0 | A | G | 0.9863 | 0.2457 | 5.95E-05 | +++?+ | 0 | 0.7498 | intergenic | |
| rs10219736 | 12 | 111788402 | 5 | 0 | 0 | T | C | 1.0613 | 0.2644 | 5.96E-05 | +-+++ | 50.8 | 0.0869 | downstream-gene | |
| rs4886162 | 13 | 60002411 | 4 | 0 | 0 | A | G | 1.0089 | 0.2513 | 5.96E-05 | +++?+ | 0 | 0.9066 | intergenic | |
| rs73159611 | 3 | 142833548 | 4 | 0 | 0 | T | C | -2.3765 | 0.5921 | 5.98E-05 | ---?- | 0 | 0.4492 | intergenic | |
| 11:30633714 | 11 | 30633714 | 3 | 0 | 0 | A | AT | -1.1633 | 0.2899 | 6.00E-05 | ---?? | 7.7 | 0.3385 | | |
| rs9408715 | 9 | 7823117 | 4 | 0 | 0 | T | G | 1.8453 | 0.4599 | 6.00E-05 | +++?+ | 23.4 | 0.2707 | intergenic | |
| rs1932742 | 13 | 60015379 | 4 | 0 | 0 | T | C | 0.9851 | 0.2455 | 6.01E-05 | +++?+ | 0 | 0.752 | intergenic | |
| rs33995001 | 4 | 47514685 | 5 | 0 | 0 | T | C | -0.8414 | 0.2097 | 6.02E-05 | ----+ | 42.5 | 0.138 | missense | ATP10D |
| rs77366533 | 8 | 86275921 | 4 | 0 | 0 | T | C | -2.0764 | 0.5175 | 6.02E-05 | ---?- | 38.4 | 0.1814 | intron | CA1 |
| rs112163423 | 17 | 47451755 | 3 | 0 | 0 | A | T | 1.2245 | 0.3052 | 6.02E-05 | +++?? | 0 | 0.8993 | intergenic | |
| 6:83842103 | 6 | 83842103 | 3 | 0 | 0 | CAA | C | -3.5609 | 0.888 | 6.07E-05 | ---?? | 0 | 0.4001 | | |
| rs2210772 | 6 | 77647626 | 4 | 0 | 0 | T | G | 1.0942 | 0.2729 | 6.09E-05 | +++?+ | 17.5 | 0.3034 | intergenic | |
| rs60127116 | 10 | 131473853 | 3 | 0 | 0 | T | C | 1.4952 | 0.373 | 6.10E-05 | +++?? | 44.8 | 0.1637 | intron | MGMT |
| rs35727081 | 6 | 77656432 | 4 | 0 | 0 | A | G | -1.0948 | 0.2732 | 6.13E-05 | ---?- | 16.1 | 0.311 | intergenic | |
| rs757554 | 17 | 47345847 | 4 | 0 | 0 | T | C | -1.1729 | 0.2927 | 6.13E-05 | ---?+ | 18.5 | 0.2979 | intergenic | |
| rs7751205 | 6 | 168579302 | 4 | 0 | 0 | T | C | -1.9829 | 0.4948 | 6.15E-05 | ---?- | 0 | 0.8435 | intergenic | |
| rs149272904 | 6 | 83874326 | 4 | 0 | 0 | A | G | -3.4969 | 0.8728 | 6.16E-05 | ---?- | 9 | 0.348 | intron | DOPEY1 |
| rs78554731 | 22 | 39400966 | 4 | 0 | 0 | T | C | -2.8034 | 0.6997 | 6.16E-05 | ---?- | 6.2 | 0.362 | intergenic | |
| rs12296488 | 12 | 43130547 | 4 | 0 | 0 | A | G | -1.6035 | 0.4002 | 6.17E-05 | ---?- | 20.7 | 0.2859 | intergenic | |
| rs4794049 | 17 | 47452076 | 3 | 0 | 0 | A | C | 1.2323 | 0.3076 | 6.18E-05 | +++?? | 0 | 0.9129 | intergenic | |
| rs17284012 | 5 | 66275800 | 3 | 0 | 0 | T | C | 2.5997 | 0.649 | 6.18E-05 | +++?? | 0 | 0.3888 | intron | MAST4 |
| rs217672 | 14 | 62361021 | 4 | 0 | 0 | A | C | 1.1554 | 0.2884 | 6.18E-05 | +++?+ | 0 | 0.5818 | intron | SYT16 |

| | | | | | | | | | | | | | | | |
|---|---|---|---|---|---|---|---|---|---|---|---|---|---|---|---|
| rs55688238 | 15 | 54348699 | 4 | 0 | 0 | A | G | 1.3493 | 0.3369 | 6.21E-05 | +++?+ | 0 | 0.8791 | intron | UNC13C |
| rs476549 | 16 | 5462094 | 4 | 0 | 0 | T | G | 1.0764 | 0.2688 | 6.22E-05 | +++?+ | 0 | 0.8833 | intron | RBFOX1 |
| rs144477493 | 6 | 83851376 | 4 | 0 | 0 | T | C | -3.5222 | 0.8796 | 6.22E-05 | ---?- | 0 | 0.4656 | intron | DOPEY1 |
| rs2447563 | 8 | 140464740 | 3 | 0 | 0 | A | T | 1.7224 | 0.4302 | 6.22E-05 | +++?? | 0 | 0.5281 | intergenic | |
| rs56409536 | 1 | 64800957 | 4 | 0 | 0 | A | C | -0.9734 | 0.2431 | 6.23E-05 | ---?- | 50.9 | 0.1063 | intergenic | |
| rs12423926 | 12 | 111781055 | 5 | 0 | 0 | A | G | -1.0621 | 0.2653 | 6.23E-05 | -+--- | 49.4 | 0.09486 | intron | CUX2 |
| rs114913055 | 3 | 159448496 | 4 | 0 | 0 | T | C | 2.7908 | 0.6971 | 6.24E-05 | +++?- | 21.3 | 0.2822 | intron | SCHIP1,IQCJ-SCHIP1 |
| rs948218 | 11 | 102593042 | 5 | 0 | 0 | A | T | -0.8358 | 0.2088 | 6.24E-05 | ----- | 0 | 0.8565 | intron | MMP8 |
| rs183322572 | 2 | 216988532 | 3 | 0 | 0 | A | G | -3.5453 | 0.8856 | 6.24E-05 | ---?? | 54.5 | 0.1109 | intron | XRCC5 |
| rs1390475 | 1 | 64800809 | 4 | 0 | 0 | T | C | 0.9733 | 0.2431 | 6.25E-05 | +++?+ | 50.9 | 0.1064 | intergenic | |
| rs1938525 | 10 | 62300929 | 4 | 0 | 0 | T | C | 2.6561 | 0.6636 | 6.26E-05 | +++?+ | 0 | 0.6858 | intron | ANK3 |
| rs1049056 | 6 | 32634369 | 5 | 0 | 1 | A | C | 1.2153 | 0.3038 | 6.33E-05 | +++++ | 38 | 0.168 | missense | HLA-DQB1 |
| rs13199575 | 6 | 99674560 | 4 | 0 | 0 | C | G | -1.0552 | 0.2638 | 6.35E-05 | ---?- | 58.4 | 0.06543 | intergenic | |
| rs28877654 | 12 | 111783179 | 5 | 0 | 0 | T | G | 1.0582 | 0.2646 | 6.35E-05 | +-+++ | 50.2 | 0.09046 | intron | CUX2 |
| rs77268645 | 5 | 82595179 | 4 | 0 | 0 | A | G | 2.5773 | 0.6444 | 6.36E-05 | +++?+ | 0 | 0.5126 | intron | XRCC4 |
| rs1843752 | 13 | 60001522 | 4 | 0 | 0 | A | G | -0.9978 | 0.2495 | 6.37E-05 | ---?- | 0 | 0.6981 | intergenic | |
| 6:77629298 | 6 | 77629298 | 3 | 0 | 0 | CT | C | 1.208 | 0.3021 | 6.37E-05 | +++?? | 49.9 | 0.1358 | | |
| rs6549413 | 3 | 71908430 | 4 | 0 | 0 | C | G | -1.0888 | 0.2723 | 6.38E-05 | ---?- | 0 | 0.8318 | intergenic | |
| rs1390480 | 1 | 64800510 | 4 | 0 | 0 | T | C | -0.972 | 0.2431 | 6.38E-05 | ---?- | 50.7 | 0.1076 | intergenic | |
| rs2186602 | 11 | 102591945 | 5 | 0 | 0 | T | C | 0.8358 | 0.2091 | 6.40E-05 | +++++ | 0 | 0.8561 | intron | MMP8 |
| rs2895476 | 15 | 98048437 | 3 | 0 | 0 | A | T | -1.1162 | 0.2792 | 6.40E-05 | ---?? | 0 | 0.5747 | intergenic | |
| rs1847170 | 3 | 2480037 | 4 | 0 | 0 | T | C | -1.1292 | 0.2825 | 6.41E-05 | ---?+ | 45.5 | 0.1385 | intron | CNTN4 |
| rs7450848 | 6 | 144760786 | 4 | 0 | 0 | A | G | -3.166 | 0.7921 | 6.42E-05 | ---?- | 0 | 0.6491 | intron | UTRN |
| rs59592557 | 8 | 23566084 | 3 | 0 | 0 | T | C | 1.3136 | 0.3287 | 6.44E-05 | +++?? | 0 | 0.6487 | upstream-gene | |
| rs77786255 | 13 | 74903448 | 4 | 0 | 0 | A | G | 3.1612 | 0.7912 | 6.46E-05 | +++?+ | 0 | 0.6732 | intergenic | |
| rs67147537 | 6 | 32634426 | 5 | 0 | 1 | A | G | 1.2135 | 0.3038 | 6.48E-05 | +++++ | 38.3 | 0.166 | 5-prime-UTR | HLA-DQB1 |
| rs598905 | 17 | 47329295 | 4 | 0 | 0 | C | G | -1.157 | 0.2897 | 6.49E-05 | ---?+ | 29.3 | 0.2361 | intron | FLJ40194 |
| rs114948848 | 6 | 32664797 | 5 | 0 | 1 | A | G | -1.5203 | 0.3807 | 6.50E-05 | ----- | 0 | 0.4153 | intergenic |  |
| rs16976165 | 18 | 40051678 | 4 | 0 | 0 | C | G | 1.2767 | 0.3197 | 6.51E-05 | +++?- | 0 | 0.4732 | intron | LINC00907 |
| rs2092408 | 22 | 37000586 | 3 | 0 | 0 | T | C | 1.8683 | 0.4679 | 6.52E-05 | +++?? | 0 | 0.9679 | intron | CACNG2 |
| rs201163813 | 6 | 77610283 | 3 | 0 | 0 | A | G | 1.1281 | 0.2825 | 6.53E-05 | +++?? | 41.4 | 0.1817 | intergenic | |
| rs67373575 | 6 | 32682043 | 4 | 0 | 1 | A | T | -1.6244 | 0.4069 | 6.54E-05 | ----? | 0 | 0.3968 | intergenic |  |
| rs12405224 | 1 | 64800134 | 4 | 0 | 0 | T | G | -0.9707 | 0.2432 | 6.55E-05 | ---?- | 50.6 | 0.108 | intergenic | |
| rs299363 | 5 | 134331822 | 4 | 0 | 0 | A | G | -1.0954 | 0.2745 | 6.60E-05 | ---?- | 0 | 0.6363 | intron | CATSPER3 |
| rs12004195 | 9 | 132026519 | 4 | 0 | 0 | T | C | 3.0175 | 0.7562 | 6.60E-05 | +++?+ | 0 | 0.8017 | intergenic | |
| rs114416888 | 4 | 20997560 | 4 | 0 | 0 | T | C | -2.8981 | 0.7263 | 6.60E-05 | ---?- | 38.3 | 0.1821 | intron | KCNIP4 |
| rs9542407 | 13 | 71143653 | 4 | 0 | 0 | T | C | 1.0123 | 0.2537 | 6.60E-05 | +++?+ | 0 | 0.8146 | intergenic | |
| 8:96219892 | 8 | 96219892 | 3 | 0 | 0 | CA | C | 1.0965 | 0.2748 | 6.60E-05 | +++?? | 0 | 0.6163 | | |
| rs35904708 | 6 | 77610014 | 3 | 0 | 0 | A | G | 1.1266 | 0.2824 | 6.60E-05 | +++?? | 41.3 | 0.1819 | intergenic | |
| rs10111532 | 8 | 76116517 | 4 | 0 | 0 | T | G | 1.0976 | 0.2751 | 6.61E-05 | +++?+ | 20.8 | 0.2852 | intergenic | |
| rs3765620 | 11 | 102595492 | 5 | 0 | 0 | A | G | -0.8327 | 0.2087 | 6.62E-05 | ----- | 0 | 0.8658 | missense | MMP8 |
| rs2210773 | 6 | 77647523 | 4 | 0 | 0 | A | C | -1.0902 | 0.2733 | 6.62E-05 | ---?- | 18.4 | 0.2988 | intergenic | |
| rs331104 | 2 | 150838972 | 4 | 0 | 0 | T | G | 1.0964 | 0.2748 | 6.63E-05 | +++?+ | 0 | 0.6985 | intergenic | |
| rs76449306 | 18 | 39935951 | 4 | 0 | 0 | T | G | 1.5859 | 0.3975 | 6.63E-05 | +++?+ | 0 | 0.8045 | intron | LINC00907 |
| rs35965935 | 7 | 105386201 | 4 | 0 | 0 | C | G | -1.0385 | 0.2604 | 6.64E-05 | ---?- | 0 | 0.9285 | intron | ATXN7L1 |
| rs17199558 | 16 | 23272014 | 2 | 0 | 0 | T | C | -5.7738 | 1.4476 | 6.65E-05 | -?-?? | 3 | 0.31 | intergenic | |
| rs1049107 | 6 | 32629936 | 5 | 0 | 1 | T | C | 1.2206 | 0.3061 | 6.67E-05 | +++++ | 39.6 | 0.1572 | missense | HLA-DQB1 |
| rs34904621 | 3 | 33259063 | 4 | 0 | 0 | T | C | 1.1326 | 0.284 | 6.67E-05 | +++?+ | 0 | 0.3962 | intron | SUSD5 |
| rs34214702 | 6 | 32666081 | 5 | 0 | 1 | A | C | 1.522 | 0.3817 | 6.68E-05 | +++++ | 12.4 | 0.3346 | intergenic | |
| rs7306529 | 12 | 111780644 | 5 | 0 | 0 | T | C | -1.0537 | 0.2643 | 6.68E-05 | -+--- | 50 | 0.09176 | intron | CUX2 |
| rs36062941 | 6 | 32666051 | 5 | 0 | 1 | T | C | 1.5218 | 0.3817 | 6.68E-05 | +++++ | 12.4 | 0.3346 | intergenic | |
| rs6590983 | 11 | 102594376 | 5 | 0 | 0 | A | C | 0.8322 | 0.2087 | 6.69E-05 | +++++ | 0 | 0.8661 | intron | MMP8 |
| rs4708298 | 6 | 77614319 | 3 | 0 | 0 | A | T | -1.1277 | 0.2829 | 6.70E-05 | ---?? | 40.3 | 0.1874 | intergenic | |
| rs11767159 | 7 | 107373258 | 3 | 0 | 0 | T | C | 1.1851 | 0.2973 | 6.71E-05 | +++?? | 50.8 | 0.1313 | intergenic | |
| 20:6009784 | 20 | 6009784 | 3 | 0 | 0 | AT | ATT | 1.0046 | 0.252 | 6.71E-05 | +++?? | 0 | 0.9528 | | |
| rs149014808 | 1 | 207302252 | 3 | 0 | 0 | T | C | -4.295 | 1.0777 | 6.73E-05 | ---?? | 0 | 0.943 | intron | C4BPA |
| rs7841496 | 8 | 76115944 | 4 | 0 | 0 | T | C | -1.097 | 0.2753 | 6.74E-05 | ---?- | 20.7 | 0.2859 | intergenic | |
| rs9273912 | 6 | 32629737 | 5 | 0 | 1 | T | C | 1.2201 | 0.3062 | 6.75E-05 | +++++ | 39.6 | 0.1571 | intron-near-splice | HLA-DQB1 |
| rs4949904 | 1 | 85909275 | 3 | 0 | 0 | T | C | 1.2739 | 0.3197 | 6.75E-05 | +++?? | 40.1 | 0.1886 | intron | DDAH1 |

| | | | | | | | | | | | | | | | |
|---|---|---|---|---|---|---|---|---|---|---|---|---|---|---|---|
| 9:90836984 | 9 | 90836984 | 3 | 0 | 0 | T | TA | 2.1198 | 0.532 | 6.76E-05 | +++?? | 0 | 0.386 | | |
| rs28866046 | 18 | 40047092 | 4 | 0 | 0 | A | G | 1.2708 | 0.3189 | 6.76E-05 | +++?- | 0 | 0.4622 | intron | LINC00907 |
| rs12739069 | 1 | 64799147 | 4 | 0 | 0 | T | C | -0.9685 | 0.2431 | 6.77E-05 | ---?- | 50.2 | 0.1107 | intergenic | |
| rs7961572 | 12 | 111797013 | 5 | 0 | 0 | A | G | -1.0014 | 0.2514 | 6.79E-05 | -+--- | 47.4 | 0.1072 | downstream-gene | |
| rs186570021 | 12 | 91425973 | 3 | 0 | 0 | A | G | 5.2523 | 1.3189 | 6.83E-05 | +++?? | 0 | 0.9422 | intergenic | |
| rs75406241 | 11 | 129311835 | 4 | 0 | 0 | A | G | 1.7544 | 0.4406 | 6.83E-05 | +++?+ | 30.8 | 0.2275 | intron | BARX2 |
| rs13082889 | 3 | 187898523 | 4 | 0 | 0 | T | C | -1.5939 | 0.4003 | 6.84E-05 | ---?- | 0 | 0.7077 | intron | LPP,FLJ42393 |
| rs117336243 | 6 | 83846821 | 4 | 0 | 0 | A | G | -3.5126 | 0.8822 | 6.84E-05 | ---?- | 0 | 0.4832 | intron | DOPEY1 |
| rs11651335 | 17 | 47446107 | 5 | 0 | 0 | T | C | -1.0234 | 0.257 | 6.85E-05 | ----+ | 44.2 | 0.127 | intergenic |  |
| rs56313004 | 16 | 5456157 | 4 | 0 | 0 | A | G | -1.1631 | 0.2921 | 6.86E-05 | ---?- | 0 | 0.7419 | intron | RBFOX1 |
| rs28778702 | 18 | 40045131 | 4 | 0 | 0 | A | G | -1.2742 | 0.3201 | 6.87E-05 | ---?+ | 0 | 0.4641 | intron | LINC00907 |
| rs28852767 | 6 | 32509655 | 5 | 0 | 1 | C | G | -1.4792 | 0.3716 | 6.88E-05 | ----- | 7.8 | 0.3619 | intergenic |  |
| rs11225395 | 11 | 102596480 | 5 | 0 | 0 | A | G | 0.8307 | 0.2087 | 6.89E-05 | +++++ | 0 | 0.866 | upstream-gene | |
| rs9310224 | 3 | 71916824 | 4 | 0 | 0 | T | C | 1.144 | 0.2875 | 6.91E-05 | +++?+ | 0 | 0.6312 | intergenic | |
| rs9860718 | 3 | 71917851 | 4 | 0 | 0 | T | C | 1.1355 | 0.2854 | 6.92E-05 | +++?+ | 0 | 0.6032 | intergenic | |
| rs9517873 | 13 | 100404491 | 4 | 0 | 0 | T | C | 1.1589 | 0.2912 | 6.92E-05 | +++?+ | 35.9 | 0.1967 | intron | CLYBL |
| rs181365231 | 9 | 136873240 | 4 | 0 | 0 | A | G | -3.5135 | 0.8831 | 6.93E-05 | ---?- | 30.5 | 0.2293 | intergenic | |
| rs4343159 | 13 | 60014647 | 4 | 0 | 0 | A | G | 0.9764 | 0.2454 | 6.94E-05 | +++?+ | 0 | 0.7596 | intergenic | |
| rs28454048 | 18 | 40049623 | 4 | 0 | 0 | A | G | -1.2698 | 0.3192 | 6.95E-05 | ---?+ | 0 | 0.4716 | intron | LINC00907 |
| rs28760027 | 6 | 32509985 | 5 | 0 | 1 | A | G | 1.4785 | 0.3717 | 6.95E-05 | +++++ | 8 | 0.3611 | intergenic | |
| rs552477 | 16 | 5454823 | 4 | 0 | 0 | C | G | -0.9827 | 0.247 | 6.96E-05 | ---?- | 0 | 0.6722 | intron | RBFOX1 |
| rs12528622 | 6 | 32658259 | 5 | 0 | 1 | A | G | 1.4849 | 0.3734 | 6.98E-05 | +++++ | 9.3 | 0.3532 | intergenic | |
| rs17427431 | 6 | 32663644 | 5 | 0 | 1 | A | G | 1.4849 | 0.3734 | 6.98E-05 | +++++ | 9.3 | 0.3532 | intergenic | |
| rs35122881 | 6 | 32663677 | 5 | 0 | 1 | T | C | 1.4849 | 0.3734 | 6.98E-05 | +++++ | 9.3 | 0.3532 | intergenic | |
| rs28889022 | 6 | 32510104 | 5 | 0 | 1 | T | C | -1.4796 | 0.372 | 6.98E-05 | ----- | 8.3 | 0.359 | intergenic |  |
| rs6953687 | 7 | 141162887 | 4 | 0 | 0 | A | G | -5.4382 | 1.3676 | 6.99E-05 | ---?- | 0 | 0.8298 | intron | TMEM178B |
| rs10750653 | 11 | 102600006 | 5 | 0 | 0 | A | G | 0.8305 | 0.2089 | 7.00E-05 | +++++ | 0 | 0.9037 | upstream-gene | |
| rs2184034 | 9 | 7822486 | 4 | 0 | 0 | T | C | 1.8627 | 0.4684 | 7.00E-05 | +++?+ | 34 | 0.2083 | intergenic | |
| 5:179966295 | 5 | 179966295 | 3 | 0 | 0 | CA | C | 1.2527 | 0.3151 | 7.03E-05 | +++?? | 32.3 | 0.2281 | | |
| rs17091783 | 12 | 43137564 | 4 | 0 | 0 | A | G | 1.6455 | 0.414 | 7.04E-05 | +++?+ | 0 | 0.3962 | intergenic | |
| rs1871230 | 2 | 218340808 | 4 | 0 | 0 | T | C | -0.9694 | 0.2439 | 7.05E-05 | ---?- | 0 | 0.6422 | intron | DIRC3 |
| rs5757429 | 22 | 39424124 | 4 | 0 | 0 | T | C | 1.6073 | 0.4044 | 7.05E-05 | +++?+ | 0 | 0.8825 | intron | APOBEC3D |
| rs818524 | 1 | 85201228 | 5 | 0 | 0 | T | C | 0.9126 | 0.2296 | 7.06E-05 | +++++ | 26.6 | 0.2439 | intergenic | |
| rs66881448 | 6 | 32508843 | 5 | 0 | 1 | A | G | 1.4799 | 0.3724 | 7.07E-05 | +++++ | 8.5 | 0.3583 | intergenic | |
| 11:19824257 | 11 | 19824257 | 3 | 0 | 0 | CA | C | 2.7124 | 0.6826 | 7.08E-05 | +++?? | 29.4 | 0.2426 | | |
| rs77916100 | 2 | 116424275 | 3 | 0 | 0 | A | C | 3.0419 | 0.7656 | 7.09E-05 | +++?? | 58.3 | 0.09099 | intron | DPP10 |
| rs3758858 | 11 | 102598457 | 5 | 0 | 0 | A | G | -0.8291 | 0.2087 | 7.10E-05 | ----- | 0 | 0.8696 | upstream-gene |  |
| rs116130911 | 2 | 122473097 | 4 | 0 | 0 | A | G | -3.2096 | 0.8079 | 7.10E-05 | ---?- | 0 | 0.8148 | intron | LOC254128 |
| rs12583031 | 13 | 60014508 | 4 | 0 | 0 | A | G | 0.9747 | 0.2454 | 7.11E-05 | +++?+ | 0 | 0.7584 | intergenic | |
| rs16976162 | 18 | 40050171 | 4 | 0 | 0 | T | C | 1.2679 | 0.3192 | 7.11E-05 | +++?- | 0 | 0.4749 | intron | LINC00907 |
| rs3758857 | 11 | 102598359 | 5 | 0 | 0 | T | C | 0.8289 | 0.2087 | 7.12E-05 | +++++ | 0 | 0.8692 | upstream-gene | |
| rs139633505 | 7 | 156413003 | 4 | 0 | 0 | T | C | 4.3309 | 1.0904 | 7.14E-05 | +++?- | 54.1 | 0.08818 | intergenic | |
| rs6891344 | 5 | 123136656 | 4 | 0 | 0 | A | G | -1.313 | 0.3306 | 7.14E-05 | ---?- | 0.4 | 0.3899 | intergenic | |
| rs1130432 | 6 | 32629129 | 5 | 0 | 0 | T | C | -1.2211 | 0.3076 | 7.18E-05 | ----- | 39.8 | 0.1559 | missense | HLA-DQB1 |
| rs7453913 | 6 | 77580164 | 3 | 0 | 0 | A | C | 1.1276 | 0.284 | 7.18E-05 | +++?? | 39.2 | 0.1928 | intergenic | |
| rs113863000 | 3 | 126464440 | 4 | 0 | 0 | A | G | 2.9326 | 0.7387 | 7.19E-05 | +++?+ | 0 | 0.999 | intron | CHCHD6 |
| rs79870097 | 6 | 32661995 | 5 | 0 | 1 | T | C | -1.5105 | 0.3805 | 7.20E-05 | ----- | 0 | 0.5021 | intergenic |  |
| 6:106184720 | 6 | 106184720 | 3 | 0 | 0 | A | C | -3.2067 | 0.8079 | 7.21E-05 | ---?? | 39.6 | 0.1908 | intergenic | |
| rs72844399 | 6 | 32634264 | 5 | 0 | 1 | T | C | 1.2078 | 0.3043 | 7.22E-05 | +++++ | 38.1 | 0.1672 | intron | HLA-DQB1 |
| rs75400359 | 6 | 32517990 | 5 | 0 | 1 | A | G | 1.4116 | 0.3556 | 7.22E-05 | +++++ | 8.9 | 0.3556 | downstream-gene | |
| 14:43248413 | 14 | 43248413 | 3 | 0 | 0 | A | AT | 1.571 | 0.3958 | 7.22E-05 | +++?? | 52.8 | 0.1201 | | |
| rs28418438 | 16 | 15068908 | 3 | 0 | 0 | C | G | -1.5443 | 0.3891 | 7.23E-05 | ---?? | 0 | 0.7725 | 5-prime-UTR | PDXDC1 |
| rs4309288 | 13 | 60015124 | 4 | 0 | 0 | A | G | -0.9751 | 0.2457 | 7.23E-05 | ---?- | 0 | 0.7539 | intergenic | |
| rs9538425 | 13 | 60014172 | 4 | 0 | 0 | A | G | 0.9732 | 0.2452 | 7.23E-05 | +++?+ | 0 | 0.7625 | intergenic | |
| 6:106184721 | 6 | 106184721 | 3 | 0 | 0 | T | TA | -3.2065 | 0.808 | 7.23E-05 | ---?? | 39.7 | 0.1907 | | |
| rs9538426 | 13 | 60014179 | 4 | 0 | 0 | A | G | 0.9732 | 0.2452 | 7.23E-05 | +++?+ | 0 | 0.7622 | intergenic | |
| rs8040917 | 15 | 75592851 | 3 | 0 | 0 | C | G | 1.9329 | 0.4871 | 7.23E-05 | +++?? | 0 | 0.9773 | downstream-gene | |
| 6:24071036 | 6 | 24071036 | 3 | 0 | 0 | CA | C | 1.759 | 0.4433 | 7.23E-05 | +++?? | 0 | 0.5978 | | |
| rs71536508 | 6 | 32518030 | 5 | 0 | 1 | A | G | -1.4115 | 0.3557 | 7.24E-05 | ----- | 8.9 | 0.3555 | downstream-gene |  |

| | | | | | | | | | | | | | | | |
|---|---|---|---|---|---|---|---|---|---|---|---|---|---|---|---|
| rs11963589 | 6 | 168584187 | 4 | 0 | 0 | T | C | -1.7879 | 0.4506 | 7.24E-05 | ---?- | 0 | 0.6125 | intergenic | |
| rs1939014 | 11 | 102597912 | 5 | 0 | 0 | A | T | 0.828 | 0.2087 | 7.25E-05 | +++++ | 0 | 0.8679 | upstream-gene | |
| 13:73819341 | 13 | 73819341 | 3 | 0 | 0 | CA | C | -1.7465 | 0.4402 | 7.25E-05 | ---?? | 0 | 0.5496 | | |
| 6:106184722 | 6 | 106184722 | 3 | 0 | 0 | T | C | -3.2061 | 0.8081 | 7.26E-05 | ---?? | 39.6 | 0.1908 | intergenic | |
| rs9462089 | 6 | 35440192 | 5 | 0 | 0 | A | G | 3.1295 | 0.7889 | 7.28E-05 | +++++ | 0 | 0.9027 | downstream-gene | |
| rs62552264 | 9 | 11814454 | 4 | 0 | 0 | A | G | 1.9909 | 0.5018 | 7.28E-05 | +++?+ | 58.5 | 0.06476 | intergenic | |
| rs4337195 | 13 | 60013930 | 4 | 0 | 0 | A | G | 0.9725 | 0.2452 | 7.28E-05 | +++?+ | 0 | 0.7654 | intergenic | |
| rs6590986 | 11 | 102598859 | 5 | 0 | 0 | T | C | -0.8281 | 0.2088 | 7.29E-05 | ----- | 0 | 0.8714 | upstream-gene |  |
| rs7583077 | 2 | 156919559 | 4 | 0 | 0 | T | C | 1.056 | 0.2662 | 7.30E-05 | +++?+ | 0 | 0.9411 | intergenic | |
| rs208966 | 15 | 85559617 | 4 | 0 | 0 | T | C | -0.9563 | 0.2411 | 7.30E-05 | ---?+ | 58.1 | 0.06702 | intron | PDE8A |
| 6:106184723 | 6 | 106184723 | 3 | 0 | 0 | A | T | 3.2052 | 0.8081 | 7.30E-05 | +++?? | 39.6 | 0.1907 | intergenic | |
| rs7148769 | 14 | 62359041 | 4 | 0 | 0 | T | C | -1.2947 | 0.3264 | 7.31E-05 | ---?- | 0 | 0.494 | intron | SYT16 |
| rs299365 | 5 | 134332469 | 4 | 0 | 0 | T | G | 1.0819 | 0.2728 | 7.31E-05 | +++?+ | 0 | 0.64 | intron | CATSPER3 |
| rs1322100 | 6 | 77740109 | 4 | 0 | 0 | A | C | -1.1612 | 0.2928 | 7.33E-05 | ---?- | 30.9 | 0.2268 | intergenic | |
| rs1390476 | 1 | 64800659 | 4 | 0 | 0 | T | C | -0.9609 | 0.2423 | 7.33E-05 | ---?- | 51.6 | 0.1023 | intergenic | |
| rs1939013 | 11 | 102597810 | 5 | 0 | 0 | T | C | 0.8274 | 0.2087 | 7.34E-05 | +++++ | 0 | 0.867 | upstream-gene | |
| rs10849937 | 12 | 111792427 | 5 | 0 | 0 | A | G | -1.042 | 0.2628 | 7.34E-05 | -+--- | 48.3 | 0.102 | downstream-gene | |
| rs1390477 | 1 | 64800647 | 4 | 0 | 0 | T | C | -0.9608 | 0.2423 | 7.35E-05 | ---?- | 51.6 | 0.1024 | intergenic | |
| rs299369 | 5 | 134336705 | 4 | 0 | 0 | T | G | 1.0734 | 0.2707 | 7.36E-05 | +++?+ | 0 | 0.614 | intron | CATSPER3 |
| rs7621924 | 3 | 65840591 | 3 | 0 | 0 | C | G | 2.3295 | 0.5876 | 7.37E-05 | +++?? | 15.4 | 0.3067 | intron | MAGI1 |
| rs12039037 | 1 | 187897389 | 4 | 0 | 0 | A | G | -1.2027 | 0.3034 | 7.38E-05 | ---?- | 0 | 0.755 | intergenic | |
| rs1390479 | 1 | 64800585 | 4 | 0 | 0 | T | C | -0.9603 | 0.2423 | 7.39E-05 | ---?- | 51.5 | 0.1028 | intergenic | |
| rs17201763 | 6 | 32185246 | 5 | 0 | 0 | T | G | 1.9336 | 0.4881 | 7.44E-05 | ++++- | 33.4 | 0.1984 | intron | NOTCH4 |
| rs142086573 | 12 | 91308646 | 3 | 0 | 0 | T | C | -5.4273 | 1.37 | 7.45E-05 | ---?? | 0 | 0.9098 | upstream-gene | |
| rs59447536 | 18 | 40047692 | 4 | 0 | 0 | A | C | -1.2636 | 0.319 | 7.46E-05 | ---?+ | 0 | 0.4675 | intron | LINC00907 |
| rs73741246 | 5 | 10301616 | 4 | 0 | 0 | C | G | 2.2375 | 0.5649 | 7.46E-05 | +++?+ | 45.9 | 0.1358 | intron | CMBL |
| rs10015016 | 4 | 176432982 | 4 | 0 | 0 | A | G | 3.0335 | 0.7658 | 7.46E-05 | +++?+ | 0 | 0.7231 | intergenic | |
| rs73217462 | 8 | 21468749 | 3 | 0 | 0 | C | G | -3.0872 | 0.7794 | 7.46E-05 | ---?? | 0 | 0.5443 | intergenic | |
| rs9621452 | 22 | 32840741 | 4 | 0 | 0 | T | C | -1.8349 | 0.4632 | 7.46E-05 | ---?- | 0 | 0.9482 | intron | BPIFC |
| rs5004245 | 1 | 64800486 | 4 | 0 | 0 | A | G | -0.9597 | 0.2423 | 7.46E-05 | ---?- | 51.4 | 0.1033 | intergenic | |
| rs5004244 | 1 | 64800456 | 4 | 0 | 0 | T | C | -0.9596 | 0.2423 | 7.49E-05 | ---?- | 51.4 | 0.1035 | intergenic | |
| rs4335683 | 13 | 60013629 | 4 | 0 | 0 | A | G | 0.9701 | 0.245 | 7.49E-05 | +++?+ | 0 | 0.7715 | intergenic | |
| rs4319644 | 13 | 60013628 | 4 | 0 | 0 | T | C | -0.97 | 0.245 | 7.51E-05 | ---?- | 0 | 0.7716 | intergenic | |
| 8:140402248 | 8 | 140402248 | 3 | 0 | 0 | A | ATACT | 1.5453 | 0.3903 | 7.52E-05 | +++?? | 0 | 0.9226 | | |
| rs61090270 | 3 | 71895116 | 4 | 0 | 0 | A | C | 1.077 | 0.272 | 7.52E-05 | +++?+ | 0 | 0.8724 | intergenic | |
| rs12854181 | 13 | 60014434 | 4 | 0 | 0 | T | C | 0.971 | 0.2453 | 7.53E-05 | +++?+ | 0 | 0.7644 | intergenic | |
| rs5004243 | 1 | 64800345 | 4 | 0 | 0 | T | C | 0.9593 | 0.2423 | 7.53E-05 | +++?+ | 51.4 | 0.1038 | intergenic | |
| rs9570129 | 13 | 60013483 | 4 | 0 | 0 | C | G | 0.9698 | 0.245 | 7.53E-05 | +++?+ | 0 | 0.7718 | intergenic | |
| rs9570128 | 13 | 60013450 | 4 | 0 | 0 | A | G | -0.9698 | 0.245 | 7.54E-05 | ---?- | 0 | 0.7718 | intergenic | |
| rs497811 | 3 | 191148875 | 4 | 0 | 0 | A | C | -1.494 | 0.3774 | 7.54E-05 | ---?+ | 37.2 | 0.1886 | intergenic | |
| rs874286 | 12 | 111799681 | 5 | 0 | 0 | T | C | -0.9968 | 0.2518 | 7.55E-05 | -+--- | 47.3 | 0.1077 | 3-prime-UTR | FAM109A |
| 7:107343129 | 7 | 107343129 | 3 | 0 | 0 | CA | C | -1.1498 | 0.2905 | 7.55E-05 | ---?? | 36.1 | 0.2089 | | |
| rs10865559 | 3 | 77406508 | 4 | 0 | 0 | T | C | 0.9817 | 0.248 | 7.57E-05 | +++?+ | 0 | 0.5594 | intron | ROBO2 |
| rs9570127 | 13 | 60013402 | 4 | 0 | 0 | A | G | 0.9695 | 0.245 | 7.58E-05 | +++?+ | 0 | 0.7722 | intergenic | |
| rs13273502 | 8 | 76112835 | 4 | 0 | 0 | T | G | 1.0927 | 0.2761 | 7.58E-05 | +++?+ | 20.3 | 0.2881 | intergenic | |
| rs12407651 | 1 | 64800026 | 4 | 0 | 0 | A | T | 0.9587 | 0.2423 | 7.60E-05 | +++?+ | 51.2 | 0.1044 | intergenic | |
| rs9570126 | 13 | 60013303 | 4 | 0 | 0 | T | C | 0.9693 | 0.245 | 7.61E-05 | +++?+ | 0 | 0.7725 | intergenic | |
| rs9570125 | 13 | 60013265 | 4 | 0 | 0 | T | C | 0.9692 | 0.245 | 7.63E-05 | +++?+ | 0 | 0.7726 | intergenic | |
| rs73469184 | 18 | 39933192 | 4 | 0 | 0 | A | T | -1.5613 | 0.3947 | 7.64E-05 | ---?- | 0 | 0.8349 | intron | LINC00907 |
| rs9570124 | 13 | 60013140 | 4 | 0 | 0 | T | C | 0.9691 | 0.245 | 7.64E-05 | +++?+ | 0 | 0.7728 | intergenic | |
| rs9570123 | 13 | 60013001 | 4 | 0 | 0 | A | G | -0.969 | 0.245 | 7.67E-05 | ---?- | 0 | 0.772 | intergenic | |
| 6:31997909 | 6 | 31997909 | 3 | 0 | 0 | A | C | 1.4783 | 0.3738 | 7.67E-05 | +++?? | 27.6 | 0.2511 | intron | C4B |
| rs350833 | 19 | 4068878 | 4 | 0 | 0 | A | C | -3.231 | 0.8171 | 7.67E-05 | ---?- | 0 | 0.9762 | upstream-gene | |
| rs10849938 | 12 | 111800059 | 5 | 0 | 0 | T | C | -0.9958 | 0.2519 | 7.70E-05 | -+--- | 47.2 | 0.1086 | 3-prime-UTR | FAM109A |
| rs1577425 | 6 | 77716404 | 4 | 0 | 0 | T | C | -1.173 | 0.2967 | 7.71E-05 | ---?- | 15 | 0.317 | intergenic | |
| rs1470637 | 1 | 205033283 | 5 | 0 | 0 | A | G | 0.8633 | 0.2184 | 7.71E-05 | +++++ | 0 | 0.8976 | intron | CNTN2 |
| rs4638458 | 13 | 60008313 | 4 | 0 | 0 | T | G | 0.9738 | 0.2463 | 7.72E-05 | +++?+ | 0 | 0.776 | intergenic | |
| rs7398343 | 12 | 111774068 | 5 | 0 | 0 | A | C | -1.032 | 0.2611 | 7.72E-05 | -+--- | 56.3 | 0.05728 | intron | CUX2 |
| rs4736097 | 8 | 140813683 | 4 | 0 | 0 | A | G | -1.4886 | 0.3766 | 7.72E-05 | ---?- | 61.6 | 0.0501 | intron | TRAPPC9 |

| | | | | | | | | | | | | | | | |
|---|---|---|---|---|---|---|---|---|---|---|---|---|---|---|---|
| rs72975533 | 2 | 235413234 | 4 | 0 | 0 | A | G | -1.5115 | 0.3824 | 7.73E-05 | ---?- | 0 | 0.7696 | intergenic | |
| rs809306 | 20 | 5682782 | 3 | 0 | 0 | A | C | -1.2162 | 0.3077 | 7.73E-05 | ---?? | 0 | 0.4862 | intergenic | |
| rs72848251 | 6 | 32601288 | 5 | 0 | 0 | A | G | 1.2164 | 0.3078 | 7.74E-05 | +++++ | 29 | 0.2284 | upstream-gene | |
| rs148788468 | 1 | 102927842 | 3 | 0 | 0 | T | C | 0.9823 | 0.2485 | 7.74E-05 | +++?? | 0 | 0.9474 | intergenic | |
| rs2890763 | 9 | 7820989 | 4 | 0 | 0 | A | G | 1.8959 | 0.4797 | 7.75E-05 | +++?+ | 2.9 | 0.3779 | intergenic | |
| rs1577426 | 6 | 77716185 | 4 | 0 | 0 | T | C | 1.1735 | 0.2969 | 7.75E-05 | +++?+ | 15 | 0.3172 | intergenic | |
| rs146459802 | 8 | 4071329 | 4 | 0 | 0 | A | T | 4.6989 | 1.189 | 7.75E-05 | +++?+ | 0 | 0.5392 | intron | CSMD1 |
| rs4713574 | 6 | 32627038 | 4 | 0 | 0 | C | G | -0.9289 | 0.2351 | 7.79E-05 | ----? | 0 | 0.4292 | downstream-gene |  |
| rs57835944 | 18 | 40045822 | 4 | 0 | 0 | A | G | -1.2603 | 0.319 | 7.79E-05 | ---?+ | 0 | 0.4652 | intron | LINC00907 |
| rs9563727 | 13 | 60012692 | 4 | 0 | 0 | A | G | 0.9689 | 0.2452 | 7.80E-05 | +++?+ | 0 | 0.7474 | intergenic | |
| rs113684515 | 6 | 83841659 | 4 | 0 | 0 | A | G | 3.4956 | 0.8849 | 7.80E-05 | +++?+ | 0 | 0.5077 | intron | DOPEY1 |
| rs9869453 | 3 | 71897788 | 4 | 0 | 0 | A | G | 1.0771 | 0.2727 | 7.82E-05 | +++?+ | 0 | 0.8795 | intergenic | |
| rs35671040 | 4 | 190023705 | 4 | 0 | 0 | T | G | 0.9515 | 0.2409 | 7.82E-05 | +++?+ | 13.1 | 0.3272 | intergenic | |
| rs9563726 | 13 | 60012691 | 4 | 0 | 0 | T | C | 0.9687 | 0.2453 | 7.82E-05 | +++?+ | 0 | 0.7476 | intergenic | |
| 5:179969208 | 5 | 179969208 | 3 | 0 | 0 | T | TAGGTT | 1.2498 | 0.3164 | 7.82E-05 | +++?? | 31.7 | 0.2314 | | |
| 2:170722660 | 2 | 170722660 | 3 | 0 | 0 | CAAAAA | C | -1.2262 | 0.3105 | 7.82E-05 | ---?? | 64.1 | 0.06174 | | |
| rs6590985 | 11 | 102598655 | 5 | 0 | 0 | T | C | 0.8247 | 0.2088 | 7.85E-05 | +++++ | 0 | 0.8718 | upstream-gene | |
| rs13096994 | 3 | 187897283 | 4 | 0 | 0 | A | G | 1.5703 | 0.3977 | 7.86E-05 | +++?+ | 0 | 0.7123 | intron | LPP,FLJ42393 |
| rs16888465 | 6 | 77741488 | 4 | 0 | 0 | T | C | 1.1604 | 0.2939 | 7.89E-05 | +++?+ | 21.9 | 0.279 | intergenic | |
| rs7115590 | 11 | 391450 | 4 | 0 | 0 | C | G | -1.2252 | 0.3104 | 7.90E-05 | ---?- | 0 | 0.5286 | upstream-gene | |
| rs2131751 | 16 | 84843207 | 3 | 0 | 0 | T | C | 1.3209 | 0.3346 | 7.91E-05 | +++?? | 0 | 0.5891 | intergenic | |
| rs146287659 | 6 | 99744936 | 4 | 0 | 0 | T | C | -3.0084 | 0.7622 | 7.91E-05 | ---?- | 0 | 0.8545 | intron | FAXC |
| rs9570120 | 13 | 60012431 | 4 | 0 | 0 | A | G | -0.9673 | 0.2451 | 7.91E-05 | ---?- | 0 | 0.7751 | intergenic | |
| rs9570119 | 13 | 60012404 | 4 | 0 | 0 | A | G | -0.9672 | 0.2451 | 7.93E-05 | ---?- | 0 | 0.7753 | intergenic | |
| rs34713614 | 6 | 77748678 | 4 | 0 | 0 | T | C | 1.173 | 0.2973 | 7.95E-05 | +++?+ | 20.2 | 0.2888 | intergenic | |
| rs12869748 | 13 | 60012648 | 4 | 0 | 0 | T | C | -0.9671 | 0.2451 | 7.95E-05 | ---?- | 0 | 0.775 | intergenic | |
| rs2184781 | 13 | 60008998 | 4 | 0 | 0 | T | C | -0.9708 | 0.246 | 7.96E-05 | ---?- | 0 | 0.7776 | intergenic | |
| rs62433220 | 6 | 77610939 | 3 | 0 | 0 | T | C | 1.114 | 0.2823 | 7.96E-05 | +++?? | 49.9 | 0.1357 | intergenic | |
| rs11553979 | 6 | 35436571 | 5 | 0 | 0 | T | C | -3.1189 | 0.7905 | 7.97E-05 | ----- | 0 | 0.863 | intron-near-splice | RPL10A |
| rs12527643 | 6 | 77742851 | 4 | 0 | 0 | A | G | 1.1616 | 0.2944 | 7.97E-05 | +++?+ | 21.3 | 0.2822 | intergenic | |
| rs116881264 | 6 | 155753110 | 5 | 0 | 0 | A | C | 3.1988 | 0.8108 | 7.97E-05 | +++++ | 44.4 | 0.1258 | intron | NOX3 |
| rs2152657 | 13 | 60009096 | 4 | 0 | 0 | A | G | -0.9705 | 0.246 | 7.98E-05 | ---?- | 0 | 0.7777 | intergenic | |
| rs1806965 | 13 | 60011518 | 4 | 0 | 0 | T | C | 0.9669 | 0.2451 | 8.00E-05 | +++?+ | 0 | 0.7765 | intergenic | |
| rs7944118 | 11 | 34359965 | 3 | 0 | 0 | T | C | -5.3847 | 1.3651 | 8.00E-05 | ---?? | 0 | 0.7425 | intron | ABTB2 |
| rs11754143 | 6 | 77725748 | 4 | 0 | 0 | T | C | -1.1657 | 0.2955 | 8.01E-05 | ---?- | 17.4 | 0.3042 | intergenic | |
| rs4713406 | 6 | 30980943 | 3 | 0 | 0 | T | G | 1.7666 | 0.4479 | 8.02E-05 | +++?? | 0 | 0.8078 | intron | MUC22 |
| rs1577427 | 6 | 77716049 | 4 | 0 | 0 | T | C | 1.1693 | 0.2965 | 8.02E-05 | +++?+ | 15 | 0.3168 | intergenic | |
| rs34770694 | 6 | 77713143 | 4 | 0 | 0 | A | G | 1.17 | 0.2967 | 8.04E-05 | +++?+ | 14.4 | 0.3201 | intergenic | |
| rs746815 | 13 | 60012238 | 4 | 0 | 0 | T | C | -0.9664 | 0.2451 | 8.04E-05 | ---?- | 0 | 0.7761 | intergenic | |
| rs9570121 | 13 | 60012602 | 4 | 0 | 0 | T | G | 0.9667 | 0.2452 | 8.05E-05 | +++?+ | 0 | 0.7765 | intergenic | |
| 20:6016187 | 20 | 6016187 | 3 | 0 | 0 | A | AT | 0.9618 | 0.2439 | 8.05E-05 | +++?? | 0 | 0.916 | | |
| rs79022314 | 13 | 71138734 | 4 | 0 | 0 | T | C | 2.7733 | 0.7035 | 8.07E-05 | +++?+ | 0 | 0.6838 | intergenic | |
| rs184221110 | 9 | 101141296 | 3 | 0 | 0 | C | G | 4.3768 | 1.1105 | 8.11E-05 | +++?? | 28.3 | 0.2477 | intron | GABBR2 |
| rs76566928 | 6 | 168583942 | 4 | 0 | 0 | T | C | -1.771 | 0.4494 | 8.11E-05 | ---?- | 0 | 0.6022 | intergenic | |
| rs746817 | 13 | 60011900 | 4 | 0 | 0 | A | G | 0.9659 | 0.2451 | 8.11E-05 | +++?+ | 0 | 0.777 | intergenic | |
| rs13216145 | 6 | 77743979 | 4 | 0 | 0 | T | C | -1.1615 | 0.2947 | 8.12E-05 | ---?- | 21.3 | 0.2828 | intergenic | |
| rs415331 | 3 | 191148436 | 4 | 0 | 0 | T | C | 1.4803 | 0.3757 | 8.13E-05 | +++?- | 36.5 | 0.1929 | intergenic | |
| rs117452990 | 15 | 56204268 | 2 | 0 | 0 | A | C | -7.9307 | 2.0128 | 8.14E-05 | ?--?? | 0 | 0.4311 | intron | NEDD4 |
| rs71565344 | 6 | 32673965 | 5 | 0 | 1 | A | G | 1.5039 | 0.3817 | 8.15E-05 | +++++ | 18.8 | 0.2951 | intergenic | |
| rs78755075 | 17 | 52329129 | 4 | 0 | 0 | T | C | 2.5808 | 0.6551 | 8.16E-05 | +++?- | 0 | 0.7866 | intergenic | |
| 16:12613075 | 16 | 12613075 | 3 | 0 | 0 | CCA | TCA | -7.9557 | 2.0195 | 8.17E-05 | ---?? | 42.1 | 0.1779 | | |
| rs72774993 | 2 | 2769662 | 4 | 0 | 0 | T | G | -2.0128 | 0.511 | 8.19E-05 | ---?- | 0 | 0.4923 | intergenic | |
| 6:77608366 | 6 | 77608366 | 3 | 0 | 0 | T | TA | -1.1082 | 0.2815 | 8.24E-05 | ---?? | 49.8 | 0.1362 | | |
| 6:77609569 | 6 | 77609569 | 3 | 0 | 0 | A | AT | -1.1093 | 0.2817 | 8.24E-05 | ---?? | 49.9 | 0.1357 | | |
| rs4496010 | 13 | 60015153 | 4 | 0 | 0 | T | C | 0.9678 | 0.2458 | 8.25E-05 | +++?+ | 0 | 0.7579 | intergenic | |
| rs4530018 | 13 | 60015132 | 4 | 0 | 0 | T | C | 0.9677 | 0.2458 | 8.25E-05 | +++?+ | 0 | 0.7581 | intergenic | |
| rs143039420 | 2 | 105050518 | 4 | 0 | 0 | A | C | -4.0443 | 1.0274 | 8.27E-05 | ---?- | 0 | 0.5244 | upstream-gene | |
| 6:77607516 | 6 | 77607516 | 3 | 0 | 0 | T | TG | 1.1075 | 0.2814 | 8.28E-05 | +++?? | 50 | 0.1352 | | |
| rs73217457 | 8 | 21457627 | 3 | 0 | 0 | A | G | -2.995 | 0.7609 | 8.28E-05 | ---?? | 0 | 0.5057 | intergenic | |

| | | | | | | | | | | | | | | | |
|---|---|---|---|---|---|---|---|---|---|---|---|---|---|---|---|
| rs79348499 | 13 | 74989267 | 4 | 0 | 0 | C | G | 3.2167 | 0.8173 | 8.29E-05 | +++?+ | 0 | 0.8698 | upstream-gene | |
| rs75686544 | 7 | 153662692 | 4 | 0 | 0 | A | G | 1.5401 | 0.3913 | 8.31E-05 | +++?+ | 0 | 0.7051 | intron | DPP6 |
| 4:100548138 | 4 | 100548138 | 3 | 0 | 0 | A | G | 1.7862 | 0.4539 | 8.32E-05 | +++?? | 54.7 | 0.11 | downstream-gene | |
| 6:77609214 | 6 | 77609214 | 3 | 0 | 0 | A | AAGAGA | -1.108 | 0.2816 | 8.33E-05 | ---?? | 49.6 | 0.1373 | | |
| rs67662266 | 6 | 77609233 | 3 | 0 | 0 | A | G | 1.1079 | 0.2816 | 8.34E-05 | +++?? | 49.6 | 0.1374 | intergenic | |
| rs4886163 | 13 | 60011075 | 4 | 0 | 0 | A | G | 0.9644 | 0.2452 | 8.36E-05 | +++?+ | 0 | 0.7793 | intergenic | |
| 6:77609878 | 6 | 77609878 | 3 | 0 | 0 | G | GAACT | 1.1075 | 0.2815 | 8.37E-05 | +++?? | 49.5 | 0.138 | | |
| rs9447898 | 6 | 77610103 | 3 | 0 | 0 | T | C | 1.1075 | 0.2816 | 8.38E-05 | +++?? | 49.4 | 0.1385 | intergenic | |
| rs13210002 | 6 | 77585881 | 3 | 0 | 0 | A | G | -1.1126 | 0.2829 | 8.39E-05 | ---?? | 42.7 | 0.1746 | intergenic | |
| rs183265686 | 19 | 17078647 | 2 | 0 | 0 | A | G | 6.1759 | 1.5706 | 8.41E-05 | ???++ | 7.3 | 0.2989 | intron | CPAMD8 |
| rs78959106 | 3 | 38537237 | 5 | 0 | 0 | A | G | 2.5941 | 0.6597 | 8.42E-05 | +++++ | 0 | 0.744 | upstream-gene | |
| rs141874684 | 6 | 77611234 | 3 | 0 | 0 | T | C | -1.1073 | 0.2816 | 8.44E-05 | ---?? | 49.2 | 0.1398 | intergenic | |
| rs189917403 | 9 | 101141297 | 3 | 0 | 0 | A | C | 4.3676 | 1.1109 | 8.44E-05 | +++?? | 27.3 | 0.2525 | intron | GABBR2 |
| rs142216966 | 18 | 40043519 | 4 | 0 | 0 | A | G | -1.2532 | 0.3188 | 8.45E-05 | ---?+ | 0 | 0.4715 | intron | LINC00907 |
| rs72958595 | 1 | 100209631 | 3 | 0 | 0 | A | G | -3.145 | 0.8 | 8.45E-05 | ---?? | 0 | 0.78 | intron | FRRS1 |
| rs1160785 | 13 | 60010326 | 4 | 0 | 0 | T | C | 0.9647 | 0.2454 | 8.46E-05 | +++?+ | 0 | 0.7808 | intergenic | |
| rs6490077 | 12 | 111791677 | 5 | 0 | 0 | A | C | 0.9837 | 0.2503 | 8.46E-05 | +-+++ | 44.8 | 0.1234 | downstream-gene | |
| 6:23872241 | 6 | 23872241 | 3 | 0 | 0 | T | TAC | 2.9068 | 0.7395 | 8.47E-05 | +++?? | 0 | 0.4833 | | |
| rs4270316 | 2 | 170619349 | 4 | 0 | 0 | A | G | 1.3117 | 0.3338 | 8.50E-05 | +++?+ | 0 | 0.6219 | intergenic | |
| 14:22504306 | 14 | 22504306 | 3 | 0 | 0 | T | TAAAG | -1.1128 | 0.2832 | 8.50E-05 | ---?? | 0 | 0.9877 | | |
| rs78338435 | 10 | 128259311 | 4 | 0 | 0 | T | C | -4.1822 | 1.0643 | 8.51E-05 | ---?- | 0 | 0.8207 | intergenic | |
| rs7139615 | 13 | 60007764 | 4 | 0 | 0 | T | C | 0.9698 | 0.2468 | 8.51E-05 | +++?+ | 0 | 0.779 | intergenic | |
| rs1160784 | 13 | 60010451 | 4 | 0 | 0 | A | C | -0.9638 | 0.2453 | 8.53E-05 | ---?- | 0 | 0.7815 | intergenic | |
| rs112247764 | 18 | 40044393 | 4 | 0 | 0 | A | G | 1.2527 | 0.3189 | 8.57E-05 | +++?- | 0 | 0.4691 | intron | LINC00907 |
| 5:179973031 | 5 | 179973031 | 3 | 0 | 0 | CT | C | 1.2398 | 0.3156 | 8.57E-05 | +++?? | 31.3 | 0.2334 | | |
| rs13217113 | 6 | 77747893 | 4 | 0 | 0 | T | C | 1.1623 | 0.2959 | 8.59E-05 | +++?+ | 20.8 | 0.2851 | intergenic | |
| rs72704202 | 9 | 11812469 | 4 | 0 | 0 | C | G | -1.9708 | 0.5019 | 8.60E-05 | ---?- | 55.2 | 0.08206 | intergenic | |
| rs117020040 | 10 | 131484799 | 4 | 0 | 0 | T | C | 1.6213 | 0.4128 | 8.60E-05 | +++?+ | 0 | 0.9053 | intron | MGMT |
| rs2322164 | 3 | 71896948 | 4 | 0 | 0 | A | C | 1.0696 | 0.2724 | 8.61E-05 | +++?+ | 0 | 0.8839 | intergenic | |
| rs11181671 | 12 | 43145517 | 4 | 0 | 0 | A | G | 1.6023 | 0.4081 | 8.62E-05 | +++?+ | 0 | 0.4472 | intergenic | |
| rs6921437 | 6 | 77615393 | 3 | 0 | 0 | A | G | -1.107 | 0.2819 | 8.63E-05 | ---?? | 48 | 0.146 | intergenic | |
| 6:77615387 | 6 | 77615387 | 3 | 0 | 0 | C | G | 1.1069 | 0.2819 | 8.63E-05 | +++?? | 48 | 0.146 | intergenic | |
| rs695885 | 3 | 191150638 | 4 | 0 | 0 | T | G | -1.4758 | 0.376 | 8.66E-05 | ---?+ | 33.3 | 0.2126 | intergenic | |
| rs72975538 | 2 | 235416711 | 4 | 0 | 0 | C | G | 1.4992 | 0.382 | 8.69E-05 | +++?+ | 0 | 0.8627 | intergenic | |
| rs9538428 | 13 | 60015219 | 4 | 0 | 0 | A | G | 0.965 | 0.2459 | 8.69E-05 | +++?+ | 0 | 0.7618 | intergenic | |
| rs7143855 | 14 | 22508676 | 4 | 0 | 0 | T | C | 1.0616 | 0.2705 | 8.69E-05 | +++?+ | 0 | 0.8089 | intergenic | |
| rs9470038 | 6 | 35444903 | 5 | 0 | 0 | A | G | 3.1299 | 0.7976 | 8.71E-05 | +++++ | 0 | 0.8727 | intron | TEAD3 |
| rs11755689 | 6 | 32586376 | 3 | 0 | 0 | A | G | -1.1264 | 0.287 | 8.71E-05 | ---?? | 0 | 0.7226 | intergenic | |
| 4:117277271 | 4 | 117277271 | 3 | 0 | 0 | CT | C | -3.8466 | 0.9803 | 8.71E-05 | ---?? | 41.1 | 0.1832 | | |
| rs116533472 | 4 | 117277274 | 3 | 0 | 0 | C | G | 3.846 | 0.9802 | 8.73E-05 | +++?? | 41 | 0.1834 | intergenic | |
| rs12679196 | 8 | 140812347 | 4 | 0 | 0 | T | C | 1.4789 | 0.377 | 8.74E-05 | +++?+ | 59.5 | 0.06009 | intron | TRAPPC9 |
| rs2305277 | 1 | 205033374 | 5 | 0 | 0 | T | C | -0.8632 | 0.22 | 8.74E-05 | ----- | 0 | 0.9708 | intron | CNTN2 |
| 12:91349851 | 12 | 91349851 | 3 | 0 | 0 | CACG | C | -5.3813 | 1.3718 | 8.75E-05 | ---?? | 0 | 0.9312 | | |
| rs201745547 | 5 | 179960341 | 4 | 0 | 0 | T | C | -1.1981 | 0.3055 | 8.77E-05 | ---?- | 0 | 0.3947 | intron | CNOT6 |
| rs17113684 | 14 | 22508144 | 4 | 0 | 0 | T | C | -1.0658 | 0.2717 | 8.78E-05 | ---?- | 0 | 0.8037 | intergenic | |
| rs4595999 | 2 | 150837420 | 4 | 0 | 0 | A | G | -1.0665 | 0.2719 | 8.78E-05 | ---?- | 0 | 0.6512 | intergenic | |
| rs117895637 | 7 | 27807521 | 4 | 0 | 0 | T | C | -2.0568 | 0.5245 | 8.80E-05 | ---?- | 0 | 0.6598 | intron | TAX1BP1 |
| rs11760092 | 6 | 77750678 | 4 | 0 | 0 | T | C | -1.1652 | 0.2972 | 8.81E-05 | ---?- | 20.5 | 0.2869 | intergenic | |
| rs77065514 | 3 | 16474995 | 4 | 0 | 0 | T | G | 2.0826 | 0.5311 | 8.81E-05 | +++?+ | 45.7 | 0.1374 | intron | RFTN1 |
| rs11785697 | 8 | 76106596 | 4 | 0 | 0 | T | C | -1.087 | 0.2772 | 8.82E-05 | ---?- | 19 | 0.2951 | intergenic | |
| rs78374008 | 17 | 9692477 | 4 | 0 | 0 | A | G | -1.5329 | 0.3909 | 8.82E-05 | ---?- | 0 | 0.926 | intron | DHRS7C |
| rs11751359 | 6 | 77750967 | 4 | 0 | 0 | T | C | -1.1643 | 0.297 | 8.85E-05 | ---?- | 20.4 | 0.2873 | intergenic | |
| 5:179970335 | 5 | 179970335 | 3 | 0 | 0 | A | AG | 1.2375 | 0.3157 | 8.85E-05 | +++?? | 31.1 | 0.2341 | | |
| rs10880359 | 12 | 43130986 | 4 | 0 | 0 | C | G | -1.5573 | 0.3974 | 8.89E-05 | ---?- | 15.9 | 0.3122 | intergenic | |
| rs12022789 | 1 | 102945678 | 3 | 0 | 0 | T | C | -0.952 | 0.2429 | 8.89E-05 | ---?? | 0 | 0.9263 | intergenic | |
| rs6453670 | 6 | 63369264 | 3 | 0 | 0 | T | G | 2.9715 | 0.7582 | 8.89E-05 | ?++?+ | 0 | 0.8117 | intergenic | |
| rs117036263 | 6 | 135680371 | 2 | 0 | 0 | T | C | 4.01 | 1.0233 | 8.91E-05 | +?+?? | 0 | 0.7702 | intron | AHI1 |
| rs12022817 | 1 | 102945785 | 3 | 0 | 0 | T | C | -0.9516 | 0.2429 | 8.92E-05 | ---?? | 0 | 0.9266 | intergenic | |
| rs12278598 | 11 | 95698325 | 4 | 0 | 0 | T | G | -1.1095 | 0.2832 | 8.92E-05 | ---?- | 0 | 0.5179 | intergenic | |

| | | | | | | | | | | | | | | | |
|---|---|---|---|---|---|---|---|---|---|---|---|---|---|---|---|
| rs111848852 | 18 | 61231105 | 4 | 0 | 0 | A | G | 2.3981 | 0.6121 | 8.93E-05 | +++?+ | 37.3 | 0.1884 | intron | SERPINB12 |
| rs6948861 | 7 | 107369949 | 5 | 0 | 0 | T | C | 0.8411 | 0.2147 | 8.94E-05 | +++++ | 34.2 | 0.1931 | intergenic | |
| rs7136440 | 12 | 43134769 | 4 | 0 | 0 | T | G | 1.6165 | 0.4126 | 8.94E-05 | +++?+ | 0 | 0.4742 | intergenic | |
| rs59738244 | 22 | 32842072 | 4 | 0 | 0 | T | C | -1.8087 | 0.4617 | 8.94E-05 | ---?- | 0 | 0.912 | intron | BPIFC |
| rs1317701 | 17 | 47444794 | 4 | 0 | 0 | T | C | 1.054 | 0.2691 | 8.95E-05 | +++?- | 32.5 | 0.2174 | upstream-gene | |
| rs13208216 | 6 | 77758706 | 4 | 0 | 0 | T | G | 1.1732 | 0.2995 | 8.96E-05 | +++?+ | 6.7 | 0.3596 | intergenic | |
| rs72633388 | 13 | 85650911 | 4 | 0 | 0 | T | G | -3.0277 | 0.773 | 8.97E-05 | ---?- | 0 | 0.8264 | intergenic | |
| rs33997707 | 6 | 32501715 | 5 | 0 | 1 | A | G | 1.4681 | 0.3748 | 8.97E-05 | +++++ | 18.5 | 0.297 | upstream-gene | |
| 10:126849636 | 10 | 126849636 | 3 | 0 | 0 | T | TGG | 1.3625 | 0.3479 | 8.98E-05 | +++?? | 33.8 | 0.2206 | | |
| rs61969743 | 14 | 22507186 | 4 | 0 | 0 | T | C | 1.0615 | 0.2711 | 9.00E-05 | +++?+ | 0 | 0.8091 | intergenic | |
| rs75162513 | 6 | 35447391 | 5 | 0 | 0 | A | G | -3.1464 | 0.8034 | 9.00E-05 | ----- | 0 | 0.865 | intron | TEAD3 |
| 2:168008281 | 2 | 168008281 | 3 | 0 | 0 | A | AG | -2.6013 | 0.6642 | 9.00E-05 | ---?? | 0 | 0.9613 | | |
| rs981725 | 15 | 98043012 | 4 | 0 | 0 | T | C | 1.0526 | 0.2688 | 9.00E-05 | +++?+ | 0 | 0.957 | intergenic | |
| rs1020088 | 2 | 150828833 | 4 | 0 | 0 | T | C | -1.0654 | 0.2721 | 9.00E-05 | ---?- | 0 | 0.6741 | intergenic | |
| rs6889859 | 5 | 123148901 | 4 | 0 | 0 | C | G | -1.2921 | 0.33 | 9.02E-05 | ---?- | 0 | 0.4745 | intergenic | |
| rs28360270 | 5 | 82588354 | 4 | 0 | 0 | A | G | 2.5073 | 0.6404 | 9.03E-05 | +++?+ | 0 | 0.5027 | intron | XRCC4 |
| rs76319806 | 17 | 52310826 | 4 | 0 | 0 | A | G | -2.4901 | 0.6361 | 9.06E-05 | ---?+ | 0 | 0.7749 | intergenic | |
| rs148966068 | 2 | 24699553 | 3 | 0 | 0 | A | C | 4.3685 | 1.116 | 9.06E-05 | +++?? | 56.6 | 0.1001 | intergenic | |
| rs16972089 | 15 | 42101694 | 5 | 0 | 0 | A | G | 1.1156 | 0.285 | 9.07E-05 | +++++ | 17.4 | 0.304 | intron | MAPKBP1 |
| rs78237592 | 5 | 76939016 | 3 | 0 | 0 | T | C | -2.8712 | 0.7336 | 9.09E-05 | ---?? | 0 | 0.9693 | upstream-gene | |
| rs76795350 | 6 | 168579820 | 4 | 0 | 0 | A | G | -1.8765 | 0.4795 | 9.11E-05 | ---?- | 0 | 0.7711 | intergenic | |
| rs35845603 | 19 | 46160458 | 5 | 0 | 0 | A | G | 0.85 | 0.2172 | 9.11E-05 | +++++ | 0 | 0.9049 | intergenic | |
| rs68081734 | 6 | 32628828 | 4 | 0 | 0 | T | C | -1.2356 | 0.3158 | 9.12E-05 | ----? | 60.6 | 0.0545 | intron | HLA-DQB1 |
| rs10149062 | 14 | 43268407 | 3 | 0 | 0 | T | G | 1.5134 | 0.3869 | 9.15E-05 | +++?? | 50.9 | 0.1306 | intergenic | |
| rs12681350 | 8 | 140812177 | 4 | 0 | 0 | A | G | -1.4741 | 0.3768 | 9.15E-05 | ---?- | 59 | 0.06238 | intron | TRAPPC9 |
| rs11708730 | 3 | 71925988 | 4 | 0 | 0 | T | C | -1.1343 | 0.29 | 9.17E-05 | ---?- | 0 | 0.6519 | intergenic | |
| 1:93498232 | 1 | 93498232 | 3 | 0 | 0 | <ins:me: | T | -1.2912 | 0.3301 | 9.18E-05 | ---?? | 17.5 | 0.2977 | | |
| rs67150904 | 6 | 32508658 | 5 | 0 | 1 | T | C | -1.4599 | 0.3733 | 9.19E-05 | ----- | 6.2 | 0.3716 | intergenic |  |
| rs9538429 | 13 | 60016739 | 4 | 0 | 0 | T | G | -0.9867 | 0.2523 | 9.20E-05 | ---?- | 0 | 0.8748 | intergenic | |
| rs8064862 | 17 | 47323380 | 4 | 0 | 0 | A | G | -1.1495 | 0.2939 | 9.20E-05 | ---?+ | 39.7 | 0.1738 | upstream-gene | |
| rs523369 | 3 | 191149390 | 4 | 0 | 0 | T | G | -1.5096 | 0.386 | 9.21E-05 | ---?+ | 40.6 | 0.1683 | intergenic | |
| rs72627228 | 18 | 52533177 | 4 | 0 | 0 | A | C | -2.2643 | 0.579 | 9.21E-05 | ---?- | 0 | 0.9868 | intron | RAB27B |
| rs9570122 | 13 | 60012845 | 4 | 0 | 0 | A | C | -0.9593 | 0.2453 | 9.22E-05 | ---?- | 0 | 0.686 | intergenic | |
| rs67202503 | 6 | 32513102 | 5 | 0 | 1 | T | C | -1.4303 | 0.3659 | 9.27E-05 | ----- | 0.7 | 0.4019 | intergenic |  |
| rs11056531 | 12 | 15659024 | 4 | 0 | 0 | T | C | -1.3147 | 0.3363 | 9.27E-05 | ---?- | 0 | 0.8424 | intron | PTPRO |
| 4:100548148 | 4 | 100548148 | 3 | 0 | 0 | A | T | 1.7725 | 0.4535 | 9.28E-05 | +++?? | 55.3 | 0.107 | downstream-gene | |
| rs67074075 | 18 | 40043454 | 4 | 0 | 0 | A | C | -1.2461 | 0.3188 | 9.28E-05 | ---?+ | 0 | 0.4716 | intron | LINC00907 |
| rs585964 | 1 | 187886683 | 4 | 0 | 0 | A | G | -1.1246 | 0.2878 | 9.31E-05 | ---?- | 0 | 0.5392 | intergenic | |
| rs9570118 | 13 | 60004432 | 4 | 0 | 0 | T | C | -0.9706 | 0.2484 | 9.32E-05 | ---?- | 0 | 0.6923 | intergenic | |
| rs145662700 | 6 | 83832806 | 4 | 0 | 0 | A | G | -3.479 | 0.8903 | 9.32E-05 | ---?- | 0 | 0.5443 | intron | DOPEY1 |
| rs9323377 | 14 | 62375656 | 4 | 0 | 0 | C | G | 1.1961 | 0.3061 | 9.33E-05 | +++?- | 0 | 0.5184 | intron | SYT16 |
| rs1502580 | 3 | 2475995 | 4 | 0 | 0 | T | C | 1.1692 | 0.2992 | 9.33E-05 | +++?+ | 0 | 0.809 | intron | CNTN4 |
| rs1317700 | 17 | 47444788 | 5 | 0 | 0 | A | G | -1.0002 | 0.256 | 9.33E-05 | ----+ | 42.2 | 0.1399 | upstream-gene |  |
| rs11965861 | 6 | 23867575 | 4 | 0 | 0 | T | G | 2.8652 | 0.7334 | 9.35E-05 | +++?- | 0 | 0.438 | intergenic | |
| rs16888501 | 6 | 77752845 | 4 | 0 | 0 | T | C | -1.1614 | 0.2973 | 9.35E-05 | ---?- | 20.2 | 0.2885 | intergenic | |
| rs2063166 | 17 | 52316824 | 4 | 0 | 0 | T | C | -2.4747 | 0.6334 | 9.36E-05 | ---?+ | 0 | 0.7751 | intergenic | |
| rs74386752 | 17 | 52318374 | 4 | 0 | 0 | A | G | -2.4728 | 0.633 | 9.36E-05 | ---?+ | 0 | 0.7745 | intergenic | |
| rs9323375 | 14 | 62375432 | 4 | 0 | 0 | A | T | -1.1952 | 0.306 | 9.36E-05 | ---?+ | 0 | 0.5187 | intron | SYT16 |
| rs7358744 | 12 | 27776230 | 4 | 0 | 0 | A | G | 1.0184 | 0.2608 | 9.40E-05 | +++?+ | 25.9 | 0.2561 | intron | PPFIBP1 |
| rs28383461 | 6 | 32609607 | 3 | 0 | 0 | A | G | 1.4769 | 0.3782 | 9.41E-05 | +++?? | 0 | 0.8274 | intron | HLA-DQA1 |
| rs17793142 | 14 | 22507126 | 4 | 0 | 0 | T | C | 1.0595 | 0.2713 | 9.41E-05 | +++?+ | 0 | 0.8101 | intergenic | |
| rs9323376 | 14 | 62375557 | 4 | 0 | 0 | C | G | 1.1951 | 0.306 | 9.42E-05 | +++?- | 0 | 0.5203 | intron | SYT16 |
| rs2023847 | 7 | 82461293 | 4 | 0 | 0 | T | G | -1.0345 | 0.2649 | 9.43E-05 | ---?- | 46.7 | 0.131 | intron | PCLO |
| rs7978701 | 12 | 111788680 | 5 | 0 | 0 | A | C | -0.977 | 0.2502 | 9.45E-05 | -+--- | 44.7 | 0.124 | downstream-gene | |
| rs17126413 | 1 | 64806201 | 4 | 0 | 0 | T | C | -1.393 | 0.3569 | 9.49E-05 | ---?- | 0 | 0.429 | intergenic | |
| rs61969296 | 13 | 104431967 | 3 | 0 | 0 | A | G | 5.0154 | 1.2851 | 9.51E-05 | +++?? | 0 | 0.5858 | intergenic | |
| rs62514368 | 8 | 113213752 | 3 | 0 | 0 | C | G | 1.0162 | 0.2604 | 9.52E-05 | +++?? | 0 | 0.9156 | intergenic | |
| rs7215163 | 17 | 52320395 | 4 | 0 | 0 | A | T | 2.4677 | 0.6324 | 9.54E-05 | +++?- | 0 | 0.7733 | intergenic | |
| rs9379639 | 6 | 24090993 | 5 | 0 | 0 | A | T | 1.5377 | 0.3941 | 9.54E-05 | ++++- | 0 | 0.5299 | intergenic | |

| | | | | | | | | | | | | | | | |
|---|---|---|---|---|---|---|---|---|---|---|---|---|---|---|---|
| rs12591949 | 15 | 92449667 | 4 | 0 | 0 | A | T | 0.9351 | 0.2397 | 9.56E-05 | +++?+ | 0 | 0.7069 | intron | SLCO3A1 |
| 5:179929700 | 5 | 179929700 | 3 | 0 | 0 | A | AT | -1.2328 | 0.316 | 9.57E-05 | ---?? | 21.5 | 0.2798 | | |
| rs192342165 | 3 | 142806441 | 3 | 0 | 0 | T | C | -2.3749 | 0.6088 | 9.58E-05 | ---?? | 23.3 | 0.2716 | intergenic | |
| rs746816 | 13 | 60012187 | 4 | 0 | 0 | T | C | -0.9571 | 0.2454 | 9.59E-05 | ---?- | 0 | 0.6892 | intergenic | |
| rs73644368 | 9 | 30877580 | 4 | 0 | 0 | A | G | -4.22 | 1.0819 | 9.60E-05 | ---?- | 11.7 | 0.3345 | intergenic | |
| rs2252230 | 8 | 113188928 | 4 | 0 | 0 | A | G | 1.0353 | 0.2654 | 9.60E-05 | +++?+ | 0 | 0.8023 | intergenic | |
| 1:102959976 | 1 | 102959976 | 3 | 0 | 0 | CATT | C | 0.9514 | 0.2439 | 9.60E-05 | +++?? | 0 | 0.9175 | | |
| rs79552826 | 17 | 52324150 | 4 | 0 | 0 | A | T | 2.4439 | 0.6266 | 9.61E-05 | +++?- | 0 | 0.771 | intergenic | |
| rs41271261 | 6 | 35437451 | 5 | 0 | 0 | T | C | 3.1018 | 0.7954 | 9.63E-05 | +++++ | 0 | 0.8731 | intron | RPL10A |
| rs55823336 | 15 | 86320164 | 3 | 0 | 0 | A | T | 1.4393 | 0.3691 | 9.63E-05 | +++?? | 0 | 0.6164 | intron | KLHL25 |
| 5:179932612 | 5 | 179932612 | 3 | 0 | 0 | A | ATC | 1.2328 | 0.3161 | 9.63E-05 | +++?? | 22.2 | 0.2764 | | |
| rs185462550 | 3 | 196491906 | 3 | 0 | 0 | A | G | 2.4649 | 0.6321 | 9.63E-05 | +++?? | 0 | 0.9783 | intron | PAK2 |
| rs368082185 | 5 | 67619936 | 3 | 0 | 0 | A | G | 2.1116 | 0.5416 | 9.67E-05 | +++?? | 0 | 0.6274 | intergenic | |
| rs145527145 | 9 | 119648204 | 3 | 0 | 0 | A | G | 3.9195 | 1.0054 | 9.68E-05 | +++?? | 61.7 | 0.07332 | intron | ASTN2 |
| rs77381455 | 8 | 85787159 | 4 | 0 | 0 | A | G | 2.3025 | 0.5907 | 9.69E-05 | +++?+ | 0 | 0.8338 | intron | RALYL |
| 8:113213749 | 8 | 113213749 | 3 | 0 | 0 | A | AG | 1.0155 | 0.2605 | 9.70E-05 | +++?? | 0 | 0.9161 | | |
| rs17126383 | 1 | 64793849 | 4 | 0 | 0 | A | G | 1.416 | 0.3633 | 9.71E-05 | +++?+ | 0 | 0.5803 | intergenic | |
| rs11049070 | 12 | 27773861 | 4 | 0 | 0 | A | G | -1.0221 | 0.2623 | 9.72E-05 | ---?- | 25.3 | 0.2597 | intron | PPFIBP1 |
| rs78261019 | 10 | 131501162 | 4 | 0 | 0 | C | G | -1.4773 | 0.3791 | 9.73E-05 | ---?- | 0 | 0.9362 | intron | MGMT |
| 22:48335193 | 22 | 48335193 | 3 | 0 | 0 | T | TA | -1.0764 | 0.2762 | 9.74E-05 | ---?? | 0 | 0.9852 | | |
| rs10149061 | 14 | 43268406 | 3 | 0 | 0 | C | G | 1.5078 | 0.3869 | 9.74E-05 | +++?? | 49.3 | 0.1389 | intergenic | |
| rs76389331 | 13 | 36253676 | 2 | 0 | 0 | T | G | 5.8597 | 1.5038 | 9.76E-05 | ?++?? | 0 | 0.4247 | intergenic | |
| rs323234 | 6 | 77669406 | 3 | 0 | 0 | A | G | 1.1051 | 0.2836 | 9.76E-05 | +++?? | 41.9 | 0.1789 | intergenic | |
| rs2229866 | 1 | 205031116 | 5 | 0 | 0 | T | C | 0.8543 | 0.2193 | 9.77E-05 | +++++ | 0 | 0.975 | missense | CNTN2 |
| 5:179952920 | 5 | 179952920 | 3 | 0 | 0 | T | TA | -1.2321 | 0.3162 | 9.77E-05 | ---?? | 25.8 | 0.2598 | | |
| rs2786046 | 1 | 187885757 | 4 | 0 | 0 | A | G | 1.1228 | 0.2882 | 9.80E-05 | +++?+ | 0 | 0.5405 | intergenic | |
| rs12056252 | 7 | 107368091 | 5 | 0 | 0 | T | C | -0.8378 | 0.2151 | 9.81E-05 | ----- | 34.1 | 0.1937 | intergenic |  |
| rs77801483 | 6 | 32673277 | 5 | 0 | 1 | C | G | -1.5535 | 0.3988 | 9.81E-05 | ----- | 18.2 | 0.299 | intergenic |  |
| rs28634221 | 14 | 22507175 | 4 | 0 | 0 | A | G | 1.0572 | 0.2715 | 9.84E-05 | +++?+ | 0 | 0.8118 | intergenic | |
| rs2568157 | 2 | 218335258 | 4 | 0 | 0 | A | G | -0.9549 | 0.2452 | 9.85E-05 | ---?- | 0 | 0.6521 | intron | DIRC3 |
| rs181440637 | 16 | 75115050 | 2 | 0 | 0 | A | G | 5.8219 | 1.495 | 9.85E-05 | +?+?? | 13 | 0.2835 | intron | ZNRF1 |
| rs12787239 | 11 | 12817798 | 4 | 0 | 0 | A | G | -1.4012 | 0.3598 | 9.87E-05 | ---?- | 34 | 0.2082 | intron | TEAD1 |
| rs66963476 | 5 | 179936846 | 3 | 0 | 0 | T | C | -1.2311 | 0.3162 | 9.87E-05 | ---?? | 22.9 | 0.2735 | intron | CNOT6 |
| rs6685698 | 1 | 64805465 | 4 | 0 | 0 | A | G | -1.3891 | 0.3568 | 9.88E-05 | ---?- | 0 | 0.4278 | intergenic | |
| rs79617549 | 10 | 131504203 | 4 | 0 | 0 | A | G | 1.4754 | 0.379 | 9.90E-05 | +++?+ | 0 | 0.9225 | intron | MGMT |
| 14:43257610 | 14 | 43257610 | 3 | 0 | 0 | A | AATAAA | -1.5309 | 0.3933 | 9.90E-05 | ---?? | 55 | 0.1082 | | |
| rs117420548 | 7 | 104775976 | 2 | 0 | 0 | A | G | 6.5435 | 1.6809 | 9.91E-05 | ?++?? | 0 | 0.3307 | intron | SRPK2 |
| rs79977771 | 18 | 52548564 | 4 | 0 | 0 | A | G | -2.425 | 0.623 | 9.92E-05 | ---?- | 0 | 0.9907 | intron | RAB27B |
| rs60099385 | 5 | 502923 | 3 | 0 | 0 | T | C | 1.7261 | 0.4436 | 9.97E-05 | +++?? | 0 | 0.883 | intron | SLC9A3 |
| rs13220775 | 6 | 80552840 | 4 | 0 | 0 | T | C | -1.0372 | 0.2665 | 9.97E-05 | ---?- | 1.9 | 0.3827 | intron | C6orf7 |
| rs75080550 | 15 | 42198244 | 4 | 0 | 0 | A | C | 3.1022 | 0.7973 | 9.98E-05 | +-+?+ | 36.7 | 0.1921 | intron | EHD4 |
| rs3828793 | 6 | 32635883 | 5 | 0 | 1 | T | C | 1.2085 | 0.3106 | 9.99E-05 | +++++ | 41.6 | 0.1439 | upstream-gene | |
| 20:31201137 | 20 | 31201137 | 3 | 0 | 0 | CATGG | C | -1.5133 | 0.389 | 9.99E-05 | ---?? | 58.8 | 0.08848 | | |
| rs323236 | 6 | 77669614 | 3 | 0 | 0 | T | C | 1.1029 | 0.2835 | 1.00E-04 | +++?? | 42.5 | 0.1757 | intergenic | |
| rs34442410 | 6 | 32442086 | 5 | 0 | 1 | C | G | -1.567 | 0.4028 | 1.00E-04 | ----- | 0 | 0.535 | intergenic |  |
| rs7398833 | 12 | 111786892 | 5 | 0 | 0 | T | C | -0.9734 | 0.2502 | 1.00E-04 | -+--- | 44.7 | 0.1242 | 3-prime-UTR | CUX2 |
| 3:196476217 | 3 | 196476217 | 3 | 0 | 0 | CA | C | 2.478 | 0.6369 | 0.0001 | +++?? | 0 | 0.9485 | | |
| rs3924427 | 15 | 92448520 | 4 | 0 | 0 | A | G | -0.9327 | 0.2398 | 0.0001001 | ---?- | 0 | 0.6723 | intron | SLCO3A1 |
| rs1110557 | 16 | 78733197 | 3 | 0 | 0 | T | C | 4.9181 | 1.2644 | 0.0001004 | +++?? | 41.9 | 0.1788 | intron | WWOX |
| rs1158403 | 13 | 60006613 | 4 | 0 | 0 | T | C | -0.9632 | 0.2476 | 0.0001004 | ---?- | 0 | 0.6968 | intergenic | |
| 11:42554002 | 11 | 42554002 | 3 | 0 | 0 | A | G | -4.2732 | 1.0987 | 0.0001005 | ---?? | 0 | 0.6659 | intergenic | |
| 11:42554003 | 11 | 42554003 | 3 | 0 | 0 | A | G | 4.273 | 1.0986 | 0.0001005 | +++?? | 0 | 0.6662 | intergenic | |
| rs144146286 | 3 | 196478575 | 3 | 0 | 0 | A | T | 2.4774 | 0.6371 | 0.0001008 | +++?? | 0 | 0.9508 | intron | PAK2 |
| rs10455366 | 6 | 80549910 | 4 | 0 | 0 | T | C | -1.0421 | 0.268 | 0.000101 | ---?- | 3 | 0.3774 | intron | C6orf7 |
| rs181556254 | 17 | 68528449 | 3 | 0 | 0 | T | C | 3.6425 | 0.9368 | 0.0001011 | +++?? | 0 | 0.7126 | intergenic | |
| rs679245 | 1 | 187887066 | 4 | 0 | 0 | T | C | 1.1155 | 0.287 | 0.0001013 | +++?+ | 0 | 0.53 | intergenic | |
| rs7323008 | 13 | 60028050 | 4 | 0 | 0 | C | G | 0.9961 | 0.2562 | 0.0001013 | +++?+ | 26 | 0.2555 | intergenic | |
| rs74634292 | 17 | 52324587 | 4 | 0 | 0 | A | G | -2.4283 | 0.6247 | 0.0001013 | ---?+ | 0 | 0.767 | intergenic | |
| rs78532069 | 4 | 21020275 | 4 | 0 | 0 | T | C | -2.5345 | 0.652 | 0.0001013 | ---?+ | 13.3 | 0.3262 | intron | KCNIP4 |

| | | | | | | | | | | | | | | | |
|---|---|---|---|---|---|---|---|---|---|---|---|---|---|---|---|
| rs78588119 | 11 | 42556215 | 3 | 0 | 0 | A | G | -4.2653 | 1.0972 | 0.0001013 | ---?? | 0 | 0.6681 | intergenic | |
| 4:59659949 | 4 | 59659949 | 3 | 0 | 0 | AT | ATT | -2.6093 | 0.6713 | 0.0001014 | ---?? | 16.9 | 0.3004 | | |
| 5:179990933 | 5 | 179990933 | 3 | 0 | 0 | A | AT | -1.233 | 0.3172 | 0.0001015 | ---?? | 31.7 | 0.2315 | | |
| rs755643 | 5 | 173136667 | 4 | 0 | 1 | A | G | -1.0728 | 0.276 | 0.0001016 | ---?- | 20.5 | 0.2868 | intergenic | |
| rs112757452 | 10 | 1006099 | 4 | 0 | 0 | T | C | -1.988 | 0.5115 | 0.0001016 | ---?- | 33.2 | 0.2132 | intergenic | |
| rs147132576 | 6 | 83829876 | 4 | 0 | 0 | A | G | 3.468 | 0.8923 | 0.0001016 | +++?+ | 0 | 0.5602 | intron | DOPEY1 |
| rs9563725 | 13 | 60010888 | 4 | 0 | 0 | T | C | -0.9534 | 0.2453 | 0.0001017 | ---?- | 0 | 0.6965 | intergenic | |
| 13:110880888 | 13 | 110880888 | 3 | 0 | 0 | G | GA | 1.2634 | 0.3251 | 0.0001017 | +++?? | 25.2 | 0.2628 | | |
| rs45489601 | 6 | 83829279 | 4 | 0 | 0 | A | G | -3.4655 | 0.8918 | 0.0001019 | ---?- | 0 | 0.5645 | intron | DOPEY1 |
| rs72748374 | 1 | 209040443 | 3 | 0 | 0 | A | C | 4.2071 | 1.0826 | 0.0001019 | ?++?+ | 0 | 0.3895 | intergenic | |
| rs10943400 | 6 | 77700022 | 4 | 0 | 0 | A | G | 1.119 | 0.288 | 0.0001023 | +++?+ | 13.7 | 0.3238 | intergenic | |
| rs73052446 | 3 | 29711526 | 4 | 0 | 0 | A | T | 4.2799 | 1.1018 | 0.0001025 | +++?+ | 25 | 0.2614 | intron | RBMS3 |
| rs12030222 | 1 | 187886858 | 4 | 0 | 0 | A | T | 1.1157 | 0.2872 | 0.0001026 | +++?+ | 0 | 0.5289 | intergenic | |
| rs680116 | 1 | 187886885 | 4 | 0 | 0 | A | C | 1.1155 | 0.2872 | 0.0001027 | +++?+ | 0 | 0.5319 | intergenic | |
| rs11580315 | 1 | 64786496 | 4 | 0 | 0 | A | G | 1.3997 | 0.3604 | 0.0001027 | +++?+ | 0 | 0.5812 | intergenic | |
| rs323266 | 6 | 77688466 | 3 | 0 | 0 | T | C | -1.1059 | 0.2847 | 0.0001028 | ---?? | 39.8 | 0.1898 | intergenic | |
| rs9863718 | 3 | 77411137 | 4 | 0 | 0 | A | G | -0.9603 | 0.2473 | 0.0001031 | ---?- | 0 | 0.5276 | intron | ROBO2 |
| rs13197584 | 6 | 77700939 | 4 | 0 | 0 | A | G | 1.1178 | 0.2879 | 0.0001031 | +++?+ | 14 | 0.3222 | intergenic | |
| rs201632070 | 11 | 30631257 | 3 | 0 | 0 | A | G | -1.0855 | 0.2795 | 0.0001031 | ---?? | 0.7 | 0.3654 | intergenic | |
| rs35656734 | 6 | 32600153 | 5 | 0 | 0 | T | C | 0.9722 | 0.2504 | 0.0001034 | +++++ | 0 | 0.4285 | intergenic | |
| rs11181672 | 12 | 43145563 | 5 | 0 | 0 | A | C | 1.0249 | 0.264 | 0.0001035 | ++++- | 0 | 0.4859 | intergenic | |
| rs11644227 | 16 | 12095738 | 4 | 0 | 0 | T | C | -0.9537 | 0.2457 | 0.0001035 | +--?- | 61.4 | 0.05079 | intron | SNX29 |
| rs62435719 | 6 | 77700510 | 4 | 0 | 0 | A | C | 1.1175 | 0.2878 | 0.0001035 | +++?+ | 13.9 | 0.3227 | intergenic | |
| rs5000880 | 13 | 60016434 | 4 | 0 | 0 | T | G | 0.9682 | 0.2494 | 0.0001037 | +++?+ | 0 | 0.8944 | intergenic | |
| 1:229165976 | 1 | 229165976 | 3 | 0 | 0 | CTAT | C | 1.4844 | 0.3824 | 0.0001037 | +++?? | 0 | 0.9785 | | |
| 6:30982124 | 6 | 30982124 | 3 | 0 | 0 | T | TGAGCA | 1.5525 | 0.4 | 0.0001038 | +++?? | 0 | 0.7977 | | |
| rs2467193 | 8 | 140478586 | 4 | 0 | 0 | A | T | 1.6165 | 0.4165 | 0.0001039 | +++?+ | 0 | 0.7587 | intergenic | |
| rs112991248 | 17 | 47321763 | 4 | 0 | 0 | T | C | -1.1732 | 0.3023 | 0.0001039 | ---?+ | 31.9 | 0.221 | upstream-gene | |
| rs17683667 | 12 | 68321776 | 3 | 0 | 0 | T | C | -4.8801 | 1.2574 | 0.000104 | ---?? | 0 | 0.7176 | intergenic | |
| rs141651313 | 15 | 95162886 | 3 | 0 | 0 | A | C | 7.5906 | 1.9561 | 0.0001042 | +-+?? | 65.4 | 0.05543 | intergenic | |
| rs7182628 | 15 | 53765494 | 4 | 0 | 0 | A | G | -0.9903 | 0.2552 | 0.0001043 | ---?- | 41.4 | 0.1633 | intergenic | |
| rs7147431 | 14 | 22506095 | 4 | 0 | 0 | A | G | 1.0565 | 0.2723 | 0.0001045 | +++?+ | 0 | 0.8121 | intergenic | |
| rs57284063 | 12 | 43146432 | 5 | 0 | 0 | T | C | -1.0251 | 0.2642 | 0.0001045 | ----+ | 0 | 0.4848 | intergenic |  |
| 11:86178723 | 11 | 86178723 | 3 | 0 | 0 | A | T | 3.8035 | 0.9804 | 0.0001046 | +++?? | 0 | 0.9303 | intron | ME3 |
| rs11976156 | 7 | 107365946 | 5 | 0 | 0 | T | C | 0.8392 | 0.2163 | 0.0001046 | +++++ | 37.5 | 0.1712 | intergenic | |
| rs9527990 | 13 | 60016002 | 4 | 0 | 0 | A | G | 0.9608 | 0.2477 | 0.0001047 | +++?+ | 0 | 0.8177 | intergenic | |
| rs17150778 | 5 | 123141745 | 4 | 0 | 0 | T | C | -1.2736 | 0.3283 | 0.0001047 | ---?- | 0 | 0.4621 | intergenic | |
| rs7702977 | 5 | 10306644 | 4 | 0 | 0 | A | G | -2.0901 | 0.5388 | 0.0001049 | ---?- | 22.6 | 0.2751 | intron | CMBL |
| rs7433362 | 3 | 77411196 | 4 | 0 | 0 | A | G | 0.9589 | 0.2472 | 0.0001051 | +++?+ | 0 | 0.5273 | intron | ROBO2 |
| rs61302145 | 7 | 141149862 | 4 | 0 | 0 | A | G | 5.8998 | 1.5212 | 0.0001052 | +++?+ | 0 | 0.8919 | intron | TMEM178B |
| rs140357870 | 12 | 27796085 | 3 | 0 | 0 | A | G | 1.1373 | 0.2933 | 0.0001053 | +++?? | 66.4 | 0.05117 | intron | PPFIBP1 |
| 1:102935521 | 1 | 102935521 | 3 | 0 | 0 | G | GT | 0.9582 | 0.2471 | 0.0001053 | +++?? | 0 | 0.9233 | | |
| rs9598011 | 13 | 60017478 | 4 | 0 | 0 | A | G | 0.9687 | 0.2498 | 0.0001055 | +++?+ | 0 | 0.8471 | intergenic | |
| rs55820429 | 3 | 187901660 | 4 | 0 | 0 | T | G | 1.5267 | 0.3937 | 0.0001055 | +++?+ | 0 | 0.777 | intron | LPP |
| 11:1117166 | 11 | 1117166 | 3 | 0 | 0 | G | GTC | 1.5825 | 0.4081 | 0.0001055 | +++?? | 54.1 | 0.1131 | | |
| rs9812557 | 3 | 77392829 | 4 | 0 | 0 | A | C | 0.9744 | 0.2513 | 0.0001056 | +++?+ | 0 | 0.4593 | intron | ROBO2 |
| rs28687254 | 6 | 32501879 | 5 | 0 | 1 | A | T | 1.344 | 0.3466 | 0.0001056 | +++++ | 28.9 | 0.2292 | upstream-gene | |
| rs78848723 | 18 | 39926597 | 4 | 0 | 0 | A | G | -1.5326 | 0.3953 | 0.0001056 | ---?- | 0 | 0.8141 | intron | LINC00907 |
| rs621793 | 1 | 187889323 | 4 | 0 | 0 | A | G | 1.1027 | 0.2844 | 0.0001059 | +++?+ | 0 | 0.4951 | intergenic | |
| rs13192505 | 6 | 32442242 | 5 | 0 | 0 | A | G | 1.4962 | 0.3859 | 0.0001059 | +++++ | 0 | 0.5048 | intergenic | |
| rs16883549 | 8 | 113409694 | 4 | 0 | 0 | A | G | -1.0985 | 0.2834 | 0.0001059 | ---?- | 0 | 0.836 | intron | CSMD3 |
| rs665059 | 1 | 187888513 | 4 | 0 | 0 | T | C | 1.1052 | 0.2851 | 0.000106 | +++?+ | 0 | 0.4996 | intergenic | |
| rs12621982 | 2 | 170651802 | 4 | 0 | 0 | A | G | -1.1036 | 0.2847 | 0.000106 | ---?- | 38 | 0.1838 | upstream-gene | |
| rs62514456 | 8 | 113409713 | 4 | 0 | 0 | A | G | 1.0984 | 0.2834 | 0.000106 | +++?+ | 0 | 0.8358 | intron | CSMD3 |
| rs9317077 | 13 | 60016892 | 4 | 0 | 0 | T | C | -0.968 | 0.2498 | 0.0001063 | ---?- | 0 | 0.8502 | intergenic | |
| rs293818 | 3 | 191152845 | 4 | 0 | 0 | A | C | -1.464 | 0.3778 | 0.0001064 | ---?+ | 37.4 | 0.1875 | intergenic | |
| 10:131925614 | 10 | 131925614 | 3 | 0 | 0 | T | TA | 1.1005 | 0.284 | 0.0001064 | +++?? | 37.9 | 0.1997 | | |
| 3:183202084 | 3 | 183202084 | 3 | 0 | 0 | CA | C | -1.7367 | 0.4481 | 0.0001065 | ---?? | 0 | 0.4657 | | |
| rs12175687 | 6 | 24089375 | 5 | 0 | 0 | A | G | -1.5046 | 0.3883 | 0.0001067 | ----+ | 0 | 0.47 | intergenic |  |

| | | | | | | | | | | | | | | | |
|---|---|---|---|---|---|---|---|---|---|---|---|---|---|---|---|
| rs36000620 | 13 | 75234405 | 4 | 0 | 0 | T | C | 3.7794 | 0.9754 | 0.0001067 | +++?+ | 0 | 0.7831 | intergenic | |
| rs180764126 | 7 | 25643353 | 3 | 0 | 0 | T | G | -3.8053 | 0.9821 | 0.0001067 | ---?? | 52.9 | 0.1198 | intergenic | |
| rs217671 | 14 | 62360464 | 4 | 0 | 0 | A | G | 1.1065 | 0.2856 | 0.0001068 | +++?+ | 0 | 0.656 | intron | SYT16 |
| rs9875924 | 3 | 71893500 | 4 | 0 | 0 | A | G | 1.0447 | 0.2696 | 0.0001068 | +++?+ | 0 | 0.8869 | intergenic | |
| rs13257461 | 8 | 96919351 | 4 | 0 | 0 | A | G | 1.2369 | 0.3192 | 0.0001068 | +++?+ | 0 | 0.9068 | intergenic | |
| rs8032291 | 15 | 85868712 | 3 | 0 | 0 | C | G | -5.8684 | 1.5147 | 0.0001069 | --??- | 0 | 0.7959 | intergenic | |
| rs77931858 | 8 | 17467215 | 4 | 0 | 0 | T | C | -3.0782 | 0.7945 | 0.0001069 | ---?- | 0 | 0.6446 | intron | PDGFRL |
| rs56677840 | 14 | 22527409 | 4 | 0 | 0 | A | G | -1.2183 | 0.3145 | 0.0001071 | ---?- | 0 | 0.98 | intergenic | |
| rs117784775 | 14 | 77134602 | 3 | 0 | 0 | A | C | -3.5998 | 0.9292 | 0.0001071 | ---?? | 0 | 0.8349 | intergenic | |
| rs115804937 | 1 | 152797307 | 3 | 0 | 0 | T | G | 4.6466 | 1.1995 | 0.0001072 | +++?? | 0 | 0.42 | upstream-gene | |
| rs9538424 | 13 | 60013510 | 4 | 0 | 0 | T | C | 0.9821 | 0.2536 | 0.0001073 | +++?+ | 0 | 0.8283 | intergenic | |
| rs2029869 | 1 | 64796824 | 4 | 0 | 0 | T | G | -0.9384 | 0.2423 | 0.0001075 | ---?- | 48.7 | 0.119 | intergenic | |
| rs293820 | 3 | 191150836 | 4 | 0 | 0 | A | G | -1.4559 | 0.3759 | 0.0001076 | ---?+ | 35.8 | 0.1972 | intergenic | |
| rs61924560 | 12 | 43148050 | 5 | 0 | 0 | A | T | 1.0248 | 0.2646 | 0.0001076 | ++++- | 0 | 0.4815 | intergenic | |
| rs9457466 | 6 | 159224561 | 4 | 0 | 0 | A | T | 0.959 | 0.2476 | 0.0001077 | +++?+ | 27.3 | 0.248 | intron | EZR |
| rs9538411 | 13 | 60000534 | 4 | 0 | 0 | T | C | 0.9875 | 0.255 | 0.0001077 | +++?+ | 0 | 0.6191 | intergenic | |
| rs41286560 | 14 | 101349454 | 4 | 0 | 0 | T | G | -2.8796 | 0.7437 | 0.0001079 | ----? | 0 | 0.6067 | missense | RTL1 |
| 6:77580161 | 6 | 77580161 | 3 | 0 | 0 | CA | C | -1.1552 | 0.2983 | 0.0001079 | ---?? | 48.9 | 0.1413 | | |
| rs117195240 | 14 | 43265011 | 3 | 0 | 0 | A | T | -1.4998 | 0.3874 | 0.000108 | ---?? | 53.5 | 0.1164 | intergenic | |
| rs11588412 | 1 | 205034723 | 5 | 0 | 0 | T | C | 0.873 | 0.2255 | 0.0001082 | +++++ | 0 | 0.9887 | intron | CNTN2 |
| 6:42760188 | 6 | 42760188 | 3 | 0 | 0 | A | T | 2.8419 | 0.7341 | 0.0001083 | +++?? | 0 | 0.9992 | intron | GLTSCR1L |
| rs9956599 | 18 | 47528242 | 4 | 0 | 0 | A | C | -0.9976 | 0.2577 | 0.0001083 | ---?- | 0 | 0.5983 | intron | MYO5B |
| 13:45628414 | 13 | 45628414 | 3 | 0 | 0 | T | TCA | 2.4798 | 0.6406 | 0.0001084 | +++?? | 0 | 0.4353 | | |
| rs16947546 | 17 | 47324353 | 4 | 0 | 0 | T | C | 1.1444 | 0.2956 | 0.0001085 | +++?- | 33 | 0.2141 | upstream-gene | |
| rs72853980 | 6 | 32511628 | 5 | 0 | 1 | A | G | 1.4235 | 0.3678 | 0.0001085 | +++++ | 5.3 | 0.3768 | intergenic | |
| rs789672 | 5 | 127630394 | 3 | 0 | 0 | A | C | -1.0589 | 0.2736 | 0.0001086 | ---?? | 62.3 | 0.07026 | intron | FBN2 |
| rs61784470 | 1 | 64796202 | 4 | 0 | 0 | T | G | -0.9376 | 0.2423 | 0.0001087 | ---?- | 48.7 | 0.119 | intergenic | |
| rs141083117 | 2 | 160170899 | 4 | 0 | 0 | C | G | 3.2493 | 0.8395 | 0.0001087 | +++?+ | 21.9 | 0.2794 | downstream-gene | |
| rs7593092 | 2 | 137010468 | 4 | 0 | 0 | C | G | 1.0211 | 0.2639 | 0.0001088 | +++?+ | 0 | 0.8499 | intergenic | |
| rs6913465 | 6 | 168626744 | 4 | 0 | 0 | T | C | 1.1279 | 0.2915 | 0.000109 | +++?+ | 0 | 0.8953 | intergenic | |
| rs11748632 | 5 | 123139333 | 4 | 0 | 0 | C | G | 1.2753 | 0.3296 | 0.000109 | +++?+ | 0 | 0.4629 | intergenic | |
| rs11181674 | 12 | 43149010 | 5 | 0 | 0 | A | T | -1.0245 | 0.2648 | 0.0001091 | ----+ | 0 | 0.4804 | intergenic |  |
| rs16876584 | 6 | 35435295 | 5 | 0 | 0 | C | G | 3.1066 | 0.8029 | 0.0001092 | +++++ | 0 | 0.8824 | upstream-gene | |
| rs11748346 | 5 | 179957781 | 3 | 0 | 0 | A | G | 1.2247 | 0.3165 | 0.0001093 | +++?? | 26.2 | 0.2579 | intron | CNOT6 |
| rs12662542 | 6 | 106216678 | 3 | 0 | 1 | T | C | -2.8019 | 0.7242 | 0.0001093 | ---?? | 62.8 | 0.06814 | intergenic | |
| rs147807608 | 15 | 85829890 | 3 | 0 | 0 | A | G | -5.9305 | 1.5328 | 0.0001093 | --??- | 0 | 0.837 | intergenic | |
| rs978776 | 6 | 109086427 | 2 | 0 | 0 | T | C | 5.4496 | 1.4086 | 0.0001094 | +?+?? | 0 | 0.7346 | intron | LINC00222 |
| rs11181675 | 12 | 43149090 | 5 | 0 | 0 | C | G | -1.0243 | 0.2648 | 0.0001096 | ----+ | 0 | 0.4803 | intergenic |  |
| rs12660988 | 6 | 106216679 | 3 | 0 | 1 | T | C | 2.8013 | 0.7242 | 0.0001096 | +++?? | 62.8 | 0.06806 | intergenic | |
| rs75764593 | 17 | 47446956 | 5 | 0 | 0 | A | G | -0.9909 | 0.2562 | 0.0001096 | ----+ | 41.1 | 0.1472 | intergenic |  |
| rs389062 | 3 | 191147791 | 4 | 0 | 0 | A | G | -1.4381 | 0.3718 | 0.0001098 | ---?+ | 31.9 | 0.2207 | intergenic | |
| rs75050966 | 7 | 36193565 | 4 | 0 | 0 | T | C | -3.5078 | 0.9069 | 0.0001098 | ---?- | 0 | 0.7448 | intron | EEPD1 |
| rs776798 | 1 | 187888876 | 4 | 0 | 0 | A | G | 1.1026 | 0.2851 | 0.0001099 | +++?+ | 0 | 0.498 | intergenic | |
| rs117574462 | 6 | 99781118 | 4 | 0 | 0 | A | G | -2.9837 | 0.7714 | 0.0001099 | ---?- | 0 | 0.6531 | intron | FAXC |
| rs945739 | 1 | 187883524 | 4 | 0 | 0 | T | C | -1.1175 | 0.289 | 0.00011 | ---?- | 0 | 0.5578 | intergenic | |
| rs11855017 | 15 | 42096146 | 5 | 0 | 0 | A | C | 1.109 | 0.2868 | 0.00011 | +++++ | 4.9 | 0.379 | intron | MAPKBP1 |
| rs144770972 | 6 | 83881993 | 4 | 0 | 0 | T | C | 3.4218 | 0.8848 | 0.00011 | +++?+ | 18.2 | 0.2999 | intron | PGM3 |
| rs11065875 | 12 | 111783802 | 5 | 0 | 0 | A | G | -0.9683 | 0.2504 | 0.0001101 | -+--- | 45.4 | 0.1196 | intron | CUX2 |
| rs56362123 | 15 | 85823779 | 3 | 0 | 0 | A | G | -6.0013 | 1.552 | 0.0001102 | --??- | 0 | 0.8259 | intergenic | |
| rs118076362 | 7 | 156396185 | 3 | 0 | 0 | T | C | 3.8766 | 1.0025 | 0.0001102 | +++?? | 0 | 0.6517 | upstream-gene | |
| rs150518446 | 6 | 83825624 | 4 | 0 | 0 | A | T | 3.4589 | 0.8944 | 0.0001102 | +++?+ | 0 | 0.5819 | intron | DOPEY1 |
| rs7724808 | 5 | 10303807 | 4 | 0 | 0 | T | C | -2.0449 | 0.5289 | 0.0001103 | ---?- | 38.5 | 0.181 | intron | CMBL |
| rs141334098 | 16 | 19846620 | 4 | 0 | 0 | A | G | 4.0644 | 1.0511 | 0.0001103 | +++?+ | 0 | 0.8249 | intron | IQCK |
| rs10834473 | 11 | 24769313 | 4 | 0 | 0 | C | G | -1.2931 | 0.3344 | 0.0001104 | +--?- | 57 | 0.07262 | intron | LUZP2 |
| 12:127112645 | 12 | 127112645 | 3 | 0 | 0 | G | GAA | 1.8711 | 0.4839 | 0.0001105 | +++?? | 54.6 | 0.1106 | | |
| rs116596433 | 15 | 85867109 | 3 | 0 | 0 | A | G | -5.8566 | 1.5148 | 0.0001106 | --??- | 0 | 0.7983 | intergenic | |
| rs12221058 | 10 | 114724501 | 3 | 0 | 0 | A | G | 1.8234 | 0.4717 | 0.0001107 | +++?? | 26.8 | 0.2549 | intron | TCF7L2 |
| rs1496015 | 1 | 64797745 | 4 | 0 | 0 | A | C | 0.9371 | 0.2424 | 0.000111 | +++?+ | 48.4 | 0.1209 | intergenic | |
| rs16827732 | 1 | 187889812 | 4 | 0 | 0 | T | C | -1.0967 | 0.2837 | 0.000111 | ---?- | 0 | 0.4909 | intergenic | |

| | | | | | | | | | | | | | | | |
|---|---|---|---|---|---|---|---|---|---|---|---|---|---|---|---|
| rs12313884 | 12 | 43149770 | 5 | 0 | 0 | A | T | -1.024 | 0.265 | 0.0001113 | ----+ | 0 | 0.4785 | intergenic |  |
| rs145067941 | 19 | 45391941 | 2 | 0 | 0 | T | C | -4.345 | 1.1243 | 0.0001113 | -?-?? | 29.3 | 0.2343 | 3-prime-UTR | PVRL2 |
| rs148440921 | 6 | 83825393 | 4 | 0 | 0 | T | G | 3.4574 | 0.8947 | 0.0001115 | +++?+ | 0 | 0.584 | intron | DOPEY1 |
| rs67793506 | 18 | 52533010 | 4 | 0 | 0 | A | G | -2.2296 | 0.577 | 0.0001116 | ---?- | 0 | 0.9924 | intron | RAB27B |
| rs11181676 | 12 | 43150251 | 5 | 0 | 0 | A | G | 1.0243 | 0.2651 | 0.0001119 | ++++- | 0 | 0.4779 | intergenic | |
| rs35182655 | 6 | 32522087 | 5 | 0 | 1 | T | C | 1.4436 | 0.3737 | 0.0001119 | +++++ | 0 | 0.4142 | intron | HLA-DRB6 |
| rs1548791 | 1 | 187884585 | 4 | 0 | 0 | T | C | -1.114 | 0.2884 | 0.0001121 | ---?- | 0 | 0.5561 | intergenic | |
| rs75644017 | 10 | 131496611 | 4 | 0 | 0 | A | G | 1.4549 | 0.3766 | 0.0001121 | +++?+ | 0 | 0.9228 | intron | MGMT |
| rs12312435 | 12 | 43149592 | 5 | 0 | 0 | A | G | -1.0206 | 0.2642 | 0.0001122 | ----+ | 0 | 0.4572 | intergenic |  |
| rs12757392 | 1 | 64790214 | 4 | 0 | 0 | C | G | -1.3995 | 0.3624 | 0.0001124 | ---?- | 0 | 0.5801 | intergenic | |
| rs12691880 | 2 | 137029511 | 4 | 0 | 0 | T | C | -1.0521 | 0.2724 | 0.0001125 | ---?- | 0 | 0.7503 | intergenic | |
| rs143012354 | 22 | 49401419 | 3 | 0 | 0 | A | G | 1.5952 | 0.4131 | 0.0001126 | +++?? | 0 | 0.501 | intergenic | |
| 5:179984905 | 5 | 179984905 | 3 | 0 | 0 | T | TTTTG | 1.2192 | 0.3157 | 0.0001126 | +++?? | 35 | 0.2149 | | |
| rs4566851 | 6 | 80551741 | 4 | 0 | 0 | T | C | -1.0328 | 0.2675 | 0.000113 | ---?- | 0.5 | 0.3892 | intron | C6orf7 |
| rs146570028 | 6 | 83882599 | 4 | 0 | 0 | A | G | 3.4347 | 0.8896 | 0.000113 | +++?+ | 17.7 | 0.3022 | intron | PGM3 |
| 3:196478567 | 3 | 196478567 | 3 | 0 | 0 | A | C | -1.9215 | 0.4978 | 0.0001132 | ---?? | 0 | 0.4316 | intron | PAK2 |
| rs868765 | 12 | 43150747 | 5 | 0 | 0 | A | G | -1.0235 | 0.2652 | 0.0001134 | ----+ | 0 | 0.477 | intergenic |  |
| rs9470027 | 6 | 35416759 | 5 | 0 | 0 | A | G | -3.0789 | 0.7976 | 0.0001134 | ----- | 0 | 0.8165 | upstream-gene |  |
| rs117931079 | 6 | 83824651 | 4 | 0 | 0 | T | C | 3.4548 | 0.895 | 0.0001134 | +++?+ | 0 | 0.5872 | intron | DOPEY1 |
| rs9638081 | 7 | 156175854 | 4 | 0 | 0 | T | C | -0.9496 | 0.246 | 0.0001135 | ---?- | 39.9 | 0.1722 | intergenic | |
| rs2145572 | 6 | 80550286 | 4 | 0 | 0 | T | C | -1.0353 | 0.2683 | 0.000114 | ---?- | 1.4 | 0.3852 | intron | C6orf7 |
| rs34093026 | 6 | 77683671 | 4 | 0 | 0 | A | C | -1.1106 | 0.2878 | 0.000114 | ---?- | 15.4 | 0.3146 | intergenic | |
| 10:74586710 | 10 | 74586710 | 2 | 0 | 0 | T | TG | -8.1709 | 2.1176 | 0.000114 | --??? | 0 | 0.6571 | | |
| rs7501971 | 17 | 1392707 | 4 | 0 | 0 | A | G | 2.228 | 0.5774 | 0.0001141 | +++?+ | 1.3 | 0.3857 | intron | MYO1C |
| rs2010066 | 15 | 85828519 | 3 | 0 | 0 | A | G | 5.9084 | 1.5313 | 0.0001142 | ++??+ | 0 | 0.8337 | intergenic | |
| rs10889476 | 1 | 64794749 | 4 | 0 | 0 | T | C | -0.9345 | 0.2422 | 0.0001142 | ---?- | 48.7 | 0.1192 | intergenic | |
| rs10969926 | 9 | 30878999 | 4 | 0 | 0 | A | T | 4.1789 | 1.0832 | 0.0001143 | +++?+ | 13.2 | 0.3265 | intergenic | |
| rs299364 | 5 | 134331921 | 4 | 0 | 0 | T | G | 1.0387 | 0.2693 | 0.0001146 | +++?+ | 0 | 0.5842 | intron | CATSPER3 |
| rs950173 | 2 | 172570419 | 4 | 0 | 0 | A | G | 1.5208 | 0.3943 | 0.0001146 | +++?+ | 0 | 0.7138 | intron | DYNC1I2 |
| rs10969913 | 9 | 30866808 | 4 | 0 | 0 | A | G | -4.1136 | 1.0665 | 0.0001148 | ---?- | 14 | 0.3222 | intergenic | |
| rs1005755 | 2 | 170650831 | 4 | 0 | 0 | A | T | -0.9612 | 0.2492 | 0.000115 | ---?- | 0 | 0.8671 | upstream-gene | |
| rs13010558 | 2 | 170650661 | 4 | 0 | 0 | A | G | -0.9613 | 0.2493 | 0.000115 | ---?- | 0 | 0.8666 | upstream-gene | |
| rs1600047 | 8 | 140459422 | 4 | 0 | 0 | T | G | -1.2072 | 0.313 | 0.0001151 | ---?- | 0 | 0.7538 | intergenic | |
| rs57951898 | 18 | 40041906 | 4 | 0 | 0 | A | G | 1.2278 | 0.3184 | 0.0001153 | +++?- | 0 | 0.4783 | intron | LINC00907 |
| rs61153228 | 12 | 41301251 | 4 | 0 | 0 | C | G | 1.2398 | 0.3215 | 0.0001153 | +++?+ | 0 | 0.602 | intron | CNTN1 |
| rs150171640 | 6 | 83883206 | 4 | 0 | 0 | T | G | -3.4417 | 0.8926 | 0.0001153 | ---?- | 17.3 | 0.3048 | intron | PGM3 |
| rs12904572 | 15 | 91589221 | 4 | 0 | 0 | A | G | -1.3756 | 0.3568 | 0.0001154 | ---?+ | 0 | 0.691 | intergenic | |
| rs56080646 | 15 | 85863030 | 3 | 0 | 0 | T | C | 5.8403 | 1.5147 | 0.0001154 | ++??+ | 0 | 0.7988 | intergenic | |
| rs6139889 | 20 | 5989838 | 3 | 0 | 0 | A | G | -0.9577 | 0.2484 | 0.0001155 | ---?? | 0 | 0.9216 | intron | CRLS1 |
| rs10890620 | 11 | 106764360 | 4 | 0 | 0 | A | G | -1.1428 | 0.2964 | 0.0001156 | ---?+ | 38.8 | 0.1793 | intron | GUCY1A2 |
| rs13204650 | 6 | 80552965 | 4 | 0 | 0 | A | G | -1.0306 | 0.2673 | 0.0001156 | ---?- | 1.7 | 0.3836 | intron | C6orf7 |
| rs11208422 | 1 | 64793422 | 4 | 0 | 0 | T | C | -0.9332 | 0.2421 | 0.0001157 | ---?- | 48.7 | 0.1194 | intergenic | |
| rs72642409 | 18 | 40042002 | 4 | 0 | 0 | A | G | -1.2277 | 0.3185 | 0.0001157 | ---?+ | 0 | 0.4783 | intron | LINC00907 |
| rs6725295 | 2 | 170599690 | 4 | 0 | 0 | A | C | -1.2662 | 0.3285 | 0.0001158 | ---?- | 0 | 0.5973 | intron | KLHL23,PHOSPHO2-KLHL23 |
| 3:196474042 | 3 | 196474042 | 3 | 0 | 0 | CT | CTTT | -2.461 | 0.6385 | 0.000116 | ---?? | 0 | 0.9478 | | |
| 5:101490557 | 5 | 101490557 | 2 | 0 | 0 | C | G | -5.6217 | 1.4586 | 0.0001161 | ?--?? | 70.7 | 0.06453 | intergenic | |
| rs670385 | 1 | 187895234 | 4 | 0 | 0 | T | C | -1.0881 | 0.2823 | 0.0001161 | ---?- | 0 | 0.4973 | intergenic | |
| rs8086183 | 18 | 40053940 | 4 | 0 | 0 | A | G | 1.2327 | 0.3198 | 0.0001161 | +++?- | 0 | 0.4833 | intron | LINC00907 |
| rs57638662 | 8 | 140807080 | 4 | 0 | 0 | T | C | -1.4469 | 0.3754 | 0.0001161 | ---?- | 59.8 | 0.05863 | intron | TRAPPC9 |
| rs73644367 | 9 | 30877456 | 4 | 0 | 0 | A | C | -4.1704 | 1.082 | 0.0001161 | ---?- | 13.5 | 0.3249 | intergenic | |
| rs117688307 | 9 | 136851850 | 4 | 0 | 0 | A | C | -3.3606 | 0.8719 | 0.0001161 | -+-?- | 39.8 | 0.1729 | intron | VAV2 |
| rs72682423 | 4 | 131499315 | 4 | 0 | 0 | T | C | -5.0301 | 1.3053 | 0.0001164 | ---?- | 0 | 0.5624 | intergenic | |
| rs685123 | 1 | 187892358 | 4 | 0 | 0 | A | G | -1.0879 | 0.2823 | 0.0001165 | ---?- | 0 | 0.4846 | intergenic | |
| rs11655704 | 17 | 47448172 | 4 | 0 | 0 | T | C | 1.038 | 0.2694 | 0.0001169 | +++?- | 32.2 | 0.2194 | intergenic | |
| rs115756139 | 2 | 122231708 | 4 | 0 | 0 | A | G | 3.4866 | 0.905 | 0.0001169 | +++?+ | 0 | 0.6138 | intron | CLASP1 |
| rs9356924 | 6 | 24057329 | 5 | 0 | 0 | T | G | -1.5653 | 0.4063 | 0.000117 | ----+ | 0 | 0.6215 | intergenic |  |
| rs4363645 | 12 | 27777195 | 4 | 0 | 0 | A | G | 1.0019 | 0.2601 | 0.0001171 | +++?+ | 22.5 | 0.2759 | intron | PPFIBP1 |
| rs12912907 | 15 | 91590587 | 4 | 0 | 0 | T | G | 1.6355 | 0.4246 | 0.0001171 | +++?+ | 0 | 0.9825 | intergenic | |
| 14:100780691 | 14 | 100780691 | 3 | 0 | 0 | CT | C | 3.361 | 0.8725 | 0.0001171 | +++?? | 0 | 0.5195 | | |

| | | | | | | | | | | | | | | | |
|---|---|---|---|---|---|---|---|---|---|---|---|---|---|---|---|
| rs13000153 | 2 | 172586067 | 4 | 0 | 0 | A | G | -1.4846 | 0.3854 | 0.0001172 | ---?- | 0 | 0.693 | intron | DYNC1I2 |
| rs9348627 | 6 | 24065658 | 5 | 0 | 0 | T | G | -1.5399 | 0.3998 | 0.0001173 | ----+ | 0 | 0.5536 | intergenic |  |
| rs3800376 | 6 | 35425990 | 5 | 0 | 0 | T | C | 3.0531 | 0.7927 | 0.0001174 | +++++ | 0 | 0.8359 | intron | FANCE |
| 1:64825652 | 1 | 64825652 | 3 | 0 | 0 | T | C | -1.9072 | 0.4952 | 0.0001176 | ---?? | 47.7 | 0.1476 | intergenic | |
| rs7550108 | 1 | 85191736 | 5 | 0 | 0 | T | C | 0.8822 | 0.2291 | 0.0001177 | +++++ | 30.4 | 0.2188 | intergenic | |
| rs148537164 | 3 | 120286638 | 3 | 0 | 0 | T | C | 4.0226 | 1.0446 | 0.0001177 | +++?? | 0 | 0.9843 | intergenic | |
| rs8028542 | 15 | 54347686 | 4 | 0 | 0 | A | G | -1.2699 | 0.3298 | 0.0001178 | ---?- | 0 | 0.9068 | intron | UNC13C |
| rs141931213 | 17 | 68603066 | 3 | 0 | 0 | A | C | 3.5654 | 0.926 | 0.0001179 | +++?? | 0 | 0.8828 | intergenic | |
| rs192854694 | 1 | 64816327 | 3 | 0 | 0 | C | G | -1.5069 | 0.3914 | 0.0001179 | ---?? | 0 | 0.5576 | intergenic | |
| rs2319365 | 8 | 96922476 | 4 | 0 | 0 | A | G | 1.2277 | 0.3189 | 0.0001182 | +++?+ | 0 | 0.9193 | intergenic | |
| rs8079668 | 17 | 52306524 | 4 | 0 | 0 | A | C | -2.2148 | 0.5753 | 0.0001182 | ---?- | 0 | 0.8206 | intergenic | |
| rs73428082 | 22 | 28847273 | 4 | 0 | 0 | A | G | 2.6837 | 0.6971 | 0.0001182 | +++?+ | 0 | 0.8373 | intron | TTC28 |
| rs1940442 | 18 | 396461 | 4 | 0 | 0 | A | G | 1.2782 | 0.332 | 0.0001183 | +++?+ | 0 | 0.4979 | intron | COLEC12 |
| rs114399632 | 3 | 155317519 | 4 | 0 | 0 | A | C | 2.5797 | 0.6702 | 0.0001184 | +++?+ | 17.3 | 0.3044 | intron | PLCH1 |
| rs144735925 | 15 | 87538221 | 4 | 0 | 0 | A | C | 2.5142 | 0.6531 | 0.0001184 | +++?+ | 0 | 0.9946 | intron | AGBL1 |
| rs10225434 | 7 | 156173333 | 4 | 0 | 0 | T | C | -0.943 | 0.245 | 0.0001186 | ---?- | 39.8 | 0.1729 | intergenic | |
| rs12673127 | 7 | 130367061 | 4 | 0 | 0 | A | G | 0.9301 | 0.2416 | 0.0001186 | +-+?+ | 51.4 | 0.1034 | intron | TSGA13 |
| rs6453953 | 6 | 77663444 | 4 | 0 | 0 | T | C | -1.1026 | 0.2865 | 0.0001187 | ---?- | 14.8 | 0.3179 | intergenic | |
| rs111692482 | 9 | 30876567 | 4 | 0 | 0 | T | C | 4.1531 | 1.0792 | 0.0001189 | +++?+ | 14 | 0.3224 | intergenic | |
| rs11753071 | 6 | 77690637 | 4 | 0 | 0 | A | G | -1.1076 | 0.2878 | 0.000119 | ---?- | 15.8 | 0.3129 | intergenic | |
| rs293819 | 3 | 191151320 | 4 | 0 | 0 | C | G | 1.4654 | 0.3808 | 0.0001191 | +++?- | 40.2 | 0.1707 | intergenic | |
| rs10464980 | 8 | 140807485 | 4 | 0 | 0 | T | C | -1.4444 | 0.3754 | 0.0001191 | ---?- | 59.8 | 0.05838 | intron | TRAPPC9 |
| 20:5988736 | 20 | 5988736 | 3 | 0 | 0 | CT | C | -1.0323 | 0.2683 | 0.0001191 | ---?? | 0 | 0.8852 | | |
| rs60929232 | 3 | 196575989 | 3 | 0 | 0 | T | C | -3.7214 | 0.9673 | 0.0001194 | ---?? | 0 | 0.9245 | intergenic | |
| rs72746758 | 15 | 85855914 | 3 | 0 | 0 | T | C | 5.8269 | 1.5145 | 0.0001194 | ++??+ | 0 | 0.7989 | intergenic | |
| 12:18864463 | 12 | 18864463 | 3 | 0 | 0 | T | TATA | -1.1121 | 0.2891 | 0.0001196 | ---?? | 0 | 0.986 | | |
| 2:79660750 | 2 | 79660750 | 3 | 0 | 0 | G | GAAAA | -1.049 | 0.2727 | 0.0001197 | ---?? | 46 | 0.157 | | |
| rs16954330 | 17 | 52314280 | 4 | 0 | 0 | A | T | 2.442 | 0.6349 | 0.0001198 | +++?- | 0 | 0.7857 | intergenic | |
| rs607013 | 1 | 187892299 | 4 | 0 | 0 | T | G | 1.0867 | 0.2825 | 0.0001199 | +++?+ | 0 | 0.4927 | intergenic | |
| rs112134389 | 9 | 30876262 | 4 | 0 | 0 | T | C | -4.155 | 1.0802 | 0.0001199 | ---?- | 14.1 | 0.3217 | intergenic | |
| 5:179981517 | 5 | 179981517 | 3 | 0 | 0 | G | GTTT | -1.2118 | 0.3151 | 0.0001199 | ---?? | 35.7 | 0.2112 | | |
| rs17162293 | 7 | 141185491 | 4 | 0 | 0 | C | G | -5.3077 | 1.38 | 0.00012 | ---?- | 0 | 0.7623 | intergenic | |
| rs11753087 | 6 | 77690787 | 4 | 0 | 0 | A | G | -1.107 | 0.2879 | 0.0001204 | ---?- | 15.9 | 0.3123 | intergenic | |
| rs56336640 | 7 | 107370821 | 5 | 0 | 0 | T | C | 0.8247 | 0.2145 | 0.0001204 | +++++ | 37.2 | 0.1731 | intergenic | |
| rs6901056 | 6 | 77689092 | 4 | 0 | 0 | A | G | 1.1066 | 0.2878 | 0.0001206 | +++?+ | 15.8 | 0.3129 | intergenic | |
| rs10969918 | 9 | 30875780 | 4 | 0 | 0 | T | C | -4.151 | 1.0797 | 0.0001208 | ---?- | 14.3 | 0.3205 | intergenic | |
| rs2897590 | 12 | 43151728 | 5 | 0 | 0 | A | G | 1.0205 | 0.2655 | 0.0001209 | ++++- | 0 | 0.4765 | intergenic | |
| rs62031614 | 16 | 9307510 | 3 | 0 | 0 | T | C | 4.6455 | 1.2084 | 0.0001209 | ?++?+ | 0 | 0.7828 | intergenic | |
| rs16954329 | 17 | 52313383 | 4 | 0 | 0 | T | C | 2.441 | 0.635 | 0.0001211 | +++?- | 0 | 0.7803 | intergenic | |
| rs71536530 | 6 | 32521970 | 5 | 0 | 1 | C | G | -1.43 | 0.372 | 0.0001211 | ----- | 0 | 0.4102 | intron | HLA-DRB6 |
| rs1437331 | 2 | 139897771 | 4 | 0 | 0 | T | G | 2.5807 | 0.6714 | 0.0001212 | +++?+ | 0 | 0.7952 | intergenic | |
| rs34927613 | 6 | 32600211 | 5 | 0 | 1 | A | G | 1.0654 | 0.2772 | 0.0001212 | +++++ | 0 | 0.6419 | upstream-gene | |
| rs7138064 | 12 | 43200941 | 5 | 0 | 0 | T | C | -1.3259 | 0.345 | 0.0001213 | ----- | 19.7 | 0.289 | intergenic |  |
| rs11181677 | 12 | 43150285 | 5 | 0 | 0 | C | G | -1.0199 | 0.2654 | 0.0001213 | ----+ | 0 | 0.4779 | intergenic |  |
| rs190528058 | 13 | 85404307 | 3 | 0 | 0 | A | G | -3.7352 | 0.9718 | 0.0001213 | -?-?+ | 16.3 | 0.3029 | intergenic | |
| rs3935712 | 2 | 235412576 | 4 | 0 | 0 | A | G | 1.4685 | 0.3821 | 0.0001214 | +++?+ | 0 | 0.7577 | intergenic | |
| rs60547794 | 4 | 20750521 | 3 | 0 | 0 | A | G | 1.0593 | 0.2756 | 0.0001214 | +++?? | 0 | 0.5381 | intron | KCNIP4 |
| rs72746771 | 15 | 85864169 | 3 | 0 | 0 | T | C | 5.822 | 1.515 | 0.0001216 | ++??+ | 0 | 0.8004 | intergenic | |
| 2:170664316 | 2 | 170664316 | 3 | 0 | 0 | G | GTGTC | 0.9887 | 0.2573 | 0.0001216 | +++?? | 60.4 | 0.0802 | | |
| 12:95866183 | 12 | 95866183 | 2 | 0 | 0 | A | ATGGAT | -4.969 | 1.2932 | 0.0001218 | -?-?? | 0 | 0.9496 | | |
| 6:83823279 | 6 | 83823279 | 3 | 0 | 0 | T | TAAG | 3.5376 | 0.9208 | 0.0001222 | +++?? | 0 | 0.4225 | | |
| rs117810414 | 6 | 155765492 | 3 | 0 | 0 | A | C | 5.2077 | 1.3557 | 0.0001223 | +++?? | 65.5 | 0.05514 | intron | NOX3 |
| rs1388962 | 8 | 113437239 | 4 | 0 | 0 | T | G | -1.0611 | 0.2763 | 0.0001227 | ---?- | 0 | 0.791 | intron | CSMD3 |
| rs150987003 | 17 | 68590321 | 3 | 0 | 0 | T | C | 3.5089 | 0.9137 | 0.0001228 | +++?? | 0 | 0.8426 | intergenic | |
| 4:89084794 | 4 | 89084794 | 3 | 0 | 0 | A | AC | 3.8823 | 1.0109 | 0.0001228 | +++?? | 1.6 | 0.3619 | | |
| 17:17770337 | 17 | 17770337 | 3 | 0 | 0 | A | C | -1.9644 | 0.5116 | 0.0001231 | ---?? | 0 | 0.4394 | intron | TOM1L2 |
| 8:64890840 | 8 | 64890840 | 3 | 0 | 0 | A | AT | -0.9648 | 0.2513 | 0.0001232 | ---?? | 0 | 0.6225 | | |
| 4:18368712 | 4 | 18368712 | 3 | 0 | 0 | CAT | C | -1.6243 | 0.4231 | 0.0001235 | ---?? | 58.9 | 0.08792 | | |
| rs913244 | 20 | 62457488 | 3 | 0 | 0 | C | G | 4.4161 | 1.1504 | 0.0001236 | +++?? | 0 | 0.7808 | intron | ZBTB46 |

| | | | | | | | | | | | | | | | |
|---|---|---|---|---|---|---|---|---|---|---|---|---|---|---|---|
| rs77552257 | 7 | 107910206 | 4 | 0 | 0 | T | G | 2.6837 | 0.6991 | 0.0001237 | +++?+ | 0 | 0.5903 | intron | NRCAM |
| rs80220949 | 17 | 56694745 | 5 | 0 | 0 | A | G | 2.6414 | 0.6881 | 0.0001237 | ++++- | 0 | 0.7131 | intron | TEX14 |
| rs343042 | 7 | 35506790 | 4 | 0 | 0 | T | C | 1.1047 | 0.2878 | 0.0001242 | +++?+ | 0 | 0.6083 | intergenic | |
| rs75068366 | 5 | 122547493 | 4 | 0 | 0 | T | C | 2.0606 | 0.5369 | 0.0001242 | +++?+ | 0 | 0.889 | intergenic | |
| rs748312 | 5 | 122551343 | 4 | 0 | 0 | A | G | 2.0668 | 0.5386 | 0.0001244 | +++?+ | 0 | 0.8789 | intergenic | |
| rs7816565 | 8 | 140806010 | 4 | 0 | 0 | A | T | -1.4415 | 0.3757 | 0.0001247 | ---?- | 60.2 | 0.05639 | intron | TRAPPC9 |
| rs146514404 | 17 | 56725263 | 5 | 0 | 0 | T | C | 2.6405 | 0.6883 | 0.0001249 | +++++ | 0 | 0.8226 | intron | TEX14 |
| rs13359926 | 5 | 123429898 | 4 | 0 | 0 | T | G | 0.9523 | 0.2482 | 0.000125 | +++?+ | 0 | 0.4238 | intergenic | |
| rs13223150 | 7 | 130369609 | 4 | 0 | 0 | A | C | 0.925 | 0.2412 | 0.0001251 | +++?+ | 49.9 | 0.1123 | intron | TSGA13 |
| rs4600177 | 11 | 106777540 | 4 | 0 | 0 | T | C | -1.1454 | 0.2986 | 0.0001253 | ---?+ | 35.8 | 0.1974 | intron | GUCY1A2 |
| rs76140389 | 17 | 56726020 | 5 | 0 | 0 | A | G | -2.64 | 0.6884 | 0.0001255 | ----- | 0 | 0.8228 | intron | TEX14 |
| rs62012919 | 15 | 63592201 | 5 | 0 | 0 | A | T | -2.0253 | 0.5281 | 0.0001256 | ----- | 21.1 | 0.2804 | intron | APH1B |
| rs1854778 | 1 | 152880723 | 4 | 0 | 0 | A | G | 2.0829 | 0.5432 | 0.0001257 | +++?+ | 0 | 0.5177 | upstream-gene | |
| rs77414312 | 10 | 4353698 | 3 | 0 | 0 | A | T | 3.0735 | 0.8015 | 0.0001258 | +++?? | 58.6 | 0.08921 | intergenic | |
| rs9264584 | 6 | 31236034 | 3 | 0 | 0 | T | C | -1.0074 | 0.2627 | 0.000126 | ---?? | 0 | 0.4885 | downstream-gene | |
| rs10493356 | 1 | 64789617 | 4 | 0 | 0 | A | T | 0.9269 | 0.2418 | 0.000126 | +++?+ | 48.9 | 0.118 | intergenic | |
| rs151129971 | 6 | 83820122 | 4 | 0 | 0 | A | G | -3.4413 | 0.8975 | 0.000126 | ---?- | 0 | 0.6123 | intron | DOPEY1 |
| 3:196469974 | 3 | 196469974 | 3 | 0 | 0 | G | GA | -2.4261 | 0.6328 | 0.000126 | ---?? | 0 | 0.9727 | | |
| rs9844958 | 3 | 71894573 | 4 | 0 | 0 | T | C | 1.0377 | 0.2707 | 0.0001261 | +++?+ | 0 | 0.8653 | intergenic | |
| rs77208224 | 17 | 52299932 | 4 | 0 | 0 | A | G | 2.473 | 0.645 | 0.0001261 | +++?- | 0 | 0.7852 | intergenic | |
| 2:238525619 | 2 | 238525619 | 2 | 0 | 0 | A | G | -4.2472 | 1.1079 | 0.0001262 | -?-?? | 0 | 0.5417 | intergenic | |
| rs28368843 | 6 | 32452587 | 5 | 0 | 1 | T | C | 1.3172 | 0.3436 | 0.0001262 | +++++ | 0 | 0.5279 | intergenic | |
| rs10493357 | 1 | 64789908 | 4 | 0 | 0 | A | G | -0.927 | 0.2418 | 0.0001264 | ---?- | 48.9 | 0.118 | intergenic | |
| 4:100547828 | 4 | 100547828 | 3 | 0 | 0 | T | TA | 1.6341 | 0.4263 | 0.0001266 | +++?? | 55.7 | 0.1048 | | |
| 12:14697908 | 12 | 14697908 | 3 | 0 | 0 | A | AC | 1.8401 | 0.4801 | 0.0001267 | +++?? | 0 | 0.6538 | | |
| rs148186333 | 17 | 56680224 | 5 | 0 | 0 | A | C | -2.6462 | 0.6904 | 0.0001268 | ----- | 0 | 0.7746 | intron | TEX14 |
| rs78371887 | 17 | 56730173 | 5 | 0 | 0 | T | C | -2.6413 | 0.6892 | 0.0001269 | ----- | 0 | 0.8213 | intron | TEX14 |
| rs138573089 | 19 | 3175543 | 3 | 0 | 0 | T | C | 1.1776 | 0.3073 | 0.0001269 | +++?? | 0 | 0.9048 | upstream-gene | |
| rs113013738 | 20 | 24566680 | 3 | 0 | 0 | A | G | -3.7129 | 0.9689 | 0.000127 | ---?? | 53.2 | 0.118 | intron | SYNDIG1 |
| 2:238525618 | 2 | 238525618 | 2 | 0 | 0 | A | ATTTC | 4.251 | 1.1094 | 0.0001272 | +?+?? | 0 | 0.5403 | | |
| 5:123515600 | 5 | 123515600 | 3 | 0 | 0 | CT | C | -0.9811 | 0.2561 | 0.0001274 | ---?? | 0 | 0.7965 | | |
| rs28762986 | 8 | 113419152 | 4 | 0 | 0 | C | G | -1.0792 | 0.2817 | 0.0001275 | ---?- | 0 | 0.7889 | intron | CSMD3 |
| rs12127472 | 1 | 64787327 | 4 | 0 | 0 | A | T | 0.9251 | 0.2415 | 0.0001276 | +++?+ | 50.2 | 0.1107 | intergenic | |
| rs11581785 | 1 | 64787206 | 4 | 0 | 0 | A | G | 0.925 | 0.2415 | 0.0001277 | +++?+ | 50.2 | 0.1107 | intergenic | |
| 8:113211760 | 8 | 113211760 | 3 | 0 | 0 | A | ACT | 0.9998 | 0.261 | 0.0001277 | +++?? | 0 | 0.9183 | | |
| rs7017899 | 8 | 113415551 | 4 | 0 | 0 | A | G | -1.081 | 0.2822 | 0.0001279 | ---?- | 0 | 0.7963 | intron | CSMD3 |
| rs60972226 | 10 | 131467850 | 3 | 0 | 0 | A | T | -1.7423 | 0.4549 | 0.0001279 | ---?? | 0 | 0.7463 | intron | MGMT |
| rs74528545 | 6 | 168600194 | 4 | 0 | 0 | A | G | 1.9211 | 0.5015 | 0.0001279 | +++?+ | 0 | 0.745 | intergenic | |
| rs144900800 | 2 | 104908002 | 4 | 0 | 0 | C | G | -3.6466 | 0.9521 | 0.0001281 | ---?- | 0 | 0.4013 | intergenic | |
| rs537930 | 5 | 134348703 | 5 | 0 | 0 | T | G | -0.9495 | 0.2479 | 0.0001282 | ----- | 21.4 | 0.2784 | downstream-gene |  |
| 13:30322795 | 13 | 30322795 | 3 | 0 | 0 | T | TCAAA | -1.6248 | 0.4242 | 0.0001282 | ---?? | 0 | 0.686 | | |
| rs744335 | 1 | 64792532 | 4 | 0 | 0 | C | G | -1.3818 | 0.3608 | 0.0001283 | ---?- | 0 | 0.5599 | intergenic | |
| rs223841 | 16 | 57457271 | 4 | 0 | 0 | A | G | 1.2621 | 0.3296 | 0.0001284 | +++?+ | 0 | 0.5039 | downstream-gene | |
| 5:173153912 | 5 | 173153912 | 3 | 0 | 0 | G | GA | 1.1002 | 0.2873 | 0.0001285 | +++?? | 0 | 0.5623 | | |
| 4:100548263 | 4 | 100548263 | 3 | 0 | 0 | A | ATCT | -1.6337 | 0.4266 | 0.0001286 | ---?? | 55.5 | 0.1055 | | |
| rs1407207 | 6 | 80555189 | 4 | 0 | 0 | A | G | 1.0401 | 0.2716 | 0.0001287 | +++?+ | 12 | 0.3329 | intron | C6orf7 |
| 10:131467848 | 10 | 131467848 | 3 | 0 | 0 | T | TA | 1.7426 | 0.4551 | 0.0001287 | +++?? | 0 | 0.745 | | |
| rs7356937 | 6 | 144749694 | 4 | 0 | 0 | A | G | -3.0636 | 0.8002 | 0.0001288 | ---?- | 0 | 0.8366 | intron | UTRN |
| rs1480079 | 12 | 98689961 | 4 | 0 | 0 | A | C | -1.7563 | 0.4587 | 0.0001289 | -+-?- | 47.2 | 0.1282 | intergenic | |
| rs56714706 | 16 | 5454657 | 4 | 0 | 0 | T | G | -1.1323 | 0.2957 | 0.0001289 | ---?- | 0 | 0.6438 | intron | RBFOX1 |
| rs75033022 | 6 | 168580484 | 4 | 0 | 0 | T | C | -1.7741 | 0.4634 | 0.000129 | ---?- | 0 | 0.7043 | intergenic | |
| rs3129882 | 6 | 32409530 | 5 | 0 | 0 | A | G | 0.8167 | 0.2133 | 0.0001291 | -++++ | 45.4 | 0.1197 | intron | HLA-DRA |
| rs2023211 | 14 | 62374092 | 4 | 0 | 0 | T | C | 1.1682 | 0.3052 | 0.0001292 | +++?- | 0 | 0.5679 | intron | SYT16 |
| rs2447562 | 8 | 140477913 | 4 | 0 | 0 | T | G | 1.5525 | 0.4056 | 0.0001292 | +++?+ | 0 | 0.8072 | intergenic | |
| rs142755670 | 15 | 85847085 | 3 | 0 | 0 | A | G | 5.7921 | 1.5131 | 0.0001292 | ++??+ | 0 | 0.7991 | intergenic | |
| rs181091901 | 5 | 29445961 | 4 | 0 | 0 | A | G | 3.9167 | 1.0232 | 0.0001293 | +++?+ | 0 | 0.7008 | intergenic | |
| rs74394962 | 12 | 43193720 | 4 | 0 | 0 | A | G | -1.7695 | 0.4623 | 0.0001294 | ---?- | 0 | 0.7672 | intergenic | |
| rs6588086 | 1 | 64792485 | 4 | 0 | 0 | C | G | 1.3806 | 0.3607 | 0.0001297 | +++?+ | 0 | 0.5591 | intergenic | |
| rs4623740 | 1 | 64792410 | 4 | 0 | 0 | A | G | -1.3804 | 0.3607 | 0.0001298 | ---?- | 0 | 0.5592 | intergenic | |

| | | | | | | | | | | | | | | | |
|---|---|---|---|---|---|---|---|---|---|---|---|---|---|---|---|
| rs35693214 | 14 | 62374319 | 4 | 0 | 0 | T | C | -1.1686 | 0.3054 | 0.0001299 | ---?+ | 0 | 0.569 | intron | SYT16 |
| rs62143234 | 2 | 43023660 | 4 | 0 | 0 | T | C | -1.3118 | 0.3428 | 0.0001299 | ---?- | 0 | 0.8706 | upstream-gene | |
| rs79294753 | 1 | 102912796 | 3 | 0 | 0 | T | C | 0.9765 | 0.2552 | 0.0001299 | +++?? | 0 | 0.8841 | intergenic | |
| rs6785980 | 3 | 77391735 | 4 | 0 | 0 | T | C | -0.964 | 0.252 | 0.0001302 | ---?- | 0 | 0.425 | intron | ROBO2 |
| rs2078385 | 11 | 86168344 | 4 | 0 | 0 | A | G | -2.478 | 0.6477 | 0.0001303 | ---?- | 0 | 0.5497 | intron | ME3 |
| rs11892927 | 2 | 170561950 | 4 | 0 | 0 | T | C | 1.3715 | 0.3585 | 0.0001303 | +++?+ | 0 | 0.9305 | intron | PHOSPHO2-KLHL23 |
| rs116357634 | 17 | 56706194 | 5 | 0 | 0 | T | C | -2.605 | 0.6809 | 0.0001303 | ----+ | 0 | 0.7672 | intron | TEX14 |
| rs144607105 | 9 | 24400113 | 4 | 0 | 0 | A | T | -3.736 | 0.9765 | 0.0001303 | ---?- | 0 | 0.8542 | intergenic | |
| rs661732 | 13 | 29528992 | 4 | 0 | 0 | T | C | 1.0034 | 0.2623 | 0.0001305 | +++?+ | 15.3 | 0.3156 | intron | MTUS2 |
| rs1197769 | 15 | 42055856 | 5 | 0 | 0 | A | G | 1.107 | 0.2894 | 0.0001305 | +++++ | 13 | 0.3314 | intron | MGA |
| rs2076536 | 6 | 32339348 | 5 | 0 | 0 | T | C | 0.8024 | 0.2097 | 0.0001305 | +++++ | 0 | 0.87 | intron | C6orf10 |
| rs75227727 | 17 | 56702064 | 5 | 0 | 0 | T | C | -2.6088 | 0.682 | 0.0001305 | ----+ | 0 | 0.7564 | intron | TEX14 |
| rs75737863 | 6 | 168588209 | 4 | 0 | 0 | A | G | -1.7738 | 0.4637 | 0.0001305 | ---?- | 0 | 0.8738 | intergenic | |
| rs1496017 | 1 | 64792347 | 4 | 0 | 0 | C | G | -1.3798 | 0.3607 | 0.0001306 | ---?- | 0 | 0.5597 | intergenic | |
| rs4951162 | 1 | 205022977 | 5 | 0 | 0 | A | G | -0.8381 | 0.2191 | 0.0001306 | ----- | 0 | 0.9453 | intron | CNTN2 |
| rs6490072 | 12 | 111784495 | 5 | 0 | 0 | T | G | 0.9572 | 0.2502 | 0.0001306 | +-+++ | 43.5 | 0.1316 | intron | CUX2 |
| rs76104108 | 2 | 51131195 | 4 | 0 | 0 | T | C | -2.4129 | 0.6308 | 0.0001306 | ---?- | 0 | 0.9644 | intron | NRXN1 |
| rs7071424 | 10 | 131474247 | 4 | 0 | 0 | A | G | -1.4968 | 0.3913 | 0.0001309 | ---?- | 0 | 0.4084 | intron | MGMT |
| rs6975897 | 7 | 107354033 | 4 | 0 | 0 | T | C | -0.9607 | 0.2512 | 0.0001311 | ---?- | 0 | 0.3995 | intron | SLC26A4 |
| rs77627426 | 2 | 131512823 | 4 | 0 | 0 | A | G | -2.2372 | 0.585 | 0.0001311 | ---?- | 0 | 0.3993 | upstream-gene | |
| rs6684121 | 1 | 64792956 | 4 | 0 | 0 | T | C | 1.3792 | 0.3607 | 0.0001312 | +++?+ | 0 | 0.558 | intergenic | |
| rs9356925 | 6 | 24058402 | 5 | 0 | 0 | A | C | -1.5549 | 0.4066 | 0.0001313 | ----+ | 0 | 0.7292 | intergenic |  |
| rs112280198 | 6 | 83886350 | 4 | 0 | 0 | A | C | -3.4615 | 0.9052 | 0.0001313 | ---?- | 14.4 | 0.3203 | intron | PGM3 |
| rs75453353 | 6 | 168580485 | 4 | 0 | 0 | T | G | -1.7695 | 0.4628 | 0.0001314 | ---?- | 0 | 0.703 | intergenic | |
| rs181033999 | 3 | 183311899 | 3 | 0 | 0 | T | C | -4.3099 | 1.1272 | 0.0001314 | ---?? | 40.4 | 0.1868 | intergenic | |
| rs78510670 | 6 | 98910539 | 3 | 0 | 0 | A | C | -2.5921 | 0.6779 | 0.0001315 | ---?? | 33.2 | 0.2236 | intergenic | |
| 6:151700772 | 6 | 151700772 | 3 | 0 | 0 | G | GAA | -1.3587 | 0.3553 | 0.0001315 | ---?? | 58.9 | 0.08794 | | |
| rs1475241 | 14 | 62373734 | 4 | 0 | 0 | A | G | -1.1661 | 0.305 | 0.0001316 | ---?+ | 0 | 0.5701 | intron | SYT16 |
| rs6053591 | 20 | 5647414 | 3 | 0 | 0 | C | G | -1.2127 | 0.3172 | 0.0001316 | ---?? | 0 | 0.4966 | intergenic | |
| rs6457822 | 6 | 35418118 | 5 | 0 | 0 | A | G | -3.0212 | 0.7902 | 0.0001316 | ----- | 0 | 0.7923 | upstream-gene |  |
| rs146754634 | 17 | 56816093 | 5 | 0 | 0 | T | G | 2.6797 | 0.701 | 0.0001319 | ++++- | 0 | 0.6451 | downstream-gene | |
| rs72746750 | 15 | 85845728 | 3 | 0 | 0 | A | G | 5.7825 | 1.5127 | 0.000132 | ++??+ | 0 | 0.7987 | intergenic | |
| rs1071505 | 3 | 172329251 | 3 | 0 | 0 | A | G | 2.134 | 0.5584 | 0.0001324 | +++?? | 0 | 0.7182 | intergenic | |
| rs114250738 | 6 | 32601442 | 5 | 0 | 1 | A | G | -1.5059 | 0.394 | 0.0001324 | ----- | 9.8 | 0.3506 | upstream-gene |  |
| rs651874 | 1 | 187895196 | 4 | 0 | 0 | A | G | -1.0793 | 0.2824 | 0.0001325 | ---?- | 0 | 0.5116 | intergenic | |
| rs552493 | 9 | 103714623 | 3 | 0 | 0 | T | G | -4.9053 | 1.2836 | 0.0001326 | ---?? | 0 | 0.6053 | intergenic | |
| rs72746748 | 15 | 85844821 | 3 | 0 | 0 | T | C | -5.7797 | 1.5124 | 0.0001326 | --??- | 0 | 0.7989 | intergenic | |
| rs116280466 | 15 | 85844991 | 3 | 0 | 0 | A | G | -5.7803 | 1.5125 | 0.0001326 | --??- | 0 | 0.7987 | intergenic | |
| rs6930893 | 6 | 144751754 | 4 | 0 | 0 | A | G | 3.0532 | 0.799 | 0.0001329 | +++?+ | 0 | 0.8254 | intron | UTRN |
| rs1412406 | 9 | 30869687 | 4 | 0 | 0 | T | C | 4.0948 | 1.0718 | 0.0001332 | +++?+ | 16.6 | 0.3085 | intergenic | |
| rs117271113 | 6 | 83807323 | 4 | 0 | 0 | T | C | 3.4361 | 0.8994 | 0.0001333 | +++?+ | 0 | 0.6568 | intron | DOPEY1 |
| rs7135571 | 12 | 98710599 | 4 | 0 | 0 | A | G | -1.8027 | 0.4719 | 0.0001334 | -+-?- | 44.8 | 0.1426 | intergenic | |
| rs13206007 | 6 | 32663442 | 5 | 0 | 1 | C | G | -1.4305 | 0.3745 | 0.0001336 | ----- | 9.2 | 0.3542 | intergenic |  |
| rs6687371 | 1 | 187895760 | 4 | 0 | 0 | A | C | -1.0769 | 0.282 | 0.0001339 | ---?- | 0 | 0.5108 | intergenic | |
| rs6986274 | 8 | 113410972 | 4 | 0 | 0 | C | G | 1.081 | 0.283 | 0.000134 | +++?+ | 0 | 0.8147 | intron | CSMD3 |
| rs10969916 | 9 | 30869372 | 4 | 0 | 0 | C | G | -4.0967 | 1.0727 | 0.000134 | ---?- | 16.7 | 0.3077 | intergenic | |
| rs35042834 | 2 | 137030136 | 4 | 0 | 0 | A | G | -1.041 | 0.2726 | 0.000134 | ---?- | 0 | 0.7804 | intergenic | |
| rs72746742 | 15 | 85827911 | 3 | 0 | 0 | A | G | 5.8128 | 1.5221 | 0.000134 | ++??+ | 0 | 0.8062 | intergenic | |
| rs9470030 | 6 | 35422674 | 5 | 0 | 0 | T | C | -3.0172 | 0.7901 | 0.0001341 | ----- | 0 | 0.813 | intron | FANCE |
| rs6139899 | 20 | 6019481 | 5 | 0 | 0 | A | G | 0.7875 | 0.2062 | 0.0001343 | ++++- | 3.8 | 0.3849 | 3-prime-UTR | CRLS1 |
| rs4932589 | 15 | 92448863 | 4 | 0 | 0 | T | G | -0.9133 | 0.2392 | 0.0001344 | ---?- | 0 | 0.7225 | intron | SLCO3A1 |
| rs116159977 | 17 | 56743960 | 5 | 0 | 0 | T | G | 2.6459 | 0.693 | 0.0001345 | +++++ | 0 | 0.7991 | intron | TEX14 |
| rs12025846 | 1 | 64787990 | 4 | 0 | 0 | A | G | -0.9224 | 0.2416 | 0.0001346 | ---?- | 49.6 | 0.1142 | intergenic | |
| rs116894890 | 11 | 59131715 | 4 | 0 | 0 | C | G | -4.0649 | 1.0647 | 0.0001346 | ---?- | 24.6 | 0.264 | upstream-gene | |
| rs16976159 | 18 | 40039846 | 4 | 0 | 0 | T | G | -1.2139 | 0.318 | 0.0001347 | ---?+ | 0 | 0.4835 | intron | LINC00907 |
| 1:30369735 | 1 | 30369735 | 3 | 0 | 0 | T | TA | -1.3858 | 0.363 | 0.0001347 | ---?? | 0 | 0.4916 | | |
| rs1480078 | 12 | 98688295 | 4 | 0 | 0 | T | C | 1.7491 | 0.4582 | 0.0001348 | +-+?+ | 47.3 | 0.1275 | intergenic | |
| rs72746743 | 15 | 85827941 | 3 | 0 | 0 | C | G | 5.8103 | 1.522 | 0.0001348 | ++??+ | 0 | 0.8055 | intergenic | |
| rs10969915 | 9 | 30868871 | 4 | 0 | 0 | A | C | -4.093 | 1.0723 | 0.000135 | ---?- | 16.9 | 0.3069 | intergenic | |

| | | | | | | | | | | | | | | | |
|---|---|---|---|---|---|---|---|---|---|---|---|---|---|---|---|
| rs17212460 | 7 | 82442273 | 4 | 0 | 0 | A | G | -0.9994 | 0.2618 | 0.000135 | ---?- | 47.4 | 0.1269 | intron | PCLO |
| rs74631766 | 6 | 72578499 | 4 | 0 | 0 | T | C | 2.0433 | 0.5353 | 0.0001351 | +++?+ | 0 | 0.4667 | intergenic | |
| rs746632 | 1 | 64791827 | 4 | 0 | 0 | A | G | -1.3755 | 0.3604 | 0.0001355 | ---?- | 0 | 0.5595 | intergenic | |
| rs8093311 | 18 | 40050428 | 4 | 0 | 0 | A | C | -1.2066 | 0.3162 | 0.0001355 | ---?+ | 0 | 0.477 | intron | LINC00907 |
| rs12452694 | 17 | 52322466 | 4 | 0 | 0 | A | C | -2.1706 | 0.5689 | 0.0001358 | ---?- | 0 | 0.8462 | intergenic | |
| rs74378461 | 2 | 197053556 | 4 | 0 | 0 | A | G | 2.6036 | 0.6823 | 0.0001358 | +++?- | 0 | 0.7201 | intergenic | |
| 19:36421693 | 19 | 36421693 | 3 | 0 | 0 | T | TA | 3.8349 | 1.005 | 0.0001358 | +++?? | 0 | 0.9869 | | |
| rs149311699 | 6 | 83816604 | 4 | 0 | 0 | T | C | 3.4296 | 0.8989 | 0.000136 | +++?+ | 0 | 0.6282 | intron | DOPEY1 |
| rs148165455 | 17 | 56735296 | 5 | 0 | 0 | A | G | -2.666 | 0.6988 | 0.0001362 | ----- | 0 | 0.7827 | intron | TEX14 |
| rs4915972 | 1 | 64791688 | 4 | 0 | 0 | T | C | -1.3746 | 0.3604 | 0.0001365 | ---?- | 0 | 0.5593 | intergenic | |
| rs9285221 | 13 | 60021610 | 4 | 0 | 0 | A | C | -0.978 | 0.2564 | 0.0001365 | ---?- | 13.7 | 0.3238 | intergenic | |
| rs4915971 | 1 | 64791678 | 4 | 0 | 0 | A | G | 1.3745 | 0.3604 | 0.0001367 | +++?+ | 0 | 0.5593 | intergenic | |
| rs74487240 | 16 | 53043626 | 4 | 0 | 0 | C | G | 3.2234 | 0.8452 | 0.0001368 | +++?+ | 0 | 0.9007 | intergenic | |
| 2:212575752 | 2 | 212575752 | 3 | 0 | 0 | T | TA | 1.5019 | 0.3938 | 0.0001368 | +++?? | 0 | 0.9967 | | |
| rs75049735 | 5 | 122542868 | 4 | 0 | 0 | T | C | 1.9988 | 0.5241 | 0.0001369 | +++?+ | 0 | 0.7713 | intergenic | |
| 13:60009592 | 13 | 60009592 | 3 | 0 | 0 | A | AT | -0.9693 | 0.2542 | 0.0001369 | ---?? | 0 | 0.501 | | |
| rs10251802 | 7 | 107377409 | 5 | 0 | 0 | T | C | 0.8115 | 0.2128 | 0.000137 | +++++ | 41.1 | 0.1475 | intergenic | |
| rs62194739 | 20 | 23839427 | 4 | 0 | 0 | T | C | 1.9124 | 0.5015 | 0.0001371 | +++?+ | 30.1 | 0.2317 | intergenic | |
| rs77875686 | 10 | 131488958 | 4 | 0 | 0 | A | G | -1.567 | 0.4109 | 0.0001371 | ---?- | 0 | 0.9061 | intron | MGMT |
| rs387027 | 16 | 12085343 | 4 | 0 | 0 | A | C | 0.9723 | 0.255 | 0.0001373 | +++?+ | 0 | 0.4199 | intron | SNX29 |
| rs9448836 | 6 | 80553853 | 4 | 0 | 0 | T | C | -1.0232 | 0.2684 | 0.0001373 | ---?- | 0 | 0.3996 | intron | C6orf7 |
| rs10465723 | 1 | 85198523 | 5 | 0 | 0 | T | C | -0.8674 | 0.2275 | 0.0001373 | ----- | 26.4 | 0.2455 | intergenic |  |
| rs13010581 | 2 | 170633492 | 4 | 0 | 0 | C | G | 0.9308 | 0.2441 | 0.0001373 | +++?+ | 0 | 0.6314 | intergenic | |
| rs28383314 | 6 | 32587213 | 5 | 0 | 0 | T | C | -0.7846 | 0.2058 | 0.0001373 | ----- | 0 | 0.9133 | intergenic |  |
| rs138544456 | 12 | 91282674 | 3 | 0 | 0 | C | G | 5.1895 | 1.361 | 0.0001373 | +++?? | 0 | 0.8878 | intergenic | |
| rs77504626 | 6 | 168627866 | 4 | 0 | 0 | A | G | 2.1604 | 0.5666 | 0.0001375 | +++?+ | 0 | 0.7437 | intergenic | |
| rs666282 | 1 | 187896205 | 4 | 0 | 0 | A | C | -1.075 | 0.282 | 0.0001376 | ---?- | 0 | 0.5161 | intergenic | |
| rs4915966 | 1 | 64789238 | 4 | 0 | 0 | A | G | -1.3725 | 0.36 | 0.0001377 | ---?- | 0 | 0.5577 | intergenic | |
| rs4915967 | 1 | 64789261 | 4 | 0 | 0 | A | G | 1.3725 | 0.36 | 0.0001377 | +++?+ | 0 | 0.5582 | intergenic | |
| rs7868409 | 9 | 30868163 | 4 | 0 | 0 | T | C | -4.0831 | 1.0711 | 0.0001378 | ---?- | 17.2 | 0.3053 | intergenic | |
| rs16943170 | 17 | 56746010 | 5 | 0 | 0 | A | C | 2.6382 | 0.6921 | 0.0001379 | +++++ | 0 | 0.8242 | intron | TEX14 |
| rs78319792 | 2 | 170564407 | 4 | 0 | 0 | A | G | -1.3661 | 0.3584 | 0.0001379 | ---?- | 0 | 0.9317 | intron | PHOSPHO2-KLHL23 |
| rs4915968 | 1 | 64789294 | 4 | 0 | 0 | A | T | 1.3722 | 0.36 | 0.0001382 | +++?+ | 0 | 0.5582 | intergenic | |
| rs2351590 | 7 | 10134884 | 4 | 0 | 0 | T | C | 1.0715 | 0.2812 | 0.0001383 | +++?+ | 54.5 | 0.08626 | intergenic | |
| rs10092138 | 8 | 113426663 | 4 | 0 | 0 | A | G | 1.0717 | 0.2812 | 0.0001386 | +++?+ | 0 | 0.7852 | intron | CSMD3 |
| rs415126 | 3 | 191148502 | 4 | 0 | 0 | T | G | 1.4476 | 0.3799 | 0.0001387 | +++?- | 39.3 | 0.1762 | intergenic | |
| rs73762566 | 6 | 109082617 | 2 | 0 | 0 | C | G | 5.481 | 1.4384 | 0.0001388 | +?+?? | 0 | 0.6602 | intron | LINC00222 |
| rs1960125 | 1 | 64791316 | 4 | 0 | 0 | T | C | -1.3726 | 0.3603 | 0.0001389 | ---?- | 0 | 0.5595 | intergenic | |
| rs138878675 | 19 | 47304401 | 4 | 0 | 0 | T | C | 3.5898 | 0.9422 | 0.0001389 | +++?- | 0 | 0.5033 | intergenic | |
| rs2888484 | 2 | 218327912 | 4 | 0 | 0 | T | G | -0.9395 | 0.2466 | 0.000139 | ---?- | 0 | 0.7443 | intron | DIRC3 |
| rs62194740 | 20 | 23841567 | 4 | 0 | 0 | T | C | 1.9108 | 0.5015 | 0.000139 | +++?+ | 31.7 | 0.222 | intergenic | |
| rs3734088 | 5 | 82577833 | 4 | 0 | 0 | A | G | -2.4234 | 0.6361 | 0.0001391 | ---?- | 0 | 0.5273 | intron | XRCC4 |
| rs111414342 | 9 | 101615757 | 4 | 0 | 0 | T | C | 2.6297 | 0.6903 | 0.0001393 | +++?+ | 56.6 | 0.0749 | downstream-gene | |
| rs7463918 | 8 | 113404601 | 4 | 0 | 0 | A | T | 1.0782 | 0.283 | 0.0001395 | +++?+ | 0 | 0.8301 | intron | CSMD3 |
| rs8082933 | 18 | 40039381 | 4 | 0 | 0 | C | G | -1.2108 | 0.3179 | 0.0001395 | ---?+ | 0 | 0.4846 | intron | LINC00907 |
| rs12654898 | 5 | 17185263 | 4 | 0 | 0 | T | G | 0.969 | 0.2544 | 0.0001395 | +++?+ | 0 | 0.7372 | intron | LOC285696 |
| rs6688061 | 1 | 90266436 | 5 | 0 | 0 | T | C | -0.7946 | 0.2086 | 0.0001397 | ----+ | 0 | 0.5839 | intergenic |  |
| rs112245252 | 22 | 39420941 | 3 | 0 | 0 | C | G | -1.5704 | 0.4124 | 0.0001399 | ---?? | 0 | 0.9003 | intron | APOBEC3D |
| rs13164729 | 5 | 10304701 | 4 | 0 | 0 | T | G | 2.0241 | 0.5315 | 0.00014 | +++?+ | 33.6 | 0.2109 | intron | CMBL |
| rs76854796 | 17 | 52322176 | 4 | 0 | 0 | A | G | -2.1671 | 0.5691 | 0.00014 | ---?- | 0 | 0.8482 | intergenic | |
| rs674464 | 5 | 134349392 | 5 | 0 | 0 | T | C | 0.8893 | 0.2335 | 0.0001401 | +++++ | 4.8 | 0.3795 | downstream-gene | |
| rs2605570 | 15 | 42059521 | 5 | 0 | 0 | A | T | 1.1015 | 0.2893 | 0.0001403 | +++++ | 14.4 | 0.3228 | 3-prime-UTR | MGA |
| rs71565305 | 6 | 32011369 | 3 | 0 | 0 | T | G | 1.6038 | 0.4212 | 0.0001405 | +++?? | 0 | 0.5849 | intron | TNXB |
| rs116922184 | 15 | 87587256 | 4 | 0 | 0 | A | C | -2.4316 | 0.6387 | 0.0001406 | ---?- | 0 | 0.965 | intergenic | |
| 17:47414304 | 17 | 47414304 | 3 | 0 | 0 | G | GTTT | -1.064 | 0.2795 | 0.0001408 | ---?? | 0 | 0.4836 | | |
| rs4939623 | 18 | 47766165 | 4 | 0 | 0 | A | T | 1.0186 | 0.2676 | 0.000141 | +++?+ | 0 | 0.8878 | intron | CCDC11 |
| rs4915662 | 1 | 64789489 | 4 | 0 | 0 | A | G | -1.3701 | 0.36 | 0.0001413 | ---?- | 0 | 0.5573 | intergenic | |
| rs117455145 | 11 | 938759 | 4 | 0 | 0 | C | G | -2.4148 | 0.6345 | 0.0001414 | ---?- | 0 | 0.7637 | intron | AP2A2 |
| rs7398796 | 12 | 111814155 | 5 | 0 | 0 | T | C | -0.9398 | 0.247 | 0.0001416 | -+--- | 44.6 | 0.1246 | intergenic | |

| | | | | | | | | | | | | | | | |
|---|---|---|---|---|---|---|---|---|---|---|---|---|---|---|---|
| 2:79691111 | 2 | 79691111 | 3 | 0 | 0 | G | GT | -1.0349 | 0.272 | 0.0001417 | ---?? | 54.3 | 0.1119 | | |
| rs6700995 | 1 | 90266925 | 5 | 0 | 0 | A | T | 0.7948 | 0.2089 | 0.0001418 | ++++- | 0 | 0.5821 | intergenic | |
| rs145366123 | 1 | 78486490 | 3 | 0 | 0 | T | G | 4.9847 | 1.3101 | 0.0001419 | +++?? | 0 | 0.5291 | downstream-gene | |
| rs1986481 | 7 | 82496683 | 4 | 0 | 0 | A | T | -1.0267 | 0.2699 | 0.0001421 | -+-?- | 51.7 | 0.102 | intron | PCLO |
| 15:32607261 | 15 | 32607261 | 2 | 0 | 0 | A | G | -7.1224 | 1.8722 | 0.0001422 | ?--?? | 0 | 0.5303 | intergenic | |
| rs4915969 | 1 | 64789758 | 4 | 0 | 0 | T | C | 1.3695 | 0.36 | 0.0001423 | +++?+ | 0 | 0.558 | intergenic | |
| rs7788732 | 7 | 16905351 | 4 | 0 | 0 | A | G | 1.8394 | 0.4835 | 0.0001424 | +++?+ | 0 | 0.4889 | intron | AGR3 |
| rs183595026 | 12 | 91278219 | 3 | 0 | 0 | T | C | -5.1883 | 1.3639 | 0.0001424 | ---?? | 0 | 0.8755 | intergenic | |
| rs79627529 | 3 | 130176733 | 3 | 0 | 0 | T | C | -3.8374 | 1.0089 | 0.0001426 | -+-?? | 52.6 | 0.1212 | intron | COL6A5 |
| 17:47450536 | 17 | 47450536 | 3 | 0 | 0 | TTCTC | TTCTCTC | -1.1468 | 0.3015 | 0.0001426 | ---?? | 0 | 0.8944 | | |
| rs11208419 | 1 | 64790062 | 4 | 0 | 0 | C | G | -1.3692 | 0.36 | 0.0001428 | ---?- | 0 | 0.5588 | intergenic | |
| rs8099219 | 18 | 40039037 | 4 | 0 | 0 | A | G | 1.2086 | 0.3178 | 0.0001431 | +++?- | 0 | 0.4854 | intron | LINC00907 |
| rs10835693 | 11 | 30660955 | 4 | 0 | 0 | A | G | 1.0359 | 0.2724 | 0.0001431 | +++?- | 37.2 | 0.1891 | intergenic | |
| 6:144649741 | 6 | 144649741 | 3 | 0 | 0 | A | ATAT | 2.0974 | 0.5515 | 0.0001431 | +++?? | 0 | 0.9822 | | |
| 3:196488940 | 3 | 196488940 | 3 | 0 | 0 | T | C | 2.7411 | 0.7209 | 0.0001433 | +++?? | 0 | 0.9564 | intron | PAK2 |
| rs186772014 | 17 | 8736883 | 3 | 0 | 0 | A | G | 3.4204 | 0.8995 | 0.0001433 | +++?? | 0 | 0.8401 | intron | PIK3R6 |
| rs9787334 | 1 | 205023316 | 5 | 0 | 0 | T | C | -0.8606 | 0.2264 | 0.0001435 | ----- | 0 | 0.9405 | intron | CNTN2 |
| rs150177745 | 17 | 56714568 | 5 | 0 | 0 | T | C | -2.6378 | 0.6939 | 0.0001439 | ----+ | 0 | 0.7215 | intron | TEX14 |
| 9:90829380 | 9 | 90829380 | 3 | 0 | 0 | G | GACAGC | 2.0364 | 0.5357 | 0.0001439 | +++?? | 0 | 0.6376 | | |
| rs343055 | 7 | 35523883 | 4 | 0 | 0 | A | G | 1.2893 | 0.3392 | 0.0001443 | -++?+ | 44 | 0.1471 | intergenic | |
| 7:48900126 | 7 | 48900126 | 3 | 0 | 0 | CAG | C | 1.186 | 0.312 | 0.0001443 | +++?? | 54.1 | 0.1132 | | |
| rs217669 | 14 | 62360075 | 4 | 0 | 0 | T | C | 1.0801 | 0.2842 | 0.0001447 | +++?+ | 0 | 0.5894 | intron | SYT16 |
| rs7799164 | 7 | 141191807 | 4 | 0 | 0 | T | C | 4.9923 | 1.3138 | 0.0001447 | +++?+ | 0 | 0.6195 | intergenic | |
| rs9358746 | 6 | 24058264 | 5 | 0 | 0 | C | G | -1.5548 | 0.4091 | 0.0001447 | ----+ | 0 | 0.7003 | intergenic |  |
| rs75821275 | 17 | 56673081 | 5 | 0 | 0 | C | G | -2.5793 | 0.6788 | 0.0001447 | ----- | 0 | 0.8359 | intron | TEX14 |
| rs35122295 | 16 | 469188 | 5 | 0 | 0 | A | C | 0.8505 | 0.2239 | 0.0001452 | +++++ | 14.4 | 0.3228 | intergenic | |
| rs1970100 | 1 | 157437040 | 3 | 0 | 0 | A | C | 0.9787 | 0.2576 | 0.0001453 | +++?? | 16.9 | 0.3003 | intergenic | |
| rs17593863 | 4 | 119023828 | 4 | 0 | 0 | T | C | -1.7165 | 0.4518 | 0.0001453 | ---?- | 0 | 0.4409 | intron | NDST3 |
| rs1197768 | 15 | 42055810 | 5 | 0 | 0 | T | G | 1.0977 | 0.289 | 0.0001454 | +++++ | 13.9 | 0.3255 | intron | MGA |
| rs76365224 | 17 | 71073582 | 4 | 0 | 0 | A | T | 2.5866 | 0.681 | 0.0001456 | +++?+ | 27.6 | 0.2465 | intron | SLC39A11 |
| rs17149140 | 10 | 10977601 | 4 | 0 | 0 | A | C | -1.2324 | 0.3245 | 0.0001457 | ---?- | 51.2 | 0.1048 | intron | CELF2,LINC00710 |
| 6:77757952 | 6 | 77757952 | 3 | 0 | 0 | CAAAAT | C | -1.1762 | 0.3097 | 0.0001458 | ---?? | 37.9 | 0.1997 | | |
| rs17113486 | 14 | 43244622 | 4 | 0 | 0 | A | T | 1.4516 | 0.3822 | 0.0001459 | +++?- | 57.5 | 0.06985 | intergenic | |
| rs674079 | 5 | 134349342 | 5 | 0 | 0 | T | G | 0.8872 | 0.2336 | 0.0001461 | +++++ | 4 | 0.3841 | downstream-gene | |
| rs4331243 | 13 | 60015053 | 3 | 0 | 0 | T | C | -0.966 | 0.2544 | 0.0001461 | ---?? | 0 | 0.557 | intergenic | |
| 5:123511711 | 5 | 123511711 | 3 | 0 | 0 | A | AT | 0.9617 | 0.2533 | 0.0001463 | +++?? | 0 | 0.8454 | | |
| rs4283121 | 13 | 60015095 | 3 | 0 | 0 | A | T | 0.966 | 0.2544 | 0.0001464 | +++?? | 0 | 0.5562 | intergenic | |
| rs4533162 | 13 | 60015061 | 3 | 0 | 0 | A | G | -0.966 | 0.2544 | 0.0001464 | ---?? | 0 | 0.557 | intergenic | |
| rs565244 | 5 | 134349385 | 5 | 0 | 0 | T | G | 0.887 | 0.2336 | 0.0001466 | +++++ | 4 | 0.3841 | downstream-gene | |
| rs2072629 | 6 | 32013500 | 3 | 0 | 0 | T | C | -1.6081 | 0.4236 | 0.0001471 | ---?? | 0 | 0.5834 | intron | TNXB |
| rs6981814 | 8 | 113410506 | 4 | 0 | 0 | A | G | 1.079 | 0.2843 | 0.0001472 | +++?+ | 0 | 0.8039 | intron | CSMD3 |
| rs7907715 | 10 | 20149813 | 4 | 0 | 0 | T | C | -0.9467 | 0.2494 | 0.0001474 | ---?- | 39.4 | 0.1755 | intron | PLXDC2 |
| 17:76044234 | 17 | 76044234 | 3 | 0 | 0 | A | AT | -2.4169 | 0.6368 | 0.0001474 | ---?? | 25.2 | 0.2625 | | |
| 1:87449722 | 1 | 87449722 | 3 | 0 | 0 | CA | C | -1.1813 | 0.3113 | 0.0001474 | ---?? | 0 | 0.4127 | | |
| rs11512202 | 11 | 30664365 | 4 | 0 | 0 | A | G | 1.0374 | 0.2733 | 0.0001475 | +++?- | 37.6 | 0.1864 | intergenic | |
| rs60873537 | 16 | 84842680 | 4 | 0 | 0 | A | G | 1.2411 | 0.327 | 0.0001476 | +++?- | 0 | 0.5834 | intergenic | |
| rs62542100 | 9 | 32938469 | 3 | 0 | 0 | A | C | 1.157 | 0.3049 | 0.0001476 | +++?? | 34.7 | 0.216 | intergenic | |
| rs146294525 | 17 | 56752010 | 5 | 0 | 0 | A | G | -2.6594 | 0.7008 | 0.0001476 | ----- | 0 | 0.786 | intron | TEX14 |
| rs725391 | 1 | 187901195 | 4 | 0 | 0 | A | G | 1.0797 | 0.2845 | 0.0001479 | +++?+ | 0 | 0.4633 | intergenic | |
| rs2311726 | 11 | 30664503 | 4 | 0 | 0 | T | C | 1.0375 | 0.2734 | 0.000148 | +++?- | 37.6 | 0.1863 | intergenic | |
| rs11148469 | 13 | 60004501 | 3 | 0 | 0 | T | G | 0.9751 | 0.257 | 0.0001484 | +++?? | 0 | 0.4791 | intergenic | |
| rs9835479 | 3 | 71904983 | 4 | 0 | 0 | T | G | 1.0344 | 0.2727 | 0.0001487 | +++?+ | 0 | 0.9121 | intergenic | |
| rs16940541 | 15 | 85830693 | 3 | 0 | 0 | A | C | 5.7174 | 1.5072 | 0.0001487 | ++??+ | 0 | 0.7961 | intergenic | |
| 12:27786750 | 12 | 27786750 | 3 | 0 | 0 | CACCAA | C | -0.9973 | 0.2629 | 0.0001488 | ---?? | 58.6 | 0.08919 | | |
| 18:50285042 | 18 | 50285042 | 3 | 0 | 0 | G | GA | -6.3757 | 1.6809 | 0.0001488 | ---?? | 0 | 0.7391 | | |
| 7:102475752 | 7 | 102475752 | 3 | 0 | 0 | G | GGA | 2.3495 | 0.6195 | 0.0001489 | +++?? | 0 | 0.6745 | | |
| rs4786286 | 16 | 2552534 | 3 | 0 | 0 | T | C | 3.3032 | 0.871 | 0.0001492 | +++?? | 31.2 | 0.234 | 3-prime-UTR | TBC1D24 |
| rs1565059 | 5 | 122555432 | 4 | 0 | 0 | A | G | -2.0563 | 0.5422 | 0.0001493 | ---?- | 0 | 0.8731 | intergenic | |
| rs1407208 | 6 | 80555339 | 4 | 0 | 0 | T | C | 1.031 | 0.2719 | 0.0001498 | +++?+ | 10.6 | 0.3398 | intron | C6orf7 |

| | | | | | | | | | | | | | | | |
|---|---|---|---|---|---|---|---|---|---|---|---|---|---|---|---|
| 6:24058190 | 6 | 24058190 | 3 | 0 | 0 | A | AT | 1.813 | 0.4782 | 0.0001499 | +++?? | 0 | 0.6964 | | |
| rs1940208 | 11 | 95694713 | 4 | 0 | 0 | A | G | 1.1333 | 0.2989 | 0.00015 | +++?+ | 0.1 | 0.391 | intergenic | |
| rs13013766 | 2 | 218829527 | 3 | 0 | 0 | C | G | 1.406 | 0.3709 | 0.0001501 | +++?? | 0 | 0.3754 | intergenic | |
| rs580990 | 3 | 191158662 | 4 | 0 | 0 | T | C | 1.4618 | 0.3856 | 0.0001502 | +++?- | 36.7 | 0.192 | intergenic | |
| rs42240 | 16 | 6538214 | 4 | 0 | 0 | T | C | 1.0053 | 0.2652 | 0.0001503 | +++?+ | 0 | 0.547 | intron | RBFOX1 |
| rs12985050 | 19 | 46156372 | 3 | 0 | 1 | C | G | 1.0233 | 0.27 | 0.0001504 | +++?? | 0 | 0.4453 | intergenic | |
| rs11600642 | 11 | 30666231 | 4 | 0 | 0 | T | C | 1.0379 | 0.2738 | 0.0001505 | +++?- | 37.8 | 0.1853 | intergenic | |
| rs2191851 | 7 | 10131856 | 4 | 0 | 0 | A | G | 1.0611 | 0.28 | 0.0001506 | +++?+ | 52.5 | 0.097 | intergenic | |
| rs35936561 | 7 | 156187274 | 4 | 0 | 0 | A | G | 0.9859 | 0.2601 | 0.0001506 | +++?+ | 18.2 | 0.2995 | intergenic | |
| rs141969119 | 5 | 9457841 | 3 | 0 | 0 | A | T | 5.2171 | 1.3765 | 0.0001506 | +++?? | 0 | 0.9546 | intron | SEMA5A |
| rs17470838 | 5 | 122555744 | 4 | 0 | 0 | A | G | -2.0566 | 0.5427 | 0.000151 | ---?- | 0 | 0.8573 | intergenic | |
| 2:199080892 | 2 | 199080892 | 3 | 0 | 0 | CA | C | 1.3504 | 0.3564 | 0.000151 | +++?? | 48.5 | 0.1433 | | |
| rs8022660 | 14 | 22535262 | 4 | 0 | 0 | A | G | 0.9918 | 0.2617 | 0.0001511 | +++?+ | 0 | 0.6109 | intergenic | |
| rs147464235 | 6 | 83812622 | 4 | 0 | 0 | A | G | 3.4144 | 0.9011 | 0.0001511 | +++?+ | 0 | 0.6531 | intron | DOPEY1 |
| rs11649878 | 17 | 47448345 | 5 | 0 | 0 | C | G | 0.9698 | 0.2559 | 0.0001512 | ++++- | 41.7 | 0.1436 | intergenic | |
| rs34287435 | 6 | 77753216 | 3 | 0 | 0 | T | C | -1.1659 | 0.3077 | 0.0001512 | ---?? | 46.6 | 0.1535 | intergenic | |
| rs150141957 | 17 | 56759658 | 5 | 0 | 0 | A | G | -2.6585 | 0.7016 | 0.0001513 | ----- | 0 | 0.7896 | intron | TEX14 |
| rs78089618 | 6 | 137548611 | 3 | 0 | 0 | T | C | 3.3513 | 0.8846 | 0.0001515 | +++?? | 0 | 0.8738 | intergenic | |
| rs11591088 | 1 | 205135883 | 4 | 0 | 0 | A | T | 2.563 | 0.6765 | 0.0001516 | +++?- | 0 | 0.5026 | intron | DSTYK |
| rs58517893 | 2 | 137029429 | 4 | 0 | 0 | T | G | -1.0497 | 0.2771 | 0.0001517 | ---?- | 0 | 0.6815 | intergenic | |
| rs6760734 | 2 | 170603399 | 4 | 0 | 0 | T | C | -1.2507 | 0.3302 | 0.0001518 | ---?- | 0 | 0.6594 | intron | KLHL23,PHOSPHO2-KLHL23 |
| 4:100548139 | 4 | 100548139 | 3 | 0 | 0 | A | G | -1.9388 | 0.5119 | 0.0001519 | ---?? | 0 | 0.4148 | downstream-gene | |
| rs368167436 | 4 | 190758396 | 3 | 0 | 0 | T | C | -1.3267 | 0.3502 | 0.0001519 | ---?? | 0 | 0.8051 | intergenic | |
| rs7759204 | 6 | 144762887 | 4 | 0 | 0 | A | G | 3.0976 | 0.8178 | 0.0001521 | +++?+ | 0 | 0.8376 | intron | UTRN |
| rs350821 | 19 | 4083969 | 4 | 0 | 0 | A | G | 2.9486 | 0.7785 | 0.0001522 | +++?+ | 0 | 0.8428 | intergenic | |
| rs16939836 | 15 | 87622943 | 4 | 0 | 0 | T | C | -2.2377 | 0.5908 | 0.0001522 | ---?- | 0 | 0.7812 | intergenic | |
| rs6005751 | 22 | 28675783 | 4 | 0 | 0 | A | G | 2.7306 | 0.721 | 0.0001524 | +++?+ | 0 | 0.7728 | intron | TTC28 |
| rs10108308 | 8 | 96929239 | 4 | 0 | 0 | T | G | 1.1959 | 0.3158 | 0.0001524 | +++?+ | 0 | 0.9288 | intergenic | |
| rs12201775 | 6 | 80556359 | 4 | 0 | 0 | A | G | -1.0382 | 0.2741 | 0.0001524 | ---?- | 17.5 | 0.3033 | intron | C6orf7 |
| rs151078485 | 6 | 83815108 | 3 | 0 | 0 | T | C | 3.5012 | 0.9245 | 0.0001524 | +++?? | 0 | 0.4675 | intron | DOPEY1 |
| rs57118960 | 1 | 3571214 | 3 | 0 | 0 | A | C | -5.9359 | 1.5675 | 0.0001525 | ---?? | 25 | 0.2638 | intron | TP73 |
| rs114508509 | 6 | 32596324 | 5 | 0 | 0 | T | G | -1.301 | 0.3436 | 0.0001525 | ----- | 48.3 | 0.1018 | intergenic |  |
| rs117506611 | 8 | 8206954 | 3 | 0 | 0 | A | G | -3.4066 | 0.8996 | 0.0001527 | ---?? | 0 | 0.4797 | intron | SGK223 |
| rs8011284 | 14 | 43242241 | 4 | 0 | 0 | T | C | -1.4442 | 0.3814 | 0.0001529 | ---?+ | 58.9 | 0.06297 | intergenic | |
| rs10835696 | 11 | 30667403 | 4 | 0 | 0 | A | G | -1.0389 | 0.2744 | 0.0001529 | ---?+ | 38 | 0.1838 | intergenic | |
| 1:222982198 | 1 | 222982198 | 3 | 0 | 0 | CT | CTT | -1.23 | 0.3248 | 0.0001529 | ---?? | 19.6 | 0.2884 | | |
| rs74906361 | 2 | 10762608 | 3 | 0 | 0 | A | G | 5.6469 | 1.4915 | 0.0001531 | +++?? | 0 | 0.6035 | intron | NOL10 |
| rs2431150 | 5 | 134348146 | 5 | 0 | 0 | C | G | -0.8856 | 0.2339 | 0.0001532 | ----- | 7.2 | 0.3654 | downstream-gene |  |
| rs11894266 | 2 | 170636642 | 4 | 0 | 0 | T | C | 0.9232 | 0.2439 | 0.0001532 | +++?+ | 0 | 0.6297 | intergenic | |
| rs12706944 | 7 | 130150705 | 4 | 0 | 0 | A | G | -0.9373 | 0.2476 | 0.0001532 | -+-?- | 58.4 | 0.06552 | intron | COPG2 |
| rs9448835 | 6 | 80553848 | 4 | 0 | 0 | A | T | -1.0173 | 0.2687 | 0.0001533 | ---?- | 0 | 0.4074 | intron | C6orf7 |
| rs72839982 | 17 | 56381729 | 5 | 0 | 0 | A | G | 2.0618 | 0.5446 | 0.0001534 | +++++ | 46.4 | 0.1131 | 3-prime-UTR | BZRAP1 |
| rs13267101 | 8 | 18579176 | 5 | 0 | 0 | T | C | 1.2798 | 0.3381 | 0.0001535 | +++++ | 0 | 0.603 | intron | PSD3 |
| rs145308121 | 6 | 83894347 | 4 | 0 | 0 | C | G | -3.5312 | 0.9329 | 0.0001535 | ---?- | 11.7 | 0.3344 | intron | PGM3 |
| rs76848572 | 17 | 52307315 | 4 | 0 | 0 | T | G | 2.1784 | 0.5755 | 0.0001537 | +++?+ | 0 | 0.8405 | intergenic | |
| rs9275493 | 6 | 32673475 | 2 | 0 | 0 | T | G | -2.4818 | 0.6557 | 0.0001538 | ???-- | 0 | 0.758 | intergenic | |
| rs35678932 | 6 | 32666863 | 2 | 0 | 0 | A | C | 2.4818 | 0.6557 | 0.0001538 | ???++ | 0 | 0.758 | intergenic | |
| rs67887332 | 6 | 32664399 | 2 | 0 | 0 | A | G | 2.4818 | 0.6557 | 0.0001538 | ???++ | 0 | 0.758 | intergenic | |
| rs9358747 | 6 | 24069549 | 5 | 0 | 0 | A | T | -1.5402 | 0.407 | 0.0001539 | ----+ | 0 | 0.5716 | intergenic |  |
| rs6846270 | 4 | 38747146 | 5 | 0 | 0 | T | G | 1.2528 | 0.331 | 0.000154 | +++++ | 0 | 0.9654 | intergenic | |
| rs11972418 | 7 | 107365851 | 5 | 0 | 0 | T | C | -0.8208 | 0.2169 | 0.000154 | ----- | 35.7 | 0.1832 | intergenic |  |
| 2:131127129 | 2 | 131127129 | 3 | 0 | 0 | T | TTG | -1.1661 | 0.3081 | 0.000154 | ---?? | 29.7 | 0.241 | | |
| rs813498 | 9 | 94188386 | 4 | 0 | 0 | A | G | -2.682 | 0.7088 | 0.0001545 | ---?- | 0 | 0.9396 | upstream-gene | |
| rs79514210 | 5 | 122568526 | 4 | 0 | 0 | T | C | -2.0746 | 0.5483 | 0.0001545 | ---?- | 0 | 0.8308 | intergenic | |
| rs75082706 | 11 | 27099113 | 4 | 0 | 0 | A | G | 5.6257 | 1.4869 | 0.0001546 | +++?+ | 0 | 0.6428 | intron | BBOX1 |
| rs2017067 | 6 | 77767170 | 4 | 0 | 0 | A | C | -1.1403 | 0.3014 | 0.0001547 | ---?- | 3.5 | 0.3749 | intergenic | |
| rs6817333 | 4 | 38746079 | 5 | 0 | 0 | A | T | -1.2458 | 0.3293 | 0.0001547 | ----- | 0 | 0.9674 | intergenic |  |
| rs35168843 | 6 | 32442408 | 5 | 0 | 1 | A | G | 1.5123 | 0.3997 | 0.0001548 | +++++ | 0 | 0.5391 | intergenic | |
| 20:5992676 | 20 | 5992676 | 3 | 0 | 0 | G | GT | 0.9695 | 0.2563 | 0.0001548 | +++?? | 0 | 0.7755 | | |

| | | | | | | | | | | | | | | | |
|---|---|---|---|---|---|---|---|---|---|---|---|---|---|---|---|
| rs299366 | 5 | 134335383 | 3 | 0 | 0 | T | G | 1.0586 | 0.2799 | 0.0001553 | +++?? | 0 | 0.3706 | intron | CATSPER3 |
| rs10835697 | 11 | 30668234 | 4 | 0 | 0 | T | C | 1.0386 | 0.2746 | 0.0001553 | +++?- | 38.2 | 0.183 | intergenic | |
| rs28986409 | 6 | 32755053 | 5 | 0 | 0 | A | C | -1.5406 | 0.4073 | 0.0001553 | ----- | 23.4 | 0.2656 | intergenic |  |
| rs11017085 | 10 | 131916809 | 4 | 0 | 0 | T | C | 1.0276 | 0.2717 | 0.0001555 | +++?+ | 0 | 0.8416 | intergenic | |
| rs141550245 | 17 | 56761748 | 5 | 0 | 0 | A | G | -2.6542 | 0.7018 | 0.0001555 | ----- | 0 | 0.7895 | intron | TEX14 |
| rs117900200 | 12 | 67559774 | 3 | 0 | 0 | C | G | -3.8701 | 1.0234 | 0.0001558 | ---?? | 0 | 0.8019 | intergenic | |
| rs13199084 | 6 | 80554986 | 4 | 0 | 0 | T | C | -1.0266 | 0.2715 | 0.0001559 | ---?- | 8.4 | 0.3514 | intron | C6orf7 |
| rs8192585 | 6 | 32188823 | 5 | 0 | 0 | A | G | -1.8586 | 0.4915 | 0.000156 | ----- | 0 | 0.4783 | missense | NOTCH4 |
| rs62161820 | 2 | 131208401 | 3 | 0 | 0 | T | G | -3.2812 | 0.8679 | 0.0001564 | ---?? | 0 | 0.5244 | intergenic | |
| rs12934679 | 16 | 6541408 | 4 | 0 | 0 | A | G | -1.009 | 0.2669 | 0.0001567 | ---?- | 0 | 0.6158 | intron | RBFOX1 |
| rs7916958 | 10 | 28227533 | 4 | 0 | 0 | T | C | -0.8927 | 0.2362 | 0.0001568 | ---?+ | 46.1 | 0.135 | intron | ARMC4 |
| rs55730260 | 14 | 22529536 | 4 | 0 | 0 | A | G | 1.1955 | 0.3163 | 0.000157 | +++?+ | 0 | 0.989 | intergenic | |
| rs17481232 | 5 | 123151676 | 4 | 0 | 0 | A | C | 1.2684 | 0.3356 | 0.0001572 | +++?+ | 0 | 0.5133 | intergenic | |
| rs6561198 | 13 | 45636682 | 4 | 0 | 0 | T | C | 2.1558 | 0.5704 | 0.0001574 | +++?- | 0 | 0.4171 | intergenic | |
| rs1500642 | 15 | 98044868 | 4 | 0 | 0 | A | T | -0.9983 | 0.2642 | 0.0001577 | ---?- | 0 | 0.9298 | intergenic | |
| rs6588087 | 1 | 64792898 | 4 | 0 | 0 | T | C | -1.3539 | 0.3584 | 0.0001581 | ---?- | 0 | 0.5415 | intergenic | |
| rs75513876 | 17 | 52327635 | 4 | 0 | 0 | T | C | 2.4428 | 0.6466 | 0.0001581 | +++?- | 0 | 0.7786 | intergenic | |
| rs11815780 | 10 | 6409333 | 4 | 0 | 0 | A | G | 2.4709 | 0.6541 | 0.0001582 | +++?- | 0 | 0.7605 | intergenic | |
| rs117373773 | 11 | 103698222 | 4 | 0 | 0 | T | C | 2.3214 | 0.6145 | 0.0001583 | +++?+ | 0 | 0.9491 | intergenic | |
| rs7200664 | 16 | 6544129 | 4 | 0 | 0 | T | C | 1.0223 | 0.2706 | 0.0001584 | +++?+ | 0 | 0.7235 | intron | RBFOX1 |
| rs72889456 | 2 | 171149024 | 4 | 0 | 0 | A | G | -4.877 | 1.2911 | 0.0001585 | ---?+ | 51.4 | 0.1037 | intron | MYO3B |
| rs150534425 | 17 | 56767432 | 5 | 0 | 0 | A | G | -2.6518 | 0.7022 | 0.000159 | ----- | 0 | 0.79 | intron | TEX14 |
| rs66837178 | 4 | 38748498 | 5 | 0 | 0 | A | G | 1.2659 | 0.3352 | 0.0001592 | +++++ | 0 | 0.9521 | intergenic | |
| rs78838894 | 19 | 57419030 | 3 | 0 | 0 | A | T | 2.8309 | 0.7496 | 0.0001592 | +++?? | 0 | 0.8603 | intergenic | |
| rs11604063 | 11 | 30647780 | 4 | 0 | 0 | T | C | 1.0231 | 0.2709 | 0.0001593 | +++?- | 33.1 | 0.2139 | intergenic | |
| rs7201579 | 16 | 6544205 | 4 | 0 | 0 | A | C | -1.022 | 0.2707 | 0.0001594 | ---?- | 0 | 0.7239 | intron | RBFOX1 |
| rs7201049 | 16 | 6544303 | 4 | 0 | 0 | T | C | 1.0221 | 0.2707 | 0.0001595 | +++?+ | 0 | 0.7242 | intron | RBFOX1 |
| 14:44913314 | 14 | 44913314 | 3 | 0 | 0 | CG | C | -1.5384 | 0.4075 | 0.0001596 | ---?? | 0 | 0.912 | | |
| rs151255005 | 20 | 62447132 | 3 | 0 | 0 | A | C | 4.3697 | 1.1574 | 0.0001597 | +++?? | 0 | 0.7593 | intron | ZBTB46 |
| rs12419170 | 11 | 12647369 | 4 | 0 | 0 | A | G | -0.9952 | 0.2636 | 0.0001598 | ---?- | 0 | 0.7938 | intergenic | |
| rs2083842 | 8 | 92881946 | 3 | 0 | 0 | A | T | 4.0131 | 1.0631 | 0.0001601 | +++?? | 0 | 0.9799 | intergenic | |
| rs56252815 | 14 | 104581950 | 4 | 0 | 0 | T | C | 2.9033 | 0.7691 | 0.0001601 | +++?- | 43 | 0.1536 | upstream-gene | |
| rs79781489 | 6 | 32507622 | 4 | 0 | 1 | T | C | 1.4578 | 0.3862 | 0.0001601 | ++++? | 30 | 0.2321 | intergenic | |
| rs79331451 | 6 | 168579450 | 4 | 0 | 0 | A | G | -1.8463 | 0.4891 | 0.0001602 | ---?- | 0 | 0.8382 | intergenic | |
| rs10955616 | 8 | 113213815 | 4 | 0 | 0 | A | G | 0.9538 | 0.2527 | 0.0001603 | +++?- | 0 | 0.7609 | intergenic | |
| rs17205135 | 6 | 32602222 | 4 | 0 | 1 | A | G | -1.0591 | 0.2806 | 0.0001603 | ----? | 0 | 0.5798 | upstream-gene |  |
| rs7333890 | 13 | 60015652 | 4 | 0 | 0 | T | C | 0.9355 | 0.2479 | 0.0001604 | +++?+ | 0 | 0.7675 | intergenic | |
| rs4961670 | 9 | 16206760 | 4 | 0 | 0 | T | G | 1.0565 | 0.2799 | 0.0001606 | +++?+ | 0 | 0.9063 | intron | C9orf92 |
| rs72800657 | 16 | 47512691 | 3 | 0 | 0 | T | C | -3.2682 | 0.8659 | 0.0001606 | ---?? | 0 | 0.8618 | intron | PHKB |
| rs116390233 | 2 | 65985296 | 3 | 0 | 0 | C | G | 4.9104 | 1.3011 | 0.0001606 | ?++?+ | 0 | 0.8768 | intergenic | |
| rs34751870 | 6 | 32673332 | 5 | 0 | 1 | T | C | 1.5447 | 0.4093 | 0.0001607 | +++++ | 0 | 0.5096 | intergenic | |
| rs10215444 | 7 | 16905498 | 4 | 0 | 0 | A | G | 1.793 | 0.4751 | 0.0001609 | +++?+ | 0 | 0.413 | intron | AGR3 |
| rs149089616 | 11 | 39237753 | 2 | 0 | 0 | A | G | -8.4146 | 2.2299 | 0.000161 | ?-??- | 0 | 0.8496 | intergenic | |
| rs1473478 | 8 | 18584440 | 5 | 0 | 0 | A | G | -1.2082 | 0.3202 | 0.0001611 | ----- | 0 | 0.7982 | intron | PSD3 |
| rs138684328 | 16 | 84272040 | 4 | 0 | 0 | T | G | 4.2269 | 1.1202 | 0.0001611 | +++?+ | 0 | 0.617 | intron | KCNG4 |
| 13:71296258 | 13 | 71296258 | 3 | 0 | 0 | CCTTT | C | 5.1268 | 1.3587 | 0.0001611 | +++?? | 0 | 0.7688 | | |
| rs10505175 | 8 | 113435406 | 4 | 0 | 0 | A | C | 1.0606 | 0.2811 | 0.0001612 | +++?+ | 0 | 0.7452 | intron | CSMD3 |
| rs7928247 | 11 | 12640489 | 4 | 0 | 0 | T | C | 1.0099 | 0.2677 | 0.0001613 | +++?+ | 0 | 0.8835 | intergenic | |
| 6:109094439 | 6 | 109094439 | 2 | 0 | 0 | T | TTAATC | 5.3245 | 1.4112 | 0.0001613 | +?+?? | 0 | 0.7547 | | |
| rs117267733 | 8 | 86262827 | 4 | 0 | 0 | A | G | -2.6745 | 0.7089 | 0.0001614 | ---?- | 2.5 | 0.38 | intron | CA1 |
| rs35433555 | 6 | 135768282 | 4 | 0 | 0 | T | C | -3.0041 | 0.7963 | 0.0001615 | ----? | 15 | 0.3168 | missense | AHI1 |
| rs9496975 | 6 | 144767524 | 4 | 0 | 0 | C | G | 3.1807 | 0.8431 | 0.0001616 | +++?+ | 0 | 0.8674 | intron | UTRN |
| rs2302826 | 7 | 130147307 | 4 | 0 | 0 | T | G | 0.9332 | 0.2474 | 0.0001619 | +-+?+ | 58.4 | 0.06523 | intron | COPG2 |
| rs114951998 | 3 | 16436460 | 4 | 0 | 0 | C | G | -2.0271 | 0.5375 | 0.0001622 | ---?- | 0 | 0.5498 | intron | RFTN1 |
| rs17741142 | 8 | 86216519 | 4 | 0 | 0 | A | G | 1.8039 | 0.4783 | 0.0001623 | +++?+ | 39.3 | 0.1758 | intergenic | |
| rs138545399 | 6 | 83768501 | 4 | 0 | 0 | T | C | 3.5761 | 0.9482 | 0.0001623 | +++?+ | 0 | 0.6837 | intron | UBE3D |
| 2:235312947 | 2 | 235312947 | 3 | 0 | 0 | T | TTTTC | 1.1665 | 0.3093 | 0.0001623 | +++?? | 0 | 0.4678 | | |
| rs7330546 | 13 | 60009634 | 4 | 0 | 0 | A | T | 0.9534 | 0.2528 | 0.0001625 | +++?+ | 0 | 0.8314 | intergenic | |
| 6:24069934 | 6 | 24069934 | 3 | 0 | 0 | T | TACTTTC | 1.7916 | 0.4751 | 0.0001625 | +++?? | 0 | 0.5872 | | |

| | | | | | | | | | | | | | | | |
|---|---|---|---|---|---|---|---|---|---|---|---|---|---|---|---|
| 12:4720646 | 12 | 4720646 | 3 | 0 | 0 | T | TTA | -3.3888 | 0.8987 | 0.0001626 | ---?? | 0 | 0.6352 | | |
| 4:128228764 | 4 | 128228764 | 3 | 0 | 0 | A | AT | 4.8009 | 1.2731 | 0.0001626 | +++?? | 0 | 0.6192 | | |
| rs538853 | 5 | 134348825 | 5 | 0 | 0 | A | G | -0.8826 | 0.2341 | 0.0001627 | ----- | 6.6 | 0.3694 | downstream-gene |  |
| rs7103179 | 11 | 103731910 | 4 | 0 | 0 | T | C | -1.7281 | 0.4583 | 0.0001628 | ---?- | 30.5 | 0.2293 | intergenic | |
| rs12472818 | 2 | 218358312 | 4 | 0 | 0 | T | C | 0.8896 | 0.2359 | 0.0001629 | +++?+ | 0 | 0.5276 | intron | DIRC3 |
| rs142620103 | 3 | 65590008 | 3 | 0 | 0 | T | C | -3.4496 | 0.9149 | 0.0001629 | ---?? | 47.7 | 0.1478 | intron | MAGI1 |
| rs2741734 | 14 | 21458989 | 3 | 0 | 0 | T | G | -1.2678 | 0.3362 | 0.000163 | ---?? | 35.5 | 0.2124 | intron | METTL17 |
| rs79874333 | 17 | 56714916 | 5 | 0 | 0 | T | C | -2.6157 | 0.6938 | 0.0001631 | ----+ | 0 | 0.709 | intron | TEX14 |
| 13:60009168 | 13 | 60009168 | 3 | 0 | 0 | A | AC | -0.9601 | 0.2547 | 0.0001632 | ---?? | 0 | 0.5846 | | |
| rs4074120 | 2 | 137026587 | 4 | 0 | 0 | T | C | -1.039 | 0.2756 | 0.0001633 | ---?- | 0 | 0.6857 | intergenic | |
| rs139412530 | 17 | 56773194 | 5 | 0 | 0 | A | G | -2.6283 | 0.6972 | 0.0001633 | ----- | 0 | 0.8268 | intron | RAD51C |
| rs79611853 | 2 | 168214638 | 4 | 0 | 0 | C | G | -2.5941 | 0.6882 | 0.0001637 | ---?- | 0 | 0.7252 | intergenic | |
| rs113577306 | 5 | 29479712 | 4 | 0 | 0 | A | G | 3.9352 | 1.0442 | 0.0001642 | +++?+ | 0 | 0.5289 | intergenic | |
| rs1512359 | 11 | 37058631 | 4 | 0 | 0 | A | C | -0.9157 | 0.243 | 0.0001643 | ---?- | 0 | 0.445 | intergenic | |
| rs6697019 | 1 | 90265407 | 5 | 0 | 0 | A | G | 0.7882 | 0.2092 | 0.0001643 | ++++- | 0 | 0.5871 | intergenic | |
| rs9960550 | 18 | 45935965 | 3 | 0 | 0 | C | G | -3.2297 | 0.8571 | 0.0001645 | ---?? | 0 | 0.8326 | upstream-gene | |
| rs112766411 | 6 | 82727148 | 4 | 0 | 0 | T | C | -2.9586 | 0.7852 | 0.0001646 | ---?- | 0 | 0.5075 | intergenic | |
| rs12932046 | 16 | 6544558 | 4 | 0 | 0 | A | G | -1.0202 | 0.2708 | 0.0001647 | ---?- | 0 | 0.7258 | intron | RBFOX1 |
| rs193283368 | 4 | 119867111 | 3 | 0 | 0 | A | G | -4.2589 | 1.1304 | 0.0001647 | ---?? | 0 | 0.9791 | intron | SYNPO2 |
| rs6005834 | 22 | 29075126 | 4 | 0 | 0 | T | C | -2.536 | 0.6731 | 0.0001648 | ---?- | 0 | 0.5232 | intron | TTC28 |
| rs78193316 | 19 | 57420124 | 3 | 0 | 0 | T | C | 2.7983 | 0.7428 | 0.0001649 | +++?? | 0 | 0.8754 | intergenic | |
| rs2605569 | 15 | 42074016 | 5 | 0 | 0 | T | C | 1.0755 | 0.2855 | 0.0001651 | +++++ | 19.7 | 0.2891 | intron | MAPKBP1 |
| rs12653191 | 5 | 17182699 | 4 | 0 | 0 | T | C | 0.9548 | 0.2535 | 0.0001654 | +++?+ | 0 | 0.7442 | intron | LOC285696 |
| rs1913589 | 1 | 64807870 | 3 | 0 | 0 | T | C | -1.4346 | 0.3809 | 0.0001655 | ---?? | 0 | 0.5233 | intergenic | |
| rs66878487 | 11 | 95460622 | 4 | 0 | 0 | A | G | 2.0895 | 0.5548 | 0.0001656 | +++?+ | 0 | 0.8121 | intergenic | |
| rs74368772 | 19 | 55238036 | 3 | 0 | 0 | A | G | 1.0426 | 0.2768 | 0.0001658 | +++?? | 18.8 | 0.292 | intron | KIR3DL3 |
| rs8055744 | 16 | 7209937 | 4 | 0 | 0 | A | G | 1.1337 | 0.301 | 0.0001659 | +++?+ | 0 | 0.8593 | intron | RBFOX1 |
| rs10141062 | 14 | 44904949 | 4 | 0 | 0 | A | T | -1.4785 | 0.3926 | 0.0001659 | ---?- | 0 | 0.7781 | intergenic | |
| rs17282763 | 7 | 82520166 | 4 | 0 | 0 | T | C | -1.0291 | 0.2733 | 0.000166 | ---?- | 15.5 | 0.3141 | intron | PCLO |
| rs1940440 | 18 | 374047 | 3 | 0 | 0 | A | T | 1.129 | 0.2998 | 0.0001661 | +++?? | 0 | 0.947 | intron | COLEC12 |
| rs72554039 | 4 | 89063354 | 4 | 0 | 0 | A | G | -4.3565 | 1.1569 | 0.0001661 | -?--- | 0 | 0.7881 | intron | ABCG2 |
| rs111396166 | 17 | 2643831 | 3 | 0 | 0 | T | C | -3.7156 | 0.9867 | 0.0001661 | ---?? | 0 | 0.8157 | intron | LOC101928991 |
| rs4815882 | 20 | 6009512 | 5 | 0 | 0 | A | C | -0.7774 | 0.2065 | 0.0001662 | ----+ | 0 | 0.4653 | intron | CRLS1 |
| rs6663300 | 1 | 90266701 | 5 | 0 | 0 | A | G | -0.7873 | 0.2091 | 0.0001663 | ----+ | 0 | 0.5884 | intergenic |  |
| rs6684610 | 1 | 90265414 | 5 | 0 | 0 | T | C | -0.7875 | 0.2092 | 0.0001664 | ----+ | 0 | 0.5932 | intergenic |  |
| rs74698326 | 6 | 23882821 | 4 | 0 | 0 | A | C | -2.7449 | 0.729 | 0.0001664 | ---?+ | 26.6 | 0.2523 | intergenic | |
| 5:134365708 | 5 | 134365708 | 3 | 0 | 0 | CTATT | C | 1.0499 | 0.2788 | 0.0001664 | +++?? | 0 | 0.5657 | | |
| rs77377473 | 4 | 89061802 | 4 | 0 | 0 | A | G | 4.3591 | 1.1578 | 0.0001666 | +?+++ | 0 | 0.7841 | intron | ABCG2 |
| rs4338876 | 18 | 39958620 | 3 | 0 | 0 | A | G | -1.2903 | 0.3427 | 0.0001668 | ---?? | 0 | 0.9233 | intron | LINC00907 |
| rs660273 | 5 | 134348564 | 5 | 0 | 0 | T | G | -0.8814 | 0.2342 | 0.000167 | ----- | 6.8 | 0.3679 | downstream-gene |  |
| rs6468472 | 8 | 96923783 | 4 | 0 | 0 | A | G | -1.198 | 0.3182 | 0.000167 | ---?- | 0 | 0.9185 | intergenic | |
| rs9393522 | 6 | 24066174 | 5 | 0 | 0 | A | G | -1.521 | 0.4041 | 0.0001671 | ----+ | 0 | 0.6245 | intergenic |  |
| rs139463886 | 17 | 52323560 | 3 | 0 | 0 | A | G | 2.1764 | 0.5782 | 0.0001674 | +++?? | 0 | 0.9715 | intergenic | |
| rs1028410 | 20 | 6009174 | 5 | 0 | 0 | A | G | -0.7772 | 0.2065 | 0.0001675 | ----+ | 0 | 0.4659 | intron | CRLS1 |
| rs6780193 | 3 | 71905876 | 4 | 0 | 0 | A | G | -1.0245 | 0.2722 | 0.0001675 | ---?- | 0 | 0.897 | intergenic | |
| rs10022328 | 4 | 38747649 | 5 | 0 | 0 | A | G | 1.2532 | 0.333 | 0.0001676 | +++++ | 0 | 0.9576 | intergenic | |
| rs35413565 | 1 | 229165924 | 4 | 0 | 0 | T | G | -1.3876 | 0.3687 | 0.0001676 | ---?- | 0 | 0.9921 | intergenic | |
| rs9496974 | 6 | 144765205 | 4 | 0 | 0 | C | G | 3.1015 | 0.8241 | 0.0001677 | +++?+ | 0 | 0.8521 | intron | UTRN |
| rs7906761 | 10 | 20149836 | 4 | 0 | 0 | A | G | -0.938 | 0.2493 | 0.0001678 | ---?- | 41.5 | 0.1625 | intron | PLXDC2 |
| rs6688572 | 1 | 90266911 | 5 | 0 | 0 | T | C | -0.7871 | 0.2092 | 0.0001679 | ----+ | 0 | 0.588 | intergenic |  |
| rs66525318 | 6 | 32504136 | 4 | 0 | 1 | A | C | 1.2978 | 0.3449 | 0.000168 | ++++? | 34 | 0.208 | intergenic | |
| rs192264321 | 5 | 179958283 | 3 | 0 | 0 | A | G | 1.1994 | 0.3187 | 0.000168 | +++?? | 20.4 | 0.2845 | intron | CNOT6 |
| rs138329492 | 3 | 175872781 | 4 | 0 | 0 | T | C | 2.7825 | 0.7396 | 0.0001684 | +++?+ | 0 | 0.5819 | intergenic | |
| rs59751637 | 2 | 40469201 | 4 | 0 | 0 | T | C | -1.565 | 0.416 | 0.0001685 | ---?- | 0 | 0.8246 | intron | SLC8A1,SLC8A1-AS1 |
| rs10022317 | 4 | 38747605 | 5 | 0 | 0 | A | G | 1.2519 | 0.3328 | 0.0001686 | +++++ | 0 | 0.9582 | intergenic | |
| 14:76943933 | 14 | 76943933 | 3 | 0 | 0 | A | ATGGCC | 3.996 | 1.0622 | 0.0001686 | +++?? | 0 | 0.9954 | | |
| rs2244132 | 8 | 140488410 | 4 | 0 | 0 | A | T | 1.5458 | 0.4109 | 0.0001687 | +++?+ | 0 | 0.7431 | intergenic | |
| rs7687495 | 4 | 176476127 | 4 | 0 | 0 | A | G | 2.6544 | 0.7057 | 0.0001688 | +++?+ | 0 | 0.6249 | intergenic | |
| rs4331318 | 15 | 91582257 | 4 | 0 | 0 | T | C | 2.0838 | 0.554 | 0.0001691 | +++?+ | 0 | 0.7199 | intergenic | |

| | | | | | | | | | | | | | | | |
|---|---|---|---|---|---|---|---|---|---|---|---|---|---|---|---|
| rs6705349 | 2 | 170528157 | 4 | 0 | 0 | T | C | 1.3378 | 0.3557 | 0.0001691 | +++?+ | 0 | 0.966 | intron | CCDC173 |
| rs57952769 | 4 | 38736246 | 5 | 0 | 0 | A | T | 1.3237 | 0.3519 | 0.0001692 | +++++ | 0 | 0.9054 | intergenic | |
| rs10014476 | 4 | 38747544 | 5 | 0 | 0 | T | C | -1.2508 | 0.3326 | 0.0001693 | ----- | 0 | 0.9586 | intergenic |  |
| rs4786873 | 16 | 6539512 | 4 | 0 | 0 | T | G | 1.0234 | 0.2721 | 0.0001694 | +++?+ | 0 | 0.6831 | intron | RBFOX1 |
| rs112709300 | 9 | 90825391 | 4 | 0 | 0 | A | T | -1.9724 | 0.5245 | 0.0001694 | ---?- | 0 | 0.7747 | intergenic | |
| rs111794383 | 9 | 90825352 | 4 | 0 | 0 | A | G | -1.9724 | 0.5245 | 0.0001695 | ---?- | 0 | 0.7749 | intergenic | |
| rs2235816 | 20 | 6018711 | 5 | 0 | 0 | A | G | -0.7737 | 0.2057 | 0.0001696 | ----+ | 0 | 0.4628 | 3-prime-UTR | CRLS1 |
| rs6663717 | 1 | 90267116 | 5 | 0 | 0 | A | G | -0.7868 | 0.2092 | 0.0001699 | ----+ | 0 | 0.5878 | intergenic |  |
| rs116579026 | 17 | 56776528 | 5 | 0 | 0 | T | C | -2.6227 | 0.6975 | 0.00017 | ----- | 0 | 0.827 | intron | RAD51C |
| 18:40432793 | 18 | 40432793 | 3 | 0 | 0 | CCTTTCT | C | 4.8879 | 1.3001 | 0.0001702 | +++?? | 0 | 0.6227 | | |
| rs139565004 | 8 | 119562006 | 4 | 0 | 0 | T | C | -3.2929 | 0.8761 | 0.0001709 | ---?- | 0 | 0.3927 | intron | SAMD12 |
| rs368186991 | 17 | 52242376 | 3 | 0 | 0 | T | C | 2.4718 | 0.6576 | 0.0001709 | +++?? | 0 | 0.8377 | intergenic | |
| rs9677195 | 2 | 137013195 | 4 | 0 | 0 | A | G | 0.9971 | 0.2653 | 0.0001712 | +++?+ | 0 | 0.8421 | intergenic | |
| rs72761114 | 16 | 1618995 | 4 | 0 | 0 | T | C | 2.7928 | 0.7432 | 0.0001712 | +++?+ | 15.4 | 0.3149 | intron | IFT140 |
| rs112013068 | 6 | 83768885 | 4 | 0 | 0 | C | G | 3.5534 | 0.9456 | 0.0001714 | +++?+ | 0 | 0.7062 | intron | UBE3D |
| rs140592005 | 6 | 83766608 | 4 | 0 | 0 | A | G | -3.5668 | 0.9492 | 0.0001714 | ---?- | 0 | 0.6906 | intron | UBE3D |
| rs2131297 | 8 | 113403781 | 4 | 0 | 0 | T | C | 1.0632 | 0.283 | 0.0001717 | +++?+ | 0 | 0.7905 | intron | CSMD3 |
| rs57802612 | 16 | 84838571 | 3 | 0 | 0 | T | C | 1.2749 | 0.3393 | 0.0001718 | +++?? | 0 | 0.8096 | intergenic | |
| rs10922727 | 1 | 90267513 | 5 | 0 | 0 | T | C | 0.7864 | 0.2093 | 0.0001721 | ++++- | 0 | 0.5874 | intergenic | |
| rs11639821 | 16 | 12092575 | 4 | 0 | 0 | T | C | -0.934 | 0.2486 | 0.0001722 | +--?- | 59.1 | 0.06182 | intron | SNX29 |
| rs76883600 | 2 | 98992488 | 4 | 0 | 0 | A | T | 1.8725 | 0.4985 | 0.0001723 | +++?- | 0 | 0.6191 | intron | CNGA3 |
| rs2602381 | 2 | 234584324 | 5 | 0 | 0 | T | C | -0.7981 | 0.2125 | 0.0001724 | ----- | 0 | 0.4913 | intron | UGT1A10,UGT1A8,UGT1A9 |
| rs145394926 | 17 | 56782387 | 5 | 0 | 0 | A | G | -2.641 | 0.7032 | 0.0001727 | ----- | 0 | 0.7938 | intron | RAD51C |
| 10:11384339 | 10 | 11384339 | 3 | 0 | 0 | CA | C | 1.2062 | 0.3212 | 0.0001728 | +++?? | 0 | 0.4703 | | |
| rs10497174 | 2 | 156926773 | 4 | 0 | 0 | T | C | -1.036 | 0.2758 | 0.0001729 | ---?- | 0 | 0.8172 | intergenic | |
| rs233087 | 1 | 85821000 | 5 | 0 | 0 | T | C | 1.0173 | 0.2709 | 0.000173 | +++++ | 35.5 | 0.1847 | intron | DDAH1 |
| 17:47328770 | 17 | 47328770 | 3 | 0 | 0 | CT | C | 1.1261 | 0.2999 | 0.0001731 | +++?? | 0 | 0.6747 | | |
| rs10922728 | 1 | 90267692 | 5 | 0 | 0 | T | C | -0.7864 | 0.2094 | 0.0001735 | ----+ | 0 | 0.5871 | intergenic |  |
| 9:121328408 | 9 | 121328408 | 3 | 0 | 0 | T | TGTAA | 3.6813 | 0.9804 | 0.0001735 | +++?? | 11.6 | 0.3225 | | |
| rs7895714 | 10 | 20149718 | 4 | 0 | 0 | A | T | -0.9364 | 0.2494 | 0.0001736 | ---?- | 41.4 | 0.163 | intron | PLXDC2 |
| rs117343210 | 17 | 56633979 | 5 | 0 | 0 | T | G | -2.6095 | 0.6951 | 0.0001738 | ----- | 0 | 0.6975 | downstream-gene |  |
| rs10146815 | 14 | 44906192 | 4 | 0 | 0 | A | G | -1.4732 | 0.3925 | 0.0001741 | ---?- | 0 | 0.782 | intergenic | |
| 22:49401406 | 22 | 49401406 | 3 | 0 | 0 | A | G | -1.5437 | 0.4113 | 0.0001744 | ---?? | 0 | 0.5336 | intergenic | |
| rs6490061 | 12 | 111773345 | 5 | 0 | 0 | T | C | -0.9961 | 0.2654 | 0.0001745 | -+--- | 43.8 | 0.1299 | intron | CUX2 |
| rs7915632 | 10 | 129759110 | 4 | 0 | 0 | T | C | -1.0578 | 0.2818 | 0.0001745 | ---?+ | 0 | 0.434 | intron | PTPRE |
| rs147495098 | 22 | 27707003 | 3 | 0 | 0 | T | C | 4.2871 | 1.1423 | 0.0001746 | +++?? | 46.7 | 0.1533 | intergenic | |
| rs10922717 | 1 | 90248248 | 5 | 0 | 0 | C | G | -0.8002 | 0.2132 | 0.0001748 | ----- | 0 | 0.9806 | intergenic |  |
| rs189023842 | 7 | 98427003 | 4 | 0 | 0 | C | G | -3.9301 | 1.0472 | 0.0001748 | ---?- | 48.8 | 0.1185 | intergenic | |
| 2:218359774 | 2 | 218359774 | 3 | 0 | 0 | T | TA | 1.141 | 0.304 | 0.0001748 | +++?? | 28.4 | 0.2472 | | |
| rs11925615 | 3 | 196566147 | 3 | 0 | 0 | A | G | -3.4625 | 0.9227 | 0.000175 | ---?? | 0 | 0.6365 | intergenic | |
| rs1597171 | 1 | 30361717 | 4 | 0 | 0 | T | C | 1.335 | 0.3558 | 0.0001751 | +++?- | 2 | 0.3821 | intergenic | |
| rs72675809 | 4 | 130694725 | 4 | 0 | 0 | A | G | 2.2241 | 0.5928 | 0.0001753 | +++?- | 0 | 0.613 | intergenic | |
| rs1492664 | 8 | 113447003 | 4 | 0 | 0 | T | G | -1.0464 | 0.2789 | 0.0001754 | ---?- | 0 | 0.7638 | intron | CSMD3 |
| rs193134444 | 4 | 89449670 | 3 | 0 | 0 | T | G | 4.3348 | 1.1554 | 0.0001756 | +++?? | 15.7 | 0.3053 | upstream-gene | |
| 22:49401409 | 22 | 49401409 | 3 | 0 | 0 | A | C | 1.5425 | 0.4112 | 0.0001757 | +++?? | 0 | 0.5399 | intergenic | |
| rs32205 | 5 | 127616518 | 5 | 0 | 0 | A | C | 0.8984 | 0.2395 | 0.0001757 | -++++ | 23.9 | 0.2618 | intron | FBN2 |
| rs941979 | 1 | 90268189 | 5 | 0 | 0 | T | C | -0.786 | 0.2095 | 0.000176 | ----+ | 0 | 0.5868 | intergenic |  |
| rs2599365 | 15 | 85552341 | 3 | 0 | 0 | A | T | 0.9548 | 0.2545 | 0.000176 | +++?? | 54.4 | 0.1117 | intron | PDE8A |
| 20:5997120 | 20 | 5997120 | 3 | 0 | 0 | T | TGTGTG | 0.9361 | 0.2496 | 0.0001761 | +++?? | 0 | 0.8741 | | |
| rs200779030 | 15 | 85552339 | 3 | 0 | 0 | A | T | 0.9547 | 0.2545 | 0.0001762 | +++?? | 54.4 | 0.1117 | intron | PDE8A |
| rs951742 | 15 | 98045563 | 4 | 0 | 0 | T | C | -0.9916 | 0.2644 | 0.0001763 | ---?- | 0 | 0.9284 | intergenic | |
| rs9542406 | 13 | 71143002 | 4 | 0 | 0 | T | G | 0.943 | 0.2514 | 0.0001763 | +++?+ | 0 | 0.6894 | intergenic | |
| rs34343213 | 15 | 85552337 | 3 | 0 | 0 | A | T | 0.9546 | 0.2545 | 0.0001763 | +++?? | 54.4 | 0.1116 | intron | PDE8A |
| rs77249848 | 1 | 95232284 | 2 | 0 | 0 | T | C | 6.498 | 1.7325 | 0.0001763 | ++??? | 0 | 0.3588 | intergenic | |
| rs9932611 | 16 | 87624787 | 4 | 0 | 0 | A | G | 2.0588 | 0.5489 | 0.0001764 | +++?- | 0 | 0.436 | intergenic | |
| rs10174909 | 2 | 79635825 | 3 | 0 | 0 | T | C | -1.0362 | 0.2763 | 0.0001764 | ---?? | 40.7 | 0.1853 | intergenic | |
| 15:75459221 | 15 | 75459221 | 3 | 0 | 0 | A | C | -2.571 | 0.6855 | 0.0001765 | ---?? | 0 | 0.5744 | intergenic | |
| 22:49401403 | 22 | 49401403 | 3 | 0 | 0 | T | C | -1.5416 | 0.411 | 0.0001765 | ---?? | 0 | 0.5385 | intergenic | |
| 22:49401405 | 22 | 49401405 | 3 | 0 | 0 | T | C | 1.5418 | 0.4111 | 0.0001765 | +++?? | 0 | 0.5388 | intergenic | |

| | | | | | | | | | | | | | | | |
|---|---|---|---|---|---|---|---|---|---|---|---|---|---|---|---|
| rs2880876 | 12 | 26798565 | 4 | 0 | 0 | T | C | 1.932 | 0.5151 | 0.0001765 | +++?+ | 9.4 | 0.3459 | intron | ITPR2 |
| rs6955309 | 7 | 107344566 | 4 | 0 | 0 | T | C | -0.9473 | 0.2526 | 0.0001765 | ---?- | 0 | 0.4597 | intron | SLC26A4 |
| rs35034857 | 7 | 107368974 | 5 | 0 | 0 | A | C | -0.9312 | 0.2483 | 0.0001766 | ---+- | 53.5 | 0.07182 | intergenic | |
| rs143705409 | 11 | 924665 | 4 | 0 | 0 | A | G | -2.3432 | 0.6248 | 0.0001767 | ---?- | 0 | 0.7604 | upstream-gene | |
| 12:48093209 | 12 | 48093209 | 3 | 0 | 0 | A | AAAG | 1.9484 | 0.5196 | 0.0001768 | +++?? | 43.3 | 0.1715 | | |
| 15:75459218 | 15 | 75459218 | 3 | 0 | 0 | G | GT | 2.5707 | 0.6855 | 0.0001768 | +++?? | 0 | 0.5743 | | |
| rs426987 | 5 | 127613791 | 5 | 0 | 0 | C | G | -0.8985 | 0.2396 | 0.000177 | +---- | 24.2 | 0.26 | intron | FBN2 |
| rs117328801 | 12 | 46006834 | 3 | 0 | 0 | T | C | 3.8986 | 1.0397 | 0.0001771 | ?++?+ | 0 | 0.8993 | intergenic | |
| rs498408 | 3 | 191157960 | 4 | 0 | 0 | A | G | 1.4582 | 0.3889 | 0.0001772 | +++?- | 39.3 | 0.1759 | intergenic | |
| rs1078095 | 1 | 90268992 | 5 | 0 | 0 | T | C | -0.7864 | 0.2097 | 0.0001773 | ----+ | 0 | 0.5813 | intergenic |  |
| rs8055897 | 16 | 7209970 | 4 | 0 | 0 | A | G | 1.1284 | 0.301 | 0.0001776 | +++?+ | 0 | 0.858 | intron | RBFOX1 |
| rs76900409 | 13 | 104570802 | 3 | 0 | 0 | T | G | 5.2845 | 1.4096 | 0.0001777 | +++?? | 0 | 0.8644 | intergenic | |
| rs17210782 | 7 | 82398970 | 4 | 0 | 0 | T | C | -1.0198 | 0.272 | 0.0001778 | -+-?- | 43.3 | 0.1519 | intron | PCLO |
| rs11049072 | 12 | 27778515 | 3 | 0 | 0 | T | G | 1.0633 | 0.2837 | 0.0001779 | +++?? | 7.4 | 0.3394 | intron | PPFIBP1 |
| rs184977907 | 8 | 8408023 | 3 | 0 | 0 | A | T | 4.6572 | 1.2424 | 0.0001779 | +++?? | 0 | 0.9817 | intergenic | |
| rs141838784 | 10 | 132819155 | 2 | 0 | 0 | T | C | -4.9976 | 1.3334 | 0.0001783 | -?-?? | 0 | 0.5872 | intergenic | |
| rs56862634 | 6 | 144769996 | 4 | 0 | 0 | C | G | 3.1339 | 0.8362 | 0.0001784 | +++?+ | 0 | 0.8567 | intron | UTRN |
| rs982663 | 7 | 107344033 | 4 | 0 | 0 | A | G | 0.9465 | 0.2526 | 0.0001785 | +++?+ | 0 | 0.4608 | intron | SLC26A4 |
| rs686574 | 1 | 187900587 | 4 | 0 | 0 | C | G | -1.0631 | 0.2837 | 0.0001786 | ---?- | 0 | 0.4663 | intergenic | |
| rs73427747 | 22 | 28582273 | 4 | 0 | 0 | A | G | 2.5912 | 0.6915 | 0.0001787 | +++?+ | 0 | 0.704 | intron | TTC28 |
| rs61885969 | 11 | 30675456 | 4 | 0 | 0 | T | C | -1.0327 | 0.2756 | 0.0001788 | ---?+ | 39.9 | 0.1725 | intergenic | |
| 4:105466574 | 4 | 105466574 | 3 | 0 | 0 | <ins:me: | T | -3.0549 | 0.8153 | 0.0001789 | ---?? | 55.4 | 0.1065 | | |
| rs117413537 | 8 | 21710032 | 3 | 0 | 0 | T | C | 4.9418 | 1.3189 | 0.0001791 | +++?? | 0 | 0.7417 | intergenic | |
| rs190450 | 5 | 127614472 | 5 | 0 | 0 | A | G | -0.8973 | 0.2395 | 0.0001792 | +---- | 23.9 | 0.2621 | synonymous | FBN2 |
| rs1352995 | 8 | 113449760 | 4 | 0 | 0 | A | T | 1.0451 | 0.2789 | 0.0001792 | +++?+ | 0 | 0.7635 | intron | CSMD3 |
| rs74570147 | 1 | 102927526 | 3 | 0 | 0 | T | C | -0.9476 | 0.2529 | 0.0001792 | ---?? | 0 | 0.7573 | intergenic | |
| rs41417552 | 5 | 10305452 | 4 | 0 | 0 | A | G | 1.6518 | 0.4409 | 0.0001793 | +++?+ | 20.9 | 0.2847 | intron | CMBL |
| rs74020233 | 16 | 60135878 | 3 | 0 | 0 | T | C | 4.9536 | 1.3222 | 0.0001793 | ?++?+ | 0 | 0.4067 | intergenic | |
| 8:113498343 | 8 | 113498343 | 3 | 0 | 0 | G | GT | -1.0583 | 0.2825 | 0.0001793 | ---?? | 0 | 0.5318 | | |
| rs77997588 | 13 | 30314265 | 3 | 0 | 0 | T | C | -1.6381 | 0.4373 | 0.0001794 | ---?? | 0 | 0.6241 | intergenic | |
| rs11607000 | 11 | 103677856 | 4 | 0 | 0 | A | G | 2.3486 | 0.6269 | 0.0001796 | +++?+ | 0 | 0.8651 | intergenic | |
| rs139984152 | 6 | 137094890 | 4 | 0 | 0 | A | T | 2.7009 | 0.7211 | 0.0001799 | +++?+ | 19.7 | 0.2912 | intron | MAP3K5 |
| rs1492675 | 8 | 113425607 | 4 | 0 | 0 | A | T | -1.0491 | 0.2801 | 0.00018 | ---?- | 0 | 0.8164 | intron | CSMD3 |
| rs1500641 | 15 | 98045655 | 4 | 0 | 0 | A | G | -0.9903 | 0.2644 | 0.00018 | ---?- | 0 | 0.9285 | intergenic | |
| rs6977338 | 7 | 130143073 | 4 | 0 | 0 | T | C | 0.9262 | 0.2473 | 0.00018 | +-+?+ | 58.3 | 0.06589 | intron | MEST |
| rs116457097 | 2 | 129158433 | 4 | 0 | 0 | A | T | -3.9103 | 1.044 | 0.0001801 | ---?- | 0 | 0.9717 | intergenic | |
| rs67109144 | 17 | 9104530 | 4 | 0 | 0 | T | C | 1.6361 | 0.4369 | 0.0001802 | +++?+ | 0 | 0.5072 | intron | NTN1 |
| rs57229781 | 16 | 60140948 | 3 | 0 | 0 | A | C | -4.9238 | 1.3147 | 0.0001803 | ?--?- | 0 | 0.402 | intergenic | |
| rs194277 | 5 | 127612979 | 5 | 0 | 0 | A | T | -0.8984 | 0.2399 | 0.0001804 | +---- | 22.9 | 0.2688 | intron | FBN2 |
| rs139683975 | 6 | 23893950 | 4 | 0 | 0 | T | C | 2.714 | 0.7247 | 0.0001804 | +++?- | 24.1 | 0.2665 | intergenic | |
| rs35205163 | 16 | 85239385 | 4 | 0 | 0 | T | G | -1.2527 | 0.3345 | 0.0001806 | ---?+ | 0 | 0.4144 | intron | GSE1 |
| rs56158443 | 6 | 118676213 | 4 | 0 | 0 | C | G | 2.5146 | 0.6715 | 0.0001806 | +++?- | 0 | 0.5468 | intergenic | |
| rs1993207 | 8 | 113450380 | 4 | 0 | 0 | T | C | -1.0446 | 0.279 | 0.0001807 | ---?- | 0 | 0.763 | intron | CSMD3 |
| rs1078096 | 1 | 90268985 | 5 | 0 | 0 | A | C | -0.7851 | 0.2097 | 0.000181 | ----+ | 0 | 0.5862 | intergenic |  |
| rs73485737 | 9 | 90816410 | 4 | 0 | 0 | T | G | 1.9606 | 0.5237 | 0.000181 | +++?+ | 0 | 0.7714 | intergenic | |
| 6:12211632 | 6 | 12211632 | 3 | 0 | 0 | CA | C | -1.6376 | 0.4374 | 0.0001811 | ---?? | 0 | 0.4926 | | |
| rs7094035 | 10 | 131918109 | 4 | 0 | 0 | T | G | -1.0225 | 0.2731 | 0.0001812 | ---?- | 0 | 0.9539 | intergenic | |
| rs17282114 | 7 | 82504017 | 4 | 0 | 0 | T | C | -1.0114 | 0.2702 | 0.0001814 | -+-?- | 46.1 | 0.1345 | intron | PCLO |
| rs57422856 | 2 | 137021394 | 4 | 0 | 0 | T | C | 1.0202 | 0.2725 | 0.0001814 | +++?+ | 0 | 0.8378 | intergenic | |
| rs7784670 | 7 | 16905115 | 4 | 0 | 0 | C | G | 1.7765 | 0.4746 | 0.0001816 | +++?+ | 0 | 0.4436 | intron | AGR3 |
| rs10922719 | 1 | 90254411 | 5 | 0 | 0 | C | G | 0.7863 | 0.2101 | 0.0001816 | ++++- | 0 | 0.6036 | intergenic | |
| rs138140369 | 6 | 134260466 | 2 | 0 | 0 | A | G | 8.7097 | 2.3268 | 0.0001816 | ++??? | 0 | 0.6885 | intergenic | |
| rs17620386 | 18 | 40062880 | 4 | 0 | 0 | T | G | 1.4459 | 0.3863 | 0.0001818 | +++?+ | 0 | 0.9889 | intron | LINC00907 |
| rs1319683 | 8 | 113428972 | 4 | 0 | 0 | T | G | 1.0458 | 0.2794 | 0.0001819 | +++?+ | 0 | 0.8101 | intron | CSMD3 |
| rs17212221 | 7 | 82433696 | 4 | 0 | 0 | C | G | 0.9795 | 0.2617 | 0.000182 | +-+?+ | 49.5 | 0.1143 | intron | PCLO |
| rs111326306 | 11 | 79039186 | 4 | 0 | 0 | A | G | -2.3301 | 0.6226 | 0.000182 | ---?- | 30.1 | 0.2314 | intron | TENM4 |
| rs143274456 | 6 | 77670551 | 3 | 0 | 0 | T | C | 1.11 | 0.2966 | 0.0001822 | +++?? | 43.3 | 0.1715 | intergenic | |
| 17:56959758 | 17 | 56959758 | 3 | 0 | 0 | A | AT | 2.8607 | 0.7644 | 0.0001822 | +++?? | 0 | 0.5383 | | |
| 10:131485636 | 10 | 131485636 | 3 | 0 | 0 | A | AAAGTT | 1.5868 | 0.424 | 0.0001823 | +++?? | 0 | 0.7794 | | |

| | | | | | | | | | | | | | | | |
|---|---|---|---|---|---|---|---|---|---|---|---|---|---|---|---|
| rs11048629 | 12 | 26798413 | 4 | 0 | 0 | T | C | 1.929 | 0.5155 | 0.0001825 | +++?+ | 5 | 0.3679 | intron | ITPR2 |
| rs142750312 | 9 | 101609006 | 4 | 0 | 0 | A | G | 3.0728 | 0.8211 | 0.0001825 | +++?+ | 12.6 | 0.3297 | intron | GALNT12 |
| rs7111952 | 11 | 106755744 | 4 | 0 | 0 | A | G | -1.1014 | 0.2944 | 0.0001827 | ---?+ | 36.8 | 0.1915 | intron | GUCY1A2 |
| rs7298086 | 12 | 27773544 | 4 | 0 | 0 | A | G | -0.9791 | 0.2617 | 0.0001827 | ---?- | 12.2 | 0.3317 | intron | PPFIBP1 |
| rs10107134 | 8 | 113429866 | 4 | 0 | 0 | T | G | 1.0449 | 0.2793 | 0.0001828 | +++?+ | 0 | 0.8086 | intron | CSMD3 |
| rs4357818 | 13 | 60014039 | 3 | 0 | 0 | T | C | 0.9566 | 0.2557 | 0.000183 | +++?? | 0 | 0.5855 | intergenic | |
| rs4338670 | 13 | 60014052 | 3 | 0 | 0 | A | G | 0.9567 | 0.2557 | 0.0001831 | +++?? | 0 | 0.5847 | intergenic | |
| 17:65477893 | 17 | 65477893 | 3 | 0 | 0 | T | TAA | 0.9972 | 0.2665 | 0.0001831 | +++?? | 0 | 0.6447 | | |
| rs10749881 | 11 | 106756696 | 4 | 0 | 0 | A | C | -1.0994 | 0.2939 | 0.0001832 | ---?+ | 36 | 0.1964 | intron | GUCY1A2 |
| rs79387179 | 6 | 77671360 | 3 | 0 | 0 | A | G | -1.1091 | 0.2965 | 0.0001833 | ---?? | 43.3 | 0.1716 | intergenic | |
| rs7997712 | 13 | 60014076 | 3 | 0 | 0 | T | C | -0.9567 | 0.2557 | 0.0001834 | ---?? | 0 | 0.5843 | intergenic | |
| rs17562397 | 7 | 130329325 | 4 | 0 | 0 | T | C | 0.9185 | 0.2455 | 0.0001834 | +-+?+ | 59 | 0.06266 | intron | COPG2 |
| rs149402168 | 6 | 83804710 | 4 | 0 | 0 | A | G | 3.3829 | 0.9043 | 0.0001834 | +++?+ | 0 | 0.6953 | intron | DOPEY1 |
| 6:135234711 | 6 | 135234711 | 3 | 0 | 0 | CT | C | 1.6965 | 0.4535 | 0.0001834 | +++?? | 1.5 | 0.3624 | | |
| rs13439841 | 8 | 113431006 | 4 | 0 | 0 | T | C | 1.0437 | 0.279 | 0.0001835 | +++?+ | 0 | 0.8051 | intron | CSMD3 |
| rs13243136 | 7 | 130323971 | 4 | 0 | 0 | T | C | 0.9204 | 0.2461 | 0.0001837 | +-+?+ | 58.7 | 0.06413 | intron | COPG2 |
| rs80049847 | 4 | 162123483 | 4 | 0 | 0 | A | T | 2.0208 | 0.5403 | 0.0001837 | +++?- | 46.9 | 0.1299 | intergenic | |
| rs148119182 | 17 | 56797357 | 5 | 0 | 0 | A | T | -2.6132 | 0.6987 | 0.000184 | ----- | 0 | 0.8309 | intron | RAD51C |
| rs11103534 | 9 | 137700749 | 4 | 0 | 0 | T | G | 1.3836 | 0.37 | 0.0001841 | +++?+ | 40.7 | 0.1673 | intron | COL5A1 |
| rs78740046 | 3 | 108546644 | 3 | 0 | 0 | T | C | -4.4036 | 1.1775 | 0.0001841 | ---?? | 0 | 0.9622 | intron | TRAT1 |
| rs78962664 | 4 | 155640297 | 4 | 0 | 0 | A | G | -3.4589 | 0.9249 | 0.0001841 | ---?- | 0 | 0.9625 | intergenic | |
| 8:113481389 | 8 | 113481389 | 3 | 0 | 0 | CAAAAA | C | -1.0129 | 0.2708 | 0.0001841 | ---?? | 0 | 0.6789 | | |
| rs741482 | 3 | 185903412 | 4 | 0 | 0 | C | G | -1.5984 | 0.4274 | 0.0001842 | ---?+ | 32.2 | 0.2192 | intron | DGKG |
| rs10147039 | 14 | 22572503 | 4 | 0 | 0 | A | G | 1.131 | 0.3024 | 0.0001842 | +++?+ | 6.4 | 0.3611 | intergenic | |
| rs10754292 | 1 | 90258821 | 5 | 0 | 0 | A | T | 0.7845 | 0.2098 | 0.0001842 | ++++- | 0 | 0.6086 | intergenic | |
| rs10922721 | 1 | 90258177 | 5 | 0 | 0 | A | G | 0.7849 | 0.2099 | 0.0001842 | ++++- | 0 | 0.6084 | intergenic | |
| rs150912885 | 5 | 9479609 | 3 | 0 | 0 | A | C | -5.9489 | 1.5908 | 0.0001843 | ?--?- | 0 | 0.8022 | intron | SEMA5A |
| rs62516467 | 8 | 113454459 | 4 | 0 | 0 | T | C | 1.0436 | 0.2791 | 0.0001844 | +++?+ | 0 | 0.76 | intron | CSMD3 |
| 2:51912409 | 2 | 51912409 | 3 | 0 | 0 | G | GTA | -3.7053 | 0.9909 | 0.0001845 | ---?? | 0 | 0.6587 | | |
| rs765205 | 7 | 130140766 | 4 | 0 | 0 | A | C | -0.9238 | 0.2471 | 0.0001846 | -+-?- | 58.4 | 0.0653 | intron | MEST |
| rs733467 | 1 | 90270146 | 5 | 0 | 0 | A | G | 0.7863 | 0.2103 | 0.0001847 | ++++- | 0 | 0.5762 | intergenic | |
| rs79715619 | 5 | 11745759 | 2 | 0 | 0 | C | G | 5.9249 | 1.5846 | 0.0001847 | ?+??+ | 0 | 0.6719 | intron | CTNND2 |
| rs74757859 | 18 | 27068256 | 4 | 0 | 0 | A | G | 3.1674 | 0.8472 | 0.0001848 | +++?+ | 60.1 | 0.05728 | intergenic | |
| rs10835700 | 11 | 30674773 | 3 | 0 | 0 | A | G | 1.0776 | 0.2882 | 0.0001849 | +++?? | 45 | 0.1622 | intergenic | |
| rs11859292 | 16 | 6541820 | 4 | 0 | 0 | A | G | 1.0353 | 0.2769 | 0.000185 | +++?+ | 0 | 0.6347 | intron | RBFOX1 |
| rs12658439 | 5 | 165467653 | 4 | 0 | 0 | A | T | 1.4438 | 0.3862 | 0.000185 | +++?+ | 0 | 0.6728 | intergenic | |
| rs6695797 | 1 | 90269347 | 5 | 0 | 0 | T | C | -0.7845 | 0.2098 | 0.0001851 | ----+ | 0 | 0.5858 | intergenic |  |
| rs11919816 | 3 | 77409297 | 4 | 0 | 0 | T | C | -0.9251 | 0.2474 | 0.0001851 | ---?- | 0 | 0.4759 | intron | ROBO2 |
| 9:123009268 | 9 | 123009268 | 3 | 0 | 0 | CA | C | -1.3527 | 0.3619 | 0.0001853 | ---?? | 2.4 | 0.3588 | | |
| rs16972136 | 15 | 42131770 | 5 | 0 | 0 | T | C | 1.0634 | 0.2845 | 0.0001855 | ++++- | 20.7 | 0.2825 | intron | JMJD7-PLA2G4B,PLA2G4B |
| rs72839978 | 17 | 56369948 | 5 | 0 | 0 | T | C | 2.0314 | 0.5435 | 0.0001857 | +++++ | 46.4 | 0.1132 | intergenic | |
| rs77267780 | 6 | 106264584 | 4 | 0 | 0 | A | G | -2.2617 | 0.6051 | 0.0001858 | ---?- | 0 | 0.576 | intergenic | |
| rs11961468 | 6 | 168580903 | 4 | 0 | 0 | A | C | -1.6751 | 0.4482 | 0.0001859 | ---?- | 0 | 0.6807 | intergenic | |
| rs342899 | 3 | 146278616 | 4 | 0 | 0 | A | C | -1.4916 | 0.3991 | 0.0001861 | ---?- | 0 | 0.6205 | intergenic | |
| rs6839269 | 4 | 38746082 | 5 | 0 | 0 | A | G | 1.2295 | 0.329 | 0.0001861 | +++++ | 0 | 0.97 | intergenic | |
| rs79865268 | 6 | 77672289 | 3 | 0 | 0 | A | G | 1.1071 | 0.2962 | 0.0001861 | +++?? | 43.2 | 0.172 | intergenic | |
| rs6531658 | 4 | 38743996 | 5 | 0 | 0 | T | C | 1.2306 | 0.3293 | 0.0001864 | +++++ | 0 | 0.961 | intergenic | |
| rs185824426 | 6 | 77669812 | 3 | 0 | 0 | A | T | 1.1063 | 0.2961 | 0.0001865 | +++?? | 43.2 | 0.1721 | intergenic | |
| rs76768530 | 12 | 17046854 | 3 | 0 | 0 | A | G | -3.6381 | 0.9738 | 0.0001868 | ---?? | 29.7 | 0.241 | intergenic | |
| rs13239136 | 7 | 130331735 | 4 | 0 | 0 | A | G | -0.9184 | 0.2458 | 0.0001871 | -+-?- | 58.6 | 0.06459 | intron | COPG2 |
| 5:165391446 | 5 | 165391446 | 3 | 0 | 0 | T | TA | -0.9341 | 0.25 | 0.0001872 | ---?? | 0 | 0.3957 | | |
| rs5001403 | 15 | 98045947 | 4 | 0 | 0 | T | C | 0.9875 | 0.2644 | 0.0001876 | +++?+ | 0 | 0.9284 | intergenic | |
| rs12536613 | 7 | 82414992 | 4 | 0 | 0 | T | C | 0.988 | 0.2645 | 0.0001876 | +-+?+ | 52.6 | 0.09652 | intron | PCLO |
| rs7108150 | 11 | 12644674 | 4 | 0 | 0 | A | G | -0.9938 | 0.2661 | 0.0001877 | ---?- | 0 | 0.8265 | intergenic | |
| rs116737693 | 1 | 31729634 | 4 | 0 | 0 | T | C | -4.4306 | 1.1862 | 0.0001877 | ---?- | 56 | 0.07808 | downstream-gene | |
| rs2312789 | 7 | 130327546 | 4 | 0 | 0 | T | C | -0.9177 | 0.2457 | 0.0001879 | -+-?- | 58.8 | 0.06363 | intron | COPG2 |
| 6:77664495 | 6 | 77664495 | 3 | 0 | 0 | CTCA | C | -1.1044 | 0.2957 | 0.000188 | ---?? | 43.1 | 0.1725 | | |
| rs6469412 | 8 | 113213542 | 4 | 0 | 0 | T | C | 0.9431 | 0.2525 | 0.0001881 | +++?- | 0 | 0.6906 | intergenic | |
| rs6924040 | 6 | 77663630 | 3 | 0 | 0 | T | C | 1.1039 | 0.2956 | 0.0001886 | +++?? | 43.1 | 0.1724 | intergenic | |

| | | | | | | | | | | | | | | | |
|---|---|---|---|---|---|---|---|---|---|---|---|---|---|---|---|
| rs3792318 | 7 | 130300270 | 4 | 0 | 0 | T | C | 0.9195 | 0.2463 | 0.0001888 | +-+?+ | 57.5 | 0.06989 | intron | COPG2 |
| rs28844095 | 12 | 111783778 | 5 | 0 | 0 | A | G | -0.9425 | 0.2524 | 0.0001889 | -+--- | 35.5 | 0.1846 | intron | CUX2 |
| rs1005171 | 7 | 130144402 | 4 | 0 | 0 | C | G | 0.9233 | 0.2473 | 0.0001892 | +-+?+ | 57.7 | 0.06909 | intron | MEST |
| rs9379634 | 6 | 24061896 | 5 | 0 | 0 | A | G | -1.5183 | 0.4067 | 0.0001892 | ----+ | 0 | 0.6257 | intergenic |  |
| rs3803167 | 12 | 111785586 | 5 | 0 | 0 | T | C | -0.9417 | 0.2523 | 0.0001893 | -+--- | 33.8 | 0.1959 | synonymous | CUX2 |
| rs585030 | 10 | 131915486 | 4 | 0 | 0 | A | G | 1.0191 | 0.273 | 0.0001896 | +++?+ | 0 | 0.9111 | intergenic | |
| rs59325484 | 16 | 57460799 | 4 | 0 | 0 | A | G | -1.6995 | 0.4554 | 0.0001899 | ---?- | 0 | 0.8038 | downstream-gene | |
| 6:24063403 | 6 | 24063403 | 3 | 0 | 0 | T | TA | 1.7481 | 0.4684 | 0.0001899 | +++?? | 0 | 0.6619 | | |
| rs907622 | 11 | 37059314 | 4 | 0 | 0 | A | C | -0.9088 | 0.2435 | 0.0001902 | ---?- | 0 | 0.4091 | intergenic | |
| rs10922725 | 1 | 90265153 | 5 | 0 | 0 | A | T | 0.7804 | 0.2091 | 0.0001902 | ++++- | 0 | 0.5777 | intergenic | |
| rs2072575 | 7 | 130148507 | 4 | 0 | 0 | A | G | -0.9249 | 0.2479 | 0.0001903 | -+-?- | 58 | 0.06762 | missense | COPG2 |
| rs8049227 | 16 | 87625263 | 4 | 0 | 0 | A | G | -2.0536 | 0.5503 | 0.0001903 | ---?+ | 0 | 0.4248 | intergenic | |
| rs733468 | 1 | 90270404 | 5 | 0 | 0 | T | C | 0.7841 | 0.2102 | 0.0001907 | ++++- | 0 | 0.5853 | intergenic | |
| rs9641881 | 7 | 130336026 | 4 | 0 | 0 | T | C | 0.9184 | 0.2461 | 0.0001907 | +-+?+ | 58 | 0.06769 | intron | COPG2 |
| 15:86320155 | 15 | 86320155 | 3 | 0 | 0 | CA | C | 1.4417 | 0.3864 | 0.0001907 | +++?? | 4.5 | 0.3508 | | |
| rs12529704 | 6 | 77687484 | 3 | 0 | 0 | T | G | 1.1081 | 0.297 | 0.0001908 | +++?? | 43.8 | 0.1687 | intergenic | |
| rs11634588 | 15 | 25109384 | 3 | 0 | 0 | A | C | 1.1888 | 0.3187 | 0.000191 | +++?? | 0 | 0.7331 | intron | SNRPN |
| rs72839979 | 17 | 56373719 | 5 | 0 | 0 | T | C | 2.0284 | 0.5437 | 0.000191 | +++++ | 45.9 | 0.1166 | downstream-gene | |
| rs982664 | 7 | 107344216 | 3 | 0 | 0 | T | C | -0.9765 | 0.2618 | 0.0001912 | ---?? | 17.7 | 0.2966 | intron | SLC26A4 |
| rs143538396 | 11 | 67497186 | 3 | 0 | 0 | T | C | -1.6491 | 0.4421 | 0.0001912 | ---?? | 0 | 0.5651 | intergenic | |
| rs12667336 | 7 | 130318972 | 4 | 0 | 0 | T | C | -0.9182 | 0.2462 | 0.0001913 | -+-?- | 58.5 | 0.06486 | intron | COPG2 |
| rs111747875 | 7 | 80338013 | 4 | 0 | 0 | T | G | 3.6635 | 0.9822 | 0.0001915 | +++?+ | 0 | 0.8869 | intergenic | |
| rs7824183 | 8 | 113460269 | 4 | 0 | 0 | T | C | 1.0414 | 0.2792 | 0.0001916 | +++?+ | 0 | 0.7564 | intron | CSMD3 |
| rs114420104 | 6 | 32596533 | 5 | 0 | 1 | T | G | 1.5717 | 0.4214 | 0.0001916 | +++++ | 0 | 0.4594 | intergenic | |
| rs144792473 | 18 | 37881050 | 4 | 0 | 0 | C | G | -2.781 | 0.7456 | 0.0001917 | ---?- | 0 | 0.6235 | intergenic | |
| rs6838865 | 4 | 38745883 | 5 | 0 | 0 | T | G | 1.2258 | 0.3287 | 0.0001918 | +++++ | 0 | 0.9594 | intergenic | |
| rs8048070 | 16 | 87624947 | 4 | 0 | 0 | T | C | 2.0517 | 0.5502 | 0.000192 | +++?- | 0 | 0.425 | intergenic | |
| rs8090196 | 18 | 40075956 | 4 | 0 | 0 | T | C | -1.2646 | 0.3391 | 0.0001921 | ---?+ | 11.6 | 0.3346 | intron | LINC00907 |
| 6:159232907 | 6 | 159232907 | 3 | 0 | 0 | CAAAAA | C | -0.952 | 0.2553 | 0.0001921 | ---?? | 0 | 0.4996 | | |
| rs7009582 | 8 | 113212913 | 4 | 0 | 0 | T | C | 0.9418 | 0.2526 | 0.0001922 | +++?- | 0 | 0.6911 | intergenic | |
| rs61972160 | 14 | 22523852 | 4 | 0 | 0 | A | G | 1.1705 | 0.3139 | 0.0001922 | +++?+ | 0 | 0.9862 | intergenic | |
| rs2072574 | 7 | 130137692 | 4 | 0 | 0 | A | G | 0.9208 | 0.2469 | 0.0001923 | +-+?+ | 58.4 | 0.06544 | intron | MEST |
| rs10003163 | 4 | 38745351 | 5 | 0 | 0 | A | G | -1.226 | 0.3288 | 0.0001923 | ----- | 0 | 0.9603 | intergenic |  |
| rs11876579 | 18 | 63871520 | 4 | 0 | 0 | T | G | -0.9251 | 0.2481 | 0.0001923 | ---?- | 0 | 0.4558 | intergenic | |
| rs45592033 | 6 | 32162559 | 5 | 0 | 0 | A | G | 1.8318 | 0.4912 | 0.0001923 | ++++- | 19.6 | 0.29 | 5-prime-UTR | GPSM3 |
| rs2269778 | 7 | 107408870 | 4 | 0 | 0 | A | G | -0.9462 | 0.2538 | 0.0001925 | ---?- | 3.4 | 0.3758 | intron | SLC26A3 |
| rs13434428 | 4 | 38745138 | 5 | 0 | 0 | T | C | 1.2263 | 0.3289 | 0.0001925 | +++++ | 0 | 0.9607 | intergenic | |
| rs149819055 | 14 | 22501337 | 4 | 0 | 0 | T | C | 1.0298 | 0.2762 | 0.0001925 | +++?+ | 0 | 0.8439 | intergenic | |
| rs13434431 | 4 | 38745161 | 5 | 0 | 0 | T | C | 1.2263 | 0.3289 | 0.0001926 | +++++ | 0 | 0.9607 | intergenic | |
| rs4605665 | 4 | 38744961 | 5 | 0 | 0 | A | G | -1.2266 | 0.329 | 0.0001927 | ----- | 0 | 0.961 | intergenic |  |
| rs6133284 | 20 | 6005634 | 5 | 0 | 0 | A | G | 0.7712 | 0.2069 | 0.0001927 | ++++- | 0 | 0.4659 | intron | CRLS1 |
| rs35416015 | 7 | 130137572 | 4 | 0 | 0 | A | G | -0.9206 | 0.2469 | 0.0001927 | -+-?- | 58.4 | 0.06545 | intron | MEST |
| 13:60015286 | 13 | 60015286 | 3 | 0 | 0 | G | GC | -0.949 | 0.2545 | 0.0001929 | ---?? | 0 | 0.4145 | | |
| rs115787626 | 3 | 47721512 | 2 | 0 | 0 | A | G | 6.8445 | 1.836 | 0.000193 | ?+??+ | 8.6 | 0.2955 | intron | SMARCC1 |
| rs4385073 | 4 | 38745026 | 5 | 0 | 0 | T | C | -1.2263 | 0.329 | 0.0001931 | ----- | 0 | 0.961 | intergenic |  |
| rs28403936 | 12 | 111783020 | 5 | 0 | 0 | A | G | -0.9409 | 0.2524 | 0.0001933 | -+--- | 35.7 | 0.1836 | intron | CUX2 |
| 9:16199842 | 9 | 16199842 | 3 | 0 | 0 | CTTTTTT | C | 1.4268 | 0.3828 | 0.0001935 | +++?? | 0 | 0.502 | | |
| rs74866176 | 9 | 112415406 | 4 | 0 | 0 | T | C | -3.4824 | 0.9343 | 0.0001936 | ---?- | 0 | 0.4337 | intron | PALM2 |
| rs1424417 | 7 | 130315598 | 4 | 0 | 0 | A | T | -0.9179 | 0.2463 | 0.0001938 | -+-?- | 58.4 | 0.0657 | intron | COPG2 |
| rs1590297 | 1 | 90259286 | 5 | 0 | 0 | T | C | -0.7817 | 0.2098 | 0.0001939 | ----+ | 0 | 0.5983 | intergenic |  |
| rs7006962 | 8 | 113473594 | 4 | 0 | 0 | A | G | -1.0612 | 0.2847 | 0.0001939 | ---?- | 0 | 0.701 | intron | CSMD3 |
| rs13241924 | 7 | 130315289 | 4 | 0 | 0 | A | G | 0.9179 | 0.2463 | 0.0001939 | +-+?+ | 58.3 | 0.06572 | intron | COPG2 |
| rs1590296 | 1 | 90259087 | 5 | 0 | 0 | T | C | 0.7817 | 0.2098 | 0.0001942 | ++++- | 0 | 0.598 | intergenic | |
| rs7158591 | 14 | 28193549 | 4 | 0 | 0 | T | C | -2.1729 | 0.5831 | 0.0001942 | ---?- | 0 | 0.991 | intergenic | |
| rs113299073 | 12 | 75502238 | 4 | 0 | 0 | C | G | -3.6613 | 0.9826 | 0.0001944 | ---?- | 0 | 0.6264 | intron | KCNC2 |
| rs17637133 | 14 | 43650399 | 4 | 0 | 0 | T | C | -1.8373 | 0.4931 | 0.0001945 | ---?+ | 0 | 0.497 | intergenic | |
| rs117613042 | 8 | 102576576 | 3 | 0 | 0 | A | G | -3.0291 | 0.813 | 0.0001945 | ---?? | 41.8 | 0.1791 | intron | GRHL2 |
| rs6987206 | 8 | 113212412 | 4 | 0 | 0 | T | C | -0.9411 | 0.2526 | 0.0001946 | ---?+ | 0 | 0.6915 | intergenic | |
| rs11903754 | 2 | 79691562 | 3 | 0 | 0 | A | C | -0.9912 | 0.266 | 0.0001946 | ---?? | 46 | 0.1572 | intergenic | |

| | | | | | | | | | | | | | | | |
|---|---|---|---|---|---|---|---|---|---|---|---|---|---|---|---|
| rs62516479 | 8 | 113475540 | 4 | 0 | 0 | A | G | 1.0557 | 0.2833 | 0.0001946 | +++?+ | 0 | 0.7509 | intron | CSMD3 |
| rs142962821 | 2 | 168029045 | 4 | 0 | 0 | A | G | 2.5267 | 0.6781 | 0.0001946 | +++?- | 0 | 0.4455 | intron | XIRP2 |
| 6:28544545 | 6 | 28544545 | 3 | 0 | 0 | CT | C | -1.3306 | 0.3571 | 0.0001947 | ---?? | 16.1 | 0.3034 | | |
| rs6853665 | 4 | 169324016 | 4 | 0 | 0 | A | G | 0.9241 | 0.2481 | 0.0001948 | +++?+ | 0 | 0.7759 | intron | DDX60L |
| rs116965761 | 20 | 41096292 | 2 | 0 | 0 | T | C | -4.5061 | 1.2096 | 0.000195 | -?-?? | 0 | 0.971 | intron | PTPRT |
| rs145282567 | 20 | 16055382 | 3 | 0 | 0 | T | C | 3.3234 | 0.8921 | 0.0001951 | +++?? | 0 | 0.4487 | intergenic | |
| rs255695 | 5 | 127613299 | 5 | 0 | 0 | T | G | 0.8924 | 0.2396 | 0.0001952 | -++++ | 24.1 | 0.2607 | intron | FBN2 |
| rs9783328 | 11 | 106756129 | 4 | 0 | 0 | T | C | 1.0948 | 0.2939 | 0.0001954 | +++?- | 32.7 | 0.2158 | intron | GUCY1A2 |
| rs4364658 | 8 | 113212297 | 4 | 0 | 0 | C | G | 0.9408 | 0.2526 | 0.0001957 | +++?- | 0 | 0.6916 | intergenic | |
| rs11736367 | 4 | 125541779 | 4 | 0 | 0 | T | C | 1.0299 | 0.2765 | 0.0001957 | +++?+ | 0 | 0.7155 | intergenic | |
| rs34186070 | 14 | 22514534 | 4 | 0 | 0 | A | G | -1.1633 | 0.3123 | 0.0001957 | ---?- | 0 | 0.8948 | intergenic | |
| rs9919551 | 11 | 30637258 | 4 | 0 | 0 | T | C | 1.0067 | 0.2703 | 0.0001958 | +++?- | 31.6 | 0.2229 | intergenic | |
| rs9625547 | 22 | 29133029 | 4 | 0 | 0 | A | G | 2.4485 | 0.6575 | 0.0001959 | +++?+ | 0 | 0.8466 | intron | CHEK2 |
| rs9866753 | 3 | 71899594 | 4 | 0 | 0 | T | C | -1.0032 | 0.2694 | 0.0001959 | ---?- | 0 | 0.8982 | intergenic | |
| rs1073138 | 11 | 95703401 | 4 | 0 | 0 | A | T | -1.04 | 0.2793 | 0.0001961 | ---?- | 17.7 | 0.3022 | intergenic | |
| rs9952112 | 18 | 47775076 | 4 | 0 | 0 | A | G | -0.9843 | 0.2643 | 0.0001961 | ---?- | 0 | 0.7152 | intron | CCDC11 |
| rs12361624 | 11 | 30680067 | 4 | 0 | 0 | A | G | -1.0268 | 0.2757 | 0.0001962 | ---?+ | 40.3 | 0.1701 | intergenic | |
| rs138272887 | 12 | 32695012 | 2 | 0 | 0 | T | G | -8.2549 | 2.2171 | 0.0001967 | ?-??- | 0 | 0.8099 | intron | FGD4 |
| rs140362750 | 17 | 56804726 | 5 | 0 | 0 | A | C | -2.6229 | 0.7045 | 0.0001968 | ----- | 0 | 0.7984 | intron | RAD51C |
| rs62194756 | 20 | 23852463 | 4 | 0 | 0 | T | C | 1.563 | 0.4198 | 0.0001969 | +++?+ | 9.5 | 0.3457 | downstream-gene | |
| rs73270582 | 6 | 168583810 | 4 | 0 | 0 | T | C | -1.6685 | 0.4482 | 0.000197 | ---?- | 0 | 0.5068 | intergenic | |
| 11:86178725 | 11 | 86178725 | 3 | 0 | 0 | A | T | 3.5468 | 0.9527 | 0.0001971 | +++?? | 0 | 0.9175 | intron | ME3 |
| rs10000466 | 4 | 38747536 | 5 | 0 | 0 | T | C | 1.2344 | 0.3316 | 0.0001971 | +++++ | 0 | 0.9828 | intergenic | |
| rs2353274 | 8 | 113212190 | 4 | 0 | 0 | A | T | -0.9403 | 0.2526 | 0.0001972 | ---?+ | 0 | 0.6918 | intergenic | |
| rs77735008 | 2 | 170561704 | 4 | 0 | 0 | T | G | -1.3378 | 0.3594 | 0.0001974 | ---?- | 0 | 0.9162 | intron | PHOSPHO2-KLHL23 |
| rs626378 | 5 | 134353501 | 5 | 0 | 0 | T | C | -0.8655 | 0.2325 | 0.0001976 | ----- | 0 | 0.5053 | intergenic |  |
| rs6114335 | 20 | 23825328 | 4 | 0 | 0 | A | C | 1.5526 | 0.4171 | 0.0001977 | +++?+ | 0 | 0.4262 | intergenic | |
| 12:43180406 | 12 | 43180406 | 3 | 0 | 0 | G | GT | -1.4952 | 0.4017 | 0.0001977 | ---?? | 7.9 | 0.3376 | | |
| rs17197692 | 3 | 36775946 | 4 | 0 | 0 | A | C | -1.7061 | 0.4584 | 0.0001978 | ---?- | 0 | 0.5986 | intron | DCLK3 |
| 6:159204431 | 6 | 159204431 | 3 | 0 | 0 | <ins:me: | T | -1.0644 | 0.286 | 0.0001978 | ---?? | 0 | 0.4786 | | |
| rs13246634 | 7 | 130311073 | 4 | 0 | 0 | A | G | 0.9167 | 0.2463 | 0.0001981 | +-+?+ | 58.1 | 0.06691 | intron | COPG2 |
| rs113707427 | 18 | 40096300 | 3 | 0 | 0 | A | G | -1.4739 | 0.3961 | 0.0001981 | ---?? | 0 | 0.9547 | intron | LINC00907 |
| rs4483563 | 11 | 106758700 | 4 | 0 | 0 | T | C | -1.0941 | 0.294 | 0.0001982 | ---?+ | 46.2 | 0.1342 | intron | GUCY1A2 |
| rs62464848 | 7 | 57276874 | 4 | 0 | 0 | C | G | 1.1595 | 0.3116 | 0.0001982 | +++?- | 0 | 0.4803 | intergenic | |
| rs74093732 | 1 | 75278726 | 4 | 0 | 0 | A | G | -2.1793 | 0.5858 | 0.000199 | ---?- | 5 | 0.3681 | intergenic | |
| rs74553399 | 6 | 168654414 | 4 | 0 | 0 | A | G | 2.0454 | 0.5499 | 0.0001994 | +++?+ | 0 | 0.6923 | intergenic | |
| rs77955561 | 20 | 23844959 | 4 | 0 | 0 | A | G | 1.5454 | 0.4155 | 0.0001994 | +++?+ | 0.4 | 0.39 | intergenic | |
| 5:134353360 | 5 | 134353360 | 3 | 0 | 0 | T | TA | 1.0497 | 0.2822 | 0.0001994 | +++?? | 0 | 0.4252 | | |
| rs6139891 | 20 | 5999437 | 5 | 0 | 0 | A | G | 0.7725 | 0.2077 | 0.0001995 | ++++- | 0 | 0.4874 | intron | CRLS1 |
| rs13357144 | 5 | 119700563 | 4 | 0 | 0 | A | G | -1.2433 | 0.3342 | 0.0001996 | ---?- | 42.9 | 0.1538 | intergenic | |
| rs11031164 | 11 | 30676301 | 4 | 0 | 0 | T | C | 1.0262 | 0.2759 | 0.0001998 | +++?- | 40.3 | 0.17 | intergenic | |
| rs11787085 | 8 | 113482086 | 4 | 0 | 0 | A | G | 1.0552 | 0.2837 | 0.0001998 | +++?+ | 0 | 0.7527 | intron | CSMD3 |
| rs1750290 | 10 | 131913871 | 4 | 0 | 0 | C | G | -1.0232 | 0.2751 | 0.0001999 | ---?- | 0 | 0.912 | upstream-gene | |
| rs7710190 | 5 | 55932140 | 4 | 0 | 0 | A | C | 2.7504 | 0.7395 | 0.0001999 | +++?- | 4.9 | 0.3685 | intergenic | |
| 2:242013702 | 2 | 242013702 | 3 | 0 | 0 | T | TC | 0.9342 | 0.2512 | 0.0001999 | +++?? | 0 | 0.3901 | | |
| rs12706947 | 7 | 130307434 | 4 | 0 | 0 | T | C | -0.916 | 0.2463 | 0.0002001 | -+-?- | 58 | 0.06756 | intron | COPG2 |
| rs148360796 | 17 | 56823312 | 5 | 0 | 0 | C | G | 2.6129 | 0.7026 | 0.0002002 | +++++ | 0 | 0.801 | intergenic | |
| rs4750788 | 10 | 131925850 | 4 | 0 | 0 | T | C | -0.8972 | 0.2413 | 0.0002003 | ---?- | 0 | 0.7064 | intergenic | |
| rs79697546 | 6 | 168581244 | 4 | 0 | 0 | T | C | -1.6542 | 0.4448 | 0.0002003 | ---?- | 0 | 0.674 | intergenic | |
| rs143346199 | 17 | 56823305 | 5 | 0 | 0 | C | G | -2.6128 | 0.7026 | 0.0002003 | ----- | 0 | 0.8011 | intergenic |  |
| rs2353272 | 8 | 113211907 | 4 | 0 | 0 | A | G | 0.9392 | 0.2526 | 0.0002006 | +++?- | 0 | 0.6921 | intergenic | |
| rs6923101 | 6 | 83804744 | 3 | 0 | 0 | T | C | 3.4549 | 0.9292 | 0.0002006 | +++?? | 0 | 0.5283 | intron | DOPEY1 |
| rs36194706 | 15 | 54351638 | 3 | 0 | 0 | A | G | 1.3134 | 0.3532 | 0.0002006 | +++?? | 0 | 0.9995 | intron | UNC13C |
| rs115005548 | 2 | 168032917 | 4 | 0 | 0 | A | T | -2.5125 | 0.6758 | 0.0002009 | ---?+ | 0 | 0.4482 | intron | XIRP2 |
| rs7535521 | 1 | 90256113 | 5 | 0 | 0 | A | G | 0.7809 | 0.2101 | 0.0002012 | ++++- | 0 | 0.595 | intergenic | |
| rs62516475 | 8 | 113465740 | 4 | 0 | 0 | A | C | 1.0387 | 0.2794 | 0.0002013 | +++?+ | 0 | 0.7557 | intron | CSMD3 |
| rs116866858 | 20 | 21486885 | 4 | 0 | 0 | A | T | -2.6369 | 0.7094 | 0.0002014 | ---?- | 46.2 | 0.1343 | downstream-gene | |
| rs2129905 | 7 | 130305479 | 4 | 0 | 0 | A | G | 0.9154 | 0.2463 | 0.0002016 | +-+?+ | 57.9 | 0.06782 | intron | COPG2 |
| rs6133281 | 20 | 5998627 | 5 | 0 | 0 | A | G | -0.7722 | 0.2077 | 0.0002017 | ----+ | 0 | 0.4878 | intron | CRLS1 |

| | | | | | | | | | | | | | | | |
|---|---|---|---|---|---|---|---|---|---|---|---|---|---|---|---|
| rs12674293 | 7 | 130341228 | 4 | 0 | 0 | T | C | -0.9161 | 0.2465 | 0.0002018 | -+-?- | 57.2 | 0.07168 | intron | COPG2 |
| rs113410424 | 14 | 32935871 | 4 | 0 | 0 | A | C | 1.4406 | 0.3876 | 0.0002018 | +++?+ | 0 | 0.748 | intron | AKAP6 |
| rs10922722 | 1 | 90258453 | 5 | 0 | 0 | A | G | -0.7799 | 0.2099 | 0.0002024 | ----+ | 0 | 0.6012 | intergenic |  |
| rs12670178 | 7 | 130304343 | 4 | 0 | 0 | T | C | -0.915 | 0.2462 | 0.0002027 | -+-?- | 57.8 | 0.06825 | intron | COPG2 |
| rs6819122 | 4 | 38749785 | 5 | 0 | 0 | T | C | 1.253 | 0.3372 | 0.0002028 | +++++ | 0 | 0.9533 | intergenic | |
| rs12435942 | 14 | 44911686 | 4 | 0 | 0 | A | C | -1.4561 | 0.3919 | 0.0002028 | ---?- | 0 | 0.8305 | intergenic | |
| rs7600311 | 2 | 51418267 | 4 | 0 | 0 | T | G | -1.7016 | 0.458 | 0.000203 | ---?- | 0 | 0.6993 | intergenic | |
| rs58882734 | 6 | 23866906 | 4 | 0 | 0 | A | C | 2.7313 | 0.7352 | 0.0002031 | +++?- | 42.5 | 0.1567 | intergenic | |
| rs116586720 | 4 | 128443767 | 4 | 0 | 0 | A | G | -3.6959 | 0.9948 | 0.0002031 | ---?- | 0 | 0.9615 | intergenic | |
| rs62194757 | 20 | 23854616 | 4 | 0 | 0 | T | C | 1.5621 | 0.4206 | 0.0002036 | +++?+ | 11.1 | 0.3373 | downstream-gene | |
| rs7894528 | 10 | 4415882 | 4 | 0 | 0 | A | G | 1.0126 | 0.2726 | 0.0002037 | +++?+ | 0 | 0.3972 | intergenic | |
| 22:29044919 | 22 | 29044919 | 3 | 0 | 0 | G | GGA | -2.6551 | 0.7148 | 0.0002038 | ---?? | 31.7 | 0.2311 | | |
| rs2871802 | 11 | 30635384 | 4 | 0 | 0 | A | G | 1.0038 | 0.2703 | 0.0002039 | +++?- | 31.7 | 0.222 | intergenic | |
| rs111316602 | 5 | 29500507 | 4 | 0 | 0 | A | G | 3.9104 | 1.0529 | 0.000204 | +++?+ | 0 | 0.4759 | intergenic | |
| rs142132502 | 4 | 169652989 | 4 | 0 | 0 | T | C | -4.6765 | 1.2592 | 0.0002041 | ---?- | 0 | 0.4368 | intron | PALLD |
| rs774226 | 9 | 94195371 | 4 | 0 | 0 | A | G | -2.5335 | 0.6822 | 0.0002042 | ---?- | 0 | 0.8049 | intergenic | |
| 6:30985692 | 6 | 30985692 | 3 | 0 | 0 | CTT | C | -1.5021 | 0.4045 | 0.0002042 | ---?? | 0 | 0.7402 | | |
| 12:2244650 | 12 | 2244650 | 2 | 0 | 0 | T | C | 7.8954 | 2.1261 | 0.0002044 | ++??? | 0 | 0.736 | intron | CACNA1C |
| rs4658140 | 1 | 90257330 | 5 | 0 | 0 | A | T | 0.7796 | 0.2099 | 0.0002044 | ++++- | 0 | 0.6002 | intergenic | |
| rs9962010 | 18 | 45836338 | 4 | 0 | 0 | C | G | 0.9368 | 0.2523 | 0.0002044 | +++?- | 0 | 0.7805 | intron | ZBTB7C |
| rs115348750 | 2 | 168022043 | 4 | 0 | 0 | T | C | -2.3162 | 0.6237 | 0.0002044 | ---?+ | 0 | 0.6517 | intron | XIRP2 |
| rs12433125 | 14 | 44911758 | 4 | 0 | 0 | T | C | -1.4553 | 0.3919 | 0.0002045 | ---?- | 0 | 0.8308 | intergenic | |
| rs73270580 | 6 | 168583792 | 4 | 0 | 0 | A | G | -1.6632 | 0.4479 | 0.0002045 | ---?- | 0 | 0.507 | intergenic | |
| rs62094326 | 18 | 63871048 | 4 | 0 | 0 | T | G | -0.9184 | 0.2473 | 0.0002047 | ---?- | 0 | 0.454 | intergenic | |
| rs75828185 | 2 | 168022920 | 4 | 0 | 0 | A | C | -2.3152 | 0.6235 | 0.0002047 | ---?+ | 0 | 0.652 | intron | XIRP2 |
| rs9654563 | 6 | 168609602 | 3 | 0 | 0 | T | C | -1.2593 | 0.3392 | 0.0002049 | ---?? | 0 | 0.8233 | intergenic | |
| rs6114339 | 20 | 23833711 | 4 | 0 | 0 | T | C | 1.5526 | 0.4182 | 0.000205 | +++?+ | 0.2 | 0.3908 | intergenic | |
| rs117270690 | 18 | 40638683 | 4 | 0 | 0 | T | C | 2.1904 | 0.59 | 0.000205 | +++?+ | 0 | 0.4406 | intron | RIT2 |
| rs6139895 | 20 | 6003893 | 5 | 0 | 0 | C | G | 0.769 | 0.2071 | 0.0002051 | ++++- | 0 | 0.4702 | intron | CRLS1 |
| rs6979027 | 7 | 141196846 | 3 | 0 | 0 | T | G | -6.6075 | 1.7797 | 0.0002051 | --??- | 43.4 | 0.1711 | intergenic | |
| rs427534 | 7 | 101886704 | 4 | 0 | 0 | A | G | 0.9445 | 0.2544 | 0.0002052 | +++?+ | 0 | 0.8064 | intron | CUX1 |
| rs7538578 | 1 | 90257041 | 5 | 0 | 0 | A | G | 0.7795 | 0.21 | 0.0002052 | ++++- | 0 | 0.6 | intergenic | |
| rs6114338 | 20 | 23833705 | 4 | 0 | 0 | T | G | 1.5525 | 0.4182 | 0.0002053 | +++?+ | 0.1 | 0.391 | intergenic | |
| rs2381667 | 9 | 7842823 | 4 | 0 | 0 | A | C | -1.1911 | 0.3209 | 0.0002054 | ---?- | 19.1 | 0.2947 | intergenic | |
| rs114159804 | 2 | 168023372 | 4 | 0 | 0 | A | G | 2.3138 | 0.6233 | 0.0002054 | +++?- | 0 | 0.6525 | intron | XIRP2 |
| rs114466899 | 2 | 35755298 | 3 | 0 | 0 | A | G | 3.5707 | 0.9619 | 0.0002054 | +++?? | 0 | 0.861 | intergenic | |
| 6:32631397 | 6 | 32631397 | 4 | 0 | 0 | A | C | -1.1632 | 0.3134 | 0.0002055 | ----? | 41.7 | 0.1615 | intron | HLA-DQB1 |
| rs13191565 | 6 | 32441577 | 5 | 0 | 1 | C | G | 1.4371 | 0.3871 | 0.0002055 | +++++ | 0 | 0.473 | intergenic | |
| 3:15950433 | 3 | 15950433 | 3 | 0 | 0 | G | GT | 2.3023 | 0.6202 | 0.0002055 | +++?? | 0 | 0.6853 | | |
| rs62031580 | 16 | 9297003 | 3 | 0 | 0 | A | G | 4.5926 | 1.2373 | 0.0002057 | ?++?+ | 0 | 0.772 | intergenic | |
| rs114567049 | 6 | 32601719 | 4 | 0 | 1 | A | G | 1.07 | 0.2883 | 0.0002058 | ++++? | 0 | 0.4461 | upstream-gene | |
| rs368639360 | 6 | 72869490 | 3 | 0 | 0 | T | C | -1.8163 | 0.4893 | 0.0002058 | ---?? | 0 | 0.9801 | intron | RIMS1 |
| rs6114341 | 20 | 23834281 | 4 | 0 | 0 | C | G | 1.5525 | 0.4183 | 0.0002059 | +++?+ | 1 | 0.387 | intergenic | |
| rs6114340 | 20 | 23834266 | 4 | 0 | 0 | A | G | 1.5525 | 0.4183 | 0.000206 | +++?+ | 0.9 | 0.3872 | intergenic | |
| rs71632791 | 4 | 190758511 | 3 | 0 | 0 | T | C | -1.2277 | 0.3308 | 0.000206 | ---?? | 0 | 0.4179 | intergenic | |
| rs12433153 | 14 | 44911852 | 4 | 0 | 0 | A | T | 1.4545 | 0.3919 | 0.0002061 | +++?+ | 0 | 0.8317 | intergenic | |
| rs181639231 | 5 | 123511933 | 3 | 0 | 0 | A | T | -0.946 | 0.2549 | 0.0002062 | ---?? | 0 | 0.8333 | intergenic | |
| 16:84842863 | 16 | 84842863 | 3 | 0 | 0 | A | AGTGAT | 1.3846 | 0.3731 | 0.0002062 | +++?? | 0 | 0.8744 | | |
| rs57899856 | 3 | 183188619 | 4 | 0 | 0 | T | C | 1.5277 | 0.4117 | 0.0002063 | +++?+ | 10.8 | 0.3391 | intergenic | |
| rs115870168 | 6 | 109099881 | 2 | 0 | 0 | C | G | 5.1726 | 1.3938 | 0.0002063 | +?+?? | 0 | 0.7739 | intergenic | |
| rs34529082 | 16 | 7208279 | 4 | 0 | 0 | A | G | 1.1409 | 0.3075 | 0.0002065 | +++?+ | 0 | 0.7814 | intron | RBFOX1 |
| rs10486616 | 7 | 33785578 | 4 | 0 | 0 | T | C | 0.9376 | 0.2527 | 0.0002066 | +++?+ | 17.8 | 0.3016 | intergenic | |
| rs17204696 | 2 | 208343421 | 4 | 0 | 0 | A | G | -1.5611 | 0.4207 | 0.0002066 | ---?- | 0 | 0.4264 | intergenic | |
| rs61374672 | 9 | 79458516 | 4 | 0 | 0 | T | C | 2.1686 | 0.5844 | 0.0002066 | +++?+ | 0 | 0.5308 | intron | PRUNE2 |
| 16:2487904 | 16 | 2487904 | 3 | 0 | 0 | T | TA | 1.2146 | 0.3273 | 0.0002066 | +++?? | 29.3 | 0.243 | | |
| 6:72869501 | 6 | 72869501 | 3 | 0 | 0 | A | T | -1.8169 | 0.4896 | 0.0002067 | ---?? | 0 | 0.9783 | intron | RIMS1 |
| rs9393521 | 6 | 24062856 | 5 | 0 | 0 | T | C | 1.5057 | 0.4058 | 0.0002067 | ++++- | 0 | 0.6182 | intergenic | |
| rs1171726 | 10 | 131910952 | 4 | 0 | 0 | A | C | 1.031 | 0.2779 | 0.0002068 | +++?+ | 0 | 0.9133 | upstream-gene | |
| rs2123489 | 8 | 113440838 | 4 | 0 | 0 | A | G | -1.028 | 0.2771 | 0.000207 | ---?- | 0 | 0.7936 | intron | CSMD3 |

| | | | | | | | | | | | | | | | |
|---|---|---|---|---|---|---|---|---|---|---|---|---|---|---|---|
| rs4233831 | 2 | 170598331 | 4 | 0 | 0 | A | G | -1.2279 | 0.3309 | 0.000207 | ---?- | 0 | 0.6395 | intron | KLHL23,PHOSPHO2-KLHL23 |
| rs2150312 | 14 | 96973712 | 3 | 0 | 0 | T | C | -1.0258 | 0.2765 | 0.0002071 | ---?? | 0 | 0.6046 | intron | PAPOLA |
| rs11048636 | 12 | 26820414 | 4 | 0 | 0 | A | G | 1.783 | 0.4806 | 0.0002073 | +-+?+ | 49.1 | 0.1168 | intron | ITPR2 |
| rs2167777 | 8 | 113441240 | 4 | 0 | 0 | T | C | -1.0279 | 0.2771 | 0.0002074 | ---?- | 0 | 0.7937 | intron | CSMD3 |
| rs55789914 | 4 | 183654596 | 3 | 0 | 0 | A | C | 1.0257 | 0.2765 | 0.0002076 | +++?? | 0 | 0.913 | intron | TENM3 |
| rs13217979 | 6 | 32441126 | 5 | 0 | 1 | T | C | 1.4359 | 0.3871 | 0.0002079 | +++++ | 0 | 0.4722 | intergenic | |
| rs12037938 | 1 | 90254973 | 5 | 0 | 0 | T | C | 0.778 | 0.2098 | 0.0002081 | ++++- | 0 | 0.5184 | intergenic | |
| rs16883689 | 8 | 113485802 | 4 | 0 | 0 | A | G | 1.0518 | 0.2836 | 0.0002081 | +++?+ | 0 | 0.7526 | intron | CSMD3 |
| rs1418297 | 6 | 82704394 | 4 | 0 | 0 | A | G | 2.5575 | 0.6896 | 0.0002082 | +++?+ | 0 | 0.8016 | intergenic | |
| rs2765489 | 10 | 131912151 | 4 | 0 | 0 | T | C | 1.0261 | 0.2767 | 0.0002082 | +++?+ | 0 | 0.9127 | upstream-gene | |
| 2:86823958 | 2 | 86823958 | 3 | 0 | 0 | CT | C | -0.9377 | 0.2528 | 0.0002082 | ---?? | 1 | 0.3642 | | |
| rs6846433 | 4 | 38747187 | 5 | 0 | 0 | A | G | 1.2282 | 0.3311 | 0.0002083 | +++++ | 0 | 0.972 | intergenic | |
| rs4658325 | 1 | 90260235 | 5 | 0 | 0 | T | C | 0.7782 | 0.2098 | 0.0002085 | ++++- | 0 | 0.607 | intergenic | |
| rs74533782 | 20 | 23813723 | 4 | 0 | 0 | A | G | 1.6278 | 0.439 | 0.0002085 | +++?+ | 0 | 0.4446 | intergenic | |
| rs4658324 | 1 | 90260133 | 5 | 0 | 0 | A | C | -0.7782 | 0.2099 | 0.0002087 | ----+ | 0 | 0.6069 | intergenic |  |
| rs11033907 | 11 | 37061128 | 4 | 0 | 0 | T | G | -0.9017 | 0.2432 | 0.0002087 | ---?- | 0 | 0.4307 | intergenic | |
| rs28707773 | 6 | 32572278 | 5 | 0 | 1 | A | C | 1.4203 | 0.383 | 0.0002087 | +++++ | 22.8 | 0.2692 | intergenic | |
| rs4846895 | 1 | 230196165 | 4 | 0 | 0 | T | C | 1.4021 | 0.3781 | 0.0002088 | +-+?+ | 54.9 | 0.08374 | intergenic | |
| rs67970757 | 6 | 32440880 | 5 | 0 | 1 | A | G | 1.4353 | 0.3871 | 0.0002091 | +++++ | 0 | 0.4718 | intergenic | |
| 17:52274715 | 17 | 52274715 | 3 | 0 | 0 | A | G | 2.562 | 0.691 | 0.0002093 | +++?? | 0 | 0.7363 | intergenic | |
| rs2588536 | 4 | 37007538 | 3 | 0 | 0 | T | C | -0.9581 | 0.2584 | 0.0002093 | ---?? | 27.2 | 0.2534 | intergenic | |
| rs9348628 | 6 | 24066519 | 5 | 0 | 0 | T | C | 1.4998 | 0.4045 | 0.0002093 | ++++- | 0 | 0.6045 | intergenic | |
| rs1866714 | 11 | 12810971 | 4 | 0 | 0 | A | G | -0.9831 | 0.2652 | 0.0002094 | +--?- | 51 | 0.106 | intron | TEAD1 |
| rs11031148 | 11 | 30633117 | 4 | 0 | 0 | A | C | 1.0009 | 0.27 | 0.0002095 | +++?- | 31.4 | 0.224 | intergenic | |
| rs28661390 | 8 | 113417273 | 3 | 0 | 0 | A | G | 1.082 | 0.2919 | 0.0002095 | +++?? | 0 | 0.5958 | intron | CSMD3 |
| rs187063742 | 2 | 51056194 | 3 | 0 | 0 | A | G | -1.2085 | 0.326 | 0.0002095 | ---?? | 0 | 0.4247 | intron | NRXN1 |
| rs2477532 | 10 | 131910301 | 4 | 0 | 0 | T | G | -1.0315 | 0.2782 | 0.0002096 | ---?- | 0 | 0.9134 | upstream-gene | |
| rs3916255 | 16 | 12093755 | 4 | 0 | 0 | C | G | -0.9189 | 0.2479 | 0.0002099 | +--?- | 60.5 | 0.0551 | intron | SNX29 |
| rs73157791 | 7 | 82407344 | 4 | 0 | 0 | A | G | -1.0166 | 0.2743 | 0.0002099 | ---?- | 20.5 | 0.2872 | intron | PCLO |
| rs184085239 | 4 | 100510073 | 3 | 0 | 0 | T | C | 1.5962 | 0.4307 | 0.0002104 | -++?? | 53.8 | 0.1148 | intron | MTTP |
| rs62194758 | 20 | 23856029 | 4 | 0 | 0 | A | C | 1.5606 | 0.4211 | 0.0002106 | +++?+ | 12.6 | 0.3298 | downstream-gene | |
| rs12706946 | 7 | 130292531 | 4 | 0 | 0 | T | C | 0.9126 | 0.2463 | 0.0002107 | +-+?+ | 57.2 | 0.07174 | intron | COPG2 |
| rs1336517 | 13 | 60034724 | 4 | 0 | 0 | T | C | 0.9529 | 0.2572 | 0.0002108 | +++?+ | 26.3 | 0.2542 | intergenic | |
| rs62288981 | 3 | 185915812 | 3 | 0 | 0 | A | G | -2.0509 | 0.5535 | 0.0002108 | ---?? | 0 | 0.6785 | intron | DGKG |
| rs5014672 | 8 | 113417475 | 3 | 0 | 0 | T | C | -1.0841 | 0.2926 | 0.0002109 | ---?? | 0 | 0.6079 | intron | CSMD3 |
| rs5014673 | 8 | 113417473 | 3 | 0 | 0 | T | C | -1.0841 | 0.2926 | 0.000211 | ---?? | 0 | 0.6083 | intron | CSMD3 |
| rs8068065 | 17 | 52279645 | 3 | 0 | 0 | C | G | 2.2109 | 0.5967 | 0.000211 | +++?? | 0 | 0.7438 | intergenic | |
| rs1392677 | 15 | 98046180 | 4 | 0 | 0 | A | C | 0.9801 | 0.2645 | 0.0002111 | +++?+ | 0 | 0.9256 | intergenic | |
| rs73509106 | 6 | 109081395 | 2 | 0 | 0 | A | G | 5.3685 | 1.449 | 0.0002113 | +?+?? | 0 | 0.5737 | intron | LINC00222 |
| rs141002583 | 5 | 109412843 | 4 | 0 | 0 | C | G | 2.1161 | 0.5711 | 0.0002113 | +++?+ | 11.7 | 0.3343 | intergenic | |
| rs72883348 | 2 | 166648027 | 4 | 0 | 0 | T | C | 2.7013 | 0.7291 | 0.0002114 | +++?- | 50.9 | 0.1065 | intron | GALNT3 |
| rs9379633 | 6 | 24057572 | 5 | 0 | 0 | A | G | -1.5137 | 0.4086 | 0.0002118 | ----+ | 0 | 0.7015 | intergenic |  |
| rs62194737 | 20 | 23838795 | 4 | 0 | 0 | T | C | 1.5509 | 0.4187 | 0.0002119 | +++?+ | 5.4 | 0.3662 | intergenic | |
| rs56407831 | 10 | 82010606 | 3 | 0 | 0 | T | C | 1.0012 | 0.2703 | 0.0002121 | +++?? | 34.8 | 0.2158 | intergenic | |
| rs145964215 | 19 | 41507057 | 3 | 0 | 0 | T | C | -3.7113 | 1.0019 | 0.0002121 | ---?? | 0 | 0.9585 | intron | CYP2B6 |
| rs71599527 | 5 | 10304213 | 4 | 0 | 0 | T | C | -1.9752 | 0.5333 | 0.0002122 | ---?- | 27 | 0.2497 | intron | CMBL |
| rs6493008 | 15 | 41996222 | 5 | 0 | 0 | C | G | 1.0513 | 0.2838 | 0.0002124 | +++++ | 18.5 | 0.2966 | intron | MGA |
| rs1354027 | 15 | 98038605 | 4 | 0 | 0 | T | G | 0.946 | 0.2554 | 0.0002126 | +++?+ | 0 | 0.9875 | intergenic | |
| 6:83802367 | 6 | 83802367 | 3 | 0 | 0 | G | GTT | -3.4598 | 0.9342 | 0.0002127 | ---?? | 0 | 0.5443 | | |
| 7:98428439 | 7 | 98428439 | 3 | 0 | 0 | T | TTA | 1.2347 | 0.3334 | 0.0002128 | +++?? | 0 | 0.7717 | | |
| rs12612349 | 2 | 40473047 | 4 | 0 | 0 | T | C | 1.3899 | 0.3753 | 0.0002131 | +++?- | 1.1 | 0.3866 | intron | SLC8A1,SLC8A1-AS1 |
| rs61900277 | 11 | 110689944 | 4 | 0 | 0 | T | G | 2.6758 | 0.7227 | 0.0002133 | +++?+ | 0 | 0.9906 | intergenic | |
| rs9379635 | 6 | 24067548 | 5 | 0 | 0 | T | C | -1.4979 | 0.4046 | 0.0002134 | ----+ | 0 | 0.5992 | intergenic |  |
| rs76422810 | 17 | 56820982 | 5 | 0 | 0 | A | G | 2.6082 | 0.7044 | 0.0002134 | +++++ | 0 | 0.8011 | intergenic | |
| 1:204386132 | 1 | 204386132 | 3 | 0 | 0 | AGG | AGGG | -1.6228 | 0.4383 | 0.0002134 | ---?? | 57.9 | 0.09279 | | |
| rs1392678 | 15 | 98046162 | 4 | 0 | 0 | T | C | 0.9795 | 0.2646 | 0.0002135 | +++?+ | 0 | 0.9265 | intergenic | |
| rs115648680 | 6 | 23846176 | 4 | 0 | 0 | T | C | 2.8259 | 0.7633 | 0.0002135 | +++?- | 34 | 0.2083 | intergenic | |
| rs12148330 | 15 | 41995199 | 5 | 0 | 0 | A | T | 1.0533 | 0.2845 | 0.0002137 | +++++ | 16.1 | 0.3118 | intron | MGA |
| rs35954087 | 6 | 32440049 | 5 | 0 | 1 | T | C | 1.433 | 0.3871 | 0.0002138 | +++++ | 0 | 0.4702 | intergenic | |

| | | | | | | | | | | | | | | | |
|---|---|---|---|---|---|---|---|---|---|---|---|---|---|---|---|
| rs12524496 | 6 | 32438513 | 5 | 0 | 1 | C | G | 1.4329 | 0.3871 | 0.0002142 | +++++ | 0 | 0.4699 | intergenic | |
| rs12525692 | 6 | 32438546 | 5 | 0 | 1 | A | T | -1.4329 | 0.3871 | 0.0002142 | ----- | 0 | 0.4699 | intergenic |  |
| rs13193827 | 6 | 32430337 | 5 | 0 | 1 | A | C | 1.4329 | 0.3871 | 0.0002142 | +++++ | 0 | 0.4699 | intergenic | |
| rs17202892 | 6 | 32434479 | 5 | 0 | 1 | A | C | 1.4329 | 0.3871 | 0.0002142 | +++++ | 0 | 0.4699 | intergenic | |
| rs28377518 | 6 | 32432650 | 5 | 0 | 1 | T | C | 1.4329 | 0.3871 | 0.0002142 | +++++ | 0 | 0.4699 | intergenic | |
| rs34200087 | 6 | 32438691 | 5 | 0 | 1 | A | G | 1.4329 | 0.3871 | 0.0002142 | +++++ | 0 | 0.4699 | intergenic | |
| rs35458464 | 6 | 32431239 | 5 | 0 | 1 | A | G | -1.4329 | 0.3871 | 0.0002142 | ----- | 0 | 0.4699 | intergenic |  |
| rs67209722 | 6 | 32439782 | 5 | 0 | 1 | T | C | -1.4329 | 0.3871 | 0.0002142 | ----- | 0 | 0.47 | intergenic |  |
| 19:58170219 | 19 | 58170219 | 3 | 0 | 0 | A | G | -2.4358 | 0.6581 | 0.0002143 | ---?? | 31.3 | 0.2333 | intergenic | |
| rs34750784 | 11 | 12837644 | 4 | 0 | 0 | A | G | -1.3034 | 0.3521 | 0.0002144 | ---?- | 5 | 0.3681 | intron | TEAD1 |
| rs2925337 | 15 | 42065422 | 5 | 0 | 0 | A | C | 1.0542 | 0.2848 | 0.0002145 | +++++ | 17.6 | 0.3025 | upstream-gene | |
| rs9379636 | 6 | 24069649 | 5 | 0 | 0 | T | G | 1.4986 | 0.4049 | 0.0002145 | ++++- | 0 | 0.5809 | intergenic | |
| rs117865215 | 12 | 43221075 | 4 | 0 | 0 | C | G | -1.7835 | 0.4819 | 0.0002146 | ---?- | 0 | 0.8413 | intergenic | |
| 7:46177065 | 7 | 46177065 | 3 | 0 | 0 | G | GTAGA | -1.3369 | 0.3613 | 0.000215 | ---?? | 12.3 | 0.3197 | | |
| rs2695169 | 15 | 42035713 | 5 | 0 | 0 | A | C | -1.0471 | 0.283 | 0.0002151 | ----- | 14.2 | 0.3236 | intron | MGA |
| rs28986347 | 6 | 32749109 | 5 | 0 | 0 | A | G | -1.5023 | 0.406 | 0.0002151 | ----- | 0 | 0.7285 | intergenic |  |
| rs28986348 | 6 | 32749122 | 5 | 0 | 0 | T | C | 1.5024 | 0.406 | 0.0002151 | +++++ | 0 | 0.7285 | intergenic | |
| rs28986349 | 6 | 32749140 | 5 | 0 | 0 | A | G | 1.5024 | 0.406 | 0.0002151 | +++++ | 0 | 0.7285 | intergenic | |
| rs28986350 | 6 | 32749175 | 5 | 0 | 0 | T | C | 1.5024 | 0.406 | 0.0002151 | +++++ | 0 | 0.7285 | intergenic | |
| rs7745167 | 6 | 144755448 | 3 | 0 | 0 | A | C | 3.0596 | 0.8268 | 0.0002152 | +++?? | 0 | 0.6118 | intron | UTRN |
| 8:113423228 | 8 | 113423228 | 3 | 0 | 0 | T | G | -1.0796 | 0.2917 | 0.0002153 | ---?? | 0 | 0.5969 | intron | CSMD3 |
| rs74520856 | 14 | 99696075 | 4 | 0 | 0 | A | G | -4.2981 | 1.1615 | 0.0002153 | ---?- | 0 | 0.8336 | intron | BCL11B |
| 8:113423229 | 8 | 113423229 | 3 | 0 | 0 | T | G | -1.0795 | 0.2917 | 0.0002154 | ---?? | 0 | 0.5967 | intron | CSMD3 |
| rs9761947 | 4 | 125533861 | 3 | 0 | 0 | A | G | 1.2359 | 0.334 | 0.0002154 | +++?? | 0 | 0.9689 | intergenic | |
| 8:113423230 | 8 | 113423230 | 3 | 0 | 0 | T | G | -1.0794 | 0.2917 | 0.0002155 | ---?? | 0 | 0.5964 | intron | CSMD3 |
| rs7168151 | 15 | 54348185 | 4 | 0 | 0 | A | G | 1.2085 | 0.3266 | 0.0002158 | +++?+ | 0 | 0.9073 | intron | UNC13C |
| rs12527323 | 6 | 32429694 | 5 | 0 | 1 | A | C | 1.4322 | 0.3871 | 0.0002158 | +++++ | 0 | 0.4681 | intergenic | |
| rs13177199 | 5 | 10303738 | 4 | 0 | 0 | A | C | -1.9691 | 0.5323 | 0.0002159 | ---?- | 28.6 | 0.2407 | intron | CMBL |
| 8:113423237 | 8 | 113423237 | 3 | 0 | 0 | A | ATT | 1.0792 | 0.2917 | 0.0002159 | +++?? | 0 | 0.5962 | | |
| rs35421268 | 6 | 32435196 | 5 | 0 | 1 | T | C | 1.4322 | 0.3871 | 0.0002161 | +++++ | 0 | 0.4684 | intergenic | |
| rs142065062 | 8 | 123595415 | 2 | 0 | 0 | T | C | 5.4574 | 1.4752 | 0.0002161 | ?++?? | 0 | 0.9967 | intergenic | |
| rs150582433 | 2 | 170421707 | 4 | 0 | 0 | T | C | -1.4878 | 0.4022 | 0.0002162 | ---?- | 0 | 0.8245 | intron | FASTKD1 |
| 9:30873589 | 9 | 30873589 | 3 | 0 | 0 | A | AT | -3.6986 | 0.9998 | 0.0002162 | ---?? | 0 | 0.6442 | | |
| rs313888 | 4 | 125543019 | 4 | 0 | 0 | A | C | -0.966 | 0.2611 | 0.0002163 | ---?- | 0 | 0.9221 | intergenic | |
| rs331128 | 2 | 150866543 | 3 | 0 | 0 | T | C | 1.0444 | 0.2824 | 0.0002164 | +++?? | 0 | 0.6088 | intergenic | |
| rs17053965 | 9 | 90831332 | 4 | 0 | 0 | T | C | -1.8261 | 0.4937 | 0.0002164 | ---?+ | 0 | 0.7772 | intergenic | |
| rs74514210 | 9 | 101131044 | 4 | 0 | 0 | A | C | 3.7299 | 1.0083 | 0.0002164 | +++?- | 43.3 | 0.1516 | intron | GABBR2 |
| rs2695164 | 15 | 42046568 | 5 | 0 | 0 | A | G | 1.0515 | 0.2843 | 0.0002165 | +++++ | 15.3 | 0.3172 | intron | MGA |
| rs9791280 | 6 | 32428994 | 5 | 0 | 1 | C | G | 1.4327 | 0.3873 | 0.0002165 | +++++ | 0 | 0.4687 | intergenic | |
| rs4306395 | 13 | 60014071 | 3 | 0 | 0 | A | G | 0.9462 | 0.2558 | 0.0002166 | +++?? | 0 | 0.4909 | intergenic | |
| rs6858786 | 4 | 189067001 | 4 | 0 | 0 | A | G | -1.2986 | 0.3511 | 0.0002166 | ---?- | 0 | 0.8346 | intron | TRIML1 |
| rs28435616 | 6 | 32519057 | 4 | 0 | 1 | A | C | 1.4212 | 0.3842 | 0.0002166 | ++++? | 23.2 | 0.2719 | downstream-gene | |
| rs62017086 | 16 | 7191371 | 4 | 0 | 0 | C | G | -1.0177 | 0.2752 | 0.0002167 | ---?- | 0 | 0.4407 | intron | RBFOX1 |
| 19:36552850 | 19 | 36552850 | 3 | 0 | 0 | CT | C | -1.4721 | 0.398 | 0.0002168 | ---?? | 19.6 | 0.2882 | | |
| 4:183654595 | 4 | 183654595 | 3 | 0 | 0 | A | AC | 1.0217 | 0.2762 | 0.0002169 | +++?? | 0 | 0.9066 | | |
| rs2162517 | 2 | 170566881 | 4 | 0 | 0 | T | C | -1.3277 | 0.359 | 0.0002173 | ---?- | 0 | 0.9198 | intron | PHOSPHO2-KLHL23 |
| rs1466510 | 12 | 26804358 | 4 | 0 | 0 | A | G | -1.8923 | 0.5117 | 0.0002174 | ---?- | 15.6 | 0.3138 | intron | ITPR2 |
| rs7167665 | 15 | 53750574 | 4 | 0 | 0 | T | C | -0.8776 | 0.2373 | 0.0002174 | ---?- | 49.3 | 0.1159 | intergenic | |
| rs148348800 | 2 | 170418867 | 4 | 0 | 0 | C | G | 1.4869 | 0.4021 | 0.0002175 | +++?+ | 0 | 0.8227 | intron | FASTKD1 |
| rs150183253 | 7 | 114644691 | 2 | 0 | 0 | T | C | -5.3244 | 1.4399 | 0.0002175 | ?--?? | 0 | 0.8816 | intron | MDFIC |
| rs34391029 | 11 | 122199277 | 4 | 0 | 0 | T | C | 2.1434 | 0.5797 | 0.0002176 | +++?+ | 0 | 0.6851 | intergenic | |
| rs36173887 | 6 | 32602086 | 5 | 0 | 1 | A | G | -1.4728 | 0.3983 | 0.0002178 | ----- | 18.8 | 0.2949 | upstream-gene |  |
| rs4460698 | 1 | 205034488 | 5 | 0 | 0 | A | G | -0.8434 | 0.2281 | 0.0002179 | ----- | 0 | 0.8541 | intron | CNTN2 |
| rs138090525 | 17 | 56826434 | 5 | 0 | 0 | T | C | -2.604 | 0.7043 | 0.0002179 | ----- | 0 | 0.8017 | intergenic |  |
| rs116270542 | 9 | 101131013 | 4 | 0 | 0 | T | C | 3.7279 | 1.0083 | 0.000218 | +++?- | 43.4 | 0.1514 | intron | GABBR2 |
| rs2028030 | 7 | 107360005 | 5 | 0 | 0 | T | C | -0.8023 | 0.217 | 0.0002181 | ----- | 34.9 | 0.1885 | downstream-gene |  |
| rs3134215 | 8 | 96559978 | 4 | 0 | 0 | T | C | 1.8121 | 0.4902 | 0.0002183 | +-+?+ | 44.4 | 0.1447 | intron | LOC100616530 |
| rs7000525 | 8 | 124704361 | 4 | 0 | 0 | T | C | -1.0102 | 0.2733 | 0.0002184 | ---?- | 0 | 0.7825 | intron | ANXA13 |
| rs4363229 | 8 | 113470609 | 4 | 0 | 0 | T | C | 1.0343 | 0.2798 | 0.0002188 | +++?+ | 0 | 0.7561 | intron | CSMD3 |

| | | | | | | | | | | | | | | | |
|---|---|---|---|---|---|---|---|---|---|---|---|---|---|---|---|
| rs35397258 | 18 | 47743810 | 4 | 0 | 0 | T | C | 1.0765 | 0.2912 | 0.0002188 | +++?+ | 16.7 | 0.3079 | intergenic | |
| rs114955471 | 2 | 168027736 | 4 | 0 | 0 | A | G | -2.2963 | 0.6213 | 0.0002188 | ---?+ | 0 | 0.6577 | intron | XIRP2 |
| rs31203 | 5 | 134351470 | 5 | 0 | 0 | T | C | 0.86 | 0.2327 | 0.0002189 | +++++ | 0 | 0.4734 | downstream-gene | |
| rs17095763 | 10 | 118920455 | 4 | 0 | 0 | A | C | 1.5317 | 0.4144 | 0.0002191 | +++?+ | 0 | 0.8847 | intergenic | |
| rs147986014 | 6 | 83761723 | 4 | 0 | 0 | C | G | 3.5183 | 0.9519 | 0.0002191 | +++?+ | 0 | 0.7202 | intron | UBE3D |
| rs12693084 | 2 | 176738510 | 4 | 0 | 0 | C | G | 0.9138 | 0.2473 | 0.0002196 | +++?+ | 0 | 0.8157 | intergenic | |
| rs1625987 | 14 | 43236006 | 4 | 0 | 0 | A | G | -1.406 | 0.3805 | 0.0002197 | ---?+ | 61.4 | 0.05082 | intergenic | |
| 7:35974808 | 7 | 35974808 | 3 | 0 | 0 | A | AACACA | 4.3178 | 1.1685 | 0.0002197 | +++?? | 0 | 0.5974 | | |
| rs6730147 | 2 | 156928123 | 4 | 0 | 0 | T | C | 1.0003 | 0.2707 | 0.0002202 | +++?+ | 0 | 0.8788 | intergenic | |
| rs72975520 | 2 | 235408509 | 4 | 0 | 0 | A | G | -1.412 | 0.3822 | 0.0002203 | ---?- | 0 | 0.7904 | upstream-gene | |
| rs3026098 | 17 | 5276611 | 5 | 0 | 0 | A | G | -2.6949 | 0.7295 | 0.0002206 | --+-- | 55.3 | 0.06251 | synonymous | RABEP1 |
| rs116996945 | 12 | 117281231 | 3 | 0 | 0 | A | G | -2.6182 | 0.7087 | 0.0002206 | ---?? | 0 | 0.5212 | intron | RNFT2 |
| rs3753036 | 1 | 203677250 | 4 | 0 | 0 | A | G | -1.1951 | 0.3235 | 0.0002207 | ---?- | 0 | 0.9542 | intron | ATP2B4 |
| rs199587127 | 1 | 6789104 | 3 | 0 | 0 | T | C | 2.7343 | 0.7402 | 0.0002207 | +++?? | 0 | 0.4787 | intergenic | |
| rs10798909 | 1 | 30364081 | 4 | 0 | 0 | T | C | -1.3188 | 0.357 | 0.0002209 | ---?+ | 7.8 | 0.354 | intergenic | |
| rs145190985 | 2 | 168027563 | 4 | 0 | 0 | T | C | -2.2949 | 0.6213 | 0.0002209 | ---?+ | 0 | 0.6584 | intron | XIRP2 |
| 5:172634939 | 5 | 172634939 | 3 | 0 | 0 | CAAACA | C | 4.0654 | 1.1006 | 0.0002209 | +++?? | 0 | 0.5413 | | |
| rs12875806 | 13 | 100413797 | 4 | 0 | 0 | T | G | -1.0502 | 0.2843 | 0.000221 | ---?- | 48.9 | 0.1182 | intron | CLYBL |
| rs13220931 | 7 | 33790575 | 3 | 0 | 0 | T | G | 0.9432 | 0.2554 | 0.0002211 | +++?? | 22.7 | 0.2744 | intergenic | |
| rs3824249 | 8 | 124709106 | 5 | 0 | 0 | C | G | -0.889 | 0.2407 | 0.0002212 | ----+ | 0 | 0.4784 | intron | ANXA13 |
| rs10268857 | 7 | 141200063 | 3 | 0 | 0 | A | C | -6.2477 | 1.6916 | 0.0002214 | --??- | 44.3 | 0.1663 | intergenic | |
| rs114614561 | 2 | 168028451 | 4 | 0 | 0 | A | C | -2.2927 | 0.6208 | 0.0002214 | ---?+ | 0 | 0.6589 | intron | XIRP2 |
| rs148494466 | 4 | 190304364 | 3 | 0 | 0 | T | C | 5.3782 | 1.4563 | 0.0002216 | +++?? | 0 | 0.8212 | intergenic | |
| rs1579929 | 1 | 30361904 | 4 | 0 | 0 | A | C | -1.3228 | 0.3582 | 0.0002217 | ---?+ | 7.9 | 0.3534 | intergenic | |
| rs79073005 | 17 | 56655482 | 5 | 0 | 0 | T | C | -2.4885 | 0.6739 | 0.0002217 | ----- | 0 | 0.8417 | intron | TEX14 |
| rs469750 | 5 | 127637634 | 5 | 0 | 0 | A | T | 0.8757 | 0.2371 | 0.0002218 | -++++ | 24.7 | 0.2566 | intron | FBN2 |
| rs55856558 | 4 | 38748027 | 5 | 0 | 0 | T | C | -1.2342 | 0.3342 | 0.0002218 | ----- | 0 | 0.9629 | intergenic |  |
| rs9366559 | 6 | 24078056 | 3 | 0 | 0 | T | C | 1.7651 | 0.478 | 0.0002219 | +++?? | 0 | 0.4503 | intergenic | |
| rs12875158 | 13 | 100413876 | 4 | 0 | 0 | A | G | 1.0499 | 0.2843 | 0.0002221 | +++?+ | 48.9 | 0.1181 | intron | CLYBL |
| 4:123801400 | 4 | 123801400 | 3 | 0 | 0 | CT | C | 1.9345 | 0.5239 | 0.0002221 | +++?? | 0 | 0.6661 | | |
| rs2695162 | 15 | 42039902 | 5 | 0 | 0 | A | G | -1.0451 | 0.283 | 0.0002222 | ----- | 14.1 | 0.3242 | intron | MGA |
| rs113201623 | 18 | 40046670 | 4 | 0 | 0 | A | T | 1.4149 | 0.3833 | 0.0002227 | +++?+ | 0 | 0.8158 | intron | LINC00907 |
| rs74586884 | 18 | 40047207 | 4 | 0 | 0 | A | C | -1.4149 | 0.3833 | 0.0002228 | ---?- | 0 | 0.8158 | intron | LINC00907 |
| rs111866406 | 18 | 40045577 | 4 | 0 | 0 | T | C | -1.4149 | 0.3833 | 0.0002228 | ---?- | 0 | 0.8158 | intron | LINC00907 |
| rs150019899 | 17 | 56836563 | 5 | 0 | 0 | A | G | -2.5968 | 0.7035 | 0.0002231 | ----- | 0 | 0.8039 | intron | PPM1E |
| rs2577950 | 15 | 42032655 | 5 | 0 | 0 | T | C | 1.0427 | 0.2825 | 0.0002233 | +++++ | 12.8 | 0.3322 | intron | MGA |
| rs9949668 | 18 | 47735555 | 4 | 0 | 0 | T | C | -1.0447 | 0.283 | 0.0002233 | ---?- | 14.6 | 0.319 | intergenic | |
| rs4912570 | 3 | 184216323 | 4 | 0 | 0 | C | G | 1.1027 | 0.2988 | 0.0002234 | +++?+ | 0 | 0.5434 | intergenic | |
| rs138326378 | 9 | 119639179 | 3 | 0 | 0 | T | C | 3.6659 | 0.9932 | 0.0002234 | +++?? | 58.3 | 0.09086 | intron | ASTN2 |
| rs1983777 | 11 | 30631366 | 4 | 0 | 0 | C | G | -0.9959 | 0.2698 | 0.0002235 | ---?+ | 30.7 | 0.2281 | intergenic | |
| rs114224242 | 2 | 168030107 | 4 | 0 | 0 | T | C | 2.2871 | 0.6197 | 0.0002235 | +++?- | 0 | 0.6603 | intron | XIRP2 |
| rs148509474 | 6 | 83798385 | 4 | 0 | 0 | T | C | 3.3476 | 0.907 | 0.0002236 | +++?+ | 0 | 0.7378 | intron | DOPEY1 |
| rs71365489 | 19 | 55238463 | 3 | 0 | 0 | A | G | -1.0157 | 0.2752 | 0.0002237 | ---?? | 14.3 | 0.3115 | intron | KIR3DL3 |
| rs11849911 | 14 | 22533591 | 4 | 0 | 0 | C | G | 0.9688 | 0.2625 | 0.0002238 | +++?+ | 0 | 0.6346 | intergenic | |
| rs17023923 | 2 | 39761367 | 4 | 0 | 0 | C | G | -1.6304 | 0.4418 | 0.0002239 | ---?- | 0 | 0.76 | intron | LOC728730 |
| rs9661104 | 1 | 229159552 | 4 | 0 | 0 | T | C | -1.3359 | 0.362 | 0.000224 | ---?- | 0 | 0.995 | intergenic | |
| rs17426565 | 6 | 32607356 | 5 | 0 | 1 | A | G | 1.5719 | 0.426 | 0.0002243 | +++++ | 9.5 | 0.3519 | intron | HLA-DQA1 |
| rs7038980 | 9 | 7846040 | 4 | 0 | 0 | A | G | -1.1809 | 0.32 | 0.0002244 | ---?- | 19.8 | 0.2911 | intergenic | |
| rs73485730 | 9 | 90812034 | 4 | 0 | 0 | T | C | 1.9313 | 0.5234 | 0.0002246 | +++?+ | 0 | 0.8077 | intergenic | |
| rs12024004 | 1 | 90265039 | 5 | 0 | 0 | T | C | -0.7715 | 0.2091 | 0.0002247 | ----+ | 0 | 0.5653 | intergenic |  |
| 6:30980562 | 6 | 30980562 | 3 | 0 | 0 | CGCCGT | C | -1.4922 | 0.4044 | 0.0002247 | ---?? | 0 | 0.7252 | | |
| rs75484580 | 5 | 38898593 | 4 | 0 | 0 | T | C | -3.2535 | 0.8819 | 0.0002248 | ---?- | 0 | 0.6524 | intron | OSMR |
| rs151002670 | 6 | 32170837 | 5 | 0 | 0 | T | C | -1.8148 | 0.4919 | 0.0002248 | ----+ | 22.5 | 0.271 | intron | NOTCH4 |
| 8:113431450 | 8 | 113431450 | 3 | 0 | 0 | G | GT | -1.0704 | 0.2901 | 0.0002249 | ---?? | 0 | 0.5725 | | |
| rs2858314 | 6 | 32663448 | 3 | 0 | 0 | A | G | 1.2312 | 0.3338 | 0.0002251 | +++?? | 25.9 | 0.2595 | intergenic | |
| rs78180536 | 5 | 122542394 | 4 | 0 | 0 | T | C | -1.9777 | 0.5361 | 0.0002252 | ---?- | 0 | 0.8357 | intergenic | |
| rs10801778 | 1 | 90264689 | 5 | 0 | 0 | T | G | -0.7716 | 0.2092 | 0.0002253 | ----+ | 0 | 0.5646 | intergenic |  |
| rs56288411 | 7 | 156177177 | 4 | 0 | 0 | T | C | 0.9106 | 0.2469 | 0.0002253 | +++?+ | 30.8 | 0.2276 | intergenic | |
| rs4128796 | 8 | 139175708 | 3 | 0 | 0 | A | C | -1.2316 | 0.3339 | 0.0002255 | ---?? | 0 | 0.6609 | intron | FAM135B |

| | | | | | | | | | | | | | | | |
|---|---|---|---|---|---|---|---|---|---|---|---|---|---|---|---|
| rs10801777 | 1 | 90264465 | 5 | 0 | 0 | A | G | 0.7716 | 0.2092 | 0.0002257 | ++++- | 0 | 0.5643 | intergenic | |
| 4:41332615 | 4 | 41332615 | 3 | 0 | 0 | T | TTTTATT | -1.5164 | 0.4112 | 0.0002258 | ---?? | 0 | 0.5852 | | |
| rs76494412 | 19 | 55238016 | 4 | 0 | 0 | T | C | 0.8646 | 0.2344 | 0.000226 | ++++? | 27.6 | 0.2461 | intron | KIR3DL3 |
| rs72750371 | 9 | 97756345 | 4 | 0 | 0 | A | G | -1.8997 | 0.5151 | 0.0002262 | ---?- | 0 | 0.8595 | intron | C9orf3 |
| 15:85499897 | 15 | 85499897 | 3 | 0 | 0 | CTT | C | 1.0583 | 0.287 | 0.0002262 | +++?? | 0 | 0.5763 | | |
| rs75709284 | 12 | 45980957 | 3 | 0 | 0 | T | G | -4.0097 | 1.0873 | 0.0002263 | ?--?- | 0 | 0.9357 | intergenic | |
| rs6751177 | 2 | 170590610 | 4 | 0 | 0 | T | C | 0.895 | 0.2427 | 0.0002264 | +++?+ | 0 | 0.5931 | intron | KLHL23,PHOSPHO2-KLHL23 |
| rs62444695 | 7 | 16903047 | 4 | 0 | 0 | T | C | -1.7872 | 0.4846 | 0.0002264 | ---?- | 0 | 0.4469 | intron | AGR3 |
| 14:43273810 | 14 | 43273810 | 3 | 0 | 0 | A | ATGCCT | 1.4575 | 0.3953 | 0.0002266 | +++?? | 44.6 | 0.1644 | | |
| rs4782729 | 16 | 82822747 | 4 | 0 | 0 | T | C | -1.3688 | 0.3712 | 0.0002268 | ---?- | 55.8 | 0.07906 | intron | CDH13 |
| rs12269969 | 11 | 86122104 | 4 | 0 | 0 | T | C | 2.6636 | 0.7224 | 0.0002268 | +++?+ | 0 | 0.4997 | intron | CCDC81 |
| 10:28230429 | 10 | 28230429 | 3 | 0 | 0 | G | GAA | 0.9016 | 0.2446 | 0.0002271 | +++?? | 45.3 | 0.1605 | | |
| rs187179647 | 11 | 123151250 | 3 | 0 | 0 | A | G | 4.8458 | 1.3144 | 0.0002273 | +++?? | 50.2 | 0.1341 | intergenic | |
| rs77671383 | 2 | 117135584 | 4 | 0 | 0 | T | C | 1.9102 | 0.5182 | 0.0002275 | +++?+ | 0 | 0.5865 | intergenic | |
| rs79065409 | 2 | 168033550 | 4 | 0 | 0 | T | C | 2.2771 | 0.6177 | 0.0002275 | +++?- | 0 | 0.6627 | intron | XIRP2 |
| rs150157625 | 13 | 22353073 | 4 | 0 | 0 | T | C | 3.1162 | 0.8453 | 0.0002275 | +++?+ | 0 | 0.4195 | intergenic | |
| rs34755442 | 5 | 124741926 | 4 | 0 | 0 | T | C | -2.8714 | 0.7789 | 0.0002276 | ---?- | 0 | 0.6048 | intergenic | |
| rs143382777 | 6 | 119050027 | 4 | 0 | 0 | T | C | 3.4092 | 0.9249 | 0.0002278 | +++?+ | 0 | 0.87 | intergenic | |
| rs1049088 | 6 | 32629868 | 4 | 0 | 1 | A | G | -1.157 | 0.3139 | 0.0002279 | ----? | 47.4 | 0.1272 | synonymous | HLA-DQB1 |
| rs9455597 | 6 | 168601002 | 4 | 0 | 0 | T | C | 1.2151 | 0.3297 | 0.0002279 | +++?+ | 0 | 0.9195 | intergenic | |
| 5:179940699 | 5 | 179940699 | 3 | 0 | 0 | T | TTTTA | -1.1503 | 0.3121 | 0.000228 | ---?? | 4 | 0.353 | | |
| rs4606820 | 18 | 70535901 | 4 | 0 | 0 | C | G | 1.4934 | 0.4052 | 0.0002281 | +++?+ | 41.9 | 0.1603 | upstream-gene | |
| rs74945390 | 12 | 127551467 | 4 | 0 | 0 | C | G | 4.9764 | 1.3503 | 0.0002283 | -++?+ | 31 | 0.2261 | intergenic | |
| rs4906137 | 14 | 43251077 | 4 | 0 | 0 | T | C | 1.4119 | 0.3832 | 0.0002286 | +++?- | 60.7 | 0.0542 | intergenic | |
| rs12135351 | 1 | 90254276 | 5 | 0 | 0 | A | G | -0.7727 | 0.2097 | 0.0002286 | ----+ | 0 | 0.546 | intergenic |  |
| rs114085989 | 9 | 101130470 | 4 | 0 | 0 | T | C | 3.7136 | 1.0078 | 0.0002287 | +++?- | 43.7 | 0.1494 | intron | GABBR2 |
| rs10111988 | 8 | 113489468 | 4 | 0 | 0 | T | G | 1.0506 | 0.2851 | 0.0002288 | +++?+ | 0 | 0.7307 | intron | CSMD3 |
| rs10461651 | 5 | 50281855 | 4 | 0 | 0 | A | G | 1.2568 | 0.3411 | 0.0002289 | +++?+ | 0 | 0.969 | intergenic | |
| rs116185178 | 2 | 170568216 | 4 | 0 | 0 | T | C | -1.3225 | 0.3589 | 0.0002289 | ---?- | 0 | 0.9232 | intron | PHOSPHO2-KLHL23 |
| rs11781694 | 8 | 113451839 | 4 | 0 | 0 | T | G | -1.022 | 0.2774 | 0.0002292 | ---?- | 0 | 0.7943 | intron | CSMD3 |
| rs7850515 | 9 | 16194652 | 4 | 0 | 0 | A | G | -1.0198 | 0.2768 | 0.0002294 | ---?- | 0 | 0.9782 | intergenic | |
| rs11852571 | 15 | 53750430 | 4 | 0 | 0 | T | C | -0.8733 | 0.2371 | 0.0002298 | ---?- | 48.8 | 0.1186 | intergenic | |
| rs6736471 | 2 | 170547746 | 4 | 0 | 0 | A | T | 1.3128 | 0.3564 | 0.00023 | +++?+ | 0 | 0.9725 | intron | CCDC173 |
| rs62020494 | 15 | 88264638 | 4 | 0 | 0 | T | G | 1.0177 | 0.2763 | 0.00023 | +++?+ | 0 | 0.5467 | intergenic | |
| rs78004704 | 2 | 170568419 | 4 | 0 | 0 | A | C | 1.3221 | 0.3589 | 0.0002303 | +++?+ | 0 | 0.9229 | intron | PHOSPHO2-KLHL23 |
| rs1027222 | 4 | 100613610 | 4 | 0 | 0 | A | C | -2.0678 | 0.5614 | 0.0002305 | +--?- | 43.9 | 0.148 | intergenic | |
| rs1880937 | 15 | 42044291 | 5 | 0 | 0 | T | C | 1.0431 | 0.2832 | 0.0002305 | +++++ | 15.8 | 0.3139 | intron | MGA |
| rs73167383 | 7 | 82420324 | 4 | 0 | 0 | A | T | 0.9657 | 0.2622 | 0.0002305 | +-+?+ | 52.6 | 0.09662 | intron | PCLO |
| rs143108728 | 17 | 56379131 | 4 | 0 | 0 | T | C | -2.7758 | 0.7537 | 0.0002305 | ---?- | 53.8 | 0.08974 | 3-prime-UTR | BZRAP1 |
| 3:172330583 | 3 | 172330583 | 3 | 0 | 0 | T | C | 2.0152 | 0.5472 | 0.0002306 | +++?? | 0 | 0.7764 | intergenic | |
| rs28986395 | 6 | 32754388 | 5 | 0 | 0 | A | G | 1.4741 | 0.4003 | 0.0002307 | +++++ | 0 | 0.669 | intergenic | |
| rs6543464 | 2 | 107676307 | 4 | 0 | 0 | T | C | 1.0459 | 0.284 | 0.0002309 | +++?+ | 0 | 0.5758 | intergenic | |
| rs79013747 | 4 | 100615309 | 4 | 0 | 0 | C | G | -2.0724 | 0.5628 | 0.000231 | +--?- | 44.1 | 0.1469 | intergenic | |
| 2:137022080 | 2 | 137022080 | 3 | 0 | 0 | CT | C | -1.0967 | 0.2978 | 0.0002312 | ---?? | 0 | 0.9881 | | |
| rs2949220 | 2 | 107676619 | 4 | 0 | 0 | C | G | -1.0448 | 0.2837 | 0.0002313 | ---?- | 0 | 0.5722 | intergenic | |
| 14:62384369 | 14 | 62384369 | 3 | 0 | 0 | CT | C | -1.1802 | 0.3205 | 0.0002313 | ---?? | 0 | 0.4287 | | |
| 7:4674157 | 7 | 4674157 | 3 | 0 | 0 | A | AT | 1.2361 | 0.3357 | 0.0002313 | +++?? | 0 | 0.5831 | | |
| rs7446001 | 5 | 161408547 | 4 | 0 | 0 | A | T | -1.1669 | 0.3169 | 0.0002317 | ---?- | 0 | 0.8696 | intergenic | |
| rs77969886 | 9 | 106369779 | 3 | 0 | 0 | A | G | -1.8928 | 0.5141 | 0.0002317 | ---?? | 0 | 0.6594 | intergenic | |
| rs113733485 | 1 | 245101256 | 4 | 0 | 0 | A | C | 1.5805 | 0.4293 | 0.0002319 | +++?- | 0 | 0.4722 | intergenic | |
| rs117964330 | 13 | 33185297 | 4 | 0 | 0 | A | G | 5.8597 | 1.5917 | 0.0002319 | +++?+ | 2.6 | 0.3794 | intron | PDS5B |
| rs145336556 | 21 | 19116941 | 4 | 0 | 0 | C | G | 2.6428 | 0.7179 | 0.0002321 | +++?- | 0 | 0.5207 | intergenic | |
| rs148124593 | 4 | 100624149 | 4 | 0 | 0 | T | G | 2.0928 | 0.5685 | 0.0002321 | -++?+ | 44.8 | 0.1423 | intergenic | |
| rs2577947 | 15 | 42037570 | 5 | 0 | 0 | T | C | -1.0427 | 0.2833 | 0.0002322 | ----- | 13.2 | 0.33 | intron | MGA |
| rs185786041 | 11 | 933097 | 4 | 0 | 0 | T | C | -2.3282 | 0.6325 | 0.0002325 | ---?- | 0 | 0.7418 | intron | AP2A2 |
| rs503035 | 5 | 134353734 | 5 | 0 | 0 | A | G | 0.8446 | 0.2295 | 0.0002326 | +++++ | 0 | 0.5035 | intergenic | |
| rs77101096 | 4 | 100626668 | 4 | 0 | 0 | A | G | -2.0974 | 0.5699 | 0.0002326 | +--?- | 45 | 0.1414 | intergenic | |
| rs11608421 | 12 | 27778920 | 4 | 0 | 0 | A | T | -0.9469 | 0.2573 | 0.0002329 | ---?+ | 45 | 0.1412 | intron | PPFIBP1 |
| rs1531605 | 6 | 168583210 | 4 | 0 | 0 | A | G | 1.6433 | 0.4465 | 0.000233 | +++?+ | 0 | 0.5055 | intergenic | |

| | | | | | | | | | | | | | | | |
|---|---|---|---|---|---|---|---|---|---|---|---|---|---|---|---|
| 5:154651411 | 5 | 154651411 | 2 | 0 | 0 | A | ATTTTTT | -1.2767 | 0.3469 | 0.0002331 | ?--?? | 48.8 | 0.1622 | | |
| rs117451605 | 6 | 148210586 | 4 | 0 | 0 | A | G | 3.9762 | 1.0805 | 0.0002332 | +++?+ | 21.4 | 0.2817 | intergenic | |
| rs10109807 | 8 | 113456944 | 4 | 0 | 0 | T | C | -1.0209 | 0.2774 | 0.0002334 | ---?- | 0 | 0.7916 | intron | CSMD3 |
| rs78173096 | 2 | 168037709 | 4 | 0 | 0 | A | G | -2.2647 | 0.6155 | 0.0002335 | ---?+ | 0 | 0.6653 | intron | XIRP2 |
| rs4713407 | 6 | 30980972 | 3 | 0 | 0 | T | C | 1.4878 | 0.4043 | 0.0002336 | +++?? | 0 | 0.719 | intron | MUC22 |
| 2:131131481 | 2 | 131131481 | 3 | 0 | 0 | T | TAGTC | 1.1318 | 0.3076 | 0.0002336 | +++?? | 0.3 | 0.3669 | | |
| rs8041315 | 15 | 98036484 | 4 | 0 | 0 | T | C | 0.944 | 0.2565 | 0.0002337 | +++?+ | 0 | 0.9904 | intergenic | |
| 18:40436577 | 18 | 40436577 | 3 | 0 | 0 | G | GACAA | -4.8788 | 1.326 | 0.0002337 | ---?? | 0 | 0.5417 | | |
| rs115748396 | 1 | 31698400 | 4 | 0 | 0 | T | C | 4.2814 | 1.1637 | 0.000234 | +++?+ | 56.7 | 0.07442 | intron | NKAIN1 |
| rs76531892 | 17 | 56852962 | 5 | 0 | 0 | T | C | 2.5924 | 0.7046 | 0.0002341 | +++++ | 0 | 0.8011 | intron | PPM1E |
| rs3134214 | 8 | 96561107 | 4 | 0 | 0 | C | G | -1.7935 | 0.4875 | 0.0002343 | -+-?- | 42.4 | 0.157 | intron | LOC100616530 |
| rs111444636 | 3 | 172329792 | 3 | 0 | 0 | A | T | -1.9979 | 0.5431 | 0.0002343 | ---?? | 0 | 0.7571 | intergenic | |
| rs2695168 | 15 | 42034002 | 5 | 0 | 0 | T | C | 1.041 | 0.283 | 0.0002344 | +++++ | 12.4 | 0.3347 | intron | MGA |
| rs114261145 | 11 | 86153550 | 4 | 0 | 0 | T | C | 2.6346 | 0.7162 | 0.0002344 | +++?+ | 0 | 0.53 | intron | ME3 |
| rs5757454 | 22 | 39449014 | 4 | 0 | 0 | T | C | -1.6344 | 0.4443 | 0.0002345 | ---?- | 0 | 0.9778 | 3-prime-UTR | APOBEC3F |
| rs62094323 | 18 | 63865640 | 4 | 0 | 0 | A | G | 0.9006 | 0.2448 | 0.0002345 | +++?+ | 0 | 0.4525 | intergenic | |
| rs144159201 | 12 | 1470973 | 3 | 0 | 0 | A | G | -4.1874 | 1.1383 | 0.0002345 | ---?? | 0 | 0.4825 | intron | ERC1 |
| rs3207555 | 6 | 31236687 | 3 | 0 | 0 | A | G | 0.9926 | 0.2698 | 0.0002349 | +++?? | 56.4 | 0.1007 | 3-prime-UTR | HLA-C |
| rs12728379 | 1 | 57886354 | 4 | 0 | 0 | T | C | 1.8654 | 0.5072 | 0.0002349 | +++?+ | 0 | 0.6208 | intron | DAB1 |
| rs2415762 | 14 | 43247915 | 4 | 0 | 0 | A | C | 1.4077 | 0.3827 | 0.000235 | +++?- | 60.9 | 0.05328 | intergenic | |
| rs7240278 | 18 | 40022868 | 4 | 0 | 0 | T | C | -1.1683 | 0.3176 | 0.000235 | ---?+ | 0 | 0.4087 | intron | LINC00907 |
| rs1761833 | 6 | 117280229 | 4 | 0 | 0 | T | C | 0.8915 | 0.2424 | 0.0002351 | +++?+ | 0 | 0.9528 | intergenic | |
| rs1632196 | 13 | 45624269 | 4 | 0 | 0 | A | T | 2.2678 | 0.6166 | 0.0002352 | +++?- | 42 | 0.1594 | intergenic | |
| rs79017656 | 4 | 100633291 | 4 | 0 | 0 | A | G | -2.1075 | 0.5731 | 0.0002354 | +--?- | 45.4 | 0.1392 | intergenic | |
| rs73157796 | 7 | 82409987 | 4 | 0 | 0 | A | G | 0.9994 | 0.2718 | 0.0002356 | +++?+ | 38 | 0.1839 | intron | PCLO |
| 6:144763759 | 6 | 144763759 | 3 | 0 | 0 | A | AG | 3.1284 | 0.8507 | 0.0002356 | +++?? | 0 | 0.6697 | | |
| 12:41352153 | 12 | 41352153 | 3 | 0 | 0 | A | ATG | 1.2073 | 0.3283 | 0.0002357 | +++?? | 0 | 0.7696 | | |
| rs76137233 | 13 | 74988373 | 4 | 0 | 0 | T | C | 3.7345 | 1.0156 | 0.0002358 | +++?+ | 0 | 0.7828 | upstream-gene | |
| rs55905885 | 1 | 90263781 | 3 | 0 | 0 | A | G | 0.9185 | 0.2498 | 0.0002359 | +++?? | 0 | 0.5638 | intergenic | |
| 3:196474999 | 3 | 196474999 | 3 | 0 | 0 | CTT | C | -2.3128 | 0.629 | 0.0002359 | ---?? | 0 | 0.9534 | | |
| rs78676416 | 2 | 168040425 | 4 | 0 | 0 | A | G | -2.257 | 0.6138 | 0.000236 | ---?+ | 0 | 0.6671 | intron | XIRP2 |
| rs183564746 | 5 | 30670764 | 3 | 0 | 0 | A | T | 2.6038 | 0.7082 | 0.0002362 | +++?? | 37.1 | 0.2039 | intergenic | |
| 20:62526696 | 20 | 62526696 | 3 | 0 | 0 | A | G | 3.6726 | 0.9989 | 0.0002363 | +++?? | 0 | 0.8849 | intron | DNAJC5 |
| rs11779960 | 8 | 113460930 | 4 | 0 | 0 | A | C | -1.02 | 0.2774 | 0.0002365 | ---?- | 0 | 0.7899 | intron | CSMD3 |
| 6:83760545 | 6 | 83760545 | 3 | 0 | 0 | G | GTAATT | -3.6177 | 0.984 | 0.0002365 | ---?? | 0 | 0.574 | | |
| rs13358089 | 5 | 174021949 | 4 | 0 | 0 | A | C | -1.607 | 0.4371 | 0.0002367 | ---?- | 0 | 0.7773 | intergenic | |
| rs13108707 | 4 | 100639912 | 4 | 0 | 0 | A | G | -2.121 | 0.577 | 0.0002368 | +--?- | 47.4 | 0.1267 | intergenic | |
| rs7750916 | 6 | 144622347 | 5 | 0 | 0 | T | C | -1.4738 | 0.4009 | 0.0002369 | ----- | 28.2 | 0.2334 | intron | UTRN |
| rs139527737 | 17 | 56855344 | 5 | 0 | 0 | A | G | 2.5901 | 0.7046 | 0.000237 | +++++ | 0 | 0.802 | intron | PPM1E |
| rs79972201 | 2 | 117133950 | 4 | 0 | 0 | A | C | 1.902 | 0.5175 | 0.0002371 | +++?+ | 0 | 0.5892 | intergenic | |
| rs945017 | 14 | 101831671 | 4 | 0 | 0 | A | G | -1.5269 | 0.4154 | 0.0002373 | ---?- | 0 | 0.4797 | intergenic | |
| 6:83760542 | 6 | 83760542 | 3 | 0 | 0 | T | TGCATT | -3.617 | 0.9841 | 0.0002373 | ---?? | 0 | 0.5747 | | |
| 7:64851852 | 7 | 64851852 | 3 | 0 | 0 | CT | C | -1.356 | 0.3689 | 0.0002374 | ---?? | 45.2 | 0.161 | | |
| rs16939837 | 15 | 87623102 | 4 | 0 | 0 | C | G | -2.1744 | 0.5916 | 0.0002375 | ---?- | 0 | 0.8336 | intergenic | |
| rs1564160 | 8 | 140488714 | 4 | 0 | 0 | A | T | 1.5135 | 0.4118 | 0.0002376 | +++?+ | 0 | 0.7971 | intergenic | |
| rs117096776 | 17 | 56856012 | 5 | 0 | 0 | T | C | -2.5894 | 0.7046 | 0.0002377 | ----- | 0 | 0.802 | intron | PPM1E |
| rs117224287 | 17 | 56856387 | 5 | 0 | 0 | T | C | -2.5894 | 0.7046 | 0.0002377 | ----- | 0 | 0.802 | intron | PPM1E |
| rs74490292 | 2 | 170569407 | 4 | 0 | 0 | T | C | -1.3185 | 0.3588 | 0.0002378 | ---?- | 0 | 0.9244 | intron | PHOSPHO2-KLHL23 |
| rs116208116 | 4 | 100636227 | 4 | 0 | 0 | T | C | 2.1108 | 0.5744 | 0.0002378 | -++?+ | 45.6 | 0.1378 | intergenic | |
| rs367703892 | 15 | 84979248 | 3 | 0 | 0 | T | G | -3.24 | 0.8816 | 0.0002378 | ---?? | 0 | 0.9078 | intergenic | |
| rs2468729 | 8 | 140487399 | 4 | 0 | 0 | C | G | 1.49 | 0.4055 | 0.000238 | +++?+ | 0 | 0.5976 | intergenic | |
| 2:122521061 | 2 | 122521061 | 3 | 0 | 0 | CTT | C | -3.7862 | 1.0304 | 0.0002381 | ---?? | 0 | 0.8212 | | |
| rs9592000 | 13 | 60035424 | 4 | 0 | 0 | T | C | 0.9239 | 0.2514 | 0.0002382 | +++?+ | 0 | 0.4423 | intergenic | |
| rs116848058 | 18 | 40719671 | 4 | 0 | 0 | T | C | 2.1594 | 0.5877 | 0.0002382 | +++?+ | 0 | 0.6844 | intergenic | |
| 6:109101921 | 6 | 109101921 | 2 | 0 | 0 | A | AG | -5.1246 | 1.3947 | 0.0002384 | -?-?? | 0 | 0.7972 | | |
| rs114843082 | 2 | 168034976 | 4 | 0 | 0 | T | C | -2.267 | 0.617 | 0.0002385 | ---?+ | 0 | 0.6629 | intron | XIRP2 |
| rs17470406 | 5 | 122541246 | 4 | 0 | 0 | T | C | -1.9686 | 0.5358 | 0.0002386 | ---?- | 0 | 0.8376 | intergenic | |
| rs12297122 | 12 | 41463680 | 4 | 0 | 0 | T | C | 1.8537 | 0.5045 | 0.0002387 | +++?+ | 0 | 0.7562 | intron | CNTN1 |
| rs28752534 | 6 | 32585068 | 3 | 0 | 0 | T | C | -1.0266 | 0.2794 | 0.0002387 | ---?? | 0 | 0.9989 | intergenic | |

| | | | | | | | | | | | | | | | |
|---|---|---|---|---|---|---|---|---|---|---|---|---|---|---|---|
| rs116272587 | 17 | 56867906 | 5 | 0 | 0 | A | G | -2.5701 | 0.6995 | 0.0002387 | ----- | 0 | 0.8364 | intron | PPM1E |
| rs4812994 | 20 | 44729089 | 5 | 0 | 0 | A | C | 0.783 | 0.2131 | 0.000239 | -++++ | 0.9 | 0.401 | intergenic | |
| rs118146691 | 10 | 81085221 | 4 | 0 | 0 | A | G | 3.1015 | 0.8442 | 0.000239 | +++?+ | 0 | 0.7213 | intergenic | |
| rs11821949 | 11 | 34368110 | 2 | 0 | 0 | T | C | -6.3701 | 1.734 | 0.0002391 | -?-?? | 0 | 0.5748 | intron | ABTB2 |
| rs10754291 | 1 | 90247139 | 5 | 0 | 0 | A | G | -0.7837 | 0.2133 | 0.0002392 | ----- | 0 | 0.9752 | intergenic |  |
| rs10767871 | 11 | 30677354 | 4 | 0 | 0 | T | C | -1.0229 | 0.2784 | 0.0002392 | ---?+ | 45.7 | 0.1371 | intergenic | |
| rs627294 | 5 | 134353743 | 5 | 0 | 0 | A | C | 0.8428 | 0.2294 | 0.0002394 | +++++ | 0 | 0.5051 | intergenic | |
| rs78643477 | 2 | 170569799 | 4 | 0 | 0 | A | G | -1.3178 | 0.3587 | 0.0002394 | ---?- | 0 | 0.9247 | intron | PHOSPHO2-KLHL23 |
| rs115922434 | 2 | 168042060 | 4 | 0 | 0 | A | G | -2.2524 | 0.6132 | 0.0002394 | ---?+ | 0 | 0.6682 | intron | XIRP2 |
| rs74412402 | 2 | 117134038 | 4 | 0 | 0 | T | C | -1.9012 | 0.5176 | 0.0002397 | ---?- | 0 | 0.5906 | intergenic | |
| rs10152128 | 15 | 65228708 | 4 | 0 | 0 | T | C | 0.9572 | 0.2606 | 0.0002403 | +++?+ | 0 | 0.4852 | intron | ANKDD1A |
| rs6852464 | 4 | 37019341 | 3 | 0 | 0 | T | C | 0.9542 | 0.2599 | 0.0002408 | +++?? | 33.5 | 0.2221 | intergenic | |
| rs72769495 | 1 | 245101788 | 4 | 0 | 0 | T | G | -1.5758 | 0.4292 | 0.0002408 | ---?+ | 0 | 0.468 | intergenic | |
| rs111616349 | 1 | 228559654 | 4 | 0 | 0 | A | G | -2.959 | 0.8059 | 0.000241 | ---?- | 0 | 0.6881 | missense | OBSCN |
| rs139293339 | 17 | 56862582 | 5 | 0 | 0 | A | G | 2.5864 | 0.7045 | 0.0002411 | +++++ | 0 | 0.803 | intron | PPM1E |
| rs1445715 | 5 | 165419631 | 4 | 0 | 0 | T | C | -0.9109 | 0.2481 | 0.0002412 | ---?- | 0 | 0.4412 | intergenic | |
| 18:63867495 | 18 | 63867495 | 3 | 0 | 0 | A | ATATGT | 0.933 | 0.2541 | 0.0002413 | +++?? | 19.9 | 0.2871 | | |
| rs61973789 | 14 | 28153243 | 4 | 0 | 0 | T | C | -2.2335 | 0.6084 | 0.0002414 | ---?- | 0 | 0.9967 | intergenic | |
| 18:63867424 | 18 | 63867424 | 3 | 0 | 0 | G | GTA | 0.9327 | 0.2541 | 0.0002417 | +++?? | 19.5 | 0.2886 | | |
| rs60701213 | 14 | 28201865 | 4 | 0 | 0 | T | C | -2.1525 | 0.5864 | 0.0002419 | ---?- | 0 | 0.9666 | intergenic | |
| rs77388514 | 14 | 28178638 | 4 | 0 | 0 | T | C | -2.1192 | 0.5774 | 0.000242 | ---?- | 0 | 0.9917 | intergenic | |
| rs7835735 | 8 | 113480276 | 4 | 0 | 0 | T | C | 1.035 | 0.282 | 0.0002422 | +++?+ | 0 | 0.7926 | intron | CSMD3 |
| rs12061474 | 1 | 204389314 | 4 | 0 | 0 | T | C | -1.5026 | 0.4094 | 0.0002422 | ---?- | 23.7 | 0.269 | downstream-gene | |
| rs113032380 | 1 | 245101546 | 4 | 0 | 0 | A | G | 1.5764 | 0.4295 | 0.0002422 | +++?- | 0 | 0.4678 | intergenic | |
| 2:131129928 | 2 | 131129928 | 3 | 0 | 0 | A | AGACGG | -1.1306 | 0.308 | 0.0002422 | ---?? | 0 | 0.3848 | | |
| rs55746965 | 6 | 72864633 | 3 | 0 | 0 | A | T | -1.6998 | 0.4631 | 0.0002424 | ---?? | 0 | 0.9956 | intron | RIMS1 |
| rs56125271 | 6 | 72864632 | 3 | 0 | 0 | A | G | 1.6997 | 0.4631 | 0.0002425 | +++?? | 0 | 0.9957 | intron | RIMS1 |
| rs78070269 | 18 | 40086783 | 4 | 0 | 0 | A | G | 1.4173 | 0.3862 | 0.0002425 | +++?+ | 0 | 0.9215 | intron | LINC00907 |
| rs1500638 | 15 | 98047622 | 4 | 0 | 0 | A | G | 0.9333 | 0.2543 | 0.0002426 | +++?+ | 0 | 0.5393 | intergenic | |
| rs28698889 | 15 | 88263572 | 4 | 0 | 0 | C | G | -1.0052 | 0.2739 | 0.0002426 | ---?- | 0 | 0.4249 | intergenic | |
| rs74480211 | 18 | 27048751 | 4 | 0 | 0 | C | G | -3.0739 | 0.8376 | 0.0002426 | ---?- | 60.9 | 0.05344 | intergenic | |
| rs114296979 | 2 | 168045326 | 4 | 0 | 0 | T | C | 2.2438 | 0.6114 | 0.0002426 | +++?- | 0 | 0.6704 | intron | XIRP2 |
| rs333804 | 11 | 37043091 | 4 | 0 | 0 | T | G | -0.9776 | 0.2664 | 0.0002427 | ---?- | 0 | 0.4866 | intergenic | |
| rs76380052 | 2 | 168043319 | 4 | 0 | 0 | T | G | 2.2477 | 0.6125 | 0.0002427 | +++?- | 0 | 0.6691 | intron | XIRP2 |
| 6:83901878 | 6 | 83901878 | 2 | 0 | 0 | A | G | -6.4704 | 1.7632 | 0.0002428 | ??-?- | 0 | 0.493 | intron | PGM3 |
| rs17621853 | 18 | 40103062 | 4 | 0 | 0 | A | G | 1.417 | 0.3862 | 0.000243 | +++?+ | 0 | 0.9223 | intron | LINC00907 |
| rs942674 | 10 | 131922572 | 4 | 0 | 0 | A | C | -0.8969 | 0.2444 | 0.0002431 | ---?- | 0 | 0.7231 | intergenic | |
| rs895792 | 1 | 30369610 | 4 | 0 | 0 | A | G | 1.2924 | 0.3522 | 0.0002434 | +++?- | 0 | 0.4625 | intergenic | |
| rs7835916 | 8 | 113480463 | 4 | 0 | 0 | A | G | -1.0346 | 0.282 | 0.0002434 | ---?- | 0 | 0.7907 | intron | CSMD3 |
| rs116181989 | 2 | 168044900 | 4 | 0 | 0 | T | C | 2.2438 | 0.6116 | 0.0002436 | +++?- | 0 | 0.6701 | intron | XIRP2 |
| rs7836340 | 8 | 113480718 | 4 | 0 | 0 | T | C | 1.0345 | 0.282 | 0.0002437 | +++?+ | 0 | 0.7933 | intron | CSMD3 |
| rs12878512 | 14 | 43253912 | 4 | 0 | 0 | A | G | 1.4042 | 0.3827 | 0.0002437 | +++?- | 60.6 | 0.05467 | intergenic | |
| rs2067748 | 7 | 141200242 | 3 | 0 | 0 | T | C | -6.2151 | 1.6943 | 0.0002443 | --??- | 49.6 | 0.1377 | intergenic | |
| rs11222720 | 11 | 131492764 | 4 | 0 | 0 | A | C | 0.8807 | 0.2401 | 0.0002444 | +++?+ | 0 | 0.7029 | intron | NTM |
| rs10083390 | 14 | 22530148 | 4 | 0 | 0 | T | C | 0.9725 | 0.2652 | 0.0002447 | +++?+ | 0 | 0.6899 | intergenic | |
| rs11617584 | 13 | 50365319 | 5 | 0 | 0 | A | C | -1.4296 | 0.3898 | 0.000245 | ----+ | 0 | 0.814 | intron | KPNA3 |
| rs16977063 | 18 | 40435751 | 3 | 0 | 0 | C | G | 4.8503 | 1.3226 | 0.000245 | +++?? | 0 | 0.5486 | intron | RIT2 |
| rs111889695 | 8 | 123491748 | 3 | 0 | 0 | A | G | -5.728 | 1.5619 | 0.000245 | --??- | 0 | 0.4108 | intergenic | |
| rs80128417 | 2 | 168046550 | 4 | 0 | 0 | C | G | 2.2406 | 0.611 | 0.0002451 | +++?- | 0 | 0.6708 | intron | XIRP2 |
| rs9861629 | 3 | 172329273 | 3 | 0 | 0 | C | G | -1.9986 | 0.545 | 0.0002453 | ---?? | 0 | 0.7469 | intergenic | |
| rs11060702 | 12 | 130591803 | 4 | 0 | 0 | T | C | 0.9004 | 0.2456 | 0.0002454 | +++?- | 57.5 | 0.07005 | intergenic | |
| rs72769494 | 1 | 245101425 | 4 | 0 | 0 | A | G | -1.577 | 0.4301 | 0.0002454 | ---?+ | 0 | 0.4673 | intergenic | |
| rs72839990 | 17 | 56390260 | 5 | 0 | 0 | C | G | -2.0361 | 0.5552 | 0.0002454 | ----- | 42.9 | 0.1357 | intron | BZRAP1 |
| 21:31407054 | 21 | 31407054 | 3 | 0 | 0 | G | GA | -1.0979 | 0.2994 | 0.0002454 | ---?? | 0 | 0.4368 | | |
| rs1212425 | 15 | 42151938 | 5 | 0 | 0 | A | G | 1.024 | 0.2793 | 0.0002456 | +++++ | 0 | 0.8232 | intron | SPTBN5 |
| rs61973825 | 14 | 28177152 | 4 | 0 | 0 | T | C | -2.1154 | 0.5769 | 0.0002457 | ---?- | 0 | 0.9918 | intergenic | |
| rs625732 | 10 | 131915104 | 4 | 0 | 0 | T | C | -0.8942 | 0.2439 | 0.0002458 | ---?- | 0 | 0.8055 | intergenic | |
| rs61973824 | 14 | 28177070 | 4 | 0 | 0 | A | T | -2.1153 | 0.5769 | 0.0002458 | ---?- | 0 | 0.9918 | intergenic | |
| 6:30983640 | 6 | 30983640 | 3 | 0 | 0 | A | AT | -1.4819 | 0.4042 | 0.0002458 | ---?? | 0 | 0.7116 | | |

| | | | | | | | | | | | | | | | |
|---|---|---|---|---|---|---|---|---|---|---|---|---|---|---|---|
| rs2045033 | 8 | 113479400 | 4 | 0 | 0 | A | C | 1.0339 | 0.282 | 0.0002461 | +++?+ | 0 | 0.7902 | intron | CSMD3 |
| rs72668218 | 14 | 28177047 | 4 | 0 | 0 | A | G | -2.115 | 0.5769 | 0.0002463 | ---?- | 0 | 0.9918 | intergenic | |
| rs61973823 | 14 | 28176869 | 4 | 0 | 0 | A | G | 2.1147 | 0.5769 | 0.0002466 | +++?+ | 0 | 0.9918 | intergenic | |
| rs9285219 | 13 | 60018618 | 4 | 0 | 0 | A | C | -0.9308 | 0.2539 | 0.0002467 | ---?- | 0 | 0.7829 | intergenic | |
| rs2151062 | 9 | 16205136 | 4 | 0 | 0 | A | G | 0.9909 | 0.2703 | 0.0002468 | +++?+ | 0 | 0.8327 | intron | C9orf92 |
| rs79622111 | 3 | 68224746 | 4 | 0 | 0 | A | G | 2.3264 | 0.6347 | 0.0002468 | +++?+ | 30.3 | 0.2303 | intron | FAM19A1 |
| rs1537581 | 1 | 232833681 | 4 | 0 | 0 | T | G | 0.9101 | 0.2483 | 0.0002469 | +++?+ | 39.3 | 0.1763 | intergenic | |
| rs1408417 | 1 | 64590697 | 4 | 0 | 0 | A | G | -1.1437 | 0.312 | 0.0002472 | ---?- | 0 | 0.8504 | intron | ROR1 |
| rs76102705 | 2 | 170551049 | 4 | 0 | 0 | A | G | 1.3211 | 0.3604 | 0.0002472 | +++?+ | 0 | 0.9286 | 5-prime-UTR | PHOSPHO2,PHOSPHO2-KLHL23 |
| rs312175 | 19 | 47337124 | 4 | 0 | 0 | A | G | -1.4239 | 0.3885 | 0.0002473 | ---?- | 0 | 0.6758 | upstream-gene | |
| rs1604172 | 15 | 98040555 | 4 | 0 | 0 | A | G | -0.9322 | 0.2544 | 0.0002473 | ---?- | 0 | 0.9562 | intergenic | |
| rs28495356 | 6 | 32512164 | 4 | 0 | 1 | T | C | 1.3877 | 0.3786 | 0.0002473 | ++++? | 28.9 | 0.2388 | intergenic | |
| rs633220 | 15 | 53768798 | 4 | 0 | 0 | T | C | 0.9436 | 0.2575 | 0.0002474 | +++?+ | 41.4 | 0.1632 | intergenic | |
| rs1387330 | 4 | 169329321 | 4 | 0 | 0 | A | G | -0.9489 | 0.2589 | 0.0002475 | ---?- | 0 | 0.4833 | intron | DDX60L |
| rs80075966 | 6 | 109099430 | 3 | 0 | 0 | A | G | -4.2123 | 1.1494 | 0.0002475 | -+-?? | 55.6 | 0.105 | intergenic | |
| rs6848080 | 4 | 100618982 | 4 | 0 | 0 | T | C | 2.0752 | 0.5663 | 0.0002476 | -++?+ | 44.3 | 0.1457 | intergenic | |
| rs2250964 | 15 | 42017542 | 5 | 0 | 0 | T | C | 1.0379 | 0.2833 | 0.0002479 | +++++ | 8.1 | 0.3601 | intron | MGA |
| rs79329147 | 1 | 64590661 | 4 | 0 | 0 | A | G | 1.1435 | 0.312 | 0.0002479 | +++?+ | 0 | 0.8507 | intron | ROR1 |
| rs949513 | 1 | 64791374 | 4 | 0 | 0 | A | G | 1.5181 | 0.4143 | 0.000248 | +++?+ | 0 | 0.8593 | intergenic | |
| rs12454579 | 18 | 40434690 | 3 | 0 | 0 | A | G | 4.8305 | 1.3182 | 0.000248 | +++?? | 0 | 0.5489 | intron | RIT2 |
| rs114881599 | 17 | 56876787 | 5 | 0 | 0 | T | C | -2.5802 | 0.7041 | 0.0002481 | ----- | 0 | 0.8053 | intron | PPM1E |
| rs4600922 | 4 | 100602754 | 4 | 0 | 0 | A | T | -2.0677 | 0.5643 | 0.0002482 | +--?- | 45.8 | 0.1365 | intergenic | |
| rs56063841 | 3 | 29749213 | 4 | 0 | 0 | A | G | -4.0178 | 1.0965 | 0.0002482 | ---?- | 20.8 | 0.2854 | intron | RBMS3 |
| rs61973821 | 14 | 28175755 | 4 | 0 | 0 | A | G | -2.1129 | 0.5767 | 0.0002484 | ---?- | 0 | 0.992 | intergenic | |
| rs9285220 | 13 | 60018644 | 4 | 0 | 0 | T | C | -0.9297 | 0.2538 | 0.0002486 | ---?- | 0 | 0.7818 | intergenic | |
| rs45535740 | 20 | 36771795 | 4 | 0 | 0 | T | G | -3.4702 | 0.9472 | 0.0002487 | ---?- | 0 | 0.5554 | intron | TGM2 |
| rs67037615 | 6 | 32453163 | 5 | 0 | 1 | T | C | -1.4176 | 0.3869 | 0.0002488 | ----- | 0 | 0.5043 | intergenic |  |
| rs148756573 | 3 | 68223695 | 4 | 0 | 0 | A | G | 2.3107 | 0.6308 | 0.000249 | +++?+ | 31 | 0.2264 | intron | FAM19A1 |
| rs32216 | 5 | 127640076 | 5 | 0 | 0 | T | C | 0.8695 | 0.2374 | 0.0002492 | -++++ | 24.6 | 0.2575 | intron | FBN2 |
| rs895795 | 1 | 30369052 | 4 | 0 | 0 | A | C | 1.2933 | 0.3531 | 0.0002492 | +++?- | 0 | 0.4647 | intergenic | |
| rs2904080 | 20 | 47157001 | 4 | 0 | 0 | A | G | 1.2063 | 0.3293 | 0.0002492 | +++?+ | 0 | 0.7367 | intergenic | |
| rs7331706 | 13 | 60018700 | 4 | 0 | 0 | T | G | 0.9295 | 0.2538 | 0.0002493 | +++?+ | 0 | 0.7812 | intergenic | |
| rs1604170 | 15 | 98040887 | 4 | 0 | 0 | T | C | -0.9275 | 0.2532 | 0.0002494 | ---?- | 0 | 0.9646 | intergenic | |
| rs76654537 | 2 | 170570823 | 4 | 0 | 0 | T | C | 1.3133 | 0.3585 | 0.0002494 | +++?+ | 0 | 0.9253 | intron | PHOSPHO2-KLHL23 |
| rs7330793 | 13 | 60018772 | 4 | 0 | 0 | A | G | -0.9295 | 0.2538 | 0.0002495 | ---?- | 0 | 0.7806 | intergenic | |
| rs9961643 | 18 | 47531478 | 4 | 0 | 0 | T | C | -0.9564 | 0.2611 | 0.0002496 | ---?- | 0 | 0.4538 | intron | MYO5B |
| rs57533195 | 7 | 130303017 | 4 | 0 | 0 | T | C | 0.9029 | 0.2465 | 0.0002496 | +-+?+ | 58.1 | 0.06722 | intron | COPG2 |
| 10:131498656 | 10 | 131498656 | 3 | 0 | 0 | G | GT | -1.4296 | 0.3903 | 0.0002497 | ---?? | 0 | 0.8515 | | |
| rs1486952 | 13 | 60016174 | 3 | 0 | 0 | T | C | 0.9448 | 0.258 | 0.0002498 | +++?? | 0 | 0.6261 | intergenic | |
| rs77627143 | 18 | 61208398 | 4 | 0 | 0 | A | G | -2.2649 | 0.6184 | 0.0002499 | ---?- | 20.3 | 0.288 | intergenic | |
| rs146110995 | 6 | 31994535 | 3 | 0 | 0 | T | C | -1.4123 | 0.3856 | 0.0002499 | ---?? | 38.1 | 0.1989 | intron | C4B |
| rs77771357 | 6 | 23898057 | 4 | 0 | 0 | C | G | 2.5941 | 0.7084 | 0.0002501 | +++?- | 26.9 | 0.2507 | intergenic | |
| 8:60386083 | 8 | 60386083 | 3 | 0 | 0 | CA | C | -1.2614 | 0.3445 | 0.0002503 | ---?? | 0 | 0.5185 | | |
| rs72668216 | 14 | 28175214 | 4 | 0 | 0 | T | C | 2.1112 | 0.5765 | 0.0002504 | +++?+ | 0 | 0.992 | intergenic | |
| rs685850 | 15 | 53751702 | 4 | 0 | 0 | A | G | -0.8733 | 0.2385 | 0.0002506 | ---?- | 48.4 | 0.1208 | intergenic | |
| 9:34440848 | 9 | 34440848 | 3 | 0 | 0 | CA | C | -1.4916 | 0.4074 | 0.0002506 | ---?? | 60.6 | 0.07881 | | |
| rs895796 | 1 | 30368883 | 4 | 0 | 0 | T | C | -1.2934 | 0.3532 | 0.0002508 | ---?+ | 0 | 0.465 | intergenic | |
| rs116029711 | 6 | 32603564 | 5 | 0 | 1 | T | C | -1.4399 | 0.3933 | 0.0002509 | ----- | 21.5 | 0.2778 | upstream-gene |  |
| rs6751587 | 2 | 170909530 | 4 | 0 | 1 | T | C | 1.0619 | 0.29 | 0.000251 | +++?+ | 32.1 | 0.2198 | intron | UBR3 |
| rs72705407 | 1 | 116337755 | 4 | 0 | 0 | A | G | -2.0618 | 0.5632 | 0.0002511 | ---?- | 34.9 | 0.2029 | intergenic | |
| rs76361729 | 5 | 122540503 | 4 | 0 | 0 | A | G | 1.9606 | 0.5355 | 0.0002511 | +++?+ | 0 | 0.8376 | intergenic | |
| rs143492821 | 6 | 83756757 | 4 | 0 | 0 | A | G | 3.4772 | 0.9498 | 0.0002512 | +++?+ | 0 | 0.7732 | intron | UBE3D |
| rs61973820 | 14 | 28173793 | 4 | 0 | 0 | T | C | -2.1097 | 0.5763 | 0.0002517 | ---?- | 0 | 0.992 | intergenic | |
| rs28986381 | 6 | 32752429 | 5 | 0 | 0 | A | G | -1.4636 | 0.3999 | 0.0002519 | ----- | 0.9 | 0.4009 | intergenic |  |
| rs28986383 | 6 | 32752436 | 5 | 0 | 0 | A | G | -1.4636 | 0.3999 | 0.0002519 | ----- | 0.9 | 0.4009 | intergenic |  |
| rs35091561 | 13 | 30313491 | 5 | 0 | 0 | A | G | -1.3665 | 0.3733 | 0.000252 | ----- | 0 | 0.6546 | intergenic |  |
| rs78651151 | 10 | 131469339 | 4 | 0 | 0 | T | C | 1.8317 | 0.5004 | 0.0002521 | +++?+ | 0 | 0.701 | intron | MGMT |
| rs1355616 | 11 | 37063507 | 4 | 0 | 0 | A | G | -0.8891 | 0.2429 | 0.0002522 | ---?- | 0 | 0.4259 | intergenic | |
| rs10774619 | 12 | 111773925 | 5 | 0 | 0 | A | G | 0.913 | 0.2495 | 0.0002523 | +-+++ | 48.4 | 0.1012 | intron | CUX2 |

| | | | | | | | | | | | | | | | |
|---|---|---|---|---|---|---|---|---|---|---|---|---|---|---|---|
| rs59220463 | 7 | 141173120 | 4 | 0 | 0 | A | G | -5.062 | 1.3831 | 0.0002523 | ---?- | 0 | 0.7905 | 3-prime-UTR | TMEM178B |
| rs72800392 | 5 | 134356061 | 5 | 0 | 0 | T | C | -0.8504 | 0.2324 | 0.0002524 | ----- | 0 | 0.4852 | intergenic |  |
| rs2993295 | 13 | 113667732 | 2 | 0 | 0 | C | G | 1.4113 | 0.3856 | 0.0002526 | ???++ | 6.9 | 0.3001 | intron | MCF2L |
| rs7604068 | 2 | 168050281 | 4 | 0 | 0 | A | G | -2.2303 | 0.6094 | 0.0002526 | ---?+ | 0 | 0.673 | intron | XIRP2 |
| rs4620705 | 11 | 12760609 | 4 | 0 | 0 | A | G | 1.4881 | 0.4067 | 0.0002528 | +++?+ | 1.3 | 0.3857 | intron | TEAD1 |
| rs61973818 | 14 | 28172301 | 4 | 0 | 0 | T | C | -2.1079 | 0.576 | 0.0002529 | ---?- | 0 | 0.9922 | intergenic | |
| rs77066079 | 6 | 72867826 | 3 | 0 | 0 | T | C | -1.6961 | 0.4635 | 0.000253 | ---?? | 0 | 0.9973 | intron | RIMS1 |
| rs12050467 | 14 | 43244785 | 4 | 0 | 0 | A | G | -1.4101 | 0.3854 | 0.0002531 | ---?+ | 61.1 | 0.05229 | intergenic | |
| rs7970181 | 12 | 112609998 | 5 | 0 | 0 | T | C | 1.3511 | 0.3693 | 0.0002532 | +-+++ | 54.6 | 0.06577 | intron | HECTD4 |
| rs17621438 | 18 | 40088617 | 4 | 0 | 0 | A | G | 1.412 | 0.3859 | 0.0002532 | +++?+ | 0 | 0.9245 | intron | LINC00907 |
| rs10276990 | 7 | 16903052 | 4 | 0 | 0 | A | C | 1.7458 | 0.4771 | 0.0002533 | +++?+ | 0 | 0.4163 | intron | AGR3 |
| rs9527989 | 13 | 60016001 | 3 | 0 | 0 | C | G | 0.9388 | 0.2566 | 0.0002534 | +++?? | 0 | 0.6628 | intergenic | |
| rs35312721 | 13 | 30313336 | 5 | 0 | 0 | A | G | -1.3662 | 0.3734 | 0.0002534 | ----- | 0 | 0.6534 | intergenic |  |
| rs141104396 | 13 | 74893847 | 4 | 0 | 0 | T | C | -2.4955 | 0.6821 | 0.0002535 | ---?- | 0 | 0.4345 | intergenic | |
| rs61973817 | 14 | 28171656 | 4 | 0 | 0 | T | C | -2.107 | 0.5759 | 0.0002536 | ---?- | 0 | 0.9922 | intergenic | |
| rs77739181 | 18 | 40026046 | 4 | 0 | 0 | A | G | 1.4029 | 0.3835 | 0.0002536 | +++?+ | 0 | 0.8265 | intron | LINC00907 |
| rs11253556 | 10 | 1040356 | 4 | 0 | 0 | A | G | -1.9016 | 0.5198 | 0.0002537 | ---?- | 35.3 | 0.2002 | intron | GTPBP4 |
| rs738597 | 22 | 49389183 | 4 | 0 | 0 | A | C | -1.6974 | 0.464 | 0.0002539 | ---?- | 15.5 | 0.3143 | intergenic | |
| rs79155198 | 9 | 79458780 | 4 | 0 | 0 | A | G | -2.1308 | 0.5825 | 0.0002539 | ---?- | 0 | 0.5403 | intron | PRUNE2 |
| rs72668214 | 14 | 28171509 | 4 | 0 | 0 | T | C | 2.1067 | 0.5759 | 0.000254 | +++?+ | 0 | 0.9922 | intergenic | |
| 6:31221116 | 6 | 31221116 | 3 | 0 | 0 | T | G | 0.8828 | 0.2413 | 0.0002541 | +++?? | 0 | 0.3768 | intergenic | |
| rs35567463 | 6 | 32633970 | 4 | 0 | 1 | A | C | 1.1458 | 0.3132 | 0.0002541 | ++++? | 44.2 | 0.1464 | intron | HLA-DQB1 |
| rs60570441 | 15 | 85815902 | 2 | 0 | 0 | A | T | -6.8362 | 1.8689 | 0.0002543 | --??? | 0 | 0.4871 | intergenic | |
| rs1550968 | 1 | 30368387 | 4 | 0 | 0 | T | C | -1.2938 | 0.3537 | 0.0002546 | ---?+ | 0 | 0.4669 | intergenic | |
| rs72842493 | 11 | 1127503 | 4 | 0 | 0 | A | C | 1.4462 | 0.3954 | 0.0002547 | +++?- | 55.1 | 0.0828 | intergenic | |
| rs147546565 | 5 | 179161563 | 2 | 0 | 0 | T | G | 5.1761 | 1.4152 | 0.0002547 | +?+?? | 0 | 0.8186 | intron | MAML1 |
| 4:125519803 | 4 | 125519803 | 3 | 0 | 0 | G | GTT | -0.9895 | 0.2705 | 0.0002547 | ---?? | 0 | 0.9314 | | |
| rs17232129 | 10 | 131921200 | 4 | 0 | 0 | T | C | 0.897 | 0.2453 | 0.0002548 | +++?+ | 0 | 0.7253 | intergenic | |
| rs55721553 | 12 | 41224229 | 3 | 0 | 0 | A | G | -1.8912 | 0.5171 | 0.0002548 | ---?? | 0 | 0.9041 | intron | CNTN1 |
| rs75445705 | 2 | 232135014 | 4 | 0 | 0 | C | G | 1.5343 | 0.4195 | 0.000255 | +++?+ | 0 | 0.7342 | intron | ARMC9 |
| rs34484108 | 13 | 30313282 | 5 | 0 | 0 | A | G | -1.3659 | 0.3735 | 0.0002551 | ----- | 0 | 0.6525 | intergenic |  |
| rs4772237 | 13 | 100403452 | 4 | 0 | 0 | A | G | -1.2897 | 0.3527 | 0.0002554 | ---?- | 32.1 | 0.2198 | intron | CLYBL |
| rs114425975 | 2 | 168051650 | 4 | 0 | 0 | T | C | 2.2258 | 0.6087 | 0.0002554 | +++?- | 0 | 0.6739 | intron | XIRP2 |
| rs72668212 | 14 | 28170737 | 4 | 0 | 0 | C | G | -2.105 | 0.5757 | 0.0002556 | ---?- | 0 | 0.9923 | intergenic | |
| 22:45954547 | 22 | 45954547 | 3 | 0 | 0 | A | AAC | -0.944 | 0.2582 | 0.0002557 | ---?? | 55.2 | 0.1072 | | |
| rs9893863 | 17 | 56883935 | 5 | 0 | 0 | A | C | 2.5737 | 0.7039 | 0.0002559 | +++++ | 0 | 0.806 | intron | PPM1E |
| rs2306550 | 12 | 26809537 | 4 | 0 | 0 | A | G | -1.8537 | 0.507 | 0.000256 | ---?- | 17.8 | 0.3017 | intron | ITPR2 |
| rs10895431 | 11 | 103324020 | 4 | 0 | 0 | A | G | 1.0239 | 0.2801 | 0.0002561 | +++?+ | 49.3 | 0.1159 | intron | DYNC2H1 |
| rs78071700 | 17 | 56885997 | 5 | 0 | 0 | A | G | 2.5735 | 0.7039 | 0.0002561 | +++++ | 0 | 0.8059 | intron | PPM1E |
| rs142819518 | 8 | 15531400 | 3 | 0 | 0 | A | G | -3.4404 | 0.941 | 0.0002561 | ---?? | 0 | 0.3902 | intron | TUSC3 |
| rs1566668 | 12 | 130594316 | 4 | 0 | 0 | A | G | -0.8681 | 0.2374 | 0.0002562 | ---?+ | 55.3 | 0.08174 | intergenic | |
| rs117560340 | 9 | 4408866 | 3 | 0 | 0 | C | G | 4.0717 | 1.1137 | 0.0002562 | +++?? | 0 | 0.5961 | intergenic | |
| rs6905581 | 6 | 109109235 | 2 | 0 | 0 | C | G | -4.9163 | 1.3448 | 0.0002563 | -?-?? | 0 | 0.9981 | intergenic | |
| rs7832171 | 8 | 113466827 | 4 | 0 | 0 | A | G | 1.0157 | 0.2778 | 0.0002563 | +++?+ | 0 | 0.7886 | intron | CSMD3 |
| rs10847413 | 12 | 128048949 | 4 | 0 | 0 | T | C | -2.0307 | 0.5555 | 0.0002563 | ---?- | 0 | 0.452 | intergenic | |
| rs28572641 | 6 | 23899574 | 4 | 0 | 0 | C | G | 2.5854 | 0.7072 | 0.0002565 | +++?- | 26.4 | 0.2535 | intergenic | |
| rs114309418 | 3 | 60615629 | 4 | 0 | 0 | T | C | 2.1455 | 0.5869 | 0.0002566 | +++?+ | 17 | 0.3063 | intron | FHIT |
| rs191620250 | 6 | 72869717 | 3 | 0 | 0 | A | T | -1.6956 | 0.4638 | 0.0002566 | ---?? | 0 | 0.9966 | intron | RIMS1 |
| rs10133742 | 14 | 43251455 | 4 | 0 | 0 | A | G | -1.4009 | 0.3832 | 0.0002567 | ---?+ | 61 | 0.05281 | intergenic | |
| rs11783041 | 8 | 113485014 | 4 | 0 | 0 | A | C | -1.0302 | 0.2818 | 0.0002568 | ---?- | 0 | 0.791 | intron | CSMD3 |
| rs13133065 | 4 | 189069680 | 4 | 0 | 0 | T | C | -1.2759 | 0.3491 | 0.0002568 | ---?- | 0 | 0.8478 | downstream-gene | |
| 6:159186947 | 6 | 159186947 | 3 | 0 | 0 | G | GCAAA | -0.9355 | 0.2559 | 0.0002568 | ---?? | 0 | 0.3728 | | |
| rs11737197 | 4 | 38708403 | 5 | 0 | 0 | C | G | -0.792 | 0.2167 | 0.000257 | ----- | 0 | 0.6167 | intergenic |  |
| rs75480510 | 12 | 112240192 | 5 | 0 | 0 | A | G | 1.3893 | 0.3801 | 0.000257 | +-+++ | 50.4 | 0.08905 | intron | ALDH2 |
| rs61973815 | 14 | 28169912 | 4 | 0 | 0 | A | C | 2.1034 | 0.5755 | 0.0002571 | +++?+ | 0 | 0.9924 | intergenic | |
| rs74645577 | 2 | 117130437 | 4 | 0 | 0 | A | G | -1.8847 | 0.5156 | 0.0002572 | ---?- | 0 | 0.5951 | intergenic | |
| rs61973814 | 14 | 28169870 | 4 | 0 | 0 | T | C | -2.1033 | 0.5755 | 0.0002573 | ---?- | 0 | 0.9924 | intergenic | |
| rs145831674 | 6 | 83748913 | 4 | 0 | 0 | A | G | 3.4955 | 0.9564 | 0.0002574 | +++?+ | 0 | 0.7627 | intron | UBE3D |
| rs10969923 | 9 | 30877317 | 4 | 0 | 0 | T | C | -3.9147 | 1.0711 | 0.0002575 | ---?- | 3.5 | 0.3751 | intergenic | |

| | | | | | | | | | | | | | | | |
|---|---|---|---|---|---|---|---|---|---|---|---|---|---|---|---|
| rs143518633 | 6 | 83744405 | 4 | 0 | 0 | A | T | -3.5024 | 0.9583 | 0.0002575 | ---?- | 0 | 0.7792 | intron | UBE3D |
| rs79270529 | 17 | 56647421 | 5 | 0 | 0 | T | C | -2.475 | 0.6772 | 0.0002576 | ----- | 0 | 0.8401 | intron | TEX14 |
| rs6858434 | 4 | 189066878 | 4 | 0 | 0 | A | G | -1.2838 | 0.3513 | 0.0002577 | ---?- | 0 | 0.8267 | intron | TRIML1 |
| rs10431056 | 11 | 106739921 | 3 | 0 | 0 | T | C | 1.0165 | 0.2781 | 0.0002577 | +++?? | 0 | 0.6164 | intron | GUCY1A2 |
| rs74412217 | 4 | 100599931 | 4 | 0 | 0 | A | G | -2.0652 | 0.5651 | 0.0002578 | +--?- | 46.4 | 0.1329 | intergenic | |
| rs333805 | 11 | 37043351 | 4 | 0 | 0 | A | G | -0.973 | 0.2663 | 0.0002582 | ---?- | 0 | 0.5122 | intergenic | |
| rs7788159 | 7 | 98445111 | 4 | 0 | 0 | A | G | 1.2464 | 0.3411 | 0.0002583 | +++?+ | 0 | 0.933 | 3-prime-UTR | TMEM130 |
| rs6812418 | 4 | 38741573 | 5 | 0 | 0 | T | C | 1.2114 | 0.3316 | 0.0002586 | +++++ | 0 | 0.9694 | intergenic | |
| rs13267095 | 8 | 18579169 | 5 | 0 | 0 | T | C | 1.1836 | 0.324 | 0.0002586 | +++++ | 0 | 0.7691 | intron | PSD3 |
| rs28359884 | 6 | 32584330 | 3 | 0 | 0 | A | C | 0.9488 | 0.2597 | 0.0002587 | +++?? | 0 | 0.8754 | intergenic | |
| rs1566669 | 12 | 130594356 | 4 | 0 | 0 | A | G | 0.8671 | 0.2373 | 0.0002588 | +++?- | 55.3 | 0.0817 | intergenic | |
| rs1392675 | 15 | 98046790 | 4 | 0 | 0 | T | G | 0.9671 | 0.2647 | 0.0002589 | +++?+ | 0 | 0.9214 | intergenic | |
| rs27713 | 5 | 127640783 | 5 | 0 | 0 | A | G | 0.8661 | 0.2371 | 0.000259 | -++++ | 17.5 | 0.3032 | intron | FBN2 |
| rs77995714 | 7 | 147536649 | 4 | 0 | 0 | T | C | 1.8343 | 0.5022 | 0.0002593 | +++?+ | 5.5 | 0.3654 | intron | CNTNAP2 |
| rs3931213 | 11 | 37070372 | 4 | 0 | 0 | A | G | -0.9063 | 0.2481 | 0.0002597 | ---?- | 0 | 0.4546 | intergenic | |
| rs4723330 | 7 | 33806117 | 4 | 0 | 0 | A | C | -0.9251 | 0.2533 | 0.0002597 | ---?- | 29.3 | 0.2366 | intergenic | |
| rs7965087 | 12 | 112548562 | 5 | 0 | 0 | A | G | 1.3468 | 0.3687 | 0.0002597 | +-+++ | 56.4 | 0.05665 | upstream-gene | |
| 6:106185906 | 6 | 106185906 | 3 | 0 | 0 | A | ACAGCA | 3.1019 | 0.8493 | 0.0002597 | +++?? | 18.9 | 0.2914 | | |
| rs61973790 | 14 | 28153387 | 4 | 0 | 0 | T | C | -2.2167 | 0.6069 | 0.0002599 | ---?- | 0 | 0.9972 | intergenic | |
| rs1880938 | 15 | 42025221 | 5 | 0 | 0 | A | G | -1.0321 | 0.2826 | 0.0002601 | ----- | 10.1 | 0.3487 | intron | MGA |
| rs9968331 | 4 | 38740056 | 5 | 0 | 0 | A | T | -1.2155 | 0.3328 | 0.0002603 | ----- | 0 | 0.9727 | intergenic |  |
| rs1959981 | 14 | 28161878 | 4 | 0 | 0 | T | C | -2.1442 | 0.5872 | 0.0002605 | ---?- | 0 | 0.9945 | intergenic | |
| rs4511427 | 13 | 60023226 | 4 | 0 | 0 | C | G | -0.9246 | 0.2532 | 0.0002605 | ---?- | 0 | 0.5786 | intergenic | |
| 3:29526872 | 3 | 29526872 | 3 | 0 | 0 | CT | C | 1.5212 | 0.4166 | 0.0002605 | +++?? | 0 | 0.8716 | | |
| rs6053592 | 20 | 5647438 | 3 | 0 | 0 | T | C | 1.1612 | 0.318 | 0.0002607 | +++?? | 0 | 0.4734 | intergenic | |
| rs817446 | 1 | 85646151 | 5 | 0 | 0 | A | G | 1.4145 | 0.3874 | 0.0002608 | +++++ | 0 | 0.9869 | intron | SYDE2 |
| rs17613816 | 18 | 40016012 | 4 | 0 | 0 | A | C | 1.3987 | 0.3831 | 0.0002608 | +++?+ | 0 | 0.8301 | intron | LINC00907 |
| rs28638783 | 6 | 32572737 | 5 | 0 | 1 | A | G | 1.3929 | 0.3815 | 0.0002608 | +++++ | 25.4 | 0.2521 | intergenic | |
| rs141575117 | 18 | 40020543 | 4 | 0 | 0 | C | G | -1.3989 | 0.3831 | 0.0002608 | ---?- | 0 | 0.83 | intron | LINC00907 |
| rs2012194 | 18 | 52527070 | 4 | 0 | 0 | T | C | -2.1668 | 0.5934 | 0.000261 | ---?- | 0 | 0.9452 | intron | RAB27B |
| rs11157306 | 14 | 43256726 | 4 | 0 | 0 | A | C | 1.392 | 0.3812 | 0.000261 | +++?- | 60.4 | 0.05551 | intergenic | |
| rs61973811 | 14 | 28167385 | 4 | 0 | 0 | T | C | -2.1015 | 0.5756 | 0.000261 | ---?- | 0 | 0.9926 | intergenic | |
| rs17613626 | 18 | 40011482 | 4 | 0 | 0 | A | G | 1.3985 | 0.383 | 0.0002611 | +++?+ | 0 | 0.8302 | intron | LINC00907 |
| rs61973812 | 14 | 28167441 | 4 | 0 | 0 | A | G | 2.1011 | 0.5755 | 0.0002612 | +++?+ | 0 | 0.9925 | intergenic | |
| rs143030234 | 20 | 48780062 | 3 | 0 | 0 | A | G | 3.7698 | 1.0325 | 0.0002612 | +++?? | 3.4 | 0.3552 | intergenic | |
| rs3104996 | 8 | 96563805 | 4 | 0 | 0 | A | T | 1.634 | 0.4476 | 0.0002613 | +-+?+ | 55.3 | 0.08158 | intron | LOC100616530 |
| rs817445 | 1 | 85646305 | 5 | 0 | 0 | A | G | 1.4143 | 0.3874 | 0.0002614 | +++++ | 0 | 0.9869 | intron | SYDE2 |
| rs10784414 | 12 | 64921819 | 4 | 0 | 0 | A | G | 0.9095 | 0.2492 | 0.0002618 | +++?+ | 0 | 0.9017 | intergenic | |
| rs61973813 | 14 | 28167643 | 4 | 0 | 0 | T | C | 2.099 | 0.575 | 0.0002618 | +++?+ | 0 | 0.9925 | intergenic | |
| rs4432647 | 3 | 68221597 | 4 | 0 | 0 | T | C | -2.2883 | 0.6269 | 0.0002619 | ---?- | 31.5 | 0.2232 | intron | FAM19A1 |
| 5:706584 | 5 | 706584 | 3 | 0 | 0 | CA | C | 1.7795 | 0.4875 | 0.0002619 | +++?? | 60.8 | 0.07791 | | |
| rs115343530 | 2 | 168053648 | 4 | 0 | 0 | A | G | -2.2237 | 0.6092 | 0.000262 | ---?+ | 0 | 0.6736 | intron | XIRP2 |
| rs10875246 | 1 | 100206801 | 4 | 0 | 0 | T | C | -1.8192 | 0.4984 | 0.0002621 | ---?- | 0 | 0.7339 | intron | FRRS1 |
| rs291938 | 3 | 173754274 | 4 | 0 | 0 | A | C | -2.8741 | 0.7874 | 0.0002622 | ---?- | 0 | 0.8678 | intron | NLGN1 |
| rs2577958 | 15 | 42024462 | 5 | 0 | 0 | T | C | 1.0313 | 0.2825 | 0.0002622 | +++++ | 9.5 | 0.3523 | intron | MGA |
| 10:11677669 | 10 | 11677669 | 3 | 0 | 0 | A | AT | -3.6086 | 0.9887 | 0.0002623 | ---?? | 0 | 0.9828 | | |
| rs60690688 | 12 | 112541983 | 5 | 0 | 0 | A | G | 1.3463 | 0.3689 | 0.0002625 | +-+++ | 56.4 | 0.05675 | intron | NAA25 |
| rs142548329 | 6 | 83746768 | 4 | 0 | 0 | A | G | 3.4944 | 0.9574 | 0.0002625 | +++?+ | 0 | 0.7701 | intron | UBE3D |
| 20:35717815 | 20 | 35717815 | 3 | 0 | 0 | A | AG | 1.2461 | 0.3414 | 0.0002626 | +++?? | 0 | 0.8196 | | |
| rs148965771 | 6 | 83757704 | 4 | 0 | 0 | C | G | 3.4765 | 0.9526 | 0.0002627 | +++?+ | 0 | 0.7407 | intron | UBE3D |
| rs188080944 | 6 | 131715232 | 2 | 0 | 0 | T | G | -7.9046 | 2.1659 | 0.0002627 | -???- | 0 | 0.909 | intergenic | |
| rs8026126 | 15 | 88264151 | 4 | 0 | 0 | T | C | -1.0004 | 0.2741 | 0.0002628 | ---?- | 0 | 0.4127 | intergenic | |
| rs9358745 | 6 | 24053389 | 5 | 0 | 0 | T | C | -1.398 | 0.3831 | 0.000263 | ----+ | 0 | 0.7864 | intergenic |  |
| rs116816620 | 2 | 167427341 | 3 | 0 | 0 | A | G | -3.8856 | 1.0648 | 0.0002631 | ---?? | 64.6 | 0.05951 | intergenic | |
| 6:32373591 | 6 | 32373591 | 3 | 0 | 0 | T | TTG | -0.9847 | 0.2699 | 0.0002632 | ---?? | 0 | 0.6242 | | |
| rs1384077 | 11 | 103322838 | 4 | 0 | 0 | T | C | 1.022 | 0.2801 | 0.0002633 | +++?+ | 50 | 0.1116 | intron | DYNC2H1 |
| rs67526112 | 1 | 235076866 | 4 | 0 | 0 | A | G | -0.8982 | 0.2461 | 0.0002633 | ---?- | 0 | 0.9373 | intergenic | |
| 6:32373595 | 6 | 32373595 | 3 | 0 | 0 | T | G | 0.9847 | 0.2699 | 0.0002634 | +++?? | 0 | 0.624 | intron | BTNL2 |
| rs2695166 | 15 | 42024579 | 5 | 0 | 0 | C | G | -1.0309 | 0.2826 | 0.0002635 | ----- | 9.8 | 0.3506 | intron | MGA |

| | | | | | | | | | | | | | | | |
|---|---|---|---|---|---|---|---|---|---|---|---|---|---|---|---|
| rs72668237 | 14 | 28200419 | 4 | 0 | 0 | T | C | -2.1384 | 0.586 | 0.0002635 | ---?- | 0 | 0.9809 | intergenic | |
| 6:32373598 | 6 | 32373598 | 3 | 0 | 0 | T | TG | -0.9847 | 0.2699 | 0.0002635 | ---?? | 0 | 0.624 | | |
| rs2274070 | 14 | 62361500 | 4 | 0 | 0 | A | G | 1.1273 | 0.309 | 0.0002637 | +++?- | 0 | 0.6399 | intron | SYT16 |
| rs1384076 | 11 | 103322991 | 4 | 0 | 0 | T | C | 1.0219 | 0.2801 | 0.0002638 | +++?+ | 50 | 0.1118 | intron | DYNC2H1 |
| rs7118634 | 11 | 103326785 | 4 | 0 | 0 | T | C | 1.0216 | 0.28 | 0.0002638 | +++?+ | 49.3 | 0.1158 | intron | DYNC2H1 |
| rs12613803 | 2 | 117132068 | 4 | 0 | 0 | T | G | 1.8857 | 0.5169 | 0.0002638 | +++?+ | 0 | 0.5883 | intergenic | |
| rs141005140 | 6 | 83750688 | 4 | 0 | 0 | T | G | 3.4871 | 0.9558 | 0.0002638 | +++?+ | 0 | 0.7602 | intron | UBE3D |
| rs11225811 | 11 | 103325204 | 4 | 0 | 0 | A | G | -1.0217 | 0.28 | 0.0002639 | ---?- | 49.5 | 0.1144 | intron | DYNC2H1 |
| rs55691906 | 6 | 24068410 | 3 | 0 | 0 | T | C | 1.7233 | 0.4724 | 0.0002639 | +++?? | 0 | 0.5727 | intergenic | |
| rs113802042 | 4 | 48050681 | 3 | 0 | 0 | A | C | -0.9694 | 0.2657 | 0.0002639 | ---?? | 0 | 0.5847 | intergenic | |
| rs144948468 | 11 | 59176837 | 4 | 0 | 0 | A | G | -3.655 | 1.002 | 0.0002644 | ---?- | 47.3 | 0.1278 | intergenic | |
| rs11225810 | 11 | 103325199 | 4 | 0 | 0 | C | G | -1.0216 | 0.28 | 0.0002645 | ---?- | 49.5 | 0.1143 | intron | DYNC2H1 |
| rs77659617 | 9 | 79458805 | 4 | 0 | 0 | A | G | -2.1159 | 0.5801 | 0.0002645 | ---?- | 0 | 0.5369 | intron | PRUNE2 |
| rs17100945 | 11 | 103323742 | 4 | 0 | 0 | A | G | 1.0216 | 0.2801 | 0.0002646 | +++?+ | 49.9 | 0.1125 | intron | DYNC2H1 |
| rs143710716 | 12 | 41280438 | 3 | 0 | 0 | A | C | -1.8339 | 0.5028 | 0.0002647 | ---?? | 0 | 0.8528 | intron | CNTN1 |
| rs147002299 | 8 | 5209397 | 2 | 0 | 0 | T | C | -4.8897 | 1.3406 | 0.0002648 | ?--?? | 0 | 0.8891 | intergenic | |
| rs2151061 | 9 | 16205165 | 4 | 0 | 0 | A | G | 0.9826 | 0.2694 | 0.0002649 | +++?+ | 0 | 0.8625 | intron | C9orf92 |
| 10:15914453 | 10 | 15914453 | 3 | 0 | 0 | G | GT | 1.0307 | 0.2826 | 0.0002649 | +++?? | 0 | 0.8195 | | |
| rs4587863 | 14 | 43257977 | 4 | 0 | 0 | T | C | -1.3901 | 0.3811 | 0.0002651 | ---?+ | 60.1 | 0.05729 | intergenic | |
| rs12982924 | 19 | 10201117 | 4 | 0 | 0 | A | G | -0.976 | 0.2676 | 0.0002651 | ---?- | 19.7 | 0.2914 | intron | C19orf66 |
| rs79326936 | 3 | 61110706 | 4 | 0 | 0 | A | T | -3.2646 | 0.8951 | 0.0002653 | ---?- | 0 | 0.84 | intron | FHIT |
| 9:133822641 | 9 | 133822641 | 3 | 0 | 0 | T | TTCAG | -4.4064 | 1.2083 | 0.0002654 | ---?? | 12.9 | 0.3171 | | |
| rs2131753 | 16 | 84843275 | 4 | 0 | 0 | A | G | -1.182 | 0.3241 | 0.0002656 | ---?+ | 0 | 0.6283 | intergenic | |
| rs10895434 | 11 | 103328121 | 4 | 0 | 0 | A | G | 1.0219 | 0.2802 | 0.0002656 | +++?+ | 48.9 | 0.1182 | intron | DYNC2H1 |
| rs117413706 | 15 | 87514282 | 4 | 0 | 0 | T | C | 2.5435 | 0.6975 | 0.0002656 | +++?+ | 0 | 0.903 | intron | AGBL1 |
| rs10872963 | 14 | 43256999 | 4 | 0 | 0 | C | G | -1.3919 | 0.3817 | 0.0002657 | ---?+ | 60.3 | 0.05592 | intergenic | |
| rs57502165 | 9 | 79458833 | 4 | 0 | 0 | A | G | -2.1131 | 0.5795 | 0.000266 | ---?- | 0 | 0.5356 | intron | PRUNE2 |
| rs76522354 | 4 | 117278403 | 4 | 0 | 0 | T | C | -3.314 | 0.9089 | 0.0002663 | ---?- | 0 | 0.8198 | intergenic | |
| rs11261008 | 1 | 18764102 | 4 | 0 | 0 | T | C | -1.0801 | 0.2963 | 0.0002664 | ---?- | 36.5 | 0.1929 | intergenic | |
| rs10280750 | 7 | 16903770 | 4 | 0 | 0 | T | C | 1.7378 | 0.4767 | 0.0002665 | +++?+ | 0 | 0.4156 | intron | AGR3 |
| rs76060979 | 12 | 26811820 | 4 | 0 | 0 | A | G | -1.8456 | 0.5062 | 0.0002665 | ---?- | 25 | 0.2613 | intron | ITPR2 |
| rs113700231 | 11 | 1130306 | 4 | 0 | 0 | C | G | -1.4418 | 0.3955 | 0.0002665 | ---?+ | 56.3 | 0.07609 | intergenic | |
| rs117763930 | 15 | 93011237 | 4 | 0 | 0 | A | G | 2.5462 | 0.6984 | 0.0002665 | +++?+ | 0 | 0.5123 | 3-prime-UTR | ST8SIA2 |
| rs148234966 | 6 | 83752311 | 4 | 0 | 0 | T | C | 3.4815 | 0.9549 | 0.0002666 | +++?+ | 0 | 0.756 | intron | UBE3D |
| 13:30325624 | 13 | 30325624 | 3 | 0 | 0 | T | TA | -1.5607 | 0.4281 | 0.0002666 | ---?? | 0 | 0.653 | | |
| rs72677035 | 4 | 119210133 | 4 | 0 | 0 | A | G | 1.6779 | 0.4602 | 0.0002667 | +++?+ | 5.1 | 0.3674 | intron | PRSS12 |
| rs114031402 | 4 | 39585489 | 4 | 0 | 0 | T | C | 3.949 | 1.0832 | 0.0002667 | +++?+ | 0 | 0.9771 | intron | SMIM14,UGDH-AS1 |
| rs183112019 | 4 | 174943269 | 3 | 0 | 0 | A | G | 3.0829 | 0.8457 | 0.0002668 | +-+?? | 64.5 | 0.05979 | intergenic | |
| rs61660159 | 6 | 109105308 | 2 | 0 | 0 | T | C | 5.0493 | 1.3851 | 0.0002669 | +?+?? | 0 | 0.8392 | intergenic | |
| rs182960130 | 2 | 167442756 | 3 | 0 | 0 | A | T | 3.8469 | 1.0553 | 0.0002669 | +++?? | 63.8 | 0.06321 | intergenic | |
| rs9659782 | 1 | 90261516 | 3 | 0 | 0 | A | G | 0.9108 | 0.2499 | 0.0002671 | +++?? | 0 | 0.5204 | intergenic | |
| rs117393454 | 16 | 62476025 | 4 | 0 | 0 | A | G | 2.2803 | 0.6255 | 0.0002671 | +++?+ | 0 | 0.5208 | intergenic | |
| rs6850040 | 4 | 38742953 | 5 | 0 | 0 | T | C | -1.2036 | 0.3302 | 0.0002674 | ----- | 0 | 0.9669 | intergenic |  |
| rs8181418 | 10 | 18982716 | 4 | 0 | 0 | C | G | 1.0841 | 0.2974 | 0.0002674 | -++?+ | 52.1 | 0.09968 | intergenic | |
| rs4772243 | 13 | 100422879 | 4 | 0 | 0 | A | G | -1.0483 | 0.2876 | 0.0002675 | ---?- | 57.1 | 0.0721 | intron | CLYBL |
| rs10907311 | 1 | 18764191 | 4 | 0 | 0 | T | C | -1.0798 | 0.2963 | 0.0002675 | ---?- | 36.7 | 0.192 | intergenic | |
| rs8058967 | 16 | 82762046 | 4 | 0 | 0 | T | C | -0.9134 | 0.2506 | 0.0002676 | ---?+ | 38.7 | 0.1798 | intron | CDH13 |
| rs193166702 | 19 | 45380521 | 2 | 0 | 0 | A | G | -4.272 | 1.1721 | 0.0002678 | -?-?? | 42.5 | 0.1874 | intron | PVRL2 |
| rs190410197 | 6 | 83757606 | 4 | 0 | 0 | C | G | -3.4721 | 0.9527 | 0.000268 | ---?- | 0 | 0.7433 | intron | UBE3D |
| rs62131757 | 19 | 55238242 | 3 | 0 | 0 | T | G | -1.0024 | 0.2751 | 0.0002682 | ---?? | 9.6 | 0.3306 | intron | KIR3DL3 |
| rs62131758 | 19 | 55238313 | 3 | 0 | 0 | T | C | -1.0024 | 0.2751 | 0.0002682 | ---?? | 9.6 | 0.3309 | intron | KIR3DL3 |
| rs72665994 | 14 | 28157095 | 4 | 0 | 0 | A | G | -2.1846 | 0.5995 | 0.0002683 | ---?- | 0 | 0.9966 | intergenic | |
| 8:113403256 | 8 | 113403256 | 3 | 0 | 0 | T | TAACTC | 1.0675 | 0.2929 | 0.0002684 | +++?? | 0 | 0.5945 | | |
| 20:54324101 | 20 | 54324101 | 3 | 0 | 0 | CA | C | -1.379 | 0.3784 | 0.0002685 | ---?? | 0 | 0.9011 | | |
| 20:6020389 | 20 | 6020389 | 3 | 0 | 0 | T | TA | -0.9313 | 0.2556 | 0.0002685 | ---?? | 0 | 0.8824 | | |
| rs4723328 | 7 | 33798880 | 4 | 0 | 0 | T | C | -0.8736 | 0.2398 | 0.0002686 | ---?- | 0 | 0.4847 | intergenic | |
| rs28693734 | 6 | 32603949 | 5 | 0 | 1 | A | G | -1.4305 | 0.3926 | 0.0002687 | ----- | 22.7 | 0.2701 | upstream-gene |  |
| rs17649886 | 4 | 169324868 | 4 | 0 | 0 | A | G | -0.9519 | 0.2613 | 0.0002688 | ---?- | 0 | 0.481 | intron | DDX60L |
| rs1772173 | 13 | 45626098 | 4 | 0 | 0 | A | T | -2.2225 | 0.61 | 0.000269 | ---?+ | 39.8 | 0.1729 | intergenic | |

| | | | | | | | | | | | | | | | |
|---|---|---|---|---|---|---|---|---|---|---|---|---|---|---|---|
| 11:7228367 | 11 | 7228367 | 3 | 0 | 0 | CAGAAA | C | -3.1611 | 0.8676 | 0.000269 | ---?? | 0 | 0.8906 | | |
| rs6133280 | 20 | 5994488 | 5 | 0 | 0 | A | G | -0.7592 | 0.2084 | 0.0002691 | ----+ | 0 | 0.5396 | intron | CRLS1 |
| rs987023 | 2 | 142402259 | 4 | 0 | 0 | T | C | -2.0713 | 0.5685 | 0.0002692 | ---?- | 0 | 0.8784 | intron | LRP1B |
| rs17127710 | 1 | 85918009 | 5 | 0 | 0 | T | C | 0.9736 | 0.2672 | 0.0002692 | +++-+ | 54.1 | 0.06838 | intron | DDAH1 |
| rs56208846 | 1 | 90263714 | 3 | 0 | 0 | T | C | 0.9091 | 0.2496 | 0.0002696 | +++?? | 0 | 0.521 | intergenic | |
| rs140764822 | 2 | 35677063 | 3 | 0 | 0 | T | C | -3.5996 | 0.9881 | 0.0002696 | ---?? | 0 | 0.6472 | intergenic | |
| rs3104994 | 8 | 96571283 | 4 | 0 | 0 | A | C | -1.5741 | 0.4321 | 0.0002697 | -+-?- | 53.9 | 0.08922 | intron | LOC100616530 |
| rs6114347 | 20 | 23842082 | 4 | 0 | 0 | T | C | 1.522 | 0.4178 | 0.0002697 | +++?+ | 0.6 | 0.3886 | intergenic | |
| rs7120664 | 11 | 59193265 | 3 | 0 | 0 | C | G | 3.6523 | 1.0026 | 0.0002697 | +?++? | 6.7 | 0.3426 | upstream-gene | |
| rs9965290 | 18 | 39998518 | 4 | 0 | 0 | A | G | 1.1765 | 0.323 | 0.0002697 | +++?- | 0 | 0.3982 | intron | LINC00907 |
| rs12889410 | 14 | 43230954 | 4 | 0 | 0 | A | T | -1.412 | 0.3876 | 0.0002697 | ---?+ | 61.6 | 0.05008 | intergenic | |
| rs146820240 | 6 | 83757577 | 4 | 0 | 0 | A | G | 3.4705 | 0.9527 | 0.0002697 | +++?+ | 0 | 0.7446 | intron | UBE3D |
| 6:35434337 | 6 | 35434337 | 3 | 0 | 0 | T | TC | -3.4853 | 0.9568 | 0.0002697 | ---?? | 0 | 0.7558 | | |
| rs1486951 | 13 | 60016356 | 3 | 0 | 0 | A | G | -0.9424 | 0.2587 | 0.0002698 | ---?? | 0 | 0.7963 | intergenic | |
| rs12684337 | 9 | 79456175 | 4 | 0 | 0 | A | G | -2.1882 | 0.6007 | 0.0002699 | ---?- | 0 | 0.4975 | intron | PRUNE2 |
| rs17283888 | 7 | 82423731 | 4 | 0 | 0 | T | C | -0.9485 | 0.2604 | 0.0002699 | -+-?- | 52.6 | 0.09684 | intron | PCLO |
| rs58087888 | 9 | 79458943 | 4 | 0 | 0 | T | C | -2.1085 | 0.5789 | 0.0002699 | ---?- | 0 | 0.5348 | intron | PRUNE2 |
| rs61973796 | 14 | 28156668 | 4 | 0 | 0 | A | G | -2.1884 | 0.6008 | 0.0002699 | ---?- | 0 | 0.9968 | intergenic | |
| rs61973795 | 14 | 28156504 | 4 | 0 | 0 | T | C | -2.1892 | 0.601 | 0.00027 | ---?- | 0 | 0.9968 | intergenic | |
| rs9302792 | 16 | 5164105 | 3 | 0 | 0 | C | G | 1.8503 | 0.508 | 0.0002701 | +++?? | 0 | 0.895 | intergenic | |
| rs189769121 | 4 | 174950152 | 4 | 0 | 0 | A | C | 3.0586 | 0.8397 | 0.0002701 | +-+?+ | 50.8 | 0.1071 | intergenic | |
| rs6931819 | 6 | 83793729 | 4 | 0 | 0 | T | G | 3.3064 | 0.9078 | 0.0002702 | +++?+ | 0 | 0.77 | intron | DOPEY1 |
| rs57136285 | 4 | 153834841 | 4 | 0 | 0 | A | T | -2.6403 | 0.7249 | 0.0002703 | ---?- | 0 | 0.5657 | downstream-gene | |
| rs6430615 | 2 | 137014907 | 4 | 0 | 0 | A | G | 0.9787 | 0.2687 | 0.0002707 | +++?+ | 0 | 0.8323 | intergenic | |
| rs12606027 | 18 | 39998860 | 4 | 0 | 0 | T | C | 1.1757 | 0.3228 | 0.0002708 | +++?- | 0 | 0.3984 | intron | LINC00907 |
| rs73125055 | 7 | 63442545 | 4 | 0 | 0 | A | C | -1.4667 | 0.4028 | 0.0002708 | ---?- | 0 | 0.8585 | intergenic | |
| rs10869722 | 9 | 78796931 | 4 | 0 | 0 | C | G | -0.8792 | 0.2414 | 0.0002709 | ---?- | 0 | 0.9993 | intron | PCSK5 |
| rs111439570 | 13 | 24965153 | 3 | 0 | 0 | A | G | 4.7056 | 1.2922 | 0.000271 | +++?? | 0 | 0.4119 | intergenic | |
| rs115713806 | 4 | 100633370 | 4 | 0 | 0 | A | G | 2.0973 | 0.5759 | 0.000271 | -++?+ | 45.4 | 0.139 | intergenic | |
| 21:21823237 | 21 | 21823237 | 3 | 0 | 0 | T | TA | -4.0388 | 1.1091 | 0.0002711 | ---?? | 17 | 0.2999 | | |
| rs2198452 | 11 | 59191240 | 3 | 0 | 0 | T | G | -3.655 | 1.0038 | 0.0002714 | -?--? | 6.7 | 0.3424 | upstream-gene | |
| rs145452949 | 17 | 56615060 | 5 | 0 | 0 | A | G | -2.4328 | 0.6681 | 0.0002714 | ----- | 0 | 0.8298 | intron | SEPTIN4 |
| rs145553917 | 16 | 57942620 | 3 | 0 | 0 | T | C | 4.2997 | 1.1809 | 0.0002714 | +++?? | 45.7 | 0.1586 | intron | CNGB1 |
| rs12302278 | 12 | 112512081 | 5 | 0 | 0 | C | G | 1.3449 | 0.3694 | 0.0002716 | +-+++ | 56.8 | 0.05494 | intron | NAA25 |
| rs28574646 | 15 | 85499722 | 3 | 0 | 0 | C | G | -0.999 | 0.2744 | 0.0002716 | ---?? | 0 | 0.5616 | intergenic | |
| rs76733895 | 2 | 137018651 | 4 | 0 | 0 | T | C | 0.9882 | 0.2714 | 0.0002717 | +++?+ | 0 | 0.8289 | intergenic | |
| rs1415266 | 1 | 162104228 | 5 | 0 | 0 | A | C | 0.9246 | 0.254 | 0.0002718 | ++++- | 39.8 | 0.156 | intron | NOS1AP |
| rs4533266 | 15 | 98048366 | 4 | 0 | 0 | T | C | -0.961 | 0.264 | 0.0002718 | ---?- | 0 | 0.8895 | intergenic | |
| rs7592856 | 2 | 137010272 | 4 | 0 | 0 | T | C | -0.9667 | 0.2655 | 0.0002718 | ---?- | 0 | 0.92 | intergenic | |
| rs12582389 | 12 | 15667104 | 4 | 0 | 0 | A | G | -1.2079 | 0.3318 | 0.0002718 | ---?- | 0 | 0.6053 | intron | PTPRO |
| rs115049330 | 2 | 170574385 | 4 | 0 | 0 | A | G | -1.3032 | 0.358 | 0.0002718 | ---?- | 0 | 0.9307 | intron | PHOSPHO2-KLHL23 |
| rs201435 | 7 | 101732865 | 4 | 0 | 0 | C | G | 0.9092 | 0.2497 | 0.0002722 | +++?+ | 0 | 0.8123 | intron | CUX1 |
| rs7575856 | 2 | 168054912 | 4 | 0 | 0 | A | G | 2.2197 | 0.6097 | 0.0002722 | +++?- | 0 | 0.6732 | intron | XIRP2 |
| rs6430614 | 2 | 137013930 | 4 | 0 | 0 | A | G | 0.9774 | 0.2685 | 0.0002723 | +++?+ | 0 | 0.8323 | intergenic | |
| rs34337585 | 11 | 12753678 | 4 | 0 | 0 | T | G | 1.4193 | 0.3899 | 0.0002723 | +++?+ | 0 | 0.4511 | intron | TEAD1 |
| 2:65972780 | 2 | 65972780 | 2 | 0 | 0 | C | G | -5.5207 | 1.5166 | 0.0002725 | ?--?? | 0 | 0.9088 | intergenic | |
| rs75647453 | 15 | 65914938 | 3 | 0 | 0 | A | G | 2.948 | 0.8099 | 0.0002726 | +++?? | 31.6 | 0.2318 | intron | SLC24A1 |
| rs1008710 | 15 | 42142392 | 5 | 0 | 0 | C | G | -0.9978 | 0.2741 | 0.0002727 | ----- | 0 | 0.5484 | intron | SPTBN5 |
| rs9570130 | 13 | 60021815 | 4 | 0 | 0 | A | G | -0.9152 | 0.2514 | 0.0002728 | ---?- | 0 | 0.543 | intergenic | |
| 1:64806450 | 1 | 64806450 | 3 | 0 | 0 | CA | C | -1.3466 | 0.37 | 0.0002728 | ---?? | 20.3 | 0.285 | | |
| 17:6687838 | 17 | 6687838 | 3 | 0 | 0 | CCAAAA | C | 3.631 | 0.9976 | 0.000273 | +++?? | 0 | 0.4504 | | |
| rs6564077 | 16 | 84840518 | 4 | 0 | 0 | C | G | 1.1656 | 0.3203 | 0.0002732 | +++?- | 0 | 0.6141 | intergenic | |
| rs6484706 | 11 | 34356694 | 3 | 0 | 0 | A | G | -4.9713 | 1.366 | 0.0002734 | ---?? | 0 | 0.6492 | intron | ABTB2 |
| 2:65972776 | 2 | 65972776 | 2 | 0 | 0 | G | GT | 5.5207 | 1.517 | 0.0002735 | ?++?? | 0 | 0.9083 | | |
| rs9317079 | 13 | 60021864 | 4 | 0 | 0 | A | G | -0.9149 | 0.2514 | 0.0002738 | ---?- | 0 | 0.5413 | intergenic | |
| rs11780166 | 8 | 113489275 | 4 | 0 | 0 | T | C | 1.0315 | 0.2835 | 0.0002738 | +++?+ | 0 | 0.7726 | intron | CSMD3 |
| rs12879026 | 14 | 43232017 | 4 | 0 | 0 | T | C | 1.4037 | 0.3858 | 0.0002739 | +++?- | 61.5 | 0.05045 | intergenic | |
| 10:51117969 | 10 | 51117969 | 2 | 0 | 0 | T | C | -6.4729 | 1.7789 | 0.0002741 | --??? | 68.4 | 0.07548 | intron | PARG |
| rs182788419 | 22 | 27617178 | 3 | 0 | 0 | T | G | 2.9043 | 0.7982 | 0.0002742 | +++?? | 0 | 0.8902 | intergenic | |

| | | | | | | | | | | | | | | | |
|---|---|---|---|---|---|---|---|---|---|---|---|---|---|---|---|
| rs6725367 | 2 | 137018214 | 4 | 0 | 0 | A | G | 0.9856 | 0.2709 | 0.0002744 | +++?+ | 0 | 0.8312 | intergenic | |
| rs992859 | 7 | 130120122 | 4 | 0 | 0 | T | C | -0.8906 | 0.2448 | 0.0002745 | -+-?- | 60.4 | 0.05582 | intergenic | |
| rs2156588 | 11 | 106774133 | 4 | 0 | 0 | T | G | -1.0845 | 0.2981 | 0.0002748 | ---?+ | 45.1 | 0.1406 | intron | GUCY1A2 |
| rs884317 | 9 | 30904898 | 3 | 0 | 0 | T | G | 3.2999 | 0.9071 | 0.000275 | +++?? | 0 | 0.6226 | intergenic | |
| rs73194416 | 13 | 50367273 | 5 | 0 | 0 | A | G | 1.4232 | 0.3912 | 0.000275 | ++++- | 0 | 0.7183 | upstream-gene | |
| rs17063064 | 9 | 79459014 | 4 | 0 | 0 | C | G | -2.1013 | 0.5776 | 0.0002751 | ---?- | 0 | 0.5325 | intron | PRUNE2 |
| rs34028389 | 11 | 12697361 | 4 | 0 | 0 | T | C | 1.3842 | 0.3805 | 0.0002752 | +++?+ | 14.6 | 0.3189 | intron | TEAD1 |
| rs73651356 | 9 | 95837930 | 5 | 0 | 0 | A | G | 2.4832 | 0.6827 | 0.0002752 | +++++ | 0 | 0.9718 | intron | SUSD3 |
| rs1818672 | 1 | 18759858 | 4 | 0 | 0 | T | C | -1.0713 | 0.2945 | 0.0002753 | ---?- | 32.6 | 0.2166 | intergenic | |
| rs201090642 | 15 | 85499703 | 3 | 0 | 0 | T | C | 1.2302 | 0.3382 | 0.0002753 | +++?? | 0 | 0.4249 | intergenic | |
| rs868025 | 9 | 30904925 | 3 | 0 | 0 | C | G | -3.2998 | 0.9072 | 0.0002754 | ---?? | 0 | 0.6226 | intergenic | |
| rs35324266 | 6 | 32593244 | 2 | 0 | 0 | A | G | 2.079 | 0.5716 | 0.0002754 | ???++ | 0 | 0.6682 | intergenic | |
| rs76805623 | 16 | 84858877 | 4 | 0 | 0 | T | G | 1.3377 | 0.3678 | 0.0002754 | +++?+ | 0 | 0.7608 | intron | CRISPLD2 |
| rs144283499 | 7 | 86840833 | 2 | 0 | 0 | A | G | -9.0197 | 2.4797 | 0.0002754 | ?-??- | 55.5 | 0.1339 | intron | TMEM243 |
| rs7959011 | 12 | 112501237 | 5 | 0 | 0 | A | G | 1.3444 | 0.3696 | 0.0002755 | +-+++ | 57.1 | 0.05367 | intron | NAA25 |
| rs72842490 | 11 | 1125618 | 4 | 0 | 0 | A | G | -1.4398 | 0.3959 | 0.0002758 | ---?+ | 54.3 | 0.08708 | intergenic | |
| rs149583738 | 6 | 35432038 | 3 | 0 | 0 | T | C | -3.4766 | 0.9559 | 0.0002758 | ---?? | 0 | 0.7498 | intron | FANCE |
| rs1336121 | 1 | 18761198 | 4 | 0 | 0 | T | C | 1.0744 | 0.2954 | 0.0002759 | +++?+ | 33.6 | 0.2106 | intergenic | |
| rs7660114 | 4 | 38732464 | 5 | 0 | 0 | T | C | -1.0511 | 0.289 | 0.0002759 | ----- | 0 | 0.897 | intergenic |  |
| rs56174638 | 1 | 204391371 | 4 | 0 | 0 | A | G | -1.5004 | 0.4126 | 0.0002759 | ---?- | 9 | 0.3481 | downstream-gene | |
| rs11066120 | 12 | 112436195 | 5 | 0 | 0 | T | C | 1.346 | 0.3701 | 0.0002761 | +-+++ | 57.5 | 0.05159 | intron | TMEM116 |
| rs12316797 | 12 | 43160744 | 5 | 0 | 0 | T | G | -1.1409 | 0.3137 | 0.0002765 | ----+ | 24.5 | 0.2578 | intergenic |  |
| rs4258520 | 14 | 43254770 | 4 | 0 | 0 | T | C | 1.3916 | 0.3827 | 0.0002766 | +++?- | 60.9 | 0.05343 | intergenic | |
| rs72842489 | 11 | 1125601 | 4 | 0 | 0 | T | C | -1.4396 | 0.3959 | 0.0002766 | ---?+ | 54.3 | 0.08722 | intergenic | |
| rs73082118 | 7 | 16904437 | 4 | 0 | 0 | T | C | -1.713 | 0.4711 | 0.0002766 | ---?- | 0 | 0.4119 | intron | AGR3 |
| rs72709093 | 15 | 45698327 | 5 | 0 | 0 | T | C | 2.1487 | 0.591 | 0.0002772 | +++++ | 0 | 0.6878 | intron | SPATA5L1 |
| rs77021556 | 15 | 65809642 | 3 | 0 | 0 | A | G | 2.9278 | 0.8053 | 0.0002773 | +++?? | 13.3 | 0.3155 | intron | DPP8 |
| rs79470844 | 6 | 106270389 | 4 | 0 | 0 | A | G | 2.1563 | 0.5931 | 0.0002773 | +++?+ | 0 | 0.5608 | intergenic | |
| rs6493010 | 15 | 42007520 | 5 | 0 | 0 | C | G | 1.0309 | 0.2836 | 0.0002774 | +++++ | 9.3 | 0.3536 | intron | MGA |
| rs7572721 | 2 | 170910733 | 4 | 0 | 1 | T | C | -1.0627 | 0.2923 | 0.0002774 | ---?- | 41.3 | 0.1637 | intron | UBR3 |
| rs16822616 | 6 | 32410083 | 5 | 0 | 1 | A | C | 1.4068 | 0.387 | 0.0002774 | +++++ | 0 | 0.4362 | intron | HLA-DRA |
| rs17496549 | 6 | 32409708 | 5 | 0 | 1 | T | C | 1.4068 | 0.387 | 0.0002774 | +++++ | 0 | 0.4362 | intron | HLA-DRA |
| rs71318346 | 3 | 183572055 | 4 | 0 | 0 | A | G | -1.7578 | 0.4835 | 0.0002774 | ---?- | 0 | 0.8437 | intron | PARL |
| rs72668247 | 14 | 28206565 | 4 | 0 | 0 | T | C | 2.1387 | 0.5883 | 0.0002775 | +++?+ | 0 | 0.976 | intergenic | |
| rs149580742 | 17 | 56638557 | 5 | 0 | 0 | A | T | 2.4979 | 0.6871 | 0.0002775 | +++++ | 0 | 0.7615 | intron | TEX14 |
| rs2767419 | 10 | 134512521 | 4 | 0 | 0 | T | C | -0.9176 | 0.2524 | 0.0002776 | ---?- | 0 | 0.6504 | intron | INPP5A |
| rs7603806 | 2 | 137019036 | 4 | 0 | 0 | T | C | 0.9877 | 0.2717 | 0.0002776 | +++?+ | 0 | 0.8319 | intergenic | |
| rs10505174 | 8 | 113485645 | 4 | 0 | 0 | T | C | 1.0253 | 0.282 | 0.0002776 | +++?+ | 0 | 0.7386 | intron | CSMD3 |
| rs10907310 | 1 | 18759952 | 4 | 0 | 0 | T | C | -1.0713 | 0.2947 | 0.0002776 | ---?- | 32.9 | 0.2146 | intergenic | |
| rs4886165 | 13 | 60034843 | 4 | 0 | 0 | A | G | -0.9164 | 0.2521 | 0.0002777 | ---?- | 0 | 0.4537 | intergenic | |
| rs13144335 | 4 | 81178916 | 3 | 0 | 0 | A | C | -2.5695 | 0.7068 | 0.0002777 | ---?? | 34.8 | 0.2155 | intergenic | |
| rs144390575 | 6 | 83735987 | 4 | 0 | 0 | A | G | 3.4922 | 0.9606 | 0.0002777 | +++?+ | 0 | 0.7944 | intron | UBE3D |
| rs7659947 | 4 | 38732398 | 5 | 0 | 0 | T | C | -1.0506 | 0.289 | 0.0002779 | ----- | 0 | 0.8966 | intergenic |  |
| rs2647074 | 6 | 32574360 | 3 | 0 | 0 | T | C | 1.0303 | 0.2834 | 0.000278 | +++?? | 0 | 0.4112 | intergenic | |
| rs34734720 | 14 | 43233274 | 4 | 0 | 0 | A | T | 1.4018 | 0.3856 | 0.000278 | +++?- | 61.6 | 0.05019 | intergenic | |
| rs61974366 | 14 | 28206905 | 4 | 0 | 0 | T | C | -2.139 | 0.5885 | 0.0002781 | ---?- | 0 | 0.9756 | intergenic | |
| 3:31814492 | 3 | 31814492 | 3 | 0 | 0 | CT | C | 1.4606 | 0.4018 | 0.0002781 | +++?? | 0 | 0.949 | | |
| rs115892942 | 2 | 168064023 | 4 | 0 | 0 | T | C | 2.4296 | 0.6684 | 0.0002782 | +++?- | 0 | 0.4618 | intron | XIRP2 |
| rs1985532 | 10 | 116563616 | 4 | 0 | 0 | A | T | -1.505 | 0.4141 | 0.0002784 | ---?- | 0 | 0.9404 | intergenic | |
| rs12915325 | 15 | 41986639 | 5 | 0 | 0 | T | C | 1.0356 | 0.2849 | 0.0002785 | +++++ | 13.8 | 0.3262 | intron | MGA |
| rs13245080 | 7 | 107345392 | 4 | 0 | 0 | A | G | -0.9195 | 0.253 | 0.0002785 | ---?- | 0 | 0.4171 | intron | SLC26A4 |
| rs144920858 | 17 | 4782937 | 5 | 0 | 0 | C | G | 1.6469 | 0.4531 | 0.0002786 | ++++- | 20 | 0.2872 | intron | MINK1 |
| 12:1388893 | 12 | 1388893 | 3 | 0 | 0 | T | C | -4.1082 | 1.1304 | 0.0002787 | ---?? | 0 | 0.3899 | intron | ERC1 |
| rs6564079 | 16 | 84840883 | 4 | 0 | 0 | T | C | 1.1639 | 0.3202 | 0.0002787 | +++?- | 0 | 0.4114 | intergenic | |
| rs6717894 | 2 | 131122276 | 3 | 0 | 0 | A | G | 1.1222 | 0.3088 | 0.0002787 | +++?? | 7 | 0.3411 | intron | PTPN18 |
| rs72630468 | 15 | 85635528 | 3 | 0 | 0 | A | T | 1.4641 | 0.4029 | 0.0002788 | +++?? | 43.7 | 0.1691 | intron | PDE8A |
| rs11055596 | 12 | 13858382 | 4 | 0 | 0 | T | C | -0.8968 | 0.2468 | 0.0002789 | ---?- | 0 | 0.6725 | intron | GRIN2B |
| rs28619515 | 16 | 84840957 | 4 | 0 | 0 | A | C | 1.1749 | 0.3233 | 0.0002789 | +++?- | 0 | 0.5963 | intergenic | |
| rs116520637 | 4 | 100592532 | 4 | 0 | 0 | C | G | 2.0593 | 0.5667 | 0.0002789 | -++?+ | 47.7 | 0.1249 | intergenic | |

| | | | | | | | | | | | | | | | |
|---|---|---|---|---|---|---|---|---|---|---|---|---|---|---|---|
| rs10087242 | 8 | 113489902 | 4 | 0 | 0 | T | G | 1.0312 | 0.2837 | 0.000279 | +++?+ | 0 | 0.7682 | intron | CSMD3 |
| rs142021399 | 10 | 28134417 | 3 | 0 | 0 | T | C | -2.0781 | 0.5718 | 0.000279 | ---?? | 0 | 0.5486 | intron | ARMC4 |
| rs11587876 | 1 | 85915183 | 5 | 0 | 0 | T | C | -0.9681 | 0.2664 | 0.0002792 | ---+- | 54.1 | 0.06879 | intron | DDAH1 |
| rs1902 | 7 | 101768791 | 4 | 0 | 0 | T | C | 0.8765 | 0.2412 | 0.0002799 | +++?+ | 0 | 0.7329 | intron | CUX1 |
| rs36006606 | 6 | 32591424 | 5 | 0 | 1 | A | G | -1.4425 | 0.397 | 0.00028 | ----- | 30.8 | 0.216 | intergenic |  |
| rs201516 | 7 | 101769507 | 4 | 0 | 0 | T | C | 0.8763 | 0.2412 | 0.0002801 | +++?+ | 0 | 0.7329 | intron | CUX1 |
| rs6992428 | 8 | 113405396 | 3 | 0 | 0 | T | C | 1.0644 | 0.293 | 0.0002801 | +++?? | 0 | 0.583 | intron | CSMD3 |
| rs7525893 | 1 | 90246906 | 5 | 0 | 0 | T | C | 0.7751 | 0.2134 | 0.0002801 | +++++ | 0 | 0.9715 | intergenic | |
| rs13032026 | 2 | 18519857 | 5 | 0 | 0 | T | C | -0.8168 | 0.2248 | 0.0002801 | ----- | 0 | 0.9834 | intergenic |  |
| 4:8678883 | 4 | 8678883 | 3 | 0 | 0 | A | AG | 3.7132 | 1.0221 | 0.0002801 | +++?? | 0 | 0.8301 | | |
| rs60586860 | 17 | 47946055 | 5 | 0 | 0 | T | C | -2.1407 | 0.5893 | 0.0002802 | ----- | 0 | 0.8983 | intergenic |  |
| rs149476073 | 1 | 85909708 | 5 | 0 | 0 | A | G | 0.9696 | 0.2669 | 0.0002803 | +++-+ | 55.1 | 0.06343 | intron | DDAH1 |
| rs77964710 | 7 | 134736969 | 4 | 0 | 0 | T | C | 4.9535 | 1.3636 | 0.0002806 | +++?+ | 35.3 | 0.2004 | intron | AGBL3 |
| rs112127105 | 6 | 83740130 | 4 | 0 | 0 | C | G | -3.4763 | 0.957 | 0.0002806 | ---?- | 0 | 0.8018 | intron | UBE3D |
| rs13389089 | 2 | 170909854 | 4 | 0 | 1 | T | C | -1.0621 | 0.2924 | 0.0002807 | ---?- | 42.9 | 0.1542 | intron | UBR3 |
| rs72687745 | 4 | 182796188 | 4 | 0 | 0 | A | G | -3.1271 | 0.8609 | 0.0002807 | ---?- | 0 | 0.8781 | intergenic | |
| 1:196473354 | 1 | 196473354 | 2 | 0 | 0 | T | C | -5.644 | 1.5538 | 0.0002808 | -?-?? | 74 | 0.05001 | intron | KCNT2 |
| rs4148153 | 4 | 89056715 | 4 | 0 | 0 | A | G | -4.1767 | 1.1498 | 0.0002808 | -?--- | 0 | 0.7237 | intron | ABCG2 |
| rs7787199 | 7 | 82425463 | 4 | 0 | 0 | T | C | 0.9426 | 0.2595 | 0.0002808 | +-+?+ | 52.1 | 0.09953 | intron | PCLO |
| rs948084 | 11 | 130631608 | 4 | 0 | 0 | A | C | 1.7783 | 0.4896 | 0.000281 | +++?+ | 12.1 | 0.3323 | intergenic | |
| rs2455788 | 8 | 140494834 | 4 | 0 | 0 | A | T | 1.5 | 0.413 | 0.000281 | +++?+ | 0 | 0.7908 | intergenic | |
| rs201514 | 7 | 101768910 | 4 | 0 | 0 | T | C | -0.8762 | 0.2412 | 0.0002812 | ---?- | 0 | 0.7328 | intron | CUX1 |
| rs201515 | 7 | 101769164 | 4 | 0 | 0 | A | G | 0.8761 | 0.2412 | 0.0002813 | +++?+ | 0 | 0.733 | intron | CUX1 |
| rs10747295 | 14 | 22530819 | 4 | 0 | 0 | A | G | -0.9513 | 0.2619 | 0.0002814 | ---?- | 0 | 0.6575 | intergenic | |
| rs11963997 | 6 | 12206709 | 4 | 0 | 0 | T | G | -3.3757 | 0.9295 | 0.0002814 | ---?- | 3.1 | 0.3771 | intergenic | |
| rs3762856 | 4 | 169327191 | 4 | 0 | 0 | T | C | -0.9485 | 0.2612 | 0.0002815 | ---?- | 0 | 0.483 | synonymous | DDX60L |
| rs2204974 | 14 | 22506703 | 4 | 0 | 0 | A | G | 0.9526 | 0.2623 | 0.0002816 | +++?+ | 0 | 0.9651 | intergenic | |
| rs79376596 | 20 | 2399550 | 3 | 0 | 0 | A | G | -3.9149 | 1.0781 | 0.0002818 | ---?? | 6.6 | 0.3429 | intron | TGM6 |
| rs72842485 | 11 | 1122277 | 4 | 0 | 0 | A | G | -1.4313 | 0.3942 | 0.0002819 | ---?+ | 58.9 | 0.06302 | intergenic | |
| rs9538421 | 13 | 60009579 | 4 | 0 | 0 | A | T | 0.9305 | 0.2562 | 0.000282 | +++?+ | 0 | 0.5357 | intergenic | |
| rs9598012 | 13 | 60019811 | 4 | 0 | 0 | A | T | -0.9179 | 0.2528 | 0.000282 | ---?- | 0 | 0.5238 | intergenic | |
| rs10498380 | 14 | 43622917 | 4 | 0 | 0 | T | C | 1.849 | 0.5092 | 0.000282 | +++?- | 0 | 0.5757 | intergenic | |
| rs7129592 | 11 | 37060502 | 4 | 0 | 0 | T | C | 0.8839 | 0.2434 | 0.0002822 | +++?+ | 8.1 | 0.3528 | intergenic | |
| rs1536087 | 10 | 131920160 | 4 | 0 | 0 | T | C | -0.8932 | 0.246 | 0.0002824 | ---?- | 0 | 0.7336 | intergenic | |
| rs10459490 | 14 | 43248147 | 4 | 0 | 0 | A | T | -1.3985 | 0.3852 | 0.0002825 | ---?+ | 60.8 | 0.05366 | intergenic | |
| rs805745 | 20 | 5685929 | 4 | 0 | 0 | A | G | -1.0472 | 0.2884 | 0.0002826 | ---?- | 0 | 0.6275 | intergenic | |
| rs6482617 | 10 | 18982965 | 4 | 0 | 0 | A | T | -1.0806 | 0.2976 | 0.0002826 | +--?- | 52.2 | 0.09895 | intergenic | |
| rs79426697 | 11 | 21168714 | 2 | 0 | 0 | A | G | 6.4373 | 1.773 | 0.0002826 | +???+ | 50.4 | 0.1554 | intron | NELL1 |
| rs201523 | 7 | 101771851 | 4 | 0 | 0 | T | C | 0.876 | 0.2413 | 0.0002832 | +++?+ | 0 | 0.7377 | intron | CUX1 |
| rs6489824 | 12 | 112468417 | 5 | 0 | 0 | T | C | 1.3414 | 0.3695 | 0.0002832 | +-+++ | 57.5 | 0.05145 | intron | NAA25 |
| rs17754389 | 8 | 1268027 | 4 | 0 | 0 | C | G | 1.1396 | 0.3139 | 0.0002834 | +++?+ | 0 | 0.4471 | intron | DLGAP2 |
| 4:130153203 | 4 | 130153203 | 3 | 0 | 0 | CA | C | -1.0756 | 0.2963 | 0.0002834 | ---?? | 38.4 | 0.1972 | | |
| rs201517 | 7 | 101769616 | 4 | 0 | 0 | T | G | 0.8756 | 0.2412 | 0.0002835 | +++?+ | 0 | 0.733 | intron | CUX1 |
| rs4772245 | 13 | 100428074 | 4 | 0 | 0 | T | C | -1.0483 | 0.2888 | 0.0002835 | ---?- | 59.2 | 0.0615 | intron | CLYBL |
| rs578475 | 5 | 134354896 | 5 | 0 | 0 | C | G | 0.8326 | 0.2294 | 0.0002836 | +++++ | 0 | 0.4802 | intergenic | |
| rs12312538 | 12 | 112489521 | 5 | 0 | 0 | T | C | -1.3413 | 0.3695 | 0.0002836 | -+--- | 57.6 | 0.05133 | intron | NAA25 |
| rs58368428 | 12 | 112462275 | 5 | 0 | 0 | A | G | 1.3412 | 0.3695 | 0.0002836 | +-+++ | 57.6 | 0.05099 | downstream-gene | |
| rs142437699 | 8 | 96558727 | 4 | 0 | 0 | A | T | -1.6697 | 0.46 | 0.0002837 | -+-?- | 55.9 | 0.07864 | intron | LOC100616530 |
| rs6929008 | 6 | 83738744 | 4 | 0 | 0 | T | C | -3.4741 | 0.9572 | 0.000284 | ---?- | 0 | 0.804 | intron | UBE3D |
| rs4344456 | 11 | 106767880 | 3 | 0 | 0 | A | T | -1.1509 | 0.3171 | 0.0002842 | ---?? | 1.6 | 0.3619 | intron | GUCY1A2 |
| rs201518 | 7 | 101770267 | 4 | 0 | 0 | T | C | 0.8752 | 0.2412 | 0.0002843 | +++?+ | 0 | 0.7336 | intron | CUX1 |
| rs77865444 | 9 | 79459080 | 4 | 0 | 0 | T | C | 2.0872 | 0.5751 | 0.0002843 | +++?+ | 0 | 0.529 | intron | PRUNE2 |
| rs78713455 | 4 | 100591628 | 4 | 0 | 0 | T | C | -2.0596 | 0.5675 | 0.0002844 | +--?- | 48 | 0.1233 | intergenic | |
| rs36214155 | 6 | 32611778 | 4 | 0 | 1 | T | C | -1.8776 | 0.5174 | 0.0002845 | ----? | 0 | 0.4199 | 3-prime-UTR | HLA-DQA1 |
| rs78387246 | 20 | 23814024 | 4 | 0 | 0 | T | G | 1.5104 | 0.4162 | 0.0002845 | +++?+ | 0 | 0.5715 | intergenic | |
| rs16852191 | 1 | 203675105 | 4 | 0 | 0 | T | C | -1.1644 | 0.3209 | 0.0002851 | ---?- | 0 | 0.9616 | intron | ATP2B4 |
| rs2950446 | 15 | 42012724 | 5 | 0 | 0 | T | C | 1.0269 | 0.283 | 0.0002852 | +++++ | 8.5 | 0.358 | intron | MGA |
| rs201519 | 7 | 101770505 | 4 | 0 | 0 | A | G | 0.875 | 0.2412 | 0.0002853 | +++?+ | 0 | 0.7341 | intron | CUX1 |
| rs6458 | 6 | 32007624 | 5 | 0 | 0 | A | G | -1.2334 | 0.34 | 0.0002854 | ---+- | 47.8 | 0.1045 | intron | CYP21A2 |

| | | | | | | | | | | | | | | | |
|---|---|---|---|---|---|---|---|---|---|---|---|---|---|---|---|
| rs28484707 | 18 | 35857546 | 4 | 0 | 0 | T | G | 1.3993 | 0.3857 | 0.0002854 | +++?+ | 0 | 0.9981 | intergenic | |
| rs78121801 | 7 | 36152873 | 4 | 0 | 0 | T | C | -2.633 | 0.7257 | 0.0002854 | ---?- | 0 | 0.9069 | intergenic | |
| rs140763142 | 17 | 69888759 | 3 | 0 | 0 | A | T | 1.2103 | 0.3336 | 0.0002855 | +++?? | 0 | 0.4733 | intergenic | |
| rs9960053 | 18 | 47734985 | 4 | 0 | 0 | C | G | 1.0343 | 0.2851 | 0.0002856 | +++?+ | 19.7 | 0.2914 | intergenic | |
| rs201520 | 7 | 101770677 | 4 | 0 | 0 | A | G | 0.8748 | 0.2411 | 0.0002857 | +++?+ | 0 | 0.7337 | intron | CUX1 |
| rs11031146 | 11 | 30630607 | 4 | 0 | 0 | A | G | 0.9574 | 0.2639 | 0.0002857 | +++?- | 32.1 | 0.2196 | intergenic | |
| rs28986410 | 6 | 32755087 | 5 | 0 | 0 | T | C | -1.4725 | 0.4059 | 0.0002857 | ----- | 0 | 0.7054 | intergenic |  |
| rs28653778 | 18 | 35857288 | 4 | 0 | 0 | A | G | -1.399 | 0.3856 | 0.0002859 | ---?- | 0 | 0.9981 | intergenic | |
| rs72846303 | 11 | 1133795 | 4 | 0 | 0 | A | G | -1.4408 | 0.3972 | 0.0002859 | ---?+ | 57.8 | 0.06849 | intergenic | |
| 12:41298235 | 12 | 41298235 | 3 | 0 | 0 | T | TTTTC | 1.82 | 0.5017 | 0.0002859 | +++?? | 0 | 0.8615 | | |
| rs28472766 | 18 | 35857502 | 4 | 0 | 0 | T | C | -1.4208 | 0.3917 | 0.0002863 | ---?- | 0 | 0.9995 | intergenic | |
| rs76654283 | 2 | 170442751 | 4 | 0 | 0 | T | C | 1.2889 | 0.3553 | 0.0002864 | +++?+ | 0 | 0.9236 | intron | PPIG |
| rs79615801 | 6 | 5495106 | 4 | 0 | 0 | A | G | 3.3736 | 0.9301 | 0.0002864 | +++?+ | 0 | 0.5943 | intron | FARS2 |
| rs2072065 | 7 | 107334661 | 4 | 0 | 0 | T | C | -0.9164 | 0.2527 | 0.0002866 | ---?- | 0 | 0.5178 | intron | SLC26A4 |
| rs7295294 | 12 | 112464893 | 5 | 0 | 0 | T | C | 1.3402 | 0.3695 | 0.0002867 | +-+++ | 57.5 | 0.05142 | 3-prime-UTR | NAA25 |
| rs73020379 | 2 | 170593234 | 4 | 0 | 0 | A | G | -1.1831 | 0.3262 | 0.0002867 | ---?- | 0 | 0.9977 | intron | KLHL23,PHOSPHO2-KLHL23 |
| rs6740242 | 2 | 167981617 | 4 | 0 | 0 | T | G | -2.3058 | 0.6358 | 0.0002869 | ---?+ | 8.7 | 0.3496 | intron | XIRP2 |
| rs2577954 | 15 | 42013977 | 5 | 0 | 0 | A | G | -1.0233 | 0.2822 | 0.000287 | ----- | 11.7 | 0.3392 | intron | MGA |
| rs60582557 | 11 | 1121615 | 4 | 0 | 0 | T | G | -1.4289 | 0.394 | 0.000287 | ---?+ | 58.6 | 0.06445 | intergenic | |
| rs4980353 | 15 | 85512767 | 3 | 0 | 0 | C | G | -0.9699 | 0.2674 | 0.0002871 | ---?? | 0 | 0.4734 | intergenic | |
| rs72668257 | 14 | 28214456 | 4 | 0 | 0 | T | C | 2.1483 | 0.5924 | 0.0002874 | +++?+ | 0 | 0.965 | intergenic | |
| rs6557136 | 6 | 151729237 | 4 | 0 | 0 | T | G | 0.9937 | 0.2741 | 0.0002878 | +++?+ | 0 | 0.5027 | intron | RMND1 |
| rs114881711 | 4 | 100594594 | 4 | 0 | 0 | T | C | 2.0586 | 0.5677 | 0.0002878 | -++?+ | 47.6 | 0.1256 | intergenic | |
| rs201521 | 7 | 101771197 | 4 | 0 | 0 | A | G | 0.8742 | 0.2411 | 0.000288 | +++?+ | 0 | 0.7341 | intron | CUX1 |
| rs12766706 | 10 | 134581520 | 4 | 0 | 0 | T | C | -1.0079 | 0.278 | 0.000288 | ---?- | 0 | 0.4887 | intron | INPP5A |
| rs4321533 | 3 | 68217843 | 4 | 0 | 0 | C | G | -2.2483 | 0.6201 | 0.0002881 | ---?- | 32 | 0.2206 | intron | FAM19A1 |
| rs117542611 | 7 | 27409314 | 4 | 0 | 0 | C | G | 1.9062 | 0.5257 | 0.0002881 | +++?+ | 0 | 0.7293 | intergenic | |
| rs7169496 | 15 | 41982203 | 5 | 0 | 0 | A | G | 1.0298 | 0.2841 | 0.0002883 | +++++ | 14.6 | 0.3214 | intron | MGA |
| rs137954053 | 10 | 1072185 | 4 | 0 | 0 | C | G | -2.0251 | 0.5586 | 0.0002883 | ---?- | 43.3 | 0.1516 | intron | IDI2-AS1 |
| rs114586213 | 4 | 100593805 | 4 | 0 | 0 | T | C | -2.0583 | 0.5678 | 0.0002885 | +--?- | 47.7 | 0.1254 | intergenic | |
| rs79723107 | 9 | 79459084 | 4 | 0 | 0 | A | G | -2.0827 | 0.5745 | 0.0002886 | ---?- | 0 | 0.5279 | intron | PRUNE2 |
| rs115555108 | 3 | 176754939 | 3 | 0 | 0 | T | C | 3.3267 | 0.9176 | 0.0002886 | +++?? | 0 | 0.6836 | intron | TBL1XR1 |
| rs73170452 | 8 | 1267895 | 4 | 0 | 0 | A | G | 1.1391 | 0.3142 | 0.0002888 | +++?+ | 0 | 0.4376 | intron | DLGAP2 |
| rs1662892 | 18 | 47731764 | 4 | 0 | 0 | A | C | -1.0464 | 0.2887 | 0.0002889 | ---?- | 0 | 0.3992 | intergenic | |
| rs111949920 | 12 | 102324197 | 4 | 0 | 0 | T | C | 2.0912 | 0.5769 | 0.000289 | +++?+ | 0 | 0.505 | intergenic | |
| rs1572822 | 1 | 102934942 | 4 | 0 | 0 | A | G | -0.8652 | 0.2387 | 0.0002891 | ---?+ | 7.4 | 0.3562 | intergenic | |
| rs7297186 | 12 | 111778178 | 5 | 0 | 0 | T | C | -0.8968 | 0.2474 | 0.0002891 | -+--- | 46.2 | 0.1148 | intron | CUX2 |
| rs61973808 | 14 | 28166805 | 4 | 0 | 0 | T | C | -2.0868 | 0.5757 | 0.0002892 | ---?- | 0 | 0.9938 | intergenic | |
| rs2695349 | 17 | 55457550 | 4 | 0 | 0 | T | C | 1.5869 | 0.4378 | 0.0002893 | +++?- | 28.5 | 0.2412 | intron | MSI2 |
| rs9366560 | 6 | 24084276 | 3 | 0 | 0 | A | G | 1.7633 | 0.4865 | 0.0002893 | +++?? | 0 | 0.4171 | intergenic | |
| rs10134949 | 14 | 43870979 | 4 | 0 | 0 | A | G | 1.9781 | 0.5457 | 0.0002894 | +++?+ | 0 | 0.8008 | intergenic | |
| rs13088984 | 3 | 183556726 | 4 | 0 | 0 | T | C | 1.7583 | 0.4851 | 0.0002894 | +++?+ | 0 | 0.8395 | intron | PARL |
| rs55690535 | 5 | 701329 | 3 | 0 | 0 | T | C | 1.398 | 0.3857 | 0.0002894 | +++?? | 0 | 0.3936 | intergenic | |
| rs1511472 | 6 | 23871249 | 4 | 0 | 0 | T | C | -2.1417 | 0.5909 | 0.0002895 | ---?+ | 39.8 | 0.1729 | intergenic | |
| rs115772710 | 2 | 168060211 | 4 | 0 | 0 | C | G | -2.2165 | 0.6115 | 0.0002896 | ---?+ | 0 | 0.6719 | intron | XIRP2 |
| rs2577953 | 15 | 42014864 | 5 | 0 | 0 | T | C | -1.0238 | 0.2825 | 0.0002899 | ----- | 8 | 0.3611 | intron | MGA |
| rs2918051 | 15 | 42018654 | 5 | 0 | 0 | T | C | 1.0224 | 0.2821 | 0.0002899 | +++++ | 7.2 | 0.3657 | intron | MGA |
| rs1185325 | 10 | 131913705 | 4 | 0 | 0 | T | C | -0.8887 | 0.2452 | 0.0002902 | ---?- | 0 | 0.796 | upstream-gene | |
| rs61973809 | 14 | 28166920 | 4 | 0 | 0 | A | G | -2.0852 | 0.5754 | 0.0002902 | ---?- | 0 | 0.9937 | intergenic | |
| rs4270151 | 15 | 41994307 | 5 | 0 | 0 | T | C | 1.029 | 0.284 | 0.0002903 | +++++ | 12.7 | 0.3332 | intron | MGA |
| rs61973810 | 14 | 28166968 | 4 | 0 | 0 | A | G | -2.0848 | 0.5753 | 0.0002904 | ---?- | 0 | 0.9937 | intergenic | |
| rs8034261 | 15 | 41994589 | 5 | 0 | 0 | T | G | 1.0289 | 0.2839 | 0.0002907 | +++++ | 12.6 | 0.3337 | intron | MGA |
| rs28613297 | 18 | 35879633 | 4 | 0 | 0 | A | G | 1.4379 | 0.3968 | 0.0002907 | +++?+ | 0 | 0.9985 | intergenic | |
| rs59169616 | 2 | 137021719 | 4 | 0 | 0 | A | G | 0.9913 | 0.2736 | 0.0002907 | +++?+ | 0 | 0.8313 | intergenic | |
| rs117960406 | 14 | 65455632 | 2 | 0 | 0 | A | G | -4.7057 | 1.2987 | 0.0002907 | -?-?? | 0 | 0.6462 | intron | FNTB,CHURC1-FNTB |
| rs10459431 | 14 | 43250337 | 4 | 0 | 0 | T | C | -1.4039 | 0.3875 | 0.0002908 | ---?+ | 61 | 0.05298 | intergenic | |
| rs115850588 | 3 | 176803134 | 3 | 0 | 0 | T | C | 3.3604 | 0.9274 | 0.0002908 | +++?? | 0 | 0.6651 | intron | TBL1XR1 |
| rs56399633 | 9 | 79459219 | 4 | 0 | 0 | A | G | 2.0794 | 0.5739 | 0.0002909 | +++?+ | 0 | 0.5272 | intron | PRUNE2 |
| rs62516466 | 8 | 113454168 | 3 | 0 | 0 | T | G | -1.046 | 0.2887 | 0.0002909 | ---?? | 0 | 0.5573 | intron | CSMD3 |

| | | | | | | | | | | | | | | | |
|---|---|---|---|---|---|---|---|---|---|---|---|---|---|---|---|
| rs72684428 | 14 | 22517913 | 4 | 0 | 0 | A | C | -1.2226 | 0.3374 | 0.0002909 | ---?- | 0 | 0.9832 | intergenic | |
| rs78865321 | 10 | 68612190 | 4 | 0 | 0 | T | C | -4.1559 | 1.147 | 0.0002909 | ---?+ | 0 | 0.5222 | intron | CTNNA3 |
| 15:87578025 | 15 | 87578025 | 3 | 0 | 0 | CATGT | C | 1.2899 | 0.356 | 0.0002909 | +++?? | 14.6 | 0.3102 | | |
| rs6474499 | 8 | 38617326 | 4 | 0 | 0 | A | C | 1.0104 | 0.2789 | 0.000291 | +++?+ | 0 | 0.938 | intron | TACC1 |
| rs3916220 | 1 | 102934974 | 4 | 0 | 0 | A | G | -0.8657 | 0.2389 | 0.0002911 | ---?+ | 7.4 | 0.3559 | intergenic | |
| 10:134483887 | 10 | 134483887 | 3 | 0 | 0 | CCT | C | 0.9142 | 0.2523 | 0.0002911 | +++?? | 0 | 0.582 | | |
| rs7169668 | 15 | 41980342 | 5 | 0 | 0 | T | C | 1.0331 | 0.2852 | 0.0002913 | +++++ | 14.2 | 0.3241 | intron | MGA |
| rs16852198 | 1 | 203676554 | 4 | 0 | 0 | C | G | -1.1654 | 0.3217 | 0.0002913 | ---?- | 0 | 0.9602 | intron | ATP2B4 |
| rs817415 | 1 | 85628317 | 5 | 0 | 0 | T | C | 1.3981 | 0.3859 | 0.0002914 | +++++ | 0 | 0.9786 | intron | SYDE2 |
| rs11161534 | 1 | 85621173 | 3 | 0 | 0 | A | T | 1.6312 | 0.4503 | 0.0002915 | +++?? | 0 | 0.8042 | downstream-gene | |
| rs303789 | 12 | 52231393 | 3 | 0 | 0 | A | G | -4.1425 | 1.1435 | 0.0002916 | ---?? | 9.6 | 0.331 | intron | FIGNL2 |
| rs7801017 | 7 | 154578128 | 4 | 0 | 0 | A | G | -1.0142 | 0.28 | 0.0002916 | ---?- | 0 | 0.7784 | intron | DPP6 |
| rs28986340 | 6 | 32748924 | 5 | 0 | 0 | C | G | 1.4599 | 0.403 | 0.0002916 | +++++ | 0 | 0.699 | intergenic | |
| rs166613 | 17 | 47943461 | 5 | 0 | 0 | A | C | 2.116 | 0.5841 | 0.0002917 | +++++ | 0 | 0.9475 | intergenic | |
| rs72665991 | 14 | 28152343 | 4 | 0 | 0 | A | C | 2.1971 | 0.6065 | 0.0002917 | +++?+ | 0 | 0.9488 | intergenic | |
| rs6051634 | 20 | 3177188 | 3 | 0 | 0 | A | G | 0.918 | 0.2534 | 0.0002922 | +++?? | 62.9 | 0.06763 | intron | DDRGK1 |
| rs114320764 | 6 | 23850101 | 4 | 0 | 0 | T | C | 2.7572 | 0.7612 | 0.0002923 | +++?- | 44.7 | 0.1431 | intergenic | |
| 20:6404783 | 20 | 6404783 | 3 | 0 | 0 | CT | C | 1.1293 | 0.3118 | 0.0002923 | +++?? | 0 | 0.7122 | | |
| rs7560183 | 2 | 142403763 | 4 | 0 | 0 | T | G | 2.0544 | 0.5672 | 0.0002926 | +++?+ | 0 | 0.8901 | intron | LRP1B |
| rs72801659 | 2 | 43033931 | 4 | 0 | 0 | T | G | -2.2106 | 0.6104 | 0.0002927 | ---?- | 0 | 0.8715 | intergenic | |
| rs75453627 | 2 | 170525469 | 4 | 0 | 0 | T | C | 1.2914 | 0.3566 | 0.0002927 | +++?+ | 0 | 0.9663 | intron | CCDC173 |
| rs79774028 | 2 | 170525764 | 4 | 0 | 0 | T | G | -1.2914 | 0.3566 | 0.0002928 | ---?- | 0 | 0.9665 | intron | CCDC173 |
| rs4658139 | 1 | 90249663 | 5 | 0 | 0 | T | C | -0.775 | 0.214 | 0.0002929 | ----- | 0 | 0.9701 | intergenic |  |
| rs6459 | 6 | 32007625 | 5 | 0 | 0 | T | C | 1.2311 | 0.34 | 0.0002931 | +++-+ | 47.7 | 0.1053 | intron | CYP21A2 |
| rs374298914 | 2 | 229290324 | 3 | 0 | 0 | A | T | -3.7706 | 1.0413 | 0.0002932 | ---?? | 0 | 0.8317 | intergenic | |
| rs10894240 | 11 | 130629933 | 4 | 0 | 0 | T | G | 1.7746 | 0.4901 | 0.0002933 | +++?+ | 14.9 | 0.3177 | intergenic | |
| rs11056185 | 12 | 14930840 | 4 | 0 | 0 | A | C | -2.158 | 0.596 | 0.0002934 | ---?- | 0 | 0.7671 | 3-prime-UTR | H2AFJ |
| rs4923908 | 15 | 41979672 | 5 | 0 | 0 | A | C | 1.0287 | 0.2841 | 0.0002936 | +++++ | 14.7 | 0.3207 | intron | MGA |
| 14:43230089 | 14 | 43230089 | 3 | 0 | 0 | A | ATG | -1.3264 | 0.3663 | 0.0002936 | ---?? | 0 | 0.7181 | | |
| rs73029706 | 2 | 168057979 | 4 | 0 | 0 | T | C | -2.2131 | 0.6112 | 0.0002937 | ---?+ | 0 | 0.6763 | intron | XIRP2 |
| rs55856335 | 2 | 170523549 | 4 | 0 | 0 | C | G | -1.291 | 0.3566 | 0.0002938 | ---?- | 0 | 0.9653 | intron | CCDC173 |
| rs672142 | 10 | 131911558 | 4 | 0 | 0 | A | G | 0.8929 | 0.2466 | 0.0002939 | +++?+ | 0 | 0.7926 | upstream-gene | |
| rs73167393 | 7 | 82426299 | 4 | 0 | 0 | A | C | 0.9374 | 0.2589 | 0.000294 | +-+?+ | 52.1 | 0.09945 | intron | PCLO |
| rs77366910 | 2 | 170519369 | 4 | 0 | 0 | T | C | -1.2906 | 0.3565 | 0.000294 | ---?- | 0 | 0.9629 | intron | CCDC173 |
| rs12591256 | 15 | 98041194 | 4 | 0 | 0 | C | G | -0.9136 | 0.2523 | 0.0002941 | ---?- | 0 | 0.947 | intergenic | |
| rs7212096 | 17 | 4779875 | 5 | 0 | 0 | T | C | -1.6396 | 0.4529 | 0.0002943 | ----+ | 20.7 | 0.2831 | intron | MINK1 |
| rs77431347 | 15 | 87930352 | 3 | 0 | 0 | A | G | -4.0441 | 1.1171 | 0.0002944 | ---?? | 14.6 | 0.3099 | intergenic | |
| rs75958843 | 8 | 25090778 | 4 | 0 | 0 | C | G | -2.7589 | 0.7621 | 0.0002945 | ---?- | 0 | 0.6654 | intron | DOCK5 |
| rs7544110 | 1 | 68860340 | 3 | 0 | 0 | A | C | -2.7674 | 0.7645 | 0.0002947 | ---?? | 0 | 0.4903 | intergenic | |
| rs13015353 | 2 | 2752308 | 4 | 0 | 0 | A | C | -1.514 | 0.4182 | 0.0002947 | ---?- | 2.5 | 0.3801 | intergenic | |
| rs60649710 | 1 | 223109883 | 3 | 0 | 0 | A | C | 1.7102 | 0.4725 | 0.0002947 | +++?? | 0 | 0.6936 | intron | DISP1 |
| rs113665854 | 2 | 170528009 | 4 | 0 | 0 | A | G | -1.2913 | 0.3567 | 0.0002947 | ---?- | 0 | 0.9661 | intron | CCDC173 |
| rs77647932 | 2 | 170521014 | 4 | 0 | 0 | T | G | 1.2905 | 0.3565 | 0.0002949 | +++?+ | 0 | 0.9641 | intron | CCDC173 |
| 5:179950452 | 5 | 179950452 | 3 | 0 | 0 | A | AT | -1.09 | 0.3011 | 0.000295 | ---?? | 0 | 0.4181 | | |
| rs79101807 | 2 | 170522148 | 4 | 0 | 0 | A | T | -1.2905 | 0.3565 | 0.0002951 | ---?- | 0 | 0.9646 | intron | CCDC173 |
| rs11766156 | 7 | 35528567 | 4 | 0 | 0 | C | G | -1.1692 | 0.323 | 0.0002952 | ---?- | 32.1 | 0.2194 | intergenic | |
| rs13100438 | 3 | 183604836 | 4 | 0 | 0 | T | C | -1.6875 | 0.4663 | 0.0002955 | ---?- | 0 | 0.9573 | upstream-gene | |
| rs7779332 | 7 | 154578186 | 4 | 0 | 0 | A | C | -1.0816 | 0.2989 | 0.0002956 | ---?- | 0 | 0.6835 | intron | DPP6 |
| rs150259582 | 2 | 170520473 | 4 | 0 | 0 | T | C | -1.2902 | 0.3565 | 0.0002956 | ---?- | 0 | 0.9639 | intron | CCDC173 |
| rs62334121 | 4 | 169328451 | 4 | 0 | 0 | T | C | -0.9398 | 0.2597 | 0.0002957 | ---?- | 0 | 0.4836 | intron | DDX60L |
| rs6432982 | 2 | 168058902 | 4 | 0 | 0 | A | C | 2.2125 | 0.6114 | 0.0002958 | +++?- | 0 | 0.6759 | intron | XIRP2 |
| rs10969954 | 9 | 30897107 | 3 | 0 | 0 | T | G | 3.4463 | 0.9523 | 0.0002958 | +++?? | 0 | 0.5088 | intergenic | |
| rs12582791 | 12 | 98974239 | 3 | 0 | 0 | A | G | -1.0486 | 0.2898 | 0.000296 | ---?? | 0 | 0.9298 | intergenic | |
| rs748030 | 20 | 61599958 | 4 | 0 | 0 | T | C | 0.9155 | 0.253 | 0.0002962 | +++?+ | 26.5 | 0.2529 | downstream-gene | |
| rs35955398 | 5 | 179952921 | 4 | 0 | 0 | A | T | 1.1062 | 0.3057 | 0.0002962 | +++?- | 41.4 | 0.1635 | intron | CNOT6 |
| rs112443141 | 6 | 83790754 | 4 | 0 | 0 | A | G | 3.2875 | 0.9085 | 0.0002963 | +++?+ | 0 | 0.7872 | intron | DOPEY1 |
| rs1912461 | 15 | 36344539 | 4 | 0 | 0 | T | C | -2.9963 | 0.8281 | 0.0002964 | ---?- | 21.4 | 0.2821 | intergenic | |
| rs73194418 | 13 | 50373278 | 5 | 0 | 0 | C | G | 1.4197 | 0.3924 | 0.0002965 | ++++- | 0 | 0.7151 | intergenic | |
| rs12978792 | 19 | 46164500 | 5 | 0 | 1 | T | G | -0.9722 | 0.2687 | 0.0002966 | ----+ | 6.8 | 0.3681 | intergenic |  |

| | | | | | | | | | | | | | | |
|---|---|---|---|---|---|---|---|---|---|---|---|---|---|---|
| rs9570133 | 13 | 60036025 | 4 | 0 | 0 | T | G | -0.9209 | 0.2545 | 0.0002967 | ---?- | 0 | 0.5748 | intergenic | |
| rs117392891 | 13 | 71335558 | 4 | 0 | 0 | T | C | -4.082 | 1.1282 | 0.0002968 | ---?- | 0 | 0.5823 | intergenic | |
| rs12742342 | 1 | 102926373 | 4 | 0 | 0 | A | T | -0.8735 | 0.2414 | 0.0002969 | ---?+ | 11.6 | 0.3347 | intergenic | |
| rs34888032 | 10 | 129678113 | 4 | 0 | 0 | A | G | -0.9031 | 0.2496 | 0.0002969 | ---?- | 0 | 0.5812 | intron | CLRN3 |
| 5:161430398 | 5 | 161430398 | 3 | 0 | 0 | A | AAGAC | 1.2458 | 0.3443 | 0.0002969 | +++?? | 0 | 0.8926 | | |
| rs77435712 | 14 | 53477762 | 2 | 0 | 0 | T | C | 4.6757 | 1.2924 | 0.0002972 | +?+?? | 0 | 0.6228 | intergenic | |
| rs12634358 | 3 | 183562662 | 4 | 0 | 0 | A | G | 1.7519 | 0.4843 | 0.0002974 | +++?+ | 0 | 0.8141 | intron | PARL |
| rs1361461 | 1 | 102925321 | 4 | 0 | 0 | T | C | -0.8764 | 0.2423 | 0.0002975 | ---?+ | 12.1 | 0.3325 | intergenic | |
| rs73167394 | 7 | 82426326 | 4 | 0 | 0 | A | C | -0.9367 | 0.2589 | 0.0002975 | -+-?- | 52.5 | 0.09698 | intron | PCLO |
| rs2012489 | 13 | 60029238 | 4 | 0 | 0 | A | T | -0.9094 | 0.2514 | 0.0002977 | ---?- | 0 | 0.452 | intergenic | |
| rs36141397 | 7 | 130261601 | 4 | 0 | 0 | A | G | -0.8941 | 0.2472 | 0.0002978 | -+-?- | 54.7 | 0.08507 | intron | COPG2 |
| rs2867751 | 11 | 59233436 | 3 | 0 | 0 | A | G | -3.5849 | 0.9911 | 0.0002981 | -?--? | 8.8 | 0.3338 | intergenic | |
| rs76384003 | 15 | 65871017 | 3 | 0 | 0 | T | C | 2.9473 | 0.8148 | 0.0002981 | +++?? | 25.8 | 0.2597 | downstream-gene | |
| 5:103139160 | 5 | 103139160 | 3 | 0 | 0 | CT | C | 1.6043 | 0.4436 | 0.0002981 | +++?? | 0 | 0.5361 | | |
| rs2045054 | 11 | 59238778 | 3 | 0 | 0 | T | C | 3.5708 | 0.9873 | 0.0002982 | +?++? | 9.5 | 0.3314 | intergenic | |
| rs112247416 | 6 | 83734416 | 4 | 0 | 0 | A | T | 3.4636 | 0.9576 | 0.0002982 | +++?+ | 0 | 0.8113 | intron | UBE3D |
| rs57614263 | 11 | 1124575 | 4 | 0 | 0 | T | C | -1.4329 | 0.3962 | 0.0002983 | ---?+ | 53.5 | 0.09167 | intergenic | |
| rs12562868 | 1 | 232832430 | 4 | 0 | 0 | A | G | 0.8997 | 0.2488 | 0.0002984 | +++?+ | 35.2 | 0.2009 | intergenic | |
| rs55887518 | 2 | 170517332 | 4 | 0 | 0 | A | G | -1.289 | 0.3564 | 0.0002986 | ---?- | 0 | 0.9619 | intron | CCDC173 |
| rs860140 | 1 | 85660811 | 5 | 0 | 0 | T | C | 1.4177 | 0.392 | 0.0002988 | +++++ | 0 | 0.9782 | intron | SYDE2 |
| rs2695170 | 15 | 42036412 | 5 | 0 | 0 | A | T | 1.0241 | 0.2832 | 0.000299 | ++++- | 15.5 | 0.316 | intron | MGA |
| rs12564255 | 1 | 232832400 | 4 | 0 | 0 | T | C | -0.8997 | 0.2488 | 0.0002992 | ---?- | 35.3 | 0.2005 | intergenic | |
| rs9944631 | 18 | 35843093 | 4 | 0 | 0 | A | G | 1.4115 | 0.3903 | 0.0002993 | +++?+ | 0 | 0.9985 | intergenic | |
| rs12633272 | 3 | 183564099 | 4 | 0 | 0 | T | C | 1.7512 | 0.4843 | 0.0002993 | +++?+ | 0 | 0.8184 | intron | PARL |
| rs115285861 | 2 | 167990975 | 4 | 0 | 0 | A | C | -2.2911 | 0.6336 | 0.0002994 | ---?+ | 4.2 | 0.3718 | intron | XIRP2 |
| 2:170759123 | 2 | 170759123 | 3 | 0 | 0 | CT | C | -1.2786 | 0.3536 | 0.0002994 | ---?? | 52.8 | 0.1204 | | |
| 17:56793466 | 17 | 56793466 | 3 | 0 | 0 | G | GA | 1.4763 | 0.4083 | 0.0002997 | +++?? | 0 | 0.6646 | | |
| rs73020383 | 2 | 170594105 | 4 | 0 | 0 | T | C | -1.1815 | 0.3268 | 0.0002999 | ---?- | 0 | 0.9972 | intron | KLHL23,PHOSPHO2-KLHL23 |
| rs930716 | 15 | 85532241 | 4 | 0 | 0 | A | G | 0.8929 | 0.247 | 0.0003 | +++?- | 57.3 | 0.0713 | intron | PDE8A |
| rs111863368 | 2 | 170590979 | 4 | 0 | 0 | A | C | 1.1805 | 0.3265 | 0.0003001 | +++?+ | 0 | 0.9978 | intron | KLHL23,PHOSPHO2-KLHL23 |
| rs6578026 | 8 | 140358700 | 4 | 0 | 0 | A | G | -1.0761 | 0.2977 | 0.0003002 | ---?- | 0 | 0.8963 | intergenic | |
| rs7126269 | 11 | 59239758 | 3 | 0 | 0 | A | C | -3.5644 | 0.986 | 0.0003003 | -?--? | 9.9 | 0.3297 | intergenic | |
| 5:52935911 | 5 | 52935911 | 3 | 0 | 0 | G | GT | 1.469 | 0.4064 | 0.0003003 | +++?? | 0 | 0.5502 | | |
| rs147862429 | 11 | 906896 | 4 | 0 | 0 | T | C | 2.2798 | 0.6307 | 0.0003004 | +++?+ | 0 | 0.9754 | intron | CHID1 |
| rs1340388 | 1 | 102939009 | 4 | 0 | 0 | T | C | 0.8601 | 0.2379 | 0.0003008 | +++?- | 6.7 | 0.3596 | intergenic | |
| rs74661315 | 2 | 170511559 | 4 | 0 | 0 | A | G | -1.2876 | 0.3562 | 0.0003008 | ---?- | 0 | 0.9593 | intron | CCDC173 |
| rs56302843 | 7 | 82386297 | 4 | 0 | 0 | T | C | -1.0795 | 0.2986 | 0.000301 | -+-?- | 45.1 | 0.1407 | 3-prime-UTR | PCLO |
| rs73471179 | 18 | 39989008 | 4 | 0 | 0 | T | C | -1.1791 | 0.3262 | 0.0003011 | ---?+ | 0 | 0.3988 | intron | LINC00907 |
| rs187625897 | 20 | 62484684 | 3 | 0 | 0 | A | G | 4.1327 | 1.1434 | 0.0003011 | +++?? | 0 | 0.7475 | intergenic | |
| 2:170541158 | 2 | 170541158 | 3 | 0 | 0 | CGTTGT | CGTTGT | 1.3275 | 0.3673 | 0.0003011 | +++?? | 0 | 0.8794 | | |
| rs138255254 | 13 | 71366666 | 4 | 0 | 0 | T | C | -3.9717 | 1.0989 | 0.0003012 | ---?- | 0 | 0.5608 | intergenic | |
| rs8019089 | 14 | 43256084 | 4 | 0 | 0 | T | C | 1.3826 | 0.3825 | 0.0003013 | +++?- | 60.9 | 0.05305 | intergenic | |
| rs17496307 | 6 | 32401036 | 5 | 0 | 1 | A | T | -1.3912 | 0.3849 | 0.0003014 | ----- | 0 | 0.5048 | intergenic |  |
| rs7991399 | 13 | 60022214 | 4 | 0 | 0 | T | C | -0.9178 | 0.254 | 0.0003015 | ---?- | 0 | 0.6248 | intergenic | |
| 1:87468973 | 1 | 87468973 | 3 | 0 | 0 | G | GA | 1.097 | 0.3036 | 0.0003015 | +++?? | 0 | 0.6976 | | |
| rs17202717 | 6 | 32397863 | 5 | 0 | 1 | T | G | 1.3911 | 0.3849 | 0.0003016 | +++++ | 0 | 0.5047 | intergenic | |
| rs115796185 | 4 | 162098821 | 4 | 0 | 0 | A | C | 1.9078 | 0.5279 | 0.0003016 | +++?- | 39.6 | 0.1743 | intergenic | |
| rs2175042 | 17 | 6688546 | 3 | 0 | 0 | A | C | 3.6177 | 1.0011 | 0.0003018 | +++?? | 0 | 0.4484 | intron | FBXO39 |
| rs5757456 | 22 | 39455105 | 4 | 0 | 0 | A | G | 1.6198 | 0.4482 | 0.0003018 | +++?+ | 0 | 0.9626 | downstream-gene | |
| rs8001020 | 13 | 60030215 | 4 | 0 | 0 | T | C | 0.9088 | 0.2515 | 0.0003018 | +++?+ | 0 | 0.4498 | intergenic | |
| rs74396071 | 3 | 36740063 | 4 | 0 | 0 | A | G | -1.6776 | 0.4642 | 0.0003018 | ---?- | 0 | 0.4595 | intergenic | |
| rs62076383 | 17 | 47321967 | 4 | 0 | 0 | T | C | -1.094 | 0.3028 | 0.000302 | ---?+ | 45.2 | 0.14 | upstream-gene | |
| rs7113590 | 11 | 37060603 | 4 | 0 | 0 | A | G | 0.8804 | 0.2437 | 0.0003022 | +++?+ | 9.1 | 0.3474 | intergenic | |
| rs7446888 | 5 | 178205533 | 4 | 0 | 0 | T | C | 1.1157 | 0.3088 | 0.0003022 | +-+?+ | 31.4 | 0.2237 | upstream-gene | |
| 2:142402354 | 2 | 142402354 | 3 | 0 | 0 | A | AAC | 2.1363 | 0.5912 | 0.0003022 | +++?? | 0 | 0.7752 | | |
| rs1361460 | 1 | 102925266 | 4 | 0 | 0 | A | C | 0.8743 | 0.242 | 0.0003023 | +++?- | 11.6 | 0.3346 | intergenic | |
| rs10152332 | 15 | 41976037 | 5 | 0 | 0 | A | C | 1.0306 | 0.2852 | 0.0003025 | +++++ | 14.4 | 0.3223 | intron | MGA |
| 9:30897193 | 9 | 30897193 | 3 | 0 | 0 | T | TTA | 3.4404 | 0.9522 | 0.0003025 | +++?? | 0 | 0.5128 | | |
| 7:107346087 | 7 | 107346087 | 3 | 0 | 0 | G | GT | 0.9488 | 0.2626 | 0.0003026 | +++?? | 16.5 | 0.3019 | | |

| | | | | | | | | | | | | | | | |
|---|---|---|---|---|---|---|---|---|---|---|---|---|---|---|---|
| 8:113316606 | 8 | 113316606 | 3 | 0 | 0 | G | GT | 1.0294 | 0.2849 | 0.0003026 | +++?? | 0 | 0.7163 | | |
| rs76487317 | 2 | 170514877 | 4 | 0 | 0 | T | C | 1.2875 | 0.3564 | 0.0003027 | +++?+ | 0 | 0.9605 | intron | CCDC173 |
| rs62446116 | 7 | 16903504 | 4 | 0 | 0 | T | C | 1.7501 | 0.4844 | 0.0003028 | +++?+ | 0 | 0.4246 | intron | AGR3 |
| rs2086892 | 3 | 196325516 | 4 | 0 | 0 | T | C | -1.0758 | 0.2978 | 0.000303 | ---?- | 0 | 0.7944 | intergenic | |
| rs10237800 | 7 | 16904068 | 4 | 0 | 0 | T | C | -1.7109 | 0.4736 | 0.0003033 | ---?- | 0 | 0.4053 | intron | AGR3 |
| rs76966099 | 2 | 170512674 | 4 | 0 | 0 | C | G | 1.2871 | 0.3563 | 0.0003035 | +++?+ | 0 | 0.9597 | intron | CCDC173 |
| rs9379612 | 6 | 23873195 | 4 | 0 | 0 | A | G | -2.133 | 0.5906 | 0.000304 | ---?+ | 39.3 | 0.1758 | intergenic | |
| rs1944417 | 18 | 40438734 | 3 | 0 | 0 | A | C | -4.8412 | 1.3404 | 0.0003041 | ---?? | 0 | 0.5246 | intron | RIT2 |
| rs817447 | 1 | 85649983 | 5 | 0 | 0 | T | C | -1.4026 | 0.3884 | 0.0003044 | ----- | 0 | 0.9858 | intron | SYDE2 |
| 1:6157040 | 1 | 6157040 | 2 | 0 | 0 | A | AC | 5.8299 | 1.6143 | 0.0003044 | ?++?? | 0 | 0.6035 | | |
| rs1934713 | 1 | 102931654 | 4 | 0 | 0 | T | C | -0.8672 | 0.2401 | 0.0003045 | ---?+ | 11.1 | 0.3376 | intergenic | |
| rs12622156 | 2 | 234701686 | 4 | 0 | 0 | A | G | 1.2383 | 0.3429 | 0.0003045 | +++?+ | 0 | 0.919 | intron | MROH2A |
| rs9542405 | 13 | 71142999 | 4 | 0 | 0 | T | G | 0.915 | 0.2534 | 0.0003046 | +++?+ | 0 | 0.5237 | intergenic | |
| rs6713210 | 2 | 170956187 | 4 | 0 | 1 | A | C | 1.0384 | 0.2875 | 0.0003048 | +++?+ | 55.2 | 0.08201 | intergenic | |
| rs62017061 | 16 | 7179455 | 4 | 0 | 0 | A | G | -1.0069 | 0.2788 | 0.0003048 | ---?- | 8.9 | 0.3487 | intron | RBFOX1 |
| 1:228228714 | 1 | 228228714 | 3 | 0 | 0 | A | ATATC | -1.0186 | 0.2821 | 0.0003048 | ---?? | 0 | 0.4962 | | |
| rs4712792 | 6 | 23856411 | 4 | 0 | 0 | T | C | 2.5956 | 0.7188 | 0.0003049 | +++?- | 54.8 | 0.08435 | intergenic | |
| rs4557308 | 4 | 183655750 | 4 | 0 | 0 | A | G | 0.9819 | 0.2719 | 0.000305 | +++?- | 0 | 0.6815 | intron | TENM3 |
| rs73020363 | 2 | 170580303 | 4 | 0 | 0 | T | C | -1.2804 | 0.3546 | 0.000305 | ---?- | 0 | 0.9378 | intron | PHOSPHO2-KLHL23 |
| 12:27784124 | 12 | 27784124 | 3 | 0 | 0 | T | TA | 1.0151 | 0.2811 | 0.000305 | +++?? | 0 | 0.4044 | | |
| rs6543469 | 2 | 107676769 | 4 | 0 | 0 | A | G | -0.9924 | 0.2748 | 0.0003051 | ---?- | 0 | 0.4559 | intergenic | |
| rs75789635 | 2 | 170512850 | 4 | 0 | 0 | A | G | 1.2866 | 0.3563 | 0.0003051 | +++?+ | 0 | 0.9597 | intron | CCDC173 |
| rs8001038 | 13 | 60030255 | 4 | 0 | 0 | T | G | 0.908 | 0.2515 | 0.0003054 | +++?+ | 0 | 0.4533 | intergenic | |
| rs58404411 | 2 | 170580349 | 4 | 0 | 0 | T | C | -1.2803 | 0.3546 | 0.0003054 | ---?- | 0 | 0.938 | intron | PHOSPHO2-KLHL23 |
| rs1880871 | 12 | 112452262 | 5 | 0 | 0 | T | C | -1.3347 | 0.3697 | 0.0003056 | -+--- | 57.7 | 0.05043 | intron | ERP29 |
| rs12374101 | 3 | 65657028 | 4 | 0 | 0 | T | C | 0.9613 | 0.2662 | 0.0003056 | +++?+ | 48.3 | 0.1215 | intron | MAGI1 |
| rs11066128 | 12 | 112450754 | 5 | 0 | 0 | T | C | -1.3345 | 0.3696 | 0.0003057 | -+--- | 57.8 | 0.05038 | intron | TMEM116 |
| rs12300496 | 12 | 112450433 | 5 | 0 | 0 | A | G | -1.3345 | 0.3696 | 0.0003057 | -+--- | 57.7 | 0.0505 | intron | TMEM116 |
| rs7316563 | 12 | 112453635 | 5 | 0 | 0 | A | C | -1.3347 | 0.3697 | 0.0003058 | -+--- | 57.7 | 0.05044 | intron | ERP29 |
| rs35929624 | 3 | 183571618 | 4 | 0 | 0 | A | G | -1.7459 | 0.4836 | 0.0003058 | ---?- | 0 | 0.8396 | intron | PARL |
| rs7313773 | 12 | 112441688 | 5 | 0 | 0 | T | C | -1.3332 | 0.3693 | 0.0003059 | -+--- | 57.8 | 0.05033 | intron | TMEM116 |
| rs28525994 | 12 | 112442586 | 5 | 0 | 0 | T | C | -1.3333 | 0.3693 | 0.0003059 | -+--- | 57.8 | 0.05026 | intron | TMEM116 |
| rs436245 | 2 | 40433420 | 4 | 0 | 0 | T | G | 1.091 | 0.3022 | 0.000306 | +++?- | 0 | 0.5517 | intron | SLC8A1,SLC8A1-AS1 |
| rs7132863 | 12 | 112448883 | 5 | 0 | 0 | A | G | -1.3342 | 0.3696 | 0.000306 | -+--- | 57.8 | 0.05041 | intron | TMEM116 |
| rs12423960 | 12 | 112442416 | 5 | 0 | 0 | A | C | 1.3332 | 0.3693 | 0.000306 | +-+++ | 57.8 | 0.05032 | intron | TMEM116 |
| rs376347530 | 4 | 39596183 | 3 | 0 | 0 | T | C | -2.9048 | 0.8046 | 0.0003061 | ---?? | 0 | 0.7185 | intron | SMIM14,UGDH-AS1 |
| rs7316650 | 12 | 112439664 | 5 | 0 | 0 | T | C | 1.3331 | 0.3693 | 0.0003062 | +-+++ | 57.8 | 0.0503 | intron | TMEM116 |
| rs10849992 | 12 | 112443017 | 5 | 0 | 0 | T | C | 1.3334 | 0.3694 | 0.0003062 | +-+++ | 57.8 | 0.05034 | intron | TMEM116 |
| rs76127956 | 11 | 129303558 | 4 | 0 | 0 | A | G | 1.5671 | 0.4341 | 0.0003062 | +++?+ | 34.4 | 0.2056 | intron | BARX2 |
| rs3104993 | 8 | 96572162 | 4 | 0 | 0 | A | G | 1.5627 | 0.4329 | 0.0003063 | +-+?+ | 50.9 | 0.1062 | intron | LOC100616530 |
| rs7312472 | 12 | 112438770 | 5 | 0 | 0 | C | G | -1.3333 | 0.3693 | 0.0003063 | -+--- | 57.7 | 0.05061 | intron | TMEM116 |
| rs7952986 | 12 | 112448162 | 5 | 0 | 0 | A | C | -1.3341 | 0.3696 | 0.0003063 | -+--- | 57.8 | 0.0504 | intron | TMEM116 |
| rs57439171 | 12 | 112439287 | 5 | 0 | 0 | A | G | 1.333 | 0.3693 | 0.0003063 | +-+++ | 57.8 | 0.05029 | intron | TMEM116 |
| rs3782304 | 12 | 26820465 | 4 | 0 | 0 | T | C | -1.8202 | 0.5042 | 0.0003065 | -+-?- | 30.5 | 0.2295 | intron | ITPR2 |
| rs2398139 | 5 | 6429667 | 4 | 0 | 0 | T | C | -0.9792 | 0.2713 | 0.0003066 | ---?- | 0 | 0.3945 | intergenic | |
| rs3759929 | 15 | 91410009 | 4 | 0 | 0 | T | G | 0.8729 | 0.2418 | 0.0003066 | +++?- | 20.1 | 0.2893 | upstream-gene | |
| rs11066127 | 12 | 112446355 | 5 | 0 | 0 | T | C | -1.3339 | 0.3695 | 0.0003066 | -+--- | 57.7 | 0.05043 | intron | TMEM116 |
| rs80082680 | 2 | 170508953 | 4 | 0 | 0 | A | G | 1.2859 | 0.3562 | 0.0003066 | +++?+ | 0 | 0.959 | intron | CCDC173 |
| rs7296841 | 12 | 112444109 | 5 | 0 | 0 | A | C | -1.3334 | 0.3694 | 0.0003067 | -+--- | 57.8 | 0.05033 | intron | TMEM116 |
| rs7310545 | 12 | 112443960 | 5 | 0 | 0 | A | G | 1.3334 | 0.3694 | 0.0003067 | +-+++ | 57.8 | 0.05033 | intron | TMEM116 |
| 1:187886312 | 1 | 187886312 | 3 | 0 | 0 | T | TAAC | 1.0703 | 0.2965 | 0.0003067 | +++?? | 8.3 | 0.3361 | | |
| rs4924571 | 15 | 41974718 | 5 | 0 | 0 | A | G | 1.0296 | 0.2853 | 0.0003068 | +++++ | 14.5 | 0.3218 | intron | MGA |
| rs7314232 | 12 | 112444517 | 5 | 0 | 0 | C | G | 1.3335 | 0.3694 | 0.0003068 | +-+++ | 57.8 | 0.05035 | intron | TMEM116 |
| 2:196462685 | 2 | 196462685 | 3 | 0 | 0 | CT | C | 1.0809 | 0.2995 | 0.0003068 | +++?? | 0 | 0.9018 | | |
| rs7958596 | 12 | 112665013 | 5 | 0 | 0 | T | C | 1.3345 | 0.3697 | 0.0003069 | +-+++ | 54.6 | 0.06611 | intron | HECTD4 |
| rs73644378 | 9 | 30897754 | 3 | 0 | 0 | A | G | 3.4265 | 0.9493 | 0.0003069 | +++?? | 0 | 0.5096 | intergenic | |
| rs148388204 | 14 | 53600630 | 2 | 0 | 0 | T | C | 4.7735 | 1.3225 | 0.0003069 | +?+?? | 0 | 0.6033 | intron | DDHD1 |
| 22:28643485 | 22 | 28643485 | 3 | 0 | 0 | CT | C | -2.7079 | 0.7502 | 0.0003069 | ---?? | 0 | 0.5332 | | |
| rs1171727 | 10 | 131908016 | 4 | 0 | 0 | T | C | -1.02 | 0.2826 | 0.000307 | ---?- | 0 | 0.9374 | intron | LINC00959 |

| | | | | | | | | | | | | | | |
|---|---|---|---|---|---|---|---|---|---|---|---|---|---|---|
| rs11936029 | 4 | 41328487 | 4 | 0 | 0 | A | G | -1.446 | 0.4006 | 0.000307 | ---?+ | 0 | 0.4267 | intergenic | |
| rs61904791 | 11 | 103322257 | 4 | 0 | 0 | T | C | -1.0092 | 0.2796 | 0.000307 | ---?- | 42.8 | 0.1548 | intron | DYNC2H1 |
| rs77365616 | 2 | 170508530 | 4 | 0 | 0 | T | C | -1.2857 | 0.3562 | 0.0003071 | ---?- | 0 | 0.9587 | intron | CCDC173 |
| rs35719907 | 4 | 110093685 | 5 | 0 | 0 | T | C | -1.4671 | 0.4065 | 0.0003072 | ----- | 0 | 0.5574 | intron | COL25A1 |
| rs8011254 | 14 | 22531910 | 4 | 0 | 0 | T | C | -0.9456 | 0.262 | 0.0003073 | ---?- | 0 | 0.6458 | intergenic | |
| rs11066119 | 12 | 112434504 | 5 | 0 | 0 | A | G | -1.3335 | 0.3695 | 0.0003074 | -+--- | 57.8 | 0.0501 | intron | TMEM116 |
| rs9317078 | 13 | 60020830 | 4 | 0 | 0 | A | G | -0.9165 | 0.2539 | 0.0003075 | ---?- | 0 | 0.632 | intergenic | |
| rs11066118 | 12 | 112434152 | 5 | 0 | 0 | T | C | 1.3332 | 0.3694 | 0.0003076 | +-+++ | 57.8 | 0.05005 | intron | TMEM116 |
| 6:83764862 | 6 | 83764862 | 3 | 0 | 0 | G | GAA | -3.6855 | 1.0213 | 0.0003077 | ---?? | 0 | 0.53 | | |
| rs72966501 | 2 | 208299204 | 4 | 0 | 0 | A | G | -1.6851 | 0.467 | 0.0003078 | ---?- | 0 | 0.7257 | intergenic | |
| rs2819800 | 14 | 85152503 | 4 | 0 | 0 | T | C | -1.1598 | 0.3214 | 0.0003079 | ---?- | 0 | 0.8722 | intergenic | |
| rs55887530 | 2 | 170454094 | 4 | 0 | 0 | T | C | -1.279 | 0.3544 | 0.0003079 | ---?- | 0 | 0.9192 | intron | PPIG |
| rs77164627 | 2 | 170534073 | 4 | 0 | 0 | A | G | 1.2883 | 0.357 | 0.0003079 | +++?+ | 0 | 0.9655 | intron | CCDC173 |
| rs11066117 | 12 | 112432709 | 5 | 0 | 0 | A | G | -1.3333 | 0.3695 | 0.000308 | -+--- | 57.8 | 0.05029 | intron | TMEM116 |
| rs34652275 | 3 | 183608316 | 4 | 0 | 0 | A | G | -1.6853 | 0.467 | 0.000308 | ---?- | 0 | 0.9576 | downstream-gene | |
| rs185944037 | 5 | 67642790 | 4 | 0 | 0 | A | G | -2.5996 | 0.7204 | 0.0003081 | ---?- | 0 | 0.605 | intergenic | |
| rs62623401 | 22 | 18021936 | 4 | 0 | 0 | A | G | -3.642 | 1.0093 | 0.0003082 | ---?- | 0 | 0.8684 | missense | CECR2 |
| rs12035863 | 1 | 102944137 | 4 | 0 | 0 | A | G | -0.8525 | 0.2363 | 0.0003083 | ---?+ | 5.8 | 0.364 | intergenic | |
| rs148165404 | 6 | 83774034 | 4 | 0 | 0 | T | C | 3.2875 | 0.9112 | 0.0003085 | +++?+ | 0 | 0.8406 | intron | UBE3D |
| rs73421065 | 12 | 112559379 | 5 | 0 | 0 | T | C | 1.3322 | 0.3692 | 0.0003086 | +-+++ | 55.2 | 0.0628 | upstream-gene | |
| rs112219670 | 17 | 56894220 | 5 | 0 | 0 | A | G | -2.5268 | 0.7003 | 0.0003086 | ----- | 0 | 0.842 | intron | PPM1E |
| rs1131015 | 6 | 31238010 | 3 | 0 | 0 | T | G | -0.9175 | 0.2543 | 0.0003087 | ---?? | 66.5 | 0.05061 | missense | HLA-C |
| rs35462761 | 1 | 102944533 | 4 | 0 | 0 | C | G | -0.8519 | 0.2361 | 0.0003088 | ---?+ | 5.7 | 0.3644 | intergenic | |
| rs116167881 | 2 | 170578134 | 4 | 0 | 0 | T | G | 2.0068 | 0.5563 | 0.0003089 | +++?+ | 0 | 0.5004 | intron | PHOSPHO2-KLHL23 |
| rs960784 | 11 | 106781077 | 4 | 0 | 0 | A | G | 1.0754 | 0.2981 | 0.000309 | +++?- | 42.8 | 0.1546 | intron | GUCY1A2 |
| rs11854420 | 15 | 88256014 | 4 | 0 | 0 | A | T | -0.9728 | 0.2696 | 0.000309 | ---?- | 10.9 | 0.3385 | intergenic | |
| rs2762840 | 1 | 64590419 | 4 | 0 | 0 | A | G | -1.1205 | 0.3106 | 0.0003091 | ---?- | 0 | 0.7589 | intron | ROR1 |
| rs10795264 | 10 | 5394397 | 4 | 0 | 0 | A | C | 1.0734 | 0.2976 | 0.0003091 | +++?+ | 0 | 0.7291 | intergenic | |
| rs56388913 | 13 | 30322804 | 3 | 0 | 0 | T | C | -1.5414 | 0.4273 | 0.0003091 | ---?? | 0 | 0.6411 | intergenic | |
| rs139181081 | 8 | 96559364 | 4 | 0 | 0 | T | G | -1.6547 | 0.4587 | 0.0003091 | -+-?- | 55.1 | 0.08267 | intron | LOC100616530 |
| rs4809437 | 20 | 61600190 | 4 | 0 | 0 | A | G | 0.9125 | 0.2529 | 0.0003093 | +++?+ | 20.9 | 0.285 | downstream-gene | |
| rs76073214 | 2 | 170445880 | 4 | 0 | 0 | T | C | 1.2808 | 0.3551 | 0.0003095 | +++?+ | 0 | 0.9146 | intron | PPIG |
| rs77340320 | 13 | 20830373 | 4 | 0 | 0 | T | C | -1.6445 | 0.4559 | 0.0003095 | ---?- | 0 | 0.9654 | intergenic | |
| 13:30322735 | 13 | 30322735 | 3 | 0 | 0 | G | GT | 1.541 | 0.4272 | 0.0003095 | +++?? | 0 | 0.6408 | | |
| rs67839461 | 16 | 5455574 | 3 | 0 | 0 | C | G | 0.9146 | 0.2536 | 0.0003098 | +++?? | 0 | 0.4762 | intron | RBFOX1 |
| rs143908827 | 2 | 167995346 | 4 | 0 | 0 | T | C | -2.2789 | 0.6318 | 0.0003099 | ---?+ | 3.6 | 0.3747 | intron | XIRP2 |
| rs78605026 | 17 | 56624411 | 5 | 0 | 0 | T | C | 2.4733 | 0.6857 | 0.00031 | +++++ | 0 | 0.7541 | upstream-gene | |
| rs73644379 | 9 | 30898351 | 3 | 0 | 0 | A | G | 3.4171 | 0.9475 | 0.0003102 | +++?? | 0 | 0.508 | intergenic | |
| rs8027390 | 15 | 41985135 | 5 | 0 | 0 | A | G | -1.0251 | 0.2842 | 0.0003103 | ----- | 14.1 | 0.3246 | intron | MGA |
| rs10216803 | 8 | 113499341 | 4 | 0 | 0 | T | C | 1.0394 | 0.2882 | 0.0003104 | +++?+ | 0 | 0.6642 | intron | CSMD3 |
| rs17040014 | 4 | 110081861 | 5 | 0 | 0 | T | C | 1.4691 | 0.4074 | 0.0003105 | +++++ | 0 | 0.5765 | intron | COL25A1 |
| rs35817460 | 6 | 32394122 | 5 | 0 | 1 | T | G | 1.3883 | 0.385 | 0.0003105 | +++++ | 0 | 0.5006 | intergenic | |
| rs140902791 | 2 | 170458918 | 4 | 0 | 0 | T | C | -1.2772 | 0.3541 | 0.0003105 | ---?- | 0 | 0.9224 | intron | PPIG |
| rs4954594 | 2 | 137028349 | 4 | 0 | 0 | A | C | -1.0095 | 0.2799 | 0.0003108 | ---?- | 0 | 0.7971 | intergenic | |
| rs73921601 | 19 | 12800597 | 3 | 0 | 0 | T | C | -4.0799 | 1.1314 | 0.0003108 | ?---? | 61.6 | 0.07392 | missense | FBXW9 |
| rs58003274 | 3 | 68215298 | 4 | 0 | 0 | A | G | 2.2177 | 0.615 | 0.0003109 | +++?+ | 32.4 | 0.2179 | intron | FAM19A1 |
| rs78610013 | 2 | 170532210 | 4 | 0 | 0 | A | C | 1.2881 | 0.3572 | 0.0003109 | +++?+ | 0 | 0.9605 | intron | CCDC173 |
| rs12486027 | 3 | 194263921 | 3 | 0 | 0 | A | C | 1.1061 | 0.3067 | 0.000311 | +++?? | 0 | 0.4405 | intergenic | |
| rs4692658 | 4 | 169329405 | 4 | 0 | 0 | A | C | 0.8954 | 0.2483 | 0.0003111 | +++?+ | 0 | 0.8492 | intron | DDX60L |
| rs2083617 | 8 | 96219799 | 3 | 0 | 0 | A | G | -1.0094 | 0.2799 | 0.0003114 | ---?? | 0 | 0.8886 | intron | C8orf69 |
| rs10792242 | 11 | 59213532 | 3 | 0 | 0 | A | G | 3.58 | 0.9929 | 0.0003114 | +?++? | 9.1 | 0.3327 | downstream-gene | |
| rs73778598 | 6 | 144612231 | 5 | 0 | 0 | T | C | 1.4558 | 0.4038 | 0.0003115 | ++++- | 44.8 | 0.1232 | intron | UTRN |
| rs116334171 | 2 | 168221678 | 4 | 0 | 0 | T | C | -2.4764 | 0.6869 | 0.0003116 | ---?- | 0 | 0.7094 | intergenic | |
| rs4658321 | 1 | 90249829 | 5 | 0 | 0 | A | G | 0.7715 | 0.214 | 0.0003117 | +++++ | 0 | 0.974 | intergenic | |
| rs6591537 | 11 | 59214258 | 3 | 0 | 0 | T | C | 3.5797 | 0.9929 | 0.0003117 | +?++? | 9.1 | 0.3326 | downstream-gene | |
| rs36127202 | 7 | 130274180 | 4 | 0 | 0 | T | C | -0.8904 | 0.247 | 0.0003118 | -+-?- | 55.5 | 0.08044 | intron | COPG2 |
| rs73752513 | 6 | 83788264 | 4 | 0 | 0 | A | G | 3.2476 | 0.9008 | 0.0003118 | +++?+ | 0 | 0.7411 | intron | DOPEY1 |
| 12:26821740 | 12 | 26821740 | 3 | 0 | 0 | CA | C | 1.0201 | 0.2829 | 0.0003118 | +++?? | 0 | 0.6268 | | |
| rs1336518 | 13 | 60044904 | 4 | 0 | 0 | A | G | -0.9444 | 0.262 | 0.0003121 | ---?- | 0 | 0.5388 | intergenic | |

| | | | | | | | | | | | | | | | |
|---|---|---|---|---|---|---|---|---|---|---|---|---|---|---|---|
| rs1413586 | 1 | 232833416 | 4 | 0 | 0 | A | C | -0.8968 | 0.2488 | 0.0003121 | ---?- | 39.5 | 0.175 | intergenic | |
| rs17403278 | 11 | 103665712 | 4 | 0 | 0 | A | G | 2.3325 | 0.647 | 0.0003121 | +++?+ | 0 | 0.8498 | intergenic | |
| rs35589082 | 1 | 102944257 | 4 | 0 | 0 | T | C | 0.8527 | 0.2365 | 0.0003121 | +++?- | 5.7 | 0.3644 | intergenic | |
| rs6432983 | 2 | 168062531 | 4 | 0 | 0 | T | G | -2.2065 | 0.6121 | 0.0003122 | ---?+ | 0 | 0.6748 | intron | XIRP2 |
| rs36153397 | 7 | 130255938 | 4 | 0 | 0 | A | G | 0.8908 | 0.2471 | 0.0003122 | +-+?+ | 54.4 | 0.08684 | intron | COPG2 |
| rs78840359 | 2 | 170577311 | 4 | 0 | 0 | T | C | -1.2885 | 0.3574 | 0.0003122 | ---?- | 0 | 0.9179 | intron | PHOSPHO2-KLHL23 |
| rs2285809 | 12 | 112708426 | 5 | 0 | 0 | T | C | -1.3336 | 0.3699 | 0.0003123 | -+--- | 54.4 | 0.06684 | intron | HECTD4 |
| rs12028006 | 1 | 87451275 | 3 | 0 | 0 | A | G | 1.1015 | 0.3056 | 0.0003123 | +++?? | 0 | 0.6975 | intron | HS2ST1 |
| rs72846309 | 11 | 1136928 | 4 | 0 | 0 | A | G | -1.4464 | 0.4012 | 0.0003123 | ---?+ | 60.4 | 0.05541 | intergenic | |
| rs374504354 | 5 | 147739144 | 2 | 0 | 0 | A | C | -5.822 | 1.6151 | 0.0003123 | --??? | 48.8 | 0.1621 | intergenic | |
| 11:1123756 | 11 | 1123756 | 4 | 0 | 0 | A | G | -1.429 | 0.3964 | 0.0003123 | ---?+ | 53.1 | 0.09384 | | |
| rs1413587 | 1 | 232833398 | 4 | 0 | 0 | A | G | 0.8968 | 0.2488 | 0.0003124 | +++?+ | 39.5 | 0.1749 | intergenic | |
| rs117095343 | 17 | 56892851 | 5 | 0 | 0 | A | G | 2.5425 | 0.7053 | 0.0003124 | +++++ | 0 | 0.8102 | intron | PPM1E |
| rs6591538 | 11 | 59218869 | 3 | 0 | 0 | T | C | -3.5789 | 0.9928 | 0.0003125 | -?--? | 9.2 | 0.3322 | intergenic | |
| rs1965012 | 12 | 111773467 | 5 | 0 | 0 | T | C | -0.9005 | 0.2498 | 0.0003127 | -+--- | 48.9 | 0.09789 | intron | CUX2 |
| rs151187951 | 18 | 25148360 | 4 | 0 | 0 | T | C | 2.9332 | 0.8138 | 0.0003127 | +++?+ | 0 | 0.5893 | intergenic | |
| rs1453543 | 11 | 59224720 | 3 | 0 | 0 | A | G | 3.5782 | 0.9928 | 0.0003132 | +?++? | 9.4 | 0.3318 | missense | OR4D6 |
| rs78652924 | 9 | 79459268 | 4 | 0 | 0 | T | C | -2.0729 | 0.5751 | 0.0003132 | ---?- | 0 | 0.5305 | intron | PRUNE2 |
| rs10750921 | 11 | 59217581 | 3 | 0 | 0 | C | G | -3.5789 | 0.993 | 0.0003134 | -?--? | 9.3 | 0.3321 | intergenic | |
| rs61992826 | 14 | 43257573 | 4 | 0 | 0 | T | C | 1.379 | 0.3826 | 0.0003134 | +++?- | 60.9 | 0.0534 | intergenic | |
| rs34730165 | 2 | 170580219 | 4 | 0 | 0 | C | G | 1.2772 | 0.3544 | 0.0003136 | +++?+ | 0 | 0.9405 | intron | PHOSPHO2-KLHL23 |
| rs7948562 | 11 | 59210480 | 3 | 0 | 0 | T | C | -3.579 | 0.9932 | 0.0003138 | -?--? | 9.2 | 0.3325 | upstream-gene | |
| 11:102630721 | 11 | 102630721 | 3 | 0 | 0 | CA | C | -1.0559 | 0.293 | 0.0003138 | ---?? | 51.2 | 0.1287 | | |
| rs28986374 | 6 | 32752199 | 5 | 0 | 0 | A | G | -1.439 | 0.3993 | 0.000314 | ----- | 0 | 0.5738 | intergenic |  |
| rs1453545 | 11 | 59223388 | 3 | 0 | 0 | T | C | -3.5776 | 0.9928 | 0.0003141 | -?--? | 9.4 | 0.3317 | upstream-gene | |
| rs17706535 | 13 | 76335109 | 4 | 0 | 0 | A | G | 1.3514 | 0.375 | 0.0003141 | +++?+ | 0 | 0.8791 | synonymous | LMO7 |
| rs6668305 | 1 | 18766063 | 4 | 0 | 0 | A | G | -1.0695 | 0.2968 | 0.0003142 | ---?- | 36.5 | 0.1931 | intergenic | |
| 3:71924035 | 3 | 71924035 | 3 | 0 | 0 | CA | C | -1.0856 | 0.3013 | 0.0003143 | ---?? | 0 | 0.4073 | | |
| rs76392976 | 2 | 167989205 | 4 | 0 | 0 | T | C | -2.2853 | 0.6343 | 0.0003145 | ---?+ | 3.7 | 0.3743 | intron | XIRP2 |
| rs76410583 | 2 | 170504412 | 4 | 0 | 0 | T | C | -1.2828 | 0.356 | 0.0003147 | ---?- | 0 | 0.9567 | intron | CCDC173 |
| rs4465584 | 15 | 41971901 | 5 | 0 | 0 | T | C | -1.0277 | 0.2853 | 0.0003148 | ----- | 14.6 | 0.3212 | intron | MGA |
| rs74973841 | 6 | 63177712 | 4 | 0 | 0 | A | T | 2.4134 | 0.6699 | 0.0003148 | +++?+ | 0 | 0.5512 | intergenic | |
| rs75900172 | 11 | 1119714 | 4 | 0 | 0 | A | G | 1.4187 | 0.3938 | 0.000315 | +++?- | 58 | 0.06757 | intergenic | |
| rs117838548 | 17 | 56615626 | 5 | 0 | 0 | A | T | 2.4688 | 0.6853 | 0.000315 | +++++ | 0 | 0.752 | intron | SEPTIN4 |
| rs61992827 | 14 | 43257578 | 4 | 0 | 0 | T | C | -1.3785 | 0.3826 | 0.0003151 | ---?+ | 60.9 | 0.05346 | intergenic | |
| rs10517816 | 4 | 63343844 | 4 | 0 | 0 | A | G | 0.9956 | 0.2764 | 0.0003152 | +++?+ | 32.4 | 0.2176 | intergenic | |
| rs904726 | 5 | 123002797 | 4 | 0 | 0 | T | C | -0.9554 | 0.2652 | 0.0003154 | ---?- | 58.1 | 0.06708 | intergenic | |
| rs11825033 | 11 | 86165659 | 4 | 0 | 0 | A | G | 2.5694 | 0.7133 | 0.0003154 | +++?+ | 0 | 0.5926 | intron | ME3 |
| 7:131558104 | 7 | 131558104 | 3 | 0 | 0 | A | AAAACA | -2.2856 | 0.6345 | 0.0003154 | ---?? | 0 | 0.6657 | | |
| rs9832564 | 3 | 33258380 | 4 | 0 | 0 | T | G | -0.9702 | 0.2693 | 0.0003155 | ---?- | 30.3 | 0.2305 | intron | SUSD5 |
| rs1336128 | 1 | 18765308 | 4 | 0 | 0 | A | G | -1.0674 | 0.2963 | 0.0003156 | ---?- | 33.4 | 0.2121 | intergenic | |
| rs1382026 | 8 | 140495818 | 4 | 0 | 0 | T | C | 1.4891 | 0.4134 | 0.0003157 | +++?+ | 0 | 0.7917 | intergenic | |
| rs145392935 | 18 | 25150796 | 4 | 0 | 0 | A | G | 2.8947 | 0.8036 | 0.0003158 | +++?+ | 0 | 0.5975 | intergenic | |
| rs1772177 | 13 | 45627290 | 4 | 0 | 0 | T | G | -2.1902 | 0.6081 | 0.000316 | ---?+ | 40.3 | 0.1699 | intergenic | |
| rs117416085 | 21 | 42586822 | 3 | 0 | 0 | A | T | -4.8125 | 1.3361 | 0.000316 | ---?? | 0 | 0.5171 | intron | BACE2 |
| rs7025506 | 9 | 7825198 | 4 | 0 | 0 | C | G | -1.209 | 0.3357 | 0.0003161 | ---?- | 8.5 | 0.3507 | intergenic | |
| rs972571 | 17 | 56899570 | 5 | 0 | 0 | A | T | 2.5212 | 0.7 | 0.0003162 | +++++ | 0 | 0.8434 | intron | PPM1E |
| rs13207229 | 6 | 165821041 | 4 | 0 | 0 | T | C | 0.9725 | 0.27 | 0.0003163 | +++?+ | 0 | 0.6618 | intron | PDE10A |
| rs114902592 | 6 | 32156895 | 5 | 0 | 0 | A | G | -1.7814 | 0.4946 | 0.0003163 | ----- | 26.9 | 0.2421 | intron | PBX2 |
| rs221764 | 19 | 29247283 | 4 | 0 | 0 | T | C | 1.3899 | 0.3859 | 0.0003164 | +++?- | 20.4 | 0.2877 | intergenic | |
| rs17162291 | 7 | 141184033 | 4 | 0 | 0 | T | C | -5.1553 | 1.4314 | 0.0003164 | ---?- | 0 | 0.7458 | downstream-gene | |
| 6:32583885 | 6 | 32583885 | 3 | 0 | 0 | G | GATGT | -0.8916 | 0.2476 | 0.0003164 | ---?? | 0 | 0.9098 | | |
| rs1020597 | 8 | 113495545 | 4 | 0 | 0 | T | C | -1.0299 | 0.286 | 0.0003165 | ---?- | 0 | 0.7391 | intron | CSMD3 |
| rs1806224 | 17 | 5290658 | 5 | 0 | 0 | T | C | -2.6595 | 0.7385 | 0.0003165 | ----- | 48.2 | 0.1022 | intron | NUP88 |
| rs4488098 | 10 | 18982031 | 4 | 0 | 0 | A | G | -1.0796 | 0.2998 | 0.0003165 | +--?- | 51.9 | 0.1008 | intergenic | |
| rs7982389 | 13 | 60027250 | 4 | 0 | 0 | T | C | 0.9022 | 0.2505 | 0.0003167 | +++?+ | 0 | 0.4831 | intergenic | |
| rs1340413 | 1 | 102952062 | 4 | 0 | 0 | T | C | -0.8426 | 0.234 | 0.0003168 | ---?+ | 0 | 0.3993 | intergenic | |
| rs9381248 | 6 | 43734606 | 4 | 0 | 0 | T | C | -1.3474 | 0.3742 | 0.000317 | ---?- | 0 | 0.678 | upstream-gene | |
| rs55770006 | 6 | 118693802 | 4 | 0 | 0 | C | G | 2.4225 | 0.6727 | 0.000317 | +++?- | 0 | 0.6528 | intergenic | |

| | | | | | | | | | | | | | | | |
|---|---|---|---|---|---|---|---|---|---|---|---|---|---|---|---|
| rs73170450 | 8 | 1267629 | 4 | 0 | 0 | A | C | 1.1188 | 0.3107 | 0.000317 | +++?+ | 20 | 0.2899 | intron | DLGAP2 |
| rs74027454 | 15 | 85815480 | 2 | 0 | 0 | A | T | -6.6731 | 1.8531 | 0.000317 | --??? | 0 | 0.4347 | intergenic | |
| rs34600111 | 3 | 183576147 | 4 | 0 | 0 | C | G | -1.7365 | 0.4822 | 0.0003171 | ---?- | 0 | 0.8548 | intron | PARL |
| rs28728149 | 4 | 183654185 | 4 | 0 | 0 | T | C | -0.962 | 0.2672 | 0.0003172 | ---?+ | 0 | 0.6984 | intron | TENM3 |
| rs6671735 | 1 | 232832958 | 4 | 0 | 0 | T | C | -0.8951 | 0.2486 | 0.0003173 | ---?- | 36.2 | 0.1951 | intergenic | |
| rs13018351 | 2 | 218829609 | 4 | 0 | 0 | A | C | 1.3022 | 0.3617 | 0.0003173 | +++?+ | 0 | 0.6046 | intergenic | |
| rs34989128 | 6 | 32391772 | 5 | 0 | 1 | A | G | 1.3868 | 0.3851 | 0.0003173 | +++++ | 0 | 0.5141 | intergenic | |
| rs35875335 | 6 | 32391991 | 5 | 0 | 1 | T | C | -1.3868 | 0.3851 | 0.0003173 | ----- | 0 | 0.514 | intergenic |  |
| rs78889410 | 10 | 4362700 | 4 | 0 | 0 | A | G | -2.6937 | 0.7481 | 0.0003173 | ---?- | 42.6 | 0.156 | intergenic | |
| rs11148472 | 13 | 60020189 | 4 | 0 | 0 | A | C | 0.9059 | 0.2516 | 0.0003174 | +++?+ | 0 | 0.5604 | intergenic | |
| rs182123754 | 11 | 624779 | 4 | 0 | 0 | A | C | 2.7612 | 0.7668 | 0.0003174 | +++?+ | 0 | 0.7269 | intron | CDHR5 |
| rs10203212 | 2 | 170917648 | 4 | 0 | 1 | A | G | 1.0438 | 0.2899 | 0.0003177 | +++?+ | 43.7 | 0.1492 | synonymous | UBR3 |
| rs17649898 | 4 | 169325886 | 4 | 0 | 0 | T | C | -0.9373 | 0.2603 | 0.0003177 | ---?- | 0 | 0.5032 | intron | DDX60L |
| rs1384075 | 11 | 103323092 | 4 | 0 | 0 | T | C | -1.0066 | 0.2796 | 0.0003178 | ---?- | 42.4 | 0.157 | intron | DYNC2H1 |
| 1:87347930 | 1 | 87347930 | 3 | 0 | 0 | CAAAAA | C | -1.3585 | 0.3773 | 0.0003178 | ---?? | 56.9 | 0.0981 | | |
| 15:87614744 | 15 | 87614744 | 3 | 0 | 0 | T | TTGTC | 0.9754 | 0.2709 | 0.0003181 | +++?? | 0 | 0.5515 | | |
| rs10174880 | 2 | 170632625 | 4 | 0 | 0 | A | C | 1.1533 | 0.3204 | 0.0003183 | +++?+ | 27.8 | 0.2452 | intergenic | |
| rs34548593 | 3 | 183605169 | 4 | 0 | 0 | A | G | -1.708 | 0.4745 | 0.0003185 | ---?- | 0 | 0.9357 | upstream-gene | |
| rs59233748 | 5 | 76073962 | 5 | 0 | 0 | T | C | 0.9643 | 0.2679 | 0.0003185 | ++++- | 0 | 0.4971 | intergenic | |
| rs16883735 | 8 | 113494316 | 4 | 0 | 0 | A | G | 1.0279 | 0.2855 | 0.0003186 | +++?+ | 0 | 0.7377 | intron | CSMD3 |
| 6:83760546 | 6 | 83760546 | 3 | 0 | 0 | C | G | 3.5602 | 0.9891 | 0.0003188 | +++?? | 0 | 0.586 | intron | UBE3D |
| rs1577676 | 13 | 60046011 | 4 | 0 | 0 | A | T | -0.9459 | 0.2628 | 0.0003189 | ---?- | 0 | 0.56 | intergenic | |
| rs2078395 | 4 | 180903364 | 4 | 0 | 0 | A | T | 0.9407 | 0.2614 | 0.000319 | +++?- | 41.1 | 0.1654 | intergenic | |
| rs7190800 | 16 | 52919377 | 4 | 0 | 0 | T | C | 0.8756 | 0.2433 | 0.000319 | +++?+ | 0 | 0.5165 | intergenic | |
| rs138406775 | 19 | 32092544 | 3 | 0 | 0 | T | G | -3.6058 | 1.0018 | 0.0003191 | ---?? | 0 | 0.9251 | intergenic | |
| rs111405629 | 6 | 144767443 | 4 | 0 | 0 | A | G | 3.2429 | 0.901 | 0.0003192 | +++?+ | 0 | 0.7922 | intron | UTRN |
| rs79104083 | 2 | 170502343 | 4 | 0 | 0 | A | T | 1.281 | 0.3559 | 0.0003194 | +++?+ | 0 | 0.9553 | 3-prime-UTR | CCDC173 |
| rs867960 | 15 | 85815585 | 2 | 0 | 0 | A | T | -6.6659 | 1.8522 | 0.0003195 | --??? | 0 | 0.4363 | intergenic | |
| rs7939127 | 11 | 30685132 | 4 | 0 | 0 | C | G | 1.0095 | 0.2805 | 0.0003197 | +++?- | 0 | 0.7009 | intergenic | |
| rs62418176 | 6 | 63218149 | 4 | 0 | 0 | T | C | -2.3713 | 0.6589 | 0.0003198 | ---?- | 0 | 0.5503 | intergenic | |
| rs145757146 | 2 | 11018067 | 3 | 0 | 0 | C | G | 4.2736 | 1.1876 | 0.0003199 | +++?? | 0 | 0.7091 | intergenic | |
| rs7107956 | 11 | 59209328 | 3 | 0 | 0 | A | T | 3.5786 | 0.9945 | 0.0003203 | +?++? | 9.4 | 0.3317 | upstream-gene | |
| rs79034995 | 2 | 170463156 | 4 | 0 | 0 | A | G | 1.2757 | 0.3545 | 0.0003204 | +++?+ | 0 | 0.9257 | intron | PPIG |
| rs116527475 | 17 | 56898401 | 5 | 0 | 0 | T | G | -2.5212 | 0.7007 | 0.0003204 | ----- | 0 | 0.8223 | intron | PPM1E |
| rs111939436 | 8 | 76172546 | 4 | 0 | 0 | A | G | 1.626 | 0.4519 | 0.0003205 | +++?+ | 0 | 0.4811 | intron | CASC9 |
| rs884213 | 15 | 85815588 | 2 | 0 | 0 | T | C | -6.6635 | 1.852 | 0.0003206 | --??? | 0 | 0.4357 | intergenic | |
| rs13175749 | 5 | 123003804 | 4 | 0 | 0 | T | G | -0.9546 | 0.2653 | 0.0003207 | ---?- | 58.2 | 0.06649 | intergenic | |
| 9:139297059 | 9 | 139297059 | 3 | 0 | 0 | G | GC | 2.831 | 0.7869 | 0.0003208 | +++?? | 0 | 0.8543 | | |
| rs7358745 | 12 | 27782021 | 4 | 0 | 0 | A | G | -0.9165 | 0.2548 | 0.0003209 | ---?+ | 53.9 | 0.08919 | intron | PPFIBP1 |
| rs6869437 | 5 | 6428642 | 4 | 0 | 0 | A | G | 0.9767 | 0.2715 | 0.000321 | +++?+ | 0.2 | 0.3906 | intergenic | |
| rs75396695 | 3 | 183601126 | 4 | 0 | 0 | C | G | -1.6954 | 0.4712 | 0.000321 | ---?- | 0 | 0.9309 | intron | PARL |
| rs12807395 | 11 | 12716675 | 4 | 0 | 0 | T | G | 1.6567 | 0.4605 | 0.0003211 | +++?+ | 0 | 0.6957 | intron | TEAD1 |
| rs75671286 | 10 | 6431765 | 4 | 0 | 0 | T | C | -2.5866 | 0.719 | 0.0003211 | ---?- | 0 | 0.5262 | downstream-gene | |
| rs111382834 | 2 | 170469174 | 4 | 0 | 0 | A | G | -1.2765 | 0.3548 | 0.0003211 | ---?- | 0 | 0.9324 | intron | PPIG |
| rs8019241 | 14 | 43256168 | 4 | 0 | 0 | T | C | 1.3772 | 0.3828 | 0.0003212 | +++?- | 60.9 | 0.05319 | intergenic | |
| rs35047344 | 3 | 183577841 | 4 | 0 | 0 | A | G | 1.733 | 0.4817 | 0.0003214 | +++?+ | 0 | 0.8577 | intron | PARL |
| rs9496922 | 6 | 144622950 | 5 | 0 | 0 | A | G | 1.4434 | 0.4012 | 0.0003215 | ++++- | 34 | 0.1944 | intron | UTRN |
| rs112654745 | 8 | 102198376 | 2 | 0 | 0 | A | G | -7.0842 | 1.9693 | 0.0003215 | --??? | 0 | 0.6796 | intergenic | |
| rs114516506 | 3 | 196479715 | 4 | 0 | 0 | A | G | 2.2195 | 0.617 | 0.0003215 | +++?- | 22.8 | 0.2741 | intron | PAK2 |
| 8:102199257 | 8 | 102199257 | 2 | 0 | 0 | A | AATG | -7.0773 | 1.9674 | 0.0003216 | --??? | 0 | 0.6718 | | |
| 1:202059492 | 1 | 202059492 | 3 | 0 | 0 | CT | C | 1.3216 | 0.3674 | 0.0003216 | +++?? | 11.4 | 0.3233 | | |
| rs62418172 | 6 | 63209780 | 4 | 0 | 0 | A | G | 2.3679 | 0.6583 | 0.0003218 | +++?+ | 0 | 0.5443 | intergenic | |
| 9:30897227 | 9 | 30897227 | 3 | 0 | 0 | A | T | 3.4625 | 0.9626 | 0.0003219 | +++?? | 0 | 0.5199 | intergenic | |
| rs4267038 | 11 | 103325335 | 4 | 0 | 0 | A | C | -1.0055 | 0.2795 | 0.0003219 | ---?- | 41.9 | 0.1601 | intron | DYNC2H1 |
| rs78769140 | 6 | 63210304 | 4 | 0 | 0 | A | G | -2.368 | 0.6583 | 0.000322 | ---?- | 0 | 0.5444 | intergenic | |
| rs11671296 | 19 | 6638982 | 4 | 0 | 0 | A | G | -0.9152 | 0.2544 | 0.0003221 | ---?- | 0 | 0.7904 | intergenic | |
| 9:30897225 | 9 | 30897225 | 3 | 0 | 0 | A | T | 3.4634 | 0.9629 | 0.0003222 | +++?? | 0 | 0.5205 | intergenic | |
| rs10797591 | 1 | 232831936 | 4 | 0 | 0 | C | G | 0.8946 | 0.2487 | 0.0003222 | +++?+ | 36.7 | 0.192 | intergenic | |
| rs1348293 | 18 | 39986819 | 4 | 0 | 0 | T | C | -1.4118 | 0.3925 | 0.0003223 | ---?- | 0 | 0.8249 | intron | LINC00907 |

| | | | | | | | | | | | | | | | |
|---|---|---|---|---|---|---|---|---|---|---|---|---|---|---|---|
| rs111397662 | 2 | 170581888 | 4 | 0 | 0 | T | G | -1.274 | 0.3542 | 0.0003223 | ---?- | 0 | 0.9449 | intron | PHOSPHO2-KLHL23 |
| rs2741727 | 14 | 21455438 | 4 | 0 | 0 | A | C | -0.9001 | 0.2503 | 0.0003224 | ---?- | 0 | 0.7965 | upstream-gene | |
| rs10077693 | 5 | 123429408 | 4 | 0 | 0 | A | C | 0.8977 | 0.2496 | 0.0003224 | +++?+ | 0 | 0.7393 | intergenic | |
| 9:30897226 | 9 | 30897226 | 3 | 0 | 0 | A | T | 3.4627 | 0.9628 | 0.0003225 | +++?? | 0 | 0.5204 | intergenic | |
| rs7943941 | 11 | 59207480 | 3 | 0 | 0 | A | G | 3.5805 | 0.9955 | 0.0003225 | +?++? | 9.4 | 0.3318 | upstream-gene | |
| rs79599650 | 2 | 170468759 | 4 | 0 | 0 | T | C | 1.2761 | 0.3548 | 0.0003225 | +++?+ | 0 | 0.9319 | intron | PPIG |
| rs2361019 | 19 | 22729613 | 4 | 0 | 0 | A | T | -0.9138 | 0.2541 | 0.0003226 | ---?- | 0 | 0.5893 | intergenic | |
| rs139346616 | 20 | 46071614 | 4 | 0 | 0 | A | G | 4.4062 | 1.2252 | 0.0003226 | +++?+ | 0 | 0.6745 | intergenic | |
| rs13145207 | 4 | 66971018 | 4 | 0 | 0 | T | C | 3.784 | 1.0522 | 0.0003227 | +++?+ | 45.8 | 0.1363 | intergenic | |
| rs60514651 | 1 | 232832131 | 4 | 0 | 0 | A | G | -0.8944 | 0.2487 | 0.0003227 | ---?- | 36.6 | 0.1926 | intergenic | |
| rs61974397 | 14 | 28223495 | 4 | 0 | 0 | T | C | -2.209 | 0.6142 | 0.0003227 | ---?- | 0 | 0.9609 | intergenic | |
| 9:106877865 | 9 | 106877865 | 3 | 0 | 0 | CA | C | -4.6496 | 1.2929 | 0.0003227 | ---?? | 0 | 0.8296 | | |
| rs72760196 | 15 | 87669825 | 4 | 0 | 0 | A | G | -3.2119 | 0.8931 | 0.0003228 | ---?- | 0 | 0.7876 | intergenic | |
| rs2231145 | 4 | 89039584 | 4 | 0 | 0 | T | C | 4.0588 | 1.1286 | 0.0003229 | +?+++ | 0 | 0.525 | intron | ABCG2 |
| rs8000074 | 13 | 60030469 | 4 | 0 | 0 | A | C | -0.9095 | 0.2529 | 0.000323 | ---?- | 0 | 0.4469 | intergenic | |
| rs7977054 | 12 | 43208500 | 5 | 0 | 0 | T | C | 0.9777 | 0.2719 | 0.0003231 | +++++ | 0 | 0.4374 | intergenic | |
| rs77796406 | 2 | 170468115 | 4 | 0 | 0 | T | C | 1.2757 | 0.3548 | 0.0003231 | +++?+ | 0 | 0.9305 | intron | PPIG |
| rs7737087 | 5 | 55930045 | 3 | 0 | 0 | T | C | 3.6961 | 1.0278 | 0.0003232 | +++?? | 0 | 0.3875 | intergenic | |
| rs140858337 | 18 | 25151208 | 4 | 0 | 0 | A | G | -2.884 | 0.802 | 0.0003232 | ---?- | 0 | 0.5969 | intergenic | |
| rs60665153 | 2 | 170649930 | 4 | 0 | 0 | T | C | -1.1077 | 0.308 | 0.0003233 | ---?- | 54.5 | 0.08601 | intergenic | |
| rs1468004 | 8 | 140377620 | 4 | 0 | 0 | T | C | 1.043 | 0.2901 | 0.0003234 | +++?+ | 0 | 0.8013 | intergenic | |
| rs10797590 | 1 | 232831866 | 4 | 0 | 0 | T | C | -0.8944 | 0.2487 | 0.0003234 | ---?- | 36.7 | 0.1919 | intergenic | |
| rs13269492 | 8 | 124710110 | 5 | 0 | 0 | T | G | 0.8326 | 0.2315 | 0.0003234 | ++++- | 21.2 | 0.2793 | intron | ANXA13 |
| rs10501379 | 11 | 59241747 | 3 | 0 | 0 | T | C | 3.5498 | 0.9872 | 0.0003235 | +?++? | 10.8 | 0.326 | upstream-gene | |
| rs7943445 | 11 | 59207104 | 3 | 0 | 0 | A | T | 3.5804 | 0.9958 | 0.0003236 | +?++? | 9.4 | 0.3317 | upstream-gene | |
| rs76504461 | 13 | 73927993 | 3 | 0 | 0 | T | C | -3.2452 | 0.9025 | 0.0003236 | ---?? | 0 | 0.9682 | intergenic | |
| 2:170556589 | 2 | 170556589 | 3 | 0 | 0 | T | TG | 1.3365 | 0.3717 | 0.0003236 | +++?? | 0 | 0.7739 | | |
| rs884214 | 15 | 85815948 | 2 | 0 | 0 | A | C | 6.655 | 1.8509 | 0.0003237 | ++??? | 0 | 0.4357 | intergenic | |
| rs6693788 | 1 | 102959894 | 4 | 0 | 0 | A | G | -0.8489 | 0.2361 | 0.0003239 | ---?+ | 0 | 0.3921 | intergenic | |
| 9:106406983 | 9 | 106406983 | 3 | 0 | 0 | A | AT | 1.7191 | 0.4781 | 0.0003239 | +++?? | 0 | 0.9691 | | |
| rs1159648 | 1 | 217988984 | 4 | 0 | 0 | A | T | 0.9518 | 0.2647 | 0.000324 | +++?+ | 0 | 0.7842 | intron | SPATA17 |
| rs116222471 | 4 | 100579903 | 4 | 0 | 0 | A | G | -2.0433 | 0.5683 | 0.000324 | +--?- | 49.8 | 0.113 | upstream-gene | |
| rs9570132 | 13 | 60033082 | 4 | 0 | 0 | T | C | 0.8989 | 0.25 | 0.0003241 | +++?+ | 0 | 0.4421 | intergenic | |
| 1:120233236 | 1 | 120233236 | 3 | 0 | 0 | CT | C | 1.3918 | 0.3871 | 0.0003241 | +++?? | 46.3 | 0.1553 | | |
| rs114224304 | 2 | 167999282 | 4 | 0 | 0 | A | G | -2.2686 | 0.631 | 0.0003243 | ---?+ | 2.9 | 0.378 | intron | XIRP2 |
| rs10161290 | 12 | 112718198 | 5 | 0 | 0 | A | G | 1.3293 | 0.3698 | 0.0003245 | +-+++ | 54.7 | 0.06545 | intron | HECTD4 |
| rs116727398 | 2 | 167992961 | 4 | 0 | 0 | A | T | 2.2777 | 0.6336 | 0.0003245 | +++?- | 3.2 | 0.3767 | intron | XIRP2 |
| rs10737711 | 1 | 90243771 | 5 | 0 | 0 | T | C | 0.7646 | 0.2127 | 0.0003247 | +++++ | 0 | 0.9915 | intergenic | |
| rs10792241 | 11 | 59206233 | 3 | 0 | 0 | T | C | -3.5815 | 0.9963 | 0.0003247 | -?--? | 9.4 | 0.3315 | upstream-gene | |
| rs12916992 | 15 | 41975127 | 5 | 0 | 0 | T | C | -1.0202 | 0.2838 | 0.0003247 | ----- | 14.7 | 0.3207 | intron | MGA |
| rs7800843 | 7 | 154577986 | 4 | 0 | 0 | C | G | -1.0083 | 0.2805 | 0.0003248 | ---?- | 0 | 0.7457 | intron | DPP6 |
| 10:2487439 | 10 | 2487439 | 3 | 0 | 0 | A | AT | -1.1213 | 0.3119 | 0.0003248 | ---?? | 0 | 0.4451 | | |
| rs1421141 | 7 | 130124927 | 4 | 0 | 0 | A | G | 0.8777 | 0.2442 | 0.000325 | +-+?+ | 59.9 | 0.05808 | upstream-gene | |
| rs11151253 | 18 | 63862928 | 4 | 0 | 0 | T | C | 0.8957 | 0.2492 | 0.000325 | +++?+ | 0 | 0.547 | intergenic | |
| rs1510263 | 1 | 217989437 | 4 | 0 | 0 | T | C | -0.9511 | 0.2646 | 0.0003251 | ---?- | 0 | 0.7846 | intron | SPATA17 |
| rs4144280 | 8 | 140371269 | 4 | 0 | 0 | A | G | -1.0428 | 0.2901 | 0.0003254 | ---?- | 0 | 0.8089 | intergenic | |
| rs6015350 | 20 | 57304777 | 3 | 0 | 0 | T | C | -1.4735 | 0.41 | 0.0003254 | ---?? | 0 | 0.5725 | intergenic | |
| rs13062365 | 3 | 129888192 | 4 | 0 | 0 | A | T | -0.9922 | 0.2761 | 0.0003254 | ---?- | 37.8 | 0.185 | intergenic | |
| rs143057839 | 16 | 7591394 | 4 | 0 | 0 | T | C | -3.894 | 1.0834 | 0.0003254 | ---?- | 0 | 0.6334 | intron | RBFOX1 |
| rs73887565 | 3 | 183613287 | 4 | 0 | 0 | A | G | -1.7159 | 0.4774 | 0.0003256 | ---?- | 0 | 0.9451 | intergenic | |
| rs2231146 | 4 | 89039500 | 4 | 0 | 0 | T | C | 4.0558 | 1.1285 | 0.0003257 | +?+++ | 0 | 0.5244 | intron | ABCG2 |
| rs2288331 | 2 | 170592709 | 4 | 0 | 0 | A | G | 1.1719 | 0.3261 | 0.0003257 | +++?+ | 0 | 0.9975 | synonymous | KLHL23,PHOSPHO2-KLHL23 |
| rs12470684 | 2 | 156862228 | 4 | 0 | 0 | C | G | -0.9385 | 0.2612 | 0.0003259 | ---?- | 0 | 0.8099 | intergenic | |
| rs28699545 | 14 | 43259298 | 4 | 0 | 0 | T | G | 1.3657 | 0.38 | 0.0003259 | +++?- | 61.5 | 0.05062 | intergenic | |
| 8:140468359 | 8 | 140468359 | 3 | 0 | 0 | G | GA | 1.4585 | 0.4059 | 0.0003261 | +++?? | 0 | 0.6548 | | |
| rs35140688 | 15 | 99614558 | 4 | 0 | 0 | A | G | -1.2493 | 0.3476 | 0.0003262 | ---?- | 10.5 | 0.3406 | intergenic | |
| 3:65664035 | 3 | 65664035 | 3 | 0 | 0 | T | TAA | -3.7525 | 1.0442 | 0.0003262 | ---?? | 6.8 | 0.3418 | | |
| rs75282167 | 3 | 143021123 | 4 | 0 | 0 | A | C | -2.3469 | 0.6531 | 0.0003263 | ---?- | 0 | 0.7815 | intron | SLC9A9 |
| rs7984999 | 13 | 60031165 | 4 | 0 | 0 | A | T | -0.9023 | 0.2511 | 0.0003264 | ---?- | 0 | 0.4503 | intergenic | |

| | | | | | | | | | | | | | | | |
|---|---|---|---|---|---|---|---|---|---|---|---|---|---|---|---|
| rs116719170 | 1 | 85621172 | 3 | 0 | 0 | A | T | 1.6543 | 0.4604 | 0.0003267 | +++?? | 0 | 0.8088 | downstream-gene | |
| rs73178517 | 12 | 102325023 | 4 | 0 | 0 | T | C | -2.0719 | 0.5766 | 0.0003268 | ---?- | 0 | 0.4987 | intergenic | |
| rs75339401 | 14 | 83295517 | 4 | 0 | 0 | C | G | 2.0716 | 0.5765 | 0.0003268 | -++?+ | 56 | 0.07796 | intergenic | |
| rs76265176 | 1 | 102947753 | 4 | 0 | 0 | A | G | 0.8443 | 0.235 | 0.0003271 | +++?- | 4.7 | 0.3695 | intergenic | |
| rs11823666 | 11 | 129303601 | 4 | 0 | 0 | A | G | -1.5121 | 0.4209 | 0.0003272 | ---?- | 36 | 0.1959 | intron | BARX2 |
| rs10506219 | 12 | 43162578 | 5 | 0 | 0 | T | C | 0.9578 | 0.2666 | 0.0003274 | ++++- | 0 | 0.4178 | intergenic | |
| rs142765517 | 6 | 83728661 | 4 | 0 | 0 | T | C | -3.4573 | 0.9623 | 0.0003274 | ---?- | 0 | 0.8007 | intron | UBE3D |
| rs7309841 | 12 | 112565053 | 5 | 0 | 0 | A | G | -1.3266 | 0.3693 | 0.0003275 | -+--- | 55.3 | 0.06215 | intron | TRAFD1 |
| rs12750240 | 1 | 92139645 | 4 | 0 | 0 | T | C | -0.905 | 0.2519 | 0.0003275 | ---?- | 0 | 0.5191 | intergenic | |
| 9:101134146 | 9 | 101134146 | 3 | 0 | 0 | CTGCTG | C | -3.5852 | 0.998 | 0.0003275 | ---?? | 27.8 | 0.2505 | | |
| rs62418206 | 6 | 63233002 | 4 | 0 | 0 | A | T | 2.3974 | 0.6673 | 0.0003276 | +++?+ | 0 | 0.5523 | intergenic | |
| rs77798972 | 12 | 26841694 | 4 | 0 | 0 | A | G | -1.6956 | 0.472 | 0.0003276 | -+-?- | 54 | 0.08896 | intron | ITPR2 |
| rs151053676 | 17 | 56902216 | 5 | 0 | 0 | T | G | -2.5313 | 0.7046 | 0.0003276 | ----- | 0 | 0.8107 | intron | PPM1E |
| rs685277 | 10 | 131911012 | 4 | 0 | 0 | A | C | -0.8865 | 0.2468 | 0.0003277 | ---?- | 0 | 0.7838 | upstream-gene | |
| rs62418203 | 6 | 63225618 | 4 | 0 | 0 | C | G | 2.3474 | 0.6534 | 0.0003278 | +++?+ | 0 | 0.5574 | intergenic | |
| rs148403760 | 14 | 77181250 | 3 | 0 | 0 | T | C | -3.6004 | 1.0023 | 0.0003278 | ---?? | 0 | 0.9571 | intergenic | |
| rs1625367 | 13 | 45628228 | 4 | 0 | 0 | T | C | 2.1869 | 0.6088 | 0.0003279 | +++?- | 39.9 | 0.1726 | intergenic | |
| rs4630768 | 2 | 122588545 | 4 | 0 | 0 | A | G | 2.0707 | 0.5764 | 0.0003279 | +++?+ | 0 | 0.4428 | intergenic | |
| rs28986328 | 6 | 32741002 | 5 | 0 | 0 | T | C | -1.4412 | 0.4012 | 0.0003279 | ----- | 0 | 0.6916 | intergenic |  |
| rs1345141 | 2 | 170496015 | 4 | 0 | 0 | T | C | -1.2772 | 0.3556 | 0.0003281 | ---?- | 0 | 0.9522 | 3-prime-UTR | PPIG |
| 13:60016207 | 13 | 60016207 | 3 | 0 | 0 | CA | C | -0.9238 | 0.2572 | 0.0003281 | ---?? | 0 | 0.6429 | | |
| rs12620397 | 2 | 170615181 | 4 | 0 | 0 | T | C | -1.2525 | 0.3487 | 0.0003282 | ---?- | 0 | 0.979 | intergenic | |
| rs6487621 | 12 | 27792774 | 4 | 0 | 0 | A | G | 0.9434 | 0.2627 | 0.0003283 | +++?- | 56 | 0.07809 | intron | PPFIBP1 |
| rs62414739 | 6 | 63173181 | 4 | 0 | 0 | T | C | -2.3826 | 0.6633 | 0.0003283 | ---?- | 0 | 0.561 | intergenic | |
| rs12590496 | 14 | 43255995 | 4 | 0 | 0 | A | G | 1.3813 | 0.3846 | 0.0003284 | +++?- | 60.8 | 0.05357 | intergenic | |
| rs62418204 | 6 | 63230069 | 4 | 0 | 0 | A | G | 2.3966 | 0.6672 | 0.0003285 | +++?+ | 0 | 0.5517 | intergenic | |
| 17:47461409 | 17 | 47461409 | 3 | 0 | 0 | T | TA | -1.1273 | 0.3139 | 0.0003285 | ---?? | 0 | 0.7964 | | |
| rs4357921 | 15 | 98048360 | 4 | 0 | 0 | T | C | -0.9486 | 0.2641 | 0.0003286 | ---?- | 0 | 0.9135 | intergenic | |
| rs7031186 | 9 | 106406981 | 3 | 0 | 0 | A | T | 1.7346 | 0.483 | 0.0003286 | +++?? | 0 | 0.8606 | intergenic | |
| rs16865043 | 2 | 178023428 | 4 | 0 | 0 | A | G | -1.7592 | 0.4898 | 0.0003286 | ---?- | 13.4 | 0.3254 | intergenic | |
| rs62418209 | 6 | 63235917 | 4 | 0 | 0 | T | C | -2.3977 | 0.6676 | 0.0003286 | ---?- | 0 | 0.5535 | intergenic | |
| rs66923335 | 2 | 170635683 | 4 | 0 | 0 | A | C | 1.1502 | 0.3202 | 0.0003286 | +++?+ | 27 | 0.25 | intergenic | |
| rs6689 | 6 | 32627700 | 5 | 0 | 0 | A | G | -1.0054 | 0.2799 | 0.0003287 | ----- | 28.1 | 0.2342 | 3-prime-UTR | HLA-DQB1 |
| rs60316007 | 3 | 196475356 | 4 | 0 | 0 | T | C | -2.2152 | 0.6168 | 0.0003287 | ---?+ | 22.7 | 0.2749 | intron | PAK2 |
| rs752606 | 20 | 62522014 | 3 | 0 | 0 | A | G | -4.0389 | 1.1246 | 0.0003288 | ---?? | 0 | 0.7668 | 3-prime-UTR | TPD52L2 |
| rs10062074 | 5 | 124691021 | 4 | 0 | 0 | A | G | 0.8906 | 0.248 | 0.0003289 | +++?+ | 0 | 0.5326 | intergenic | |
| 1:80729508 | 1 | 80729508 | 3 | 0 | 0 | G | GA | -0.9776 | 0.2722 | 0.000329 | ---?? | 58.6 | 0.08911 | | |
| rs1499502 | 3 | 65658054 | 4 | 0 | 0 | T | G | 0.9574 | 0.2666 | 0.0003292 | +++?+ | 47.3 | 0.1274 | intron | MAGI1 |
| rs1648826 | 15 | 42147206 | 5 | 0 | 0 | T | C | 1.2147 | 0.3382 | 0.0003292 | +++++ | 16.5 | 0.3093 | intron | SPTBN5 |
| rs7294753 | 12 | 112567450 | 5 | 0 | 0 | T | G | -1.3263 | 0.3693 | 0.0003293 | -+--- | 55.4 | 0.06191 | intron | TRAFD1 |
| rs663699 | 15 | 53766542 | 4 | 0 | 0 | A | G | 0.904 | 0.2517 | 0.0003294 | +++?+ | 35.2 | 0.2012 | intergenic | |
| rs9264218 | 6 | 31221895 | 3 | 0 | 0 | A | T | -0.8919 | 0.2484 | 0.0003294 | ---?? | 0 | 0.3824 | intergenic | |
| rs12445450 | 16 | 85211373 | 4 | 0 | 0 | T | C | 1.0613 | 0.2956 | 0.0003294 | +++?+ | 0 | 0.5109 | intron | GSE1 |
| rs28620921 | 4 | 57869527 | 4 | 0 | 0 | T | C | -0.8586 | 0.2391 | 0.0003294 | ---?- | 40.3 | 0.17 | intron | POLR2B |
| 6:83807562 | 6 | 83807562 | 3 | 0 | 0 | CTATTT | C | -3.4619 | 0.9641 | 0.0003295 | ---?? | 0 | 0.4163 | | |
| rs72687725 | 4 | 182757735 | 4 | 0 | 0 | T | C | 3.2205 | 0.8968 | 0.0003296 | +++?+ | 0 | 0.8607 | intergenic | |
| rs1026318 | 11 | 59204920 | 3 | 0 | 0 | A | G | 3.58 | 0.997 | 0.0003297 | +?++? | 9.6 | 0.331 | intergenic | |
| rs4931208 | 12 | 27782898 | 4 | 0 | 0 | C | G | -0.9118 | 0.2539 | 0.0003297 | -+-?+ | 54.5 | 0.0859 | intron | PPFIBP1 |
| 1:152779715 | 1 | 152779715 | 3 | 0 | 0 | T | TTGTCA | -2.2619 | 0.6299 | 0.0003297 | ---?? | 40.8 | 0.1845 | | |
| rs4930866 | 12 | 27782920 | 4 | 0 | 0 | T | G | 0.9116 | 0.2539 | 0.0003298 | +-+?- | 54.6 | 0.08569 | intron | PPFIBP1 |
| rs1545064 | 12 | 41513751 | 3 | 0 | 0 | A | G | -3.8261 | 1.0656 | 0.0003299 | ---?? | 0 | 0.4332 | intergenic | |
| rs4908249 | 1 | 102950021 | 4 | 0 | 0 | T | C | -0.8389 | 0.2336 | 0.0003299 | ---?+ | 4.1 | 0.3724 | intergenic | |
| rs4930867 | 12 | 27783057 | 4 | 0 | 0 | A | G | -0.9113 | 0.2538 | 0.0003301 | -+-?+ | 54.6 | 0.08559 | intron | PPFIBP1 |
| rs72846316 | 11 | 1139039 | 4 | 0 | 0 | A | T | -1.4342 | 0.3995 | 0.0003301 | ---?+ | 59.8 | 0.05839 | upstream-gene | |
| rs79110568 | 2 | 170541884 | 4 | 0 | 0 | A | T | 1.2827 | 0.3573 | 0.0003302 | +++?+ | 0 | 0.9638 | intron | CCDC173 |
| rs3843392 | 3 | 188549655 | 4 | 0 | 0 | A | T | 0.8995 | 0.2505 | 0.0003303 | +++?+ | 0 | 0.7141 | intron | LPP |
| rs141003011 | 6 | 148250504 | 4 | 0 | 0 | T | C | 3.7984 | 1.0579 | 0.0003303 | +++?+ | 26.7 | 0.2515 | intergenic | |
| rs10086605 | 8 | 140364275 | 4 | 0 | 0 | T | C | 1.042 | 0.2902 | 0.0003306 | +++?+ | 0 | 0.8079 | intergenic | |
| rs62418210 | 6 | 63237293 | 4 | 0 | 0 | A | G | -2.3976 | 0.6678 | 0.0003306 | ---?- | 0 | 0.5535 | intergenic | |

| | | | | | | | | | | | | | | | |
|---|---|---|---|---|---|---|---|---|---|---|---|---|---|---|---|
| rs1020609 | 11 | 59203603 | 3 | 0 | 0 | A | G | -3.5825 | 0.9979 | 0.0003308 | -?--? | 9.6 | 0.3309 | intergenic | |
| rs7645396 | 3 | 196476209 | 4 | 0 | 0 | A | G | 2.2154 | 0.6171 | 0.0003308 | +++?- | 23 | 0.273 | intron | PAK2 |
| rs10095578 | 8 | 140367700 | 4 | 0 | 0 | T | C | -1.0419 | 0.2902 | 0.0003308 | ---?- | 0 | 0.8087 | intergenic | |
| rs78302551 | 2 | 62723722 | 4 | 0 | 0 | A | G | 3.3863 | 0.9433 | 0.0003308 | +++?+ | 52.8 | 0.09558 | downstream-gene | |
| rs4931209 | 12 | 27783079 | 4 | 0 | 0 | A | G | 0.9109 | 0.2538 | 0.0003309 | +-+?- | 54.6 | 0.08552 | intron | PPFIBP1 |
| rs7017126 | 8 | 140376925 | 4 | 0 | 0 | C | G | -1.0414 | 0.2901 | 0.0003309 | ---?- | 0 | 0.7993 | intergenic | |
| rs248545 | 5 | 165392865 | 3 | 0 | 0 | T | C | -0.8998 | 0.2507 | 0.000331 | ---?? | 14.7 | 0.3095 | intergenic | |
| rs62414747 | 6 | 63175338 | 4 | 0 | 0 | A | G | 2.3738 | 0.6613 | 0.0003312 | +++?+ | 0 | 0.5572 | intergenic | |
| rs116495888 | 3 | 143021205 | 4 | 0 | 0 | A | G | 2.3419 | 0.6524 | 0.0003312 | +++?+ | 0 | 0.7807 | intron | SLC9A9 |
| rs1026317 | 11 | 59204904 | 3 | 0 | 0 | T | C | -3.5789 | 0.9971 | 0.0003313 | -?--? | 9.6 | 0.3307 | intergenic | |
| rs11070347 | 15 | 41976289 | 5 | 0 | 0 | A | G | -1.021 | 0.2844 | 0.0003313 | ----- | 14.6 | 0.3216 | intron | MGA |
| rs74986046 | 9 | 133795946 | 3 | 0 | 0 | C | G | -3.2337 | 0.9009 | 0.0003313 | ---?? | 65.5 | 0.05519 | intron | FIBCD1 |
| rs2468672 | 8 | 140497279 | 4 | 0 | 0 | T | G | -1.4858 | 0.4139 | 0.0003314 | ---?- | 0 | 0.7837 | intergenic | |
| rs13094830 | 3 | 183615272 | 4 | 0 | 0 | T | C | 1.7169 | 0.4783 | 0.0003314 | +++?+ | 0 | 0.9455 | intergenic | |
| rs62418173 | 6 | 63210152 | 4 | 0 | 0 | T | C | -2.3413 | 0.6523 | 0.0003315 | ---?- | 0 | 0.5485 | intergenic | |
| rs6552732 | 4 | 169324777 | 4 | 0 | 0 | A | G | -0.8924 | 0.2486 | 0.0003316 | ---?- | 0 | 0.8446 | intron | DDX60L |
| rs62414749 | 6 | 63177028 | 4 | 0 | 0 | A | G | -2.3695 | 0.6601 | 0.0003316 | ---?- | 0 | 0.5556 | intergenic | |
| 16:77959733 | 16 | 77959733 | 2 | 0 | 0 | T | TC | -5.1208 | 1.4267 | 0.0003316 | ?--?? | 68.9 | 0.07313 | | |
| rs9276715 | 6 | 32760723 | 5 | 0 | 0 | A | G | 1.4235 | 0.3966 | 0.0003318 | +++++ | 0 | 0.617 | intergenic | |
| rs8017291 | 14 | 22503062 | 4 | 0 | 0 | T | C | -0.9436 | 0.2629 | 0.0003319 | ---?- | 0 | 0.9441 | intergenic | |
| rs76084741 | 6 | 63231026 | 4 | 0 | 0 | T | C | -2.3944 | 0.6671 | 0.0003319 | ---?- | 0 | 0.5543 | intergenic | |
| rs2498499 | 6 | 163467189 | 4 | 0 | 0 | C | G | 2.3839 | 0.6642 | 0.000332 | +++?+ | 0 | 0.6889 | intron | PACRG |
| rs370710514 | 6 | 32562498 | 3 | 0 | 0 | T | G | -1.199 | 0.3341 | 0.0003321 | ---?? | 0 | 0.6489 | upstream-gene | |
| rs12564280 | 1 | 232832633 | 4 | 0 | 0 | T | C | -0.8922 | 0.2486 | 0.0003322 | ---?- | 37.8 | 0.1853 | intergenic | |
| rs79025737 | 5 | 122537007 | 4 | 0 | 0 | C | G | -1.911 | 0.5325 | 0.0003322 | ---?- | 0 | 0.845 | intergenic | |
| rs2807950 | 1 | 87557999 | 4 | 0 | 0 | A | G | -1.1043 | 0.3077 | 0.0003326 | ---?- | 0 | 0.6673 | intron | HS2ST1 |
| rs79332534 | 3 | 18142124 | 4 | 0 | 0 | A | T | -2.2424 | 0.6249 | 0.0003326 | ---?- | 0 | 0.8947 | intron | LOC339862 |
| rs115541968 | 3 | 15698228 | 4 | 0 | 0 | A | T | 2.5784 | 0.7185 | 0.0003326 | +++?+ | 0 | 0.7524 | intergenic | |
| rs148503580 | 18 | 25154711 | 4 | 0 | 0 | A | G | 2.8335 | 0.7896 | 0.0003326 | +++?+ | 0 | 0.6043 | intergenic | |
| rs10152364 | 15 | 41975667 | 5 | 0 | 0 | T | C | 1.0207 | 0.2844 | 0.0003327 | +++++ | 14.6 | 0.3213 | intron | MGA |
| rs7754074 | 6 | 63216701 | 4 | 0 | 0 | A | G | 2.3421 | 0.6527 | 0.0003328 | +++?+ | 0 | 0.5577 | intergenic | |
| rs17073596 | 6 | 144621908 | 5 | 0 | 0 | A | T | 1.4269 | 0.3976 | 0.0003328 | ++++- | 39.5 | 0.158 | intron | UTRN |
| rs57115621 | 18 | 55754878 | 4 | 0 | 0 | A | G | 1.6347 | 0.4556 | 0.0003328 | +++?+ | 0 | 0.9691 | intron | NEDD4L |
| rs77782673 | 4 | 100582399 | 4 | 0 | 0 | A | T | -2.0386 | 0.5681 | 0.0003328 | +--?- | 49.2 | 0.1162 | intergenic | |
| rs1964314 | 12 | 27783162 | 4 | 0 | 0 | A | T | -0.9096 | 0.2535 | 0.0003331 | -+-?+ | 54.7 | 0.08489 | intron | PPFIBP1 |
| rs10836662 | 11 | 37046429 | 4 | 0 | 0 | T | G | 0.8988 | 0.2505 | 0.0003331 | +++?+ | 0 | 0.4237 | intergenic | |
| rs78090129 | 6 | 63209316 | 4 | 0 | 0 | A | G | -2.3401 | 0.6522 | 0.0003331 | ---?- | 0 | 0.548 | intergenic | |
| rs201438 | 7 | 101735236 | 4 | 0 | 0 | A | G | 0.892 | 0.2486 | 0.0003332 | +++?+ | 0 | 0.8382 | intron | CUX1 |
| rs12029730 | 1 | 232831207 | 4 | 0 | 0 | A | T | 0.8928 | 0.2488 | 0.0003335 | +++?+ | 37.2 | 0.189 | intergenic | |
| rs180894532 | 14 | 80987328 | 3 | 0 | 0 | A | T | -2.2574 | 0.6292 | 0.0003336 | ---?? | 0 | 0.8619 | intron | CEP128 |
| rs10246481 | 7 | 156184748 | 4 | 0 | 0 | A | G | -0.9076 | 0.253 | 0.0003338 | ---?- | 8.8 | 0.3491 | intergenic | |
| rs111369835 | 4 | 184807591 | 2 | 0 | 0 | T | C | 5.4835 | 1.5285 | 0.0003339 | +?+?? | 0 | 0.4139 | intergenic | |
| rs147723591 | 17 | 56588345 | 5 | 0 | 0 | A | G | -2.4612 | 0.686 | 0.0003339 | ----- | 0 | 0.7364 | intron | MTMR4 |
| 11:20152754 | 11 | 20152754 | 3 | 0 | 0 | T | TA | 1.7607 | 0.4908 | 0.000334 | +++?? | 0 | 0.813 | | |
| rs62418213 | 6 | 63244155 | 4 | 0 | 0 | T | C | -2.4018 | 0.6695 | 0.0003342 | ---?- | 0 | 0.5555 | intergenic | |
| rs115420353 | 6 | 63206177 | 4 | 0 | 0 | A | C | 2.3381 | 0.6518 | 0.0003342 | +++?+ | 0 | 0.5456 | intergenic | |
| 5:179942256 | 5 | 179942256 | 3 | 0 | 0 | CA | C | -1.0802 | 0.3011 | 0.0003342 | ---?? | 0 | 0.4021 | | |
| rs62418167 | 6 | 63206272 | 4 | 0 | 0 | C | G | 2.3381 | 0.6518 | 0.0003343 | +++?+ | 0 | 0.5457 | intergenic | |
| rs7736130 | 5 | 55930153 | 3 | 0 | 0 | A | G | 3.6704 | 1.0232 | 0.0003344 | +++?? | 1.2 | 0.3634 | intergenic | |
| rs7114559 | 11 | 59202250 | 3 | 0 | 0 | C | G | -3.583 | 0.9989 | 0.0003345 | -?--? | 9.6 | 0.3306 | intergenic | |
| rs16883746 | 8 | 113499408 | 4 | 0 | 0 | A | T | 0.9261 | 0.2582 | 0.0003345 | +++?+ | 0 | 0.6696 | intron | CSMD3 |
| rs73020374 | 2 | 170591040 | 4 | 0 | 0 | A | G | -1.1749 | 0.3275 | 0.0003345 | ---?- | 0 | 0.998 | intron | KLHL23,PHOSPHO2-KLHL23 |
| rs117264323 | 17 | 34455874 | 2 | 0 | 0 | C | G | 6.7233 | 1.8744 | 0.0003347 | +?+?? | 0 | 0.3309 | intergenic | |
| rs62416857 | 6 | 63201943 | 4 | 0 | 0 | T | C | 2.3356 | 0.6512 | 0.0003348 | +++?+ | 0 | 0.5439 | intergenic | |
| rs7812523 | 8 | 140367142 | 4 | 0 | 0 | T | C | 1.041 | 0.2902 | 0.0003349 | +++?+ | 0 | 0.809 | intergenic | |
| rs1499503 | 3 | 65658302 | 4 | 0 | 0 | T | C | 0.9564 | 0.2667 | 0.0003351 | +++?+ | 47.1 | 0.1289 | intron | MAGI1 |
| rs7834006 | 8 | 140367089 | 4 | 0 | 0 | T | G | -1.0408 | 0.2902 | 0.0003353 | ---?- | 0 | 0.8089 | intergenic | |
| rs62418215 | 6 | 63252328 | 4 | 0 | 0 | T | C | 2.4298 | 0.6775 | 0.0003353 | +++?+ | 0 | 0.5526 | intergenic | |
| rs56174601 | 3 | 183597731 | 4 | 0 | 0 | C | G | 1.6931 | 0.4721 | 0.0003354 | +++?+ | 0 | 0.92 | intron | PARL |

| | | | | | | | | | | | | | | |
|---|---|---|---|---|---|---|---|---|---|---|---|---|---|---|
| rs61595241 | 6 | 109107919 | 2 | 0 | 0 | A | G | -4.9645 | 1.3843 | 0.0003356 | -?-?? | 0 | 0.8785 | intergenic | |
| rs76835051 | 9 | 90849840 | 3 | 0 | 0 | A | G | 3.377 | 0.9417 | 0.0003356 | +++?? | 1.6 | 0.3619 | intergenic | |
| rs1964315 | 12 | 27783444 | 4 | 0 | 0 | A | G | 0.9077 | 0.2531 | 0.0003357 | +-+?- | 54.9 | 0.0838 | intron | PPFIBP1 |
| rs7330487 | 13 | 60018768 | 4 | 0 | 0 | T | C | -0.917 | 0.2557 | 0.0003357 | ---?- | 0 | 0.6209 | intergenic | |
| rs1646988 | 7 | 35526024 | 4 | 0 | 0 | A | T | 1.1504 | 0.3208 | 0.0003358 | +++?+ | 36.1 | 0.1958 | intergenic | |
| rs62416859 | 6 | 63202318 | 4 | 0 | 0 | A | G | -2.3351 | 0.6512 | 0.0003359 | ---?- | 0 | 0.5429 | intergenic | |
| rs1646989 | 7 | 35526022 | 4 | 0 | 0 | T | G | -1.1504 | 0.3208 | 0.0003361 | ---?- | 36 | 0.1962 | intergenic | |
| rs1382022 | 8 | 140497775 | 4 | 0 | 0 | A | G | -1.4844 | 0.414 | 0.0003363 | ---?- | 0 | 0.7806 | intergenic | |
| rs1959500 | 14 | 45058892 | 3 | 0 | 0 | T | C | 1.514 | 0.4223 | 0.0003363 | +++?? | 0 | 0.5611 | intergenic | |
| rs12319782 | 12 | 43160201 | 5 | 0 | 0 | A | G | 0.9558 | 0.2666 | 0.0003363 | ++++- | 3.3 | 0.3879 | intergenic | |
| rs201440 | 7 | 101735599 | 4 | 0 | 0 | T | C | 0.8887 | 0.2479 | 0.0003365 | +++?+ | 0 | 0.8386 | intron | CUX1 |
| rs12297983 | 12 | 43165721 | 5 | 0 | 0 | A | G | -1.2304 | 0.3432 | 0.0003365 | ----- | 23.1 | 0.2674 | intergenic |  |
| 13:20802825 | 13 | 20802825 | 3 | 0 | 0 | G | GTGTTA( | 1.0943 | 0.3052 | 0.0003365 | +++?? | 0 | 0.8501 | | |
| rs666317 | 15 | 53765980 | 4 | 0 | 0 | T | G | 0.8977 | 0.2504 | 0.0003366 | +++?+ | 33 | 0.2144 | intergenic | |
| rs139966869 | 2 | 122637446 | 4 | 0 | 0 | T | C | 3.2088 | 0.895 | 0.0003366 | +++?+ | 0 | 0.9849 | intergenic | |
| rs4143626 | 8 | 113279478 | 4 | 0 | 0 | T | C | 1.0218 | 0.285 | 0.0003367 | +++?+ | 0 | 0.8098 | intron | CSMD3 |
| rs11041251 | 11 | 7235856 | 4 | 0 | 0 | A | G | 3.0136 | 0.8405 | 0.0003367 | +++?- | 0 | 0.7929 | intergenic | |
| rs7769348 | 6 | 63181615 | 4 | 0 | 0 | T | C | 2.3537 | 0.6565 | 0.0003368 | +++?+ | 0 | 0.5503 | intergenic | |
| rs13201932 | 6 | 32756950 | 5 | 0 | 0 | C | G | 1.4395 | 0.4015 | 0.0003368 | +++++ | 0 | 0.6868 | intergenic | |
| rs75821709 | 2 | 170480465 | 4 | 0 | 0 | T | C | -1.2731 | 0.3551 | 0.0003368 | ---?- | 0 | 0.9411 | intron | PPIG |
| rs35618016 | 5 | 51556387 | 4 | 0 | 0 | A | G | -2.8101 | 0.7838 | 0.0003369 | ---?- | 35.7 | 0.1982 | intergenic | |
| rs11488254 | 1 | 64667085 | 3 | 0 | 0 | A | G | -0.955 | 0.2664 | 0.0003371 | ---?? | 25.5 | 0.2615 | upstream-gene | |
| rs13217824 | 6 | 32756953 | 5 | 0 | 0 | T | G | -1.4392 | 0.4015 | 0.0003373 | ----- | 0 | 0.6868 | intergenic |  |
| rs62418216 | 6 | 63253634 | 4 | 0 | 0 | A | G | -2.4308 | 0.6781 | 0.0003373 | ---?- | 0 | 0.5534 | intergenic | |
| rs75233893 | 6 | 135110090 | 3 | 0 | 0 | T | C | 2.808 | 0.7833 | 0.0003373 | +++?? | 0 | 0.5657 | intergenic | |
| rs140254592 | 6 | 83725811 | 4 | 0 | 0 | A | G | -3.4477 | 0.9617 | 0.0003373 | ---?- | 0 | 0.8041 | intron | UBE3D |
| rs115626688 | 17 | 56910227 | 5 | 0 | 0 | T | C | 2.5058 | 0.699 | 0.0003375 | +++++ | 0 | 0.8454 | intron | PPM1E |
| rs12904651 | 15 | 98041875 | 4 | 0 | 0 | T | C | 0.9018 | 0.2516 | 0.0003377 | +++?+ | 0 | 0.9324 | intergenic | |
| rs62416852 | 6 | 63197149 | 4 | 0 | 0 | C | G | 2.3318 | 0.6505 | 0.0003377 | +++?+ | 0 | 0.5394 | intergenic | |
| rs1334727 | 6 | 63195923 | 4 | 0 | 0 | T | C | -2.3313 | 0.6504 | 0.0003378 | ---?- | 0 | 0.5392 | intergenic | |
| rs6981195 | 8 | 140380087 | 4 | 0 | 0 | T | C | -1.0425 | 0.2908 | 0.0003378 | ---?- | 0 | 0.7994 | intergenic | |
| 22:31527907 | 22 | 31527907 | 3 | 0 | 0 | CAG | C | 3.6823 | 1.0273 | 0.0003378 | +++?? | 0 | 0.4298 | | |
| rs4243509 | 8 | 135304218 | 4 | 0 | 0 | T | C | -1.3087 | 0.3651 | 0.000338 | ---?- | 0 | 0.8449 | intergenic | |
| rs1453532 | 11 | 59199917 | 3 | 0 | 0 | T | C | -3.5833 | 0.9998 | 0.0003382 | -?--? | 9.7 | 0.3305 | intergenic | |
| rs139459590 | 19 | 2076412 | 4 | 0 | 0 | C | G | 2.8206 | 0.787 | 0.0003382 | +++?+ | 0 | 0.8343 | intron | MOB3A |
| rs17099333 | 14 | 62376495 | 4 | 0 | 0 | A | T | -1.0792 | 0.3011 | 0.0003383 | ---?+ | 0 | 0.4629 | intron | SYT16 |
| rs7718808 | 5 | 55930154 | 3 | 0 | 0 | A | T | -3.6679 | 1.0234 | 0.0003384 | ---?? | 1.1 | 0.3636 | intergenic | |
| rs117187408 | 7 | 122919880 | 4 | 0 | 0 | A | C | -3.5492 | 0.9903 | 0.0003384 | ---?- | 58.5 | 0.06486 | intergenic | |
| rs73131633 | 5 | 76072070 | 5 | 0 | 0 | A | G | 0.9546 | 0.2664 | 0.0003387 | ++++- | 0 | 0.5153 | intergenic | |
| 8:113468740 | 8 | 113468740 | 3 | 0 | 0 | G | GA | -1.037 | 0.2894 | 0.0003388 | ---?? | 0 | 0.5521 | | |
| rs59403859 | 4 | 38736272 | 5 | 0 | 0 | A | G | -1.1784 | 0.3288 | 0.0003389 | ----- | 0 | 0.9529 | intergenic |  |
| rs1387332 | 4 | 169332232 | 4 | 0 | 0 | T | C | 0.8896 | 0.2482 | 0.000339 | +++?+ | 0 | 0.8632 | intron | DDX60L |
| rs9988824 | 11 | 59189405 | 3 | 0 | 0 | T | C | 3.6013 | 1.005 | 0.000339 | +?++? | 9.3 | 0.3321 | downstream-gene | |
| rs7981195 | 13 | 60027109 | 4 | 0 | 0 | A | G | -0.9039 | 0.2522 | 0.0003391 | ---?- | 0 | 0.4842 | intergenic | |
| rs62416851 | 6 | 63194835 | 4 | 0 | 0 | T | C | 2.3305 | 0.6504 | 0.0003392 | +++?+ | 0 | 0.54 | intergenic | |
| rs62418214 | 6 | 63251070 | 4 | 0 | 0 | A | G | -2.4071 | 0.6717 | 0.0003392 | ---?- | 0 | 0.5561 | intergenic | |
| rs62418217 | 6 | 63257165 | 4 | 0 | 0 | A | C | -2.433 | 0.679 | 0.0003392 | ---?- | 0 | 0.5536 | intergenic | |
| rs1499501 | 3 | 65657837 | 4 | 0 | 0 | A | G | 0.9554 | 0.2666 | 0.0003393 | +++?+ | 47 | 0.1294 | intron | MAGI1 |
| rs116676664 | 2 | 170474689 | 4 | 0 | 0 | A | C | 1.2722 | 0.355 | 0.0003393 | +++?+ | 0 | 0.9375 | intron | PPIG |
| rs6851367 | 4 | 180905243 | 4 | 0 | 0 | C | G | -0.9386 | 0.2619 | 0.0003394 | ---?+ | 42.3 | 0.1579 | intergenic | |
| rs12424641 | 12 | 112716037 | 5 | 0 | 0 | A | G | 1.3265 | 0.3702 | 0.0003394 | +-+++ | 54.4 | 0.06697 | intron | HECTD4 |
| rs115846403 | 17 | 56913430 | 5 | 0 | 0 | T | G | -2.5039 | 0.6988 | 0.0003395 | ----- | 0 | 0.8441 | intron | PPM1E |
| rs34039405 | 3 | 183596387 | 4 | 0 | 0 | A | G | -1.6923 | 0.4723 | 0.0003396 | ---?- | 0 | 0.9166 | intron | PARL |
| rs116913119 | 2 | 168005245 | 4 | 0 | 0 | A | T | -2.2562 | 0.6297 | 0.0003396 | ---?+ | 2.1 | 0.3816 | intron | XIRP2 |
| rs4949511 | 1 | 30367470 | 4 | 0 | 0 | A | C | -1.2668 | 0.3535 | 0.0003397 | ---?+ | 14.5 | 0.3198 | intergenic | |
| rs4961665 | 9 | 16191258 | 4 | 0 | 0 | T | G | -0.9908 | 0.2765 | 0.0003398 | ---?- | 0 | 0.9422 | intergenic | |
| rs343056 | 7 | 35526390 | 4 | 0 | 0 | A | G | -1.1468 | 0.3201 | 0.0003399 | ---?- | 35.8 | 0.1972 | intergenic | |
| rs200124554 | 8 | 113454162 | 3 | 0 | 0 | A | T | 1.0393 | 0.2901 | 0.0003399 | +++?? | 0 | 0.557 | intron | CSMD3 |
| 1:245107672 | 1 | 245107672 | 3 | 0 | 0 | CA | C | -1.5784 | 0.4405 | 0.0003399 | ---?? | 5.9 | 0.3454 | | |

| | | | | | | | | | | | | | | | |
|---|---|---|---|---|---|---|---|---|---|---|---|---|---|---|---|
| 22:49409335 | 22 | 49409335 | 3 | 0 | 0 | A | AATT | -1.5515 | 0.4331 | 0.0003399 | ---?? | 20 | 0.2863 | | |
| 8:113454157 | 8 | 113454157 | 3 | 0 | 0 | A | AT | 1.0393 | 0.2901 | 0.0003399 | +++?? | 0 | 0.5571 | | |
| rs4434601 | 8 | 140363316 | 4 | 0 | 0 | A | C | -1.0395 | 0.2902 | 0.0003402 | ---?- | 0 | 0.8077 | intergenic | |
| rs62416815 | 6 | 63184506 | 4 | 0 | 0 | T | C | -2.3466 | 0.655 | 0.0003403 | ---?- | 0 | 0.5489 | intergenic | |
| 5:67795642 | 5 | 67795642 | 3 | 0 | 0 | CA | C | -1.9589 | 0.5468 | 0.0003403 | ---?? | 0 | 0.3885 | | |
| rs74323155 | 3 | 36740194 | 4 | 0 | 0 | A | G | 1.6636 | 0.4644 | 0.0003405 | +++?+ | 0 | 0.4474 | intergenic | |
| 9:139090465 | 9 | 139090465 | 3 | 0 | 0 | T | TC | 1.0071 | 0.2811 | 0.0003408 | +++?? | 0 | 0.7205 | | |
| rs60962282 | 3 | 196474029 | 4 | 0 | 0 | T | G | -2.2076 | 0.6163 | 0.0003409 | ---?+ | 22.3 | 0.2771 | intron | PAK2 |
| rs117777892 | 14 | 65455086 | 2 | 0 | 0 | T | C | 4.651 | 1.2984 | 0.000341 | +?+?? | 0 | 0.62 | intron | FNTB,CHURC1-FNTB |
| rs141444542 | 18 | 40019915 | 3 | 0 | 0 | T | C | -1.4171 | 0.3956 | 0.0003411 | ---?? | 0 | 0.6454 | intron | LINC00907 |
| rs7970397 | 12 | 112580071 | 5 | 0 | 0 | T | C | 1.3238 | 0.3696 | 0.0003413 | +-+++ | 55.3 | 0.06221 | synonymous | TRAFD1 |
| rs181741151 | 5 | 21985551 | 3 | 0 | 0 | T | C | -2.6483 | 0.7394 | 0.0003413 | ---?? | 0 | 0.5769 | intron | CDH12 |
| rs76844899 | 3 | 36739897 | 4 | 0 | 0 | C | G | 1.661 | 0.4638 | 0.0003417 | +++?+ | 0 | 0.4513 | intergenic | |
| rs7959454 | 12 | 112732608 | 5 | 0 | 0 | A | T | 1.3239 | 0.3697 | 0.0003419 | +-+++ | 54.4 | 0.06686 | intron | HECTD4 |
| rs11697360 | 20 | 26305178 | 2 | 0 | 0 | A | G | 2.8169 | 0.7866 | 0.0003419 | +???+ | 32.9 | 0.2222 | intergenic | |
| rs10733009 | 1 | 30366086 | 4 | 0 | 0 | T | G | -1.277 | 0.3566 | 0.000342 | ---?+ | 9.4 | 0.3461 | intergenic | |
| rs80025411 | 4 | 100570403 | 4 | 0 | 0 | T | C | -2.0359 | 0.5685 | 0.0003422 | +--?- | 51.5 | 0.103 | 3-prime-UTR | LOC285556 |
| rs140147587 | 5 | 35718573 | 3 | 0 | 0 | A | C | 3.1936 | 0.8918 | 0.0003422 | +++?? | 0 | 0.5606 | intron | SPEF2 |
| rs7010421 | 8 | 140375854 | 4 | 0 | 0 | A | G | 1.0401 | 0.2905 | 0.0003423 | +++?+ | 0 | 0.798 | intergenic | |
| rs7522292 | 1 | 102953488 | 4 | 0 | 0 | T | C | -0.8415 | 0.235 | 0.0003423 | ---?+ | 0 | 0.4011 | intergenic | |
| rs113503805 | 12 | 5547233 | 4 | 0 | 0 | A | C | -2.5718 | 0.7182 | 0.0003423 | ---?- | 0 | 0.4997 | intron | NTF3 |
| rs147400384 | 6 | 63184702 | 4 | 0 | 0 | A | G | -2.3441 | 0.6546 | 0.0003425 | ---?- | 0 | 0.5484 | intergenic | |
| rs148966266 | 6 | 148195998 | 4 | 0 | 0 | A | G | 4.2097 | 1.1756 | 0.0003425 | -++?+ | 11.4 | 0.3358 | intergenic | |
| rs4457961 | 15 | 22960447 | 4 | 0 | 0 | T | G | 0.9102 | 0.2542 | 0.0003426 | +++?+ | 1 | 0.3871 | intron | CYFIP1 |
| rs11744162 | 5 | 124141265 | 4 | 0 | 0 | T | C | 1.0501 | 0.2932 | 0.0003426 | +++?+ | 19.7 | 0.2912 | intergenic | |
| rs61712778 | 12 | 112756009 | 5 | 0 | 0 | T | C | -1.3275 | 0.3707 | 0.0003427 | -+--- | 54.1 | 0.06857 | intron | HECTD4 |
| 9:108228973 | 9 | 108228973 | 3 | 0 | 0 | CA | C | 1.061 | 0.2963 | 0.0003427 | +++?? | 0 | 0.9953 | | |
| rs9990928 | 4 | 8679493 | 3 | 0 | 0 | A | G | 3.6048 | 1.0068 | 0.0003429 | +++?? | 0 | 0.8361 | intergenic | |
| rs6489837 | 12 | 112758105 | 5 | 0 | 0 | A | T | 1.3276 | 0.3708 | 0.0003431 | +-+++ | 54.1 | 0.0687 | intron | HECTD4 |
| rs5004568 | 8 | 113470469 | 3 | 0 | 0 | T | G | -1.0368 | 0.2896 | 0.0003432 | ---?? | 0 | 0.5527 | intron | CSMD3 |
| rs6735719 | 2 | 137031718 | 4 | 0 | 0 | A | G | 0.9962 | 0.2783 | 0.0003432 | +++?+ | 0 | 0.7959 | intergenic | |
| rs113455436 | 17 | 77129778 | 4 | 0 | 0 | A | G | 2.2199 | 0.62 | 0.0003432 | +++?- | 1.7 | 0.3835 | intron | RBFOX3 |
| rs62080922 | 18 | 25206550 | 4 | 0 | 0 | A | G | 2.5571 | 0.7142 | 0.0003433 | +++?+ | 0 | 0.3998 | intergenic | |
| rs79489686 | 2 | 170486445 | 4 | 0 | 0 | T | C | 1.2715 | 0.3551 | 0.0003433 | +++?+ | 0 | 0.947 | intron | PPIG |
| rs149270358 | 15 | 83881299 | 4 | 0 | 0 | T | C | 3.895 | 1.0879 | 0.0003433 | +++?+ | 0 | 0.9739 | upstream-gene | |
| 3:77392235 | 3 | 77392235 | 3 | 0 | 0 | G | GAA | -0.9355 | 0.2613 | 0.0003434 | ---?? | 0 | 0.524 | | |
| rs72842479 | 11 | 1117707 | 4 | 0 | 0 | T | C | -1.4093 | 0.3937 | 0.0003435 | ---?+ | 57.4 | 0.07039 | intergenic | |
| rs1453553 | 11 | 59196518 | 3 | 0 | 0 | A | G | 3.5846 | 1.0013 | 0.0003437 | +?++? | 9.8 | 0.3299 | intergenic | |
| rs1377558 | 11 | 59196954 | 3 | 0 | 0 | A | G | -3.5831 | 1.0009 | 0.0003439 | -?--? | 9.9 | 0.3297 | intergenic | |
| rs114248729 | 2 | 170582116 | 4 | 0 | 0 | A | G | 1.2715 | 0.3552 | 0.0003439 | +++?+ | 0 | 0.9367 | intron | PHOSPHO2-KLHL23 |
| rs143461772 | 8 | 142141485 | 4 | 0 | 0 | A | G | -3.266 | 0.9124 | 0.0003439 | ---?- | 0 | 0.492 | intron | DENND3 |
| rs6931088 | 6 | 63191121 | 4 | 0 | 0 | T | C | -2.328 | 0.6503 | 0.000344 | ---?- | 0 | 0.5421 | intergenic | |
| rs7167085 | 15 | 91406188 | 4 | 0 | 0 | A | G | 0.8529 | 0.2383 | 0.000344 | +++?- | 8.9 | 0.3485 | intergenic | |
| rs114767899 | 2 | 168182656 | 4 | 0 | 0 | A | G | 2.4227 | 0.6768 | 0.000344 | +++?- | 0 | 0.5699 | intergenic | |
| rs28986403 | 6 | 32754869 | 5 | 0 | 0 | A | G | 1.4389 | 0.402 | 0.0003441 | +++++ | 0 | 0.6851 | intergenic | |
| rs117842363 | 10 | 851177 | 4 | 0 | 0 | A | C | -2.4092 | 0.6731 | 0.0003442 | ---?- | 0 | 0.4932 | downstream-gene | |
| rs76426403 | 3 | 196485127 | 4 | 0 | 0 | A | G | 2.2081 | 0.6169 | 0.0003443 | +++?- | 22.4 | 0.2763 | intron | PAK2 |
| 7:130122722 | 7 | 130122722 | 3 | 0 | 0 | A | AT | 0.9356 | 0.2614 | 0.0003443 | +++?? | 58 | 0.09231 | | |
| rs7965261 | 12 | 112754449 | 5 | 0 | 0 | A | G | 1.3266 | 0.3706 | 0.0003444 | +-+++ | 54.1 | 0.06846 | intron | HECTD4 |
| rs17111354 | 11 | 110394313 | 4 | 0 | 0 | T | C | 3.4785 | 0.9718 | 0.0003445 | +++?+ | 0 | 0.9581 | intergenic | |
| 11:119383758 | 11 | 119383758 | 3 | 0 | 0 | T | TAAG | 2.3869 | 0.6669 | 0.0003445 | +++?? | 44.3 | 0.166 | | |
| rs13087953 | 3 | 183595425 | 4 | 0 | 0 | T | C | -1.6912 | 0.4725 | 0.0003446 | ---?- | 0 | 0.9129 | intron | PARL |
| rs817422 | 1 | 85632596 | 5 | 0 | 0 | T | G | -1.3773 | 0.3848 | 0.0003448 | ----- | 0 | 0.9734 | intron | SYDE2 |
| rs7693489 | 4 | 41330010 | 4 | 0 | 0 | C | G | -1.432 | 0.4001 | 0.0003448 | ---?+ | 0 | 0.458 | intergenic | |
| rs140291329 | 2 | 170582329 | 4 | 0 | 0 | T | C | 1.2711 | 0.3551 | 0.0003448 | +++?+ | 0 | 0.9378 | intron | PHOSPHO2-KLHL23 |
| 16:7178371 | 16 | 7178371 | 3 | 0 | 0 | A | AATGAG | 1.0253 | 0.2865 | 0.0003451 | +++?? | 37 | 0.2044 | | |
| rs62418218 | 6 | 63257371 | 4 | 0 | 0 | A | G | -2.4111 | 0.6737 | 0.0003452 | ---?- | 0 | 0.5573 | intergenic | |
| rs184193367 | 6 | 83762373 | 4 | 0 | 0 | T | G | -3.2815 | 0.917 | 0.0003454 | ---?- | 0 | 0.8125 | intron | UBE3D |
| rs11066260 | 12 | 112766116 | 5 | 0 | 0 | A | C | 1.3291 | 0.3714 | 0.0003455 | +-+++ | 53.8 | 0.07022 | intron | HECTD4 |

| | | | | | | | | | | | | | | | | |
|---|---|---|---|---|---|---|---|---|---|---|---|---|---|---|---|---|
| rs117195876 | 4 | 89041399 | 4 | 0 | 0 | T | C | 4.0364 | 1.1279 | 0.0003455 | +?+++ | 0 | 0.5196 | intron | | ABCG2 |
| rs11927118 | 3 | 196485416 | 4 | 0 | 0 | A | G | -2.2074 | 0.6169 | 0.0003457 | ---?+ | 22.4 | 0.2764 | intron | | PAK2 |
| rs192526735 | 3 | 147430343 | 3 | 0 | 0 | T | C | -4.3459 | 1.2145 | 0.0003458 | ---?? | 0 | 0.9679 | intergenic | | |
| rs28986404 | 6 | 32754921 | 5 | 0 | 0 | T | G | -1.4378 | 0.4018 | 0.0003459 | ----- | 0 | 0.6846 | intergenic |  |  |
| rs77502429 | 12 | 112767851 | 5 | 0 | 0 | T | C | -1.3294 | 0.3715 | 0.0003459 | -+--- | 53.8 | 0.07019 | intron | | HECTD4 |
| rs1934712 | 1 | 102954001 | 4 | 0 | 0 | A | G | -0.8412 | 0.2351 | 0.000346 | ---?+ | 0 | 0.401 | intergenic | | |
| rs62416850 | 6 | 63194389 | 4 | 0 | 0 | A | G | -2.3166 | 0.6474 | 0.000346 | ---?- | 0 | 0.4616 | intergenic | | |
| rs79587808 | 4 | 157478923 | 4 | 0 | 0 | A | G | 2.903 | 0.8113 | 0.000346 | +++?+ | 0 | 0.6224 | intergenic | | |
| rs817423 | 1 | 85632660 | 5 | 0 | 0 | A | G | 1.3769 | 0.3848 | 0.0003461 | +++++ | 0 | 0.9735 | intron | | SYDE2 |
| rs56073653 | 8 | 76052265 | 5 | 0 | 0 | A | C | 0.986 | 0.2756 | 0.0003461 | +++++ | 19.5 | 0.2907 | intergenic | | |
| rs11927473 | 3 | 196482833 | 4 | 0 | 0 | A | G | 2.2083 | 0.6172 | 0.0003462 | +++?- | 22.6 | 0.2754 | intron | | PAK2 |
| rs28986405 | 6 | 32754949 | 5 | 0 | 0 | A | G | 1.4375 | 0.4018 | 0.0003464 | +++++ | 0 | 0.6845 | intergenic | | |
| rs4559814 | 13 | 60045877 | 4 | 0 | 0 | A | G | 0.9346 | 0.2612 | 0.0003465 | +++?+ | 0 | 0.5594 | intergenic | | |
| 22:50830769 | 22 | 50830769 | 3 | 0 | 0 | G | GAA | 1.0194 | 0.2849 | 0.0003465 | +++?? | 52.6 | 0.1214 | | | |
| rs1598807 | 5 | 123479815 | 4 | 0 | 0 | A | G | -0.9313 | 0.2603 | 0.0003466 | ---?- | 0 | 0.5305 | intergenic | | |
| rs10800813 | 1 | 202055955 | 4 | 0 | 0 | T | C | 0.9065 | 0.2534 | 0.0003466 | +++?+ | 0 | 0.947 | intergenic | | |
| rs6489836 | 12 | 112736504 | 5 | 0 | 0 | T | C | 1.3224 | 0.3696 | 0.0003467 | +-+++ | 54.3 | 0.06737 | intron | | HECTD4 |
| rs73834866 | 3 | 68210542 | 4 | 0 | 0 | A | G | -2.1737 | 0.6076 | 0.0003467 | ---?- | 33 | 0.2145 | intron | | FAM19A1 |
| rs139961030 | 16 | 5558505 | 4 | 0 | 0 | T | C | 3.5525 | 0.993 | 0.0003467 | +++?+ | 0 | 0.4206 | intron | | RBFOX1 |
| rs73834865 | 3 | 68210305 | 4 | 0 | 0 | C | G | -2.175 | 0.608 | 0.0003468 | ---?- | 33 | 0.2144 | intron | | FAM19A1 |
| rs75473190 | 16 | 62485091 | 4 | 0 | 0 | A | G | 2.183 | 0.6102 | 0.0003469 | +++?+ | 0 | 0.4916 | intergenic | | |
| rs73834864 | 3 | 68209227 | 4 | 0 | 0 | A | G | 2.1803 | 0.6095 | 0.0003471 | +++?+ | 32.9 | 0.2146 | intron | | FAM19A1 |
| rs11774739 | 8 | 3987971 | 4 | 0 | 0 | T | C | 1.1662 | 0.326 | 0.0003472 | +++?+ | 42.3 | 0.1579 | intron | | CSMD1 |
| rs57196059 | 12 | 112764543 | 5 | 0 | 0 | T | C | 1.3282 | 0.3713 | 0.0003472 | +-+++ | 53.8 | 0.07037 | intron | | HECTD4 |
| rs143600244 | 11 | 112923269 | 2 | 0 | 0 | A | T | -7.3476 | 2.054 | 0.0003473 | ?--?? | 0 | 0.7776 | intron | | NCAM1 |
| rs1382020 | 8 | 140499374 | 4 | 0 | 0 | A | G | -1.4897 | 0.4165 | 0.0003474 | ---?- | 0 | 0.7576 | intergenic | | |
| rs817425 | 1 | 85633294 | 5 | 0 | 0 | T | C | 1.3765 | 0.3848 | 0.0003475 | +++++ | 0 | 0.9737 | intron | | SYDE2 |
| rs4766504 | 12 | 111760814 | 5 | 0 | 0 | A | G | -0.9657 | 0.27 | 0.0003476 | ----- | 0 | 0.4673 | intron |  | CUX2 |
| rs71474858 | 15 | 54344134 | 3 | 0 | 0 | A | G | -1.2327 | 0.3446 | 0.0003476 | ---?? | 0 | 0.9748 | intron | | UNC13C |
| rs76166785 | 5 | 50260095 | 4 | 0 | 0 | T | C | 1.2837 | 0.3589 | 0.0003477 | +++?+ | 0 | 0.6345 | intergenic | | |
| rs4658138 | 1 | 90248302 | 5 | 0 | 0 | A | C | -0.763 | 0.2133 | 0.0003482 | ----- | 0 | 0.9709 | intergenic |  |  |
| rs17761664 | 2 | 85266776 | 5 | 0 | 0 | A | G | -3.109 | 0.8693 | 0.0003483 | ----+ | 0 | 0.9428 | intron |  | KCMF1 |
| rs12525623 | 6 | 151731479 | 4 | 0 | 0 | T | C | -1.1762 | 0.3289 | 0.0003484 | ---?- | 57.7 | 0.06913 | intron | | RMND1 |
| rs115999222 | 9 | 95836793 | 5 | 0 | 0 | A | G | 2.4507 | 0.6852 | 0.0003484 | +++++ | 0 | 0.9715 | intron | | SUSD3 |
| rs148590874 | 18 | 25170528 | 4 | 0 | 0 | A | T | -2.6752 | 0.748 | 0.0003485 | ---?- | 0 | 0.6256 | intergenic | | |
| rs17073578 | 6 | 144618513 | 5 | 0 | 0 | A | G | -1.4277 | 0.3992 | 0.0003487 | ----+ | 47.8 | 0.1048 | intron |  | UTRN |
| 7:112705840 | 7 | 112705840 | 3 | 0 | 0 | TTA | TTT | 1.2568 | 0.3514 | 0.0003488 | +++?? | 31.5 | 0.2325 | | | |
| rs7763219 | 6 | 63185535 | 4 | 0 | 0 | A | C | -2.3386 | 0.654 | 0.0003489 | ---?- | 0 | 0.5502 | intergenic | | |
| rs147678167 | 17 | 56915328 | 5 | 0 | 0 | T | C | 2.5157 | 0.7035 | 0.0003489 | +++++ | 0 | 0.8124 | intron | | PPM1E |
| rs28986399 | 6 | 32754593 | 5 | 0 | 0 | T | G | -1.4352 | 0.4014 | 0.000349 | ----- | 0 | 0.6838 | intergenic |  |  |
| rs28986411 | 6 | 32755275 | 5 | 0 | 0 | A | G | -1.4353 | 0.4014 | 0.000349 | ----- | 0 | 0.6837 | intergenic |  |  |
| rs7763047 | 6 | 63185435 | 4 | 0 | 0 | A | C | -2.3389 | 0.6541 | 0.0003491 | ---?- | 0 | 0.5504 | intergenic | | |
| rs28986412 | 6 | 32755327 | 5 | 0 | 0 | A | T | -1.4351 | 0.4013 | 0.0003491 | ----- | 0 | 0.6836 | intergenic |  |  |
| rs2551652 | 2 | 208362080 | 4 | 0 | 0 | T | C | -1.5121 | 0.4229 | 0.0003492 | ---?- | 0 | 0.5909 | intergenic | | |
| rs28533694 | 6 | 32587235 | 5 | 0 | 0 | T | C | 0.8233 | 0.2302 | 0.0003492 | +++++ | 0 | 0.5461 | intergenic | | |
| rs12884462 | 14 | 43263743 | 4 | 0 | 0 | A | C | 1.3513 | 0.3779 | 0.0003494 | +++?- | 61.4 | 0.0508 | intergenic | | |
| rs28409699 | 6 | 32571295 | 5 | 0 | 1 | T | C | 1.3902 | 0.3888 | 0.0003494 | +++++ | 6.1 | 0.3721 | intergenic | | |
| rs60551991 | 3 | 68208715 | 4 | 0 | 0 | T | C | 2.1816 | 0.6101 | 0.0003494 | +++?+ | 32.9 | 0.2148 | intron | | FAM19A1 |
| rs147410307 | 19 | 47180963 | 4 | 0 | 0 | T | C | 2.8634 | 0.8008 | 0.0003494 | +++?- | 0 | 0.436 | intron | | PRKD2 |
| rs6021639 | 20 | 50644475 | 4 | 0 | 0 | T | C | 1.0929 | 0.3057 | 0.0003495 | +++?- | 0 | 0.4597 | intergenic | | |
| rs7973349 | 12 | 112738985 | 5 | 0 | 0 | T | C | 1.3216 | 0.3696 | 0.0003495 | +-+++ | 54.3 | 0.06753 | intron | | HECTD4 |
| rs10226548 | 7 | 67409119 | 4 | 0 | 0 | A | G | -1.8617 | 0.5207 | 0.0003495 | ---?+ | 0 | 0.4269 | intergenic | | |
| rs28986413 | 6 | 32755375 | 5 | 0 | 0 | T | G | -1.4348 | 0.4013 | 0.0003497 | ----- | 0 | 0.6835 | intergenic |  |  |
| rs6489830 | 12 | 112582540 | 5 | 0 | 0 | A | G | -1.3213 | 0.3696 | 0.0003498 | -+--- | 55.5 | 0.06163 | intron | | TRAFD1 |
| rs28986415 | 6 | 32755465 | 5 | 0 | 0 | T | C | -1.4345 | 0.4012 | 0.0003498 | ----- | 0 | 0.6834 | intergenic |  |  |
| rs143699556 | 6 | 63261131 | 4 | 0 | 0 | A | T | -2.4135 | 0.6751 | 0.0003498 | ---?- | 0 | 0.5562 | intergenic | | |
| rs28986414 | 6 | 32755460 | 5 | 0 | 0 | A | G | -1.4345 | 0.4013 | 0.0003501 | ----- | 0 | 0.6834 | intergenic |  |  |
| rs34330585 | 6 | 32390703 | 5 | 0 | 1 | C | G | 1.3768 | 0.3851 | 0.0003501 | +++++ | 0 | 0.5564 | intergenic | | |
| rs28986416 | 6 | 32755506 | 5 | 0 | 0 | C | G | -1.4343 | 0.4012 | 0.0003502 | ----- | 0 | 0.6833 | intergenic |  |  |

| | | | | | | | | | | | | | | | |
|---|---|---|---|---|---|---|---|---|---|---|---|---|---|---|---|
| rs12052761 | 2 | 69065841 | 4 | 0 | 0 | A | G | -0.8687 | 0.243 | 0.0003503 | ---?- | 0 | 0.4075 | intergenic | |
| rs34001756 | 1 | 102954618 | 4 | 0 | 0 | T | C | -0.8413 | 0.2353 | 0.0003504 | ---?+ | 0 | 0.4012 | intergenic | |
| rs1340389 | 1 | 102960346 | 4 | 0 | 0 | T | C | 0.8466 | 0.2368 | 0.0003505 | +++?- | 0 | 0.395 | intergenic | |
| rs28986398 | 6 | 32754516 | 5 | 0 | 0 | A | G | 1.4339 | 0.4011 | 0.0003505 | +++++ | 0 | 0.6833 | intergenic | |
| 5:65168618 | 5 | 65168618 | 3 | 0 | 0 | CT | C | -0.9964 | 0.2787 | 0.0003505 | ---?? | 0 | 0.8727 | | |
| rs28986417 | 6 | 32755535 | 5 | 0 | 0 | T | C | -1.4341 | 0.4012 | 0.0003506 | ----- | 0 | 0.6833 | intergenic |  |
| 8:72276115 | 8 | 72276115 | 3 | 0 | 0 | A | AAG | 4.8693 | 1.3621 | 0.0003506 | +++?? | 41.1 | 0.1831 | | |
| rs28986418 | 6 | 32755536 | 5 | 0 | 0 | T | C | -1.4339 | 0.4011 | 0.0003507 | ----- | 0 | 0.6832 | intergenic |  |
| rs7718652 | 5 | 55930051 | 3 | 0 | 0 | A | G | -3.6522 | 1.0217 | 0.0003509 | ---?? | 0.3 | 0.3668 | intergenic | |
| rs12547267 | 8 | 65036628 | 4 | 0 | 0 | A | G | -0.8767 | 0.2453 | 0.0003512 | ---?- | 0 | 0.9489 | intergenic | |
| rs79162651 | 5 | 103177710 | 4 | 0 | 0 | T | C | 1.6294 | 0.4559 | 0.0003512 | +++?+ | 0 | 0.9045 | intergenic | |
| rs34874825 | 3 | 183628271 | 4 | 0 | 0 | T | C | -1.8439 | 0.5159 | 0.0003514 | ---?- | 0 | 0.9151 | intergenic | |
| rs28986397 | 6 | 32754458 | 5 | 0 | 0 | T | C | -1.4332 | 0.401 | 0.0003515 | ----- | 0 | 0.683 | intergenic |  |
| rs12366012 | 11 | 90862963 | 3 | 0 | 0 | T | C | -4.456 | 1.2468 | 0.0003516 | ---?? | 0 | 0.8928 | intergenic | |
| rs145680698 | 6 | 32755712 | 5 | 0 | 0 | A | G | -1.4331 | 0.401 | 0.0003517 | ----- | 0 | 0.6828 | intergenic |  |
| rs8094704 | 18 | 63863227 | 4 | 0 | 0 | T | C | -0.8904 | 0.2492 | 0.0003518 | ---?- | 0 | 0.537 | intergenic | |
| rs12422864 | 12 | 112584201 | 5 | 0 | 0 | A | G | -1.3207 | 0.3696 | 0.0003518 | -+--- | 55.5 | 0.06149 | intron | TRAFD1 |
| rs28986419 | 6 | 32755633 | 5 | 0 | 0 | T | G | -1.4331 | 0.401 | 0.0003518 | ----- | 0 | 0.6829 | intergenic |  |
| rs115781591 | 2 | 168009775 | 4 | 0 | 0 | T | C | -2.2447 | 0.6281 | 0.0003518 | ---?+ | 1.4 | 0.385 | intron | XIRP2 |
| 22:50789273 | 22 | 50789273 | 3 | 0 | 0 | CA | C | -1.0112 | 0.2829 | 0.000352 | ---?? | 0 | 0.6392 | | |
| rs28986396 | 6 | 32754430 | 5 | 0 | 0 | A | G | 1.4327 | 0.4009 | 0.0003521 | +++++ | 0 | 0.6828 | intergenic | |
| rs28893562 | 6 | 32755918 | 5 | 0 | 0 | A | G | -1.4326 | 0.4009 | 0.0003522 | ----- | 0 | 0.6826 | intergenic |  |
| rs114518174 | 2 | 168009790 | 4 | 0 | 0 | C | G | -2.2444 | 0.6281 | 0.0003522 | ---?+ | 1.4 | 0.3851 | intron | XIRP2 |
| rs11049075 | 12 | 27781040 | 4 | 0 | 0 | A | G | 0.9133 | 0.2556 | 0.0003523 | +++?- | 52.8 | 0.09564 | intron | PPFIBP1 |
| rs11962764 | 6 | 144616425 | 5 | 0 | 0 | T | C | 1.4266 | 0.3992 | 0.0003524 | ++++- | 47.9 | 0.1043 | intron | UTRN |
| rs1013110 | 11 | 95705574 | 4 | 0 | 0 | A | G | 1.0299 | 0.2882 | 0.0003525 | +-+?+ | 47.1 | 0.1289 | intergenic | |
| rs2881467 | 12 | 27781162 | 4 | 0 | 0 | A | G | 0.9129 | 0.2555 | 0.0003525 | +++?- | 53 | 0.09422 | intron | PPFIBP1 |
| rs2346750 | 12 | 27781153 | 4 | 0 | 0 | A | G | 0.9131 | 0.2555 | 0.0003526 | +++?- | 53 | 0.09441 | intron | PPFIBP1 |
| rs11833302 | 12 | 43205919 | 5 | 0 | 0 | T | C | -0.9708 | 0.2717 | 0.0003526 | ----+ | 2.5 | 0.3926 | intergenic |  |
| rs62330354 | 5 | 706557 | 3 | 0 | 0 | T | C | 1.4749 | 0.4128 | 0.0003526 | +++?? | 53.1 | 0.1183 | downstream-gene | |
| rs74365378 | 2 | 40481188 | 4 | 0 | 0 | A | G | 1.2666 | 0.3545 | 0.0003526 | +++?- | 39.1 | 0.1774 | intron | SLC8A1,SLC8A1-AS1 |
| rs117375939 | 18 | 25169815 | 4 | 0 | 0 | T | C | -2.6781 | 0.7495 | 0.0003526 | ---?- | 0 | 0.6243 | intergenic | |
| rs62330353 | 5 | 706555 | 3 | 0 | 0 | A | G | 1.4747 | 0.4127 | 0.0003527 | +++?? | 53.2 | 0.1182 | downstream-gene | |
| rs117573560 | 18 | 25169732 | 4 | 0 | 0 | A | G | 2.6788 | 0.7497 | 0.0003527 | +++?+ | 0 | 0.6241 | intergenic | |
| rs28755999 | 4 | 38737532 | 5 | 0 | 0 | C | G | -1.2207 | 0.3416 | 0.0003528 | ----- | 0 | 0.8343 | intergenic |  |
| rs117156096 | 20 | 22449417 | 4 | 0 | 0 | T | C | -2.14 | 0.5989 | 0.0003528 | ---?+ | 0 | 0.4089 | intergenic | |
| rs28752515 | 6 | 32583603 | 3 | 0 | 0 | A | G | 0.8855 | 0.2478 | 0.0003531 | +++?? | 0 | 0.917 | intergenic | |
| rs56403824 | 7 | 107331612 | 4 | 0 | 0 | T | C | -0.9029 | 0.2527 | 0.0003533 | ---?- | 0 | 0.5173 | intron | SLC26A4 |
| 7:107862386 | 7 | 107862386 | 3 | 0 | 0 | CA | C | 1.485 | 0.4157 | 0.0003533 | +++?? | 47 | 0.1515 | | |
| rs146779118 | 3 | 29705412 | 4 | 0 | 0 | T | C | 3.7833 | 1.0589 | 0.0003534 | +++?+ | 34.6 | 0.2048 | intron | RBMS3 |
| 6:72862831 | 6 | 72862831 | 3 | 0 | 0 | A | ATTCCC | 1.7222 | 0.4821 | 0.0003534 | +++?? | 0 | 0.8606 | | |
| rs76687058 | 2 | 170548771 | 4 | 0 | 0 | T | C | 1.2764 | 0.3573 | 0.0003536 | +++?+ | 0 | 0.9622 | intron | CCDC173 |
| 22:45969862 | 22 | 45969862 | 3 | 0 | 0 | T | TCTC | -1.0357 | 0.2899 | 0.0003536 | ---?? | 0 | 0.6918 | | |
| 6:72862840 | 6 | 72862840 | 3 | 0 | 0 | G | GTA | 1.7222 | 0.4821 | 0.0003536 | +++?? | 0 | 0.8604 | | |
| rs62514389 | 8 | 113288068 | 4 | 0 | 0 | C | G | 1.0172 | 0.2847 | 0.0003537 | +++?+ | 0 | 0.8063 | intron | CSMD3 |
| rs148517925 | 19 | 47262197 | 4 | 0 | 0 | T | C | 3.0984 | 0.8673 | 0.0003539 | +++?- | 0 | 0.4156 | downstream-gene | |
| rs7260557 | 19 | 22729821 | 4 | 0 | 0 | T | G | 0.9665 | 0.2706 | 0.0003541 | +++?+ | 18.8 | 0.2965 | intergenic | |
| rs35152189 | 16 | 85238972 | 4 | 0 | 0 | A | C | 1.2291 | 0.3441 | 0.0003541 | +++?- | 0 | 0.4415 | intron | GSE1 |
| rs16853036 | 2 | 168001470 | 4 | 0 | 0 | T | C | 2.2434 | 0.628 | 0.0003543 | +++?- | 0 | 0.5045 | intron | XIRP2 |
| rs112996392 | 12 | 102326863 | 4 | 0 | 0 | A | G | 2.0585 | 0.5763 | 0.0003544 | +++?+ | 0 | 0.4947 | intergenic | |
| rs182033683 | 2 | 168016684 | 4 | 0 | 0 | A | T | 2.4422 | 0.6838 | 0.0003547 | +++?- | 32.8 | 0.2156 | intron | XIRP2 |
| rs13197727 | 6 | 32756292 | 5 | 0 | 0 | A | G | -1.4308 | 0.4006 | 0.0003548 | ----- | 0 | 0.6819 | intergenic |  |
| rs189259344 | 10 | 74321569 | 2 | 0 | 0 | T | C | -7.5419 | 2.1116 | 0.0003548 | --??? | 0 | 0.5302 | intron | MICU1 |
| 4:54383089 | 4 | 54383089 | 3 | 0 | 0 | CAG | C | -2.4517 | 0.6865 | 0.0003548 | ---?? | 22.5 | 0.2753 | | |
| rs7973819 | 12 | 27781274 | 4 | 0 | 0 | A | T | 0.9122 | 0.2554 | 0.0003549 | +++?- | 53.1 | 0.0941 | intron | PPFIBP1 |
| rs28506828 | 1 | 211759378 | 4 | 0 | 0 | T | C | 2.8935 | 0.8102 | 0.0003549 | +++?- | 0 | 0.7455 | intergenic | |
| rs28383230 | 6 | 32584088 | 3 | 0 | 0 | T | C | -0.8832 | 0.2473 | 0.000355 | ---?? | 0 | 0.912 | intergenic | |
| rs28383231 | 6 | 32584099 | 3 | 0 | 0 | T | C | 0.8832 | 0.2473 | 0.0003551 | +++?? | 0 | 0.912 | intergenic | |
| rs117815474 | 4 | 89041878 | 4 | 0 | 0 | T | C | -4.0279 | 1.1278 | 0.0003551 | -?--- | 0 | 0.5184 | intron | ABCG2 |

| | | | | | | | | | | | | | | | |
|---|---|---|---|---|---|---|---|---|---|---|---|---|---|---|---|
| rs184311710 | 10 | 74321568 | 2 | 0 | 0 | T | C | -7.5403 | 2.1113 | 0.0003551 | --??? | 0 | 0.5296 | intron | MICU1 |
| rs10036119 | 5 | 94801296 | 2 | 0 | 0 | A | C | -1.8646 | 0.5221 | 0.0003553 | ???-- | 0 | 0.8424 | intron | TTC37 |
| rs13197854 | 6 | 32756385 | 5 | 0 | 0 | A | C | -1.4303 | 0.4005 | 0.0003554 | ----- | 0 | 0.6818 | intergenic |  |
| 15:45989264 | 15 | 45989264 | 3 | 0 | 0 | CT | C | -1.3516 | 0.3785 | 0.0003555 | ---?? | 0 | 0.9651 | | |
| rs28383223 | 6 | 32583477 | 3 | 0 | 0 | T | C | 0.8853 | 0.2479 | 0.0003556 | +++?? | 0 | 0.9131 | intergenic | |
| rs333801 | 11 | 37042494 | 3 | 0 | 0 | A | C | 0.9959 | 0.2789 | 0.0003557 | +++?? | 3.7 | 0.3541 | intergenic | |
| rs6962631 | 7 | 67411817 | 4 | 0 | 0 | A | C | -1.7657 | 0.4945 | 0.0003557 | ---?+ | 0 | 0.5185 | intergenic | |
| rs11783519 | 8 | 113288338 | 4 | 0 | 0 | T | C | -1.0164 | 0.2846 | 0.0003557 | ---?- | 0 | 0.8054 | intron | CSMD3 |
| rs9356923 | 6 | 24050912 | 5 | 0 | 0 | T | C | 1.3775 | 0.3857 | 0.0003558 | ++++- | 0 | 0.7373 | intergenic | |
| rs13197975 | 6 | 32756480 | 5 | 0 | 0 | A | G | -1.4298 | 0.4004 | 0.0003558 | ----- | 0 | 0.6816 | intergenic |  |
| rs13198082 | 6 | 32756457 | 5 | 0 | 0 | T | C | 1.4299 | 0.4004 | 0.0003558 | +++++ | 0 | 0.6816 | intergenic | |
| rs62158790 | 2 | 137012170 | 4 | 0 | 0 | T | C | 0.9639 | 0.2699 | 0.0003558 | +++?+ | 0 | 0.8451 | intergenic | |
| rs183694510 | 5 | 150343874 | 4 | 0 | 0 | T | C | -4.4408 | 1.2436 | 0.0003559 | ---?- | 0 | 0.5059 | intergenic | |
| rs10453088 | 8 | 65016969 | 4 | 0 | 0 | A | C | 0.8851 | 0.2479 | 0.000356 | +++?+ | 0 | 0.9463 | intergenic | |
| rs13198307 | 6 | 32756486 | 5 | 0 | 0 | A | G | 1.4296 | 0.4004 | 0.0003561 | +++++ | 0 | 0.6815 | intergenic | |
| rs72908403 | 6 | 88489052 | 4 | 0 | 0 | T | C | -2.2348 | 0.6259 | 0.0003561 | ---?- | 0 | 0.8746 | intergenic | |
| 15:45989286 | 15 | 45989286 | 3 | 0 | 0 | CTCTT | C | -1.3516 | 0.3786 | 0.0003563 | ---?? | 0 | 0.9649 | | |
| rs10910596 | 1 | 232834067 | 4 | 0 | 0 | T | G | -0.8744 | 0.2449 | 0.0003566 | ---?- | 12.2 | 0.3319 | intergenic | |
| 2:139909308 | 2 | 139909308 | 3 | 0 | 0 | CTT | CTTT | -1.3575 | 0.3802 | 0.0003566 | ---?? | 0 | 0.9236 | | |
| rs10872974 | 14 | 22530493 | 4 | 0 | 0 | A | G | -0.9355 | 0.262 | 0.0003567 | ---?- | 0 | 0.7053 | intergenic | |
| rs13198109 | 6 | 32756575 | 5 | 0 | 0 | A | G | -1.4292 | 0.4003 | 0.0003567 | ----- | 0 | 0.6814 | intergenic |  |
| rs28986394 | 6 | 32754342 | 5 | 0 | 0 | A | G | 1.4292 | 0.4003 | 0.0003567 | +++++ | 0 | 0.6815 | intergenic | |
| rs964045 | 13 | 60028921 | 4 | 0 | 0 | A | G | -0.8979 | 0.2515 | 0.0003568 | ---?- | 0 | 0.4529 | intergenic | |
| rs9276717 | 6 | 32761398 | 5 | 0 | 0 | A | C | 1.4158 | 0.3966 | 0.0003569 | +++++ | 0 | 0.6388 | intergenic | |
| rs11729139 | 4 | 63338769 | 4 | 0 | 0 | T | C | -0.9575 | 0.2682 | 0.0003569 | ---?- | 25.6 | 0.258 | intergenic | |
| 18:40054080 | 18 | 40054080 | 3 | 0 | 0 | <Cn0> | C | -1.4138 | 0.396 | 0.0003569 | ---?? | 0 | 0.6547 | | |
| rs58894129 | 9 | 16196387 | 4 | 0 | 0 | T | C | -1.2756 | 0.3573 | 0.000357 | ---?- | 0 | 0.5988 | intergenic | |
| rs4598707 | 12 | 27781484 | 4 | 0 | 0 | T | C | 0.9109 | 0.2552 | 0.0003571 | +++?- | 53.1 | 0.09358 | intron | PPFIBP1 |
| rs77245836 | 6 | 63264916 | 4 | 0 | 0 | T | G | -2.4159 | 0.6767 | 0.0003572 | ---?- | 0 | 0.5571 | intergenic | |
| rs79448298 | 3 | 183609735 | 4 | 0 | 0 | T | C | -1.7096 | 0.4789 | 0.0003572 | ---?- | 0 | 0.9594 | intergenic | |
| 14:102776426 | 14 | 102776426 | 3 | 0 | 0 | CAA | C | 1.7006 | 0.4764 | 0.0003572 | +++?? | 0 | 0.9514 | | |
| rs79618830 | 2 | 170439091 | 4 | 0 | 0 | C | G | -1.2682 | 0.3553 | 0.0003573 | ---?- | 0 | 0.9671 | upstream-gene | |
| rs28986393 | 6 | 32754340 | 5 | 0 | 0 | A | G | 1.4285 | 0.4002 | 0.0003577 | +++++ | 0 | 0.6812 | intergenic | |
| rs376714549 | 8 | 96564550 | 3 | 0 | 0 | T | C | 1.7599 | 0.493 | 0.0003577 | +-+?? | 61.7 | 0.07337 | intron | LOC100616530 |
| rs10083378 | 14 | 22530369 | 4 | 0 | 0 | A | G | 0.9352 | 0.262 | 0.0003578 | +++?+ | 0 | 0.7063 | intergenic | |
| rs11048652 | 12 | 26842486 | 4 | 0 | 0 | T | C | 1.7656 | 0.4947 | 0.0003578 | +-+?+ | 41.5 | 0.1625 | intron | ITPR2 |
| 11:78875109 | 11 | 78875109 | 3 | 0 | 0 | A | AGT | -1.1105 | 0.3111 | 0.0003578 | ---?? | 28.3 | 0.2479 | | |
| rs6662967 | 1 | 244884573 | 4 | 0 | 0 | A | G | -1.2244 | 0.343 | 0.0003579 | ---?- | 0 | 0.436 | intergenic | |
| rs62418220 | 6 | 63265279 | 4 | 0 | 0 | T | C | -2.4162 | 0.6769 | 0.000358 | ---?- | 0 | 0.5575 | intergenic | |
| rs75348876 | 2 | 168013767 | 4 | 0 | 0 | T | C | -2.2355 | 0.6263 | 0.0003581 | ---?+ | 0.9 | 0.3875 | intron | XIRP2 |
| rs114316022 | 1 | 167437715 | 3 | 0 | 0 | A | T | -5.1937 | 1.4552 | 0.0003581 | ---?? | 0 | 0.8596 | intron | CD247 |
| rs1128982 | 11 | 400040 | 4 | 0 | 0 | A | G | 1.3177 | 0.3692 | 0.0003583 | +++?+ | 0 | 0.7017 | synonymous | PKP3 |
| rs28383236 | 6 | 32584188 | 3 | 0 | 0 | T | G | -0.8831 | 0.2474 | 0.0003583 | ---?? | 0 | 0.9118 | intergenic | |
| rs61861879 | 10 | 126126519 | 4 | 0 | 0 | T | C | -1.5494 | 0.4341 | 0.0003583 | ---?- | 0 | 0.6092 | intergenic | |
| rs192786227 | 12 | 17392607 | 3 | 0 | 0 | T | C | -4.7029 | 1.3177 | 0.0003583 | ---?? | 0 | 0.5172 | intergenic | |
| rs112634937 | 6 | 63265400 | 4 | 0 | 0 | T | C | 2.4163 | 0.677 | 0.0003584 | +++?+ | 0 | 0.5577 | intergenic | |
| rs13059476 | 3 | 183592521 | 4 | 0 | 0 | T | C | -1.6879 | 0.4729 | 0.0003585 | ---?- | 0 | 0.9025 | intron | PARL |
| rs62621446 | 2 | 170550963 | 4 | 0 | 0 | A | G | -1.2749 | 0.3572 | 0.0003586 | ---?- | 0 | 0.9615 | 5-prime-UTR | PHOSPHO2 |
| rs76196397 | 17 | 57166232 | 5 | 0 | 0 | T | G | 2.4275 | 0.6802 | 0.0003586 | +++++ | 0 | 0.8361 | intron | TRIM37 |
| 18:67218013 | 18 | 67218013 | 3 | 0 | 0 | CTTTT | C | 1.3945 | 0.3908 | 0.0003586 | +++?? | 14.1 | 0.312 | | |
| rs11011648 | 10 | 20151090 | 4 | 0 | 0 | A | G | -0.894 | 0.2505 | 0.0003587 | ---?- | 0 | 0.4564 | intron | PLXDC2 |
| rs115263034 | 17 | 56921416 | 5 | 0 | 0 | A | G | 2.4935 | 0.6987 | 0.0003587 | +++++ | 0 | 0.8242 | intron | PPM1E |
| rs116490666 | 18 | 25177797 | 4 | 0 | 0 | T | G | -2.6304 | 0.7371 | 0.0003588 | ---?- | 0 | 0.6264 | intergenic | |
| rs78475275 | 1 | 223121349 | 4 | 0 | 0 | A | G | -3.1156 | 0.8731 | 0.0003589 | ---?- | 50.8 | 0.107 | intron | DISP1 |
| rs13201537 | 6 | 32756857 | 5 | 0 | 0 | A | C | -1.4275 | 0.4 | 0.0003591 | ----- | 0 | 0.6807 | intergenic |  |
| rs61973856 | 14 | 28192034 | 4 | 0 | 0 | A | G | -2.0177 | 0.5654 | 0.0003591 | ---?- | 0 | 0.8619 | intergenic | |
| rs10148273 | 14 | 43264623 | 4 | 0 | 0 | A | T | 1.3478 | 0.3777 | 0.0003592 | +++?- | 61.4 | 0.05108 | intergenic | |
| rs76801468 | 2 | 168015745 | 4 | 0 | 0 | T | C | -2.2333 | 0.6259 | 0.0003592 | ---?+ | 0.8 | 0.388 | intron | XIRP2 |
| rs4658137 | 1 | 90245736 | 5 | 0 | 0 | A | G | -0.7595 | 0.2128 | 0.0003594 | ----- | 0 | 0.9833 | intergenic |  |

| | | | | | | | | | | | | | | | |
|---|---|---|---|---|---|---|---|---|---|---|---|---|---|---|---|
| rs76089248 | 14 | 43264687 | 4 | 0 | 0 | C | G | -1.3476 | 0.3777 | 0.0003594 | ---?+ | 61.4 | 0.05106 | intergenic | |
| rs2083996 | 8 | 65014961 | 4 | 0 | 0 | A | C | -0.8873 | 0.2487 | 0.0003595 | ---?- | 0 | 0.9473 | intergenic | |
| rs7974256 | 12 | 27781327 | 4 | 0 | 0 | A | G | -0.9112 | 0.2554 | 0.0003595 | ---?+ | 53.3 | 0.09292 | intron | PPFIBP1 |
| rs12549416 | 8 | 38736820 | 3 | 0 | 0 | A | T | -1.2882 | 0.361 | 0.0003596 | ---?? | 0 | 0.3939 | intergenic | |
| rs13251387 | 8 | 65014043 | 4 | 0 | 0 | A | G | 0.886 | 0.2483 | 0.0003598 | +++?+ | 0 | 0.9482 | intergenic | |
| rs149207630 | 15 | 54190989 | 4 | 0 | 0 | A | T | -4.2602 | 1.194 | 0.0003599 | ---?- | 24.4 | 0.2647 | intergenic | |
| 18:40439944 | 18 | 40439944 | 3 | 0 | 0 | TA | TAA | 4.746 | 1.3302 | 0.00036 | +++?? | 0 | 0.3844 | | |
| rs12278188 | 11 | 86114240 | 4 | 0 | 0 | A | G | -2.5864 | 0.7249 | 0.0003601 | ---?- | 0 | 0.5528 | intron | CCDC81 |
| rs28986392 | 6 | 32754322 | 5 | 0 | 0 | A | T | -1.4269 | 0.4 | 0.0003602 | ----- | 0 | 0.6805 | intergenic |  |
| rs1483627 | 3 | 176027750 | 4 | 0 | 0 | T | C | 1.3292 | 0.3726 | 0.0003604 | +++?+ | 24.4 | 0.2649 | intergenic | |
| rs57453103 | 3 | 33262069 | 4 | 0 | 0 | T | C | 0.9696 | 0.2718 | 0.0003606 | +++?+ | 1.8 | 0.3831 | upstream-gene | |
| rs28986343 | 6 | 32748953 | 5 | 0 | 0 | T | C | 1.4249 | 0.3995 | 0.0003608 | +++++ | 0 | 0.6807 | intergenic | |
| rs8017061 | 14 | 22502880 | 4 | 0 | 0 | T | C | -0.9434 | 0.2645 | 0.0003609 | ---?- | 0 | 0.9588 | intergenic | |
| 6:72838109 | 6 | 72838109 | 3 | 0 | 0 | T | TA | 1.7046 | 0.4779 | 0.000361 | +++?? | 0 | 0.9231 | | |
| rs10478657 | 5 | 124691852 | 4 | 0 | 0 | A | C | 0.8861 | 0.2484 | 0.0003611 | +++?+ | 0 | 0.5444 | intergenic | |
| rs11005662 | 10 | 58906151 | 4 | 0 | 0 | A | C | -2.9956 | 0.8398 | 0.0003612 | ---?- | 0 | 0.5042 | intergenic | |
| rs12404964 | 1 | 232834256 | 4 | 0 | 0 | A | G | -0.8741 | 0.2451 | 0.0003614 | ---?- | 12.3 | 0.3314 | intergenic | |
| rs35589320 | 8 | 65015921 | 4 | 0 | 0 | C | G | 0.8842 | 0.2479 | 0.0003615 | +++?+ | 0 | 0.9486 | intergenic | |
| rs1405869 | 2 | 156869592 | 4 | 0 | 0 | A | G | 0.9291 | 0.2605 | 0.0003617 | +++?+ | 0 | 0.8316 | intergenic | |
| rs9273058 | 6 | 32612110 | 3 | 0 | 0 | T | C | -1.2502 | 0.3505 | 0.0003618 | ---?? | 0 | 0.5665 | 3-prime-UTR | HLA-DQA1 |
| rs8092022 | 18 | 63859634 | 4 | 0 | 0 | C | G | -0.8893 | 0.2493 | 0.0003619 | ---?- | 0 | 0.5279 | intergenic | |
| rs28986391 | 6 | 32754294 | 5 | 0 | 0 | T | C | -1.4256 | 0.3997 | 0.0003619 | ----- | 0 | 0.68 | intergenic |  |
| rs28986420 | 6 | 32757468 | 5 | 0 | 0 | T | C | 1.4241 | 0.3993 | 0.0003619 | +++++ | 0 | 0.6785 | intergenic | |
| rs149788850 | 18 | 25183871 | 4 | 0 | 0 | T | G | 2.6351 | 0.7389 | 0.0003619 | +++?+ | 0 | 0.6571 | intergenic | |
| rs11778262 | 8 | 113276821 | 4 | 0 | 0 | A | G | 1.0173 | 0.2853 | 0.0003625 | +++?+ | 0 | 0.8092 | intron | CSMD3 |
| rs141070545 | 6 | 63267402 | 4 | 0 | 0 | A | G | -2.4181 | 0.6781 | 0.0003625 | ---?- | 0 | 0.5579 | intergenic | |
| rs6679502 | 1 | 90244083 | 5 | 0 | 0 | T | C | 0.7583 | 0.2127 | 0.0003627 | +++++ | 0 | 0.9832 | intergenic | |
| rs62418242 | 6 | 63268737 | 4 | 0 | 0 | A | T | 2.4205 | 0.6788 | 0.0003627 | +++?+ | 0 | 0.5583 | intergenic | |
| rs76427911 | 11 | 129294294 | 4 | 0 | 0 | A | G | 1.541 | 0.4322 | 0.0003627 | +++?+ | 18 | 0.301 | intron | BARX2 |
| rs147506504 | 14 | 27771503 | 4 | 0 | 0 | T | G | -3.451 | 0.9678 | 0.0003628 | ---?- | 0 | 0.4685 | intergenic | |
| rs4721746 | 7 | 19153843 | 4 | 0 | 0 | A | C | -2.4367 | 0.6834 | 0.0003629 | ---?- | 0 | 0.7305 | intron | TWIST1 |
| rs28986421 | 6 | 32757598 | 5 | 0 | 0 | A | G | -1.4225 | 0.399 | 0.0003629 | ----- | 0 | 0.6771 | intergenic |  |
| rs6139888 | 20 | 5987634 | 5 | 0 | 0 | A | C | 0.7479 | 0.2098 | 0.0003631 | ++++- | 0 | 0.5073 | intron | CRLS1 |
| rs28986422 | 6 | 32757724 | 5 | 0 | 0 | T | C | -1.422 | 0.3988 | 0.0003631 | ----- | 0 | 0.6765 | intergenic |  |
| rs35292277 | 9 | 37422276 | 4 | 0 | 0 | T | C | -2.9283 | 0.8213 | 0.0003631 | ---?- | 0 | 0.6885 | upstream-gene | |
| rs16883325 | 8 | 113276635 | 4 | 0 | 0 | A | G | -1.0161 | 0.285 | 0.0003632 | ---?- | 0 | 0.8112 | intron | CSMD3 |
| rs10411774 | 19 | 11872011 | 3 | 0 | 0 | T | G | -5.1399 | 1.4416 | 0.0003633 | ---?? | 0 | 0.762 | intergenic | |
| rs115343878 | 18 | 25178309 | 4 | 0 | 0 | A | T | -2.6225 | 0.7355 | 0.0003633 | ---?- | 0 | 0.6276 | intergenic | |
| 6:75124255 | 6 | 75124255 | 3 | 0 | 0 | T | C | -6.5946 | 1.8497 | 0.0003635 | ---?? | 0 | 0.5353 | intergenic | |
| rs118168385 | 18 | 25161211 | 4 | 0 | 0 | A | G | 2.7403 | 0.7686 | 0.0003635 | +++?+ | 0 | 0.6118 | intergenic | |
| rs140913877 | 15 | 87553155 | 4 | 0 | 0 | T | C | 2.0697 | 0.5805 | 0.0003636 | +++?+ | 0 | 0.9888 | intron | AGBL1 |
| rs201436 | 7 | 101733576 | 4 | 0 | 0 | T | C | -0.8885 | 0.2492 | 0.0003639 | ---?- | 0 | 0.8375 | intron | CUX1 |
| rs10458387 | 1 | 162093507 | 5 | 0 | 0 | T | C | -0.8952 | 0.2511 | 0.0003639 | ----+ | 40.6 | 0.1509 | intron | NOS1AP |
| rs12478330 | 2 | 137011741 | 4 | 0 | 0 | A | G | -0.9623 | 0.2699 | 0.0003639 | ---?- | 0 | 0.8435 | intergenic | |
| rs10228820 | 7 | 156154582 | 4 | 0 | 0 | A | G | 0.9369 | 0.2628 | 0.000364 | +++?+ | 0 | 0.4313 | intergenic | |
| rs685244 | 6 | 32584769 | 3 | 0 | 0 | A | G | -0.8834 | 0.2478 | 0.0003643 | ---?? | 0 | 0.9138 | intergenic | |
| rs142317702 | 17 | 57009596 | 5 | 0 | 0 | A | G | -2.4597 | 0.69 | 0.0003643 | ----- | 0 | 0.8296 | intron | PPM1E |
| rs4473089 | 14 | 43264961 | 4 | 0 | 0 | T | C | 1.3457 | 0.3775 | 0.0003644 | +++?- | 61.4 | 0.05112 | intergenic | |
| rs28594446 | 18 | 63860846 | 4 | 0 | 0 | T | C | 0.8885 | 0.2493 | 0.0003644 | +++?+ | 0 | 0.5708 | intergenic | |
| rs140410252 | 5 | 67730105 | 2 | 0 | 0 | T | C | 5.2377 | 1.4693 | 0.0003644 | +?+?? | 0 | 0.9359 | intergenic | |
| rs4965201 | 15 | 98048483 | 4 | 0 | 0 | A | T | 0.9577 | 0.2687 | 0.0003645 | +++?+ | 0 | 0.826 | intergenic | |
| rs35216590 | 6 | 32593246 | 2 | 0 | 0 | C | G | -2.039 | 0.572 | 0.0003645 | ???-- | 0 | 0.6927 | intergenic | |
| rs967748 | 11 | 37049352 | 4 | 0 | 0 | T | C | 0.9439 | 0.2648 | 0.0003647 | +++?+ | 0 | 0.4602 | intergenic | |
| rs28752527 | 6 | 32584928 | 3 | 0 | 0 | T | C | 0.8823 | 0.2475 | 0.0003647 | +++?? | 0 | 0.9135 | intergenic | |
| rs958798 | 1 | 110770223 | 5 | 0 | 0 | T | G | -0.9684 | 0.2717 | 0.0003648 | ----+ | 0 | 0.5122 | intron | KCNC4 |
| rs139467994 | 22 | 29084385 | 3 | 0 | 0 | A | G | 2.4932 | 0.6995 | 0.000365 | +++?? | 14.6 | 0.3099 | intron | CHEK2 |
| rs370575452 | 6 | 83777675 | 4 | 0 | 0 | T | G | 3.2501 | 0.9119 | 0.0003652 | +++?+ | 0 | 0.8426 | intron | DOPEY1 |
| rs74925869 | 8 | 5605500 | 2 | 0 | 0 | C | G | 6.8735 | 1.9288 | 0.0003658 | ?+??+ | 0 | 0.4702 | intergenic | |
| rs77403528 | 18 | 25178743 | 4 | 0 | 0 | T | C | -2.6187 | 0.7349 | 0.0003659 | ---?- | 0 | 0.6286 | intergenic | |

| | | | | | | | | | | | | | | | |
|---|---|---|---|---|---|---|---|---|---|---|---|---|---|---|---|
| rs146556037 | 3 | 65613283 | 3 | 0 | 0 | C | G | -3.4402 | 0.9654 | 0.000366 | ---?? | 55.3 | 0.1067 | intron | MAGI1 |
| rs2116413 | 18 | 39980604 | 4 | 0 | 0 | A | G | 1.1679 | 0.3278 | 0.0003665 | +++?- | 0 | 0.401 | intron | LINC00907 |
| rs4531123 | 9 | 137701553 | 4 | 0 | 0 | A | G | -1.5215 | 0.427 | 0.0003665 | ---?- | 31.5 | 0.2234 | intron | COL5A1 |
| rs28752514 | 6 | 32583576 | 3 | 0 | 0 | T | C | -0.8847 | 0.2483 | 0.0003665 | ---?? | 0 | 0.9175 | intergenic | |
| rs12662462 | 6 | 32753433 | 5 | 0 | 0 | A | G | 1.4221 | 0.3991 | 0.0003666 | +++++ | 0 | 0.6786 | intergenic | |
| rs12663317 | 6 | 32742827 | 5 | 0 | 0 | A | C | -1.4221 | 0.3991 | 0.0003666 | ----- | 0 | 0.6786 | intergenic |  |
| rs12663979 | 6 | 32753654 | 5 | 0 | 0 | T | C | -1.4221 | 0.3991 | 0.0003666 | ----- | 0 | 0.6786 | intergenic |  |
| rs12665315 | 6 | 32753403 | 5 | 0 | 0 | A | G | -1.4221 | 0.3991 | 0.0003666 | ----- | 0 | 0.6786 | intergenic |  |
| rs13203578 | 6 | 32752540 | 5 | 0 | 0 | A | G | -1.4221 | 0.3991 | 0.0003666 | ----- | 0 | 0.6786 | intergenic |  |
| rs13203581 | 6 | 32752546 | 5 | 0 | 0 | A | G | -1.4221 | 0.3991 | 0.0003666 | ----- | 0 | 0.6786 | intergenic |  |
| rs13203642 | 6 | 32752477 | 5 | 0 | 0 | T | C | 1.4221 | 0.3991 | 0.0003666 | +++++ | 0 | 0.6786 | intergenic | |
| rs13203652 | 6 | 32752504 | 5 | 0 | 0 | T | C | 1.4221 | 0.3991 | 0.0003666 | +++++ | 0 | 0.6786 | intergenic | |
| rs13203691 | 6 | 32752647 | 5 | 0 | 0 | A | G | -1.4221 | 0.3991 | 0.0003666 | ----- | 0 | 0.6786 | intergenic |  |
| rs13203859 | 6 | 32752492 | 5 | 0 | 0 | A | G | 1.4221 | 0.3991 | 0.0003666 | +++++ | 0 | 0.6786 | intergenic | |
| rs13204139 | 6 | 32752685 | 5 | 0 | 0 | A | G | 1.4221 | 0.3991 | 0.0003666 | +++++ | 0 | 0.6786 | intergenic | |
| rs13215244 | 6 | 32753053 | 5 | 0 | 0 | A | G | 1.4221 | 0.3991 | 0.0003666 | +++++ | 0 | 0.6786 | intergenic | |
| rs13219960 | 6 | 32752641 | 5 | 0 | 0 | T | C | -1.4221 | 0.3991 | 0.0003666 | ----- | 0 | 0.6786 | intergenic |  |
| rs13219964 | 6 | 32752649 | 5 | 0 | 0 | T | C | -1.4221 | 0.3991 | 0.0003666 | ----- | 0 | 0.6786 | intergenic |  |
| rs28986363 | 6 | 32751555 | 5 | 0 | 0 | A | C | -1.4221 | 0.3991 | 0.0003666 | ----- | 0 | 0.6786 | intergenic |  |
| rs28986364 | 6 | 32751691 | 5 | 0 | 0 | A | C | -1.4221 | 0.3991 | 0.0003666 | ----- | 0 | 0.6786 | intergenic |  |
| rs28986366 | 6 | 32751751 | 5 | 0 | 0 | A | G | 1.4221 | 0.3991 | 0.0003666 | +++++ | 0 | 0.6786 | intergenic | |
| rs28986367 | 6 | 32751800 | 5 | 0 | 0 | T | C | 1.4221 | 0.3991 | 0.0003666 | +++++ | 0 | 0.6786 | intergenic | |
| rs28986378 | 6 | 32752242 | 5 | 0 | 0 | A | C | 1.4221 | 0.3991 | 0.0003666 | +++++ | 0 | 0.6786 | intergenic | |
| rs28986379 | 6 | 32752357 | 5 | 0 | 0 | T | G | 1.4221 | 0.3991 | 0.0003666 | +++++ | 0 | 0.6786 | intergenic | |
| rs28986380 | 6 | 32752392 | 5 | 0 | 0 | A | G | -1.4221 | 0.3991 | 0.0003666 | ----- | 0 | 0.6786 | intergenic |  |
| rs28986386 | 6 | 32753145 | 5 | 0 | 0 | T | C | 1.4221 | 0.3991 | 0.0003666 | +++++ | 0 | 0.6786 | intergenic | |
| rs28986388 | 6 | 32753150 | 5 | 0 | 0 | A | G | -1.4221 | 0.3991 | 0.0003666 | ----- | 0 | 0.6786 | intergenic |  |
| rs35902780 | 1 | 160030743 | 4 | 0 | 0 | A | G | -0.9925 | 0.2785 | 0.0003666 | ---?+ | 9.2 | 0.3472 | intron | KCNJ10 |
| rs28986359 | 6 | 32751349 | 5 | 0 | 0 | C | G | 1.422 | 0.3991 | 0.0003667 | +++++ | 0 | 0.6786 | intergenic | |
| rs28986360 | 6 | 32751370 | 5 | 0 | 0 | A | C | 1.422 | 0.3991 | 0.0003667 | +++++ | 0 | 0.6786 | intergenic | |
| rs28986361 | 6 | 32751384 | 5 | 0 | 0 | A | G | 1.4221 | 0.3991 | 0.0003667 | +++++ | 0 | 0.6786 | intergenic | |
| rs28986362 | 6 | 32751451 | 5 | 0 | 0 | C | G | -1.4221 | 0.3991 | 0.0003667 | ----- | 0 | 0.6786 | intergenic |  |
| rs28986365 | 6 | 32751746 | 5 | 0 | 0 | T | C | -1.4221 | 0.3991 | 0.0003667 | ----- | 0 | 0.6786 | intergenic |  |
| rs28986373 | 6 | 32752150 | 5 | 0 | 0 | A | C | 1.4221 | 0.3991 | 0.0003667 | +++++ | 0 | 0.6786 | intergenic | |
| rs57253991 | 12 | 112781852 | 5 | 0 | 0 | T | C | 1.3258 | 0.3721 | 0.0003667 | +-+++ | 53.5 | 0.07178 | intron | HECTD4 |
| rs12660504 | 6 | 32743138 | 5 | 0 | 0 | A | G | 1.4223 | 0.3992 | 0.0003668 | +++++ | 0 | 0.6787 | intergenic | |
| rs60424767 | 18 | 63859048 | 4 | 0 | 0 | T | C | -0.8886 | 0.2494 | 0.0003668 | ---?- | 0 | 0.5252 | intergenic | |
| rs73020380 | 2 | 170593293 | 4 | 0 | 0 | A | C | -1.2537 | 0.3519 | 0.0003668 | ---?- | 0 | 0.9636 | intron | KLHL23,PHOSPHO2-KLHL23 |
| 6:32603442 | 6 | 32603442 | 3 | 0 | 0 | CAA | C | -1.2525 | 0.3515 | 0.0003668 | ---?? | 0 | 0.7359 | | |
| rs28986357 | 6 | 32751192 | 5 | 0 | 0 | A | G | 1.422 | 0.3991 | 0.0003669 | +++++ | 0 | 0.6786 | intergenic | |
| rs28986358 | 6 | 32751228 | 5 | 0 | 0 | A | T | -1.422 | 0.3991 | 0.0003669 | ----- | 0 | 0.6786 | intergenic |  |
| rs28383241 | 6 | 32584319 | 3 | 0 | 0 | A | C | 0.8856 | 0.2486 | 0.000367 | +++?? | 0 | 0.907 | intergenic | |
| rs28893556 | 6 | 32750675 | 5 | 0 | 0 | T | G | -1.422 | 0.3991 | 0.000367 | ----- | 0 | 0.6787 | intergenic |  |
| rs28893559 | 6 | 32750938 | 5 | 0 | 0 | T | C | 1.422 | 0.3991 | 0.000367 | +++++ | 0 | 0.6787 | intergenic | |
| rs28986356 | 6 | 32751185 | 5 | 0 | 0 | C | G | 1.422 | 0.3991 | 0.000367 | +++++ | 0 | 0.6786 | intergenic | |
| rs61740646 | 11 | 65168284 | 5 | 0 | 0 | T | C | 3.2488 | 0.9119 | 0.000367 | +++++ | 0 | 0.4919 | synonymous | FRMD8 |
| rs28383242 | 6 | 32584325 | 3 | 0 | 0 | A | G | -0.8856 | 0.2486 | 0.0003671 | ---?? | 0 | 0.9071 | intergenic | |
| rs28893554 | 6 | 32750504 | 5 | 0 | 0 | T | C | 1.422 | 0.3991 | 0.0003671 | +++++ | 0 | 0.6787 | intergenic | |
| rs28893555 | 6 | 32750573 | 5 | 0 | 0 | A | G | -1.422 | 0.3991 | 0.0003671 | ----- | 0 | 0.6787 | intergenic |  |
| rs28986351 | 6 | 32750650 | 5 | 0 | 0 | T | C | 1.422 | 0.3991 | 0.0003671 | +++++ | 0 | 0.6787 | intergenic | |
| rs28986352 | 6 | 32750656 | 5 | 0 | 0 | A | G | -1.422 | 0.3991 | 0.0003671 | ----- | 0 | 0.6787 | intergenic |  |
| rs28986354 | 6 | 32750983 | 5 | 0 | 0 | A | T | -1.422 | 0.3991 | 0.0003671 | ----- | 0 | 0.6786 | intergenic |  |
| rs28986355 | 6 | 32751105 | 5 | 0 | 0 | A | G | -1.422 | 0.3991 | 0.0003671 | ----- | 0 | 0.6786 | intergenic |  |
| rs79555028 | 3 | 196489822 | 4 | 0 | 0 | A | C | -2.1975 | 0.6168 | 0.0003671 | ---?+ | 22.1 | 0.2778 | intron | PAK2 |
| rs28744289 | 6 | 32774920 | 5 | 0 | 0 | T | G | 1.4056 | 0.3945 | 0.0003672 | +++++ | 0 | 0.6455 | intergenic | |
| rs28893560 | 6 | 32750940 | 5 | 0 | 0 | T | C | 1.422 | 0.3991 | 0.0003672 | +++++ | 0 | 0.6786 | intergenic | |
| rs28986370 | 6 | 32751960 | 5 | 0 | 0 | A | G | 1.4365 | 0.4032 | 0.0003672 | +++++ | 0 | 0.6862 | intergenic | |
| rs28893551 | 6 | 32750369 | 5 | 0 | 0 | T | C | -1.422 | 0.3992 | 0.0003673 | ----- | 0 | 0.6787 | intergenic |  |
| rs28986368 | 6 | 32751942 | 5 | 0 | 0 | T | C | 1.4365 | 0.4032 | 0.0003673 | +++++ | 0 | 0.6862 | intergenic | |

| | | | | | | | | | | | | | | | |
|---|---|---|---|---|---|---|---|---|---|---|---|---|---|---|---|
| rs28986369 | 6 | 32751946 | 5 | 0 | 0 | A | G | -1.4365 | 0.4032 | 0.0003673 | ----- | 0 | 0.6862 | intergenic |  |
| rs75626083 | 11 | 129292261 | 4 | 0 | 0 | A | G | 1.564 | 0.439 | 0.0003673 | +++?+ | 29.9 | 0.233 | intron | BARX2 |
| rs7132314 | 12 | 27790370 | 4 | 0 | 0 | A | G | 0.9117 | 0.2559 | 0.0003674 | +-+?- | 56.3 | 0.07622 | intron | PPFIBP1 |
| rs12985669 | 19 | 46163605 | 5 | 0 | 1 | A | G | -0.9471 | 0.2659 | 0.0003674 | ----+ | 25 | 0.2551 | intergenic |  |
| rs28893545 | 6 | 32750134 | 5 | 0 | 0 | A | C | 1.422 | 0.3992 | 0.0003675 | +++++ | 0 | 0.6787 | intergenic | |
| rs28893546 | 6 | 32750156 | 5 | 0 | 0 | A | G | -1.422 | 0.3992 | 0.0003675 | ----- | 0 | 0.6787 | intergenic |  |
| 1:243774467 | 1 | 243774467 | 3 | 0 | 0 | C | G | -5.0416 | 1.4153 | 0.0003676 | ---?? | 0 | 0.5497 | intron | AKT3 |
| rs11066264 | 12 | 112781117 | 5 | 0 | 0 | A | C | 1.3254 | 0.3721 | 0.0003676 | +-+++ | 53.5 | 0.07173 | intron | HECTD4 |
| rs12674813 | 8 | 140360355 | 4 | 0 | 0 | T | C | -1.0349 | 0.2905 | 0.0003677 | ---?- | 0 | 0.7913 | intergenic | |
| rs112062732 | 6 | 32453936 | 4 | 0 | 0 | T | C | -1.0986 | 0.3084 | 0.0003677 | ---?- | 5.4 | 0.3658 | intergenic | |
| 10:117605212 | 10 | 117605212 | 3 | 0 | 0 | G | GT | -1.0158 | 0.2852 | 0.0003677 | ---?? | 0 | 0.4495 | | |
| rs1887462 | 10 | 129681306 | 4 | 0 | 0 | A | G | 1.5388 | 0.432 | 0.0003678 | +++?- | 0 | 0.5691 | intron | CLRN3 |
| rs28986335 | 6 | 32748137 | 5 | 0 | 0 | A | T | -1.4218 | 0.3991 | 0.0003678 | ----- | 0 | 0.6788 | intergenic |  |
| rs28986338 | 6 | 32748810 | 5 | 0 | 0 | T | C | 1.4218 | 0.3991 | 0.0003678 | +++++ | 0 | 0.6788 | intergenic | |
| rs3796531 | 4 | 57850846 | 4 | 0 | 0 | A | G | 0.8543 | 0.2398 | 0.0003679 | +++?+ | 33.7 | 0.2103 | intron | POLR2B |
| rs13217231 | 6 | 32748690 | 5 | 0 | 0 | T | C | 1.4218 | 0.3991 | 0.0003679 | +++++ | 0 | 0.6788 | intergenic | |
| rs28986337 | 6 | 32748398 | 5 | 0 | 0 | T | C | 1.4218 | 0.3991 | 0.0003679 | +++++ | 0 | 0.6788 | intergenic | |
| rs28893527 | 6 | 32747592 | 5 | 0 | 0 | T | C | 1.4217 | 0.3991 | 0.000368 | +++++ | 0 | 0.6788 | intergenic | |
| rs28893532 | 6 | 32747817 | 5 | 0 | 0 | T | C | 1.4217 | 0.3991 | 0.000368 | +++++ | 0 | 0.6788 | intergenic | |
| rs28986336 | 6 | 32748313 | 5 | 0 | 0 | T | C | -1.4218 | 0.3991 | 0.000368 | ----- | 0 | 0.6788 | intergenic |  |
| 6:32584192 | 6 | 32584192 | 3 | 0 | 0 | CAA | C | 0.8835 | 0.248 | 0.000368 | +++?? | 0 | 0.9137 | | |
| rs28893533 | 6 | 32747827 | 5 | 0 | 0 | T | G | 1.4217 | 0.3991 | 0.0003681 | +++++ | 0 | 0.6788 | intergenic | |
| 4:99247071 | 4 | 99247071 | 3 | 0 | 0 | T | TA | 1.1728 | 0.3293 | 0.0003681 | +++?? | 0 | 0.886 | | |
| rs28986333 | 6 | 32747727 | 5 | 0 | 0 | A | C | -1.4217 | 0.3991 | 0.0003682 | ----- | 0 | 0.6788 | intergenic |  |
| rs28986334 | 6 | 32748021 | 5 | 0 | 0 | T | C | -1.4217 | 0.3991 | 0.0003682 | ----- | 0 | 0.6788 | intergenic |  |
| rs74649950 | 3 | 196490322 | 4 | 0 | 0 | A | G | 2.1967 | 0.6167 | 0.0003682 | +++?- | 22.1 | 0.2779 | intron | PAK2 |
| rs71352282 | 19 | 46162482 | 5 | 0 | 1 | T | C | 0.9473 | 0.266 | 0.0003683 | ++++- | 25.5 | 0.2517 | intergenic | |
| rs1586128 | 11 | 103662703 | 4 | 0 | 0 | A | G | -2.2891 | 0.6427 | 0.0003684 | ---?- | 0 | 0.9576 | intergenic | |
| rs28893525 | 6 | 32747262 | 5 | 0 | 0 | A | G | 1.422 | 0.3992 | 0.0003684 | +++++ | 0 | 0.679 | intergenic | |
| rs149001409 | 6 | 32176308 | 5 | 0 | 0 | A | G | 2.7563 | 0.7739 | 0.0003684 | +++++ | 22.6 | 0.2705 | intron | NOTCH4 |
| rs7423627 | 2 | 40463896 | 3 | 0 | 0 | A | G | -1.124 | 0.3156 | 0.0003685 | ---?? | 0 | 0.6168 | intron | SLC8A1,SLC8A1-AS1 |
| rs4954591 | 2 | 137010302 | 4 | 0 | 0 | A | G | -0.9588 | 0.2692 | 0.0003686 | ---?- | 0 | 0.8408 | intergenic | |
| rs73942938 | 19 | 46162346 | 5 | 0 | 1 | T | C | 0.9473 | 0.266 | 0.0003686 | ++++- | 25.5 | 0.2512 | intergenic | |
| 12:70659783 | 12 | 70659783 | 3 | 0 | 0 | CT | C | 0.983 | 0.276 | 0.0003686 | +-+?? | 64.2 | 0.06099 | | |
| rs12971691 | 19 | 46163561 | 5 | 0 | 1 | T | C | 0.9466 | 0.2658 | 0.0003687 | ++++- | 25 | 0.2548 | intergenic | |
| rs10458392 | 1 | 162092110 | 5 | 0 | 0 | T | G | 0.8931 | 0.2508 | 0.0003688 | ++++- | 40.5 | 0.1514 | intron | NOS1AP |
| rs13212165 | 6 | 32747029 | 5 | 0 | 0 | A | G | 1.4224 | 0.3994 | 0.0003689 | +++++ | 0 | 0.6791 | intergenic | |
| 12:128051957 | 12 | 128051957 | 3 | 0 | 0 | CAAA | C | 2.211 | 0.6208 | 0.0003689 | +++?? | 0 | 0.3965 | | |
| rs2288333 | 2 | 170593191 | 4 | 0 | 0 | C | G | 1.2532 | 0.3519 | 0.000369 | +++?+ | 0 | 0.9643 | intron | KLHL23,PHOSPHO2-KLHL23 |
| rs71352281 | 19 | 46161878 | 5 | 0 | 1 | T | C | -0.9474 | 0.266 | 0.0003691 | ----+ | 25.8 | 0.2494 | intergenic |  |
| rs79216324 | 2 | 122694586 | 4 | 0 | 0 | A | G | 2.6611 | 0.7472 | 0.0003691 | +++?+ | 0 | 0.8791 | intergenic | |
| rs4768321 | 12 | 41304969 | 3 | 0 | 0 | A | G | -1.806 | 0.5071 | 0.0003692 | ---?? | 0 | 0.9106 | intron | CNTN1 |
| rs115441677 | 3 | 1778569 | 4 | 0 | 0 | T | G | 2.6274 | 0.7378 | 0.0003692 | +++?- | 58 | 0.06737 | intergenic | |
| rs1879053 | 4 | 111396948 | 4 | 0 | 0 | T | C | -0.861 | 0.2418 | 0.0003693 | ---?- | 0 | 0.998 | upstream-gene | |
| rs74997859 | 2 | 170509231 | 4 | 0 | 0 | T | C | -1.2662 | 0.3556 | 0.0003693 | ---?- | 0 | 0.9812 | intron | CCDC173 |
| rs12983068 | 19 | 46161612 | 5 | 0 | 1 | T | C | -0.9473 | 0.266 | 0.0003696 | ----+ | 25.9 | 0.2488 | intergenic |  |
| rs72777541 | 16 | 5457174 | 4 | 0 | 0 | T | C | 1.0615 | 0.2981 | 0.0003696 | +++?+ | 0 | 0.7299 | intron | RBFOX1 |
| rs7178205 | 15 | 41971213 | 5 | 0 | 0 | A | G | 1.0157 | 0.2852 | 0.0003697 | +++++ | 11.5 | 0.3401 | intron | MGA |
| rs72750256 | 1 | 232834596 | 4 | 0 | 0 | A | G | -0.8736 | 0.2453 | 0.0003698 | ---?- | 12.2 | 0.3318 | intergenic | |
| rs117471661 | 12 | 5521519 | 4 | 0 | 0 | T | C | 2.635 | 0.74 | 0.0003698 | +++?+ | 0 | 0.5056 | intergenic | |
| rs9893925 | 17 | 4738909 | 5 | 0 | 0 | A | G | -1.6388 | 0.4603 | 0.00037 | ----+ | 23.4 | 0.265 | intron | MINK1 |
| rs77947625 | 18 | 25179415 | 4 | 0 | 0 | A | G | 2.6153 | 0.7345 | 0.00037 | +++?+ | 0 | 0.6299 | intergenic | |
| 11:109797718 | 11 | 109797718 | 3 | 0 | 0 | T | TGAA | 1.018 | 0.2859 | 0.00037 | +++?? | 0 | 0.9989 | | |
| rs79882652 | 6 | 63276874 | 4 | 0 | 0 | A | C | -2.4429 | 0.6861 | 0.0003701 | ---?- | 0 | 0.5576 | intergenic | |
| rs77259451 | 7 | 7788634 | 4 | 0 | 0 | T | C | 1.6759 | 0.4707 | 0.0003703 | +++?+ | 0 | 0.9115 | intron | RPA3-AS1 |
| 14:44017357 | 14 | 44017357 | 3 | 0 | 0 | T | TATGA | 2.1062 | 0.5916 | 0.0003704 | +++?? | 0 | 0.9486 | | |
| rs62418243 | 6 | 63276430 | 4 | 0 | 0 | A | G | 2.4246 | 0.681 | 0.0003705 | +++?+ | 0 | 0.5606 | intergenic | |
| rs223848 | 16 | 57460493 | 4 | 0 | 0 | T | C | 1.342 | 0.377 | 0.0003707 | +++?+ | 0 | 0.8244 | downstream-gene | |
| rs74172499 | 19 | 46164415 | 5 | 0 | 1 | A | G | 0.9458 | 0.2657 | 0.0003707 | ++++- | 24.4 | 0.259 | intergenic | |

| | | | | | | | | | | | | | | | |
|---|---|---|---|---|---|---|---|---|---|---|---|---|---|---|---|
| 6:32631313 | 6 | 32631313 | 4 | 0 | 0 | A | G | 1.1413 | 0.3206 | 0.0003708 | ++++? | 39.4 | 0.1754 | intron | HLA-DQB1 |
| rs28893536 | 6 | 32749659 | 5 | 0 | 0 | T | C | 1.4282 | 0.4012 | 0.0003709 | +++++ | 0 | 0.679 | intergenic | |
| rs145449914 | 8 | 92963619 | 3 | 0 | 0 | T | C | 3.6377 | 1.0218 | 0.0003709 | +++?? | 0 | 0.9116 | downstream-gene | |
| rs111373319 | 19 | 46165067 | 5 | 0 | 1 | T | C | -0.9452 | 0.2655 | 0.000371 | ----+ | 23.8 | 0.2628 | intergenic |  |
| rs149564747 | 6 | 83719896 | 4 | 0 | 0 | A | G | 3.4169 | 0.9599 | 0.0003712 | +++?+ | 0 | 0.8127 | intron | UBE3D |
| 15:87616327 | 15 | 87616327 | 3 | 0 | 0 | CAAAAA | C | -1.0724 | 0.3012 | 0.0003712 | ---?? | 0 | 0.7063 | | |
| rs11822939 | 11 | 129285091 | 4 | 0 | 0 | T | C | 1.5616 | 0.4387 | 0.0003714 | +++?+ | 38.8 | 0.1793 | intron | BARX2 |
| rs72765243 | 15 | 98032739 | 4 | 0 | 0 | T | C | 1.1387 | 0.3199 | 0.0003714 | +++?- | 0 | 0.5057 | intergenic | |
| rs139776315 | 5 | 132983696 | 4 | 0 | 0 | A | C | -3.1382 | 0.8816 | 0.0003715 | ---?- | 0 | 0.7512 | intergenic | |
| rs7969932 | 12 | 64922187 | 4 | 0 | 0 | T | C | -0.8866 | 0.2491 | 0.0003716 | ---?- | 0 | 0.8908 | intergenic | |
| rs991912 | 3 | 120352900 | 4 | 0 | 0 | C | G | -1.012 | 0.2843 | 0.0003717 | ---?- | 61.5 | 0.05037 | intron | HGD |
| rs11947311 | 4 | 169335657 | 4 | 0 | 0 | A | G | -0.8835 | 0.2482 | 0.0003717 | ---?- | 0 | 0.8782 | intron | DDX60L |
| rs12023188 | 1 | 232834832 | 4 | 0 | 0 | C | G | 0.8739 | 0.2455 | 0.0003717 | +++?+ | 12.3 | 0.3312 | intergenic | |
| rs35121749 | 19 | 46165770 | 5 | 0 | 1 | A | C | 0.9442 | 0.2653 | 0.0003718 | ++++- | 23.3 | 0.2661 | intergenic | |
| rs61286727 | 2 | 170593586 | 4 | 0 | 0 | A | G | 1.2528 | 0.352 | 0.0003718 | +++?+ | 0 | 0.9629 | intron | KLHL23,PHOSPHO2-KLHL23 |
| rs78921312 | 8 | 25095982 | 4 | 0 | 0 | A | G | 2.7411 | 0.7701 | 0.0003719 | +++?+ | 0 | 0.7498 | intron | DOCK5 |
| rs28986331 | 6 | 32744892 | 5 | 0 | 0 | T | C | 1.4238 | 0.4 | 0.000372 | +++++ | 0 | 0.6793 | intergenic | |
| rs12661326 | 6 | 32745435 | 5 | 0 | 0 | T | C | 1.4237 | 0.4 | 0.0003723 | +++++ | 0 | 0.6793 | intergenic | |
| rs12661352 | 6 | 32745527 | 5 | 0 | 0 | T | G | 1.4237 | 0.4 | 0.0003723 | +++++ | 0 | 0.6792 | intergenic | |
| rs56109734 | 4 | 1388379 | 3 | 0 | 0 | A | G | 1.5843 | 0.4452 | 0.0003723 | +++?? | 0 | 0.5738 | missense | CRIPAK |
| rs76360526 | 2 | 170481727 | 4 | 0 | 0 | A | G | -1.1729 | 0.3296 | 0.0003724 | ---?- | 0 | 0.9834 | intron | PPIG |
| rs77241604 | 17 | 57117822 | 5 | 0 | 0 | T | C | -2.3792 | 0.6685 | 0.0003724 | ----- | 0 | 0.8281 | intron | TRIM37 |
| rs12220969 | 10 | 122489133 | 4 | 0 | 0 | A | G | 1.0026 | 0.2817 | 0.0003725 | +++?+ | 13.4 | 0.3254 | intergenic | |
| rs74443608 | 9 | 101129729 | 4 | 0 | 0 | A | G | 3.5735 | 1.0041 | 0.0003725 | +++?- | 49.4 | 0.1154 | intron | GABBR2 |
| rs28986371 | 6 | 32752006 | 5 | 0 | 0 | T | C | 1.4199 | 0.399 | 0.0003727 | +++++ | 0 | 0.6767 | intergenic | |
| rs35016841 | 19 | 46166617 | 5 | 0 | 1 | T | C | -0.9432 | 0.265 | 0.0003727 | ----+ | 22.7 | 0.2696 | upstream-gene |  |
| rs9295601 | 6 | 23868354 | 4 | 0 | 0 | T | C | 2.5601 | 0.7194 | 0.0003728 | +++?- | 53 | 0.09427 | intergenic | |
| rs35922055 | 19 | 46165979 | 5 | 0 | 1 | T | C | -0.9437 | 0.2652 | 0.000373 | ----+ | 23.2 | 0.2669 | intergenic |  |
| rs9358743 | 6 | 24047987 | 5 | 0 | 0 | T | C | 1.3745 | 0.3863 | 0.0003733 | ++++- | 0 | 0.7385 | intergenic | |
| rs7978571 | 12 | 27780899 | 4 | 0 | 0 | A | T | -0.9104 | 0.2559 | 0.0003734 | ---?+ | 52.5 | 0.09719 | intron | PPFIBP1 |
| rs28893558 | 6 | 32750865 | 5 | 0 | 0 | A | G | -1.4223 | 0.3997 | 0.0003735 | ----- | 0 | 0.6791 | intergenic |  |
| rs150197842 | 18 | 25163103 | 4 | 0 | 0 | C | G | -2.7066 | 0.7607 | 0.0003735 | ---?- | 0 | 0.6125 | intergenic | |
| rs12137740 | 1 | 209659731 | 4 | 0 | 0 | A | C | -2.7735 | 0.7795 | 0.0003736 | ---?- | 0 | 0.5467 | intergenic | |
| rs61973845 | 14 | 28182304 | 4 | 0 | 0 | A | G | -1.9977 | 0.5614 | 0.0003736 | ---?- | 0 | 0.8793 | intergenic | |
| rs62418244 | 6 | 63278275 | 4 | 0 | 0 | A | C | 2.4252 | 0.6816 | 0.0003736 | +++?+ | 0 | 0.56 | intergenic | |
| rs77728018 | 2 | 170551132 | 4 | 0 | 0 | A | G | -1.8546 | 0.5212 | 0.0003737 | +--?- | 47.5 | 0.1265 | 5-prime-UTR | PHOSPHO2,PHOSPHO2-KLHL23 |
| rs201573624 | 4 | 24511333 | 2 | 0 | 0 | A | T | 3.5763 | 1.0051 | 0.0003737 | ??+?+ | 0 | 0.7209 | intergenic | |
| rs28893557 | 6 | 32750859 | 5 | 0 | 0 | T | C | 1.4222 | 0.3997 | 0.0003738 | +++++ | 0 | 0.6791 | intergenic | |
| rs7533409 | 1 | 203674637 | 4 | 0 | 0 | T | C | -1.1439 | 0.3215 | 0.000374 | ---?- | 0 | 0.9575 | intron | ATP2B4 |
| rs77657330 | 3 | 196472869 | 4 | 0 | 0 | T | C | -2.1941 | 0.6167 | 0.0003741 | ---?+ | 22.1 | 0.2778 | intron | PAK2 |
| rs140573327 | 6 | 88488082 | 4 | 0 | 0 | T | C | -2.2272 | 0.626 | 0.0003741 | ---?- | 0 | 0.8841 | intergenic | |
| rs11048647 | 12 | 26834792 | 4 | 0 | 0 | A | C | 1.7689 | 0.4972 | 0.0003742 | +-+?+ | 37.6 | 0.1864 | intron | ITPR2 |
| rs57946278 | 14 | 28181577 | 4 | 0 | 0 | C | G | 1.9963 | 0.5611 | 0.0003743 | +++?+ | 0 | 0.88 | intergenic | |
| rs58548812 | 14 | 28186167 | 4 | 0 | 0 | A | G | -2.0037 | 0.5632 | 0.0003744 | ---?- | 0 | 0.8748 | intergenic | |
| rs80040850 | 2 | 170477033 | 4 | 0 | 0 | A | G | 1.2634 | 0.3551 | 0.0003745 | +++?+ | 0 | 0.9458 | intron | PPIG |
| rs13065559 | 3 | 155127210 | 4 | 0 | 0 | A | T | 1.0484 | 0.2947 | 0.0003746 | +++?+ | 0 | 0.5627 | intergenic | |
| rs60756607 | 14 | 28181789 | 4 | 0 | 0 | T | G | -1.9964 | 0.5612 | 0.0003748 | ---?- | 0 | 0.8799 | intergenic | |
| rs113542180 | 10 | 14434737 | 4 | 0 | 0 | A | T | -4.9853 | 1.4014 | 0.0003748 | ---?- | 0 | 0.8707 | intergenic | |
| rs12323878 | 14 | 44912392 | 4 | 0 | 0 | A | G | -1.3863 | 0.3897 | 0.0003749 | ---?- | 0 | 0.8298 | intergenic | |
| 10:117610869 | 10 | 117610869 | 3 | 0 | 0 | A | AT | 1.0079 | 0.2834 | 0.0003749 | +++?? | 0 | 0.3996 | | |
| rs56118374 | 14 | 28181804 | 4 | 0 | 0 | T | G | -1.9964 | 0.5612 | 0.000375 | ---?- | 0 | 0.8799 | intergenic | |
| rs11049074 | 12 | 27780901 | 4 | 0 | 0 | T | C | 0.91 | 0.2558 | 0.0003751 | +++?- | 52.5 | 0.097 | intron | PPFIBP1 |
| rs62514386 | 8 | 113275493 | 4 | 0 | 0 | T | G | 1.0136 | 0.285 | 0.0003751 | +++?+ | 0 | 0.8536 | intron | CSMD3 |
| rs16933625 | 8 | 68661890 | 3 | 0 | 0 | T | C | 5.7189 | 1.6079 | 0.0003754 | +?+?+ | 5.2 | 0.3483 | upstream-gene | |
| rs9689019 | 6 | 63280143 | 4 | 0 | 0 | A | G | -2.4263 | 0.6822 | 0.0003755 | ---?- | 0 | 0.5608 | intergenic | |
| rs77962087 | 2 | 170615580 | 4 | 0 | 0 | A | T | 1.2401 | 0.3487 | 0.0003755 | +++?+ | 0 | 0.9749 | intergenic | |
| rs1871426 | 6 | 168619182 | 4 | 0 | 0 | A | G | -0.9355 | 0.263 | 0.0003757 | ---?- | 0 | 0.8029 | intergenic | |
| rs77019683 | 5 | 122534395 | 4 | 0 | 0 | A | G | -1.8867 | 0.5305 | 0.0003758 | ---?- | 0 | 0.8398 | downstream-gene | |
| 16:84846612 | 16 | 84846612 | 3 | 0 | 0 | T | TTCACTC | 1.2448 | 0.35 | 0.0003759 | +++?? | 0 | 0.8019 | | |

| | | | | | | | | | | | | | | | |
|---|---|---|---|---|---|---|---|---|---|---|---|---|---|---|---|
| rs112105676 | 12 | 41304975 | 3 | 0 | 0 | T | G | 1.8037 | 0.5072 | 0.000376 | +++?? | 0 | 0.91 | intron | CNTN1 |
| rs11874009 | 18 | 5913575 | 4 | 0 | 0 | T | C | 1.2641 | 0.3554 | 0.0003761 | +++?+ | 0 | 0.4076 | intergenic | |
| 6:30982139 | 6 | 30982139 | 4 | 0 | 0 | A | G | 1.4108 | 0.3967 | 0.0003762 | +++?- | 0 | 0.7052 | intron | MUC22 |
| rs7706338 | 5 | 170926547 | 3 | 0 | 0 | T | G | -0.8705 | 0.2448 | 0.0003762 | ---?? | 0 | 0.661 | intergenic | |
| rs201648223 | 6 | 32004917 | 3 | 0 | 0 | T | G | 1.3752 | 0.3867 | 0.0003762 | +++?? | 17.1 | 0.2995 | upstream-gene | |
| rs9469084 | 6 | 32080383 | 5 | 0 | 0 | T | C | 1.3977 | 0.393 | 0.0003764 | +++-+ | 45.3 | 0.1204 | upstream-gene | |
| rs145663802 | 3 | 47697657 | 2 | 0 | 0 | T | C | 6.6803 | 1.8786 | 0.0003765 | ?+??+ | 1.6 | 0.3134 | intron | SMARCC1 |
| 13:82985894 | 13 | 82985894 | 3 | 0 | 0 | T | TAC | -1.0527 | 0.296 | 0.0003765 | ---?? | 0 | 0.7991 | | |
| rs9689459 | 6 | 63283633 | 4 | 0 | 0 | A | G | -2.4461 | 0.6879 | 0.0003767 | ---?- | 0 | 0.5588 | intergenic | |
| rs61904575 | 11 | 95706680 | 4 | 0 | 0 | A | G | -0.9794 | 0.2754 | 0.0003767 | ---?- | 30.6 | 0.2288 | downstream-gene | |
| 12:41304977 | 12 | 41304977 | 3 | 0 | 0 | G | GA | 1.8036 | 0.5072 | 0.0003767 | +++?? | 0 | 0.91 | | |
| rs76541820 | 4 | 15219264 | 4 | 0 | 0 | A | G | 1.9697 | 0.5539 | 0.0003768 | +++?+ | 5.1 | 0.3673 | intergenic | |
| 1:42986649 | 1 | 42986649 | 3 | 0 | 0 | CT | C | 1.2389 | 0.3484 | 0.0003768 | +++?? | 15.4 | 0.3066 | | |
| rs10469726 | 2 | 185452799 | 4 | 0 | 0 | A | G | -0.9778 | 0.275 | 0.000377 | ---?- | 0 | 0.7044 | intergenic | |
| rs62076829 | 18 | 5913148 | 4 | 0 | 0 | A | G | -1.2642 | 0.3556 | 0.000377 | ---?- | 0 | 0.4085 | intergenic | |
| rs9448837 | 6 | 80553896 | 3 | 0 | 0 | C | G | 0.986 | 0.2773 | 0.0003771 | +++?? | 26 | 0.259 | intron | C6orf7 |
| rs35477410 | 3 | 183628509 | 4 | 0 | 0 | C | G | 1.8309 | 0.5149 | 0.0003771 | +++?+ | 0 | 0.937 | intergenic | |
| rs1577771 | 1 | 162078088 | 5 | 0 | 0 | A | G | -0.891 | 0.2506 | 0.0003772 | ----+ | 38.9 | 0.1617 | intron | NOS1AP |
| rs1339374 | 9 | 35660990 | 5 | 0 | 0 | A | G | 0.9483 | 0.2667 | 0.0003773 | ++++- | 14.2 | 0.3239 | missense | ARHGEF39,CCDC107 |
| rs1607251 | 12 | 43195140 | 5 | 0 | 0 | A | T | 0.9523 | 0.2678 | 0.0003774 | ++++- | 0 | 0.4612 | intergenic | |
| rs78956672 | 5 | 122534378 | 4 | 0 | 0 | A | G | -1.8859 | 0.5304 | 0.0003774 | ---?- | 0 | 0.8394 | downstream-gene | |
| rs58894228 | 14 | 28186436 | 4 | 0 | 0 | T | C | -2.0026 | 0.5633 | 0.0003775 | ---?- | 0 | 0.8739 | intergenic | |
| rs12322102 | 12 | 112775474 | 5 | 0 | 0 | T | C | 1.328 | 0.3735 | 0.0003776 | +-+++ | 53.6 | 0.07144 | intron | HECTD4 |
| rs143129850 | 13 | 50393787 | 5 | 0 | 0 | A | G | -1.4414 | 0.4054 | 0.0003779 | ----+ | 0 | 0.6817 | intergenic |  |
| rs79960326 | 1 | 152766672 | 3 | 0 | 0 | T | C | 2.27 | 0.6385 | 0.000378 | +++?? | 38.9 | 0.1945 | upstream-gene | |
| rs139743512 | 17 | 56843771 | 4 | 0 | 0 | T | C | -2.5847 | 0.7271 | 0.000378 | ----? | 0 | 0.7495 | intron | PPM1E |
| rs12706948 | 7 | 130362027 | 4 | 0 | 0 | C | G | 0.8831 | 0.2484 | 0.0003781 | +-+?+ | 54.9 | 0.08409 | intron | TSGA13 |
| rs61973847 | 14 | 28185907 | 4 | 0 | 0 | T | C | -2.0016 | 0.563 | 0.0003781 | ---?- | 0 | 0.8748 | intergenic | |
| rs2390796 | 1 | 90240317 | 5 | 0 | 0 | C | G | 0.7572 | 0.213 | 0.0003783 | +++++ | 0 | 0.9779 | intergenic | |
| rs368175987 | 7 | 29846240 | 4 | 0 | 0 | A | G | 4.5065 | 1.2678 | 0.0003784 | +++?+ | 0 | 0.4828 | 5-prime-UTR | WIPF3 |
| rs7160256 | 14 | 28188438 | 4 | 0 | 0 | T | C | -2.0053 | 0.5641 | 0.0003785 | ---?- | 0 | 0.8716 | intergenic | |
| rs11132611 | 4 | 190019261 | 4 | 0 | 0 | T | C | -0.8901 | 0.2504 | 0.0003785 | ---?- | 0 | 0.5491 | intergenic | |
| rs8011867 | 14 | 101833831 | 4 | 0 | 0 | T | G | 1.4316 | 0.4028 | 0.0003788 | +++?+ | 0 | 0.398 | intergenic | |
| rs12618211 | 2 | 185445333 | 4 | 0 | 0 | A | G | 0.9641 | 0.2712 | 0.0003788 | +++?+ | 0 | 0.7728 | intergenic | |
| rs62418249 | 6 | 63285382 | 4 | 0 | 0 | T | G | -2.4464 | 0.6883 | 0.0003788 | ---?- | 0 | 0.5597 | intergenic | |
| rs150061704 | 1 | 102909744 | 4 | 0 | 0 | A | G | 0.8872 | 0.2496 | 0.0003789 | +++?- | 0 | 0.415 | intergenic | |
| rs3922407 | 8 | 113290548 | 4 | 0 | 0 | A | C | -1.0113 | 0.2845 | 0.000379 | ---?- | 0 | 0.7977 | intron | CSMD3 |
| rs61973851 | 14 | 28188737 | 4 | 0 | 0 | A | C | -2.0054 | 0.5642 | 0.000379 | ---?- | 0 | 0.871 | intergenic | |
| rs114803358 | 6 | 32601708 | 4 | 0 | 1 | A | G | 1.0242 | 0.2882 | 0.0003791 | ++++? | 0 | 0.5718 | upstream-gene | |
| rs4547579 | 2 | 137008308 | 4 | 0 | 0 | T | C | 0.9483 | 0.2668 | 0.0003793 | +++?+ | 0 | 0.8297 | intergenic | |
| 2:40454398 | 2 | 40454398 | 3 | 0 | 0 | CCTAT | C | 1.1125 | 0.313 | 0.0003794 | +++?? | 0 | 0.6248 | | |
| rs11600148 | 11 | 12708614 | 4 | 0 | 0 | A | G | -1.4242 | 0.4007 | 0.0003795 | ---?- | 0 | 0.5361 | intron | TEAD1 |
| 14:38489810 | 14 | 38489810 | 3 | 0 | 0 | A | AGAGG( | -1.118 | 0.3146 | 0.0003795 | ---?? | 0 | 0.9328 | | |
| rs12409856 | 1 | 185390063 | 3 | 0 | 0 | A | G | -0.975 | 0.2743 | 0.0003796 | ---?? | 0 | 0.8433 | intergenic | |
| rs10920327 | 1 | 202061947 | 4 | 0 | 0 | T | C | -0.9243 | 0.2601 | 0.0003797 | ---?- | 0 | 0.9438 | intergenic | |
| rs1849343 | 11 | 37068238 | 4 | 0 | 0 | T | C | -0.8787 | 0.2473 | 0.0003798 | ---?- | 10 | 0.3429 | intergenic | |
| rs7957169 | 12 | 111772770 | 5 | 0 | 0 | A | C | -0.9019 | 0.2538 | 0.0003799 | -+--- | 46.3 | 0.1143 | intron | CUX2 |
| rs113800430 | 6 | 106277440 | 4 | 0 | 0 | A | G | 1.9999 | 0.5628 | 0.0003799 | +++?+ | 0 | 0.4134 | intergenic | |
| 3:77402253 | 3 | 77402253 | 3 | 0 | 0 | T | TG | -0.912 | 0.2566 | 0.0003799 | ---?? | 0 | 0.7859 | | |
| rs6581575 | 12 | 64923107 | 4 | 0 | 0 | A | G | -0.8855 | 0.2492 | 0.00038 | ---?- | 0 | 0.8918 | intergenic | |
| rs11924950 | 3 | 196472468 | 4 | 0 | 0 | T | G | -2.192 | 0.6168 | 0.00038 | ---?+ | 22.1 | 0.2781 | intron | PAK2 |
| rs72919041 | 2 | 199075207 | 4 | 0 | 0 | T | C | -1.1922 | 0.3355 | 0.0003801 | ---?+ | 39.7 | 0.1735 | intergenic | |
| 8:64913550 | 8 | 64913550 | 3 | 0 | 0 | A | AT | -0.9019 | 0.2538 | 0.0003802 | ---?? | 0 | 0.5163 | | |
| rs9276714 | 6 | 32760318 | 5 | 0 | 0 | A | G | -1.4107 | 0.397 | 0.0003803 | ----- | 0 | 0.6149 | intergenic |  |
| rs13096198 | 3 | 183619177 | 4 | 0 | 0 | A | C | -1.7074 | 0.4805 | 0.0003803 | ---?- | 0 | 0.9509 | intergenic | |
| rs4750922 | 10 | 129673226 | 4 | 0 | 0 | T | C | 0.9181 | 0.2584 | 0.0003804 | +++?+ | 4 | 0.3728 | downstream-gene | |
| rs7973898 | 12 | 112616167 | 5 | 0 | 0 | T | C | -1.3123 | 0.3693 | 0.0003805 | -+--- | 55.3 | 0.06249 | intron | HECTD4 |
| rs116964922 | 15 | 87486755 | 4 | 0 | 0 | A | G | 2.374 | 0.6681 | 0.0003805 | +++?+ | 0 | 0.956 | intron | AGBL1 |
| 2:40446174 | 2 | 40446174 | 3 | 0 | 0 | T | TCTACC | 1.1226 | 0.3159 | 0.0003806 | +++?? | 0 | 0.5919 | | |

| | | | | | | | | | | | | | | | |
|---|---|---|---|---|---|---|---|---|---|---|---|---|---|---|---|
| rs79014413 | 17 | 56939859 | 5 | 0 | 0 | T | G | -2.4737 | 0.6962 | 0.0003807 | ----- | 0 | 0.8462 | intron | PPM1E |
| 6:154516925 | 6 | 154516925 | 3 | 0 | 0 | A | AT | -1.038 | 0.2921 | 0.0003808 | ---?? | 0 | 0.4069 | | |
| rs35274284 | 2 | 82993127 | 4 | 0 | 0 | A | G | 2.4905 | 0.701 | 0.0003809 | +++?+ | 0 | 0.5695 | intergenic | |
| 6:104711931 | 6 | 104711931 | 3 | 0 | 0 | A | AT | -4.3727 | 1.2308 | 0.000381 | ---?? | 0 | 0.4296 | | |
| rs10478656 | 5 | 124691789 | 4 | 0 | 0 | T | G | 0.8829 | 0.2485 | 0.0003811 | +++?+ | 0 | 0.5668 | intergenic | |
| rs17427599 | 6 | 32667364 | 5 | 0 | 0 | T | C | 1.0087 | 0.2839 | 0.0003811 | +++++ | 0 | 0.845 | intergenic | |
| rs6658062 | 1 | 102910651 | 4 | 0 | 0 | T | C | -0.8846 | 0.249 | 0.0003813 | ---?+ | 0 | 0.4159 | intergenic | |
| rs6756711 | 2 | 40448560 | 3 | 0 | 0 | A | C | -1.1205 | 0.3154 | 0.0003813 | ---?? | 0 | 0.6028 | intron | SLC8A1,SLC8A1-AS1 |
| rs12566660 | 1 | 185390570 | 4 | 0 | 0 | T | G | 0.9446 | 0.2659 | 0.0003813 | +++?+ | 0 | 0.9189 | intergenic | |
| 20:37731813 | 20 | 37731813 | 3 | 0 | 0 | T | TA | 2.9208 | 0.8221 | 0.0003813 | +++?? | 0 | 0.4866 | | |
| rs73214528 | 5 | 107571902 | 4 | 0 | 0 | T | C | 1.1617 | 0.327 | 0.0003814 | +++?+ | 0 | 0.6057 | intron | FBXL17 |
| rs11634025 | 15 | 98036632 | 4 | 0 | 0 | A | C | 1.1241 | 0.3164 | 0.0003815 | +++?- | 0 | 0.4813 | intergenic | |
| rs139371393 | 18 | 40021298 | 3 | 0 | 0 | A | G | -1.3974 | 0.3934 | 0.0003815 | ---?? | 0 | 0.6431 | intron | LINC00907 |
| rs71633416 | 5 | 51514855 | 4 | 0 | 0 | A | C | -2.77 | 0.7798 | 0.0003819 | ---?- | 24.2 | 0.2663 | intergenic | |
| rs1028098 | 1 | 64774778 | 4 | 0 | 0 | T | C | -0.988 | 0.2781 | 0.000382 | ---?- | 0 | 0.4174 | intergenic | |
| rs16977068 | 18 | 40440609 | 3 | 0 | 0 | T | C | 4.7757 | 1.3444 | 0.000382 | +++?? | 0 | 0.5119 | intron | RIT2 |
| rs61973855 | 14 | 28191614 | 4 | 0 | 0 | T | C | -2.0079 | 0.5652 | 0.000382 | ---?- | 0 | 0.8673 | intergenic | |
| rs3811306 | 14 | 22534497 | 4 | 0 | 0 | T | G | -0.9312 | 0.2622 | 0.0003821 | ---?- | 0 | 0.644 | intergenic | |
| rs6606812 | 15 | 22963272 | 4 | 0 | 0 | T | C | -0.8911 | 0.2509 | 0.0003822 | ---?- | 0 | 0.5203 | intron | CYFIP1 |
| rs12221835 | 11 | 95706507 | 4 | 0 | 0 | T | C | -0.9767 | 0.275 | 0.0003822 | -+-?- | 31 | 0.2262 | downstream-gene | |
| rs140006486 | 11 | 91054290 | 3 | 0 | 0 | A | C | 3.5589 | 1.0019 | 0.0003823 | +++?? | 0 | 0.4499 | intergenic | |
| rs10155453 | 4 | 190017685 | 4 | 0 | 0 | A | C | 0.8969 | 0.2525 | 0.0003824 | +++?+ | 0 | 0.6389 | intergenic | |
| 19:49721419 | 19 | 49721419 | 3 | 0 | 0 | T | C | -1.2234 | 0.3444 | 0.0003825 | ---?? | 0 | 0.6836 | intergenic | |
| rs10836663 | 11 | 37048379 | 4 | 0 | 0 | T | C | 0.9387 | 0.2643 | 0.0003826 | +++?+ | 0 | 0.4507 | intergenic | |
| rs61973854 | 14 | 28191299 | 4 | 0 | 0 | A | G | -2.0071 | 0.5651 | 0.0003826 | ---?- | 0 | 0.8673 | intergenic | |
| rs75052360 | 12 | 112594017 | 5 | 0 | 0 | A | C | 1.3121 | 0.3694 | 0.0003827 | +-+++ | 56.6 | 0.05588 | downstream-gene | |
| 11:12661420 | 11 | 12661420 | 3 | 0 | 0 | A | AACT | -1.3756 | 0.3873 | 0.0003827 | ---?? | 59.8 | 0.08324 | | |
| rs59504640 | 18 | 5915709 | 4 | 0 | 0 | A | G | -1.2597 | 0.3547 | 0.0003829 | ---?- | 0 | 0.4042 | intergenic | |
| rs62400276 | 6 | 24040373 | 5 | 0 | 0 | A | G | 1.379 | 0.3883 | 0.000383 | ++++- | 0 | 0.7349 | intergenic | |
| 3:77402256 | 3 | 77402256 | 3 | 0 | 0 | A | ATCAGT | -0.9116 | 0.2567 | 0.000383 | ---?? | 0 | 0.7867 | | |
| 5:172102132 | 5 | 172102132 | 2 | 0 | 0 | T | G | -5.2203 | 1.4699 | 0.0003831 | -?-?? | 0 | 0.6407 | intron | NEURL1B |
| rs61973857 | 14 | 28192331 | 4 | 0 | 0 | A | C | 2.0081 | 0.5654 | 0.0003832 | +++?+ | 0 | 0.8662 | intergenic | |
| rs142394371 | 4 | 178478769 | 4 | 0 | 0 | A | G | -3.7196 | 1.0474 | 0.0003833 | ---?- | 0 | 0.4286 | intron | LOC285500 |
| 1:87423476 | 1 | 87423476 | 3 | 0 | 0 | A | AT | 1.1361 | 0.3199 | 0.0003833 | +++?? | 0 | 0.5731 | | |
| rs115004527 | 3 | 173594998 | 4 | 0 | 0 | T | C | -2.725 | 0.7673 | 0.0003834 | ---?- | 0 | 0.4913 | intron | NLGN1 |
| rs143491386 | 2 | 168065048 | 4 | 0 | 0 | T | G | -2.3568 | 0.6637 | 0.0003835 | ---?+ | 0 | 0.4616 | intron | XIRP2 |
| rs7930663 | 11 | 130621232 | 4 | 0 | 0 | T | C | 1.7277 | 0.4865 | 0.0003837 | +++?+ | 30.5 | 0.2294 | intergenic | |
| rs3776082 | 5 | 149544045 | 4 | 0 | 0 | A | G | 0.8515 | 0.2398 | 0.0003842 | +++?+ | 0 | 0.9596 | upstream-gene | |
| rs57699736 | 18 | 5916161 | 4 | 0 | 0 | T | C | -1.2586 | 0.3545 | 0.0003842 | ---?- | 0 | 0.4036 | intergenic | |
| rs73043807 | 19 | 39383238 | 4 | 0 | 0 | C | G | 1.244 | 0.3503 | 0.0003842 | +++?+ | 0 | 0.5474 | intron | SIRT2 |
| rs4853773 | 2 | 2763135 | 4 | 0 | 0 | T | C | -1.3533 | 0.3812 | 0.0003843 | ---?- | 0 | 0.8807 | intergenic | |
| rs35970251 | 7 | 130362423 | 4 | 0 | 0 | A | C | -0.8823 | 0.2485 | 0.0003845 | -+-?- | 54.9 | 0.08378 | intron | TSGA13 |
| rs74656478 | 17 | 56937939 | 5 | 0 | 0 | T | C | -2.4749 | 0.697 | 0.0003845 | ----- | 0 | 0.8248 | intron | PPM1E |
| rs115444818 | 3 | 156749560 | 4 | 0 | 0 | A | G | 3.0287 | 0.853 | 0.0003846 | +++?+ | 9.3 | 0.3468 | intron | LEKR1 |
| rs7159849 | 14 | 22500911 | 4 | 0 | 0 | A | C | -0.9366 | 0.2638 | 0.0003847 | ---?- | 0 | 0.9288 | intergenic | |
| rs9667239 | 11 | 1143101 | 4 | 0 | 0 | T | C | -1.4192 | 0.3997 | 0.0003848 | +--?+ | 61.3 | 0.05139 | intron | MUC5AC |
| rs114575065 | 18 | 25181147 | 4 | 0 | 0 | T | C | -2.6005 | 0.7325 | 0.0003848 | ---?- | 0 | 0.634 | intergenic | |
| rs115423105 | 17 | 56600490 | 5 | 0 | 0 | T | C | 2.3843 | 0.6716 | 0.0003848 | +++++ | 0 | 0.8419 | intron | SEPTIN4 |
| rs61973858 | 14 | 28192407 | 4 | 0 | 0 | A | G | -2.0076 | 0.5655 | 0.000385 | ---?- | 0 | 0.8659 | intergenic | |
| rs73834885 | 3 | 68243722 | 4 | 0 | 0 | T | C | -2.3888 | 0.6729 | 0.000385 | ---?- | 23 | 0.2729 | intron | FAM19A1 |
| rs876076 | 3 | 183616241 | 4 | 0 | 0 | T | G | -1.7057 | 0.4805 | 0.0003851 | ---?- | 0 | 0.9524 | intergenic | |
| rs5997367 | 22 | 28842447 | 4 | 0 | 0 | T | C | -2.5477 | 0.7177 | 0.0003851 | ---?- | 0 | 0.723 | intron | TTC28 |
| rs10780067 | 4 | 169325683 | 4 | 0 | 0 | T | C | -0.8868 | 0.2498 | 0.0003851 | ---?- | 0 | 0.8518 | intron | DDX60L |
| rs12026041 | 1 | 232834967 | 4 | 0 | 0 | T | C | 0.8717 | 0.2456 | 0.0003852 | +++?+ | 12.6 | 0.3299 | intergenic | |
| rs72668232 | 14 | 28194985 | 4 | 0 | 0 | T | C | 2.0103 | 0.5663 | 0.0003857 | +++?+ | 0 | 0.863 | intergenic | |
| rs2272308 | 12 | 112601250 | 5 | 0 | 0 | A | G | 1.3125 | 0.3698 | 0.0003861 | +-+++ | 55.4 | 0.0617 | intron | HECTD4 |
| rs7349241 | 2 | 170889997 | 4 | 0 | 1 | T | G | 1.0242 | 0.2885 | 0.0003861 | +++?+ | 43.4 | 0.1509 | intron | UBR3 |
| rs189407431 | 17 | 56974563 | 5 | 0 | 0 | T | C | -2.4753 | 0.6974 | 0.0003861 | ----- | 0 | 0.7992 | intron | PPM1E |
| rs2143009 | 6 | 144630226 | 5 | 0 | 0 | T | C | -1.4522 | 0.4091 | 0.0003862 | ----+ | 34.3 | 0.1923 | intron | UTRN |

| | | | | | | | | | | | | | | | |
|---|---|---|---|---|---|---|---|---|---|---|---|---|---|---|---|
| rs2331491 | 14 | 22522428 | 4 | 0 | 0 | T | C | 0.9262 | 0.2609 | 0.0003862 | +++?+ | 0 | 0.6993 | intergenic | |
| rs7969049 | 12 | 27783723 | 4 | 0 | 0 | T | C | 0.8966 | 0.2526 | 0.0003864 | +-+?- | 54.1 | 0.08852 | intron | PPFIBP1 |
| rs67753607 | 14 | 28195163 | 4 | 0 | 0 | T | C | -2.0103 | 0.5664 | 0.0003864 | ---?- | 0 | 0.8625 | intergenic | |
| 3:172328799 | 3 | 172328799 | 3 | 0 | 0 | G | GAAAAA | -1.941 | 0.5469 | 0.0003868 | ---?? | 0 | 0.6612 | | |
| rs12775208 | 10 | 134597157 | 4 | 0 | 0 | A | C | 0.9571 | 0.2697 | 0.0003873 | +++?+ | 0 | 0.5945 | downstream-gene | |
| rs72799522 | 16 | 84842438 | 4 | 0 | 0 | C | G | 1.2927 | 0.3643 | 0.0003874 | +++?- | 0 | 0.5102 | intergenic | |
| rs141899940 | 17 | 56953537 | 5 | 0 | 0 | T | C | 2.501 | 0.7048 | 0.0003874 | +++++ | 0 | 0.8209 | intron | PPM1E |
| rs12056383 | 8 | 130711839 | 4 | 0 | 0 | C | G | 1.654 | 0.4661 | 0.0003875 | +++?+ | 0 | 0.8129 | intergenic | |
| 2:170594129 | 2 | 170594129 | 3 | 0 | 0 | CTTTTTC | C | -1.1967 | 0.3372 | 0.0003875 | ---?? | 0 | 0.9889 | | |
| rs28408170 | 8 | 113503869 | 4 | 0 | 0 | T | C | 1.028 | 0.2897 | 0.0003876 | +++?+ | 0 | 0.6308 | intron | CSMD3 |
| rs114673658 | 2 | 50990348 | 4 | 0 | 0 | T | C | -2.4921 | 0.7023 | 0.0003877 | ---?- | 0 | 0.5973 | intron | NRXN1 |
| rs117885771 | 8 | 113257249 | 4 | 0 | 0 | A | T | 1.0185 | 0.2871 | 0.0003877 | +++?+ | 0 | 0.945 | intron | CSMD3 |
| rs141482113 | 17 | 56975015 | 5 | 0 | 0 | T | G | -2.474 | 0.6972 | 0.0003877 | ----- | 0 | 0.7998 | intron | PPM1E |
| rs28399486 | 19 | 41512500 | 2 | 0 | 0 | T | G | 3.7832 | 1.0662 | 0.0003878 | +?+?? | 0 | 0.7073 | intron | CYP2B6 |
| rs936462 | 11 | 636690 | 3 | 0 | 0 | A | G | -1.6987 | 0.4788 | 0.0003881 | ---?? | 40.5 | 0.1862 | upstream-gene | |
| rs2102080 | 15 | 98048649 | 4 | 0 | 0 | T | C | 0.9499 | 0.2677 | 0.0003882 | +++?+ | 0 | 0.8069 | intergenic | |
| rs6591510 | 11 | 58995669 | 5 | 0 | 0 | T | C | -0.764 | 0.2153 | 0.0003882 | ----+ | 10.7 | 0.3452 | intergenic |  |
| rs10904240 | 10 | 4416232 | 4 | 0 | 0 | A | G | -0.9747 | 0.2747 | 0.0003882 | ---?- | 0 | 0.3968 | intergenic | |
| rs62177279 | 2 | 180015523 | 3 | 0 | 0 | A | G | -6.7274 | 1.8962 | 0.0003883 | ---?? | 13.9 | 0.3132 | intron | SESTD1 |
| 17:52167365 | 17 | 52167365 | 3 | 0 | 0 | T | TC | -2.3502 | 0.6624 | 0.0003883 | ---?? | 0 | 0.8882 | | |
| rs116786842 | 9 | 101128951 | 4 | 0 | 0 | A | C | -3.5591 | 1.0032 | 0.0003884 | ---?+ | 49.6 | 0.1139 | intron | GABBR2 |
| rs16932246 | 12 | 27784629 | 4 | 0 | 0 | A | G | 0.9732 | 0.2743 | 0.0003885 | +++?- | 0 | 0.4344 | intron | PPFIBP1 |
| rs114941331 | 2 | 168092600 | 4 | 0 | 0 | T | C | 2.3571 | 0.6644 | 0.0003885 | +++?- | 0 | 0.5917 | intron | XIRP2 |
| rs139026785 | 15 | 83872205 | 4 | 0 | 0 | A | C | 3.7873 | 1.0676 | 0.0003888 | +++?+ | 0 | 0.9787 | intron | HDGFRP3 |
| rs59900917 | 19 | 56223486 | 4 | 0 | 0 | T | G | -1.2321 | 0.3473 | 0.000389 | ---?- | 0 | 0.8126 | intron | NLRP9 |
| 1:117573025 | 1 | 117573025 | 2 | 0 | 0 | CAAAG | C | -5.3749 | 1.5151 | 0.000389 | -?-?? | 0 | 0.8695 | | |
| rs1953975 | 1 | 102955981 | 4 | 0 | 0 | T | C | 0.8357 | 0.2356 | 0.0003892 | +++?- | 0 | 0.3964 | intergenic | |
| rs4486432 | 1 | 90240982 | 5 | 0 | 0 | A | C | -0.7559 | 0.2131 | 0.0003892 | ----- | 0 | 0.9877 | intergenic |  |
| rs74531234 | 12 | 112619761 | 5 | 0 | 0 | T | C | 1.3097 | 0.3692 | 0.0003894 | +-+++ | 55.2 | 0.0628 | intron | HECTD4 |
| rs4908007 | 1 | 100198799 | 4 | 0 | 0 | T | C | 1.7914 | 0.505 | 0.0003895 | +++?+ | 0 | 0.6707 | intron | FRRS1 |
| rs7512106 | 1 | 162081822 | 5 | 0 | 0 | T | C | -0.8854 | 0.2496 | 0.0003895 | ----+ | 38.1 | 0.1675 | intron | NOS1AP |
| rs146062758 | 2 | 25935851 | 2 | 0 | 0 | A | C | 5.0271 | 1.4172 | 0.0003895 | ?++?? | 0 | 0.4591 | intergenic | |
| rs11927524 | 3 | 196489527 | 4 | 0 | 0 | T | C | 2.1875 | 0.6167 | 0.0003897 | +++?- | 21.5 | 0.2814 | intron | PAK2 |
| rs7938152 | 11 | 95706254 | 4 | 0 | 0 | A | G | -0.9737 | 0.2745 | 0.00039 | -+-?- | 31.6 | 0.2225 | intergenic | |
| 12:103256724 | 12 | 103256724 | 3 | 0 | 0 | G | GA | -0.9251 | 0.2608 | 0.0003903 | ---?? | 0 | 0.521 | | |
| rs12902548 | 15 | 41915809 | 5 | 0 | 0 | A | G | 1.0195 | 0.2875 | 0.0003905 | +++++ | 12 | 0.3374 | intron | MGA |
| rs73020372 | 2 | 170588737 | 4 | 0 | 0 | A | G | -1.2484 | 0.352 | 0.0003906 | ---?- | 0 | 0.967 | intron | PHOSPHO2-KLHL23 |
| rs4133728 | 12 | 26836395 | 4 | 0 | 0 | C | G | -1.7608 | 0.4965 | 0.0003907 | -+-?- | 38.1 | 0.1837 | intron | ITPR2 |
| rs62148799 | 2 | 40447142 | 3 | 0 | 0 | C | G | 1.1185 | 0.3154 | 0.0003907 | +++?? | 0 | 0.5975 | intron | SLC8A1,SLC8A1-AS1 |
| rs12023295 | 1 | 232835224 | 4 | 0 | 0 | T | G | 0.8726 | 0.2461 | 0.0003908 | +++?+ | 17.4 | 0.3039 | intergenic | |
| rs16853108 | 4 | 41330982 | 4 | 0 | 0 | T | C | 1.4222 | 0.401 | 0.0003909 | +++?- | 0 | 0.4342 | intergenic | |
| rs1747109 | 13 | 45630008 | 5 | 0 | 0 | A | G | -1.2442 | 0.3509 | 0.0003911 | ----+ | 0 | 0.508 | intergenic |  |
| rs9689376 | 6 | 63283341 | 4 | 0 | 0 | A | G | -2.422 | 0.683 | 0.0003912 | ---?- | 0 | 0.5588 | intergenic | |
| 16:59132614 | 16 | 59132614 | 3 | 0 | 0 | A | C | -3.9175 | 1.1048 | 0.0003913 | ---?? | 0 | 0.9186 | intergenic | |
| rs369251 | 2 | 40476117 | 4 | 0 | 0 | T | G | -1.2077 | 0.3406 | 0.0003915 | ---?+ | 34.2 | 0.2071 | intron | SLC8A1,SLC8A1-AS1 |
| rs114991517 | 1 | 152871305 | 3 | 0 | 0 | T | C | -4.0682 | 1.1473 | 0.0003915 | ---?? | 15.1 | 0.308 | intergenic | |
| rs11870867 | 17 | 55459682 | 4 | 0 | 0 | T | G | 1.1169 | 0.315 | 0.0003917 | +++?- | 15.4 | 0.3146 | intron | MSI2 |
| rs62418253 | 6 | 63289416 | 4 | 0 | 0 | T | C | 2.4277 | 0.6847 | 0.0003918 | +++?+ | 0 | 0.5632 | intergenic | |
| rs17552785 | 18 | 35864561 | 4 | 0 | 0 | A | T | -1.3697 | 0.3863 | 0.0003921 | ---?- | 0 | 0.9896 | intergenic | |
| rs34874721 | 2 | 139896412 | 4 | 0 | 0 | A | C | 1.354 | 0.3819 | 0.0003922 | +++?+ | 0 | 0.9928 | intergenic | |
| rs10489481 | 1 | 185391167 | 4 | 0 | 0 | A | G | 0.942 | 0.2657 | 0.0003924 | +++?+ | 0 | 0.9198 | intergenic | |
| rs72701248 | 14 | 83268567 | 4 | 0 | 0 | T | G | -1.9577 | 0.5522 | 0.0003925 | +--?- | 57.3 | 0.07117 | intergenic | |
| rs55816821 | 10 | 117074621 | 4 | 0 | 0 | C | G | 3.8131 | 1.0756 | 0.0003926 | +++?- | 0 | 0.4047 | intron | ATRNL1 |
| rs56023747 | 10 | 117074623 | 4 | 0 | 0 | A | C | -3.8131 | 1.0756 | 0.0003926 | ---?+ | 0 | 0.4047 | intron | ATRNL1 |
| rs4448132 | 6 | 32548122 | 5 | 0 | 1 | C | G | -1.3605 | 0.3838 | 0.0003927 | ----- | 12.8 | 0.3321 | intron | HLA-DRB1 |
| rs12311080 | 12 | 43174747 | 5 | 0 | 0 | A | G | 0.9438 | 0.2662 | 0.0003927 | ++++- | 0 | 0.4062 | intergenic | |
| rs72842486 | 11 | 1123100 | 4 | 0 | 0 | T | C | -1.4059 | 0.3966 | 0.0003927 | ---?+ | 54.3 | 0.08741 | intergenic | |
| rs6925769 | 6 | 83721902 | 4 | 0 | 0 | A | G | 3.3933 | 0.9572 | 0.0003928 | +++?+ | 0 | 0.8213 | intron | UBE3D |
| rs56303484 | 12 | 81423653 | 2 | 0 | 0 | A | G | 6.7802 | 1.9127 | 0.0003928 | +???+ | 37.9 | 0.2045 | intergenic | |

| | | | | | | | | | | | | | | | |
|---|---|---|---|---|---|---|---|---|---|---|---|---|---|---|---|
| rs140966179 | 3 | 125883584 | 4 | 0 | 0 | T | C | 3.7949 | 1.0705 | 0.0003928 | +++?+ | 0 | 0.4764 | intron | ALDH1L1 |
| rs825202 | 1 | 64773433 | 4 | 0 | 0 | T | C | 0.9852 | 0.2779 | 0.0003929 | +++?+ | 0 | 0.4248 | intergenic | |
| rs77753889 | 8 | 130711599 | 4 | 0 | 0 | A | G | 1.6482 | 0.4649 | 0.0003929 | +++?+ | 0 | 0.8013 | intergenic | |
| rs7974644 | 12 | 130593579 | 4 | 0 | 0 | A | G | -0.847 | 0.239 | 0.0003931 | ---?+ | 19.6 | 0.2922 | intergenic | |
| rs74586224 | 17 | 57128625 | 5 | 0 | 0 | A | G | 2.358 | 0.6652 | 0.0003931 | +++++ | 0 | 0.8348 | synonymous | TRIM37 |
| rs6836454 | 4 | 169333585 | 4 | 0 | 0 | C | G | 0.8804 | 0.2484 | 0.0003932 | +++?+ | 0 | 0.8758 | intron | DDX60L |
| rs11060711 | 12 | 130600785 | 4 | 0 | 0 | A | G | 0.8679 | 0.2449 | 0.0003932 | +++?- | 48.3 | 0.1215 | intergenic | |
| rs74783722 | 6 | 109106048 | 2 | 0 | 0 | T | G | -4.9569 | 1.3985 | 0.0003933 | -?-?? | 0 | 0.8432 | intergenic | |
| rs1038638 | 7 | 130361092 | 4 | 0 | 0 | A | G | -0.8801 | 0.2483 | 0.0003935 | -+-?- | 54.7 | 0.08512 | intron | TSGA13 |
| rs77808663 | 1 | 85180950 | 4 | 0 | 0 | T | C | 2.3604 | 0.6659 | 0.0003935 | +++?+ | 0 | 0.8348 | intergenic | |
| rs75468753 | 16 | 84840856 | 4 | 0 | 0 | T | C | 1.2782 | 0.3606 | 0.0003936 | +++?- | 20.5 | 0.2868 | intergenic | |
| rs7333078 | 13 | 60033271 | 4 | 0 | 0 | C | G | 0.8866 | 0.2501 | 0.0003937 | +++?+ | 0 | 0.4416 | intergenic | |
| rs12325235 | 16 | 7178972 | 4 | 0 | 0 | A | C | -0.9825 | 0.2772 | 0.0003937 | ---?- | 11.6 | 0.3346 | intron | RBFOX1 |
| rs2148606 | 10 | 4418869 | 4 | 0 | 0 | T | C | -0.9734 | 0.2746 | 0.0003938 | ---?- | 6 | 0.3632 | intergenic | |
| rs6533524 | 4 | 111407451 | 4 | 0 | 0 | A | G | 0.8564 | 0.2416 | 0.0003939 | +++?+ | 0 | 0.9934 | intron | ENPEP |
| rs114961696 | 4 | 89120118 | 4 | 0 | 0 | A | G | -3.1388 | 0.8857 | 0.000394 | ---?+ | 38.7 | 0.1799 | intron | ABCG2 |
| rs7690142 | 4 | 41742722 | 4 | 0 | 0 | C | G | 1.5054 | 0.4248 | 0.0003941 | +++?+ | 29.4 | 0.2357 | downstream-gene | |
| rs4678420 | 3 | 138185860 | 2 | 0 | 0 | A | G | 1.7914 | 0.5055 | 0.0003943 | +???+ | 0 | 0.5146 | intron | ESYT3 |
| rs35880697 | 6 | 32433759 | 5 | 0 | 1 | T | C | 1.3023 | 0.3675 | 0.0003943 | +++++ | 0 | 0.454 | intergenic | |
| rs62418254 | 6 | 63292002 | 4 | 0 | 0 | T | C | 2.429 | 0.6854 | 0.0003943 | +++?+ | 0 | 0.5637 | intergenic | |
| rs143633080 | 8 | 86885175 | 4 | 0 | 0 | T | G | 1.9615 | 0.5535 | 0.0003943 | +++?+ | 0 | 0.507 | intergenic | |
| rs4924568 | 15 | 41948046 | 5 | 0 | 0 | A | G | 1.0038 | 0.2833 | 0.0003945 | +++++ | 14.9 | 0.3197 | intron | MGA |
| rs817456 | 1 | 85666327 | 5 | 0 | 0 | T | C | -1.3902 | 0.3923 | 0.0003946 | ----- | 0 | 0.9793 | missense | SYDE2 |
| 1:181881058 | 1 | 181881058 | 3 | 0 | 0 | A | ATT | -3.4479 | 0.973 | 0.0003947 | ---?? | 0.7 | 0.3652 | | |
| rs1474630 | 6 | 144619627 | 5 | 0 | 0 | A | C | 1.4147 | 0.3992 | 0.0003948 | ++++- | 46.8 | 0.1106 | intron | UTRN |
| rs2123495 | 8 | 113512349 | 4 | 0 | 0 | A | G | -0.922 | 0.2602 | 0.0003949 | ---?- | 0 | 0.7172 | intron | CSMD3 |
| 17:65481486 | 17 | 65481486 | 3 | 0 | 0 | A | ATT | -0.9632 | 0.2718 | 0.000395 | ---?? | 0 | 0.5497 | | |
| rs1940162 | 11 | 95707290 | 4 | 0 | 0 | T | C | 0.9781 | 0.276 | 0.0003951 | +++?+ | 26.8 | 0.2511 | downstream-gene | |
| 10:117133936 | 10 | 117133936 | 3 | 0 | 0 | A | ATTCTC( | -3.9018 | 1.1012 | 0.0003951 | ---?? | 0 | 0.7904 | | |
| rs10875555 | 5 | 149543790 | 4 | 0 | 0 | A | C | 0.8473 | 0.2391 | 0.0003953 | +++?+ | 0 | 0.926 | upstream-gene | |
| rs141789230 | 6 | 121418306 | 4 | 0 | 0 | A | C | 2.6401 | 0.7451 | 0.0003953 | +++?+ | 21.7 | 0.2803 | intron | TBC1D32 |
| rs7850564 | 9 | 79462008 | 4 | 0 | 0 | T | C | -2.1226 | 0.5991 | 0.0003954 | ---?- | 0 | 0.6087 | intron | PRUNE2 |
| rs72752288 | 1 | 234998104 | 3 | 0 | 0 | A | G | 3.674 | 1.037 | 0.0003955 | +++?? | 0 | 0.7182 | intergenic | |
| rs12499156 | 4 | 169334418 | 4 | 0 | 0 | T | G | 0.8796 | 0.2483 | 0.0003958 | +++?+ | 0 | 0.8786 | intron | DDX60L |
| rs62173308 | 2 | 170576381 | 4 | 0 | 0 | A | C | 1.0029 | 0.2831 | 0.0003958 | +++?+ | 0 | 0.9658 | intron | PHOSPHO2-KLHL23 |
| 9:128885037 | 9 | 128885037 | 3 | 0 | 0 | G | GA | -3.5611 | 1.0051 | 0.0003958 | ---?? | 0 | 0.7421 | | |
| rs146259605 | 1 | 230982645 | 3 | 0 | 0 | T | C | 1.6494 | 0.4656 | 0.0003961 | +++?? | 0 | 0.5909 | intron | C1orf198 |
| rs9914357 | 17 | 4801125 | 5 | 0 | 0 | C | G | 1.5902 | 0.4489 | 0.0003964 | ++++- | 9.8 | 0.3503 | 3-prime-UTR | CHRNE,MINK1 |
| rs12760421 | 1 | 229220081 | 4 | 0 | 0 | A | G | -1.2925 | 0.3649 | 0.0003968 | ---?- | 0 | 0.8983 | intergenic | |
| rs62418255 | 6 | 63292819 | 4 | 0 | 0 | A | C | 2.4286 | 0.6856 | 0.0003969 | +++?+ | 0 | 0.5637 | intergenic | |
| rs34130265 | 9 | 4923110 | 4 | 0 | 0 | T | C | 2.0608 | 0.5819 | 0.0003973 | +++?+ | 0 | 0.444 | intergenic | |
| 12:41305782 | 12 | 41305782 | 3 | 0 | 0 | CT | C | 1.801 | 0.5085 | 0.0003975 | +++?? | 0 | 0.9111 | | |
| rs6895633 | 5 | 55638314 | 2 | 0 | 0 | T | C | 5.9021 | 1.6665 | 0.0003976 | +?+?? | 0 | 0.6039 | intergenic | |
| rs66942203 | 14 | 28201044 | 4 | 0 | 0 | T | C | -2.0118 | 0.568 | 0.0003976 | ---?- | 0 | 0.854 | intergenic | |
| rs145063256 | 16 | 7599784 | 4 | 0 | 0 | A | G | 3.8846 | 1.0968 | 0.0003976 | +++?+ | 0 | 0.7054 | intron | RBFOX1 |
| rs2282850 | 6 | 33267103 | 5 | 0 | 0 | T | C | 0.9783 | 0.2762 | 0.0003977 | +++++ | 0 | 0.5375 | 5-prime-UTR | RGL2 |
| 5:173159720 | 5 | 173159720 | 3 | 0 | 0 | CG | C | 1.0487 | 0.2961 | 0.0003977 | +++?? | 0 | 0.7548 | | |
| rs837990 | 9 | 110907870 | 2 | 0 | 0 | T | C | 5.6047 | 1.5826 | 0.0003978 | ++??? | 56.3 | 0.1303 | intergenic | |
| rs185387557 | 10 | 93966454 | 2 | 0 | 0 | T | C | 5.9314 | 1.6748 | 0.0003978 | +?+?? | 0 | 0.7314 | intron | CPEB3 |
| rs11632100 | 15 | 41946620 | 5 | 0 | 0 | A | G | -1.0029 | 0.2832 | 0.0003979 | ----- | 14.9 | 0.3193 | intron | MGA |
| rs4494523 | 16 | 85232249 | 4 | 0 | 0 | T | C | 1.301 | 0.3674 | 0.0003981 | +++?+ | 0 | 0.7362 | intron | GSE1 |
| rs145566024 | 6 | 63296603 | 4 | 0 | 0 | T | C | -2.4485 | 0.6914 | 0.0003981 | ---?- | 0 | 0.5615 | intergenic | |
| rs78469685 | 10 | 86716336 | 4 | 0 | 0 | A | C | -1.7927 | 0.5062 | 0.0003983 | ---?- | 0 | 0.842 | intergenic | |
| rs4357832 | 14 | 43267432 | 4 | 0 | 0 | T | C | 1.3321 | 0.3762 | 0.0003985 | +++?- | 61.3 | 0.05151 | intergenic | |
| rs2043544 | 2 | 2761141 | 4 | 0 | 0 | T | C | -1.3604 | 0.3842 | 0.0003988 | ---?- | 0 | 0.9034 | intergenic | |
| rs12233121 | 2 | 170895402 | 4 | 0 | 1 | T | C | 1.0305 | 0.291 | 0.0003988 | +++?+ | 42.7 | 0.1555 | intron | UBR3 |
| rs7990366 | 13 | 45631100 | 4 | 0 | 0 | A | G | 2.1639 | 0.6111 | 0.0003989 | +++?- | 38.5 | 0.1808 | intergenic | |
| rs4963184 | 11 | 524397 | 5 | 0 | 0 | T | C | -0.8142 | 0.23 | 0.000399 | ----- | 0 | 0.473 | upstream-gene |  |
| rs66973094 | 1 | 232837334 | 4 | 0 | 0 | A | G | 0.8867 | 0.2504 | 0.0003991 | +++?+ | 40.9 | 0.1665 | intergenic | |

| | | | | | | | | | | | | | | | |
|---|---|---|---|---|---|---|---|---|---|---|---|---|---|---|---|
| rs7137412 | 12 | 43175633 | 5 | 0 | 0 | C | G | -0.9426 | 0.2662 | 0.0003992 | ----+ | 0 | 0.4244 | intergenic |  |
| rs11513656 | 12 | 15600169 | 4 | 0 | 0 | A | G | -1.193 | 0.3369 | 0.0003993 | ---?- | 0 | 0.9321 | intron | PTPRO |
| rs7497418 | 15 | 91407994 | 4 | 0 | 0 | A | G | 0.8509 | 0.2403 | 0.0003995 | +++?- | 11.6 | 0.3349 | upstream-gene | |
| rs116448128 | 2 | 155685653 | 4 | 0 | 0 | T | C | -4.0017 | 1.1303 | 0.0003995 | ---?- | 36.1 | 0.1956 | intron | KCNJ3 |
| rs72672874 | 8 | 113291242 | 4 | 0 | 0 | A | C | -1.0074 | 0.2845 | 0.0003996 | ---?- | 0 | 0.7925 | intron | CSMD3 |
| rs72799520 | 16 | 84841172 | 4 | 0 | 0 | T | C | 1.2768 | 0.3607 | 0.0003998 | +++?- | 20.3 | 0.2882 | intergenic | |
| rs149641549 | 17 | 57147504 | 5 | 0 | 0 | T | C | 2.3536 | 0.6648 | 0.0003998 | +++++ | 0 | 0.8431 | intron | TRIM37 |
| rs113061159 | 2 | 199082794 | 4 | 0 | 0 | C | G | -1.1819 | 0.3339 | 0.0003999 | ---?+ | 40.9 | 0.1664 | intergenic | |
| rs13044967 | 20 | 669301 | 4 | 0 | 0 | A | G | -1.0346 | 0.2923 | 0.0004 | ---?- | 0 | 0.8031 | intergenic | |
| 5:148724594 | 5 | 148724594 | 3 | 0 | 0 | T | TTC | 1.5735 | 0.4445 | 0.0004002 | +++?? | 0 | 0.8052 | | |
| rs1617492 | 2 | 40472469 | 4 | 0 | 0 | T | C | -1.2018 | 0.3395 | 0.0004003 | ---?+ | 40 | 0.1716 | intron | SLC8A1,SLC8A1-AS1 |
| rs13071135 | 3 | 183556902 | 4 | 0 | 0 | A | G | -1.7236 | 0.4869 | 0.0004004 | ---?- | 0 | 0.799 | intron | PARL |
| rs12783367 | 10 | 134587334 | 4 | 0 | 0 | T | C | -0.9807 | 0.277 | 0.0004005 | ---?- | 0 | 0.4885 | intron | INPP5A |
| rs11864173 | 16 | 7176978 | 4 | 0 | 0 | C | G | 0.98 | 0.2769 | 0.0004008 | +++?+ | 11.2 | 0.3369 | intron | RBFOX1 |
| rs73080066 | 4 | 8656926 | 3 | 0 | 0 | C | G | -3.5647 | 1.0071 | 0.0004008 | ---?? | 0 | 0.8193 | intergenic | |
| rs115581902 | 2 | 3060917 | 4 | 0 | 0 | T | C | -3.0981 | 0.8753 | 0.0004008 | ---?- | 0 | 0.7093 | intergenic | |
| rs76232919 | 17 | 57162196 | 5 | 0 | 0 | A | G | -2.3948 | 0.6766 | 0.0004009 | ----- | 0 | 0.8233 | intron | TRIM37 |
| rs79394728 | 11 | 86129819 | 4 | 0 | 0 | T | G | -2.5814 | 0.7293 | 0.0004009 | ---?- | 0 | 0.5346 | intron | CCDC81 |
| rs7176052 | 15 | 85707257 | 4 | 0 | 0 | C | G | -1.6396 | 0.4632 | 0.0004011 | ---?+ | 44 | 0.1474 | intergenic | |
| rs72730588 | 15 | 75361683 | 3 | 0 | 0 | A | G | -3.2614 | 0.9215 | 0.0004011 | ---?? | 0 | 0.7123 | intergenic | |
| 2:206985954 | 2 | 206985954 | 3 | 0 | 0 | T | TC | -4.0727 | 1.1507 | 0.0004012 | ---?? | 0 | 0.8563 | | |
| rs6820060 | 4 | 169333837 | 4 | 0 | 0 | T | G | -0.8789 | 0.2483 | 0.0004014 | ---?- | 0 | 0.8772 | intron | DDX60L |
| rs12664558 | 6 | 32758677 | 5 | 0 | 0 | A | G | 1.409 | 0.3981 | 0.0004015 | +++++ | 0 | 0.6634 | intergenic | |
| rs10475003 | 5 | 14032887 | 4 | 0 | 0 | A | G | 0.926 | 0.2617 | 0.0004016 | +++?+ | 0 | 0.5544 | intergenic | |
| rs116176885 | 1 | 160491074 | 2 | 0 | 0 | A | T | -6.1311 | 1.7324 | 0.0004016 | -?-?? | 0 | 0.5913 | intron | SLAMF6 |
| rs62418256 | 6 | 63294927 | 4 | 0 | 0 | C | G | 2.4287 | 0.6863 | 0.0004017 | +++?+ | 0 | 0.5647 | intergenic | |
| rs78314191 | 6 | 63294453 | 4 | 0 | 0 | A | T | 2.4284 | 0.6862 | 0.0004017 | +++?+ | 0 | 0.5643 | intergenic | |
| rs7954424 | 12 | 43191701 | 5 | 0 | 0 | T | C | 0.9453 | 0.2671 | 0.0004018 | ++++- | 7 | 0.3667 | intergenic | |
| rs201365886 | 6 | 31981428 | 3 | 0 | 0 | A | G | 1.3521 | 0.3821 | 0.0004019 | +++?? | 34.1 | 0.2194 | upstream-gene | |
| rs57255502 | 10 | 86712757 | 4 | 0 | 0 | A | G | 1.7867 | 0.5049 | 0.000402 | +++?+ | 0 | 0.8401 | intergenic | |
| rs59279338 | 9 | 79462094 | 4 | 0 | 0 | T | C | 2.1211 | 0.5994 | 0.000402 | +++?+ | 0 | 0.6106 | intron | PRUNE2 |
| rs12033159 | 1 | 162077218 | 5 | 0 | 0 | A | G | -0.8875 | 0.2508 | 0.0004023 | ----+ | 38.6 | 0.1636 | intron | NOS1AP |
| rs57168159 | 12 | 112626845 | 5 | 0 | 0 | T | C | -1.3058 | 0.369 | 0.0004023 | -+--- | 55.1 | 0.06323 | intron | HECTD4 |
| rs78247600 | 7 | 11743554 | 3 | 0 | 0 | A | T | -4.4628 | 1.2612 | 0.0004024 | ---?? | 21.5 | 0.2795 | intron | THSD7A |
| rs114383657 | 6 | 32595636 | 5 | 0 | 1 | T | G | -1.2467 | 0.3523 | 0.0004024 | ----- | 45.2 | 0.1212 | intergenic |  |
| rs9407717 | 9 | 16190735 | 4 | 0 | 0 | T | G | 0.9674 | 0.2734 | 0.0004025 | +++?+ | 0 | 0.9304 | intergenic | |
| rs4861134 | 4 | 41740375 | 4 | 0 | 0 | T | G | -1.4975 | 0.4232 | 0.0004026 | ---?- | 23.8 | 0.2684 | intergenic | |
| rs12796054 | 11 | 37069042 | 4 | 0 | 0 | C | G | 0.8751 | 0.2473 | 0.0004026 | +++?+ | 5.2 | 0.3668 | intergenic | |
| rs145378804 | 8 | 96554215 | 4 | 0 | 0 | C | G | -1.771 | 0.5005 | 0.0004026 | -+-?- | 37 | 0.1898 | intron | LOC100616530 |
| rs13168740 | 5 | 56881455 | 4 | 0 | 0 | C | G | -0.8884 | 0.2511 | 0.0004027 | ---?- | 0 | 0.717 | intergenic | |
| rs12986533 | 2 | 139901993 | 4 | 0 | 0 | A | T | 1.3655 | 0.3859 | 0.0004028 | +++?+ | 0 | 0.9923 | intergenic | |
| rs79007940 | 2 | 129307377 | 3 | 0 | 0 | T | C | 3.0627 | 0.8656 | 0.0004028 | +++?? | 0 | 0.8696 | intergenic | |
| rs6812955 | 4 | 111407479 | 4 | 0 | 0 | T | C | 0.855 | 0.2417 | 0.0004032 | +++?+ | 0 | 0.9983 | intron | ENPEP |
| rs2331478 | 14 | 22503398 | 4 | 0 | 0 | C | G | -0.9358 | 0.2645 | 0.0004035 | ---?- | 0 | 0.9606 | intergenic | |
| rs113799213 | 21 | 36777301 | 4 | 0 | 0 | T | C | 1.6585 | 0.4688 | 0.0004036 | +++?+ | 0 | 0.5307 | intron | LOC100506403 |
| rs142966111 | 18 | 25183087 | 4 | 0 | 0 | T | C | -2.588 | 0.7315 | 0.0004036 | ---?- | 0 | 0.642 | intergenic | |
| rs146499587 | 6 | 83717622 | 4 | 0 | 0 | C | G | -3.3812 | 0.9558 | 0.0004037 | ---?- | 0 | 0.812 | intron | UBE3D |
| 14:43229277 | 14 | 43229277 | 3 | 0 | 0 | CA | C | 1.3227 | 0.3739 | 0.0004039 | +++?? | 0 | 0.7644 | | |
| 22:28725359 | 22 | 28725359 | 3 | 0 | 0 | T | TTAAA | 2.7986 | 0.7911 | 0.0004039 | +++?? | 0 | 0.7227 | | |
| rs111644677 | 6 | 83720013 | 4 | 0 | 0 | A | C | -3.3824 | 0.9562 | 0.000404 | ---?- | 0 | 0.8239 | intron | UBE3D |
| rs142671052 | 5 | 124691375 | 4 | 0 | 0 | A | G | -4.0888 | 1.1559 | 0.0004041 | ---?- | 19.5 | 0.2926 | intergenic | |
| rs73834891 | 3 | 68246368 | 4 | 0 | 0 | A | G | 2.4222 | 0.6848 | 0.0004043 | +++?+ | 21.1 | 0.2836 | intron | FAM19A1 |
| rs113473833 | 17 | 56744387 | 5 | 0 | 0 | T | C | -2.4992 | 0.7065 | 0.0004043 | ----- | 0 | 0.8337 | intron | TEX14 |
| 12:121073087 | 12 | 121073087 | 3 | 0 | 0 | CT | C | 1.2898 | 0.3646 | 0.0004043 | +++?? | 0 | 0.9264 | | |
| rs8102704 | 19 | 46384459 | 4 | 0 | 0 | T | C | -1.1069 | 0.3129 | 0.0004044 | ---?- | 0 | 0.7564 | downstream-gene | |
| rs59487183 | 14 | 28201900 | 4 | 0 | 0 | A | C | 2.0108 | 0.5685 | 0.0004044 | +++?+ | 0 | 0.8514 | intergenic | |
| rs62154212 | 2 | 107674235 | 4 | 0 | 0 | A | G | 1.0175 | 0.2877 | 0.0004045 | +++?+ | 0 | 0.5837 | intergenic | |
| rs11989521 | 8 | 113252097 | 4 | 0 | 0 | T | C | 1.0137 | 0.2866 | 0.0004046 | +++?+ | 0 | 0.949 | intron | CSMD3 |
| 6:140644758 | 6 | 140644758 | 2 | 0 | 0 | A | T | -4.9169 | 1.3902 | 0.0004048 | ?--?? | 40.6 | 0.1946 | intergenic | |

| | | | | | | | | | | | | | | | |
|---|---|---|---|---|---|---|---|---|---|---|---|---|---|---|---|
| rs10798014 | 1 | 185394834 | 4 | 0 | 0 | A | G | -0.9373 | 0.265 | 0.0004048 | ---?- | 0 | 0.9291 | intergenic | |
| rs9502950 | 6 | 1490104 | 4 | 0 | 0 | T | G | -0.8605 | 0.2433 | 0.000405 | ---?- | 0.5 | 0.3895 | intergenic | |
| rs71440146 | 13 | 85614011 | 4 | 0 | 0 | C | G | 2.5778 | 0.7289 | 0.000405 | +++?+ | 42.4 | 0.1574 | intergenic | |
| rs2331477 | 14 | 22503247 | 4 | 0 | 0 | A | T | -0.9357 | 0.2646 | 0.0004052 | ---?- | 0 | 0.9606 | intergenic | |
| rs13140461 | 4 | 41331251 | 4 | 0 | 0 | T | C | 1.4185 | 0.4011 | 0.0004052 | +++?- | 0 | 0.4352 | intergenic | |
| 16:57454611 | 16 | 57454611 | 3 | 0 | 0 | A | AAAGG | -1.1952 | 0.338 | 0.0004053 | ---?? | 6 | 0.3452 | | |
| rs148166179 | 6 | 63297326 | 4 | 0 | 0 | C | G | 2.4338 | 0.6882 | 0.0004054 | +++?+ | 0 | 0.564 | intergenic | |
| rs201530 | 7 | 101777382 | 4 | 0 | 0 | A | G | -0.8559 | 0.242 | 0.0004055 | ---?- | 0 | 0.7795 | intron | CUX1 |
| rs2889270 | 1 | 102959308 | 4 | 0 | 0 | A | G | -0.8384 | 0.2371 | 0.0004055 | ---?+ | 0 | 0.4067 | intergenic | |
| rs12716066 | 4 | 59628770 | 3 | 0 | 0 | T | C | 2.4874 | 0.7034 | 0.0004055 | +++?? | 0 | 0.8055 | intergenic | |
| rs140893623 | 17 | 57176338 | 5 | 0 | 0 | C | G | -2.4305 | 0.6873 | 0.0004055 | ----- | 0 | 0.8275 | intron | TRIM37 |
| rs189815723 | 15 | 25438001 | 3 | 0 | 0 | A | G | -3.9367 | 1.1132 | 0.0004056 | ---?? | 0 | 0.5286 | upstream-gene | |
| rs2634444 | 8 | 18438866 | 4 | 0 | 0 | A | G | -0.8638 | 0.2443 | 0.0004057 | ---?- | 0 | 0.7873 | intron | PSD3 |
| rs28716813 | 8 | 113283172 | 4 | 0 | 0 | A | G | 1.0076 | 0.2849 | 0.0004057 | +++?+ | 0 | 0.7957 | intron | CSMD3 |
| rs10459260 | 12 | 64926338 | 4 | 0 | 0 | T | C | 0.8806 | 0.249 | 0.0004058 | +++?+ | 0 | 0.9004 | intergenic | |
| rs148364501 | 17 | 56984507 | 5 | 0 | 0 | T | C | 2.4595 | 0.6955 | 0.0004058 | +++++ | 0 | 0.8005 | intron | PPM1E |
| rs12059922 | 1 | 204388180 | 4 | 0 | 0 | T | C | -1.4535 | 0.411 | 0.0004059 | ---?- | 18.9 | 0.2957 | downstream-gene | |
| rs2035332 | 11 | 37071950 | 4 | 0 | 0 | A | G | 0.8827 | 0.2496 | 0.000406 | +++?+ | 0.6 | 0.3887 | intergenic | |
| rs6918317 | 6 | 32381210 | 5 | 0 | 0 | T | C | -0.7462 | 0.211 | 0.0004061 | ----- | 0 | 0.865 | intergenic |  |
| rs62419063 | 6 | 63300972 | 4 | 0 | 0 | C | G | 2.4491 | 0.6926 | 0.0004061 | +++?+ | 0 | 0.5625 | intergenic | |
| rs12034640 | 1 | 64667959 | 3 | 0 | 0 | A | T | -0.9531 | 0.2696 | 0.0004062 | ---?? | 35 | 0.2145 | upstream-gene | |
| rs61630054 | 14 | 28202063 | 4 | 0 | 0 | A | C | -2.0105 | 0.5686 | 0.0004062 | ---?- | 0 | 0.8508 | intergenic | |
| rs78736381 | 3 | 196472700 | 4 | 0 | 0 | A | G | -2.1817 | 0.617 | 0.0004062 | ---?+ | 22 | 0.2788 | intron | PAK2 |
| rs4277845 | 4 | 111407832 | 4 | 0 | 0 | A | T | -0.8544 | 0.2416 | 0.0004064 | ---?- | 0 | 0.9985 | intron | ENPEP |
| rs3026124 | 17 | 5292033 | 5 | 0 | 0 | A | T | -2.5944 | 0.7338 | 0.0004066 | --+-- | 51.3 | 0.08402 | intron | NUP88 |
| rs7235962 | 18 | 68668966 | 4 | 0 | 0 | T | C | -1.6512 | 0.467 | 0.0004066 | ---?- | 0 | 0.9907 | intergenic | |
| rs62076828 | 18 | 5910986 | 4 | 0 | 0 | T | C | -1.2604 | 0.3565 | 0.0004066 | ---?- | 0 | 0.4251 | intergenic | |
| rs201441 | 7 | 101737327 | 4 | 0 | 0 | T | G | -0.8729 | 0.2469 | 0.0004067 | ---?- | 0 | 0.8092 | intron | CUX1 |
| rs35207157 | 11 | 13713421 | 4 | 0 | 0 | T | G | -1.1361 | 0.3213 | 0.0004067 | ---?- | 0 | 0.5843 | intron | FAR1 |
| rs59534399 | 8 | 113294853 | 4 | 0 | 0 | A | G | 1.0064 | 0.2847 | 0.0004068 | +++?+ | 0 | 0.7887 | intron | CSMD3 |
| 11:12772832 | 11 | 12772832 | 3 | 0 | 0 | CA | C | 0.9964 | 0.2818 | 0.0004068 | +++?? | 14.2 | 0.3117 | | |
| rs72754517 | 15 | 86005108 | 3 | 0 | 0 | C | G | -5.2935 | 1.4973 | 0.0004071 | --??- | 0 | 0.9507 | intron | AKAP13 |
| rs112120464 | 16 | 84842158 | 4 | 0 | 0 | A | C | 1.2872 | 0.3641 | 0.0004072 | +++?- | 0 | 0.5146 | intergenic | |
| rs201443 | 7 | 101737688 | 4 | 0 | 0 | A | T | -0.8704 | 0.2462 | 0.0004073 | ---?- | 0 | 0.8096 | intron | CUX1 |
| rs4936113 | 11 | 130622957 | 4 | 0 | 0 | A | G | 1.7196 | 0.4864 | 0.0004073 | +++?+ | 30 | 0.2322 | intergenic | |
| rs77212474 | 1 | 75287584 | 4 | 0 | 0 | A | G | 2.5251 | 0.7143 | 0.0004076 | +++?- | 23.3 | 0.271 | intergenic | |
| rs142205146 | 6 | 83717344 | 4 | 0 | 0 | A | G | -3.3781 | 0.9556 | 0.0004076 | ---?- | 0 | 0.8124 | intron | UBE3D |
| rs11654339 | 17 | 47456829 | 4 | 0 | 0 | C | G | -1.0652 | 0.3013 | 0.0004077 | ---?+ | 47.5 | 0.1266 | intergenic | |
| rs62076827 | 18 | 5909942 | 4 | 0 | 0 | T | G | -1.2609 | 0.3567 | 0.0004077 | ---?- | 0 | 0.4275 | intergenic | |
| rs11775143 | 8 | 113283680 | 4 | 0 | 0 | A | G | 1.0072 | 0.2849 | 0.0004078 | +++?+ | 0 | 0.7951 | intron | CSMD3 |
| rs13198822 | 6 | 165806518 | 4 | 0 | 0 | T | C | -0.8997 | 0.2545 | 0.0004078 | ---?- | 0 | 0.6245 | intron | PDE10A |
| rs180991008 | 17 | 9030403 | 3 | 0 | 0 | C | G | -4.6729 | 1.3219 | 0.0004078 | ---?? | 0 | 0.6015 | intron | NTN1 |
| rs11885797 | 2 | 137005737 | 4 | 0 | 0 | A | G | 0.9434 | 0.2669 | 0.0004079 | +++?+ | 0 | 0.8221 | intergenic | |
| rs16883360 | 8 | 113294967 | 4 | 0 | 0 | A | G | -1.0062 | 0.2846 | 0.0004079 | ---?- | 0 | 0.7884 | intron | CSMD3 |
| rs6711711 | 2 | 123442086 | 4 | 0 | 0 | C | G | -1.6861 | 0.477 | 0.0004081 | ---?- | 56.5 | 0.07527 | intergenic | |
| rs13106719 | 4 | 41331266 | 4 | 0 | 0 | A | G | -1.4177 | 0.4011 | 0.0004081 | ---?+ | 0 | 0.4355 | intergenic | |
| rs78186717 | 5 | 179019868 | 4 | 0 | 0 | A | G | -1.9224 | 0.5438 | 0.0004081 | ---?+ | 0 | 0.5067 | intron | RUFY1 |
| rs6139887 | 20 | 5986234 | 5 | 0 | 0 | T | C | 0.7446 | 0.2107 | 0.0004083 | ++++- | 0 | 0.425 | upstream-gene | |
| rs144669603 | 6 | 83716701 | 4 | 0 | 0 | A | G | 3.3761 | 0.9551 | 0.0004083 | +++?+ | 0 | 0.8123 | intron | UBE3D |
| 2:235346620 | 2 | 235346620 | 3 | 0 | 0 | CTT | C | 2.1961 | 0.6213 | 0.0004083 | +++?? | 0 | 0.8029 | | |
| 1:64666321 | 1 | 64666321 | 3 | 0 | 0 | T | C | 0.9553 | 0.2703 | 0.0004084 | +++?? | 4 | 0.3529 | upstream-gene | |
| rs10754290 | 1 | 90241470 | 5 | 0 | 0 | T | C | -0.7529 | 0.213 | 0.0004084 | ----- | 0 | 0.9866 | intergenic |  |
| rs990616 | 8 | 113274853 | 4 | 0 | 0 | A | G | -1.0071 | 0.2849 | 0.0004085 | ---?- | 0 | 0.8486 | intron | CSMD3 |
| rs9443316 | 6 | 77611307 | 3 | 0 | 0 | A | C | -1.1073 | 0.3133 | 0.0004086 | ---?? | 56.8 | 0.0989 | intergenic | |
| rs71494241 | 1 | 64666323 | 3 | 0 | 0 | T | C | 0.9552 | 0.2703 | 0.0004086 | +++?? | 4.1 | 0.3524 | upstream-gene | |
| rs717602 | 6 | 165807107 | 4 | 0 | 0 | T | C | 0.8913 | 0.2522 | 0.0004087 | +++?+ | 0 | 0.6148 | intron | PDE10A |
| rs9447899 | 6 | 77611300 | 3 | 0 | 0 | A | G | -1.1073 | 0.3133 | 0.0004087 | ---?? | 56.8 | 0.09881 | intergenic | |
| rs6774972 | 3 | 183617044 | 4 | 0 | 0 | A | G | -1.6868 | 0.4773 | 0.0004088 | ---?- | 0 | 0.9458 | intergenic | |
| rs9471209 | 6 | 12211354 | 4 | 0 | 0 | T | C | -3.2337 | 0.915 | 0.0004089 | ---?- | 15.1 | 0.3162 | intergenic | |

| | | | | | | | | | | | | | | | |
|---|---|---|---|---|---|---|---|---|---|---|---|---|---|---|---|
| rs9538437 | 13 | 60045413 | 4 | 0 | 0 | A | G | 0.9221 | 0.2609 | 0.0004089 | +++?+ | 33.7 | 0.2102 | intergenic | |
| rs12151330 | 19 | 53797421 | 4 | 0 | 0 | A | G | -1.2685 | 0.3589 | 0.000409 | ---?- | 1.2 | 0.386 | upstream-gene | |
| rs6682083 | 1 | 232835935 | 4 | 0 | 0 | T | C | -0.8696 | 0.246 | 0.0004091 | ---?- | 16.3 | 0.3101 | intergenic | |
| rs80190945 | 17 | 54082965 | 3 | 0 | 0 | T | G | -0.9864 | 0.2791 | 0.0004091 | ---?? | 20.1 | 0.2862 | intergenic | |
| rs72672877 | 8 | 113294856 | 4 | 0 | 0 | C | G | -1.0054 | 0.2845 | 0.0004092 | ---?- | 0 | 0.7899 | intron | CSMD3 |
| rs7314031 | 12 | 112830579 | 5 | 0 | 0 | T | G | -1.322 | 0.3741 | 0.0004093 | -+--- | 52.6 | 0.07703 | intergenic | |
| rs74519491 | 4 | 157636909 | 4 | 0 | 0 | C | G | -3.3919 | 0.9598 | 0.0004094 | ---?- | 0 | 0.8542 | intergenic | |
| rs141872975 | 6 | 63299212 | 4 | 0 | 0 | A | C | -2.4297 | 0.6875 | 0.0004096 | ---?- | 0 | 0.5651 | intergenic | |
| rs6757136 | 2 | 40448676 | 3 | 0 | 0 | A | G | -1.1148 | 0.3155 | 0.0004097 | ---?? | 0 | 0.6546 | intron | SLC8A1,SLC8A1-AS1 |
| rs150563915 | 6 | 63299214 | 4 | 0 | 0 | T | C | -2.4296 | 0.6876 | 0.0004098 | ---?- | 0 | 0.5651 | intergenic | |
| rs77854194 | 9 | 101127438 | 4 | 0 | 0 | A | G | 3.5326 | 0.9997 | 0.0004099 | +++?- | 48.7 | 0.119 | intron | GABBR2 |
| rs116244064 | 5 | 111786073 | 4 | 0 | 0 | A | G | 3.5098 | 0.9933 | 0.00041 | +++?+ | 0 | 0.5422 | intergenic | |
| rs16883359 | 8 | 113294891 | 4 | 0 | 0 | T | C | 1.0052 | 0.2845 | 0.0004101 | +++?+ | 0 | 0.7897 | intron | CSMD3 |
| rs1778011 | 1 | 87567780 | 4 | 0 | 0 | A | G | -1.0837 | 0.3067 | 0.0004102 | ---?- | 0 | 0.6576 | intron | HS2ST1 |
| rs57162756 | 14 | 28204644 | 4 | 0 | 0 | A | C | 2.0118 | 0.5694 | 0.0004103 | +++?+ | 0 | 0.8462 | intergenic | |
| rs73173316 | 3 | 172329696 | 4 | 0 | 0 | T | C | 1.9058 | 0.5394 | 0.0004103 | +++?- | 10.6 | 0.3402 | intergenic | |
| rs11786836 | 8 | 113248109 | 4 | 0 | 0 | A | G | -1.0132 | 0.2868 | 0.0004104 | ---?- | 0 | 0.9507 | intron | CSMD3 |
| rs191336922 | 2 | 118076827 | 3 | 0 | 0 | A | G | -3.0602 | 0.8661 | 0.0004105 | ---?? | 42.6 | 0.1752 | intergenic | |
| rs7150901 | 14 | 22571660 | 4 | 0 | 0 | T | G | 0.9047 | 0.2561 | 0.0004106 | +++?+ | 43.6 | 0.1499 | intergenic | |
| 19:36562104 | 19 | 36562104 | 3 | 0 | 0 | CAAAAA | C | -1.5917 | 0.4505 | 0.0004106 | -+-?? | 61.8 | 0.07277 | | |
| rs55899575 | 1 | 102915151 | 4 | 0 | 0 | A | G | -0.8705 | 0.2464 | 0.0004107 | ---?+ | 0 | 0.4175 | intergenic | |
| rs144891972 | 2 | 199066822 | 3 | 0 | 0 | A | G | -1.2071 | 0.3417 | 0.0004107 | ---?? | 47.7 | 0.1478 | intergenic | |
| rs9837287 | 3 | 29526264 | 4 | 0 | 0 | T | C | 1.4356 | 0.4063 | 0.0004108 | +++?- | 0 | 0.6977 | intron | RBMS3 |
| rs72625292 | 2 | 185441009 | 4 | 0 | 0 | A | G | 0.9679 | 0.274 | 0.0004108 | +++?+ | 0 | 0.7522 | intergenic | |
| rs17039980 | 4 | 110076212 | 5 | 0 | 0 | A | G | -1.4425 | 0.4083 | 0.0004109 | ----- | 0 | 0.5643 | intron | COL25A1 |
| rs114278864 | 3 | 196496401 | 4 | 0 | 0 | A | G | 2.1756 | 0.6158 | 0.0004112 | +++?- | 21.4 | 0.2818 | intron | PAK2 |
| rs7753825 | 6 | 168621944 | 4 | 0 | 0 | T | C | -0.932 | 0.2638 | 0.0004115 | ---?- | 0 | 0.8223 | intergenic | |
| rs7782033 | 7 | 107457403 | 4 | 0 | 0 | T | G | -1.0951 | 0.31 | 0.0004117 | ---?- | 7.3 | 0.3564 | intergenic | |
| rs10150077 | 14 | 43891689 | 4 | 0 | 0 | A | C | 1.9232 | 0.5444 | 0.0004117 | +++?+ | 0 | 0.8403 | intergenic | |
| rs34566568 | 5 | 83025710 | 4 | 0 | 0 | A | C | 1.3684 | 0.3874 | 0.0004117 | +++?- | 0 | 0.6172 | intergenic | |
| rs74938865 | 20 | 21517472 | 4 | 0 | 0 | T | G | -2.4605 | 0.6966 | 0.0004121 | ---?- | 21.6 | 0.2811 | intergenic | |
| rs185302143 | 1 | 63702267 | 2 | 0 | 0 | T | C | 4.7221 | 1.3369 | 0.0004121 | +?+?? | 43.3 | 0.1843 | intron | LINC00466 |
| rs16883753 | 8 | 113501308 | 4 | 0 | 0 | T | C | 1.0159 | 0.2876 | 0.0004122 | +++?+ | 0 | 0.6908 | intron | CSMD3 |
| rs79382248 | 3 | 196481901 | 4 | 0 | 0 | T | C | 2.1965 | 0.6219 | 0.0004122 | +++?- | 23.3 | 0.2713 | intron | PAK2 |
| rs11919061 | 3 | 183617617 | 4 | 0 | 0 | T | C | -1.6866 | 0.4775 | 0.0004123 | ---?- | 0 | 0.9455 | intergenic | |
| rs116106466 | 4 | 117282174 | 4 | 0 | 0 | A | T | -3.2062 | 0.9078 | 0.0004124 | ---?- | 0 | 0.7948 | intergenic | |
| rs4863098 | 4 | 190018899 | 4 | 0 | 0 | T | C | -0.8845 | 0.2504 | 0.0004125 | ---?- | 0 | 0.5518 | intergenic | |
| rs117055607 | 15 | 65830876 | 3 | 0 | 0 | A | G | 2.9691 | 0.8406 | 0.0004125 | +++?? | 12.1 | 0.3205 | intron | PTPLAD1 |
| rs3799168 | 6 | 165805454 | 4 | 0 | 0 | A | G | -0.9017 | 0.2553 | 0.0004127 | ---?- | 0 | 0.6341 | intron | PDE10A |
| rs71612099 | 3 | 31710915 | 2 | 0 | 0 | A | G | -5.263 | 1.4902 | 0.0004127 | ?--?? | 0 | 0.9353 | intron | OSBPL10 |
| rs184460437 | 8 | 92904692 | 3 | 0 | 0 | A | G | 3.6416 | 1.0311 | 0.0004127 | +++?? | 0 | 0.8976 | intergenic | |
| rs72763573 | 16 | 7178205 | 4 | 0 | 0 | T | G | -0.9819 | 0.278 | 0.0004128 | ---?- | 13.1 | 0.3272 | intron | RBFOX1 |
| rs115203376 | 17 | 54080344 | 3 | 0 | 0 | A | G | -0.9841 | 0.2786 | 0.0004128 | ---?? | 24.9 | 0.2639 | intergenic | |
| rs117607915 | 4 | 89046617 | 4 | 0 | 0 | A | G | -3.945 | 1.117 | 0.000413 | -?--- | 0 | 0.6 | intron | ABCG2 |
| rs2141141 | 15 | 41969474 | 5 | 0 | 0 | C | G | 1.0043 | 0.2844 | 0.0004131 | +++++ | 11.8 | 0.3384 | intron | MGA |
| rs1355457 | 6 | 23871607 | 4 | 0 | 0 | C | G | 2.5349 | 0.7178 | 0.0004133 | +++?- | 52.2 | 0.09887 | intergenic | |
| rs4711173 | 6 | 28514583 | 5 | 0 | 0 | T | C | -0.9364 | 0.2652 | 0.0004133 | ----- | 0 | 0.858 | intergenic |  |
| rs303790 | 12 | 52230194 | 4 | 0 | 0 | T | C | 3.4648 | 0.9812 | 0.0004134 | +++?- | 26.8 | 0.2512 | intron | FIGNL2 |
| rs12036268 | 1 | 162075736 | 5 | 0 | 0 | A | G | 0.8872 | 0.2512 | 0.0004134 | ++++- | 38.8 | 0.1626 | intron | NOS1AP |
| rs7296050 | 12 | 112811783 | 5 | 0 | 0 | A | G | -1.3196 | 0.3737 | 0.0004138 | -+--- | 52.9 | 0.07488 | intron | HECTD4 |
| 19:36571206 | 19 | 36571206 | 3 | 0 | 0 | CG | C | -1.5696 | 0.4445 | 0.0004138 | -+-?? | 62.1 | 0.07138 | | |
| rs201445 | 7 | 101737921 | 4 | 0 | 0 | T | G | 0.8689 | 0.2461 | 0.0004139 | +++?+ | 0 | 0.8085 | intron | CUX1 |
| rs12767469 | 10 | 134587261 | 4 | 0 | 0 | C | G | -0.9793 | 0.2773 | 0.0004139 | ---?- | 0 | 0.4872 | intron | INPP5A |
| 5:124741463 | 5 | 124741463 | 3 | 0 | 0 | T | TC | -1.3327 | 0.3774 | 0.0004139 | ---?? | 26.8 | 0.2553 | | |
| rs28551113 | 12 | 112814745 | 5 | 0 | 0 | A | G | -1.3198 | 0.3738 | 0.000414 | -+--- | 52.9 | 0.07536 | intron | HECTD4 |
| rs11848869 | 14 | 21335266 | 3 | 0 | 0 | T | C | 4.4378 | 1.2568 | 0.0004142 | +++?? | 0 | 0.6062 | intergenic | |
| rs825208 | 1 | 64771723 | 4 | 0 | 0 | A | T | 0.9813 | 0.2779 | 0.0004143 | +++?+ | 0 | 0.4341 | intergenic | |
| rs55691798 | 8 | 113273852 | 4 | 0 | 0 | C | G | 1.0062 | 0.285 | 0.0004146 | +++?+ | 0 | 0.8519 | intron | CSMD3 |
| rs78052154 | 4 | 117279368 | 4 | 0 | 0 | T | C | 3.7492 | 1.0619 | 0.0004146 | +++?+ | 58.6 | 0.06433 | intergenic | |

| | | | | | | | | | | | | | | | |
|---|---|---|---|---|---|---|---|---|---|---|---|---|---|---|---|
| rs113108104 | 16 | 58416479 | 4 | 0 | 0 | A | G | 1.6967 | 0.4806 | 0.0004146 | +++?- | 0 | 0.4757 | intergenic | |
| rs950983 | 8 | 113255323 | 4 | 0 | 0 | A | T | 1.0123 | 0.2867 | 0.0004148 | +++?+ | 0 | 0.9459 | intron | CSMD3 |
| rs11928605 | 3 | 183591014 | 4 | 0 | 0 | T | C | -1.6693 | 0.4728 | 0.0004149 | ---?- | 0 | 0.8971 | intron | PARL |
| 17:56956225 | 17 | 56956225 | 5 | 0 | 0 | T | C | -2.4683 | 0.6992 | 0.000415 | ----- | 0 | 0.8152 | intron | PPM1E |
| rs73161596 | 22 | 39526303 | 4 | 0 | 0 | A | G | -2.8906 | 0.8188 | 0.000415 | ---?- | 26.8 | 0.2508 | downstream-gene | |
| rs60258199 | 12 | 112785582 | 5 | 0 | 0 | T | C | -1.3152 | 0.3725 | 0.0004152 | -+--- | 53.6 | 0.07135 | intron | HECTD4 |
| rs201112889 | 5 | 124741465 | 3 | 0 | 0 | C | G | 1.3325 | 0.3775 | 0.0004152 | +++?? | 26.7 | 0.2555 | intergenic | |
| rs57617581 | 12 | 112793560 | 5 | 0 | 0 | T | C | -1.3166 | 0.3729 | 0.0004153 | -+--- | 53.4 | 0.07228 | intron | HECTD4 |
| 18:55810067 | 18 | 55810067 | 3 | 0 | 0 | A | AT | 1.0878 | 0.3081 | 0.0004153 | +++?? | 0 | 0.8108 | | |
| rs1899821 | 15 | 41956695 | 5 | 0 | 0 | T | G | 1.0025 | 0.284 | 0.0004154 | +++++ | 14.6 | 0.321 | intron | MGA |
| rs885869 | 6 | 12208093 | 4 | 0 | 0 | A | G | -3.2733 | 0.9273 | 0.0004156 | ---?- | 4.8 | 0.369 | intergenic | |
| 17:51152012 | 17 | 51152012 | 3 | 0 | 0 | A | ATGATT | 1.4616 | 0.4141 | 0.0004156 | +++?? | 0 | 0.4191 | | |
| rs323249 | 6 | 77672401 | 3 | 0 | 0 | A | G | 1.0068 | 0.2852 | 0.0004159 | +++?? | 13.7 | 0.3138 | intergenic | |
| rs7323592 | 13 | 45530194 | 3 | 0 | 0 | T | C | -2.3046 | 0.6529 | 0.0004159 | ---?? | 0 | 0.8004 | intron | NUFIP1 |
| rs112320667 | 6 | 83722318 | 4 | 0 | 0 | A | G | 3.3785 | 0.9571 | 0.0004159 | +++?+ | 0 | 0.8152 | intron | UBE3D |
| rs9912393 | 17 | 6695656 | 3 | 0 | 0 | C | G | 3.7185 | 1.0535 | 0.0004161 | +++?? | 0 | 0.5217 | downstream-gene | |
| rs17036313 | 2 | 47385622 | 4 | 0 | 0 | T | C | 1.3762 | 0.3899 | 0.0004161 | +++?+ | 0 | 0.9647 | upstream-gene | |
| rs142794369 | 6 | 63301322 | 4 | 0 | 0 | C | G | -2.4328 | 0.6893 | 0.0004163 | ---?- | 0 | 0.5649 | intergenic | |
| rs7111111 | 11 | 131486558 | 4 | 0 | 0 | C | G | -0.8595 | 0.2435 | 0.0004164 | ---?- | 0 | 0.7567 | intron | NTM |
| rs10035746 | 5 | 165420160 | 4 | 0 | 0 | T | C | -0.8771 | 0.2485 | 0.0004164 | ---?- | 0 | 0.5497 | intergenic | |
| rs10516635 | 4 | 118079907 | 4 | 0 | 0 | A | G | 1.2547 | 0.3555 | 0.0004164 | +++?+ | 0 | 0.9815 | intergenic | |
| rs60569935 | 12 | 112755835 | 5 | 0 | 0 | T | C | 1.3096 | 0.371 | 0.0004164 | +-+++ | 53.6 | 0.07119 | intron | HECTD4 |
| rs183574507 | 3 | 59807483 | 3 | 0 | 0 | A | T | 1.4292 | 0.4049 | 0.0004164 | +++?? | 0 | 0.8433 | intron | FHIT |
| rs817434 | 1 | 85637979 | 5 | 0 | 0 | A | G | -1.3578 | 0.3847 | 0.0004165 | ----- | 0 | 0.9674 | intron | SYDE2 |
| rs6887526 | 5 | 149543812 | 4 | 0 | 0 | T | C | -0.8444 | 0.2392 | 0.0004166 | ---?- | 0 | 0.9172 | upstream-gene | |
| rs11488192 | 1 | 64666836 | 3 | 0 | 0 | T | C | -0.9394 | 0.2662 | 0.0004166 | ---?? | 28.7 | 0.2461 | upstream-gene | |
| rs11635205 | 15 | 23688555 | 2 | 0 | 0 | A | G | -6.615 | 1.8743 | 0.0004166 | ?--?? | 0 | 0.7854 | intron | GOLGA6L2 |
| rs117327491 | 17 | 56989329 | 5 | 0 | 0 | T | G | 2.4513 | 0.6945 | 0.0004166 | +++++ | 0 | 0.8005 | intron | PPM1E |
| rs11017093 | 10 | 131926121 | 4 | 0 | 0 | T | C | 0.9157 | 0.2595 | 0.0004168 | +++?+ | 0 | 0.8333 | intergenic | |
| rs76715371 | 12 | 27789261 | 3 | 0 | 0 | T | C | 0.9798 | 0.2776 | 0.0004168 | +++?? | 24.4 | 0.2662 | intron | PPFIBP1 |
| rs4772419 | 13 | 102538018 | 4 | 0 | 0 | T | C | 1.0025 | 0.2841 | 0.000417 | +++?+ | 2.3 | 0.381 | intron | FGF14 |
| rs12036529 | 1 | 162070706 | 5 | 0 | 0 | A | T | 0.8907 | 0.2524 | 0.0004173 | ++++- | 40 | 0.1547 | intron | NOS1AP |
| rs16883320 | 8 | 113274464 | 4 | 0 | 0 | A | C | 1.0056 | 0.285 | 0.0004173 | +++?+ | 0 | 0.851 | intron | CSMD3 |
| rs73971638 | 2 | 170443035 | 4 | 0 | 0 | A | C | 1.2519 | 0.3548 | 0.0004173 | +++?+ | 0 | 0.9288 | intron | PPIG |
| rs7963631 | 12 | 43187465 | 5 | 0 | 0 | C | G | -0.9399 | 0.2664 | 0.0004174 | ----+ | 21.3 | 0.2786 | intergenic |  |
| rs66982543 | 3 | 142414603 | 2 | 0 | 0 | T | C | 1.4354 | 0.4068 | 0.0004174 | ???++ | 0 | 0.8765 | intron | PLS1 |
| rs6489839 | 12 | 112791373 | 5 | 0 | 0 | T | G | 1.3156 | 0.3728 | 0.0004176 | +-+++ | 53.4 | 0.07259 | intron | HECTD4 |
| rs7969739 | 12 | 27787244 | 4 | 0 | 0 | C | G | -0.8926 | 0.253 | 0.0004179 | ---?+ | 52.6 | 0.0969 | intron | PPFIBP1 |
| rs16915236 | 9 | 101126718 | 4 | 0 | 0 | T | C | -3.522 | 0.9982 | 0.0004179 | ---?+ | 48.3 | 0.1216 | intron | GABBR2 |
| rs6746027 | 2 | 199087090 | 4 | 0 | 0 | T | C | 1.1741 | 0.3328 | 0.000418 | +++?- | 41.1 | 0.1649 | intergenic | |
| rs13206337 | 6 | 165807717 | 4 | 0 | 0 | T | C | -0.8906 | 0.2524 | 0.000418 | ---?- | 0 | 0.6113 | intron | PDE10A |
| rs10148796 | 14 | 43268288 | 4 | 0 | 0 | T | C | 1.325 | 0.3755 | 0.0004182 | +++?- | 61.2 | 0.05182 | intergenic | |
| rs77186014 | 6 | 63302678 | 4 | 0 | 0 | T | C | 2.4293 | 0.6885 | 0.0004182 | +++?+ | 0 | 0.5661 | intergenic | |
| rs7654060 | 4 | 190020998 | 4 | 0 | 0 | A | G | 0.881 | 0.2497 | 0.0004184 | +++?+ | 0 | 0.5485 | intergenic | |
| 8:114886273 | 8 | 114886273 | 3 | 0 | 0 | T | TCCTACT | -2.0036 | 0.5679 | 0.0004184 | ---?? | 0 | 0.4672 | | |
| rs12039094 | 1 | 185391626 | 4 | 0 | 0 | T | G | -0.9371 | 0.2656 | 0.0004185 | ---?- | 0 | 0.9166 | intergenic | |
| rs12606859 | 18 | 40443383 | 3 | 0 | 0 | A | G | -4.7885 | 1.3572 | 0.0004185 | ---?? | 0 | 0.4829 | intron | RIT2 |
| rs1674175 | 1 | 85654128 | 5 | 0 | 0 | C | G | -1.371 | 0.3886 | 0.0004186 | ----- | 0 | 0.9748 | intron | SYDE2 |
| rs7993781 | 13 | 60045118 | 4 | 0 | 0 | T | C | 0.9315 | 0.264 | 0.0004186 | +++?+ | 0 | 0.574 | intergenic | |
| 19:46161130 | 19 | 46161130 | 3 | 0 | 0 | A | AC | 0.9451 | 0.2679 | 0.0004186 | +++?? | 0 | 0.6356 | | |
| rs6697463 | 1 | 187858115 | 4 | 0 | 0 | T | C | -1.177 | 0.3336 | 0.0004187 | ---?- | 0 | 0.8799 | intergenic | |
| rs28744288 | 6 | 32774318 | 5 | 0 | 0 | A | G | 1.3973 | 0.3961 | 0.0004189 | +++++ | 0 | 0.6656 | intergenic | |
| rs11658873 | 17 | 56423750 | 5 | 0 | 0 | C | G | -1.1304 | 0.3204 | 0.0004191 | ----- | 33.6 | 0.1971 | intron | SUPT4H1,BZRAP1-AS1 |
| rs3829312 | 12 | 27795866 | 4 | 0 | 0 | A | G | 0.9602 | 0.2722 | 0.0004192 | +-+?- | 58.2 | 0.06646 | intron | PPFIBP1 |
| rs9276729 | 6 | 32764246 | 5 | 0 | 0 | A | G | 1.3969 | 0.396 | 0.0004193 | +++++ | 0 | 0.6653 | intergenic | |
| rs13212256 | 6 | 32768171 | 5 | 0 | 0 | A | T | 1.3971 | 0.396 | 0.0004193 | +++++ | 0 | 0.6654 | intergenic | |
| rs76854676 | 6 | 63303193 | 4 | 0 | 0 | A | G | 2.4292 | 0.6886 | 0.0004193 | +++?+ | 0 | 0.566 | intergenic | |
| rs4484958 | 1 | 64667314 | 3 | 0 | 0 | T | C | -0.9423 | 0.2671 | 0.0004194 | ---?? | 30.5 | 0.2373 | upstream-gene | |
| rs9276723 | 6 | 32763405 | 5 | 0 | 0 | A | G | -1.3969 | 0.396 | 0.0004194 | ----- | 0 | 0.6653 | intergenic |  |

| | | | | | | | | | | | | | | | |
|---|---|---|---|---|---|---|---|---|---|---|---|---|---|---|---|
| rs9276724 | 6 | 32763635 | 5 | 0 | 0 | T | C | 1.3969 | 0.396 | 0.0004194 | +++++ | 0 | 0.6653 | intergenic | |
| rs9276728 | 6 | 32764042 | 5 | 0 | 0 | T | C | 1.3969 | 0.396 | 0.0004194 | +++++ | 0 | 0.6653 | intergenic | |
| rs9276722 | 6 | 32762391 | 5 | 0 | 0 | A | G | -1.3969 | 0.396 | 0.0004195 | ----- | 0 | 0.6653 | intergenic |  |
| rs9276725 | 6 | 32763818 | 5 | 0 | 0 | T | C | -1.3969 | 0.396 | 0.0004195 | ----- | 0 | 0.6653 | intergenic |  |
| rs28361064 | 6 | 32783295 | 5 | 0 | 0 | T | G | -1.3969 | 0.396 | 0.0004195 | ----- | 0 | 0.6653 | intron | HLA-DOB |
| rs35888696 | 8 | 65024549 | 4 | 0 | 0 | T | C | 0.8701 | 0.2466 | 0.0004195 | +++?+ | 0 | 0.9476 | intergenic | |
| rs2042483 | 2 | 208365344 | 4 | 0 | 0 | A | G | -1.4952 | 0.4239 | 0.0004196 | ---?- | 0 | 0.5993 | intergenic | |
| rs78690211 | 3 | 134815187 | 4 | 0 | 0 | A | T | 1.5417 | 0.4371 | 0.0004198 | +++?+ | 43.9 | 0.1479 | intron | EPHB1 |
| 1:90270016 | 1 | 90270016 | 3 | 0 | 0 | CA | C | -0.8962 | 0.2541 | 0.0004199 | ---?? | 8.7 | 0.3346 | | |
| rs1019351 | 8 | 113296811 | 4 | 0 | 0 | A | T | 1.0039 | 0.2846 | 0.00042 | +++?+ | 0 | 0.7854 | intron | CSMD3 |
| rs11048650 | 12 | 26841194 | 4 | 0 | 0 | C | G | -1.747 | 0.4953 | 0.00042 | -+-?- | 38.6 | 0.1804 | intron | ITPR2 |
| rs12094596 | 1 | 172848303 | 3 | 0 | 0 | A | G | 0.9054 | 0.2567 | 0.0004201 | +++?? | 0 | 0.4019 | intergenic | |
| rs13438706 | 7 | 67409478 | 4 | 0 | 0 | A | T | -1.8559 | 0.5262 | 0.0004202 | ---?+ | 0 | 0.4177 | intergenic | |
| rs10916411 | 1 | 229169328 | 4 | 0 | 0 | A | G | 1.3642 | 0.3868 | 0.0004203 | +++?+ | 0 | 0.9598 | intergenic | |
| rs966935 | 10 | 4419830 | 4 | 0 | 0 | A | G | 0.9684 | 0.2746 | 0.0004204 | +++?+ | 3.5 | 0.375 | intergenic | |
| rs201442 | 7 | 101737627 | 4 | 0 | 0 | A | G | -0.8699 | 0.2467 | 0.0004207 | ---?- | 0 | 0.8071 | intron | CUX1 |
| rs888254 | 8 | 113275266 | 4 | 0 | 0 | C | G | -1.005 | 0.285 | 0.0004207 | ---?- | 0 | 0.85 | intron | CSMD3 |
| rs9276721 | 6 | 32762309 | 5 | 0 | 0 | T | C | 1.3966 | 0.396 | 0.0004208 | +++++ | 0 | 0.665 | intergenic | |
| rs147405800 | 5 | 158412406 | 4 | 0 | 0 | T | G | 3.5263 | 0.9999 | 0.0004209 | +++?+ | 12.2 | 0.3317 | intron | EBF1 |
| rs1547179 | 10 | 4421518 | 4 | 0 | 0 | T | G | -0.9358 | 0.2654 | 0.0004211 | ---?- | 3.9 | 0.3733 | intergenic | |
| rs303791 | 12 | 52229859 | 4 | 0 | 0 | T | C | 3.4637 | 0.9823 | 0.0004214 | +++?- | 26.7 | 0.2517 | intron | FIGNL2 |
| rs7969216 | 12 | 27787114 | 4 | 0 | 0 | A | G | 0.892 | 0.2529 | 0.0004214 | +++?- | 52.7 | 0.0961 | intron | PPFIBP1 |
| rs35454635 | 8 | 65024997 | 4 | 0 | 0 | A | C | -0.8696 | 0.2466 | 0.0004215 | ---?- | 0 | 0.9477 | intergenic | |
| rs12660559 | 6 | 32758774 | 5 | 0 | 0 | T | C | -1.4033 | 0.398 | 0.0004219 | ----- | 0 | 0.6576 | intergenic |  |
| rs139366813 | 17 | 57139319 | 5 | 0 | 0 | A | T | -2.331 | 0.6611 | 0.0004219 | ----- | 0 | 0.842 | intron | TRIM37 |
| rs149197511 | 17 | 56959910 | 5 | 0 | 0 | T | C | -2.4629 | 0.6985 | 0.0004219 | ----- | 0 | 0.8148 | intron | PPM1E |
| rs73160523 | 13 | 22348058 | 4 | 0 | 0 | A | T | -2.9877 | 0.8474 | 0.0004224 | ---?- | 0 | 0.4765 | intergenic | |
| rs451367 | 19 | 16412842 | 4 | 0 | 0 | T | C | 1.1014 | 0.3124 | 0.0004225 | +++?+ | 43 | 0.1533 | intergenic | |
| rs10490594 | 2 | 208357762 | 4 | 0 | 0 | A | G | 1.461 | 0.4144 | 0.0004225 | +++?+ | 0 | 0.4108 | intergenic | |
| rs368897 | 5 | 746657 | 3 | 0 | 0 | A | G | 1.6055 | 0.4554 | 0.0004226 | +++?? | 0 | 0.5872 | intron | ZDHHC11B |
| rs201491353 | 11 | 85733815 | 3 | 0 | 0 | T | C | -4.6387 | 1.3158 | 0.0004227 | ---?? | 0 | 0.5542 | intron | PICALM |
| rs118059879 | 18 | 25186115 | 4 | 0 | 0 | A | G | -2.5703 | 0.7291 | 0.0004228 | ---?- | 0 | 0.6467 | intergenic | |
| 2:206961362 | 2 | 206961362 | 3 | 0 | 0 | CA | C | -3.9897 | 1.1317 | 0.0004228 | ---?? | 0 | 0.8002 | | |
| rs355570 | 2 | 142418428 | 4 | 0 | 0 | T | C | 1.9696 | 0.5587 | 0.0004229 | +++?+ | 0 | 0.9219 | intron | LRP1B |
| 1:117947973 | 1 | 117947973 | 3 | 0 | 0 | A | AT | 2.2374 | 0.6347 | 0.0004229 | +++?? | 0 | 0.6207 | | |
| rs75930837 | 2 | 170542248 | 3 | 0 | 0 | T | C | -1.3001 | 0.3688 | 0.000423 | ---?? | 0 | 0.9083 | intron | CCDC173 |
| rs11781364 | 8 | 113285886 | 4 | 0 | 0 | T | C | -1.0044 | 0.2849 | 0.0004231 | ---?- | 0 | 0.7848 | intron | CSMD3 |
| rs1890177 | 10 | 134491348 | 4 | 0 | 0 | A | C | -0.8569 | 0.2431 | 0.0004232 | ---?- | 0 | 0.7733 | intron | INPP5A |
| rs7897273 | 10 | 123504920 | 4 | 0 | 0 | T | C | 0.8633 | 0.2449 | 0.0004232 | +++?+ | 0 | 0.8025 | intron | ATE1 |
| 14:95896800 | 14 | 95896800 | 3 | 0 | 0 | CT | C | 1.4318 | 0.4062 | 0.0004232 | +++?? | 42.1 | 0.1776 | | |
| rs734476 | 2 | 2913914 | 4 | 0 | 0 | C | G | -2.6264 | 0.7451 | 0.0004233 | ---?- | 43.4 | 0.151 | intergenic | |
| rs35355256 | 1 | 229171692 | 4 | 0 | 0 | T | C | -1.3644 | 0.387 | 0.0004233 | ---?- | 0 | 0.9579 | intergenic | |
| rs145324030 | 17 | 57140015 | 5 | 0 | 0 | A | G | -2.3298 | 0.6609 | 0.0004233 | ----- | 0 | 0.8425 | intron-near-splice | TRIM37 |
| rs7540690 | 1 | 162114217 | 5 | 0 | 0 | A | G | -0.8979 | 0.2547 | 0.0004236 | ----+ | 37.9 | 0.1687 | intron | NOS1AP |
| rs10461311 | 4 | 190021478 | 4 | 0 | 0 | A | T | 0.8798 | 0.2496 | 0.0004236 | +++?+ | 0 | 0.5487 | intergenic | |
| rs75251303 | 4 | 4753308 | 4 | 0 | 0 | A | G | 2.4314 | 0.6898 | 0.0004236 | +++?+ | 0 | 0.5277 | intergenic | |
| rs74009292 | 16 | 14093598 | 4 | 0 | 0 | T | C | 2.1364 | 0.6061 | 0.0004238 | +++?+ | 15.2 | 0.3156 | intergenic | |
| rs11928606 | 3 | 183591031 | 4 | 0 | 0 | T | C | -1.665 | 0.4724 | 0.000424 | ---?- | 0 | 0.8653 | intron | PARL |
| rs11964375 | 6 | 109110541 | 2 | 0 | 0 | A | G | -4.8666 | 1.3807 | 0.000424 | -?-?? | 0 | 0.9496 | intergenic | |
| rs193291442 | 17 | 56959951 | 5 | 0 | 0 | T | C | -2.4468 | 0.6942 | 0.000424 | ----- | 0 | 0.8262 | intron | PPM1E |
| 2:170570322 | 2 | 170570322 | 3 | 0 | 0 | T | TTTTG | -1.3092 | 0.3714 | 0.0004241 | ---?? | 0 | 0.8148 | | |
| rs144807019 | 7 | 114299758 | 3 | 0 | 0 | C | G | 3.98 | 1.1293 | 0.0004244 | +++?? | 0 | 0.5973 | intron | FOXP2 |
| rs78779116 | 4 | 63517115 | 4 | 0 | 0 | T | C | -2.993 | 0.8492 | 0.0004245 | ---?- | 13.9 | 0.323 | intergenic | |
| rs10775123 | 15 | 41953420 | 5 | 0 | 0 | A | T | 1.0002 | 0.2838 | 0.0004246 | +++++ | 14.7 | 0.3205 | intron | MGA |
| rs10775124 | 15 | 41953702 | 5 | 0 | 0 | T | C | 1.0003 | 0.2838 | 0.0004248 | +++++ | 14.8 | 0.3201 | intron | MGA |
| rs185153112 | 17 | 56959952 | 5 | 0 | 0 | A | G | 2.4463 | 0.6942 | 0.0004248 | +++++ | 0 | 0.8263 | intron | PPM1E |
| rs6114326 | 20 | 23815299 | 4 | 0 | 0 | T | C | 1.469 | 0.4169 | 0.000425 | +++?+ | 0 | 0.5543 | intergenic | |
| rs79688075 | 1 | 230948206 | 4 | 0 | 0 | T | C | -1.8688 | 0.5303 | 0.0004252 | ---?- | 0 | 0.5069 | intergenic | |
| rs1021438 | 8 | 113502325 | 4 | 0 | 0 | A | T | 1.0148 | 0.288 | 0.0004253 | +++?+ | 0 | 0.6826 | intron | CSMD3 |

| | | | | | | | | | | | | | | |
|---|---|---|---|---|---|---|---|---|---|---|---|---|---|---|
| 20:16323229 | 20 | 16323229 | 3 | 0 | 0 | T | TA | 3.2273 | 0.9158 | 0.0004253 | +++?? | 0 | 0.8254 | |
| rs74009291 | 16 | 14093584 | 4 | 0 | 0 | T | C | -2.1189 | 0.6013 | 0.0004254 | ---?- | 19.1 | 0.2948 | intergenic | |
| rs79873431 | 5 | 132991927 | 4 | 0 | 0 | T | C | 3.0374 | 0.862 | 0.0004254 | +++?+ | 0 | 0.7656 | intergenic | |
| rs12508695 | 4 | 110109456 | 5 | 0 | 0 | T | G | 1.4412 | 0.409 | 0.0004255 | +++++ | 0 | 0.6175 | intron | COL25A1 |
| rs78620996 | 8 | 86865022 | 4 | 0 | 0 | T | C | -1.9612 | 0.5566 | 0.0004255 | ---?- | 0 | 0.6318 | intergenic | |
| 4:26165742 | 4 | 26165742 | 3 | 0 | 0 | T | TAC | 1.3242 | 0.3758 | 0.0004255 | +++?? | 0 | 0.5329 | | |
| rs775194 | 11 | 109440253 | 3 | 0 | 0 | A | T | -1.0148 | 0.288 | 0.0004256 | -+-?? | 59.1 | 0.08661 | intergenic | |
| rs10461312 | 4 | 190021682 | 4 | 0 | 0 | A | T | -0.8794 | 0.2496 | 0.0004256 | ---?- | 0 | 0.5485 | intergenic | |
| 9:30897223 | 9 | 30897223 | 3 | 0 | 0 | A | AT | 3.3051 | 0.938 | 0.0004256 | +++?? | 0 | 0.5109 | | |
| rs113737480 | 18 | 25174486 | 4 | 0 | 0 | T | C | -2.6076 | 0.74 | 0.0004257 | ---?- | 0 | 0.6463 | intergenic | |
| rs6982539 | 8 | 113285772 | 4 | 0 | 0 | T | G | 1.0037 | 0.2848 | 0.0004258 | +++?+ | 0 | 0.7883 | intron | CSMD3 |
| rs115677860 | 17 | 56947727 | 5 | 0 | 0 | T | G | -2.4499 | 0.6953 | 0.0004258 | ----- | 0 | 0.8314 | intron | PPM1E |
| rs7155956 | 14 | 43268599 | 4 | 0 | 0 | A | C | 1.3223 | 0.3753 | 0.0004259 | +++?- | 61.2 | 0.05193 | intergenic | |
| rs117348664 | 8 | 21483268 | 3 | 0 | 0 | A | G | 3.3061 | 0.9383 | 0.0004259 | +++?? | 0 | 0.6857 | intergenic | |
| rs28527338 | 19 | 56222998 | 4 | 0 | 0 | T | C | -1.2139 | 0.3445 | 0.000426 | ---?- | 0 | 0.7644 | intron | NLRP9 |
| rs10904241 | 10 | 4419524 | 4 | 0 | 0 | A | G | -0.9678 | 0.2747 | 0.0004261 | ---?- | 5.2 | 0.3667 | intergenic | |
| rs2111426 | 8 | 113298153 | 4 | 0 | 0 | A | T | 1.0027 | 0.2846 | 0.0004262 | +++?+ | 0 | 0.7843 | intron | CSMD3 |
| 2:170544370 | 2 | 170544370 | 3 | 0 | 0 | CTGCCT( | C | 1.2999 | 0.3689 | 0.0004262 | +++?? | 0 | 0.9068 | | |
| rs1019350 | 8 | 113297131 | 4 | 0 | 0 | T | G | -1.0028 | 0.2846 | 0.0004263 | ---?- | 0 | 0.7843 | intron | CSMD3 |
| rs2111427 | 8 | 113298122 | 4 | 0 | 0 | A | G | 1.0027 | 0.2846 | 0.0004264 | +++?+ | 0 | 0.7841 | intron | CSMD3 |
| 7:38145351 | 7 | 38145351 | 3 | 0 | 0 | CAAA | C | -1.2821 | 0.3639 | 0.0004264 | ---?? | 24.2 | 0.2676 | | |
| rs2052339 | 8 | 113297893 | 4 | 0 | 0 | T | C | -1.0017 | 0.2843 | 0.0004265 | ---?- | 0 | 0.7853 | intron | CSMD3 |
| rs76132719 | 5 | 10299180 | 4 | 0 | 0 | T | C | -1.8943 | 0.5377 | 0.0004266 | ---?- | 56.3 | 0.07635 | intron | CMBL |
| 7:156160508 | 7 | 156160508 | 3 | 0 | 0 | CA | C | 0.9446 | 0.2681 | 0.0004267 | +++?? | 0 | 0.7023 | | |
| rs11735180 | 4 | 57918564 | 4 | 0 | 0 | T | G | -1.0043 | 0.2851 | 0.0004269 | ---?- | 0 | 0.7081 | intron | IGFBP7 |
| rs114082943 | 6 | 23886266 | 4 | 0 | 0 | A | G | 2.6985 | 0.766 | 0.0004269 | +++?- | 24.3 | 0.2655 | intergenic | |
| rs117685986 | 11 | 97073870 | 4 | 0 | 0 | C | G | 2.9614 | 0.8406 | 0.0004269 | +++?+ | 0 | 0.756 | intergenic | |
| 19:36559647 | 19 | 36559647 | 3 | 0 | 0 | A | G | -1.5962 | 0.4531 | 0.000427 | -+-?? | 61.4 | 0.07498 | intron | WDR62 |
| rs2432709 | 2 | 208365719 | 4 | 0 | 0 | T | C | -1.4933 | 0.4239 | 0.000427 | ---?- | 0 | 0.5989 | intergenic | |
| rs73971648 | 2 | 170459391 | 4 | 0 | 0 | A | G | 1.2452 | 0.3535 | 0.000427 | +++?+ | 0 | 0.9378 | intron | PPIG |
| rs72898666 | 18 | 40743591 | 4 | 0 | 0 | A | G | 2.0142 | 0.5718 | 0.0004271 | +++?+ | 0 | 0.7268 | intergenic | |
| rs77568953 | 5 | 43399355 | 3 | 0 | 0 | A | G | 3.1936 | 0.9066 | 0.0004271 | +++?? | 0 | 0.6101 | intron | CCL28 |
| rs118141397 | 11 | 39300938 | 2 | 0 | 0 | A | T | 7.6791 | 2.1799 | 0.0004273 | ?+??+ | 0 | 0.7104 | intergenic | |
| rs145338825 | 4 | 141334521 | 3 | 0 | 0 | T | C | -5.223 | 1.4827 | 0.0004274 | ?--?+ | 58.9 | 0.08773 | intron | CLGN |
| rs10490209 | 2 | 40476866 | 4 | 0 | 0 | T | G | -1.2243 | 0.3476 | 0.0004275 | ---?+ | 39.5 | 0.1747 | intron | SLC8A1,SLC8A1-AS1 |
| rs112933880 | 17 | 57158388 | 5 | 0 | 0 | T | C | 2.3703 | 0.6729 | 0.0004275 | +++++ | 0 | 0.823 | intron | TRIM37 |
| rs343053 | 7 | 35522553 | 4 | 0 | 0 | A | G | -1.2019 | 0.3412 | 0.0004277 | ---?- | 34.4 | 0.2061 | intergenic | |
| 1:160043844 | 1 | 160043844 | 3 | 0 | 0 | CT | C | -0.9727 | 0.2761 | 0.0004277 | ---?? | 0 | 0.6488 | | |
| rs4667617 | 2 | 170968414 | 4 | 0 | 1 | T | G | -0.8981 | 0.255 | 0.0004279 | ---?- | 38.7 | 0.18 | intergenic | |
| rs114507829 | 2 | 170396272 | 4 | 0 | 0 | A | T | -1.2773 | 0.3627 | 0.000428 | ---?- | 0 | 0.9979 | intron | FASTKD1 |
| rs17885011 | 6 | 32552127 | 4 | 0 | 1 | T | C | 1.5818 | 0.4491 | 0.0004281 | ++++? | 20.6 | 0.2866 | synonymous | HLA-DRB1 |
| rs61974368 | 14 | 28209685 | 4 | 0 | 0 | A | T | -2.0122 | 0.5713 | 0.0004281 | ---?- | 0 | 0.8331 | intergenic | |
| 11:111531073 | 11 | 111531073 | 3 | 0 | 0 | CTCT | C | -4.2177 | 1.1975 | 0.0004281 | ---?? | 0 | 0.8101 | | |
| 2:170570862 | 2 | 170570862 | 3 | 0 | 0 | A | AGTGCT | -1.3051 | 0.3705 | 0.0004281 | ---?? | 0 | 0.793 | | |
| rs116326235 | 11 | 86138154 | 4 | 0 | 0 | T | C | -2.5482 | 0.7235 | 0.0004282 | ---?- | 0 | 0.5638 | downstream-gene | |
| rs11048651 | 12 | 26842301 | 4 | 0 | 0 | A | G | 1.7433 | 0.495 | 0.0004285 | +-+?+ | 39 | 0.178 | intron | ITPR2 |
| rs72898662 | 18 | 40740904 | 4 | 0 | 0 | A | G | 2.0075 | 0.5701 | 0.0004288 | +++?+ | 0 | 0.7273 | intergenic | |
| rs62419099 | 6 | 63305215 | 4 | 0 | 0 | C | G | -2.4302 | 0.6901 | 0.0004291 | ---?- | 0 | 0.5664 | intergenic | |
| rs7644525 | 3 | 61096225 | 4 | 0 | 0 | A | G | -3.2158 | 0.9132 | 0.0004292 | ---?- | 0 | 0.9086 | intron | FHIT |
| rs79557429 | 17 | 57116114 | 5 | 0 | 0 | T | C | 2.3561 | 0.6691 | 0.0004292 | +++++ | 0 | 0.8118 | intron | TRIM37 |
| rs10953358 | 7 | 101738226 | 4 | 0 | 0 | T | C | 0.8656 | 0.2458 | 0.0004293 | +++?+ | 0 | 0.8061 | intron | CUX1 |
| rs57956200 | 4 | 12223811 | 4 | 0 | 0 | A | C | -2.7648 | 0.7851 | 0.0004293 | ---?- | 0 | 0.7258 | intergenic | |
| rs12606847 | 18 | 40443555 | 3 | 0 | 0 | A | T | 4.7809 | 1.3577 | 0.0004295 | +++?? | 0 | 0.4822 | intron | RIT2 |
| rs705568 | 1 | 64770542 | 4 | 0 | 0 | A | G | -0.9784 | 0.2779 | 0.0004297 | ---?- | 0 | 0.4409 | intergenic | |
| rs77768584 | 3 | 196498060 | 4 | 0 | 0 | A | G | 2.1835 | 0.6201 | 0.00043 | +++?- | 22.7 | 0.2749 | intron | PAK2 |
| rs13256664 | 8 | 65026660 | 4 | 0 | 0 | A | G | 0.8676 | 0.2464 | 0.0004302 | +++?+ | 0 | 0.9476 | intergenic | |
| rs116209043 | 2 | 170379531 | 4 | 0 | 0 | A | C | -1.3695 | 0.389 | 0.0004302 | ---?- | 0 | 0.8438 | intron | KLHL41 |
| rs56059071 | 3 | 68247293 | 4 | 0 | 0 | A | T | -2.3867 | 0.6779 | 0.0004303 | ---?- | 21 | 0.2842 | intron | FAM19A1 |
| rs78758811 | 6 | 106286466 | 4 | 0 | 0 | T | C | -2.1024 | 0.5971 | 0.0004303 | ---?- | 0 | 0.4615 | intergenic | |

| | | | | | | | | | | | | | | | |
|---|---|---|---|---|---|---|---|---|---|---|---|---|---|---|---|
| rs79868284 | 11 | 86137125 | 4 | 0 | 0 | T | C | 2.5489 | 0.724 | 0.0004304 | +++?+ | 0 | 0.5635 | downstream-gene | |
| rs200549089 | 11 | 39129460 | 3 | 0 | 0 | T | C | 1.1913 | 0.3384 | 0.0004304 | +++?? | 0 | 0.7259 | intergenic | |
| rs6759667 | 2 | 170463679 | 4 | 0 | 0 | A | G | 1.2455 | 0.3538 | 0.0004305 | +++?+ | 0 | 0.9415 | intron | PPIG |
| rs7953873 | 12 | 43188454 | 5 | 0 | 0 | A | G | -0.9367 | 0.2661 | 0.0004307 | ----+ | 21.4 | 0.2781 | intergenic |  |
| rs68027833 | 6 | 32630028 | 4 | 0 | 0 | A | G | 1.1025 | 0.3132 | 0.0004307 | ++++? | 31.8 | 0.2218 | intron-near-splice | HLA-DQB1 |
| rs7965657 | 12 | 27786415 | 4 | 0 | 0 | A | G | 0.8903 | 0.2529 | 0.000431 | +++?- | 53.2 | 0.09352 | intron | PPFIBP1 |
| rs34598665 | 1 | 160039278 | 4 | 0 | 0 | T | G | -0.928 | 0.2636 | 0.000431 | ---?- | 0 | 0.7302 | intron | KCNJ10 |
| rs16857073 | 2 | 170469839 | 4 | 0 | 0 | A | G | 1.2474 | 0.3544 | 0.0004312 | +++?+ | 0 | 0.9468 | intron | PPIG |
| rs219986 | 6 | 12208819 | 4 | 0 | 0 | T | G | 0.841 | 0.2389 | 0.0004313 | +++?+ | 0 | 0.7018 | intergenic | |
| rs10092988 | 8 | 130624661 | 4 | 0 | 0 | A | G | 0.8479 | 0.2409 | 0.0004318 | +++?+ | 9 | 0.3482 | intergenic | |
| 6:83716688 | 6 | 83716688 | 3 | 0 | 0 | GA | GAA | -3.4684 | 0.9854 | 0.0004318 | ---?? | 0 | 0.6686 | | |
| rs12809496 | 12 | 13857909 | 4 | 0 | 0 | A | G | -0.8933 | 0.2538 | 0.0004319 | ---?- | 28.3 | 0.2422 | intron | GRIN2B |
| rs62419100 | 6 | 63306209 | 4 | 0 | 0 | A | G | -2.4279 | 0.6898 | 0.0004319 | ---?- | 0 | 0.5664 | intergenic | |
| rs184686929 | 15 | 98676225 | 3 | 0 | 0 | T | C | 5.1649 | 1.4674 | 0.0004319 | ?++?- | 33.5 | 0.2224 | intergenic | |
| rs74562870 | 7 | 121273208 | 3 | 0 | 0 | T | C | -2.1466 | 0.6099 | 0.000432 | ---?? | 44 | 0.1677 | intergenic | |
| rs118019940 | 17 | 57143357 | 5 | 0 | 0 | T | C | 2.3224 | 0.6598 | 0.000432 | +++++ | 0 | 0.8441 | intron | TRIM37 |
| rs7128181 | 11 | 106748529 | 4 | 0 | 0 | T | G | -1.0289 | 0.2923 | 0.0004326 | ---?+ | 38.8 | 0.1792 | intron | GUCY1A2 |
| rs10789549 | 11 | 106743516 | 4 | 0 | 0 | A | T | 1.0293 | 0.2925 | 0.0004327 | +++?- | 37 | 0.1903 | intron | GUCY1A2 |
| rs11784683 | 8 | 113286111 | 4 | 0 | 0 | A | G | -1.0023 | 0.2848 | 0.0004327 | ---?- | 0 | 0.7863 | intron | CSMD3 |
| rs62201019 | 20 | 46537054 | 4 | 0 | 0 | T | C | -1.791 | 0.5089 | 0.000433 | ---?- | 10 | 0.3428 | intergenic | |
| rs28680981 | 6 | 32606579 | 3 | 0 | 0 | A | G | 1.2932 | 0.3675 | 0.0004331 | +++?? | 0 | 0.3842 | intron | HLA-DQA1 |
| rs11580741 | 1 | 64783615 | 4 | 0 | 0 | T | G | -1.2406 | 0.3526 | 0.0004332 | ---?- | 0 | 0.5365 | intergenic | |
| rs1867417 | 15 | 41964134 | 5 | 0 | 0 | A | G | -1.0003 | 0.2843 | 0.0004333 | ----- | 12 | 0.3369 | intron | MGA |
| rs1867492 | 8 | 65025766 | 4 | 0 | 0 | T | C | -0.8674 | 0.2465 | 0.0004334 | ---?- | 0 | 0.9487 | intergenic | |
| rs62419101 | 6 | 63307052 | 4 | 0 | 0 | A | G | -2.4278 | 0.69 | 0.0004335 | ---?- | 0 | 0.5662 | intergenic | |
| rs62519731 | 8 | 65027287 | 4 | 0 | 0 | T | C | -0.8668 | 0.2463 | 0.0004335 | ---?- | 0 | 0.9475 | intergenic | |
| rs73173317 | 3 | 172331005 | 4 | 0 | 0 | T | C | -1.9222 | 0.5463 | 0.0004335 | ---?+ | 9.4 | 0.3463 | intergenic | |
| 19:3329359 | 19 | 3329359 | 3 | 0 | 0 | T | TA | 4.3966 | 1.2495 | 0.0004335 | +++?? | 0 | 0.3896 | | |
| rs10890618 | 11 | 106747093 | 4 | 0 | 0 | T | C | -1.0288 | 0.2924 | 0.0004336 | ---?+ | 36.5 | 0.1932 | intron | GUCY1A2 |
| rs1061354 | 15 | 42140166 | 5 | 0 | 0 | T | G | -0.9567 | 0.2719 | 0.0004337 | ----- | 0 | 0.539 | 3-prime-UTR | JMJD7-PLA2G4B,PLA2G4B |
| 6:159210662 | 6 | 159210662 | 3 | 0 | 0 | T | TTTTC | -0.8981 | 0.2552 | 0.0004338 | ---?? | 4.7 | 0.35 | | |
| rs112439040 | 2 | 170461307 | 4 | 0 | 0 | C | G | 1.2444 | 0.3537 | 0.000434 | +++?+ | 0 | 0.9392 | intron | PPIG |
| rs35535980 | 6 | 165817058 | 4 | 0 | 0 | C | G | 0.8901 | 0.253 | 0.0004343 | +++?+ | 0 | 0.6667 | intron | PDE10A |
| rs72696282 | 9 | 11315721 | 4 | 0 | 0 | A | G | -1.8581 | 0.5281 | 0.0004344 | ---?- | 29.2 | 0.237 | intergenic | |
| rs10097465 | 8 | 113510766 | 4 | 0 | 0 | A | C | 0.9149 | 0.26 | 0.0004345 | +++?+ | 0 | 0.7069 | intron | CSMD3 |
| rs62518601 | 8 | 124709275 | 5 | 0 | 0 | A | G | 0.8194 | 0.2329 | 0.0004345 | ++++- | 19.7 | 0.2892 | intron | ANXA13 |
| 7:88636875 | 7 | 88636875 | 3 | 0 | 0 | T | TATTAT | 0.9247 | 0.2628 | 0.0004348 | +++?? | 44.8 | 0.1635 | | |
| rs4939264 | 11 | 59003722 | 5 | 0 | 0 | A | C | 0.7582 | 0.2155 | 0.0004349 | ++++- | 16.6 | 0.3089 | intergenic | |
| 22:28884508 | 22 | 28884508 | 3 | 0 | 0 | A | AT | -2.4051 | 0.6837 | 0.0004349 | ---?? | 0 | 0.5145 | | |
| rs56141483 | 5 | 723000 | 4 | 0 | 0 | T | C | 1.4399 | 0.4093 | 0.000435 | +++?+ | 58.2 | 0.06657 | intron | ZDHHC11B |
| rs138513618 | 17 | 73249830 | 3 | 0 | 0 | A | C | 4.9647 | 1.4113 | 0.000435 | +++?? | 0 | 0.6474 | intron | GGA3 |
| rs817451 | 1 | 85656602 | 5 | 0 | 0 | T | C | -1.3698 | 0.3894 | 0.0004351 | ----- | 0 | 0.9757 | intron | SYDE2 |
| rs1867416 | 15 | 41963640 | 5 | 0 | 0 | A | G | -1 | 0.2843 | 0.0004351 | ----- | 12.1 | 0.3366 | intron | MGA |
| rs6730530 | 2 | 75647440 | 3 | 0 | 0 | T | C | -1.0145 | 0.2884 | 0.0004351 | ---?? | 0 | 0.4648 | intergenic | |
| rs6753328 | 2 | 2780239 | 4 | 0 | 0 | T | C | -0.938 | 0.2667 | 0.0004354 | ---?- | 35.4 | 0.1998 | intergenic | |
| 1:87480731 | 1 | 87480731 | 3 | 0 | 0 | T | TTTTG | -1.0747 | 0.3055 | 0.0004354 | ---?? | 0 | 0.7281 | | |
| rs9276720 | 6 | 32761939 | 5 | 0 | 0 | C | G | -1.3935 | 0.3962 | 0.0004358 | ----- | 0 | 0.6614 | intergenic |  |
| rs62419102 | 6 | 63307925 | 4 | 0 | 0 | A | G | -2.4273 | 0.6901 | 0.000436 | ---?- | 0 | 0.5677 | intergenic | |
| rs34267806 | 6 | 165817089 | 4 | 0 | 0 | A | G | -0.8898 | 0.253 | 0.0004362 | ---?- | 0 | 0.6662 | intron | PDE10A |
| 2:167691046 | 2 | 167691046 | 3 | 0 | 0 | CTTTAT | C | 2.5177 | 0.7158 | 0.0004362 | +++?? | 0 | 0.9154 | | |
| rs11998556 | 8 | 140379919 | 4 | 0 | 0 | T | C | 1.0163 | 0.289 | 0.0004363 | +++?+ | 0 | 0.9415 | intergenic | |
| 19:36552803 | 19 | 36552803 | 3 | 0 | 0 | T | TATAC | -1.607 | 0.4569 | 0.0004363 | -+-?? | 60.8 | 0.0778 | | |
| rs4924572 | 15 | 42004962 | 5 | 0 | 0 | C | G | -1.0003 | 0.2844 | 0.0004364 | ----+ | 16.4 | 0.3104 | intron | MGA |
| rs149278920 | 15 | 86341406 | 4 | 0 | 0 | A | G | -3.6495 | 1.0377 | 0.0004364 | ---?- | 28.6 | 0.2407 | upstream-gene | |
| rs7562374 | 2 | 131115424 | 3 | 0 | 0 | A | G | 1.0815 | 0.3075 | 0.0004365 | +++?? | 0 | 0.4606 | intron | PTPN18 |
| rs74009293 | 16 | 14094837 | 4 | 0 | 0 | C | G | 2.1389 | 0.6082 | 0.0004365 | +++?+ | 17.7 | 0.3022 | intergenic | |
| rs4078788 | 15 | 42140998 | 5 | 0 | 0 | C | G | -0.9559 | 0.2718 | 0.0004367 | ----- | 0 | 0.5383 | intron | SPTBN5 |
| rs56346755 | 2 | 170466527 | 4 | 0 | 0 | A | T | 1.245 | 0.354 | 0.0004367 | +++?+ | 0 | 0.9447 | intron | PPIG |
| rs141018652 | 10 | 109272074 | 4 | 0 | 0 | T | C | -2.4527 | 0.6974 | 0.0004368 | ---?- | 0 | 0.9854 | intergenic | |

| | | | | | | | | | | | | | | | |
|---|---|---|---|---|---|---|---|---|---|---|---|---|---|---|---|
| rs149701693 | 3 | 46612493 | 4 | 0 | 0 | A | G | 3.1331 | 0.8909 | 0.0004368 | +++?- | 31.9 | 0.2208 | intron | LRRC2 |
| rs116805392 | 17 | 57119113 | 5 | 0 | 0 | T | C | -2.3502 | 0.6683 | 0.0004369 | ----- | 0 | 0.8139 | intron | TRIM37 |
| 3:115914565 | 3 | 115914565 | 3 | 0 | 0 | G | GA | 1.0416 | 0.2962 | 0.0004369 | +++?? | 0 | 0.5028 | | |
| rs809568 | 9 | 94199485 | 4 | 0 | 0 | A | G | 2.31 | 0.6569 | 0.000437 | +++?+ | 0 | 0.8186 | intergenic | |
| rs817436 | 1 | 85639827 | 5 | 0 | 0 | T | C | -1.3555 | 0.3855 | 0.000437 | ----- | 0 | 0.9673 | intron | SYDE2 |
| rs1859246 | 12 | 112740315 | 5 | 0 | 0 | A | G | 1.2983 | 0.3692 | 0.000437 | +-+++ | 52.5 | 0.07728 | intron | HECTD4 |
| rs73008494 | 2 | 170620649 | 4 | 0 | 0 | A | G | 1.2239 | 0.348 | 0.0004372 | +++?+ | 0 | 0.9692 | intergenic | |
| rs2098455 | 8 | 113264960 | 4 | 0 | 0 | T | C | 1.0091 | 0.287 | 0.0004373 | +++?+ | 0 | 0.9333 | intron | CSMD3 |
| 2:137509074 | 2 | 137509074 | 3 | 0 | 0 | CT | C | 3.54 | 1.0067 | 0.0004374 | +++?? | 65.7 | 0.05412 | | |
| rs201446 | 7 | 101738323 | 4 | 0 | 0 | C | G | -0.864 | 0.2457 | 0.0004375 | ---?- | 0 | 0.8049 | intron | CUX1 |
| rs1810910 | 9 | 16199886 | 4 | 0 | 0 | T | C | 1.2152 | 0.3456 | 0.0004376 | +++?+ | 0 | 0.9535 | downstream-gene | |
| rs116603501 | 17 | 57167603 | 5 | 0 | 0 | A | T | -2.3964 | 0.6815 | 0.0004377 | ----- | 0 | 0.8173 | intron | TRIM37 |
| 6:56012163 | 6 | 56012163 | 3 | 0 | 0 | CT | C | -3.0773 | 0.8752 | 0.0004379 | ---?? | 31.4 | 0.2329 | | |
| rs11914437 | 3 | 183592432 | 4 | 0 | 0 | A | G | -1.6595 | 0.472 | 0.000438 | ---?- | 0 | 0.898 | intron | PARL |
| rs76592655 | 5 | 56336901 | 2 | 0 | 0 | A | G | -5.8189 | 1.6549 | 0.000438 | ?--?? | 0 | 0.4625 | intergenic | |
| rs817411 | 1 | 85623859 | 5 | 0 | 0 | A | G | 1.3605 | 0.3869 | 0.0004381 | +++++ | 0 | 0.9526 | 3-prime-UTR | SYDE2 |
| rs4861135 | 4 | 41740605 | 4 | 0 | 0 | A | G | -1.4873 | 0.423 | 0.0004382 | ---?- | 26.2 | 0.2548 | intergenic | |
| rs6577129 | 1 | 100204302 | 4 | 0 | 0 | T | C | -1.7669 | 0.5025 | 0.0004382 | ---?- | 0 | 0.7077 | intron | FRRS1 |
| rs3830123 | 6 | 32548646 | 5 | 0 | 1 | T | G | 1.3488 | 0.3836 | 0.0004383 | +++++ | 14.2 | 0.3236 | intron | HLA-DRB1 |
| rs8027806 | 15 | 42146632 | 5 | 0 | 0 | C | G | 1.0257 | 0.2917 | 0.0004384 | +-+++ | 26 | 0.2479 | intron | SPTBN5 |
| rs66869205 | 14 | 28214550 | 4 | 0 | 0 | T | C | -2.0182 | 0.5741 | 0.0004385 | ---?- | 0 | 0.8169 | intergenic | |
| rs2771334 | 14 | 21452603 | 4 | 0 | 0 | A | G | 0.8727 | 0.2482 | 0.0004386 | +++?+ | 0 | 0.8549 | upstream-gene | |
| rs79111787 | 3 | 47715545 | 2 | 0 | 0 | T | C | -6.6592 | 1.8941 | 0.0004386 | ?-??- | 0.9 | 0.3151 | intron | SMARCC1 |
| rs268502 | 5 | 9501082 | 2 | 0 | 0 | A | G | -7.685 | 2.186 | 0.0004388 | ?--?? | 0 | 0.7298 | intron | SEMA5A |
| rs3129961 | 6 | 32378940 | 5 | 0 | 0 | T | C | 0.7515 | 0.2138 | 0.0004388 | -++++ | 42.2 | 0.1404 | upstream-gene | |
| rs3751231 | 12 | 27786115 | 4 | 0 | 0 | T | C | -0.8903 | 0.2533 | 0.0004391 | ---?+ | 53.3 | 0.09263 | intron | PPFIBP1 |
| rs17427521 | 8 | 3987036 | 4 | 0 | 0 | T | C | -1.181 | 0.336 | 0.0004391 | +--?- | 56.3 | 0.07619 | intron | CSMD1 |
| rs16963330 | 16 | 14091340 | 4 | 0 | 0 | A | G | 2.1413 | 0.6091 | 0.0004392 | +++?+ | 16.3 | 0.3101 | intergenic | |
| rs6878843 | 5 | 103167572 | 4 | 0 | 0 | T | G | -1.5828 | 0.4503 | 0.0004394 | ---?- | 0 | 0.7437 | intergenic | |
| rs117400276 | 7 | 4140699 | 4 | 0 | 0 | A | G | 2.7981 | 0.796 | 0.0004394 | +++?- | 39.1 | 0.1771 | intron | SDK1 |
| rs10836670 | 11 | 37056349 | 4 | 0 | 0 | A | G | -0.8711 | 0.2478 | 0.0004395 | ---?- | 0 | 0.3919 | intergenic | |
| rs11741852 | 5 | 39186351 | 4 | 0 | 0 | T | C | 2.6963 | 0.7671 | 0.0004397 | +++?+ | 0 | 0.6428 | intron | FYB |
| 13:30321181 | 13 | 30321181 | 3 | 0 | 0 | CT | C | 1.1878 | 0.3379 | 0.0004397 | +++?? | 0 | 0.8736 | | |
| rs28611286 | 4 | 111393438 | 4 | 0 | 0 | T | C | -0.9246 | 0.2631 | 0.0004398 | ---?- | 0 | 0.8198 | upstream-gene | |
| rs12758997 | 1 | 245104596 | 4 | 0 | 0 | T | C | 1.1741 | 0.334 | 0.0004399 | +++?+ | 8.4 | 0.3511 | intergenic | |
| rs35487354 | 11 | 65366882 | 5 | 0 | 0 | T | C | 2.9122 | 0.8285 | 0.0004399 | ++++- | 22.9 | 0.2687 | missense | MAP3K11 |
| rs114478971 | 4 | 105324175 | 4 | 0 | 0 | A | G | 2.5386 | 0.7222 | 0.0004399 | +++?+ | 0 | 0.64 | intergenic | |
| rs1414978 | 13 | 60042376 | 4 | 0 | 0 | A | T | -0.9257 | 0.2634 | 0.00044 | ---?- | 0 | 0.5515 | intergenic | |
| rs115740540 | 12 | 27791020 | 3 | 0 | 0 | A | G | -0.9819 | 0.2794 | 0.00044 | ---?? | 25.5 | 0.2613 | intron | PPFIBP1 |
| rs188466358 | 2 | 13556955 | 4 | 0 | 0 | T | G | -4.1512 | 1.1811 | 0.0004401 | ---?- | 35.5 | 0.1993 | intergenic | |
| rs9288556 | 2 | 220771575 | 4 | 0 | 0 | T | C | 0.9307 | 0.2648 | 0.0004402 | +++?+ | 0 | 0.6233 | upstream-gene | |
| 17:55448911 | 17 | 55448911 | 3 | 0 | 0 | T | TACACA | -1.1639 | 0.3311 | 0.0004402 | ---?? | 0 | 0.6772 | | |
| rs10798013 | 1 | 185394729 | 4 | 0 | 0 | A | C | -0.93 | 0.2646 | 0.0004403 | ---?- | 0 | 0.9237 | intergenic | |
| rs77935032 | 8 | 140813230 | 3 | 0 | 0 | T | C | 1.365 | 0.3884 | 0.0004403 | +++?? | 66.2 | 0.05185 | intron | TRAPPC9 |
| 2:170732594 | 2 | 170732594 | 3 | 0 | 0 | CT | C | 1.5012 | 0.4271 | 0.0004403 | +++?? | 0 | 0.944 | | |
| rs817453 | 1 | 85657310 | 5 | 0 | 0 | A | G | -1.3696 | 0.3897 | 0.0004404 | ----- | 0 | 0.976 | intron | SYDE2 |
| rs1333476 | 13 | 45520889 | 4 | 0 | 0 | T | G | -2.2613 | 0.6434 | 0.0004404 | ---?+ | 23.5 | 0.2703 | intron | NUFIP1 |
| rs73077560 | 3 | 46657202 | 3 | 0 | 0 | T | C | -3.6306 | 1.033 | 0.0004405 | ---?? | 0 | 0.4293 | intron | LOC100132146 |
| rs112579857 | 18 | 40149534 | 4 | 0 | 0 | T | C | 1.4 | 0.3984 | 0.0004406 | +++?+ | 0 | 0.9412 | intron | LINC00907 |
| rs74351555 | 2 | 170467630 | 4 | 0 | 0 | T | G | -1.2447 | 0.3542 | 0.0004407 | ---?- | 0 | 0.9462 | intron | PPIG |
| 16:74394379 | 16 | 74394379 | 3 | 0 | 0 | T | TA | -1.4082 | 0.4007 | 0.0004407 | ---?? | 0.4 | 0.3663 | | |
| rs1002422 | 12 | 27788474 | 4 | 0 | 0 | A | G | 0.8891 | 0.253 | 0.0004409 | +++?- | 52.8 | 0.09546 | intron | PPFIBP1 |
| rs76193383 | 3 | 60585595 | 4 | 0 | 0 | A | G | 1.7554 | 0.4995 | 0.0004409 | +++?+ | 24.7 | 0.2631 | intron | FHIT |
| rs1589977 | 2 | 156897347 | 4 | 0 | 0 | A | C | 0.9241 | 0.263 | 0.000441 | +++?+ | 0 | 0.7072 | intergenic | |
| rs3120939 | 1 | 90245167 | 5 | 0 | 0 | A | G | 0.7491 | 0.2132 | 0.0004411 | +++++ | 0 | 0.9889 | intergenic | |
| rs62419103 | 6 | 63308798 | 4 | 0 | 0 | T | C | 2.4245 | 0.6899 | 0.0004412 | +++?+ | 0 | 0.5706 | intergenic | |
| rs76791478 | 3 | 143070313 | 4 | 0 | 0 | A | G | 2.1306 | 0.6063 | 0.0004412 | +++?+ | 0 | 0.6955 | intron | SLC9A9 |
| rs5997365 | 22 | 28826994 | 4 | 0 | 0 | T | C | 2.5294 | 0.7198 | 0.0004413 | +++?+ | 0 | 0.7321 | intron | TTC28 |
| rs74703510 | 8 | 4010605 | 4 | 0 | 0 | T | C | -3.8733 | 1.1022 | 0.0004413 | ---?- | 0 | 0.6406 | intron | CSMD1 |

| | | | | | | | | | | | | | | | |
|---|---|---|---|---|---|---|---|---|---|---|---|---|---|---|---|
| rs61974404 | 14 | 28234822 | 4 | 0 | 0 | T | C | -2.1016 | 0.5981 | 0.0004414 | ---?- | 0 | 0.8532 | intergenic | |
| rs62514388 | 8 | 113287158 | 4 | 0 | 0 | T | C | 1.0008 | 0.2848 | 0.0004414 | +++?+ | 0 | 0.7836 | intron | CSMD3 |
| 18:6547546 | 18 | 6547546 | 3 | 0 | 0 | CA | C | -0.973 | 0.2769 | 0.0004414 | ---?? | 35.2 | 0.2139 | | |
| rs76712087 | 7 | 8884722 | 3 | 0 | 0 | T | C | -2.3608 | 0.6718 | 0.0004416 | ---?? | 0 | 0.7399 | intergenic | |
| rs149678265 | 11 | 24890453 | 3 | 0 | 0 | T | C | 4.2447 | 1.208 | 0.0004417 | ?++?+ | 36.4 | 0.2076 | intron | LUZP2 |
| rs2015030 | 15 | 85611292 | 4 | 0 | 0 | A | T | 0.8443 | 0.2403 | 0.0004418 | +++?- | 59.6 | 0.0596 | intron | PDE8A |
| 19:5561075 | 19 | 5561075 | 3 | 0 | 0 | A | G | 5.3795 | 1.531 | 0.0004419 | +++?? | 0 | 0.7823 | non-coding-exon | TINCR |
| rs55947663 | 9 | 16188797 | 3 | 0 | 0 | T | C | -0.9889 | 0.2815 | 0.000442 | ---?? | 0 | 0.9108 | intergenic | |
| 2:170788586 | 2 | 170788586 | 3 | 0 | 0 | A | AT | -0.9138 | 0.2601 | 0.000442 | ---?? | 2.5 | 0.3586 | | |
| rs4431223 | 4 | 20741370 | 4 | 0 | 0 | T | C | 0.9164 | 0.2608 | 0.0004421 | +++?+ | 0 | 0.8847 | intron | KCNIP4 |
| rs7073234 | 10 | 129677153 | 4 | 0 | 0 | C | G | -0.8807 | 0.2507 | 0.0004423 | ---?- | 0 | 0.5436 | intron | CLRN3 |
| rs77422707 | 12 | 112244053 | 5 | 0 | 0 | A | G | 1.3363 | 0.3803 | 0.0004423 | +-+++ | 46.9 | 0.11 | intron | ALDH2 |
| rs72839991 | 17 | 56392938 | 5 | 0 | 0 | T | C | 1.9537 | 0.5561 | 0.0004425 | +++++ | 48.6 | 0.1 | intron | BZRAP1 |
| rs75231029 | 1 | 85725032 | 5 | 0 | 0 | T | G | -1.6297 | 0.4639 | 0.0004425 | ----- | 13.7 | 0.3266 | intron | C1orf52 |
| rs7177911 | 15 | 41948157 | 5 | 0 | 0 | T | C | -0.996 | 0.2835 | 0.0004426 | ----- | 14.9 | 0.3193 | intron | MGA |
| rs12049531 | 1 | 187858866 | 4 | 0 | 0 | A | T | -1.1679 | 0.3324 | 0.0004426 | ---?- | 0 | 0.879 | intergenic | |
| rs77690229 | 17 | 57120996 | 5 | 0 | 0 | A | C | -2.3459 | 0.6677 | 0.0004426 | ----- | 0 | 0.8153 | intron | TRIM37 |
| rs2890764 | 9 | 7847266 | 4 | 0 | 0 | A | T | 1.1633 | 0.3311 | 0.0004427 | +++?+ | 24.1 | 0.2665 | intergenic | |
| rs3129948 | 6 | 32354644 | 5 | 0 | 0 | A | C | 0.7503 | 0.2136 | 0.0004427 | -++++ | 38.7 | 0.1632 | upstream-gene | |
| rs115479198 | 2 | 79463655 | 4 | 0 | 0 | A | C | 1.9505 | 0.5552 | 0.0004429 | +++?+ | 0 | 0.8828 | intergenic | |
| rs1995029 | 1 | 64666102 | 3 | 0 | 0 | T | C | -0.935 | 0.2662 | 0.000443 | ---?? | 28.7 | 0.2459 | upstream-gene | |
| rs2279859 | 2 | 98994134 | 4 | 0 | 0 | A | G | -1.4672 | 0.4176 | 0.000443 | ---?- | 0 | 0.8377 | intron | CNGA3 |
| rs202174933 | 1 | 98600672 | 3 | 0 | 0 | A | T | 1.4163 | 0.4032 | 0.000443 | +++?? | 50.6 | 0.1322 | intergenic | |
| 12:96800169 | 12 | 96800169 | 3 | 0 | 0 | CT | C | 1.7424 | 0.496 | 0.000443 | +++?? | 0 | 0.7248 | | |
| rs971072 | 3 | 65659641 | 4 | 0 | 0 | A | G | 0.9383 | 0.2671 | 0.0004431 | +-+?+ | 47.9 | 0.1237 | intron | MAGI1 |
| rs1414976 | 13 | 60031893 | 4 | 0 | 0 | A | G | 0.8777 | 0.2498 | 0.0004433 | +++?+ | 0 | 0.4577 | intergenic | |
| rs13423574 | 2 | 170923296 | 4 | 0 | 1 | A | T | 1.0188 | 0.29 | 0.0004433 | +++?+ | 46.3 | 0.1337 | intron | UBR3 |
| rs4836064 | 5 | 123515729 | 4 | 0 | 0 | A | G | -0.8536 | 0.243 | 0.0004435 | ---?+ | 10.3 | 0.3416 | intergenic | |
| rs140091303 | 17 | 57088145 | 5 | 0 | 0 | T | C | -2.3689 | 0.6744 | 0.0004436 | ----+ | 0 | 0.6833 | intron | TRIM37 |
| rs6534464 | 4 | 80556161 | 4 | 0 | 0 | T | G | 1.4413 | 0.4103 | 0.0004438 | +++?+ | 41.1 | 0.165 | intergenic | |
| rs8104137 | 19 | 14748260 | 4 | 0 | 0 | T | C | 0.9248 | 0.2633 | 0.0004438 | +++?- | 47.6 | 0.1256 | intron | EMR3 |
| rs971073 | 3 | 65659656 | 4 | 0 | 0 | A | G | 0.9383 | 0.2671 | 0.0004439 | +-+?+ | 47.9 | 0.1239 | intron | MAGI1 |
| rs6598014 | 11 | 524919 | 5 | 0 | 0 | T | C | -0.8088 | 0.2303 | 0.0004439 | ----- | 0 | 0.445 | upstream-gene |  |
| rs57461927 | 2 | 137007096 | 4 | 0 | 0 | A | G | -0.9403 | 0.2677 | 0.000444 | ---?- | 0 | 0.8156 | intergenic | |
| rs16883454 | 8 | 113342457 | 4 | 0 | 0 | C | G | 0.9981 | 0.2842 | 0.0004442 | +++?+ | 0 | 0.7961 | intron | CSMD3 |
| 4:36680575 | 4 | 36680575 | 3 | 0 | 0 | CCTAAA( | C | 1.3171 | 0.375 | 0.0004442 | +++?? | 0 | 0.9018 | | |
| rs9563733 | 13 | 60040286 | 4 | 0 | 0 | T | C | -0.9221 | 0.2625 | 0.0004443 | ---?- | 0 | 0.6057 | intergenic | |
| rs7192280 | 16 | 84838639 | 4 | 0 | 0 | A | C | 1.1618 | 0.3308 | 0.0004445 | +++?- | 0 | 0.4354 | intergenic | |
| rs12916248 | 15 | 98043537 | 4 | 0 | 0 | T | C | -0.8778 | 0.2499 | 0.0004446 | ---?- | 0 | 0.9376 | intergenic | |
| rs56019797 | 1 | 90977104 | 3 | 0 | 0 | A | T | 1.7222 | 0.4904 | 0.0004447 | +++?? | 0 | 0.4214 | intergenic | |
| rs138145505 | 9 | 139516136 | 4 | 0 | 0 | A | G | 3.3235 | 0.9463 | 0.0004447 | +++?+ | 0 | 0.8156 | intergenic | |
| rs1385252 | 8 | 96994618 | 4 | 0 | 0 | T | C | 1.4215 | 0.4048 | 0.0004448 | +++?+ | 0 | 0.9628 | intergenic | |
| rs38421 | 7 | 30546193 | 3 | 0 | 0 | T | C | 1.0923 | 0.311 | 0.0004449 | +++?? | 45.4 | 0.1602 | upstream-gene | |
| rs201447 | 7 | 101738384 | 4 | 0 | 0 | A | C | 0.8623 | 0.2455 | 0.000445 | +++?+ | 0 | 0.8038 | intron | CUX1 |
| rs117955927 | 17 | 57091092 | 5 | 0 | 0 | T | C | 2.3664 | 0.6738 | 0.0004451 | ++++- | 0 | 0.6854 | intron | TRIM37 |
| rs72888283 | 2 | 47744260 | 4 | 0 | 0 | T | G | -1.5745 | 0.4484 | 0.0004452 | ---?- | 38.4 | 0.1815 | downstream-gene | |
| rs72655733 | 4 | 88639362 | 4 | 0 | 0 | T | C | 3.1524 | 0.8977 | 0.0004456 | +++?- | 0 | 0.6067 | intergenic | |
| rs148790149 | 6 | 31976899 | 3 | 0 | 0 | T | C | 1.3073 | 0.3723 | 0.0004456 | +++?? | 66 | 0.05284 | non-coding-exon | TNXA |
| rs151075319 | 6 | 8054710 | 2 | 0 | 0 | A | C | 4.4521 | 1.2679 | 0.0004456 | +?+?? | 0 | 0.8095 | intron | BLOC1S5,BLOC1S5-TXNDC5,EEF1E1- |
| rs113934011 | 18 | 25160997 | 4 | 0 | 0 | A | G | -2.6873 | 0.7653 | 0.0004457 | ---?- | 0 | 0.6301 | intergenic | |
| 7:64566808 | 7 | 64566808 | 3 | 0 | 0 | A | ATTCTTT | -1.326 | 0.3776 | 0.0004457 | ---?? | 0 | 0.6735 | | |
| rs12643049 | 4 | 175017856 | 4 | 0 | 0 | A | G | -2.8695 | 0.8172 | 0.0004458 | ---?- | 0 | 0.9575 | intergenic | |
| rs79860095 | 13 | 65189008 | 4 | 0 | 0 | A | G | -2.0135 | 0.5734 | 0.0004458 | ---?- | 0 | 0.3953 | intergenic | |
| rs9276719 | 6 | 32761690 | 5 | 0 | 0 | A | C | -1.3913 | 0.3963 | 0.000446 | ----- | 0 | 0.6588 | intergenic |  |
| rs72717429 | 1 | 177769090 | 4 | 0 | 0 | A | G | -3.0516 | 0.8691 | 0.0004461 | ---?+ | 0 | 0.5205 | intergenic | |
| rs113800707 | 13 | 56123878 | 2 | 0 | 0 | A | C | 7.3072 | 2.0811 | 0.0004461 | ?+??+ | 2.9 | 0.3101 | intergenic | |
| rs11783495 | 8 | 113288316 | 4 | 0 | 0 | T | C | -0.9999 | 0.2848 | 0.0004463 | ---?- | 0 | 0.7993 | intron | CSMD3 |
| rs74327757 | 2 | 79223870 | 4 | 0 | 0 | T | C | 1.805 | 0.5141 | 0.0004463 | +++?+ | 0 | 0.807 | intergenic | |
| rs111260948 | 6 | 109111386 | 2 | 0 | 0 | A | G | -4.7192 | 1.3441 | 0.0004464 | -?-?? | 0 | 0.924 | intergenic | |

| | | | | | | | | | | | | | | | |
|---|---|---|---|---|---|---|---|---|---|---|---|---|---|---|---|
| 4:166250726 | 4 | 166250726 | 3 | 0 | 0 | CA | C | -1.2882 | 0.3669 | 0.0004464 | ---?? | 0 | 0.4665 | | |
| rs11018383 | 10 | 129677797 | 4 | 0 | 0 | T | C | 0.8777 | 0.25 | 0.0004465 | +++?+ | 0 | 0.5457 | intron | CLRN3 |
| 1:64666320 | 1 | 64666320 | 3 | 0 | 0 | A | G | -0.9343 | 0.2661 | 0.0004467 | ---?? | 29 | 0.2447 | upstream-gene | |
| rs140141827 | 17 | 4742378 | 5 | 0 | 0 | A | G | 1.6081 | 0.458 | 0.0004467 | ++++- | 20.7 | 0.283 | intron | MINK1 |
| rs219985 | 6 | 12208665 | 4 | 0 | 0 | A | T | -0.8403 | 0.2394 | 0.0004473 | ---?- | 0 | 0.7012 | intergenic | |
| rs13340827 | 12 | 41460426 | 4 | 0 | 0 | A | G | 1.8004 | 0.5129 | 0.0004474 | +++?+ | 0 | 0.6229 | intron | CNTN1 |
| rs55724920 | 15 | 98037802 | 4 | 0 | 0 | A | C | -1.1078 | 0.3156 | 0.0004474 | ---?+ | 0 | 0.4738 | intergenic | |
| rs423358 | 6 | 1251437 | 4 | 0 | 0 | T | C | 0.8665 | 0.2469 | 0.0004475 | +-+?+ | 38.4 | 0.1818 | intergenic | |
| rs9276718 | 6 | 32761674 | 5 | 0 | 0 | C | G | 1.391 | 0.3963 | 0.0004475 | +++++ | 0 | 0.6584 | intergenic | |
| rs11639249 | 15 | 41914597 | 5 | 0 | 0 | C | G | -1.0046 | 0.2862 | 0.0004475 | ----- | 7.5 | 0.3642 | intron | MGA |
| rs11754677 | 6 | 165813070 | 4 | 0 | 0 | C | G | 0.8876 | 0.2528 | 0.0004475 | +++?+ | 0 | 0.7038 | intron | PDE10A |
| 3:1801213 | 3 | 1801213 | 3 | 0 | 0 | T | TA | -1.1228 | 0.3199 | 0.0004476 | ---?? | 40.7 | 0.185 | | |
| 4:140923203 | 4 | 140923203 | 3 | 0 | 0 | A | T | 1.6469 | 0.4692 | 0.0004477 | +++?? | 10.1 | 0.3288 | intron | MAML3 |
| rs1511747 | 6 | 165817390 | 4 | 0 | 0 | T | C | -0.8901 | 0.2536 | 0.0004477 | ---?- | 0 | 0.6612 | intron | PDE10A |
| rs28986423 | 6 | 32758843 | 5 | 0 | 0 | A | G | 1.3967 | 0.3979 | 0.0004477 | +++++ | 0 | 0.6509 | intergenic | |
| rs76360871 | 20 | 2567327 | 4 | 0 | 0 | A | G | 3.1402 | 0.8946 | 0.0004477 | +++?+ | 0 | 0.794 | intron | TMC2 |
| rs80129023 | 12 | 5534794 | 4 | 0 | 0 | T | C | -2.6951 | 0.7679 | 0.0004485 | ---?- | 0 | 0.5057 | intergenic | |
| rs138416979 | 17 | 57004648 | 5 | 0 | 0 | T | C | -2.4274 | 0.6916 | 0.0004486 | ----- | 0 | 0.8011 | intron | PPM1E |
| rs1861755 | 8 | 113301548 | 4 | 0 | 0 | A | G | -0.9973 | 0.2842 | 0.0004489 | ---?- | 0 | 0.7795 | intron | CSMD3 |
| rs12470875 | 2 | 19535518 | 4 | 0 | 0 | A | C | 0.8497 | 0.2421 | 0.000449 | +++?+ | 0 | 0.5955 | intergenic | |
| 16:84789401 | 16 | 84789401 | 3 | 0 | 0 | CAA | C | 1.0702 | 0.305 | 0.0004492 | +++?? | 0 | 0.8221 | | |
| 1:54915296 | 1 | 54915296 | 3 | 0 | 0 | A | AATTTT | -0.9243 | 0.2634 | 0.0004493 | ---?? | 38.1 | 0.199 | | |
| rs6456584 | 6 | 23876191 | 4 | 0 | 0 | T | G | -2.513 | 0.7161 | 0.0004494 | ---?+ | 51.3 | 0.1042 | intergenic | |
| rs16883301 | 8 | 113262032 | 4 | 0 | 0 | T | C | -1.0076 | 0.2871 | 0.0004495 | ---?- | 0 | 0.9369 | intron | CSMD3 |
| rs55993388 | 9 | 104028710 | 4 | 0 | 0 | T | G | -0.9515 | 0.2711 | 0.0004497 | +--?- | 55.2 | 0.08249 | intron | LPPR1 |
| rs75000529 | 10 | 109283511 | 4 | 0 | 0 | T | C | -2.453 | 0.6991 | 0.0004499 | ---?- | 0 | 0.9802 | intergenic | |
| rs79146657 | 9 | 74205358 | 3 | 0 | 0 | A | C | 3.679 | 1.0485 | 0.0004499 | ?++?- | 33.5 | 0.2225 | intergenic | |
| rs13140211 | 4 | 63341773 | 4 | 0 | 0 | T | C | 0.9583 | 0.2731 | 0.0004502 | +++?+ | 34.9 | 0.2028 | intergenic | |
| 19:46159035 | 19 | 46159035 | 3 | 0 | 0 | T | TTTTGTT | 0.936 | 0.2668 | 0.0004502 | +++?? | 0 | 0.6315 | | |
| rs117539473 | 17 | 57064021 | 5 | 0 | 0 | T | C | -2.3972 | 0.6832 | 0.0004503 | ----- | 0 | 0.8159 | intron | TRIM37 |
| 10:99143837 | 10 | 99143837 | 2 | 0 | 0 | A | G | 8.1653 | 2.3273 | 0.0004506 | ?++?? | 0 | 0.361 | intron | RRP12 |
| rs2499072 | 10 | 19564421 | 4 | 0 | 0 | A | C | 0.94 | 0.2679 | 0.0004507 | +++?+ | 0 | 0.9519 | intron | C10orf112 |
| rs12878625 | 14 | 95896843 | 4 | 0 | 0 | A | G | -1.3209 | 0.3765 | 0.0004509 | ---?- | 0 | 0.4598 | intron | SYNE3 |
| 6:159230257 | 6 | 159230257 | 3 | 0 | 0 | CACAG | C | -0.8903 | 0.2538 | 0.0004509 | ---?? | 13.5 | 0.3145 | | |
| rs3117102 | 6 | 32350107 | 5 | 0 | 0 | T | C | 0.7493 | 0.2136 | 0.000451 | -++++ | 39.1 | 0.1608 | intergenic | |
| rs7545591 | 1 | 185395846 | 4 | 0 | 0 | C | G | -0.9255 | 0.2638 | 0.000451 | ---?- | 0 | 0.9225 | intergenic | |
| rs3784282 | 15 | 42141200 | 5 | 0 | 0 | C | G | 0.9532 | 0.2717 | 0.0004511 | +++++ | 0 | 0.5421 | intron | SPTBN5 |
| rs11580562 | 1 | 64737389 | 4 | 0 | 0 | A | G | 1.2505 | 0.3564 | 0.0004511 | +++?+ | 0 | 0.4602 | downstream-gene | |
| rs72886873 | 18 | 33710132 | 3 | 0 | 0 | A | G | 3.7115 | 1.058 | 0.0004513 | +++?? | 0 | 0.6613 | intron | ELP2 |
| rs115656500 | 3 | 159272819 | 4 | 0 | 0 | A | G | 2.4789 | 0.7066 | 0.0004513 | +++?+ | 43.1 | 0.1527 | intron | SCHIP1,IQCJ-SCHIP1 |
| rs10742108 | 11 | 25664199 | 4 | 0 | 0 | T | C | -0.8951 | 0.2551 | 0.0004514 | ---?- | 0 | 0.6107 | intergenic | |
| rs11070345 | 15 | 41958907 | 5 | 0 | 0 | A | G | -0.9967 | 0.2841 | 0.0004517 | ----- | 12.3 | 0.3356 | intron | MGA |
| rs35754451 | 3 | 33266421 | 4 | 0 | 0 | T | C | -0.9353 | 0.2666 | 0.0004517 | ---?- | 9.9 | 0.3438 | intergenic | |
| rs6415459 | 8 | 113301936 | 4 | 0 | 0 | A | T | 0.9968 | 0.2842 | 0.0004518 | +++?+ | 0 | 0.7789 | intron | CSMD3 |
| rs139602069 | 3 | 143070599 | 4 | 0 | 0 | A | G | -2.1264 | 0.6062 | 0.0004518 | ---?- | 0 | 0.7134 | intron | SLC9A9 |
| rs77771142 | 13 | 73928262 | 3 | 0 | 0 | T | G | -3.1745 | 0.905 | 0.0004519 | ---?? | 0 | 0.935 | intergenic | |
| 8:96942324 | 8 | 96942324 | 3 | 0 | 0 | A | AG | -1.1107 | 0.3167 | 0.0004521 | ---?? | 0 | 0.765 | | |
| rs11226013 | 11 | 103659661 | 4 | 0 | 0 | T | G | 1.5101 | 0.4305 | 0.0004523 | +++?+ | 0 | 0.5183 | intergenic | |
| 22:50775538 | 22 | 50775538 | 3 | 0 | 0 | A | AGT | -0.9919 | 0.2828 | 0.0004523 | ---?? | 0 | 0.9691 | | |
| 15:85567249 | 15 | 85567249 | 3 | 0 | 0 | CAAA | C | -0.8879 | 0.2532 | 0.0004525 | ---?? | 65.8 | 0.05386 | | |
| rs3129946 | 6 | 32346772 | 5 | 0 | 0 | A | G | -0.7489 | 0.2135 | 0.0004526 | +---- | 39.3 | 0.1594 | intergenic |  |
| rs147670765 | 8 | 65772858 | 4 | 0 | 0 | A | G | 3.8186 | 1.0887 | 0.0004526 | +++?+ | 0 | 0.78 | intergenic | |
| rs929686 | 8 | 113302023 | 4 | 0 | 0 | A | G | -0.9977 | 0.2845 | 0.0004527 | ---?- | 0 | 0.7773 | intron | CSMD3 |
| rs72839992 | 17 | 56393128 | 5 | 0 | 0 | A | G | 1.9501 | 0.556 | 0.0004527 | +++++ | 48.7 | 0.09921 | intron | BZRAP1 |
| rs139925737 | 11 | 59005239 | 5 | 0 | 0 | A | G | -0.7563 | 0.2157 | 0.000453 | ----+ | 17 | 0.3063 | intergenic |  |
| rs763110 | 1 | 172627498 | 4 | 0 | 0 | T | C | 0.9344 | 0.2664 | 0.0004531 | +++?+ | 32.3 | 0.2186 | upstream-gene | |
| rs375436029 | 2 | 28797411 | 3 | 0 | 0 | A | G | 4.1707 | 1.1892 | 0.0004531 | +++?? | 0 | 0.6488 | intron | PLB1 |
| rs4974287 | 3 | 70445283 | 4 | 0 | 0 | T | C | 1.0263 | 0.2927 | 0.0004534 | +++?+ | 4.6 | 0.37 | intergenic | |
| rs915510 | 13 | 60041767 | 4 | 0 | 0 | T | G | -0.9228 | 0.2632 | 0.0004537 | ---?- | 0 | 0.62 | intergenic | |

| | | | | | | | | | | | | | | | |
|---|---|---|---|---|---|---|---|---|---|---|---|---|---|---|---|
| rs3117104 | 6 | 32348309 | 5 | 0 | 0 | A | T | 0.749 | 0.2136 | 0.0004537 | -++++ | 39.2 | 0.1599 | intergenic | |
| rs201182972 | 6 | 31223685 | 3 | 0 | 0 | T | C | -0.9877 | 0.2817 | 0.0004538 | ---?? | 18.9 | 0.2914 | intergenic | |
| rs117158628 | 22 | 48485449 | 4 | 0 | 0 | A | G | -1.7199 | 0.4905 | 0.0004539 | ---?+ | 61.3 | 0.05149 | intergenic | |
| rs4530903 | 6 | 32581889 | 5 | 0 | 1 | T | C | 1.3619 | 0.3884 | 0.000454 | +++++ | 12.9 | 0.3317 | intergenic | |
| rs4981401 | 14 | 22498482 | 4 | 0 | 0 | A | T | -0.9245 | 0.2637 | 0.0004541 | ---?- | 0 | 0.9317 | intergenic | |
| rs7834691 | 8 | 113240538 | 4 | 0 | 0 | T | C | -0.9968 | 0.2843 | 0.0004541 | ---?- | 0 | 0.9654 | intron | CSMD3 |
| rs7108052 | 11 | 59242976 | 3 | 0 | 0 | A | G | -3.469 | 0.9894 | 0.0004544 | -?--? | 15.9 | 0.3047 | upstream-gene | |
| rs117867565 | 9 | 115435679 | 4 | 0 | 0 | A | G | 2.6639 | 0.7598 | 0.0004545 | +++?+ | 0 | 0.6562 | intergenic | |
| 5:106810477 | 5 | 106810477 | 3 | 0 | 0 | T | TGCTGG | -1.4077 | 0.4015 | 0.0004545 | ---?? | 65.7 | 0.05424 | | |
| rs4915661 | 1 | 64747984 | 4 | 0 | 0 | T | G | 1.2411 | 0.354 | 0.0004549 | +++?+ | 0 | 0.4973 | intergenic | |
| rs76440435 | 9 | 95838450 | 5 | 0 | 0 | T | C | -2.3984 | 0.6841 | 0.000455 | ----- | 0 | 0.9775 | intron | SUSD3 |
| rs73337077 | 17 | 47947919 | 5 | 0 | 0 | T | C | -2.0588 | 0.5872 | 0.0004551 | ----- | 0 | 0.9739 | intergenic |  |
| 6:149328251 | 6 | 149328251 | 3 | 0 | 0 | CTTTT | C | -1.0236 | 0.292 | 0.0004551 | ---?? | 26 | 0.259 | | |
| rs3114670 | 1 | 90244009 | 5 | 0 | 0 | A | G | 0.7464 | 0.2129 | 0.0004553 | +++++ | 0 | 0.987 | intergenic | |
| rs7799309 | 7 | 141176988 | 4 | 0 | 0 | T | C | -4.7734 | 1.3616 | 0.0004555 | ---?- | 0 | 0.8253 | 3-prime-UTR | TMEM178B |
| rs61974395 | 14 | 28218023 | 4 | 0 | 0 | T | C | -2.0164 | 0.5752 | 0.0004556 | ---?- | 0 | 0.8048 | intergenic | |
| 9:21570274 | 9 | 21570274 | 3 | 0 | 0 | A | AT | 1.6722 | 0.477 | 0.0004556 | +++?? | 0 | 0.9295 | | |
| rs4324488 | 3 | 195931868 | 2 | 0 | 0 | T | C | 1.6898 | 0.482 | 0.0004557 | ???++ | 33.1 | 0.2216 | intron | ZDHHC19 |
| rs17155896 | 5 | 103172273 | 4 | 0 | 0 | A | G | -1.5583 | 0.4445 | 0.0004557 | ---?- | 0 | 0.9016 | intergenic | |
| rs1160073 | 6 | 99687655 | 4 | 0 | 0 | T | C | -0.8619 | 0.2459 | 0.0004559 | ---?- | 55.6 | 0.08019 | intergenic | |
| rs1861753 | 8 | 113267771 | 4 | 0 | 0 | T | C | -1.0063 | 0.2871 | 0.000456 | ---?- | 0 | 0.9289 | intron | CSMD3 |
| rs1008411 | 3 | 70445977 | 4 | 0 | 0 | T | C | -1.1542 | 0.3293 | 0.0004563 | ---?- | 0 | 0.5342 | intergenic | |
| rs10789550 | 11 | 106754354 | 4 | 0 | 0 | T | C | 1.0234 | 0.292 | 0.0004564 | +++?- | 35.7 | 0.198 | intron | GUCY1A2 |
| rs12644782 | 4 | 20738401 | 4 | 0 | 0 | A | G | 0.9124 | 0.2603 | 0.0004565 | +++?+ | 0 | 0.8265 | intron | KCNIP4 |
| rs35484767 | 15 | 41945568 | 5 | 0 | 0 | A | G | -0.9921 | 0.283 | 0.0004565 | ----- | 14.2 | 0.3236 | intron | MGA |
| rs189720659 | 13 | 89516376 | 3 | 0 | 0 | A | G | 3.2305 | 0.9217 | 0.0004567 | +++?? | 0 | 0.4005 | intergenic | |
| rs3751230 | 12 | 27786025 | 4 | 0 | 0 | T | G | -0.888 | 0.2534 | 0.000457 | ---?+ | 53.5 | 0.09158 | intron | PPFIBP1 |
| rs11071019 | 15 | 54351589 | 3 | 0 | 0 | A | G | -1.2105 | 0.3454 | 0.000457 | ---?? | 0 | 0.9824 | intron | UNC13C |
| rs73080274 | 12 | 27774693 | 4 | 0 | 0 | A | G | -0.9779 | 0.279 | 0.0004572 | ---?+ | 0 | 0.4274 | intron | PPFIBP1 |
| rs13192471 | 6 | 32671103 | 5 | 0 | 1 | T | C | -1.1322 | 0.3231 | 0.0004574 | ----- | 18.6 | 0.296 | intergenic |  |
| rs17206147 | 6 | 32665311 | 5 | 0 | 1 | T | C | -1.1322 | 0.3231 | 0.0004574 | ----- | 18.6 | 0.296 | intergenic |  |
| rs35120848 | 6 | 32670495 | 5 | 0 | 1 | T | C | 1.1322 | 0.3231 | 0.0004574 | +++++ | 18.6 | 0.296 | intergenic | |
| rs66709408 | 6 | 32670414 | 5 | 0 | 1 | T | C | 1.1322 | 0.3231 | 0.0004574 | +++++ | 18.6 | 0.296 | intergenic | |
| rs545925 | 1 | 90243692 | 5 | 0 | 0 | A | G | 0.7461 | 0.2129 | 0.0004575 | +++++ | 0 | 0.9871 | intergenic | |
| 22:49402144 | 22 | 49402144 | 3 | 0 | 0 | A | AGTGTG | -1.1623 | 0.3317 | 0.0004575 | ---?? | 44.2 | 0.1664 | | |
| rs2882044 | 7 | 135500933 | 4 | 0 | 0 | A | G | -1.9771 | 0.5642 | 0.0004577 | ---?+ | 0 | 0.7361 | intergenic | |
| rs1049100 | 6 | 32629963 | 4 | 0 | 1 | T | C | 1.1016 | 0.3144 | 0.0004578 | ++++? | 32.8 | 0.2156 | missense | HLA-DQB1 |
| rs11226010 | 11 | 103651279 | 4 | 0 | 0 | A | C | -1.5145 | 0.4322 | 0.0004578 | ---?- | 0 | 0.5351 | intergenic | |
| rs148507186 | 17 | 57018515 | 5 | 0 | 0 | T | C | -2.4152 | 0.6893 | 0.0004581 | ----- | 0 | 0.8033 | intron | PPM1E |
| rs35040284 | 6 | 32586068 | 5 | 0 | 1 | T | C | -1.3488 | 0.3849 | 0.0004583 | ----- | 16.7 | 0.3084 | intergenic |  |
| rs200888561 | 4 | 12247942 | 3 | 0 | 0 | A | G | -1.2784 | 0.3648 | 0.0004584 | ---?? | 0 | 0.4678 | intergenic | |
| rs17451397 | 5 | 107563200 | 4 | 0 | 0 | A | G | 1.3971 | 0.3987 | 0.0004586 | +++?+ | 0 | 0.9949 | intron | FBXL17 |
| rs2568164 | 2 | 218328738 | 4 | 0 | 0 | A | C | -0.8683 | 0.2478 | 0.0004587 | ---?- | 17 | 0.3061 | intron | DIRC3 |
| rs7693669 | 4 | 14433990 | 4 | 0 | 0 | A | G | -0.8953 | 0.2555 | 0.0004587 | ---?- | 0 | 0.69 | intergenic | |
| rs11603584 | 11 | 12802158 | 4 | 0 | 0 | A | T | 1.4562 | 0.4156 | 0.0004589 | +++?+ | 0 | 0.6075 | intron | TEAD1 |
| 20:56984956 | 20 | 56984956 | 3 | 0 | 0 | CAT | C | -1.6356 | 0.4668 | 0.0004589 | ---?? | 0 | 0.4053 | | |
| rs142362232 | 18 | 52482894 | 3 | 0 | 0 | T | C | -2.1078 | 0.6016 | 0.000459 | ---?? | 0 | 0.5574 | intron | RAB27B |
| rs4884338 | 13 | 60031815 | 4 | 0 | 0 | T | C | 0.8736 | 0.2494 | 0.0004592 | +++?+ | 0 | 0.4547 | intergenic | |
| rs10916417 | 1 | 229221155 | 4 | 0 | 0 | T | C | 1.281 | 0.3656 | 0.0004592 | +++?+ | 0 | 0.9013 | intergenic | |
| rs73780509 | 6 | 144623043 | 5 | 0 | 0 | T | C | 1.4189 | 0.405 | 0.0004596 | ++++- | 40.4 | 0.1519 | intron | UTRN |
| rs7322235 | 13 | 45530211 | 3 | 0 | 0 | A | G | -2.2914 | 0.6541 | 0.0004597 | ---?? | 0 | 0.8184 | intron | NUFIP1 |
| rs35406540 | 6 | 32670368 | 5 | 0 | 1 | A | G | -1.1318 | 0.3231 | 0.0004598 | ----- | 18.6 | 0.2961 | intergenic |  |
| rs187662999 | 2 | 35664260 | 3 | 0 | 0 | A | G | -3.565 | 1.0177 | 0.00046 | ---?? | 0 | 0.6547 | intergenic | |
| rs61744121 | 17 | 56572347 | 5 | 0 | 0 | A | G | -2.4204 | 0.6909 | 0.0004601 | ----- | 0 | 0.7092 | synonymous | MTMR4 |
| rs3930234 | 3 | 184216130 | 5 | 0 | 0 | T | C | -0.9483 | 0.2707 | 0.0004602 | ----- | 0 | 0.7521 | intergenic |  |
| rs10895525 | 11 | 103663377 | 4 | 0 | 0 | A | G | -1.5043 | 0.4295 | 0.0004602 | ---?- | 0 | 0.5081 | intergenic | |
| 1:102960063 | 1 | 102960063 | 3 | 0 | 0 | A | ATT | 0.889 | 0.2538 | 0.0004603 | +++?? | 0 | 0.9985 | | |
| 1:78444764 | 1 | 78444764 | 3 | 0 | 0 | CGGCCG | C | 1.5412 | 0.44 | 0.0004604 | +++?? | 0 | 0.5536 | | |
| rs9671298 | 14 | 22501595 | 4 | 0 | 0 | T | G | -0.9295 | 0.2654 | 0.0004605 | ---?- | 0 | 0.9544 | intergenic | |

| | | | | | | | | | | | | | | | |
|---|---|---|---|---|---|---|---|---|---|---|---|---|---|---|---|
| rs12732786 | 1 | 29022560 | 4 | 0 | 0 | A | G | -1.0654 | 0.3042 | 0.0004605 | ---?- | 0 | 0.7252 | intron | GMEB1 |
| rs2882045 | 7 | 135500931 | 4 | 0 | 0 | A | G | -1.9761 | 0.5642 | 0.0004608 | ---?+ | 0 | 0.7363 | intergenic | |
| rs9743943 | 14 | 43268067 | 4 | 0 | 0 | A | G | 1.3151 | 0.3755 | 0.0004608 | +++?- | 61.6 | 0.05024 | intergenic | |
| rs16943321 | 17 | 57049095 | 5 | 0 | 0 | A | G | 2.3847 | 0.6809 | 0.0004609 | +++++ | 0 | 0.8451 | intron | PPM1E |
| rs60302127 | 17 | 5865114 | 2 | 0 | 0 | A | G | 4.3107 | 1.2308 | 0.0004609 | ??++? | 0 | 0.9414 | intergenic | |
| rs79529635 | 2 | 68140768 | 4 | 0 | 0 | A | G | -4.1204 | 1.1764 | 0.0004609 | ---?- | 0 | 0.7443 | intergenic | |
| rs76958973 | 7 | 25604123 | 4 | 0 | 0 | T | C | 1.6537 | 0.4722 | 0.0004611 | +++?- | 50.2 | 0.1103 | intergenic | |
| rs114432555 | 18 | 40776877 | 3 | 0 | 0 | T | C | 2.2318 | 0.6372 | 0.0004611 | +++?? | 0 | 0.6498 | intergenic | |
| rs1158628 | 8 | 140406286 | 4 | 0 | 0 | A | T | -1.0402 | 0.297 | 0.0004612 | ---?- | 0 | 0.8853 | intergenic | |
| rs115418659 | 6 | 32167532 | 5 | 0 | 0 | A | G | -2.7932 | 0.7975 | 0.0004612 | ----- | 24.4 | 0.259 | intron | NOTCH4 |
| rs7324221 | 13 | 60044372 | 4 | 0 | 0 | A | C | 0.9191 | 0.2624 | 0.0004613 | +++?+ | 0 | 0.5711 | intergenic | |
| rs72746602 | 9 | 97615467 | 4 | 0 | 0 | T | C | 1.7611 | 0.5029 | 0.0004614 | +++?+ | 0 | 0.9355 | intron | C9orf3 |
| rs147002250 | 4 | 26170990 | 3 | 0 | 0 | A | G | -1.2547 | 0.3583 | 0.0004614 | ---?? | 41.4 | 0.1816 | intron | RBPJ |
| rs67879832 | 5 | 39600099 | 4 | 0 | 0 | T | G | 1.031 | 0.2944 | 0.0004615 | +++?+ | 0 | 0.8318 | intergenic | |
| 17:65478217 | 17 | 65478217 | 3 | 0 | 0 | A | AC | 0.9316 | 0.266 | 0.0004616 | +++?? | 0 | 0.621 | | |
| rs115272165 | 2 | 232137162 | 4 | 0 | 0 | T | C | 1.5464 | 0.4416 | 0.0004619 | +++?+ | 0 | 0.7114 | intron | ARMC9 |
| rs8111925 | 19 | 14748113 | 4 | 0 | 0 | T | C | -0.9221 | 0.2633 | 0.000462 | ---?+ | 47.8 | 0.1245 | intron | EMR3 |
| rs10875247 | 1 | 100206896 | 4 | 0 | 0 | A | C | 1.7625 | 0.5033 | 0.000462 | +++?+ | 0 | 0.7176 | intron | FRRS1 |
| rs79215842 | 17 | 57036018 | 5 | 0 | 0 | A | T | 2.4064 | 0.6872 | 0.0004622 | +++++ | 0 | 0.8057 | intron | PPM1E |
| rs12002003 | 9 | 16196717 | 4 | 0 | 0 | A | G | -1.2497 | 0.3569 | 0.0004625 | ---?- | 0 | 0.5804 | intergenic | |
| rs11242508 | 5 | 103160351 | 4 | 0 | 0 | T | C | -1.5828 | 0.452 | 0.0004627 | ---?- | 0 | 0.7982 | intergenic | |
| rs12131316 | 1 | 217984483 | 4 | 0 | 0 | T | C | 1.1378 | 0.325 | 0.0004627 | +++?+ | 0 | 0.957 | intron | SPATA17 |
| rs73412716 | 12 | 112545197 | 5 | 0 | 0 | T | G | 1.4483 | 0.4136 | 0.0004627 | +-+++ | 33.7 | 0.1967 | intron | NAA25 |
| rs9317080 | 13 | 60041410 | 4 | 0 | 0 | A | C | -0.9218 | 0.2633 | 0.0004628 | ---?- | 0 | 0.6043 | intergenic | |
| rs12120342 | 1 | 217984553 | 4 | 0 | 0 | A | T | 1.1379 | 0.325 | 0.0004628 | +++?+ | 0 | 0.9573 | intron | SPATA17 |
| rs79383531 | 14 | 43748358 | 4 | 0 | 0 | T | C | -1.7496 | 0.4997 | 0.0004629 | ---?+ | 0 | 0.5967 | intergenic | |
| rs13146801 | 4 | 148023828 | 3 | 0 | 0 | T | C | -0.8798 | 0.2513 | 0.0004631 | ---?? | 55.3 | 0.1066 | intergenic | |
| rs77192360 | 9 | 36951451 | 4 | 0 | 0 | A | G | 1.8144 | 0.5182 | 0.0004631 | +++?+ | 0 | 0.5419 | intron | PAX5 |
| 6:99921261 | 6 | 99921261 | 3 | 0 | 0 | CGAA | C | -1.8171 | 0.519 | 0.0004631 | ---?? | 0 | 0.651 | | |
| rs2465879 | 8 | 18438022 | 4 | 0 | 0 | A | T | 0.853 | 0.2436 | 0.0004633 | +++?+ | 0 | 0.8079 | intron | PSD3 |
| rs7647686 | 3 | 29528415 | 4 | 0 | 0 | T | C | -1.4409 | 0.4116 | 0.0004633 | ---?+ | 0 | 0.823 | intron | RBMS3 |
| rs144304610 | 17 | 57050603 | 5 | 0 | 0 | T | C | -2.4014 | 0.6859 | 0.0004633 | ----- | 0 | 0.8106 | intron | PPM1E |
| rs62222174 | 22 | 20022746 | 4 | 0 | 0 | T | C | 1.5198 | 0.4341 | 0.0004635 | +++?- | 0 | 0.5453 | intron | TANGO2 |
| rs1899819 | 15 | 41913668 | 5 | 0 | 0 | T | C | 1.0077 | 0.2878 | 0.0004636 | +++++ | 8 | 0.3612 | intron | MGA |
| rs193054699 | 12 | 132052467 | 4 | 0 | 0 | A | G | -3.1438 | 0.898 | 0.0004636 | ---?- | 0 | 0.9863 | intergenic | |
| rs140938064 | 17 | 57020302 | 5 | 0 | 0 | T | C | 2.4122 | 0.689 | 0.0004637 | +++++ | 0 | 0.8031 | intron | PPM1E |
| rs188721333 | 19 | 33452970 | 2 | 0 | 0 | A | G | -5.0634 | 1.4463 | 0.0004637 | --??? | 0 | 0.6597 | intron | CEP89 |
| rs36071900 | 4 | 190021448 | 4 | 0 | 0 | T | C | -0.8723 | 0.2492 | 0.0004639 | ---?- | 0 | 0.5545 | intergenic | |
| rs79848013 | 1 | 208186917 | 4 | 0 | 0 | T | C | -2.1907 | 0.6258 | 0.0004639 | ---?- | 0 | 0.8624 | intergenic | |
| rs7549625 | 1 | 90251996 | 5 | 0 | 0 | A | G | -0.7492 | 0.214 | 0.000464 | ----- | 0 | 0.9302 | intergenic |  |
| rs1416220 | 6 | 83713276 | 4 | 0 | 0 | A | G | -3.1726 | 0.9063 | 0.0004641 | ---?- | 0 | 0.8888 | intron | UBE3D |
| rs7123567 | 11 | 37057699 | 4 | 0 | 0 | A | C | 0.9277 | 0.265 | 0.0004642 | +++?+ | 0 | 0.4744 | intergenic | |
| rs142252418 | 20 | 62404871 | 4 | 0 | 0 | A | C | 3.9193 | 1.1196 | 0.0004644 | +++?- | 0 | 0.6341 | intron | ZBTB46 |
| rs185282488 | 12 | 132051466 | 4 | 0 | 0 | A | G | -3.1435 | 0.898 | 0.0004644 | ---?- | 0 | 0.9863 | intergenic | |
| rs10463558 | 5 | 103147801 | 4 | 0 | 0 | A | T | 1.5533 | 0.4437 | 0.0004645 | +++?+ | 0 | 0.7432 | intergenic | |
| rs3117098 | 6 | 32358513 | 5 | 0 | 0 | A | G | 0.7476 | 0.2136 | 0.0004648 | -++++ | 39.6 | 0.1574 | non-coding-exon | HCG23 |
| 2:170494851 | 2 | 170494851 | 3 | 0 | 0 | T | TA | 1.1994 | 0.3427 | 0.0004649 | +++?? | 0 | 0.9433 | | |
| rs200281315 | 11 | 37057342 | 4 | 0 | 0 | T | G | -0.9271 | 0.2649 | 0.0004656 | ---?- | 0 | 0.4751 | intergenic | |
| 6:72885850 | 6 | 72885850 | 3 | 0 | 0 | CTTTAG | C | -1.6872 | 0.4821 | 0.0004657 | ---?? | 0 | 0.8676 | | |
| rs117999099 | 13 | 73927839 | 3 | 0 | 0 | A | T | -3.17 | 0.9059 | 0.0004661 | ---?? | 0 | 0.9346 | intergenic | |
| rs143157978 | 17 | 57022199 | 5 | 0 | 0 | T | G | -2.4103 | 0.6888 | 0.0004661 | ----- | 0 | 0.8034 | intron | PPM1E |
| rs2347617 | 2 | 47386395 | 4 | 0 | 0 | C | G | 1.3708 | 0.3917 | 0.0004663 | +++?+ | 0 | 0.964 | upstream-gene | |
| rs138888809 | 8 | 4004756 | 4 | 0 | 0 | T | G | -3.8871 | 1.1108 | 0.0004665 | ---?- | 0 | 0.7445 | intron | CSMD1 |
| rs62516487 | 8 | 113508984 | 4 | 0 | 0 | C | G | -0.9091 | 0.2598 | 0.0004666 | ---?- | 0 | 0.7034 | intron | CSMD3 |
| rs117260136 | 6 | 72563892 | 4 | 0 | 0 | T | C | -1.7483 | 0.4996 | 0.0004666 | ---?- | 0 | 0.6013 | intergenic | |
| rs10107418 | 8 | 113508993 | 4 | 0 | 0 | T | G | -0.9091 | 0.2598 | 0.0004667 | ---?- | 0 | 0.7034 | intron | CSMD3 |
| rs6447977 | 4 | 20736741 | 4 | 0 | 0 | A | C | -0.9102 | 0.2601 | 0.0004669 | ---?- | 0 | 0.8281 | intron | KCNIP4 |
| rs2771335 | 14 | 21453373 | 4 | 0 | 0 | C | G | -0.8709 | 0.2489 | 0.000467 | ---?- | 0 | 0.8587 | upstream-gene | |
| rs201448 | 7 | 101738860 | 4 | 0 | 0 | T | C | -0.8578 | 0.2452 | 0.0004673 | ---?- | 0 | 0.8004 | intron | CUX1 |

| | | | | | | | | | | | | | | | |
|---|---|---|---|---|---|---|---|---|---|---|---|---|---|---|---|
| rs6664162 | 1 | 192462574 | 4 | 0 | 0 | A | G | -1.1035 | 0.3154 | 0.0004673 | ---?+ | 33.9 | 0.2092 | intergenic | |
| rs73839213 | 3 | 61066311 | 4 | 0 | 0 | A | C | 3.3391 | 0.9543 | 0.0004673 | +++?- | 0 | 0.6456 | intron | FHIT |
| rs76362302 | 2 | 238522083 | 3 | 0 | 0 | A | C | -4.476 | 1.2793 | 0.0004673 | -?-?- | 0 | 0.4908 | intergenic | |
| rs11161572 | 1 | 85722829 | 5 | 0 | 0 | A | T | 1.6228 | 0.4638 | 0.0004675 | +++++ | 13.9 | 0.3256 | intron | C1orf52 |
| rs7009195 | 8 | 113304534 | 4 | 0 | 0 | T | C | -0.9938 | 0.2841 | 0.0004676 | ---?- | 0 | 0.775 | intron | CSMD3 |
| rs72900958 | 2 | 68127630 | 4 | 0 | 0 | A | G | -4.0655 | 1.162 | 0.0004677 | ---?- | 0 | 0.5577 | intergenic | |
| rs11925380 | 3 | 61066520 | 4 | 0 | 0 | T | C | 3.338 | 0.9542 | 0.0004683 | +++?- | 0 | 0.6449 | intron | FHIT |
| rs139281924 | 11 | 85451543 | 2 | 0 | 0 | A | G | 6.4274 | 1.8373 | 0.0004683 | +???+ | 0 | 0.7681 | intron | SYTL2 |
| rs113028686 | 15 | 91537820 | 4 | 0 | 0 | A | G | 2.7442 | 0.7844 | 0.0004684 | +++?+ | 0 | 0.5934 | 5-prime-UTR | PRC1 |
| rs145548106 | 10 | 11696266 | 3 | 0 | 0 | T | G | 3.5874 | 1.0255 | 0.0004684 | +++?? | 0 | 0.9596 | intergenic | |
| 8:114891449 | 8 | 114891449 | 3 | 0 | 0 | CACAA | C | -2.0143 | 0.5758 | 0.0004684 | ---?? | 0 | 0.4782 | | |
| rs113392659 | 15 | 86349364 | 3 | 0 | 0 | A | G | 1.6348 | 0.4673 | 0.0004685 | +++?? | 0 | 0.6253 | intergenic | |
| rs34075028 | 13 | 20832915 | 4 | 0 | 0 | A | G | -1.0454 | 0.2989 | 0.0004688 | ---?- | 0 | 0.9144 | intergenic | |
| rs1773883 | 10 | 29183994 | 4 | 0 | 0 | A | G | 0.9221 | 0.2636 | 0.000469 | +++?+ | 0 | 0.468 | intergenic | |
| rs11925426 | 3 | 61066648 | 4 | 0 | 0 | T | C | 3.3371 | 0.9541 | 0.000469 | +++?- | 0 | 0.6449 | intron | FHIT |
| rs60606703 | 10 | 86700022 | 4 | 0 | 0 | A | G | -1.7599 | 0.5031 | 0.0004691 | ---?- | 0 | 0.849 | intergenic | |
| 4:36716729 | 4 | 36716729 | 3 | 0 | 0 | AT | ATT | 1.2872 | 0.368 | 0.0004695 | +++?? | 0 | 0.9207 | | |
| rs35900428 | 6 | 32591755 | 5 | 0 | 1 | A | G | 1.3529 | 0.3868 | 0.0004696 | +++++ | 14.2 | 0.3236 | intergenic | |
| rs55649307 | 7 | 82412950 | 4 | 0 | 0 | A | C | 0.9071 | 0.2593 | 0.0004696 | +-+?+ | 46.1 | 0.1346 | intron | PCLO |
| rs193299383 | 17 | 16325340 | 2 | 0 | 0 | T | C | -6.3164 | 1.806 | 0.0004696 | --??? | 71 | 0.06329 | intron | TRPV2 |
| rs7467823 | 9 | 103872681 | 4 | 0 | 0 | A | G | 1.0522 | 0.3009 | 0.0004699 | +++?+ | 59.3 | 0.061 | intron | LPPR1 |
| rs62524607 | 8 | 88834747 | 4 | 0 | 0 | T | C | 1.4916 | 0.4265 | 0.0004701 | +++?+ | 0 | 0.6759 | intergenic | |
| rs71624487 | 4 | 41332427 | 4 | 0 | 0 | A | C | -1.4029 | 0.4012 | 0.0004706 | ---?+ | 0 | 0.4403 | intergenic | |
| rs117391892 | 11 | 1489458 | 4 | 0 | 0 | A | G | 2.4356 | 0.6965 | 0.0004706 | +++?+ | 0 | 0.7734 | downstream-gene | |
| 4:148024074 | 4 | 148024074 | 3 | 0 | 0 | CA | C | 0.8883 | 0.254 | 0.0004706 | +++?? | 40.6 | 0.1859 | | |
| rs10921171 | 1 | 192462941 | 4 | 0 | 0 | T | G | -1.1082 | 0.3169 | 0.0004709 | ---?+ | 34.1 | 0.2077 | intergenic | |
| rs112513636 | 2 | 47385434 | 4 | 0 | 0 | T | C | 1.3681 | 0.3913 | 0.0004709 | +++?+ | 0 | 0.9634 | upstream-gene | |
| rs6573766 | 14 | 24852046 | 5 | 0 | 0 | T | G | 0.9152 | 0.2617 | 0.0004711 | +++++ | 0 | 0.9373 | downstream-gene | |
| rs7935490 | 11 | 103669593 | 4 | 0 | 0 | T | C | 1.4914 | 0.4265 | 0.0004712 | +++?+ | 0 | 0.4916 | intergenic | |
| 1:85615105 | 1 | 85615105 | 3 | 0 | 0 | GA | GAAA | 1.5359 | 0.4392 | 0.0004712 | +++?? | 0 | 0.7742 | | |
| rs3117108 | 6 | 32342822 | 5 | 0 | 0 | C | G | -0.7471 | 0.2137 | 0.0004714 | +---- | 39.5 | 0.1581 | upstream-gene |  |
| rs17403397 | 11 | 103667245 | 4 | 0 | 0 | A | G | -1.4929 | 0.427 | 0.0004715 | ---?- | 0 | 0.4945 | intergenic | |
| rs7834235 | 8 | 118538683 | 3 | 0 | 0 | T | G | 1.0691 | 0.3058 | 0.0004716 | ?++?+ | 0 | 0.7883 | intron | MED30 |
| rs140322127 | 3 | 59879149 | 3 | 0 | 0 | A | C | 4.6748 | 1.337 | 0.0004716 | +?+?+ | 28.4 | 0.2474 | intron | FHIT |
| rs17809667 | 2 | 208285598 | 4 | 0 | 0 | C | G | -1.6202 | 0.4634 | 0.0004717 | ---?- | 0 | 0.7097 | intergenic | |
| rs56260470 | 3 | 61067457 | 4 | 0 | 0 | T | G | 3.3346 | 0.9537 | 0.0004717 | +++?- | 0 | 0.644 | intron | FHIT |
| rs147928696 | 17 | 57132519 | 5 | 0 | 0 | A | G | -2.3218 | 0.6641 | 0.0004717 | ----- | 0 | 0.8215 | intron | TRIM37 |
| rs10098507 | 8 | 113512207 | 4 | 0 | 0 | T | G | -0.9098 | 0.2602 | 0.0004718 | ---?- | 0 | 0.6995 | intron | CSMD3 |
| 1:20068235 | 1 | 20068235 | 3 | 0 | 0 | CT | C | 1.4944 | 0.4274 | 0.0004718 | +++?? | 1.9 | 0.3608 | | |
| rs1770701 | 1 | 85606661 | 4 | 0 | 0 | T | C | -1.2447 | 0.356 | 0.000472 | ---?- | 0 | 0.9985 | intergenic | |
| rs62521370 | 8 | 76209295 | 4 | 0 | 0 | A | G | 1.5684 | 0.4486 | 0.000472 | +++?+ | 0 | 0.6079 | intergenic | |
| rs116245601 | 4 | 157975647 | 3 | 0 | 0 | T | G | -4.2412 | 1.2131 | 0.000472 | ?--?- | 0 | 0.7307 | intergenic | |
| rs891988 | 5 | 141467283 | 5 | 0 | 0 | T | G | 0.8279 | 0.2368 | 0.0004721 | -++++ | 30.9 | 0.2154 | intergenic | |
| rs34772037 | 1 | 64778490 | 4 | 0 | 0 | A | G | 1.2318 | 0.3523 | 0.0004723 | +++?+ | 0 | 0.5223 | intergenic | |
| rs56240052 | 6 | 118580405 | 4 | 0 | 0 | A | G | -2.4372 | 0.6971 | 0.0004723 | ---?- | 0 | 0.9988 | intron | SLC35F1 |
| rs115032105 | 17 | 57174257 | 5 | 0 | 0 | T | C | -2.3978 | 0.6859 | 0.0004724 | ----- | 0 | 0.8117 | intron | TRIM37 |
| rs77592522 | 1 | 85144317 | 4 | 0 | 0 | A | G | -2.3416 | 0.6698 | 0.0004725 | ---?- | 0 | 0.7898 | intron | SSX2IP |
| rs6489833 | 12 | 112709566 | 5 | 0 | 0 | T | G | -1.3033 | 0.3728 | 0.0004726 | -+--- | 55.4 | 0.06214 | intron | HECTD4 |
| rs4934011 | 10 | 82022878 | 3 | 0 | 0 | T | G | -1.0018 | 0.2866 | 0.0004727 | ---?? | 0 | 0.6275 | intergenic | |
| rs3799162 | 6 | 165799614 | 4 | 0 | 0 | A | G | -0.9152 | 0.2618 | 0.0004729 | ---?- | 0 | 0.8734 | intron | PDE10A |
| rs12363457 | 11 | 103665923 | 4 | 0 | 0 | A | C | 1.4979 | 0.4285 | 0.0004729 | +++?+ | 0 | 0.4958 | intergenic | |
| rs74382543 | 5 | 122531550 | 4 | 0 | 0 | T | G | -1.8417 | 0.5269 | 0.0004729 | ---?- | 0 | 0.8221 | downstream-gene | |
| 7:4608261 | 7 | 4608261 | 3 | 0 | 0 | CT | C | 1.0089 | 0.2886 | 0.0004729 | +++?? | 36.8 | 0.2056 | | |
| rs1878976 | 1 | 18758343 | 4 | 0 | 0 | T | C | -1.0636 | 0.3043 | 0.000473 | ---?- | 46 | 0.1352 | intergenic | |
| rs11809033 | 1 | 203689116 | 4 | 0 | 0 | T | C | 1.1305 | 0.3234 | 0.0004731 | +++?+ | 0 | 0.9308 | intron | ATP2B4 |
| rs45622432 | 9 | 35659107 | 5 | 0 | 0 | T | C | 0.9456 | 0.2705 | 0.0004731 | ++++- | 29 | 0.2284 | intron | CCDC107 |
| rs1899820 | 15 | 41951572 | 5 | 0 | 0 | A | C | 1.0019 | 0.2866 | 0.0004733 | +++++ | 12.5 | 0.3343 | intron | MGA |
| rs77764116 | 17 | 47452864 | 4 | 0 | 0 | A | G | -1.0459 | 0.2992 | 0.0004734 | ---?+ | 54.5 | 0.08611 | intergenic | |
| rs60690718 | 2 | 123823909 | 4 | 0 | 0 | A | T | 0.923 | 0.2641 | 0.0004737 | +++?+ | 0 | 0.9284 | intergenic | |

| | | | | | | | | | | | | | | | |
|---|---|---|---|---|---|---|---|---|---|---|---|---|---|---|---|
| rs74362234 | 11 | 125786163 | 4 | 0 | 0 | T | C | -1.8321 | 0.5242 | 0.0004737 | ---?- | 0 | 0.4451 | intron | DDX25 |
| rs190709101 | 17 | 57186996 | 5 | 0 | 0 | A | G | 2.4258 | 0.6941 | 0.0004737 | +++++ | 0 | 0.815 | upstream-gene | |
| rs6482992 | 10 | 129678674 | 4 | 0 | 0 | T | C | 0.8686 | 0.2485 | 0.0004738 | +++?- | 1.1 | 0.3865 | intron | CLRN3 |
| rs60776684 | 18 | 25184580 | 4 | 0 | 0 | A | G | -2.5407 | 0.7269 | 0.0004738 | ---?- | 0 | 0.6242 | intergenic | |
| 19:36538739 | 19 | 36538739 | 3 | 0 | 0 | A | AAAAG | 1.6197 | 0.4634 | 0.0004738 | +-+?? | 59.5 | 0.08448 | | |
| rs2555658 | 4 | 175489046 | 4 | 0 | 0 | A | C | -0.9151 | 0.2618 | 0.0004739 | ---?- | 0 | 0.4623 | intergenic | |
| rs10489818 | 1 | 118099243 | 4 | 0 | 0 | A | G | 0.9576 | 0.274 | 0.0004739 | +++?+ | 0 | 0.4875 | intergenic | |
| rs531948 | 5 | 103139693 | 4 | 0 | 0 | T | C | -1.5488 | 0.4431 | 0.000474 | ---?- | 0 | 0.705 | intergenic | |
| rs1849206 | 6 | 165818537 | 4 | 0 | 0 | A | G | 0.8784 | 0.2513 | 0.0004742 | +++?+ | 0 | 0.6519 | intron | PDE10A |
| rs28393315 | 6 | 32603177 | 3 | 0 | 0 | A | G | -1.1532 | 0.33 | 0.0004743 | ---?? | 0 | 0.4128 | upstream-gene | |
| rs75226964 | 4 | 110083339 | 5 | 0 | 0 | T | C | 1.4211 | 0.4066 | 0.0004743 | +++++ | 0 | 0.6538 | intron | COL25A1 |
| rs201449 | 7 | 101739054 | 4 | 0 | 0 | A | G | -0.8564 | 0.245 | 0.0004744 | ---?- | 0 | 0.7996 | intron | CUX1 |
| rs6450379 | 5 | 55635039 | 3 | 0 | 0 | A | G | -5.1855 | 1.4838 | 0.0004744 | ---?? | 0 | 0.5875 | intergenic | |
| rs9853310 | 3 | 2493161 | 4 | 0 | 0 | A | G | -0.9404 | 0.2691 | 0.0004749 | ---?+ | 9.9 | 0.3434 | intron | CNTN4 |
| rs11966018 | 6 | 12209228 | 4 | 0 | 0 | C | G | -3.2319 | 0.9249 | 0.000475 | ---?- | 7.3 | 0.3567 | intergenic | |
| rs113082967 | 22 | 28688462 | 4 | 0 | 0 | T | C | 2.5716 | 0.7359 | 0.0004751 | +++?+ | 0 | 0.742 | intron | TTC28 |
| rs192338329 | 7 | 127406084 | 2 | 0 | 0 | T | C | 6.6108 | 1.8918 | 0.0004751 | ++??? | 63.2 | 0.09915 | intron | SND1 |
| rs6009354 | 22 | 49414516 | 4 | 0 | 0 | A | G | 1.4349 | 0.4106 | 0.0004752 | +++?+ | 0 | 0.3971 | intergenic | |
| rs78631540 | 13 | 104583110 | 3 | 0 | 0 | A | G | 5.5037 | 1.575 | 0.0004752 | +++?? | 0 | 0.4195 | intergenic | |
| rs1861754 | 8 | 113267916 | 4 | 0 | 0 | T | G | 1.0037 | 0.2873 | 0.0004754 | +++?+ | 0 | 0.9285 | intron | CSMD3 |
| rs11691444 | 2 | 185439634 | 4 | 0 | 0 | T | C | -0.9582 | 0.2742 | 0.0004757 | ---?- | 0 | 0.7649 | intergenic | |
| rs7990995 | 13 | 60049691 | 4 | 0 | 0 | T | C | 0.914 | 0.2616 | 0.0004758 | +++?+ | 0 | 0.7666 | intergenic | |
| rs9512507 | 13 | 27606554 | 5 | 0 | 0 | A | C | 1.0579 | 0.3028 | 0.0004758 | +++-+ | 36.5 | 0.1782 | intergenic | |
| 7:65481608 | 7 | 65481608 | 3 | 0 | 0 | A | AT | -1.6177 | 0.463 | 0.0004758 | ---?? | 0 | 0.6774 | | |
| rs117963807 | 8 | 17464395 | 3 | 0 | 0 | T | G | -1.2407 | 0.3551 | 0.0004759 | ---?? | 0 | 0.4803 | intron | PDGFRL |
| rs6868659 | 5 | 165429829 | 4 | 0 | 0 | A | G | -0.878 | 0.2513 | 0.000476 | ---?- | 3.4 | 0.3755 | intergenic | |
| rs11229899 | 11 | 59029526 | 5 | 0 | 0 | T | G | -0.7625 | 0.2182 | 0.000476 | -+--+ | 10.1 | 0.3485 | intergenic | |
| rs116584264 | 3 | 68195411 | 3 | 0 | 0 | T | C | 4.1664 | 1.1925 | 0.000476 | +++?? | 0 | 0.6268 | intron | FAM19A1 |
| rs116717924 | 4 | 133559200 | 3 | 0 | 0 | T | G | -4.6038 | 1.3177 | 0.0004761 | ?--?- | 0 | 0.8536 | intergenic | |
| rs184069664 | 2 | 168024009 | 4 | 0 | 0 | A | T | -2.6143 | 0.7483 | 0.0004762 | ---?+ | 30.7 | 0.228 | intron | XIRP2 |
| rs115752667 | 3 | 71318158 | 3 | 0 | 0 | T | C | 4.6204 | 1.3225 | 0.0004765 | +++?? | 8.3 | 0.3359 | intron | FOXP1 |
| 19:46158702 | 19 | 46158702 | 3 | 0 | 0 | CATTTA1 | C | 0.9314 | 0.2666 | 0.0004765 | +++?? | 0 | 0.6332 | | |
| rs73839214 | 3 | 61067848 | 4 | 0 | 0 | A | C | 3.3306 | 0.9534 | 0.0004767 | +++?- | 0 | 0.6442 | intron | FHIT |
| rs147057003 | 12 | 68455947 | 4 | 0 | 0 | A | G | 3.7454 | 1.0721 | 0.0004767 | +++?+ | 0 | 0.6167 | intergenic | |
| rs3117105 | 6 | 32343714 | 5 | 0 | 0 | T | C | -0.7462 | 0.2136 | 0.0004769 | +---- | 40.5 | 0.1515 | upstream-gene |  |
| rs1778012 | 1 | 87554556 | 4 | 0 | 0 | A | G | 1.0558 | 0.3022 | 0.000477 | +++?+ | 0.5 | 0.3892 | intron | HS2ST1 |
| rs2163932 | 4 | 21098882 | 4 | 0 | 0 | A | G | 1.8571 | 0.5316 | 0.0004773 | +++?+ | 0 | 0.9283 | intron | KCNIP4 |
| rs9956614 | 18 | 39960608 | 4 | 0 | 0 | A | G | 1.1654 | 0.3336 | 0.0004774 | +++?- | 0 | 0.4058 | intron | LINC00907 |
| rs111951658 | 7 | 7793764 | 4 | 0 | 0 | C | G | -1.645 | 0.4709 | 0.0004774 | ---?- | 0 | 0.9402 | intron | RPA3-AS1 |
| rs369112445 | 14 | 28214352 | 3 | 0 | 0 | A | C | -2.1553 | 0.617 | 0.0004777 | ---?? | 0 | 0.8874 | intergenic | |
| 18:55740110 | 18 | 55740110 | 3 | 0 | 0 | A | G | -1.7524 | 0.5017 | 0.0004778 | ---?? | 0 | 0.8718 | intron | NEDD4L |
| rs75913369 | 11 | 103670026 | 4 | 0 | 0 | A | G | -1.491 | 0.4269 | 0.0004779 | ---?- | 0 | 0.4921 | intergenic | |
| rs17036311 | 2 | 47385059 | 4 | 0 | 0 | C | G | 1.3658 | 0.3911 | 0.0004783 | +++?+ | 0 | 0.963 | upstream-gene | |
| rs17599077 | 6 | 32591058 | 5 | 0 | 1 | A | C | -1.3453 | 0.3852 | 0.0004784 | ----- | 14.8 | 0.32 | intergenic |  |
| rs12428774 | 13 | 65198145 | 4 | 0 | 0 | T | C | 1.9364 | 0.5544 | 0.0004785 | +++?+ | 11.1 | 0.3376 | intergenic | |
| rs112050805 | 2 | 199089079 | 4 | 0 | 0 | A | T | 1.3956 | 0.3996 | 0.0004785 | +++?+ | 49.9 | 0.1119 | intergenic | |
| 5:95802278 | 5 | 95802278 | 3 | 0 | 0 | T | G | -5.8079 | 1.663 | 0.0004786 | ---?? | 48.6 | 0.1427 | intergenic | |
| rs2635336 | 4 | 118460715 | 4 | 0 | 0 | A | G | -2.327 | 0.6663 | 0.0004786 | ---?- | 0 | 0.5454 | intergenic | |
| rs8106861 | 19 | 46158156 | 3 | 0 | 1 | A | G | 0.9345 | 0.2676 | 0.0004786 | +++?? | 0 | 0.6967 | intergenic | |
| rs146358837 | 17 | 57148758 | 5 | 0 | 0 | T | C | 2.3267 | 0.6662 | 0.0004789 | +++++ | 0 | 0.8234 | intron | TRIM37 |
| rs35826015 | 6 | 32590669 | 5 | 0 | 1 | A | T | 1.3451 | 0.3852 | 0.000479 | +++++ | 14.8 | 0.3201 | intergenic | |
| rs12517593 | 5 | 103160773 | 4 | 0 | 0 | A | G | -1.5731 | 0.4504 | 0.0004791 | ---?- | 0 | 0.8045 | intergenic | |
| rs3898728 | 11 | 109407405 | 4 | 0 | 0 | A | C | 0.8964 | 0.2567 | 0.0004792 | +-+?+ | 49.2 | 0.1161 | intergenic | |
| rs10842956 | 12 | 27785926 | 4 | 0 | 0 | T | C | 0.8833 | 0.2529 | 0.0004792 | +-+?- | 55.6 | 0.08018 | intron | PPFIBP1 |
| rs13007438 | 2 | 18528390 | 3 | 0 | 0 | T | C | -1.8634 | 0.5336 | 0.0004792 | ---?? | 0 | 0.5231 | intergenic | |
| rs142395446 | 6 | 72833978 | 3 | 0 | 0 | T | G | -1.6733 | 0.4792 | 0.0004794 | ---?? | 0 | 0.9179 | intron | RIMS1 |
| rs34053446 | 6 | 32586809 | 5 | 0 | 1 | A | G | -1.3525 | 0.3873 | 0.0004795 | ----- | 15.3 | 0.317 | intergenic |  |
| 2:170474511 | 2 | 170474511 | 3 | 0 | 0 | G | GTTC | -1.409 | 0.4035 | 0.0004796 | ---?? | 12.5 | 0.3188 | | |
| rs17036316 | 2 | 47385726 | 4 | 0 | 0 | A | T | 1.3667 | 0.3914 | 0.0004797 | +++?+ | 0 | 0.9635 | upstream-gene | |

| | | | | | | | | | | | | | | | |
|---|---|---|---|---|---|---|---|---|---|---|---|---|---|---|---|
| rs3910844 | 8 | 64901019 | 4 | 0 | 0 | T | C | -0.8743 | 0.2504 | 0.0004799 | ---?- | 0 | 0.9521 | intergenic | |
| 5:103151722 | 5 | 103151722 | 3 | 0 | 0 | G | GTTTTC | -1.622 | 0.4645 | 0.00048 | ---?? | 0 | 0.6672 | | |
| 6:159232962 | 6 | 159232962 | 3 | 0 | 0 | T | TAAAAG | 0.8902 | 0.255 | 0.0004801 | +++?? | 11.1 | 0.3247 | | |
| rs34642101 | 19 | 46163990 | 5 | 0 | 1 | T | C | 0.9254 | 0.265 | 0.0004802 | ++++- | 26.4 | 0.2457 | intergenic | |
| rs11229898 | 11 | 59029485 | 5 | 0 | 0 | T | C | 0.762 | 0.2182 | 0.0004803 | +-++- | 10.2 | 0.3482 | intergenic | |
| rs2422456 | 11 | 37051648 | 4 | 0 | 0 | T | C | 0.9224 | 0.2642 | 0.0004804 | +++?+ | 0 | 0.4402 | intergenic | |
| rs185221278 | 12 | 132063810 | 4 | 0 | 0 | T | C | -3.139 | 0.8991 | 0.0004804 | ---?- | 0 | 0.986 | intergenic | |
| rs535456 | 1 | 90246069 | 5 | 0 | 0 | A | G | 0.7443 | 0.2132 | 0.0004808 | +++++ | 0 | 0.9878 | intergenic | |
| rs1419131 | 1 | 87555165 | 4 | 0 | 0 | A | G | -1.055 | 0.3022 | 0.0004808 | ---?- | 1 | 0.387 | intron | HS2ST1 |
| rs147679359 | 4 | 124308243 | 4 | 0 | 0 | A | T | -4.545 | 1.3018 | 0.0004808 | -+-?- | 47.2 | 0.128 | intergenic | |
| rs62181733 | 2 | 220759653 | 3 | 0 | 0 | T | C | 5.5407 | 1.587 | 0.0004809 | +?+?+ | 53.4 | 0.1168 | intergenic | |
| rs28430893 | 7 | 159006661 | 4 | 0 | 0 | A | C | 1.883 | 0.5394 | 0.000481 | +++?+ | 44 | 0.1474 | intergenic | |
| rs3812948 | 16 | 88716534 | 5 | 0 | 0 | A | G | -0.8662 | 0.2481 | 0.0004811 | ----- | 0 | 0.8606 | intron | CYBA |
| 5:103152458 | 5 | 103152458 | 3 | 0 | 0 | G | GA | -1.6153 | 0.4627 | 0.0004811 | ---?? | 0 | 0.5981 | | |
| rs57634784 | 2 | 159095326 | 4 | 0 | 0 | T | C | -0.8685 | 0.2488 | 0.0004812 | ---?- | 0 | 0.5187 | intron | CCDC148 |
| rs9929260 | 16 | 52918182 | 4 | 0 | 0 | A | G | 0.873 | 0.2501 | 0.0004814 | +++?+ | 0 | 0.7312 | intergenic | |
| rs10196002 | 2 | 170949144 | 4 | 0 | 1 | T | G | 0.9996 | 0.2863 | 0.0004814 | +++?+ | 58.1 | 0.0669 | intergenic | |
| rs28872589 | 6 | 32584989 | 5 | 0 | 1 | T | C | -1.3432 | 0.3848 | 0.0004814 | ----- | 15.2 | 0.3175 | intergenic |  |
| rs79411636 | 13 | 73927269 | 3 | 0 | 0 | T | C | 3.1649 | 0.9066 | 0.0004814 | +++?? | 0 | 0.9335 | intergenic | |
| rs233128 | 1 | 85794413 | 5 | 0 | 0 | C | G | -0.821 | 0.2352 | 0.0004815 | ----- | 0 | 0.9817 | intron | DDAH1 |
| rs959668 | 2 | 208357838 | 4 | 0 | 0 | T | C | -1.4361 | 0.4114 | 0.0004815 | ---?- | 0 | 0.3977 | intergenic | |
| rs3857923 | 8 | 140375503 | 4 | 0 | 0 | A | G | -1.0058 | 0.2881 | 0.0004816 | ---?- | 0 | 0.9384 | intergenic | |
| rs62174940 | 2 | 185439307 | 4 | 0 | 0 | A | G | 0.9575 | 0.2743 | 0.0004818 | +++?+ | 0 | 0.7654 | intergenic | |
| rs11616510 | 13 | 112829964 | 4 | 0 | 0 | T | C | 0.9602 | 0.2751 | 0.000482 | +++?+ | 8.7 | 0.3494 | intergenic | |
| rs28493768 | 5 | 76081093 | 5 | 0 | 0 | T | C | 0.9399 | 0.2693 | 0.000482 | ++++- | 0 | 0.4932 | intergenic | |
| rs75758800 | 13 | 65194796 | 4 | 0 | 0 | A | G | -1.9616 | 0.562 | 0.000482 | ---?- | 8.7 | 0.3496 | intergenic | |
| rs111665670 | 6 | 32567065 | 5 | 0 | 1 | A | G | 1.3438 | 0.385 | 0.000482 | +++++ | 15.8 | 0.314 | intergenic | |
| rs1928560 | 12 | 41241645 | 4 | 0 | 0 | T | C | 1.732 | 0.4962 | 0.0004821 | +++?- | 0 | 0.6211 | intron | CNTN1 |
| 4:100487454 | 4 | 100487454 | 3 | 0 | 0 | G | GAAAAA | -1.1924 | 0.3416 | 0.0004822 | ---?? | 45.1 | 0.1616 | | |
| 20:50649303 | 20 | 50649303 | 3 | 0 | 0 | CA | C | -1.1825 | 0.3388 | 0.0004823 | ---?? | 51.1 | 0.1294 | | |
| rs7685481 | 4 | 20740583 | 4 | 0 | 0 | T | G | 0.9116 | 0.2612 | 0.0004824 | +++?+ | 0 | 0.8232 | intron | KCNIP4 |
| rs182650214 | 22 | 29082204 | 3 | 0 | 0 | A | G | -2.5523 | 0.7312 | 0.0004824 | ---?? | 0 | 0.5536 | downstream-gene | |
| rs12705420 | 7 | 107383133 | 5 | 0 | 0 | T | C | 0.8658 | 0.2481 | 0.0004826 | +++-+ | 53.5 | 0.0717 | upstream-gene | |
| rs73315138 | 17 | 56977012 | 5 | 0 | 0 | A | G | -2.3901 | 0.6848 | 0.0004826 | ----- | 0 | 0.7684 | intron | PPM1E |
| rs74637534 | 11 | 103702613 | 4 | 0 | 0 | A | G | -1.4871 | 0.4261 | 0.0004827 | ---?- | 0 | 0.4428 | intergenic | |
| rs73362199 | 8 | 140813232 | 3 | 0 | 0 | C | G | 1.3549 | 0.3882 | 0.000483 | +++?? | 65.8 | 0.05376 | intron | TRAPPC9 |
| rs137862911 | 18 | 25191997 | 4 | 0 | 0 | A | C | 2.5196 | 0.722 | 0.0004831 | +++?+ | 0 | 0.6568 | intergenic | |
| rs11518504 | 11 | 103705692 | 4 | 0 | 0 | A | C | -1.4856 | 0.4257 | 0.0004836 | ---?- | 0 | 0.4502 | intergenic | |
| rs12769959 | 10 | 134579026 | 4 | 0 | 0 | A | C | -0.9709 | 0.2782 | 0.0004838 | ---?- | 0 | 0.4733 | intron | INPP5A |
| rs138123890 | 2 | 199062271 | 4 | 0 | 0 | C | G | 1.1627 | 0.3332 | 0.0004838 | +++?+ | 31.2 | 0.2254 | intergenic | |
| rs142701669 | 2 | 199062272 | 4 | 0 | 0 | A | T | -1.1627 | 0.3332 | 0.0004839 | ---?- | 31.2 | 0.2253 | intergenic | |
| rs139263557 | 17 | 57135177 | 5 | 0 | 0 | A | G | -2.3125 | 0.6627 | 0.000484 | ----- | 0 | 0.8242 | intron | TRIM37 |
| rs11968981 | 6 | 23907278 | 4 | 0 | 0 | T | C | 2.6695 | 0.765 | 0.0004841 | +++?- | 43.2 | 0.1524 | intergenic | |
| rs72690354 | 4 | 101264897 | 4 | 0 | 0 | A | T | 0.9563 | 0.2741 | 0.0004842 | +++?+ | 0 | 0.9686 | intergenic | |
| rs7970242 | 12 | 112838209 | 5 | 0 | 0 | A | G | 1.3062 | 0.3743 | 0.0004843 | +-+++ | 53 | 0.07469 | downstream-gene | |
| rs12001383 | 9 | 16195440 | 4 | 0 | 0 | C | G | -1.2549 | 0.3597 | 0.0004845 | ---?- | 0 | 0.5752 | intergenic | |
| rs140427694 | 17 | 57195089 | 5 | 0 | 0 | T | C | 2.4339 | 0.6976 | 0.0004846 | +++++ | 0 | 0.7955 | intron | SKA2 |
| rs142038422 | 15 | 87656549 | 3 | 0 | 0 | A | C | 4.0973 | 1.1743 | 0.0004846 | +++?? | 0 | 0.8872 | intergenic | |
| rs10202446 | 2 | 170929880 | 4 | 0 | 1 | A | G | -1.0064 | 0.2884 | 0.0004847 | ---?- | 47.9 | 0.1242 | intron | UBR3 |
| rs62340525 | 4 | 190010558 | 4 | 0 | 0 | T | C | 0.8758 | 0.251 | 0.0004847 | +++?+ | 29.5 | 0.2353 | intergenic | |
| 6:168584307 | 6 | 168584307 | 3 | 0 | 0 | A | AT | 1.6334 | 0.4682 | 0.0004849 | +++?? | 0 | 0.7964 | | |
| rs146065243 | 2 | 170578818 | 3 | 0 | 0 | T | C | -1.2798 | 0.3668 | 0.0004851 | ---?? | 0 | 0.8805 | intron | PHOSPHO2-KLHL23 |
| rs28510673 | 6 | 32569506 | 5 | 0 | 1 | T | G | 1.3405 | 0.3842 | 0.0004852 | +++++ | 16.1 | 0.3119 | intergenic | |
| rs28567460 | 6 | 32569513 | 5 | 0 | 1 | C | G | 1.3405 | 0.3842 | 0.0004852 | +++++ | 16.1 | 0.3119 | intergenic | |
| rs2577952 | 15 | 42073402 | 5 | 0 | 0 | T | C | 0.9563 | 0.2741 | 0.0004854 | +++++ | 0 | 0.5045 | intron | MAPKBP1 |
| rs547460 | 1 | 90238738 | 5 | 0 | 0 | A | G | -0.7444 | 0.2134 | 0.0004855 | ----- | 0 | 0.985 | intergenic |  |
| rs4509992 | 15 | 92185715 | 3 | 0 | 0 | A | G | 1.4139 | 0.4053 | 0.0004855 | +++?? | 0 | 0.3794 | intergenic | |
| 6:32045874 | 6 | 32045874 | 3 | 0 | 0 | A | AAG | 1.4649 | 0.4199 | 0.0004857 | +++?? | 0 | 0.4888 | | |
| rs6433161 | 2 | 170833222 | 4 | 0 | 1 | A | G | 1.0114 | 0.2899 | 0.0004862 | +++?+ | 44.7 | 0.1432 | intron | UBR3 |

| | | | | | | | | | | | | | | | |
|---|---|---|---|---|---|---|---|---|---|---|---|---|---|---|---|
| rs11033911 | 11 | 37065402 | 4 | 0 | 0 | A | G | -0.9349 | 0.268 | 0.0004863 | ---?- | 0 | 0.4899 | intergenic | |
| rs3931608 | 13 | 108408959 | 4 | 0 | 0 | T | C | -1.2556 | 0.36 | 0.0004864 | ---?- | 11 | 0.338 | intron | FAM155A |
| rs10053555 | 5 | 123444620 | 4 | 0 | 0 | C | G | 0.8889 | 0.2548 | 0.0004864 | +++?+ | 0 | 0.6298 | intergenic | |
| rs61523045 | 6 | 32585762 | 5 | 0 | 1 | A | G | -1.342 | 0.3847 | 0.0004864 | ----- | 15 | 0.3186 | intergenic |  |
| rs9297476 | 8 | 113505364 | 4 | 0 | 0 | A | G | 1.0085 | 0.2891 | 0.0004865 | +++?+ | 0 | 0.6631 | intron | CSMD3 |
| rs35150023 | 6 | 32586754 | 5 | 0 | 1 | A | T | 1.342 | 0.3847 | 0.0004865 | +++++ | 15.7 | 0.3142 | intergenic | |
| rs34032759 | 6 | 32585775 | 5 | 0 | 1 | A | G | 1.342 | 0.3847 | 0.0004866 | +++++ | 15 | 0.3188 | intergenic | |
| rs35637654 | 6 | 32586141 | 5 | 0 | 1 | A | C | 1.342 | 0.3847 | 0.0004866 | +++++ | 15 | 0.3191 | intergenic | |
| rs12221838 | 11 | 95706538 | 4 | 0 | 0 | A | C | -0.9615 | 0.2757 | 0.0004867 | -+-?- | 30.1 | 0.2314 | downstream-gene | |
| 7:16064474 | 7 | 16064474 | 3 | 0 | 0 | T | TA | -1.0217 | 0.2929 | 0.0004867 | ---?? | 0 | 0.3826 | | |
| rs10491081 | 17 | 5866285 | 2 | 0 | 0 | A | G | 4.2717 | 1.2247 | 0.000487 | ??++? | 0 | 0.9198 | intergenic | |
| rs36233210 | 6 | 32577417 | 5 | 0 | 1 | T | G | -1.3419 | 0.3847 | 0.000487 | ----- | 16 | 0.3127 | intergenic |  |
| rs7256688 | 19 | 14747455 | 4 | 0 | 0 | C | G | -0.9189 | 0.2635 | 0.0004871 | ---?+ | 48.1 | 0.1226 | intron | EMR3 |
| rs28833508 | 6 | 32584794 | 5 | 0 | 1 | T | C | -1.3418 | 0.3847 | 0.0004872 | ----- | 15.6 | 0.3151 | intergenic |  |
| rs116096899 | 2 | 47380844 | 4 | 0 | 0 | A | G | 1.3561 | 0.3888 | 0.0004872 | +++?+ | 0 | 0.9595 | intron | C2orf61 |
| 22:28758396 | 22 | 28758396 | 3 | 0 | 0 | CTCTT | C | -2.6057 | 0.7471 | 0.0004872 | ---?? | 0 | 0.6476 | | |
| rs5803960 | 13 | 60057209 | 3 | 0 | 0 | A | C | -1.0462 | 0.3 | 0.0004873 | ---?? | 0 | 0.471 | intergenic | |
| rs6716901 | 2 | 172673453 | 4 | 0 | 0 | A | G | 1.3337 | 0.3824 | 0.0004874 | +++?+ | 0 | 0.899 | intron | SLC25A12 |
| rs12021941 | 1 | 162443355 | 4 | 0 | 0 | T | C | 1.5154 | 0.4345 | 0.0004875 | +++?+ | 0 | 0.9217 | intergenic | |
| rs28587541 | 6 | 32584652 | 5 | 0 | 1 | A | G | -1.3418 | 0.3847 | 0.0004875 | ----- | 15.7 | 0.3146 | intergenic |  |
| rs1081022 | 15 | 42075123 | 5 | 0 | 0 | T | G | 0.9573 | 0.2745 | 0.0004877 | +++++ | 0 | 0.5135 | intron | MAPKBP1 |
| rs28986353 | 6 | 32750777 | 5 | 0 | 0 | T | C | 1.3817 | 0.3962 | 0.0004877 | +++++ | 0 | 0.586 | intergenic | |
| rs72672894 | 8 | 113319943 | 4 | 0 | 0 | C | G | 0.9898 | 0.2838 | 0.0004877 | +++?+ | 0 | 0.7556 | intron | CSMD3 |
| rs77977170 | 8 | 113307827 | 4 | 0 | 0 | T | C | -0.9901 | 0.2839 | 0.0004877 | ---?- | 0 | 0.7706 | intron | CSMD3 |
| rs113711404 | 6 | 32583757 | 5 | 0 | 1 | A | G | 1.3417 | 0.3847 | 0.0004877 | +++++ | 15.9 | 0.313 | intergenic | |
| rs9277988 | 6 | 33306235 | 5 | 0 | 0 | T | C | -0.9505 | 0.2726 | 0.0004878 | ----- | 0 | 0.961 | intergenic |  |
| rs55873471 | 6 | 32581712 | 5 | 0 | 1 | A | G | -1.3416 | 0.3847 | 0.0004878 | ----- | 15.9 | 0.3129 | intergenic |  |
| rs34471348 | 6 | 32581175 | 5 | 0 | 1 | T | C | -1.3416 | 0.3847 | 0.0004879 | ----- | 16 | 0.3129 | intergenic |  |
| rs72844253 | 6 | 32556263 | 5 | 0 | 1 | T | G | -1.3416 | 0.3847 | 0.0004879 | ----- | 16 | 0.3129 | intron | HLA-DRB1 |
| 14:28153481 | 14 | 28153481 | 3 | 0 | 0 | G | GC | 2.1862 | 0.6269 | 0.0004879 | +++?? | 0 | 0.9933 | | |
| 1:7127478 | 1 | 7127478 | 3 | 0 | 0 | A | ATTC | -1.0736 | 0.3079 | 0.0004879 | ---?? | 16.5 | 0.3021 | | |
| rs17211356 | 6 | 32578426 | 5 | 0 | 1 | A | G | -1.3416 | 0.3847 | 0.000488 | ----- | 16 | 0.3129 | intergenic |  |
| rs34694416 | 6 | 32584371 | 5 | 0 | 1 | T | C | 1.3416 | 0.3847 | 0.000488 | +++++ | 15.8 | 0.3138 | intergenic | |
| rs72847591 | 6 | 32570494 | 5 | 0 | 1 | T | C | 1.3416 | 0.3847 | 0.000488 | +++++ | 16 | 0.3129 | intergenic | |
| rs112270441 | 6 | 32582980 | 5 | 0 | 1 | T | C | 1.3416 | 0.3847 | 0.000488 | +++++ | 16 | 0.3129 | intergenic | |
| rs113195871 | 6 | 32580640 | 5 | 0 | 1 | A | T | 1.3416 | 0.3847 | 0.000488 | +++++ | 16 | 0.3129 | intergenic | |
| rs113979555 | 6 | 32583188 | 5 | 0 | 1 | A | G | 1.3416 | 0.3847 | 0.000488 | +++++ | 16 | 0.3129 | intergenic | |
| rs4504507 | 6 | 32582288 | 5 | 0 | 1 | T | C | 1.3416 | 0.3847 | 0.0004881 | +++++ | 16 | 0.3129 | intergenic | |
| rs56245106 | 6 | 32577846 | 5 | 0 | 1 | T | C | -1.3416 | 0.3847 | 0.0004882 | ----- | 16 | 0.3129 | intergenic |  |
| rs71564270 | 6 | 99920118 | 3 | 0 | 0 | A | G | -1.8109 | 0.5193 | 0.0004882 | ---?? | 0 | 0.6389 | intron | USP45 |
| rs117505823 | 12 | 41252791 | 4 | 0 | 0 | T | G | 1.7171 | 0.4924 | 0.0004882 | +++?- | 0 | 0.6209 | intron | CNTN1 |
| rs9886391 | 8 | 96989116 | 4 | 0 | 0 | A | G | 1.4135 | 0.4054 | 0.0004883 | +++?+ | 0 | 0.9518 | intergenic | |
| rs35515288 | 6 | 32576356 | 5 | 0 | 1 | T | C | -1.3415 | 0.3847 | 0.0004883 | ----- | 16 | 0.3128 | intergenic |  |
| rs35899011 | 6 | 32577245 | 5 | 0 | 1 | T | C | 1.3415 | 0.3847 | 0.0004883 | +++++ | 16 | 0.3129 | intergenic | |
| rs36202959 | 6 | 32576324 | 5 | 0 | 1 | C | G | -1.3415 | 0.3847 | 0.0004883 | ----- | 16 | 0.3128 | intergenic |  |
| rs55978013 | 15 | 42145049 | 5 | 0 | 0 | T | C | 1.2287 | 0.3524 | 0.0004883 | +++++ | 12.9 | 0.332 | synonymous | SPTBN5 |
| rs76851726 | 12 | 96628037 | 4 | 0 | 0 | T | C | -2.6086 | 0.7481 | 0.0004883 | ---?- | 0 | 0.4669 | intron | ELK3 |
| rs11639146 | 15 | 41914592 | 5 | 0 | 0 | A | G | 0.9991 | 0.2865 | 0.0004884 | +++++ | 7.6 | 0.3633 | intron | MGA |
| rs36202958 | 6 | 32576368 | 5 | 0 | 1 | T | C | 1.3415 | 0.3847 | 0.0004884 | +++++ | 16 | 0.3128 | intergenic | |
| rs36233212 | 6 | 32575820 | 5 | 0 | 1 | T | C | -1.3415 | 0.3847 | 0.0004884 | ----- | 16 | 0.3128 | intergenic |  |
| rs7605205 | 2 | 170832805 | 4 | 0 | 1 | A | T | -1.011 | 0.2899 | 0.0004885 | ---?- | 44.6 | 0.1436 | intron | UBR3 |
| rs9500775 | 6 | 62117494 | 3 | 0 | 0 | A | C | -1.3542 | 0.3883 | 0.0004885 | ---?? | 0 | 0.9862 | intergenic | |
| rs117184243 | 12 | 41252627 | 4 | 0 | 0 | T | C | 1.717 | 0.4924 | 0.0004885 | +++?- | 0 | 0.6211 | intron | CNTN1 |
| rs4912571 | 3 | 184216615 | 4 | 0 | 0 | T | C | -1.0316 | 0.2958 | 0.0004886 | ---?- | 0 | 0.718 | intergenic | |
| rs35946908 | 8 | 65030591 | 4 | 0 | 0 | A | G | 0.8583 | 0.2461 | 0.0004886 | +++?+ | 0 | 0.9474 | intergenic | |
| rs61670843 | 6 | 32574453 | 5 | 0 | 1 | T | C | 1.3416 | 0.3847 | 0.0004886 | +++++ | 16.1 | 0.3122 | intergenic | |
| rs343044 | 7 | 35508859 | 4 | 0 | 0 | A | G | -1.1148 | 0.3197 | 0.0004887 | ---?- | 12 | 0.3328 | intergenic | |
| rs7035277 | 9 | 16167502 | 4 | 0 | 0 | A | G | 0.9388 | 0.2693 | 0.0004887 | +++?+ | 0 | 0.8882 | intergenic | |
| rs144015605 | 9 | 79457923 | 4 | 0 | 0 | T | C | 2.0346 | 0.5835 | 0.0004887 | +++?+ | 0 | 0.5581 | intron | PRUNE2 |

| | | | | | | | | | | | | | | | |
|---|---|---|---|---|---|---|---|---|---|---|---|---|---|---|---|
| rs150763002 | 11 | 39328472 | 2 | 0 | 0 | A | T | -7.565 | 2.1696 | 0.0004887 | ?-??- | 0 | 0.6898 | intergenic | |
| rs4382332 | 6 | 32572155 | 5 | 0 | 1 | T | C | -1.3418 | 0.3848 | 0.0004889 | ----- | 15.8 | 0.3137 | intergenic |  |
| rs34113780 | 6 | 32575258 | 5 | 0 | 1 | A | G | -1.3415 | 0.3847 | 0.000489 | ----- | 16 | 0.3126 | intergenic |  |
| rs72850021 | 6 | 32573895 | 5 | 0 | 1 | A | G | -1.3414 | 0.3847 | 0.000489 | ----- | 16 | 0.3126 | intergenic |  |
| rs9500776 | 6 | 62117497 | 3 | 0 | 0 | T | C | 1.354 | 0.3883 | 0.0004891 | +++?? | 0 | 0.9862 | intergenic | |
| rs35488182 | 6 | 32574048 | 5 | 0 | 1 | A | G | 1.3414 | 0.3847 | 0.0004891 | +++++ | 16 | 0.3126 | intergenic | |
| rs79355658 | 12 | 4355889 | 3 | 0 | 0 | A | G | -3.1758 | 0.9109 | 0.0004892 | ---?? | 0 | 0.9912 | intergenic | |
| rs1420857 | 8 | 113321012 | 4 | 0 | 0 | A | G | 0.9884 | 0.2835 | 0.0004893 | +++?+ | 0 | 0.7553 | intron | CSMD3 |
| rs34822594 | 6 | 32587062 | 5 | 0 | 1 | T | C | -1.3414 | 0.3847 | 0.0004893 | ----- | 16 | 0.3126 | intergenic |  |
| rs35752949 | 6 | 32574329 | 5 | 0 | 1 | T | C | 1.3414 | 0.3847 | 0.0004893 | +++++ | 16 | 0.3126 | intergenic | |
| rs59108952 | 6 | 32574244 | 5 | 0 | 1 | T | C | 1.3414 | 0.3847 | 0.0004893 | +++++ | 16 | 0.3126 | intergenic | |
| rs72810419 | 2 | 47381677 | 4 | 0 | 0 | T | C | 1.3581 | 0.3895 | 0.0004895 | +++?+ | 0 | 0.9611 | intron | C2orf61 |
| rs192490354 | 16 | 5871424 | 3 | 0 | 0 | C | G | -3.8868 | 1.1149 | 0.0004897 | -?-?- | 30.6 | 0.2365 | intron | RBFOX1 |
| rs115195914 | 2 | 199058338 | 4 | 0 | 0 | A | C | 1.1639 | 0.3338 | 0.0004898 | +++?+ | 30.1 | 0.2318 | intergenic | |
| rs10269491 | 7 | 82414119 | 4 | 0 | 0 | A | G | -0.9009 | 0.2584 | 0.0004899 | -+-?- | 46.3 | 0.1334 | intron | PCLO |
| rs17128925 | 12 | 41301806 | 4 | 0 | 0 | T | C | -1.7275 | 0.4955 | 0.00049 | ---?+ | 0 | 0.6321 | intron | CNTN1 |
| 8:91112061 | 8 | 91112061 | 3 | 0 | 0 | A | C | 2.2663 | 0.6501 | 0.0004902 | +++?? | 0 | 0.9963 | intergenic | |
| rs1096223 | 5 | 165397086 | 5 | 0 | 0 | T | C | 0.7467 | 0.2142 | 0.0004904 | +++++ | 28.8 | 0.2295 | intergenic | |
| 1:100229424 | 1 | 100229424 | 3 | 0 | 0 | CA | C | -1.622 | 0.4653 | 0.0004906 | ---?? | 0 | 0.4999 | | |
| rs10029167 | 4 | 18856773 | 4 | 0 | 0 | A | G | 1.7488 | 0.5017 | 0.000491 | +++?+ | 0 | 0.976 | intergenic | |
| rs62423829 | 6 | 120475016 | 4 | 0 | 0 | T | C | 2.7889 | 0.8001 | 0.000491 | +++?+ | 35.6 | 0.1988 | intergenic | |
| rs76036093 | 11 | 103677956 | 4 | 0 | 0 | T | C | -1.4852 | 0.4261 | 0.0004912 | ---?- | 0 | 0.4812 | intergenic | |
| rs76693806 | 6 | 63309761 | 4 | 0 | 0 | A | G | -2.4379 | 0.6994 | 0.0004912 | ---?- | 0 | 0.5083 | intergenic | |
| 14:28189324 | 14 | 28189324 | 3 | 0 | 0 | CT | C | 2.0167 | 0.5786 | 0.0004912 | +++?? | 0 | 0.7249 | | |
| rs1354241 | 3 | 2462577 | 4 | 0 | 0 | C | G | -0.8497 | 0.2438 | 0.0004913 | ---?- | 0 | 0.3927 | intron | CNTN4 |
| rs16934205 | 9 | 16195381 | 4 | 0 | 0 | T | C | -1.2532 | 0.3596 | 0.0004913 | ---?- | 0 | 0.5752 | intergenic | |
| rs751699 | 2 | 208284570 | 4 | 0 | 0 | C | G | 1.6141 | 0.4631 | 0.0004914 | +++?+ | 0 | 0.7065 | intergenic | |
| rs4937562 | 11 | 130621675 | 4 | 0 | 0 | T | C | 1.6857 | 0.4837 | 0.0004915 | +++?+ | 31.8 | 0.2214 | intergenic | |
| rs62419105 | 6 | 63311195 | 4 | 0 | 0 | A | G | 2.4547 | 0.7043 | 0.0004916 | +++?+ | 0 | 0.5087 | intergenic | |
| rs189745421 | 9 | 36919714 | 4 | 0 | 0 | T | G | 2.3549 | 0.6757 | 0.0004916 | -++?+ | 48.6 | 0.1195 | intron | PAX5 |
| rs1442185 | 12 | 41253935 | 4 | 0 | 0 | A | G | -1.7162 | 0.4924 | 0.0004917 | ---?+ | 0 | 0.6201 | intron | CNTN1 |
| rs4767993 | 12 | 41299621 | 4 | 0 | 0 | T | C | -1.7097 | 0.4906 | 0.0004917 | ---?+ | 0 | 0.6433 | intron | CNTN1 |
| rs7700119 | 4 | 190020650 | 4 | 0 | 0 | A | G | 0.8683 | 0.2491 | 0.0004918 | +++?+ | 0 | 0.5347 | intergenic | |
| rs73839215 | 3 | 61070288 | 4 | 0 | 0 | A | G | 3.3174 | 0.9518 | 0.0004918 | +++?- | 0 | 0.6421 | intron | FHIT |
| rs1891672 | 13 | 36573162 | 4 | 0 | 0 | A | C | 1.043 | 0.2993 | 0.0004919 | +++?- | 0 | 0.5604 | intron | DCLK1 |
| rs9945067 | 18 | 63822382 | 4 | 0 | 0 | A | C | -0.8597 | 0.2467 | 0.0004919 | ---?- | 9.5 | 0.3455 | intergenic | |
| 6:159198945 | 6 | 159198945 | 3 | 0 | 0 | T | TA | -0.8917 | 0.2559 | 0.0004919 | ---?? | 7.8 | 0.338 | | |
| rs6705754 | 2 | 170831308 | 4 | 0 | 1 | A | G | 1.0106 | 0.29 | 0.000492 | +++?+ | 44.5 | 0.1441 | intron | UBR3 |
| rs36022997 | 6 | 32573361 | 5 | 0 | 1 | A | C | -1.3408 | 0.3847 | 0.000492 | ----- | 16.1 | 0.3118 | intergenic |  |
| rs144513061 | 17 | 57082670 | 5 | 0 | 0 | T | C | -2.3579 | 0.6766 | 0.0004923 | ----+ | 0 | 0.657 | intron | TRIM37 |
| rs2176753 | 3 | 148841194 | 4 | 0 | 0 | T | G | 1.1819 | 0.3392 | 0.0004926 | +++?+ | 3.2 | 0.3768 | intergenic | |
| rs12900295 | 15 | 41927449 | 5 | 0 | 0 | T | C | -0.9856 | 0.2828 | 0.0004928 | ----- | 10.6 | 0.3456 | intron | MGA |
| rs74388756 | 13 | 73926986 | 3 | 0 | 0 | T | C | 3.1611 | 0.9071 | 0.0004928 | +++?? | 0 | 0.9326 | intergenic | |
| rs76206226 | 4 | 127978725 | 4 | 0 | 0 | T | G | 2.9674 | 0.8516 | 0.0004928 | +++?+ | 0 | 0.7161 | intergenic | |
| rs142641056 | 16 | 62117384 | 3 | 0 | 0 | A | C | 5.0056 | 1.4365 | 0.0004928 | +++?? | 0 | 0.4403 | intergenic | |
| rs35172561 | 3 | 33265987 | 4 | 0 | 0 | A | G | 0.9314 | 0.2673 | 0.0004929 | +++?+ | 3.3 | 0.376 | intergenic | |
| rs181308749 | 5 | 751589 | 3 | 0 | 0 | T | C | -1.5939 | 0.4574 | 0.0004929 | ---?? | 63.3 | 0.06546 | intron | ZDHHC11B |
| 15:87561879 | 15 | 87561879 | 3 | 0 | 0 | AT | ATT | -2.1384 | 0.6137 | 0.000493 | ---?? | 0 | 0.9347 | | |
| rs2744671 | 1 | 14195122 | 4 | 0 | 0 | T | C | 0.896 | 0.2572 | 0.0004931 | +++?+ | 41.7 | 0.1616 | intergenic | |
| rs9276713 | 6 | 32759771 | 5 | 0 | 0 | A | G | 1.3843 | 0.3973 | 0.0004932 | +++++ | 0 | 0.6335 | intergenic | |
| rs9310945 | 3 | 30825860 | 4 | 0 | 0 | A | G | 2.9144 | 0.8364 | 0.0004934 | +++?+ | 0 | 0.6655 | intron | GADL1 |
| rs115386102 | 18 | 25188093 | 4 | 0 | 0 | A | G | -2.5191 | 0.723 | 0.0004935 | ---?- | 0 | 0.635 | intergenic | |
| rs77953280 | 1 | 152698223 | 4 | 0 | 0 | A | T | -2.5807 | 0.7407 | 0.0004937 | ---?- | 20.3 | 0.2883 | intergenic | |
| 2:213547305 | 2 | 213547305 | 3 | 0 | 0 | A | AT | -0.8836 | 0.2536 | 0.0004937 | ---?? | 2 | 0.3604 | | |
| rs117374334 | 13 | 29579449 | 4 | 0 | 0 | A | G | 3.7373 | 1.0727 | 0.0004938 | +++?+ | 28.5 | 0.2409 | intron | MTUS2 |
| rs10495490 | 2 | 2778929 | 4 | 0 | 0 | A | G | -0.93 | 0.2669 | 0.0004939 | ---?- | 35.1 | 0.2015 | intergenic | |
| rs7927598 | 11 | 129284072 | 4 | 0 | 0 | A | G | 1.5294 | 0.439 | 0.000494 | +++?+ | 30.4 | 0.2298 | intron | BARX2 |
| rs73552615 | 8 | 21545403 | 3 | 0 | 0 | A | G | -4.276 | 1.2273 | 0.000494 | ?--?- | 0 | 0.9289 | downstream-gene | |
| rs111553658 | 6 | 32583417 | 5 | 0 | 1 | T | C | 1.3409 | 0.3849 | 0.000494 | +++++ | 16.1 | 0.3122 | intergenic | |

| | | | | | | | | | | | | | | | |
|---|---|---|---|---|---|---|---|---|---|---|---|---|---|---|---|
| rs41288889 | 6 | 32712009 | 5 | 0 | 0 | A | C | -1.5135 | 0.4344 | 0.0004941 | ----- | 0 | 0.7015 | intron | HLA-DQA2 |
| rs2354443 | 1 | 146566245 | 4 | 0 | 0 | A | G | -1.1873 | 0.3408 | 0.0004942 | ---?- | 0 | 0.4506 | downstream-gene | |
| rs2850535 | 18 | 71697714 | 4 | 0 | 0 | A | G | 0.8781 | 0.252 | 0.0004943 | +++?+ | 0 | 0.4674 | intergenic | |
| 2:137512977 | 2 | 137512977 | 3 | 0 | 0 | A | AT | 3.6429 | 1.0457 | 0.0004944 | +++?? | 65.5 | 0.0552 | | |
| rs17069085 | 8 | 3987816 | 4 | 0 | 0 | T | C | 1.0882 | 0.3124 | 0.0004945 | -++?+ | 43.9 | 0.148 | intron | CSMD1 |
| 7:33772554 | 7 | 33772554 | 3 | 0 | 0 | CA | C | 0.9593 | 0.2754 | 0.0004946 | +++?? | 51.4 | 0.1279 | | |
| rs62410159 | 4 | 20739031 | 4 | 0 | 0 | T | C | 0.91 | 0.2612 | 0.0004947 | +++?+ | 0 | 0.8121 | intron | KCNIP4 |
| rs801975 | 1 | 233709522 | 4 | 0 | 0 | T | C | 0.9722 | 0.2791 | 0.0004948 | +++?+ | 0 | 0.5971 | intergenic | |
| rs3105849 | 2 | 235347523 | 4 | 0 | 0 | T | C | -1.3969 | 0.401 | 0.0004948 | ---?- | 0 | 0.8297 | intergenic | |
| rs2224651 | 1 | 90252187 | 5 | 0 | 0 | T | C | -0.7454 | 0.214 | 0.0004949 | ----- | 0 | 0.9324 | intergenic |  |
| rs11033908 | 11 | 37061204 | 4 | 0 | 0 | T | C | 0.8642 | 0.2481 | 0.0004949 | +++?+ | 0 | 0.3927 | intergenic | |
| rs57785510 | 12 | 112829868 | 5 | 0 | 0 | T | C | -1.3052 | 0.3747 | 0.0004949 | -+--- | 53 | 0.07435 | intergenic | |
| rs58427200 | 22 | 28862474 | 4 | 0 | 0 | C | G | -2.3648 | 0.6788 | 0.0004949 | ---?- | 0 | 0.6122 | intron | TTC28 |
| rs191122358 | 10 | 93744679 | 2 | 0 | 0 | A | G | -5.1315 | 1.4731 | 0.0004949 | -?-?? | 0 | 0.422 | intron | BTAF1 |
| rs1197666 | 15 | 42141484 | 5 | 0 | 0 | T | G | -0.9477 | 0.2721 | 0.000495 | ----- | 0 | 0.5538 | intron | SPTBN5 |
| rs56220527 | 3 | 176029989 | 4 | 0 | 0 | T | C | 1.3013 | 0.3736 | 0.000495 | +++?+ | 24.9 | 0.262 | intergenic | |
| rs113399573 | 6 | 32559159 | 3 | 0 | 0 | T | C | 1.3604 | 0.3905 | 0.0004951 | +++?? | 0 | 0.3769 | upstream-gene | |
| rs4658323 | 1 | 90251626 | 5 | 0 | 0 | A | G | 0.7453 | 0.214 | 0.0004952 | +++++ | 0 | 0.9313 | intergenic | |
| 22:29900468 | 22 | 29900468 | 3 | 0 | 0 | T | C | 5.1368 | 1.4748 | 0.0004955 | +++?? | 0 | 0.408 | downstream-gene | |
| rs12894805 | 14 | 62371427 | 4 | 0 | 0 | A | G | 1.0491 | 0.3012 | 0.0004955 | +++?- | 0 | 0.4894 | intron | SYT16 |
| rs62514394 | 8 | 113308857 | 4 | 0 | 0 | A | G | -0.9886 | 0.2838 | 0.0004955 | ---?- | 0 | 0.7688 | intron | CSMD3 |
| rs56132609 | 8 | 65031387 | 3 | 0 | 0 | T | G | 1.27 | 0.3646 | 0.0004956 | +++?? | 3.7 | 0.3542 | intergenic | |
| rs10180260 | 2 | 170830481 | 4 | 0 | 1 | A | G | 1.0101 | 0.29 | 0.0004957 | +++?+ | 44.5 | 0.1446 | intron | UBR3 |
| rs343045 | 7 | 35509024 | 4 | 0 | 0 | A | C | 1.1133 | 0.3196 | 0.0004958 | +++?+ | 12.6 | 0.3296 | intergenic | |
| rs36004380 | 10 | 20137048 | 4 | 0 | 0 | A | G | 1.1054 | 0.3174 | 0.0004958 | +++?+ | 0 | 0.8237 | intron | PLXDC2 |
| rs76543013 | 8 | 113309267 | 4 | 0 | 0 | A | G | 0.9885 | 0.2838 | 0.0004959 | +++?+ | 0 | 0.7685 | intron | CSMD3 |
| rs74514780 | 2 | 156807959 | 4 | 0 | 0 | A | G | -1.131 | 0.3247 | 0.000496 | +--?- | 54.1 | 0.08808 | intergenic | |
| rs78836863 | 6 | 63312161 | 4 | 0 | 0 | A | T | -2.4379 | 0.7 | 0.0004961 | ---?- | 0 | 0.5104 | intergenic | |
| rs12916525 | 15 | 41900710 | 5 | 0 | 0 | T | C | 1.0098 | 0.2899 | 0.0004962 | +++++ | 0.4 | 0.4039 | intergenic | |
| rs35585151 | 6 | 32600046 | 4 | 0 | 0 | A | G | 0.8903 | 0.2556 | 0.0004962 | ++++? | 0 | 0.5893 | intergenic | |
| rs72775756 | 9 | 139275006 | 3 | 0 | 0 | T | C | 3.0928 | 0.888 | 0.0004963 | +++?? | 0 | 0.8326 | intron | SNAPC4 |
| rs58847767 | 3 | 61070643 | 4 | 0 | 0 | C | G | 3.3137 | 0.9515 | 0.0004964 | +++?- | 0 | 0.642 | intron | FHIT |
| 4:992571 | 4 | 992571 | 3 | 0 | 0 | A | AC | -1.1587 | 0.3327 | 0.0004967 | -+-?? | 64.8 | 0.0586 | | |
| rs3768234 | 1 | 85740812 | 5 | 0 | 0 | A | G | -1.6315 | 0.4685 | 0.000497 | ----- | 0 | 0.4065 | intron | BCL10 |
| rs4976974 | 8 | 143309627 | 4 | 0 | 0 | A | G | -2.2636 | 0.65 | 0.0004971 | ---?- | 0 | 0.9736 | intron | TSNARE1 |
| rs7821329 | 8 | 64897046 | 4 | 0 | 0 | A | T | -0.8712 | 0.2502 | 0.0004974 | ---?- | 0 | 0.9525 | intergenic | |
| rs146998757 | 9 | 79457529 | 4 | 0 | 0 | T | C | -2.041 | 0.5862 | 0.0004975 | ---?- | 0 | 0.56 | intron | PRUNE2 |
| rs59971851 | 3 | 61070616 | 4 | 0 | 0 | T | C | 3.3136 | 0.9516 | 0.0004976 | +++?- | 0 | 0.6418 | intron | FHIT |
| rs1013109 | 11 | 95705590 | 4 | 0 | 0 | C | G | -0.9415 | 0.2704 | 0.0004978 | ---?- | 25.8 | 0.2571 | intergenic | |
| rs7822258 | 8 | 64897415 | 4 | 0 | 0 | A | G | 0.871 | 0.2502 | 0.000498 | +++?+ | 0 | 0.952 | intergenic | |
| rs1483254 | 11 | 103681813 | 4 | 0 | 0 | T | C | 1.4821 | 0.4257 | 0.0004982 | +++?+ | 0 | 0.4779 | intergenic | |
| rs2030191 | 1 | 103113403 | 4 | 0 | 0 | A | G | 0.848 | 0.2436 | 0.0004982 | +++?- | 0 | 0.4836 | intergenic | |
| rs473400 | 2 | 40433184 | 4 | 0 | 0 | A | G | -1.0777 | 0.3095 | 0.0004983 | ---?+ | 0 | 0.5586 | intron | SLC8A1,SLC8A1-AS1 |
| rs201519051 | 3 | 14969454 | 3 | 0 | 0 | A | T | 2.2088 | 0.6344 | 0.0004983 | +++?? | 0 | 0.6301 | intron | FGD5 |
| rs62514395 | 8 | 113309628 | 4 | 0 | 0 | A | G | 0.9881 | 0.2838 | 0.0004984 | +++?+ | 0 | 0.768 | intron | CSMD3 |
| rs2020304 | 1 | 64782409 | 4 | 0 | 0 | A | G | 1.2216 | 0.3509 | 0.0004985 | +++?+ | 0 | 0.5174 | intergenic | |
| rs13283078 | 9 | 21580031 | 4 | 0 | 0 | T | C | 1.6072 | 0.4616 | 0.0004986 | +++?+ | 0 | 0.946 | intergenic | |
| rs140816968 | 2 | 171142059 | 4 | 0 | 0 | A | G | 4.2073 | 1.2085 | 0.0004987 | +++?- | 30.8 | 0.2276 | intron | MYO3B |
| rs2741729 | 14 | 21456092 | 4 | 0 | 0 | A | C | 0.9139 | 0.2625 | 0.0004988 | +++?+ | 0 | 0.9885 | upstream-gene | |
| rs6490060 | 12 | 111773172 | 5 | 0 | 0 | A | G | -0.8733 | 0.2509 | 0.0004988 | -+--- | 51.6 | 0.08258 | intron | CUX2 |
| 1:102955385 | 1 | 102955385 | 3 | 0 | 0 | CA | C | -0.8801 | 0.2528 | 0.0004988 | ---?? | 0 | 0.9974 | | |
| 14:104632135 | 14 | 104632135 | 3 | 0 | 0 | CAT | C | -4.1479 | 1.1914 | 0.0004988 | ---?? | 0 | 0.9538 | | |
| rs147481461 | 18 | 25188555 | 4 | 0 | 0 | T | C | -2.516 | 0.7227 | 0.0004989 | ---?- | 0 | 0.6363 | intergenic | |
| rs36305 | 5 | 120320117 | 4 | 0 | 0 | A | C | 1.2926 | 0.3713 | 0.000499 | +++?+ | 0 | 0.9169 | intergenic | |
| rs115750212 | 3 | 132287727 | 4 | 0 | 0 | T | C | -2.1708 | 0.6236 | 0.000499 | ---?- | 0 | 0.7164 | intron | ACAD11,NPHP3-ACAD11 |
| rs1603510 | 4 | 189997650 | 4 | 0 | 0 | T | C | 0.8881 | 0.2551 | 0.0004994 | +++?+ | 30 | 0.2323 | intergenic | |
| rs77073692 | 9 | 16188572 | 3 | 0 | 0 | A | G | -0.9721 | 0.2793 | 0.0004996 | ---?? | 0 | 0.8453 | intergenic | |
| rs2949219 | 2 | 107674422 | 4 | 0 | 0 | A | G | -0.9642 | 0.277 | 0.0004998 | ---?- | 0 | 0.4023 | intergenic | |
| rs1609234 | 7 | 145186070 | 4 | 0 | 0 | A | G | -1.0053 | 0.2888 | 0.0005 | ---?- | 35.3 | 0.2005 | intergenic | |

| | | | | | | | | | | | | | | | |
|---|---|---|---|---|---|---|---|---|---|---|---|---|---|---|---|
| rs11712650 | 3 | 155101136 | 4 | 0 | 0 | A | G | -1.0243 | 0.2943 | 0.0005 | ---?- | 9.8 | 0.3439 | intergenic | |
| rs11778532 | 8 | 113322357 | 4 | 0 | 0 | T | C | 0.9855 | 0.2831 | 0.0005 | +++?+ | 0 | 0.754 | intron | CSMD3 |
| rs115752743 | 6 | 32559163 | 3 | 0 | 0 | T | C | 1.3585 | 0.3903 | 0.0005 | +++?? | 0 | 0.3774 | upstream-gene | |
| rs62514396 | 8 | 113309730 | 4 | 0 | 0 | A | G | -0.9878 | 0.2838 | 0.0005001 | ---?- | 0 | 0.7675 | intron | CSMD3 |
| rs12705418 | 7 | 107349285 | 4 | 0 | 0 | T | G | -0.8796 | 0.2527 | 0.0005002 | ---?- | 0 | 0.5799 | intron | SLC26A4 |
| rs187291837 | 11 | 92637202 | 3 | 0 | 0 | A | T | 4.8763 | 1.401 | 0.0005002 | +++?? | 0 | 0.5167 | intergenic | |
| rs9985234 | 3 | 188546826 | 4 | 0 | 0 | A | G | -0.9659 | 0.2775 | 0.0005003 | ---?- | 11 | 0.3381 | intron | LPP |
| rs72799524 | 16 | 84843430 | 4 | 0 | 0 | A | G | 1.2646 | 0.3633 | 0.0005004 | +++?- | 0 | 0.5369 | intergenic | |
| 13:65193320 | 13 | 65193320 | 3 | 0 | 0 | T | TG | 2.0339 | 0.5844 | 0.0005004 | +++?? | 22.8 | 0.274 | | |
| 5:55633904 | 5 | 55633904 | 3 | 0 | 0 | T | TTC | -5.172 | 1.486 | 0.0005004 | ---?? | 0 | 0.5778 | | |
| rs62039817 | 16 | 64361647 | 4 | 0 | 0 | A | G | 2.7152 | 0.7801 | 0.0005005 | +++?+ | 40.5 | 0.1685 | intergenic | |
| rs113420674 | 12 | 114246794 | 4 | 0 | 0 | T | C | -2.3965 | 0.6886 | 0.0005005 | ---?- | 0 | 0.8876 | intergenic | |
| rs148377938 | 2 | 232098425 | 4 | 0 | 0 | T | G | 1.5605 | 0.4484 | 0.0005007 | +++?+ | 0 | 0.6736 | intron | ARMC9 |
| 8:113292830 | 8 | 113292830 | 3 | 0 | 0 | CTATT | C | -1.0205 | 0.2932 | 0.0005008 | ---?? | 0 | 0.6062 | | |
| rs76138634 | 17 | 47453156 | 4 | 0 | 0 | A | T | 1.0425 | 0.2996 | 0.000501 | +++?- | 54.4 | 0.08657 | intergenic | |
| rs34967517 | 3 | 115081563 | 4 | 0 | 0 | A | C | -0.9604 | 0.276 | 0.0005011 | ---?- | 0 | 0.9672 | intergenic | |
| rs72929131 | 2 | 199089265 | 4 | 0 | 0 | T | C | -1.3926 | 0.4002 | 0.0005012 | ---?- | 49.5 | 0.1148 | intergenic | |
| rs1281290 | 1 | 181924155 | 3 | 0 | 0 | C | G | -0.9001 | 0.2586 | 0.0005014 | ---?? | 0 | 0.5098 | intergenic | |
| rs7621652 | 3 | 116036079 | 4 | 0 | 0 | T | C | -1.0861 | 0.3121 | 0.0005014 | ---?- | 0 | 0.5856 | intron | LSAMP |
| rs62276916 | 3 | 155100847 | 4 | 0 | 0 | T | G | 1.0241 | 0.2943 | 0.0005014 | +++?+ | 9.7 | 0.3448 | intergenic | |
| rs6782488 | 3 | 16696226 | 4 | 0 | 0 | T | C | 1.4639 | 0.4207 | 0.0005015 | +++?+ | 0 | 0.5216 | intergenic | |
| rs11826066 | 11 | 86172176 | 4 | 0 | 0 | T | C | 2.4752 | 0.7113 | 0.0005016 | +++?+ | 0 | 0.8388 | intron | ME3 |
| rs78068702 | 18 | 25204531 | 4 | 0 | 0 | T | C | -2.5659 | 0.7373 | 0.0005016 | ---?- | 0 | 0.6442 | intergenic | |
| rs148065815 | 11 | 67497233 | 3 | 0 | 0 | T | C | -1.5039 | 0.4322 | 0.0005016 | ---?? | 0 | 0.6628 | intergenic | |
| 2:119062844 | 2 | 119062844 | 3 | 0 | 0 | CAT | C | -2.9045 | 0.8346 | 0.0005016 | ---?? | 0 | 0.5855 | | |
| rs13274344 | 8 | 65047796 | 4 | 0 | 0 | T | C | -0.8464 | 0.2432 | 0.0005017 | ---?- | 0 | 0.9435 | intergenic | |
| 1:181876969 | 1 | 181876969 | 3 | 0 | 0 | A | ACAC | 3.6122 | 1.038 | 0.0005017 | +++?? | 22.1 | 0.2769 | | |
| rs62419106 | 6 | 63311321 | 4 | 0 | 0 | A | G | 2.4348 | 0.6997 | 0.0005018 | +++?+ | 0 | 0.5116 | intergenic | |
| rs1197772 | 15 | 42064541 | 5 | 0 | 0 | A | G | -0.953 | 0.2739 | 0.000502 | ----- | 0 | 0.5386 | upstream-gene |  |
| rs141175348 | 6 | 157223851 | 3 | 0 | 0 | T | C | -3.8481 | 1.1059 | 0.000502 | ---?? | 0 | 0.6386 | intron | ARID1B |
| 1:87538182 | 1 | 87538182 | 3 | 0 | 0 | T | TTG | -1.0791 | 0.3101 | 0.0005022 | ---?? | 24.9 | 0.2642 | | |
| 4:166588202 | 4 | 166588202 | 2 | 0 | 0 | CAGGAA | C | 2.6676 | 0.7667 | 0.0005025 | ?++?? | 0 | 0.8144 | | |
| rs17022368 | 2 | 82979403 | 4 | 0 | 0 | A | G | -2.503 | 0.7194 | 0.0005028 | ---?- | 0 | 0.5967 | intergenic | |
| 6:159217682 | 6 | 159217682 | 3 | 0 | 0 | T | TG | 0.8864 | 0.2548 | 0.0005028 | +++?? | 10.3 | 0.3278 | | |
| 6:159219285 | 6 | 159219285 | 3 | 0 | 0 | T | TG | 0.8859 | 0.2546 | 0.0005028 | +++?? | 10.4 | 0.3274 | | |
| rs4076698 | 9 | 16192115 | 4 | 0 | 0 | C | G | 1.2524 | 0.36 | 0.0005029 | +++?+ | 0 | 0.6101 | intergenic | |
| rs7939252 | 11 | 37052054 | 4 | 0 | 0 | T | G | 0.9191 | 0.2642 | 0.0005029 | +++?+ | 0 | 0.4639 | intergenic | |
| rs56239199 | 5 | 737422 | 3 | 0 | 0 | A | T | 1.5656 | 0.45 | 0.0005029 | +++?? | 32.8 | 0.226 | intron | ZDHHC11B |
| 6:159223768 | 6 | 159223768 | 3 | 0 | 0 | T | TA | -0.8886 | 0.2554 | 0.000503 | ---?? | 3.4 | 0.3552 | | |
| rs2035664 | 7 | 130393537 | 4 | 0 | 0 | T | C | -0.8426 | 0.2422 | 0.0005031 | -+-?- | 53.6 | 0.09103 | intergenic | |
| 8:113287398 | 8 | 113287398 | 3 | 0 | 0 | T | TA | -1.0245 | 0.2945 | 0.0005031 | ---?? | 0 | 0.6201 | | |
| rs72701247 | 14 | 83268537 | 4 | 0 | 0 | A | G | 1.9295 | 0.5546 | 0.0005032 | -++?+ | 56.2 | 0.07672 | intergenic | |
| rs201450 | 7 | 101739441 | 4 | 0 | 0 | A | T | -0.8509 | 0.2446 | 0.0005033 | ---?- | 0 | 0.7952 | intron | CUX1 |
| rs13210961 | 6 | 32789388 | 5 | 0 | 0 | A | G | 1.3773 | 0.3959 | 0.0005034 | +++++ | 0 | 0.6754 | upstream-gene | |
| rs78124565 | 3 | 183160070 | 3 | 0 | 0 | C | G | -1.7345 | 0.4986 | 0.0005034 | ---?? | 0 | 0.6168 | intergenic | |
| rs4442530 | 11 | 37061671 | 4 | 0 | 0 | T | G | -0.8626 | 0.248 | 0.0005035 | ---?- | 0.4 | 0.3896 | intergenic | |
| rs115783327 | 5 | 44643475 | 3 | 0 | 0 | T | C | -3.6146 | 1.0391 | 0.0005039 | -?-?- | 0 | 0.8752 | intergenic | |
| 1:100328506 | 1 | 100328506 | 3 | 0 | 0 | A | AT | 2.8122 | 0.8084 | 0.0005039 | +++?? | 0 | 0.5037 | | |
| rs5023474 | 11 | 58997505 | 5 | 0 | 0 | T | C | -0.7482 | 0.2151 | 0.000504 | ----+ | 7.2 | 0.3657 | intergenic |  |
| rs62419104 | 6 | 63311107 | 4 | 0 | 0 | A | G | 2.4332 | 0.6995 | 0.000504 | +++?+ | 0 | 0.5108 | intergenic | |
| rs77891994 | 7 | 13252242 | 4 | 0 | 0 | A | G | 1.3659 | 0.3927 | 0.000504 | +++?+ | 0 | 0.5582 | intergenic | |
| rs116610937 | 18 | 25198695 | 4 | 0 | 0 | T | C | 2.5085 | 0.7211 | 0.000504 | +++?+ | 0 | 0.6195 | intergenic | |
| 10:91148551 | 10 | 91148551 | 3 | 0 | 0 | T | TA | -1.1374 | 0.327 | 0.000504 | ---?? | 66.5 | 0.05061 | | |
| rs17126327 | 1 | 64781794 | 4 | 0 | 0 | C | G | -1.2205 | 0.3509 | 0.0005043 | ---?- | 0 | 0.5166 | intergenic | |
| rs148029528 | 2 | 118027385 | 3 | 0 | 0 | T | C | -3.0494 | 0.8767 | 0.0005044 | ---?? | 43.3 | 0.1715 | intergenic | |
| rs9821959 | 3 | 2479136 | 4 | 0 | 0 | T | C | 0.9369 | 0.2693 | 0.0005046 | +++?- | 24.9 | 0.2622 | intron | CNTN4 |
| rs11158366 | 14 | 62366535 | 4 | 0 | 0 | A | G | 1.0571 | 0.3039 | 0.0005046 | +++?+ | 0 | 0.7394 | intron | SYT16 |
| rs74172501 | 19 | 46164609 | 5 | 0 | 1 | T | C | 0.926 | 0.2662 | 0.0005047 | ++++- | 14.5 | 0.3221 | intergenic | |
| 2:170542228 | 2 | 170542228 | 3 | 0 | 0 | <ins:me: | T | -1.5851 | 0.4557 | 0.0005047 | ---?? | 0 | 0.7754 | | |

| | | | | | | | | | | | | | | | |
|---|---|---|---|---|---|---|---|---|---|---|---|---|---|---|---|
| rs117334070 | 8 | 4701064 | 4 | 0 | 0 | T | C | -3.2959 | 0.9476 | 0.0005048 | ---?- | 0 | 0.9258 | intron | CSMD1 |
| 9:136651657 | 9 | 136651657 | 3 | 0 | 0 | G | GC | 1.9701 | 0.5664 | 0.0005048 | +++?? | 0 | 0.8952 | | |
| 22:29900469 | 22 | 29900469 | 3 | 0 | 0 | A | G | -5.1295 | 1.4748 | 0.000505 | ---?? | 0 | 0.408 | downstream-gene | |
| rs117341173 | 19 | 46164600 | 5 | 0 | 1 | A | G | -0.926 | 0.2662 | 0.000505 | ----+ | 14.5 | 0.322 | intergenic |  |
| rs34136830 | 7 | 98432711 | 4 | 0 | 0 | A | G | -1.1585 | 0.3331 | 0.0005052 | ---?- | 0 | 0.9389 | intergenic | |
| rs74172500 | 19 | 46164591 | 5 | 0 | 1 | C | G | -0.926 | 0.2662 | 0.0005052 | ----+ | 14.5 | 0.3218 | intergenic |  |
| rs116971894 | 16 | 2516417 | 3 | 0 | 0 | C | G | 3.3257 | 0.9562 | 0.0005052 | +++?? | 26.3 | 0.2574 | downstream-gene | |
| rs10775122 | 15 | 41898711 | 5 | 0 | 0 | T | C | -1.0093 | 0.2902 | 0.0005053 | ----- | 0 | 0.4068 | intergenic |  |
| rs11017087 | 10 | 131917244 | 3 | 0 | 0 | A | G | 0.9782 | 0.2813 | 0.0005053 | +++?? | 0 | 0.9963 | intergenic | |
| rs118069429 | 18 | 60704256 | 2 | 0 | 0 | A | G | 4.8208 | 1.3861 | 0.0005053 | ?++?? | 0 | 0.8577 | intergenic | |
| rs116025419 | 14 | 43839764 | 4 | 0 | 0 | T | C | 1.8542 | 0.5332 | 0.0005058 | +++?- | 0 | 0.4506 | intergenic | |
| rs114600612 | 12 | 27791019 | 3 | 0 | 0 | T | C | 0.9727 | 0.2797 | 0.0005062 | +++?? | 25.5 | 0.2614 | intron | PPFIBP1 |
| rs5023475 | 11 | 58997523 | 5 | 0 | 0 | T | C | -0.748 | 0.2151 | 0.0005064 | ----+ | 7.3 | 0.3648 | intergenic |  |
| rs12553738 | 9 | 78461766 | 4 | 0 | 0 | C | G | -1.284 | 0.3692 | 0.0005064 | ---?- | 54 | 0.08872 | intergenic | |
| rs3102998 | 6 | 159212764 | 3 | 0 | 0 | A | G | 0.8876 | 0.2552 | 0.0005066 | +++?? | 10.4 | 0.3277 | intron | EZR |
| rs10200017 | 2 | 170819774 | 4 | 0 | 1 | T | C | 1.0086 | 0.29 | 0.0005066 | +++?+ | 43.8 | 0.1486 | intron | UBR3 |
| 6:159213316 | 6 | 159213316 | 3 | 0 | 0 | G | GCT | 0.8874 | 0.2552 | 0.0005068 | +++?? | 10.4 | 0.3277 | | |
| rs16884556 | 5 | 10309877 | 4 | 0 | 0 | A | T | -1.9313 | 0.5554 | 0.0005069 | ---?- | 14.7 | 0.3184 | upstream-gene | |
| rs147520972 | 10 | 11583094 | 4 | 0 | 0 | A | G | -3.2457 | 0.9334 | 0.0005069 | ---?- | 0 | 0.9163 | intron | USP6NL |
| 19:36585458 | 19 | 36585458 | 3 | 0 | 0 | A | AAAT | -1.6803 | 0.4833 | 0.0005069 | -+-?? | 66 | 0.053 | | |
| 19:14748205 | 19 | 14748205 | 3 | 0 | 0 | CT | C | 0.9812 | 0.2822 | 0.000507 | +++?? | 50.8 | 0.1311 | | |
| rs149166828 | 17 | 57142304 | 5 | 0 | 0 | T | C | 2.2961 | 0.6604 | 0.0005072 | +++++ | 0 | 0.8274 | intron | TRIM37 |
| rs16986656 | 22 | 29132458 | 4 | 0 | 0 | T | C | -2.2971 | 0.6607 | 0.0005074 | ---?- | 0 | 0.8131 | intron | CHEK2 |
| rs77572620 | 17 | 57087246 | 5 | 0 | 0 | A | T | 2.3444 | 0.6743 | 0.0005076 | ++++- | 0 | 0.6684 | intron | TRIM37 |
| 3:183406579 | 3 | 183406579 | 3 | 0 | 0 | G | GC | -1.007 | 0.2896 | 0.0005078 | ---?? | 0 | 0.9557 | | |
| 3:183564558 | 3 | 183564558 | 3 | 0 | 0 | T | TA | -1.7462 | 0.5023 | 0.0005081 | ---?? | 0 | 0.6295 | | |
| rs73072117 | 3 | 45535867 | 4 | 0 | 0 | T | C | -2.4766 | 0.7124 | 0.0005083 | ---?- | 0 | 0.78 | intron | LARS2,LARS2-AS1 |
| rs140330649 | 5 | 716207 | 3 | 0 | 0 | A | T | -1.5302 | 0.4402 | 0.0005087 | ---?? | 66.5 | 0.05032 | intron | ZDHHC11B |
| rs76415080 | 2 | 129197489 | 4 | 0 | 0 | T | C | -2.8254 | 0.8128 | 0.0005088 | ---?- | 0 | 0.9641 | intergenic | |
| rs9276712 | 6 | 32759252 | 5 | 0 | 0 | A | C | -1.381 | 0.3973 | 0.000509 | ----- | 0 | 0.633 | intergenic |  |
| 22:45914936 | 22 | 45914936 | 3 | 0 | 0 | T | TGCAGC | -1.5674 | 0.451 | 0.0005093 | ---?? | 65.3 | 0.05612 | | |
| rs1123952 | 3 | 16697146 | 4 | 0 | 0 | A | G | -1.4634 | 0.4211 | 0.0005096 | ---?- | 0 | 0.5259 | intergenic | |
| 13:30320988 | 13 | 30320988 | 3 | 0 | 0 | CA | C | -1.5381 | 0.4426 | 0.0005099 | ---?? | 0 | 0.6193 | | |
| rs71624488 | 4 | 41332876 | 4 | 0 | 0 | T | C | -1.3941 | 0.4012 | 0.0005102 | ---?+ | 0 | 0.443 | intergenic | |
| 22:40544666 | 22 | 40544666 | 3 | 0 | 0 | G | GTCTGC | 0.9101 | 0.2619 | 0.0005103 | +++?? | 30.9 | 0.2352 | | |
| rs11132612 | 4 | 190021957 | 4 | 0 | 0 | T | C | -0.8685 | 0.2499 | 0.0005104 | ---?- | 0 | 0.5385 | intergenic | |
| rs7302587 | 12 | 102338853 | 4 | 0 | 0 | A | C | 0.8722 | 0.251 | 0.0005105 | +++?+ | 0 | 0.6121 | intergenic | |
| rs62514397 | 8 | 113311221 | 4 | 0 | 0 | A | G | -0.9859 | 0.2837 | 0.0005105 | ---?- | 0 | 0.7654 | intron | CSMD3 |
| rs201451 | 7 | 101739550 | 4 | 0 | 0 | A | G | 0.8496 | 0.2445 | 0.0005107 | +++?+ | 0 | 0.7944 | intron | CUX1 |
| rs4081413 | 11 | 120371653 | 4 | 0 | 0 | A | C | -0.9616 | 0.2767 | 0.0005108 | ---?- | 0 | 0.9997 | intergenic | |
| rs11033913 | 11 | 37072634 | 4 | 0 | 0 | T | G | 0.9479 | 0.2728 | 0.0005108 | +++?+ | 0 | 0.4122 | intergenic | |
| rs12318506 | 12 | 75718423 | 4 | 0 | 0 | T | G | 2.0489 | 0.5896 | 0.0005108 | +++?+ | 0 | 0.7823 | intron | CAPS2 |
| rs59086942 | 5 | 719581 | 4 | 0 | 0 | C | G | -1.432 | 0.4121 | 0.0005109 | ---?- | 54.5 | 0.0859 | intron | ZDHHC11B |
| rs6447978 | 4 | 20736742 | 4 | 0 | 0 | T | C | -0.9069 | 0.261 | 0.0005111 | ---?- | 0 | 0.8132 | intron | KCNIP4 |
| rs78804403 | 22 | 28777895 | 4 | 0 | 0 | T | C | 2.5214 | 0.7257 | 0.0005115 | +++?+ | 0 | 0.7419 | intron | TTC28 |
| rs80352209 | 18 | 40141030 | 4 | 0 | 0 | T | C | -1.3879 | 0.3994 | 0.0005115 | ---?- | 0 | 0.8936 | intron | LINC00907 |
| rs76748975 | 2 | 170476679 | 4 | 0 | 0 | A | G | 1.2322 | 0.3546 | 0.0005117 | +++?+ | 0 | 0.9618 | intron | PPIG |
| rs7805644 | 7 | 51834666 | 4 | 0 | 0 | T | C | 2.0895 | 0.6014 | 0.0005118 | +++?+ | 0 | 0.9336 | intergenic | |
| rs55835449 | 16 | 48094415 | 3 | 0 | 0 | A | G | 3.1632 | 0.9104 | 0.0005119 | +++?? | 0 | 0.4909 | intergenic | |
| rs60520388 | 19 | 46158936 | 5 | 0 | 1 | A | C | 0.9257 | 0.2665 | 0.0005121 | ++++- | 22.4 | 0.2719 | intergenic | |
| rs66861863 | 3 | 1945253 | 4 | 0 | 0 | T | C | 0.9625 | 0.277 | 0.0005122 | +++?+ | 14.7 | 0.3184 | intergenic | |
| rs475636 | 1 | 90237804 | 5 | 0 | 0 | T | C | 0.7414 | 0.2134 | 0.0005124 | +++++ | 0 | 0.9858 | intergenic | |
| rs333583 | 15 | 66322702 | 2 | 0 | 0 | T | G | -8.0697 | 2.3228 | 0.0005126 | ?-??- | 0 | 0.4716 | intron | MEGF11 |
| rs73453214 | 9 | 30819908 | 4 | 0 | 0 | A | G | -3.3093 | 0.9526 | 0.0005126 | ---?- | 0 | 0.4622 | intergenic | |
| rs146238393 | 17 | 57096805 | 5 | 0 | 0 | T | C | 2.3358 | 0.6724 | 0.0005126 | ++++- | 0 | 0.6752 | intron | TRIM37 |
| rs1419132 | 1 | 87555358 | 4 | 0 | 0 | T | C | -1.0584 | 0.3047 | 0.0005128 | ---?- | 1.8 | 0.3834 | intron | HS2ST1 |
| rs3114669 | 1 | 90242988 | 5 | 0 | 0 | T | G | 0.74 | 0.213 | 0.000513 | +++++ | 0 | 0.9889 | intergenic | |
| rs7588899 | 2 | 184043396 | 4 | 0 | 0 | A | T | 1.1699 | 0.3368 | 0.0005131 | +++?+ | 0 | 0.791 | intergenic | |
| rs56143451 | 1 | 30355908 | 4 | 0 | 0 | T | C | 1.3228 | 0.3808 | 0.0005132 | +++?- | 0 | 0.5438 | intergenic | |

| | | | | | | | | | | | | | | | |
|---|---|---|---|---|---|---|---|---|---|---|---|---|---|---|---|
| rs7646887 | 3 | 61072434 | 4 | 0 | 0 | C | G | 3.3009 | 0.9503 | 0.0005133 | +++?- | 0 | 0.642 | intron | FHIT |
| 8:113294734 | 8 | 113294734 | 3 | 0 | 0 | A | ATCTT | -1.0192 | 0.2934 | 0.0005136 | ---?? | 0 | 0.6007 | | |
| 1:64791255 | 1 | 64791255 | 3 | 0 | 0 | G | GCT | 1.2927 | 0.3722 | 0.0005138 | +++?? | 0 | 0.5104 | | |
| rs114014201 | 17 | 39389205 | 3 | 0 | 0 | A | C | -3.8034 | 1.095 | 0.0005139 | ---?? | 0 | 0.9674 | missense | KRTAP9-3 |
| rs11609636 | 12 | 41482984 | 4 | 0 | 0 | A | G | 1.6893 | 0.4864 | 0.0005141 | +++?+ | 8.7 | 0.3496 | intergenic | |
| rs1548986 | 8 | 113324432 | 4 | 0 | 0 | T | C | -0.9829 | 0.283 | 0.0005142 | ---?- | 0 | 0.7508 | intron | CSMD3 |
| rs1958513 | 14 | 44900268 | 4 | 0 | 0 | A | G | 1.3621 | 0.3922 | 0.0005143 | +++?- | 0 | 0.6078 | intergenic | |
| rs113522861 | 11 | 129281178 | 4 | 0 | 0 | A | G | -1.5326 | 0.4413 | 0.0005143 | ---?- | 30.1 | 0.2314 | intron | BARX2 |
| rs148706118 | 17 | 57096981 | 5 | 0 | 0 | A | G | 2.3349 | 0.6723 | 0.0005143 | ++++- | 0 | 0.6757 | intron | TRIM37 |
| rs60886194 | 8 | 3986820 | 4 | 0 | 0 | A | G | 1.1694 | 0.3367 | 0.0005144 | -++?+ | 55.6 | 0.0798 | intron | CSMD1 |
| rs1433450 | 15 | 87541062 | 4 | 0 | 0 | A | G | 1.9664 | 0.5662 | 0.0005149 | +++?+ | 0 | 0.9935 | intron | AGBL1 |
| rs7240523 | 18 | 68662810 | 4 | 0 | 0 | A | G | 1.6249 | 0.4679 | 0.0005149 | +++?+ | 0 | 0.9845 | intergenic | |
| rs10918738 | 1 | 162110568 | 5 | 0 | 0 | T | G | -0.885 | 0.2548 | 0.000515 | ----+ | 33.9 | 0.1955 | intron | NOS1AP |
| rs28878173 | 15 | 32182828 | 4 | 0 | 0 | T | C | -2.542 | 0.732 | 0.000515 | ---?- | 0 | 0.8047 | intergenic | |
| rs111605897 | 17 | 76910346 | 3 | 0 | 0 | T | C | 3.2705 | 0.9417 | 0.000515 | +++?? | 0 | 0.7282 | intron | TIMP2 |
| rs2365435 | 12 | 74416562 | 4 | 0 | 0 | A | T | 0.8858 | 0.2551 | 0.0005154 | +++?+ | 0 | 0.6834 | intergenic | |
| rs57476558 | 18 | 55757297 | 4 | 0 | 0 | C | G | -1.5827 | 0.4558 | 0.0005154 | ---?- | 0 | 0.9745 | intron | NEDD4L |
| rs58184590 | 18 | 52575875 | 4 | 0 | 0 | A | T | -2.0252 | 0.5832 | 0.0005156 | ---?- | 0 | 0.9219 | intron | CCDC68 |
| rs10836665 | 11 | 37052608 | 4 | 0 | 0 | T | C | -0.9172 | 0.2641 | 0.0005158 | ---?- | 0 | 0.4623 | intergenic | |
| rs17153461 | 11 | 58989046 | 5 | 0 | 0 | A | G | -0.748 | 0.2154 | 0.0005162 | ----+ | 12.9 | 0.3319 | intergenic |  |
| rs17793153 | 2 | 199055155 | 4 | 0 | 0 | A | G | 1.1518 | 0.3317 | 0.0005162 | +++?+ | 31.5 | 0.2236 | intergenic | |
| 1:203688453 | 1 | 203688453 | 3 | 0 | 0 | A | ATTGCC | 1.1746 | 0.3383 | 0.0005162 | +++?? | 0 | 0.786 | | |
| 2:150349955 | 2 | 150349955 | 3 | 0 | 0 | CA | C | 0.8965 | 0.2582 | 0.0005164 | +++?? | 41.2 | 0.1825 | | |
| rs958883 | 5 | 122998955 | 4 | 0 | 0 | T | C | -0.9187 | 0.2646 | 0.0005169 | ---?- | 54.5 | 0.0858 | intergenic | |
| rs1841656 | 2 | 170816628 | 4 | 0 | 1 | A | G | -1.0069 | 0.29 | 0.0005171 | ---?- | 43.8 | 0.1489 | intron | UBR3 |
| rs145360440 | 3 | 143027790 | 4 | 0 | 0 | C | G | -2.2128 | 0.6374 | 0.0005172 | ---?- | 0 | 0.7877 | intron | SLC9A9 |
| rs13153874 | 5 | 122999649 | 4 | 0 | 0 | T | C | -0.9189 | 0.2647 | 0.0005173 | ---?- | 54.8 | 0.08431 | intergenic | |
| rs71318345 | 3 | 183566276 | 4 | 0 | 0 | A | G | -1.8383 | 0.5295 | 0.0005174 | ---?- | 0 | 0.9396 | intron | PARL |
| rs62419062 | 6 | 63300513 | 4 | 0 | 0 | C | G | -2.43 | 0.7 | 0.0005177 | ---?- | 0 | 0.5459 | intergenic | |
| rs28451714 | 6 | 32671601 | 5 | 0 | 1 | T | G | 1.1226 | 0.3234 | 0.000518 | +++++ | 20.7 | 0.283 | intergenic | |
| rs6656617 | 1 | 87435442 | 3 | 0 | 0 | T | G | -1.0702 | 0.3083 | 0.0005181 | ---?? | 0 | 0.7536 | intron | HS2ST1 |
| rs116944085 | 7 | 27397513 | 4 | 0 | 0 | T | G | 1.8554 | 0.5345 | 0.0005181 | +++?+ | 0 | 0.8071 | intergenic | |
| 17:54096393 | 17 | 54096393 | 3 | 0 | 0 | A | AT | 0.9574 | 0.2758 | 0.0005183 | +++?? | 24.1 | 0.2676 | | |
| rs6728423 | 2 | 231406529 | 4 | 0 | 0 | T | C | -1.1086 | 0.3194 | 0.0005184 | ---?- | 0 | 0.5915 | intron | SP100 |
| rs6876704 | 5 | 123001882 | 4 | 0 | 0 | A | T | -0.9202 | 0.2651 | 0.0005184 | ---?- | 55.1 | 0.08278 | intergenic | |
| rs56260739 | 15 | 65456723 | 4 | 0 | 0 | T | G | -3.3693 | 0.9707 | 0.0005184 | ---?- | 0 | 0.8646 | intron | CLPX |
| rs17205170 | 6 | 32602483 | 5 | 0 | 1 | T | G | 1.067 | 0.3074 | 0.0005185 | -++++ | 43.9 | 0.1289 | upstream-gene | |
| rs113600224 | 7 | 139185162 | 3 | 0 | 0 | C | G | -2.5051 | 0.7217 | 0.0005185 | ---?? | 0 | 0.5966 | intergenic | |
| rs12373600 | 2 | 150853048 | 4 | 0 | 0 | C | G | -1.0384 | 0.2992 | 0.0005186 | ---?- | 0 | 0.7584 | intergenic | |
| rs117540823 | 11 | 127114795 | 4 | 0 | 0 | A | G | 3.9727 | 1.1445 | 0.0005186 | +++?+ | 0 | 0.7677 | intergenic | |
| rs10150304 | 14 | 43520246 | 4 | 0 | 0 | A | G | -1.6628 | 0.4791 | 0.0005187 | ---?+ | 0 | 0.5988 | intergenic | |
| rs1727128 | 7 | 99809921 | 2 | 0 | 0 | C | G | 1.3841 | 0.3988 | 0.0005188 | ???++ | 39 | 0.2004 | intron | STAG3,GATS |
| rs35109842 | 5 | 118699537 | 3 | 0 | 0 | A | G | 3.3565 | 0.967 | 0.0005188 | +++?? | 0 | 0.9367 | intron | TNFAIP8 |
| rs4485702 | 3 | 5389179 | 4 | 0 | 0 | T | C | -1.4098 | 0.4062 | 0.0005189 | ---?- | 25.2 | 0.2605 | intergenic | |
| rs113588564 | 9 | 79457947 | 4 | 0 | 0 | A | G | 2.0219 | 0.5825 | 0.0005189 | +++?+ | 0 | 0.5554 | intron | PRUNE2 |
| rs56223457 | 9 | 79458102 | 4 | 0 | 0 | A | G | -2.0188 | 0.5817 | 0.000519 | ---?- | 0 | 0.5529 | intron | PRUNE2 |
| rs62048629 | 16 | 85199643 | 4 | 0 | 0 | T | C | 1.6925 | 0.4877 | 0.000519 | +++?+ | 0 | 0.9788 | upstream-gene | |
| rs933305 | 12 | 111738287 | 5 | 0 | 0 | A | G | 0.9392 | 0.2706 | 0.0005192 | +++++ | 7.2 | 0.3657 | intron | CUX2 |
| rs3750579 | 10 | 134597211 | 4 | 0 | 0 | A | G | -0.9415 | 0.2713 | 0.0005192 | ---?- | 0 | 0.5888 | downstream-gene | |
| rs716979 | 8 | 86946269 | 4 | 0 | 0 | A | T | -1.9449 | 0.5604 | 0.0005195 | ---?- | 0 | 0.5872 | intergenic | |
| rs74761450 | 9 | 126859585 | 4 | 0 | 0 | A | G | -2.2411 | 0.6458 | 0.0005196 | ---?- | 44.4 | 0.1452 | intergenic | |
| rs10895331 | 11 | 102511736 | 5 | 0 | 0 | A | G | 0.879 | 0.2533 | 0.0005197 | -++++ | 19.1 | 0.2933 | intergenic | |
| rs61689139 | 11 | 59023025 | 5 | 0 | 0 | A | G | -0.7533 | 0.2171 | 0.0005197 | -+--+ | 11.8 | 0.3385 | intergenic | |
| rs10007184 | 4 | 18873165 | 4 | 0 | 0 | T | C | -1.8116 | 0.522 | 0.0005199 | ---?- | 0 | 0.9768 | intergenic | |
| rs7660006 | 4 | 20740339 | 4 | 0 | 0 | A | G | 0.9087 | 0.2618 | 0.00052 | +++?+ | 0 | 0.8097 | intron | KCNIP4 |
| rs11853525 | 15 | 88262136 | 4 | 0 | 0 | A | G | -0.9522 | 0.2744 | 0.0005202 | ---?- | 0 | 0.5004 | intergenic | |
| rs112280486 | 4 | 20744552 | 4 | 0 | 0 | A | G | 0.9112 | 0.2626 | 0.0005202 | +++?+ | 0 | 0.8039 | intron | KCNIP4 |
| rs148032365 | 4 | 20743661 | 4 | 0 | 0 | A | G | 0.9103 | 0.2623 | 0.0005202 | +++?+ | 0 | 0.8727 | intron | KCNIP4 |
| 2:104691441 | 2 | 104691441 | 3 | 0 | 0 | G | GA | 2.8486 | 0.8209 | 0.0005202 | +++?? | 0 | 0.5164 | | |

| | | | | | | | | | | | | | | | |
|---|---|---|---|---|---|---|---|---|---|---|---|---|---|---|---|
| 16:6542589 | 16 | 6542589 | 3 | 0 | 0 | T | TTAAA | 0.9731 | 0.2804 | 0.0005203 | +++?? | 0 | 0.6717 | | |
| rs817457 | 1 | 85666886 | 5 | 0 | 0 | T | C | -1.3589 | 0.3916 | 0.0005205 | ----- | 0 | 0.9673 | missense | SYDE2 |
| 3:183606994 | 3 | 183606994 | 3 | 0 | 0 | CA | C | -1.6868 | 0.4861 | 0.0005205 | ---?? | 0 | 0.858 | | |
| 4:41746024 | 4 | 41746024 | 3 | 0 | 0 | TA | TAA | -1.4709 | 0.4239 | 0.0005205 | ---?? | 50.1 | 0.1347 | | |
| rs8093214 | 18 | 45824508 | 4 | 0 | 0 | A | G | -0.9164 | 0.2641 | 0.0005207 | ---?+ | 0 | 0.5088 | intron | ZBTB7C |
| rs77419648 | 1 | 111849088 | 4 | 0 | 0 | A | G | 1.9626 | 0.5656 | 0.0005207 | +++?+ | 0 | 0.6823 | intron | CHIA |
| rs193094033 | 8 | 113471590 | 3 | 0 | 0 | T | G | 1.0429 | 0.3006 | 0.0005207 | +++?? | 0 | 0.8978 | intron | CSMD3 |
| rs10834247 | 11 | 23963682 | 4 | 0 | 0 | T | G | 1.2668 | 0.3651 | 0.000521 | +++?+ | 0 | 0.7293 | intergenic | |
| rs6830723 | 4 | 41333410 | 4 | 0 | 0 | T | C | -1.3919 | 0.4012 | 0.0005211 | ---?+ | 0 | 0.4438 | intergenic | |
| rs72954873 | 6 | 118906089 | 4 | 0 | 0 | A | G | 2.4577 | 0.7083 | 0.0005211 | +++?- | 0 | 0.5098 | intron | CEP85L |
| 1:64790602 | 1 | 64790602 | 3 | 0 | 0 | CT | C | -1.2908 | 0.372 | 0.0005211 | ---?? | 0 | 0.5109 | | |
| rs74484761 | 9 | 79457956 | 4 | 0 | 0 | T | C | -2.0197 | 0.5821 | 0.0005212 | ---?- | 0 | 0.5553 | intron | PRUNE2 |
| rs1199679 | 1 | 87564757 | 4 | 0 | 0 | A | T | 1.0428 | 0.3005 | 0.0005213 | +++?+ | 9.4 | 0.3461 | intron | HS2ST1 |
| rs10465169 | 9 | 78450030 | 4 | 0 | 0 | T | C | 1.2684 | 0.3656 | 0.0005214 | +++?+ | 58.2 | 0.0667 | intergenic | |
| rs13253431 | 8 | 65031560 | 4 | 0 | 0 | A | G | 0.8524 | 0.2457 | 0.0005216 | +++?+ | 0 | 0.9499 | intergenic | |
| rs149976621 | 4 | 38413474 | 2 | 0 | 0 | C | G | 5.2502 | 1.5133 | 0.0005217 | ?++?? | 0 | 0.3463 | intergenic | |
| 2:137509372 | 2 | 137509372 | 3 | 0 | 0 | G | GA | -3.6118 | 1.0411 | 0.0005217 | ---?? | 66 | 0.05285 | | |
| rs233111 | 1 | 85785655 | 5 | 0 | 0 | T | C | 0.8124 | 0.2342 | 0.0005218 | +++++ | 0 | 0.9768 | 3-prime-UTR | DDAH1 |
| rs62174937 | 2 | 185433550 | 4 | 0 | 0 | T | C | -0.9506 | 0.274 | 0.0005218 | ---?- | 0 | 0.7816 | intergenic | |
| rs7109866 | 11 | 58997684 | 5 | 0 | 0 | A | G | -0.7466 | 0.2152 | 0.0005219 | ----+ | 7.1 | 0.3662 | intergenic |  |
| rs73143041 | 7 | 78492596 | 4 | 0 | 0 | A | G | -2.1965 | 0.6331 | 0.0005219 | ---?- | 0 | 0.4938 | intron | MAGI2 |
| rs7177232 | 15 | 41910538 | 5 | 0 | 0 | A | G | 1.0022 | 0.2889 | 0.0005222 | +++++ | 4.3 | 0.3821 | upstream-gene | |
| rs8092030 | 18 | 63859697 | 4 | 0 | 0 | A | G | -0.865 | 0.2493 | 0.0005222 | ---?- | 0 | 0.5844 | intergenic | |
| rs11161436 | 1 | 85206477 | 5 | 0 | 0 | T | C | 0.8059 | 0.2323 | 0.0005222 | +++++ | 0 | 0.7672 | intergenic | |
| rs116847425 | 7 | 25608659 | 4 | 0 | 0 | A | G | -1.788 | 0.5154 | 0.0005222 | ---?- | 19.6 | 0.2917 | intergenic | |
| rs6005758 | 22 | 28719080 | 4 | 0 | 0 | A | G | 2.5314 | 0.7297 | 0.0005224 | +++?+ | 0 | 0.7314 | intron | TTC28 |
| rs12611647 | 2 | 139894894 | 4 | 0 | 0 | A | C | 1.3169 | 0.3796 | 0.0005224 | +++?+ | 0 | 0.9869 | intergenic | |
| rs116440583 | 18 | 25200946 | 4 | 0 | 0 | T | C | 2.5215 | 0.7269 | 0.0005226 | +++?+ | 0 | 0.6236 | intergenic | |
| rs34520993 | 12 | 27789780 | 4 | 0 | 0 | C | G | -0.9398 | 0.2709 | 0.0005227 | ---?- | 6.1 | 0.3625 | intron | PPFIBP1 |
| rs9506225 | 13 | 30323830 | 3 | 0 | 0 | A | T | -1.4947 | 0.4309 | 0.0005228 | ---?? | 0 | 0.5918 | intergenic | |
| rs11657381 | 17 | 47450133 | 5 | 0 | 0 | A | G | 0.8827 | 0.2545 | 0.0005228 | ++++- | 49.9 | 0.09202 | intergenic | |
| rs117018131 | 8 | 54302802 | 4 | 0 | 0 | T | C | -2.4014 | 0.6923 | 0.0005229 | ---?- | 55.1 | 0.08262 | intergenic | |
| rs9864602 | 3 | 123534758 | 4 | 0 | 0 | A | G | -2.619 | 0.7551 | 0.0005232 | ---?- | 0 | 0.9767 | intron | MYLK |
| rs7810268 | 7 | 156189327 | 4 | 0 | 0 | A | C | 0.8856 | 0.2553 | 0.0005236 | +++?+ | 21.8 | 0.2795 | intergenic | |
| rs12520743 | 5 | 103157759 | 4 | 0 | 0 | A | C | -1.5633 | 0.4507 | 0.0005236 | ---?- | 0 | 0.7592 | intergenic | |
| 1:85863673 | 1 | 85863673 | 3 | 0 | 0 | CTT | C | -1.5701 | 0.4527 | 0.0005236 | ---?? | 0 | 0.8467 | | |
| rs10508596 | 10 | 20151577 | 4 | 0 | 0 | A | T | -0.8706 | 0.251 | 0.0005237 | ---?- | 13 | 0.3277 | intron | PLXDC2 |
| rs61334489 | 4 | 190023789 | 4 | 0 | 0 | T | C | -0.8666 | 0.2499 | 0.0005237 | ---?- | 0 | 0.4342 | intergenic | |
| rs73116513 | 3 | 5389290 | 4 | 0 | 0 | T | C | 1.4081 | 0.406 | 0.0005237 | +++?+ | 25.1 | 0.2607 | intergenic | |
| rs151006853 | 13 | 85085443 | 4 | 0 | 0 | A | G | -3.3109 | 0.9546 | 0.0005237 | ---?- | 0 | 0.8269 | intron | LINC00333 |
| 6:31121874 | 6 | 31121874 | 3 | 0 | 0 | CAA | C | -0.8525 | 0.2458 | 0.0005237 | ---?? | 48.9 | 0.1415 | | |
| rs11844872 | 14 | 21335191 | 3 | 0 | 0 | A | G | -4.3993 | 1.2685 | 0.0005238 | ---?? | 0 | 0.6685 | intergenic | |
| rs56216434 | 3 | 29473054 | 4 | 0 | 0 | T | C | -1.7983 | 0.5185 | 0.0005238 | ---?- | 0 | 0.5373 | intron | RBMS3 |
| rs73552611 | 8 | 21544791 | 3 | 0 | 0 | A | T | 4.253 | 1.2262 | 0.0005238 | ?++?+ | 0 | 0.9252 | downstream-gene | |
| rs6781815 | 3 | 155102234 | 4 | 0 | 0 | T | G | 1.0218 | 0.2946 | 0.000524 | +++?+ | 9.3 | 0.3465 | intergenic | |
| rs79362739 | 4 | 110065442 | 5 | 0 | 0 | A | G | -1.4169 | 0.4086 | 0.0005243 | ----- | 0 | 0.5698 | intron | COL25A1 |
| rs75381868 | 6 | 32501629 | 2 | 0 | 0 | T | C | 2.1339 | 0.6153 | 0.0005244 | ???++ | 0 | 0.8592 | upstream-gene | |
| rs6689803 | 1 | 102931983 | 3 | 0 | 0 | A | T | -0.8854 | 0.2553 | 0.0005247 | ---?? | 0 | 0.8412 | intergenic | |
| rs11512511 | 11 | 103706538 | 4 | 0 | 0 | T | C | 1.4758 | 0.4256 | 0.0005248 | +++?+ | 0 | 0.4399 | intergenic | |
| rs2529094 | 7 | 101739617 | 4 | 0 | 0 | T | C | 0.8477 | 0.2444 | 0.0005249 | +++?+ | 0 | 0.7955 | intron | CUX1 |
| rs6699779 | 1 | 203688785 | 4 | 0 | 0 | A | G | -1.1245 | 0.3243 | 0.0005249 | ---?- | 0 | 0.9097 | intron | ATP2B4 |
| rs6844145 | 4 | 41319184 | 4 | 0 | 0 | A | G | 1.4961 | 0.4314 | 0.0005251 | +++?- | 0 | 0.6919 | intergenic | |
| 2:170520306 | 2 | 170520306 | 3 | 0 | 0 | T | TA | 1.282 | 0.3697 | 0.0005253 | +++?? | 0 | 0.8832 | | |
| rs2219002 | 8 | 113509584 | 4 | 0 | 0 | A | C | 1.0058 | 0.2901 | 0.0005256 | +++?+ | 0 | 0.6604 | intron | CSMD3 |
| 6:159233087 | 6 | 159233087 | 3 | 0 | 0 | G | GCT | 0.8833 | 0.2548 | 0.0005256 | +++?? | 12.3 | 0.3199 | | |
| rs2031467 | 10 | 131923470 | 4 | 0 | 0 | T | C | 0.902 | 0.2602 | 0.0005258 | +++?+ | 0 | 0.858 | intergenic | |
| rs2038751 | 1 | 64576126 | 4 | 0 | 0 | A | G | 1.0707 | 0.3088 | 0.000526 | +++?+ | 0 | 0.8472 | intron | ROR1 |
| rs35663295 | 3 | 120342843 | 4 | 0 | 0 | T | C | 1.0065 | 0.2903 | 0.000526 | +++?+ | 61.4 | 0.05089 | downstream-gene | |
| rs78976052 | 1 | 100196380 | 4 | 0 | 0 | A | G | 1.9245 | 0.5551 | 0.000526 | +++?+ | 0 | 0.472 | intron | FRRS1 |

| | | | | | | | | | | | | | | | |
|---|---|---|---|---|---|---|---|---|---|---|---|---|---|---|---|
| rs79718248 | 11 | 103707398 | 4 | 0 | 0 | A | G | 1.4755 | 0.4256 | 0.000526 | +++?+ | 0 | 0.4387 | intergenic | |
| rs13149237 | 4 | 189999482 | 4 | 0 | 0 | T | C | 0.8822 | 0.2545 | 0.0005262 | +++?+ | 29.3 | 0.2364 | intergenic | |
| rs2353271 | 8 | 113325575 | 4 | 0 | 0 | T | C | 0.9805 | 0.2828 | 0.0005264 | +++?+ | 0 | 0.7476 | intron | CSMD3 |
| 14:21453947 | 14 | 21453947 | 3 | 0 | 0 | T | TA | 0.9402 | 0.2712 | 0.0005264 | +++?? | 38.4 | 0.1974 | | |
| rs9958338 | 18 | 377105 | 4 | 0 | 0 | T | C | 0.8974 | 0.2589 | 0.0005267 | +++?+ | 0 | 0.5505 | intron | COLEC12 |
| rs4471754 | 18 | 45936862 | 4 | 0 | 0 | C | G | -2.7037 | 0.7799 | 0.0005268 | ---?+ | 26.3 | 0.2542 | upstream-gene | |
| rs78419816 | 6 | 106059742 | 4 | 0 | 0 | T | C | 2.1773 | 0.628 | 0.0005268 | +++?+ | 0 | 0.9692 | intergenic | |
| 6:95159430 | 6 | 95159430 | 3 | 0 | 0 | T | TAC | -0.8807 | 0.2541 | 0.0005268 | ---?? | 0 | 0.7241 | | |
| 16:84115343 | 16 | 84115343 | 3 | 0 | 0 | T | G | -1.6042 | 0.4627 | 0.0005269 | ---?? | 0 | 0.9346 | intron | MBTPS1 |
| rs3099835 | 5 | 3067377 | 3 | 0 | 0 | T | C | 1.658 | 0.4783 | 0.0005269 | +++?? | 14.4 | 0.3109 | intergenic | |
| rs4666613 | 2 | 184016667 | 3 | 0 | 0 | A | T | -1.2247 | 0.3533 | 0.000527 | ---?? | 0 | 0.7384 | intron | NUP35 |
| rs11877953 | 18 | 63806606 | 4 | 0 | 0 | A | G | -0.855 | 0.2466 | 0.000527 | ---?- | 7.5 | 0.3554 | intergenic | |
| rs12448392 | 16 | 5440346 | 4 | 0 | 0 | A | G | -0.8681 | 0.2504 | 0.000527 | ---?- | 0 | 0.7892 | intron | RBFOX1 |
| rs12972158 | 19 | 46158473 | 5 | 0 | 1 | T | C | -0.9252 | 0.2669 | 0.000527 | ----+ | 23.7 | 0.2633 | intergenic |  |
| 22:28787535 | 22 | 28787535 | 3 | 0 | 0 | A | T | -2.7749 | 0.8005 | 0.0005271 | ---?? | 0 | 0.7398 | intron | TTC28 |
| 1:181876400 | 1 | 181876400 | 3 | 0 | 0 | T | TAG | -3.6037 | 1.0396 | 0.0005271 | ---?? | 21.4 | 0.2802 | | |
| rs7820425 | 8 | 113325684 | 4 | 0 | 0 | A | T | 0.9803 | 0.2828 | 0.0005274 | +++?+ | 0 | 0.7476 | intron | CSMD3 |
| rs11070350 | 15 | 42127918 | 5 | 0 | 0 | A | G | -0.952 | 0.2746 | 0.0005274 | ----- | 0 | 0.5728 | intron | JMJD7-PLA2G4B,JMJD7 |
| rs146157869 | 12 | 68464073 | 4 | 0 | 0 | T | C | -3.6731 | 1.0596 | 0.0005276 | ---?- | 0 | 0.6312 | intergenic | |
| rs35688602 | 1 | 87465380 | 4 | 0 | 0 | A | C | -1.0019 | 0.289 | 0.0005277 | ---?- | 0 | 0.6004 | intron | HS2ST1 |
| rs80110170 | 6 | 63320028 | 4 | 0 | 0 | A | G | 2.4482 | 0.7063 | 0.0005278 | +++?+ | 0 | 0.5133 | intergenic | |
| rs4951107 | 1 | 203687154 | 4 | 0 | 0 | T | G | 1.125 | 0.3246 | 0.000528 | +++?+ | 0 | 0.8947 | intron | ATP2B4 |
| rs61089532 | 17 | 77316244 | 4 | 0 | 0 | C | G | 1.3149 | 0.3794 | 0.000528 | +++?- | 18.1 | 0.3 | intron | RBFOX3 |
| rs6812543 | 4 | 41333429 | 4 | 0 | 0 | T | G | 1.3904 | 0.4012 | 0.0005281 | +++?- | 0 | 0.4441 | intergenic | |
| 8:113296246 | 8 | 113296246 | 3 | 0 | 0 | T | TA | 1.0169 | 0.2934 | 0.0005281 | +++?? | 0 | 0.5969 | | |
| rs2031468 | 10 | 131923448 | 4 | 0 | 0 | T | C | 0.9018 | 0.2602 | 0.0005282 | +++?+ | 0 | 0.8585 | intergenic | |
| rs201452 | 7 | 101739663 | 4 | 0 | 0 | T | C | 0.8465 | 0.2442 | 0.0005284 | +++?+ | 0 | 0.7924 | intron | CUX1 |
| rs343048 | 7 | 35515295 | 4 | 0 | 0 | A | G | 1.1096 | 0.3201 | 0.0005285 | +++?+ | 21.1 | 0.2833 | intergenic | |
| rs28383311 | 6 | 32587011 | 3 | 0 | 0 | A | T | -1.134 | 0.3272 | 0.0005286 | ---?? | 0 | 0.4331 | intergenic | |
| rs62178690 | 2 | 217036351 | 3 | 0 | 0 | T | C | -3.7664 | 1.0867 | 0.0005286 | -?-?- | 0 | 0.8056 | intron | XRCC5 |
| 11:125821884 | 11 | 125821884 | 3 | 0 | 0 | CAAG | C | -4.1208 | 1.189 | 0.0005286 | ---?? | 0 | 0.4341 | | |
| rs75754615 | 18 | 40141604 | 4 | 0 | 0 | T | C | -1.383 | 0.3991 | 0.0005287 | ---?- | 0 | 0.9005 | intron | LINC00907 |
| rs12305437 | 12 | 41502813 | 4 | 0 | 0 | T | C | 1.8861 | 0.5442 | 0.0005288 | +++?+ | 0 | 0.4976 | intergenic | |
| rs80139511 | 12 | 41302108 | 4 | 0 | 0 | C | G | -1.6992 | 0.4903 | 0.0005288 | ---?+ | 0 | 0.639 | intron | CNTN1 |
| rs9287928 | 2 | 170944471 | 4 | 0 | 1 | A | G | -0.9901 | 0.2857 | 0.0005292 | ---?- | 60 | 0.05761 | downstream-gene | |
| rs113668614 | 11 | 103708198 | 4 | 0 | 0 | T | C | 1.4748 | 0.4256 | 0.0005292 | +++?+ | 0 | 0.4374 | intergenic | |
| rs66692444 | 8 | 98186294 | 5 | 0 | 0 | A | C | 1.46 | 0.4213 | 0.0005294 | +++++ | 8.9 | 0.3557 | intergenic | |
| rs145467050 | 6 | 106275116 | 4 | 0 | 0 | A | C | 1.9508 | 0.5629 | 0.0005294 | +++?+ | 0 | 0.424 | intergenic | |
| rs114418931 | 3 | 47776704 | 2 | 0 | 0 | T | C | -6.6336 | 1.9144 | 0.0005301 | ?-??- | 0.1 | 0.3171 | intron | SMARCC1 |
| rs4244585 | 15 | 41941111 | 5 | 0 | 0 | A | G | -0.9808 | 0.2831 | 0.0005305 | ----- | 12.6 | 0.3334 | intron | MGA |
| rs12706940 | 7 | 130128793 | 4 | 0 | 0 | T | C | -0.8521 | 0.2459 | 0.0005305 | -+-?- | 60.5 | 0.0554 | intron | MEST,MESTIT1 |
| 2:24666774 | 2 | 24666774 | 3 | 0 | 0 | A | AT | -4.5047 | 1.3001 | 0.0005305 | ---?? | 55 | 0.1086 | | |
| 3:45341730 | 3 | 45341730 | 3 | 0 | 0 | G | GAT | -1.2883 | 0.3718 | 0.0005305 | ---?? | 19.7 | 0.2877 | | |
| rs513743 | 1 | 90242428 | 5 | 0 | 0 | A | T | 0.7385 | 0.2131 | 0.0005307 | +++++ | 0 | 0.9897 | intergenic | |
| rs6888738 | 5 | 123000271 | 4 | 0 | 0 | T | C | -0.9174 | 0.2648 | 0.0005308 | ---?- | 54.7 | 0.08472 | intergenic | |
| rs5997369 | 22 | 28885620 | 4 | 0 | 0 | C | G | -2.2935 | 0.662 | 0.0005309 | ---?- | 0 | 0.5825 | intron | TTC28 |
| rs6841534 | 4 | 190013351 | 4 | 0 | 0 | A | G | -0.8791 | 0.2537 | 0.0005312 | ---?- | 0 | 0.5529 | intergenic | |
| rs11977793 | 7 | 82413911 | 4 | 0 | 0 | T | C | 0.899 | 0.2595 | 0.0005312 | +-+?+ | 47.4 | 0.1272 | intron | PCLO |
| rs59586663 | 9 | 137747703 | 4 | 0 | 0 | T | G | -3.4843 | 1.0057 | 0.0005312 | ---?- | 0 | 0.4854 | intergenic | |
| rs149938743 | 9 | 11335337 | 4 | 0 | 0 | C | G | 2.6704 | 0.7708 | 0.0005312 | +++?+ | 10.2 | 0.3419 | intergenic | |
| rs12157518 | 22 | 28754042 | 4 | 0 | 0 | A | T | 2.5153 | 0.726 | 0.0005315 | +++?+ | 0 | 0.7508 | intron | TTC28 |
| 12:96643890 | 12 | 96643890 | 3 | 0 | 0 | A | AAGG | -1.713 | 0.4945 | 0.0005315 | ---?? | 0 | 0.447 | | |
| rs76464945 | 3 | 68250723 | 4 | 0 | 0 | A | G | 2.3703 | 0.6842 | 0.0005316 | +++?+ | 21.8 | 0.2797 | intron | FAM19A1 |
| rs114879185 | 3 | 143035468 | 4 | 0 | 0 | T | C | -2.0784 | 0.5999 | 0.0005316 | ---?- | 0 | 0.8127 | intron | SLC9A9 |
| 8:113296420 | 8 | 113296420 | 3 | 0 | 0 | CTCAGT | C | -1.0154 | 0.2931 | 0.0005318 | ---?? | 0 | 0.5978 | | |
| rs6889036 | 5 | 123000403 | 4 | 0 | 0 | A | C | -0.9173 | 0.2648 | 0.0005319 | ---?- | 54.8 | 0.08462 | intergenic | |
| rs12422294 | 12 | 27787619 | 4 | 0 | 0 | A | G | 0.9423 | 0.272 | 0.0005319 | +++?+ | 5.9 | 0.3634 | intron | PPFIBP1 |
| rs117475884 | 10 | 131476071 | 4 | 0 | 0 | A | G | -1.291 | 0.3727 | 0.000532 | ---?- | 0 | 0.8264 | intron | MGMT |
| rs7175554 | 15 | 54350824 | 3 | 0 | 0 | C | G | 1.1899 | 0.3435 | 0.0005322 | +++?? | 0 | 0.9856 | intron | UNC13C |

| | | | | | | | | | | | | | | | |
|---|---|---|---|---|---|---|---|---|---|---|---|---|---|---|---|
| rs192699549 | 7 | 36581487 | 3 | 0 | 0 | A | G | -4.6168 | 1.3328 | 0.0005322 | ?--?- | 35.9 | 0.2103 | intron | AOAH |
| rs55985982 | 8 | 65032910 | 4 | 0 | 0 | T | C | -0.8503 | 0.2455 | 0.0005324 | ---?- | 0 | 0.9499 | intergenic | |
| rs4658322 | 1 | 90251584 | 5 | 0 | 0 | T | C | -0.7416 | 0.2141 | 0.0005325 | ----- | 0 | 0.931 | intergenic |  |
| rs11982147 | 7 | 82414164 | 4 | 0 | 0 | T | C | -0.8972 | 0.259 | 0.0005325 | -+-?- | 47.6 | 0.1258 | intron | PCLO |
| rs35836546 | 8 | 65033058 | 4 | 0 | 0 | T | C | -0.8501 | 0.2454 | 0.0005325 | ---?- | 0 | 0.9499 | intergenic | |
| 8:89040568 | 8 | 89040568 | 3 | 0 | 0 | CA | C | -1.2444 | 0.3593 | 0.0005326 | ---?? | 47.5 | 0.1489 | | |
| rs76137647 | 12 | 17008386 | 3 | 0 | 0 | A | G | 3.4781 | 1.0041 | 0.0005327 | +++?? | 42.5 | 0.1755 | intergenic | |
| rs114534594 | 17 | 57104386 | 5 | 0 | 0 | A | T | -2.3236 | 0.6708 | 0.0005327 | ----+ | 0 | 0.681 | intron | TRIM37 |
| rs117749892 | 18 | 68766298 | 4 | 0 | 0 | T | C | -2.6413 | 0.7625 | 0.0005327 | ---?- | 0 | 0.6091 | intergenic | |
| rs141793176 | 12 | 103252065 | 3 | 0 | 0 | A | G | 0.8745 | 0.2525 | 0.0005328 | +++?? | 0 | 0.5918 | intron | PAH |
| rs173009 | 11 | 37042300 | 3 | 0 | 0 | T | C | 0.9 | 0.2598 | 0.000533 | +++?? | 30.3 | 0.2381 | intergenic | |
| rs34337233 | 8 | 65033069 | 4 | 0 | 0 | T | C | -0.8501 | 0.2454 | 0.000533 | ---?- | 0 | 0.9499 | intergenic | |
| rs6773774 | 3 | 196503053 | 4 | 0 | 0 | A | G | 2.1272 | 0.6142 | 0.0005331 | +++?- | 19.9 | 0.2902 | intron | PAK2 |
| rs1363350 | 5 | 123000895 | 4 | 0 | 0 | A | G | -0.9173 | 0.2649 | 0.0005333 | ---?- | 54.8 | 0.08468 | intergenic | |
| rs7170637 | 15 | 22969232 | 5 | 0 | 0 | A | G | -0.9747 | 0.2814 | 0.0005333 | ----- | 39.3 | 0.1592 | missense | CYFIP1 |
| rs2720230 | 3 | 127194308 | 3 | 0 | 0 | A | C | -1.0295 | 0.2972 | 0.0005334 | ---?? | 39.7 | 0.1907 | intergenic | |
| rs112923461 | 12 | 26210817 | 2 | 0 | 0 | A | G | 5.3882 | 1.5558 | 0.0005334 | ++??? | 48 | 0.1655 | intron | RASSF8 |
| rs9542404 | 13 | 71142214 | 4 | 0 | 0 | A | G | -0.8705 | 0.2513 | 0.0005335 | ---?- | 1.2 | 0.3858 | intergenic | |
| rs58840465 | 9 | 79458114 | 4 | 0 | 0 | T | C | -2.0109 | 0.5806 | 0.0005335 | ---?- | 0 | 0.5526 | intron | PRUNE2 |
| rs2422455 | 11 | 37051498 | 4 | 0 | 0 | A | G | 0.8626 | 0.2491 | 0.0005336 | +++?+ | 5.8 | 0.3641 | intergenic | |
| rs4752460 | 10 | 122500375 | 4 | 0 | 0 | T | C | -1.0088 | 0.2913 | 0.0005336 | ---?- | 12.2 | 0.3315 | intergenic | |
| rs11823076 | 11 | 129286380 | 4 | 0 | 0 | T | C | -1.5002 | 0.4332 | 0.0005337 | ---?- | 47.4 | 0.1268 | intron | BARX2 |
| 12:94936027 | 12 | 94936027 | 3 | 0 | 0 | CA | C | 0.9444 | 0.2727 | 0.000534 | +++?? | 52.6 | 0.1214 | | |
| rs17374025 | 4 | 155611649 | 4 | 0 | 0 | C | G | 1.436 | 0.4147 | 0.0005343 | +++?+ | 0 | 0.6457 | intergenic | |
| 2:184016937 | 2 | 184016937 | 3 | 0 | 0 | CAG | C | 1.2233 | 0.3532 | 0.0005343 | +++?? | 0 | 0.7373 | | |
| rs6839421 | 4 | 111404410 | 4 | 0 | 0 | T | C | -0.8358 | 0.2414 | 0.0005349 | ---?- | 0 | 0.9989 | intron | ENPEP |
| rs1199680 | 1 | 87565676 | 4 | 0 | 0 | T | C | -1.0402 | 0.3004 | 0.0005352 | ---?- | 8.7 | 0.3494 | intron | HS2ST1 |
| rs140208030 | 17 | 57105706 | 5 | 0 | 0 | T | C | 2.3215 | 0.6705 | 0.0005352 | ++++- | 0 | 0.6815 | intron | TRIM37 |
| rs4768318 | 12 | 41295547 | 4 | 0 | 0 | A | G | -1.697 | 0.4901 | 0.0005353 | ---?+ | 0 | 0.5962 | intron | CNTN1 |
| rs144376615 | 6 | 77601141 | 3 | 0 | 0 | A | G | -1.2057 | 0.3482 | 0.0005353 | ---?? | 6.7 | 0.3423 | intergenic | |
| rs1882441 | 2 | 184046474 | 4 | 0 | 0 | A | G | 1.164 | 0.3362 | 0.0005354 | +++?+ | 0 | 0.7917 | intergenic | |
| rs932818 | 10 | 134498564 | 4 | 0 | 0 | A | T | -0.8848 | 0.2556 | 0.0005356 | ---?- | 0 | 0.7645 | intron | INPP5A |
| rs1515664 | 1 | 235076299 | 4 | 0 | 0 | C | G | -0.8654 | 0.25 | 0.0005356 | ---?- | 0 | 0.8678 | intergenic | |
| rs2294014 | 8 | 124710277 | 5 | 0 | 0 | A | G | -0.8012 | 0.2314 | 0.0005357 | ----+ | 21.2 | 0.2797 | intron | ANXA13 |
| rs62276914 | 3 | 155091218 | 4 | 0 | 0 | A | G | 1.0405 | 0.3005 | 0.0005358 | +++?+ | 6.4 | 0.3613 | intergenic | |
| rs7536241 | 1 | 87566713 | 4 | 0 | 0 | T | G | 1.0395 | 0.3003 | 0.0005359 | +++?+ | 9.2 | 0.347 | intron | HS2ST1 |
| rs958881 | 5 | 122999412 | 4 | 0 | 0 | A | G | 0.9163 | 0.2647 | 0.000536 | +++?+ | 54.7 | 0.08482 | intergenic | |
| rs6813719 | 4 | 111404413 | 4 | 0 | 0 | A | G | -0.8357 | 0.2414 | 0.0005361 | ---?- | 0 | 0.9989 | intron | ENPEP |
| rs11157397 | 14 | 44889891 | 4 | 0 | 0 | T | C | -1.3634 | 0.3938 | 0.0005361 | ---?+ | 0 | 0.4839 | intergenic | |
| 6:75630288 | 6 | 75630288 | 3 | 0 | 0 | CT | C | 1.8176 | 0.525 | 0.0005361 | +++?? | 0 | 0.6585 | | |
| 7:64918121 | 7 | 64918121 | 3 | 0 | 0 | A | AT | -1.3422 | 0.3877 | 0.0005361 | ---?? | 0 | 0.5644 | | |
| rs333772 | 11 | 37052785 | 4 | 0 | 0 | A | G | 0.8618 | 0.2489 | 0.0005364 | +++?+ | 7.1 | 0.3575 | intergenic | |
| rs78117862 | 2 | 117123431 | 4 | 0 | 0 | T | C | 1.7657 | 0.51 | 0.0005364 | +++?+ | 0 | 0.6717 | intergenic | |
| rs6476237 | 9 | 30885773 | 3 | 0 | 0 | A | G | 4.0662 | 1.1746 | 0.0005366 | +++?? | 0 | 0.5763 | intergenic | |
| rs7176474 | 15 | 41931506 | 5 | 0 | 0 | A | G | -0.9755 | 0.2818 | 0.0005369 | ----- | 13.8 | 0.3262 | intron | MGA |
| rs147677499 | 22 | 29076344 | 4 | 0 | 0 | A | G | 2.3548 | 0.6803 | 0.0005369 | +++?+ | 0 | 0.4069 | upstream-gene | |
| rs34832499 | 16 | 5156583 | 3 | 0 | 0 | A | G | -1.8308 | 0.5289 | 0.0005372 | ---?? | 0 | 0.9561 | intergenic | |
| rs1849205 | 6 | 165818395 | 4 | 0 | 0 | A | G | 0.8702 | 0.2514 | 0.0005373 | +++?+ | 0 | 0.7053 | intron | PDE10A |
| rs138555714 | 17 | 57203250 | 5 | 0 | 0 | C | G | 2.4234 | 0.7001 | 0.0005373 | +++++ | 0 | 0.7853 | intron | SKA2 |
| rs6509989 | 19 | 56815923 | 4 | 0 | 0 | A | G | 1.5721 | 0.4542 | 0.0005377 | +++?+ | 0 | 0.4522 | intron | ZSCAN5A,LOC100506374 |
| rs7721201 | 5 | 55638989 | 2 | 0 | 0 | T | C | -5.6935 | 1.645 | 0.0005377 | -?-?? | 0 | 0.461 | intergenic | |
| rs17128913 | 12 | 41291809 | 4 | 0 | 0 | T | C | -1.697 | 0.4903 | 0.0005377 | ---?+ | 0 | 0.5965 | intron | CNTN1 |
| rs1665149 | 11 | 44917578 | 4 | 0 | 0 | C | G | -1.0135 | 0.2928 | 0.000538 | ---?- | 0 | 0.7156 | intron | TSPAN18 |
| rs113971128 | 13 | 24009060 | 3 | 0 | 0 | C | G | 3.715 | 1.0734 | 0.000538 | +++?? | 0 | 0.6528 | upstream-gene | |
| rs201453 | 7 | 101739788 | 4 | 0 | 0 | T | C | -0.8446 | 0.244 | 0.0005382 | ---?- | 0 | 0.791 | intron | CUX1 |
| rs7349323 | 2 | 170970668 | 4 | 0 | 1 | A | G | -0.887 | 0.2563 | 0.0005382 | ---?- | 40.5 | 0.1684 | intergenic | |
| 3:30825887 | 3 | 30825887 | 3 | 0 | 0 | T | TTGTC | 2.9448 | 0.8508 | 0.0005382 | +++?? | 0 | 0.4636 | | |
| rs75970534 | 12 | 41302424 | 4 | 0 | 0 | A | G | -1.6973 | 0.4904 | 0.0005385 | ---?+ | 0 | 0.6391 | intron | CNTN1 |
| 22:41833209 | 22 | 41833209 | 2 | 0 | 0 | T | C | -4.1794 | 1.2076 | 0.0005386 | ?--?? | 64.4 | 0.09381 | synonymous | TOB2 |

| | | | | | | | | | | | | | | | |
|---|---|---|---|---|---|---|---|---|---|---|---|---|---|---|---|
| rs2771340 | 14 | 21455760 | 4 | 0 | 0 | T | C | -0.9071 | 0.2621 | 0.0005386 | ---?- | 0 | 0.9454 | upstream-gene | |
| rs7515864 | 1 | 192461456 | 4 | 0 | 0 | A | G | -1.1004 | 0.318 | 0.0005386 | ---?+ | 34.1 | 0.2077 | intergenic | |
| rs16883401 | 8 | 113327247 | 4 | 0 | 0 | T | G | -0.9783 | 0.2827 | 0.0005386 | ---?- | 0 | 0.7454 | intron | CSMD3 |
| rs61692854 | 15 | 91582496 | 4 | 0 | 0 | T | C | 1.8968 | 0.5481 | 0.0005388 | +++?+ | 0 | 0.8664 | intergenic | |
| rs116174973 | 1 | 85117083 | 4 | 0 | 0 | T | C | -2.3505 | 0.6792 | 0.000539 | ---?- | 0 | 0.8248 | intron | SSX2IP |
| rs1372540 | 12 | 41325471 | 4 | 0 | 0 | A | T | -1.4788 | 0.4273 | 0.0005392 | ---?- | 0 | 0.6237 | intron | CNTN1 |
| rs16883404 | 8 | 113327282 | 4 | 0 | 0 | A | T | 0.9782 | 0.2827 | 0.0005392 | +++?+ | 0 | 0.7451 | intron | CSMD3 |
| rs57560641 | 7 | 141196465 | 2 | 0 | 0 | C | G | 6.4969 | 1.8774 | 0.0005392 | ++??? | 71.3 | 0.06196 | intergenic | |
| 2:137508228 | 2 | 137508228 | 3 | 0 | 0 | G | GTGTC | 3.5975 | 1.0396 | 0.0005392 | +++?? | 66.3 | 0.05143 | | |
| rs43215 | 5 | 35086969 | 4 | 0 | 0 | T | C | 1.2649 | 0.3655 | 0.0005393 | +++?- | 48.2 | 0.1221 | intron | PRLR |
| rs6831336 | 4 | 41333674 | 4 | 0 | 0 | C | G | 1.3882 | 0.4012 | 0.0005394 | +++?- | 0 | 0.4449 | intergenic | |
| rs17128908 | 12 | 41288366 | 4 | 0 | 0 | T | C | -1.6988 | 0.4909 | 0.0005394 | ---?+ | 0 | 0.6019 | intron | CNTN1 |
| rs148323943 | 3 | 143065416 | 4 | 0 | 0 | A | G | 2.0366 | 0.5886 | 0.0005394 | +++?+ | 0 | 0.6301 | intron | SLC9A9,SLC9A9-AS1 |
| rs6596827 | 6 | 1489300 | 4 | 0 | 0 | C | G | -0.8426 | 0.2435 | 0.0005396 | ---?- | 0 | 0.4186 | intergenic | |
| rs61981114 | 14 | 32887583 | 4 | 0 | 0 | T | G | -0.8739 | 0.2525 | 0.0005396 | -+-?- | 46.6 | 0.132 | intron | AKAP6 |
| rs10070162 | 5 | 123001509 | 4 | 0 | 0 | C | G | -0.9166 | 0.2649 | 0.0005397 | ---?- | 54.9 | 0.084 | intergenic | |
| rs17036320 | 2 | 47387187 | 4 | 0 | 0 | T | C | 1.3554 | 0.3917 | 0.0005398 | +++?+ | 0 | 0.9619 | upstream-gene | |
| rs142938427 | 3 | 143068728 | 4 | 0 | 0 | A | G | 2.0431 | 0.5905 | 0.0005399 | +++?+ | 0 | 0.6136 | intron | SLC9A9 |
| rs289389 | 15 | 85575044 | 4 | 0 | 0 | A | G | -0.8288 | 0.2395 | 0.00054 | +--?+ | 56 | 0.07799 | intron | PDE8A |
| rs79329889 | 4 | 177264338 | 4 | 0 | 0 | C | G | -2.8857 | 0.834 | 0.00054 | ---?- | 0 | 0.6781 | intergenic | |
| rs10010429 | 4 | 122361381 | 4 | 0 | 0 | T | C | 1.0733 | 0.3102 | 0.0005401 | +++?+ | 0 | 0.6468 | intergenic | |
| 16:7597606 | 16 | 7597606 | 3 | 0 | 0 | T | TAGAAT | 3.8944 | 1.1255 | 0.0005401 | +++?? | 0 | 0.4774 | | |
| rs2053645 | 8 | 65011693 | 4 | 0 | 0 | T | C | -0.859 | 0.2483 | 0.0005404 | ---?- | 0 | 0.9274 | intergenic | |
| rs75369211 | 2 | 117123327 | 4 | 0 | 0 | T | C | -1.7643 | 0.5099 | 0.0005405 | ---?- | 0 | 0.6719 | intergenic | |
| rs55829616 | 1 | 204387463 | 4 | 0 | 0 | A | C | -1.4194 | 0.4103 | 0.0005406 | ---?- | 0 | 0.4073 | downstream-gene | |
| rs7916315 | 10 | 129680261 | 4 | 0 | 0 | A | G | 0.8613 | 0.249 | 0.0005408 | +++?+ | 0 | 0.7749 | intron | CLRN3 |
| rs147778951 | 4 | 166231305 | 4 | 0 | 0 | A | G | -3.7761 | 1.0915 | 0.0005409 | ---?- | 0 | 0.9573 | intron | KLHL2 |
| rs60587813 | 9 | 79458231 | 4 | 0 | 0 | A | G | -2.0036 | 0.5791 | 0.000541 | ---?- | 0 | 0.5506 | intron | PRUNE2 |
| 19:46158113 | 19 | 46158113 | 3 | 0 | 0 | A | AAAAAG | 0.9245 | 0.2672 | 0.0005411 | +++?? | 0 | 0.7472 | | |
| rs28730515 | 15 | 98043663 | 4 | 0 | 0 | T | C | 0.8598 | 0.2485 | 0.0005412 | +++?+ | 0 | 0.9009 | intergenic | |
| rs138199167 | 6 | 23731799 | 4 | 0 | 0 | T | C | -3.198 | 0.9244 | 0.0005412 | ---?+ | 25.8 | 0.2567 | intergenic | |
| 6:106273898 | 6 | 106273898 | 3 | 0 | 0 | G | GA | 2.0003 | 0.5782 | 0.0005415 | +++?? | 24.4 | 0.2666 | | |
| rs145334823 | 8 | 85103861 | 4 | 0 | 0 | A | G | -4.2883 | 1.2397 | 0.0005416 | ---?- | 0 | 0.8008 | intron | RALYL |
| 9:135303561 | 9 | 135303561 | 3 | 0 | 0 | G | GT | 1.2771 | 0.3692 | 0.0005417 | +-+?? | 54.8 | 0.1096 | | |
| 9:16161124 | 9 | 16161124 | 3 | 0 | 0 | G | GCA | -1.3568 | 0.3922 | 0.0005417 | ---?? | 54.4 | 0.1117 | | |
| rs2276153 | 11 | 58407740 | 5 | 0 | 0 | C | G | 0.8498 | 0.2457 | 0.0005419 | +++++ | 0 | 0.9765 | intergenic | |
| rs77463648 | 2 | 125035415 | 3 | 0 | 0 | T | C | -2.9685 | 0.8582 | 0.0005419 | ---?? | 0 | 0.953 | intron | CNTNAP5 |
| rs10955618 | 8 | 113328308 | 4 | 0 | 0 | T | G | -0.9797 | 0.2832 | 0.000542 | ---?- | 0 | 0.7412 | intron | CSMD3 |
| rs17128906 | 12 | 41287466 | 4 | 0 | 0 | A | G | 1.6983 | 0.491 | 0.000542 | +++?- | 0 | 0.6028 | intron | CNTN1 |
| 1:102941479 | 1 | 102941479 | 3 | 0 | 0 | A | AC | 0.8799 | 0.2544 | 0.000542 | +++?? | 0 | 0.8372 | | |
| rs76604079 | 4 | 177263964 | 4 | 0 | 0 | T | C | -2.8843 | 0.8338 | 0.0005421 | ---?- | 0 | 0.6778 | intergenic | |
| rs2273404 | 6 | 165808448 | 4 | 0 | 0 | A | G | 0.8779 | 0.2538 | 0.0005422 | +++?+ | 0 | 0.6464 | intron | PDE10A |
| rs34835838 | 11 | 74300010 | 4 | 0 | 0 | T | C | 1.259 | 0.364 | 0.0005422 | +++?+ | 0 | 0.6921 | upstream-gene | |
| rs60999840 | 9 | 79458183 | 4 | 0 | 0 | T | C | 2.005 | 0.5796 | 0.0005422 | +++?+ | 0 | 0.5513 | intron | PRUNE2 |
| rs117452698 | 8 | 139905950 | 3 | 0 | 0 | T | C | 3.2478 | 0.9389 | 0.0005422 | +++?? | 14.7 | 0.3098 | intron | COL22A1 |
| rs144074900 | 2 | 191094804 | 4 | 0 | 0 | T | C | -3.6618 | 1.0586 | 0.0005423 | ---?- | 0 | 0.5198 | intron | HIBCH |
| rs1156211 | 4 | 38737461 | 5 | 0 | 0 | A | G | -0.7652 | 0.2212 | 0.0005425 | ----- | 0 | 0.9112 | intergenic |  |
| rs12539454 | 7 | 49469540 | 4 | 0 | 0 | T | C | -0.97 | 0.2804 | 0.0005425 | ---?+ | 24.4 | 0.2651 | intergenic | |
| rs62182221 | 2 | 218831041 | 3 | 0 | 0 | T | C | 1.4828 | 0.4287 | 0.0005427 | +++?? | 0 | 0.7101 | intergenic | |
| rs356672 | 9 | 97952689 | 4 | 0 | 0 | T | C | -1.696 | 0.4904 | 0.0005428 | ---?- | 0 | 0.6212 | intron | FANCC |
| rs142155153 | 10 | 86690873 | 4 | 0 | 0 | T | C | -1.7588 | 0.5085 | 0.0005428 | ---?- | 0 | 0.8248 | intergenic | |
| rs146024120 | 4 | 157715981 | 4 | 0 | 0 | A | G | -3.2492 | 0.9395 | 0.0005429 | ---?- | 0 | 0.9222 | intron | PDGFC |
| 1:117847289 | 1 | 117847289 | 3 | 0 | 0 | A | AAATT | 2.8656 | 0.8286 | 0.000543 | +++?? | 20.2 | 0.2857 | | |
| 11:87629821 | 11 | 87629821 | 2 | 0 | 0 | T | TA | 6.209 | 1.7953 | 0.0005432 | +?+?? | 0 | 0.7385 | | |
| 2:173371566 | 2 | 173371566 | 3 | 0 | 0 | T | C | 4.5539 | 1.3168 | 0.0005437 | +++?? | 0 | 0.9556 | downstream-gene | |
| rs1985239 | 12 | 27790632 | 4 | 0 | 0 | A | G | 0.9361 | 0.2707 | 0.0005438 | +++?+ | 5.7 | 0.3643 | intron | PPFIBP1 |
| rs117909295 | 19 | 58318570 | 4 | 0 | 0 | A | G | -2.2338 | 0.646 | 0.0005439 | ---?+ | 54.3 | 0.08706 | 3-prime-UTR | ZNF552 |
| rs76212147 | 2 | 117123229 | 4 | 0 | 0 | A | G | -1.7629 | 0.5098 | 0.000544 | ---?- | 0 | 0.6717 | intergenic | |
| rs62514400 | 8 | 113315839 | 4 | 0 | 0 | A | C | 0.9815 | 0.2838 | 0.0005442 | +++?+ | 0 | 0.7567 | intron | CSMD3 |

| | | | | | | | | | | | | | | | |
|---|---|---|---|---|---|---|---|---|---|---|---|---|---|---|---|
| rs57568306 | 8 | 65034483 | 4 | 0 | 0 | T | C | 0.8479 | 0.2452 | 0.0005443 | +++?+ | 0 | 0.9501 | intergenic | |
| rs17150184 | 5 | 122530459 | 4 | 0 | 0 | A | T | -1.8158 | 0.5251 | 0.0005444 | ---?- | 0 | 0.8134 | downstream-gene | |
| rs8105523 | 19 | 46154609 | 5 | 0 | 1 | A | G | 0.9606 | 0.2778 | 0.0005445 | +-++- | 53.3 | 0.07305 | intergenic | |
| rs62514401 | 8 | 113315924 | 4 | 0 | 0 | T | C | 0.9814 | 0.2838 | 0.0005446 | +++?+ | 0 | 0.7565 | intron | CSMD3 |
| rs10156195 | 7 | 7794799 | 4 | 0 | 0 | A | G | -1.6297 | 0.4713 | 0.000545 | ---?- | 0 | 0.9477 | intron | RPA3-AS1 |
| rs148254818 | 3 | 189477641 | 3 | 0 | 0 | A | G | 4.2209 | 1.2207 | 0.000545 | +++?? | 0 | 0.4932 | intron | TP63 |
| rs4602017 | 15 | 92185737 | 3 | 0 | 0 | T | C | 1.4029 | 0.4058 | 0.0005451 | +++?? | 0 | 0.4669 | intergenic | |
| rs17716530 | 21 | 29022751 | 4 | 0 | 0 | T | C | 2.7235 | 0.7877 | 0.0005451 | +++?+ | 0 | 0.6946 | intergenic | |
| rs4733950 | 8 | 38658394 | 4 | 0 | 0 | A | C | 1.2913 | 0.3735 | 0.0005453 | +++?+ | 19.3 | 0.2938 | intron | TACC1 |
| rs1352254 | 1 | 98597553 | 4 | 0 | 0 | A | G | -1.3514 | 0.3909 | 0.0005454 | ---?- | 18.8 | 0.2964 | intergenic | |
| rs78423415 | 12 | 41303388 | 4 | 0 | 0 | T | G | -1.6961 | 0.4906 | 0.0005455 | ---?+ | 0 | 0.639 | intron | CNTN1 |
| 19:56073115 | 19 | 56073115 | 3 | 0 | 0 | A | T | 1.6696 | 0.4829 | 0.0005457 | +++?? | 0 | 0.4727 | intergenic | |
| rs28383224 | 6 | 32583653 | 5 | 0 | 0 | A | G | -0.7193 | 0.208 | 0.0005457 | ----- | 0 | 0.7194 | intergenic |  |
| rs4542372 | 11 | 95706776 | 4 | 0 | 0 | A | G | 0.9362 | 0.2708 | 0.0005461 | +++?+ | 24.1 | 0.2666 | downstream-gene | |
| 6:159187877 | 6 | 159187877 | 3 | 0 | 0 | G | GAC | -0.8849 | 0.256 | 0.0005461 | ---?? | 42.6 | 0.1749 | | |
| rs10102788 | 8 | 96970499 | 4 | 0 | 0 | T | C | 0.991 | 0.2867 | 0.0005462 | +++?+ | 0 | 0.87 | intergenic | |
| rs62077690 | 17 | 47450644 | 5 | 0 | 0 | T | C | -0.8819 | 0.2551 | 0.0005463 | ----+ | 49.9 | 0.09223 | intergenic |  |
| rs1106743 | 17 | 77125223 | 4 | 0 | 0 | A | C | -0.9503 | 0.2749 | 0.0005465 | ---?- | 0 | 0.5737 | intron | RBFOX3 |
| rs71318344 | 3 | 183559085 | 4 | 0 | 0 | T | G | -1.8307 | 0.5296 | 0.0005465 | ---?- | 0 | 0.9355 | intron | PARL |
| rs13209654 | 6 | 32792659 | 5 | 0 | 0 | T | C | -1.3698 | 0.3963 | 0.0005467 | ----- | 0 | 0.6661 | intron | TAP2 |
| rs10505185 | 8 | 113234556 | 4 | 0 | 0 | A | G | -0.9832 | 0.2845 | 0.000547 | ---?- | 0 | 0.9693 | downstream-gene | |
| rs148168003 | 6 | 155700439 | 2 | 0 | 0 | T | G | 6.845 | 1.9803 | 0.000547 | ++??? | 0 | 0.3958 | intergenic | |
| 3:185789508 | 3 | 185789508 | 3 | 0 | 0 | T | C | -3.0697 | 0.8881 | 0.0005471 | ---?? | 0 | 0.8 | intron | ETV5 |
| rs11541295 | 17 | 57184046 | 5 | 0 | 0 | T | C | 2.3931 | 0.6923 | 0.0005472 | +++++ | 0 | 0.7997 | 5-prime-UTR | TRIM37 |
| rs78298569 | 3 | 143068304 | 4 | 0 | 0 | T | C | 2.0389 | 0.5899 | 0.0005472 | +++?+ | 0 | 0.6116 | intron | SLC9A9 |
| rs3767297 | 1 | 205036826 | 5 | 0 | 0 | A | G | -0.9761 | 0.2824 | 0.0005473 | ----- | 0 | 0.6624 | intron | CNTN2 |
| rs6817475 | 4 | 20761511 | 4 | 0 | 0 | T | G | 0.911 | 0.2636 | 0.0005473 | +++?+ | 0 | 0.7986 | intron | KCNIP4 |
| rs7937620 | 11 | 23963902 | 4 | 0 | 0 | T | C | -1.2537 | 0.3627 | 0.0005474 | ---?- | 0 | 0.7263 | intergenic | |
| rs61542621 | 9 | 16191889 | 4 | 0 | 0 | A | G | -1.2431 | 0.3597 | 0.0005475 | ---?- | 0 | 0.6048 | intergenic | |
| rs9655918 | 7 | 82416315 | 4 | 0 | 0 | A | G | 0.8923 | 0.2582 | 0.0005477 | +-+?+ | 46.4 | 0.1332 | intron | PCLO |
| rs4906150 | 14 | 43270942 | 4 | 0 | 0 | A | T | -1.295 | 0.3747 | 0.0005478 | ---?+ | 61.3 | 0.05128 | intergenic | |
| 8:113250908 | 8 | 113250908 | 3 | 0 | 0 | T | TTTG | 1.0209 | 0.2954 | 0.000548 | +++?? | 0 | 0.8434 | | |
| 3:185789510 | 3 | 185789510 | 3 | 0 | 0 | A | G | 3.0692 | 0.8881 | 0.0005481 | +++?? | 0 | 0.7999 | intron | ETV5 |
| rs28383233 | 6 | 32584153 | 5 | 0 | 0 | A | G | 0.719 | 0.208 | 0.0005481 | +++++ | 0 | 0.7195 | intergenic | |
| rs115316908 | 3 | 143028638 | 4 | 0 | 0 | C | G | -2.1969 | 0.6357 | 0.0005481 | ---?- | 0 | 0.7877 | intron | SLC9A9 |
| rs79990996 | 9 | 112533872 | 5 | 0 | 0 | T | C | -2.698 | 0.7807 | 0.0005482 | ----- | 7.5 | 0.364 | intron | PALM2 |
| rs10097216 | 8 | 113510549 | 4 | 0 | 0 | T | C | 1.0032 | 0.2903 | 0.0005485 | +++?+ | 0 | 0.6543 | intron | CSMD3 |
| rs10775241 | 15 | 87608399 | 4 | 0 | 0 | C | G | -0.8912 | 0.2579 | 0.0005485 | ---?- | 0 | 0.6213 | intergenic | |
| rs11785270 | 8 | 113328833 | 4 | 0 | 0 | T | G | 0.9764 | 0.2825 | 0.0005485 | +++?+ | 0 | 0.7425 | intron | CSMD3 |
| 5:105653672 | 5 | 105653672 | 2 | 0 | 0 | T | TTGTG | 1.3227 | 0.3828 | 0.0005485 | ?++?? | 0 | 0.7085 | | |
| rs1979960 | 11 | 7249257 | 4 | 0 | 0 | T | C | -2.9802 | 0.8624 | 0.000549 | ---?+ | 0 | 0.8232 | intergenic | |
| rs28383221 | 6 | 32583212 | 5 | 0 | 0 | A | G | -0.7195 | 0.2082 | 0.000549 | ----- | 0 | 0.7183 | intergenic |  |
| 13:51755914 | 13 | 51755914 | 3 | 0 | 0 | A | ATAAC | 2.234 | 0.6465 | 0.000549 | +++?? | 0 | 0.6824 | | |
| rs72678820 | 8 | 97244456 | 4 | 0 | 0 | T | G | 1.872 | 0.5417 | 0.0005491 | +++?+ | 0 | 0.7555 | intron | UQCRB |
| rs34346046 | 7 | 36679928 | 4 | 0 | 0 | A | G | 1.1484 | 0.3323 | 0.0005493 | +++?+ | 0 | 0.5716 | intron | AOAH |
| rs148932605 | 3 | 143067955 | 4 | 0 | 0 | A | C | -2.0372 | 0.5896 | 0.0005496 | ---?- | 0 | 0.6104 | intron | SLC9A9 |
| 6:71455485 | 6 | 71455485 | 2 | 0 | 0 | A | G | -5.0793 | 1.47 | 0.0005498 | ?--?? | 0 | 0.6351 | intron | SMAP1 |
| rs11241706 | 5 | 123001867 | 4 | 0 | 0 | T | C | 0.9156 | 0.265 | 0.0005498 | +++?+ | 55.2 | 0.08219 | intergenic | |
| rs7825635 | 8 | 113330037 | 4 | 0 | 0 | T | C | 0.977 | 0.2828 | 0.0005499 | +++?+ | 0 | 0.7453 | intron | CSMD3 |
| rs145193796 | 17 | 56394484 | 5 | 0 | 0 | A | G | -2.7511 | 0.7962 | 0.0005499 | ----- | 38.4 | 0.1652 | intron | BZRAP1 |
| rs2232778 | 19 | 8370120 | 4 | 0 | 0 | T | C | 0.9293 | 0.269 | 0.00055 | +++?+ | 0 | 0.8092 | intron | CD320 |
| rs77172798 | 1 | 98438757 | 3 | 0 | 0 | T | G | -1.0292 | 0.2979 | 0.0005501 | ---?? | 0 | 0.7397 | intergenic | |
| rs12725099 | 1 | 237795118 | 3 | 0 | 0 | T | C | -1.3597 | 0.3935 | 0.0005502 | ---?? | 52.7 | 0.1209 | intron | RYR2 |
| rs62077691 | 17 | 47451117 | 5 | 0 | 0 | T | C | -0.8819 | 0.2553 | 0.0005502 | ----+ | 49.9 | 0.0923 | intergenic |  |
| rs112321201 | 12 | 130974561 | 3 | 0 | 0 | T | C | 3.4171 | 0.989 | 0.0005502 | +++?? | 57.6 | 0.09459 | intron | RIMBP2 |
| rs12495003 | 3 | 155092978 | 4 | 0 | 0 | A | C | 1.0226 | 0.296 | 0.0005503 | +++?+ | 32.4 | 0.2178 | intergenic | |
| rs114884516 | 6 | 10962637 | 4 | 0 | 0 | T | C | 2.8381 | 0.8215 | 0.0005504 | +-+?+ | 47.6 | 0.1257 | intron | SYCP2L |
| rs72783777 | 2 | 7229619 | 3 | 0 | 0 | A | C | -2.175 | 0.6296 | 0.0005506 | ---?? | 0 | 0.6179 | intergenic | |
| rs7826270 | 8 | 113330490 | 4 | 0 | 0 | T | C | 0.9771 | 0.2828 | 0.0005507 | +++?+ | 0 | 0.7464 | intron | CSMD3 |

| | | | | | | | | | | | | | | | |
|---|---|---|---|---|---|---|---|---|---|---|---|---|---|---|---|
| rs6841400 | 4 | 190013426 | 4 | 0 | 0 | T | C | -0.8762 | 0.2536 | 0.0005508 | ---?- | 0 | 0.4972 | intergenic | |
| rs11236666 | 11 | 75974052 | 4 | 0 | 0 | A | C | 1.0677 | 0.3091 | 0.0005509 | +++?+ | 0 | 0.9736 | intergenic | |
| rs80335813 | 19 | 46153804 | 5 | 0 | 0 | T | G | -0.9608 | 0.2781 | 0.000551 | -+--+ | 54.6 | 0.06586 | intergenic | |
| rs6005735 | 22 | 28579487 | 4 | 0 | 0 | T | C | 2.4962 | 0.7226 | 0.0005513 | +++?+ | 0 | 0.7336 | intron | TTC28 |
| rs78983883 | 2 | 117103544 | 4 | 0 | 0 | T | C | -2.7396 | 0.7931 | 0.0005515 | ---?- | 24.7 | 0.2631 | intergenic | |
| rs1538278 | 13 | 112829112 | 4 | 0 | 0 | A | G | 0.9512 | 0.2754 | 0.0005519 | +++?+ | 12.9 | 0.3279 | intergenic | |
| 3:172330585 | 3 | 172330585 | 3 | 0 | 0 | G | GAACTC | 1.8812 | 0.5446 | 0.0005519 | +++?? | 0 | 0.7837 | | |
| rs75241650 | 14 | 53504851 | 2 | 0 | 0 | A | G | -4.165 | 1.2058 | 0.000552 | -?-?? | 0 | 0.7585 | 3-prime-UTR | DDHD1 |
| 3:172330578 | 3 | 172330578 | 3 | 0 | 0 | CAAG | C | -1.8789 | 0.544 | 0.000552 | ---?? | 0 | 0.7816 | | |
| rs395755 | 2 | 40432937 | 4 | 0 | 0 | A | G | -1.0726 | 0.3105 | 0.0005522 | ---?+ | 0 | 0.5418 | intron | SLC8A1,SLC8A1-AS1 |
| 7:43358992 | 7 | 43358992 | 3 | 0 | 0 | CAAAA | C | -3.5656 | 1.0323 | 0.0005522 | ---?? | 43.2 | 0.1719 | | |
| rs7931091 | 11 | 528575 | 5 | 0 | 0 | A | G | -0.8072 | 0.2337 | 0.0005524 | ----- | 0 | 0.4697 | intron | LOC101059906 |
| rs201454 | 7 | 101740056 | 4 | 0 | 0 | A | G | -0.8422 | 0.2438 | 0.0005526 | ---?- | 0 | 0.7889 | intron | CUX1 |
| rs147638801 | 3 | 143067446 | 4 | 0 | 0 | A | T | 2.0346 | 0.5891 | 0.0005526 | +++?+ | 0 | 0.6097 | intron | SLC9A9 |
| rs17126319 | 1 | 64777977 | 4 | 0 | 0 | C | G | -1.2114 | 0.3508 | 0.0005528 | ---?- | 0 | 0.507 | intergenic | |
| rs36066467 | 3 | 183596010 | 4 | 0 | 0 | T | C | -1.6221 | 0.4697 | 0.0005528 | ---?- | 0 | 0.9479 | intron | PARL |
| 1:7125714 | 1 | 7125714 | 3 | 0 | 0 | CTTG | C | 0.9116 | 0.264 | 0.0005528 | +++?? | 57.2 | 0.09673 | | |
| rs7165230 | 15 | 42129168 | 5 | 0 | 0 | A | T | -0.9478 | 0.2744 | 0.0005529 | ----- | 0 | 0.5756 | intron | JMJD7-PLA2G4B,JMJD7 |
| rs113106530 | 3 | 23807709 | 4 | 0 | 0 | T | C | 3.6157 | 1.0469 | 0.0005531 | +++?+ | 0 | 0.4675 | intergenic | |
| rs180892194 | 16 | 47591324 | 3 | 0 | 0 | A | C | 3.2514 | 0.9415 | 0.0005533 | +++?? | 0 | 0.7555 | intron | PHKB |
| 13:60025745 | 13 | 60025745 | 3 | 0 | 0 | A | AG | -0.9069 | 0.2626 | 0.0005533 | ---?? | 12 | 0.321 | | |
| rs11689614 | 2 | 18519734 | 4 | 0 | 0 | A | G | 0.8907 | 0.2579 | 0.0005535 | +++?+ | 0 | 0.9196 | intergenic | |
| rs7031123 | 9 | 78456701 | 4 | 0 | 0 | A | G | 1.2566 | 0.3639 | 0.0005536 | +++?+ | 56.2 | 0.07672 | intergenic | |
| rs55994245 | 3 | 176022695 | 4 | 0 | 0 | A | G | -1.2908 | 0.3738 | 0.0005536 | ---?- | 0 | 0.3971 | intergenic | |
| rs1938593 | 11 | 58984717 | 5 | 0 | 0 | A | G | -0.7437 | 0.2154 | 0.000554 | -+--+ | 18.1 | 0.2993 | upstream-gene | |
| rs10836666 | 11 | 37052706 | 4 | 0 | 0 | A | G | -0.9123 | 0.2642 | 0.000554 | ---?- | 0 | 0.4554 | intergenic | |
| rs12002637 | 9 | 137745126 | 4 | 0 | 0 | A | C | -3.4062 | 0.9864 | 0.000554 | ---?- | 0 | 0.6152 | upstream-gene | |
| rs445300 | 5 | 743547 | 3 | 0 | 0 | A | C | -1.5823 | 0.4582 | 0.0005543 | ---?? | 0 | 0.4951 | intron | ZDHHC11B |
| rs74544369 | 1 | 64576649 | 4 | 0 | 0 | C | G | 1.0653 | 0.3085 | 0.0005543 | +++?+ | 0 | 0.8471 | intron | ROR1 |
| rs37389 | 5 | 35085180 | 4 | 0 | 0 | A | G | 1.2565 | 0.3639 | 0.0005545 | +++?- | 50.2 | 0.1102 | intron | PRLR |
| rs7844829 | 8 | 113330755 | 4 | 0 | 0 | T | C | -0.9764 | 0.2828 | 0.0005546 | ---?- | 0 | 0.7461 | intron | CSMD3 |
| rs57498313 | 15 | 42131965 | 5 | 0 | 0 | A | G | -0.9465 | 0.2741 | 0.0005546 | ----- | 0 | 0.5825 | intron | JMJD7-PLA2G4B,PLA2G4B |
| rs1674162 | 1 | 85673871 | 5 | 0 | 0 | T | C | 1.3607 | 0.3941 | 0.0005548 | +++++ | 0 | 0.978 | intergenic | |
| rs817403 | 1 | 85617351 | 4 | 0 | 0 | T | C | 1.5703 | 0.4548 | 0.0005551 | +++?+ | 0 | 0.9849 | intergenic | |
| rs3893283 | 17 | 4770094 | 5 | 0 | 0 | T | C | -1.5673 | 0.454 | 0.0005555 | ----+ | 17 | 0.3062 | intron | MINK1 |
| rs72751894 | 15 | 86324557 | 4 | 0 | 0 | C | G | 3.5423 | 1.026 | 0.0005555 | +++?+ | 33.2 | 0.213 | intron | KLHL25 |
| rs1878410 | 13 | 60037723 | 4 | 0 | 0 | T | C | -0.9036 | 0.2617 | 0.0005556 | ---?- | 0 | 0.644 | intergenic | |
| rs10504958 | 8 | 96959819 | 4 | 0 | 0 | T | C | -1.0014 | 0.2901 | 0.0005557 | ---?- | 0 | 0.9224 | non-coding-exon | LOC100500773 |
| 15:87597346 | 15 | 87597346 | 3 | 0 | 0 | T | TACAC | 1.2214 | 0.3538 | 0.0005557 | +++?? | 0 | 0.6795 | | |
| rs2880064 | 16 | 5456593 | 4 | 0 | 0 | A | C | -0.9768 | 0.283 | 0.0005559 | ---?- | 0 | 0.6792 | intron | RBFOX1 |
| rs10107977 | 8 | 140521282 | 4 | 0 | 0 | C | G | 0.8862 | 0.2567 | 0.000556 | +++?+ | 0 | 0.4671 | intergenic | |
| rs6490056 | 12 | 111769292 | 5 | 0 | 0 | T | C | -0.8644 | 0.2504 | 0.0005561 | -+--- | 48.8 | 0.09867 | intron | CUX2 |
| rs17618064 | 7 | 127903879 | 2 | 0 | 0 | T | C | -5.2141 | 1.5104 | 0.0005561 | -?-?? | 0 | 0.4319 | intergenic | |
| 5:706797 | 5 | 706797 | 3 | 0 | 0 | A | C | 1.559 | 0.4516 | 0.0005562 | +++?? | 47.4 | 0.1494 | | |
| rs28383310 | 6 | 32586930 | 3 | 0 | 0 | A | G | 1.1302 | 0.3274 | 0.0005563 | +++?? | 0 | 0.4293 | intergenic | |
| rs41533445 | 8 | 87212048 | 3 | 0 | 0 | A | T | 4.5137 | 1.3076 | 0.0005564 | ?++?+ | 0 | 0.999 | intergenic | |
| rs145224985 | 10 | 75586395 | 2 | 0 | 0 | A | C | 6.6544 | 1.9277 | 0.0005564 | ??+?+ | 0 | 0.8938 | intron | CAMK2G |
| rs11033910 | 11 | 37065382 | 4 | 0 | 0 | A | G | 0.9248 | 0.2679 | 0.0005568 | +++?+ | 0 | 0.4756 | intergenic | |
| rs78806125 | 10 | 86693865 | 4 | 0 | 0 | T | C | -1.7508 | 0.5072 | 0.0005568 | ---?- | 0 | 0.8258 | intergenic | |
| rs192745420 | 12 | 60413593 | 2 | 0 | 0 | A | G | -7.7097 | 2.2335 | 0.0005568 | ?-??- | 0 | 0.5098 | intergenic | |
| rs72672901 | 8 | 113332510 | 4 | 0 | 0 | A | G | 0.9768 | 0.283 | 0.000557 | +++?+ | 0 | 0.7494 | intron | CSMD3 |
| rs7937981 | 11 | 23964034 | 4 | 0 | 0 | C | G | -1.251 | 0.3624 | 0.0005572 | ---?- | 0 | 0.7256 | intergenic | |
| rs187076 | 10 | 55228462 | 4 | 0 | 0 | A | G | 0.9112 | 0.264 | 0.0005573 | +++?+ | 23.7 | 0.2689 | intergenic | |
| rs35883088 | 19 | 46154442 | 5 | 0 | 1 | T | C | -0.9593 | 0.2779 | 0.0005574 | -+--+ | 53.8 | 0.0703 | intergenic | |
| rs12101909 | 15 | 92205107 | 4 | 0 | 0 | A | G | 1.3385 | 0.3878 | 0.0005575 | +++?+ | 0 | 0.75 | intergenic | |
| 11:86178122 | 11 | 86178122 | 3 | 0 | 0 | T | TG | -1.6666 | 0.4829 | 0.0005575 | ---?? | 6.8 | 0.3421 | | |
| rs11066196 | 12 | 112655122 | 3 | 0 | 0 | T | C | -1.4649 | 0.4244 | 0.0005576 | -+-?? | 65.5 | 0.05506 | intron | HECTD4 |
| rs11991589 | 8 | 113231424 | 4 | 0 | 0 | A | G | -0.9819 | 0.2845 | 0.0005576 | ---?- | 0 | 0.9694 | downstream-gene | |
| 1:107962245 | 1 | 107962245 | 3 | 0 | 0 | CA | C | -1.1904 | 0.3449 | 0.0005577 | ---?? | 0 | 0.7934 | | |

| | | | | | | | | | | | | | | | |
|---|---|---|---|---|---|---|---|---|---|---|---|---|---|---|---|
| rs12297355 | 12 | 41481613 | 4 | 0 | 0 | A | G | 1.6795 | 0.4866 | 0.0005578 | +++?+ | 14.3 | 0.3209 | intergenic | |
| rs78645394 | 2 | 170793660 | 4 | 0 | 1 | A | C | 0.998 | 0.2892 | 0.0005579 | +++?+ | 43.6 | 0.1498 | intron | UBR3 |
| rs9382617 | 6 | 56175482 | 4 | 0 | 0 | A | T | -1.2749 | 0.3694 | 0.0005582 | ---?+ | 27.9 | 0.2447 | intergenic | |
| rs12319487 | 12 | 41481630 | 4 | 0 | 0 | T | G | -1.6795 | 0.4867 | 0.0005582 | ---?- | 14.2 | 0.321 | intergenic | |
| rs78383841 | 5 | 10301075 | 4 | 0 | 0 | A | G | 1.9418 | 0.5626 | 0.0005582 | +++?+ | 42.6 | 0.1561 | intron | CMBL |
| rs10923344 | 1 | 118096628 | 4 | 0 | 0 | A | G | -0.9289 | 0.2692 | 0.0005583 | ---?- | 0 | 0.5393 | intergenic | |
| rs147553880 | 2 | 184040619 | 4 | 0 | 0 | T | C | 1.1633 | 0.3371 | 0.0005583 | +++?+ | 0 | 0.7974 | intergenic | |
| rs140029 | 22 | 36950740 | 3 | 0 | 0 | T | C | -1.1391 | 0.3301 | 0.0005586 | ---?? | 37.2 | 0.2032 | intergenic | |
| rs1462536 | 2 | 184058178 | 4 | 0 | 0 | T | C | 1.1769 | 0.341 | 0.0005586 | +++?+ | 0 | 0.7225 | intergenic | |
| rs2468728 | 8 | 140437534 | 4 | 0 | 0 | A | G | -1.2849 | 0.3723 | 0.0005586 | ---?- | 0 | 0.7666 | intergenic | |
| rs12543669 | 8 | 65071623 | 4 | 0 | 0 | A | T | -0.8462 | 0.2452 | 0.0005586 | ---?- | 0 | 0.9381 | intergenic | |
| rs17312403 | 8 | 65069907 | 4 | 0 | 0 | T | C | 0.8451 | 0.2449 | 0.0005586 | +++?+ | 0 | 0.9408 | intergenic | |
| rs113176001 | 6 | 32571483 | 3 | 0 | 0 | C | G | 1.2398 | 0.3593 | 0.0005587 | +++?? | 8.1 | 0.337 | intergenic | |
| rs4767991 | 12 | 41219126 | 4 | 0 | 0 | T | C | -1.7429 | 0.5051 | 0.0005588 | ---?+ | 0 | 0.6132 | intron | CNTN1 |
| 2:185438605 | 2 | 185438605 | 3 | 0 | 0 | A | AC | 0.9708 | 0.2813 | 0.0005588 | +++?? | 0 | 0.6032 | | |
| rs9271573 | 6 | 32590501 | 5 | 0 | 0 | A | C | -0.7147 | 0.2071 | 0.000559 | ----- | 0 | 0.665 | intergenic |  |
| rs1749448 | 1 | 87567793 | 4 | 0 | 0 | A | T | -1.0379 | 0.3008 | 0.0005591 | ---?- | 6.7 | 0.3596 | intron | HS2ST1 |
| rs72674703 | 8 | 113333124 | 4 | 0 | 0 | T | C | -0.9773 | 0.2832 | 0.0005593 | ---?- | 0 | 0.7519 | intron | CSMD3 |
| rs56385303 | 5 | 165387243 | 5 | 0 | 0 | A | C | 0.7542 | 0.2186 | 0.0005596 | +++++ | 36.6 | 0.1771 | intergenic | |
| rs4816448 | 21 | 34507631 | 4 | 0 | 0 | T | C | 1.0029 | 0.2907 | 0.0005597 | +++?- | 3.7 | 0.3743 | intergenic | |
| rs17297879 | 10 | 131921229 | 4 | 0 | 0 | C | G | 0.8976 | 0.2601 | 0.0005597 | +++?+ | 0 | 0.8597 | intergenic | |
| rs1462535 | 2 | 184058307 | 4 | 0 | 0 | A | C | 1.1769 | 0.3411 | 0.00056 | +++?+ | 0 | 0.7216 | intergenic | |
| rs185892800 | 4 | 166207579 | 4 | 0 | 0 | A | C | -3.8832 | 1.1255 | 0.00056 | ---?- | 0 | 0.9083 | intron | KLHL2 |
| rs145217032 | 7 | 103928926 | 3 | 0 | 0 | A | G | 4.6515 | 1.3482 | 0.0005601 | +++?? | 0 | 0.7041 | intergenic | |
| rs763098 | 6 | 118481215 | 4 | 0 | 0 | A | G | -1.0974 | 0.3181 | 0.0005602 | ---?- | 0 | 0.4065 | intron | SLC35F1 |
| rs9656532 | 7 | 82414933 | 4 | 0 | 0 | A | G | 0.8943 | 0.2592 | 0.0005604 | +-+?+ | 47.2 | 0.128 | intron | PCLO |
| rs2134356 | 3 | 2487786 | 4 | 0 | 0 | T | C | 0.9101 | 0.2638 | 0.0005605 | +++?- | 0 | 0.4038 | intron | CNTN4 |
| rs77519740 | 4 | 38443737 | 3 | 0 | 0 | T | C | 4.8622 | 1.4093 | 0.0005605 | ?++?+ | 0 | 0.5186 | intergenic | |
| rs1933948 | 10 | 86694529 | 4 | 0 | 0 | A | G | 1.7486 | 0.5068 | 0.0005607 | +++?+ | 0 | 0.8259 | intergenic | |
| rs13207328 | 6 | 165808386 | 4 | 0 | 0 | A | C | -0.8752 | 0.2537 | 0.0005607 | ---?- | 0 | 0.645 | intron | PDE10A |
| rs1420861 | 8 | 113230122 | 4 | 0 | 0 | A | G | 0.9815 | 0.2845 | 0.0005609 | +++?+ | 0 | 0.9694 | intergenic | |
| rs10111382 | 8 | 113511997 | 4 | 0 | 0 | A | C | 1.0049 | 0.2913 | 0.0005609 | +++?+ | 0 | 0.5816 | intron | CSMD3 |
| rs9828932 | 3 | 70446735 | 4 | 0 | 0 | T | C | -1.0276 | 0.2979 | 0.0005611 | ---?- | 40.5 | 0.169 | intergenic | |
| rs13195853 | 6 | 165808391 | 4 | 0 | 0 | T | C | 0.8752 | 0.2537 | 0.0005612 | +++?+ | 0 | 0.6447 | intron | PDE10A |
| 14:28227276 | 14 | 28227276 | 3 | 0 | 0 | T | TA | -2.0953 | 0.6074 | 0.0005612 | ---?? | 0 | 0.8226 | | |
| 20:34176844 | 20 | 34176844 | 3 | 0 | 0 | G | GA | 1.5502 | 0.4494 | 0.0005612 | +++?? | 8.7 | 0.3346 | | |
| rs11877248 | 18 | 5903391 | 4 | 0 | 0 | A | G | -1.0228 | 0.2965 | 0.0005613 | ---?- | 0 | 0.5074 | intergenic | |
| rs111842857 | 16 | 5455531 | 4 | 0 | 0 | C | G | -0.9896 | 0.2869 | 0.0005613 | ---?- | 0 | 0.8359 | intron | RBFOX1 |
| 4:38622478 | 4 | 38622478 | 3 | 0 | 0 | A | AGCATA | 0.9693 | 0.281 | 0.0005615 | +++?? | 21.2 | 0.2813 | | |
| rs58998183 | 7 | 46183837 | 4 | 0 | 0 | A | C | 1.9768 | 0.5731 | 0.0005616 | +++?+ | 0 | 0.9435 | intergenic | |
| rs2281415 | 1 | 185392845 | 4 | 0 | 0 | T | C | 0.9165 | 0.2657 | 0.0005618 | +++?+ | 0 | 0.9348 | intergenic | |
| rs3818177 | 6 | 165808429 | 4 | 0 | 0 | T | C | -0.8751 | 0.2537 | 0.0005618 | ---?- | 0 | 0.6445 | intron | PDE10A |
| rs201455 | 7 | 101740212 | 4 | 0 | 0 | T | C | 0.8404 | 0.2436 | 0.0005619 | +++?+ | 0 | 0.788 | intron | CUX1 |
| rs4296226 | 15 | 85580547 | 4 | 0 | 0 | T | C | -0.8262 | 0.2395 | 0.0005619 | +--?+ | 56.4 | 0.07604 | intron | PDE8A |
| rs2752978 | 6 | 63408727 | 4 | 0 | 0 | T | C | -2.1137 | 0.6128 | 0.0005621 | +--?- | 23.5 | 0.2698 | intergenic | |
| rs4392919 | 8 | 3986246 | 4 | 0 | 0 | T | G | -1.165 | 0.3378 | 0.0005622 | +--?- | 55.2 | 0.08207 | intron | CSMD1 |
| rs12542914 | 8 | 65042779 | 4 | 0 | 0 | T | C | 0.8414 | 0.2439 | 0.0005622 | +++?+ | 0 | 0.9493 | intergenic | |
| rs80312119 | 18 | 47048113 | 4 | 0 | 0 | T | C | 1.9845 | 0.5753 | 0.0005622 | +++?+ | 0 | 0.9252 | intergenic | |
| rs13414991 | 2 | 170933176 | 4 | 0 | 1 | A | C | -0.9935 | 0.288 | 0.0005623 | ---?- | 47.8 | 0.1247 | intron | UBR3 |
| 8:113261587 | 8 | 113261587 | 3 | 0 | 0 | A | AACAGT | 1.019 | 0.2954 | 0.0005625 | +++?? | 0 | 0.8205 | | |
| rs2715297 | 18 | 63723550 | 4 | 0 | 0 | A | C | -0.8643 | 0.2506 | 0.0005626 | ---?- | 44.4 | 0.1451 | intergenic | |
| 12:74343924 | 12 | 74343924 | 3 | 0 | 0 | CA | C | 0.9981 | 0.2894 | 0.0005626 | +++?? | 48.5 | 0.1437 | | |
| rs17036325 | 2 | 47390058 | 4 | 0 | 0 | T | C | 1.347 | 0.3906 | 0.0005627 | +++?+ | 0 | 0.9582 | intron | CALM2 |
| rs1385251 | 8 | 96994590 | 4 | 0 | 0 | T | C | -1.42 | 0.4117 | 0.0005629 | ---?- | 0 | 0.9591 | intergenic | |
| rs369365368 | 7 | 56883409 | 3 | 0 | 0 | C | G | 3.5524 | 1.03 | 0.0005629 | +++?? | 0 | 0.421 | upstream-gene | |
| rs4939260 | 11 | 58982694 | 5 | 0 | 0 | A | G | 0.7426 | 0.2153 | 0.0005633 | +-++- | 19.6 | 0.2898 | upstream-gene | |
| rs9570135 | 13 | 60038560 | 4 | 0 | 0 | T | C | 0.8981 | 0.2604 | 0.0005633 | +++?+ | 0 | 0.6237 | intergenic | |
| rs62337111 | 4 | 183667541 | 4 | 0 | 0 | T | G | 0.9482 | 0.275 | 0.0005633 | +++?- | 0 | 0.6545 | intron | TENM3 |
| rs12508139 | 4 | 169322975 | 4 | 0 | 0 | C | G | 0.8545 | 0.2478 | 0.0005634 | +++?+ | 0 | 0.9053 | intron | DDX60L |

| | | | | | | | | | | | | | | | |
|---|---|---|---|---|---|---|---|---|---|---|---|---|---|---|---|
| rs376005 | 21 | 31401556 | 4 | 0 | 0 | T | C | 0.976 | 0.283 | 0.0005635 | +++?- | 15.6 | 0.314 | intergenic | |
| rs114353093 | 4 | 2799465 | 5 | 0 | 0 | T | C | 2.2582 | 0.6548 | 0.0005635 | +++++ | 0 | 0.6835 | intron | SH3BP2 |
| rs6005779 | 22 | 28807277 | 4 | 0 | 0 | T | C | -2.2795 | 0.661 | 0.000564 | ---?- | 0 | 0.7223 | intron | TTC28 |
| rs115817471 | 2 | 73653835 | 2 | 0 | 0 | A | G | 4.8007 | 1.3922 | 0.0005642 | +?+?? | 0 | 0.746 | intron | ALMS1 |
| rs114798663 | 17 | 56879913 | 5 | 0 | 0 | T | G | -2.394 | 0.6943 | 0.0005644 | ----- | 0 | 0.8541 | intron | PPM1E |
| rs7285141 | 22 | 50840783 | 5 | 0 | 0 | T | C | -0.7998 | 0.2319 | 0.0005645 | ----+ | 0 | 0.5782 | intron | PPP6R2 |
| rs6713257 | 2 | 47378943 | 4 | 0 | 0 | A | G | 1.3376 | 0.3879 | 0.0005646 | +++?+ | 0 | 0.9551 | intron | C2orf61 |
| rs145265776 | 10 | 6421626 | 4 | 0 | 0 | A | G | -2.6548 | 0.77 | 0.0005647 | ---?- | 0 | 0.803 | intergenic | |
| rs11778155 | 8 | 113334425 | 4 | 0 | 0 | A | G | -0.9773 | 0.2834 | 0.0005649 | ---?- | 0 | 0.7552 | intron | CSMD3 |
| rs58036270 | 9 | 79458380 | 4 | 0 | 0 | A | G | -1.9863 | 0.5761 | 0.0005649 | ---?- | 0 | 0.5459 | intron | PRUNE2 |
| rs77604539 | 4 | 158176114 | 3 | 0 | 0 | A | G | -2.3639 | 0.6856 | 0.0005649 | ---?? | 0 | 0.9012 | intron | GRIA2 |
| rs139224754 | 9 | 44331413 | 3 | 0 | 0 | T | C | -3.3098 | 0.96 | 0.0005649 | ---?? | 0 | 0.8377 | intergenic | |
| rs190539388 | 16 | 52370738 | 2 | 0 | 0 | T | C | -5.2649 | 1.527 | 0.0005649 | --??? | 0 | 0.4912 | intergenic | |
| rs17452998 | 18 | 386203 | 4 | 0 | 0 | T | C | 1.184 | 0.3434 | 0.0005652 | +++?+ | 0 | 0.9049 | intron | COLEC12 |
| rs1875568 | 11 | 8700167 | 5 | 0 | 0 | T | C | 0.7965 | 0.231 | 0.0005656 | ++++- | 0 | 0.8664 | intron | TRIM66 |
| rs17460042 | 7 | 114894984 | 4 | 0 | 0 | A | G | -2.7746 | 0.8048 | 0.0005656 | ---?- | 18.3 | 0.2989 | intergenic | |
| rs6798777 | 3 | 196502817 | 4 | 0 | 0 | C | G | -2.1172 | 0.6141 | 0.0005657 | ---?+ | 19.6 | 0.2919 | intron | PAK2 |
| rs3746687 | 20 | 2463592 | 4 | 0 | 0 | T | C | 1.097 | 0.3182 | 0.000566 | -++?+ | 60.8 | 0.05397 | 3-prime-UTR | ZNF343 |
| rs17128926 | 12 | 41304649 | 4 | 0 | 0 | A | G | -1.6923 | 0.4909 | 0.000566 | ---?+ | 0 | 0.6403 | intron | CNTN1 |
| rs143582335 | 9 | 112531359 | 5 | 0 | 0 | A | G | 2.9836 | 0.8655 | 0.0005661 | +++++ | 7.1 | 0.3663 | intron | PALM2 |
| rs74850961 | 2 | 15572788 | 4 | 0 | 0 | T | C | -1.7466 | 0.5067 | 0.0005662 | ---?- | 0 | 0.6208 | intron | NBAS |
| rs146036609 | 4 | 166208621 | 4 | 0 | 0 | A | T | 3.8746 | 1.124 | 0.0005663 | +++?+ | 0 | 0.9112 | intron | KLHL2 |
| rs28470301 | 4 | 116909584 | 4 | 0 | 0 | A | G | 2.8887 | 0.838 | 0.0005664 | +-+?+ | 50.9 | 0.1062 | intergenic | |
| 6:99932254 | 6 | 99932254 | 3 | 0 | 0 | CTT | C | 1.7939 | 0.5204 | 0.0005664 | +++?? | 0 | 0.6239 | | |
| rs1419130 | 1 | 87576481 | 4 | 0 | 0 | A | G | -1.047 | 0.3037 | 0.0005666 | ---?- | 0 | 0.5231 | downstream-gene | |
| rs141300970 | 15 | 59530960 | 3 | 0 | 0 | A | G | 4.8062 | 1.3943 | 0.0005668 | ?++?+ | 30.4 | 0.2377 | intron | MYO1E |
| 1:203688368 | 1 | 203688368 | 3 | 0 | 0 | A | AAAAAA | -1.1405 | 0.3309 | 0.0005668 | ---?? | 0 | 0.9784 | | |
| rs150971903 | 18 | 40417374 | 4 | 0 | 0 | T | C | -3.9407 | 1.1433 | 0.000567 | ---?+ | 53.6 | 0.09117 | intron | RIT2 |
| rs80323386 | 9 | 11576708 | 3 | 0 | 0 | T | G | 2.876 | 0.8344 | 0.0005672 | +++?? | 61.7 | 0.07367 | intergenic | |
| rs1665685 | 10 | 73416482 | 4 | 0 | 0 | A | G | -1.8746 | 0.5439 | 0.0005673 | ---?- | 0 | 0.9958 | intron | CDH23 |
| rs12579954 | 12 | 112250158 | 5 | 0 | 0 | A | G | 1.3088 | 0.3797 | 0.0005673 | +-+++ | 48.9 | 0.09799 | downstream-gene | |
| rs450450 | 2 | 40434239 | 4 | 0 | 0 | T | C | -1.0669 | 0.3096 | 0.0005674 | ---?+ | 0 | 0.5551 | intron | SLC8A1,SLC8A1-AS1 |
| rs2201203 | 8 | 113336210 | 4 | 0 | 0 | A | G | -0.9795 | 0.2842 | 0.0005678 | ---?- | 0 | 0.7546 | intron | CSMD3 |
| rs10869660 | 9 | 78462617 | 4 | 0 | 0 | T | C | -1.2742 | 0.3697 | 0.0005678 | ---?- | 55.7 | 0.07976 | intergenic | |
| rs743928 | 6 | 118481811 | 4 | 0 | 0 | T | C | -1.0965 | 0.3181 | 0.0005679 | ---?- | 0 | 0.4056 | intron | SLC35F1 |
| rs28406237 | 6 | 93902701 | 4 | 0 | 0 | T | C | 2.0931 | 0.6073 | 0.0005681 | +++?+ | 0 | 0.5771 | intergenic | |
| rs1994796 | 8 | 113333701 | 4 | 0 | 0 | A | G | 0.9783 | 0.2839 | 0.0005682 | +++?+ | 0 | 0.7492 | intron | CSMD3 |
| rs17142721 | 16 | 7206700 | 4 | 0 | 0 | T | C | -1.0555 | 0.3063 | 0.0005682 | ---?- | 0 | 0.9536 | intron | RBFOX1 |
| rs11111408 | 12 | 103255228 | 3 | 0 | 0 | A | C | 0.8642 | 0.2507 | 0.0005683 | +++?? | 0 | 0.6222 | intron | PAH |
| rs77647964 | 5 | 97125367 | 3 | 0 | 0 | T | C | 1.2006 | 0.3484 | 0.0005683 | +++?? | 39.1 | 0.1934 | intergenic | |
| 6:12213303 | 6 | 12213303 | 3 | 0 | 0 | A | AAG | 1.3806 | 0.4006 | 0.0005684 | +++?? | 28.4 | 0.2475 | | |
| rs3129954 | 6 | 32365580 | 5 | 0 | 0 | A | G | -0.7362 | 0.2136 | 0.0005685 | +---- | 43.1 | 0.1346 | intron | BTNL2 |
| rs79350915 | 7 | 46196046 | 4 | 0 | 0 | C | G | 1.9789 | 0.5742 | 0.0005687 | +++?+ | 0 | 0.9505 | intergenic | |
| 8:113511066 | 8 | 113511066 | 3 | 0 | 0 | T | TA | -0.925 | 0.2684 | 0.0005687 | ---?? | 0 | 0.5008 | | |
| rs9285222 | 13 | 60039368 | 4 | 0 | 0 | A | G | -0.8988 | 0.2608 | 0.0005689 | ---?- | 0 | 0.645 | intergenic | |
| rs17128929 | 12 | 41304672 | 4 | 0 | 0 | A | G | 1.6916 | 0.4909 | 0.0005689 | +++?- | 0 | 0.6398 | intron | CNTN1 |
| rs3991306 | 5 | 119821638 | 4 | 0 | 0 | A | T | 1.0058 | 0.2919 | 0.0005691 | +++?- | 40.5 | 0.1686 | intron | PRR16 |
| 11:86182228 | 11 | 86182228 | 3 | 0 | 0 | CTG | C | -1.6664 | 0.4836 | 0.0005691 | ---?? | 0 | 0.3704 | | |
| rs9407716 | 9 | 16167434 | 4 | 0 | 0 | C | G | 0.9252 | 0.2685 | 0.0005693 | +++?+ | 0 | 0.8886 | intergenic | |
| rs187988604 | 3 | 28570942 | 4 | 0 | 0 | T | G | -2.7909 | 0.81 | 0.0005694 | ---?+ | 56.9 | 0.07299 | downstream-gene | |
| rs74585232 | 1 | 192463740 | 4 | 0 | 0 | T | C | -1.1528 | 0.3346 | 0.0005695 | ---?+ | 0 | 0.3998 | intergenic | |
| rs112244376 | 2 | 64244890 | 2 | 0 | 0 | T | C | 7.3362 | 2.129 | 0.0005695 | ++??? | 0 | 0.8707 | intron | VPS54 |
| rs62521160 | 8 | 65060001 | 4 | 0 | 0 | T | G | -0.8342 | 0.2421 | 0.0005696 | ---?- | 0 | 0.9478 | intergenic | |
| rs185817908 | 17 | 5240719 | 5 | 0 | 0 | A | G | -2.5245 | 0.7327 | 0.0005696 | ----- | 55.6 | 0.06075 | intron | RABEP1 |
| rs11921110 | 3 | 196561599 | 3 | 0 | 0 | T | G | -2.7145 | 0.7878 | 0.0005697 | ---?? | 0 | 0.7117 | downstream-gene | |
| rs148952682 | 18 | 40417254 | 4 | 0 | 0 | T | C | 3.9377 | 1.1428 | 0.0005697 | +++?- | 53.6 | 0.09106 | intron | RIT2 |
| rs78317300 | 18 | 47049476 | 4 | 0 | 0 | A | C | -1.9868 | 0.5766 | 0.0005698 | ---?- | 0 | 0.924 | intergenic | |
| rs77234249 | 19 | 14746814 | 3 | 0 | 0 | T | C | 1.4206 | 0.4123 | 0.00057 | +++?? | 0 | 0.4893 | intron | EMR3 |
| rs11893450 | 2 | 184059063 | 4 | 0 | 0 | A | G | 1.1763 | 0.3414 | 0.0005702 | +++?+ | 0 | 0.7143 | intergenic | |

| | | | | | | | | | | | | | | | |
|---|---|---|---|---|---|---|---|---|---|---|---|---|---|---|---|
| rs192439830 | 10 | 111081499 | 3 | 0 | 0 | T | C | -5.9068 | 1.7144 | 0.0005703 | ---?? | 0 | 0.9759 | intergenic | |
| 6:31135732 | 6 | 31135732 | 3 | 0 | 0 | CC | C | 0.8567 | 0.2487 | 0.0005703 | +++?? | 44.8 | 0.1632 | | |
| rs72810415 | 2 | 47377833 | 4 | 0 | 0 | A | G | -1.3306 | 0.3862 | 0.0005704 | ---?- | 0 | 0.9728 | intron | C2orf61 |
| rs6802122 | 3 | 143531170 | 4 | 0 | 0 | A | G | 0.826 | 0.2397 | 0.0005705 | +++?+ | 0 | 0.572 | intron | SLC9A9 |
| rs248546 | 5 | 165390149 | 5 | 0 | 0 | A | T | -0.7397 | 0.2147 | 0.0005706 | ---+- | 37.6 | 0.1704 | intergenic | |
| rs143643209 | 16 | 47839022 | 4 | 0 | 0 | T | G | -3.0221 | 0.8772 | 0.0005708 | ---?- | 0 | 0.6337 | intergenic | |
| rs12902574 | 15 | 92204973 | 3 | 0 | 0 | A | G | 1.3824 | 0.4013 | 0.0005709 | +++?? | 0 | 0.5959 | intergenic | |
| rs149247179 | 4 | 166210145 | 4 | 0 | 0 | A | G | -3.8655 | 1.122 | 0.0005709 | ---?- | 0 | 0.9146 | intron | KLHL2 |
| rs28416724 | 18 | 386309 | 4 | 0 | 0 | A | T | -1.1829 | 0.3433 | 0.000571 | ---?- | 0 | 0.9028 | intron | COLEC12 |
| 2:170514204 | 2 | 170514204 | 3 | 0 | 0 | A | AG | -1.2701 | 0.3687 | 0.000571 | ---?? | 0 | 0.8789 | | |
| rs2053644 | 8 | 65063298 | 4 | 0 | 0 | A | T | 0.838 | 0.2432 | 0.0005711 | +++?+ | 0 | 0.9402 | intergenic | |
| rs58902245 | 3 | 33267346 | 4 | 0 | 0 | C | G | -0.9155 | 0.2657 | 0.0005711 | ---?- | 0.2 | 0.3907 | intergenic | |
| 17:47319331 | 17 | 47319331 | 3 | 0 | 0 | T | TACAC | 1.256 | 0.3646 | 0.0005715 | +++?? | 0 | 0.451 | | |
| rs141490325 | 5 | 150162847 | 4 | 0 | 0 | A | G | 4.0638 | 1.1797 | 0.0005716 | +++?+ | 0 | 0.7918 | intron | SMIM3 |
| rs2619778 | 5 | 82380733 | 4 | 0 | 0 | T | C | 2.1775 | 0.6321 | 0.0005717 | +++?+ | 0 | 0.4906 | intron | XRCC4 |
| rs3129955 | 6 | 32365840 | 5 | 0 | 0 | T | C | -0.7357 | 0.2136 | 0.0005719 | +---- | 43.1 | 0.1345 | intron | BTNL2 |
| rs2717717 | 8 | 18437206 | 4 | 0 | 0 | A | G | -0.8361 | 0.2427 | 0.000572 | ---?- | 0 | 0.7869 | intron | PSD3 |
| rs6456824 | 6 | 28513135 | 5 | 0 | 0 | T | C | -1.0346 | 0.3004 | 0.0005721 | ----- | 14.4 | 0.3225 | intergenic |  |
| rs34118615 | 13 | 30312184 | 5 | 0 | 0 | T | C | -1.2824 | 0.3723 | 0.0005725 | ----- | 0 | 0.5501 | intergenic |  |
| rs75385514 | 6 | 12210560 | 4 | 0 | 0 | A | C | 3.176 | 0.9221 | 0.0005725 | +++?+ | 2.6 | 0.3794 | intergenic | |
| rs78129949 | 18 | 25195847 | 4 | 0 | 0 | T | C | 2.4707 | 0.7173 | 0.0005727 | +++?+ | 0 | 0.65 | intergenic | |
| rs2166094 | 4 | 36982008 | 4 | 0 | 0 | A | G | 0.8685 | 0.2522 | 0.0005728 | +++?- | 30.9 | 0.2267 | intergenic | |
| rs10123717 | 9 | 16168242 | 4 | 0 | 0 | T | C | 0.9056 | 0.2629 | 0.0005731 | +++?- | 0 | 0.538 | intergenic | |
| rs74547705 | 3 | 143904151 | 3 | 0 | 0 | T | C | 4.3594 | 1.2658 | 0.0005735 | +++?? | 0 | 0.4095 | intergenic | |
| 7:23129485 | 7 | 23129485 | 3 | 0 | 0 | CAAA | C | -0.9119 | 0.2648 | 0.0005735 | ---?? | 11.4 | 0.3236 | | |
| rs13380897 | 18 | 25196184 | 4 | 0 | 0 | T | G | 2.4731 | 0.7181 | 0.0005736 | +++?+ | 0 | 0.6508 | intergenic | |
| 2:218831644 | 2 | 218831644 | 3 | 0 | 0 | T | TG | 1.3893 | 0.4034 | 0.0005737 | +++?? | 0 | 0.8521 | | |
| rs79455345 | 7 | 46193546 | 4 | 0 | 0 | A | T | 1.9775 | 0.5742 | 0.0005738 | +++?+ | 0 | 0.95 | intergenic | |
| rs12041494 | 1 | 187857590 | 4 | 0 | 0 | A | G | 1.1485 | 0.3335 | 0.0005739 | +++?+ | 0 | 0.8908 | intergenic | |
| rs62514402 | 8 | 113319754 | 4 | 0 | 0 | T | C | 0.9754 | 0.2832 | 0.000574 | +++?+ | 0 | 0.7546 | intron | CSMD3 |
| rs72777540 | 16 | 5456802 | 4 | 0 | 0 | A | G | -0.9693 | 0.2815 | 0.000574 | ---?- | 0 | 0.7688 | intron | RBFOX1 |
| rs66608459 | 16 | 57456698 | 4 | 0 | 0 | A | G | 1.5311 | 0.4446 | 0.0005741 | +++?+ | 0 | 0.8746 | intergenic | |
| 1:246412833 | 1 | 246412833 | 3 | 0 | 0 | A | ATT | -1.7138 | 0.4977 | 0.0005741 | ---?? | 55.2 | 0.1075 | | |
| rs11179797 | 12 | 41488497 | 4 | 0 | 0 | T | C | -1.6827 | 0.4887 | 0.0005745 | ---?- | 12.8 | 0.3285 | intergenic | |
| rs16883432 | 8 | 113337565 | 4 | 0 | 0 | T | C | 0.9769 | 0.2837 | 0.0005745 | +++?+ | 0 | 0.7598 | intron | CSMD3 |
| rs35939681 | 3 | 175669636 | 4 | 0 | 0 | T | C | 1.8229 | 0.5294 | 0.0005745 | +++?+ | 0 | 0.9633 | intergenic | |
| rs139695521 | 2 | 216088675 | 3 | 0 | 0 | A | G | -2.6135 | 0.759 | 0.0005746 | ---?? | 0 | 0.6496 | intergenic | |
| 11:20152764 | 11 | 20152764 | 3 | 0 | 0 | T | TA | 1.6815 | 0.4884 | 0.0005751 | +++?? | 0 | 0.711 | | |
| 3:183597150 | 3 | 183597150 | 3 | 0 | 0 | CT | C | 1.6831 | 0.4888 | 0.0005751 | +++?? | 0 | 0.7799 | | |
| rs4836063 | 5 | 123515303 | 4 | 0 | 0 | T | C | -0.8378 | 0.2434 | 0.0005755 | ---?+ | 27.1 | 0.2492 | intergenic | |
| rs72888749 | 18 | 33747714 | 3 | 0 | 0 | A | G | 3.6876 | 1.0711 | 0.0005755 | +++?? | 0 | 0.647 | intron | ELP2 |
| rs11250255 | 10 | 1132215 | 4 | 0 | 0 | T | C | -1.8843 | 0.5473 | 0.0005756 | ---?- | 31.6 | 0.2225 | intron | WDR37 |
| rs76280953 | 16 | 9195327 | 3 | 0 | 0 | T | C | -2.772 | 0.8051 | 0.0005756 | ---?? | 0 | 0.9069 | intron | C16orf72 |
| rs6924633 | 6 | 63323064 | 4 | 0 | 0 | A | T | -2.4227 | 0.7037 | 0.0005758 | ---?- | 0 | 0.5083 | intergenic | |
| 5:51478435 | 5 | 51478435 | 3 | 0 | 0 | G | GA | -1.0154 | 0.2949 | 0.0005759 | ---?? | 34.5 | 0.2173 | | |
| rs62402381 | 6 | 28531538 | 5 | 0 | 0 | C | G | 1.1517 | 0.3345 | 0.000576 | +++++ | 0 | 0.5128 | intergenic | |
| rs57091586 | 16 | 12138097 | 4 | 0 | 0 | A | G | 0.9101 | 0.2644 | 0.0005761 | -++?+ | 56.4 | 0.07597 | intron | SNX29 |
| rs145943094 | 3 | 47881942 | 2 | 0 | 0 | T | C | 6.1238 | 1.7788 | 0.0005761 | ?+??+ | 56 | 0.1319 | intron | DHX30 |
| rs62276915 | 3 | 155096878 | 4 | 0 | 0 | A | C | -1.0182 | 0.2958 | 0.0005762 | ---?- | 12.7 | 0.329 | intergenic | |
| 7:16656204 | 7 | 16656204 | 3 | 0 | 0 | C | G | -4.637 | 1.347 | 0.0005764 | ---?? | 0 | 0.5364 | intron | ANKMY2 |
| rs12742456 | 1 | 64787868 | 3 | 0 | 0 | A | G | 1.2809 | 0.3721 | 0.0005764 | +++?? | 0 | 0.4762 | intergenic | |
| rs1994795 | 8 | 113338239 | 4 | 0 | 0 | C | G | -0.9791 | 0.2844 | 0.0005766 | ---?- | 0 | 0.7573 | intron | CSMD3 |
| rs4676808 | 3 | 120338500 | 4 | 0 | 0 | T | C | -1.0006 | 0.2907 | 0.0005766 | ---?- | 59.5 | 0.06018 | intergenic | |
| rs11245225 | 10 | 128599021 | 3 | 0 | 0 | C | G | 4.4142 | 1.2823 | 0.0005767 | +++?? | 29.1 | 0.244 | intergenic | |
| rs148476806 | 17 | 57193540 | 5 | 0 | 0 | T | C | 2.3997 | 0.6971 | 0.0005767 | +++++ | 0 | 0.7778 | intron | SKA2 |
| 22:29108082 | 22 | 29108082 | 3 | 0 | 0 | G | GTATTA[illegible] | -2.3242 | 0.6752 | 0.0005769 | ---?? | 0 | 0.4171 | | |
| rs950199 | 9 | 16184708 | 4 | 0 | 0 | A | G | -0.9391 | 0.2728 | 0.000577 | ---?- | 0 | 0.9387 | intergenic | |
| rs57398776 | 18 | 40128279 | 4 | 0 | 0 | A | C | -1.2002 | 0.3487 | 0.0005774 | ---?+ | 25.1 | 0.2611 | intron | LINC00907 |
| rs74722520 | 5 | 10300463 | 4 | 0 | 0 | A | T | -1.937 | 0.5628 | 0.0005774 | ---?- | 42.8 | 0.1548 | intron | CMBL |

| | | | | | | | | | | | | | | | |
|---|---|---|---|---|---|---|---|---|---|---|---|---|---|---|---|
| rs11724874 | 4 | 189998442 | 4 | 0 | 0 | A | T | -0.8775 | 0.2549 | 0.0005779 | ---?- | 33.3 | 0.2126 | intergenic | |
| 6:18473227 | 6 | 18473227 | 3 | 0 | 0 | CA | C | -1.7837 | 0.5183 | 0.000578 | ---?? | 0 | 0.9347 | | |
| rs6948464 | 7 | 82417332 | 4 | 0 | 0 | T | C | 0.8849 | 0.2571 | 0.0005781 | +-+?+ | 46.2 | 0.1342 | intron | PCLO |
| rs76032993 | 15 | 87472210 | 4 | 0 | 0 | A | C | 2.41 | 0.7002 | 0.0005782 | +++?+ | 0 | 0.9716 | intron | AGBL1 |
| rs145352381 | 14 | 104133690 | 4 | 0 | 0 | T | C | 4.0104 | 1.1653 | 0.0005782 | +++?+ | 0 | 0.4237 | intron | KLC1 |
| 16:26814427 | 16 | 26814427 | 3 | 0 | 0 | A | G | 1.4582 | 0.4237 | 0.0005783 | +++?? | 5.9 | 0.3456 | intergenic | |
| rs6481496 | 10 | 28235153 | 4 | 0 | 0 | A | C | -1.6838 | 0.4893 | 0.0005786 | ---?- | 20.6 | 0.2862 | intron | ARMC4 |
| rs16883437 | 8 | 113338786 | 4 | 0 | 0 | T | C | 0.9767 | 0.2838 | 0.0005786 | +++?+ | 0 | 0.7614 | intron | CSMD3 |
| 8:113302842 | 8 | 113302842 | 3 | 0 | 0 | T | TC | 1.0079 | 0.2929 | 0.0005786 | +++?? | 0 | 0.586 | | |
| rs16986377 | 22 | 28799312 | 4 | 0 | 0 | T | C | -2.2784 | 0.6621 | 0.0005787 | ---?- | 0 | 0.7262 | intron | TTC28 |
| rs60039318 | 5 | 123116001 | 4 | 0 | 0 | A | G | -0.9853 | 0.2863 | 0.0005787 | ---?- | 0 | 0.4049 | intergenic | |
| 2:170466989 | 2 | 170466989 | 3 | 0 | 0 | A | ATT | 1.2654 | 0.3677 | 0.0005787 | +++?? | 0 | 0.8133 | | |
| rs13280898 | 8 | 65039823 | 4 | 0 | 0 | A | G | 0.841 | 0.2444 | 0.0005788 | +++?+ | 0 | 0.95 | intergenic | |
| rs113180452 | 17 | 56425704 | 5 | 0 | 0 | A | G | -1.1035 | 0.3207 | 0.0005788 | ----- | 37.3 | 0.1724 | intron | SUPT4H1,BZRAP1-AS1 |
| 6:72807992 | 6 | 72807992 | 3 | 0 | 0 | CT | C | -0.9113 | 0.2648 | 0.000579 | ---?? | 0 | 0.972 | | |
| rs10262970 | 7 | 82417078 | 4 | 0 | 0 | A | T | 0.8848 | 0.2571 | 0.0005792 | +-+?+ | 46.2 | 0.1344 | intron | PCLO |
| rs1831037 | 9 | 78458935 | 4 | 0 | 0 | T | C | 1.2482 | 0.3627 | 0.0005793 | +++?+ | 56 | 0.07784 | intergenic | |
| rs9538440 | 13 | 60068406 | 4 | 0 | 0 | A | G | 0.9393 | 0.273 | 0.0005793 | +++?+ | 1.6 | 0.3842 | intergenic | |
| 16:26814426 | 16 | 26814426 | 3 | 0 | 0 | A | G | 1.4582 | 0.4238 | 0.0005794 | +++?? | 5.8 | 0.3461 | intergenic | |
| rs11048656 | 12 | 26845060 | 4 | 0 | 0 | C | G | 1.7029 | 0.4949 | 0.0005794 | +-+?+ | 35.6 | 0.1984 | intron | ITPR2 |
| rs61050506 | 5 | 123116030 | 4 | 0 | 0 | A | G | 0.9853 | 0.2863 | 0.0005794 | +++?+ | 0 | 0.406 | intergenic | |
| rs12685478 | 9 | 30896810 | 3 | 0 | 0 | C | G | -3.3674 | 0.9786 | 0.0005795 | ---?? | 0 | 0.457 | intergenic | |
| rs4906138 | 14 | 43251153 | 4 | 0 | 0 | T | G | -1.2142 | 0.3529 | 0.0005797 | ---?- | 0 | 0.6269 | intergenic | |
| rs36026993 | 8 | 65039572 | 4 | 0 | 0 | T | C | -0.8409 | 0.2444 | 0.0005797 | ---?- | 0 | 0.95 | intergenic | |
| rs12524664 | 6 | 32029431 | 5 | 0 | 0 | A | G | 1.6754 | 0.4869 | 0.0005798 | ++++- | 42.8 | 0.1364 | missense | TNXB |
| rs2695163 | 15 | 42041712 | 5 | 0 | 0 | A | G | -0.9379 | 0.2726 | 0.00058 | ----- | 0 | 0.5533 | synonymous | MGA |
| rs4711175 | 6 | 28570416 | 5 | 0 | 0 | A | G | -1.1489 | 0.3339 | 0.00058 | ----- | 0 | 0.5254 | intergenic |  |
| rs140203858 | 3 | 143066327 | 4 | 0 | 0 | T | G | -2.019 | 0.5868 | 0.00058 | ---?- | 0 | 0.6148 | intron | SLC9A9 |
| rs10734347 | 11 | 23968245 | 4 | 0 | 0 | A | C | 1.2406 | 0.3606 | 0.0005801 | +++?+ | 0 | 0.7285 | intergenic | |
| rs12295895 | 11 | 109492771 | 3 | 0 | 0 | C | G | -0.9151 | 0.266 | 0.0005801 | -+-?? | 59.7 | 0.08373 | intergenic | |
| rs11808009 | 1 | 203689216 | 4 | 0 | 0 | T | C | -1.1161 | 0.3244 | 0.0005803 | ---?- | 0 | 0.9066 | intron | ATP2B4 |
| rs17064433 | 3 | 61074705 | 4 | 0 | 0 | T | C | -3.263 | 0.9484 | 0.0005804 | ---?+ | 0 | 0.6384 | intron | FHIT |
| rs7993572 | 13 | 45636684 | 4 | 0 | 0 | A | T | -2.2202 | 0.6453 | 0.0005805 | ---?+ | 23.3 | 0.2714 | intergenic | |
| rs16986640 | 22 | 29108112 | 4 | 0 | 0 | A | C | -2.2418 | 0.6516 | 0.0005806 | ---?- | 0 | 0.5791 | intron | CHEK2 |
| rs75334189 | 10 | 86668797 | 4 | 0 | 0 | T | C | 1.7814 | 0.5178 | 0.0005808 | +++?+ | 0 | 0.7774 | intergenic | |
| rs71434755 | 13 | 30321657 | 5 | 0 | 0 | A | C | 1.2794 | 0.3719 | 0.0005809 | +++++ | 0 | 0.5573 | intergenic | |
| rs12280107 | 11 | 59025022 | 5 | 0 | 0 | T | C | -0.7466 | 0.217 | 0.0005811 | -+--+ | 10.8 | 0.3443 | intergenic | |
| rs13174873 | 5 | 123003548 | 4 | 0 | 0 | A | C | 0.9121 | 0.2651 | 0.0005811 | +++?+ | 55.5 | 0.08074 | intergenic | |
| rs343052 | 7 | 35519519 | 4 | 0 | 0 | T | G | 1.1004 | 0.3199 | 0.0005812 | +++?+ | 25.1 | 0.2609 | intergenic | |
| rs37387 | 5 | 35085600 | 4 | 0 | 0 | T | C | 1.2495 | 0.3632 | 0.0005813 | +++?- | 49.7 | 0.1133 | intron | PRLR |
| rs75482448 | 9 | 137742408 | 3 | 0 | 0 | C | G | -3.6449 | 1.0595 | 0.0005813 | ---?? | 31.9 | 0.2301 | upstream-gene | |
| rs248547 | 5 | 165390351 | 5 | 0 | 0 | A | T | -0.7316 | 0.2127 | 0.0005815 | ---+- | 39.1 | 0.1604 | intergenic | |
| rs12405255 | 1 | 203667797 | 4 | 0 | 0 | A | C | 1.1651 | 0.3387 | 0.0005815 | +++?+ | 0 | 0.9368 | intron | ATP2B4 |
| rs74903468 | 2 | 85284578 | 5 | 0 | 0 | A | G | -2.8816 | 0.8376 | 0.0005815 | ----+ | 0 | 0.8319 | 3-prime-UTR | KCMF1 |
| rs825213 | 1 | 64762226 | 4 | 0 | 0 | A | G | -0.9526 | 0.2769 | 0.0005817 | ---?- | 0 | 0.4962 | intergenic | |
| rs77831665 | 16 | 48187570 | 3 | 0 | 0 | T | C | 3.2421 | 0.9425 | 0.0005817 | +++?? | 0 | 0.5056 | intergenic | |
| rs2089917 | 11 | 37063291 | 4 | 0 | 0 | T | C | 0.852 | 0.2477 | 0.0005818 | +++?+ | 0.6 | 0.3888 | intergenic | |
| rs10869661 | 9 | 78462976 | 4 | 0 | 0 | T | C | -1.2726 | 0.3699 | 0.0005818 | ---?- | 55.2 | 0.08235 | intergenic | |
| rs117667649 | 14 | 92867411 | 3 | 0 | 0 | T | C | 3.3675 | 0.9789 | 0.0005818 | +++?? | 2.1 | 0.3602 | intron | SLC24A4 |
| rs10931063 | 2 | 184030555 | 4 | 0 | 0 | A | G | 1.167 | 0.3392 | 0.0005819 | +++?+ | 0 | 0.8091 | downstream-gene | |
| rs3830135 | 6 | 32548464 | 5 | 0 | 0 | A | G | 1.3352 | 0.3881 | 0.000582 | +++++ | 5.3 | 0.3764 | intron | HLA-DRB1 |
| rs1366351 | 5 | 123516908 | 4 | 0 | 0 | A | T | 0.8383 | 0.2437 | 0.0005821 | +++?- | 4 | 0.3727 | intergenic | |
| rs76770617 | 6 | 106283037 | 4 | 0 | 0 | T | C | -1.9344 | 0.5624 | 0.0005821 | ---?- | 0 | 0.4079 | intergenic | |
| 12:41306204 | 12 | 41306204 | 3 | 0 | 0 | G | GAA | 1.7479 | 0.5081 | 0.0005821 | +++?? | 0 | 0.9301 | | |
| rs1778007 | 1 | 87567091 | 4 | 0 | 0 | T | C | -1.0323 | 0.3001 | 0.0005822 | ---?- | 11.1 | 0.3374 | intron | HS2ST1 |
| rs10836669 | 11 | 37055016 | 4 | 0 | 0 | T | C | -0.908 | 0.264 | 0.0005822 | ---?- | 0 | 0.4535 | intergenic | |
| rs111427464 | 4 | 148938833 | 3 | 0 | 0 | A | G | 1.5903 | 0.4623 | 0.0005822 | +++?? | 36.4 | 0.2075 | intron | ARHGAP10 |
| rs35603491 | 4 | 100652297 | 4 | 0 | 0 | A | G | -2.0426 | 0.5938 | 0.0005823 | +--?- | 50.6 | 0.108 | intergenic | |
| 13:69250915 | 13 | 69250915 | 3 | 0 | 0 | CCCCCT | C | 1.0926 | 0.3177 | 0.0005823 | +++?? | 61.2 | 0.07616 | | |

| | | | | | | | | | | | | | | | |
|---|---|---|---|---|---|---|---|---|---|---|---|---|---|---|---|
| rs191080170 | 15 | 86326577 | 4 | 0 | 0 | T | G | 3.5361 | 1.0281 | 0.0005825 | +++?+ | 33 | 0.2143 | intron | KLHL25 |
| rs12186497 | 5 | 123515763 | 4 | 0 | 0 | T | G | 0.8365 | 0.2432 | 0.0005826 | +++?- | 26.8 | 0.2511 | intergenic | |
| rs3921861 | 3 | 176029464 | 4 | 0 | 0 | T | C | 1.2898 | 0.375 | 0.0005827 | +++?+ | 23.6 | 0.2697 | intergenic | |
| rs35165052 | 11 | 74289625 | 4 | 0 | 0 | T | G | -1.2687 | 0.3689 | 0.0005827 | ---?- | 0 | 0.8389 | intergenic | |
| rs187877887 | 4 | 166213995 | 4 | 0 | 0 | T | C | 3.8423 | 1.1171 | 0.0005827 | +++?+ | 0 | 0.9225 | intron | KLHL2 |
| rs11773142 | 7 | 49471189 | 4 | 0 | 0 | T | G | -0.9674 | 0.2813 | 0.000583 | ---?+ | 25.3 | 0.2596 | intergenic | |
| rs12788170 | 11 | 635569 | 3 | 0 | 0 | A | C | -1.0117 | 0.2942 | 0.000583 | ---?? | 0 | 0.8892 | upstream-gene | |
| rs139422331 | 3 | 196554926 | 4 | 0 | 0 | T | C | -3.9211 | 1.1401 | 0.0005833 | ---?- | 0 | 0.8514 | intron | PAK2 |
| rs185160495 | 16 | 87310916 | 2 | 0 | 0 | T | C | -4.0615 | 1.1809 | 0.0005833 | -?-?? | 0 | 0.8971 | intergenic | |
| rs144250015 | 12 | 5640229 | 2 | 0 | 0 | T | G | 5.0975 | 1.4822 | 0.0005834 | +?+?? | 0 | 0.3636 | intergenic | |
| rs4932566 | 15 | 92180972 | 4 | 0 | 0 | C | G | -1.3416 | 0.3901 | 0.0005837 | ---?- | 0 | 0.5621 | intergenic | |
| rs6005833 | 22 | 29068539 | 4 | 0 | 0 | T | C | 2.3515 | 0.6838 | 0.0005837 | +++?+ | 0 | 0.4224 | intron | TTC28 |
| rs2365053 | 7 | 49462209 | 4 | 0 | 0 | A | G | 0.9578 | 0.2785 | 0.000584 | +++?- | 21.8 | 0.2797 | intergenic | |
| rs10230606 | 7 | 49471064 | 4 | 0 | 0 | C | G | 0.967 | 0.2812 | 0.0005841 | +++?- | 25.3 | 0.2598 | intergenic | |
| rs1778008 | 1 | 87567346 | 4 | 0 | 0 | T | C | -1.0318 | 0.3 | 0.0005843 | ---?- | 11 | 0.3381 | intron | HS2ST1 |
| rs3739123 | 2 | 131131424 | 5 | 0 | 0 | C | G | -0.8903 | 0.2589 | 0.0005845 | ----- | 34.2 | 0.193 | 3-prime-UTR | PTPN18 |
| rs35426010 | 14 | 21445441 | 3 | 0 | 0 | C | G | -0.9471 | 0.2754 | 0.0005847 | ---?? | 24.7 | 0.2652 | intergenic | |
| rs56138634 | 5 | 169554202 | 5 | 0 | 0 | A | C | -1.626 | 0.4729 | 0.0005847 | ----- | 0 | 0.7384 | intergenic |  |
| rs72902882 | 18 | 37918556 | 4 | 0 | 0 | A | C | -2.6207 | 0.7622 | 0.0005847 | ---?- | 0 | 0.7049 | intergenic | |
| 5:109132294 | 5 | 109132294 | 3 | 0 | 0 | T | TA | -0.8795 | 0.2558 | 0.0005847 | ---?? | 0 | 0.8477 | | |
| rs4680169 | 3 | 155094465 | 4 | 0 | 0 | T | C | 1.0198 | 0.2966 | 0.0005848 | +++?+ | 13.4 | 0.3253 | intergenic | |
| rs11180678 | 12 | 76175019 | 4 | 0 | 0 | T | C | 2.1803 | 0.6341 | 0.0005848 | +++?+ | 0 | 0.9201 | intergenic | |
| 6:118430816 | 6 | 118430816 | 3 | 0 | 0 | G | GA | 1.176 | 0.342 | 0.0005851 | +++?? | 0 | 0.6182 | | |
| rs11936334 | 4 | 169322336 | 4 | 0 | 0 | A | C | -0.8512 | 0.2476 | 0.0005853 | ---?- | 0 | 0.9164 | intron | DDX60L |
| rs34060848 | 11 | 74311667 | 4 | 0 | 0 | A | G | -1.2636 | 0.3675 | 0.0005853 | ---?- | 0 | 0.5897 | intron | POLD3 |
| rs143246593 | 19 | 50706619 | 4 | 0 | 0 | T | G | -2.6279 | 0.7643 | 0.0005854 | ---?- | 13 | 0.3275 | upstream-gene | |
| rs56363430 | 15 | 56220607 | 3 | 0 | 0 | A | G | 5.8045 | 1.6883 | 0.0005857 | -++?? | 57.6 | 0.0944 | intron | NEDD4 |
| rs114252443 | 4 | 2811737 | 5 | 0 | 0 | A | G | 2.2593 | 0.6571 | 0.0005858 | +++++ | 0 | 0.6722 | intron | SH3BP2 |
| rs2345714 | 22 | 28508025 | 4 | 0 | 0 | T | C | 2.5413 | 0.7392 | 0.0005861 | +++?+ | 0 | 0.759 | intron | TTC28 |
| rs248548 | 5 | 165390453 | 5 | 0 | 0 | T | G | 0.7309 | 0.2126 | 0.0005862 | +++-+ | 39 | 0.161 | intergenic | |
| rs73082283 | 12 | 27787023 | 4 | 0 | 0 | T | C | 0.9346 | 0.2719 | 0.0005863 | +++?+ | 3.1 | 0.3771 | intron | PPFIBP1 |
| rs12567725 | 1 | 98595058 | 4 | 0 | 0 | A | G | -1.3485 | 0.3923 | 0.0005864 | ---?- | 19.5 | 0.2927 | intergenic | |
| rs57503016 | 16 | 7206439 | 4 | 0 | 0 | C | G | 1.0526 | 0.3062 | 0.0005868 | +++?+ | 0 | 0.9541 | intron | RBFOX1 |
| rs59234435 | 8 | 65056927 | 4 | 0 | 0 | T | C | -0.8324 | 0.2421 | 0.0005869 | ---?- | 0 | 0.9493 | intergenic | |
| rs4142468 | 6 | 78085506 | 4 | 0 | 0 | A | T | 0.9021 | 0.2624 | 0.000587 | +++?+ | 0 | 0.7252 | intergenic | |
| rs138796605 | 8 | 140041109 | 3 | 0 | 0 | A | C | -3.848 | 1.1194 | 0.000587 | ---?? | 0 | 0.6356 | intergenic | |
| 6:109114094 | 6 | 109114094 | 2 | 0 | 0 | A | T | -4.7536 | 1.3829 | 0.0005871 | -?-?? | 0 | 0.975 | intergenic | |
| rs78525734 | 5 | 82396896 | 4 | 0 | 0 | T | C | -2.1616 | 0.6288 | 0.0005872 | ---?- | 0 | 0.496 | intron | XRCC4 |
| rs185994038 | 2 | 62845992 | 4 | 0 | 0 | T | C | -3.5691 | 1.0383 | 0.0005872 | ---?- | 0 | 0.4291 | intergenic | |
| rs6927891 | 6 | 1479768 | 4 | 0 | 0 | A | G | 0.8803 | 0.2561 | 0.0005873 | +++?+ | 0 | 0.5486 | intergenic | |
| rs6979685 | 7 | 67408091 | 4 | 0 | 0 | T | C | -1.8145 | 0.5279 | 0.0005874 | ---?+ | 0 | 0.3981 | intergenic | |
| rs7928370 | 11 | 8725268 | 5 | 0 | 0 | A | G | -0.8036 | 0.2338 | 0.0005874 | ----- | 0 | 0.926 | intron | ST5 |
| rs6675783 | 1 | 102916806 | 4 | 0 | 0 | T | C | 0.8453 | 0.2459 | 0.0005875 | +++?- | 0 | 0.5077 | intergenic | |
| rs75278585 | 10 | 90282687 | 4 | 0 | 0 | T | C | 2.0793 | 0.6049 | 0.0005875 | +++?+ | 0 | 0.8274 | intron | RNLS |
| rs4742416 | 9 | 7851742 | 4 | 0 | 0 | T | C | -1.0472 | 0.3047 | 0.0005876 | -+-?- | 53.5 | 0.09161 | intergenic | |
| rs9343608 | 6 | 78086656 | 4 | 0 | 0 | T | C | 0.902 | 0.2624 | 0.0005876 | +++?+ | 0 | 0.7249 | intergenic | |
| rs148405566 | 15 | 54349215 | 3 | 0 | 0 | A | G | 1.1666 | 0.3394 | 0.0005876 | +++?? | 0 | 0.9781 | intron | UNC13C |
| rs1342635 | 6 | 78084035 | 4 | 0 | 0 | T | C | -0.9022 | 0.2625 | 0.0005879 | ---?- | 0 | 0.7249 | intergenic | |
| rs1833194 | 16 | 52917922 | 4 | 0 | 0 | A | C | 0.8558 | 0.249 | 0.000588 | +++?+ | 0 | 0.6341 | intergenic | |
| rs28517561 | 22 | 39422762 | 3 | 0 | 0 | T | G | -0.9218 | 0.2682 | 0.000588 | ---?? | 0 | 0.9787 | intron | APOBEC3D |
| rs11229896 | 11 | 59027039 | 5 | 0 | 0 | A | T | -0.7472 | 0.2174 | 0.0005882 | -+--+ | 11.1 | 0.3428 | intergenic | |
| rs142977187 | 3 | 143065577 | 4 | 0 | 0 | A | G | 2.0145 | 0.5861 | 0.0005882 | +++?+ | 0 | 0.6125 | intron | SLC9A9,SLC9A9-AS1 |
| rs73234751 | 7 | 129039300 | 3 | 0 | 0 | A | G | 1.3705 | 0.3988 | 0.0005883 | +++?? | 0 | 0.5161 | intron | AHCYL2 |
| rs75810822 | 4 | 171473359 | 3 | 0 | 0 | T | C | 5.6144 | 1.6335 | 0.0005883 | ++??- | 18.4 | 0.2936 | intergenic | |
| 1:103347728 | 1 | 103347728 | 3 | 0 | 0 | G | GTATT | -0.879 | 0.2557 | 0.0005883 | ---?? | 0 | 0.8464 | | |
| rs62419109 | 6 | 63324512 | 4 | 0 | 0 | T | C | -2.425 | 0.7056 | 0.0005884 | ---?- | 0 | 0.5059 | intergenic | |
| rs146635818 | 18 | 68724063 | 4 | 0 | 0 | T | C | 2.5141 | 0.7315 | 0.0005885 | +++?+ | 0 | 0.6692 | intergenic | |
| rs1562098 | 4 | 36995289 | 4 | 0 | 0 | T | C | 0.866 | 0.252 | 0.0005886 | +++?- | 44.7 | 0.1435 | intergenic | |
| rs17142713 | 16 | 7205858 | 4 | 0 | 0 | A | G | -1.0493 | 0.3053 | 0.0005886 | ---?- | 0 | 0.9561 | intron | RBFOX1 |

| | | | | | | | | | | | | | | | |
|---|---|---|---|---|---|---|---|---|---|---|---|---|---|---|---|
| rs73644369 | 9 | 30877811 | 4 | 0 | 0 | A | T | 2.748 | 0.7996 | 0.0005887 | +++?+ | 0 | 0.5027 | intergenic | |
| rs112403136 | 22 | 29031126 | 3 | 0 | 0 | A | G | 2.4632 | 0.7167 | 0.0005887 | +++?? | 0.5 | 0.3659 | intron | TTC28 |
| rs61858412 | 10 | 55853850 | 4 | 0 | 0 | T | C | -2.9689 | 0.8639 | 0.0005889 | ---?+ | 27.5 | 0.247 | intron | PCDH15 |
| 3:123718428 | 3 | 123718428 | 3 | 0 | 0 | CT | C | -3.2134 | 0.935 | 0.0005889 | ---?? | 0 | 0.7528 | | |
| rs35003102 | 1 | 64576889 | 4 | 0 | 0 | A | G | 1.0589 | 0.3081 | 0.000589 | +++?+ | 0 | 0.847 | intron | ROR1 |
| 22:41366743 | 22 | 41366743 | 3 | 0 | 0 | T | TA | 1.6963 | 0.4936 | 0.000589 | +++?? | 0 | 0.5146 | | |
| rs630482 | 8 | 12555445 | 4 | 0 | 0 | T | C | 1.1159 | 0.3247 | 0.0005892 | +++?+ | 45.2 | 0.1404 | intergenic | |
| rs12139634 | 1 | 87431586 | 4 | 0 | 0 | T | C | 1.2312 | 0.3583 | 0.0005893 | +++?- | 8.2 | 0.3523 | intron | HS2ST1 |
| rs142354534 | 11 | 20157338 | 4 | 0 | 0 | C | G | 1.6756 | 0.4876 | 0.0005893 | +++?+ | 0 | 0.7554 | intergenic | |
| rs117147243 | 18 | 68652672 | 4 | 0 | 0 | A | C | -1.5797 | 0.4597 | 0.0005894 | ---?- | 0 | 0.9753 | intergenic | |
| rs117174082 | 18 | 68700227 | 4 | 0 | 0 | T | C | 2.4534 | 0.7139 | 0.0005894 | +++?+ | 0 | 0.4171 | intergenic | |
| rs190046901 | 12 | 60965474 | 2 | 0 | 0 | T | C | 7.4614 | 2.1713 | 0.0005894 | ?+??+ | 0 | 0.4753 | intergenic | |
| rs12685479 | 9 | 30896865 | 3 | 0 | 0 | A | C | 3.3663 | 0.9796 | 0.0005898 | +++?? | 0 | 0.4586 | intergenic | |
| rs75485436 | 3 | 47935009 | 2 | 0 | 0 | A | G | 6.1292 | 1.7837 | 0.0005898 | ?+??+ | 56.1 | 0.1312 | intron | MAP4 |
| rs1778009 | 1 | 87567415 | 4 | 0 | 0 | A | G | -1.0308 | 0.3 | 0.0005899 | ---?- | 10.8 | 0.3388 | intron | HS2ST1 |
| rs143616153 | 18 | 73328106 | 4 | 0 | 0 | A | C | 2.4305 | 0.7073 | 0.00059 | +++?+ | 0 | 0.7147 | intergenic | |
| 6:159223446 | 6 | 159223446 | 3 | 0 | 0 | T | TTA | 0.8736 | 0.2542 | 0.00059 | +++?? | 12.1 | 0.3204 | | |
| 2:170454031 | 2 | 170454031 | 3 | 0 | 0 | CAG | C | 1.2598 | 0.3666 | 0.0005901 | +++?? | 0 | 0.7959 | | |
| rs3830133 | 6 | 32548469 | 5 | 0 | 0 | A | G | -1.3189 | 0.3838 | 0.0005902 | ----- | 6.5 | 0.3696 | intron | HLA-DRB1 |
| rs150771819 | 6 | 63324680 | 4 | 0 | 0 | T | C | 2.4246 | 0.7056 | 0.0005904 | +++?+ | 0 | 0.5052 | intergenic | |
| rs9967795 | 2 | 131131866 | 5 | 0 | 0 | A | G | 0.8904 | 0.2592 | 0.0005907 | +++++ | 32.1 | 0.2074 | 3-prime-UTR | PTPN18 |
| rs140558787 | 2 | 199064496 | 4 | 0 | 0 | A | T | -1.1431 | 0.3327 | 0.0005907 | ---?+ | 35.8 | 0.1976 | intergenic | |
| rs10955056 | 8 | 96993697 | 4 | 0 | 0 | T | C | -1.4148 | 0.4118 | 0.0005909 | ---?- | 0 | 0.9601 | intergenic | |
| rs143958297 | 2 | 168600921 | 2 | 0 | 0 | A | G | -6.6195 | 1.9266 | 0.0005909 | --??? | 37.8 | 0.2049 | intron | B3GALT1 |
| rs17069103 | 8 | 3988689 | 4 | 0 | 0 | A | G | -1.0946 | 0.3186 | 0.0005912 | +--?- | 58 | 0.06723 | intron | CSMD1 |
| rs10263065 | 7 | 49461592 | 4 | 0 | 0 | T | G | -0.9561 | 0.2783 | 0.0005913 | ---?+ | 22.4 | 0.2763 | intergenic | |
| rs60505450 | 12 | 112258579 | 5 | 0 | 0 | A | G | -1.3007 | 0.3786 | 0.0005914 | -+--- | 49.7 | 0.09341 | intergenic | |
| rs7129035 | 11 | 58982071 | 5 | 0 | 0 | T | C | -0.7395 | 0.2153 | 0.0005915 | -+--+ | 21.1 | 0.2804 | upstream-gene | |
| rs1342633 | 6 | 78082678 | 4 | 0 | 0 | T | C | -0.9016 | 0.2624 | 0.0005916 | ---?- | 0 | 0.7239 | intergenic | |
| rs7187508 | 16 | 6533549 | 4 | 0 | 0 | A | C | -0.8752 | 0.2547 | 0.0005916 | ---?- | 0 | 0.8023 | intron | RBFOX1 |
| rs144732945 | 4 | 166215822 | 4 | 0 | 0 | A | T | 3.8275 | 1.1141 | 0.0005916 | +++?+ | 0 | 0.9259 | intron | KLHL2 |
| rs190676802 | 10 | 16761882 | 2 | 0 | 0 | T | C | 3.557 | 1.0354 | 0.0005918 | +?+?? | 0 | 0.7658 | intron | RSU1 |
| rs1483628 | 3 | 176027779 | 4 | 0 | 0 | T | C | -1.2866 | 0.3745 | 0.0005919 | ---?- | 17.5 | 0.3037 | intergenic | |
| rs12683426 | 9 | 30896876 | 3 | 0 | 0 | A | G | -3.3649 | 0.9795 | 0.0005919 | ---?? | 0 | 0.4582 | intergenic | |
| rs34499829 | 12 | 734001 | 3 | 0 | 0 | A | G | -3.825 | 1.1135 | 0.0005921 | ---?? | 48.1 | 0.1457 | intron | NINJ2 |
| rs10769940 | 11 | 8702469 | 5 | 0 | 0 | A | G | -0.7957 | 0.2316 | 0.0005924 | ----+ | 0 | 0.9101 | intron | TRIM66 |
| rs184492725 | 5 | 100594993 | 2 | 0 | 0 | A | G | 5.6885 | 1.656 | 0.0005924 | +?+?? | 0 | 0.5059 | intergenic | |
| rs984015 | 13 | 60039626 | 4 | 0 | 0 | T | C | -0.8949 | 0.2605 | 0.0005925 | ---?- | 15.3 | 0.3152 | intergenic | |
| rs12046836 | 1 | 98598745 | 4 | 0 | 0 | A | G | -1.3467 | 0.3921 | 0.0005925 | ---?- | 15.8 | 0.3129 | intergenic | |
| rs56768329 | 16 | 7205434 | 4 | 0 | 0 | C | G | 1.0469 | 0.3048 | 0.0005927 | +++?+ | 0 | 0.9569 | intron | RBFOX1 |
| rs2577951 | 15 | 42030783 | 5 | 0 | 0 | A | G | 0.9342 | 0.272 | 0.0005928 | +++++ | 0 | 0.5746 | intron | MGA |
| rs13123573 | 4 | 174904309 | 4 | 0 | 0 | T | C | 2.0662 | 0.6015 | 0.0005929 | +++?+ | 0 | 0.5493 | intergenic | |
| rs55676791 | 2 | 62783449 | 4 | 0 | 0 | T | G | -3.1954 | 0.9303 | 0.0005929 | ---?- | 49.1 | 0.1168 | intergenic | |
| 14:101835379 | 14 | 101835379 | 3 | 0 | 0 | CT | C | -1.4182 | 0.4129 | 0.0005929 | ---?? | 19.9 | 0.2868 | | |
| rs4768322 | 12 | 41305222 | 4 | 0 | 0 | C | G | 1.7011 | 0.4953 | 0.0005931 | +++?- | 0 | 0.7971 | intron | CNTN1 |
| rs149681550 | 22 | 39593405 | 3 | 0 | 0 | A | G | -2.6962 | 0.785 | 0.0005931 | ---?? | 23.7 | 0.2695 | intergenic | |
| 1:102932003 | 1 | 102932003 | 3 | 0 | 0 | A | ACAT | -0.8783 | 0.2557 | 0.0005931 | ---?? | 0 | 0.7685 | | |
| rs6696364 | 1 | 18753647 | 4 | 0 | 0 | T | C | -1.0451 | 0.3043 | 0.0005932 | ---?- | 49.1 | 0.117 | intergenic | |
| rs10149275 | 14 | 43268640 | 4 | 0 | 0 | A | G | -1.3953 | 0.4062 | 0.0005932 | ---?+ | 53.1 | 0.09391 | intergenic | |
| rs34103704 | 19 | 46153512 | 5 | 0 | 1 | A | C | -0.9555 | 0.2782 | 0.0005932 | -+--+ | 55.6 | 0.06091 | upstream-gene | |
| rs76901156 | 2 | 116711775 | 3 | 0 | 0 | A | G | 2.4909 | 0.7252 | 0.0005934 | +++?? | 59 | 0.08746 | intergenic | |
| rs4768323 | 12 | 41305249 | 4 | 0 | 0 | A | G | 1.7009 | 0.4953 | 0.0005942 | +++?- | 0 | 0.7971 | intron | CNTN1 |
| rs35835288 | 1 | 222982059 | 4 | 0 | 0 | T | C | -1.0091 | 0.2938 | 0.0005943 | ---?+ | 57.9 | 0.06812 | intergenic | |
| rs6872047 | 5 | 123004718 | 4 | 0 | 0 | T | C | -0.9106 | 0.2652 | 0.0005944 | ---?- | 55.5 | 0.0808 | intergenic | |
| rs357910 | 18 | 46562289 | 4 | 0 | 0 | T | C | -1.0073 | 0.2933 | 0.0005946 | ---?- | 7.5 | 0.3557 | intergenic | |
| rs369931 | 21 | 31401942 | 4 | 0 | 0 | A | T | 0.9752 | 0.284 | 0.0005946 | +++?- | 6.2 | 0.362 | intergenic | |
| rs6735968 | 2 | 184045757 | 4 | 0 | 0 | A | G | 1.1554 | 0.3365 | 0.0005947 | +++?+ | 0 | 0.8331 | intergenic | |
| 6:156029094 | 6 | 156029094 | 3 | 0 | 0 | T | TTA | -0.9918 | 0.2888 | 0.0005948 | ---?? | 0 | 0.7692 | | |
| rs10102048 | 8 | 137677221 | 3 | 0 | 0 | T | C | 1.3394 | 0.39 | 0.0005949 | +++?? | 0 | 0.5061 | intergenic | |

| | | | | | | | | | | | | | | | |
|---|---|---|---|---|---|---|---|---|---|---|---|---|---|---|---|
| rs57740160 | 4 | 20760644 | 4 | 0 | 0 | A | G | 0.9055 | 0.2637 | 0.0005951 | +++?+ | 0 | 0.786 | intron | KCNIP4 |
| rs79928580 | 3 | 143838425 | 3 | 0 | 0 | A | G | 4.0396 | 1.1764 | 0.0005951 | +++?? | 0 | 0.3903 | intergenic | |
| rs937476 | 12 | 103247564 | 4 | 0 | 0 | A | T | 0.8296 | 0.2416 | 0.0005953 | +++?+ | 0 | 0.6648 | intron | PAH |
| rs6490055 | 12 | 111768973 | 5 | 0 | 0 | A | G | -0.8541 | 0.2488 | 0.0005955 | -+--- | 52.4 | 0.07791 | intron | CUX2 |
| rs7790587 | 7 | 82418795 | 4 | 0 | 0 | A | G | 0.8826 | 0.257 | 0.0005955 | +-+?+ | 46.4 | 0.1331 | intron | PCLO |
| rs74497197 | 10 | 86673380 | 4 | 0 | 0 | A | G | -1.7734 | 0.5165 | 0.0005955 | ---?- | 0 | 0.7802 | intergenic | |
| rs12943935 | 17 | 59317457 | 4 | 0 | 0 | A | G | -1.0397 | 0.3028 | 0.0005957 | ---?- | 0 | 0.5793 | intron | BCAS3 |
| 1:87465376 | 1 | 87465376 | 3 | 0 | 0 | T | TC | -1.0161 | 0.2959 | 0.0005957 | ---?? | 0 | 0.4034 | | |
| rs9967796 | 2 | 131131869 | 5 | 0 | 0 | A | G | 0.8898 | 0.2592 | 0.0005959 | +++++ | 32.4 | 0.2054 | 3-prime-UTR | PTPN18 |
| rs10895432 | 11 | 103328079 | 4 | 0 | 0 | A | T | -0.9397 | 0.2737 | 0.000596 | ---?- | 49.6 | 0.1139 | intron | DYNC2H1 |
| rs10895433 | 11 | 103328081 | 4 | 0 | 0 | A | C | -0.9397 | 0.2737 | 0.000596 | ---?- | 49.6 | 0.114 | intron | DYNC2H1 |
| rs11809431 | 1 | 98595242 | 4 | 0 | 0 | C | G | -1.3467 | 0.3922 | 0.000596 | ---?- | 19.6 | 0.2918 | intergenic | |
| rs3136645 | 19 | 39398649 | 4 | 0 | 0 | T | C | -1.0544 | 0.3071 | 0.0005961 | ---?- | 11.9 | 0.3332 | intron | NFKBIB |
| rs79881617 | 9 | 11834891 | 4 | 0 | 0 | A | C | -1.981 | 0.577 | 0.0005963 | ---?- | 47.5 | 0.1263 | intergenic | |
| rs74637529 | 4 | 127746848 | 4 | 0 | 0 | T | C | 3.4881 | 1.016 | 0.0005967 | +++?+ | 0 | 0.9863 | intergenic | |
| rs78094547 | 6 | 12210721 | 4 | 0 | 0 | A | G | 3.1645 | 0.9218 | 0.0005968 | +++?+ | 5.2 | 0.367 | intergenic | |
| rs142485458 | 15 | 54349227 | 3 | 0 | 0 | T | G | -1.166 | 0.3397 | 0.0005969 | ---?? | 0 | 0.9802 | intron | UNC13C |
| rs66593336 | 5 | 747535 | 3 | 0 | 0 | T | C | 1.5668 | 0.4564 | 0.000597 | +++?? | 0 | 0.5064 | intron | ZDHHC11B |
| rs1487301 | 1 | 98595815 | 4 | 0 | 0 | C | G | 1.3464 | 0.3922 | 0.0005972 | +++?+ | 19.7 | 0.2916 | intergenic | |
| rs2701369 | 15 | 87605554 | 4 | 0 | 0 | T | C | -0.8825 | 0.2571 | 0.0005972 | ---?- | 0 | 0.4744 | intergenic | |
| rs4711174 | 6 | 28532960 | 5 | 0 | 0 | A | G | 1.1476 | 0.3343 | 0.0005973 | +++++ | 0 | 0.4935 | intergenic | |
| rs6509990 | 19 | 56815963 | 4 | 0 | 0 | A | G | 1.5607 | 0.4547 | 0.0005973 | +++?+ | 0 | 0.4444 | intron | ZSCAN5A,LOC100506374 |
| rs3739122 | 2 | 131131525 | 5 | 0 | 0 | T | C | -0.8886 | 0.2589 | 0.0005977 | ----- | 34.9 | 0.1888 | 3-prime-UTR | PTPN18 |
| rs6898834 | 5 | 80526113 | 4 | 0 | 0 | T | C | -1.7192 | 0.5008 | 0.0005977 | ---?- | 41.9 | 0.1603 | upstream-gene | |
| rs4810093 | 20 | 56220444 | 4 | 0 | 0 | A | G | 1.435 | 0.4181 | 0.000598 | +++?+ | 0 | 0.9616 | downstream-gene | |
| rs7544563 | 1 | 185400748 | 4 | 0 | 0 | T | C | -0.8989 | 0.2619 | 0.000598 | ---?- | 0 | 0.9063 | intergenic | |
| rs80293647 | 5 | 82332254 | 4 | 0 | 0 | A | G | 2.52 | 0.7342 | 0.0005981 | +++?+ | 0 | 0.6875 | intergenic | |
| rs7663644 | 4 | 41335150 | 4 | 0 | 0 | A | C | 1.3786 | 0.4016 | 0.0005982 | +++?- | 0 | 0.4488 | intergenic | |
| rs6436173 | 2 | 220766768 | 4 | 0 | 0 | T | C | 0.9498 | 0.2767 | 0.0005983 | +++?+ | 0 | 0.5734 | downstream-gene | |
| 13:20810339 | 13 | 20810339 | 3 | 0 | 0 | CA | C | -1.1541 | 0.3362 | 0.0005984 | ---?? | 0 | 0.3816 | | |
| 19:46889230 | 19 | 46889230 | 3 | 0 | 0 | T | TG | -1.1065 | 0.3224 | 0.0005986 | ---?? | 33.7 | 0.2215 | | |
| rs12545053 | 8 | 65073605 | 4 | 0 | 0 | A | G | -0.8432 | 0.2457 | 0.0005987 | ---?- | 0 | 0.9195 | intergenic | |
| rs73428096 | 22 | 28870469 | 4 | 0 | 0 | A | G | 2.2798 | 0.6642 | 0.0005987 | +++?+ | 0 | 0.6132 | intron | TTC28 |
| rs76484592 | 8 | 53132284 | 2 | 0 | 0 | T | C | -4.3456 | 1.2662 | 0.000599 | -?-?? | 0 | 0.641 | intron | ST18 |
| 5:10305363 | 5 | 10305363 | 3 | 0 | 0 | G | GCCA | -1.8909 | 0.551 | 0.000599 | ---?? | 29.1 | 0.2441 | | |
| rs72914798 | 2 | 199016124 | 4 | 0 | 0 | T | C | 1.136 | 0.331 | 0.0005992 | +++?+ | 25.8 | 0.2571 | downstream-gene | |
| rs4938915 | 11 | 59022865 | 5 | 0 | 0 | A | G | -0.7443 | 0.2169 | 0.0005993 | -+--+ | 8.8 | 0.3564 | intergenic | |
| rs34446915 | 2 | 184046847 | 4 | 0 | 0 | A | G | 1.157 | 0.3371 | 0.0005993 | +++?+ | 0 | 0.827 | intergenic | |
| rs138109345 | 8 | 131420396 | 3 | 0 | 0 | T | C | -4.8903 | 1.425 | 0.0005996 | ---?? | 0 | 0.7885 | intron | ASAP1 |
| rs145248145 | 8 | 139896075 | 3 | 0 | 0 | A | G | -3.217 | 0.9374 | 0.0005996 | ---?? | 26.9 | 0.2546 | intron | COL22A1 |
| rs261556 | 8 | 97001222 | 4 | 0 | 0 | A | G | -1.4128 | 0.4117 | 0.0005998 | ---?- | 0 | 0.9384 | intergenic | |
| rs693940 | 15 | 53753766 | 4 | 0 | 0 | A | C | 0.8227 | 0.2397 | 0.0005998 | +++?+ | 51.4 | 0.1035 | intergenic | |
| rs7124836 | 11 | 12650458 | 4 | 0 | 0 | A | G | 1.0596 | 0.3088 | 0.0005999 | +++?+ | 0 | 0.7713 | intergenic | |
| 10:131916901 | 10 | 131916901 | 3 | 0 | 0 | G | GGTTT | -0.8637 | 0.2517 | 0.0006 | ---?? | 0 | 0.6841 | | |
| rs13254721 | 8 | 65066904 | 4 | 0 | 0 | C | G | 0.8379 | 0.2442 | 0.0006001 | +++?+ | 0 | 0.9367 | intergenic | |
| rs115057399 | 2 | 14450340 | 2 | 0 | 0 | A | G | 7.9872 | 2.3276 | 0.0006002 | ?+??+ | 0 | 0.8891 | intron | LINC00276 |
| rs13137795 | 4 | 10051506 | 5 | 0 | 0 | A | G | 0.7227 | 0.2106 | 0.0006003 | ++++- | 12.2 | 0.3361 | intergenic | |
| rs35312619 | 8 | 64930684 | 4 | 0 | 0 | A | T | -0.9186 | 0.2677 | 0.0006003 | ---?- | 0 | 0.8382 | intergenic | |
| 2:185433514 | 2 | 185433514 | 3 | 0 | 0 | CAT | C | -0.9679 | 0.2821 | 0.0006003 | ---?? | 0 | 0.6036 | | |
| 12:13858018 | 12 | 13858018 | 3 | 0 | 0 | CT | C | -0.873 | 0.2544 | 0.0006004 | ---?? | 0 | 0.6008 | | |
| rs148529031 | 4 | 122992232 | 2 | 0 | 0 | A | G | 5.9014 | 1.7198 | 0.0006006 | ++??? | 0 | 0.4441 | intergenic | |
| rs12401542 | 1 | 98589828 | 4 | 0 | 0 | T | C | -1.3459 | 0.3923 | 0.0006009 | ---?- | 17.8 | 0.3021 | intergenic | |
| rs113222491 | 4 | 95072369 | 4 | 0 | 0 | C | G | 3.4436 | 1.0036 | 0.0006009 | +++?+ | 0 | 0.6172 | intergenic | |
| 4:1347834 | 4 | 1347834 | 3 | 0 | 0 | CTT | C | 2.4234 | 0.7063 | 0.0006012 | +++?? | 26.9 | 0.2547 | | |
| rs117113898 | 16 | 27981530 | 4 | 0 | 0 | A | G | -1.9761 | 0.576 | 0.0006014 | ---?- | 0 | 0.9659 | intron | GSG1L |
| rs10986044 | 9 | 101135387 | 4 | 0 | 0 | C | G | 3.2847 | 0.9574 | 0.0006016 | +++?- | 28.4 | 0.2419 | intron | GABBR2 |
| rs26217 | 5 | 14534994 | 4 | 0 | 0 | A | G | -1.187 | 0.346 | 0.0006017 | ---?- | 0 | 0.6316 | intergenic | |
| rs66830107 | 8 | 64921986 | 4 | 0 | 0 | T | C | 0.918 | 0.2676 | 0.0006018 | +++?+ | 0 | 0.8423 | intergenic | |
| rs113732400 | 5 | 152505645 | 4 | 0 | 0 | A | T | -1.5219 | 0.4436 | 0.0006018 | ---?- | 0 | 0.5857 | intergenic | |

| | | | | | | | | | | | | | | | |
|---|---|---|---|---|---|---|---|---|---|---|---|---|---|---|---|
| rs145638726 | 22 | 28783036 | 4 | 0 | 0 | A | G | 2.6415 | 0.7699 | 0.0006018 | +++?+ | 0 | 0.8109 | intron | TTC28 |
| rs28651717 | 7 | 1196394 | 4 | 0 | 0 | T | G | 1.505 | 0.4387 | 0.000602 | +++?+ | 0 | 0.8762 | intron | ZFAND2A |
| rs62334870 | 4 | 160952965 | 4 | 0 | 0 | A | T | -3.1266 | 0.9114 | 0.0006022 | ---?- | 0 | 0.7242 | intergenic | |
| 3:196467076 | 3 | 196467076 | 4 | 0 | 0 | A | C | 2.2665 | 0.6607 | 0.0006023 | +++?- | 0 | 0.3986 | intron | PAK2 |
| rs62419110 | 6 | 63325684 | 4 | 0 | 0 | T | C | 2.4182 | 0.7049 | 0.0006024 | +++?+ | 0 | 0.5046 | intergenic | |
| rs409133 | 21 | 31400829 | 4 | 0 | 0 | A | C | -0.9706 | 0.2829 | 0.0006025 | ---?+ | 8.1 | 0.3525 | intergenic | |
| rs2952359 | 2 | 184007278 | 4 | 0 | 0 | T | C | -1.1773 | 0.3432 | 0.0006029 | ---?- | 0 | 0.8334 | intron | NUP35 |
| rs11785513 | 8 | 113225710 | 4 | 0 | 0 | T | C | -0.9769 | 0.2848 | 0.0006029 | ---?- | 0 | 0.9694 | intergenic | |
| rs112985139 | 3 | 23811415 | 4 | 0 | 0 | A | G | -3.5645 | 1.0391 | 0.0006029 | ---?- | 0 | 0.6222 | intergenic | |
| rs59861187 | 7 | 49478057 | 4 | 0 | 0 | T | C | -0.9735 | 0.2838 | 0.000603 | ---?+ | 26.8 | 0.251 | intergenic | |
| rs6886731 | 5 | 103163206 | 3 | 0 | 0 | A | C | -1.6101 | 0.4694 | 0.0006032 | ---?? | 0 | 0.6099 | intergenic | |
| rs117652943 | 11 | 20157483 | 4 | 0 | 0 | A | G | 1.6713 | 0.4872 | 0.0006032 | +++?+ | 0 | 0.7548 | intergenic | |
| rs80218654 | 5 | 122528152 | 4 | 0 | 0 | A | G | -1.795 | 0.5233 | 0.0006033 | ---?- | 0 | 0.7749 | 3-prime-UTR | PRDM6 |
| rs150906309 | 6 | 32093029 | 3 | 0 | 0 | T | C | -3.6451 | 1.0627 | 0.0006033 | ?---? | 23.4 | 0.2713 | intron | ATF6B |
| rs7048896 | 9 | 7848988 | 4 | 0 | 0 | T | G | -1.0891 | 0.3175 | 0.0006035 | ---?- | 47.9 | 0.1242 | intergenic | |
| rs11737937 | 5 | 123117002 | 4 | 0 | 0 | A | C | 0.9826 | 0.2865 | 0.0006035 | +++?+ | 0 | 0.4063 | intergenic | |
| rs233080 | 1 | 85814552 | 5 | 0 | 0 | A | G | 0.8047 | 0.2346 | 0.0006036 | +++++ | 0 | 0.9811 | intron | DDAH1 |
| rs1945593 | 11 | 59025429 | 5 | 0 | 0 | T | C | -0.7448 | 0.2172 | 0.0006037 | -+--+ | 10.5 | 0.3462 | intergenic | |
| rs112627226 | 6 | 32582969 | 5 | 0 | 1 | T | C | -1.6329 | 0.4761 | 0.0006039 | ----- | 0 | 0.5303 | intergenic |  |
| rs140097699 | 3 | 143026579 | 4 | 0 | 0 | T | C | -2.1935 | 0.6396 | 0.0006043 | ---?- | 0 | 0.7763 | intron | SLC9A9 |
| rs149619649 | 13 | 63867347 | 4 | 0 | 0 | A | T | 3.9293 | 1.1457 | 0.0006044 | +++?- | 12.2 | 0.3317 | intergenic | |
| rs11590380 | 1 | 185400939 | 4 | 0 | 0 | T | C | 0.8997 | 0.2623 | 0.0006045 | +++?+ | 0 | 0.9053 | intergenic | |
| rs10264137 | 7 | 82418234 | 4 | 0 | 0 | A | C | -0.8803 | 0.2567 | 0.0006046 | -+-?- | 46.1 | 0.1348 | intron | PCLO |
| rs72656146 | 13 | 113154019 | 2 | 0 | 0 | A | C | 4.7583 | 1.3874 | 0.0006046 | +?+?? | 0 | 0.4336 | intron | TUBGCP3 |
| rs2290273 | 15 | 85544050 | 4 | 0 | 0 | T | C | -0.8208 | 0.2393 | 0.0006047 | +--?+ | 56 | 0.07778 | intron | PDE8A |
| 6:163543281 | 6 | 163543281 | 3 | 0 | 0 | CAT | C | -1.3621 | 0.3972 | 0.0006047 | ---?? | 0 | 0.792 | | |
| rs2356339 | 8 | 65065863 | 4 | 0 | 0 | T | C | 0.8363 | 0.2439 | 0.0006048 | +++?+ | 0 | 0.9386 | intergenic | |
| rs79305846 | 1 | 208187889 | 4 | 0 | 0 | T | C | -2.0991 | 0.6121 | 0.0006049 | ---?- | 0 | 0.8672 | intergenic | |
| 3:196467075 | 3 | 196467075 | 4 | 0 | 0 | A | T | 2.2656 | 0.6607 | 0.0006051 | +++?- | 0 | 0.3989 | intron | PAK2 |
| rs10508597 | 10 | 20151881 | 4 | 0 | 0 | C | G | -0.8586 | 0.2504 | 0.0006051 | ---?- | 10.2 | 0.3419 | intron | PLXDC2 |
| rs825214 | 1 | 64760886 | 4 | 0 | 0 | A | T | 0.949 | 0.2767 | 0.0006052 | +++?+ | 0 | 0.5035 | intergenic | |
| rs2834098 | 21 | 34506306 | 4 | 0 | 0 | T | C | 1.0098 | 0.2945 | 0.0006053 | +++?- | 0 | 0.6475 | intergenic | |
| rs4768324 | 12 | 41305430 | 4 | 0 | 0 | A | G | 1.6987 | 0.4954 | 0.0006054 | +++?- | 0 | 0.7974 | intron | CNTN1 |
| 2:170499431 | 2 | 170499431 | 3 | 0 | 0 | T | TAAATA | -1.2658 | 0.3691 | 0.0006054 | ---?? | 0 | 0.8658 | | |
| rs2883908 | 8 | 65065872 | 4 | 0 | 0 | T | C | -0.8362 | 0.2439 | 0.0006055 | ---?- | 0 | 0.9387 | intergenic | |
| rs1920605 | 12 | 114691384 | 3 | 0 | 0 | T | C | 1.8001 | 0.525 | 0.0006056 | +++?? | 0 | 0.4116 | intergenic | |
| rs7128566 | 11 | 12641391 | 4 | 0 | 0 | T | G | -1.0466 | 0.3052 | 0.0006056 | ---?- | 0 | 0.8012 | intergenic | |
| rs12120723 | 1 | 102954040 | 4 | 0 | 0 | T | C | 0.8156 | 0.2378 | 0.0006057 | +++?- | 3.3 | 0.376 | intergenic | |
| rs12056484 | 8 | 65040149 | 4 | 0 | 0 | T | G | -0.84 | 0.245 | 0.0006058 | ---?- | 0 | 0.9397 | intergenic | |
| rs77911294 | 18 | 2417865 | 4 | 0 | 0 | A | T | 3.5676 | 1.0404 | 0.0006058 | +++?+ | 0 | 0.4695 | intergenic | |
| rs56183165 | 15 | 45956106 | 4 | 0 | 0 | T | C | 1.3815 | 0.4029 | 0.0006059 | +++?+ | 44.1 | 0.1466 | intron | SQRDL |
| rs12903546 | 15 | 92179005 | 4 | 0 | 0 | T | G | -1.3432 | 0.3917 | 0.000606 | ---?- | 0 | 0.4934 | intergenic | |
| 17:47392921 | 17 | 47392921 | 3 | 0 | 0 | T | TAA | -1.3643 | 0.3979 | 0.000606 | ---?? | 0 | 0.4022 | | |
| rs10233248 | 7 | 49474666 | 4 | 0 | 0 | A | T | -0.9697 | 0.2828 | 0.0006061 | ---?+ | 26.3 | 0.2537 | intergenic | |
| 10:18982786 | 10 | 18982786 | 3 | 0 | 0 | A | AAAGAA | -1.0562 | 0.308 | 0.0006061 | +--?? | 65.8 | 0.05353 | | |
| 20:34127336 | 20 | 34127336 | 3 | 0 | 0 | A | AT | -1.6765 | 0.4889 | 0.0006061 | ---?? | 0 | 0.4217 | | |
| rs58433811 | 5 | 123105721 | 4 | 0 | 0 | A | G | 0.9775 | 0.2851 | 0.0006062 | +++?+ | 12.7 | 0.3289 | intergenic | |
| rs7375941 | 4 | 189049361 | 4 | 0 | 0 | C | G | 1.2453 | 0.3632 | 0.0006064 | +++?+ | 0 | 0.8741 | intergenic | |
| rs79174153 | 6 | 106286430 | 4 | 0 | 0 | A | G | 1.9256 | 0.5616 | 0.0006065 | +++?+ | 0 | 0.4021 | intergenic | |
| rs10875257 | 1 | 100242048 | 4 | 0 | 0 | A | C | -1.7068 | 0.4978 | 0.0006066 | ---?- | 0 | 0.8738 | intergenic | |
| rs57502306 | 12 | 112290443 | 5 | 0 | 0 | A | G | 1.2895 | 0.3761 | 0.0006067 | +-+++ | 48.9 | 0.09833 | intron | MAPKAPK5 |
| rs3130506 | 6 | 31145451 | 5 | 0 | 0 | A | G | 0.7315 | 0.2134 | 0.0006068 | +++++ | 28.7 | 0.2301 | non-coding-exon | PSORS1C3 |
| rs3131018 | 6 | 31143582 | 5 | 0 | 0 | A | C | -0.7315 | 0.2134 | 0.0006069 | ----- | 28.7 | 0.23 | intron | PSORS1C3 |
| rs1648831 | 15 | 42135541 | 5 | 0 | 0 | T | G | -0.9403 | 0.2743 | 0.0006072 | ----- | 0 | 0.5775 | intron | JMJD7-PLA2G4B,PLA2G4B |
| rs1485510 | 8 | 65059823 | 4 | 0 | 0 | T | C | 0.8289 | 0.2418 | 0.0006074 | +++?+ | 0 | 0.9508 | intergenic | |
| rs17128977 | 12 | 41377851 | 4 | 0 | 0 | T | C | 1.4564 | 0.4248 | 0.0006075 | +++?+ | 0 | 0.5005 | intron | CNTN1 |
| rs147743438 | 18 | 14278172 | 3 | 0 | 0 | A | G | 3.6131 | 1.0539 | 0.0006075 | +++?? | 60 | 0.08192 | intergenic | |
| rs77101935 | 5 | 152449259 | 4 | 0 | 0 | T | C | 1.3223 | 0.3857 | 0.0006076 | +++?+ | 0 | 0.8513 | intergenic | |
| rs7642143 | 3 | 61096082 | 4 | 0 | 0 | A | G | -3.2113 | 0.9368 | 0.0006078 | ---?- | 0 | 0.7112 | intron | FHIT |

| | | | | | | | | | | | | | | | |
|---|---|---|---|---|---|---|---|---|---|---|---|---|---|---|---|
| rs17133288 | 7 | 49473723 | 4 | 0 | 0 | T | C | -0.9687 | 0.2826 | 0.0006078 | ---?+ | 26.2 | 0.2543 | intergenic | |
| rs73143047 | 7 | 78497411 | 4 | 0 | 0 | T | C | 2.1642 | 0.6313 | 0.0006079 | +++?+ | 0 | 0.4849 | intron | MAGI2 |
| rs74927567 | 6 | 32597064 | 5 | 0 | 0 | A | G | 0.7141 | 0.2083 | 0.0006079 | +++++ | 0 | 0.6846 | intergenic | |
| rs116957931 | 16 | 47804858 | 4 | 0 | 0 | A | G | -3.0055 | 0.8767 | 0.0006079 | ---?- | 0 | 0.6625 | intergenic | |
| rs12079395 | 1 | 203687572 | 4 | 0 | 0 | A | G | -1.1085 | 0.3234 | 0.0006083 | ---?- | 0 | 0.9064 | intron | ATP2B4 |
| rs56136802 | 2 | 170504700 | 3 | 0 | 0 | A | T | 1.2628 | 0.3684 | 0.0006085 | +++?? | 0 | 0.8688 | intron | CCDC173 |
| rs483846 | 6 | 12626572 | 4 | 0 | 0 | A | G | -0.9405 | 0.2744 | 0.0006086 | ---?- | 0 | 0.8828 | intergenic | |
| rs13256238 | 8 | 65060422 | 4 | 0 | 0 | A | C | -0.8299 | 0.2421 | 0.0006087 | ---?- | 0 | 0.9496 | intergenic | |
| rs1648834 | 15 | 42134371 | 5 | 0 | 0 | C | G | -0.9402 | 0.2743 | 0.0006088 | ----- | 0 | 0.5943 | intron | JMJD7-PLA2G4B,PLA2G4B |
| rs12989026 | 2 | 116587692 | 4 | 0 | 0 | A | G | 1.2961 | 0.3781 | 0.0006089 | +++?+ | 10.1 | 0.3424 | intron | DPP10 |
| rs117307218 | 8 | 131424594 | 3 | 0 | 0 | T | C | 4.8804 | 1.4239 | 0.0006093 | +++?? | 0 | 0.7928 | intron | ASAP1 |
| rs13261202 | 8 | 98208085 | 5 | 0 | 0 | T | C | -1.4995 | 0.4375 | 0.0006096 | ----- | 9.4 | 0.3525 | intergenic |  |
| rs1281286 | 1 | 181926744 | 3 | 0 | 0 | T | C | -0.8849 | 0.2582 | 0.0006098 | ---?? | 0 | 0.5829 | intergenic | |
| rs77184128 | 6 | 32672135 | 5 | 0 | 0 | T | C | 1.1115 | 0.3243 | 0.00061 | +++++ | 23 | 0.2678 | intergenic | |
| rs138265770 | 15 | 86379985 | 4 | 0 | 0 | T | G | -1.9569 | 0.571 | 0.0006101 | ---?- | 0 | 0.9951 | intergenic | |
| rs201466 | 7 | 101742140 | 4 | 0 | 0 | T | C | 0.8331 | 0.2431 | 0.0006102 | +++?+ | 0 | 0.7794 | intron | CUX1 |
| rs12424677 | 12 | 27790064 | 4 | 0 | 0 | T | C | -0.9283 | 0.2709 | 0.0006105 | ---?- | 1.8 | 0.3833 | intron | PPFIBP1 |
| rs148208644 | 8 | 65018192 | 3 | 0 | 0 | T | C | -0.8731 | 0.2548 | 0.0006105 | ---?? | 0 | 0.8385 | intergenic | |
| 3:29511155 | 3 | 29511155 | 2 | 0 | 0 | CAT | C | -1.9032 | 0.5554 | 0.0006105 | ?--?? | 66.2 | 0.08522 | | |
| rs2317914 | 13 | 112827235 | 4 | 0 | 0 | T | C | 0.9463 | 0.2761 | 0.0006106 | +++?+ | 13 | 0.3273 | intergenic | |
| rs10836671 | 11 | 37056381 | 4 | 0 | 0 | A | C | 0.9042 | 0.2638 | 0.0006106 | +++?+ | 0 | 0.4483 | intergenic | |
| rs1937833 | 10 | 122505327 | 4 | 0 | 0 | A | G | 1.0007 | 0.292 | 0.0006107 | +++?+ | 31.6 | 0.2229 | intergenic | |
| rs191534655 | 5 | 51618874 | 2 | 0 | 0 | T | C | -5.3819 | 1.5705 | 0.0006107 | --??? | 40.5 | 0.1949 | intergenic | |
| rs71504796 | 9 | 21586484 | 4 | 0 | 0 | T | C | -1.5934 | 0.465 | 0.0006109 | ---?- | 0 | 0.9464 | intergenic | |
| rs78845406 | 18 | 8302561 | 4 | 0 | 0 | A | G | -1.6415 | 0.479 | 0.000611 | ---?- | 15.9 | 0.3122 | intron | PTPRM |
| rs140816862 | 9 | 94076353 | 3 | 0 | 0 | A | G | 3.7288 | 1.0882 | 0.0006115 | +++?? | 49.9 | 0.136 | intron | AUH |
| rs4915962 | 1 | 64751610 | 4 | 0 | 0 | A | G | 1.2049 | 0.3517 | 0.0006117 | +++?+ | 0 | 0.5088 | intergenic | |
| rs2675085 | 2 | 184012311 | 3 | 0 | 0 | T | C | 1.219 | 0.3558 | 0.0006118 | +++?? | 0 | 0.7915 | intron | NUP35 |
| rs10239613 | 7 | 29827106 | 3 | 0 | 0 | A | G | -0.8916 | 0.2602 | 0.0006119 | ---?? | 0 | 0.9642 | intergenic | |
| rs55870311 | 8 | 64936334 | 4 | 0 | 0 | A | G | 0.9171 | 0.2677 | 0.000612 | +++?+ | 0 | 0.8229 | intergenic | |
| rs3850664 | 7 | 49465665 | 4 | 0 | 0 | A | G | 0.9543 | 0.2785 | 0.0006121 | +++?- | 23.9 | 0.2676 | intergenic | |
| rs6489181 | 12 | 128051831 | 4 | 0 | 0 | A | G | 1.9511 | 0.5695 | 0.0006122 | +++?+ | 5 | 0.3681 | intergenic | |
| rs7364556 | 1 | 185401229 | 4 | 0 | 0 | T | C | 0.9006 | 0.2629 | 0.0006123 | +++?+ | 0 | 0.9046 | intergenic | |
| rs7761954 | 6 | 137742990 | 4 | 0 | 0 | A | G | 1.1385 | 0.3323 | 0.0006123 | +++?+ | 0 | 0.7164 | intergenic | |
| rs17128930 | 12 | 41305737 | 4 | 0 | 0 | A | G | -1.7001 | 0.4962 | 0.0006123 | ---?+ | 0 | 0.798 | intron | CNTN1 |
| rs75862345 | 4 | 128752634 | 4 | 0 | 0 | A | G | 3.2793 | 0.9572 | 0.0006123 | +++?+ | 0 | 0.7124 | intron | HSPA4L |
| rs1637257 | 7 | 101740606 | 4 | 0 | 0 | T | G | -0.8324 | 0.243 | 0.0006124 | ---?- | 0 | 0.7811 | intron | CUX1 |
| 13:60030690 | 13 | 60030690 | 3 | 0 | 0 | CT | C | 0.9112 | 0.2659 | 0.0006124 | +++?? | 44.9 | 0.163 | | |
| rs10268748 | 7 | 29827111 | 3 | 0 | 0 | A | G | 0.8916 | 0.2602 | 0.0006129 | +++?? | 0 | 0.9641 | intergenic | |
| rs183885762 | 11 | 65210838 | 5 | 0 | 0 | T | G | 3.4325 | 1.0019 | 0.0006129 | +++++ | 0 | 0.8548 | upstream-gene | |
| 1:87502816 | 1 | 87502816 | 3 | 0 | 0 | CT | C | -1.0367 | 0.3026 | 0.0006129 | ---?? | 0 | 0.3746 | | |
| rs75307378 | 1 | 192463324 | 4 | 0 | 0 | A | T | 1.151 | 0.336 | 0.000613 | +++?- | 0 | 0.4056 | intergenic | |
| rs142650345 | 16 | 47382902 | 3 | 0 | 0 | A | T | 2.9796 | 0.8698 | 0.0006133 | +++?? | 0 | 0.63 | intron | ITFG1 |
| rs11018388 | 10 | 129680772 | 4 | 0 | 0 | A | G | -0.8558 | 0.2498 | 0.0006137 | ---?- | 0 | 0.718 | intron | CLRN3 |
| rs12777745 | 10 | 131914341 | 3 | 0 | 0 | A | C | -0.8636 | 0.2521 | 0.0006139 | ---?? | 0 | 0.7008 | intergenic | |
| rs113370581 | 12 | 41305902 | 4 | 0 | 0 | A | G | 1.6998 | 0.4962 | 0.0006141 | +++?- | 0 | 0.798 | intron | CNTN1 |
| rs13380794 | 16 | 7205228 | 4 | 0 | 0 | A | C | -1.0453 | 0.3052 | 0.0006142 | ---?- | 0 | 0.9574 | intron | RBFOX1 |
| 5:165010817 | 5 | 165010817 | 3 | 0 | 0 | G | GA | -1.4873 | 0.4342 | 0.0006144 | ---?? | 0 | 0.9417 | | |
| rs16943335 | 17 | 57059257 | 5 | 0 | 0 | A | G | 2.3204 | 0.6775 | 0.0006145 | +++++ | 0 | 0.8679 | 3-prime-UTR | PPM1E |
| rs34659453 | 11 | 12664053 | 4 | 0 | 0 | T | C | 1.2602 | 0.3679 | 0.0006145 | +++?- | 53.6 | 0.09116 | intergenic | |
| rs1621247 | 1 | 87570398 | 4 | 0 | 0 | A | G | -1.0267 | 0.2998 | 0.0006146 | ---?- | 9 | 0.3479 | 3-prime-UTR | HS2ST1 |
| 8:113514901 | 8 | 113514901 | 3 | 0 | 0 | A | AT | -0.9216 | 0.2691 | 0.0006146 | ---?? | 0 | 0.4926 | | |
| rs440202 | 21 | 31402735 | 4 | 0 | 0 | A | G | -0.9754 | 0.2848 | 0.0006148 | ---?+ | 7 | 0.3583 | intergenic | |
| rs12936438 | 17 | 54085778 | 3 | 0 | 0 | T | C | -0.9248 | 0.27 | 0.0006148 | ---?? | 21 | 0.2821 | intergenic | |
| 12:103259861 | 12 | 103259861 | 3 | 0 | 0 | G | GAAAA | -0.8695 | 0.2539 | 0.0006148 | ---?? | 0 | 0.5405 | | |
| rs7910200 | 10 | 15981388 | 4 | 0 | 0 | T | C | 0.8712 | 0.2544 | 0.0006149 | +++?+ | 35.2 | 0.2009 | intergenic | |
| 10:131914340 | 10 | 131914340 | 3 | 0 | 0 | A | AG | -0.8636 | 0.2522 | 0.0006151 | ---?? | 0 | 0.7009 | | |
| rs2251799 | 12 | 103249952 | 4 | 0 | 0 | T | C | 0.8283 | 0.2419 | 0.0006152 | +++?+ | 0 | 0.6679 | intron | PAH |
| rs7174762 | 15 | 85533539 | 4 | 0 | 0 | A | G | 0.8465 | 0.2472 | 0.0006152 | +++?- | 54.1 | 0.08844 | intron | PDE8A |

| | | | | | | | | | | | | | | | |
|---|---|---|---|---|---|---|---|---|---|---|---|---|---|---|---|
| rs56833097 | 2 | 185428312 | 4 | 0 | 0 | A | G | 0.9428 | 0.2753 | 0.0006152 | +++?+ | 0 | 0.8028 | intergenic | |
| rs16856425 | 3 | 171233756 | 4 | 0 | 0 | A | G | 2.721 | 0.7945 | 0.0006153 | +++?+ | 0 | 0.7203 | intergenic | |
| rs2294489 | 1 | 1664124 | 5 | 0 | 0 | T | C | -1.4403 | 0.4206 | 0.0006157 | ----- | 30.4 | 0.2188 | intron | SLC35E2 |
| rs62397281 | 5 | 179023990 | 4 | 0 | 0 | A | G | 1.6185 | 0.4726 | 0.0006159 | +++?- | 0 | 0.4142 | intron | RUFY1 |
| rs4659226 | 1 | 120224972 | 4 | 0 | 0 | A | G | -0.8454 | 0.2469 | 0.000616 | ---?- | 0 | 0.8052 | intergenic | |
| rs1734879 | 7 | 101740610 | 4 | 0 | 0 | A | C | 0.8318 | 0.2429 | 0.0006161 | +++?+ | 0 | 0.7805 | intron | CUX1 |
| rs7603532 | 2 | 170850214 | 4 | 0 | 1 | A | T | -0.9976 | 0.2913 | 0.0006162 | ---?- | 48.3 | 0.1218 | intron | UBR3 |
| rs72800702 | 16 | 47796888 | 4 | 0 | 0 | T | C | -3.0006 | 0.8763 | 0.0006163 | ---?- | 0 | 0.6705 | intergenic | |
| rs116807308 | 3 | 143064368 | 4 | 0 | 0 | T | C | -2.0003 | 0.5842 | 0.0006165 | ---?- | 0 | 0.608 | intron | SLC9A9,SLC9A9-AS1 |
| rs2573614 | 15 | 100458481 | 4 | 0 | 0 | C | G | -0.8454 | 0.2469 | 0.0006168 | -+-?- | 37.7 | 0.186 | intergenic | |
| rs9341664 | 6 | 78085186 | 4 | 0 | 0 | A | C | -0.8859 | 0.2587 | 0.0006168 | ---?- | 0 | 0.4289 | intergenic | |
| rs10742018 | 11 | 23967985 | 4 | 0 | 0 | A | G | -1.236 | 0.361 | 0.0006168 | ---?- | 0 | 0.7219 | intergenic | |
| rs182910263 | 20 | 5830377 | 3 | 0 | 0 | A | G | -1.8348 | 0.5358 | 0.0006168 | ---?? | 0 | 0.5032 | intron | C20orf196 |
| rs9570136 | 13 | 60039063 | 4 | 0 | 0 | A | G | 0.892 | 0.2605 | 0.0006169 | +++?+ | 0 | 0.6331 | intergenic | |
| rs11165951 | 1 | 98593653 | 4 | 0 | 0 | A | C | 1.343 | 0.3922 | 0.0006169 | +++?+ | 18.5 | 0.2979 | intergenic | |
| rs73592522 | 10 | 21651292 | 2 | 0 | 0 | A | T | 7.9281 | 2.3155 | 0.0006174 | +?+?? | 0 | 0.5776 | intergenic | |
| rs79990351 | 10 | 86679556 | 4 | 0 | 0 | T | C | -1.7621 | 0.5146 | 0.0006174 | ---?- | 0 | 0.7866 | intergenic | |
| rs138124058 | 17 | 54080610 | 3 | 0 | 0 | T | C | -0.9478 | 0.2768 | 0.0006174 | ---?? | 24.6 | 0.2656 | intergenic | |
| rs440730 | 21 | 31401798 | 4 | 0 | 0 | A | G | 0.9715 | 0.2837 | 0.0006175 | +++?- | 9 | 0.3483 | intergenic | |
| rs76164043 | 12 | 64922746 | 4 | 0 | 0 | C | G | -1.6997 | 0.4964 | 0.0006175 | ---?- | 12.8 | 0.3288 | intergenic | |
| rs1616090 | 16 | 12136323 | 4 | 0 | 0 | T | C | 0.9068 | 0.2648 | 0.0006178 | -++?+ | 56.4 | 0.07555 | intron | SNX29 |
| rs10885602 | 10 | 116506525 | 4 | 0 | 0 | A | G | -1.3555 | 0.3959 | 0.0006178 | ---?- | 0 | 0.8566 | intergenic | |
| rs6540063 | 16 | 87626215 | 4 | 0 | 0 | A | G | 0.8686 | 0.2537 | 0.0006179 | +++?+ | 44.5 | 0.1447 | intergenic | |
| rs12586065 | 13 | 45518586 | 4 | 0 | 0 | A | G | 2.1771 | 0.6359 | 0.0006179 | +++?- | 37.8 | 0.1851 | intron | NUFIP1 |
| rs17117759 | 1 | 98600618 | 4 | 0 | 0 | T | C | -1.3425 | 0.3921 | 0.000618 | ---?- | 19.9 | 0.2904 | intergenic | |
| rs149011071 | 1 | 213632676 | 3 | 0 | 0 | T | G | 3.9356 | 1.1496 | 0.000618 | ?++?+ | 59 | 0.0875 | intergenic | |
| rs115951138 | 4 | 141055791 | 4 | 0 | 0 | A | G | -2.8638 | 0.8365 | 0.0006182 | ---?- | 0 | 0.9924 | intron | MAML3 |
| rs1197670 | 15 | 42138321 | 5 | 0 | 0 | A | G | 0.939 | 0.2743 | 0.0006183 | +++++ | 0 | 0.5879 | intron | JMJD7-PLA2G4B,PLA2G4B |
| rs12577722 | 11 | 12641918 | 4 | 0 | 0 | A | G | 1.0369 | 0.3029 | 0.0006184 | +++?+ | 0 | 0.8037 | intergenic | |
| 1:87395788 | 1 | 87395788 | 3 | 0 | 0 | A | AT | -1.0623 | 0.3103 | 0.0006185 | ---?? | 0 | 0.7338 | | |
| rs7978923 | 12 | 111780259 | 5 | 0 | 0 | T | G | 0.8262 | 0.2413 | 0.0006186 | +-+++ | 35 | 0.1879 | intron | CUX2 |
| rs72699015 | 1 | 112985836 | 4 | 0 | 0 | A | G | 1.537 | 0.449 | 0.0006186 | +++?+ | 21.2 | 0.2832 | intron | CTTNBP2NL |
| rs76582422 | 7 | 159051849 | 4 | 0 | 0 | A | G | 1.7914 | 0.5233 | 0.0006187 | +++?+ | 25.2 | 0.2601 | intergenic | |
| rs765400 | 9 | 107923457 | 4 | 0 | 0 | T | G | -1.2408 | 0.3625 | 0.0006188 | ---?- | 0 | 0.6637 | intergenic | |
| rs1342634 | 6 | 78083971 | 4 | 0 | 0 | T | C | 0.8858 | 0.2588 | 0.0006189 | +++?+ | 0 | 0.4294 | intergenic | |
| rs56249221 | 1 | 90263598 | 3 | 0 | 0 | A | G | -0.8636 | 0.2523 | 0.0006189 | ---?? | 0 | 0.5449 | intergenic | |
| rs7685300 | 4 | 41335413 | 4 | 0 | 0 | A | G | -1.3755 | 0.4018 | 0.0006191 | ---?+ | 0 | 0.4485 | intergenic | |
| rs80327850 | 2 | 150828285 | 4 | 0 | 0 | A | G | 1.0249 | 0.2994 | 0.0006191 | +++?+ | 0 | 0.8294 | intergenic | |
| rs17100555 | 14 | 63408564 | 4 | 0 | 0 | A | G | -2.113 | 0.6173 | 0.0006192 | ---?- | 47.1 | 0.1289 | intron | KCNH5 |
| rs4999618 | 1 | 227135503 | 3 | 0 | 0 | A | T | -0.9274 | 0.2709 | 0.0006195 | ---?? | 49.9 | 0.1358 | intron | ADCK3 |
| rs4793923 | 17 | 55975204 | 3 | 0 | 0 | A | T | -1.3581 | 0.3968 | 0.0006196 | ---?? | 60.7 | 0.07844 | intron | CUEDC1 |
| 4:129699995 | 4 | 129699995 | 3 | 0 | 0 | CA | C | 1.3024 | 0.3805 | 0.0006197 | +++?? | 34.5 | 0.2171 | | |
| rs7071570 | 10 | 28214488 | 4 | 0 | 0 | T | C | -1.6619 | 0.4855 | 0.0006198 | ---?- | 22.3 | 0.2769 | intron | ARMC4 |
| rs3024987 | 6 | 43740840 | 4 | 0 | 0 | T | C | -1.2513 | 0.3656 | 0.0006199 | ---?- | 0 | 0.8001 | intron | VEGFA |
| rs28396409 | 9 | 44267749 | 3 | 0 | 0 | T | C | -1.7465 | 0.5103 | 0.0006199 | ---?? | 0 | 0.8473 | intergenic | |
| rs115064525 | 4 | 133738635 | 3 | 0 | 0 | T | C | -4.2503 | 1.2418 | 0.0006199 | ?--?- | 0 | 0.7196 | intergenic | |
| rs9402894 | 6 | 137741905 | 4 | 0 | 0 | A | T | -1.1359 | 0.3319 | 0.0006201 | ---?- | 0 | 0.7172 | intergenic | |
| rs2161724 | 16 | 84240607 | 4 | 0 | 0 | A | G | -0.8291 | 0.2422 | 0.0006202 | ---?- | 0 | 0.5558 | intergenic | |
| 2:60602579 | 2 | 60602579 | 2 | 0 | 0 | G | GCACAG | 5.0596 | 1.4783 | 0.0006204 | ?++?? | 15.8 | 0.2757 | | |
| rs142423992 | 17 | 57059069 | 5 | 0 | 0 | A | G | 2.3355 | 0.6824 | 0.0006205 | +++++ | 0 | 0.8413 | 3-prime-UTR | PPM1E |
| rs76464068 | 18 | 52667092 | 4 | 0 | 0 | A | G | 2.0816 | 0.6082 | 0.0006206 | +++?+ | 0 | 0.8493 | intergenic | |
| rs817392 | 1 | 85612739 | 4 | 0 | 0 | A | G | 1.5462 | 0.4518 | 0.0006207 | +++?+ | 0 | 0.983 | intergenic | |
| rs10944332 | 6 | 88492526 | 4 | 0 | 0 | A | G | 1.3464 | 0.3934 | 0.0006207 | +++?+ | 0 | 0.8377 | intergenic | |
| rs74473416 | 15 | 92199454 | 4 | 0 | 0 | T | C | -1.3281 | 0.3881 | 0.0006207 | ---?- | 0 | 0.7309 | intergenic | |
| 3:45481951 | 3 | 45481951 | 3 | 0 | 0 | T | TG | -2.5488 | 0.7447 | 0.0006207 | ---?? | 0 | 0.5219 | | |
| rs201461 | 7 | 101741625 | 4 | 0 | 0 | T | C | -0.8313 | 0.2429 | 0.0006208 | ---?- | 0 | 0.7794 | intron | CUX1 |
| rs111770053 | 9 | 79462917 | 4 | 0 | 0 | T | C | -2.0503 | 0.5991 | 0.0006209 | ---?- | 0 | 0.6249 | intron | PRUNE2 |
| rs17084873 | 5 | 94861118 | 4 | 0 | 0 | C | G | 0.966 | 0.2823 | 0.000621 | ++++? | 44.2 | 0.1461 | missense | TTC37 |
| rs146911911 | 12 | 42889835 | 3 | 0 | 0 | T | G | -5.1595 | 1.5076 | 0.000621 | --??- | 0 | 0.7839 | intron | PRICKLE1 |

| | | | | | | | | | | | | | | | |
|---|---|---|---|---|---|---|---|---|---|---|---|---|---|---|---|
| rs817393 | 1 | 85612760 | 4 | 0 | 0 | A | G | 1.5461 | 0.4518 | 0.0006212 | +++?+ | 0 | 0.983 | intergenic | |
| rs1500643 | 15 | 98042731 | 4 | 0 | 0 | A | G | 1.1175 | 0.3265 | 0.0006213 | +++?- | 0 | 0.678 | intergenic | |
| rs945500 | 1 | 18753139 | 4 | 0 | 0 | T | G | -1.0409 | 0.3042 | 0.0006214 | ---?- | 49.4 | 0.1151 | intergenic | |
| rs10873689 | 1 | 85605914 | 4 | 0 | 0 | A | T | -1.2396 | 0.3622 | 0.0006214 | ---?- | 0 | 0.9931 | intergenic | |
| rs62419111 | 6 | 63326592 | 4 | 0 | 0 | T | C | -2.4175 | 0.7064 | 0.0006214 | ---?- | 0 | 0.5057 | intergenic | |
| rs77549377 | 12 | 91510355 | 4 | 0 | 0 | A | G | -2.4956 | 0.7293 | 0.0006214 | ---?+ | 59.2 | 0.06152 | upstream-gene | |
| rs7858165 | 9 | 30744605 | 4 | 0 | 0 | A | G | 3.4928 | 1.0207 | 0.0006217 | +++?+ | 0 | 0.4225 | intergenic | |
| rs13132853 | 4 | 38680015 | 5 | 0 | 0 | A | G | 0.8066 | 0.2357 | 0.000622 | +++++ | 20.2 | 0.286 | intron | KLF3 |
| rs13150928 | 4 | 10052961 | 5 | 0 | 0 | T | G | 0.7209 | 0.2107 | 0.0006222 | ++++- | 13.9 | 0.3254 | intergenic | |
| rs12545274 | 8 | 113222995 | 4 | 0 | 0 | A | G | 0.9741 | 0.2847 | 0.0006223 | +++?+ | 0 | 0.9695 | intergenic | |
| rs41495354 | 3 | 61097010 | 4 | 0 | 0 | T | C | -3.2045 | 0.9365 | 0.0006224 | ---?- | 0 | 0.7058 | intron | FHIT |
| rs78970277 | 17 | 33218289 | 2 | 0 | 0 | T | C | -5.4405 | 1.59 | 0.0006224 | ?--?? | 0 | 0.4228 | intergenic | |
| rs10860346 | 12 | 98973753 | 4 | 0 | 0 | T | C | -0.9783 | 0.2859 | 0.0006226 | ---?- | 0 | 0.9697 | intergenic | |
| rs73839217 | 3 | 61097771 | 4 | 0 | 0 | A | G | -3.2041 | 0.9364 | 0.0006226 | ---?- | 0 | 0.707 | intron | FHIT |
| rs114155199 | 3 | 143042850 | 4 | 0 | 0 | T | G | -1.9975 | 0.5838 | 0.0006227 | ---?- | 0 | 0.6161 | intron | SLC9A9 |
| 17:31967792 | 17 | 31967792 | 3 | 0 | 0 | CTA | C | 1.1797 | 0.3448 | 0.0006227 | +++?? | 18.1 | 0.295 | | |
| rs1158895 | 8 | 113348699 | 4 | 0 | 0 | A | G | -0.9747 | 0.2849 | 0.0006228 | ---?- | 0 | 0.7752 | intron | CSMD3 |
| rs201459 | 7 | 101741502 | 4 | 0 | 0 | A | G | 0.8309 | 0.2429 | 0.0006229 | +++?+ | 0 | 0.7793 | intron | CUX1 |
| rs62017092 | 16 | 7204441 | 4 | 0 | 0 | A | G | -1.0406 | 0.3041 | 0.000623 | ---?- | 0 | 0.9578 | intron | RBFOX1 |
| rs17878936 | 22 | 29132338 | 4 | 0 | 0 | T | C | 2.2579 | 0.66 | 0.0006231 | +++?+ | 0 | 0.8178 | intron | CHEK2 |
| rs817399 | 1 | 85613551 | 4 | 0 | 0 | A | G | 1.5462 | 0.4519 | 0.0006232 | +++?+ | 0 | 0.9827 | intergenic | |
| rs200520887 | 1 | 90263430 | 3 | 0 | 0 | A | G | 0.8627 | 0.2522 | 0.0006232 | +++?? | 0 | 0.4242 | intergenic | |
| 9:30645845 | 9 | 30645845 | 3 | 0 | 0 | G | GTGTTG | -2.4026 | 0.7022 | 0.0006232 | ---?? | 58.7 | 0.08903 | | |
| rs7669153 | 4 | 41335420 | 4 | 0 | 0 | T | G | 1.3743 | 0.4017 | 0.0006235 | +++?- | 0 | 0.4491 | intergenic | |
| rs61109891 | 15 | 70305117 | 4 | 0 | 0 | T | C | -1.7102 | 0.4999 | 0.0006235 | ---?- | 0 | 0.612 | intergenic | |
| rs113733553 | 7 | 107331959 | 3 | 0 | 0 | T | C | 0.9266 | 0.2708 | 0.0006235 | +++?? | 0 | 0.7107 | intron | SLC26A4 |
| rs1722381 | 12 | 103247836 | 4 | 0 | 0 | A | G | -0.8372 | 0.2447 | 0.0006237 | ---?- | 0 | 0.659 | intron | PAH |
| rs112150089 | 9 | 90803949 | 3 | 0 | 0 | A | G | 3.1986 | 0.935 | 0.0006237 | +-+?? | 32.1 | 0.2294 | intergenic | |
| 4:188131892 | 4 | 188131892 | 2 | 0 | 0 | T | C | -6.9728 | 2.0383 | 0.000624 | -???- | 28.3 | 0.2375 | intergenic | |
| rs817394 | 1 | 85613163 | 4 | 0 | 0 | A | C | -1.5456 | 0.4518 | 0.0006241 | ---?- | 0 | 0.9831 | intergenic | |
| rs139790369 | 3 | 42540043 | 4 | 0 | 0 | A | G | -2.9351 | 0.858 | 0.0006244 | ---?- | 0 | 0.8599 | intron | VIPR1 |
| rs635798 | 11 | 125818297 | 5 | 0 | 0 | T | C | -0.7703 | 0.2252 | 0.0006245 | ----- | 0 | 0.5089 | synonymous | LOC338667 |
| rs876055 | 2 | 170731122 | 4 | 0 | 1 | A | T | 0.9935 | 0.2904 | 0.0006245 | +++?+ | 47.7 | 0.125 | intron | UBR3 |
| rs6700749 | 1 | 85739685 | 5 | 0 | 0 | A | G | -1.6174 | 0.4729 | 0.000625 | ----- | 0 | 0.4083 | intron | BCL10 |
| rs79438518 | 9 | 18625666 | 4 | 0 | 0 | A | T | 4.4547 | 1.3023 | 0.000625 | +++?+ | 48.2 | 0.122 | intron | ADAMTSL1 |
| 6:151755082 | 6 | 151755082 | 3 | 0 | 0 | CCG | C | 1.5105 | 0.4416 | 0.000625 | +++?? | 7 | 0.3413 | | |
| rs2118812 | 3 | 61098350 | 4 | 0 | 0 | A | G | -3.203 | 0.9364 | 0.0006251 | ---?- | 0 | 0.7066 | intron | FHIT |
| rs6692579 | 1 | 187878097 | 4 | 0 | 0 | A | G | -0.9971 | 0.2915 | 0.0006251 | ---?- | 0 | 0.6218 | intergenic | |
| rs61741210 | 17 | 56386732 | 5 | 0 | 0 | T | C | -2.0857 | 0.6098 | 0.0006251 | ----+ | 21 | 0.281 | missense | BZRAP1 |
| 6:49558990 | 6 | 49558990 | 3 | 0 | 0 | T | TAAAC | 1.2298 | 0.3595 | 0.0006252 | +++?? | 3.6 | 0.3543 | | |
| rs8096634 | 18 | 13880125 | 3 | 0 | 0 | A | G | 1.0366 | 0.3031 | 0.0006253 | +++?? | 52 | 0.1246 | downstream-gene | |
| rs28494517 | 9 | 44271050 | 3 | 0 | 0 | T | C | 1.7411 | 0.509 | 0.0006254 | +++?? | 0 | 0.8521 | intergenic | |
| rs78890790 | 1 | 98601644 | 4 | 0 | 0 | A | G | 1.3409 | 0.392 | 0.0006255 | +++?+ | 20 | 0.2897 | intergenic | |
| rs191368549 | 12 | 14858866 | 3 | 0 | 0 | T | G | -1.8257 | 0.5338 | 0.0006255 | ---?? | 15.4 | 0.3065 | intergenic | |
| rs76033224 | 10 | 86681207 | 4 | 0 | 0 | A | G | -1.7583 | 0.5141 | 0.0006256 | ---?- | 0 | 0.7882 | intergenic | |
| rs113479363 | 6 | 8927693 | 4 | 0 | 0 | T | C | -2.5637 | 0.7495 | 0.0006256 | ---?- | 56.6 | 0.0749 | intergenic | |
| rs201458 | 7 | 101741439 | 4 | 0 | 0 | T | C | -0.8304 | 0.2428 | 0.0006257 | ---?- | 0 | 0.779 | intron | CUX1 |
| rs117192952 | 20 | 22591941 | 3 | 0 | 0 | T | C | -3.9942 | 1.1678 | 0.0006257 | ---?? | 61.6 | 0.0738 | intergenic | |
| rs35662477 | 12 | 112313208 | 5 | 0 | 0 | A | T | -1.2817 | 0.3748 | 0.0006261 | -+--- | 48.5 | 0.1006 | intron | MAPKAPK5 |
| rs10840113 | 11 | 8719615 | 5 | 0 | 0 | C | G | -0.7899 | 0.231 | 0.0006262 | ----+ | 0 | 0.8327 | intron | ST5 |
| rs10239605 | 7 | 29827097 | 3 | 0 | 0 | A | G | -0.8902 | 0.2603 | 0.0006263 | ---?? | 0 | 0.963 | intergenic | |
| rs11737526 | 4 | 189998281 | 4 | 0 | 0 | T | C | -0.8719 | 0.2549 | 0.0006263 | ---?- | 32.7 | 0.2159 | intergenic | |
| rs201457 | 7 | 101741338 | 4 | 0 | 0 | A | C | 0.8303 | 0.2428 | 0.0006264 | +++?+ | 0 | 0.7791 | intron | CUX1 |
| 22:28843233 | 22 | 28843233 | 3 | 0 | 0 | T | TA | 2.4676 | 0.7216 | 0.0006266 | +++?? | 0 | 0.836 | | |
| rs7792186 | 7 | 82419141 | 4 | 0 | 0 | T | C | 0.878 | 0.2567 | 0.0006268 | +-+?+ | 45.8 | 0.1366 | intron | PCLO |
| rs73096186 | 4 | 12221624 | 4 | 0 | 0 | T | C | 2.7179 | 0.7948 | 0.0006269 | +++?+ | 0 | 0.7075 | intergenic | |
| rs2050188 | 6 | 32339897 | 5 | 0 | 0 | T | C | 0.7242 | 0.2118 | 0.000627 | +++++ | 41.2 | 0.1468 | upstream-gene | |
| rs3843905 | 5 | 165427280 | 4 | 0 | 0 | T | C | 0.8602 | 0.2515 | 0.0006271 | +++?+ | 0 | 0.4619 | intergenic | |
| rs201456 | 7 | 101741059 | 4 | 0 | 0 | C | G | -0.8301 | 0.2428 | 0.0006272 | ---?- | 0 | 0.7791 | intron | CUX1 |

| | | | | | | | | | | | | | | | |
|---|---|---|---|---|---|---|---|---|---|---|---|---|---|---|---|
| rs4668162 | 2 | 170638901 | 4 | 0 | 0 | A | G | 0.8384 | 0.2452 | 0.0006275 | +++?+ | 0 | 0.7553 | intergenic | |
| rs817391 | 1 | 85612615 | 4 | 0 | 0 | A | G | 1.5458 | 0.4521 | 0.0006276 | +++?+ | 0 | 0.983 | intergenic | |
| rs79018332 | 2 | 83951526 | 4 | 0 | 0 | A | G | 3.2264 | 0.9436 | 0.0006276 | +++?+ | 55.7 | 0.0796 | intergenic | |
| 16:52918165 | 16 | 52918165 | 3 | 0 | 0 | T | TGG | -0.8793 | 0.2571 | 0.0006276 | ---?? | 0 | 0.4365 | | |
| 1:87386774 | 1 | 87386774 | 3 | 0 | 0 | T | TG | -1.0588 | 0.3097 | 0.0006276 | ---?? | 0 | 0.7269 | | |
| rs7758287 | 6 | 62042253 | 4 | 0 | 0 | A | C | -0.9879 | 0.2889 | 0.0006277 | ---?- | 0 | 0.569 | intergenic | |
| rs7019540 | 9 | 97598150 | 4 | 0 | 0 | T | C | -1.6982 | 0.4967 | 0.0006278 | ---?- | 0 | 0.9588 | intron | C9orf3 |
| 7:1429784 | 7 | 1429784 | 3 | 0 | 0 | G | GGGAA | 1.158 | 0.3387 | 0.0006279 | +++?? | 52.5 | 0.1218 | | |
| 3:45312379 | 3 | 45312379 | 3 | 0 | 0 | G | GA | 1.3718 | 0.4012 | 0.000628 | +++?? | 51.3 | 0.1285 | | |
| rs2647071 | 6 | 32573910 | 3 | 0 | 0 | T | C | 1.115 | 0.3261 | 0.0006281 | +++?? | 0 | 0.3974 | intergenic | |
| rs34080245 | 9 | 35657063 | 5 | 0 | 0 | T | C | 0.9349 | 0.2734 | 0.0006281 | ++++- | 28.6 | 0.2312 | upstream-gene | |
| rs12005826 | 9 | 16196674 | 4 | 0 | 0 | A | T | 1.2251 | 0.3583 | 0.0006282 | +++?+ | 0 | 0.6282 | intergenic | |
| 3:138741558 | 3 | 138741558 | 3 | 0 | 0 | G | GT | -2.177 | 0.6367 | 0.0006283 | ---?? | 23.9 | 0.2689 | | |
| rs7018185 | 8 | 113475721 | 4 | 0 | 0 | T | G | 0.9042 | 0.2645 | 0.0006284 | +++?+ | 0 | 0.7147 | intron | CSMD3 |
| rs72908425 | 6 | 88493474 | 4 | 0 | 0 | A | G | -2.0017 | 0.5855 | 0.0006286 | ---?- | 0 | 0.8379 | intergenic | |
| rs817398 | 1 | 85613315 | 4 | 0 | 0 | T | C | 1.545 | 0.4519 | 0.0006288 | +++?+ | 0 | 0.986 | intergenic | |
| rs11807935 | 1 | 98595170 | 4 | 0 | 0 | A | T | 1.3409 | 0.3922 | 0.0006289 | +++?+ | 18.6 | 0.2974 | intergenic | |
| rs12077945 | 1 | 203686504 | 4 | 0 | 0 | A | G | -1.1094 | 0.3245 | 0.0006291 | ---?- | 0 | 0.8863 | intron | ATP2B4 |
| rs78378925 | 9 | 139059318 | 4 | 0 | 0 | A | G | 1.7081 | 0.4996 | 0.0006291 | +++?+ | 0 | 0.7061 | intergenic | |
| rs1672465 | 15 | 42134839 | 5 | 0 | 0 | T | C | 0.938 | 0.2744 | 0.0006292 | +++++ | 0 | 0.5916 | intron | JMJD7-PLA2G4B,PLA2G4B |
| 9:30656642 | 9 | 30656642 | 3 | 0 | 0 | A | AT | -2.401 | 0.7023 | 0.0006292 | ---?? | 59 | 0.08708 | | |
| rs1932209 | 6 | 77659682 | 4 | 0 | 0 | A | G | -1.1464 | 0.3353 | 0.0006293 | ---?- | 0 | 0.5545 | intergenic | |
| rs2675099 | 2 | 184006027 | 4 | 0 | 0 | T | C | -1.1717 | 0.3427 | 0.0006293 | ---?- | 0 | 0.8411 | intron | NUP35 |
| rs7531123 | 1 | 18742827 | 4 | 0 | 0 | T | C | -1.0626 | 0.3108 | 0.0006293 | ---?- | 51.7 | 0.1017 | intergenic | |
| rs1648836 | 15 | 42133886 | 5 | 0 | 0 | A | G | -0.9374 | 0.2742 | 0.0006296 | ----- | 0 | 0.5894 | intron | JMJD7-PLA2G4B,PLA2G4B |
| rs74988866 | 3 | 61098835 | 4 | 0 | 0 | A | G | 3.2013 | 0.9365 | 0.0006296 | +++?+ | 0 | 0.7059 | intron | FHIT |
| rs1648835 | 15 | 42134301 | 5 | 0 | 0 | A | C | 0.9374 | 0.2742 | 0.0006299 | +++++ | 0 | 0.5899 | intron | JMJD7-PLA2G4B,PLA2G4B |
| rs79132188 | 10 | 131478225 | 4 | 0 | 0 | A | G | 1.2708 | 0.3718 | 0.0006299 | +++?+ | 0 | 0.8139 | intron | MGMT |
| rs59640908 | 2 | 184064771 | 4 | 0 | 0 | A | G | 1.1737 | 0.3434 | 0.00063 | +++?+ | 0 | 0.6643 | intergenic | |
| rs16943331 | 17 | 57057113 | 5 | 0 | 0 | T | C | 2.3205 | 0.6788 | 0.0006301 | +++++ | 0 | 0.8674 | intron | PPM1E |
| rs139108720 | 18 | 52725597 | 4 | 0 | 0 | A | G | -4.4172 | 1.2922 | 0.0006302 | ---?- | 0 | 0.845 | intergenic | |
| rs146267868 | 3 | 143048744 | 4 | 0 | 0 | T | C | -1.9924 | 0.5829 | 0.0006302 | ---?- | 0 | 0.6064 | intron | SLC9A9 |
| 2:231347412 | 2 | 231347412 | 3 | 0 | 0 | CATAT | C | 1.4966 | 0.4378 | 0.0006302 | +++?? | 43 | 0.173 | | |
| rs11056489 | 12 | 15592643 | 4 | 0 | 0 | A | G | -1.1428 | 0.3343 | 0.0006303 | ---?- | 0 | 0.9205 | intron | PTPRO |
| rs141431628 | 3 | 143053013 | 4 | 0 | 0 | T | C | -1.9923 | 0.5829 | 0.0006303 | ---?- | 0 | 0.6058 | intron | SLC9A9 |
| rs147477070 | 3 | 143053179 | 4 | 0 | 0 | A | T | 1.9923 | 0.5829 | 0.0006303 | +++?+ | 0 | 0.6058 | intron | SLC9A9 |
| rs436709 | 7 | 101878968 | 3 | 0 | 0 | T | C | -0.9043 | 0.2646 | 0.0006304 | ---?? | 0 | 0.6718 | intron | CUX1 |
| rs1002311 | 15 | 65188424 | 4 | 0 | 0 | C | G | 0.9173 | 0.2684 | 0.0006304 | ++++? | 0 | 0.7536 | intergenic | |
| rs7791668 | 7 | 20488976 | 4 | 0 | 0 | A | T | 0.9262 | 0.271 | 0.0006304 | +++?+ | 0 | 0.9105 | intergenic | |
| 2:199095451 | 2 | 199095451 | 3 | 0 | 0 | A | AAAC | -1.4147 | 0.4139 | 0.0006304 | ---?? | 64.3 | 0.06094 | | |
| rs115331795 | 3 | 143051541 | 4 | 0 | 0 | A | G | 1.9923 | 0.5829 | 0.0006305 | +++?+ | 0 | 0.606 | intron | SLC9A9 |
| rs138943876 | 3 | 143044931 | 4 | 0 | 0 | T | C | -1.9922 | 0.5828 | 0.0006305 | ---?- | 0 | 0.6065 | intron | SLC9A9 |
| rs139698202 | 3 | 143049362 | 4 | 0 | 0 | T | C | 1.9923 | 0.5829 | 0.0006305 | +++?+ | 0 | 0.6064 | intron | SLC9A9 |
| rs78528748 | 3 | 143051209 | 4 | 0 | 0 | T | C | 1.9923 | 0.5829 | 0.0006306 | +++?+ | 0 | 0.6061 | intron | SLC9A9 |
| rs2675097 | 2 | 184005357 | 4 | 0 | 0 | T | C | -1.1717 | 0.3428 | 0.0006307 | ---?- | 0 | 0.8422 | intron | NUP35 |
| rs76439962 | 2 | 237854361 | 4 | 0 | 0 | A | G | -2.0248 | 0.5924 | 0.0006308 | ---?- | 0 | 0.8241 | intergenic | |
| rs1918314 | 15 | 42026764 | 5 | 0 | 0 | T | C | -0.9294 | 0.2719 | 0.0006309 | ----- | 0 | 0.5887 | synonymous | MGA |
| rs11787146 | 8 | 8538102 | 4 | 0 | 0 | C | G | 0.7492 | 0.2192 | 0.0006312 | ++++? | 0 | 0.4249 | intergenic | |
| rs13268622 | 8 | 64924306 | 4 | 0 | 0 | T | C | -0.915 | 0.2677 | 0.0006312 | ---?- | 0 | 0.8491 | intergenic | |
| rs147383582 | 3 | 143043203 | 4 | 0 | 0 | A | G | -1.9919 | 0.5828 | 0.0006313 | ---?- | 0 | 0.6069 | intron | SLC9A9 |
| rs149939727 | 3 | 143043431 | 4 | 0 | 0 | A | T | -1.9919 | 0.5828 | 0.0006313 | ---?- | 0 | 0.6069 | intron | SLC9A9 |
| rs147062384 | 8 | 65019913 | 3 | 0 | 0 | A | G | -0.8699 | 0.2545 | 0.0006314 | ---?? | 0 | 0.8387 | intergenic | |
| 22:29126347 | 22 | 29126347 | 3 | 0 | 0 | G | GAAAG[illegible] | 2.3289 | 0.6814 | 0.0006315 | +++?? | 0 | 0.7031 | | |
| rs4908596 | 1 | 7127054 | 4 | 0 | 0 | A | T | -0.8761 | 0.2563 | 0.0006317 | ---?- | 40.4 | 0.1692 | intron | CAMTA1 |
| rs6958698 | 7 | 67405642 | 4 | 0 | 0 | A | C | -1.7706 | 0.5181 | 0.0006317 | ---?+ | 3.5 | 0.375 | intergenic | |
| 4:12246895 | 4 | 12246895 | 3 | 0 | 0 | G | GAA | -1.2309 | 0.3602 | 0.0006318 | ---?? | 0 | 0.4161 | | |
| 6:159204277 | 6 | 159204277 | 3 | 0 | 0 | G | GC | -0.8797 | 0.2574 | 0.0006318 | ---?? | 26.7 | 0.2557 | | |
| 9:19010476 | 9 | 19010476 | 3 | 0 | 0 | G | GAA | 1.6451 | 0.4814 | 0.0006318 | +++?? | 58.4 | 0.09056 | | |
| rs10168942 | 2 | 170940845 | 4 | 0 | 1 | T | C | 0.976 | 0.2856 | 0.0006319 | +++?+ | 60.3 | 0.05589 | downstream-gene | |

| | | | | | | | | | | | | | | | |
|---|---|---|---|---|---|---|---|---|---|---|---|---|---|---|---|
| rs73096188 | 4 | 12221739 | 4 | 0 | 0 | C | G | 2.7157 | 0.7946 | 0.000632 | +++?+ | 0 | 0.709 | intergenic | |
| rs7964647 | 12 | 112839366 | 5 | 0 | 0 | A | T | 1.3026 | 0.3812 | 0.0006321 | +-+++ | 56.7 | 0.05566 | downstream-gene | |
| rs35501905 | 4 | 10053431 | 5 | 0 | 0 | T | C | 0.7206 | 0.2108 | 0.0006321 | ++++- | 15.5 | 0.3157 | intergenic | |
| rs10765980 | 11 | 12639979 | 4 | 0 | 0 | T | C | 1.057 | 0.3093 | 0.0006322 | +++?+ | 0 | 0.8058 | intergenic | |
| rs2705759 | 2 | 184005260 | 4 | 0 | 0 | A | G | 1.1716 | 0.3428 | 0.0006323 | +++?+ | 0 | 0.8427 | intron | NUP35 |
| rs7623637 | 3 | 71884247 | 4 | 0 | 0 | C | G | -0.9345 | 0.2735 | 0.0006324 | ---?- | 0 | 0.7107 | intergenic | |
| rs72820086 | 10 | 90321949 | 4 | 0 | 0 | T | C | -2.0044 | 0.5865 | 0.0006324 | ---?- | 0 | 0.8568 | intron | RNLS |
| rs1751433 | 9 | 97527855 | 4 | 0 | 0 | A | G | -1.7951 | 0.5253 | 0.0006325 | ---?- | 0 | 0.8718 | intron | C9orf3 |
| rs3844072 | 8 | 64921157 | 4 | 0 | 0 | T | G | 0.8457 | 0.2475 | 0.0006325 | +++?- | 0 | 0.5025 | intergenic | |
| rs2927707 | 19 | 8370921 | 4 | 0 | 0 | T | C | -0.9157 | 0.268 | 0.0006326 | ---?- | 0 | 0.8146 | intron | CD320 |
| rs13108944 | 4 | 189996319 | 4 | 0 | 0 | A | C | 0.8736 | 0.2557 | 0.0006326 | +++?+ | 36.6 | 0.1928 | intergenic | |
| rs12455810 | 18 | 8375601 | 2 | 0 | 0 | A | G | 7.2467 | 2.1207 | 0.0006328 | ?+??+ | 0 | 0.7098 | intron | PTPRM |
| rs6483998 | 11 | 23966159 | 4 | 0 | 0 | T | C | -1.2316 | 0.3604 | 0.0006331 | ---?- | 0 | 0.7308 | intergenic | |
| rs148140512 | 18 | 68460478 | 3 | 0 | 0 | T | C | -4.6619 | 1.3643 | 0.0006331 | ?--?- | 29.1 | 0.2442 | intergenic | |
| rs10771293 | 12 | 26843918 | 4 | 0 | 0 | T | C | -0.8357 | 0.2446 | 0.0006333 | ---?+ | 0 | 0.4153 | intron | ITPR2 |
| 2:184056407 | 2 | 184056407 | 3 | 0 | 0 | G | GTTT | -1.2011 | 0.3515 | 0.0006333 | ---?? | 0 | 0.6225 | | |
| rs11939585 | 4 | 37016697 | 4 | 0 | 0 | A | G | 0.8519 | 0.2493 | 0.0006334 | +++?- | 42.9 | 0.1542 | intergenic | |
| rs12177624 | 6 | 159201212 | 3 | 0 | 0 | A | C | 0.8736 | 0.2557 | 0.0006337 | +++?? | 8.1 | 0.3367 | intron | EZR |
| rs1648833 | 15 | 42134523 | 5 | 0 | 0 | T | C | -0.9369 | 0.2742 | 0.000634 | ----- | 0 | 0.5897 | intron | JMJD7-PLA2G4B,PLA2G4B |
| rs72890612 | 18 | 33781695 | 3 | 0 | 0 | T | C | -3.6434 | 1.0664 | 0.000634 | ---?? | 0 | 0.6502 | intron | MOCOS |
| rs7978137 | 12 | 98944324 | 4 | 0 | 0 | A | G | 0.9498 | 0.278 | 0.0006341 | +++?+ | 0 | 0.957 | downstream-gene | |
| rs139060682 | 3 | 143062891 | 4 | 0 | 0 | T | C | -1.9914 | 0.5829 | 0.0006341 | ---?- | 0 | 0.6047 | intron | SLC9A9,SLC9A9-AS1 |
| rs6070200 | 20 | 56214292 | 4 | 0 | 0 | T | G | -1.1738 | 0.3436 | 0.0006342 | ---?- | 0 | 0.8456 | intergenic | |
| 9:81793764 | 9 | 81793764 | 3 | 0 | 0 | A | G | -2.83 | 0.8283 | 0.0006343 | ---?? | 0 | 0.8991 | intergenic | |
| rs1536088 | 10 | 131920290 | 4 | 0 | 0 | T | C | 0.8893 | 0.2603 | 0.0006343 | +++?+ | 0 | 0.8662 | intergenic | |
| 6:24056076 | 6 | 24056076 | 3 | 0 | 0 | G | GTGAA | -1.3821 | 0.4045 | 0.0006345 | ---?? | 0 | 0.9765 | | |
| rs11165957 | 1 | 98602415 | 4 | 0 | 0 | A | T | -1.339 | 0.3919 | 0.0006347 | ---?- | 20.3 | 0.288 | intergenic | |
| rs117472323 | 22 | 28739173 | 4 | 0 | 0 | T | C | 2.6404 | 0.7729 | 0.0006348 | +++?+ | 0 | 0.8098 | intron | TTC28 |
| rs10921172 | 1 | 192463190 | 4 | 0 | 0 | A | G | 1.0952 | 0.3206 | 0.0006349 | +++?- | 33.4 | 0.2117 | intergenic | |
| rs1648837 | 15 | 42133885 | 5 | 0 | 0 | T | C | 0.9371 | 0.2743 | 0.000635 | +++++ | 0 | 0.594 | intron | JMJD7-PLA2G4B,PLA2G4B |
| rs148540863 | 18 | 27032484 | 4 | 0 | 0 | T | C | 2.8528 | 0.8351 | 0.0006352 | -++?+ | 37.8 | 0.1851 | intergenic | |
| rs145154133 | 3 | 179950887 | 3 | 0 | 0 | C | G | 4.4739 | 1.3097 | 0.0006353 | +++?? | 0 | 0.8112 | intergenic | |
| rs11227637 | 11 | 66749824 | 4 | 0 | 0 | C | G | 1.6313 | 0.4775 | 0.0006354 | -++?+ | 55.6 | 0.0801 | intergenic | |
| rs112611171 | 9 | 44271408 | 3 | 0 | 0 | A | G | -1.7448 | 0.5108 | 0.0006355 | ---?? | 0 | 0.838 | intergenic | |
| rs142493084 | 17 | 57053300 | 5 | 0 | 0 | A | T | 2.3372 | 0.6842 | 0.0006355 | +++++ | 0 | 0.8396 | intron | PPM1E |
| rs28507575 | 9 | 44263893 | 3 | 0 | 0 | T | C | -1.7429 | 0.5102 | 0.0006356 | ---?? | 0 | 0.8192 | intergenic | |
| rs75182964 | 6 | 109095163 | 3 | 0 | 0 | A | G | -4.1094 | 1.203 | 0.0006357 | -+-?? | 57.8 | 0.09336 | downstream-gene | |
| rs76384559 | 21 | 36817202 | 4 | 0 | 0 | T | G | 1.522 | 0.4456 | 0.0006357 | +++?+ | 0 | 0.5085 | intron | LOC100506403 |
| rs6490042 | 12 | 111719060 | 5 | 0 | 0 | A | T | 0.8568 | 0.2508 | 0.0006358 | +++++ | 0 | 0.471 | intron | CUX2 |
| rs9264492 | 6 | 31232676 | 3 | 0 | 0 | T | G | -0.9696 | 0.2839 | 0.0006358 | ---?? | 23.2 | 0.2719 | downstream-gene | |
| rs11165958 | 1 | 98602431 | 4 | 0 | 0 | T | C | -1.3388 | 0.3919 | 0.0006358 | ---?- | 20.3 | 0.2879 | intergenic | |
| rs35479996 | 3 | 65660688 | 4 | 0 | 0 | A | G | 0.9108 | 0.2666 | 0.0006359 | +++?+ | 46.6 | 0.1318 | intron | MAGI1 |
| 1:59389806 | 1 | 59389806 | 3 | 0 | 0 | A | T | 4.8326 | 1.4148 | 0.000636 | +++?? | 16.3 | 0.3028 | intergenic | |
| rs76582776 | 17 | 54083012 | 3 | 0 | 0 | T | C | 0.9506 | 0.2783 | 0.0006362 | +++?? | 28.3 | 0.2481 | intergenic | |
| 13:90902013 | 13 | 90902013 | 2 | 0 | 0 | A | G | -7.438 | 2.1776 | 0.0006363 | --??? | 0 | 0.5805 | intergenic | |
| rs1392676 | 15 | 98046783 | 4 | 0 | 0 | A | G | -0.8498 | 0.2488 | 0.0006364 | ---?- | 0 | 0.8622 | intergenic | |
| rs7609788 | 3 | 61099264 | 4 | 0 | 0 | A | G | -3.1988 | 0.9365 | 0.0006364 | ---?- | 0 | 0.7064 | intron | FHIT |
| rs9298746 | 9 | 16168116 | 4 | 0 | 0 | A | G | 0.9191 | 0.2691 | 0.0006364 | +++?+ | 0 | 0.8852 | intergenic | |
| rs817401 | 1 | 85615880 | 4 | 0 | 0 | A | G | -1.5441 | 0.4521 | 0.0006365 | ---?- | 0 | 0.9831 | intergenic | |
| rs1648830 | 15 | 42136199 | 5 | 0 | 0 | A | G | 0.9368 | 0.2743 | 0.0006365 | +++++ | 0 | 0.5905 | intron | JMJD7-PLA2G4B,PLA2G4B |
| rs7331020 | 13 | 112826632 | 4 | 0 | 0 | A | G | 0.9432 | 0.2762 | 0.0006365 | +++?+ | 13.3 | 0.3259 | intergenic | |
| rs78130108 | 3 | 118977553 | 4 | 0 | 0 | A | T | -2.0584 | 0.6027 | 0.0006365 | ---?- | 50.1 | 0.1108 | intergenic | |
| rs17150734 | 5 | 123115534 | 4 | 0 | 0 | A | G | -0.978 | 0.2863 | 0.0006366 | ---?- | 0 | 0.4003 | intergenic | |
| rs817390 | 1 | 85612492 | 4 | 0 | 0 | C | G | -1.5452 | 0.4524 | 0.0006369 | ---?- | 0 | 0.983 | intergenic | |
| rs66543347 | 2 | 2760049 | 4 | 0 | 0 | T | C | -1.3173 | 0.3857 | 0.0006373 | ---?- | 0 | 0.942 | intergenic | |
| rs6692079 | 1 | 18742198 | 4 | 0 | 0 | T | C | -1.063 | 0.3113 | 0.0006374 | ---?- | 51.9 | 0.1005 | intergenic | |
| rs6531659 | 4 | 38744030 | 5 | 0 | 0 | A | G | -0.75 | 0.2196 | 0.0006375 | ----- | 0 | 0.8112 | intergenic |  |
| rs75535992 | 5 | 123104498 | 4 | 0 | 0 | T | C | 0.9731 | 0.285 | 0.0006375 | +++?+ | 13.5 | 0.3249 | intergenic | |
| 5:115784451 | 5 | 115784451 | 3 | 0 | 0 | CA | C | 1.9342 | 0.5664 | 0.0006375 | +++?? | 0 | 0.8742 | | |

| | | | | | | | | | | | | | | | |
|---|---|---|---|---|---|---|---|---|---|---|---|---|---|---|---|
| 19:3687557 | 19 | 3687557 | 3 | 0 | 0 | ACA | GCA | -5.661 | 1.6576 | 0.0006376 | ---?? | 0 | 0.9616 | | |
| rs6434018 | 2 | 184044339 | 4 | 0 | 0 | T | C | 1.1539 | 0.3379 | 0.0006379 | +++?+ | 0 | 0.8308 | intergenic | |
| rs2577959 | 15 | 42026004 | 5 | 0 | 0 | A | G | 0.9285 | 0.2719 | 0.0006381 | +++++ | 0 | 0.5907 | intron | MGA |
| rs2675096 | 2 | 184005321 | 4 | 0 | 0 | A | G | -1.1708 | 0.3429 | 0.0006381 | ---?- | 0 | 0.8425 | intron | NUP35 |
| rs28429242 | 9 | 44274255 | 3 | 0 | 0 | T | C | -1.7809 | 0.5215 | 0.0006381 | ---?? | 0 | 0.8377 | intergenic | |
| rs10930377 | 2 | 170638324 | 4 | 0 | 0 | A | G | 0.8322 | 0.2437 | 0.0006382 | +++?+ | 0 | 0.8506 | intergenic | |
| rs3801938 | 7 | 107428573 | 4 | 0 | 0 | C | G | -1.0049 | 0.2943 | 0.0006383 | ---?- | 0 | 0.7415 | intron | SLC26A3 |
| 6:43240634 | 6 | 43240634 | 3 | 0 | 0 | CG | C | -2.3421 | 0.6859 | 0.0006383 | ---?? | 36.2 | 0.2086 | | |
| rs1487302 | 1 | 98595853 | 4 | 0 | 0 | A | C | -1.3392 | 0.3922 | 0.0006384 | ---?- | 18.8 | 0.2966 | intergenic | |
| rs75711592 | 12 | 43118607 | 4 | 0 | 0 | T | C | 1.7582 | 0.5149 | 0.0006384 | +++?+ | 0 | 0.7954 | intergenic | |
| rs139843377 | 16 | 85238926 | 4 | 0 | 0 | T | C | 1.2392 | 0.3629 | 0.0006384 | +++?+ | 0 | 0.6586 | intron | GSE1 |
| rs9600759 | 13 | 77266350 | 4 | 0 | 0 | T | C | 2.5809 | 0.7558 | 0.0006387 | +++?+ | 0 | 0.8004 | intergenic | |
| rs10264600 | 7 | 49462933 | 4 | 0 | 0 | A | G | -0.9457 | 0.2769 | 0.0006389 | ---?+ | 22.9 | 0.2734 | intergenic | |
| rs12650007 | 4 | 20737411 | 4 | 0 | 0 | A | G | -0.8983 | 0.2631 | 0.0006389 | ---?- | 0 | 0.8314 | intron | KCNIP4 |
| rs34722008 | 4 | 38659594 | 5 | 0 | 0 | A | G | -0.7997 | 0.2342 | 0.000639 | ----- | 3.6 | 0.386 | intron | FLJ13197 |
| rs72919027 | 2 | 199069966 | 4 | 0 | 0 | A | C | 1.1323 | 0.3316 | 0.000639 | +++?- | 34.9 | 0.203 | intergenic | |
| 11:15856491 | 11 | 15856491 | 3 | 0 | 0 | T | TC | 2.4511 | 0.7179 | 0.0006392 | +++?? | 0 | 0.4903 | | |
| rs443829 | 21 | 31402972 | 4 | 0 | 0 | T | C | 0.9737 | 0.2852 | 0.0006393 | +++?- | 7.9 | 0.3537 | intergenic | |
| rs78267120 | 3 | 196503856 | 4 | 0 | 0 | A | C | -2.0916 | 0.6126 | 0.0006393 | ---?+ | 18.9 | 0.2958 | intron | PAK2 |
| rs11179880 | 12 | 74423511 | 4 | 0 | 0 | T | C | -0.8697 | 0.2547 | 0.0006394 | ---?- | 0 | 0.6709 | intergenic | |
| rs1672466 | 15 | 42135988 | 5 | 0 | 0 | T | C | -0.9361 | 0.2742 | 0.0006395 | ----- | 0 | 0.5897 | synonymous | JMJD7-PLA2G4B,PLA2G4B |
| rs4915942 | 1 | 64581219 | 4 | 0 | 0 | A | G | 1.0499 | 0.3075 | 0.0006395 | +++?+ | 0 | 0.8413 | intron | ROR1 |
| rs4667257 | 2 | 189880810 | 4 | 0 | 0 | A | G | 0.9697 | 0.284 | 0.0006396 | +++?+ | 43.5 | 0.1506 | downstream-gene | |
| rs10165827 | 2 | 208336308 | 4 | 0 | 0 | T | C | -1.3852 | 0.4057 | 0.0006398 | ---?- | 0 | 0.477 | intergenic | |
| rs57618647 | 16 | 85241528 | 4 | 0 | 0 | C | G | -1.2301 | 0.3603 | 0.0006398 | ---?- | 0 | 0.7099 | intron | GSE1 |
| rs4950096 | 1 | 98597309 | 4 | 0 | 0 | A | G | 1.3387 | 0.3921 | 0.00064 | +++?+ | 19.2 | 0.2943 | intergenic | |
| rs146984456 | 20 | 55446839 | 3 | 0 | 0 | T | C | -4.0732 | 1.1931 | 0.0006401 | ---?? | 32.4 | 0.2278 | intergenic | |
| rs4642112 | 3 | 188521929 | 4 | 0 | 0 | T | C | 0.8561 | 0.2508 | 0.0006402 | +++?+ | 0 | 0.6832 | intron | LPP |
| rs56314211 | 9 | 16194370 | 4 | 0 | 0 | T | C | -1.2312 | 0.3606 | 0.0006402 | ---?- | 0 | 0.5977 | intergenic | |
| rs1517111 | 11 | 103717688 | 4 | 0 | 0 | A | G | 1.6117 | 0.4721 | 0.0006403 | +++?+ | 0 | 0.4565 | upstream-gene | |
| rs28485331 | 9 | 44266761 | 3 | 0 | 0 | T | C | -1.743 | 0.5106 | 0.0006404 | ---?? | 0 | 0.8244 | intergenic | |
| rs1829127 | 12 | 74429110 | 4 | 0 | 0 | A | G | 0.87 | 0.2548 | 0.0006406 | +++?+ | 0 | 0.6798 | intergenic | |
| rs74785798 | 9 | 116143275 | 4 | 0 | 0 | A | T | -2.5263 | 0.7401 | 0.0006409 | ---?- | 0 | 0.9231 | upstream-gene | |
| rs11665772 | 19 | 56221878 | 4 | 0 | 0 | A | T | 1.1752 | 0.3443 | 0.0006413 | +++?+ | 0 | 0.7863 | intron | NLRP9 |
| rs72826891 | 10 | 117542884 | 4 | 0 | 0 | A | G | 3.0254 | 0.8863 | 0.0006415 | +++?+ | 0 | 0.7115 | intron | ATRNL1 |
| 20:39823175 | 20 | 39823175 | 3 | 0 | 0 | CT | C | -1.0606 | 0.3107 | 0.0006417 | ---?? | 0 | 0.9777 | | |
| rs12664109 | 6 | 120394813 | 4 | 0 | 0 | A | G | 2.3836 | 0.6983 | 0.0006419 | +++?+ | 47.3 | 0.1274 | intergenic | |
| rs1131118 | 6 | 31239108 | 3 | 0 | 0 | A | T | 1.0098 | 0.2959 | 0.0006421 | +++?? | 0 | 0.6875 | missense | HLA-C |
| rs76107995 | 12 | 75905055 | 4 | 0 | 0 | C | G | -2.9832 | 0.874 | 0.0006421 | ---?+ | 57.8 | 0.06874 | intron | KRR1 |
| rs7168951 | 15 | 91406146 | 4 | 0 | 0 | A | C | -0.8185 | 0.2398 | 0.0006422 | ---?+ | 23.1 | 0.2723 | intergenic | |
| rs143904538 | 20 | 35641617 | 3 | 0 | 0 | A | T | 1.2419 | 0.3639 | 0.0006422 | +++?? | 0 | 0.875 | intron | RBL1 |
| rs77234902 | 2 | 184055988 | 4 | 0 | 0 | A | G | 1.1661 | 0.3417 | 0.0006425 | +++?+ | 0 | 0.7773 | intergenic | |
| rs152226 | 5 | 142970450 | 4 | 0 | 0 | C | G | -1.8137 | 0.5314 | 0.0006426 | -+-?- | 60.6 | 0.05463 | intron | NR3C1 |
| 9:44257671 | 9 | 44257671 | 3 | 0 | 0 | T | TA | -1.7848 | 0.5229 | 0.0006426 | ---?? | 0 | 0.8567 | | |
| rs12041998 | 1 | 101865012 | 4 | 0 | 0 | T | C | 1.4716 | 0.4312 | 0.0006428 | +++?+ | 0 | 0.6755 | intergenic | |
| rs79115430 | 3 | 60618443 | 4 | 0 | 0 | T | C | 1.6526 | 0.4842 | 0.0006428 | +++?+ | 23.4 | 0.2705 | intron | FHIT |
| rs114484479 | 17 | 68743570 | 4 | 0 | 0 | A | G | -1.3534 | 0.3966 | 0.0006429 | ---?- | 0 | 0.7381 | intergenic | |
| rs150257209 | 15 | 86379743 | 4 | 0 | 0 | T | C | -1.9467 | 0.5704 | 0.0006429 | ---?- | 0 | 0.9944 | intergenic | |
| rs1992617 | 4 | 37005546 | 4 | 0 | 0 | A | G | -0.8557 | 0.2507 | 0.000643 | ---?+ | 46.2 | 0.1344 | intergenic | |
| rs6577067 | 13 | 112826806 | 4 | 0 | 0 | T | C | -0.9428 | 0.2763 | 0.000643 | ---?- | 14.8 | 0.3179 | intergenic | |
| rs970059 | 12 | 98624406 | 4 | 0 | 0 | A | G | -1.5187 | 0.445 | 0.0006431 | -+-?- | 58.6 | 0.06461 | intergenic | |
| rs7011371 | 8 | 3985130 | 4 | 0 | 0 | A | G | 1.1591 | 0.3397 | 0.0006433 | -++?+ | 54.5 | 0.08619 | intron | CSMD1 |
| rs74131299 | 13 | 112825250 | 4 | 0 | 0 | A | T | -0.9446 | 0.2768 | 0.0006433 | ---?- | 10.7 | 0.3397 | intergenic | |
| 1:15859198 | 1 | 15859198 | 3 | 0 | 0 | G | GT | 1.0151 | 0.2975 | 0.0006434 | +++?? | 54.6 | 0.1108 | | |
| 3:195549882 | 3 | 195549882 | 2 | 0 | 0 | CTTTTTT | C | -5.4654 | 1.6015 | 0.0006436 | ?--?? | 0 | 0.3224 | | |
| rs78398422 | 10 | 4344038 | 4 | 0 | 0 | T | C | -2.7065 | 0.7931 | 0.0006437 | ---?- | 51.4 | 0.1033 | intergenic | |
| rs75412544 | 19 | 50698559 | 4 | 0 | 0 | A | G | 2.6801 | 0.7854 | 0.000644 | +++?+ | 5.1 | 0.3675 | intergenic | |
| rs140128850 | 22 | 28927879 | 3 | 0 | 0 | C | G | -2.3917 | 0.7009 | 0.000644 | ---?? | 0 | 0.4452 | intron | TTC28 |
| rs13136504 | 4 | 169321009 | 4 | 0 | 0 | T | G | -0.8357 | 0.2449 | 0.0006442 | ---?- | 0 | 0.5246 | intron | DDX60L |

| | | | | | | | | | | | | | | | |
|---|---|---|---|---|---|---|---|---|---|---|---|---|---|---|---|
| 3:183591316 | 3 | 183591316 | 3 | 0 | 0 | G | GAC | -1.6844 | 0.4936 | 0.0006442 | ---?? | 0 | 0.8113 | | |
| rs3104969 | 8 | 96592975 | 4 | 0 | 0 | T | G | 1.3642 | 0.3998 | 0.0006445 | +-+?+ | 35.4 | 0.1999 | intron | LOC100616530 |
| rs111551930 | 11 | 541961 | 3 | 0 | 0 | A | G | -1.0085 | 0.2956 | 0.0006446 | ---?? | 0 | 0.4706 | intron | LRRC56 |
| 13:42637220 | 13 | 42637220 | 2 | 0 | 0 | T | TA | -5.8793 | 1.7231 | 0.0006447 | --??? | 0 | 0.772 | | |
| rs8002385 | 13 | 27578256 | 4 | 0 | 0 | T | C | -1.4356 | 0.4208 | 0.0006448 | ---?- | 0 | 0.9465 | intergenic | |
| rs12468028 | 2 | 170779905 | 4 | 0 | 1 | A | T | -0.9848 | 0.2886 | 0.0006449 | ---?- | 43.4 | 0.1509 | intron | UBR3 |
| rs6751043 | 2 | 208352656 | 3 | 0 | 0 | T | C | -1.4495 | 0.4248 | 0.000645 | ---?? | 37.8 | 0.2003 | intergenic | |
| rs12468029 | 2 | 170779906 | 4 | 0 | 1 | A | T | -0.9848 | 0.2886 | 0.0006451 | ---?- | 43.4 | 0.151 | intron | UBR3 |
| rs35254973 | 21 | 42187213 | 3 | 0 | 0 | T | C | 4.7068 | 1.3795 | 0.0006451 | +++?? | 0 | 0.846 | intron | DSCAM |
| 19:3599325 | 19 | 3599325 | 3 | 0 | 0 | T | TTATTTT | -0.893 | 0.2617 | 0.0006451 | ---?? | 0 | 0.4918 | | |
| rs12775990 | 10 | 134575836 | 4 | 0 | 0 | T | C | 0.9624 | 0.2821 | 0.0006453 | +++?+ | 0 | 0.4553 | intron | INPP5A |
| rs6730980 | 2 | 170777579 | 4 | 0 | 1 | A | T | 0.9846 | 0.2886 | 0.0006455 | +++?+ | 43.9 | 0.148 | intron | UBR3 |
| 8:113310316 | 8 | 113310316 | 3 | 0 | 0 | A | AAAAT | 0.9986 | 0.2927 | 0.0006455 | +++?? | 0 | 0.573 | | |
| rs115765185 | 9 | 44274916 | 3 | 0 | 0 | A | C | -1.79 | 0.5247 | 0.0006457 | ---?? | 0 | 0.8379 | intergenic | |
| rs36009794 | 19 | 46152715 | 5 | 0 | 1 | T | C | 0.9501 | 0.2785 | 0.0006458 | +-++- | 56.2 | 0.05798 | upstream-gene | |
| rs117869545 | 8 | 76211596 | 4 | 0 | 0 | T | G | -2.6453 | 0.7754 | 0.0006458 | ---?+ | 31.3 | 0.2244 | intergenic | |
| rs72797686 | 10 | 27992837 | 4 | 0 | 0 | A | G | -2.3696 | 0.6946 | 0.0006461 | ---?- | 0 | 0.7385 | intron | MKX |
| rs76133128 | 17 | 4796271 | 5 | 0 | 0 | T | C | -1.5579 | 0.4567 | 0.0006462 | ----+ | 12.6 | 0.3338 | intron-near-splice | MINK1 |
| rs72839993 | 17 | 56395293 | 5 | 0 | 0 | T | C | 1.8944 | 0.5553 | 0.0006466 | +++++ | 50.5 | 0.08866 | intron | BZRAP1 |
| rs112757554 | 3 | 62364138 | 2 | 0 | 0 | C | G | -3.6256 | 1.0628 | 0.0006467 | ?--?? | 0 | 0.5756 | upstream-gene | |
| rs112976408 | 22 | 29126770 | 4 | 0 | 0 | T | C | 2.2415 | 0.6571 | 0.0006468 | +++?+ | 0 | 0.8131 | intron | CHEK2 |
| rs181003402 | 5 | 693446 | 3 | 0 | 0 | A | C | 1.4537 | 0.4262 | 0.0006468 | +++?? | 0 | 0.7259 | 5-prime-UTR | TPPP |
| rs9513656 | 13 | 100403988 | 4 | 0 | 0 | A | G | -0.9836 | 0.2884 | 0.000647 | ---?- | 31.8 | 0.2214 | intron | CLYBL |
| rs31210 | 5 | 134361020 | 5 | 0 | 0 | A | G | -0.8268 | 0.2424 | 0.0006471 | ---+- | 18.1 | 0.2995 | downstream-gene | |
| rs3129849 | 6 | 32397309 | 5 | 0 | 0 | A | G | -0.7867 | 0.2306 | 0.0006471 | ----- | 33.6 | 0.1974 | intergenic |  |
| rs72914779 | 2 | 198985004 | 4 | 0 | 0 | T | C | -1.6038 | 0.4702 | 0.0006471 | ---?- | 10.3 | 0.3414 | intron | PLCL1 |
| rs4504509 | 6 | 32399159 | 5 | 0 | 0 | T | C | -0.7867 | 0.2306 | 0.0006472 | ----- | 33.6 | 0.1974 | intergenic |  |
| rs9268633 | 6 | 32406473 | 5 | 0 | 0 | A | G | -0.7867 | 0.2306 | 0.0006472 | ----- | 33.6 | 0.1974 | upstream-gene |  |
| rs7755363 | 6 | 137747267 | 4 | 0 | 0 | A | G | 1.1391 | 0.334 | 0.0006473 | +++?+ | 0 | 0.7639 | intergenic | |
| rs7918950 | 10 | 71008705 | 4 | 0 | 0 | A | G | 1.0162 | 0.2979 | 0.0006473 | +++?- | 44.1 | 0.1469 | intron | HKDC1 |
| rs28501426 | 9 | 44259781 | 3 | 0 | 0 | A | C | 1.7391 | 0.5098 | 0.0006473 | +++?? | 0 | 0.8154 | intergenic | |
| rs150632910 | 8 | 110710517 | 4 | 0 | 0 | T | C | 3.5741 | 1.0478 | 0.0006473 | +++?+ | 0 | 0.7645 | intergenic | |
| rs1648827 | 15 | 42137581 | 5 | 0 | 0 | A | C | 0.9348 | 0.2741 | 0.0006474 | +++++ | 0 | 0.5884 | intron | JMJD7-PLA2G4B,PLA2G4B |
| rs8009726 | 14 | 43272103 | 4 | 0 | 0 | A | C | 1.2911 | 0.3785 | 0.0006474 | +++?- | 60.3 | 0.05629 | intergenic | |
| rs3763321 | 6 | 32406704 | 5 | 0 | 0 | T | G | -0.7867 | 0.2306 | 0.0006475 | ----- | 33.6 | 0.1973 | upstream-gene |  |
| rs7927966 | 11 | 58376276 | 5 | 0 | 0 | T | C | -0.8202 | 0.2405 | 0.0006475 | ----- | 0 | 0.9816 | intron | ZFP91,ZFP91-CNTF |
| 16:84804374 | 16 | 84804374 | 3 | 0 | 0 | T | TC | -1.0693 | 0.3135 | 0.0006476 | ---?? | 0 | 0.3997 | | |
| rs114498847 | 9 | 44275172 | 3 | 0 | 0 | T | C | 1.7928 | 0.5256 | 0.0006477 | +++?? | 0 | 0.8378 | intergenic | |
| rs877674 | 1 | 32242206 | 4 | 0 | 0 | A | G | -1.2219 | 0.3582 | 0.0006478 | ---?+ | 41.9 | 0.1599 | intergenic | |
| rs1978573 | 2 | 185459544 | 4 | 0 | 0 | T | C | 0.9395 | 0.2755 | 0.0006479 | +++?+ | 0 | 0.6337 | upstream-gene | |
| rs28756462 | 9 | 44260434 | 3 | 0 | 0 | T | C | -1.7391 | 0.5099 | 0.0006479 | ---?? | 0 | 0.8176 | intergenic | |
| rs2322688 | 4 | 20751278 | 4 | 0 | 0 | A | G | 0.902 | 0.2645 | 0.000648 | +++?+ | 0 | 0.8985 | intron-near-splice | KCNIP4 |
| rs12812622 | 12 | 718750 | 3 | 0 | 0 | A | G | 3.7589 | 1.1021 | 0.000648 | +++?? | 44.7 | 0.164 | intron | NINJ2 |
| rs7027513 | 9 | 78456306 | 4 | 0 | 0 | A | G | 1.2417 | 0.3641 | 0.0006481 | +++?+ | 56.5 | 0.07541 | intergenic | |
| rs16954312 | 17 | 52287259 | 4 | 0 | 0 | T | C | -2.0893 | 0.6126 | 0.0006481 | ---?+ | 35.2 | 0.2008 | intergenic | |
| rs117717080 | 17 | 70043095 | 4 | 0 | 0 | T | C | 3.056 | 0.896 | 0.0006481 | +++?+ | 0 | 0.8618 | intergenic | |
| rs2275993 | 11 | 58379987 | 5 | 0 | 0 | A | G | 0.8212 | 0.2408 | 0.0006482 | +++++ | 0 | 0.9815 | intron | ZFP91,ZFP91-CNTF |
| rs9736056 | 11 | 130626242 | 3 | 0 | 0 | A | C | 1.7495 | 0.513 | 0.0006482 | +++?? | 51.9 | 0.1251 | intergenic | |
| rs1800837 | 6 | 33272626 | 5 | 0 | 0 | T | C | 0.9209 | 0.27 | 0.0006483 | +++++ | 0 | 0.9217 | intron | TAPBP |
| rs3135334 | 6 | 32402686 | 5 | 0 | 0 | T | C | 0.7866 | 0.2306 | 0.0006483 | +++++ | 33.6 | 0.1973 | upstream-gene | |
| rs6750479 | 2 | 170774719 | 4 | 0 | 1 | C | G | -0.984 | 0.2885 | 0.0006483 | ---?- | 44.6 | 0.1435 | intron | UBR3 |
| rs77178060 | 17 | 52285781 | 4 | 0 | 0 | C | G | -2.0861 | 0.6117 | 0.0006483 | ---?+ | 35.3 | 0.2006 | intergenic | |
| rs4972927 | 2 | 230915945 | 4 | 0 | 0 | T | C | 1.4465 | 0.4241 | 0.0006484 | +++?+ | 13.2 | 0.3266 | intron | SLC16A14 |
| rs77972902 | 1 | 180131941 | 4 | 0 | 0 | C | G | -2.2123 | 0.6487 | 0.0006484 | ---?+ | 37.6 | 0.1863 | intron | QSOX1 |
| rs1648829 | 15 | 42137282 | 5 | 0 | 0 | A | G | -0.9347 | 0.2741 | 0.0006485 | ----- | 0 | 0.5885 | intron | JMJD7-PLA2G4B,PLA2G4B |
| rs10097310 | 8 | 96217666 | 4 | 0 | 0 | T | G | 0.8508 | 0.2495 | 0.0006485 | +++?+ | 0 | 0.8739 | downstream-gene | |
| rs10931066 | 2 | 184042680 | 4 | 0 | 0 | T | C | 1.1544 | 0.3385 | 0.0006485 | +++?+ | 0 | 0.8309 | intergenic | |
| rs111725927 | 9 | 44260003 | 3 | 0 | 0 | T | C | -1.7388 | 0.5098 | 0.0006485 | ---?? | 0 | 0.8169 | intergenic | |
| rs10056109 | 5 | 103164240 | 4 | 0 | 0 | T | C | -1.5397 | 0.4515 | 0.0006486 | ---?- | 0 | 0.7209 | intergenic | |

| | | | | | | | | | | | | | | |
|---|---|---|---|---|---|---|---|---|---|---|---|---|---|---|
| rs11684034 | 2 | 184042603 | 4 | 0 | 0 | T | C | -1.1545 | 0.3385 | 0.0006487 | ---?- | 0 | 0.8308 | intergenic | |
| rs62285964 | 3 | 184182619 | 4 | 0 | 0 | A | G | -1.8922 | 0.5548 | 0.0006487 | ---?- | 0 | 0.7673 | intergenic | |
| rs11161435 | 1 | 85206106 | 5 | 0 | 0 | A | T | 0.7934 | 0.2327 | 0.0006488 | +++++ | 0 | 0.7496 | intergenic | |
| rs140753044 | 7 | 141210513 | 3 | 0 | 0 | A | G | 6.3793 | 1.8706 | 0.0006488 | ++??+ | 32.3 | 0.2285 | intergenic | |
| rs659711 | 15 | 53759512 | 4 | 0 | 0 | A | G | -0.8417 | 0.2468 | 0.0006489 | ---?- | 47.9 | 0.1237 | intergenic | |
| rs1457691 | 5 | 123501288 | 4 | 0 | 0 | C | G | -0.8478 | 0.2486 | 0.0006489 | ---?+ | 20.6 | 0.2864 | intergenic | |
| rs6747591 | 2 | 170768891 | 4 | 0 | 1 | A | C | -0.9835 | 0.2884 | 0.0006489 | ---?- | 45.6 | 0.1377 | intron | UBR3 |
| rs10955617 | 8 | 113241438 | 4 | 0 | 0 | T | C | -0.968 | 0.2838 | 0.0006489 | ---?- | 0 | 0.9688 | intron | CSMD3 |
| rs7102510 | 11 | 58371283 | 5 | 0 | 0 | T | G | 0.818 | 0.2399 | 0.000649 | +++++ | 0 | 0.9817 | intron | ZFP91,ZFP91-CNTF |
| rs139317622 | 1 | 102970712 | 3 | 0 | 0 | T | C | -0.8644 | 0.2535 | 0.000649 | ---?? | 0 | 0.6209 | intergenic | |
| rs141465091 | 22 | 28629663 | 4 | 0 | 0 | A | C | 2.641 | 0.7744 | 0.000649 | +++?+ | 0 | 0.7835 | intron | TTC28 |
| rs1648828 | 15 | 42137576 | 5 | 0 | 0 | T | C | -0.9346 | 0.274 | 0.0006491 | ----- | 0 | 0.5881 | intron | JMJD7-PLA2G4B,PLA2G4B |
| rs28594668 | 2 | 170773127 | 4 | 0 | 1 | T | G | 0.9839 | 0.2885 | 0.0006491 | +++?+ | 44.9 | 0.1423 | intron | UBR3 |
| rs7096347 | 10 | 10091918 | 2 | 0 | 0 | A | C | -6.869 | 2.0143 | 0.0006492 | ?--?? | 0 | 0.8333 | intergenic | |
| rs3129852 | 6 | 32397784 | 5 | 0 | 0 | A | G | 0.7864 | 0.2306 | 0.0006493 | +++++ | 33.4 | 0.1984 | intergenic | |
| rs3135341 | 6 | 32398748 | 5 | 0 | 0 | A | C | 0.7864 | 0.2306 | 0.0006493 | +++++ | 33.4 | 0.1984 | intergenic | |
| rs11199523 | 10 | 122494215 | 4 | 0 | 0 | A | G | -0.9524 | 0.2793 | 0.0006493 | ---?- | 28.7 | 0.24 | intergenic | |
| rs1672464 | 15 | 42134652 | 5 | 0 | 0 | A | G | -0.935 | 0.2742 | 0.0006494 | ----- | 0 | 0.592 | intron | JMJD7-PLA2G4B,PLA2G4B |
| rs116707672 | 17 | 52285718 | 4 | 0 | 0 | T | C | -2.0858 | 0.6116 | 0.0006494 | ---?+ | 35.3 | 0.2006 | intergenic | |
| rs75045680 | 6 | 164988405 | 4 | 0 | 0 | A | T | -2.5219 | 0.7395 | 0.0006495 | ---?- | 0 | 0.431 | intergenic | |
| rs114957141 | 4 | 95063320 | 4 | 0 | 0 | A | T | -3.2084 | 0.9409 | 0.0006495 | ---?- | 6.9 | 0.3587 | intergenic | |
| rs115343618 | 6 | 31980773 | 3 | 0 | 0 | A | G | -1.2983 | 0.3807 | 0.0006495 | ---?? | 29.6 | 0.2415 | non-coding-exon | TNXA |
| rs16934204 | 9 | 16194182 | 4 | 0 | 0 | A | T | 1.2293 | 0.3605 | 0.0006496 | +++?+ | 0 | 0.5943 | intergenic | |
| rs73288629 | 10 | 74640812 | 4 | 0 | 0 | C | G | 2.7869 | 0.8173 | 0.0006497 | +++?+ | 20 | 0.29 | intron | MCU |
| rs10267372 | 7 | 49462604 | 4 | 0 | 0 | T | C | -0.9468 | 0.2777 | 0.0006498 | ---?+ | 22.7 | 0.2744 | intergenic | |
| rs28583884 | 9 | 44259611 | 3 | 0 | 0 | C | G | -1.7373 | 0.5095 | 0.0006498 | ---?? | 0 | 0.8154 | intergenic | |
| rs7515049 | 1 | 18743016 | 4 | 0 | 0 | T | G | 1.0598 | 0.3108 | 0.00065 | +++?+ | 51.2 | 0.1047 | intergenic | |
| rs8373 | 11 | 58378424 | 5 | 0 | 0 | A | G | 0.8207 | 0.2407 | 0.0006501 | +++++ | 0 | 0.9815 | splice-acceptor | ZFP91,ZFP91-CNTF |
| rs9734285 | 11 | 130626240 | 3 | 0 | 0 | A | G | -1.7491 | 0.513 | 0.0006502 | ---?? | 51.9 | 0.1249 | intergenic | |
| rs6732048 | 2 | 170765215 | 4 | 0 | 1 | T | C | -0.9833 | 0.2884 | 0.0006503 | ---?- | 46.3 | 0.1337 | intron | UBR3 |
| rs10231902 | 7 | 49466193 | 4 | 0 | 0 | C | G | 0.9519 | 0.2792 | 0.0006503 | +++?- | 24.2 | 0.2662 | intergenic | |
| rs10144987 | 14 | 43272811 | 4 | 0 | 0 | A | G | -1.2999 | 0.3812 | 0.0006504 | ---?+ | 58.8 | 0.06321 | intergenic | |
| rs3168135 | 11 | 58386177 | 5 | 0 | 0 | A | G | -0.8227 | 0.2413 | 0.0006506 | ----- | 0 | 0.9811 | 3-prime-UTR | ZFP91,ZFP91-CNTF |
| rs10200004 | 2 | 170767137 | 4 | 0 | 1 | T | C | 0.9834 | 0.2884 | 0.0006506 | +++?+ | 46 | 0.1355 | intron | UBR3 |
| rs11229543 | 11 | 58370698 | 5 | 0 | 0 | A | G | -0.8176 | 0.2398 | 0.0006506 | ----- | 0 | 0.9818 | intron | ZFP91,ZFP91-CNTF |
| rs11229545 | 11 | 58386650 | 5 | 0 | 0 | A | G | 0.8228 | 0.2413 | 0.0006506 | +++++ | 0 | 0.9811 | 3-prime-UTR | ZFP91,ZFP91-CNTF |
| rs7217340 | 17 | 51120278 | 3 | 0 | 0 | T | C | -1.5597 | 0.4575 | 0.0006507 | ---?? | 27.4 | 0.2522 | intergenic | |
| rs10197717 | 2 | 170770224 | 4 | 0 | 1 | T | C | -0.9837 | 0.2885 | 0.0006507 | ---?- | 45.4 | 0.1389 | intron | UBR3 |
| 1:182009553 | 1 | 182009553 | 3 | 0 | 0 | A | AAAAAC | 2.9507 | 0.8654 | 0.0006508 | +++?? | 0 | 0.8829 | | |
| rs12411065 | 1 | 32241767 | 4 | 0 | 0 | T | C | 1.2201 | 0.3579 | 0.0006509 | +++?- | 41.8 | 0.1606 | intergenic | |
| rs62174934 | 2 | 185426237 | 4 | 0 | 0 | A | G | 0.9439 | 0.2769 | 0.0006509 | +++?+ | 0 | 0.7948 | intergenic | |
| rs10849935 | 12 | 111767891 | 5 | 0 | 0 | A | G | 0.8523 | 0.25 | 0.000651 | +-+++ | 50 | 0.09178 | intron | CUX2 |
| rs890508 | 15 | 42139038 | 5 | 0 | 0 | A | G | -0.9337 | 0.2739 | 0.0006512 | ----- | 0 | 0.5854 | intron-near-splice | JMJD7-PLA2G4B,PLA2G4B |
| rs4568494 | 6 | 32672361 | 5 | 0 | 1 | A | G | -1.1067 | 0.3246 | 0.0006512 | ----- | 24 | 0.2614 | intergenic |  |
| rs7201543 | 16 | 84748541 | 3 | 0 | 0 | A | T | 1.0778 | 0.3161 | 0.0006513 | +++?? | 37.4 | 0.2024 | intron | USP10 |
| 13:20812713 | 13 | 20812713 | 3 | 0 | 0 | CG | C | -1.153 | 0.3382 | 0.0006513 | ---?? | 0 | 0.3749 | | |
| rs34863361 | 1 | 245106985 | 4 | 0 | 0 | C | G | 1.4355 | 0.4211 | 0.0006517 | +++?- | 15.4 | 0.3148 | intergenic | |
| rs76557791 | 11 | 86174538 | 4 | 0 | 0 | T | C | 2.4295 | 0.7127 | 0.0006518 | +++?+ | 0 | 0.8661 | intron | ME3 |
| 6:159189422 | 6 | 159189422 | 3 | 0 | 0 | G | GT | -0.8717 | 0.2557 | 0.0006518 | ---?? | 43.5 | 0.1703 | | |
| rs113402437 | 1 | 97404322 | 2 | 0 | 0 | A | C | -7.3662 | 2.1608 | 0.000652 | -???- | 24 | 0.2514 | intergenic | |
| rs1587018 | 3 | 87375276 | 4 | 0 | 0 | T | C | 1.1604 | 0.3404 | 0.0006521 | +++?+ | 0 | 0.832 | intergenic | |
| rs2373067 | 2 | 218360362 | 4 | 0 | 0 | A | G | -0.8227 | 0.2413 | 0.0006523 | ---?- | 0 | 0.9285 | intron | DIRC3 |
| rs116371541 | 4 | 39584720 | 4 | 0 | 0 | T | C | -3.6118 | 1.0595 | 0.0006524 | ---?- | 0 | 0.965 | intron | SMIM14,UGDH-AS1 |
| rs10191174 | 2 | 170761262 | 4 | 0 | 1 | A | G | 0.9828 | 0.2883 | 0.0006525 | +++?+ | 46.8 | 0.1305 | intron | UBR3 |
| rs11165952 | 1 | 98598635 | 4 | 0 | 0 | A | G | -1.3364 | 0.3921 | 0.0006526 | ---?- | 19.4 | 0.2929 | intergenic | |
| rs62402376 | 6 | 28507446 | 5 | 0 | 0 | A | C | -0.9091 | 0.2667 | 0.0006527 | ----- | 0 | 0.9277 | downstream-gene |  |
| rs55747410 | 14 | 96975299 | 4 | 0 | 0 | C | G | 0.9074 | 0.2662 | 0.0006529 | +++?- | 0 | 0.5403 | intron | PAPOLA |
| 6:106272100 | 6 | 106272100 | 3 | 0 | 0 | CAT | C | -1.9715 | 0.5784 | 0.0006529 | ---?? | 28 | 0.2496 | | |
| rs377715374 | 3 | 111872939 | 4 | 0 | 0 | T | C | -3.7127 | 1.0892 | 0.0006532 | ---?- | 0 | 0.7146 | intron | SLC9C1 |

| | | | | | | | | | | | | | | | |
|---|---|---|---|---|---|---|---|---|---|---|---|---|---|---|---|
| rs9631 | 2 | 170939679 | 4 | 0 | 1 | T | C | 0.9748 | 0.286 | 0.0006535 | +++?+ | 58 | 0.06757 | 3-prime-UTR | UBR3 |
| rs78830128 | 10 | 86689829 | 4 | 0 | 0 | A | G | -1.7443 | 0.5118 | 0.0006536 | ---?- | 0 | 0.7915 | intergenic | |
| rs9379629 | 6 | 24025720 | 5 | 0 | 0 | A | G | -1.3265 | 0.3892 | 0.0006537 | ----+ | 0 | 0.8109 | intergenic |  |
| rs10896795 | 11 | 58366561 | 5 | 0 | 0 | A | G | 0.8161 | 0.2395 | 0.0006538 | +++++ | 0 | 0.9819 | intron | ZFP91,ZFP91-CNTF |
| 3:15822911 | 3 | 15822911 | 3 | 0 | 0 | G | GA | -2.3136 | 0.6788 | 0.0006539 | ---?? | 0 | 0.5785 | | |
| rs1711773 | 13 | 60052825 | 4 | 0 | 0 | A | C | -0.8914 | 0.2615 | 0.000654 | ---?- | 43.2 | 0.1523 | intergenic | |
| 13:60029348 | 13 | 60029348 | 3 | 0 | 0 | A | AT | 0.8882 | 0.2606 | 0.000654 | +++?? | 23.5 | 0.2708 | | |
| rs17612860 | 15 | 85640282 | 4 | 0 | 0 | A | C | -1.1606 | 0.3406 | 0.0006545 | ---?+ | 40.9 | 0.1664 | intron | PDE8A |
| 2:184009316 | 2 | 184009316 | 3 | 0 | 0 | T | TAATAA | 1.2165 | 0.357 | 0.0006545 | +++?? | 0 | 0.7948 | | |
| 4:144565353 | 4 | 144565353 | 3 | 0 | 0 | T | C | 2.0283 | 0.5952 | 0.0006546 | +++?? | 18.2 | 0.2946 | intron | FREM3 |
| rs6447976 | 4 | 20736090 | 4 | 0 | 0 | T | C | -0.8865 | 0.2601 | 0.0006546 | ---?- | 0 | 0.8454 | intron | KCNIP4 |
| rs13380598 | 16 | 7205098 | 4 | 0 | 0 | C | G | -1.0401 | 0.3052 | 0.0006546 | ---?- | 0 | 0.9568 | intron | RBFOX1 |
| rs1372543 | 12 | 41312147 | 4 | 0 | 0 | A | G | 1.6921 | 0.4965 | 0.000655 | +++?+ | 0 | 0.8645 | intron | CNTN1 |
| rs11644621 | 16 | 85212930 | 4 | 0 | 0 | A | G | 0.92 | 0.27 | 0.000655 | +++?+ | 0 | 0.4594 | intron | GSE1 |
| rs4688243 | 3 | 65659336 | 4 | 0 | 0 | C | G | -0.9062 | 0.2659 | 0.0006551 | -+-?- | 32.8 | 0.2153 | intron | MAGI1 |
| rs28377659 | 9 | 44280136 | 3 | 0 | 0 | A | C | 1.8708 | 0.549 | 0.0006552 | +++?? | 0 | 0.8339 | intergenic | |
| 22:28583474 | 22 | 28583474 | 3 | 0 | 0 | A | ATACT | -2.4611 | 0.7222 | 0.0006553 | ---?? | 0 | 0.4983 | | |
| rs17637448 | 14 | 43720236 | 4 | 0 | 0 | A | G | 1.6111 | 0.4728 | 0.0006555 | +++?- | 0 | 0.5619 | intergenic | |
| rs9268520 | 6 | 32380860 | 5 | 0 | 0 | A | C | 0.7228 | 0.2121 | 0.0006557 | +++++ | 0 | 0.6553 | intergenic | |
| rs142373973 | 9 | 112731067 | 4 | 0 | 0 | A | G | -2.899 | 0.8508 | 0.0006558 | ---?- | 0 | 0.9637 | intron | PALM2-AKAP2 |
| rs10106754 | 8 | 96991901 | 4 | 0 | 0 | A | G | -1.4035 | 0.4119 | 0.0006559 | ---?- | 0 | 0.9613 | intergenic | |
| rs58540150 | 7 | 49477889 | 4 | 0 | 0 | A | G | -0.9672 | 0.2839 | 0.0006559 | ---?+ | 27.2 | 0.2486 | intergenic | |
| rs190881411 | 4 | 117432222 | 3 | 0 | 0 | A | G | 3.9115 | 1.148 | 0.000656 | +++?? | 66.3 | 0.05132 | intergenic | |
| rs1500639 | 15 | 98047533 | 4 | 0 | 0 | A | G | 0.8472 | 0.2486 | 0.0006563 | +++?+ | 0 | 0.8295 | intergenic | |
| rs12291827 | 11 | 95698414 | 4 | 0 | 0 | T | G | 0.9066 | 0.2661 | 0.0006563 | +++?+ | 0 | 0.646 | intergenic | |
| 17:39655591 | 17 | 39655591 | 3 | 0 | 0 | A | ACCCC | -2.0915 | 0.6139 | 0.0006564 | ---?? | 0 | 0.5851 | | |
| rs3829132 | 10 | 134579044 | 4 | 0 | 0 | A | C | -0.9609 | 0.282 | 0.0006566 | ---?- | 0 | 0.4609 | intron | INPP5A |
| rs7357155 | 7 | 49465437 | 4 | 0 | 0 | A | G | 0.9495 | 0.2787 | 0.0006566 | +++?- | 23.9 | 0.2677 | intergenic | |
| 17:51144713 | 17 | 51144713 | 3 | 0 | 0 | CT | C | 1.4537 | 0.4267 | 0.0006568 | +++?? | 36.7 | 0.2059 | | |
| rs112621128 | 1 | 192468550 | 4 | 0 | 0 | A | C | -1.1331 | 0.3326 | 0.0006569 | ---?+ | 42 | 0.1596 | intergenic | |
| rs114649924 | 1 | 217306713 | 4 | 0 | 0 | T | G | 1.7734 | 0.5205 | 0.0006571 | -++?- | 59.1 | 0.06215 | intron | ESRRG |
| rs1980160 | 2 | 183998237 | 4 | 0 | 0 | T | C | -1.1765 | 0.3453 | 0.0006573 | ---?- | 0 | 0.8509 | intron | NUP35 |
| rs12044342 | 1 | 98599144 | 4 | 0 | 0 | A | G | 1.3355 | 0.392 | 0.0006575 | +++?+ | 19.5 | 0.2926 | intergenic | |
| rs56821450 | 4 | 190023756 | 4 | 0 | 0 | A | G | -0.8501 | 0.2496 | 0.0006579 | ---?- | 0 | 0.4289 | intergenic | |
| rs11179885 | 12 | 74433655 | 4 | 0 | 0 | T | C | 0.8693 | 0.2552 | 0.000658 | +++?+ | 0 | 0.6865 | intergenic | |
| rs112578012 | 3 | 23806633 | 4 | 0 | 0 | T | C | -3.6456 | 1.0702 | 0.0006582 | ---?- | 0 | 0.7849 | intergenic | |
| 2:184022776 | 2 | 184022776 | 3 | 0 | 0 | CATT | C | 1.2147 | 0.3566 | 0.0006582 | +++?? | 0 | 0.7532 | | |
| rs144399508 | 4 | 166223704 | 4 | 0 | 0 | T | C | 3.7598 | 1.1037 | 0.0006583 | +++?+ | 0 | 0.9374 | intron | KLHL2 |
| rs6748654 | 2 | 230914766 | 4 | 0 | 0 | T | C | -1.4499 | 0.4257 | 0.0006585 | ---?- | 13.6 | 0.3245 | intron | SLC16A14 |
| rs13361535 | 5 | 119700340 | 4 | 0 | 0 | A | G | 1.1514 | 0.338 | 0.0006585 | +-+?+ | 51.9 | 0.1009 | intergenic | |
| 9:33430209 | 9 | 33430209 | 3 | 0 | 0 | T | C | -2.8375 | 0.833 | 0.0006586 | ---?? | 0 | 0.771 | intergenic | |
| rs10741592 | 11 | 12492188 | 4 | 0 | 0 | A | C | 3.0685 | 0.9008 | 0.0006586 | +++?+ | 43.7 | 0.149 | intron | PARVA |
| rs11961231 | 6 | 163542049 | 3 | 0 | 0 | A | T | -1.4302 | 0.4199 | 0.0006586 | ---?? | 0 | 0.8337 | intron | PACRG |
| rs71434749 | 13 | 30312455 | 5 | 0 | 0 | T | C | 1.271 | 0.3731 | 0.0006586 | +++++ | 0 | 0.5575 | intergenic | |
| rs71419908 | 14 | 32900776 | 4 | 0 | 0 | A | C | 0.8449 | 0.2481 | 0.0006588 | +++?+ | 0 | 0.8298 | intron | AKAP6 |
| rs149882883 | 3 | 143040210 | 4 | 0 | 0 | A | G | -2.0046 | 0.5885 | 0.0006588 | ---?- | 0 | 0.6972 | intron | SLC9A9 |
| rs144203347 | 22 | 40773310 | 3 | 0 | 0 | C | G | -3.8828 | 1.14 | 0.0006589 | ---?? | 8.8 | 0.3339 | intron | SGSM3 |
| rs4712125 | 6 | 56168802 | 4 | 0 | 0 | T | C | 1.3102 | 0.3847 | 0.0006591 | +++?+ | 4.1 | 0.3725 | intergenic | |
| rs4932770 | 19 | 22730114 | 4 | 0 | 0 | T | C | -0.8763 | 0.2573 | 0.0006593 | ---?- | 0 | 0.9779 | intergenic | |
| rs138123479 | 9 | 44263905 | 3 | 0 | 0 | A | C | 1.7471 | 0.513 | 0.0006595 | +++?? | 0 | 0.8169 | intergenic | |
| rs144042960 | 18 | 52761810 | 4 | 0 | 0 | A | G | 4.191 | 1.2305 | 0.0006595 | +++?+ | 0 | 0.8516 | intergenic | |
| rs817404 | 1 | 85618630 | 4 | 0 | 0 | T | C | -1.5408 | 0.4524 | 0.0006596 | ---?- | 0 | 0.9837 | downstream-gene | |
| rs12258248 | 10 | 134573767 | 4 | 0 | 0 | A | G | -0.9603 | 0.282 | 0.0006596 | ---?- | 0 | 0.4624 | intron | INPP5A |
| rs12797431 | 11 | 521119 | 5 | 0 | 0 | A | G | 0.7786 | 0.2286 | 0.0006599 | +++++ | 5.7 | 0.3743 | intergenic | |
| rs13157333 | 5 | 10308440 | 4 | 0 | 0 | T | G | 1.8521 | 0.5438 | 0.0006599 | +++?+ | 19.5 | 0.2927 | upstream-gene | |
| 4:143393477 | 4 | 143393477 | 3 | 0 | 0 | G | GA | -1.1313 | 0.3322 | 0.00066 | ---?? | 0 | 0.5237 | | |
| rs28515692 | 9 | 44265793 | 3 | 0 | 0 | T | G | 1.7476 | 0.5132 | 0.0006603 | +++?? | 0 | 0.8201 | intergenic | |
| 5:97159279 | 5 | 97159279 | 3 | 0 | 0 | CCT | C | 1.1833 | 0.3475 | 0.0006603 | +++?? | 50.5 | 0.1327 | | |
| rs11011649 | 10 | 20151213 | 4 | 0 | 0 | T | C | 0.8755 | 0.2571 | 0.0006604 | +++?+ | 11.4 | 0.3356 | intron | PLXDC2 |

| | | | | | | | | | | | | | | | |
|---|---|---|---|---|---|---|---|---|---|---|---|---|---|---|---|
| 12:70492559 | 12 | 70492559 | 3 | 0 | 0 | G | GAGAGT | 0.8752 | 0.257 | 0.0006604 | +++?? | 14.9 | 0.3088 | | |
| 4:88396443 | 4 | 88396443 | 2 | 0 | 0 | A | ACTT | -3.8839 | 1.1405 | 0.0006605 | -?-?? | 0 | 0.6027 | | |
| rs7868819 | 9 | 90816092 | 3 | 0 | 0 | A | G | 1.6851 | 0.4948 | 0.0006606 | +++?? | 0 | 0.8842 | intergenic | |
| rs145930075 | 18 | 52761829 | 4 | 0 | 0 | A | G | -4.1905 | 1.2305 | 0.0006606 | ---?- | 0 | 0.8516 | intergenic | |
| rs1487289 | 1 | 98603170 | 4 | 0 | 0 | A | G | -1.3338 | 0.3917 | 0.0006607 | ---?- | 21.3 | 0.2826 | intergenic | |
| rs9277976 | 6 | 33292745 | 5 | 0 | 0 | T | C | -1.3576 | 0.3987 | 0.0006607 | ----- | 6.8 | 0.3678 | upstream-gene |  |
| rs11890854 | 2 | 137017060 | 3 | 0 | 0 | A | C | -0.9549 | 0.2804 | 0.0006607 | ---?? | 0 | 0.7686 | intergenic | |
| rs73445897 | 15 | 86436918 | 4 | 0 | 0 | A | G | 2.202 | 0.6466 | 0.0006607 | +++?+ | 0 | 0.7319 | intergenic | |
| rs11808688 | 1 | 203689314 | 4 | 0 | 0 | A | G | 1.1128 | 0.3268 | 0.0006608 | +++?+ | 0 | 0.8903 | intron | ATP2B4 |
| rs147559682 | 12 | 48085655 | 4 | 0 | 0 | A | C | -1.7108 | 0.5024 | 0.0006608 | ---?+ | 53.7 | 0.09075 | intron | RPAP3 |
| rs852965 | 6 | 72134958 | 4 | 0 | 0 | T | C | 0.8835 | 0.2595 | 0.0006609 | +++?+ | 1 | 0.3871 | upstream-gene | |
| rs60591831 | 1 | 32240968 | 4 | 0 | 0 | T | C | -1.2182 | 0.3577 | 0.0006609 | ---?+ | 41.6 | 0.1618 | intergenic | |
| rs6689602 | 1 | 180132688 | 4 | 0 | 0 | A | G | 2.2027 | 0.6469 | 0.0006613 | +++?- | 37.5 | 0.187 | intron | QSOX1 |
| rs112254014 | 18 | 47052853 | 4 | 0 | 0 | A | G | -1.9763 | 0.5804 | 0.0006615 | ---?- | 0 | 0.9186 | intergenic | |
| rs9937047 | 16 | 58145585 | 4 | 0 | 0 | A | G | -0.9083 | 0.2667 | 0.0006616 | ---?- | 3.9 | 0.3733 | downstream-gene | |
| rs34517439 | 1 | 78450517 | 4 | 0 | 0 | A | C | 1.4486 | 0.4254 | 0.0006618 | +++?- | 0 | 0.5067 | intron | DNAJB4 |
| rs10056035 | 5 | 743206 | 3 | 0 | 0 | C | G | -1.5459 | 0.454 | 0.0006619 | ---?? | 0 | 0.4132 | intron | ZDHHC11B |
| rs117177345 | 8 | 3665476 | 3 | 0 | 0 | C | G | -3.5818 | 1.0519 | 0.0006619 | ---?? | 29 | 0.2447 | intron | CSMD1 |
| rs11467 | 8 | 38644914 | 4 | 0 | 0 | A | G | 0.8815 | 0.2589 | 0.0006621 | +++?+ | 0.2 | 0.3908 | 5-prime-UTR | TACC1 |
| rs3849743 | 5 | 165434873 | 4 | 0 | 0 | A | G | -0.8598 | 0.2525 | 0.0006622 | ---?- | 0 | 0.4421 | intergenic | |
| rs12607685 | 18 | 73330298 | 4 | 0 | 0 | T | C | 2.3938 | 0.7031 | 0.0006622 | +++?+ | 0 | 0.6989 | intergenic | |
| rs78763644 | 17 | 52280662 | 4 | 0 | 0 | A | G | 2.0839 | 0.6121 | 0.0006624 | +++?- | 35.4 | 0.1995 | intergenic | |
| rs56169804 | 4 | 190023778 | 4 | 0 | 0 | T | C | 0.8495 | 0.2495 | 0.0006627 | +++?+ | 0 | 0.4289 | intergenic | |
| rs140965658 | 2 | 165686129 | 4 | 0 | 0 | A | G | 3.3519 | 0.9845 | 0.0006627 | +-+?+ | 23.5 | 0.2701 | intron | COBLL1 |
| rs917074 | 7 | 49472674 | 4 | 0 | 0 | A | T | -0.9613 | 0.2824 | 0.0006628 | ---?+ | 26.6 | 0.2525 | intergenic | |
| rs9303955 | 18 | 457867 | 4 | 0 | 0 | A | G | -0.8435 | 0.2478 | 0.0006628 | ---?- | 31.8 | 0.2216 | intron | COLEC12 |
| rs12808544 | 11 | 58373221 | 5 | 0 | 0 | A | C | -0.8175 | 0.2401 | 0.0006628 | ----- | 0 | 0.9831 | intron | ZFP91,ZFP91-CNTF |
| rs250512 | 19 | 8370959 | 4 | 0 | 0 | C | G | -1.639 | 0.4814 | 0.0006629 | ---?- | 43.1 | 0.153 | intron | CD320 |
| rs9268519 | 6 | 32380826 | 5 | 0 | 0 | A | G | 0.7221 | 0.2121 | 0.0006629 | +++++ | 0 | 0.6566 | intergenic | |
| rs11719892 | 3 | 143526418 | 4 | 0 | 0 | T | C | 0.8278 | 0.2431 | 0.000663 | +++?+ | 0 | 0.5808 | intron | SLC9A9 |
| rs61928861 | 12 | 11443303 | 4 | 0 | 0 | A | G | 1.5672 | 0.4603 | 0.0006632 | +++?+ | 33 | 0.214 | intergenic | |
| rs75330487 | 18 | 7198875 | 4 | 0 | 0 | A | C | 2.1504 | 0.6317 | 0.0006632 | +-+?+ | 24.9 | 0.2621 | intergenic | |
| rs1032077 | 3 | 176025504 | 4 | 0 | 0 | T | G | 1.2759 | 0.3748 | 0.0006634 | +++?+ | 13.7 | 0.3237 | intergenic | |
| rs78355072 | 11 | 86176540 | 4 | 0 | 0 | T | C | 2.4259 | 0.7126 | 0.0006636 | +++?+ | 0 | 0.8774 | intron | ME3 |
| rs12741042 | 1 | 64786970 | 3 | 0 | 0 | T | G | 1.266 | 0.3719 | 0.0006637 | +++?? | 0 | 0.5233 | intergenic | |
| rs6956904 | 7 | 49472204 | 4 | 0 | 0 | C | G | 0.9603 | 0.2821 | 0.000664 | +++?- | 26.4 | 0.2535 | intergenic | |
| rs182157089 | 2 | 170431900 | 3 | 0 | 0 | T | C | -1.2925 | 0.3797 | 0.0006641 | ---?? | 0 | 0.9996 | upstream-gene | |
| rs11165953 | 1 | 98599780 | 4 | 0 | 0 | T | C | -1.3343 | 0.392 | 0.0006642 | ---?- | 19.6 | 0.2919 | intergenic | |
| 20:42406461 | 20 | 42406461 | 3 | 0 | 0 | A | AT | 4.0357 | 1.1856 | 0.0006645 | +++?? | 0 | 0.6171 | | |
| rs3445 | 6 | 33246518 | 5 | 0 | 0 | A | G | 1.0399 | 0.3055 | 0.0006646 | +++++ | 21.8 | 0.2755 | 3-prime-UTR | B3GALT4 |
| 10:28213917 | 10 | 28213917 | 3 | 0 | 0 | G | GA | 0.8881 | 0.2609 | 0.0006646 | +++?? | 14.2 | 0.3119 | | |
| 1:182013610 | 1 | 182013610 | 3 | 0 | 0 | A | AATAG | -2.9308 | 0.861 | 0.0006646 | ---?? | 0 | 0.8861 | | |
| 1:72168409 | 1 | 72168409 | 3 | 0 | 0 | T | TA | -1.1027 | 0.324 | 0.0006646 | ---?? | 0 | 0.9134 | | |
| 12:25545721 | 12 | 25545721 | 2 | 0 | 0 | A | AGAGT | -4.9537 | 1.4554 | 0.0006647 | -?-?? | 41.4 | 0.1913 | | |
| rs16954302 | 17 | 52278314 | 4 | 0 | 0 | T | C | 2.0895 | 0.6139 | 0.0006648 | +++?- | 35.5 | 0.1993 | intergenic | |
| rs189156520 | 3 | 146129360 | 3 | 0 | 0 | T | C | 5.2111 | 1.531 | 0.0006648 | +?+?+ | 36.4 | 0.2077 | intron | PLSCR2 |
| rs1032196 | 1 | 98602249 | 4 | 0 | 0 | A | G | 1.3337 | 0.3918 | 0.000665 | +++?+ | 20.4 | 0.2873 | intergenic | |
| rs9268518 | 6 | 32380818 | 5 | 0 | 0 | A | G | -0.7219 | 0.2121 | 0.0006652 | ----- | 0 | 0.6571 | intergenic |  |
| rs6811505 | 4 | 20733963 | 4 | 0 | 0 | C | G | 0.8867 | 0.2605 | 0.0006653 | +++?+ | 0 | 0.8469 | intron | KCNIP4 |
| rs11165954 | 1 | 98600110 | 4 | 0 | 0 | A | G | -1.334 | 0.392 | 0.0006654 | ---?- | 19.7 | 0.2914 | intergenic | |
| rs36118801 | 13 | 30312359 | 5 | 0 | 0 | T | G | -1.2699 | 0.3731 | 0.0006658 | ----- | 0 | 0.561 | intergenic |  |
| rs115007184 | 4 | 127996574 | 4 | 0 | 0 | T | C | 2.7423 | 0.8058 | 0.000666 | +++?+ | 0 | 0.5157 | intergenic | |
| rs72620505 | 19 | 1641922 | 4 | 0 | 0 | T | G | -2.7134 | 0.7973 | 0.0006661 | ---?- | 0 | 0.8216 | intron | TCF3 |
| rs6806591 | 3 | 196517566 | 4 | 0 | 0 | A | G | 1.6347 | 0.4804 | 0.0006662 | +++?- | 42 | 0.1597 | intron | PAK2 |
| rs11185569 | 7 | 49471861 | 4 | 0 | 0 | T | C | -0.9595 | 0.2819 | 0.0006662 | ---?+ | 26.3 | 0.2541 | intergenic | |
| rs62332113 | 5 | 746877 | 3 | 0 | 0 | A | G | 1.5496 | 0.4553 | 0.0006662 | +++?? | 0 | 0.4778 | intron | ZDHHC11B |
| rs4544392 | 2 | 184038693 | 4 | 0 | 0 | T | C | 1.1551 | 0.3394 | 0.0006663 | +++?+ | 0 | 0.8308 | intergenic | |
| rs5997372 | 22 | 28909942 | 4 | 0 | 0 | A | G | -2.2907 | 0.6731 | 0.0006663 | ---?- | 0 | 0.5689 | intron | TTC28 |
| rs7644694 | 3 | 61096355 | 4 | 0 | 0 | A | G | -3.1925 | 0.9381 | 0.0006665 | ---?- | 0 | 0.7115 | intron | FHIT |

| | | | | | | | | | | | | | | | |
|---|---|---|---|---|---|---|---|---|---|---|---|---|---|---|---|
| rs56974443 | 9 | 36216426 | 3 | 0 | 0 | A | G | -0.8714 | 0.2561 | 0.0006666 | ---?? | 0 | 0.9036 | 3-prime-UTR | GNE |
| rs201869387 | 9 | 44284389 | 3 | 0 | 0 | T | C | -1.9043 | 0.5596 | 0.0006667 | ---?? | 0 | 0.8311 | intergenic | |
| rs111377128 | 3 | 77350289 | 4 | 0 | 0 | T | G | 2.1938 | 0.6447 | 0.000667 | +++?+ | 54.2 | 0.08755 | intron | ROBO2 |
| rs2179704 | 20 | 56210224 | 4 | 0 | 0 | T | C | -1.191 | 0.35 | 0.0006671 | ---?- | 0 | 0.967 | intergenic | |
| rs2675044 | 2 | 184000167 | 4 | 0 | 0 | A | G | -1.1697 | 0.3438 | 0.0006672 | ---?- | 0 | 0.8519 | intron | NUP35 |
| 1:87469392 | 1 | 87469392 | 3 | 0 | 0 | CT | C | 1.0119 | 0.2974 | 0.0006672 | +++?? | 0.9 | 0.3646 | | |
| rs28550821 | 9 | 44260853 | 3 | 0 | 0 | A | G | 1.7443 | 0.5126 | 0.0006674 | +++?? | 0 | 0.8142 | intergenic | |
| rs144408238 | 3 | 143041115 | 4 | 0 | 0 | T | C | -1.9963 | 0.5867 | 0.0006674 | ---?- | 0 | 0.6767 | intron | SLC9A9 |
| rs115879342 | 2 | 184043213 | 4 | 0 | 0 | A | G | 1.1583 | 0.3404 | 0.0006677 | +++?+ | 0 | 0.8303 | intergenic | |
| rs35484347 | 13 | 30312175 | 5 | 0 | 0 | T | G | 1.2701 | 0.3733 | 0.0006678 | +++++ | 0 | 0.5557 | intergenic | |
| rs1974988 | 2 | 170647837 | 3 | 0 | 0 | T | C | 0.8738 | 0.2568 | 0.0006681 | +++?? | 0.2 | 0.3672 | intergenic | |
| rs11165955 | 1 | 98600210 | 4 | 0 | 0 | T | C | -1.3335 | 0.3919 | 0.0006682 | ---?- | 19.7 | 0.2912 | intergenic | |
| 16:57454295 | 16 | 57454295 | 3 | 0 | 0 | G | GA | 1.2959 | 0.3809 | 0.0006684 | +++?? | 0 | 0.8628 | | |
| rs11165956 | 1 | 98600355 | 4 | 0 | 0 | T | C | 1.3334 | 0.3919 | 0.0006686 | +++?+ | 19.7 | 0.2912 | intergenic | |
| rs34166160 | 12 | 732505 | 4 | 0 | 0 | A | C | -3.6882 | 1.0841 | 0.0006689 | ---?- | 26.4 | 0.2536 | intron | NINJ2 |
| rs17228585 | 15 | 32985305 | 4 | 0 | 0 | A | T | 2.3059 | 0.6778 | 0.000669 | +++?+ | 0 | 0.6048 | intron | SCG5 |
| rs11783119 | 8 | 113214505 | 4 | 0 | 0 | A | G | -0.9666 | 0.2841 | 0.0006691 | ---?- | 0 | 0.9757 | intergenic | |
| rs28372665 | 9 | 44268312 | 3 | 0 | 0 | A | G | 1.7538 | 0.5155 | 0.0006692 | +++?? | 0 | 0.8286 | intergenic | |
| 10:114137985 | 10 | 114137985 | 3 | 0 | 0 | G | GTTT | 0.9775 | 0.2873 | 0.0006693 | +++?? | 34.1 | 0.2191 | | |
| rs28528169 | 9 | 44260872 | 3 | 0 | 0 | T | C | 1.744 | 0.5127 | 0.0006694 | +++?? | 0 | 0.8146 | intergenic | |
| rs12043365 | 1 | 98598019 | 4 | 0 | 0 | A | T | -1.3337 | 0.3921 | 0.0006699 | ---?- | 19.1 | 0.2947 | intergenic | |
| rs2570127 | 15 | 87612901 | 4 | 0 | 0 | A | C | -0.912 | 0.2681 | 0.00067 | ---?- | 0 | 0.6473 | intergenic | |
| rs968154 | 6 | 32380785 | 5 | 0 | 0 | A | G | 0.7214 | 0.2121 | 0.0006701 | +++++ | 0 | 0.658 | intergenic | |
| rs1751582 | 1 | 208187670 | 4 | 0 | 0 | A | G | -2.0624 | 0.6063 | 0.0006702 | ---?- | 0 | 0.6863 | intergenic | |
| rs2114473 | 4 | 20733310 | 4 | 0 | 0 | T | G | 0.8865 | 0.2606 | 0.0006702 | +++?+ | 0 | 0.8468 | intron | KCNIP4 |
| rs16954306 | 17 | 52281378 | 4 | 0 | 0 | T | G | 2.0807 | 0.6117 | 0.0006702 | +++?- | 35.4 | 0.2001 | intergenic | |
| rs12254034 | 10 | 134571007 | 4 | 0 | 0 | T | C | -0.9584 | 0.2818 | 0.0006703 | ---?- | 0 | 0.4593 | intron | INPP5A |
| rs2701414 | 15 | 87612902 | 4 | 0 | 0 | T | C | 0.912 | 0.2681 | 0.0006705 | +++?+ | 0 | 0.6471 | intergenic | |
| rs61883647 | 11 | 14781646 | 4 | 0 | 0 | A | G | 0.9417 | 0.2769 | 0.0006707 | +++?+ | 39.1 | 0.1773 | intron | PDE3B |
| rs62050673 | 16 | 85241278 | 4 | 0 | 0 | T | C | 1.2252 | 0.3602 | 0.0006707 | +++?+ | 0 | 0.7047 | intron | GSE1 |
| rs73414942 | 18 | 25187462 | 4 | 0 | 0 | A | G | 2.4694 | 0.726 | 0.0006707 | +++?+ | 0 | 0.629 | intergenic | |
| rs7927532 | 11 | 12640551 | 4 | 0 | 0 | A | G | 1.0441 | 0.307 | 0.0006708 | +++?+ | 0 | 0.796 | intergenic | |
| rs138901704 | 3 | 29716072 | 4 | 0 | 0 | A | G | 4.1291 | 1.214 | 0.0006709 | +++?+ | 12.8 | 0.3287 | intron | RBMS3 |
| rs12120811 | 1 | 157454010 | 4 | 0 | 0 | A | G | 0.9383 | 0.2759 | 0.000671 | +++?- | 31.7 | 0.2218 | intergenic | |
| rs16823091 | 2 | 170670059 | 5 | 0 | 0 | A | G | 0.8012 | 0.2356 | 0.000671 | +++++ | 48.8 | 0.09874 | intron | METTL5 |
| rs76255363 | 19 | 46169474 | 5 | 0 | 1 | A | G | 0.8926 | 0.2625 | 0.0006711 | +-++- | 27.1 | 0.2408 | upstream-gene | |
| 6:165844094 | 6 | 165844094 | 3 | 0 | 0 | A | AACATC | -1.3413 | 0.3944 | 0.0006711 | ---?? | 66.4 | 0.05119 | | |
| rs10885600 | 10 | 116485847 | 4 | 0 | 0 | T | C | 1.3756 | 0.4045 | 0.0006713 | +++?+ | 0 | 0.869 | intergenic | |
| rs56121565 | 2 | 170670163 | 5 | 0 | 0 | A | G | -0.8013 | 0.2356 | 0.0006713 | ----- | 48.6 | 0.09964 | intron | METTL5 |
| rs7858447 | 9 | 35677245 | 5 | 0 | 0 | A | G | -0.9387 | 0.276 | 0.0006716 | ----+ | 30.9 | 0.2157 | intron | CA9 |
| rs10910557 | 1 | 232692293 | 4 | 0 | 0 | A | G | 0.9924 | 0.2918 | 0.0006718 | +++?+ | 10 | 0.3429 | intron | SIPA1L2 |
| rs146783327 | 4 | 20968119 | 3 | 0 | 0 | A | G | 3.2107 | 0.9441 | 0.0006718 | +++?? | 0 | 0.5786 | intron | KCNIP4 |
| rs1442178 | 12 | 41313189 | 4 | 0 | 0 | A | G | -1.6889 | 0.4966 | 0.000672 | ---?- | 0 | 0.8646 | intron | CNTN1 |
| rs79766819 | 2 | 218831043 | 4 | 0 | 0 | T | C | 1.3536 | 0.398 | 0.0006722 | +++?- | 0 | 0.7369 | intergenic | |
| rs34279346 | 2 | 138963333 | 4 | 0 | 0 | A | G | 1.6451 | 0.4838 | 0.0006723 | +++?+ | 0 | 0.8324 | intergenic | |
| rs150968661 | 1 | 152806778 | 3 | 0 | 0 | A | C | 1.0625 | 0.3124 | 0.0006723 | +++?? | 0 | 0.5499 | intergenic | |
| rs7932515 | 11 | 23966043 | 4 | 0 | 0 | T | C | 1.2265 | 0.3607 | 0.0006724 | +++?+ | 0 | 0.7325 | intergenic | |
| 12:10870397 | 12 | 10870397 | 3 | 0 | 0 | T | TC | -3.6273 | 1.0667 | 0.0006726 | ---?? | 0 | 0.8481 | | |
| rs117024557 | 22 | 40743813 | 3 | 0 | 0 | A | G | 3.8652 | 1.1366 | 0.0006727 | +++?? | 10.3 | 0.3282 | intron | ADSL |
| 3:173480755 | 3 | 173480755 | 3 | 0 | 0 | T | TAC | 1.2378 | 0.364 | 0.0006727 | +++?? | 0 | 0.5958 | | |
| rs61922202 | 12 | 29786823 | 4 | 0 | 0 | T | C | -1.4656 | 0.431 | 0.0006729 | ---?- | 0 | 0.7499 | intron | TMTC1 |
| 14:21456633 | 14 | 21456633 | 3 | 0 | 0 | CAATTT | C | -0.9221 | 0.2712 | 0.0006729 | ---?? | 17.8 | 0.2961 | | |
| rs10234478 | 7 | 10156712 | 4 | 0 | 0 | A | T | -0.9679 | 0.2847 | 0.000673 | ---?- | 45.7 | 0.137 | intergenic | |
| 18:40413105 | 18 | 40413105 | 3 | 0 | 0 | T | TC | 4.073 | 1.1978 | 0.000673 | +++?? | 0 | 0.7846 | | |
| rs150142020 | 11 | 67513187 | 3 | 0 | 0 | T | C | -2.0912 | 0.615 | 0.0006731 | ---?? | 64.3 | 0.06097 | intergenic | |
| rs78270617 | 10 | 33794579 | 4 | 0 | 0 | A | T | -3.4226 | 1.0066 | 0.0006732 | ---?- | 55.1 | 0.0829 | intergenic | |
| rs73201092 | 21 | 36314277 | 4 | 0 | 0 | A | C | -1.102 | 0.3241 | 0.0006733 | ---?- | 0 | 0.9347 | intron | RUNX1 |
| rs78167629 | 10 | 109285700 | 4 | 0 | 0 | A | C | 2.3875 | 0.7022 | 0.0006735 | +++?+ | 0 | 0.9733 | intergenic | |
| rs6729378 | 2 | 231410520 | 4 | 0 | 0 | T | C | 1.1189 | 0.3291 | 0.0006736 | +++?+ | 0 | 0.5025 | downstream-gene | |

| | | | | | | | | | | | | | | | |
|---|---|---|---|---|---|---|---|---|---|---|---|---|---|---|---|
| rs1457687 | 5 | 123490628 | 4 | 0 | 0 | T | G | 0.8876 | 0.261 | 0.0006737 | +++?+ | 0 | 0.7447 | intergenic | |
| rs28697924 | 9 | 44257192 | 3 | 0 | 0 | A | T | -1.7283 | 0.5083 | 0.0006739 | ---?? | 0 | 0.8001 | intergenic | |
| rs10501168 | 11 | 37056944 | 4 | 0 | 0 | C | G | -0.8445 | 0.2484 | 0.000674 | ---?- | 6 | 0.3631 | intergenic | |
| rs114632657 | 5 | 38997170 | 4 | 0 | 0 | A | G | 3.0253 | 0.8898 | 0.0006741 | +++?+ | 0 | 0.5552 | intron | RICTOR |
| rs28438551 | 9 | 44256735 | 3 | 0 | 0 | T | G | -1.7282 | 0.5083 | 0.0006743 | ---?? | 0 | 0.7996 | intergenic | |
| rs139678865 | 17 | 76877067 | 3 | 0 | 0 | A | G | -3.2246 | 0.9485 | 0.0006745 | ---?? | 0 | 0.6902 | intron | TIMP2 |
| 6:159235577 | 6 | 159235577 | 3 | 0 | 0 | CTT | C | 0.9435 | 0.2775 | 0.0006746 | +++?? | 0 | 0.8549 | | |
| rs4670034 | 2 | 5271928 | 4 | 0 | 0 | T | C | -0.9606 | 0.2825 | 0.0006747 | ---?- | 0 | 0.9691 | intergenic | |
| rs805749 | 20 | 5682567 | 3 | 0 | 0 | A | G | -0.9177 | 0.27 | 0.0006754 | ---?? | 0 | 0.9164 | intergenic | |
| rs2717716 | 8 | 18437200 | 4 | 0 | 0 | A | G | 0.8271 | 0.2433 | 0.0006754 | +++?+ | 0 | 0.7202 | intron | PSD3 |
| 10:72468242 | 10 | 72468242 | 3 | 0 | 0 | A | G | -4.7535 | 1.3984 | 0.0006755 | ---?? | 0 | 0.9967 | intron | ADAMTS14 |
| rs72672825 | 8 | 113217832 | 4 | 0 | 0 | A | G | 0.9669 | 0.2844 | 0.0006756 | +++?+ | 0 | 0.9737 | intergenic | |
| rs1764507 | 1 | 208187652 | 4 | 0 | 0 | T | G | 2.0674 | 0.6082 | 0.0006759 | +++?+ | 0 | 0.6819 | intergenic | |
| rs8081007 | 17 | 77989580 | 4 | 0 | 0 | T | C | -0.8708 | 0.2562 | 0.0006759 | ---?- | 0 | 0.8124 | intron | TBC1D16 |
| rs12139888 | 1 | 117879399 | 4 | 0 | 0 | A | C | 2.7233 | 0.8012 | 0.0006759 | +++?- | 29.6 | 0.2348 | intergenic | |
| rs2028732 | 4 | 37004281 | 4 | 0 | 0 | T | G | -0.8521 | 0.2507 | 0.000676 | ---?+ | 46.2 | 0.1339 | intergenic | |
| rs117807021 | 12 | 91770919 | 3 | 0 | 0 | A | G | 4.3957 | 1.2932 | 0.000676 | +++?? | 0 | 0.7017 | intergenic | |
| rs28406482 | 9 | 44264363 | 3 | 0 | 0 | T | C | 1.7526 | 0.5156 | 0.0006762 | +++?? | 0 | 0.8143 | intergenic | |
| rs71645580 | 1 | 64786472 | 4 | 0 | 0 | T | C | -1.0379 | 0.3054 | 0.0006765 | ---?- | 0 | 0.4568 | intergenic | |
| rs4458518 | 4 | 111396971 | 4 | 0 | 0 | A | T | -0.8877 | 0.2612 | 0.0006767 | ---?- | 0 | 0.8072 | upstream-gene | |
| rs72888282 | 2 | 47744130 | 4 | 0 | 0 | T | C | 1.5204 | 0.4473 | 0.0006767 | +++?+ | 37 | 0.1902 | downstream-gene | |
| rs80050931 | 8 | 113314439 | 3 | 0 | 0 | T | C | -0.9932 | 0.2922 | 0.0006767 | ---?? | 0 | 0.5648 | intron | CSMD3 |
| rs191791016 | 6 | 32055163 | 4 | 0 | 0 | A | C | 2.798 | 0.8232 | 0.0006767 | ++++? | 18.2 | 0.2998 | intron | TNXB |
| rs1359830 | 7 | 49470255 | 4 | 0 | 0 | A | G | -0.9556 | 0.2811 | 0.0006769 | ---?+ | 25.9 | 0.2564 | intergenic | |
| rs2588538 | 4 | 37004743 | 4 | 0 | 0 | A | G | 0.8519 | 0.2507 | 0.0006769 | +++?- | 46.2 | 0.1339 | intergenic | |
| 6:31105078 | 6 | 31105078 | 3 | 0 | 0 | CAAA | C | 0.8984 | 0.2643 | 0.0006769 | +++?? | 0 | 0.646 | | |
| rs35209562 | 11 | 12647771 | 4 | 0 | 0 | T | C | 1.0399 | 0.306 | 0.000677 | +++?+ | 0 | 0.8111 | intergenic | |
| rs17117764 | 1 | 98601002 | 4 | 0 | 0 | A | G | 1.3318 | 0.3919 | 0.0006771 | +++?+ | 19.9 | 0.2905 | intergenic | |
| rs4666615 | 2 | 184016917 | 4 | 0 | 0 | A | G | 1.1686 | 0.3438 | 0.0006772 | +++?+ | 0 | 0.8524 | intron | NUP35 |
| rs115280199 | 3 | 47868235 | 2 | 0 | 0 | T | C | 6.5838 | 1.9372 | 0.0006772 | ?+??+ | 0 | 0.3221 | intron | DHX30 |
| rs116695008 | 17 | 57075501 | 5 | 0 | 0 | A | C | 2.3092 | 0.6794 | 0.0006772 | ++++- | 0 | 0.6873 | intron | TRIM37 |
| rs2108814 | 2 | 158823617 | 4 | 0 | 0 | T | G | 0.8275 | 0.2435 | 0.0006776 | +++?+ | 57.2 | 0.07149 | intron | UPP2 |
| 16:57453685 | 16 | 57453685 | 3 | 0 | 0 | G | GC | 1.2949 | 0.381 | 0.0006777 | +++?? | 0 | 0.8639 | | |
| rs62510572 | 8 | 49812579 | 4 | 0 | 0 | A | G | -3.0583 | 0.8999 | 0.000678 | ---?+ | 0 | 0.4086 | intergenic | |
| rs73573239 | 19 | 46169446 | 5 | 0 | 1 | A | T | -0.8921 | 0.2625 | 0.000678 | -+--+ | 27.2 | 0.2404 | upstream-gene | |
| rs17117766 | 1 | 98601353 | 4 | 0 | 0 | A | T | 1.3315 | 0.3918 | 0.0006785 | +++?+ | 20 | 0.2897 | intergenic | |
| rs201634389 | 9 | 44285696 | 3 | 0 | 0 | A | G | -1.8857 | 0.5549 | 0.0006787 | ---?? | 0 | 0.8321 | intergenic | |
| rs7810097 | 7 | 10157264 | 4 | 0 | 0 | T | C | -0.9673 | 0.2847 | 0.0006788 | ---?- | 46.2 | 0.1341 | intergenic | |
| rs2588537 | 4 | 37005939 | 4 | 0 | 0 | A | G | 0.8514 | 0.2506 | 0.0006792 | +++?- | 46.3 | 0.1337 | intergenic | |
| rs74844716 | 2 | 208392215 | 4 | 0 | 0 | T | C | 1.4102 | 0.415 | 0.0006792 | +++?+ | 0 | 0.7897 | upstream-gene | |
| rs4244510 | 11 | 23974818 | 4 | 0 | 0 | C | G | 1.217 | 0.3582 | 0.0006795 | +++?+ | 0 | 0.7237 | intergenic | |
| 9:107790574 | 9 | 107790574 | 3 | 0 | 0 | T | TA | 3.3818 | 0.9953 | 0.0006797 | +++?? | 37.3 | 0.203 | | |
| rs2701415 | 15 | 87612828 | 4 | 0 | 0 | T | C | -0.9102 | 0.2679 | 0.0006799 | ---?- | 0 | 0.65 | intergenic | |
| rs74130658 | 1 | 192460535 | 4 | 0 | 0 | T | G | -1.1338 | 0.3337 | 0.0006799 | ---?+ | 0 | 0.4134 | intergenic | |
| rs7291143 | 22 | 41395588 | 4 | 0 | 0 | T | C | -1.5743 | 0.4634 | 0.0006801 | ---?+ | 0 | 0.4693 | intergenic | |
| rs72723727 | 1 | 100183357 | 4 | 0 | 0 | A | C | 2.1602 | 0.6358 | 0.0006801 | +++?+ | 0 | 0.4641 | intron | FRRS1 |
| rs4244511 | 11 | 23974839 | 4 | 0 | 0 | A | C | 1.2169 | 0.3582 | 0.0006802 | +++?+ | 0 | 0.7236 | intergenic | |
| rs112076601 | 8 | 102198249 | 3 | 0 | 0 | T | C | -6.0478 | 1.7801 | 0.0006803 | --??- | 0 | 0.4028 | intergenic | |
| rs729767 | 19 | 12829463 | 3 | 0 | 0 | C | G | -3.7217 | 1.0955 | 0.0006804 | ?---? | 64 | 0.06203 | intron | TNPO2 |
| rs2108813 | 2 | 158823641 | 4 | 0 | 0 | A | T | -0.8278 | 0.2437 | 0.0006804 | ---?- | 57.2 | 0.07184 | intron | UPP2 |
| rs78595522 | 11 | 20153625 | 4 | 0 | 0 | T | G | 1.735 | 0.5107 | 0.0006805 | +++?+ | 0 | 0.7049 | intergenic | |
| rs10896796 | 11 | 58394256 | 5 | 0 | 0 | A | G | 0.8224 | 0.2421 | 0.0006807 | +++++ | 0 | 0.9811 | downstream-gene | |
| 6:129207440 | 6 | 129207440 | 3 | 0 | 0 | T | TA | 3.6478 | 1.0738 | 0.0006807 | +++?? | 65 | 0.05732 | | |
| rs75410527 | 7 | 49466542 | 4 | 0 | 0 | T | C | 0.9495 | 0.2795 | 0.0006808 | +++?- | 25.1 | 0.261 | intergenic | |
| rs2570128 | 15 | 87612741 | 4 | 0 | 0 | A | G | 0.91 | 0.2679 | 0.0006809 | +++?+ | 0 | 0.6502 | intergenic | |
| rs6511837 | 19 | 12829697 | 3 | 0 | 0 | A | G | 3.7212 | 1.0954 | 0.000681 | ?+++? | 64 | 0.06196 | intron | TNPO2 |
| rs75477470 | 1 | 42704957 | 4 | 0 | 0 | C | G | 1.8163 | 0.5347 | 0.0006811 | +++?+ | 0 | 0.5688 | intron | FOXJ3 |
| rs35750364 | 4 | 10049049 | 5 | 0 | 0 | T | C | -0.7147 | 0.2104 | 0.0006812 | ----+ | 8.7 | 0.3566 | intergenic |  |
| rs66519675 | 2 | 47377795 | 4 | 0 | 0 | T | C | 1.3085 | 0.3852 | 0.0006814 | +++?+ | 0 | 0.9728 | intron | C2orf61 |

| | | | | | | | | | | | | | | | |
|---|---|---|---|---|---|---|---|---|---|---|---|---|---|---|---|
| rs146675635 | 14 | 33288052 | 4 | 0 | 0 | T | G | -3.8575 | 1.1356 | 0.0006814 | ---?- | 0 | 0.9404 | intron | AKAP6 |
| rs4880275 | 10 | 134576552 | 4 | 0 | 0 | T | C | 0.9585 | 0.2822 | 0.0006815 | +++?+ | 0 | 0.4639 | intron | INPP5A |
| rs10769941 | 11 | 8710244 | 5 | 0 | 0 | T | G | 0.7721 | 0.2273 | 0.0006816 | ++++- | 0 | 0.8438 | 3-prime-UTR | RPL27A |
| rs28457799 | 9 | 44286238 | 3 | 0 | 0 | A | G | 1.8936 | 0.5575 | 0.0006816 | +++?? | 0 | 0.8323 | intergenic | |
| rs79418941 | 6 | 71750514 | 4 | 0 | 0 | A | G | -1.3975 | 0.4114 | 0.0006818 | ---?+ | 0 | 0.4661 | intergenic | |
| 15:57675496 | 15 | 57675496 | 3 | 0 | 0 | T | TTTTTTT | -1.2534 | 0.369 | 0.0006818 | -+-?? | 49.9 | 0.1359 | | |
| 1:87947641 | 1 | 87947641 | 3 | 0 | 0 | CA | C | -1.2226 | 0.3599 | 0.0006819 | ---?? | 26.1 | 0.2585 | | |
| rs7259207 | 19 | 12828375 | 3 | 0 | 0 | T | C | -3.7246 | 1.0965 | 0.000682 | ?---? | 63.9 | 0.06263 | intron | TNPO2 |
| rs6907622 | 6 | 31235610 | 5 | 0 | 0 | A | C | 0.8113 | 0.2388 | 0.0006821 | +++++ | 0 | 0.484 | downstream-gene | |
| rs145043902 | 1 | 40407659 | 3 | 0 | 0 | T | G | 3.7213 | 1.0956 | 0.0006822 | +++?? | 0 | 0.3761 | intergenic | |
| rs142589589 | 16 | 27998516 | 4 | 0 | 0 | T | C | 2.0157 | 0.5935 | 0.0006823 | +++?+ | 0 | 0.9271 | intron | GSG1L |
| rs28497748 | 9 | 16187335 | 4 | 0 | 0 | T | C | -0.9118 | 0.2684 | 0.0006824 | ---?- | 0 | 0.9265 | intergenic | |
| rs2176231 | 2 | 183999281 | 4 | 0 | 0 | A | G | -1.168 | 0.3439 | 0.0006825 | ---?- | 0 | 0.8554 | intron | NUP35 |
| rs10501718 | 11 | 90022286 | 3 | 0 | 0 | C | G | 3.4153 | 1.0055 | 0.0006825 | +++?? | 0 | 0.4535 | intergenic | |
| rs10927352 | 1 | 245106683 | 4 | 0 | 0 | T | G | -1.1552 | 0.3401 | 0.0006825 | ---?- | 0.1 | 0.3913 | intergenic | |
| rs72898506 | 2 | 184057386 | 4 | 0 | 0 | A | G | 1.1654 | 0.3431 | 0.0006828 | +++?+ | 0 | 0.7671 | intergenic | |
| rs139868625 | 12 | 103250733 | 4 | 0 | 0 | T | C | 0.831 | 0.2447 | 0.0006828 | +++?+ | 0 | 0.652 | intron | PAH |
| rs2760338 | 4 | 37006559 | 4 | 0 | 0 | A | T | 0.8507 | 0.2505 | 0.0006831 | +++?- | 46.3 | 0.1336 | intergenic | |
| 17:51135954 | 17 | 51135954 | 3 | 0 | 0 | CTTTAA | C | 1.4506 | 0.4271 | 0.0006834 | +++?? | 0 | 0.3732 | | |
| rs2678140 | 12 | 43200916 | 5 | 0 | 0 | T | G | -0.7624 | 0.2245 | 0.0006836 | ----- | 0 | 0.6677 | intergenic |  |
| 1:203684397 | 1 | 203684397 | 3 | 0 | 0 | T | C | 1.1475 | 0.3379 | 0.0006837 | +++?? | 0 | 0.7356 | intron | ATP2B4 |
| rs67398881 | 2 | 185422672 | 4 | 0 | 0 | T | G | -0.9384 | 0.2763 | 0.0006837 | ---?- | 0 | 0.8125 | intergenic | |
| rs2228564 | 1 | 40773149 | 5 | 0 | 0 | T | C | -0.8696 | 0.2561 | 0.0006842 | ----- | 31.8 | 0.2096 | missense | COL9A2 |
| 16:52875388 | 16 | 52875388 | 3 | 0 | 0 | T | TA | 0.8938 | 0.2632 | 0.0006842 | +++?? | 0 | 0.8724 | | |
| rs4768320 | 12 | 41304909 | 4 | 0 | 0 | A | G | 1.682 | 0.4953 | 0.0006843 | +++?- | 0 | 0.684 | intron | CNTN1 |
| rs57879729 | 19 | 1636162 | 4 | 0 | 0 | A | T | 2.5852 | 0.7613 | 0.0006845 | +++?+ | 0 | 0.6794 | intron | TCF3 |
| rs60939466 | 4 | 190023424 | 4 | 0 | 0 | A | G | -0.8469 | 0.2494 | 0.0006845 | ---?- | 0 | 0.4036 | intergenic | |
| rs72889449 | 2 | 171147816 | 4 | 0 | 0 | T | C | 4.4625 | 1.3142 | 0.0006845 | +++?- | 40.7 | 0.1678 | intron | MYO3B |
| rs10520648 | 15 | 87618593 | 4 | 0 | 0 | T | C | 0.9812 | 0.289 | 0.0006846 | +++?- | 6.2 | 0.3621 | intergenic | |
| rs1442181 | 12 | 41310159 | 4 | 0 | 0 | A | T | 1.6869 | 0.4968 | 0.0006849 | +++?- | 0 | 0.7991 | intron | CNTN1 |
| rs7078409 | 10 | 131914331 | 3 | 0 | 0 | A | G | 0.8554 | 0.2519 | 0.000685 | +++?? | 0 | 0.6594 | intergenic | |
| 7:130365345 | 7 | 130365345 | 3 | 0 | 0 | G | GA | -0.8821 | 0.2598 | 0.0006853 | -+-?? | 62.9 | 0.06764 | | |
| rs28637181 | 4 | 20749624 | 4 | 0 | 0 | T | C | -0.8999 | 0.265 | 0.0006855 | ---?- | 0 | 0.89 | intron | KCNIP4 |
| rs150833766 | 2 | 35800098 | 3 | 0 | 0 | T | G | 3.7219 | 1.0962 | 0.0006855 | +++?? | 0 | 0.8261 | intergenic | |
| 8:95921576 | 8 | 95921576 | 3 | 0 | 0 | CAAAAA | C | -1.7392 | 0.5122 | 0.0006855 | ---?? | 52.5 | 0.1218 | | |
| rs2904994 | 8 | 12558496 | 4 | 0 | 0 | A | C | 1.0808 | 0.3183 | 0.0006857 | +++?+ | 43.6 | 0.15 | intergenic | |
| rs1427387 | 2 | 170639647 | 4 | 0 | 0 | T | C | -0.8325 | 0.2452 | 0.0006858 | ---?- | 0 | 0.7833 | intergenic | |
| rs12360957 | 11 | 134611417 | 4 | 0 | 0 | A | T | -0.8959 | 0.2639 | 0.0006858 | ---?- | 0 | 0.6016 | intergenic | |
| rs117713769 | 6 | 164933046 | 4 | 0 | 0 | A | G | -2.4345 | 0.717 | 0.0006858 | ---?- | 20.8 | 0.2855 | intergenic | |
| rs2705762 | 2 | 184009676 | 4 | 0 | 0 | A | G | 1.171 | 0.3449 | 0.0006859 | +++?+ | 0 | 0.8586 | intron | NUP35 |
| rs67128820 | 3 | 143527157 | 4 | 0 | 0 | A | G | -0.8224 | 0.2422 | 0.000686 | ---?- | 0 | 0.5842 | intron | SLC9A9 |
| 13:60016563 | 13 | 60016563 | 3 | 0 | 0 | A | AAC | 0.9022 | 0.2657 | 0.0006862 | +++?? | 30.1 | 0.2392 | | |
| rs28698122 | 9 | 44256022 | 3 | 0 | 0 | A | T | -1.726 | 0.5084 | 0.0006863 | ---?? | 0 | 0.8051 | intergenic | |
| rs17220928 | 8 | 65066584 | 4 | 0 | 0 | T | C | 0.8303 | 0.2446 | 0.0006866 | +++?+ | 0 | 0.9085 | intergenic | |
| rs12092787 | 1 | 229166965 | 4 | 0 | 0 | T | C | 1.3074 | 0.3851 | 0.000687 | +++?+ | 0 | 0.9625 | intergenic | |
| rs145015776 | 3 | 143041963 | 4 | 0 | 0 | T | C | 1.9851 | 0.5848 | 0.000687 | +++?+ | 0 | 0.6595 | intron | SLC9A9 |
| rs187601735 | 3 | 11020591 | 3 | 0 | 0 | A | G | 3.8113 | 1.1227 | 0.000687 | +++?? | 0 | 0.5622 | intergenic | |
| rs333802 | 11 | 37042544 | 3 | 0 | 0 | A | G | 0.9319 | 0.2745 | 0.0006871 | +++?? | 0 | 0.408 | intergenic | |
| rs2246149 | 2 | 170668106 | 5 | 0 | 0 | A | G | 0.8015 | 0.2361 | 0.0006871 | +++++ | 49.4 | 0.09539 | intron | SSB |
| rs12154325 | 7 | 20236203 | 4 | 0 | 0 | T | C | 1.4675 | 0.4323 | 0.0006872 | +++?+ | 32.5 | 0.2174 | intron | MACC1 |
| rs114720932 | 10 | 132822002 | 2 | 0 | 0 | A | T | -4.7151 | 1.389 | 0.0006872 | -?-?? | 0 | 0.4914 | intergenic | |
| rs28403240 | 7 | 82421634 | 4 | 0 | 0 | A | G | 0.8682 | 0.2558 | 0.0006873 | +-+?+ | 45.7 | 0.1371 | intron | PCLO |
| rs9358744 | 6 | 24053169 | 5 | 0 | 0 | A | G | -1.2082 | 0.3559 | 0.0006874 | ----+ | 0 | 0.8341 | intergenic |  |
| rs9788089 | 12 | 43205273 | 5 | 0 | 0 | A | G | -0.8341 | 0.2457 | 0.0006876 | ----- | 0 | 0.6591 | intergenic |  |
| rs56123464 | 19 | 46169794 | 5 | 0 | 1 | C | G | 0.8937 | 0.2633 | 0.0006878 | ++++- | 25.7 | 0.2502 | upstream-gene | |
| rs73260699 | 8 | 64925971 | 3 | 0 | 0 | C | G | 0.8661 | 0.2551 | 0.0006878 | +++?? | 0 | 0.4601 | intergenic | |
| rs76936644 | 13 | 82103611 | 2 | 0 | 0 | A | C | 8.0014 | 2.3573 | 0.0006878 | ?+??+ | 0 | 0.3707 | intergenic | |
| rs9325212 | 12 | 43205187 | 5 | 0 | 0 | A | G | 0.8341 | 0.2457 | 0.000688 | +++++ | 0 | 0.6591 | intergenic | |
| rs9393526 | 6 | 24072458 | 5 | 0 | 0 | T | C | 1.2921 | 0.3807 | 0.000688 | ++++- | 25.9 | 0.2487 | intergenic | |

| | | | | | | | | | | | | | | | |
|---|---|---|---|---|---|---|---|---|---|---|---|---|---|---|---|
| rs116605210 | 2 | 167806871 | 4 | 0 | 0 | A | G | -3.4153 | 1.0062 | 0.000688 | ---?- | 23.8 | 0.2683 | intron | XIRP2 |
| rs142987262 | 2 | 117118854 | 4 | 0 | 0 | T | C | -1.7172 | 0.5059 | 0.000688 | ---?- | 0 | 0.6766 | intergenic | |
| rs1049074 | 6 | 32632711 | 4 | 0 | 1 | T | C | 1.3588 | 0.4003 | 0.0006881 | ++++? | 0 | 0.4805 | synonymous | HLA-DQB1 |
| rs61641071 | 5 | 123105434 | 4 | 0 | 0 | T | C | 0.9668 | 0.2848 | 0.0006881 | +++?+ | 2.1 | 0.3818 | intergenic | |
| rs61939789 | 12 | 119412202 | 4 | 0 | 0 | A | G | -1.6607 | 0.4893 | 0.0006881 | ---?- | 20.1 | 0.2893 | intergenic | |
| 17:39389597 | 17 | 39389597 | 3 | 0 | 0 | C | G | 3.7528 | 1.1056 | 0.0006882 | +++?? | 0 | 0.988 | 3-prime-UTR | KRTAP9-3 |
| rs111262662 | 9 | 71981671 | 4 | 0 | 0 | T | C | 1.8335 | 0.5402 | 0.0006883 | +++?+ | 0 | 0.7351 | intron | FAM189A2 |
| rs10767869 | 11 | 30650649 | 4 | 0 | 0 | A | T | 0.8974 | 0.2644 | 0.0006884 | +++?- | 61.3 | 0.05136 | intergenic | |
| rs77105473 | 5 | 146980603 | 2 | 0 | 0 | A | T | 4.1459 | 1.2215 | 0.0006884 | +?+?? | 0 | 0.4871 | intron | JAKMIP2,JAKMIP2-AS1 |
| 8:64925938 | 8 | 64925938 | 3 | 0 | 0 | CTTCT | C | 0.866 | 0.2551 | 0.0006884 | +++?? | 0 | 0.4602 | | |
| rs1852915 | 7 | 112707313 | 4 | 0 | 0 | T | C | -0.9716 | 0.2863 | 0.0006885 | ---?- | 0 | 0.5937 | intergenic | |
| rs75176010 | 10 | 86706986 | 4 | 0 | 0 | C | G | -1.7167 | 0.5058 | 0.0006888 | ---?- | 0 | 0.797 | intergenic | |
| rs1975498 | 4 | 37008129 | 4 | 0 | 0 | A | G | 0.8491 | 0.2502 | 0.0006889 | +++?- | 46.3 | 0.1333 | intergenic | |
| 15:87556764 | 15 | 87556764 | 3 | 0 | 0 | G | GTT | 0.9392 | 0.2767 | 0.0006889 | +++?? | 24.8 | 0.2645 | | |
| rs4389295 | 2 | 185421477 | 4 | 0 | 0 | T | C | -0.9378 | 0.2763 | 0.0006895 | ---?- | 0 | 0.8086 | intergenic | |
| rs12770291 | 10 | 134586400 | 3 | 0 | 0 | A | G | -0.9928 | 0.2925 | 0.0006895 | ---?? | 6.3 | 0.3441 | intron | INPP5A |
| rs4348720 | 1 | 111806066 | 4 | 0 | 0 | A | T | -1.9124 | 0.5635 | 0.0006896 | ---?- | 0 | 0.541 | intergenic | |
| rs79248845 | 11 | 86178083 | 4 | 0 | 0 | A | G | -1.5906 | 0.4687 | 0.0006896 | ---?- | 0 | 0.463 | intron | ME3 |
| 1:87432579 | 1 | 87432579 | 3 | 0 | 0 | CT | C | -1.0969 | 0.3232 | 0.0006898 | ---?? | 0 | 0.7702 | | |
| rs7787792 | 7 | 10157295 | 4 | 0 | 0 | A | C | 0.9663 | 0.2847 | 0.00069 | +++?+ | 46.4 | 0.1329 | intergenic | |
| rs9913377 | 17 | 4765906 | 5 | 0 | 0 | A | G | -1.5377 | 0.4531 | 0.0006901 | ----+ | 17.2 | 0.3054 | intron | MINK1 |
| rs11783361 | 8 | 113357209 | 4 | 0 | 0 | T | C | 0.9693 | 0.2856 | 0.0006901 | +++?+ | 0 | 0.7972 | intron | CSMD3 |
| rs11828773 | 11 | 86176674 | 4 | 0 | 0 | T | G | 1.5892 | 0.4683 | 0.0006902 | +++?+ | 0 | 0.4574 | intron | ME3 |
| rs150204991 | 2 | 168180827 | 4 | 0 | 0 | A | G | 2.4603 | 0.725 | 0.0006902 | +++?+ | 0 | 0.5435 | intergenic | |
| rs2248788 | 4 | 37008595 | 4 | 0 | 0 | A | G | -0.8487 | 0.2501 | 0.0006905 | ---?+ | 46.3 | 0.1333 | intergenic | |
| rs6553157 | 4 | 189046697 | 4 | 0 | 0 | T | C | -1.2291 | 0.3622 | 0.0006906 | ---?- | 0 | 0.8647 | intergenic | |
| rs8084348 | 18 | 25363418 | 4 | 0 | 0 | T | C | 0.9655 | 0.2845 | 0.0006906 | +++?+ | 6.8 | 0.3593 | intergenic | |
| rs12312616 | 12 | 41503140 | 4 | 0 | 0 | A | G | -1.8513 | 0.5456 | 0.0006906 | ---?- | 0 | 0.4805 | intergenic | |
| rs75749778 | 11 | 86176878 | 4 | 0 | 0 | T | C | 1.5892 | 0.4683 | 0.0006907 | +++?+ | 0 | 0.4584 | intron | ME3 |
| 15:73700137 | 15 | 73700137 | 3 | 0 | 0 | CA | C | 2.3707 | 0.6987 | 0.0006907 | +++?? | 0 | 0.459 | | |
| rs2043380 | 15 | 87612368 | 4 | 0 | 0 | A | G | 0.9085 | 0.2678 | 0.0006908 | +++?+ | 0 | 0.6503 | intergenic | |
| rs78379542 | 4 | 39570110 | 4 | 0 | 0 | T | G | -2.6519 | 0.7815 | 0.0006908 | ---?- | 0 | 0.565 | intron | SMIM14,UGDH-AS1 |
| rs1627883 | 13 | 45624805 | 3 | 0 | 0 | A | T | 2.281 | 0.6723 | 0.0006912 | +++?? | 0 | 0.6469 | intergenic | |
| rs10767868 | 11 | 30650634 | 4 | 0 | 0 | T | G | 0.8971 | 0.2644 | 0.0006912 | +++?- | 61.3 | 0.05141 | intergenic | |
| rs149910861 | 5 | 120076746 | 2 | 0 | 0 | A | G | 5.9714 | 1.7599 | 0.0006912 | ?++?? | 0 | 0.485 | intergenic | |
| rs6938337 | 6 | 32381041 | 5 | 0 | 0 | C | G | -0.7183 | 0.2117 | 0.0006914 | ----- | 0 | 0.6443 | intergenic |  |
| rs17828460 | 6 | 130949362 | 4 | 0 | 0 | T | C | -1.4399 | 0.4244 | 0.0006914 | ---?- | 0 | 0.7076 | intergenic | |
| rs4579569 | 8 | 64913762 | 4 | 0 | 0 | A | G | 0.8366 | 0.2466 | 0.0006915 | +++?- | 0 | 0.4706 | intergenic | |
| rs373273 | 2 | 40434662 | 4 | 0 | 0 | T | C | 1.0513 | 0.3098 | 0.0006917 | +++?- | 0 | 0.4132 | intron | SLC8A1,SLC8A1-AS1 |
| rs66991667 | 3 | 172329736 | 4 | 0 | 0 | T | C | 1.7866 | 0.5266 | 0.0006919 | +++?- | 3 | 0.3775 | intergenic | |
| rs28409734 | 9 | 44255591 | 3 | 0 | 0 | A | G | 1.7246 | 0.5083 | 0.0006922 | +++?? | 0 | 0.8093 | intergenic | |
| rs2949222 | 2 | 107677489 | 4 | 0 | 0 | A | C | -0.8663 | 0.2553 | 0.0006923 | ---?- | 0 | 0.6981 | intergenic | |
| 15:99927410 | 15 | 99927410 | 3 | 0 | 0 | T | C | 3.8322 | 1.1296 | 0.0006925 | +++?? | 0 | 0.9741 | intron | LRRC28 |
| rs185486 | 8 | 97002215 | 4 | 0 | 0 | A | G | 1.3965 | 0.4116 | 0.0006926 | +++?+ | 0 | 0.9551 | intergenic | |
| rs12107202 | 3 | 183572790 | 4 | 0 | 0 | T | C | -1.0907 | 0.3215 | 0.0006926 | ---?- | 0 | 0.7137 | intron | PARL |
| rs57707806 | 10 | 86696840 | 4 | 0 | 0 | C | G | 1.722 | 0.5076 | 0.0006926 | +++?+ | 0 | 0.7946 | intergenic | |
| rs12167378 | 22 | 29100889 | 4 | 0 | 0 | A | C | 2.1815 | 0.6431 | 0.0006927 | +++?+ | 0 | 0.4986 | intron | CHEK2 |
| rs62194738 | 20 | 23839394 | 3 | 0 | 0 | T | C | 1.4743 | 0.4346 | 0.0006927 | +++?? | 18.5 | 0.2933 | intergenic | |
| 1:182008573 | 1 | 182008573 | 3 | 0 | 0 | A | AT | -2.9339 | 0.8648 | 0.0006927 | ---?? | 0 | 0.8753 | | |
| rs57142217 | 13 | 103286989 | 3 | 0 | 0 | C | G | 1.534 | 0.4522 | 0.0006928 | ?++?+ | 27.4 | 0.2524 | intron | TPP2 |
| rs1171725 | 10 | 131913142 | 3 | 0 | 0 | T | C | -0.8586 | 0.2531 | 0.0006929 | ---?? | 0 | 0.6873 | upstream-gene | |
| 17:76787211 | 17 | 76787211 | 3 | 0 | 0 | CTTTTTT | C | 1.0012 | 0.2951 | 0.000693 | +++?? | 0 | 0.7126 | | |
| rs351791 | 10 | 55209376 | 4 | 0 | 0 | A | G | -0.8747 | 0.2579 | 0.0006933 | ---?+ | 38.6 | 0.1805 | intergenic | |
| rs2588532 | 4 | 37009932 | 4 | 0 | 0 | A | T | 0.8479 | 0.25 | 0.0006933 | +++?- | 46.4 | 0.133 | intergenic | |
| rs1528890 | 18 | 68738292 | 4 | 0 | 0 | A | T | 0.9654 | 0.2846 | 0.0006936 | +++?- | 21.7 | 0.2803 | intergenic | |
| rs3739751 | 9 | 98498387 | 4 | 0 | 0 | T | C | 1.3872 | 0.409 | 0.0006936 | +++?+ | 0 | 0.666 | intergenic | |
| rs4668163 | 2 | 170639070 | 4 | 0 | 0 | T | C | 0.8314 | 0.2451 | 0.0006936 | +++?+ | 0 | 0.7814 | intergenic | |
| rs9872860 | 3 | 116035402 | 4 | 0 | 0 | A | G | 1.0541 | 0.3108 | 0.0006936 | +++?+ | 0 | 0.5199 | intron | LSAMP |
| rs117436165 | 6 | 163431868 | 4 | 0 | 0 | A | G | -1.8658 | 0.5501 | 0.0006936 | ---?- | 0 | 0.9787 | intron | PACRG |

| | | | | | | | | | | | | | | | |
|---|---|---|---|---|---|---|---|---|---|---|---|---|---|---|---|
| rs368013375 | 1 | 31140384 | 3 | 0 | 0 | A | T | -1.9709 | 0.581 | 0.0006936 | ---?? | 0 | 0.941 | intergenic | |
| 8:86223442 | 8 | 86223442 | 3 | 0 | 0 | CT | C | -3.4771 | 1.0251 | 0.0006936 | ---?? | 0 | 0.8902 | | |
| rs1571449 | 9 | 103871031 | 4 | 0 | 0 | A | G | -0.8498 | 0.2505 | 0.0006937 | ---?- | 44.5 | 0.1442 | intron | LPPR1 |
| rs11776386 | 8 | 113357053 | 4 | 0 | 0 | A | G | -0.9684 | 0.2855 | 0.0006939 | ---?- | 0 | 0.7965 | intron | CSMD3 |
| rs73116958 | 12 | 41381970 | 4 | 0 | 0 | A | G | 1.443 | 0.4254 | 0.0006939 | +++?+ | 0 | 0.6045 | intron | CNTN1 |
| rs9913346 | 17 | 4765864 | 5 | 0 | 0 | A | G | -1.5369 | 0.4531 | 0.000694 | ----+ | 17.1 | 0.3054 | intron | MINK1 |
| rs783640 | 6 | 117295140 | 4 | 0 | 0 | T | C | -0.7993 | 0.2357 | 0.0006942 | ---?- | 0 | 0.6917 | intergenic | |
| rs1360917 | 1 | 18750770 | 4 | 0 | 0 | A | G | -1.033 | 0.3046 | 0.0006944 | ---?- | 51.5 | 0.1029 | intergenic | |
| rs9631774 | 4 | 169323348 | 4 | 0 | 0 | A | G | 0.8863 | 0.2613 | 0.0006944 | +++?+ | 0 | 0.4808 | intron | DDX60L |
| rs17162784 | 7 | 10250272 | 4 | 0 | 0 | T | C | -1.5131 | 0.4461 | 0.0006947 | ---?- | 25.7 | 0.2575 | intergenic | |
| rs2224720 | 4 | 37018478 | 4 | 0 | 0 | T | C | -0.8444 | 0.249 | 0.0006948 | ---?+ | 46.7 | 0.1309 | intergenic | |
| rs6483999 | 11 | 23969457 | 4 | 0 | 0 | T | C | 1.2145 | 0.3581 | 0.0006949 | +++?+ | 0 | 0.7307 | intergenic | |
| rs9518471 | 13 | 102301284 | 4 | 0 | 0 | A | C | 0.9957 | 0.2936 | 0.0006949 | +++?+ | 0 | 0.467 | intron | ITGBL1 |
| rs192884282 | 4 | 166227250 | 4 | 0 | 0 | T | C | -3.7252 | 1.0984 | 0.0006951 | ---?- | 0 | 0.9412 | intron | KLHL2 |
| rs55861928 | 7 | 49467389 | 4 | 0 | 0 | A | G | 0.9485 | 0.2797 | 0.0006954 | +++?- | 25.1 | 0.2606 | intergenic | |
| rs141950136 | 4 | 166226581 | 4 | 0 | 0 | T | C | 3.7267 | 1.0989 | 0.0006955 | +++?+ | 0 | 0.9407 | intron | KLHL2 |
| 11:3246567 | 11 | 3246567 | 3 | 0 | 0 | AG | CG | 0.9204 | 0.2714 | 0.0006955 | +++?? | 0 | 0.5587 | | |
| 3:183616886 | 3 | 183616886 | 3 | 0 | 0 | T | TTAAAA | -1.6764 | 0.4943 | 0.0006955 | ---?? | 0 | 0.832 | | |
| rs76435080 | 17 | 52275561 | 4 | 0 | 0 | T | C | 2.0875 | 0.6156 | 0.0006959 | +++?- | 35.3 | 0.2002 | intergenic | |
| rs1933947 | 10 | 86694492 | 4 | 0 | 0 | T | C | -1.7239 | 0.5083 | 0.000696 | ---?- | 0 | 0.7955 | intergenic | |
| rs4640668 | 4 | 20741331 | 4 | 0 | 0 | A | G | -0.8805 | 0.2596 | 0.0006962 | ---?- | 0 | 0.9671 | intron | KCNIP4 |
| rs59604187 | 5 | 103171712 | 4 | 0 | 0 | T | C | -1.4981 | 0.4418 | 0.0006962 | ---?- | 0 | 0.8122 | intergenic | |
| rs148666045 | 6 | 165000686 | 4 | 0 | 0 | T | G | -2.5309 | 0.7463 | 0.0006962 | ---?- | 0 | 0.5002 | intergenic | |
| rs1010010 | 4 | 37011104 | 4 | 0 | 0 | T | C | -0.8471 | 0.2498 | 0.0006965 | ---?+ | 46.4 | 0.1329 | intergenic | |
| rs7143812 | 14 | 28234681 | 4 | 0 | 0 | T | G | 1.9662 | 0.5798 | 0.0006965 | +++?+ | 0 | 0.6483 | intergenic | |
| rs10960431 | 9 | 12003841 | 4 | 0 | 0 | A | G | 1.8546 | 0.5469 | 0.0006966 | +++?+ | 0 | 0.4702 | intergenic | |
| 10:122109126 | 10 | 122109126 | 3 | 0 | 0 | T | TG | 0.8759 | 0.2583 | 0.0006966 | +++?? | 0 | 0.7416 | | |
| rs825589 | 16 | 87669371 | 4 | 0 | 0 | C | G | -0.9386 | 0.2768 | 0.0006967 | ---?- | 0 | 0.9533 | intron | JPH3 |
| rs28445893 | 9 | 44272885 | 3 | 0 | 0 | A | T | 1.784 | 0.5261 | 0.0006969 | +++?? | 0 | 0.8265 | intergenic | |
| rs73834878 | 3 | 68233409 | 4 | 0 | 0 | T | C | 2.175 | 0.6414 | 0.0006969 | +++?+ | 8.1 | 0.3527 | intron | FAM19A1 |
| rs75709983 | 22 | 28475427 | 4 | 0 | 0 | A | G | 2.5448 | 0.7505 | 0.000697 | +++?+ | 0 | 0.7619 | intron | TTC28 |
| rs140619432 | 18 | 25175175 | 3 | 0 | 0 | T | C | -3.894 | 1.1484 | 0.000697 | ---?? | 0 | 0.8971 | intergenic | |
| rs10894233 | 11 | 130623433 | 4 | 0 | 0 | T | C | 1.6603 | 0.4896 | 0.0006971 | +++?+ | 17 | 0.3062 | intergenic | |
| 20:12002524 | 20 | 12002524 | 3 | 0 | 0 | A | ACTAT | -0.9791 | 0.2887 | 0.0006971 | ---?? | 0 | 0.4219 | | |
| rs79326429 | 1 | 42801012 | 4 | 0 | 0 | T | G | 1.8676 | 0.5508 | 0.0006974 | +++?+ | 0 | 0.5791 | intron | FOXJ3 |
| rs28383282 | 6 | 32586375 | 3 | 0 | 0 | T | C | 1.3158 | 0.3881 | 0.0006975 | +++?? | 0 | 0.4028 | intergenic | |
| rs117635845 | 13 | 48771056 | 4 | 0 | 0 | C | G | -2.5221 | 0.7439 | 0.0006976 | ---?- | 18.6 | 0.2973 | intergenic | |
| 2:69065544 | 2 | 69065544 | 3 | 0 | 0 | CTATTAT | C | -0.913 | 0.2693 | 0.0006976 | ---?? | 0 | 0.3986 | | |
| rs6897605 | 5 | 744428 | 3 | 0 | 0 | T | C | -1.5337 | 0.4524 | 0.0006977 | ---?? | 0 | 0.4459 | intron | ZDHHC11B |
| rs4680178 | 3 | 155151059 | 4 | 0 | 0 | A | T | 0.8807 | 0.2598 | 0.0006982 | +++?+ | 0 | 0.6514 | intergenic | |
| rs142361046 | 10 | 129714780 | 3 | 0 | 0 | A | G | -2.8982 | 0.8549 | 0.0006985 | ---?? | 10.4 | 0.3276 | intron | PTPRE |
| rs17117765 | 1 | 98601247 | 4 | 0 | 0 | T | C | 1.3283 | 0.3918 | 0.0006986 | +++?+ | 19.8 | 0.2911 | intergenic | |
| rs2771342 | 14 | 21457048 | 4 | 0 | 0 | T | C | 0.8763 | 0.2585 | 0.0006987 | +++?+ | 0 | 0.9515 | upstream-gene | |
| 20:60657941 | 20 | 60657941 | 3 | 0 | 0 | A | G | -3.0189 | 0.8905 | 0.0006988 | ---?? | 0 | 0.9697 | intergenic | |
| rs116723633 | 9 | 44271490 | 3 | 0 | 0 | T | G | -1.7658 | 0.5209 | 0.0006991 | ---?? | 0 | 0.8273 | intergenic | |
| rs817389 | 1 | 85611061 | 4 | 0 | 0 | C | G | 1.5399 | 0.4543 | 0.0006992 | +++?+ | 0 | 0.9833 | intergenic | |
| rs77389857 | 3 | 15707596 | 4 | 0 | 0 | T | C | -2.2419 | 0.6614 | 0.0006995 | ---?- | 0 | 0.5011 | downstream-gene | |
| rs55712837 | 11 | 14799072 | 4 | 0 | 0 | T | C | -0.9395 | 0.2772 | 0.0006999 | ---?- | 36.9 | 0.191 | intron | PDE3B |
| rs78443590 | 5 | 109421626 | 4 | 0 | 0 | A | G | -1.9378 | 0.5717 | 0.0006999 | ---?+ | 0 | 0.4574 | intergenic | |
| 14:89914408 | 14 | 89914408 | 3 | 0 | 0 | C | G | -2.0401 | 0.6019 | 0.0007002 | ---?? | 0 | 0.4045 | intron | FOXN3 |
| rs12787765 | 11 | 58400686 | 5 | 0 | 0 | A | G | -0.8258 | 0.2436 | 0.0007002 | ----- | 0 | 0.983 | intergenic |  |
| rs150011196 | 10 | 80608473 | 3 | 0 | 0 | T | C | 4.7395 | 1.3983 | 0.0007004 | ?++?- | 10.2 | 0.3285 | intergenic | |
| 10:126824090 | 10 | 126824090 | 3 | 0 | 0 | CT | C | -1.5816 | 0.4666 | 0.0007006 | ---?? | 0 | 0.9935 | | |
| rs11724386 | 4 | 119047969 | 4 | 0 | 0 | T | C | 1.6395 | 0.4837 | 0.0007007 | +++?+ | 0 | 0.4414 | intron | NDST3 |
| rs4748615 | 10 | 20150185 | 4 | 0 | 0 | A | G | -0.8708 | 0.2569 | 0.0007009 | ---?- | 42.4 | 0.1572 | intron | PLXDC2 |
| 2:184013338 | 2 | 184013338 | 3 | 0 | 0 | A | ATTCAT | -1.2155 | 0.3586 | 0.0007009 | ---?? | 0 | 0.7994 | | |
| rs72900295 | 11 | 19321297 | 2 | 0 | 0 | T | C | -6.0211 | 1.7766 | 0.0007011 | ?-??- | 0 | 0.6411 | intergenic | |
| rs148435998 | 3 | 143045881 | 4 | 0 | 0 | A | G | -1.9728 | 0.5821 | 0.0007011 | ---?- | 0 | 0.6341 | intron | SLC9A9 |
| 5:180002892 | 5 | 180002892 | 3 | 0 | 0 | G | GT | 1.0477 | 0.3091 | 0.0007011 | +++?? | 0 | 0.6386 | | |

| | | | | | | | | | | | | | | | |
|---|---|---|---|---|---|---|---|---|---|---|---|---|---|---|---|
| rs825200 | 1 | 64773973 | 4 | 0 | 0 | T | C | 0.9409 | 0.2776 | 0.0007012 | +++?+ | 0 | 0.448 | intergenic | |
| rs141249909 | 3 | 39931439 | 4 | 0 | 0 | A | G | -3.1787 | 0.9379 | 0.0007012 | ---?- | 25.4 | 0.2593 | intron | MYRIP |
| 8:90883445 | 8 | 90883445 | 3 | 0 | 0 | G | GA | -2.6654 | 0.7865 | 0.0007014 | ---?? | 19.6 | 0.2883 | | |
| rs2034544 | 2 | 2759551 | 4 | 0 | 0 | C | G | 1.3207 | 0.3897 | 0.0007015 | +++?+ | 0 | 0.9547 | intergenic | |
| rs79944685 | 2 | 168275353 | 4 | 0 | 0 | A | G | -2.3457 | 0.6922 | 0.0007015 | ---?- | 0 | 0.6666 | intergenic | |
| rs117512776 | 20 | 62505223 | 2 | 0 | 0 | T | C | 5.5596 | 1.6405 | 0.0007015 | ?++?? | 0 | 0.618 | intron | TPD52L2 |
| rs28535317 | 6 | 32589065 | 5 | 0 | 1 | A | G | -1.6171 | 0.4772 | 0.0007016 | ----- | 0 | 0.5583 | intergenic |  |
| rs7585172 | 2 | 184017978 | 4 | 0 | 0 | A | G | -1.171 | 0.3455 | 0.0007017 | ---?- | 0 | 0.7799 | intron | NUP35 |
| 12:74292556 | 12 | 74292556 | 3 | 0 | 0 | CT | C | -0.8997 | 0.2655 | 0.0007018 | ---?? | 5 | 0.3489 | | |
| rs116924548 | 15 | 39290682 | 3 | 0 | 0 | C | G | -2.074 | 0.612 | 0.0007019 | ---?? | 0 | 0.3805 | intergenic | |
| rs2577955 | 15 | 42012178 | 5 | 0 | 0 | T | C | -0.9234 | 0.2725 | 0.000702 | ----- | 0 | 0.6051 | intron | MGA |
| 9:44249865 | 9 | 44249865 | 3 | 0 | 0 | T | TA | -1.7827 | 0.5261 | 0.0007022 | ---?? | 0 | 0.9727 | | |
| rs6747405 | 2 | 212574618 | 4 | 0 | 0 | A | C | 1.2897 | 0.3806 | 0.0007024 | +++?- | 0 | 0.7608 | intron | ERBB4 |
| 14:37536508 | 14 | 37536508 | 3 | 0 | 0 | G | GAAGTA | -4.1367 | 1.2208 | 0.0007024 | ---?? | 16.1 | 0.3038 | | |
| rs115078509 | 3 | 143054596 | 4 | 0 | 0 | A | G | -1.9726 | 0.5821 | 0.0007025 | ---?- | 0 | 0.634 | intron | SLC9A9 |
| rs112859941 | 9 | 44269978 | 3 | 0 | 0 | T | C | -1.7603 | 0.5195 | 0.0007026 | ---?? | 0 | 0.825 | intergenic | |
| rs201037449 | 4 | 38735305 | 3 | 0 | 0 | A | G | -1.4574 | 0.4301 | 0.0007027 | ---?? | 5.8 | 0.3461 | intergenic | |
| rs7092082 | 10 | 119516757 | 4 | 0 | 0 | T | C | 1.5674 | 0.4626 | 0.0007028 | +++?+ | 0 | 0.8188 | intergenic | |
| rs7105348 | 11 | 23970654 | 4 | 0 | 0 | T | C | 1.2116 | 0.3576 | 0.0007028 | +++?+ | 0 | 0.7293 | intergenic | |
| rs78147978 | 6 | 120431032 | 4 | 0 | 0 | T | C | -2.7859 | 0.8222 | 0.0007028 | ---?- | 45.5 | 0.1384 | intergenic | |
| rs6005829 | 22 | 29043969 | 4 | 0 | 0 | T | C | -2.3356 | 0.6893 | 0.0007029 | ---?- | 0 | 0.449 | intron | TTC28 |
| rs117242407 | 14 | 21295683 | 3 | 0 | 0 | A | G | 2.8045 | 0.8277 | 0.0007029 | +++?? | 5.2 | 0.3482 | intergenic | |
| 5:39602723 | 5 | 39602723 | 3 | 0 | 0 | A | AG | -1.0432 | 0.3079 | 0.0007029 | ---?? | 0 | 0.9437 | | |
| 8:60300480 | 8 | 60300480 | 3 | 0 | 0 | CGTGTG | C | -1.471 | 0.4341 | 0.0007029 | ---?? | 0 | 0.6733 | | |
| rs1868183 | 3 | 143526685 | 4 | 0 | 0 | C | G | 0.8223 | 0.2427 | 0.000703 | +++?+ | 0 | 0.5832 | intron | SLC9A9 |
| rs28555341 | 9 | 44257533 | 3 | 0 | 0 | A | G | -1.7419 | 0.5141 | 0.000703 | ---?? | 0 | 0.7913 | intergenic | |
| rs78458917 | 6 | 72700667 | 4 | 0 | 0 | A | T | 1.3349 | 0.394 | 0.000703 | +++?+ | 48.6 | 0.12 | intron | RIMS1 |
| rs75226449 | 6 | 120431033 | 4 | 0 | 0 | T | C | -2.7858 | 0.8222 | 0.0007031 | ---?- | 45.5 | 0.1385 | intergenic | |
| rs58067470 | 20 | 37490445 | 2 | 0 | 0 | A | G | -6.3687 | 1.8796 | 0.0007033 | ??-?- | 10 | 0.2917 | intron | PPP1R16B |
| rs147867534 | 5 | 103163469 | 4 | 0 | 0 | A | G | -1.5257 | 0.4503 | 0.0007033 | ---?- | 0 | 0.6839 | intergenic | |
| rs7874278 | 9 | 16205719 | 4 | 0 | 0 | C | G | -1.1669 | 0.3444 | 0.0007034 | ---?- | 0 | 0.9177 | intron | C9orf92 |
| rs13248395 | 8 | 64909495 | 4 | 0 | 0 | T | G | 0.836 | 0.2467 | 0.0007037 | +++?- | 0 | 0.4795 | intergenic | |
| rs55660658 | 6 | 32581500 | 4 | 0 | 1 | A | G | -1.3407 | 0.3957 | 0.0007037 | ----? | 38.9 | 0.1784 | intergenic |  |
| rs112294537 | 9 | 44272178 | 3 | 0 | 0 | A | G | 1.7754 | 0.524 | 0.0007038 | +++?? | 0 | 0.8283 | intergenic | |
| 15:75355071 | 15 | 75355071 | 2 | 0 | 0 | CAG | C | -5.9211 | 1.7477 | 0.000704 | -?-?? | 0 | 0.8348 | | |
| 8:64961708 | 8 | 64961708 | 3 | 0 | 0 | G | GA | -0.8661 | 0.2556 | 0.000704 | ---?? | 0 | 0.4999 | | |
| rs140504890 | 10 | 1741328 | 3 | 0 | 0 | A | G | -4.4497 | 1.3134 | 0.0007041 | ---?? | 0 | 0.7072 | intron | ADARB2 |
| 14:21453280 | 14 | 21453280 | 3 | 0 | 0 | CTG | C | 2.4459 | 0.7219 | 0.0007042 | +++?? | 12.6 | 0.3184 | | |
| 16:58441868 | 16 | 58441868 | 3 | 0 | 0 | CT | C | -1.59 | 0.4693 | 0.0007042 | ---?? | 47.9 | 0.1468 | | |
| 7:101859933 | 7 | 101859933 | 3 | 0 | 0 | A | ACT | -0.8755 | 0.2584 | 0.0007044 | ---?? | 0 | 0.8118 | | |
| rs2556352 | 2 | 170665108 | 5 | 0 | 0 | A | G | 0.8022 | 0.2368 | 0.0007045 | +++++ | 50 | 0.0915 | intron | SSB |
| 1:49165743 | 1 | 49165743 | 3 | 0 | 0 | A | AC | 4.0679 | 1.2007 | 0.0007045 | +++?? | 24 | 0.2683 | | |
| rs75121681 | 1 | 180134075 | 4 | 0 | 0 | A | G | -2.1712 | 0.6409 | 0.0007046 | ---?+ | 37 | 0.1902 | intron | QSOX1 |
| rs35250962 | 4 | 10048271 | 5 | 0 | 0 | T | C | 0.7134 | 0.2106 | 0.0007047 | ++++- | 10.2 | 0.3481 | intergenic | |
| rs113584265 | 5 | 39610806 | 3 | 0 | 0 | C | G | 1.0487 | 0.3095 | 0.0007047 | +++?? | 0 | 0.9672 | intergenic | |
| rs112324104 | 11 | 90024181 | 3 | 0 | 0 | A | G | -3.4197 | 1.0095 | 0.0007048 | ---?? | 0 | 0.4684 | intergenic | |
| rs114458886 | 2 | 116687834 | 3 | 0 | 0 | T | C | 2.5134 | 0.7419 | 0.0007048 | +++?? | 55.5 | 0.1055 | intergenic | |
| rs4798462 | 18 | 6547519 | 3 | 0 | 0 | A | T | 0.9522 | 0.2811 | 0.0007049 | +++?? | 40.9 | 0.1842 | intron | LOC100130480 |
| rs112984162 | 17 | 78096922 | 5 | 0 | 0 | C | G | -1.3202 | 0.3897 | 0.0007049 | ----+ | 0 | 0.6961 | downstream-gene |  |
| rs7237973 | 18 | 47514403 | 4 | 0 | 0 | A | C | 1.0228 | 0.3019 | 0.0007052 | +++?+ | 0 | 0.952 | intron | MYO5B |
| rs74732096 | 8 | 21035558 | 4 | 0 | 0 | A | G | -1.7904 | 0.5285 | 0.0007053 | ---?+ | 0 | 0.6303 | intergenic | |
| rs11980441 | 7 | 82422080 | 4 | 0 | 0 | T | C | 0.8654 | 0.2555 | 0.0007054 | +-+?+ | 45.7 | 0.1372 | intron | PCLO |
| rs139378959 | 18 | 39869846 | 4 | 0 | 0 | T | G | 2.5541 | 0.754 | 0.0007055 | +++?+ | 23.8 | 0.2682 | intron | LINC00907 |
| rs28422091 | 17 | 68608228 | 3 | 0 | 0 | A | G | -0.9425 | 0.2782 | 0.0007056 | ---?? | 0 | 0.9934 | intergenic | |
| rs11603890 | 11 | 58402251 | 5 | 0 | 0 | T | C | 0.8267 | 0.2441 | 0.0007057 | +++++ | 0 | 0.9834 | intergenic | |
| rs8068233 | 17 | 55450074 | 4 | 0 | 0 | A | G | 1.0853 | 0.3204 | 0.0007058 | +++?- | 0 | 0.562 | intron | MSI2 |
| rs75984940 | 11 | 1474637 | 4 | 0 | 0 | T | C | -2.4037 | 0.7096 | 0.0007058 | ---?- | 0 | 0.4496 | intron | BRSK2 |
| rs28505561 | 9 | 44274370 | 3 | 0 | 0 | C | G | 1.8045 | 0.5327 | 0.0007059 | +++?? | 0 | 0.827 | intergenic | |
| rs12228343 | 12 | 96736570 | 4 | 0 | 0 | T | C | -2.5726 | 0.7595 | 0.000706 | ---?- | 0 | 0.4386 | intron | CDK17 |

| | | | | | | | | | | | | | | | |
|---|---|---|---|---|---|---|---|---|---|---|---|---|---|---|---|
| rs7580321 | 2 | 230905265 | 4 | 0 | 0 | A | T | 1.4902 | 0.44 | 0.0007061 | +++?+ | 26.4 | 0.2531 | intron | SLC16A14 |
| rs10752435 | 1 | 18749949 | 4 | 0 | 0 | A | C | -1.0353 | 0.3057 | 0.0007061 | ---?- | 52 | 0.1 | intergenic | |
| rs112887643 | 9 | 79463110 | 4 | 0 | 0 | A | G | 2.0153 | 0.595 | 0.0007061 | +++?+ | 0 | 0.5701 | intron | PRUNE2 |
| rs333803 | 11 | 37042823 | 3 | 0 | 0 | T | C | 0.9296 | 0.2744 | 0.0007062 | +++?? | 0 | 0.4067 | intergenic | |
| rs6984541 | 8 | 136705975 | 4 | 0 | 0 | A | G | 0.9222 | 0.2723 | 0.0007062 | +++?+ | 53.5 | 0.09168 | intergenic | |
| rs7251103 | 19 | 12796028 | 3 | 0 | 0 | T | C | 3.8167 | 1.1268 | 0.0007062 | ?+++? | 66 | 0.05272 | upstream-gene | |
| rs2024447 | 16 | 52855494 | 4 | 0 | 0 | A | C | -0.8196 | 0.242 | 0.0007064 | ---?- | 0 | 0.8026 | intergenic | |
| rs61651400 | 14 | 28229918 | 4 | 0 | 0 | A | C | -1.4011 | 0.4137 | 0.0007064 | ---?- | 12.9 | 0.328 | intergenic | |
| rs11179780 | 12 | 41486580 | 4 | 0 | 0 | T | G | -1.6528 | 0.488 | 0.0007069 | ---?- | 14.2 | 0.3212 | intergenic | |
| rs56000503 | 1 | 30352294 | 4 | 0 | 0 | A | G | 1.2976 | 0.3831 | 0.0007069 | +++?- | 0 | 0.4129 | intergenic | |
| rs1267860 | 6 | 6351620 | 3 | 0 | 0 | A | C | 1.5379 | 0.4541 | 0.000707 | +++?? | 32.3 | 0.2284 | intron | LY86-AS1 |
| 7:67401776 | 7 | 67401776 | 3 | 0 | 0 | A | AAAG | -1.8608 | 0.5494 | 0.000707 | ---?? | 32.5 | 0.2271 | | |
| rs1496931 | 4 | 190023934 | 4 | 0 | 0 | T | C | 0.8441 | 0.2492 | 0.0007072 | +++?+ | 0 | 0.4054 | intergenic | |
| rs3793681 | 10 | 134564451 | 4 | 0 | 0 | T | C | -0.9514 | 0.2809 | 0.0007072 | ---?- | 0 | 0.4521 | intron | INPP5A |
| rs6583330 | 3 | 196329468 | 4 | 0 | 0 | A | C | 0.9609 | 0.2837 | 0.0007073 | +++?+ | 0 | 0.9003 | intergenic | |
| rs75366971 | 7 | 10251207 | 4 | 0 | 0 | A | T | 1.5088 | 0.4455 | 0.0007073 | +++?+ | 23.5 | 0.2701 | intergenic | |
| rs7727384 | 5 | 105736353 | 4 | 0 | 0 | T | C | -1.732 | 0.5114 | 0.0007074 | ---?- | 0 | 0.8031 | intergenic | |
| rs12085512 | 1 | 203687524 | 4 | 0 | 0 | A | T | 1.0992 | 0.3246 | 0.0007075 | +++?+ | 0 | 0.9164 | intron | ATP2B4 |
| rs75114484 | 22 | 36974532 | 4 | 0 | 0 | A | G | -1.3919 | 0.411 | 0.0007076 | ---?+ | 13.5 | 0.325 | intron | CACNG2 |
| rs56024390 | 3 | 176022983 | 4 | 0 | 0 | A | C | 1.2637 | 0.3732 | 0.0007077 | +++?+ | 3.5 | 0.3753 | intergenic | |
| rs113447643 | 3 | 23806742 | 4 | 0 | 0 | T | C | -3.6183 | 1.0685 | 0.0007079 | ---?- | 0 | 0.7759 | intergenic | |
| 16:84803189 | 16 | 84803189 | 3 | 0 | 0 | T | TTGGC | 1.0479 | 0.3094 | 0.000708 | +++?? | 0.9 | 0.3647 | | |
| rs10031879 | 4 | 183669589 | 4 | 0 | 0 | A | T | 0.9381 | 0.277 | 0.0007081 | +++?- | 0 | 0.741 | intron | TENM3 |
| rs76333139 | 10 | 21332920 | 3 | 0 | 0 | A | G | -3.2465 | 0.9587 | 0.0007081 | ---?? | 0 | 0.9427 | intron | NEBL |
| rs2701413 | 15 | 87613837 | 3 | 0 | 0 | T | C | -0.9402 | 0.2777 | 0.0007082 | ---?? | 0 | 0.4579 | intergenic | |
| rs12627226 | 21 | 19081619 | 4 | 0 | 0 | T | C | -1.3378 | 0.3951 | 0.0007082 | ---?- | 0 | 0.7972 | intergenic | |
| rs13336084 | 16 | 52937615 | 4 | 0 | 0 | T | C | 0.8374 | 0.2473 | 0.0007083 | +++?+ | 0 | 0.4556 | intergenic | |
| rs72929153 | 2 | 199094550 | 4 | 0 | 0 | A | G | -1.3646 | 0.403 | 0.0007083 | ---?- | 49.4 | 0.1148 | intergenic | |
| rs62419112 | 6 | 63329420 | 4 | 0 | 0 | A | G | 2.4073 | 0.7109 | 0.0007086 | +++?+ | 0 | 0.4899 | intergenic | |
| rs77404150 | 6 | 130953004 | 4 | 0 | 0 | A | C | -1.4385 | 0.4248 | 0.0007087 | ---?- | 0 | 0.7016 | intergenic | |
| 5:39589369 | 5 | 39589369 | 3 | 0 | 0 | T | TTAG | -0.8777 | 0.2592 | 0.0007087 | ---?? | 0 | 0.5235 | | |
| 6:159235419 | 6 | 159235419 | 3 | 0 | 0 | A | ACTGCA | 0.8648 | 0.2554 | 0.0007087 | +++?? | 13.5 | 0.3147 | | |
| rs1923449 | 10 | 123934456 | 4 | 0 | 0 | A | C | -1.4812 | 0.4374 | 0.0007088 | ---?- | 0 | 0.9947 | intron | TACC2 |
| rs151237826 | 17 | 68796257 | 3 | 0 | 0 | A | C | -3.476 | 1.0265 | 0.0007088 | ---?? | 0 | 0.4764 | intergenic | |
| rs2288388 | 5 | 175310713 | 4 | 0 | 0 | T | C | 1.1776 | 0.3478 | 0.0007089 | +++?+ | 0 | 0.711 | 3-prime-UTR | CPLX2 |
| rs60980548 | 19 | 12827395 | 4 | 0 | 0 | A | G | 3.303 | 0.9755 | 0.0007091 | ++++? | 53 | 0.0943 | intron | TNPO2 |
| rs73056807 | 7 | 10251831 | 4 | 0 | 0 | A | G | 1.5067 | 0.445 | 0.0007091 | +++?+ | 23.2 | 0.2718 | intergenic | |
| rs34359137 | 13 | 20808487 | 4 | 0 | 0 | T | C | 1.1262 | 0.3326 | 0.0007095 | +++?+ | 0 | 0.5234 | upstream-gene | |
| rs74132557 | 1 | 180134220 | 4 | 0 | 0 | A | G | -2.166 | 0.6397 | 0.0007095 | ---?+ | 36.9 | 0.1907 | intron | QSOX1 |
| rs6005697 | 22 | 28423935 | 4 | 0 | 0 | T | C | 2.7134 | 0.8014 | 0.0007096 | +++?+ | 0 | 0.7975 | intron | TTC28 |
| 5:39616698 | 5 | 39616698 | 3 | 0 | 0 | CT | CTT | 1.1145 | 0.3292 | 0.0007096 | +++?? | 0 | 0.9783 | | |
| rs76684929 | 5 | 120085776 | 2 | 0 | 0 | T | G | 5.9586 | 1.7599 | 0.0007098 | ?++?? | 0 | 0.4853 | intergenic | |
| rs77311365 | 18 | 39928868 | 4 | 0 | 0 | T | C | -1.3824 | 0.4083 | 0.0007098 | ---?- | 0 | 0.7898 | intron | LINC00907 |
| rs75939173 | 11 | 86184272 | 4 | 0 | 0 | A | C | 1.5895 | 0.4695 | 0.0007099 | +++?+ | 0 | 0.5111 | intron | ME3 |
| rs2731866 | 5 | 82547566 | 4 | 0 | 0 | A | G | -2.0449 | 0.604 | 0.0007101 | ---?- | 0 | 0.5355 | intron | XRCC4 |
| rs7793401 | 7 | 10175504 | 4 | 0 | 0 | A | C | 0.966 | 0.2853 | 0.0007101 | +++?+ | 48.2 | 0.1223 | intergenic | |
| rs10511867 | 9 | 30814128 | 4 | 0 | 0 | C | G | -3.2166 | 0.9501 | 0.0007102 | ---?- | 2.1 | 0.382 | intergenic | |
| 12:43209837 | 12 | 43209837 | 3 | 0 | 0 | CTG | C | 0.9844 | 0.2908 | 0.0007103 | +++?? | 0 | 0.7891 | | |
| rs113149638 | 9 | 44274893 | 3 | 0 | 0 | A | G | 1.8084 | 0.5342 | 0.0007104 | +++?? | 0 | 0.8277 | intergenic | |
| rs11838404 | 13 | 112823350 | 4 | 0 | 0 | T | C | 0.9358 | 0.2764 | 0.0007108 | +++?+ | 13.5 | 0.3251 | intergenic | |
| rs2541918 | 7 | 35475322 | 4 | 0 | 0 | A | T | -0.9717 | 0.287 | 0.0007111 | ---?- | 5.8 | 0.3641 | intergenic | |
| rs10249218 | 7 | 49462699 | 4 | 0 | 0 | T | C | 0.94 | 0.2777 | 0.0007111 | +++?- | 23.6 | 0.2696 | intergenic | |
| 1:213099171 | 1 | 213099171 | 3 | 0 | 0 | A | AAG | 1.3293 | 0.3927 | 0.0007112 | +++?? | 0 | 0.4077 | | |
| rs9927048 | 16 | 52918002 | 4 | 0 | 0 | A | G | 0.8398 | 0.2481 | 0.0007113 | +++?+ | 0 | 0.6644 | intergenic | |
| rs57425824 | 6 | 32586011 | 5 | 0 | 1 | T | C | 1.6135 | 0.4767 | 0.0007115 | +++++ | 0 | 0.5621 | intergenic | |
| rs12626851 | 21 | 19081705 | 4 | 0 | 0 | T | C | 1.337 | 0.395 | 0.0007118 | +++?+ | 0 | 0.7974 | intergenic | |
| rs6811030 | 4 | 20730601 | 4 | 0 | 0 | T | C | 0.8864 | 0.2619 | 0.000712 | +++?+ | 0 | 0.8498 | 3-prime-UTR | KCNIP4 |
| rs151278309 | 6 | 31846544 | 4 | 0 | 0 | A | T | 3.3284 | 0.9833 | 0.0007121 | ++++? | 29.6 | 0.2349 | intron | SLC44A4 |
| rs807104 | 17 | 51246492 | 5 | 0 | 0 | T | G | 1.0586 | 0.3127 | 0.0007124 | ++++- | 2 | 0.395 | intergenic | |

| | | | | | | | | | | | | | | | |
|---|---|---|---|---|---|---|---|---|---|---|---|---|---|---|---|
| rs111673045 | 9 | 44271437 | 3 | 0 | 0 | A | C | 1.771 | 0.5232 | 0.0007126 | +++?? | 0 | 0.8243 | intergenic | |
| rs6962352 | 7 | 75189471 | 5 | 0 | 0 | A | G | 0.7582 | 0.224 | 0.0007127 | +++++ | 0 | 0.4564 | intron | HIP1 |
| 8:65030446 | 8 | 65030446 | 3 | 0 | 0 | T | TA | 0.8568 | 0.2531 | 0.0007128 | +++?? | 0 | 0.8395 | | |
| rs2298608 | 11 | 58404467 | 5 | 0 | 0 | T | C | -0.8277 | 0.2446 | 0.0007131 | ----- | 0 | 0.9841 | intergenic |  |
| rs62421003 | 6 | 137739360 | 4 | 0 | 0 | A | G | 1.116 | 0.3298 | 0.0007131 | +++?+ | 0 | 0.7319 | intergenic | |
| rs1080291 | 6 | 95160757 | 4 | 0 | 0 | A | G | -0.8339 | 0.2464 | 0.0007133 | ---?- | 0 | 0.7565 | intergenic | |
| rs12531190 | 7 | 49464971 | 4 | 0 | 0 | C | G | 0.9425 | 0.2785 | 0.0007133 | +++?- | 24.4 | 0.265 | intergenic | |
| rs7019003 | 9 | 103870462 | 4 | 0 | 0 | A | T | 0.8466 | 0.2501 | 0.0007136 | +++?+ | 44.9 | 0.1418 | intron | LPPR1 |
| rs76025597 | 11 | 86230008 | 4 | 0 | 0 | T | C | 2.5131 | 0.7426 | 0.0007136 | +++?+ | 0 | 0.8381 | intron | ME3 |
| rs1025667 | 8 | 96935794 | 4 | 0 | 0 | T | C | -0.9039 | 0.2671 | 0.0007138 | ---?- | 0 | 0.7776 | intergenic | |
| rs1814264 | 16 | 52857809 | 4 | 0 | 0 | T | C | -0.8204 | 0.2424 | 0.0007138 | ---?- | 0 | 0.793 | intergenic | |
| rs28750405 | 15 | 32183126 | 4 | 0 | 0 | A | C | 2.4949 | 0.7372 | 0.0007139 | +++?+ | 0 | 0.8339 | intergenic | |
| rs12118621 | 1 | 100197059 | 4 | 0 | 0 | T | C | -1.7991 | 0.5316 | 0.000714 | ---?- | 0 | 0.5781 | intron | FRRS1 |
| rs12647270 | 4 | 96510901 | 4 | 0 | 0 | T | C | -1.2752 | 0.3768 | 0.000714 | ---?- | 26.6 | 0.252 | intergenic | |
| rs61820438 | 1 | 187855454 | 4 | 0 | 0 | T | C | 1.138 | 0.3363 | 0.0007141 | +++?+ | 0 | 0.8719 | intergenic | |
| rs112058906 | 9 | 44271431 | 3 | 0 | 0 | A | C | -1.77 | 0.5231 | 0.0007145 | ---?? | 0 | 0.8236 | intergenic | |
| rs10986041 | 9 | 101134187 | 4 | 0 | 0 | T | C | 3.257 | 0.9625 | 0.0007148 | +++?- | 30.6 | 0.2288 | intron | GABBR2 |
| rs11164516 | 1 | 102903311 | 4 | 0 | 0 | T | C | -0.8507 | 0.2514 | 0.000715 | ---?+ | 0 | 0.5846 | intergenic | |
| rs28479973 | 9 | 44271486 | 3 | 0 | 0 | A | T | -1.7721 | 0.5237 | 0.000715 | ---?? | 0 | 0.8243 | intergenic | |
| rs7820539 | 8 | 113487537 | 4 | 0 | 0 | T | G | 0.8909 | 0.2633 | 0.0007151 | +++?+ | 0 | 0.7866 | intron | CSMD3 |
| rs72854962 | 2 | 131118610 | 5 | 0 | 0 | A | G | 0.8819 | 0.2606 | 0.0007151 | +++++ | 25.5 | 0.2515 | intron | PTPN18 |
| 6:139335986 | 6 | 139335986 | 3 | 0 | 0 | T | TCAAA | 1.4512 | 0.4289 | 0.0007151 | +++?? | 0 | 0.9767 | | |
| rs896001 | 13 | 103342039 | 4 | 0 | 0 | T | C | -0.9456 | 0.2795 | 0.0007152 | ---?- | 0 | 0.7956 | intron | METTL21C |
| rs1938598 | 11 | 58413910 | 5 | 0 | 0 | T | C | 0.8337 | 0.2464 | 0.0007153 | +++++ | 0 | 0.9875 | intergenic | |
| rs71401691 | 15 | 75602162 | 5 | 0 | 0 | A | G | -1.5227 | 0.45 | 0.0007153 | ----+ | 0 | 0.7182 | intergenic |  |
| rs141501030 | 5 | 707472 | 3 | 0 | 0 | A | G | 1.4883 | 0.4399 | 0.0007156 | +++?? | 60.9 | 0.07766 | downstream-gene | |
| 1:202046518 | 1 | 202046518 | 3 | 0 | 0 | A | C | 1.1927 | 0.3525 | 0.0007157 | +++?? | 32.2 | 0.2287 | intergenic | |
| rs1326914 | 9 | 78438434 | 4 | 0 | 0 | T | C | 1.2327 | 0.3643 | 0.0007158 | +++?+ | 58.3 | 0.06577 | intergenic | |
| rs116865693 | 16 | 58872564 | 4 | 0 | 0 | A | G | 3.9266 | 1.1605 | 0.0007158 | +++?+ | 0 | 0.8553 | intergenic | |
| rs1487300 | 1 | 98595814 | 3 | 0 | 0 | A | G | -1.368 | 0.4043 | 0.0007161 | ---?? | 45.7 | 0.1583 | intergenic | |
| rs10874617 | 1 | 102902818 | 4 | 0 | 0 | T | C | 0.851 | 0.2515 | 0.0007161 | +++?- | 0 | 0.5838 | intergenic | |
| rs61862917 | 10 | 112572856 | 4 | 0 | 0 | A | G | 2.2832 | 0.6748 | 0.0007161 | +++?+ | 0 | 0.9809 | intron | RBM20 |
| rs184110624 | 15 | 25201817 | 3 | 0 | 0 | A | T | 4.0476 | 1.1963 | 0.0007162 | +++?? | 0 | 0.5209 | intron | SNRPN,SNURF |
| rs567792 | 18 | 57849587 | 4 | 0 | 0 | C | G | 0.823 | 0.2433 | 0.0007163 | +-+?+ | 53.3 | 0.09284 | intergenic | |
| rs12131728 | 1 | 117891913 | 4 | 0 | 0 | A | G | -2.7577 | 0.8151 | 0.0007164 | ---?+ | 28.2 | 0.2431 | intergenic | |
| rs17100574 | 14 | 63424825 | 4 | 0 | 0 | A | C | -2.0356 | 0.6017 | 0.0007165 | ---?+ | 51.1 | 0.1054 | intron | KCNH5 |
| rs539498 | 18 | 57850355 | 4 | 0 | 0 | A | G | -0.8228 | 0.2432 | 0.0007168 | -+-?- | 53.2 | 0.09303 | intergenic | |
| rs10108081 | 8 | 113488660 | 4 | 0 | 0 | A | G | -0.892 | 0.2637 | 0.0007168 | ---?- | 0 | 0.7883 | intron | CSMD3 |
| 14:89914363 | 14 | 89914363 | 3 | 0 | 0 | A | G | -2.0357 | 0.6018 | 0.0007172 | ---?? | 0 | 0.4041 | intron | FOXN3 |
| rs10986039 | 9 | 101133931 | 4 | 0 | 0 | A | G | 3.255 | 0.9623 | 0.0007178 | +++?- | 30.5 | 0.2295 | intron | GABBR2 |
| rs77575406 | 19 | 49154373 | 4 | 0 | 0 | T | C | -1.8812 | 0.5561 | 0.0007178 | ---?+ | 0 | 0.7383 | intron | SEC1P |
| rs367568966 | 19 | 56071709 | 3 | 0 | 0 | A | T | 1.625 | 0.4804 | 0.0007178 | +++?? | 0 | 0.4657 | intergenic | |
| rs115746499 | 1 | 42800913 | 4 | 0 | 0 | T | C | 1.8655 | 0.5515 | 0.0007179 | +++?+ | 0 | 0.58 | intron | FOXJ3 |
| rs7662115 | 4 | 190016749 | 4 | 0 | 0 | C | G | 0.8502 | 0.2513 | 0.0007182 | +++?+ | 25.4 | 0.2591 | intergenic | |
| rs13204631 | 6 | 137739024 | 4 | 0 | 0 | T | C | 1.1148 | 0.3296 | 0.0007183 | +++?+ | 0 | 0.7322 | intergenic | |
| rs11075649 | 16 | 52859323 | 4 | 0 | 0 | A | G | -0.8208 | 0.2427 | 0.0007184 | ---?- | 0 | 0.7862 | intergenic | |
| rs114510568 | 1 | 42800394 | 4 | 0 | 0 | A | G | 1.8645 | 0.5512 | 0.0007184 | +++?+ | 0 | 0.5798 | intron | FOXJ3 |
| 3:45345471 | 3 | 45345471 | 3 | 0 | 0 | T | TTAA | -1.3487 | 0.3987 | 0.0007185 | ---?? | 48.7 | 0.1425 | | |
| rs28523819 | 9 | 44265521 | 3 | 0 | 0 | A | T | -1.7678 | 0.5227 | 0.0007186 | ---?? | 0 | 0.806 | intergenic | |
| rs116632574 | 9 | 44276038 | 3 | 0 | 0 | T | C | 1.8288 | 0.5407 | 0.0007189 | +++?? | 0 | 0.8264 | intergenic | |
| rs62547350 | 9 | 16208483 | 3 | 0 | 0 | A | T | 0.9674 | 0.286 | 0.0007192 | +++?? | 0 | 0.9908 | intron | C9orf92 |
| rs143288512 | 11 | 86227647 | 4 | 0 | 0 | C | G | 2.5082 | 0.7416 | 0.0007192 | +++?+ | 0 | 0.8348 | intron | ME3 |
| rs150911833 | 1 | 85621171 | 3 | 0 | 0 | A | T | 1.6809 | 0.497 | 0.0007192 | +++?? | 0 | 0.9496 | downstream-gene | |
| 10:117602197 | 10 | 117602197 | 3 | 0 | 0 | T | TAGTC | -0.9546 | 0.2822 | 0.0007193 | ---?? | 0 | 0.5596 | | |
| 1:203684252 | 1 | 203684252 | 3 | 0 | 0 | <Cn0> | T | -1.1314 | 0.3345 | 0.0007193 | ---?? | 0 | 0.7513 | | |
| rs13085587 | 3 | 33269181 | 4 | 0 | 0 | T | C | -0.89 | 0.2631 | 0.0007194 | ---?- | 13.6 | 0.3244 | intergenic | |
| rs7080744 | 10 | 122498087 | 4 | 0 | 0 | T | C | 0.9492 | 0.2807 | 0.0007195 | +++?+ | 37.7 | 0.1856 | intergenic | |
| rs9277965 | 6 | 33256397 | 5 | 0 | 0 | A | G | 0.986 | 0.2915 | 0.0007195 | +++++ | 0 | 0.678 | synonymous | WDR46 |
| rs7727455 | 5 | 94798315 | 2 | 0 | 0 | A | G | 1.7741 | 0.5246 | 0.0007196 | ???++ | 0 | 0.8993 | downstream-gene | |

| | | | | | | | | | | | | | | | |
|---|---|---|---|---|---|---|---|---|---|---|---|---|---|---|---|
| rs11075659 | 16 | 52918219 | 4 | 0 | 0 | A | G | -0.878 | 0.2596 | 0.0007197 | ---?- | 0 | 0.6985 | intergenic | |
| rs940527 | 12 | 103243601 | 4 | 0 | 0 | A | G | 0.8317 | 0.2459 | 0.0007198 | +++?+ | 0 | 0.6621 | intron | PAH |
| rs1932040 | 6 | 45696788 | 4 | 0 | 0 | A | G | -1.0848 | 0.3208 | 0.0007201 | ---?+ | 58.6 | 0.06434 | intergenic | |
| rs72819904 | 10 | 106980349 | 4 | 0 | 0 | A | G | 1.4723 | 0.4354 | 0.0007202 | +++?- | 0 | 0.7248 | intron | SORCS3 |
| rs929026 | 22 | 35346625 | 4 | 0 | 0 | A | G | 1.1233 | 0.3322 | 0.0007203 | +++?+ | 0 | 0.9537 | intergenic | |
| rs28572754 | 9 | 44272389 | 3 | 0 | 0 | A | T | -1.7833 | 0.5273 | 0.0007203 | ---?? | 0 | 0.824 | intergenic | |
| rs10911009 | 1 | 182021804 | 4 | 0 | 0 | T | C | -1.0998 | 0.3252 | 0.0007204 | ---?- | 9.4 | 0.3463 | downstream-gene | |
| rs148343688 | 9 | 44280257 | 3 | 0 | 0 | T | G | -1.9143 | 0.5661 | 0.0007204 | ---?? | 0 | 0.8149 | intergenic | |
| rs16883708 | 8 | 113487450 | 4 | 0 | 0 | C | G | 0.89 | 0.2632 | 0.0007205 | +++?+ | 0 | 0.7862 | intron | CSMD3 |
| rs76886335 | 12 | 104876132 | 4 | 0 | 0 | C | G | 1.0769 | 0.3185 | 0.0007208 | +++?+ | 0 | 0.5113 | intron | CHST11 |
| rs11011647 | 10 | 20150688 | 4 | 0 | 0 | T | C | -0.8627 | 0.2551 | 0.0007209 | ---?- | 10.4 | 0.3411 | intron | PLXDC2 |
| rs10165565 | 2 | 170785484 | 4 | 0 | 1 | A | G | -0.9774 | 0.289 | 0.000721 | ---?- | 43.9 | 0.148 | intron | UBR3 |
| rs142998346 | 12 | 96653280 | 4 | 0 | 0 | T | C | -2.6206 | 0.775 | 0.000721 | ---?- | 0 | 0.3955 | intron | ELK3 |
| 18:39952119 | 18 | 39952119 | 3 | 0 | 0 | T | TTG | -1.4066 | 0.416 | 0.0007211 | ---?? | 0 | 0.6165 | | |
| rs17369151 | 1 | 42798311 | 4 | 0 | 0 | T | C | -1.8619 | 0.5506 | 0.0007214 | ---?- | 0 | 0.5781 | intron | FOXJ3 |
| rs145912985 | 2 | 170425520 | 3 | 0 | 0 | A | C | 1.2766 | 0.3775 | 0.0007214 | +++?? | 0 | 0.9968 | intron | FASTKD1 |
| rs1908635 | 1 | 64805503 | 4 | 0 | 0 | A | C | -1.0411 | 0.3079 | 0.0007216 | ---?- | 0 | 0.5538 | intergenic | |
| rs11734229 | 4 | 14376177 | 4 | 0 | 0 | A | G | -0.9069 | 0.2682 | 0.0007217 | -+-?- | 51.7 | 0.1016 | intergenic | |
| rs4973248 | 2 | 230916403 | 4 | 0 | 0 | A | G | -1.4313 | 0.4233 | 0.0007218 | ---?- | 21.6 | 0.2811 | intron | SLC16A14 |
| rs7585433 | 2 | 184066737 | 4 | 0 | 0 | T | C | 1.1646 | 0.3445 | 0.000722 | +++?+ | 0 | 0.6609 | intergenic | |
| rs57137336 | 22 | 23940458 | 3 | 0 | 0 | T | C | 4.6978 | 1.3894 | 0.000722 | ?++?- | 41.3 | 0.1823 | intergenic | |
| rs4666616 | 2 | 184017124 | 4 | 0 | 0 | T | G | 1.1649 | 0.3445 | 0.0007221 | +++?+ | 0 | 0.8518 | intron | NUP35 |
| rs9653385 | 2 | 158822406 | 4 | 0 | 0 | T | C | 0.8206 | 0.2427 | 0.0007221 | +++?+ | 56.6 | 0.07466 | intron | UPP2 |
| rs13301369 | 9 | 21592386 | 4 | 0 | 0 | A | T | 1.0757 | 0.3182 | 0.0007222 | +++?+ | 0 | 0.8688 | intergenic | |
| rs62378384 | 5 | 148247866 | 4 | 0 | 0 | A | G | 2.082 | 0.6158 | 0.0007223 | +++?+ | 0 | 0.9255 | intron | ADRB2 |
| rs10421327 | 19 | 14716909 | 4 | 0 | 0 | T | C | -1.1693 | 0.3458 | 0.0007224 | ---?- | 26.5 | 0.2529 | intron | CLEC17A |
| rs76977067 | 7 | 147487758 | 4 | 0 | 0 | T | C | -1.6485 | 0.4876 | 0.0007224 | ---?- | 12.6 | 0.3294 | intron | CNTNAP2 |
| rs11077507 | 17 | 68599873 | 4 | 0 | 0 | A | G | 0.9109 | 0.2694 | 0.0007225 | +++?+ | 0 | 0.9677 | intergenic | |
| 9:16208482 | 9 | 16208482 | 3 | 0 | 0 | CCT | C | -0.9685 | 0.2865 | 0.0007225 | ---?? | 0 | 0.985 | | |
| 7:10236886 | 7 | 10236886 | 3 | 0 | 0 | G | GA | 0.9294 | 0.2749 | 0.0007227 | +++?? | 27.5 | 0.2518 | | |
| 3:18056428 | 3 | 18056428 | 3 | 0 | 0 | T | TTAAA | -2.3481 | 0.6946 | 0.0007229 | ---?? | 0 | 0.5459 | | |
| rs2887233 | 2 | 184015089 | 4 | 0 | 0 | T | G | 1.1658 | 0.3448 | 0.000723 | +++?+ | 0 | 0.8538 | intron | NUP35 |
| rs4961664 | 9 | 16161233 | 4 | 0 | 0 | T | C | 0.9028 | 0.2671 | 0.000723 | +++?- | 0 | 0.6561 | intergenic | |
| rs4957107 | 5 | 699647 | 3 | 0 | 0 | A | G | -1.3456 | 0.398 | 0.0007233 | ---?? | 0 | 0.735 | intergenic | |
| rs1522306 | 12 | 103240630 | 4 | 0 | 0 | A | C | 0.8347 | 0.2469 | 0.0007236 | +++?+ | 0 | 0.6804 | intron | PAH |
| rs10742020 | 11 | 23975873 | 4 | 0 | 0 | A | G | 1.2106 | 0.3581 | 0.0007237 | +++?+ | 0 | 0.7869 | intergenic | |
| rs11734375 | 4 | 10046298 | 5 | 0 | 0 | A | G | -0.7115 | 0.2105 | 0.0007238 | ----+ | 9.3 | 0.3532 | upstream-gene |  |
| rs113509411 | 22 | 20048447 | 4 | 0 | 0 | A | G | -2.1426 | 0.6338 | 0.0007239 | ---?- | 16.2 | 0.3108 | intron | TANGO2 |
| rs28605404 | 6 | 32569687 | 5 | 0 | 1 | A | G | -1.2307 | 0.3641 | 0.0007242 | ----- | 27.3 | 0.2393 | intergenic |  |
| rs17073943 | 4 | 183657059 | 4 | 0 | 0 | A | G | 0.8963 | 0.2651 | 0.0007243 | +++?+ | 0 | 0.9515 | intron | TENM3 |
| rs11607177 | 11 | 102513324 | 5 | 0 | 0 | T | C | -0.8586 | 0.254 | 0.0007244 | +---- | 21.3 | 0.2788 | intergenic |  |
| rs1496930 | 4 | 190023992 | 4 | 0 | 0 | T | C | 0.8413 | 0.2489 | 0.0007245 | +++?+ | 0 | 0.4043 | intergenic | |
| 15:87572737 | 15 | 87572737 | 3 | 0 | 0 | CAAT | C | -0.9251 | 0.2737 | 0.0007247 | ---?? | 19.8 | 0.2872 | | |
| rs28438690 | 9 | 44271139 | 3 | 0 | 0 | A | T | 1.765 | 0.5222 | 0.0007249 | +++?? | 0 | 0.8238 | intergenic | |
| rs140769288 | 16 | 62238226 | 3 | 0 | 0 | T | C | 4.6485 | 1.3753 | 0.0007249 | +++?? | 12.1 | 0.3206 | intergenic | |
| 19:7191939 | 19 | 7191939 | 3 | 0 | 0 | G | GGGAA | -0.9202 | 0.2722 | 0.0007251 | ---?? | 0 | 0.7103 | | |
| rs4748616 | 10 | 20150263 | 4 | 0 | 0 | C | G | -0.8683 | 0.2569 | 0.0007253 | ---?- | 42.3 | 0.1579 | intron | PLXDC2 |
| rs918844 | 2 | 208355781 | 3 | 0 | 0 | T | C | 1.4417 | 0.4265 | 0.0007254 | +++?? | 34.3 | 0.2181 | intergenic | |
| rs11889063 | 2 | 75666640 | 4 | 0 | 0 | T | C | -0.8897 | 0.2633 | 0.0007255 | ---?- | 0 | 0.9899 | intergenic | |
| rs34687248 | 3 | 33265607 | 4 | 0 | 0 | A | G | 0.8672 | 0.2566 | 0.0007255 | +++?+ | 22.7 | 0.2745 | upstream-gene | |
| rs2701412 | 15 | 87613838 | 3 | 0 | 0 | T | C | 0.9361 | 0.277 | 0.0007258 | +++?? | 0 | 0.4596 | intergenic | |
| rs9995074 | 4 | 91495454 | 3 | 0 | 0 | A | C | 0.9338 | 0.2763 | 0.0007258 | +++?? | 0 | 0.6628 | intron | CCSER1 |
| rs41373051 | 18 | 8564183 | 3 | 0 | 0 | A | G | -4.3712 | 1.2934 | 0.000726 | ---?? | 0 | 0.4447 | intergenic | |
| rs151316944 | 6 | 118841862 | 4 | 0 | 0 | T | C | -3.7428 | 1.1075 | 0.000726 | ---?- | 0 | 0.7423 | intron | CEP85L |
| rs28391166 | 7 | 82421177 | 4 | 0 | 0 | A | G | 0.8671 | 0.2566 | 0.0007261 | +-+?+ | 47 | 0.1295 | intron | PCLO |
| rs61922200 | 12 | 29785292 | 4 | 0 | 0 | C | G | 1.4554 | 0.4307 | 0.0007262 | +++?+ | 0 | 0.7412 | intron | TMTC1 |
| 3:59807484 | 3 | 59807484 | 3 | 0 | 0 | A | T | 1.473 | 0.4359 | 0.0007264 | +++?? | 0 | 0.5887 | intron | FHIT |
| rs113409670 | 9 | 44275068 | 3 | 0 | 0 | A | T | 1.8167 | 0.5376 | 0.0007264 | +++?? | 0 | 0.824 | intergenic | |
| 5:10306101 | 5 | 10306101 | 3 | 0 | 0 | T | TACA | -1.8731 | 0.5543 | 0.0007265 | ---?? | 17 | 0.2998 | | |

| | | | | | | | | | | | | | | | |
|---|---|---|---|---|---|---|---|---|---|---|---|---|---|---|---|
| rs139437925 | 9 | 44275243 | 3 | 0 | 0 | A | G | -1.8204 | 0.5387 | 0.0007269 | ---?? | 0 | 0.8248 | intergenic | |
| rs56087177 | 12 | 5514425 | 4 | 0 | 0 | A | G | -2.421 | 0.7164 | 0.000727 | ---?- | 0 | 0.49 | intergenic | |
| rs72810414 | 2 | 47377714 | 4 | 0 | 0 | A | G | 1.3048 | 0.3861 | 0.000727 | +++?+ | 0 | 0.9761 | intron | C2orf61 |
| rs10869651 | 9 | 78441553 | 4 | 0 | 0 | T | G | 1.2299 | 0.364 | 0.0007271 | +++?+ | 57.9 | 0.06824 | intergenic | |
| rs77070969 | 5 | 122559358 | 4 | 0 | 0 | T | C | -1.6335 | 0.4834 | 0.0007273 | ---?- | 0 | 0.9116 | intergenic | |
| rs10054364 | 5 | 103163820 | 4 | 0 | 0 | T | C | -1.5231 | 0.4508 | 0.0007274 | ---?- | 0 | 0.7131 | intergenic | |
| 9:19007682 | 9 | 19007682 | 3 | 0 | 0 | T | TCAGCT | -1.6323 | 0.4831 | 0.0007275 | ---?? | 58.5 | 0.09001 | | |
| rs4643758 | 4 | 176496244 | 4 | 0 | 0 | A | T | -2.5776 | 0.7628 | 0.0007276 | ---?- | 0 | 0.7805 | intergenic | |
| rs138954315 | 22 | 28433902 | 4 | 0 | 0 | T | C | 2.6976 | 0.7983 | 0.0007276 | +++?+ | 0 | 0.8013 | intron | TTC28 |
| rs2675081 | 2 | 184011979 | 4 | 0 | 0 | T | C | -1.1668 | 0.3453 | 0.000728 | ---?- | 0 | 0.8569 | intron | NUP35 |
| 2:239252287 | 2 | 239252287 | 3 | 0 | 0 | CT | C | 0.9435 | 0.2792 | 0.0007283 | +-+?? | 56.8 | 0.0988 | | |
| rs10205516 | 2 | 170865588 | 4 | 0 | 1 | T | G | -0.9918 | 0.2935 | 0.0007284 | ---?- | 47.6 | 0.1256 | intron | UBR3 |
| rs72753620 | 15 | 86340958 | 4 | 0 | 0 | A | G | 3.5168 | 1.0409 | 0.0007284 | +++?+ | 27.6 | 0.2461 | upstream-gene | |
| rs12997351 | 2 | 116519568 | 4 | 0 | 0 | A | C | -1.2147 | 0.3595 | 0.0007286 | ---?- | 7.3 | 0.3565 | intron | DPP10 |
| rs2024446 | 16 | 52853507 | 4 | 0 | 0 | A | G | 0.8153 | 0.2413 | 0.0007289 | +++?+ | 0 | 0.8121 | intergenic | |
| rs4847021 | 1 | 230959298 | 4 | 0 | 0 | T | C | 1.7054 | 0.5048 | 0.0007291 | +++?+ | 0 | 0.5808 | intergenic | |
| rs72962242 | 1 | 42795757 | 4 | 0 | 0 | A | G | 1.8551 | 0.5491 | 0.0007291 | +++?+ | 0 | 0.5774 | intron | FOXJ3 |
| rs3129964 | 6 | 32380717 | 4 | 0 | 0 | C | G | -0.7438 | 0.2202 | 0.0007292 | +---? | 57.9 | 0.06778 | intergenic | |
| rs17635482 | 17 | 51189763 | 4 | 0 | 0 | A | T | 1.021 | 0.3022 | 0.0007295 | +++?+ | 38.4 | 0.1814 | intergenic | |
| rs2459749 | 19 | 56071753 | 3 | 0 | 0 | T | C | 1.6213 | 0.4799 | 0.0007296 | +++?? | 0 | 0.4615 | intergenic | |
| rs112565313 | 17 | 71066896 | 4 | 0 | 0 | A | G | 2.2462 | 0.6649 | 0.0007296 | +++?+ | 28.3 | 0.2422 | intron | SLC39A11 |
| rs597697 | 6 | 10538945 | 4 | 0 | 0 | A | G | 0.8251 | 0.2443 | 0.0007297 | -++?+ | 54.4 | 0.08655 | intron | GCNT2 |
| rs17706332 | 13 | 76323507 | 4 | 0 | 0 | A | G | -1.2615 | 0.3734 | 0.0007298 | ---?- | 0 | 0.8796 | intron | LMO7 |
| rs4766502 | 12 | 111769641 | 5 | 0 | 0 | T | G | 0.8509 | 0.2519 | 0.00073 | +-+++ | 47.5 | 0.1064 | intron | CUX2 |
| rs8022710 | 14 | 22496029 | 4 | 0 | 0 | A | C | 0.8628 | 0.2554 | 0.00073 | +++?+ | 0 | 0.9453 | intergenic | |
| rs12784446 | 10 | 134589094 | 4 | 0 | 0 | T | G | -0.9355 | 0.277 | 0.0007301 | ---?- | 0 | 0.5583 | intron | INPP5A |
| rs6593901 | 1 | 203688090 | 4 | 0 | 0 | T | C | -1.0998 | 0.3256 | 0.0007302 | ---?- | 0 | 0.8276 | intron | ATP2B4 |
| 22:50760507 | 22 | 50760507 | 3 | 0 | 0 | A | ATT | 0.8883 | 0.263 | 0.0007306 | +++?? | 0 | 0.782 | | |
| rs411337 | 6 | 32077380 | 5 | 0 | 0 | T | C | 1.3294 | 0.3936 | 0.0007308 | +++-+ | 40.8 | 0.1494 | upstream-gene | |
| 1:80748197 | 1 | 80748197 | 3 | 0 | 0 | CATAA | C | 2.7961 | 0.8278 | 0.0007308 | +++?? | 0 | 0.9017 | | |
| rs940528 | 12 | 103243355 | 4 | 0 | 0 | T | C | -0.8309 | 0.246 | 0.0007309 | ---?- | 0 | 0.6626 | intron | PAH |
| rs10505173 | 8 | 113486216 | 4 | 0 | 0 | T | C | -0.8864 | 0.2624 | 0.0007309 | ---?- | 0 | 0.7823 | intron | CSMD3 |
| 4:20730541 | 4 | 20730541 | 3 | 0 | 0 | A | ATGAT | 0.9202 | 0.2724 | 0.0007309 | +++?? | 0 | 0.7169 | | |
| rs35877939 | 13 | 100407309 | 4 | 0 | 0 | A | G | 0.9555 | 0.2829 | 0.0007311 | +++?+ | 32.3 | 0.2187 | intron | CLYBL |
| 6:63191060 | 6 | 63191060 | 3 | 0 | 0 | T | TAG | -2.2991 | 0.6807 | 0.0007313 | ---?? | 3.1 | 0.3562 | | |
| rs35635311 | 12 | 75712220 | 4 | 0 | 0 | T | C | 2.0289 | 0.6007 | 0.0007318 | +++?+ | 0 | 0.7231 | intron | CAPS2 |
| rs150297366 | 13 | 61294132 | 2 | 0 | 0 | A | T | -5.5081 | 1.6309 | 0.000732 | --??? | 0 | 0.788 | intergenic | |
| rs9923327 | 16 | 12785655 | 3 | 0 | 0 | A | G | -1.0787 | 0.3194 | 0.0007321 | ---?? | 0 | 0.9064 | intron | CPPED1 |
| rs184088360 | 9 | 16188142 | 3 | 0 | 0 | T | G | 0.9308 | 0.2756 | 0.0007321 | +++?? | 0 | 0.9177 | intergenic | |
| rs2116865 | 19 | 22711952 | 4 | 0 | 0 | T | C | 0.8667 | 0.2566 | 0.0007322 | +++?+ | 0 | 0.7547 | intergenic | |
| rs11832384 | 12 | 41369322 | 4 | 0 | 0 | A | C | -1.4309 | 0.4237 | 0.0007322 | ---?- | 0 | 0.5144 | intron | CNTN1 |
| rs115949773 | 10 | 132820072 | 2 | 0 | 0 | A | G | -4.6685 | 1.3824 | 0.0007323 | -?-?? | 0 | 0.4941 | intergenic | |
| rs145879016 | 1 | 42794455 | 4 | 0 | 0 | A | C | -1.8563 | 0.5497 | 0.0007324 | ---?- | 0 | 0.5762 | intron | FOXJ3 |
| 7:135690864 | 7 | 135690864 | 3 | 0 | 0 | CA | C | -0.9978 | 0.2955 | 0.0007324 | ---?? | 0 | 0.5365 | | |
| rs1428417 | 5 | 123517721 | 4 | 0 | 0 | T | C | 0.8238 | 0.2439 | 0.0007325 | +++?- | 23.2 | 0.2716 | intergenic | |
| rs7818544 | 8 | 113483171 | 4 | 0 | 0 | T | G | -0.8871 | 0.2627 | 0.0007325 | ---?- | 0 | 0.7742 | intron | CSMD3 |
| rs7955574 | 12 | 30822414 | 4 | 0 | 0 | C | G | 0.8055 | 0.2385 | 0.0007325 | ?++++ | 0 | 0.8039 | intron | IPO8 |
| rs2675100 | 2 | 184006844 | 4 | 0 | 0 | T | C | 1.1681 | 0.3459 | 0.0007327 | +++?+ | 0 | 0.8613 | intron | NUP35 |
| rs34281679 | 6 | 157162456 | 4 | 0 | 0 | A | G | 1.6461 | 0.4874 | 0.0007327 | +++?+ | 0 | 0.7731 | intron | ARID1B |
| rs2123487 | 8 | 113481629 | 4 | 0 | 0 | C | G | 0.888 | 0.263 | 0.0007329 | +++?+ | 0 | 0.7715 | intron | CSMD3 |
| rs149261546 | 9 | 44283171 | 3 | 0 | 0 | A | C | 1.9744 | 0.5847 | 0.0007332 | +++?? | 0 | 0.8251 | intergenic | |
| rs2426744 | 20 | 56207927 | 4 | 0 | 0 | T | C | -1.2084 | 0.3579 | 0.0007333 | ---?- | 0 | 0.9495 | intergenic | |
| rs2798684 | 1 | 87574217 | 4 | 0 | 0 | A | G | 1.0075 | 0.2984 | 0.0007333 | +++?+ | 21.4 | 0.2821 | 3-prime-UTR | HS2ST1 |
| rs72936947 | 6 | 72856261 | 4 | 0 | 0 | A | C | 1.5752 | 0.4665 | 0.0007333 | +++?- | 0 | 0.5796 | intron | RIMS1 |
| rs114441788 | 1 | 42791705 | 4 | 0 | 0 | T | G | -1.8534 | 0.5489 | 0.0007333 | ---?- | 0 | 0.5734 | intron | FOXJ3 |
| rs71484661 | 10 | 122929721 | 3 | 0 | 0 | T | C | 3.5357 | 1.0471 | 0.0007334 | +++?? | 0 | 0.8283 | intergenic | |
| rs12160232 | 22 | 28399915 | 4 | 0 | 0 | T | C | 2.746 | 0.8132 | 0.0007336 | +++?+ | 0 | 0.795 | intron | TTC28 |
| rs117047314 | 18 | 68729752 | 4 | 0 | 0 | A | G | 2.4518 | 0.7261 | 0.000734 | +++?+ | 0 | 0.6749 | intergenic | |
| rs1538829 | 1 | 102942170 | 4 | 0 | 0 | A | T | -0.8235 | 0.2439 | 0.0007342 | ---?+ | 0 | 0.4105 | intergenic | |

| | | | | | | | | | | | | | | |
|---|---|---|---|---|---|---|---|---|---|---|---|---|---|---|
| rs2705761 | 2 | 184005859 | 4 | 0 | 0 | A | T | -1.1681 | 0.346 | 0.0007342 | ---?- | 0 | 0.8618 | intron | NUP35 |
| rs7105248 | 11 | 8722325 | 5 | 0 | 0 | C | G | 0.7869 | 0.2331 | 0.0007342 | ++++- | 0 | 0.8841 | intron | ST5 |
| rs145088754 | 15 | 83767799 | 4 | 0 | 0 | T | C | 3.3478 | 0.9915 | 0.0007342 | +++?+ | 0 | 0.9208 | intergenic | |
| rs6447975 | 4 | 20735769 | 4 | 0 | 0 | T | G | -0.8822 | 0.2613 | 0.0007346 | ---?- | 0 | 0.8322 | intron | KCNIP4 |
| rs6831516 | 4 | 20735519 | 4 | 0 | 0 | T | C | -0.8824 | 0.2613 | 0.0007347 | ---?- | 0 | 0.8322 | intron | KCNIP4 |
| 11:12877494 | 11 | 12877494 | 3 | 0 | 0 | T | TA | 1.0369 | 0.3071 | 0.0007347 | +++?? | 0 | 0.8506 | | |
| rs11048659 | 12 | 26851607 | 4 | 0 | 0 | C | G | -1.6752 | 0.4962 | 0.0007348 | -+-?- | 35.9 | 0.1965 | intron | ITPR2 |
| rs1907802 | 8 | 113492766 | 4 | 0 | 0 | A | C | -0.8962 | 0.2654 | 0.0007349 | ---?- | 0 | 0.7905 | intron | CSMD3 |
| rs10021016 | 4 | 183661413 | 4 | 0 | 0 | A | G | 0.9158 | 0.2713 | 0.0007349 | +++?- | 0 | 0.7185 | intron | TENM3 |
| rs850137 | 7 | 29794434 | 4 | 0 | 0 | T | C | 0.8746 | 0.2591 | 0.0007351 | +++?+ | 0 | 0.6525 | intergenic | |
| rs10493105 | 1 | 42792005 | 4 | 0 | 0 | A | G | -1.8533 | 0.5489 | 0.0007351 | ---?- | 0 | 0.5734 | intron | FOXJ3 |
| rs10493104 | 1 | 42791484 | 4 | 0 | 0 | A | G | -1.8529 | 0.5488 | 0.0007352 | ---?- | 0 | 0.5732 | intron | FOXJ3 |
| rs2705726 | 2 | 183996495 | 4 | 0 | 0 | A | G | 1.178 | 0.3489 | 0.0007354 | +++?+ | 0 | 0.8827 | intron | NUP35 |
| 3:185789495 | 3 | 185789495 | 3 | 0 | 0 | A | G | -2.9502 | 0.8739 | 0.0007357 | ---?? | 0 | 0.8296 | intron | ETV5 |
| rs6831295 | 4 | 20735408 | 4 | 0 | 0 | C | G | 0.8823 | 0.2613 | 0.0007357 | +++?+ | 0 | 0.8322 | intron | KCNIP4 |
| rs11888068 | 2 | 185460295 | 4 | 0 | 0 | A | C | 0.9291 | 0.2752 | 0.0007357 | +++?+ | 0 | 0.5213 | upstream-gene | |
| rs16827910 | 1 | 187959236 | 4 | 0 | 0 | A | T | -1.3703 | 0.4059 | 0.0007358 | ---?- | 47.3 | 0.1274 | intergenic | |
| 3:185789494 | 3 | 185789494 | 3 | 0 | 0 | A | C | 2.9503 | 0.8739 | 0.0007359 | +++?? | 0 | 0.8295 | intron | ETV5 |
| rs9878431 | 3 | 2476848 | 4 | 0 | 0 | A | G | 0.9097 | 0.2695 | 0.0007359 | +++?- | 38.4 | 0.1815 | intron | CNTN4 |
| rs9888667 | 15 | 57665702 | 4 | 0 | 0 | A | T | 1.0739 | 0.3181 | 0.0007359 | +-+?+ | 41.6 | 0.1622 | upstream-gene | |
| rs78556852 | 6 | 161858768 | 3 | 0 | 0 | C | G | 3.6409 | 1.0786 | 0.0007362 | +++?? | 0 | 0.6554 | intron | PARK2 |
| rs12130344 | 1 | 217990918 | 4 | 0 | 0 | A | G | -1.1045 | 0.3272 | 0.0007363 | ---?- | 0 | 0.9709 | intron | SPATA17 |
| rs181627680 | 15 | 30326555 | 3 | 0 | 0 | A | G | -4.3824 | 1.2982 | 0.0007364 | ---?? | 0 | 0.3903 | intergenic | |
| rs12799215 | 11 | 58405674 | 5 | 0 | 0 | T | C | -0.8269 | 0.245 | 0.0007369 | ----- | 0 | 0.9847 | intergenic |  |
| rs151028799 | 18 | 25892961 | 3 | 0 | 0 | T | C | -5.6118 | 1.6625 | 0.0007371 | --??- | 0 | 0.6152 | intergenic | |
| 4:36702006 | 4 | 36702006 | 3 | 0 | 0 | CT | C | 1.2227 | 0.3622 | 0.0007371 | +++?? | 0 | 0.8484 | | |
| rs76098034 | 3 | 196512144 | 4 | 0 | 0 | T | C | 2.0577 | 0.6096 | 0.0007373 | +++?- | 18.7 | 0.2967 | intron | PAK2 |
| rs12903530 | 15 | 91405684 | 4 | 0 | 0 | A | G | -0.9744 | 0.2887 | 0.0007374 | ---?- | 29.6 | 0.2348 | intergenic | |
| rs4353890 | 4 | 8311951 | 2 | 0 | 0 | A | C | -6.938 | 2.0556 | 0.0007375 | -???- | 0 | 0.5845 | downstream-gene | |
| rs62191759 | 2 | 230916715 | 4 | 0 | 0 | T | C | -1.4349 | 0.4251 | 0.0007375 | ---?- | 22.2 | 0.2777 | intron | SLC16A14 |
| rs1563064 | 8 | 113476586 | 4 | 0 | 0 | A | G | 0.8884 | 0.2632 | 0.0007376 | +++?+ | 0 | 0.7567 | intron | CSMD3 |
| rs6911225 | 6 | 10538394 | 4 | 0 | 0 | A | G | 0.8326 | 0.2467 | 0.0007377 | -++?+ | 57.9 | 0.06778 | intron | GCNT2 |
| rs80340154 | 6 | 63332895 | 4 | 0 | 0 | T | C | -2.4016 | 0.7115 | 0.0007377 | ---?- | 0 | 0.4872 | intergenic | |
| 4:40189926 | 4 | 40189926 | 3 | 0 | 0 | T | TCA | 1.3681 | 0.4053 | 0.0007377 | +++?? | 8 | 0.3371 | | |
| 7:130381056 | 7 | 130381056 | 3 | 0 | 0 | CT | C | -0.8616 | 0.2553 | 0.0007377 | -+-?? | 60.4 | 0.07993 | | |
| rs2132836 | 4 | 91498411 | 3 | 0 | 0 | T | C | -0.926 | 0.2744 | 0.0007378 | ---?? | 0 | 0.6684 | intron | CCSER1 |
| rs1948796 | 3 | 2485134 | 4 | 0 | 0 | T | C | 0.9501 | 0.2815 | 0.0007379 | +++?+ | 0 | 0.9175 | intron | CNTN4 |
| 6:80281827 | 6 | 80281827 | 2 | 0 | 0 | CTA | C | -5.1755 | 1.5334 | 0.0007379 | -?-?? | 54.1 | 0.14 | | |
| rs11120454 | 1 | 215132091 | 4 | 0 | 0 | A | G | -1.0809 | 0.3203 | 0.0007383 | ---?- | 0 | 0.658 | intergenic | |
| rs4768325 | 12 | 41316848 | 4 | 0 | 0 | T | G | 1.6755 | 0.4965 | 0.0007386 | +++?+ | 0 | 0.8595 | intron | CNTN1 |
| rs11240274 | 1 | 203690852 | 4 | 0 | 0 | T | G | -1.1028 | 0.3268 | 0.0007387 | ---?- | 0 | 0.8634 | intron | ATP2B4 |
| rs6848627 | 4 | 190014411 | 4 | 0 | 0 | A | G | -0.8489 | 0.2515 | 0.0007391 | ---?- | 29.7 | 0.2342 | intergenic | |
| 2:40471190 | 2 | 40471190 | 3 | 0 | 0 | T | TA | 1.2168 | 0.3606 | 0.0007395 | +++?? | 0 | 0.753 | | |
| 8:64963167 | 8 | 64963167 | 3 | 0 | 0 | A | ATCTC | -0.978 | 0.2898 | 0.0007396 | ---?? | 0 | 0.914 | | |
| rs75377718 | 7 | 46184240 | 4 | 0 | 0 | A | T | 1.9277 | 0.5713 | 0.0007398 | +++?+ | 0 | 0.967 | intergenic | |
| rs17115 | 17 | 51246147 | 5 | 0 | 0 | T | C | 1.0542 | 0.3124 | 0.0007399 | ++++- | 6 | 0.3727 | intergenic | |
| rs10106343 | 8 | 113493173 | 4 | 0 | 0 | A | T | -0.896 | 0.2655 | 0.0007399 | ---?- | 0 | 0.7856 | intron | CSMD3 |
| 14:95898487 | 14 | 95898487 | 3 | 0 | 0 | AA | G | -1.4218 | 0.4213 | 0.0007399 | ---?? | 33.4 | 0.2229 | | |
| rs74664350 | 9 | 20723320 | 4 | 0 | 0 | C | G | -1.7017 | 0.5043 | 0.0007401 | ---?- | 0 | 0.767 | intron | FOCAD |
| rs2705725 | 2 | 183995536 | 4 | 0 | 0 | T | C | -1.179 | 0.3494 | 0.0007404 | ---?- | 0 | 0.8847 | intron | NUP35 |
| rs4859055 | 3 | 87373894 | 4 | 0 | 0 | T | C | -1.0729 | 0.318 | 0.0007404 | ---?- | 0 | 0.9976 | intergenic | |
| rs10272463 | 7 | 82428904 | 4 | 0 | 0 | C | G | -0.8536 | 0.253 | 0.0007404 | -+-?- | 43.6 | 0.1499 | intron | PCLO |
| rs10943451 | 6 | 78081840 | 4 | 0 | 0 | A | G | -0.8865 | 0.2627 | 0.0007404 | ---?- | 0 | 0.7549 | intergenic | |
| rs117739035 | 11 | 408174 | 5 | 0 | 0 | T | G | -2.0364 | 0.6035 | 0.0007404 | ----+ | 0 | 0.8027 | missense | SIGIRR |
| rs10742021 | 11 | 23976050 | 4 | 0 | 0 | T | C | -1.2088 | 0.3583 | 0.0007405 | ---?- | 0 | 0.791 | intergenic | |
| rs117803663 | 16 | 87310136 | 2 | 0 | 0 | T | C | 3.8693 | 1.1468 | 0.0007412 | +?+?? | 0 | 0.7392 | intergenic | |
| rs145913976 | 15 | 85728155 | 4 | 0 | 0 | A | G | 1.9615 | 0.5814 | 0.0007413 | +++?+ | 41.3 | 0.1637 | intergenic | |
| rs180926195 | 5 | 120054832 | 2 | 0 | 0 | A | C | 5.925 | 1.7562 | 0.0007415 | ?++?? | 0 | 0.5017 | intergenic | |
| 1:53111094 | 1 | 53111094 | 3 | 0 | 0 | G | GTTTTT | 2.6276 | 0.7789 | 0.0007418 | +++?? | 0 | 0.6871 | | |

| | | | | | | | | | | | | | | | |
|---|---|---|---|---|---|---|---|---|---|---|---|---|---|---|---|
| rs2026763 | 20 | 56199626 | 4 | 0 | 0 | T | C | -1.1182 | 0.3315 | 0.0007419 | ---?- | 0 | 0.7936 | upstream-gene | |
| rs62516481 | 8 | 113477764 | 4 | 0 | 0 | A | G | -0.8874 | 0.2631 | 0.000742 | ---?- | 0 | 0.7565 | intron | CSMD3 |
| 17:51141777 | 17 | 51141777 | 3 | 0 | 0 | T | TTA | -1.4221 | 0.4215 | 0.000742 | ---?? | 0 | 0.3838 | | |
| rs6708552 | 2 | 170754071 | 4 | 0 | 1 | T | C | -0.9735 | 0.2886 | 0.0007422 | ---?- | 49.1 | 0.1167 | intron | UBR3 |
| rs79973871 | 11 | 86190639 | 4 | 0 | 0 | A | T | -2.3943 | 0.7098 | 0.0007424 | ---?- | 0 | 0.9459 | intron | ME3 |
| rs7784031 | 7 | 82430153 | 4 | 0 | 0 | A | G | 0.8559 | 0.2537 | 0.0007425 | +-+?+ | 44.8 | 0.1425 | intron | PCLO |
| rs73769681 | 6 | 120395416 | 4 | 0 | 0 | A | G | -2.3341 | 0.6919 | 0.0007426 | ---?- | 48.1 | 0.1229 | intergenic | |
| rs2212522 | 18 | 5900773 | 4 | 0 | 0 | T | C | 0.976 | 0.2893 | 0.0007427 | +++?+ | 0 | 0.4169 | intergenic | |
| rs7542928 | 1 | 182002021 | 4 | 0 | 0 | T | C | -1.131 | 0.3353 | 0.0007428 | ---?- | 0 | 0.4389 | intergenic | |
| rs73380893 | 7 | 67403435 | 4 | 0 | 0 | A | G | -1.7927 | 0.5314 | 0.0007428 | ---?+ | 16.3 | 0.3099 | intergenic | |
| rs78528486 | 7 | 46184413 | 4 | 0 | 0 | C | G | 1.9273 | 0.5714 | 0.0007428 | +++?+ | 0 | 0.9675 | intergenic | |
| rs6812257 | 4 | 20734171 | 4 | 0 | 0 | A | G | -0.8824 | 0.2616 | 0.000743 | ---?- | 0 | 0.8329 | intron | KCNIP4 |
| rs191171792 | 7 | 70485743 | 3 | 0 | 0 | T | G | 4.0168 | 1.1908 | 0.0007431 | +++?? | 5.7 | 0.3462 | intergenic | |
| rs66908195 | 15 | 85725733 | 4 | 0 | 0 | T | G | 1.9581 | 0.5805 | 0.0007432 | +++?+ | 40.5 | 0.169 | intergenic | |
| rs11055601 | 12 | 13869312 | 4 | 0 | 0 | A | T | 0.9002 | 0.2669 | 0.0007434 | +++?+ | 18.4 | 0.2987 | intron | GRIN2B |
| rs9804649 | 11 | 23976380 | 4 | 0 | 0 | T | G | 1.2081 | 0.3582 | 0.0007435 | +++?+ | 0 | 0.7814 | intergenic | |
| rs11048658 | 12 | 26850168 | 4 | 0 | 0 | T | G | 1.6663 | 0.494 | 0.0007435 | +-+?+ | 33.4 | 0.2122 | intron | ITPR2 |
| rs12322831 | 12 | 41512705 | 4 | 0 | 0 | A | T | 1.843 | 0.5464 | 0.0007435 | +++?+ | 0 | 0.489 | intergenic | |
| rs8051660 | 16 | 7203953 | 4 | 0 | 0 | A | G | 1.0249 | 0.3039 | 0.0007436 | +++?+ | 0 | 0.9751 | intron | RBFOX1 |
| rs2134309 | 11 | 8717607 | 5 | 0 | 0 | C | G | 0.7815 | 0.2317 | 0.0007437 | ++++- | 0 | 0.8446 | intron | ST5 |
| rs9378123 | 6 | 32196864 | 5 | 0 | 0 | A | G | 0.9736 | 0.2887 | 0.0007439 | +++++ | 0 | 0.7092 | intergenic | |
| 17:11223985 | 17 | 11223985 | 3 | 0 | 0 | A | AAAGAC | 0.9748 | 0.289 | 0.0007439 | +++?? | 0 | 0.5385 | | |
| rs11117918 | 1 | 217908761 | 4 | 0 | 0 | A | C | 0.8582 | 0.2545 | 0.000744 | +++?+ | 28 | 0.2438 | intron | SPATA17 |
| rs111999865 | 15 | 73170791 | 4 | 0 | 0 | T | G | -2.9355 | 0.8703 | 0.000744 | ---?- | 46 | 0.1355 | intergenic | |
| rs17113511 | 14 | 43272727 | 4 | 0 | 0 | T | C | -1.3018 | 0.386 | 0.0007441 | ---?+ | 59.5 | 0.06019 | intergenic | |
| rs16986462 | 22 | 28863019 | 4 | 0 | 0 | T | C | -2.3147 | 0.6863 | 0.0007443 | ---?- | 0 | 0.5849 | intron | TTC28 |
| rs9436255 | 1 | 64733378 | 4 | 0 | 0 | T | C | 1.1925 | 0.3536 | 0.0007444 | +++?+ | 6.3 | 0.3614 | downstream-gene | |
| 17:51144999 | 17 | 51144999 | 3 | 0 | 0 | T | TATTA | 1.4145 | 0.4194 | 0.0007448 | +++?? | 0.1 | 0.3674 | | |
| rs35273246 | 2 | 116643333 | 4 | 0 | 0 | A | G | -1.3212 | 0.3917 | 0.0007449 | ---?- | 38.7 | 0.18 | intergenic | |
| rs4553453 | 13 | 27607690 | 5 | 0 | 0 | T | C | 1.0351 | 0.3069 | 0.000745 | +++-+ | 39 | 0.1613 | intergenic | |
| 20:5489563 | 20 | 5489563 | 3 | 0 | 0 | CA | CAA | -2.2257 | 0.66 | 0.000745 | ---?? | 48.6 | 0.1427 | | |
| rs2381668 | 9 | 7847180 | 4 | 0 | 0 | T | C | 1.0227 | 0.3033 | 0.0007451 | +++?+ | 23.2 | 0.272 | intergenic | |
| rs11616286 | 13 | 112823895 | 4 | 0 | 0 | T | C | 0.9344 | 0.2771 | 0.0007454 | +++?+ | 9.6 | 0.3449 | intergenic | |
| rs11683911 | 2 | 158822102 | 4 | 0 | 0 | T | C | -0.8167 | 0.2422 | 0.0007454 | ---?- | 40.7 | 0.1676 | intron | UPP2 |
| rs9264481 | 6 | 31232318 | 4 | 0 | 0 | T | C | -0.8248 | 0.2446 | 0.0007459 | ----? | 8 | 0.3531 | downstream-gene |  |
| rs16933794 | 9 | 36956418 | 4 | 0 | 0 | T | G | 1.7454 | 0.5176 | 0.0007459 | +++?+ | 0 | 0.5243 | intron | PAX5 |
| rs80282384 | 16 | 6005391 | 3 | 0 | 0 | A | G | -0.9008 | 0.2671 | 0.0007462 | ---?? | 0 | 0.912 | intron | RBFOX1 |
| rs140312969 | 11 | 24780693 | 3 | 0 | 0 | A | G | 3.2802 | 0.9728 | 0.0007464 | +++?? | 15.5 | 0.3062 | intron | LUZP2 |
| rs56970199 | 6 | 109120932 | 2 | 0 | 0 | A | G | -4.6934 | 1.3919 | 0.0007465 | -?-?? | 0 | 0.7984 | intergenic | |
| rs76547283 | 16 | 6005386 | 3 | 0 | 0 | A | G | -0.9008 | 0.2672 | 0.0007465 | ---?? | 0 | 0.9123 | intron | RBFOX1 |
| rs4404434 | 3 | 2478578 | 4 | 0 | 0 | T | C | 0.9087 | 0.2695 | 0.0007466 | +++?- | 33.8 | 0.2094 | intron | CNTN4 |
| rs4790009 | 17 | 77318220 | 4 | 0 | 0 | A | G | -1.2982 | 0.385 | 0.0007469 | ---?+ | 15.2 | 0.3158 | intron | RBFOX3 |
| rs12133263 | 1 | 85712792 | 5 | 0 | 0 | T | G | -1.5644 | 0.464 | 0.0007469 | ----- | 0 | 0.4511 | downstream-gene |  |
| 2:170380756 | 2 | 170380756 | 3 | 0 | 0 | CA | C | 1.2613 | 0.3741 | 0.0007469 | +++?? | 0 | 0.9816 | | |
| rs7709025 | 5 | 123489092 | 4 | 0 | 0 | A | T | -0.8786 | 0.2606 | 0.0007471 | ---?- | 0 | 0.7339 | intergenic | |
| rs10513966 | 18 | 67229281 | 4 | 0 | 0 | T | G | 1.2399 | 0.3677 | 0.0007473 | +++?+ | 59 | 0.0626 | intron | DOK6 |
| rs11229555 | 11 | 58408687 | 5 | 0 | 0 | T | G | -0.8281 | 0.2456 | 0.0007474 | ----- | 0 | 0.9854 | intergenic |  |
| rs59831782 | 3 | 68231025 | 4 | 0 | 0 | A | G | -2.1423 | 0.6354 | 0.0007474 | ---?- | 10.1 | 0.3424 | intron | FAM19A1 |
| rs72802511 | 16 | 47818273 | 4 | 0 | 0 | A | G | 2.9268 | 0.8682 | 0.0007484 | +++?+ | 0 | 0.6466 | intergenic | |
| 8:64911737 | 8 | 64911737 | 3 | 0 | 0 | T | TG | 0.9302 | 0.2759 | 0.0007486 | +++?? | 0 | 0.6871 | | |
| rs11144637 | 9 | 78440473 | 4 | 0 | 0 | T | C | -1.2363 | 0.3668 | 0.0007489 | ---?- | 57 | 0.07286 | intergenic | |
| rs28413746 | 9 | 44280071 | 3 | 0 | 0 | T | C | 1.9199 | 0.5695 | 0.0007489 | +++?? | 0 | 0.8113 | intergenic | |
| rs79357102 | 8 | 253700 | 3 | 0 | 0 | C | G | 1.0371 | 0.3077 | 0.0007492 | +++?? | 61.8 | 0.07299 | intergenic | |
| rs35976740 | 12 | 63257810 | 3 | 0 | 0 | T | C | 3.5366 | 1.0492 | 0.0007493 | +++?? | 0 | 0.5881 | intron | PPM1H |
| rs112774341 | 4 | 21114444 | 4 | 0 | 0 | T | C | -2.1467 | 0.6369 | 0.0007496 | ---?+ | 8.2 | 0.3523 | intron | KCNIP4 |
| rs184791507 | 11 | 65310451 | 3 | 0 | 0 | A | G | 3.4958 | 1.0371 | 0.0007497 | +++?? | 0 | 0.9656 | intron | LTBP3 |
| rs59415749 | 2 | 47377300 | 4 | 0 | 0 | A | G | -1.3006 | 0.3859 | 0.0007499 | ---?- | 0 | 0.9752 | intron | C2orf61 |
| rs12174203 | 6 | 151727582 | 4 | 0 | 0 | T | C | 1.0633 | 0.3155 | 0.00075 | +++?+ | 41.6 | 0.1619 | intron | RMND1 |
| 4:182943683 | 4 | 182943683 | 3 | 0 | 0 | T | TA | 1.8016 | 0.5345 | 0.00075 | +++?? | 0 | 0.7281 | | |

| | | | | | | | | | | | | | | | |
|---|---|---|---|---|---|---|---|---|---|---|---|---|---|---|---|
| rs12781152 | 10 | 134590257 | 4 | 0 | 0 | C | G | 0.9298 | 0.2759 | 0.0007501 | +++?+ | 0 | 0.5027 | intron | INPP5A |
| rs6926737 | 6 | 32375745 | 5 | 0 | 0 | A | G | 0.7114 | 0.2111 | 0.0007503 | +++++ | 0 | 0.702 | upstream-gene | |
| rs5007264 | 6 | 32378877 | 5 | 0 | 0 | T | C | 0.712 | 0.2113 | 0.0007505 | +++++ | 0 | 0.6783 | upstream-gene | |
| rs117554162 | 8 | 81864743 | 3 | 0 | 0 | T | G | 4.5151 | 1.3396 | 0.0007505 | ?++?+ | 0 | 0.4214 | intergenic | |
| rs6938611 | 6 | 130911839 | 4 | 0 | 0 | A | C | 0.8838 | 0.2622 | 0.0007507 | +++?+ | 0 | 0.9629 | intergenic | |
| 2:170406883 | 2 | 170406883 | 3 | 0 | 0 | G | GA | 1.2675 | 0.3761 | 0.0007507 | +++?? | 0 | 0.9938 | | |
| rs10153262 | 17 | 8735003 | 4 | 0 | 0 | A | G | 0.9596 | 0.2847 | 0.0007509 | +++?+ | 0 | 0.4455 | intron | PIK3R6 |
| rs3915557 | 1 | 182016728 | 4 | 0 | 0 | T | G | -1.127 | 0.3344 | 0.0007517 | ---?- | 0 | 0.4604 | intergenic | |
| 3:59903943 | 3 | 59903943 | 3 | 0 | 0 | G | GT | -3.0683 | 0.9105 | 0.0007517 | ---?? | 14.8 | 0.3094 | | |
| rs1508019 | 17 | 52295016 | 4 | 0 | 0 | T | G | 1.965 | 0.5831 | 0.0007518 | +++?- | 29 | 0.2382 | intergenic | |
| rs10742019 | 11 | 23973175 | 4 | 0 | 0 | A | G | -1.1971 | 0.3552 | 0.0007518 | ---?- | 0 | 0.7183 | intergenic | |
| rs146619362 | 8 | 113210935 | 4 | 0 | 0 | T | C | 0.958 | 0.2843 | 0.0007518 | +++?+ | 0 | 0.9765 | intergenic | |
| 4:26167305 | 4 | 26167305 | 3 | 0 | 0 | G | GA | 1.2621 | 0.3745 | 0.0007518 | +++?? | 0 | 0.4618 | | |
| rs2034648 | 2 | 236957822 | 5 | 0 | 0 | A | G | 0.7058 | 0.2094 | 0.0007519 | +++++ | 45.3 | 0.12 | missense | AGAP1 |
| rs12712685 | 2 | 23111137 | 4 | 0 | 0 | A | G | 0.8875 | 0.2633 | 0.0007519 | +++?+ | 0 | 0.6114 | intergenic | |
| rs1357407 | 2 | 138954903 | 4 | 0 | 0 | T | C | -1.6271 | 0.4828 | 0.000752 | ---?- | 0 | 0.8206 | intergenic | |
| rs8058648 | 16 | 25828907 | 4 | 0 | 0 | C | G | -0.8362 | 0.2482 | 0.0007521 | ---?- | 0 | 0.6731 | intron | HS3ST4 |
| rs28668517 | 9 | 44254366 | 3 | 0 | 0 | A | C | -1.7398 | 0.5163 | 0.0007521 | ---?? | 0 | 0.839 | intergenic | |
| rs62133248 | 19 | 57260625 | 4 | 0 | 0 | T | C | 0.878 | 0.2605 | 0.0007521 | +++?+ | 0 | 0.464 | intergenic | |
| rs3741127 | 11 | 74346933 | 4 | 0 | 0 | A | G | -1.2351 | 0.3665 | 0.0007522 | ---?- | 0 | 0.6664 | intron | POLD3 |
| 18:73317863 | 18 | 73317863 | 3 | 0 | 0 | A | AT | 2.512 | 0.7454 | 0.0007523 | +++?? | 0 | 0.534 | | |
| rs6886414 | 5 | 103163072 | 4 | 0 | 0 | T | C | -1.5131 | 0.449 | 0.0007525 | ---?- | 0 | 0.6838 | intergenic | |
| rs67774070 | 10 | 116495139 | 4 | 0 | 0 | A | T | 1.349 | 0.4004 | 0.0007531 | +++?+ | 0 | 0.8596 | intergenic | |
| rs76019796 | 16 | 12107270 | 4 | 0 | 0 | T | G | 0.7979 | 0.2368 | 0.0007534 | -++?+ | 60.6 | 0.05491 | intron | SNX29 |
| rs61806392 | 1 | 182008282 | 4 | 0 | 0 | T | C | -1.1295 | 0.3352 | 0.0007535 | ---?- | 0 | 0.4469 | intergenic | |
| rs4299948 | 7 | 82429908 | 4 | 0 | 0 | T | G | 0.8546 | 0.2536 | 0.0007539 | +-+?+ | 44.9 | 0.1418 | intron | PCLO |
| rs4646988 | 12 | 103253157 | 4 | 0 | 0 | A | G | 0.8264 | 0.2453 | 0.000754 | +++?+ | 0 | 0.6601 | intron | PAH |
| rs10502334 | 18 | 5903698 | 4 | 0 | 0 | A | G | 0.9879 | 0.2932 | 0.000754 | +++?+ | 0 | 0.636 | intergenic | |
| rs75924914 | 17 | 56429585 | 5 | 0 | 0 | C | G | 1.0828 | 0.3214 | 0.0007542 | +++++ | 35.7 | 0.1834 | 5-prime-UTR | SUPT4H1,BZRAP1-AS1 |
| rs144735785 | 13 | 103456935 | 4 | 0 | 0 | A | G | 2.3514 | 0.698 | 0.0007542 | +++?+ | 0 | 0.9463 | intron | BIVM |
| rs74880187 | 15 | 100426460 | 4 | 0 | 0 | T | C | 0.9475 | 0.2812 | 0.0007543 | +++?+ | 41.9 | 0.1601 | intergenic | |
| rs416819 | 5 | 141908924 | 3 | 0 | 0 | T | G | 2.1816 | 0.6476 | 0.0007544 | +++?? | 0 | 0.7027 | intergenic | |
| rs10054363 | 5 | 103163817 | 4 | 0 | 0 | T | C | -1.5185 | 0.4507 | 0.0007545 | ---?- | 0 | 0.7168 | intergenic | |
| rs73062405 | 19 | 56220504 | 4 | 0 | 0 | T | C | 1.1602 | 0.3444 | 0.0007545 | +++?+ | 0 | 0.8017 | intron | NLRP9 |
| rs12644069 | 4 | 94736617 | 3 | 0 | 0 | A | G | 3.0905 | 0.9174 | 0.0007546 | +++?? | 45 | 0.1621 | intergenic | |
| 6:63256378 | 6 | 63256378 | 3 | 0 | 0 | A | ATT | -2.3775 | 0.7057 | 0.0007548 | ---?? | 0.5 | 0.3659 | | |
| rs12565870 | 1 | 182008478 | 4 | 0 | 0 | A | G | -1.1292 | 0.3352 | 0.0007551 | ---?- | 0 | 0.4469 | intergenic | |
| 8:113223251 | 8 | 113223251 | 3 | 0 | 0 | CTGTT | C | -0.9874 | 0.2931 | 0.0007551 | ---?? | 0 | 0.8987 | | |
| rs1326242 | 10 | 20150639 | 4 | 0 | 0 | A | G | -0.8601 | 0.2553 | 0.0007552 | ---?- | 10.9 | 0.3385 | intron | PLXDC2 |
| rs62419113 | 6 | 63334565 | 4 | 0 | 0 | A | G | -2.3855 | 0.7081 | 0.0007552 | ---?- | 0 | 0.4859 | intergenic | |
| rs186210766 | 11 | 86232434 | 4 | 0 | 0 | C | G | -2.5836 | 0.767 | 0.0007553 | ---?- | 4.6 | 0.3697 | intron | ME3 |
| rs11601792 | 11 | 12839976 | 4 | 0 | 0 | A | G | 1.4239 | 0.4227 | 0.0007554 | +++?+ | 0 | 0.6485 | intron | TEAD1 |
| rs2806542 | 1 | 64612756 | 4 | 0 | 0 | A | G | 0.8648 | 0.2567 | 0.0007555 | +++?+ | 0 | 0.4836 | intron | ROR1 |
| rs186301560 | 22 | 39426281 | 3 | 0 | 0 | T | C | -0.8698 | 0.2582 | 0.0007555 | ---?? | 0 | 0.9655 | intron | APOBEC3D |
| rs7767677 | 6 | 151715975 | 4 | 0 | 0 | T | G | -1.0487 | 0.3113 | 0.0007557 | ---?- | 45.6 | 0.1379 | upstream-gene | |
| rs78686508 | 3 | 143043173 | 4 | 0 | 0 | A | G | 1.9591 | 0.5816 | 0.0007557 | +++?+ | 0 | 0.5607 | intron | SLC9A9 |
| 4:188131906 | 4 | 188131906 | 2 | 0 | 0 | T | C | -6.7344 | 1.9993 | 0.0007559 | -???- | 42.1 | 0.1886 | intergenic | |
| rs10912204 | 1 | 187853046 | 4 | 0 | 0 | A | G | -1.1404 | 0.3385 | 0.0007559 | ---?- | 0 | 0.7962 | intergenic | |
| rs12406981 | 1 | 182009035 | 4 | 0 | 0 | A | G | -1.129 | 0.3352 | 0.0007559 | ---?- | 0 | 0.448 | intergenic | |
| rs144723316 | 4 | 140541160 | 4 | 0 | 0 | C | G | 3.1089 | 0.923 | 0.000756 | +++?+ | 41.9 | 0.1601 | intergenic | |
| rs438239 | 6 | 155734754 | 5 | 0 | 0 | T | C | 0.7273 | 0.2159 | 0.0007564 | -++++ | 49.3 | 0.0957 | intron | NOX3 |
| rs8056587 | 16 | 85205255 | 4 | 0 | 0 | T | C | 0.9068 | 0.2692 | 0.0007564 | +++?+ | 0 | 0.4438 | synonymous | GSE1 |
| rs12409451 | 1 | 182009171 | 4 | 0 | 0 | A | G | 1.1289 | 0.3352 | 0.0007564 | +++?+ | 0 | 0.4482 | intergenic | |
| rs17785193 | 9 | 79460804 | 4 | 0 | 0 | T | C | 1.8777 | 0.5575 | 0.0007566 | +++?+ | 0 | 0.4324 | intron | PRUNE2 |
| rs7546388 | 1 | 182003433 | 4 | 0 | 0 | T | C | -1.1295 | 0.3354 | 0.0007568 | ---?- | 0 | 0.4396 | intergenic | |
| 7:67404999 | 7 | 67404999 | 3 | 0 | 0 | T | TA | 1.8394 | 0.5461 | 0.0007568 | +++?? | 0 | 0.4301 | | |
| rs4916526 | 3 | 196330156 | 4 | 0 | 0 | T | C | -0.959 | 0.2847 | 0.0007569 | ---?- | 0 | 0.8973 | intergenic | |
| rs10962315 | 9 | 16176912 | 4 | 0 | 0 | T | C | -1.2434 | 0.3692 | 0.000757 | ---?- | 0 | 0.6217 | intergenic | |
| rs61869110 | 10 | 116495377 | 4 | 0 | 0 | T | C | 1.348 | 0.4003 | 0.0007573 | +++?+ | 0 | 0.8593 | intergenic | |

| | | | | | | | | | | | | | | | |
|---|---|---|---|---|---|---|---|---|---|---|---|---|---|---|---|
| rs641459 | 7 | 114722787 | 4 | 0 | 0 | T | G | 0.8274 | 0.2457 | 0.0007575 | +++?+ | 2.9 | 0.3781 | intergenic | |
| rs6886374 | 5 | 103162999 | 4 | 0 | 0 | T | C | -1.5115 | 0.4488 | 0.0007575 | ---?- | 0 | 0.6841 | intergenic | |
| rs6932542 | 6 | 32380262 | 5 | 0 | 0 | A | G | -0.7125 | 0.2116 | 0.0007576 | ----- | 0 | 0.654 | intergenic |  |
| rs825209 | 1 | 64771492 | 4 | 0 | 0 | T | C | 0.9346 | 0.2775 | 0.0007577 | +++?+ | 0 | 0.4629 | intergenic | |
| rs151125265 | 3 | 68223771 | 4 | 0 | 0 | A | G | 2.0875 | 0.6199 | 0.0007578 | +++?+ | 10.7 | 0.3395 | intron | FAM19A1 |
| 15:87612561 | 15 | 87612561 | 3 | 0 | 0 | CT | C | -0.9321 | 0.2768 | 0.0007579 | ---?? | 0 | 0.4637 | | |
| rs589606 | 7 | 114722698 | 4 | 0 | 0 | A | T | -0.8274 | 0.2457 | 0.0007581 | ---?- | 2.9 | 0.378 | intergenic | |
| rs144549680 | 4 | 188131875 | 2 | 0 | 0 | A | G | 6.7293 | 1.9983 | 0.0007583 | +???+ | 42 | 0.1892 | intergenic | |
| rs28986401 | 6 | 32754767 | 5 | 0 | 0 | A | G | 1.4272 | 0.4238 | 0.0007589 | +++++ | 0 | 0.6496 | intergenic | |
| rs181231870 | 9 | 83275425 | 3 | 0 | 0 | C | G | 3.5018 | 1.0399 | 0.0007589 | +++?? | 0 | 0.48 | intergenic | |
| rs9997989 | 4 | 190015318 | 4 | 0 | 0 | T | C | -0.8468 | 0.2515 | 0.000759 | ---?- | 29.3 | 0.2363 | intergenic | |
| rs55896799 | 5 | 55923915 | 4 | 0 | 0 | A | C | 3.5469 | 1.0534 | 0.0007595 | +++?- | 19.8 | 0.2907 | intergenic | |
| rs77283968 | 1 | 202063456 | 4 | 0 | 0 | A | G | -1.1562 | 0.3434 | 0.0007595 | ---?+ | 7 | 0.3583 | intergenic | |
| rs143587838 | 8 | 113210566 | 4 | 0 | 0 | A | G | 0.9571 | 0.2842 | 0.0007595 | +++?+ | 0 | 0.9766 | intergenic | |
| 4:38749791 | 4 | 38749791 | 3 | 0 | 0 | G | GT | -1.3664 | 0.4058 | 0.0007595 | ---?? | 0 | 0.7795 | | |
| rs2339719 | 12 | 111726850 | 5 | 0 | 0 | A | T | -0.9951 | 0.2955 | 0.0007596 | ----- | 0 | 0.565 | intron | CUX2 |
| rs4973247 | 2 | 230913033 | 4 | 0 | 0 | A | C | 1.4446 | 0.429 | 0.0007598 | +++?+ | 23.2 | 0.2718 | intron | SLC16A14 |
| rs10912208 | 1 | 187865354 | 4 | 0 | 0 | T | C | 1.106 | 0.3285 | 0.0007598 | +++?+ | 0 | 0.9395 | intergenic | |
| rs76528106 | 14 | 48564030 | 4 | 0 | 0 | T | G | 2.1665 | 0.6434 | 0.0007598 | +++?+ | 0 | 0.5081 | intergenic | |
| rs150360581 | 9 | 114122492 | 4 | 0 | 0 | T | C | 4.5205 | 1.3426 | 0.00076 | +++?+ | 25.1 | 0.2612 | downstream-gene | |
| rs204888 | 6 | 32089142 | 5 | 0 | 0 | T | G | 1.3269 | 0.3941 | 0.0007601 | +++-+ | 40.3 | 0.1527 | intron | ATF6B |
| 9:27621239 | 9 | 27621239 | 3 | 0 | 0 | CAAAT | C | -0.9643 | 0.2864 | 0.0007602 | ---?? | 39.9 | 0.1896 | | |
| rs2288308 | 4 | 20731047 | 4 | 0 | 0 | T | C | 0.8834 | 0.2624 | 0.0007603 | +++?+ | 0 | 0.835 | 3-prime-UTR | KCNIP4 |
| rs77892116 | 5 | 80304524 | 2 | 0 | 0 | T | C | 6.905 | 2.0509 | 0.0007603 | ??+?+ | 0 | 0.6521 | intron | RASGRF2 |
| rs139094111 | 11 | 119373544 | 4 | 0 | 0 | T | C | 2.9458 | 0.875 | 0.0007604 | +++?+ | 0 | 0.526 | downstream-gene | |
| 11:13694210 | 11 | 13694210 | 3 | 0 | 0 | A | AG | 1.1062 | 0.3286 | 0.0007605 | +++?? | 0 | 0.4044 | | |
| rs12683842 | 9 | 4352621 | 2 | 0 | 0 | A | G | -4.8015 | 1.4262 | 0.0007607 | ?--?? | 0 | 0.7599 | intergenic | |
| rs140008794 | 2 | 3689214 | 3 | 0 | 0 | A | G | 5.3741 | 1.5963 | 0.0007608 | +++?? | 0 | 0.7105 | intron | COLEC11 |
| rs4482199 | 14 | 43273646 | 4 | 0 | 0 | C | G | -1.2933 | 0.3842 | 0.0007609 | ---?+ | 57.3 | 0.07121 | intergenic | |
| rs62419114 | 6 | 63334762 | 4 | 0 | 0 | C | G | 2.384 | 0.7082 | 0.0007613 | +++?+ | 0 | 0.4848 | intergenic | |
| rs74929712 | 10 | 68809676 | 4 | 0 | 0 | C | G | -2.2392 | 0.6651 | 0.0007614 | ---?- | 0 | 0.9837 | intron | CTNNA3,LRRTM3 |
| rs6495530 | 15 | 81179725 | 4 | 0 | 0 | A | G | -0.7973 | 0.2368 | 0.0007615 | ---?+ | 17.8 | 0.3016 | intron | KIAA1199 |
| rs76763773 | 9 | 30554411 | 4 | 0 | 0 | C | G | -3.5531 | 1.0555 | 0.0007616 | ---?- | 0 | 0.4393 | intergenic | |
| rs7231131 | 18 | 56546022 | 4 | 0 | 0 | A | G | -1.1834 | 0.3516 | 0.0007618 | ---?- | 0 | 0.788 | intron | ZNF532 |
| rs34941618 | 13 | 20810296 | 4 | 0 | 0 | T | C | -1.1033 | 0.3278 | 0.0007618 | ---?- | 0 | 0.4939 | upstream-gene | |
| rs55927654 | 12 | 41379657 | 4 | 0 | 0 | T | G | 1.4326 | 0.4256 | 0.0007618 | +++?+ | 0 | 0.59 | intron | CNTN1 |
| rs1158175 | 1 | 217994138 | 4 | 0 | 0 | T | C | -1.0969 | 0.3259 | 0.0007623 | ---?- | 0 | 0.9808 | intron | SPATA17 |
| 6:63240991 | 6 | 63240991 | 3 | 0 | 0 | G | GAATAT | -2.3369 | 0.6942 | 0.0007623 | ---?? | 0 | 0.3681 | | |
| rs2038294 | 6 | 95161946 | 4 | 0 | 0 | A | G | 0.8305 | 0.2467 | 0.0007624 | +++?+ | 0 | 0.7504 | intergenic | |
| rs7531336 | 1 | 22329523 | 2 | 0 | 0 | A | G | 8.1016 | 2.4068 | 0.0007624 | ++??? | 0 | 0.404 | missense | CELA3A |
| rs59439814 | 10 | 133355893 | 3 | 0 | 0 | A | T | -1.2928 | 0.3841 | 0.0007626 | ---?? | 0 | 0.7665 | intergenic | |
| rs6899678 | 6 | 130893264 | 4 | 0 | 0 | T | C | 0.8405 | 0.2497 | 0.000763 | +++?+ | 0 | 0.7198 | intergenic | |
| rs7799607 | 7 | 82423571 | 4 | 0 | 0 | T | C | -0.8556 | 0.2542 | 0.000763 | -+-?- | 45.9 | 0.136 | intron | PCLO |
| 5:124668597 | 5 | 124668597 | 3 | 0 | 0 | G | GA | -1.1771 | 0.3497 | 0.000763 | ---?? | 0 | 0.6626 | | |
| rs355536 | 2 | 142423065 | 4 | 0 | 0 | A | G | 2.4148 | 0.7175 | 0.0007634 | +++?+ | 4.7 | 0.3695 | intron | LRP1B |
| rs34765230 | 16 | 20200700 | 4 | 0 | 0 | T | G | 0.9915 | 0.2946 | 0.0007634 | +++?+ | 5.3 | 0.3662 | intergenic | |
| rs10953518 | 7 | 106399882 | 4 | 0 | 0 | A | G | 1.3208 | 0.3924 | 0.0007636 | +++?+ | 0 | 0.5659 | intergenic | |
| rs5007266 | 6 | 32378834 | 5 | 0 | 0 | T | C | -0.7109 | 0.2112 | 0.0007637 | ----- | 0 | 0.682 | upstream-gene |  |
| rs6005733 | 22 | 28564931 | 4 | 0 | 0 | T | C | 2.4736 | 0.735 | 0.0007637 | +++?+ | 0 | 0.7276 | intron | TTC28 |
| rs6932810 | 6 | 32380190 | 5 | 0 | 0 | A | G | 0.7119 | 0.2115 | 0.0007637 | +++++ | 0 | 0.6557 | intergenic | |
| rs72929156 | 2 | 199095911 | 4 | 0 | 0 | T | C | 1.3575 | 0.4033 | 0.0007639 | +++?+ | 49.3 | 0.1158 | intergenic | |
| rs2620895 | 10 | 50023801 | 4 | 0 | 0 | A | G | -3.4757 | 1.0327 | 0.000764 | +--?- | 14.3 | 0.3206 | intron | WDFY4 |
| rs144377865 | 19 | 2066066 | 4 | 0 | 0 | A | G | -2.1247 | 0.6313 | 0.0007641 | ---?- | 0 | 0.9388 | downstream-gene | |
| rs2798682 | 1 | 87573963 | 4 | 0 | 0 | T | C | 1.0051 | 0.2987 | 0.0007646 | +++?+ | 21.2 | 0.2828 | 3-prime-UTR | HS2ST1 |
| rs62440966 | 7 | 5489653 | 3 | 0 | 0 | A | G | 5.7519 | 1.7092 | 0.0007646 | ++??+ | 0 | 0.5476 | intergenic | |
| rs377132059 | 15 | 59812310 | 3 | 0 | 0 | A | T | -3.8543 | 1.1453 | 0.0007647 | ---?? | 0 | 0.5434 | intron | FAM81A |
| rs6005843 | 22 | 29108444 | 4 | 0 | 0 | T | C | -1.8203 | 0.5409 | 0.0007648 | ---?- | 0 | 0.9837 | intron | CHEK2 |
| rs12956748 | 18 | 37415724 | 4 | 0 | 0 | T | C | 0.9068 | 0.2695 | 0.000765 | +++?+ | 0 | 0.5317 | intergenic | |
| rs28986402 | 6 | 32754781 | 5 | 0 | 0 | T | C | 1.4265 | 0.4239 | 0.000765 | +++++ | 0 | 0.6483 | intergenic | |

| | | | | | | | | | | | | | | | |
|---|---|---|---|---|---|---|---|---|---|---|---|---|---|---|---|
| rs187293856 | 10 | 9315466 | 3 | 0 | 0 | A | G | -1.7668 | 0.525 | 0.000765 | ---?? | 0 | 0.4934 | intergenic | |
| rs3776980 | 5 | 80527477 | 4 | 0 | 0 | T | C | -1.6879 | 0.5016 | 0.0007652 | ---?- | 43.3 | 0.1518 | upstream-gene | |
| rs7753601 | 6 | 120387048 | 4 | 0 | 0 | T | C | 2.3167 | 0.6885 | 0.0007656 | +++?+ | 51.2 | 0.1048 | intergenic | |
| rs7635335 | 3 | 71884576 | 4 | 0 | 0 | A | G | -0.9136 | 0.2715 | 0.0007657 | ---?- | 0 | 0.7003 | intergenic | |
| rs10808170 | 7 | 112695484 | 4 | 0 | 0 | T | C | 0.9476 | 0.2816 | 0.0007657 | +++?+ | 0 | 0.6808 | intergenic | |
| rs12407644 | 1 | 182010860 | 4 | 0 | 0 | A | G | -1.1272 | 0.335 | 0.0007659 | ---?- | 0 | 0.4506 | intergenic | |
| rs4763672 | 12 | 11446612 | 4 | 0 | 0 | A | G | -0.8473 | 0.2518 | 0.000766 | ---?- | 38 | 0.1842 | intergenic | |
| rs150426486 | 5 | 109168557 | 3 | 0 | 0 | C | G | -0.863 | 0.2565 | 0.000766 | ---?? | 0 | 0.8171 | intron | MAN2A1 |
| rs6687373 | 1 | 185398983 | 4 | 0 | 0 | A | C | -0.8854 | 0.2631 | 0.0007661 | ---?- | 0 | 0.9173 | intergenic | |
| rs55972184 | 4 | 183664946 | 4 | 0 | 0 | A | G | -0.9022 | 0.2681 | 0.0007661 | ---?- | 0 | 0.9325 | intron | TENM3 |
| 2:124501953 | 2 | 124501953 | 3 | 0 | 0 | T | TTATA | 4.7503 | 1.4118 | 0.0007661 | +++?? | 0 | 0.6003 | | |
| rs10880364 | 12 | 43210572 | 5 | 0 | 0 | A | G | -0.8315 | 0.2471 | 0.0007663 | ----- | 0 | 0.6832 | intergenic |  |
| rs73834875 | 3 | 68228629 | 4 | 0 | 0 | T | G | -2.127 | 0.6321 | 0.0007663 | ---?- | 9.5 | 0.3458 | intron | FAM19A1 |
| rs12732882 | 1 | 64805059 | 4 | 0 | 0 | A | G | 1.0311 | 0.3065 | 0.0007664 | +++?+ | 0 | 0.5423 | intergenic | |
| rs79523952 | 15 | 87613107 | 4 | 0 | 0 | A | G | 1.4984 | 0.4453 | 0.0007664 | +++?+ | 0 | 0.5485 | intergenic | |
| rs141362408 | 5 | 109168556 | 3 | 0 | 0 | A | G | -0.863 | 0.2565 | 0.0007664 | ---?? | 0 | 0.817 | intron | MAN2A1 |
| rs73156577 | 7 | 143786235 | 3 | 0 | 0 | A | T | -3.9694 | 1.1798 | 0.0007667 | ---?? | 36 | 0.2096 | intergenic | |
| rs138243526 | 5 | 109168558 | 3 | 0 | 0 | A | G | -0.8629 | 0.2565 | 0.0007667 | ---?? | 0 | 0.8173 | intron | MAN2A1 |
| rs6911383 | 6 | 32379682 | 5 | 0 | 0 | T | C | -0.7113 | 0.2114 | 0.0007668 | ----- | 0 | 0.6648 | upstream-gene |  |
| rs12402682 | 1 | 182011172 | 4 | 0 | 0 | T | C | -1.127 | 0.335 | 0.000767 | ---?- | 0 | 0.4503 | intergenic | |
| rs9349326 | 6 | 45695564 | 4 | 0 | 0 | A | G | -1.1435 | 0.3399 | 0.0007671 | ---?+ | 57.3 | 0.07124 | intergenic | |
| rs79019275 | 7 | 84846613 | 4 | 0 | 0 | A | C | 3.7309 | 1.109 | 0.0007673 | +++?+ | 8.4 | 0.3509 | intergenic | |
| rs7159088 | 14 | 21451913 | 4 | 0 | 0 | A | C | -0.8808 | 0.2618 | 0.000768 | ---?- | 19.5 | 0.2923 | intergenic | |
| rs1616017 | 16 | 12136356 | 4 | 0 | 0 | T | C | -0.9048 | 0.269 | 0.0007681 | +--?- | 49.4 | 0.1152 | intron | SNX29 |
| rs78598167 | 18 | 6110865 | 4 | 0 | 0 | T | C | 2.6679 | 0.7931 | 0.0007681 | +++?- | 0.7 | 0.3886 | intron | L3MBTL4 |
| rs9343611 | 6 | 78095134 | 4 | 0 | 0 | T | C | -0.8753 | 0.2602 | 0.0007683 | ---?- | 0 | 0.4941 | intergenic | |
| rs16986290 | 22 | 28705589 | 4 | 0 | 0 | T | C | 2.3599 | 0.7015 | 0.0007683 | +++?+ | 0 | 0.5895 | intron | TTC28 |
| rs73282980 | 5 | 122555913 | 4 | 0 | 0 | A | G | -1.5789 | 0.4693 | 0.0007684 | ---?- | 0 | 0.822 | intergenic | |
| rs147110482 | 6 | 145307800 | 3 | 0 | 0 | A | C | 4.2262 | 1.2564 | 0.0007686 | +++?? | 0 | 0.732 | intergenic | |
| rs930887 | 5 | 123462991 | 4 | 0 | 0 | T | C | -0.8885 | 0.2641 | 0.0007687 | ---?- | 0 | 0.9489 | intergenic | |
| rs6905090 | 6 | 130961459 | 4 | 0 | 0 | C | G | 1.4275 | 0.4244 | 0.0007689 | +++?+ | 0 | 0.69 | intergenic | |
| rs1703493 | 16 | 12136526 | 4 | 0 | 0 | A | G | 0.9047 | 0.269 | 0.000769 | -++?+ | 49.4 | 0.1149 | intron | SNX29 |
| rs7626279 | 3 | 196331714 | 4 | 0 | 0 | C | G | -0.952 | 0.283 | 0.000769 | ---?- | 0 | 0.9057 | intergenic | |
| 8:113223335 | 8 | 113223335 | 3 | 0 | 0 | T | TA | -0.9868 | 0.2934 | 0.000769 | ---?? | 0 | 0.9003 | | |
| rs705570 | 1 | 64771013 | 4 | 0 | 0 | T | C | -0.9333 | 0.2775 | 0.0007692 | ---?- | 0 | 0.466 | intergenic | |
| rs5007258 | 6 | 32379239 | 5 | 0 | 0 | A | G | -0.7104 | 0.2112 | 0.0007692 | ----- | 0 | 0.6812 | upstream-gene |  |
| rs5007262 | 6 | 32379011 | 5 | 0 | 0 | A | G | -0.7104 | 0.2112 | 0.0007692 | ----- | 0 | 0.6812 | upstream-gene |  |
| rs5007263 | 6 | 32378982 | 5 | 0 | 0 | A | G | -0.7104 | 0.2112 | 0.0007692 | ----- | 0 | 0.6812 | upstream-gene |  |
| rs5007265 | 6 | 32378866 | 5 | 0 | 0 | T | G | -0.7104 | 0.2112 | 0.0007692 | ----- | 0 | 0.6812 | upstream-gene |  |
| rs9268506 | 6 | 32377470 | 5 | 0 | 0 | T | C | 0.7104 | 0.2112 | 0.0007692 | +++++ | 0 | 0.6812 | upstream-gene | |
| rs9268507 | 6 | 32377539 | 5 | 0 | 0 | A | G | -0.7104 | 0.2112 | 0.0007692 | ----- | 0 | 0.6812 | upstream-gene |  |
| rs9268513 | 6 | 32378787 | 5 | 0 | 0 | T | C | -0.7104 | 0.2112 | 0.0007692 | ----- | 0 | 0.6812 | upstream-gene |  |
| rs354288 | 5 | 14237265 | 4 | 0 | 0 | A | G | 1.0625 | 0.3159 | 0.0007693 | +++?+ | 0 | 0.7256 | intron | TRIO |
| rs79786600 | 12 | 30976057 | 4 | 0 | 0 | T | C | 2.8763 | 0.8551 | 0.0007693 | +++?+ | 9.2 | 0.347 | intergenic | |
| 9:107922808 | 9 | 107922808 | 3 | 0 | 0 | G | GA | -1.4448 | 0.4295 | 0.0007693 | ---?? | 0 | 0.7477 | | |
| rs12907779 | 15 | 91405752 | 4 | 0 | 0 | A | T | 0.9706 | 0.2886 | 0.0007694 | +++?+ | 28.9 | 0.2386 | intergenic | |
| rs17567260 | 11 | 116250056 | 4 | 0 | 0 | T | C | -1.7351 | 0.5158 | 0.0007694 | ---?- | 4.5 | 0.3702 | intergenic | |
| rs4543822 | 1 | 7124926 | 4 | 0 | 0 | T | C | -0.858 | 0.2551 | 0.0007696 | ---?- | 42.4 | 0.1574 | intron | CAMTA1 |
| rs5007260 | 6 | 32379047 | 5 | 0 | 0 | A | G | 0.7103 | 0.2112 | 0.0007696 | +++++ | 0 | 0.6816 | upstream-gene | |
| rs5007261 | 6 | 32379031 | 5 | 0 | 0 | A | G | 0.7103 | 0.2112 | 0.0007696 | +++++ | 0 | 0.6816 | upstream-gene | |
| rs3010791 | 1 | 213211858 | 4 | 0 | 0 | T | C | 2.0978 | 0.6237 | 0.0007697 | +++?+ | 21.3 | 0.2825 | intergenic | |
| rs75173335 | 1 | 118223693 | 4 | 0 | 0 | A | G | -1.4523 | 0.4318 | 0.0007699 | ---?+ | 60.3 | 0.05628 | intergenic | |
| rs541845 | 9 | 44277687 | 3 | 0 | 0 | T | C | 2.1144 | 0.6287 | 0.0007703 | +++?? | 0 | 0.8248 | intergenic | |
| rs34715053 | 13 | 20810537 | 4 | 0 | 0 | T | C | 1.1038 | 0.3282 | 0.0007706 | +++?+ | 0 | 0.4929 | upstream-gene | |
| rs149023127 | 17 | 41947181 | 3 | 0 | 0 | A | G | 4.4268 | 1.3163 | 0.0007706 | +++?? | 37.8 | 0.2002 | intergenic | |
| rs113732694 | 6 | 63257678 | 3 | 0 | 0 | A | G | -2.3548 | 0.7002 | 0.0007707 | ---?? | 0 | 0.3706 | intergenic | |
| rs5008218 | 1 | 232833427 | 3 | 0 | 0 | A | G | -0.858 | 0.2551 | 0.0007711 | ---?? | 53 | 0.1189 | intergenic | |
| rs6906730 | 6 | 32379445 | 5 | 0 | 0 | T | C | -0.7106 | 0.2113 | 0.0007711 | ----- | 0 | 0.6733 | upstream-gene |  |
| rs73062410 | 19 | 56224778 | 4 | 0 | 0 | A | G | -1.2008 | 0.3571 | 0.0007712 | ---?- | 0 | 0.6588 | intron | NLRP9 |

| | | | | | | | | | | | | | | | |
|---|---|---|---|---|---|---|---|---|---|---|---|---|---|---|---|
| rs10084802 | 4 | 20746450 | 4 | 0 | 0 | A | G | -0.8682 | 0.2582 | 0.0007713 | ---?- | 0 | 0.9624 | intron | KCNIP4 |
| 10:93166934 | 10 | 93166934 | 3 | 0 | 0 | A | AATTT | 1.0572 | 0.3144 | 0.0007714 | +++?? | 0 | 0.4174 | | |
| rs9268504 | 6 | 32377116 | 5 | 0 | 0 | C | G | -0.7102 | 0.2112 | 0.0007716 | ----- | 0 | 0.6821 | upstream-gene |  |
| rs13018687 | 2 | 218829678 | 4 | 0 | 0 | A | G | 1.1781 | 0.3503 | 0.0007717 | +++?+ | 0 | 0.8188 | intergenic | |
| rs145257751 | 4 | 166228467 | 4 | 0 | 0 | A | G | -3.6956 | 1.099 | 0.0007717 | ---?- | 0 | 0.9445 | intron | KLHL2 |
| 5:159351202 | 5 | 159351202 | 3 | 0 | 0 | <ins:me: | T | -1.5905 | 0.473 | 0.0007717 | ---?? | 0 | 0.8278 | | |
| rs35016710 | 4 | 190013728 | 4 | 0 | 0 | A | G | -0.8447 | 0.2512 | 0.0007718 | ---?- | 29.4 | 0.2359 | intergenic | |
| 10:112238279 | 10 | 112238279 | 3 | 0 | 0 | G | GA | 1.0928 | 0.325 | 0.0007718 | +++?? | 0 | 0.523 | | |
| rs12723416 | 1 | 64803741 | 4 | 0 | 0 | A | C | -1.0274 | 0.3055 | 0.0007719 | ---?- | 0 | 0.5425 | intergenic | |
| 7:116170262 | 7 | 116170262 | 3 | 0 | 0 | CTTT | C | 0.9817 | 0.2919 | 0.000772 | +++?? | 45.6 | 0.1591 | | |
| rs2090029 | 2 | 124501976 | 3 | 0 | 0 | T | C | -4.7471 | 1.4118 | 0.0007723 | ---?? | 0 | 0.6003 | intergenic | |
| rs4141975 | 20 | 61611400 | 4 | 0 | 0 | C | G | -0.8211 | 0.2442 | 0.0007724 | ---?- | 0.5 | 0.3895 | intergenic | |
| rs8111428 | 19 | 46168101 | 5 | 0 | 1 | A | G | 0.879 | 0.2614 | 0.0007725 | +-++- | 26.2 | 0.2469 | upstream-gene | |
| rs10835691 | 11 | 30646177 | 4 | 0 | 0 | A | G | 0.8871 | 0.2638 | 0.0007725 | +++?- | 60.8 | 0.05369 | intergenic | |
| rs58590144 | 17 | 56882237 | 5 | 0 | 0 | A | T | -2.6535 | 0.7892 | 0.0007725 | ----- | 0 | 0.9292 | intron | PPM1E |
| rs13022979 | 2 | 218841301 | 4 | 0 | 0 | A | G | 1.3677 | 0.4068 | 0.0007726 | +++?- | 0 | 0.8127 | intergenic | |
| rs2616503 | 8 | 89008059 | 4 | 0 | 0 | T | C | -1.0929 | 0.325 | 0.0007729 | ---?+ | 12.8 | 0.3284 | intergenic | |
| rs11723840 | 4 | 176735253 | 4 | 0 | 0 | T | C | 2.8858 | 0.8583 | 0.000773 | +++?+ | 0 | 0.3939 | intron | GPM6A |
| 6:159241284 | 6 | 159241284 | 3 | 0 | 0 | T | TG | -0.863 | 0.2567 | 0.000773 | ---?? | 7.3 | 0.3401 | | |
| rs12525554 | 6 | 77663087 | 4 | 0 | 0 | T | C | 0.9586 | 0.2851 | 0.0007731 | +++?+ | 0 | 0.4366 | intergenic | |
| rs35030589 | 6 | 32672903 | 5 | 0 | 1 | A | G | 1.0936 | 0.3253 | 0.0007736 | +++++ | 26.4 | 0.2453 | intergenic | |
| rs34639600 | 13 | 20810602 | 4 | 0 | 0 | A | G | 1.1039 | 0.3283 | 0.0007737 | +++?+ | 0 | 0.4925 | upstream-gene | |
| rs117606710 | 6 | 80309407 | 2 | 0 | 0 | A | G | -5.1125 | 1.5207 | 0.0007737 | -?-?? | 50 | 0.1575 | intergenic | |
| rs10939669 | 4 | 10045827 | 5 | 0 | 0 | A | G | -0.7078 | 0.2105 | 0.0007738 | ----+ | 12.3 | 0.3353 | upstream-gene |  |
| rs61998187 | 10 | 131476700 | 4 | 0 | 0 | T | C | 1.2526 | 0.3726 | 0.0007739 | +++?+ | 0 | 0.8203 | intron | MGMT |
| 2:79681829 | 2 | 79681829 | 3 | 0 | 0 | T | TCTCAT/ | 0.9074 | 0.2699 | 0.0007739 | +++?? | 40.2 | 0.1879 | | |
| rs16994566 | 4 | 38465646 | 3 | 0 | 0 | T | C | -4.6331 | 1.3781 | 0.000774 | ?--?- | 0 | 0.5432 | intergenic | |
| rs4638785 | 2 | 230917177 | 4 | 0 | 0 | A | G | -1.4271 | 0.4245 | 0.0007742 | ---?- | 21 | 0.2844 | intron | SLC16A14 |
| rs7159682 | 14 | 100085637 | 4 | 0 | 0 | A | C | -0.9576 | 0.2849 | 0.0007742 | ---?- | 0 | 0.9119 | intergenic | |
| rs6665692 | 1 | 245107400 | 4 | 0 | 0 | C | G | 1.1485 | 0.3416 | 0.0007744 | +++?+ | 0 | 0.4 | intergenic | |
| rs76614496 | 2 | 104649498 | 3 | 0 | 0 | A | T | 2.6725 | 0.795 | 0.0007744 | +++?? | 0 | 0.6781 | intergenic | |
| rs12744084 | 1 | 87251746 | 4 | 0 | 0 | T | G | 1.1938 | 0.3551 | 0.0007745 | +++?- | 0 | 0.5078 | intergenic | |
| rs17027391 | 1 | 111852792 | 4 | 0 | 0 | T | C | -1.893 | 0.5631 | 0.0007745 | ---?- | 0 | 0.6917 | intron | CHIA |
| rs7553335 | 1 | 7126617 | 4 | 0 | 0 | C | G | -0.862 | 0.2564 | 0.0007746 | ---?- | 40.5 | 0.169 | intron | CAMTA1 |
| rs115876338 | 5 | 6167892 | 4 | 0 | 0 | T | C | 2.6282 | 0.7818 | 0.0007746 | +++?+ | 24.7 | 0.2629 | intergenic | |
| 20:7909706 | 20 | 7909706 | 3 | 0 | 0 | T | TTTTG | 1.0336 | 0.3075 | 0.0007747 | +-+?? | 65.8 | 0.05391 | | |
| rs16933796 | 9 | 36956733 | 4 | 0 | 0 | A | G | -1.7399 | 0.5176 | 0.0007748 | ---?- | 0 | 0.5276 | intron | PAX5 |
| rs111724290 | 22 | 29101636 | 4 | 0 | 0 | A | G | 2.1592 | 0.6423 | 0.0007748 | +++?+ | 0 | 0.5109 | intron | CHEK2 |
| rs62419115 | 6 | 63335597 | 4 | 0 | 0 | A | G | 2.3936 | 0.7121 | 0.000775 | +++?+ | 0 | 0.4841 | intergenic | |
| 1:232833431 | 1 | 232833431 | 3 | 0 | 0 | T | TG | -0.8578 | 0.2552 | 0.0007751 | ---?? | 52.9 | 0.1199 | | |
| 9:44253804 | 9 | 44253804 | 3 | 0 | 0 | T | TAC | -1.9788 | 0.5887 | 0.0007751 | ---?? | 0 | 0.8685 | | |
| rs13030534 | 2 | 138953990 | 4 | 0 | 0 | A | G | 1.6223 | 0.4826 | 0.0007752 | +++?+ | 0 | 0.8189 | intergenic | |
| rs367560998 | 9 | 44281235 | 3 | 0 | 0 | A | G | 2.1844 | 0.6498 | 0.0007752 | +++?? | 0 | 0.8109 | intergenic | |
| rs71198965 | 13 | 20810561 | 4 | 0 | 0 | T | G | -1.1042 | 0.3285 | 0.0007753 | ---?- | 0 | 0.495 | upstream-gene | |
| 21:19069257 | 21 | 19069257 | 3 | 0 | 0 | CA | C | -2.0091 | 0.5977 | 0.0007753 | ---?? | 47.6 | 0.1485 | | |
| rs12524441 | 6 | 32024234 | 4 | 0 | 0 | A | G | 2.7747 | 0.8254 | 0.0007755 | ++++? | 18.8 | 0.2965 | intron | TNXB |
| rs9268503 | 6 | 32377061 | 5 | 0 | 0 | T | C | 0.71 | 0.2112 | 0.0007756 | +++++ | 0 | 0.6794 | upstream-gene | |
| rs10259627 | 7 | 82416922 | 4 | 0 | 0 | A | G | -0.8668 | 0.2579 | 0.0007758 | -+-?- | 46.9 | 0.1298 | intron | PCLO |
| rs57499049 | 19 | 6639040 | 4 | 0 | 0 | T | C | -0.8831 | 0.2627 | 0.0007759 | ---?- | 0 | 0.5975 | intergenic | |
| rs35916937 | 1 | 64786412 | 4 | 0 | 0 | T | C | -1.0268 | 0.3055 | 0.000776 | ---?- | 0 | 0.4847 | intergenic | |
| rs112894535 | 15 | 87612920 | 4 | 0 | 0 | A | G | -1.4948 | 0.4447 | 0.000776 | ---?- | 0 | 0.5484 | intergenic | |
| rs173571 | 5 | 165390936 | 5 | 0 | 0 | T | C | -0.7074 | 0.2105 | 0.0007762 | ---+- | 45 | 0.1219 | intergenic | |
| rs147191051 | 6 | 32126977 | 4 | 0 | 0 | T | C | 2.7322 | 0.8129 | 0.0007762 | ++++? | 49.6 | 0.1139 | intron | PPT2,PPT2-EGFL8 |
| rs12632491 | 3 | 193748506 | 4 | 0 | 0 | A | G | 2.2858 | 0.6801 | 0.0007763 | +++?+ | 0 | 0.917 | intergenic | |
| rs73363855 | 17 | 51252607 | 5 | 0 | 0 | T | C | -1.0539 | 0.3135 | 0.0007765 | ----+ | 0 | 0.414 | intergenic |  |
| rs2241591 | 19 | 8386367 | 4 | 0 | 0 | T | C | 1.5969 | 0.4751 | 0.0007767 | +++?+ | 42.6 | 0.1561 | upstream-gene | |
| 14:93178874 | 14 | 93178874 | 2 | 0 | 0 | A | ATGGAG | 5.955 | 1.7718 | 0.0007768 | ?++?? | 4.3 | 0.3066 | | |
| rs261555 | 8 | 97003139 | 4 | 0 | 0 | A | G | 0.9882 | 0.294 | 0.0007769 | +++?+ | 0 | 0.7879 | intergenic | |
| rs140676113 | 4 | 128834250 | 4 | 0 | 0 | T | C | 3.6375 | 1.0823 | 0.0007769 | +++?+ | 0 | 0.8547 | downstream-gene | |

| | | | | | | | | | | | | | | | |
|---|---|---|---|---|---|---|---|---|---|---|---|---|---|---|---|
| rs79563157 | 2 | 131535579 | 4 | 0 | 0 | T | G | 2.0782 | 0.6184 | 0.000777 | +++?+ | 19.7 | 0.2913 | intergenic | |
| rs6129778 | 20 | 39870578 | 3 | 0 | 0 | A | C | 1.0826 | 0.3221 | 0.0007771 | +++?? | 40.1 | 0.1883 | intron | ZHX3 |
| rs28840500 | 9 | 44253019 | 3 | 0 | 0 | A | T | 1.7242 | 0.513 | 0.0007771 | +++?? | 0 | 0.8567 | intergenic | |
| 15:85684280 | 15 | 85684280 | 3 | 0 | 0 | CAA | C | -1.1707 | 0.3483 | 0.0007771 | ---?? | 1.1 | 0.3638 | | |
| 5:119816849 | 5 | 119816849 | 3 | 0 | 0 | A | AT | -0.9891 | 0.2943 | 0.0007771 | ---?? | 23.1 | 0.2727 | | |
| 1:164259953 | 1 | 164259953 | 3 | 0 | 0 | T | C | 1.7611 | 0.524 | 0.0007772 | +++?? | 0 | 0.9371 | intergenic | |
| rs7017435 | 8 | 114880103 | 4 | 0 | 0 | A | G | 1.8071 | 0.5377 | 0.0007773 | +++?- | 0.9 | 0.3876 | intergenic | |
| rs62010504 | 15 | 33909648 | 4 | 0 | 0 | A | G | -1.1724 | 0.3489 | 0.0007773 | ---?- | 25 | 0.2614 | intron | RYR3 |
| rs7995742 | 13 | 20810103 | 4 | 0 | 0 | A | G | 1.1057 | 0.329 | 0.0007774 | +++?+ | 0 | 0.5029 | upstream-gene | |
| rs67929979 | 3 | 71886184 | 4 | 0 | 0 | T | C | 0.9129 | 0.2716 | 0.0007775 | +++?+ | 0 | 0.807 | intergenic | |
| rs6901175 | 6 | 32198527 | 5 | 0 | 0 | A | G | -0.9728 | 0.2895 | 0.0007776 | ----- | 0 | 0.6703 | intergenic |  |
| rs2945931 | 9 | 44277696 | 3 | 0 | 0 | C | G | 2.1135 | 0.6289 | 0.0007778 | +++?? | 0 | 0.825 | intergenic | |
| rs42385 | 16 | 6536933 | 4 | 0 | 0 | A | C | 0.9116 | 0.2713 | 0.0007779 | +++?+ | 0 | 0.6899 | intron | RBFOX1 |
| rs35583396 | 1 | 117856355 | 4 | 0 | 0 | T | C | 2.6453 | 0.7872 | 0.0007781 | +++?- | 34 | 0.208 | intergenic | |
| rs3021090 | 6 | 32021976 | 5 | 0 | 0 | A | G | -1.6375 | 0.4873 | 0.0007782 | ----+ | 41.4 | 0.1454 | intron | TNXB |
| rs1806573 | 12 | 43210052 | 5 | 0 | 0 | A | G | -0.8295 | 0.2469 | 0.0007789 | ----- | 0 | 0.6826 | intergenic |  |
| rs6774890 | 3 | 61069187 | 4 | 0 | 0 | T | C | 3.2003 | 0.9524 | 0.0007789 | +++?- | 0 | 0.6812 | intron | FHIT |
| rs73334983 | 8 | 114880666 | 4 | 0 | 0 | T | G | -1.8101 | 0.5387 | 0.000779 | ---?+ | 0.6 | 0.3887 | intergenic | |
| rs117997300 | 15 | 102061671 | 4 | 0 | 0 | A | C | -2.069 | 0.6157 | 0.000779 | ---?- | 0 | 0.6283 | intergenic | |
| rs9268502 | 6 | 32376932 | 5 | 0 | 0 | T | C | 0.7096 | 0.2112 | 0.0007793 | +++++ | 0 | 0.6837 | upstream-gene | |
| 10:69185202 | 10 | 69185202 | 3 | 0 | 0 | A | AATTAT | 2.7213 | 0.8099 | 0.0007794 | +++?? | 0 | 0.8452 | | |
| rs1641835 | 16 | 12136584 | 4 | 0 | 0 | T | C | 0.9035 | 0.2689 | 0.0007795 | -++?+ | 49.5 | 0.1143 | intron | SNX29 |
| rs11075042 | 16 | 12122639 | 4 | 0 | 0 | T | C | -0.8038 | 0.2392 | 0.0007795 | +--?- | 56.6 | 0.07448 | intron | SNX29 |
| rs7102330 | 11 | 46364604 | 4 | 0 | 0 | T | G | -2.4683 | 0.7346 | 0.0007796 | ---?- | 0 | 0.8619 | intron | DGKZ |
| rs77704819 | 9 | 118324574 | 4 | 0 | 0 | T | G | 2.184 | 0.65 | 0.0007796 | +++?+ | 0 | 0.6432 | intergenic | |
| 15:22828332 | 15 | 22828332 | 3 | 0 | 0 | A | G | -4.3182 | 1.2852 | 0.0007797 | ---?? | 20.3 | 0.285 | intergenic | |
| rs3091794 | 20 | 44701765 | 4 | 0 | 0 | T | C | -0.8634 | 0.257 | 0.0007799 | +--?- | 37.7 | 0.1861 | intron | NCOA5 |
| rs75607798 | 10 | 131477440 | 4 | 0 | 0 | A | G | -1.2501 | 0.3721 | 0.0007799 | ---?- | 0 | 0.8158 | intron | MGMT |
| rs17169080 | 7 | 15833678 | 4 | 0 | 0 | C | G | 1.8244 | 0.543 | 0.00078 | +++?- | 58 | 0.0677 | intergenic | |
| rs17828682 | 6 | 130964273 | 4 | 0 | 0 | T | C | 1.4262 | 0.4245 | 0.00078 | +++?+ | 0 | 0.6851 | intergenic | |
| rs10136731 | 14 | 43272995 | 4 | 0 | 0 | T | C | -1.2824 | 0.3817 | 0.0007802 | ---?+ | 59.1 | 0.06184 | intergenic | |
| rs77925086 | 6 | 130954698 | 4 | 0 | 0 | A | G | -1.427 | 0.4247 | 0.0007802 | ---?- | 0 | 0.7394 | intergenic | |
| rs140905715 | 18 | 8576035 | 3 | 0 | 0 | T | G | 4.5517 | 1.3548 | 0.0007802 | +++?? | 0 | 0.4351 | intergenic | |
| 7:4599338 | 7 | 4599338 | 3 | 0 | 0 | T | TA | -0.8425 | 0.2508 | 0.0007803 | ---?? | 19.1 | 0.2905 | | |
| rs7641225 | 3 | 87373667 | 4 | 0 | 0 | A | T | 1.069 | 0.3182 | 0.0007804 | +++?+ | 0 | 0.9981 | intergenic | |
| rs2895614 | 6 | 106053771 | 4 | 0 | 0 | A | C | -0.8456 | 0.2517 | 0.0007806 | ---?- | 0 | 0.9992 | intergenic | |
| rs11918956 | 3 | 61092405 | 4 | 0 | 0 | T | C | -3.1819 | 0.9471 | 0.0007807 | ---?- | 0 | 0.6935 | intron | FHIT |
| rs4920092 | 21 | 43781114 | 5 | 0 | 0 | A | G | -0.7731 | 0.2301 | 0.000781 | ----- | 0 | 0.8373 | downstream-gene |  |
| rs11969759 | 6 | 32021130 | 5 | 0 | 0 | T | C | 1.6374 | 0.4874 | 0.0007811 | ++++- | 41.4 | 0.1455 | intron | TNXB |
| rs62055322 | 16 | 68681201 | 5 | 0 | 0 | T | C | 2.3089 | 0.6873 | 0.0007811 | +++++ | 3.7 | 0.3857 | intron | CDH3 |
| rs75307819 | 11 | 35314384 | 3 | 0 | 0 | A | T | -3.372 | 1.0038 | 0.0007811 | ---?? | 0 | 0.7985 | intron | SLC1A2 |
| rs142736923 | 4 | 38488585 | 3 | 0 | 0 | T | G | 4.4365 | 1.3206 | 0.0007811 | ?++?+ | 0 | 0.5215 | intergenic | |
| 10:131475037 | 10 | 131475037 | 3 | 0 | 0 | A | ATACAT | 1.2977 | 0.3863 | 0.0007813 | +++?? | 0 | 0.6525 | | |
| rs9268501 | 6 | 32376882 | 5 | 0 | 0 | T | G | -0.7095 | 0.2112 | 0.0007814 | ----- | 0 | 0.6846 | upstream-gene |  |
| rs12866227 | 13 | 20810737 | 4 | 0 | 0 | A | C | -1.104 | 0.3287 | 0.0007814 | ---?- | 0 | 0.4945 | upstream-gene | |
| rs17006127 | 4 | 123692125 | 4 | 0 | 0 | A | G | -2.1344 | 0.6354 | 0.0007814 | ---?- | 60.7 | 0.05399 | intergenic | |
| rs429701 | 2 | 40436561 | 4 | 0 | 0 | T | G | 1.0438 | 0.3107 | 0.0007815 | +++?- | 0 | 0.422 | intron | SLC8A1,SLC8A1-AS1 |
| rs4686663 | 3 | 193747979 | 4 | 0 | 0 | A | G | -2.2847 | 0.6801 | 0.0007815 | ---?- | 0 | 0.9169 | intergenic | |
| 9:96568060 | 9 | 96568060 | 3 | 0 | 0 | G | GA | -0.9549 | 0.2843 | 0.0007816 | -+-?? | 65 | 0.05732 | | |
| rs11004040 | 10 | 55830753 | 4 | 0 | 0 | T | C | -2.134 | 0.6353 | 0.0007817 | ---?+ | 0 | 0.4199 | intron | PCDH15 |
| rs9812198 | 3 | 71879327 | 4 | 0 | 0 | A | G | -0.8415 | 0.2505 | 0.0007818 | ---?- | 0 | 0.7503 | intergenic | |
| rs11588217 | 1 | 64727239 | 4 | 0 | 0 | A | C | 1.2168 | 0.3622 | 0.0007823 | +++?+ | 0 | 0.4409 | intron | UBE2U |
| rs7533776 | 1 | 22329525 | 2 | 0 | 0 | T | C | 8.0907 | 2.4088 | 0.0007827 | ++??? | 0 | 0.3733 | missense | CELA3A |
| rs74822731 | 10 | 131477508 | 4 | 0 | 0 | A | G | 1.2497 | 0.3721 | 0.0007827 | +++?+ | 0 | 0.8159 | intron | MGMT |
| rs460568 | 6 | 33232025 | 5 | 0 | 0 | T | C | 0.938 | 0.2793 | 0.0007828 | +++++ | 0 | 0.7972 | intron | VPS52 |
| rs56296536 | 4 | 186806882 | 4 | 0 | 0 | A | T | -2.5859 | 0.7699 | 0.0007828 | ---?- | 0 | 0.5967 | intron | SORBS2 |
| rs76555996 | 2 | 23019848 | 4 | 0 | 0 | A | C | -2.9615 | 0.8817 | 0.0007828 | ---?- | 0 | 0.8668 | intergenic | |
| rs59705399 | 19 | 49095805 | 4 | 0 | 0 | C | G | -1.586 | 0.4722 | 0.0007829 | ---?- | 0 | 0.8947 | intron | SULT2B1 |
| rs115029566 | 1 | 182014592 | 4 | 0 | 0 | T | C | 1.1239 | 0.3346 | 0.0007829 | +++?+ | 0 | 0.4538 | intergenic | |

| | | | | | | | | | | | | | | | |
|---|---|---|---|---|---|---|---|---|---|---|---|---|---|---|---|
| rs74771174 | 6 | 130955025 | 4 | 0 | 0 | A | G | 1.4267 | 0.4248 | 0.0007831 | +++?+ | 0 | 0.7383 | intergenic | |
| 12:116848653 | 12 | 116848653 | 3 | 0 | 0 | CGT | C | -1.4396 | 0.4286 | 0.0007832 | ---?? | 0 | 0.5267 | | |
| rs2075763 | 17 | 4802685 | 5 | 0 | 0 | A | G | -1.5155 | 0.4512 | 0.0007835 | ----+ | 9 | 0.3554 | intron-near-splice | CHRNE |
| rs705566 | 1 | 64769980 | 4 | 0 | 0 | T | G | -0.9321 | 0.2775 | 0.0007837 | ---?- | 0 | 0.4758 | intergenic | |
| rs2033897 | 4 | 47834151 | 5 | 0 | 0 | A | G | 1.6438 | 0.4894 | 0.0007838 | +++++ | 11.9 | 0.3378 | intron | CORIN |
| 10:56705084 | 10 | 56705084 | 3 | 0 | 0 | G | GGATA | -0.9649 | 0.2873 | 0.0007838 | ---?? | 0 | 0.615 | | |
| rs12456464 | 18 | 3832168 | 4 | 0 | 0 | T | C | 0.8549 | 0.2546 | 0.0007839 | +++?+ | 0 | 0.4713 | intron | DLGAP1 |
| 20:12000715 | 20 | 12000715 | 3 | 0 | 0 | A | ATC | -0.9647 | 0.2873 | 0.0007839 | ---?? | 0 | 0.4535 | | |
| rs2407116 | 12 | 43209766 | 5 | 0 | 0 | T | C | -0.8286 | 0.2467 | 0.0007842 | ----- | 0 | 0.6822 | intergenic |  |
| rs1878582 | 12 | 43209772 | 5 | 0 | 0 | C | G | 0.8288 | 0.2468 | 0.0007843 | +++++ | 0 | 0.6834 | intergenic | |
| rs16880565 | 6 | 18348630 | 4 | 0 | 0 | T | G | -2.0457 | 0.6091 | 0.0007843 | ---?- | 20.5 | 0.2869 | intron | RNF144B |
| rs62340526 | 4 | 190014276 | 4 | 0 | 0 | T | C | 0.8434 | 0.2512 | 0.0007843 | +++?+ | 29.2 | 0.2371 | intergenic | |
| 15:87561281 | 15 | 87561281 | 3 | 0 | 0 | A | AAAT | -0.925 | 0.2754 | 0.0007844 | ---?? | 26.5 | 0.2567 | | |
| rs2131606 | 17 | 71059408 | 4 | 0 | 0 | A | G | -2.2895 | 0.6818 | 0.0007845 | ---?- | 26.1 | 0.2549 | intron | SLC39A11 |
| rs6737988 | 2 | 184027598 | 4 | 0 | 0 | T | C | -1.1759 | 0.3502 | 0.0007846 | ---?- | 0 | 0.8033 | downstream-gene | |
| rs73334978 | 8 | 114879988 | 4 | 0 | 0 | A | T | -1.8065 | 0.538 | 0.0007848 | ---?+ | 1.3 | 0.3854 | intergenic | |
| rs2188869 | 7 | 20497113 | 4 | 0 | 0 | T | C | -0.8199 | 0.2442 | 0.0007852 | ---?- | 36 | 0.196 | intergenic | |
| rs75930458 | 10 | 131477560 | 4 | 0 | 0 | T | G | -1.2494 | 0.3721 | 0.0007852 | ---?- | 0 | 0.8161 | intron | MGMT |
| rs111572871 | 17 | 56427077 | 5 | 0 | 0 | A | G | -1.0735 | 0.3197 | 0.0007853 | ----- | 36.2 | 0.1798 | intron | SUPT4H1,BZRAP1-AS1 |
| rs10947230 | 6 | 32024395 | 5 | 0 | 0 | T | C | 1.6358 | 0.4871 | 0.0007854 | ++++- | 41.4 | 0.1454 | missense | TNXB |
| rs78522389 | 17 | 56428283 | 5 | 0 | 0 | A | T | -1.0735 | 0.3197 | 0.0007854 | ----- | 36.2 | 0.1799 | intron | SUPT4H1,BZRAP1-AS1 |
| rs6005726 | 22 | 28547846 | 4 | 0 | 0 | A | G | 2.4403 | 0.7268 | 0.0007856 | +++?+ | 0 | 0.7457 | intron | TTC28 |
| rs35654349 | 10 | 20140666 | 4 | 0 | 0 | A | C | -0.9122 | 0.2717 | 0.0007856 | ---?- | 0 | 0.7571 | intron | PLXDC2 |
| rs10186203 | 2 | 116957966 | 3 | 0 | 0 | C | G | -1.0605 | 0.3159 | 0.0007858 | ---?? | 0 | 0.6905 | intergenic | |
| rs75136052 | 3 | 185769644 | 3 | 0 | 0 | A | G | -3.6506 | 1.0872 | 0.0007858 | ---?? | 0 | 0.6286 | intron | ETV5 |
| rs3763314 | 6 | 32376618 | 5 | 0 | 0 | T | C | 0.7092 | 0.2112 | 0.000786 | +++++ | 0 | 0.6905 | upstream-gene | |
| rs12157749 | 22 | 26256918 | 3 | 0 | 0 | T | C | -4.7153 | 1.4043 | 0.000786 | ---?? | 19.4 | 0.2891 | intron | MYO18B |
| 9:29940725 | 9 | 29940725 | 3 | 0 | 0 | CATATA[illegible] | C | -3.691 | 1.0993 | 0.0007862 | ---?? | 52.9 | 0.1196 | | |
| rs3827908 | 14 | 21451743 | 4 | 0 | 0 | A | G | -0.8792 | 0.2619 | 0.0007863 | ---?- | 18.9 | 0.2956 | intergenic | |
| rs7669549 | 4 | 108290300 | 4 | 0 | 0 | A | C | -0.8462 | 0.252 | 0.0007864 | ---?- | 0 | 0.3996 | intergenic | |
| rs12429088 | 13 | 65199689 | 4 | 0 | 0 | C | G | -1.8787 | 0.5596 | 0.0007864 | ---?- | 28.3 | 0.2424 | intergenic | |
| rs11734805 | 4 | 111398252 | 4 | 0 | 0 | A | C | -0.8743 | 0.2604 | 0.0007865 | ---?- | 0 | 0.8089 | intron | ENPEP |
| rs78965241 | 1 | 42771238 | 4 | 0 | 0 | T | C | 1.8243 | 0.5434 | 0.0007866 | +++?+ | 0 | 0.5406 | intron | FOXJ3 |
| rs4404800 | 6 | 32581913 | 4 | 0 | 1 | A | G | -1.3279 | 0.3955 | 0.0007869 | ----? | 36.7 | 0.1921 | intergenic |  |
| rs9264524 | 6 | 31233845 | 5 | 0 | 0 | T | G | -0.8017 | 0.2388 | 0.0007869 | ----- | 0 | 0.4742 | downstream-gene |  |
| rs11061314 | 12 | 131563328 | 4 | 0 | 0 | T | C | 0.8495 | 0.253 | 0.0007869 | +++?+ | 0 | 0.9656 | intron | GPR133 |
| rs61145885 | 3 | 30827071 | 3 | 0 | 0 | A | G | 3.0377 | 0.9048 | 0.0007869 | +++?? | 0 | 0.5306 | intron | GADL1 |
| rs8111430 | 19 | 46168066 | 5 | 0 | 1 | T | C | -0.8788 | 0.2618 | 0.000787 | -+--+ | 25.9 | 0.2487 | upstream-gene | |
| rs4636067 | 6 | 32578621 | 4 | 0 | 1 | T | G | 1.3279 | 0.3955 | 0.0007872 | ++++? | 36.7 | 0.1921 | intergenic | |
| rs12726916 | 1 | 157401437 | 4 | 0 | 0 | T | G | 1.106 | 0.3294 | 0.0007872 | +++?+ | 61.3 | 0.05165 | intergenic | |
| rs138991200 | 14 | 44854926 | 4 | 0 | 0 | T | C | 1.3235 | 0.3942 | 0.0007873 | +++?+ | 0 | 0.8166 | intergenic | |
| rs28653355 | 15 | 27585034 | 4 | 0 | 0 | A | G | -0.8313 | 0.2476 | 0.0007874 | ---?- | 0 | 0.9309 | intron | GABRG3 |
| 10:131911333 | 10 | 131911333 | 3 | 0 | 0 | T | TA | -0.8534 | 0.2542 | 0.0007874 | ---?? | 0 | 0.6787 | | |
| rs16869940 | 4 | 20767378 | 4 | 0 | 0 | A | G | -1.0751 | 0.3203 | 0.0007876 | ---?- | 37.6 | 0.1863 | intron | KCNIP4 |
| rs16976223 | 17 | 68746950 | 4 | 0 | 0 | T | C | -1.3104 | 0.3903 | 0.0007876 | ---?- | 0 | 0.7504 | intergenic | |
| rs58587654 | 17 | 52292601 | 4 | 0 | 0 | A | G | 1.9548 | 0.5823 | 0.0007876 | +++?- | 28.3 | 0.2424 | intergenic | |
| rs12409709 | 1 | 182015366 | 4 | 0 | 0 | A | G | 1.1228 | 0.3344 | 0.0007877 | +++?+ | 0 | 0.4548 | intergenic | |
| rs12751821 | 1 | 157401352 | 4 | 0 | 0 | A | T | 1.1058 | 0.3294 | 0.0007877 | +++?+ | 61.1 | 0.05216 | intergenic | |
| rs2454837 | 10 | 2991774 | 4 | 0 | 0 | T | C | 1.2327 | 0.3672 | 0.000788 | +++?+ | 0 | 0.8077 | intergenic | |
| rs13167234 | 5 | 60554362 | 5 | 0 | 0 | A | G | -0.961 | 0.2863 | 0.0007881 | ----+ | 27.3 | 0.2392 | intergenic |  |
| rs61806398 | 1 | 182016344 | 4 | 0 | 0 | T | C | -1.1224 | 0.3343 | 0.0007881 | ---?- | 0 | 0.4554 | intergenic | |
| rs139807715 | 3 | 111888461 | 4 | 0 | 0 | T | C | -3.6862 | 1.0981 | 0.0007881 | ---?- | 0 | 0.7185 | intron | SLC9C1 |
| rs2889182 | 2 | 138952478 | 4 | 0 | 0 | T | C | -1.619 | 0.4823 | 0.0007882 | ---?- | 0 | 0.816 | intergenic | |
| rs79040109 | 12 | 51909909 | 3 | 0 | 0 | C | G | 2.9406 | 0.876 | 0.0007882 | +++?? | 35.5 | 0.2123 | downstream-gene | |
| rs10413249 | 19 | 12830209 | 4 | 0 | 0 | T | C | -3.4736 | 1.0348 | 0.0007884 | ----? | 52.5 | 0.09734 | intron | TNPO2 |
| rs140366827 | 3 | 111957808 | 4 | 0 | 0 | A | G | -4.0173 | 1.1968 | 0.0007885 | ---?- | 0 | 0.7876 | intron | SLC9C1 |
| rs145634562 | 14 | 36652410 | 2 | 0 | 0 | A | G | -6.5027 | 1.9372 | 0.0007888 | -???- | 0 | 0.6611 | intergenic | |
| 10:117721149 | 10 | 117721149 | 3 | 0 | 0 | CTG | C | 0.9222 | 0.2747 | 0.0007888 | +++?? | 0 | 0.8988 | | |
| 2:86591238 | 2 | 86591238 | 3 | 0 | 0 | A | AT | 2.9706 | 0.885 | 0.0007891 | +++?? | 0.5 | 0.3661 | | |

| | | | | | | | | | | | | | | | |
|---|---|---|---|---|---|---|---|---|---|---|---|---|---|---|---|
| rs148808086 | 5 | 140961364 | 5 | 0 | 0 | C | G | -1.2585 | 0.3749 | 0.0007893 | ----- | 0 | 0.4124 | intron | DIAPH1 |
| rs11165961 | 1 | 98621065 | 4 | 0 | 0 | A | C | 1.3069 | 0.3894 | 0.0007894 | +++?+ | 23.9 | 0.2678 | intergenic | |
| rs1879230 | 11 | 23982573 | 4 | 0 | 0 | T | C | 1.2074 | 0.3597 | 0.0007895 | +++?+ | 0 | 0.785 | intergenic | |
| rs2037905 | 7 | 158972018 | 4 | 0 | 0 | T | C | -1.8414 | 0.5486 | 0.0007895 | ---?- | 0 | 0.5601 | intergenic | |
| rs55740958 | 6 | 32073700 | 5 | 0 | 0 | T | C | 1.3412 | 0.3996 | 0.0007896 | +++-+ | 50.5 | 0.08861 | intron | TNXB |
| rs11656427 | 17 | 4723541 | 5 | 0 | 0 | A | G | -1.5252 | 0.4544 | 0.0007898 | ----- | 21.5 | 0.2775 | intron | PLD2 |
| rs149020771 | 6 | 63267094 | 3 | 0 | 0 | T | C | 2.3847 | 0.7105 | 0.0007898 | +++?? | 0.3 | 0.3667 | intergenic | |
| rs78649963 | 7 | 46178703 | 4 | 0 | 0 | T | C | 1.9109 | 0.5693 | 0.0007901 | +++?+ | 0 | 0.9688 | intergenic | |
| rs817428 | 1 | 85634286 | 4 | 0 | 0 | C | G | 1.5186 | 0.4525 | 0.0007904 | +++?+ | 0 | 0.9875 | intron | SYDE2 |
| rs114836315 | 3 | 155402741 | 4 | 0 | 0 | T | C | 2.2578 | 0.6727 | 0.0007904 | +++?+ | 13.8 | 0.3232 | intron | PLCH1 |
| rs6005724 | 22 | 28544093 | 4 | 0 | 0 | C | G | 2.443 | 0.7279 | 0.0007905 | +++?+ | 0 | 0.7443 | intron | TTC28 |
| rs7189943 | 16 | 57534258 | 4 | 0 | 0 | A | C | 1.5378 | 0.4582 | 0.0007905 | +++?+ | 0 | 0.9083 | intergenic | |
| rs12441249 | 15 | 41984538 | 5 | 0 | 0 | A | G | -0.9194 | 0.2739 | 0.0007906 | ----- | 0 | 0.5927 | intron | MGA |
| rs76494695 | 4 | 100561204 | 4 | 0 | 0 | T | C | 1.7924 | 0.5341 | 0.0007906 | -++?+ | 43.2 | 0.1525 | 3-prime-UTR | LOC285556 |
| rs142534596 | 4 | 166234897 | 4 | 0 | 0 | T | C | 3.6362 | 1.0835 | 0.0007906 | +++?+ | 0 | 0.9534 | intron | KLHL2 |
| rs146413733 | 12 | 117516254 | 4 | 0 | 0 | A | G | -2.6839 | 0.7997 | 0.0007906 | ---?- | 0 | 0.5547 | intron | TESC |
| rs3768875 | 2 | 225645937 | 2 | 0 | 0 | T | G | -1.5091 | 0.4497 | 0.0007907 | -???- | 0.8 | 0.3153 | intron | DOCK10 |
| 14:21455793 | 14 | 21455793 | 3 | 0 | 0 | G | GA | -0.9154 | 0.2728 | 0.0007907 | ---?? | 14.5 | 0.3105 | | |
| rs12038214 | 1 | 217999341 | 4 | 0 | 0 | A | G | -1.116 | 0.3325 | 0.0007909 | ---?- | 0 | 0.9442 | intron | SPATA17 |
| rs36043260 | 1 | 117919159 | 4 | 0 | 0 | T | C | 2.7712 | 0.8258 | 0.0007909 | +++?- | 21.9 | 0.2793 | intron | MAN1A2 |
| 12:112207947 | 12 | 112207947 | 3 | 0 | 0 | CA | C | 1.157 | 0.3448 | 0.0007911 | +-+?? | 65.2 | 0.05648 | | |
| 2:184027882 | 2 | 184027882 | 3 | 0 | 0 | T | TAA | -1.0821 | 0.3224 | 0.0007911 | ---?? | 0 | 0.9212 | | |
| rs10225534 | 7 | 82428636 | 4 | 0 | 0 | A | G | -0.8504 | 0.2534 | 0.0007912 | -+-?- | 45.3 | 0.1396 | intron | PCLO |
| rs11226009 | 11 | 103643692 | 3 | 0 | 0 | C | G | -1.6087 | 0.4794 | 0.0007913 | ?--?- | 28.7 | 0.2462 | intergenic | |
| rs7530993 | 1 | 18739603 | 4 | 0 | 0 | C | G | -1.0493 | 0.3127 | 0.0007916 | ---?- | 50.6 | 0.1079 | intergenic | |
| rs35303461 | 2 | 218830626 | 4 | 0 | 0 | T | C | 1.3066 | 0.3894 | 0.0007916 | +++?- | 0 | 0.7965 | intergenic | |
| rs1583808 | 4 | 47836096 | 5 | 0 | 0 | T | C | 1.6425 | 0.4895 | 0.0007917 | +++++ | 12 | 0.3374 | intron | CORIN |
| rs2678142 | 12 | 43193602 | 5 | 0 | 0 | T | C | 0.7489 | 0.2232 | 0.0007917 | +++++ | 0 | 0.6618 | intergenic | |
| rs7258212 | 19 | 12802481 | 4 | 0 | 0 | A | C | -3.527 | 1.0511 | 0.0007918 | ----? | 48.2 | 0.122 | intron | FBXW9 |
| rs4862070 | 4 | 183392126 | 4 | 0 | 0 | A | C | 1.0231 | 0.3049 | 0.0007921 | +++?+ | 0 | 0.4793 | intron | TENM3 |
| rs34813208 | 2 | 40588614 | 3 | 0 | 0 | T | G | -3.1677 | 0.944 | 0.0007923 | ---?? | 0 | 0.56 | intron | SLC8A1 |
| rs4907890 | 1 | 100244236 | 4 | 0 | 0 | A | C | -1.683 | 0.5016 | 0.0007929 | ---?- | 0 | 0.9028 | intergenic | |
| rs75402183 | 8 | 2862189 | 4 | 0 | 0 | C | G | 3.5924 | 1.0707 | 0.000793 | +++?+ | 0 | 0.9815 | intron | CSMD1 |
| 1:98602825 | 1 | 98602825 | 3 | 0 | 0 | G | GT | 1.3547 | 0.4038 | 0.000793 | +++?? | 46.9 | 0.1522 | | |
| rs17692820 | 15 | 91579417 | 4 | 0 | 0 | A | G | -1.5668 | 0.467 | 0.0007935 | ---?- | 0 | 0.6546 | intergenic | |
| rs35090855 | 2 | 218830592 | 4 | 0 | 0 | T | C | 1.3062 | 0.3893 | 0.0007935 | +++?- | 0 | 0.7965 | intergenic | |
| rs58100540 | 15 | 70304324 | 4 | 0 | 0 | A | C | 1.6769 | 0.4998 | 0.0007937 | +++?+ | 0 | 0.6929 | intergenic | |
| 2:86817050 | 2 | 86817050 | 3 | 0 | 0 | A | AT | -0.8327 | 0.2482 | 0.0007937 | ---?? | 0 | 0.6325 | | |
| rs5997341 | 22 | 28549292 | 4 | 0 | 0 | A | T | 2.4348 | 0.7257 | 0.0007938 | +++?+ | 0 | 0.7448 | intron | TTC28 |
| rs67044368 | 2 | 212573895 | 4 | 0 | 0 | T | C | 1.2464 | 0.3715 | 0.0007938 | +++?+ | 0 | 0.9314 | intron | ERBB4 |
| rs55659927 | 18 | 58958094 | 4 | 0 | 0 | T | G | -2.0527 | 0.6118 | 0.0007939 | ---?- | 0 | 0.8666 | intergenic | |
| rs186301413 | 8 | 30348006 | 4 | 0 | 0 | T | C | 3.5055 | 1.0449 | 0.000794 | +-+?+ | 49.6 | 0.1137 | intron | RBPMS |
| rs9884519 | 4 | 176485606 | 4 | 0 | 0 | T | C | 2.5377 | 0.7564 | 0.0007942 | +++?+ | 0 | 0.811 | intergenic | |
| rs145696092 | 6 | 144779211 | 4 | 0 | 0 | A | G | 3.7023 | 1.1036 | 0.0007942 | +++?+ | 0 | 0.696 | intron | UTRN |
| rs17117881 | 1 | 98649772 | 4 | 0 | 0 | T | C | -1.3502 | 0.4025 | 0.0007944 | ---?- | 0 | 0.54 | intergenic | |
| rs35848199 | 7 | 112699789 | 4 | 0 | 0 | T | G | 1.1383 | 0.3393 | 0.0007944 | +++?- | 14.8 | 0.3182 | intergenic | |
| rs7724634 | 5 | 744666 | 3 | 0 | 0 | T | C | 1.5188 | 0.4527 | 0.0007945 | +++?? | 0 | 0.4381 | intron | ZDHHC11B |
| rs114182581 | 4 | 92746736 | 4 | 0 | 0 | T | G | 2.3995 | 0.7153 | 0.0007946 | +++?+ | 0 | 0.8687 | intergenic | |
| rs11601197 | 11 | 12835130 | 4 | 0 | 0 | C | G | -1.414 | 0.4215 | 0.0007947 | ---?- | 0 | 0.6294 | intron | TEAD1 |
| rs78834684 | 7 | 46178262 | 4 | 0 | 0 | A | T | 1.9095 | 0.5692 | 0.0007947 | +++?+ | 0 | 0.9693 | intergenic | |
| 3:5389734 | 3 | 5389734 | 4 | 0 | 0 | A | G | 1.348 | 0.4018 | 0.0007948 | +++?+ | 2.3 | 0.3807 | | |
| rs76045334 | 7 | 46178307 | 4 | 0 | 0 | A | G | 1.9095 | 0.5692 | 0.0007949 | +++?+ | 0 | 0.9693 | intergenic | |
| 6:168596570 | 6 | 168596570 | 3 | 0 | 0 | A | AG | 0.9755 | 0.2908 | 0.0007949 | +++?? | 0 | 0.9775 | | |
| rs7553743 | 1 | 182017678 | 4 | 0 | 0 | T | C | 1.1211 | 0.3342 | 0.000795 | +++?+ | 0 | 0.4556 | intergenic | |
| 1:153157815 | 1 | 153157815 | 3 | 0 | 0 | CT | C | -0.9288 | 0.2769 | 0.0007951 | ---?? | 65.4 | 0.05542 | | |
| rs16880181 | 7 | 40340186 | 4 | 0 | 0 | A | G | -3.0942 | 0.9224 | 0.0007952 | ---?- | 0 | 0.9088 | intron | C7orf10 |
| rs10853677 | 18 | 25372386 | 4 | 0 | 0 | A | G | -0.8541 | 0.2546 | 0.0007953 | ---?- | 0 | 0.5081 | intergenic | |
| rs28632030 | 19 | 46168273 | 5 | 0 | 1 | A | G | -0.8805 | 0.2625 | 0.0007953 | -+--+ | 24.8 | 0.2559 | upstream-gene | |
| rs185027431 | 15 | 23727113 | 3 | 0 | 0 | T | C | 4.9224 | 1.4675 | 0.0007955 | +++?? | 0 | 0.6584 | intergenic | |

| | | | | | | | | | | | | | | | | |
|---|---|---|---|---|---|---|---|---|---|---|---|---|---|---|---|---|
| rs77588619 | 9 | 118320433 | 4 | 0 | 0 | T | C | 2.1792 | 0.6497 | 0.0007956 | +++?+ | 0 | 0.6424 | intergenic | | |
| 1:202056430 | 1 | 202056430 | 3 | 0 | 0 | G | GGGAT | 0.88 | 0.2623 | 0.0007956 | +++?? | 0 | 0.8987 | | | |
| rs7137346 | 12 | 130596396 | 4 | 0 | 0 | A | G | -0.7985 | 0.238 | 0.0007957 | ---?+ | 20.3 | 0.2883 | intergenic | | |
| rs55836759 | 5 | 737506 | 3 | 0 | 0 | T | G | 1.5083 | 0.4497 | 0.0007958 | +++?? | 5.4 | 0.3475 | intron | | ZDHHC11B |
| rs2136686 | 1 | 217994792 | 4 | 0 | 0 | A | G | -1.0937 | 0.3261 | 0.0007964 | ---?- | 0 | 0.9803 | intron | | SPATA17 |
| rs12933772 | 16 | 6560430 | 4 | 0 | 0 | C | G | -0.9266 | 0.2763 | 0.0007968 | ---?- | 0 | 0.6666 | intron | | RBFOX1 |
| rs147190717 | 14 | 104149666 | 4 | 0 | 0 | T | G | 3.7868 | 1.1291 | 0.0007969 | +++?+ | 0 | 0.5989 | intron | | KLC1 |
| 1:232830699 | 1 | 232830699 | 3 | 0 | 0 | CT | C | 1.0281 | 0.3065 | 0.0007971 | +++?? | 57.8 | 0.09359 | | | |
| rs75361139 | 6 | 164375334 | 4 | 0 | 0 | A | G | 1.5437 | 0.4603 | 0.0007972 | +++?+ | 0 | 0.862 | intergenic | | |
| rs1914898 | 1 | 18744658 | 4 | 0 | 0 | C | G | -1.0419 | 0.3107 | 0.0007975 | ---?- | 52.2 | 0.09919 | intergenic | | |
| 17:11219477 | 17 | 11219477 | 3 | 0 | 0 | T | TTTA | -0.9631 | 0.2872 | 0.0007976 | ---?? | 0 | 0.5129 | | | |
| rs28853331 | 9 | 44253241 | 3 | 0 | 0 | A | G | -1.7322 | 0.5165 | 0.0007977 | ---?? | 0 | 0.8496 | intergenic | | |
| rs35976116 | 19 | 3597031 | 3 | 0 | 0 | C | G | -0.8883 | 0.2649 | 0.0007977 | ---?? | 0 | 0.6069 | intron | | TBXA2R |
| rs11756394 | 6 | 63337004 | 4 | 0 | 0 | A | G | -2.376 | 0.7085 | 0.0007979 | ---?- | 0 | 0.4845 | intergenic | | |
| rs61753649 | 4 | 153874666 | 5 | 0 | 0 | A | G | 2.6185 | 0.7808 | 0.0007983 | -++++ | 38.1 | 0.1672 | missense | | FHDC1 |
| rs117249145 | 7 | 40342521 | 4 | 0 | 0 | T | C | 3.0919 | 0.922 | 0.0007983 | +++?+ | 0 | 0.909 | intron | | C7orf10 |
| rs7686643 | 4 | 190016610 | 4 | 0 | 0 | A | T | -0.8428 | 0.2513 | 0.0007984 | ---?- | 28.7 | 0.2399 | intergenic | | |
| rs12604549 | 18 | 26522562 | 3 | 0 | 0 | C | G | 1.4155 | 0.4221 | 0.0007984 | +++?? | 0 | 0.5151 | intergenic | | |
| 8:14498158 | 8 | 14498158 | 3 | 0 | 0 | G | GGTGTG | 0.8594 | 0.2563 | 0.0007984 | +++?? | 0 | 0.4729 | | | |
| rs4794416 | 17 | 51198961 | 5 | 0 | 0 | T | C | -1.0342 | 0.3084 | 0.0007985 | ----+ | 0 | 0.6056 | intergenic |  |  |
| rs9508549 | 13 | 30323668 | 5 | 0 | 0 | T | G | -1.2253 | 0.3654 | 0.0007985 | ----- | 0 | 0.5548 | intergenic |  |  |
| rs28550631 | 15 | 42199780 | 3 | 0 | 0 | T | C | 2.8192 | 0.8407 | 0.0007985 | +-+?? | 46.4 | 0.1547 | intron | | EHD4 |
| rs72900978 | 2 | 68139187 | 4 | 0 | 0 | A | G | 4.0676 | 1.2131 | 0.000799 | +++?+ | 0 | 0.4624 | intergenic | | |
| rs771767 | 3 | 101748638 | 5 | 0 | 0 | A | G | -0.766 | 0.2285 | 0.0007993 | ----- | 0 | 0.9234 | intergenic |  |  |
| rs62419116 | 6 | 63336321 | 4 | 0 | 0 | A | G | 2.3742 | 0.7081 | 0.0007994 | +++?+ | 0 | 0.4876 | intergenic | | |
| rs79306889 | 4 | 36410125 | 4 | 0 | 0 | T | C | -1.5836 | 0.4723 | 0.0007995 | ---?+ | 0 | 0.572 | intergenic | | |
| rs3893305 | 16 | 5559527 | 4 | 0 | 0 | T | G | -3.1019 | 0.9251 | 0.0007996 | -+-?- | 12.5 | 0.3301 | intron | | RBFOX1 |
| rs11590269 | 1 | 64771845 | 4 | 0 | 0 | A | G | 1.1727 | 0.3498 | 0.0007996 | +++?+ | 0 | 0.469 | intergenic | | |
| rs77316248 | 2 | 46276930 | 4 | 0 | 0 | C | G | -3.9702 | 1.1841 | 0.0007997 | ---?- | 0 | 0.8917 | intron | | PRKCE |
| rs10491392 | 5 | 106813852 | 4 | 0 | 0 | C | G | 1.2881 | 0.3842 | 0.0007998 | +++?- | 57.8 | 0.06827 | intron | | EFNA5 |
| rs80187804 | 3 | 29325798 | 4 | 0 | 0 | T | G | 1.1741 | 0.3502 | 0.0008001 | +++?+ | 2 | 0.382 | intron | | RBMS3 |
| 12:18862177 | 12 | 18862177 | 3 | 0 | 0 | G | GT | -0.9391 | 0.2801 | 0.0008002 | ---?? | 0 | 0.9378 | | | |
| rs2425753 | 20 | 44705624 | 4 | 0 | 0 | A | G | 0.8586 | 0.2561 | 0.0008003 | -++?+ | 36.7 | 0.192 | intron | | NCOA5 |
| rs11158732 | 14 | 68825191 | 4 | 0 | 0 | A | C | -0.8417 | 0.251 | 0.0008003 | ---?- | 0 | 0.6798 | intron | | RAD51B |
| rs186538556 | 5 | 2933657 | 2 | 0 | 0 | A | G | -6.7397 | 2.0102 | 0.0008004 | ?--?? | 0 | 0.4917 | intergenic | | |
| rs38428 | 7 | 30549074 | 5 | 0 | 0 | C | G | 0.8718 | 0.26 | 0.0008005 | +++++ | 20 | 0.2873 | upstream-gene | | |
| rs1751581 | 1 | 208187836 | 4 | 0 | 0 | A | G | 1.9407 | 0.5788 | 0.0008005 | +++?+ | 0 | 0.6979 | intergenic | | |
| rs71633532 | 1 | 205062466 | 5 | 0 | 0 | A | G | 1.501 | 0.4477 | 0.0008005 | +++-+ | 22.4 | 0.2718 | intron | | RBBP5 |
| 17:11221178 | 17 | 11221178 | 3 | 0 | 0 | CT | C | -0.959 | 0.2861 | 0.0008005 | ---?? | 0 | 0.4656 | | | |
| rs78579111 | 12 | 96653965 | 4 | 0 | 0 | T | C | 2.636 | 0.7863 | 0.0008008 | +++?+ | 0 | 0.4041 | intron | | ELK3 |
| rs17050837 | 4 | 140546424 | 4 | 0 | 0 | T | C | -3.0016 | 0.8953 | 0.0008009 | ---?- | 22.9 | 0.2733 | intergenic | | |
| rs61876327 | 11 | 471923 | 3 | 0 | 0 | A | G | -0.8963 | 0.2674 | 0.000801 | ---?? | 0 | 0.4688 | intron | | PTDSS2 |
| rs59806423 | 2 | 68139107 | 4 | 0 | 0 | A | G | -4.0661 | 1.2129 | 0.0008011 | ---?- | 0 | 0.4629 | intergenic | | |
| rs7190381 | 16 | 84841750 | 4 | 0 | 0 | T | C | -1.2075 | 0.3602 | 0.0008012 | ---?+ | 3.2 | 0.3765 | intergenic | | |
| rs57146051 | 2 | 68139111 | 4 | 0 | 0 | A | G | 4.0663 | 1.213 | 0.0008012 | +++?+ | 0 | 0.4628 | intergenic | | |
| rs116863593 | 11 | 59225382 | 3 | 0 | 0 | T | G | 3.4461 | 1.028 | 0.0008013 | +?++? | 23.3 | 0.2717 | downstream-gene | | |
| rs17153732 | 11 | 59210795 | 3 | 0 | 0 | A | G | -3.4461 | 1.028 | 0.0008014 | -?--? | 23.3 | 0.2717 | missense | | OR5A1 |
| rs74453752 | 11 | 59216279 | 3 | 0 | 0 | C | G | 3.4461 | 1.028 | 0.0008014 | +?++? | 23.3 | 0.2717 | downstream-gene | | |
| rs168776 | 5 | 141450326 | 5 | 0 | 0 | A | G | -0.995 | 0.2968 | 0.0008015 | ----- | 0 | 0.9641 | intergenic |  |  |
| rs146579370 | 8 | 21802686 | 4 | 0 | 0 | T | C | 1.5724 | 0.4691 | 0.0008017 | +++?+ | 0 | 0.6859 | intron | | XPO7 |
| rs6997512 | 8 | 95873541 | 3 | 0 | 0 | C | G | 0.8859 | 0.2643 | 0.0008021 | +++?? | 0 | 0.4488 | intron | | INTS8 |
| rs111553113 | 5 | 38805153 | 3 | 0 | 0 | A | C | 6.05 | 1.8049 | 0.0008022 | +++?? | 0 | 0.9843 | intergenic | | |
| rs2570114 | 15 | 87561720 | 3 | 0 | 0 | A | G | 0.923 | 0.2754 | 0.0008024 | +++?? | 26.9 | 0.2546 | intron | | AGBL1 |
| rs7000126 | 8 | 114880541 | 4 | 0 | 0 | T | C | -1.8268 | 0.545 | 0.0008024 | ---?+ | 3.1 | 0.3768 | intergenic | | |
| rs80055345 | 4 | 100557051 | 4 | 0 | 0 | T | C | 1.7878 | 0.5334 | 0.0008029 | -++?+ | 43.5 | 0.1505 | downstream-gene | | |
| rs1019706 | 2 | 185455629 | 4 | 0 | 0 | A | G | -0.9296 | 0.2774 | 0.0008033 | ---?- | 0 | 0.5064 | intergenic | | |
| rs2304818 | 12 | 41316381 | 4 | 0 | 0 | A | C | -1.6654 | 0.4969 | 0.0008033 | ---?+ | 0 | 0.7984 | intron | | CNTN1 |
| rs1960124 | 1 | 64791446 | 4 | 0 | 0 | A | G | 1.0242 | 0.3056 | 0.0008034 | +++?+ | 0 | 0.5265 | intergenic | | |
| rs4656130 | 1 | 87450654 | 5 | 0 | 0 | A | G | 1.0285 | 0.3069 | 0.0008034 | +++++ | 0 | 0.9258 | intron | | HS2ST1 |

| | | | | | | | | | | | | | | | |
|---|---|---|---|---|---|---|---|---|---|---|---|---|---|---|---|
| rs61771745 | 1 | 87451542 | 5 | 0 | 0 | A | G | -1.0283 | 0.3068 | 0.0008038 | ----- | 0 | 0.9258 | intron | HS2ST1 |
| 7:129463874 | 7 | 129463874 | 3 | 0 | 0 | CAAAAA | C | -0.9255 | 0.2761 | 0.0008038 | ---?? | 0 | 0.925 | | |
| rs2295440 | 20 | 57035076 | 4 | 0 | 0 | T | C | -0.9395 | 0.2803 | 0.0008043 | -+-?- | 40.9 | 0.1666 | 3-prime-UTR | APCDD1L |
| rs75599511 | 7 | 46177237 | 4 | 0 | 0 | T | C | 1.9041 | 0.5682 | 0.0008043 | +++?+ | 0 | 0.9693 | intergenic | |
| rs80224102 | 2 | 170327146 | 4 | 0 | 0 | A | G | -1.3855 | 0.4134 | 0.0008043 | ---?- | 0 | 0.9191 | intergenic | |
| rs5747208 | 22 | 17984458 | 4 | 0 | 0 | T | C | 3.2858 | 0.9805 | 0.0008044 | +++?+ | 0 | 0.7516 | intron | CECR2 |
| 10:134572870 | 10 | 134572870 | 3 | 0 | 0 | CATGGG | C | 0.9744 | 0.2908 | 0.0008044 | +++?? | 21.3 | 0.2808 | | |
| rs77705384 | 10 | 131477739 | 4 | 0 | 0 | T | C | 1.2466 | 0.372 | 0.0008046 | +++?+ | 0 | 0.8174 | intron | MGMT |
| rs138170488 | 3 | 176002804 | 4 | 0 | 0 | T | C | -2.3403 | 0.6984 | 0.0008047 | ---?- | 0 | 0.4243 | intergenic | |
| rs149389761 | 2 | 183735118 | 2 | 0 | 0 | A | G | 5.1865 | 1.5477 | 0.0008048 | ++??? | 0 | 0.706 | upstream-gene | |
| 4:162209564 | 4 | 162209564 | 2 | 0 | 0 | A | AT | -6.1491 | 1.8349 | 0.0008049 | ?--?? | 0 | 0.5784 | | |
| rs74345732 | 4 | 100558037 | 4 | 0 | 0 | A | G | 1.7879 | 0.5335 | 0.0008053 | -++?+ | 43.5 | 0.1508 | 3-prime-UTR | LOC285556 |
| rs149922127 | 4 | 47848465 | 5 | 0 | 0 | T | C | 1.6428 | 0.4903 | 0.0008053 | +++++ | 12.3 | 0.3355 | downstream-gene | |
| rs34230758 | 6 | 32633558 | 5 | 0 | 1 | A | G | -1.0456 | 0.312 | 0.0008054 | ----- | 0 | 0.5085 | intron | HLA-DQB1 |
| rs76231354 | 9 | 90841360 | 4 | 0 | 0 | A | G | -1.8855 | 0.5627 | 0.0008054 | ---?- | 0 | 0.7943 | intergenic | |
| rs113135111 | 2 | 98972714 | 4 | 0 | 0 | T | C | -1.601 | 0.4778 | 0.0008056 | ---?+ | 0 | 0.6167 | intron | CNGA3 |
| rs145922975 | 4 | 47847686 | 5 | 0 | 0 | A | G | -1.6424 | 0.4902 | 0.0008057 | ----- | 12.2 | 0.3357 | downstream-gene |  |
| rs116687432 | 4 | 100555700 | 4 | 0 | 0 | A | G | -1.7863 | 0.5331 | 0.000806 | +--?- | 43.6 | 0.1497 | downstream-gene | |
| rs9272400 | 6 | 32604898 | 3 | 0 | 0 | A | G | 0.9776 | 0.2918 | 0.0008065 | +++?? | 0 | 0.4302 | upstream-gene | |
| rs11694770 | 2 | 80914954 | 4 | 0 | 0 | A | T | -1.8011 | 0.5376 | 0.0008068 | ---?- | 0 | 0.6289 | intergenic | |
| rs112708953 | 2 | 98972687 | 4 | 0 | 0 | C | G | 1.6002 | 0.4776 | 0.000807 | +++?- | 0 | 0.6165 | intron | CNGA3 |
| rs76964538 | 9 | 101136647 | 4 | 0 | 0 | A | T | 3.2045 | 0.9565 | 0.0008071 | +++?- | 26.1 | 0.2552 | intron | GABBR2 |
| rs111993171 | 8 | 54249090 | 4 | 0 | 0 | A | G | 1.6343 | 0.4878 | 0.0008071 | +++?+ | 45.1 | 0.141 | intergenic | |
| rs142264100 | 6 | 63267915 | 3 | 0 | 0 | A | C | -2.3623 | 0.7051 | 0.0008071 | ---?? | 0 | 0.3704 | intergenic | |
| rs3792723 | 5 | 670706 | 4 | 0 | 0 | A | G | 0.9514 | 0.284 | 0.0008072 | +++?+ | 0 | 0.5417 | intron | TPPP |
| rs34761192 | 9 | 32914907 | 4 | 0 | 0 | A | T | -2.2833 | 0.6815 | 0.0008072 | ---?- | 54.1 | 0.08849 | intergenic | |
| rs11066085 | 12 | 112355949 | 5 | 0 | 0 | T | G | 0.7703 | 0.2299 | 0.0008073 | +-+++ | 56.7 | 0.05562 | intergenic | |
| rs7770203 | 6 | 116526409 | 5 | 0 | 0 | T | C | 0.794 | 0.237 | 0.0008074 | +++++ | 0 | 0.562 | intron | NT5DC1 |
| rs4588030 | 17 | 6717883 | 3 | 0 | 0 | C | G | 3.9778 | 1.1873 | 0.0008075 | +++?? | 0 | 0.6647 | intron | TEKT1 |
| rs71568147 | 6 | 103819901 | 4 | 0 | 0 | T | C | 1.4067 | 0.4199 | 0.0008075 | +++?+ | 0 | 0.8595 | intergenic | |
| 1:118095687 | 1 | 118095687 | 3 | 0 | 0 | CAAAAA | C | -1.0626 | 0.3172 | 0.0008076 | ---?? | 0 | 0.4723 | | |
| rs76186109 | 19 | 56073000 | 3 | 0 | 0 | A | G | 1.5991 | 0.4773 | 0.0008077 | +++?? | 0 | 0.4425 | intergenic | |
| rs6866655 | 5 | 165455512 | 3 | 0 | 0 | A | T | 2.7008 | 0.8062 | 0.0008078 | +++?? | 62.1 | 0.07126 | intergenic | |
| rs17034612 | 2 | 68138416 | 4 | 0 | 0 | A | G | -4.0325 | 1.2037 | 0.0008078 | ---?- | 0 | 0.4266 | intergenic | |
| rs77265144 | 15 | 66312194 | 2 | 0 | 0 | A | T | -7.4869 | 2.2349 | 0.0008081 | ?-??- | 0 | 0.3901 | intron | MEGF11 |
| 17:51139867 | 17 | 51139867 | 3 | 0 | 0 | T | TTTC | 1.4231 | 0.4248 | 0.0008084 | +++?? | 3.1 | 0.3561 | | |
| 18:37405828 | 18 | 37405828 | 3 | 0 | 0 | T | TTA | -0.9393 | 0.2804 | 0.0008085 | ---?? | 36.7 | 0.2061 | | |
| rs6601839 | 10 | 4390790 | 4 | 0 | 0 | T | C | -0.9765 | 0.2915 | 0.0008087 | ---?- | 0 | 0.7612 | intergenic | |
| rs2570130 | 15 | 87608945 | 4 | 0 | 0 | T | C | 0.8851 | 0.2642 | 0.0008091 | +++?+ | 0 | 0.642 | intergenic | |
| rs147630753 | 6 | 63268377 | 3 | 0 | 0 | A | G | -2.3631 | 0.7055 | 0.0008092 | ---?? | 0 | 0.3708 | intergenic | |
| rs3088113 | 12 | 106633569 | 3 | 0 | 0 | C | G | -1.8641 | 0.5565 | 0.0008095 | ---?? | 0 | 0.7146 | missense | CKAP4 |
| rs9393519 | 6 | 24056489 | 5 | 0 | 0 | T | G | -1.2016 | 0.3587 | 0.0008096 | ----+ | 0 | 0.8369 | intergenic |  |
| rs78105443 | 4 | 100554436 | 4 | 0 | 0 | A | G | 1.785 | 0.5329 | 0.0008096 | -++?+ | 43.7 | 0.1492 | downstream-gene | |
| 1:182028845 | 1 | 182028845 | 3 | 0 | 0 | CA | C | -2.9906 | 0.8929 | 0.0008096 | ---?? | 0 | 0.8691 | | |
| rs2782374 | 6 | 63337882 | 4 | 0 | 0 | T | G | -2.2956 | 0.6854 | 0.0008097 | ---?- | 0 | 0.4055 | intergenic | |
| rs11006919 | 10 | 28498112 | 4 | 0 | 0 | A | G | 1.9422 | 0.5798 | 0.0008097 | +++?- | 22.5 | 0.2755 | intron | MPP7 |
| rs2116727 | 5 | 158193049 | 4 | 0 | 0 | A | G | 2.3986 | 0.7161 | 0.0008098 | +++?+ | 0 | 0.97 | intron | EBF1 |
| rs60469982 | 19 | 12826710 | 4 | 0 | 0 | A | G | -3.4882 | 1.0414 | 0.0008098 | ----? | 51.6 | 0.1024 | intron | TNPO2 |
| 4:63342873 | 4 | 63342873 | 3 | 0 | 0 | CT | C | 0.9541 | 0.2848 | 0.0008098 | +++?? | 42.1 | 0.1778 | | |
| 9:106455642 | 9 | 106455642 | 3 | 0 | 0 | A | AT | -1.5476 | 0.462 | 0.0008098 | ---?? | 0 | 0.7696 | | |
| rs4695286 | 4 | 47852625 | 5 | 0 | 0 | T | C | -1.6433 | 0.4906 | 0.0008099 | ----- | 12.4 | 0.3346 | intron | NFXL1 |
| rs3129951 | 6 | 32358286 | 5 | 0 | 0 | T | C | -0.7752 | 0.2314 | 0.0008101 | +---- | 39.4 | 0.1586 | upstream-gene |  |
| rs57423571 | 2 | 68138529 | 4 | 0 | 0 | A | C | 4.0316 | 1.2037 | 0.0008101 | +++?+ | 0 | 0.4272 | intergenic | |
| rs79590084 | 1 | 117861595 | 4 | 0 | 0 | A | C | 1.8288 | 0.546 | 0.0008103 | +++?+ | 0 | 0.7064 | intergenic | |
| 2:79635416 | 2 | 79635416 | 3 | 0 | 0 | T | TTTA | -0.9216 | 0.2752 | 0.0008104 | ---?? | 50.3 | 0.1339 | | |
| rs28879456 | 9 | 44252675 | 3 | 0 | 0 | A | G | 1.7265 | 0.5155 | 0.0008105 | +++?? | 0 | 0.8573 | intergenic | |
| rs116972220 | 13 | 67125356 | 4 | 0 | 0 | T | C | -3.437 | 1.0262 | 0.0008106 | ---?- | 0 | 0.5784 | intron | PCDH9 |
| 3:119581371 | 3 | 119581371 | 3 | 0 | 0 | T | TA | -0.973 | 0.2905 | 0.0008107 | ---?? | 56.8 | 0.09858 | | |
| rs6474563 | 9 | 10615489 | 4 | 0 | 0 | A | G | 1.3853 | 0.4137 | 0.0008114 | +++?- | 0 | 0.602 | upstream-gene | |

| | | | | | | | | | | | | | | | |
|---|---|---|---|---|---|---|---|---|---|---|---|---|---|---|---|
| rs76267749 | 13 | 37303811 | 4 | 0 | 0 | T | C | 2.2426 | 0.6697 | 0.0008117 | +++?+ | 0 | 0.6515 | intergenic | |
| rs73334977 | 8 | 114878680 | 4 | 0 | 0 | T | C | 1.8031 | 0.5385 | 0.0008121 | +++?- | 2.2 | 0.3813 | intergenic | |
| rs55770649 | 19 | 12826863 | 4 | 0 | 0 | T | C | -3.4871 | 1.0414 | 0.0008122 | ----? | 51.7 | 0.1019 | intron | TNPO2 |
| rs11776799 | 8 | 123877155 | 4 | 0 | 0 | A | G | 1.0683 | 0.319 | 0.0008124 | +++?+ | 17.3 | 0.3049 | intron | ZHX2 |
| rs114981407 | 5 | 39029140 | 4 | 0 | 0 | A | T | -3.144 | 0.9389 | 0.0008124 | ---?- | 0 | 0.7626 | intron | RICTOR |
| rs7632824 | 3 | 183647081 | 4 | 0 | 0 | T | C | 1.0723 | 0.3202 | 0.0008125 | +-+?+ | 48.1 | 0.123 | intron | ABCC5 |
| rs76002428 | 10 | 131477903 | 4 | 0 | 0 | A | G | 1.2454 | 0.3719 | 0.0008126 | +++?+ | 0 | 0.8181 | intron | MGMT |
| rs11715619 | 3 | 196334416 | 4 | 0 | 0 | A | G | 0.9415 | 0.2812 | 0.0008128 | +++?+ | 0 | 0.8837 | intergenic | |
| rs7205113 | 16 | 84847823 | 4 | 0 | 0 | T | C | 1.2065 | 0.3603 | 0.000813 | +++?- | 0 | 0.5808 | intergenic | |
| rs114051008 | 9 | 72170060 | 4 | 0 | 0 | T | C | 2.871 | 0.8575 | 0.0008131 | +++?+ | 0 | 0.9961 | intron | APBA1 |
| rs6673004 | 1 | 30370065 | 4 | 0 | 0 | A | G | 1.1315 | 0.3379 | 0.0008133 | +++?+ | 0 | 0.7219 | intergenic | |
| rs117246429 | 21 | 28344968 | 2 | 0 | 0 | T | C | -6.1106 | 1.825 | 0.0008133 | --??? | 0 | 0.4468 | intergenic | |
| 15:27241768 | 15 | 27241768 | 3 | 0 | 0 | CTTATTT | CTTATTT | -2.6398 | 0.7885 | 0.0008139 | -+-?? | 63.1 | 0.06635 | | |
| 16:52918098 | 16 | 52918098 | 3 | 0 | 0 | A | AG | 0.8699 | 0.2598 | 0.0008139 | +++?? | 0 | 0.5682 | | |
| rs8113527 | 19 | 12824447 | 4 | 0 | 0 | A | G | 3.4934 | 1.0434 | 0.000814 | ++++? | 51.1 | 0.1051 | intron | TNPO2 |
| rs10019243 | 4 | 171627133 | 4 | 0 | 0 | A | G | -0.8812 | 0.2632 | 0.000814 | ---?- | 48.6 | 0.1201 | intergenic | |
| rs10231679 | 7 | 82428803 | 4 | 0 | 0 | T | C | -0.847 | 0.253 | 0.000814 | -+-?- | 45.1 | 0.1409 | intron | PCLO |
| rs6499108 | 16 | 52847573 | 4 | 0 | 0 | T | C | -0.8178 | 0.2443 | 0.0008141 | ---?- | 0 | 0.7948 | intergenic | |
| rs6511835 | 19 | 12824853 | 4 | 0 | 0 | A | G | -3.4923 | 1.0431 | 0.0008141 | ----? | 51.2 | 0.1048 | intron | TNPO2 |
| rs10834241 | 11 | 23958837 | 4 | 0 | 0 | A | G | -1.2266 | 0.3664 | 0.0008141 | ---?- | 0 | 0.8768 | intergenic | |
| rs56299514 | 15 | 87562620 | 3 | 0 | 0 | A | G | 0.9219 | 0.2754 | 0.0008143 | +++?? | 26.9 | 0.2547 | intron | AGBL1 |
| rs140865787 | 4 | 100552764 | 4 | 0 | 0 | T | C | -1.7825 | 0.5324 | 0.0008143 | +--?- | 43.7 | 0.149 | downstream-gene | |
| 1:98600662 | 1 | 98600662 | 3 | 0 | 0 | A | AT | 1.3525 | 0.404 | 0.0008143 | +++?? | 46 | 0.1571 | | |
| rs185768634 | 17 | 31986847 | 4 | 0 | 0 | T | C | -1.7536 | 0.5238 | 0.0008146 | ---?- | 0 | 0.4207 | intron | ASIC2 |
| rs76876166 | 16 | 1486598 | 4 | 0 | 0 | A | G | 1.2253 | 0.366 | 0.0008147 | +++?+ | 2.3 | 0.381 | intron | CCDC154 |
| rs59861013 | 15 | 87562630 | 3 | 0 | 0 | T | G | 0.9219 | 0.2754 | 0.0008148 | +++?? | 26.9 | 0.2548 | intron | AGBL1 |
| rs10269313 | 7 | 82428754 | 4 | 0 | 0 | C | G | -0.8469 | 0.253 | 0.0008149 | -+-?- | 45.1 | 0.1408 | intron | PCLO |
| rs13379414 | 14 | 43502084 | 4 | 0 | 0 | A | G | -1.6822 | 0.5025 | 0.0008149 | ---?+ | 0 | 0.6124 | intergenic | |
| rs62451211 | 7 | 40667551 | 3 | 0 | 0 | A | G | 4.2972 | 1.2836 | 0.0008149 | ?++?+ | 42.7 | 0.1744 | intron | C7orf10 |
| 16:27980925 | 16 | 27980925 | 3 | 0 | 0 | T | TA | 1.9665 | 0.5874 | 0.0008152 | +++?? | 0 | 0.8995 | | |
| rs76376003 | 1 | 192465280 | 4 | 0 | 0 | T | C | -1.1326 | 0.3383 | 0.0008153 | ---?+ | 21.3 | 0.2824 | intergenic | |
| rs189197630 | 1 | 184556349 | 2 | 0 | 0 | A | G | -5.5433 | 1.6559 | 0.0008153 | --??? | 0 | 0.4382 | intron | C1orf21 |
| 17:51142108 | 17 | 51142108 | 3 | 0 | 0 | T | TAC | -1.352 | 0.4039 | 0.0008153 | ---?? | 13.3 | 0.3157 | | |
| rs79125032 | 16 | 6005381 | 3 | 0 | 0 | A | G | -0.8949 | 0.2673 | 0.0008157 | ---?? | 0 | 0.9167 | intron | RBFOX1 |
| 1:182015285 | 1 | 182015285 | 3 | 0 | 0 | A | AT | 1.1545 | 0.3449 | 0.0008157 | +++?? | 19.2 | 0.2899 | | |
| rs56379111 | 3 | 61118262 | 4 | 0 | 0 | T | C | -3.1049 | 0.9276 | 0.0008159 | ---?- | 0 | 0.771 | intron | FHIT |
| rs4469231 | 5 | 737564 | 3 | 0 | 0 | T | C | 1.5039 | 0.4493 | 0.0008161 | +++?? | 4.5 | 0.3508 | intron | ZDHHC11B |
| rs2009535 | 13 | 30324744 | 5 | 0 | 0 | A | G | 1.2237 | 0.3656 | 0.0008164 | +++++ | 0 | 0.5561 | intergenic | |
| rs12990532 | 2 | 230915321 | 4 | 0 | 0 | T | C | 1.4228 | 0.4251 | 0.0008164 | +++?+ | 18.4 | 0.2988 | intron | SLC16A14 |
| rs34852239 | 13 | 20811820 | 4 | 0 | 0 | A | G | -1.102 | 0.3292 | 0.0008164 | ---?- | 0 | 0.4886 | intergenic | |
| rs6466178 | 7 | 107347796 | 4 | 0 | 0 | A | T | 0.8486 | 0.2535 | 0.0008165 | +++?+ | 0 | 0.5886 | intron | SLC26A4 |
| rs67785287 | 4 | 67905347 | 4 | 0 | 0 | A | G | 1.3337 | 0.3985 | 0.0008165 | +++?+ | 59.8 | 0.05831 | intergenic | |
| rs2182776 | 13 | 30322970 | 5 | 0 | 0 | A | G | 1.2226 | 0.3653 | 0.0008166 | +++++ | 0 | 0.5554 | intergenic | |
| rs1107154 | 5 | 165387313 | 5 | 0 | 0 | A | G | 0.7321 | 0.2187 | 0.0008167 | +++-+ | 40.7 | 0.1503 | intergenic | |
| rs9994314 | 4 | 190013831 | 4 | 0 | 0 | A | C | -0.8505 | 0.2541 | 0.000817 | ---?- | 0 | 0.5042 | intergenic | |
| rs144224547 | 8 | 134966338 | 2 | 0 | 0 | T | G | -6.3711 | 1.9036 | 0.0008172 | -?-?? | 35.3 | 0.2139 | intergenic | |
| rs1026156 | 4 | 123694013 | 4 | 0 | 0 | T | C | 2.1199 | 0.6334 | 0.0008173 | +++?+ | 60.6 | 0.0547 | intergenic | |
| rs4367043 | 3 | 5390506 | 4 | 0 | 0 | A | T | 1.3461 | 0.4022 | 0.0008175 | +++?+ | 2.2 | 0.3814 | intergenic | |
| rs11956982 | 5 | 39586352 | 4 | 0 | 0 | T | C | -0.996 | 0.2976 | 0.0008175 | ---?+ | 0 | 0.8024 | intergenic | |
| rs17211712 | 14 | 21454062 | 4 | 0 | 0 | T | C | 0.877 | 0.262 | 0.0008176 | +++?+ | 25.5 | 0.2586 | upstream-gene | |
| rs140426418 | 2 | 63578617 | 3 | 0 | 0 | A | G | 3.7816 | 1.1299 | 0.0008176 | +?+?+ | 56.4 | 0.1012 | intron | WDPCP |
| rs4798111 | 18 | 377746 | 4 | 0 | 0 | T | G | 1.0588 | 0.3164 | 0.0008177 | +++?+ | 0 | 0.9213 | intron | COLEC12 |
| 14:35338122 | 14 | 35338122 | 3 | 0 | 0 | G | GTA | 0.9623 | 0.2875 | 0.0008177 | +++?? | 0 | 0.4334 | | |
| rs2182775 | 13 | 30323020 | 5 | 0 | 0 | C | G | -1.2225 | 0.3653 | 0.0008178 | ----- | 0 | 0.5555 | intergenic |  |
| rs62065309 | 16 | 58699365 | 3 | 0 | 0 | T | C | 4.4762 | 1.3375 | 0.0008178 | +?+?+ | 0 | 0.6771 | downstream-gene | |
| rs77858651 | 5 | 132977783 | 4 | 0 | 0 | A | G | -3.0423 | 0.909 | 0.0008178 | ---?- | 0 | 0.8409 | intergenic | |
| rs145622340 | 3 | 130368337 | 2 | 0 | 0 | T | C | 7.2642 | 2.1706 | 0.0008178 | ?+??+ | 36.7 | 0.2088 | synonymous | COL6A6 |
| rs186681177 | 1 | 98600673 | 3 | 0 | 0 | A | T | 1.3519 | 0.404 | 0.0008179 | +++?? | 46 | 0.157 | intergenic | |
| rs2083074 | 15 | 61348432 | 4 | 0 | 0 | T | C | -0.8521 | 0.2546 | 0.000818 | ---?- | 0 | 0.9033 | intron | RORA |

| | | | | | | | | | | | | | | | |
|---|---|---|---|---|---|---|---|---|---|---|---|---|---|---|---|
| rs3850737 | 8 | 64921643 | 4 | 0 | 0 | T | C | -0.8297 | 0.2479 | 0.0008181 | ---?+ | 0 | 0.4887 | intergenic | |
| rs4239467 | 18 | 52742732 | 4 | 0 | 0 | C | G | 4.1945 | 1.2534 | 0.0008182 | +++?+ | 0 | 0.8662 | intergenic | |
| rs4589643 | 18 | 52753240 | 4 | 0 | 0 | T | C | 4.1317 | 1.2346 | 0.0008182 | +++?+ | 0 | 0.8662 | intergenic | |
| 15:87563292 | 15 | 87563292 | 3 | 0 | 0 | G | GA | -0.9217 | 0.2754 | 0.0008182 | ---?? | 26.8 | 0.2551 | | |
| rs188300214 | 4 | 166238319 | 4 | 0 | 0 | T | C | -3.6141 | 1.08 | 0.0008183 | ---?- | 0 | 0.9523 | intron | KLHL2 |
| rs1496937 | 4 | 190004805 | 4 | 0 | 0 | A | G | -0.8408 | 0.2513 | 0.0008184 | ---?- | 33.2 | 0.2133 | intergenic | |
| rs4769023 | 13 | 30322833 | 5 | 0 | 0 | A | G | 1.2222 | 0.3652 | 0.0008187 | +++++ | 0 | 0.5557 | intergenic | |
| rs34959638 | 16 | 12070207 | 4 | 0 | 0 | A | G | 0.8876 | 0.2652 | 0.0008187 | +++?+ | 15.4 | 0.3149 | upstream-gene | |
| rs73244433 | 4 | 47871842 | 5 | 0 | 0 | T | C | 1.6453 | 0.4917 | 0.0008188 | +++++ | 12.5 | 0.3342 | intron | NFXL1 |
| rs142429045 | 22 | 28786875 | 3 | 0 | 0 | T | C | -2.346 | 0.7011 | 0.0008188 | ---?? | 0 | 0.482 | intron | TTC28 |
| rs114002582 | 4 | 100553091 | 4 | 0 | 0 | A | G | 1.7821 | 0.5325 | 0.0008189 | -++?+ | 43.8 | 0.1484 | downstream-gene | |
| 2:170469930 | 2 | 170469930 | 3 | 0 | 0 | G | GT | 1.2265 | 0.3665 | 0.0008189 | +++?? | 0 | 0.8543 | | |
| rs144782417 | 9 | 118381465 | 4 | 0 | 0 | T | C | 2.2393 | 0.6692 | 0.000819 | +++?+ | 0 | 0.6984 | intergenic | |
| rs202247462 | 12 | 2244448 | 3 | 0 | 0 | T | C | -1.8506 | 0.553 | 0.000819 | ---?? | 59.3 | 0.08556 | intron | CACNA1C |
| rs75687346 | 16 | 4277580 | 2 | 0 | 0 | C | G | 6.4124 | 1.9163 | 0.0008191 | ?+??+ | 0 | 0.5697 | intron | SRL |
| rs4239468 | 18 | 52742748 | 4 | 0 | 0 | A | T | -4.1938 | 1.2533 | 0.0008194 | ---?- | 0 | 0.8662 | intergenic | |
| rs13001908 | 2 | 107701621 | 4 | 0 | 0 | A | G | -1.1003 | 0.3288 | 0.0008194 | ---?- | 0 | 0.6818 | intergenic | |
| rs188505954 | 11 | 7247873 | 3 | 0 | 0 | T | C | -3.0098 | 0.8995 | 0.0008196 | ---?? | 0 | 0.9708 | intergenic | |
| rs458434 | 6 | 33242825 | 5 | 0 | 0 | A | C | 1.0236 | 0.3059 | 0.0008197 | +++++ | 20.3 | 0.2853 | intron | RPS18 |
| rs458679 | 6 | 33242492 | 5 | 0 | 0 | C | G | -1.0236 | 0.3059 | 0.0008198 | ----- | 20.3 | 0.2853 | intron | RPS18 |
| rs466886 | 6 | 33240699 | 5 | 0 | 0 | T | G | 1.0236 | 0.3059 | 0.0008198 | +++++ | 20.3 | 0.2853 | intron | RPS18 |
| rs4291041 | 5 | 38723715 | 4 | 0 | 0 | A | T | -1.2302 | 0.3676 | 0.0008198 | ---?- | 53.8 | 0.08994 | intergenic | |
| rs79860514 | 14 | 46602646 | 4 | 0 | 0 | T | C | -1.0245 | 0.3062 | 0.0008198 | ---?- | 37 | 0.19 | intron | LINC00871 |
| rs147813227 | 6 | 31224488 | 3 | 0 | 0 | T | C | 0.9351 | 0.2795 | 0.00082 | +++?? | 41.7 | 0.1799 | intergenic | |
| rs12601665 | 17 | 47322264 | 5 | 0 | 0 | C | G | 0.7119 | 0.2128 | 0.0008202 | +++++ | 0 | 0.8455 | upstream-gene | |
| rs945366 | 13 | 20801844 | 4 | 0 | 0 | A | G | -1.1139 | 0.3329 | 0.0008203 | ---?- | 0 | 0.961 | intron | GJB6 |
| rs144714392 | 6 | 31224494 | 3 | 0 | 0 | T | C | 0.935 | 0.2795 | 0.0008204 | +++?? | 41.7 | 0.18 | intergenic | |
| 15:87578009 | 15 | 87578009 | 3 | 0 | 0 | T | C | 1.2526 | 0.3744 | 0.0008206 | +++?? | 0 | 0.7006 | intergenic | |
| rs76667108 | 14 | 90049406 | 4 | 0 | 0 | T | C | 3.7323 | 1.1155 | 0.0008206 | +++?+ | 0 | 0.5608 | intron | FOXN3 |
| rs72979352 | 4 | 162210511 | 2 | 0 | 0 | T | C | -6.0586 | 1.8109 | 0.0008207 | ?--?? | 0 | 0.582 | intergenic | |
| rs140722727 | 6 | 31224495 | 3 | 0 | 0 | T | G | -0.935 | 0.2795 | 0.0008207 | ---?? | 41.7 | 0.1801 | intergenic | |
| rs1586360 | 7 | 46176155 | 4 | 0 | 0 | T | C | -1.8985 | 0.5675 | 0.0008209 | ---?- | 0 | 0.9702 | intergenic | |
| rs181813800 | 21 | 46398632 | 2 | 0 | 0 | T | C | -6.7268 | 2.0106 | 0.0008209 | --??? | 0 | 0.6783 | downstream-gene | |
| 2:231876435 | 2 | 231876435 | 3 | 0 | 0 | CAG | C | 1.734 | 0.5183 | 0.0008209 | +++?? | 0 | 0.9179 | | |
| rs11586134 | 1 | 64730659 | 4 | 0 | 0 | A | T | -1.2146 | 0.363 | 0.000821 | ---?- | 0 | 0.4346 | intron | UBE2U |
| rs11883224 | 19 | 12799686 | 4 | 0 | 0 | A | G | 3.5024 | 1.0469 | 0.0008214 | ++++? | 49.4 | 0.1152 | downstream-gene | |
| rs73834877 | 3 | 68232284 | 4 | 0 | 0 | A | G | 2.1404 | 0.6398 | 0.0008214 | +++?+ | 6.6 | 0.36 | intron | FAM19A1 |
| rs116504256 | 2 | 35840684 | 3 | 0 | 0 | A | G | -3.648 | 1.0904 | 0.0008215 | ---?? | 0 | 0.633 | intergenic | |
| rs13145872 | 4 | 190015291 | 4 | 0 | 0 | T | G | -0.8399 | 0.2511 | 0.0008216 | ---?- | 28.8 | 0.239 | intergenic | |
| 13:60033826 | 13 | 60033826 | 3 | 0 | 0 | G | GT | 0.8678 | 0.2594 | 0.0008216 | +++?? | 23.2 | 0.272 | | |
| rs1115280 | 22 | 48436543 | 4 | 0 | 0 | A | G | 1.2671 | 0.3788 | 0.0008219 | +++?+ | 30.7 | 0.2282 | intergenic | |
| rs2968164 | 3 | 25765252 | 5 | 0 | 0 | A | T | 0.9509 | 0.2843 | 0.0008222 | -++++ | 11.2 | 0.3419 | intron | NGLY1 |
| rs4433886 | 18 | 52743556 | 4 | 0 | 0 | T | C | 4.1887 | 1.2521 | 0.0008222 | +++?+ | 0 | 0.8661 | intergenic | |
| rs464921 | 6 | 33240506 | 5 | 0 | 0 | A | G | -1.0234 | 0.3059 | 0.0008223 | ----- | 20.3 | 0.2854 | intron-near-splice | RPS18 |
| rs2782375 | 6 | 63339097 | 4 | 0 | 0 | T | C | 2.2927 | 0.6854 | 0.0008224 | +++?+ | 0 | 0.4001 | intergenic | |
| rs142071896 | 2 | 168299949 | 4 | 0 | 0 | A | G | -2.3154 | 0.6922 | 0.0008225 | ---?- | 0 | 0.6548 | intergenic | |
| 5:108621263 | 5 | 108621263 | 3 | 0 | 0 | CA | C | -1.4119 | 0.4221 | 0.0008225 | ---?? | 0 | 0.9853 | | |
| rs71424092 | 13 | 20811907 | 4 | 0 | 0 | T | C | -1.1023 | 0.3295 | 0.0008226 | ---?- | 0 | 0.4904 | intergenic | |
| rs465658 | 6 | 33239905 | 5 | 0 | 0 | A | G | -1.0234 | 0.3059 | 0.0008228 | ----- | 20.2 | 0.2862 | intron-near-splice | RPS18 |
| rs79690264 | 6 | 31224098 | 3 | 0 | 0 | T | G | -0.9364 | 0.2799 | 0.0008228 | ---?? | 39.8 | 0.19 | intergenic | |
| 10:9315579 | 10 | 9315579 | 3 | 0 | 0 | T | TGGGC | -1.777 | 0.5313 | 0.0008229 | ---?? | 0 | 0.6731 | | |
| rs11667628 | 19 | 6638837 | 4 | 0 | 0 | A | G | 0.8412 | 0.2515 | 0.0008231 | +++?+ | 0 | 0.7689 | intergenic | |
| 2:103455906 | 2 | 103455906 | 3 | 0 | 0 | CTA | C | 1.3306 | 0.3978 | 0.0008231 | +++?? | 0 | 0.6718 | | |
| rs1621340 | 6 | 117298959 | 4 | 0 | 0 | A | G | 0.7889 | 0.2359 | 0.0008233 | +++?+ | 0 | 0.7199 | intergenic | |
| rs79510007 | 6 | 95454311 | 4 | 0 | 0 | A | C | -2.6034 | 0.7783 | 0.0008233 | ---?- | 0 | 0.8513 | intergenic | |
| rs415748 | 6 | 33238824 | 5 | 0 | 0 | A | G | -1.0234 | 0.306 | 0.0008234 | ----- | 20 | 0.287 | intron | VPS52 |
| rs1641453 | 2 | 40448015 | 4 | 0 | 0 | A | C | -1.0296 | 0.3078 | 0.0008235 | ---?+ | 1.5 | 0.3846 | intron | SLC8A1,SLC8A1-AS1 |
| 5:82609737 | 5 | 82609737 | 3 | 0 | 0 | T | TAA | -2.3764 | 0.7105 | 0.0008236 | ---?? | 0 | 0.3999 | | |
| rs7522545 | 1 | 7126671 | 4 | 0 | 0 | T | C | -0.8495 | 0.254 | 0.000824 | ---?- | 40.8 | 0.1671 | intron | CAMTA1 |

| | | | | | | | | | | | | | | | |
|---|---|---|---|---|---|---|---|---|---|---|---|---|---|---|---|
| rs60499412 | 19 | 12824015 | 4 | 0 | 0 | A | G | 3.4913 | 1.0439 | 0.000824 | ++++? | 51.1 | 0.1053 | intron | TNPO2 |
| rs62065305 | 16 | 58684165 | 3 | 0 | 0 | T | C | 4.4384 | 1.327 | 0.0008241 | +?+?+ | 0 | 0.7031 | intergenic | |
| rs392317 | 2 | 40473297 | 4 | 0 | 0 | A | G | -1.1878 | 0.3551 | 0.0008242 | ---?- | 0 | 0.7942 | intron | SLC8A1,SLC8A1-AS1 |
| rs6000298 | 22 | 36923480 | 4 | 0 | 0 | C | G | -0.856 | 0.256 | 0.0008243 | ---?- | 0 | 0.8895 | intron | EIF3D |
| rs10268057 | 7 | 141199472 | 4 | 0 | 0 | A | T | -3.6898 | 1.1032 | 0.0008243 | ---?- | 20.1 | 0.2893 | intergenic | |
| rs58730238 | 12 | 66018252 | 3 | 0 | 0 | T | C | -3.8678 | 1.1565 | 0.0008243 | ---?? | 0 | 0.6445 | intergenic | |
| rs72966488 | 2 | 208280779 | 4 | 0 | 0 | T | C | -1.4998 | 0.4484 | 0.0008245 | ---?- | 0 | 0.477 | intergenic | |
| rs13015861 | 2 | 222485359 | 4 | 0 | 0 | A | T | -1.8405 | 0.5503 | 0.0008248 | ---?- | 0 | 0.9039 | intergenic | |
| rs34448371 | 2 | 138942607 | 4 | 0 | 0 | C | G | -1.6059 | 0.4802 | 0.0008249 | ---?- | 0 | 0.7885 | intergenic | |
| rs375585740 | 7 | 153662029 | 3 | 0 | 0 | T | C | 2.3983 | 0.7172 | 0.000825 | +++?? | 0 | 0.7102 | intron | DPP6 |
| rs28542780 | 4 | 8663924 | 4 | 0 | 0 | A | G | 3.2321 | 0.9665 | 0.0008251 | +++?- | 50.7 | 0.1074 | intergenic | |
| rs13110980 | 4 | 63337758 | 4 | 0 | 0 | T | C | 0.9101 | 0.2721 | 0.0008252 | +++?+ | 44.1 | 0.1468 | intergenic | |
| rs78865813 | 12 | 109982025 | 3 | 0 | 0 | T | C | 2.5148 | 0.752 | 0.0008253 | +++?? | 0 | 0.7616 | intergenic | |
| rs80286802 | 4 | 153874309 | 5 | 0 | 0 | T | C | 2.6141 | 0.7817 | 0.0008254 | -++++ | 37.9 | 0.1684 | intron | FHDC1 |
| rs990206 | 4 | 20758408 | 4 | 0 | 0 | A | T | -0.8834 | 0.2642 | 0.0008257 | ---?- | 0 | 0.8198 | intron | KCNIP4 |
| 1:107993730 | 1 | 107993730 | 3 | 0 | 0 | CT | C | 0.8526 | 0.255 | 0.0008257 | +++?? | 0 | 0.6906 | | |
| rs55969561 | 2 | 170367783 | 4 | 0 | 0 | T | C | -1.1759 | 0.3516 | 0.0008259 | ---?- | 0 | 0.9919 | intron | KLHL41 |
| rs3769771 | 2 | 170368438 | 4 | 0 | 0 | T | C | -1.1752 | 0.3514 | 0.0008261 | ---?- | 0 | 0.9917 | intron | KLHL41 |
| rs7655727 | 4 | 11424957 | 4 | 0 | 0 | A | T | 0.8275 | 0.2475 | 0.0008263 | +++?+ | 39.3 | 0.1762 | intron | HS3ST1 |
| rs8013773 | 14 | 104636584 | 3 | 0 | 0 | A | G | 1.0127 | 0.3029 | 0.0008263 | +++?? | 0 | 0.5752 | intron | KIF26A |
| rs380590 | 2 | 40473347 | 4 | 0 | 0 | T | G | 1.1878 | 0.3552 | 0.0008264 | +++?+ | 0 | 0.7953 | intron | SLC8A1,SLC8A1-AS1 |
| rs11253452 | 10 | 858033 | 4 | 0 | 0 | A | G | -2.4015 | 0.7182 | 0.0008264 | ---?- | 0 | 0.3955 | 3-prime-UTR | LARP4B |
| rs13346030 | 19 | 12797589 | 4 | 0 | 0 | T | C | 3.51 | 1.0497 | 0.0008264 | ++++? | 49.9 | 0.1123 | upstream-gene | |
| 8:64907394 | 8 | 64907394 | 3 | 0 | 0 | A | AT | -0.9596 | 0.287 | 0.0008265 | ---?? | 0 | 0.9054 | | |
| rs1349881 | 2 | 52949270 | 4 | 0 | 0 | A | G | 1.2903 | 0.3859 | 0.0008267 | +++?+ | 0 | 0.7897 | intergenic | |
| rs55836157 | 1 | 101864228 | 4 | 0 | 0 | A | G | -1.4814 | 0.443 | 0.0008267 | ---?- | 0 | 0.7691 | intergenic | |
| rs7254207 | 19 | 12801715 | 4 | 0 | 0 | A | C | -3.4964 | 1.0457 | 0.0008268 | ----? | 49.2 | 0.1161 | intron | FBXW9 |
| rs111683084 | 5 | 29439556 | 3 | 0 | 0 | A | G | -3.4203 | 1.0229 | 0.0008269 | ---?? | 17.5 | 0.2978 | intergenic | |
| rs76901383 | 6 | 32669817 | 5 | 0 | 1 | T | C | -1.1002 | 0.3291 | 0.000827 | ----- | 0.4 | 0.4041 | intergenic |  |
| rs78280604 | 5 | 158245189 | 4 | 0 | 0 | T | C | 2.4873 | 0.7439 | 0.000827 | +++?+ | 0 | 0.958 | intron | EBF1 |
| rs78103518 | 11 | 106802349 | 3 | 0 | 0 | T | G | -1.3743 | 0.411 | 0.0008271 | ---?? | 0 | 0.5464 | intron | GUCY1A2 |
| rs74908205 | 11 | 106802348 | 3 | 0 | 0 | A | G | 1.3743 | 0.411 | 0.0008273 | +++?? | 0 | 0.546 | intron | GUCY1A2 |
| rs180996633 | 10 | 134885068 | 2 | 0 | 0 | A | G | 7.4391 | 2.2249 | 0.0008273 | +???+ | 53.9 | 0.1409 | intron | GPR123 |
| 1:172866786 | 1 | 172866786 | 3 | 0 | 0 | CT | C | -0.8768 | 0.2623 | 0.0008273 | ---?? | 0 | 0.4237 | | |
| rs72620504 | 19 | 1641296 | 4 | 0 | 0 | A | G | 2.658 | 0.795 | 0.0008275 | +++?+ | 0 | 0.8122 | intron | TCF3 |
| 2:19515395 | 2 | 19515395 | 3 | 0 | 0 | CAA | C | 0.8624 | 0.2579 | 0.0008275 | +++?? | 0 | 0.7977 | | |
| rs135428 | 22 | 44657259 | 4 | 0 | 0 | T | C | -0.9062 | 0.271 | 0.0008276 | ---?+ | 48.8 | 0.1189 | intron | KIAA1644 |
| rs6705938 | 2 | 68137125 | 4 | 0 | 0 | T | C | 4.0188 | 1.202 | 0.0008276 | +++?+ | 0 | 0.4195 | intergenic | |
| rs114700122 | 3 | 109741385 | 4 | 0 | 0 | A | G | -2.4766 | 0.7407 | 0.0008276 | ---?- | 0 | 0.5548 | intergenic | |
| rs62096989 | 18 | 71751072 | 4 | 0 | 0 | T | C | 1.3991 | 0.4185 | 0.0008278 | +++?+ | 21.4 | 0.2818 | intron | FBXO15 |
| rs6538848 | 12 | 98628305 | 4 | 0 | 0 | A | G | 1.4794 | 0.4425 | 0.000828 | +-+?+ | 60.4 | 0.05579 | intergenic | |
| rs1620521 | 6 | 117299048 | 4 | 0 | 0 | T | C | 0.7886 | 0.2359 | 0.0008281 | +++?+ | 0 | 0.7204 | intergenic | |
| rs7258853 | 19 | 12799245 | 4 | 0 | 0 | C | G | 3.5022 | 1.0476 | 0.0008283 | ++++? | 49.5 | 0.1146 | downstream-gene | |
| rs79700555 | 1 | 42672413 | 4 | 0 | 0 | T | C | 1.7579 | 0.5258 | 0.0008284 | +++?+ | 0 | 0.5392 | intron | FOXJ3 |
| rs11020136 | 11 | 92728050 | 3 | 0 | 0 | T | G | 1.1262 | 0.3369 | 0.0008285 | +++?? | 0 | 0.4381 | intergenic | |
| rs113959729 | 18 | 40108351 | 4 | 0 | 0 | T | C | 1.3122 | 0.3925 | 0.0008286 | +++?+ | 0 | 0.9153 | intron | LINC00907 |
| rs4401687 | 6 | 144831355 | 4 | 0 | 0 | A | C | -0.9876 | 0.2954 | 0.0008287 | ---?- | 0 | 0.566 | intron | UTRN |
| rs79615241 | 5 | 103162693 | 4 | 0 | 0 | A | G | 1.4933 | 0.4467 | 0.0008291 | +++?+ | 0 | 0.6862 | intergenic | |
| rs7328044 | 13 | 20812563 | 4 | 0 | 0 | T | C | 1.1017 | 0.3296 | 0.0008294 | +++?+ | 0 | 0.4877 | intergenic | |
| rs34553151 | 2 | 138942217 | 4 | 0 | 0 | T | C | -1.6049 | 0.4801 | 0.0008294 | ---?- | 0 | 0.7881 | intergenic | |
| rs62030662 | 16 | 12785726 | 3 | 0 | 0 | T | C | 1.0685 | 0.3197 | 0.0008295 | +++?? | 0 | 0.8714 | intron | CPPED1 |
| rs12603271 | 17 | 71056200 | 4 | 0 | 0 | T | C | -2.2141 | 0.6624 | 0.0008297 | ---?- | 21.6 | 0.2808 | intron | SLC39A11 |
| 2:220494604 | 2 | 220494604 | 3 | 0 | 0 | A | AG | -1.3941 | 0.4171 | 0.0008297 | ---?? | 0 | 0.4457 | | |
| rs2225982 | 20 | 26272824 | 4 | 0 | 0 | C | G | 1.3262 | 0.3968 | 0.0008302 | +++?+ | 60.4 | 0.05555 | intergenic | |
| 13:45550259 | 13 | 45550259 | 3 | 0 | 0 | A | AG | -2.1271 | 0.6364 | 0.0008303 | ---?? | 0 | 0.7195 | | |
| rs114136736 | 2 | 35841795 | 3 | 0 | 0 | T | C | 3.6432 | 1.09 | 0.0008304 | +++?? | 0 | 0.6276 | intergenic | |
| rs35592480 | 7 | 112699069 | 4 | 0 | 0 | A | G | 1.1329 | 0.3389 | 0.0008306 | +++?- | 13.4 | 0.3256 | intergenic | |
| rs4610542 | 6 | 63362450 | 4 | 0 | 0 | T | G | -2.1869 | 0.6543 | 0.000831 | ---?- | 0 | 0.4377 | intergenic | |
| rs463749 | 6 | 33238962 | 5 | 0 | 0 | A | T | -1.0226 | 0.306 | 0.0008311 | ----- | 20 | 0.2876 | intron | VPS52 |

| | | | | | | | | | | | | | | | |
|---|---|---|---|---|---|---|---|---|---|---|---|---|---|---|---|
| rs78203755 | 15 | 86341697 | 4 | 0 | 0 | T | C | 1.3481 | 0.4034 | 0.0008311 | +++?+ | 0 | 0.7795 | upstream-gene | |
| rs3851005 | 16 | 12071213 | 4 | 0 | 0 | T | C | 0.8588 | 0.257 | 0.0008316 | +++?+ | 16.6 | 0.3084 | intron | SNX29 |
| rs72898559 | 2 | 184066507 | 4 | 0 | 0 | A | C | 1.1596 | 0.347 | 0.0008316 | +++?+ | 0 | 0.6982 | intergenic | |
| rs6005783 | 22 | 28838013 | 4 | 0 | 0 | T | C | 2.3353 | 0.6988 | 0.0008318 | +++?+ | 0 | 0.6432 | intron | TTC28 |
| rs147138607 | 1 | 181855562 | 3 | 0 | 0 | C | G | -3.6934 | 1.1051 | 0.0008318 | ---?? | 19.1 | 0.2904 | intergenic | |
| rs3742230 | 13 | 113509919 | 5 | 0 | 0 | A | G | -0.8008 | 0.2396 | 0.0008321 | ----- | 0 | 0.9286 | intron | ATP11A |
| rs13344177 | 19 | 12798027 | 4 | 0 | 0 | A | G | -3.5065 | 1.0493 | 0.0008322 | ----? | 49.8 | 0.1128 | upstream-gene |  |
| rs9965327 | 18 | 52738476 | 4 | 0 | 0 | T | C | 4.2133 | 1.2608 | 0.0008324 | +++?+ | 0 | 0.8647 | intergenic | |
| rs111635263 | 4 | 15217358 | 3 | 0 | 0 | T | C | 1.8327 | 0.5484 | 0.0008327 | +++?? | 15.7 | 0.3054 | intergenic | |
| rs2182620 | 10 | 134590715 | 4 | 0 | 0 | T | C | 0.9218 | 0.2758 | 0.0008328 | +++?+ | 0 | 0.5116 | intron | INPP5A |
| rs76739112 | 7 | 48549947 | 4 | 0 | 0 | A | G | -2.4601 | 0.7362 | 0.0008329 | ---?- | 0 | 0.7305 | intron | ABCA13 |
| 1:64738094 | 1 | 64738094 | 3 | 0 | 0 | CAAA | C | -0.8919 | 0.2669 | 0.0008332 | ---?? | 63.9 | 0.0628 | | |
| rs12831319 | 12 | 98932707 | 4 | 0 | 0 | A | G | -0.9258 | 0.2771 | 0.0008333 | ---?- | 0 | 0.9309 | intron | TMPO |
| rs13344580 | 19 | 12797836 | 4 | 0 | 0 | T | C | -3.5067 | 1.0494 | 0.0008333 | ----? | 49.9 | 0.1125 | upstream-gene |  |
| 1:87429366 | 1 | 87429366 | 3 | 0 | 0 | CG | C | 1.0357 | 0.31 | 0.0008336 | +++?? | 0 | 0.6876 | | |
| rs595748 | 19 | 50715092 | 3 | 0 | 0 | A | T | 1.1493 | 0.344 | 0.0008337 | +++?? | 12.1 | 0.3204 | intron | MYH14 |
| rs56168069 | 8 | 252880 | 4 | 0 | 0 | A | G | 0.9802 | 0.2933 | 0.0008337 | +-+?+ | 50.1 | 0.1109 | intergenic | |
| rs62293415 | 4 | 10043886 | 5 | 0 | 0 | A | C | -0.7032 | 0.2105 | 0.0008337 | ----+ | 13.8 | 0.3261 | upstream-gene |  |
| rs11117946 | 1 | 218003361 | 4 | 0 | 0 | A | C | -0.8972 | 0.2685 | 0.000834 | ---?- | 0 | 0.8058 | intron | SPATA17 |
| rs1893057 | 11 | 95701595 | 4 | 0 | 0 | T | G | 0.8856 | 0.2651 | 0.0008343 | +++?+ | 0 | 0.4678 | intergenic | |
| rs807108 | 17 | 51240423 | 5 | 0 | 0 | A | G | 1.0397 | 0.3112 | 0.0008344 | +-++- | 6.3 | 0.3706 | intergenic | |
| rs11539247 | 19 | 11664422 | 3 | 0 | 0 | A | C | 2.4215 | 0.7248 | 0.0008344 | +++?? | 58 | 0.09233 | 3-prime-UTR | ELOF1 |
| rs17486311 | 2 | 2901761 | 4 | 0 | 0 | A | G | 1.3325 | 0.3988 | 0.0008344 | +++?+ | 0 | 0.5067 | intergenic | |
| rs187795064 | 2 | 170578931 | 3 | 0 | 0 | T | C | 1.0749 | 0.3217 | 0.0008344 | +++?? | 0 | 0.9302 | intron | PHOSPHO2-KLHL23 |
| rs55672351 | 7 | 4607963 | 3 | 0 | 0 | C | G | 0.9873 | 0.2955 | 0.0008345 | +++?? | 0 | 0.467 | intergenic | |
| rs514618 | 22 | 27646755 | 4 | 0 | 0 | C | G | 1.1625 | 0.3479 | 0.0008347 | +++?- | 32.7 | 0.216 | intergenic | |
| rs78215794 | 9 | 118316056 | 4 | 0 | 0 | T | C | 2.1804 | 0.6526 | 0.000835 | +++?+ | 0 | 0.6416 | intergenic | |
| rs10834242 | 11 | 23958901 | 4 | 0 | 0 | A | G | -1.2221 | 0.3658 | 0.0008351 | ---?- | 0 | 0.8749 | intergenic | |
| rs140981723 | 1 | 231421937 | 3 | 0 | 0 | T | C | -4.9896 | 1.4935 | 0.0008351 | --??- | 0 | 0.6018 | intergenic | |
| rs2133298 | 12 | 103260634 | 4 | 0 | 0 | T | C | -0.8177 | 0.2448 | 0.0008353 | ---?- | 0 | 0.6173 | intron | PAH |
| rs56190903 | 4 | 154923685 | 4 | 0 | 0 | T | C | 0.8429 | 0.2523 | 0.0008357 | +++?+ | 0 | 0.9101 | intergenic | |
| 10:9315140 | 10 | 9315140 | 3 | 0 | 0 | A | G | 1.7494 | 0.5237 | 0.0008358 | +++?? | 0 | 0.5184 | intergenic | |
| rs9998081 | 4 | 123725449 | 4 | 0 | 0 | A | G | -2.2044 | 0.6599 | 0.0008361 | ---?- | 53.4 | 0.09195 | intergenic | |
| rs72802536 | 16 | 47991323 | 4 | 0 | 0 | T | C | 2.8316 | 0.8477 | 0.0008361 | +++?+ | 0 | 0.6721 | intergenic | |
| rs147458182 | 6 | 109132983 | 2 | 0 | 0 | T | C | 4.7149 | 1.4114 | 0.0008361 | +?+?? | 0 | 0.686 | intergenic | |
| rs78348121 | 6 | 109127762 | 2 | 0 | 0 | T | C | 4.7277 | 1.4153 | 0.0008362 | +?+?? | 0 | 0.7208 | intergenic | |
| rs4880426 | 10 | 134590610 | 4 | 0 | 0 | A | G | 0.9219 | 0.276 | 0.0008363 | +++?+ | 0 | 0.5112 | intron | INPP5A |
| rs1368350 | 5 | 145231050 | 4 | 0 | 0 | A | G | -2.3271 | 0.6966 | 0.0008364 | ---?- | 35.3 | 0.2004 | intergenic | |
| rs943561 | 13 | 30324946 | 5 | 0 | 0 | T | C | 1.2183 | 0.3647 | 0.0008365 | +++++ | 0 | 0.5606 | intergenic | |
| rs16954225 | 17 | 52228865 | 4 | 0 | 0 | T | C | 2.1537 | 0.6447 | 0.0008366 | +++?- | 35.6 | 0.1987 | intergenic | |
| rs117095766 | 14 | 93486038 | 4 | 0 | 0 | T | C | -2.6035 | 0.7794 | 0.0008366 | ---?- | 0 | 0.871 | intron | ITPK1 |
| 2:47388764 | 2 | 47388764 | 3 | 0 | 0 | T | TG | 1.347 | 0.4033 | 0.0008367 | +++?? | 0 | 0.9942 | | |
| 8:89060661 | 8 | 89060661 | 3 | 0 | 0 | T | TG | -1.1953 | 0.3578 | 0.0008368 | ---?? | 57.5 | 0.095 | | |
| rs4676844 | 3 | 71883565 | 4 | 0 | 0 | A | G | 0.9075 | 0.2717 | 0.000837 | +++?+ | 0 | 0.7286 | intergenic | |
| rs60981622 | 7 | 67860916 | 3 | 0 | 0 | A | G | -2.4184 | 0.724 | 0.000837 | ---?? | 0 | 0.4556 | intergenic | |
| rs4521247 | 3 | 20847171 | 3 | 0 | 0 | A | T | 1.7513 | 0.5243 | 0.0008372 | +++?? | 0 | 0.9704 | intergenic | |
| rs354287 | 5 | 14236440 | 4 | 0 | 0 | T | C | -1.0593 | 0.3172 | 0.0008373 | ---?- | 0 | 0.7241 | intron | TRIO |
| rs4695285 | 4 | 47851944 | 5 | 0 | 0 | T | C | 1.6385 | 0.4905 | 0.0008373 | +++++ | 12.3 | 0.3353 | intron | NFXL1 |
| rs807109 | 17 | 51239725 | 5 | 0 | 0 | T | C | 1.0388 | 0.311 | 0.0008377 | +-++- | 6.2 | 0.3714 | intergenic | |
| 8:20442293 | 8 | 20442293 | 3 | 0 | 0 | G | GT | -1.8661 | 0.5587 | 0.0008378 | ---?? | 23.7 | 0.2699 | | |
| rs28607086 | 7 | 141200745 | 4 | 0 | 0 | A | G | 3.6562 | 1.0947 | 0.0008379 | +++?+ | 28.1 | 0.2434 | intergenic | |
| 18:59178565 | 18 | 59178565 | 3 | 0 | 0 | CT | C | -4.1913 | 1.2549 | 0.0008379 | ---?? | 0 | 0.644 | | |
| rs1926815 | 10 | 4379965 | 4 | 0 | 0 | A | C | -1.0004 | 0.2995 | 0.0008382 | ---?- | 0 | 0.6719 | intergenic | |
| rs7245489 | 19 | 12823575 | 4 | 0 | 0 | T | C | -3.4853 | 1.0435 | 0.0008382 | ----? | 51.2 | 0.1045 | intron | TNPO2 |
| rs74778840 | 10 | 131489527 | 4 | 0 | 0 | C | G | -1.2304 | 0.3684 | 0.0008382 | ---?- | 0 | 0.8047 | intron | MGMT |
| rs1507997 | 17 | 52230469 | 4 | 0 | 0 | A | C | -2.1519 | 0.6444 | 0.0008392 | ---?+ | 35.7 | 0.1979 | intergenic | |
| rs10110127 | 8 | 109123400 | 4 | 0 | 0 | A | G | -1.4459 | 0.433 | 0.0008392 | ---?+ | 60.2 | 0.05668 | intergenic | |
| rs191304959 | 2 | 183986879 | 3 | 0 | 0 | A | G | 1.3599 | 0.4072 | 0.0008394 | +++?? | 0 | 0.6617 | intron | NUP35 |
| rs686250 | 6 | 32585055 | 3 | 0 | 0 | A | G | -0.8344 | 0.2499 | 0.0008396 | ---?? | 0 | 0.9952 | intergenic | |

| | | | | | | | | | | | | | | |
|---|---|---|---|---|---|---|---|---|---|---|---|---|---|---|
| rs1859176 | 7 | 82428058 | 4 | 0 | 0 | T | C | 0.8442 | 0.2528 | 0.0008397 | +-+?+ | 45.3 | 0.1393 intron | PCLO |
| rs151055402 | 2 | 168221664 | 4 | 0 | 0 | T | C | -2.4649 | 0.7382 | 0.0008401 | ---?- | 0 | 0.5705 intergenic | |
| rs7976188 | 12 | 117170368 | 5 | 0 | 0 | C | G | 1.2394 | 0.3712 | 0.0008402 | +++++ | 0 | 0.836 intron | C12orf49 |
| rs7180383 | 15 | 86023905 | 3 | 0 | 0 | A | G | -5.0117 | 1.5009 | 0.0008404 | --??- | 0 | 0.9415 intron | AKAP13 |
| rs10235074 | 7 | 82425679 | 4 | 0 | 0 | A | T | -0.8451 | 0.2531 | 0.0008404 | -+-?- | 45.8 | 0.1366 intron | PCLO |
| rs11601521 | 11 | 12645699 | 4 | 0 | 0 | A | G | 1.0207 | 0.3057 | 0.0008404 | +++?+ | 0 | 0.809 intergenic | |
| rs4414951 | 4 | 20743161 | 3 | 0 | 0 | T | C | -0.906 | 0.2713 | 0.0008408 | ---?? | 0 | 0.7058 intron | KCNIP4 |
| rs10220444 | 14 | 43273561 | 4 | 0 | 0 | A | T | 1.2815 | 0.3838 | 0.0008408 | +++?- | 58.4 | 0.06559 intergenic | |
| rs12776110 | 10 | 122114196 | 3 | 0 | 0 | A | T | -0.9271 | 0.2777 | 0.0008409 | ---?? | 0 | 0.7852 intergenic | |
| rs17393990 | 1 | 64789374 | 4 | 0 | 0 | C | G | -1.0211 | 0.3058 | 0.0008409 | ---?- | 0 | 0.5195 intergenic | |
| rs28766141 | 9 | 44252265 | 3 | 0 | 0 | A | G | -1.7291 | 0.5179 | 0.0008409 | ---?? | 0 | 0.8604 intergenic | |
| rs9264453 | 6 | 31231234 | 3 | 0 | 0 | A | G | 0.9499 | 0.2845 | 0.000841 | +++?? | 26.5 | 0.2565 intergenic | |
| rs9863528 | 3 | 30826658 | 3 | 0 | 0 | A | G | 3.0192 | 0.9043 | 0.0008412 | +++?? | 0 | 0.521 intron | GADL1 |
| 20:4369866 | 20 | 4369866 | 3 | 0 | 0 | T | G | 5.8287 | 1.7457 | 0.0008413 | +++?? | 0 | 0.7264 intergenic | |
| rs250511 | 19 | 8367504 | 4 | 0 | 0 | A | T | 1.6609 | 0.4974 | 0.0008413 | +++?+ | 46.2 | 0.1339 intron | CD320 |
| rs4976261 | 5 | 134374108 | 5 | 0 | 0 | C | G | -0.8069 | 0.2417 | 0.0008414 | ----- | 10.4 | 0.347 intron | LOC100996485 |
| rs11033909 | 11 | 37064503 | 4 | 0 | 0 | T | G | 0.8346 | 0.25 | 0.0008416 | +++?+ | 8.7 | 0.3496 intergenic | |
| rs74919572 | 16 | 26133990 | 3 | 0 | 0 | A | G | 4.4366 | 1.3288 | 0.0008416 | +++?? | 61.5 | 0.07458 intron | HS3ST4 |
| rs28714128 | 4 | 20757220 | 4 | 0 | 0 | A | G | 0.8824 | 0.2643 | 0.0008418 | +++?+ | 0 | 0.8256 intron | KCNIP4 |
| rs28699005 | 19 | 12820910 | 4 | 0 | 0 | C | G | -3.4998 | 1.0483 | 0.0008419 | ----? | 50.6 | 0.1083 intron | TNPO2 |
| rs1030430 | 2 | 185412099 | 4 | 0 | 0 | A | G | -0.9273 | 0.2777 | 0.0008421 | ---?- | 0 | 0.8535 intergenic | |
| rs1940163 | 11 | 95707042 | 4 | 0 | 0 | A | C | -0.8721 | 0.2612 | 0.0008422 | ---?- | 0 | 0.5213 downstream-gene | |
| rs9264450 | 6 | 31231193 | 3 | 0 | 0 | A | G | -0.9498 | 0.2845 | 0.0008422 | ---?? | 26.6 | 0.2561 intergenic | |
| rs62152019 | 2 | 80871004 | 4 | 0 | 0 | A | G | 1.8928 | 0.5669 | 0.0008422 | +++?+ | 0 | 0.7831 intron | CTNNA2 |
| rs2550409 | 16 | 84801482 | 3 | 0 | 0 | T | C | -1.0231 | 0.3064 | 0.0008424 | ---?? | 3.7 | 0.354 intron | USP10 |
| rs1204841 | 6 | 116519640 | 5 | 0 | 0 | A | C | 0.7925 | 0.2374 | 0.0008425 | +++++ | 0 | 0.5779 intron | NT5DC1 |
| rs1204842 | 6 | 116520348 | 5 | 0 | 0 | A | C | -0.7923 | 0.2373 | 0.0008429 | ----- | 0 | 0.5763 intron | NT5DC1 |
| rs8080654 | 17 | 51256940 | 5 | 0 | 0 | T | C | 1.044 | 0.3127 | 0.000843 | ++++- | 6.3 | 0.3706 intergenic | |
| rs183813 | 4 | 96527094 | 4 | 0 | 0 | A | G | 1.2537 | 0.3755 | 0.0008432 | +++?+ | 26.2 | 0.2542 intergenic | |
| 3:183619001 | 3 | 183619001 | 3 | 0 | 0 | CAA | C | -1.6935 | 0.5073 | 0.0008433 | ---?? | 0 | 0.9314 | |
| rs143291247 | 17 | 79779423 | 3 | 0 | 0 | C | G | 1.5398 | 0.4613 | 0.0008434 | +++?? | 0 | 0.8694 downstream-gene | |
| rs1410063 | 10 | 129679650 | 4 | 0 | 0 | A | G | 0.8313 | 0.249 | 0.0008435 | +++?+ | 0 | 0.6968 intron | CLRN3 |
| rs56753619 | 2 | 68138679 | 4 | 0 | 0 | T | C | -4.0182 | 1.2037 | 0.0008436 | ---?- | 0 | 0.4279 intergenic | |
| rs73323454 | 10 | 95245023 | 3 | 0 | 0 | T | C | 4.7212 | 1.4144 | 0.0008436 | ?++?+ | 0 | 0.9191 upstream-gene | |
| 15:87564735 | 15 | 87564735 | 3 | 0 | 0 | T | TTAAAC | -0.9195 | 0.2755 | 0.0008437 | ---?? | 26.7 | 0.2556 | |
| rs10939665 | 4 | 10037628 | 5 | 0 | 0 | T | C | -0.7019 | 0.2103 | 0.0008438 | ----+ | 13.7 | 0.3268 intron | SLC2A9 |
| rs79714331 | 12 | 85261005 | 4 | 0 | 0 | T | C | -1.7997 | 0.5391 | 0.0008438 | ---?- | 0 | 0.8508 intron | SLC6A15 |
| 3:45357945 | 3 | 45357945 | 3 | 0 | 0 | CAAA | CAAAAA | 1.3497 | 0.4044 | 0.0008438 | +++?? | 47.7 | 0.1479 | |
| rs72900971 | 2 | 68135266 | 4 | 0 | 0 | A | G | 4.0072 | 1.2005 | 0.000844 | +++?+ | 0 | 0.4146 intergenic | |
| rs144196980 | 18 | 52736463 | 4 | 0 | 0 | T | C | 4.2196 | 1.2641 | 0.0008442 | +++?+ | 0 | 0.8626 intergenic | |
| 8:113214968 | 8 | 113214968 | 3 | 0 | 0 | T | TA | 0.9772 | 0.2928 | 0.0008442 | +++?? | 0 | 0.9125 | |
| rs12810174 | 12 | 51472461 | 5 | 0 | 0 | A | T | -0.9288 | 0.2783 | 0.0008445 | ----- | 0 | 0.5174 intron | CSRNP2 |
| rs1204843 | 6 | 116520985 | 5 | 0 | 0 | A | G | -0.7921 | 0.2373 | 0.0008449 | ----- | 0 | 0.5759 intron | NT5DC1 |
| rs16885881 | 5 | 55934534 | 4 | 0 | 0 | T | C | -2.4385 | 0.7306 | 0.0008449 | ---?+ | 28.9 | 0.2387 intergenic | |
| rs7574489 | 2 | 53853156 | 4 | 0 | 0 | T | G | 0.8144 | 0.244 | 0.0008452 | +++?+ | 25.9 | 0.2565 intergenic | |
| rs2701378 | 15 | 87561723 | 3 | 0 | 0 | A | T | -0.9217 | 0.2762 | 0.0008453 | ---?? | 30.7 | 0.2363 intron | AGBL1 |
| rs150271987 | 9 | 44250705 | 3 | 0 | 0 | T | C | 1.7086 | 0.5119 | 0.0008454 | +++?? | 0 | 0.8778 intergenic | |
| rs28548350 | 19 | 12819542 | 4 | 0 | 0 | A | G | -3.4906 | 1.0459 | 0.0008457 | ----? | 50.5 | 0.1086 intron | TNPO2 |
| rs35059864 | 13 | 20812759 | 4 | 0 | 0 | A | G | 1.1007 | 0.3298 | 0.0008461 | +++?+ | 0 | 0.487 intergenic | |
| rs13185378 | 5 | 55819789 | 4 | 0 | 0 | A | T | 2.0455 | 0.6129 | 0.0008463 | +++?+ | 27.4 | 0.2477 intergenic | |
| rs9429528 | 1 | 114887458 | 4 | 0 | 0 | A | C | -1.3761 | 0.4124 | 0.0008464 | ---?- | 0 | 0.7197 intergenic | |
| rs6553275 | 4 | 190001080 | 4 | 0 | 0 | C | G | 0.848 | 0.2541 | 0.0008468 | +++?+ | 35.1 | 0.2014 intergenic | |
| rs6713511 | 2 | 131121149 | 5 | 0 | 0 | A | G | 0.8695 | 0.2606 | 0.0008469 | +++++ | 26.4 | 0.2456 intron | PTPN18 |
| rs10799548 | 1 | 229832654 | 4 | 0 | 0 | T | C | -2.7213 | 0.8155 | 0.0008469 | ---?- | 43.7 | 0.1493 intergenic | |
| rs78866102 | 4 | 39437905 | 4 | 0 | 0 | A | G | 2.584 | 0.7744 | 0.000847 | +-+?+ | 42.7 | 0.1553 intron | KLB |
| rs12602669 | 17 | 71055604 | 4 | 0 | 0 | A | G | -2.2027 | 0.6601 | 0.0008471 | ---?- | 25.1 | 0.2608 intron | SLC39A11 |
| 15:79731601 | 15 | 79731601 | 3 | 0 | 0 | T | TAGTA | 1.4321 | 0.4292 | 0.0008471 | +++?? | 0 | 0.7167 | |
| rs36194872 | 5 | 51463037 | 3 | 0 | 0 | A | T | 2.4529 | 0.7351 | 0.0008473 | +++?? | 47.7 | 0.1476 intergenic | |
| rs56084652 | 5 | 706437 | 4 | 0 | 0 | T | C | -1.2564 | 0.3765 | 0.0008476 | ---?- | 28.3 | 0.2425 downstream-gene | |

| | | | | | | | | | | | | | | | |
|---|---|---|---|---|---|---|---|---|---|---|---|---|---|---|---|
| 1:103048896 | 1 | 103048896 | 3 | 0 | 0 | A | ATC | 0.895 | 0.2682 | 0.0008476 | +++?? | 26 | 0.2591 | | |
| 20:3033479 | 20 | 3033479 | 3 | 0 | 0 | T | TA | -1.0152 | 0.3043 | 0.0008477 | ---?? | 0 | 0.3856 | | |
| rs16986103 | 22 | 28559595 | 4 | 0 | 0 | C | G | 2.4061 | 0.7211 | 0.0008478 | +++?+ | 0 | 0.7478 | intron-near-splice | TTC28 |
| rs13137343 | 4 | 10043028 | 5 | 0 | 0 | A | C | 0.702 | 0.2104 | 0.000848 | ++++- | 14.1 | 0.3246 | upstream-gene | |
| rs28623709 | 19 | 12821816 | 4 | 0 | 0 | T | C | -3.4854 | 1.0446 | 0.000848 | ----? | 51 | 0.1058 | intron | TNPO2 |
| rs113105683 | 4 | 91563901 | 4 | 0 | 0 | A | G | -3.4285 | 1.0276 | 0.0008482 | ---?- | 0 | 0.4273 | intron | CCSER1 |
| rs11833022 | 12 | 51465785 | 5 | 0 | 0 | T | C | -0.9302 | 0.2788 | 0.0008483 | ----- | 0 | 0.4633 | intron | CSRNP2 |
| rs111539503 | 17 | 56434481 | 5 | 0 | 0 | C | G | 1.0703 | 0.3208 | 0.0008483 | +++++ | 36.7 | 0.1766 | intron | RNF43 |
| rs114346141 | 6 | 30034289 | 5 | 0 | 0 | T | C | -1.4796 | 0.4435 | 0.0008484 | ----- | 0 | 0.7631 | upstream-gene |  |
| 11:131659622 | 11 | 131659622 | 3 | 0 | 0 | A | G | -4.4162 | 1.3237 | 0.0008488 | ---?? | 0 | 0.5991 | intron | NTM |
| rs77368744 | 10 | 37512116 | 3 | 0 | 0 | T | C | 4.2272 | 1.267 | 0.0008488 | +?+?+ | 0 | 0.4935 | intron | ANKRD30A |
| 20:34185344 | 20 | 34185344 | 3 | 0 | 0 | T | TC | -1.4843 | 0.4449 | 0.0008488 | ---?? | 1.4 | 0.3626 | | |
| rs8108290 | 19 | 12819840 | 4 | 0 | 0 | C | G | 3.4889 | 1.0458 | 0.0008493 | ++++? | 50.6 | 0.1079 | intron | TNPO2 |
| rs28478248 | 19 | 12819452 | 4 | 0 | 0 | A | G | -3.4893 | 1.0459 | 0.0008496 | ----? | 50.5 | 0.1086 | intron | TNPO2 |
| rs149168659 | 5 | 35751415 | 3 | 0 | 0 | A | G | 3.103 | 0.9301 | 0.0008496 | +++?? | 0 | 0.518 | intron | SPEF2 |
| rs12876903 | 13 | 20812795 | 4 | 0 | 0 | A | G | 1.1005 | 0.3299 | 0.0008497 | +++?+ | 0 | 0.4867 | intergenic | |
| rs61934011 | 12 | 98947524 | 4 | 0 | 0 | A | G | -0.9251 | 0.2773 | 0.00085 | ---?- | 0 | 0.9404 | downstream-gene | |
| rs4252251 | 11 | 117859629 | 4 | 0 | 0 | A | C | 1.3674 | 0.4099 | 0.0008501 | +++?+ | 0 | 0.7244 | intron | IL10RA |
| rs184415474 | 1 | 100292473 | 3 | 0 | 0 | A | G | -2.7964 | 0.8383 | 0.0008501 | ---?? | 0 | 0.5088 | intergenic | |
| rs185080705 | 5 | 35748576 | 3 | 0 | 0 | A | G | 3.1028 | 0.9301 | 0.0008502 | +++?? | 0 | 0.5168 | intron | SPEF2 |
| rs4350391 | 11 | 97090996 | 4 | 0 | 0 | C | G | 2.0081 | 0.602 | 0.0008503 | +++?+ | 0 | 0.4978 | intergenic | |
| rs6670810 | 1 | 182019480 | 4 | 0 | 0 | A | C | 1.1143 | 0.334 | 0.0008504 | +++?+ | 0 | 0.447 | downstream-gene | |
| rs9393515 | 6 | 24024396 | 5 | 0 | 0 | A | G | 1.3042 | 0.391 | 0.0008505 | ++++- | 0 | 0.8113 | intergenic | |
| rs12978299 | 19 | 52154398 | 4 | 0 | 0 | A | G | 0.9763 | 0.2927 | 0.0008505 | +++?+ | 0 | 0.8928 | upstream-gene | |
| rs140202846 | 9 | 11346429 | 3 | 0 | 0 | A | G | -2.9845 | 0.8947 | 0.0008505 | ---?? | 59.9 | 0.08268 | intergenic | |
| rs2079139 | 7 | 82424231 | 4 | 0 | 0 | T | C | 0.8488 | 0.2544 | 0.0008507 | +-+?+ | 47.1 | 0.1286 | intron | PCLO |
| rs143751757 | 1 | 227087994 | 3 | 0 | 0 | T | G | 4.2359 | 1.2699 | 0.0008509 | ?++?+ | 0 | 0.9419 | intron | ADCK3 |
| rs75898556 | 11 | 59245593 | 3 | 0 | 0 | T | G | 3.3206 | 0.9955 | 0.0008511 | +?++? | 46.2 | 0.1561 | stop-gained | OR4D10 |
| rs12877035 | 13 | 20812865 | 4 | 0 | 0 | A | G | 1.1005 | 0.3299 | 0.0008514 | +++?+ | 0 | 0.4865 | intergenic | |
| rs115771581 | 9 | 44250579 | 3 | 0 | 0 | A | C | -1.7072 | 0.5118 | 0.0008514 | ---?? | 0 | 0.8788 | intergenic | |
| rs7583684 | 2 | 47376995 | 4 | 0 | 0 | T | C | -1.287 | 0.3859 | 0.0008516 | ---?- | 0 | 0.976 | intron | C2orf61 |
| rs6670536 | 1 | 182019252 | 4 | 0 | 0 | A | G | 1.1142 | 0.3341 | 0.0008517 | +++?+ | 0 | 0.447 | downstream-gene | |
| 5:83027531 | 5 | 83027531 | 3 | 0 | 0 | G | GT | 1.4759 | 0.4425 | 0.0008517 | +++?? | 0 | 0.3888 | | |
| rs11731110 | 4 | 10037346 | 5 | 0 | 0 | T | C | -0.7013 | 0.2103 | 0.0008518 | ----+ | 13.8 | 0.3265 | intron | SLC2A9 |
| rs147436376 | 18 | 52730178 | 4 | 0 | 0 | A | T | -4.2502 | 1.2743 | 0.0008518 | ---?- | 0 | 0.8584 | intergenic | |
| rs8104187 | 19 | 12818888 | 4 | 0 | 0 | A | G | 3.4891 | 1.0461 | 0.0008519 | ++++? | 50.4 | 0.1091 | intron | TNPO2 |
| rs11208392 | 1 | 64726065 | 4 | 0 | 0 | T | G | -1.2094 | 0.3626 | 0.000852 | ---?- | 0 | 0.4185 | intron | UBE2U |
| rs1566584 | 1 | 42797765 | 4 | 0 | 0 | A | G | 1.8205 | 0.5458 | 0.0008522 | +++?+ | 0 | 0.5621 | intron | FOXJ3 |
| rs75056841 | 12 | 124882604 | 3 | 0 | 0 | A | G | 2.6333 | 0.7895 | 0.0008522 | +++?? | 8.7 | 0.3344 | intron | NCOR2 |
| rs9809878 | 3 | 116032441 | 4 | 0 | 0 | C | G | 1.037 | 0.3109 | 0.0008523 | +++?+ | 0 | 0.5066 | intron | LSAMP |
| 3:45345041 | 3 | 45345041 | 3 | 0 | 0 | T | TAGTAT | -1.4515 | 0.4352 | 0.0008523 | ---?? | 50.6 | 0.132 | | |
| rs7029730 | 9 | 16178816 | 4 | 0 | 0 | A | C | -0.9154 | 0.2745 | 0.0008526 | ---?- | 0 | 0.9036 | intergenic | |
| rs9736134 | 11 | 130627110 | 3 | 0 | 0 | A | G | 1.7132 | 0.5137 | 0.0008527 | +++?? | 45.1 | 0.1617 | intergenic | |
| rs1204820 | 6 | 116562914 | 5 | 0 | 0 | A | G | -0.786 | 0.2357 | 0.0008528 | +---- | 0 | 0.4715 | intron | NT5DC1 |
| 17:223304 | 17 | 223304 | 2 | 0 | 0 | G | GA | 6.9988 | 2.0986 | 0.0008529 | ?++?? | 0 | 0.621 | | |
| rs2120834 | 4 | 95896344 | 3 | 0 | 0 | C | G | -1.1283 | 0.3383 | 0.000853 | ---?? | 0 | 0.9954 | intron | BMPR1B |
| rs13145442 | 4 | 10037261 | 5 | 0 | 0 | A | C | 0.7012 | 0.2103 | 0.0008532 | ++++- | 13.8 | 0.3262 | intron | SLC2A9 |
| rs75188369 | 2 | 47575056 | 4 | 0 | 0 | A | G | 2.7048 | 0.8111 | 0.0008532 | +++?+ | 0 | 0.4628 | intergenic | |
| rs118037283 | 22 | 29067122 | 4 | 0 | 0 | A | G | 2.5362 | 0.7605 | 0.0008532 | +++?+ | 0 | 0.7737 | intron | TTC28 |
| rs7733193 | 5 | 165435775 | 4 | 0 | 0 | T | C | -0.8431 | 0.2528 | 0.0008534 | ---?- | 0 | 0.3971 | intergenic | |
| rs742930 | 6 | 116522615 | 5 | 0 | 0 | A | G | -0.791 | 0.2372 | 0.0008535 | ----- | 0 | 0.572 | intron | NT5DC1 |
| rs4335843 | 18 | 52731414 | 4 | 0 | 0 | C | G | 4.2442 | 1.2727 | 0.0008536 | +++?+ | 0 | 0.8601 | intergenic | |
| rs186584575 | 9 | 11323311 | 3 | 0 | 0 | T | G | -1.8475 | 0.554 | 0.0008536 | ---?? | 4 | 0.3527 | intergenic | |
| 9:11323310 | 9 | 11323310 | 3 | 0 | 0 | A | AC | 1.8475 | 0.554 | 0.0008536 | +++?? | 4.1 | 0.3527 | | |
| rs6442729 | 3 | 2485509 | 4 | 0 | 0 | T | C | 2.4922 | 0.7474 | 0.0008539 | +++?+ | 0 | 0.9121 | intron | CNTN4 |
| 2:117126715 | 2 | 117126715 | 3 | 0 | 0 | T | TTTTTTT | 1.7925 | 0.5375 | 0.000854 | +++?? | 0 | 0.5161 | | |
| rs1204821 | 6 | 116563263 | 5 | 0 | 0 | A | G | 0.7859 | 0.2357 | 0.0008541 | -++++ | 0 | 0.4712 | intron | NT5DC1 |
| rs1038373 | 15 | 86018148 | 3 | 0 | 0 | A | T | -5.0159 | 1.5042 | 0.0008542 | --??- | 0 | 0.9426 | intron | AKAP13 |
| rs530048 | 2 | 40450454 | 4 | 0 | 0 | A | G | 1.025 | 0.3074 | 0.0008543 | +++?- | 0 | 0.4068 | intron | SLC8A1,SLC8A1-AS1 |

| | | | | | | | | | | | | | | | |
|---|---|---|---|---|---|---|---|---|---|---|---|---|---|---|---|
| rs719281 | 2 | 185411685 | 4 | 0 | 0 | A | T | -0.9263 | 0.2778 | 0.0008543 | ---?- | 0 | 0.8548 | intergenic | |
| rs6659717 | 1 | 182019762 | 4 | 0 | 0 | T | C | 1.1137 | 0.334 | 0.0008543 | +++?+ | 0 | 0.4477 | downstream-gene | |
| rs873961 | 15 | 61348204 | 4 | 0 | 0 | T | C | -0.8512 | 0.2553 | 0.0008544 | ---?- | 0 | 0.7509 | intron | RORA |
| rs4908016 | 1 | 100214634 | 4 | 0 | 0 | T | C | 1.707 | 0.5119 | 0.0008544 | +++?+ | 0 | 0.8021 | intron | FRRS1 |
| rs4462690 | 18 | 52729228 | 4 | 0 | 0 | T | C | 4.2537 | 1.2757 | 0.0008545 | +++?+ | 0 | 0.8577 | intergenic | |
| rs35682092 | 16 | 6536593 | 4 | 0 | 0 | A | G | -0.9061 | 0.2717 | 0.0008546 | ---?- | 0 | 0.6953 | intron | RBFOX1 |
| 2:170413338 | 2 | 170413338 | 3 | 0 | 0 | A | AAAAC | 1.164 | 0.3491 | 0.0008546 | +++?? | 0 | 0.8193 | | |
| rs1188710 | 1 | 228528563 | 4 | 0 | 0 | C | G | -0.717 | 0.215 | 0.0008549 | ----? | 58.3 | 0.0659 | missense | OBSCN |
| rs1321365 | 6 | 117301614 | 4 | 0 | 0 | T | C | 0.7867 | 0.2359 | 0.0008549 | +++?+ | 0 | 0.7204 | intergenic | |
| rs4666768 | 2 | 189880752 | 4 | 0 | 0 | A | G | 0.9463 | 0.2838 | 0.0008552 | +++?+ | 40.6 | 0.1682 | downstream-gene | |
| rs57719132 | 3 | 127194310 | 3 | 0 | 0 | A | C | -1.0984 | 0.3294 | 0.0008552 | ---?? | 11.6 | 0.3225 | intergenic | |
| rs1204807 | 6 | 116549281 | 5 | 0 | 0 | A | C | 0.7857 | 0.2357 | 0.0008554 | -++++ | 0 | 0.4776 | intron | NT5DC1 |
| rs12506090 | 4 | 190013274 | 4 | 0 | 0 | A | G | -0.8235 | 0.247 | 0.0008554 | ---?- | 0 | 0.83 | intergenic | |
| rs9359266 | 6 | 78099258 | 4 | 0 | 0 | T | C | -0.8699 | 0.2609 | 0.0008555 | ---?- | 0 | 0.5035 | intergenic | |
| rs12784407 | 10 | 134589631 | 4 | 0 | 0 | A | G | -0.9214 | 0.2763 | 0.0008555 | ---?- | 0 | 0.4959 | intron | INPP5A |
| rs76624346 | 2 | 170779785 | 3 | 0 | 0 | T | C | 3.9813 | 1.1941 | 0.0008555 | ?++?+ | 0 | 0.9151 | intron | UBR3 |
| rs78238685 | 4 | 140546557 | 4 | 0 | 0 | A | G | 3.0077 | 0.9021 | 0.0008557 | +++?+ | 28.7 | 0.2401 | intergenic | |
| rs75162665 | 8 | 135834376 | 2 | 0 | 0 | A | T | 4.6791 | 1.4034 | 0.0008559 | +?+?? | 0 | 0.6327 | intergenic | |
| rs7526852 | 1 | 182007527 | 4 | 0 | 0 | A | G | -2.7741 | 0.8321 | 0.0008561 | ---?- | 0 | 0.7126 | intergenic | |
| rs79478914 | 4 | 162211813 | 2 | 0 | 0 | A | G | -6.003 | 1.8005 | 0.0008561 | ?--?? | 0 | 0.6315 | intergenic | |
| rs12566221 | 1 | 64728736 | 4 | 0 | 0 | A | G | -1.2093 | 0.3627 | 0.0008564 | ---?- | 0 | 0.43 | intron | UBE2U |
| 6:141765107 | 6 | 141765107 | 3 | 0 | 0 | T | TA | -1.9398 | 0.5819 | 0.0008567 | ---?? | 0 | 0.7231 | | |
| rs16829253 | 1 | 42723017 | 4 | 0 | 0 | T | C | -1.7761 | 0.5328 | 0.0008568 | ---?- | 0 | 0.5357 | intron | FOXJ3 |
| rs1172981 | 1 | 100230197 | 4 | 0 | 0 | T | C | 1.6554 | 0.4966 | 0.0008569 | +++?+ | 0 | 0.882 | intron | FRRS1 |
| rs4097591 | 9 | 44250853 | 3 | 0 | 0 | T | C | -1.7156 | 0.5146 | 0.0008569 | ---?? | 0 | 0.8734 | intergenic | |
| rs1204811 | 6 | 116552103 | 5 | 0 | 0 | T | C | -0.7857 | 0.2357 | 0.0008571 | +---- | 0 | 0.477 | intron | NT5DC1 |
| rs7484588 | 12 | 18892448 | 3 | 0 | 0 | A | T | -0.9409 | 0.2822 | 0.0008571 | ---?? | 0 | 0.6777 | upstream-gene | |
| rs75909488 | 12 | 29789330 | 4 | 0 | 0 | A | G | -1.4419 | 0.4325 | 0.0008572 | ---?- | 0 | 0.8308 | intron | TMTC1 |
| rs13101785 | 4 | 10042915 | 5 | 0 | 0 | A | T | 0.7015 | 0.2105 | 0.0008577 | ++++- | 14.1 | 0.3247 | upstream-gene | |
| rs28752516 | 6 | 32583611 | 3 | 0 | 0 | A | G | -0.8303 | 0.2491 | 0.0008577 | ---?? | 0 | 0.4531 | intergenic | |
| rs2245604 | 6 | 88498604 | 4 | 0 | 0 | T | G | 1.2107 | 0.3632 | 0.0008578 | +++?+ | 0 | 0.8251 | intergenic | |
| rs1204848 | 6 | 116524263 | 5 | 0 | 0 | A | G | 0.7903 | 0.2371 | 0.000858 | +++++ | 0 | 0.5692 | intron | NT5DC1 |
| rs72979361 | 4 | 162210863 | 2 | 0 | 0 | A | C | -6.0149 | 1.8045 | 0.0008581 | ?--?? | 0 | 0.6173 | intergenic | |
| rs2204979 | 14 | 22534990 | 4 | 0 | 0 | T | C | -0.9223 | 0.2767 | 0.0008585 | ---?- | 0 | 0.9312 | intergenic | |
| rs76746010 | 8 | 94158975 | 4 | 0 | 0 | A | G | 2.8888 | 0.8667 | 0.0008585 | +++?+ | 18.9 | 0.2959 | intron | C8orf87 |
| rs7915610 | 10 | 133803156 | 4 | 0 | 0 | T | C | -0.894 | 0.2682 | 0.0008586 | ---?- | 45.6 | 0.1378 | intergenic | |
| rs72882990 | 2 | 164433524 | 3 | 0 | 0 | A | G | 2.8905 | 0.8672 | 0.0008587 | +++?? | 10.1 | 0.3288 | intergenic | |
| rs9883767 | 3 | 122292124 | 4 | 0 | 0 | A | G | -1.2524 | 0.3758 | 0.0008588 | ---?- | 0 | 0.5989 | 3-prime-UTR | DTX3L |
| rs75344409 | 11 | 86302513 | 4 | 0 | 0 | A | G | -2.5244 | 0.7574 | 0.0008588 | ---?- | 0 | 0.505 | intron | ME3 |
| rs5007259 | 6 | 32379101 | 5 | 0 | 0 | T | C | -0.7038 | 0.2112 | 0.000859 | ----- | 0 | 0.6689 | upstream-gene |  |
| rs8113087 | 19 | 12820568 | 4 | 0 | 0 | A | G | -3.4846 | 1.0455 | 0.000859 | ----? | 50.8 | 0.1067 | intron | TNPO2 |
| rs6499105 | 16 | 52846375 | 4 | 0 | 0 | T | C | -0.8154 | 0.2446 | 0.0008592 | ---?- | 0 | 0.7956 | intergenic | |
| rs17131108 | 1 | 91091581 | 4 | 0 | 0 | A | G | -2.3112 | 0.6934 | 0.0008592 | ---?- | 0 | 0.4099 | intergenic | |
| rs8110510 | 19 | 12829183 | 4 | 0 | 0 | A | G | -3.5095 | 1.053 | 0.0008594 | +---? | 51.9 | 0.1009 | intron | TNPO2 |
| rs117898979 | 7 | 90553506 | 4 | 0 | 0 | A | C | 2.8289 | 0.8488 | 0.0008594 | +++?+ | 0 | 0.5081 | intron | CDK14 |
| rs7696163 | 4 | 119233946 | 4 | 0 | 0 | T | C | -1.5965 | 0.479 | 0.0008596 | ---?- | 0 | 0.5819 | intron | PRSS12 |
| rs74751081 | 4 | 186760139 | 4 | 0 | 0 | A | C | 1.5529 | 0.466 | 0.0008597 | +++?- | 6.2 | 0.3619 | intron | SORBS2 |
| rs10410839 | 19 | 12818475 | 4 | 0 | 0 | A | G | -3.4864 | 1.0461 | 0.0008599 | ----? | 50.4 | 0.1092 | intron | TNPO2 |
| rs10767867 | 11 | 30645229 | 4 | 0 | 0 | A | G | -0.888 | 0.2665 | 0.0008599 | ---?+ | 60.9 | 0.05323 | intergenic | |
| rs62130626 | 19 | 3711322 | 3 | 0 | 0 | A | G | -1.0009 | 0.3003 | 0.00086 | ---?? | 18.7 | 0.2922 | intron | TJP3 |
| rs7654255 | 4 | 183664522 | 4 | 0 | 0 | T | C | 0.8918 | 0.2676 | 0.0008601 | +++?+ | 0 | 0.932 | synonymous | TENM3 |
| rs116940565 | 9 | 112760741 | 4 | 0 | 0 | C | G | -2.8639 | 0.8593 | 0.0008601 | ---?- | 0 | 0.9729 | intron | PALM2-AKAP2 |
| 6:32008924 | 6 | 32008924 | 4 | 0 | 0 | A | G | 2.7455 | 0.8238 | 0.0008602 | ++++? | 20.7 | 0.2859 | 3-prime-UTR | CYP21A2 |
| rs3753805 | 1 | 180138890 | 4 | 0 | 0 | A | G | -2.0709 | 0.6214 | 0.0008602 | ---?+ | 35.2 | 0.2012 | intron | QSOX1 |
| rs13137074 | 4 | 10042842 | 5 | 0 | 0 | T | C | 0.7013 | 0.2104 | 0.0008603 | ++++- | 14.1 | 0.3247 | upstream-gene | |
| rs1210355 | 6 | 116526917 | 5 | 0 | 0 | A | T | -0.7894 | 0.2369 | 0.0008604 | ----- | 0 | 0.5618 | intron | NT5DC1 |
| rs4811890 | 20 | 56194732 | 4 | 0 | 0 | T | C | 1.1919 | 0.3576 | 0.0008604 | +++?+ | 0 | 0.8462 | intron | ZBP1 |
| rs6466432 | 7 | 112700531 | 4 | 0 | 0 | A | G | 0.9477 | 0.2844 | 0.0008605 | +++?+ | 0 | 0.6515 | intergenic | |
| rs62532369 | 8 | 127971458 | 2 | 0 | 0 | T | C | 6.9672 | 2.0907 | 0.0008605 | +???+ | 0 | 0.3934 | intergenic | |

| | | | | | | | | | | | | | | | |
|---|---|---|---|---|---|---|---|---|---|---|---|---|---|---|---|
| rs202162332 | 12 | 18892450 | 3 | 0 | 0 | A | T | -0.9407 | 0.2823 | 0.0008605 | ---?? | 0 | 0.6781 | upstream-gene | |
| 4:105217727 | 4 | 105217727 | 3 | 0 | 0 | T | TA | 3.4945 | 1.0486 | 0.0008605 | +++?? | 0 | 0.7335 | | |
| rs76839556 | 2 | 40462207 | 3 | 0 | 0 | T | G | -1.0345 | 0.3104 | 0.0008607 | ---?? | 2.3 | 0.3595 | intron | SLC8A1,SLC8A1-AS1 |
| rs706268 | 5 | 14030243 | 4 | 0 | 0 | T | G | -0.8604 | 0.2582 | 0.0008609 | ---?- | 0 | 0.6657 | intergenic | |
| rs1412408 | 9 | 30815421 | 4 | 0 | 0 | A | G | 3.1865 | 0.9563 | 0.0008615 | +++?+ | 5.2 | 0.3668 | intergenic | |
| rs6511833 | 19 | 12807187 | 4 | 0 | 0 | T | C | 3.4802 | 1.0444 | 0.0008615 | ++++? | 49.5 | 0.1148 | missense | FBXW9 |
| rs116662727 | 14 | 57216498 | 4 | 0 | 0 | C | G | -1.9988 | 0.5998 | 0.0008615 | ---?- | 0 | 0.4366 | intergenic | |
| rs817438 | 1 | 85641119 | 4 | 0 | 0 | T | G | 1.5144 | 0.4545 | 0.0008617 | +++?+ | 0 | 0.9889 | intron | SYDE2 |
| rs17376651 | 1 | 42638971 | 4 | 0 | 0 | T | C | 1.7043 | 0.5115 | 0.0008617 | +++?+ | 0 | 0.5125 | downstream-gene | |
| rs807142 | 17 | 51215712 | 5 | 0 | 0 | A | G | 1.0357 | 0.3108 | 0.0008621 | ++++- | 0 | 0.4077 | intergenic | |
| rs9990935 | 4 | 123726646 | 4 | 0 | 0 | T | C | -2.2026 | 0.661 | 0.0008621 | ---?- | 53.2 | 0.09338 | intergenic | |
| rs926829 | 6 | 116555878 | 5 | 0 | 0 | T | C | 0.7854 | 0.2357 | 0.0008622 | -++++ | 0 | 0.4756 | intron | NT5DC1 |
| rs2123978 | 2 | 52949987 | 4 | 0 | 0 | T | G | 1.2833 | 0.3851 | 0.0008623 | +++?+ | 0 | 0.7889 | intergenic | |
| 20:35955925 | 20 | 35955925 | 3 | 0 | 0 | T | TA | -1.7638 | 0.5294 | 0.0008623 | ---?? | 0 | 0.6504 | | |
| rs354285 | 5 | 14235150 | 4 | 0 | 0 | A | G | -1.0577 | 0.3174 | 0.0008624 | ---?- | 0 | 0.7227 | intron | TRIO |
| rs12650014 | 4 | 108272864 | 4 | 0 | 0 | A | G | -0.8375 | 0.2513 | 0.0008624 | ---?- | 0 | 0.4249 | intergenic | |
| rs1641838 | 16 | 12140484 | 4 | 0 | 0 | T | G | -0.8738 | 0.2623 | 0.0008628 | +--?- | 52.2 | 0.09878 | intron | SNX29 |
| rs12583719 | 13 | 20831995 | 4 | 0 | 0 | T | C | -1.0427 | 0.3129 | 0.0008628 | ---?- | 0 | 0.5157 | intergenic | |
| rs72712355 | 1 | 88003219 | 3 | 0 | 0 | A | G | 2.5636 | 0.7694 | 0.0008629 | +++?? | 0 | 0.3782 | intergenic | |
| rs115492559 | 5 | 105986224 | 4 | 0 | 0 | A | G | -2.6559 | 0.7971 | 0.0008629 | ---?+ | 0 | 0.4014 | intergenic | |
| rs118089939 | 16 | 9317093 | 4 | 0 | 0 | T | G | -2.9427 | 0.8832 | 0.0008632 | ---?- | 20.8 | 0.2855 | intergenic | |
| rs11066353 | 12 | 113012293 | 4 | 0 | 0 | A | G | 1.1149 | 0.3346 | 0.0008633 | +-+?+ | 44.5 | 0.1446 | intergenic | |
| rs61451669 | 11 | 117853666 | 4 | 0 | 0 | T | C | -1.3021 | 0.3908 | 0.0008633 | ---?- | 0 | 0.8318 | upstream-gene | |
| 15:52708518 | 15 | 52708518 | 3 | 0 | 0 | A | T | 5.2951 | 1.5893 | 0.0008634 | +++?? | 45.2 | 0.1615 | intron | MYO5A |
| rs73334976 | 8 | 114877887 | 4 | 0 | 0 | A | G | -1.7962 | 0.5391 | 0.0008635 | ---?+ | 2.8 | 0.3786 | intergenic | |
| 1:222689631 | 1 | 222689631 | 3 | 0 | 0 | A | C | 3.215 | 0.965 | 0.0008636 | +++?? | 7.5 | 0.3391 | intergenic | |
| rs7806071 | 7 | 82427324 | 4 | 0 | 0 | A | C | 0.8417 | 0.2526 | 0.000864 | +-+?+ | 45.5 | 0.1381 | intron | PCLO |
| rs146778462 | 4 | 65240286 | 2 | 0 | 0 | T | C | -4.4541 | 1.337 | 0.0008641 | ?--?? | 0 | 0.3293 | intron | TECRL |
| 3:30817064 | 3 | 30817064 | 3 | 0 | 0 | A | AT | 3.0205 | 0.9067 | 0.0008641 | +++?? | 0 | 0.6889 | | |
| rs7961442 | 12 | 98952808 | 4 | 0 | 0 | A | G | -0.9233 | 0.2772 | 0.0008642 | ---?- | 0 | 0.9436 | intergenic | |
| rs10735357 | 12 | 98627571 | 4 | 0 | 0 | T | C | 1.4758 | 0.443 | 0.0008643 | +-+?+ | 60.3 | 0.0562 | intergenic | |
| 19:40932078 | 19 | 40932078 | 3 | 0 | 0 | G | GA | -2.8021 | 0.8411 | 0.0008643 | ---?? | 0 | 0.7737 | | |
| rs6134439 | 20 | 12003484 | 4 | 0 | 0 | A | G | 0.945 | 0.2837 | 0.0008644 | +++?+ | 0 | 0.5457 | intergenic | |
| rs2588541 | 4 | 37002380 | 4 | 0 | 0 | A | G | -0.8358 | 0.2509 | 0.0008645 | ---?+ | 47.1 | 0.1288 | intergenic | |
| rs117263366 | 13 | 85445096 | 4 | 0 | 0 | T | C | -1.9659 | 0.5901 | 0.0008645 | ---?- | 29.9 | 0.2327 | intergenic | |
| rs113607963 | 3 | 23807262 | 4 | 0 | 0 | T | C | 3.574 | 1.0729 | 0.0008648 | +++?+ | 0 | 0.8069 | intergenic | |
| rs6134438 | 20 | 12003410 | 4 | 0 | 0 | A | G | 0.9442 | 0.2834 | 0.0008649 | +++?+ | 0 | 0.5469 | intergenic | |
| rs150235261 | 4 | 65240361 | 2 | 0 | 0 | A | T | -4.4536 | 1.337 | 0.0008649 | ?--?? | 0 | 0.3291 | intron | TECRL |
| rs1577636 | 6 | 62042385 | 4 | 0 | 0 | A | C | 0.9745 | 0.2926 | 0.000865 | +++?+ | 0 | 0.5962 | intergenic | |
| 13:109326232 | 13 | 109326232 | 3 | 0 | 0 | G | GT | 0.915 | 0.2747 | 0.000865 | +++?? | 0 | 0.4856 | | |
| rs3733584 | 4 | 10036506 | 5 | 0 | 0 | T | C | -0.7003 | 0.2102 | 0.0008652 | ----+ | 13.9 | 0.3259 | intron | SLC2A9 |
| rs11831810 | 12 | 51464251 | 5 | 0 | 0 | T | C | -0.9289 | 0.2789 | 0.0008653 | ----- | 0 | 0.458 | intron | CSRNP2 |
| 2:156929186 | 2 | 156929186 | 3 | 0 | 0 | G | GGT | -0.9082 | 0.2727 | 0.0008653 | ---?? | 0 | 0.6989 | | |
| rs6134437 | 20 | 12003383 | 4 | 0 | 0 | T | C | -0.9429 | 0.2831 | 0.0008654 | ---?- | 0 | 0.546 | intergenic | |
| rs160455 | 8 | 90667179 | 4 | 0 | 0 | A | G | 3.2107 | 0.9639 | 0.0008656 | +++?+ | 0 | 0.8716 | intergenic | |
| rs115236529 | 2 | 84094398 | 4 | 0 | 0 | A | G | -3.0015 | 0.9011 | 0.0008662 | -+-?- | 58.6 | 0.06469 | intergenic | |
| rs117148287 | 4 | 125480795 | 3 | 0 | 0 | A | G | 2.1144 | 0.6348 | 0.0008662 | +++?? | 0 | 0.9832 | intergenic | |
| rs4700415 | 5 | 60554858 | 5 | 0 | 0 | T | C | -0.9521 | 0.2859 | 0.0008664 | ----+ | 24.2 | 0.2598 | intergenic |  |
| rs114036883 | 5 | 132975398 | 4 | 0 | 0 | T | C | -3.0266 | 0.9087 | 0.0008664 | ---?- | 0 | 0.8283 | intergenic | |
| 8:124345649 | 8 | 124345649 | 3 | 0 | 0 | CAAA | C | 0.9594 | 0.2881 | 0.0008665 | +++?? | 0 | 0.4979 | | |
| rs67753410 | 2 | 2762080 | 4 | 0 | 0 | T | C | 1.2769 | 0.3834 | 0.0008666 | +++?+ | 47.3 | 0.1274 | intergenic | |
| rs807146 | 17 | 51212785 | 5 | 0 | 0 | A | C | -1.036 | 0.3111 | 0.0008668 | ----+ | 0 | 0.4171 | intergenic |  |
| rs419125 | 2 | 40441229 | 4 | 0 | 0 | T | C | 1.0309 | 0.3095 | 0.0008669 | +++?- | 5.6 | 0.365 | intron | SLC8A1,SLC8A1-AS1 |
| rs116358021 | 3 | 65907486 | 4 | 0 | 0 | A | G | -2.6673 | 0.8009 | 0.000867 | ---?- | 23.3 | 0.2711 | intron | MAGI1,MAGI1-AS1 |
| rs1355678 | 17 | 51154612 | 3 | 0 | 0 | T | C | 1.3529 | 0.4062 | 0.0008671 | +++?? | 3.8 | 0.3537 | intergenic | |
| rs11847646 | 14 | 64339727 | 5 | 0 | 0 | C | G | -2.7047 | 0.8121 | 0.0008671 | ----- | 0 | 0.9141 | intron | SYNE2 |
| 6:31234516 | 6 | 31234516 | 3 | 0 | 0 | CAT | C | -0.9485 | 0.2848 | 0.0008671 | ---?? | 25.4 | 0.2618 | | |
| 5:105742673 | 5 | 105742673 | 3 | 0 | 0 | T | TTAA | 2.2887 | 0.6872 | 0.0008672 | +++?? | 0 | 0.8444 | | |
| rs2745945 | 1 | 208188196 | 4 | 0 | 0 | T | C | 1.8916 | 0.568 | 0.0008673 | +++?+ | 0 | 0.7383 | intergenic | |

| | | | | | | | | | | | | | | | |
|---|---|---|---|---|---|---|---|---|---|---|---|---|---|---|---|
| rs12335923 | 9 | 30746043 | 4 | 0 | 0 | C | G | 3.336 | 1.0017 | 0.0008673 | +++?+ | 1.2 | 0.3862 | intergenic | |
| rs71394240 | 16 | 75835678 | 3 | 0 | 0 | T | C | 4.6777 | 1.4046 | 0.0008673 | ?++?+ | 0 | 0.722 | intergenic | |
| rs10990402 | 9 | 105734158 | 4 | 0 | 0 | T | G | 1.3102 | 0.3934 | 0.0008677 | +++?+ | 0 | 0.8772 | intergenic | |
| rs7548919 | 1 | 101864021 | 4 | 0 | 0 | C | G | 1.4682 | 0.4409 | 0.0008678 | +++?+ | 0 | 0.7763 | intergenic | |
| rs79501380 | 14 | 22460287 | 2 | 0 | 0 | A | G | 7.17 | 2.153 | 0.0008679 | ++??? | 0 | 0.98 | intergenic | |
| rs72979324 | 4 | 162205750 | 2 | 0 | 0 | T | C | -6.0895 | 1.8286 | 0.0008682 | ?--?? | 0 | 0.5107 | intergenic | |
| rs73449050 | 9 | 30746226 | 4 | 0 | 0 | A | C | -3.3358 | 1.0017 | 0.0008682 | ---?- | 1.1 | 0.3866 | intergenic | |
| rs2771338 | 14 | 21455159 | 4 | 0 | 0 | T | G | -0.8678 | 0.2606 | 0.0008683 | ---?- | 30.7 | 0.2278 | upstream-gene | |
| 7:82410512 | 7 | 82410512 | 3 | 0 | 0 | T | TAA | 0.9216 | 0.2767 | 0.0008683 | +++?? | 44.7 | 0.1638 | | |
| rs79914459 | 1 | 42721126 | 4 | 0 | 0 | T | C | 1.7742 | 0.5328 | 0.0008684 | +++?+ | 0 | 0.5376 | intron | FOXJ3 |
| 1:42797053 | 1 | 42797053 | 3 | 0 | 0 | A | AAAT | -1.8785 | 0.5641 | 0.0008685 | ---?? | 0 | 0.378 | | |
| rs10014579 | 4 | 46390250 | 4 | 0 | 0 | A | G | -0.865 | 0.2598 | 0.0008686 | -+-?- | 59.6 | 0.05965 | intron | GABRA2 |
| rs7193103 | 16 | 12785398 | 3 | 0 | 0 | T | C | 1.0968 | 0.3294 | 0.0008687 | +++?? | 0 | 0.8506 | intron | CPPED1 |
| rs56255108 | 16 | 5455223 | 4 | 0 | 0 | T | G | 0.9577 | 0.2876 | 0.0008687 | +++?+ | 0 | 0.7979 | intron | RBFOX1 |
| rs74924350 | 6 | 106291335 | 4 | 0 | 0 | T | C | 2.0526 | 0.6164 | 0.0008687 | +++?+ | 0 | 0.5154 | intergenic | |
| rs376303 | 2 | 40440793 | 4 | 0 | 0 | C | G | -1.0309 | 0.3096 | 0.0008688 | ---?+ | 5.5 | 0.3653 | intron | SLC8A1,SLC8A1-AS1 |
| rs78535563 | 6 | 12397127 | 4 | 0 | 0 | T | C | -2.1895 | 0.6575 | 0.0008688 | ---?- | 0 | 0.9706 | intergenic | |
| rs78265297 | 6 | 118476737 | 4 | 0 | 0 | T | C | -1.4935 | 0.4485 | 0.0008689 | ---?- | 0 | 0.5278 | intron | SLC35F1 |
| rs112869384 | 9 | 107894451 | 4 | 0 | 0 | T | G | 2.2531 | 0.6766 | 0.0008689 | +++?+ | 26.5 | 0.2529 | intergenic | |
| rs10237836 | 7 | 46631327 | 4 | 0 | 0 | A | G | 2.129 | 0.6394 | 0.000869 | +++?- | 31.7 | 0.2222 | intergenic | |
| 10:100057146 | 10 | 100057146 | 3 | 0 | 0 | A | AG | -3.1221 | 0.9376 | 0.000869 | ---?? | 0 | 0.5689 | | |
| rs401690 | 9 | 106485957 | 3 | 0 | 0 | T | C | -1.7043 | 0.5118 | 0.0008692 | ---?? | 0 | 0.6962 | intergenic | |
| rs2439636 | 8 | 96212069 | 4 | 0 | 0 | A | G | -0.8503 | 0.2554 | 0.0008692 | ---?- | 0 | 0.9658 | intergenic | |
| rs7852086 | 9 | 16189987 | 4 | 0 | 0 | T | G | -0.9197 | 0.2762 | 0.0008693 | ---?- | 0 | 0.9369 | intergenic | |
| rs62295210 | 4 | 34808662 | 3 | 0 | 0 | T | C | -1.8435 | 0.5537 | 0.0008694 | ---?? | 0 | 0.7076 | intergenic | |
| rs148289531 | 3 | 169699061 | 3 | 0 | 0 | A | G | 2.8066 | 0.8429 | 0.0008694 | +++?? | 0 | 0.9152 | intron | SEC62 |
| rs38415 | 7 | 30544747 | 5 | 0 | 0 | C | G | -0.8231 | 0.2472 | 0.0008695 | ----- | 27.7 | 0.2369 | upstream-gene |  |
| rs2151166 | 1 | 157405430 | 4 | 0 | 0 | C | G | 1.0985 | 0.3299 | 0.0008698 | +++?+ | 60.2 | 0.0564 | intergenic | |
| rs7792340 | 7 | 82427120 | 4 | 0 | 0 | T | C | -0.841 | 0.2526 | 0.0008698 | -+-?- | 45.6 | 0.1376 | intron | PCLO |
| rs10402101 | 19 | 12817973 | 4 | 0 | 0 | A | G | 3.4831 | 1.0461 | 0.0008703 | ++++? | 50.4 | 0.1093 | intron | TNPO2 |
| rs1347234 | 15 | 87555882 | 4 | 0 | 0 | A | C | -0.8826 | 0.2651 | 0.0008704 | ---?- | 0 | 0.424 | intron | AGBL1 |
| rs3733585 | 4 | 10036339 | 5 | 0 | 0 | A | G | -0.6999 | 0.2102 | 0.0008704 | ----+ | 13.9 | 0.3256 | intron | SLC2A9 |
| rs10424461 | 19 | 12805380 | 4 | 0 | 0 | T | C | -3.4777 | 1.0445 | 0.0008704 | ----? | 49.5 | 0.1143 | intron | FBXW9 |
| rs10401842 | 19 | 12817844 | 4 | 0 | 0 | A | G | 3.483 | 1.0461 | 0.0008706 | ++++? | 50.4 | 0.1094 | intron | TNPO2 |
| rs145848491 | 4 | 166242523 | 4 | 0 | 0 | A | G | 3.5796 | 1.0752 | 0.0008706 | +++?+ | 0 | 0.9501 | intron | KLHL2 |
| rs7970732 | 12 | 86243199 | 4 | 0 | 0 | A | C | 0.9273 | 0.2785 | 0.0008707 | +-+?+ | 18 | 0.3009 | intergenic | |
| rs76752455 | 6 | 109132156 | 2 | 0 | 0 | A | G | 4.6989 | 1.4114 | 0.0008707 | +?+?? | 0 | 0.6791 | intergenic | |
| rs7572986 | 2 | 170910812 | 3 | 0 | 1 | A | G | -0.9976 | 0.2997 | 0.0008708 | ---?? | 60.2 | 0.08131 | intron | UBR3 |
| rs4853877 | 2 | 2757028 | 4 | 0 | 0 | T | C | -1.2965 | 0.3894 | 0.000871 | ---?- | 0 | 0.9268 | intergenic | |
| 5:90878716 | 5 | 90878716 | 2 | 0 | 0 | T | C | -4.9759 | 1.4946 | 0.0008711 | ?--?? | 0 | 0.7911 | intergenic | |
| rs347548 | 18 | 49969813 | 4 | 0 | 0 | T | G | -1.8116 | 0.5442 | 0.0008711 | -+-?- | 24.6 | 0.264 | intron | DCC |
| rs1204851 | 6 | 116527120 | 5 | 0 | 0 | A | G | 0.7885 | 0.2368 | 0.0008712 | +++++ | 0 | 0.5604 | intron | NT5DC1 |
| rs61006093 | 15 | 87562628 | 3 | 0 | 0 | A | T | -0.9181 | 0.2758 | 0.0008712 | ---?? | 24.1 | 0.268 | intron | AGBL1 |
| rs379604 | 18 | 39942410 | 4 | 0 | 0 | T | C | -0.8599 | 0.2583 | 0.0008714 | ---?- | 0 | 0.4595 | intron | LINC00907 |
| rs7804652 | 7 | 82427102 | 4 | 0 | 0 | A | G | 0.8408 | 0.2526 | 0.0008714 | +-+?+ | 45.6 | 0.1375 | intron | PCLO |
| rs72936951 | 6 | 72860252 | 4 | 0 | 0 | T | C | -1.5428 | 0.4634 | 0.0008715 | ---?+ | 0 | 0.5937 | intron | RIMS1 |
| 18:39992060 | 18 | 39992060 | 3 | 0 | 0 | CT | C | 0.8876 | 0.2666 | 0.0008715 | +++?? | 0 | 0.376 | | |
| rs73203013 | 7 | 111025938 | 4 | 0 | 0 | A | G | 1.3514 | 0.4059 | 0.0008716 | +++?+ | 29.6 | 0.2347 | intron | IMMP2L |
| rs9264578 | 6 | 31235708 | 5 | 0 | 0 | C | G | -0.7947 | 0.2387 | 0.0008719 | ----- | 0 | 0.4696 | downstream-gene |  |
| rs13237692 | 7 | 32347546 | 4 | 0 | 0 | C | G | -1.3451 | 0.4041 | 0.0008719 | ---?- | 0 | 0.9121 | intergenic | |
| 15:87562624 | 15 | 87562624 | 3 | 0 | 0 | A | ATGT | -0.918 | 0.2758 | 0.0008719 | ---?? | 24.1 | 0.2679 | | |
| rs1940164 | 11 | 95702726 | 4 | 0 | 0 | A | C | -0.916 | 0.2752 | 0.000872 | ---?- | 9.4 | 0.3459 | intergenic | |
| rs73334975 | 8 | 114877577 | 4 | 0 | 0 | T | C | 1.7952 | 0.5393 | 0.000872 | +++?- | 3.1 | 0.3769 | intergenic | |
| 6:33580689 | 6 | 33580689 | 3 | 0 | 0 | T | C | 3.2577 | 0.9786 | 0.0008722 | +++?? | 28.8 | 0.2455 | intergenic | |
| 4:20766304 | 4 | 20766304 | 3 | 0 | 0 | G | GACAC | -0.9658 | 0.2901 | 0.0008722 | ---?? | 0 | 0.4681 | | |
| rs895668 | 18 | 46551318 | 3 | 0 | 0 | A | G | 1.9452 | 0.5844 | 0.0008723 | +++?? | 0 | 0.7213 | intergenic | |
| rs6580784 | 12 | 51468311 | 5 | 0 | 0 | A | G | -0.9275 | 0.2786 | 0.0008725 | ----- | 0 | 0.5128 | intron | CSRNP2 |
| rs7129003 | 11 | 23958122 | 4 | 0 | 0 | T | C | 1.2207 | 0.3667 | 0.0008725 | +++?+ | 0 | 0.8715 | intergenic | |
| rs150803637 | 3 | 111854002 | 4 | 0 | 0 | T | C | 3.4932 | 1.0494 | 0.0008725 | +++?+ | 0 | 0.6035 | upstream-gene | |

| | | | | | | | | | | | | | | | |
|---|---|---|---|---|---|---|---|---|---|---|---|---|---|---|---|
| rs7257128 | 19 | 12802621 | 4 | 0 | 0 | A | G | 3.4789 | 1.0451 | 0.0008727 | ++++? | 49.7 | 0.1135 | intron | FBXW9 |
| rs7132709 | 12 | 80447400 | 4 | 0 | 0 | T | C | 2.3258 | 0.6987 | 0.0008728 | +++?+ | 24.7 | 0.2631 | intergenic | |
| rs142747784 | 9 | 25003674 | 2 | 0 | 0 | T | C | 5.4137 | 1.6264 | 0.0008728 | +?+?? | 0 | 0.911 | intergenic | |
| rs4897440 | 6 | 130903604 | 3 | 0 | 0 | A | T | -0.9302 | 0.2795 | 0.0008733 | ---?? | 0 | 0.5512 | intergenic | |
| rs7754202 | 6 | 120401382 | 4 | 0 | 0 | A | G | 2.2568 | 0.678 | 0.0008733 | +++?+ | 50.4 | 0.109 | intergenic | |
| rs807137 | 17 | 51223215 | 5 | 0 | 0 | T | G | 1.0242 | 0.3077 | 0.0008734 | ++++- | 4.3 | 0.382 | intergenic | |
| rs6129729 | 20 | 39648662 | 4 | 0 | 0 | A | T | 1.1895 | 0.3574 | 0.0008734 | +++?+ | 0 | 0.4998 | intergenic | |
| rs12350524 | 9 | 30743422 | 4 | 0 | 0 | A | G | 3.3335 | 1.0015 | 0.0008735 | +++?+ | 2.3 | 0.3808 | intergenic | |
| rs7104067 | 11 | 12643870 | 4 | 0 | 0 | A | G | -1.0149 | 0.3049 | 0.0008737 | ---?- | 0 | 0.8941 | intergenic | |
| rs113116817 | 3 | 8561613 | 3 | 0 | 0 | A | T | 3.1873 | 0.9576 | 0.0008739 | +++?? | 0 | 0.5714 | intron | LMCD1 |
| rs1204780 | 6 | 116529114 | 5 | 0 | 0 | T | C | 0.7878 | 0.2367 | 0.0008741 | +++++ | 0 | 0.5558 | intron | NT5DC1 |
| rs10416965 | 19 | 12805525 | 4 | 0 | 0 | C | G | 3.4761 | 1.0445 | 0.0008741 | ++++? | 49.6 | 0.114 | synonymous | FBXW9 |
| rs117047646 | 15 | 86377573 | 4 | 0 | 0 | T | C | -1.889 | 0.5676 | 0.0008741 | ---?- | 0 | 0.9932 | intergenic | |
| rs2705724 | 2 | 183994454 | 4 | 0 | 0 | T | C | -1.1572 | 0.3477 | 0.0008744 | ---?- | 0 | 0.8235 | intron | NUP35 |
| rs11766118 | 7 | 98438576 | 4 | 0 | 0 | A | G | 1.1274 | 0.3388 | 0.0008744 | +++?+ | 0 | 0.9175 | intergenic | |
| rs6033274 | 20 | 12001554 | 4 | 0 | 0 | T | C | 0.9349 | 0.2809 | 0.0008745 | +++?+ | 0 | 0.5601 | intergenic | |
| rs60251885 | 3 | 61083737 | 4 | 0 | 0 | C | G | -3.1364 | 0.9424 | 0.0008745 | ---?+ | 0 | 0.6297 | intron | FHIT |
| rs1209221 | 6 | 116528979 | 5 | 0 | 0 | T | C | -0.7878 | 0.2367 | 0.0008747 | ----- | 0 | 0.5568 | intron | NT5DC1 |
| 2:156775881 | 2 | 156775881 | 3 | 0 | 0 | A | AGCCTG | -1.1898 | 0.3575 | 0.0008747 | +--?? | 66 | 0.05261 | | |
| rs150851337 | 4 | 119291047 | 4 | 0 | 0 | T | C | 1.6311 | 0.4901 | 0.0008749 | +++?+ | 0 | 0.7137 | intergenic | |
| 1:42795963 | 1 | 42795963 | 3 | 0 | 0 | G | GAACA | 1.8762 | 0.5638 | 0.0008753 | +++?? | 0 | 0.3757 | | |
| rs55639531 | 18 | 394341 | 4 | 0 | 0 | C | G | -0.8834 | 0.2655 | 0.0008754 | ---?- | 48.6 | 0.1201 | intron | COLEC12 |
| rs80174548 | 17 | 51112130 | 4 | 0 | 0 | A | G | 1.0316 | 0.31 | 0.0008754 | +-+?+ | 39.6 | 0.174 | intergenic | |
| rs10258433 | 7 | 156157399 | 4 | 0 | 0 | A | G | -0.8426 | 0.2532 | 0.0008755 | ---?- | 38.3 | 0.1824 | intergenic | |
| rs35464923 | 14 | 100098571 | 4 | 0 | 0 | T | C | -0.8635 | 0.2595 | 0.0008755 | ---?+ | 0 | 0.4841 | intergenic | |
| 8:105286346 | 8 | 105286346 | 3 | 0 | 0 | G | GT | 0.8947 | 0.2689 | 0.0008755 | +++?? | 0 | 0.9834 | | |
| rs116834474 | 4 | 140546376 | 4 | 0 | 0 | T | C | -3.0017 | 0.902 | 0.0008758 | ---?- | 23.7 | 0.269 | intergenic | |
| rs184270427 | 7 | 10607892 | 4 | 0 | 0 | A | G | -3.4224 | 1.0285 | 0.0008758 | ---?- | 0 | 0.4001 | intergenic | |
| rs509210 | 1 | 90234055 | 5 | 0 | 0 | A | C | 0.7099 | 0.2133 | 0.000876 | +++++ | 0 | 0.9951 | intergenic | |
| rs12538159 | 7 | 16836642 | 4 | 0 | 0 | A | T | -2.0349 | 0.6115 | 0.000876 | ---?- | 0 | 0.9456 | intron | AGR2 |
| rs35949864 | 3 | 120329134 | 4 | 0 | 0 | T | G | -0.8968 | 0.2695 | 0.000876 | ---?- | 51.5 | 0.1032 | intergenic | |
| 8:140369529 | 8 | 140369529 | 3 | 0 | 0 | G | GA | -0.9875 | 0.2968 | 0.000876 | ---?? | 0 | 0.9131 | | |
| rs75682852 | 6 | 124316871 | 3 | 0 | 0 | A | G | -3.0175 | 0.9068 | 0.0008761 | ---?? | 24.2 | 0.2673 | intron | NKAIN2 |
| rs12870711 | 13 | 20802844 | 3 | 0 | 0 | T | C | -1.0943 | 0.3289 | 0.0008763 | ---?? | 0 | 0.7659 | intron | GJB6 |
| rs601955 | 6 | 165937338 | 5 | 0 | 0 | C | G | 0.7814 | 0.2348 | 0.0008764 | +++++ | 52.4 | 0.07804 | intron | PDE10A |
| rs150976309 | 16 | 77954091 | 3 | 0 | 0 | A | G | -5.8828 | 1.768 | 0.0008764 | ---?? | 0 | 0.8673 | intron | VAT1L |
| 5:42459773 | 5 | 42459773 | 3 | 0 | 0 | G | GA | -1.4413 | 0.4332 | 0.0008765 | ---?? | 0 | 0.5357 | | |
| 22:36927058 | 22 | 36927058 | 3 | 0 | 0 | CT | C | -1.1224 | 0.3373 | 0.0008767 | ---?? | 0 | 0.998 | | |
| rs760087 | 20 | 61467800 | 3 | 0 | 0 | A | G | 1.4819 | 0.4454 | 0.0008769 | +++?? | 0 | 0.4125 | intron | COL9A3 |
| rs78755797 | 13 | 98367042 | 4 | 0 | 0 | A | G | -1.6679 | 0.5013 | 0.000877 | ---?- | 0 | 0.6289 | intergenic | |
| rs145767873 | 1 | 246404454 | 3 | 0 | 0 | A | G | 1.6301 | 0.4899 | 0.0008771 | +++?? | 63.6 | 0.06402 | intron | SMYD3 |
| rs1592216 | 6 | 78099695 | 4 | 0 | 0 | T | C | 0.8682 | 0.2609 | 0.0008773 | +++?+ | 0 | 0.5027 | intergenic | |
| rs2248682 | 21 | 27461329 | 4 | 0 | 0 | A | T | 0.9371 | 0.2817 | 0.0008773 | +++?+ | 44.7 | 0.1435 | intron | APP |
| rs10013356 | 4 | 96524664 | 4 | 0 | 0 | A | T | -1.2507 | 0.3759 | 0.0008773 | ---?- | 27.4 | 0.2476 | intergenic | |
| rs73373407 | 12 | 93992424 | 4 | 0 | 0 | T | G | 2.4646 | 0.7407 | 0.0008773 | +++?+ | 25.7 | 0.2574 | intergenic | |
| rs6812216 | 4 | 38651939 | 5 | 0 | 0 | A | G | -0.7798 | 0.2344 | 0.0008776 | ----+ | 0 | 0.4168 | intron | FLJ13197 |
| rs10072362 | 5 | 134342720 | 4 | 0 | 0 | A | C | 0.8054 | 0.2421 | 0.0008776 | +++?+ | 0 | 0.7565 | intron | CATSPER3 |
| rs12111254 | 6 | 117305572 | 4 | 0 | 0 | T | C | -0.7851 | 0.236 | 0.0008776 | ---?- | 0 | 0.7194 | intergenic | |
| rs4915950 | 1 | 64667955 | 3 | 0 | 0 | A | T | -0.9063 | 0.2724 | 0.0008778 | ---?? | 42.4 | 0.1764 | upstream-gene | |
| rs7679916 | 4 | 10042160 | 5 | 0 | 0 | T | C | 0.7 | 0.2104 | 0.0008778 | ++++- | 14.2 | 0.3236 | upstream-gene | |
| rs12293545 | 11 | 86115657 | 4 | 0 | 0 | T | G | 2.1867 | 0.6573 | 0.000878 | +++?+ | 24 | 0.2669 | intron | CCDC81 |
| rs1120172 | 5 | 39598316 | 4 | 0 | 0 | T | G | -0.9899 | 0.2976 | 0.0008781 | ---?- | 0 | 0.8837 | intergenic | |
| rs4131538 | 7 | 130154197 | 4 | 0 | 0 | T | G | 0.8423 | 0.2532 | 0.0008781 | +-+?+ | 54.5 | 0.08622 | intron | COPG2 |
| rs779189 | 3 | 22142591 | 4 | 0 | 0 | T | C | -0.8593 | 0.2583 | 0.0008782 | ---?- | 0 | 0.7503 | intron | ZNF385D |
| rs6813852 | 4 | 38651997 | 5 | 0 | 0 | T | C | -0.7798 | 0.2344 | 0.0008783 | ----+ | 0 | 0.4165 | intron | FLJ13197 |
| rs368822756 | 1 | 32170249 | 3 | 0 | 0 | A | C | -1.3 | 0.3908 | 0.0008783 | +--?? | 58.1 | 0.09176 | upstream-gene | |
| rs11759843 | 6 | 117310608 | 4 | 0 | 0 | A | G | -0.7849 | 0.2359 | 0.0008784 | ---?- | 0 | 0.7162 | intergenic | |
| rs2705723 | 2 | 183993665 | 4 | 0 | 0 | A | G | 1.1678 | 0.351 | 0.0008785 | +++?+ | 0 | 0.8439 | intron | NUP35 |
| rs4957485 | 5 | 39589159 | 4 | 0 | 0 | A | T | -0.9892 | 0.2973 | 0.0008787 | ---?+ | 0 | 0.8107 | intergenic | |

| | | | | | | | | | | | | | | | |
|---|---|---|---|---|---|---|---|---|---|---|---|---|---|---|---|
| rs10417846 | 19 | 12805948 | 4 | 0 | 0 | A | G | 3.4747 | 1.0445 | 0.0008787 | ++++? | 49.6 | 0.1138 | intron | FBXW9 |
| rs34933499 | 9 | 137741993 | 3 | 0 | 0 | A | G | -2.6383 | 0.7931 | 0.0008787 | ---?? | 0 | 0.4038 | non-coding-exon | MIR3689B |
| rs2090855 | 3 | 114876972 | 4 | 0 | 0 | T | C | 1.785 | 0.5366 | 0.0008789 | +++?+ | 0 | 0.4876 | intergenic | |
| rs9691334 | 7 | 98435152 | 4 | 0 | 0 | A | G | 1.1285 | 0.3392 | 0.0008789 | +++?+ | 0 | 0.895 | intergenic | |
| rs2843039 | 6 | 63342653 | 4 | 0 | 0 | A | G | -2.2807 | 0.6856 | 0.000879 | ---?- | 0 | 0.3971 | intergenic | |
| rs151297784 | 9 | 37071932 | 3 | 0 | 0 | C | G | -2.9234 | 0.8788 | 0.000879 | ---?? | 55.8 | 0.1043 | intergenic | |
| rs9679384 | 2 | 40460805 | 3 | 0 | 0 | T | C | 1.0521 | 0.3163 | 0.0008791 | +++?? | 0 | 0.6018 | intron | SLC8A1,SLC8A1-AS1 |
| rs339456 | 12 | 117168067 | 5 | 0 | 0 | A | G | 1.2297 | 0.3697 | 0.0008792 | +++++ | 0 | 0.8483 | intron | C12orf49 |
| 5:127635617 | 5 | 127635617 | 3 | 0 | 0 | CTAGA | C | 0.9603 | 0.2887 | 0.0008792 | -++?? | 63.6 | 0.06434 | | |
| rs947890 | 11 | 117874493 | 4 | 0 | 0 | A | G | 1.2069 | 0.3628 | 0.0008794 | +++?+ | 0 | 0.5084 | downstream-gene | |
| rs2322795 | 4 | 143427874 | 4 | 0 | 0 | C | G | 1.1251 | 0.3382 | 0.0008794 | +++?+ | 0 | 0.9501 | intron | INPP4B |
| rs72764522 | 15 | 85819215 | 2 | 0 | 0 | A | G | 6.871 | 2.0655 | 0.0008794 | ?+??+ | 0 | 0.7229 | intergenic | |
| rs6871729 | 5 | 122027048 | 4 | 0 | 0 | T | C | -1.6087 | 0.4836 | 0.0008798 | ---?- | 0 | 0.9358 | intergenic | |
| rs59092585 | 4 | 162213481 | 2 | 0 | 0 | T | C | 6.2718 | 1.8855 | 0.0008798 | ?++?? | 0 | 0.6312 | intergenic | |
| rs11761897 | 7 | 57272599 | 3 | 0 | 0 | T | C | -0.9694 | 0.2914 | 0.0008799 | ---?? | 0 | 0.9186 | intergenic | |
| rs112099596 | 13 | 37306316 | 4 | 0 | 0 | T | C | 2.2805 | 0.6856 | 0.0008799 | +++?+ | 0 | 0.8731 | intergenic | |
| 15:85573935 | 15 | 85573935 | 3 | 0 | 0 | CTT | C | 0.8347 | 0.2509 | 0.00088 | -++?? | 63.7 | 0.06377 | | |
| rs13241771 | 7 | 98432607 | 4 | 0 | 0 | A | G | 1.1312 | 0.3401 | 0.0008801 | +++?+ | 0 | 0.8351 | intergenic | |
| rs73122682 | 3 | 79085246 | 5 | 0 | 0 | T | C | -1.9871 | 0.5974 | 0.0008801 | ----- | 54.8 | 0.06472 | intron | ROBO1 |
| rs9264581 | 6 | 31235786 | 5 | 0 | 0 | T | C | 0.7941 | 0.2387 | 0.0008802 | +++++ | 0 | 0.4684 | downstream-gene | |
| rs807147 | 17 | 51212612 | 5 | 0 | 0 | A | G | 1.035 | 0.3112 | 0.0008803 | ++++- | 0 | 0.4183 | intergenic | |
| rs6711360 | 2 | 52950475 | 4 | 0 | 0 | T | C | 1.2814 | 0.3852 | 0.0008803 | +++?+ | 0 | 0.7563 | intergenic | |
| rs59299234 | 19 | 12806172 | 4 | 0 | 0 | A | G | -3.4742 | 1.0445 | 0.0008803 | ----? | 49.6 | 0.1137 | intron | FBXW9 |
| rs201493 | 7 | 101754865 | 4 | 0 | 0 | T | C | -0.8153 | 0.2451 | 0.0008804 | ---?- | 0 | 0.7069 | intron | CUX1 |
| rs3910845 | 8 | 64918476 | 4 | 0 | 0 | T | C | 0.8252 | 0.2481 | 0.0008804 | +++?- | 0 | 0.4885 | intergenic | |
| rs1204781 | 6 | 116530201 | 5 | 0 | 0 | C | G | -0.787 | 0.2366 | 0.0008806 | ----- | 0 | 0.5531 | intron | NT5DC1 |
| rs79250382 | 5 | 141463515 | 5 | 0 | 0 | A | G | -0.9596 | 0.2885 | 0.0008807 | +---- | 0 | 0.5864 | intergenic |  |
| 2:184031753 | 2 | 184031753 | 3 | 0 | 0 | T | TACTG | -1.1836 | 0.3558 | 0.0008807 | ---?? | 0 | 0.7992 | | |
| rs3753804 | 1 | 180138988 | 4 | 0 | 0 | A | T | 2.0627 | 0.6202 | 0.0008809 | +++?- | 35 | 0.2021 | intron | QSOX1 |
| rs117645909 | 15 | 102073408 | 4 | 0 | 0 | A | T | 2.0958 | 0.6301 | 0.000881 | +++?- | 4.8 | 0.3688 | intergenic | |
| rs772896 | 12 | 103237793 | 4 | 0 | 0 | A | G | 0.8246 | 0.2479 | 0.0008811 | +++?+ | 0 | 0.6741 | intron | PAH |
| rs118093614 | 16 | 85242361 | 4 | 0 | 0 | T | C | 1.1894 | 0.3576 | 0.0008811 | +++?+ | 0 | 0.6961 | intron | GSE1 |
| rs7800433 | 7 | 82426565 | 4 | 0 | 0 | A | G | 0.8399 | 0.2525 | 0.0008813 | +-+?+ | 45.8 | 0.1367 | intron | PCLO |
| rs117526932 | 19 | 3387152 | 4 | 0 | 0 | A | G | -2.6825 | 0.8065 | 0.0008814 | ---?- | 0 | 0.5943 | intron | NFIC |
| rs12812650 | 12 | 51463697 | 5 | 0 | 0 | A | G | 0.928 | 0.279 | 0.0008816 | +++++ | 0 | 0.4546 | intron | CSRNP2 |
| rs73430108 | 22 | 28890561 | 4 | 0 | 0 | T | G | 2.2576 | 0.6788 | 0.0008817 | +++?+ | 0 | 0.568 | intron | TTC28 |
| rs7804206 | 7 | 82426805 | 4 | 0 | 0 | C | G | 0.8397 | 0.2525 | 0.0008818 | +-+?+ | 45.7 | 0.137 | intron | PCLO |
| rs12503725 | 4 | 42768857 | 3 | 0 | 0 | T | C | -1.1534 | 0.3468 | 0.0008819 | ---?? | 0 | 0.5586 | intergenic | |
| rs13108399 | 4 | 190017030 | 4 | 0 | 0 | T | C | -0.8341 | 0.2508 | 0.0008819 | ---?- | 27.5 | 0.2468 | intergenic | |
| rs17726447 | 16 | 84124359 | 4 | 0 | 0 | T | C | 3.1867 | 0.9582 | 0.000882 | +++?+ | 0 | 0.4109 | intron | MBTPS1 |
| rs223831 | 16 | 57453897 | 4 | 0 | 0 | A | G | -1.2317 | 0.3704 | 0.0008821 | ---?- | 0 | 0.8433 | downstream-gene | |
| rs6074015 | 20 | 44712175 | 4 | 0 | 0 | T | C | 0.8579 | 0.258 | 0.0008821 | +++?+ | 32.8 | 0.2157 | intron | NCOA5 |
| rs75483607 | 13 | 75252699 | 4 | 0 | 0 | A | C | 1.6339 | 0.4913 | 0.0008822 | +++?+ | 22.8 | 0.274 | intergenic | |
| rs9846777 | 3 | 196332734 | 4 | 0 | 0 | A | G | 0.9409 | 0.2829 | 0.0008824 | +++?+ | 0 | 0.885 | intergenic | |
| 5:10155839 | 5 | 10155839 | 3 | 0 | 0 | A | AG | -0.9101 | 0.2737 | 0.0008824 | ---?? | 36.9 | 0.2051 | | |
| rs13162906 | 5 | 124617261 | 4 | 0 | 0 | A | C | -2.009 | 0.6041 | 0.0008826 | ---?- | 26.7 | 0.2515 | intergenic | |
| rs55719957 | 11 | 74365045 | 4 | 0 | 0 | C | G | 1.2464 | 0.3748 | 0.0008826 | +++?+ | 0 | 0.6144 | intron | POLD3 |
| rs78745584 | 11 | 97029941 | 3 | 0 | 0 | A | G | -2.6086 | 0.7844 | 0.0008827 | ---?? | 25.4 | 0.2617 | intergenic | |
| rs7297455 | 12 | 30864924 | 5 | 0 | 0 | A | G | 0.702 | 0.2111 | 0.000883 | +++++ | 0 | 0.7372 | intron | CAPRIN2 |
| rs11722930 | 4 | 10035454 | 5 | 0 | 0 | A | G | 0.699 | 0.2102 | 0.000883 | ++++- | 14.1 | 0.3242 | intron | SLC2A9 |
| rs28799386 | 9 | 30741749 | 4 | 0 | 0 | A | G | 3.3297 | 1.0013 | 0.000883 | +++?+ | 2.6 | 0.3795 | intergenic | |
| rs78066335 | 4 | 63506722 | 4 | 0 | 0 | A | G | 2.9019 | 0.8726 | 0.000883 | +++?+ | 0 | 0.6681 | intergenic | |
| 10:93057933 | 10 | 93057933 | 3 | 0 | 0 | G | GT | 0.9446 | 0.2841 | 0.000883 | +++?? | 0 | 0.6703 | | |
| rs1204782 | 6 | 116530767 | 5 | 0 | 0 | T | C | 0.7866 | 0.2366 | 0.0008831 | +++++ | 0 | 0.5518 | intron | NT5DC1 |
| rs3850738 | 8 | 64921894 | 4 | 0 | 0 | A | T | 0.8253 | 0.2482 | 0.0008831 | +++?- | 0 | 0.4869 | intergenic | |
| rs339455 | 12 | 117168476 | 5 | 0 | 0 | C | G | 1.23 | 0.3699 | 0.0008832 | +++++ | 0 | 0.8475 | intron | C12orf49 |
| rs13167685 | 5 | 60554610 | 5 | 0 | 0 | T | C | 0.9506 | 0.2859 | 0.0008832 | ++++- | 23.9 | 0.2619 | intergenic | |
| 18:32867839 | 18 | 32867839 | 3 | 0 | 0 | CT | C | 0.9195 | 0.2765 | 0.0008832 | +++?? | 0 | 0.8291 | | |
| rs9956259 | 18 | 37411265 | 4 | 0 | 0 | T | C | 0.9005 | 0.2708 | 0.0008834 | +++?+ | 0 | 0.6257 | intergenic | |

| | | | | | | | | | | | | | | | |
|---|---|---|---|---|---|---|---|---|---|---|---|---|---|---|---|
| rs55722303 | 4 | 94711288 | 4 | 0 | 0 | A | G | 0.9021 | 0.2713 | 0.0008834 | +++?- | 0 | 0.7461 | intergenic | |
| rs141798189 | 12 | 127184468 | 3 | 0 | 0 | A | G | 2.2475 | 0.6759 | 0.0008834 | +++?? | 0 | 0.4522 | intergenic | |
| rs1204783 | 6 | 116530795 | 5 | 0 | 0 | A | G | 0.7866 | 0.2366 | 0.0008836 | +++++ | 0 | 0.5515 | intron | NT5DC1 |
| rs1491283 | 3 | 163458069 | 3 | 0 | 0 | T | C | -1.4588 | 0.4387 | 0.0008836 | ?--?- | 0 | 0.7601 | intergenic | |
| rs67354673 | 12 | 15589309 | 4 | 0 | 0 | T | C | 1.1065 | 0.3328 | 0.0008836 | +++?+ | 0 | 0.9414 | intron | PTPRO |
| rs339458 | 12 | 117165833 | 5 | 0 | 0 | T | C | 1.2241 | 0.3681 | 0.0008837 | +++++ | 0 | 0.8535 | intron | C12orf49 |
| rs3914506 | 9 | 30742141 | 4 | 0 | 0 | T | C | -3.3296 | 1.0014 | 0.0008838 | ---?- | 2.4 | 0.3805 | intergenic | |
| rs3916917 | 13 | 102560610 | 4 | 0 | 0 | T | C | 0.9918 | 0.2983 | 0.000884 | +++?+ | 17 | 0.306 | intron | FGF14 |
| rs79233179 | 11 | 86181023 | 4 | 0 | 0 | T | C | 1.5628 | 0.47 | 0.000884 | +++?+ | 0 | 0.4533 | intron | ME3 |
| rs7784058 | 7 | 82426607 | 4 | 0 | 0 | A | G | -0.8395 | 0.2525 | 0.0008841 | -+-?- | 45.8 | 0.1368 | intron | PCLO |
| rs1355681 | 17 | 51127870 | 4 | 0 | 0 | A | C | 1.4188 | 0.4267 | 0.0008843 | +++?- | 15.9 | 0.3123 | intergenic | |
| rs7745661 | 6 | 130913854 | 4 | 0 | 0 | A | G | 0.8631 | 0.2596 | 0.0008843 | +++?+ | 0 | 0.9595 | intergenic | |
| 9:94273105 | 9 | 94273105 | 2 | 0 | 0 | <ins:me: | C | -5.6469 | 1.6984 | 0.0008844 | --??? | 0 | 0.3406 | | |
| rs825536 | 2 | 19505964 | 3 | 0 | 0 | A | G | 0.8447 | 0.254 | 0.0008845 | +++?? | 0 | 0.8324 | intergenic | |
| rs11865810 | 16 | 62315105 | 4 | 0 | 0 | A | T | 2.1519 | 0.6472 | 0.0008848 | +++?+ | 12.9 | 0.3282 | intergenic | |
| rs75960714 | 13 | 106177600 | 4 | 0 | 0 | T | C | 1.7293 | 0.5201 | 0.0008848 | +++?+ | 0 | 0.757 | intergenic | |
| rs201418979 | 11 | 25667294 | 3 | 0 | 0 | A | G | 0.8683 | 0.2612 | 0.0008849 | +++?? | 0 | 0.4224 | intergenic | |
| rs7377536 | 4 | 189064509 | 4 | 0 | 0 | A | G | 1.2333 | 0.371 | 0.0008852 | +++?+ | 0 | 0.9244 | intron | TRIML1 |
| rs57858781 | 19 | 1639684 | 4 | 0 | 0 | C | G | -2.6167 | 0.7871 | 0.0008852 | ---?- | 0 | 0.8114 | intron | TCF3 |
| rs8034410 | 15 | 53758807 | 4 | 0 | 0 | T | C | -0.8133 | 0.2446 | 0.0008855 | ---?- | 45.9 | 0.136 | intergenic | |
| rs28827536 | 9 | 30741767 | 4 | 0 | 0 | T | G | 3.3289 | 1.0013 | 0.0008856 | +++?+ | 2.5 | 0.3801 | intergenic | |
| rs12432292 | 14 | 95897037 | 4 | 0 | 0 | T | G | 1.2441 | 0.3742 | 0.0008858 | +++?+ | 0 | 0.455 | intron | SYNE3 |
| rs11734786 | 4 | 10041499 | 5 | 0 | 0 | A | C | -0.6994 | 0.2104 | 0.0008859 | ----+ | 14.3 | 0.3229 | intron | SLC2A9 |
| rs12494652 | 3 | 116028380 | 4 | 0 | 0 | C | G | -1.0305 | 0.31 | 0.0008861 | ---?- | 0 | 0.4839 | intron | LSAMP |
| rs35544666 | 2 | 125003188 | 4 | 0 | 0 | A | T | 2.5617 | 0.7706 | 0.0008861 | +++?- | 0 | 0.5743 | intron | CNTNAP5 |
| rs10409749 | 19 | 12816794 | 4 | 0 | 0 | A | G | -3.4778 | 1.0462 | 0.0008862 | ----? | 50.4 | 0.1094 | intron | TNPO2 |
| rs372766190 | 1 | 32170250 | 3 | 0 | 0 | A | C | -1.299 | 0.3907 | 0.0008862 | +--?? | 58.2 | 0.09151 | upstream-gene | |
| rs10415321 | 19 | 12816790 | 4 | 0 | 0 | T | C | -3.4778 | 1.0462 | 0.0008863 | ----? | 50.4 | 0.1095 | intron | TNPO2 |
| rs12663033 | 6 | 24024083 | 5 | 0 | 0 | T | C | -1.2991 | 0.3908 | 0.0008864 | ----+ | 0 | 0.8127 | intergenic |  |
| rs448591 | 7 | 101873585 | 4 | 0 | 0 | A | G | -0.8405 | 0.2528 | 0.0008865 | ---?- | 0 | 0.8567 | intron | CUX1 |
| rs116084968 | 1 | 42713564 | 4 | 0 | 0 | T | C | -1.7695 | 0.5323 | 0.0008865 | ---?- | 0 | 0.5474 | intron | FOXJ3 |
| rs17546500 | 7 | 7711468 | 4 | 0 | 0 | T | G | -1.7839 | 0.5366 | 0.0008869 | ---?- | 48.5 | 0.1206 | intron | RPA3,RPA3-AS1 |
| rs17677605 | 10 | 20138208 | 4 | 0 | 0 | A | G | -0.909 | 0.2735 | 0.000887 | ---?- | 0 | 0.955 | intron | PLXDC2 |
| rs807136 | 17 | 51223965 | 5 | 0 | 0 | T | C | 1.0228 | 0.3077 | 0.0008871 | ++++- | 5.4 | 0.3761 | intergenic | |
| rs11828585 | 11 | 86181711 | 4 | 0 | 0 | A | G | 1.5627 | 0.4701 | 0.0008871 | +++?+ | 0 | 0.4588 | intron | ME3 |
| rs17088878 | 18 | 71823945 | 4 | 0 | 0 | T | G | -1.4097 | 0.4241 | 0.0008871 | +--?- | 50.3 | 0.1098 | intron | TIMM21 |
| rs6532564 | 4 | 96464931 | 3 | 0 | 0 | C | G | 1.0492 | 0.3156 | 0.0008873 | +++?? | 3.2 | 0.3558 | intron | UNC5C |
| rs134821 | 22 | 45970029 | 4 | 0 | 0 | T | C | 0.9369 | 0.2819 | 0.0008875 | +++?- | 4.2 | 0.3717 | intron | FBLN1 |
| rs61796994 | 1 | 109588530 | 4 | 0 | 0 | T | C | -0.9726 | 0.2926 | 0.0008876 | ---?- | 56.2 | 0.07681 | upstream-gene | |
| rs4876223 | 8 | 2065092 | 4 | 0 | 0 | T | G | -0.8954 | 0.2694 | 0.0008878 | ---?- | 0 | 0.5886 | intron | MYOM2 |
| rs10834510 | 11 | 24883203 | 2 | 0 | 0 | T | C | 4.1868 | 1.2596 | 0.0008878 | ?++?? | 66.3 | 0.08518 | intron | LUZP2 |
| rs181716200 | 5 | 21963310 | 3 | 0 | 0 | T | C | -4.051 | 1.2188 | 0.0008878 | +--?? | 58.5 | 0.08979 | intron | CDH12 |
| rs223832 | 16 | 57454356 | 4 | 0 | 0 | T | C | 1.2315 | 0.3705 | 0.0008879 | +++?+ | 0 | 0.844 | downstream-gene | |
| rs639132 | 16 | 57466539 | 4 | 0 | 0 | A | G | 1.2661 | 0.3809 | 0.0008879 | +++?+ | 0 | 0.8856 | intron | CIAPIN1 |
| rs1431311 | 17 | 54115205 | 4 | 0 | 0 | A | C | -0.9792 | 0.2946 | 0.0008879 | ---?+ | 40.3 | 0.1701 | intergenic | |
| rs11196898 | 10 | 116508805 | 4 | 0 | 0 | T | G | -1.3173 | 0.3963 | 0.000888 | ---?- | 0 | 0.8395 | intergenic | |
| rs807135 | 17 | 51224241 | 5 | 0 | 0 | T | C | -1.0201 | 0.3069 | 0.0008882 | ----+ | 6.4 | 0.3701 | intergenic |  |
| rs12152652 | 4 | 20753490 | 4 | 0 | 0 | T | C | -0.8776 | 0.2641 | 0.0008885 | ---?- | 0 | 0.7475 | intron | KCNIP4 |
| rs4814058 | 20 | 12002704 | 4 | 0 | 0 | T | C | 0.9368 | 0.2819 | 0.0008886 | +++?+ | 0 | 0.5482 | intergenic | |
| rs6068695 | 20 | 52480241 | 4 | 0 | 0 | A | G | 0.8104 | 0.2438 | 0.0008886 | +++?+ | 32.8 | 0.2154 | intergenic | |
| rs112225687 | 6 | 135008361 | 3 | 0 | 0 | T | C | 2.2512 | 0.6773 | 0.0008886 | +++?? | 0 | 0.5406 | intergenic | |
| rs1204806 | 6 | 116548944 | 5 | 0 | 0 | C | G | -0.8433 | 0.2537 | 0.0008887 | ----- | 0 | 0.833 | intron | NT5DC1 |
| rs36108077 | 4 | 190017202 | 4 | 0 | 0 | T | C | 0.8337 | 0.2508 | 0.0008887 | +++?+ | 27.5 | 0.2467 | intergenic | |
| rs75065042 | 6 | 109102424 | 3 | 0 | 0 | A | G | -3.961 | 1.1918 | 0.0008887 | -+-?? | 55.1 | 0.108 | intergenic | |
| rs9264531 | 6 | 31234239 | 5 | 0 | 0 | A | T | -0.8064 | 0.2426 | 0.0008888 | ----- | 0 | 0.9415 | downstream-gene |  |
| rs639463 | 2 | 20716757 | 3 | 0 | 0 | T | C | 1.5786 | 0.475 | 0.0008891 | +++?? | 0 | 0.5835 | intergenic | |
| rs9761637 | 4 | 38750185 | 5 | 0 | 0 | A | G | -0.754 | 0.2269 | 0.0008892 | ----- | 0 | 0.9401 | intergenic |  |
| rs4973249 | 2 | 230918326 | 4 | 0 | 0 | A | G | 1.4147 | 0.4257 | 0.0008893 | +++?+ | 21.9 | 0.2789 | intron | SLC16A14 |
| rs1204805 | 6 | 116548891 | 5 | 0 | 0 | T | C | 0.8432 | 0.2537 | 0.0008894 | +++++ | 0 | 0.833 | intron | NT5DC1 |

| | | | | | | | | | | | | | | | |
|---|---|---|---|---|---|---|---|---|---|---|---|---|---|---|---|
| rs1334684 | 6 | 117312352 | 4 | 0 | 0 | T | C | 0.7839 | 0.2359 | 0.0008894 | +++?+ | 0 | 0.7162 | intergenic | |
| rs6511832 | 19 | 12807066 | 4 | 0 | 0 | C | G | 3.4713 | 1.0445 | 0.0008894 | ++++? | 49.7 | 0.1132 | synonymous | FBXW9 |
| 10:82305802 | 10 | 82305802 | 3 | 0 | 0 | CT | C | -1.3762 | 0.4141 | 0.0008894 | ---?? | 0 | 0.8787 | | |
| rs6531657 | 4 | 38718732 | 5 | 0 | 0 | T | C | -0.7504 | 0.2258 | 0.0008895 | ----- | 0 | 0.9027 | intergenic |  |
| 5:1717551 | 5 | 1717551 | 4 | 0 | 0 | C | G | -2.3838 | 0.7173 | 0.0008895 | ---?- | 44.3 | 0.1456 | | |
| rs143282921 | 4 | 14723587 | 3 | 0 | 0 | A | C | 2.1069 | 0.634 | 0.0008896 | +++?? | 47.7 | 0.1475 | intergenic | |
| rs13148371 | 4 | 10032561 | 5 | 0 | 0 | T | C | 0.6986 | 0.2102 | 0.0008897 | ++++- | 14.8 | 0.3199 | intron | SLC2A9 |
| rs72900970 | 2 | 68134824 | 4 | 0 | 0 | T | C | -4.0357 | 1.2144 | 0.0008897 | ---?- | 0 | 0.4335 | intergenic | |
| rs28749936 | 9 | 44252408 | 3 | 0 | 0 | A | T | -1.7435 | 0.5247 | 0.0008898 | ---?? | 0 | 0.8484 | intergenic | |
| rs10843465 | 12 | 29805354 | 4 | 0 | 0 | T | C | -0.8947 | 0.2692 | 0.0008906 | ---?- | 0 | 0.9364 | intron | TMTC1 |
| 1:42784893 | 1 | 42784893 | 3 | 0 | 0 | CTGTTT | C | -1.8618 | 0.5603 | 0.0008908 | ---?? | 0.4 | 0.3665 | | |
| rs61696770 | 2 | 117108325 | 4 | 0 | 0 | T | C | 1.6597 | 0.4995 | 0.000891 | +++?+ | 0 | 0.6607 | intergenic | |
| rs7248726 | 19 | 12816559 | 4 | 0 | 0 | T | G | 3.476 | 1.0461 | 0.0008912 | ++++? | 50.4 | 0.1095 | synonymous | TNPO2 |
| rs12000926 | 9 | 16189079 | 4 | 0 | 0 | A | G | -0.9102 | 0.2739 | 0.0008914 | ---?- | 0 | 0.9294 | intergenic | |
| rs16998686 | 21 | 41218070 | 4 | 0 | 0 | T | C | 2.2294 | 0.671 | 0.0008914 | +++?+ | 36.5 | 0.1933 | intergenic | |
| rs6032644 | 20 | 44693296 | 4 | 0 | 0 | A | T | -0.8605 | 0.259 | 0.0008915 | +--?- | 41.4 | 0.1632 | intron | NCOA5 |
| rs117142879 | 19 | 45305584 | 3 | 0 | 0 | T | C | 2.9412 | 0.8852 | 0.0008915 | +++?? | 33.8 | 0.2208 | downstream-gene | |
| 8:65038761 | 8 | 65038761 | 3 | 0 | 0 | T | TA | 0.8365 | 0.2518 | 0.0008915 | +++?? | 0 | 0.8479 | | |
| rs113536508 | 8 | 1041555 | 4 | 0 | 0 | A | G | 2.5854 | 0.7781 | 0.0008918 | +++?+ | 0 | 0.5696 | intron | DLGAP2,ERICH1-AS1 |
| rs7189422 | 16 | 84841883 | 4 | 0 | 0 | T | C | 1.1968 | 0.3602 | 0.0008919 | +++?- | 1.6 | 0.3841 | intergenic | |
| 2:170490173 | 2 | 170490173 | 3 | 0 | 0 | T | TA | 1.2193 | 0.367 | 0.0008919 | +++?? | 0 | 0.89 | | |
| rs74737216 | 13 | 48760626 | 4 | 0 | 0 | A | G | -2.4843 | 0.7477 | 0.0008921 | ---?- | 5.9 | 0.3636 | intergenic | |
| rs13167244 | 5 | 60554377 | 3 | 0 | 0 | A | G | -1.1283 | 0.3396 | 0.0008924 | ---?? | 46.1 | 0.1562 | intergenic | |
| rs192214348 | 20 | 22466816 | 2 | 0 | 0 | T | C | -5.2638 | 1.5843 | 0.0008924 | -?-?? | 0 | 0.3667 | intergenic | |
| rs11179652 | 12 | 41453822 | 4 | 0 | 0 | T | G | 1.7204 | 0.5178 | 0.0008929 | +++?+ | 0 | 0.6258 | intron | CNTN1 |
| rs10775210 | 15 | 33931265 | 4 | 0 | 0 | A | C | 1.1703 | 0.3523 | 0.0008932 | +++?+ | 55.1 | 0.08286 | intron | RYR3 |
| rs60001648 | 3 | 30829361 | 4 | 0 | 0 | A | C | 2.7725 | 0.8345 | 0.0008932 | +++?- | 9.5 | 0.3458 | intron | GADL1 |
| rs365308 | 5 | 176510140 | 3 | 0 | 0 | A | T | 0.9341 | 0.2812 | 0.0008933 | +++?? | 0 | 0.7906 | upstream-gene | |
| rs182021935 | 4 | 89057338 | 2 | 0 | 0 | T | C | -3.9462 | 1.1878 | 0.0008933 | -?-?? | 0 | 0.4594 | intron | ABCG2 |
| rs1393589 | 4 | 108242111 | 4 | 0 | 0 | A | G | -0.815 | 0.2453 | 0.0008935 | ---?- | 0 | 0.5602 | intergenic | |
| rs12208867 | 6 | 30026236 | 5 | 0 | 0 | A | G | 1.4685 | 0.4421 | 0.0008935 | +++++ | 0 | 0.7553 | intron | ZNRD1-AS1 |
| rs377210569 | 1 | 32170251 | 3 | 0 | 0 | T | G | 1.298 | 0.3907 | 0.0008935 | -++?? | 58.2 | 0.09132 | upstream-gene | |
| rs667380 | 16 | 57466549 | 4 | 0 | 0 | T | C | -1.2657 | 0.381 | 0.0008939 | ---?- | 0 | 0.8856 | intron | CIAPIN1 |
| rs13167246 | 5 | 60554383 | 3 | 0 | 0 | A | G | -1.13 | 0.3402 | 0.0008939 | ---?? | 47.5 | 0.149 | intergenic | |
| rs1413751 | 6 | 117315210 | 4 | 0 | 0 | A | G | -0.7837 | 0.2359 | 0.0008941 | ---?- | 0 | 0.7159 | intergenic | |
| rs5005997 | 6 | 118474562 | 4 | 0 | 0 | T | C | -0.8415 | 0.2533 | 0.0008943 | ---?- | 0 | 0.8305 | intron | SLC35F1 |
| rs80285458 | 18 | 4202503 | 4 | 0 | 0 | A | T | -3.9251 | 1.1816 | 0.0008943 | ---?- | 0 | 0.7376 | intron | DLGAP1 |
| rs7117762 | 11 | 12643601 | 4 | 0 | 0 | C | G | 1.0127 | 0.3049 | 0.0008944 | +++?+ | 0 | 0.8948 | intergenic | |
| rs56222853 | 9 | 16188771 | 3 | 0 | 0 | A | G | -0.9363 | 0.2819 | 0.0008944 | ---?? | 0 | 0.8262 | intergenic | |
| rs111497785 | 12 | 6303905 | 2 | 0 | 0 | A | C | 6.8263 | 2.055 | 0.0008944 | +???+ | 0 | 0.781 | intergenic | |
| rs16934192 | 9 | 16185090 | 4 | 0 | 0 | T | G | -1.2064 | 0.3632 | 0.0008945 | ---?- | 0 | 0.6186 | intergenic | |
| rs6543468 | 2 | 107676592 | 4 | 0 | 0 | A | G | 0.8901 | 0.268 | 0.0008948 | +++?+ | 34.9 | 0.2027 | intergenic | |
| rs7781603 | 7 | 107348372 | 4 | 0 | 0 | A | C | -0.8423 | 0.2536 | 0.0008948 | ---?- | 0 | 0.5502 | intron | SLC26A4 |
| rs9883507 | 3 | 131424084 | 4 | 0 | 0 | A | G | 2.0433 | 0.6152 | 0.0008951 | +++?+ | 0 | 0.6294 | intron | CPNE4 |
| rs60746804 | 4 | 158729043 | 4 | 0 | 0 | A | T | 0.8233 | 0.2479 | 0.0008951 | +++?+ | 48.5 | 0.1207 | intergenic | |
| rs72819936 | 10 | 107029061 | 4 | 0 | 0 | T | C | 1.6557 | 0.4985 | 0.0008952 | +++?+ | 0 | 0.8386 | downstream-gene | |
| rs7113773 | 11 | 134473947 | 4 | 0 | 0 | A | T | 0.99 | 0.2981 | 0.0008954 | +++?+ | 0 | 0.9671 | intergenic | |
| rs13289323 | 9 | 85592091 | 4 | 0 | 0 | A | C | 2.3659 | 0.7123 | 0.0008956 | +++?- | 0 | 0.8765 | downstream-gene | |
| rs4667256 | 2 | 189880726 | 4 | 0 | 0 | A | G | -0.9437 | 0.2841 | 0.0008957 | ---?- | 40.8 | 0.1672 | downstream-gene | |
| rs825212 | 1 | 64766900 | 4 | 0 | 0 | A | C | 0.9203 | 0.2771 | 0.000896 | +++?+ | 0 | 0.4955 | intergenic | |
| rs11856240 | 15 | 46001514 | 3 | 0 | 0 | C | G | -1.2978 | 0.3908 | 0.0008961 | ---?? | 0 | 0.9398 | intergenic | |
| rs4964010 | 12 | 26798389 | 4 | 0 | 0 | T | G | -0.8541 | 0.2572 | 0.0008962 | ---?+ | 29.2 | 0.2368 | intron | ITPR2 |
| rs1997893 | 14 | 85976907 | 4 | 0 | 0 | A | C | 2.172 | 0.654 | 0.0008963 | +++?+ | 34.7 | 0.2041 | intergenic | |
| rs12508991 | 4 | 10041104 | 5 | 0 | 0 | T | C | 0.6987 | 0.2104 | 0.0008963 | ++++- | 14.4 | 0.3225 | intron | SLC2A9 |
| rs12779673 | 10 | 5508434 | 3 | 0 | 0 | C | G | 1.1362 | 0.3421 | 0.0008963 | +++?? | 0 | 0.5403 | intergenic | |
| 1:168852863 | 1 | 168852863 | 3 | 0 | 0 | T | TTTA | 1.2737 | 0.3835 | 0.0008964 | +++?? | 24.7 | 0.2651 | | |
| rs8050519 | 16 | 82822618 | 4 | 0 | 0 | A | C | 1.6572 | 0.499 | 0.0008966 | +++?+ | 39.9 | 0.1724 | intron | CDH13 |
| rs7112309 | 11 | 86175078 | 4 | 0 | 0 | A | G | 1.5573 | 0.4689 | 0.0008967 | +++?+ | 0 | 0.4251 | intron | ME3 |
| rs9295943 | 6 | 30989188 | 3 | 0 | 0 | A | C | 1.3655 | 0.4112 | 0.0008969 | +++?? | 0 | 0.8486 | intron | MUC22 |

| | | | | | | | | | | | | | | | |
|---|---|---|---|---|---|---|---|---|---|---|---|---|---|---|---|
| rs4352461 | 4 | 26169972 | 3 | 0 | 0 | A | G | -1.2746 | 0.3838 | 0.000897 | ---?? | 0 | 0.4794 | intron | RBPJ |
| rs2594330 | 15 | 87539733 | 4 | 0 | 0 | T | C | -0.8587 | 0.2586 | 0.0008972 | ---?- | 0 | 0.4459 | intron | AGBL1 |
| rs4273126 | 18 | 25374914 | 4 | 0 | 0 | T | C | -0.8312 | 0.2503 | 0.0008972 | ---?- | 4.7 | 0.3693 | intergenic | |
| rs4907025 | 1 | 85608721 | 4 | 0 | 0 | C | G | -1.5062 | 0.4536 | 0.0008972 | ---?- | 0 | 0.9883 | intergenic | |
| rs11999291 | 9 | 16186212 | 4 | 0 | 0 | A | G | -0.8977 | 0.2703 | 0.0008972 | ---?- | 0 | 0.9257 | intergenic | |
| rs79532807 | 2 | 237849999 | 4 | 0 | 0 | A | G | 1.8973 | 0.5713 | 0.0008972 | +++?+ | 0 | 0.4171 | intergenic | |
| rs72936931 | 6 | 72840385 | 4 | 0 | 0 | T | C | -1.5428 | 0.4646 | 0.0008974 | ---?+ | 0 | 0.5939 | intron | RIMS1 |
| rs113808930 | 12 | 6304790 | 2 | 0 | 0 | T | G | 6.8206 | 2.0538 | 0.0008974 | +???+ | 0 | 0.7821 | upstream-gene | |
| rs73334973 | 8 | 114877361 | 4 | 0 | 0 | T | C | -1.8122 | 0.5457 | 0.0008977 | ---?+ | 5.4 | 0.3659 | intergenic | |
| rs7042604 | 9 | 7852471 | 4 | 0 | 0 | A | G | -1.0097 | 0.3041 | 0.0008978 | -+-?- | 45.8 | 0.1368 | intergenic | |
| rs12791577 | 11 | 95535510 | 4 | 0 | 0 | C | G | -2.0286 | 0.6109 | 0.0008978 | ---?- | 0 | 0.8021 | intron | CEP57 |
| rs4145648 | 5 | 103176234 | 4 | 0 | 0 | T | C | 1.4633 | 0.4407 | 0.0008979 | +++?+ | 0 | 0.8107 | intergenic | |
| rs4695240 | 4 | 47516526 | 5 | 0 | 0 | A | G | 0.7475 | 0.2251 | 0.0008979 | ++++- | 0 | 0.4696 | intron | ATP10D |
| rs6820188 | 4 | 10051655 | 5 | 0 | 0 | T | C | 0.7077 | 0.2131 | 0.0008979 | ++++- | 27.3 | 0.2393 | intergenic | |
| rs143647552 | 1 | 246405138 | 3 | 0 | 0 | A | G | 1.6325 | 0.4916 | 0.0008979 | +++?? | 63.8 | 0.06303 | intron | SMYD3 |
| rs35368771 | 11 | 95535415 | 4 | 0 | 0 | T | C | 2.0285 | 0.6109 | 0.000898 | +++?+ | 0 | 0.8019 | intron | CEP57 |
| rs35634995 | 4 | 47514967 | 5 | 0 | 0 | A | C | 0.7453 | 0.2245 | 0.0008985 | ++++- | 0 | 0.4697 | intron | ATP10D |
| rs72936929 | 6 | 72840182 | 4 | 0 | 0 | T | C | -1.5427 | 0.4646 | 0.0008988 | ---?+ | 0 | 0.594 | intron | RIMS1 |
| rs2367481 | 3 | 188520543 | 4 | 0 | 0 | A | G | -0.8523 | 0.2567 | 0.000899 | ---?- | 26.1 | 0.2554 | intron | LPP |
| rs37388 | 5 | 35085562 | 4 | 0 | 0 | T | C | 1.2126 | 0.3652 | 0.0008992 | -++?- | 54.1 | 0.08843 | intron | PRLR |
| rs147468633 | 16 | 20532904 | 3 | 0 | 0 | C | G | 3.8433 | 1.1575 | 0.0008992 | +++?? | 46.4 | 0.1545 | intergenic | |
| rs66528019 | 19 | 56830757 | 4 | 0 | 0 | T | C | 2.3277 | 0.7011 | 0.0008995 | +-+?+ | 42.9 | 0.1538 | upstream-gene | |
| rs79655983 | 1 | 112163327 | 4 | 0 | 0 | T | C | 2.4824 | 0.7477 | 0.0008995 | +++?- | 26.8 | 0.251 | intron | RAP1A |
| rs11213688 | 11 | 110853756 | 4 | 0 | 0 | A | G | -0.9342 | 0.2814 | 0.0008998 | ---?- | 0 | 0.4444 | intergenic | |
| rs807130 | 17 | 51229953 | 5 | 0 | 0 | A | G | -1.0259 | 0.309 | 0.0008999 | ----+ | 5.2 | 0.377 | intergenic |  |
| 13:71704904 | 13 | 71704904 | 3 | 0 | 0 | T | TA | -0.9178 | 0.2764 | 0.0009001 | ---?? | 20.9 | 0.2826 | | |
| rs62418252 | 6 | 63288410 | 3 | 0 | 0 | T | C | -2.3641 | 0.7121 | 0.0009004 | ---?? | 0.4 | 0.3664 | intergenic | |
| rs75230005 | 11 | 86182560 | 4 | 0 | 0 | A | G | 1.5612 | 0.4703 | 0.0009004 | +++?+ | 0 | 0.4688 | intron | ME3 |
| rs9556766 | 13 | 98265720 | 3 | 0 | 0 | T | G | -4.6583 | 1.4031 | 0.0009005 | ---?? | 0 | 0.5316 | intergenic | |
| rs1502570 | 3 | 2477066 | 4 | 0 | 0 | A | G | -0.8032 | 0.2419 | 0.0009009 | ---?- | 0 | 0.4219 | intron | CNTN4 |
| rs8040401 | 15 | 98049176 | 4 | 0 | 0 | T | C | 0.8481 | 0.2555 | 0.0009009 | +++?+ | 0 | 0.8582 | intergenic | |
| 9:30866629 | 9 | 30866629 | 3 | 0 | 0 | CATT | C | -3.6541 | 1.1007 | 0.0009011 | ---?? | 0 | 0.5971 | | |
| rs142816451 | 3 | 169648513 | 3 | 0 | 0 | A | G | -2.7971 | 0.8426 | 0.0009012 | ---?? | 0 | 0.9708 | intron | SAMD7 |
| rs369530185 | 1 | 32170252 | 3 | 0 | 0 | C | G | 1.2969 | 0.3907 | 0.0009013 | -++?? | 58.3 | 0.09115 | upstream-gene | |
| rs200534587 | 1 | 184001001 | 3 | 0 | 0 | A | C | -2.253 | 0.6787 | 0.0009014 | ---?? | 0 | 0.9784 | intron | COLGALT2 |
| rs118185280 | 12 | 60874473 | 2 | 0 | 0 | A | G | 6.4383 | 1.9395 | 0.0009017 | ?+??+ | 11.4 | 0.288 | intergenic | |
| rs13202963 | 6 | 117308972 | 4 | 0 | 0 | C | G | 0.7833 | 0.236 | 0.0009018 | +++?+ | 0 | 0.7186 | intergenic | |
| 17:47447808 | 17 | 47447808 | 3 | 0 | 0 | A | AAG | 0.9795 | 0.2951 | 0.0009018 | +++?? | 0 | 0.6477 | | |
| rs4848145 | 2 | 121937956 | 4 | 0 | 0 | A | G | 1.0741 | 0.3236 | 0.0009023 | +++?+ | 0 | 0.8005 | intergenic | |
| rs12506455 | 4 | 10031569 | 5 | 0 | 0 | A | T | 0.6979 | 0.2103 | 0.0009023 | ++++- | 15.1 | 0.3185 | intron | SLC2A9 |
| rs7037253 | 9 | 103870616 | 4 | 0 | 0 | T | C | -0.8298 | 0.25 | 0.0009027 | ---?- | 42.8 | 0.1547 | intron | LPPR1 |
| rs115580594 | 6 | 32673569 | 5 | 0 | 0 | A | G | -1.0804 | 0.3255 | 0.0009027 | ----- | 28.7 | 0.2304 | intergenic |  |
| rs10872964 | 14 | 43273386 | 4 | 0 | 0 | A | G | 1.2833 | 0.3866 | 0.0009028 | +++?- | 59.1 | 0.06207 | intergenic | |
| rs12000929 | 9 | 16189225 | 4 | 0 | 0 | A | C | -0.9103 | 0.2742 | 0.0009029 | ---?- | 0 | 0.9438 | intergenic | |
| rs55832820 | 10 | 28467350 | 4 | 0 | 0 | T | C | -2.5637 | 0.7724 | 0.0009029 | ---?+ | 0 | 0.7277 | intron | MPP7 |
| 12:19587923 | 12 | 19587923 | 3 | 0 | 0 | CT | C | -2.6757 | 0.8061 | 0.0009029 | ---?? | 0 | 0.9692 | | |
| 1:181852442 | 1 | 181852442 | 3 | 0 | 0 | T | TCTCCTC | 3.6944 | 1.1131 | 0.000903 | +++?? | 19.3 | 0.2898 | | |
| rs803805 | 13 | 71599180 | 4 | 0 | 0 | T | G | -0.9321 | 0.2808 | 0.0009033 | ---?- | 3.5 | 0.3752 | intergenic | |
| rs12432294 | 14 | 95897053 | 4 | 0 | 0 | T | C | 1.2426 | 0.3744 | 0.0009034 | +++?+ | 0 | 0.454 | intron | SYNE3 |
| rs147100484 | 2 | 101631390 | 2 | 0 | 0 | T | G | -7.563 | 2.2787 | 0.0009035 | ?-??- | 16.4 | 0.2741 | intron | RPL31,TBC1D8 |
| rs35141512 | 13 | 20807828 | 4 | 0 | 0 | T | G | -1.1074 | 0.3337 | 0.0009036 | ---?- | 0 | 0.5686 | upstream-gene | |
| rs72979327 | 4 | 162206936 | 2 | 0 | 0 | T | C | 6.0483 | 1.8223 | 0.0009036 | ?++?? | 0 | 0.5448 | intergenic | |
| 2:184067166 | 2 | 184067166 | 3 | 0 | 0 | CT | CTT | -0.9261 | 0.279 | 0.0009036 | ---?? | 15.4 | 0.3066 | | |
| rs9446589 | 6 | 72888264 | 4 | 0 | 0 | T | G | -1.5453 | 0.4656 | 0.0009037 | ---?+ | 0 | 0.5723 | intron | RIMS1 |
| rs62330359 | 5 | 708980 | 3 | 0 | 0 | T | C | -1.4816 | 0.4464 | 0.0009037 | ---?? | 65.4 | 0.05547 | downstream-gene | |
| rs2570117 | 15 | 87553090 | 4 | 0 | 0 | C | G | -0.8764 | 0.2641 | 0.0009038 | ---?- | 0 | 0.4153 | intron | AGBL1 |
| 15:87578187 | 15 | 87578187 | 3 | 0 | 0 | T | TAATA | 0.9074 | 0.2734 | 0.0009038 | +++?? | 17.7 | 0.2967 | | |
| rs9264528 | 6 | 31234058 | 3 | 0 | 0 | A | G | -0.9439 | 0.2844 | 0.000904 | ---?? | 22.6 | 0.2745 | downstream-gene | |
| rs10024436 | 4 | 96517899 | 4 | 0 | 0 | A | G | -1.2507 | 0.3769 | 0.000904 | ---?- | 28.2 | 0.2427 | intergenic | |

| | | | | | | | | | | | | | | | |
|---|---|---|---|---|---|---|---|---|---|---|---|---|---|---|---|
| rs12796605 | 11 | 630029 | 5 | 0 | 0 | A | G | 0.805 | 0.2426 | 0.0009041 | +++++ | 0 | 0.4168 | upstream-gene | |
| 5:709572 | 5 | 709572 | 3 | 0 | 0 | A | G | -1.5151 | 0.4565 | 0.0009043 | ---?? | 63.6 | 0.06392 | | |
| rs78869065 | 2 | 52950169 | 4 | 0 | 0 | A | C | -1.2808 | 0.386 | 0.0009045 | ---?- | 0 | 0.7515 | intergenic | |
| rs75064711 | 11 | 86174335 | 4 | 0 | 0 | A | G | 1.5551 | 0.4686 | 0.0009046 | +++?+ | 0 | 0.4207 | intron | ME3 |
| rs7774197 | 6 | 32046275 | 5 | 0 | 0 | A | C | -1.6077 | 0.4844 | 0.0009048 | ----+ | 35 | 0.1877 | intron | TNXB |
| rs313806 | 7 | 65499211 | 4 | 0 | 0 | A | C | -1.4866 | 0.448 | 0.000905 | ---?+ | 0 | 0.5361 | intergenic | |
| rs3990202 | 5 | 123097858 | 4 | 0 | 0 | T | C | -0.943 | 0.2842 | 0.000905 | ---?- | 6.5 | 0.3605 | intergenic | |
| rs9264500 | 6 | 31232825 | 5 | 0 | 0 | A | G | 0.7921 | 0.2387 | 0.000905 | +++++ | 0 | 0.4505 | downstream-gene | |
| rs1183161 | 6 | 95521291 | 4 | 0 | 0 | T | G | 1.6817 | 0.5068 | 0.0009051 | +++?+ | 0 | 0.6897 | intergenic | |
| rs6543465 | 2 | 107676353 | 4 | 0 | 0 | T | C | -0.8897 | 0.2681 | 0.0009051 | ---?- | 34.9 | 0.2027 | intergenic | |
| rs12626988 | 21 | 19085866 | 4 | 0 | 0 | A | G | 1.4226 | 0.4287 | 0.0009051 | +++?+ | 0 | 0.7316 | intergenic | |
| rs4380750 | 6 | 56104833 | 3 | 0 | 0 | A | G | 1.3429 | 0.4047 | 0.0009052 | +++?? | 0 | 0.8611 | intron | COL21A1 |
| 20:12000933 | 20 | 12000933 | 3 | 0 | 0 | CAAA | C | -0.9453 | 0.2849 | 0.0009053 | ---?? | 0 | 0.5618 | | |
| rs114176938 | 8 | 3153997 | 4 | 0 | 0 | T | C | 3.5984 | 1.0844 | 0.0009054 | +++?+ | 0 | 0.746 | intron | CSMD1 |
| rs12500810 | 4 | 10031068 | 5 | 0 | 0 | A | G | 0.698 | 0.2103 | 0.0009055 | ++++- | 14.6 | 0.3213 | intron | SLC2A9 |
| rs73334971 | 8 | 114877235 | 4 | 0 | 0 | T | C | 1.8112 | 0.5458 | 0.0009057 | +++?- | 5.6 | 0.365 | intergenic | |
| rs117846280 | 10 | 65907697 | 4 | 0 | 0 | T | C | -2.3156 | 0.6978 | 0.0009057 | ---?- | 0 | 0.8735 | intergenic | |
| 10:131476538 | 10 | 131476538 | 3 | 0 | 0 | G | GTTTCT( | 1.2779 | 0.3851 | 0.0009058 | +++?? | 0 | 0.6531 | | |
| 5:124273733 | 5 | 124273733 | 3 | 0 | 0 | G | GTATAT/ | -0.9634 | 0.2903 | 0.0009059 | ---?? | 0 | 0.7689 | | |
| rs10847204 | 12 | 127179160 | 4 | 0 | 0 | A | G | 1.1492 | 0.3463 | 0.000906 | +++?+ | 31.2 | 0.2248 | intergenic | |
| rs2402350 | 5 | 17172790 | 4 | 0 | 0 | T | C | 0.8098 | 0.244 | 0.0009063 | +++?+ | 0 | 0.7811 | intron | LOC285696 |
| rs112245017 | 5 | 1687238 | 3 | 0 | 0 | A | G | 4.4376 | 1.3374 | 0.0009067 | +++?? | 0 | 0.8506 | intergenic | |
| rs1204814 | 6 | 116553574 | 5 | 0 | 0 | A | T | -0.7821 | 0.2357 | 0.0009068 | +---- | 0 | 0.4607 | intron | NT5DC1 |
| rs3845451 | 1 | 182017018 | 4 | 0 | 0 | C | G | -1.1082 | 0.334 | 0.0009068 | ---?- | 0 | 0.4947 | intergenic | |
| rs60931903 | 7 | 32346695 | 4 | 0 | 0 | A | C | 1.3415 | 0.4043 | 0.0009069 | +++?+ | 0 | 0.9062 | intergenic | |
| rs9264529 | 6 | 31234074 | 3 | 0 | 0 | A | G | 0.9436 | 0.2844 | 0.000907 | +++?? | 22.6 | 0.2747 | downstream-gene | |
| rs144855060 | 6 | 63297240 | 3 | 0 | 0 | T | C | -2.3722 | 0.715 | 0.0009073 | ---?? | 0 | 0.3767 | intergenic | |
| rs7441586 | 4 | 189059078 | 4 | 0 | 0 | T | C | 1.2311 | 0.3711 | 0.0009074 | +++?+ | 0 | 0.9509 | upstream-gene | |
| 4:12266458 | 4 | 12266458 | 2 | 0 | 0 | T | C | -5.4378 | 1.639 | 0.0009076 | --??? | 0 | 0.6099 | intergenic | |
| rs2066168 | 1 | 102941373 | 4 | 0 | 0 | T | C | 0.8108 | 0.2444 | 0.0009077 | +++?- | 34.6 | 0.2046 | intergenic | |
| rs2877398 | 17 | 51112697 | 4 | 0 | 0 | T | C | -1.0337 | 0.3116 | 0.0009077 | -+-?- | 40.4 | 0.1693 | intergenic | |
| 2:29682430 | 2 | 29682430 | 3 | 0 | 0 | A | AT | 1.2089 | 0.3644 | 0.0009077 | +++?? | 0 | 0.5758 | | |
| rs7510988 | 22 | 28580269 | 4 | 0 | 0 | T | C | -2.3144 | 0.6976 | 0.000908 | ---?- | 0 | 0.5818 | intron | TTC28 |
| rs66624704 | 6 | 31232343 | 3 | 0 | 0 | A | C | -0.9435 | 0.2844 | 0.0009082 | ---?? | 22.9 | 0.2733 | downstream-gene | |
| rs2843037 | 6 | 63344536 | 4 | 0 | 0 | A | C | -2.2746 | 0.6857 | 0.000909 | ---?- | 0 | 0.396 | intergenic | |
| rs373728107 | 1 | 32170253 | 3 | 0 | 0 | A | C | -1.2958 | 0.3906 | 0.0009094 | +--?? | 58.3 | 0.09082 | upstream-gene | |
| 10:11651049 | 10 | 11651049 | 3 | 0 | 0 | GCTTTT | GCTTTT( | 1.2033 | 0.3628 | 0.0009095 | +++?? | 0 | 0.9255 | | |
| rs9559625 | 13 | 110339480 | 3 | 0 | 0 | T | C | 1.1705 | 0.3529 | 0.0009099 | +++?? | 58.6 | 0.08925 | intergenic | |
| rs77694532 | 3 | 119007475 | 5 | 0 | 0 | A | G | 1.6179 | 0.4878 | 0.0009101 | +++++ | 32.4 | 0.2053 | intergenic | |
| rs79483040 | 11 | 86173808 | 4 | 0 | 0 | A | G | 1.5537 | 0.4684 | 0.0009101 | +++?+ | 0 | 0.4185 | intron | ME3 |
| rs184852236 | 3 | 47967917 | 2 | 0 | 0 | A | G | 6.3791 | 1.9232 | 0.0009101 | ?+??+ | 55.5 | 0.1338 | intron | MAP4 |
| rs34735738 | 12 | 51462922 | 5 | 0 | 0 | T | C | -0.9255 | 0.279 | 0.0009102 | ----- | 0 | 0.4464 | intron | CSRNP2 |
| rs187735226 | 10 | 93580321 | 3 | 0 | 0 | A | G | 3.9467 | 1.1899 | 0.0009102 | +-+?? | 49.4 | 0.1386 | intron | TNKS2 |
| rs8084343 | 18 | 37415264 | 4 | 0 | 0 | T | C | -0.9 | 0.2713 | 0.0009104 | ---?- | 0 | 0.6306 | intergenic | |
| rs138478706 | 15 | 26085081 | 4 | 0 | 0 | T | C | 2.2627 | 0.6822 | 0.0009105 | +++?+ | 0 | 0.8639 | intron | ATP10A |
| rs140318960 | 4 | 166225096 | 4 | 0 | 0 | T | C | 3.6233 | 1.0924 | 0.0009106 | +++?+ | 0 | 0.925 | intron | KLHL2 |
| 2:170647325 | 2 | 170647325 | 3 | 0 | 0 | A | AG | 0.8604 | 0.2594 | 0.0009106 | +++?? | 0 | 0.4466 | | |
| rs79224634 | 1 | 42705017 | 4 | 0 | 0 | T | C | -1.7626 | 0.5314 | 0.0009111 | ---?- | 0 | 0.5568 | intron | FOXJ3 |
| rs5028779 | 2 | 230918514 | 4 | 0 | 0 | T | C | 1.4124 | 0.4259 | 0.0009114 | +++?+ | 21.7 | 0.2803 | intron | SLC16A14 |
| rs11942333 | 4 | 46389486 | 4 | 0 | 0 | A | G | 0.8607 | 0.2595 | 0.0009114 | +-+?+ | 59.5 | 0.06011 | intron | GABRA2 |
| rs78546351 | 10 | 101256797 | 3 | 0 | 0 | A | G | -2.9914 | 0.902 | 0.0009118 | ---?? | 0 | 0.8669 | intergenic | |
| rs12701215 | 7 | 32371676 | 4 | 0 | 0 | A | C | 1.3495 | 0.4069 | 0.000912 | +++?+ | 0 | 0.9356 | intergenic | |
| rs7549764 | 1 | 208186003 | 4 | 0 | 0 | A | G | -1.9921 | 0.6007 | 0.0009122 | ---?- | 0 | 0.7069 | intergenic | |
| rs72979325 | 4 | 162205882 | 2 | 0 | 0 | C | G | -6.0515 | 1.8248 | 0.0009122 | ?--?? | 0 | 0.5347 | intergenic | |
| rs116741554 | 3 | 23811049 | 4 | 0 | 0 | A | T | 3.4752 | 1.048 | 0.0009128 | +++?+ | 0 | 0.5477 | intergenic | |
| 9:44247029 | 9 | 44247029 | 3 | 0 | 0 | A | ATGCTA( | -1.7001 | 0.5127 | 0.0009128 | ---?? | 0 | 0.9331 | | |
| rs148926269 | 11 | 101564072 | 3 | 0 | 0 | T | C | 2.9096 | 0.8774 | 0.0009129 | +++?? | 0 | 0.954 | intergenic | |
| rs150224165 | 3 | 18439917 | 3 | 0 | 0 | T | C | -3.2166 | 0.97 | 0.0009135 | -+-?? | 56.3 | 0.1016 | intron | SATB1 |
| rs138877763 | 2 | 232168899 | 4 | 0 | 0 | T | C | 1.5644 | 0.4718 | 0.0009136 | +++?+ | 0 | 0.8614 | intron | ARMC9 |

| | | | | | | | | | | | | | | | |
|---|---|---|---|---|---|---|---|---|---|---|---|---|---|---|---|
| 1:164313892 | 1 | 164313892 | 3 | 0 | 0 | G | GA | 0.8726 | 0.2632 | 0.0009136 | +++?? | 0 | 0.5524 | | |
| rs28811897 | 1 | 109592473 | 4 | 0 | 0 | T | G | -0.9634 | 0.2905 | 0.0009137 | ---?- | 54.2 | 0.08772 | intergenic | |
| rs62130627 | 19 | 3711334 | 3 | 0 | 0 | T | C | -0.9955 | 0.3002 | 0.0009138 | ---?? | 18.1 | 0.2947 | intron | TJP3 |
| rs121570 | 16 | 57455846 | 4 | 0 | 0 | T | C | -1.2311 | 0.3713 | 0.0009139 | ---?- | 0 | 0.8472 | intergenic | |
| rs2617852 | 5 | 82430588 | 4 | 0 | 0 | T | C | -1.8404 | 0.5551 | 0.0009139 | ---?- | 0 | 0.8231 | intron | XRCC4 |
| rs79551517 | 12 | 114501664 | 3 | 0 | 0 | T | C | -1.0547 | 0.3181 | 0.000914 | ---?? | 41 | 0.1839 | intergenic | |
| rs74110446 | 12 | 75564951 | 4 | 0 | 0 | A | G | -2.2142 | 0.6678 | 0.0009141 | ---?- | 0 | 0.5301 | intron | KCNC2 |
| rs34737022 | 12 | 772656 | 4 | 0 | 0 | A | T | 3.2175 | 0.9704 | 0.0009142 | +++?+ | 0 | 0.4091 | synonymous | NINJ2 |
| 4:91488458 | 4 | 91488458 | 3 | 0 | 0 | A | AT | 0.9947 | 0.3 | 0.0009142 | +++?? | 0 | 0.6715 | | |
| rs191625896 | 17 | 56814495 | 5 | 0 | 0 | T | C | 2.6038 | 0.7853 | 0.0009144 | +++++ | 0 | 0.9222 | downstream-gene | |
| rs107861 | 6 | 33218868 | 5 | 0 | 0 | C | G | -1.0123 | 0.3053 | 0.0009146 | ----- | 24 | 0.2612 | intron | VPS52,HCG25 |
| rs1496008 | 1 | 64814887 | 4 | 0 | 0 | A | T | -1.0461 | 0.3155 | 0.0009146 | ---?- | 0 | 0.5803 | intergenic | |
| rs17401789 | 7 | 90629203 | 4 | 0 | 0 | T | C | -2.7538 | 0.8306 | 0.0009146 | ---?- | 13.6 | 0.3246 | intron | CDK14 |
| rs79774278 | 14 | 85874744 | 4 | 0 | 0 | C | G | 2.0097 | 0.6061 | 0.0009146 | +++?+ | 19 | 0.2953 | intron | LINC00911 |
| rs1715862 | 10 | 126856255 | 4 | 0 | 0 | T | C | -0.7953 | 0.2399 | 0.0009147 | ---?- | 36.5 | 0.1934 | intergenic | |
| rs117783203 | 17 | 79107632 | 3 | 0 | 0 | A | G | 3.2 | 0.9652 | 0.0009147 | +++?? | 0 | 0.792 | intron | AATK |
| rs147154522 | 6 | 124350045 | 3 | 0 | 0 | T | C | -2.8552 | 0.8612 | 0.0009147 | ---?? | 0 | 0.4237 | intron | NKAIN2 |
| 4:162205297 | 4 | 162205297 | 2 | 0 | 0 | T | TA | 6.0508 | 1.825 | 0.0009148 | ?++?? | 0 | 0.5233 | | |
| rs1204786 | 6 | 116534246 | 5 | 0 | 0 | A | G | 0.7839 | 0.2364 | 0.000915 | +++++ | 0 | 0.5444 | intron | NT5DC1 |
| rs28594536 | 8 | 143308691 | 4 | 0 | 0 | T | C | -2.2665 | 0.6836 | 0.000915 | ---?- | 0 | 0.8646 | intron | TSNARE1 |
| rs13007218 | 2 | 170903703 | 3 | 0 | 0 | C | G | 0.9006 | 0.2716 | 0.0009151 | +++?? | 63.2 | 0.06597 | intron | UBR3 |
| rs28673515 | 4 | 181526409 | 4 | 0 | 0 | C | G | 0.94 | 0.2835 | 0.0009151 | +++?+ | 0 | 0.4722 | intergenic | |
| rs72871735 | 3 | 45321453 | 4 | 0 | 0 | C | G | -1.2767 | 0.3851 | 0.0009153 | ---?- | 25.9 | 0.2564 | intergenic | |
| rs3104970 | 8 | 96592356 | 4 | 0 | 0 | T | G | -1.2588 | 0.3797 | 0.0009155 | -+-?- | 42.1 | 0.1592 | intron | LOC100616530 |
| rs75571761 | 15 | 36630616 | 4 | 0 | 0 | T | C | 1.634 | 0.4929 | 0.0009157 | +++?+ | 0 | 0.5915 | intergenic | |
| rs6860707 | 5 | 134342295 | 4 | 0 | 0 | T | G | 0.8045 | 0.2427 | 0.0009159 | +++?+ | 0 | 0.7573 | intron | CATSPER3 |
| rs17666023 | 21 | 29168645 | 4 | 0 | 0 | A | G | 3.1176 | 0.9404 | 0.0009162 | +++?+ | 0 | 0.7211 | intergenic | |
| rs2003748 | 22 | 29102645 | 4 | 0 | 0 | A | G | 2.1412 | 0.6459 | 0.0009165 | +++?+ | 0 | 0.585 | intron | CHEK2 |
| rs9554989 | 13 | 104379789 | 4 | 0 | 0 | A | G | 1.2864 | 0.3881 | 0.0009165 | +++?+ | 0 | 0.7341 | intergenic | |
| rs10927351 | 1 | 245105192 | 4 | 0 | 0 | T | C | -1.1482 | 0.3464 | 0.0009166 | ---?- | 9.3 | 0.3463 | intergenic | |
| rs118027248 | 16 | 60131445 | 2 | 0 | 0 | T | G | -5.5425 | 1.672 | 0.0009166 | ??-?- | 33.4 | 0.2205 | intergenic | |
| rs2705722 | 2 | 183993289 | 4 | 0 | 0 | T | C | -1.1547 | 0.3483 | 0.0009169 | ---?- | 0 | 0.8287 | intron | NUP35 |
| rs66991463 | 8 | 142046080 | 3 | 0 | 0 | T | C | 1.4333 | 0.4324 | 0.000917 | +++?? | 0 | 0.9263 | intergenic | |
| rs12547990 | 8 | 252583 | 4 | 0 | 0 | A | G | 0.9789 | 0.2953 | 0.0009172 | +-+?+ | 51.1 | 0.1052 | intergenic | |
| rs6483468 | 11 | 95706031 | 4 | 0 | 0 | C | G | 0.8642 | 0.2607 | 0.0009175 | +++?+ | 0 | 0.5213 | intergenic | |
| 1:85612648 | 1 | 85612648 | 3 | 0 | 0 | G | GT | 1.5598 | 0.4706 | 0.0009175 | +++?? | 0 | 0.9258 | | |
| rs12021635 | 1 | 98625675 | 4 | 0 | 0 | A | G | 1.2881 | 0.3886 | 0.0009176 | +++?+ | 24.1 | 0.2667 | intergenic | |
| rs28863518 | 1 | 109592261 | 4 | 0 | 0 | A | G | 0.9634 | 0.2907 | 0.0009177 | +++?+ | 54.2 | 0.08763 | intergenic | |
| rs1080022 | 6 | 144664731 | 4 | 0 | 0 | T | C | -1.7247 | 0.5203 | 0.0009178 | ---?+ | 6.3 | 0.3615 | intron | UTRN |
| rs4653751 | 1 | 226741284 | 4 | 0 | 0 | A | T | 1.3083 | 0.3947 | 0.000918 | +++?+ | 24.3 | 0.2656 | intron | C1orf95 |
| rs8084011 | 18 | 37415011 | 4 | 0 | 0 | T | C | -0.9002 | 0.2716 | 0.000918 | ---?- | 0 | 0.6278 | intergenic | |
| rs28478985 | 16 | 5180666 | 3 | 0 | 0 | A | G | 0.8525 | 0.2572 | 0.000918 | +++?? | 0 | 0.9879 | intergenic | |
| rs80348820 | 20 | 21484763 | 4 | 0 | 0 | T | C | 2.0744 | 0.6259 | 0.000918 | +++?+ | 0 | 0.6356 | intergenic | |
| rs1492649 | 8 | 96402339 | 4 | 0 | 0 | A | C | -1.1159 | 0.3367 | 0.0009181 | ---?- | 0 | 0.6998 | intron | LOC100616530 |
| rs2247836 | 12 | 103260608 | 4 | 0 | 0 | T | C | -0.8156 | 0.2461 | 0.0009181 | ---?- | 0 | 0.6212 | intron | PAH |
| rs150371334 | 12 | 14917997 | 4 | 0 | 0 | A | G | -1.9794 | 0.5972 | 0.0009181 | ---?- | 0 | 0.6196 | downstream-gene | |
| rs73552610 | 8 | 21544627 | 3 | 0 | 0 | T | C | -3.9633 | 1.1958 | 0.0009183 | ?--?- | 0 | 0.9367 | downstream-gene | |
| rs10407626 | 19 | 3599535 | 3 | 0 | 0 | A | G | -0.8639 | 0.2607 | 0.0009185 | ---?? | 0 | 0.5197 | intron | TBXA2R |
| rs9264549 | 6 | 31234774 | 3 | 0 | 0 | C | G | -0.9432 | 0.2846 | 0.0009186 | ---?? | 22.6 | 0.2749 | downstream-gene | |
| rs62222057 | 21 | 21370331 | 4 | 0 | 0 | A | G | -3.2506 | 0.9808 | 0.0009186 | ---?- | 0 | 0.8937 | intergenic | |
| rs12974183 | 19 | 46169005 | 5 | 0 | 1 | A | C | 0.8717 | 0.263 | 0.0009187 | +-++- | 26.3 | 0.2462 | upstream-gene | |
| rs28515780 | 1 | 109590798 | 4 | 0 | 0 | A | G | -0.9661 | 0.2915 | 0.0009188 | ---?- | 54.6 | 0.08525 | intergenic | |
| 10:6230791 | 10 | 6230791 | 3 | 0 | 0 | A | ATTTT | -1.1709 | 0.3533 | 0.000919 | ---?? | 0 | 0.5587 | | |
| rs2931473 | 8 | 21168676 | 3 | 0 | 0 | A | G | 1.4639 | 0.4417 | 0.0009191 | +++?? | 0 | 0.4055 | intergenic | |
| rs2297200 | 20 | 44684526 | 4 | 0 | 0 | A | T | 0.875 | 0.264 | 0.0009192 | -++?+ | 47.3 | 0.1276 | intron | SLC12A5 |
| rs6848675 | 4 | 96522533 | 4 | 0 | 0 | A | G | -1.2477 | 0.3765 | 0.0009193 | ---?- | 27.1 | 0.2492 | intergenic | |
| rs9264550 | 6 | 31234786 | 3 | 0 | 0 | T | C | -0.9431 | 0.2846 | 0.0009195 | ---?? | 22.5 | 0.2751 | downstream-gene | |
| rs61876664 | 11 | 635073 | 5 | 0 | 0 | A | G | 0.8119 | 0.245 | 0.0009195 | +++++ | 0 | 0.4887 | upstream-gene | |
| rs28628461 | 9 | 44246927 | 3 | 0 | 0 | T | C | -1.7835 | 0.5382 | 0.0009196 | ---?? | 0 | 0.924 | intergenic | |

| | | | | | | | | | | | | | | | |
|---|---|---|---|---|---|---|---|---|---|---|---|---|---|---|---|
| rs6998082 | 8 | 64916288 | 4 | 0 | 0 | A | T | -0.8217 | 0.2479 | 0.0009197 | ---?+ | 0 | 0.4723 | intergenic | |
| rs72877054 | 2 | 47377096 | 4 | 0 | 0 | A | G | 1.2788 | 0.3859 | 0.0009197 | +++?+ | 0 | 0.9756 | intron | C2orf61 |
| rs73530236 | 13 | 85124610 | 3 | 0 | 0 | C | G | -1.1331 | 0.3419 | 0.0009197 | ---?? | 0 | 0.8342 | intron | LINC00333 |
| rs115981015 | 6 | 109121362 | 2 | 0 | 0 | T | G | -4.6544 | 1.4045 | 0.0009197 | -?-?? | 0 | 0.7769 | intergenic | |
| rs372402375 | 15 | 83775259 | 3 | 0 | 0 | A | T | 3.4807 | 1.0503 | 0.0009197 | +++?? | 0 | 0.8193 | upstream-gene | |
| rs4666893 | 2 | 184023884 | 4 | 0 | 0 | T | C | 1.1443 | 0.3453 | 0.0009198 | +++?+ | 0 | 0.794 | intron | NUP35 |
| rs6602353 | 10 | 9309987 | 4 | 0 | 0 | T | C | -1.6836 | 0.508 | 0.0009201 | ---?- | 0 | 0.6519 | intergenic | |
| rs2468727 | 8 | 140435235 | 4 | 0 | 0 | A | G | -0.9493 | 0.2865 | 0.0009202 | ---?- | 0 | 0.5982 | intergenic | |
| rs787363 | 4 | 67906156 | 4 | 0 | 0 | A | G | 1.3117 | 0.3958 | 0.0009203 | +++?+ | 57.1 | 0.07191 | intergenic | |
| 1:87409707 | 1 | 87409707 | 3 | 0 | 0 | CA | C | 1.0233 | 0.3088 | 0.0009205 | +++?? | 4.6 | 0.3505 | | |
| rs6051285 | 20 | 26286219 | 4 | 0 | 0 | A | G | -1.4696 | 0.4435 | 0.0009209 | ---?- | 59.8 | 0.05854 | intergenic | |
| rs5021855 | 4 | 143418682 | 3 | 0 | 0 | A | G | 1.2097 | 0.3651 | 0.0009211 | +++?? | 0 | 0.8771 | intron | INPP4B |
| rs6076416 | 20 | 26286170 | 4 | 0 | 0 | T | C | -1.4687 | 0.4432 | 0.0009212 | ---?- | 59.9 | 0.05817 | intergenic | |
| rs13272262 | 8 | 6145581 | 4 | 0 | 0 | A | C | -1.3572 | 0.4096 | 0.0009214 | ---?- | 0 | 0.8612 | intergenic | |
| rs66905872 | 6 | 144835040 | 4 | 0 | 0 | T | C | -0.9925 | 0.2995 | 0.0009214 | ---?- | 0 | 0.6685 | intron-near-splice | UTRN |
| rs79999788 | 16 | 82753625 | 4 | 0 | 0 | A | G | -1.5527 | 0.4686 | 0.0009214 | ---?- | 56.4 | 0.07603 | intron | CDH13 |
| rs144012442 | 15 | 74585206 | 3 | 0 | 0 | T | C | 2.9842 | 0.9006 | 0.0009215 | +++?? | 0 | 0.6255 | intron | CCDC33 |
| rs7596514 | 2 | 189878769 | 4 | 0 | 0 | A | G | 0.9471 | 0.2858 | 0.0009216 | +++?+ | 40.8 | 0.1667 | downstream-gene | |
| rs17629979 | 13 | 29099266 | 4 | 0 | 0 | T | G | -1.612 | 0.4865 | 0.0009218 | ---?- | 20.4 | 0.2874 | intergenic | |
| rs12618495 | 2 | 18048949 | 4 | 0 | 0 | A | G | -0.8279 | 0.2499 | 0.0009219 | ---?- | 9.4 | 0.3459 | intergenic | |
| rs141448390 | 5 | 90921494 | 2 | 0 | 0 | A | G | -4.9548 | 1.4954 | 0.0009221 | ?--?? | 0 | 0.8485 | intergenic | |
| rs817396 | 1 | 85613292 | 3 | 0 | 0 | A | T | 1.559 | 0.4705 | 0.0009223 | +++?? | 0 | 0.9263 | intergenic | |
| rs13186867 | 5 | 39600815 | 4 | 0 | 0 | A | G | -0.9872 | 0.298 | 0.0009223 | ---?- | 0 | 0.8841 | intergenic | |
| rs1204789 | 6 | 116536612 | 5 | 0 | 0 | A | T | -0.7829 | 0.2363 | 0.0009225 | ----- | 0 | 0.5242 | intron | NT5DC1 |
| rs386984 | 2 | 40439952 | 4 | 0 | 0 | A | G | -1.0285 | 0.3104 | 0.0009229 | ---?+ | 5.5 | 0.3653 | intron | SLC8A1,SLC8A1-AS1 |
| rs9501344 | 6 | 30989537 | 3 | 0 | 0 | T | C | 1.3624 | 0.4112 | 0.000923 | +++?? | 0 | 0.8442 | intron | MUC22 |
| rs10409244 | 19 | 3599529 | 3 | 0 | 0 | T | C | -0.8638 | 0.2607 | 0.000923 | ---?? | 0 | 0.52 | intron | TBXA2R |
| rs11180677 | 12 | 76174859 | 4 | 0 | 0 | T | G | -2.0895 | 0.6307 | 0.000923 | ---?+ | 0 | 0.7325 | intergenic | |
| rs13186060 | 5 | 113207613 | 4 | 0 | 0 | T | C | 0.9434 | 0.2848 | 0.0009231 | +++?+ | 0 | 0.8236 | intergenic | |
| rs145206552 | 3 | 183163864 | 4 | 0 | 0 | A | T | -1.6213 | 0.4894 | 0.0009233 | ---?- | 0 | 0.7529 | upstream-gene | |
| rs2594321 | 15 | 87527583 | 4 | 0 | 0 | T | C | -0.8489 | 0.2562 | 0.0009234 | ---?- | 0 | 0.419 | intron | AGBL1 |
| rs16888503 | 6 | 24022898 | 5 | 0 | 0 | T | C | -1.2883 | 0.3889 | 0.0009234 | ----+ | 0 | 0.819 | intergenic |  |
| rs34732626 | 15 | 67422638 | 4 | 0 | 0 | C | G | -1.6388 | 0.4947 | 0.0009236 | ---?- | 0 | 0.8579 | intron | SMAD3 |
| rs5021853 | 4 | 143418684 | 3 | 0 | 0 | T | C | -1.2098 | 0.3652 | 0.0009237 | ---?? | 0 | 0.8776 | intron | INPP4B |
| rs5021854 | 4 | 143418683 | 3 | 0 | 0 | T | G | 1.2095 | 0.3651 | 0.0009238 | +++?? | 0 | 0.8773 | intron | INPP4B |
| rs2256677 | 14 | 21443154 | 4 | 0 | 0 | A | G | 0.8509 | 0.2569 | 0.0009243 | +++?+ | 0 | 0.957 | intergenic | |
| rs8177055 | 10 | 30731348 | 3 | 0 | 0 | A | C | -2.6115 | 0.7883 | 0.0009243 | ---?? | 56.6 | 0.09982 | intron | MAP3K8 |
| rs5021852 | 4 | 143418685 | 3 | 0 | 0 | T | G | 1.2102 | 0.3653 | 0.0009245 | +++?? | 0 | 0.8777 | intron | INPP4B |
| 6:32608589 | 6 | 32608589 | 3 | 0 | 0 | T | TA | -1.2667 | 0.3824 | 0.0009249 | ---?? | 39.7 | 0.1905 | | |
| rs200260167 | 5 | 123553005 | 3 | 0 | 0 | C | G | -0.9775 | 0.2951 | 0.000925 | ---?? | 0 | 0.9195 | intergenic | |
| rs1204788 | 6 | 116536251 | 5 | 0 | 0 | A | G | 0.7824 | 0.2362 | 0.0009251 | +++++ | 0 | 0.5332 | intron | NT5DC1 |
| rs12824225 | 12 | 51462145 | 5 | 0 | 0 | C | G | -0.9247 | 0.2792 | 0.0009254 | ----- | 0 | 0.4428 | intron | CSRNP2 |
| 1:27950883 | 1 | 27950883 | 2 | 0 | 0 | CTA | C | -3.5781 | 1.0803 | 0.0009254 | ?--?? | 0 | 0.6741 | | |
| rs4760255 | 12 | 57855180 | 5 | 0 | 0 | A | G | -0.7308 | 0.2206 | 0.0009255 | ----- | 0 | 0.9785 | intron | GLI1 |
| rs180305 | 12 | 117164747 | 5 | 0 | 0 | T | C | -1.2216 | 0.3688 | 0.0009257 | ----- | 0 | 0.8486 | intron | C12orf49 |
| rs12588852 | 14 | 43271889 | 4 | 0 | 0 | C | G | -1.3919 | 0.4202 | 0.0009259 | ---?+ | 45.5 | 0.1384 | intergenic | |
| rs6906908 | 6 | 31235254 | 3 | 0 | 0 | A | G | 0.9421 | 0.2845 | 0.000926 | +++?? | 22.1 | 0.2771 | downstream-gene | |
| rs139208600 | 17 | 52242931 | 4 | 0 | 0 | T | C | -1.9395 | 0.5856 | 0.000926 | ---?+ | 29.8 | 0.2334 | intergenic | |
| 4:162205222 | 4 | 162205222 | 2 | 0 | 0 | T | TATG | 6.0496 | 1.8265 | 0.0009261 | ?++?? | 0 | 0.5246 | | |
| rs35492781 | 4 | 180911535 | 4 | 0 | 0 | T | C | -0.8967 | 0.2708 | 0.0009262 | ---?- | 0 | 0.6278 | intergenic | |
| rs61806396 | 1 | 182014335 | 3 | 0 | 0 | A | C | 1.1478 | 0.3466 | 0.0009262 | +++?? | 19.2 | 0.2902 | intergenic | |
| rs13071226 | 3 | 188520947 | 4 | 0 | 0 | T | C | -0.8506 | 0.2568 | 0.0009264 | ---?- | 23.8 | 0.2685 | intron | LPP |
| rs79060386 | 4 | 4739364 | 4 | 0 | 0 | A | C | 2.3159 | 0.6992 | 0.0009264 | +++?+ | 0 | 0.3957 | intergenic | |
| rs74609155 | 7 | 32349013 | 4 | 0 | 0 | T | C | -1.3395 | 0.4044 | 0.0009266 | ---?- | 0 | 0.9131 | intergenic | |
| rs4533254 | 15 | 87527198 | 4 | 0 | 0 | T | G | -0.8488 | 0.2563 | 0.0009268 | ---?- | 0 | 0.4197 | intron | AGBL1 |
| rs74856816 | 17 | 52285092 | 3 | 0 | 0 | A | G | -1.6655 | 0.5029 | 0.0009269 | ---?? | 0 | 0.4833 | intergenic | |
| rs1061801 | 6 | 33282338 | 5 | 0 | 0 | A | G | 0.9213 | 0.2782 | 0.0009275 | +++++ | 0 | 0.5861 | 3-prime-UTR | ZBTB22 |
| rs232 | 4 | 185284364 | 4 | 0 | 0 | A | G | -1.1471 | 0.3464 | 0.0009276 | ---?- | 0 | 0.9815 | intergenic | |
| rs9264506 | 6 | 31233143 | 5 | 0 | 0 | T | C | 0.7904 | 0.2387 | 0.0009276 | +++++ | 0 | 0.4563 | downstream-gene | |

| | | | | | | | | | | | | | | | |
|---|---|---|---|---|---|---|---|---|---|---|---|---|---|---|---|
| rs11196916 | 10 | 116551764 | 4 | 0 | 0 | A | G | 1.317 | 0.3977 | 0.0009276 | +++?+ | 0 | 0.8393 | intergenic | |
| rs4252270 | 11 | 117863846 | 4 | 0 | 0 | T | C | 1.3056 | 0.3942 | 0.0009277 | +++?+ | 0 | 0.7472 | intron | IL10RA |
| rs2717715 | 8 | 18436242 | 3 | 0 | 0 | T | C | -0.8224 | 0.2484 | 0.0009278 | ---?? | 0 | 0.7635 | intron | PSD3 |
| rs9810721 | 3 | 30818057 | 3 | 0 | 0 | T | C | -3.0002 | 0.906 | 0.0009281 | ---?? | 0 | 0.6714 | intron | GADL1 |
| rs117458179 | 12 | 43098413 | 4 | 0 | 0 | A | G | 1.8676 | 0.564 | 0.0009281 | +++?+ | 0 | 0.604 | intergenic | |
| 17:51146662 | 17 | 51146662 | 3 | 0 | 0 | A | AC | 1.3778 | 0.4161 | 0.0009281 | +++?? | 0.1 | 0.3676 | | |
| rs4915955 | 1 | 64709216 | 4 | 0 | 0 | T | C | 1.2233 | 0.3694 | 0.0009282 | +++?+ | 0 | 0.5115 | intron | UBE2U |
| rs142521315 | 5 | 21083268 | 4 | 0 | 0 | T | C | -4.6996 | 1.4192 | 0.0009282 | ---?- | 0 | 0.9262 | intergenic | |
| rs9264555 | 6 | 31234911 | 3 | 0 | 0 | A | G | -0.942 | 0.2845 | 0.0009287 | ---?? | 22.3 | 0.2763 | downstream-gene | |
| rs17822733 | 13 | 28287105 | 3 | 0 | 0 | T | C | 2.9008 | 0.876 | 0.0009288 | +++?? | 11.3 | 0.324 | intergenic | |
| rs76501190 | 2 | 236715201 | 3 | 0 | 0 | A | T | -1.7419 | 0.526 | 0.0009288 | -+-?? | 46.7 | 0.1533 | intron | AGAP1 |
| rs12504232 | 4 | 105911875 | 4 | 0 | 0 | A | G | -0.8895 | 0.2686 | 0.0009289 | ---?+ | 13.9 | 0.3226 | intergenic | |
| rs112339677 | 16 | 20175488 | 3 | 0 | 0 | T | G | -1.9559 | 0.5907 | 0.0009289 | ---?? | 47.8 | 0.1472 | intergenic | |
| rs11828425 | 11 | 86172006 | 4 | 0 | 0 | T | C | -1.5485 | 0.4677 | 0.000929 | ---?- | 0 | 0.4109 | intron | ME3 |
| rs2594335 | 15 | 87560304 | 4 | 0 | 0 | A | G | -0.8838 | 0.2669 | 0.0009292 | ---?+ | 9.8 | 0.3442 | intron | AGBL1 |
| rs1926814 | 10 | 4392256 | 4 | 0 | 0 | T | G | -0.9662 | 0.2918 | 0.0009293 | ---?- | 0 | 0.7652 | intergenic | |
| rs6434006 | 2 | 184024702 | 4 | 0 | 0 | T | C | -1.1427 | 0.3451 | 0.0009293 | ---?- | 0 | 0.7934 | intron | NUP35 |
| rs9264445 | 6 | 31231114 | 3 | 0 | 0 | T | C | 0.943 | 0.2848 | 0.0009296 | +++?? | 21.6 | 0.2791 | intergenic | |
| rs6543467 | 2 | 107676552 | 4 | 0 | 0 | T | C | -0.8884 | 0.2683 | 0.0009298 | ---?- | 35.1 | 0.2018 | intergenic | |
| rs9264455 | 6 | 31231382 | 3 | 0 | 0 | A | T | 0.9417 | 0.2844 | 0.0009298 | +++?? | 23.5 | 0.2705 | intergenic | |
| rs11251843 | 10 | 3339644 | 3 | 0 | 0 | A | T | -1.8517 | 0.5593 | 0.0009299 | ---?? | 0 | 0.8589 | intergenic | |
| 10:95222495 | 10 | 95222495 | 3 | 0 | 0 | CA | C | 4.4163 | 1.3339 | 0.0009302 | +++?? | 24.5 | 0.2659 | | |
| rs6845302 | 4 | 38743114 | 5 | 0 | 0 | A | G | -0.7282 | 0.22 | 0.0009304 | ----- | 0 | 0.863 | intergenic |  |
| rs462618 | 6 | 33222163 | 5 | 0 | 0 | T | G | 1.0111 | 0.3054 | 0.0009306 | +++++ | 23 | 0.2676 | intron | VPS52,HCG25 |
| rs461338 | 6 | 33218180 | 5 | 0 | 0 | T | C | -1.0111 | 0.3054 | 0.0009307 | ----- | 23 | 0.2676 | 3-prime-UTR | VPS52,HCG25 |
| rs148564520 | 5 | 4917578 | 4 | 0 | 0 | T | C | 2.6213 | 0.7918 | 0.0009307 | +++?+ | 0 | 0.5986 | intergenic | |
| rs74429448 | 2 | 238522913 | 4 | 0 | 0 | A | G | -3.6652 | 1.1071 | 0.0009308 | ---?- | 0 | 0.4898 | intergenic | |
| rs2616531 | 3 | 5113306 | 4 | 0 | 0 | A | G | -0.9428 | 0.2848 | 0.0009309 | ---?- | 0 | 0.4484 | intergenic | |
| rs6829727 | 4 | 10051672 | 5 | 0 | 0 | A | C | -0.7054 | 0.2131 | 0.0009312 | ----+ | 27.7 | 0.2369 | intergenic |  |
| rs6936854 | 6 | 130946962 | 4 | 0 | 0 | A | G | -1.3811 | 0.4172 | 0.0009312 | ---?- | 0 | 0.7394 | intergenic | |
| rs75098407 | 16 | 9117005 | 3 | 0 | 0 | C | G | 4.1392 | 1.2503 | 0.0009312 | +++?? | 0 | 0.5589 | intergenic | |
| rs12075337 | 1 | 180145545 | 4 | 0 | 0 | A | G | -2.0175 | 0.6094 | 0.0009313 | ---?+ | 32.6 | 0.2167 | intron | QSOX1 |
| rs115852905 | 5 | 44404464 | 4 | 0 | 0 | A | C | -3.7113 | 1.1211 | 0.0009313 | ---?- | 0 | 0.7049 | intergenic | |
| rs346450 | 18 | 39942161 | 4 | 0 | 0 | T | C | -0.8552 | 0.2583 | 0.0009314 | ---?- | 0 | 0.438 | intron | LINC00907 |
| rs2767569 | 1 | 208188125 | 4 | 0 | 0 | T | C | 1.8896 | 0.5708 | 0.0009314 | +++?+ | 0 | 0.7439 | intergenic | |
| rs28410185 | 9 | 44249662 | 3 | 0 | 0 | T | G | 1.708 | 0.5159 | 0.0009314 | +++?? | 0 | 0.8882 | intergenic | |
| rs169494 | 6 | 32097876 | 5 | 0 | 0 | A | G | 1.3081 | 0.3952 | 0.0009315 | +++-+ | 42.7 | 0.137 | 5-prime-UTR | FKBPL |
| rs9264526 | 6 | 31233988 | 5 | 0 | 0 | T | G | 0.7902 | 0.2387 | 0.0009316 | +++++ | 0 | 0.4572 | downstream-gene | |
| rs77614584 | 10 | 11351678 | 4 | 0 | 0 | T | C | -1.4192 | 0.4287 | 0.0009317 | ---?- | 0 | 0.716 | intron | CELF2 |
| rs6128112 | 20 | 56211853 | 4 | 0 | 0 | T | C | 1.0839 | 0.3274 | 0.0009318 | +++?+ | 0 | 0.9135 | intergenic | |
| rs10467873 | 14 | 96995877 | 3 | 0 | 0 | T | G | -1.063 | 0.3211 | 0.0009318 | ---?? | 0 | 0.8082 | intron | PAPOLA |
| rs190934677 | 11 | 16291075 | 2 | 0 | 0 | A | C | -6.3411 | 1.9156 | 0.0009321 | ?--?? | 0 | 0.6765 | intron | SOX6 |
| rs3850736 | 8 | 64912021 | 4 | 0 | 0 | C | G | -0.8214 | 0.2481 | 0.0009324 | ---?+ | 0 | 0.483 | intergenic | |
| rs4502931 | 6 | 32380782 | 4 | 0 | 0 | A | T | 0.7185 | 0.2171 | 0.0009324 | ++++? | 0 | 0.4889 | intergenic | |
| rs9264527 | 6 | 31233992 | 5 | 0 | 0 | T | C | 0.7901 | 0.2387 | 0.0009324 | +++++ | 0 | 0.4573 | downstream-gene | |
| rs1159304 | 3 | 30825421 | 3 | 0 | 0 | T | C | -2.989 | 0.903 | 0.0009325 | ---?? | 0 | 0.5066 | intron | GADL1 |
| rs28463264 | 17 | 7067549 | 3 | 0 | 0 | A | G | 4.1813 | 1.2632 | 0.0009326 | +++?? | 0 | 0.7491 | intergenic | |
| rs4670283 | 2 | 39755436 | 4 | 0 | 0 | T | C | -1.3001 | 0.3928 | 0.0009328 | ---?- | 0 | 0.7446 | intron | LOC728730 |
| rs12588853 | 14 | 43271891 | 4 | 0 | 0 | A | G | -1.3916 | 0.4204 | 0.0009328 | ---?+ | 45.4 | 0.1387 | intergenic | |
| rs807133 | 17 | 51228816 | 5 | 0 | 0 | T | C | 1.023 | 0.3091 | 0.000933 | ++++- | 5.1 | 0.3778 | intergenic | |
| rs2785651 | 1 | 208188103 | 4 | 0 | 0 | T | C | 1.8919 | 0.5716 | 0.0009331 | +++?+ | 0 | 0.7417 | intergenic | |
| rs12622845 | 2 | 117103110 | 4 | 0 | 0 | C | G | 1.6631 | 0.5024 | 0.0009331 | +++?+ | 0 | 0.673 | intergenic | |
| rs62553724 | 9 | 11518595 | 4 | 0 | 0 | A | T | -1.7242 | 0.5209 | 0.0009331 | ---?- | 56 | 0.07778 | intergenic | |
| rs2420169 | 7 | 65655918 | 3 | 0 | 0 | C | G | 1.5719 | 0.4749 | 0.0009333 | +++?? | 0 | 0.7131 | intergenic | |
| rs1010419 | 15 | 87536101 | 4 | 0 | 0 | A | G | -0.851 | 0.2571 | 0.0009334 | ---?- | 0 | 0.4497 | intron | AGBL1 |
| 4:59660124 | 4 | 59660124 | 3 | 0 | 0 | A | AT | -2.3598 | 0.713 | 0.0009335 | ---?? | 0 | 0.7686 | | |
| rs1905852 | 4 | 96520014 | 4 | 0 | 0 | A | G | 1.2452 | 0.3762 | 0.0009337 | +++?+ | 28.1 | 0.2432 | intergenic | |
| rs61450873 | 19 | 12813206 | 4 | 0 | 0 | T | G | -3.4611 | 1.0457 | 0.0009337 | ----? | 50.4 | 0.1094 | intron | TNPO2 |
| rs10425615 | 19 | 12808326 | 4 | 0 | 0 | A | G | 3.4563 | 1.0443 | 0.0009338 | ++++? | 50.2 | 0.1104 | upstream-gene | |

| | | | | | | | | | | | | | | | |
|---|---|---|---|---|---|---|---|---|---|---|---|---|---|---|---|
| rs185008697 | 8 | 90893314 | 4 | 0 | 0 | A | T | -2.4897 | 0.7522 | 0.0009338 | ---?- | 0 | 0.9693 | intergenic | |
| rs1557119 | 10 | 29244183 | 4 | 0 | 0 | T | C | -0.8539 | 0.258 | 0.0009341 | ---?- | 0 | 0.7493 | intergenic | |
| rs190425904 | 3 | 143800136 | 2 | 0 | 0 | T | G | 4.8295 | 1.4592 | 0.0009341 | +?+?? | 62.4 | 0.103 | intergenic | |
| rs996664 | 1 | 187868951 | 4 | 0 | 0 | T | C | 0.9875 | 0.2984 | 0.0009342 | +++?+ | 0 | 0.6612 | intergenic | |
| 6:31096562 | 6 | 31096562 | 3 | 0 | 0 | T | TC | 1.9556 | 0.5909 | 0.0009344 | +++?? | 0 | 0.8875 | | |
| rs9264483 | 6 | 31232446 | 5 | 0 | 0 | T | C | -0.7899 | 0.2387 | 0.0009347 | ----- | 0 | 0.4562 | downstream-gene |  |
| rs146945912 | 12 | 43107063 | 4 | 0 | 0 | C | G | -1.7019 | 0.5143 | 0.0009347 | ---?- | 0 | 0.8258 | intergenic | |
| rs2468720 | 8 | 140415969 | 4 | 0 | 0 | A | T | 0.978 | 0.2955 | 0.0009348 | +++?+ | 0 | 0.4559 | intergenic | |
| rs12506932 | 4 | 10039528 | 5 | 0 | 0 | A | C | 0.6961 | 0.2103 | 0.0009349 | ++++- | 14.7 | 0.3207 | intron | SLC2A9 |
| rs76697635 | 4 | 41314694 | 4 | 0 | 0 | A | G | -1.5368 | 0.4644 | 0.0009349 | ---?+ | 0 | 0.6501 | intergenic | |
| rs6917425 | 6 | 130947351 | 4 | 0 | 0 | T | G | -1.3808 | 0.4172 | 0.000935 | ---?- | 0 | 0.7388 | intergenic | |
| rs2570131 | 15 | 87607311 | 4 | 0 | 0 | A | G | 0.8538 | 0.258 | 0.0009352 | +++?+ | 0 | 0.6826 | intergenic | |
| rs4791464 | 17 | 11217860 | 3 | 0 | 0 | T | G | -0.9549 | 0.2886 | 0.0009352 | ---?? | 0 | 0.525 | intron | SHISA6 |
| rs6447982 | 4 | 20760238 | 4 | 0 | 0 | A | C | -0.8785 | 0.2655 | 0.0009352 | ---?- | 0 | 0.7914 | intron | KCNIP4 |
| rs56296358 | 11 | 134478665 | 4 | 0 | 0 | T | C | -1.1037 | 0.3335 | 0.0009353 | ---?- | 0 | 0.9918 | intergenic | |
| rs55757636 | 19 | 56829207 | 4 | 0 | 0 | A | T | 2.3245 | 0.7024 | 0.0009356 | +-+?+ | 41.3 | 0.1637 | upstream-gene | |
| rs117976266 | 20 | 1517215 | 5 | 0 | 0 | A | G | 1.9052 | 0.5757 | 0.0009356 | ++++- | 35 | 0.1881 | intron | SIRPD |
| rs4068735 | 2 | 69122839 | 4 | 0 | 0 | T | C | 0.8255 | 0.2494 | 0.0009357 | +++?+ | 0 | 0.7475 | intergenic | |
| rs11081113 | 18 | 452104 | 4 | 0 | 0 | T | C | -0.7942 | 0.24 | 0.0009357 | ---?- | 32.4 | 0.2179 | intron | COLEC12 |
| 6:31231965 | 6 | 31231965 | 3 | 0 | 0 | T | TC | 0.9411 | 0.2844 | 0.0009358 | +++?? | 22.9 | 0.2731 | | |
| 3:194649490 | 3 | 194649490 | 3 | 0 | 0 | T | G | 3.0564 | 0.9236 | 0.0009359 | +++?? | 0 | 0.9627 | intergenic | |
| rs148924071 | 12 | 61478341 | 2 | 0 | 0 | A | G | 5.7252 | 1.7301 | 0.0009359 | ?+??+ | 66 | 0.08644 | intergenic | |
| rs13216467 | 6 | 159182358 | 3 | 0 | 0 | A | G | 0.8554 | 0.2585 | 0.0009361 | +++?? | 0 | 0.6413 | intron | SYTL3 |
| 4:38745238 | 4 | 38745238 | 3 | 0 | 0 | CA | C | -1.1917 | 0.3601 | 0.0009361 | ---?? | 0 | 0.6428 | | |
| rs11779170 | 8 | 135204911 | 3 | 0 | 0 | T | C | 3.1123 | 0.9406 | 0.0009362 | +++?? | 0 | 0.9201 | intergenic | |
| rs6076417 | 20 | 26287724 | 4 | 0 | 0 | T | C | -1.49 | 0.4503 | 0.0009366 | ---?- | 59.7 | 0.05886 | intergenic | |
| rs76915951 | 17 | 4378105 | 4 | 0 | 0 | C | G | 4.021 | 1.2152 | 0.0009366 | +++?+ | 2.6 | 0.3793 | intron | SPNS3 |
| rs976635 | 1 | 217997362 | 4 | 0 | 0 | A | T | -1.0827 | 0.3272 | 0.0009367 | ---?- | 0 | 0.9785 | intron | SPATA17 |
| rs9625544 | 22 | 29115035 | 4 | 0 | 0 | T | C | 2.1287 | 0.6433 | 0.0009367 | +++?+ | 0 | 0.6131 | intron | CHEK2 |
| rs12104166 | 19 | 12808502 | 4 | 0 | 0 | T | C | -3.4552 | 1.0442 | 0.0009367 | ----? | 50.2 | 0.1102 | upstream-gene |  |
| rs35731009 | 22 | 28987988 | 4 | 0 | 0 | A | G | 2.3808 | 0.7195 | 0.0009367 | +++?+ | 0 | 0.4839 | intron | TTC28 |
| rs116697544 | 11 | 86192213 | 4 | 0 | 0 | A | G | -2.3457 | 0.7089 | 0.0009367 | ---?- | 0 | 0.9621 | intron | ME3 |
| rs2045856 | 8 | 140409588 | 4 | 0 | 0 | A | C | 0.9978 | 0.3016 | 0.0009369 | +++?+ | 0 | 0.8441 | intergenic | |
| rs7743762 | 6 | 164372151 | 4 | 0 | 0 | T | C | 1.2436 | 0.3759 | 0.0009369 | +++?+ | 0 | 0.9812 | intergenic | |
| rs74924524 | 2 | 52950830 | 4 | 0 | 0 | A | G | 1.2769 | 0.3859 | 0.0009369 | +++?+ | 0 | 0.7622 | intergenic | |
| rs4252267 | 11 | 117863292 | 4 | 0 | 0 | A | G | 1.3017 | 0.3934 | 0.000937 | +++?+ | 0 | 0.757 | intron | IL10RA |
| rs75996500 | 1 | 152737593 | 4 | 0 | 0 | T | C | -2.0806 | 0.6288 | 0.000937 | ---?- | 0 | 0.451 | downstream-gene | |
| rs144605585 | 1 | 26961670 | 4 | 0 | 0 | T | C | 1.7974 | 0.5432 | 0.000937 | +++?+ | 0 | 0.7568 | intergenic | |
| rs72759171 | 5 | 65905163 | 4 | 0 | 0 | T | G | 1.1651 | 0.3521 | 0.0009372 | +++?+ | 0 | 0.8156 | intron | MAST4 |
| 1:85616116 | 1 | 85616116 | 3 | 0 | 0 | G | GATAAA | 1.5579 | 0.4708 | 0.0009372 | +++?? | 0 | 0.9264 | | |
| rs2345917 | 4 | 42764552 | 4 | 0 | 0 | T | G | 1.1063 | 0.3344 | 0.0009373 | +++?+ | 0 | 0.6125 | intergenic | |
| rs17142816 | 7 | 103058720 | 5 | 0 | 0 | T | C | 0.9091 | 0.2748 | 0.0009377 | ++++- | 10.4 | 0.3471 | intron | SLC26A5 |
| rs3758877 | 11 | 86171405 | 4 | 0 | 0 | T | G | 1.546 | 0.4673 | 0.0009379 | +++?+ | 0 | 0.4081 | intron | ME3 |
| rs79658777 | 18 | 74931364 | 4 | 0 | 0 | C | G | 2.2276 | 0.6733 | 0.0009379 | +++?- | 0 | 0.4564 | intergenic | |
| rs2102081 | 15 | 98048628 | 4 | 0 | 0 | A | G | -0.834 | 0.2521 | 0.000938 | ---?- | 0 | 0.8754 | intergenic | |
| rs10765982 | 11 | 12652308 | 4 | 0 | 0 | T | C | 1.0876 | 0.3287 | 0.0009382 | +++?+ | 0 | 0.8292 | intergenic | |
| rs17059337 | 6 | 130947684 | 4 | 0 | 0 | T | C | 1.3807 | 0.4173 | 0.0009384 | +++?+ | 0 | 0.7384 | intergenic | |
| rs1034156 | 9 | 30808739 | 4 | 0 | 0 | A | G | -3.1698 | 0.9581 | 0.0009385 | ---?- | 0.1 | 0.3913 | intergenic | |
| 5:170931820 | 5 | 170931820 | 3 | 0 | 0 | A | AG | -0.8288 | 0.2505 | 0.0009386 | ---?? | 0 | 0.981 | | |
| rs190943161 | 6 | 72862541 | 3 | 0 | 0 | A | G | -1.243 | 0.3757 | 0.0009387 | ---?? | 45.2 | 0.1611 | intron | RIMS1 |
| rs28875291 | 9 | 44250995 | 3 | 0 | 0 | A | G | 1.7335 | 0.524 | 0.0009389 | +++?? | 0 | 0.8606 | intergenic | |
| rs138727547 | 2 | 35077576 | 3 | 0 | 0 | T | C | -3.4987 | 1.0576 | 0.000939 | ---?? | 42.7 | 0.1747 | intergenic | |
| rs138102338 | 2 | 136631443 | 3 | 0 | 0 | A | G | -3.8456 | 1.1625 | 0.0009391 | ---?? | 0 | 0.6155 | intron | MCM6 |
| rs62404200 | 6 | 19873870 | 5 | 0 | 0 | T | C | 0.9254 | 0.2797 | 0.0009392 | +++++ | 0 | 0.9271 | intergenic | |
| rs2764541 | 1 | 111811430 | 4 | 0 | 0 | C | G | -1.8745 | 0.5667 | 0.0009394 | ---?+ | 0 | 0.4595 | intergenic | |
| rs3758878 | 11 | 86171003 | 4 | 0 | 0 | C | G | 1.5454 | 0.4672 | 0.0009395 | +++?+ | 0 | 0.407 | intron | ME3 |
| rs11180676 | 12 | 76174818 | 4 | 0 | 0 | T | C | 2.0855 | 0.6304 | 0.0009395 | +++?- | 0 | 0.7331 | intergenic | |
| rs139161675 | 17 | 80026978 | 3 | 0 | 0 | A | G | -5.1558 | 1.5586 | 0.0009395 | --??- | 0 | 0.8758 | upstream-gene | |
| rs9264464 | 6 | 31231622 | 3 | 0 | 0 | C | G | 0.9409 | 0.2844 | 0.0009397 | +++?? | 22.4 | 0.2755 | downstream-gene | |

| | | | | | | | | | | | | | | | |
|---|---|---|---|---|---|---|---|---|---|---|---|---|---|---|---|
| 3:87378820 | 3 | 87378820 | 3 | 0 | 0 | CT | CTT | 1.1667 | 0.3527 | 0.0009399 | +++?? | 0 | 0.7012 | | |
| rs6005780 | 22 | 28807540 | 4 | 0 | 0 | T | C | 2.3293 | 0.7042 | 0.0009402 | +++?+ | 0 | 0.6395 | intron | TTC28 |
| rs71419569 | 2 | 138944797 | 4 | 0 | 0 | T | C | -1.5886 | 0.4803 | 0.0009402 | ---?- | 0 | 0.789 | intergenic | |
| rs182456270 | 6 | 109140281 | 2 | 0 | 0 | A | G | 4.6505 | 1.4059 | 0.0009402 | +?+?? | 0 | 0.749 | intergenic | |
| 7:4585684 | 7 | 4585684 | 3 | 0 | 0 | A | AC | 1.017 | 0.3074 | 0.0009402 | +++?? | 0 | 0.5063 | | |
| 9:101508864 | 9 | 101508864 | 3 | 0 | 0 | CT | C | 1.2893 | 0.3898 | 0.0009402 | +++?? | 7 | 0.3411 | | |
| rs10114437 | 9 | 139059275 | 4 | 0 | 0 | T | C | 0.8742 | 0.2643 | 0.0009403 | +++?+ | 46.9 | 0.1299 | intergenic | |
| rs77716993 | 11 | 86170938 | 4 | 0 | 0 | A | T | -1.5451 | 0.4671 | 0.0009404 | ---?- | 0 | 0.4068 | intron | ME3 |
| rs10910532 | 1 | 235082127 | 4 | 0 | 0 | A | G | -0.8082 | 0.2443 | 0.0009405 | ---?- | 0 | 0.8888 | intergenic | |
| rs181600309 | 1 | 42743149 | 3 | 0 | 0 | T | C | 1.81 | 0.5472 | 0.0009405 | +++?? | 13.6 | 0.3141 | intron | FOXJ3 |
| rs2118842 | 2 | 196507984 | 4 | 0 | 0 | A | G | 1.069 | 0.3232 | 0.0009406 | +++?+ | 0 | 0.9758 | intron | SLC39A10 |
| rs2248705 | 2 | 184010492 | 4 | 0 | 0 | A | C | 1.1446 | 0.346 | 0.0009408 | +++?+ | 0 | 0.834 | intron | NUP35 |
| rs11942930 | 4 | 140543648 | 4 | 0 | 0 | T | C | -2.9607 | 0.8951 | 0.0009408 | ---?- | 20.2 | 0.2889 | intergenic | |
| rs13201420 | 6 | 30989847 | 3 | 0 | 0 | T | C | -1.3621 | 0.4118 | 0.0009408 | ---?? | 0 | 0.8412 | intron | MUC22 |
| rs145759519 | 16 | 6005316 | 3 | 0 | 0 | A | G | -0.9199 | 0.2781 | 0.0009409 | ---?? | 0 | 0.628 | intron | RBFOX1 |
| rs2926574 | 1 | 69810770 | 4 | 0 | 0 | A | G | -0.8511 | 0.2573 | 0.0009413 | ---?- | 0 | 0.8308 | intergenic | |
| rs12999790 | 2 | 117822459 | 3 | 0 | 0 | A | T | 0.8996 | 0.272 | 0.0009413 | +++?? | 1.5 | 0.3622 | intergenic | |
| 16:13768108 | 16 | 13768108 | 3 | 0 | 0 | CA | C | 2.5596 | 0.7739 | 0.0009416 | +++?? | 0 | 0.9837 | | |
| 8:118493477 | 8 | 118493477 | 3 | 0 | 0 | CA | C | -0.9057 | 0.2738 | 0.0009416 | +--?? | 65.8 | 0.05382 | | |
| rs369305 | 18 | 39942414 | 4 | 0 | 0 | C | G | 0.8543 | 0.2583 | 0.0009417 | +++?+ | 0 | 0.4406 | intron | LINC00907 |
| rs78439417 | 2 | 47400734 | 4 | 0 | 0 | T | G | 1.3778 | 0.4166 | 0.0009418 | +++?+ | 0 | 0.6351 | intron | CALM2 |
| rs12552109 | 9 | 14639150 | 4 | 0 | 0 | T | C | -1.6491 | 0.4986 | 0.0009419 | ---?- | 0 | 0.8215 | intron | ZDHHC21 |
| rs4252287 | 11 | 117868638 | 4 | 0 | 0 | A | G | 1.3003 | 0.3932 | 0.0009421 | +++?+ | 0 | 0.7122 | intron | IL10RA |
| rs5011787 | 6 | 74387935 | 3 | 0 | 0 | A | T | -1.106 | 0.3344 | 0.0009421 | ---?? | 0 | 0.7552 | intergenic | |
| rs10261162 | 7 | 22132118 | 4 | 0 | 0 | C | G | 1.8516 | 0.5599 | 0.0009421 | +++?+ | 0 | 0.7769 | intergenic | |
| rs17256536 | 8 | 124699175 | 4 | 0 | 0 | A | G | 0.8168 | 0.247 | 0.0009422 | +++?+ | 0 | 0.8691 | intron | ANXA13 |
| rs6803806 | 3 | 188508478 | 4 | 0 | 0 | T | C | 0.86 | 0.26 | 0.0009423 | +++?+ | 15.9 | 0.3121 | intron | LPP |
| rs10033935 | 4 | 123724257 | 4 | 0 | 0 | A | G | 2.1373 | 0.6463 | 0.0009423 | +++?+ | 58.8 | 0.06332 | intergenic | |
| rs10189689 | 2 | 131130678 | 5 | 0 | 0 | T | C | 0.8587 | 0.2596 | 0.0009423 | +++++ | 16.5 | 0.3095 | intron | PTPN18 |
| rs11097945 | 4 | 108214738 | 4 | 0 | 0 | A | G | -0.8087 | 0.2445 | 0.0009426 | ---?- | 0 | 0.5537 | intergenic | |
| rs12165352 | 22 | 29117308 | 4 | 0 | 0 | A | C | 2.1311 | 0.6444 | 0.0009426 | +++?+ | 0 | 0.615 | intron | CHEK2 |
| rs12564589 | 1 | 64706579 | 4 | 0 | 0 | A | G | -1.2224 | 0.3696 | 0.0009428 | ---?- | 0 | 0.4555 | intron | UBE2U |
| rs118175016 | 21 | 46953849 | 3 | 0 | 0 | A | G | 2.5679 | 0.7765 | 0.0009429 | +++?? | 0 | 0.9808 | intron | SLC19A1 |
| rs143878674 | 11 | 26295799 | 3 | 0 | 0 | T | C | 4.26 | 1.2882 | 0.0009429 | +++?? | 59.5 | 0.08469 | intron | ANO3 |
| rs747021 | 9 | 90850936 | 4 | 0 | 0 | T | C | -1.864 | 0.5637 | 0.000943 | ---?- | 0 | 0.7434 | intergenic | |
| rs11208387 | 1 | 64706743 | 4 | 0 | 0 | T | C | -1.2222 | 0.3696 | 0.0009431 | ---?- | 0 | 0.4548 | intron | UBE2U |
| rs59736952 | 12 | 132336799 | 3 | 0 | 0 | A | C | 4.6648 | 1.4106 | 0.0009431 | +?+?+ | 0 | 0.9587 | downstream-gene | |
| 2:167399927 | 2 | 167399927 | 3 | 0 | 0 | A | AAG | 1.5908 | 0.481 | 0.0009433 | +++?? | 0 | 0.7831 | | |
| rs78866296 | 4 | 162212644 | 2 | 0 | 0 | A | T | 6.0479 | 1.8289 | 0.0009434 | ?++?? | 0 | 0.6594 | intergenic | |
| rs6076415 | 20 | 26285399 | 4 | 0 | 0 | A | G | 1.4549 | 0.4399 | 0.0009435 | +++?+ | 59.5 | 0.06009 | intergenic | |
| rs2203602 | 2 | 189877911 | 4 | 0 | 0 | A | T | -0.9452 | 0.2858 | 0.0009436 | ---?- | 42.2 | 0.1584 | downstream-gene | |
| rs116462335 | 5 | 120177871 | 2 | 0 | 0 | A | G | -5.3189 | 1.6085 | 0.0009436 | ?--?? | 0 | 0.715 | intergenic | |
| rs2594334 | 15 | 87553198 | 4 | 0 | 0 | A | C | 0.8733 | 0.2641 | 0.0009437 | +++?+ | 0 | 0.4241 | intron | AGBL1 |
| rs7091418 | 10 | 86634920 | 4 | 0 | 0 | A | G | 1.5118 | 0.4572 | 0.0009439 | +++?+ | 4.5 | 0.3704 | intergenic | |
| rs9423904 | 10 | 5718794 | 2 | 0 | 0 | A | T | -5.5167 | 1.6683 | 0.0009439 | --??? | 0 | 0.8159 | intergenic | |
| rs11825064 | 11 | 134477854 | 4 | 0 | 0 | C | G | -1.1153 | 0.3373 | 0.0009439 | ---?- | 0 | 0.9881 | intergenic | |
| rs74464606 | 17 | 42271881 | 3 | 0 | 0 | A | G | 3.8473 | 1.1635 | 0.0009439 | +-+?? | 58.5 | 0.08966 | intron-near-splice | ATXN7L3 |
| rs76401566 | 11 | 86170722 | 4 | 0 | 0 | T | C | 1.544 | 0.4669 | 0.0009439 | +++?+ | 0 | 0.4054 | intron | ME3 |
| 1:194302202 | 1 | 194302202 | 3 | 0 | 0 | T | TA | -2.5148 | 0.7605 | 0.000944 | ---?? | 0 | 0.945 | | |
| 12:95821249 | 12 | 95821249 | 3 | 0 | 0 | CA | C | 1.9351 | 0.5852 | 0.000944 | +++?? | 19.1 | 0.2905 | | |
| rs117392224 | 17 | 76957170 | 4 | 0 | 0 | A | G | 2.2223 | 0.6721 | 0.0009444 | +++?- | 53.4 | 0.09229 | intergenic | |
| rs6079982 | 20 | 15820353 | 2 | 0 | 0 | A | G | 1.5246 | 0.4611 | 0.0009446 | ???++ | 0 | 0.4632 | intron | MACROD2 |
| rs2644692 | 12 | 117159954 | 5 | 0 | 0 | A | G | 1.2664 | 0.383 | 0.0009448 | +++++ | 0 | 0.784 | intron | C12orf49 |
| rs10921369 | 1 | 193475831 | 5 | 0 | 0 | A | G | 2.7132 | 0.8206 | 0.0009449 | -++++ | 0.3 | 0.4042 | intergenic | |
| rs10773842 | 12 | 131562094 | 4 | 0 | 0 | A | G | -0.8418 | 0.2546 | 0.0009451 | ---?- | 0 | 0.9613 | intron | GPR133 |
| rs11828283 | 11 | 86110521 | 4 | 0 | 0 | A | C | 2.1688 | 0.6559 | 0.0009452 | +++?+ | 21.6 | 0.281 | intron | CCDC81 |
| rs12165475 | 22 | 29117216 | 4 | 0 | 0 | A | T | -2.1304 | 0.6443 | 0.0009452 | ---?- | 0 | 0.615 | intron | CHEK2 |
| rs3130543 | 6 | 31232147 | 5 | 0 | 0 | T | C | 0.7892 | 0.2387 | 0.0009453 | +++++ | 0 | 0.4571 | downstream-gene | |
| rs346446 | 18 | 39942821 | 4 | 0 | 0 | A | C | 0.854 | 0.2583 | 0.0009457 | +++?+ | 0 | 0.4423 | intron | LINC00907 |

| | | | | | | | | | | | | | | | |
|---|---|---|---|---|---|---|---|---|---|---|---|---|---|---|---|
| rs75463542 | 22 | 31646982 | 4 | 0 | 0 | A | G | 3.6685 | 1.1096 | 0.0009459 | +++?+ | 0 | 0.7391 | intron | LIMK2 |
| rs112740321 | 7 | 48553678 | 4 | 0 | 0 | A | G | 2.5378 | 0.7676 | 0.0009459 | +++?+ | 0 | 0.6949 | intron | ABCA13 |
| 8:30281340 | 8 | 30281340 | 3 | 0 | 0 | G | GTTTT | 0.8678 | 0.2625 | 0.0009459 | +++?? | 63.6 | 0.06389 | | |
| rs79475932 | 1 | 98631848 | 4 | 0 | 0 | A | C | 1.4326 | 0.4333 | 0.0009461 | +++?+ | 0 | 0.4291 | intergenic | |
| rs11586998 | 1 | 64714704 | 4 | 0 | 0 | T | C | 1.2139 | 0.3672 | 0.0009463 | +++?+ | 0 | 0.4756 | intron | UBE2U |
| rs80085001 | 4 | 100552473 | 4 | 0 | 0 | T | C | 1.7553 | 0.5309 | 0.0009463 | -++?+ | 51.8 | 0.1011 | intergenic | |
| rs140702813 | 6 | 83902972 | 4 | 0 | 0 | T | C | -2.7419 | 0.8294 | 0.0009463 | ---?- | 0 | 0.4906 | intron | PGM3 |
| rs7027525 | 9 | 25616027 | 3 | 0 | 0 | T | G | 0.9705 | 0.2936 | 0.0009464 | +++?? | 62.8 | 0.06806 | intergenic | |
| rs2298206 | 1 | 180145088 | 4 | 0 | 0 | A | G | -2.0135 | 0.6091 | 0.0009465 | ---?+ | 32.5 | 0.2171 | synonymous | QSOX1 |
| rs7739307 | 6 | 130948135 | 4 | 0 | 0 | A | G | 1.3801 | 0.4175 | 0.0009466 | +++?+ | 0 | 0.7366 | intergenic | |
| rs12630163 | 3 | 5387267 | 4 | 0 | 0 | A | G | 1.3364 | 0.4042 | 0.0009467 | +++?+ | 5.5 | 0.3653 | intergenic | |
| rs17270127 | 6 | 157314607 | 4 | 0 | 0 | T | G | -1.4247 | 0.431 | 0.0009468 | ---?+ | 3.7 | 0.3744 | intron | ARID1B |
| rs4666617 | 2 | 184019721 | 4 | 0 | 0 | C | G | 1.1439 | 0.346 | 0.0009469 | +++?+ | 0 | 0.8004 | intron | NUP35 |
| rs2468718 | 8 | 140416137 | 4 | 0 | 0 | T | G | -0.9768 | 0.2955 | 0.000947 | ---?- | 0 | 0.4555 | intergenic | |
| rs4974266 | 3 | 70432819 | 4 | 0 | 0 | A | G | 0.9386 | 0.2839 | 0.000947 | +++?+ | 0 | 0.5583 | intergenic | |
| rs10888115 | 8 | 12568750 | 4 | 0 | 0 | A | G | 0.9445 | 0.2857 | 0.000947 | +++?+ | 48.6 | 0.1198 | intergenic | |
| rs77046295 | 3 | 1676853 | 4 | 0 | 0 | A | T | 1.5963 | 0.4829 | 0.0009473 | +++?+ | 0 | 0.9565 | intergenic | |
| rs12593787 | 15 | 92449426 | 4 | 0 | 0 | T | C | -0.8242 | 0.2493 | 0.0009474 | ---?- | 0 | 0.527 | intron | SLCO3A1 |
| rs10990394 | 9 | 105725266 | 4 | 0 | 0 | T | C | 1.3069 | 0.3953 | 0.0009475 | +++?+ | 0 | 0.8775 | intergenic | |
| rs2594328 | 15 | 87534285 | 4 | 0 | 0 | A | G | -0.8492 | 0.2569 | 0.0009476 | ---?- | 0 | 0.4462 | intron | AGBL1 |
| rs12536248 | 7 | 32369296 | 4 | 0 | 0 | A | G | 1.3431 | 0.4063 | 0.0009476 | +++?+ | 0 | 0.9373 | intergenic | |
| rs17612346 | 1 | 152737144 | 4 | 0 | 0 | A | G | 2.0782 | 0.6287 | 0.000948 | +++?+ | 0 | 0.4525 | downstream-gene | |
| rs5028780 | 2 | 230918496 | 4 | 0 | 0 | A | C | -1.4058 | 0.4253 | 0.0009481 | ---?- | 16.3 | 0.3102 | intron | SLC16A14 |
| rs11611439 | 12 | 131561902 | 4 | 0 | 0 | T | C | 0.8422 | 0.2548 | 0.0009481 | +++?+ | 0 | 0.9615 | intron | GPR133 |
| rs75367913 | 2 | 131114042 | 5 | 0 | 0 | T | G | -0.8596 | 0.26 | 0.0009481 | ----- | 21 | 0.2809 | intron | PTPN18 |
| rs28868927 | 9 | 44248134 | 3 | 0 | 0 | A | G | 1.7009 | 0.5146 | 0.0009483 | +++?? | 0 | 0.894 | intergenic | |
| rs59578882 | 12 | 132336621 | 3 | 0 | 0 | T | C | 4.6572 | 1.409 | 0.0009485 | +?+?+ | 0 | 0.9582 | downstream-gene | |
| rs6005841 | 22 | 29103453 | 4 | 0 | 0 | A | C | 2.1144 | 0.6397 | 0.0009486 | +++?+ | 0 | 0.5929 | intron | CHEK2 |
| rs78060822 | 1 | 14205141 | 4 | 0 | 0 | T | C | 1.8185 | 0.5502 | 0.0009487 | +-+?+ | 50.4 | 0.1092 | intergenic | |
| rs141251706 | 20 | 35632692 | 4 | 0 | 0 | T | C | -1.1752 | 0.3555 | 0.0009488 | ---?- | 0 | 0.9698 | intron | RBL1 |
| rs2004300 | 15 | 96668810 | 4 | 0 | 0 | A | T | -0.8599 | 0.2602 | 0.0009491 | ---?- | 0 | 0.6517 | intergenic | |
| rs79234252 | 17 | 52247118 | 4 | 0 | 0 | T | C | 2.1036 | 0.6364 | 0.0009491 | +++?- | 35.2 | 0.2008 | intergenic | |
| rs1535867 | 10 | 9305165 | 4 | 0 | 0 | T | G | 1.6799 | 0.5083 | 0.0009492 | +++?+ | 0 | 0.663 | intergenic | |
| rs774297 | 16 | 84804409 | 4 | 0 | 0 | C | G | -1.0129 | 0.3065 | 0.0009495 | ---?+ | 0 | 0.465 | intron | USP10 |
| rs11693165 | 2 | 59980995 | 4 | 0 | 0 | A | G | -1.0612 | 0.3211 | 0.0009495 | ---?- | 0 | 0.6896 | intergenic | |
| 1:85625679 | 1 | 85625679 | 3 | 0 | 0 | A | ACAGAG | 2.5841 | 0.7819 | 0.0009501 | +++?? | 0 | 0.6141 | | |
| 15:87604748 | 15 | 87604748 | 3 | 0 | 0 | T | TA | -0.8882 | 0.2688 | 0.0009502 | ---?? | 10.2 | 0.3284 | | |
| 19:14718362 | 19 | 14718362 | 3 | 0 | 0 | A | ATTAT | -1.0737 | 0.3249 | 0.0009503 | ---?? | 12.3 | 0.3199 | | |
| rs7908103 | 10 | 9306064 | 4 | 0 | 0 | A | T | -1.68 | 0.5083 | 0.0009507 | ---?- | 0 | 0.6612 | intergenic | |
| rs13226085 | 7 | 92628043 | 2 | 0 | 0 | A | G | 5.3317 | 1.6134 | 0.0009507 | ??+?+ | 0 | 0.8037 | intergenic | |
| rs12292316 | 11 | 86109900 | 4 | 0 | 0 | A | C | 2.168 | 0.656 | 0.0009508 | +++?+ | 21.6 | 0.2808 | intron | CCDC81 |
| rs72701767 | 4 | 185276008 | 4 | 0 | 0 | T | C | -1.1422 | 0.3456 | 0.0009508 | ---?- | 0 | 0.9724 | upstream-gene | |
| rs146665713 | 4 | 166246988 | 4 | 0 | 0 | A | G | 3.5332 | 1.0692 | 0.0009508 | +++?+ | 0 | 0.951 | upstream-gene | |
| rs390145 | 2 | 40457049 | 4 | 0 | 0 | A | C | 1.1685 | 0.3536 | 0.0009509 | +++?+ | 0 | 0.7891 | intron | SLC8A1,SLC8A1-AS1 |
| rs2701406 | 15 | 87539694 | 4 | 0 | 0 | A | C | 0.8558 | 0.259 | 0.0009509 | +++?+ | 0 | 0.4419 | intron | AGBL1 |
| rs10409345 | 19 | 12809159 | 4 | 0 | 0 | A | T | 3.4501 | 1.044 | 0.000951 | ++++? | 50.4 | 0.1093 | upstream-gene | |
| rs11789032 | 9 | 19011586 | 4 | 0 | 0 | C | G | -1.546 | 0.4678 | 0.0009511 | ---?- | 44.7 | 0.1432 | intron | FAM154A |
| rs7238560 | 18 | 399245 | 4 | 0 | 0 | A | G | 0.8357 | 0.2529 | 0.0009512 | +++?+ | 0 | 0.5087 | intron | COLEC12 |
| rs7684213 | 4 | 101283387 | 4 | 0 | 0 | T | C | 0.8681 | 0.2627 | 0.0009514 | +++?+ | 0 | 0.492 | intergenic | |
| rs11789009 | 9 | 19011593 | 4 | 0 | 0 | T | C | -1.5459 | 0.4678 | 0.0009514 | ---?- | 44.7 | 0.1431 | intron | FAM154A |
| rs72730594 | 15 | 75394798 | 3 | 0 | 0 | T | C | -3.1982 | 0.9678 | 0.0009514 | ---?? | 0 | 0.8077 | intergenic | |
| rs16950635 | 15 | 67418205 | 4 | 0 | 0 | A | G | 1.6491 | 0.499 | 0.0009515 | +++?+ | 0 | 0.8035 | 5-prime-UTR | SMAD3 |
| rs6902922 | 6 | 130943526 | 4 | 0 | 0 | C | G | -1.3726 | 0.4154 | 0.0009516 | ---?- | 0 | 0.7399 | intergenic | |
| 12:95821253 | 12 | 95821253 | 3 | 0 | 0 | A | C | 1.9338 | 0.5852 | 0.0009519 | +++?? | 19.2 | 0.29 | intergenic | |
| rs3853275 | 9 | 30443903 | 4 | 0 | 0 | A | G | 1.1786 | 0.3567 | 0.0009519 | +++?+ | 0 | 0.6683 | intergenic | |
| 6:124350031 | 6 | 124350031 | 3 | 0 | 0 | T | G | -2.8439 | 0.8607 | 0.0009523 | ---?? | 0 | 0.4226 | intron | NKAIN2 |
| rs7428050 | 3 | 77403911 | 4 | 0 | 0 | A | G | 0.8281 | 0.2506 | 0.0009523 | +++?+ | 0 | 0.4039 | intron | ROBO2 |
| rs9475522 | 6 | 55895491 | 4 | 0 | 0 | A | T | -2.2178 | 0.6712 | 0.0009524 | ---?- | 0 | 0.8012 | intergenic | |
| rs16885254 | 8 | 114886519 | 4 | 0 | 0 | A | G | -1.8136 | 0.5489 | 0.0009525 | ---?+ | 6.4 | 0.3609 | intergenic | |

| | | | | | | | | | | | | | | | |
|---|---|---|---|---|---|---|---|---|---|---|---|---|---|---|---|
| rs6869413 | 5 | 39606335 | 4 | 0 | 0 | A | G | -0.9865 | 0.2986 | 0.0009526 | ---?- | 0 | 0.8859 | intergenic | |
| rs71428627 | 2 | 83097948 | 4 | 0 | 0 | A | C | 2.2598 | 0.6839 | 0.0009529 | +++?+ | 0 | 0.5145 | intergenic | |
| rs2004301 | 15 | 96668796 | 4 | 0 | 0 | A | G | -0.8595 | 0.2601 | 0.0009532 | ---?- | 0 | 0.6515 | intergenic | |
| rs12891137 | 14 | 79623717 | 5 | 0 | 0 | T | C | 1.1694 | 0.3539 | 0.0009534 | +++++ | 0 | 0.5697 | intron | NRXN3 |
| rs1385677 | 4 | 96508701 | 4 | 0 | 0 | A | G | 1.2452 | 0.3769 | 0.0009535 | +++?+ | 27.4 | 0.2473 | intergenic | |
| rs6940272 | 6 | 118471856 | 4 | 0 | 0 | T | C | -0.8364 | 0.2532 | 0.0009535 | ---?- | 0 | 0.8161 | intron | SLC35F1 |
| rs10051886 | 5 | 67620863 | 5 | 0 | 0 | A | G | -0.8668 | 0.2623 | 0.0009535 | ----+ | 0 | 0.4808 | intergenic |  |
| rs1862576 | 5 | 37754628 | 4 | 0 | 0 | A | T | -1.0012 | 0.303 | 0.0009537 | ---?- | 0 | 0.7814 | downstream-gene | |
| rs12429521 | 13 | 45522180 | 4 | 0 | 0 | T | C | 2.2063 | 0.6678 | 0.0009537 | +++?- | 22.8 | 0.2743 | intron | NUFIP1 |
| rs74643884 | 11 | 90024410 | 3 | 0 | 0 | T | C | -3.3215 | 1.0054 | 0.0009538 | ---?? | 0 | 0.5466 | intergenic | |
| rs1116224 | 4 | 123714231 | 4 | 0 | 0 | A | G | -2.1394 | 0.6476 | 0.000954 | ---?- | 53.2 | 0.09349 | intergenic | |
| rs6902528 | 6 | 130943338 | 4 | 0 | 0 | T | C | 1.3723 | 0.4154 | 0.000954 | +++?+ | 0 | 0.7397 | intergenic | |
| rs7255525 | 19 | 3388149 | 4 | 0 | 0 | A | G | -2.4905 | 0.7538 | 0.000954 | ---?- | 0 | 0.6981 | intron | NFIC |
| rs77879634 | 6 | 130948689 | 4 | 0 | 0 | T | C | 1.3797 | 0.4176 | 0.0009541 | +++?+ | 0 | 0.7352 | intergenic | |
| rs112085846 | 19 | 56828351 | 4 | 0 | 0 | A | G | 2.3227 | 0.7031 | 0.0009544 | +-+?+ | 41 | 0.1657 | upstream-gene | |
| rs17047218 | 3 | 68204352 | 4 | 0 | 0 | A | G | 1.9984 | 0.6049 | 0.0009546 | +++?+ | 14.4 | 0.32 | intron | FAM19A1 |
| rs145698372 | 7 | 44723531 | 4 | 0 | 0 | A | G | -2.9241 | 0.8851 | 0.0009546 | ---?- | 0 | 0.8084 | intron | OGDH |
| rs2482548 | 9 | 30812044 | 4 | 0 | 0 | A | G | -3.1603 | 0.9567 | 0.000955 | ---?- | 1.9 | 0.3827 | intergenic | |
| rs16885238 | 8 | 114875815 | 4 | 0 | 0 | T | G | 1.8054 | 0.5465 | 0.000955 | +++?- | 7 | 0.3581 | intergenic | |
| rs4995902 | 6 | 31230645 | 3 | 0 | 0 | T | C | 0.9394 | 0.2844 | 0.0009553 | +++?? | 23.1 | 0.2722 | intergenic | |
| 4:59737471 | 4 | 59737471 | 3 | 0 | 0 | G | GA | 2.6118 | 0.7906 | 0.0009553 | +++?? | 0 | 0.6334 | | |
| rs1882443 | 2 | 184046119 | 4 | 0 | 0 | T | C | -1.1245 | 0.3404 | 0.0009554 | ---?- | 0 | 0.7796 | intergenic | |
| rs10834248 | 11 | 23965566 | 4 | 0 | 0 | T | G | -1.1906 | 0.3604 | 0.0009555 | ---?- | 0 | 0.8863 | intergenic | |
| rs16978630 | 19 | 12813683 | 4 | 0 | 0 | T | C | -3.4518 | 1.045 | 0.0009556 | ----? | 50.7 | 0.1074 | synonymous | TNPO2 |
| 12:95821256 | 12 | 95821256 | 3 | 0 | 0 | A | C | 1.933 | 0.5852 | 0.0009559 | +++?? | 19.2 | 0.2901 | intergenic | |
| rs3863624 | 8 | 251173 | 4 | 0 | 0 | A | C | -0.9384 | 0.2841 | 0.0009562 | -+-?- | 61.2 | 0.05179 | intergenic | |
| rs79005415 | 10 | 21340772 | 3 | 0 | 0 | A | T | -3.2583 | 0.9864 | 0.0009562 | ---?? | 0 | 0.8329 | intron | NEBL |
| rs461512 | 6 | 33215280 | 5 | 0 | 0 | T | G | 1.0108 | 0.306 | 0.0009567 | +++++ | 25.4 | 0.2521 | upstream-gene | |
| rs1447981 | 3 | 60502641 | 4 | 0 | 0 | T | C | -0.9073 | 0.2747 | 0.0009567 | ---?- | 0 | 0.9737 | intron | FHIT |
| rs138697696 | 1 | 5862189 | 3 | 0 | 0 | A | G | 4.8704 | 1.4746 | 0.0009568 | +++?? | 0 | 0.7361 | intergenic | |
| rs6051275 | 20 | 26278200 | 4 | 0 | 0 | A | G | -1.3563 | 0.4106 | 0.0009569 | ---?- | 61.3 | 0.05127 | intergenic | |
| rs77789747 | 9 | 19010620 | 4 | 0 | 0 | T | C | 1.5476 | 0.4686 | 0.0009571 | +++?+ | 44.5 | 0.1445 | intron | FAM154A |
| 15:45961376 | 15 | 45961376 | 3 | 0 | 0 | A | AT | 1.0623 | 0.3216 | 0.0009571 | +++?? | 0 | 0.8621 | | |
| rs11792398 | 9 | 19010681 | 4 | 0 | 0 | A | T | 1.5467 | 0.4683 | 0.0009572 | +++?+ | 44.5 | 0.1446 | intron | FAM154A |
| rs62560429 | 9 | 19010686 | 4 | 0 | 0 | T | C | -1.5466 | 0.4683 | 0.0009575 | ---?- | 44.5 | 0.1444 | intron | FAM154A |
| rs9475523 | 6 | 55895569 | 4 | 0 | 0 | T | C | -2.2168 | 0.6712 | 0.0009576 | ---?- | 0 | 0.8008 | intergenic | |
| rs1355679 | 17 | 51154613 | 3 | 0 | 0 | A | G | -1.344 | 0.4069 | 0.0009578 | ---?? | 0 | 0.3927 | intergenic | |
| rs73834859 | 3 | 68204400 | 4 | 0 | 0 | A | G | -1.9972 | 0.6047 | 0.0009578 | ---?- | 14.3 | 0.3206 | intron | FAM19A1 |
| rs142772045 | 5 | 140986124 | 5 | 0 | 0 | T | G | -1.2456 | 0.3772 | 0.0009578 | +---- | 0 | 0.4253 | intron | DIAPH1 |
| rs11004054 | 10 | 55844165 | 4 | 0 | 0 | A | G | -2.1428 | 0.6488 | 0.0009579 | ---?+ | 0 | 0.4188 | intron | PCDH15 |
| rs17799440 | 2 | 199089918 | 4 | 0 | 0 | T | C | -1.3241 | 0.4009 | 0.0009579 | ---?- | 48.4 | 0.1209 | intergenic | |
| rs66507056 | 9 | 19010621 | 4 | 0 | 0 | A | G | -1.5474 | 0.4685 | 0.0009579 | ---?- | 44.5 | 0.1445 | intron | FAM154A |
| 12:95821259 | 12 | 95821259 | 3 | 0 | 0 | A | C | 1.9326 | 0.5852 | 0.000958 | +++?? | 19.2 | 0.29 | intergenic | |
| rs4379924 | 13 | 30314486 | 5 | 0 | 0 | A | G | 1.2115 | 0.3668 | 0.0009581 | +++++ | 0 | 0.5453 | intergenic | |
| rs72712353 | 1 | 87995645 | 3 | 0 | 0 | A | G | 2.474 | 0.7491 | 0.0009582 | -++?? | 0 | 0.4055 | intergenic | |
| rs75945975 | 21 | 24221837 | 4 | 0 | 0 | T | C | 3.796 | 1.1494 | 0.0009583 | +++?- | 0 | 0.4171 | intergenic | |
| rs12445385 | 16 | 6549160 | 4 | 0 | 0 | A | C | 0.883 | 0.2674 | 0.0009586 | +++?+ | 0 | 0.7112 | intron | RBFOX1 |
| rs149435450 | 3 | 29365954 | 2 | 0 | 0 | T | C | -6.1555 | 1.8639 | 0.0009586 | ??-?- | 61.9 | 0.1053 | intron | RBMS3 |
| rs6005789 | 22 | 28874313 | 4 | 0 | 0 | T | C | 2.2469 | 0.6804 | 0.0009587 | +++?+ | 0 | 0.5669 | intron | TTC28 |
| rs6496387 | 15 | 87562109 | 4 | 0 | 0 | T | C | 0.8809 | 0.2668 | 0.0009588 | +++?- | 8.5 | 0.3506 | intron | AGBL1 |
| rs11748851 | 5 | 174026875 | 4 | 0 | 0 | T | G | -1.2143 | 0.3677 | 0.0009588 | ---?- | 0 | 0.6869 | intergenic | |
| rs1329785 | 1 | 64770200 | 4 | 0 | 0 | A | T | 1.1486 | 0.3478 | 0.0009589 | +++?+ | 0 | 0.4515 | intergenic | |
| 2:86628604 | 2 | 86628604 | 3 | 0 | 0 | CTG | C | 2.9463 | 0.8922 | 0.0009591 | +++?? | 14.3 | 0.3115 | | |
| rs113506289 | 5 | 134875938 | 4 | 0 | 0 | A | C | 4.0448 | 1.2249 | 0.0009592 | +++?+ | 0 | 0.7649 | upstream-gene | |
| rs965619 | 4 | 183079082 | 4 | 0 | 0 | T | C | -0.8449 | 0.2559 | 0.0009594 | ---?- | 0 | 0.4066 | intron | TENM3 |
| rs3856919 | 3 | 188520252 | 4 | 0 | 0 | T | C | -0.8462 | 0.2563 | 0.0009595 | ---?- | 28.1 | 0.2436 | intron | LPP |
| rs34387057 | 9 | 19010406 | 4 | 0 | 0 | T | C | 1.5473 | 0.4686 | 0.0009595 | +++?+ | 44.5 | 0.1446 | intron | FAM154A |
| rs2570115 | 15 | 87555339 | 4 | 0 | 0 | A | G | -0.8742 | 0.2647 | 0.0009598 | ---?+ | 0 | 0.4012 | intron | AGBL1 |
| 6:159245489 | 6 | 159245489 | 3 | 0 | 0 | CT | C | -0.85 | 0.2574 | 0.0009599 | ---?? | 5.8 | 0.3459 | | |

| | | | | | | | | | | | | | | | |
|---|---|---|---|---|---|---|---|---|---|---|---|---|---|---|---|
| 2:47390577 | 2 | 47390577 | 3 | 0 | 0 | G | GT | 1.3267 | 0.4018 | 0.00096 | +++?? | 0 | 0.9401 | | |
| rs35151631 | 11 | 97688223 | 2 | 0 | 0 | T | C | -4.941 | 1.4964 | 0.0009601 | ??-?+ | 71.7 | 0.06013 | intergenic | |
| rs4126016 | 3 | 188520256 | 4 | 0 | 0 | A | G | 0.8466 | 0.2564 | 0.0009603 | +++?+ | 27.8 | 0.2454 | intron | LPP |
| rs76701220 | 2 | 117112240 | 4 | 0 | 0 | T | C | -1.6495 | 0.4996 | 0.0009603 | ---?- | 0 | 0.6797 | intergenic | |
| rs9999191 | 4 | 108275465 | 4 | 0 | 0 | T | C | 0.8287 | 0.251 | 0.0009604 | +++?+ | 0 | 0.403 | intergenic | |
| 2:170697881 | 2 | 170697881 | 3 | 0 | 0 | CA | C | -1.8417 | 0.5578 | 0.0009604 | ---?? | 0 | 0.9992 | | |
| rs2116671 | 7 | 32384945 | 4 | 0 | 0 | A | C | -1.4645 | 0.4435 | 0.0009606 | ---?- | 0 | 0.8324 | intergenic | |
| 1:48633111 | 1 | 48633111 | 2 | 0 | 0 | CATTATT | C | 1.4717 | 0.4457 | 0.0009606 | ?++?? | 48.5 | 0.1633 | | |
| 4:59658998 | 4 | 59658998 | 3 | 0 | 0 | T | TTTTTTG | 2.3548 | 0.7132 | 0.0009606 | +++?? | 0 | 0.7689 | | |
| rs7874465 | 9 | 19011246 | 4 | 0 | 0 | T | C | -1.5453 | 0.468 | 0.0009607 | ---?- | 44.5 | 0.1447 | intron | FAM154A |
| rs16937266 | 9 | 19010558 | 4 | 0 | 0 | C | G | -1.5467 | 0.4685 | 0.0009608 | ---?- | 44.5 | 0.1445 | intron | FAM154A |
| 2:170639545 | 2 | 170639545 | 3 | 0 | 0 | G | GGTT | 0.8355 | 0.2531 | 0.0009608 | +++?? | 0 | 0.5843 | | |
| rs17203578 | 2 | 208270827 | 4 | 0 | 0 | T | C | 1.4732 | 0.4462 | 0.0009609 | +++?+ | 0 | 0.4671 | intergenic | |
| rs9475524 | 6 | 55895876 | 4 | 0 | 0 | T | C | 2.2155 | 0.671 | 0.0009611 | +++?+ | 0 | 0.801 | intergenic | |
| rs78012716 | 2 | 204516802 | 4 | 0 | 0 | A | G | 1.4015 | 0.4245 | 0.0009611 | +++?+ | 59.4 | 0.06033 | intergenic | |
| rs9506223 | 13 | 30314766 | 5 | 0 | 0 | A | T | 1.2114 | 0.3669 | 0.0009612 | +++++ | 0 | 0.5485 | intergenic | |
| rs80299584 | 2 | 51417519 | 4 | 0 | 0 | T | C | 2.3136 | 0.7007 | 0.0009612 | +++?+ | 0 | 0.9284 | intergenic | |
| rs150366857 | 4 | 26171406 | 3 | 0 | 0 | A | G | 1.2796 | 0.3876 | 0.0009612 | +++?? | 0 | 0.4699 | intron | RBPJ |
| rs576879 | 1 | 90228377 | 5 | 0 | 0 | A | C | 0.7047 | 0.2135 | 0.0009615 | +++++ | 0 | 0.9969 | intergenic | |
| rs6051274 | 20 | 26278121 | 4 | 0 | 0 | A | G | -1.3544 | 0.4102 | 0.0009617 | ---?- | 61.4 | 0.05081 | intergenic | |
| rs73552608 | 8 | 21542205 | 3 | 0 | 0 | A | C | -3.9277 | 1.1897 | 0.0009618 | ?--?- | 0 | 0.9343 | intergenic | |
| rs6005842 | 22 | 29103606 | 4 | 0 | 0 | A | T | -2.1113 | 0.6395 | 0.0009619 | ---?- | 0 | 0.5979 | intron | CHEK2 |
| rs11764380 | 7 | 38586366 | 4 | 0 | 0 | T | C | 0.8645 | 0.2618 | 0.000962 | +++?+ | 0 | 0.6123 | intron | AMPH |
| rs73102123 | 12 | 48031236 | 4 | 0 | 0 | A | G | 1.7188 | 0.5206 | 0.0009622 | +++?- | 56.4 | 0.07595 | intergenic | |
| rs1976374 | 16 | 5182110 | 3 | 0 | 0 | T | G | -0.8497 | 0.2574 | 0.0009623 | ---?? | 0 | 0.9701 | intergenic | |
| rs62560428 | 9 | 19010282 | 4 | 0 | 0 | T | C | 1.5472 | 0.4687 | 0.0009623 | +++?+ | 44.5 | 0.1447 | intron | FAM154A |
| rs72620503 | 19 | 1638933 | 4 | 0 | 0 | C | G | 2.5808 | 0.7818 | 0.0009623 | +++?+ | 0 | 0.8059 | intron | TCF3 |
| rs72979307 | 4 | 162200698 | 2 | 0 | 0 | A | G | -6.0564 | 1.8346 | 0.0009624 | ?--?? | 0 | 0.4746 | intergenic | |
| rs59206581 | 1 | 235082292 | 4 | 0 | 0 | T | G | 0.808 | 0.2448 | 0.0009627 | +++?+ | 0 | 0.8993 | intergenic | |
| rs141945510 | 1 | 111842864 | 4 | 0 | 0 | A | C | 1.9312 | 0.585 | 0.0009629 | +++?+ | 0 | 0.5474 | intron | CHIA |
| rs1969207 | 12 | 127179444 | 4 | 0 | 0 | A | G | -1.1433 | 0.3463 | 0.000963 | ---?- | 29.4 | 0.2358 | intergenic | |
| rs4607231 | 4 | 123713381 | 4 | 0 | 0 | C | G | 2.13 | 0.6452 | 0.000963 | +++?+ | 52.8 | 0.09578 | intergenic | |
| rs2782378 | 6 | 63347987 | 4 | 0 | 0 | T | C | -2.2649 | 0.6861 | 0.0009632 | ---?- | 0 | 0.393 | intergenic | |
| rs201289222 | 7 | 3133070 | 4 | 0 | 0 | A | C | 0.8721 | 0.2642 | 0.0009633 | +++?+ | 0 | 0.9623 | intergenic | |
| rs4130775 | 6 | 55796190 | 4 | 0 | 0 | T | C | 0.8618 | 0.2611 | 0.0009634 | +++?+ | 0 | 0.4944 | intergenic | |
| rs807134 | 17 | 51228624 | 5 | 0 | 0 | A | T | -1.0168 | 0.308 | 0.0009635 | ----+ | 7.3 | 0.3648 | intergenic |  |
| rs2138861 | 12 | 43166458 | 5 | 0 | 0 | A | T | -0.7345 | 0.2225 | 0.0009635 | ----- | 0 | 0.6484 | intergenic |  |
| rs807149 | 17 | 51211383 | 5 | 0 | 0 | A | G | -1.0238 | 0.3102 | 0.0009637 | ----+ | 0 | 0.4494 | intergenic |  |
| rs56013076 | 6 | 72873985 | 4 | 0 | 0 | T | C | 1.5358 | 0.4653 | 0.0009639 | +++?- | 0 | 0.5911 | intron | RIMS1 |
| rs143710780 | 1 | 111839962 | 4 | 0 | 0 | T | C | -1.9265 | 0.5836 | 0.000964 | ---?- | 0 | 0.5661 | intron | CHIA |
| 1:194308648 | 1 | 194308648 | 3 | 0 | 0 | A | AT | 2.5079 | 0.7598 | 0.0009645 | +++?? | 0 | 0.9439 | | |
| rs3753797 | 1 | 180145978 | 4 | 0 | 0 | A | G | -2.011 | 0.6093 | 0.0009647 | ---?+ | 32.3 | 0.2183 | intron | QSOX1 |
| rs78360656 | 1 | 42689161 | 4 | 0 | 0 | T | C | 1.7453 | 0.5288 | 0.0009647 | +++?+ | 0 | 0.5744 | intron | FOXJ3 |
| rs141688006 | 4 | 180006008 | 2 | 0 | 0 | A | G | -6.7461 | 2.0439 | 0.0009647 | ?--?? | 0 | 0.5006 | intergenic | |
| rs6433151 | 2 | 170700579 | 5 | 0 | 1 | T | C | -0.8306 | 0.2516 | 0.0009649 | ----- | 48.7 | 0.09923 | intron | UBR3 |
| rs62560426 | 9 | 19009920 | 4 | 0 | 0 | A | T | 1.5477 | 0.4689 | 0.0009649 | +++?+ | 44.4 | 0.1449 | intron | FAM154A |
| 7:23118371 | 7 | 23118371 | 3 | 0 | 0 | CT | C | -0.8916 | 0.2701 | 0.0009649 | ---?? | 33.7 | 0.2213 | | |
| rs6070196 | 20 | 56212531 | 4 | 0 | 0 | A | G | 1.0794 | 0.327 | 0.0009651 | +++?+ | 0 | 0.9094 | intergenic | |
| rs412609 | 16 | 84493515 | 2 | 0 | 0 | T | C | -6.0301 | 1.8271 | 0.0009655 | --??? | 12.6 | 0.2848 | intron | ATP2C2 |
| rs17194188 | 6 | 30064702 | 5 | 0 | 0 | T | C | 1.4625 | 0.4431 | 0.0009655 | +++++ | 0 | 0.717 | intergenic | |
| rs607730 | 3 | 59851920 | 4 | 0 | 0 | T | G | 0.8011 | 0.2427 | 0.0009656 | +++?+ | 0 | 0.9475 | intron | FHIT |
| rs9264429 | 6 | 31230498 | 3 | 0 | 0 | A | G | 0.9387 | 0.2844 | 0.0009658 | +++?? | 23 | 0.2728 | intergenic | |
| rs6496386 | 15 | 87562020 | 4 | 0 | 0 | A | T | -0.8806 | 0.2668 | 0.0009659 | ---?+ | 9 | 0.3484 | intron | AGBL1 |
| rs31211 | 5 | 134363145 | 5 | 0 | 0 | A | G | -0.782 | 0.2369 | 0.000966 | ----- | 15.5 | 0.3158 | downstream-gene |  |
| rs35030005 | 4 | 190006378 | 4 | 0 | 0 | T | G | 0.8289 | 0.2512 | 0.0009661 | +++?+ | 31.9 | 0.2208 | intergenic | |
| rs55920634 | 10 | 127952177 | 4 | 0 | 0 | T | C | 1.4707 | 0.4456 | 0.0009661 | +++?+ | 0 | 0.9784 | intron | ADAM12 |
| rs4908019 | 1 | 100239486 | 4 | 0 | 0 | A | C | 1.6483 | 0.4994 | 0.0009662 | +++?+ | 0 | 0.8925 | intergenic | |
| rs10879708 | 12 | 74321820 | 4 | 0 | 0 | T | C | -0.8141 | 0.2467 | 0.0009665 | ---?- | 0 | 0.5215 | intergenic | |
| 1:80349541 | 1 | 80349541 | 2 | 0 | 0 | G | GA | 4.4715 | 1.355 | 0.0009668 | +?+?? | 0 | 0.9205 | | |

| | | | | | | | | | | | | | | | |
|---|---|---|---|---|---|---|---|---|---|---|---|---|---|---|---|
| rs1297621 | 7 | 101890901 | 4 | 0 | 0 | T | C | -0.8369 | 0.2536 | 0.0009669 | ---?- | 0 | 0.7773 | intron | CUX1 |
| rs61820441 | 1 | 187866251 | 4 | 0 | 0 | A | G | 1 | 0.303 | 0.0009669 | +++?+ | 0 | 0.6325 | intergenic | |
| rs60750232 | 3 | 68202124 | 4 | 0 | 0 | T | C | -2.0081 | 0.6085 | 0.000967 | ---?- | 13.2 | 0.3263 | intron | FAM19A1 |
| rs140123563 | 22 | 49386930 | 3 | 0 | 0 | T | G | 2.7426 | 0.8311 | 0.000967 | +++?? | 0 | 0.9854 | intergenic | |
| rs57856931 | 1 | 182014924 | 4 | 0 | 0 | A | C | 2.7608 | 0.8366 | 0.0009671 | +++?+ | 0 | 0.7242 | intergenic | |
| rs55808005 | 15 | 87578513 | 3 | 0 | 0 | A | C | 0.9008 | 0.273 | 0.0009673 | +++?? | 18 | 0.2952 | intergenic | |
| rs16986548 | 22 | 29001380 | 4 | 0 | 0 | T | C | -2.2965 | 0.696 | 0.0009674 | ---?- | 0 | 0.4867 | intron | TTC28 |
| rs73933998 | 2 | 68074934 | 3 | 0 | 0 | T | C | 2.8165 | 0.8535 | 0.0009675 | +++?? | 0 | 0.8473 | intergenic | |
| rs2065156 | 1 | 90231738 | 5 | 0 | 0 | A | G | 0.7035 | 0.2132 | 0.0009677 | +++++ | 0 | 0.9947 | intergenic | |
| rs4413424 | 4 | 123713190 | 4 | 0 | 0 | T | C | -2.128 | 0.6449 | 0.0009679 | ---?- | 52.7 | 0.0959 | intergenic | |
| rs779177 | 3 | 22148612 | 4 | 0 | 0 | A | G | 0.891 | 0.27 | 0.0009683 | +++?+ | 0 | 0.8208 | intron | ZNF385D |
| rs17781768 | 6 | 72875138 | 4 | 0 | 0 | A | G | -1.5356 | 0.4654 | 0.0009684 | ---?+ | 0 | 0.5906 | intron | RIMS1 |
| rs10805219 | 4 | 20754630 | 4 | 0 | 0 | T | C | -0.8705 | 0.2638 | 0.0009686 | ---?- | 0 | 0.7408 | intron | KCNIP4 |
| rs67844957 | 9 | 19009308 | 4 | 0 | 0 | A | C | -1.5486 | 0.4694 | 0.0009686 | ---?- | 44.4 | 0.1448 | intron | FAM154A |
| rs76920721 | 2 | 117111901 | 4 | 0 | 0 | T | C | 1.6477 | 0.4994 | 0.0009687 | +++?+ | 0 | 0.6796 | intergenic | |
| 1:100298988 | 1 | 100298988 | 3 | 0 | 0 | A | G | 1.9636 | 0.5951 | 0.0009688 | +++?? | 0 | 0.9639 | intergenic | |
| rs6001840 | 22 | 40604131 | 3 | 0 | 0 | A | C | 0.9136 | 0.2769 | 0.0009688 | +++?? | 40.4 | 0.1867 | intron | TNRC6B |
| rs10939672 | 4 | 10052136 | 5 | 0 | 0 | A | G | -0.7024 | 0.2129 | 0.0009689 | ----+ | 27.8 | 0.2359 | intergenic |  |
| rs72826507 | 6 | 18306822 | 4 | 0 | 0 | T | C | -1.9992 | 0.6059 | 0.0009689 | ---?- | 24.4 | 0.2646 | intron | RNF144B |
| rs376880074 | 2 | 223012653 | 3 | 0 | 0 | A | C | 1.2046 | 0.3651 | 0.000969 | -++?? | 63.2 | 0.06585 | intergenic | |
| rs807128 | 17 | 51231444 | 5 | 0 | 0 | A | T | 1.0206 | 0.3093 | 0.0009691 | +-++- | 5.3 | 0.3764 | intergenic | |
| rs10492134 | 12 | 13858882 | 4 | 0 | 0 | C | G | -0.8543 | 0.2589 | 0.0009692 | ---?- | 15.5 | 0.3144 | intron | GRIN2B |
| rs150172415 | 18 | 62125113 | 4 | 0 | 0 | T | C | -2.016 | 0.611 | 0.0009694 | ---?+ | 51.6 | 0.1025 | intergenic | |
| rs73799758 | 5 | 123096916 | 4 | 0 | 0 | T | C | 0.937 | 0.284 | 0.0009695 | +++?+ | 7.2 | 0.3574 | intergenic | |
| rs66474121 | 21 | 19080953 | 4 | 0 | 0 | T | C | 1.3773 | 0.4175 | 0.0009698 | +++?+ | 0 | 0.744 | intergenic | |
| rs9475525 | 6 | 55896060 | 4 | 0 | 0 | A | G | -2.2137 | 0.671 | 0.0009699 | ---?- | 0 | 0.8017 | intergenic | |
| 8:113219572 | 8 | 113219572 | 3 | 0 | 0 | T | TAA | -0.9752 | 0.2956 | 0.0009702 | ---?? | 0 | 0.8679 | | |
| rs34365814 | 7 | 159075276 | 4 | 0 | 0 | A | C | -1.7011 | 0.5157 | 0.0009703 | ---?- | 21.8 | 0.2798 | intergenic | |
| rs2894198 | 6 | 31230990 | 5 | 0 | 0 | A | G | 0.7875 | 0.2387 | 0.0009704 | +++++ | 0 | 0.457 | intergenic | |
| rs10190260 | 2 | 170713214 | 3 | 0 | 1 | T | C | -0.9948 | 0.3015 | 0.0009705 | ---?? | 65.8 | 0.05357 | intron | UBR3 |
| rs144435319 | 3 | 45669479 | 3 | 0 | 0 | T | G | -2.2797 | 0.691 | 0.0009705 | ---?? | 14.5 | 0.3107 | intron | LIMD1 |
| rs4637645 | 6 | 55811672 | 4 | 0 | 0 | A | T | -0.8605 | 0.2608 | 0.0009706 | ---?- | 0 | 0.4611 | intergenic | |
| rs78608089 | 8 | 86540417 | 4 | 0 | 0 | T | C | 1.5804 | 0.4791 | 0.0009706 | +++?+ | 0.9 | 0.3874 | intergenic | |
| rs9264463 | 6 | 31231591 | 3 | 0 | 0 | T | C | 0.9384 | 0.2845 | 0.0009707 | +++?? | 21 | 0.2821 | downstream-gene | |
| rs13417081 | 2 | 9770050 | 3 | 0 | 0 | A | G | 1.1015 | 0.3339 | 0.0009707 | +++?? | 0 | 0.6627 | intron | YWHAQ |
| 12:15588492 | 12 | 15588492 | 3 | 0 | 0 | CT | C | -1.1235 | 0.3406 | 0.0009709 | ---?? | 0 | 0.952 | | |
| rs60671196 | 4 | 162200614 | 2 | 0 | 0 | T | C | -6.0523 | 1.8347 | 0.000971 | ?--?? | 0 | 0.4724 | intergenic | |
| rs62162778 | 2 | 85111444 | 3 | 0 | 0 | T | C | 3.1121 | 0.9434 | 0.000971 | +++?? | 0 | 0.5694 | upstream-gene | |
| rs12828349 | 12 | 51459675 | 5 | 0 | 0 | A | G | 0.9208 | 0.2791 | 0.0009711 | +++++ | 0 | 0.4302 | intron | CSRNP2 |
| rs73184562 | 12 | 86245024 | 4 | 0 | 0 | A | T | -0.9201 | 0.2789 | 0.0009711 | -+-?- | 17 | 0.306 | intergenic | |
| rs12751730 | 1 | 235077568 | 4 | 0 | 0 | A | G | 0.7857 | 0.2382 | 0.0009712 | +++?+ | 0 | 0.9455 | intergenic | |
| rs7571767 | 2 | 184039594 | 4 | 0 | 0 | A | C | -1.127 | 0.3416 | 0.0009713 | ---?- | 0 | 0.7809 | intergenic | |
| rs72889498 | 2 | 171168178 | 4 | 0 | 0 | A | G | -4.1047 | 1.2443 | 0.0009714 | ---?+ | 44.8 | 0.1428 | intron | MYO3B |
| rs373984563 | 2 | 223012651 | 3 | 0 | 0 | A | T | -1.2044 | 0.3651 | 0.0009714 | +--?? | 63.2 | 0.06584 | intergenic | |
| rs58953222 | 2 | 184065035 | 3 | 0 | 0 | A | G | 1.1846 | 0.3591 | 0.0009715 | +++?? | 0 | 0.5761 | intergenic | |
| rs144584607 | 6 | 31223823 | 3 | 0 | 0 | A | G | 0.9257 | 0.2806 | 0.0009715 | +++?? | 33 | 0.2247 | intergenic | |
| rs9264426 | 6 | 31230319 | 3 | 0 | 0 | A | C | 0.941 | 0.2853 | 0.0009716 | +++?? | 19.7 | 0.2876 | intergenic | |
| rs7597111 | 2 | 121938362 | 4 | 0 | 0 | A | G | -1.0653 | 0.3229 | 0.0009717 | ---?- | 0 | 0.7958 | intergenic | |
| rs62560424 | 9 | 19008466 | 4 | 0 | 0 | A | G | 1.5497 | 0.4698 | 0.0009718 | +++?+ | 44.4 | 0.1452 | intron | FAM154A |
| rs114097391 | 4 | 157731617 | 4 | 0 | 0 | T | C | 2.5873 | 0.7844 | 0.0009718 | +++?+ | 0 | 0.5105 | intron | PDGFC |
| rs4521002 | 2 | 184039724 | 4 | 0 | 0 | T | C | -1.1269 | 0.3416 | 0.0009719 | ---?- | 0 | 0.781 | intergenic | |
| rs9464329 | 6 | 55896218 | 4 | 0 | 0 | A | T | 2.2131 | 0.6709 | 0.0009719 | +++?+ | 0 | 0.8019 | intergenic | |
| rs75097147 | 2 | 15608123 | 4 | 0 | 0 | T | C | -1.5766 | 0.478 | 0.0009719 | ---?- | 0 | 0.8621 | intron | NBAS |
| rs7823277 | 8 | 113199716 | 3 | 0 | 0 | A | C | 0.8782 | 0.2662 | 0.0009722 | +++?? | 0 | 0.8516 | intergenic | |
| rs9264425 | 6 | 31230318 | 3 | 0 | 0 | A | G | -0.941 | 0.2853 | 0.0009722 | ---?? | 19.7 | 0.2877 | intergenic | |
| rs62560425 | 9 | 19008595 | 4 | 0 | 0 | C | G | -1.5492 | 0.4697 | 0.0009722 | ---?- | 44.4 | 0.145 | intron | FAM154A |
| rs4666895 | 2 | 184035368 | 4 | 0 | 0 | T | G | -1.1304 | 0.3427 | 0.0009724 | ---?- | 0 | 0.7823 | intergenic | |
| rs7540079 | 1 | 90231341 | 5 | 0 | 0 | C | G | 0.7032 | 0.2132 | 0.0009724 | +++++ | 0 | 0.9947 | intergenic | |
| rs2323248 | 3 | 5389519 | 4 | 0 | 0 | A | T | 1.3206 | 0.4004 | 0.0009729 | +++?+ | 0 | 0.4573 | intergenic | |

| | | | | | | | | | | | | | | | |
|---|---|---|---|---|---|---|---|---|---|---|---|---|---|---|---|
| rs13140892 | 4 | 46386896 | 4 | 0 | 0 | T | C | 0.8541 | 0.259 | 0.000973 | +-+?+ | 59 | 0.06226 | intron | GABRA2 |
| rs139420342 | 7 | 44759169 | 4 | 0 | 0 | T | C | 2.7892 | 0.8457 | 0.0009732 | +++?+ | 0 | 0.4948 | intergenic | |
| rs147523712 | 2 | 109976671 | 2 | 0 | 0 | A | T | -5.5413 | 1.6801 | 0.0009732 | --??? | 17.2 | 0.2719 | intron | SH3RF3 |
| rs76701585 | 2 | 138931645 | 4 | 0 | 0 | T | C | 1.5841 | 0.4803 | 0.0009733 | +++?+ | 0 | 0.7816 | intergenic | |
| rs28890275 | 4 | 123707945 | 4 | 0 | 0 | A | G | 2.1024 | 0.6375 | 0.0009734 | +++?+ | 51.9 | 0.1007 | intergenic | |
| rs36084205 | 4 | 10028678 | 5 | 0 | 0 | A | C | -0.6933 | 0.2102 | 0.0009734 | ----+ | 13 | 0.3311 | intron | SLC2A9 |
| rs7231012 | 18 | 413686 | 4 | 0 | 0 | A | G | 0.8444 | 0.256 | 0.0009735 | +++?+ | 0 | 0.8165 | intron | COLEC12 |
| rs10415831 | 19 | 12809726 | 4 | 0 | 0 | T | C | -3.4428 | 1.0439 | 0.0009735 | ----? | 50.6 | 0.1079 | upstream-gene |  |
| rs55792405 | 8 | 103721202 | 4 | 0 | 0 | A | G | -1.0995 | 0.3334 | 0.0009735 | ---?+ | 5.4 | 0.366 | intergenic | |
| rs9961290 | 18 | 63857388 | 4 | 0 | 0 | C | G | 0.8097 | 0.2455 | 0.0009737 | +++?+ | 0 | 0.4377 | intergenic | |
| rs12212039 | 6 | 30068195 | 5 | 0 | 0 | T | C | 1.4695 | 0.4456 | 0.0009738 | +++++ | 0 | 0.733 | downstream-gene | |
| rs17480154 | 5 | 123122388 | 4 | 0 | 0 | T | C | -1.2065 | 0.3658 | 0.0009739 | ---?- | 39.5 | 0.1749 | intergenic | |
| 6:12701451 | 6 | 12701451 | 3 | 0 | 0 | A | ACC | -1.8473 | 0.5601 | 0.0009741 | ---?? | 0 | 0.51 | | |
| rs4331705 | 3 | 129911911 | 3 | 0 | 0 | T | C | 0.9522 | 0.2887 | 0.0009744 | +++?? | 38.1 | 0.1989 | intergenic | |
| rs9940817 | 16 | 6535942 | 4 | 0 | 0 | T | C | -0.8991 | 0.2726 | 0.0009745 | ---?- | 0 | 0.7019 | intron | RBFOX1 |
| rs112679444 | 2 | 170686323 | 3 | 0 | 0 | A | G | -1.8473 | 0.5602 | 0.0009749 | ---?? | 0 | 0.9985 | intron | UBR3 |
| 12:68674717 | 12 | 68674717 | 3 | 0 | 0 | G | GTTGT | -3.2287 | 0.9791 | 0.0009749 | ---?? | 0 | 0.6127 | | |
| rs9475526 | 6 | 55896317 | 4 | 0 | 0 | A | G | -2.2124 | 0.6709 | 0.0009753 | ---?- | 0 | 0.8017 | intergenic | |
| rs190265382 | 1 | 80334756 | 2 | 0 | 0 | A | G | -4.4519 | 1.3501 | 0.0009754 | -?-?? | 0 | 0.9143 | intergenic | |
| rs270776 | 19 | 55222807 | 4 | 0 | 0 | T | G | -0.8408 | 0.255 | 0.0009755 | ---?- | 0 | 0.7926 | intron | LILRP2 |
| rs483452 | 1 | 208191383 | 3 | 0 | 0 | T | C | 1.9575 | 0.5936 | 0.0009755 | +++?? | 0 | 0.7134 | downstream-gene | |
| rs1069699 | 1 | 64764403 | 4 | 0 | 0 | T | C | -0.9127 | 0.2768 | 0.0009755 | ---?- | 0 | 0.5128 | intergenic | |
| rs57235722 | 19 | 12812748 | 4 | 0 | 0 | T | C | -3.4868 | 1.0574 | 0.0009756 | ----? | 55.1 | 0.08273 | intron | TNPO2 |
| rs71605582 | 4 | 59656971 | 3 | 0 | 0 | A | T | -2.353 | 0.7136 | 0.000976 | ---?? | 0 | 0.7681 | intergenic | |
| rs2570137 | 15 | 87596850 | 4 | 0 | 0 | A | C | -0.8541 | 0.259 | 0.0009761 | ---?- | 0 | 0.4426 | intergenic | |
| rs2691046 | 12 | 103254342 | 3 | 0 | 0 | A | C | 0.8538 | 0.2589 | 0.0009761 | +++?? | 0 | 0.4449 | intron | PAH |
| rs7009994 | 8 | 38606564 | 4 | 0 | 0 | A | C | 1.0128 | 0.3072 | 0.0009761 | +++?+ | 0 | 0.5653 | intron | TACC1 |
| rs13120348 | 4 | 10053155 | 5 | 0 | 0 | C | G | -0.7018 | 0.2128 | 0.0009761 | ----+ | 28.2 | 0.2337 | intergenic |  |
| 6:63241017 | 6 | 63241017 | 3 | 0 | 0 | A | G | 3.3749 | 1.0235 | 0.0009763 | +++?? | 0 | 0.603 | intergenic | |
| 4:59656961 | 4 | 59656961 | 3 | 0 | 0 | GT | GTTT | -2.353 | 0.7136 | 0.0009766 | ---?? | 0 | 0.7681 | | |
| rs7069362 | 10 | 86628822 | 4 | 0 | 0 | A | G | -1.51 | 0.458 | 0.0009768 | ---?- | 4.4 | 0.3709 | intergenic | |
| rs6751275 | 2 | 184027625 | 4 | 0 | 0 | A | T | -1.162 | 0.3524 | 0.0009769 | ---?- | 0 | 0.8352 | downstream-gene | |
| rs116257403 | 2 | 51422654 | 4 | 0 | 0 | C | G | -2.3125 | 0.7014 | 0.0009769 | ---?- | 0 | 0.9201 | intergenic | |
| rs7244572 | 18 | 458383 | 4 | 0 | 0 | A | G | -0.8435 | 0.2558 | 0.000977 | ---?- | 0 | 0.8321 | intron | COLEC12 |
| rs9464330 | 6 | 55896887 | 4 | 0 | 0 | T | G | 2.2118 | 0.6708 | 0.0009771 | +++?+ | 0 | 0.8022 | intergenic | |
| 13:45604140 | 13 | 45604140 | 3 | 0 | 0 | A | AACTTT | 2.1585 | 0.6547 | 0.0009773 | +++?? | 0 | 0.483 | | |
| rs2626386 | 2 | 52944632 | 4 | 0 | 0 | T | C | 1.2943 | 0.3926 | 0.0009774 | +++?+ | 0 | 0.7064 | intergenic | |
| rs2594338 | 15 | 87561537 | 4 | 0 | 0 | T | C | 0.879 | 0.2666 | 0.0009778 | +++?- | 8.3 | 0.3516 | intron | AGBL1 |
| rs2902997 | 20 | 56213126 | 4 | 0 | 0 | A | G | -1.0894 | 0.3304 | 0.0009778 | ---?- | 0 | 0.9105 | intergenic | |
| rs115382951 | 4 | 188131741 | 2 | 0 | 0 | T | C | -6.4648 | 1.9609 | 0.0009778 | -???- | 52.9 | 0.1451 | intergenic | |
| rs16860658 | 4 | 47821397 | 5 | 0 | 0 | T | G | -1.3926 | 0.4224 | 0.0009779 | -+--- | 15.7 | 0.3145 | intron | CORIN |
| rs12159502 | 22 | 28923924 | 4 | 0 | 0 | A | G | 2.3918 | 0.7255 | 0.000978 | +++?+ | 0 | 0.679 | intron | TTC28 |
| 2:114362665 | 2 | 114362665 | 3 | 0 | 0 | CT | C | 1.0958 | 0.3324 | 0.000978 | +++?? | 50.2 | 0.1341 | | |
| rs4812992 | 20 | 44721594 | 4 | 0 | 0 | A | C | -0.8604 | 0.261 | 0.0009781 | ---?- | 31.5 | 0.2231 | upstream-gene | |
| rs9997378 | 4 | 123704311 | 4 | 0 | 0 | A | G | -2.0844 | 0.6323 | 0.0009781 | ---?- | 48.2 | 0.1221 | intergenic | |
| rs77636116 | 5 | 173113535 | 3 | 0 | 0 | A | G | -4.5051 | 1.3665 | 0.0009781 | --??- | 60 | 0.08215 | intergenic | |
| rs7249929 | 19 | 12825211 | 4 | 0 | 0 | C | G | -3.4289 | 1.0401 | 0.0009783 | ----? | 53.7 | 0.09037 | intron | TNPO2 |
| rs11073692 | 15 | 87603927 | 4 | 0 | 0 | A | G | 0.848 | 0.2572 | 0.0009783 | +++?+ | 0 | 0.4644 | intergenic | |
| rs71440087 | 13 | 111550069 | 5 | 0 | 0 | A | G | 2.1253 | 0.6447 | 0.0009783 | +-+++ | 28.2 | 0.2337 | intron | ANKRD10 |
| rs200464597 | 9 | 115891675 | 3 | 0 | 0 | T | C | -3.7054 | 1.124 | 0.0009785 | ---?? | 0 | 0.4769 | intergenic | |
| rs34709913 | 4 | 10028955 | 5 | 0 | 0 | T | C | -0.6929 | 0.2102 | 0.0009787 | ----+ | 12.6 | 0.3333 | intron | SLC2A9 |
| rs2782380 | 6 | 63349180 | 4 | 0 | 0 | A | T | 2.2619 | 0.6861 | 0.0009788 | +++?+ | 0 | 0.3928 | intergenic | |
| rs73102114 | 12 | 48029185 | 4 | 0 | 0 | T | G | 1.7192 | 0.5215 | 0.0009789 | +++?- | 56.5 | 0.07551 | intergenic | |
| rs2122439 | 4 | 183079434 | 4 | 0 | 0 | T | G | -0.8422 | 0.2555 | 0.000979 | ---?- | 1.3 | 0.3854 | intron | TENM3 |
| rs138797400 | 13 | 70866762 | 4 | 0 | 0 | C | G | 3.4574 | 1.0488 | 0.000979 | +++?- | 53.2 | 0.09325 | intergenic | |
| 3:183151223 | 3 | 183151223 | 3 | 0 | 0 | G | GA | 1.5447 | 0.4686 | 0.000979 | +++?? | 0.7 | 0.3652 | | |
| rs11694381 | 2 | 59981123 | 4 | 0 | 0 | A | G | -1.0566 | 0.3205 | 0.0009791 | ---?- | 0 | 0.7042 | intergenic | |
| rs117145251 | 8 | 94167381 | 4 | 0 | 0 | T | C | 2.8554 | 0.8662 | 0.0009791 | +++?+ | 3 | 0.3774 | intron | C8orf87 |
| rs182869640 | 13 | 82503957 | 3 | 0 | 0 | T | C | -4.4404 | 1.347 | 0.0009791 | ?--?- | 0 | 0.3774 | intergenic | |

| | | | | | | | | | | | | | | | |
|---|---|---|---|---|---|---|---|---|---|---|---|---|---|---|---|
| rs35424673 | 3 | 155123709 | 4 | 0 | 0 | T | C | 0.8181 | 0.2482 | 0.0009794 | +++?+ | 0 | 0.4611 | intergenic | |
| rs4513028 | 14 | 95896632 | 4 | 0 | 0 | C | G | 1.224 | 0.3714 | 0.0009804 | +++?+ | 0 | 0.4937 | intron | SYNE3 |
| 15:87571267 | 15 | 87571267 | 3 | 0 | 0 | T | TTAAAA | 0.9054 | 0.2747 | 0.0009805 | +++?? | 19.1 | 0.2904 | | |
| rs2977379 | 8 | 59066811 | 4 | 0 | 0 | T | C | 1.0802 | 0.3277 | 0.0009808 | +++?+ | 20.7 | 0.2861 | downstream-gene | |
| 1:203667577 | 1 | 203667577 | 3 | 0 | 0 | T | TA | -1.157 | 0.351 | 0.0009808 | ---?? | 0 | 0.8017 | | |
| rs7666624 | 4 | 123706512 | 4 | 0 | 0 | A | T | 2.0947 | 0.6355 | 0.000981 | +++?+ | 51.7 | 0.1018 | intergenic | |
| rs4315093 | 11 | 134475682 | 4 | 0 | 0 | T | C | 0.9754 | 0.296 | 0.0009811 | +++?+ | 0 | 0.9851 | intergenic | |
| rs2967634 | 19 | 8849265 | 4 | 0 | 0 | C | G | 0.8667 | 0.263 | 0.0009812 | +++?+ | 6.1 | 0.3623 | intergenic | |
| rs4995903 | 6 | 31230605 | 5 | 0 | 0 | T | C | 0.7868 | 0.2387 | 0.0009812 | +++++ | 0 | 0.4576 | intergenic | |
| rs13141233 | 4 | 10028729 | 5 | 0 | 0 | T | C | -0.6927 | 0.2102 | 0.0009813 | ----+ | 12.7 | 0.3331 | intron | SLC2A9 |
| rs6662466 | 1 | 111837922 | 4 | 0 | 0 | C | G | 1.9189 | 0.5822 | 0.0009814 | +++?+ | 0 | 0.5819 | intron | CHIA |
| rs142442619 | 13 | 36091229 | 2 | 0 | 0 | A | G | 5.3564 | 1.6252 | 0.0009814 | ?++?? | 0 | 0.5626 | intron | NBEA |
| rs935114 | 9 | 32918314 | 4 | 0 | 0 | T | C | 2.2661 | 0.6876 | 0.0009815 | +++?+ | 52.8 | 0.0955 | intergenic | |
| rs11240275 | 1 | 203690904 | 4 | 0 | 0 | A | C | -1.0795 | 0.3275 | 0.0009817 | ---?- | 0 | 0.7717 | intron | ATP2B4 |
| rs9475529 | 6 | 55896944 | 4 | 0 | 0 | A | G | 2.2108 | 0.6708 | 0.0009818 | +++?+ | 0 | 0.8019 | intergenic | |
| rs12536117 | 7 | 32345872 | 4 | 0 | 0 | T | C | -1.3339 | 0.4047 | 0.0009818 | ---?- | 0 | 0.8979 | intergenic | |
| 5:127620898 | 5 | 127620898 | 3 | 0 | 0 | A | AGTTT | -0.9703 | 0.2944 | 0.0009818 | +--?? | 57.7 | 0.09399 | | |
| 22:29113143 | 22 | 29113143 | 3 | 0 | 0 | G | GATC | 2.1908 | 0.6648 | 0.0009819 | +++?? | 0 | 0.4451 | | |
| 7:64927848 | 7 | 64927848 | 3 | 0 | 0 | A | ATG | -1.0623 | 0.3223 | 0.0009819 | ---?? | 5.3 | 0.3478 | | |
| rs7182264 | 15 | 87588003 | 4 | 0 | 0 | A | G | 0.8631 | 0.2619 | 0.0009821 | +++?+ | 0 | 0.4186 | intergenic | |
| rs4559278 | 8 | 65017530 | 3 | 0 | 0 | A | G | -0.8431 | 0.2558 | 0.0009823 | ---?? | 0 | 0.7218 | intergenic | |
| rs4343208 | 14 | 95896536 | 4 | 0 | 0 | A | C | 1.2244 | 0.3715 | 0.0009825 | +++?+ | 0 | 0.4857 | intron | SYNE3 |
| rs67854134 | 6 | 36900103 | 4 | 0 | 0 | A | G | 1.2043 | 0.3654 | 0.0009825 | +++?+ | 44.6 | 0.144 | downstream-gene | |
| rs143793884 | 15 | 85890746 | 2 | 0 | 0 | A | T | -6.7284 | 2.0417 | 0.0009826 | ?-??- | 0 | 0.6976 | intergenic | |
| rs32527 | 5 | 14479991 | 2 | 0 | 0 | T | C | -4.9039 | 1.4881 | 0.0009827 | -?-?? | 0 | 0.7254 | intron | TRIO |
| rs7780974 | 7 | 21798233 | 5 | 0 | 0 | A | G | 0.7278 | 0.2209 | 0.0009828 | ++++- | 27.3 | 0.2397 | intron | DNAH11 |
| rs72814167 | 5 | 172478593 | 5 | 0 | 0 | T | C | -1.0406 | 0.3158 | 0.0009828 | ----- | 0 | 0.9403 | upstream-gene |  |
| rs8026531 | 15 | 65160874 | 5 | 0 | 0 | T | G | -0.8919 | 0.2706 | 0.0009829 | ----+ | 0 | 0.8209 | downstream-gene |  |
| rs6005793 | 22 | 28914573 | 4 | 0 | 0 | A | G | 2.257 | 0.6849 | 0.000983 | +++?+ | 0 | 0.5358 | intron | TTC28 |
| rs13397323 | 2 | 42711038 | 4 | 0 | 0 | T | C | -1.6689 | 0.5064 | 0.0009831 | ---?- | 47.1 | 0.1285 | intron | KCNG3 |
| rs7659887 | 4 | 119285459 | 4 | 0 | 0 | T | G | 1.6095 | 0.4884 | 0.0009832 | +++?+ | 0 | 0.6995 | intergenic | |
| rs144767684 | 18 | 40022677 | 4 | 0 | 0 | A | T | 2.5432 | 0.7718 | 0.0009834 | +++?+ | 0 | 0.6401 | intron | LINC00907 |
| rs7644142 | 3 | 45362447 | 4 | 0 | 0 | T | C | -1.2895 | 0.3913 | 0.0009836 | ---?- | 27.1 | 0.2491 | intergenic | |
| rs77297528 | 4 | 5770125 | 4 | 0 | 0 | T | G | -1.1261 | 0.3418 | 0.0009838 | ---?- | 0 | 0.729 | intron | EVC |
| rs2782377 | 6 | 63347849 | 4 | 0 | 0 | A | C | -2.2592 | 0.6856 | 0.000984 | ---?- | 0 | 0.3949 | intergenic | |
| rs6827414 | 4 | 38650464 | 5 | 0 | 0 | T | C | -0.7718 | 0.2342 | 0.0009841 | ----+ | 0 | 0.4128 | intron | FLJ13197 |
| rs56861485 | 2 | 29588261 | 4 | 0 | 0 | T | G | 0.7884 | 0.2393 | 0.0009841 | +++?+ | 0 | 0.7067 | intron | ALK |
| rs79907108 | 2 | 170360577 | 4 | 0 | 0 | A | C | -1.1874 | 0.3604 | 0.0009842 | ---?- | 0 | 0.9947 | intron | BBS5 |
| rs113547809 | 3 | 155472721 | 4 | 0 | 0 | A | G | 2.1865 | 0.6636 | 0.0009843 | +++?+ | 0 | 0.4751 | intergenic | |
| rs10802133 | 1 | 120247855 | 4 | 0 | 0 | T | C | 0.838 | 0.2543 | 0.0009847 | +++?+ | 0 | 0.9514 | intergenic | |
| rs72696420 | 9 | 19007775 | 4 | 0 | 0 | A | T | -1.5492 | 0.4702 | 0.0009847 | ---?- | 44.2 | 0.1459 | intron | FAM154A |
| 4:180874904 | 4 | 180874904 | 3 | 0 | 0 | G | GT | -0.9266 | 0.2812 | 0.0009847 | ---?? | 2.3 | 0.3595 | | |
| 6:170231793 | 6 | 170231793 | 3 | 0 | 0 | G | GC | 1.0581 | 0.3211 | 0.0009848 | +++?? | 24.5 | 0.2659 | | |
| rs62518599 | 8 | 124699329 | 4 | 0 | 0 | A | G | -0.8163 | 0.2477 | 0.0009849 | ---?- | 0 | 0.8615 | intron | ANXA13 |
| rs34563071 | 12 | 27910879 | 4 | 0 | 0 | A | G | -1.9056 | 0.5784 | 0.0009851 | ---?- | 1.4 | 0.385 | downstream-gene | |
| rs117760782 | 11 | 129349195 | 4 | 0 | 0 | A | T | 3.8939 | 1.1819 | 0.0009853 | +++?+ | 22.4 | 0.2764 | intergenic | |
| rs149592376 | 6 | 31223738 | 3 | 0 | 0 | T | C | -0.9505 | 0.2885 | 0.0009854 | ---?? | 0 | 0.7816 | intergenic | |
| rs12867718 | 13 | 20813572 | 4 | 0 | 0 | T | C | 1.1012 | 0.3342 | 0.0009855 | +++?+ | 0 | 0.4884 | intergenic | |
| 7:117466880 | 7 | 117466880 | 3 | 0 | 0 | CATT | C | 1.1205 | 0.3401 | 0.0009857 | +++?? | 65.4 | 0.05576 | | |
| 22:19174454 | 22 | 19174454 | 3 | 0 | 0 | CT | C | -0.979 | 0.2972 | 0.0009858 | ---?? | 30.5 | 0.2373 | | |
| rs79174869 | 8 | 113196925 | 4 | 0 | 0 | A | G | -0.9348 | 0.2838 | 0.0009859 | ---?- | 0 | 0.9794 | intergenic | |
| 1:90252836 | 1 | 90252836 | 3 | 0 | 0 | G | GT | -0.8435 | 0.256 | 0.0009859 | ---?? | 0 | 0.8496 | | |
| rs117040538 | 10 | 35938664 | 4 | 0 | 0 | A | G | 4.2257 | 1.2827 | 0.000986 | +++?+ | 8.1 | 0.3525 | intergenic | |
| rs185256459 | 12 | 102083333 | 3 | 0 | 0 | A | C | -1.8942 | 0.575 | 0.0009861 | ---?? | 0 | 0.3988 | downstream-gene | |
| rs7680512 | 4 | 123712072 | 4 | 0 | 0 | T | C | -2.1201 | 0.6435 | 0.0009862 | ---?- | 52.4 | 0.09764 | intergenic | |
| rs2344498 | 12 | 26798755 | 4 | 0 | 0 | A | G | 0.8467 | 0.257 | 0.0009863 | +++?- | 29.4 | 0.2357 | intron | ITPR2 |
| rs140288420 | 7 | 146314892 | 3 | 0 | 0 | T | C | 4.3774 | 1.3287 | 0.0009864 | +?+?- | 13.7 | 0.3137 | intron | CNTNAP2 |
| rs77247238 | 6 | 130940119 | 4 | 0 | 0 | A | G | 1.3852 | 0.4205 | 0.0009867 | +++?+ | 0 | 0.7416 | intergenic | |
| rs6421618 | 11 | 134475781 | 4 | 0 | 0 | T | C | -0.9748 | 0.2959 | 0.0009868 | ---?- | 0 | 0.9852 | intergenic | |

| | | | | | | | | | | | | | | | |
|---|---|---|---|---|---|---|---|---|---|---|---|---|---|---|---|
| rs6987049 | 8 | 113198661 | 4 | 0 | 0 | A | G | -0.8501 | 0.258 | 0.0009868 | ---?- | 0 | 0.9211 | intergenic | |
| rs77265705 | 19 | 57428735 | 3 | 0 | 0 | A | T | -3.1104 | 0.9442 | 0.0009869 | ---?? | 0 | 0.9102 | intergenic | |
| rs117610027 | 13 | 67075367 | 4 | 0 | 0 | T | C | -1.4998 | 0.4553 | 0.000987 | ---?- | 61.3 | 0.05159 | intron | PCDH9 |
| rs10415389 | 19 | 3599428 | 3 | 0 | 0 | T | C | 0.8589 | 0.2607 | 0.0009871 | +++?? | 0 | 0.5678 | intron | TBXA2R |
| rs62508609 | 8 | 49638219 | 3 | 0 | 0 | T | C | -3.2788 | 0.9953 | 0.0009872 | ---?? | 30.3 | 0.238 | intron | EFCAB1 |
| rs807131 | 17 | 51229932 | 5 | 0 | 0 | T | G | -1.0155 | 0.3083 | 0.0009873 | ----+ | 7.5 | 0.3636 | intergenic |  |
| rs116273226 | 4 | 59661319 | 3 | 0 | 0 | T | G | 2.3516 | 0.7139 | 0.0009874 | +++?? | 0 | 0.7668 | intergenic | |
| 6:159211720 | 6 | 159211720 | 3 | 0 | 0 | A | AACA | 0.841 | 0.2553 | 0.0009874 | +++?? | 39.9 | 0.1892 | | |
| rs10734791 | 12 | 30863909 | 5 | 0 | 0 | T | G | -0.6952 | 0.2111 | 0.0009877 | ----- | 0 | 0.7323 | intron | CAPRIN2 |
| rs3764451 | 5 | 42398465 | 4 | 0 | 0 | T | C | 1.2145 | 0.3687 | 0.0009879 | +++?+ | 0 | 0.5684 | intergenic | |
| rs55813192 | 2 | 170359943 | 4 | 0 | 0 | A | G | -1.188 | 0.3607 | 0.0009882 | ---?- | 0 | 0.9948 | intron | BBS5 |
| rs118002892 | 15 | 86376426 | 4 | 0 | 0 | A | G | -1.8653 | 0.5663 | 0.0009882 | ---?- | 0 | 0.9926 | intergenic | |
| 1:23567667 | 1 | 23567667 | 3 | 0 | 0 | CG | C | -1.1085 | 0.3365 | 0.0009882 | ---?? | 0 | 0.8171 | | |
| rs10149421 | 14 | 43268943 | 4 | 0 | 0 | C | G | -1.1523 | 0.3498 | 0.0009883 | ---?+ | 5.7 | 0.3643 | intergenic | |
| rs11746044 | 5 | 123092028 | 4 | 0 | 0 | T | C | -0.9333 | 0.2834 | 0.0009884 | ---?- | 19.7 | 0.2913 | intergenic | |
| rs59035627 | 10 | 74713046 | 4 | 0 | 0 | T | G | 2.8267 | 0.8582 | 0.0009886 | +++?+ | 37.1 | 0.1894 | intron | PLA2G12B |
| 1:107983662 | 1 | 107983662 | 3 | 0 | 0 | G | GA | 0.9361 | 0.2842 | 0.0009886 | +++?? | 0 | 0.9013 | | |
| rs9264493 | 6 | 31232697 | 3 | 0 | 0 | A | G | -0.9383 | 0.2849 | 0.0009887 | ---?? | 21.8 | 0.2784 | downstream-gene | |
| rs9654082 | 4 | 47777626 | 5 | 0 | 0 | T | C | -1.265 | 0.3841 | 0.0009887 | ----- | 0 | 0.5577 | intron | CORIN |
| rs34905798 | 13 | 111551366 | 5 | 0 | 0 | A | G | -2.1248 | 0.6451 | 0.0009887 | -+--- | 28.3 | 0.2331 | intron | ANKRD10 |
| 11:134683331 | 11 | 134683331 | 3 | 0 | 0 | T | G | -2.7928 | 0.8479 | 0.0009888 | ---?? | 0 | 0.8263 | intergenic | |
| rs4836037 | 5 | 123091599 | 4 | 0 | 0 | T | C | 0.9327 | 0.2832 | 0.0009888 | +++?+ | 20 | 0.2899 | intergenic | |
| rs1204810 | 6 | 116551638 | 5 | 0 | 0 | A | C | -0.7144 | 0.2169 | 0.000989 | ----- | 0 | 0.612 | intron | NT5DC1 |
| rs3909469 | 5 | 39424010 | 4 | 0 | 0 | A | G | -0.9061 | 0.2751 | 0.0009891 | ---?- | 0 | 0.4093 | intron | DAB2 |
| rs73102111 | 12 | 48028501 | 4 | 0 | 0 | A | G | 1.7216 | 0.5227 | 0.0009891 | +++?- | 56.9 | 0.07327 | intergenic | |
| rs144375974 | 3 | 1066095 | 2 | 0 | 0 | T | C | -6.9603 | 2.1133 | 0.0009891 | ?-??- | 0 | 0.3982 | intergenic | |
| rs146660442 | 4 | 190537947 | 3 | 0 | 0 | A | G | 1.7702 | 0.5375 | 0.0009891 | +++?? | 0 | 0.6706 | intergenic | |
| 6:18687899 | 6 | 18687899 | 3 | 0 | 0 | T | TTTTTCT | 0.8764 | 0.2661 | 0.0009894 | +++?? | 41 | 0.1835 | | |
| rs9425267 | 1 | 181987479 | 4 | 0 | 0 | T | C | 1.0758 | 0.3266 | 0.0009897 | +++?+ | 0 | 0.5068 | intergenic | |
| rs34164540 | 12 | 78680545 | 4 | 0 | 0 | A | G | -1.7613 | 0.5348 | 0.0009897 | ---?+ | 0 | 0.4251 | intergenic | |
| 6:31230003 | 6 | 31230003 | 3 | 0 | 0 | T | TG | 0.9366 | 0.2844 | 0.0009897 | +++?? | 22.6 | 0.2746 | | |
| 11:110195880 | 11 | 110195880 | 2 | 0 | 0 | T | C | 4.7867 | 1.4534 | 0.0009899 | ?++?? | 50.9 | 0.1534 | intergenic | |
| rs72854965 | 2 | 131134511 | 5 | 0 | 0 | A | G | 0.8644 | 0.2625 | 0.00099 | +++++ | 35.7 | 0.1835 | downstream-gene | |
| 11:130626264 | 11 | 130626264 | 3 | 0 | 0 | T | TG | -1.6856 | 0.5118 | 0.00099 | ---?? | 49.2 | 0.1394 | | |
| rs921301 | 4 | 123705387 | 4 | 0 | 0 | T | C | 2.0875 | 0.6339 | 0.0009901 | +++?+ | 51.5 | 0.1028 | intergenic | |
| rs62514362 | 8 | 113202366 | 4 | 0 | 0 | T | C | 0.9334 | 0.2834 | 0.0009902 | +++?+ | 0 | 0.9741 | intergenic | |
| 15:32272268 | 15 | 32272268 | 2 | 0 | 0 | T | G | 5.0615 | 1.5369 | 0.0009903 | ?++?? | 0 | 0.9519 | intergenic | |
| rs479710 | 6 | 12624669 | 4 | 0 | 0 | A | G | 0.8891 | 0.27 | 0.0009903 | +++?+ | 0 | 0.769 | intergenic | |
| 15:83905586 | 15 | 83905586 | 3 | 0 | 0 | T | G | 1.2313 | 0.3739 | 0.0009906 | +++?? | 51.2 | 0.1288 | intergenic | |
| rs11727199 | 4 | 10036190 | 5 | 0 | 0 | T | C | 0.692 | 0.2101 | 0.0009906 | ++++- | 15.2 | 0.3177 | intron | SLC2A9 |
| rs77978666 | 2 | 3773068 | 3 | 0 | 0 | A | G | -1.1657 | 0.354 | 0.0009906 | ---?? | 0 | 0.5961 | intron | DCDC2C |
| rs12143319 | 1 | 85727335 | 5 | 0 | 0 | A | G | -1.2732 | 0.3866 | 0.0009908 | ----- | 0 | 0.8052 | upstream-gene |  |
| 6:159212738 | 6 | 159212738 | 3 | 0 | 0 | A | ATTTTT | 0.8449 | 0.2566 | 0.0009908 | +++?? | 0 | 0.3745 | | |
| rs9848198 | 3 | 77404621 | 4 | 0 | 0 | C | G | -0.8259 | 0.2508 | 0.0009909 | ---?- | 0 | 0.4027 | intron | ROBO2 |
| rs372664112 | 15 | 83905583 | 3 | 0 | 0 | A | G | 1.2312 | 0.3739 | 0.0009909 | +++?? | 51.2 | 0.1289 | intergenic | |
| rs3849552 | 3 | 21881441 | 4 | 0 | 0 | A | C | -0.9922 | 0.3013 | 0.000991 | ---?- | 0 | 0.7037 | intron | ZNF385D |
| rs146450130 | 5 | 152680753 | 4 | 0 | 0 | T | G | -3.042 | 0.9238 | 0.0009911 | ---?- | 0 | 0.5479 | intergenic | |
| rs56293740 | 2 | 170359758 | 4 | 0 | 0 | C | G | -1.1882 | 0.3608 | 0.0009912 | ---?- | 0 | 0.9948 | intron | BBS5 |
| rs3742183 | 13 | 111551952 | 5 | 0 | 0 | T | C | -2.1251 | 0.6454 | 0.0009913 | -+--- | 28.3 | 0.2327 | intron | ANKRD10 |
| rs173665 | 19 | 8366914 | 4 | 0 | 0 | A | G | 1.6189 | 0.4916 | 0.0009914 | +++?+ | 40.6 | 0.168 | downstream-gene | |
| rs921300 | 4 | 123705169 | 4 | 0 | 0 | T | C | -2.0857 | 0.6334 | 0.0009914 | ---?- | 51.5 | 0.1031 | intergenic | |
| rs4252279 | 11 | 117867187 | 4 | 0 | 0 | T | C | 1.3102 | 0.3979 | 0.0009915 | +++?+ | 0 | 0.7057 | intron | IL10RA |
| rs73424702 | 18 | 32643044 | 4 | 0 | 0 | A | G | 2.8549 | 0.867 | 0.0009915 | +++?+ | 0 | 0.6812 | intron | MAPRE2 |
| rs118021549 | 18 | 40420881 | 3 | 0 | 0 | C | G | -1.9768 | 0.6003 | 0.0009916 | ---?? | 0 | 0.7273 | intron | RIT2 |
| rs58281223 | 2 | 184074989 | 4 | 0 | 0 | T | C | 1.1934 | 0.3624 | 0.0009917 | +++?+ | 0 | 0.7347 | intergenic | |
| rs71530587 | 7 | 32361901 | 4 | 0 | 0 | T | G | 1.3396 | 0.4068 | 0.0009918 | +++?+ | 0 | 0.9567 | intergenic | |
| rs2351125 | 6 | 74387936 | 3 | 0 | 0 | A | T | -0.9653 | 0.2932 | 0.0009919 | ---?? | 0 | 0.5632 | intergenic | |
| rs71398296 | 15 | 67419315 | 4 | 0 | 0 | T | C | 1.642 | 0.4986 | 0.0009919 | +++?+ | 0 | 0.8293 | intron | SMAD3 |
| rs887565 | 4 | 14349798 | 4 | 0 | 0 | A | T | 0.8439 | 0.2563 | 0.000992 | +++?+ | 0 | 0.6531 | intergenic | |

| | | | | | | | | | | | | | | | |
|---|---|---|---|---|---|---|---|---|---|---|---|---|---|---|---|
| rs75450037 | 12 | 78831475 | 4 | 0 | 0 | A | C | 1.5739 | 0.478 | 0.000992 | +++?+ | 0 | 0.5168 | intergenic | |
| rs7732928 | 5 | 123425663 | 4 | 0 | 0 | T | C | -0.8046 | 0.2444 | 0.0009921 | ---?- | 5.6 | 0.365 | intergenic | |
| rs58357998 | 17 | 71050300 | 4 | 0 | 0 | A | G | -2.2257 | 0.6759 | 0.0009921 | ---?- | 0 | 0.453 | intron | SLC39A11 |
| rs2782381 | 6 | 63349898 | 4 | 0 | 0 | A | G | -2.2593 | 0.6861 | 0.0009922 | ---?- | 0 | 0.3925 | intergenic | |
| rs9975484 | 21 | 28707627 | 4 | 0 | 0 | A | T | 1.4881 | 0.452 | 0.0009922 | +++?+ | 26.8 | 0.2508 | intergenic | |
| rs7127461 | 11 | 75032518 | 4 | 0 | 0 | C | G | 0.9392 | 0.2852 | 0.0009929 | +++?+ | 0 | 0.4124 | intron | ARRB1 |
| 4:47988991 | 4 | 47988991 | 3 | 0 | 0 | A | ATTGTG | 1.5455 | 0.4694 | 0.000993 | +++?? | 0 | 0.5218 | | |
| rs62182224 | 2 | 218843880 | 4 | 0 | 0 | A | T | -1.3264 | 0.4028 | 0.0009931 | ---?+ | 0 | 0.7604 | intergenic | |
| rs2843034 | 6 | 63349954 | 4 | 0 | 0 | A | G | -2.2591 | 0.6862 | 0.0009932 | ---?- | 0 | 0.3928 | intergenic | |
| rs72842401 | 6 | 24021534 | 5 | 0 | 0 | A | C | 1.2858 | 0.3905 | 0.0009932 | ++++- | 0 | 0.8195 | intergenic | |
| rs2253303 | 3 | 59849717 | 4 | 0 | 0 | A | T | -0.7965 | 0.2419 | 0.0009933 | ---?- | 0 | 0.9465 | intron | FHIT |
| rs113874861 | 13 | 31108216 | 4 | 0 | 0 | T | G | 1.491 | 0.4528 | 0.0009934 | +++?+ | 35.7 | 0.1982 | intron | HMGB1 |
| rs1502766 | 4 | 108276296 | 4 | 0 | 0 | T | C | -0.8278 | 0.2514 | 0.0009935 | ---?- | 0 | 0.424 | intergenic | |
| rs12662262 | 6 | 118466410 | 4 | 0 | 0 | T | C | 0.8335 | 0.2532 | 0.0009935 | +++?+ | 0 | 0.8122 | intron | SLC35F1 |
| rs1204793 | 6 | 116540842 | 5 | 0 | 0 | A | G | 0.7762 | 0.2358 | 0.0009937 | +++++ | 0 | 0.5083 | intron | NT5DC1 |
| rs190807712 | 2 | 170616067 | 4 | 0 | 0 | A | G | -3.665 | 1.1132 | 0.0009938 | ---?- | 10.1 | 0.3426 | intergenic | |
| rs9975935 | 21 | 28707316 | 4 | 0 | 0 | T | C | 1.4905 | 0.4527 | 0.0009939 | +++?+ | 27.3 | 0.2483 | intergenic | |
| rs1204785 | 6 | 116532265 | 5 | 0 | 0 | A | C | -0.8381 | 0.2546 | 0.0009942 | ----- | 0 | 0.854 | intron | NT5DC1 |
| rs4730558 | 7 | 112308347 | 4 | 0 | 0 | T | G | 1.271 | 0.3861 | 0.0009942 | -++?+ | 26.8 | 0.2508 | intergenic | |
| rs1442191 | 12 | 41370616 | 4 | 0 | 0 | T | C | 1.3991 | 0.425 | 0.0009944 | +++?+ | 0 | 0.6055 | intron | CNTN1 |
| rs62133246 | 19 | 57259821 | 4 | 0 | 0 | A | G | -0.8948 | 0.2718 | 0.0009944 | ---?- | 0 | 0.9475 | intergenic | |
| rs17296192 | 12 | 51524459 | 5 | 0 | 0 | A | T | 0.8951 | 0.2719 | 0.0009945 | +++++ | 0 | 0.7023 | intron | TFCP2 |
| rs28558512 | 4 | 46386157 | 4 | 0 | 0 | A | G | -0.8518 | 0.2587 | 0.0009945 | -+-?- | 58.9 | 0.06297 | intron | GABRA2 |
| rs1605133 | 2 | 117111221 | 4 | 0 | 0 | T | C | -1.6426 | 0.499 | 0.0009946 | ---?- | 0 | 0.6798 | intergenic | |
| rs526896 | 5 | 134356705 | 5 | 0 | 0 | T | G | 0.7676 | 0.2332 | 0.0009947 | +++++ | 0 | 0.4545 | intergenic | |
| 3:123541511 | 3 | 123541511 | 3 | 0 | 0 | A | AAG | 2.54 | 0.7716 | 0.0009947 | +++?? | 0 | 0.9664 | | |
| rs35185382 | 5 | 39608726 | 4 | 0 | 0 | T | C | -0.9843 | 0.299 | 0.0009951 | ---?- | 0 | 0.8893 | intergenic | |
| 2:165723888 | 2 | 165723888 | 3 | 0 | 0 | T | C | 3.2168 | 0.9772 | 0.0009952 | +-+?? | 65.2 | 0.05641 | intergenic | |
| rs2843033 | 6 | 63350053 | 4 | 0 | 0 | A | G | -2.2587 | 0.6862 | 0.0009952 | ---?- | 0 | 0.3927 | intergenic | |
| rs111485498 | 13 | 31108218 | 4 | 0 | 0 | A | C | 1.4908 | 0.4529 | 0.0009952 | +++?+ | 35.8 | 0.1977 | intron | HMGB1 |
| rs7355427 | 2 | 184024026 | 4 | 0 | 0 | T | G | -1.1465 | 0.3483 | 0.0009953 | ---?- | 0 | 0.818 | intron | NUP35 |
| 6:144940937 | 6 | 144940937 | 3 | 0 | 0 | G | GC | 1.1129 | 0.3381 | 0.0009953 | +++?? | 2.8 | 0.3575 | | |
| rs62343778 | 4 | 174557499 | 4 | 0 | 0 | T | C | 0.8082 | 0.2455 | 0.0009954 | +++?+ | 0 | 0.6261 | intergenic | |
| rs62423695 | 6 | 118466219 | 4 | 0 | 0 | T | C | 0.8333 | 0.2531 | 0.0009955 | +++?+ | 0 | 0.8116 | intron | SLC35F1 |
| rs79478110 | 8 | 21030476 | 3 | 0 | 0 | C | G | 3.9126 | 1.1886 | 0.0009955 | +?+?- | 51.9 | 0.1253 | intergenic | |
| rs2254867 | 2 | 183991048 | 4 | 0 | 0 | A | G | 1.161 | 0.3527 | 0.0009957 | +++?+ | 0 | 0.8544 | intron | NUP35 |
| rs56893636 | 19 | 12797314 | 4 | 0 | 0 | T | C | -3.4692 | 1.0539 | 0.0009957 | ----? | 50.6 | 0.1083 | upstream-gene |  |
| rs11693598 | 2 | 82973341 | 4 | 0 | 0 | A | G | -2.4949 | 0.7579 | 0.0009958 | ---?- | 0 | 0.4694 | intergenic | |
| rs12281645 | 11 | 86151931 | 4 | 0 | 0 | A | T | -2.1016 | 0.6384 | 0.0009959 | ---?- | 0 | 0.4033 | downstream-gene | |
| rs34321532 | 16 | 12118260 | 4 | 0 | 0 | C | G | -0.7903 | 0.2401 | 0.0009959 | +--?- | 58.8 | 0.06341 | intron | SNX29 |
| rs7221521 | 17 | 77125461 | 3 | 0 | 0 | T | C | -0.9183 | 0.279 | 0.0009961 | ---?? | 8.5 | 0.3352 | intron | RBFOX3 |
| rs787367 | 4 | 67907853 | 4 | 0 | 0 | T | C | 1.3036 | 0.396 | 0.0009966 | +++?- | 57.1 | 0.0719 | intergenic | |
| rs118176228 | 10 | 99696771 | 3 | 0 | 0 | T | C | 3.0451 | 0.9251 | 0.0009966 | +++?? | 0 | 0.4471 | intron | CRTAC1 |
| rs1109009 | 20 | 2457788 | 4 | 0 | 0 | T | G | -0.9789 | 0.2974 | 0.0009967 | +--?- | 51.3 | 0.104 | downstream-gene | |
| rs10892202 | 11 | 117857338 | 4 | 0 | 0 | C | G | 1.2892 | 0.3917 | 0.0009967 | +++?+ | 0 | 0.823 | intron | IL10RA |
| 1:64585905 | 1 | 64585905 | 3 | 0 | 0 | T | TG | -1.0565 | 0.321 | 0.0009967 | ---?? | 0 | 0.8553 | | |
| rs1199667 | 1 | 87581167 | 4 | 0 | 0 | A | G | -0.9742 | 0.296 | 0.000997 | ---?- | 26.3 | 0.2537 | intergenic | |
| rs9551734 | 13 | 30320185 | 5 | 0 | 0 | T | C | -1.2125 | 0.3684 | 0.000997 | ----- | 0 | 0.5443 | intergenic |  |
| rs11664017 | 18 | 10723068 | 4 | 0 | 0 | C | G | -0.8373 | 0.2544 | 0.000997 | ---?+ | 35.8 | 0.1972 | intron | PIEZO2 |
| rs12777794 | 10 | 122969810 | 3 | 0 | 0 | T | C | 3.3805 | 1.0271 | 0.0009971 | +++?? | 0 | 0.7696 | intergenic | |
| rs187208564 | 22 | 28824880 | 4 | 0 | 0 | A | G | 2.6554 | 0.8068 | 0.0009971 | +++?+ | 0 | 0.8855 | intron | TTC28 |
| rs6500789 | 16 | 6534217 | 4 | 0 | 0 | A | G | -0.8377 | 0.2545 | 0.0009972 | ---?- | 0 | 0.7602 | intron | RBFOX1 |
| 14:21440332 | 14 | 21440332 | 3 | 0 | 0 | A | ATG | -0.869 | 0.264 | 0.0009972 | ---?? | 0 | 0.5067 | | |
| rs9475531 | 6 | 55897373 | 4 | 0 | 0 | A | G | -2.2071 | 0.6706 | 0.0009974 | ---?- | 0 | 0.803 | intergenic | |
| rs6562708 | 13 | 73049071 | 3 | 0 | 0 | T | C | 0.9175 | 0.2788 | 0.0009975 | +++?? | 57.2 | 0.09647 | intergenic | |
| rs115623743 | 1 | 42677647 | 4 | 0 | 0 | A | G | 1.7343 | 0.5269 | 0.0009975 | +++?+ | 0 | 0.5832 | intron | FOXJ3 |
| rs117811283 | 17 | 69946546 | 4 | 0 | 0 | A | T | 2.607 | 0.7921 | 0.0009976 | +++?+ | 4.5 | 0.3705 | intergenic | |
| 1:85657440 | 1 | 85657440 | 3 | 0 | 0 | CA | C | 1.1708 | 0.3557 | 0.0009976 | +++?? | 0 | 0.8898 | | |
| 4:38728970 | 4 | 38728970 | 3 | 0 | 0 | CTT | C | -0.9354 | 0.2842 | 0.0009978 | ---?? | 0 | 0.6152 | | |

| | | | | | | | | | | | | | | | |
|---|---|---|---|---|---|---|---|---|---|---|---|---|---|---|---|
| rs75018710 | 3 | 119007096 | 5 | 0 | 0 | A | T | -1.5135 | 0.4599 | 0.0009979 | ----- | 15.8 | 0.3138 | intergenic |  |
| rs74543541 | 16 | 14173036 | 3 | 0 | 0 | A | G | -1.9868 | 0.6037 | 0.000998 | ---?? | 0 | 0.9136 | intron | MKL2 |
| rs3850739 | 8 | 64939772 | 4 | 0 | 0 | A | C | 0.8166 | 0.2481 | 0.0009983 | +++?- | 0 | 0.5097 | intergenic | |
| rs7551490 | 1 | 182045043 | 4 | 0 | 0 | T | C | 2.3124 | 0.7027 | 0.0009986 | +++?+ | 0 | 0.8933 | intergenic | |
| rs73337636 | 8 | 114889984 | 4 | 0 | 0 | A | C | -1.8096 | 0.5499 | 0.0009986 | ---?+ | 4.1 | 0.3724 | intergenic | |
| rs9475532 | 6 | 55897432 | 4 | 0 | 0 | A | G | -2.2067 | 0.6706 | 0.0009988 | ---?- | 0 | 0.8031 | intergenic | |
| rs55959894 | 4 | 10026580 | 5 | 0 | 0 | A | G | 0.6917 | 0.2102 | 0.0009988 | ++++- | 12.9 | 0.3317 | intron | SLC2A9 |
| rs62216867 | 21 | 20402131 | 4 | 0 | 0 | A | T | -1.5798 | 0.4801 | 0.0009988 | ---?- | 0 | 0.6034 | intergenic | |
| rs1961377 | 17 | 65480264 | 4 | 0 | 0 | A | C | -0.8591 | 0.2611 | 0.000999 | ---?+ | 3.8 | 0.3739 | intron | PITPNC1 |
| rs5997386 | 22 | 29103911 | 3 | 0 | 0 | A | G | 2.1767 | 0.6615 | 0.0009991 | +++?? | 0 | 0.4274 | intron | CHEK2 |
| 20:35645344 | 20 | 35645344 | 3 | 0 | 0 | A | AG | -1.2028 | 0.3655 | 0.0009991 | ---?? | 0 | 0.9346 | | |
| rs751911 | 11 | 86194654 | 4 | 0 | 0 | A | C | -2.331 | 0.7084 | 0.0009994 | ---?- | 0 | 0.9706 | intron | ME3 |
| rs75530201 | 3 | 119007073 | 5 | 0 | 0 | T | C | 1.5132 | 0.4598 | 0.0009994 | +++++ | 15.8 | 0.3137 | intergenic | |
| rs79292354 | 3 | 119007095 | 5 | 0 | 0 | T | G | 1.5133 | 0.4599 | 0.0009994 | +++++ | 15.8 | 0.3137 | intergenic | |
| rs113433323 | 6 | 109118980 | 2 | 0 | 0 | A | G | 4.5904 | 1.395 | 0.0009994 | +?+?? | 0 | 0.826 | intergenic | |
| rs7202791 | 16 | 52855926 | 4 | 0 | 0 | A | G | 0.8079 | 0.2455 | 0.0009996 | +++?+ | 0 | 0.7842 | intergenic | |
| rs1452137 | 11 | 86151015 | 4 | 0 | 0 | A | G | -2.1011 | 0.6385 | 0.001 | ---?- | 0 | 0.4014 | downstream-gene | |
| rs9684523 | 4 | 108209364 | 4 | 0 | 0 | T | C | -0.8052 | 0.2447 | 0.001 | ---?- | 0 | 0.4679 | intergenic | |
| rs10100514 | 8 | 96985292 | 4 | 0 | 0 | T | C | 1.3279 | 0.4036 | 0.001 | +++?+ | 0 | 0.9827 | intergenic | |
| rs11754433 | 6 | 118465941 | 4 | 0 | 0 | A | G | 0.8329 | 0.2531 | 0.001 | +++?+ | 0 | 0.8112 | intron | SLC35F1 |
| rs11781755 | 8 | 113195249 | 4 | 0 | 0 | T | C | 0.9337 | 0.2838 | 0.001 | +++?+ | 0 | 0.9794 | intergenic | |
| rs12206897 | 6 | 859664 | 4 | 0 | 0 | T | C | -2.2858 | 0.6947 | 0.001 | ---?- | 0 | 0.8747 | intergenic | |
| rs72940742 | 6 | 72880825 | 4 | 0 | 0 | A | G | 1.5323 | 0.4657 | 0.001 | +++?- | 0 | 0.5896 | intron | RIMS1 |
| rs73143032 | 7 | 78482523 | 4 | 0 | 0 | A | G | 2.1555 | 0.6551 | 0.001 | +++?+ | 0 | 0.484 | intron | MAGI2 |
| rs150401628 | 11 | 25669058 | 3 | 0 | 0 | C | G | -0.8576 | 0.2606 | 0.001 | ---?? | 0 | 0.4151 | intergenic | |
| 14:21439972 | 14 | 21439972 | 3 | 0 | 0 | G | GT | 0.8687 | 0.264 | 0.001 | +++?? | 0 | 0.5051 | | |

MUTED
