## Supplementary Table 4 for "A higher burden of multiple sclerosis genetic risk confers an earlier onset"

**Supplementary Table 4. Results from MAGMA Gene-Based Test**

| Column Header | Key |
|---|---|
| **Gene** | Ensemble Gene Id |
| **CHR** | Chromosome of gene |
| **START** | Gene start position (hg19) |
| **STOP** | Gene end posiion (hg19) |
| **NSNPS** | Number of SNPs annotated to the gene |
| **NPARAM** | Number of SNPs available and used in the model |
| **Z** | The z-score for the specified gene based on permutation p-value |
| **P** | P-value for the specified gene using an asymptotic sampling distribution |
| **SYMBOL** | Official Gene Symbol |
| **P_0kb** | Significance level for exact gene position |
| **P_10kb** | Significance level for 10kb window around gene |
| **P_25kb** | Significance level for 25kb window around gene |

| GENE | CHR | START | STOP | NSNPS | NPARAM | Z | P | SYMBOL | P_0kb | P_10kb | P_25kb |
|---|---|---|---|---|---|---|---|---|---|---|---|
| ENSG00000138385 | 2 | 170648443 | 170668574 | 63 | 16 | 4.0493 | 0.000025689 | SSB | 0.000025689 | 0.00027952 | 0.000024291 |
| ENSG00000135148 | 12 | 112563305 | 112591407 | 23 | 9 | 4.0402 | 0.000026705 | TRAFD1 | 0.000026705 | 0.000044708 | 0.000049794 |
| ENSG00000173064 | 12 | 112597992 | 112819896 | 225 | 29 | 4.0308 | 0.00002779 | HECTD4 | 0.00002779 | 0.000028863 | 0.00002498 |
| ENSG00000118113 | 11 | 102582526 | 102597781 | 85 | 18 | 4.0246 | 0.000028542 | MMP8 | 0.000028542 | 0.000010425 | 0.000032219 |
| ENSG00000111300 | 12 | 112464500 | 112546826 | 78 | 18 | 3.9162 | 0.000044975 | NAA25 | 0.000044975 | 0.000051595 | 0.000023699 |
| ENSG00000144357 | 2 | 170683968 | 170940641 | 1091 | 28 | 3.9114 | 0.000045881 | UBR3 | 0.000045881 | 0.000040312 | 0.000013429 |
| ENSG00000206077 | 5 | 710475 | 767067 | 123 | 17 | 3.8718 | 0.000054022 | ZDHHC11B | 0.000054022 | 0.00018555 | 0.000064936 |
| ENSG00000147526 | 8 | 38585704 | 38710546 | 278 | 23 | 3.8671 | 0.000055073 | TACC1 | 0.000055073 | 0.00024257 | 0.000076099 |
| ENSG00000089248 | 12 | 112451120 | 112461255 | 15 | 6 | 3.8421 | 0.000060987 | ERP29 | 0.000060987 | 0.000041054 | 0.000059898 |
| ENSG00000169126 | 10 | 28064115 | 28287977 | 541 | 30 | 3.8054 | 0.000070796 | ARMC4 | 0.000070796 | 0.000095611 | 0.00025746 |
| ENSG00000105879 | 7 | 107384142 | 107402112 | 45 | 6 | 3.7814 | 0.000077979 | CBLL1 | 0.000077979 | 0.00010198 | 0.000039504 |
| ENSG00000213160 | 2 | 170550998 | 170633499 | 468 | 33 | 3.7648 | 0.000083327 | KLHL23 | 0.000083327 | 0.000107 | 0.000059012 |
| ENSG00000224389 | 6 | 31982539 | 32003195 | 8 | 4 | 3.7508 | 0.000088123 | C4B | 0.000088123 | 0.000044586 | 0.000051417 |
| ENSG00000242019 | 19 | 55235984 | 55248003 | 60 | 12 | 3.7505 | 0.000088232 | KIR3DL3 | 0.000088232 | 0.00053114 | 0.00019896 |
| ENSG00000138382 | 2 | 170666591 | 170681441 | 59 | 12 | 3.7451 | 0.000090168 | METTL5 | 0.000090168 | 0.000019994 | 2.9195E-06 |
| ENSG00000180745 | 10 | 129676105 | 129691211 | 62 | 20 | 3.7132 | 0.00010233 | CLRN3 | 0.00010233 | 0.00094675 | 0.006322 |
| ENSG00000198270 | 12 | 112369086 | 112450970 | 75 | 10 | 3.6931 | 0.00011075 | TMEM116 | 0.00011075 | 0.00013026 | 0.000066642 |
| ENSG00000091137 | 7 | 107301080 | 107358254 | 112 | 22 | 3.6463 | 0.000133 | SLC26A4 | 0.000133 | 0.000074692 | 0.00028719 |
| ENSG00000165792 | 14 | 21457929 | 21465189 | 37 | 11 | 3.6448 | 0.0001338 | METTL17 | 0.0001338 | 0.000046065 | 0.00018784 |
| ENSG00000121742 | 13 | 20796110 | 20806534 | 51 | 15 | 3.6291 | 0.00014218 | GJB6 | 0.00014218 | 0.000047181 | 0.00070654 |
| ENSG00000198740 | 17 | 47366568 | 47439835 | 204 | 18 | 3.5477 | 0.00019429 | ZNF652 | 0.00019429 | 0.00013135 | 0.000060969 |
| ENSG00000213246 | 17 | 56422539 | 56430454 | 20 | 6 | 3.52 | 0.00021577 | SUPT4H1 | 0.00021577 | 0.0033225 | 0.009759 |
| ENSG00000198324 | 12 | 111798455 | 111806925 | 10 | 4 | 3.4249 | 0.00030756 | FAM109A | 0.00030756 | 0.00001006 | 9.0352E-06 |
| ENSG00000072135 | 2 | 131113580 | 131132982 | 34 | 4 | 3.4066 | 0.00032892 | PTPN18 | 0.00032892 | 0.00032067 | 0.00028702 |
| ENSG00000152705 | 5 | 134303596 | 134347392 | 94 | 22 | 3.3892 | 0.00035046 | CATSPER3 | 0.00035046 | 0.00006334 | 0.00025199 |
| ENSG00000187079 | 11 | 12695969 | 12966298 | 610 | 47 | 3.3803 | 0.00036197 | TEAD1 | 0.00036197 | 0.00057183 | 0.0022972 |
| ENSG00000179295 | 12 | 112856155 | 112947717 | 98 | 20 | 3.3497 | 0.00040442 | PTPN11 | 0.00040442 | 0.00041096 | 0.00031911 |
| ENSG00000173467 | 7 | 16899029 | 16921611 | 95 | 21 | 3.3472 | 0.00040812 | AGR3 | 0.00040812 | 0.029643 | 0.032951 |
| ENSG00000148123 | 9 | 103790991 | 104087417 | 1157 | 68 | 3.341 | 0.00041731 | LPPR1 | 0.00041731 | 0.00066271 | 0.000941 |
| ENSG00000257767 | 12 | 112191694 | 112229222 | 10 | 6 | 3.3135 | 0.00046072 | RP11-162P23.2 | 0.00046072 | 0.01231 | 0.0012739 |
| ENSG00000188004 | 1 | 159804264 | 159825137 | 43 | 16 | 3.296 | 0.00049039 | C1orf204 | 0.00049039 | 0.0012634 | 0.012588 |
| ENSG00000137675 | 11 | 102562218 | 102576537 | 70 | 11 | 3.2935 | 0.00049468 | MMP27 | 0.00049468 | 0.00024579 | 0.000028193 |
| ENSG00000175175 | 17 | 56833230 | 57058983 | 309 | 25 | 3.2815 | 0.00051633 | PPM1E | 0.00051633 | 0.00052486 | 0.00048058 |
| ENSG00000204873 | 17 | 39388715 | 39389705 | 6 | 3 | 3.2782 | 0.00052233 | KRTAP9-3 | 0.00052233 | 0.57542 | 0.54302 |
| ENSG00000204525 | 6 | 31236526 | 31239907 | 175 | 15 | 3.261 | 0.00055503 | HLA-C | 0.00055503 | 0.00030231 | 0.00028478 |
| ENSG00000243649 | 6 | 31895475 | 31919861 | 61 | 25 | 3.2563 | 0.00056428 | CFB | 0.00056428 | 0.0006642 | 0.00030926 |
| ENSG00000106484 | 7 | 130126012 | 130146133 | 37 | 11 | 3.2443 | 0.00058878 | MEST | 0.00058878 | 0.00073167 | 0.00059485 |
| ENSG00000161328 | 11 | 537527 | 554916 | 109 | 18 | 3.2388 | 0.00060012 | LRRC56 | 0.00060012 | 0.00084508 | 0.00050106 |
| ENSG00000090060 | 14 | 96967770 | 97033448 | 126 | 17 | 3.2368 | 0.00060435 | PAPOLA | 0.00060435 | 0.00064831 | 0.00066264 |
| ENSG00000244255 | 6 | 31895475 | 31919825 | 60 | 26 | 3.234 | 0.00061042 | CFB | 0.00061042 | 0.0006642 | 0.00030926 |
| ENSG00000271949 | 1 | 90098699 | 90398809 | 662 | 47 | 3.2315 | 0.00061579 | RP11-302M6.4 | 0.00061579 | 0.00059664 | 0.00080758 |
| ENSG00000173200 | 3 | 122296449 | 122357894 | 252 | 27 | 3.2285 | 0.00062219 | PARP15 | 0.00062219 | 0.00053434 | 0.0037191 |
| ENSG00000268632 | 15 | 42073346 | 42076229 | 11 | 4 | 3.2182 | 0.00064507 | AC073657.1 | 0.00064507 | 0.0038412 | 0.0038709 |
| ENSG00000163138 | 4 | 20697905 | 20754530 | 247 | 20 | 3.2156 | 0.00065087 | PACRGL | 0.00065087 | 0.00044946 | 0.0005177 |
| ENSG00000243708 | 15 | 42120283 | 42140345 | 74 | 11 | 3.2151 | 0.00065196 | PLA2G4B | 0.00065196 | 0.0010688 | 0.0014996 |
| ENSG00000168970 | 15 | 42120293 | 42140353 | 74 | 11 | 3.2151 | 0.00065196 | JMJD7-PLA2G4B | 0.00065196 | 0.0010688 | 0.0014996 |
| ENSG00000144791 | 3 | 45596886 | 45727830 | 555 | 29 | 3.2142 | 0.00065407 | LIMD1 | 0.00065407 | 0.0010897 | 0.0027207 |
| ENSG00000111249 | 12 | 111471828 | 111788358 | 509 | 37 | 3.1714 | 0.00075866 | CUX2 | 0.00075866 | 0.00061942 | 0.00041363 |
| ENSG00000196312 | 9 | 99660348 | 99775862 | 145 | 22 | 3.155 | 0.00080254 | HIATL2 | 0.00080254 | 0.0008808 | 0.00090107 |
| ENSG00000184144 | 1 | 205012325 | 205047627 | 155 | 19 | 3.1488 | 0.00081967 | CNTN2 | 0.00081967 | 0.0010615 | 0.0013694 |
| ENSG00000254466 | 11 | 59244846 | 59245869 | 1 | 1 | 3.1378 | 0.0008511 | OR4D10 | 0.0008511 | 0.055244 | 0.14139 |
| ENSG00000006638 | 19 | 3594504 | 3606838 | 71 | 11 | 3.1192 | 0.00090683 | TBXA2R | 0.00090683 | 0.0035304 | 0.0025274 |
| ENSG00000111252 | 12 | 111843752 | 111889427 | 33 | 8 | 3.1178 | 0.00091113 | SH2B3 | 0.00091113 | 0.00091273 | 0.00039173 |
| ENSG00000231852 | 6 | 32006042 | 32009447 | 33 | 16 | 3.1146 | 0.00092098 | CYP21A2 | 0.00092098 | 0.000059846 | 0.00013681 |
| ENSG00000088766 | 20 | 5986736 | 6020699 | 121 | 14 | 3.113 | 0.00092586 | CRLS1 | 0.00092586 | 0.0016436 | 0.0065771 |
| ENSG00000092621 | 1 | 120202421 | 120286838 | 235 | 30 | 3.1106 | 0.00093347 | PHGDH | 0.00093347 | 0.001662 | 0.0018314 |

| | | | | | | | | | | | |
|---|---|---|---|---|---|---|---|---|---|---|---|
| ENSG00000133704 | 12 | 30781922 | 30848920 | 238 | 15 | 3.1101 | 0.00093512 | IPO8 | 0.00093512 | 0.0011262 | 0.0009685 |
| ENSG00000112039 | 6 | 35420138 | 35434880 | 42 | 6 | 3.1044 | 0.00095324 | FANCE | 0.00095324 | 0.00082459 | 0.0065832 |
| ENSG00000239713 | 22 | 39436924 | 39483748 | 133 | 17 | 3.0663 | 0.0010837 | APOBEC3G | 0.0010837 | 0.0012555 | 0.0055546 |
| ENSG00000105576 | 19 | 12810008 | 12834825 | 70 | 5 | 3.053 | 0.0011329 | TNPO2 | 0.0011329 | 0.0011469 | 0.00095061 |
| ENSG00000166278 | 6 | 31865562 | 31913449 | 128 | 37 | 3.0499 | 0.0011446 | C2 | 0.0011446 | 0.00058354 | 0.00029409 |
| ENSG00000188517 | 4 | 109731877 | 110223813 | 1764 | 81 | 3.0446 | 0.0011649 | COL25A1 | 0.0011649 | 0.0012709 | 0.0010484 |
| ENSG00000205593 | 22 | 50747459 | 50765489 | 87 | 13 | 3.0268 | 0.0012357 | DENND6B | 0.0012357 | 0.0011131 | 0.001172 |
| ENSG00000204344 | 6 | 31938868 | 31950598 | 38 | 15 | 3.0228 | 0.0012524 | STK19 | 0.0012524 | 0.00090602 | 0.0002857 |
| ENSG00000104213 | 8 | 17433942 | 17501580 | 603 | 54 | 2.9943 | 0.0013752 | PDGFRL | 0.0013752 | 0.0014658 | 0.0023925 |
| ENSG00000204371 | 6 | 31847536 | 31865464 | 54 | 19 | 2.9843 | 0.0014213 | EHMT2 | 0.0014213 | 0.00222 | 0.0021012 |
| ENSG00000006625 | 7 | 30536237 | 30591095 | 165 | 17 | 2.9757 | 0.0014617 | GGCT | 0.0014617 | 0.0031548 | 0.0059902 |
| ENSG00000174915 | 11 | 448268 | 491393 | 159 | 18 | 2.9595 | 0.0015408 | PTDSS2 | 0.0015408 | 0.0018935 | 0.0043513 |
| ENSG00000171368 | 5 | 660883 | 693510 | 200 | 24 | 2.9561 | 0.0015579 | TPPP | 0.0015579 | 0.0012893 | 0.00054357 |
| ENSG00000107159 | 9 | 35673853 | 35681156 | 20 | 7 | 2.9551 | 0.0015627 | CA9 | 0.0015627 | 0.0017675 | 0.0034667 |
| ENSG00000244731 | 6 | 31949801 | 31970458 | 5 | 3 | 2.953 | 0.0015736 | C4A | 0.0015736 | 0.0012617 | 0.00037287 |
| ENSG00000268434 | 19 | 46282695 | 46289231 | 10 | 4 | 2.9505 | 0.0015864 | AC011530.4 | 0.0015864 | 0.0048086 | 0.0015354 |
| ENSG00000198356 | 19 | 12847953 | 12859137 | 9 | 4 | 2.9465 | 0.0016072 | ASNA1 | 0.0016072 | 0.012424 | 0.0013627 |
| ENSG00000158623 | 7 | 130146089 | 130353598 | 264 | 20 | 2.9451 | 0.0016142 | COPG2 | 0.0016142 | 0.001574 | 0.0010651 |
| ENSG00000174197 | 15 | 41913422 | 42062141 | 355 | 22 | 2.9423 | 0.0016288 | MGA | 0.0016288 | 0.002735 | 0.002878 |
| ENSG00000153936 | 1 | 87380331 | 87602334 | 664 | 24 | 2.9404 | 0.0016389 | HS2ST1 | 0.0016389 | 0.0021554 | 0.0022098 |
| ENSG00000132004 | 19 | 12798867 | 12807457 | 28 | 2 | 2.9334 | 0.0016764 | FBXW9 | 0.0016764 | 0.0010422 | 0.00089774 |
| ENSG00000090534 | 3 | 184089723 | 184095932 | 25 | 8 | 2.9233 | 0.0017317 | THPO | 0.0017317 | 0.0028399 | 0.024263 |
| ENSG00000237452 | 19 | 46236509 | 46267792 | 47 | 8 | 2.9165 | 0.00177 | AC074212.3 | 0.00177 | 0.002349 | 0.0018362 |
| ENSG00000137135 | 9 | 35658872 | 35675863 | 35 | 8 | 2.8997 | 0.0018673 | ARHGEF39 | 0.0018673 | 0.002496 | 0.0098429 |
| ENSG00000129194 | 17 | 7491496 | 7493488 | 5 | 3 | 2.8978 | 0.001879 | SOX15 | 0.001879 | 0.023024 | 0.015907 |
| ENSG00000078687 | 17 | 76000249 | 76104916 | 260 | 23 | 2.8974 | 0.0018812 | TNRC6C | 0.0018812 | 0.0013552 | 0.0034935 |
| ENSG00000109787 | 4 | 38665817 | 38702663 | 102 | 25 | 2.8973 | 0.0018822 | KLF3 | 0.0018822 | 0.00038617 | 0.0006242 |
| ENSG00000128394 | 22 | 39436609 | 39449915 | 46 | 4 | 2.8955 | 0.001893 | APOBEC3F | 0.001893 | 0.001629 | 0.0014196 |
| ENSG00000097096 | 1 | 85622556 | 85666729 | 159 | 17 | 2.894 | 0.0019016 | SYDE2 | 0.0019016 | 0.0010491 | 0.0020329 |
| ENSG00000168477 | 6 | 32008931 | 32083111 | 200 | 43 | 2.8854 | 0.0019547 | TNXB | 0.0019547 | 0.00086861 | 0.00074298 |
| ENSG00000163053 | 2 | 230899698 | 230933715 | 138 | 19 | 2.885 | 0.0019571 | SLC16A14 | 0.0019571 | 0.0038208 | 0.0019683 |
| ENSG00000196735 | 6 | 32595956 | 32614839 | 1813 | 26 | 2.8821 | 0.0019749 | HLA-DQA1 | 0.0019749 | 0.0028464 | 0.0064216 |
| ENSG00000092820 | 6 | 159186773 | 159240444 | 187 | 24 | 2.8756 | 0.002016 | EZR | 0.002016 | 0.0032181 | 0.0092985 |
| ENSG00000183291 | 1 | 87328132 | 87380107 | 205 | 16 | 2.864 | 0.0020915 | SELENOF | 0.0020915 | 0.0028177 | 0.0030967 |
| ENSG00000100099 | 22 | 26839389 | 26879803 | 155 | 21 | 2.8604 | 0.0021158 | HPS4 | 0.0021158 | 0.0029296 | 0.0030391 |
| ENSG00000159884 | 9 | 35658301 | 35661508 | 10 | 3 | 2.8579 | 0.0021322 | CCDC107 | 0.0021322 | 0.0066703 | 0.0084688 |
| ENSG00000081386 | 9 | 99518147 | 99540411 | 24 | 11 | 2.8542 | 0.0021571 | ZNF510 | 0.0021571 | 0.0042127 | 0.0032814 |
| ENSG00000166441 | 11 | 8703958 | 8736306 | 94 | 18 | 2.8517 | 0.0021741 | RPL27A | 0.0021741 | 0.0031308 | 0.018489 |
| ENSG00000180233 | 7 | 30323923 | 30452118 | 230 | 39 | 2.8487 | 0.0021947 | ZNRF2 | 0.0021947 | 0.002235 | 0.0044105 |
| ENSG00000163093 | 2 | 170335688 | 170382432 | 160 | 23 | 2.8472 | 0.0022051 | BBS5 | 0.0022051 | 0.0019351 | 0.010699 |
| ENSG00000251569 | 2 | 170335990 | 170382432 | 160 | 23 | 2.8472 | 0.0022051 | RP11-724O16.1 | 0.0022051 | 0.00199 | 0.009635 |
| ENSG00000239474 | 2 | 170366212 | 170382772 | 42 | 13 | 2.837 | 0.0022771 | KLHL41 | 0.0022771 | 0.00042975 | 0.0028044 |
| ENSG00000204287 | 6 | 32407619 | 32412823 | 71 | 12 | 2.8365 | 0.0022808 | HLA-DRA | 0.0022808 | 0.0015958 | 0.0048256 |
| ENSG00000089009 | 12 | 112842994 | 112856642 | 13 | 6 | 2.8321 | 0.0023122 | RPL6 | 0.0023122 | 0.00085024 | 0.00059934 |
| ENSG00000111087 | 12 | 57853918 | 57866045 | 31 | 5 | 2.8297 | 0.0023293 | GLI1 | 0.0023293 | 0.0014663 | 0.0011293 |
| ENSG00000100888 | 14 | 21853353 | 21924285 | 131 | 16 | 2.8128 | 0.0024555 | CHD8 | 0.0024555 | 0.0027536 | 0.014484 |
| ENSG00000213265 | 7 | 130353488 | 130372268 | 47 | 11 | 2.8101 | 0.0024767 | TSGA13 | 0.0024767 | 0.004946 | 0.0027891 |
| ENSG00000170430 | 10 | 131265448 | 131566271 | 1437 | 63 | 2.7954 | 0.0025921 | MGMT | 0.0025921 | 0.0068456 | 0.0096197 |
| ENSG00000197457 | 20 | 62271061 | 62284780 | 54 | 11 | 2.7902 | 0.0026339 | STMN3 | 0.0026339 | 0.0093977 | 0.034842 |
| ENSG00000176198 | 14 | 20710898 | 20711979 | 3 | 2 | 2.7827 | 0.0026952 | OR11H4 | 0.0026952 | 0.0029083 | 0.0079797 |
| ENSG00000179930 | 1 | 182023705 | 182030847 | 8 | 2 | 2.7754 | 0.0027567 | ZNF648 | 0.0027567 | 0.00011689 | 0.00019088 |
| ENSG00000180176 | 11 | 2185159 | 2193107 | 14 | 7 | 2.7713 | 0.0027919 | TH | 0.0027919 | 0.16631 | 0.49432 |
| ENSG00000166508 | 7 | 99690351 | 99699563 | 28 | 6 | 2.7671 | 0.0028277 | MCM7 | 0.0028277 | 0.0052031 | 0.0082252 |
| ENSG00000101194 | 20 | 61584052 | 61599949 | 66 | 16 | 2.7604 | 0.0028862 | SLC17A9 | 0.0028862 | 0.0028609 | 0.0069685 |
| ENSG00000136048 | 12 | 102271129 | 102405908 | 621 | 59 | 2.7575 | 0.0029122 | DRAM1 | 0.0029122 | 0.0038557 | 0.0031737 |
| ENSG00000178425 | 6 | 116422012 | 116570660 | 412 | 28 | 2.7557 | 0.0029286 | NT5DC1 | 0.0029286 | 0.002763 | 0.0038618 |
| ENSG00000104774 | 19 | 12757325 | 12777556 | 40 | 10 | 2.754 | 0.0029437 | MAN2B1 | 0.0029437 | 0.0029736 | 0.00091199 |
| ENSG00000111412 | 12 | 117153593 | 117175875 | 83 | 18 | 2.7496 | 0.0029832 | C12orf49 | 0.0029832 | 0.022564 | 0.011806 |
| ENSG00000188818 | 5 | 710471 | 851101 | 441 | 34 | 2.7449 | 0.003026 | ZDHHC11 | 0.003026 | 0.0029114 | 0.000836 |
| ENSG00000163882 | 3 | 184079506 | 184086384 | 21 | 6 | 2.7358 | 0.0031115 | POLR2H | 0.0031115 | 0.0016564 | 0.028405 |
| ENSG00000162063 | 16 | 2479395 | 2508855 | 103 | 6 | 2.7284 | 0.0031825 | CCNF | 0.0031825 | 0.0034646 | 0.0035306 |
| ENSG00000144362 | 2 | 170550975 | 170558218 | 16 | 4 | 2.7201 | 0.0032627 | PHOSPHO2 | 0.0032627 | 0.0022475 | 0.00085413 |
| ENSG00000119335 | 9 | 131445703 | 131458679 | 10 | 5 | 2.72 | 0.0032643 | SET | 0.0032643 | 0.063009 | 0.1257 |
| ENSG00000152818 | 6 | 144606837 | 145174170 | 1393 | 84 | 2.7155 | 0.0033085 | UTRN | 0.0033085 | 0.0042276 | 0.0034391 |
| ENSG00000121101 | 17 | 56634039 | 56769416 | 290 | 22 | 2.7072 | 0.0033929 | TEX14 | 0.0033929 | 0.003009 | 0.0033175 |
| ENSG00000188152 | 9 | 99690592 | 99702535 | 13 | 5 | 2.7055 | 0.0034099 | NUTM2G | 0.0034099 | 0.00097461 | 0.00073797 |
| ENSG00000177414 | 1 | 64669310 | 64733051 | 183 | 18 | 2.701 | 0.0034562 | UBE2U | 0.0034562 | 0.00047875 | 0.00082752 |
| ENSG00000163453 | 4 | 57896939 | 57976551 | 493 | 48 | 2.6923 | 0.0035476 | IGFBP7 | 0.0035476 | 0.0030019 | 0.0020549 |
| ENSG00000111596 | 12 | 70636774 | 70748773 | 353 | 21 | 2.6895 | 0.0035775 | CNOT2 | 0.0035775 | 0.00567 | 0.0065002 |
| ENSG00000177045 | 19 | 46268043 | 46272484 | 6 | 3 | 2.6837 | 0.0036405 | SIX5 | 0.0036405 | 0.0036268 | 0.0028085 |
| ENSG00000130813 | 19 | 10196798 | 10203928 | 16 | 7 | 2.678 | 0.0037034 | C19orf66 | 0.0037034 | 0.014077 | 0.094702 |
| ENSG00000154479 | 2 | 170501935 | 170550943 | 247 | 15 | 2.6764 | 0.0037211 | CCDC173 | 0.0037211 | 0.0032035 | 0.0024103 |
| ENSG00000235156 | 3 | 99904531 | 99913035 | 24 | 5 | 2.6737 | 0.0037512 | TMEM30C | 0.0037512 | 0.0065811 | 0.019465 |

| | | | | | | | | | | | |
|---|---|---|---|---|---|---|---|---|---|---|---|
| ENSG00000151466 | 4 | 129786076 | 130014764 | 549 | 24 | 2.673 | 0.0037591 | SCLT1 | 0.0037591 | 0.003266 | 0.0040204 |
| ENSG00000084623 | 1 | 32687529 | 32697205 | 14 | 3 | 2.672 | 0.00377 | EIF3I | 0.00377 | 0.0039651 | 0.003244 |
| ENSG00000164758 | 8 | 118532952 | 118552501 | 54 | 11 | 2.6715 | 0.0037757 | MED30 | 0.0037757 | 0.0098794 | 0.016006 |
| ENSG00000137877 | 15 | 42140345 | 42186275 | 142 | 27 | 2.6672 | 0.003824 | SPTBN5 | 0.003824 | 0.0017971 | 0.0069966 |
| ENSG00000164951 | 8 | 94870035 | 94938294 | 217 | 19 | 2.6621 | 0.0038832 | PDP1 | 0.0038832 | 0.0040486 | 0.014689 |
| ENSG00000167858 | 17 | 6692452 | 6735080 | 213 | 24 | 2.6618 | 0.0038865 | TEKT1 | 0.0038865 | 0.0033431 | 0.012486 |
| ENSG00000171488 | 1 | 90098631 | 90235462 | 295 | 16 | 2.6607 | 0.0038986 | LRRC8C | 0.0038986 | 0.0020247 | 0.0011231 |
| ENSG00000269242 | 19 | 12754645 | 12759211 | 8 | 3 | 2.6567 | 0.0039454 | CTD-2192J16.22 | 0.0039454 | 0.011763 | 0.0056278 |
| ENSG00000066117 | 12 | 50478755 | 50494495 | 25 | 5 | 2.653 | 0.0039892 | SMARCD1 | 0.0039892 | 0.0082864 | 0.035531 |
| ENSG00000108395 | 17 | 57059999 | 57184282 | 180 | 23 | 2.6522 | 0.0039987 | TRIM37 | 0.0039987 | 0.0022949 | 0.0014135 |
| ENSG00000047315 | 4 | 57843888 | 57897334 | 186 | 28 | 2.6504 | 0.0040199 | POLR2B | 0.0040199 | 0.0021061 | 0.0048983 |
| ENSG00000144644 | 3 | 30767692 | 30936257 | 766 | 41 | 2.6461 | 0.0040708 | GADL1 | 0.0040708 | 0.0052275 | 0.0043665 |
| ENSG00000164796 | 8 | 113235157 | 114449328 | 3469 | 72 | 2.6412 | 0.0041304 | CSMD3 | 0.0041304 | 0.0059588 | 0.0046849 |
| ENSG00000204310 | 6 | 32135989 | 32145873 | 19 | 8 | 2.6329 | 0.004233 | AGPAT1 | 0.004233 | 0.070036 | 0.034056 |
| ENSG00000181733 | 19 | 8841316 | 8842371 | 2 | 1 | 2.623 | 0.0043583 | OR2Z1 | 0.0043583 | 0.00033398 | 0.00016386 |
| ENSG00000151882 | 5 | 43376747 | 43412493 | 78 | 10 | 2.6201 | 0.0043955 | CCL28 | 0.0043955 | 0.016752 | 0.05036 |
| ENSG00000111785 | 12 | 107168373 | 107283090 | 193 | 26 | 2.618 | 0.0044218 | RIC8B | 0.0044218 | 0.0054154 | 0.0081265 |
| ENSG00000197142 | 10 | 114133776 | 114188138 | 69 | 15 | 2.6137 | 0.0044781 | ACSL5 | 0.0044781 | 0.0035026 | 0.0095989 |
| ENSG00000227839 | 1 | 50459990 | 50461874 | 4 | 2 | 2.6124 | 0.0044955 | AL645730.2 | 0.0044955 | 0.033927 | 0.050385 |
| ENSG00000100104 | 22 | 26879843 | 26890624 | 28 | 8 | 2.611 | 0.0045141 | SRRD | 0.0045141 | 0.0024009 | 0.0038974 |
| ENSG00000176407 | 2 | 85198216 | 85286595 | 196 | 19 | 2.6073 | 0.0045631 | KCMF1 | 0.0045631 | 0.0079433 | 0.011661 |
| ENSG00000151353 | 2 | 667335 | 677439 | 26 | 5 | 2.6048 | 0.0045968 | TMEM18 | 0.0045968 | 0.0034569 | 0.0082528 |
| ENSG00000188909 | 11 | 122848278 | 122852428 | 10 | 3 | 2.6024 | 0.0046293 | BSX | 0.0046293 | 0.0021629 | 0.003977 |
| ENSG00000161956 | 17 | 7465192 | 7475287 | 26 | 9 | 2.602 | 0.0046338 | SENP3 | 0.0046338 | 0.0029599 | 0.018744 |
| ENSG00000174083 | 17 | 8706041 | 8770994 | 183 | 29 | 2.5986 | 0.0046804 | PIK3R6 | 0.0046804 | 0.045843 | 0.09404 |
| ENSG00000124164 | 20 | 56964178 | 57026157 | 216 | 22 | 2.5982 | 0.0046854 | VAPB | 0.0046854 | 0.009035 | 0.0066591 |
| ENSG00000257923 | 7 | 101458959 | 101927249 | 1427 | 76 | 2.5919 | 0.004773 | CUX1 | 0.004773 | 0.0043323 | 0.004073 |
| ENSG00000175193 | 3 | 183547173 | 183602721 | 141 | 20 | 2.5901 | 0.004798 | PARL | 0.004798 | 0.0022348 | 0.0020393 |
| ENSG00000204740 | 10 | 19492779 | 20079330 | 2653 | 95 | 2.588 | 0.0048267 | MALRD1 | 0.0048267 | 0.0044663 | 0.0051052 |
| ENSG00000196597 | 9 | 99578754 | 99637905 | 70 | 19 | 2.587 | 0.0048403 | ZNF782 | 0.0048403 | 0.004446 | 0.0021144 |
| ENSG00000215790 | 1 | 1656277 | 1677431 | 102 | 12 | 2.581 | 0.0049257 | SLC35E2 | 0.0049257 | 0.012349 | 0.025645 |
| ENSG00000113300 | 5 | 179921412 | 180005405 | 425 | 19 | 2.5806 | 0.0049312 | CNOT6 | 0.0049312 | 0.0076913 | 0.0082407 |
| ENSG00000083635 | 13 | 45513384 | 45563618 | 76 | 15 | 2.5796 | 0.0049464 | NUFIP1 | 0.0049464 | 0.018328 | 0.072492 |
| ENSG00000089127 | 12 | 113344582 | 113369990 | 111 | 4 | 2.5791 | 0.0049528 | OAS1 | 0.0049528 | 0.0051015 | 0.0054683 |
| ENSG00000138823 | 4 | 100484918 | 100545156 | 254 | 26 | 2.5787 | 0.0049585 | MTTP | 0.0049585 | 0.0012888 | 0.0020228 |
| ENSG00000065491 | 6 | 37225548 | 37300746 | 174 | 14 | 2.578 | 0.0049683 | TBC1D22B | 0.0049683 | 0.0030711 | 0.0023115 |
| ENSG00000121775 | 1 | 32537632 | 32568467 | 52 | 7 | 2.5779 | 0.0049707 | TMEM39B | 0.0049707 | 0.0049864 | 0.0077667 |
| ENSG00000108387 | 17 | 56597611 | 56618179 | 27 | 9 | 2.5749 | 0.0050131 | SEPTIN4 | 0.0050131 | 0.0059095 | 0.0096082 |
| ENSG00000089022 | 12 | 112279782 | 112334343 | 37 | 10 | 2.5688 | 0.0051028 | MAPKAPK5 | 0.0051028 | 0.0040876 | 0.0034212 |
| ENSG00000144119 | 2 | 119913819 | 119916465 | 10 | 3 | 2.5669 | 0.0051307 | C1QL2 | 0.0051307 | 0.0029561 | 0.0037127 |
| ENSG00000204351 | 6 | 31926857 | 31937532 | 28 | 12 | 2.5648 | 0.0051619 | SKIV2L | 0.0051619 | 0.00066322 | 0.0002551 |
| ENSG00000111275 | 12 | 112204691 | 112247782 | 25 | 11 | 2.5585 | 0.0052561 | ALDH2 | 0.0052561 | 0.0027629 | 0.0056101 |
| ENSG00000243811 | 22 | 39410368 | 39429281 | 69 | 11 | 2.5581 | 0.0052624 | APOBEC3D | 0.0052624 | 0.0021011 | 0.0014232 |
| ENSG00000143376 | 1 | 151584541 | 151671567 | 169 | 18 | 2.5574 | 0.0052733 | SNX27 | 0.0052733 | 0.0052072 | 0.0057869 |
| ENSG00000133627 | 7 | 152456834 | 152552463 | 246 | 25 | 2.5563 | 0.0052894 | ACTR3B | 0.0052894 | 0.0055858 | 0.0039336 |
| ENSG00000181804 | 3 | 142984064 | 143567373 | 2162 | 162 | 2.556 | 0.0052947 | SLC9A9 | 0.0052947 | 0.0048103 | 0.0056016 |
| ENSG00000267561 | 1 | 87458692 | 87634884 | 491 | 33 | 2.5515 | 0.0053629 | RP5-1052I5.2 | 0.0053629 | 0.0033056 | 0.0032833 |
| ENSG00000184162 | 19 | 19312225 | 19314233 | 1 | 1 | 2.5497 | 0.005391 | NR2C2AP | 0.005391 | 0.8018 | 0.84553 |
| ENSG00000254106 | 5 | 109218883 | 109220452 | 10 | 5 | 2.5461 | 0.0054467 | AC011366.3 | 0.0054467 | 0.025663 | 0.016525 |
| ENSG00000224130 | 11 | 58672897 | 58676071 | 10 | 1 | 2.5439 | 0.0054813 | AP001652.1 | 0.0054813 | 0.0044431 | 0.025302 |
| ENSG00000171720 | 5 | 141000443 | 141016437 | 34 | 10 | 2.5424 | 0.0055045 | HDAC3 | 0.0055045 | 0.017298 | 0.070474 |
| ENSG00000158270 | 18 | 319361 | 500722 | 1030 | 78 | 2.5411 | 0.0055246 | COLEC12 | 0.0055246 | 0.0087594 | 0.01248 |
| ENSG00000198900 | 20 | 39657458 | 39753127 | 170 | 28 | 2.5391 | 0.0055563 | TOP1 | 0.0055563 | 0.0087931 | 0.016974 |
| ENSG00000070610 | 9 | 35736863 | 35749983 | 10 | 5 | 2.5361 | 0.0056049 | GBA2 | 0.0056049 | 0.0068663 | 0.01005 |
| ENSG00000183023 | 2 | 40324410 | 40838193 | 2181 | 118 | 2.5349 | 0.0056234 | SLC8A1 | 0.0056234 | 0.0051721 | 0.0066152 |
| ENSG00000163564 | 1 | 158900586 | 158946844 | 140 | 17 | 2.533 | 0.0056548 | PYHIN1 | 0.0056548 | 0.028771 | 0.16583 |
| ENSG00000217236 | 2 | 175199674 | 175203220 | 16 | 7 | 2.5262 | 0.0057645 | SP9 | 0.0057645 | 0.046436 | 0.060732 |
| ENSG00000198515 | 4 | 47937994 | 48018689 | 355 | 19 | 2.5239 | 0.0058025 | CNGA1 | 0.0058025 | 0.010315 | 0.016282 |
| ENSG00000105371 | 19 | 10397643 | 10399198 | 1 | 1 | 2.5223 | 0.00583 | ICAM4 | 0.00583 | 0.12627 | 0.083381 |
| ENSG00000136527 | 3 | 185633694 | 185655924 | 30 | 7 | 2.5159 | 0.0059359 | TRA2B | 0.0059359 | 0.0059831 | 0.0053126 |
| ENSG00000172738 | 6 | 37179956 | 37225931 | 67 | 14 | 2.5128 | 0.0059882 | TMEM217 | 0.0059882 | 0.0041112 | 0.0049897 |
| ENSG00000179979 | 4 | 1385340 | 1389780 | 57 | 12 | 2.5103 | 0.0060309 | CRIPAK | 0.0060309 | 0.02563 | 0.060579 |
| ENSG00000213676 | 6 | 32065953 | 32096030 | 82 | 28 | 2.5057 | 0.0061107 | ATF6B | 0.0061107 | 0.0092144 | 0.0075256 |
| ENSG00000197283 | 6 | 33387847 | 33421466 | 69 | 19 | 2.5042 | 0.0061372 | SYNGAP1 | 0.0061372 | 0.01487 | 0.04881 |
| ENSG00000119185 | 2 | 9543604 | 9563676 | 60 | 9 | 2.5039 | 0.0061412 | ITGB1BP1 | 0.0061412 | 0.026989 | 0.025088 |
| ENSG00000159387 | 16 | 55357672 | 55364672 | 40 | 7 | 2.5033 | 0.0061519 | IRX6 | 0.0061519 | 0.0091091 | 0.0097156 |
| ENSG00000125872 | 20 | 6021424 | 6034695 | 52 | 8 | 2.4997 | 0.006215 | LRRN4 | 0.006215 | 0.0061631 | 0.009598 |
| ENSG00000091138 | 7 | 107405912 | 107443670 | 113 | 25 | 2.4989 | 0.0062283 | SLC26A3 | 0.0062283 | 0.0035635 | 0.000074897 |
| ENSG00000108384 | 17 | 56769934 | 56811703 | 91 | 15 | 2.4974 | 0.0062554 | RAD51C | 0.0062554 | 0.0026556 | 0.0026516 |
| ENSG00000211460 | 2 | 122494679 | 122525429 | 51 | 10 | 2.4936 | 0.0063221 | TSN | 0.0063221 | 0.016258 | 0.058867 |
| ENSG00000267885 | 1 | 33439268 | 33439642 | 1 | 1 | 2.4913 | 0.006363 | FKSG48 | 0.006363 | 0.027788 | 0.078539 |
| ENSG00000139687 | 13 | 48877887 | 49056122 | 430 | 21 | 2.4913 | 0.0063639 | RB1 | 0.0063639 | 0.0070318 | 0.0067999 |
| ENSG00000005379 | 17 | 56378592 | 56406152 | 81 | 15 | 2.4908 | 0.0063726 | BZRAP1 | 0.0063726 | 0.020035 | 0.02152 |

| | | | | | | | | | | | |
|---|---|---|---|---|---|---|---|---|---|---|---|
| ENSG00000160051 | 1 | 32671236 | 32674288 | 5 | 1 | 2.4875 | 0.0064326 | IQCC | 0.0064326 | 0.004649 | 0.0037846 |
| ENSG00000167658 | 19 | 3976054 | 3985467 | 30 | 9 | 2.4873 | 0.0064365 | EEF2 | 0.0064365 | 0.0067092 | 0.021949 |
| ENSG00000234972 | 17 | 34580902 | 34757050 | 40 | 13 | 2.4741 | 0.0066794 | TBC1D3C | 0.0066794 | 0.0029423 | 0.0087708 |
| ENSG00000186660 | 11 | 58346584 | 58388515 | 95 | 15 | 2.4681 | 0.0067918 | ZFP91 | 0.0067918 | 0.0053031 | 0.0056252 |
| ENSG00000070047 | 11 | 576486 | 612222 | 129 | 18 | 2.4677 | 0.0067991 | PHRF1 | 0.0067991 | 0.014946 | 0.0040425 |
| ENSG00000239305 | 2 | 86830516 | 86850989 | 57 | 11 | 2.4641 | 0.0068677 | RNF103 | 0.0068677 | 0.003983 | 0.009258 |
| ENSG00000130812 | 19 | 10203014 | 10213472 | 21 | 8 | 2.4628 | 0.0068921 | ANGPTL6 | 0.0068921 | 0.001968 | 0.080441 |
| ENSG00000269700 | 10 | 52152766 | 52154321 | 5 | 3 | 2.4588 | 0.0069693 | AC069547.2 | 0.0069693 | 0.20129 | 0.086728 |
| ENSG00000163683 | 4 | 39547950 | 39640710 | 299 | 28 | 2.458 | 0.0069855 | SMIM14 | 0.0069855 | 0.010567 | 0.005331 |
| ENSG00000138050 | 2 | 39963200 | 40006407 | 123 | 11 | 2.4514 | 0.0071152 | THUMPD2 | 0.0071152 | 0.0047551 | 0.0066623 |
| ENSG00000185522 | 11 | 554855 | 560779 | 32 | 10 | 2.4494 | 0.0071542 | C11orf35 | 0.0071542 | 0.001285 | 0.00079131 |
| ENSG00000129244 | 17 | 7549945 | 7561086 | 40 | 7 | 2.4463 | 0.0072162 | ATP1B2 | 0.0072162 | 0.019295 | 0.049624 |
| ENSG00000084652 | 1 | 32645287 | 32663886 | 38 | 9 | 2.4459 | 0.0072254 | TXLNA | 0.0072254 | 0.0042793 | 0.0046726 |
| ENSG00000117155 | 1 | 85109390 | 85156486 | 118 | 15 | 2.4453 | 0.0072367 | SSX2IP | 0.0072367 | 0.012202 | 0.012332 |
| ENSG00000123329 | 12 | 57866038 | 57882597 | 43 | 10 | 2.444 | 0.0072634 | ARHGAP9 | 0.0072634 | 0.0024287 | 0.0055322 |
| ENSG00000204356 | 6 | 31919864 | 31926887 | 21 | 11 | 2.4431 | 0.0072806 | NELFE | 0.0072806 | 0.00041162 | 0.00032154 |
| ENSG00000138398 | 2 | 170440850 | 170497916 | 261 | 15 | 2.4341 | 0.0074645 | PPIG | 0.0074645 | 0.0077147 | 0.0049931 |
| ENSG00000183655 | 15 | 86302554 | 86338261 | 151 | 26 | 2.4279 | 0.0075928 | KLHL25 | 0.0075928 | 0.0044309 | 0.0068161 |
| ENSG00000110324 | 11 | 117857063 | 117872196 | 42 | 8 | 2.4268 | 0.0076169 | IL10RA | 0.0076169 | 0.040773 | 0.18396 |
| ENSG00000198026 | 20 | 44577292 | 44600833 | 82 | 18 | 2.4266 | 0.0076201 | ZNF335 | 0.0076201 | 0.014188 | 0.010039 |
| ENSG00000255073 | 11 | 58346645 | 58392112 | 102 | 15 | 2.4232 | 0.0076916 | ZFP91-CNTF | 0.0076916 | 0.0055051 | 0.0062894 |
| ENSG00000119866 | 2 | 60678302 | 60780702 | 189 | 38 | 2.4165 | 0.0078352 | BCL11A | 0.0078352 | 0.010367 | 0.017297 |
| ENSG00000011028 | 17 | 60704762 | 60770958 | 139 | 22 | 2.4139 | 0.0078923 | MRC2 | 0.0078923 | 0.0094816 | 0.014173 |
| ENSG00000125551 | 2 | 88047606 | 88058312 | 1 | 1 | 2.4133 | 0.007905 | PLGLB2 | 0.007905 | 0.013582 | 0.019232 |
| ENSG00000134640 | 11 | 92702886 | 92718232 | 46 | 9 | 2.4104 | 0.0079673 | MTNR1B | 0.0079673 | 0.0043737 | 0.02566 |
| ENSG00000176219 | 14 | 20691791 | 20692964 | 9 | 2 | 2.4096 | 0.0079843 | OR11H6 | 0.0079843 | 0.014069 | 0.015137 |
| ENSG00000133636 | 12 | 86268073 | 86276770 | 32 | 4 | 2.4074 | 0.0080322 | NTS | 0.0080322 | 0.0081041 | 0.0050258 |
| ENSG00000111783 | 12 | 106976685 | 107156581 | 403 | 32 | 2.4072 | 0.0080368 | RFX4 | 0.0080368 | 0.011034 | 0.0087231 |
| ENSG00000121743 | 13 | 20712394 | 20735188 | 101 | 17 | 2.4065 | 0.0080538 | GJA3 | 0.0080538 | 0.013778 | 0.081223 |
| ENSG00000213420 | 7 | 99767229 | 99774995 | 14 | 6 | 2.4054 | 0.0080774 | GPC2 | 0.0080774 | 0.027445 | 0.039544 |
| ENSG00000068650 | 13 | 113344643 | 113541482 | 769 | 57 | 2.4042 | 0.0081047 | ATP11A | 0.0081047 | 0.012037 | 0.0071896 |
| ENSG00000107317 | 9 | 139871956 | 139879887 | 22 | 6 | 2.4027 | 0.0081368 | PTGDS | 0.0081368 | 0.012251 | 0.016324 |
| ENSG00000000938 | 1 | 27938575 | 27961788 | 55 | 11 | 2.4018 | 0.0081575 | FGR | 0.0081575 | 0.01388 | 0.055868 |
| ENSG00000177938 | 12 | 18891045 | 18892121 | 1 | 1 | 2.4007 | 0.008181 | CAPZA3 | 0.008181 | 0.052836 | 0.019122 |
| ENSG00000111670 | 12 | 102139275 | 102224716 | 188 | 27 | 2.3996 | 0.0082064 | GNPTAB | 0.0082064 | 0.0065175 | 0.006472 |
| ENSG00000128578 | 7 | 129074274 | 129128240 | 134 | 14 | 2.3995 | 0.0082086 | STRIP2 | 0.0082086 | 0.007981 | 0.016698 |
| ENSG00000105963 | 7 | 937540 | 995043 | 354 | 50 | 2.398 | 0.0082432 | ADAP1 | 0.0082432 | 0.0092532 | 0.013835 |
| ENSG00000156970 | 15 | 40453224 | 40513337 | 107 | 11 | 2.3949 | 0.0083117 | BUB1B | 0.0083117 | 0.0039213 | 0.012461 |
| ENSG00000105205 | 19 | 40221890 | 40228668 | 63 | 4 | 2.3926 | 0.008365 | CLC | 0.008365 | 0.015741 | 0.040645 |
| ENSG00000168481 | 8 | 22004338 | 22014597 | 25 | 4 | 2.3926 | 0.0083653 | LGI3 | 0.0083653 | 0.033284 | 0.17787 |
| ENSG00000198774 | 12 | 86198331 | 86230348 | 117 | 13 | 2.3922 | 0.0083749 | RASSF9 | 0.0083749 | 0.014481 | 0.0089605 |
| ENSG00000175274 | 11 | 44907454 | 44972840 | 202 | 44 | 2.3915 | 0.0083887 | TP53I11 | 0.0083887 | 0.0094408 | 0.0524 |
| ENSG00000107623 | 10 | 48425815 | 48438976 | 40 | 10 | 2.3908 | 0.0084069 | GDF10 | 0.0084069 | 0.031163 | 0.021618 |
| ENSG00000185800 | 19 | 46286205 | 46296060 | 20 | 9 | 2.3902 | 0.0084191 | DMWD | 0.0084191 | 0.0019014 | 0.002238 |
| ENSG00000214402 | 9 | 139876356 | 139880862 | 14 | 3 | 2.3897 | 0.0084303 | LCNL1 | 0.0084303 | 0.0064273 | 0.01532 |
| ENSG00000068383 | 10 | 134351324 | 134596979 | 903 | 38 | 2.3885 | 0.0084578 | INPP5A | 0.0084578 | 0.0058017 | 0.0035527 |
| ENSG00000107929 | 10 | 855484 | 977564 | 326 | 16 | 2.3811 | 0.0086304 | LARP4B | 0.0086304 | 0.0097775 | 0.0072937 |
| ENSG00000113732 | 5 | 172410760 | 172462448 | 124 | 26 | 2.3804 | 0.008648 | ATP6V0E1 | 0.008648 | 0.029889 | 0.028045 |
| ENSG00000139354 | 12 | 100967461 | 101022064 | 143 | 18 | 2.3791 | 0.0086769 | GAS2L3 | 0.0086769 | 0.011867 | 0.030923 |
| ENSG00000164938 | 8 | 95938200 | 95961639 | 75 | 11 | 2.379 | 0.0086799 | TP53INP1 | 0.0086799 | 0.0098753 | 0.016338 |
| ENSG00000150433 | 11 | 124966398 | 124981659 | 65 | 15 | 2.3766 | 0.0087374 | TMEM218 | 0.0087374 | 0.027805 | 0.063571 |
| ENSG00000136161 | 13 | 49063095 | 49107369 | 86 | 12 | 2.3765 | 0.00874 | RCBTB2 | 0.00874 | 0.006898 | 0.0075165 |
| ENSG00000151702 | 11 | 128556430 | 128683162 | 454 | 67 | 2.3763 | 0.0087447 | FLI1 | 0.0087447 | 0.024759 | 0.014202 |
| ENSG00000101180 | 20 | 60790026 | 60795323 | 8 | 5 | 2.3753 | 0.008767 | HRH3 | 0.008767 | 0.47106 | 0.36681 |
| ENSG00000188042 | 2 | 235401685 | 235405697 | 13 | 4 | 2.3747 | 0.0087826 | ARL4C | 0.0087826 | 0.014785 | 0.10883 |
| ENSG00000196954 | 11 | 104813593 | 104840163 | 71 | 13 | 2.3734 | 0.0088137 | CASP4 | 0.0088137 | 0.011248 | 0.01694 |
| ENSG00000141485 | 17 | 6588032 | 6616886 | 140 | 28 | 2.3706 | 0.0088798 | SLC13A5 | 0.0088798 | 0.014017 | 0.018472 |
| ENSG00000123154 | 19 | 12777614 | 12786646 | 16 | 8 | 2.3699 | 0.008897 | WDR83 | 0.008897 | 0.00065558 | 0.00020229 |
| ENSG00000111331 | 12 | 113376157 | 113411054 | 122 | 13 | 2.3688 | 0.0089225 | OAS3 | 0.0089225 | 0.0060195 | 0.0055964 |
| ENSG00000161960 | 17 | 7476024 | 7482323 | 21 | 7 | 2.368 | 0.0089414 | EIF4A1 | 0.0089414 | 0.0052021 | 0.010735 |
| ENSG00000011485 | 19 | 46850251 | 46896238 | 105 | 17 | 2.3668 | 0.0089705 | PPP5C | 0.0089705 | 0.013803 | 0.079805 |
| ENSG00000169129 | 10 | 116054583 | 116164515 | 320 | 54 | 2.3665 | 0.0089795 | AFAP1L2 | 0.0089795 | 0.018222 | 0.041866 |
| ENSG00000145012 | 3 | 187871072 | 188608460 | 2501 | 226 | 2.3635 | 0.0090519 | LPP | 0.0090519 | 0.031028 | 0.066978 |
| ENSG00000005194 | 16 | 57462081 | 57481440 | 77 | 16 | 2.3617 | 0.0090947 | CIAPIN1 | 0.0090947 | 0.0012788 | 0.0069595 |
| ENSG00000104936 | 19 | 46272975 | 46285810 | 30 | 4 | 2.3591 | 0.0091589 | DMPK | 0.0091589 | 0.0032777 | 0.0014621 |
| ENSG00000268885 | 5 | 667759 | 668839 | 18 | 7 | 2.3541 | 0.0092849 | AC026740.1 | 0.0092849 | 0.0012702 | 0.0017826 |
| ENSG00000242384 | 17 | 34581086 | 34808061 | 67 | 24 | 2.35 | 0.009386 | TBC1D3H | 0.009386 | 0.054596 | 0.1192 |
| ENSG00000198755 | 6 | 35436185 | 35438562 | 9 | 3 | 2.3498 | 0.0093919 | RPL10A | 0.0093919 | 0.00042965 | 0.0007271 |
| ENSG00000089737 | 14 | 94517266 | 94547591 | 107 | 9 | 2.3491 | 0.0094106 | DDX24 | 0.0094106 | 0.014175 | 0.043924 |
| ENSG00000151092 | 3 | 25760435 | 25831530 | 140 | 9 | 2.3488 | 0.0094162 | NGLY1 | 0.0094162 | 0.011148 | 0.011965 |
| ENSG00000138660 | 4 | 113152893 | 113191203 | 71 | 11 | 2.3461 | 0.0094861 | AP1AR | 0.0094861 | 0.015415 | 0.023617 |
| ENSG00000262039 | 17 | 47448102 | 47554350 | 355 | 51 | 2.3423 | 0.0095827 | RP11-81K2.1 | 0.0095827 | 0.0018632 | 0.0017358 |
| ENSG00000143436 | 1 | 151732119 | 151736040 | 13 | 3 | 2.3413 | 0.009609 | MRPL9 | 0.009609 | 0.0071051 | 0.023706 |

| | | | | | | | | | | | |
|---|---|---|---|---|---|---|---|---|---|---|---|
| ENSG00000130427 | 7 | 100318423 | 100321323 | 7 | 4 | 2.34 | 0.0096427 | EPO | 0.0096427 | 0.19289 | 0.2399 |
| ENSG00000240230 | 7 | 938415 | 1015235 | 475 | 63 | 2.338 | 0.0096925 | COX19 | 0.0096925 | 0.0095116 | 0.015723 |
| ENSG00000121073 | 17 | 47778305 | 47786376 | 13 | 6 | 2.3348 | 0.0097771 | SLC35B1 | 0.0097771 | 0.0081537 | 0.036742 |
| ENSG00000066230 | 5 | 473425 | 524447 | 335 | 40 | 2.3339 | 0.0098006 | SLC9A3 | 0.0098006 | 0.0060466 | 0.0078528 |
| ENSG00000151468 | 10 | 12938627 | 13141652 | 1100 | 91 | 2.3332 | 0.0098195 | CCDC3 | 0.0098195 | 0.0096743 | 0.0072415 |
| ENSG00000083097 | 6 | 83777385 | 83881069 | 184 | 15 | 2.3325 | 0.0098374 | DOPEY1 | 0.0098374 | 0.0077031 | 0.0047028 |
| ENSG00000068078 | 4 | 1795034 | 1810599 | 56 | 13 | 2.3278 | 0.0099602 | FGFR3 | 0.0099602 | 0.011143 | 0.015789 |
| ENSG00000171759 | 12 | 103230663 | 103352188 | 469 | 30 | 2.3261 | 0.010006 | PAH | 0.010006 | 0.025224 | 0.032149 |
| ENSG00000177051 | 19 | 46213887 | 46234162 | 19 | 8 | 2.3246 | 0.010047 | FBXO46 | 0.010047 | 0.014716 | 0.0046881 |
| ENSG00000214866 | 2 | 3751453 | 3836122 | 381 | 54 | 2.323 | 0.010089 | DCDC2C | 0.010089 | 0.0045301 | 0.0049609 |
| ENSG00000070761 | 16 | 58147496 | 58163354 | 55 | 7 | 2.3222 | 0.01011 | C16orf80 | 0.01011 | 0.0058363 | 0.0065058 |
| ENSG00000196950 | 2 | 196440701 | 196602426 | 431 | 33 | 2.3167 | 0.01026 | SLC39A10 | 0.01026 | 0.006915 | 0.013179 |
| ENSG00000160055 | 1 | 32680073 | 32687972 | 14 | 6 | 2.3137 | 0.010342 | TMEM234 | 0.010342 | 0.0038197 | 0.0038054 |
| ENSG00000073417 | 15 | 85523671 | 85682376 | 473 | 29 | 2.3118 | 0.010395 | PDE8A | 0.010395 | 0.0057998 | 0.0023784 |
| ENSG00000127989 | 7 | 91321323 | 91510034 | 367 | 22 | 2.3114 | 0.010406 | MTERF | 0.010406 | 0.013976 | 0.013568 |
| ENSG00000023191 | 11 | 494512 | 507300 | 82 | 11 | 2.3096 | 0.010456 | RNH1 | 0.010456 | 0.0053491 | 0.0019924 |
| ENSG00000159289 | 15 | 74362198 | 74374891 | 10 | 6 | 2.3066 | 0.010537 | GOLGA6A | 0.010537 | 0.12694 | 0.21231 |
| ENSG00000267467 | 19 | 45445495 | 45452820 | 33 | 6 | 2.3055 | 0.010569 | APOC4 | 0.010569 | 0.0091086 | 0.0086725 |
| ENSG00000224916 | 19 | 45445495 | 45452822 | 33 | 6 | 2.3055 | 0.010569 | APOC4-APOC2 | 0.010569 | 0.0091086 | 0.0086725 |
| ENSG00000204291 | 9 | 101705461 | 101833069 | 383 | 40 | 2.3045 | 0.010597 | COL15A1 | 0.010597 | 0.0069605 | 0.0078859 |
| ENSG00000111142 | 12 | 95867296 | 95909615 | 164 | 18 | 2.3039 | 0.010615 | METAP2 | 0.010615 | 0.015645 | 0.03966 |
| ENSG00000196355 | 4 | 38628029 | 38666430 | 105 | 21 | 2.3034 | 0.010628 | AC021860.1 | 0.010628 | 0.016113 | 0.012326 |
| ENSG00000136997 | 8 | 128747680 | 128753674 | 22 | 9 | 2.3019 | 0.010669 | MYC | 0.010669 | 0.42291 | 0.86466 |
| ENSG00000122042 | 13 | 30338508 | 30424821 | 258 | 13 | 2.3015 | 0.010681 | UBL3 | 0.010681 | 0.010424 | 0.0049809 |
| ENSG00000114796 | 3 | 183353356 | 183402265 | 234 | 3 | 2.3011 | 0.010693 | KLHL24 | 0.010693 | 0.0074733 | 0.0042936 |
| ENSG00000139874 | 14 | 38677204 | 38682272 | 10 | 3 | 2.3007 | 0.010704 | SSTR1 | 0.010704 | 0.018857 | 0.04083 |
| ENSG00000106804 | 9 | 123714616 | 123812554 | 260 | 19 | 2.3007 | 0.010705 | C5 | 0.010705 | 0.01249 | 0.022841 |
| ENSG00000139372 | 12 | 104359582 | 104382652 | 116 | 20 | 2.2997 | 0.010733 | TDG | 0.010733 | 0.018553 | 0.046568 |
| ENSG00000111335 | 12 | 113416200 | 113449528 | 105 | 15 | 2.2996 | 0.010735 | OAS2 | 0.010735 | 0.040749 | 0.03236 |
| ENSG00000140015 | 14 | 63173287 | 63568755 | 1253 | 69 | 2.2959 | 0.010841 | KCNH5 | 0.010841 | 0.013298 | 0.013879 |
| ENSG00000156150 | 1 | 110602616 | 110613322 | 21 | 3 | 2.2948 | 0.010872 | ALX3 | 0.010872 | 0.010971 | 0.018676 |
| ENSG00000182628 | 17 | 57187312 | 57232630 | 57 | 13 | 2.293 | 0.010924 | SKA2 | 0.010924 | 0.01427 | 0.014872 |
| ENSG00000080166 | 13 | 95089558 | 95131936 | 165 | 20 | 2.2921 | 0.010949 | DCT | 0.010949 | 0.013227 | 0.029518 |
| ENSG00000025800 | 1 | 32573639 | 32642169 | 114 | 12 | 2.2901 | 0.011008 | KPNA6 | 0.011008 | 0.0084885 | 0.0070012 |
| ENSG00000163964 | 3 | 196366646 | 196462878 | 453 | 75 | 2.289 | 0.011038 | PIGX | 0.011038 | 0.0014743 | 0.0040884 |
| ENSG00000168899 | 2 | 85811531 | 85820535 | 23 | 6 | 2.2878 | 0.011075 | VAMP5 | 0.011075 | 0.02575 | 0.033958 |
| ENSG00000198162 | 1 | 117910071 | 118071494 | 525 | 21 | 2.2849 | 0.01116 | MAN1A2 | 0.01116 | 0.010704 | 0.0077778 |
| ENSG00000127951 | 7 | 76822688 | 76829143 | 6 | 3 | 2.2841 | 0.011183 | FGL2 | 0.011183 | 0.21353 | 0.29332 |
| ENSG00000150627 | 4 | 176986985 | 177103978 | 483 | 22 | 2.2818 | 0.01125 | WDR17 | 0.01125 | 0.022997 | 0.035915 |
| ENSG00000102978 | 16 | 57496299 | 57505922 | 28 | 10 | 2.2795 | 0.011318 | POLR2C | 0.011318 | 0.022622 | 0.014709 |
| ENSG00000187902 | 19 | 55940107 | 55954230 | 52 | 12 | 2.279 | 0.011335 | SHISA7 | 0.011335 | 0.025243 | 0.087686 |
| ENSG00000007001 | 2 | 158733214 | 158992666 | 906 | 68 | 2.2761 | 0.011421 | UPP2 | 0.011421 | 0.011739 | 0.016209 |
| ENSG00000181371 | 11 | 56257825 | 56258871 | 1 | 1 | 2.2721 | 0.01154 | OR5M8 | 0.01154 | 0.057074 | 0.063728 |
| ENSG00000143093 | 1 | 110574199 | 110617263 | 74 | 11 | 2.2719 | 0.011546 | STRIP1 | 0.011546 | 0.011598 | 0.022685 |
| ENSG00000234631 | 10 | 115674530 | 115676949 | 2 | 1 | 2.2714 | 0.011561 | AL162407.1 | 0.011561 | 0.22224 | 0.1181 |
| ENSG00000105501 | 19 | 52114781 | 52150151 | 142 | 33 | 2.2689 | 0.011637 | SIGLEC5 | 0.011637 | 0.0084627 | 0.0099615 |
| ENSG00000104888 | 19 | 49932658 | 49945617 | 30 | 9 | 2.2667 | 0.011704 | SLC17A7 | 0.011704 | 0.015931 | 0.036868 |
| ENSG00000128191 | 22 | 20067755 | 20099400 | 122 | 15 | 2.2652 | 0.011751 | DGCR8 | 0.011751 | 0.0086711 | 0.021288 |
| ENSG00000100109 | 22 | 26887191 | 26908471 | 78 | 19 | 2.2645 | 0.011773 | TFIP11 | 0.011773 | 0.016674 | 0.017551 |
| ENSG00000198797 | 1 | 177140633 | 177251558 | 241 | 33 | 2.2641 | 0.011784 | BRINP2 | 0.011784 | 0.017847 | 0.020354 |
| ENSG00000117560 | 1 | 172628154 | 172636014 | 12 | 4 | 2.2631 | 0.011815 | FASLG | 0.011815 | 0.056248 | 0.1634 |
| ENSG00000234906 | 19 | 45449243 | 45452822 | 21 | 4 | 2.2596 | 0.011922 | APOC2 | 0.011922 | 0.0094909 | 0.0098709 |
| ENSG00000100239 | 22 | 50781733 | 50883514 | 487 | 32 | 2.2581 | 0.011971 | PPP6R2 | 0.011971 | 0.011878 | 0.0051547 |
| ENSG00000163083 | 2 | 121103719 | 121109384 | 10 | 5 | 2.2554 | 0.012055 | INHBB | 0.012055 | 0.28429 | 0.5404 |
| ENSG00000107282 | 9 | 72042446 | 72287222 | 706 | 38 | 2.2542 | 0.01209 | APBA1 | 0.01209 | 0.013037 | 0.013038 |
| ENSG00000204311 | 2 | 179316163 | 179326117 | 17 | 6 | 2.2537 | 0.012108 | DFNB59 | 0.012108 | 0.034546 | 0.030388 |
| ENSG00000198856 | 4 | 109571740 | 109588976 | 61 | 4 | 2.2464 | 0.012338 | OSTC | 0.012338 | 0.011086 | 0.0078585 |
| ENSG00000227011 | 17 | 51062880 | 51065012 | 4 | 2 | 2.246 | 0.012351 | C17orf112 | 0.012351 | 0.011358 | 0.010076 |
| ENSG00000196684 | 19 | 16244838 | 16269386 | 82 | 21 | 2.2459 | 0.012356 | HSH2D | 0.012356 | 0.062885 | 0.050748 |
| ENSG00000023041 | 10 | 114190058 | 114206672 | 25 | 6 | 2.2438 | 0.012423 | ZDHHC6 | 0.012423 | 0.036593 | 0.062938 |
| ENSG00000138615 | 15 | 65488337 | 65503826 | 30 | 5 | 2.2436 | 0.012428 | CILP | 0.012428 | 0.0051887 | 0.014519 |
| ENSG00000116786 | 1 | 16010827 | 16061264 | 185 | 16 | 2.2436 | 0.01243 | PLEKHM2 | 0.01243 | 0.010366 | 0.010646 |
| ENSG00000189403 | 13 | 31032884 | 31191734 | 437 | 75 | 2.2424 | 0.012467 | HMGB1 | 0.012467 | 0.015273 | 0.00914 |
| ENSG00000174775 | 11 | 532242 | 537287 | 30 | 7 | 2.2419 | 0.012482 | HRAS | 0.012482 | 0.0015791 | 0.00032556 |
| ENSG00000204385 | 6 | 31830969 | 31846823 | 77 | 16 | 2.2416 | 0.012494 | SLC44A4 | 0.012494 | 0.0046104 | 0.010255 |
| ENSG00000151458 | 4 | 125585207 | 125633887 | 98 | 15 | 2.2405 | 0.01253 | ANKRD50 | 0.01253 | 0.014258 | 0.020248 |
| ENSG00000240510 | 1 | 151682909 | 151702070 | 39 | 11 | 2.2391 | 0.012576 | AL589765.1 | 0.012576 | 0.0029127 | 0.0031161 |
| ENSG00000178796 | 1 | 151682909 | 151702281 | 39 | 11 | 2.2391 | 0.012576 | RIIAD1 | 0.012576 | 0.0027115 | 0.0031161 |
| ENSG00000132199 | 18 | 670324 | 712676 | 249 | 20 | 2.239 | 0.012579 | ENOSF1 | 0.012579 | 0.020116 | 0.036003 |
| ENSG00000184838 | 5 | 119799973 | 120023027 | 1001 | 58 | 2.2348 | 0.012715 | PRR16 | 0.012715 | 0.0075846 | 0.012287 |
| ENSG00000249240 | 15 | 65134082 | 65223118 | 297 | 20 | 2.2346 | 0.01272 | AC069368.3 | 0.01272 | 0.015214 | 0.034005 |
| ENSG00000225899 | 10 | 135437399 | 135440299 | 4 | 2 | 2.2342 | 0.012736 | FRG2B | 0.012736 | 0.2432 | 0.27168 |
| ENSG00000099834 | 11 | 616565 | 626078 | 46 | 11 | 2.2318 | 0.012813 | CDHR5 | 0.012813 | 0.0094489 | 0.00363 |

| | | | | | | | | | | | |
|---|---|---|---|---|---|---|---|---|---|---|---|
| ENSG00000177294 | 17 | 6679542 | 6700420 | 113 | 19 | 2.2318 | 0.012813 | FBXO39 | 0.012813 | 0.0081311 | 0.008509 |
| ENSG00000158806 | 8 | 21881636 | 21894408 | 35 | 12 | 2.2283 | 0.012929 | NPM2 | 0.012929 | 0.023454 | 0.024842 |
| ENSG00000160352 | 19 | 21264965 | 21308073 | 203 | 14 | 2.228 | 0.012941 | ZNF714 | 0.012941 | 0.020397 | 0.021029 |
| ENSG00000186472 | 7 | 82383329 | 82792246 | 1409 | 49 | 2.2233 | 0.013098 | PCLO | 0.013098 | 0.01487 | 0.014581 |
| ENSG00000244242 | 11 | 1753640 | 1771821 | 64 | 7 | 2.2225 | 0.013123 | IFITM10 | 0.013123 | 0.014156 | 0.019147 |
| ENSG00000183718 | 5 | 180681417 | 180688119 | 18 | 7 | 2.2224 | 0.013127 | TRIM52 | 0.013127 | 0.049173 | 0.048292 |
| ENSG00000269563 | 2 | 47799601 | 47801998 | 8 | 2 | 2.2222 | 0.013136 | AC138655.1 | 0.013136 | 0.070964 | 0.48867 |
| ENSG00000183018 | 17 | 4402133 | 4442330 | 174 | 34 | 2.2173 | 0.013301 | SPNS2 | 0.013301 | 0.029764 | 0.037186 |
| ENSG00000172154 | 11 | 55860753 | 55861762 | 8 | 2 | 2.2169 | 0.013314 | OR8I2 | 0.013314 | 0.018891 | 0.020801 |
| ENSG00000110888 | 12 | 30862486 | 30907885 | 154 | 16 | 2.2166 | 0.013324 | CAPRIN2 | 0.013324 | 0.010136 | 0.0056111 |
| ENSG00000187173 | 1 | 152670840 | 152671918 | 2 | 1 | 2.2161 | 0.013344 | LCE2A | 0.013344 | 0.091086 | 0.14472 |
| ENSG00000142186 | 11 | 65292548 | 65306175 | 21 | 8 | 2.215 | 0.01338 | SCYL1 | 0.01338 | 0.019162 | 0.041274 |
| ENSG00000189013 | 19 | 55236002 | 55325972 | 208 | 27 | 2.2119 | 0.013488 | KIR2DL4 | 0.013488 | 0.012956 | 0.0049429 |
| ENSG00000160094 | 1 | 33722146 | 33766320 | 99 | 14 | 2.2115 | 0.013502 | ZNF362 | 0.013502 | 0.012753 | 0.027356 |
| ENSG00000168887 | 2 | 85833777 | 85839189 | 9 | 6 | 2.209 | 0.013587 | C2orf68 | 0.013587 | 0.29106 | 0.088149 |
| ENSG00000167230 | 17 | 35732928 | 35749662 | 47 | 20 | 2.2077 | 0.013631 | C17orf78 | 0.013631 | 0.039027 | 0.035769 |
| ENSG00000142867 | 1 | 85731931 | 85742773 | 39 | 12 | 2.2073 | 0.013648 | BCL10 | 0.013648 | 0.0090166 | 0.015328 |
| ENSG00000141542 | 17 | 80612849 | 80656604 | 201 | 38 | 2.2051 | 0.013724 | RAB40B | 0.013724 | 0.041043 | 0.16911 |
| ENSG00000116991 | 1 | 232533711 | 232697304 | 549 | 41 | 2.2033 | 0.013786 | SIPA1L2 | 0.013786 | 0.016254 | 0.009074 |
| ENSG00000151292 | 5 | 122847793 | 122952739 | 129 | 18 | 2.203 | 0.013799 | CSNK1G3 | 0.013799 | 0.0076048 | 0.011229 |
| ENSG00000197629 | 11 | 58975983 | 58980424 | 10 | 4 | 2.2017 | 0.013845 | MPEG1 | 0.013845 | 0.016928 | 0.0078169 |
| ENSG00000173875 | 19 | 12721732 | 12742735 | 84 | 8 | 2.1996 | 0.013919 | ZNF791 | 0.013919 | 0.015019 | 0.016745 |
| ENSG00000268500 | 19 | 52115344 | 52150142 | 139 | 32 | 2.1982 | 0.013967 | SIGLEC5 | 0.013967 | 0.0084627 | 0.0099615 |
| ENSG00000165661 | 9 | 139098179 | 139137687 | 140 | 22 | 2.1964 | 0.014032 | QSOX2 | 0.014032 | 0.010268 | 0.010164 |
| ENSG00000019991 | 7 | 81328322 | 81399754 | 149 | 13 | 2.1961 | 0.014042 | HGF | 0.014042 | 0.020968 | 0.032361 |
| ENSG00000159409 | 1 | 151674880 | 151689290 | 28 | 6 | 2.1959 | 0.014048 | CELF3 | 0.014048 | 0.0048077 | 0.0030257 |
| ENSG00000103150 | 16 | 83932731 | 83949787 | 129 | 12 | 2.1959 | 0.014048 | MLYCD | 0.014048 | 0.020607 | 0.050627 |
| ENSG00000143933 | 2 | 47387221 | 47403740 | 54 | 12 | 2.1946 | 0.014097 | CALM2 | 0.014097 | 0.006534 | 0.015742 |
| ENSG00000204842 | 12 | 111890018 | 112037480 | 80 | 29 | 2.1937 | 0.01413 | ATXN2 | 0.01413 | 0.0029514 | 0.0037651 |
| ENSG00000172365 | 11 | 58189738 | 58190786 | 3 | 1 | 2.1914 | 0.014212 | OR5B2 | 0.014212 | 0.014057 | 0.014846 |
| ENSG00000152402 | 11 | 106544738 | 106889250 | 1249 | 65 | 2.1912 | 0.014218 | GUCY1A2 | 0.014218 | 0.01367 | 0.013906 |
| ENSG00000150269 | 11 | 56229945 | 56230877 | 8 | 2 | 2.1909 | 0.014229 | OR5M9 | 0.014229 | 0.016634 | 0.014078 |
| ENSG00000148840 | 10 | 103892787 | 103910082 | 26 | 9 | 2.1893 | 0.014288 | PPRC1 | 0.014288 | 0.0073862 | 0.015104 |
| ENSG00000129566 | 14 | 20833826 | 20881588 | 185 | 36 | 2.185 | 0.014445 | TEP1 | 0.014445 | 0.014085 | 0.071028 |
| ENSG00000103194 | 16 | 84733584 | 84813528 | 385 | 27 | 2.1841 | 0.014477 | USP10 | 0.014477 | 0.01052 | 0.0079523 |
| ENSG00000120279 | 6 | 153019030 | 153045702 | 105 | 12 | 2.1804 | 0.014616 | MYCT1 | 0.014616 | 0.051328 | 0.037348 |
| ENSG00000131355 | 19 | 14729929 | 14800839 | 312 | 34 | 2.1754 | 0.014798 | EMR3 | 0.014798 | 0.012097 | 0.015454 |
| ENSG00000111665 | 12 | 6953957 | 6961230 | 13 | 4 | 2.1736 | 0.014869 | CDCA3 | 0.014869 | 0.079889 | 0.0854 |
| ENSG00000074054 | 2 | 122095352 | 122407163 | 844 | 15 | 2.1724 | 0.014912 | CLASP1 | 0.014912 | 0.015889 | 0.016711 |
| ENSG00000138685 | 4 | 123747863 | 123819391 | 325 | 33 | 2.1719 | 0.014933 | FGF2 | 0.014933 | 0.012586 | 0.011509 |
| ENSG00000185774 | 4 | 20730239 | 21950422 | 5049 | 203 | 2.1712 | 0.014959 | KCNIP4 | 0.014959 | 0.01502 | 0.015256 |
| ENSG00000137802 | 15 | 42066632 | 42120053 | 107 | 17 | 2.1708 | 0.014972 | MAPKBP1 | 0.014972 | 0.015454 | 0.0018411 |
| ENSG00000189292 | 2 | 279558 | 288851 | 23 | 5 | 2.1691 | 0.015038 | FAM150B | 0.015038 | 0.033008 | 0.073231 |
| ENSG00000070770 | 16 | 58191811 | 58231824 | 155 | 14 | 2.1683 | 0.015069 | CSNK2A2 | 0.015069 | 0.021068 | 0.030648 |
| ENSG00000164237 | 5 | 10275987 | 10308138 | 135 | 17 | 2.168 | 0.015081 | CMBL | 0.015081 | 0.032983 | 0.060161 |
| ENSG00000166884 | 11 | 59224411 | 59225464 | 8 | 3 | 2.1674 | 0.015102 | OR4D6 | 0.015102 | 0.17874 | 0.042958 |
| ENSG00000060762 | 6 | 166778407 | 166796486 | 36 | 5 | 2.1672 | 0.015109 | MPC1 | 0.015109 | 0.017258 | 0.012163 |
| ENSG00000164151 | 5 | 5420777 | 5490347 | 137 | 14 | 2.1661 | 0.01515 | KIAA0947 | 0.01515 | 0.006636 | 0.0037783 |
| ENSG00000166250 | 11 | 122943035 | 123065989 | 606 | 43 | 2.164 | 0.015232 | CLMP | 0.015232 | 0.014896 | 0.016044 |
| ENSG00000267114 | 19 | 45453301 | 45457264 | 32 | 2 | 2.1624 | 0.015295 | CTB-129P6.11 | 0.015295 | 0.011256 | 0.0095876 |
| ENSG00000132824 | 20 | 43124862 | 43150750 | 45 | 7 | 2.1596 | 0.015402 | SERINC3 | 0.015402 | 0.022098 | 0.027719 |
| ENSG00000115561 | 2 | 86730554 | 86948245 | 532 | 31 | 2.1575 | 0.015483 | CHMP3 | 0.015483 | 0.015738 | 0.021101 |
| ENSG00000163519 | 3 | 108541545 | 108573852 | 94 | 17 | 2.1574 | 0.015489 | TRAT1 | 0.015489 | 0.0052298 | 0.0099512 |
| ENSG00000071539 | 5 | 892758 | 919472 | 81 | 5 | 2.1563 | 0.015531 | TRIP13 | 0.015531 | 0.015419 | 0.013558 |
| ENSG00000174749 | 4 | 113066553 | 113116412 | 126 | 15 | 2.1529 | 0.015662 | C4orf32 | 0.015662 | 0.017998 | 0.029513 |
| ENSG00000258388 | 6 | 32121622 | 32139755 | 51 | 16 | 2.1519 | 0.015702 | PPT2-EGFL8 | 0.015702 | 0.046577 | 0.029407 |
| ENSG00000134765 | 18 | 28709199 | 28742819 | 176 | 14 | 2.1512 | 0.015732 | DSC1 | 0.015732 | 0.034459 | 0.081157 |
| ENSG00000170836 | 17 | 58677544 | 58741849 | 23 | 12 | 2.151 | 0.015737 | PPM1D | 0.015737 | 0.025339 | 0.0055096 |
| ENSG00000163293 | 4 | 47916159 | 48042188 | 508 | 21 | 2.1507 | 0.015749 | NIPAL1 | 0.015749 | 0.014305 | 0.015385 |
| ENSG00000115641 | 2 | 105974169 | 106054970 | 269 | 48 | 2.1505 | 0.015757 | FHL2 | 0.015757 | 0.017558 | 0.0077381 |
| ENSG00000163464 | 2 | 219027568 | 219031718 | 5 | 2 | 2.1482 | 0.015849 | CXCR1 | 0.015849 | 0.027049 | 0.05037 |
| ENSG00000118620 | 19 | 21203426 | 21242856 | 107 | 10 | 2.1475 | 0.015877 | ZNF430 | 0.015877 | 0.018127 | 0.017756 |
| ENSG00000159516 | 1 | 153122058 | 153123345 | 4 | 1 | 2.1474 | 0.015882 | SPRR2G | 0.015882 | 0.015358 | 0.011542 |
| ENSG00000182183 | 1 | 53099016 | 53135355 | 103 | 23 | 2.1471 | 0.015891 | FAM159A | 0.015891 | 0.028261 | 0.021623 |
| ENSG00000157423 | 16 | 70841281 | 71264625 | 719 | 49 | 2.1461 | 0.015933 | HYDIN | 0.015933 | 0.015426 | 0.020914 |
| ENSG00000163002 | 2 | 183982241 | 184026408 | 322 | 21 | 2.1451 | 0.015971 | NUP35 | 0.015971 | 0.014403 | 0.01086 |
| ENSG00000241945 | 21 | 45527171 | 45551063 | 62 | 6 | 2.1435 | 0.016036 | PWP2 | 0.016036 | 0.015353 | 0.017475 |
| ENSG00000204104 | 2 | 239229082 | 239309541 | 272 | 18 | 2.1431 | 0.016054 | TRAF3IP1 | 0.016054 | 0.012461 | 0.011681 |
| ENSG00000101323 | 20 | 7863628 | 7921121 | 190 | 30 | 2.1409 | 0.016143 | HAO1 | 0.016143 | 0.020666 | 0.02953 |
| ENSG00000095059 | 19 | 12786531 | 12792716 | 11 | 6 | 2.1385 | 0.016239 | DHPS | 0.016239 | 0.00039573 | 0.00024445 |
| ENSG00000179820 | 19 | 54369477 | 54379691 | 30 | 6 | 2.1376 | 0.016275 | MYADM | 0.016275 | 0.023868 | 0.060056 |
| ENSG00000198704 | 6 | 28471073 | 28495992 | 94 | 18 | 2.1374 | 0.016283 | GPX6 | 0.016283 | 0.011429 | 0.011416 |
| ENSG00000132906 | 1 | 15817327 | 15853029 | 168 | 10 | 2.1367 | 0.01631 | CASP9 | 0.01631 | 0.015775 | 0.021582 |

| | | | | | | | | | | | |
|---|---|---|---|---|---|---|---|---|---|---|---|
| ENSG00000108950 | 17 | 66531254 | 66597530 | 182 | 30 | 2.1353 | 0.016367 | FAM20A | 0.016367 | 0.020694 | 0.046827 |
| ENSG00000249884 | 2 | 86732791 | 86948245 | 526 | 31 | 2.1352 | 0.016372 | RNF103-CHMP3 | 0.016372 | 0.015395 | 0.020896 |
| ENSG00000155265 | 10 | 99609996 | 99631337 | 46 | 8 | 2.1349 | 0.016387 | GOLGA7B | 0.016387 | 0.021613 | 0.022216 |
| ENSG00000124939 | 11 | 61976140 | 61981408 | 20 | 4 | 2.1312 | 0.016536 | SCGB2A1 | 0.016536 | 0.049865 | 0.087188 |
| ENSG00000174004 | 3 | 196366557 | 196388875 | 126 | 38 | 2.1299 | 0.016591 | NRROS | 0.016591 | 0.0027175 | 0.0034223 |
| ENSG00000133056 | 1 | 204391756 | 204463852 | 221 | 23 | 2.1283 | 0.016654 | PIK3C2B | 0.016654 | 0.0082803 | 0.0079709 |
| ENSG00000149679 | 20 | 60963688 | 60982341 | 103 | 14 | 2.128 | 0.01667 | CABLES2 | 0.01667 | 0.018085 | 0.045953 |
| ENSG00000166546 | 16 | 66461200 | 66527432 | 147 | 27 | 2.1266 | 0.016726 | BEAN1 | 0.016726 | 0.026717 | 0.035097 |
| ENSG00000163703 | 3 | 9975506 | 9987097 | 35 | 7 | 2.1245 | 0.016815 | CRELD1 | 0.016815 | 0.0066228 | 0.023588 |
| ENSG00000104941 | 19 | 46298968 | 46318577 | 70 | 13 | 2.1241 | 0.016829 | RSPH6A | 0.016829 | 0.025833 | 0.02272 |
| ENSG00000104853 | 19 | 45457842 | 45496599 | 135 | 17 | 2.1236 | 0.016853 | CLPTM1 | 0.016853 | 0.0087482 | 0.012388 |
| ENSG00000204010 | 10 | 91137813 | 91144962 | 17 | 5 | 2.1226 | 0.016893 | IFIT1B | 0.016893 | 0.012528 | 0.014479 |
| ENSG00000151376 | 11 | 86152150 | 86383678 | 1128 | 56 | 2.1218 | 0.016928 | ME3 | 0.016928 | 0.010921 | 0.0081491 |
| ENSG00000174332 | 1 | 53971910 | 54199877 | 694 | 49 | 2.1189 | 0.017051 | GLIS1 | 0.017051 | 0.012933 | 0.020659 |
| ENSG00000157870 | 1 | 2517930 | 2522908 | 10 | 4 | 2.1161 | 0.017169 | FAM213B | 0.017169 | 0.025426 | 0.034232 |
| ENSG00000145246 | 4 | 47487305 | 47595503 | 427 | 42 | 2.1153 | 0.017202 | ATP10D | 0.017202 | 0.020012 | 0.064045 |
| ENSG00000129038 | 15 | 74218330 | 74244478 | 132 | 19 | 2.1151 | 0.017209 | LOXL1 | 0.017209 | 0.029071 | 0.064269 |
| ENSG00000273269 | 2 | 47293080 | 47403650 | 569 | 54 | 2.1144 | 0.017242 | RP11-761B3.1 | 0.017242 | 0.021587 | 0.046376 |
| ENSG00000256980 | 6 | 73933267 | 73935493 | 5 | 2 | 2.1121 | 0.017338 | KHDC1L | 0.017338 | 0.099557 | 0.2106 |
| ENSG00000185745 | 10 | 91152303 | 91163745 | 39 | 9 | 2.1107 | 0.017399 | IFIT1 | 0.017399 | 0.016133 | 0.010304 |
| ENSG00000166197 | 10 | 103911933 | 103923627 | 17 | 5 | 2.1092 | 0.017465 | NOLC1 | 0.017465 | 0.024345 | 0.024393 |
| ENSG00000164253 | 5 | 76721795 | 76916436 | 884 | 42 | 2.1077 | 0.017528 | WDR41 | 0.017528 | 0.014541 | 0.02044 |
| ENSG00000111816 | 6 | 116252312 | 116381921 | 276 | 25 | 2.1059 | 0.017607 | FRK | 0.017607 | 0.019692 | 0.02754 |
| ENSG00000174502 | 1 | 205882176 | 205912588 | 130 | 20 | 2.104 | 0.017688 | SLC26A9 | 0.017688 | 0.035446 | 0.051441 |
| ENSG00000204003 | 9 | 139632619 | 139642905 | 30 | 5 | 2.1026 | 0.017751 | LCN6 | 0.017751 | 0.036914 | 0.22606 |
| ENSG00000137449 | 4 | 15004298 | 15071777 | 200 | 24 | 2.1025 | 0.017757 | CPEB2 | 0.017757 | 0.016423 | 0.018064 |
| ENSG00000138433 | 2 | 175212750 | 175260443 | 124 | 17 | 2.1001 | 0.017859 | CIR1 | 0.017859 | 0.022839 | 0.036488 |
| ENSG00000176182 | 19 | 46393278 | 46405862 | 62 | 13 | 2.0987 | 0.017922 | MYPOP | 0.017922 | 0.014524 | 0.025135 |
| ENSG00000241106 | 6 | 32780540 | 32784825 | 39 | 10 | 2.0973 | 0.017985 | HLA-DOB | 0.017985 | 0.11383 | 0.017476 |
| ENSG00000036672 | 11 | 119225925 | 119252436 | 68 | 16 | 2.0971 | 0.017994 | USP2 | 0.017994 | 0.0034437 | 0.0071805 |
| ENSG00000155229 | 10 | 99218081 | 99258551 | 86 | 14 | 2.0958 | 0.018049 | MMS19 | 0.018049 | 0.021011 | 0.019807 |
| ENSG00000101448 | 20 | 44165625 | 44176391 | 12 | 6 | 2.0941 | 0.018125 | EPPIN | 0.018125 | 0.46712 | 0.5542 |
| ENSG00000196417 | 19 | 53893046 | 53930574 | 261 | 28 | 2.0941 | 0.018127 | ZNF765 | 0.018127 | 0.07368 | 0.17883 |
| ENSG00000143590 | 1 | 155036224 | 155060014 | 44 | 9 | 2.0914 | 0.018246 | EFNA3 | 0.018246 | 0.015187 | 0.010169 |
| ENSG00000119227 | 3 | 196673214 | 196695931 | 117 | 25 | 2.0913 | 0.018252 | PIGZ | 0.018252 | 0.048851 | 0.091395 |
| ENSG00000163217 | 2 | 69092613 | 69098649 | 14 | 4 | 2.0894 | 0.018337 | BMP10 | 0.018337 | 0.17357 | 0.030583 |
| ENSG00000269169 | 19 | 999601 | 999952 | 3 | 1 | 2.0882 | 0.018392 | AC004528.1 | 0.018392 | 0.10244 | 0.27704 |
| ENSG00000226979 | 6 | 31539831 | 31542101 | 13 | 5 | 2.0881 | 0.018394 | LTA | 0.018394 | 0.02969 | 0.10779 |
| ENSG00000139679 | 13 | 48963707 | 49018840 | 120 | 14 | 2.0878 | 0.018409 | LPAR6 | 0.018409 | 0.019008 | 0.009605 |
| ENSG00000169876 | 7 | 100663353 | 100702020 | 111 | 15 | 2.0867 | 0.018458 | MUC17 | 0.018458 | 0.01127 | 0.0086524 |
| ENSG00000028310 | 5 | 850406 | 892939 | 143 | 12 | 2.0862 | 0.018479 | BRD9 | 0.018479 | 0.015439 | 0.016395 |
| ENSG00000038382 | 5 | 14143811 | 14532235 | 1078 | 49 | 2.0797 | 0.018778 | TRIO | 0.018778 | 0.012896 | 0.01238 |
| ENSG00000205496 | 11 | 4976002 | 4976943 | 13 | 4 | 2.0789 | 0.018814 | OR51A2 | 0.018814 | 0.3895 | 0.42886 |
| ENSG00000188033 | 19 | 12688775 | 12750912 | 199 | 15 | 2.0772 | 0.018892 | ZNF490 | 0.018892 | 0.021269 | 0.01393 |
| ENSG00000146776 | 7 | 105245514 | 105517050 | 867 | 119 | 2.0746 | 0.019013 | ATXN7L1 | 0.019013 | 0.023204 | 0.050729 |
| ENSG00000139351 | 12 | 102122426 | 102133250 | 17 | 6 | 2.0742 | 0.01903 | SYCP3 | 0.01903 | 0.016703 | 0.023862 |
| ENSG00000187922 | 9 | 139632619 | 139637808 | 16 | 2 | 2.0737 | 0.019055 | LCN10 | 0.019055 | 0.01949 | 0.19932 |
| ENSG00000013375 | 6 | 83870869 | 83903655 | 49 | 8 | 2.0719 | 0.019136 | PGM3 | 0.019136 | 0.0013039 | 0.00038545 |
| ENSG00000198728 | 10 | 103867317 | 103880210 | 23 | 9 | 2.0713 | 0.019165 | LDB1 | 0.019165 | 0.0091684 | 0.02571 |
| ENSG00000152778 | 10 | 91174343 | 91180758 | 27 | 8 | 2.071 | 0.019179 | IFIT5 | 0.019179 | 0.016078 | 0.011849 |
| ENSG00000268509 | 3 | 5229227 | 5229982 | 1 | 1 | 2.0695 | 0.01925 | AC026202.1 | 0.01925 | 0.42546 | 0.3568 |
| ENSG00000197483 | 19 | 55987699 | 55995854 | 40 | 11 | 2.066 | 0.019414 | ZNF628 | 0.019414 | 0.13609 | 0.1654 |
| ENSG00000068903 | 19 | 39369197 | 39390502 | 56 | 18 | 2.0626 | 0.019575 | SIRT2 | 0.019575 | 0.026669 | 0.076754 |
| ENSG00000139515 | 13 | 28494157 | 28500368 | 18 | 4 | 2.0608 | 0.019662 | PDX1 | 0.019662 | 0.030791 | 0.27246 |
| ENSG00000115414 | 2 | 216225163 | 216300895 | 287 | 21 | 2.0607 | 0.019665 | FN1 | 0.019665 | 0.026062 | 0.0333 |
| ENSG00000161326 | 17 | 35849937 | 35873603 | 75 | 13 | 2.0571 | 0.019837 | DUSP14 | 0.019837 | 0.042625 | 0.037768 |
| ENSG00000174937 | 11 | 56236964 | 56238014 | 3 | 1 | 2.0567 | 0.019859 | OR5M3 | 0.019859 | 0.028496 | 0.017803 |
| ENSG00000256566 | 20 | 2447858 | 2489553 | 120 | 20 | 2.0544 | 0.01997 | RP4-734P14.4 | 0.01997 | 0.018549 | 0.070447 |
| ENSG00000129255 | 17 | 7486847 | 7496107 | 19 | 7 | 2.0533 | 0.020022 | MPDU1 | 0.020022 | 0.030126 | 0.018891 |
| ENSG00000158714 | 1 | 159796540 | 159807039 | 22 | 13 | 2.0523 | 0.020072 | SLAMF8 | 0.020072 | 0.03113 | 0.014404 |
| ENSG00000066032 | 2 | 79412357 | 80875905 | 5236 | 302 | 2.0513 | 0.02012 | CTNNA2 | 0.02012 | 0.025019 | 0.025999 |
| ENSG00000172081 | 19 | 2071037 | 2096672 | 56 | 15 | 2.0509 | 0.020137 | MOB3A | 0.020137 | 0.038897 | 0.11668 |
| ENSG00000162461 | 1 | 16062900 | 16067891 | 9 | 4 | 2.0493 | 0.020217 | SLC25A34 | 0.020217 | 0.024541 | 0.0030305 |
| ENSG00000139289 | 12 | 76419227 | 76427712 | 30 | 12 | 2.0491 | 0.020228 | PHLDA1 | 0.020228 | 0.29787 | 0.5503 |
| ENSG00000179253 | 20 | 60292562 | 60294804 | 22 | 10 | 2.0489 | 0.020234 | RP11-429E11.3 | 0.020234 | 0.13128 | 0.18852 |
| ENSG00000100292 | 22 | 35776354 | 35790207 | 63 | 8 | 2.0487 | 0.020248 | HMOX1 | 0.020248 | 0.025283 | 0.051451 |
| ENSG00000125170 | 16 | 57505863 | 57521239 | 53 | 14 | 2.0481 | 0.020277 | DOK4 | 0.020277 | 0.02034 | 0.0034325 |
| ENSG00000146267 | 6 | 99719045 | 99797938 | 298 | 23 | 2.048 | 0.020282 | FAXC | 0.020282 | 0.034457 | 0.027417 |
| ENSG00000116138 | 1 | 15853308 | 15918874 | 255 | 13 | 2.0474 | 0.020309 | DNAJC16 | 0.020309 | 0.024904 | 0.021346 |
| ENSG00000181761 | 11 | 55889849 | 55890787 | 14 | 3 | 2.047 | 0.02033 | OR8H3 | 0.02033 | 0.019233 | 0.019547 |
| ENSG00000148362 | 9 | 139886870 | 139888436 | 2 | 1 | 2.0462 | 0.020366 | C9orf142 | 0.020366 | 0.0053794 | 0.0071849 |
| ENSG00000168298 | 6 | 26156559 | 26157343 | 2 | 1 | 2.0441 | 0.02047 | HIST1H1E | 0.02047 | 0.73608 | 0.6364 |
| ENSG00000112096 | 6 | 160090089 | 160183561 | 375 | 24 | 2.0429 | 0.020531 | SOD2 | 0.020531 | 0.023143 | 0.02422 |

| | | | | | | | | | | | |
|---|---|---|---|---|---|---|---|---|---|---|---|
| ENSG00000197217 | 8 | 23243296 | 23315208 | 229 | 28 | 2.0429 | 0.020531 | ENTPD4 | 0.020531 | 0.029194 | 0.037116 |
| ENSG00000005175 | 12 | 48057070 | 48099844 | 82 | 17 | 2.0417 | 0.020591 | RPAP3 | 0.020591 | 0.049569 | 0.091246 |
| ENSG00000138308 | 10 | 74694928 | 74714536 | 16 | 8 | 2.0413 | 0.020612 | PLA2G12B | 0.020612 | 0.072256 | 0.51136 |
| ENSG00000124160 | 20 | 44689624 | 44718591 | 69 | 10 | 2.0411 | 0.020621 | NCOA5 | 0.020621 | 0.025119 | 0.015179 |
| ENSG00000164509 | 5 | 55147207 | 55218678 | 227 | 31 | 2.0407 | 0.02064 | IL31RA | 0.02064 | 0.024779 | 0.024213 |
| ENSG00000167470 | 19 | 1248552 | 1259142 | 32 | 8 | 2.0405 | 0.020648 | MIDN | 0.020648 | 0.05115 | 0.26997 |
| ENSG00000136504 | 17 | 47865917 | 47906458 | 42 | 15 | 2.0405 | 0.020651 | KAT7 | 0.020651 | 0.019323 | 0.01557 |
| ENSG00000147571 | 8 | 67088620 | 67090960 | 2 | 1 | 2.0396 | 0.020696 | CRH | 0.020696 | 0.85692 | 0.8186 |
| ENSG00000169131 | 5 | 178138593 | 178157703 | 94 | 6 | 2.0372 | 0.020813 | ZNF354A | 0.020813 | 0.031826 | 0.075475 |
| ENSG00000109452 | 4 | 142944313 | 143768585 | 2664 | 99 | 2.0362 | 0.020864 | INPP4B | 0.020864 | 0.022549 | 0.031773 |
| ENSG00000261210 | 16 | 19297061 | 19333467 | 149 | 18 | 2.035 | 0.020925 | CLEC19A | 0.020925 | 0.039265 | 0.029366 |
| ENSG00000110841 | 12 | 27676364 | 27848497 | 703 | 53 | 2.0346 | 0.020948 | PPFIBP1 | 0.020948 | 0.019063 | 0.03985 |
| ENSG00000236104 | 6 | 33282183 | 33285719 | 7 | 3 | 2.0332 | 0.021014 | ZBTB22 | 0.021014 | 0.055747 | 0.047168 |
| ENSG00000170604 | 19 | 46386866 | 46389376 | 6 | 3 | 2.0331 | 0.02102 | IRF2BP1 | 0.02102 | 0.0119 | 0.0084834 |
| ENSG00000159921 | 9 | 36214438 | 36277053 | 183 | 17 | 2.0321 | 0.021071 | GNE | 0.021071 | 0.026537 | 0.037075 |
| ENSG00000171863 | 2 | 3622795 | 3628509 | 24 | 5 | 2.032 | 0.021076 | RPS7 | 0.021076 | 0.047851 | 0.10276 |
| ENSG00000099984 | 22 | 24322339 | 24326106 | 20 | 6 | 2.0311 | 0.021123 | GSTT2 | 0.021123 | 0.046194 | 0.017854 |
| ENSG00000184956 | 11 | 1012821 | 1036706 | 78 | 21 | 2.0306 | 0.021146 | MUC6 | 0.021146 | 0.053982 | 0.04828 |
| ENSG00000181472 | 6 | 151685252 | 151712683 | 72 | 12 | 2.0294 | 0.02121 | ZBTB2 | 0.02121 | 0.0059046 | 0.0047323 |
| ENSG00000155906 | 6 | 151725989 | 151773259 | 235 | 24 | 2.0291 | 0.021222 | RMND1 | 0.021222 | 0.0072789 | 0.0045935 |
| ENSG00000147443 | 8 | 21766384 | 21771371 | 13 | 6 | 2.0274 | 0.021309 | DOK2 | 0.021309 | 0.062157 | 0.055309 |
| ENSG00000113722 | 5 | 149546358 | 149564120 | 35 | 8 | 2.0265 | 0.021358 | CDX1 | 0.021358 | 0.019872 | 0.018803 |
| ENSG00000173812 | 17 | 39845137 | 39848920 | 6 | 3 | 2.0261 | 0.021375 | EIF1 | 0.021375 | 0.15464 | 0.16325 |
| ENSG00000095002 | 2 | 47630108 | 47789450 | 543 | 32 | 2.0256 | 0.0214 | MSH2 | 0.0214 | 0.018284 | 0.023424 |
| ENSG00000172489 | 11 | 56019676 | 56020698 | 1 | 1 | 2.0255 | 0.02141 | OR5T3 | 0.02141 | 0.0273 | 0.031067 |
| ENSG00000257545 | 12 | 106889736 | 107168696 | 683 | 56 | 2.0244 | 0.021464 | RP11-144F15.1 | 0.021464 | 0.021289 | 0.020914 |
| ENSG00000136436 | 17 | 46908350 | 46943884 | 115 | 10 | 2.0228 | 0.021545 | CALCOCO2 | 0.021545 | 0.022976 | 0.016069 |
| ENSG00000139684 | 13 | 47345391 | 47371367 | 55 | 9 | 2.022 | 0.021588 | ESD | 0.021588 | 0.061444 | 0.14948 |
| ENSG00000018236 | 12 | 41086244 | 41466220 | 1514 | 55 | 2.0196 | 0.021714 | CNTN1 | 0.021714 | 0.021093 | 0.016224 |
| ENSG00000047936 | 6 | 117609463 | 117747018 | 488 | 48 | 2.0177 | 0.021809 | ROS1 | 0.021809 | 0.026863 | 0.040467 |
| ENSG00000204531 | 6 | 31132119 | 31148508 | 184 | 30 | 2.0173 | 0.02183 | POU5F1 | 0.02183 | 0.070322 | 0.12464 |
| ENSG00000188910 | 1 | 35246790 | 35251970 | 14 | 6 | 2.0155 | 0.021924 | GJB3 | 0.021924 | 0.02433 | 0.15415 |
| ENSG00000260272 | 16 | 2546033 | 2570219 | 22 | 12 | 2.0135 | 0.022031 | RP11-20I23.1 | 0.022031 | 0.024228 | 0.071093 |
| ENSG00000107185 | 9 | 35749203 | 35758572 | 12 | 3 | 2.0126 | 0.022077 | RGP1 | 0.022077 | 0.0092162 | 0.022537 |
| ENSG00000206560 | 3 | 15708743 | 15901278 | 437 | 18 | 2.0123 | 0.022095 | ANKRD28 | 0.022095 | 0.024966 | 0.018498 |
| ENSG00000239697 | 17 | 7452208 | 7464925 | 46 | 10 | 2.0109 | 0.02217 | TNFSF12 | 0.02217 | 0.039754 | 0.050626 |
| ENSG00000248871 | 17 | 7452416 | 7464918 | 46 | 10 | 2.0109 | 0.02217 | TNFSF12-TNFSF13 | 0.02217 | 0.040745 | 0.049773 |
| ENSG00000187240 | 11 | 102980160 | 103350591 | 1927 | 32 | 2.0106 | 0.022186 | DYNC2H1 | 0.022186 | 0.022045 | 0.027539 |
| ENSG00000156467 | 8 | 97238148 | 97247862 | 23 | 4 | 2.0104 | 0.022195 | UQCRB | 0.022195 | 0.039377 | 0.074378 |
| ENSG00000225556 | 1 | 151810295 | 151813033 | 5 | 2 | 2.0099 | 0.02222 | C2CD4D | 0.02222 | 0.046191 | 0.21269 |
| ENSG00000266956 | 17 | 34310692 | 34317306 | 20 | 3 | 2.0093 | 0.022251 | CTB-186H2.3 | 0.022251 | 0.27893 | 0.39766 |
| ENSG00000183765 | 22 | 29083731 | 29138410 | 181 | 28 | 2.0064 | 0.022408 | CHEK2 | 0.022408 | 0.033226 | 0.063945 |
| ENSG00000130433 | 19 | 54495542 | 54515923 | 98 | 16 | 2.0061 | 0.022424 | CACNG6 | 0.022424 | 0.035754 | 0.068259 |
| ENSG00000180305 | 20 | 44258165 | 44259835 | 9 | 2 | 2.0035 | 0.02256 | WFDC10A | 0.02256 | 0.05621 | 0.040474 |
| ENSG00000096654 | 6 | 27418522 | 27440897 | 102 | 7 | 2.0021 | 0.022635 | ZNF184 | 0.022635 | 0.022137 | 0.029125 |
| ENSG00000075429 | 17 | 64831235 | 64881603 | 211 | 47 | 2.002 | 0.022645 | CACNG5 | 0.022645 | 0.040293 | 0.087745 |
| ENSG00000251246 | 1 | 155036224 | 155059283 | 42 | 9 | 2.0019 | 0.022648 | EFNA3 | 0.022648 | 0.015187 | 0.010169 |
| ENSG00000162434 | 1 | 65298912 | 65432187 | 428 | 34 | 2.0015 | 0.022667 | JAK1 | 0.022667 | 0.030546 | 0.053222 |
| ENSG00000092067 | 14 | 23586513 | 23588825 | 9 | 4 | 2.0013 | 0.022677 | CEBPE | 0.022677 | 0.018421 | 0.070034 |
| ENSG00000126460 | 19 | 50083903 | 50094272 | 23 | 5 | 2.001 | 0.022696 | PRRG2 | 0.022696 | 0.045452 | 0.079963 |
| ENSG00000185792 | 19 | 56219798 | 56249768 | 128 | 30 | 1.9997 | 0.022768 | NLRP9 | 0.022768 | 0.034861 | 0.10442 |
| ENSG00000168515 | 11 | 61957688 | 61961011 | 11 | 3 | 1.9992 | 0.022794 | SCGB1D1 | 0.022794 | 0.13219 | 0.046502 |
| ENSG00000135334 | 6 | 88384790 | 88411927 | 94 | 15 | 1.9983 | 0.022845 | AKIRIN2 | 0.022845 | 0.012692 | 0.016815 |
| ENSG00000162814 | 1 | 217804666 | 218045038 | 803 | 34 | 1.9978 | 0.022869 | SPATA17 | 0.022869 | 0.023372 | 0.041963 |
| ENSG00000111817 | 6 | 116575336 | 116762424 | 497 | 45 | 1.9976 | 0.022882 | DSE | 0.022882 | 0.042014 | 0.024406 |
| ENSG00000105643 | 19 | 18111941 | 18124911 | 61 | 11 | 1.9971 | 0.022907 | ARRDC2 | 0.022907 | 0.06016 | 0.12827 |
| ENSG00000101199 | 20 | 61904137 | 61921142 | 47 | 5 | 1.9961 | 0.022963 | ARFGAP1 | 0.022963 | 0.033252 | 0.10267 |
| ENSG00000136235 | 7 | 23275586 | 23314727 | 51 | 11 | 1.9955 | 0.022996 | GPNMB | 0.022996 | 0.020085 | 0.025973 |
| ENSG00000125037 | 3 | 10004221 | 10052800 | 168 | 7 | 1.9946 | 0.023043 | EMC3 | 0.023043 | 0.023467 | 0.019349 |
| ENSG00000133433 | 22 | 24299601 | 24303373 | 22 | 6 | 1.9939 | 0.023081 | GSTT2B | 0.023081 | 0.018488 | 0.034039 |
| ENSG00000180398 | 2 | 47129009 | 47168994 | 202 | 22 | 1.9924 | 0.023164 | MCFD2 | 0.023164 | 0.026616 | 0.041481 |
| ENSG00000149742 | 11 | 63137261 | 63177766 | 133 | 23 | 1.9915 | 0.023214 | SLC22A9 | 0.023214 | 0.033631 | 0.035253 |
| ENSG00000100982 | 20 | 44563267 | 44576662 | 31 | 10 | 1.9903 | 0.023277 | PCIF1 | 0.023277 | 0.0054433 | 0.029577 |
| ENSG00000112893 | 5 | 109025067 | 109205326 | 712 | 28 | 1.99 | 0.023297 | MAN2A1 | 0.023297 | 0.024982 | 0.02321 |
| ENSG00000204540 | 6 | 31082527 | 31107869 | 351 | 26 | 1.9893 | 0.023337 | PSORS1C1 | 0.023337 | 0.087906 | 0.11385 |
| ENSG00000166002 | 11 | 93211638 | 93276674 | 247 | 28 | 1.9881 | 0.023401 | SMCO4 | 0.023401 | 0.026193 | 0.037753 |
| ENSG00000118972 | 12 | 4477393 | 4488894 | 39 | 7 | 1.9865 | 0.023486 | FGF23 | 0.023486 | 0.018484 | 0.081885 |
| ENSG00000189283 | 3 | 59735036 | 61237133 | 7323 | 377 | 1.9835 | 0.023658 | FHIT | 0.023658 | 0.027846 | 0.030062 |
| ENSG00000255730 | 19 | 41856758 | 41930906 | 143 | 16 | 1.9827 | 0.0237 | CTC-435M10.3 | 0.0237 | 0.024105 | 0.042728 |
| ENSG00000105341 | 19 | 41937223 | 41946622 | 20 | 3 | 1.9823 | 0.023723 | ATP5SL | 0.023723 | 0.054823 | 0.093709 |
| ENSG00000106541 | 7 | 16831435 | 16873057 | 253 | 20 | 1.9821 | 0.023732 | AGR2 | 0.023732 | 0.024728 | 0.052962 |
| ENSG00000153904 | 1 | 85784164 | 86043933 | 911 | 69 | 1.9815 | 0.023768 | DDAH1 | 0.023768 | 0.023452 | 0.050087 |
| ENSG00000231500 | 6 | 33239787 | 33244287 | 22 | 5 | 1.9811 | 0.023792 | RPS18 | 0.023792 | 0.044274 | 0.035975 |

| | | | | | | | | | | | |
|---|---|---|---|---|---|---|---|---|---|---|---|
| ENSG00000109667 | 4 | 9772777 | 10056560 | 1388 | 32 | 1.9804 | 0.023831 | SLC2A9 | 0.023831 | 0.024582 | 0.021206 |
| ENSG00000135457 | 12 | 51487446 | 51566926 | 272 | 21 | 1.9804 | 0.023832 | TFCP2 | 0.023832 | 0.022579 | 0.027852 |
| ENSG00000182931 | 20 | 44313292 | 44333658 | 80 | 9 | 1.9803 | 0.023835 | WFDC10B | 0.023835 | 0.025037 | 0.046773 |
| ENSG00000175449 | 5 | 94982458 | 95020477 | 96 | 16 | 1.9802 | 0.023838 | RFESD | 0.023838 | 0.040891 | 0.086156 |
| ENSG00000257327 | 12 | 104235229 | 104238442 | 18 | 6 | 1.98 | 0.023852 | RP11-650K20.3 | 0.023852 | 0.043302 | 0.029392 |
| ENSG00000103888 | 15 | 81071684 | 81244117 | 422 | 80 | 1.9756 | 0.024099 | KIAA1199 | 0.024099 | 0.04467 | 0.29203 |
| ENSG00000107872 | 10 | 104178946 | 104182893 | 3 | 2 | 1.9741 | 0.024187 | FBXL15 | 0.024187 | 0.17762 | 0.22692 |
| ENSG00000181752 | 11 | 55926870 | 55927793 | 1 | 1 | 1.9703 | 0.0244 | OR8K5 | 0.0244 | 0.035647 | 0.029822 |
| ENSG00000114942 | 2 | 207024309 | 207027652 | 7 | 2 | 1.9682 | 0.024522 | EEF1B2 | 0.024522 | 0.051929 | 0.061703 |
| ENSG00000010626 | 12 | 6982733 | 7023407 | 160 | 6 | 1.9678 | 0.024545 | LRRC23 | 0.024545 | 0.024116 | 0.021548 |
| ENSG00000185567 | 14 | 105403581 | 105444694 | 256 | 22 | 1.9672 | 0.024582 | AHNAK2 | 0.024582 | 0.053164 | 0.070973 |
| ENSG00000152193 | 13 | 79188426 | 79233314 | 120 | 9 | 1.967 | 0.02459 | RNF219 | 0.02459 | 0.032952 | 0.033342 |
| ENSG00000173349 | 2 | 128458597 | 128461385 | 9 | 4 | 1.9665 | 0.024623 | SFT2D3 | 0.024623 | 0.27165 | 0.4155 |
| ENSG00000149451 | 20 | 3648612 | 3662893 | 72 | 25 | 1.9661 | 0.024646 | ADAM33 | 0.024646 | 0.06203 | 0.23662 |
| ENSG00000058668 | 1 | 203595689 | 203713209 | 304 | 35 | 1.9654 | 0.024685 | ATP2B4 | 0.024685 | 0.02884 | 0.041706 |
| ENSG00000248098 | 19 | 41884215 | 41930910 | 78 | 8 | 1.9652 | 0.024693 | BCKDHA | 0.024693 | 0.018233 | 0.023343 |
| ENSG00000151470 | 4 | 130014472 | 130037795 | 100 | 12 | 1.9651 | 0.024702 | C4orf33 | 0.024702 | 0.012283 | 0.016425 |
| ENSG00000125877 | 20 | 3189514 | 3204516 | 77 | 11 | 1.9648 | 0.024719 | ITPA | 0.024719 | 0.020511 | 0.020465 |
| ENSG00000255223 | 11 | 56309746 | 56310757 | 5 | 2 | 1.9647 | 0.024723 | OR5M11 | 0.024723 | 0.11533 | 0.075822 |
| ENSG00000130675 | 7 | 156786745 | 156803345 | 67 | 20 | 1.9629 | 0.024827 | MNX1 | 0.024827 | 0.082501 | 0.35383 |
| ENSG00000162078 | 16 | 2880170 | 2888967 | 21 | 3 | 1.9626 | 0.024847 | ZG16B | 0.024847 | 0.034508 | 0.16945 |
| ENSG00000197595 | 13 | 113301358 | 113338811 | 171 | 28 | 1.9622 | 0.024868 | C13orf35 | 0.024868 | 0.017581 | 0.023484 |
| ENSG00000135502 | 12 | 58013310 | 58019934 | 6 | 3 | 1.9602 | 0.024984 | SLC26A10 | 0.024984 | 0.21149 | 0.24071 |
| ENSG00000105289 | 19 | 3708107 | 3750811 | 207 | 36 | 1.9593 | 0.02504 | TJP3 | 0.02504 | 0.0049531 | 0.0044338 |
| ENSG00000136861 | 9 | 123151147 | 123342448 | 312 | 27 | 1.9588 | 0.025067 | CDK5RAP2 | 0.025067 | 0.035027 | 0.042892 |
| ENSG00000189275 | 10 | 133604734 | 133622535 | 81 | 8 | 1.9579 | 0.025123 | AL450307.1 | 0.025123 | 0.048911 | 0.055106 |
| ENSG00000213132 | 3 | 187896331 | 187898598 | 13 | 4 | 1.9569 | 0.02518 | AC022498.1 | 0.02518 | 0.038961 | 0.075934 |
| ENSG00000125944 | 1 | 23630264 | 23670829 | 54 | 10 | 1.9547 | 0.02531 | HNRNPR | 0.02531 | 0.024501 | 0.023753 |
| ENSG00000124181 | 20 | 39765600 | 39811629 | 70 | 18 | 1.954 | 0.025351 | PLCG1 | 0.025351 | 0.044097 | 0.028242 |
| ENSG00000132746 | 11 | 67429633 | 67448671 | 62 | 7 | 1.9539 | 0.025357 | ALDH3B2 | 0.025357 | 0.026146 | 0.027411 |
| ENSG00000185177 | 7 | 57187321 | 57207571 | 90 | 8 | 1.9528 | 0.025422 | ZNF479 | 0.025422 | 0.016027 | 0.019857 |
| ENSG00000102468 | 13 | 47405685 | 47471169 | 263 | 35 | 1.9523 | 0.025452 | HTR2A | 0.025452 | 0.026582 | 0.053831 |
| ENSG00000140481 | 15 | 74509613 | 74628813 | 349 | 35 | 1.9523 | 0.025452 | CCDC33 | 0.025452 | 0.028399 | 0.038591 |
| ENSG00000104267 | 8 | 86376081 | 86393722 | 37 | 9 | 1.9518 | 0.02548 | CA2 | 0.02548 | 0.067729 | 0.1369 |
| ENSG00000173137 | 8 | 145596790 | 145618457 | 30 | 7 | 1.9518 | 0.025481 | ADCK5 | 0.025481 | 0.04962 | 0.055297 |
| ENSG00000213139 | 3 | 186256230 | 186264491 | 10 | 4 | 1.9518 | 0.025483 | CRYGS | 0.025483 | 0.11772 | 0.35184 |
| ENSG00000267772 | 17 | 58641908 | 58663984 | 32 | 7 | 1.9483 | 0.02569 | RP11-15E18.4 | 0.02569 | 0.039092 | 0.067229 |
| ENSG00000005513 | 16 | 1031808 | 1036979 | 26 | 7 | 1.9473 | 0.025748 | SOX8 | 0.025748 | 0.15407 | 0.50094 |
| ENSG00000197471 | 16 | 29674300 | 29682187 | 21 | 7 | 1.9453 | 0.025869 | SPN | 0.025869 | 0.080817 | 0.080434 |
| ENSG00000151422 | 5 | 108083523 | 108532542 | 1949 | 50 | 1.9449 | 0.025892 | FER | 0.025892 | 0.027794 | 0.033348 |
| ENSG00000120802 | 12 | 98909290 | 98944157 | 113 | 18 | 1.9442 | 0.025936 | TMPO | 0.025936 | 0.0089733 | 0.014063 |
| ENSG00000135318 | 6 | 86159809 | 86205500 | 74 | 13 | 1.9442 | 0.025938 | NT5E | 0.025938 | 0.012458 | 0.0095773 |
| ENSG00000259649 | 15 | 81188532 | 81202118 | 18 | 9 | 1.9411 | 0.026122 | RP11-351M8.1 | 0.026122 | 0.017342 | 0.0062854 |
| ENSG00000170579 | 18 | 3496030 | 4455335 | 3984 | 328 | 1.9406 | 0.026152 | DLGAP1 | 0.026152 | 0.041777 | 0.054312 |
| ENSG00000198929 | 1 | 162039564 | 162353321 | 1160 | 72 | 1.9392 | 0.026239 | NOS1AP | 0.026239 | 0.0166 | 0.013775 |
| ENSG00000204628 | 5 | 180663909 | 180675096 | 51 | 8 | 1.9388 | 0.02626 | GNB2L1 | 0.02626 | 0.088523 | 0.14408 |
| ENSG00000196670 | 5 | 180274611 | 180288286 | 56 | 8 | 1.9387 | 0.026271 | ZFP62 | 0.026271 | 0.011574 | 0.039234 |
| ENSG00000148814 | 10 | 134145614 | 134195010 | 249 | 29 | 1.9384 | 0.026285 | LRRC27 | 0.026285 | 0.03898 | 0.042292 |
| ENSG00000160221 | 21 | 45553487 | 45565605 | 37 | 5 | 1.9384 | 0.026285 | C21orf33 | 0.026285 | 0.021057 | 0.027404 |
| ENSG00000175029 | 10 | 126676421 | 126849739 | 772 | 75 | 1.9372 | 0.026361 | CTBP2 | 0.026361 | 0.011352 | 0.0080035 |
| ENSG00000221988 | 6 | 32121218 | 32134011 | 36 | 12 | 1.9372 | 0.026363 | PPT2 | 0.026363 | 0.023811 | 0.034092 |
| ENSG00000119599 | 14 | 73393040 | 73426411 | 163 | 23 | 1.9367 | 0.026388 | DCAF4 | 0.026388 | 0.042103 | 0.067089 |
| ENSG00000022567 | 8 | 142217265 | 142318404 | 418 | 49 | 1.9364 | 0.026409 | SLC45A4 | 0.026409 | 0.045214 | 0.066673 |
| ENSG00000215906 | 1 | 23279536 | 23291831 | 40 | 11 | 1.9356 | 0.026458 | LACTBL1 | 0.026458 | 0.26221 | 0.42505 |
| ENSG00000008438 | 19 | 46522411 | 46526323 | 9 | 3 | 1.9356 | 0.026459 | PGLYRP1 | 0.026459 | 0.22737 | 0.38857 |
| ENSG00000182473 | 17 | 74077087 | 74117657 | 154 | 13 | 1.9344 | 0.026533 | EXOC7 | 0.026533 | 0.054695 | 0.062382 |
| ENSG00000130368 | 6 | 160327974 | 160329108 | 1 | 1 | 1.933 | 0.02662 | MAS1 | 0.02662 | 0.11759 | 0.060369 |
| ENSG00000177596 | 13 | 74987094 | 74993252 | 27 | 9 | 1.9325 | 0.026649 | AL355390.1 | 0.026649 | 0.15267 | 0.329 |
| ENSG00000164068 | 3 | 49726932 | 49758962 | 58 | 11 | 1.9323 | 0.02666 | RNF123 | 0.02666 | 0.03701 | 0.064878 |
| ENSG00000004478 | 12 | 2904119 | 2914576 | 14 | 5 | 1.9317 | 0.026698 | FKBP4 | 0.026698 | 0.055384 | 0.093629 |
| ENSG00000188404 | 1 | 169659808 | 169680839 | 100 | 15 | 1.9269 | 0.026995 | SELL | 0.026995 | 0.050585 | 0.067049 |
| ENSG00000118420 | 6 | 83602117 | 83775560 | 521 | 20 | 1.9265 | 0.027022 | UBE3D | 0.027022 | 0.037646 | 0.049951 |
| ENSG00000174307 | 1 | 201434620 | 201438365 | 14 | 5 | 1.9255 | 0.027084 | PHLDA3 | 0.027084 | 0.0071078 | 0.026616 |
| ENSG00000196189 | 1 | 156117157 | 156147543 | 67 | 13 | 1.9247 | 0.027131 | SEMA4A | 0.027131 | 0.15048 | 0.30351 |
| ENSG00000243284 | 1 | 159824106 | 159832447 | 34 | 10 | 1.9245 | 0.027147 | VSIG8 | 0.027147 | 0.018835 | 0.021745 |
| ENSG00000184304 | 14 | 30045687 | 30661104 | 1739 | 118 | 1.9241 | 0.027168 | PRKD1 | 0.027168 | 0.030322 | 0.026141 |
| ENSG00000203813 | 6 | 27777842 | 27778314 | 1 | 1 | 1.924 | 0.02718 | HIST1H3H | 0.02718 | 0.69893 | 0.72191 |
| ENSG00000135063 | 9 | 71939488 | 72007371 | 294 | 28 | 1.9234 | 0.027217 | FAM189A2 | 0.027217 | 0.047536 | 0.052756 |
| ENSG00000144410 | 2 | 207804278 | 207834198 | 94 | 17 | 1.9214 | 0.02734 | CPO | 0.02734 | 0.013888 | 0.030275 |
| ENSG00000166006 | 12 | 75433857 | 75603648 | 516 | 34 | 1.9193 | 0.027473 | KCNC2 | 0.027473 | 0.023759 | 0.024023 |
| ENSG00000185002 | 6 | 117198375 | 117253326 | 224 | 20 | 1.9169 | 0.027626 | RFX6 | 0.027626 | 0.051273 | 0.086433 |
| ENSG00000077943 | 10 | 15555948 | 15762124 | 626 | 41 | 1.9155 | 0.027713 | ITGA8 | 0.027713 | 0.027383 | 0.041641 |
| ENSG00000146826 | 7 | 99752043 | 99756338 | 3 | 2 | 1.9129 | 0.02788 | C7orf43 | 0.02788 | 0.050193 | 0.01991 |

| | | | | | | | | | | | |
|---|---|---|---|---|---|---|---|---|---|---|---|
| ENSG00000123500 | 6 | 116440086 | 116479910 | 94 | 18 | 1.9118 | 0.027949 | COL10A1 | 0.027949 | 0.01992 | 0.022948 |
| ENSG00000125746 | 19 | 46110252 | 46148887 | 91 | 16 | 1.9093 | 0.028112 | EML2 | 0.028112 | 0.000023209 | 0.000014706 |
| ENSG00000101608 | 18 | 3247479 | 3256234 | 49 | 6 | 1.9083 | 0.028175 | MYL12A | 0.028175 | 0.085355 | 0.097412 |
| ENSG00000222046 | 1 | 32674695 | 32681797 | 13 | 6 | 1.9068 | 0.028272 | DCDC2B | 0.028272 | 0.0035344 | 0.0035052 |
| ENSG00000177191 | 19 | 41931264 | 41934635 | 8 | 3 | 1.9068 | 0.028274 | B3GNT8 | 0.028274 | 0.024205 | 0.047781 |
| ENSG00000215474 | 18 | 44746293 | 44775554 | 61 | 10 | 1.9061 | 0.028319 | SKOR2 | 0.028319 | 0.029128 | 0.033968 |
| ENSG00000177946 | 16 | 90036206 | 90038942 | 9 | 4 | 1.9024 | 0.028558 | CENPBD1 | 0.028558 | 0.062248 | 0.094238 |
| ENSG00000075426 | 2 | 28615315 | 28640179 | 52 | 9 | 1.9018 | 0.028599 | FOSL2 | 0.028599 | 0.04269 | 0.11596 |
| ENSG00000156222 | 15 | 85427885 | 85518876 | 476 | 59 | 1.9017 | 0.028608 | SLC28A1 | 0.028608 | 0.020652 | 0.0084488 |
| ENSG00000145365 | 4 | 113195698 | 113207059 | 49 | 3 | 1.9014 | 0.028628 | TIFA | 0.028628 | 0.046593 | 0.031667 |
| ENSG00000138663 | 4 | 83955600 | 83996971 | 72 | 13 | 1.9002 | 0.028706 | COPS4 | 0.028706 | 0.028412 | 0.072542 |
| ENSG00000262601 | 16 | 85204882 | 85218969 | 54 | 17 | 1.8989 | 0.028788 | CTC-786C10.1 | 0.028788 | 0.0050535 | 0.0042399 |
| ENSG00000160712 | 1 | 154377669 | 154441926 | 202 | 22 | 1.8984 | 0.028819 | IL6R | 0.028819 | 0.031145 | 0.056195 |
| ENSG00000160218 | 21 | 45432200 | 45526433 | 222 | 13 | 1.8965 | 0.028944 | TRAPPC10 | 0.028944 | 0.02839 | 0.029869 |
| ENSG00000134986 | 5 | 110998318 | 111333161 | 1085 | 58 | 1.8958 | 0.028994 | NREP | 0.028994 | 0.030113 | 0.034696 |
| ENSG00000137486 | 11 | 74975226 | 75062873 | 217 | 44 | 1.8956 | 0.029008 | ARRB1 | 0.029008 | 0.049047 | 0.087849 |
| ENSG00000268635 | 11 | 77184416 | 77188074 | 10 | 4 | 1.8949 | 0.029055 | DKFZP434E1119 | 0.029055 | 0.15689 | 0.22575 |
| ENSG00000249139 | 20 | 44165625 | 44176065 | 11 | 6 | 1.8928 | 0.029193 | EPPIN-WFDC6 | 0.029193 | 0.44185 | 0.5542 |
| ENSG00000085998 | 1 | 46654354 | 46685977 | 56 | 11 | 1.8922 | 0.029234 | POMGNT1 | 0.029234 | 0.025075 | 0.042575 |
| ENSG00000080839 | 20 | 35624752 | 35724398 | 151 | 23 | 1.8905 | 0.029348 | RBL1 | 0.029348 | 0.015235 | 0.050699 |
| ENSG00000182326 | 12 | 7096351 | 7178336 | 177 | 18 | 1.8869 | 0.029586 | C1S | 0.029586 | 0.022733 | 0.022461 |
| ENSG00000143355 | 1 | 197881037 | 197904608 | 70 | 16 | 1.885 | 0.029717 | LHX9 | 0.029717 | 0.03882 | 0.056945 |
| ENSG00000152154 | 2 | 39892122 | 39945103 | 79 | 8 | 1.8842 | 0.029766 | TMEM178A | 0.029766 | 0.031909 | 0.013503 |
| ENSG00000111846 | 6 | 10492456 | 10629601 | 565 | 49 | 1.8839 | 0.029792 | GCNT2 | 0.029792 | 0.046678 | 0.036788 |
| ENSG00000101158 | 20 | 57556263 | 57570188 | 37 | 7 | 1.8838 | 0.029796 | NELFCD | 0.029796 | 0.10486 | 0.14076 |
| ENSG00000172005 | 2 | 95691422 | 95719737 | 31 | 10 | 1.8824 | 0.029893 | MAL | 0.029893 | 0.066138 | 0.12911 |
| ENSG00000151490 | 12 | 15475331 | 15750333 | 739 | 62 | 1.882 | 0.029916 | PTPRO | 0.029916 | 0.029336 | 0.028928 |
| ENSG00000168269 | 5 | 169532901 | 169536727 | 21 | 8 | 1.8818 | 0.029932 | FOXI1 | 0.029932 | 0.12675 | 0.18804 |
| ENSG00000176732 | 2 | 24338241 | 24346347 | 31 | 5 | 1.8812 | 0.02997 | PFN4 | 0.02997 | 0.036988 | 0.065435 |
| ENSG00000257955 | 12 | 48413554 | 48419165 | 33 | 6 | 1.8785 | 0.030159 | RP1-228P16.5 | 0.030159 | 0.15836 | 0.42619 |
| ENSG00000205108 | 9 | 34723052 | 34729464 | 30 | 7 | 1.8773 | 0.030237 | FAM205A | 0.030237 | 0.030399 | 0.022244 |
| ENSG00000134013 | 8 | 23154702 | 23282841 | 771 | 37 | 1.8764 | 0.0303 | LOXL2 | 0.0303 | 0.028832 | 0.030833 |
| ENSG00000185885 | 11 | 313506 | 315272 | 5 | 2 | 1.8763 | 0.030307 | IFITM1 | 0.030307 | 0.40706 | 0.52304 |
| ENSG00000258818 | 14 | 21152259 | 21168761 | 78 | 14 | 1.8763 | 0.030309 | RNASE4 | 0.030309 | 0.073834 | 0.28223 |
| ENSG00000124120 | 20 | 43104526 | 43123244 | 40 | 8 | 1.8762 | 0.030311 | TTPAL | 0.030311 | 0.026215 | 0.036515 |
| ENSG00000167524 | 17 | 26934982 | 26941218 | 4 | 2 | 1.8757 | 0.030351 | SGK494 | 0.030351 | 0.18027 | 0.17761 |
| ENSG00000115282 | 2 | 74710200 | 74722013 | 5 | 3 | 1.8756 | 0.030354 | TTC31 | 0.030354 | 0.38519 | 0.41877 |
| ENSG00000167716 | 17 | 1619817 | 1641893 | 74 | 12 | 1.8754 | 0.03037 | WDR81 | 0.03037 | 0.055315 | 0.078507 |
| ENSG00000235608 | 4 | 1396720 | 1400119 | 5 | 2 | 1.8722 | 0.03059 | NKX1-1 | 0.03059 | 0.035355 | 0.046378 |
| ENSG00000131386 | 3 | 16216156 | 16273499 | 238 | 37 | 1.8721 | 0.030595 | GALNT15 | 0.030595 | 0.063559 | 0.1357 |
| ENSG00000204306 | 11 | 120382511 | 120385732 | 9 | 3 | 1.8719 | 0.030611 | AP002348.1 | 0.030611 | 0.059736 | 0.05374 |
| ENSG00000119688 | 14 | 74752126 | 74769759 | 41 | 9 | 1.871 | 0.030674 | ABCD4 | 0.030674 | 0.10592 | 0.12076 |
| ENSG00000204859 | 1 | 6640061 | 6649340 | 13 | 5 | 1.871 | 0.030676 | ZBTB48 | 0.030676 | 0.099339 | 0.096072 |
| ENSG00000222031 | 2 | 151857734 | 151905288 | 221 | 24 | 1.8709 | 0.030677 | AC023469.1 | 0.030677 | 0.045522 | 0.033897 |
| ENSG00000065243 | 1 | 89149905 | 89301938 | 479 | 15 | 1.8709 | 0.030678 | PKN2 | 0.030678 | 0.027433 | 0.02475 |
| ENSG00000204304 | 6 | 32152512 | 32157963 | 19 | 9 | 1.87 | 0.03074 | PBX2 | 0.03074 | 0.046139 | 0.052387 |
| ENSG00000125945 | 1 | 23685941 | 23695935 | 33 | 7 | 1.8673 | 0.030929 | ZNF436 | 0.030929 | 0.051509 | 0.12486 |
| ENSG00000181767 | 11 | 55872519 | 55873457 | 9 | 2 | 1.8669 | 0.030961 | OR8H2 | 0.030961 | 0.018481 | 0.019292 |
| ENSG00000074657 | 18 | 56529832 | 56653712 | 368 | 31 | 1.8657 | 0.031042 | ZNF532 | 0.031042 | 0.033924 | 0.012586 |
| ENSG00000125910 | 19 | 3172344 | 3180329 | 68 | 19 | 1.8646 | 0.031122 | S1PR4 | 0.031122 | 0.15876 | 0.29181 |
| ENSG00000268484 | 15 | 96904487 | 96904900 | 1 | 1 | 1.8643 | 0.03114 | AC087477.1 | 0.03114 | 0.29486 | 0.22388 |
| ENSG00000260851 | 16 | 66503715 | 66551215 | 83 | 10 | 1.8643 | 0.031143 | RP11-403P17.5 | 0.031143 | 0.02317 | 0.017481 |
| ENSG00000137563 | 8 | 63927638 | 63951730 | 82 | 12 | 1.8636 | 0.031191 | GGH | 0.031191 | 0.023413 | 0.023386 |
| ENSG00000149133 | 11 | 55761157 | 55762101 | 6 | 2 | 1.8618 | 0.031315 | OR5F1 | 0.031315 | 0.028628 | 0.024498 |
| ENSG00000081800 | 7 | 122753585 | 122840040 | 212 | 28 | 1.8615 | 0.031339 | SLC13A1 | 0.031339 | 0.05138 | 0.075856 |
| ENSG00000177807 | 1 | 160007257 | 160040038 | 73 | 16 | 1.8615 | 0.03134 | KCNJ10 | 0.03134 | 0.021878 | 0.08441 |
| ENSG00000108264 | 17 | 35766965 | 35839835 | 247 | 22 | 1.8614 | 0.031346 | TADA2A | 0.031346 | 0.033329 | 0.028025 |
| ENSG00000185896 | 13 | 113951556 | 113977987 | 23 | 14 | 1.8614 | 0.031347 | LAMP1 | 0.031347 | 0.0050471 | 0.11985 |
| ENSG00000137809 | 15 | 68594050 | 68724501 | 513 | 52 | 1.8607 | 0.031392 | ITGA11 | 0.031392 | 0.034303 | 0.030794 |
| ENSG00000179348 | 3 | 128198270 | 128212028 | 46 | 15 | 1.8597 | 0.031461 | GATA2 | 0.031461 | 0.042874 | 0.04434 |
| ENSG00000154359 | 8 | 12579403 | 12613582 | 111 | 12 | 1.8562 | 0.031714 | LONRF1 | 0.031714 | 0.016991 | 0.015483 |
| ENSG00000179855 | 19 | 3585551 | 3593539 | 27 | 8 | 1.856 | 0.031725 | GIPC3 | 0.031725 | 0.0020386 | 0.0023876 |
| ENSG00000178462 | 10 | 5435061 | 5446793 | 44 | 6 | 1.8559 | 0.031737 | TUBAL3 | 0.031737 | 0.030475 | 0.12722 |
| ENSG00000124772 | 6 | 36708552 | 36807778 | 398 | 60 | 1.8558 | 0.03174 | CPNE5 | 0.03174 | 0.030653 | 0.08026 |
| ENSG00000189308 | 4 | 83831126 | 83934079 | 276 | 11 | 1.8555 | 0.031765 | LIN54 | 0.031765 | 0.034364 | 0.031476 |
| ENSG00000260240 | 16 | 59743897 | 59788752 | 137 | 25 | 1.8554 | 0.03177 | RP11-105C20.2 | 0.03177 | 0.037987 | 0.056555 |
| ENSG00000181609 | 11 | 5509915 | 5510979 | 9 | 3 | 1.8521 | 0.032004 | OR52D1 | 0.032004 | 0.062653 | 0.19615 |
| ENSG00000182580 | 3 | 184279572 | 184300197 | 70 | 16 | 1.8515 | 0.032051 | EPHB3 | 0.032051 | 0.071718 | 0.077463 |
| ENSG00000258227 | 7 | 141627157 | 141646807 | 58 | 7 | 1.8494 | 0.032203 | CLEC5A | 0.032203 | 0.025479 | 0.039187 |
| ENSG00000230657 | 12 | 11460017 | 11463369 | 14 | 3 | 1.8483 | 0.032281 | PRB4 | 0.032281 | 0.037159 | 0.052701 |
| ENSG00000146250 | 6 | 84222194 | 84235423 | 38 | 5 | 1.8472 | 0.032357 | PRSS35 | 0.032357 | 0.08163 | 0.039091 |
| ENSG00000187492 | 3 | 49828165 | 49837268 | 19 | 7 | 1.8464 | 0.032417 | CDHR4 | 0.032417 | 0.036378 | 0.047123 |
| ENSG00000257704 | 19 | 47778142 | 47778979 | 3 | 1 | 1.8453 | 0.032496 | PRR24 | 0.032496 | 0.093297 | 0.078703 |

| | | | | | | | | | | | |
|---|---|---|---|---|---|---|---|---|---|---|---|
| ENSG00000111348 | 12 | 15094951 | 15114662 | 55 | 10 | 1.8453 | 0.032497 | ARHGDIB | 0.032497 | 0.18069 | 0.28142 |
| ENSG00000259171 | 14 | 21152746 | 21168735 | 71 | 15 | 1.8448 | 0.032533 | AL163636.6 | 0.032533 | 0.071276 | 0.26839 |
| ENSG00000112877 | 5 | 612387 | 667283 | 276 | 20 | 1.8446 | 0.03255 | CEP72 | 0.03255 | 0.01064 | 0.0058083 |
| ENSG00000099949 | 22 | 21333751 | 21353327 | 65 | 21 | 1.8441 | 0.032582 | LZTR1 | 0.032582 | 0.13038 | 0.25208 |
| ENSG00000111271 | 12 | 112123857 | 112194903 | 38 | 19 | 1.8441 | 0.032587 | ACAD10 | 0.032587 | 0.011626 | 0.010906 |
| ENSG00000162873 | 1 | 205305220 | 205326218 | 107 | 17 | 1.844 | 0.032588 | KLHDC8A | 0.032588 | 0.025712 | 0.016828 |
| ENSG00000153814 | 7 | 27870192 | 28220362 | 1008 | 92 | 1.8439 | 0.032595 | JAZF1 | 0.032595 | 0.041553 | 0.044278 |
| ENSG00000160999 | 7 | 101928405 | 101962178 | 94 | 21 | 1.8439 | 0.032599 | SH2B2 | 0.032599 | 0.062262 | 0.12585 |
| ENSG00000170471 | 20 | 37101459 | 37207504 | 197 | 18 | 1.8434 | 0.032639 | RALGAPB | 0.032639 | 0.029115 | 0.04304 |
| ENSG00000167775 | 19 | 8367011 | 8373240 | 46 | 5 | 1.8429 | 0.032674 | CD320 | 0.032674 | 0.046765 | 0.074941 |
| ENSG00000066735 | 14 | 104605060 | 104647231 | 247 | 43 | 1.8423 | 0.032717 | KIF26A | 0.032717 | 0.034571 | 0.063996 |
| ENSG00000214309 | 7 | 99724317 | 99726118 | 3 | 1 | 1.8419 | 0.032748 | MBLAC1 | 0.032748 | 0.025068 | 0.031981 |
| ENSG00000108819 | 17 | 48211104 | 48227991 | 25 | 9 | 1.8416 | 0.032767 | PPP1R9B | 0.032767 | 0.088706 | 0.42134 |
| ENSG00000111671 | 12 | 6980099 | 6998522 | 58 | 6 | 1.8409 | 0.032821 | SPSB2 | 0.032821 | 0.02305 | 0.023139 |
| ENSG00000169242 | 1 | 155099936 | 155107333 | 14 | 4 | 1.8397 | 0.032903 | EFNA1 | 0.032903 | 0.045817 | 0.037878 |
| ENSG00000145107 | 3 | 196046213 | 196065374 | 91 | 13 | 1.8397 | 0.032908 | TM4SF19 | 0.032908 | 0.038268 | 0.072532 |
| ENSG00000135736 | 16 | 57546090 | 57570511 | 95 | 16 | 1.8395 | 0.032921 | CCDC102A | 0.032921 | 0.0036806 | 0.0020542 |
| ENSG00000106346 | 7 | 6144515 | 6201195 | 192 | 24 | 1.8388 | 0.032971 | USP42 | 0.032971 | 0.024047 | 0.020786 |
| ENSG00000112541 | 6 | 165740776 | 166400091 | 2890 | 221 | 1.837 | 0.033103 | PDE10A | 0.033103 | 0.03312 | 0.05764 |
| ENSG00000162384 | 1 | 53679771 | 53686289 | 11 | 3 | 1.8369 | 0.033111 | C1orf123 | 0.033111 | 0.036532 | 0.026133 |
| ENSG00000181785 | 11 | 55797895 | 55798869 | 7 | 2 | 1.8363 | 0.033157 | OR5AS1 | 0.033157 | 0.021153 | 0.021111 |
| ENSG00000137941 | 1 | 84330711 | 84464833 | 442 | 18 | 1.8341 | 0.033316 | TTLL7 | 0.033316 | 0.03374 | 0.047256 |
| ENSG00000254834 | 11 | 56344212 | 56345263 | 3 | 2 | 1.8323 | 0.033457 | OR5M10 | 0.033457 | 0.014007 | 0.020431 |
| ENSG00000166471 | 11 | 9302201 | 9336327 | 139 | 15 | 1.8322 | 0.03346 | TMEM41B | 0.03346 | 0.044708 | 0.068133 |
| ENSG00000077380 | 2 | 172543919 | 172604930 | 146 | 20 | 1.8315 | 0.033512 | DYNC1I2 | 0.033512 | 0.046861 | 0.062826 |
| ENSG00000174243 | 12 | 49223547 | 49246625 | 48 | 8 | 1.8312 | 0.033536 | DDX23 | 0.033536 | 0.051183 | 0.079673 |
| ENSG00000115239 | 2 | 53759810 | 54087170 | 1237 | 74 | 1.8304 | 0.033595 | GPR75-ASB3 | 0.033595 | 0.029925 | 0.037711 |
| ENSG00000174306 | 20 | 39807088 | 39946312 | 299 | 19 | 1.8297 | 0.033646 | ZHX3 | 0.033646 | 0.035823 | 0.038149 |
| ENSG00000168152 | 4 | 83821837 | 83841122 | 47 | 5 | 1.8294 | 0.033667 | THAP9 | 0.033667 | 0.033521 | 0.043401 |
| ENSG00000174970 | 11 | 55734975 | 55735990 | 11 | 1 | 1.8293 | 0.03368 | OR10AG1 | 0.03368 | 0.027355 | 0.029688 |
| ENSG00000106819 | 9 | 95218487 | 95244788 | 61 | 10 | 1.8282 | 0.033756 | ASPN | 0.033756 | 0.049048 | 0.050874 |
| ENSG00000099849 | 11 | 560404 | 564021 | 18 | 8 | 1.8281 | 0.033768 | RASSF7 | 0.033768 | 0.0097249 | 0.001513 |
| ENSG00000178726 | 20 | 23026270 | 23030378 | 7 | 4 | 1.8281 | 0.033768 | THBD | 0.033768 | 0.20667 | 0.36909 |
| ENSG00000255804 | 14 | 23102673 | 23103716 | 15 | 3 | 1.8265 | 0.033884 | OR6J1 | 0.033884 | 0.010744 | 0.0067861 |
| ENSG00000179639 | 1 | 159259504 | 159278014 | 64 | 13 | 1.8256 | 0.033959 | FCER1A | 0.033959 | 0.050263 | 0.14275 |
| ENSG00000173581 | 19 | 56152428 | 56164527 | 34 | 8 | 1.8253 | 0.033979 | CCDC106 | 0.033979 | 0.082254 | 0.26729 |
| ENSG00000149201 | 11 | 86085778 | 86134151 | 192 | 31 | 1.8244 | 0.034049 | CCDC81 | 0.034049 | 0.0059763 | 0.014221 |
| ENSG00000140022 | 14 | 81727000 | 81902809 | 548 | 38 | 1.8236 | 0.034105 | STON2 | 0.034105 | 0.051655 | 0.075435 |
| ENSG00000185565 | 3 | 115521235 | 117716095 | 7694 | 327 | 1.8231 | 0.034145 | LSAMP | 0.034145 | 0.033966 | 0.031815 |
| ENSG00000166503 | 15 | 83784320 | 83876770 | 168 | 13 | 1.8227 | 0.034177 | HDGFRP3 | 0.034177 | 0.026975 | 0.032211 |
| ENSG00000102575 | 19 | 11685475 | 11689823 | 10 | 3 | 1.8221 | 0.034221 | ACP5 | 0.034221 | 0.035293 | 0.05333 |
| ENSG00000258512 | 14 | 102196774 | 102198859 | 12 | 1 | 1.8205 | 0.034338 | RP11-796G6.2 | 0.034338 | 0.043143 | 0.053593 |
| ENSG00000148429 | 10 | 11495945 | 11653753 | 458 | 31 | 1.8194 | 0.034427 | USP6NL | 0.034427 | 0.029533 | 0.031602 |
| ENSG00000121542 | 3 | 122920774 | 122992977 | 320 | 7 | 1.8178 | 0.03455 | SEC22A | 0.03455 | 0.033138 | 0.032091 |
| ENSG00000122705 | 9 | 36190853 | 36304778 | 342 | 25 | 1.8168 | 0.034627 | CLTA | 0.034627 | 0.07081 | 0.1206 |
| ENSG00000240386 | 1 | 152748848 | 152749445 | 3 | 1 | 1.8163 | 0.034661 | LCE1F | 0.034661 | 0.13542 | 0.11884 |
| ENSG00000136541 | 2 | 158175137 | 158184225 | 7 | 3 | 1.8162 | 0.034672 | ERMN | 0.034672 | 0.11309 | 0.62604 |
| ENSG00000104738 | 8 | 48872745 | 48890720 | 14 | 5 | 1.8148 | 0.034775 | MCM4 | 0.034775 | 0.045676 | 0.044797 |
| ENSG00000145075 | 3 | 180320646 | 180588793 | 750 | 49 | 1.8147 | 0.034785 | CCDC39 | 0.034785 | 0.033124 | 0.037553 |
| ENSG00000130024 | 6 | 170104001 | 170124151 | 37 | 10 | 1.8147 | 0.034785 | PHF10 | 0.034785 | 0.040548 | 0.029656 |
| ENSG00000154556 | 4 | 186506598 | 186877806 | 1803 | 157 | 1.8145 | 0.034801 | SORBS2 | 0.034801 | 0.039247 | 0.02445 |
| ENSG00000111676 | 12 | 7033626 | 7051484 | 32 | 14 | 1.8139 | 0.034845 | ATN1 | 0.034845 | 0.011613 | 0.012322 |
| ENSG00000258713 | 20 | 2795633 | 2796479 | 4 | 2 | 1.8139 | 0.034849 | C20orf141 | 0.034849 | 0.40119 | 0.19467 |
| ENSG00000071127 | 4 | 10075963 | 10118573 | 181 | 18 | 1.8132 | 0.034902 | WDR1 | 0.034902 | 0.040768 | 0.073393 |
| ENSG00000139899 | 14 | 24895738 | 24900160 | 20 | 5 | 1.8127 | 0.03494 | CBLN3 | 0.03494 | 0.10327 | 0.18796 |
| ENSG00000124613 | 6 | 27342394 | 27371683 | 181 | 7 | 1.8125 | 0.034955 | ZNF391 | 0.034955 | 0.028519 | 0.027806 |
| ENSG00000069011 | 5 | 134363425 | 134370503 | 29 | 4 | 1.812 | 0.034993 | PITX1 | 0.034993 | 0.0079785 | 0.00033906 |
| ENSG00000168214 | 4 | 26165077 | 26436541 | 711 | 63 | 1.8115 | 0.035032 | RBPJ | 0.035032 | 0.050907 | 0.050065 |
| ENSG00000181693 | 11 | 56057529 | 56058566 | 5 | 2 | 1.8113 | 0.03505 | OR8H1 | 0.03505 | 0.019454 | 0.025655 |
| ENSG00000154832 | 18 | 47808713 | 47814674 | 10 | 4 | 1.8105 | 0.035112 | CXXC1 | 0.035112 | 0.054664 | 0.14746 |
| ENSG00000144554 | 3 | 10068098 | 10143614 | 270 | 6 | 1.8095 | 0.035188 | FANCD2 | 0.035188 | 0.034599 | 0.032481 |
| ENSG00000182348 | 7 | 88388682 | 88966346 | 2334 | 90 | 1.8093 | 0.035205 | ZNF804B | 0.035205 | 0.03981 | 0.04503 |
| ENSG00000170448 | 4 | 47849257 | 47916653 | 201 | 16 | 1.8091 | 0.035219 | NFXL1 | 0.035219 | 0.057532 | 0.069697 |
| ENSG00000013441 | 2 | 201717732 | 201729422 | 37 | 14 | 1.8086 | 0.035257 | CLK1 | 0.035257 | 0.1642 | 0.24654 |
| ENSG00000249049 | 4 | 92240185 | 92246457 | 28 | 3 | 1.8085 | 0.035265 | RP11-763F8.1 | 0.035265 | 0.049406 | 0.058225 |
| ENSG00000173928 | 19 | 11485361 | 11487627 | 2 | 1 | 1.8085 | 0.035268 | SWSAP1 | 0.035268 | 0.63921 | 0.68051 |
| ENSG00000008130 | 1 | 1682671 | 1711896 | 87 | 12 | 1.8083 | 0.03528 | NADK | 0.03528 | 0.034447 | 0.013632 |
| ENSG00000254656 | 14 | 101346992 | 101351184 | 11 | 5 | 1.808 | 0.035301 | RTL1 | 0.035301 | 0.34181 | 0.29169 |
| ENSG00000121753 | 1 | 32192718 | 32230494 | 48 | 10 | 1.8073 | 0.035357 | BAI2 | 0.035357 | 0.046251 | 0.0024236 |
| ENSG00000160049 | 1 | 10516579 | 10532583 | 32 | 7 | 1.8062 | 0.035441 | DFFA | 0.035441 | 0.047154 | 0.052536 |
| ENSG00000007866 | 6 | 35441374 | 35464853 | 43 | 15 | 1.8054 | 0.035504 | TEAD3 | 0.035504 | 0.001572 | 0.0024051 |
| ENSG00000171487 | 19 | 56511092 | 56573179 | 580 | 68 | 1.805 | 0.035536 | NLRP5 | 0.035536 | 0.069126 | 0.084976 |
| ENSG00000114124 | 3 | 141496990 | 141537332 | 137 | 21 | 1.8043 | 0.035589 | GRK7 | 0.035589 | 0.035714 | 0.04304 |

| | | | | | | | | | | | |
|---|---|---|---|---|---|---|---|---|---|---|---|
| ENSG00000138621 | 15 | 75315896 | 75409803 | 295 | 18 | 1.8042 | 0.0356 | PPCDC | 0.0356 | 0.034683 | 0.033881 |
| ENSG00000184584 | 5 | 138855119 | 138862520 | 13 | 3 | 1.8032 | 0.035682 | TMEM173 | 0.035682 | 0.14549 | 0.19119 |
| ENSG00000185966 | 1 | 152538130 | 152539248 | 1 | 1 | 1.8024 | 0.03574 | LCE3E | 0.03574 | 0.03877 | 0.11133 |
| ENSG00000069696 | 11 | 637293 | 640706 | 19 | 7 | 1.8022 | 0.035757 | DRD4 | 0.035757 | 0.0025761 | 0.0097487 |
| ENSG00000273331 | 3 | 196042953 | 196065244 | 101 | 14 | 1.8018 | 0.035791 | TM4SF19 | 0.035791 | 0.038268 | 0.072532 |
| ENSG00000106823 | 9 | 95256365 | 95298937 | 129 | 16 | 1.8018 | 0.035792 | ECM2 | 0.035792 | 0.034414 | 0.034163 |
| ENSG00000088836 | 20 | 3208063 | 3219836 | 59 | 13 | 1.8015 | 0.03581 | SLC4A11 | 0.03581 | 0.029099 | 0.019618 |
| ENSG00000136155 | 13 | 78109809 | 78219398 | 305 | 28 | 1.8014 | 0.035821 | SCEL | 0.035821 | 0.052921 | 0.10644 |
| ENSG00000176095 | 3 | 49761727 | 49823975 | 147 | 14 | 1.8008 | 0.03587 | IP6K1 | 0.03587 | 0.036271 | 0.031018 |
| ENSG00000175866 | 17 | 79008948 | 79091232 | 452 | 57 | 1.7989 | 0.036017 | BAIAP2 | 0.036017 | 0.022073 | 0.073546 |
| ENSG00000162642 | 1 | 85715639 | 85725355 | 39 | 10 | 1.7988 | 0.036028 | C1orf52 | 0.036028 | 0.0085377 | 0.012532 |
| ENSG00000157193 | 1 | 53711217 | 53793742 | 208 | 31 | 1.7982 | 0.03607 | LRP8 | 0.03607 | 0.016804 | 0.0076109 |
| ENSG00000185614 | 3 | 49840687 | 49842463 | 2 | 1 | 1.7966 | 0.036202 | FAM212A | 0.036202 | 0.027691 | 0.042993 |
| ENSG00000169683 | 17 | 79981178 | 79989027 | 11 | 5 | 1.7964 | 0.036213 | LRRC45 | 0.036213 | 0.4992 | 0.68485 |
| ENSG00000171314 | 10 | 99185917 | 99193198 | 14 | 5 | 1.7964 | 0.036216 | PGAM1 | 0.036216 | 0.10104 | 0.033939 |
| ENSG00000221838 | 7 | 99699172 | 99707968 | 26 | 6 | 1.7959 | 0.036253 | AP4M1 | 0.036253 | 0.0063998 | 0.0077499 |
| ENSG00000268822 | 2 | 241624478 | 241624705 | 3 | 1 | 1.7957 | 0.03627 | AC011298.1 | 0.03627 | 0.36121 | 0.4759 |
| ENSG00000164576 | 5 | 153825517 | 153840614 | 33 | 15 | 1.7953 | 0.036301 | SAP30L | 0.036301 | 0.13576 | 0.10634 |
| ENSG00000102760 | 13 | 42031695 | 42045018 | 39 | 11 | 1.7952 | 0.036314 | RGCC | 0.036314 | 0.026947 | 0.0049545 |
| ENSG00000077348 | 19 | 41892279 | 41903384 | 18 | 3 | 1.7945 | 0.036365 | EXOSC5 | 0.036365 | 0.022252 | 0.017796 |
| ENSG00000149418 | 11 | 130029457 | 130080271 | 189 | 33 | 1.7943 | 0.036381 | ST14 | 0.036381 | 0.034893 | 0.028045 |
| ENSG00000110031 | 11 | 58294344 | 58345693 | 99 | 12 | 1.7942 | 0.036389 | LPXN | 0.036389 | 0.018453 | 0.01568 |
| ENSG00000268821 | 2 | 172560844 | 172567314 | 14 | 5 | 1.7941 | 0.036397 | AC068039.1 | 0.036397 | 0.045986 | 0.036631 |
| ENSG00000182199 | 12 | 57623110 | 57628718 | 7 | 4 | 1.7938 | 0.036422 | SHMT2 | 0.036422 | 0.30928 | 0.33846 |
| ENSG00000178522 | 4 | 71457973 | 71473005 | 69 | 17 | 1.7923 | 0.036543 | AMBN | 0.036543 | 0.17769 | 0.37351 |
| ENSG00000165275 | 9 | 37753804 | 37778969 | 78 | 14 | 1.7922 | 0.036554 | TRMT10B | 0.036554 | 0.02724 | 0.037515 |
| ENSG00000177106 | 11 | 694438 | 727727 | 159 | 36 | 1.7918 | 0.036584 | EPS8L2 | 0.036584 | 0.087354 | 0.11172 |
| ENSG00000172785 | 9 | 121041 | 188979 | 253 | 20 | 1.7904 | 0.036693 | CBWD1 | 0.036693 | 0.045181 | 0.077162 |
| ENSG00000105929 | 7 | 138391040 | 138484305 | 484 | 65 | 1.7901 | 0.036715 | ATP6V0A4 | 0.036715 | 0.087223 | 0.086029 |
| ENSG00000198276 | 20 | 62571186 | 62587769 | 36 | 19 | 1.7899 | 0.036735 | UCKL1 | 0.036735 | 0.062729 | 0.19649 |
| ENSG00000196172 | 19 | 23921997 | 23941693 | 48 | 13 | 1.7897 | 0.036748 | ZNF681 | 0.036748 | 0.02764 | 0.029239 |
| ENSG00000175189 | 12 | 57828543 | 57844611 | 39 | 9 | 1.7892 | 0.036788 | INHBC | 0.036788 | 0.026315 | 0.0059353 |
| ENSG00000010310 | 19 | 46171502 | 46186982 | 46 | 11 | 1.7892 | 0.03679 | GIPR | 0.03679 | 0.00022113 | 0.000029926 |
| ENSG00000171307 | 10 | 99205927 | 99217127 | 21 | 5 | 1.7891 | 0.036797 | ZDHHC16 | 0.036797 | 0.017135 | 0.023031 |
| ENSG00000161091 | 19 | 3538259 | 3574288 | 130 | 33 | 1.7888 | 0.036821 | MFSD12 | 0.036821 | 0.026031 | 0.0025002 |
| ENSG00000189181 | 1 | 248844628 | 248845629 | 5 | 1 | 1.7884 | 0.036855 | OR14I1 | 0.036855 | 0.030379 | 0.035908 |
| ENSG00000168542 | 2 | 189839046 | 189877472 | 106 | 16 | 1.7883 | 0.036865 | COL3A1 | 0.036865 | 0.023782 | 0.034794 |
| ENSG00000166845 | 18 | 51884287 | 51911588 | 115 | 7 | 1.788 | 0.036887 | C18orf54 | 0.036887 | 0.033489 | 0.033698 |
| ENSG00000173692 | 2 | 231921578 | 232037541 | 225 | 24 | 1.7876 | 0.036924 | PSMD1 | 0.036924 | 0.036106 | 0.04816 |
| ENSG00000183463 | 13 | 28551851 | 28562791 | 31 | 6 | 1.7865 | 0.03701 | URAD | 0.03701 | 0.048677 | 0.065258 |
| ENSG00000112232 | 6 | 62389865 | 62996132 | 1698 | 39 | 1.7864 | 0.037017 | KHDRBS2 | 0.037017 | 0.041327 | 0.039889 |
| ENSG00000224586 | 6 | 28493702 | 28502729 | 29 | 7 | 1.7861 | 0.037045 | GPX5 | 0.037045 | 0.035788 | 0.0077296 |
| ENSG00000113088 | 5 | 54320081 | 54330398 | 33 | 9 | 1.786 | 0.037047 | GZMK | 0.037047 | 0.044397 | 0.057623 |
| ENSG00000163705 | 3 | 10123001 | 10149915 | 76 | 5 | 1.786 | 0.037051 | FANCD2OS | 0.037051 | 0.036611 | 0.047514 |
| ENSG00000162949 | 2 | 30945637 | 31043408 | 480 | 69 | 1.7851 | 0.037124 | CAPN13 | 0.037124 | 0.03657 | 0.040864 |
| ENSG00000112511 | 6 | 33378176 | 33384230 | 16 | 7 | 1.7842 | 0.037198 | PHF1 | 0.037198 | 0.025293 | 0.043241 |
| ENSG00000119559 | 19 | 1461142 | 1479555 | 34 | 6 | 1.7835 | 0.037255 | C19orf25 | 0.037255 | 0.063092 | 0.17944 |
| ENSG00000185231 | 18 | 13882043 | 13915706 | 153 | 12 | 1.7832 | 0.03728 | MC2R | 0.03728 | 0.034628 | 0.06396 |
| ENSG00000258256 | 8 | 86134675 | 86157946 | 34 | 7 | 1.7828 | 0.037308 | RP11-219B4.5 | 0.037308 | 0.12663 | 0.13215 |
| ENSG00000187513 | 1 | 35258599 | 35261348 | 13 | 4 | 1.7808 | 0.037474 | GJA4 | 0.037474 | 0.055732 | 0.16703 |
| ENSG00000198574 | 1 | 162365056 | 162381928 | 68 | 9 | 1.7802 | 0.037526 | SH2D1B | 0.037526 | 0.049465 | 0.074636 |
| ENSG00000007038 | 16 | 2867164 | 2876305 | 18 | 4 | 1.7796 | 0.037567 | PRSS21 | 0.037567 | 0.017212 | 0.055963 |
| ENSG00000197226 | 5 | 179289066 | 179334859 | 41 | 14 | 1.7793 | 0.037598 | TBC1D9B | 0.037598 | 0.15341 | 0.17748 |
| ENSG00000267206 | 9 | 139638463 | 139642980 | 13 | 5 | 1.7791 | 0.037613 | LCN6 | 0.037613 | 0.036914 | 0.22606 |
| ENSG00000181961 | 11 | 55110627 | 55111707 | 13 | 4 | 1.7789 | 0.037629 | OR4A16 | 0.037629 | 0.078279 | 0.082266 |
| ENSG00000163013 | 2 | 73481810 | 73511559 | 60 | 18 | 1.778 | 0.0377 | FBXO41 | 0.0377 | 0.12592 | 0.16943 |
| ENSG00000021852 | 1 | 57394883 | 57431813 | 119 | 22 | 1.7767 | 0.037805 | C8B | 0.037805 | 0.047476 | 0.12902 |
| ENSG00000179796 | 3 | 26664297 | 26752267 | 280 | 30 | 1.7753 | 0.037924 | LRRC3B | 0.037924 | 0.047057 | 0.070295 |
| ENSG00000166840 | 11 | 58672871 | 58811000 | 453 | 20 | 1.7753 | 0.037926 | GLYATL1 | 0.037926 | 0.036732 | 0.045115 |
| ENSG00000204103 | 20 | 39314488 | 39317880 | 5 | 1 | 1.7748 | 0.037963 | MAFB | 0.037963 | 0.031458 | 0.012703 |
| ENSG00000125835 | 20 | 2442280 | 2451499 | 15 | 3 | 1.7742 | 0.038015 | SNRPB | 0.038015 | 0.021949 | 0.042619 |
| ENSG00000120436 | 6 | 167569759 | 167571817 | 4 | 2 | 1.7724 | 0.038167 | GPR31 | 0.038167 | 0.035566 | 0.10193 |
| ENSG00000111554 | 12 | 68666223 | 68726161 | 59 | 20 | 1.7716 | 0.038228 | MDM1 | 0.038228 | 0.069992 | 0.051447 |
| ENSG00000196715 | 7 | 65338254 | 65424550 | 220 | 16 | 1.7715 | 0.03824 | VKORC1L1 | 0.03824 | 0.04572 | 0.047722 |
| ENSG00000214274 | 14 | 21152336 | 21167130 | 74 | 14 | 1.7714 | 0.038248 | ANG | 0.038248 | 0.072644 | 0.28122 |
| ENSG00000105135 | 19 | 15225795 | 15236596 | 22 | 9 | 1.7713 | 0.038251 | ILVBL | 0.038251 | 0.19444 | 0.34646 |
| ENSG00000205978 | 14 | 24867992 | 24888494 | 66 | 17 | 1.7713 | 0.038256 | NYNRIN | 0.038256 | 0.010187 | 0.01202 |
| ENSG00000168056 | 11 | 65306276 | 65326401 | 40 | 11 | 1.7711 | 0.038271 | LTBP3 | 0.038271 | 0.049158 | 0.050904 |
| ENSG00000164022 | 4 | 107236701 | 107270383 | 79 | 12 | 1.7702 | 0.038347 | AIMP1 | 0.038347 | 0.038777 | 0.032165 |
| ENSG00000100065 | 22 | 37886400 | 37915549 | 117 | 19 | 1.77 | 0.038365 | CARD10 | 0.038365 | 0.10921 | 0.14407 |
| ENSG00000101017 | 20 | 44746911 | 44758502 | 36 | 14 | 1.7693 | 0.038422 | CD40 | 0.038422 | 0.12789 | 0.030848 |
| ENSG00000105321 | 19 | 47759237 | 47775210 | 62 | 6 | 1.7679 | 0.03854 | CCDC9 | 0.03854 | 0.041927 | 0.026371 |
| ENSG00000166407 | 11 | 8245851 | 8290263 | 228 | 43 | 1.7671 | 0.038608 | LMO1 | 0.038608 | 0.043166 | 0.099097 |

| | | | | | | | | | | | |
|---|---|---|---|---|---|---|---|---|---|---|---|
| ENSG00000111669 | 12 | 6976283 | 6980112 | 5 | 1 | 1.7657 | 0.038724 | TPI1 | 0.038724 | 0.019366 | 0.024824 |
| ENSG00000146352 | 6 | 123317116 | 123394072 | 202 | 21 | 1.7651 | 0.03877 | CLVS2 | 0.03877 | 0.047609 | 0.060812 |
| ENSG00000183571 | 15 | 99511459 | 99551024 | 159 | 25 | 1.7644 | 0.038829 | PGPEP1L | 0.038829 | 0.061164 | 0.051694 |
| ENSG00000167553 | 12 | 49582519 | 49667114 | 186 | 35 | 1.7644 | 0.038835 | TUBA1C | 0.038835 | 0.043657 | 0.030563 |
| ENSG00000131055 | 20 | 30225691 | 30232809 | 18 | 4 | 1.7641 | 0.038859 | COX4I2 | 0.038859 | 0.074506 | 0.087645 |
| ENSG00000068912 | 2 | 54014181 | 54045956 | 91 | 16 | 1.764 | 0.038867 | ERLEC1 | 0.038867 | 0.051699 | 0.046132 |
| ENSG00000143375 | 1 | 151482986 | 151511168 | 49 | 12 | 1.7634 | 0.038916 | CGN | 0.038916 | 0.063735 | 0.12004 |
| ENSG00000060069 | 18 | 77439801 | 77514510 | 324 | 21 | 1.7628 | 0.038963 | CTDP1 | 0.038963 | 0.040615 | 0.077996 |
| ENSG00000076258 | 1 | 171283347 | 171311223 | 45 | 10 | 1.7628 | 0.038969 | FMO4 | 0.038969 | 0.063434 | 0.06254 |
| ENSG00000188124 | 11 | 6789218 | 6790286 | 4 | 2 | 1.7616 | 0.039069 | OR2AG2 | 0.039069 | 0.020171 | 0.033323 |
| ENSG00000137310 | 6 | 31126319 | 31134936 | 46 | 11 | 1.7608 | 0.039135 | TCF19 | 0.039135 | 0.03013 | 0.062487 |
| ENSG00000104332 | 8 | 41119481 | 41167016 | 145 | 19 | 1.7601 | 0.0392 | SFRP1 | 0.0392 | 0.05918 | 0.15281 |
| ENSG00000136141 | 13 | 47127303 | 47327175 | 647 | 33 | 1.7595 | 0.039244 | LRCH1 | 0.039244 | 0.039521 | 0.044516 |
| ENSG00000198178 | 12 | 7882011 | 7904201 | 131 | 14 | 1.7595 | 0.03925 | CLEC4C | 0.03925 | 0.069962 | 0.095965 |
| ENSG00000097021 | 1 | 6324329 | 6454451 | 534 | 28 | 1.759 | 0.039285 | ACOT7 | 0.039285 | 0.062409 | 0.077053 |
| ENSG00000197705 | 18 | 30252634 | 30353025 | 285 | 12 | 1.7589 | 0.039293 | KLHL14 | 0.039293 | 0.033882 | 0.030432 |
| ENSG00000111674 | 12 | 7022909 | 7032861 | 17 | 4 | 1.7588 | 0.039303 | ENO2 | 0.039303 | 0.015048 | 0.020573 |
| ENSG00000163376 | 3 | 67048727 | 67061634 | 17 | 5 | 1.7587 | 0.03931 | KBTBD8 | 0.03931 | 0.027704 | 0.043637 |
| ENSG00000170683 | 11 | 7960081 | 7961141 | 1 | 1 | 1.7585 | 0.03933 | OR10A3 | 0.03933 | 0.18019 | 0.22568 |
| ENSG00000205718 | 19 | 7037759 | 7040190 | 1 | 1 | 1.7583 | 0.03935 | MBD3L4 | 0.03935 | 0.071809 | 0.15366 |
| ENSG00000113070 | 5 | 139712428 | 139726216 | 36 | 9 | 1.7574 | 0.039429 | HBEGF | 0.039429 | 0.081684 | 0.11974 |
| ENSG00000002726 | 7 | 150521715 | 150558592 | 203 | 19 | 1.7569 | 0.039469 | AOC1 | 0.039469 | 0.04962 | 0.051891 |
| ENSG00000177272 | 1 | 111214310 | 111217655 | 3 | 1 | 1.7558 | 0.039559 | KCNA3 | 0.039559 | 0.080023 | 0.12427 |
| ENSG00000140506 | 15 | 75105057 | 75118099 | 28 | 7 | 1.7549 | 0.039637 | LMAN1L | 0.039637 | 0.078245 | 0.082258 |
| ENSG00000256206 | 11 | 14515329 | 14541890 | 54 | 11 | 1.7537 | 0.039745 | PSMA1 | 0.039745 | 0.044757 | 0.060243 |
| ENSG00000101200 | 20 | 3063202 | 3065370 | 3 | 1 | 1.7536 | 0.039751 | AVP | 0.039751 | 0.058522 | 0.063901 |
| ENSG00000268324 | 3 | 46598888 | 46601178 | 14 | 3 | 1.7534 | 0.039766 | LUZPP1 | 0.039766 | 0.046229 | 0.0484 |
| ENSG00000172139 | 3 | 111859734 | 112013105 | 663 | 17 | 1.7534 | 0.039769 | SLC9C1 | 0.039769 | 0.045962 | 0.071254 |
| ENSG00000100271 | 22 | 43435522 | 43485434 | 227 | 30 | 1.7531 | 0.039796 | TTLL1 | 0.039796 | 0.081668 | 0.063051 |
| ENSG00000127481 | 1 | 19401000 | 19536770 | 251 | 27 | 1.7521 | 0.039878 | UBR4 | 0.039878 | 0.055505 | 0.062635 |
| ENSG00000215912 | 1 | 2567415 | 2718286 | 348 | 24 | 1.752 | 0.03989 | TTC34 | 0.03989 | 0.048886 | 0.061673 |
| ENSG00000070081 | 11 | 17229700 | 17371521 | 328 | 25 | 1.751 | 0.039974 | NUCB2 | 0.039974 | 0.045758 | 0.07714 |
| ENSG00000197054 | 19 | 12035890 | 12091196 | 194 | 10 | 1.7508 | 0.039986 | ZNF763 | 0.039986 | 0.046788 | 0.05579 |
| ENSG00000268790 | 19 | 16688453 | 16770926 | 204 | 28 | 1.7499 | 0.040064 | CTC-429P9.4 | 0.040064 | 0.045045 | 0.063597 |
| ENSG00000110925 | 12 | 51454990 | 51477447 | 73 | 13 | 1.7494 | 0.040114 | CSRNP2 | 0.040114 | 0.026812 | 0.052339 |
| ENSG00000188283 | 19 | 37708828 | 37734828 | 47 | 9 | 1.7472 | 0.040299 | ZNF383 | 0.040299 | 0.061671 | 0.12896 |
| ENSG00000131127 | 4 | 331603 | 378653 | 200 | 20 | 1.7457 | 0.040433 | ZNF141 | 0.040433 | 0.046443 | 0.049185 |
| ENSG00000112282 | 6 | 131895106 | 131949369 | 72 | 14 | 1.7455 | 0.040448 | MED23 | 0.040448 | 0.05357 | 0.072314 |
| ENSG00000132002 | 19 | 14625582 | 14640582 | 32 | 7 | 1.7455 | 0.040448 | DNAJB1 | 0.040448 | 0.023286 | 0.051333 |
| ENSG00000111666 | 12 | 102090725 | 102137918 | 114 | 24 | 1.7451 | 0.040485 | CHPT1 | 0.040485 | 0.043439 | 0.06309 |
| ENSG00000145244 | 4 | 47596015 | 47840123 | 713 | 40 | 1.7442 | 0.04056 | CORIN | 0.04056 | 0.049649 | 0.049298 |
| ENSG00000267179 | 19 | 12035890 | 12090105 | 184 | 10 | 1.7436 | 0.040613 | ZNF763 | 0.040613 | 0.046788 | 0.05579 |
| ENSG00000183617 | 19 | 3762662 | 3768573 | 24 | 6 | 1.743 | 0.040665 | MRPL54 | 0.040665 | 0.049326 | 0.23932 |
| ENSG00000269605 | 20 | 43234346 | 43248126 | 37 | 12 | 1.7417 | 0.040776 | Z97053.1 | 0.040776 | 0.087506 | 0.095649 |
| ENSG00000258474 | 14 | 32414059 | 32419588 | 45 | 6 | 1.7417 | 0.040781 | RP11-187E13.1 | 0.040781 | 0.12375 | 0.20204 |
| ENSG00000151093 | 3 | 25824408 | 25836025 | 15 | 4 | 1.7409 | 0.040849 | OXSM | 0.040849 | 0.041691 | 0.034524 |
| ENSG00000114098 | 3 | 137906109 | 138017231 | 105 | 31 | 1.7409 | 0.040851 | ARMC8 | 0.040851 | 0.075814 | 0.15237 |
| ENSG00000186111 | 19 | 3630181 | 3700477 | 226 | 32 | 1.7396 | 0.040969 | PIP5K1C | 0.040969 | 0.0073774 | 0.0019595 |
| ENSG00000160753 | 1 | 155290687 | 155300905 | 5 | 3 | 1.7381 | 0.0411 | RUSC1 | 0.0411 | 0.16247 | 0.20122 |
| ENSG00000185634 | 15 | 49115932 | 49255641 | 399 | 51 | 1.7374 | 0.041162 | SHC4 | 0.041162 | 0.056817 | 0.11178 |
| ENSG00000043039 | 11 | 129245835 | 129322171 | 291 | 47 | 1.737 | 0.041196 | BARX2 | 0.041196 | 0.084364 | 0.089829 |
| ENSG00000127241 | 3 | 186935942 | 187009810 | 255 | 43 | 1.7368 | 0.04121 | MASP1 | 0.04121 | 0.081891 | 0.092587 |
| ENSG00000168090 | 7 | 99686577 | 99689823 | 6 | 2 | 1.7362 | 0.041262 | COPS6 | 0.041262 | 0.0039161 | 0.0076514 |
| ENSG00000130023 | 6 | 170151718 | 170181680 | 117 | 15 | 1.7356 | 0.041318 | ERMARD | 0.041318 | 0.031045 | 0.020214 |
| ENSG00000114251 | 3 | 55499743 | 55523973 | 62 | 15 | 1.7328 | 0.04157 | WNT5A | 0.04157 | 0.026431 | 0.02475 |
| ENSG00000167825 | 11 | 55702932 | 55703876 | 8 | 2 | 1.7326 | 0.041584 | OR5I1 | 0.041584 | 0.036783 | 0.032645 |
| ENSG00000269549 | 20 | 44074071 | 44077220 | 16 | 7 | 1.7312 | 0.041711 | AL031663.2 | 0.041711 | 0.45315 | 0.42276 |
| ENSG00000198551 | 19 | 11670189 | 11729950 | 220 | 26 | 1.7306 | 0.041764 | ZNF627 | 0.041764 | 0.04158 | 0.044649 |
| ENSG00000134207 | 1 | 114631913 | 114696541 | 277 | 54 | 1.73 | 0.041819 | SYT6 | 0.041819 | 0.091826 | 0.2113 |
| ENSG00000257230 | 12 | 59917059 | 59917965 | 2 | 1 | 1.7295 | 0.041858 | RP11-272B17.2 | 0.041858 | 0.03897 | 0.15782 |
| ENSG00000243364 | 1 | 155036207 | 155042029 | 7 | 3 | 1.7294 | 0.041865 | EFNA4 | 0.041865 | 0.025593 | 0.0084783 |
| ENSG00000230522 | 19 | 7049332 | 7051746 | 4 | 2 | 1.7284 | 0.04196 | MBD3L2 | 0.04196 | 0.074927 | 0.14504 |
| ENSG00000269590 | 19 | 12775642 | 12780211 | 5 | 3 | 1.7274 | 0.042047 | CTD-2192J16.24 | 0.042047 | 0.0014194 | 0.0003621 |
| ENSG00000184560 | 17 | 7328934 | 7330887 | 4 | 2 | 1.7266 | 0.042116 | C17orf74 | 0.042116 | 0.23531 | 0.38994 |
| ENSG00000114062 | 15 | 25582381 | 25684128 | 274 | 20 | 1.7261 | 0.042169 | UBE3A | 0.042169 | 0.047122 | 0.048496 |
| ENSG00000180083 | 20 | 44277202 | 44298909 | 90 | 7 | 1.7255 | 0.042217 | WFDC11 | 0.042217 | 0.032313 | 0.033969 |
| ENSG00000167216 | 18 | 44497455 | 44627658 | 388 | 29 | 1.7253 | 0.042238 | KATNAL2 | 0.042238 | 0.044528 | 0.045622 |
| ENSG00000041353 | 18 | 52385091 | 52562747 | 627 | 29 | 1.7241 | 0.042343 | RAB27B | 0.042343 | 0.046586 | 0.043756 |
| ENSG00000086991 | 11 | 89057524 | 89322779 | 594 | 42 | 1.7235 | 0.042395 | NOX4 | 0.042395 | 0.042008 | 0.048704 |
| ENSG00000105427 | 19 | 42891171 | 42894439 | 7 | 2 | 1.7231 | 0.042435 | CNFN | 0.042435 | 0.073894 | 0.13338 |
| ENSG00000079841 | 6 | 72596406 | 73112845 | 1616 | 48 | 1.721 | 0.042622 | RIMS1 | 0.042622 | 0.04477 | 0.046146 |
| ENSG00000149089 | 11 | 34874641 | 34938046 | 369 | 15 | 1.7206 | 0.042665 | APIP | 0.042665 | 0.03263 | 0.035096 |
| ENSG00000107187 | 9 | 139088096 | 139096955 | 29 | 9 | 1.7201 | 0.042707 | LHX3 | 0.042707 | 0.0091992 | 0.00098735 |

| | | | | | | | | | | | |
|---|---|---|---|---|---|---|---|---|---|---|---|
| ENSG00000135899 | 2 | 231032009 | 231090444 | 253 | 16 | 1.7192 | 0.042786 | SP110 | 0.042786 | 0.077395 | 0.07916 |
| ENSG00000167220 | 18 | 44633774 | 44676891 | 155 | 20 | 1.7191 | 0.042796 | HDHD2 | 0.042796 | 0.044098 | 0.054395 |
| ENSG00000037757 | 19 | 13875346 | 13885096 | 29 | 6 | 1.7189 | 0.042813 | MRI1 | 0.042813 | 0.26303 | 0.36801 |
| ENSG00000007314 | 17 | 62015914 | 62050278 | 99 | 18 | 1.7181 | 0.042885 | SCN4A | 0.042885 | 0.064352 | 0.08321 |
| ENSG00000167765 | 19 | 52095036 | 52097630 | 32 | 10 | 1.7172 | 0.042969 | AC018755.1 | 0.042969 | 0.28538 | 0.075499 |
| ENSG00000171298 | 17 | 78075355 | 78093678 | 114 | 11 | 1.7165 | 0.043031 | GAA | 0.043031 | 0.097353 | 0.084766 |
| ENSG00000170703 | 17 | 46839597 | 46894576 | 180 | 19 | 1.7164 | 0.043043 | TTLL6 | 0.043043 | 0.02893 | 0.018923 |
| ENSG00000103485 | 16 | 29674600 | 29710020 | 77 | 16 | 1.7161 | 0.043076 | QPRT | 0.043076 | 0.039709 | 0.024654 |
| ENSG00000188897 | 16 | 11453702 | 11617444 | 1173 | 79 | 1.7157 | 0.043112 | CTD-3088G3.8 | 0.043112 | 0.068297 | 0.059311 |
| ENSG00000118242 | 2 | 216809213 | 216898819 | 404 | 68 | 1.7153 | 0.043142 | MREG | 0.043142 | 0.040176 | 0.10343 |
| ENSG00000151247 | 4 | 99792835 | 99851788 | 219 | 25 | 1.7149 | 0.043181 | EIF4E | 0.043181 | 0.028482 | 0.027471 |
| ENSG00000007168 | 17 | 2496504 | 2588909 | 244 | 17 | 1.7147 | 0.043201 | PAFAH1B1 | 0.043201 | 0.04747 | 0.046525 |
| ENSG00000106336 | 7 | 100181605 | 100198740 | 19 | 8 | 1.7142 | 0.043242 | FBXO24 | 0.043242 | 0.10346 | 0.18953 |
| ENSG00000196427 | 1 | 108765963 | 108786689 | 16 | 5 | 1.7139 | 0.043271 | NBPF4 | 0.043271 | 0.17476 | 0.18339 |
| ENSG00000163634 | 3 | 63819546 | 63849579 | 62 | 18 | 1.7132 | 0.043338 | THOC7 | 0.043338 | 0.031086 | 0.093614 |
| ENSG00000150054 | 10 | 28339922 | 28623415 | 1303 | 46 | 1.713 | 0.043354 | MPP7 | 0.043354 | 0.061917 | 0.059014 |
| ENSG00000177943 | 9 | 139745395 | 139755249 | 10 | 4 | 1.713 | 0.043359 | MAMDC4 | 0.043359 | 0.078906 | 0.12033 |
| ENSG00000115128 | 2 | 24290454 | 24299313 | 48 | 9 | 1.7121 | 0.043436 | SF3B14 | 0.043436 | 0.059354 | 0.1139 |
| ENSG00000135446 | 12 | 58141510 | 58149796 | 16 | 9 | 1.7121 | 0.04344 | CDK4 | 0.04344 | 0.22485 | 0.52037 |
| ENSG00000121691 | 11 | 34460472 | 34493609 | 99 | 13 | 1.712 | 0.04345 | CAT | 0.04345 | 0.086382 | 0.10402 |
| ENSG00000126705 | 1 | 27860546 | 27930942 | 91 | 13 | 1.7117 | 0.043473 | AHDC1 | 0.043473 | 0.046154 | 0.019711 |
| ENSG00000196436 | 16 | 74411776 | 74425978 | 20 | 10 | 1.7117 | 0.043479 | NPIPB15 | 0.043479 | 0.33076 | 0.20461 |
| ENSG00000105298 | 19 | 3610639 | 3626813 | 72 | 10 | 1.7107 | 0.043569 | CACTIN | 0.043569 | 0.019668 | 0.0065265 |
| ENSG00000152256 | 2 | 173420101 | 173489823 | 166 | 29 | 1.7107 | 0.04357 | PDK1 | 0.04357 | 0.03725 | 0.039055 |
| ENSG00000214106 | 7 | 154720208 | 154741618 | 55 | 5 | 1.7101 | 0.043623 | PAXIP1-AS2 | 0.043623 | 0.060976 | 0.078215 |
| ENSG00000247270 | 16 | 66516775 | 66519747 | 4 | 1 | 1.7096 | 0.043673 | AC132186.1 | 0.043673 | 0.04092 | 0.016979 |
| ENSG00000181938 | 16 | 58328984 | 58440048 | 478 | 51 | 1.7094 | 0.043689 | GINS3 | 0.043689 | 0.037245 | 0.071522 |
| ENSG00000141425 | 18 | 33564350 | 33647539 | 206 | 17 | 1.7089 | 0.043739 | RPRD1A | 0.043739 | 0.040565 | 0.039951 |
| ENSG00000125816 | 20 | 21376005 | 21378666 | 2 | 1 | 1.7068 | 0.043926 | NKX2-4 | 0.043926 | 0.056791 | 0.083724 |
| ENSG00000173442 | 11 | 65343509 | 65360121 | 29 | 9 | 1.7067 | 0.043936 | EHBP1L1 | 0.043936 | 0.072808 | 0.092381 |
| ENSG00000242612 | 16 | 451826 | 462487 | 29 | 5 | 1.7065 | 0.043953 | DECR2 | 0.043953 | 0.077164 | 0.063305 |
| ENSG00000165699 | 9 | 135766735 | 135820020 | 126 | 16 | 1.7054 | 0.044055 | TSC1 | 0.044055 | 0.047874 | 0.10373 |
| ENSG00000153823 | 2 | 229715242 | 230136001 | 1513 | 106 | 1.7045 | 0.04414 | PID1 | 0.04414 | 0.042208 | 0.049481 |
| ENSG00000180205 | 20 | 44236578 | 44259907 | 99 | 11 | 1.704 | 0.04419 | WFDC9 | 0.04419 | 0.073692 | 0.12974 |
| ENSG00000134308 | 2 | 9724101 | 9771143 | 108 | 14 | 1.7037 | 0.044222 | YWHAQ | 0.044222 | 0.020998 | 0.016283 |
| ENSG00000172410 | 1 | 67263424 | 67266939 | 6 | 3 | 1.7022 | 0.04436 | INSL5 | 0.04436 | 0.37987 | 0.48336 |
| ENSG00000109944 | 11 | 122753391 | 122830506 | 253 | 32 | 1.7016 | 0.044411 | C11orf63 | 0.044411 | 0.039002 | 0.016534 |
| ENSG00000185269 | 17 | 79910388 | 79919716 | 30 | 4 | 1.701 | 0.044474 | NOTUM | 0.044474 | 0.067746 | 0.070156 |
| ENSG00000168671 | 5 | 36035119 | 36071460 | 143 | 19 | 1.7005 | 0.044519 | UGT3A2 | 0.044519 | 0.086412 | 0.16744 |
| ENSG00000099954 | 22 | 17840837 | 18037850 | 1032 | 101 | 1.7005 | 0.044519 | CECR2 | 0.044519 | 0.043512 | 0.099746 |
| ENSG00000136718 | 2 | 131099798 | 131105383 | 10 | 4 | 1.699 | 0.044655 | IMP4 | 0.044655 | 0.0047103 | 0.0035635 |
| ENSG00000256683 | 19 | 52467596 | 52490109 | 85 | 12 | 1.6987 | 0.044691 | ZNF350 | 0.044691 | 0.051635 | 0.09361 |
| ENSG00000166997 | 7 | 99717236 | 99723134 | 13 | 5 | 1.6967 | 0.044873 | CNPY4 | 0.044873 | 0.033899 | 0.01407 |
| ENSG00000174672 | 11 | 1411129 | 1483919 | 368 | 38 | 1.6964 | 0.044904 | BRSK2 | 0.044904 | 0.024685 | 0.03633 |
| ENSG00000174914 | 11 | 56467864 | 56468781 | 7 | 3 | 1.6964 | 0.044904 | OR9G1 | 0.044904 | 0.027475 | 0.023417 |
| ENSG00000139151 | 12 | 18836103 | 18890991 | 234 | 29 | 1.6964 | 0.044904 | PLCZ1 | 0.044904 | 0.051163 | 0.029764 |
| ENSG00000181381 | 4 | 169277886 | 169458937 | 852 | 57 | 1.6964 | 0.044906 | DDX60L | 0.044906 | 0.048972 | 0.059716 |
| ENSG00000161649 | 17 | 41924516 | 41940997 | 27 | 5 | 1.6963 | 0.044911 | CD300LG | 0.044911 | 0.090186 | 0.12586 |
| ENSG00000136404 | 15 | 83776159 | 83813606 | 55 | 11 | 1.6953 | 0.045013 | TM6SF1 | 0.045013 | 0.033636 | 0.062128 |
| ENSG00000106069 | 7 | 29161890 | 29553944 | 1573 | 133 | 1.694 | 0.045129 | CHN2 | 0.045129 | 0.047831 | 0.05586 |
| ENSG00000253958 | 8 | 8559448 | 8561616 | 17 | 2 | 1.6902 | 0.045495 | CLDN23 | 0.045495 | 0.02632 | 0.023277 |
| ENSG00000131467 | 17 | 40976402 | 40995774 | 9 | 6 | 1.6893 | 0.045584 | PSME3 | 0.045584 | 0.19714 | 0.35624 |
| ENSG00000168944 | 5 | 122680579 | 122759286 | 266 | 11 | 1.6887 | 0.045639 | CEP120 | 0.045639 | 0.047002 | 0.044068 |
| ENSG00000183010 | 17 | 79890260 | 79900288 | 16 | 4 | 1.6885 | 0.045659 | PYCR1 | 0.045659 | 0.046242 | 0.056419 |
| ENSG00000136327 | 14 | 37049784 | 37051812 | 3 | 1 | 1.6883 | 0.045674 | NKX2-8 | 0.045674 | 0.63893 | 0.70688 |
| ENSG00000102804 | 13 | 45007655 | 45151283 | 378 | 18 | 1.687 | 0.045805 | TSC22D1 | 0.045805 | 0.048334 | 0.049775 |
| ENSG00000166986 | 12 | 57869228 | 57911352 | 88 | 9 | 1.6867 | 0.045826 | MARS | 0.045826 | 0.026799 | 0.017977 |
| ENSG00000162636 | 1 | 109102711 | 109187522 | 182 | 10 | 1.6865 | 0.045851 | FAM102B | 0.045851 | 0.051078 | 0.056281 |
| ENSG00000254415 | 19 | 52145806 | 52150054 | 11 | 5 | 1.686 | 0.045897 | SIGLEC14 | 0.045897 | 0.024459 | 0.016982 |
| ENSG00000154310 | 3 | 170779128 | 171178197 | 1286 | 121 | 1.6859 | 0.045911 | TNIK | 0.045911 | 0.053913 | 0.065885 |
| ENSG00000100664 | 14 | 103799881 | 103811362 | 27 | 7 | 1.6852 | 0.045975 | EIF5 | 0.045975 | 0.13265 | 0.28564 |
| ENSG00000196409 | 9 | 40760700 | 40836415 | 83 | 10 | 1.6847 | 0.046024 | ZNF658 | 0.046024 | 0.05137 | 0.048697 |
| ENSG00000005448 | 2 | 74648805 | 74652882 | 3 | 2 | 1.6846 | 0.046034 | WDR54 | 0.046034 | 0.26801 | 0.35284 |
| ENSG00000155592 | 16 | 25247322 | 25269252 | 56 | 9 | 1.6839 | 0.046105 | ZKSCAN2 | 0.046105 | 0.024938 | 0.17608 |
| ENSG00000175931 | 17 | 74385532 | 74449288 | 194 | 25 | 1.6824 | 0.046244 | UBE2O | 0.046244 | 0.057533 | 0.14338 |
| ENSG00000197888 | 4 | 69402902 | 69434245 | 81 | 7 | 1.6805 | 0.046434 | UGT2B17 | 0.046434 | 0.062094 | 0.086078 |
| ENSG00000213654 | 6 | 32158543 | 32163300 | 11 | 5 | 1.6802 | 0.046456 | GPSM3 | 0.046456 | 0.04079 | 0.076744 |
| ENSG00000164483 | 6 | 130465460 | 130686570 | 1226 | 35 | 1.6799 | 0.046486 | SAMD3 | 0.046486 | 0.04362 | 0.044814 |
| ENSG00000137947 | 1 | 89318615 | 89357627 | 138 | 10 | 1.6792 | 0.046557 | GTF2B | 0.046557 | 0.0586 | 0.075571 |
| ENSG00000267041 | 19 | 37205285 | 37263727 | 146 | 15 | 1.6784 | 0.04663 | ZNF850 | 0.04663 | 0.079191 | 0.1086 |
| ENSG00000099974 | 22 | 24309089 | 24314721 | 14 | 4 | 1.6783 | 0.046641 | DDTL | 0.046641 | 0.011776 | 0.019956 |
| ENSG00000075702 | 19 | 36545783 | 36596008 | 151 | 21 | 1.678 | 0.04667 | WDR62 | 0.04667 | 0.056751 | 0.089841 |
| ENSG00000157184 | 1 | 53662101 | 53679869 | 32 | 7 | 1.6779 | 0.046681 | CPT2 | 0.046681 | 0.034635 | 0.080004 |

| | | | | | | | | | | | |
|---|---|---|---|---|---|---|---|---|---|---|---|
| ENSG00000116771 | 1 | 15898848 | 15911605 | 66 | 6 | 1.677 | 0.046775 | AGMAT | 0.046775 | 0.028142 | 0.038046 |
| ENSG00000163918 | 3 | 186507669 | 186524847 | 35 | 6 | 1.676 | 0.046872 | RFC4 | 0.046872 | 0.05522 | 0.070823 |
| ENSG00000132570 | 5 | 134240596 | 134343649 | 190 | 29 | 1.6748 | 0.046984 | PCBD2 | 0.046984 | 0.0022095 | 0.00088661 |
| ENSG00000146872 | 17 | 60536019 | 60692842 | 248 | 22 | 1.6742 | 0.047045 | TLK2 | 0.047045 | 0.047685 | 0.038987 |
| ENSG00000227057 | 6 | 33246885 | 33257304 | 44 | 12 | 1.6732 | 0.047148 | WDR46 | 0.047148 | 0.02288 | 0.038792 |
| ENSG00000187912 | 19 | 14693896 | 14721969 | 127 | 21 | 1.6713 | 0.047334 | CLEC17A | 0.047334 | 0.042963 | 0.003924 |
| ENSG00000151224 | 10 | 82031576 | 82049440 | 52 | 14 | 1.671 | 0.047359 | MAT1A | 0.047359 | 0.016676 | 0.049962 |
| ENSG00000204316 | 17 | 73894724 | 73905899 | 28 | 8 | 1.6698 | 0.047484 | MRPL38 | 0.047484 | 0.16002 | 0.21611 |
| ENSG00000164266 | 5 | 147204131 | 147211349 | 8 | 4 | 1.6692 | 0.047537 | SPINK1 | 0.047537 | 0.22654 | 0.34179 |
| ENSG00000178226 | 16 | 31150246 | 31161415 | 18 | 9 | 1.6686 | 0.047603 | PRSS36 | 0.047603 | 0.24931 | 0.49682 |
| ENSG00000100625 | 14 | 61176246 | 61191066 | 14 | 5 | 1.6681 | 0.047646 | SIX4 | 0.047646 | 0.046526 | 0.01854 |
| ENSG00000146701 | 7 | 75677369 | 75696826 | 53 | 7 | 1.6673 | 0.047727 | MDH2 | 0.047727 | 0.064315 | 0.10765 |
| ENSG00000128581 | 7 | 100956975 | 100965104 | 44 | 9 | 1.6671 | 0.047744 | RABL5 | 0.047744 | 0.048437 | 0.040619 |
| ENSG00000159403 | 12 | 7187513 | 7245203 | 18 | 9 | 1.6668 | 0.04778 | C1R | 0.04778 | 0.062102 | 0.13367 |
| ENSG00000188373 | 10 | 85933494 | 85945050 | 47 | 11 | 1.6666 | 0.047793 | C10orf99 | 0.047793 | 0.051844 | 0.095368 |
| ENSG00000105085 | 19 | 16685718 | 16739873 | 113 | 14 | 1.666 | 0.047853 | MED26 | 0.047853 | 0.05944 | 0.045253 |
| ENSG00000100191 | 22 | 32614465 | 32651328 | 150 | 27 | 1.6659 | 0.047864 | SLC5A4 | 0.047864 | 0.10735 | 0.20202 |
| ENSG00000152192 | 13 | 79172497 | 79177673 | 11 | 5 | 1.6657 | 0.047884 | POU4F1 | 0.047884 | 0.034368 | 0.029862 |
| ENSG00000173890 | 3 | 169755717 | 169803191 | 120 | 11 | 1.6657 | 0.047888 | GPR160 | 0.047888 | 0.023849 | 0.008002 |
| ENSG00000163081 | 2 | 223162866 | 223169936 | 19 | 9 | 1.6647 | 0.047983 | CCDC140 | 0.047983 | 0.17947 | 0.42816 |
| ENSG00000148459 | 10 | 26986588 | 27035727 | 174 | 24 | 1.6637 | 0.048086 | PDSS1 | 0.048086 | 0.112 | 0.14271 |
| ENSG00000074771 | 6 | 155716504 | 155777037 | 227 | 55 | 1.6626 | 0.0482 | NOX3 | 0.0482 | 0.12348 | 0.17327 |
| ENSG00000107798 | 10 | 90973326 | 91174314 | 629 | 60 | 1.6616 | 0.048293 | LIPA | 0.048293 | 0.045614 | 0.048078 |
| ENSG00000149635 | 20 | 45169585 | 45179213 | 32 | 11 | 1.6614 | 0.048315 | OCSTAMP | 0.048315 | 0.016268 | 0.026364 |
| ENSG00000140398 | 15 | 75639296 | 75647592 | 11 | 5 | 1.6609 | 0.048362 | NEIL1 | 0.048362 | 0.07542 | 0.026396 |
| ENSG00000189057 | 11 | 58874658 | 58894883 | 30 | 7 | 1.6601 | 0.048442 | FAM111B | 0.048442 | 0.079738 | 0.082168 |
| ENSG00000198417 | 16 | 56691606 | 56694610 | 6 | 3 | 1.6585 | 0.048609 | MT1F | 0.048609 | 0.40314 | 0.49566 |
| ENSG00000243789 | 15 | 42120283 | 42129779 | 36 | 7 | 1.6585 | 0.048613 | JMJD7 | 0.048613 | 0.0023865 | 0.0014335 |
| ENSG00000130717 | 9 | 134399188 | 134406655 | 25 | 7 | 1.6583 | 0.048624 | UCK1 | 0.048624 | 0.087492 | 0.36203 |
| ENSG00000241839 | 15 | 65134088 | 65160206 | 91 | 11 | 1.6566 | 0.048802 | PLEKHO2 | 0.048802 | 0.05897 | 0.065763 |
| ENSG00000116095 | 2 | 179345195 | 179381189 | 80 | 14 | 1.6565 | 0.048813 | PLEKHA3 | 0.048813 | 0.036554 | 0.02563 |
| ENSG00000165861 | 14 | 73436159 | 73493920 | 300 | 20 | 1.6562 | 0.048844 | ZFYVE1 | 0.048844 | 0.062968 | 0.062758 |
| ENSG00000163734 | 4 | 74902306 | 74904524 | 5 | 3 | 1.6555 | 0.048907 | CXCL3 | 0.048907 | 0.077161 | 0.15271 |
| ENSG00000156384 | 10 | 105881816 | 105886143 | 13 | 3 | 1.655 | 0.048966 | SFR1 | 0.048966 | 0.14776 | 0.13476 |
| ENSG00000078898 | 20 | 31595406 | 31611515 | 43 | 11 | 1.655 | 0.048967 | BPIFB2 | 0.048967 | 0.27959 | 0.60119 |
| ENSG00000155363 | 1 | 113215763 | 113243368 | 68 | 12 | 1.653 | 0.049164 | MOV10 | 0.049164 | 0.10384 | 0.19165 |
| ENSG00000121104 | 17 | 47787694 | 47866542 | 100 | 27 | 1.653 | 0.04917 | FAM117A | 0.04917 | 0.021805 | 0.019825 |
| ENSG00000166529 | 7 | 99647390 | 99662661 | 29 | 5 | 1.6525 | 0.049218 | ZSCAN21 | 0.049218 | 0.063367 | 0.065109 |
| ENSG00000023228 | 2 | 206979541 | 207024327 | 150 | 13 | 1.6523 | 0.049236 | NDUFS1 | 0.049236 | 0.04501 | 0.055761 |
| ENSG00000178105 | 11 | 108535752 | 108811657 | 582 | 20 | 1.6518 | 0.049291 | DDX10 | 0.049291 | 0.054216 | 0.0508 |
| ENSG00000118514 | 6 | 135238528 | 135271260 | 43 | 8 | 1.6507 | 0.0494 | ALDH8A1 | 0.0494 | 0.033531 | 0.052086 |
| ENSG00000042088 | 14 | 90421283 | 90511106 | 278 | 13 | 1.6505 | 0.049421 | TDP1 | 0.049421 | 0.052844 | 0.054102 |
| ENSG00000137076 | 9 | 35696945 | 35732392 | 49 | 13 | 1.6505 | 0.049422 | TLN1 | 0.049422 | 0.041539 | 0.0067655 |
| ENSG00000127191 | 9 | 139776364 | 139821059 | 132 | 8 | 1.6501 | 0.049462 | TRAF2 | 0.049462 | 0.043199 | 0.049142 |
| ENSG00000144550 | 3 | 9745491 | 9771592 | 70 | 28 | 1.6491 | 0.049565 | CPNE9 | 0.049565 | 0.078861 | 0.09807 |
| ENSG00000123159 | 19 | 14588572 | 14606944 | 89 | 19 | 1.649 | 0.049577 | GIPC1 | 0.049577 | 0.079944 | 0.026152 |
| ENSG00000011566 | 2 | 39476407 | 39664453 | 516 | 24 | 1.6488 | 0.049597 | MAP4K3 | 0.049597 | 0.044897 | 0.048603 |
| ENSG00000177169 | 12 | 132379196 | 132407712 | 81 | 17 | 1.6485 | 0.049624 | ULK1 | 0.049624 | 0.060234 | 0.064091 |
| ENSG00000155508 | 5 | 154237113 | 154256353 | 33 | 5 | 1.6485 | 0.049629 | CNOT8 | 0.049629 | 0.054428 | 0.075631 |
| ENSG00000165527 | 14 | 50359810 | 50361490 | 4 | 2 | 1.6483 | 0.049645 | ARF6 | 0.049645 | 0.30445 | 0.40085 |
| ENSG00000162065 | 16 | 2525147 | 2555735 | 27 | 15 | 1.648 | 0.049676 | TBC1D24 | 0.049676 | 0.0079624 | 0.00058038 |
| ENSG00000167780 | 12 | 53497302 | 53518322 | 123 | 12 | 1.6478 | 0.049697 | SOAT2 | 0.049697 | 0.043129 | 0.054567 |
| ENSG00000105137 | 19 | 15218214 | 15225799 | 9 | 3 | 1.6478 | 0.049702 | SYDE1 | 0.049702 | 0.13429 | 0.19979 |
| ENSG00000230891 | 3 | 25900024 | 25915191 | 41 | 6 | 1.6475 | 0.049729 | LINC00692 | 0.049729 | 0.029321 | 0.025314 |
| ENSG00000169641 | 1 | 23410516 | 23504301 | 224 | 25 | 1.6474 | 0.049737 | LUZP1 | 0.049737 | 0.05733 | 0.056154 |
| ENSG00000101134 | 20 | 53092136 | 53267710 | 513 | 32 | 1.647 | 0.049784 | DOK5 | 0.049784 | 0.034661 | 0.042605 |
| ENSG00000179046 | 4 | 189012427 | 189030757 | 133 | 20 | 1.6467 | 0.049805 | TRIML2 | 0.049805 | 0.054626 | 0.061282 |
| ENSG00000120438 | 6 | 160199530 | 160210781 | 41 | 8 | 1.6466 | 0.049821 | TCP1 | 0.049821 | 0.076924 | 0.077907 |
| ENSG00000188312 | 9 | 95087766 | 95382815 | 893 | 25 | 1.6465 | 0.049827 | CENPP | 0.049827 | 0.055066 | 0.04945 |
| ENSG00000221944 | 2 | 233412779 | 233415226 | 9 | 2 | 1.6454 | 0.049941 | TIGD1 | 0.049941 | 0.084989 | 0.13307 |
| ENSG00000165879 | 10 | 99079022 | 99081672 | 6 | 2 | 1.6454 | 0.049947 | FRAT1 | 0.049947 | 0.023434 | 0.024766 |
| ENSG00000164385 | 6 | 2622147 | 2634837 | 50 | 10 | 1.6452 | 0.049962 | C6orf195 | 0.049962 | 0.12575 | 0.17024 |
| ENSG00000204209 | 6 | 33286335 | 33297046 | 33 | 9 | 1.6433 | 0.050156 | DAXX | 0.050156 | 0.045668 | 0.048935 |
| ENSG00000197496 | 20 | 45338126 | 45364965 | 80 | 12 | 1.6423 | 0.050267 | SLC2A10 | 0.050267 | 0.086937 | 0.16516 |
| ENSG00000154027 | 1 | 77747736 | 78025651 | 1303 | 83 | 1.6422 | 0.050272 | AK5 | 0.050272 | 0.057156 | 0.053982 |
| ENSG00000105281 | 19 | 47278140 | 47291851 | 44 | 11 | 1.6418 | 0.050317 | SLC1A5 | 0.050317 | 0.20695 | 0.17423 |
| ENSG00000185811 | 7 | 50343720 | 50472799 | 237 | 47 | 1.6413 | 0.050369 | IKZF1 | 0.050369 | 0.055523 | 0.069668 |
| ENSG00000187045 | 22 | 37461476 | 37505603 | 174 | 20 | 1.6412 | 0.050383 | TMPRSS6 | 0.050383 | 0.050682 | 0.070805 |
| ENSG00000136630 | 1 | 221051699 | 221058401 | 27 | 7 | 1.641 | 0.050394 | HLX | 0.050394 | 0.32486 | 0.54374 |
| ENSG00000171858 | 20 | 60962172 | 60963576 | 4 | 2 | 1.6405 | 0.050453 | RPS21 | 0.050453 | 0.027871 | 0.05273 |
| ENSG00000182325 | 8 | 145579091 | 145583036 | 10 | 2 | 1.64 | 0.050506 | FBXL6 | 0.050506 | 0.094231 | 0.081551 |
| ENSG00000164169 | 4 | 148558936 | 148605381 | 39 | 11 | 1.6397 | 0.050531 | PRMT10 | 0.050531 | 0.065708 | 0.069772 |
| ENSG00000137054 | 9 | 37485932 | 37503694 | 46 | 10 | 1.6395 | 0.050551 | POLR1E | 0.050551 | 0.12445 | 0.2162 |

| | | | | | | | | | | | |
|---|---|---|---|---|---|---|---|---|---|---|---|
| ENSG00000142046 | 19 | 41856816 | 41889988 | 81 | 12 | 1.6394 | 0.050569 | TMEM91 | 0.050569 | 0.046813 | 0.043528 |
| ENSG00000101365 | 20 | 2639041 | 2644865 | 16 | 4 | 1.6391 | 0.050595 | IDH3B | 0.050595 | 0.046321 | 0.18773 |
| ENSG00000198049 | 1 | 206223976 | 206231639 | 48 | 7 | 1.6389 | 0.050621 | AVPR1B | 0.050621 | 0.083621 | 0.10329 |
| ENSG00000182179 | 3 | 49842640 | 49851379 | 10 | 3 | 1.6386 | 0.050648 | UBA7 | 0.050648 | 0.038205 | 0.043433 |
| ENSG00000168071 | 11 | 64107695 | 64125006 | 47 | 9 | 1.6385 | 0.05066 | CCDC88B | 0.05066 | 0.061156 | 0.10937 |
| ENSG00000119408 | 9 | 127019885 | 127115586 | 287 | 23 | 1.6384 | 0.050665 | NEK6 | 0.050665 | 0.037593 | 0.046139 |
| ENSG00000188389 | 2 | 242792033 | 242801060 | 24 | 6 | 1.6382 | 0.050695 | PDCD1 | 0.050695 | 0.12904 | 0.17222 |
| ENSG00000244274 | 20 | 44034697 | 44039250 | 4 | 1 | 1.6379 | 0.050718 | DBNDD2 | 0.050718 | 0.11148 | 0.096523 |
| ENSG00000157212 | 7 | 154735397 | 154794794 | 125 | 10 | 1.6377 | 0.050743 | PAXIP1 | 0.050743 | 0.036479 | 0.034761 |
| ENSG00000204227 | 6 | 33176272 | 33180499 | 11 | 6 | 1.6377 | 0.050746 | RING1 | 0.050746 | 0.082437 | 0.11701 |
| ENSG00000107897 | 10 | 27484146 | 27531059 | 345 | 15 | 1.6375 | 0.050766 | ACBD5 | 0.050766 | 0.037635 | 0.03115 |
| ENSG00000164167 | 4 | 147096837 | 147121152 | 87 | 6 | 1.6366 | 0.050858 | LSM6 | 0.050858 | 0.087716 | 0.11597 |
| ENSG00000132330 | 2 | 238969530 | 239008054 | 159 | 15 | 1.6364 | 0.050883 | SCLY | 0.050883 | 0.068335 | 0.082637 |
| ENSG00000104872 | 19 | 49949555 | 49956754 | 28 | 4 | 1.6357 | 0.050948 | PIH1D1 | 0.050948 | 0.030134 | 0.034814 |
| ENSG00000197992 | 12 | 10183276 | 10218565 | 64 | 10 | 1.6353 | 0.050993 | CLEC9A | 0.050993 | 0.0084249 | 0.01266 |
| ENSG00000143942 | 2 | 53994929 | 54002333 | 24 | 8 | 1.6352 | 0.051005 | CHAC2 | 0.051005 | 0.060185 | 0.059617 |
| ENSG00000145975 | 6 | 4049668 | 4087578 | 148 | 16 | 1.6346 | 0.051064 | FAM217A | 0.051064 | 0.067349 | 0.10863 |
| ENSG00000176567 | 11 | 48285413 | 48286330 | 6 | 2 | 1.6344 | 0.051087 | OR4X1 | 0.051087 | 0.04468 | 0.046067 |
| ENSG00000131482 | 17 | 41052814 | 41065386 | 25 | 6 | 1.6344 | 0.051091 | G6PC | 0.051091 | 0.14383 | 0.16409 |
| ENSG00000169340 | 16 | 20370492 | 20416059 | 168 | 23 | 1.6333 | 0.0512 | PDILT | 0.0512 | 0.076348 | 0.13786 |
| ENSG00000128342 | 22 | 30636436 | 30642840 | 14 | 5 | 1.633 | 0.05123 | LIF | 0.05123 | 0.1386 | 0.10857 |
| ENSG00000166126 | 14 | 103388993 | 103399933 | 26 | 8 | 1.6326 | 0.051278 | AMN | 0.051278 | 0.096902 | 0.19646 |
| ENSG00000182898 | 1 | 152056620 | 152061540 | 3 | 2 | 1.6325 | 0.05129 | TCHHL1 | 0.05129 | 0.71856 | 0.83297 |
| ENSG00000271698 | 8 | 145576888 | 145579269 | 8 | 3 | 1.6317 | 0.051375 | GS1-393G12.13 | 0.051375 | 0.10576 | 0.075447 |
| ENSG00000156469 | 8 | 97251626 | 97273838 | 49 | 10 | 1.6313 | 0.051415 | MTERFD1 | 0.051415 | 0.050079 | 0.060614 |
| ENSG00000088854 | 20 | 3229951 | 3388272 | 709 | 20 | 1.6313 | 0.051416 | C20orf194 | 0.051416 | 0.055043 | 0.0533 |
| ENSG00000165025 | 9 | 93564069 | 93660831 | 468 | 60 | 1.6308 | 0.051471 | SYK | 0.051471 | 0.077621 | 0.12127 |
| ENSG00000117318 | 1 | 23884409 | 23886285 | 8 | 3 | 1.6301 | 0.051545 | ID3 | 0.051545 | 0.098826 | 0.11888 |
| ENSG00000181458 | 3 | 100211463 | 100296288 | 264 | 14 | 1.6298 | 0.051574 | TMEM45A | 0.051574 | 0.061849 | 0.085694 |
| ENSG00000244165 | 19 | 10222214 | 10226048 | 14 | 4 | 1.6284 | 0.051715 | P2RY11 | 0.051715 | 0.21052 | 0.05988 |
| ENSG00000100024 | 22 | 24863206 | 24924358 | 168 | 19 | 1.628 | 0.051766 | UPB1 | 0.051766 | 0.05761 | 0.059633 |
| ENSG00000145632 | 5 | 57749809 | 57756087 | 25 | 5 | 1.6279 | 0.051774 | PLK2 | 0.051774 | 0.24424 | 0.35073 |
| ENSG00000127318 | 12 | 68642022 | 68647387 | 14 | 5 | 1.6274 | 0.051831 | IL22 | 0.051831 | 0.11935 | 0.1389 |
| ENSG00000183813 | 3 | 32993066 | 32997841 | 11 | 2 | 1.627 | 0.051873 | CCR4 | 0.051873 | 0.16749 | 0.3794 |
| ENSG00000106333 | 7 | 100199800 | 100205798 | 5 | 3 | 1.6269 | 0.05188 | PCOLCE | 0.05188 | 0.22407 | 0.32655 |
| ENSG00000198648 | 2 | 168810530 | 169104651 | 1023 | 50 | 1.6265 | 0.051921 | STK39 | 0.051921 | 0.059961 | 0.068294 |
| ENSG00000161016 | 8 | 146015150 | 146017972 | 8 | 4 | 1.6265 | 0.051924 | RPL8 | 0.051924 | 0.036573 | 0.075744 |
| ENSG00000181958 | 11 | 55135360 | 55136394 | 5 | 2 | 1.6259 | 0.051986 | OR4A15 | 0.051986 | 0.098376 | 0.077985 |
| ENSG00000107175 | 9 | 35732332 | 35737001 | 5 | 3 | 1.6256 | 0.052018 | CREB3 | 0.052018 | 0.01281 | 0.012549 |
| ENSG00000114770 | 3 | 183637722 | 183735803 | 255 | 17 | 1.625 | 0.052078 | ABCC5 | 0.052078 | 0.069462 | 0.024908 |
| ENSG00000103381 | 16 | 12756919 | 12897874 | 619 | 52 | 1.624 | 0.05219 | CPPED1 | 0.05219 | 0.059644 | 0.17366 |
| ENSG00000125246 | 13 | 100258919 | 100549387 | 1014 | 88 | 1.6238 | 0.052211 | CLYBL | 0.052211 | 0.06838 | 0.089924 |
| ENSG00000164251 | 5 | 76114758 | 76131140 | 51 | 8 | 1.6237 | 0.052216 | F2RL1 | 0.052216 | 0.049419 | 0.056166 |
| ENSG00000078304 | 14 | 102228135 | 102394326 | 530 | 45 | 1.6234 | 0.052252 | PPP2R5C | 0.052252 | 0.036197 | 0.049014 |
| ENSG00000196378 | 8 | 145998499 | 146012730 | 43 | 6 | 1.6232 | 0.052271 | ZNF34 | 0.052271 | 0.083542 | 0.0933 |
| ENSG00000197093 | 7 | 99756867 | 99766373 | 16 | 4 | 1.6226 | 0.052342 | GAL3ST4 | 0.052342 | 0.022849 | 0.030709 |
| ENSG00000103495 | 16 | 29817427 | 29823649 | 4 | 2 | 1.6222 | 0.052375 | MAZ | 0.052375 | 0.17848 | 0.19175 |
| ENSG00000141428 | 18 | 33552046 | 33559241 | 16 | 3 | 1.6214 | 0.052467 | C18orf21 | 0.052467 | 0.033793 | 0.036562 |
| ENSG00000220201 | 19 | 10415479 | 10420556 | 15 | 6 | 1.6209 | 0.052518 | ZGLP1 | 0.052518 | 0.088001 | 0.11702 |
| ENSG00000110665 | 11 | 2316875 | 2324279 | 55 | 9 | 1.6205 | 0.052563 | C11orf21 | 0.052563 | 0.14744 | 0.53268 |
| ENSG00000142606 | 1 | 2522078 | 2564481 | 193 | 10 | 1.6186 | 0.052771 | MMEL1 | 0.052771 | 0.042797 | 0.031386 |
| ENSG00000099977 | 22 | 24313554 | 24322660 | 4 | 2 | 1.6185 | 0.05278 | DDT | 0.05278 | 0.042206 | 0.015567 |
| ENSG00000259120 | 17 | 73642323 | 73644057 | 2 | 1 | 1.6184 | 0.052792 | SMIM6 | 0.052792 | 0.52318 | 0.43287 |
| ENSG00000162408 | 1 | 6581407 | 6614595 | 158 | 6 | 1.6167 | 0.052975 | NOL9 | 0.052975 | 0.054928 | 0.068079 |
| ENSG00000233232 | 16 | 28467693 | 28481868 | 4 | 2 | 1.616 | 0.053046 | NPIPB7 | 0.053046 | 0.84869 | 0.86291 |
| ENSG00000184108 | 4 | 189060573 | 189068897 | 74 | 15 | 1.6153 | 0.053126 | TRIML1 | 0.053126 | 0.022587 | 0.017053 |
| ENSG00000166526 | 7 | 99661656 | 99680171 | 48 | 13 | 1.6151 | 0.053149 | ZNF3 | 0.053149 | 0.019806 | 0.012389 |
| ENSG00000170634 | 2 | 54197975 | 54532437 | 1119 | 47 | 1.6139 | 0.053275 | ACYP2 | 0.053275 | 0.057698 | 0.063476 |
| ENSG00000269018 | 11 | 65358665 | 65359947 | 4 | 2 | 1.6137 | 0.053292 | AP001362.1 | 0.053292 | 0.083681 | 0.094993 |
| ENSG00000167487 | 19 | 18747775 | 18781309 | 95 | 16 | 1.6129 | 0.05338 | KLHL26 | 0.05338 | 0.072061 | 0.13625 |
| ENSG00000196757 | 19 | 12035883 | 12061588 | 71 | 9 | 1.6129 | 0.053382 | ZNF700 | 0.053382 | 0.052564 | 0.054711 |
| ENSG00000179776 | 16 | 66400533 | 66438686 | 112 | 24 | 1.6123 | 0.053454 | CDH5 | 0.053454 | 0.042511 | 0.123 |
| ENSG00000136319 | 14 | 20724717 | 20774153 | 222 | 35 | 1.6119 | 0.053488 | TTC5 | 0.053488 | 0.099552 | 0.0068937 |
| ENSG00000099617 | 19 | 1286153 | 1301430 | 38 | 6 | 1.6108 | 0.053613 | EFNA2 | 0.053613 | 0.18513 | 0.33558 |
| ENSG00000181004 | 4 | 123653857 | 123666098 | 38 | 6 | 1.6094 | 0.053766 | BBS12 | 0.053766 | 0.15443 | 0.11032 |
| ENSG00000135931 | 2 | 232063260 | 232239548 | 552 | 39 | 1.6094 | 0.053767 | ARMC9 | 0.053767 | 0.058718 | 0.064738 |
| ENSG00000260970 | 9 | 122257653 | 122286403 | 71 | 15 | 1.6093 | 0.053775 | RP11-295D22.1 | 0.053775 | 0.15859 | 0.18267 |
| ENSG00000168454 | 18 | 9885763 | 9889272 | 23 | 8 | 1.6092 | 0.053788 | TXNDC2 | 0.053788 | 0.014717 | 0.12217 |
| ENSG00000134375 | 1 | 201924619 | 201939789 | 53 | 10 | 1.6089 | 0.053822 | TIMM17A | 0.053822 | 0.23052 | 0.44923 |
| ENSG00000044012 | 1 | 42619092 | 42621495 | 9 | 4 | 1.6088 | 0.053832 | GUCA2B | 0.053832 | 0.034751 | 0.026469 |
| ENSG00000125630 | 2 | 113299492 | 113334635 | 94 | 12 | 1.6082 | 0.053893 | POLR1B | 0.053893 | 0.099593 | 0.15365 |
| ENSG00000124253 | 20 | 56136136 | 56141513 | 30 | 10 | 1.6082 | 0.053894 | PCK1 | 0.053894 | 0.10735 | 0.23636 |
| ENSG00000172320 | 11 | 59210617 | 59211667 | 6 | 2 | 1.6077 | 0.05395 | OR5A1 | 0.05395 | 0.018662 | 0.063748 |

| | | | | | | | | | | | |
|---|---|---|---|---|---|---|---|---|---|---|---|
| ENSG00000164168 | 4 | 148538534 | 148593195 | 57 | 16 | 1.6075 | 0.053977 | TMEM184C | 0.053977 | 0.028936 | 0.21513 |
| ENSG00000154957 | 17 | 11880756 | 11900827 | 38 | 9 | 1.6067 | 0.054058 | ZNF18 | 0.054058 | 0.10652 | 0.11974 |
| ENSG00000079150 | 2 | 179328391 | 179343327 | 23 | 7 | 1.6061 | 0.054122 | FKBP7 | 0.054122 | 0.029032 | 0.040788 |
| ENSG00000167600 | 19 | 41698954 | 41713444 | 45 | 12 | 1.6057 | 0.054171 | CYP2S1 | 0.054171 | 0.28661 | 0.12534 |
| ENSG00000106701 | 9 | 108210077 | 108314714 | 301 | 24 | 1.6054 | 0.054198 | FSD1L | 0.054198 | 0.046979 | 0.03034 |
| ENSG00000186496 | 18 | 32946661 | 32957301 | 32 | 8 | 1.6051 | 0.05424 | ZNF396 | 0.05424 | 0.10862 | 0.15629 |
| ENSG00000111664 | 12 | 6949118 | 6956557 | 14 | 5 | 1.6049 | 0.054254 | GNB3 | 0.054254 | 0.08622 | 0.21736 |
| ENSG00000144035 | 2 | 73867959 | 73869520 | 4 | 2 | 1.6046 | 0.054288 | NAT8 | 0.054288 | 0.35822 | 0.40828 |
| ENSG00000203392 | 15 | 75970672 | 75973093 | 4 | 1 | 1.6042 | 0.054331 | AC105020.1 | 0.054331 | 0.11754 | 0.17748 |
| ENSG00000100997 | 20 | 25275379 | 25371619 | 294 | 10 | 1.604 | 0.05436 | ABHD12 | 0.05436 | 0.053433 | 0.053874 |
| ENSG00000267281 | 12 | 53900472 | 54019763 | 237 | 19 | 1.6034 | 0.054421 | RP11-793H13.10 | 0.054421 | 0.062319 | 0.046046 |
| ENSG00000134709 | 1 | 60280458 | 60342050 | 137 | 19 | 1.6029 | 0.054475 | HOOK1 | 0.054475 | 0.056797 | 0.058486 |
| ENSG00000162999 | 2 | 183943287 | 183964733 | 74 | 15 | 1.6024 | 0.054531 | DUSP19 | 0.054531 | 0.054641 | 0.068035 |
| ENSG00000170803 | 11 | 6806248 | 6807245 | 4 | 1 | 1.602 | 0.05458 | OR2AG1 | 0.05458 | 0.060028 | 0.036642 |
| ENSG00000139324 | 12 | 88536073 | 88593664 | 110 | 8 | 1.6016 | 0.054624 | TMTC3 | 0.054624 | 0.042998 | 0.042478 |
| ENSG00000269783 | 10 | 99627889 | 99631294 | 13 | 2 | 1.6007 | 0.054723 | GOLGA7B | 0.054723 | 0.021613 | 0.022216 |
| ENSG00000184384 | 11 | 95709762 | 96076344 | 1383 | 136 | 1.6005 | 0.054741 | MAML2 | 0.054741 | 0.02033 | 0.010466 |
| ENSG00000179772 | 20 | 30432103 | 30433420 | 2 | 1 | 1.6003 | 0.054766 | FOXS1 | 0.054766 | 0.065974 | 0.098759 |
| ENSG00000148604 | 10 | 86004809 | 86019716 | 70 | 14 | 1.5995 | 0.054856 | RGR | 0.054856 | 0.061389 | 0.10723 |
| ENSG00000101294 | 20 | 30102231 | 30157370 | 144 | 8 | 1.5994 | 0.054865 | HM13 | 0.054865 | 0.054889 | 0.048032 |
| ENSG00000170653 | 12 | 53901640 | 54020199 | 236 | 19 | 1.5992 | 0.054883 | ATF7 | 0.054883 | 0.057761 | 0.051483 |
| ENSG00000122692 | 9 | 33041762 | 33076665 | 136 | 16 | 1.5989 | 0.054924 | SMU1 | 0.054924 | 0.074801 | 0.087494 |
| ENSG00000085982 | 2 | 234384166 | 234475428 | 203 | 10 | 1.5974 | 0.055084 | USP40 | 0.055084 | 0.055834 | 0.058122 |
| ENSG00000030110 | 6 | 33540329 | 33548019 | 28 | 8 | 1.5958 | 0.055267 | BAK1 | 0.055267 | 0.033397 | 0.17017 |
| ENSG00000125998 | 20 | 33873534 | 33880204 | 12 | 2 | 1.5953 | 0.055322 | FAM83C | 0.055322 | 0.061956 | 0.095116 |
| ENSG00000169594 | 15 | 83924655 | 83953466 | 45 | 9 | 1.5942 | 0.055446 | BNC1 | 0.055446 | 0.060752 | 0.036235 |
| ENSG00000171885 | 18 | 24432002 | 24445782 | 38 | 9 | 1.5934 | 0.055536 | AQP4 | 0.055536 | 0.16132 | 0.24879 |
| ENSG00000074621 | 15 | 65903704 | 65953333 | 96 | 7 | 1.5933 | 0.055548 | SLC24A1 | 0.055548 | 0.063096 | 0.077186 |
| ENSG00000062038 | 16 | 68670092 | 68756519 | 216 | 27 | 1.5931 | 0.055565 | CDH3 | 0.055565 | 0.080043 | 0.091604 |
| ENSG00000112992 | 5 | 43602794 | 43707507 | 326 | 12 | 1.593 | 0.055585 | NNT | 0.055585 | 0.057545 | 0.05516 |
| ENSG00000112514 | 6 | 33384219 | 33386094 | 4 | 2 | 1.5912 | 0.05578 | CUTA | 0.05578 | 0.027742 | 0.024513 |
| ENSG00000111684 | 12 | 7085348 | 7125814 | 74 | 10 | 1.5905 | 0.055863 | LPCAT3 | 0.055863 | 0.068007 | 0.05515 |
| ENSG00000163632 | 3 | 63805038 | 63834312 | 49 | 14 | 1.59 | 0.055919 | C3orf49 | 0.055919 | 0.14475 | 0.087555 |
| ENSG00000108389 | 17 | 56566898 | 56595266 | 30 | 7 | 1.5899 | 0.055931 | MTMR4 | 0.055931 | 0.046745 | 0.055051 |
| ENSG00000065518 | 3 | 120315156 | 120321347 | 21 | 7 | 1.5892 | 0.056012 | NDUFB4 | 0.056012 | 0.10888 | 0.082278 |
| ENSG00000123178 | 13 | 50486842 | 50510626 | 58 | 10 | 1.5888 | 0.056053 | SPRYD7 | 0.056053 | 0.060673 | 0.039472 |
| ENSG00000166984 | 6 | 167559902 | 167610401 | 197 | 33 | 1.5877 | 0.056177 | TCP10L2 | 0.056177 | 0.20389 | 0.33063 |
| ENSG00000104432 | 8 | 79587978 | 79717758 | 353 | 24 | 1.5867 | 0.056291 | IL7 | 0.056291 | 0.089831 | 0.12807 |
| ENSG00000168159 | 1 | 228674762 | 228683467 | 27 | 9 | 1.586 | 0.056368 | RNF187 | 0.056368 | 0.040188 | 0.047096 |
| ENSG00000242372 | 20 | 33866714 | 33872788 | 23 | 3 | 1.585 | 0.056478 | EIF6 | 0.056478 | 0.07839 | 0.10486 |
| ENSG00000064652 | 5 | 122179134 | 122365049 | 715 | 23 | 1.5849 | 0.056492 | SNX24 | 0.056492 | 0.057304 | 0.064805 |
| ENSG00000175928 | 3 | 3841121 | 3889387 | 178 | 25 | 1.5848 | 0.056505 | LRRN1 | 0.056505 | 0.077859 | 0.15499 |
| ENSG00000162688 | 1 | 100315640 | 100389579 | 292 | 20 | 1.5843 | 0.056563 | AGL | 0.056563 | 0.049212 | 0.022678 |
| ENSG00000137880 | 15 | 41056218 | 41059906 | 4 | 2 | 1.5839 | 0.056605 | GCHFR | 0.056605 | 0.30997 | 0.53869 |
| ENSG00000164258 | 5 | 52856463 | 52979168 | 434 | 26 | 1.5838 | 0.056616 | NDUFS4 | 0.056616 | 0.044205 | 0.035806 |
| ENSG00000164346 | 5 | 74062817 | 74072737 | 22 | 4 | 1.5833 | 0.056679 | NSA2 | 0.056679 | 0.047052 | 0.045438 |
| ENSG00000166548 | 16 | 66541906 | 66586447 | 101 | 7 | 1.5798 | 0.057079 | TK2 | 0.057079 | 0.055117 | 0.064904 |
| ENSG00000015475 | 22 | 18216906 | 18257536 | 205 | 40 | 1.5792 | 0.057148 | BID | 0.057148 | 0.14118 | 0.23895 |
| ENSG00000166224 | 10 | 72575717 | 72640930 | 207 | 24 | 1.5786 | 0.05721 | SGPL1 | 0.05721 | 0.10346 | 0.13318 |
| ENSG00000181718 | 11 | 55999490 | 56000737 | 12 | 4 | 1.5781 | 0.057277 | OR5T2 | 0.057277 | 0.032252 | 0.027631 |
| ENSG00000169758 | 15 | 76352178 | 76521462 | 474 | 49 | 1.5778 | 0.057305 | C15orf27 | 0.057305 | 0.073793 | 0.078927 |
| ENSG00000214026 | 11 | 1968508 | 2005752 | 163 | 24 | 1.5772 | 0.057369 | MRPL23 | 0.057369 | 0.046384 | 0.019357 |
| ENSG00000176160 | 17 | 56497528 | 56565745 | 103 | 14 | 1.5765 | 0.05746 | HSF5 | 0.05746 | 0.062591 | 0.064631 |
| ENSG00000253729 | 8 | 48685669 | 48872743 | 179 | 26 | 1.5755 | 0.057568 | PRKDC | 0.057568 | 0.054449 | 0.051431 |
| ENSG00000079308 | 2 | 218664512 | 218867718 | 629 | 96 | 1.5749 | 0.057637 | TNS1 | 0.057637 | 0.087819 | 0.11948 |
| ENSG00000189143 | 7 | 73213872 | 73247014 | 61 | 10 | 1.5743 | 0.057707 | CLDN4 | 0.057707 | 0.044627 | 0.073769 |
| ENSG00000176230 | 14 | 20585566 | 20586597 | 4 | 2 | 1.5738 | 0.05777 | OR4K17 | 0.05777 | 0.047714 | 0.0057296 |
| ENSG00000077044 | 2 | 234263153 | 234380750 | 262 | 23 | 1.5735 | 0.057799 | DGKD | 0.057799 | 0.067374 | 0.073829 |
| ENSG00000149269 | 11 | 77032752 | 77185680 | 427 | 22 | 1.5731 | 0.057849 | PAK1 | 0.057849 | 0.052009 | 0.076304 |
| ENSG00000155506 | 5 | 154092462 | 154197167 | 235 | 29 | 1.5728 | 0.057879 | LARP1 | 0.057879 | 0.086195 | 0.1553 |
| ENSG00000170006 | 4 | 153539784 | 153601317 | 206 | 36 | 1.5728 | 0.057888 | TMEM154 | 0.057888 | 0.077254 | 0.12932 |
| ENSG00000167608 | 19 | 54663846 | 54676944 | 48 | 12 | 1.5715 | 0.058037 | TMC4 | 0.058037 | 0.14515 | 0.18664 |
| ENSG00000204511 | 6 | 31496494 | 31498009 | 23 | 8 | 1.5714 | 0.058048 | MCCD1 | 0.058048 | 0.14178 | 0.24647 |
| ENSG00000095739 | 10 | 28966271 | 28971868 | 18 | 5 | 1.5705 | 0.058152 | BAMBI | 0.058152 | 0.056054 | 0.13304 |
| ENSG00000177576 | 18 | 47008028 | 47013622 | 19 | 7 | 1.57 | 0.058206 | C18orf32 | 0.058206 | 0.027204 | 0.020905 |
| ENSG00000182636 | 15 | 23930565 | 23932450 | 2 | 1 | 1.5698 | 0.058226 | NDN | 0.058226 | 0.43865 | 0.78958 |
| ENSG00000198467 | 9 | 35681989 | 35691017 | 24 | 6 | 1.5693 | 0.058286 | TPM2 | 0.058286 | 0.013819 | 0.0033158 |
| ENSG00000117394 | 1 | 43391052 | 43424530 | 101 | 14 | 1.5691 | 0.05831 | SLC2A1 | 0.05831 | 0.047143 | 0.074157 |
| ENSG00000150261 | 11 | 56113421 | 56114507 | 4 | 2 | 1.5679 | 0.058448 | OR8K1 | 0.058448 | 0.021708 | 0.017127 |
| ENSG00000176020 | 3 | 49754267 | 49761349 | 6 | 3 | 1.5672 | 0.058531 | AMIGO3 | 0.058531 | 0.028972 | 0.034175 |
| ENSG00000173540 | 3 | 49754277 | 49761384 | 6 | 3 | 1.5672 | 0.058531 | GMPPB | 0.058531 | 0.028972 | 0.034175 |
| ENSG00000125652 | 19 | 6372444 | 6375042 | 6 | 2 | 1.5666 | 0.058603 | ALKBH7 | 0.058603 | 0.12866 | 0.17885 |
| ENSG00000185803 | 8 | 145577795 | 145584932 | 18 | 3 | 1.5665 | 0.058617 | SLC52A2 | 0.058617 | 0.10561 | 0.080459 |

| | | | | | | | | | | | |
|---|---|---|---|---|---|---|---|---|---|---|---|
| ENSG00000212768 | 15 | 35270552 | 35272268 | 6 | 2 | 1.5663 | 0.058644 | AC114546.1 | 0.058644 | 0.14664 | 0.19364 |
| ENSG00000162385 | 1 | 53692564 | 53704282 | 24 | 6 | 1.5637 | 0.058944 | MAGOH | 0.058944 | 0.0234 | 0.024272 |
| ENSG00000140365 | 15 | 75628232 | 75634268 | 14 | 7 | 1.5637 | 0.05895 | COMMD4 | 0.05895 | 0.029168 | 0.02143 |
| ENSG00000164879 | 8 | 86285665 | 86361269 | 139 | 17 | 1.5634 | 0.058979 | CA3 | 0.058979 | 0.052692 | 0.053221 |
| ENSG00000255582 | 14 | 22101992 | 22103096 | 11 | 4 | 1.563 | 0.059024 | OR10G2 | 0.059024 | 0.01332 | 0.010721 |
| ENSG00000054611 | 22 | 47158518 | 47571336 | 1730 | 95 | 1.5627 | 0.059056 | TBC1D22A | 0.059056 | 0.059308 | 0.074137 |
| ENSG00000101162 | 20 | 57594309 | 57601709 | 18 | 8 | 1.562 | 0.059146 | TUBB1 | 0.059146 | 0.14181 | 0.095908 |
| ENSG00000169208 | 14 | 22037934 | 22038875 | 8 | 5 | 1.562 | 0.05915 | OR10G3 | 0.05915 | 0.35641 | 0.326 |
| ENSG00000161618 | 19 | 49956426 | 49974305 | 77 | 9 | 1.5618 | 0.059163 | ALDH16A1 | 0.059163 | 0.05159 | 0.053669 |
| ENSG00000249624 | 21 | 34619079 | 34655517 | 101 | 24 | 1.5616 | 0.059197 | AP000295.9 | 0.059197 | 0.10612 | 0.18955 |
| ENSG00000182040 | 17 | 72912176 | 72919351 | 6 | 3 | 1.5609 | 0.059272 | USH1G | 0.059272 | 0.12459 | 0.55652 |
| ENSG00000144228 | 2 | 139259371 | 139331117 | 103 | 19 | 1.5608 | 0.059283 | SPOPL | 0.059283 | 0.049169 | 0.053478 |
| ENSG00000198707 | 12 | 88442793 | 88535993 | 133 | 17 | 1.5604 | 0.059338 | CEP290 | 0.059338 | 0.0658 | 0.069366 |
| ENSG00000103342 | 16 | 11961985 | 12009939 | 147 | 17 | 1.5602 | 0.05935 | GSPT1 | 0.05935 | 0.049155 | 0.045902 |
| ENSG00000087995 | 17 | 60501228 | 60527454 | 27 | 7 | 1.5602 | 0.05936 | METTL2A | 0.05936 | 0.045159 | 0.028923 |
| ENSG00000110881 | 12 | 50451331 | 50477394 | 52 | 8 | 1.5599 | 0.059387 | ASIC1 | 0.059387 | 0.030204 | 0.047197 |
| ENSG00000111241 | 12 | 4537321 | 4554780 | 37 | 6 | 1.5599 | 0.059395 | FGF6 | 0.059395 | 0.15773 | 0.40909 |
| ENSG00000169994 | 2 | 128293378 | 128395304 | 294 | 28 | 1.5596 | 0.059424 | MYO7B | 0.059424 | 0.066842 | 0.052753 |
| ENSG00000271271 | 4 | 70454135 | 70505360 | 328 | 21 | 1.5588 | 0.059528 | UGT2A2 | 0.059528 | 0.060299 | 0.066073 |
| ENSG00000171105 | 19 | 7112266 | 7294045 | 996 | 107 | 1.557 | 0.059735 | INSR | 0.059735 | 0.035584 | 0.04818 |
| ENSG00000125975 | 20 | 34111014 | 34117481 | 12 | 4 | 1.5568 | 0.059764 | C20orf173 | 0.059764 | 0.077584 | 0.098807 |
| ENSG00000167994 | 11 | 61664773 | 61687741 | 51 | 15 | 1.5567 | 0.059777 | RAB3IL1 | 0.059777 | 0.18018 | 0.15322 |
| ENSG00000270898 | 2 | 53897430 | 54087297 | 642 | 32 | 1.5563 | 0.059816 | ASB3 | 0.059816 | 0.057493 | 0.04877 |
| ENSG00000120437 | 6 | 160181360 | 160200144 | 83 | 12 | 1.5561 | 0.059839 | ACAT2 | 0.059839 | 0.057743 | 0.076242 |
| ENSG00000095574 | 10 | 124750322 | 124768333 | 56 | 11 | 1.556 | 0.059858 | IKZF5 | 0.059858 | 0.10744 | 0.26253 |
| ENSG00000167807 | 19 | 10416103 | 10426685 | 43 | 10 | 1.5557 | 0.05989 | CTD-2369P2.10 | 0.05989 | 0.084791 | 0.16716 |
| ENSG00000267673 | 19 | 10416103 | 10426691 | 43 | 10 | 1.5557 | 0.05989 | FDX1L | 0.05989 | 0.084791 | 0.16716 |
| ENSG00000118640 | 2 | 85788685 | 85809154 | 85 | 11 | 1.5556 | 0.059898 | VAMP8 | 0.059898 | 0.038531 | 0.041618 |
| ENSG00000149256 | 11 | 78363876 | 79151992 | 2557 | 231 | 1.5556 | 0.0599 | TENM4 | 0.0599 | 0.068625 | 0.071202 |
| ENSG00000137996 | 1 | 100731763 | 100758325 | 78 | 8 | 1.5553 | 0.059936 | RTCA | 0.059936 | 0.080545 | 0.080462 |
| ENSG00000198663 | 6 | 36839646 | 36896740 | 148 | 23 | 1.5546 | 0.060016 | C6orf89 | 0.060016 | 0.05755 | 0.18721 |
| ENSG00000068793 | 15 | 22892005 | 23006016 | 533 | 65 | 1.5546 | 0.060016 | CYFIP1 | 0.060016 | 0.060915 | 0.054673 |
| ENSG00000089234 | 12 | 112079950 | 112123790 | 25 | 13 | 1.5541 | 0.060085 | BRAP | 0.060085 | 0.086206 | 0.047075 |
| ENSG00000035862 | 17 | 76849059 | 76921469 | 340 | 29 | 1.5539 | 0.060105 | TIMP2 | 0.060105 | 0.064794 | 0.078318 |
| ENSG00000153944 | 17 | 55333212 | 55762046 | 1368 | 179 | 1.5538 | 0.060111 | MSI2 | 0.060111 | 0.081664 | 0.11463 |
| ENSG00000129226 | 17 | 7482785 | 7485429 | 7 | 4 | 1.5535 | 0.060152 | CD68 | 0.060152 | 0.0082407 | 0.0098693 |
| ENSG00000130950 | 9 | 97077605 | 97090926 | 52 | 13 | 1.5531 | 0.060198 | NUTM2F | 0.060198 | 0.14383 | 0.14242 |
| ENSG00000119938 | 10 | 93388199 | 93392811 | 6 | 4 | 1.553 | 0.060213 | PPP1R3C | 0.060213 | 0.41336 | 0.7086 |
| ENSG00000116157 | 1 | 53068044 | 53074723 | 21 | 5 | 1.5521 | 0.060313 | GPX7 | 0.060313 | 0.030453 | 0.010004 |
| ENSG00000107669 | 10 | 123499939 | 123688316 | 576 | 42 | 1.552 | 0.060331 | ATE1 | 0.060331 | 0.051702 | 0.025999 |
| ENSG00000175619 | 11 | 48238344 | 48239314 | 6 | 3 | 1.5512 | 0.060421 | OR4B1 | 0.060421 | 0.065789 | 0.04962 |
| ENSG00000174501 | 2 | 96514587 | 96657541 | 139 | 30 | 1.5512 | 0.06043 | ANKRD36C | 0.06043 | 0.060613 | 0.053498 |
| ENSG00000070601 | 9 | 37650997 | 37746901 | 404 | 45 | 1.551 | 0.060453 | FRMPD1 | 0.060453 | 0.041235 | 0.045487 |
| ENSG00000150867 | 10 | 22823778 | 23003484 | 671 | 33 | 1.5503 | 0.060541 | PIP4K2A | 0.060541 | 0.076286 | 0.11043 |
| ENSG00000127946 | 7 | 75162621 | 75368280 | 850 | 67 | 1.5497 | 0.060604 | HIP1 | 0.060604 | 0.067433 | 0.093835 |
| ENSG00000023572 | 1 | 193065598 | 193075244 | 14 | 5 | 1.5497 | 0.060606 | GLRX2 | 0.060606 | 0.21458 | 0.29834 |
| ENSG00000130309 | 19 | 17666403 | 17693971 | 89 | 19 | 1.5487 | 0.060727 | COLGALT1 | 0.060727 | 0.026049 | 0.063127 |
| ENSG00000178917 | 3 | 44540462 | 44552128 | 14 | 6 | 1.5485 | 0.060755 | ZNF852 | 0.060755 | 0.14604 | 0.2677 |
| ENSG00000183763 | 3 | 49866034 | 49894007 | 51 | 8 | 1.5481 | 0.060805 | TRAIP | 0.060805 | 0.061904 | 0.05913 |
| ENSG00000133742 | 8 | 86239837 | 86291243 | 111 | 19 | 1.5473 | 0.06089 | CA1 | 0.06089 | 0.068524 | 0.062369 |
| ENSG00000214597 | 8 | 145575878 | 145578574 | 8 | 2 | 1.5472 | 0.060906 | TMEM249 | 0.060906 | 0.10106 | 0.077616 |
| ENSG00000107957 | 10 | 105348285 | 105615301 | 774 | 69 | 1.5471 | 0.060922 | SH3PXD2A | 0.060922 | 0.067824 | 0.065572 |
| ENSG00000104221 | 8 | 37700786 | 37707422 | 9 | 2 | 1.5451 | 0.061166 | BRF2 | 0.061166 | 0.18937 | 0.54538 |
| ENSG00000140564 | 15 | 91411822 | 91426688 | 30 | 6 | 1.5444 | 0.061248 | FURIN | 0.061248 | 0.020717 | 0.03336 |
| ENSG00000156983 | 3 | 9773413 | 9789702 | 33 | 13 | 1.5444 | 0.061252 | BRPF1 | 0.061252 | 0.023002 | 0.049787 |
| ENSG00000156976 | 3 | 186500994 | 186507689 | 14 | 7 | 1.5433 | 0.06138 | EIF4A2 | 0.06138 | 0.044084 | 0.078346 |
| ENSG00000198128 | 1 | 248223984 | 248224922 | 10 | 2 | 1.5428 | 0.061444 | OR2L3 | 0.061444 | 0.027292 | 0.019083 |
| ENSG00000197837 | 12 | 14920933 | 14924065 | 11 | 4 | 1.5423 | 0.061502 | HIST4H4 | 0.061502 | 0.3582 | 0.50942 |
| ENSG00000155016 | 4 | 108852525 | 108874613 | 87 | 8 | 1.5414 | 0.061608 | CYP2U1 | 0.061608 | 0.060222 | 0.06005 |
| ENSG00000119899 | 6 | 74303102 | 74363878 | 433 | 13 | 1.5409 | 0.061673 | SLC17A5 | 0.061673 | 0.064569 | 0.053722 |
| ENSG00000164520 | 6 | 150204511 | 150219238 | 72 | 4 | 1.5405 | 0.061723 | RAET1E | 0.061723 | 0.065842 | 0.091076 |
| ENSG00000168002 | 11 | 62529016 | 62534182 | 12 | 7 | 1.5403 | 0.061745 | POLR2G | 0.061745 | 0.50762 | 0.57599 |
| ENSG00000085872 | 19 | 16628700 | 16653341 | 65 | 14 | 1.5399 | 0.06179 | CHERP | 0.06179 | 0.128 | 0.18534 |
| ENSG00000174529 | 1 | 205052258 | 205053645 | 6 | 3 | 1.5381 | 0.062011 | TMEM81 | 0.062011 | 0.025223 | 0.0089371 |
| ENSG00000101335 | 20 | 35169887 | 35178228 | 11 | 6 | 1.5376 | 0.062079 | MYL9 | 0.062079 | 0.14194 | 0.37885 |
| ENSG00000162302 | 11 | 64126620 | 64139687 | 32 | 4 | 1.537 | 0.062145 | RPS6KA4 | 0.062145 | 0.13387 | 0.13579 |
| ENSG00000085999 | 1 | 46713360 | 46744145 | 44 | 13 | 1.5359 | 0.06228 | RAD54L | 0.06228 | 0.047809 | 0.056802 |
| ENSG00000169919 | 7 | 65425671 | 65447301 | 63 | 10 | 1.5355 | 0.062333 | GUSB | 0.062333 | 0.044453 | 0.047354 |
| ENSG00000162972 | 2 | 200820040 | 200873263 | 110 | 18 | 1.5351 | 0.062385 | C2orf47 | 0.062385 | 0.11085 | 0.11168 |
| ENSG00000171357 | 1 | 46669006 | 46686933 | 31 | 10 | 1.5342 | 0.062491 | LURAP1 | 0.062491 | 0.013929 | 0.022725 |
| ENSG00000138792 | 4 | 111286889 | 111486441 | 673 | 51 | 1.5337 | 0.062551 | ENPEP | 0.062551 | 0.07706 | 0.15297 |
| ENSG00000182271 | 17 | 28643351 | 28661077 | 48 | 8 | 1.5335 | 0.062577 | TMIGD1 | 0.062577 | 0.08881 | 0.14393 |
| ENSG00000197863 | 19 | 37308330 | 37341689 | 55 | 10 | 1.533 | 0.06264 | ZNF790 | 0.06264 | 0.057085 | 0.056028 |

| | | | | | | | | | | | |
|---|---|---|---|---|---|---|---|---|---|---|---|
| ENSG00000116525 | 1 | 33611003 | 33647660 | 102 | 8 | 1.5325 | 0.062695 | TRIM62 | 0.062695 | 0.059066 | 0.063445 |
| ENSG00000106290 | 7 | 99704693 | 99717464 | 26 | 7 | 1.5321 | 0.062751 | TAF6 | 0.062751 | 0.015194 | 0.0094652 |
| ENSG00000081721 | 1 | 161719548 | 161727028 | 33 | 7 | 1.5319 | 0.06277 | DUSP12 | 0.06277 | 0.12636 | 0.23548 |
| ENSG00000170075 | 1 | 202091986 | 202102720 | 51 | 14 | 1.5315 | 0.062822 | GPR37L1 | 0.062822 | 0.13834 | 0.14066 |
| ENSG00000007923 | 1 | 6694228 | 6761984 | 171 | 19 | 1.5313 | 0.06285 | DNAJC11 | 0.06285 | 0.057756 | 0.083266 |
| ENSG00000123143 | 19 | 14543865 | 14582679 | 131 | 29 | 1.5309 | 0.062891 | PKN1 | 0.062891 | 0.19494 | 0.11471 |
| ENSG00000182545 | 14 | 20973696 | 20979328 | 14 | 7 | 1.5309 | 0.062896 | RNASE10 | 0.062896 | 0.063442 | 0.4287 |
| ENSG00000163827 | 3 | 46556913 | 46621589 | 284 | 21 | 1.5309 | 0.062901 | LRRC2 | 0.062901 | 0.073919 | 0.087263 |
| ENSG00000163114 | 4 | 96761239 | 96762625 | 2 | 1 | 1.5301 | 0.062991 | PDHA2 | 0.062991 | 0.44797 | 0.4159 |
| ENSG00000091622 | 17 | 6354584 | 6459814 | 426 | 36 | 1.53 | 0.063004 | PITPNM3 | 0.063004 | 0.11376 | 0.21825 |
| ENSG00000204882 | 8 | 142366600 | 142377367 | 45 | 9 | 1.53 | 0.063013 | GPR20 | 0.063013 | 0.11454 | 0.23074 |
| ENSG00000127080 | 9 | 95375466 | 95432547 | 180 | 20 | 1.5296 | 0.06306 | IPPK | 0.06306 | 0.067465 | 0.081666 |
| ENSG00000167525 | 17 | 27030215 | 27038872 | 13 | 5 | 1.5291 | 0.063118 | PROCA1 | 0.063118 | 0.079932 | 0.1074 |
| ENSG00000181274 | 10 | 99092255 | 99094458 | 3 | 2 | 1.5288 | 0.063156 | FRAT2 | 0.063156 | 0.0093914 | 0.018685 |
| ENSG00000269657 | 17 | 66539369 | 66542970 | 9 | 4 | 1.5284 | 0.063207 | AC079210.1 | 0.063207 | 0.0082653 | 0.015474 |
| ENSG00000123407 | 12 | 54348618 | 54352740 | 16 | 4 | 1.5282 | 0.063234 | HOXC12 | 0.063234 | 0.13491 | 0.11197 |
| ENSG00000116106 | 2 | 222282747 | 222438922 | 496 | 44 | 1.5274 | 0.063331 | EPHA4 | 0.063331 | 0.091424 | 0.11626 |
| ENSG00000117862 | 1 | 52485803 | 52521843 | 84 | 10 | 1.5271 | 0.063367 | TXNDC12 | 0.063367 | 0.04702 | 0.065631 |
| ENSG00000196811 | 2 | 233404437 | 233411113 | 30 | 7 | 1.5266 | 0.063426 | CHRNG | 0.063426 | 0.16492 | 0.13507 |
| ENSG00000107371 | 9 | 37766975 | 37801434 | 114 | 14 | 1.5266 | 0.06343 | EXOSC3 | 0.06343 | 0.081853 | 0.073143 |
| ENSG00000185905 | 16 | 29753784 | 29757327 | 5 | 2 | 1.5264 | 0.063456 | C16orf54 | 0.063456 | 0.11707 | 0.054888 |
| ENSG00000185561 | 17 | 1611064 | 1613732 | 7 | 3 | 1.5234 | 0.063828 | TLCD2 | 0.063828 | 0.21308 | 0.060502 |
| ENSG00000119514 | 9 | 101569981 | 101612363 | 114 | 29 | 1.5233 | 0.063841 | GALNT12 | 0.063841 | 0.10508 | 0.24138 |
| ENSG00000205246 | 19 | 23945807 | 24010937 | 135 | 22 | 1.5226 | 0.063931 | RPSAP58 | 0.063931 | 0.064534 | 0.049716 |
| ENSG00000162290 | 3 | 53317447 | 53381654 | 132 | 18 | 1.5226 | 0.063932 | DCP1A | 0.063932 | 0.055714 | 0.057252 |
| ENSG00000051523 | 16 | 88709691 | 88717560 | 40 | 13 | 1.5221 | 0.063993 | CYBA | 0.063993 | 0.15845 | 0.25275 |
| ENSG00000099901 | 22 | 20103461 | 20114878 | 41 | 11 | 1.5219 | 0.064016 | RANBP1 | 0.064016 | 0.11918 | 0.058815 |
| ENSG00000163704 | 3 | 9987226 | 9994078 | 13 | 7 | 1.5218 | 0.064029 | PRRT3 | 0.064029 | 0.024628 | 0.012071 |
| ENSG00000172362 | 11 | 58206593 | 58207646 | 3 | 2 | 1.5213 | 0.064086 | OR5B12 | 0.064086 | 0.015048 | 0.02075 |
| ENSG00000167555 | 19 | 52901102 | 52921665 | 40 | 6 | 1.5209 | 0.064138 | ZNF528 | 0.064138 | 0.093369 | 0.15555 |
| ENSG00000115392 | 2 | 58386378 | 58468507 | 225 | 27 | 1.5204 | 0.064207 | FANCL | 0.064207 | 0.11799 | 0.19969 |
| ENSG00000121211 | 4 | 154265801 | 154336270 | 257 | 20 | 1.5196 | 0.064306 | MND1 | 0.064306 | 0.063764 | 0.070558 |
| ENSG00000008256 | 7 | 6201407 | 6312275 | 320 | 26 | 1.519 | 0.064386 | CYTH3 | 0.064386 | 0.049971 | 0.034473 |
| ENSG00000126777 | 14 | 56025790 | 56168244 | 502 | 14 | 1.5188 | 0.064405 | KTN1 | 0.064405 | 0.059233 | 0.068168 |
| ENSG00000205549 | 9 | 16203933 | 16276311 | 356 | 46 | 1.5178 | 0.064532 | C9orf92 | 0.064532 | 0.006005 | 0.00079955 |
| ENSG00000040608 | 22 | 20228938 | 20270769 | 104 | 22 | 1.5168 | 0.064659 | RTN4R | 0.064659 | 0.070569 | 0.12833 |
| ENSG00000146083 | 5 | 175953698 | 175965026 | 37 | 13 | 1.5163 | 0.064723 | RNF44 | 0.064723 | 0.4378 | 0.2737 |
| ENSG00000164291 | 5 | 94890778 | 94940768 | 85 | 10 | 1.5161 | 0.064747 | ARSK | 0.064747 | 0.10651 | 0.12721 |
| ENSG00000151883 | 5 | 49961733 | 50142356 | 358 | 18 | 1.5151 | 0.064877 | PARP8 | 0.064877 | 0.063682 | 0.06774 |
| ENSG00000213085 | 1 | 159842154 | 159869953 | 96 | 16 | 1.5148 | 0.064917 | CCDC19 | 0.064917 | 0.10791 | 0.062756 |
| ENSG00000074696 | 15 | 65822756 | 65870687 | 121 | 11 | 1.5147 | 0.064922 | PTPLAD1 | 0.064922 | 0.063618 | 0.060685 |
| ENSG00000173699 | 2 | 231860836 | 231889770 | 194 | 16 | 1.5142 | 0.064983 | SPATA3 | 0.064983 | 0.071982 | 0.06369 |
| ENSG00000158528 | 7 | 94536514 | 94925727 | 795 | 32 | 1.5141 | 0.065 | PPP1R9A | 0.065 | 0.064661 | 0.078647 |
| ENSG00000196834 | 2 | 131217028 | 131267239 | 45 | 5 | 1.5136 | 0.065069 | POTEI | 0.065069 | 0.068376 | 0.073019 |
| ENSG00000154099 | 16 | 84178865 | 84212373 | 160 | 25 | 1.5134 | 0.065085 | DNAAF1 | 0.065085 | 0.1065 | 0.15842 |
| ENSG00000137460 | 4 | 153857504 | 153900848 | 140 | 34 | 1.5125 | 0.065201 | FHDC1 | 0.065201 | 0.044787 | 0.050361 |
| ENSG00000005981 | 7 | 95107756 | 95169544 | 193 | 31 | 1.5122 | 0.065239 | ASB4 | 0.065239 | 0.11959 | 0.21836 |
| ENSG00000007908 | 1 | 169691781 | 169733846 | 295 | 12 | 1.5116 | 0.065321 | SELE | 0.065321 | 0.060757 | 0.066803 |
| ENSG00000151789 | 3 | 21459915 | 22414812 | 5159 | 247 | 1.5115 | 0.065337 | ZNF385D | 0.065337 | 0.064441 | 0.070483 |
| ENSG00000212928 | 14 | 57671888 | 57673291 | 6 | 2 | 1.5112 | 0.065363 | AL391152.1 | 0.065363 | 0.079309 | 0.08885 |
| ENSG00000094975 | 1 | 172501489 | 172580971 | 137 | 19 | 1.5112 | 0.065372 | SUCO | 0.065372 | 0.079172 | 0.104 |
| ENSG00000143578 | 1 | 153940010 | 153946839 | 11 | 3 | 1.51 | 0.065519 | CREB3L4 | 0.065519 | 0.22087 | 0.19595 |
| ENSG00000204228 | 6 | 33172419 | 33174608 | 7 | 3 | 1.5095 | 0.065586 | HSD17B8 | 0.065586 | 0.077544 | 0.1888 |
| ENSG00000166578 | 12 | 113633246 | 113658899 | 60 | 5 | 1.5095 | 0.06559 | IQCD | 0.06559 | 0.068878 | 0.073775 |
| ENSG00000111667 | 12 | 6961292 | 6975796 | 13 | 5 | 1.5094 | 0.065594 | USP5 | 0.065594 | 0.03944 | 0.021918 |
| ENSG00000088682 | 16 | 57481337 | 57495187 | 34 | 10 | 1.5088 | 0.065674 | COQ9 | 0.065674 | 0.011493 | 0.0063955 |
| ENSG00000128886 | 15 | 44064798 | 44069741 | 6 | 2 | 1.5083 | 0.065735 | ELL3 | 0.065735 | 0.23305 | 0.20614 |
| ENSG00000258366 | 20 | 62289163 | 62328416 | 186 | 21 | 1.5075 | 0.065839 | RTEL1 | 0.065839 | 0.046642 | 0.050893 |
| ENSG00000198794 | 15 | 75249560 | 75313837 | 187 | 20 | 1.5067 | 0.065944 | SCAMP5 | 0.065944 | 0.061025 | 0.065935 |
| ENSG00000171840 | 12 | 673462 | 772945 | 517 | 34 | 1.5064 | 0.065977 | NINJ2 | 0.065977 | 0.076673 | 0.046811 |
| ENSG00000130695 | 1 | 26560691 | 26605299 | 120 | 15 | 1.5064 | 0.065979 | CEP85 | 0.065979 | 0.090072 | 0.1201 |
| ENSG00000181039 | 1 | 145470508 | 145475646 | 7 | 4 | 1.5064 | 0.065985 | ANKRD34A | 0.065985 | 0.21819 | 0.39274 |
| ENSG00000146476 | 6 | 151773422 | 151791236 | 59 | 10 | 1.5056 | 0.066084 | C6orf211 | 0.066084 | 0.1454 | 0.12911 |
| ENSG00000134020 | 8 | 22570769 | 22857513 | 1052 | 110 | 1.5054 | 0.066111 | PEBP4 | 0.066111 | 0.085746 | 0.11615 |
| ENSG00000172188 | 11 | 55370830 | 55371874 | 4 | 2 | 1.5052 | 0.066131 | OR4C11 | 0.066131 | 0.057874 | 0.068195 |
| ENSG00000151651 | 10 | 135075907 | 135090372 | 55 | 7 | 1.505 | 0.066163 | ADAM8 | 0.066163 | 0.039587 | 0.047513 |
| ENSG00000182782 | 12 | 123185840 | 123187890 | 8 | 3 | 1.5039 | 0.066302 | HCAR2 | 0.066302 | 0.16469 | 0.19173 |
| ENSG00000134061 | 5 | 66478103 | 66492627 | 36 | 9 | 1.5031 | 0.066407 | CD180 | 0.066407 | 0.060877 | 0.064291 |
| ENSG00000115718 | 2 | 128176003 | 128186822 | 27 | 5 | 1.5028 | 0.066444 | PROC | 0.066444 | 0.11249 | 0.25856 |
| ENSG00000258053 | 12 | 71441182 | 71512027 | 227 | 16 | 1.5027 | 0.066458 | CTD-2021H9.3 | 0.066458 | 0.056371 | 0.043069 |
| ENSG00000100151 | 22 | 38452318 | 38471708 | 32 | 6 | 1.5027 | 0.066458 | PICK1 | 0.066458 | 0.099194 | 0.16945 |
| ENSG00000134824 | 11 | 61560452 | 61634826 | 196 | 21 | 1.502 | 0.066549 | FADS2 | 0.066549 | 0.064212 | 0.084624 |
| ENSG00000153896 | 19 | 35248981 | 35264134 | 29 | 9 | 1.502 | 0.066555 | ZNF599 | 0.066555 | 0.12563 | 0.13084 |

| | | | | | | | | | | | |
|---|---|---|---|---|---|---|---|---|---|---|---|
| ENSG00000100360 | 22 | 37154246 | 37172300 | 46 | 6 | 1.5013 | 0.066639 | IFT27 | 0.066639 | 0.13836 | 0.18473 |
| ENSG00000153993 | 7 | 84624869 | 84816171 | 309 | 33 | 1.5013 | 0.066643 | SEMA3D | 0.066643 | 0.061398 | 0.046208 |
| ENSG00000180917 | 16 | 71315292 | 71323618 | 24 | 7 | 1.5002 | 0.066778 | CMTR2 | 0.066778 | 0.083677 | 0.079415 |
| ENSG00000167970 | 16 | 2303738 | 2312343 | 12 | 5 | 1.4992 | 0.066905 | AC009065.1 | 0.066905 | 0.2436 | 0.44768 |
| ENSG00000177103 | 11 | 117298489 | 117688240 | 1571 | 128 | 1.4981 | 0.067059 | DSCAML1 | 0.067059 | 0.062755 | 0.062508 |
| ENSG00000171984 | 20 | 5731039 | 5844558 | 493 | 67 | 1.4974 | 0.067141 | C20orf196 | 0.067141 | 0.10633 | 0.16338 |
| ENSG00000034053 | 15 | 29129629 | 29410518 | 706 | 33 | 1.4972 | 0.067166 | APBA2 | 0.067166 | 0.106 | 0.093504 |
| ENSG00000105464 | 19 | 48898132 | 48948188 | 147 | 31 | 1.4972 | 0.067173 | GRIN2D | 0.067173 | 0.11858 | 0.16814 |
| ENSG00000186326 | 19 | 33166313 | 33169206 | 16 | 5 | 1.4968 | 0.067223 | RGS9BP | 0.067223 | 0.048024 | 0.08936 |
| ENSG00000126458 | 19 | 50138549 | 50143458 | 9 | 2 | 1.4968 | 0.067226 | RRAS | 0.067226 | 0.1195 | 0.14689 |
| ENSG00000183876 | 5 | 149675906 | 149718870 | 112 | 19 | 1.4965 | 0.067261 | ARSI | 0.067261 | 0.09413 | 0.10758 |
| ENSG00000151360 | 2 | 3705785 | 3750261 | 191 | 23 | 1.4964 | 0.06728 | ALLC | 0.06728 | 0.049095 | 0.023892 |
| ENSG00000173610 | 4 | 70454135 | 70518965 | 391 | 25 | 1.496 | 0.067321 | UGT2A1 | 0.067321 | 0.085248 | 0.083888 |
| ENSG00000119778 | 2 | 23971534 | 24149984 | 627 | 25 | 1.496 | 0.067328 | ATAD2B | 0.067328 | 0.071627 | 0.076993 |
| ENSG00000270386 | 4 | 70454912 | 70518967 | 388 | 25 | 1.4953 | 0.067413 | UGT2A1 | 0.067413 | 0.085248 | 0.083888 |
| ENSG00000162631 | 1 | 107682629 | 108026080 | 876 | 75 | 1.4953 | 0.067419 | NTNG1 | 0.067419 | 0.059829 | 0.06314 |
| ENSG00000155893 | 3 | 140947568 | 141013748 | 303 | 24 | 1.4952 | 0.067427 | ACPL2 | 0.067427 | 0.081137 | 0.061326 |
| ENSG00000174448 | 18 | 51850728 | 51884334 | 99 | 8 | 1.4947 | 0.067499 | STARD6 | 0.067499 | 0.055351 | 0.050658 |
| ENSG00000132854 | 1 | 62702651 | 62785085 | 411 | 28 | 1.4942 | 0.067561 | KANK4 | 0.067561 | 0.067293 | 0.10689 |
| ENSG00000167601 | 19 | 41725108 | 41767671 | 121 | 19 | 1.4941 | 0.067575 | AXL | 0.067575 | 0.051456 | 0.05283 |
| ENSG00000110436 | 11 | 35272753 | 35441610 | 592 | 37 | 1.4938 | 0.067619 | SLC1A2 | 0.067619 | 0.074907 | 0.085354 |
| ENSG00000159556 | 15 | 76629065 | 76634817 | 12 | 5 | 1.4937 | 0.067626 | ISL2 | 0.067626 | 0.080406 | 0.11919 |
| ENSG00000125650 | 19 | 6375159 | 6379069 | 26 | 5 | 1.4937 | 0.067632 | PSPN | 0.067632 | 0.11206 | 0.15578 |
| ENSG00000142549 | 19 | 51815102 | 51833608 | 64 | 11 | 1.4932 | 0.067688 | IGLON5 | 0.067688 | 0.10743 | 0.21759 |
| ENSG00000127415 | 4 | 980785 | 998316 | 86 | 21 | 1.4929 | 0.067732 | IDUA | 0.067732 | 0.12421 | 0.12222 |
| ENSG00000134198 | 1 | 115590632 | 115632121 | 179 | 25 | 1.4929 | 0.067734 | TSPAN2 | 0.067734 | 0.080709 | 0.077428 |
| ENSG00000157657 | 9 | 116638562 | 116818871 | 617 | 44 | 1.4925 | 0.067779 | ZNF618 | 0.067779 | 0.072956 | 0.096817 |
| ENSG00000110315 | 11 | 10533225 | 10562777 | 129 | 16 | 1.4924 | 0.0678 | RNF141 | 0.0678 | 0.10378 | 0.17326 |
| ENSG00000069482 | 11 | 68451247 | 68458643 | 17 | 5 | 1.4921 | 0.06783 | GAL | 0.06783 | 0.032307 | 0.023718 |
| ENSG00000127419 | 4 | 926175 | 952444 | 90 | 20 | 1.4918 | 0.06787 | TMEM175 | 0.06787 | 0.051024 | 0.10177 |
| ENSG00000181264 | 11 | 120195838 | 120204391 | 23 | 6 | 1.4902 | 0.068082 | TMEM136 | 0.068082 | 0.18382 | 0.22928 |
| ENSG00000186231 | 6 | 97372605 | 97588630 | 825 | 41 | 1.4892 | 0.068216 | KLHL32 | 0.068216 | 0.072724 | 0.092489 |
| ENSG00000109158 | 4 | 46920917 | 46996424 | 255 | 20 | 1.4889 | 0.068256 | GABRA4 | 0.068256 | 0.059849 | 0.05075 |
| ENSG00000152213 | 13 | 50202435 | 50208008 | 18 | 7 | 1.4888 | 0.068268 | ARL11 | 0.068268 | 0.18936 | 0.29705 |
| ENSG00000170289 | 8 | 87566205 | 87755903 | 672 | 53 | 1.4883 | 0.068335 | CNGB3 | 0.068335 | 0.073617 | 0.13631 |
| ENSG00000203863 | 6 | 89674246 | 89676685 | 9 | 5 | 1.4879 | 0.06839 | AL079342.1 | 0.06839 | 0.49452 | 0.78996 |
| ENSG00000070814 | 5 | 149737202 | 149779871 | 94 | 15 | 1.4868 | 0.068532 | TCOF1 | 0.068532 | 0.078602 | 0.076784 |
| ENSG00000168509 | 1 | 145413095 | 145417545 | 3 | 2 | 1.4867 | 0.06855 | HFE2 | 0.06855 | 0.66058 | 0.46697 |
| ENSG00000100427 | 22 | 50497820 | 50524331 | 212 | 17 | 1.4866 | 0.068566 | MLC1 | 0.068566 | 0.10077 | 0.19281 |
| ENSG00000171936 | 19 | 15852203 | 15853153 | 7 | 3 | 1.4859 | 0.068655 | OR10H3 | 0.068655 | 0.030571 | 0.093698 |
| ENSG00000159069 | 9 | 139834887 | 139839148 | 10 | 4 | 1.4858 | 0.068664 | FBXW5 | 0.068664 | 0.079843 | 0.057672 |
| ENSG00000161714 | 17 | 43186335 | 43210721 | 79 | 21 | 1.4853 | 0.068727 | PLCD3 | 0.068727 | 0.13886 | 0.32549 |
| ENSG00000070886 | 1 | 22890057 | 22930087 | 205 | 30 | 1.4851 | 0.068763 | EPHA8 | 0.068763 | 0.035096 | 0.069475 |
| ENSG00000071054 | 2 | 102313312 | 102511149 | 426 | 30 | 1.485 | 0.068776 | MAP4K4 | 0.068776 | 0.076773 | 0.077764 |
| ENSG00000146457 | 6 | 160146617 | 160177351 | 96 | 14 | 1.4849 | 0.068786 | WTAP | 0.068786 | 0.048966 | 0.034998 |
| ENSG00000143450 | 1 | 151735445 | 151743808 | 28 | 4 | 1.4843 | 0.068871 | OAZ3 | 0.068871 | 0.024706 | 0.036279 |
| ENSG00000154319 | 8 | 11278972 | 11332224 | 327 | 26 | 1.4836 | 0.068951 | FAM167A | 0.068951 | 0.095254 | 0.1854 |
| ENSG00000102753 | 13 | 50273447 | 50367057 | 365 | 26 | 1.4833 | 0.068997 | KPNA3 | 0.068997 | 0.051402 | 0.056101 |
| ENSG00000185730 | 8 | 144371846 | 144380231 | 41 | 4 | 1.4829 | 0.069047 | ZNF696 | 0.069047 | 0.059529 | 0.067134 |
| ENSG00000204314 | 6 | 32116136 | 32122150 | 7 | 3 | 1.4819 | 0.069177 | PRRT1 | 0.069177 | 0.025421 | 0.030307 |
| ENSG00000234949 | 2 | 238499663 | 238504624 | 38 | 11 | 1.4816 | 0.069218 | AC104667.3 | 0.069218 | 0.34395 | 0.26394 |
| ENSG00000171450 | 2 | 219824377 | 219826876 | 6 | 1 | 1.4815 | 0.069241 | CDK5R2 | 0.069241 | 0.12 | 0.072648 |
| ENSG00000145217 | 4 | 972861 | 987228 | 40 | 15 | 1.4814 | 0.069247 | SLC26A1 | 0.069247 | 0.12455 | 0.10895 |
| ENSG00000155875 | 9 | 18927656 | 19033251 | 558 | 29 | 1.4803 | 0.069392 | FAM154A | 0.069392 | 0.064652 | 0.083796 |
| ENSG00000105880 | 7 | 96649704 | 96654409 | 14 | 8 | 1.4802 | 0.069407 | DLX5 | 0.069407 | 0.093289 | 0.071402 |
| ENSG00000128512 | 7 | 111366166 | 111846466 | 1335 | 147 | 1.4802 | 0.069409 | DOCK4 | 0.069409 | 0.061096 | 0.055423 |
| ENSG00000204315 | 6 | 32096484 | 32098068 | 6 | 3 | 1.4801 | 0.069425 | FKBPL | 0.069425 | 0.016447 | 0.011677 |
| ENSG00000169814 | 3 | 15642848 | 15687329 | 94 | 16 | 1.48 | 0.069435 | BTD | 0.069435 | 0.075119 | 0.050105 |
| ENSG00000183978 | 17 | 40947165 | 40950722 | 2 | 1 | 1.4791 | 0.069561 | COA3 | 0.069561 | 0.33822 | 0.53947 |
| ENSG00000105583 | 19 | 12778885 | 12782170 | 6 | 3 | 1.4785 | 0.069636 | WDR83OS | 0.069636 | 0.0019052 | 0.00021025 |
| ENSG00000143067 | 1 | 120162045 | 120190396 | 76 | 10 | 1.4768 | 0.069867 | ZNF697 | 0.069867 | 0.17113 | 0.079486 |
| ENSG00000124172 | 20 | 57600522 | 57607437 | 9 | 4 | 1.4762 | 0.069939 | ATP5E | 0.069939 | 0.079113 | 0.093645 |
| ENSG00000135406 | 12 | 49687035 | 49692465 | 13 | 5 | 1.476 | 0.069978 | PRPH | 0.069978 | 0.051065 | 0.057261 |
| ENSG00000171532 | 17 | 37759789 | 37766030 | 9 | 4 | 1.4757 | 0.070011 | NEUROD2 | 0.070011 | 0.15978 | 0.27955 |
| ENSG00000162783 | 1 | 181057638 | 181059977 | 7 | 3 | 1.4756 | 0.070029 | IER5 | 0.070029 | 0.036157 | 0.20422 |
| ENSG00000188803 | 17 | 11144580 | 11467380 | 1292 | 118 | 1.4751 | 0.070088 | SHISA6 | 0.070088 | 0.083298 | 0.10419 |
| ENSG00000196391 | 15 | 90895477 | 90909324 | 61 | 7 | 1.4745 | 0.07018 | ZNF774 | 0.07018 | 0.1046 | 0.069171 |
| ENSG00000170185 | 4 | 144106070 | 144144983 | 89 | 8 | 1.474 | 0.070243 | USP38 | 0.070243 | 0.081078 | 0.078987 |
| ENSG00000223501 | 6 | 33218049 | 33239824 | 76 | 12 | 1.4729 | 0.070391 | VPS52 | 0.070391 | 0.044062 | 0.03646 |
| ENSG00000183347 | 1 | 89829617 | 89852020 | 84 | 16 | 1.4729 | 0.070392 | GBP6 | 0.070392 | 0.27283 | 0.25121 |
| ENSG00000163624 | 4 | 85504132 | 85572491 | 126 | 17 | 1.4727 | 0.070417 | CDS1 | 0.070417 | 0.063852 | 0.061284 |
| ENSG00000186710 | 12 | 113587663 | 113597081 | 20 | 2 | 1.4726 | 0.070426 | CCDC42B | 0.070426 | 0.13632 | 0.094849 |
| ENSG00000118520 | 6 | 131894284 | 131905472 | 27 | 9 | 1.4723 | 0.070476 | ARG1 | 0.070476 | 0.11432 | 0.11001 |

| | | | | | | | | | | | |
|---|---|---|---|---|---|---|---|---|---|---|---|
| ENSG00000178404 | 17 | 76866992 | 76899299 | 180 | 23 | 1.4722 | 0.070481 | DDC8 | 0.070481 | 0.057377 | 0.069326 |
| ENSG00000181273 | 11 | 56756347 | 56757342 | 4 | 2 | 1.4709 | 0.070656 | OR5AK2 | 0.070656 | 0.39936 | 0.46505 |
| ENSG00000198363 | 8 | 62413116 | 62627155 | 604 | 41 | 1.4702 | 0.070761 | ASPH | 0.070761 | 0.073706 | 0.083161 |
| ENSG00000088325 | 20 | 30327074 | 30389608 | 133 | 13 | 1.4701 | 0.070761 | TPX2 | 0.070761 | 0.078077 | 0.0796 |
| ENSG00000249087 | 1 | 23695490 | 23698332 | 3 | 2 | 1.4701 | 0.07077 | C1orf213 | 0.07077 | 0.067859 | 0.18304 |
| ENSG00000119862 | 2 | 64681103 | 64688515 | 20 | 9 | 1.4695 | 0.070852 | LGALSL | 0.070852 | 0.12885 | 0.15277 |
| ENSG00000248713 | 4 | 100557686 | 100575805 | 73 | 17 | 1.4685 | 0.070988 | RP11-766F14.2 | 0.070988 | 0.00075573 | 0.00035314 |
| ENSG00000267426 | 17 | 73894726 | 73926210 | 67 | 13 | 1.4672 | 0.07116 | RP11-552F3.12 | 0.07116 | 0.11958 | 0.16917 |
| ENSG00000009724 | 1 | 11086580 | 11107290 | 78 | 11 | 1.4669 | 0.071204 | MASP2 | 0.071204 | 0.092133 | 0.16998 |
| ENSG00000164347 | 5 | 74017029 | 74063196 | 95 | 9 | 1.4668 | 0.071222 | GFM2 | 0.071222 | 0.060303 | 0.043005 |
| ENSG00000168924 | 4 | 1813206 | 1857974 | 49 | 17 | 1.4654 | 0.071409 | LETM1 | 0.071409 | 0.072959 | 0.020634 |
| ENSG00000166557 | 15 | 79603404 | 79704334 | 458 | 27 | 1.4649 | 0.071471 | TMED3 | 0.071471 | 0.072532 | 0.065875 |
| ENSG00000188916 | 10 | 128933694 | 128994422 | 162 | 19 | 1.464 | 0.071592 | FAM196A | 0.071592 | 0.07389 | 0.11544 |
| ENSG00000198780 | 5 | 74073399 | 74162776 | 189 | 12 | 1.464 | 0.071594 | FAM169A | 0.071594 | 0.07097 | 0.068299 |
| ENSG00000169554 | 2 | 145141648 | 145282147 | 238 | 46 | 1.4629 | 0.071752 | ZEB2 | 0.071752 | 0.10153 | 0.18075 |
| ENSG00000100697 | 14 | 95552565 | 95624347 | 125 | 21 | 1.4619 | 0.071883 | DICER1 | 0.071883 | 0.068473 | 0.10915 |
| ENSG00000215472 | 18 | 47008051 | 47017956 | 39 | 9 | 1.4618 | 0.071901 | RPL17-C18orf32 | 0.071901 | 0.02784 | 0.026372 |
| ENSG00000114125 | 3 | 141457046 | 141466402 | 26 | 3 | 1.4617 | 0.071916 | RNF7 | 0.071916 | 0.045796 | 0.01723 |
| ENSG00000120948 | 1 | 11072414 | 11085796 | 35 | 4 | 1.4615 | 0.07194 | TARDBP | 0.07194 | 0.098933 | 0.1405 |
| ENSG00000056972 | 6 | 111877657 | 111927481 | 166 | 22 | 1.4612 | 0.071985 | TRAF3IP2 | 0.071985 | 0.08162 | 0.04676 |
| ENSG00000170178 | 2 | 176964458 | 176966408 | 4 | 2 | 1.4607 | 0.072052 | HOXD12 | 0.072052 | 0.15017 | 0.33089 |
| ENSG00000250571 | 8 | 144349603 | 144359101 | 37 | 5 | 1.4604 | 0.072086 | GLI4 | 0.072086 | 0.10621 | 0.076863 |
| ENSG00000075043 | 20 | 62037542 | 62103993 | 318 | 30 | 1.459 | 0.072289 | KCNQ2 | 0.072289 | 0.034885 | 0.042411 |
| ENSG00000139517 | 13 | 28120050 | 28194541 | 230 | 26 | 1.458 | 0.072421 | LNX2 | 0.072421 | 0.078645 | 0.14314 |
| ENSG00000164604 | 7 | 112718386 | 112727833 | 27 | 11 | 1.4579 | 0.072438 | GPR85 | 0.072438 | 0.0060885 | 0.0058936 |
| ENSG00000197601 | 11 | 13690217 | 13753893 | 134 | 22 | 1.4575 | 0.072483 | FAR1 | 0.072483 | 0.069481 | 0.16155 |
| ENSG00000172349 | 15 | 81451916 | 81605104 | 532 | 35 | 1.4574 | 0.072504 | IL16 | 0.072504 | 0.076109 | 0.07031 |
| ENSG00000184925 | 9 | 139844003 | 139849949 | 32 | 8 | 1.4574 | 0.072507 | LCN12 | 0.072507 | 0.17453 | 0.051326 |
| ENSG00000161277 | 19 | 36525887 | 36545664 | 57 | 12 | 1.4573 | 0.072515 | THAP8 | 0.072515 | 0.077202 | 0.069099 |
| ENSG00000101003 | 20 | 25388363 | 25433264 | 141 | 12 | 1.4571 | 0.072548 | GINS1 | 0.072548 | 0.07682 | 0.076155 |
| ENSG00000127666 | 19 | 4815944 | 4831716 | 74 | 15 | 1.4558 | 0.072728 | TICAM1 | 0.072728 | 0.16778 | 0.12917 |
| ENSG00000250254 | 4 | 37962056 | 37962746 | 3 | 2 | 1.4556 | 0.072757 | PTTG2 | 0.072757 | 0.50245 | 0.54296 |
| ENSG00000188186 | 7 | 99746530 | 99753567 | 9 | 3 | 1.4554 | 0.07278 | LAMTOR4 | 0.07278 | 0.065464 | 0.017416 |
| ENSG00000189051 | 17 | 8294022 | 8301144 | 24 | 7 | 1.4553 | 0.072788 | RNF222 | 0.072788 | 0.23711 | 0.22899 |
| ENSG00000172774 | 11 | 57982217 | 57983195 | 9 | 2 | 1.455 | 0.072837 | OR1S1 | 0.072837 | 0.060305 | 0.14265 |
| ENSG00000163682 | 4 | 39455744 | 39460568 | 25 | 7 | 1.4546 | 0.072888 | RPL9 | 0.072888 | 0.045416 | 0.17974 |
| ENSG00000129472 | 14 | 21927179 | 21945132 | 63 | 8 | 1.4542 | 0.072943 | RAB2B | 0.072943 | 0.064543 | 0.029238 |
| ENSG00000088926 | 4 | 187187099 | 187210835 | 76 | 17 | 1.4537 | 0.07301 | F11 | 0.07301 | 0.096226 | 0.19326 |
| ENSG00000134452 | 10 | 5931535 | 5979556 | 169 | 20 | 1.4537 | 0.073012 | FBXO18 | 0.073012 | 0.11944 | 0.13432 |
| ENSG00000223443 | 8 | 11994677 | 11996586 | 2 | 1 | 1.4522 | 0.073221 | USP17L2 | 0.073221 | 0.037449 | 0.040823 |
| ENSG00000115266 | 19 | 1446300 | 1473243 | 85 | 18 | 1.4515 | 0.073315 | APC2 | 0.073315 | 0.098766 | 0.25003 |
| ENSG00000184459 | 22 | 32809834 | 32860471 | 253 | 44 | 1.4512 | 0.073357 | BPIFC | 0.073357 | 0.15765 | 0.36193 |
| ENSG00000123836 | 1 | 207222801 | 207254369 | 76 | 15 | 1.451 | 0.073394 | PFKFB2 | 0.073394 | 0.10129 | 0.13938 |
| ENSG00000153207 | 1 | 247002400 | 247095280 | 290 | 27 | 1.4499 | 0.073539 | AHCTF1 | 0.073539 | 0.070824 | 0.080357 |
| ENSG00000174942 | 11 | 56184734 | 56185708 | 8 | 3 | 1.4499 | 0.073539 | OR5R1 | 0.073539 | 0.13996 | 0.054983 |
| ENSG00000155980 | 12 | 57943781 | 57980415 | 65 | 16 | 1.4498 | 0.073554 | KIF5A | 0.073554 | 0.050331 | 0.077873 |
| ENSG00000214999 | 17 | 7982800 | 7984883 | 7 | 4 | 1.4491 | 0.073659 | AC129492.6 | 0.073659 | 0.21933 | 0.14195 |
| ENSG00000120727 | 5 | 138677276 | 138705406 | 35 | 8 | 1.449 | 0.073664 | PAIP2 | 0.073664 | 0.092225 | 0.17625 |
| ENSG00000124942 | 11 | 62201016 | 62323707 | 300 | 51 | 1.4483 | 0.073764 | AHNAK | 0.073764 | 0.11369 | 0.1289 |
| ENSG00000125304 | 13 | 100153671 | 100216260 | 166 | 23 | 1.4476 | 0.07387 | TM9SF2 | 0.07387 | 0.071924 | 0.096212 |
| ENSG00000129219 | 17 | 4710391 | 4726729 | 61 | 15 | 1.4472 | 0.07392 | PLD2 | 0.07392 | 0.10516 | 0.16429 |
| ENSG00000158815 | 8 | 21899909 | 21906320 | 16 | 8 | 1.4468 | 0.073982 | FGF17 | 0.073982 | 0.022424 | 0.02008 |
| ENSG00000071564 | 19 | 1609291 | 1652604 | 120 | 27 | 1.4461 | 0.074074 | TCF3 | 0.074074 | 0.15024 | 0.23118 |
| ENSG00000132749 | 11 | 68474908 | 68519032 | 99 | 10 | 1.446 | 0.074095 | MTL5 | 0.074095 | 0.081388 | 0.081557 |
| ENSG00000138166 | 10 | 112257596 | 112271302 | 27 | 5 | 1.4456 | 0.074148 | DUSP5 | 0.074148 | 0.043354 | 0.0059811 |
| ENSG00000197110 | 19 | 39734246 | 39735646 | 8 | 2 | 1.4454 | 0.074172 | IFNL3 | 0.074172 | 0.19375 | 0.22359 |
| ENSG00000131737 | 17 | 39533902 | 39538655 | 11 | 4 | 1.4452 | 0.074195 | KRT34 | 0.074195 | 0.14635 | 0.20191 |
| ENSG00000124900 | 11 | 55650773 | 55659286 | 25 | 8 | 1.4452 | 0.074207 | TRIM51 | 0.074207 | 0.03162 | 0.042586 |
| ENSG00000140470 | 15 | 100511794 | 100882210 | 2093 | 141 | 1.4442 | 0.074342 | ADAMTS17 | 0.074342 | 0.087698 | 0.16411 |
| ENSG00000181698 | 11 | 56043029 | 56044146 | 3 | 2 | 1.4439 | 0.074385 | OR5T1 | 0.074385 | 0.024927 | 0.025822 |
| ENSG00000136710 | 2 | 131095814 | 131099922 | 4 | 2 | 1.4438 | 0.0744 | CCDC115 | 0.0744 | 0.017139 | 0.0078387 |
| ENSG00000126432 | 11 | 64085560 | 64089283 | 5 | 3 | 1.4436 | 0.074424 | PRDX5 | 0.074424 | 0.34743 | 0.17266 |
| ENSG00000136250 | 7 | 36552456 | 36764154 | 806 | 68 | 1.4434 | 0.074448 | AOAH | 0.074448 | 0.092722 | 0.15672 |
| ENSG00000132275 | 11 | 6616305 | 6624850 | 22 | 10 | 1.4432 | 0.074477 | RRP8 | 0.074477 | 0.0657 | 0.082408 |
| ENSG00000169359 | 3 | 155538813 | 155572218 | 108 | 14 | 1.4416 | 0.074711 | SLC33A1 | 0.074711 | 0.09963 | 0.18876 |
| ENSG00000152689 | 2 | 33661391 | 33789817 | 501 | 73 | 1.4414 | 0.07473 | RASGRP3 | 0.07473 | 0.13218 | 0.20855 |
| ENSG00000138650 | 4 | 134070470 | 134129356 | 115 | 22 | 1.44 | 0.074939 | PCDH10 | 0.074939 | 0.011757 | 0.0083647 |
| ENSG00000197273 | 1 | 42628362 | 42630389 | 4 | 2 | 1.4399 | 0.074953 | GUCA2A | 0.074953 | 0.010458 | 0.039366 |
| ENSG00000089041 | 12 | 121570622 | 121623876 | 308 | 28 | 1.4387 | 0.075122 | P2RX7 | 0.075122 | 0.061447 | 0.085498 |
| ENSG00000138614 | 15 | 65871091 | 65903627 | 80 | 10 | 1.4387 | 0.075124 | VWA9 | 0.075124 | 0.066432 | 0.059962 |
| ENSG00000112584 | 6 | 170599791 | 170716153 | 351 | 21 | 1.4386 | 0.075139 | FAM120B | 0.075139 | 0.11295 | 0.1446 |
| ENSG00000140105 | 14 | 100800125 | 100843142 | 156 | 9 | 1.4383 | 0.075174 | WARS | 0.075174 | 0.062888 | 0.042967 |
| ENSG00000072401 | 10 | 60094735 | 60130513 | 73 | 15 | 1.438 | 0.075215 | UBE2D1 | 0.075215 | 0.043316 | 0.035089 |

| | | | | | | | | | | | |
|---|---|---|---|---|---|---|---|---|---|---|---|
| ENSG00000125741 | 19 | 46030685 | 46105470 | 265 | 33 | 1.4378 | 0.075239 | OPA3 | 0.075239 | 0.076472 | 0.063556 |
| ENSG00000108518 | 17 | 4848947 | 4852356 | 13 | 6 | 1.4377 | 0.075256 | PFN1 | 0.075256 | 0.27501 | 0.19559 |
| ENSG00000050438 | 12 | 51785101 | 51902980 | 313 | 22 | 1.4375 | 0.075283 | SLC4A8 | 0.075283 | 0.073138 | 0.07247 |
| ENSG00000134461 | 10 | 5903580 | 5931869 | 111 | 20 | 1.4372 | 0.075328 | ANKRD16 | 0.075328 | 0.03621 | 0.039638 |
| ENSG00000133961 | 14 | 73741815 | 73930348 | 527 | 26 | 1.4371 | 0.075349 | NUMB | 0.075349 | 0.085615 | 0.099248 |
| ENSG00000127526 | 19 | 16660642 | 16683193 | 57 | 15 | 1.4364 | 0.075451 | SLC35E1 | 0.075451 | 0.094424 | 0.059188 |
| ENSG00000141543 | 17 | 78109013 | 78120982 | 57 | 14 | 1.4359 | 0.075514 | EIF4A3 | 0.075514 | 0.10216 | 0.076327 |
| ENSG00000066654 | 16 | 20744986 | 20753406 | 27 | 6 | 1.4359 | 0.075516 | THUMPD1 | 0.075516 | 0.12722 | 0.15204 |
| ENSG00000118004 | 2 | 3642426 | 3692048 | 241 | 24 | 1.4358 | 0.075526 | COLEC11 | 0.075526 | 0.054083 | 0.022059 |
| ENSG00000148344 | 9 | 132500610 | 132515326 | 75 | 7 | 1.4358 | 0.075531 | PTGES | 0.075531 | 0.026787 | 0.028196 |
| ENSG00000264668 | 8 | 144329280 | 144358573 | 101 | 9 | 1.4356 | 0.075551 | ZFP41 | 0.075551 | 0.075832 | 0.073558 |
| ENSG00000171425 | 19 | 56146825 | 56156988 | 36 | 10 | 1.4348 | 0.075679 | ZNF581 | 0.075679 | 0.099925 | 0.20618 |
| ENSG00000095015 | 5 | 56111401 | 56191979 | 206 | 23 | 1.4347 | 0.075689 | MAP3K1 | 0.075689 | 0.071697 | 0.064713 |
| ENSG00000132623 | 20 | 9966774 | 10037410 | 315 | 30 | 1.4347 | 0.07569 | ANKEF1 | 0.07569 | 0.083724 | 0.015035 |
| ENSG00000184828 | 18 | 45553044 | 45937123 | 1622 | 166 | 1.4345 | 0.07572 | ZBTB7C | 0.07572 | 0.075007 | 0.065811 |
| ENSG00000140459 | 15 | 74630100 | 74660081 | 61 | 13 | 1.4343 | 0.075746 | CYP11A1 | 0.075746 | 0.072379 | 0.070605 |
| ENSG00000105695 | 19 | 35783028 | 35804707 | 61 | 15 | 1.4339 | 0.075801 | MAG | 0.075801 | 0.22888 | 0.27303 |
| ENSG00000164615 | 5 | 134074191 | 134087847 | 13 | 5 | 1.4339 | 0.075806 | CAMLG | 0.075806 | 0.35194 | 0.38023 |
| ENSG00000182870 | 12 | 132680924 | 132905935 | 658 | 69 | 1.4332 | 0.075893 | GALNT9 | 0.075893 | 0.069387 | 0.083391 |
| ENSG00000149262 | 11 | 77589766 | 77705724 | 285 | 13 | 1.4332 | 0.075905 | INTS4 | 0.075905 | 0.061248 | 0.06047 |
| ENSG00000172538 | 10 | 50339199 | 50342053 | 16 | 4 | 1.4331 | 0.075918 | FAM170B | 0.075918 | 0.085349 | 0.13878 |
| ENSG00000164556 | 7 | 38724946 | 38726637 | 6 | 3 | 1.4327 | 0.075965 | FAM183B | 0.075965 | 0.24575 | 0.23582 |
| ENSG00000158296 | 20 | 45186463 | 45304714 | 361 | 62 | 1.4327 | 0.075965 | SLC13A3 | 0.075965 | 0.039192 | 0.04581 |
| ENSG00000127922 | 7 | 96110938 | 96339203 | 421 | 40 | 1.4319 | 0.07608 | SHFM1 | 0.07608 | 0.08607 | 0.083104 |
| ENSG00000184261 | 2 | 47743720 | 47798078 | 148 | 38 | 1.4316 | 0.076132 | KCNK12 | 0.076132 | 0.01943 | 0.070545 |
| ENSG00000268870 | 19 | 12502763 | 12551896 | 354 | 9 | 1.4315 | 0.076147 | CTD-3105H18.16 | 0.076147 | 0.11416 | 0.13129 |
| ENSG00000255470 | 11 | 18161733 | 18211042 | 285 | 9 | 1.4315 | 0.076148 | RP11-113D6.6 | 0.076148 | 0.068516 | 0.093384 |
| ENSG00000071051 | 2 | 106361354 | 106510730 | 522 | 29 | 1.4305 | 0.07628 | NCK2 | 0.07628 | 0.070444 | 0.093184 |
| ENSG00000124155 | 20 | 44044717 | 44054884 | 20 | 7 | 1.4302 | 0.076336 | PIGT | 0.076336 | 0.06606 | 0.070512 |
| ENSG00000125651 | 19 | 6379580 | 6393992 | 72 | 10 | 1.43 | 0.076352 | GTF2F1 | 0.076352 | 0.082135 | 0.20163 |
| ENSG00000125848 | 20 | 14303634 | 14318262 | 36 | 12 | 1.43 | 0.076364 | FLRT3 | 0.076364 | 0.13134 | 0.1719 |
| ENSG00000122585 | 7 | 24323782 | 24331484 | 40 | 10 | 1.4299 | 0.076375 | NPY | 0.076375 | 0.096661 | 0.101 |
| ENSG00000145220 | 4 | 4269428 | 4291896 | 125 | 15 | 1.4299 | 0.076379 | LYAR | 0.076379 | 0.22868 | 0.38072 |
| ENSG00000269693 | 19 | 12636534 | 12691789 | 123 | 22 | 1.4295 | 0.076436 | CTD-2192J16.20 | 0.076436 | 0.046344 | 0.10112 |
| ENSG00000140575 | 15 | 90931450 | 91045475 | 398 | 18 | 1.4294 | 0.076445 | IQGAP1 | 0.076445 | 0.080932 | 0.087289 |
| ENSG00000163219 | 2 | 68906733 | 69053965 | 645 | 49 | 1.4293 | 0.076464 | ARHGAP25 | 0.076464 | 0.10417 | 0.058602 |
| ENSG00000198113 | 9 | 140172201 | 140177093 | 10 | 6 | 1.4291 | 0.076481 | TOR4A | 0.076481 | 0.17445 | 0.37908 |
| ENSG00000160949 | 8 | 145654165 | 145669827 | 54 | 3 | 1.429 | 0.076509 | TONSL | 0.076509 | 0.07021 | 0.077822 |
| ENSG00000167178 | 15 | 74392652 | 74430881 | 35 | 16 | 1.4285 | 0.076578 | ISLR2 | 0.076578 | 0.052196 | 0.053271 |
| ENSG00000163357 | 1 | 155006300 | 155023406 | 29 | 7 | 1.4282 | 0.07662 | DCST1 | 0.07662 | 0.10195 | 0.062713 |
| ENSG00000156564 | 6 | 40359325 | 40555204 | 824 | 53 | 1.4275 | 0.076713 | LRFN2 | 0.076713 | 0.082557 | 0.083873 |
| ENSG00000166979 | 21 | 33784314 | 33887707 | 302 | 22 | 1.4274 | 0.076733 | EVA1C | 0.076733 | 0.075759 | 0.087432 |
| ENSG00000125991 | 20 | 34129770 | 34145405 | 38 | 7 | 1.4272 | 0.076763 | ERGIC3 | 0.076763 | 0.05755 | 0.058795 |
| ENSG00000239605 | 2 | 47272677 | 47382517 | 578 | 54 | 1.427 | 0.076791 | C2orf61 | 0.076791 | 0.037978 | 0.038655 |
| ENSG00000133454 | 22 | 26138111 | 26427007 | 1199 | 126 | 1.4267 | 0.076829 | MYO18B | 0.076829 | 0.10797 | 0.090227 |
| ENSG00000091409 | 2 | 173292082 | 173371181 | 359 | 29 | 1.4266 | 0.076854 | ITGA6 | 0.076854 | 0.050214 | 0.060047 |
| ENSG00000138347 | 10 | 69865912 | 69971774 | 522 | 37 | 1.4262 | 0.07691 | MYPN | 0.07691 | 0.099868 | 0.10436 |
| ENSG00000197147 | 1 | 89990395 | 90063423 | 149 | 19 | 1.4256 | 0.076998 | LRRC8B | 0.076998 | 0.061027 | 0.078118 |
| ENSG00000078902 | 11 | 1295601 | 1330884 | 134 | 27 | 1.4254 | 0.077021 | TOLLIP | 0.077021 | 0.061598 | 0.085411 |
| ENSG00000213390 | 10 | 98978996 | 99052413 | 276 | 17 | 1.4251 | 0.077059 | ARHGAP19 | 0.077059 | 0.054167 | 0.038248 |
| ENSG00000175877 | 7 | 73275489 | 73280223 | 14 | 4 | 1.4249 | 0.077095 | WBSCR28 | 0.077095 | 0.032883 | 0.023733 |
| ENSG00000167550 | 12 | 49458468 | 49463808 | 9 | 3 | 1.4244 | 0.077163 | RHEBL1 | 0.077163 | 0.3933 | 0.31239 |
| ENSG00000172238 | 4 | 94750042 | 94751221 | 1 | 1 | 1.424 | 0.07723 | ATOH1 | 0.07723 | 0.050954 | 0.034444 |
| ENSG00000104967 | 19 | 46436992 | 46476804 | 89 | 19 | 1.4239 | 0.077231 | NOVA2 | 0.077231 | 0.12953 | 0.41138 |
| ENSG00000185482 | 12 | 57637236 | 57644976 | 11 | 5 | 1.4235 | 0.077297 | STAC3 | 0.077297 | 0.12505 | 0.20507 |
| ENSG00000087111 | 17 | 26880401 | 26898890 | 20 | 8 | 1.4235 | 0.077302 | PIGS | 0.077302 | 0.069082 | 0.20186 |
| ENSG00000157873 | 1 | 2487078 | 2496821 | 24 | 5 | 1.4219 | 0.077532 | TNFRSF14 | 0.077532 | 0.066044 | 0.039949 |
| ENSG00000154277 | 4 | 41258430 | 41270472 | 49 | 9 | 1.4218 | 0.077539 | UCHL1 | 0.077539 | 0.1148 | 0.18206 |
| ENSG00000124749 | 6 | 55921388 | 56258892 | 1589 | 55 | 1.4209 | 0.077677 | COL21A1 | 0.077677 | 0.077911 | 0.032546 |
| ENSG00000186564 | 1 | 47901689 | 47906363 | 10 | 3 | 1.4209 | 0.077679 | FOXD2 | 0.077679 | 0.12686 | 0.12036 |
| ENSG00000121570 | 3 | 109044988 | 109056419 | 45 | 7 | 1.4207 | 0.077706 | DPPA4 | 0.077706 | 0.12472 | 0.14657 |
| ENSG00000162438 | 1 | 15764935 | 15775737 | 30 | 9 | 1.4199 | 0.077824 | CTRC | 0.077824 | 0.047527 | 0.097849 |
| ENSG00000198146 | 15 | 35270542 | 35280488 | 22 | 5 | 1.4192 | 0.077922 | ZNF770 | 0.077922 | 0.20697 | 0.21785 |
| ENSG00000116514 | 1 | 33402046 | 33430286 | 72 | 9 | 1.418 | 0.078093 | RNF19B | 0.078093 | 0.053539 | 0.070865 |
| ENSG00000170608 | 19 | 46367247 | 46377055 | 28 | 9 | 1.418 | 0.078101 | FOXA3 | 0.078101 | 0.042533 | 0.012188 |
| ENSG00000123612 | 2 | 158383279 | 158485517 | 168 | 28 | 1.4177 | 0.078143 | ACVR1C | 0.078143 | 0.047454 | 0.038672 |
| ENSG00000047932 | 6 | 117639374 | 117923691 | 1086 | 49 | 1.4177 | 0.078144 | GOPC | 0.078144 | 0.08992 | 0.091431 |
| ENSG00000178896 | 8 | 145133529 | 145135550 | 4 | 2 | 1.4173 | 0.078203 | EXOSC4 | 0.078203 | 0.095347 | 0.11166 |
| ENSG00000177042 | 11 | 695428 | 705028 | 62 | 17 | 1.4171 | 0.078227 | TMEM80 | 0.078227 | 0.029775 | 0.038694 |
| ENSG00000179271 | 19 | 13064972 | 13068050 | 4 | 2 | 1.416 | 0.078389 | GADD45GIP1 | 0.078389 | 0.70784 | 0.62007 |
| ENSG00000172819 | 12 | 53604354 | 53626764 | 46 | 11 | 1.4151 | 0.078517 | RARG | 0.078517 | 0.25549 | 0.41513 |
| ENSG00000265096 | 17 | 77015291 | 77023737 | 19 | 8 | 1.4151 | 0.078522 | C1QTNF1-AS1 | 0.078522 | 0.22075 | 0.43866 |
| ENSG00000163515 | 3 | 108462271 | 108476231 | 46 | 9 | 1.4146 | 0.078589 | RETNLB | 0.078589 | 0.093559 | 0.07779 |

| | | | | | | | | | | | |
|---|---|---|---|---|---|---|---|---|---|---|---|
| ENSG00000144381 | 2 | 198351305 | 198381461 | 59 | 10 | 1.4142 | 0.078653 | HSPD1 | 0.078653 | 0.076546 | 0.1223 |
| ENSG00000138378 | 2 | 191894302 | 192016322 | 270 | 28 | 1.4133 | 0.078786 | STAT4 | 0.078786 | 0.10163 | 0.11998 |
| ENSG00000156170 | 8 | 95907995 | 96128683 | 809 | 33 | 1.4132 | 0.078799 | NDUFAF6 | 0.078799 | 0.073138 | 0.069761 |
| ENSG00000196126 | 6 | 32546546 | 32557625 | 375 | 15 | 1.4131 | 0.078817 | HLA-DRB1 | 0.078817 | 0.069701 | 0.098808 |
| ENSG00000179817 | 11 | 18194384 | 18195827 | 8 | 2 | 1.413 | 0.078829 | MRGPRX4 | 0.078829 | 0.0742 | 0.081382 |
| ENSG00000164414 | 6 | 88180341 | 88222054 | 164 | 17 | 1.4123 | 0.078934 | SLC35A1 | 0.078934 | 0.11264 | 0.16348 |
| ENSG00000244752 | 22 | 25615489 | 25627836 | 69 | 18 | 1.4115 | 0.079048 | CRYBB2 | 0.079048 | 0.27203 | 0.72383 |
| ENSG00000115290 | 2 | 165349322 | 165478358 | 250 | 36 | 1.4102 | 0.079233 | GRB14 | 0.079233 | 0.11216 | 0.14788 |
| ENSG00000175054 | 3 | 142168077 | 142297668 | 354 | 16 | 1.4102 | 0.079235 | ATR | 0.079235 | 0.078377 | 0.10475 |
| ENSG00000135452 | 12 | 58131796 | 58143994 | 23 | 14 | 1.4101 | 0.079257 | TSPAN31 | 0.079257 | 0.28762 | 0.43854 |
| ENSG00000006659 | 19 | 40194946 | 40200084 | 13 | 6 | 1.4099 | 0.079287 | LGALS14 | 0.079287 | 0.078386 | 0.02811 |
| ENSG00000203985 | 1 | 54474412 | 54483856 | 27 | 5 | 1.4095 | 0.079338 | LDLRAD1 | 0.079338 | 0.14164 | 0.21371 |
| ENSG00000242107 | 3 | 159733811 | 159749210 | 77 | 11 | 1.4092 | 0.079388 | LINC01100 | 0.079388 | 0.096698 | 0.10919 |
| ENSG00000132773 | 1 | 45805342 | 45809647 | 4 | 2 | 1.4092 | 0.079395 | TOE1 | 0.079395 | 0.21323 | 0.31456 |
| ENSG00000140391 | 15 | 77336359 | 77376326 | 35 | 18 | 1.4091 | 0.0794 | TSPAN3 | 0.0794 | 0.089117 | 0.13378 |
| ENSG00000154438 | 7 | 117003276 | 117068177 | 108 | 16 | 1.4088 | 0.079446 | ASZ1 | 0.079446 | 0.076932 | 0.055335 |
| ENSG00000187189 | 6 | 116571151 | 116575261 | 13 | 4 | 1.4086 | 0.079474 | TSPYL4 | 0.079474 | 0.02743 | 0.020016 |
| ENSG00000188878 | 17 | 73905655 | 73937221 | 64 | 12 | 1.4074 | 0.079658 | FBF1 | 0.079658 | 0.058248 | 0.15946 |
| ENSG00000162618 | 1 | 79355449 | 79472403 | 544 | 25 | 1.4073 | 0.079666 | ELTD1 | 0.079666 | 0.11921 | 0.17318 |
| ENSG00000138639 | 4 | 86396267 | 86923823 | 1890 | 80 | 1.4068 | 0.079736 | ARHGAP24 | 0.079736 | 0.076907 | 0.047133 |
| ENSG00000176473 | 14 | 100842755 | 100996640 | 270 | 30 | 1.4068 | 0.079739 | WDR25 | 0.079739 | 0.066362 | 0.060039 |
| ENSG00000189060 | 22 | 38201114 | 38203442 | 6 | 2 | 1.4061 | 0.079844 | H1F0 | 0.079844 | 0.19064 | 0.26732 |
| ENSG00000123064 | 12 | 113594979 | 113623284 | 61 | 7 | 1.4053 | 0.079965 | DDX54 | 0.079965 | 0.081956 | 0.076559 |
| ENSG00000006377 | 7 | 96634860 | 96640351 | 9 | 3 | 1.4046 | 0.080071 | DLX6 | 0.080071 | 0.037012 | 0.047016 |
| ENSG00000163463 | 1 | 155141884 | 155145951 | 12 | 4 | 1.4041 | 0.080149 | KRTCAP2 | 0.080149 | 0.091546 | 0.073177 |
| ENSG00000166908 | 12 | 57984957 | 57997198 | 22 | 5 | 1.4036 | 0.080214 | PIP4K2C | 0.080214 | 0.14344 | 0.071861 |
| ENSG00000054148 | 9 | 139743176 | 139745488 | 8 | 4 | 1.4036 | 0.080217 | PHPT1 | 0.080217 | 0.099477 | 0.1982 |
| ENSG00000050426 | 12 | 51441745 | 51454207 | 42 | 7 | 1.4029 | 0.080325 | LETMD1 | 0.080325 | 0.092125 | 0.062797 |
| ENSG00000104833 | 19 | 6494330 | 6502859 | 30 | 13 | 1.4029 | 0.080328 | TUBB4A | 0.080328 | 0.17274 | 0.18765 |
| ENSG00000173960 | 2 | 24150155 | 24227779 | 299 | 22 | 1.4027 | 0.080354 | UBXN2A | 0.080354 | 0.086017 | 0.089574 |
| ENSG00000186815 | 12 | 113658855 | 113736390 | 167 | 9 | 1.4024 | 0.080393 | TPCN1 | 0.080393 | 0.077474 | 0.085123 |
| ENSG00000113396 | 5 | 127873706 | 128369335 | 2411 | 47 | 1.4021 | 0.080438 | SLC27A6 | 0.080438 | 0.082513 | 0.10019 |
| ENSG00000121058 | 17 | 55015548 | 55038415 | 77 | 21 | 1.402 | 0.080463 | COIL | 0.080463 | 0.14681 | 0.24333 |
| ENSG00000130303 | 19 | 17513748 | 17516457 | 10 | 4 | 1.4013 | 0.080557 | BST2 | 0.080557 | 0.257 | 0.2321 |
| ENSG00000187658 | 5 | 157098561 | 157107151 | 34 | 6 | 1.4012 | 0.080581 | C5orf52 | 0.080581 | 0.24384 | 0.39553 |
| ENSG00000170967 | 11 | 103907308 | 103909922 | 5 | 2 | 1.4011 | 0.080589 | DDI1 | 0.080589 | 0.12298 | 0.039804 |
| ENSG00000100994 | 20 | 25228705 | 25278650 | 206 | 15 | 1.4011 | 0.080595 | PYGB | 0.080595 | 0.073988 | 0.066614 |
| ENSG00000129484 | 14 | 20811741 | 20826064 | 70 | 10 | 1.4 | 0.080759 | PARP2 | 0.080759 | 0.078539 | 0.062417 |
| ENSG00000134548 | 12 | 21679241 | 21690311 | 35 | 7 | 1.3994 | 0.080842 | C12orf39 | 0.080842 | 0.1436 | 0.19487 |
| ENSG00000159182 | 17 | 46799084 | 46799884 | 3 | 2 | 1.399 | 0.080905 | PRAC1 | 0.080905 | 0.34369 | 0.41898 |
| ENSG00000167515 | 16 | 88922628 | 88929094 | 33 | 6 | 1.3986 | 0.080962 | TRAPPC2L | 0.080962 | 0.18227 | 0.43255 |
| ENSG00000159618 | 16 | 57576333 | 57625593 | 189 | 22 | 1.3981 | 0.081046 | GPR114 | 0.081046 | 0.077493 | 0.080018 |
| ENSG00000163702 | 3 | 9958758 | 9975314 | 55 | 19 | 1.398 | 0.081063 | IL17RC | 0.081063 | 0.10682 | 0.11853 |
| ENSG00000130770 | 1 | 28562620 | 28573417 | 46 | 4 | 1.3975 | 0.081129 | ATPIF1 | 0.081129 | 0.10598 | 0.14681 |
| ENSG00000152583 | 4 | 88394487 | 88452213 | 231 | 18 | 1.3975 | 0.081138 | SPARCL1 | 0.081138 | 0.14382 | 0.14733 |
| ENSG00000204590 | 6 | 30509154 | 30524951 | 29 | 9 | 1.3971 | 0.081188 | GNL1 | 0.081188 | 0.074231 | 0.077722 |
| ENSG00000109685 | 4 | 1873151 | 1983934 | 98 | 40 | 1.396 | 0.081359 | WHSC1 | 0.081359 | 0.14916 | 0.35085 |
| ENSG00000166902 | 11 | 59573608 | 59578345 | 10 | 5 | 1.3959 | 0.081377 | MRPL16 | 0.081377 | 0.096808 | 0.19254 |
| ENSG00000186792 | 3 | 50330262 | 50336899 | 6 | 4 | 1.3958 | 0.08139 | HYAL3 | 0.08139 | 0.27406 | 0.24208 |
| ENSG00000138399 | 2 | 170386259 | 170430385 | 161 | 26 | 1.3958 | 0.081393 | FASTKD1 | 0.081393 | 0.036634 | 0.0080615 |
| ENSG00000143614 | 1 | 153777201 | 153895451 | 243 | 22 | 1.3958 | 0.081394 | GATAD2B | 0.081394 | 0.087394 | 0.094622 |
| ENSG00000180855 | 19 | 12540521 | 12551926 | 130 | 8 | 1.3948 | 0.081535 | ZNF443 | 0.081535 | 0.1481 | 0.12152 |
| ENSG00000074603 | 15 | 65734801 | 65810042 | 222 | 19 | 1.3942 | 0.081622 | DPP8 | 0.081622 | 0.096332 | 0.099539 |
| ENSG00000163104 | 4 | 95128762 | 95212443 | 223 | 14 | 1.3941 | 0.081638 | SMARCAD1 | 0.081638 | 0.081975 | 0.11002 |
| ENSG00000182851 | 8 | 144295068 | 144299044 | 28 | 7 | 1.3915 | 0.082044 | GPIHBP1 | 0.082044 | 0.047937 | 0.043222 |
| ENSG00000107779 | 10 | 88516407 | 88692595 | 539 | 27 | 1.3912 | 0.082082 | BMPR1A | 0.082082 | 0.072815 | 0.053889 |
| ENSG00000022976 | 14 | 102783714 | 102809044 | 58 | 18 | 1.391 | 0.082113 | ZNF839 | 0.082113 | 0.033754 | 0.064913 |
| ENSG00000163673 | 3 | 36753913 | 36781352 | 62 | 16 | 1.3898 | 0.082299 | DCLK3 | 0.082299 | 0.2405 | 0.059752 |
| ENSG00000103653 | 15 | 75074398 | 75095539 | 39 | 8 | 1.3894 | 0.082354 | CSK | 0.082354 | 0.066532 | 0.079368 |
| ENSG00000265681 | 18 | 47014851 | 47018906 | 15 | 4 | 1.3882 | 0.082536 | RPL17 | 0.082536 | 0.045456 | 0.026767 |
| ENSG00000166363 | 11 | 6866883 | 6867936 | 2 | 1 | 1.3874 | 0.08266 | OR10A5 | 0.08266 | 0.039659 | 0.080338 |
| ENSG00000268781 | 4 | 3511523 | 3511678 | 1 | 1 | 1.3865 | 0.0828 | AL590235.1 | 0.0828 | 0.74905 | 0.7478 |
| ENSG00000136750 | 10 | 26505236 | 26593487 | 263 | 13 | 1.3864 | 0.082819 | GAD2 | 0.082819 | 0.10455 | 0.10912 |
| ENSG00000269526 | 19 | 53517174 | 53519375 | 3 | 2 | 1.3851 | 0.083008 | ERVV-1 | 0.083008 | 0.19081 | 0.18777 |
| ENSG00000198246 | 10 | 73079015 | 73123142 | 166 | 25 | 1.3832 | 0.083294 | SLC29A3 | 0.083294 | 0.099668 | 0.19064 |
| ENSG00000125089 | 4 | 8183799 | 8243530 | 221 | 42 | 1.3829 | 0.083342 | SH3TC1 | 0.083342 | 0.078547 | 0.059831 |
| ENSG00000206260 | 3 | 138722804 | 138725110 | 4 | 2 | 1.3827 | 0.083374 | PRR23A | 0.083374 | 0.071758 | 0.051079 |
| ENSG00000173404 | 20 | 20348765 | 20351590 | 1 | 1 | 1.3826 | 0.08339 | INSM1 | 0.08339 | 0.27204 | 0.36832 |
| ENSG00000126749 | 12 | 7072408 | 7105520 | 61 | 11 | 1.3813 | 0.083589 | EMG1 | 0.083589 | 0.077734 | 0.047714 |
| ENSG00000204296 | 6 | 32256303 | 32339684 | 901 | 25 | 1.3812 | 0.083602 | C6orf10 | 0.083602 | 0.048799 | 0.041758 |
| ENSG00000168734 | 20 | 43160426 | 43252888 | 202 | 23 | 1.3809 | 0.083651 | PKIG | 0.083651 | 0.078439 | 0.080606 |
| ENSG00000038532 | 16 | 11038345 | 11276046 | 803 | 28 | 1.3808 | 0.083667 | CLEC16A | 0.083667 | 0.078964 | 0.071129 |
| ENSG00000197256 | 19 | 11274943 | 11308467 | 138 | 14 | 1.3807 | 0.08369 | KANK2 | 0.08369 | 0.081123 | 0.15043 |

| | | | | | | | | | | | |
|---|---|---|---|---|---|---|---|---|---|---|---|
| ENSG00000163617 | 3 | 113682984 | 113775460 | 156 | 24 | 1.3802 | 0.083767 | KIAA1407 | 0.083767 | 0.097385 | 0.12397 |
| ENSG00000088876 | 20 | 2462463 | 2505348 | 138 | 25 | 1.3797 | 0.083838 | ZNF343 | 0.083838 | 0.092467 | 0.079758 |
| ENSG00000135097 | 12 | 120779133 | 120806983 | 73 | 11 | 1.3789 | 0.083969 | MSI1 | 0.083969 | 0.095995 | 0.097467 |
| ENSG00000187583 | 1 | 901877 | 911245 | 44 | 17 | 1.3782 | 0.08407 | PLEKHN1 | 0.08407 | 0.27243 | 0.44092 |
| ENSG00000157653 | 9 | 116172334 | 116191964 | 62 | 7 | 1.3779 | 0.084113 | C9orf43 | 0.084113 | 0.095847 | 0.096027 |
| ENSG00000187612 | 11 | 55681126 | 55682058 | 9 | 3 | 1.3779 | 0.084119 | OR5W2 | 0.084119 | 0.03869 | 0.034054 |
| ENSG00000115317 | 2 | 74756504 | 74760472 | 3 | 2 | 1.3773 | 0.084206 | HTRA2 | 0.084206 | 0.61868 | 0.26587 |
| ENSG00000168124 | 16 | 3254247 | 3255188 | 2 | 1 | 1.3767 | 0.084298 | OR1F1 | 0.084298 | 0.222 | 0.4309 |
| ENSG00000232941 | 8 | 52730143 | 52730969 | 1 | 1 | 1.3766 | 0.08432 | AC090186.1 | 0.08432 | 0.15827 | 0.13764 |
| ENSG00000169330 | 15 | 79724858 | 79764632 | 172 | 25 | 1.3763 | 0.084368 | KIAA1024 | 0.084368 | 0.12286 | 0.10944 |
| ENSG00000026036 | 20 | 62290653 | 62330037 | 192 | 21 | 1.375 | 0.084573 | RTEL1-TNFRSF6B | 0.084573 | 0.051951 | 0.055553 |
| ENSG00000134717 | 1 | 52521797 | 52556388 | 67 | 13 | 1.3749 | 0.084587 | BTF3L4 | 0.084587 | 0.098167 | 0.09048 |
| ENSG00000127528 | 19 | 16435628 | 16438685 | 8 | 4 | 1.3746 | 0.084631 | KLF2 | 0.084631 | 0.35714 | 0.4828 |
| ENSG00000162062 | 16 | 2510081 | 2514964 | 5 | 3 | 1.3743 | 0.084681 | C16orf59 | 0.084681 | 0.00069212 | 0.0021461 |
| ENSG00000035115 | 2 | 217730 | 266398 | 113 | 11 | 1.3739 | 0.084734 | SH3YL1 | 0.084734 | 0.10142 | 0.087881 |
| ENSG00000166188 | 16 | 58028572 | 58034357 | 10 | 3 | 1.3736 | 0.084777 | ZNF319 | 0.084777 | 0.091051 | 0.093126 |
| ENSG00000127870 | 13 | 26706253 | 26796791 | 422 | 43 | 1.3735 | 0.084791 | RNF6 | 0.084791 | 0.082432 | 0.13485 |
| ENSG00000115421 | 2 | 60983365 | 61029220 | 131 | 18 | 1.3728 | 0.08491 | PAPOLG | 0.08491 | 0.096399 | 0.09321 |
| ENSG00000143727 | 2 | 264140 | 278283 | 38 | 7 | 1.3724 | 0.084971 | ACP1 | 0.084971 | 0.049137 | 0.05167 |
| ENSG00000181927 | 11 | 55405834 | 55406772 | 2 | 1 | 1.3722 | 0.084997 | OR4P4 | 0.084997 | 0.17464 | 0.11737 |
| ENSG00000179632 | 8 | 145159402 | 145162514 | 4 | 1 | 1.372 | 0.085038 | MAF1 | 0.085038 | 0.20205 | 0.17727 |
| ENSG00000042445 | 2 | 85569211 | 85581743 | 42 | 12 | 1.3717 | 0.085072 | RETSAT | 0.085072 | 0.073024 | 0.12958 |
| ENSG00000204850 | 19 | 46997765 | 46999755 | 3 | 2 | 1.3717 | 0.085072 | AC011484.1 | 0.085072 | 0.58218 | 0.43377 |
| ENSG00000137819 | 15 | 69591286 | 69700119 | 392 | 24 | 1.3716 | 0.085099 | PAQR5 | 0.085099 | 0.080803 | 0.096443 |
| ENSG00000139648 | 12 | 52937693 | 52946931 | 44 | 6 | 1.3712 | 0.085159 | KRT71 | 0.085159 | 0.24103 | 0.60658 |
| ENSG00000214189 | 19 | 12203078 | 12248050 | 95 | 13 | 1.3711 | 0.085174 | ZNF788 | 0.085174 | 0.11113 | 0.16094 |
| ENSG00000188996 | 6 | 655939 | 656963 | 5 | 2 | 1.371 | 0.085191 | HUS1B | 0.085191 | 0.43284 | 0.68353 |
| ENSG00000267303 | 19 | 10426147 | 10431354 | 21 | 7 | 1.3707 | 0.085228 | CTD-2369P2.12 | 0.085228 | 0.10836 | 0.24968 |
| ENSG00000106100 | 7 | 30464143 | 30518400 | 125 | 18 | 1.3707 | 0.08523 | NOD1 | 0.08523 | 0.067706 | 0.046571 |
| ENSG00000105755 | 19 | 44010871 | 44031396 | 77 | 11 | 1.3706 | 0.08525 | ETHE1 | 0.08525 | 0.10155 | 0.19395 |
| ENSG00000113494 | 5 | 35048861 | 35230794 | 652 | 42 | 1.3705 | 0.085265 | PRLR | 0.085265 | 0.073583 | 0.080689 |
| ENSG00000164035 | 4 | 101316498 | 101801283 | 1833 | 55 | 1.3701 | 0.085329 | EMCN | 0.085329 | 0.088155 | 0.079787 |
| ENSG00000170775 | 7 | 124386051 | 124405681 | 52 | 10 | 1.3698 | 0.085374 | GPR37 | 0.085374 | 0.15607 | 0.14247 |
| ENSG00000206052 | 18 | 67068291 | 67516323 | 1851 | 131 | 1.3695 | 0.085425 | DOK6 | 0.085425 | 0.090028 | 0.11996 |
| ENSG00000113734 | 5 | 172571445 | 172591390 | 69 | 13 | 1.3694 | 0.085434 | BNIP1 | 0.085434 | 0.11143 | 0.0988 |
| ENSG00000095110 | 11 | 114392437 | 114430617 | 130 | 8 | 1.3694 | 0.085444 | NXPE1 | 0.085444 | 0.070162 | 0.066425 |
| ENSG00000163060 | 2 | 95537178 | 95542574 | 16 | 5 | 1.3687 | 0.08554 | TEKT4 | 0.08554 | 0.070604 | 0.073236 |
| ENSG00000164182 | 5 | 60240956 | 60448853 | 599 | 21 | 1.3679 | 0.085669 | NDUFAF2 | 0.085669 | 0.087328 | 0.088035 |
| ENSG00000108946 | 17 | 66507921 | 66547460 | 89 | 14 | 1.3672 | 0.085776 | PRKAR1A | 0.085776 | 0.017507 | 0.015017 |
| ENSG00000164941 | 8 | 95825539 | 95893974 | 256 | 17 | 1.367 | 0.085808 | INTS8 | 0.085808 | 0.062797 | 0.042222 |
| ENSG00000102755 | 13 | 28874489 | 29069265 | 537 | 45 | 1.367 | 0.085818 | FLT1 | 0.085818 | 0.073734 | 0.037905 |
| ENSG00000163638 | 3 | 64501333 | 64673676 | 545 | 46 | 1.3669 | 0.085822 | ADAMTS9 | 0.085822 | 0.11387 | 0.15861 |
| ENSG00000011304 | 19 | 797075 | 812327 | 63 | 14 | 1.3669 | 0.085831 | PTBP1 | 0.085831 | 0.10728 | 0.38731 |
| ENSG00000132394 | 3 | 127872297 | 128127485 | 486 | 22 | 1.3668 | 0.085839 | EEFSEC | 0.085839 | 0.051386 | 0.03678 |
| ENSG00000171729 | 1 | 15479028 | 15546976 | 353 | 28 | 1.3664 | 0.085899 | TMEM51 | 0.085899 | 0.13614 | 0.20122 |
| ENSG00000129521 | 14 | 34393437 | 34931980 | 2316 | 199 | 1.3661 | 0.085956 | EGLN3 | 0.085956 | 0.084552 | 0.084141 |
| ENSG00000102996 | 16 | 58059470 | 58080805 | 31 | 7 | 1.3656 | 0.086029 | MMP15 | 0.086029 | 0.065032 | 0.029336 |
| ENSG00000250264 | 6 | 32781544 | 32806599 | 247 | 33 | 1.3652 | 0.086101 | TAP2 | 0.086101 | 0.11743 | 0.038556 |
| ENSG00000203301 | 1 | 2143360 | 2145620 | 7 | 4 | 1.365 | 0.08612 | AL590822.1 | 0.08612 | 0.25619 | 0.39936 |
| ENSG00000170791 | 8 | 57124245 | 57131357 | 8 | 5 | 1.365 | 0.08613 | CHCHD7 | 0.08613 | 0.43948 | 0.65166 |
| ENSG00000162004 | 16 | 772582 | 776954 | 14 | 3 | 1.3645 | 0.08621 | CCDC78 | 0.08621 | 0.2793 | 0.36244 |
| ENSG00000088002 | 19 | 49055332 | 49102682 | 350 | 27 | 1.3641 | 0.086263 | SULT2B1 | 0.086263 | 0.070946 | 0.022879 |
| ENSG00000157211 | 1 | 54598747 | 54619443 | 115 | 22 | 1.364 | 0.086288 | CDCP2 | 0.086288 | 0.11243 | 0.16178 |
| ENSG00000197780 | 1 | 109605108 | 109618624 | 58 | 14 | 1.3629 | 0.086457 | TAF13 | 0.086457 | 0.072928 | 0.065795 |
| ENSG00000148680 | 10 | 92500578 | 92617671 | 365 | 20 | 1.3624 | 0.086533 | HTR7 | 0.086533 | 0.073993 | 0.073987 |
| ENSG00000165953 | 14 | 94953611 | 94984181 | 157 | 30 | 1.3624 | 0.086542 | SERPINA12 | 0.086542 | 0.11072 | 0.16991 |
| ENSG00000114805 | 3 | 155093369 | 155462856 | 1201 | 36 | 1.3619 | 0.086613 | PLCH1 | 0.086613 | 0.070768 | 0.063459 |
| ENSG00000147874 | 9 | 19053141 | 19103117 | 198 | 22 | 1.3615 | 0.086686 | HAUS6 | 0.086686 | 0.094104 | 0.066028 |
| ENSG00000167531 | 12 | 48961467 | 48963849 | 3 | 2 | 1.3613 | 0.086706 | LALBA | 0.086706 | 0.32303 | 0.71033 |
| ENSG00000140505 | 15 | 75041185 | 75048543 | 16 | 6 | 1.3613 | 0.086716 | CYP1A2 | 0.086716 | 0.075467 | 0.072572 |
| ENSG00000182035 | 20 | 37209838 | 37217106 | 15 | 7 | 1.3607 | 0.086802 | ADIG | 0.086802 | 0.25057 | 0.19544 |
| ENSG00000184557 | 17 | 76352864 | 76356158 | 5 | 3 | 1.3602 | 0.086888 | SOCS3 | 0.086888 | 0.18186 | 0.15157 |
| ENSG00000188763 | 7 | 72848109 | 72850450 | 2 | 1 | 1.3599 | 0.08693 | FZD9 | 0.08693 | 0.65013 | 0.53855 |
| ENSG00000131379 | 3 | 14716606 | 14814541 | 287 | 37 | 1.3588 | 0.087109 | C3orf20 | 0.087109 | 0.12346 | 0.13721 |
| ENSG00000141965 | 19 | 4791728 | 4795571 | 5 | 2 | 1.3586 | 0.08714 | FEM1A | 0.08714 | 0.074278 | 0.056113 |
| ENSG00000267880 | 19 | 51377552 | 51378040 | 2 | 1 | 1.3578 | 0.08727 | AC037199.1 | 0.08727 | 0.48743 | 0.78824 |
| ENSG00000146426 | 6 | 155153831 | 155578857 | 1732 | 165 | 1.3576 | 0.0873 | TIAM2 | 0.0873 | 0.098142 | 0.12957 |
| ENSG00000163377 | 3 | 68780917 | 68981761 | 647 | 27 | 1.3573 | 0.087347 | FAM19A4 | 0.087347 | 0.098087 | 0.11182 |
| ENSG00000196150 | 8 | 146076632 | 146127553 | 70 | 10 | 1.356 | 0.087544 | ZNF250 | 0.087544 | 0.11556 | 0.14573 |
| ENSG00000015520 | 7 | 44552134 | 44580914 | 59 | 10 | 1.3558 | 0.087579 | NPC1L1 | 0.087579 | 0.079515 | 0.10548 |
| ENSG00000104825 | 19 | 39390340 | 39399533 | 29 | 7 | 1.3551 | 0.087695 | NFKBIB | 0.087695 | 0.066546 | 0.054304 |
| ENSG00000214215 | 12 | 93096619 | 93102325 | 7 | 5 | 1.355 | 0.087705 | C12orf74 | 0.087705 | 0.14772 | 0.26591 |
| ENSG00000173264 | 11 | 64037534 | 64056972 | 36 | 14 | 1.3549 | 0.087728 | GPR137 | 0.087728 | 0.18173 | 0.28642 |

| | | | | | | | | | | | |
|---|---|---|---|---|---|---|---|---|---|---|---|
| ENSG00000187766 | 21 | 46031996 | 46032871 | 2 | 1 | 1.3541 | 0.087846 | KRTAP10-8 | 0.087846 | 0.13154 | 0.25995 |
| ENSG00000177685 | 11 | 826144 | 831991 | 19 | 7 | 1.3537 | 0.087911 | EFCAB4A | 0.087911 | 0.65627 | 0.72588 |
| ENSG00000224877 | 17 | 79213039 | 79215081 | 8 | 5 | 1.3537 | 0.087917 | C17orf89 | 0.087917 | 0.32386 | 0.51904 |
| ENSG00000137634 | 11 | 114441313 | 114466484 | 57 | 14 | 1.3535 | 0.087951 | NXPE4 | 0.087951 | 0.12516 | 0.16217 |
| ENSG00000185015 | 8 | 86132816 | 86196302 | 103 | 10 | 1.3524 | 0.088118 | CA13 | 0.088118 | 0.08394 | 0.11483 |
| ENSG00000197786 | 11 | 58125597 | 58126542 | 3 | 1 | 1.3523 | 0.088132 | OR5B17 | 0.088132 | 0.11276 | 0.098436 |
| ENSG00000167645 | 19 | 38795753 | 38807913 | 39 | 9 | 1.3521 | 0.088177 | YIF1B | 0.088177 | 0.19549 | 0.33865 |
| ENSG00000136167 | 13 | 46700055 | 46786006 | 290 | 40 | 1.3509 | 0.088365 | LCP1 | 0.088365 | 0.12584 | 0.11534 |
| ENSG00000119203 | 2 | 9563697 | 9613230 | 116 | 10 | 1.3506 | 0.088413 | CPSF3 | 0.088413 | 0.062924 | 0.051417 |
| ENSG00000187554 | 1 | 223282748 | 223316624 | 124 | 13 | 1.3505 | 0.088424 | TLR5 | 0.088424 | 0.097784 | 0.093826 |
| ENSG00000146857 | 7 | 134916731 | 134943244 | 57 | 8 | 1.3504 | 0.088448 | STRA8 | 0.088448 | 0.088648 | 0.11026 |
| ENSG00000176153 | 14 | 65405870 | 65409531 | 12 | 4 | 1.3493 | 0.088623 | GPX2 | 0.088623 | 0.18102 | 0.20915 |
| ENSG00000073067 | 7 | 1022835 | 1029276 | 24 | 9 | 1.3488 | 0.088703 | CYP2W1 | 0.088703 | 0.10016 | 0.13394 |
| ENSG00000204038 | 10 | 82009466 | 82013395 | 38 | 5 | 1.3487 | 0.088717 | AL359195.1 | 0.088717 | 0.12371 | 0.045946 |
| ENSG00000268397 | 5 | 180682720 | 180684578 | 2 | 1 | 1.3465 | 0.089078 | AC008443.1 | 0.089078 | 0.040904 | 0.04367 |
| ENSG00000178342 | 18 | 77623668 | 77660184 | 133 | 19 | 1.346 | 0.089159 | KCNG2 | 0.089159 | 0.083231 | 0.084808 |
| ENSG00000152270 | 11 | 14665269 | 14892350 | 359 | 24 | 1.3458 | 0.089183 | PDE3B | 0.089183 | 0.094987 | 0.091703 |
| ENSG00000138674 | 4 | 83739814 | 83822319 | 246 | 15 | 1.3457 | 0.089198 | SEC31A | 0.089198 | 0.070179 | 0.064263 |
| ENSG00000130701 | 20 | 60985293 | 61002589 | 66 | 15 | 1.3454 | 0.089248 | RBBP8NL | 0.089248 | 0.071282 | 0.051266 |
| ENSG00000213015 | 19 | 56146382 | 56154835 | 32 | 9 | 1.3452 | 0.089279 | ZNF580 | 0.089279 | 0.092326 | 0.18345 |
| ENSG00000137942 | 1 | 93913688 | 94020218 | 147 | 23 | 1.3449 | 0.089331 | FNBP1L | 0.089331 | 0.099296 | 0.14229 |
| ENSG00000163012 | 2 | 187692562 | 187713935 | 114 | 5 | 1.3447 | 0.089353 | ZSWIM2 | 0.089353 | 0.10589 | 0.10467 |
| ENSG00000169598 | 1 | 3773845 | 3801993 | 186 | 25 | 1.3447 | 0.089361 | DFFB | 0.089361 | 0.1878 | 0.15731 |
| ENSG00000078295 | 5 | 7396321 | 7830194 | 1160 | 92 | 1.3442 | 0.089436 | ADCY2 | 0.089436 | 0.10923 | 0.14108 |
| ENSG00000182993 | 12 | 14956506 | 15059520 | 292 | 15 | 1.3436 | 0.089535 | C12orf60 | 0.089535 | 0.097271 | 0.12685 |
| ENSG00000143552 | 1 | 153965161 | 154127592 | 311 | 20 | 1.3432 | 0.08961 | NUP210L | 0.08961 | 0.11251 | 0.12916 |
| ENSG00000061938 | 3 | 195590235 | 195638816 | 176 | 31 | 1.343 | 0.08964 | TNK2 | 0.08964 | 0.12303 | 0.13138 |
| ENSG00000150048 | 12 | 10222153 | 10264226 | 195 | 10 | 1.3428 | 0.089663 | CLEC1A | 0.089663 | 0.13858 | 0.15254 |
| ENSG00000104883 | 19 | 7541761 | 7562335 | 107 | 15 | 1.3425 | 0.08971 | PEX11G | 0.08971 | 0.17393 | 0.21613 |
| ENSG00000143178 | 1 | 168250278 | 168283664 | 100 | 19 | 1.3425 | 0.089716 | TBX19 | 0.089716 | 0.13191 | 0.13548 |
| ENSG00000254550 | 11 | 76813886 | 76814377 | 2 | 1 | 1.3423 | 0.089746 | OMP | 0.089746 | 0.030714 | 0.10587 |
| ENSG00000198765 | 1 | 115397424 | 115537991 | 388 | 24 | 1.3421 | 0.089774 | SYCP1 | 0.089774 | 0.093602 | 0.09336 |
| ENSG00000124490 | 6 | 49660073 | 49681274 | 58 | 13 | 1.3419 | 0.08981 | CRISP2 | 0.08981 | 0.096837 | 0.11537 |
| ENSG00000166801 | 11 | 58910221 | 58922512 | 20 | 5 | 1.3413 | 0.089906 | FAM111A | 0.089906 | 0.055769 | 0.057598 |
| ENSG00000260415 | 12 | 52203789 | 52206636 | 5 | 2 | 1.3413 | 0.089916 | AC068987.1 | 0.089916 | 0.40067 | 0.26646 |
| ENSG00000159267 | 21 | 38123189 | 38362536 | 880 | 44 | 1.3409 | 0.08997 | HLCS | 0.08997 | 0.098185 | 0.10971 |
| ENSG00000171735 | 1 | 6845384 | 7829766 | 3378 | 229 | 1.3403 | 0.090078 | CAMTA1 | 0.090078 | 0.085482 | 0.10337 |
| ENSG00000149016 | 11 | 62342517 | 62359649 | 22 | 7 | 1.3402 | 0.09009 | TUT1 | 0.09009 | 0.1367 | 0.2763 |
| ENSG00000150403 | 13 | 114145310 | 114204542 | 371 | 18 | 1.3399 | 0.09014 | TMCO3 | 0.09014 | 0.10902 | 0.11896 |
| ENSG00000147789 | 8 | 146052849 | 146072894 | 61 | 11 | 1.3398 | 0.090155 | ZNF7 | 0.090155 | 0.073743 | 0.089153 |
| ENSG00000111432 | 12 | 130647004 | 130650285 | 11 | 5 | 1.3393 | 0.090235 | FZD10 | 0.090235 | 0.04092 | 0.17703 |
| ENSG00000148848 | 10 | 127700950 | 128077024 | 1575 | 114 | 1.3393 | 0.09024 | ADAM12 | 0.09024 | 0.098286 | 0.10875 |
| ENSG00000198912 | 1 | 3805689 | 3816857 | 53 | 9 | 1.339 | 0.090286 | C1orf174 | 0.090286 | 0.06783 | 0.02492 |
| ENSG00000124610 | 6 | 26017260 | 26018040 | 4 | 2 | 1.339 | 0.090287 | HIST1H1A | 0.090287 | 0.4739 | 0.34723 |
| ENSG00000135643 | 12 | 70760056 | 70828072 | 277 | 27 | 1.3389 | 0.090307 | KCNMB4 | 0.090307 | 0.082521 | 0.073173 |
| ENSG00000189195 | 1 | 92545862 | 92613393 | 179 | 14 | 1.3387 | 0.090338 | BTBD8 | 0.090338 | 0.087104 | 0.15453 |
| ENSG00000129250 | 17 | 4901243 | 4931696 | 87 | 17 | 1.3386 | 0.090353 | KIF1C | 0.090353 | 0.11171 | 0.17993 |
| ENSG00000197841 | 19 | 35225060 | 35233777 | 18 | 7 | 1.3386 | 0.090357 | ZNF181 | 0.090357 | 0.22577 | 0.21586 |
| ENSG00000167483 | 19 | 17634110 | 17664648 | 98 | 23 | 1.3385 | 0.090361 | FAM129C | 0.090361 | 0.081927 | 0.1009 |
| ENSG00000145506 | 5 | 1008944 | 1039058 | 146 | 22 | 1.3381 | 0.09044 | NKD2 | 0.09044 | 0.069161 | 0.08355 |
| ENSG00000115919 | 2 | 143635067 | 143799890 | 507 | 31 | 1.3374 | 0.090545 | KYNU | 0.090545 | 0.077676 | 0.053073 |
| ENSG00000005339 | 16 | 3775055 | 3930727 | 184 | 45 | 1.3372 | 0.090577 | CREBBP | 0.090577 | 0.12127 | 0.18242 |
| ENSG00000110042 | 11 | 58938903 | 58976060 | 71 | 12 | 1.3371 | 0.090603 | DTX4 | 0.090603 | 0.019531 | 0.0096325 |
| ENSG00000101542 | 18 | 59000815 | 59223006 | 664 | 115 | 1.3356 | 0.090848 | CDH20 | 0.090848 | 0.10299 | 0.13362 |
| ENSG00000126522 | 7 | 65540785 | 65558545 | 47 | 8 | 1.3354 | 0.090871 | ASL | 0.090871 | 0.094656 | 0.10741 |
| ENSG00000149507 | 11 | 59807748 | 59815517 | 18 | 4 | 1.3353 | 0.090888 | OOSP2 | 0.090888 | 0.10198 | 0.11436 |
| ENSG00000186566 | 17 | 42472652 | 42580798 | 225 | 15 | 1.3351 | 0.090919 | GPATCH8 | 0.090919 | 0.095905 | 0.12065 |
| ENSG00000186281 | 2 | 96687694 | 96705199 | 9 | 4 | 1.3336 | 0.091161 | GPAT2 | 0.091161 | 0.10475 | 0.12421 |
| ENSG00000106524 | 7 | 16639401 | 16685442 | 205 | 27 | 1.3328 | 0.091303 | ANKMY2 | 0.091303 | 0.098689 | 0.17766 |
| ENSG00000219626 | 2 | 24299396 | 24392509 | 379 | 26 | 1.3325 | 0.091348 | FAM228B | 0.091348 | 0.11116 | 0.12109 |
| ENSG00000269210 | 2 | 39186764 | 39187483 | 4 | 2 | 1.3319 | 0.091444 | AC019171.1 | 0.091444 | 0.16797 | 0.24993 |
| ENSG00000170776 | 15 | 85923802 | 86292586 | 1456 | 41 | 1.3303 | 0.091715 | AKAP13 | 0.091715 | 0.056342 | 0.027838 |
| ENSG00000019582 | 5 | 149781200 | 149792492 | 27 | 9 | 1.3299 | 0.091783 | CD74 | 0.091783 | 0.31156 | 0.27077 |
| ENSG00000170381 | 7 | 82993222 | 83278326 | 1154 | 44 | 1.3294 | 0.091857 | SEMA3E | 0.091857 | 0.091712 | 0.095276 |
| ENSG00000107331 | 9 | 139901686 | 139923367 | 44 | 12 | 1.3292 | 0.091888 | ABCA2 | 0.091888 | 0.13417 | 0.065405 |
| ENSG00000002834 | 17 | 37026112 | 37078023 | 196 | 39 | 1.3291 | 0.091915 | LASP1 | 0.091915 | 0.18467 | 0.29497 |
| ENSG00000142669 | 1 | 26605667 | 26608007 | 5 | 3 | 1.3279 | 0.092105 | SH3BGRL3 | 0.092105 | 0.088509 | 0.12006 |
| ENSG00000102781 | 13 | 30776767 | 30881621 | 444 | 39 | 1.3277 | 0.092147 | KATNAL1 | 0.092147 | 0.12511 | 0.18587 |
| ENSG00000163508 | 3 | 27757440 | 27764206 | 12 | 5 | 1.3276 | 0.092159 | EOMES | 0.092159 | 0.048495 | 0.089218 |
| ENSG00000177030 | 11 | 644233 | 706715 | 312 | 22 | 1.3275 | 0.092164 | DEAF1 | 0.092164 | 0.031653 | 0.0099063 |
| ENSG00000169402 | 7 | 6793740 | 6838996 | 47 | 19 | 1.3271 | 0.09224 | RSPH10B2 | 0.09224 | 0.26605 | 0.63874 |
| ENSG00000152207 | 13 | 49280951 | 49283498 | 3 | 2 | 1.327 | 0.092262 | CYSLTR2 | 0.092262 | 0.0094967 | 0.039874 |
| ENSG00000258984 | 2 | 238875656 | 239008053 | 465 | 17 | 1.3269 | 0.092278 | UBE2F-SCLY | 0.092278 | 0.10326 | 0.12365 |

| | | | | | | | | | | | |
|---|---|---|---|---|---|---|---|---|---|---|---|
| ENSG00000135914 | 2 | 231972944 | 231989832 | 19 | 6 | 1.3263 | 0.092375 | HTR2B | 0.092375 | 0.053705 | 0.052784 |
| ENSG00000090402 | 3 | 164696686 | 164796283 | 244 | 14 | 1.3256 | 0.092491 | SI | 0.092491 | 0.088007 | 0.088999 |
| ENSG00000181013 | 17 | 56618948 | 56621729 | 4 | 2 | 1.3255 | 0.092503 | C17orf47 | 0.092503 | 0.0018613 | 0.0054803 |
| ENSG00000129084 | 11 | 14515329 | 14665181 | 211 | 36 | 1.3248 | 0.092611 | PSMA1 | 0.092611 | 0.044757 | 0.060243 |
| ENSG00000067082 | 10 | 3818188 | 3827473 | 27 | 9 | 1.3248 | 0.092626 | KLF6 | 0.092626 | 0.15406 | 0.083664 |
| ENSG00000106052 | 7 | 27778950 | 27884183 | 258 | 25 | 1.3245 | 0.092671 | TAX1BP1 | 0.092671 | 0.076558 | 0.081287 |
| ENSG00000169564 | 2 | 70314585 | 70316332 | 2 | 1 | 1.3244 | 0.092692 | PCBP1 | 0.092692 | 0.29109 | 0.30984 |
| ENSG00000100665 | 14 | 95027428 | 95036250 | 37 | 7 | 1.3241 | 0.092727 | SERPINA4 | 0.092727 | 0.066328 | 0.13288 |
| ENSG00000101557 | 18 | 158383 | 214629 | 158 | 21 | 1.3233 | 0.092863 | USP14 | 0.092863 | 0.094131 | 0.080034 |
| ENSG00000179409 | 17 | 647654 | 657239 | 40 | 10 | 1.322 | 0.093087 | GEMIN4 | 0.093087 | 0.17926 | 0.23152 |
| ENSG00000188095 | 15 | 90303822 | 90321982 | 58 | 9 | 1.3217 | 0.093136 | MESP2 | 0.093136 | 0.10416 | 0.14666 |
| ENSG00000141376 | 17 | 58754814 | 59470199 | 1275 | 58 | 1.3216 | 0.093157 | BCAS3 | 0.093157 | 0.12138 | 0.13496 |
| ENSG00000033100 | 7 | 150929575 | 150935908 | 15 | 3 | 1.3206 | 0.093324 | CHPF2 | 0.093324 | 0.20307 | 0.27448 |
| ENSG00000184349 | 5 | 106712590 | 107006596 | 829 | 101 | 1.3197 | 0.093472 | EFNA5 | 0.093472 | 0.081997 | 0.066724 |
| ENSG00000171552 | 20 | 30252255 | 30311792 | 97 | 8 | 1.3188 | 0.093616 | BCL2L1 | 0.093616 | 0.090624 | 0.094294 |
| ENSG00000088320 | 20 | 30063096 | 30072708 | 18 | 6 | 1.3188 | 0.093617 | REM1 | 0.093617 | 0.057013 | 0.054058 |
| ENSG00000082515 | 5 | 154320630 | 154348971 | 69 | 12 | 1.3185 | 0.09367 | MRPL22 | 0.09367 | 0.07221 | 0.076415 |
| ENSG00000239886 | 17 | 39382900 | 39383904 | 4 | 2 | 1.3184 | 0.093692 | KRTAP9-2 | 0.093692 | 0.17797 | 0.52183 |
| ENSG00000145191 | 3 | 183852826 | 184402546 | 2193 | 181 | 1.3177 | 0.093798 | EIF2B5 | 0.093798 | 0.13198 | 0.14513 |
| ENSG00000173175 | 3 | 123001143 | 123168605 | 440 | 36 | 1.3175 | 0.093839 | ADCY5 | 0.093839 | 0.072565 | 0.039983 |
| ENSG00000129437 | 19 | 51580752 | 51587502 | 32 | 7 | 1.3162 | 0.094047 | KLK14 | 0.094047 | 0.37929 | 0.4416 |
| ENSG00000184897 | 3 | 129033615 | 129035120 | 6 | 2 | 1.3161 | 0.094072 | H1FX | 0.094072 | 0.3415 | 0.40947 |
| ENSG00000144659 | 3 | 39424839 | 39438842 | 43 | 7 | 1.3159 | 0.094112 | SLC25A38 | 0.094112 | 0.10336 | 0.14906 |
| ENSG00000111339 | 12 | 14978503 | 14996429 | 55 | 8 | 1.3154 | 0.094191 | ART4 | 0.094191 | 0.091344 | 0.082056 |
| ENSG00000122136 | 9 | 138437985 | 138441815 | 57 | 7 | 1.3151 | 0.09423 | OBP2A | 0.09423 | 0.12803 | 0.22662 |
| ENSG00000177721 | 5 | 43039335 | 43043272 | 7 | 3 | 1.315 | 0.094256 | ANXA2R | 0.094256 | 0.24529 | 0.51772 |
| ENSG00000264364 | 17 | 56160776 | 56172897 | 24 | 6 | 1.3147 | 0.094314 | DYNLL2 | 0.094314 | 0.066441 | 0.15354 |
| ENSG00000100121 | 22 | 22988780 | 22990368 | 8 | 3 | 1.3143 | 0.094365 | GGTLC2 | 0.094365 | 0.030423 | 0.073984 |
| ENSG00000132010 | 19 | 12203658 | 12251222 | 103 | 14 | 1.3142 | 0.094382 | ZNF20 | 0.094382 | 0.13397 | 0.18732 |
| ENSG00000172878 | 2 | 172864490 | 172947158 | 307 | 27 | 1.314 | 0.094425 | METAP1D | 0.094425 | 0.095944 | 0.081504 |
| ENSG00000109113 | 17 | 27041299 | 27045447 | 2 | 1 | 1.3138 | 0.094463 | RAB34 | 0.094463 | 0.096293 | 0.11301 |
| ENSG00000009790 | 1 | 209929377 | 209955668 | 110 | 14 | 1.3135 | 0.09451 | TRAF3IP3 | 0.09451 | 0.11163 | 0.14146 |
| ENSG00000106344 | 7 | 127937738 | 127983962 | 87 | 17 | 1.3134 | 0.094522 | RBM28 | 0.094522 | 0.1422 | 0.13547 |
| ENSG00000107223 | 9 | 139756571 | 139760738 | 9 | 4 | 1.3126 | 0.094663 | EDF1 | 0.094663 | 0.090345 | 0.097531 |
| ENSG00000269738 | 9 | 125148344 | 125148928 | 3 | 2 | 1.3125 | 0.094672 | AL162424.1 | 0.094672 | 0.2494 | 0.71869 |
| ENSG00000130182 | 16 | 3138891 | 3149318 | 37 | 9 | 1.3125 | 0.094679 | ZSCAN10 | 0.094679 | 0.062555 | 0.10863 |
| ENSG00000157303 | 9 | 95820989 | 95847420 | 107 | 15 | 1.3119 | 0.094769 | SUSD3 | 0.094769 | 0.16379 | 0.2499 |
| ENSG00000178187 | 5 | 178368192 | 178393434 | 102 | 17 | 1.3118 | 0.094796 | ZNF454 | 0.094796 | 0.036508 | 0.059782 |
| ENSG00000109971 | 11 | 122928197 | 122933938 | 41 | 10 | 1.3112 | 0.094893 | HSPA8 | 0.094893 | 0.054448 | 0.046026 |
| ENSG00000157110 | 8 | 30241944 | 30429778 | 405 | 27 | 1.3108 | 0.094958 | RBPMS | 0.094958 | 0.096729 | 0.14767 |
| ENSG00000170290 | 11 | 107578104 | 107590419 | 33 | 7 | 1.3106 | 0.094994 | SLN | 0.094994 | 0.12052 | 0.30473 |
| ENSG00000243207 | 19 | 10216899 | 10225456 | 30 | 9 | 1.3099 | 0.095115 | PPAN-P2RY11 | 0.095115 | 0.067168 | 0.052068 |
| ENSG00000130810 | 19 | 10216965 | 10225414 | 30 | 9 | 1.3099 | 0.095115 | PPAN | 0.095115 | 0.065392 | 0.052068 |
| ENSG00000198553 | 13 | 50589390 | 50595058 | 16 | 4 | 1.3099 | 0.095123 | KCNRG | 0.095123 | 0.10559 | 0.079084 |
| ENSG00000162545 | 1 | 20808884 | 20812713 | 1 | 1 | 1.3092 | 0.09524 | CAMK2N1 | 0.09524 | 0.21805 | 0.24882 |
| ENSG00000164144 | 4 | 153701089 | 153839615 | 374 | 22 | 1.3092 | 0.09524 | ARFIP1 | 0.09524 | 0.10894 | 0.097887 |
| ENSG00000118526 | 6 | 134210276 | 134216691 | 20 | 4 | 1.3091 | 0.095248 | TCF21 | 0.095248 | 0.22052 | 0.25396 |
| ENSG00000178338 | 5 | 178286954 | 178315123 | 237 | 9 | 1.3084 | 0.095369 | ZNF354B | 0.095369 | 0.06867 | 0.13911 |
| ENSG00000017373 | 17 | 36686251 | 36762183 | 176 | 24 | 1.3081 | 0.095419 | SRCIN1 | 0.095419 | 0.038536 | 0.049163 |
| ENSG00000105327 | 19 | 47724081 | 47736023 | 13 | 7 | 1.3078 | 0.095477 | BBC3 | 0.095477 | 0.033759 | 0.016598 |
| ENSG00000076554 | 8 | 80870571 | 81143467 | 925 | 36 | 1.3075 | 0.095524 | TPD52 | 0.095524 | 0.099957 | 0.090567 |
| ENSG00000140374 | 15 | 76507696 | 76603813 | 241 | 16 | 1.3074 | 0.095531 | ETFA | 0.095531 | 0.10392 | 0.10254 |
| ENSG00000205022 | 16 | 88928034 | 88933068 | 28 | 9 | 1.3055 | 0.095863 | PABPN1L | 0.095863 | 0.27204 | 0.3108 |
| ENSG00000178445 | 9 | 6532464 | 6645650 | 495 | 54 | 1.3051 | 0.095932 | GLDC | 0.095932 | 0.10579 | 0.25392 |
| ENSG00000178172 | 5 | 147582357 | 147594700 | 40 | 10 | 1.3049 | 0.095961 | SPINK6 | 0.095961 | 0.44137 | 0.27976 |
| ENSG00000161638 | 12 | 54789045 | 54813244 | 50 | 8 | 1.3048 | 0.095985 | ITGA5 | 0.095985 | 0.10193 | 0.20611 |
| ENSG00000164935 | 8 | 105351315 | 105368917 | 64 | 14 | 1.3046 | 0.096021 | DCSTAMP | 0.096021 | 0.19619 | 0.21741 |
| ENSG00000136731 | 2 | 128848774 | 128953251 | 237 | 21 | 1.3038 | 0.096149 | UGGT1 | 0.096149 | 0.1127 | 0.19526 |
| ENSG00000184886 | 17 | 34890847 | 34895159 | 11 | 4 | 1.3032 | 0.096246 | PIGW | 0.096246 | 0.16604 | 0.2967 |
| ENSG00000154945 | 17 | 48770551 | 48785285 | 35 | 9 | 1.3028 | 0.096317 | ANKRD40 | 0.096317 | 0.088116 | 0.19334 |
| ENSG00000155087 | 8 | 103563800 | 103573245 | 64 | 10 | 1.3026 | 0.096363 | ODF1 | 0.096363 | 0.16304 | 0.15244 |
| ENSG00000180616 | 17 | 71161151 | 71167185 | 14 | 3 | 1.3025 | 0.096366 | SSTR2 | 0.096366 | 0.32141 | 0.47741 |
| ENSG00000132026 | 19 | 12936292 | 12946242 | 20 | 5 | 1.3024 | 0.096387 | RTBDN | 0.096387 | 0.18491 | 0.43929 |
| ENSG00000139722 | 12 | 123349882 | 123380991 | 43 | 12 | 1.3024 | 0.096394 | VPS37B | 0.096394 | 0.10036 | 0.25616 |
| ENSG00000104427 | 8 | 79578282 | 79632000 | 159 | 15 | 1.3017 | 0.096516 | ZC2HC1A | 0.096516 | 0.093581 | 0.083715 |
| ENSG00000059915 | 10 | 104162376 | 104181296 | 25 | 10 | 1.3015 | 0.096539 | PSD | 0.096539 | 0.17352 | 0.24205 |
| ENSG00000132635 | 20 | 2815960 | 2821836 | 9 | 5 | 1.3013 | 0.096583 | PCED1A | 0.096583 | 0.37295 | 0.38553 |
| ENSG00000059691 | 4 | 152591656 | 152682175 | 250 | 13 | 1.2998 | 0.096842 | PET112 | 0.096842 | 0.088552 | 0.080159 |
| ENSG00000215612 | 4 | 8847802 | 8873543 | 3 | 1 | 1.2988 | 0.097008 | HMX1 | 0.097008 | 0.096241 | 0.096241 |
| ENSG00000196071 | 1 | 248100493 | 248264224 | 650 | 29 | 1.2979 | 0.097154 | OR2L13 | 0.097154 | 0.086461 | 0.11764 |
| ENSG00000184903 | 7 | 110303110 | 111202573 | 2133 | 83 | 1.2977 | 0.097201 | IMMP2L | 0.097201 | 0.10597 | 0.1144 |
| ENSG00000084092 | 4 | 57829536 | 57844989 | 44 | 14 | 1.2975 | 0.097236 | NOA1 | 0.097236 | 0.020905 | 0.019848 |
| ENSG00000184206 | 15 | 84904525 | 84914120 | 7 | 4 | 1.2974 | 0.09724 | GOLGA6L4 | 0.09724 | 0.17327 | 0.16367 |

| | | | | | | | | | | | |
|---|---|---|---|---|---|---|---|---|---|---|---|
| ENSG00000186952 | 5 | 109624934 | 110074657 | 1365 | 29 | 1.2974 | 0.097253 | TMEM232 | 0.097253 | 0.1149 | 0.13139 |
| ENSG00000178690 | 18 | 52254988 | 52266724 | 25 | 8 | 1.2973 | 0.097258 | DYNAP | 0.097258 | 0.057897 | 0.066396 |
| ENSG00000106554 | 7 | 132469629 | 132766848 | 649 | 25 | 1.2973 | 0.097259 | CHCHD3 | 0.097259 | 0.089293 | 0.075759 |
| ENSG00000181291 | 17 | 32907768 | 32966337 | 149 | 33 | 1.297 | 0.097324 | TMEM132E | 0.097324 | 0.20738 | 0.24161 |
| ENSG00000164542 | 7 | 36363830 | 36429734 | 178 | 28 | 1.2968 | 0.097343 | KIAA0895 | 0.097343 | 0.12291 | 0.2051 |
| ENSG00000198792 | 22 | 38615298 | 38669040 | 140 | 20 | 1.2968 | 0.097352 | TMEM184B | 0.097352 | 0.17554 | 0.25603 |
| ENSG00000134014 | 8 | 27947190 | 28048673 | 248 | 19 | 1.2965 | 0.097397 | ELP3 | 0.097397 | 0.074025 | 0.081765 |
| ENSG00000144040 | 2 | 73169165 | 73302747 | 288 | 20 | 1.2956 | 0.097553 | SFXN5 | 0.097553 | 0.11509 | 0.1152 |
| ENSG00000108666 | 17 | 30651584 | 30677042 | 32 | 10 | 1.2953 | 0.097608 | C17orf75 | 0.097608 | 0.080355 | 0.11729 |
| ENSG00000164463 | 5 | 172483355 | 172566291 | 266 | 18 | 1.2952 | 0.097632 | CREBRF | 0.097632 | 0.08233 | 0.081672 |
| ENSG00000135835 | 1 | 180882290 | 180920750 | 115 | 29 | 1.2949 | 0.097675 | KIAA1614 | 0.097675 | 0.15965 | 0.36928 |
| ENSG00000058866 | 3 | 185823457 | 186080026 | 722 | 108 | 1.2934 | 0.09794 | DGKG | 0.09794 | 0.080157 | 0.12166 |
| ENSG00000139793 | 13 | 97873688 | 98046374 | 330 | 36 | 1.2932 | 0.097965 | MBNL2 | 0.097965 | 0.11114 | 0.13784 |
| ENSG00000110880 | 12 | 109038885 | 109125372 | 245 | 25 | 1.2928 | 0.098041 | CORO1C | 0.098041 | 0.099425 | 0.13837 |
| ENSG00000241119 | 2 | 234580499 | 234681946 | 408 | 23 | 1.2928 | 0.098045 | UGT1A9 | 0.098045 | 0.10975 | 0.10003 |
| ENSG00000137038 | 9 | 7796490 | 7888380 | 381 | 41 | 1.2918 | 0.098205 | TMEM261 | 0.098205 | 0.11536 | 0.14847 |
| ENSG00000166535 | 12 | 8975068 | 9039597 | 248 | 29 | 1.2915 | 0.098263 | A2ML1 | 0.098263 | 0.15345 | 0.26929 |
| ENSG00000117625 | 1 | 211431719 | 211489727 | 206 | 25 | 1.2914 | 0.098282 | RCOR3 | 0.098282 | 0.16516 | 0.19797 |
| ENSG00000213760 | 6 | 31512239 | 31516204 | 19 | 8 | 1.2914 | 0.098286 | ATP6V1G2 | 0.098286 | 0.24827 | 0.078387 |
| ENSG00000250173 | 5 | 979480 | 981415 | 5 | 3 | 1.29 | 0.098527 | RP11-661C8.3 | 0.098527 | 0.050561 | 0.0086863 |
| ENSG00000204536 | 6 | 31110216 | 31126015 | 161 | 21 | 1.2898 | 0.098557 | CCHCR1 | 0.098557 | 0.03254 | 0.022232 |
| ENSG00000183508 | 1 | 118148556 | 118170994 | 71 | 8 | 1.2895 | 0.098606 | FAM46C | 0.098606 | 0.093235 | 0.068982 |
| ENSG00000164850 | 7 | 1121844 | 1133451 | 52 | 13 | 1.2895 | 0.098615 | GPER1 | 0.098615 | 0.21214 | 0.27557 |
| ENSG00000082701 | 3 | 119540170 | 119813264 | 526 | 25 | 1.2889 | 0.098718 | GSK3B | 0.098718 | 0.09455 | 0.068487 |
| ENSG00000268194 | 2 | 121221911 | 121223697 | 3 | 1 | 1.2886 | 0.098769 | LINC01101 | 0.098769 | 0.039337 | 0.28425 |
| ENSG00000129083 | 11 | 14464986 | 14521573 | 114 | 14 | 1.2884 | 0.098799 | COPB1 | 0.098799 | 0.11356 | 0.12424 |
| ENSG00000091640 | 17 | 4862521 | 4871167 | 25 | 11 | 1.2869 | 0.099072 | SPAG7 | 0.099072 | 0.31785 | 0.29408 |
| ENSG00000106415 | 7 | 8008425 | 8133902 | 450 | 26 | 1.2868 | 0.099083 | GLCCI1 | 0.099083 | 0.10382 | 0.12847 |
| ENSG00000174844 | 3 | 57327727 | 57530071 | 916 | 26 | 1.2866 | 0.099113 | DNAH12 | 0.099113 | 0.095431 | 0.097854 |
| ENSG00000071282 | 3 | 8543393 | 8609805 | 265 | 21 | 1.2864 | 0.09915 | LMCD1 | 0.09915 | 0.12157 | 0.16925 |
| ENSG00000204977 | 13 | 50570024 | 50594617 | 65 | 9 | 1.2864 | 0.099151 | TRIM13 | 0.099151 | 0.099517 | 0.077106 |
| ENSG00000143537 | 1 | 155023042 | 155035252 | 21 | 7 | 1.2863 | 0.099172 | ADAM15 | 0.099172 | 0.011036 | 0.013716 |
| ENSG00000109255 | 4 | 56461396 | 56502865 | 177 | 21 | 1.2857 | 0.099275 | NMU | 0.099275 | 0.15266 | 0.30871 |
| ENSG00000196743 | 5 | 150591711 | 150650001 | 210 | 29 | 1.2857 | 0.099275 | GM2A | 0.099275 | 0.11889 | 0.26155 |
| ENSG00000176540 | 11 | 48387037 | 48388017 | 2 | 1 | 1.2856 | 0.099287 | OR4C5 | 0.099287 | 0.1773 | 0.21138 |
| ENSG00000269948 | 10 | 97709725 | 97744785 | 121 | 19 | 1.2852 | 0.099362 | RP11-248J23.6 | 0.099362 | 0.26363 | 0.39615 |
| ENSG00000133111 | 13 | 37393361 | 37403241 | 15 | 4 | 1.2846 | 0.099469 | RFXAP | 0.099469 | 0.18915 | 0.31988 |
| ENSG00000172315 | 20 | 45313004 | 45318418 | 13 | 7 | 1.2845 | 0.099491 | TP53RK | 0.099491 | 0.23558 | 0.17294 |
| ENSG00000164588 | 5 | 45259349 | 45696253 | 613 | 32 | 1.2841 | 0.099549 | HCN1 | 0.099549 | 0.092522 | 0.094488 |
| ENSG00000203871 | 6 | 88106840 | 88109467 | 13 | 2 | 1.2836 | 0.099634 | C6orf164 | 0.099634 | 0.10026 | 0.11269 |
| ENSG00000006114 | 17 | 35874900 | 35969544 | 167 | 28 | 1.2835 | 0.099653 | SYNRG | 0.099653 | 0.069307 | 0.056146 |
| ENSG00000155876 | 9 | 19049372 | 19051019 | 2 | 1 | 1.2835 | 0.09966 | RRAGA | 0.09966 | 0.078777 | 0.03563 |
| ENSG00000143740 | 1 | 227916240 | 227968927 | 144 | 19 | 1.2829 | 0.099755 | SNAP47 | 0.099755 | 0.082308 | 0.062854 |
| ENSG00000173546 | 15 | 75966663 | 76005189 | 69 | 10 | 1.2829 | 0.099759 | CSPG4 | 0.099759 | 0.1699 | 0.21296 |
| ENSG00000157131 | 1 | 57320479 | 57383894 | 261 | 26 | 1.2823 | 0.099877 | C8A | 0.099877 | 0.082462 | 0.056372 |
| ENSG00000185658 | 21 | 40556102 | 40693485 | 462 | 19 | 1.282 | 0.099913 | BRWD1 | 0.099913 | 0.095175 | 0.1002 |
| ENSG00000135747 | 1 | 247108849 | 247242113 | 684 | 30 | 1.2814 | 0.10003 | ZNF670 | 0.10003 | 0.085413 | 0.083351 |
| ENSG00000169618 | 2 | 68870721 | 68882708 | 50 | 18 | 1.2812 | 0.10005 | PROKR1 | 0.10005 | 0.067307 | 0.23846 |
| ENSG00000137507 | 11 | 76368568 | 76381791 | 33 | 10 | 1.2812 | 0.10006 | LRRC32 | 0.10006 | 0.10022 | 0.26841 |
| ENSG00000112335 | 6 | 108532426 | 108582464 | 109 | 18 | 1.281 | 0.10009 | SNX3 | 0.10009 | 0.1541 | 0.14989 |
| ENSG00000104915 | 19 | 13254872 | 13261197 | 8 | 3 | 1.2808 | 0.10014 | STX10 | 0.10014 | 0.41411 | 0.34382 |
| ENSG00000110066 | 11 | 67922330 | 67981295 | 121 | 15 | 1.2807 | 0.10016 | SUV420H1 | 0.10016 | 0.1147 | 0.116 |
| ENSG00000197406 | 14 | 102027688 | 102029789 | 1 | 1 | 1.2804 | 0.1002 | DIO3 | 0.1002 | 0.19481 | 0.08876 |
| ENSG00000143333 | 1 | 182567758 | 182573543 | 9 | 3 | 1.2801 | 0.10025 | RGS16 | 0.10025 | 0.35773 | 0.49791 |
| ENSG00000142655 | 1 | 10532345 | 10690815 | 447 | 31 | 1.2797 | 0.10033 | PEX14 | 0.10033 | 0.09613 | 0.099028 |
| ENSG00000198171 | 20 | 3170996 | 3185331 | 82 | 18 | 1.2796 | 0.10034 | DDRGK1 | 0.10034 | 0.031419 | 0.040031 |
| ENSG00000137628 | 4 | 169137444 | 169239958 | 379 | 25 | 1.2793 | 0.10039 | DDX60 | 0.10039 | 0.094666 | 0.083536 |
| ENSG00000140107 | 14 | 100789674 | 100796715 | 25 | 7 | 1.279 | 0.10045 | SLC25A47 | 0.10045 | 0.027904 | 0.057531 |
| ENSG00000007968 | 1 | 23832922 | 23857712 | 68 | 9 | 1.2788 | 0.10049 | E2F2 | 0.10049 | 0.18755 | 0.089933 |
| ENSG00000143653 | 1 | 246887349 | 246931439 | 105 | 17 | 1.2786 | 0.10052 | SCCPDH | 0.10052 | 0.034352 | 0.029235 |
| ENSG00000171551 | 2 | 233344537 | 233352538 | 32 | 14 | 1.2781 | 0.10061 | ECEL1 | 0.10061 | 0.24769 | 0.35481 |
| ENSG00000076351 | 17 | 26721661 | 26734215 | 20 | 5 | 1.2779 | 0.10064 | SLC46A1 | 0.10064 | 0.19302 | 0.38021 |
| ENSG00000122490 | 18 | 77662420 | 77711664 | 188 | 27 | 1.2779 | 0.10065 | PQLC1 | 0.10065 | 0.077598 | 0.05011 |
| ENSG00000176555 | 11 | 48327775 | 48328704 | 4 | 2 | 1.2777 | 0.10068 | OR4S1 | 0.10068 | 0.039155 | 0.050266 |
| ENSG00000198832 | 22 | 31500758 | 31516055 | 65 | 12 | 1.2777 | 0.10068 | SELM | 0.10068 | 0.16243 | 0.11148 |
| ENSG00000065802 | 2 | 239335383 | 239360891 | 122 | 14 | 1.2775 | 0.1007 | ASB1 | 0.1007 | 0.083551 | 0.052866 |
| ENSG00000226209 | 10 | 13350145 | 13350447 | 1 | 1 | 1.2776 | 0.1007 | AL138764.1 | 0.1007 | 0.29205 | 0.53876 |
| ENSG00000198964 | 10 | 52065360 | 52384923 | 952 | 46 | 1.2774 | 0.10072 | SGMS1 | 0.10072 | 0.104 | 0.11944 |
| ENSG00000165997 | 10 | 18948334 | 18970568 | 79 | 15 | 1.277 | 0.1008 | ARL5B | 0.1008 | 0.042769 | 0.01722 |
| ENSG00000184999 | 11 | 62905339 | 63137190 | 792 | 23 | 1.2769 | 0.10082 | SLC22A10 | 0.10082 | 0.11244 | 0.11449 |
| ENSG00000166821 | 15 | 90220995 | 90234014 | 30 | 6 | 1.2765 | 0.10089 | PEX11A | 0.10089 | 0.29302 | 0.34661 |
| ENSG00000052749 | 10 | 99116115 | 99161127 | 219 | 9 | 1.276 | 0.10097 | RRP12 | 0.10097 | 0.10005 | 0.070747 |
| ENSG00000141380 | 18 | 23596578 | 23671181 | 171 | 17 | 1.2756 | 0.10104 | SS18 | 0.10104 | 0.12877 | 0.15575 |

| | | | | | | | | | | | |
|---|---|---|---|---|---|---|---|---|---|---|---|
| ENSG00000125531 | 20 | 62184373 | 62188061 | 13 | 5 | 1.275 | 0.10116 | C20orf195 | 0.10116 | 0.15126 | 0.32385 |
| ENSG00000187239 | 9 | 132649466 | 132805473 | 635 | 33 | 1.2742 | 0.10129 | FNBP1 | 0.10129 | 0.099357 | 0.13092 |
| ENSG00000140577 | 15 | 91073157 | 91188577 | 418 | 24 | 1.2743 | 0.10129 | CRTC3 | 0.10129 | 0.090104 | 0.06407 |
| ENSG00000173269 | 10 | 88695297 | 88729238 | 117 | 17 | 1.2737 | 0.10138 | MMRN2 | 0.10138 | 0.11239 | 0.12351 |
| ENSG00000105492 | 19 | 52022779 | 52035110 | 73 | 9 | 1.2737 | 0.10139 | SIGLEC6 | 0.10139 | 0.093639 | 0.051799 |
| ENSG00000158553 | 6 | 27253682 | 27279949 | 112 | 21 | 1.2726 | 0.10158 | POM121L2 | 0.10158 | 0.038614 | 0.072527 |
| ENSG00000132334 | 10 | 129705325 | 129884119 | 796 | 101 | 1.272 | 0.10169 | PTPRE | 0.10169 | 0.1318 | 0.15763 |
| ENSG00000100354 | 22 | 40440821 | 40731812 | 355 | 31 | 1.272 | 0.10169 | TNRC6B | 0.10169 | 0.086181 | 0.092753 |
| ENSG00000152315 | 14 | 90528109 | 90652201 | 557 | 43 | 1.2716 | 0.10176 | KCNK13 | 0.10176 | 0.078388 | 0.052289 |
| ENSG00000271786 | 10 | 18931405 | 18944123 | 50 | 8 | 1.2706 | 0.10194 | RP11-139J15.7 | 0.10194 | 0.11145 | 0.08878 |
| ENSG00000146285 | 6 | 108025308 | 108145521 | 424 | 32 | 1.2705 | 0.10195 | SCML4 | 0.10195 | 0.090152 | 0.096721 |
| ENSG00000121577 | 3 | 119355304 | 119384171 | 127 | 15 | 1.2705 | 0.10196 | POPDC2 | 0.10196 | 0.097129 | 0.12486 |
| ENSG00000257127 | 12 | 92815307 | 92824778 | 36 | 6 | 1.2704 | 0.10198 | CLLU1 | 0.10198 | 0.096653 | 0.11331 |
| ENSG00000203684 | 1 | 228351787 | 228353213 | 1 | 1 | 1.2702 | 0.102 | IBA57-AS1 | 0.102 | 0.49058 | 0.36523 |
| ENSG00000011132 | 19 | 3750817 | 3761697 | 41 | 10 | 1.27 | 0.10205 | APBA3 | 0.10205 | 0.056008 | 0.30321 |
| ENSG00000132326 | 2 | 239152679 | 239198743 | 121 | 22 | 1.2694 | 0.10214 | PER2 | 0.10214 | 0.12581 | 0.072993 |
| ENSG00000243156 | 22 | 18270415 | 18507325 | 1114 | 68 | 1.2693 | 0.10217 | MICAL3 | 0.10217 | 0.12187 | 0.15255 |
| ENSG00000258847 | 14 | 66424015 | 66471241 | 155 | 30 | 1.2684 | 0.10232 | CTD-2014B16.3 | 0.10232 | 0.16083 | 0.11726 |
| ENSG00000138675 | 4 | 81187753 | 81257834 | 182 | 28 | 1.2682 | 0.10236 | FGF5 | 0.10236 | 0.11725 | 0.21768 |
| ENSG00000163599 | 2 | 204732509 | 204738683 | 4 | 2 | 1.2671 | 0.10255 | CTLA4 | 0.10255 | 0.57735 | 0.71347 |
| ENSG00000180667 | 1 | 207217194 | 207226325 | 15 | 4 | 1.2671 | 0.10256 | YOD1 | 0.10256 | 0.084654 | 0.095721 |
| ENSG00000198087 | 6 | 47445525 | 47594999 | 446 | 17 | 1.2669 | 0.10259 | CD2AP | 0.10259 | 0.10726 | 0.13274 |
| ENSG00000150275 | 10 | 55562531 | 57387702 | 7689 | 219 | 1.2667 | 0.10264 | PCDH15 | 0.10264 | 0.12179 | 0.088109 |
| ENSG00000155130 | 6 | 114178541 | 114184648 | 9 | 5 | 1.2666 | 0.10265 | MARCKS | 0.10265 | 0.052408 | 0.073151 |
| ENSG00000130948 | 9 | 98997588 | 99064434 | 208 | 31 | 1.2665 | 0.10267 | HSD17B3 | 0.10267 | 0.29299 | 0.38747 |
| ENSG00000113580 | 5 | 142657496 | 142815077 | 264 | 22 | 1.2662 | 0.10272 | NR3C1 | 0.10272 | 0.072999 | 0.064475 |
| ENSG00000118816 | 4 | 77968311 | 77997158 | 68 | 15 | 1.2656 | 0.10283 | CCNI | 0.10283 | 0.10478 | 0.16201 |
| ENSG00000175868 | 11 | 14926543 | 15103888 | 371 | 14 | 1.2656 | 0.10283 | CALCB | 0.10283 | 0.085721 | 0.086027 |
| ENSG00000146453 | 6 | 160221298 | 160241736 | 73 | 14 | 1.265 | 0.10293 | PNLDC1 | 0.10293 | 0.060145 | 0.070335 |
| ENSG00000267477 | 19 | 11577055 | 11616513 | 111 | 12 | 1.2648 | 0.10298 | CTC-398G3.6 | 0.10298 | 0.13156 | 0.1853 |
| ENSG00000250673 | 4 | 147145709 | 147150208 | 5 | 3 | 1.2647 | 0.10299 | RP11-6L6.2 | 0.10299 | 0.25355 | 0.090734 |
| ENSG00000011083 | 5 | 149569520 | 149602351 | 154 | 24 | 1.2645 | 0.10302 | SLC6A7 | 0.10302 | 0.10985 | 0.065445 |
| ENSG00000183657 | 3 | 171509580 | 171527714 | 60 | 8 | 1.2642 | 0.10308 | PP13439 | 0.10308 | 0.080894 | 0.18686 |
| ENSG00000241837 | 21 | 35275757 | 35288284 | 46 | 11 | 1.2633 | 0.10324 | ATP5O | 0.10324 | 0.0795 | 0.098627 |
| ENSG00000186130 | 9 | 125670335 | 125675609 | 11 | 4 | 1.2625 | 0.10339 | ZBTB6 | 0.10339 | 0.2907 | 0.3491 |
| ENSG00000158042 | 11 | 6702013 | 6704632 | 15 | 2 | 1.2623 | 0.10341 | MRPL17 | 0.10341 | 0.20307 | 0.42209 |
| ENSG00000115944 | 2 | 42560686 | 42652228 | 244 | 25 | 1.262 | 0.10348 | COX7A2L | 0.10348 | 0.12475 | 0.13862 |
| ENSG00000069712 | 1 | 92632542 | 92650280 | 42 | 5 | 1.2618 | 0.10351 | KIAA1107 | 0.10351 | 0.13065 | 0.11877 |
| ENSG00000183695 | 11 | 19076002 | 19082228 | 42 | 6 | 1.2618 | 0.10351 | MRGPRX2 | 0.10351 | 0.01319 | 0.020175 |
| ENSG00000126464 | 19 | 50094900 | 50129696 | 100 | 13 | 1.2616 | 0.10354 | PRR12 | 0.10354 | 0.087576 | 0.087436 |
| ENSG00000141505 | 17 | 7076750 | 7082883 | 50 | 16 | 1.2612 | 0.10362 | ASGR1 | 0.10362 | 0.13215 | 0.10472 |
| ENSG00000150244 | 11 | 55029658 | 55038595 | 40 | 6 | 1.2608 | 0.10368 | TRIM48 | 0.10368 | 0.11656 | 0.098091 |
| ENSG00000108055 | 10 | 112327449 | 112364394 | 125 | 11 | 1.2607 | 0.1037 | SMC3 | 0.1037 | 0.11544 | 0.11327 |
| ENSG00000162542 | 1 | 20008706 | 20126438 | 423 | 55 | 1.2606 | 0.10373 | TMCO4 | 0.10373 | 0.11693 | 0.10481 |
| ENSG00000185926 | 11 | 51515282 | 51516211 | 9 | 3 | 1.2604 | 0.10375 | OR4C46 | 0.10375 | 0.16002 | 0.2412 |
| ENSG00000134987 | 5 | 110427414 | 110466200 | 112 | 13 | 1.2603 | 0.10379 | WDR36 | 0.10379 | 0.13691 | 0.15948 |
| ENSG00000087152 | 17 | 42269173 | 42277481 | 13 | 9 | 1.2591 | 0.10399 | ATXN7L3 | 0.10399 | 0.82066 | 0.89699 |
| ENSG00000166748 | 15 | 86685227 | 87572283 | 4237 | 164 | 1.259 | 0.10401 | AGBL1 | 0.10401 | 0.064316 | 0.047753 |
| ENSG00000172731 | 10 | 72058729 | 72142382 | 375 | 35 | 1.2586 | 0.10408 | LRRC20 | 0.10408 | 0.084855 | 0.10041 |
| ENSG00000157856 | 2 | 26624784 | 26679579 | 151 | 24 | 1.2583 | 0.10415 | DRC1 | 0.10415 | 0.1226 | 0.14452 |
| ENSG00000101203 | 20 | 61924538 | 61966203 | 147 | 14 | 1.2572 | 0.10433 | COL20A1 | 0.10433 | 0.060641 | 0.076373 |
| ENSG00000171311 | 10 | 99195899 | 99205774 | 24 | 7 | 1.2565 | 0.10446 | EXOSC1 | 0.10446 | 0.030892 | 0.018865 |
| ENSG00000112038 | 6 | 154331631 | 154568001 | 805 | 54 | 1.2561 | 0.10455 | OPRM1 | 0.10455 | 0.11404 | 0.13424 |
| ENSG00000095139 | 11 | 118443105 | 118473748 | 74 | 6 | 1.2553 | 0.10468 | ARCN1 | 0.10468 | 0.14094 | 0.16873 |
| ENSG00000010244 | 17 | 30677136 | 30708905 | 50 | 13 | 1.2551 | 0.10472 | ZNF207 | 0.10472 | 0.1156 | 0.10512 |
| ENSG00000183638 | 8 | 10463859 | 10569697 | 349 | 50 | 1.2549 | 0.10476 | RP1L1 | 0.10476 | 0.10214 | 0.1131 |
| ENSG00000135469 | 12 | 56660642 | 56664750 | 4 | 2 | 1.2548 | 0.10478 | COQ10A | 0.10478 | 0.19564 | 0.21377 |
| ENSG00000204952 | 17 | 37092685 | 37123655 | 51 | 13 | 1.2548 | 0.10478 | FBXO47 | 0.10478 | 0.19945 | 0.3871 |
| ENSG00000115274 | 2 | 74682150 | 74688011 | 2 | 1 | 1.2542 | 0.10489 | INO80B | 0.10489 | 0.66883 | 0.7309 |
| ENSG00000124145 | 20 | 43953928 | 43977064 | 88 | 11 | 1.2542 | 0.10489 | SDC4 | 0.10489 | 0.025003 | 0.04502 |
| ENSG00000204301 | 6 | 32162620 | 32191844 | 171 | 30 | 1.2535 | 0.10502 | NOTCH4 | 0.10502 | 0.034404 | 0.022066 |
| ENSG00000228594 | 1 | 1533392 | 1535476 | 2 | 1 | 1.2528 | 0.10515 | C1orf233 | 0.10515 | 0.60178 | 0.77504 |
| ENSG00000181625 | 16 | 29465822 | 29469540 | 1 | 1 | 1.2525 | 0.1052 | SLX1B | 0.1052 | 0.1971 | 0.12737 |
| ENSG00000215704 | 1 | 15792404 | 15817895 | 96 | 15 | 1.2518 | 0.10533 | CELA2B | 0.10533 | 0.058532 | 0.027754 |
| ENSG00000143171 | 1 | 165370159 | 165414433 | 175 | 26 | 1.2517 | 0.10533 | RXRG | 0.10533 | 0.088682 | 0.079626 |
| ENSG00000187144 | 1 | 16713612 | 16763919 | 157 | 25 | 1.2502 | 0.10562 | SPATA21 | 0.10562 | 0.14262 | 0.17268 |
| ENSG00000238244 | 15 | 90890819 | 90892669 | 13 | 2 | 1.25 | 0.10564 | GABARAPL3 | 0.10564 | 0.086166 | 0.063596 |
| ENSG00000175065 | 18 | 28956740 | 28994875 | 93 | 11 | 1.25 | 0.10565 | DSG4 | 0.10565 | 0.11173 | 0.13276 |
| ENSG00000119888 | 2 | 47572297 | 47614740 | 158 | 27 | 1.2497 | 0.1057 | EPCAM | 0.1057 | 0.18157 | 0.2191 |
| ENSG00000243480 | 1 | 104159999 | 104168402 | 3 | 2 | 1.2493 | 0.10578 | AMY2A | 0.10578 | 0.046924 | 0.049887 |
| ENSG00000196600 | 11 | 62931248 | 62997124 | 256 | 12 | 1.2491 | 0.10581 | SLC22A25 | 0.10581 | 0.1099 | 0.10512 |
| ENSG00000227471 | 7 | 134233888 | 134264627 | 104 | 11 | 1.2488 | 0.10586 | AKR1B15 | 0.10586 | 0.069094 | 0.1435 |
| ENSG00000172487 | 11 | 56127691 | 56128764 | 9 | 4 | 1.248 | 0.10601 | OR8J1 | 0.10601 | 0.025004 | 0.03661 |

| | | | | | | | | | | | |
|---|---|---|---|---|---|---|---|---|---|---|---|
| ENSG00000156802 | 8 | 124332090 | 124428590 | 142 | 40 | 1.2476 | 0.10609 | ATAD2 | 0.10609 | 0.25734 | 0.24916 |
| ENSG00000196588 | 22 | 40806285 | 41032706 | 473 | 19 | 1.2473 | 0.10614 | MKL1 | 0.10614 | 0.11544 | 0.12421 |
| ENSG00000254852 | 16 | 14841923 | 14859270 | 1 | 1 | 1.247 | 0.1062 | NPIPA2 | 0.1062 | 0.68877 | 0.81944 |
| ENSG00000179912 | 12 | 57643392 | 57824788 | 318 | 16 | 1.2469 | 0.10622 | R3HDM2 | 0.10622 | 0.093783 | 0.084601 |
| ENSG00000158863 | 8 | 21946670 | 21962409 | 44 | 10 | 1.2468 | 0.10623 | FAM160B2 | 0.10623 | 0.14617 | 0.21678 |
| ENSG00000152520 | 13 | 28712643 | 28869475 | 348 | 20 | 1.2468 | 0.10624 | PAN3 | 0.10624 | 0.11826 | 0.076913 |
| ENSG00000138777 | 4 | 106290234 | 106395238 | 504 | 20 | 1.2465 | 0.1063 | PPA2 | 0.1063 | 0.13978 | 0.17283 |
| ENSG00000122367 | 10 | 88428206 | 88495825 | 262 | 35 | 1.246 | 0.10638 | LDB3 | 0.10638 | 0.11973 | 0.13066 |
| ENSG00000174236 | 12 | 27849428 | 27850566 | 2 | 1 | 1.2459 | 0.1064 | REP15 | 0.1064 | 0.19583 | 0.33918 |
| ENSG00000164305 | 4 | 185548850 | 185570663 | 66 | 11 | 1.2454 | 0.10648 | CASP3 | 0.10648 | 0.11288 | 0.16762 |
| ENSG00000188620 | 10 | 124895478 | 124897257 | 1 | 1 | 1.2454 | 0.1065 | HMX3 | 0.1065 | 0.037056 | 0.11202 |
| ENSG00000183889 | 16 | 16411301 | 16444447 | 3 | 2 | 1.2451 | 0.10655 | NPIPA7 | 0.10655 | 0.073767 | 0.28034 |
| ENSG00000116906 | 1 | 231376953 | 231413719 | 99 | 9 | 1.2448 | 0.1066 | GNPAT | 0.1066 | 0.14358 | 0.33269 |
| ENSG00000160877 | 19 | 13228917 | 13251955 | 41 | 9 | 1.2448 | 0.10661 | NACC1 | 0.10661 | 0.10955 | 0.17164 |
| ENSG00000179774 | 10 | 69990386 | 69991871 | 4 | 2 | 1.2446 | 0.10664 | ATOH7 | 0.10664 | 0.19142 | 0.19242 |
| ENSG00000179344 | 6 | 32627244 | 32636160 | 831 | 21 | 1.2445 | 0.10665 | HLA-DQB1 | 0.10665 | 0.11149 | 0.018059 |
| ENSG00000268923 | 6 | 31973945 | 31974881 | 12 | 6 | 1.2443 | 0.1067 | AL645922.1 | 0.1067 | 0.0025701 | 0.00027782 |
| ENSG00000159086 | 21 | 34106210 | 34144169 | 56 | 13 | 1.2443 | 0.1067 | PAXBP1 | 0.1067 | 0.18586 | 0.16448 |
| ENSG00000213927 | 9 | 34661877 | 34664045 | 6 | 2 | 1.2438 | 0.10678 | CCL27 | 0.10678 | 0.13645 | 0.16581 |
| ENSG00000189253 | 11 | 89602450 | 89609185 | 6 | 3 | 1.2435 | 0.10685 | TRIM64B | 0.10685 | 0.22635 | 0.4352 |
| ENSG00000155660 | 7 | 148700154 | 148725733 | 86 | 8 | 1.2433 | 0.10687 | PDIA4 | 0.10687 | 0.20889 | 0.32017 |
| ENSG00000198150 | 17 | 8261731 | 8263538 | 2 | 1 | 1.2432 | 0.1069 | AC135178.1 | 0.1069 | 0.5221 | 0.48336 |
| ENSG00000228008 | 3 | 49941278 | 49954370 | 13 | 3 | 1.2431 | 0.10691 | CTD-2330K9.3 | 0.10691 | 0.18028 | 0.1841 |
| ENSG00000122870 | 10 | 60272900 | 60591195 | 813 | 37 | 1.2426 | 0.10701 | BICC1 | 0.10701 | 0.10696 | 0.14193 |
| ENSG00000133048 | 1 | 203148059 | 203155877 | 28 | 5 | 1.2425 | 0.10703 | CHI3L1 | 0.10703 | 0.30749 | 0.37204 |
| ENSG00000269403 | 19 | 51853627 | 51871353 | 85 | 13 | 1.2418 | 0.10716 | CTD-2616J11.11 | 0.10716 | 0.23945 | 0.50455 |
| ENSG00000197826 | 4 | 81256874 | 81884910 | 1637 | 39 | 1.2416 | 0.10719 | C4orf22 | 0.10719 | 0.099662 | 0.10626 |
| ENSG00000178950 | 4 | 843064 | 926161 | 430 | 38 | 1.2416 | 0.1072 | GAK | 0.1072 | 0.087402 | 0.14407 |
| ENSG00000205029 | 11 | 55606228 | 55607214 | 5 | 3 | 1.2403 | 0.10743 | OR5D16 | 0.10743 | 0.10524 | 0.21872 |
| ENSG00000101353 | 20 | 35729629 | 35807991 | 151 | 23 | 1.2397 | 0.10754 | MROH8 | 0.10754 | 0.11129 | 0.11501 |
| ENSG00000148826 | 10 | 134598297 | 134599556 | 1 | 1 | 1.2394 | 0.1076 | NKX6-2 | 0.1076 | 0.00016749 | 0.0019258 |
| ENSG00000179840 | 1 | 9712668 | 9714644 | 5 | 2 | 1.2392 | 0.10764 | C1orf200 | 0.10764 | 0.21431 | 0.29188 |
| ENSG00000221900 | 7 | 53103349 | 53104617 | 9 | 2 | 1.2392 | 0.10764 | POM121L12 | 0.10764 | 0.2104 | 0.25711 |
| ENSG00000186451 | 3 | 57094469 | 57109460 | 49 | 10 | 1.2391 | 0.10765 | SPATA12 | 0.10765 | 0.23779 | 0.37209 |
| ENSG00000258572 | 14 | 95982473 | 95984248 | 11 | 4 | 1.2389 | 0.1077 | RP11-1070N10.3 | 0.1077 | 0.22428 | 0.46857 |
| ENSG00000170619 | 8 | 146066427 | 146079121 | 44 | 5 | 1.2387 | 0.10773 | COMMD5 | 0.10773 | 0.083083 | 0.079905 |
| ENSG00000082516 | 5 | 154266976 | 154317769 | 147 | 16 | 1.2374 | 0.10796 | GEMIN5 | 0.10796 | 0.077196 | 0.073781 |
| ENSG00000166450 | 15 | 55903744 | 56035288 | 431 | 35 | 1.2371 | 0.10803 | PRTG | 0.10803 | 0.11574 | 0.21054 |
| ENSG00000128309 | 22 | 37415676 | 37425863 | 32 | 8 | 1.2371 | 0.10803 | MPST | 0.10803 | 0.20886 | 0.25172 |
| ENSG00000109511 | 4 | 169013666 | 169108841 | 265 | 20 | 1.2368 | 0.10807 | ANXA10 | 0.10807 | 0.11475 | 0.14058 |
| ENSG00000104064 | 15 | 50569389 | 50647605 | 254 | 12 | 1.2369 | 0.10807 | GABPB1 | 0.10807 | 0.15045 | 0.18755 |
| ENSG00000213029 | 1 | 229440129 | 229441248 | 2 | 1 | 1.2368 | 0.10808 | SPHAR | 0.10808 | 0.32172 | 0.46708 |
| ENSG00000067066 | 2 | 231280657 | 231408805 | 397 | 45 | 1.2364 | 0.10816 | SP100 | 0.10816 | 0.11343 | 0.17345 |
| ENSG00000206557 | 3 | 32859510 | 32939318 | 256 | 31 | 1.2361 | 0.1082 | TRIM71 | 0.1082 | 0.13611 | 0.1825 |
| ENSG00000138744 | 4 | 76831809 | 76862204 | 161 | 15 | 1.2362 | 0.1082 | NAAA | 0.1082 | 0.10491 | 0.17339 |
| ENSG00000183324 | 15 | 73735499 | 73852355 | 280 | 20 | 1.2361 | 0.10821 | C15orf60 | 0.10821 | 0.10186 | 0.07882 |
| ENSG00000109111 | 17 | 26989109 | 27029697 | 45 | 10 | 1.2356 | 0.10831 | SUPT6H | 0.10831 | 0.10472 | 0.12803 |
| ENSG00000122863 | 10 | 73724123 | 73773322 | 178 | 16 | 1.2349 | 0.10843 | CHST3 | 0.10843 | 0.20951 | 0.28522 |
| ENSG00000203805 | 10 | 122216466 | 122349367 | 554 | 45 | 1.2348 | 0.10846 | PPAPDC1A | 0.10846 | 0.13524 | 0.20385 |
| ENSG00000268788 | 4 | 76871883 | 76878526 | 13 | 5 | 1.2345 | 0.10851 | AC110615.1 | 0.10851 | 0.11051 | 0.18463 |
| ENSG00000145431 | 4 | 157681606 | 157892546 | 391 | 32 | 1.2335 | 0.1087 | PDGFC | 0.1087 | 0.11289 | 0.13632 |
| ENSG00000213638 | 19 | 1905377 | 1913444 | 37 | 6 | 1.2333 | 0.10873 | ADAT3 | 0.10873 | 0.14536 | 0.36942 |
| ENSG00000234745 | 6 | 31321649 | 31324965 | 205 | 42 | 1.2324 | 0.1089 | HLA-B | 0.1089 | 0.13581 | 0.21425 |
| ENSG00000106772 | 9 | 79226292 | 79521003 | 1163 | 92 | 1.2321 | 0.10895 | PRUNE2 | 0.10895 | 0.13673 | 0.15339 |
| ENSG00000136869 | 9 | 120466610 | 120479149 | 29 | 9 | 1.2318 | 0.10901 | TLR4 | 0.10901 | 0.29393 | 0.47293 |
| ENSG00000218416 | 2 | 241388732 | 241396131 | 30 | 8 | 1.2318 | 0.10902 | AC110619.2 | 0.10902 | 0.41153 | 0.68908 |
| ENSG00000154485 | 10 | 127455022 | 127464390 | 11 | 5 | 1.2317 | 0.10903 | MMP21 | 0.10903 | 0.37908 | 0.53403 |
| ENSG00000139405 | 12 | 113623331 | 113630173 | 17 | 3 | 1.2316 | 0.10905 | RITA1 | 0.10905 | 0.070898 | 0.075425 |
| ENSG00000175463 | 11 | 67171386 | 67177560 | 15 | 5 | 1.2309 | 0.10918 | TBC1D10C | 0.10918 | 0.33601 | 0.52524 |
| ENSG00000169393 | 19 | 48497908 | 48528410 | 221 | 21 | 1.2304 | 0.10927 | ELSPBP1 | 0.10927 | 0.10125 | 0.1625 |
| ENSG00000125817 | 20 | 3764498 | 3767337 | 2 | 1 | 1.2304 | 0.10927 | CENPB | 0.10927 | 0.73477 | 0.76427 |
| ENSG00000135766 | 1 | 231499497 | 231560790 | 365 | 9 | 1.23 | 0.10934 | EGLN1 | 0.10934 | 0.11551 | 0.15173 |
| ENSG00000089723 | 14 | 94492675 | 94515276 | 96 | 11 | 1.2292 | 0.1095 | OTUB2 | 0.1095 | 0.072081 | 0.061354 |
| ENSG00000051596 | 5 | 175344876 | 175461683 | 125 | 26 | 1.2292 | 0.10951 | THOC3 | 0.10951 | 0.16634 | 0.19864 |
| ENSG00000231925 | 6 | 33267471 | 33282164 | 51 | 10 | 1.2286 | 0.10961 | TAPBP | 0.10961 | 0.067039 | 0.033311 |
| ENSG00000242515 | 2 | 234545100 | 234681951 | 500 | 31 | 1.2285 | 0.10962 | UGT1A10 | 0.10962 | 0.13095 | 0.12769 |
| ENSG00000174444 | 15 | 66790355 | 66816870 | 87 | 16 | 1.228 | 0.10973 | RPL4 | 0.10973 | 0.19025 | 0.1698 |
| ENSG00000166938 | 15 | 66585555 | 66626236 | 145 | 25 | 1.2277 | 0.10978 | DIS3L | 0.10978 | 0.12824 | 0.17388 |
| ENSG00000106351 | 7 | 100136834 | 100165842 | 42 | 8 | 1.2274 | 0.10984 | AGFG2 | 0.10984 | 0.16311 | 0.18763 |
| ENSG00000171490 | 16 | 11929056 | 11945442 | 47 | 8 | 1.2271 | 0.10989 | RSL1D1 | 0.10989 | 0.0447 | 0.10656 |
| ENSG00000205838 | 5 | 34838938 | 34899561 | 189 | 32 | 1.2268 | 0.10995 | TTC23L | 0.10995 | 0.21831 | 0.3662 |
| ENSG00000196376 | 6 | 118228689 | 118638839 | 1207 | 72 | 1.2265 | 0.11001 | SLC35F1 | 0.11001 | 0.082825 | 0.073717 |
| ENSG00000261739 | 15 | 23599895 | 23612378 | 4 | 1 | 1.2264 | 0.11003 | GOLGA8S | 0.11003 | 0.11066 | 0.184 |

| | | | | | | | | | | | |
|---|---|---|---|---|---|---|---|---|---|---|---|
| ENSG00000232859 | 17 | 26205340 | 26221778 | 17 | 4 | 1.2256 | 0.11018 | LYRM9 | 0.11018 | 0.096525 | 0.14472 |
| ENSG00000123104 | 12 | 26490342 | 26986131 | 1814 | 100 | 1.2255 | 0.11019 | ITPR2 | 0.11019 | 0.10848 | 0.084426 |
| ENSG00000118596 | 12 | 59989848 | 60176395 | 555 | 20 | 1.2255 | 0.11019 | SLC16A7 | 0.11019 | 0.047995 | 0.022159 |
| ENSG00000135709 | 16 | 85061375 | 85127836 | 378 | 32 | 1.2252 | 0.11026 | KIAA0513 | 0.11026 | 0.11364 | 0.13498 |
| ENSG00000176358 | 17 | 47915671 | 47925379 | 11 | 6 | 1.2247 | 0.11034 | TAC4 | 0.11034 | 0.0052414 | 0.0066005 |
| ENSG00000185245 | 17 | 4835592 | 4838325 | 9 | 5 | 1.2247 | 0.11035 | GP1BA | 0.11035 | 0.16245 | 0.27192 |
| ENSG00000215218 | 5 | 6448736 | 6495022 | 184 | 38 | 1.2246 | 0.11037 | UBE2QL1 | 0.11037 | 0.05937 | 0.016989 |
| ENSG00000143320 | 1 | 156669398 | 156675608 | 13 | 5 | 1.2245 | 0.11039 | CRABP2 | 0.11039 | 0.065314 | 0.080098 |
| ENSG00000138623 | 15 | 74701630 | 74726808 | 65 | 14 | 1.2244 | 0.1104 | SEMA7A | 0.1104 | 0.15982 | 0.17896 |
| ENSG00000115486 | 2 | 85771846 | 85788670 | 65 | 13 | 1.2243 | 0.11043 | GGCX | 0.11043 | 0.064877 | 0.027662 |
| ENSG00000001617 | 3 | 50192478 | 50226508 | 65 | 11 | 1.2242 | 0.11045 | SEMA3F | 0.11045 | 0.15991 | 0.165 |
| ENSG00000149516 | 11 | 59824060 | 59838601 | 54 | 8 | 1.224 | 0.11048 | MS4A3 | 0.11048 | 0.064432 | 0.048136 |
| ENSG00000138495 | 3 | 119373360 | 119396301 | 110 | 10 | 1.223 | 0.11067 | COX17 | 0.11067 | 0.11313 | 0.10523 |
| ENSG00000063176 | 19 | 49122548 | 49133974 | 20 | 6 | 1.2229 | 0.11068 | SPHK2 | 0.11068 | 0.19974 | 0.030671 |
| ENSG00000226777 | 14 | 106383838 | 106391825 | 8 | 1 | 1.2224 | 0.11078 | KIAA0125 | 0.11078 | 0.17699 | 0.20365 |
| ENSG00000178814 | 8 | 145106167 | 145118735 | 40 | 7 | 1.2218 | 0.11089 | OPLAH | 0.11089 | 0.053435 | 0.10537 |
| ENSG00000169018 | 15 | 68570141 | 68588203 | 33 | 7 | 1.2217 | 0.11091 | FEM1B | 0.11091 | 0.0893 | 0.10314 |
| ENSG00000130332 | 19 | 2321516 | 2328619 | 37 | 8 | 1.2213 | 0.11099 | LSM7 | 0.11099 | 0.1389 | 0.15213 |
| ENSG00000125657 | 19 | 6531010 | 6535931 | 23 | 9 | 1.2208 | 0.11107 | TNFSF9 | 0.11107 | 0.21844 | 0.41385 |
| ENSG00000129245 | 17 | 7494548 | 7518189 | 54 | 13 | 1.2207 | 0.1111 | FXR2 | 0.1111 | 0.081771 | 0.01797 |
| ENSG00000145348 | 4 | 106962756 | 107242652 | 679 | 23 | 1.2202 | 0.1112 | TBCK | 0.1112 | 0.11443 | 0.1225 |
| ENSG00000268488 | 15 | 44075041 | 44075231 | 2 | 1 | 1.22 | 0.11123 | AC018512.1 | 0.11123 | 0.22662 | 0.20341 |
| ENSG00000181638 | 8 | 144328991 | 144344875 | 50 | 7 | 1.2195 | 0.11132 | ZFP41 | 0.11132 | 0.075832 | 0.073558 |
| ENSG00000106003 | 7 | 2552163 | 2568811 | 56 | 18 | 1.2194 | 0.11134 | LFNG | 0.11134 | 0.05477 | 0.067167 |
| ENSG00000100092 | 22 | 38030661 | 38062939 | 131 | 15 | 1.2192 | 0.11138 | SH3BP1 | 0.11138 | 0.14836 | 0.13396 |
| ENSG00000176349 | 7 | 1878222 | 1889567 | 80 | 9 | 1.219 | 0.11142 | AC110781.3 | 0.11142 | 0.092828 | 0.12169 |
| ENSG00000168634 | 20 | 44330655 | 44337456 | 21 | 7 | 1.2185 | 0.11152 | WFDC13 | 0.11152 | 0.028422 | 0.041724 |
| ENSG00000053524 | 3 | 182895831 | 183146566 | 843 | 59 | 1.2184 | 0.11153 | MCF2L2 | 0.11153 | 0.10181 | 0.10752 |
| ENSG00000049089 | 1 | 40766159 | 40783488 | 64 | 11 | 1.2182 | 0.11157 | COL9A2 | 0.11157 | 0.21405 | 0.2789 |
| ENSG00000065609 | 6 | 84262599 | 84419410 | 343 | 22 | 1.2178 | 0.11165 | SNAP91 | 0.11165 | 0.11399 | 0.11409 |
| ENSG00000107593 | 10 | 102047903 | 102090243 | 144 | 24 | 1.2169 | 0.11181 | PKD2L1 | 0.11181 | 0.12505 | 0.251 |
| ENSG00000111713 | 12 | 21689123 | 21757781 | 323 | 22 | 1.2169 | 0.11182 | GYS2 | 0.11182 | 0.11083 | 0.1323 |
| ENSG00000177181 | 1 | 42846466 | 42888249 | 184 | 11 | 1.2166 | 0.11188 | RIMKLA | 0.11188 | 0.11054 | 0.10213 |
| ENSG00000136068 | 3 | 57994127 | 58157982 | 465 | 35 | 1.2165 | 0.1119 | FLNB | 0.1119 | 0.076912 | 0.068003 |
| ENSG00000165462 | 11 | 71950121 | 71956708 | 10 | 4 | 1.2161 | 0.11197 | PHOX2A | 0.11197 | 0.14764 | 0.18418 |
| ENSG00000186007 | 1 | 205350506 | 205425082 | 272 | 31 | 1.216 | 0.112 | LEMD1 | 0.112 | 0.058743 | 0.036451 |
| ENSG00000148688 | 10 | 92631473 | 92668312 | 89 | 12 | 1.2147 | 0.11224 | RPP30 | 0.11224 | 0.086304 | 0.054453 |
| ENSG00000204682 | 10 | 21781587 | 21786191 | 9 | 4 | 1.2145 | 0.11228 | CASC10 | 0.11228 | 0.19638 | 0.16163 |
| ENSG00000173614 | 1 | 10003486 | 10045559 | 97 | 14 | 1.2144 | 0.11229 | NMNAT1 | 0.11229 | 0.14127 | 0.28222 |
| ENSG00000153575 | 15 | 22833395 | 22873892 | 149 | 15 | 1.2139 | 0.11238 | TUBGCP5 | 0.11238 | 0.089693 | 0.07231 |
| ENSG00000198039 | 7 | 64330550 | 64391344 | 291 | 12 | 1.2139 | 0.1124 | ZNF273 | 0.1124 | 0.086105 | 0.078709 |
| ENSG00000214324 | 3 | 126911974 | 126917025 | 11 | 5 | 1.2135 | 0.11246 | C3orf56 | 0.11246 | 0.16068 | 0.23606 |
| ENSG00000163518 | 1 | 157543539 | 157567870 | 72 | 12 | 1.2135 | 0.11247 | FCRL4 | 0.11247 | 0.14344 | 0.21646 |
| ENSG00000183963 | 22 | 31460091 | 31500743 | 170 | 21 | 1.2134 | 0.11248 | SMTN | 0.11248 | 0.10457 | 0.12446 |
| ENSG00000142611 | 1 | 2985732 | 3355185 | 1797 | 180 | 1.2133 | 0.1125 | PRDM16 | 0.1125 | 0.14748 | 0.21722 |
| ENSG00000187609 | 9 | 140201348 | 140317714 | 418 | 67 | 1.2133 | 0.11251 | EXD3 | 0.11251 | 0.18247 | 0.18899 |
| ENSG00000243477 | 3 | 50333833 | 50336852 | 3 | 2 | 1.2131 | 0.11254 | NAT6 | 0.11254 | 0.27825 | 0.2374 |
| ENSG00000135537 | 6 | 108616098 | 108847999 | 433 | 33 | 1.2131 | 0.11255 | LACE1 | 0.11255 | 0.11655 | 0.14039 |
| ENSG00000186226 | 1 | 152758690 | 152760902 | 17 | 5 | 1.2129 | 0.11258 | LCE1E | 0.11258 | 0.15614 | 0.14218 |
| ENSG00000186777 | 4 | 264464 | 299110 | 160 | 16 | 1.2127 | 0.11263 | ZNF732 | 0.11263 | 0.12124 | 0.11123 |
| ENSG00000189233 | 8 | 27879481 | 27941388 | 156 | 30 | 1.2125 | 0.11266 | NUGGC | 0.11266 | 0.084234 | 0.061032 |
| ENSG00000148229 | 9 | 116169515 | 116172952 | 7 | 2 | 1.2124 | 0.11268 | POLE3 | 0.11268 | 0.10982 | 0.11106 |
| ENSG00000133641 | 12 | 88427623 | 88443937 | 15 | 5 | 1.2123 | 0.11269 | C12orf29 | 0.11269 | 0.13579 | 0.35895 |
| ENSG00000103710 | 15 | 65345679 | 65369028 | 55 | 9 | 1.2122 | 0.11271 | RASL12 | 0.11271 | 0.16673 | 0.29542 |
| ENSG00000198743 | 21 | 35445870 | 35478559 | 45 | 16 | 1.2116 | 0.11284 | SLC5A3 | 0.11284 | 0.12298 | 0.12673 |
| ENSG00000272962 | 21 | 35445870 | 35478561 | 45 | 16 | 1.2116 | 0.11284 | SLC5A3 | 0.11284 | 0.12298 | 0.12673 |
| ENSG00000149260 | 11 | 76777979 | 76837201 | 186 | 34 | 1.2114 | 0.11287 | CAPN5 | 0.11287 | 0.26599 | 0.4649 |
| ENSG00000115289 | 2 | 74732170 | 74735707 | 2 | 1 | 1.2109 | 0.11297 | PCGF1 | 0.11297 | 0.45886 | 0.33822 |
| ENSG00000078725 | 9 | 121915736 | 122131745 | 586 | 49 | 1.2108 | 0.11299 | BRINP1 | 0.11299 | 0.14616 | 0.19075 |
| ENSG00000268367 | 12 | 57810198 | 57810536 | 1 | 1 | 1.2102 | 0.1131 | AC126614.1 | 0.1131 | 0.1685 | 0.10365 |
| ENSG00000162910 | 1 | 228294380 | 228297013 | 4 | 2 | 1.2102 | 0.11311 | MRPL55 | 0.11311 | 0.67576 | 0.35674 |
| ENSG00000155463 | 14 | 23235731 | 23241007 | 14 | 5 | 1.2097 | 0.1132 | OXA1L | 0.1132 | 0.070188 | 0.056557 |
| ENSG00000188474 | 19 | 12203078 | 12225491 | 49 | 8 | 1.2096 | 0.11322 | ZNF788 | 0.11322 | 0.11113 | 0.16094 |
| ENSG00000174238 | 17 | 1421012 | 1466110 | 170 | 17 | 1.209 | 0.11334 | PITPNA | 0.11334 | 0.094991 | 0.11009 |
| ENSG00000109103 | 17 | 26873725 | 26879686 | 2 | 1 | 1.2087 | 0.11339 | UNC119 | 0.11339 | 0.030045 | 0.45079 |
| ENSG00000150471 | 4 | 62066976 | 62944053 | 2413 | 136 | 1.2079 | 0.11355 | LPHN3 | 0.11355 | 0.12123 | 0.10652 |
| ENSG00000162728 | 1 | 160051360 | 160060353 | 23 | 6 | 1.2074 | 0.11364 | KCNJ9 | 0.11364 | 0.037788 | 0.066594 |
| ENSG00000114861 | 3 | 71003844 | 71633140 | 1575 | 136 | 1.2073 | 0.11365 | FOXP1 | 0.11365 | 0.10234 | 0.078671 |
| ENSG00000160224 | 21 | 45705721 | 45718531 | 34 | 15 | 1.2072 | 0.11367 | AIRE | 0.11367 | 0.33158 | 0.33735 |
| ENSG00000115425 | 2 | 216861052 | 216947678 | 341 | 34 | 1.2069 | 0.11374 | PECR | 0.11374 | 0.098683 | 0.13364 |
| ENSG00000170385 | 1 | 211744910 | 211752084 | 16 | 5 | 1.2058 | 0.11394 | SLC30A1 | 0.11394 | 0.056534 | 0.054136 |
| ENSG00000104177 | 15 | 48431625 | 48470714 | 7 | 4 | 1.2058 | 0.11395 | MYEF2 | 0.11395 | 0.040085 | 0.094762 |
| ENSG00000135378 | 11 | 32851489 | 32879669 | 102 | 15 | 1.2056 | 0.11399 | PRRG4 | 0.11399 | 0.10782 | 0.15579 |

| | | | | | | | | | | | |
|---|---|---|---|---|---|---|---|---|---|---|---|
| ENSG00000142303 | 19 | 8645126 | 8675620 | 94 | 28 | 1.2053 | 0.11405 | ADAMTS10 | 0.11405 | 0.072695 | 0.04758 |
| ENSG00000180329 | 17 | 42750437 | 42767147 | 16 | 7 | 1.2045 | 0.1142 | CCDC43 | 0.1142 | 0.25028 | 0.4088 |
| ENSG00000144591 | 2 | 220363589 | 220371710 | 11 | 5 | 1.2045 | 0.11421 | GMPPA | 0.11421 | 0.37713 | 0.34994 |
| ENSG00000171522 | 5 | 40679600 | 40693837 | 23 | 6 | 1.2039 | 0.11431 | PTGER4 | 0.11431 | 0.27131 | 0.26 |
| ENSG00000053702 | 12 | 2934514 | 2944710 | 33 | 6 | 1.2038 | 0.11434 | NRIP2 | 0.11434 | 0.14938 | 0.16103 |
| ENSG00000139410 | 12 | 113860042 | 113876081 | 44 | 5 | 1.2034 | 0.1144 | SDSL | 0.1144 | 0.16525 | 0.23597 |
| ENSG00000156395 | 10 | 106400859 | 107024993 | 2107 | 54 | 1.2034 | 0.11441 | SORCS3 | 0.11441 | 0.096625 | 0.10723 |
| ENSG00000232423 | 1 | 12998181 | 13117751 | 12 | 2 | 1.2031 | 0.11446 | PRAMEF6 | 0.11446 | 0.11302 | 0.25394 |
| ENSG00000181722 | 3 | 114056941 | 114866118 | 1683 | 115 | 1.2031 | 0.11447 | ZBTB20 | 0.11447 | 0.091421 | 0.069892 |
| ENSG00000180071 | 9 | 38540566 | 38620657 | 318 | 50 | 1.203 | 0.11449 | ANKRD18A | 0.11449 | 0.11195 | 0.21149 |
| ENSG00000179331 | 11 | 107799229 | 107834208 | 131 | 22 | 1.203 | 0.11449 | RAB39A | 0.11449 | 0.1389 | 0.11391 |
| ENSG00000151116 | 11 | 18551156 | 18610294 | 216 | 25 | 1.2028 | 0.11452 | UEVLD | 0.11452 | 0.092431 | 0.11359 |
| ENSG00000268069 | 12 | 48178706 | 48179787 | 10 | 3 | 1.2026 | 0.11456 | AC004466.1 | 0.11456 | 0.43923 | 0.34808 |
| ENSG00000141437 | 18 | 29339525 | 29340843 | 2 | 1 | 1.2025 | 0.11458 | SLC25A52 | 0.11458 | 0.63972 | 0.47859 |
| ENSG00000100362 | 22 | 37196728 | 37215523 | 84 | 15 | 1.2024 | 0.1146 | PVALB | 0.1146 | 0.14469 | 0.17473 |
| ENSG00000140025 | 14 | 90261013 | 90421121 | 530 | 36 | 1.2021 | 0.11467 | EFCAB11 | 0.11467 | 0.08548 | 0.046359 |
| ENSG00000039987 | 19 | 12862516 | 12869272 | 7 | 5 | 1.202 | 0.11467 | BEST2 | 0.11467 | 0.03673 | 0.072055 |
| ENSG00000152137 | 12 | 119616447 | 119658936 | 130 | 27 | 1.2015 | 0.11478 | HSPB8 | 0.11478 | 0.15647 | 0.097542 |
| ENSG00000214967 | 16 | 16472912 | 16487811 | 1 | 1 | 1.2014 | 0.1148 | NPIPA7 | 0.1148 | 0.073767 | 0.28034 |
| ENSG00000115325 | 2 | 74776153 | 74784681 | 9 | 3 | 1.2013 | 0.11483 | DOK1 | 0.11483 | 0.24209 | 0.24952 |
| ENSG00000163945 | 4 | 1341054 | 1381837 | 194 | 21 | 1.2004 | 0.115 | UVSSA | 0.115 | 0.055745 | 0.065385 |
| ENSG00000085465 | 1 | 111956936 | 111970399 | 53 | 10 | 1.1997 | 0.11512 | OVGP1 | 0.11512 | 0.13028 | 0.097669 |
| ENSG00000102763 | 13 | 42140973 | 42535256 | 1204 | 52 | 1.1994 | 0.11519 | VWA8 | 0.11519 | 0.11294 | 0.11362 |
| ENSG00000145794 | 5 | 126626523 | 126801429 | 632 | 29 | 1.1994 | 0.1152 | MEGF10 | 0.1152 | 0.12666 | 0.13729 |
| ENSG00000131791 | 1 | 146626685 | 146644129 | 53 | 9 | 1.1991 | 0.11524 | PRKAB2 | 0.11524 | 0.17372 | 0.21143 |
| ENSG00000237521 | 19 | 9361606 | 9362820 | 6 | 3 | 1.199 | 0.11526 | OR7E24 | 0.11526 | 0.41148 | 0.46061 |
| ENSG00000159388 | 1 | 203274619 | 203278730 | 13 | 6 | 1.1985 | 0.11536 | BTG2 | 0.11536 | 0.31736 | 0.60202 |
| ENSG00000185201 | 11 | 307631 | 315272 | 37 | 10 | 1.1982 | 0.11542 | IFITM2 | 0.11542 | 0.44569 | 0.58 |
| ENSG00000203786 | 1 | 152730506 | 152734529 | 9 | 3 | 1.1978 | 0.11549 | KPRP | 0.11549 | 0.099753 | 0.11558 |
| ENSG00000185313 | 3 | 38738293 | 38835501 | 365 | 33 | 1.1979 | 0.11549 | SCN10A | 0.11549 | 0.14078 | 0.19519 |
| ENSG00000159648 | 16 | 58010339 | 58022020 | 39 | 7 | 1.1975 | 0.11555 | TEPP | 0.11555 | 0.10186 | 0.17236 |
| ENSG00000213171 | 1 | 151772740 | 151778630 | 9 | 3 | 1.1973 | 0.11559 | LINGO4 | 0.11559 | 0.15581 | 0.22002 |
| ENSG00000100814 | 14 | 20779527 | 20801471 | 79 | 13 | 1.1971 | 0.11564 | CCNB1IP1 | 0.11564 | 0.10612 | 0.069901 |
| ENSG00000166333 | 11 | 6624961 | 6632102 | 34 | 5 | 1.1969 | 0.11568 | ILK | 0.11568 | 0.096002 | 0.065349 |
| ENSG00000141052 | 17 | 12569207 | 12672266 | 378 | 47 | 1.1956 | 0.11592 | MYOCD | 0.11592 | 0.087617 | 0.074719 |
| ENSG00000125965 | 20 | 34021145 | 34042568 | 40 | 9 | 1.1955 | 0.11596 | GDF5 | 0.11596 | 0.17409 | 0.1966 |
| ENSG00000101166 | 20 | 57608200 | 57617964 | 19 | 7 | 1.195 | 0.11605 | SLMO2 | 0.11605 | 0.084388 | 0.060499 |
| ENSG00000169914 | 1 | 20209006 | 20239438 | 91 | 15 | 1.1948 | 0.11608 | OTUD3 | 0.11608 | 0.15654 | 0.14975 |
| ENSG00000268942 | 5 | 61808070 | 61808309 | 1 | 1 | 1.1947 | 0.1161 | CKS1B | 0.1161 | 0.44668 | 0.61149 |
| ENSG00000179941 | 12 | 76738254 | 76742222 | 5 | 3 | 1.1947 | 0.11611 | BBS10 | 0.11611 | 0.51631 | 0.36738 |
| ENSG00000162188 | 11 | 62475130 | 62476673 | 3 | 2 | 1.1944 | 0.11616 | GNG3 | 0.11616 | 0.3157 | 0.40228 |
| ENSG00000162869 | 2 | 48667737 | 48742525 | 270 | 23 | 1.1935 | 0.11633 | PPP1R21 | 0.11633 | 0.12388 | 0.11626 |
| ENSG00000185105 | 17 | 79897521 | 79905109 | 29 | 4 | 1.1935 | 0.11634 | MYADML2 | 0.11634 | 0.04476 | 0.055767 |
| ENSG00000093072 | 22 | 17660194 | 17702879 | 247 | 51 | 1.193 | 0.11644 | CECR1 | 0.11644 | 0.17531 | 0.23402 |
| ENSG00000071462 | 7 | 73097355 | 73119491 | 72 | 10 | 1.1927 | 0.11649 | WBSCR22 | 0.11649 | 0.11746 | 0.17524 |
| ENSG00000104371 | 8 | 42231586 | 42234750 | 9 | 3 | 1.1926 | 0.11651 | DKK4 | 0.11651 | 0.19581 | 0.26296 |
| ENSG00000125875 | 20 | 416124 | 443197 | 90 | 12 | 1.1923 | 0.11658 | TBC1D20 | 0.11658 | 0.23847 | 0.30543 |
| ENSG00000173276 | 21 | 43406940 | 43430496 | 67 | 12 | 1.1914 | 0.11675 | ZBTB21 | 0.11675 | 0.093597 | 0.047127 |
| ENSG00000171772 | 10 | 135367404 | 135382876 | 136 | 5 | 1.191 | 0.11682 | SYCE1 | 0.11682 | 0.11332 | 0.1127 |
| ENSG00000137267 | 6 | 3153903 | 3157760 | 6 | 2 | 1.191 | 0.11683 | TUBB2A | 0.11683 | 0.37425 | 0.43921 |
| ENSG00000206181 | 18 | 44558943 | 44561988 | 16 | 6 | 1.1905 | 0.11692 | TCEB3B | 0.11692 | 0.13825 | 0.083626 |
| ENSG00000260300 | 16 | 83941738 | 83985058 | 307 | 35 | 1.1904 | 0.11694 | RP11-505K9.4 | 0.11694 | 0.082776 | 0.098971 |
| ENSG00000135316 | 6 | 86318053 | 86353510 | 69 | 15 | 1.19 | 0.11701 | SYNCRIP | 0.11701 | 0.11598 | 0.14671 |
| ENSG00000143140 | 1 | 147228332 | 147245484 | 54 | 10 | 1.1899 | 0.11703 | GJA5 | 0.11703 | 0.13159 | 0.19397 |
| ENSG00000164039 | 4 | 104000592 | 104021040 | 54 | 7 | 1.1899 | 0.11705 | BDH2 | 0.11705 | 0.12566 | 0.12527 |
| ENSG00000182004 | 1 | 203830731 | 203839678 | 44 | 6 | 1.1897 | 0.11707 | SNRPE | 0.11707 | 0.10764 | 0.10159 |
| ENSG00000145757 | 5 | 94987885 | 95034415 | 148 | 18 | 1.1897 | 0.11708 | SPATA9 | 0.11708 | 0.12218 | 0.078916 |
| ENSG00000100241 | 22 | 50883429 | 50913454 | 113 | 24 | 1.1896 | 0.11709 | SBF1 | 0.11709 | 0.052777 | 0.26166 |
| ENSG00000137331 | 6 | 30710976 | 30712331 | 6 | 2 | 1.1895 | 0.11712 | IER3 | 0.11712 | 0.24895 | 0.42554 |
| ENSG00000105261 | 19 | 36602104 | 36604613 | 6 | 2 | 1.1894 | 0.11714 | OVOL3 | 0.11714 | 0.35106 | 0.23962 |
| ENSG00000176410 | 7 | 73095299 | 73097783 | 14 | 4 | 1.1892 | 0.11718 | DNAJC30 | 0.11718 | 0.13653 | 0.16237 |
| ENSG00000227500 | 19 | 1905213 | 1926016 | 94 | 17 | 1.189 | 0.11723 | SCAMP4 | 0.11723 | 0.22557 | 0.47895 |
| ENSG00000165171 | 7 | 73248920 | 73256865 | 10 | 5 | 1.1887 | 0.11727 | WBSCR27 | 0.11727 | 0.036799 | 0.040324 |
| ENSG00000131504 | 5 | 140894583 | 140998622 | 206 | 30 | 1.1877 | 0.11747 | DIAPH1 | 0.11747 | 0.10057 | 0.096755 |
| ENSG00000185947 | 16 | 31885079 | 31928668 | 75 | 8 | 1.1873 | 0.11755 | ZNF267 | 0.11755 | 0.08419 | 0.068444 |
| ENSG00000146276 | 6 | 89887220 | 89940997 | 262 | 25 | 1.1868 | 0.11765 | GABRR1 | 0.11765 | 0.17669 | 0.28567 |
| ENSG00000023516 | 13 | 42846289 | 42897396 | 154 | 19 | 1.1868 | 0.11766 | AKAP11 | 0.11766 | 0.13316 | 0.16436 |
| ENSG00000073536 | 17 | 33455772 | 33469334 | 13 | 6 | 1.1865 | 0.11771 | NLE1 | 0.11771 | 0.36079 | 0.50738 |
| ENSG00000215252 | 15 | 34817408 | 34880704 | 179 | 21 | 1.1859 | 0.11783 | GOLGA8B | 0.11783 | 0.11174 | 0.12079 |
| ENSG00000106105 | 7 | 30634297 | 30673649 | 113 | 16 | 1.1854 | 0.11792 | GARS | 0.11792 | 0.094025 | 0.082038 |
| ENSG00000000460 | 1 | 169631245 | 169823221 | 981 | 35 | 1.1852 | 0.11797 | C1orf112 | 0.11797 | 0.12849 | 0.13529 |
| ENSG00000218891 | 19 | 56088899 | 56092211 | 11 | 3 | 1.185 | 0.118 | ZNF579 | 0.118 | 0.11106 | 0.0082052 |
| ENSG00000180785 | 11 | 4664650 | 4676718 | 59 | 10 | 1.1843 | 0.11814 | OR51E1 | 0.11814 | 0.20881 | 0.27723 |

| | | | | | | | | | | | |
|---|---|---|---|---|---|---|---|---|---|---|---|
| ENSG00000135119 | 12 | 117176096 | 117291436 | 392 | 33 | 1.1838 | 0.11824 | RNFT2 | 0.11824 | 0.074782 | 0.038468 |
| ENSG00000104313 | 8 | 72109668 | 72274467 | 430 | 59 | 1.1834 | 0.11833 | EYA1 | 0.11833 | 0.072948 | 0.1016 |
| ENSG00000115524 | 2 | 198254508 | 198299815 | 63 | 11 | 1.1824 | 0.11852 | SF3B1 | 0.11852 | 0.10834 | 0.13027 |
| ENSG00000167083 | 17 | 47280153 | 47287936 | 27 | 8 | 1.1824 | 0.11852 | GNGT2 | 0.11852 | 0.087949 | 0.14203 |
| ENSG00000106853 | 9 | 114312002 | 114362135 | 218 | 38 | 1.1823 | 0.11855 | PTGR1 | 0.11855 | 0.13869 | 0.1902 |
| ENSG00000205445 | 21 | 45970240 | 45971388 | 10 | 2 | 1.1823 | 0.11855 | KRTAP10-2 | 0.11855 | 0.11499 | 0.2017 |
| ENSG00000162129 | 11 | 72003469 | 72145692 | 249 | 18 | 1.182 | 0.1186 | CLPB | 0.1186 | 0.12544 | 0.13105 |
| ENSG00000074319 | 11 | 18489883 | 18548779 | 197 | 18 | 1.1819 | 0.11863 | TSG101 | 0.11863 | 0.17181 | 0.2155 |
| ENSG00000134200 | 1 | 115572415 | 115576941 | 25 | 4 | 1.1817 | 0.11867 | TSHB | 0.11867 | 0.087356 | 0.11803 |
| ENSG00000188305 | 19 | 2274631 | 2282181 | 48 | 9 | 1.1815 | 0.1187 | C19orf35 | 0.1187 | 0.26048 | 0.59024 |
| ENSG00000189091 | 16 | 70557691 | 70608820 | 105 | 10 | 1.1814 | 0.11872 | SF3B3 | 0.11872 | 0.12268 | 0.17196 |
| ENSG00000152086 | 2 | 130949318 | 130956034 | 15 | 2 | 1.1813 | 0.11874 | TUBA3E | 0.11874 | 0.12064 | 0.13045 |
| ENSG00000161914 | 19 | 11594242 | 11616738 | 76 | 7 | 1.1811 | 0.11877 | ZNF653 | 0.11877 | 0.090946 | 0.13285 |
| ENSG00000198939 | 5 | 178322895 | 178360213 | 196 | 18 | 1.1806 | 0.11887 | ZFP2 | 0.11887 | 0.077434 | 0.071449 |
| ENSG00000112695 | 6 | 75947391 | 75960039 | 33 | 6 | 1.1802 | 0.11897 | COX7A2 | 0.11897 | 0.11884 | 0.11163 |
| ENSG00000196666 | 11 | 47608198 | 47610746 | 4 | 2 | 1.1798 | 0.11905 | FAM180B | 0.11905 | 0.53776 | 0.672 |
| ENSG00000007933 | 1 | 171060018 | 171086959 | 139 | 17 | 1.1796 | 0.11907 | FMO3 | 0.11907 | 0.13028 | 0.18871 |
| ENSG00000127515 | 19 | 14951760 | 14952689 | 5 | 2 | 1.1796 | 0.11907 | OR7A10 | 0.11907 | 0.052118 | 0.14147 |
| ENSG00000267987 | 9 | 124103625 | 124110410 | 14 | 6 | 1.1796 | 0.11908 | AL161784.1 | 0.11908 | 0.27286 | 0.21828 |
| ENSG00000076382 | 17 | 26904588 | 26926297 | 15 | 9 | 1.1792 | 0.11916 | SPAG5 | 0.11916 | 0.17801 | 0.084487 |
| ENSG00000118496 | 6 | 146114638 | 146135889 | 44 | 8 | 1.1792 | 0.11917 | FBXO30 | 0.11917 | 0.13424 | 0.23414 |
| ENSG00000154217 | 17 | 65373575 | 65693372 | 1123 | 133 | 1.179 | 0.11919 | PITPNC1 | 0.11919 | 0.12096 | 0.079412 |
| ENSG00000152670 | 5 | 55033845 | 55112985 | 262 | 23 | 1.1789 | 0.11922 | DDX4 | 0.11922 | 0.096323 | 0.080182 |
| ENSG00000183662 | 3 | 68053359 | 68594776 | 2118 | 92 | 1.1788 | 0.11923 | FAM19A1 | 0.11923 | 0.1305 | 0.15898 |
| ENSG00000165606 | 10 | 50572237 | 50603497 | 60 | 7 | 1.1785 | 0.1193 | DRGX | 0.1193 | 0.18197 | 0.20122 |
| ENSG00000143278 | 1 | 197008321 | 197036397 | 61 | 7 | 1.1784 | 0.11932 | F13B | 0.11932 | 0.18907 | 0.22062 |
| ENSG00000250424 | 7 | 30893010 | 30963427 | 205 | 46 | 1.1781 | 0.11939 | AQP1 | 0.11939 | 0.15354 | 0.22896 |
| ENSG00000151576 | 3 | 113724680 | 113807269 | 156 | 20 | 1.1779 | 0.11942 | QTRTD1 | 0.11942 | 0.11316 | 0.084766 |
| ENSG00000128203 | 22 | 26825239 | 26840981 | 71 | 14 | 1.1777 | 0.11946 | ASPHD2 | 0.11946 | 0.025116 | 0.0088405 |
| ENSG00000205857 | 12 | 7917812 | 7926717 | 16 | 6 | 1.1776 | 0.11949 | NANOGNB | 0.11949 | 0.066961 | 0.13053 |
| ENSG00000163513 | 3 | 30647994 | 30735634 | 325 | 45 | 1.1773 | 0.11954 | TGFBR2 | 0.11954 | 0.072675 | 0.11531 |
| ENSG00000164283 | 5 | 54273692 | 54318499 | 165 | 21 | 1.1769 | 0.11961 | ESM1 | 0.11961 | 0.10666 | 0.10581 |
| ENSG00000075391 | 1 | 178062864 | 178448644 | 715 | 24 | 1.1767 | 0.11966 | RASAL2 | 0.11966 | 0.088348 | 0.075003 |
| ENSG00000162944 | 2 | 198432948 | 198540769 | 231 | 16 | 1.1761 | 0.11978 | RFTN2 | 0.11978 | 0.11868 | 0.12134 |
| ENSG00000102802 | 13 | 31480328 | 31499709 | 64 | 16 | 1.1759 | 0.11983 | MEDAG | 0.11983 | 0.23748 | 0.29457 |
| ENSG00000131043 | 20 | 34824381 | 34858840 | 49 | 16 | 1.1754 | 0.11991 | AAR2 | 0.11991 | 0.14245 | 0.29323 |
| ENSG00000148110 | 9 | 97136833 | 97223324 | 351 | 19 | 1.1753 | 0.11993 | HIATL1 | 0.11993 | 0.10266 | 0.092479 |
| ENSG00000117595 | 1 | 209959036 | 209979465 | 65 | 8 | 1.1752 | 0.11996 | IRF6 | 0.11996 | 0.21841 | 0.16716 |
| ENSG00000203942 | 10 | 99349450 | 99350690 | 3 | 1 | 1.1752 | 0.11996 | C10orf62 | 0.11996 | 0.40486 | 0.71542 |
| ENSG00000149257 | 11 | 75273101 | 75283828 | 39 | 5 | 1.175 | 0.11999 | SERPINH1 | 0.11999 | 0.2013 | 0.62144 |
| ENSG00000110435 | 11 | 34937376 | 35042138 | 513 | 18 | 1.1744 | 0.12012 | PDHX | 0.12012 | 0.077794 | 0.064538 |
| ENSG00000254521 | 19 | 51994611 | 52005043 | 57 | 16 | 1.1744 | 0.12012 | SIGLEC12 | 0.12012 | 0.099974 | 0.12972 |
| ENSG00000212743 | 10 | 6392278 | 6394723 | 5 | 2 | 1.1741 | 0.12018 | DKFZP667F0711 | 0.12018 | 0.4634 | 0.15134 |
| ENSG00000159455 | 1 | 152647771 | 152659877 | 39 | 6 | 1.1739 | 0.12021 | LCE2B | 0.12021 | 0.12907 | 0.1537 |
| ENSG00000144488 | 2 | 239008798 | 239041928 | 206 | 25 | 1.1737 | 0.12025 | ESPNL | 0.12025 | 0.093954 | 0.10324 |
| ENSG00000166261 | 11 | 123594885 | 123612383 | 69 | 7 | 1.1722 | 0.12056 | ZNF202 | 0.12056 | 0.064485 | 0.059262 |
| ENSG00000105251 | 19 | 4278598 | 4290721 | 70 | 12 | 1.1717 | 0.12066 | SHD | 0.12066 | 0.096485 | 0.16018 |
| ENSG00000173757 | 17 | 40351186 | 40428725 | 128 | 13 | 1.1711 | 0.12079 | STAT5B | 0.12079 | 0.12801 | 0.13834 |
| ENSG00000111341 | 12 | 15034115 | 15038860 | 8 | 1 | 1.1708 | 0.12085 | MGP | 0.12085 | 0.10794 | 0.095065 |
| ENSG00000130311 | 19 | 17420327 | 17434110 | 81 | 13 | 1.1707 | 0.12086 | DDA1 | 0.12086 | 0.23239 | 0.19687 |
| ENSG00000137692 | 11 | 102932805 | 102962944 | 52 | 8 | 1.1704 | 0.12091 | DCUN1D5 | 0.12091 | 0.1094 | 0.049872 |
| ENSG00000137168 | 6 | 36822603 | 36842800 | 75 | 14 | 1.1703 | 0.12095 | PPIL1 | 0.12095 | 0.16162 | 0.04702 |
| ENSG00000268241 | 2 | 175200601 | 175202151 | 2 | 1 | 1.1701 | 0.12098 | AC018470.1 | 0.12098 | 0.038105 | 0.059129 |
| ENSG00000182809 | 14 | 105939299 | 105946499 | 4 | 2 | 1.1699 | 0.12101 | CRIP2 | 0.12101 | 0.14245 | 0.326 |
| ENSG00000259159 | 11 | 119209652 | 119217368 | 20 | 10 | 1.1695 | 0.1211 | MFRP | 0.1211 | 0.0050927 | 0.0045278 |
| ENSG00000235718 | 11 | 119209652 | 119217383 | 20 | 10 | 1.1695 | 0.1211 | MFRP | 0.1211 | 0.0050927 | 0.0045278 |
| ENSG00000223953 | 11 | 119209652 | 119217383 | 20 | 10 | 1.1695 | 0.1211 | C1QTNF5 | 0.1211 | 0.0050927 | 0.0045278 |
| ENSG00000181803 | 14 | 21108855 | 21109850 | 8 | 2 | 1.1692 | 0.12117 | OR6S1 | 0.12117 | 0.28979 | 0.34989 |
| ENSG00000148677 | 10 | 92671853 | 92681033 | 21 | 7 | 1.169 | 0.12119 | ANKRD1 | 0.12119 | 0.097122 | 0.069308 |
| ENSG00000143761 | 1 | 228270361 | 228286912 | 27 | 7 | 1.169 | 0.1212 | ARF1 | 0.1212 | 0.16762 | 0.35765 |
| ENSG00000116824 | 1 | 117297007 | 117311850 | 38 | 9 | 1.1687 | 0.12126 | CD2 | 0.12126 | 0.038568 | 0.0084962 |
| ENSG00000180228 | 2 | 179296141 | 179316239 | 38 | 11 | 1.1686 | 0.12127 | PRKRA | 0.12127 | 0.039358 | 0.012681 |
| ENSG00000248167 | 6 | 30297359 | 30314631 | 60 | 16 | 1.1686 | 0.12128 | TRIM39-RPP21 | 0.12128 | 0.16175 | 0.2477 |
| ENSG00000120949 | 1 | 12123434 | 12204264 | 216 | 42 | 1.1684 | 0.12132 | TNFRSF8 | 0.12132 | 0.022378 | 0.025982 |
| ENSG00000181788 | 3 | 150458914 | 150481264 | 59 | 5 | 1.1684 | 0.12133 | SIAH2 | 0.12133 | 0.13147 | 0.06637 |
| ENSG00000086504 | 16 | 417384 | 420527 | 23 | 8 | 1.1683 | 0.12134 | MRPL28 | 0.12134 | 0.23971 | 0.2608 |
| ENSG00000222014 | 2 | 130737235 | 130740311 | 13 | 3 | 1.168 | 0.12141 | RAB6C | 0.12141 | 0.084477 | 0.11642 |
| ENSG00000173848 | 10 | 5454514 | 5500426 | 129 | 11 | 1.1671 | 0.12159 | NET1 | 0.12159 | 0.088857 | 0.069091 |
| ENSG00000168010 | 11 | 72525353 | 72554719 | 35 | 14 | 1.1671 | 0.12159 | ATG16L2 | 0.12159 | 0.23734 | 0.32795 |
| ENSG00000090432 | 1 | 20825943 | 20834654 | 23 | 8 | 1.1664 | 0.12172 | MUL1 | 0.12172 | 0.147 | 0.14341 |
| ENSG00000053328 | 6 | 110567131 | 110679475 | 345 | 26 | 1.166 | 0.1218 | METTL24 | 0.1218 | 0.08517 | 0.10637 |
| ENSG00000185504 | 17 | 79506911 | 79520987 | 53 | 10 | 1.1658 | 0.12185 | C17orf70 | 0.12185 | 0.3528 | 0.65788 |
| ENSG00000122035 | 13 | 27844464 | 27847827 | 6 | 2 | 1.1657 | 0.12186 | RASL11A | 0.12186 | 0.36169 | 0.36903 |

| | | | | | | | | | | | |
|---|---|---|---|---|---|---|---|---|---|---|---|
| ENSG00000162753 | 1 | 173469603 | 173572233 | 250 | 13 | 1.1657 | 0.12187 | SLC9C2 | 0.12187 | 0.14331 | 0.17135 |
| ENSG00000197619 | 19 | 52494585 | 52511483 | 78 | 7 | 1.1654 | 0.12193 | ZNF615 | 0.12193 | 0.11342 | 0.097689 |
| ENSG00000162391 | 1 | 55074855 | 55089229 | 57 | 17 | 1.1652 | 0.12198 | FAM151A | 0.12198 | 0.38877 | 0.25789 |
| ENSG00000129214 | 17 | 7517382 | 7536700 | 58 | 15 | 1.1652 | 0.12198 | SHBG | 0.12198 | 0.016383 | 0.018206 |
| ENSG00000176547 | 11 | 48346472 | 48347507 | 2 | 1 | 1.1646 | 0.12209 | OR4C3 | 0.12209 | 0.070162 | 0.057319 |
| ENSG00000242173 | 16 | 318726 | 333003 | 71 | 23 | 1.1642 | 0.12218 | ARHGDIG | 0.12218 | 0.25955 | 0.37437 |
| ENSG00000175857 | 5 | 57787262 | 57792917 | 11 | 5 | 1.1641 | 0.12219 | GAPT | 0.12219 | 0.25256 | 0.29729 |
| ENSG00000074855 | 19 | 17434032 | 17445638 | 35 | 9 | 1.1637 | 0.12228 | ANO8 | 0.12228 | 0.11459 | 0.22003 |
| ENSG00000146678 | 7 | 45927956 | 45933267 | 16 | 5 | 1.1634 | 0.12234 | IGFBP1 | 0.12234 | 0.19619 | 0.35024 |
| ENSG00000101306 | 20 | 30407111 | 30422492 | 43 | 7 | 1.163 | 0.12241 | MYLK2 | 0.12241 | 0.11852 | 0.084273 |
| ENSG00000149124 | 11 | 58407899 | 58499447 | 402 | 30 | 1.1629 | 0.12244 | GLYAT | 0.12244 | 0.096651 | 0.16336 |
| ENSG00000212901 | 17 | 39164773 | 39175576 | 30 | 4 | 1.1614 | 0.12273 | KRTAP3-1 | 0.12273 | 0.32165 | 0.45696 |
| ENSG00000182208 | 11 | 1490687 | 1522477 | 106 | 11 | 1.16 | 0.12302 | MOB2 | 0.12302 | 0.1129 | 0.054285 |
| ENSG00000037637 | 1 | 16573334 | 16678949 | 286 | 13 | 1.1595 | 0.12312 | FBXO42 | 0.12312 | 0.12335 | 0.1309 |
| ENSG00000164438 | 5 | 170736288 | 170739138 | 2 | 1 | 1.1586 | 0.12331 | TLX3 | 0.12331 | 0.10155 | 0.075093 |
| ENSG00000121057 | 17 | 55162453 | 55198710 | 133 | 25 | 1.1585 | 0.12332 | AKAP1 | 0.12332 | 0.20055 | 0.33264 |
| ENSG00000258555 | 22 | 24666866 | 24838324 | 419 | 26 | 1.1585 | 0.12333 | SPECC1L-ADORA2A | 0.12333 | 0.11474 | 0.098845 |
| ENSG00000109132 | 4 | 41746099 | 41750987 | 13 | 4 | 1.1583 | 0.12338 | PHOX2B | 0.12338 | 0.0061709 | 0.035817 |
| ENSG00000067113 | 5 | 54720682 | 54830878 | 359 | 13 | 1.1577 | 0.12349 | PPAP2A | 0.12349 | 0.13008 | 0.12718 |
| ENSG00000134250 | 1 | 120454176 | 120612240 | 187 | 15 | 1.1572 | 0.1236 | NOTCH2 | 0.1236 | 0.13382 | 0.14246 |
| ENSG00000241360 | 22 | 38054734 | 38062941 | 24 | 4 | 1.1568 | 0.12367 | PDXP | 0.12367 | 0.15523 | 0.14312 |
| ENSG00000124578 | 6 | 26246886 | 26247259 | 1 | 1 | 1.1567 | 0.1237 | HIST1H4G | 0.1237 | 0.40308 | 0.33089 |
| ENSG00000121897 | 4 | 39460620 | 39479273 | 86 | 14 | 1.1561 | 0.12383 | LIAS | 0.12383 | 0.17467 | 0.18723 |
| ENSG00000156096 | 4 | 70345883 | 70391732 | 239 | 13 | 1.1555 | 0.12395 | UGT2B4 | 0.12395 | 0.083373 | 0.14152 |
| ENSG00000010932 | 1 | 171217638 | 171255117 | 119 | 21 | 1.155 | 0.12404 | FMO1 | 0.12404 | 0.18612 | 0.26316 |
| ENSG00000235863 | 6 | 33244917 | 33252609 | 29 | 9 | 1.1549 | 0.12406 | B3GALT4 | 0.12406 | 0.020219 | 0.033535 |
| ENSG00000163536 | 3 | 167453031 | 167543356 | 338 | 22 | 1.1548 | 0.12408 | SERPINI1 | 0.12408 | 0.16037 | 0.20398 |
| ENSG00000128250 | 22 | 29834572 | 29838444 | 18 | 4 | 1.1548 | 0.12409 | RFPL1 | 0.12409 | 0.17374 | 0.28706 |
| ENSG00000125968 | 20 | 30193086 | 30194318 | 5 | 2 | 1.1547 | 0.1241 | ID1 | 0.1241 | 0.073768 | 0.069476 |
| ENSG00000174992 | 16 | 29789561 | 29793096 | 2 | 1 | 1.1547 | 0.12411 | ZG16 | 0.12411 | 0.05557 | 0.14699 |
| ENSG00000002587 | 4 | 11394774 | 11431389 | 125 | 39 | 1.1545 | 0.12415 | HS3ST1 | 0.12415 | 0.28237 | 0.54937 |
| ENSG00000240583 | 7 | 30893010 | 30965131 | 210 | 47 | 1.1544 | 0.12417 | AQP1 | 0.12417 | 0.15354 | 0.22896 |
| ENSG00000198028 | 19 | 9577183 | 9609283 | 131 | 5 | 1.1544 | 0.12417 | ZNF560 | 0.12417 | 0.11597 | 0.06929 |
| ENSG00000196660 | 1 | 219858769 | 220131989 | 1142 | 51 | 1.1543 | 0.12419 | SLC30A10 | 0.12419 | 0.10424 | 0.11058 |
| ENSG00000170271 | 5 | 154198051 | 154238812 | 82 | 13 | 1.1536 | 0.12433 | FAXDC2 | 0.12433 | 0.090671 | 0.059728 |
| ENSG00000032219 | 14 | 58765103 | 58840605 | 122 | 15 | 1.1535 | 0.12436 | ARID4A | 0.12436 | 0.15475 | 0.15675 |
| ENSG00000112110 | 6 | 160210844 | 160219468 | 59 | 6 | 1.153 | 0.12445 | MRPL18 | 0.12445 | 0.092974 | 0.074508 |
| ENSG00000186377 | 1 | 47427036 | 47516423 | 231 | 23 | 1.153 | 0.12446 | CYP4X1 | 0.12446 | 0.13116 | 0.15151 |
| ENSG00000173727 | 11 | 65222728 | 65234028 | 40 | 5 | 1.1529 | 0.12448 | AP000769.1 | 0.12448 | 0.12627 | 0.19345 |
| ENSG00000116455 | 1 | 111982512 | 111991998 | 18 | 5 | 1.1528 | 0.1245 | WDR77 | 0.1245 | 0.20748 | 0.15987 |
| ENSG00000269415 | 3 | 59957215 | 59957583 | 4 | 2 | 1.1521 | 0.12464 | NPCDR1 | 0.12464 | 0.5687 | 0.50028 |
| ENSG00000125166 | 16 | 58741035 | 58768261 | 131 | 15 | 1.152 | 0.12466 | GOT2 | 0.12466 | 0.12688 | 0.1883 |
| ENSG00000214530 | 11 | 72465774 | 72504726 | 87 | 12 | 1.1515 | 0.12476 | STARD10 | 0.12476 | 0.15225 | 0.23179 |
| ENSG00000163497 | 2 | 219845809 | 219850379 | 2 | 1 | 1.1514 | 0.12477 | FEV | 0.12477 | 0.13867 | 0.067228 |
| ENSG00000154611 | 18 | 23713816 | 23773319 | 122 | 12 | 1.1513 | 0.12481 | PSMA8 | 0.12481 | 0.14194 | 0.12922 |
| ENSG00000245017 | 12 | 98879322 | 98897633 | 78 | 14 | 1.1511 | 0.12484 | RP11-181C3.1 | 0.12484 | 0.053618 | 0.055081 |
| ENSG00000099204 | 10 | 116190872 | 116444762 | 836 | 64 | 1.1507 | 0.12492 | ABLIM1 | 0.12492 | 0.12142 | 0.076686 |
| ENSG00000198502 | 6 | 32485120 | 32498064 | 32 | 7 | 1.1488 | 0.12532 | HLA-DRB5 | 0.12532 | 0.018605 | 0.029657 |
| ENSG00000225781 | 7 | 142749417 | 142750444 | 5 | 2 | 1.1486 | 0.12535 | OR6V1 | 0.12535 | 0.2678 | 0.20245 |
| ENSG00000250644 | 11 | 1768897 | 1780281 | 45 | 12 | 1.1484 | 0.1254 | RP11-295K3.1 | 0.1254 | 0.04348 | 0.02523 |
| ENSG00000182324 | 19 | 48958766 | 48969367 | 18 | 5 | 1.1484 | 0.12541 | KCNJ14 | 0.12541 | 0.16309 | 0.15573 |
| ENSG00000103202 | 16 | 446725 | 460367 | 55 | 7 | 1.1482 | 0.12544 | NME4 | 0.12544 | 0.074334 | 0.07452 |
| ENSG00000141979 | 19 | 16589878 | 16739015 | 407 | 30 | 1.1482 | 0.12544 | CTD-3222D19.2 | 0.12544 | 0.1239 | 0.11453 |
| ENSG00000136709 | 2 | 128458596 | 128568761 | 339 | 29 | 1.1477 | 0.12555 | WDR33 | 0.12555 | 0.15206 | 0.19469 |
| ENSG00000162888 | 1 | 206664449 | 206671061 | 31 | 7 | 1.1475 | 0.12558 | C1orf147 | 0.12558 | 0.093346 | 0.11894 |
| ENSG00000069493 | 12 | 9817565 | 9848413 | 136 | 16 | 1.1474 | 0.1256 | CLEC2D | 0.1256 | 0.13629 | 0.14639 |
| ENSG00000174842 | 1 | 92711959 | 92764544 | 78 | 11 | 1.1471 | 0.12567 | GLMN | 0.12567 | 0.13203 | 0.14685 |
| ENSG00000166501 | 16 | 23847322 | 24231932 | 1415 | 105 | 1.1468 | 0.12573 | PRKCB | 0.12573 | 0.15927 | 0.17621 |
| ENSG00000173486 | 11 | 64008475 | 64011604 | 3 | 2 | 1.1468 | 0.12574 | FKBP2 | 0.12574 | 0.25539 | 0.42968 |
| ENSG00000197748 | 10 | 105889646 | 105992120 | 270 | 21 | 1.1466 | 0.12577 | WDR96 | 0.12577 | 0.12768 | 0.13727 |
| ENSG00000168411 | 16 | 74655292 | 74700779 | 199 | 20 | 1.1462 | 0.12585 | RFWD3 | 0.12585 | 0.10691 | 0.15533 |
| ENSG00000267855 | 19 | 8373490 | 8386280 | 44 | 6 | 1.1462 | 0.12585 | NDUFA7 | 0.12585 | 0.059813 | 0.087246 |
| ENSG00000181939 | 11 | 55321783 | 55322895 | 10 | 2 | 1.146 | 0.1259 | OR4C15 | 0.1259 | 0.093168 | 0.092143 |
| ENSG00000204237 | 17 | 79632066 | 79633665 | 3 | 2 | 1.1454 | 0.12602 | OXLD1 | 0.12602 | 0.2668 | 0.47772 |
| ENSG00000257103 | 19 | 34663409 | 34720420 | 195 | 16 | 1.1454 | 0.12602 | LSM14A | 0.12602 | 0.14698 | 0.18698 |
| ENSG00000196337 | 19 | 49557531 | 49562117 | 17 | 8 | 1.1447 | 0.12617 | CGB7 | 0.12617 | 0.36536 | 0.31417 |
| ENSG00000246705 | 12 | 14927270 | 14930936 | 13 | 7 | 1.144 | 0.12632 | H2AFJ | 0.12632 | 0.10679 | 0.50274 |
| ENSG00000101361 | 20 | 2632791 | 2639039 | 30 | 9 | 1.1425 | 0.12661 | NOP56 | 0.12661 | 0.1636 | 0.18235 |
| ENSG00000134575 | 11 | 47260853 | 47270457 | 28 | 6 | 1.1424 | 0.12665 | ACP2 | 0.12665 | 0.14154 | 0.22356 |
| ENSG00000186205 | 1 | 220960101 | 220987735 | 141 | 32 | 1.1421 | 0.12671 | MTARC1 | 0.12671 | 0.18133 | 0.30778 |
| ENSG00000004766 | 7 | 92861653 | 92988338 | 215 | 32 | 1.142 | 0.12672 | CCDC132 | 0.12672 | 0.13288 | 0.18518 |
| ENSG00000160233 | 21 | 45875369 | 45878739 | 9 | 4 | 1.142 | 0.12672 | LRRC3 | 0.12672 | 0.38559 | 0.54166 |
| ENSG00000132589 | 17 | 27206353 | 27224697 | 30 | 6 | 1.1418 | 0.12676 | FLOT2 | 0.12676 | 0.16681 | 0.16481 |

| | | | | | | | | | | | |
|---|---|---|---|---|---|---|---|---|---|---|---|
| ENSG00000205057 | 12 | 92813870 | 92821924 | 24 | 5 | 1.1411 | 0.12692 | CLLU1OS | 0.12692 | 0.084243 | 0.12847 |
| ENSG00000089199 | 20 | 5892076 | 5906007 | 47 | 8 | 1.1408 | 0.12699 | CHGB | 0.12699 | 0.1664 | 0.35416 |
| ENSG00000204290 | 6 | 32361740 | 32374905 | 223 | 17 | 1.1407 | 0.12701 | BTNL2 | 0.12701 | 0.01196 | 0.0097782 |
| ENSG00000130598 | 11 | 1860219 | 1862910 | 7 | 4 | 1.1399 | 0.12717 | TNNI2 | 0.12717 | 0.85912 | 0.82641 |
| ENSG00000139132 | 12 | 32552463 | 32798984 | 1130 | 56 | 1.1396 | 0.12723 | FGD4 | 0.12723 | 0.12678 | 0.19989 |
| ENSG00000196476 | 20 | 251504 | 271390 | 164 | 23 | 1.1395 | 0.12724 | C20orf96 | 0.12724 | 0.18563 | 0.38103 |
| ENSG00000243772 | 19 | 55249980 | 55295776 | 77 | 7 | 1.1395 | 0.12725 | KIR2DL3 | 0.12725 | 0.084156 | 0.0028985 |
| ENSG00000198805 | 14 | 20937113 | 20945253 | 27 | 6 | 1.1393 | 0.12729 | PNP | 0.12729 | 0.4085 | 0.22148 |
| ENSG00000108244 | 17 | 39078948 | 39093886 | 69 | 10 | 1.1392 | 0.12731 | KRT23 | 0.12731 | 0.31248 | 0.49231 |
| ENSG00000249209 | 21 | 34956993 | 35284635 | 696 | 75 | 1.1387 | 0.12741 | AP000304.12 | 0.12741 | 0.13468 | 0.10905 |
| ENSG00000249319 | 7 | 65552756 | 65617228 | 221 | 18 | 1.1379 | 0.12758 | AC068533.7 | 0.12758 | 0.13891 | 0.13983 |
| ENSG00000184363 | 11 | 392614 | 404908 | 52 | 13 | 1.1378 | 0.1276 | PKP3 | 0.1276 | 0.2677 | 0.34918 |
| ENSG00000135338 | 6 | 80194708 | 80247175 | 129 | 18 | 1.1378 | 0.12761 | LCA5 | 0.12761 | 0.12206 | 0.13213 |
| ENSG00000183207 | 19 | 49496705 | 49519252 | 70 | 7 | 1.1369 | 0.1278 | RUVBL2 | 0.1278 | 0.24429 | 0.27887 |
| ENSG00000242689 | 11 | 58390146 | 58393198 | 6 | 3 | 1.1366 | 0.12785 | CNTF | 0.12785 | 0.0069739 | 0.0076298 |
| ENSG00000123179 | 13 | 50234859 | 50265611 | 179 | 22 | 1.1361 | 0.12796 | EBPL | 0.12796 | 0.095232 | 0.12861 |
| ENSG00000050405 | 12 | 50569571 | 50677329 | 197 | 16 | 1.136 | 0.12799 | LIMA1 | 0.12799 | 0.14142 | 0.14598 |
| ENSG00000092421 | 5 | 115779312 | 115910630 | 513 | 76 | 1.1359 | 0.128 | SEMA6A | 0.128 | 0.13229 | 0.13705 |
| ENSG00000166157 | 21 | 10906201 | 11029719 | 3 | 2 | 1.1354 | 0.1281 | TPTE | 0.1281 | 0.1281 | 0.10775 |
| ENSG00000233502 | 2 | 10223908 | 10226228 | 9 | 4 | 1.1349 | 0.1282 | AC104794.4 | 0.1282 | 0.13203 | 0.23061 |
| ENSG00000147677 | 8 | 117654369 | 117779164 | 311 | 20 | 1.1348 | 0.12822 | EIF3H | 0.12822 | 0.13421 | 0.17847 |
| ENSG00000256029 | 1 | 159827690 | 159842843 | 63 | 9 | 1.1348 | 0.12824 | RP11-190A12.7 | 0.12824 | 0.047279 | 0.024361 |
| ENSG00000076944 | 19 | 7701767 | 7712759 | 29 | 7 | 1.1344 | 0.12832 | STXBP2 | 0.12832 | 0.29111 | 0.511 |
| ENSG00000108947 | 17 | 7608520 | 7614696 | 11 | 4 | 1.1341 | 0.12838 | EFNB3 | 0.12838 | 0.30286 | 0.51465 |
| ENSG00000100968 | 14 | 24834879 | 24848810 | 40 | 11 | 1.1336 | 0.12847 | NFATC4 | 0.12847 | 0.031835 | 0.0084845 |
| ENSG00000134313 | 2 | 8865408 | 8977760 | 217 | 24 | 1.1334 | 0.12852 | KIDINS220 | 0.12852 | 0.11549 | 0.17674 |
| ENSG00000155657 | 2 | 179390716 | 179695529 | 612 | 38 | 1.1326 | 0.12869 | TTN | 0.12869 | 0.10743 | 0.097089 |
| ENSG00000258231 | 12 | 58937907 | 59206842 | 775 | 62 | 1.1326 | 0.1287 | RP11-362K2.2 | 0.1287 | 0.048236 | 0.035197 |
| ENSG00000170615 | 7 | 102993177 | 103086624 | 248 | 32 | 1.1325 | 0.12871 | SLC26A5 | 0.12871 | 0.11338 | 0.15754 |
| ENSG00000180834 | 3 | 183533664 | 183543382 | 32 | 6 | 1.1321 | 0.1288 | MAP6D1 | 0.1288 | 0.029451 | 0.019284 |
| ENSG00000152926 | 7 | 64432154 | 64467062 | 140 | 10 | 1.1319 | 0.12884 | ZNF117 | 0.12884 | 0.13861 | 0.13033 |
| ENSG00000233828 | 5 | 86512423 | 86534822 | 31 | 12 | 1.1316 | 0.1289 | AC008394.1 | 0.1289 | 0.07824 | 0.11403 |
| ENSG00000204538 | 6 | 31105313 | 31107127 | 22 | 11 | 1.1312 | 0.12898 | PSORS1C2 | 0.12898 | 0.013282 | 0.032708 |
| ENSG00000197265 | 8 | 30435835 | 30515768 | 185 | 19 | 1.1312 | 0.12899 | GTF2E2 | 0.12899 | 0.11487 | 0.1841 |
| ENSG00000145743 | 5 | 107194736 | 107717799 | 1454 | 47 | 1.1306 | 0.12912 | FBXL17 | 0.12912 | 0.14274 | 0.14367 |
| ENSG00000172201 | 6 | 19837617 | 19840915 | 19 | 8 | 1.1304 | 0.12916 | ID4 | 0.12916 | 0.086002 | 0.15876 |
| ENSG00000168779 | 3 | 157814948 | 157824292 | 35 | 4 | 1.1298 | 0.12929 | SHOX2 | 0.12929 | 0.11636 | 0.19062 |
| ENSG00000167619 | 19 | 42817477 | 42829214 | 11 | 3 | 1.1297 | 0.1293 | TMEM145 | 0.1293 | 0.17214 | 0.21861 |
| ENSG00000090339 | 19 | 10381511 | 10397291 | 55 | 17 | 1.1295 | 0.12934 | ICAM1 | 0.12934 | 0.10187 | 0.10906 |
| ENSG00000261221 | 19 | 56116771 | 56128635 | 27 | 11 | 1.1295 | 0.12934 | ZNF865 | 0.12934 | 0.19394 | 0.15079 |
| ENSG00000100014 | 22 | 24666786 | 24813708 | 363 | 21 | 1.1294 | 0.12937 | SPECC1L | 0.12937 | 0.12399 | 0.11712 |
| ENSG00000123810 | 19 | 41860326 | 41870078 | 27 | 6 | 1.1292 | 0.12941 | B9D2 | 0.12941 | 0.13341 | 0.063998 |
| ENSG00000177733 | 5 | 137087075 | 137090039 | 5 | 3 | 1.1289 | 0.12946 | HNRNPA0 | 0.12946 | 0.55176 | 0.71267 |
| ENSG00000135956 | 2 | 96914254 | 96931732 | 17 | 8 | 1.1288 | 0.12949 | TMEM127 | 0.12949 | 0.10576 | 0.086111 |
| ENSG00000129673 | 17 | 74449433 | 74466199 | 68 | 16 | 1.1288 | 0.12949 | AANAT | 0.12949 | 0.17118 | 0.18442 |
| ENSG00000184182 | 2 | 238875469 | 238951236 | 258 | 11 | 1.1286 | 0.12954 | UBE2F | 0.12954 | 0.14312 | 0.1772 |
| ENSG00000131848 | 19 | 56732681 | 56879752 | 591 | 80 | 1.1283 | 0.12959 | ZSCAN5A | 0.12959 | 0.16339 | 0.22894 |
| ENSG00000135390 | 12 | 54026510 | 54071192 | 153 | 13 | 1.1282 | 0.12961 | ATP5G2 | 0.12961 | 0.098152 | 0.10879 |
| ENSG00000176165 | 14 | 29235050 | 29238870 | 9 | 5 | 1.1281 | 0.12964 | FOXG1 | 0.12964 | 0.27206 | 0.34173 |
| ENSG00000105939 | 7 | 138728266 | 138794465 | 244 | 29 | 1.1276 | 0.12975 | ZC3HAV1 | 0.12975 | 0.19293 | 0.41467 |
| ENSG00000168496 | 11 | 61560109 | 61564716 | 4 | 2 | 1.1272 | 0.12983 | FEN1 | 0.12983 | 0.13869 | 0.17524 |
| ENSG00000162723 | 1 | 159921282 | 159924044 | 11 | 3 | 1.1263 | 0.13002 | SLAMF9 | 0.13002 | 0.34994 | 0.67516 |
| ENSG00000166337 | 11 | 6627526 | 6633898 | 19 | 4 | 1.1262 | 0.13003 | TAF10 | 0.13003 | 0.098966 | 0.055328 |
| ENSG00000104412 | 8 | 109455830 | 109499145 | 132 | 10 | 1.1261 | 0.13007 | EMC2 | 0.13007 | 0.15573 | 0.16651 |
| ENSG00000110944 | 12 | 56732663 | 56734193 | 1 | 1 | 1.1259 | 0.1301 | IL23A | 0.1301 | 0.2122 | 0.2096 |
| ENSG00000213064 | 1 | 168195176 | 168222263 | 57 | 7 | 1.1252 | 0.13025 | SFT2D2 | 0.13025 | 0.1018 | 0.1183 |
| ENSG00000059588 | 1 | 234527059 | 234614849 | 476 | 48 | 1.1242 | 0.13046 | TARBP1 | 0.13046 | 0.129 | 0.17127 |
| ENSG00000205791 | 12 | 12508342 | 12510001 | 6 | 3 | 1.1241 | 0.1305 | LOH12CR2 | 0.1305 | 0.7495 | 0.59301 |
| ENSG00000117262 | 1 | 145764411 | 145827103 | 11 | 5 | 1.1239 | 0.13052 | GPR89A | 0.13052 | 0.14573 | 0.21524 |
| ENSG00000072849 | 17 | 5374571 | 5390131 | 50 | 15 | 1.1238 | 0.13054 | DERL2 | 0.13054 | 0.33513 | 0.26643 |
| ENSG00000174450 | 15 | 23684645 | 23692381 | 40 | 4 | 1.1236 | 0.13059 | GOLGA6L2 | 0.13059 | 0.19589 | 0.2978 |
| ENSG00000217555 | 16 | 66586466 | 66600154 | 27 | 4 | 1.1235 | 0.13062 | CKLF | 0.13062 | 0.10083 | 0.10228 |
| ENSG00000110046 | 11 | 64662007 | 64684722 | 22 | 7 | 1.1233 | 0.13065 | ATG2A | 0.13065 | 0.23012 | 0.44143 |
| ENSG00000167633 | 19 | 55235969 | 55378448 | 706 | 41 | 1.1231 | 0.13069 | KIR3DL1 | 0.13069 | 0.0889 | 0.030884 |
| ENSG00000065029 | 6 | 35226686 | 35263762 | 99 | 11 | 1.1228 | 0.13077 | ZNF76 | 0.13077 | 0.13722 | 0.14094 |
| ENSG00000167674 | 19 | 4472284 | 4502223 | 156 | 17 | 1.1227 | 0.13079 | HDGFRP2 | 0.13079 | 0.17854 | 0.23692 |
| ENSG00000243501 | 6 | 73919469 | 73972919 | 191 | 26 | 1.1211 | 0.13113 | RP11-257K9.8 | 0.13113 | 0.25895 | 0.25235 |
| ENSG00000125861 | 20 | 3639939 | 3644046 | 23 | 8 | 1.1209 | 0.13116 | GFRA4 | 0.13116 | 0.29098 | 0.16206 |
| ENSG00000180383 | 20 | 30053309 | 30064560 | 28 | 8 | 1.1209 | 0.13116 | DEFB124 | 0.13116 | 0.11758 | 0.10028 |
| ENSG00000074416 | 3 | 127407909 | 127542051 | 441 | 59 | 1.1208 | 0.13119 | MGLL | 0.13119 | 0.1826 | 0.25943 |
| ENSG00000106261 | 7 | 99613204 | 99639312 | 55 | 13 | 1.1206 | 0.13122 | ZKSCAN1 | 0.13122 | 0.12835 | 0.096531 |
| ENSG00000167774 | 19 | 8373167 | 8386263 | 45 | 6 | 1.1202 | 0.1313 | NDUFA7 | 0.1313 | 0.059813 | 0.087246 |
| ENSG00000075975 | 3 | 12598513 | 12625212 | 96 | 17 | 1.1201 | 0.13134 | MKRN2 | 0.13134 | 0.078303 | 0.12611 |

| | | | | | | | | | | | |
|---|---|---|---|---|---|---|---|---|---|---|---|
| ENSG00000179010 | 4 | 6641818 | 6644472 | 10 | 4 | 1.1198 | 0.13139 | MRFAP1 | 0.13139 | 0.38338 | 0.27842 |
| ENSG00000135974 | 2 | 105953816 | 105965668 | 36 | 9 | 1.1195 | 0.13146 | C2orf49 | 0.13146 | 0.14059 | 0.12689 |
| ENSG00000270394 | 4 | 117220016 | 117221520 | 3 | 2 | 1.1195 | 0.13147 | MTRNR2L13 | 0.13147 | 0.092362 | 0.064421 |
| ENSG00000241258 | 7 | 65579591 | 65619555 | 140 | 15 | 1.1193 | 0.13152 | CRCP | 0.13152 | 0.14006 | 0.1528 |
| ENSG00000113231 | 5 | 76506274 | 76725632 | 636 | 46 | 1.1189 | 0.13158 | PDE8B | 0.13158 | 0.085002 | 0.04837 |
| ENSG00000096006 | 6 | 49695097 | 49712150 | 48 | 9 | 1.1188 | 0.1316 | CRISP3 | 0.1316 | 0.14299 | 0.18431 |
| ENSG00000134955 | 11 | 124932963 | 124959131 | 120 | 29 | 1.1186 | 0.13165 | SLC37A2 | 0.13165 | 0.08132 | 0.072761 |
| ENSG00000139428 | 12 | 109991542 | 110011679 | 96 | 14 | 1.1184 | 0.13169 | MMAB | 0.13169 | 0.11394 | 0.13691 |
| ENSG00000139865 | 14 | 38065052 | 38510647 | 1186 | 67 | 1.1183 | 0.13172 | TTC6 | 0.13172 | 0.13873 | 0.14474 |
| ENSG00000164111 | 4 | 122589110 | 122618268 | 141 | 16 | 1.1182 | 0.13173 | ANXA5 | 0.13173 | 0.13163 | 0.18174 |
| ENSG00000167195 | 15 | 75550940 | 75562540 | 6 | 3 | 1.1177 | 0.13185 | GOLGA6C | 0.13185 | 0.010401 | 0.020588 |
| ENSG00000154144 | 11 | 124492732 | 124505287 | 27 | 11 | 1.1174 | 0.1319 | TBRG1 | 0.1319 | 0.12993 | 0.088699 |
| ENSG00000173801 | 17 | 39775692 | 39943183 | 698 | 41 | 1.1174 | 0.13191 | JUP | 0.13191 | 0.12298 | 0.22869 |
| ENSG00000164299 | 5 | 79615444 | 79617661 | 10 | 5 | 1.1173 | 0.13194 | SPZ1 | 0.13194 | 0.10411 | 0.10675 |
| ENSG00000256574 | 10 | 45798102 | 45811056 | 51 | 4 | 1.1172 | 0.13195 | OR13A1 | 0.13195 | 0.12349 | 0.10963 |
| ENSG00000141504 | 17 | 7529552 | 7531194 | 4 | 2 | 1.117 | 0.132 | SAT2 | 0.132 | 0.050607 | 0.015454 |
| ENSG00000181323 | 17 | 7323679 | 7324951 | 3 | 1 | 1.1169 | 0.13202 | SPEM1 | 0.13202 | 0.2251 | 0.35166 |
| ENSG00000138778 | 4 | 104026963 | 104119566 | 209 | 14 | 1.1164 | 0.13212 | CENPE | 0.13212 | 0.1246 | 0.12667 |
| ENSG00000104880 | 19 | 7459999 | 7537363 | 352 | 19 | 1.1163 | 0.13214 | ARHGEF18 | 0.13214 | 0.11559 | 0.12599 |
| ENSG00000172273 | 11 | 118992297 | 119006752 | 42 | 4 | 1.116 | 0.13221 | HINFP | 0.13221 | 0.2294 | 0.32071 |
| ENSG00000162621 | 1 | 74935562 | 74978298 | 76 | 8 | 1.115 | 0.13242 | LRRC53 | 0.13242 | 0.052393 | 0.13529 |
| ENSG00000151834 | 4 | 46250444 | 46477247 | 602 | 29 | 1.115 | 0.13242 | GABRA2 | 0.13242 | 0.14197 | 0.18717 |
| ENSG00000120594 | 10 | 20105168 | 20578785 | 2129 | 100 | 1.115 | 0.13242 | PLXDC2 | 0.13242 | 0.13166 | 0.18397 |
| ENSG00000179943 | 19 | 56102746 | 56113336 | 33 | 9 | 1.1147 | 0.13248 | FIZ1 | 0.13248 | 0.13547 | 0.097928 |
| ENSG00000105507 | 19 | 48533210 | 48547310 | 119 | 8 | 1.1147 | 0.13249 | CABP5 | 0.13249 | 0.1321 | 0.089476 |
| ENSG00000196151 | 2 | 160092304 | 160143310 | 274 | 12 | 1.1143 | 0.13257 | WDSUB1 | 0.13257 | 0.13154 | 0.14489 |
| ENSG00000126870 | 7 | 158649269 | 158749438 | 421 | 26 | 1.1142 | 0.13259 | WDR60 | 0.13259 | 0.19102 | 0.2844 |
| ENSG00000136099 | 13 | 53418109 | 53422775 | 9 | 3 | 1.1135 | 0.13276 | PCDH8 | 0.13276 | 0.16552 | 0.094308 |
| ENSG00000115539 | 2 | 101179152 | 101193197 | 58 | 7 | 1.1134 | 0.13277 | PDCL3 | 0.13277 | 0.12436 | 0.15794 |
| ENSG00000204599 | 6 | 30294256 | 30311506 | 47 | 14 | 1.1134 | 0.13277 | TRIM39 | 0.13277 | 0.14522 | 0.23762 |
| ENSG00000171067 | 11 | 68028803 | 68039469 | 28 | 8 | 1.1129 | 0.13287 | C11orf24 | 0.13287 | 0.24184 | 0.22706 |
| ENSG00000054598 | 6 | 1610681 | 1614127 | 3 | 2 | 1.1128 | 0.1329 | FOXC1 | 0.1329 | 0.020476 | 0.1096 |
| ENSG00000168028 | 3 | 39448180 | 39454033 | 40 | 4 | 1.1126 | 0.13293 | RPSA | 0.13293 | 0.12176 | 0.13767 |
| ENSG00000198945 | 6 | 130334844 | 130462594 | 278 | 25 | 1.1126 | 0.13293 | L3MBTL3 | 0.13293 | 0.12419 | 0.15703 |
| ENSG00000168538 | 4 | 184580420 | 184634745 | 241 | 23 | 1.1125 | 0.13297 | TRAPPC11 | 0.13297 | 0.17734 | 0.22112 |
| ENSG00000110274 | 11 | 117185273 | 117283984 | 247 | 28 | 1.112 | 0.13307 | CEP164 | 0.13307 | 0.1155 | 0.1347 |
| ENSG00000116254 | 1 | 6161853 | 6240183 | 355 | 39 | 1.1119 | 0.13309 | CHD5 | 0.13309 | 0.081823 | 0.097722 |
| ENSG00000164808 | 8 | 48173167 | 48648868 | 474 | 32 | 1.1119 | 0.13309 | SPIDR | 0.13309 | 0.13202 | 0.12992 |
| ENSG00000153237 | 2 | 159027593 | 159313265 | 1495 | 50 | 1.1117 | 0.13313 | CCDC148 | 0.13313 | 0.098739 | 0.10691 |
| ENSG00000198464 | 19 | 52800430 | 52829175 | 94 | 14 | 1.1116 | 0.13316 | ZNF480 | 0.13316 | 0.20161 | 0.17829 |
| ENSG00000171385 | 1 | 112313284 | 112531777 | 765 | 91 | 1.1109 | 0.13331 | KCND3 | 0.13331 | 0.19447 | 0.30787 |
| ENSG00000140090 | 14 | 92788925 | 92962596 | 792 | 59 | 1.1109 | 0.13331 | SLC24A4 | 0.13331 | 0.13409 | 0.14885 |
| ENSG00000135744 | 1 | 230838269 | 230850043 | 52 | 11 | 1.1105 | 0.13338 | AGT | 0.13338 | 0.17003 | 0.2657 |
| ENSG00000213588 | 6 | 33422356 | 33425325 | 6 | 4 | 1.1104 | 0.13342 | ZBTB9 | 0.13342 | 0.05632 | 0.10652 |
| ENSG00000146676 | 7 | 44915896 | 44924960 | 13 | 6 | 1.1103 | 0.13344 | PURB | 0.13344 | 0.22481 | 0.19141 |
| ENSG00000140943 | 16 | 84087368 | 84150511 | 319 | 28 | 1.1101 | 0.13347 | MBTPS1 | 0.13347 | 0.14371 | 0.16772 |
| ENSG00000075415 | 12 | 98987369 | 98995946 | 19 | 8 | 1.1101 | 0.13348 | SLC25A3 | 0.13348 | 0.3998 | 0.14478 |
| ENSG00000146197 | 6 | 35182190 | 35220856 | 66 | 11 | 1.1098 | 0.13354 | SCUBE3 | 0.13354 | 0.11335 | 0.13679 |
| ENSG00000131591 | 1 | 1017198 | 1051741 | 133 | 15 | 1.1088 | 0.13376 | C1orf159 | 0.13376 | 0.15512 | 0.19235 |
| ENSG00000115828 | 2 | 37571717 | 37600465 | 132 | 28 | 1.1088 | 0.13376 | QPCT | 0.13376 | 0.26587 | 0.22395 |
| ENSG00000167085 | 17 | 47481414 | 47492246 | 36 | 13 | 1.1086 | 0.1338 | PHB | 0.1338 | 0.056985 | 0.0086862 |
| ENSG00000218537 | 22 | 24236613 | 24241117 | 26 | 7 | 1.1083 | 0.13388 | AP000350.4 | 0.13388 | 0.069272 | 0.094035 |
| ENSG00000112624 | 6 | 42749765 | 42836298 | 183 | 22 | 1.1081 | 0.13391 | GLTSCR1L | 0.13391 | 0.16258 | 0.15696 |
| ENSG00000186509 | 11 | 57791353 | 57949088 | 613 | 27 | 1.1078 | 0.13397 | OR9Q1 | 0.13397 | 0.15381 | 0.17748 |
| ENSG00000140368 | 15 | 77285700 | 77329673 | 106 | 20 | 1.1068 | 0.13419 | PSTPIP1 | 0.13419 | 0.16164 | 0.20773 |
| ENSG00000167822 | 11 | 55904247 | 55905194 | 5 | 2 | 1.1065 | 0.13425 | OR8J3 | 0.13425 | 0.027081 | 0.01996 |
| ENSG00000204366 | 6 | 31867384 | 31869769 | 9 | 5 | 1.1062 | 0.13432 | ZBTB12 | 0.13432 | 0.0028247 | 0.00059259 |
| ENSG00000214046 | 19 | 16741562 | 16771253 | 87 | 15 | 1.1062 | 0.13432 | SMIM7 | 0.13432 | 0.093493 | 0.075531 |
| ENSG00000187109 | 12 | 76438670 | 76478813 | 79 | 10 | 1.1053 | 0.13451 | NAP1L1 | 0.13451 | 0.12734 | 0.14277 |
| ENSG00000241635 | 2 | 234526291 | 234681956 | 545 | 30 | 1.1052 | 0.13453 | UGT1A8 | 0.13453 | 0.18913 | 0.18215 |
| ENSG00000100299 | 22 | 51061182 | 51066607 | 36 | 12 | 1.1049 | 0.13459 | ARSA | 0.13459 | 0.35169 | 0.47018 |
| ENSG00000181935 | 11 | 55339604 | 55340536 | 12 | 4 | 1.1046 | 0.13466 | OR4C16 | 0.13466 | 0.086384 | 0.079351 |
| ENSG00000213088 | 1 | 159173097 | 159176290 | 16 | 6 | 1.1039 | 0.13482 | DARC | 0.13482 | 0.37453 | 0.15237 |
| ENSG00000078369 | 1 | 1716729 | 1822495 | 200 | 24 | 1.1035 | 0.1349 | GNB1 | 0.1349 | 0.13323 | 0.14766 |
| ENSG00000095397 | 9 | 117164360 | 117267730 | 392 | 28 | 1.1035 | 0.1349 | DFNB31 | 0.1349 | 0.13261 | 0.12629 |
| ENSG00000135045 | 9 | 77561497 | 77567802 | 8 | 3 | 1.1033 | 0.13495 | C9orf40 | 0.13495 | 0.06207 | 0.047015 |
| ENSG00000166589 | 16 | 66942025 | 66952887 | 17 | 6 | 1.1029 | 0.13503 | CDH16 | 0.13503 | 0.24388 | 0.41475 |
| ENSG00000092758 | 20 | 61447596 | 61472511 | 126 | 29 | 1.1024 | 0.13514 | COL9A3 | 0.13514 | 0.18697 | 0.30782 |
| ENSG00000185220 | 1 | 249200395 | 249214145 | 31 | 8 | 1.1018 | 0.13527 | PGBD2 | 0.13527 | 0.19043 | 0.30079 |
| ENSG00000272617 | 16 | 69362547 | 69368774 | 15 | 4 | 1.1017 | 0.13529 | RP11-343C2.12 | 0.13529 | 0.25593 | 0.2921 |
| ENSG00000173988 | 13 | 46786083 | 46851501 | 271 | 18 | 1.1016 | 0.13533 | LRRC63 | 0.13533 | 0.13286 | 0.17673 |
| ENSG00000127083 | 9 | 95176527 | 95186743 | 24 | 7 | 1.1015 | 0.13534 | OMD | 0.13534 | 0.067124 | 0.086436 |
| ENSG00000138767 | 4 | 78634541 | 78740769 | 162 | 23 | 1.1015 | 0.13535 | CNOT6L | 0.13535 | 0.1133 | 0.11037 |

| | | | | | | | | | | | |
|---|---|---|---|---|---|---|---|---|---|---|---|
| ENSG00000214711 | 2 | 31395924 | 31456724 | 452 | 31 | 1.1012 | 0.1354 | CAPN14 | 0.1354 | 0.14306 | 0.19177 |
| ENSG00000176928 | 5 | 74323289 | 74326724 | 12 | 4 | 1.101 | 0.13544 | GCNT4 | 0.13544 | 0.10816 | 0.03882 |
| ENSG00000112697 | 6 | 75962640 | 75994684 | 62 | 7 | 1.1011 | 0.13544 | TMEM30A | 0.13544 | 0.12969 | 0.17804 |
| ENSG00000168938 | 5 | 122358945 | 122372436 | 43 | 11 | 1.1009 | 0.13547 | PPIC | 0.13547 | 0.076054 | 0.0687 |
| ENSG00000163386 | 1 | 145289772 | 145370303 | 13 | 4 | 1.1007 | 0.13551 | NBPF10 | 0.13551 | 0.21214 | 0.2824 |
| ENSG00000174442 | 15 | 66797297 | 66842115 | 138 | 19 | 1.1007 | 0.13551 | ZWILCH | 0.13551 | 0.15079 | 0.19339 |
| ENSG00000188566 | 9 | 140100147 | 140111461 | 34 | 8 | 1.1006 | 0.13554 | NDOR1 | 0.13554 | 0.37742 | 0.6349 |
| ENSG00000003756 | 3 | 50126341 | 50156454 | 40 | 8 | 1.1004 | 0.13559 | RBM5 | 0.13559 | 0.13369 | 0.14082 |
| ENSG00000185467 | 7 | 98771197 | 98805129 | 203 | 13 | 1.0999 | 0.13568 | KPNA7 | 0.13568 | 0.2057 | 0.24359 |
| ENSG00000161055 | 5 | 180017103 | 180018540 | 3 | 2 | 1.0998 | 0.13571 | SCGB3A1 | 0.13571 | 0.30776 | 0.12892 |
| ENSG00000151692 | 2 | 7057523 | 7208417 | 515 | 50 | 1.0995 | 0.13577 | RNF144A | 0.13577 | 0.16369 | 0.18493 |
| ENSG00000262003 | 17 | 812872 | 814452 | 9 | 3 | 1.0995 | 0.13578 | RP11-676J12.7 | 0.13578 | 0.54063 | 0.58976 |
| ENSG00000080644 | 15 | 78885394 | 78913637 | 119 | 12 | 1.0993 | 0.13581 | CHRNA3 | 0.13581 | 0.18022 | 0.16259 |
| ENSG00000137561 | 8 | 63961112 | 63998612 | 96 | 14 | 1.099 | 0.13589 | TTPA | 0.13589 | 0.10918 | 0.080872 |
| ENSG00000132688 | 1 | 156638555 | 156647189 | 19 | 4 | 1.0986 | 0.13597 | NES | 0.13597 | 0.09008 | 0.050619 |
| ENSG00000133401 | 5 | 31639517 | 32111037 | 2317 | 241 | 1.0986 | 0.13598 | PDZD2 | 0.13598 | 0.1833 | 0.24702 |
| ENSG00000196576 | 22 | 50713408 | 50746056 | 122 | 24 | 1.0985 | 0.13599 | PLXNB2 | 0.13599 | 0.027852 | 0.011356 |
| ENSG00000267893 | 17 | 70036164 | 70036872 | 5 | 2 | 1.0985 | 0.136 | AC007461.1 | 0.136 | 0.21146 | 0.27122 |
| ENSG00000176895 | 11 | 4928600 | 4929538 | 2 | 1 | 1.0976 | 0.1362 | OR51A7 | 0.1362 | 0.3078 | 0.35553 |
| ENSG00000102879 | 16 | 30194148 | 30200397 | 5 | 3 | 1.0976 | 0.1362 | CORO1A | 0.1362 | 0.59836 | 0.70791 |
| ENSG00000132376 | 17 | 1397865 | 1420182 | 109 | 16 | 1.0975 | 0.13622 | INPP5K | 0.13622 | 0.15303 | 0.13238 |
| ENSG00000105325 | 19 | 3506271 | 3538328 | 77 | 10 | 1.0973 | 0.13625 | FZR1 | 0.13625 | 0.087925 | 0.071209 |
| ENSG00000171016 | 15 | 55831088 | 55881145 | 257 | 17 | 1.0971 | 0.13629 | PYGO1 | 0.13629 | 0.12816 | 0.13101 |
| ENSG00000166387 | 11 | 7534529 | 7678358 | 653 | 65 | 1.0967 | 0.13638 | PPFIBP2 | 0.13638 | 0.14523 | 0.17824 |
| ENSG00000169021 | 19 | 29698173 | 29704448 | 26 | 6 | 1.0967 | 0.13639 | UQCRFS1 | 0.13639 | 0.11906 | 0.089315 |
| ENSG00000139625 | 12 | 53874274 | 53893847 | 18 | 9 | 1.0965 | 0.13642 | MAP3K12 | 0.13642 | 0.086559 | 0.084434 |
| ENSG00000120322 | 5 | 140557371 | 140560081 | 20 | 7 | 1.0963 | 0.13647 | PCDHB8 | 0.13647 | 0.41775 | 0.45481 |
| ENSG00000257726 | 12 | 115800817 | 115802550 | 5 | 3 | 1.0958 | 0.13659 | RP11-116D17.1 | 0.13659 | 0.35892 | 0.80507 |
| ENSG00000251493 | 5 | 72740654 | 72744352 | 7 | 4 | 1.0955 | 0.13665 | FOXD1 | 0.13665 | 0.1333 | 0.30112 |
| ENSG00000156642 | 15 | 73852355 | 73926475 | 152 | 16 | 1.0953 | 0.1367 | NPTN | 0.1367 | 0.11421 | 0.11539 |
| ENSG00000237649 | 6 | 33359313 | 33377701 | 64 | 14 | 1.095 | 0.13675 | KIFC1 | 0.13675 | 0.093213 | 0.076843 |
| ENSG00000144746 | 3 | 69134095 | 69155217 | 76 | 12 | 1.095 | 0.13677 | ARL6IP5 | 0.13677 | 0.19969 | 0.25489 |
| ENSG00000183527 | 21 | 40546695 | 40555777 | 37 | 5 | 1.0949 | 0.13678 | PSMG1 | 0.13678 | 0.083173 | 0.10824 |
| ENSG00000175264 | 11 | 45670427 | 45687172 | 48 | 7 | 1.0949 | 0.13679 | CHST1 | 0.13679 | 0.14068 | 0.13335 |
| ENSG00000080293 | 2 | 120197419 | 120282070 | 329 | 25 | 1.0945 | 0.13687 | SCTR | 0.13687 | 0.16288 | 0.18728 |
| ENSG00000094661 | 19 | 15197791 | 15199032 | 12 | 3 | 1.094 | 0.13698 | OR1I1 | 0.13698 | 0.20472 | 0.29197 |
| ENSG00000164113 | 4 | 123300121 | 123350957 | 108 | 16 | 1.0935 | 0.13709 | ADAD1 | 0.13709 | 0.12039 | 0.15262 |
| ENSG00000104755 | 8 | 39601254 | 39695808 | 227 | 28 | 1.0929 | 0.13721 | ADAM2 | 0.13721 | 0.17384 | 0.23624 |
| ENSG00000172361 | 18 | 47753563 | 47792892 | 111 | 11 | 1.0927 | 0.13725 | CCDC11 | 0.13725 | 0.098263 | 0.052595 |
| ENSG00000068120 | 17 | 40713485 | 40718295 | 6 | 2 | 1.0926 | 0.13728 | COASY | 0.13728 | 0.20802 | 0.29008 |
| ENSG00000160469 | 19 | 55793440 | 55823901 | 112 | 18 | 1.0926 | 0.1373 | BRSK1 | 0.1373 | 0.27608 | 0.40174 |
| ENSG00000163931 | 3 | 53258723 | 53290068 | 167 | 14 | 1.0918 | 0.13747 | TKT | 0.13747 | 0.15355 | 0.14647 |
| ENSG00000127903 | 19 | 57174020 | 57183151 | 56 | 8 | 1.0916 | 0.13751 | ZNF835 | 0.13751 | 0.2245 | 0.3977 |
| ENSG00000105523 | 19 | 49104067 | 49118111 | 49 | 11 | 1.0915 | 0.13752 | FAM83E | 0.13752 | 0.0078871 | 0.031588 |
| ENSG00000184724 | 21 | 31985750 | 31986249 | 1 | 1 | 1.0912 | 0.1376 | KRTAP6-1 | 0.1376 | 0.8097 | 0.77163 |
| ENSG00000269891 | 10 | 98757795 | 99052413 | 976 | 86 | 1.0905 | 0.13774 | ARHGAP19-SLIT1 | 0.13774 | 0.15898 | 0.15081 |
| ENSG00000164815 | 7 | 103766788 | 103848495 | 287 | 15 | 1.0904 | 0.13776 | ORC5 | 0.13776 | 0.13716 | 0.15334 |
| ENSG00000068878 | 2 | 54091204 | 54197977 | 376 | 25 | 1.0903 | 0.13779 | PSME4 | 0.13779 | 0.15864 | 0.16609 |
| ENSG00000186395 | 17 | 38974369 | 38978847 | 12 | 5 | 1.09 | 0.13786 | KRT10 | 0.13786 | 0.70986 | 0.74597 |
| ENSG00000065000 | 19 | 2100988 | 2164464 | 332 | 27 | 1.0895 | 0.13796 | AP3D1 | 0.13796 | 0.14498 | 0.14054 |
| ENSG00000111678 | 12 | 7052141 | 7055166 | 12 | 7 | 1.0888 | 0.13813 | C12orf57 | 0.13813 | 0.043784 | 0.0098928 |
| ENSG00000059758 | 12 | 96672039 | 96794338 | 322 | 27 | 1.0887 | 0.13813 | CDK17 | 0.13813 | 0.12452 | 0.15498 |
| ENSG00000077549 | 1 | 19665267 | 19812066 | 582 | 28 | 1.088 | 0.1383 | CAPZB | 0.1383 | 0.11571 | 0.11451 |
| ENSG00000106089 | 7 | 73113536 | 73134002 | 43 | 13 | 1.0877 | 0.13837 | STX1A | 0.13837 | 0.1722 | 0.13724 |
| ENSG00000112619 | 6 | 42664340 | 42690312 | 118 | 13 | 1.0876 | 0.13839 | PRPH2 | 0.13839 | 0.14264 | 0.16531 |
| ENSG00000140400 | 15 | 75648133 | 75660971 | 11 | 5 | 1.087 | 0.13852 | MAN2C1 | 0.13852 | 0.052298 | 0.061653 |
| ENSG00000214306 | 11 | 107643129 | 107647344 | 20 | 6 | 1.0869 | 0.13854 | AP001024.2 | 0.13854 | 0.12556 | 0.29735 |
| ENSG00000138036 | 2 | 44001178 | 44037149 | 155 | 15 | 1.0869 | 0.13855 | DYNC2LI1 | 0.13855 | 0.17874 | 0.2832 |
| ENSG00000161944 | 17 | 7004641 | 7019019 | 91 | 14 | 1.0865 | 0.13864 | ASGR2 | 0.13864 | 0.10844 | 0.18469 |
| ENSG00000133114 | 13 | 45563687 | 45607746 | 124 | 16 | 1.0862 | 0.13871 | GPALPP1 | 0.13871 | 0.14921 | 0.03744 |
| ENSG00000050344 | 7 | 26191860 | 26226745 | 118 | 16 | 1.0857 | 0.1388 | NFE2L3 | 0.1388 | 0.21018 | 0.32209 |
| ENSG00000268964 | 19 | 53547991 | 53554329 | 30 | 5 | 1.0857 | 0.1388 | ERVV-2 | 0.1388 | 0.30558 | 0.47675 |
| ENSG00000106809 | 9 | 95146249 | 95166978 | 57 | 7 | 1.0856 | 0.13882 | OGN | 0.13882 | 0.13976 | 0.14739 |
| ENSG00000257727 | 12 | 56703626 | 56710120 | 11 | 4 | 1.0844 | 0.1391 | CNPY2 | 0.1391 | 0.15291 | 0.17728 |
| ENSG00000152291 | 2 | 85545147 | 85555548 | 42 | 5 | 1.0841 | 0.13915 | TGOLN2 | 0.13915 | 0.17422 | 0.18643 |
| ENSG00000105379 | 19 | 51848423 | 51869672 | 120 | 13 | 1.0841 | 0.13915 | ETFB | 0.13915 | 0.24759 | 0.41529 |
| ENSG00000184698 | 11 | 5410607 | 5411664 | 7 | 1 | 1.0839 | 0.1392 | OR51M1 | 0.1392 | 0.32949 | 0.52796 |
| ENSG00000222009 | 1 | 45274154 | 45281257 | 20 | 5 | 1.0838 | 0.13922 | BTBD19 | 0.13922 | 0.17331 | 0.26959 |
| ENSG00000167711 | 17 | 1646130 | 1658562 | 55 | 9 | 1.0838 | 0.13922 | SERPINF2 | 0.13922 | 0.1269 | 0.10024 |
| ENSG00000113578 | 5 | 141971743 | 142077617 | 405 | 61 | 1.0838 | 0.13923 | FGF1 | 0.13923 | 0.035356 | 0.010645 |
| ENSG00000086544 | 19 | 41223008 | 41246765 | 77 | 13 | 1.0838 | 0.13923 | ITPKC | 0.13923 | 0.18227 | 0.22253 |
| ENSG00000112210 | 6 | 57053607 | 57087078 | 43 | 10 | 1.0835 | 0.1393 | RAB23 | 0.1393 | 0.27486 | 0.28711 |
| ENSG00000137871 | 15 | 56922379 | 57210769 | 624 | 27 | 1.0834 | 0.13932 | ZNF280D | 0.13932 | 0.14487 | 0.13466 |

| | | | | | | | | | | | |
|---|---|---|---|---|---|---|---|---|---|---|---|
| ENSG00000172155 | 1 | 152769227 | 152770657 | 3 | 1 | 1.0829 | 0.13943 | LCE1D | 0.13943 | 0.24577 | 0.1737 |
| ENSG00000197958 | 9 | 130209953 | 130213684 | 8 | 2 | 1.0828 | 0.13945 | RPL12 | 0.13945 | 0.21232 | 0.23851 |
| ENSG00000174123 | 4 | 38773860 | 38784611 | 91 | 10 | 1.0824 | 0.13953 | TLR10 | 0.13953 | 0.14709 | 0.12616 |
| ENSG00000254019 | 8 | 142264664 | 142266916 | 7 | 3 | 1.0825 | 0.13953 | RP11-10J21.3 | 0.13953 | 0.090872 | 0.038478 |
| ENSG00000116161 | 1 | 174968300 | 174980851 | 23 | 7 | 1.0823 | 0.13956 | CACYBP | 0.13956 | 0.24206 | 0.24628 |
| ENSG00000242252 | 1 | 156211753 | 156213112 | 1 | 1 | 1.0821 | 0.1396 | BGLAP | 0.1396 | 0.71729 | 0.6762 |
| ENSG00000130640 | 10 | 135093135 | 135125841 | 157 | 27 | 1.082 | 0.13962 | TUBGCP2 | 0.13962 | 0.12076 | 0.12771 |
| ENSG00000104361 | 8 | 99202061 | 99306621 | 278 | 38 | 1.0817 | 0.13969 | NIPAL2 | 0.13969 | 0.14526 | 0.11375 |
| ENSG00000198203 | 2 | 108905095 | 108926371 | 77 | 14 | 1.0817 | 0.1397 | SULT1C2 | 0.1397 | 0.2193 | 0.43487 |
| ENSG00000109743 | 4 | 15704573 | 15739936 | 185 | 18 | 1.0816 | 0.13973 | BST1 | 0.13973 | 0.14685 | 0.20622 |
| ENSG00000158163 | 3 | 137780832 | 137834660 | 71 | 14 | 1.0813 | 0.13979 | DZIP1L | 0.13979 | 0.13658 | 0.23515 |
| ENSG00000104537 | 8 | 124693034 | 124749647 | 205 | 32 | 1.0812 | 0.1398 | ANXA13 | 0.1398 | 0.1034 | 0.18127 |
| ENSG00000041988 | 1 | 6684926 | 6695646 | 37 | 7 | 1.0811 | 0.13983 | THAP3 | 0.13983 | 0.07569 | 0.11477 |
| ENSG00000168772 | 4 | 105389469 | 105416058 | 57 | 11 | 1.0804 | 0.13997 | CXXC4 | 0.13997 | 0.20298 | 0.14002 |
| ENSG00000163605 | 3 | 73045936 | 73118350 | 353 | 11 | 1.0804 | 0.13998 | PPP4R2 | 0.13998 | 0.16384 | 0.20737 |
| ENSG00000144026 | 2 | 95813075 | 95831158 | 18 | 7 | 1.0803 | 0.14001 | ZNF514 | 0.14001 | 0.31376 | 0.39181 |
| ENSG00000123485 | 2 | 234742062 | 234763212 | 89 | 12 | 1.08 | 0.14008 | HJURP | 0.14008 | 0.050373 | 0.071511 |
| ENSG00000138101 | 2 | 25600067 | 25896503 | 578 | 51 | 1.0798 | 0.14011 | DTNB | 0.14011 | 0.18609 | 0.29934 |
| ENSG00000204954 | 12 | 104343980 | 104359486 | 89 | 16 | 1.0796 | 0.14015 | C12orf73 | 0.14015 | 0.075524 | 0.044448 |
| ENSG00000167136 | 9 | 131580753 | 131584956 | 11 | 3 | 1.0795 | 0.14018 | ENDOG | 0.14018 | 0.25414 | 0.26295 |
| ENSG00000088930 | 20 | 21283942 | 21370463 | 209 | 18 | 1.079 | 0.14028 | XRN2 | 0.14028 | 0.1363 | 0.16475 |
| ENSG00000143322 | 1 | 179068462 | 179198819 | 367 | 20 | 1.0789 | 0.14032 | ABL2 | 0.14032 | 0.16442 | 0.1876 |
| ENSG00000179151 | 15 | 74922899 | 74988633 | 76 | 16 | 1.0785 | 0.1404 | EDC3 | 0.1404 | 0.1937 | 0.21736 |
| ENSG00000268317 | 16 | 56662971 | 56666150 | 11 | 5 | 1.0783 | 0.14045 | AC026461.1 | 0.14045 | 0.3418 | 0.46173 |
| ENSG00000130299 | 19 | 17445729 | 17453544 | 32 | 7 | 1.0783 | 0.14045 | GTPBP3 | 0.14045 | 0.11986 | 0.13122 |
| ENSG00000257028 | 15 | 80215507 | 80217194 | 13 | 5 | 1.0773 | 0.14066 | C15ORF37 | 0.14066 | 0.50801 | 0.48226 |
| ENSG00000166922 | 15 | 32933877 | 32989299 | 181 | 47 | 1.0773 | 0.14068 | SCG5 | 0.14068 | 0.13561 | 0.17071 |
| ENSG00000140839 | 16 | 74442529 | 74455368 | 18 | 10 | 1.0772 | 0.14069 | CLEC18B | 0.14069 | 0.10083 | 0.27675 |
| ENSG00000161958 | 17 | 7341592 | 7348256 | 14 | 4 | 1.0767 | 0.14081 | FGF11 | 0.14081 | 0.40319 | 0.40028 |
| ENSG00000163872 | 3 | 183415606 | 183530413 | 261 | 16 | 1.0766 | 0.14082 | YEATS2 | 0.14082 | 0.11951 | 0.018923 |
| ENSG00000183615 | 1 | 32712834 | 32714457 | 2 | 1 | 1.0764 | 0.14088 | FAM167B | 0.14088 | 0.0072151 | 0.0058577 |
| ENSG00000169241 | 1 | 155107820 | 155111329 | 3 | 2 | 1.0758 | 0.141 | SLC50A1 | 0.141 | 0.021628 | 0.046539 |
| ENSG00000123473 | 1 | 47715811 | 47779819 | 195 | 13 | 1.0756 | 0.14105 | STIL | 0.14105 | 0.15577 | 0.15333 |
| ENSG00000241058 | 10 | 18834490 | 18940551 | 548 | 20 | 1.0755 | 0.14106 | NSUN6 | 0.14106 | 0.1254 | 0.11705 |
| ENSG00000143799 | 1 | 226548392 | 226595780 | 97 | 12 | 1.0753 | 0.14112 | PARP1 | 0.14112 | 0.067018 | 0.039333 |
| ENSG00000156374 | 10 | 105062553 | 105110891 | 107 | 18 | 1.0751 | 0.14117 | PCGF6 | 0.14117 | 0.13349 | 0.16697 |
| ENSG00000125734 | 19 | 6729925 | 6737614 | 31 | 10 | 1.0751 | 0.14117 | GPR108 | 0.14117 | 0.22199 | 0.2163 |
| ENSG00000179593 | 17 | 7942335 | 7952452 | 55 | 13 | 1.0742 | 0.14137 | ALOX15B | 0.14137 | 0.075023 | 0.14296 |
| ENSG00000134291 | 12 | 48357352 | 48362661 | 18 | 6 | 1.074 | 0.14142 | TMEM106C | 0.14142 | 0.17727 | 0.41626 |
| ENSG00000131470 | 17 | 40724333 | 40729849 | 13 | 6 | 1.0733 | 0.14156 | PSMC3IP | 0.14156 | 0.18768 | 0.25053 |
| ENSG00000143036 | 1 | 95285898 | 95360802 | 317 | 50 | 1.0733 | 0.14157 | SLC44A3 | 0.14157 | 0.2165 | 0.36617 |
| ENSG00000171246 | 17 | 78440948 | 78451643 | 36 | 10 | 1.0721 | 0.14183 | NPTX1 | 0.14183 | 0.17737 | 0.38317 |
| ENSG00000257962 | 8 | 86158524 | 86162077 | 12 | 4 | 1.071 | 0.14207 | RP11-219B4.6 | 0.14207 | 0.15062 | 0.066975 |
| ENSG00000228075 | 18 | 54814293 | 54817531 | 14 | 4 | 1.0711 | 0.14207 | BOD1L2 | 0.14207 | 0.48681 | 0.44667 |
| ENSG00000155545 | 5 | 56215429 | 56267502 | 198 | 22 | 1.071 | 0.14208 | MIER3 | 0.14208 | 0.14822 | 0.12302 |
| ENSG00000233280 | 3 | 97540884 | 97594132 | 82 | 10 | 1.071 | 0.14209 | CRYBG3 | 0.14209 | 0.15265 | 0.17518 |
| ENSG00000169371 | 15 | 75890424 | 75918810 | 50 | 13 | 1.0706 | 0.14218 | SNUPN | 0.14218 | 0.14908 | 0.17998 |
| ENSG00000112305 | 6 | 71377479 | 71571718 | 516 | 21 | 1.0703 | 0.14224 | SMAP1 | 0.14224 | 0.1281 | 0.14859 |
| ENSG00000164707 | 7 | 135365985 | 135414006 | 180 | 21 | 1.07 | 0.14231 | SLC13A4 | 0.14231 | 0.19781 | 0.30225 |
| ENSG00000213204 | 6 | 88117730 | 88221352 | 395 | 26 | 1.0698 | 0.14236 | C6orf165 | 0.14236 | 0.14369 | 0.14678 |
| ENSG00000176495 | 11 | 59131885 | 59132921 | 2 | 1 | 1.0697 | 0.14237 | OR5AN1 | 0.14237 | 0.32533 | 0.40049 |
| ENSG00000086300 | 7 | 26331541 | 26413949 | 295 | 27 | 1.0695 | 0.14243 | SNX10 | 0.14243 | 0.22037 | 0.29666 |
| ENSG00000129055 | 3 | 134196548 | 134205558 | 25 | 3 | 1.0694 | 0.14245 | ANAPC13 | 0.14245 | 0.12644 | 0.1329 |
| ENSG00000167632 | 8 | 140742586 | 141468678 | 2472 | 183 | 1.0693 | 0.14247 | TRAPPC9 | 0.14247 | 0.091481 | 0.08915 |
| ENSG00000119917 | 10 | 91087651 | 91100728 | 68 | 9 | 1.0691 | 0.14252 | IFIT3 | 0.14252 | 0.10714 | 0.094526 |
| ENSG00000077942 | 22 | 45898118 | 45997015 | 475 | 63 | 1.069 | 0.14253 | FBLN1 | 0.14253 | 0.086848 | 0.15976 |
| ENSG00000118939 | 13 | 76123619 | 76180085 | 197 | 19 | 1.0676 | 0.14285 | UCHL3 | 0.14285 | 0.1432 | 0.15666 |
| ENSG00000128567 | 7 | 131185021 | 131242976 | 217 | 45 | 1.067 | 0.14298 | PODXL | 0.14298 | 0.11605 | 0.18844 |
| ENSG00000133115 | 13 | 39540062 | 39565203 | 89 | 16 | 1.0669 | 0.14302 | STOML3 | 0.14302 | 0.53605 | 0.339 |
| ENSG00000185742 | 11 | 109292846 | 109299840 | 6 | 3 | 1.0657 | 0.14328 | C11orf87 | 0.14328 | 0.098047 | 0.075156 |
| ENSG00000186090 | 3 | 183749332 | 183757157 | 31 | 8 | 1.0655 | 0.14333 | HTR3D | 0.14333 | 0.47613 | 0.41499 |
| ENSG00000103723 | 15 | 83328033 | 83378666 | 144 | 24 | 1.0653 | 0.14336 | AP3B2 | 0.14336 | 0.17935 | 0.24868 |
| ENSG00000141627 | 18 | 46570039 | 46987717 | 1350 | 29 | 1.0651 | 0.14341 | DYM | 0.14341 | 0.13677 | 0.099676 |
| ENSG00000107771 | 10 | 86088342 | 86278273 | 682 | 25 | 1.0644 | 0.14357 | CCSER2 | 0.14357 | 0.1402 | 0.15184 |
| ENSG00000172769 | 11 | 58169937 | 58170882 | 9 | 1 | 1.0643 | 0.14361 | OR5B3 | 0.14361 | 0.037294 | 0.027399 |
| ENSG00000183665 | 8 | 125463048 | 125474391 | 37 | 7 | 1.064 | 0.14367 | TRMT12 | 0.14367 | 0.18475 | 0.14106 |
| ENSG00000110911 | 12 | 51373184 | 51422349 | 160 | 15 | 1.064 | 0.14367 | SLC11A2 | 0.14367 | 0.11859 | 0.14453 |
| ENSG00000063241 | 19 | 55964352 | 55973710 | 51 | 17 | 1.0632 | 0.14384 | ISOC2 | 0.14384 | 0.1634 | 0.013356 |
| ENSG00000115129 | 2 | 24300303 | 24308731 | 22 | 5 | 1.0627 | 0.14396 | TP53I3 | 0.14396 | 0.091679 | 0.10552 |
| ENSG00000151148 | 12 | 109915207 | 109974507 | 179 | 21 | 1.0624 | 0.14402 | UBE3B | 0.14402 | 0.15926 | 0.16721 |
| ENSG00000163430 | 3 | 120111140 | 120170100 | 200 | 25 | 1.0623 | 0.14406 | FSTL1 | 0.14406 | 0.18207 | 0.22758 |
| ENSG00000105058 | 19 | 16296191 | 16302857 | 18 | 5 | 1.0622 | 0.14407 | FAM32A | 0.14407 | 0.049599 | 0.092034 |
| ENSG00000196914 | 11 | 120207787 | 120360645 | 316 | 17 | 1.062 | 0.14412 | ARHGEF12 | 0.14412 | 0.094415 | 0.078018 |

| | | | | | | | | | | | |
|---|---|---|---|---|---|---|---|---|---|---|---|
| ENSG00000130159 | 19 | 11616731 | 11639989 | 63 | 10 | 1.0616 | 0.1442 | ECSIT | 0.1442 | 0.13084 | 0.11891 |
| ENSG00000140009 | 14 | 64550950 | 64804830 | 781 | 44 | 1.0615 | 0.14423 | ESR2 | 0.14423 | 0.151 | 0.16501 |
| ENSG00000031691 | 6 | 49431091 | 49460820 | 71 | 9 | 1.0612 | 0.1443 | CENPQ | 0.1443 | 0.22798 | 0.3524 |
| ENSG00000066382 | 11 | 30406040 | 30608419 | 576 | 39 | 1.0608 | 0.14438 | MPPED2 | 0.14438 | 0.14915 | 0.091925 |
| ENSG00000215306 | 14 | 23286463 | 23291750 | 27 | 10 | 1.0606 | 0.14444 | AL135998.1 | 0.14444 | 0.44172 | 0.56391 |
| ENSG00000133026 | 17 | 8377523 | 8534079 | 297 | 20 | 1.0606 | 0.14444 | MYH10 | 0.14444 | 0.13921 | 0.11813 |
| ENSG00000183072 | 5 | 172659112 | 172662360 | 7 | 3 | 1.0605 | 0.14446 | NKX2-5 | 0.14446 | 0.031706 | 0.05474 |
| ENSG00000156171 | 1 | 111659955 | 111682838 | 59 | 8 | 1.0602 | 0.14452 | DRAM2 | 0.14452 | 0.10391 | 0.09431 |
| ENSG00000103254 | 16 | 770581 | 772601 | 8 | 1 | 1.0601 | 0.14455 | FAM173A | 0.14455 | 0.31061 | 0.3526 |
| ENSG00000234719 | 16 | 12021230 | 12070500 | 213 | 24 | 1.0599 | 0.14458 | RP11-166B2.1 | 0.14458 | 0.071256 | 0.033512 |
| ENSG00000101464 | 20 | 33148346 | 33264910 | 198 | 15 | 1.0597 | 0.14464 | PIGU | 0.14464 | 0.16238 | 0.17693 |
| ENSG00000168275 | 1 | 234509202 | 234519795 | 52 | 5 | 1.0593 | 0.14473 | COA6 | 0.14473 | 0.12814 | 0.13884 |
| ENSG00000196313 | 7 | 72349936 | 72421979 | 88 | 26 | 1.0592 | 0.14476 | POM121 | 0.14476 | 0.19258 | 0.2488 |
| ENSG00000185386 | 22 | 50702142 | 50709196 | 22 | 6 | 1.0587 | 0.14486 | MAPK11 | 0.14486 | 0.14251 | 0.16098 |
| ENSG00000131203 | 8 | 39759794 | 39785963 | 54 | 8 | 1.0587 | 0.14487 | IDO1 | 0.14487 | 0.16572 | 0.1912 |
| ENSG00000187054 | 4 | 68775103 | 68829858 | 166 | 23 | 1.0586 | 0.1449 | TMPRSS11A | 0.1449 | 0.18518 | 0.25595 |
| ENSG00000170893 | 3 | 129693148 | 129696781 | 9 | 4 | 1.0583 | 0.14496 | TRH | 0.14496 | 0.23951 | 0.35746 |
| ENSG00000118777 | 4 | 89011416 | 89152474 | 542 | 45 | 1.0583 | 0.14496 | ABCG2 | 0.14496 | 0.15715 | 0.16472 |
| ENSG00000092203 | 14 | 21944756 | 21967319 | 94 | 10 | 1.0583 | 0.14497 | TOX4 | 0.14497 | 0.18102 | 0.13078 |
| ENSG00000111615 | 12 | 75890684 | 75905416 | 48 | 8 | 1.0581 | 0.145 | KRR1 | 0.145 | 0.2117 | 0.32306 |
| ENSG00000150991 | 12 | 125396150 | 125401914 | 19 | 6 | 1.058 | 0.14502 | UBC | 0.14502 | 0.18692 | 0.48166 |
| ENSG00000170322 | 11 | 129733531 | 129765485 | 72 | 11 | 1.058 | 0.14503 | NFRKB | 0.14503 | 0.19345 | 0.18842 |
| ENSG00000092470 | 15 | 44119161 | 44160617 | 86 | 9 | 1.0577 | 0.14509 | WDR76 | 0.14509 | 0.16518 | 0.15688 |
| ENSG00000204128 | 2 | 231902205 | 231914434 | 44 | 9 | 1.0575 | 0.14513 | C2orf72 | 0.14513 | 0.099179 | 0.2078 |
| ENSG00000182308 | 4 | 41983713 | 41988476 | 16 | 4 | 1.0575 | 0.14515 | DCAF4L1 | 0.14515 | 0.24699 | 0.21202 |
| ENSG00000162878 | 2 | 42275160 | 42285668 | 40 | 6 | 1.0574 | 0.14516 | PKDCC | 0.14516 | 0.24867 | 0.4314 |
| ENSG00000159399 | 2 | 75061108 | 75120486 | 214 | 33 | 1.0572 | 0.1452 | HK2 | 0.1452 | 0.26852 | 0.28928 |
| ENSG00000152234 | 18 | 43664110 | 43684300 | 102 | 9 | 1.0572 | 0.14521 | ATP5A1 | 0.14521 | 0.15493 | 0.14086 |
| ENSG00000179934 | 3 | 39371197 | 39375171 | 6 | 3 | 1.0571 | 0.14522 | CCR8 | 0.14522 | 0.47287 | 0.51668 |
| ENSG00000067064 | 10 | 1085848 | 1095110 | 24 | 8 | 1.0571 | 0.14523 | IDI1 | 0.14523 | 0.112 | 0.13034 |
| ENSG00000183340 | 11 | 96123153 | 96240738 | 332 | 24 | 1.057 | 0.14527 | JRKL | 0.14527 | 0.13945 | 0.17116 |
| ENSG00000258830 | 12 | 57643392 | 57690267 | 77 | 16 | 1.0567 | 0.14532 | RP11-123K3.4 | 0.14532 | 0.11883 | 0.098648 |
| ENSG00000253276 | 7 | 106297211 | 106301442 | 10 | 4 | 1.0566 | 0.14534 | CCDC71L | 0.14534 | 0.11931 | 0.14867 |
| ENSG00000212722 | 17 | 39261584 | 39262740 | 13 | 4 | 1.0565 | 0.14538 | KRTAP4-9 | 0.14538 | 0.2977 | 0.36461 |
| ENSG00000151846 | 13 | 25670300 | 25673389 | 13 | 5 | 1.0558 | 0.14552 | PABPC3 | 0.14552 | 0.044978 | 0.25042 |
| ENSG00000186020 | 19 | 37025676 | 37096178 | 173 | 18 | 1.0557 | 0.14555 | ZNF529 | 0.14555 | 0.19314 | 0.26602 |
| ENSG00000110680 | 11 | 14988214 | 14993900 | 10 | 5 | 1.0556 | 0.14557 | CALCA | 0.14557 | 0.16962 | 0.077771 |
| ENSG00000162068 | 16 | 2521500 | 2524146 | 1 | 1 | 1.0555 | 0.1456 | NTN3 | 0.1456 | 0.037452 | 0.0013098 |
| ENSG00000145414 | 4 | 164031225 | 164088073 | 138 | 20 | 1.0554 | 0.14563 | NAF1 | 0.14563 | 0.16447 | 0.1932 |
| ENSG00000198818 | 6 | 166733216 | 166756079 | 60 | 7 | 1.0546 | 0.1458 | SFT2D1 | 0.1458 | 0.28845 | 0.38887 |
| ENSG00000089558 | 17 | 40308909 | 40333296 | 43 | 11 | 1.0546 | 0.1458 | KCNH4 | 0.1458 | 0.088641 | 0.24728 |
| ENSG00000146085 | 6 | 49398073 | 49430904 | 84 | 10 | 1.0543 | 0.14587 | MUT | 0.14587 | 0.16073 | 0.13155 |
| ENSG00000177464 | 19 | 46093022 | 46105466 | 43 | 8 | 1.0543 | 0.14587 | GPR4 | 0.14587 | 0.18243 | 0.031922 |
| ENSG00000112077 | 6 | 49572871 | 49604552 | 127 | 15 | 1.0542 | 0.14589 | RHAG | 0.14589 | 0.28034 | 0.24599 |
| ENSG00000182687 | 17 | 74070875 | 74073622 | 2 | 1 | 1.054 | 0.14594 | GALR2 | 0.14594 | 0.59046 | 0.58643 |
| ENSG00000214360 | 5 | 171621176 | 171630456 | 51 | 13 | 1.0539 | 0.14596 | EFCAB9 | 0.14596 | 0.20942 | 0.069543 |
| ENSG00000114354 | 3 | 100428205 | 100467810 | 63 | 9 | 1.0535 | 0.14606 | TFG | 0.14606 | 0.19301 | 0.26825 |
| ENSG00000003989 | 8 | 17354597 | 17428082 | 471 | 71 | 1.0535 | 0.14606 | SLC7A2 | 0.14606 | 0.20184 | 0.12936 |
| ENSG00000198443 | 17 | 39340354 | 39341594 | 13 | 2 | 1.0535 | 0.14606 | KRTAP4-1 | 0.14606 | 0.2099 | 0.24533 |
| ENSG00000188581 | 17 | 39196811 | 39197713 | 6 | 2 | 1.0533 | 0.14611 | KRTAP1-1 | 0.14611 | 0.67262 | 0.60339 |
| ENSG00000163975 | 3 | 196715492 | 196756687 | 159 | 32 | 1.053 | 0.14616 | MFI2 | 0.14616 | 0.11613 | 0.17116 |
| ENSG00000214290 | 11 | 111169565 | 111179460 | 33 | 7 | 1.053 | 0.14617 | COLCA2 | 0.14617 | 0.25216 | 0.28621 |
| ENSG00000196865 | 10 | 115614420 | 115676953 | 61 | 14 | 1.0529 | 0.14618 | NHLRC2 | 0.14618 | 0.20912 | 0.21431 |
| ENSG00000182923 | 3 | 134204585 | 134293859 | 252 | 7 | 1.0529 | 0.14619 | CEP63 | 0.14619 | 0.14755 | 0.15095 |
| ENSG00000254788 | 16 | 66586490 | 66613038 | 54 | 4 | 1.0528 | 0.14622 | CKLF-CMTM1 | 0.14622 | 0.11962 | 0.11856 |
| ENSG00000131381 | 3 | 15111580 | 15140670 | 70 | 5 | 1.0526 | 0.14626 | ZFYVE20 | 0.14626 | 0.13294 | 0.12734 |
| ENSG00000109819 | 4 | 23756664 | 23905712 | 569 | 74 | 1.0526 | 0.14626 | PPARGC1A | 0.14626 | 0.13579 | 0.16819 |
| ENSG00000213462 | 7 | 64451187 | 64467121 | 52 | 9 | 1.0512 | 0.14657 | ERV3-1 | 0.14657 | 0.13813 | 0.13031 |
| ENSG00000174652 | 19 | 9523272 | 9546254 | 103 | 3 | 1.051 | 0.14662 | ZNF266 | 0.14662 | 0.096726 | 0.064742 |
| ENSG00000196782 | 4 | 140637907 | 141075338 | 1340 | 124 | 1.0505 | 0.14674 | MAML3 | 0.14674 | 0.16251 | 0.18148 |
| ENSG00000100429 | 22 | 50683612 | 50689834 | 21 | 5 | 1.0502 | 0.14682 | HDAC10 | 0.14682 | 0.19237 | 0.21299 |
| ENSG00000237388 | 11 | 48510269 | 48511332 | 5 | 3 | 1.05 | 0.14685 | OR4A47 | 0.14685 | 0.19037 | 0.20886 |
| ENSG00000125755 | 19 | 46318668 | 46366548 | 205 | 13 | 1.05 | 0.14685 | SYMPK | 0.14685 | 0.10749 | 0.05113 |
| ENSG00000172421 | 17 | 60447579 | 60493837 | 87 | 14 | 1.0499 | 0.14689 | EFCAB3 | 0.14689 | 0.25212 | 0.4615 |
| ENSG00000255423 | 3 | 73110810 | 73112488 | 11 | 3 | 1.0497 | 0.14693 | EBLN2 | 0.14693 | 0.1168 | 0.1581 |
| ENSG00000153721 | 6 | 154708638 | 154831793 | 663 | 73 | 1.0496 | 0.14696 | CNKSR3 | 0.14696 | 0.147 | 0.13961 |
| ENSG00000122257 | 16 | 24549014 | 24584184 | 102 | 10 | 1.0495 | 0.14698 | RBBP6 | 0.14698 | 0.14237 | 0.22025 |
| ENSG00000183397 | 19 | 3539152 | 3544028 | 21 | 7 | 1.0495 | 0.14698 | C19orf71 | 0.14698 | 0.058841 | 0.061369 |
| ENSG00000249709 | 19 | 12636184 | 12662356 | 60 | 14 | 1.0494 | 0.147 | ZNF564 | 0.147 | 0.040095 | 0.14447 |
| ENSG00000178358 | 11 | 6942205 | 6943264 | 9 | 4 | 1.0492 | 0.14705 | OR2D3 | 0.14705 | 0.33043 | 0.3193 |
| ENSG00000181036 | 1 | 159770301 | 159786041 | 64 | 11 | 1.0485 | 0.1472 | FCRL6 | 0.1472 | 0.097962 | 0.14758 |
| ENSG00000136867 | 9 | 115913222 | 115926417 | 79 | 14 | 1.047 | 0.14756 | SLC31A2 | 0.14756 | 0.1396 | 0.19846 |
| ENSG00000064961 | 19 | 3572775 | 3579086 | 12 | 5 | 1.0465 | 0.14765 | HMG20B | 0.14765 | 0.077659 | 0.00075739 |

| | | | | | | | | | | | |
|---|---|---|---|---|---|---|---|---|---|---|---|
| ENSG00000173809 | 19 | 33210659 | 33320483 | 326 | 18 | 1.0464 | 0.14768 | TDRD12 | 0.14768 | 0.15992 | 0.17746 |
| ENSG00000255508 | 11 | 62327075 | 62359003 | 50 | 8 | 1.0464 | 0.14769 | MIR3654 | 0.14769 | 0.19455 | 0.21253 |
| ENSG00000182240 | 21 | 42539728 | 42654445 | 366 | 66 | 1.046 | 0.14777 | BACE2 | 0.14777 | 0.18614 | 0.24271 |
| ENSG00000254806 | 20 | 43991840 | 44039250 | 114 | 8 | 1.0454 | 0.14793 | SYS1-DBNDD2 | 0.14793 | 0.13113 | 0.079721 |
| ENSG00000175305 | 8 | 95891998 | 95908906 | 62 | 13 | 1.0453 | 0.14795 | CCNE2 | 0.14795 | 0.035587 | 0.010312 |
| ENSG00000100721 | 14 | 96176304 | 96180533 | 14 | 4 | 1.0452 | 0.14795 | TCL1A | 0.14795 | 0.42456 | 0.56361 |
| ENSG00000254999 | 3 | 10157276 | 10168874 | 67 | 5 | 1.045 | 0.14801 | BRK1 | 0.14801 | 0.051074 | 0.039102 |
| ENSG00000111875 | 6 | 119215384 | 119230332 | 50 | 10 | 1.0449 | 0.14803 | ASF1A | 0.14803 | 0.21517 | 0.21695 |
| ENSG00000142546 | 19 | 50058968 | 50093519 | 72 | 15 | 1.0449 | 0.14804 | NOSIP | 0.14804 | 0.1642 | 0.19696 |
| ENSG00000248050 | 8 | 110656344 | 110660313 | 16 | 6 | 1.0446 | 0.1481 | RP11-422N16.3 | 0.1481 | 0.22475 | 0.39199 |
| ENSG00000175664 | 13 | 31506840 | 31549639 | 157 | 22 | 1.0443 | 0.14817 | TEX26 | 0.14817 | 0.14687 | 0.20404 |
| ENSG00000138867 | 22 | 24936406 | 24951903 | 45 | 7 | 1.0442 | 0.1482 | GUCD1 | 0.1482 | 0.15531 | 0.11713 |
| ENSG00000187048 | 1 | 47394849 | 47407156 | 51 | 7 | 1.0441 | 0.14821 | CYP4A11 | 0.14821 | 0.15908 | 0.15054 |
| ENSG00000125901 | 20 | 3026591 | 3028900 | 6 | 3 | 1.0438 | 0.14828 | MRPS26 | 0.14828 | 0.094122 | 0.10417 |
| ENSG00000106789 | 9 | 100883257 | 100954922 | 289 | 44 | 1.0437 | 0.14831 | CORO2A | 0.14831 | 0.21557 | 0.18061 |
| ENSG00000113369 | 5 | 90664541 | 90679176 | 21 | 8 | 1.0436 | 0.14833 | ARRDC3 | 0.14833 | 0.17666 | 0.27494 |
| ENSG00000162571 | 1 | 1109264 | 1133315 | 154 | 19 | 1.0433 | 0.1484 | TTLL10 | 0.1484 | 0.22448 | 0.12705 |
| ENSG00000106459 | 7 | 129251555 | 129396922 | 451 | 34 | 1.0432 | 0.14843 | NRF1 | 0.14843 | 0.17378 | 0.17543 |
| ENSG00000140955 | 16 | 84224744 | 84230774 | 37 | 10 | 1.0432 | 0.14843 | ADAD2 | 0.14843 | 0.25018 | 0.18518 |
| ENSG00000101346 | 20 | 30795683 | 30826470 | 27 | 8 | 1.043 | 0.14848 | POFUT1 | 0.14848 | 0.29564 | 0.21467 |
| ENSG00000174839 | 3 | 57611184 | 57678816 | 184 | 22 | 1.0429 | 0.1485 | DENND6A | 0.1485 | 0.14899 | 0.17509 |
| ENSG00000102974 | 16 | 67596310 | 67673086 | 142 | 22 | 1.0418 | 0.14875 | CTCF | 0.14875 | 0.15591 | 0.21878 |
| ENSG00000108960 | 17 | 53469974 | 53499353 | 69 | 10 | 1.0416 | 0.14881 | MMD | 0.14881 | 0.1895 | 0.35529 |
| ENSG00000140262 | 15 | 57210821 | 57591479 | 942 | 29 | 1.0415 | 0.14882 | TCF12 | 0.14882 | 0.13514 | 0.13152 |
| ENSG00000180370 | 3 | 196466728 | 196559518 | 440 | 38 | 1.0415 | 0.14883 | PAK2 | 0.14883 | 0.093743 | 0.094053 |
| ENSG00000100764 | 14 | 90722839 | 90738968 | 80 | 3 | 1.0414 | 0.14884 | PSMC1 | 0.14884 | 0.15565 | 0.17052 |
| ENSG00000081041 | 4 | 74962752 | 74965010 | 3 | 2 | 1.0412 | 0.1489 | CXCL2 | 0.1489 | 0.27061 | 0.43187 |
| ENSG00000164484 | 6 | 130686879 | 130764208 | 292 | 18 | 1.0411 | 0.1489 | TMEM200A | 0.1489 | 0.14817 | 0.17919 |
| ENSG00000124251 | 20 | 44002526 | 44036529 | 89 | 8 | 1.0404 | 0.14908 | TP53TG5 | 0.14908 | 0.14149 | 0.10275 |
| ENSG00000106066 | 7 | 29034847 | 29235067 | 849 | 77 | 1.04 | 0.14916 | CPVL | 0.14916 | 0.15713 | 0.096547 |
| ENSG00000115548 | 2 | 86667770 | 86719839 | 135 | 17 | 1.0399 | 0.1492 | KDM3A | 0.1492 | 0.083785 | 0.07308 |
| ENSG00000270757 | 2 | 198365137 | 198415450 | 80 | 10 | 1.0394 | 0.14931 | HSPE1-MOB4 | 0.14931 | 0.10351 | 0.11631 |
| ENSG00000136930 | 9 | 127115745 | 127177723 | 125 | 14 | 1.0394 | 0.14932 | PSMB7 | 0.14932 | 0.11272 | 0.090854 |
| ENSG00000251503 | 1 | 10490159 | 10512210 | 72 | 7 | 1.0388 | 0.14945 | APITD1-CORT | 0.14945 | 0.16633 | 0.12392 |
| ENSG00000175279 | 1 | 10490159 | 10512210 | 72 | 7 | 1.0388 | 0.14945 | APITD1 | 0.14945 | 0.16633 | 0.12392 |
| ENSG00000104938 | 19 | 7828035 | 7834491 | 34 | 7 | 1.0385 | 0.14952 | CLEC4M | 0.14952 | 0.10239 | 0.16177 |
| ENSG00000268861 | 19 | 7445850 | 7535131 | 391 | 21 | 1.0385 | 0.14953 | CTD-2207O23.3 | 0.14953 | 0.16835 | 0.14886 |
| ENSG00000138095 | 2 | 44113647 | 44223144 | 355 | 21 | 1.0384 | 0.14955 | LRPPRC | 0.14955 | 0.1915 | 0.2734 |
| ENSG00000188277 | 15 | 41062159 | 41064643 | 4 | 2 | 1.0383 | 0.14956 | C15orf62 | 0.14956 | 0.3689 | 0.54415 |
| ENSG00000175874 | 2 | 101962013 | 102004057 | 140 | 14 | 1.0381 | 0.1496 | CREG2 | 0.1496 | 0.12022 | 0.16699 |
| ENSG00000122691 | 7 | 19060614 | 19157295 | 325 | 20 | 1.0381 | 0.14962 | TWIST1 | 0.14962 | 0.092281 | 0.058078 |
| ENSG00000112333 | 6 | 108487262 | 108510013 | 30 | 10 | 1.0377 | 0.14972 | NR2E1 | 0.14972 | 0.13438 | 0.054495 |
| ENSG00000100079 | 22 | 37966255 | 37978623 | 71 | 12 | 1.0376 | 0.14973 | LGALS2 | 0.14973 | 0.23473 | 0.31719 |
| ENSG00000178996 | 5 | 53813589 | 53842415 | 137 | 20 | 1.0375 | 0.14975 | SNX18 | 0.14975 | 0.18454 | 0.20646 |
| ENSG00000221947 | 8 | 71581600 | 71702606 | 580 | 15 | 1.0374 | 0.14977 | XKR9 | 0.14977 | 0.15024 | 0.15765 |
| ENSG00000112115 | 6 | 52051185 | 52055436 | 9 | 3 | 1.037 | 0.14987 | IL17A | 0.14987 | 0.67502 | 0.73425 |
| ENSG00000168356 | 3 | 38887260 | 38992052 | 312 | 25 | 1.0368 | 0.14992 | SCN11A | 0.14992 | 0.14559 | 0.15493 |
| ENSG00000080815 | 14 | 73603126 | 73690399 | 207 | 23 | 1.0367 | 0.14994 | PSEN1 | 0.14994 | 0.15167 | 0.15286 |
| ENSG00000197262 | 17 | 34639793 | 34641846 | 2 | 1 | 1.0366 | 0.14996 | CCL4L2 | 0.14996 | 0.095536 | 0.019144 |
| ENSG00000178409 | 6 | 107386386 | 107436473 | 156 | 25 | 1.0364 | 0.15 | BEND3 | 0.15 | 0.10369 | 0.098158 |
| ENSG00000108021 | 10 | 5726801 | 5805703 | 230 | 28 | 1.0353 | 0.15026 | FAM208B | 0.15026 | 0.11487 | 0.12911 |
| ENSG00000167613 | 19 | 54865362 | 54882165 | 63 | 17 | 1.0352 | 0.15028 | LAIR1 | 0.15028 | 0.28436 | 0.38028 |
| ENSG00000152611 | 5 | 35904397 | 35938881 | 142 | 16 | 1.0351 | 0.15032 | CAPSL | 0.15032 | 0.17357 | 0.1659 |
| ENSG00000091583 | 17 | 64208151 | 64252643 | 183 | 22 | 1.035 | 0.15034 | APOH | 0.15034 | 0.16974 | 0.20875 |
| ENSG00000091542 | 17 | 18086392 | 18113268 | 88 | 12 | 1.0349 | 0.15036 | ALKBH5 | 0.15036 | 0.12443 | 0.16907 |
| ENSG00000197608 | 19 | 52567719 | 52599018 | 122 | 11 | 1.0344 | 0.15047 | ZNF841 | 0.15047 | 0.22388 | 0.25606 |
| ENSG00000217442 | 22 | 50989541 | 51001334 | 108 | 7 | 1.0339 | 0.15059 | SYCE3 | 0.15059 | 0.36392 | 0.48048 |
| ENSG00000255284 | 11 | 777578 | 784297 | 19 | 6 | 1.0329 | 0.15083 | AP006621.5 | 0.15083 | 0.27024 | 0.4248 |
| ENSG00000164081 | 3 | 51696709 | 51738339 | 47 | 19 | 1.0328 | 0.15084 | TEX264 | 0.15084 | 0.27514 | 0.35448 |
| ENSG00000141527 | 17 | 78143791 | 78183130 | 179 | 38 | 1.0324 | 0.15095 | CARD14 | 0.15095 | 0.20246 | 0.15582 |
| ENSG00000099797 | 19 | 14627897 | 14676792 | 185 | 29 | 1.0319 | 0.15105 | TECR | 0.15105 | 0.10946 | 0.095333 |
| ENSG00000165646 | 10 | 119000604 | 119038941 | 153 | 29 | 1.031 | 0.15127 | SLC18A2 | 0.15127 | 0.19895 | 0.31777 |
| ENSG00000132631 | 20 | 18794370 | 18795033 | 3 | 1 | 1.031 | 0.15127 | SCP2D1 | 0.15127 | 0.086942 | 0.11256 |
| ENSG00000221840 | 11 | 51411378 | 51412448 | 7 | 3 | 1.0309 | 0.15129 | OR4A5 | 0.15129 | 0.093546 | 0.11152 |
| ENSG00000135966 | 2 | 105880871 | 105946491 | 228 | 28 | 1.0308 | 0.15132 | TGFBRAP1 | 0.15132 | 0.13089 | 0.18244 |
| ENSG00000157045 | 16 | 15131710 | 15149921 | 57 | 6 | 1.0303 | 0.15144 | NTAN1 | 0.15144 | 0.19561 | 0.25772 |
| ENSG00000126698 | 1 | 28525967 | 28559536 | 100 | 12 | 1.0297 | 0.15157 | DNAJC8 | 0.15157 | 0.11866 | 0.13142 |
| ENSG00000249008 | 4 | 178163693 | 178169927 | 32 | 9 | 1.0292 | 0.1517 | RP11-487E13.1 | 0.1517 | 0.12373 | 0.36124 |
| ENSG00000179172 | 1 | 12907261 | 12908578 | 7 | 3 | 1.0271 | 0.15218 | HNRNPCL1 | 0.15218 | 0.18124 | 0.19756 |
| ENSG00000156017 | 9 | 77595936 | 77643339 | 123 | 20 | 1.0271 | 0.1522 | C9orf41 | 0.1522 | 0.10664 | 0.096485 |
| ENSG00000169629 | 2 | 113127669 | 113192065 | 37 | 10 | 1.027 | 0.15222 | RGPD8 | 0.15222 | 0.17978 | 0.19857 |
| ENSG00000140853 | 16 | 57023397 | 57117443 | 449 | 77 | 1.0264 | 0.15235 | NLRC5 | 0.15235 | 0.085058 | 0.16199 |
| ENSG00000084733 | 2 | 26256976 | 26360323 | 245 | 11 | 1.0262 | 0.15239 | RAB10 | 0.15239 | 0.15171 | 0.15328 |

| | | | | | | | | | | | |
|---|---|---|---|---|---|---|---|---|---|---|---|
| ENSG00000104903 | 19 | 13209847 | 13213975 | 17 | 7 | 1.0259 | 0.15247 | LYL1 | 0.15247 | 0.16158 | 0.073779 |
| ENSG00000146066 | 5 | 175815748 | 175816772 | 5 | 2 | 1.0257 | 0.15252 | HIGD2A | 0.15252 | 0.18625 | 0.38601 |
| ENSG00000229676 | 19 | 22817126 | 22850472 | 252 | 6 | 1.0255 | 0.15255 | ZNF492 | 0.15255 | 0.15604 | 0.1646 |
| ENSG00000152422 | 5 | 82373317 | 82649606 | 760 | 33 | 1.0253 | 0.1526 | XRCC4 | 0.1526 | 0.1606 | 0.17344 |
| ENSG00000116260 | 1 | 180123969 | 180173165 | 276 | 22 | 1.0253 | 0.15262 | QSOX1 | 0.15262 | 0.12273 | 0.16115 |
| ENSG00000002330 | 11 | 64037302 | 64052176 | 28 | 11 | 1.0241 | 0.15289 | BAD | 0.15289 | 0.23083 | 0.26916 |
| ENSG00000120053 | 10 | 101156627 | 101190381 | 73 | 14 | 1.0237 | 0.153 | GOT1 | 0.153 | 0.20676 | 0.17769 |
| ENSG00000167080 | 17 | 47209822 | 47247351 | 136 | 24 | 1.0234 | 0.15306 | B4GALNT2 | 0.15306 | 0.14351 | 0.13093 |
| ENSG00000214325 | 16 | 75728367 | 75734089 | 18 | 5 | 1.0227 | 0.15322 | AC025287.1 | 0.15322 | 0.22462 | 0.33435 |
| ENSG00000196460 | 2 | 102013823 | 102091165 | 397 | 18 | 1.0226 | 0.15325 | RFX8 | 0.15325 | 0.17633 | 0.20846 |
| ENSG00000188938 | 9 | 96208776 | 96215874 | 25 | 8 | 1.0225 | 0.15327 | FAM120AOS | 0.15327 | 0.28217 | 0.2902 |
| ENSG00000205423 | 16 | 50058321 | 50070999 | 29 | 5 | 1.0223 | 0.15332 | CNEP1R1 | 0.15332 | 0.16496 | 0.2405 |
| ENSG00000104979 | 19 | 13884982 | 13889276 | 23 | 3 | 1.0217 | 0.15345 | C19orf53 | 0.15345 | 0.17609 | 0.371 |
| ENSG00000171045 | 8 | 143293441 | 143484601 | 879 | 43 | 1.0216 | 0.15347 | TSNARE1 | 0.15347 | 0.092319 | 0.058608 |
| ENSG00000107742 | 10 | 73818793 | 73848790 | 95 | 11 | 1.0215 | 0.15351 | SPOCK2 | 0.15351 | 0.18655 | 0.20395 |
| ENSG00000132016 | 19 | 13993161 | 14017265 | 81 | 4 | 1.0215 | 0.15351 | C19orf57 | 0.15351 | 0.19997 | 0.30017 |
| ENSG00000174667 | 19 | 9324526 | 9325542 | 4 | 2 | 1.0214 | 0.15354 | OR7D4 | 0.15354 | 0.78262 | 0.55953 |
| ENSG00000149443 | 20 | 18790372 | 18810797 | 73 | 17 | 1.0212 | 0.15359 | C20orf78 | 0.15359 | 0.089196 | 0.04764 |
| ENSG00000267228 | 18 | 44661027 | 44702652 | 175 | 16 | 1.0208 | 0.15367 | IER3IP1 | 0.15367 | 0.082593 | 0.083522 |
| ENSG00000269121 | 16 | 12062000 | 12062333 | 1 | 1 | 1.0207 | 0.1537 | AC007216.2 | 0.1537 | 0.09275 | 0.057499 |
| ENSG00000146067 | 5 | 176946789 | 176981542 | 64 | 9 | 1.0206 | 0.15373 | FAM193B | 0.15373 | 0.16244 | 0.18494 |
| ENSG00000099899 | 22 | 20099389 | 20104915 | 21 | 8 | 1.0203 | 0.1538 | TRMT2A | 0.1538 | 0.036392 | 0.046466 |
| ENSG00000068781 | 2 | 48757064 | 49003654 | 1081 | 65 | 1.0202 | 0.15381 | STON1-GTF2A1L | 0.15381 | 0.14732 | 0.13779 |
| ENSG00000118096 | 11 | 118415243 | 118443685 | 76 | 7 | 1.0201 | 0.15385 | IFT46 | 0.15385 | 0.13381 | 0.13846 |
| ENSG00000163444 | 1 | 202976514 | 202993976 | 35 | 7 | 1.0195 | 0.15399 | TMEM183A | 0.15399 | 0.12769 | 0.24029 |
| ENSG00000182568 | 3 | 18386879 | 18487080 | 176 | 37 | 1.0194 | 0.15402 | SATB1 | 0.15402 | 0.2821 | 0.38084 |
| ENSG00000023892 | 6 | 35265595 | 35289548 | 56 | 9 | 1.0193 | 0.15404 | DEF6 | 0.15404 | 0.16375 | 0.21144 |
| ENSG00000144306 | 2 | 175260458 | 175294303 | 125 | 8 | 1.0187 | 0.15418 | SCRN3 | 0.15418 | 0.16986 | 0.15895 |
| ENSG00000237289 | 15 | 43885252 | 43897099 | 20 | 2 | 1.0185 | 0.15423 | CKMT1B | 0.15423 | 0.18242 | 0.33295 |
| ENSG00000114812 | 3 | 42530791 | 42579059 | 145 | 30 | 1.0182 | 0.15429 | VIPR1 | 0.15429 | 0.287 | 0.43103 |
| ENSG00000159110 | 21 | 34602206 | 34637980 | 133 | 14 | 1.0181 | 0.15431 | IFNAR2 | 0.15431 | 0.19232 | 0.072485 |
| ENSG00000105339 | 8 | 142127377 | 142205907 | 211 | 43 | 1.0181 | 0.15432 | DENND3 | 0.15432 | 0.24983 | 0.28416 |
| ENSG00000180016 | 17 | 3300398 | 3301704 | 3 | 2 | 1.0181 | 0.15432 | OR1E1 | 0.15432 | 0.3266 | 0.15999 |
| ENSG00000157570 | 11 | 44748015 | 44953972 | 806 | 80 | 1.018 | 0.15434 | TSPAN18 | 0.15434 | 0.16803 | 0.25834 |
| ENSG00000105136 | 19 | 57999079 | 58006048 | 34 | 5 | 1.0179 | 0.15435 | ZNF419 | 0.15435 | 0.2629 | 0.45763 |
| ENSG00000138315 | 10 | 74653339 | 74692791 | 41 | 7 | 1.0175 | 0.15445 | OIT3 | 0.15445 | 0.17094 | 0.098163 |
| ENSG00000040487 | 1 | 19638820 | 19655794 | 105 | 15 | 1.0173 | 0.1545 | PQLC2 | 0.1545 | 0.10246 | 0.11847 |
| ENSG00000116857 | 1 | 201103900 | 201140702 | 153 | 27 | 1.0164 | 0.15471 | TMEM9 | 0.15471 | 0.17832 | 0.33113 |
| ENSG00000163050 | 1 | 227085237 | 227175246 | 261 | 43 | 1.016 | 0.15481 | ADCK3 | 0.15481 | 0.24265 | 0.25212 |
| ENSG00000120910 | 8 | 22298332 | 22398652 | 227 | 13 | 1.0158 | 0.15486 | PPP3CC | 0.15486 | 0.13758 | 0.11467 |
| ENSG00000231880 | 11 | 47599277 | 47599823 | 1 | 1 | 1.0156 | 0.1549 | KBTBD4 | 0.1549 | 0.61116 | 0.63419 |
| ENSG00000164078 | 3 | 49924435 | 49941299 | 27 | 6 | 1.0156 | 0.15491 | MST1R | 0.15491 | 0.12918 | 0.16817 |
| ENSG00000042813 | 7 | 49890017 | 50160925 | 781 | 22 | 1.0156 | 0.15492 | ZPBP | 0.15492 | 0.1763 | 0.19467 |
| ENSG00000172995 | 3 | 35680437 | 35835988 | 366 | 33 | 1.0152 | 0.155 | ARPP21 | 0.155 | 0.19114 | 0.21977 |
| ENSG00000197191 | 9 | 140119087 | 140120763 | 4 | 2 | 1.0151 | 0.15503 | C9orf169 | 0.15503 | 0.54121 | 0.45041 |
| ENSG00000105698 | 19 | 35759881 | 35770724 | 38 | 7 | 1.015 | 0.15505 | USF2 | 0.15505 | 0.21752 | 0.18084 |
| ENSG00000116983 | 1 | 40144320 | 40157361 | 78 | 13 | 1.0142 | 0.15524 | HPCAL4 | 0.15524 | 0.19595 | 0.31957 |
| ENSG00000174567 | 1 | 204167288 | 204183220 | 43 | 12 | 1.0138 | 0.15533 | GOLT1A | 0.15533 | 0.42346 | 0.50963 |
| ENSG00000116251 | 1 | 6241329 | 6269449 | 80 | 15 | 1.0137 | 0.15536 | RPL22 | 0.15536 | 0.11915 | 0.20155 |
| ENSG00000271303 | 20 | 627259 | 634014 | 27 | 9 | 1.0136 | 0.15539 | SRXN1 | 0.15539 | 0.48756 | 0.71451 |
| ENSG00000168300 | 8 | 52730140 | 52811735 | 380 | 16 | 1.0127 | 0.15559 | PCMTD1 | 0.15559 | 0.16023 | 0.17237 |
| ENSG00000188368 | 19 | 42806250 | 42814973 | 8 | 2 | 1.0128 | 0.15559 | PRR19 | 0.15559 | 0.205 | 0.19215 |
| ENSG00000130165 | 19 | 11661962 | 11670051 | 24 | 8 | 1.0122 | 0.15572 | ELOF1 | 0.15572 | 0.23869 | 0.086932 |
| ENSG00000113194 | 5 | 175874629 | 175937075 | 161 | 20 | 1.0122 | 0.15573 | FAF2 | 0.15573 | 0.22518 | 0.21739 |
| ENSG00000172461 | 6 | 96463860 | 96663488 | 813 | 38 | 1.0121 | 0.15574 | FUT9 | 0.15574 | 0.11791 | 0.12075 |
| ENSG00000105229 | 19 | 4007644 | 4039384 | 150 | 19 | 1.0115 | 0.15589 | PIAS4 | 0.15589 | 0.14656 | 0.12864 |
| ENSG00000105708 | 19 | 19821280 | 19843906 | 56 | 12 | 1.0111 | 0.15598 | ZNF14 | 0.15598 | 0.12608 | 0.12193 |
| ENSG00000169583 | 9 | 139889087 | 139891255 | 3 | 2 | 1.0109 | 0.15604 | CLIC3 | 0.15604 | 0.0055179 | 0.0073028 |
| ENSG00000106113 | 7 | 30692200 | 30739745 | 139 | 26 | 1.0103 | 0.15616 | CRHR2 | 0.15616 | 0.12524 | 0.099298 |
| ENSG00000086666 | 15 | 80351910 | 80430735 | 233 | 24 | 1.01 | 0.15624 | ZFAND6 | 0.15624 | 0.16038 | 0.16011 |
| ENSG00000188394 | 9 | 125796806 | 125797975 | 1 | 1 | 1.0094 | 0.1564 | GPR21 | 0.1564 | 0.3943 | 0.33076 |
| ENSG00000226742 | 18 | 77724561 | 77730822 | 26 | 8 | 1.0092 | 0.15643 | HSBP1L1 | 0.15643 | 0.0088121 | 0.031694 |
| ENSG00000256087 | 19 | 52534632 | 52598991 | 245 | 18 | 1.0092 | 0.15644 | ZNF432 | 0.15644 | 0.22061 | 0.27225 |
| ENSG00000197312 | 1 | 15943995 | 15995539 | 142 | 7 | 1.0086 | 0.15659 | DDI2 | 0.15659 | 0.11982 | 0.09893 |
| ENSG00000197125 | 11 | 124310022 | 124311054 | 1 | 1 | 1.0085 | 0.1566 | OR8B8 | 0.1566 | 0.14487 | 0.33768 |
| ENSG00000149636 | 20 | 35380194 | 35402221 | 20 | 12 | 1.0085 | 0.1566 | DSN1 | 0.1566 | 0.28362 | 0.26191 |
| ENSG00000184860 | 16 | 82031221 | 82045093 | 27 | 15 | 1.0085 | 0.15661 | SDR42E1 | 0.15661 | 0.31039 | 0.70542 |
| ENSG00000188215 | 16 | 20866247 | 20911706 | 90 | 13 | 1.0083 | 0.15665 | DCUN1D3 | 0.15665 | 0.19996 | 0.17655 |
| ENSG00000204463 | 6 | 31606805 | 31620482 | 55 | 16 | 1.0082 | 0.15667 | BAG6 | 0.15667 | 0.28181 | 0.30185 |
| ENSG00000184226 | 13 | 66876967 | 67804468 | 2502 | 139 | 1.0082 | 0.15669 | PCDH9 | 0.15669 | 0.15857 | 0.16112 |
| ENSG00000166839 | 15 | 65204101 | 65251042 | 173 | 30 | 1.0079 | 0.15675 | ANKDD1A | 0.15675 | 0.088402 | 0.02483 |
| ENSG00000273079 | 12 | 13693165 | 14133053 | 1580 | 93 | 1.0077 | 0.15679 | GRIN2B | 0.15679 | 0.16193 | 0.20573 |
| ENSG00000149599 | 20 | 30435440 | 30458550 | 61 | 19 | 1.0077 | 0.1568 | DUSP15 | 0.1568 | 0.19922 | 0.2311 |

| | | | | | | | | | | | |
|---|---|---|---|---|---|---|---|---|---|---|---|
| ENSG00000144730 | 3 | 57124010 | 57204334 | 243 | 26 | 1.0074 | 0.15686 | IL17RD | 0.15686 | 0.20151 | 0.21939 |
| ENSG00000117222 | 1 | 205055270 | 205091143 | 119 | 15 | 1.0072 | 0.15691 | RBBP5 | 0.15691 | 0.13063 | 0.027321 |
| ENSG00000172987 | 10 | 100218875 | 100995619 | 2380 | 57 | 1.0072 | 0.15693 | HPSE2 | 0.15693 | 0.16431 | 0.20365 |
| ENSG00000135930 | 2 | 233414762 | 233448354 | 90 | 10 | 1.0068 | 0.15701 | EIF4E2 | 0.15701 | 0.11518 | 0.26123 |
| ENSG00000187699 | 2 | 190744335 | 191068210 | 885 | 72 | 1.0062 | 0.15717 | C2orf88 | 0.15717 | 0.19295 | 0.26151 |
| ENSG00000158062 | 1 | 26607819 | 26644854 | 121 | 12 | 1.006 | 0.15721 | UBXN11 | 0.15721 | 0.13869 | 0.12882 |
| ENSG00000213494 | 17 | 34310327 | 34329100 | 90 | 12 | 1.006 | 0.15721 | CCL14 | 0.15721 | 0.44489 | 0.39857 |
| ENSG00000071894 | 8 | 145618444 | 145634753 | 25 | 9 | 1.0059 | 0.15723 | CPSF1 | 0.15723 | 0.04597 | 0.038469 |
| ENSG00000105419 | 19 | 47906381 | 47922780 | 31 | 9 | 1.0055 | 0.15734 | MEIS3 | 0.15734 | 0.54208 | 0.61525 |
| ENSG00000158079 | 9 | 96793076 | 96872138 | 149 | 22 | 1.0054 | 0.15735 | PTPDC1 | 0.15735 | 0.13273 | 0.0862 |
| ENSG00000232434 | 9 | 139738867 | 139741797 | 2 | 1 | 1.0052 | 0.1574 | C9orf172 | 0.1574 | 0.30342 | 0.25468 |
| ENSG00000138688 | 4 | 123073488 | 123283913 | 383 | 27 | 1.0052 | 0.15741 | KIAA1109 | 0.15741 | 0.16876 | 0.16096 |
| ENSG00000182903 | 4 | 419604 | 492945 | 292 | 15 | 1.005 | 0.15745 | ZNF721 | 0.15745 | 0.16717 | 0.18152 |
| ENSG00000182132 | 5 | 169780491 | 170163636 | 1422 | 149 | 1.0046 | 0.15754 | KCNIP1 | 0.15754 | 0.18548 | 0.28137 |
| ENSG00000125743 | 19 | 46190712 | 46195827 | 9 | 3 | 1.0046 | 0.15754 | SNRPD2 | 0.15754 | 0.068485 | 0.0029263 |
| ENSG00000083812 | 19 | 58978423 | 58984945 | 12 | 7 | 1.0045 | 0.15756 | ZNF324 | 0.15756 | 0.58492 | 0.72614 |
| ENSG00000183807 | 6 | 117073363 | 117086886 | 52 | 12 | 1.0045 | 0.15758 | FAM162B | 0.15758 | 0.17498 | 0.2546 |
| ENSG00000141480 | 17 | 4613784 | 4624794 | 30 | 7 | 1.0044 | 0.1576 | ARRB2 | 0.1576 | 0.32251 | 0.4822 |
| ENSG00000269679 | 18 | 77275812 | 77276057 | 1 | 1 | 1.0044 | 0.1576 | AC018445.1 | 0.1576 | 0.19978 | 0.26764 |
| ENSG00000145911 | 5 | 177540444 | 177553088 | 28 | 8 | 1.0039 | 0.15771 | N4BP3 | 0.15771 | 0.067595 | 0.14801 |
| ENSG00000023909 | 1 | 94350761 | 94374966 | 82 | 12 | 1.0038 | 0.15774 | GCLM | 0.15774 | 0.16476 | 0.20455 |
| ENSG00000222017 | 2 | 198557830 | 198639547 | 151 | 12 | 1.0037 | 0.15776 | AC011997.1 | 0.15776 | 0.16805 | 0.16708 |
| ENSG00000152784 | 4 | 81105033 | 81125483 | 64 | 11 | 1.0035 | 0.15781 | PRDM8 | 0.15781 | 0.24182 | 0.16268 |
| ENSG00000185483 | 1 | 64239693 | 64647181 | 1151 | 77 | 1.0023 | 0.15811 | ROR1 | 0.15811 | 0.12501 | 0.052994 |
| ENSG00000198625 | 1 | 204485511 | 204542871 | 193 | 17 | 1.0021 | 0.15814 | MDM4 | 0.15814 | 0.1704 | 0.20148 |
| ENSG00000198841 | 1 | 52497775 | 52499488 | 7 | 2 | 1.002 | 0.15817 | KTI12 | 0.15817 | 0.055608 | 0.054763 |
| ENSG00000166833 | 11 | 19372271 | 20143144 | 2540 | 243 | 1.002 | 0.15818 | NAV2 | 0.15818 | 0.12561 | 0.11293 |
| ENSG00000149050 | 11 | 7020549 | 7041599 | 95 | 8 | 1.0019 | 0.1582 | ZNF214 | 0.1582 | 0.19513 | 0.30694 |
| ENSG00000144785 | 12 | 56679700 | 56709843 | 66 | 6 | 1.0019 | 0.15821 | RP11-977G19.10 | 0.15821 | 0.19072 | 0.19003 |
| ENSG00000104299 | 8 | 28625178 | 28747759 | 264 | 20 | 1.0017 | 0.15825 | INTS9 | 0.15825 | 0.185 | 0.2386 |
| ENSG00000096696 | 6 | 7541808 | 7586950 | 163 | 28 | 1.0016 | 0.15826 | DSP | 0.15826 | 0.13592 | 0.27088 |
| ENSG00000186086 | 1 | 108918421 | 109013624 | 40 | 17 | 1.0013 | 0.15834 | NBPF6 | 0.15834 | 0.19587 | 0.19192 |
| ENSG00000166598 | 12 | 104323885 | 104347423 | 126 | 22 | 1.0011 | 0.15838 | HSP90B1 | 0.15838 | 0.14565 | 0.092078 |
| ENSG00000166439 | 11 | 74459913 | 74553458 | 200 | 19 | 1.0006 | 0.1585 | RNF169 | 0.1585 | 0.14815 | 0.12499 |
| ENSG00000155542 | 5 | 56205087 | 56221359 | 51 | 6 | 1.0006 | 0.15851 | SETD9 | 0.15851 | 0.20471 | 0.163 |
| ENSG00000256222 | 20 | 55933496 | 55934878 | 8 | 1 | 1.0002 | 0.1586 | MTRNR2L3 | 0.1586 | 0.18949 | 0.18706 |
| ENSG00000141696 | 17 | 39958199 | 39968856 | 25 | 6 | 1.0001 | 0.15863 | LEPREL4 | 0.15863 | 0.18857 | 0.37853 |
| ENSG00000100266 | 22 | 43231418 | 43411151 | 673 | 17 | 1 | 0.15865 | PACSIN2 | 0.15865 | 0.17449 | 0.18199 |
| ENSG00000268898 | 2 | 40488147 | 40490117 | 10 | 4 | 0.9998 | 0.1587 | AC007377.1 | 0.1587 | 0.13561 | 0.030474 |
| ENSG00000227164 | 20 | 52105495 | 52106164 | 1 | 1 | 0.99982 | 0.1587 | AL354993.1 | 0.1587 | 0.52312 | 0.51531 |
| ENSG00000022355 | 5 | 161274197 | 161326975 | 163 | 19 | 0.9994 | 0.1588 | GABRA1 | 0.1588 | 0.16865 | 0.22482 |
| ENSG00000148180 | 9 | 123970072 | 124095121 | 343 | 23 | 0.99934 | 0.15882 | GSN | 0.15882 | 0.16926 | 0.19099 |
| ENSG00000144136 | 2 | 113403434 | 113421404 | 46 | 7 | 0.99928 | 0.15883 | SLC20A1 | 0.15883 | 0.19998 | 0.2673 |
| ENSG00000259642 | 15 | 80215113 | 80217194 | 15 | 5 | 0.99907 | 0.15888 | C15orf37 | 0.15888 | 0.50801 | 0.48226 |
| ENSG00000169599 | 2 | 69622882 | 69664760 | 139 | 9 | 0.99895 | 0.15891 | NFU1 | 0.15891 | 0.17676 | 0.16653 |
| ENSG00000203663 | 1 | 248201474 | 248202607 | 1 | 1 | 0.99858 | 0.159 | OR2L2 | 0.159 | 0.11066 | 0.10199 |
| ENSG00000154229 | 17 | 64298754 | 64806861 | 1989 | 124 | 0.99845 | 0.15903 | PRKCA | 0.15903 | 0.15837 | 0.15981 |
| ENSG00000167302 | 17 | 79202077 | 79212891 | 39 | 11 | 0.99815 | 0.1591 | ENTHD2 | 0.1591 | 0.3236 | 0.56288 |
| ENSG00000198612 | 2 | 237993955 | 238009109 | 42 | 8 | 0.99786 | 0.15917 | COPS8 | 0.15917 | 0.11646 | 0.11284 |
| ENSG00000119431 | 9 | 116135699 | 116139279 | 30 | 7 | 0.99741 | 0.15928 | HDHD3 | 0.15928 | 0.26704 | 0.35166 |
| ENSG00000128271 | 22 | 24813847 | 24838328 | 56 | 14 | 0.99736 | 0.1593 | ADORA2A | 0.1593 | 0.12937 | 0.10697 |
| ENSG00000104093 | 15 | 51739908 | 51915030 | 501 | 20 | 0.99727 | 0.15932 | DMXL2 | 0.15932 | 0.15789 | 0.15961 |
| ENSG00000132382 | 17 | 4442192 | 4458926 | 64 | 22 | 0.99718 | 0.15934 | MYBBP1A | 0.15934 | 0.20695 | 0.081212 |
| ENSG00000179407 | 3 | 128181282 | 128186091 | 21 | 6 | 0.99679 | 0.15943 | DNAJB8 | 0.15943 | 0.051947 | 0.036611 |
| ENSG00000115540 | 2 | 198380295 | 198418423 | 59 | 8 | 0.99652 | 0.1595 | MOB4 | 0.1595 | 0.11161 | 0.12019 |
| ENSG00000149582 | 11 | 118401756 | 118417995 | 52 | 3 | 0.9963 | 0.15955 | TMEM25 | 0.15955 | 0.16384 | 0.15062 |
| ENSG00000162757 | 1 | 209955661 | 209957904 | 3 | 2 | 0.9962 | 0.15958 | C1orf74 | 0.15958 | 0.13886 | 0.067034 |
| ENSG00000182983 | 3 | 42947223 | 42960825 | 24 | 4 | 0.99591 | 0.15965 | ZNF662 | 0.15965 | 0.17354 | 0.25645 |
| ENSG00000196411 | 7 | 100400187 | 100425121 | 75 | 12 | 0.9959 | 0.15965 | EPHB4 | 0.15965 | 0.10294 | 0.07606 |
| ENSG00000180481 | 12 | 75784850 | 75826468 | 143 | 9 | 0.99586 | 0.15966 | GLIPR1L2 | 0.15966 | 0.20092 | 0.2229 |
| ENSG00000077463 | 19 | 4174106 | 4182601 | 18 | 6 | 0.99584 | 0.15966 | SIRT6 | 0.15966 | 0.28203 | 0.54876 |
| ENSG00000122378 | 10 | 82167585 | 82192753 | 69 | 13 | 0.9957 | 0.1597 | FAM213A | 0.1597 | 0.20982 | 0.22609 |
| ENSG00000103544 | 16 | 19566562 | 19718115 | 573 | 35 | 0.99567 | 0.15971 | C16orf62 | 0.15971 | 0.19128 | 0.23183 |
| ENSG00000130723 | 9 | 134269480 | 134375584 | 280 | 16 | 0.9953 | 0.1598 | PRRC2B | 0.1598 | 0.14629 | 0.1913 |
| ENSG00000154222 | 1 | 52811395 | 52831865 | 32 | 11 | 0.99504 | 0.15986 | CC2D1B | 0.15986 | 0.25196 | 0.22732 |
| ENSG00000075131 | 15 | 66628544 | 66679084 | 186 | 31 | 0.99496 | 0.15988 | TIPIN | 0.15988 | 0.16079 | 0.1596 |
| ENSG00000169136 | 19 | 50431959 | 50437192 | 29 | 8 | 0.99481 | 0.15991 | ATF5 | 0.15991 | 0.34229 | 0.24574 |
| ENSG00000204377 | 1 | 16555368 | 16556038 | 3 | 1 | 0.99477 | 0.15992 | C1orf134 | 0.15992 | 0.16466 | 0.15419 |
| ENSG00000250127 | 5 | 57837394 | 57854070 | 49 | 11 | 0.99463 | 0.15996 | CTD-2117L12.1 | 0.15996 | 0.34324 | 0.30093 |
| ENSG00000198851 | 11 | 118175260 | 118186890 | 35 | 7 | 0.99451 | 0.15999 | CD3E | 0.15999 | 0.16331 | 0.16094 |
| ENSG00000178115 | 15 | 30844183 | 30854704 | 1 | 1 | 0.99446 | 0.16 | GOLGA8Q | 0.16 | 0.58734 | 0.73828 |
| ENSG00000100300 | 22 | 43547520 | 43559248 | 45 | 7 | 0.99443 | 0.16001 | TSPO | 0.16001 | 0.24771 | 0.38311 |
| ENSG00000116266 | 1 | 109289296 | 109352148 | 265 | 11 | 0.99402 | 0.16011 | STXBP3 | 0.16011 | 0.16816 | 0.17551 |

| | | | | | | | | | | | |
|---|---|---|---|---|---|---|---|---|---|---|---|
| ENSG00000138039 | 2 | 48859428 | 48982880 | 534 | 46 | 0.99342 | 0.16025 | LHCGR | 0.16025 | 0.16814 | 0.13044 |
| ENSG00000141759 | 18 | 77732867 | 77793949 | 179 | 18 | 0.99341 | 0.16025 | TXNL4A | 0.16025 | 0.15128 | 0.090543 |
| ENSG00000160948 | 8 | 145649000 | 145653931 | 15 | 6 | 0.99331 | 0.16028 | VPS28 | 0.16028 | 0.085591 | 0.063334 |
| ENSG00000198000 | 9 | 95059640 | 95087918 | 70 | 9 | 0.99275 | 0.16042 | NOL8 | 0.16042 | 0.1422 | 0.1482 |
| ENSG00000184786 | 6 | 170140210 | 170151655 | 22 | 10 | 0.99257 | 0.16046 | TCTE3 | 0.16046 | 0.021085 | 0.036914 |
| ENSG00000161574 | 17 | 34310692 | 34328660 | 86 | 11 | 0.99256 | 0.16046 | CCL15-CCL14 | 0.16046 | 0.44425 | 0.40097 |
| ENSG00000169026 | 4 | 675618 | 683230 | 19 | 7 | 0.99252 | 0.16047 | MFSD7 | 0.16047 | 0.26511 | 0.40238 |
| ENSG00000158856 | 8 | 21906506 | 21940038 | 126 | 14 | 0.99244 | 0.16049 | DMTN | 0.16049 | 0.12482 | 0.04921 |
| ENSG00000181965 | 5 | 134869991 | 134871639 | 1 | 1 | 0.99199 | 0.1606 | NEUROG1 | 0.1606 | 0.42206 | 0.49813 |
| ENSG00000187372 | 5 | 140593509 | 140596993 | 10 | 4 | 0.99187 | 0.16063 | PCDHB13 | 0.16063 | 0.27856 | 0.15461 |
| ENSG00000115541 | 2 | 198364718 | 198368181 | 4 | 2 | 0.99181 | 0.16065 | HSPE1 | 0.16065 | 0.087115 | 0.080405 |
| ENSG00000150456 | 13 | 21302870 | 21348097 | 190 | 17 | 0.99149 | 0.16072 | N6AMT2 | 0.16072 | 0.19434 | 0.20118 |
| ENSG00000092054 | 14 | 23881947 | 23904927 | 68 | 12 | 0.99142 | 0.16074 | MYH7 | 0.16074 | 0.2134 | 0.49721 |
| ENSG00000231672 | 2 | 218148742 | 218621316 | 1578 | 73 | 0.9909 | 0.16087 | DIRC3 | 0.16087 | 0.1773 | 0.15246 |
| ENSG00000215431 | 18 | 60491423 | 60492724 | 1 | 1 | 0.99036 | 0.161 | AC015989.1 | 0.161 | 0.3305 | 0.48229 |
| ENSG00000251247 | 19 | 37341263 | 37403931 | 168 | 16 | 0.99029 | 0.16102 | ZNF345 | 0.16102 | 0.20648 | 0.17383 |
| ENSG00000185306 | 12 | 64660217 | 64784972 | 431 | 29 | 0.99013 | 0.16106 | C12orf56 | 0.16106 | 0.15394 | 0.20408 |
| ENSG00000124212 | 20 | 48120411 | 48184683 | 146 | 41 | 0.98972 | 0.16116 | PTGIS | 0.16116 | 0.11254 | 0.15736 |
| ENSG00000152082 | 2 | 130939310 | 130948302 | 40 | 6 | 0.98914 | 0.1613 | MZT2B | 0.1613 | 0.16242 | 0.13712 |
| ENSG00000143674 | 1 | 233463514 | 233520894 | 240 | 25 | 0.98895 | 0.16134 | MLK4 | 0.16134 | 0.20538 | 0.20445 |
| ENSG00000149485 | 11 | 61567099 | 61596790 | 44 | 5 | 0.98888 | 0.16136 | FADS1 | 0.16136 | 0.15286 | 0.16026 |
| ENSG00000122386 | 16 | 3162561 | 3170518 | 15 | 3 | 0.98881 | 0.16138 | ZNF205 | 0.16138 | 0.085402 | 0.12108 |
| ENSG00000143499 | 1 | 214454445 | 214510474 | 195 | 23 | 0.98862 | 0.16142 | SMYD2 | 0.16142 | 0.20224 | 0.17323 |
| ENSG00000160602 | 17 | 27052915 | 27070473 | 30 | 8 | 0.9886 | 0.16143 | NEK8 | 0.16143 | 0.20177 | 0.18377 |
| ENSG00000064309 | 11 | 125825691 | 125933230 | 419 | 44 | 0.98842 | 0.16147 | CDON | 0.16147 | 0.091532 | 0.084747 |
| ENSG00000151379 | 2 | 17997763 | 17998368 | 3 | 1 | 0.98783 | 0.16162 | MSGN1 | 0.16162 | 0.23691 | 0.18875 |
| ENSG00000182866 | 1 | 32716840 | 32751766 | 50 | 19 | 0.98749 | 0.1617 | LCK | 0.1617 | 0.093859 | 0.0094081 |
| ENSG00000137869 | 15 | 51500254 | 51630807 | 393 | 29 | 0.98751 | 0.1617 | CYP19A1 | 0.1617 | 0.22244 | 0.24502 |
| ENSG00000111203 | 12 | 2921788 | 2968957 | 130 | 10 | 0.98696 | 0.16183 | ITFG2 | 0.16183 | 0.15826 | 0.16573 |
| ENSG00000164307 | 5 | 96096521 | 96143803 | 297 | 19 | 0.98685 | 0.16186 | ERAP1 | 0.16186 | 0.15146 | 0.20722 |
| ENSG00000187800 | 1 | 156863490 | 156886226 | 85 | 12 | 0.98658 | 0.16192 | PEAR1 | 0.16192 | 0.18113 | 0.29206 |
| ENSG00000257335 | 7 | 141607613 | 141806547 | 742 | 84 | 0.9865 | 0.16194 | MGAM | 0.16194 | 0.17188 | 0.16785 |
| ENSG00000148308 | 9 | 135906076 | 135933890 | 56 | 6 | 0.98646 | 0.16195 | GTF3C5 | 0.16195 | 0.17851 | 0.18407 |
| ENSG00000148908 | 10 | 121259340 | 121302220 | 145 | 25 | 0.98642 | 0.16196 | RGS10 | 0.16196 | 0.23815 | 0.39879 |
| ENSG00000135451 | 12 | 49717019 | 49725514 | 20 | 6 | 0.98608 | 0.16205 | TROAP | 0.16205 | 0.11708 | 0.093883 |
| ENSG00000186635 | 11 | 72396114 | 72504644 | 296 | 23 | 0.98577 | 0.16212 | ARAP1 | 0.16212 | 0.20647 | 0.21509 |
| ENSG00000135925 | 2 | 219745085 | 219764303 | 22 | 5 | 0.9847 | 0.16239 | WNT10A | 0.16239 | 0.23652 | 0.17044 |
| ENSG00000164403 | 5 | 132157833 | 132166590 | 10 | 3 | 0.9845 | 0.16243 | SHROOM1 | 0.16243 | 0.64308 | 0.81142 |
| ENSG00000158966 | 1 | 64936428 | 65158741 | 570 | 66 | 0.98438 | 0.16246 | CACHD1 | 0.16246 | 0.18942 | 0.23773 |
| ENSG00000101638 | 18 | 44259081 | 44339220 | 335 | 52 | 0.98367 | 0.16264 | ST8SIA5 | 0.16264 | 0.16318 | 0.25099 |
| ENSG00000213030 | 19 | 49550895 | 49552400 | 2 | 1 | 0.98313 | 0.16277 | CGB8 | 0.16277 | 0.13226 | 0.36272 |
| ENSG00000198478 | 6 | 80341000 | 80413372 | 201 | 27 | 0.98278 | 0.16286 | SH3BGRL2 | 0.16286 | 0.10693 | 0.031119 |
| ENSG00000253633 | 8 | 103540968 | 103550896 | 64 | 11 | 0.9826 | 0.1629 | KB-1980E6.3 | 0.1629 | 0.1052 | 0.051528 |
| ENSG00000111405 | 12 | 48103517 | 48119350 | 41 | 10 | 0.98257 | 0.16291 | ENDOU | 0.16291 | 0.23529 | 0.11727 |
| ENSG00000067248 | 5 | 54552073 | 54603550 | 114 | 15 | 0.98246 | 0.16294 | DHX29 | 0.16294 | 0.1523 | 0.17624 |
| ENSG00000143819 | 1 | 225997794 | 226033260 | 141 | 39 | 0.98212 | 0.16302 | EPHX1 | 0.16302 | 0.40199 | 0.44599 |
| ENSG00000089505 | 16 | 66600294 | 66613040 | 27 | 3 | 0.9816 | 0.16315 | CMTM1 | 0.16315 | 0.1798 | 0.14134 |
| ENSG00000136153 | 13 | 76194570 | 76434004 | 808 | 59 | 0.98128 | 0.16323 | LMO7 | 0.16323 | 0.17241 | 0.20584 |
| ENSG00000156928 | 7 | 23338358 | 23351348 | 25 | 5 | 0.98113 | 0.16326 | MALSU1 | 0.16326 | 0.12252 | 0.06835 |
| ENSG00000187134 | 10 | 4934796 | 5025475 | 507 | 32 | 0.98116 | 0.16326 | AKR1C1 | 0.16326 | 0.14132 | 0.2344 |
| ENSG00000212710 | 18 | 19993564 | 19997878 | 10 | 7 | 0.98076 | 0.16335 | CTAGE1 | 0.16335 | 0.12041 | 0.1202 |
| ENSG00000256892 | 10 | 37890366 | 37891859 | 8 | 2 | 0.98071 | 0.16337 | MTRNR2L7 | 0.16337 | 0.22175 | 0.23979 |
| ENSG00000221990 | 5 | 441645 | 443258 | 13 | 3 | 0.98065 | 0.16338 | C5orf55 | 0.16338 | 0.17995 | 0.13594 |
| ENSG00000153187 | 1 | 245014468 | 245027844 | 23 | 7 | 0.97988 | 0.16357 | HNRNPU | 0.16357 | 0.43188 | 0.63761 |
| ENSG00000180219 | 12 | 100041531 | 100043824 | 7 | 2 | 0.97945 | 0.16368 | FAM71C | 0.16368 | 0.21985 | 0.39785 |
| ENSG00000100359 | 22 | 40766595 | 40806293 | 64 | 9 | 0.97848 | 0.16392 | SGSM3 | 0.16392 | 0.28026 | 0.21623 |
| ENSG00000162639 | 1 | 109190912 | 109204148 | 27 | 11 | 0.97773 | 0.1641 | HENMT1 | 0.1641 | 0.092984 | 0.087834 |
| ENSG00000105877 | 7 | 21582833 | 21941457 | 1738 | 127 | 0.97752 | 0.16416 | DNAH11 | 0.16416 | 0.15955 | 0.1812 |
| ENSG00000120057 | 10 | 99526508 | 99531709 | 11 | 3 | 0.97744 | 0.16417 | SFRP5 | 0.16417 | 0.14912 | 0.27318 |
| ENSG00000104490 | 8 | 102698771 | 103137135 | 1438 | 81 | 0.97744 | 0.16418 | NCALD | 0.16418 | 0.16073 | 0.19975 |
| ENSG00000245680 | 19 | 37672481 | 37709055 | 75 | 14 | 0.97744 | 0.16418 | ZNF585B | 0.16418 | 0.133 | 0.13518 |
| ENSG00000127952 | 7 | 75625656 | 75677322 | 144 | 21 | 0.97735 | 0.1642 | STYXL1 | 0.1642 | 0.15672 | 0.17809 |
| ENSG00000198346 | 19 | 53970989 | 54006950 | 283 | 38 | 0.9768 | 0.16433 | ZNF813 | 0.16433 | 0.22671 | 0.067508 |
| ENSG00000109758 | 4 | 3443614 | 3451211 | 35 | 13 | 0.97661 | 0.16438 | HGFAC | 0.16438 | 0.13781 | 0.39148 |
| ENSG00000152214 | 18 | 40323192 | 40695657 | 995 | 62 | 0.97649 | 0.16441 | RIT2 | 0.16441 | 0.19046 | 0.22834 |
| ENSG00000177679 | 7 | 75831216 | 75916605 | 181 | 21 | 0.976 | 0.16453 | SRRM3 | 0.16453 | 0.15797 | 0.19416 |
| ENSG00000197905 | 12 | 3068496 | 3149839 | 358 | 37 | 0.97597 | 0.16454 | TEAD4 | 0.16454 | 0.14703 | 0.13157 |
| ENSG00000143199 | 1 | 167778625 | 167883453 | 419 | 56 | 0.97593 | 0.16455 | ADCY10 | 0.16455 | 0.1397 | 0.17499 |
| ENSG00000120329 | 5 | 140682196 | 140683612 | 5 | 2 | 0.97534 | 0.1647 | SLC25A2 | 0.1647 | 0.22127 | 0.29152 |
| ENSG00000163932 | 3 | 53190025 | 53226733 | 94 | 15 | 0.9748 | 0.16483 | PRKCD | 0.16483 | 0.16024 | 0.25606 |
| ENSG00000163202 | 1 | 152551857 | 152552980 | 3 | 2 | 0.97409 | 0.16501 | LCE3D | 0.16501 | 0.028692 | 0.042441 |
| ENSG00000184436 | 22 | 21353393 | 21356485 | 9 | 5 | 0.97377 | 0.16509 | THAP7 | 0.16509 | 0.34598 | 0.35472 |
| ENSG00000163681 | 3 | 57741177 | 57914895 | 288 | 21 | 0.9737 | 0.1651 | SLMAP | 0.1651 | 0.19129 | 0.19784 |

| | | | | | | | | | | | |
|---|---|---|---|---|---|---|---|---|---|---|---|
| ENSG00000166579 | 17 | 8316449 | 8393729 | 149 | 24 | 0.9736 | 0.16513 | NDEL1 | 0.16513 | 0.18125 | 0.17333 |
| ENSG00000102897 | 16 | 20911190 | 20936328 | 48 | 9 | 0.97343 | 0.16517 | LYRM1 | 0.16517 | 0.17784 | 0.17957 |
| ENSG00000130449 | 5 | 60628100 | 60841997 | 362 | 22 | 0.97329 | 0.1652 | ZSWIM6 | 0.1652 | 0.1751 | 0.18559 |
| ENSG00000163746 | 3 | 146109208 | 146213778 | 375 | 16 | 0.97309 | 0.16525 | PLSCR2 | 0.16525 | 0.16482 | 0.1471 |
| ENSG00000100298 | 22 | 39493229 | 39500072 | 53 | 4 | 0.97296 | 0.16529 | APOBEC3H | 0.16529 | 0.1987 | 0.11462 |
| ENSG00000163995 | 4 | 7967039 | 8160559 | 925 | 123 | 0.97273 | 0.16534 | ABLIM2 | 0.16534 | 0.14602 | 0.1015 |
| ENSG00000169777 | 5 | 9629109 | 9712490 | 248 | 37 | 0.97265 | 0.16536 | TAS2R1 | 0.16536 | 0.18384 | 0.3274 |
| ENSG00000164076 | 3 | 49895421 | 49907655 | 20 | 5 | 0.97259 | 0.16538 | CAMKV | 0.16538 | 0.13108 | 0.087878 |
| ENSG00000177954 | 1 | 153963235 | 153964626 | 4 | 2 | 0.97203 | 0.16552 | RPS27 | 0.16552 | 0.19323 | 0.20726 |
| ENSG00000180447 | 9 | 89559279 | 89562104 | 4 | 2 | 0.97172 | 0.1656 | GAS1 | 0.1656 | 0.61606 | 0.64657 |
| ENSG00000196224 | 11 | 1628795 | 1629693 | 6 | 3 | 0.97155 | 0.16564 | KRTAP5-3 | 0.16564 | 0.47572 | 0.66872 |
| ENSG00000124256 | 20 | 56178902 | 56195632 | 54 | 19 | 0.97153 | 0.16564 | ZBP1 | 0.16564 | 0.0083142 | 0.0030207 |
| ENSG00000162365 | 1 | 47603107 | 47615413 | 54 | 7 | 0.97147 | 0.16566 | CYP4A22 | 0.16566 | 0.19471 | 0.23243 |
| ENSG00000187950 | 12 | 29565407 | 29650619 | 348 | 22 | 0.97147 | 0.16566 | OVCH1 | 0.16566 | 0.19678 | 0.24757 |
| ENSG00000167281 | 17 | 77085427 | 77613550 | 2297 | 262 | 0.97102 | 0.16577 | RBFOX3 | 0.16577 | 0.15126 | 0.18197 |
| ENSG00000163755 | 3 | 148847371 | 148891519 | 212 | 30 | 0.97098 | 0.16578 | HPS3 | 0.16578 | 0.17115 | 0.12301 |
| ENSG00000239900 | 22 | 40742507 | 40786467 | 66 | 12 | 0.97028 | 0.16595 | ADSL | 0.16595 | 0.1679 | 0.16634 |
| ENSG00000138134 | 10 | 90639491 | 90734910 | 282 | 22 | 0.97 | 0.16602 | STAMBPL1 | 0.16602 | 0.2135 | 0.258 |
| ENSG00000023734 | 12 | 16035325 | 16056412 | 15 | 5 | 0.9693 | 0.1662 | STRAP | 0.1662 | 0.096763 | 0.10904 |
| ENSG00000160752 | 1 | 155278539 | 155290457 | 13 | 2 | 0.96898 | 0.16628 | FDPS | 0.16628 | 0.10822 | 0.15092 |
| ENSG00000099624 | 19 | 1241749 | 1244824 | 4 | 2 | 0.96868 | 0.16635 | ATP5D | 0.16635 | 0.084153 | 0.16237 |
| ENSG00000230055 | 17 | 36886488 | 36891297 | 17 | 6 | 0.96846 | 0.16641 | CISD3 | 0.16641 | 0.23251 | 0.43369 |
| ENSG00000132703 | 1 | 159557615 | 159558655 | 2 | 1 | 0.9681 | 0.1665 | APCS | 0.1665 | 0.47884 | 0.45052 |
| ENSG00000218336 | 4 | 183065140 | 183724177 | 2837 | 249 | 0.96778 | 0.16658 | TENM3 | 0.16658 | 0.18063 | 0.12985 |
| ENSG00000049167 | 5 | 60169658 | 60240900 | 221 | 23 | 0.96759 | 0.16662 | ERCC8 | 0.16662 | 0.15948 | 0.15695 |
| ENSG00000159248 | 15 | 35043233 | 35047166 | 9 | 4 | 0.96746 | 0.16666 | GJD2 | 0.16666 | 0.24102 | 0.41996 |
| ENSG00000171060 | 8 | 10530147 | 10558103 | 50 | 14 | 0.96697 | 0.16678 | C8orf74 | 0.16678 | 0.0067678 | 0.012272 |
| ENSG00000168890 | 2 | 85825671 | 85830319 | 5 | 2 | 0.96681 | 0.16682 | TMEM150A | 0.16682 | 0.04847 | 0.040208 |
| ENSG00000205441 | 21 | 46020497 | 46022091 | 18 | 2 | 0.96625 | 0.16696 | KRTAP10-7 | 0.16696 | 0.13539 | 0.26231 |
| ENSG00000130649 | 10 | 135333910 | 135374724 | 252 | 16 | 0.96566 | 0.16711 | CYP2E1 | 0.16711 | 0.19159 | 0.19299 |
| ENSG00000124406 | 4 | 42410390 | 42659122 | 851 | 34 | 0.96558 | 0.16713 | ATP8A1 | 0.16713 | 0.17439 | 0.29528 |
| ENSG00000159199 | 17 | 46970127 | 46973233 | 7 | 2 | 0.96558 | 0.16713 | ATP5G1 | 0.16713 | 0.22251 | 0.15949 |
| ENSG00000203630 | 20 | 1316228 | 1317555 | 2 | 1 | 0.9655 | 0.16715 | AL136531.1 | 0.16715 | 0.036965 | 0.087471 |
| ENSG00000106771 | 9 | 111777432 | 111882225 | 458 | 30 | 0.96508 | 0.16725 | TMEM245 | 0.16725 | 0.19672 | 0.26362 |
| ENSG00000184863 | 7 | 155437145 | 155574179 | 455 | 15 | 0.96446 | 0.16741 | RBM33 | 0.16741 | 0.20066 | 0.22848 |
| ENSG00000187186 | 9 | 34664160 | 34666109 | 5 | 3 | 0.96442 | 0.16742 | RP11-195F19.5 | 0.16742 | 0.11238 | 0.16058 |
| ENSG00000180537 | 6 | 13924677 | 13980533 | 107 | 14 | 0.96345 | 0.16766 | RNF182 | 0.16766 | 0.16635 | 0.20938 |
| ENSG00000154736 | 21 | 28290231 | 28338832 | 168 | 20 | 0.96339 | 0.16768 | ADAMTS5 | 0.16768 | 0.1179 | 0.1029 |
| ENSG00000178597 | 4 | 7432022 | 7436700 | 42 | 9 | 0.96277 | 0.16783 | PSAPL1 | 0.16783 | 0.12644 | 0.44933 |
| ENSG00000221961 | 2 | 240981230 | 240982399 | 47 | 7 | 0.96271 | 0.16785 | PRR21 | 0.16785 | 0.063774 | 0.058329 |
| ENSG00000176438 | 14 | 95883831 | 95942173 | 341 | 47 | 0.9627 | 0.16785 | SYNE3 | 0.16785 | 0.2515 | 0.64668 |
| ENSG00000066923 | 7 | 99775186 | 99819111 | 124 | 10 | 0.96227 | 0.16796 | STAG3 | 0.16796 | 0.1575 | 0.11798 |
| ENSG00000183742 | 7 | 20174278 | 20257027 | 254 | 19 | 0.96218 | 0.16798 | MACC1 | 0.16798 | 0.20152 | 0.24601 |
| ENSG00000184451 | 17 | 40830907 | 40835935 | 8 | 1 | 0.96214 | 0.16799 | CCR10 | 0.16799 | 0.39606 | 0.41078 |
| ENSG00000108798 | 17 | 47287589 | 47300587 | 44 | 11 | 0.96213 | 0.16799 | ABI3 | 0.16799 | 0.10621 | 0.026582 |
| ENSG00000145321 | 4 | 72607410 | 72669758 | 179 | 25 | 0.96189 | 0.16805 | GC | 0.16805 | 0.15851 | 0.23019 |
| ENSG00000132514 | 17 | 6977856 | 6983626 | 25 | 7 | 0.96185 | 0.16806 | CLEC10A | 0.16806 | 0.17973 | 0.23503 |
| ENSG00000112739 | 6 | 4021501 | 4065217 | 160 | 14 | 0.96167 | 0.16811 | PRPF4B | 0.16811 | 0.12768 | 0.13708 |
| ENSG00000110079 | 11 | 60048014 | 60076445 | 64 | 13 | 0.96113 | 0.16824 | MS4A4A | 0.16824 | 0.084313 | 0.12976 |
| ENSG00000166710 | 15 | 45003675 | 45011075 | 17 | 5 | 0.96107 | 0.16826 | B2M | 0.16826 | 0.34035 | 0.38385 |
| ENSG00000204070 | 20 | 43990577 | 44005438 | 33 | 3 | 0.96105 | 0.16826 | SYS1 | 0.16826 | 0.14417 | 0.088473 |
| ENSG00000254995 | 20 | 57226490 | 57290466 | 231 | 29 | 0.96085 | 0.16831 | STX16-NPEPL1 | 0.16831 | 0.18176 | 0.15296 |
| ENSG00000136874 | 9 | 102668915 | 102732618 | 138 | 12 | 0.96077 | 0.16833 | STX17 | 0.16833 | 0.20141 | 0.16646 |
| ENSG00000007392 | 16 | 238968 | 279462 | 87 | 14 | 0.96063 | 0.16837 | LUC7L | 0.16837 | 0.14907 | 0.22946 |
| ENSG00000198783 | 17 | 33288549 | 33290205 | 5 | 3 | 0.96064 | 0.16837 | ZNF830 | 0.16837 | 0.23014 | 0.40224 |
| ENSG00000148498 | 10 | 34398488 | 35104253 | 1714 | 70 | 0.96025 | 0.16846 | PARD3 | 0.16846 | 0.11516 | 0.1011 |
| ENSG00000111911 | 6 | 126277927 | 126301387 | 41 | 4 | 0.9602 | 0.16848 | HINT3 | 0.16848 | 0.094853 | 0.057241 |
| ENSG00000169220 | 5 | 176784838 | 176799602 | 21 | 7 | 0.96015 | 0.16849 | RGS14 | 0.16849 | 0.22093 | 0.33825 |
| ENSG00000108175 | 10 | 80828792 | 81076276 | 900 | 111 | 0.95978 | 0.16858 | ZMIZ1 | 0.16858 | 0.12457 | 0.099474 |
| ENSG00000115507 | 2 | 63277192 | 63284971 | 8 | 4 | 0.95975 | 0.16859 | OTX1 | 0.16859 | 0.1704 | 0.59785 |
| ENSG00000092931 | 17 | 74731947 | 74777531 | 50 | 22 | 0.95972 | 0.1686 | MFSD11 | 0.1686 | 0.072382 | 0.085728 |
| ENSG00000236338 | 7 | 135611542 | 135613214 | 2 | 1 | 0.95966 | 0.16861 | AC015987.2 | 0.16861 | 0.26227 | 0.25508 |
| ENSG00000001626 | 7 | 117105838 | 117356025 | 546 | 28 | 0.95951 | 0.16865 | CFTR | 0.16865 | 0.14615 | 0.13969 |
| ENSG00000143367 | 1 | 151512781 | 151556059 | 115 | 18 | 0.95926 | 0.16871 | TUFT1 | 0.16871 | 0.089145 | 0.049325 |
| ENSG00000229292 | 19 | 56280507 | 56284545 | 25 | 6 | 0.95929 | 0.16871 | RFPL4AL1 | 0.16871 | 0.13413 | 0.51726 |
| ENSG00000070214 | 9 | 108006903 | 108201452 | 537 | 31 | 0.95902 | 0.16877 | SLC44A1 | 0.16877 | 0.15334 | 0.14796 |
| ENSG00000173660 | 1 | 46769303 | 46782448 | 32 | 10 | 0.95872 | 0.16885 | UQCRH | 0.16885 | 0.22724 | 0.36192 |
| ENSG00000197016 | 19 | 57078880 | 57100279 | 80 | 9 | 0.95856 | 0.16889 | ZNF470 | 0.16889 | 0.20661 | 0.28716 |
| ENSG00000162419 | 1 | 28995244 | 29045865 | 96 | 9 | 0.95854 | 0.1689 | GMEB1 | 0.1689 | 0.18977 | 0.22169 |
| ENSG00000105509 | 19 | 52216365 | 52227247 | 47 | 10 | 0.95846 | 0.16891 | HAS1 | 0.16891 | 0.20744 | 0.29258 |
| ENSG00000197891 | 11 | 64358113 | 64369820 | 49 | 7 | 0.9573 | 0.16921 | SLC22A12 | 0.16921 | 0.17709 | 0.17484 |
| ENSG00000181240 | 19 | 6426048 | 6433790 | 29 | 8 | 0.95708 | 0.16926 | SLC25A41 | 0.16926 | 0.20849 | 0.35036 |
| ENSG00000124177 | 20 | 40030741 | 40247133 | 599 | 26 | 0.95711 | 0.16926 | CHD6 | 0.16926 | 0.17452 | 0.22676 |

| | | | | | | | | | | | |
|---|---|---|---|---|---|---|---|---|---|---|---|
| ENSG00000146216 | 6 | 43211418 | 43255997 | 139 | 20 | 0.95664 | 0.16937 | TTBK1 | 0.16937 | 0.2616 | 0.35269 |
| ENSG00000137259 | 6 | 26033320 | 26033796 | 2 | 1 | 0.95646 | 0.16942 | HIST1H2AB | 0.16942 | 0.31474 | 0.404 |
| ENSG00000171126 | 2 | 42669157 | 42721237 | 182 | 30 | 0.95644 | 0.16943 | KCNG3 | 0.16943 | 0.09614 | 0.1429 |
| ENSG00000100156 | 22 | 38474141 | 38480100 | 12 | 6 | 0.95591 | 0.16956 | SLC16A8 | 0.16956 | 0.44003 | 0.13702 |
| ENSG00000168350 | 14 | 100612756 | 100626500 | 44 | 9 | 0.95572 | 0.16961 | DEGS2 | 0.16961 | 0.064447 | 0.1203 |
| ENSG00000127580 | 16 | 734622 | 740444 | 13 | 3 | 0.95549 | 0.16967 | WDR24 | 0.16967 | 0.15532 | 0.41383 |
| ENSG00000160293 | 9 | 136627016 | 136857726 | 1308 | 102 | 0.95484 | 0.16983 | VAV2 | 0.16983 | 0.24145 | 0.34419 |
| ENSG00000181027 | 19 | 47249303 | 47280245 | 120 | 17 | 0.95467 | 0.16987 | FKRP | 0.16987 | 0.099568 | 0.18663 |
| ENSG00000087302 | 14 | 52456193 | 52471420 | 52 | 13 | 0.95463 | 0.16988 | C14orf166 | 0.16988 | 0.18392 | 0.25806 |
| ENSG00000126217 | 13 | 113548692 | 113754053 | 930 | 89 | 0.95446 | 0.16992 | MCF2L | 0.16992 | 0.16731 | 0.11444 |
| ENSG00000125899 | 20 | 11008408 | 11010014 | 5 | 2 | 0.95421 | 0.16999 | C20orf187 | 0.16999 | 0.16857 | 0.14916 |
| ENSG00000241644 | 7 | 30737601 | 30797218 | 182 | 35 | 0.95402 | 0.17004 | INMT | 0.17004 | 0.18042 | 0.17386 |
| ENSG00000206190 | 15 | 25922420 | 26110317 | 958 | 95 | 0.95384 | 0.17008 | ATP10A | 0.17008 | 0.16618 | 0.070684 |
| ENSG00000123066 | 12 | 116395711 | 116715143 | 642 | 30 | 0.95382 | 0.17009 | MED13L | 0.17009 | 0.16892 | 0.14418 |
| ENSG00000198967 | 1 | 158576229 | 158577170 | 2 | 1 | 0.95364 | 0.17013 | OR10Z1 | 0.17013 | 0.71454 | 0.72458 |
| ENSG00000249861 | 19 | 40146534 | 40151287 | 9 | 3 | 0.95309 | 0.17027 | LGALS16 | 0.17027 | 0.36025 | 0.4127 |
| ENSG00000187080 | 1 | 248128535 | 248129641 | 5 | 2 | 0.95288 | 0.17033 | OR2AK2 | 0.17033 | 0.41421 | 0.36348 |
| ENSG00000135960 | 2 | 109510927 | 109605828 | 303 | 36 | 0.95246 | 0.17043 | EDAR | 0.17043 | 0.19535 | 0.21332 |
| ENSG00000152591 | 4 | 88529681 | 88538062 | 65 | 17 | 0.95211 | 0.17052 | DSPP | 0.17052 | 0.16446 | 0.18573 |
| ENSG00000171115 | 7 | 150147718 | 150176480 | 20 | 8 | 0.95202 | 0.17054 | GIMAP8 | 0.17054 | 0.1462 | 0.30999 |
| ENSG00000244094 | 1 | 153084590 | 153085991 | 3 | 1 | 0.95189 | 0.17058 | SPRR2F | 0.17058 | 0.18777 | 0.11939 |
| ENSG00000105664 | 19 | 18893583 | 18902123 | 16 | 7 | 0.95152 | 0.17067 | COMP | 0.17067 | 0.44237 | 0.60296 |
| ENSG00000243244 | 2 | 48756522 | 48826025 | 284 | 24 | 0.95119 | 0.17075 | STON1 | 0.17075 | 0.24057 | 0.33766 |
| ENSG00000086205 | 11 | 49168187 | 49230222 | 242 | 20 | 0.95022 | 0.171 | FOLH1 | 0.171 | 0.20008 | 0.25527 |
| ENSG00000113196 | 5 | 153854532 | 153857824 | 9 | 5 | 0.9499 | 0.17108 | HAND1 | 0.17108 | 0.19079 | 0.35333 |
| ENSG00000164440 | 6 | 139561198 | 139613276 | 161 | 15 | 0.94972 | 0.17113 | TXLNB | 0.17113 | 0.17386 | 0.23039 |
| ENSG00000101160 | 20 | 57570240 | 57582302 | 58 | 10 | 0.94947 | 0.17119 | CTSZ | 0.17119 | 0.1199 | 0.096216 |
| ENSG00000087086 | 19 | 49468558 | 49470135 | 3 | 1 | 0.94932 | 0.17123 | FTL | 0.17123 | 0.51037 | 0.75647 |
| ENSG00000130725 | 19 | 59067079 | 59070327 | 2 | 1 | 0.94895 | 0.17132 | UBE2M | 0.17132 | 0.88621 | 0.94601 |
| ENSG00000108417 | 17 | 39577005 | 39580775 | 19 | 3 | 0.94814 | 0.17153 | KRT37 | 0.17153 | 0.26576 | 0.40539 |
| ENSG00000148841 | 10 | 106071894 | 106098162 | 63 | 12 | 0.94758 | 0.17167 | ITPRIP | 0.17167 | 0.22954 | 0.20558 |
| ENSG00000196678 | 16 | 20791515 | 20911671 | 214 | 22 | 0.94731 | 0.17174 | ERI2 | 0.17174 | 0.21227 | 0.21052 |
| ENSG00000145882 | 5 | 148737570 | 148749216 | 15 | 4 | 0.9471 | 0.17179 | PCYOX1L | 0.17179 | 0.1944 | 0.17041 |
| ENSG00000164338 | 5 | 72861268 | 72877794 | 49 | 6 | 0.94682 | 0.17186 | UTP15 | 0.17186 | 0.25333 | 0.33844 |
| ENSG00000184293 | 12 | 9868456 | 9885895 | 112 | 11 | 0.94663 | 0.17191 | CLECL1 | 0.17191 | 0.22197 | 0.14755 |
| ENSG00000184545 | 11 | 1575274 | 1593150 | 68 | 8 | 0.94656 | 0.17193 | DUSP8 | 0.17193 | 0.22591 | 0.19202 |
| ENSG00000147488 | 8 | 53023399 | 53373519 | 953 | 75 | 0.94641 | 0.17197 | ST18 | 0.17197 | 0.13135 | 0.12443 |
| ENSG00000134330 | 2 | 9613787 | 9636672 | 79 | 9 | 0.94611 | 0.17205 | IAH1 | 0.17205 | 0.191 | 0.236 |
| ENSG00000205726 | 21 | 35014706 | 35272165 | 556 | 58 | 0.94598 | 0.17208 | ITSN1 | 0.17208 | 0.12619 | 0.12597 |
| ENSG00000186532 | 17 | 1682779 | 1733928 | 235 | 20 | 0.94535 | 0.17224 | SMYD4 | 0.17224 | 0.19767 | 0.17329 |
| ENSG00000184434 | 9 | 26993134 | 27005691 | 39 | 11 | 0.9453 | 0.17225 | LRRC19 | 0.17225 | 0.26072 | 0.35352 |
| ENSG00000233198 | 9 | 140122018 | 140124090 | 7 | 3 | 0.945 | 0.17233 | RNF224 | 0.17233 | 0.61026 | 0.41737 |
| ENSG00000162951 | 2 | 80515483 | 80531874 | 31 | 9 | 0.94439 | 0.17249 | LRRTM1 | 0.17249 | 0.46985 | 0.48512 |
| ENSG00000186572 | 8 | 7669242 | 7673238 | 1 | 1 | 0.94433 | 0.1725 | DEFB107A | 0.1725 | 0.1725 | 0.36425 |
| ENSG00000254585 | 15 | 23888691 | 23891175 | 1 | 1 | 0.94433 | 0.1725 | MAGEL2 | 0.1725 | 0.50953 | 0.5907 |
| ENSG00000163825 | 3 | 46538981 | 46542439 | 12 | 3 | 0.94377 | 0.17264 | RTP3 | 0.17264 | 0.32435 | 0.43147 |
| ENSG00000213923 | 22 | 38686697 | 38794527 | 255 | 35 | 0.94371 | 0.17266 | CSNK1E | 0.17266 | 0.21947 | 0.24345 |
| ENSG00000081692 | 1 | 227918126 | 227923112 | 13 | 6 | 0.94329 | 0.17277 | JMJD4 | 0.17277 | 0.18656 | 0.12638 |
| ENSG00000144354 | 2 | 174219548 | 174233725 | 54 | 11 | 0.94318 | 0.17279 | CDCA7 | 0.17279 | 0.19168 | 0.37589 |
| ENSG00000073861 | 17 | 45810610 | 45823485 | 47 | 11 | 0.94307 | 0.17282 | TBX21 | 0.17282 | 0.14182 | 0.18476 |
| ENSG00000145214 | 4 | 952675 | 980683 | 87 | 18 | 0.94281 | 0.17289 | DGKQ | 0.17289 | 0.10967 | 0.074407 |
| ENSG00000108559 | 17 | 5264258 | 5323480 | 321 | 12 | 0.94244 | 0.17298 | NUP88 | 0.17298 | 0.20053 | 0.20138 |
| ENSG00000109536 | 4 | 190861943 | 190884359 | 36 | 2 | 0.94228 | 0.17302 | FRG1 | 0.17302 | 0.16966 | 0.18643 |
| ENSG00000104974 | 19 | 55105047 | 55113555 | 68 | 7 | 0.94197 | 0.1731 | LILRA1 | 0.1731 | 0.21029 | 0.27552 |
| ENSG00000118503 | 6 | 138188351 | 138204449 | 32 | 5 | 0.94185 | 0.17313 | TNFAIP3 | 0.17313 | 0.14675 | 0.14891 |
| ENSG00000169509 | 1 | 152486978 | 152488486 | 5 | 2 | 0.94157 | 0.17321 | CRCT1 | 0.17321 | 0.3411 | 0.23764 |
| ENSG00000188396 | 1 | 45271585 | 45272957 | 5 | 3 | 0.94147 | 0.17323 | TCTEX1D4 | 0.17323 | 0.24343 | 0.22148 |
| ENSG00000171611 | 6 | 42883727 | 42893573 | 43 | 9 | 0.94113 | 0.17332 | PTCRA | 0.17332 | 0.1317 | 0.10938 |
| ENSG00000105219 | 19 | 40728115 | 40732597 | 11 | 6 | 0.94103 | 0.17334 | CNTD2 | 0.17334 | 0.20056 | 0.22372 |
| ENSG00000169981 | 3 | 44690219 | 44702283 | 17 | 8 | 0.9401 | 0.17358 | ZNF35 | 0.17358 | 0.10638 | 0.34465 |
| ENSG00000171956 | 15 | 60296421 | 60353929 | 206 | 24 | 0.94 | 0.17361 | FOXB1 | 0.17361 | 0.26095 | 0.29858 |
| ENSG00000220205 | 17 | 8062467 | 8066864 | 13 | 3 | 0.93962 | 0.17371 | VAMP2 | 0.17371 | 0.33724 | 0.5099 |
| ENSG00000121621 | 11 | 28042167 | 28129855 | 147 | 19 | 0.93939 | 0.17376 | KIF18A | 0.17376 | 0.18983 | 0.24144 |
| ENSG00000065534 | 3 | 123328896 | 123603178 | 596 | 62 | 0.93934 | 0.17378 | MYLK | 0.17378 | 0.21813 | 0.29446 |
| ENSG00000111254 | 12 | 4724674 | 4758213 | 173 | 18 | 0.93928 | 0.17379 | AKAP3 | 0.17379 | 0.23952 | 0.21358 |
| ENSG00000116670 | 1 | 11734537 | 11751707 | 77 | 16 | 0.93883 | 0.17391 | MAD2L2 | 0.17391 | 0.38853 | 0.58851 |
| ENSG00000138135 | 10 | 90965694 | 90967071 | 2 | 1 | 0.93804 | 0.17411 | CH25H | 0.17411 | 0.85439 | 0.89714 |
| ENSG00000142235 | 19 | 48988528 | 49016446 | 76 | 11 | 0.93746 | 0.17426 | LMTK3 | 0.17426 | 0.20788 | 0.23033 |
| ENSG00000179826 | 11 | 18142502 | 18160027 | 52 | 7 | 0.93742 | 0.17427 | MRGPRX3 | 0.17427 | 0.15901 | 0.10775 |
| ENSG00000162882 | 2 | 42994229 | 43019733 | 150 | 13 | 0.93667 | 0.17446 | HAAO | 0.17446 | 0.20405 | 0.20873 |
| ENSG00000129965 | 11 | 2153768 | 2182439 | 83 | 19 | 0.93651 | 0.17451 | INS-IGF2 | 0.17451 | 0.061769 | 0.13856 |
| ENSG00000189001 | 19 | 36014269 | 36019253 | 24 | 8 | 0.93647 | 0.17452 | SBSN | 0.17452 | 0.22777 | 0.4002 |
| ENSG00000132128 | 1 | 46726868 | 46769280 | 47 | 13 | 0.9362 | 0.17459 | LRRC41 | 0.17459 | 0.11346 | 0.085303 |

| | | | | | | | | | | | |
|---|---|---|---|---|---|---|---|---|---|---|---|
| ENSG00000174982 | 11 | 55418380 | 55419315 | 3 | 2 | 0.93586 | 0.17467 | OR4S2 | 0.17467 | 0.11184 | 0.12494 |
| ENSG00000178395 | 1 | 223566715 | 223568812 | 8 | 2 | 0.93545 | 0.17478 | C1orf65 | 0.17478 | 0.24407 | 0.30564 |
| ENSG00000103021 | 16 | 58265061 | 58317740 | 230 | 21 | 0.93532 | 0.17481 | CCDC113 | 0.17481 | 0.27893 | 0.30521 |
| ENSG00000151967 | 3 | 159557650 | 159615149 | 94 | 22 | 0.93508 | 0.17487 | SCHIP1 | 0.17487 | 0.3073 | 0.39072 |
| ENSG00000136014 | 12 | 95910336 | 95945266 | 118 | 25 | 0.93473 | 0.17496 | USP44 | 0.17496 | 0.055165 | 0.016249 |
| ENSG00000150361 | 13 | 70274726 | 70682591 | 2074 | 73 | 0.93457 | 0.175 | KLHL1 | 0.175 | 0.1878 | 0.21829 |
| ENSG00000177570 | 8 | 119201698 | 119634234 | 1374 | 89 | 0.93454 | 0.17501 | SAMD12 | 0.17501 | 0.22015 | 0.3203 |
| ENSG00000144485 | 2 | 239146908 | 239149303 | 10 | 3 | 0.93433 | 0.17507 | HES6 | 0.17507 | 0.37866 | 0.27295 |
| ENSG00000218739 | 2 | 37423631 | 37443336 | 60 | 15 | 0.9342 | 0.1751 | CEBPZ-AS1 | 0.1751 | 0.28488 | 0.42158 |
| ENSG00000173992 | 11 | 66360292 | 66373490 | 17 | 6 | 0.93419 | 0.1751 | CCS | 0.1751 | 0.22116 | 0.30177 |
| ENSG00000147655 | 8 | 108911544 | 109095913 | 561 | 39 | 0.93323 | 0.17535 | RSPO2 | 0.17535 | 0.23671 | 0.25958 |
| ENSG00000215021 | 12 | 7074490 | 7079988 | 7 | 4 | 0.9331 | 0.17539 | PHB2 | 0.17539 | 0.13156 | 0.051367 |
| ENSG00000100373 | 22 | 45680863 | 45691755 | 52 | 12 | 0.93303 | 0.1754 | UPK3A | 0.1754 | 0.13178 | 0.33769 |
| ENSG00000148734 | 10 | 72007085 | 72043432 | 155 | 29 | 0.93286 | 0.17545 | NPFFR1 | 0.17545 | 0.23803 | 0.14165 |
| ENSG00000253313 | 1 | 43747554 | 43751288 | 7 | 4 | 0.93254 | 0.17553 | C1orf210 | 0.17553 | 0.40697 | 0.70232 |
| ENSG00000078269 | 6 | 158402888 | 158520208 | 440 | 45 | 0.93222 | 0.17561 | SYNJ2 | 0.17561 | 0.27303 | 0.25375 |
| ENSG00000148331 | 9 | 132399171 | 132404444 | 13 | 5 | 0.93223 | 0.17561 | ASB6 | 0.17561 | 0.26103 | 0.27262 |
| ENSG00000173559 | 2 | 192542794 | 192561385 | 25 | 9 | 0.93201 | 0.17566 | NABP1 | 0.17566 | 0.089011 | 0.10778 |
| ENSG00000163254 | 2 | 208992861 | 208994554 | 4 | 2 | 0.93175 | 0.17573 | CRYGC | 0.17573 | 0.70054 | 0.59354 |
| ENSG00000105968 | 7 | 44866390 | 44887682 | 71 | 11 | 0.93164 | 0.17576 | H2AFV | 0.17576 | 0.1807 | 0.1956 |
| ENSG00000124466 | 19 | 43964939 | 43969812 | 14 | 5 | 0.93147 | 0.17581 | LYPD3 | 0.17581 | 0.34015 | 0.63078 |
| ENSG00000120963 | 8 | 102190106 | 102218421 | 40 | 8 | 0.9314 | 0.17582 | ZNF706 | 0.17582 | 0.34179 | 0.46243 |
| ENSG00000133083 | 13 | 36345478 | 36705443 | 1243 | 105 | 0.93135 | 0.17584 | DCLK1 | 0.17584 | 0.13417 | 0.11232 |
| ENSG00000037241 | 5 | 172385732 | 172396774 | 51 | 9 | 0.93124 | 0.17586 | RPL26L1 | 0.17586 | 0.18344 | 0.14127 |
| ENSG00000125851 | 20 | 17206752 | 17465223 | 1089 | 56 | 0.93112 | 0.17589 | PCSK2 | 0.17589 | 0.21262 | 0.27107 |
| ENSG00000142615 | 1 | 15783223 | 15798586 | 68 | 10 | 0.93109 | 0.1759 | CELA2A | 0.1759 | 0.17335 | 0.033096 |
| ENSG00000099139 | 9 | 78505560 | 78977255 | 1938 | 225 | 0.9309 | 0.17595 | PCSK5 | 0.17595 | 0.17826 | 0.30864 |
| ENSG00000100567 | 14 | 58711549 | 58738730 | 59 | 10 | 0.93078 | 0.17598 | PSMA3 | 0.17598 | 0.23928 | 0.31697 |
| ENSG00000173662 | 1 | 6615241 | 6639817 | 69 | 12 | 0.93064 | 0.17602 | TAS1R1 | 0.17602 | 0.069062 | 0.050631 |
| ENSG00000173976 | 19 | 3769087 | 3772233 | 14 | 4 | 0.93001 | 0.17618 | RAX2 | 0.17618 | 0.091792 | 0.14979 |
| ENSG00000066651 | 6 | 126307576 | 126360422 | 55 | 6 | 0.93 | 0.17619 | TRMT11 | 0.17619 | 0.15615 | 0.14737 |
| ENSG00000249992 | 3 | 45265958 | 45267770 | 5 | 2 | 0.92976 | 0.17625 | TMEM158 | 0.17625 | 0.23092 | 0.2954 |
| ENSG00000197122 | 20 | 35973088 | 36034453 | 175 | 21 | 0.9296 | 0.17629 | SRC | 0.17629 | 0.17526 | 0.21727 |
| ENSG00000151577 | 3 | 113847499 | 113918254 | 227 | 29 | 0.92926 | 0.17638 | DRD3 | 0.17638 | 0.18356 | 0.17527 |
| ENSG00000214642 | 6 | 49936390 | 49937338 | 1 | 1 | 0.92917 | 0.1764 | DEFB113 | 0.1764 | 0.66206 | 0.76809 |
| ENSG00000204007 | 9 | 138515502 | 138531386 | 89 | 13 | 0.92892 | 0.17646 | GLT6D1 | 0.17646 | 0.26079 | 0.31358 |
| ENSG00000198301 | 4 | 76862103 | 76912115 | 126 | 19 | 0.92795 | 0.17672 | SDAD1 | 0.17672 | 0.18566 | 0.14979 |
| ENSG00000144959 | 3 | 172348039 | 172429008 | 305 | 42 | 0.92767 | 0.17679 | NCEH1 | 0.17679 | 0.14845 | 0.03206 |
| ENSG00000186222 | 4 | 6717842 | 6719387 | 1 | 1 | 0.92724 | 0.1769 | BLOC1S4 | 0.1769 | 0.51966 | 0.29642 |
| ENSG00000042286 | 10 | 71857979 | 71892690 | 127 | 30 | 0.92702 | 0.17696 | AIFM2 | 0.17696 | 0.21927 | 0.2761 |
| ENSG00000138035 | 2 | 55861400 | 55921045 | 260 | 21 | 0.92685 | 0.177 | PNPT1 | 0.177 | 0.15826 | 0.16592 |
| ENSG00000153885 | 19 | 34286838 | 34306668 | 63 | 19 | 0.92686 | 0.177 | KCTD15 | 0.177 | 0.44847 | 0.5642 |
| ENSG00000171811 | 10 | 134621896 | 134756327 | 892 | 41 | 0.92683 | 0.17701 | TTC40 | 0.17701 | 0.24954 | 0.24575 |
| ENSG00000165060 | 9 | 71650175 | 71715094 | 283 | 18 | 0.92676 | 0.17703 | FXN | 0.17703 | 0.22584 | 0.31194 |
| ENSG00000058673 | 1 | 203764782 | 203823252 | 195 | 14 | 0.9266 | 0.17707 | ZC3H11A | 0.17707 | 0.22926 | 0.20504 |
| ENSG00000162763 | 1 | 165513244 | 165533198 | 88 | 15 | 0.92653 | 0.17708 | LRRC52 | 0.17708 | 0.201 | 0.20093 |
| ENSG00000186329 | 3 | 171561139 | 171656505 | 350 | 27 | 0.92636 | 0.17713 | TMEM212 | 0.17713 | 0.21332 | 0.25762 |
| ENSG00000119397 | 9 | 123837141 | 123939888 | 300 | 19 | 0.92577 | 0.17728 | CNTRL | 0.17728 | 0.19482 | 0.20483 |
| ENSG00000099904 | 22 | 20116979 | 20135530 | 69 | 8 | 0.92475 | 0.17755 | ZDHHC8 | 0.17755 | 0.14239 | 0.20609 |
| ENSG00000095596 | 10 | 94833232 | 94837647 | 7 | 2 | 0.92464 | 0.17758 | CYP26A1 | 0.17758 | 0.5419 | 0.52482 |
| ENSG00000139926 | 14 | 51955818 | 52197445 | 960 | 96 | 0.92441 | 0.17764 | FRMD6 | 0.17764 | 0.25142 | 0.30157 |
| ENSG00000251380 | 5 | 134779905 | 134783038 | 16 | 6 | 0.92416 | 0.1777 | C5orf20 | 0.1777 | 0.15223 | 0.19832 |
| ENSG00000178394 | 5 | 63256183 | 63258334 | 2 | 1 | 0.92388 | 0.17777 | HTR1A | 0.17777 | 0.33915 | 0.3478 |
| ENSG00000149639 | 20 | 35405845 | 35492089 | 136 | 27 | 0.92389 | 0.17777 | SOGA1 | 0.17777 | 0.14995 | 0.1578 |
| ENSG00000177963 | 11 | 207511 | 215113 | 22 | 7 | 0.92382 | 0.17779 | RIC8A | 0.17779 | 0.19089 | 0.32185 |
| ENSG00000163754 | 3 | 148709128 | 148745419 | 99 | 19 | 0.92255 | 0.17812 | GYG1 | 0.17812 | 0.18326 | 0.19825 |
| ENSG00000166012 | 11 | 93463114 | 93517557 | 154 | 9 | 0.92203 | 0.17826 | TAF1D | 0.17826 | 0.1771 | 0.20003 |
| ENSG00000127328 | 12 | 70132461 | 70216984 | 284 | 21 | 0.92196 | 0.17827 | RAB3IP | 0.17827 | 0.19007 | 0.19879 |
| ENSG00000273088 | 1 | 155141885 | 155159748 | 34 | 9 | 0.92182 | 0.17831 | RP11-201K10.3 | 0.17831 | 0.10321 | 0.085147 |
| ENSG00000143924 | 2 | 42396490 | 42559688 | 565 | 21 | 0.92182 | 0.17831 | EML4 | 0.17831 | 0.17639 | 0.17911 |
| ENSG00000173369 | 1 | 22979255 | 22988031 | 28 | 6 | 0.92178 | 0.17832 | C1QB | 0.17832 | 0.33951 | 0.49941 |
| ENSG00000065559 | 17 | 11924141 | 12047147 | 197 | 20 | 0.92131 | 0.17844 | MAP2K4 | 0.17844 | 0.22382 | 0.14199 |
| ENSG00000150401 | 13 | 114110134 | 114145267 | 91 | 16 | 0.9212 | 0.17847 | DCUN1D2 | 0.17847 | 0.10365 | 0.10889 |
| ENSG00000105887 | 7 | 135611509 | 135662101 | 221 | 29 | 0.921 | 0.17852 | MTPN | 0.17852 | 0.15073 | 0.1046 |
| ENSG00000160791 | 3 | 46411633 | 46417697 | 20 | 4 | 0.92098 | 0.17853 | CCR5 | 0.17853 | 0.37461 | 0.50139 |
| ENSG00000204099 | 2 | 242749920 | 242758739 | 56 | 12 | 0.92075 | 0.17859 | NEU4 | 0.17859 | 0.27845 | 0.41511 |
| ENSG00000205929 | 21 | 34162985 | 34186053 | 64 | 4 | 0.92039 | 0.17868 | C21orf62 | 0.17868 | 0.18792 | 0.1836 |
| ENSG00000160781 | 1 | 156213206 | 156217881 | 10 | 5 | 0.92036 | 0.17869 | PAQR6 | 0.17869 | 0.65869 | 0.65125 |
| ENSG00000179361 | 15 | 74833518 | 74890472 | 101 | 11 | 0.91994 | 0.1788 | ARID3B | 0.1788 | 0.23777 | 0.2323 |
| ENSG00000158497 | 5 | 143191726 | 143200284 | 22 | 7 | 0.91953 | 0.17891 | HMHB1 | 0.17891 | 0.16592 | 0.27006 |
| ENSG00000214686 | 3 | 51812580 | 51813009 | 2 | 1 | 0.9194 | 0.17894 | IQCF6 | 0.17894 | 0.45868 | 0.62109 |
| ENSG00000166819 | 15 | 90207596 | 90222658 | 41 | 13 | 0.91932 | 0.17896 | PLIN1 | 0.17896 | 0.14834 | 0.33487 |
| ENSG00000138760 | 4 | 77079890 | 77135046 | 207 | 18 | 0.9189 | 0.17907 | SCARB2 | 0.17907 | 0.19406 | 0.21978 |

| | | | | | | | | | | | |
|---|---|---|---|---|---|---|---|---|---|---|---|
| ENSG00000154040 | 18 | 21718942 | 21741567 | 57 | 10 | 0.91866 | 0.17914 | CABYR | 0.17914 | 0.22226 | 0.2173 |
| ENSG00000010327 | 3 | 52529354 | 52558511 | 54 | 15 | 0.9181 | 0.17928 | STAB1 | 0.17928 | 0.21609 | 0.21286 |
| ENSG00000060718 | 1 | 103342023 | 103574052 | 1140 | 32 | 0.91792 | 0.17933 | COL11A1 | 0.17933 | 0.17315 | 0.19782 |
| ENSG00000140600 | 15 | 84115980 | 84287495 | 463 | 67 | 0.91766 | 0.1794 | SH3GL3 | 0.1794 | 0.17715 | 0.15854 |
| ENSG00000243749 | 1 | 35447136 | 35450954 | 11 | 2 | 0.91733 | 0.17948 | ZMYM6NB | 0.17948 | 0.21459 | 0.21786 |
| ENSG00000198131 | 19 | 58739960 | 58788815 | 193 | 20 | 0.91705 | 0.17956 | ZNF544 | 0.17956 | 0.27654 | 0.30288 |
| ENSG00000249242 | 4 | 83404323 | 83483510 | 207 | 37 | 0.9169 | 0.1796 | TMEM150C | 0.1796 | 0.26339 | 0.43222 |
| ENSG00000180694 | 8 | 91634223 | 91803860 | 385 | 16 | 0.91675 | 0.17964 | TMEM64 | 0.17964 | 0.19659 | 0.25127 |
| ENSG00000196711 | 8 | 53446597 | 53478067 | 71 | 11 | 0.91668 | 0.17966 | FAM150A | 0.17966 | 0.26176 | 0.3018 |
| ENSG00000143110 | 1 | 112016414 | 112021134 | 10 | 3 | 0.91647 | 0.17971 | C1orf162 | 0.17971 | 0.18938 | 0.295 |
| ENSG00000204252 | 6 | 32971955 | 32977389 | 45 | 13 | 0.91643 | 0.17972 | HLA-DOA | 0.17972 | 0.29723 | 0.50289 |
| ENSG00000147416 | 8 | 20054878 | 20084330 | 47 | 12 | 0.91643 | 0.17972 | ATP6V1B2 | 0.17972 | 0.21313 | 0.51827 |
| ENSG00000242028 | 15 | 44088340 | 44095241 | 7 | 3 | 0.91645 | 0.17972 | HYPK | 0.17972 | 0.15495 | 0.20134 |
| ENSG00000130818 | 19 | 9638667 | 9649303 | 26 | 4 | 0.9162 | 0.17978 | ZNF426 | 0.17978 | 0.25596 | 0.25575 |
| ENSG00000141503 | 17 | 4736683 | 4801356 | 216 | 22 | 0.91592 | 0.17985 | MINK1 | 0.17985 | 0.25877 | 0.23399 |
| ENSG00000172992 | 17 | 43100708 | 43138473 | 144 | 18 | 0.91555 | 0.17995 | DCAKD | 0.17995 | 0.15606 | 0.22841 |
| ENSG00000233927 | 19 | 8386042 | 8388224 | 12 | 5 | 0.91553 | 0.17996 | RPS28 | 0.17996 | 0.24751 | 0.075783 |
| ENSG00000269295 | 9 | 71869331 | 71869642 | 1 | 1 | 0.91537 | 0.18 | AL358113.1 | 0.18 | 0.37229 | 0.57606 |
| ENSG00000056558 | 9 | 123664671 | 123691451 | 99 | 10 | 0.91501 | 0.18009 | TRAF1 | 0.18009 | 0.1911 | 0.17866 |
| ENSG00000187026 | 21 | 32119112 | 32119551 | 1 | 1 | 0.9146 | 0.1802 | KRTAP21-2 | 0.1802 | 0.65705 | 0.60836 |
| ENSG00000143815 | 1 | 225589204 | 225616627 | 76 | 14 | 0.91452 | 0.18022 | LBR | 0.18022 | 0.3423 | 0.49414 |
| ENSG00000100739 | 14 | 96722161 | 96735304 | 68 | 14 | 0.91428 | 0.18028 | BDKRB1 | 0.18028 | 0.23042 | 0.33446 |
| ENSG00000161647 | 17 | 41878167 | 41910538 | 52 | 6 | 0.91398 | 0.18036 | MPP3 | 0.18036 | 0.14206 | 0.16392 |
| ENSG00000197302 | 16 | 31724550 | 31806190 | 207 | 13 | 0.91394 | 0.18037 | ZNF720 | 0.18037 | 0.18573 | 0.16822 |
| ENSG00000166736 | 11 | 113845603 | 113861035 | 53 | 11 | 0.91377 | 0.18042 | HTR3A | 0.18042 | 0.31303 | 0.4165 |
| ENSG00000162729 | 1 | 160061130 | 160068733 | 14 | 4 | 0.91304 | 0.18061 | IGSF8 | 0.18061 | 0.24607 | 0.076307 |
| ENSG00000268602 | 11 | 107582998 | 107595135 | 34 | 8 | 0.91284 | 0.18066 | AP002353.1 | 0.18066 | 0.14079 | 0.30396 |
| ENSG00000151320 | 14 | 32798479 | 33300567 | 1896 | 151 | 0.91285 | 0.18066 | AKAP6 | 0.18066 | 0.2245 | 0.25185 |
| ENSG00000119707 | 14 | 73525144 | 73588122 | 162 | 27 | 0.91253 | 0.18075 | RBM25 | 0.18075 | 0.18852 | 0.18945 |
| ENSG00000204267 | 6 | 32789610 | 32806557 | 170 | 26 | 0.91208 | 0.18086 | TAP2 | 0.18086 | 0.11743 | 0.038556 |
| ENSG00000103253 | 16 | 776936 | 785525 | 29 | 5 | 0.912 | 0.18088 | HAGHL | 0.18088 | 0.23347 | 0.41008 |
| ENSG00000182950 | 15 | 76016318 | 76020029 | 2 | 1 | 0.91172 | 0.18096 | ODF3L1 | 0.18096 | 0.2648 | 0.28791 |
| ENSG00000099337 | 19 | 38810484 | 38819660 | 37 | 8 | 0.91164 | 0.18098 | KCNK6 | 0.18098 | 0.28379 | 0.2816 |
| ENSG00000147804 | 8 | 145635126 | 145642279 | 16 | 8 | 0.91156 | 0.181 | SLC39A4 | 0.181 | 0.057688 | 0.048918 |
| ENSG00000054796 | 20 | 55904815 | 55919050 | 44 | 4 | 0.91156 | 0.181 | SPO11 | 0.181 | 0.18063 | 0.19479 |
| ENSG00000151657 | 10 | 7792925 | 7829990 | 158 | 22 | 0.91044 | 0.1813 | KIN | 0.1813 | 0.27843 | 0.42837 |
| ENSG00000122176 | 1 | 203309756 | 203320617 | 36 | 10 | 0.90995 | 0.18142 | FMOD | 0.18142 | 0.29177 | 0.33257 |
| ENSG00000155846 | 5 | 149109861 | 149234585 | 475 | 50 | 0.90974 | 0.18148 | PPARGC1B | 0.18148 | 0.23709 | 0.32251 |
| ENSG00000165702 | 9 | 135820932 | 135867083 | 221 | 53 | 0.90922 | 0.18162 | GFI1B | 0.18162 | 0.28947 | 0.37136 |
| ENSG00000241186 | 3 | 46616045 | 46668033 | 248 | 31 | 0.90898 | 0.18168 | TDGF1 | 0.18168 | 0.20622 | 0.14779 |
| ENSG00000166349 | 11 | 36532259 | 36614706 | 136 | 21 | 0.90892 | 0.1817 | RAG1 | 0.1817 | 0.20429 | 0.27105 |
| ENSG00000145604 | 5 | 36152091 | 36184421 | 112 | 15 | 0.90867 | 0.18176 | SKP2 | 0.18176 | 0.24535 | 0.2899 |
| ENSG00000142319 | 5 | 1392909 | 1445545 | 177 | 35 | 0.90855 | 0.18179 | SLC6A3 | 0.18179 | 0.27795 | 0.49425 |
| ENSG00000269446 | 7 | 156902674 | 156903764 | 9 | 2 | 0.90766 | 0.18203 | AC006967.1 | 0.18203 | 0.25906 | 0.23743 |
| ENSG00000204335 | 2 | 171571861 | 171574588 | 5 | 2 | 0.90737 | 0.18211 | SP5 | 0.18211 | 0.11844 | 0.25149 |
| ENSG00000140694 | 16 | 14529558 | 14726585 | 374 | 24 | 0.9069 | 0.18223 | PARN | 0.18223 | 0.20971 | 0.24191 |
| ENSG00000100075 | 22 | 19163095 | 19166343 | 7 | 3 | 0.9069 | 0.18223 | SLC25A1 | 0.18223 | 0.021284 | 0.0073074 |
| ENSG00000138802 | 4 | 110354928 | 110462052 | 255 | 11 | 0.90636 | 0.18237 | SEC24B | 0.18237 | 0.16179 | 0.13656 |
| ENSG00000112715 | 6 | 43737921 | 43754224 | 47 | 14 | 0.90602 | 0.18246 | VEGFA | 0.18246 | 0.17924 | 0.29234 |
| ENSG00000072958 | 19 | 16308389 | 16346160 | 155 | 9 | 0.90518 | 0.18268 | AP1M1 | 0.18268 | 0.14727 | 0.10983 |
| ENSG00000177628 | 1 | 155204243 | 155214490 | 25 | 4 | 0.90517 | 0.18269 | GBA | 0.18269 | 0.17495 | 0.16295 |
| ENSG00000204001 | 9 | 139648838 | 139652678 | 18 | 3 | 0.90507 | 0.18271 | LCN8 | 0.18271 | 0.31388 | 0.32273 |
| ENSG00000164897 | 7 | 150778167 | 150780633 | 4 | 2 | 0.90502 | 0.18273 | TMUB1 | 0.18273 | 0.89261 | 0.87164 |
| ENSG00000058335 | 15 | 79252289 | 79383115 | 448 | 74 | 0.90457 | 0.18285 | RASGRF1 | 0.18285 | 0.094574 | 0.084586 |
| ENSG00000047056 | 10 | 1095478 | 1178237 | 306 | 29 | 0.9044 | 0.18289 | WDR37 | 0.18289 | 0.18845 | 0.17475 |
| ENSG00000255872 | 9 | 37588410 | 38068684 | 1617 | 62 | 0.90413 | 0.18296 | RP11-613M10.9 | 0.18296 | 0.20399 | 0.26411 |
| ENSG00000159784 | 7 | 143050493 | 143059863 | 15 | 6 | 0.90374 | 0.18307 | FAM131B | 0.18307 | 0.47144 | 0.59087 |
| ENSG00000113716 | 5 | 149379884 | 149432386 | 148 | 12 | 0.90352 | 0.18312 | HMGXB3 | 0.18312 | 0.21131 | 0.23695 |
| ENSG00000172348 | 6 | 46188475 | 46459709 | 440 | 39 | 0.90343 | 0.18315 | RCAN2 | 0.18315 | 0.19479 | 0.23652 |
| ENSG00000038295 | 4 | 166794410 | 167025047 | 592 | 65 | 0.90332 | 0.18318 | TLL1 | 0.18318 | 0.22031 | 0.29962 |
| ENSG00000258052 | 12 | 70195449 | 70249143 | 198 | 23 | 0.90317 | 0.18322 | AC025263.3 | 0.18322 | 0.18878 | 0.23344 |
| ENSG00000084093 | 4 | 57774075 | 57802010 | 69 | 19 | 0.90288 | 0.1833 | REST | 0.1833 | 0.15161 | 0.33353 |
| ENSG00000131507 | 5 | 141488070 | 141534005 | 204 | 17 | 0.9027 | 0.18334 | NDFIP1 | 0.18334 | 0.1832 | 0.15409 |
| ENSG00000186409 | 1 | 42929001 | 43120335 | 703 | 22 | 0.90263 | 0.18336 | CCDC30 | 0.18336 | 0.19404 | 0.21083 |
| ENSG00000124217 | 20 | 49575363 | 49577820 | 3 | 2 | 0.90261 | 0.18337 | MOCS3 | 0.18337 | 0.72301 | 0.92694 |
| ENSG00000137691 | 11 | 101918174 | 101955291 | 78 | 10 | 0.90235 | 0.18344 | C11orf70 | 0.18344 | 0.19121 | 0.22207 |
| ENSG00000182185 | 14 | 68286496 | 69196935 | 2493 | 117 | 0.90232 | 0.18344 | RAD51B | 0.18344 | 0.14682 | 0.1536 |
| ENSG00000120539 | 10 | 27443753 | 27475853 | 147 | 15 | 0.90226 | 0.18346 | MASTL | 0.18346 | 0.14111 | 0.14511 |
| ENSG00000129932 | 19 | 3490819 | 3500938 | 26 | 5 | 0.90226 | 0.18346 | DOHH | 0.18346 | 0.11943 | 0.19129 |
| ENSG00000233816 | 9 | 21367371 | 21368075 | 1 | 1 | 0.90173 | 0.1836 | IFNA13 | 0.1836 | 0.23615 | 0.23832 |
| ENSG00000142408 | 19 | 54466294 | 54493469 | 112 | 17 | 0.90144 | 0.18368 | CACNG8 | 0.18368 | 0.1846 | 0.021731 |
| ENSG00000197879 | 17 | 1367392 | 1396106 | 115 | 38 | 0.90058 | 0.1839 | MYO1C | 0.1839 | 0.12979 | 0.15117 |
| ENSG00000100083 | 22 | 38004481 | 38029571 | 51 | 10 | 0.9002 | 0.18401 | GGA1 | 0.18401 | 0.2056 | 0.19198 |

| | | | | | | | | | | | |
|---|---|---|---|---|---|---|---|---|---|---|---|
| ENSG00000172766 | 13 | 41885341 | 41951166 | 195 | 13 | 0.89999 | 0.18406 | NAA16 | 0.18406 | 0.18772 | 0.17665 |
| ENSG00000173457 | 11 | 64011956 | 64014413 | 6 | 2 | 0.8999 | 0.18409 | PPP1R14B | 0.18409 | 0.24808 | 0.44868 |
| ENSG00000184305 | 4 | 91048686 | 92523064 | 5671 | 175 | 0.89958 | 0.18417 | CCSER1 | 0.18417 | 0.17266 | 0.16401 |
| ENSG00000166396 | 18 | 61420169 | 61472604 | 183 | 21 | 0.8994 | 0.18422 | SERPINB7 | 0.18422 | 0.21822 | 0.36501 |
| ENSG00000134108 | 3 | 5163905 | 5222596 | 175 | 11 | 0.89936 | 0.18423 | ARL8B | 0.18423 | 0.20079 | 0.092611 |
| ENSG00000178235 | 13 | 84451344 | 84456528 | 7 | 3 | 0.89929 | 0.18425 | SLITRK1 | 0.18425 | 0.34152 | 0.5173 |
| ENSG00000168267 | 10 | 23481256 | 23483181 | 3 | 1 | 0.89913 | 0.18429 | PTF1A | 0.18429 | 0.34399 | 0.26721 |
| ENSG00000167291 | 17 | 77906142 | 78009647 | 469 | 102 | 0.89866 | 0.18442 | TBC1D16 | 0.18442 | 0.1997 | 0.20901 |
| ENSG00000136108 | 13 | 53029564 | 53050763 | 64 | 8 | 0.89842 | 0.18448 | CKAP2 | 0.18448 | 0.20411 | 0.21763 |
| ENSG00000181689 | 11 | 56085783 | 56086721 | 2 | 1 | 0.89796 | 0.1846 | OR8K3 | 0.1846 | 0.015089 | 0.019748 |
| ENSG00000120306 | 5 | 139554227 | 139661637 | 252 | 11 | 0.89793 | 0.18461 | CYSTM1 | 0.18461 | 0.17323 | 0.143 |
| ENSG00000198720 | 17 | 27916787 | 27941779 | 36 | 11 | 0.89792 | 0.18462 | ANKRD13B | 0.18462 | 0.22988 | 0.33583 |
| ENSG00000139574 | 12 | 53900474 | 53901422 | 1 | 1 | 0.8976 | 0.1847 | NPFF | 0.1847 | 0.065137 | 0.025373 |
| ENSG00000224383 | 17 | 62075711 | 62081664 | 17 | 6 | 0.89699 | 0.18486 | C17orf72 | 0.18486 | 0.1479 | 0.33306 |
| ENSG00000126067 | 1 | 36067185 | 36107445 | 31 | 7 | 0.89668 | 0.18494 | PSMB2 | 0.18494 | 0.17692 | 0.19556 |
| ENSG00000185347 | 14 | 105956192 | 105965912 | 19 | 8 | 0.89646 | 0.185 | C14orf80 | 0.185 | 0.38895 | 0.41204 |
| ENSG00000161692 | 17 | 42785976 | 42829632 | 96 | 20 | 0.896 | 0.18513 | DBF4B | 0.18513 | 0.2299 | 0.26032 |
| ENSG00000011478 | 19 | 46195741 | 46207247 | 29 | 10 | 0.89591 | 0.18515 | QPCTL | 0.18515 | 0.063639 | 0.022459 |
| ENSG00000198431 | 12 | 104609557 | 104744061 | 535 | 30 | 0.89577 | 0.18519 | TXNRD1 | 0.18519 | 0.16121 | 0.20813 |
| ENSG00000014824 | 4 | 41992489 | 42092474 | 415 | 10 | 0.89568 | 0.18521 | SLC30A9 | 0.18521 | 0.19009 | 0.22676 |
| ENSG00000167552 | 12 | 49578579 | 49583107 | 8 | 2 | 0.89552 | 0.18525 | TUBA1A | 0.18525 | 0.099121 | 0.12802 |
| ENSG00000125266 | 13 | 107142079 | 107187462 | 119 | 24 | 0.89507 | 0.18538 | EFNB2 | 0.18538 | 0.30642 | 0.40493 |
| ENSG00000205784 | 19 | 4890449 | 4902879 | 80 | 14 | 0.89475 | 0.18546 | ARRDC5 | 0.18546 | 0.15778 | 0.083991 |
| ENSG00000158467 | 7 | 128864864 | 129070052 | 494 | 29 | 0.89452 | 0.18552 | AHCYL2 | 0.18552 | 0.1486 | 0.11595 |
| ENSG00000161970 | 17 | 8280838 | 8286531 | 19 | 4 | 0.89419 | 0.18561 | RPL26 | 0.18561 | 0.21576 | 0.33633 |
| ENSG00000124785 | 6 | 5998232 | 6007200 | 19 | 8 | 0.89388 | 0.18569 | NRN1 | 0.18569 | 0.17465 | 0.1031 |
| ENSG00000139780 | 13 | 103338094 | 103346854 | 49 | 17 | 0.89378 | 0.18572 | METTL21C | 0.18572 | 0.15556 | 0.34291 |
| ENSG00000120458 | 11 | 124636394 | 124670569 | 72 | 13 | 0.89349 | 0.1858 | MSANTD2 | 0.1858 | 0.18173 | 0.42747 |
| ENSG00000152766 | 10 | 90581889 | 90611575 | 120 | 20 | 0.89298 | 0.18593 | ANKRD22 | 0.18593 | 0.23752 | 0.33122 |
| ENSG00000155761 | 1 | 118496484 | 118727846 | 483 | 23 | 0.89295 | 0.18594 | SPAG17 | 0.18594 | 0.20553 | 0.2291 |
| ENSG00000115232 | 2 | 182321929 | 182400914 | 264 | 26 | 0.89287 | 0.18596 | ITGA4 | 0.18596 | 0.16704 | 0.16619 |
| ENSG00000205111 | 2 | 39402787 | 39456729 | 160 | 14 | 0.89281 | 0.18598 | CDKL4 | 0.18598 | 0.19911 | 0.18816 |
| ENSG00000196805 | 1 | 153042700 | 153113927 | 273 | 24 | 0.8927 | 0.18601 | SPRR2B | 0.18601 | 0.13707 | 0.064817 |
| ENSG00000204533 | 19 | 56662314 | 56663250 | 2 | 1 | 0.89249 | 0.18606 | AC024580.1 | 0.18606 | 0.65282 | 0.80841 |
| ENSG00000115816 | 2 | 37428755 | 37458856 | 98 | 20 | 0.89196 | 0.18621 | CEBPZ | 0.18621 | 0.25391 | 0.45904 |
| ENSG00000115318 | 2 | 74759541 | 74782817 | 22 | 6 | 0.89178 | 0.18626 | LOXL3 | 0.18626 | 0.29379 | 0.30493 |
| ENSG00000197472 | 1 | 247108849 | 247171395 | 255 | 24 | 0.89164 | 0.18629 | ZNF695 | 0.18629 | 0.1808 | 0.12755 |
| ENSG00000006611 | 11 | 17515442 | 17565963 | 165 | 28 | 0.89138 | 0.18636 | USH1C | 0.18636 | 0.21984 | 0.32548 |
| ENSG00000268332 | 1 | 6100630 | 6101399 | 4 | 2 | 0.89085 | 0.18651 | AL035406.1 | 0.18651 | 0.61564 | 0.67393 |
| ENSG00000127463 | 1 | 19542158 | 19578046 | 89 | 12 | 0.89062 | 0.18657 | EMC1 | 0.18657 | 0.36483 | 0.39675 |
| ENSG00000186306 | 1 | 158368312 | 158369256 | 8 | 3 | 0.88965 | 0.18683 | OR10T2 | 0.18683 | 0.55721 | 0.39767 |
| ENSG00000214237 | 3 | 150588832 | 150612089 | 72 | 13 | 0.88965 | 0.18683 | FAM188B2 | 0.18683 | 0.20162 | 0.22371 |
| ENSG00000265944 | 18 | 6511415 | 6590652 | 334 | 51 | 0.88948 | 0.18687 | C18orf64 | 0.18687 | 0.24001 | 0.2479 |
| ENSG00000177453 | 5 | 43192173 | 43280952 | 267 | 34 | 0.88945 | 0.18688 | NIM1K | 0.18688 | 0.18701 | 0.25204 |
| ENSG00000140259 | 15 | 44096690 | 44117000 | 35 | 5 | 0.88908 | 0.18698 | MFAP1 | 0.18698 | 0.16234 | 0.16495 |
| ENSG00000189046 | 12 | 109525996 | 109531436 | 13 | 5 | 0.88902 | 0.187 | ALKBH2 | 0.187 | 0.26874 | 0.51748 |
| ENSG00000054356 | 2 | 220154345 | 220174370 | 54 | 11 | 0.88876 | 0.18707 | PTPRN | 0.18707 | 0.29085 | 0.48169 |
| ENSG00000197363 | 8 | 146024261 | 146036554 | 52 | 6 | 0.88861 | 0.18711 | ZNF517 | 0.18711 | 0.095095 | 0.059793 |
| ENSG00000141424 | 18 | 33688495 | 33709348 | 50 | 4 | 0.88855 | 0.18712 | SLC39A6 | 0.18712 | 0.35021 | 0.35549 |
| ENSG00000170264 | 2 | 62051989 | 62081278 | 96 | 14 | 0.88839 | 0.18717 | FAM161A | 0.18717 | 0.23655 | 0.29906 |
| ENSG00000107672 | 10 | 123716603 | 123734732 | 85 | 9 | 0.88823 | 0.18721 | NSMCE4A | 0.18721 | 0.19448 | 0.17753 |
| ENSG00000166136 | 10 | 102267203 | 102289757 | 15 | 7 | 0.88788 | 0.1873 | NDUFB8 | 0.1873 | 0.14079 | 0.29712 |
| ENSG00000228259 | 5 | 179078298 | 179079445 | 4 | 2 | 0.88764 | 0.18737 | AC136604.1 | 0.18737 | 0.43854 | 0.42043 |
| ENSG00000106031 | 7 | 27233122 | 27239725 | 15 | 4 | 0.88722 | 0.18748 | HOXA13 | 0.18748 | 0.080731 | 0.19629 |
| ENSG00000203668 | 1 | 241792155 | 241799232 | 20 | 9 | 0.8871 | 0.18751 | CHML | 0.18751 | 0.19659 | 0.1321 |
| ENSG00000113946 | 3 | 190040330 | 190129932 | 482 | 33 | 0.88705 | 0.18753 | CLDN16 | 0.18753 | 0.23516 | 0.32228 |
| ENSG00000058404 | 7 | 44256749 | 44374176 | 333 | 29 | 0.88698 | 0.18754 | CAMK2B | 0.18754 | 0.19728 | 0.26842 |
| ENSG00000163960 | 3 | 196074533 | 196159345 | 185 | 20 | 0.88685 | 0.18758 | UBXN7 | 0.18758 | 0.095262 | 0.093051 |
| ENSG00000172716 | 17 | 33677324 | 33700720 | 70 | 17 | 0.88635 | 0.18771 | SLFN11 | 0.18771 | 0.45592 | 0.67892 |
| ENSG00000103460 | 16 | 52471917 | 52581714 | 237 | 46 | 0.88631 | 0.18772 | TOX3 | 0.18772 | 0.28488 | 0.34463 |
| ENSG00000010072 | 1 | 231472850 | 231490769 | 53 | 6 | 0.88619 | 0.18776 | SPRTN | 0.18776 | 0.22487 | 0.22158 |
| ENSG00000111145 | 12 | 96588160 | 96663613 | 213 | 43 | 0.88616 | 0.18777 | ELK3 | 0.18777 | 0.17164 | 0.10196 |
| ENSG00000206069 | 22 | 25331077 | 25342662 | 27 | 8 | 0.88576 | 0.18787 | TMEM211 | 0.18787 | 0.30823 | 0.7058 |
| ENSG00000135317 | 6 | 86215214 | 86303874 | 178 | 22 | 0.88497 | 0.18809 | SNX14 | 0.18809 | 0.1374 | 0.14309 |
| ENSG00000118733 | 1 | 102268130 | 102462586 | 810 | 74 | 0.88483 | 0.18812 | OLFM3 | 0.18812 | 0.20509 | 0.25087 |
| ENSG00000196227 | 20 | 58508819 | 58523735 | 13 | 7 | 0.88431 | 0.18826 | FAM217B | 0.18826 | 0.21333 | 0.19997 |
| ENSG00000196428 | 3 | 150126122 | 150184218 | 192 | 21 | 0.88416 | 0.1883 | TSC22D2 | 0.1883 | 0.17487 | 0.16692 |
| ENSG00000198965 | 1 | 158449668 | 158450675 | 5 | 2 | 0.88413 | 0.18831 | OR10R2 | 0.18831 | 0.40277 | 0.30225 |
| ENSG00000182968 | 13 | 112721913 | 112726020 | 12 | 3 | 0.88412 | 0.18832 | SOX1 | 0.18832 | 0.042894 | 0.010363 |
| ENSG00000169519 | 11 | 28129795 | 28355054 | 394 | 21 | 0.88386 | 0.18839 | METTL15 | 0.18839 | 0.19175 | 0.1814 |
| ENSG00000180891 | 17 | 55938604 | 56032684 | 299 | 46 | 0.88357 | 0.18846 | CUEDC1 | 0.18846 | 0.24666 | 0.32728 |
| ENSG00000088280 | 1 | 23755056 | 23811061 | 143 | 17 | 0.8835 | 0.18848 | ASAP3 | 0.18848 | 0.13235 | 0.18992 |
| ENSG00000170364 | 3 | 4344988 | 4359251 | 34 | 10 | 0.88348 | 0.18849 | SETMAR | 0.18849 | 0.42565 | 0.3775 |

| | | | | | | | | | | | |
|---|---|---|---|---|---|---|---|---|---|---|---|
| ENSG00000205899 | 17 | 1173853 | 1174754 | 1 | 1 | 0.88344 | 0.1885 | BHLHA9 | 0.1885 | 0.92873 | 0.99427 |
| ENSG00000196787 | 6 | 27100832 | 27103070 | 2 | 1 | 0.88314 | 0.18858 | HIST1H2AG | 0.18858 | 0.45318 | 0.40026 |
| ENSG00000108370 | 17 | 63133549 | 63223821 | 339 | 29 | 0.88314 | 0.18858 | RGS9 | 0.18858 | 0.17922 | 0.15908 |
| ENSG00000168994 | 6 | 3722848 | 3752260 | 89 | 18 | 0.88285 | 0.18866 | PXDC1 | 0.18866 | 0.31315 | 0.41393 |
| ENSG00000142507 | 17 | 4699439 | 4701790 | 11 | 6 | 0.88265 | 0.18871 | PSMB6 | 0.18871 | 0.15764 | 0.15295 |
| ENSG00000151835 | 13 | 23902965 | 24007841 | 381 | 40 | 0.88257 | 0.18873 | SACS | 0.18873 | 0.20953 | 0.054797 |
| ENSG00000106868 | 9 | 114803065 | 114937688 | 385 | 43 | 0.88231 | 0.1888 | SUSD1 | 0.1888 | 0.28833 | 0.3636 |
| ENSG00000110777 | 11 | 111222977 | 111326355 | 466 | 29 | 0.88215 | 0.18885 | POU2AF1 | 0.18885 | 0.2279 | 0.28351 |
| ENSG00000170122 | 9 | 116237 | 118417 | 8 | 3 | 0.88211 | 0.18886 | FOXD4 | 0.18886 | 0.0524 | 0.090776 |
| ENSG00000063245 | 19 | 56186592 | 56221224 | 149 | 24 | 0.88199 | 0.18889 | EPN1 | 0.18889 | 0.1014 | 0.063614 |
| ENSG00000048162 | 5 | 175810949 | 175815976 | 14 | 3 | 0.88189 | 0.18892 | NOP16 | 0.18892 | 0.28698 | 0.34179 |
| ENSG00000131668 | 9 | 96713905 | 96717654 | 16 | 5 | 0.88132 | 0.18907 | BARX1 | 0.18907 | 0.02929 | 0.10752 |
| ENSG00000172379 | 15 | 80696692 | 80890278 | 606 | 65 | 0.88125 | 0.18909 | ARNT2 | 0.18909 | 0.17068 | 0.13329 |
| ENSG00000123610 | 2 | 152214106 | 152236560 | 72 | 9 | 0.88117 | 0.18911 | TNFAIP6 | 0.18911 | 0.18217 | 0.28638 |
| ENSG00000255398 | 12 | 123199303 | 123201439 | 15 | 3 | 0.8809 | 0.18919 | HCAR3 | 0.18919 | 0.26027 | 0.19068 |
| ENSG00000117448 | 1 | 46016215 | 46035721 | 28 | 9 | 0.88086 | 0.1892 | AKR1A1 | 0.1892 | 0.19746 | 0.344 |
| ENSG00000131378 | 3 | 16355081 | 16555533 | 783 | 73 | 0.88045 | 0.18931 | RFTN1 | 0.18931 | 0.18318 | 0.15568 |
| ENSG00000105662 | 19 | 18794487 | 18893004 | 248 | 35 | 0.87993 | 0.18945 | CRTC1 | 0.18945 | 0.18453 | 0.20986 |
| ENSG00000102524 | 13 | 108903588 | 108960832 | 197 | 47 | 0.87972 | 0.18951 | TNFSF13B | 0.18951 | 0.2658 | 0.50743 |
| ENSG00000066827 | 8 | 135490031 | 135725292 | 806 | 46 | 0.87962 | 0.18953 | ZFAT | 0.18953 | 0.19237 | 0.19541 |
| ENSG00000130222 | 9 | 92219928 | 92221470 | 8 | 3 | 0.87939 | 0.1896 | GADD45G | 0.1896 | 0.2833 | 0.25499 |
| ENSG00000198815 | 1 | 42642210 | 42801548 | 360 | 15 | 0.87898 | 0.18971 | FOXJ3 | 0.18971 | 0.17313 | 0.13606 |
| ENSG00000261667 | 8 | 144161874 | 144164418 | 10 | 4 | 0.8782 | 0.18992 | RP11-520P18.5 | 0.18992 | 0.63334 | 0.6651 |
| ENSG00000160284 | 21 | 47581062 | 47604390 | 189 | 16 | 0.87804 | 0.18996 | SPATC1L | 0.18996 | 0.19619 | 0.22136 |
| ENSG00000268427 | 22 | 50714103 | 50714350 | 1 | 1 | 0.8779 | 0.19 | AL022328.1 | 0.19 | 0.118 | 0.31023 |
| ENSG00000239857 | 7 | 916189 | 936073 | 100 | 20 | 0.87713 | 0.19021 | GET4 | 0.19021 | 0.16998 | 0.0385 |
| ENSG00000089163 | 12 | 120740119 | 120751052 | 24 | 3 | 0.87709 | 0.19022 | SIRT4 | 0.19022 | 0.22617 | 0.24085 |
| ENSG00000105650 | 19 | 18318771 | 18366229 | 168 | 19 | 0.87707 | 0.19022 | PDE4C | 0.19022 | 0.21716 | 0.28403 |
| ENSG00000168268 | 3 | 52558386 | 52569070 | 31 | 6 | 0.87654 | 0.19037 | NT5DC2 | 0.19037 | 0.16244 | 0.22177 |
| ENSG00000182134 | 1 | 151742583 | 151763892 | 65 | 5 | 0.87584 | 0.19056 | TDRKH | 0.19056 | 0.1091 | 0.037524 |
| ENSG00000172519 | 19 | 15904761 | 15905892 | 14 | 5 | 0.87566 | 0.19061 | OR10H5 | 0.19061 | 0.33993 | 0.18473 |
| ENSG00000179938 | 15 | 30375158 | 30385702 | 6 | 3 | 0.8755 | 0.19065 | GOLGA8J | 0.19065 | 0.75768 | 0.74793 |
| ENSG00000147679 | 8 | 117778742 | 117861702 | 223 | 37 | 0.87545 | 0.19067 | UTP23 | 0.19067 | 0.21432 | 0.22808 |
| ENSG00000093009 | 22 | 19466982 | 19508135 | 125 | 21 | 0.87527 | 0.19071 | CDC45 | 0.19071 | 0.24498 | 0.33911 |
| ENSG00000249016 | 5 | 54252063 | 54253674 | 12 | 2 | 0.87483 | 0.19083 | RP11-45H22.3 | 0.19083 | 0.29192 | 0.42704 |
| ENSG00000186275 | 1 | 146373546 | 146467744 | 9 | 3 | 0.87474 | 0.19086 | NBPF12 | 0.19086 | 0.27501 | 0.3549 |
| ENSG00000092094 | 14 | 20914570 | 20923264 | 47 | 6 | 0.87475 | 0.19086 | OSGEP | 0.19086 | 0.33642 | 0.38369 |
| ENSG00000134371 | 1 | 193091147 | 193223031 | 257 | 18 | 0.87453 | 0.19091 | CDC73 | 0.19091 | 0.20914 | 0.20865 |
| ENSG00000188324 | 12 | 55688072 | 55689016 | 2 | 1 | 0.87403 | 0.19105 | OR6C6 | 0.19105 | 0.95903 | 0.95698 |
| ENSG00000169567 | 5 | 130494720 | 130507428 | 51 | 4 | 0.87362 | 0.19116 | HINT1 | 0.19116 | 0.22517 | 0.28713 |
| ENSG00000229314 | 9 | 117085336 | 117088755 | 19 | 7 | 0.87338 | 0.19123 | ORM1 | 0.19123 | 0.056059 | 0.092751 |
| ENSG00000149651 | 20 | 44350990 | 44354469 | 25 | 2 | 0.87301 | 0.19133 | SPINT4 | 0.19133 | 0.19757 | 0.16497 |
| ENSG00000197008 | 7 | 64254766 | 64294054 | 177 | 16 | 0.87288 | 0.19136 | ZNF138 | 0.19136 | 0.20312 | 0.14241 |
| ENSG00000082684 | 3 | 122628041 | 122747452 | 422 | 57 | 0.87285 | 0.19137 | SEMA5B | 0.19137 | 0.17885 | 0.26332 |
| ENSG00000132383 | 17 | 1732996 | 1803376 | 245 | 23 | 0.87284 | 0.19138 | RPA1 | 0.19138 | 0.17121 | 0.19351 |
| ENSG00000170581 | 12 | 56735381 | 56753939 | 37 | 4 | 0.8727 | 0.19141 | STAT2 | 0.19141 | 0.20692 | 0.24352 |
| ENSG00000079931 | 6 | 132617194 | 132722684 | 319 | 31 | 0.8726 | 0.19144 | MOXD1 | 0.19144 | 0.28052 | 0.29734 |
| ENSG00000108375 | 17 | 56429861 | 56494956 | 95 | 16 | 0.87227 | 0.19153 | RNF43 | 0.19153 | 0.034541 | 0.018058 |
| ENSG00000109906 | 11 | 113930315 | 114121398 | 684 | 85 | 0.87205 | 0.19159 | ZBTB16 | 0.19159 | 0.24215 | 0.33761 |
| ENSG00000173876 | 10 | 92828 | 120103 | 107 | 16 | 0.87194 | 0.19162 | TUBB8 | 0.19162 | 0.18753 | 0.19557 |
| ENSG00000042062 | 20 | 49202645 | 49308065 | 389 | 56 | 0.87186 | 0.19164 | FAM65C | 0.19164 | 0.2478 | 0.34019 |
| ENSG00000115155 | 2 | 26680071 | 26781566 | 314 | 40 | 0.87161 | 0.19171 | OTOF | 0.19171 | 0.23865 | 0.32602 |
| ENSG00000100749 | 14 | 97263641 | 97398059 | 391 | 28 | 0.87099 | 0.19188 | VRK1 | 0.19188 | 0.18618 | 0.1849 |
| ENSG00000175393 | 11 | 7949180 | 7950209 | 7 | 3 | 0.87096 | 0.19189 | OR10A6 | 0.19189 | 0.14147 | 0.20473 |
| ENSG00000175793 | 1 | 27189633 | 27190947 | 2 | 1 | 0.87062 | 0.19198 | SFN | 0.19198 | 0.71974 | 0.77269 |
| ENSG00000140859 | 16 | 57792129 | 57896957 | 400 | 44 | 0.8703 | 0.19207 | KIFC3 | 0.19207 | 0.22542 | 0.17963 |
| ENSG00000090661 | 19 | 8271620 | 8327305 | 301 | 44 | 0.87003 | 0.19214 | CERS4 | 0.19214 | 0.44462 | 0.49772 |
| ENSG00000183251 | 11 | 5322244 | 5323226 | 3 | 2 | 0.86893 | 0.19244 | OR51B4 | 0.19244 | 0.38129 | 0.58095 |
| ENSG00000090316 | 4 | 1283639 | 1333935 | 345 | 22 | 0.86874 | 0.19249 | MAEA | 0.19249 | 0.1163 | 0.11168 |
| ENSG00000115665 | 2 | 108602979 | 108630450 | 88 | 26 | 0.86853 | 0.19255 | SLC5A7 | 0.19255 | 0.30906 | 0.36381 |
| ENSG00000107618 | 10 | 48381487 | 48390991 | 11 | 5 | 0.86841 | 0.19258 | RBP3 | 0.19258 | 0.23254 | 0.28733 |
| ENSG00000176444 | 1 | 155232659 | 155248282 | 26 | 4 | 0.86784 | 0.19274 | CLK2 | 0.19274 | 0.20654 | 0.18173 |
| ENSG00000130635 | 9 | 137533620 | 137736686 | 1132 | 90 | 0.86736 | 0.19287 | COL5A1 | 0.19287 | 0.22852 | 0.30229 |
| ENSG00000105675 | 19 | 36040945 | 36054560 | 33 | 11 | 0.86716 | 0.19293 | ATP4A | 0.19293 | 0.50344 | 0.52191 |
| ENSG00000123815 | 19 | 41197434 | 41224112 | 72 | 13 | 0.86704 | 0.19296 | ADCK4 | 0.19296 | 0.14928 | 0.20908 |
| ENSG00000204386 | 6 | 31825436 | 31830683 | 8 | 4 | 0.86657 | 0.19309 | NEU1 | 0.19309 | 0.03966 | 0.016993 |
| ENSG00000227868 | 1 | 23337327 | 23342343 | 10 | 3 | 0.86588 | 0.19328 | C1orf234 | 0.19328 | 0.2443 | 0.21219 |
| ENSG00000064787 | 20 | 52553316 | 52687304 | 758 | 50 | 0.86564 | 0.19334 | BCAS1 | 0.19334 | 0.19888 | 0.224 |
| ENSG00000173327 | 11 | 65365226 | 65382853 | 33 | 8 | 0.86555 | 0.19337 | MAP3K11 | 0.19337 | 0.1803 | 0.15589 |
| ENSG00000128602 | 7 | 128828713 | 128853386 | 94 | 15 | 0.86538 | 0.19341 | SMO | 0.19341 | 0.16266 | 0.13017 |
| ENSG00000112769 | 6 | 112429963 | 112576141 | 391 | 42 | 0.86511 | 0.19349 | LAMA4 | 0.19349 | 0.29966 | 0.24262 |
| ENSG00000147606 | 8 | 92221722 | 92410378 | 477 | 41 | 0.86494 | 0.19354 | SLC26A7 | 0.19354 | 0.2088 | 0.236 |
| ENSG00000180532 | 19 | 58180303 | 58190520 | 42 | 6 | 0.86464 | 0.19362 | ZSCAN4 | 0.19362 | 0.20642 | 0.30637 |

| | | | | | | | | | | | |
|---|---|---|---|---|---|---|---|---|---|---|---|
| ENSG00000116213 | 1 | 3547331 | 3569325 | 68 | 17 | 0.86451 | 0.19365 | WRAP73 | 0.19365 | 0.25252 | 0.2914 |
| ENSG00000269468 | 1 | 17655507 | 17676070 | 137 | 15 | 0.86434 | 0.1937 | AC004824.2 | 0.1937 | 0.24861 | 0.46309 |
| ENSG00000104219 | 8 | 17013538 | 17082308 | 241 | 39 | 0.8637 | 0.19388 | ZDHHC2 | 0.19388 | 0.19178 | 0.1953 |
| ENSG00000100889 | 14 | 24563262 | 24579807 | 32 | 11 | 0.86364 | 0.19389 | PCK2 | 0.19389 | 0.39364 | 0.64746 |
| ENSG00000188991 | 12 | 16341419 | 16430619 | 416 | 17 | 0.86358 | 0.19391 | SLC15A5 | 0.19391 | 0.18978 | 0.2243 |
| ENSG00000144452 | 2 | 215796266 | 216003151 | 556 | 57 | 0.86356 | 0.19392 | ABCA12 | 0.19392 | 0.23697 | 0.26178 |
| ENSG00000173208 | 12 | 39943835 | 40013553 | 167 | 23 | 0.86343 | 0.19395 | ABCD2 | 0.19395 | 0.23792 | 0.2662 |
| ENSG00000177551 | 1 | 116378998 | 116386538 | 26 | 7 | 0.86337 | 0.19397 | NHLH2 | 0.19397 | 0.1214 | 0.14586 |
| ENSG00000185982 | 20 | 168494 | 170322 | 7 | 3 | 0.86318 | 0.19402 | DEFB128 | 0.19402 | 0.1611 | 0.17581 |
| ENSG00000140416 | 15 | 63334831 | 63364114 | 121 | 17 | 0.86312 | 0.19403 | TPM1 | 0.19403 | 0.26935 | 0.21948 |
| ENSG00000182919 | 11 | 93474757 | 93497915 | 73 | 6 | 0.86292 | 0.19409 | C11orf54 | 0.19409 | 0.17761 | 0.18697 |
| ENSG00000184716 | 15 | 44086360 | 44092419 | 6 | 3 | 0.86265 | 0.19417 | SERINC4 | 0.19417 | 0.17937 | 0.21235 |
| ENSG00000141316 | 17 | 31297394 | 31324895 | 134 | 29 | 0.86264 | 0.19417 | SPACA3 | 0.19417 | 0.10464 | 0.11142 |
| ENSG00000142959 | 1 | 45249257 | 45253377 | 2 | 1 | 0.86212 | 0.19431 | BEST4 | 0.19431 | 0.33146 | 0.32809 |
| ENSG00000268107 | 19 | 57999086 | 58016749 | 112 | 7 | 0.8618 | 0.1944 | AC003005.4 | 0.1944 | 0.33711 | 0.42215 |
| ENSG00000170382 | 1 | 204586298 | 204654861 | 202 | 30 | 0.86146 | 0.19449 | LRRN2 | 0.19449 | 0.2555 | 0.32103 |
| ENSG00000145850 | 5 | 156346293 | 156390266 | 135 | 16 | 0.86116 | 0.19458 | TIMD4 | 0.19458 | 0.2223 | 0.48378 |
| ENSG00000119401 | 9 | 119449581 | 119463579 | 31 | 7 | 0.86112 | 0.19459 | TRIM32 | 0.19459 | 0.11482 | 0.15347 |
| ENSG00000117507 | 1 | 171106879 | 171130707 | 111 | 11 | 0.861 | 0.19462 | FMO6P | 0.19462 | 0.17743 | 0.11586 |
| ENSG00000169032 | 15 | 66679155 | 66784650 | 353 | 27 | 0.86038 | 0.19479 | MAP2K1 | 0.19479 | 0.20772 | 0.1832 |
| ENSG00000177947 | 11 | 196738 | 200261 | 14 | 6 | 0.86035 | 0.1948 | ODF3 | 0.1948 | 0.26284 | 0.24424 |
| ENSG00000179526 | 8 | 145153536 | 145163027 | 21 | 4 | 0.85989 | 0.19492 | SHARPIN | 0.19492 | 0.16679 | 0.16242 |
| ENSG00000176979 | 4 | 165953105 | 165962896 | 17 | 6 | 0.85964 | 0.19499 | TRIM60 | 0.19499 | 0.74014 | 0.84435 |
| ENSG00000103257 | 16 | 87863629 | 87903094 | 209 | 30 | 0.8596 | 0.195 | SLC7A5 | 0.195 | 0.59151 | 0.57296 |
| ENSG00000180340 | 17 | 42634925 | 42636907 | 1 | 1 | 0.85962 | 0.195 | FZD2 | 0.195 | 0.25954 | 0.45697 |
| ENSG00000078043 | 18 | 44388353 | 44500123 | 326 | 18 | 0.85949 | 0.19504 | PIAS2 | 0.19504 | 0.19091 | 0.18213 |
| ENSG00000165731 | 10 | 43572475 | 43625799 | 245 | 27 | 0.85943 | 0.19505 | RET | 0.19505 | 0.23674 | 0.30916 |
| ENSG00000258289 | 14 | 65381079 | 65411309 | 136 | 13 | 0.85868 | 0.19526 | CHURC1 | 0.19526 | 0.20418 | 0.21721 |
| ENSG00000172459 | 11 | 56431138 | 56432163 | 1 | 1 | 0.85853 | 0.1953 | OR5AR1 | 0.1953 | 0.014845 | 0.025629 |
| ENSG00000101004 | 20 | 25433341 | 25566153 | 402 | 23 | 0.85827 | 0.19537 | NINL | 0.19537 | 0.19202 | 0.20216 |
| ENSG00000160199 | 21 | 44394620 | 44453691 | 368 | 20 | 0.8581 | 0.19542 | PKNOX1 | 0.19542 | 0.25152 | 0.33194 |
| ENSG00000082512 | 1 | 211499957 | 211548288 | 108 | 20 | 0.85803 | 0.19544 | TRAF5 | 0.19544 | 0.29703 | 0.2339 |
| ENSG00000064726 | 15 | 83685174 | 83736106 | 169 | 23 | 0.85793 | 0.19546 | BTBD1 | 0.19546 | 0.18236 | 0.25539 |
| ENSG00000186204 | 19 | 15783567 | 15807984 | 217 | 12 | 0.85796 | 0.19546 | CYP4F12 | 0.19546 | 0.1909 | 0.27983 |
| ENSG00000204592 | 6 | 30457244 | 30461982 | 9 | 4 | 0.85784 | 0.19549 | HLA-E | 0.19549 | 0.1333 | 0.15247 |
| ENSG00000267260 | 19 | 37264055 | 37267980 | 11 | 4 | 0.85783 | 0.19549 | CTD-2162K18.4 | 0.19549 | 0.20984 | 0.10619 |
| ENSG00000090971 | 19 | 55996371 | 55998935 | 9 | 5 | 0.85753 | 0.19558 | NAT14 | 0.19558 | 0.18898 | 0.32716 |
| ENSG00000100246 | 22 | 39174513 | 39190203 | 36 | 11 | 0.85738 | 0.19562 | DNAL4 | 0.19562 | 0.41923 | 0.34847 |
| ENSG00000130037 | 12 | 5153085 | 5155949 | 3 | 2 | 0.85732 | 0.19564 | KCNA5 | 0.19564 | 0.84928 | 0.64076 |
| ENSG00000131738 | 17 | 39519746 | 39526052 | 11 | 4 | 0.85727 | 0.19565 | KRT33B | 0.19565 | 0.17276 | 0.18431 |
| ENSG00000196388 | 17 | 4891425 | 4900905 | 28 | 7 | 0.8572 | 0.19567 | INCA1 | 0.19567 | 0.28947 | 0.20559 |
| ENSG00000108671 | 17 | 30771279 | 30810336 | 38 | 10 | 0.85686 | 0.19576 | PSMD11 | 0.19576 | 0.19671 | 0.14641 |
| ENSG00000106692 | 9 | 108320411 | 108403399 | 209 | 29 | 0.85635 | 0.1959 | FKTN | 0.1959 | 0.19801 | 0.10843 |
| ENSG00000138755 | 4 | 76922428 | 76928641 | 9 | 4 | 0.85632 | 0.19591 | CXCL9 | 0.19591 | 0.18006 | 0.35889 |
| ENSG00000105447 | 19 | 48949030 | 48960279 | 42 | 6 | 0.85624 | 0.19593 | GRWD1 | 0.19593 | 0.13623 | 0.11846 |
| ENSG00000198513 | 14 | 50999227 | 51099786 | 227 | 23 | 0.85603 | 0.19599 | ATL1 | 0.19599 | 0.21096 | 0.22994 |
| ENSG00000075618 | 7 | 5632439 | 5646286 | 80 | 16 | 0.85573 | 0.19607 | FSCN1 | 0.19607 | 0.28849 | 0.31871 |
| ENSG00000176542 | 3 | 113367232 | 113415493 | 68 | 7 | 0.85532 | 0.19619 | KIAA2018 | 0.19619 | 0.22257 | 0.23931 |
| ENSG00000131023 | 6 | 149979289 | 150039392 | 104 | 6 | 0.85482 | 0.19632 | LATS1 | 0.19632 | 0.21474 | 0.2077 |
| ENSG00000142920 | 1 | 33546705 | 33586131 | 72 | 15 | 0.85469 | 0.19636 | ADC | 0.19636 | 0.33396 | 0.21313 |
| ENSG00000160606 | 17 | 27051366 | 27054953 | 8 | 4 | 0.85469 | 0.19636 | TLCD1 | 0.19636 | 0.14687 | 0.14359 |
| ENSG00000203784 | 1 | 153175919 | 153177596 | 2 | 1 | 0.85427 | 0.19648 | LELP1 | 0.19648 | 0.83741 | 0.64978 |
| ENSG00000205758 | 21 | 34961647 | 35016232 | 89 | 22 | 0.85425 | 0.19648 | CRYZL1 | 0.19648 | 0.23848 | 0.26331 |
| ENSG00000229833 | 19 | 7694623 | 7696842 | 5 | 1 | 0.8541 | 0.19653 | PET100 | 0.19653 | 0.32541 | 0.39796 |
| ENSG00000155719 | 16 | 21689835 | 21772050 | 164 | 26 | 0.85351 | 0.19669 | OTOA | 0.19669 | 0.14567 | 0.18133 |
| ENSG00000122779 | 7 | 138145079 | 138274738 | 264 | 23 | 0.853 | 0.19683 | TRIM24 | 0.19683 | 0.13413 | 0.21733 |
| ENSG00000178449 | 12 | 50505762 | 50514240 | 31 | 5 | 0.85249 | 0.19697 | COX14 | 0.19697 | 0.2119 | 0.11855 |
| ENSG00000170890 | 12 | 120759914 | 120765592 | 13 | 4 | 0.85231 | 0.19702 | PLA2G1B | 0.19702 | 0.13293 | 0.1463 |
| ENSG00000135999 | 2 | 149402009 | 149545130 | 299 | 13 | 0.85227 | 0.19703 | EPC2 | 0.19703 | 0.18794 | 0.18856 |
| ENSG00000102189 | 12 | 93164413 | 93323107 | 422 | 20 | 0.85211 | 0.19708 | EEA1 | 0.19708 | 0.20605 | 0.18557 |
| ENSG00000213648 | 16 | 29467127 | 29476300 | 2 | 1 | 0.85202 | 0.1971 | SULT1A4 | 0.1971 | 0.28008 | 0.095976 |
| ENSG00000122515 | 7 | 44788180 | 44809477 | 41 | 10 | 0.85191 | 0.19713 | ZMIZ2 | 0.19713 | 0.12528 | 0.26884 |
| ENSG00000206044 | 12 | 2958397 | 2966213 | 22 | 4 | 0.85178 | 0.19717 | AC005841.1 | 0.19717 | 0.1866 | 0.18016 |
| ENSG00000079337 | 12 | 48128455 | 48164823 | 139 | 25 | 0.85177 | 0.19717 | RAPGEF3 | 0.19717 | 0.26109 | 0.27276 |
| ENSG00000163221 | 1 | 153346184 | 153348125 | 2 | 1 | 0.85165 | 0.19721 | S100A12 | 0.19721 | 0.092227 | 0.22975 |
| ENSG00000146038 | 6 | 24171984 | 24358280 | 869 | 49 | 0.85163 | 0.19721 | DCDC2 | 0.19721 | 0.20713 | 0.17907 |
| ENSG00000108270 | 17 | 35306175 | 35414171 | 185 | 15 | 0.85125 | 0.19732 | AATF | 0.19732 | 0.22972 | 0.20621 |
| ENSG00000198242 | 17 | 27046411 | 27051377 | 7 | 3 | 0.8509 | 0.19741 | RPL23A | 0.19741 | 0.13208 | 0.14405 |
| ENSG00000196611 | 11 | 102660651 | 102668891 | 34 | 11 | 0.85034 | 0.19757 | MMP1 | 0.19757 | 0.081597 | 0.26643 |
| ENSG00000181450 | 1 | 227751244 | 227865144 | 261 | 15 | 0.85004 | 0.19765 | ZNF678 | 0.19765 | 0.18203 | 0.15763 |
| ENSG00000151414 | 1 | 198126093 | 198291550 | 628 | 25 | 0.84978 | 0.19772 | NEK7 | 0.19772 | 0.22042 | 0.22615 |
| ENSG00000079616 | 16 | 29802040 | 29816706 | 6 | 4 | 0.84973 | 0.19774 | KIF22 | 0.19774 | 0.063455 | 0.13702 |
| ENSG00000101146 | 20 | 55926066 | 55954267 | 117 | 9 | 0.84943 | 0.19782 | RAE1 | 0.19782 | 0.19741 | 0.19916 |

| | | | | | | | | | | | |
|---|---|---|---|---|---|---|---|---|---|---|---|
| ENSG00000205581 | 21 | 40714241 | 40721573 | 34 | 6 | 0.84941 | 0.19783 | HMGN1 | 0.19783 | 0.15994 | 0.13879 |
| ENSG00000138750 | 4 | 77035812 | 77069668 | 278 | 7 | 0.84899 | 0.19794 | NUP54 | 0.19794 | 0.20085 | 0.19805 |
| ENSG00000128011 | 19 | 39797208 | 39805976 | 12 | 4 | 0.849 | 0.19794 | LRFN1 | 0.19794 | 0.10503 | 0.15679 |
| ENSG00000140932 | 16 | 66613351 | 66622178 | 29 | 5 | 0.84894 | 0.19796 | CMTM2 | 0.19796 | 0.21407 | 0.21165 |
| ENSG00000108813 | 17 | 48046334 | 48052321 | 13 | 4 | 0.84878 | 0.198 | DLX4 | 0.198 | 0.29533 | 0.62837 |
| ENSG00000143891 | 2 | 38893052 | 38968379 | 268 | 26 | 0.84867 | 0.19803 | GALM | 0.19803 | 0.27517 | 0.34842 |
| ENSG00000136110 | 13 | 53277399 | 53313947 | 114 | 12 | 0.84864 | 0.19804 | LECT1 | 0.19804 | 0.18552 | 0.071051 |
| ENSG00000198208 | 14 | 75370657 | 75390099 | 57 | 10 | 0.84824 | 0.19815 | RPS6KL1 | 0.19815 | 0.35362 | 0.35625 |
| ENSG00000188868 | 19 | 12428291 | 12444534 | 38 | 4 | 0.84809 | 0.19819 | ZNF563 | 0.19819 | 0.15459 | 0.2318 |
| ENSG00000177098 | 11 | 118004092 | 118023603 | 51 | 8 | 0.84805 | 0.1982 | SCN4B | 0.1982 | 0.37703 | 0.48689 |
| ENSG00000221978 | 1 | 1321091 | 1334708 | 25 | 6 | 0.84777 | 0.19828 | CCNL2 | 0.19828 | 0.35389 | 0.25885 |
| ENSG00000167194 | 16 | 30034655 | 30039057 | 8 | 4 | 0.84769 | 0.1983 | C16orf92 | 0.1983 | 0.78581 | 0.77476 |
| ENSG00000143546 | 1 | 153362508 | 153363664 | 2 | 1 | 0.84758 | 0.19834 | S100A8 | 0.19834 | 0.13191 | 0.16755 |
| ENSG00000204869 | 19 | 46543006 | 46580376 | 131 | 19 | 0.84737 | 0.19839 | IGFL4 | 0.19839 | 0.36549 | 0.22419 |
| ENSG00000084453 | 12 | 21417534 | 21572528 | 701 | 43 | 0.84731 | 0.19841 | SLCO1A2 | 0.19841 | 0.20029 | 0.31175 |
| ENSG00000135336 | 6 | 88299839 | 88377169 | 244 | 21 | 0.84728 | 0.19842 | ORC3 | 0.19842 | 0.14889 | 0.11687 |
| ENSG00000198398 | 3 | 190146444 | 190167665 | 98 | 20 | 0.84673 | 0.19857 | TMEM207 | 0.19857 | 0.21182 | 0.30769 |
| ENSG00000106688 | 9 | 4490444 | 4587469 | 409 | 48 | 0.84651 | 0.19863 | SLC1A1 | 0.19863 | 0.25984 | 0.28964 |
| ENSG00000138472 | 3 | 108626639 | 108672742 | 205 | 27 | 0.84636 | 0.19868 | GUCA1C | 0.19868 | 0.22806 | 0.44444 |
| ENSG00000165556 | 13 | 28536274 | 28545276 | 18 | 4 | 0.84616 | 0.19873 | CDX2 | 0.19873 | 0.17458 | 0.28006 |
| ENSG00000165643 | 9 | 138585253 | 138591374 | 33 | 6 | 0.84592 | 0.1988 | SOHLH1 | 0.1988 | 0.39555 | 0.55659 |
| ENSG00000018280 | 2 | 219246752 | 219261617 | 26 | 6 | 0.84574 | 0.19885 | SLC11A1 | 0.19885 | 0.20835 | 0.62325 |
| ENSG00000118680 | 18 | 3261907 | 3278282 | 101 | 7 | 0.84575 | 0.19885 | MYL12B | 0.19885 | 0.14302 | 0.090485 |
| ENSG00000131323 | 14 | 103243813 | 103377837 | 271 | 18 | 0.84565 | 0.19887 | TRAF3 | 0.19887 | 0.24793 | 0.26154 |
| ENSG00000113504 | 5 | 1050499 | 1112150 | 258 | 32 | 0.8454 | 0.19894 | SLC12A7 | 0.19894 | 0.26056 | 0.33981 |
| ENSG00000164674 | 6 | 159071046 | 159185908 | 459 | 50 | 0.84538 | 0.19895 | SYTL3 | 0.19895 | 0.12308 | 0.084844 |
| ENSG00000234560 | 11 | 123900297 | 123901341 | 2 | 1 | 0.84522 | 0.19899 | OR10G8 | 0.19899 | 0.44402 | 0.42364 |
| ENSG00000101444 | 20 | 32868074 | 32899608 | 39 | 4 | 0.84493 | 0.19907 | AHCY | 0.19907 | 0.18437 | 0.17082 |
| ENSG00000112996 | 5 | 44809027 | 44820530 | 21 | 3 | 0.84491 | 0.19908 | MRPS30 | 0.19908 | 0.21172 | 0.23444 |
| ENSG00000186897 | 12 | 49726200 | 49730971 | 16 | 5 | 0.8447 | 0.19914 | C1QL4 | 0.19914 | 0.18239 | 0.20122 |
| ENSG00000204624 | 1 | 11539223 | 11597641 | 128 | 18 | 0.84459 | 0.19917 | PTCHD2 | 0.19917 | 0.1575 | 0.24407 |
| ENSG00000269545 | 19 | 58740831 | 58798173 | 226 | 22 | 0.84451 | 0.19919 | CTD-3138B18.4 | 0.19919 | 0.30493 | 0.33187 |
| ENSG00000163424 | 3 | 118864997 | 118878889 | 53 | 12 | 0.84429 | 0.19925 | C3orf30 | 0.19925 | 0.27513 | 0.27566 |
| ENSG00000120563 | 10 | 29577990 | 29607257 | 223 | 31 | 0.84375 | 0.19941 | LYZL1 | 0.19941 | 0.081441 | 0.047192 |
| ENSG00000175728 | 11 | 130542851 | 130587247 | 94 | 22 | 0.84366 | 0.19943 | C11orf44 | 0.19943 | 0.07631 | 0.016321 |
| ENSG00000153922 | 5 | 98190908 | 98262240 | 186 | 19 | 0.8435 | 0.19947 | CHD1 | 0.19947 | 0.20635 | 0.24005 |
| ENSG00000130283 | 19 | 18979361 | 19006905 | 73 | 18 | 0.84352 | 0.19947 | GDF1 | 0.19947 | 0.17845 | 0.27068 |
| ENSG00000223802 | 19 | 18979361 | 19007536 | 73 | 18 | 0.84352 | 0.19947 | CERS1 | 0.19947 | 0.18427 | 0.27314 |
| ENSG00000117724 | 1 | 214776538 | 214837931 | 235 | 21 | 0.8426 | 0.19973 | CENPF | 0.19973 | 0.26393 | 0.29801 |
| ENSG00000267631 | 19 | 49538831 | 49552006 | 46 | 9 | 0.84186 | 0.19993 | CGB1 | 0.19993 | 0.16656 | 0.39362 |
| ENSG00000176049 | 5 | 146967990 | 147162338 | 523 | 25 | 0.84183 | 0.19994 | JAKMIP2 | 0.19994 | 0.21349 | 0.22752 |
| ENSG00000172466 | 18 | 32912176 | 32925446 | 33 | 7 | 0.84166 | 0.19999 | ZNF24 | 0.19999 | 0.12468 | 0.10732 |
| ENSG00000168374 | 3 | 57557090 | 57583947 | 97 | 14 | 0.84163 | 0.2 | ARF4 | 0.2 | 0.18786 | 0.18514 |
| ENSG00000143630 | 1 | 155247374 | 155259639 | 26 | 3 | 0.84152 | 0.20003 | HCN3 | 0.20003 | 0.21125 | 0.18152 |
| ENSG00000177225 | 11 | 767220 | 777488 | 32 | 6 | 0.84134 | 0.20008 | PDDC1 | 0.20008 | 0.22693 | 0.35105 |
| ENSG00000183310 | 1 | 248737022 | 248738080 | 6 | 2 | 0.84131 | 0.20009 | OR2T34 | 0.20009 | 0.37063 | 0.36748 |
| ENSG00000116819 | 1 | 36038971 | 36060929 | 21 | 10 | 0.84127 | 0.2001 | TFAP2E | 0.2001 | 0.29593 | 0.27563 |
| ENSG00000103051 | 16 | 70514471 | 70557468 | 75 | 12 | 0.84124 | 0.20011 | COG4 | 0.20011 | 0.21348 | 0.22026 |
| ENSG00000203910 | 1 | 92683497 | 92711370 | 56 | 11 | 0.84117 | 0.20013 | C1orf146 | 0.20013 | 0.15911 | 0.1522 |
| ENSG00000154143 | 11 | 124481386 | 124490252 | 16 | 6 | 0.84114 | 0.20014 | PANX3 | 0.20014 | 0.048328 | 0.062461 |
| ENSG00000205126 | 11 | 44069531 | 44081527 | 25 | 9 | 0.84107 | 0.20016 | ACCSL | 0.20016 | 0.4782 | 0.55671 |
| ENSG00000172890 | 11 | 71164155 | 71235153 | 289 | 13 | 0.84104 | 0.20016 | NADSYN1 | 0.20016 | 0.25477 | 0.34765 |
| ENSG00000096088 | 6 | 41704449 | 41721847 | 66 | 21 | 0.84092 | 0.2002 | PGC | 0.2002 | 0.41282 | 0.59545 |
| ENSG00000179598 | 17 | 17104309 | 17109629 | 19 | 6 | 0.84069 | 0.20026 | PLD6 | 0.20026 | 0.30876 | 0.44144 |
| ENSG00000148377 | 10 | 1064847 | 1071799 | 35 | 8 | 0.8405 | 0.20031 | IDI2 | 0.20031 | 0.22567 | 0.19413 |
| ENSG00000270299 | 20 | 627270 | 656444 | 84 | 26 | 0.8404 | 0.20034 | RP5-850E9.3 | 0.20034 | 0.48992 | 0.61915 |
| ENSG00000139624 | 12 | 50523575 | 50561288 | 89 | 9 | 0.84029 | 0.20037 | CERS5 | 0.20037 | 0.19743 | 0.19155 |
| ENSG00000198854 | 1 | 152691998 | 152692905 | 7 | 2 | 0.84025 | 0.20038 | C1orf68 | 0.20038 | 0.24725 | 0.1806 |
| ENSG00000178074 | 2 | 200775979 | 200820658 | 130 | 19 | 0.83987 | 0.20049 | C2orf69 | 0.20049 | 0.15249 | 0.084996 |
| ENSG00000180535 | 7 | 97840739 | 97842291 | 7 | 3 | 0.83964 | 0.20056 | BHLHA15 | 0.20056 | 0.53771 | 0.47636 |
| ENSG00000182504 | 3 | 101442769 | 101489406 | 108 | 12 | 0.83926 | 0.20066 | CEP97 | 0.20066 | 0.2168 | 0.22583 |
| ENSG00000163803 | 2 | 28680012 | 28866654 | 780 | 84 | 0.83896 | 0.20075 | PLB1 | 0.20075 | 0.22933 | 0.27835 |
| ENSG00000187170 | 1 | 152680636 | 152681967 | 5 | 3 | 0.83887 | 0.20077 | LCE4A | 0.20077 | 0.17926 | 0.15015 |
| ENSG00000106399 | 7 | 7676149 | 7758238 | 356 | 22 | 0.83879 | 0.20079 | RPA3 | 0.20079 | 0.19159 | 0.17305 |
| ENSG00000183111 | 5 | 148931510 | 149014531 | 292 | 20 | 0.83833 | 0.20092 | ARHGEF37 | 0.20092 | 0.18798 | 0.17924 |
| ENSG00000204365 | 10 | 29135337 | 29170827 | 206 | 20 | 0.8379 | 0.20104 | C10orf126 | 0.20104 | 0.13698 | 0.038384 |
| ENSG00000198837 | 1 | 153901977 | 153919172 | 23 | 5 | 0.83765 | 0.20111 | DENND4B | 0.20111 | 0.24738 | 0.16595 |
| ENSG00000125868 | 20 | 17550508 | 17590564 | 118 | 15 | 0.83752 | 0.20115 | DSTN | 0.20115 | 0.22949 | 0.31723 |
| ENSG00000120756 | 3 | 142315229 | 142432506 | 322 | 21 | 0.83727 | 0.20122 | PLS1 | 0.20122 | 0.16393 | 0.10076 |
| ENSG00000110848 | 12 | 9905082 | 9913497 | 38 | 9 | 0.83717 | 0.20125 | CD69 | 0.20125 | 0.13706 | 0.19849 |
| ENSG00000177186 | 1 | 248486894 | 248487899 | 8 | 2 | 0.83686 | 0.20133 | OR2M7 | 0.20133 | 0.20325 | 0.28126 |
| ENSG00000185507 | 11 | 612553 | 615999 | 21 | 3 | 0.83685 | 0.20134 | IRF7 | 0.20134 | 0.01682 | 0.0065172 |
| ENSG00000165478 | 11 | 124789089 | 124806308 | 49 | 16 | 0.8367 | 0.20138 | HEPACAM | 0.20138 | 0.18706 | 0.16992 |

| | | | | | | | | | | | |
|---|---|---|---|---|---|---|---|---|---|---|---|
| ENSG00000146521 | 6 | 168185217 | 168197539 | 70 | 14 | 0.83648 | 0.20144 | C6orf123 | 0.20144 | 0.053486 | 0.064283 |
| ENSG00000125999 | 20 | 31861286 | 31897684 | 102 | 18 | 0.8357 | 0.20166 | BPIFB1 | 0.20166 | 0.14997 | 0.16859 |
| ENSG00000255800 | 9 | 107360650 | 107361788 | 14 | 3 | 0.83549 | 0.20172 | OR13C5 | 0.20172 | 0.25071 | 0.29016 |
| ENSG00000204843 | 2 | 74588281 | 74619214 | 24 | 9 | 0.835 | 0.20186 | DCTN1 | 0.20186 | 0.35639 | 0.45625 |
| ENSG00000169047 | 2 | 227596033 | 227664475 | 194 | 18 | 0.83477 | 0.20192 | IRS1 | 0.20192 | 0.22506 | 0.18706 |
| ENSG00000215440 | 20 | 57264187 | 57294294 | 142 | 20 | 0.83453 | 0.20199 | NPEPL1 | 0.20199 | 0.092303 | 0.071844 |
| ENSG00000127399 | 7 | 150019728 | 150035239 | 43 | 3 | 0.83443 | 0.20202 | LRRC61 | 0.20202 | 0.23066 | 0.38172 |
| ENSG00000144935 | 3 | 142442916 | 142526730 | 182 | 15 | 0.83425 | 0.20207 | TRPC1 | 0.20207 | 0.19755 | 0.14155 |
| ENSG00000174206 | 12 | 64580096 | 64616076 | 68 | 10 | 0.83404 | 0.20213 | C12orf66 | 0.20213 | 0.25465 | 0.28894 |
| ENSG00000122435 | 1 | 100598706 | 100616053 | 42 | 9 | 0.83385 | 0.20218 | TRMT13 | 0.20218 | 0.22046 | 0.25522 |
| ENSG00000254772 | 11 | 62327073 | 62342401 | 28 | 5 | 0.83356 | 0.20227 | EEF1G | 0.20227 | 0.23539 | 0.13938 |
| ENSG00000042753 | 19 | 47341393 | 47354249 | 17 | 9 | 0.8334 | 0.20231 | AP2S1 | 0.20231 | 0.34447 | 0.58649 |
| ENSG00000188175 | 7 | 92817899 | 92855837 | 71 | 18 | 0.83283 | 0.20247 | HEPACAM2 | 0.20247 | 0.25038 | 0.16317 |
| ENSG00000081138 | 18 | 63417488 | 63548638 | 392 | 18 | 0.83271 | 0.2025 | CDH7 | 0.2025 | 0.21209 | 0.24745 |
| ENSG00000129911 | 19 | 1852399 | 1863567 | 36 | 7 | 0.83222 | 0.20264 | KLF16 | 0.20264 | 0.193 | 0.31297 |
| ENSG00000186648 | 14 | 24521206 | 24538937 | 61 | 20 | 0.83215 | 0.20266 | LRRC16B | 0.20266 | 0.32534 | 0.49456 |
| ENSG00000170953 | 11 | 124412578 | 124413575 | 1 | 1 | 0.83202 | 0.2027 | OR8B12 | 0.2027 | 0.27577 | 0.29529 |
| ENSG00000109458 | 4 | 144257915 | 144395721 | 281 | 32 | 0.83181 | 0.20276 | GAB1 | 0.20276 | 0.20576 | 0.24136 |
| ENSG00000161180 | 22 | 21987005 | 21991616 | 12 | 5 | 0.83168 | 0.2028 | CCDC116 | 0.2028 | 0.58899 | 0.76855 |
| ENSG00000266118 | 2 | 24347236 | 24413305 | 262 | 22 | 0.83127 | 0.20291 | RP11-507M3.1 | 0.20291 | 0.15368 | 0.15291 |
| ENSG00000221821 | 6 | 42858005 | 42858554 | 2 | 1 | 0.83127 | 0.20291 | C6orf226 | 0.20291 | 0.08364 | 0.14166 |
| ENSG00000197238 | 6 | 27791884 | 27792257 | 2 | 1 | 0.83098 | 0.20299 | HIST1H4J | 0.20299 | 0.71745 | 0.68593 |
| ENSG00000244122 | 2 | 234590584 | 234681945 | 354 | 21 | 0.83088 | 0.20302 | UGT1A7 | 0.20302 | 0.11366 | 0.092012 |
| ENSG00000096264 | 6 | 41303393 | 41318625 | 73 | 11 | 0.83077 | 0.20305 | NCR2 | 0.20305 | 0.61553 | 0.68659 |
| ENSG00000101350 | 20 | 30865467 | 30922814 | 115 | 11 | 0.8305 | 0.20313 | KIF3B | 0.20313 | 0.19087 | 0.19133 |
| ENSG00000120054 | 10 | 101801950 | 101841634 | 147 | 23 | 0.82983 | 0.20332 | CPN1 | 0.20332 | 0.17579 | 0.17375 |
| ENSG00000182566 | 19 | 7793843 | 7798792 | 21 | 5 | 0.82962 | 0.20338 | CLEC4G | 0.20338 | 0.32284 | 0.35032 |
| ENSG00000124357 | 2 | 71291474 | 71306935 | 64 | 10 | 0.82919 | 0.2035 | NAGK | 0.2035 | 0.51266 | 0.36725 |
| ENSG00000136383 | 15 | 85359911 | 85416713 | 156 | 30 | 0.82916 | 0.20351 | ALPK3 | 0.20351 | 0.23137 | 0.25147 |
| ENSG00000145681 | 5 | 82933624 | 83017432 | 262 | 38 | 0.82898 | 0.20356 | HAPLN1 | 0.20356 | 0.17439 | 0.16063 |
| ENSG00000141040 | 17 | 16454701 | 16472520 | 23 | 4 | 0.82891 | 0.20358 | ZNF287 | 0.20358 | 0.24235 | 0.16726 |
| ENSG00000119915 | 10 | 103986085 | 103989346 | 4 | 1 | 0.82836 | 0.20373 | ELOVL3 | 0.20373 | 0.19375 | 0.20644 |
| ENSG00000167004 | 15 | 44038590 | 44065477 | 51 | 9 | 0.82835 | 0.20373 | PDIA3 | 0.20373 | 0.21729 | 0.21541 |
| ENSG00000126583 | 19 | 54382444 | 54410906 | 91 | 23 | 0.82808 | 0.20381 | PRKCG | 0.20381 | 0.14044 | 0.084565 |
| ENSG00000183578 | 15 | 51348795 | 51397473 | 132 | 16 | 0.82805 | 0.20382 | TNFAIP8L3 | 0.20382 | 0.24633 | 0.28888 |
| ENSG00000255302 | 15 | 49170083 | 49172380 | 1 | 1 | 0.82777 | 0.2039 | EID1 | 0.2039 | 0.16704 | 0.15883 |
| ENSG00000039068 | 16 | 68771128 | 68869451 | 268 | 33 | 0.82724 | 0.20405 | CDH1 | 0.20405 | 0.25639 | 0.19072 |
| ENSG00000175197 | 12 | 57910371 | 57914300 | 4 | 1 | 0.82706 | 0.2041 | DDIT3 | 0.2041 | 0.27658 | 0.15277 |
| ENSG00000204574 | 6 | 30539153 | 30564956 | 84 | 11 | 0.82687 | 0.20415 | ABCF1 | 0.20415 | 0.22314 | 0.20792 |
| ENSG00000184014 | 11 | 9160372 | 9286937 | 346 | 22 | 0.82682 | 0.20417 | DENND5A | 0.20417 | 0.20998 | 0.16364 |
| ENSG00000160844 | 7 | 99798283 | 99869855 | 160 | 18 | 0.82667 | 0.20421 | GATS | 0.20421 | 0.17253 | 0.15487 |
| ENSG00000185038 | 2 | 234684370 | 234742069 | 259 | 44 | 0.82646 | 0.20427 | MROH2A | 0.20427 | 0.26438 | 0.1611 |
| ENSG00000179630 | 13 | 44453420 | 44468068 | 62 | 11 | 0.82646 | 0.20427 | LACC1 | 0.20427 | 0.17106 | 0.1967 |
| ENSG00000183313 | 11 | 6007122 | 6008215 | 11 | 4 | 0.82613 | 0.20436 | OR52L1 | 0.20436 | 0.32421 | 0.43572 |
| ENSG00000011347 | 11 | 61282785 | 61348620 | 48 | 8 | 0.826 | 0.2044 | SYT7 | 0.2044 | 0.36483 | 0.40195 |
| ENSG00000258429 | 16 | 69362524 | 69364498 | 5 | 2 | 0.82591 | 0.20443 | PDF | 0.20443 | 0.23157 | 0.29534 |
| ENSG00000116815 | 1 | 117057157 | 117113661 | 65 | 14 | 0.82559 | 0.20452 | CD58 | 0.20452 | 0.20757 | 0.36454 |
| ENSG00000128739 | 15 | 25068794 | 25223870 | 752 | 111 | 0.82558 | 0.20452 | SNRPN | 0.20452 | 0.17186 | 0.21208 |
| ENSG00000157404 | 4 | 55524085 | 55606881 | 234 | 32 | 0.82549 | 0.20455 | KIT | 0.20455 | 0.16343 | 0.1992 |
| ENSG00000161583 | 17 | 34745936 | 34808104 | 28 | 11 | 0.82506 | 0.20467 | TBC1D3G | 0.20467 | 0.34156 | 0.41027 |
| ENSG00000141644 | 18 | 47793252 | 47808144 | 30 | 8 | 0.82498 | 0.20469 | MBD1 | 0.20469 | 0.16621 | 0.11566 |
| ENSG00000187185 | 16 | 57832098 | 57850850 | 52 | 14 | 0.82492 | 0.20471 | CTD-2600O9.1 | 0.20471 | 0.12649 | 0.14387 |
| ENSG00000061794 | 12 | 27863706 | 27909228 | 123 | 16 | 0.82489 | 0.20472 | MRPS35 | 0.20472 | 0.2233 | 0.24285 |
| ENSG00000256229 | 19 | 20278037 | 20311299 | 136 | 9 | 0.82485 | 0.20473 | ZNF486 | 0.20473 | 0.24545 | 0.34298 |
| ENSG00000130176 | 19 | 11649532 | 11661138 | 43 | 8 | 0.8247 | 0.20477 | CNN1 | 0.20477 | 0.17234 | 0.17952 |
| ENSG00000139146 | 12 | 31433518 | 31479992 | 60 | 16 | 0.82462 | 0.20479 | FAM60A | 0.20479 | 0.25173 | 0.2985 |
| ENSG00000101266 | 20 | 459116 | 524465 | 137 | 22 | 0.8246 | 0.2048 | CSNK2A1 | 0.2048 | 0.27471 | 0.19259 |
| ENSG00000054654 | 14 | 64319683 | 64693165 | 1105 | 53 | 0.82423 | 0.2049 | SYNE2 | 0.2049 | 0.19375 | 0.18517 |
| ENSG00000077514 | 11 | 74204896 | 74380162 | 692 | 35 | 0.8239 | 0.205 | POLD3 | 0.205 | 0.26489 | 0.29434 |
| ENSG00000165568 | 10 | 4828821 | 4890254 | 488 | 44 | 0.82385 | 0.20501 | AKR1E2 | 0.20501 | 0.21848 | 0.25592 |
| ENSG00000257019 | 9 | 107366924 | 107367951 | 6 | 1 | 0.82369 | 0.20506 | OR13C2 | 0.20506 | 0.25868 | 0.3196 |
| ENSG00000110514 | 11 | 47290712 | 47351582 | 178 | 18 | 0.82363 | 0.20507 | MADD | 0.20507 | 0.22913 | 0.20766 |
| ENSG00000162624 | 1 | 75594119 | 75627218 | 77 | 13 | 0.82345 | 0.20512 | LHX8 | 0.20512 | 0.363 | 0.54185 |
| ENSG00000100347 | 22 | 44351301 | 44406411 | 232 | 30 | 0.82315 | 0.20521 | SAMM50 | 0.20521 | 0.22714 | 0.072991 |
| ENSG00000154309 | 1 | 222988406 | 223179337 | 604 | 34 | 0.82262 | 0.20536 | DISP1 | 0.20536 | 0.15176 | 0.13142 |
| ENSG00000118762 | 4 | 88928820 | 88998929 | 181 | 29 | 0.8222 | 0.20548 | PKD2 | 0.20548 | 0.14641 | 0.15581 |
| ENSG00000146263 | 6 | 97590037 | 97731093 | 339 | 17 | 0.82202 | 0.20553 | MMS22L | 0.20553 | 0.20816 | 0.20954 |
| ENSG00000075420 | 3 | 171757418 | 172119455 | 1127 | 103 | 0.822 | 0.20554 | FNDC3B | 0.20554 | 0.26515 | 0.45312 |
| ENSG00000216921 | 2 | 242836136 | 242844702 | 56 | 10 | 0.82191 | 0.20556 | AC131097.4 | 0.20556 | 0.20393 | 0.21447 |
| ENSG00000180806 | 12 | 54388679 | 54397121 | 20 | 6 | 0.82168 | 0.20563 | HOXC9 | 0.20563 | 0.22028 | 0.19534 |
| ENSG00000073169 | 22 | 50639408 | 50656045 | 70 | 13 | 0.82106 | 0.20581 | SELO | 0.20581 | 0.24431 | 0.23132 |
| ENSG00000243646 | 21 | 34638663 | 34669539 | 119 | 13 | 0.82085 | 0.20587 | IL10RB | 0.20587 | 0.18548 | 0.26004 |
| ENSG00000271793 | 6 | 86267696 | 86324952 | 98 | 18 | 0.82081 | 0.20588 | RP11-321N4.5 | 0.20588 | 0.19397 | 0.16746 |

| | | | | | | | | | | | |
|---|---|---|---|---|---|---|---|---|---|---|---|
| ENSG00000269699 | 19 | 57285920 | 57352097 | 96 | 20 | 0.82049 | 0.20597 | ZIM2 | 0.20597 | 0.15055 | 0.093503 |
| ENSG00000173578 | 3 | 46058516 | 46069234 | 29 | 3 | 0.82016 | 0.20606 | XCR1 | 0.20606 | 0.21504 | 0.29913 |
| ENSG00000122484 | 1 | 92764522 | 92867613 | 191 | 22 | 0.82007 | 0.20609 | RPAP2 | 0.20609 | 0.21629 | 0.21263 |
| ENSG00000188782 | 1 | 26517052 | 26529459 | 75 | 12 | 0.81953 | 0.20624 | CATSPER4 | 0.20624 | 0.13426 | 0.16181 |
| ENSG00000148704 | 10 | 118888032 | 118897812 | 15 | 5 | 0.81943 | 0.20627 | VAX1 | 0.20627 | 0.47565 | 0.38531 |
| ENSG00000254531 | 4 | 102268937 | 102270040 | 2 | 1 | 0.81941 | 0.20628 | AP001816.1 | 0.20628 | 0.40911 | 0.48036 |
| ENSG00000083444 | 1 | 11994262 | 12035595 | 144 | 16 | 0.81896 | 0.20641 | PLOD1 | 0.20641 | 0.24675 | 0.36714 |
| ENSG00000122674 | 7 | 5938356 | 5965605 | 169 | 34 | 0.81892 | 0.20641 | CCZ1 | 0.20641 | 0.27405 | 0.32558 |
| ENSG00000184702 | 22 | 19701987 | 19712295 | 7 | 4 | 0.81851 | 0.20653 | SEPTIN5 | 0.20653 | 0.29657 | 0.57232 |
| ENSG00000204475 | 6 | 31556672 | 31560762 | 26 | 12 | 0.81831 | 0.20659 | NCR3 | 0.20659 | 0.33989 | 0.07329 |
| ENSG00000100865 | 14 | 102808956 | 102829253 | 33 | 11 | 0.81825 | 0.20661 | CINP | 0.20661 | 0.28726 | 0.37753 |
| ENSG00000100321 | 22 | 39745930 | 39781593 | 129 | 17 | 0.81825 | 0.20661 | SYNGR1 | 0.20661 | 0.27178 | 0.33316 |
| ENSG00000176605 | 14 | 99177950 | 99184098 | 18 | 7 | 0.81808 | 0.20666 | C14orf177 | 0.20666 | 0.46777 | 0.43303 |
| ENSG00000188770 | 1 | 203463271 | 203477992 | 56 | 10 | 0.81773 | 0.20676 | OPTC | 0.20676 | 0.1766 | 0.13869 |
| ENSG00000258945 | 14 | 85994943 | 85995730 | 3 | 1 | 0.81755 | 0.20681 | RP11-497E19.2 | 0.20681 | 0.17483 | 0.28826 |
| ENSG00000111879 | 6 | 119280928 | 119470552 | 690 | 49 | 0.8175 | 0.20682 | FAM184A | 0.20682 | 0.15779 | 0.1107 |
| ENSG00000134245 | 1 | 113009163 | 113072787 | 199 | 36 | 0.81674 | 0.20704 | WNT2B | 0.20704 | 0.23565 | 0.33727 |
| ENSG00000134317 | 2 | 10085341 | 10142411 | 354 | 25 | 0.8167 | 0.20705 | GRHL1 | 0.20705 | 0.22807 | 0.24042 |
| ENSG00000173436 | 1 | 19923477 | 19956314 | 92 | 16 | 0.81624 | 0.20718 | MINOS1 | 0.20718 | 0.24025 | 0.35317 |
| ENSG00000169684 | 15 | 78857862 | 78887611 | 88 | 11 | 0.81595 | 0.20726 | CHRNA5 | 0.20726 | 0.17889 | 0.17579 |
| ENSG00000163545 | 1 | 205271187 | 205290883 | 60 | 15 | 0.8159 | 0.20728 | NUAK2 | 0.20728 | 0.089481 | 0.14336 |
| ENSG00000122550 | 7 | 23145353 | 23217533 | 135 | 11 | 0.81584 | 0.2073 | KLHL7 | 0.2073 | 0.17609 | 0.07629 |
| ENSG00000141977 | 19 | 16272179 | 16284336 | 46 | 5 | 0.81582 | 0.2073 | CIB3 | 0.2073 | 0.039803 | 0.024157 |
| ENSG00000183048 | 17 | 79670404 | 79688042 | 90 | 12 | 0.81579 | 0.20731 | SLC25A10 | 0.20731 | 0.25243 | 0.31987 |
| ENSG00000130669 | 19 | 39616410 | 39673456 | 193 | 22 | 0.81567 | 0.20734 | PAK4 | 0.20734 | 0.086673 | 0.16383 |
| ENSG00000197063 | 17 | 79876146 | 79885590 | 4 | 2 | 0.8156 | 0.20736 | MAFG | 0.20736 | 0.11192 | 0.055824 |
| ENSG00000244005 | 20 | 34255977 | 34287281 | 68 | 10 | 0.81559 | 0.20737 | NFS1 | 0.20737 | 0.24847 | 0.34543 |
| ENSG00000133195 | 17 | 70642088 | 71088851 | 1920 | 170 | 0.81555 | 0.20738 | SLC39A11 | 0.20738 | 0.41676 | 0.2417 |
| ENSG00000016082 | 5 | 50678921 | 50690564 | 25 | 8 | 0.81545 | 0.20741 | ISL1 | 0.20741 | 0.49672 | 0.51532 |
| ENSG00000112149 | 6 | 14117872 | 14137149 | 59 | 14 | 0.81534 | 0.20744 | CD83 | 0.20744 | 0.23991 | 0.3638 |
| ENSG00000072954 | 19 | 16771938 | 16800840 | 105 | 17 | 0.81532 | 0.20745 | TMEM38A | 0.20745 | 0.13964 | 0.15439 |
| ENSG00000135773 | 1 | 230883130 | 230937749 | 280 | 29 | 0.81524 | 0.20747 | CAPN9 | 0.20747 | 0.2238 | 0.13895 |
| ENSG00000173567 | 2 | 26531041 | 26569685 | 122 | 23 | 0.81514 | 0.2075 | GPR113 | 0.2075 | 0.15449 | 0.17051 |
| ENSG00000070371 | 22 | 19166986 | 19279239 | 379 | 18 | 0.81509 | 0.20751 | CLTCL1 | 0.20751 | 0.19519 | 0.080281 |
| ENSG00000173214 | 6 | 111580551 | 111592370 | 35 | 10 | 0.81496 | 0.20755 | KIAA1919 | 0.20755 | 0.19807 | 0.25204 |
| ENSG00000136100 | 13 | 52986737 | 53024763 | 105 | 11 | 0.8148 | 0.20759 | VPS36 | 0.20759 | 0.22205 | 0.25157 |
| ENSG00000111877 | 6 | 119134605 | 119256327 | 436 | 23 | 0.81474 | 0.20761 | MCM9 | 0.20761 | 0.18806 | 0.16885 |
| ENSG00000054118 | 1 | 36690017 | 36770958 | 142 | 15 | 0.81469 | 0.20763 | THRAP3 | 0.20763 | 0.23537 | 0.31864 |
| ENSG00000076864 | 1 | 21922708 | 21995801 | 288 | 43 | 0.81423 | 0.20776 | RAP1GAP | 0.20776 | 0.25022 | 0.36101 |
| ENSG00000169375 | 15 | 75661720 | 75748183 | 107 | 11 | 0.81423 | 0.20776 | SIN3A | 0.20776 | 0.19761 | 0.18171 |
| ENSG00000182749 | 1 | 26187701 | 26197744 | 11 | 4 | 0.81401 | 0.20782 | PAQR7 | 0.20782 | 0.61079 | 0.66933 |
| ENSG00000165669 | 10 | 120065401 | 120101840 | 69 | 17 | 0.81402 | 0.20782 | FAM204A | 0.20782 | 0.12235 | 0.099893 |
| ENSG00000169688 | 16 | 56685811 | 56687116 | 9 | 2 | 0.81399 | 0.20782 | MT1B | 0.20782 | 0.31311 | 0.41686 |
| ENSG00000151838 | 14 | 59971257 | 60043549 | 269 | 14 | 0.81397 | 0.20783 | CCDC175 | 0.20783 | 0.22877 | 0.25123 |
| ENSG00000130758 | 19 | 40697651 | 40721482 | 42 | 10 | 0.81389 | 0.20785 | MAP3K10 | 0.20785 | 0.1252 | 0.14786 |
| ENSG00000176428 | 7 | 73082155 | 73086442 | 6 | 2 | 0.81371 | 0.2079 | VPS37D | 0.2079 | 0.23279 | 0.22265 |
| ENSG00000135722 | 16 | 67193834 | 67198473 | 4 | 2 | 0.81365 | 0.20792 | FBXL8 | 0.20792 | 0.89817 | 0.915 |
| ENSG00000113384 | 5 | 32124810 | 32174456 | 134 | 15 | 0.81359 | 0.20794 | GOLPH3 | 0.20794 | 0.26453 | 0.388 |
| ENSG00000149218 | 11 | 94822974 | 94865809 | 184 | 32 | 0.81358 | 0.20794 | ENDOD1 | 0.20794 | 0.35816 | 0.39562 |
| ENSG00000164284 | 5 | 148724993 | 148734146 | 19 | 7 | 0.81309 | 0.20808 | GRPEL2 | 0.20808 | 0.12291 | 0.16729 |
| ENSG00000253548 | 3 | 191178952 | 191179245 | 1 | 1 | 0.81303 | 0.2081 | PYDC2 | 0.2081 | 0.11792 | 0.091503 |
| ENSG00000170866 | 19 | 54799854 | 54809952 | 62 | 14 | 0.81299 | 0.20811 | LILRA3 | 0.20811 | 0.65315 | 0.66456 |
| ENSG00000267534 | 19 | 10334520 | 10341948 | 19 | 5 | 0.81248 | 0.20826 | S1PR2 | 0.20826 | 0.5139 | 0.63288 |
| ENSG00000261884 | 16 | 67963517 | 67969920 | 10 | 6 | 0.81241 | 0.20828 | CTC-479C5.12 | 0.20828 | 0.5453 | 0.69044 |
| ENSG00000197446 | 19 | 41620337 | 41634271 | 78 | 11 | 0.81239 | 0.20828 | CYP2F1 | 0.20828 | 0.19752 | 0.27231 |
| ENSG00000095970 | 6 | 41126244 | 41130924 | 4 | 2 | 0.81227 | 0.20832 | TREM2 | 0.20832 | 0.34819 | 0.43288 |
| ENSG00000254827 | 11 | 2909010 | 2924970 | 51 | 17 | 0.81209 | 0.20837 | SLC22A18AS | 0.20837 | 0.58891 | 0.65087 |
| ENSG00000196475 | 4 | 80327508 | 80329372 | 1 | 1 | 0.81199 | 0.2084 | GK2 | 0.2084 | 0.28061 | 0.36313 |
| ENSG00000138604 | 15 | 69452923 | 69564556 | 223 | 17 | 0.81184 | 0.20844 | GLCE | 0.20844 | 0.23971 | 0.22799 |
| ENSG00000156284 | 21 | 31586324 | 31588391 | 4 | 2 | 0.81177 | 0.20846 | CLDN8 | 0.20846 | 0.48008 | 0.33682 |
| ENSG00000060237 | 12 | 861759 | 1020618 | 407 | 31 | 0.81101 | 0.20868 | WNK1 | 0.20868 | 0.2479 | 0.27569 |
| ENSG00000100867 | 14 | 24099324 | 24114848 | 8 | 4 | 0.81017 | 0.20892 | DHRS2 | 0.20892 | 0.71645 | 0.88402 |
| ENSG00000148798 | 10 | 105036920 | 105050108 | 35 | 7 | 0.81008 | 0.20895 | INA | 0.20895 | 0.21366 | 0.096128 |
| ENSG00000114923 | 2 | 220492049 | 220506702 | 20 | 7 | 0.80963 | 0.20908 | SLC4A3 | 0.20908 | 0.26993 | 0.31529 |
| ENSG00000235568 | 22 | 42776416 | 42828401 | 213 | 27 | 0.80957 | 0.20909 | NFAM1 | 0.20909 | 0.23426 | 0.24195 |
| ENSG00000162413 | 1 | 6650784 | 6674667 | 47 | 10 | 0.80849 | 0.2094 | KLHL21 | 0.2094 | 0.14621 | 0.14084 |
| ENSG00000070367 | 14 | 57670518 | 57735726 | 115 | 12 | 0.8084 | 0.20943 | EXOC5 | 0.20943 | 0.14479 | 0.12356 |
| ENSG00000167447 | 17 | 57286761 | 57292608 | 6 | 3 | 0.80822 | 0.20948 | SMG8 | 0.20948 | 0.37464 | 0.39929 |
| ENSG00000039600 | 5 | 157052687 | 157098488 | 130 | 18 | 0.80821 | 0.20949 | SOX30 | 0.20949 | 0.15981 | 0.35776 |
| ENSG00000169410 | 15 | 75759462 | 75871630 | 181 | 16 | 0.8082 | 0.20949 | PTPN9 | 0.20949 | 0.21306 | 0.20189 |
| ENSG00000204183 | 20 | 34020827 | 34023248 | 6 | 2 | 0.80807 | 0.20953 | GDF5OS | 0.20953 | 0.1436 | 0.15548 |
| ENSG00000243056 | 5 | 139927251 | 139929163 | 2 | 1 | 0.80766 | 0.20964 | EIF4EBP3 | 0.20964 | 0.36701 | 0.42265 |
| ENSG00000176893 | 11 | 4935900 | 4936922 | 2 | 1 | 0.80714 | 0.20979 | OR51G2 | 0.20979 | 0.26383 | 0.46505 |

| | | | | | | | | | | | |
|---|---|---|---|---|---|---|---|---|---|---|---|
| ENSG00000103245 | 16 | 779753 | 791329 | 33 | 6 | 0.80711 | 0.2098 | NARFL | 0.2098 | 0.17488 | 0.46789 |
| ENSG00000170854 | 3 | 97660662 | 97691301 | 42 | 10 | 0.80675 | 0.20991 | MINA | 0.20991 | 0.334 | 0.44227 |
| ENSG00000124143 | 20 | 37230577 | 37279678 | 151 | 29 | 0.80674 | 0.20991 | ARHGAP40 | 0.20991 | 0.31504 | 0.24991 |
| ENSG00000113248 | 5 | 140625147 | 140627799 | 14 | 6 | 0.80665 | 0.20994 | PCDHB15 | 0.20994 | 0.05021 | 0.045605 |
| ENSG00000166509 | 16 | 78056412 | 78100658 | 396 | 17 | 0.8064 | 0.21001 | CLEC3A | 0.21001 | 0.25128 | 0.27961 |
| ENSG00000155974 | 12 | 66741211 | 67197966 | 1718 | 117 | 0.80636 | 0.21002 | GRIP1 | 0.21002 | 0.22081 | 0.27563 |
| ENSG00000248540 | 15 | 74418256 | 74421619 | 5 | 2 | 0.80635 | 0.21002 | RP11-247C2.2 | 0.21002 | 0.34925 | 0.047947 |
| ENSG00000005189 | 16 | 20817751 | 20860987 | 71 | 14 | 0.80635 | 0.21002 | AC004381.6 | 0.21002 | 0.21347 | 0.18533 |
| ENSG00000050165 | 11 | 11984653 | 12031316 | 204 | 53 | 0.80629 | 0.21004 | DKK3 | 0.21004 | 0.3677 | 0.37073 |
| ENSG00000185928 | 16 | 29827285 | 29841948 | 25 | 4 | 0.80608 | 0.2101 | PAGR1 | 0.2101 | 0.19711 | 0.24322 |
| ENSG00000121314 | 12 | 10958650 | 10959892 | 4 | 2 | 0.80578 | 0.21018 | TAS2R8 | 0.21018 | 0.67702 | 0.58785 |
| ENSG00000080200 | 3 | 97595819 | 97663810 | 111 | 10 | 0.80563 | 0.21023 | CRYBG3 | 0.21023 | 0.15265 | 0.17518 |
| ENSG00000110900 | 12 | 31079362 | 31149534 | 316 | 42 | 0.80535 | 0.21031 | TSPAN11 | 0.21031 | 0.26631 | 0.24086 |
| ENSG00000142937 | 1 | 45240923 | 45244451 | 6 | 3 | 0.8053 | 0.21032 | RPS8 | 0.21032 | 0.47484 | 0.44141 |
| ENSG00000183748 | 10 | 17851362 | 17953178 | 86 | 23 | 0.8053 | 0.21032 | MRC1L1 | 0.21032 | 0.17458 | 0.18617 |
| ENSG00000172772 | 11 | 58034264 | 58035732 | 4 | 1 | 0.8052 | 0.21035 | OR10W1 | 0.21035 | 0.21594 | 0.20112 |
| ENSG00000132361 | 17 | 2592680 | 2615957 | 55 | 10 | 0.80519 | 0.21036 | CLUH | 0.21036 | 0.29492 | 0.30851 |
| ENSG00000100095 | 22 | 26565440 | 26779562 | 867 | 84 | 0.80509 | 0.21038 | SEZ6L | 0.21038 | 0.27885 | 0.35289 |
| ENSG00000096070 | 6 | 36164521 | 36200567 | 42 | 10 | 0.80503 | 0.2104 | BRPF3 | 0.2104 | 0.32317 | 0.33431 |
| ENSG00000071575 | 2 | 12857015 | 12882860 | 76 | 13 | 0.80498 | 0.21041 | TRIB2 | 0.21041 | 0.12857 | 0.12944 |
| ENSG00000100078 | 22 | 31530795 | 31536593 | 20 | 11 | 0.80416 | 0.21065 | PLA2G3 | 0.21065 | 0.55241 | 0.35798 |
| ENSG00000185551 | 15 | 96869167 | 96883492 | 21 | 6 | 0.804 | 0.2107 | NR2F2 | 0.2107 | 0.4938 | 0.32483 |
| ENSG00000145388 | 4 | 119606523 | 119636588 | 80 | 11 | 0.80384 | 0.21074 | METTL14 | 0.21074 | 0.22385 | 0.21589 |
| ENSG00000184148 | 1 | 152943142 | 152945050 | 4 | 2 | 0.80367 | 0.21079 | SPRR4 | 0.21079 | 0.25976 | 0.56787 |
| ENSG00000163029 | 2 | 17845079 | 17981509 | 358 | 24 | 0.80343 | 0.21086 | SMC6 | 0.21086 | 0.22685 | 0.23415 |
| ENSG00000081019 | 1 | 114304454 | 114355098 | 79 | 13 | 0.80282 | 0.21104 | RSBN1 | 0.21104 | 0.22368 | 0.23813 |
| ENSG00000140718 | 16 | 53737875 | 54155853 | 1554 | 119 | 0.80264 | 0.21109 | FTO | 0.21109 | 0.28821 | 0.37763 |
| ENSG00000198090 | 17 | 39295685 | 39296739 | 7 | 4 | 0.80242 | 0.21115 | KRTAP4-6 | 0.21115 | 0.31264 | 0.23061 |
| ENSG00000125772 | 20 | 5525085 | 5591672 | 228 | 26 | 0.80232 | 0.21118 | GPCPD1 | 0.21118 | 0.23655 | 0.28168 |
| ENSG00000007312 | 17 | 62006100 | 62009714 | 15 | 3 | 0.80179 | 0.21134 | CD79B | 0.21134 | 0.31598 | 0.18797 |
| ENSG00000166341 | 11 | 6642556 | 6677085 | 92 | 20 | 0.80153 | 0.21141 | DCHS1 | 0.21141 | 0.32213 | 0.2363 |
| ENSG00000136689 | 2 | 113864791 | 113891593 | 134 | 12 | 0.80104 | 0.21155 | IL1RN | 0.21155 | 0.24571 | 0.40348 |
| ENSG00000117597 | 1 | 210001352 | 210030910 | 97 | 13 | 0.80102 | 0.21156 | DIEXF | 0.21156 | 0.27768 | 0.31117 |
| ENSG00000204539 | 6 | 31082867 | 31088223 | 83 | 9 | 0.80101 | 0.21156 | CDSN | 0.21156 | 0.21858 | 0.13977 |
| ENSG00000169442 | 1 | 26644448 | 26647014 | 12 | 3 | 0.80088 | 0.2116 | CD52 | 0.2116 | 0.10638 | 0.15825 |
| ENSG00000164715 | 7 | 97736197 | 97838945 | 309 | 21 | 0.80078 | 0.21163 | LMTK2 | 0.21163 | 0.27766 | 0.37524 |
| ENSG00000197927 | 2 | 132479948 | 132524973 | 143 | 12 | 0.80022 | 0.21179 | C2orf27A | 0.21179 | 0.16519 | 0.27213 |
| ENSG00000177156 | 11 | 747329 | 765024 | 33 | 8 | 0.80024 | 0.21179 | TALDO1 | 0.21179 | 0.33715 | 0.37676 |
| ENSG00000244734 | 11 | 5246694 | 5250625 | 18 | 9 | 0.80014 | 0.21181 | HBB | 0.21181 | 0.68244 | 0.93882 |
| ENSG00000184022 | 1 | 248756052 | 248757100 | 5 | 1 | 0.79993 | 0.21188 | OR2T10 | 0.21188 | 0.13738 | 0.075553 |
| ENSG00000011198 | 3 | 43731605 | 43775863 | 123 | 11 | 0.79945 | 0.21201 | ABHD5 | 0.21201 | 0.168 | 0.19629 |
| ENSG00000009950 | 7 | 73007524 | 73038873 | 85 | 11 | 0.79945 | 0.21201 | MLXIPL | 0.21201 | 0.22687 | 0.20977 |
| ENSG00000148584 | 10 | 52559169 | 52645435 | 141 | 25 | 0.79927 | 0.21207 | A1CF | 0.21207 | 0.31991 | 0.3812 |
| ENSG00000084234 | 11 | 129939732 | 130014699 | 160 | 15 | 0.79846 | 0.2123 | APLP2 | 0.2123 | 0.14107 | 0.091164 |
| ENSG00000269755 | 19 | 12543917 | 12595630 | 305 | 11 | 0.79824 | 0.21237 | CTD-3105H18.18 | 0.21237 | 0.17576 | 0.18372 |
| ENSG00000121680 | 11 | 45931220 | 45940363 | 23 | 5 | 0.79778 | 0.2125 | PEX16 | 0.2125 | 0.21603 | 0.45892 |
| ENSG00000039123 | 5 | 54603588 | 54721409 | 305 | 10 | 0.79773 | 0.21251 | SKIV2L2 | 0.21251 | 0.20339 | 0.20424 |
| ENSG00000243566 | 7 | 76139745 | 76648340 | 1788 | 75 | 0.79771 | 0.21252 | UPK3B | 0.21252 | 0.22821 | 0.23504 |
| ENSG00000154370 | 1 | 228581374 | 228594541 | 16 | 5 | 0.79653 | 0.21286 | TRIM11 | 0.21286 | 0.30804 | 0.14608 |
| ENSG00000168246 | 5 | 171636644 | 171711075 | 294 | 12 | 0.79655 | 0.21286 | UBTD2 | 0.21286 | 0.16582 | 0.17631 |
| ENSG00000197961 | 19 | 9671029 | 9695209 | 74 | 6 | 0.79647 | 0.21288 | ZNF121 | 0.21288 | 0.25259 | 0.29999 |
| ENSG00000173821 | 17 | 78234665 | 78372586 | 497 | 58 | 0.79644 | 0.21289 | RNF213 | 0.21289 | 0.25505 | 0.31105 |
| ENSG00000118263 | 2 | 207938861 | 208031991 | 284 | 38 | 0.7964 | 0.2129 | KLF7 | 0.2129 | 0.26177 | 0.23456 |
| ENSG00000174485 | 15 | 65950384 | 66084631 | 266 | 19 | 0.79624 | 0.21295 | DENND4A | 0.21295 | 0.17538 | 0.13779 |
| ENSG00000152582 | 5 | 35617946 | 35814713 | 683 | 37 | 0.79605 | 0.213 | SPEF2 | 0.213 | 0.21045 | 0.19619 |
| ENSG00000204876 | 7 | 155755326 | 155759037 | 9 | 4 | 0.79563 | 0.21312 | AC021218.2 | 0.21312 | 0.22728 | 0.44964 |
| ENSG00000164694 | 6 | 159590429 | 159693141 | 436 | 20 | 0.79524 | 0.21324 | FNDC1 | 0.21324 | 0.22683 | 0.24466 |
| ENSG00000105223 | 19 | 40854363 | 40886346 | 87 | 21 | 0.79511 | 0.21328 | PLD3 | 0.21328 | 0.23204 | 0.30465 |
| ENSG00000108479 | 17 | 73747675 | 73761792 | 35 | 13 | 0.79492 | 0.21333 | GALK1 | 0.21333 | 0.3035 | 0.61244 |
| ENSG00000162772 | 1 | 212738676 | 212794119 | 252 | 33 | 0.79487 | 0.21334 | ATF3 | 0.21334 | 0.23301 | 0.17776 |
| ENSG00000173020 | 11 | 67033881 | 67054027 | 16 | 6 | 0.79468 | 0.2134 | ADRBK1 | 0.2134 | 0.59587 | 0.77875 |
| ENSG00000153446 | 16 | 5094123 | 5116111 | 63 | 16 | 0.79464 | 0.21341 | C16orf89 | 0.21341 | 0.34183 | 0.5539 |
| ENSG00000055147 | 5 | 153369688 | 153418496 | 127 | 7 | 0.79442 | 0.21347 | FAM114A2 | 0.21347 | 0.20942 | 0.21976 |
| ENSG00000198129 | 8 | 7353368 | 7366833 | 2 | 1 | 0.79432 | 0.2135 | DEFB107B | 0.2135 | 0.2135 | 0.2135 |
| ENSG00000135679 | 12 | 69201956 | 69239214 | 109 | 15 | 0.79409 | 0.21357 | MDM2 | 0.21357 | 0.26669 | 0.35753 |
| ENSG00000173114 | 7 | 110731062 | 110765510 | 64 | 9 | 0.79396 | 0.21361 | LRRN3 | 0.21361 | 0.22424 | 0.29582 |
| ENSG00000075142 | 7 | 87834433 | 87856308 | 43 | 9 | 0.79394 | 0.21362 | SRI | 0.21362 | 0.27912 | 0.41049 |
| ENSG00000063046 | 12 | 53399942 | 53435993 | 139 | 9 | 0.79368 | 0.21369 | EIF4B | 0.21369 | 0.2123 | 0.33474 |
| ENSG00000155438 | 2 | 122484521 | 122494499 | 30 | 4 | 0.7934 | 0.21377 | NIFK | 0.21377 | 0.31216 | 0.20083 |
| ENSG00000224186 | 5 | 134368970 | 134691744 | 881 | 108 | 0.79325 | 0.21382 | C5orf66 | 0.21382 | 0.12935 | 0.029888 |
| ENSG00000002822 | 7 | 1855429 | 2272878 | 2025 | 34 | 0.79325 | 0.21382 | MAD1L1 | 0.21382 | 0.21381 | 0.21664 |
| ENSG00000103005 | 16 | 58033450 | 58055522 | 41 | 7 | 0.79316 | 0.21384 | USB1 | 0.21384 | 0.14802 | 0.061931 |
| ENSG00000132517 | 17 | 4935895 | 4955304 | 74 | 11 | 0.79316 | 0.21384 | SLC52A1 | 0.21384 | 0.38344 | 0.27417 |

| | | | | | | | | | | | |
|---|---|---|---|---|---|---|---|---|---|---|---|
| ENSG00000215066 | 9 | 6467618 | 6470375 | 14 | 4 | 0.79301 | 0.21389 | C9orf38 | 0.21389 | 0.37862 | 0.40813 |
| ENSG00000104205 | 8 | 67624653 | 67774257 | 125 | 30 | 0.79285 | 0.21393 | SGK3 | 0.21393 | 0.32321 | 0.35989 |
| ENSG00000172260 | 1 | 71861623 | 72748417 | 2179 | 101 | 0.7928 | 0.21395 | NEGR1 | 0.21395 | 0.21999 | 0.21818 |
| ENSG00000087258 | 16 | 56225302 | 56391356 | 498 | 55 | 0.79278 | 0.21395 | GNAO1 | 0.21395 | 0.152 | 0.1355 |
| ENSG00000175787 | 9 | 97021593 | 97063736 | 212 | 21 | 0.7927 | 0.21398 | ZNF169 | 0.21398 | 0.19272 | 0.15559 |
| ENSG00000143156 | 1 | 169101769 | 169337205 | 1081 | 26 | 0.79185 | 0.21422 | NME7 | 0.21422 | 0.23019 | 0.25129 |
| ENSG00000164683 | 8 | 80676245 | 80680098 | 10 | 4 | 0.79175 | 0.21425 | HEY1 | 0.21425 | 0.2575 | 0.45962 |
| ENSG00000155254 | 10 | 99472930 | 99477907 | 6 | 2 | 0.79155 | 0.21431 | MARVELD1 | 0.21431 | 0.69468 | 0.73604 |
| ENSG00000217825 | 7 | 154988273 | 154990165 | 15 | 4 | 0.79142 | 0.21435 | AC099552.4 | 0.21435 | 0.61823 | 0.56053 |
| ENSG00000218672 | 7 | 155174771 | 155189051 | 122 | 22 | 0.79101 | 0.21447 | AC008060.7 | 0.21447 | 0.10529 | 0.38424 |
| ENSG00000267950 | 11 | 1295809 | 1296033 | 1 | 1 | 0.7909 | 0.2145 | AC136297.1 | 0.2145 | 0.14219 | 0.14687 |
| ENSG00000120008 | 10 | 122610687 | 122669036 | 276 | 11 | 0.79076 | 0.21454 | WDR11 | 0.21454 | 0.21702 | 0.22128 |
| ENSG00000259288 | 15 | 40509629 | 40568597 | 171 | 41 | 0.79022 | 0.2147 | RP11-133K1.2 | 0.2147 | 0.11999 | 0.057786 |
| ENSG00000146535 | 7 | 2767746 | 2883958 | 423 | 25 | 0.79011 | 0.21473 | GNA12 | 0.21473 | 0.24013 | 0.23033 |
| ENSG00000023171 | 11 | 123396344 | 123498482 | 407 | 55 | 0.79002 | 0.21476 | GRAMD1B | 0.21476 | 0.25729 | 0.45146 |
| ENSG00000113269 | 5 | 179338651 | 179499118 | 472 | 40 | 0.78984 | 0.21481 | RNF130 | 0.21481 | 0.22456 | 0.23799 |
| ENSG00000127324 | 12 | 71518865 | 71835678 | 1108 | 62 | 0.78985 | 0.21481 | TSPAN8 | 0.21481 | 0.21831 | 0.22767 |
| ENSG00000112320 | 6 | 107811162 | 107981357 | 339 | 47 | 0.78979 | 0.21482 | SOBP | 0.21482 | 0.2096 | 0.29491 |
| ENSG00000100368 | 22 | 37309670 | 37336491 | 135 | 14 | 0.78975 | 0.21484 | CSF2RB | 0.21484 | 0.16709 | 0.23297 |
| ENSG00000188716 | 10 | 76797594 | 76818272 | 65 | 14 | 0.78964 | 0.21487 | DUPD1 | 0.21487 | 0.28727 | 0.2892 |
| ENSG00000069812 | 1 | 6472478 | 6484730 | 54 | 10 | 0.7875 | 0.2155 | HES2 | 0.2155 | 0.24996 | 0.26324 |
| ENSG00000186442 | 12 | 53183469 | 53189901 | 19 | 6 | 0.78734 | 0.21554 | KRT3 | 0.21554 | 0.3614 | 0.53339 |
| ENSG00000154330 | 9 | 70943224 | 71145977 | 612 | 39 | 0.78703 | 0.21563 | PGM5 | 0.21563 | 0.22576 | 0.25358 |
| ENSG00000132436 | 7 | 50511831 | 50518088 | 10 | 3 | 0.78674 | 0.21572 | FIGNL1 | 0.21572 | 0.11814 | 0.27906 |
| ENSG00000162694 | 1 | 101337943 | 101361554 | 31 | 10 | 0.78639 | 0.21582 | EXTL2 | 0.21582 | 0.19864 | 0.24646 |
| ENSG00000163867 | 1 | 35449523 | 35497569 | 91 | 3 | 0.78607 | 0.21591 | ZMYM6 | 0.21591 | 0.22371 | 0.23562 |
| ENSG00000054392 | 1 | 210501596 | 210849638 | 1576 | 116 | 0.78609 | 0.21591 | HHAT | 0.21591 | 0.20964 | 0.26795 |
| ENSG00000198380 | 2 | 69546905 | 69614382 | 171 | 11 | 0.78588 | 0.21597 | GFPT1 | 0.21597 | 0.20917 | 0.24143 |
| ENSG00000099290 | 10 | 51827648 | 51893269 | 68 | 10 | 0.78574 | 0.21601 | FAM21A | 0.21601 | 0.067413 | 0.055718 |
| ENSG00000179915 | 2 | 50145643 | 51259674 | 3781 | 140 | 0.78556 | 0.21606 | NRXN1 | 0.21606 | 0.20413 | 0.13312 |
| ENSG00000214753 | 11 | 62480102 | 62494821 | 23 | 5 | 0.78516 | 0.21618 | HNRNPUL2 | 0.21618 | 0.36153 | 0.44671 |
| ENSG00000143466 | 1 | 206643791 | 206670223 | 94 | 22 | 0.7851 | 0.2162 | IKBKE | 0.2162 | 0.17963 | 0.19744 |
| ENSG00000109771 | 4 | 186285032 | 186317053 | 117 | 21 | 0.78497 | 0.21624 | LRP2BP | 0.21624 | 0.17637 | 0.12987 |
| ENSG00000132819 | 20 | 55966463 | 55984389 | 48 | 12 | 0.78475 | 0.2163 | RBM38 | 0.2163 | 0.2245 | 0.34651 |
| ENSG00000158195 | 1 | 27730730 | 27816669 | 167 | 11 | 0.78462 | 0.21634 | WASF2 | 0.21634 | 0.20568 | 0.20648 |
| ENSG00000162627 | 1 | 99127236 | 99226056 | 374 | 22 | 0.78463 | 0.21634 | SNX7 | 0.21634 | 0.23899 | 0.26196 |
| ENSG00000134940 | 11 | 125541417 | 125551018 | 28 | 6 | 0.78429 | 0.21644 | ACRV1 | 0.21644 | 0.1996 | 0.3196 |
| ENSG00000163736 | 4 | 74852755 | 74853914 | 2 | 1 | 0.7839 | 0.21655 | PPBP | 0.21655 | 0.21474 | 0.37451 |
| ENSG00000164761 | 8 | 119935796 | 119964439 | 133 | 19 | 0.7839 | 0.21655 | TNFRSF11B | 0.21655 | 0.14834 | 0.20333 |
| ENSG00000137976 | 1 | 84864215 | 84880701 | 68 | 10 | 0.78341 | 0.21669 | DNASE2B | 0.21669 | 0.19132 | 0.12401 |
| ENSG00000175018 | 10 | 127265091 | 127371713 | 263 | 25 | 0.78343 | 0.21669 | TEX36 | 0.21669 | 0.23802 | 0.33431 |
| ENSG00000181090 | 9 | 140513444 | 140764468 | 883 | 50 | 0.78338 | 0.2167 | EHMT1 | 0.2167 | 0.21966 | 0.21949 |
| ENSG00000160282 | 21 | 47556176 | 47575481 | 115 | 31 | 0.78324 | 0.21674 | FTCD | 0.21674 | 0.3338 | 0.17735 |
| ENSG00000164306 | 4 | 185570767 | 185616117 | 217 | 11 | 0.78249 | 0.21696 | PRIMPOL | 0.21696 | 0.19138 | 0.18608 |
| ENSG00000111775 | 12 | 120875893 | 120878545 | 8 | 3 | 0.78249 | 0.21696 | COX6A1 | 0.21696 | 0.28356 | 0.42359 |
| ENSG00000143627 | 1 | 155259630 | 155271225 | 17 | 3 | 0.78245 | 0.21698 | PKLR | 0.21698 | 0.16551 | 0.16802 |
| ENSG00000156599 | 11 | 57435219 | 57468659 | 64 | 12 | 0.78238 | 0.217 | ZDHHC5 | 0.217 | 0.30051 | 0.41799 |
| ENSG00000124299 | 19 | 33877856 | 34012700 | 650 | 41 | 0.78236 | 0.217 | PEPD | 0.217 | 0.23154 | 0.27468 |
| ENSG00000137343 | 6 | 30594619 | 30614600 | 61 | 13 | 0.78229 | 0.21702 | ATAT1 | 0.21702 | 0.20873 | 0.25936 |
| ENSG00000106603 | 7 | 43648055 | 43769316 | 422 | 16 | 0.78215 | 0.21706 | COA1 | 0.21706 | 0.2125 | 0.22014 |
| ENSG00000146757 | 7 | 64838712 | 64866038 | 107 | 10 | 0.78161 | 0.21722 | ZNF92 | 0.21722 | 0.23908 | 0.19417 |
| ENSG00000244405 | 3 | 185764097 | 185828107 | 111 | 15 | 0.78155 | 0.21724 | ETV5 | 0.21724 | 0.19107 | 0.050457 |
| ENSG00000179364 | 14 | 105766900 | 105864484 | 60 | 27 | 0.78138 | 0.21729 | PACS2 | 0.21729 | 0.36188 | 0.45442 |
| ENSG00000095951 | 6 | 12008995 | 12165232 | 496 | 49 | 0.78122 | 0.21734 | HIVEP1 | 0.21734 | 0.20835 | 0.22234 |
| ENSG00000117523 | 1 | 171454651 | 171562650 | 307 | 18 | 0.78118 | 0.21735 | PRRC2C | 0.21735 | 0.22687 | 0.31673 |
| ENSG00000006715 | 7 | 38762563 | 38971994 | 816 | 31 | 0.78117 | 0.21735 | VPS41 | 0.21735 | 0.20904 | 0.21814 |
| ENSG00000120256 | 6 | 150139934 | 150186199 | 125 | 10 | 0.78112 | 0.21737 | LRP11 | 0.21737 | 0.1893 | 0.11492 |
| ENSG00000137760 | 11 | 107373452 | 107436472 | 188 | 19 | 0.78109 | 0.21737 | ALKBH8 | 0.21737 | 0.2168 | 0.16359 |
| ENSG00000091527 | 3 | 133292574 | 133309105 | 46 | 8 | 0.78098 | 0.21741 | CDV3 | 0.21741 | 0.35017 | 0.52335 |
| ENSG00000130396 | 6 | 168227602 | 168372703 | 478 | 22 | 0.78094 | 0.21742 | MLLT4 | 0.21742 | 0.18995 | 0.22429 |
| ENSG00000256188 | 12 | 11285557 | 11287243 | 7 | 2 | 0.78087 | 0.21744 | TAS2R30 | 0.21744 | 0.21198 | 0.22732 |
| ENSG00000228273 | 7 | 100547257 | 100550424 | 12 | 4 | 0.78077 | 0.21747 | MUC3A | 0.21747 | 0.42635 | 0.5218 |
| ENSG00000186625 | 6 | 149916009 | 149970108 | 138 | 6 | 0.78072 | 0.21748 | KATNA1 | 0.21748 | 0.22491 | 0.23849 |
| ENSG00000153266 | 3 | 62355356 | 62359999 | 6 | 4 | 0.78014 | 0.21765 | FEZF2 | 0.21765 | 0.10225 | 0.42031 |
| ENSG00000130382 | 19 | 6212966 | 6279959 | 220 | 17 | 0.78013 | 0.21766 | MLLT1 | 0.21766 | 0.22425 | 0.41142 |
| ENSG00000160271 | 9 | 135973107 | 136039301 | 201 | 31 | 0.78006 | 0.21768 | RALGDS | 0.21768 | 0.31371 | 0.1666 |
| ENSG00000271741 | 1 | 35447134 | 35497342 | 99 | 4 | 0.77998 | 0.2177 | RP11-244H3.4 | 0.2177 | 0.22479 | 0.23678 |
| ENSG00000196152 | 9 | 130186661 | 130207651 | 65 | 5 | 0.77991 | 0.21772 | ZNF79 | 0.21772 | 0.22978 | 0.23418 |
| ENSG00000215183 | 9 | 35752987 | 35756610 | 3 | 2 | 0.77951 | 0.21784 | MSMP | 0.21784 | 0.0112 | 0.018847 |
| ENSG00000258365 | 12 | 94671534 | 94676620 | 24 | 5 | 0.77931 | 0.2179 | RP11-1105G2.3 | 0.2179 | 0.3033 | 0.38782 |
| ENSG00000009413 | 6 | 111620234 | 111804918 | 473 | 28 | 0.77916 | 0.21794 | REV3L | 0.21794 | 0.20854 | 0.22873 |
| ENSG00000152430 | 2 | 198591603 | 198651486 | 90 | 11 | 0.77884 | 0.21804 | BOLL | 0.21804 | 0.25014 | 0.21647 |
| ENSG00000144057 | 2 | 107418056 | 107503564 | 341 | 22 | 0.77863 | 0.2181 | ST6GAL2 | 0.2181 | 0.20133 | 0.20557 |

| | | | | | | | | | | | |
|---|---|---|---|---|---|---|---|---|---|---|---|
| ENSG00000162368 | 1 | 47799469 | 47844511 | 148 | 13 | 0.77822 | 0.21822 | CMPK1 | 0.21822 | 0.18968 | 0.18515 |
| ENSG00000090013 | 19 | 40953696 | 40971747 | 75 | 13 | 0.7781 | 0.21825 | BLVRB | 0.21825 | 0.20974 | 0.20909 |
| ENSG00000169836 | 4 | 104507188 | 104640973 | 539 | 31 | 0.77792 | 0.21831 | TACR3 | 0.21831 | 0.25255 | 0.29227 |
| ENSG00000183729 | 8 | 53850991 | 53853677 | 11 | 5 | 0.77791 | 0.21831 | NPBWR1 | 0.21831 | 0.34237 | 0.42659 |
| ENSG00000204188 | 6 | 33551515 | 33556803 | 22 | 8 | 0.7775 | 0.21843 | GGNBP1 | 0.21843 | 0.071398 | 0.10234 |
| ENSG00000268400 | 19 | 7694682 | 7707934 | 48 | 4 | 0.77741 | 0.21846 | CTD-3214H19.4 | 0.21846 | 0.30553 | 0.49183 |
| ENSG00000065485 | 3 | 122785909 | 122944074 | 557 | 45 | 0.77734 | 0.21848 | PDIA5 | 0.21848 | 0.18263 | 0.12805 |
| ENSG00000239590 | 9 | 125281420 | 125282361 | 1 | 1 | 0.77727 | 0.2185 | OR1J4 | 0.2185 | 0.75153 | 0.76336 |
| ENSG00000174950 | 1 | 27705666 | 27709870 | 4 | 1 | 0.77724 | 0.21851 | CD164L2 | 0.21851 | 0.2425 | 0.16286 |
| ENSG00000165819 | 14 | 21966277 | 21979517 | 45 | 8 | 0.77721 | 0.21852 | METTL3 | 0.21852 | 0.24082 | 0.29835 |
| ENSG00000141431 | 18 | 31158579 | 31331156 | 473 | 33 | 0.77711 | 0.21855 | ASXL3 | 0.21855 | 0.28017 | 0.36323 |
| ENSG00000203443 | 3 | 172034218 | 172039513 | 11 | 5 | 0.77687 | 0.21862 | AC092964.1 | 0.21862 | 0.043367 | 0.080032 |
| ENSG00000126562 | 17 | 40932696 | 40948954 | 10 | 7 | 0.77678 | 0.21864 | WNK4 | 0.21864 | 0.53176 | 0.65853 |
| ENSG00000166435 | 11 | 74518784 | 74660245 | 293 | 23 | 0.77676 | 0.21865 | XRRA1 | 0.21865 | 0.22603 | 0.23897 |
| ENSG00000188313 | 3 | 146232967 | 146262651 | 148 | 17 | 0.77643 | 0.21875 | PLSCR1 | 0.21875 | 0.23735 | 0.18226 |
| ENSG00000182263 | 2 | 164449906 | 164592522 | 313 | 39 | 0.77612 | 0.21884 | FIGN | 0.21884 | 0.15785 | 0.52341 |
| ENSG00000181016 | 7 | 112120908 | 112131117 | 28 | 9 | 0.77603 | 0.21887 | LSMEM1 | 0.21887 | 0.062452 | 0.10159 |
| ENSG00000019505 | 11 | 45261852 | 45307870 | 132 | 16 | 0.77544 | 0.21904 | SYT13 | 0.21904 | 0.26953 | 0.35007 |
| ENSG00000164309 | 5 | 78985700 | 79096063 | 386 | 51 | 0.77521 | 0.21911 | CMYA5 | 0.21911 | 0.20626 | 0.23011 |
| ENSG00000188243 | 13 | 76099350 | 76123575 | 95 | 15 | 0.77521 | 0.21911 | COMMD6 | 0.21911 | 0.22666 | 0.16008 |
| ENSG00000041357 | 15 | 78832747 | 78841604 | 36 | 5 | 0.77444 | 0.21934 | PSMA4 | 0.21934 | 0.24151 | 0.2157 |
| ENSG00000152455 | 10 | 14920819 | 14946314 | 57 | 15 | 0.77434 | 0.21937 | SUV39H2 | 0.21937 | 0.38684 | 0.3322 |
| ENSG00000127540 | 19 | 1597171 | 1605480 | 14 | 6 | 0.77428 | 0.21938 | UQCR11 | 0.21938 | 0.42987 | 0.50656 |
| ENSG00000144820 | 3 | 100328433 | 100414323 | 184 | 14 | 0.77413 | 0.21943 | GPR128 | 0.21943 | 0.21433 | 0.21112 |
| ENSG00000141750 | 17 | 37366789 | 37382125 | 38 | 10 | 0.77374 | 0.21954 | STAC2 | 0.21954 | 0.25984 | 0.3988 |
| ENSG00000198642 | 9 | 21329670 | 21335379 | 13 | 3 | 0.77372 | 0.21955 | KLHL9 | 0.21955 | 0.29634 | 0.31111 |
| ENSG00000135473 | 12 | 56710007 | 56727837 | 28 | 5 | 0.77358 | 0.21959 | PAN2 | 0.21959 | 0.20399 | 0.18263 |
| ENSG00000188686 | 8 | 145490549 | 145492470 | 1 | 1 | 0.77354 | 0.2196 | SCXA | 0.2196 | 0.7477 | 0.7433 |
| ENSG00000229387 | 11 | 22646860 | 22647369 | 2 | 1 | 0.77355 | 0.2196 | AC103801.2 | 0.2196 | 0.26915 | 0.30717 |
| ENSG00000185053 | 8 | 13947373 | 15095848 | 7054 | 191 | 0.7735 | 0.21961 | SGCZ | 0.21961 | 0.22006 | 0.21648 |
| ENSG00000262560 | 15 | 44064815 | 44091331 | 47 | 7 | 0.77305 | 0.21975 | RP11-296A16.1 | 0.21975 | 0.20153 | 0.2029 |
| ENSG00000162437 | 1 | 65210778 | 65298915 | 199 | 24 | 0.77241 | 0.21994 | RAVER2 | 0.21994 | 0.23741 | 0.31887 |
| ENSG00000161281 | 19 | 36641824 | 36643771 | 5 | 2 | 0.77226 | 0.21998 | COX7A1 | 0.21998 | 0.14582 | 0.32995 |
| ENSG00000157103 | 3 | 11034410 | 11080933 | 175 | 44 | 0.77195 | 0.22007 | SLC6A1 | 0.22007 | 0.11859 | 0.039394 |
| ENSG00000106733 | 9 | 77675489 | 77703133 | 138 | 21 | 0.77186 | 0.2201 | NMRK1 | 0.2201 | 0.25691 | 0.22779 |
| ENSG00000169750 | 17 | 79989500 | 79992080 | 7 | 3 | 0.77179 | 0.22012 | RAC3 | 0.22012 | 0.57955 | 0.69755 |
| ENSG00000112799 | 6 | 6588341 | 6655216 | 374 | 54 | 0.77159 | 0.22018 | LY86 | 0.22018 | 0.26434 | 0.22937 |
| ENSG00000116273 | 1 | 6673745 | 6684093 | 25 | 6 | 0.77145 | 0.22022 | PHF13 | 0.22022 | 0.15096 | 0.093935 |
| ENSG00000161980 | 16 | 96407 | 103628 | 33 | 5 | 0.7713 | 0.22026 | POLR3K | 0.22026 | 0.31213 | 0.48806 |
| ENSG00000134444 | 18 | 59854491 | 59974355 | 266 | 20 | 0.77123 | 0.22028 | KIAA1468 | 0.22028 | 0.24602 | 0.27177 |
| ENSG00000128917 | 15 | 41221538 | 41231237 | 15 | 6 | 0.77084 | 0.2204 | DLL4 | 0.2204 | 0.40917 | 0.59735 |
| ENSG00000188050 | 7 | 122337766 | 122339210 | 1 | 1 | 0.77051 | 0.2205 | RNF133 | 0.2205 | 0.47782 | 0.54805 |
| ENSG00000231171 | 4 | 178649911 | 178911904 | 1014 | 57 | 0.77024 | 0.22058 | LINC01098 | 0.22058 | 0.23755 | 0.24986 |
| ENSG00000137312 | 6 | 30695486 | 30710510 | 104 | 7 | 0.76967 | 0.22075 | FLOT1 | 0.22075 | 0.27047 | 0.48473 |
| ENSG00000128713 | 2 | 176968944 | 176974722 | 16 | 5 | 0.76934 | 0.22084 | HOXD11 | 0.22084 | 0.19802 | 0.30543 |
| ENSG00000078053 | 7 | 38423305 | 38671167 | 1304 | 69 | 0.76903 | 0.22094 | AMPH | 0.22094 | 0.23045 | 0.22677 |
| ENSG00000186272 | 19 | 57922531 | 57933307 | 24 | 6 | 0.769 | 0.22095 | ZNF17 | 0.22095 | 0.23713 | 0.36881 |
| ENSG00000173714 | 17 | 48912011 | 48919709 | 25 | 4 | 0.76894 | 0.22096 | WFIKKN2 | 0.22096 | 0.20068 | 0.22279 |
| ENSG00000219435 | 11 | 64067863 | 64072242 | 7 | 3 | 0.76893 | 0.22097 | TEX40 | 0.22097 | 0.47802 | 0.21735 |
| ENSG00000104951 | 19 | 50392911 | 50432796 | 138 | 28 | 0.76893 | 0.22097 | IL4I1 | 0.22097 | 0.2805 | 0.24549 |
| ENSG00000148655 | 10 | 77360998 | 78319925 | 2400 | 158 | 0.7685 | 0.22109 | C10orf11 | 0.22109 | 0.21249 | 0.21246 |
| ENSG00000215187 | 9 | 35561828 | 35563896 | 7 | 3 | 0.76849 | 0.2211 | FAM166B | 0.2211 | 0.21862 | 0.19972 |
| ENSG00000174607 | 4 | 115519611 | 115599380 | 178 | 23 | 0.76843 | 0.22112 | UGT8 | 0.22112 | 0.37617 | 0.49793 |
| ENSG00000171903 | 19 | 16023177 | 16045677 | 136 | 13 | 0.76817 | 0.22119 | CYP4F11 | 0.22119 | 0.42378 | 0.61083 |
| ENSG00000128285 | 22 | 41074754 | 41078818 | 9 | 3 | 0.76813 | 0.2212 | MCHR1 | 0.2212 | 0.45881 | 0.51585 |
| ENSG00000136271 | 7 | 44605016 | 44614650 | 36 | 8 | 0.76748 | 0.2214 | DDX56 | 0.2214 | 0.22441 | 0.18747 |
| ENSG00000185946 | 1 | 104068313 | 104097861 | 53 | 12 | 0.76718 | 0.22149 | RNPC3 | 0.22149 | 0.24312 | 0.34431 |
| ENSG00000184933 | 11 | 6815756 | 6817128 | 4 | 2 | 0.76682 | 0.2216 | OR6A2 | 0.2216 | 0.10489 | 0.078227 |
| ENSG00000169413 | 14 | 21249210 | 21250626 | 2 | 1 | 0.76674 | 0.22162 | RNASE6 | 0.22162 | 0.72581 | 0.90975 |
| ENSG00000103316 | 16 | 21250195 | 21314404 | 99 | 26 | 0.7664 | 0.22172 | CRYM | 0.22172 | 0.29783 | 0.18923 |
| ENSG00000240542 | 17 | 39344697 | 39346891 | 7 | 4 | 0.76608 | 0.22181 | KRTAP9-1 | 0.22181 | 0.20212 | 0.23746 |
| ENSG00000048462 | 16 | 12058964 | 12061925 | 14 | 4 | 0.76602 | 0.22183 | TNFRSF17 | 0.22183 | 0.11427 | 0.059376 |
| ENSG00000156869 | 1 | 100174259 | 100232187 | 285 | 10 | 0.76585 | 0.22188 | FRRS1 | 0.22188 | 0.15059 | 0.11334 |
| ENSG00000181513 | 17 | 43209967 | 43221548 | 21 | 7 | 0.76587 | 0.22188 | ACBD4 | 0.22188 | 0.16441 | 0.19824 |
| ENSG00000020426 | 14 | 61201460 | 61436671 | 340 | 35 | 0.76581 | 0.2219 | MNAT1 | 0.2219 | 0.2329 | 0.22266 |
| ENSG00000133107 | 13 | 38210773 | 38444562 | 923 | 70 | 0.76555 | 0.22197 | TRPC4 | 0.22197 | 0.25038 | 0.2092 |
| ENSG00000124116 | 20 | 44376583 | 44420571 | 247 | 17 | 0.76548 | 0.22199 | WFDC3 | 0.22199 | 0.22179 | 0.23412 |
| ENSG00000261272 | 6 | 30978251 | 31003179 | 470 | 37 | 0.7653 | 0.22205 | MUC22 | 0.22205 | 0.32547 | 0.4567 |
| ENSG00000170458 | 5 | 140011313 | 140013286 | 3 | 1 | 0.7652 | 0.22208 | CD14 | 0.22208 | 0.33912 | 0.50097 |
| ENSG00000088256 | 19 | 3094408 | 3124002 | 171 | 26 | 0.7652 | 0.22208 | GNA11 | 0.22208 | 0.21394 | 0.50583 |
| ENSG00000113638 | 5 | 40714577 | 40756077 | 100 | 14 | 0.76506 | 0.22212 | TTC33 | 0.22212 | 0.25462 | 0.31524 |
| ENSG00000187498 | 13 | 110801318 | 110959496 | 790 | 85 | 0.76476 | 0.22221 | COL4A1 | 0.22221 | 0.14509 | 0.12628 |
| ENSG00000050130 | 14 | 59951161 | 59972128 | 82 | 8 | 0.76434 | 0.22233 | JKAMP | 0.22233 | 0.21657 | 0.22738 |

| | | | | | | | | | | | |
|---|---|---|---|---|---|---|---|---|---|---|---|
| ENSG00000178821 | 1 | 1849029 | 1850712 | 8 | 3 | 0.76383 | 0.22248 | TMEM52 | 0.22248 | 0.20337 | 0.2157 |
| ENSG00000006606 | 7 | 75398851 | 75419214 | 117 | 16 | 0.76367 | 0.22253 | CCL26 | 0.22253 | 0.14707 | 0.18288 |
| ENSG00000188488 | 14 | 95027779 | 95059457 | 191 | 24 | 0.76351 | 0.22258 | SERPINA5 | 0.22258 | 0.23 | 0.22112 |
| ENSG00000186399 | 15 | 30695943 | 30706463 | 1 | 1 | 0.76344 | 0.2226 | GOLGA8R | 0.2226 | 0.70481 | 0.70481 |
| ENSG00000100284 | 22 | 35695268 | 35743985 | 165 | 17 | 0.76323 | 0.22266 | TOM1 | 0.22266 | 0.23157 | 0.17199 |
| ENSG00000125386 | 4 | 2626988 | 2734292 | 352 | 27 | 0.76307 | 0.22271 | FAM193A | 0.22271 | 0.23012 | 0.2374 |
| ENSG00000163092 | 2 | 167744997 | 168116263 | 1498 | 42 | 0.76254 | 0.22287 | XIRP2 | 0.22287 | 0.24039 | 0.24153 |
| ENSG00000164929 | 8 | 104152938 | 104242533 | 358 | 38 | 0.76241 | 0.22291 | BAALC | 0.22291 | 0.22694 | 0.31651 |
| ENSG00000166428 | 14 | 105391153 | 105399574 | 35 | 6 | 0.76238 | 0.22292 | PLD4 | 0.22292 | 0.14197 | 0.031752 |
| ENSG00000169885 | 1 | 1846266 | 1848735 | 11 | 3 | 0.76221 | 0.22297 | CALML6 | 0.22297 | 0.22192 | 0.19147 |
| ENSG00000130208 | 19 | 45417504 | 45422606 | 15 | 6 | 0.7622 | 0.22297 | APOC1 | 0.22297 | 0.36736 | 0.064635 |
| ENSG00000139269 | 12 | 57846106 | 57853063 | 11 | 4 | 0.76203 | 0.22302 | INHBE | 0.22302 | 0.0010634 | 0.0014783 |
| ENSG00000032742 | 13 | 21140585 | 21265503 | 409 | 19 | 0.76165 | 0.22313 | IFT88 | 0.22313 | 0.23738 | 0.25107 |
| ENSG00000101342 | 20 | 35504524 | 35522638 | 48 | 14 | 0.76167 | 0.22313 | TLDC2 | 0.22313 | 0.31417 | 0.3676 |
| ENSG00000167986 | 11 | 61066923 | 61110068 | 27 | 16 | 0.76134 | 0.22323 | DDB1 | 0.22323 | 0.25799 | 0.46565 |
| ENSG00000106605 | 7 | 43798279 | 43846939 | 105 | 13 | 0.76127 | 0.22325 | BLVRA | 0.22325 | 0.18567 | 0.19114 |
| ENSG00000119333 | 9 | 131395940 | 131419066 | 68 | 9 | 0.76127 | 0.22325 | WDR34 | 0.22325 | 0.24074 | 0.19826 |
| ENSG00000123364 | 12 | 54332535 | 54340328 | 15 | 3 | 0.76097 | 0.22334 | HOXC13 | 0.22334 | 0.14817 | 0.098115 |
| ENSG00000173258 | 9 | 114287439 | 114340124 | 184 | 27 | 0.76014 | 0.22359 | ZNF483 | 0.22359 | 0.21451 | 0.18747 |
| ENSG00000065413 | 2 | 197831741 | 198175897 | 1142 | 63 | 0.75988 | 0.22366 | ANKRD44 | 0.22366 | 0.17854 | 0.095072 |
| ENSG00000181903 | 11 | 55432614 | 55433671 | 2 | 1 | 0.75987 | 0.22366 | OR4C6 | 0.22366 | 0.1224 | 0.14498 |
| ENSG00000196263 | 19 | 57019212 | 57041590 | 67 | 8 | 0.75978 | 0.22369 | ZNF471 | 0.22369 | 0.20624 | 0.34752 |
| ENSG00000147437 | 8 | 25276776 | 25282170 | 9 | 3 | 0.75943 | 0.2238 | GNRH1 | 0.2238 | 0.41708 | 0.41188 |
| ENSG00000213380 | 16 | 69354043 | 69373570 | 51 | 8 | 0.75935 | 0.22382 | COG8 | 0.22382 | 0.27014 | 0.31805 |
| ENSG00000064763 | 12 | 29302036 | 29493913 | 759 | 29 | 0.7593 | 0.22384 | FAR2 | 0.22384 | 0.20672 | 0.25927 |
| ENSG00000171942 | 19 | 15838834 | 15839862 | 5 | 2 | 0.75922 | 0.22386 | OR10H2 | 0.22386 | 0.30387 | 0.13858 |
| ENSG00000132792 | 20 | 36322408 | 36500531 | 247 | 30 | 0.75898 | 0.22393 | CTNNBL1 | 0.22393 | 0.26954 | 0.154 |
| ENSG00000170989 | 1 | 101702444 | 101707074 | 17 | 6 | 0.75896 | 0.22394 | S1PR1 | 0.22394 | 0.54118 | 0.53799 |
| ENSG00000167193 | 17 | 1323983 | 1366456 | 153 | 24 | 0.75893 | 0.22395 | CRK | 0.22395 | 0.16618 | 0.20862 |
| ENSG00000221931 | 11 | 123624196 | 123625253 | 1 | 1 | 0.75842 | 0.2241 | OR6X1 | 0.2241 | 0.040082 | 0.056806 |
| ENSG00000128016 | 19 | 39897453 | 39900052 | 2 | 1 | 0.75819 | 0.22417 | ZFP36 | 0.22417 | 0.67138 | 0.81463 |
| ENSG00000129951 | 19 | 812518 | 821967 | 56 | 9 | 0.75815 | 0.22418 | LPPR3 | 0.22418 | 0.28333 | 0.30024 |
| ENSG00000205129 | 4 | 186347394 | 186370980 | 111 | 12 | 0.75786 | 0.22427 | C4orf47 | 0.22427 | 0.25913 | 0.31273 |
| ENSG00000178297 | 19 | 2389769 | 2426237 | 266 | 23 | 0.75783 | 0.22428 | TMPRSS9 | 0.22428 | 0.17327 | 0.11222 |
| ENSG00000170745 | 2 | 18059114 | 18542882 | 2299 | 143 | 0.75777 | 0.22429 | KCNS3 | 0.22429 | 0.21811 | 0.14671 |
| ENSG00000107099 | 9 | 214854 | 465259 | 1457 | 78 | 0.75757 | 0.22436 | DOCK8 | 0.22436 | 0.21845 | 0.23486 |
| ENSG00000144647 | 3 | 43120724 | 43147568 | 53 | 10 | 0.75751 | 0.22437 | POMGNT2 | 0.22437 | 0.2786 | 0.31086 |
| ENSG00000150750 | 11 | 111126707 | 111157126 | 85 | 12 | 0.75747 | 0.22438 | C11orf53 | 0.22438 | 0.39672 | 0.34469 |
| ENSG00000131183 | 5 | 176806236 | 176825849 | 39 | 8 | 0.75716 | 0.22448 | SLC34A1 | 0.22448 | 0.22557 | 0.25019 |
| ENSG00000127585 | 16 | 742500 | 755829 | 27 | 4 | 0.75682 | 0.22458 | FBXL16 | 0.22458 | 0.39403 | 0.30425 |
| ENSG00000116726 | 1 | 12834984 | 12838049 | 13 | 4 | 0.75658 | 0.22465 | PRAMEF12 | 0.22465 | 0.096975 | 0.2793 |
| ENSG00000171864 | 20 | 4702556 | 4709106 | 34 | 12 | 0.75659 | 0.22465 | PRND | 0.22465 | 0.46505 | 0.58106 |
| ENSG00000132938 | 13 | 29598748 | 30077892 | 1926 | 94 | 0.75655 | 0.22466 | MTUS2 | 0.22466 | 0.23892 | 0.17349 |
| ENSG00000146282 | 6 | 88224096 | 88299721 | 312 | 15 | 0.75644 | 0.22469 | RARS2 | 0.22469 | 0.20864 | 0.17189 |
| ENSG00000078804 | 20 | 33292094 | 33301243 | 15 | 5 | 0.75624 | 0.22475 | TP53INP2 | 0.22475 | 0.30941 | 0.23327 |
| ENSG00000167733 | 19 | 5680615 | 5688533 | 12 | 5 | 0.75604 | 0.22481 | HSD11B1L | 0.22481 | 0.42489 | 0.55276 |
| ENSG00000139438 | 12 | 110152033 | 110208312 | 157 | 25 | 0.75581 | 0.22488 | FAM222A | 0.22488 | 0.37461 | 0.39552 |
| ENSG00000179698 | 8 | 145162629 | 145173218 | 26 | 5 | 0.7554 | 0.225 | KIAA1875 | 0.225 | 0.17163 | 0.17838 |
| ENSG00000198189 | 4 | 88257762 | 88312538 | 237 | 22 | 0.75524 | 0.22505 | HSD17B11 | 0.22505 | 0.26507 | 0.32003 |
| ENSG00000153820 | 2 | 228844666 | 229046361 | 700 | 36 | 0.75492 | 0.22515 | SPHKAP | 0.22515 | 0.26888 | 0.32606 |
| ENSG00000226524 | 6 | 88032323 | 88109459 | 342 | 18 | 0.75483 | 0.22518 | RP1-102H19.8 | 0.22518 | 0.20853 | 0.26079 |
| ENSG00000132294 | 8 | 132916335 | 133025889 | 343 | 27 | 0.75481 | 0.22518 | EFR3A | 0.22518 | 0.26747 | 0.25974 |
| ENSG00000269179 | 19 | 50412758 | 50461648 | 185 | 34 | 0.75447 | 0.22528 | CTC-326K19.6 | 0.22528 | 0.20137 | 0.20209 |
| ENSG00000214946 | 17 | 15635561 | 15649476 | 36 | 8 | 0.75402 | 0.22542 | TBC1D26 | 0.22542 | 0.49101 | 0.61654 |
| ENSG00000086288 | 7 | 37888199 | 37940003 | 273 | 27 | 0.75397 | 0.22543 | NME8 | 0.22543 | 0.25271 | 0.28831 |
| ENSG00000214891 | 11 | 49075266 | 49080664 | 15 | 4 | 0.75399 | 0.22543 | TRIM64C | 0.22543 | 0.29669 | 0.29761 |
| ENSG00000249109 | 5 | 177209198 | 177221397 | 45 | 6 | 0.75342 | 0.2256 | RP11-1026M7.2 | 0.2256 | 0.26323 | 0.32683 |
| ENSG00000160767 | 1 | 155216996 | 155225274 | 25 | 2 | 0.75338 | 0.22561 | FAM189B | 0.22561 | 0.19117 | 0.17543 |
| ENSG00000179091 | 8 | 145149930 | 145152428 | 3 | 1 | 0.75325 | 0.22565 | CYC1 | 0.22565 | 0.1735 | 0.13731 |
| ENSG00000100097 | 22 | 38071615 | 38075813 | 11 | 4 | 0.7529 | 0.22576 | LGALS1 | 0.22576 | 0.27653 | 0.19804 |
| ENSG00000005073 | 7 | 27221129 | 27224842 | 8 | 4 | 0.75277 | 0.22579 | HOXA11 | 0.22579 | 0.29962 | 0.24839 |
| ENSG00000099800 | 19 | 2425622 | 2427892 | 17 | 5 | 0.75278 | 0.22579 | TIMM13 | 0.22579 | 0.28655 | 0.22256 |
| ENSG00000163563 | 1 | 158801107 | 158819296 | 50 | 11 | 0.75267 | 0.22582 | MNDA | 0.22582 | 0.26017 | 0.24779 |
| ENSG00000144891 | 3 | 148415571 | 148460795 | 129 | 17 | 0.75268 | 0.22582 | AGTR1 | 0.22582 | 0.37139 | 0.26037 |
| ENSG00000171456 | 20 | 30946155 | 31027122 | 150 | 11 | 0.75224 | 0.22595 | ASXL1 | 0.22595 | 0.22907 | 0.24613 |
| ENSG00000137843 | 15 | 40509629 | 40569688 | 176 | 42 | 0.75204 | 0.22601 | PAK6 | 0.22601 | 0.11005 | 0.057786 |
| ENSG00000186860 | 17 | 39471173 | 39471947 | 7 | 2 | 0.75201 | 0.22602 | KRTAP17-1 | 0.22602 | 0.23315 | 0.22286 |
| ENSG00000249222 | 22 | 43035809 | 43036607 | 4 | 1 | 0.75202 | 0.22602 | ATP5L2 | 0.22602 | 0.32761 | 0.3908 |
| ENSG00000184428 | 8 | 144386554 | 144442149 | 259 | 16 | 0.75184 | 0.22607 | TOP1MT | 0.22607 | 0.19977 | 0.20394 |
| ENSG00000188811 | 13 | 39612443 | 39624246 | 22 | 8 | 0.7515 | 0.22617 | NHLRC3 | 0.22617 | 0.099868 | 0.13393 |
| ENSG00000162458 | 1 | 16083102 | 16113089 | 82 | 13 | 0.75138 | 0.22621 | FBLIM1 | 0.22621 | 0.30909 | 0.18078 |
| ENSG00000132471 | 17 | 73841780 | 73852588 | 35 | 6 | 0.75108 | 0.2263 | WBP2 | 0.2263 | 0.25984 | 0.37746 |
| ENSG00000041880 | 3 | 51976361 | 51982883 | 12 | 4 | 0.7509 | 0.22636 | PARP3 | 0.22636 | 0.7271 | 0.73161 |

| | | | | | | | | | | | |
|---|---|---|---|---|---|---|---|---|---|---|---|
| ENSG00000119973 | 10 | 120352916 | 120355160 | 11 | 5 | 0.75083 | 0.22638 | PRLHR | 0.22638 | 0.16197 | 0.12992 |
| ENSG00000100441 | 14 | 24898492 | 24910540 | 74 | 12 | 0.75077 | 0.2264 | KHNYN | 0.2264 | 0.41687 | 0.41483 |
| ENSG00000146574 | 7 | 6833765 | 6866401 | 78 | 17 | 0.7502 | 0.22657 | CCZ1B | 0.22657 | 0.24094 | 0.19978 |
| ENSG00000106211 | 7 | 75931861 | 75933612 | 4 | 2 | 0.75007 | 0.22661 | HSPB1 | 0.22661 | 0.85574 | 0.78349 |
| ENSG00000267882 | 20 | 45947246 | 45949467 | 4 | 2 | 0.74939 | 0.22681 | AL031666.2 | 0.22681 | 0.75861 | 0.84881 |
| ENSG00000128052 | 4 | 55944644 | 55991756 | 138 | 30 | 0.74931 | 0.22683 | KDR | 0.22683 | 0.42068 | 0.72018 |
| ENSG00000197694 | 9 | 131314866 | 131395941 | 120 | 9 | 0.74904 | 0.22692 | SPTAN1 | 0.22692 | 0.24838 | 0.2635 |
| ENSG00000173209 | 2 | 61404553 | 61418338 | 35 | 10 | 0.749 | 0.22693 | AHSA2 | 0.22693 | 0.42825 | 0.3994 |
| ENSG00000184465 | 6 | 169857307 | 170102159 | 871 | 34 | 0.74896 | 0.22694 | WDR27 | 0.22694 | 0.21765 | 0.22475 |
| ENSG00000103269 | 16 | 725666 | 728268 | 2 | 1 | 0.74852 | 0.22707 | RHBDL1 | 0.22707 | 0.30907 | 0.27326 |
| ENSG00000113263 | 5 | 156569944 | 156682201 | 478 | 46 | 0.74848 | 0.22709 | ITK | 0.22709 | 0.23081 | 0.24029 |
| ENSG00000215012 | 22 | 19833661 | 19842419 | 34 | 6 | 0.74838 | 0.22712 | C22orf29 | 0.22712 | 0.12863 | 0.092178 |
| ENSG00000095752 | 19 | 55875757 | 55881831 | 26 | 8 | 0.74773 | 0.22731 | IL11 | 0.22731 | 0.42231 | 0.62304 |
| ENSG00000177058 | 5 | 54921673 | 55069022 | 784 | 26 | 0.74768 | 0.22733 | SLC38A9 | 0.22733 | 0.20763 | 0.18299 |
| ENSG00000113448 | 5 | 58264865 | 59817947 | 4503 | 176 | 0.74767 | 0.22733 | PDE4D | 0.22733 | 0.22255 | 0.23075 |
| ENSG00000123405 | 12 | 54685895 | 54694905 | 8 | 3 | 0.74742 | 0.22741 | NFE2 | 0.22741 | 0.62029 | 0.50392 |
| ENSG00000101680 | 18 | 6941743 | 7117813 | 751 | 121 | 0.74739 | 0.22742 | LAMA1 | 0.22742 | 0.36877 | 0.38121 |
| ENSG00000213973 | 19 | 22939007 | 22966909 | 167 | 9 | 0.74711 | 0.2275 | ZNF99 | 0.2275 | 0.22301 | 0.24242 |
| ENSG00000179889 | 16 | 15068448 | 15233196 | 438 | 43 | 0.7467 | 0.22762 | PDXDC1 | 0.22762 | 0.21125 | 0.17107 |
| ENSG00000153339 | 18 | 29409136 | 29533099 | 363 | 21 | 0.74648 | 0.22769 | TRAPPC8 | 0.22769 | 0.28923 | 0.41781 |
| ENSG00000154118 | 16 | 87635441 | 87731762 | 585 | 63 | 0.74609 | 0.22781 | JPH3 | 0.22781 | 0.18361 | 0.1026 |
| ENSG00000145592 | 5 | 40825364 | 40835437 | 31 | 9 | 0.74586 | 0.22788 | RPL37 | 0.22788 | 0.382 | 0.54826 |
| ENSG00000189042 | 19 | 37178514 | 37218603 | 120 | 11 | 0.74565 | 0.22794 | ZNF567 | 0.22794 | 0.19675 | 0.1411 |
| ENSG00000175203 | 12 | 57923885 | 57941114 | 31 | 5 | 0.74542 | 0.22801 | DCTN2 | 0.22801 | 0.18816 | 0.15422 |
| ENSG00000162711 | 1 | 247579458 | 247612410 | 178 | 41 | 0.74532 | 0.22804 | NLRP3 | 0.22804 | 0.59212 | 0.65836 |
| ENSG00000164458 | 6 | 166571144 | 166582188 | 52 | 9 | 0.74521 | 0.22807 | T | 0.22807 | 0.28922 | 0.34343 |
| ENSG00000087128 | 4 | 69313167 | 69363322 | 215 | 17 | 0.74452 | 0.22828 | TMPRSS11E | 0.22828 | 0.24715 | 0.29418 |
| ENSG00000115084 | 2 | 114462588 | 114514400 | 197 | 19 | 0.74448 | 0.22829 | SLC35F5 | 0.22829 | 0.25307 | 0.42438 |
| ENSG00000149547 | 11 | 125439112 | 125454575 | 41 | 8 | 0.74431 | 0.22834 | EI24 | 0.22834 | 0.25108 | 0.27708 |
| ENSG00000236287 | 11 | 10833621 | 10880343 | 153 | 15 | 0.74386 | 0.22848 | ZBED5 | 0.22848 | 0.20016 | 0.11735 |
| ENSG00000151725 | 4 | 185615772 | 185655287 | 317 | 6 | 0.74365 | 0.22854 | CENPU | 0.22854 | 0.2497 | 0.25873 |
| ENSG00000174953 | 3 | 153990335 | 154042286 | 113 | 16 | 0.74346 | 0.2286 | DHX36 | 0.2286 | 0.26811 | 0.30209 |
| ENSG00000241370 | 6 | 30312908 | 30314661 | 14 | 5 | 0.7434 | 0.22862 | RPP21 | 0.22862 | 0.32232 | 0.29028 |
| ENSG00000214305 | 11 | 107650219 | 107652316 | 12 | 5 | 0.74311 | 0.22871 | AP001024.1 | 0.22871 | 0.31067 | 0.29861 |
| ENSG00000079156 | 2 | 179059208 | 179264160 | 585 | 49 | 0.74204 | 0.22903 | OSBPL6 | 0.22903 | 0.19422 | 0.14768 |
| ENSG00000206177 | 16 | 203891 | 216767 | 52 | 11 | 0.74201 | 0.22904 | HBM | 0.22904 | 0.30966 | 0.32594 |
| ENSG00000172724 | 9 | 34689564 | 34691274 | 6 | 3 | 0.74195 | 0.22906 | CCL19 | 0.22906 | 0.051622 | 0.017514 |
| ENSG00000203993 | 9 | 140509784 | 140513358 | 2 | 1 | 0.74181 | 0.2291 | C9orf37 | 0.2291 | 0.30961 | 0.33476 |
| ENSG00000108242 | 10 | 96443251 | 96495947 | 169 | 17 | 0.74182 | 0.2291 | CYP2C18 | 0.2291 | 0.28187 | 0.27829 |
| ENSG00000116521 | 1 | 155225770 | 155232221 | 15 | 3 | 0.74158 | 0.22917 | SCAMP3 | 0.22917 | 0.18975 | 0.19173 |
| ENSG00000254667 | 11 | 123301058 | 123306427 | 22 | 6 | 0.74148 | 0.2292 | AP000783.1 | 0.2292 | 0.27656 | 0.067049 |
| ENSG00000130811 | 19 | 10225693 | 10230596 | 15 | 4 | 0.74146 | 0.22921 | EIF3G | 0.22921 | 0.21146 | 0.1343 |
| ENSG00000140263 | 15 | 45315302 | 45369383 | 203 | 25 | 0.74133 | 0.22925 | SORD | 0.22925 | 0.22417 | 0.19113 |
| ENSG00000121931 | 1 | 111489807 | 111506701 | 25 | 9 | 0.74127 | 0.22927 | LRIF1 | 0.22927 | 0.016215 | 0.015134 |
| ENSG00000204231 | 6 | 33161365 | 33168630 | 14 | 5 | 0.74112 | 0.22931 | RXRB | 0.22931 | 0.10591 | 0.30226 |
| ENSG00000104907 | 19 | 13215716 | 13228381 | 28 | 14 | 0.74069 | 0.22944 | TRMT1 | 0.22944 | 0.079958 | 0.092238 |
| ENSG00000168894 | 2 | 85822848 | 85824736 | 10 | 4 | 0.74053 | 0.22949 | RNF181 | 0.22949 | 0.071255 | 0.038493 |
| ENSG00000083312 | 5 | 72112139 | 72212560 | 209 | 19 | 0.74022 | 0.22958 | TNPO1 | 0.22958 | 0.2375 | 0.2388 |
| ENSG00000152661 | 6 | 121756838 | 121770873 | 50 | 10 | 0.73962 | 0.22977 | GJA1 | 0.22977 | 0.016203 | 0.020165 |
| ENSG00000268412 | 2 | 26251074 | 26251481 | 1 | 1 | 0.73918 | 0.2299 | AC013449.1 | 0.2299 | 0.18915 | 0.08223 |
| ENSG00000041515 | 13 | 109248500 | 109860355 | 2831 | 180 | 0.73915 | 0.22991 | MYO16 | 0.22991 | 0.27758 | 0.36194 |
| ENSG00000101442 | 20 | 37377085 | 37400834 | 116 | 13 | 0.73906 | 0.22993 | ACTR5 | 0.22993 | 0.25097 | 0.23697 |
| ENSG00000212721 | 17 | 39273433 | 39274606 | 14 | 4 | 0.73893 | 0.22998 | KRTAP4-11 | 0.22998 | 0.20397 | 0.271 |
| ENSG00000165471 | 10 | 54525140 | 54531460 | 31 | 7 | 0.73865 | 0.23006 | MBL2 | 0.23006 | 0.2491 | 0.38275 |
| ENSG00000095383 | 9 | 100961311 | 101017915 | 284 | 26 | 0.73756 | 0.23039 | TBC1D2 | 0.23039 | 0.2057 | 0.21986 |
| ENSG00000151694 | 2 | 9628615 | 9695921 | 170 | 14 | 0.73708 | 0.23054 | ADAM17 | 0.23054 | 0.20636 | 0.15315 |
| ENSG00000198589 | 4 | 151185594 | 151936879 | 1742 | 41 | 0.73678 | 0.23063 | LRBA | 0.23063 | 0.23675 | 0.24139 |
| ENSG00000146540 | 7 | 1036622 | 1177896 | 945 | 37 | 0.73666 | 0.23066 | C7orf50 | 0.23066 | 0.21798 | 0.22247 |
| ENSG00000197070 | 9 | 140500106 | 140509812 | 22 | 4 | 0.73664 | 0.23067 | ARRDC1 | 0.23067 | 0.26966 | 0.33113 |
| ENSG00000105429 | 19 | 42829761 | 42882921 | 62 | 6 | 0.73657 | 0.23069 | MEGF8 | 0.23069 | 0.17949 | 0.15524 |
| ENSG00000188295 | 1 | 247263264 | 247267674 | 13 | 3 | 0.73632 | 0.23077 | ZNF669 | 0.23077 | 0.40673 | 0.29381 |
| ENSG00000169436 | 8 | 139600478 | 139926249 | 1335 | 124 | 0.73617 | 0.23081 | COL22A1 | 0.23081 | 0.23951 | 0.25867 |
| ENSG00000163394 | 4 | 26483022 | 26492084 | 27 | 6 | 0.73532 | 0.23107 | CCKAR | 0.23107 | 0.70359 | 0.60757 |
| ENSG00000015479 | 5 | 138609441 | 138667360 | 51 | 8 | 0.73513 | 0.23113 | MATR3 | 0.23113 | 0.19508 | 0.19835 |
| ENSG00000116675 | 1 | 65713902 | 65881552 | 551 | 32 | 0.73499 | 0.23117 | DNAJC6 | 0.23117 | 0.2166 | 0.26892 |
| ENSG00000197081 | 6 | 160390131 | 160534539 | 387 | 49 | 0.73495 | 0.23119 | IGF2R | 0.23119 | 0.15486 | 0.11508 |
| ENSG00000254870 | 6 | 31497996 | 31514385 | 114 | 22 | 0.73469 | 0.23126 | ATP6V1G2-DDX39B | 0.23126 | 0.1748 | 0.14515 |
| ENSG00000229453 | 3 | 48348336 | 48369831 | 53 | 3 | 0.73465 | 0.23128 | SPINK8 | 0.23128 | 0.24522 | 0.1977 |
| ENSG00000134574 | 11 | 47236493 | 47260767 | 69 | 11 | 0.73465 | 0.23128 | DDB2 | 0.23128 | 0.2726 | 0.30128 |
| ENSG00000064995 | 6 | 34845555 | 34855866 | 34 | 8 | 0.7344 | 0.23135 | TAF11 | 0.23135 | 0.30123 | 0.31033 |
| ENSG00000165633 | 10 | 50222290 | 50323554 | 410 | 22 | 0.73432 | 0.23138 | VSTM4 | 0.23138 | 0.2695 | 0.2224 |
| ENSG00000149231 | 11 | 96085933 | 96123087 | 97 | 16 | 0.73425 | 0.2314 | CCDC82 | 0.2314 | 0.20025 | 0.17381 |
| ENSG00000182197 | 8 | 118806729 | 119124092 | 890 | 58 | 0.73398 | 0.23148 | EXT1 | 0.23148 | 0.30238 | 0.35489 |

| | | | | | | | | | | | |
|---|---|---|---|---|---|---|---|---|---|---|---|
| ENSG00000101019 | 20 | 33890369 | 33999944 | 218 | 12 | 0.73378 | 0.23154 | UQCC1 | 0.23154 | 0.20321 | 0.13283 |
| ENSG00000111536 | 12 | 68595131 | 68619601 | 48 | 10 | 0.73342 | 0.23165 | IL26 | 0.23165 | 0.088194 | 0.010261 |
| ENSG00000075340 | 2 | 70834750 | 70995357 | 748 | 43 | 0.73341 | 0.23166 | ADD2 | 0.23166 | 0.26219 | 0.32674 |
| ENSG00000135638 | 2 | 73143389 | 73162020 | 23 | 6 | 0.73324 | 0.23171 | EMX1 | 0.23171 | 0.39996 | 0.26286 |
| ENSG00000107021 | 9 | 131549483 | 131572711 | 87 | 10 | 0.73257 | 0.23191 | TBC1D13 | 0.23191 | 0.22161 | 0.25821 |
| ENSG00000158125 | 2 | 31557187 | 31637581 | 328 | 42 | 0.73232 | 0.23199 | XDH | 0.23199 | 0.22474 | 0.24327 |
| ENSG00000138629 | 15 | 74738318 | 74753523 | 30 | 5 | 0.73226 | 0.232 | UBL7 | 0.232 | 0.22624 | 0.17078 |
| ENSG00000087157 | 17 | 76374721 | 76421195 | 236 | 21 | 0.73213 | 0.23204 | PGS1 | 0.23204 | 0.36738 | 0.36157 |
| ENSG00000177311 | 3 | 141043055 | 141168634 | 303 | 28 | 0.732 | 0.23208 | ZBTB38 | 0.23208 | 0.14171 | 0.15081 |
| ENSG00000182118 | 1 | 231154704 | 231175992 | 127 | 24 | 0.73196 | 0.2321 | FAM89A | 0.2321 | 0.19049 | 0.2006 |
| ENSG00000150477 | 18 | 34409069 | 34812135 | 896 | 19 | 0.73171 | 0.23217 | KIAA1328 | 0.23217 | 0.23478 | 0.23626 |
| ENSG00000145113 | 3 | 195473636 | 195539148 | 535 | 37 | 0.73159 | 0.23221 | MUC4 | 0.23221 | 0.21644 | 0.21201 |
| ENSG00000134716 | 1 | 60358980 | 60392462 | 71 | 12 | 0.73135 | 0.23228 | CYP2J2 | 0.23228 | 0.14129 | 0.12534 |
| ENSG00000221836 | 7 | 143747461 | 143748473 | 2 | 1 | 0.73085 | 0.23243 | OR2A5 | 0.23243 | 0.43877 | 0.5244 |
| ENSG00000163728 | 3 | 180319918 | 180335616 | 18 | 6 | 0.73079 | 0.23245 | TTC14 | 0.23245 | 0.26316 | 0.19723 |
| ENSG00000204566 | 10 | 23492745 | 23528745 | 51 | 11 | 0.73074 | 0.23247 | C10orf115 | 0.23247 | 0.20716 | 0.1908 |
| ENSG00000188573 | 5 | 167956121 | 167957639 | 9 | 3 | 0.73057 | 0.23252 | FBLL1 | 0.23252 | 0.37403 | 0.3307 |
| ENSG00000204519 | 19 | 58193337 | 58228669 | 94 | 16 | 0.73037 | 0.23258 | ZNF551 | 0.23258 | 0.22585 | 0.27476 |
| ENSG00000168036 | 3 | 41236328 | 41301587 | 123 | 14 | 0.73023 | 0.23262 | CTNNB1 | 0.23262 | 0.23935 | 0.29782 |
| ENSG00000119125 | 9 | 74729511 | 74872109 | 473 | 35 | 0.73026 | 0.23262 | GDA | 0.23262 | 0.24064 | 0.30267 |
| ENSG00000101190 | 20 | 61472467 | 61493115 | 95 | 10 | 0.72989 | 0.23273 | TCFL5 | 0.23273 | 0.20981 | 0.26642 |
| ENSG00000105409 | 19 | 42470734 | 42501649 | 34 | 16 | 0.72955 | 0.23283 | ATP1A3 | 0.23283 | 0.29804 | 0.45168 |
| ENSG00000183114 | 1 | 20878932 | 20881512 | 1 | 1 | 0.72933 | 0.2329 | FAM43B | 0.2329 | 0.81507 | 0.84236 |
| ENSG00000164077 | 3 | 49946302 | 49967606 | 32 | 9 | 0.72892 | 0.23303 | MON1A | 0.23303 | 0.21888 | 0.2014 |
| ENSG00000178573 | 16 | 79619740 | 79634611 | 70 | 21 | 0.72852 | 0.23315 | MAF | 0.23315 | 0.46154 | 0.48661 |
| ENSG00000272921 | 12 | 4714110 | 4771809 | 264 | 19 | 0.72819 | 0.23325 | RP11-500M8.7 | 0.23325 | 0.25785 | 0.25856 |
| ENSG00000127743 | 5 | 148750887 | 148783765 | 104 | 16 | 0.72808 | 0.23328 | IL17B | 0.23328 | 0.29426 | 0.35186 |
| ENSG00000198821 | 1 | 167399877 | 167487847 | 366 | 69 | 0.72807 | 0.23329 | CD247 | 0.23329 | 0.29107 | 0.27265 |
| ENSG00000173678 | 7 | 102290772 | 102301847 | 3 | 1 | 0.72789 | 0.23334 | SPDYE2B | 0.23334 | 0.45885 | 0.28013 |
| ENSG00000111780 | 12 | 120876001 | 120897809 | 88 | 16 | 0.7274 | 0.23349 | AL021546.6 | 0.23349 | 0.40623 | 0.48609 |
| ENSG00000269469 | 19 | 49966009 | 49989488 | 89 | 14 | 0.72724 | 0.23354 | CTD-3148I10.9 | 0.23354 | 0.14961 | 0.10309 |
| ENSG00000186807 | 10 | 47746936 | 47763041 | 10 | 3 | 0.72687 | 0.23365 | ANXA8L2 | 0.23365 | 0.68299 | 0.71658 |
| ENSG00000215033 | 10 | 47746962 | 47770871 | 10 | 3 | 0.72687 | 0.23365 | AL603965.1 | 0.23365 | 0.68299 | 0.71658 |
| ENSG00000110921 | 12 | 110011060 | 110035067 | 82 | 15 | 0.72684 | 0.23366 | MVK | 0.23366 | 0.21074 | 0.15971 |
| ENSG00000005100 | 17 | 5344232 | 5372380 | 90 | 20 | 0.72683 | 0.23367 | DHX33 | 0.23367 | 0.22586 | 0.33041 |
| ENSG00000151748 | 14 | 51098776 | 51135049 | 114 | 13 | 0.7263 | 0.23383 | SAV1 | 0.23383 | 0.2607 | 0.28493 |
| ENSG00000168906 | 2 | 85766288 | 85772403 | 10 | 5 | 0.72615 | 0.23387 | MAT2A | 0.23387 | 0.029393 | 0.03099 |
| ENSG00000247626 | 2 | 198570087 | 198573113 | 2 | 1 | 0.72581 | 0.23398 | MARS2 | 0.23398 | 0.092517 | 0.14059 |
| ENSG00000099998 | 22 | 24615622 | 24641110 | 123 | 24 | 0.72552 | 0.23407 | GGT5 | 0.23407 | 0.40869 | 0.30821 |
| ENSG00000103534 | 16 | 19421818 | 19510435 | 432 | 32 | 0.72507 | 0.2342 | TMC5 | 0.2342 | 0.2979 | 0.29392 |
| ENSG00000070759 | 1 | 45809555 | 45956872 | 292 | 14 | 0.7249 | 0.23426 | TESK2 | 0.23426 | 0.25191 | 0.2991 |
| ENSG00000122741 | 9 | 37800499 | 37867663 | 235 | 8 | 0.7248 | 0.23429 | DCAF10 | 0.23429 | 0.21586 | 0.18518 |
| ENSG00000134897 | 13 | 103451399 | 103493885 | 151 | 16 | 0.72455 | 0.23436 | BIVM | 0.23436 | 0.33184 | 0.37251 |
| ENSG00000120253 | 6 | 150045451 | 150070801 | 52 | 5 | 0.72449 | 0.23438 | NUP43 | 0.23438 | 0.20922 | 0.22757 |
| ENSG00000138829 | 5 | 127593601 | 127994878 | 1384 | 37 | 0.72444 | 0.2344 | FBN2 | 0.2344 | 0.22882 | 0.28247 |
| ENSG00000101000 | 20 | 33759876 | 33765165 | 14 | 4 | 0.7242 | 0.23447 | PROCR | 0.23447 | 0.35908 | 0.4057 |
| ENSG00000163026 | 2 | 24252210 | 24272445 | 61 | 14 | 0.72411 | 0.2345 | C2orf44 | 0.2345 | 0.44399 | 0.11873 |
| ENSG00000164654 | 7 | 7606503 | 7648560 | 184 | 19 | 0.72381 | 0.23459 | MIOS | 0.23459 | 0.21769 | 0.187 |
| ENSG00000121579 | 3 | 113435307 | 113465147 | 44 | 6 | 0.72364 | 0.23464 | NAA50 | 0.23464 | 0.20401 | 0.30962 |
| ENSG00000169575 | 22 | 22599087 | 22599927 | 8 | 4 | 0.72346 | 0.2347 | VPREB1 | 0.2347 | 0.83049 | 0.86896 |
| ENSG00000113262 | 5 | 178405328 | 178423207 | 111 | 13 | 0.7232 | 0.23478 | GRM6 | 0.23478 | 0.21886 | 0.25907 |
| ENSG00000064199 | 11 | 124543694 | 124567414 | 37 | 6 | 0.72314 | 0.2348 | SPA17 | 0.2348 | 0.17041 | 0.34843 |
| ENSG00000171094 | 2 | 29415640 | 30144432 | 2927 | 160 | 0.72309 | 0.23481 | ALK | 0.23481 | 0.23829 | 0.27033 |
| ENSG00000165383 | 10 | 50117528 | 50143242 | 158 | 23 | 0.72308 | 0.23481 | LRRC18 | 0.23481 | 0.17155 | 0.22076 |
| ENSG00000100129 | 22 | 38244875 | 38285414 | 56 | 8 | 0.72297 | 0.23485 | EIF3L | 0.23485 | 0.23972 | 0.2635 |
| ENSG00000141551 | 17 | 80196899 | 80231607 | 38 | 19 | 0.72252 | 0.23499 | CSNK1D | 0.23499 | 0.44996 | 0.69002 |
| ENSG00000198399 | 2 | 24425733 | 24583583 | 393 | 19 | 0.72198 | 0.23515 | ITSN2 | 0.23515 | 0.23061 | 0.26027 |
| ENSG00000105176 | 19 | 30414551 | 30506611 | 184 | 18 | 0.72184 | 0.2352 | URI1 | 0.2352 | 0.30264 | 0.24105 |
| ENSG00000196187 | 1 | 226033237 | 226070069 | 131 | 15 | 0.72135 | 0.23535 | TMEM63A | 0.23535 | 0.27471 | 0.22994 |
| ENSG00000198836 | 3 | 193310933 | 193415612 | 303 | 14 | 0.72089 | 0.23549 | OPA1 | 0.23549 | 0.25841 | 0.28177 |
| ENSG00000078900 | 1 | 3569084 | 3652765 | 367 | 61 | 0.7207 | 0.23555 | TP73 | 0.23555 | 0.29146 | 0.27158 |
| ENSG00000179564 | 3 | 50316458 | 50325545 | 4 | 1 | 0.72051 | 0.23561 | LSMEM2 | 0.23561 | 0.22607 | 0.24512 |
| ENSG00000075399 | 16 | 89773542 | 89787394 | 54 | 18 | 0.72004 | 0.23575 | VPS9D1 | 0.23575 | 0.76936 | 0.74955 |
| ENSG00000215595 | 20 | 1184098 | 1188918 | 12 | 4 | 0.71956 | 0.2359 | C20orf202 | 0.2359 | 0.16834 | 0.14699 |
| ENSG00000003056 | 12 | 9092959 | 9102551 | 27 | 7 | 0.71932 | 0.23597 | M6PR | 0.23597 | 0.41381 | 0.38885 |
| ENSG00000188707 | 7 | 150026938 | 150029808 | 4 | 1 | 0.71909 | 0.23604 | ZBED6CL | 0.23604 | 0.22745 | 0.34084 |
| ENSG00000157693 | 9 | 117373486 | 117408702 | 176 | 16 | 0.71909 | 0.23604 | C9orf91 | 0.23604 | 0.26638 | 0.30201 |
| ENSG00000214944 | 5 | 72921983 | 73237818 | 1136 | 71 | 0.71905 | 0.23605 | ARHGEF28 | 0.23605 | 0.24287 | 0.30286 |
| ENSG00000158874 | 1 | 161192082 | 161193421 | 1 | 1 | 0.7189 | 0.2361 | APOA2 | 0.2361 | 0.7209 | 0.83906 |
| ENSG00000140497 | 15 | 75136071 | 75165706 | 44 | 12 | 0.71845 | 0.23624 | SCAMP2 | 0.23624 | 0.31562 | 0.34195 |
| ENSG00000116745 | 1 | 68894505 | 68915642 | 76 | 14 | 0.71835 | 0.23627 | RPE65 | 0.23627 | 0.28071 | 0.34741 |
| ENSG00000142973 | 1 | 47223510 | 47285085 | 191 | 29 | 0.7183 | 0.23628 | CYP4B1 | 0.23628 | 0.1948 | 0.10429 |
| ENSG00000164181 | 5 | 60047618 | 60140216 | 312 | 25 | 0.71827 | 0.23629 | ELOVL7 | 0.23629 | 0.27797 | 0.28405 |

| | | | | | | | | | | | |
|---|---|---|---|---|---|---|---|---|---|---|---|
| ENSG00000144407 | 2 | 209224438 | 209719227 | 1741 | 72 | 0.71827 | 0.2363 | PTH2R | 0.2363 | 0.27507 | 0.29716 |
| ENSG00000213928 | 14 | 24630262 | 24635774 | 7 | 4 | 0.71821 | 0.23631 | IRF9 | 0.23631 | 0.51103 | 0.66542 |
| ENSG00000255339 | 10 | 102265385 | 102289638 | 16 | 7 | 0.71793 | 0.2364 | NDUFB8 | 0.2364 | 0.14079 | 0.29712 |
| ENSG00000119801 | 2 | 30369807 | 30383399 | 30 | 11 | 0.71747 | 0.23654 | YPEL5 | 0.23654 | 0.10564 | 0.23462 |
| ENSG00000131462 | 17 | 40761694 | 40767252 | 4 | 1 | 0.71744 | 0.23655 | TUBG1 | 0.23655 | 0.2648 | 0.26803 |
| ENSG00000154262 | 17 | 67074843 | 67138029 | 137 | 13 | 0.71614 | 0.23695 | ABCA6 | 0.23695 | 0.24261 | 0.29094 |
| ENSG00000169016 | 2 | 11584501 | 11606297 | 76 | 7 | 0.71589 | 0.23703 | E2F6 | 0.23703 | 0.20669 | 0.16674 |
| ENSG00000062485 | 12 | 56665483 | 56694176 | 60 | 7 | 0.71585 | 0.23704 | CS | 0.23704 | 0.19505 | 0.18878 |
| ENSG00000142794 | 1 | 21766621 | 21811498 | 215 | 19 | 0.71538 | 0.23719 | NBPF3 | 0.23719 | 0.24174 | 0.35011 |
| ENSG00000123892 | 11 | 87846410 | 87908635 | 259 | 25 | 0.7153 | 0.23721 | RAB38 | 0.23721 | 0.19708 | 0.17796 |
| ENSG00000158406 | 6 | 26281283 | 26285762 | 16 | 5 | 0.71499 | 0.23731 | HIST1H4H | 0.23731 | 0.35131 | 0.2838 |
| ENSG00000136653 | 1 | 206680879 | 206762616 | 228 | 34 | 0.71482 | 0.23736 | RASSF5 | 0.23736 | 0.23165 | 0.19794 |
| ENSG00000113600 | 5 | 39284364 | 39424970 | 456 | 40 | 0.71477 | 0.23737 | C9 | 0.23737 | 0.27997 | 0.23975 |
| ENSG00000141699 | 17 | 40731531 | 40762641 | 64 | 8 | 0.71474 | 0.23739 | FAM134C | 0.23739 | 0.23223 | 0.23718 |
| ENSG00000163041 | 1 | 226249552 | 226259702 | 19 | 6 | 0.71464 | 0.23742 | H3F3A | 0.23742 | 0.40617 | 0.52443 |
| ENSG00000172113 | 3 | 48334754 | 48343175 | 17 | 2 | 0.71447 | 0.23747 | NME6 | 0.23747 | 0.27834 | 0.27454 |
| ENSG00000253350 | 8 | 105602961 | 106431488 | 2415 | 122 | 0.71441 | 0.23749 | RP11-127H5.1 | 0.23749 | 0.24869 | 0.28325 |
| ENSG00000164296 | 5 | 149372681 | 149380730 | 20 | 5 | 0.71433 | 0.23751 | TIGD6 | 0.23751 | 0.22989 | 0.21432 |
| ENSG00000262660 | 17 | 79670401 | 79687569 | 82 | 11 | 0.71424 | 0.23754 | SLC25A10 | 0.23754 | 0.25243 | 0.31987 |
| ENSG00000102466 | 13 | 102372134 | 103054124 | 2207 | 122 | 0.71408 | 0.23759 | FGF14 | 0.23759 | 0.13584 | 0.1323 |
| ENSG00000176788 | 5 | 17065707 | 17276943 | 825 | 80 | 0.71373 | 0.2377 | BASP1 | 0.2377 | 0.27683 | 0.35323 |
| ENSG00000099804 | 19 | 531714 | 542097 | 79 | 12 | 0.71369 | 0.23771 | CDC34 | 0.23771 | 0.19474 | 0.28575 |
| ENSG00000180658 | 6 | 132021520 | 132022635 | 3 | 1 | 0.71341 | 0.2378 | OR2A4 | 0.2378 | 0.48692 | 0.43481 |
| ENSG00000164543 | 7 | 43622357 | 43666385 | 165 | 13 | 0.71324 | 0.23785 | STK17A | 0.23785 | 0.21105 | 0.22498 |
| ENSG00000139154 | 12 | 19556979 | 19873735 | 1771 | 66 | 0.71318 | 0.23787 | AEBP2 | 0.23787 | 0.2641 | 0.31732 |
| ENSG00000139053 | 12 | 15125956 | 15134799 | 35 | 8 | 0.71308 | 0.2379 | PDE6H | 0.2379 | 0.30528 | 0.18562 |
| ENSG00000116793 | 1 | 114239453 | 114302111 | 124 | 14 | 0.71278 | 0.23799 | PHTF1 | 0.23799 | 0.22296 | 0.19724 |
| ENSG00000172171 | 17 | 29224354 | 29233838 | 33 | 4 | 0.71241 | 0.23811 | TEFM | 0.23811 | 0.27304 | 0.30389 |
| ENSG00000147654 | 8 | 110551940 | 110578225 | 54 | 13 | 0.71199 | 0.23824 | EBAG9 | 0.23824 | 0.39423 | 0.57132 |
| ENSG00000122642 | 7 | 32997017 | 33046543 | 198 | 33 | 0.71176 | 0.23831 | FKBP9 | 0.23831 | 0.33956 | 0.41077 |
| ENSG00000177951 | 11 | 167784 | 207428 | 73 | 20 | 0.71133 | 0.23844 | BET1L | 0.23844 | 0.19956 | 0.30747 |
| ENSG00000041982 | 9 | 117782806 | 117880536 | 357 | 36 | 0.71121 | 0.23848 | TNC | 0.23848 | 0.21652 | 0.21315 |
| ENSG00000145777 | 5 | 110405760 | 110413722 | 15 | 5 | 0.71114 | 0.2385 | TSLP | 0.2385 | 0.21008 | 0.29444 |
| ENSG00000157150 | 3 | 12194551 | 12200851 | 24 | 7 | 0.71097 | 0.23855 | TIMP4 | 0.23855 | 0.081843 | 0.087224 |
| ENSG00000125434 | 17 | 8191081 | 8198661 | 21 | 8 | 0.71083 | 0.2386 | SLC25A35 | 0.2386 | 0.56943 | 0.52655 |
| ENSG00000176170 | 17 | 74372665 | 74383941 | 46 | 16 | 0.71073 | 0.23862 | SPHK1 | 0.23862 | 0.55211 | 0.59419 |
| ENSG00000115380 | 2 | 56093102 | 56151274 | 182 | 33 | 0.71065 | 0.23865 | EFEMP1 | 0.23865 | 0.27673 | 0.14151 |
| ENSG00000074966 | 4 | 48068410 | 48136273 | 200 | 23 | 0.71062 | 0.23866 | TXK | 0.23866 | 0.2582 | 0.23936 |
| ENSG00000198092 | 4 | 68918916 | 68995598 | 187 | 13 | 0.71057 | 0.23868 | TMPRSS11F | 0.23868 | 0.31556 | 0.37952 |
| ENSG00000128652 | 2 | 177001340 | 177037830 | 90 | 15 | 0.7104 | 0.23873 | HOXD3 | 0.23873 | 0.20284 | 0.30203 |
| ENSG00000173171 | 1 | 155178490 | 155183615 | 7 | 3 | 0.71024 | 0.23878 | MTX1 | 0.23878 | 0.23045 | 0.19705 |
| ENSG00000090447 | 16 | 4307187 | 4323076 | 32 | 7 | 0.7102 | 0.23879 | TFAP4 | 0.23879 | 0.28925 | 0.21604 |
| ENSG00000137767 | 15 | 45923346 | 45983492 | 287 | 37 | 0.71 | 0.23885 | SQRDL | 0.23885 | 0.38465 | 0.29757 |
| ENSG00000188501 | 15 | 66839517 | 66858317 | 45 | 12 | 0.70988 | 0.23889 | LCTL | 0.23889 | 0.18132 | 0.28298 |
| ENSG00000164330 | 5 | 158122928 | 158526769 | 890 | 65 | 0.70976 | 0.23893 | EBF1 | 0.23893 | 0.22474 | 0.221 |
| ENSG00000142731 | 4 | 128802016 | 128820350 | 28 | 6 | 0.70965 | 0.23896 | PLK4 | 0.23896 | 0.25091 | 0.18992 |
| ENSG00000175497 | 2 | 115199876 | 116603328 | 3820 | 136 | 0.70947 | 0.23902 | DPP10 | 0.23902 | 0.23547 | 0.23123 |
| ENSG00000196119 | 11 | 124439893 | 124441037 | 2 | 1 | 0.70945 | 0.23902 | OR8A1 | 0.23902 | 0.35171 | 0.28608 |
| ENSG00000004534 | 3 | 49977440 | 50137478 | 391 | 19 | 0.70918 | 0.23911 | RBM6 | 0.23911 | 0.24028 | 0.23396 |
| ENSG00000167157 | 9 | 132427920 | 132484875 | 247 | 34 | 0.70911 | 0.23913 | PRRX2 | 0.23913 | 0.21725 | 0.19198 |
| ENSG00000115956 | 2 | 68592305 | 68624585 | 185 | 16 | 0.7089 | 0.23919 | PLEK | 0.23919 | 0.29643 | 0.3756 |
| ENSG00000183644 | 11 | 111385510 | 111407756 | 38 | 7 | 0.70889 | 0.23919 | C11orf88 | 0.23919 | 0.397 | 0.42797 |
| ENSG00000090487 | 15 | 65255362 | 65282648 | 72 | 9 | 0.70884 | 0.23921 | SPG21 | 0.23921 | 0.34442 | 0.40201 |
| ENSG00000173409 | 1 | 231114727 | 231136341 | 82 | 12 | 0.70867 | 0.23926 | ARV1 | 0.23926 | 0.32898 | 0.25447 |
| ENSG00000132781 | 1 | 45794835 | 45806142 | 18 | 6 | 0.70794 | 0.23949 | MUTYH | 0.23949 | 0.27803 | 0.31494 |
| ENSG00000204510 | 1 | 12976450 | 12980569 | 1 | 1 | 0.70791 | 0.2395 | PRAMEF7 | 0.2395 | 0.34609 | 0.75563 |
| ENSG00000168843 | 4 | 162305049 | 163085187 | 3172 | 113 | 0.70775 | 0.23955 | FSTL5 | 0.23955 | 0.24802 | 0.22251 |
| ENSG00000133665 | 10 | 82104501 | 82127829 | 83 | 12 | 0.70761 | 0.23959 | DYDC2 | 0.23959 | 0.28094 | 0.18812 |
| ENSG00000169857 | 15 | 34158428 | 34331377 | 873 | 47 | 0.7076 | 0.2396 | AVEN | 0.2396 | 0.27615 | 0.25957 |
| ENSG00000105355 | 19 | 4838353 | 4867780 | 162 | 29 | 0.70749 | 0.23963 | PLIN3 | 0.23963 | 0.15615 | 0.056278 |
| ENSG00000153930 | 17 | 54188319 | 54589020 | 1001 | 57 | 0.70745 | 0.23964 | ANKFN1 | 0.23964 | 0.1975 | 0.21297 |
| ENSG00000155755 | 2 | 202484907 | 202508293 | 40 | 11 | 0.70704 | 0.23977 | TMEM237 | 0.23977 | 0.39867 | 0.51522 |
| ENSG00000133619 | 7 | 149411872 | 149431664 | 61 | 7 | 0.70697 | 0.23979 | KRBA1 | 0.23979 | 0.29638 | 0.12093 |
| ENSG00000188869 | 15 | 81623558 | 81666554 | 179 | 17 | 0.7067 | 0.23988 | TMC3 | 0.23988 | 0.25329 | 0.19044 |
| ENSG00000163840 | 3 | 122283085 | 122294050 | 33 | 16 | 0.70653 | 0.23993 | DTX3L | 0.23993 | 0.13127 | 0.040899 |
| ENSG00000256391 | 6 | 166307330 | 166309725 | 16 | 3 | 0.70623 | 0.24002 | SDIM1 | 0.24002 | 0.21535 | 0.36257 |
| ENSG00000103546 | 16 | 55689516 | 55740104 | 188 | 35 | 0.70616 | 0.24004 | SLC6A2 | 0.24004 | 0.30468 | 0.22652 |
| ENSG00000222011 | 7 | 102389418 | 102449672 | 183 | 23 | 0.70604 | 0.24008 | FAM185A | 0.24008 | 0.37928 | 0.36235 |
| ENSG00000150961 | 4 | 119643978 | 119759838 | 452 | 24 | 0.70578 | 0.24016 | SEC24D | 0.24016 | 0.29492 | 0.30625 |
| ENSG00000205869 | 11 | 1605572 | 1606513 | 2 | 1 | 0.70571 | 0.24018 | KRTAP5-1 | 0.24018 | 0.36399 | 0.25718 |
| ENSG00000134825 | 11 | 61535973 | 61560274 | 37 | 8 | 0.70532 | 0.24031 | TMEM258 | 0.24031 | 0.29486 | 0.44548 |
| ENSG00000266728 | 17 | 25950390 | 25967803 | 44 | 5 | 0.70504 | 0.24039 | AC015688.3 | 0.24039 | 0.3228 | 0.34451 |
| ENSG00000176125 | 7 | 100486346 | 100487339 | 1 | 1 | 0.70502 | 0.2404 | UFSP1 | 0.2404 | 0.43749 | 0.37464 |

| | | | | | | | | | | | |
|---|---|---|---|---|---|---|---|---|---|---|---|
| ENSG00000186567 | 19 | 45165545 | 45187631 | 35 | 11 | 0.70456 | 0.24054 | CEACAM19 | 0.24054 | 0.40982 | 0.50691 |
| ENSG00000137509 | 11 | 82534544 | 82681626 | 450 | 25 | 0.70427 | 0.24063 | PRCP | 0.24063 | 0.23456 | 0.24573 |
| ENSG00000161298 | 19 | 37095719 | 37119499 | 46 | 7 | 0.70425 | 0.24064 | ZNF382 | 0.24064 | 0.16927 | 0.15885 |
| ENSG00000149806 | 11 | 64888100 | 64889945 | 4 | 2 | 0.70419 | 0.24066 | FAU | 0.24066 | 0.4996 | 0.67606 |
| ENSG00000167657 | 19 | 3958451 | 3971121 | 29 | 6 | 0.70416 | 0.24067 | DAPK3 | 0.24067 | 0.052749 | 0.039252 |
| ENSG00000118922 | 13 | 74260226 | 74708394 | 1561 | 139 | 0.70393 | 0.24074 | KLF12 | 0.24074 | 0.22183 | 0.23219 |
| ENSG00000130227 | 8 | 21777180 | 21864096 | 221 | 17 | 0.70384 | 0.24077 | XPO7 | 0.24077 | 0.24294 | 0.10476 |
| ENSG00000257062 | 12 | 21200113 | 21329829 | 585 | 28 | 0.70382 | 0.24077 | RP11-125O5.2 | 0.24077 | 0.23598 | 0.24393 |
| ENSG00000133980 | 14 | 74769772 | 74826711 | 150 | 25 | 0.70377 | 0.24079 | VRTN | 0.24079 | 0.23143 | 0.27021 |
| ENSG00000129925 | 16 | 420773 | 437113 | 116 | 16 | 0.7034 | 0.2409 | TMEM8A | 0.2409 | 0.16112 | 0.16695 |
| ENSG00000124827 | 6 | 10873456 | 10882174 | 20 | 7 | 0.70319 | 0.24097 | GCM2 | 0.24097 | 0.20328 | 0.30558 |
| ENSG00000197380 | 19 | 47150869 | 47164395 | 29 | 8 | 0.70298 | 0.24103 | DACT3 | 0.24103 | 0.49269 | 0.64256 |
| ENSG00000100116 | 22 | 38203912 | 38213183 | 28 | 5 | 0.70277 | 0.2411 | GCAT | 0.2411 | 0.16525 | 0.27794 |
| ENSG00000122133 | 9 | 138453602 | 138458801 | 22 | 5 | 0.70222 | 0.24127 | PAEP | 0.24127 | 0.16817 | 0.235 |
| ENSG00000239282 | 22 | 30681106 | 30685616 | 10 | 5 | 0.70213 | 0.2413 | GATSL3 | 0.2413 | 0.5749 | 0.69208 |
| ENSG00000138131 | 10 | 100007447 | 100028007 | 55 | 8 | 0.70125 | 0.24157 | LOXL4 | 0.24157 | 0.29724 | 0.19162 |
| ENSG00000197808 | 19 | 37128094 | 37157755 | 71 | 11 | 0.70094 | 0.24167 | ZNF461 | 0.24167 | 0.19058 | 0.21633 |
| ENSG00000196167 | 11 | 111164114 | 111175770 | 42 | 8 | 0.70092 | 0.24168 | COLCA1 | 0.24168 | 0.20919 | 0.23185 |
| ENSG00000204308 | 6 | 32146131 | 32151930 | 18 | 5 | 0.70048 | 0.24182 | RNF5 | 0.24182 | 0.053163 | 0.03145 |
| ENSG00000173285 | 1 | 158435272 | 158436344 | 3 | 1 | 0.70024 | 0.24189 | OR10K1 | 0.24189 | 0.20023 | 0.19196 |
| ENSG00000198742 | 7 | 98625061 | 98741723 | 264 | 39 | 0.70002 | 0.24196 | SMURF1 | 0.24196 | 0.39166 | 0.50701 |
| ENSG00000149531 | 20 | 29611857 | 29634010 | 24 | 5 | 0.69999 | 0.24197 | FRG1B | 0.24197 | 0.19745 | 0.19048 |
| ENSG00000169231 | 1 | 155165379 | 155178842 | 20 | 6 | 0.69993 | 0.24198 | THBS3 | 0.24198 | 0.26639 | 0.14956 |
| ENSG00000127418 | 4 | 1003724 | 1020685 | 76 | 15 | 0.69978 | 0.24203 | FGFRL1 | 0.24203 | 0.2041 | 0.22537 |
| ENSG00000110651 | 11 | 2397407 | 2418649 | 54 | 9 | 0.69976 | 0.24204 | CD81 | 0.24204 | 0.35858 | 0.58223 |
| ENSG00000182511 | 15 | 91426925 | 91439006 | 37 | 7 | 0.69977 | 0.24204 | FES | 0.24204 | 0.31099 | 0.10529 |
| ENSG00000185666 | 22 | 32908539 | 33454358 | 1979 | 129 | 0.69965 | 0.24207 | SYN3 | 0.24207 | 0.27738 | 0.29042 |
| ENSG00000173566 | 8 | 21964383 | 21966932 | 8 | 3 | 0.69942 | 0.24214 | NUDT18 | 0.24214 | 0.33602 | 0.31023 |
| ENSG00000109943 | 11 | 122709208 | 122743347 | 111 | 27 | 0.69912 | 0.24224 | CRTAM | 0.24224 | 0.24357 | 0.33441 |
| ENSG00000142733 | 1 | 27681675 | 27693383 | 31 | 10 | 0.69902 | 0.24227 | MAP3K6 | 0.24227 | 0.15282 | 0.27692 |
| ENSG00000174255 | 3 | 113953483 | 113956425 | 18 | 7 | 0.69901 | 0.24227 | ZNF80 | 0.24227 | 0.22158 | 0.16437 |
| ENSG00000104835 | 19 | 39405906 | 39440495 | 122 | 13 | 0.6989 | 0.24231 | SARS2 | 0.24231 | 0.25469 | 0.26014 |
| ENSG00000269547 | 19 | 39406044 | 39440495 | 122 | 13 | 0.6989 | 0.24231 | CTC-360G5.8 | 0.24231 | 0.25469 | 0.26014 |
| ENSG00000144649 | 3 | 43020759 | 43101703 | 250 | 23 | 0.69877 | 0.24235 | FAM198A | 0.24235 | 0.26711 | 0.26799 |
| ENSG00000146049 | 6 | 24357131 | 24358512 | 4 | 2 | 0.69865 | 0.24238 | KAAG1 | 0.24238 | 0.15238 | 0.21206 |
| ENSG00000154342 | 1 | 228194752 | 228248961 | 119 | 15 | 0.6983 | 0.24249 | WNT3A | 0.24249 | 0.2406 | 0.24308 |
| ENSG00000059728 | 2 | 70124820 | 70170077 | 102 | 8 | 0.69833 | 0.24249 | MXD1 | 0.24249 | 0.26585 | 0.33156 |
| ENSG00000152034 | 6 | 100367786 | 100442123 | 218 | 32 | 0.69818 | 0.24253 | MCHR2 | 0.24253 | 0.16378 | 0.14198 |
| ENSG00000101290 | 20 | 5107432 | 5178533 | 222 | 24 | 0.69818 | 0.24253 | CDS2 | 0.24253 | 0.1774 | 0.14006 |
| ENSG00000093010 | 22 | 19929130 | 19957498 | 145 | 28 | 0.698 | 0.24259 | COMT | 0.24259 | 0.23595 | 0.40348 |
| ENSG00000125492 | 9 | 135457572 | 135465653 | 14 | 8 | 0.69781 | 0.24265 | BARHL1 | 0.24265 | 0.2521 | 0.34453 |
| ENSG00000196866 | 6 | 26199079 | 26199471 | 2 | 1 | 0.69769 | 0.24268 | HIST1H2AD | 0.24268 | 0.52444 | 0.51867 |
| ENSG00000186350 | 9 | 137208944 | 137332431 | 427 | 41 | 0.69767 | 0.24269 | RXRA | 0.24269 | 0.23315 | 0.23586 |
| ENSG00000171121 | 3 | 178957530 | 178984790 | 145 | 19 | 0.69765 | 0.2427 | KCNMB3 | 0.2427 | 0.28122 | 0.47799 |
| ENSG00000171431 | 17 | 39032193 | 39041479 | 31 | 8 | 0.69763 | 0.2427 | KRT20 | 0.2427 | 0.42502 | 0.76551 |
| ENSG00000114739 | 3 | 38495342 | 38534633 | 90 | 7 | 0.69747 | 0.24275 | ACVR2B | 0.24275 | 0.20176 | 0.24013 |
| ENSG00000184060 | 17 | 29233362 | 29286340 | 91 | 16 | 0.69713 | 0.24286 | ADAP2 | 0.24286 | 0.2489 | 0.28306 |
| ENSG00000037965 | 12 | 54402832 | 54407570 | 11 | 5 | 0.6971 | 0.24287 | HOXC8 | 0.24287 | 0.40701 | 0.40729 |
| ENSG00000254858 | 19 | 18303992 | 18307758 | 16 | 4 | 0.69705 | 0.24288 | MPV17L2 | 0.24288 | 0.34044 | 0.19491 |
| ENSG00000112195 | 6 | 41158015 | 41168932 | 44 | 10 | 0.69628 | 0.24313 | TREML2 | 0.24313 | 0.29308 | 0.23457 |
| ENSG00000172016 | 2 | 79384132 | 79386879 | 4 | 2 | 0.69595 | 0.24323 | REG3A | 0.24323 | 0.63375 | 0.48958 |
| ENSG00000181409 | 17 | 79091095 | 79139877 | 282 | 40 | 0.69592 | 0.24324 | AATK | 0.24324 | 0.13002 | 0.03449 |
| ENSG00000075651 | 3 | 171318195 | 171528740 | 618 | 21 | 0.69583 | 0.24327 | PLD1 | 0.24327 | 0.26367 | 0.33206 |
| ENSG00000174611 | 3 | 134321980 | 134370478 | 128 | 8 | 0.69541 | 0.2434 | KY | 0.2434 | 0.24384 | 0.26799 |
| ENSG00000149716 | 11 | 69467844 | 69490184 | 52 | 11 | 0.69534 | 0.24342 | ORAOV1 | 0.24342 | 0.33566 | 0.5109 |
| ENSG00000203618 | 22 | 19710468 | 19712294 | 1 | 1 | 0.69509 | 0.2435 | GP1BB | 0.2435 | 0.54603 | 0.50689 |
| ENSG00000149476 | 11 | 61100682 | 61120767 | 15 | 9 | 0.69491 | 0.24355 | DAK | 0.24355 | 0.22467 | 0.30359 |
| ENSG00000188629 | 19 | 9435021 | 9493293 | 175 | 11 | 0.69472 | 0.24362 | ZNF177 | 0.24362 | 0.25441 | 0.24759 |
| ENSG00000082068 | 5 | 37379314 | 37753537 | 1028 | 28 | 0.69438 | 0.24372 | WDR70 | 0.24372 | 0.24445 | 0.26462 |
| ENSG00000171303 | 2 | 26915619 | 26956288 | 94 | 14 | 0.69398 | 0.24385 | KCNK3 | 0.24385 | 0.30054 | 0.36165 |
| ENSG00000103647 | 15 | 68871308 | 69020145 | 497 | 59 | 0.69365 | 0.24395 | CORO2B | 0.24395 | 0.27422 | 0.38415 |
| ENSG00000135315 | 6 | 84833960 | 84937353 | 246 | 19 | 0.69358 | 0.24397 | KIAA1009 | 0.24397 | 0.23425 | 0.26406 |
| ENSG00000138175 | 10 | 104433488 | 104474164 | 86 | 15 | 0.69343 | 0.24402 | ARL3 | 0.24402 | 0.31655 | 0.45468 |
| ENSG00000162630 | 1 | 193148175 | 193155784 | 10 | 3 | 0.69304 | 0.24414 | B3GALT2 | 0.24414 | 0.13849 | 0.1504 |
| ENSG00000127311 | 12 | 66696325 | 66737423 | 58 | 15 | 0.6929 | 0.24419 | HELB | 0.24419 | 0.10319 | 0.08534 |
| ENSG00000188886 | 2 | 96789589 | 96804175 | 11 | 6 | 0.69253 | 0.2443 | ASTL | 0.2443 | 0.22896 | 0.21641 |
| ENSG00000168404 | 16 | 74705753 | 74734858 | 100 | 28 | 0.69233 | 0.24436 | MLKL | 0.24436 | 0.30443 | 0.29709 |
| ENSG00000108788 | 17 | 40719086 | 40725257 | 13 | 3 | 0.69228 | 0.24438 | MLX | 0.24438 | 0.20328 | 0.29576 |
| ENSG00000068028 | 3 | 50367219 | 50378411 | 17 | 6 | 0.69222 | 0.2444 | RASSF1 | 0.2444 | 0.36227 | 0.37187 |
| ENSG00000152779 | 10 | 91190051 | 91316398 | 399 | 49 | 0.69221 | 0.2444 | SLC16A12 | 0.2444 | 0.16129 | 0.09412 |
| ENSG00000176136 | 18 | 13824154 | 13826861 | 23 | 4 | 0.69206 | 0.24445 | MC5R | 0.24445 | 0.32172 | 0.37256 |
| ENSG00000114127 | 3 | 142025449 | 142166904 | 241 | 22 | 0.69202 | 0.24446 | XRN1 | 0.24446 | 0.22426 | 0.20631 |
| ENSG00000142230 | 19 | 47616531 | 47713886 | 114 | 21 | 0.69186 | 0.24451 | SAE1 | 0.24451 | 0.23925 | 0.14718 |

| | | | | | | | | | | | |
|---|---|---|---|---|---|---|---|---|---|---|---|
| ENSG00000170962 | 11 | 103777914 | 104035107 | 1095 | 67 | 0.69179 | 0.24454 | PDGFD | 0.24454 | 0.24293 | 0.24973 |
| ENSG00000213416 | 17 | 39279343 | 39280419 | 5 | 2 | 0.69113 | 0.24474 | KRTAP4-12 | 0.24474 | 0.18996 | 0.22072 |
| ENSG00000101751 | 18 | 51795774 | 51847636 | 142 | 12 | 0.69107 | 0.24476 | POLI | 0.24476 | 0.2166 | 0.19868 |
| ENSG00000100060 | 22 | 37865101 | 37882439 | 32 | 10 | 0.69077 | 0.24485 | MFNG | 0.24485 | 0.13705 | 0.044328 |
| ENSG00000270011 | 19 | 9434850 | 9493293 | 177 | 11 | 0.69055 | 0.24492 | ZNF559-ZNF177 | 0.24492 | 0.2553 | 0.24759 |
| ENSG00000120820 | 12 | 104382762 | 104457961 | 273 | 32 | 0.69043 | 0.24496 | GLT8D2 | 0.24496 | 0.21751 | 0.094345 |
| ENSG00000182053 | 11 | 49050504 | 49059579 | 56 | 7 | 0.69039 | 0.24497 | TRIM49B | 0.24497 | 0.28141 | 0.26365 |
| ENSG00000107951 | 10 | 30598730 | 30663377 | 242 | 24 | 0.6901 | 0.24507 | MTPAP | 0.24507 | 0.34105 | 0.24033 |
| ENSG00000188163 | 9 | 140138036 | 140142222 | 8 | 3 | 0.68993 | 0.24512 | FAM166A | 0.24512 | 0.59145 | 0.44377 |
| ENSG00000234602 | 5 | 54515442 | 54523143 | 17 | 4 | 0.68984 | 0.24515 | MCIDAS | 0.24515 | 0.26542 | 0.23718 |
| ENSG00000232263 | 21 | 31661463 | 31661832 | 2 | 1 | 0.6896 | 0.24522 | KRTAP25-1 | 0.24522 | 0.26264 | 0.30949 |
| ENSG00000166619 | 20 | 36120874 | 36156333 | 54 | 7 | 0.68945 | 0.24527 | BLCAP | 0.24527 | 0.2021 | 0.22421 |
| ENSG00000253117 | 8 | 133036467 | 133071627 | 141 | 16 | 0.68917 | 0.24536 | OC90 | 0.24536 | 0.26562 | 0.30812 |
| ENSG00000253457 | 8 | 30496117 | 30503581 | 13 | 3 | 0.68901 | 0.24541 | SMIM18 | 0.24541 | 0.17208 | 0.15386 |
| ENSG00000170367 | 20 | 23856572 | 23860387 | 16 | 3 | 0.68896 | 0.24543 | CST5 | 0.24543 | 0.19019 | 0.14097 |
| ENSG00000092330 | 14 | 24708849 | 24711880 | 1 | 1 | 0.68872 | 0.2455 | TINF2 | 0.2455 | 0.88943 | 0.91187 |
| ENSG00000189227 | 15 | 67813406 | 67819628 | 9 | 3 | 0.68868 | 0.24551 | C15orf61 | 0.24551 | 0.2036 | 0.16061 |
| ENSG00000197933 | 19 | 11832080 | 11849824 | 68 | 8 | 0.68858 | 0.24554 | ZNF823 | 0.24554 | 0.48975 | 0.48406 |
| ENSG00000159239 | 2 | 74641304 | 74648718 | 5 | 3 | 0.68852 | 0.24556 | C2orf81 | 0.24556 | 0.18796 | 0.43059 |
| ENSG00000140264 | 15 | 44069285 | 44094787 | 46 | 7 | 0.6884 | 0.2456 | SERF2 | 0.2456 | 0.17563 | 0.19549 |
| ENSG00000173915 | 10 | 105148798 | 105156223 | 9 | 3 | 0.68808 | 0.2457 | USMG5 | 0.2457 | 0.37742 | 0.26183 |
| ENSG00000178741 | 15 | 75212132 | 75230509 | 47 | 7 | 0.68798 | 0.24573 | COX5A | 0.24573 | 0.20432 | 0.28369 |
| ENSG00000113643 | 5 | 167913450 | 167946304 | 182 | 13 | 0.68797 | 0.24574 | RARS | 0.24574 | 0.24398 | 0.27636 |
| ENSG00000151746 | 12 | 32259769 | 32536567 | 1396 | 60 | 0.68742 | 0.24591 | BICD1 | 0.24591 | 0.24534 | 0.21638 |
| ENSG00000152527 | 2 | 43864412 | 43995126 | 539 | 40 | 0.68739 | 0.24592 | PLEKHH2 | 0.24592 | 0.20785 | 0.19741 |
| ENSG00000244482 | 19 | 54720737 | 54746649 | 241 | 35 | 0.68724 | 0.24596 | LILRA6 | 0.24596 | 0.30335 | 0.36291 |
| ENSG00000145817 | 5 | 143537723 | 143550278 | 56 | 9 | 0.68723 | 0.24597 | YIPF5 | 0.24597 | 0.25094 | 0.21986 |
| ENSG00000156603 | 11 | 57471186 | 57479693 | 11 | 5 | 0.68707 | 0.24602 | MED19 | 0.24602 | 0.38691 | 0.32306 |
| ENSG00000139631 | 12 | 53551447 | 53575135 | 42 | 6 | 0.6869 | 0.24607 | CSAD | 0.24607 | 0.29127 | 0.28113 |
| ENSG00000237541 | 6 | 32709119 | 32714992 | 97 | 11 | 0.68657 | 0.24618 | HLA-DQA2 | 0.24618 | 0.19135 | 0.15238 |
| ENSG00000131969 | 14 | 51338878 | 51371688 | 93 | 15 | 0.68624 | 0.24628 | ABHD12B | 0.24628 | 0.23107 | 0.2799 |
| ENSG00000104312 | 8 | 90769975 | 90803291 | 69 | 12 | 0.68609 | 0.24633 | RIPK2 | 0.24633 | 0.30831 | 0.41413 |
| ENSG00000204576 | 6 | 30524663 | 30531500 | 21 | 9 | 0.68592 | 0.24638 | PRR3 | 0.24638 | 0.10343 | 0.10769 |
| ENSG00000167695 | 17 | 635652 | 646206 | 51 | 12 | 0.68583 | 0.24641 | FAM57A | 0.24641 | 0.23732 | 0.24039 |
| ENSG00000133226 | 1 | 24958207 | 24999758 | 55 | 10 | 0.68567 | 0.24646 | SRRM1 | 0.24646 | 0.31492 | 0.23208 |
| ENSG00000197887 | 11 | 57970674 | 57971653 | 4 | 2 | 0.68536 | 0.24656 | OR1S2 | 0.24656 | 0.097684 | 0.099412 |
| ENSG00000106638 | 7 | 72983262 | 72993121 | 27 | 8 | 0.68526 | 0.24659 | TBL2 | 0.24659 | 0.22446 | 0.27808 |
| ENSG00000266086 | 17 | 56066399 | 56082614 | 32 | 6 | 0.68497 | 0.24668 | RP11-159D12.5 | 0.24668 | 0.18474 | 0.2148 |
| ENSG00000115607 | 2 | 103035149 | 103069025 | 158 | 16 | 0.68458 | 0.2468 | IL18RAP | 0.2468 | 0.23809 | 0.21903 |
| ENSG00000139370 | 12 | 129277739 | 129308528 | 134 | 10 | 0.68455 | 0.24681 | SLC15A4 | 0.24681 | 0.096828 | 0.052846 |
| ENSG00000136628 | 1 | 220141943 | 220220000 | 360 | 15 | 0.68436 | 0.24687 | EPRS | 0.24687 | 0.22844 | 0.25233 |
| ENSG00000152503 | 5 | 114460459 | 114516243 | 151 | 21 | 0.68437 | 0.24687 | TRIM36 | 0.24687 | 0.18482 | 0.24964 |
| ENSG00000100767 | 14 | 73704205 | 73741348 | 135 | 28 | 0.68416 | 0.24694 | PAPLN | 0.24694 | 0.15929 | 0.070966 |
| ENSG00000237435 | 1 | 96403457 | 96488436 | 318 | 23 | 0.68335 | 0.24719 | RP11-147C23.1 | 0.24719 | 0.20287 | 0.186 |
| ENSG00000168594 | 4 | 175750819 | 175899331 | 765 | 25 | 0.68333 | 0.2472 | ADAM29 | 0.2472 | 0.25227 | 0.26628 |
| ENSG00000165494 | 11 | 82868030 | 82898493 | 60 | 7 | 0.68251 | 0.24746 | PCF11 | 0.24746 | 0.36665 | 0.34879 |
| ENSG00000070019 | 12 | 14765576 | 14849519 | 126 | 26 | 0.6824 | 0.24749 | GUCY2C | 0.24749 | 0.20628 | 0.23972 |
| ENSG00000177354 | 10 | 50507187 | 50535537 | 92 | 17 | 0.68239 | 0.2475 | C10orf71 | 0.2475 | 0.28835 | 0.19467 |
| ENSG00000168685 | 5 | 35852797 | 35879705 | 107 | 10 | 0.68233 | 0.24752 | IL7R | 0.24752 | 0.20027 | 0.1888 |
| ENSG00000070540 | 17 | 66417089 | 66453654 | 125 | 22 | 0.68232 | 0.24752 | WIPI1 | 0.24752 | 0.21996 | 0.14041 |
| ENSG00000182187 | 2 | 209007297 | 209010892 | 23 | 4 | 0.68228 | 0.24753 | CRYGB | 0.24753 | 0.67482 | 0.72164 |
| ENSG00000162971 | 2 | 200794698 | 200820459 | 61 | 12 | 0.68206 | 0.2476 | TYW5 | 0.2476 | 0.16761 | 0.11618 |
| ENSG00000197437 | 1 | 247835320 | 247836365 | 7 | 2 | 0.68188 | 0.24766 | OR13G1 | 0.24766 | 0.30166 | 0.37312 |
| ENSG00000186479 | 5 | 63802084 | 63908139 | 238 | 33 | 0.68172 | 0.24771 | RGS7BP | 0.24771 | 0.18961 | 0.20184 |
| ENSG00000115827 | 2 | 172290727 | 172341562 | 168 | 10 | 0.68161 | 0.24774 | DCAF17 | 0.24774 | 0.31423 | 0.34641 |
| ENSG00000161847 | 19 | 10426888 | 10444316 | 71 | 13 | 0.68089 | 0.24797 | RAVER1 | 0.24797 | 0.25616 | 0.28109 |
| ENSG00000162374 | 1 | 50513686 | 50669458 | 235 | 40 | 0.68053 | 0.24809 | ELAVL4 | 0.24809 | 0.26257 | 0.22415 |
| ENSG00000096092 | 6 | 52535907 | 52551386 | 74 | 10 | 0.68052 | 0.24809 | TMEM14A | 0.24809 | 0.20858 | 0.11107 |
| ENSG00000214491 | 22 | 30918786 | 30942669 | 110 | 13 | 0.68025 | 0.24817 | SEC14L6 | 0.24817 | 0.29874 | 0.36299 |
| ENSG00000143632 | 1 | 229566992 | 229569845 | 9 | 3 | 0.68024 | 0.24818 | ACTA1 | 0.24818 | 0.41729 | 0.32109 |
| ENSG00000169302 | 5 | 146614526 | 146767415 | 685 | 43 | 0.68014 | 0.24821 | STK32A | 0.24821 | 0.24089 | 0.26376 |
| ENSG00000152240 | 18 | 43684298 | 43708299 | 91 | 11 | 0.67983 | 0.24831 | HAUS1 | 0.24831 | 0.17587 | 0.10798 |
| ENSG00000186113 | 11 | 55563032 | 55563976 | 6 | 3 | 0.67965 | 0.24836 | OR5D14 | 0.24836 | 0.37551 | 0.42653 |
| ENSG00000118231 | 2 | 208986331 | 208989225 | 8 | 3 | 0.67925 | 0.24849 | CRYGD | 0.24849 | 0.37372 | 0.48295 |
| ENSG00000186367 | 5 | 129083772 | 129102425 | 72 | 11 | 0.67869 | 0.24867 | KIAA1024L | 0.24867 | 0.23307 | 0.27921 |
| ENSG00000107290 | 9 | 135136743 | 135230372 | 417 | 17 | 0.67859 | 0.2487 | SETX | 0.2487 | 0.20195 | 0.19355 |
| ENSG00000162460 | 1 | 16068917 | 16074477 | 29 | 7 | 0.67803 | 0.24888 | TMEM82 | 0.24888 | 0.096646 | 0.010197 |
| ENSG00000143570 | 1 | 153931575 | 153940188 | 14 | 5 | 0.67801 | 0.24888 | SLC39A1 | 0.24888 | 0.20854 | 0.24479 |
| ENSG00000112599 | 6 | 42152139 | 42162654 | 68 | 6 | 0.67788 | 0.24892 | GUCA1B | 0.24892 | 0.28668 | 0.35804 |
| ENSG00000182359 | 11 | 105921825 | 105948492 | 64 | 13 | 0.67781 | 0.24895 | KBTBD3 | 0.24895 | 0.40607 | 0.42229 |
| ENSG00000197919 | 9 | 21440440 | 21441315 | 4 | 2 | 0.67758 | 0.24902 | IFNA1 | 0.24902 | 0.55709 | 0.56659 |
| ENSG00000119508 | 9 | 102584137 | 102629173 | 53 | 11 | 0.67754 | 0.24903 | NR4A3 | 0.24903 | 0.22648 | 0.23273 |
| ENSG00000187994 | 19 | 39358470 | 39368919 | 26 | 6 | 0.67756 | 0.24903 | RINL | 0.24903 | 0.23311 | 0.17043 |

| | | | | | | | | | | | |
|---|---|---|---|---|---|---|---|---|---|---|---|
| ENSG00000205937 | 16 | 2303117 | 2318413 | 26 | 6 | 0.67722 | 0.24913 | RNPS1 | 0.24913 | 0.27058 | 0.41176 |
| ENSG00000107554 | 10 | 101635334 | 101769676 | 563 | 23 | 0.67691 | 0.24923 | DNMBP | 0.24923 | 0.28237 | 0.29336 |
| ENSG00000144021 | 2 | 96931870 | 96939087 | 8 | 4 | 0.67687 | 0.24925 | CIAO1 | 0.24925 | 0.21391 | 0.16411 |
| ENSG00000132530 | 17 | 6658766 | 6678966 | 62 | 14 | 0.67668 | 0.24931 | XAF1 | 0.24931 | 0.089391 | 0.026072 |
| ENSG00000178467 | 3 | 49027319 | 49044587 | 14 | 4 | 0.67659 | 0.24933 | P4HTM | 0.24933 | 0.37477 | 0.54771 |
| ENSG00000225526 | 3 | 12556433 | 12602558 | 192 | 19 | 0.67621 | 0.24945 | C3orf83 | 0.24945 | 0.17427 | 0.25044 |
| ENSG00000158711 | 1 | 205577071 | 205601090 | 65 | 11 | 0.67599 | 0.24952 | ELK4 | 0.24952 | 0.62683 | 0.86975 |
| ENSG00000125384 | 14 | 52781023 | 52795324 | 39 | 6 | 0.67597 | 0.24953 | PTGER2 | 0.24953 | 0.20817 | 0.34528 |
| ENSG00000171634 | 17 | 65821640 | 65980494 | 342 | 12 | 0.67574 | 0.2496 | BPTF | 0.2496 | 0.27635 | 0.29389 |
| ENSG00000197182 | 22 | 46449749 | 46509808 | 154 | 31 | 0.6753 | 0.24974 | FLJ27365 | 0.24974 | 0.22887 | 0.44123 |
| ENSG00000153253 | 2 | 165944032 | 166060577 | 320 | 22 | 0.67523 | 0.24977 | SCN3A | 0.24977 | 0.227 | 0.22823 |
| ENSG00000006788 | 17 | 10201401 | 10276447 | 300 | 31 | 0.67506 | 0.24982 | MYH13 | 0.24982 | 0.32376 | 0.23193 |
| ENSG00000113163 | 5 | 74664311 | 74807963 | 295 | 19 | 0.67472 | 0.24993 | COL4A3BP | 0.24993 | 0.28841 | 0.32692 |
| ENSG00000267710 | 19 | 56784138 | 56821819 | 136 | 20 | 0.67414 | 0.25011 | AC006116.20 | 0.25011 | 0.20456 | 0.083799 |
| ENSG00000172322 | 12 | 10103915 | 10148293 | 251 | 22 | 0.67405 | 0.25014 | CLEC12A | 0.25014 | 0.32541 | 0.26315 |
| ENSG00000103126 | 16 | 337440 | 402673 | 307 | 25 | 0.67391 | 0.25018 | AXIN1 | 0.25018 | 0.2558 | 0.2608 |
| ENSG00000100311 | 22 | 39619364 | 39640756 | 82 | 10 | 0.67384 | 0.25021 | PDGFB | 0.25021 | 0.27885 | 0.29691 |
| ENSG00000157036 | 3 | 38537618 | 38583652 | 109 | 12 | 0.67375 | 0.25023 | EXOG | 0.25023 | 0.2865 | 0.33056 |
| ENSG00000104826 | 19 | 49519237 | 49520338 | 11 | 3 | 0.67368 | 0.25026 | LHB | 0.25026 | 0.38848 | 0.19558 |
| ENSG00000113368 | 5 | 126112315 | 126172712 | 230 | 30 | 0.67362 | 0.25028 | LMNB1 | 0.25028 | 0.21537 | 0.24707 |
| ENSG00000112343 | 6 | 25963030 | 25987384 | 53 | 16 | 0.67333 | 0.25037 | TRIM38 | 0.25037 | 0.39574 | 0.44488 |
| ENSG00000171135 | 3 | 9932238 | 9936033 | 15 | 3 | 0.67325 | 0.2504 | JAGN1 | 0.2504 | 0.52104 | 0.6112 |
| ENSG00000189144 | 19 | 38226734 | 38307940 | 203 | 25 | 0.6731 | 0.25044 | ZNF573 | 0.25044 | 0.29688 | 0.3162 |
| ENSG00000147669 | 8 | 101162812 | 101166230 | 6 | 2 | 0.67303 | 0.25046 | POLR2K | 0.25046 | 0.53516 | 0.53248 |
| ENSG00000205176 | 8 | 86568695 | 86575726 | 4 | 2 | 0.67302 | 0.25047 | REXO1L1P | 0.25047 | 0.43682 | 0.6369 |
| ENSG00000198736 | 16 | 1988211 | 1993327 | 10 | 3 | 0.67291 | 0.2505 | MSRB1 | 0.2505 | 0.48669 | 0.90554 |
| ENSG00000135346 | 6 | 87795216 | 87804824 | 41 | 7 | 0.67258 | 0.25061 | CGA | 0.25061 | 0.35398 | 0.30495 |
| ENSG00000135314 | 6 | 73951037 | 74020088 | 304 | 14 | 0.67228 | 0.2507 | KHDC1 | 0.2507 | 0.23758 | 0.18583 |
| ENSG00000070756 | 8 | 101698044 | 101735037 | 95 | 12 | 0.67229 | 0.2507 | PABPC1 | 0.2507 | 0.27386 | 0.30053 |
| ENSG00000088038 | 19 | 54641444 | 54659419 | 61 | 13 | 0.67212 | 0.25075 | CNOT3 | 0.25075 | 0.1674 | 0.23673 |
| ENSG00000152804 | 10 | 94447945 | 94455403 | 14 | 4 | 0.67183 | 0.25084 | HHEX | 0.25084 | 0.28971 | 0.13742 |
| ENSG00000115596 | 2 | 219724544 | 219738955 | 22 | 6 | 0.67182 | 0.25085 | WNT6 | 0.25085 | 0.36983 | 0.45988 |
| ENSG00000243927 | 21 | 35445524 | 35515334 | 93 | 30 | 0.67147 | 0.25096 | MRPS6 | 0.25096 | 0.30931 | 0.46978 |
| ENSG00000186094 | 1 | 48998527 | 50489585 | 2530 | 56 | 0.67115 | 0.25106 | AGBL4 | 0.25106 | 0.25605 | 0.26214 |
| ENSG00000139540 | 12 | 56623833 | 56631630 | 11 | 2 | 0.67114 | 0.25106 | SLC39A5 | 0.25106 | 0.4306 | 0.39916 |
| ENSG00000132881 | 1 | 16558195 | 16563657 | 11 | 4 | 0.67109 | 0.25108 | RSG1 | 0.25108 | 0.13192 | 0.118 |
| ENSG00000182974 | 15 | 22368478 | 22369561 | 1 | 1 | 0.67103 | 0.2511 | OR4M2 | 0.2511 | 0.045756 | 0.032875 |
| ENSG00000173088 | 10 | 97667360 | 97698480 | 124 | 21 | 0.67025 | 0.25135 | C10orf131 | 0.25135 | 0.27295 | 0.2862 |
| ENSG00000101745 | 18 | 9136226 | 9285983 | 492 | 26 | 0.67024 | 0.25135 | ANKRD12 | 0.25135 | 0.28601 | 0.31072 |
| ENSG00000196176 | 6 | 26021907 | 26022278 | 1 | 1 | 0.67009 | 0.2514 | HIST1H4A | 0.2514 | 0.46169 | 0.3677 |
| ENSG00000176531 | 19 | 43979106 | 44009087 | 99 | 18 | 0.66988 | 0.25147 | PHLDB3 | 0.25147 | 0.18607 | 0.18545 |
| ENSG00000114019 | 3 | 134074716 | 134094321 | 43 | 8 | 0.66967 | 0.25154 | AMOTL2 | 0.25154 | 0.17601 | 0.13748 |
| ENSG00000162139 | 11 | 74699179 | 74729938 | 57 | 11 | 0.66964 | 0.25154 | NEU3 | 0.25154 | 0.19516 | 0.30996 |
| ENSG00000161217 | 3 | 195941093 | 196014828 | 237 | 22 | 0.66946 | 0.2516 | PCYT1A | 0.2516 | 0.31989 | 0.35364 |
| ENSG00000111886 | 6 | 89966927 | 90025018 | 209 | 45 | 0.66938 | 0.25163 | GABRR2 | 0.25163 | 0.50997 | 0.49456 |
| ENSG00000175040 | 3 | 142838173 | 142841800 | 4 | 2 | 0.66911 | 0.25171 | CHST2 | 0.25171 | 0.67328 | 0.81031 |
| ENSG00000146755 | 7 | 72726535 | 72742085 | 22 | 10 | 0.6688 | 0.25181 | TRIM50 | 0.25181 | 0.40361 | 0.50863 |
| ENSG00000226364 | 17 | 50939481 | 50976948 | 209 | 23 | 0.6688 | 0.25181 | AC102948.2 | 0.25181 | 0.38968 | 0.39727 |
| ENSG00000188730 | 7 | 49813257 | 49961546 | 387 | 36 | 0.66859 | 0.25188 | VWC2 | 0.25188 | 0.28453 | 0.23418 |
| ENSG00000237412 | 2 | 233385173 | 233390422 | 12 | 5 | 0.66856 | 0.25189 | PRSS56 | 0.25189 | 0.49352 | 0.20228 |
| ENSG00000099284 | 10 | 71812552 | 71872036 | 202 | 35 | 0.66857 | 0.25189 | H2AFY2 | 0.25189 | 0.2408 | 0.15035 |
| ENSG00000152683 | 2 | 32390933 | 32449448 | 179 | 20 | 0.66785 | 0.25212 | SLC30A6 | 0.25212 | 0.21301 | 0.27229 |
| ENSG00000122565 | 7 | 26240782 | 26252976 | 32 | 8 | 0.66765 | 0.25218 | CBX3 | 0.25218 | 0.28117 | 0.18587 |
| ENSG00000188558 | 1 | 248684916 | 248685964 | 1 | 1 | 0.66758 | 0.2522 | OR2G6 | 0.2522 | 0.83525 | 0.93488 |
| ENSG00000077454 | 7 | 100169855 | 100183776 | 37 | 8 | 0.66754 | 0.25221 | LRCH4 | 0.25221 | 0.15752 | 0.12162 |
| ENSG00000110013 | 11 | 124503009 | 124565603 | 117 | 9 | 0.66743 | 0.25225 | SIAE | 0.25225 | 0.2311 | 0.27148 |
| ENSG00000204577 | 19 | 54720147 | 54746602 | 238 | 34 | 0.66725 | 0.25231 | LILRB3 | 0.25231 | 0.30367 | 0.36433 |
| ENSG00000183161 | 11 | 22644079 | 22647387 | 6 | 4 | 0.6671 | 0.25235 | FANCF | 0.25235 | 0.22473 | 0.30955 |
| ENSG00000132740 | 11 | 68671310 | 68708070 | 118 | 14 | 0.66705 | 0.25237 | IGHMBP2 | 0.25237 | 0.21199 | 0.25311 |
| ENSG00000110768 | 11 | 18343842 | 18388591 | 113 | 13 | 0.66699 | 0.25239 | GTF2H1 | 0.25239 | 0.29488 | 0.49786 |
| ENSG00000145779 | 5 | 118604387 | 118735383 | 545 | 42 | 0.66697 | 0.2524 | TNFAIP8 | 0.2524 | 0.27023 | 0.2747 |
| ENSG00000214575 | 15 | 83211951 | 83317612 | 382 | 27 | 0.66666 | 0.25249 | CPEB1 | 0.25249 | 0.24572 | 0.23338 |
| ENSG00000134183 | 1 | 110145889 | 110155679 | 26 | 9 | 0.66644 | 0.25257 | GNAT2 | 0.25257 | 0.39317 | 0.45915 |
| ENSG00000110076 | 11 | 64373646 | 64490660 | 182 | 17 | 0.66608 | 0.25268 | NRXN2 | 0.25268 | 0.25776 | 0.24142 |
| ENSG00000263136 | 16 | 29827832 | 29831438 | 5 | 1 | 0.66608 | 0.25268 | PAGR1 | 0.25268 | 0.19711 | 0.24322 |
| ENSG00000112941 | 5 | 6714718 | 6757161 | 187 | 20 | 0.66597 | 0.25272 | PAPD7 | 0.25272 | 0.27583 | 0.27122 |
| ENSG00000168484 | 8 | 22014426 | 22021992 | 34 | 14 | 0.6657 | 0.2528 | SFTPC | 0.2528 | 0.036931 | 0.19529 |
| ENSG00000130270 | 19 | 1782074 | 1812275 | 103 | 24 | 0.66546 | 0.25288 | ATP8B3 | 0.25288 | 0.39763 | 0.37478 |
| ENSG00000119614 | 14 | 74706175 | 74729441 | 99 | 12 | 0.66539 | 0.2529 | VSX2 | 0.2529 | 0.33051 | 0.36689 |
| ENSG00000154889 | 18 | 11882621 | 11909222 | 161 | 26 | 0.66538 | 0.2529 | MPPE1 | 0.2529 | 0.5653 | 0.72262 |
| ENSG00000095261 | 9 | 123577774 | 123605262 | 45 | 8 | 0.66534 | 0.25292 | PSMD5 | 0.25292 | 0.20846 | 0.21715 |
| ENSG00000175893 | 9 | 14611069 | 14693469 | 404 | 33 | 0.6653 | 0.25293 | ZDHHC21 | 0.25293 | 0.21336 | 0.13092 |
| ENSG00000145996 | 6 | 20534688 | 21232635 | 2590 | 89 | 0.66482 | 0.25308 | CDKAL1 | 0.25308 | 0.258 | 0.3035 |

| | | | | | | | | | | | |
|---|---|---|---|---|---|---|---|---|---|---|---|
| ENSG00000004948 | 7 | 93053799 | 93204042 | 440 | 33 | 0.66483 | 0.25308 | CALCR | 0.25308 | 0.21954 | 0.20509 |
| ENSG00000165322 | 10 | 32094365 | 32217742 | 280 | 14 | 0.66457 | 0.25316 | ARHGAP12 | 0.25316 | 0.29637 | 0.40302 |
| ENSG00000205364 | 16 | 56666145 | 56667898 | 9 | 4 | 0.66458 | 0.25316 | MT1M | 0.25316 | 0.27997 | 0.46517 |
| ENSG00000188820 | 6 | 116782533 | 116784934 | 8 | 4 | 0.66455 | 0.25317 | FAM26F | 0.25317 | 0.45992 | 0.56976 |
| ENSG00000196305 | 9 | 94972489 | 95056038 | 259 | 18 | 0.66451 | 0.25318 | IARS | 0.25318 | 0.22014 | 0.20044 |
| ENSG00000186119 | 11 | 55587086 | 55588073 | 3 | 2 | 0.66428 | 0.25326 | OR5D18 | 0.25326 | 0.58886 | 0.37463 |
| ENSG00000171928 | 17 | 18684308 | 18710027 | 90 | 14 | 0.66416 | 0.25329 | TVP23B | 0.25329 | 0.28198 | 0.29867 |
| ENSG00000256660 | 12 | 10163226 | 10171218 | 21 | 8 | 0.66399 | 0.25335 | CLEC12B | 0.25335 | 0.29545 | 0.19423 |
| ENSG00000187475 | 6 | 26107640 | 26108364 | 1 | 1 | 0.66383 | 0.2534 | HIST1H1T | 0.2534 | 0.72935 | 0.58444 |
| ENSG00000100324 | 22 | 39795746 | 39833065 | 69 | 8 | 0.66341 | 0.25354 | TAB1 | 0.25354 | 0.25666 | 0.24859 |
| ENSG00000170647 | 11 | 100862811 | 100864663 | 11 | 3 | 0.66316 | 0.25361 | TMEM133 | 0.25361 | 0.2752 | 0.55942 |
| ENSG00000204464 | 1 | 15490840 | 15497813 | 27 | 7 | 0.66305 | 0.25365 | C1orf195 | 0.25365 | 0.34544 | 0.1292 |
| ENSG00000066583 | 5 | 128430444 | 128449721 | 76 | 14 | 0.66304 | 0.25365 | ISOC1 | 0.25365 | 0.22759 | 0.32171 |
| ENSG00000135605 | 4 | 48137800 | 48271881 | 583 | 38 | 0.66296 | 0.25368 | TEC | 0.25368 | 0.2973 | 0.36745 |
| ENSG00000254614 | 11 | 64945077 | 64949305 | 14 | 6 | 0.66293 | 0.25369 | AP003068.23 | 0.25369 | 0.38257 | 0.52096 |
| ENSG00000120265 | 6 | 150070579 | 150132556 | 194 | 6 | 0.66283 | 0.25372 | PCMT1 | 0.25372 | 0.24102 | 0.24103 |
| ENSG00000124194 | 20 | 42875887 | 42909013 | 93 | 11 | 0.66257 | 0.2538 | GDAP1L1 | 0.2538 | 0.34466 | 0.31574 |
| ENSG00000160396 | 19 | 40885178 | 40896094 | 22 | 7 | 0.66256 | 0.25381 | HIPK4 | 0.25381 | 0.33562 | 0.26575 |
| ENSG00000101276 | 20 | 740724 | 749131 | 53 | 14 | 0.66256 | 0.25381 | SLC52A3 | 0.25381 | 0.36331 | 0.51979 |
| ENSG00000158113 | 12 | 122652285 | 122688018 | 80 | 16 | 0.66251 | 0.25382 | LRRC43 | 0.25382 | 0.23674 | 0.27716 |
| ENSG00000053371 | 1 | 19630459 | 19638640 | 29 | 12 | 0.66242 | 0.25385 | AKR7A2 | 0.25385 | 0.21202 | 0.19814 |
| ENSG00000006432 | 14 | 71189243 | 71276251 | 373 | 28 | 0.66241 | 0.25385 | MAP3K9 | 0.25385 | 0.21704 | 0.22047 |
| ENSG00000183117 | 8 | 2792875 | 4852494 | 18162 | 611 | 0.66239 | 0.25386 | CSMD1 | 0.25386 | 0.26169 | 0.24061 |
| ENSG00000160678 | 1 | 153600402 | 153604513 | 11 | 3 | 0.66227 | 0.2539 | S100A1 | 0.2539 | 0.42732 | 0.40694 |
| ENSG00000170369 | 20 | 23804406 | 23807368 | 22 | 6 | 0.66226 | 0.2539 | CST2 | 0.2539 | 0.064818 | 0.15885 |
| ENSG00000100281 | 22 | 35653445 | 35691800 | 98 | 5 | 0.6622 | 0.25392 | HMGXB4 | 0.25392 | 0.2402 | 0.29155 |
| ENSG00000184507 | 15 | 34635516 | 34649938 | 49 | 13 | 0.66147 | 0.25415 | NUTM1 | 0.25415 | 0.3742 | 0.46926 |
| ENSG00000168653 | 1 | 39491990 | 39500308 | 42 | 5 | 0.66147 | 0.25416 | NDUFS5 | 0.25416 | 0.16476 | 0.18632 |
| ENSG00000187486 | 11 | 17407406 | 17410878 | 12 | 4 | 0.66094 | 0.25432 | KCNJ11 | 0.25432 | 0.45674 | 0.47214 |
| ENSG00000164620 | 5 | 141016517 | 141020644 | 12 | 5 | 0.66093 | 0.25433 | RELL2 | 0.25433 | 0.093722 | 0.067988 |
| ENSG00000170373 | 20 | 23728190 | 23731905 | 31 | 7 | 0.66081 | 0.25437 | CST1 | 0.25437 | 0.34236 | 0.18802 |
| ENSG00000141194 | 17 | 56232494 | 56233517 | 4 | 2 | 0.66077 | 0.25438 | OR4D1 | 0.25438 | 0.21916 | 0.28127 |
| ENSG00000185176 | 2 | 241615835 | 241622323 | 24 | 9 | 0.66057 | 0.25444 | AQP12B | 0.25444 | 0.37917 | 0.49029 |
| ENSG00000268864 | 19 | 54102882 | 54106751 | 11 | 6 | 0.66054 | 0.25445 | CTB-167G5.5 | 0.25445 | 0.091176 | 0.16168 |
| ENSG00000090020 | 1 | 27425306 | 27493472 | 92 | 9 | 0.66053 | 0.25446 | SLC9A1 | 0.25446 | 0.27262 | 0.26717 |
| ENSG00000167964 | 16 | 2190804 | 2204166 | 28 | 6 | 0.66015 | 0.25458 | RAB26 | 0.25458 | 0.48209 | 0.47829 |
| ENSG00000035403 | 10 | 75757872 | 75879918 | 249 | 32 | 0.66012 | 0.25459 | VCL | 0.25459 | 0.29606 | 0.21861 |
| ENSG00000197360 | 19 | 22573821 | 22715287 | 597 | 39 | 0.65991 | 0.25466 | ZNF98 | 0.25466 | 0.25245 | 0.14785 |
| ENSG00000128310 | 22 | 38219389 | 38221502 | 4 | 2 | 0.65963 | 0.25475 | GALR3 | 0.25475 | 0.16573 | 0.19005 |
| ENSG00000174015 | 13 | 46276446 | 46288694 | 48 | 11 | 0.65936 | 0.25483 | SPERT | 0.25483 | 0.42136 | 0.51281 |
| ENSG00000105127 | 19 | 15464196 | 15490609 | 63 | 10 | 0.65926 | 0.25486 | AKAP8 | 0.25486 | 0.25997 | 0.33211 |
| ENSG00000186814 | 18 | 32831032 | 32870196 | 95 | 16 | 0.65892 | 0.25497 | ZSCAN30 | 0.25497 | 0.24264 | 0.2221 |
| ENSG00000270181 | 13 | 103459704 | 103524748 | 195 | 21 | 0.65891 | 0.25498 | BIVM-ERCC5 | 0.25498 | 0.36801 | 0.42138 |
| ENSG00000128881 | 15 | 43030932 | 43213007 | 260 | 28 | 0.65891 | 0.25498 | TTBK2 | 0.25498 | 0.30865 | 0.32954 |
| ENSG00000205045 | 17 | 33791268 | 33864880 | 236 | 42 | 0.6589 | 0.25498 | SLFN12L | 0.25498 | 0.32546 | 0.48334 |
| ENSG00000144320 | 2 | 176788620 | 176867567 | 222 | 27 | 0.65875 | 0.25503 | KIAA1715 | 0.25503 | 0.28004 | 0.27095 |
| ENSG00000154080 | 18 | 24495595 | 24765281 | 901 | 45 | 0.65853 | 0.2551 | CHST9 | 0.2551 | 0.2826 | 0.25477 |
| ENSG00000146618 | 7 | 19184405 | 19185044 | 1 | 1 | 0.65821 | 0.2552 | FERD3L | 0.2552 | 0.026229 | 0.038556 |
| ENSG00000109339 | 4 | 86936276 | 87515284 | 1905 | 55 | 0.65783 | 0.25532 | MAPK10 | 0.25532 | 0.27082 | 0.32903 |
| ENSG00000103942 | 15 | 83509838 | 83654661 | 383 | 31 | 0.65783 | 0.25532 | HOMER2 | 0.25532 | 0.27118 | 0.31404 |
| ENSG00000039650 | 19 | 50364461 | 50371166 | 19 | 6 | 0.6578 | 0.25533 | PNKP | 0.25533 | 0.6298 | 0.54022 |
| ENSG00000159339 | 1 | 17634690 | 17690499 | 327 | 32 | 0.65777 | 0.25534 | PADI4 | 0.25534 | 0.53857 | 0.65514 |
| ENSG00000168884 | 4 | 2743375 | 2758103 | 67 | 12 | 0.65771 | 0.25536 | TNIP2 | 0.25536 | 0.27287 | 0.32849 |
| ENSG00000086061 | 9 | 33025209 | 33039905 | 76 | 10 | 0.65755 | 0.25541 | DNAJA1 | 0.25541 | 0.21878 | 0.18169 |
| ENSG00000197858 | 8 | 145137493 | 145141119 | 7 | 2 | 0.65737 | 0.25547 | GPAA1 | 0.25547 | 0.12691 | 0.13763 |
| ENSG00000155090 | 8 | 103661007 | 103668130 | 18 | 7 | 0.65724 | 0.25551 | KLF10 | 0.25551 | 0.029558 | 0.05685 |
| ENSG00000119402 | 9 | 123514256 | 123555690 | 72 | 10 | 0.65726 | 0.25551 | FBXW2 | 0.25551 | 0.2358 | 0.32696 |
| ENSG00000138029 | 2 | 26466038 | 26513336 | 63 | 20 | 0.65721 | 0.25552 | HADHB | 0.25552 | 0.24159 | 0.12986 |
| ENSG00000091656 | 8 | 77593454 | 77779521 | 410 | 34 | 0.65716 | 0.25554 | ZFHX4 | 0.25554 | 0.26443 | 0.29969 |
| ENSG00000070010 | 22 | 19437433 | 19466738 | 88 | 13 | 0.65691 | 0.25562 | UFD1L | 0.25562 | 0.26538 | 0.33271 |
| ENSG00000182500 | 12 | 122064455 | 122080583 | 29 | 9 | 0.65646 | 0.25576 | ORAI1 | 0.25576 | 0.18242 | 0.18269 |
| ENSG00000175077 | 3 | 186915274 | 186919253 | 9 | 4 | 0.65629 | 0.25582 | RTP1 | 0.25582 | 0.26826 | 0.30541 |
| ENSG00000156876 | 1 | 100549119 | 100598511 | 143 | 17 | 0.65604 | 0.2559 | SASS6 | 0.2559 | 0.24779 | 0.24779 |
| ENSG00000179335 | 15 | 74890841 | 74932057 | 61 | 10 | 0.65594 | 0.25593 | CLK3 | 0.25593 | 0.19769 | 0.18126 |
| ENSG00000204822 | 2 | 74699085 | 74700449 | 3 | 2 | 0.6559 | 0.25594 | MRPL53 | 0.25594 | 0.78607 | 0.4113 |
| ENSG00000164070 | 4 | 128702976 | 128761888 | 88 | 12 | 0.65576 | 0.25599 | HSPA4L | 0.25599 | 0.28873 | 0.29731 |
| ENSG00000170777 | 9 | 6328349 | 6331900 | 12 | 4 | 0.65575 | 0.25599 | TPD52L3 | 0.25599 | 0.34286 | 0.36128 |
| ENSG00000069667 | 15 | 60780483 | 61521518 | 2427 | 245 | 0.65559 | 0.25604 | RORA | 0.25604 | 0.28345 | 0.23207 |
| ENSG00000143001 | 1 | 55446465 | 55457966 | 45 | 3 | 0.65558 | 0.25605 | TMEM61 | 0.25605 | 0.41051 | 0.7117 |
| ENSG00000173281 | 8 | 8993765 | 9009084 | 45 | 11 | 0.65557 | 0.25605 | PPP1R3B | 0.25605 | 0.45165 | 0.65188 |
| ENSG00000205930 | 21 | 34144411 | 34266043 | 312 | 24 | 0.65526 | 0.25615 | C21orf49 | 0.25615 | 0.24877 | 0.27987 |
| ENSG00000163810 | 3 | 44916100 | 44956482 | 204 | 18 | 0.65508 | 0.25621 | TGM4 | 0.25621 | 0.32251 | 0.35445 |
| ENSG00000188321 | 19 | 9434448 | 9461838 | 88 | 8 | 0.65503 | 0.25623 | ZNF559 | 0.25623 | 0.29756 | 0.2629 |

| | | | | | | | | | | | |
|---|---|---|---|---|---|---|---|---|---|---|---|
| ENSG00000159147 | 21 | 34931848 | 34961014 | 36 | 10 | 0.65405 | 0.25654 | DONSON | 0.25654 | 0.30082 | 0.25729 |
| ENSG00000142347 | 19 | 8585674 | 8642461 | 276 | 34 | 0.65353 | 0.25671 | MYO1F | 0.25671 | 0.27058 | 0.30416 |
| ENSG00000099937 | 22 | 21128167 | 21142008 | 54 | 8 | 0.65352 | 0.25671 | SERPIND1 | 0.25671 | 0.29044 | 0.34035 |
| ENSG00000138813 | 4 | 100432161 | 100463460 | 176 | 26 | 0.65336 | 0.25676 | C4orf17 | 0.25676 | 0.25281 | 0.25899 |
| ENSG00000104043 | 15 | 50150435 | 50475014 | 1172 | 62 | 0.65336 | 0.25676 | ATP8B4 | 0.25676 | 0.33673 | 0.36497 |
| ENSG00000203667 | 1 | 244998624 | 245008359 | 31 | 5 | 0.65314 | 0.25683 | COX20 | 0.25683 | 0.40185 | 0.53108 |
| ENSG00000138080 | 2 | 27301435 | 27309271 | 8 | 3 | 0.65313 | 0.25684 | EMILIN1 | 0.25684 | 0.57538 | 0.65869 |
| ENSG00000100815 | 14 | 92432335 | 92507240 | 202 | 20 | 0.65309 | 0.25685 | TRIP11 | 0.25685 | 0.21479 | 0.2675 |
| ENSG00000187714 | 10 | 50818347 | 50820765 | 5 | 3 | 0.65298 | 0.25689 | SLC18A3 | 0.25689 | 0.37164 | 0.28257 |
| ENSG00000170917 | 4 | 123809852 | 123844123 | 160 | 21 | 0.65267 | 0.25698 | NUDT6 | 0.25698 | 0.2563 | 0.22862 |
| ENSG00000132825 | 20 | 58511894 | 58515352 | 2 | 1 | 0.65255 | 0.25702 | PPP1R3D | 0.25702 | 0.19288 | 0.22215 |
| ENSG00000179921 | 2 | 219124219 | 219128582 | 8 | 2 | 0.65253 | 0.25703 | GPBAR1 | 0.25703 | 0.30085 | 0.43939 |
| ENSG00000132141 | 17 | 33254878 | 33308097 | 87 | 20 | 0.65254 | 0.25703 | CCT6B | 0.25703 | 0.35777 | 0.3822 |
| ENSG00000072952 | 11 | 10594638 | 10715535 | 465 | 50 | 0.65236 | 0.25709 | MRVI1 | 0.25709 | 0.2511 | 0.27003 |
| ENSG00000086062 | 9 | 33104080 | 33167354 | 305 | 25 | 0.65218 | 0.25714 | B4GALT1 | 0.25714 | 0.18446 | 0.1272 |
| ENSG00000079462 | 19 | 42801185 | 42807698 | 6 | 2 | 0.65218 | 0.25714 | PAFAH1B3 | 0.25714 | 0.2244 | 0.18739 |
| ENSG00000118181 | 11 | 118886422 | 118889401 | 16 | 3 | 0.65201 | 0.2572 | RPS25 | 0.2572 | 0.50093 | 0.71033 |
| ENSG00000114353 | 3 | 50263724 | 50296787 | 46 | 13 | 0.6519 | 0.25723 | GNAI2 | 0.25723 | 0.22491 | 0.16828 |
| ENSG00000169989 | 4 | 153690506 | 153700916 | 22 | 6 | 0.65179 | 0.25727 | TIGD4 | 0.25727 | 0.24277 | 0.13911 |
| ENSG00000166228 | 10 | 72642037 | 72648541 | 24 | 9 | 0.65164 | 0.25732 | PCBD1 | 0.25732 | 0.16965 | 0.056105 |
| ENSG00000105835 | 7 | 105888731 | 105926772 | 111 | 16 | 0.65134 | 0.25741 | NAMPT | 0.25741 | 0.28762 | 0.35154 |
| ENSG00000259458 | 15 | 62929371 | 62937490 | 28 | 10 | 0.6513 | 0.25743 | RP11-625H11.1 | 0.25743 | 0.31712 | 0.34489 |
| ENSG00000136877 | 9 | 130556876 | 130576606 | 61 | 9 | 0.65119 | 0.25746 | FPGS | 0.25746 | 0.37068 | 0.42661 |
| ENSG00000103043 | 16 | 70721342 | 70835064 | 259 | 28 | 0.65111 | 0.25749 | VAC14 | 0.25749 | 0.25226 | 0.34331 |
| ENSG00000133962 | 14 | 92047040 | 92247051 | 752 | 72 | 0.65052 | 0.25768 | CATSPERB | 0.25768 | 0.21982 | 0.24153 |
| ENSG00000089094 | 12 | 121866900 | 122018920 | 344 | 40 | 0.65032 | 0.25774 | KDM2B | 0.25774 | 0.28982 | 0.31298 |
| ENSG00000099864 | 19 | 708953 | 748329 | 253 | 40 | 0.65006 | 0.25783 | PALM | 0.25783 | 0.26484 | 0.4905 |
| ENSG00000151715 | 11 | 129685714 | 129729898 | 188 | 24 | 0.64955 | 0.25799 | TMEM45B | 0.25799 | 0.24748 | 0.2284 |
| ENSG00000112339 | 6 | 135281516 | 135424194 | 369 | 15 | 0.64907 | 0.25815 | HBS1L | 0.25815 | 0.2483 | 0.24819 |
| ENSG00000064102 | 12 | 27058114 | 27091259 | 93 | 18 | 0.64873 | 0.25826 | ASUN | 0.25826 | 0.21079 | 0.26399 |
| ENSG00000226761 | 12 | 11213964 | 11214893 | 3 | 1 | 0.64865 | 0.25828 | TAS2R46 | 0.25828 | 0.4569 | 0.60141 |
| ENSG00000085511 | 6 | 161412759 | 161538417 | 432 | 29 | 0.64851 | 0.25833 | MAP3K4 | 0.25833 | 0.21214 | 0.29519 |
| ENSG00000181827 | 15 | 56379478 | 56535483 | 478 | 21 | 0.64798 | 0.2585 | RFX7 | 0.2585 | 0.23997 | 0.22258 |
| ENSG00000068724 | 2 | 47143296 | 47303276 | 801 | 59 | 0.64794 | 0.25851 | TTC7A | 0.25851 | 0.21111 | 0.14849 |
| ENSG00000114349 | 3 | 50229045 | 50233949 | 3 | 1 | 0.64755 | 0.25864 | GNAT1 | 0.25864 | 0.27542 | 0.14566 |
| ENSG00000168758 | 2 | 97525453 | 97536494 | 7 | 3 | 0.64742 | 0.25868 | SEMA4C | 0.25868 | 0.38554 | 0.73584 |
| ENSG00000116688 | 1 | 12040238 | 12073571 | 108 | 10 | 0.64733 | 0.25871 | MFN2 | 0.25871 | 0.38609 | 0.37054 |
| ENSG00000118961 | 2 | 20883788 | 21022882 | 343 | 31 | 0.64676 | 0.25889 | C2orf43 | 0.25889 | 0.35452 | 0.41032 |
| ENSG00000108691 | 17 | 32582304 | 32584222 | 5 | 2 | 0.64668 | 0.25892 | CCL2 | 0.25892 | 0.36713 | 0.43772 |
| ENSG00000164142 | 4 | 152330368 | 152584784 | 641 | 15 | 0.64646 | 0.25899 | FAM160A1 | 0.25899 | 0.22643 | 0.20475 |
| ENSG00000064218 | 9 | 976964 | 991731 | 63 | 11 | 0.64641 | 0.25901 | DMRT3 | 0.25901 | 0.44972 | 0.69651 |
| ENSG00000206432 | 18 | 5882071 | 5895954 | 47 | 12 | 0.6464 | 0.25901 | TMEM200C | 0.25901 | 0.03021 | 0.014671 |
| ENSG00000108064 | 10 | 60144782 | 60158981 | 50 | 11 | 0.64619 | 0.25908 | TFAM | 0.25908 | 0.08151 | 0.049701 |
| ENSG00000118600 | 12 | 64173583 | 64203338 | 65 | 14 | 0.64606 | 0.25912 | TMEM5 | 0.25912 | 0.22343 | 0.40206 |
| ENSG00000115484 | 2 | 62095224 | 62115939 | 92 | 15 | 0.64584 | 0.25919 | CCT4 | 0.25919 | 0.24541 | 0.22009 |
| ENSG00000107859 | 10 | 103989943 | 104001231 | 21 | 3 | 0.64564 | 0.25925 | PITX3 | 0.25925 | 0.21869 | 0.23321 |
| ENSG00000114554 | 3 | 126707437 | 126756235 | 131 | 17 | 0.64556 | 0.25928 | PLXNA1 | 0.25928 | 0.24389 | 0.21874 |
| ENSG00000124614 | 6 | 34385231 | 34393902 | 20 | 6 | 0.64555 | 0.25929 | RPS10 | 0.25929 | 0.39381 | 0.58344 |
| ENSG00000136636 | 1 | 215740735 | 215795149 | 35 | 15 | 0.64546 | 0.25931 | KCTD3 | 0.25931 | 0.3392 | 0.37394 |
| ENSG00000135750 | 1 | 233749750 | 233808258 | 232 | 47 | 0.64529 | 0.25937 | KCNK1 | 0.25937 | 0.38941 | 0.17534 |
| ENSG00000034152 | 17 | 21187984 | 21218552 | 90 | 18 | 0.64518 | 0.2594 | MAP2K3 | 0.2594 | 0.25424 | 0.22154 |
| ENSG00000100312 | 22 | 51176624 | 51183762 | 35 | 9 | 0.64516 | 0.25941 | ACR | 0.25941 | 0.21558 | 0.21693 |
| ENSG00000091128 | 7 | 107663993 | 107770801 | 213 | 26 | 0.64481 | 0.25953 | LAMB4 | 0.25953 | 0.27401 | 0.16403 |
| ENSG00000222004 | 7 | 26677490 | 26686889 | 35 | 4 | 0.64439 | 0.25966 | C7orf71 | 0.25966 | 0.32531 | 0.43063 |
| ENSG00000088448 | 13 | 111530887 | 111567416 | 137 | 12 | 0.64415 | 0.25974 | ANKRD10 | 0.25974 | 0.29154 | 0.30154 |
| ENSG00000058272 | 12 | 80167343 | 80329240 | 321 | 14 | 0.64411 | 0.25975 | PPP1R12A | 0.25975 | 0.26431 | 0.25955 |
| ENSG00000004487 | 1 | 23345941 | 23410182 | 97 | 12 | 0.64407 | 0.25976 | KDM1A | 0.25976 | 0.26582 | 0.24344 |
| ENSG00000152076 | 2 | 130896860 | 130902707 | 28 | 7 | 0.64309 | 0.26008 | CCDC74B | 0.26008 | 0.38354 | 0.38811 |
| ENSG00000156787 | 8 | 124054208 | 124164393 | 310 | 31 | 0.6428 | 0.26018 | TBC1D31 | 0.26018 | 0.29858 | 0.32788 |
| ENSG00000166987 | 12 | 57914493 | 57923931 | 13 | 3 | 0.64276 | 0.26019 | MBD6 | 0.26019 | 0.28754 | 0.16997 |
| ENSG00000079432 | 19 | 42772689 | 42799949 | 27 | 4 | 0.64255 | 0.26026 | CIC | 0.26026 | 0.39328 | 0.42314 |
| ENSG00000124615 | 6 | 39867354 | 39902290 | 113 | 18 | 0.64236 | 0.26032 | MOCS1 | 0.26032 | 0.16302 | 0.22751 |
| ENSG00000136560 | 2 | 161993419 | 162092732 | 171 | 15 | 0.64138 | 0.26064 | TANK | 0.26064 | 0.29242 | 0.3595 |
| ENSG00000120333 | 1 | 174979925 | 174992561 | 30 | 8 | 0.64102 | 0.26075 | MRPS14 | 0.26075 | 0.2229 | 0.22666 |
| ENSG00000204390 | 6 | 31777396 | 31783437 | 19 | 8 | 0.64086 | 0.26081 | HSPA1L | 0.26081 | 0.39211 | 0.40199 |
| ENSG00000010219 | 12 | 4671370 | 4723325 | 180 | 27 | 0.64074 | 0.26084 | DYRK4 | 0.26084 | 0.21191 | 0.23588 |
| ENSG00000142494 | 17 | 19398698 | 19482347 | 182 | 29 | 0.64059 | 0.2609 | SLC47A1 | 0.2609 | 0.51041 | 0.67378 |
| ENSG00000144579 | 2 | 219262979 | 219270664 | 19 | 7 | 0.64041 | 0.26095 | CTDSP1 | 0.26095 | 0.26147 | 0.38751 |
| ENSG00000155886 | 9 | 19507450 | 19786926 | 1016 | 66 | 0.64028 | 0.26099 | SLC24A2 | 0.26099 | 0.26832 | 0.29343 |
| ENSG00000138018 | 2 | 26531415 | 26618759 | 260 | 23 | 0.63962 | 0.26121 | EPT1 | 0.26121 | 0.20398 | 0.16512 |
| ENSG00000177519 | 2 | 154333852 | 154335322 | 2 | 1 | 0.63961 | 0.26121 | RPRM | 0.26121 | 0.66321 | 0.79332 |
| ENSG00000087250 | 16 | 56622986 | 56625000 | 7 | 3 | 0.63962 | 0.26121 | MT3 | 0.26121 | 0.1061 | 0.23336 |
| ENSG00000268706 | 18 | 61602042 | 61609601 | 25 | 4 | 0.63929 | 0.26132 | AC009802.1 | 0.26132 | 0.2447 | 0.40712 |

| | | | | | | | | | | | |
|---|---|---|---|---|---|---|---|---|---|---|---|
| ENSG00000168333 | 8 | 49966870 | 49988649 | 40 | 6 | 0.63923 | 0.26134 | C8orf22 | 0.26134 | 0.21707 | 0.22534 |
| ENSG00000106571 | 7 | 42000548 | 42277469 | 952 | 84 | 0.63905 | 0.2614 | GLI3 | 0.2614 | 0.29224 | 0.38657 |
| ENSG00000104635 | 8 | 22224762 | 22291642 | 287 | 16 | 0.63901 | 0.26141 | SLC39A14 | 0.26141 | 0.27447 | 0.30182 |
| ENSG00000104341 | 8 | 98787285 | 98865241 | 469 | 15 | 0.63877 | 0.26149 | LAPTM4B | 0.26149 | 0.32203 | 0.44723 |
| ENSG00000176102 | 11 | 33098734 | 33183917 | 266 | 15 | 0.63861 | 0.26154 | CSTF3 | 0.26154 | 0.23215 | 0.2929 |
| ENSG00000141756 | 17 | 39968932 | 39979465 | 28 | 6 | 0.63844 | 0.26159 | FKBP10 | 0.26159 | 0.25612 | 0.37224 |
| ENSG00000197372 | 19 | 23708433 | 23870022 | 520 | 24 | 0.63807 | 0.26171 | ZNF675 | 0.26171 | 0.26079 | 0.25174 |
| ENSG00000105198 | 19 | 40093164 | 40098116 | 10 | 6 | 0.63786 | 0.26178 | LGALS13 | 0.26178 | 0.64533 | 0.7515 |
| ENSG00000155324 | 5 | 125695824 | 125832186 | 466 | 69 | 0.63747 | 0.26191 | GRAMD3 | 0.26191 | 0.43077 | 0.52996 |
| ENSG00000100906 | 14 | 35870717 | 35873955 | 19 | 8 | 0.63748 | 0.26191 | NFKBIA | 0.26191 | 0.45369 | 0.72568 |
| ENSG00000196081 | 19 | 23404401 | 23433192 | 89 | 8 | 0.63745 | 0.26192 | ZNF724P | 0.26192 | 0.2512 | 0.3083 |
| ENSG00000175229 | 11 | 65809423 | 65816651 | 15 | 4 | 0.63714 | 0.26202 | GAL3ST3 | 0.26202 | 0.28814 | 0.33928 |
| ENSG00000127125 | 1 | 42921788 | 42939056 | 39 | 5 | 0.6369 | 0.2621 | PPCS | 0.2621 | 0.30693 | 0.27133 |
| ENSG00000197043 | 5 | 150480273 | 150537443 | 244 | 46 | 0.63665 | 0.26218 | ANXA6 | 0.26218 | 0.48021 | 0.38745 |
| ENSG00000178028 | 1 | 44679127 | 44686353 | 13 | 5 | 0.63639 | 0.26226 | DMAP1 | 0.26226 | 0.32664 | 0.47465 |
| ENSG00000145626 | 5 | 35951112 | 36001130 | 140 | 18 | 0.63632 | 0.26228 | UGT3A1 | 0.26228 | 0.3014 | 0.39944 |
| ENSG00000251209 | 15 | 98285448 | 98417780 | 708 | 48 | 0.63609 | 0.26236 | LINC00923 | 0.26236 | 0.33367 | 0.40981 |
| ENSG00000198885 | 2 | 96991069 | 96994091 | 4 | 2 | 0.63605 | 0.26237 | ITPRIPL1 | 0.26237 | 0.43857 | 0.37674 |
| ENSG00000189241 | 6 | 116597741 | 116601066 | 9 | 3 | 0.63581 | 0.26245 | TSPYL1 | 0.26245 | 0.2003 | 0.1487 |
| ENSG00000173085 | 4 | 84182689 | 84206067 | 45 | 9 | 0.63541 | 0.26258 | COQ2 | 0.26258 | 0.36586 | 0.51365 |
| ENSG00000069869 | 15 | 56119120 | 56285944 | 776 | 23 | 0.6352 | 0.26265 | NEDD4 | 0.26265 | 0.25225 | 0.33616 |
| ENSG00000172318 | 2 | 168675182 | 168730551 | 111 | 24 | 0.63513 | 0.26267 | B3GALT1 | 0.26267 | 0.43654 | 0.40438 |
| ENSG00000169139 | 8 | 48920960 | 48977268 | 50 | 13 | 0.63492 | 0.26274 | UBE2V2 | 0.26274 | 0.24402 | 0.2762 |
| ENSG00000152894 | 6 | 128289924 | 128841870 | 1158 | 74 | 0.63485 | 0.26276 | PTPRK | 0.26276 | 0.28671 | 0.22057 |
| ENSG00000204790 | 9 | 69204538 | 69269662 | 4 | 1 | 0.63483 | 0.26277 | CBWD6 | 0.26277 | 0.26277 | 0.26277 |
| ENSG00000234224 | 7 | 123670973 | 123673523 | 5 | 2 | 0.63463 | 0.26283 | TMEM229A | 0.26283 | 0.50152 | 0.66142 |
| ENSG00000049249 | 1 | 7979907 | 8000926 | 51 | 8 | 0.63448 | 0.26289 | TNFRSF9 | 0.26289 | 0.19613 | 0.30033 |
| ENSG00000109814 | 4 | 39500375 | 39529931 | 147 | 19 | 0.63439 | 0.26291 | UGDH | 0.26291 | 0.12705 | 0.12295 |
| ENSG00000092853 | 1 | 36185819 | 36235568 | 73 | 10 | 0.63418 | 0.26298 | CLSPN | 0.26298 | 0.29243 | 0.32079 |
| ENSG00000137965 | 1 | 79115481 | 79129763 | 23 | 5 | 0.63405 | 0.26302 | IFI44 | 0.26302 | 0.23004 | 0.47543 |
| ENSG00000148834 | 10 | 105995114 | 106027217 | 74 | 16 | 0.63407 | 0.26302 | GSTO1 | 0.26302 | 0.30484 | 0.3024 |
| ENSG00000240771 | 12 | 58003963 | 58013162 | 19 | 6 | 0.6339 | 0.26307 | ARHGEF25 | 0.26307 | 0.18776 | 0.13543 |
| ENSG00000175792 | 3 | 127783621 | 127872757 | 217 | 13 | 0.63385 | 0.26309 | RUVBL1 | 0.26309 | 0.24399 | 0.19746 |
| ENSG00000116489 | 1 | 113161795 | 113214241 | 112 | 16 | 0.6338 | 0.26311 | CAPZA1 | 0.26311 | 0.19487 | 0.17426 |
| ENSG00000143801 | 1 | 227057885 | 227083806 | 77 | 9 | 0.63334 | 0.26326 | PSEN2 | 0.26326 | 0.29895 | 0.37827 |
| ENSG00000249852 | 7 | 227554 | 229557 | 16 | 4 | 0.63328 | 0.26327 | AC145676.2 | 0.26327 | 0.17049 | 0.27663 |
| ENSG00000183426 | 16 | 15016659 | 15045915 | 15 | 7 | 0.63329 | 0.26327 | NPIPA1 | 0.26327 | 0.099921 | 0.038495 |
| ENSG00000196923 | 5 | 176910395 | 176924607 | 22 | 7 | 0.63319 | 0.2633 | PDLIM7 | 0.2633 | 0.49863 | 0.33538 |
| ENSG00000188761 | 1 | 114420790 | 114430169 | 31 | 8 | 0.63305 | 0.26335 | BCL2L15 | 0.26335 | 0.2847 | 0.41877 |
| ENSG00000077782 | 8 | 38268656 | 38326352 | 106 | 21 | 0.63275 | 0.26345 | FGFR1 | 0.26345 | 0.40896 | 0.44016 |
| ENSG00000067533 | 1 | 218458629 | 218511325 | 99 | 7 | 0.63264 | 0.26349 | RRP15 | 0.26349 | 0.33918 | 0.49461 |
| ENSG00000136830 | 9 | 130267618 | 130341268 | 248 | 34 | 0.63253 | 0.26352 | FAM129B | 0.26352 | 0.32538 | 0.41084 |
| ENSG00000197928 | 19 | 53731577 | 53758151 | 130 | 10 | 0.63242 | 0.26356 | ZNF677 | 0.26356 | 0.29534 | 0.30911 |
| ENSG00000167664 | 19 | 4292229 | 4302428 | 31 | 10 | 0.63235 | 0.26358 | TMIGD2 | 0.26358 | 0.074152 | 0.11738 |
| ENSG00000183715 | 11 | 132284871 | 133402414 | 4246 | 308 | 0.63233 | 0.26359 | OPCML | 0.26359 | 0.23494 | 0.21844 |
| ENSG00000088882 | 20 | 2774715 | 2781283 | 23 | 7 | 0.63232 | 0.26359 | CPXM1 | 0.26359 | 0.23911 | 0.26862 |
| ENSG00000120251 | 4 | 158125334 | 158287227 | 305 | 23 | 0.63205 | 0.26368 | GRIA2 | 0.26368 | 0.25311 | 0.31231 |
| ENSG00000030304 | 9 | 113431051 | 113563859 | 426 | 47 | 0.63187 | 0.26374 | MUSK | 0.26374 | 0.26991 | 0.18922 |
| ENSG00000187855 | 12 | 108168162 | 108170421 | 7 | 5 | 0.63175 | 0.26377 | ASCL4 | 0.26377 | 0.73834 | 0.57527 |
| ENSG00000198937 | 6 | 37450696 | 37467698 | 33 | 9 | 0.6317 | 0.26379 | CCDC167 | 0.26379 | 0.33632 | 0.57935 |
| ENSG00000132386 | 17 | 1665253 | 1680868 | 69 | 11 | 0.63126 | 0.26393 | SERPINF1 | 0.26393 | 0.40154 | 0.23096 |
| ENSG00000221890 | 22 | 39214457 | 39239987 | 92 | 13 | 0.6312 | 0.26395 | NPTXR | 0.26395 | 0.28446 | 0.29494 |
| ENSG00000164687 | 8 | 82192598 | 82197012 | 4 | 2 | 0.63056 | 0.26417 | FABP5 | 0.26417 | 0.32835 | 0.37943 |
| ENSG00000159961 | 17 | 3323862 | 3324827 | 4 | 2 | 0.63039 | 0.26422 | OR3A3 | 0.26422 | 0.31493 | 0.4305 |
| ENSG00000107018 | 9 | 5334969 | 5339873 | 16 | 4 | 0.6303 | 0.26425 | RLN1 | 0.26425 | 0.22146 | 0.2786 |
| ENSG00000153498 | 13 | 113030633 | 113089003 | 284 | 22 | 0.63018 | 0.26429 | SPACA7 | 0.26429 | 0.25812 | 0.46641 |
| ENSG00000109466 | 4 | 166128770 | 166244308 | 366 | 22 | 0.63014 | 0.2643 | KLHL2 | 0.2643 | 0.24983 | 0.33151 |
| ENSG00000180155 | 8 | 143845752 | 143859640 | 53 | 12 | 0.63003 | 0.26434 | LYNX1 | 0.26434 | 0.67962 | 0.72517 |
| ENSG00000167543 | 17 | 27893070 | 27900175 | 14 | 5 | 0.63 | 0.26435 | TP53I13 | 0.26435 | 0.35285 | 0.37457 |
| ENSG00000140403 | 15 | 78556428 | 78574538 | 76 | 15 | 0.62988 | 0.26439 | DNAJA4 | 0.26439 | 0.19793 | 0.23873 |
| ENSG00000125148 | 16 | 56642111 | 56643409 | 5 | 3 | 0.62962 | 0.26447 | MT2A | 0.26447 | 0.53953 | 0.2691 |
| ENSG00000204305 | 6 | 32148745 | 32152101 | 8 | 3 | 0.62922 | 0.2646 | AGER | 0.2646 | 0.088671 | 0.036356 |
| ENSG00000161888 | 19 | 11242196 | 11266484 | 157 | 25 | 0.62923 | 0.2646 | SPC24 | 0.2646 | 0.24781 | 0.27832 |
| ENSG00000169154 | 8 | 37791799 | 37797647 | 9 | 3 | 0.62914 | 0.26463 | GOT1L1 | 0.26463 | 0.57838 | 0.57798 |
| ENSG00000025434 | 11 | 47269851 | 47290396 | 62 | 9 | 0.62912 | 0.26464 | NR1H3 | 0.26464 | 0.18502 | 0.21642 |
| ENSG00000140519 | 15 | 89998680 | 90039844 | 80 | 12 | 0.62894 | 0.26469 | RHCG | 0.26469 | 0.29724 | 0.57595 |
| ENSG00000160410 | 19 | 41082757 | 41097305 | 56 | 12 | 0.62889 | 0.26471 | SHKBP1 | 0.26471 | 0.35963 | 0.48919 |
| ENSG00000223572 | 15 | 43985084 | 43991420 | 6 | 2 | 0.6284 | 0.26487 | CKMT1A | 0.26487 | 0.43186 | 0.326 |
| ENSG00000113327 | 5 | 161494546 | 161582542 | 200 | 33 | 0.6282 | 0.26494 | GABRG2 | 0.26494 | 0.23296 | 0.12033 |
| ENSG00000141140 | 17 | 34851477 | 34899284 | 105 | 13 | 0.62819 | 0.26494 | MYO19 | 0.26494 | 0.32077 | 0.31948 |
| ENSG00000140950 | 16 | 84511681 | 84587639 | 597 | 46 | 0.62778 | 0.26507 | TLDC1 | 0.26507 | 0.34582 | 0.3917 |
| ENSG00000105982 | 7 | 156432975 | 156469824 | 103 | 12 | 0.62763 | 0.26512 | RNF32 | 0.26512 | 0.16169 | 0.10745 |
| ENSG00000119231 | 3 | 196594727 | 196661585 | 221 | 20 | 0.62729 | 0.26523 | SENP5 | 0.26523 | 0.29467 | 0.23895 |

| | | | | | | | | | | | |
|---|---|---|---|---|---|---|---|---|---|---|---|
| ENSG00000103175 | 16 | 84328252 | 84363450 | 236 | 50 | 0.62709 | 0.2653 | WFDC1 | 0.2653 | 0.46522 | 0.68468 |
| ENSG00000127884 | 10 | 135175984 | 135187193 | 34 | 11 | 0.62678 | 0.2654 | ECHS1 | 0.2654 | 0.6225 | 0.66863 |
| ENSG00000174840 | 3 | 57542003 | 57552571 | 28 | 7 | 0.6266 | 0.26546 | PDE12 | 0.26546 | 0.24617 | 0.1763 |
| ENSG00000141994 | 19 | 5784843 | 5791249 | 31 | 6 | 0.62657 | 0.26547 | DUS3L | 0.26547 | 0.36599 | 0.49306 |
| ENSG00000238227 | 9 | 139006427 | 139010731 | 11 | 5 | 0.62638 | 0.26553 | C9orf69 | 0.26553 | 0.52892 | 0.25164 |
| ENSG00000024862 | 6 | 139094657 | 139114456 | 88 | 13 | 0.62632 | 0.26555 | CCDC28A | 0.26555 | 0.21272 | 0.19619 |
| ENSG00000167173 | 15 | 75487984 | 75504510 | 19 | 6 | 0.62625 | 0.26558 | C15orf39 | 0.26558 | 0.32715 | 0.097603 |
| ENSG00000139645 | 12 | 56631591 | 56652175 | 29 | 5 | 0.62602 | 0.26565 | ANKRD52 | 0.26565 | 0.23393 | 0.30996 |
| ENSG00000140623 | 16 | 4827670 | 4838522 | 46 | 6 | 0.62576 | 0.26574 | SEPTIN12 | 0.26574 | 0.32271 | 0.5706 |
| ENSG00000256294 | 19 | 44616334 | 44637027 | 42 | 6 | 0.62556 | 0.2658 | ZNF225 | 0.2658 | 0.3112 | 0.31664 |
| ENSG00000124588 | 6 | 2988221 | 3019996 | 227 | 25 | 0.62545 | 0.26584 | NQO2 | 0.26584 | 0.44517 | 0.62607 |
| ENSG00000157764 | 7 | 140419127 | 140624564 | 429 | 16 | 0.62539 | 0.26586 | BRAF | 0.26586 | 0.27332 | 0.29089 |
| ENSG00000184840 | 5 | 177019159 | 177023125 | 14 | 7 | 0.62503 | 0.26598 | TMED9 | 0.26598 | 0.34348 | 0.27502 |
| ENSG00000114698 | 3 | 145910126 | 145968966 | 277 | 16 | 0.62417 | 0.26626 | PLSCR4 | 0.26626 | 0.32401 | 0.41332 |
| ENSG00000180573 | 6 | 26124373 | 26139344 | 45 | 10 | 0.62408 | 0.26629 | HIST1H2AC | 0.26629 | 0.39823 | 0.55345 |
| ENSG00000171631 | 11 | 72975550 | 73009662 | 89 | 9 | 0.62408 | 0.26629 | P2RY6 | 0.26629 | 0.37323 | 0.58243 |
| ENSG00000147509 | 8 | 54764368 | 54871863 | 198 | 32 | 0.62401 | 0.26631 | RGS20 | 0.26631 | 0.35346 | 0.44533 |
| ENSG00000159527 | 1 | 153270338 | 153283194 | 33 | 8 | 0.62375 | 0.2664 | PGLYRP3 | 0.2664 | 0.36118 | 0.44431 |
| ENSG00000057704 | 12 | 94960900 | 95044338 | 395 | 47 | 0.62369 | 0.26642 | TMCC3 | 0.26642 | 0.19687 | 0.12801 |
| ENSG00000215717 | 1 | 109632425 | 109639556 | 24 | 6 | 0.62328 | 0.26655 | TMEM167B | 0.26655 | 0.34332 | 0.28732 |
| ENSG00000204569 | 6 | 30568177 | 30586389 | 62 | 6 | 0.62324 | 0.26656 | PPP1R10 | 0.26656 | 0.27803 | 0.25935 |
| ENSG00000108187 | 10 | 70042417 | 70092806 | 223 | 14 | 0.62316 | 0.26659 | PBLD | 0.26659 | 0.26413 | 0.23086 |
| ENSG00000241563 | 1 | 10509776 | 10511959 | 5 | 2 | 0.62287 | 0.26668 | CORT | 0.26668 | 0.35629 | 0.10145 |
| ENSG00000263020 | 6 | 31633879 | 31641323 | 26 | 10 | 0.62253 | 0.2668 | CSNK2B-LY6G5B-1181 | 0.2668 | 0.47704 | 0.38624 |
| ENSG00000174136 | 5 | 98104354 | 98134347 | 84 | 13 | 0.62233 | 0.26686 | RGMB | 0.26686 | 0.30997 | 0.24615 |
| ENSG00000106366 | 7 | 100770370 | 100782547 | 58 | 15 | 0.62138 | 0.26717 | SERPINE1 | 0.26717 | 0.53875 | 0.32132 |
| ENSG00000126003 | 20 | 30780306 | 30795594 | 6 | 2 | 0.62125 | 0.26722 | PLAGL2 | 0.26722 | 0.23054 | 0.24364 |
| ENSG00000159335 | 12 | 6874682 | 6880116 | 11 | 3 | 0.62105 | 0.26728 | PTMS | 0.26728 | 0.64714 | 0.88771 |
| ENSG00000139726 | 12 | 123237321 | 123255611 | 40 | 4 | 0.62054 | 0.26745 | DENR | 0.26745 | 0.35512 | 0.40563 |
| ENSG00000146828 | 7 | 100424442 | 100464631 | 99 | 20 | 0.62045 | 0.26748 | SLC12A9 | 0.26748 | 0.2639 | 0.25625 |
| ENSG00000136888 | 9 | 117350026 | 117360653 | 29 | 10 | 0.62044 | 0.26748 | ATP6V1G1 | 0.26748 | 0.18251 | 0.23894 |
| ENSG00000151812 | 14 | 58030640 | 58448912 | 2054 | 90 | 0.62041 | 0.26749 | SLC35F4 | 0.26749 | 0.27873 | 0.3223 |
| ENSG00000135374 | 11 | 34500340 | 34535352 | 157 | 17 | 0.62032 | 0.26752 | ELF5 | 0.26752 | 0.19457 | 0.1851 |
| ENSG00000136546 | 2 | 167260083 | 167350757 | 279 | 21 | 0.62029 | 0.26753 | SCN7A | 0.26753 | 0.26045 | 0.2913 |
| ENSG00000090989 | 4 | 56719782 | 56771200 | 145 | 23 | 0.62021 | 0.26756 | EXOC1 | 0.26756 | 0.20274 | 0.20531 |
| ENSG00000055070 | 1 | 16679070 | 16724640 | 128 | 13 | 0.61996 | 0.26764 | SZRD1 | 0.26764 | 0.3303 | 0.15666 |
| ENSG00000197442 | 6 | 136878185 | 137113656 | 509 | 38 | 0.61963 | 0.26775 | MAP3K5 | 0.26775 | 0.28698 | 0.29665 |
| ENSG00000180660 | 13 | 36047926 | 36050832 | 3 | 2 | 0.61874 | 0.26804 | MAB21L1 | 0.26804 | 0.2038 | 0.33597 |
| ENSG00000095637 | 10 | 97071528 | 97321171 | 975 | 70 | 0.61859 | 0.26809 | SORBS1 | 0.26809 | 0.29169 | 0.36477 |
| ENSG00000204398 | 13 | 112240548 | 112324955 | 360 | 30 | 0.61839 | 0.26816 | RP11-65D24.2 | 0.26816 | 0.24797 | 0.18829 |
| ENSG00000233670 | 17 | 10725792 | 10741762 | 43 | 10 | 0.61817 | 0.26823 | PIRT | 0.26823 | 0.10901 | 0.042942 |
| ENSG00000108176 | 10 | 69556427 | 69597924 | 99 | 18 | 0.61778 | 0.26836 | DNAJC12 | 0.26836 | 0.34911 | 0.46278 |
| ENSG00000197430 | 10 | 98102973 | 98119092 | 62 | 10 | 0.61759 | 0.26842 | OPALIN | 0.26842 | 0.35252 | 0.5597 |
| ENSG00000106477 | 7 | 130033612 | 130082274 | 140 | 22 | 0.61753 | 0.26844 | CEP41 | 0.26844 | 0.32927 | 0.34575 |
| ENSG00000198874 | 7 | 66460160 | 66704501 | 1710 | 32 | 0.61725 | 0.26853 | TYW1 | 0.26853 | 0.27699 | 0.28344 |
| ENSG00000174788 | 19 | 7696497 | 7698634 | 11 | 2 | 0.61724 | 0.26854 | PCP2 | 0.26854 | 0.27674 | 0.43808 |
| ENSG00000163950 | 4 | 1694527 | 1714282 | 73 | 15 | 0.61711 | 0.26858 | SLBP | 0.26858 | 0.45144 | 0.46215 |
| ENSG00000147905 | 9 | 37120536 | 37358146 | 367 | 21 | 0.61698 | 0.26862 | ZCCHC7 | 0.26862 | 0.25969 | 0.28316 |
| ENSG00000188266 | 15 | 78799906 | 78829714 | 97 | 9 | 0.61699 | 0.26862 | HYKK | 0.26862 | 0.24147 | 0.26706 |
| ENSG00000119396 | 9 | 123940415 | 123985292 | 114 | 19 | 0.61694 | 0.26864 | RAB14 | 0.26864 | 0.27361 | 0.26131 |
| ENSG00000137547 | 8 | 55047770 | 55060461 | 31 | 5 | 0.61669 | 0.26872 | MRPL15 | 0.26872 | 0.2962 | 0.39835 |
| ENSG00000170835 | 9 | 135937365 | 135947248 | 11 | 5 | 0.61648 | 0.26879 | CEL | 0.26879 | 0.037331 | 0.072258 |
| ENSG00000178021 | 2 | 54480315 | 54483409 | 11 | 4 | 0.61625 | 0.26886 | TSPYL6 | 0.26886 | 0.075305 | 0.08849 |
| ENSG00000233493 | 19 | 55890613 | 55895966 | 18 | 5 | 0.61617 | 0.26889 | TMEM238 | 0.26889 | 0.75192 | 0.71481 |
| ENSG00000134438 | 18 | 56934267 | 56941318 | 20 | 4 | 0.61612 | 0.26891 | RAX | 0.26891 | 0.51082 | 0.60877 |
| ENSG00000122180 | 1 | 203052260 | 203055164 | 8 | 4 | 0.61605 | 0.26893 | MYOG | 0.26893 | 0.37542 | 0.34878 |
| ENSG00000131368 | 3 | 15083967 | 15106842 | 46 | 13 | 0.61588 | 0.26899 | MRPS25 | 0.26899 | 0.27572 | 0.20781 |
| ENSG00000104047 | 15 | 49913177 | 49937333 | 87 | 8 | 0.61575 | 0.26903 | DTWD1 | 0.26903 | 0.31235 | 0.47416 |
| ENSG00000198754 | 1 | 40235195 | 40237020 | 2 | 1 | 0.6156 | 0.26908 | OXCT2 | 0.26908 | 0.59663 | 0.28777 |
| ENSG00000054803 | 20 | 54572496 | 54580528 | 11 | 5 | 0.6154 | 0.26914 | CBLN4 | 0.26914 | 0.26471 | 0.23718 |
| ENSG00000159082 | 21 | 34001069 | 34100359 | 155 | 22 | 0.61539 | 0.26915 | SYNJ1 | 0.26915 | 0.28156 | 0.28167 |
| ENSG00000132429 | 6 | 105606155 | 105627870 | 74 | 8 | 0.6152 | 0.26921 | POPDC3 | 0.26921 | 0.35226 | 0.37333 |
| ENSG00000132622 | 20 | 3713314 | 3733758 | 59 | 15 | 0.61509 | 0.26925 | HSPA12B | 0.26925 | 0.55998 | 0.65733 |
| ENSG00000075891 | 10 | 102495360 | 102589698 | 264 | 26 | 0.61473 | 0.26937 | PAX2 | 0.26937 | 0.24477 | 0.28858 |
| ENSG00000165533 | 14 | 89290497 | 89344335 | 123 | 16 | 0.61441 | 0.26947 | TTC8 | 0.26947 | 0.35439 | 0.40938 |
| ENSG00000107518 | 10 | 116853124 | 117708503 | 1768 | 35 | 0.61434 | 0.26949 | ATRNL1 | 0.26949 | 0.2238 | 0.2036 |
| ENSG00000149091 | 11 | 46354455 | 46402104 | 66 | 13 | 0.61432 | 0.2695 | DGKZ | 0.2695 | 0.33422 | 0.37532 |
| ENSG00000135355 | 6 | 90604188 | 90605839 | 1 | 1 | 0.61402 | 0.2696 | GJA10 | 0.2696 | 0.57739 | 0.40934 |
| ENSG00000198156 | 16 | 28353876 | 28374829 | 18 | 6 | 0.61383 | 0.26966 | NPIPB6 | 0.26966 | 0.54191 | 0.55859 |
| ENSG00000216895 | 7 | 155403925 | 155437899 | 108 | 13 | 0.61336 | 0.26982 | AC009403.2 | 0.26982 | 0.37342 | 0.33995 |
| ENSG00000008517 | 16 | 3115298 | 3131908 | 86 | 13 | 0.61309 | 0.26991 | IL32 | 0.26991 | 0.41224 | 0.274 |
| ENSG00000179431 | 11 | 35639735 | 35642419 | 5 | 2 | 0.6129 | 0.26997 | FJX1 | 0.26997 | 0.66011 | 0.66436 |
| ENSG00000108561 | 17 | 5336097 | 5352150 | 59 | 13 | 0.61289 | 0.26997 | C1QBP | 0.26997 | 0.28654 | 0.24628 |

| | | | | | | | | | | | |
|---|---|---|---|---|---|---|---|---|---|---|---|
| ENSG00000062194 | 5 | 56469775 | 56560506 | 166 | 30 | 0.61278 | 0.27001 | GPBP1 | 0.27001 | 0.25778 | 0.28171 |
| ENSG00000144792 | 3 | 44619715 | 44641186 | 46 | 10 | 0.61274 | 0.27002 | ZNF660 | 0.27002 | 0.52334 | 0.51984 |
| ENSG00000188386 | 9 | 104353897 | 104357283 | 10 | 5 | 0.61276 | 0.27002 | PPP3R2 | 0.27002 | 0.42868 | 0.39279 |
| ENSG00000253110 | 5 | 171200656 | 171201743 | 7 | 3 | 0.61263 | 0.27006 | CTB-78H18.1 | 0.27006 | 0.18776 | 0.24895 |
| ENSG00000042429 | 11 | 93517393 | 93547861 | 50 | 7 | 0.61154 | 0.27042 | MED17 | 0.27042 | 0.23701 | 0.23668 |
| ENSG00000204532 | 19 | 56713670 | 56720821 | 64 | 15 | 0.61145 | 0.27045 | ZSCAN5C | 0.27045 | 0.54471 | 0.67878 |
| ENSG00000166106 | 11 | 130318869 | 130346532 | 82 | 17 | 0.61068 | 0.27071 | ADAMTS15 | 0.27071 | 0.065647 | 0.16646 |
| ENSG00000163106 | 4 | 95219686 | 95264027 | 71 | 10 | 0.61052 | 0.27076 | HPGDS | 0.27076 | 0.13045 | 0.11977 |
| ENSG00000174808 | 4 | 75669969 | 75719896 | 242 | 17 | 0.61039 | 0.2708 | BTC | 0.2708 | 0.31822 | 0.2787 |
| ENSG00000167548 | 12 | 49412758 | 49453557 | 46 | 10 | 0.6102 | 0.27087 | KMT2D | 0.27087 | 0.30518 | 0.35232 |
| ENSG00000132823 | 20 | 42825136 | 42839431 | 57 | 9 | 0.61019 | 0.27087 | OSER1 | 0.27087 | 0.31939 | 0.30903 |
| ENSG00000168878 | 2 | 85884437 | 85895864 | 21 | 9 | 0.61015 | 0.27088 | SFTPB | 0.27088 | 0.40939 | 0.46026 |
| ENSG00000156575 | 11 | 57144242 | 57148623 | 13 | 6 | 0.61002 | 0.27092 | PRG3 | 0.27092 | 0.70748 | 0.86784 |
| ENSG00000166888 | 12 | 57489191 | 57525922 | 80 | 13 | 0.61004 | 0.27092 | STAT6 | 0.27092 | 0.21776 | 0.25356 |
| ENSG00000118705 | 20 | 35806813 | 35870022 | 218 | 12 | 0.60996 | 0.27094 | RPN2 | 0.27094 | 0.23641 | 0.23294 |
| ENSG00000116574 | 1 | 228870824 | 228882416 | 47 | 7 | 0.60968 | 0.27104 | RHOU | 0.27104 | 0.22475 | 0.18324 |
| ENSG00000173868 | 17 | 47300724 | 47308128 | 22 | 7 | 0.60932 | 0.27116 | PHOSPHO1 | 0.27116 | 0.2742 | 0.0034426 |
| ENSG00000174776 | 3 | 167196472 | 167371771 | 381 | 43 | 0.60927 | 0.27117 | WDR49 | 0.27117 | 0.28918 | 0.32121 |
| ENSG00000144642 | 3 | 29322473 | 30051886 | 3238 | 180 | 0.60875 | 0.27134 | RBMS3 | 0.27134 | 0.21111 | 0.16171 |
| ENSG00000259494 | 15 | 89002707 | 89010650 | 7 | 3 | 0.6086 | 0.27139 | MRPL46 | 0.27139 | 0.42381 | 0.25439 |
| ENSG00000164970 | 9 | 34398182 | 34458568 | 168 | 22 | 0.6084 | 0.27146 | FAM219A | 0.27146 | 0.2811 | 0.30814 |
| ENSG00000214900 | 14 | 50448430 | 50474238 | 95 | 17 | 0.6082 | 0.27153 | C14orf182 | 0.27153 | 0.41634 | 0.59643 |
| ENSG00000122359 | 10 | 81910645 | 81965328 | 191 | 20 | 0.6081 | 0.27156 | ANXA11 | 0.27156 | 0.21937 | 0.19664 |
| ENSG00000065268 | 19 | 983174 | 994569 | 97 | 12 | 0.6081 | 0.27156 | WDR18 | 0.27156 | 0.10051 | 0.27829 |
| ENSG00000172733 | 8 | 30853321 | 30891231 | 66 | 13 | 0.60801 | 0.27159 | PURG | 0.27159 | 0.14833 | 0.13516 |
| ENSG00000198691 | 1 | 94458393 | 94586688 | 481 | 75 | 0.60799 | 0.2716 | ABCA4 | 0.2716 | 0.30032 | 0.26614 |
| ENSG00000163166 | 2 | 128193783 | 128284462 | 204 | 23 | 0.60796 | 0.27161 | IWS1 | 0.27161 | 0.2648 | 0.23753 |
| ENSG00000185164 | 16 | 18511182 | 18573533 | 7 | 4 | 0.60781 | 0.27166 | NOMO2 | 0.27166 | 0.16663 | 0.13485 |
| ENSG00000054282 | 1 | 243419320 | 243663394 | 522 | 36 | 0.60752 | 0.27175 | SDCCAG8 | 0.27175 | 0.32235 | 0.36439 |
| ENSG00000116957 | 1 | 235530675 | 235612283 | 312 | 21 | 0.60738 | 0.2718 | TBCE | 0.2718 | 0.23094 | 0.23277 |
| ENSG00000158109 | 1 | 3541566 | 3546691 | 16 | 4 | 0.60687 | 0.27197 | TPRG1L | 0.27197 | 0.44387 | 0.41924 |
| ENSG00000250844 | 4 | 9250356 | 9251948 | 1 | 1 | 0.60678 | 0.272 | USP17L18 | 0.272 | 0.49474 | 0.46667 |
| ENSG00000198488 | 11 | 76745385 | 76753096 | 25 | 5 | 0.60672 | 0.27202 | B3GNT6 | 0.27202 | 0.54601 | 0.68743 |
| ENSG00000204147 | 10 | 52499078 | 52576255 | 297 | 13 | 0.60665 | 0.27204 | ASAH2B | 0.27204 | 0.31032 | 0.34435 |
| ENSG00000240505 | 17 | 16832849 | 16875432 | 209 | 26 | 0.6065 | 0.27209 | TNFRSF13B | 0.27209 | 0.35294 | 0.30653 |
| ENSG00000124089 | 20 | 54823788 | 54824871 | 2 | 1 | 0.6064 | 0.27212 | MC3R | 0.27212 | 0.77379 | 0.62555 |
| ENSG00000162641 | 1 | 109358520 | 109506106 | 605 | 39 | 0.60629 | 0.27216 | AKNAD1 | 0.27216 | 0.26985 | 0.27866 |
| ENSG00000162733 | 1 | 162601163 | 162757190 | 436 | 30 | 0.60619 | 0.27219 | DDR2 | 0.27219 | 0.29219 | 0.33027 |
| ENSG00000250565 | 2 | 46717889 | 46769696 | 201 | 27 | 0.60602 | 0.27225 | ATP6V1E2 | 0.27225 | 0.33198 | 0.42854 |
| ENSG00000125834 | 20 | 2082257 | 2157684 | 246 | 26 | 0.60593 | 0.27228 | STK35 | 0.27228 | 0.40823 | 0.40638 |
| ENSG00000168828 | 9 | 35869375 | 35870461 | 1 | 1 | 0.60587 | 0.2723 | OR13J1 | 0.2723 | 0.78816 | 0.84021 |
| ENSG00000142623 | 1 | 17531621 | 17572500 | 175 | 48 | 0.60534 | 0.27248 | PADI1 | 0.27248 | 0.4107 | 0.43886 |
| ENSG00000142686 | 1 | 36179476 | 36185073 | 6 | 2 | 0.60531 | 0.27249 | C1orf216 | 0.27249 | 0.31311 | 0.31564 |
| ENSG00000156463 | 5 | 145316142 | 145461354 | 392 | 15 | 0.60506 | 0.27257 | SH3RF2 | 0.27257 | 0.28379 | 0.29632 |
| ENSG00000196248 | 11 | 123847368 | 123848488 | 5 | 2 | 0.60505 | 0.27257 | OR10S1 | 0.27257 | 0.32658 | 0.35108 |
| ENSG00000269155 | 6 | 168196210 | 168198927 | 13 | 5 | 0.60468 | 0.27269 | AL009178.1 | 0.27269 | 0.22885 | 0.053303 |
| ENSG00000187990 | 6 | 26216428 | 26216872 | 2 | 1 | 0.60436 | 0.2728 | HIST1H2BG | 0.2728 | 0.53204 | 0.50691 |
| ENSG00000162585 | 1 | 2115903 | 2144159 | 60 | 12 | 0.60398 | 0.27293 | C1orf86 | 0.27293 | 0.3173 | 0.43224 |
| ENSG00000148795 | 10 | 104590288 | 104597290 | 20 | 4 | 0.60365 | 0.27304 | CYP17A1 | 0.27304 | 0.4159 | 0.52542 |
| ENSG00000063169 | 19 | 48111453 | 48206533 | 284 | 25 | 0.6033 | 0.27315 | GLTSCR1 | 0.27315 | 0.38681 | 0.55179 |
| ENSG00000153551 | 3 | 32433163 | 32524559 | 423 | 44 | 0.6031 | 0.27322 | CMTM7 | 0.27322 | 0.38492 | 0.36249 |
| ENSG00000152104 | 1 | 214522039 | 214725792 | 799 | 56 | 0.60302 | 0.27325 | PTPN14 | 0.27325 | 0.31515 | 0.30287 |
| ENSG00000160445 | 9 | 131492065 | 131534693 | 62 | 11 | 0.60298 | 0.27326 | ZER1 | 0.27326 | 0.29834 | 0.27636 |
| ENSG00000180596 | 6 | 26115101 | 26124154 | 25 | 8 | 0.60277 | 0.27333 | HIST1H2BC | 0.27333 | 0.45108 | 0.60001 |
| ENSG00000120328 | 5 | 140588269 | 140591696 | 7 | 4 | 0.60274 | 0.27334 | PCDHB12 | 0.27334 | 0.37013 | 0.22418 |
| ENSG00000235376 | 10 | 105005644 | 105007773 | 6 | 3 | 0.6027 | 0.27335 | RPEL1 | 0.27335 | 0.067308 | 0.052176 |
| ENSG00000168505 | 2 | 237073879 | 237077012 | 6 | 4 | 0.60261 | 0.27338 | GBX2 | 0.27338 | 0.81885 | 0.8416 |
| ENSG00000014138 | 11 | 65029233 | 65073060 | 43 | 18 | 0.60262 | 0.27338 | POLA2 | 0.27338 | 0.22547 | 0.30801 |
| ENSG00000255692 | 12 | 120137573 | 120138816 | 1 | 1 | 0.60256 | 0.2734 | RP1-127H14.3 | 0.2734 | 0.77257 | 0.84058 |
| ENSG00000112238 | 6 | 100054606 | 100063454 | 17 | 7 | 0.60239 | 0.27346 | PRDM13 | 0.27346 | 0.14265 | 0.19483 |
| ENSG00000106268 | 7 | 2281857 | 2290781 | 56 | 12 | 0.60207 | 0.27356 | NUDT1 | 0.27356 | 0.3198 | 0.46167 |
| ENSG00000117472 | 1 | 46640745 | 46651630 | 15 | 6 | 0.60185 | 0.27364 | TSPAN1 | 0.27364 | 0.16989 | 0.17163 |
| ENSG00000153879 | 19 | 33864236 | 33873592 | 25 | 9 | 0.60157 | 0.27373 | CEBPG | 0.27373 | 0.36622 | 0.089742 |
| ENSG00000196156 | 17 | 39323484 | 39324424 | 1 | 1 | 0.60136 | 0.2738 | KRTAP4-3 | 0.2738 | 0.35723 | 0.2516 |
| ENSG00000137100 | 9 | 34613548 | 34620515 | 7 | 3 | 0.60099 | 0.27392 | DCTN3 | 0.27392 | 0.36367 | 0.31618 |
| ENSG00000156711 | 6 | 36095586 | 36107842 | 31 | 10 | 0.60016 | 0.2742 | MAPK13 | 0.2742 | 0.23348 | 0.39442 |
| ENSG00000188039 | 19 | 16830787 | 16928774 | 385 | 38 | 0.60005 | 0.27423 | NWD1 | 0.27423 | 0.3629 | 0.28523 |
| ENSG00000126785 | 14 | 63670832 | 63759937 | 261 | 17 | 0.59981 | 0.27432 | RHOJ | 0.27432 | 0.32347 | 0.41095 |
| ENSG00000153786 | 16 | 85007787 | 85045141 | 227 | 21 | 0.59971 | 0.27435 | ZDHHC7 | 0.27435 | 0.24186 | 0.16242 |
| ENSG00000172893 | 11 | 71139239 | 71163914 | 70 | 8 | 0.59963 | 0.27438 | DHCR7 | 0.27438 | 0.2549 | 0.31855 |
| ENSG00000164488 | 6 | 168693510 | 168720434 | 143 | 37 | 0.5994 | 0.27445 | DACT2 | 0.27445 | 0.27672 | 0.31641 |
| ENSG00000112902 | 5 | 9035138 | 9546187 | 1992 | 166 | 0.59923 | 0.27451 | SEMA5A | 0.27451 | 0.37951 | 0.40811 |
| ENSG00000182256 | 15 | 27216429 | 27778373 | 1907 | 185 | 0.59903 | 0.27458 | GABRG3 | 0.27458 | 0.30078 | 0.45682 |

| | | | | | | | | | | | |
|---|---|---|---|---|---|---|---|---|---|---|---|
| ENSG00000128849 | 15 | 57668165 | 57842925 | 811 | 78 | 0.59803 | 0.27491 | CGNL1 | 0.27491 | 0.2672 | 0.085754 |
| ENSG00000184281 | 11 | 2421718 | 2425106 | 13 | 7 | 0.59779 | 0.27499 | TSSC4 | 0.27499 | 0.086429 | 0.67938 |
| ENSG00000165389 | 14 | 34901995 | 34931562 | 125 | 12 | 0.59764 | 0.27504 | SPTSSA | 0.27504 | 0.27779 | 0.17173 |
| ENSG00000134982 | 5 | 112043195 | 112181936 | 387 | 21 | 0.59762 | 0.27505 | APC | 0.27505 | 0.2671 | 0.28894 |
| ENSG00000103319 | 16 | 22217603 | 22298554 | 193 | 25 | 0.59727 | 0.27516 | EEF2K | 0.27516 | 0.32549 | 0.37015 |
| ENSG00000213965 | 19 | 33182867 | 33204702 | 55 | 8 | 0.59702 | 0.27525 | NUDT19 | 0.27525 | 0.20908 | 0.091632 |
| ENSG00000156140 | 4 | 73146686 | 73434516 | 626 | 39 | 0.59699 | 0.27526 | ADAMTS3 | 0.27526 | 0.2247 | 0.099678 |
| ENSG00000150636 | 18 | 66382446 | 66722426 | 1308 | 69 | 0.59697 | 0.27526 | CCDC102B | 0.27526 | 0.3305 | 0.54429 |
| ENSG00000094804 | 17 | 38443885 | 38459171 | 16 | 4 | 0.59682 | 0.27531 | CDC6 | 0.27531 | 0.50667 | 0.66167 |
| ENSG00000132142 | 17 | 35441923 | 35766909 | 657 | 41 | 0.5968 | 0.27532 | ACACA | 0.27532 | 0.20529 | 0.18598 |
| ENSG00000020256 | 20 | 50668202 | 50820847 | 734 | 57 | 0.59652 | 0.27541 | ZFP64 | 0.27541 | 0.30565 | 0.27517 |
| ENSG00000163273 | 2 | 232786530 | 232791113 | 8 | 3 | 0.59638 | 0.27546 | NPPC | 0.27546 | 0.5139 | 0.72914 |
| ENSG00000089116 | 12 | 113899839 | 113910085 | 25 | 7 | 0.59637 | 0.27546 | LHX5 | 0.27546 | 0.30763 | 0.33954 |
| ENSG00000139329 | 12 | 91496406 | 91505608 | 43 | 6 | 0.59634 | 0.27547 | LUM | 0.27547 | 0.4364 | 0.63156 |
| ENSG00000140538 | 15 | 88418230 | 88799999 | 867 | 72 | 0.59636 | 0.27547 | NTRK3 | 0.27547 | 0.30945 | 0.33718 |
| ENSG00000244509 | 22 | 39410088 | 39416357 | 26 | 6 | 0.59629 | 0.27549 | APOBEC3C | 0.27549 | 0.0075799 | 0.0022866 |
| ENSG00000188641 | 1 | 97543299 | 98386605 | 2461 | 128 | 0.59626 | 0.2755 | DPYD | 0.2755 | 0.28484 | 0.27759 |
| ENSG00000008441 | 19 | 13106422 | 13209610 | 182 | 21 | 0.59619 | 0.27552 | NFIX | 0.27552 | 0.28559 | 0.23132 |
| ENSG00000269000 | 2 | 107099728 | 107103848 | 16 | 4 | 0.5961 | 0.27555 | AC108868.1 | 0.27555 | 0.40712 | 0.41381 |
| ENSG00000130300 | 19 | 17462257 | 17488159 | 134 | 28 | 0.59606 | 0.27557 | PLVAP | 0.27557 | 0.15857 | 0.12588 |
| ENSG00000232013 | 3 | 145782358 | 145784156 | 2 | 1 | 0.59562 | 0.27572 | AC107021.1 | 0.27572 | 0.39014 | 0.57043 |
| ENSG00000164180 | 5 | 87485450 | 87565293 | 158 | 22 | 0.59553 | 0.27575 | TMEM161B | 0.27575 | 0.1934 | 0.16195 |
| ENSG00000145247 | 4 | 48887036 | 48908954 | 40 | 8 | 0.59522 | 0.27585 | OCIAD2 | 0.27585 | 0.32324 | 0.43105 |
| ENSG00000197429 | 1 | 46159996 | 46216322 | 99 | 10 | 0.5949 | 0.27595 | IPP | 0.27595 | 0.30858 | 0.34325 |
| ENSG00000147457 | 8 | 23101150 | 23119512 | 82 | 19 | 0.59462 | 0.27605 | CHMP7 | 0.27605 | 0.2471 | 0.33254 |
| ENSG00000171861 | 17 | 685513 | 695749 | 39 | 11 | 0.59458 | 0.27606 | RNMTL1 | 0.27606 | 0.29779 | 0.47157 |
| ENSG00000135047 | 9 | 90340434 | 90346308 | 17 | 7 | 0.59455 | 0.27607 | CTSL | 0.27607 | 0.30614 | 0.40187 |
| ENSG00000240764 | 5 | 140868808 | 140892546 | 74 | 16 | 0.59432 | 0.27615 | PCDHGC5 | 0.27615 | 0.41473 | 0.33284 |
| ENSG00000114331 | 3 | 194995465 | 195163807 | 616 | 15 | 0.59427 | 0.27616 | ACAP2 | 0.27616 | 0.27286 | 0.27277 |
| ENSG00000111962 | 6 | 149068464 | 149398126 | 1203 | 116 | 0.59421 | 0.27619 | UST | 0.27619 | 0.21201 | 0.2752 |
| ENSG00000152969 | 4 | 6027926 | 6202318 | 665 | 115 | 0.59416 | 0.2762 | JAKMIP1 | 0.2762 | 0.2665 | 0.45175 |
| ENSG00000116996 | 1 | 238045705 | 238054094 | 26 | 9 | 0.59404 | 0.27624 | ZP4 | 0.27624 | 0.30951 | 0.28065 |
| ENSG00000143162 | 1 | 167498914 | 167523004 | 129 | 30 | 0.59389 | 0.27629 | CREG1 | 0.27629 | 0.10604 | 0.082496 |
| ENSG00000240344 | 2 | 201735630 | 201754026 | 47 | 13 | 0.59386 | 0.2763 | PPIL3 | 0.2763 | 0.31068 | 0.20092 |
| ENSG00000157916 | 1 | 2323267 | 2336883 | 42 | 12 | 0.59365 | 0.27637 | RER1 | 0.27637 | 0.4228 | 0.76661 |
| ENSG00000179520 | 12 | 100750857 | 100815837 | 196 | 30 | 0.59361 | 0.27639 | SLC17A8 | 0.27639 | 0.2918 | 0.34251 |
| ENSG00000188921 | 9 | 20995306 | 21031635 | 156 | 17 | 0.59354 | 0.27641 | PTPLAD2 | 0.27641 | 0.34999 | 0.4379 |
| ENSG00000135569 | 6 | 132909731 | 132910877 | 3 | 2 | 0.59346 | 0.27644 | TAAR5 | 0.27644 | 0.40092 | 0.3004 |
| ENSG00000170454 | 12 | 52817854 | 52828309 | 31 | 6 | 0.59291 | 0.27662 | KRT75 | 0.27662 | 0.29095 | 0.3526 |
| ENSG00000186638 | 9 | 34252379 | 34329198 | 215 | 20 | 0.59276 | 0.27667 | KIF24 | 0.27667 | 0.29854 | 0.34332 |
| ENSG00000212657 | 17 | 39463952 | 39465505 | 8 | 2 | 0.59266 | 0.27671 | KRTAP16-1 | 0.27671 | 0.20491 | 0.24606 |
| ENSG00000090006 | 19 | 41098789 | 41135725 | 117 | 13 | 0.59252 | 0.27675 | LTBP4 | 0.27675 | 0.25369 | 0.33985 |
| ENSG00000232810 | 6 | 31543344 | 31546113 | 8 | 4 | 0.59229 | 0.27683 | TNF | 0.27683 | 0.070683 | 0.0995 |
| ENSG00000105568 | 19 | 52693292 | 52730687 | 150 | 17 | 0.5922 | 0.27686 | PPP2R1A | 0.27686 | 0.23692 | 0.27243 |
| ENSG00000124813 | 6 | 45295894 | 45632086 | 987 | 71 | 0.59209 | 0.2769 | RUNX2 | 0.2769 | 0.19009 | 0.20023 |
| ENSG00000255150 | 12 | 104697517 | 104698980 | 4 | 2 | 0.59204 | 0.27691 | EID3 | 0.27691 | 0.067741 | 0.072727 |
| ENSG00000212725 | 17 | 39202793 | 39203568 | 11 | 4 | 0.592 | 0.27692 | KRTAP2-1 | 0.27692 | 0.64236 | 0.69357 |
| ENSG00000133059 | 1 | 205111632 | 205180727 | 191 | 14 | 0.592 | 0.27693 | DSTYK | 0.27693 | 0.21621 | 0.21263 |
| ENSG00000122026 | 13 | 27825446 | 27830828 | 19 | 8 | 0.59172 | 0.27702 | RPL21 | 0.27702 | 0.32958 | 0.29475 |
| ENSG00000129048 | 3 | 132316081 | 132337811 | 45 | 11 | 0.59151 | 0.27709 | ACKR4 | 0.27709 | 0.33087 | 0.3465 |
| ENSG00000183060 | 15 | 100255902 | 100273766 | 65 | 11 | 0.59146 | 0.27711 | LYSMD4 | 0.27711 | 0.16343 | 0.073745 |
| ENSG00000183034 | 17 | 72920370 | 72930007 | 8 | 4 | 0.59078 | 0.27734 | OTOP2 | 0.27734 | 0.44761 | 0.46451 |
| ENSG00000186994 | 19 | 8387468 | 8408146 | 137 | 13 | 0.59027 | 0.2775 | KANK3 | 0.2775 | 0.16802 | 0.10161 |
| ENSG00000149084 | 11 | 43577986 | 43878167 | 811 | 30 | 0.5902 | 0.27753 | HSD17B12 | 0.27753 | 0.29557 | 0.33464 |
| ENSG00000148136 | 9 | 107288534 | 107289490 | 2 | 1 | 0.58988 | 0.27763 | OR13C4 | 0.27763 | 0.56502 | 0.77238 |
| ENSG00000188092 | 1 | 147400506 | 147465753 | 55 | 4 | 0.5891 | 0.2779 | GPR89B | 0.2779 | 0.29419 | 0.31607 |
| ENSG00000111490 | 12 | 65174589 | 65274812 | 166 | 26 | 0.58906 | 0.27791 | TBC1D30 | 0.27791 | 0.23459 | 0.22025 |
| ENSG00000160654 | 11 | 118215059 | 118225876 | 28 | 7 | 0.58891 | 0.27796 | CD3G | 0.27796 | 0.53161 | 0.6884 |
| ENSG00000169085 | 8 | 67372238 | 67430759 | 204 | 30 | 0.58886 | 0.27798 | C8orf46 | 0.27798 | 0.30027 | 0.31766 |
| ENSG00000100216 | 22 | 39077953 | 39080818 | 7 | 2 | 0.5887 | 0.27803 | TOMM22 | 0.27803 | 0.42404 | 0.44187 |
| ENSG00000173267 | 10 | 88718375 | 88723017 | 22 | 5 | 0.58854 | 0.27809 | SNCG | 0.27809 | 0.17404 | 0.18312 |
| ENSG00000136098 | 13 | 52706775 | 52733996 | 86 | 12 | 0.58837 | 0.27814 | NEK3 | 0.27814 | 0.23929 | 0.23586 |
| ENSG00000169689 | 17 | 79976578 | 79981983 | 22 | 5 | 0.5883 | 0.27816 | STRA13 | 0.27816 | 0.37835 | 0.55116 |
| ENSG00000140307 | 15 | 59930261 | 59949740 | 74 | 7 | 0.58829 | 0.27817 | GTF2A2 | 0.27817 | 0.31338 | 0.37288 |
| ENSG00000101327 | 20 | 1959403 | 1974732 | 37 | 7 | 0.58811 | 0.27823 | PDYN | 0.27823 | 0.23606 | 0.096182 |
| ENSG00000132017 | 19 | 14063304 | 14072254 | 15 | 6 | 0.58792 | 0.27829 | DCAF15 | 0.27829 | 0.42715 | 0.53104 |
| ENSG00000101074 | 20 | 42965626 | 42979875 | 63 | 5 | 0.58794 | 0.27829 | R3HDML | 0.27829 | 0.37709 | 0.4456 |
| ENSG00000163157 | 1 | 151142462 | 151148760 | 8 | 4 | 0.58774 | 0.27835 | TMOD4 | 0.27835 | 0.90806 | 0.92419 |
| ENSG00000154165 | 3 | 98250743 | 98251960 | 2 | 1 | 0.58727 | 0.27851 | GPR15 | 0.27851 | 0.58415 | 0.59743 |
| ENSG00000005381 | 17 | 56347217 | 56358296 | 17 | 8 | 0.58695 | 0.27862 | MPO | 0.27862 | 0.34403 | 0.27393 |
| ENSG00000170421 | 12 | 53290977 | 53343738 | 186 | 23 | 0.58676 | 0.27868 | KRT8 | 0.27868 | 0.38543 | 0.35694 |
| ENSG00000132603 | 16 | 69373333 | 69377014 | 16 | 6 | 0.58609 | 0.27891 | NIP7 | 0.27891 | 0.28245 | 0.30327 |
| ENSG00000139193 | 12 | 6554033 | 6560884 | 27 | 14 | 0.58568 | 0.27904 | CD27 | 0.27904 | 0.12352 | 0.15944 |

| | | | | | | | | | | | |
|---|---|---|---|---|---|---|---|---|---|---|---|
| ENSG00000125888 | 20 | 17674317 | 17716516 | 158 | 20 | 0.58559 | 0.27908 | BANF2 | 0.27908 | 0.26181 | 0.29754 |
| ENSG00000172354 | 7 | 100271154 | 100276797 | 11 | 6 | 0.58555 | 0.27909 | GNB2 | 0.27909 | 0.46642 | 0.54129 |
| ENSG00000114790 | 3 | 153838792 | 153975616 | 416 | 25 | 0.58518 | 0.27921 | ARHGEF26 | 0.27921 | 0.28903 | 0.30995 |
| ENSG00000123416 | 12 | 49521565 | 49525180 | 10 | 2 | 0.58466 | 0.27939 | TUBA1B | 0.27939 | 0.26885 | 0.25052 |
| ENSG00000130363 | 6 | 159393903 | 159421219 | 199 | 8 | 0.58459 | 0.27941 | RSPH3 | 0.27941 | 0.28906 | 0.30432 |
| ENSG00000132205 | 18 | 2847028 | 2915991 | 292 | 55 | 0.58452 | 0.27944 | EMILIN2 | 0.27944 | 0.29663 | 0.44465 |
| ENSG00000168883 | 2 | 85829979 | 85876403 | 103 | 22 | 0.58425 | 0.27953 | USP39 | 0.27953 | 0.2834 | 0.097816 |
| ENSG00000174358 | 5 | 1201710 | 1225232 | 140 | 33 | 0.58409 | 0.27958 | SLC6A19 | 0.27958 | 0.30743 | 0.34482 |
| ENSG00000059378 | 7 | 139723544 | 139763521 | 167 | 23 | 0.58409 | 0.27958 | PARP12 | 0.27958 | 0.34742 | 0.33278 |
| ENSG00000145375 | 4 | 123844229 | 124240605 | 1584 | 31 | 0.58385 | 0.27966 | SPATA5 | 0.27966 | 0.28383 | 0.28586 |
| ENSG00000180104 | 5 | 443273 | 472052 | 155 | 17 | 0.58375 | 0.27969 | EXOC3 | 0.27969 | 0.3062 | 0.22304 |
| ENSG00000071794 | 3 | 148747914 | 148804341 | 145 | 19 | 0.58371 | 0.27971 | HLTF | 0.27971 | 0.21808 | 0.20458 |
| ENSG00000198917 | 9 | 131581930 | 131592100 | 40 | 6 | 0.58365 | 0.27973 | C9orf114 | 0.27973 | 0.27628 | 0.26308 |
| ENSG00000196110 | 19 | 9404951 | 9420514 | 34 | 6 | 0.58358 | 0.27975 | ZNF699 | 0.27975 | 0.25025 | 0.29484 |
| ENSG00000225614 | 16 | 88493879 | 88507165 | 40 | 15 | 0.58347 | 0.27979 | ZNF469 | 0.27979 | 0.53092 | 0.43175 |
| ENSG00000116852 | 1 | 200938520 | 200992828 | 148 | 16 | 0.58342 | 0.27981 | KIF21B | 0.27981 | 0.32041 | 0.44535 |
| ENSG00000105699 | 19 | 35739233 | 35758867 | 61 | 11 | 0.58304 | 0.27993 | LSR | 0.27993 | 0.34714 | 0.40323 |
| ENSG00000215375 | 4 | 667369 | 675822 | 30 | 10 | 0.58285 | 0.28 | MYL5 | 0.28 | 0.30444 | 0.32231 |
| ENSG00000061936 | 12 | 132195626 | 132284282 | 201 | 25 | 0.58274 | 0.28003 | SFSWAP | 0.28003 | 0.29196 | 0.36958 |
| ENSG00000263620 | 17 | 8054134 | 8066293 | 36 | 7 | 0.58274 | 0.28003 | RP11-599B13.6 | 0.28003 | 0.41529 | 0.50431 |
| ENSG00000111452 | 12 | 131438452 | 131626014 | 979 | 71 | 0.5823 | 0.28018 | GPR133 | 0.28018 | 0.29987 | 0.33268 |
| ENSG00000070731 | 17 | 74559792 | 74583038 | 138 | 16 | 0.58223 | 0.28021 | ST6GALNAC2 | 0.28021 | 0.30365 | 0.11456 |
| ENSG00000141141 | 17 | 35969787 | 36003493 | 57 | 9 | 0.58194 | 0.2803 | DDX52 | 0.2803 | 0.23862 | 0.14505 |
| ENSG00000148602 | 10 | 85991349 | 86001217 | 38 | 6 | 0.58188 | 0.28032 | LRIT1 | 0.28032 | 0.065527 | 0.11021 |
| ENSG00000145241 | 4 | 68337521 | 68411324 | 167 | 26 | 0.58187 | 0.28033 | CENPC | 0.28033 | 0.29718 | 0.33103 |
| ENSG00000182791 | 11 | 66357640 | 66360554 | 3 | 2 | 0.58172 | 0.28038 | CCDC87 | 0.28038 | 0.1002 | 0.31686 |
| ENSG00000167644 | 19 | 38794801 | 38795649 | 7 | 3 | 0.58171 | 0.28038 | C19orf33 | 0.28038 | 0.18634 | 0.27923 |
| ENSG00000140465 | 15 | 75011883 | 75017951 | 5 | 3 | 0.58162 | 0.28041 | CYP1A1 | 0.28041 | 0.40003 | 0.10563 |
| ENSG00000142765 | 1 | 27668513 | 27680421 | 26 | 9 | 0.58145 | 0.28047 | SYTL1 | 0.28047 | 0.37363 | 0.37953 |
| ENSG00000132535 | 17 | 7093209 | 7123021 | 57 | 13 | 0.58116 | 0.28057 | DLG4 | 0.28057 | 0.26894 | 0.25583 |
| ENSG00000164023 | 4 | 108745719 | 108836203 | 237 | 47 | 0.58096 | 0.28063 | SGMS2 | 0.28063 | 0.097747 | 0.038788 |
| ENSG00000183569 | 22 | 42949623 | 42970388 | 93 | 15 | 0.58096 | 0.28063 | SERHL2 | 0.28063 | 0.32577 | 0.31121 |
| ENSG00000079313 | 19 | 1815248 | 1848452 | 87 | 11 | 0.58058 | 0.28076 | REXO1 | 0.28076 | 0.29134 | 0.16895 |
| ENSG00000079134 | 18 | 214520 | 268050 | 106 | 14 | 0.58037 | 0.28083 | THOC1 | 0.28083 | 0.15064 | 0.094102 |
| ENSG00000113068 | 5 | 139624624 | 139682706 | 117 | 11 | 0.58009 | 0.28093 | PFDN1 | 0.28093 | 0.21519 | 0.18934 |
| ENSG00000099308 | 19 | 18208603 | 18262502 | 235 | 21 | 0.57994 | 0.28098 | MAST3 | 0.28098 | 0.31635 | 0.36637 |
| ENSG00000187918 | 11 | 5474638 | 5475707 | 6 | 3 | 0.57985 | 0.28101 | OR51I2 | 0.28101 | 0.64462 | 0.72508 |
| ENSG00000124374 | 2 | 71409869 | 71454213 | 203 | 23 | 0.57961 | 0.28109 | PAIP2B | 0.28109 | 0.4065 | 0.37332 |
| ENSG00000174948 | 3 | 154055461 | 154147504 | 288 | 23 | 0.57957 | 0.2811 | GPR149 | 0.2811 | 0.25029 | 0.21981 |
| ENSG00000168264 | 1 | 234740015 | 234745271 | 13 | 5 | 0.57955 | 0.28111 | IRF2BP2 | 0.28111 | 0.36289 | 0.30989 |
| ENSG00000133103 | 13 | 40229764 | 40365802 | 386 | 22 | 0.5792 | 0.28123 | COG6 | 0.28123 | 0.27526 | 0.32249 |
| ENSG00000135503 | 12 | 52345451 | 52390862 | 97 | 16 | 0.57917 | 0.28124 | ACVR1B | 0.28124 | 0.2348 | 0.53209 |
| ENSG00000152558 | 11 | 102267063 | 102341115 | 169 | 20 | 0.57887 | 0.28134 | TMEM123 | 0.28134 | 0.25518 | 0.29933 |
| ENSG00000167566 | 12 | 50184929 | 50222533 | 141 | 15 | 0.57873 | 0.28139 | NCKAP5L | 0.28139 | 0.25521 | 0.30147 |
| ENSG00000128159 | 22 | 50656118 | 50683421 | 128 | 21 | 0.5785 | 0.28146 | TUBGCP6 | 0.28146 | 0.27147 | 0.23172 |
| ENSG00000272514 | 6 | 88117701 | 88174183 | 222 | 18 | 0.57834 | 0.28152 | C6ORF165 | 0.28152 | 0.14369 | 0.14678 |
| ENSG00000129277 | 17 | 34430983 | 34433014 | 3 | 1 | 0.57797 | 0.28164 | CCL4 | 0.28164 | 0.46109 | 0.16925 |
| ENSG00000187037 | 7 | 37723399 | 37873390 | 500 | 37 | 0.57761 | 0.28176 | GPR141 | 0.28176 | 0.27382 | 0.2438 |
| ENSG00000113583 | 5 | 133291201 | 133304478 | 24 | 7 | 0.5775 | 0.2818 | C5orf15 | 0.2818 | 0.60092 | 0.72674 |
| ENSG00000180611 | 3 | 192514604 | 192635950 | 460 | 33 | 0.57745 | 0.28182 | MB21D2 | 0.28182 | 0.3688 | 0.42413 |
| ENSG00000105376 | 19 | 10400657 | 10407454 | 21 | 5 | 0.57712 | 0.28193 | ICAM5 | 0.28193 | 0.19675 | 0.068289 |
| ENSG00000174469 | 7 | 145813453 | 148118090 | 9850 | 345 | 0.57687 | 0.28201 | CNTNAP2 | 0.28201 | 0.27531 | 0.29192 |
| ENSG00000167705 | 17 | 1549444 | 1553371 | 8 | 3 | 0.57688 | 0.28201 | RILP | 0.28201 | 0.29414 | 0.43027 |
| ENSG00000153989 | 6 | 117996665 | 118031803 | 102 | 19 | 0.57686 | 0.28202 | NUS1 | 0.28202 | 0.30898 | 0.33032 |
| ENSG00000204348 | 6 | 31937587 | 31940069 | 9 | 5 | 0.57616 | 0.28225 | DXO | 0.28225 | 0.0012119 | 0.00020024 |
| ENSG00000205639 | 2 | 24232951 | 24286191 | 161 | 27 | 0.57601 | 0.2823 | MFSD2B | 0.2823 | 0.092574 | 0.082875 |
| ENSG00000107937 | 10 | 1034338 | 1065876 | 133 | 17 | 0.57596 | 0.28232 | GTPBP4 | 0.28232 | 0.2274 | 0.21422 |
| ENSG00000204560 | 6 | 30620896 | 30640814 | 46 | 12 | 0.57541 | 0.28251 | DHX16 | 0.28251 | 0.39723 | 0.43679 |
| ENSG00000167699 | 17 | 660337 | 685581 | 73 | 20 | 0.57532 | 0.28254 | GLOD4 | 0.28254 | 0.17239 | 0.21861 |
| ENSG00000248727 | 5 | 55753622 | 55777773 | 137 | 22 | 0.57499 | 0.28265 | CTC-236F12.4 | 0.28265 | 0.30739 | 0.15743 |
| ENSG00000075073 | 10 | 71163659 | 71176623 | 53 | 7 | 0.57475 | 0.28273 | TACR2 | 0.28273 | 0.55515 | 0.41735 |
| ENSG00000185359 | 17 | 79650356 | 79670168 | 97 | 12 | 0.57466 | 0.28276 | HGS | 0.28276 | 0.29346 | 0.26223 |
| ENSG00000170955 | 11 | 6340176 | 6341877 | 8 | 4 | 0.57409 | 0.28295 | PRKCDBP | 0.28295 | 0.41934 | 0.25773 |
| ENSG00000100433 | 14 | 88649113 | 88793251 | 617 | 57 | 0.57399 | 0.28299 | KCNK10 | 0.28299 | 0.33849 | 0.29408 |
| ENSG00000212124 | 12 | 11174218 | 11175219 | 8 | 4 | 0.57387 | 0.28303 | TAS2R19 | 0.28303 | 0.41843 | 0.45515 |
| ENSG00000125107 | 16 | 58553855 | 58663790 | 418 | 18 | 0.57379 | 0.28305 | CNOT1 | 0.28305 | 0.27755 | 0.2978 |
| ENSG00000130204 | 19 | 45393826 | 45406946 | 66 | 15 | 0.5737 | 0.28309 | TOMM40 | 0.28309 | 0.21063 | 0.3542 |
| ENSG00000153815 | 16 | 81478775 | 81745367 | 1204 | 120 | 0.57355 | 0.28314 | CMIP | 0.28314 | 0.29093 | 0.36143 |
| ENSG00000137074 | 9 | 32972604 | 33025166 | 285 | 8 | 0.57345 | 0.28317 | APTX | 0.28317 | 0.28962 | 0.25169 |
| ENSG00000161640 | 19 | 50452242 | 50464429 | 37 | 10 | 0.57345 | 0.28317 | SIGLEC11 | 0.28317 | 0.13183 | 0.2041 |
| ENSG00000135094 | 12 | 113830250 | 113864106 | 90 | 7 | 0.57341 | 0.28318 | SDS | 0.28318 | 0.24346 | 0.27317 |
| ENSG00000180424 | 20 | 30028322 | 30038060 | 29 | 6 | 0.57291 | 0.28335 | DEFB123 | 0.28335 | 0.42592 | 0.39109 |
| ENSG00000198812 | 12 | 70002351 | 70004942 | 4 | 2 | 0.57288 | 0.28336 | LRRC10 | 0.28336 | 0.38317 | 0.40574 |

| | | | | | | | | | | | |
|---|---|---|---|---|---|---|---|---|---|---|---|
| ENSG00000170891 | 4 | 5016313 | 5021199 | 29 | 5 | 0.57262 | 0.28345 | CYTL1 | 0.28345 | 0.25215 | 0.22339 |
| ENSG00000173612 | 6 | 117113248 | 117150200 | 118 | 17 | 0.57246 | 0.2835 | GPRC6A | 0.2835 | 0.30148 | 0.28831 |
| ENSG00000204406 | 2 | 148778580 | 149275805 | 1175 | 48 | 0.57236 | 0.28354 | MBD5 | 0.28354 | 0.30823 | 0.3219 |
| ENSG00000221977 | 14 | 22133297 | 22134238 | 4 | 2 | 0.57234 | 0.28355 | OR4E2 | 0.28355 | 0.24832 | 0.13952 |
| ENSG00000095713 | 10 | 99624757 | 99790585 | 356 | 46 | 0.57207 | 0.28364 | CRTAC1 | 0.28364 | 0.29554 | 0.17397 |
| ENSG00000006534 | 11 | 67776048 | 67796744 | 57 | 7 | 0.57201 | 0.28366 | ALDH3B1 | 0.28366 | 0.39201 | 0.43897 |
| ENSG00000135074 | 5 | 156822542 | 157002783 | 739 | 37 | 0.57183 | 0.28372 | ADAM19 | 0.28372 | 0.267 | 0.26378 |
| ENSG00000185070 | 14 | 85996488 | 86095034 | 394 | 42 | 0.57173 | 0.28375 | FLRT2 | 0.28375 | 0.23973 | 0.30716 |
| ENSG00000163110 | 4 | 95373037 | 95589377 | 546 | 24 | 0.57164 | 0.28378 | PDLIM5 | 0.28378 | 0.2758 | 0.28476 |
| ENSG00000215018 | 7 | 7395834 | 7575484 | 712 | 29 | 0.57155 | 0.28381 | COL28A1 | 0.28381 | 0.25914 | 0.23467 |
| ENSG00000115520 | 2 | 198318147 | 198340032 | 33 | 7 | 0.57115 | 0.28395 | COQ10B | 0.28395 | 0.30455 | 0.16424 |
| ENSG00000173926 | 5 | 126203406 | 126366500 | 495 | 11 | 0.57113 | 0.28396 | MARCHF3 | 0.28396 | 0.29064 | 0.27909 |
| ENSG00000164610 | 7 | 33134409 | 33149013 | 42 | 6 | 0.57107 | 0.28398 | RP9 | 0.28398 | 0.30247 | 0.39678 |
| ENSG00000186967 | 21 | 31869142 | 31869451 | 1 | 1 | 0.571 | 0.284 | KRTAP19-4 | 0.284 | 0.49628 | 0.53806 |
| ENSG00000143149 | 1 | 165631453 | 165668100 | 168 | 18 | 0.57093 | 0.28402 | ALDH9A1 | 0.28402 | 0.29437 | 0.36889 |
| ENSG00000257365 | 14 | 65381203 | 65529368 | 406 | 45 | 0.57052 | 0.28416 | FNTB | 0.28416 | 0.28231 | 0.25928 |
| ENSG00000171492 | 1 | 90286573 | 90402170 | 211 | 44 | 0.57006 | 0.28432 | LRRC8D | 0.28432 | 0.095466 | 0.00096302 |
| ENSG00000179119 | 11 | 18627948 | 18656338 | 104 | 16 | 0.57002 | 0.28433 | SPTY2D1 | 0.28433 | 0.20384 | 0.14047 |
| ENSG00000131143 | 16 | 85832239 | 85840650 | 10 | 6 | 0.57 | 0.28434 | COX4I1 | 0.28434 | 0.69433 | 0.78577 |
| ENSG00000103042 | 16 | 58699013 | 58719008 | 89 | 15 | 0.56978 | 0.28441 | SLC38A7 | 0.28441 | 0.43384 | 0.37958 |
| ENSG00000185594 | 15 | 97326619 | 97328845 | 6 | 3 | 0.56963 | 0.28446 | SPATA8 | 0.28446 | 0.41978 | 0.47564 |
| ENSG00000245317 | 5 | 179286013 | 179288283 | 6 | 2 | 0.56947 | 0.28452 | CTC-241N9.1 | 0.28452 | 0.60547 | 0.63554 |
| ENSG00000137055 | 9 | 26904081 | 26947461 | 187 | 18 | 0.56946 | 0.28452 | PLAA | 0.28452 | 0.31593 | 0.35049 |
| ENSG00000228919 | 4 | 7940728 | 7942022 | 9 | 2 | 0.56936 | 0.28456 | AC097381.1 | 0.28456 | 0.14511 | 0.15192 |
| ENSG00000269531 | 12 | 80198742 | 80202321 | 6 | 2 | 0.56906 | 0.28466 | AC073569.1 | 0.28466 | 0.34022 | 0.25889 |
| ENSG00000136021 | 12 | 100660918 | 100735502 | 151 | 18 | 0.56906 | 0.28466 | SCYL2 | 0.28466 | 0.31267 | 0.32678 |
| ENSG00000115904 | 2 | 39208537 | 39351486 | 372 | 14 | 0.56897 | 0.28469 | SOS1 | 0.28469 | 0.28145 | 0.2548 |
| ENSG00000198963 | 9 | 77112281 | 77308093 | 404 | 45 | 0.56877 | 0.28476 | RORB | 0.28476 | 0.36364 | 0.50186 |
| ENSG00000100380 | 22 | 41220539 | 41253026 | 52 | 9 | 0.56877 | 0.28476 | ST13 | 0.28476 | 0.27124 | 0.41563 |
| ENSG00000149480 | 11 | 62360686 | 62369312 | 13 | 3 | 0.56825 | 0.28493 | MTA2 | 0.28493 | 0.3089 | 0.31085 |
| ENSG00000178381 | 7 | 1191707 | 1200395 | 77 | 14 | 0.56818 | 0.28496 | ZFAND2A | 0.28496 | 0.38965 | 0.36959 |
| ENSG00000197562 | 16 | 639357 | 679272 | 141 | 10 | 0.56808 | 0.28499 | RAB40C | 0.28499 | 0.42394 | 0.41504 |
| ENSG00000146205 | 2 | 242127924 | 242164792 | 137 | 29 | 0.56757 | 0.28516 | ANO7 | 0.28516 | 0.54544 | 0.67098 |
| ENSG00000122584 | 7 | 8473585 | 8792593 | 1491 | 78 | 0.56745 | 0.28521 | NXPH1 | 0.28521 | 0.31701 | 0.36822 |
| ENSG00000114391 | 3 | 101399935 | 101405626 | 13 | 5 | 0.56702 | 0.28535 | RPL24 | 0.28535 | 0.32585 | 0.32032 |
| ENSG00000182902 | 22 | 18043139 | 18073760 | 136 | 28 | 0.56703 | 0.28535 | SLC25A18 | 0.28535 | 0.040496 | 0.15161 |
| ENSG00000162980 | 2 | 152645498 | 152685006 | 90 | 14 | 0.567 | 0.28536 | ARL5A | 0.28536 | 0.28778 | 0.22657 |
| ENSG00000115419 | 2 | 191745553 | 191830278 | 141 | 18 | 0.56689 | 0.2854 | GLS | 0.2854 | 0.38626 | 0.50086 |
| ENSG00000183486 | 21 | 42733870 | 42781317 | 172 | 38 | 0.56684 | 0.28541 | MX2 | 0.28541 | 0.26109 | 0.28369 |
| ENSG00000109063 | 17 | 10531843 | 10560626 | 111 | 8 | 0.56682 | 0.28542 | MYH3 | 0.28542 | 0.30655 | 0.48623 |
| ENSG00000188227 | 19 | 37997841 | 38039664 | 86 | 19 | 0.56618 | 0.28563 | ZNF793 | 0.28563 | 0.31405 | 0.33689 |
| ENSG00000155629 | 10 | 98353069 | 98480271 | 402 | 52 | 0.56618 | 0.28564 | PIK3AP1 | 0.28564 | 0.32164 | 0.33055 |
| ENSG00000186104 | 11 | 14899553 | 14913798 | 13 | 5 | 0.56613 | 0.28565 | CYP2R1 | 0.28565 | 0.055101 | 0.034295 |
| ENSG00000104763 | 8 | 17913934 | 17942494 | 199 | 16 | 0.56594 | 0.28572 | ASAH1 | 0.28572 | 0.37026 | 0.48178 |
| ENSG00000049759 | 18 | 55711599 | 56068772 | 1298 | 102 | 0.56594 | 0.28572 | NEDD4L | 0.28572 | 0.2723 | 0.29612 |
| ENSG00000167165 | 2 | 234600253 | 234681946 | 305 | 21 | 0.56552 | 0.28586 | UGT1A6 | 0.28586 | 0.22673 | 0.088743 |
| ENSG00000203909 | 6 | 74062785 | 74064018 | 1 | 1 | 0.5654 | 0.2859 | DPPA5 | 0.2859 | 0.63393 | 0.59617 |
| ENSG00000164849 | 7 | 1084212 | 1098897 | 104 | 14 | 0.56524 | 0.28595 | GPR146 | 0.28595 | 0.27577 | 0.24731 |
| ENSG00000156510 | 10 | 70980059 | 71027315 | 207 | 37 | 0.56526 | 0.28595 | HKDC1 | 0.28595 | 0.42266 | 0.51962 |
| ENSG00000109089 | 17 | 72983727 | 73001895 | 53 | 15 | 0.56495 | 0.28605 | CDR2L | 0.28605 | 0.74936 | 0.69009 |
| ENSG00000187955 | 8 | 121072019 | 121384275 | 960 | 59 | 0.56488 | 0.28608 | COL14A1 | 0.28608 | 0.34339 | 0.38157 |
| ENSG00000104814 | 19 | 39078281 | 39109522 | 99 | 16 | 0.56486 | 0.28608 | MAP4K1 | 0.28608 | 0.32429 | 0.39871 |
| ENSG00000134242 | 1 | 114356433 | 114414381 | 136 | 17 | 0.56464 | 0.28616 | PTPN22 | 0.28616 | 0.23706 | 0.26222 |
| ENSG00000258987 | 14 | 94405901 | 94410475 | 18 | 6 | 0.56435 | 0.28626 | RP11-131H24.4 | 0.28626 | 0.40868 | 0.85505 |
| ENSG00000132612 | 16 | 69345259 | 69358949 | 42 | 8 | 0.56398 | 0.28638 | VPS4A | 0.28638 | 0.2755 | 0.30278 |
| ENSG00000162066 | 16 | 2570358 | 2581423 | 8 | 5 | 0.56389 | 0.28641 | AMDHD2 | 0.28641 | 0.35443 | 0.059435 |
| ENSG00000113240 | 5 | 178029665 | 178057616 | 60 | 12 | 0.56362 | 0.2865 | CLK4 | 0.2865 | 0.23201 | 0.24497 |
| ENSG00000148053 | 9 | 87283466 | 87638505 | 1082 | 68 | 0.56357 | 0.28652 | NTRK2 | 0.28652 | 0.25153 | 0.2183 |
| ENSG00000255833 | 5 | 134779908 | 134788089 | 32 | 9 | 0.56346 | 0.28656 | TIFAB | 0.28656 | 0.15434 | 0.20682 |
| ENSG00000108861 | 17 | 41843489 | 41856356 | 36 | 5 | 0.56308 | 0.28669 | DUSP3 | 0.28669 | 0.32743 | 0.31938 |
| ENSG00000176092 | 1 | 26648350 | 26680621 | 113 | 17 | 0.56298 | 0.28672 | AIM1L | 0.28672 | 0.33452 | 0.32794 |
| ENSG00000103769 | 15 | 66018392 | 66184329 | 376 | 31 | 0.56285 | 0.28677 | RAB11A | 0.28677 | 0.28543 | 0.32909 |
| ENSG00000111913 | 6 | 24797601 | 25042238 | 1036 | 75 | 0.56254 | 0.28687 | FAM65B | 0.28687 | 0.37241 | 0.41307 |
| ENSG00000170759 | 10 | 32297938 | 32345359 | 129 | 14 | 0.56253 | 0.28688 | KIF5B | 0.28688 | 0.34207 | 0.33714 |
| ENSG00000131730 | 5 | 80529104 | 80562216 | 103 | 13 | 0.56223 | 0.28698 | CKMT2 | 0.28698 | 0.25072 | 0.24575 |
| ENSG00000196092 | 9 | 36833272 | 37034103 | 735 | 73 | 0.56198 | 0.28707 | PAX5 | 0.28707 | 0.25943 | 0.22345 |
| ENSG00000125954 | 14 | 65381203 | 65528521 | 404 | 46 | 0.56189 | 0.28709 | CHURC1-FNTB | 0.28709 | 0.27931 | 0.25955 |
| ENSG00000128604 | 7 | 128577666 | 128590089 | 46 | 13 | 0.56183 | 0.28712 | IRF5 | 0.28712 | 0.21172 | 0.28435 |
| ENSG00000139546 | 12 | 53894705 | 53900215 | 1 | 1 | 0.56158 | 0.2872 | TARBP2 | 0.2872 | 0.07248 | 0.022484 |
| ENSG00000171169 | 9 | 130823512 | 130830485 | 10 | 3 | 0.56151 | 0.28722 | NAIF1 | 0.28722 | 0.55606 | 0.66449 |
| ENSG00000114999 | 2 | 113239731 | 113299316 | 157 | 16 | 0.56143 | 0.28725 | TTL | 0.28725 | 0.2249 | 0.16964 |
| ENSG00000163492 | 2 | 179694484 | 179914813 | 770 | 51 | 0.56131 | 0.28729 | CCDC141 | 0.28729 | 0.20865 | 0.13763 |
| ENSG00000188467 | 15 | 48413169 | 48434869 | 7 | 3 | 0.56133 | 0.28729 | SLC24A5 | 0.28729 | 0.10498 | 0.11755 |

| | | | | | | | | | | | |
|---|---|---|---|---|---|---|---|---|---|---|---|
| ENSG00000130844 | 19 | 54024235 | 54083523 | 390 | 37 | 0.56125 | 0.28731 | ZNF331 | 0.28731 | 0.087853 | 0.046038 |
| ENSG00000163625 | 4 | 85590704 | 85887544 | 407 | 25 | 0.56123 | 0.28732 | WDFY3 | 0.28732 | 0.28428 | 0.26242 |
| ENSG00000149554 | 11 | 125495036 | 125546150 | 157 | 17 | 0.56116 | 0.28734 | CHEK1 | 0.28734 | 0.31235 | 0.32133 |
| ENSG00000213215 | 7 | 143657027 | 143658108 | 3 | 1 | 0.56112 | 0.28736 | OR2F1 | 0.28736 | 0.44416 | 0.60107 |
| ENSG00000088305 | 20 | 31350191 | 31397162 | 165 | 12 | 0.56109 | 0.28737 | DNMT3B | 0.28737 | 0.3365 | 0.43053 |
| ENSG00000118058 | 11 | 118307205 | 118397539 | 107 | 14 | 0.56073 | 0.28749 | KMT2A | 0.28749 | 0.29765 | 0.25029 |
| ENSG00000174599 | 4 | 118004718 | 118006736 | 1 | 1 | 0.56041 | 0.2876 | TRAM1L1 | 0.2876 | 0.29276 | 0.41838 |
| ENSG00000164972 | 9 | 34379017 | 34397830 | 46 | 11 | 0.56007 | 0.28772 | C9orf24 | 0.28772 | 0.24099 | 0.28506 |
| ENSG00000170525 | 10 | 6186881 | 6277495 | 515 | 103 | 0.55987 | 0.28778 | PFKFB3 | 0.28778 | 0.5859 | 0.60082 |
| ENSG00000166152 | 16 | 49407734 | 49433342 | 86 | 11 | 0.55986 | 0.28779 | C16orf78 | 0.28779 | 0.33386 | 0.45651 |
| ENSG00000172199 | 11 | 56143100 | 56144029 | 15 | 5 | 0.55954 | 0.2879 | OR8U1 | 0.2879 | 0.15655 | 0.076986 |
| ENSG00000224474 | 10 | 99179228 | 99185831 | 17 | 7 | 0.55949 | 0.28791 | AL355490.1 | 0.28791 | 0.046132 | 0.045136 |
| ENSG00000184674 | 22 | 24376133 | 24384680 | 9 | 2 | 0.55935 | 0.28796 | GSTT1 | 0.28796 | 0.16168 | 0.15684 |
| ENSG00000132359 | 17 | 2680350 | 2941033 | 1228 | 168 | 0.55931 | 0.28797 | RAP1GAP2 | 0.28797 | 0.2758 | 0.091143 |
| ENSG00000257599 | 12 | 29542227 | 29640421 | 457 | 23 | 0.55904 | 0.28807 | OVCH1-AS1 | 0.28807 | 0.26458 | 0.26168 |
| ENSG00000176105 | 18 | 721588 | 812547 | 280 | 22 | 0.55902 | 0.28807 | YES1 | 0.28807 | 0.30521 | 0.18192 |
| ENSG00000121060 | 17 | 54965270 | 54991399 | 46 | 12 | 0.55883 | 0.28814 | TRIM25 | 0.28814 | 0.35887 | 0.36781 |
| ENSG00000183091 | 2 | 152341850 | 152591001 | 461 | 41 | 0.55878 | 0.28816 | NEB | 0.28816 | 0.28395 | 0.3299 |
| ENSG00000131781 | 1 | 146646930 | 146714700 | 429 | 22 | 0.55835 | 0.2883 | FMO5 | 0.2883 | 0.29726 | 0.28129 |
| ENSG00000241043 | 9 | 32566787 | 32568619 | 6 | 3 | 0.55828 | 0.28833 | GVQW1 | 0.28833 | 0.38604 | 0.43792 |
| ENSG00000152443 | 19 | 58258164 | 58269527 | 25 | 7 | 0.55825 | 0.28834 | ZNF776 | 0.28834 | 0.3869 | 0.43953 |
| ENSG00000113360 | 5 | 31400604 | 31532303 | 306 | 26 | 0.55814 | 0.28837 | DROSHA | 0.28837 | 0.3131 | 0.33972 |
| ENSG00000198870 | 9 | 136243117 | 136271220 | 125 | 10 | 0.5581 | 0.28839 | C9orf96 | 0.28839 | 0.31655 | 0.39369 |
| ENSG00000196876 | 12 | 51984050 | 52206648 | 461 | 33 | 0.55795 | 0.28844 | SCN8A | 0.28844 | 0.25594 | 0.20359 |
| ENSG00000171791 | 18 | 60790579 | 60987361 | 633 | 98 | 0.55795 | 0.28844 | BCL2 | 0.28844 | 0.27372 | 0.32367 |
| ENSG00000013364 | 16 | 29831715 | 29859355 | 37 | 3 | 0.55769 | 0.28853 | MVP | 0.28853 | 0.28724 | 0.35385 |
| ENSG00000065183 | 1 | 118472343 | 118509165 | 81 | 7 | 0.55742 | 0.28862 | WDR3 | 0.28862 | 0.27985 | 0.24842 |
| ENSG00000117000 | 1 | 40627045 | 40706593 | 124 | 21 | 0.55739 | 0.28863 | RLF | 0.28863 | 0.33052 | 0.34692 |
| ENSG00000145649 | 5 | 54398476 | 54406080 | 12 | 3 | 0.55736 | 0.28864 | GZMA | 0.28864 | 0.40875 | 0.34983 |
| ENSG00000143971 | 2 | 67624451 | 67637677 | 41 | 6 | 0.55728 | 0.28867 | ETAA1 | 0.28867 | 0.18305 | 0.15932 |
| ENSG00000259518 | 15 | 82380935 | 82390035 | 40 | 15 | 0.55692 | 0.28879 | RP11-597K23.2 | 0.28879 | 0.57272 | 0.59926 |
| ENSG00000181786 | 19 | 8807751 | 8809172 | 4 | 2 | 0.5568 | 0.28883 | ACTL9 | 0.28883 | 0.24077 | 0.063586 |
| ENSG00000174130 | 4 | 38825336 | 38858438 | 183 | 15 | 0.5565 | 0.28894 | TLR6 | 0.28894 | 0.27023 | 0.313 |
| ENSG00000126226 | 13 | 113831891 | 113863029 | 40 | 13 | 0.55635 | 0.28899 | PCID2 | 0.28899 | 0.48445 | 0.66154 |
| ENSG00000166900 | 11 | 59480929 | 59573354 | 330 | 10 | 0.55622 | 0.28903 | STX3 | 0.28903 | 0.24725 | 0.22898 |
| ENSG00000183208 | 15 | 90777040 | 90785315 | 29 | 8 | 0.5562 | 0.28904 | GDPGP1 | 0.28904 | 0.37006 | 0.44124 |
| ENSG00000182158 | 7 | 137559725 | 137686813 | 358 | 44 | 0.55604 | 0.28909 | CREB3L2 | 0.28909 | 0.3555 | 0.34591 |
| ENSG00000181315 | 6 | 26636518 | 26659980 | 56 | 12 | 0.55589 | 0.28914 | ZNF322 | 0.28914 | 0.25843 | 0.31387 |
| ENSG00000142675 | 1 | 26503894 | 26516377 | 33 | 10 | 0.55571 | 0.28921 | CNKSR1 | 0.28921 | 0.30488 | 0.20172 |
| ENSG00000168792 | 17 | 27887565 | 27894155 | 12 | 7 | 0.55567 | 0.28922 | ABHD15 | 0.28922 | 0.27348 | 0.43025 |
| ENSG00000180818 | 12 | 54378849 | 54384063 | 7 | 4 | 0.55558 | 0.28925 | HOXC10 | 0.28925 | 0.095625 | 0.15011 |
| ENSG00000198498 | 4 | 164415594 | 164441691 | 57 | 5 | 0.55516 | 0.28939 | TMA16 | 0.28939 | 0.30035 | 0.34208 |
| ENSG00000180259 | 20 | 4711928 | 4721314 | 53 | 7 | 0.55494 | 0.28947 | PRNT | 0.28947 | 0.3553 | 0.61597 |
| ENSG00000151726 | 4 | 185676749 | 185747972 | 278 | 27 | 0.55488 | 0.28949 | ACSL1 | 0.28949 | 0.41788 | 0.47741 |
| ENSG00000140931 | 16 | 66637777 | 66647795 | 30 | 7 | 0.55487 | 0.28949 | CMTM3 | 0.28949 | 0.36853 | 0.35388 |
| ENSG00000148335 | 9 | 132371163 | 132398209 | 85 | 14 | 0.55464 | 0.28957 | NTMT1 | 0.28957 | 0.30678 | 0.20692 |
| ENSG00000112812 | 6 | 27215480 | 27224403 | 18 | 6 | 0.55407 | 0.28977 | PRSS16 | 0.28977 | 0.38946 | 0.32562 |
| ENSG00000183747 | 16 | 20462783 | 20498991 | 168 | 12 | 0.55403 | 0.28978 | ACSM2A | 0.28978 | 0.29747 | 0.46189 |
| ENSG00000128040 | 4 | 57676026 | 57687908 | 59 | 15 | 0.55362 | 0.28992 | SPINK2 | 0.28992 | 0.3306 | 0.35152 |
| ENSG00000122008 | 5 | 74807581 | 74896969 | 151 | 18 | 0.5536 | 0.28993 | POLK | 0.28993 | 0.2703 | 0.26695 |
| ENSG00000104408 | 8 | 109213445 | 109447562 | 785 | 26 | 0.55358 | 0.28993 | EIF3E | 0.28993 | 0.29497 | 0.30657 |
| ENSG00000137674 | 11 | 102447566 | 102496063 | 235 | 19 | 0.55335 | 0.29001 | MMP20 | 0.29001 | 0.19345 | 0.058208 |
| ENSG00000054983 | 14 | 88304164 | 88460009 | 809 | 31 | 0.55327 | 0.29004 | GALC | 0.29004 | 0.26098 | 0.17537 |
| ENSG00000023318 | 9 | 102741461 | 102861322 | 260 | 14 | 0.55312 | 0.29009 | ERP44 | 0.29009 | 0.31362 | 0.29071 |
| ENSG00000157978 | 1 | 25870071 | 25895377 | 82 | 12 | 0.55306 | 0.29011 | LDLRAP1 | 0.29011 | 0.34013 | 0.30975 |
| ENSG00000164051 | 3 | 48473574 | 48481866 | 14 | 3 | 0.55297 | 0.29014 | CCDC51 | 0.29014 | 0.44438 | 0.39794 |
| ENSG00000214285 | 10 | 129347613 | 129350935 | 36 | 8 | 0.55285 | 0.29018 | NPS | 0.29018 | 0.34476 | 0.4163 |
| ENSG00000184371 | 1 | 110452864 | 110473614 | 47 | 14 | 0.5528 | 0.2902 | CSF1 | 0.2902 | 0.32628 | 0.32336 |
| ENSG00000144677 | 3 | 37903451 | 38025960 | 332 | 29 | 0.55275 | 0.29022 | CTDSPL | 0.29022 | 0.35799 | 0.44062 |
| ENSG00000213922 | 19 | 39930212 | 39932082 | 8 | 2 | 0.55271 | 0.29023 | AC011500.1 | 0.29023 | 0.33133 | 0.35364 |
| ENSG00000157613 | 11 | 46299212 | 46342972 | 102 | 20 | 0.5526 | 0.29027 | CREB3L1 | 0.29027 | 0.2367 | 0.20184 |
| ENSG00000182557 | 17 | 4336983 | 4391503 | 375 | 39 | 0.55258 | 0.29027 | SPNS3 | 0.29027 | 0.43191 | 0.3659 |
| ENSG00000146463 | 1 | 35734568 | 35887659 | 258 | 19 | 0.55219 | 0.29041 | ZMYM4 | 0.29041 | 0.29013 | 0.31564 |
| ENSG00000079332 | 10 | 71907045 | 71930279 | 98 | 14 | 0.5518 | 0.29054 | SAR1A | 0.29054 | 0.41137 | 0.34823 |
| ENSG00000267981 | 11 | 129936836 | 129944012 | 20 | 4 | 0.55156 | 0.29063 | AP003041.2 | 0.29063 | 0.23403 | 0.28001 |
| ENSG00000146094 | 5 | 176928908 | 176938275 | 24 | 9 | 0.55116 | 0.29076 | DOK3 | 0.29076 | 0.32567 | 0.26616 |
| ENSG00000133935 | 14 | 76116134 | 76127532 | 20 | 7 | 0.55107 | 0.29079 | C14orf1 | 0.29079 | 0.33417 | 0.48172 |
| ENSG00000204568 | 6 | 30585486 | 30594172 | 21 | 4 | 0.55062 | 0.29095 | MRPS18B | 0.29095 | 0.30409 | 0.26794 |
| ENSG00000166596 | 17 | 9479944 | 9546776 | 145 | 27 | 0.5506 | 0.29095 | WDR16 | 0.29095 | 0.46733 | 0.59371 |
| ENSG00000088756 | 18 | 6729717 | 6915715 | 573 | 55 | 0.55055 | 0.29097 | ARHGAP28 | 0.29097 | 0.33699 | 0.4239 |
| ENSG00000167861 | 17 | 72946838 | 72969261 | 67 | 16 | 0.55054 | 0.29098 | HID1 | 0.29098 | 0.39525 | 0.35144 |
| ENSG00000163380 | 3 | 69156023 | 69172183 | 57 | 10 | 0.55046 | 0.291 | LMOD3 | 0.291 | 0.29933 | 0.26997 |
| ENSG00000254349 | 8 | 75512010 | 75735548 | 587 | 45 | 0.55016 | 0.2911 | RP11-758M4.1 | 0.2911 | 0.26154 | 0.24315 |

| | | | | | | | | | | | |
|---|---|---|---|---|---|---|---|---|---|---|---|
| ENSG00000168826 | 4 | 4291924 | 4323513 | 143 | 15 | 0.54991 | 0.29119 | ZBTB49 | 0.29119 | 0.10094 | 0.10501 |
| ENSG00000196358 | 9 | 135037334 | 135119921 | 236 | 33 | 0.54993 | 0.29119 | NTNG2 | 0.29119 | 0.37769 | 0.55582 |
| ENSG00000138413 | 2 | 209100951 | 209130798 | 72 | 14 | 0.54952 | 0.29133 | IDH1 | 0.29133 | 0.21829 | 0.31433 |
| ENSG00000082898 | 2 | 61704984 | 61765761 | 136 | 13 | 0.54922 | 0.29143 | XPO1 | 0.29143 | 0.32418 | 0.33847 |
| ENSG00000185523 | 1 | 213003483 | 213020991 | 105 | 16 | 0.54919 | 0.29144 | C1orf227 | 0.29144 | 0.23103 | 0.32144 |
| ENSG00000115750 | 2 | 9983483 | 10074545 | 339 | 20 | 0.54873 | 0.2916 | TAF1B | 0.2916 | 0.24904 | 0.15874 |
| ENSG00000187747 | 11 | 5602107 | 5603114 | 7 | 2 | 0.54867 | 0.29162 | OR52B6 | 0.29162 | 0.12747 | 0.17929 |
| ENSG00000165762 | 14 | 20344391 | 20345468 | 2 | 1 | 0.5486 | 0.29164 | OR4K2 | 0.29164 | 0.18645 | 0.21846 |
| ENSG00000164066 | 4 | 128544426 | 128647892 | 202 | 18 | 0.54848 | 0.29168 | INTU | 0.29168 | 0.29642 | 0.33311 |
| ENSG00000253626 | 10 | 81272357 | 81276188 | 19 | 5 | 0.5484 | 0.29171 | EIF5AL1 | 0.29171 | 0.49943 | 0.53508 |
| ENSG00000188783 | 1 | 203444956 | 203460480 | 55 | 10 | 0.54821 | 0.29177 | PRELP | 0.29177 | 0.2362 | 0.30919 |
| ENSG00000153822 | 17 | 68049570 | 68131749 | 338 | 20 | 0.54817 | 0.29179 | KCNJ16 | 0.29179 | 0.37099 | 0.47674 |
| ENSG00000170234 | 5 | 159488808 | 159546430 | 121 | 15 | 0.54724 | 0.29211 | PWWP2A | 0.29211 | 0.26278 | 0.28948 |
| ENSG00000106077 | 7 | 73150424 | 73153197 | 6 | 2 | 0.54706 | 0.29217 | ABHD11 | 0.29217 | 0.35794 | 0.41328 |
| ENSG00000109775 | 4 | 186320694 | 186347139 | 86 | 13 | 0.54693 | 0.29221 | UFSP2 | 0.29221 | 0.27702 | 0.25844 |
| ENSG00000070423 | 19 | 647526 | 663277 | 49 | 17 | 0.54693 | 0.29221 | RNF126 | 0.29221 | 0.6588 | 0.82779 |
| ENSG00000114859 | 3 | 184063973 | 184079439 | 45 | 13 | 0.54668 | 0.2923 | CLCN2 | 0.2923 | 0.061493 | 0.050298 |
| ENSG00000116212 | 1 | 54411750 | 54433841 | 35 | 9 | 0.54652 | 0.29235 | LRRC42 | 0.29235 | 0.20185 | 0.32339 |
| ENSG00000146856 | 7 | 134671259 | 134832715 | 442 | 13 | 0.54652 | 0.29235 | AGBL3 | 0.29235 | 0.31052 | 0.33122 |
| ENSG00000168619 | 8 | 39442008 | 39587583 | 303 | 24 | 0.54636 | 0.29241 | ADAM18 | 0.29241 | 0.25278 | 0.19122 |
| ENSG00000183128 | 10 | 105232561 | 105238997 | 12 | 4 | 0.54627 | 0.29244 | CALHM3 | 0.29244 | 0.43631 | 0.45727 |
| ENSG00000186009 | 13 | 114303119 | 114312501 | 22 | 9 | 0.54602 | 0.29253 | ATP4B | 0.29253 | 0.39675 | 0.41017 |
| ENSG00000160972 | 8 | 145703352 | 145727504 | 36 | 7 | 0.54573 | 0.29263 | PPP1R16A | 0.29263 | 0.37943 | 0.48299 |
| ENSG00000198756 | 1 | 183898796 | 184006863 | 403 | 38 | 0.54557 | 0.29268 | COLGALT2 | 0.29268 | 0.32351 | 0.3929 |
| ENSG00000134249 | 1 | 120436156 | 120439118 | 4 | 1 | 0.54552 | 0.2927 | ADAM30 | 0.2927 | 0.26746 | 0.30174 |
| ENSG00000087266 | 4 | 2794750 | 2842825 | 226 | 35 | 0.54509 | 0.29285 | SH3BP2 | 0.29285 | 0.35551 | 0.41776 |
| ENSG00000205923 | 16 | 2577081 | 2581420 | 4 | 2 | 0.54485 | 0.29293 | CEMP1 | 0.29293 | 0.38784 | 0.033594 |
| ENSG00000134954 | 11 | 128328656 | 128457453 | 312 | 45 | 0.54417 | 0.29316 | ETS1 | 0.29316 | 0.32772 | 0.38234 |
| ENSG00000144118 | 2 | 120997640 | 121052289 | 200 | 11 | 0.5441 | 0.29319 | RALB | 0.29319 | 0.2956 | 0.3574 |
| ENSG00000101773 | 18 | 20378224 | 20606451 | 590 | 21 | 0.54363 | 0.29335 | RBBP8 | 0.29335 | 0.29664 | 0.32775 |
| ENSG00000188508 | 19 | 35978226 | 35986460 | 25 | 6 | 0.54349 | 0.2934 | KRTDAP | 0.2934 | 0.39562 | 0.59054 |
| ENSG00000126216 | 13 | 113139325 | 113242481 | 302 | 18 | 0.54334 | 0.29345 | TUBGCP3 | 0.29345 | 0.31423 | 0.40384 |
| ENSG00000253207 | 8 | 114612539 | 114629187 | 44 | 9 | 0.54331 | 0.29346 | RP11-67H2.1 | 0.29346 | 0.25747 | 0.28477 |
| ENSG00000179041 | 8 | 67341263 | 67342966 | 7 | 4 | 0.54302 | 0.29356 | RRS1 | 0.29356 | 0.47755 | 0.31315 |
| ENSG00000120875 | 8 | 29190581 | 29208185 | 59 | 11 | 0.54278 | 0.29364 | DUSP4 | 0.29364 | 0.31948 | 0.32019 |
| ENSG00000175414 | 5 | 175792471 | 175828866 | 103 | 22 | 0.54245 | 0.29376 | ARL10 | 0.29376 | 0.37597 | 0.41196 |
| ENSG00000176386 | 9 | 116018115 | 116037869 | 69 | 15 | 0.54239 | 0.29377 | CDC26 | 0.29377 | 0.31184 | 0.33662 |
| ENSG00000085433 | 1 | 109512836 | 109584850 | 229 | 22 | 0.54232 | 0.2938 | WDR47 | 0.2938 | 0.19481 | 0.16156 |
| ENSG00000169964 | 3 | 44903361 | 44907162 | 3 | 2 | 0.54215 | 0.29386 | TMEM42 | 0.29386 | 0.37363 | 0.27134 |
| ENSG00000136699 | 2 | 130908981 | 130940323 | 54 | 4 | 0.54203 | 0.2939 | SMPD4 | 0.2939 | 0.22573 | 0.23633 |
| ENSG00000162551 | 1 | 21835858 | 21904905 | 359 | 34 | 0.54177 | 0.29399 | ALPL | 0.29399 | 0.3522 | 0.32931 |
| ENSG00000272268 | 2 | 6111712 | 6114374 | 9 | 3 | 0.54173 | 0.294 | DKFZP761K2322 | 0.294 | 0.13656 | 0.2412 |
| ENSG00000233050 | 8 | 11921898 | 11929256 | 1 | 1 | 0.54174 | 0.294 | DEFB130 | 0.294 | 0.23201 | 0.56833 |
| ENSG00000226321 | 2 | 241858202 | 241932728 | 252 | 27 | 0.54158 | 0.29405 | AC104809.3 | 0.29405 | 0.3461 | 0.52579 |
| ENSG00000121964 | 2 | 144695635 | 145090135 | 611 | 39 | 0.54037 | 0.29447 | GTDC1 | 0.29447 | 0.31284 | 0.35388 |
| ENSG00000218823 | 7 | 4897364 | 4901625 | 15 | 4 | 0.54021 | 0.29453 | PAPOLB | 0.29453 | 0.61064 | 0.85159 |
| ENSG00000081026 | 1 | 113933371 | 114228545 | 722 | 32 | 0.54005 | 0.29458 | MAGI3 | 0.29458 | 0.30613 | 0.32793 |
| ENSG00000135002 | 9 | 79000433 | 79009433 | 26 | 6 | 0.54007 | 0.29458 | RFK | 0.29458 | 0.31196 | 0.28301 |
| ENSG00000157483 | 15 | 59427113 | 59665099 | 869 | 66 | 0.54001 | 0.2946 | MYO1E | 0.2946 | 0.32141 | 0.31611 |
| ENSG00000268465 | 19 | 48969094 | 48975688 | 9 | 3 | 0.5397 | 0.2947 | CTC-273B12.7 | 0.2947 | 0.20193 | 0.11541 |
| ENSG00000167208 | 16 | 50700211 | 50715264 | 31 | 12 | 0.53953 | 0.29476 | SNX20 | 0.29476 | 0.24416 | 0.43487 |
| ENSG00000131196 | 18 | 77155856 | 77289325 | 718 | 56 | 0.5394 | 0.29481 | NFATC1 | 0.29481 | 0.20706 | 0.36603 |
| ENSG00000267270 | 18 | 77905807 | 77936315 | 131 | 17 | 0.53915 | 0.29489 | AC139100.2 | 0.29489 | 0.29178 | 0.32922 |
| ENSG00000160182 | 21 | 43782391 | 43786703 | 24 | 9 | 0.5391 | 0.29491 | TFF1 | 0.29491 | 0.064289 | 0.25509 |
| ENSG00000141905 | 19 | 3359561 | 3469215 | 473 | 62 | 0.53907 | 0.29492 | NFIC | 0.29492 | 0.30291 | 0.36844 |
| ENSG00000130749 | 19 | 47567444 | 47617009 | 82 | 11 | 0.53864 | 0.29507 | ZC3H4 | 0.29507 | 0.34048 | 0.37764 |
| ENSG00000130584 | 20 | 62375019 | 62462597 | 305 | 21 | 0.53858 | 0.29509 | ZBTB46 | 0.29509 | 0.3117 | 0.26945 |
| ENSG00000154447 | 4 | 170015407 | 170192256 | 410 | 33 | 0.53803 | 0.29528 | SH3RF1 | 0.29528 | 0.31354 | 0.43194 |
| ENSG00000242419 | 5 | 140864741 | 140892546 | 79 | 16 | 0.53759 | 0.29543 | PCDHGC4 | 0.29543 | 0.44371 | 0.38069 |
| ENSG00000143158 | 1 | 167885967 | 167906278 | 30 | 6 | 0.53748 | 0.29547 | MPC2 | 0.29547 | 0.35388 | 0.36139 |
| ENSG00000205309 | 17 | 17206649 | 17250977 | 162 | 20 | 0.5372 | 0.29557 | NT5M | 0.29557 | 0.26602 | 0.3538 |
| ENSG00000146143 | 6 | 57179603 | 57513375 | 622 | 28 | 0.53704 | 0.29562 | PRIM2 | 0.29562 | 0.30355 | 0.30085 |
| ENSG00000108679 | 17 | 76967320 | 76976191 | 23 | 9 | 0.53692 | 0.29566 | LGALS3BP | 0.29566 | 0.388 | 0.40889 |
| ENSG00000115902 | 2 | 65215611 | 65250999 | 126 | 29 | 0.53684 | 0.29569 | SLC1A4 | 0.29569 | 0.30528 | 0.48318 |
| ENSG00000116285 | 1 | 8064464 | 8086368 | 45 | 9 | 0.53664 | 0.29576 | ERRFI1 | 0.29576 | 0.39053 | 0.38371 |
| ENSG00000235478 | 22 | 17602476 | 17612994 | 57 | 13 | 0.53657 | 0.29578 | AC006946.15 | 0.29578 | 0.18688 | 0.32423 |
| ENSG00000183691 | 17 | 54671060 | 54672951 | 1 | 1 | 0.53652 | 0.2958 | NOG | 0.2958 | 0.86017 | 0.93912 |
| ENSG00000178229 | 19 | 57831877 | 57842144 | 49 | 12 | 0.53621 | 0.29591 | ZNF543 | 0.29591 | 0.37977 | 0.43585 |
| ENSG00000144815 | 3 | 101498046 | 101547073 | 103 | 12 | 0.536 | 0.29598 | NXPE3 | 0.29598 | 0.31009 | 0.3415 |
| ENSG00000177483 | 2 | 238707032 | 238751451 | 118 | 17 | 0.53584 | 0.29604 | RBM44 | 0.29604 | 0.32954 | 0.63677 |
| ENSG00000111364 | 12 | 124086624 | 124105488 | 83 | 10 | 0.53581 | 0.29605 | DDX55 | 0.29605 | 0.38558 | 0.38738 |
| ENSG00000163807 | 3 | 44779153 | 44803154 | 38 | 9 | 0.53572 | 0.29608 | KIAA1143 | 0.29608 | 0.40107 | 0.52047 |
| ENSG00000180044 | 3 | 159943423 | 159945999 | 6 | 3 | 0.5355 | 0.29615 | C3orf80 | 0.29615 | 0.60731 | 0.8763 |

| | | | | | | | | | | | |
|---|---|---|---|---|---|---|---|---|---|---|---|
| ENSG00000136856 | 9 | 130159421 | 130170703 | 27 | 5 | 0.53545 | 0.29617 | SLC2A8 | 0.29617 | 0.38954 | 0.35086 |
| ENSG00000186136 | 12 | 11338599 | 11339543 | 8 | 2 | 0.53514 | 0.29628 | TAS2R42 | 0.29628 | 0.29963 | 0.26135 |
| ENSG00000138768 | 4 | 76649777 | 76735382 | 257 | 14 | 0.535 | 0.29633 | USO1 | 0.29633 | 0.33053 | 0.37932 |
| ENSG00000161551 | 19 | 52359055 | 52394203 | 231 | 15 | 0.53478 | 0.2964 | ZNF577 | 0.2964 | 0.31128 | 0.33925 |
| ENSG00000196431 | 22 | 27017928 | 27026636 | 41 | 7 | 0.53475 | 0.29641 | CRYBA4 | 0.29641 | 0.31532 | 0.45787 |
| ENSG00000162402 | 1 | 55532032 | 55680786 | 345 | 20 | 0.53454 | 0.29648 | USP24 | 0.29648 | 0.31951 | 0.3296 |
| ENSG00000139629 | 12 | 51745031 | 51786651 | 111 | 32 | 0.53369 | 0.29678 | GALNT6 | 0.29678 | 0.096182 | 0.14514 |
| ENSG00000164363 | 5 | 1225470 | 1246304 | 111 | 25 | 0.53299 | 0.29702 | SLC6A18 | 0.29702 | 0.34071 | 0.34898 |
| ENSG00000130173 | 19 | 11348178 | 11352619 | 6 | 2 | 0.53287 | 0.29706 | C19orf80 | 0.29706 | 0.28521 | 0.62882 |
| ENSG00000169692 | 9 | 139567595 | 139581875 | 43 | 11 | 0.53281 | 0.29708 | AGPAT2 | 0.29708 | 0.27739 | 0.46071 |
| ENSG00000157322 | 16 | 69984810 | 69998141 | 32 | 9 | 0.53281 | 0.29708 | CLEC18A | 0.29708 | 0.34108 | 0.38581 |
| ENSG00000205277 | 7 | 100612904 | 100662230 | 151 | 21 | 0.53259 | 0.29716 | MUC12 | 0.29716 | 0.18527 | 0.071475 |
| ENSG00000141552 | 17 | 79848666 | 79858867 | 9 | 6 | 0.5324 | 0.29722 | ANAPC11 | 0.29722 | 0.43918 | 0.58378 |
| ENSG00000267531 | 19 | 55851254 | 55856561 | 23 | 5 | 0.53241 | 0.29722 | AC020922.1 | 0.29722 | 0.42553 | 0.53566 |
| ENSG00000136305 | 14 | 24774302 | 24780636 | 14 | 4 | 0.53228 | 0.29726 | CIDEB | 0.29726 | 0.56797 | 0.64735 |
| ENSG00000164743 | 8 | 13424352 | 13425796 | 3 | 1 | 0.53191 | 0.29739 | C8orf48 | 0.29739 | 0.31485 | 0.25675 |
| ENSG00000198624 | 5 | 150560613 | 150603706 | 189 | 35 | 0.53159 | 0.2975 | CCDC69 | 0.2975 | 0.10152 | 0.12903 |
| ENSG00000163864 | 3 | 139279022 | 139396859 | 321 | 34 | 0.53151 | 0.29753 | NMNAT3 | 0.29753 | 0.23195 | 0.20192 |
| ENSG00000073008 | 19 | 45147098 | 45166850 | 49 | 11 | 0.53122 | 0.29763 | PVR | 0.29763 | 0.27429 | 0.53107 |
| ENSG00000117676 | 1 | 26856252 | 26901521 | 107 | 14 | 0.53117 | 0.29765 | RPS6KA1 | 0.29765 | 0.37411 | 0.15176 |
| ENSG00000188778 | 8 | 37820516 | 37824483 | 5 | 1 | 0.53103 | 0.2977 | ADRB3 | 0.2977 | 0.4778 | 0.52588 |
| ENSG00000166444 | 11 | 8714898 | 8932498 | 681 | 34 | 0.53056 | 0.29786 | ST5 | 0.29786 | 0.18779 | 0.16379 |
| ENSG00000185958 | 12 | 50720013 | 50790405 | 177 | 16 | 0.53048 | 0.29789 | FAM186A | 0.29789 | 0.34618 | 0.35122 |
| ENSG00000146192 | 6 | 36973422 | 36996846 | 78 | 20 | 0.53022 | 0.29798 | FGD2 | 0.29798 | 0.655 | 0.76208 |
| ENSG00000135083 | 5 | 159678659 | 159772403 | 305 | 31 | 0.53013 | 0.29801 | CCNJL | 0.29801 | 0.30763 | 0.40155 |
| ENSG00000130706 | 20 | 60877149 | 60883918 | 13 | 5 | 0.52928 | 0.2983 | ADRM1 | 0.2983 | 0.35643 | 0.25311 |
| ENSG00000159479 | 1 | 43849588 | 43855479 | 4 | 2 | 0.52921 | 0.29833 | MED8 | 0.29833 | 0.4187 | 0.54696 |
| ENSG00000242247 | 22 | 43192508 | 43254112 | 248 | 16 | 0.52914 | 0.29835 | ARFGAP3 | 0.29835 | 0.27071 | 0.29215 |
| ENSG00000144635 | 3 | 32567463 | 32612366 | 112 | 19 | 0.52897 | 0.29841 | DYNC1LI1 | 0.29841 | 0.42441 | 0.50647 |
| ENSG00000183671 | 2 | 207040040 | 207082771 | 126 | 20 | 0.52892 | 0.29843 | GPR1 | 0.29843 | 0.23593 | 0.1268 |
| ENSG00000151491 | 12 | 15773092 | 16035263 | 277 | 43 | 0.52892 | 0.29843 | EPS8 | 0.29843 | 0.31538 | 0.26716 |
| ENSG00000176124 | 13 | 50656307 | 51297372 | 1892 | 103 | 0.5286 | 0.29854 | DLEU1 | 0.29854 | 0.29102 | 0.28506 |
| ENSG00000215343 | 8 | 11961898 | 11973025 | 1 | 1 | 0.52843 | 0.2986 | ZNF705D | 0.2986 | 0.10679 | 0.075805 |
| ENSG00000169403 | 1 | 28473677 | 28520447 | 62 | 14 | 0.52831 | 0.29864 | PTAFR | 0.29864 | 0.23137 | 0.2868 |
| ENSG00000255298 | 11 | 124134723 | 124135763 | 9 | 3 | 0.52783 | 0.29881 | OR8G5 | 0.29881 | 0.21033 | 0.24665 |
| ENSG00000196646 | 19 | 12273879 | 12300064 | 56 | 12 | 0.52745 | 0.29894 | ZNF136 | 0.29894 | 0.34159 | 0.36262 |
| ENSG00000148835 | 10 | 105127724 | 105148822 | 31 | 8 | 0.52742 | 0.29895 | TAF5 | 0.29895 | 0.2325 | 0.21792 |
| ENSG00000130518 | 19 | 18367908 | 18385319 | 114 | 13 | 0.52737 | 0.29897 | KIAA1683 | 0.29897 | 0.45187 | 0.55443 |
| ENSG00000126790 | 14 | 59927081 | 59951148 | 81 | 10 | 0.52713 | 0.29905 | L3HYPDH | 0.29905 | 0.33518 | 0.27806 |
| ENSG00000140474 | 15 | 75128457 | 75135687 | 16 | 4 | 0.52704 | 0.29908 | ULK3 | 0.29908 | 0.27171 | 0.14346 |
| ENSG00000129535 | 14 | 24549316 | 24584223 | 60 | 18 | 0.52651 | 0.29927 | NRL | 0.29927 | 0.53003 | 0.53048 |
| ENSG00000166823 | 15 | 90291892 | 90294541 | 10 | 4 | 0.52631 | 0.29934 | MESP1 | 0.29934 | 0.1145 | 0.11405 |
| ENSG00000196661 | 11 | 124266278 | 124267264 | 4 | 1 | 0.52628 | 0.29935 | OR8B3 | 0.29935 | 0.74962 | 0.76821 |
| ENSG00000101230 | 20 | 13202418 | 13281298 | 377 | 50 | 0.52621 | 0.29937 | ISM1 | 0.29937 | 0.38683 | 0.40172 |
| ENSG00000109083 | 17 | 26655352 | 26662515 | 7 | 3 | 0.52609 | 0.29941 | IFT20 | 0.29941 | 0.47514 | 0.37669 |
| ENSG00000148175 | 9 | 124101355 | 124132531 | 91 | 17 | 0.52566 | 0.29956 | STOM | 0.29956 | 0.27303 | 0.24922 |
| ENSG00000067715 | 12 | 79257773 | 79845788 | 1702 | 74 | 0.52549 | 0.29962 | SYT1 | 0.29962 | 0.29686 | 0.33851 |
| ENSG00000102572 | 13 | 99102455 | 99230194 | 432 | 30 | 0.5255 | 0.29962 | STK24 | 0.29962 | 0.34036 | 0.41581 |
| ENSG00000103310 | 16 | 21208773 | 21225831 | 45 | 13 | 0.52531 | 0.29968 | ZP2 | 0.29968 | 0.24056 | 0.33244 |
| ENSG00000007341 | 1 | 113066140 | 113163447 | 268 | 17 | 0.5252 | 0.29972 | ST7L | 0.29972 | 0.29794 | 0.29838 |
| ENSG00000133027 | 17 | 17408877 | 17495022 | 280 | 19 | 0.52517 | 0.29973 | PEMT | 0.29973 | 0.37133 | 0.47398 |
| ENSG00000239998 | 19 | 55084387 | 55099027 | 58 | 9 | 0.52511 | 0.29975 | LILRA2 | 0.29975 | 0.22156 | 0.28103 |
| ENSG00000148358 | 9 | 132815705 | 132902448 | 370 | 25 | 0.5251 | 0.29976 | GPR107 | 0.29976 | 0.4161 | 0.25446 |
| ENSG00000070808 | 5 | 149599054 | 149669854 | 176 | 31 | 0.52473 | 0.29988 | CAMK2A | 0.29988 | 0.54456 | 0.22861 |
| ENSG00000146477 | 6 | 160769300 | 160876014 | 237 | 19 | 0.52414 | 0.30009 | SLC22A3 | 0.30009 | 0.36985 | 0.47732 |
| ENSG00000213024 | 19 | 50410082 | 50433020 | 92 | 18 | 0.52391 | 0.30017 | NUP62 | 0.30017 | 0.31465 | 0.20752 |
| ENSG00000121289 | 19 | 33369902 | 33462897 | 339 | 19 | 0.52352 | 0.30031 | CEP89 | 0.30031 | 0.31754 | 0.39566 |
| ENSG00000120324 | 5 | 140571942 | 140575215 | 16 | 8 | 0.52341 | 0.30034 | PCDHB10 | 0.30034 | 0.45426 | 0.41842 |
| ENSG00000123200 | 13 | 46528600 | 46626894 | 209 | 12 | 0.52337 | 0.30036 | ZC3H13 | 0.30036 | 0.3214 | 0.41211 |
| ENSG00000185880 | 15 | 45021186 | 45060027 | 174 | 14 | 0.52336 | 0.30036 | TRIM69 | 0.30036 | 0.29591 | 0.28925 |
| ENSG00000258804 | 14 | 87372122 | 87389099 | 63 | 8 | 0.52335 | 0.30037 | RP11-322L20.1 | 0.30037 | 0.2065 | 0.25594 |
| ENSG00000135108 | 12 | 117581146 | 117628336 | 198 | 11 | 0.52312 | 0.30045 | FBXO21 | 0.30045 | 0.3077 | 0.38556 |
| ENSG00000182986 | 19 | 53367043 | 53400946 | 214 | 23 | 0.52286 | 0.30054 | ZNF320 | 0.30054 | 0.30987 | 0.46299 |
| ENSG00000178343 | 4 | 42399856 | 42404504 | 17 | 6 | 0.52246 | 0.30067 | SHISA3 | 0.30067 | 0.49181 | 0.56823 |
| ENSG00000137462 | 4 | 154622652 | 154626851 | 9 | 5 | 0.52239 | 0.3007 | TLR2 | 0.3007 | 0.41204 | 0.48203 |
| ENSG00000179546 | 1 | 23516993 | 23521222 | 15 | 3 | 0.52217 | 0.30078 | HTR1D | 0.30078 | 0.065178 | 0.057045 |
| ENSG00000164325 | 5 | 72469022 | 72470970 | 5 | 2 | 0.52201 | 0.30083 | TMEM174 | 0.30083 | 0.31162 | 0.73525 |
| ENSG00000172292 | 2 | 169312372 | 169631644 | 1093 | 78 | 0.52195 | 0.30085 | CERS6 | 0.30085 | 0.3184 | 0.3236 |
| ENSG00000173320 | 4 | 184774584 | 184944679 | 617 | 79 | 0.52195 | 0.30085 | STOX2 | 0.30085 | 0.3892 | 0.50908 |
| ENSG00000186513 | 11 | 57957906 | 57958990 | 4 | 1 | 0.52138 | 0.30105 | OR9Q2 | 0.30105 | 0.19036 | 0.091058 |
| ENSG00000140157 | 15 | 23004684 | 23034427 | 183 | 18 | 0.52009 | 0.3015 | NIPA2 | 0.3015 | 0.4637 | 0.48047 |
| ENSG00000100387 | 22 | 41347351 | 41369313 | 25 | 7 | 0.51958 | 0.30168 | RBX1 | 0.30168 | 0.38631 | 0.32309 |
| ENSG00000164125 | 4 | 159045626 | 159094470 | 127 | 9 | 0.51913 | 0.30183 | FAM198B | 0.30183 | 0.28842 | 0.26908 |

| | | | | | | | | | | | |
|---|---|---|---|---|---|---|---|---|---|---|---|
| ENSG00000138757 | 4 | 76567966 | 76649709 | 260 | 17 | 0.51859 | 0.30202 | G3BP2 | 0.30202 | 0.2885 | 0.29681 |
| ENSG00000106009 | 7 | 2577511 | 2595361 | 92 | 13 | 0.51855 | 0.30204 | BRAT1 | 0.30204 | 0.27353 | 0.34708 |
| ENSG00000148600 | 10 | 85954410 | 85979377 | 94 | 16 | 0.51855 | 0.30204 | CDHR1 | 0.30204 | 0.20628 | 0.10147 |
| ENSG00000178125 | 8 | 67876334 | 67968839 | 49 | 28 | 0.51851 | 0.30205 | PPP1R42 | 0.30205 | 0.36406 | 0.44492 |
| ENSG00000203730 | 1 | 182367252 | 182369751 | 7 | 2 | 0.51822 | 0.30215 | TEDDM1 | 0.30215 | 0.31882 | 0.39232 |
| ENSG00000124831 | 2 | 238536219 | 238722325 | 797 | 46 | 0.51751 | 0.3024 | LRRFIP1 | 0.3024 | 0.32168 | 0.23854 |
| ENSG00000056736 | 3 | 53880607 | 53899827 | 65 | 16 | 0.51745 | 0.30242 | IL17RB | 0.30242 | 0.36019 | 0.53148 |
| ENSG00000181873 | 1 | 228353516 | 228369958 | 25 | 7 | 0.51738 | 0.30245 | IBA57 | 0.30245 | 0.36445 | 0.38841 |
| ENSG00000181751 | 5 | 102594403 | 102614361 | 55 | 5 | 0.51698 | 0.30258 | C5orf30 | 0.30258 | 0.37985 | 0.43673 |
| ENSG00000099194 | 10 | 102106881 | 102124591 | 41 | 8 | 0.51683 | 0.30264 | SCD | 0.30264 | 0.20328 | 0.14938 |
| ENSG00000136824 | 9 | 106856541 | 106903698 | 110 | 10 | 0.51656 | 0.30273 | SMC2 | 0.30273 | 0.24056 | 0.12991 |
| ENSG00000070526 | 17 | 74620843 | 74639920 | 90 | 12 | 0.51651 | 0.30275 | ST6GALNAC1 | 0.30275 | 0.34152 | 0.17348 |
| ENSG00000129654 | 17 | 74132414 | 74137380 | 9 | 3 | 0.51637 | 0.3028 | FOXJ1 | 0.3028 | 0.23054 | 0.025972 |
| ENSG00000269093 | 19 | 32895915 | 32896466 | 2 | 1 | 0.51623 | 0.30285 | AC007773.3 | 0.30285 | 0.51467 | 0.52598 |
| ENSG00000160951 | 19 | 14583278 | 14586174 | 10 | 4 | 0.51551 | 0.3031 | PTGER1 | 0.3031 | 0.06831 | 0.049959 |
| ENSG00000140464 | 15 | 74287014 | 74340153 | 162 | 17 | 0.51539 | 0.30314 | PML | 0.30314 | 0.33499 | 0.31934 |
| ENSG00000148296 | 9 | 136197552 | 136203235 | 15 | 4 | 0.51531 | 0.30317 | SURF6 | 0.30317 | 0.38781 | 0.58543 |
| ENSG00000145920 | 5 | 175223313 | 175311023 | 263 | 32 | 0.51506 | 0.30326 | CPLX2 | 0.30326 | 0.35088 | 0.36794 |
| ENSG00000183775 | 5 | 143550396 | 143865249 | 655 | 26 | 0.5145 | 0.30345 | KCTD16 | 0.30345 | 0.28896 | 0.31373 |
| ENSG00000188000 | 19 | 9296279 | 9299493 | 12 | 4 | 0.51435 | 0.3035 | OR7D2 | 0.3035 | 0.27208 | 0.51606 |
| ENSG00000197948 | 5 | 141018869 | 141030986 | 54 | 13 | 0.51376 | 0.30371 | FCHSD1 | 0.30371 | 0.21944 | 0.10247 |
| ENSG00000120889 | 8 | 22877646 | 22926692 | 200 | 35 | 0.51365 | 0.30375 | TNFRSF10B | 0.30375 | 0.33991 | 0.45128 |
| ENSG00000272414 | 4 | 77172874 | 77232282 | 283 | 18 | 0.51351 | 0.3038 | FAM47E-STBD1 | 0.3038 | 0.3152 | 0.3827 |
| ENSG00000186352 | 4 | 186317175 | 186321782 | 16 | 4 | 0.51343 | 0.30383 | ANKRD37 | 0.30383 | 0.33127 | 0.26258 |
| ENSG00000069702 | 1 | 92145902 | 92371892 | 849 | 65 | 0.51339 | 0.30384 | TGFBR3 | 0.30384 | 0.16951 | 0.18013 |
| ENSG00000121440 | 3 | 73431584 | 73674091 | 723 | 81 | 0.51338 | 0.30384 | PDZRN3 | 0.30384 | 0.28049 | 0.3813 |
| ENSG00000148408 | 9 | 140772241 | 141019076 | 451 | 41 | 0.51326 | 0.30388 | CACNA1B | 0.30388 | 0.28602 | 0.17194 |
| ENSG00000187017 | 1 | 6484848 | 6521430 | 136 | 15 | 0.51293 | 0.304 | ESPN | 0.304 | 0.31823 | 0.34045 |
| ENSG00000153395 | 5 | 1456595 | 1524092 | 312 | 45 | 0.5124 | 0.30419 | LPCAT1 | 0.30419 | 0.33267 | 0.41029 |
| ENSG00000131459 | 5 | 179727690 | 179780387 | 259 | 26 | 0.51222 | 0.30425 | GFPT2 | 0.30425 | 0.3455 | 0.44239 |
| ENSG00000056291 | 4 | 72897521 | 73013784 | 399 | 14 | 0.5119 | 0.30436 | NPFFR2 | 0.30436 | 0.29127 | 0.16794 |
| ENSG00000165905 | 11 | 45943172 | 45950647 | 19 | 6 | 0.51173 | 0.30442 | GYLTL1B | 0.30442 | 0.30133 | 0.37068 |
| ENSG00000138160 | 10 | 94353043 | 94415150 | 294 | 19 | 0.51159 | 0.30447 | KIF11 | 0.30447 | 0.29051 | 0.31743 |
| ENSG00000169347 | 16 | 20320894 | 20339130 | 60 | 12 | 0.51158 | 0.30447 | GP2 | 0.30447 | 0.30637 | 0.38324 |
| ENSG00000134802 | 11 | 57174427 | 57195053 | 40 | 12 | 0.51153 | 0.30449 | SLC43A3 | 0.30449 | 0.49203 | 0.56051 |
| ENSG00000172264 | 20 | 13976015 | 16033842 | 7442 | 348 | 0.51154 | 0.30449 | MACROD2 | 0.30449 | 0.33035 | 0.37729 |
| ENSG00000150625 | 4 | 176554085 | 176923815 | 1373 | 100 | 0.51121 | 0.3046 | GPM6A | 0.3046 | 0.32394 | 0.36484 |
| ENSG00000144589 | 2 | 220462582 | 220481173 | 22 | 9 | 0.5112 | 0.30461 | STK11IP | 0.30461 | 0.19273 | 0.24949 |
| ENSG00000081479 | 2 | 169983619 | 170219195 | 873 | 78 | 0.51087 | 0.30472 | LRP2 | 0.30472 | 0.26639 | 0.3005 |
| ENSG00000077147 | 10 | 98277866 | 98347209 | 200 | 15 | 0.51077 | 0.30476 | TM9SF3 | 0.30476 | 0.30775 | 0.26675 |
| ENSG00000150676 | 11 | 85566144 | 85631064 | 229 | 19 | 0.50978 | 0.3051 | CCDC83 | 0.3051 | 0.27206 | 0.28998 |
| ENSG00000137474 | 11 | 76839310 | 76926284 | 318 | 32 | 0.50909 | 0.30535 | MYO7A | 0.30535 | 0.30233 | 0.29824 |
| ENSG00000116898 | 1 | 36921319 | 36930038 | 41 | 6 | 0.50865 | 0.3055 | MRPS15 | 0.3055 | 0.34577 | 0.41408 |
| ENSG00000117289 | 1 | 145438469 | 145442635 | 5 | 1 | 0.50856 | 0.30553 | TXNIP | 0.30553 | 0.41085 | 0.49968 |
| ENSG00000133800 | 11 | 10578513 | 10633236 | 181 | 26 | 0.50857 | 0.30553 | LYVE1 | 0.30553 | 0.28658 | 0.20255 |
| ENSG00000097007 | 9 | 133589333 | 133763062 | 572 | 33 | 0.50836 | 0.3056 | ABL1 | 0.3056 | 0.36571 | 0.46487 |
| ENSG00000269048 | 6 | 166945369 | 166947709 | 6 | 4 | 0.50814 | 0.30568 | Z98049.1 | 0.30568 | 0.19274 | 0.16295 |
| ENSG00000175356 | 11 | 9041071 | 9159661 | 396 | 52 | 0.50796 | 0.30574 | SCUBE2 | 0.30574 | 0.30072 | 0.38605 |
| ENSG00000118804 | 4 | 77172886 | 77232752 | 285 | 17 | 0.50788 | 0.30577 | FAM47E-STBD1 | 0.30577 | 0.3152 | 0.3827 |
| ENSG00000177119 | 12 | 45609770 | 45834187 | 550 | 22 | 0.50769 | 0.30583 | ANO6 | 0.30583 | 0.31519 | 0.34788 |
| ENSG00000117362 | 1 | 150237804 | 150241980 | 3 | 2 | 0.50768 | 0.30584 | APH1A | 0.30584 | 0.76301 | 0.90107 |
| ENSG00000196867 | 19 | 57050317 | 57068169 | 49 | 11 | 0.50763 | 0.30586 | ZFP28 | 0.30586 | 0.15023 | 0.15866 |
| ENSG00000165325 | 11 | 93063137 | 93171653 | 414 | 23 | 0.5074 | 0.30594 | CCDC67 | 0.30594 | 0.26034 | 0.25287 |
| ENSG00000139194 | 12 | 7276280 | 7281538 | 12 | 4 | 0.50727 | 0.30598 | RBP5 | 0.30598 | 0.36951 | 0.56604 |
| ENSG00000168079 | 8 | 27727399 | 27850244 | 575 | 54 | 0.50726 | 0.30599 | SCARA5 | 0.30599 | 0.30998 | 0.37369 |
| ENSG00000214109 | 19 | 13890884 | 13900972 | 69 | 9 | 0.50713 | 0.30603 | AC008686.1 | 0.30603 | 0.21674 | 0.42275 |
| ENSG00000165046 | 8 | 38243725 | 38267045 | 42 | 6 | 0.50696 | 0.30609 | LETM2 | 0.30609 | 0.40314 | 0.53998 |
| ENSG00000177283 | 10 | 35927177 | 35930362 | 4 | 2 | 0.50686 | 0.30613 | FZD8 | 0.30613 | 0.082547 | 0.053782 |
| ENSG00000082293 | 6 | 70576463 | 70919679 | 997 | 65 | 0.50616 | 0.30637 | COL19A1 | 0.30637 | 0.29631 | 0.27818 |
| ENSG00000263809 | 17 | 8271956 | 8286505 | 41 | 7 | 0.50583 | 0.30649 | RP11-849F2.7 | 0.30649 | 0.23321 | 0.39773 |
| ENSG00000221923 | 19 | 52873170 | 52889048 | 69 | 8 | 0.50565 | 0.30655 | ZNF880 | 0.30655 | 0.38149 | 0.34745 |
| ENSG00000181523 | 17 | 78180515 | 78194722 | 64 | 8 | 0.50551 | 0.3066 | SGSH | 0.3066 | 0.26662 | 0.21833 |
| ENSG00000168491 | 4 | 186366336 | 186392913 | 125 | 27 | 0.50531 | 0.30667 | CCDC110 | 0.30667 | 0.39211 | 0.49629 |
| ENSG00000133597 | 7 | 140372953 | 140396061 | 50 | 8 | 0.50484 | 0.30683 | ADCK2 | 0.30683 | 0.17198 | 0.22691 |
| ENSG00000163206 | 1 | 152850793 | 152857523 | 6 | 3 | 0.5048 | 0.30685 | SMCP | 0.30685 | 0.60203 | 0.43551 |
| ENSG00000159184 | 17 | 46802125 | 46806540 | 9 | 4 | 0.50438 | 0.307 | HOXB13 | 0.307 | 0.32213 | 0.42429 |
| ENSG00000167588 | 12 | 50497602 | 50505102 | 10 | 5 | 0.50432 | 0.30702 | GPD1 | 0.30702 | 0.071937 | 0.083805 |
| ENSG00000169508 | 13 | 99946784 | 99959659 | 28 | 10 | 0.50422 | 0.30705 | GPR183 | 0.30705 | 0.47031 | 0.5815 |
| ENSG00000136950 | 9 | 127624409 | 127640003 | 25 | 6 | 0.50345 | 0.30732 | ARPC5L | 0.30732 | 0.48962 | 0.5942 |
| ENSG00000198668 | 14 | 90862846 | 90874605 | 33 | 9 | 0.50342 | 0.30734 | CALM1 | 0.30734 | 0.27179 | 0.25093 |
| ENSG00000145916 | 5 | 177557997 | 177577566 | 69 | 10 | 0.50274 | 0.30757 | RMND5B | 0.30757 | 0.19476 | 0.12637 |
| ENSG00000177600 | 11 | 809647 | 812880 | 6 | 2 | 0.50264 | 0.30761 | RPLP2 | 0.30761 | 0.76236 | 0.72015 |
| ENSG00000061455 | 5 | 122424816 | 122529960 | 459 | 33 | 0.50251 | 0.30765 | PRDM6 | 0.30765 | 0.24301 | 0.16747 |

| | | | | | | | | | | | |
|---|---|---|---|---|---|---|---|---|---|---|---|
| ENSG00000137766 | 15 | 54305101 | 54920806 | 2854 | 108 | 0.50235 | 0.30771 | UNC13C | 0.30771 | 0.31317 | 0.39815 |
| ENSG00000261833 | 16 | 76587314 | 76669520 | 268 | 47 | 0.50193 | 0.30786 | RP11-58C22.1 | 0.30786 | 0.40907 | 0.4409 |
| ENSG00000124839 | 2 | 238482965 | 238510257 | 182 | 25 | 0.50179 | 0.30791 | RAB17 | 0.30791 | 0.55865 | 0.30079 |
| ENSG00000185272 | 21 | 15588451 | 15600693 | 74 | 12 | 0.50141 | 0.30804 | RBM11 | 0.30804 | 0.070109 | 0.077121 |
| ENSG00000158636 | 11 | 76155967 | 76264069 | 250 | 28 | 0.50123 | 0.30811 | C11orf30 | 0.30811 | 0.3298 | 0.34598 |
| ENSG00000114416 | 3 | 180585929 | 180700541 | 327 | 19 | 0.50093 | 0.30821 | FXR1 | 0.30821 | 0.30934 | 0.34502 |
| ENSG00000006116 | 16 | 24266874 | 24374122 | 393 | 61 | 0.50078 | 0.30826 | CACNG3 | 0.30826 | 0.38473 | 0.34968 |
| ENSG00000169760 | 3 | 173114074 | 174004434 | 2954 | 148 | 0.50028 | 0.30844 | NLGN1 | 0.30844 | 0.32516 | 0.40761 |
| ENSG00000188155 | 21 | 46011149 | 46012386 | 9 | 4 | 0.50009 | 0.30851 | KRTAP10-6 | 0.30851 | 0.26614 | 0.28644 |
| ENSG00000125255 | 13 | 103696350 | 103719196 | 170 | 25 | 0.49944 | 0.30873 | SLC10A2 | 0.30873 | 0.32981 | 0.47637 |
| ENSG00000163874 | 1 | 37940153 | 37949980 | 15 | 6 | 0.49934 | 0.30877 | ZC3H12A | 0.30877 | 0.49754 | 0.63867 |
| ENSG00000165724 | 9 | 140476531 | 140484942 | 20 | 3 | 0.49922 | 0.30881 | ZMYND19 | 0.30881 | 0.27993 | 0.28033 |
| ENSG00000012822 | 12 | 54104903 | 54121529 | 45 | 10 | 0.49895 | 0.30891 | CALCOCO1 | 0.30891 | 0.2445 | 0.19452 |
| ENSG00000076685 | 10 | 104845940 | 104953056 | 311 | 18 | 0.49866 | 0.30901 | NT5C2 | 0.30901 | 0.29525 | 0.21533 |
| ENSG00000146223 | 6 | 42847356 | 42857663 | 40 | 8 | 0.49831 | 0.30913 | RPL7L1 | 0.30913 | 0.17842 | 0.12273 |
| ENSG00000049860 | 5 | 73935848 | 74018472 | 292 | 46 | 0.49822 | 0.30917 | HEXB | 0.30917 | 0.25578 | 0.25387 |
| ENSG00000176472 | 19 | 44029649 | 44040282 | 29 | 8 | 0.49804 | 0.30923 | ZNF575 | 0.30923 | 0.1823 | 0.17522 |
| ENSG00000073150 | 22 | 50609160 | 50618723 | 31 | 12 | 0.49768 | 0.30936 | PANX2 | 0.30936 | 0.091233 | 0.12563 |
| ENSG00000181518 | 11 | 123777109 | 123778183 | 6 | 2 | 0.49747 | 0.30943 | OR8D4 | 0.30943 | 0.57695 | 0.45984 |
| ENSG00000089101 | 20 | 20033158 | 20341346 | 1126 | 63 | 0.49744 | 0.30944 | C20orf26 | 0.30944 | 0.31891 | 0.34772 |
| ENSG00000112972 | 5 | 43289497 | 43313614 | 58 | 5 | 0.49717 | 0.30954 | HMGCS1 | 0.30954 | 0.27039 | 0.25329 |
| ENSG00000203785 | 1 | 153065611 | 153078660 | 43 | 7 | 0.49699 | 0.3096 | SPRR2E | 0.3096 | 0.23494 | 0.24915 |
| ENSG00000094796 | 17 | 39549976 | 39553844 | 15 | 4 | 0.49667 | 0.30971 | KRT31 | 0.30971 | 0.30183 | 0.23517 |
| ENSG00000164494 | 6 | 107473761 | 107780768 | 813 | 46 | 0.49638 | 0.30981 | PDSS2 | 0.30981 | 0.30865 | 0.29361 |
| ENSG00000162456 | 1 | 47011316 | 47017199 | 12 | 3 | 0.49621 | 0.30987 | KNCN | 0.30987 | 0.41279 | 0.54929 |
| ENSG00000141279 | 17 | 45600308 | 45700642 | 231 | 19 | 0.49623 | 0.30987 | NPEPPS | 0.30987 | 0.31586 | 0.33251 |
| ENSG00000176731 | 8 | 86126311 | 86132650 | 10 | 2 | 0.49608 | 0.30992 | C8orf59 | 0.30992 | 0.07768 | 0.14423 |
| ENSG00000173673 | 1 | 6304252 | 6305638 | 5 | 2 | 0.49586 | 0.31 | HES3 | 0.31 | 0.74196 | 0.77519 |
| ENSG00000123552 | 6 | 99880190 | 99969604 | 295 | 18 | 0.49586 | 0.31 | USP45 | 0.31 | 0.3524 | 0.43233 |
| ENSG00000137571 | 8 | 70579282 | 70747299 | 510 | 43 | 0.49571 | 0.31005 | SLCO5A1 | 0.31005 | 0.37695 | 0.49694 |
| ENSG00000175643 | 16 | 11343476 | 11445619 | 535 | 23 | 0.49572 | 0.31005 | RMI2 | 0.31005 | 0.31159 | 0.35001 |
| ENSG00000196705 | 19 | 21324827 | 21373034 | 111 | 14 | 0.4957 | 0.31005 | ZNF431 | 0.31005 | 0.31857 | 0.21323 |
| ENSG00000182793 | 6 | 52696453 | 52710893 | 44 | 8 | 0.49567 | 0.31006 | GSTA5 | 0.31006 | 0.34054 | 0.37769 |
| ENSG00000120699 | 13 | 37572953 | 37583750 | 19 | 4 | 0.49548 | 0.31013 | EXOSC8 | 0.31013 | 0.32865 | 0.35795 |
| ENSG00000061273 | 12 | 48176505 | 48226915 | 194 | 38 | 0.49522 | 0.31022 | HDAC7 | 0.31022 | 0.40625 | 0.4982 |
| ENSG00000103852 | 15 | 99676528 | 99791428 | 379 | 31 | 0.49506 | 0.31028 | TTC23 | 0.31028 | 0.30841 | 0.45756 |
| ENSG00000203722 | 6 | 150238014 | 150244257 | 20 | 5 | 0.49494 | 0.31032 | RAET1G | 0.31032 | 0.51755 | 0.40384 |
| ENSG00000241553 | 3 | 9834179 | 9849410 | 33 | 10 | 0.49465 | 0.31043 | ARPC4 | 0.31043 | 0.43946 | 0.50065 |
| ENSG00000091262 | 16 | 16242785 | 16317379 | 316 | 34 | 0.49443 | 0.3105 | ABCC6 | 0.3105 | 0.31737 | 0.27234 |
| ENSG00000134668 | 1 | 32256023 | 32281652 | 47 | 11 | 0.49419 | 0.31059 | SPOCD1 | 0.31059 | 0.2241 | 0.15105 |
| ENSG00000118518 | 6 | 127587755 | 127609712 | 55 | 10 | 0.49405 | 0.31064 | RNF146 | 0.31064 | 0.22515 | 0.25289 |
| ENSG00000163283 | 2 | 233243244 | 233247599 | 20 | 6 | 0.49401 | 0.31065 | ALPP | 0.31065 | 0.15871 | 0.069545 |
| ENSG00000141665 | 18 | 71740588 | 71815100 | 260 | 11 | 0.49386 | 0.3107 | FBXO15 | 0.3107 | 0.3604 | 0.37935 |
| ENSG00000163006 | 2 | 109403213 | 109501933 | 240 | 19 | 0.49383 | 0.31071 | CCDC138 | 0.31071 | 0.32006 | 0.30108 |
| ENSG00000142409 | 19 | 56598732 | 56632649 | 160 | 16 | 0.49373 | 0.31075 | ZNF787 | 0.31075 | 0.38143 | 0.47494 |
| ENSG00000136295 | 7 | 2671585 | 2704436 | 99 | 23 | 0.49364 | 0.31078 | TTYH3 | 0.31078 | 0.33043 | 0.31905 |
| ENSG00000268533 | 19 | 57905551 | 57949825 | 83 | 14 | 0.49361 | 0.31079 | AC004076.7 | 0.31079 | 0.38051 | 0.56033 |
| ENSG00000158604 | 7 | 44617493 | 44621886 | 15 | 5 | 0.49337 | 0.31087 | TMED4 | 0.31087 | 0.19725 | 0.21932 |
| ENSG00000106536 | 7 | 39017598 | 39532694 | 1658 | 68 | 0.49281 | 0.31107 | POU6F2 | 0.31107 | 0.29861 | 0.26661 |
| ENSG00000146039 | 6 | 25754927 | 25781419 | 115 | 16 | 0.4928 | 0.31108 | SLC17A4 | 0.31108 | 0.3956 | 0.52623 |
| ENSG00000258386 | 14 | 32476108 | 32487891 | 24 | 6 | 0.49264 | 0.31114 | RP11-187E13.2 | 0.31114 | 0.31575 | 0.30517 |
| ENSG00000167785 | 19 | 8920380 | 8943004 | 73 | 10 | 0.49229 | 0.31126 | ZNF558 | 0.31126 | 0.24066 | 0.18742 |
| ENSG00000185043 | 15 | 90773207 | 90777279 | 18 | 3 | 0.49212 | 0.31132 | CIB1 | 0.31132 | 0.21181 | 0.46224 |
| ENSG00000010282 | 3 | 42734155 | 42744319 | 29 | 8 | 0.49201 | 0.31136 | HHATL | 0.31136 | 0.42989 | 0.43122 |
| ENSG00000158077 | 11 | 7041677 | 7092539 | 231 | 20 | 0.49199 | 0.31136 | NLRP14 | 0.31136 | 0.26819 | 0.26517 |
| ENSG00000165495 | 11 | 125034583 | 125303285 | 897 | 96 | 0.49137 | 0.31158 | PKNOX2 | 0.31158 | 0.32201 | 0.44282 |
| ENSG00000214078 | 20 | 34213953 | 34252878 | 96 | 9 | 0.49134 | 0.31159 | CPNE1 | 0.31159 | 0.30586 | 0.28915 |
| ENSG00000095303 | 9 | 125132824 | 125157982 | 88 | 14 | 0.49133 | 0.3116 | PTGS1 | 0.3116 | 0.67888 | 0.81613 |
| ENSG00000198771 | 1 | 167599330 | 167675486 | 340 | 55 | 0.49123 | 0.31163 | RCSD1 | 0.31163 | 0.41076 | 0.66626 |
| ENSG00000090565 | 16 | 475619 | 573011 | 356 | 27 | 0.49119 | 0.31164 | RAB11FIP3 | 0.31164 | 0.24448 | 0.1924 |
| ENSG00000224470 | 16 | 71879894 | 71919171 | 93 | 16 | 0.49121 | 0.31164 | ATXN1L | 0.31164 | 0.31398 | 0.34052 |
| ENSG00000179058 | 9 | 132374504 | 132383055 | 33 | 7 | 0.49093 | 0.31174 | C9orf50 | 0.31174 | 0.2883 | 0.20764 |
| ENSG00000244537 | 17 | 39333698 | 39334460 | 5 | 2 | 0.49093 | 0.31174 | KRTAP4-2 | 0.31174 | 0.20666 | 0.2506 |
| ENSG00000170426 | 12 | 57316938 | 57328189 | 35 | 6 | 0.49087 | 0.31176 | SDR9C7 | 0.31176 | 0.70585 | 0.55999 |
| ENSG00000171295 | 19 | 11925099 | 11946016 | 84 | 5 | 0.49084 | 0.31177 | ZNF440 | 0.31177 | 0.28478 | 0.37453 |
| ENSG00000120549 | 10 | 23983675 | 24836772 | 2900 | 165 | 0.49057 | 0.31186 | KIAA1217 | 0.31186 | 0.32455 | 0.26433 |
| ENSG00000122477 | 1 | 100614409 | 100643771 | 74 | 15 | 0.49057 | 0.31187 | LRRC39 | 0.31187 | 0.33188 | 0.32608 |
| ENSG00000182077 | 10 | 27687116 | 27703297 | 112 | 15 | 0.49052 | 0.31188 | PTCHD3 | 0.31188 | 0.29293 | 0.26001 |
| ENSG00000172339 | 1 | 95439963 | 95538501 | 290 | 25 | 0.49042 | 0.31192 | ALG14 | 0.31192 | 0.31044 | 0.2496 |
| ENSG00000082213 | 5 | 31532373 | 31555165 | 57 | 16 | 0.49011 | 0.31203 | C5orf22 | 0.31203 | 0.11311 | 0.1241 |
| ENSG00000117983 | 11 | 1244296 | 1283406 | 292 | 17 | 0.49005 | 0.31205 | MUC5B | 0.31205 | 0.28853 | 0.33604 |
| ENSG00000169826 | 10 | 43633934 | 43680756 | 123 | 15 | 0.48994 | 0.31209 | CSGALNACT2 | 0.31209 | 0.32061 | 0.31166 |
| ENSG00000224940 | 7 | 127990379 | 128001739 | 32 | 9 | 0.48954 | 0.31223 | PRRT4 | 0.31223 | 0.015487 | 0.0064604 |

| | | | | | | | | | | | |
|---|---|---|---|---|---|---|---|---|---|---|---|
| ENSG00000187258 | 7 | 34697851 | 34917944 | 898 | 49 | 0.48946 | 0.31226 | NPSR1 | 0.31226 | 0.27515 | 0.26544 |
| ENSG00000149633 | 20 | 36838890 | 36889174 | 155 | 20 | 0.48934 | 0.3123 | KIAA1755 | 0.3123 | 0.35848 | 0.41646 |
| ENSG00000178295 | 2 | 17935125 | 17966632 | 109 | 17 | 0.48923 | 0.31234 | GEN1 | 0.31234 | 0.38741 | 0.41296 |
| ENSG00000203782 | 1 | 153232176 | 153234598 | 9 | 4 | 0.48919 | 0.31235 | LOR | 0.31235 | 0.80996 | 0.88896 |
| ENSG00000169908 | 3 | 149086809 | 149095652 | 19 | 3 | 0.48914 | 0.31237 | TM4SF1 | 0.31237 | 0.31974 | 0.25359 |
| ENSG00000153531 | 13 | 114076260 | 114107839 | 129 | 30 | 0.48911 | 0.31238 | ADPRHL1 | 0.31238 | 0.41302 | 0.33641 |
| ENSG00000065621 | 10 | 106028631 | 106064703 | 64 | 13 | 0.48904 | 0.31241 | GSTO2 | 0.31241 | 0.41331 | 0.34458 |
| ENSG00000171557 | 4 | 155525286 | 155534119 | 7 | 4 | 0.48875 | 0.31251 | FGG | 0.31251 | 0.51782 | 0.72046 |
| ENSG00000144028 | 2 | 96940074 | 96971297 | 35 | 7 | 0.48873 | 0.31252 | SNRNP200 | 0.31252 | 0.27963 | 0.29029 |
| ENSG00000166394 | 11 | 7686331 | 7698453 | 71 | 16 | 0.48849 | 0.3126 | CYB5R2 | 0.3126 | 0.43497 | 0.33857 |
| ENSG00000137673 | 11 | 102391239 | 102401484 | 45 | 9 | 0.48801 | 0.31277 | MMP7 | 0.31277 | 0.60217 | 0.46999 |
| ENSG00000118432 | 6 | 88849583 | 88876078 | 65 | 20 | 0.48791 | 0.31281 | CNR1 | 0.31281 | 0.2418 | 0.12438 |
| ENSG00000006007 | 16 | 19513015 | 19533467 | 78 | 10 | 0.48732 | 0.31302 | GDE1 | 0.31302 | 0.36851 | 0.40035 |
| ENSG00000134255 | 1 | 111682249 | 111727724 | 71 | 14 | 0.48697 | 0.31314 | CEPT1 | 0.31314 | 0.26101 | 0.33314 |
| ENSG00000089597 | 11 | 62392298 | 62414104 | 29 | 6 | 0.48691 | 0.31316 | GANAB | 0.31316 | 0.48817 | 0.5511 |
| ENSG00000198673 | 12 | 62102040 | 62672931 | 1970 | 107 | 0.48677 | 0.31321 | FAM19A2 | 0.31321 | 0.31117 | 0.32673 |
| ENSG00000135917 | 2 | 228549926 | 228582728 | 142 | 23 | 0.48671 | 0.31323 | SLC19A3 | 0.31323 | 0.55682 | 0.41369 |
| ENSG00000234438 | 15 | 65369154 | 65372276 | 10 | 3 | 0.48665 | 0.31325 | KBTBD13 | 0.31325 | 0.21763 | 0.20674 |
| ENSG00000185385 | 19 | 14991138 | 14992264 | 9 | 3 | 0.48666 | 0.31325 | OR7A17 | 0.31325 | 0.012527 | 0.049538 |
| ENSG00000137133 | 9 | 35812957 | 35815351 | 1 | 1 | 0.48652 | 0.3133 | HINT2 | 0.3133 | 0.76872 | 0.7303 |
| ENSG00000052850 | 11 | 44281994 | 44331716 | 214 | 40 | 0.48633 | 0.31337 | ALX4 | 0.31337 | 0.35388 | 0.42342 |
| ENSG00000186862 | 10 | 102767440 | 102790890 | 103 | 32 | 0.4862 | 0.31341 | PDZD7 | 0.31341 | 0.58326 | 0.7381 |
| ENSG00000124557 | 6 | 26501449 | 26510650 | 27 | 5 | 0.48611 | 0.31345 | BTN1A1 | 0.31345 | 0.4681 | 0.50679 |
| ENSG00000152229 | 18 | 43563502 | 43652238 | 279 | 18 | 0.48587 | 0.31353 | PSTPIP2 | 0.31353 | 0.27743 | 0.25456 |
| ENSG00000039523 | 16 | 67552321 | 67580691 | 40 | 11 | 0.48585 | 0.31354 | FAM65A | 0.31354 | 0.32427 | 0.26428 |
| ENSG00000112242 | 6 | 20402398 | 20493941 | 250 | 40 | 0.48574 | 0.31358 | E2F3 | 0.31358 | 0.32717 | 0.24748 |
| ENSG00000197417 | 17 | 3511556 | 3539616 | 134 | 25 | 0.48557 | 0.31364 | SHPK | 0.31364 | 0.44399 | 0.62358 |
| ENSG00000134571 | 11 | 47352957 | 47374253 | 55 | 16 | 0.48549 | 0.31366 | MYBPC3 | 0.31366 | 0.23439 | 0.2215 |
| ENSG00000105642 | 19 | 18062102 | 18110889 | 152 | 23 | 0.48522 | 0.31376 | KCNN1 | 0.31376 | 0.24509 | 0.17214 |
| ENSG00000027847 | 5 | 177027101 | 177037348 | 35 | 8 | 0.48467 | 0.31395 | B4GALT7 | 0.31395 | 0.32982 | 0.31325 |
| ENSG00000162804 | 2 | 241938255 | 242034983 | 253 | 40 | 0.48459 | 0.31398 | SNED1 | 0.31398 | 0.43262 | 0.63573 |
| ENSG00000185298 | 17 | 79633394 | 79640934 | 51 | 7 | 0.48459 | 0.31398 | CCDC137 | 0.31398 | 0.29304 | 0.47246 |
| ENSG00000212938 | 21 | 31964759 | 31965394 | 1 | 1 | 0.48454 | 0.314 | KRTAP6-3 | 0.314 | 0.7191 | 0.68881 |
| ENSG00000131069 | 20 | 33459949 | 33515769 | 112 | 8 | 0.48426 | 0.3141 | ACSS2 | 0.3141 | 0.33159 | 0.41921 |
| ENSG00000107201 | 9 | 32455300 | 32526322 | 215 | 19 | 0.48419 | 0.31413 | DDX58 | 0.31413 | 0.33781 | 0.38693 |
| ENSG00000142687 | 1 | 35899091 | 36023551 | 154 | 25 | 0.48383 | 0.31425 | KIAA0319L | 0.31425 | 0.34174 | 0.37175 |
| ENSG00000151014 | 4 | 139936943 | 139966906 | 66 | 16 | 0.4838 | 0.31427 | CCRN4L | 0.31427 | 0.52736 | 0.57897 |
| ENSG00000143776 | 1 | 227177566 | 227506175 | 1973 | 24 | 0.48373 | 0.31429 | CDC42BPA | 0.31429 | 0.31033 | 0.29673 |
| ENSG00000152669 | 5 | 54526980 | 54529508 | 6 | 3 | 0.48369 | 0.3143 | CCNO | 0.3143 | 0.20618 | 0.18345 |
| ENSG00000156471 | 8 | 97273943 | 97349223 | 186 | 25 | 0.48367 | 0.31431 | PTDSS1 | 0.31431 | 0.29773 | 0.31578 |
| ENSG00000167419 | 17 | 56295909 | 56345879 | 78 | 18 | 0.48346 | 0.31438 | LPO | 0.31438 | 0.32285 | 0.39087 |
| ENSG00000142330 | 2 | 241526133 | 241557122 | 170 | 21 | 0.48328 | 0.31445 | CAPN10 | 0.31445 | 0.31379 | 0.33421 |
| ENSG00000170516 | 4 | 46736844 | 46911252 | 452 | 28 | 0.48311 | 0.31451 | COX7B2 | 0.31451 | 0.28796 | 0.26383 |
| ENSG00000085491 | 1 | 108676658 | 108743471 | 266 | 17 | 0.48293 | 0.31457 | SLC25A24 | 0.31457 | 0.32855 | 0.40918 |
| ENSG00000162747 | 1 | 161592986 | 161601753 | 68 | 9 | 0.4829 | 0.31458 | FCGR3B | 0.31458 | 0.38119 | 0.51428 |
| ENSG00000115839 | 2 | 135809835 | 135933964 | 244 | 19 | 0.48288 | 0.31459 | RAB3GAP1 | 0.31459 | 0.36744 | 0.41428 |
| ENSG00000158402 | 5 | 137620954 | 137674044 | 123 | 15 | 0.48284 | 0.3146 | CDC25C | 0.3146 | 0.34237 | 0.41674 |
| ENSG00000138294 | 10 | 51549498 | 51562517 | 38 | 5 | 0.48256 | 0.3147 | MSMB | 0.3147 | 0.54149 | 0.54135 |
| ENSG00000204220 | 6 | 33257079 | 33266178 | 17 | 5 | 0.48222 | 0.31483 | PFDN6 | 0.31483 | 0.064393 | 0.031227 |
| ENSG00000137707 | 11 | 111338251 | 111383079 | 106 | 14 | 0.48193 | 0.31493 | BTG4 | 0.31493 | 0.33345 | 0.31352 |
| ENSG00000142327 | 2 | 241505221 | 241520789 | 50 | 17 | 0.48179 | 0.31498 | RNPEPL1 | 0.31498 | 0.35213 | 0.46429 |
| ENSG00000112294 | 6 | 24495080 | 24537435 | 214 | 26 | 0.48175 | 0.31499 | ALDH5A1 | 0.31499 | 0.35607 | 0.33298 |
| ENSG00000082014 | 7 | 150935850 | 150974982 | 124 | 22 | 0.48147 | 0.31509 | SMARCD3 | 0.31509 | 0.21881 | 0.26561 |
| ENSG00000011454 | 9 | 125703112 | 125867145 | 362 | 20 | 0.48124 | 0.31517 | RABGAP1 | 0.31517 | 0.32414 | 0.32325 |
| ENSG00000148356 | 9 | 130213765 | 130265780 | 163 | 12 | 0.4811 | 0.31522 | LRSAM1 | 0.31522 | 0.30882 | 0.29395 |
| ENSG00000142698 | 1 | 34632484 | 34684732 | 206 | 30 | 0.48103 | 0.31525 | C1orf94 | 0.31525 | 0.32999 | 0.22559 |
| ENSG00000143479 | 1 | 206808881 | 206857764 | 144 | 20 | 0.48089 | 0.3153 | DYRK3 | 0.3153 | 0.37692 | 0.52866 |
| ENSG00000196177 | 10 | 124768495 | 124817827 | 198 | 28 | 0.48043 | 0.31546 | ACADSB | 0.31546 | 0.28729 | 0.28788 |
| ENSG00000134121 | 3 | 238279 | 451090 | 999 | 75 | 0.48016 | 0.31556 | CHL1 | 0.31556 | 0.37069 | 0.3498 |
| ENSG00000024048 | 6 | 42531800 | 42661242 | 298 | 21 | 0.48003 | 0.3156 | UBR2 | 0.3156 | 0.22978 | 0.26668 |
| ENSG00000222040 | 2 | 96778707 | 96781984 | 4 | 2 | 0.47986 | 0.31566 | ADRA2B | 0.31566 | 0.29259 | 0.32946 |
| ENSG00000047249 | 8 | 54628117 | 54756118 | 128 | 15 | 0.47969 | 0.31572 | ATP6V1H | 0.31572 | 0.33796 | 0.40975 |
| ENSG00000168930 | 11 | 89530823 | 89541743 | 21 | 11 | 0.47971 | 0.31572 | TRIM49 | 0.31572 | 0.42705 | 0.38872 |
| ENSG00000166860 | 12 | 57392618 | 57400230 | 14 | 5 | 0.47965 | 0.31574 | ZBTB39 | 0.31574 | 0.36773 | 0.56007 |
| ENSG00000128268 | 22 | 39853349 | 39888199 | 58 | 16 | 0.47957 | 0.31577 | MGAT3 | 0.31577 | 0.28771 | 0.2369 |
| ENSG00000185267 | 10 | 14861249 | 14880574 | 57 | 14 | 0.47944 | 0.31581 | CDNF | 0.31581 | 0.19514 | 0.30389 |
| ENSG00000188234 | 10 | 46321042 | 46349323 | 3 | 1 | 0.47938 | 0.31583 | AGAP4 | 0.31583 | 0.30014 | 0.30014 |
| ENSG00000103044 | 16 | 69139467 | 69152622 | 23 | 8 | 0.4793 | 0.31586 | HAS3 | 0.31586 | 0.29368 | 0.24982 |
| ENSG00000145912 | 5 | 177576461 | 177580968 | 11 | 3 | 0.4792 | 0.3159 | NHP2 | 0.3159 | 0.17963 | 0.12624 |
| ENSG00000162946 | 1 | 231762561 | 232177018 | 1307 | 116 | 0.47901 | 0.31597 | DISC1 | 0.31597 | 0.33659 | 0.40487 |
| ENSG00000078328 | 16 | 6069095 | 7763340 | 11488 | 490 | 0.47876 | 0.31606 | RBFOX1 | 0.31606 | 0.251 | 0.29777 |
| ENSG00000197299 | 15 | 91260558 | 91358859 | 353 | 23 | 0.47873 | 0.31607 | BLM | 0.31607 | 0.3309 | 0.40058 |
| ENSG00000140382 | 15 | 77712754 | 77777949 | 132 | 14 | 0.47857 | 0.31612 | HMG20A | 0.31612 | 0.28411 | 0.32648 |

| | | | | | | | | | | | |
|---|---|---|---|---|---|---|---|---|---|---|---|
| ENSG00000122783 | 7 | 134777115 | 134855547 | 240 | 11 | 0.47849 | 0.31615 | C7orf49 | 0.31615 | 0.37865 | 0.39931 |
| ENSG00000258417 | 8 | 133036467 | 133097902 | 213 | 29 | 0.47823 | 0.31624 | OC90 | 0.31624 | 0.26562 | 0.30812 |
| ENSG00000204345 | 17 | 72575504 | 72588422 | 42 | 8 | 0.4781 | 0.31629 | CD300LD | 0.31629 | 0.38826 | 0.42205 |
| ENSG00000129657 | 17 | 75082798 | 75213179 | 541 | 44 | 0.4778 | 0.3164 | SEC14L1 | 0.3164 | 0.39341 | 0.41836 |
| ENSG00000137411 | 6 | 30876019 | 30894236 | 98 | 10 | 0.4776 | 0.31647 | VARS2 | 0.31647 | 0.31134 | 0.33405 |
| ENSG00000163472 | 1 | 156252726 | 156262976 | 21 | 5 | 0.47744 | 0.31653 | TMEM79 | 0.31653 | 0.42477 | 0.42643 |
| ENSG00000111679 | 12 | 7055631 | 7070479 | 30 | 10 | 0.47737 | 0.31655 | PTPN6 | 0.31655 | 0.032197 | 0.026442 |
| ENSG00000123358 | 12 | 52416616 | 52453291 | 115 | 26 | 0.47734 | 0.31656 | NR4A1 | 0.31656 | 0.46714 | 0.456 |
| ENSG00000140983 | 16 | 718086 | 724174 | 25 | 6 | 0.47719 | 0.31661 | RHOT2 | 0.31661 | 0.41872 | 0.28805 |
| ENSG00000205740 | 10 | 1017097 | 1034281 | 60 | 11 | 0.47707 | 0.31666 | AL359878.1 | 0.31666 | 0.23169 | 0.14843 |
| ENSG00000137478 | 11 | 72547790 | 72853306 | 343 | 38 | 0.47668 | 0.31679 | FCHSD2 | 0.31679 | 0.32555 | 0.3565 |
| ENSG00000141338 | 17 | 66863433 | 66951533 | 237 | 26 | 0.47637 | 0.31691 | ABCA8 | 0.31691 | 0.30274 | 0.38909 |
| ENSG00000212123 | 19 | 5782971 | 5784776 | 7 | 2 | 0.47637 | 0.31691 | PRR22 | 0.31691 | 0.34952 | 0.44801 |
| ENSG00000060656 | 1 | 29563028 | 29653325 | 244 | 37 | 0.4762 | 0.31697 | PTPRU | 0.31697 | 0.38426 | 0.36176 |
| ENSG00000182022 | 10 | 125767184 | 125853206 | 211 | 37 | 0.47613 | 0.31699 | CHST15 | 0.31699 | 0.31847 | 0.58999 |
| ENSG00000078668 | 8 | 42249142 | 42263415 | 21 | 4 | 0.476 | 0.31704 | VDAC3 | 0.31704 | 0.31936 | 0.25165 |
| ENSG00000141076 | 16 | 69165194 | 69265033 | 229 | 29 | 0.47548 | 0.31722 | CIRH1A | 0.31722 | 0.33989 | 0.34348 |
| ENSG00000244414 | 1 | 196788887 | 196801319 | 93 | 5 | 0.47526 | 0.3173 | CFHR1 | 0.3173 | 0.40503 | 0.59988 |
| ENSG00000143157 | 1 | 166808681 | 166825581 | 31 | 6 | 0.47517 | 0.31733 | POGK | 0.31733 | 0.31913 | 0.26032 |
| ENSG00000081177 | 14 | 69658228 | 69709075 | 122 | 13 | 0.47511 | 0.31735 | EXD2 | 0.31735 | 0.34113 | 0.41415 |
| ENSG00000042980 | 8 | 24151553 | 24216531 | 177 | 15 | 0.47509 | 0.31736 | ADAM28 | 0.31736 | 0.38233 | 0.459 |
| ENSG00000119900 | 6 | 71998506 | 72018653 | 81 | 13 | 0.47506 | 0.31737 | OGFRL1 | 0.31737 | 0.33005 | 0.31852 |
| ENSG00000215695 | 1 | 15986364 | 15988217 | 3 | 2 | 0.47504 | 0.31738 | RSC1A1 | 0.31738 | 0.12187 | 0.086013 |
| ENSG00000172247 | 11 | 47611216 | 47616211 | 6 | 4 | 0.47488 | 0.31744 | C1QTNF4 | 0.31744 | 0.5976 | 0.75682 |
| ENSG00000134698 | 1 | 36273773 | 36323491 | 74 | 14 | 0.47476 | 0.31748 | AGO4 | 0.31748 | 0.33004 | 0.37764 |
| ENSG00000121634 | 1 | 147374946 | 147381393 | 31 | 8 | 0.47456 | 0.31755 | GJA8 | 0.31755 | 0.28215 | 0.19147 |
| ENSG00000152785 | 4 | 81952119 | 81978685 | 65 | 10 | 0.47452 | 0.31756 | BMP3 | 0.31756 | 0.26778 | 0.298 |
| ENSG00000033050 | 7 | 150904923 | 150924316 | 73 | 11 | 0.47429 | 0.31765 | ABCF2 | 0.31765 | 0.35031 | 0.43884 |
| ENSG00000134910 | 11 | 125461607 | 125495110 | 166 | 12 | 0.47399 | 0.31775 | STT3A | 0.31775 | 0.33706 | 0.30688 |
| ENSG00000258989 | 14 | 61995846 | 62124682 | 363 | 61 | 0.47375 | 0.31784 | RP11-47I22.4 | 0.31784 | 0.35391 | 0.51515 |
| ENSG00000198182 | 19 | 38187264 | 38210691 | 61 | 10 | 0.4736 | 0.31789 | ZNF607 | 0.31789 | 0.35146 | 0.38839 |
| ENSG00000171872 | 1 | 44584522 | 44600812 | 62 | 14 | 0.47358 | 0.3179 | KLF17 | 0.3179 | 0.44301 | 0.6569 |
| ENSG00000138587 | 15 | 56713742 | 56757335 | 117 | 17 | 0.47355 | 0.31791 | MNS1 | 0.31791 | 0.39664 | 0.33809 |
| ENSG00000183833 | 3 | 119421869 | 119485949 | 135 | 17 | 0.47308 | 0.31808 | MAATS1 | 0.31808 | 0.28499 | 0.34486 |
| ENSG00000110330 | 11 | 102217942 | 102249401 | 70 | 9 | 0.47301 | 0.3181 | BIRC2 | 0.3181 | 0.31331 | 0.30103 |
| ENSG00000134864 | 13 | 101183810 | 101241782 | 115 | 15 | 0.47249 | 0.31829 | GGACT | 0.31829 | 0.2875 | 0.32568 |
| ENSG00000129170 | 11 | 19203578 | 19232120 | 115 | 25 | 0.47235 | 0.31834 | CSRP3 | 0.31834 | 0.24247 | 0.29977 |
| ENSG00000108349 | 17 | 38296571 | 38328436 | 67 | 8 | 0.47234 | 0.31834 | CASC3 | 0.31834 | 0.28248 | 0.33374 |
| ENSG00000182247 | 3 | 23244511 | 23633284 | 1508 | 25 | 0.47226 | 0.31837 | UBE2E2 | 0.31837 | 0.32747 | 0.32991 |
| ENSG00000236761 | 6 | 132029581 | 132032206 | 5 | 2 | 0.47219 | 0.31839 | CTAGE9 | 0.31839 | 0.31235 | 0.34614 |
| ENSG00000164821 | 8 | 6793344 | 6795860 | 14 | 6 | 0.4722 | 0.31839 | DEFA4 | 0.31839 | 0.26507 | 0.036165 |
| ENSG00000175097 | 11 | 36597124 | 36619829 | 50 | 10 | 0.47176 | 0.31855 | RAG2 | 0.31855 | 0.43392 | 0.30814 |
| ENSG00000197971 | 18 | 74690783 | 74845639 | 853 | 113 | 0.47158 | 0.31861 | MBP | 0.31861 | 0.24993 | 0.39404 |
| ENSG00000225828 | 1 | 32826871 | 32829879 | 1 | 1 | 0.47134 | 0.3187 | FAM229A | 0.3187 | 0.37559 | 0.73699 |
| ENSG00000169962 | 1 | 1266694 | 1270686 | 16 | 5 | 0.4713 | 0.31871 | TAS1R3 | 0.31871 | 0.46613 | 0.48377 |
| ENSG00000118508 | 6 | 146864829 | 146876101 | 23 | 5 | 0.47124 | 0.31873 | RAB32 | 0.31873 | 0.24098 | 0.25968 |
| ENSG00000197245 | 1 | 26485511 | 26489119 | 11 | 5 | 0.47097 | 0.31883 | FAM110D | 0.31883 | 0.34515 | 0.35631 |
| ENSG00000104972 | 19 | 55085346 | 55148979 | 463 | 23 | 0.47077 | 0.3189 | LILRB1 | 0.3189 | 0.26907 | 0.31197 |
| ENSG00000171130 | 7 | 149570057 | 149577784 | 39 | 4 | 0.47072 | 0.31892 | ATP6V0E2 | 0.31892 | 0.30874 | 0.3999 |
| ENSG00000213626 | 2 | 30454397 | 30546596 | 382 | 38 | 0.47044 | 0.31902 | LBH | 0.31902 | 0.33073 | 0.34059 |
| ENSG00000175768 | 9 | 37582643 | 37592639 | 30 | 7 | 0.47036 | 0.31905 | TOMM5 | 0.31905 | 0.26809 | 0.43894 |
| ENSG00000186106 | 8 | 101521980 | 101572012 | 146 | 19 | 0.47027 | 0.31908 | ANKRD46 | 0.31908 | 0.30915 | 0.26597 |
| ENSG00000118894 | 16 | 5134305 | 5147809 | 90 | 22 | 0.47013 | 0.31913 | FAM86A | 0.31913 | 0.18012 | 0.11475 |
| ENSG00000136451 | 17 | 56048910 | 56065620 | 29 | 7 | 0.47 | 0.31918 | VEZF1 | 0.31918 | 0.3271 | 0.29564 |
| ENSG00000048707 | 1 | 12290124 | 12572099 | 467 | 37 | 0.46993 | 0.3192 | VPS13D | 0.3192 | 0.20612 | 0.11945 |
| ENSG00000196911 | 6 | 117002350 | 117063029 | 126 | 14 | 0.4699 | 0.31921 | KPNA5 | 0.31921 | 0.36499 | 0.35332 |
| ENSG00000073282 | 3 | 189349205 | 189615068 | 1161 | 139 | 0.46983 | 0.31924 | TP63 | 0.31924 | 0.27449 | 0.3036 |
| ENSG00000170949 | 19 | 53569859 | 53606689 | 219 | 33 | 0.46953 | 0.31935 | ZNF160 | 0.31935 | 0.25691 | 0.27781 |
| ENSG00000182117 | 15 | 34633917 | 34635378 | 14 | 4 | 0.46925 | 0.31945 | NOP10 | 0.31945 | 0.38519 | 0.37074 |
| ENSG00000128951 | 15 | 48623208 | 48635570 | 10 | 3 | 0.46913 | 0.31949 | DUT | 0.31949 | 0.55871 | 0.62046 |
| ENSG00000186446 | 3 | 44771088 | 44778575 | 16 | 4 | 0.46885 | 0.31959 | ZNF501 | 0.31959 | 0.53791 | 0.57356 |
| ENSG00000165264 | 9 | 32552997 | 32573160 | 60 | 8 | 0.46882 | 0.3196 | NDUFB6 | 0.3196 | 0.39581 | 0.43115 |
| ENSG00000185499 | 1 | 155158300 | 155162707 | 8 | 5 | 0.46853 | 0.3197 | MUC1 | 0.3197 | 0.24909 | 0.13663 |
| ENSG00000169213 | 1 | 52373628 | 52456436 | 205 | 26 | 0.46831 | 0.31978 | RAB3B | 0.31978 | 0.30609 | 0.29463 |
| ENSG00000185689 | 6 | 4079440 | 4131185 | 185 | 18 | 0.46815 | 0.31984 | C6orf201 | 0.31984 | 0.24453 | 0.24718 |
| ENSG00000128573 | 7 | 113726382 | 114333827 | 1011 | 62 | 0.46774 | 0.31998 | FOXP2 | 0.31998 | 0.31758 | 0.32162 |
| ENSG00000138162 | 10 | 123748689 | 124014060 | 1282 | 127 | 0.46758 | 0.32004 | TACC2 | 0.32004 | 0.27598 | 0.40017 |
| ENSG00000105821 | 7 | 102952921 | 102985320 | 65 | 14 | 0.4675 | 0.32007 | DNAJC2 | 0.32007 | 0.35172 | 0.35986 |
| ENSG00000173511 | 11 | 64002010 | 64006259 | 13 | 4 | 0.46727 | 0.32015 | VEGFB | 0.32015 | 0.38576 | 0.44827 |
| ENSG00000135624 | 2 | 73460548 | 73480149 | 54 | 10 | 0.46724 | 0.32016 | CCT7 | 0.32016 | 0.49775 | 0.44832 |
| ENSG00000269103 | 22 | 21358030 | 21360736 | 22 | 7 | 0.46726 | 0.32016 | AC002472.1 | 0.32016 | 0.42396 | 0.41314 |
| ENSG00000136872 | 9 | 104182860 | 104198105 | 51 | 10 | 0.46721 | 0.32017 | ALDOB | 0.32017 | 0.46022 | 0.64491 |
| ENSG00000140798 | 16 | 48116884 | 48189929 | 61 | 11 | 0.46718 | 0.32018 | ABCC12 | 0.32018 | 0.29961 | 0.32432 |

| | | | | | | | | | | | |
|---|---|---|---|---|---|---|---|---|---|---|---|
| ENSG00000138769 | 4 | 76503215 | 76555900 | 159 | 15 | 0.46704 | 0.32024 | CDKL2 | 0.32024 | 0.29493 | 0.3335 |
| ENSG00000203705 | 1 | 212965170 | 212989968 | 137 | 11 | 0.46691 | 0.32028 | TATDN3 | 0.32028 | 0.36242 | 0.4066 |
| ENSG00000168421 | 4 | 40192673 | 40248587 | 198 | 32 | 0.4668 | 0.32032 | RHOH | 0.32032 | 0.19737 | 0.18644 |
| ENSG00000001084 | 6 | 53362139 | 53481768 | 473 | 65 | 0.46628 | 0.32051 | GCLC | 0.32051 | 0.41076 | 0.45563 |
| ENSG00000110321 | 11 | 10818597 | 10830657 | 33 | 8 | 0.46603 | 0.3206 | EIF4G2 | 0.3206 | 0.24405 | 0.27622 |
| ENSG00000134086 | 3 | 10182692 | 10193904 | 51 | 8 | 0.46573 | 0.32071 | VHL | 0.32071 | 0.05515 | 0.090692 |
| ENSG00000106682 | 7 | 73588575 | 73611431 | 31 | 11 | 0.46557 | 0.32076 | EIF4H | 0.32076 | 0.69882 | 0.78188 |
| ENSG00000175084 | 2 | 220283099 | 220291461 | 19 | 5 | 0.4654 | 0.32082 | DES | 0.32082 | 0.61885 | 0.64697 |
| ENSG00000163098 | 19 | 53792856 | 53794875 | 12 | 5 | 0.46511 | 0.32093 | BIRC8 | 0.32093 | 0.47213 | 0.37866 |
| ENSG00000188130 | 22 | 50683879 | 50700254 | 60 | 12 | 0.46504 | 0.32095 | MAPK12 | 0.32095 | 0.16902 | 0.20238 |
| ENSG00000066279 | 1 | 197053258 | 197115824 | 115 | 9 | 0.4641 | 0.32129 | ASPM | 0.32129 | 0.30093 | 0.29562 |
| ENSG00000081051 | 4 | 74296855 | 74321891 | 54 | 9 | 0.46366 | 0.32145 | AFP | 0.32145 | 0.43512 | 0.52638 |
| ENSG00000135324 | 6 | 84743475 | 84800600 | 167 | 17 | 0.46364 | 0.32145 | MRAP2 | 0.32145 | 0.28328 | 0.32673 |
| ENSG00000157617 | 21 | 43305221 | 43373999 | 370 | 66 | 0.46319 | 0.32162 | C2CD2 | 0.32162 | 0.26183 | 0.21115 |
| ENSG00000213079 | 6 | 155054459 | 155155192 | 252 | 16 | 0.46309 | 0.32165 | SCAF8 | 0.32165 | 0.32065 | 0.2886 |
| ENSG00000211584 | 12 | 48147699 | 48176536 | 101 | 16 | 0.46279 | 0.32176 | SLC48A1 | 0.32176 | 0.27657 | 0.269 |
| ENSG00000266733 | 17 | 28884130 | 28890511 | 11 | 4 | 0.46262 | 0.32182 | TBC1D29 | 0.32182 | 0.23819 | 0.4763 |
| ENSG00000167081 | 9 | 128509624 | 128729656 | 383 | 21 | 0.46231 | 0.32193 | PBX3 | 0.32193 | 0.29576 | 0.33765 |
| ENSG00000198715 | 1 | 156259880 | 156265463 | 14 | 5 | 0.46221 | 0.32197 | C1orf85 | 0.32197 | 0.41226 | 0.45578 |
| ENSG00000198125 | 22 | 36002811 | 36033998 | 114 | 15 | 0.46161 | 0.32218 | MB | 0.32218 | 0.06716 | 0.12069 |
| ENSG00000171777 | 19 | 38899695 | 38916945 | 60 | 15 | 0.46149 | 0.32222 | RASGRP4 | 0.32222 | 0.58433 | 0.69639 |
| ENSG00000145439 | 4 | 169784921 | 169931426 | 323 | 15 | 0.46137 | 0.32227 | CBR4 | 0.32227 | 0.31798 | 0.31173 |
| ENSG00000105640 | 19 | 17970685 | 17974962 | 14 | 3 | 0.46107 | 0.32237 | RPL18A | 0.32237 | 0.20501 | 0.32738 |
| ENSG00000181467 | 3 | 152880029 | 152886265 | 9 | 5 | 0.46105 | 0.32238 | RAP2B | 0.32238 | 0.35418 | 0.24843 |
| ENSG00000141506 | 17 | 8782233 | 8869029 | 234 | 39 | 0.46101 | 0.3224 | PIK3R5 | 0.3224 | 0.28572 | 0.22018 |
| ENSG00000142192 | 21 | 27252861 | 27543446 | 947 | 70 | 0.46096 | 0.32241 | APP | 0.32241 | 0.31512 | 0.29467 |
| ENSG00000121152 | 2 | 97001525 | 97039583 | 47 | 9 | 0.46086 | 0.32245 | NCAPH | 0.32245 | 0.4226 | 0.34995 |
| ENSG00000074706 | 6 | 154475631 | 154677926 | 884 | 70 | 0.46084 | 0.32246 | IPCEF1 | 0.32246 | 0.31609 | 0.29538 |
| ENSG00000268856 | 21 | 46492796 | 46492927 | 1 | 1 | 0.46072 | 0.3225 | AP001579.1 | 0.3225 | 0.63915 | 0.70395 |
| ENSG00000175538 | 11 | 74165886 | 74178774 | 46 | 11 | 0.46065 | 0.32253 | KCNE3 | 0.32253 | 0.46731 | 0.79934 |
| ENSG00000254402 | 8 | 145747761 | 145752416 | 7 | 3 | 0.46015 | 0.3227 | LRRC24 | 0.3227 | 0.62715 | 0.61303 |
| ENSG00000164074 | 4 | 128886461 | 128960866 | 133 | 13 | 0.46012 | 0.32272 | C4orf29 | 0.32272 | 0.33608 | 0.35077 |
| ENSG00000105851 | 7 | 106505723 | 106547590 | 90 | 18 | 0.46001 | 0.32275 | PIK3CG | 0.32275 | 0.44034 | 0.50418 |
| ENSG00000173531 | 3 | 49721380 | 49726934 | 10 | 4 | 0.45974 | 0.32285 | MST1 | 0.32285 | 0.22319 | 0.12016 |
| ENSG00000108639 | 17 | 76164639 | 76169608 | 25 | 7 | 0.45965 | 0.32288 | SYNGR2 | 0.32288 | 0.70829 | 0.84279 |
| ENSG00000151650 | 10 | 135050908 | 135055433 | 28 | 6 | 0.45954 | 0.32292 | VENTX | 0.32292 | 0.32412 | 0.069226 |
| ENSG00000240563 | 1 | 62660503 | 62678000 | 142 | 21 | 0.45908 | 0.32309 | L1TD1 | 0.32309 | 0.3379 | 0.37878 |
| ENSG00000232671 | 1 | 151252500 | 151254405 | 4 | 2 | 0.45875 | 0.32321 | RP11-126K1.2 | 0.32321 | 0.70874 | 0.86228 |
| ENSG00000130595 | 11 | 1940792 | 1959936 | 90 | 20 | 0.45869 | 0.32323 | TNNT3 | 0.32323 | 0.1251 | 0.044151 |
| ENSG00000205143 | 9 | 34621376 | 34628104 | 15 | 5 | 0.45866 | 0.32324 | ARID3C | 0.32324 | 0.30018 | 0.3801 |
| ENSG00000182667 | 11 | 131240373 | 132206716 | 3386 | 253 | 0.45848 | 0.3233 | NTM | 0.3233 | 0.28624 | 0.25647 |
| ENSG00000169035 | 19 | 51479729 | 51487355 | 40 | 9 | 0.45832 | 0.32336 | KLK7 | 0.32336 | 0.67813 | 0.74283 |
| ENSG00000104856 | 19 | 45504688 | 45541452 | 121 | 28 | 0.45827 | 0.32338 | RELB | 0.32338 | 0.1361 | 0.076658 |
| ENSG00000187094 | 3 | 42299317 | 42307699 | 25 | 6 | 0.45825 | 0.32339 | CCK | 0.32339 | 0.488 | 0.68011 |
| ENSG00000136933 | 9 | 127962821 | 127996437 | 106 | 13 | 0.45816 | 0.32342 | RABEPK | 0.32342 | 0.28816 | 0.34877 |
| ENSG00000267552 | 19 | 38180766 | 38210089 | 71 | 10 | 0.45814 | 0.32343 | CTD-2528L19.4 | 0.32343 | 0.38396 | 0.389 |
| ENSG00000100249 | 22 | 29454660 | 29457832 | 9 | 6 | 0.458 | 0.32348 | C22orf31 | 0.32348 | 0.88199 | 0.80406 |
| ENSG00000121380 | 12 | 12202778 | 12364018 | 721 | 81 | 0.45758 | 0.32363 | BCL2L14 | 0.32363 | 0.41206 | 0.46803 |
| ENSG00000164465 | 6 | 117774980 | 117891021 | 449 | 33 | 0.45746 | 0.32367 | DCBLD1 | 0.32367 | 0.3125 | 0.32007 |
| ENSG00000126266 | 19 | 35842445 | 35843367 | 1 | 1 | 0.45738 | 0.3237 | FFAR1 | 0.3237 | 0.67662 | 0.84026 |
| ENSG00000138041 | 2 | 55774428 | 55846015 | 258 | 21 | 0.45717 | 0.32377 | SMEK2 | 0.32377 | 0.28556 | 0.30389 |
| ENSG00000166855 | 15 | 65440557 | 65477680 | 41 | 11 | 0.45681 | 0.3239 | CLPX | 0.3239 | 0.21082 | 0.040643 |
| ENSG00000205426 | 12 | 52679697 | 52685318 | 25 | 6 | 0.45676 | 0.32392 | KRT81 | 0.32392 | 0.53478 | 0.62179 |
| ENSG00000226650 | 5 | 154393260 | 154397685 | 7 | 3 | 0.45654 | 0.324 | KIF4B | 0.324 | 0.15741 | 0.093682 |
| ENSG00000118017 | 3 | 137842560 | 137851229 | 14 | 3 | 0.45593 | 0.32422 | A4GNT | 0.32422 | 0.31493 | 0.38093 |
| ENSG00000054938 | 11 | 74407474 | 74442430 | 105 | 27 | 0.45565 | 0.32432 | CHRDL2 | 0.32432 | 0.22508 | 0.15554 |
| ENSG00000152348 | 5 | 81267844 | 81572676 | 487 | 39 | 0.45559 | 0.32434 | ATG10 | 0.32434 | 0.30093 | 0.30606 |
| ENSG00000197540 | 19 | 544034 | 549919 | 56 | 13 | 0.45555 | 0.32436 | GZMM | 0.32436 | 0.20821 | 0.26555 |
| ENSG00000148399 | 9 | 140449356 | 140473387 | 38 | 8 | 0.45527 | 0.32446 | DPH7 | 0.32446 | 0.32475 | 0.3075 |
| ENSG00000128346 | 22 | 38339528 | 38349676 | 12 | 6 | 0.45523 | 0.32447 | C22orf23 | 0.32447 | 0.47799 | 0.4923 |
| ENSG00000174226 | 8 | 101585116 | 101675643 | 452 | 41 | 0.45514 | 0.3245 | SNX31 | 0.3245 | 0.30023 | 0.28659 |
| ENSG00000110851 | 12 | 108126643 | 108155049 | 84 | 8 | 0.45484 | 0.32461 | PRDM4 | 0.32461 | 0.35675 | 0.45194 |
| ENSG00000100243 | 22 | 43013846 | 43045574 | 129 | 17 | 0.4548 | 0.32463 | CYB5R3 | 0.32463 | 0.36548 | 0.47719 |
| ENSG00000197415 | 3 | 156977531 | 157251408 | 885 | 42 | 0.4546 | 0.3247 | VEPH1 | 0.3247 | 0.32232 | 0.34199 |
| ENSG00000128908 | 15 | 41271078 | 41408552 | 301 | 20 | 0.45461 | 0.3247 | INO80 | 0.3247 | 0.36407 | 0.40543 |
| ENSG00000175104 | 11 | 36508577 | 36531822 | 53 | 9 | 0.45448 | 0.32474 | TRAF6 | 0.32474 | 0.29495 | 0.27025 |
| ENSG00000185340 | 22 | 29702572 | 29708774 | 15 | 6 | 0.4544 | 0.32477 | GAS2L1 | 0.32477 | 0.25446 | 0.29143 |
| ENSG00000093183 | 3 | 42589461 | 42642572 | 157 | 17 | 0.45431 | 0.3248 | SEC22C | 0.3248 | 0.41318 | 0.38883 |
| ENSG00000196381 | 19 | 38158652 | 38183223 | 69 | 14 | 0.45425 | 0.32482 | ZNF781 | 0.32482 | 0.28385 | 0.36139 |
| ENSG00000101440 | 20 | 32782375 | 32857150 | 140 | 8 | 0.45418 | 0.32485 | ASIP | 0.32485 | 0.32302 | 0.31464 |
| ENSG00000196659 | 2 | 178413726 | 178417742 | 6 | 2 | 0.45402 | 0.32491 | TTC30B | 0.32491 | 0.54744 | 0.60566 |
| ENSG00000169116 | 4 | 75858305 | 75975325 | 214 | 26 | 0.45362 | 0.32505 | PARM1 | 0.32505 | 0.3555 | 0.40378 |
| ENSG00000163959 | 3 | 195938358 | 195970049 | 111 | 19 | 0.4536 | 0.32506 | SLC51A | 0.32506 | 0.22996 | 0.2637 |

| | | | | | | | | | | | |
|---|---|---|---|---|---|---|---|---|---|---|---|
| ENSG00000171346 | 17 | 39669995 | 39678781 | 8 | 3 | 0.45341 | 0.32513 | KRT15 | 0.32513 | 0.76313 | 0.84909 |
| ENSG00000114503 | 3 | 196662273 | 196669468 | 32 | 7 | 0.4533 | 0.32517 | NCBP2 | 0.32517 | 0.42197 | 0.081953 |
| ENSG00000161955 | 17 | 7461609 | 7464925 | 7 | 4 | 0.45313 | 0.32523 | TNFSF13 | 0.32523 | 0.0098741 | 0.024389 |
| ENSG00000131100 | 22 | 18074902 | 18111584 | 210 | 16 | 0.45296 | 0.32529 | ATP6V1E1 | 0.32529 | 0.32388 | 0.37168 |
| ENSG00000107821 | 10 | 102821598 | 102827888 | 12 | 5 | 0.45214 | 0.32558 | KAZALD1 | 0.32558 | 0.69181 | 0.74924 |
| ENSG00000130764 | 1 | 3696784 | 3713068 | 89 | 17 | 0.45212 | 0.32559 | LRRC47 | 0.32559 | 0.51277 | 0.70187 |
| ENSG00000165935 | 12 | 27619743 | 27655118 | 230 | 15 | 0.45195 | 0.32565 | SMCO2 | 0.32565 | 0.36303 | 0.28444 |
| ENSG00000083845 | 19 | 58897767 | 58906173 | 29 | 4 | 0.4519 | 0.32567 | RPS5 | 0.32567 | 0.56313 | 0.6106 |
| ENSG00000256436 | 12 | 11182986 | 11184006 | 11 | 4 | 0.45144 | 0.32584 | TAS2R31 | 0.32584 | 0.45322 | 0.42588 |
| ENSG00000149474 | 20 | 18118762 | 18169031 | 191 | 35 | 0.45141 | 0.32585 | CSRP2BP | 0.32585 | 0.41661 | 0.53107 |
| ENSG00000168040 | 11 | 70049269 | 70053496 | 8 | 2 | 0.45135 | 0.32587 | FADD | 0.32587 | 0.22063 | 0.13614 |
| ENSG00000182578 | 5 | 149432854 | 149492935 | 201 | 38 | 0.45132 | 0.32588 | CSF1R | 0.32588 | 0.35021 | 0.37492 |
| ENSG00000144015 | 2 | 96257766 | 96265526 | 4 | 2 | 0.45103 | 0.32599 | TRIM43 | 0.32599 | 0.52835 | 0.46812 |
| ENSG00000168959 | 11 | 88237744 | 88799113 | 2108 | 45 | 0.45088 | 0.32604 | GRM5 | 0.32604 | 0.33345 | 0.35831 |
| ENSG00000167384 | 19 | 44979854 | 45004576 | 158 | 15 | 0.45079 | 0.32607 | ZNF180 | 0.32607 | 0.26767 | 0.29031 |
| ENSG00000125733 | 19 | 6737936 | 6751537 | 38 | 12 | 0.45072 | 0.3261 | TRIP10 | 0.3261 | 0.20671 | 0.37737 |
| ENSG00000178585 | 1 | 9908334 | 9970394 | 81 | 16 | 0.45069 | 0.32611 | CTNNBIP1 | 0.32611 | 0.40633 | 0.64076 |
| ENSG00000134153 | 15 | 34376218 | 34394149 | 52 | 9 | 0.4503 | 0.32625 | EMC7 | 0.32625 | 0.36215 | 0.41717 |
| ENSG00000204389 | 6 | 31783291 | 31785723 | 5 | 3 | 0.45027 | 0.32626 | HSPA1A | 0.32626 | 0.25492 | 0.39232 |
| ENSG00000174574 | 1 | 39456895 | 39471731 | 58 | 7 | 0.45006 | 0.32633 | AKIRIN1 | 0.32633 | 0.30211 | 0.22501 |
| ENSG00000121481 | 1 | 185014496 | 185071740 | 62 | 16 | 0.44996 | 0.32637 | RNF2 | 0.32637 | 0.48103 | 0.63048 |
| ENSG00000197579 | 9 | 32540542 | 32552551 | 24 | 7 | 0.44986 | 0.32641 | TOPORS | 0.32641 | 0.29828 | 0.37426 |
| ENSG00000196505 | 1 | 118406107 | 118472253 | 147 | 6 | 0.44972 | 0.32646 | GDAP2 | 0.32646 | 0.31805 | 0.31575 |
| ENSG00000087303 | 14 | 52471521 | 52535712 | 387 | 27 | 0.44959 | 0.3265 | NID2 | 0.3265 | 0.34992 | 0.39611 |
| ENSG00000132680 | 1 | 155882834 | 155904191 | 36 | 7 | 0.44955 | 0.32652 | KIAA0907 | 0.32652 | 0.39278 | 0.42772 |
| ENSG00000196396 | 20 | 49126891 | 49201299 | 169 | 23 | 0.44944 | 0.32656 | PTPN1 | 0.32656 | 0.28819 | 0.25117 |
| ENSG00000139998 | 14 | 65412532 | 65439494 | 61 | 14 | 0.4491 | 0.32668 | RAB15 | 0.32668 | 0.21444 | 0.1635 |
| ENSG00000130699 | 20 | 60528525 | 60640866 | 398 | 25 | 0.44903 | 0.32671 | TAF4 | 0.32671 | 0.34319 | 0.35583 |
| ENSG00000181817 | 1 | 36856839 | 36863493 | 43 | 6 | 0.44896 | 0.32673 | LSM10 | 0.32673 | 0.29977 | 0.22288 |
| ENSG00000108848 | 17 | 48796905 | 48833574 | 117 | 19 | 0.44883 | 0.32678 | LUC7L3 | 0.32678 | 0.27945 | 0.31832 |
| ENSG00000177174 | 1 | 248512077 | 248513015 | 6 | 2 | 0.44853 | 0.32688 | OR14C36 | 0.32688 | 0.51676 | 0.51789 |
| ENSG00000138646 | 4 | 89378268 | 89427314 | 52 | 18 | 0.44842 | 0.32693 | HERC5 | 0.32693 | 0.29367 | 0.26776 |
| ENSG00000166135 | 10 | 102288829 | 102319755 | 43 | 10 | 0.4483 | 0.32697 | HIF1AN | 0.32697 | 0.21705 | 0.20553 |
| ENSG00000185897 | 19 | 35849362 | 35851387 | 5 | 3 | 0.44815 | 0.32702 | FFAR3 | 0.32702 | 0.53388 | 0.88562 |
| ENSG00000198961 | 5 | 108670410 | 108745695 | 337 | 16 | 0.44799 | 0.32708 | PJA2 | 0.32708 | 0.30119 | 0.21769 |
| ENSG00000130175 | 19 | 11546109 | 11561783 | 32 | 8 | 0.44765 | 0.3272 | PRKCSH | 0.3272 | 0.4338 | 0.44552 |
| ENSG00000155011 | 4 | 107842959 | 108204963 | 1129 | 97 | 0.44739 | 0.3273 | DKK2 | 0.3273 | 0.14784 | 0.077077 |
| ENSG00000072110 | 14 | 69340860 | 69446157 | 398 | 43 | 0.44714 | 0.32739 | ACTN1 | 0.32739 | 0.1673 | 0.19945 |
| ENSG00000162706 | 1 | 159141399 | 159173103 | 68 | 15 | 0.447 | 0.32744 | CADM3 | 0.32744 | 0.30816 | 0.16703 |
| ENSG00000008128 | 1 | 1634169 | 1655766 | 102 | 10 | 0.44687 | 0.32748 | CDK11A | 0.32748 | 0.1236 | 0.052698 |
| ENSG00000178171 | 2 | 131513008 | 131525707 | 18 | 7 | 0.44619 | 0.32773 | AMER3 | 0.32773 | 0.1595 | 0.36736 |
| ENSG00000164816 | 8 | 6912831 | 6914256 | 8 | 4 | 0.4461 | 0.32776 | DEFA5 | 0.32776 | 0.85711 | 0.8837 |
| ENSG00000180438 | 3 | 13978756 | 14124311 | 414 | 19 | 0.44604 | 0.32778 | TPRXL | 0.32778 | 0.33131 | 0.34479 |
| ENSG00000151388 | 5 | 33523640 | 33892297 | 1431 | 97 | 0.4458 | 0.32787 | ADAMTS12 | 0.32787 | 0.36308 | 0.4385 |
| ENSG00000243709 | 1 | 226073982 | 226099082 | 104 | 13 | 0.44564 | 0.32793 | LEFTY1 | 0.32793 | 0.2903 | 0.2813 |
| ENSG00000122574 | 7 | 29846102 | 29956682 | 418 | 38 | 0.44541 | 0.32801 | WIPF3 | 0.32801 | 0.25583 | 0.15034 |
| ENSG00000101251 | 20 | 13829893 | 13977089 | 514 | 40 | 0.44539 | 0.32802 | SEL1L2 | 0.32802 | 0.35432 | 0.4107 |
| ENSG00000077684 | 4 | 129730779 | 129796379 | 98 | 15 | 0.44524 | 0.32807 | JADE1 | 0.32807 | 0.39207 | 0.12452 |
| ENSG00000137392 | 6 | 35762759 | 35765088 | 17 | 7 | 0.44518 | 0.32809 | CLPS | 0.32809 | 0.35603 | 0.43119 |
| ENSG00000198570 | 1 | 211649864 | 211666259 | 52 | 13 | 0.44517 | 0.3281 | RD3 | 0.3281 | 0.18197 | 0.33484 |
| ENSG00000197502 | 14 | 50300810 | 50311552 | 21 | 8 | 0.44515 | 0.32811 | AL627171.1 | 0.32811 | 0.36514 | 0.47273 |
| ENSG00000168566 | 6 | 7590432 | 7612200 | 72 | 11 | 0.44512 | 0.32812 | SNRNP48 | 0.32812 | 0.40971 | 0.44131 |
| ENSG00000175711 | 17 | 80900031 | 81009686 | 673 | 46 | 0.44496 | 0.32817 | B3GNTL1 | 0.32817 | 0.34973 | 0.34629 |
| ENSG00000115350 | 2 | 75185619 | 75197255 | 37 | 10 | 0.44493 | 0.32819 | POLE4 | 0.32819 | 0.28668 | 0.26629 |
| ENSG00000133243 | 19 | 1985447 | 2034880 | 281 | 38 | 0.44488 | 0.3282 | BTBD2 | 0.3282 | 0.35564 | 0.38315 |
| ENSG00000077809 | 7 | 74071994 | 74175026 | 155 | 22 | 0.44469 | 0.32827 | GTF2I | 0.32827 | 0.35466 | 0.42291 |
| ENSG00000164329 | 5 | 78907943 | 78982471 | 253 | 9 | 0.44467 | 0.32828 | PAPD4 | 0.32828 | 0.26034 | 0.18922 |
| ENSG00000128709 | 2 | 176987088 | 176989853 | 8 | 3 | 0.44452 | 0.32833 | HOXD9 | 0.32833 | 0.39401 | 0.19889 |
| ENSG00000132744 | 11 | 67410026 | 67418130 | 32 | 6 | 0.44443 | 0.32837 | ACY3 | 0.32837 | 0.21251 | 0.13463 |
| ENSG00000126878 | 9 | 133971863 | 133998539 | 81 | 14 | 0.44435 | 0.3284 | AIF1L | 0.3284 | 0.3665 | 0.36383 |
| ENSG00000066336 | 11 | 47376411 | 47400127 | 102 | 11 | 0.44432 | 0.32841 | SPI1 | 0.32841 | 0.35801 | 0.37458 |
| ENSG00000138696 | 4 | 95679119 | 96079599 | 1635 | 56 | 0.44416 | 0.32846 | BMPR1B | 0.32846 | 0.32969 | 0.34009 |
| ENSG00000188352 | 9 | 20658308 | 20995954 | 978 | 44 | 0.44401 | 0.32852 | FOCAD | 0.32852 | 0.35523 | 0.33577 |
| ENSG00000154269 | 6 | 131949582 | 132068553 | 296 | 23 | 0.44396 | 0.32854 | ENPP3 | 0.32854 | 0.20739 | 0.060324 |
| ENSG00000143867 | 2 | 19551246 | 19558414 | 15 | 6 | 0.44379 | 0.3286 | OSR1 | 0.3286 | 0.4689 | 0.29117 |
| ENSG00000163462 | 1 | 155145873 | 155157447 | 18 | 6 | 0.44337 | 0.32875 | TRIM46 | 0.32875 | 0.10964 | 0.092783 |
| ENSG00000213199 | 7 | 150745379 | 150749843 | 7 | 4 | 0.44328 | 0.32878 | ASIC3 | 0.32878 | 0.67568 | 0.81552 |
| ENSG00000240891 | 3 | 111393523 | 111565294 | 705 | 50 | 0.44312 | 0.32884 | PLCXD2 | 0.32884 | 0.34047 | 0.32245 |
| ENSG00000231274 | 19 | 56052023 | 56056909 | 21 | 9 | 0.44308 | 0.32885 | SBK3 | 0.32885 | 0.59406 | 0.056964 |
| ENSG00000167244 | 11 | 2150342 | 2170833 | 67 | 17 | 0.44267 | 0.329 | IGF2 | 0.329 | 0.066766 | 0.064866 |
| ENSG00000175470 | 10 | 133747955 | 133773331 | 90 | 9 | 0.44266 | 0.32901 | PPP2R2D | 0.32901 | 0.35508 | 0.37064 |
| ENSG00000036448 | 8 | 1993155 | 2113475 | 1196 | 79 | 0.44262 | 0.32902 | MYOM2 | 0.32902 | 0.27471 | 0.15517 |
| ENSG00000074356 | 17 | 3714460 | 3749545 | 89 | 18 | 0.44256 | 0.32904 | C17orf85 | 0.32904 | 0.40284 | 0.30366 |

| | | | | | | | | | | | |
|---|---|---|---|---|---|---|---|---|---|---|---|
| ENSG00000172531 | 11 | 67165654 | 67188654 | 47 | 8 | 0.4424 | 0.3291 | PPP1CA | 0.3291 | 0.43641 | 0.62571 |
| ENSG00000119673 | 14 | 74034324 | 74042357 | 45 | 13 | 0.44205 | 0.32923 | ACOT2 | 0.32923 | 0.45796 | 0.5624 |
| ENSG00000176208 | 17 | 29158988 | 29222887 | 195 | 11 | 0.44191 | 0.32928 | ATAD5 | 0.32928 | 0.33927 | 0.3331 |
| ENSG00000066427 | 14 | 92524896 | 92572965 | 300 | 8 | 0.44182 | 0.32931 | ATXN3 | 0.32931 | 0.31574 | 0.41216 |
| ENSG00000104760 | 8 | 17721889 | 17767874 | 385 | 19 | 0.44159 | 0.32939 | FGL1 | 0.32939 | 0.37508 | 0.37586 |
| ENSG00000123213 | 5 | 65018023 | 65167553 | 555 | 40 | 0.44144 | 0.32945 | NLN | 0.32945 | 0.1803 | 0.12994 |
| ENSG00000101187 | 20 | 61273797 | 61317137 | 157 | 35 | 0.44136 | 0.32947 | SLCO4A1 | 0.32947 | 0.42364 | 0.40391 |
| ENSG00000177565 | 3 | 176737143 | 176915261 | 602 | 40 | 0.44135 | 0.32948 | TBL1XR1 | 0.32948 | 0.32711 | 0.33825 |
| ENSG00000184967 | 12 | 132628993 | 132637013 | 24 | 13 | 0.4411 | 0.32957 | NOC4L | 0.32957 | 0.59791 | 0.58032 |
| ENSG00000100154 | 22 | 28374004 | 29075853 | 1486 | 43 | 0.44105 | 0.32959 | TTC28 | 0.32959 | 0.31062 | 0.30943 |
| ENSG00000053254 | 14 | 89591215 | 90085493 | 1678 | 184 | 0.44092 | 0.32964 | FOXN3 | 0.32964 | 0.3378 | 0.55686 |
| ENSG00000136840 | 9 | 130670165 | 130679317 | 29 | 13 | 0.44073 | 0.3297 | ST6GALNAC4 | 0.3297 | 0.55961 | 0.61096 |
| ENSG00000163820 | 3 | 45959396 | 46037316 | 241 | 20 | 0.4405 | 0.32979 | FYCO1 | 0.32979 | 0.40055 | 0.38741 |
| ENSG00000143196 | 1 | 168664697 | 168698502 | 151 | 21 | 0.4399 | 0.33 | DPT | 0.33 | 0.33173 | 0.39956 |
| ENSG00000188316 | 10 | 118609023 | 118671299 | 95 | 18 | 0.43988 | 0.33001 | ENO4 | 0.33001 | 0.34532 | 0.39221 |
| ENSG00000188984 | 1 | 12776118 | 12788726 | 53 | 8 | 0.43979 | 0.33005 | AADACL3 | 0.33005 | 0.22115 | 0.20457 |
| ENSG00000111640 | 12 | 6643093 | 6647537 | 16 | 3 | 0.43974 | 0.33006 | GAPDH | 0.33006 | 0.28361 | 0.46304 |
| ENSG00000134028 | 8 | 24241798 | 24263526 | 33 | 8 | 0.43963 | 0.3301 | ADAMDEC1 | 0.3301 | 0.40994 | 0.56358 |
| ENSG00000219607 | 6 | 5085720 | 5087455 | 7 | 2 | 0.43934 | 0.33021 | PPP1R3G | 0.33021 | 0.3676 | 0.43331 |
| ENSG00000237441 | 6 | 33259431 | 33267101 | 14 | 4 | 0.43933 | 0.33021 | RGL2 | 0.33021 | 0.046445 | 0.027747 |
| ENSG00000120235 | 9 | 21349834 | 21351377 | 4 | 1 | 0.43889 | 0.33037 | IFNA6 | 0.33037 | 0.22772 | 0.24212 |
| ENSG00000184761 | 2 | 131296668 | 131307456 | 2 | 1 | 0.43843 | 0.33054 | AC013269.5 | 0.33054 | 0.33054 | 0.33054 |
| ENSG00000109436 | 4 | 141541919 | 141677274 | 416 | 38 | 0.43839 | 0.33055 | TBC1D9 | 0.33055 | 0.37753 | 0.48511 |
| ENSG00000196936 | 1 | 248112160 | 248113098 | 10 | 3 | 0.43823 | 0.33061 | OR2L8 | 0.33061 | 0.60225 | 0.48781 |
| ENSG00000213413 | 7 | 99815864 | 99819113 | 15 | 4 | 0.43806 | 0.33067 | PVRIG | 0.33067 | 0.26028 | 0.22114 |
| ENSG00000272897 | 20 | 34220262 | 34262466 | 92 | 10 | 0.43779 | 0.33077 | RP1-309K20.6 | 0.33077 | 0.30211 | 0.25947 |
| ENSG00000224051 | 1 | 1260136 | 1264277 | 11 | 2 | 0.43742 | 0.3309 | GLTPD1 | 0.3309 | 0.40343 | 0.45704 |
| ENSG00000186187 | 16 | 75032928 | 75144892 | 280 | 20 | 0.4374 | 0.33091 | ZNRF1 | 0.33091 | 0.29167 | 0.43844 |
| ENSG00000152822 | 6 | 146348782 | 146758734 | 870 | 43 | 0.4373 | 0.33095 | GRM1 | 0.33095 | 0.33593 | 0.34658 |
| ENSG00000080511 | 19 | 10123925 | 10132955 | 56 | 9 | 0.4373 | 0.33095 | RDH8 | 0.33095 | 0.77878 | 0.86744 |
| ENSG00000261052 | 16 | 30205754 | 30215631 | 2 | 1 | 0.43725 | 0.33096 | SULT1A3 | 0.33096 | 0.26974 | 0.67602 |
| ENSG00000056586 | 9 | 125606835 | 125667620 | 134 | 18 | 0.43675 | 0.33115 | RC3H2 | 0.33115 | 0.24575 | 0.2442 |
| ENSG00000213906 | 14 | 24774940 | 24781259 | 13 | 4 | 0.43674 | 0.33115 | LTB4R2 | 0.33115 | 0.54878 | 0.65034 |
| ENSG00000162337 | 11 | 68080077 | 68216743 | 401 | 33 | 0.43657 | 0.33121 | LRP5 | 0.33121 | 0.30295 | 0.30337 |
| ENSG00000008952 | 3 | 169684423 | 169716161 | 62 | 6 | 0.43649 | 0.33124 | SEC62 | 0.33124 | 0.22962 | 0.060283 |
| ENSG00000159173 | 1 | 201373625 | 201398994 | 66 | 20 | 0.43631 | 0.33131 | TNNI1 | 0.33131 | 0.34866 | 0.35182 |
| ENSG00000146966 | 7 | 140218220 | 140373793 | 399 | 29 | 0.4362 | 0.33135 | DENND2A | 0.33135 | 0.3424 | 0.35734 |
| ENSG00000159140 | 21 | 34914924 | 34949812 | 41 | 9 | 0.43566 | 0.33154 | SON | 0.33154 | 0.40294 | 0.55468 |
| ENSG00000127362 | 7 | 141463897 | 141464997 | 2 | 1 | 0.43533 | 0.33166 | TAS2R3 | 0.33166 | 0.37354 | 0.34476 |
| ENSG00000176946 | 2 | 242523820 | 242576864 | 149 | 12 | 0.43522 | 0.3317 | THAP4 | 0.3317 | 0.3744 | 0.50159 |
| ENSG00000166634 | 18 | 61223393 | 61234244 | 26 | 6 | 0.43477 | 0.33186 | SERPINB12 | 0.33186 | 0.458 | 0.49428 |
| ENSG00000164574 | 5 | 153570290 | 153800544 | 898 | 57 | 0.43467 | 0.3319 | GALNT10 | 0.3319 | 0.32257 | 0.28385 |
| ENSG00000106723 | 9 | 91003334 | 91093609 | 205 | 6 | 0.43461 | 0.33192 | SPIN1 | 0.33192 | 0.35607 | 0.39042 |
| ENSG00000160993 | 7 | 102096685 | 102105323 | 38 | 10 | 0.43454 | 0.33195 | ALKBH4 | 0.33195 | 0.43559 | 0.4737 |
| ENSG00000073146 | 22 | 50528308 | 50600119 | 474 | 18 | 0.43451 | 0.33196 | MOV10L1 | 0.33196 | 0.29209 | 0.17185 |
| ENSG00000132432 | 7 | 54819943 | 54827667 | 35 | 7 | 0.43416 | 0.33209 | SEC61G | 0.33209 | 0.31473 | 0.41215 |
| ENSG00000112936 | 5 | 40909354 | 40983041 | 345 | 28 | 0.43401 | 0.33214 | C7 | 0.33214 | 0.37622 | 0.51349 |
| ENSG00000110080 | 11 | 126225535 | 126310239 | 259 | 38 | 0.43393 | 0.33217 | ST3GAL4 | 0.33217 | 0.41415 | 0.35896 |
| ENSG00000138835 | 9 | 116207011 | 116360018 | 461 | 16 | 0.4336 | 0.33229 | RGS3 | 0.33229 | 0.30826 | 0.29627 |
| ENSG00000106538 | 7 | 150035408 | 150038763 | 14 | 3 | 0.43346 | 0.33234 | RARRES2 | 0.33234 | 0.33588 | 0.42805 |
| ENSG00000204580 | 6 | 30844198 | 30867933 | 91 | 15 | 0.43345 | 0.33235 | DDR1 | 0.33235 | 0.31298 | 0.25927 |
| ENSG00000101337 | 20 | 30697309 | 30755061 | 138 | 9 | 0.43334 | 0.33238 | TM9SF4 | 0.33238 | 0.31545 | 0.29543 |
| ENSG00000172725 | 11 | 67205519 | 67211292 | 4 | 2 | 0.43306 | 0.33248 | CORO1B | 0.33248 | 0.72317 | 0.6582 |
| ENSG00000203740 | 1 | 170115142 | 170136931 | 77 | 13 | 0.43289 | 0.33255 | METTL11B | 0.33255 | 0.35066 | 0.49422 |
| ENSG00000112473 | 6 | 33168222 | 33172216 | 4 | 2 | 0.43286 | 0.33256 | SLC39A7 | 0.33256 | 0.071838 | 0.21471 |
| ENSG00000139116 | 12 | 39687030 | 39837192 | 301 | 19 | 0.43216 | 0.33281 | KIF21A | 0.33281 | 0.32936 | 0.34486 |
| ENSG00000160209 | 21 | 45138975 | 45182188 | 170 | 33 | 0.43185 | 0.33293 | PDXK | 0.33293 | 0.50922 | 0.604 |
| ENSG00000101336 | 20 | 30639991 | 30689659 | 111 | 17 | 0.43146 | 0.33307 | HCK | 0.33307 | 0.33107 | 0.35036 |
| ENSG00000205856 | 22 | 32544993 | 32555309 | 63 | 12 | 0.43122 | 0.33315 | C22orf42 | 0.33315 | 0.36726 | 0.39377 |
| ENSG00000182836 | 5 | 41307056 | 41510730 | 604 | 37 | 0.43063 | 0.33337 | PLCXD3 | 0.33337 | 0.32113 | 0.31153 |
| ENSG00000114982 | 2 | 97258907 | 97308524 | 50 | 13 | 0.43053 | 0.33341 | KANSL3 | 0.33341 | 0.34547 | 0.30004 |
| ENSG00000120129 | 5 | 172195093 | 172198198 | 10 | 3 | 0.43046 | 0.33343 | DUSP1 | 0.33343 | 0.078965 | 0.097863 |
| ENSG00000115041 | 2 | 95963052 | 96051825 | 111 | 10 | 0.43039 | 0.33346 | KCNIP3 | 0.33346 | 0.33574 | 0.40584 |
| ENSG00000148090 | 9 | 93976097 | 94124195 | 408 | 24 | 0.43036 | 0.33347 | AUH | 0.33347 | 0.33232 | 0.34425 |
| ENSG00000205821 | 17 | 2310279 | 2318731 | 40 | 7 | 0.4303 | 0.33349 | AC006435.1 | 0.33349 | 0.32666 | 0.55642 |
| ENSG00000122543 | 7 | 5919458 | 5925993 | 45 | 12 | 0.43017 | 0.33354 | OCM | 0.33354 | 0.29838 | 0.20343 |
| ENSG00000182645 | 10 | 118083940 | 118139541 | 187 | 23 | 0.43005 | 0.33358 | CCDC172 | 0.33358 | 0.35758 | 0.35313 |
| ENSG00000130703 | 20 | 60813580 | 60871268 | 171 | 16 | 0.43001 | 0.3336 | OSBPL2 | 0.3336 | 0.28085 | 0.28621 |
| ENSG00000113312 | 5 | 159436120 | 159492550 | 154 | 17 | 0.4299 | 0.33363 | TTC1 | 0.33363 | 0.3772 | 0.36989 |
| ENSG00000166272 | 10 | 104503727 | 104576021 | 259 | 21 | 0.42991 | 0.33363 | WBP1L | 0.33363 | 0.36393 | 0.37479 |
| ENSG00000187688 | 17 | 16318856 | 16340317 | 74 | 13 | 0.42987 | 0.33364 | TRPV2 | 0.33364 | 0.31116 | 0.20077 |
| ENSG00000056050 | 4 | 170650616 | 170679104 | 75 | 13 | 0.4297 | 0.33371 | C4orf27 | 0.33371 | 0.54058 | 0.69965 |
| ENSG00000157181 | 1 | 186344890 | 186390510 | 88 | 15 | 0.42968 | 0.33372 | C1orf27 | 0.33372 | 0.36041 | 0.36766 |

| | | | | | | | | | | | |
|---|---|---|---|---|---|---|---|---|---|---|---|
| ENSG00000114378 | 3 | 50337320 | 50349812 | 26 | 9 | 0.42967 | 0.33372 | HYAL1 | 0.33372 | 0.3648 | 0.27522 |
| ENSG00000198597 | 19 | 30719197 | 31204445 | 1258 | 136 | 0.42967 | 0.33372 | ZNF536 | 0.33372 | 0.37566 | 0.3888 |
| ENSG00000131019 | 6 | 150384286 | 150390231 | 32 | 9 | 0.42927 | 0.33386 | ULBP3 | 0.33386 | 0.46729 | 0.48533 |
| ENSG00000239887 | 1 | 162336698 | 162356600 | 76 | 23 | 0.42924 | 0.33387 | C1orf226 | 0.33387 | 0.24631 | 0.14153 |
| ENSG00000143486 | 1 | 206744620 | 206785904 | 126 | 22 | 0.4291 | 0.33393 | EIF2D | 0.33393 | 0.4101 | 0.53027 |
| ENSG00000152266 | 11 | 13513602 | 13517728 | 9 | 2 | 0.42901 | 0.33396 | PTH | 0.33396 | 0.33818 | 0.37201 |
| ENSG00000269074 | 2 | 11603927 | 11606274 | 10 | 3 | 0.42874 | 0.33406 | AC099344.1 | 0.33406 | 0.2561 | 0.16087 |
| ENSG00000119321 | 9 | 115923286 | 115983641 | 279 | 19 | 0.42859 | 0.33411 | FKBP15 | 0.33411 | 0.29663 | 0.30134 |
| ENSG00000166734 | 15 | 44580927 | 44707956 | 215 | 10 | 0.4286 | 0.33411 | CASC4 | 0.33411 | 0.34551 | 0.37483 |
| ENSG00000242180 | 11 | 5363744 | 5364809 | 11 | 3 | 0.42847 | 0.33416 | OR51B5 | 0.33416 | 0.20279 | 0.32951 |
| ENSG00000103187 | 16 | 84599200 | 84651683 | 313 | 48 | 0.42845 | 0.33416 | COTL1 | 0.33416 | 0.58769 | 0.89143 |
| ENSG00000164096 | 4 | 120218207 | 120225600 | 21 | 7 | 0.42838 | 0.33419 | C4orf3 | 0.33419 | 0.38648 | 0.53727 |
| ENSG00000164896 | 7 | 150773711 | 150777953 | 5 | 2 | 0.42822 | 0.33424 | FASTK | 0.33424 | 0.86442 | 0.86113 |
| ENSG00000110693 | 11 | 15987995 | 16761138 | 1707 | 64 | 0.42824 | 0.33424 | SOX6 | 0.33424 | 0.34344 | 0.33097 |
| ENSG00000011426 | 7 | 36429415 | 36493400 | 222 | 32 | 0.42804 | 0.33431 | ANLN | 0.33431 | 0.30687 | 0.26114 |
| ENSG00000073614 | 12 | 389295 | 498620 | 372 | 18 | 0.428 | 0.33433 | KDM5A | 0.33433 | 0.30803 | 0.27344 |
| ENSG00000099260 | 1 | 100111499 | 100160097 | 148 | 21 | 0.42793 | 0.33435 | PALMD | 0.33435 | 0.17349 | 0.069134 |
| ENSG00000203711 | 6 | 159290971 | 159331385 | 202 | 35 | 0.42789 | 0.33436 | C6orf99 | 0.33436 | 0.38822 | 0.27925 |
| ENSG00000214511 | 12 | 51347705 | 51364289 | 113 | 12 | 0.42754 | 0.33449 | HIGD1C | 0.33449 | 0.2821 | 0.26384 |
| ENSG00000175309 | 5 | 177635498 | 177659792 | 120 | 21 | 0.42753 | 0.3345 | PHYKPL | 0.3345 | 0.15759 | 0.14267 |
| ENSG00000174276 | 11 | 64883875 | 64885170 | 1 | 1 | 0.42752 | 0.3345 | ZNHIT2 | 0.3345 | 0.70395 | 0.7134 |
| ENSG00000268798 | 19 | 1440838 | 1441937 | 3 | 1 | 0.42747 | 0.33452 | AC027307.3 | 0.33452 | 0.64845 | 0.39807 |
| ENSG00000196262 | 7 | 44836279 | 44864163 | 112 | 16 | 0.4274 | 0.33454 | PPIA | 0.33454 | 0.2901 | 0.25603 |
| ENSG00000142634 | 1 | 15736391 | 15756839 | 76 | 23 | 0.42733 | 0.33457 | EFHD2 | 0.33457 | 0.16169 | 0.086083 |
| ENSG00000187118 | 3 | 28283075 | 28366633 | 196 | 19 | 0.42716 | 0.33463 | CMC1 | 0.33463 | 0.38101 | 0.36473 |
| ENSG00000205571 | 5 | 69345350 | 69374349 | 3 | 2 | 0.42712 | 0.33465 | SMN2 | 0.33465 | 0.62115 | 0.79544 |
| ENSG00000139973 | 14 | 62453803 | 62568431 | 564 | 53 | 0.42709 | 0.33466 | SYT16 | 0.33466 | 0.32727 | 0.21041 |
| ENSG00000176463 | 15 | 92396925 | 92715665 | 1222 | 92 | 0.42704 | 0.33467 | SLCO3A1 | 0.33467 | 0.33475 | 0.32988 |
| ENSG00000198563 | 6 | 31497996 | 31510225 | 92 | 20 | 0.42682 | 0.33475 | DDX39B | 0.33475 | 0.13409 | 0.16442 |
| ENSG00000174125 | 4 | 38792298 | 38858438 | 329 | 18 | 0.42681 | 0.33476 | TLR1 | 0.33476 | 0.30294 | 0.23483 |
| ENSG00000198454 | 9 | 139863695 | 139866442 | 18 | 2 | 0.4268 | 0.33476 | C9orf141 | 0.33476 | 0.084135 | 0.041734 |
| ENSG00000019995 | 10 | 126630692 | 126676758 | 128 | 13 | 0.42681 | 0.33476 | ZRANB1 | 0.33476 | 0.32814 | 0.3475 |
| ENSG00000124237 | 20 | 56725960 | 56736206 | 18 | 5 | 0.42672 | 0.33479 | C20orf85 | 0.33479 | 0.73133 | 0.7143 |
| ENSG00000166682 | 11 | 113558272 | 113577095 | 74 | 14 | 0.42638 | 0.33492 | TMPRSS5 | 0.33492 | 0.50009 | 0.47908 |
| ENSG00000197603 | 5 | 37106330 | 37249530 | 205 | 19 | 0.42628 | 0.33495 | C5orf42 | 0.33495 | 0.35209 | 0.3531 |
| ENSG00000267127 | 18 | 77794346 | 77854968 | 312 | 19 | 0.42558 | 0.33521 | RP11-795F19.5 | 0.33521 | 0.37422 | 0.36538 |
| ENSG00000006042 | 17 | 31254928 | 31272124 | 64 | 10 | 0.42546 | 0.33525 | TMEM98 | 0.33525 | 0.16311 | 0.087418 |
| ENSG00000136026 | 12 | 106631655 | 106698057 | 212 | 49 | 0.42523 | 0.33533 | CKAP4 | 0.33533 | 0.43458 | 0.61613 |
| ENSG00000108984 | 17 | 67410839 | 67539472 | 379 | 51 | 0.42502 | 0.33541 | MAP2K6 | 0.33541 | 0.32637 | 0.36819 |
| ENSG00000204262 | 2 | 189896622 | 190044605 | 477 | 17 | 0.42464 | 0.33555 | COL5A2 | 0.33555 | 0.35005 | 0.25662 |
| ENSG00000272104 | 3 | 50388323 | 50405354 | 14 | 7 | 0.42446 | 0.33562 | XXcos-LUCA11.5 | 0.33562 | 0.43095 | 0.48348 |
| ENSG00000160213 | 21 | 45192393 | 45196326 | 10 | 5 | 0.42431 | 0.33567 | CSTB | 0.33567 | 0.55222 | 0.65411 |
| ENSG00000144671 | 3 | 38323785 | 38360066 | 196 | 9 | 0.42419 | 0.33572 | SLC22A14 | 0.33572 | 0.29634 | 0.38309 |
| ENSG00000148303 | 9 | 136215069 | 136218281 | 9 | 2 | 0.42395 | 0.3358 | RPL7A | 0.3358 | 0.54163 | 0.44593 |
| ENSG00000167549 | 17 | 27941774 | 27949925 | 9 | 4 | 0.42382 | 0.33585 | CORO6 | 0.33585 | 0.34701 | 0.32295 |
| ENSG00000174576 | 11 | 66188475 | 66194178 | 8 | 3 | 0.42368 | 0.3359 | NPAS4 | 0.3359 | 0.67588 | 0.77404 |
| ENSG00000136940 | 9 | 125560668 | 125590910 | 73 | 17 | 0.42354 | 0.33595 | PDCL | 0.33595 | 0.28388 | 0.56645 |
| ENSG00000182768 | 15 | 90808891 | 90816463 | 9 | 4 | 0.42354 | 0.33595 | NGRN | 0.33595 | 0.62099 | 0.49273 |
| ENSG00000010438 | 9 | 33750515 | 33799230 | 167 | 25 | 0.42343 | 0.33599 | PRSS3 | 0.33599 | 0.3575 | 0.31434 |
| ENSG00000184470 | 22 | 19863040 | 19929341 | 334 | 34 | 0.42343 | 0.33599 | TXNRD2 | 0.33599 | 0.17435 | 0.13208 |
| ENSG00000182257 | 22 | 46445358 | 46450024 | 15 | 4 | 0.42337 | 0.33601 | C22orf26 | 0.33601 | 0.4269 | 0.55352 |
| ENSG00000186487 | 2 | 1792885 | 2335032 | 1953 | 97 | 0.42335 | 0.33602 | MYT1L | 0.33602 | 0.38705 | 0.3998 |
| ENSG00000067596 | 17 | 41561233 | 41621831 | 135 | 9 | 0.4233 | 0.33604 | DHX8 | 0.33604 | 0.35147 | 0.39043 |
| ENSG00000099812 | 19 | 751126 | 764319 | 53 | 18 | 0.42316 | 0.33609 | MISP | 0.33609 | 0.47218 | 0.58121 |
| ENSG00000224870 | 1 | 1334902 | 1337426 | 10 | 3 | 0.42301 | 0.33614 | RP4-758J18.2 | 0.33614 | 0.51669 | 0.64039 |
| ENSG00000049618 | 6 | 157099063 | 157531913 | 1184 | 88 | 0.42279 | 0.33622 | ARID1B | 0.33622 | 0.3203 | 0.30354 |
| ENSG00000163466 | 2 | 219081817 | 219119079 | 86 | 10 | 0.42267 | 0.33627 | ARPC2 | 0.33627 | 0.3301 | 0.40522 |
| ENSG00000112308 | 6 | 24705089 | 24721064 | 58 | 16 | 0.42264 | 0.33628 | C6orf62 | 0.33628 | 0.23289 | 0.19823 |
| ENSG00000246922 | 15 | 65385338 | 65407535 | 25 | 7 | 0.42169 | 0.33663 | UBAP1L | 0.33663 | 0.53324 | 0.29964 |
| ENSG00000169682 | 16 | 28985542 | 28995869 | 26 | 8 | 0.42153 | 0.33668 | SPNS1 | 0.33668 | 0.51072 | 0.59146 |
| ENSG00000119782 | 2 | 24272571 | 24286551 | 40 | 8 | 0.42127 | 0.33678 | FKBP1B | 0.33678 | 0.13225 | 0.10838 |
| ENSG00000077150 | 10 | 104153867 | 104162281 | 17 | 8 | 0.42126 | 0.33678 | NFKB2 | 0.33678 | 0.33303 | 0.21532 |
| ENSG00000177971 | 15 | 75931426 | 75941047 | 14 | 6 | 0.42114 | 0.33683 | IMP3 | 0.33683 | 0.44656 | 0.56101 |
| ENSG00000196214 | 19 | 52772824 | 52795977 | 140 | 16 | 0.42112 | 0.33683 | ZNF766 | 0.33683 | 0.31309 | 0.23193 |
| ENSG00000112144 | 6 | 52866077 | 52926600 | 122 | 20 | 0.42097 | 0.33689 | ICK | 0.33689 | 0.33384 | 0.44191 |
| ENSG00000164116 | 4 | 156587863 | 156653501 | 205 | 31 | 0.42093 | 0.3369 | GUCY1A3 | 0.3369 | 0.46573 | 0.57747 |
| ENSG00000015676 | 7 | 44418720 | 44530479 | 246 | 10 | 0.42079 | 0.33696 | NUDCD3 | 0.33696 | 0.32742 | 0.31656 |
| ENSG00000171497 | 4 | 159630286 | 159644548 | 23 | 4 | 0.42052 | 0.33705 | PPID | 0.33705 | 0.43775 | 0.61788 |
| ENSG00000145103 | 3 | 121706170 | 121741051 | 81 | 15 | 0.42041 | 0.33709 | ILDR1 | 0.33709 | 0.26824 | 0.24557 |
| ENSG00000109610 | 4 | 24791534 | 24802464 | 26 | 7 | 0.42042 | 0.33709 | SOD3 | 0.33709 | 0.45716 | 0.61039 |
| ENSG00000108292 | 17 | 36861795 | 36886056 | 22 | 10 | 0.42025 | 0.33715 | MLLT6 | 0.33715 | 0.16729 | 0.20223 |
| ENSG00000174231 | 17 | 1553923 | 1588176 | 132 | 13 | 0.41966 | 0.33737 | PRPF8 | 0.33737 | 0.34265 | 0.41443 |
| ENSG00000085644 | 16 | 3179778 | 3192806 | 25 | 6 | 0.41948 | 0.33743 | ZNF213 | 0.33743 | 0.51613 | 0.40601 |

| | | | | | | | | | | | |
|---|---|---|---|---|---|---|---|---|---|---|---|
| ENSG00000115170 | 2 | 158592958 | 158732374 | 279 | 40 | 0.41942 | 0.33745 | ACVR1 | 0.33745 | 0.39454 | 0.44965 |
| ENSG00000163635 | 3 | 63850233 | 63989138 | 332 | 29 | 0.41929 | 0.3375 | ATXN7 | 0.3375 | 0.28593 | 0.28339 |
| ENSG00000180423 | 11 | 46624411 | 46639459 | 17 | 5 | 0.41927 | 0.33751 | HARBI1 | 0.33751 | 0.50271 | 0.50303 |
| ENSG00000135069 | 9 | 80912059 | 80945009 | 157 | 20 | 0.419 | 0.33761 | PSAT1 | 0.33761 | 0.51937 | 0.59922 |
| ENSG00000135297 | 6 | 74171301 | 74218959 | 169 | 20 | 0.41894 | 0.33763 | MTO1 | 0.33763 | 0.39503 | 0.4163 |
| ENSG00000134755 | 18 | 28645940 | 28682378 | 63 | 15 | 0.41877 | 0.33769 | DSC2 | 0.33769 | 0.37811 | 0.60074 |
| ENSG00000185219 | 3 | 44481262 | 44519162 | 81 | 15 | 0.41786 | 0.33803 | ZNF445 | 0.33803 | 0.35019 | 0.31279 |
| ENSG00000152904 | 1 | 235490665 | 235507847 | 49 | 8 | 0.4177 | 0.33808 | GGPS1 | 0.33808 | 0.32349 | 0.25286 |
| ENSG00000102984 | 16 | 71893583 | 71929239 | 86 | 15 | 0.41763 | 0.33811 | ZNF821 | 0.33811 | 0.39743 | 0.33431 |
| ENSG00000125900 | 20 | 1514897 | 1539489 | 86 | 15 | 0.41756 | 0.33814 | SIRPD | 0.33814 | 0.49017 | 0.55387 |
| ENSG00000203685 | 1 | 226736501 | 226796915 | 184 | 30 | 0.4171 | 0.3383 | C1orf95 | 0.3383 | 0.42541 | 0.52257 |
| ENSG00000165621 | 13 | 97637973 | 97646984 | 21 | 6 | 0.41701 | 0.33833 | OXGR1 | 0.33833 | 0.34385 | 0.20176 |
| ENSG00000005961 | 17 | 42449548 | 42466873 | 25 | 6 | 0.41692 | 0.33837 | ITGA2B | 0.33837 | 0.34995 | 0.31896 |
| ENSG00000175604 | 16 | 12180603 | 12184159 | 21 | 3 | 0.41688 | 0.33838 | RP11-276H1.3 | 0.33838 | 0.23636 | 0.32186 |
| ENSG00000152484 | 13 | 27640293 | 27746033 | 359 | 28 | 0.41678 | 0.33842 | USP12 | 0.33842 | 0.32459 | 0.32853 |
| ENSG00000182054 | 15 | 90626277 | 90645736 | 92 | 17 | 0.41677 | 0.33842 | IDH2 | 0.33842 | 0.56877 | 0.56915 |
| ENSG00000182481 | 17 | 66031635 | 66042958 | 44 | 5 | 0.41672 | 0.33844 | KPNA2 | 0.33844 | 0.39164 | 0.48954 |
| ENSG00000104824 | 19 | 39327028 | 39342987 | 43 | 10 | 0.41624 | 0.33862 | HNRNPL | 0.33862 | 0.46246 | 0.39821 |
| ENSG00000172458 | 13 | 21276266 | 21297237 | 66 | 8 | 0.41612 | 0.33866 | IL17D | 0.33866 | 0.25192 | 0.17713 |
| ENSG00000142227 | 19 | 48824766 | 48833810 | 46 | 11 | 0.41597 | 0.33872 | EMP3 | 0.33872 | 0.35893 | 0.59434 |
| ENSG00000196826 | 19 | 12572373 | 12662288 | 322 | 21 | 0.41594 | 0.33873 | ZNF709 | 0.33873 | 0.30367 | 0.23926 |
| ENSG00000111906 | 6 | 125541108 | 125623282 | 286 | 21 | 0.4158 | 0.33878 | HDDC2 | 0.33878 | 0.36157 | 0.42036 |
| ENSG00000268328 | 2 | 70129482 | 70129841 | 1 | 1 | 0.41574 | 0.3388 | AC019206.1 | 0.3388 | 0.28852 | 0.30584 |
| ENSG00000129810 | 3 | 20202085 | 20227784 | 107 | 14 | 0.4156 | 0.33885 | SGOL1 | 0.33885 | 0.46101 | 0.52015 |
| ENSG00000130529 | 19 | 49660998 | 49715093 | 263 | 19 | 0.41529 | 0.33897 | TRPM4 | 0.33897 | 0.27149 | 0.33745 |
| ENSG00000066422 | 3 | 101367733 | 101396339 | 65 | 11 | 0.41525 | 0.33898 | ZBTB11 | 0.33898 | 0.3526 | 0.36465 |
| ENSG00000166128 | 15 | 63481668 | 63559981 | 202 | 25 | 0.41502 | 0.33906 | RAB8B | 0.33906 | 0.33008 | 0.32325 |
| ENSG00000258691 | 14 | 96671181 | 96730266 | 283 | 36 | 0.4147 | 0.33918 | RP11-404P21.8 | 0.33918 | 0.41172 | 0.58391 |
| ENSG00000085514 | 7 | 99965153 | 99997719 | 62 | 10 | 0.41454 | 0.33924 | PILRA | 0.33924 | 0.44735 | 0.50741 |
| ENSG00000206532 | 3 | 110607231 | 110612323 | 17 | 7 | 0.41435 | 0.33931 | RP11-553A10.1 | 0.33931 | 0.31053 | 0.28304 |
| ENSG00000117505 | 1 | 93811445 | 93835055 | 49 | 6 | 0.41432 | 0.33932 | DR1 | 0.33932 | 0.40557 | 0.39543 |
| ENSG00000128641 | 2 | 192109911 | 192290115 | 415 | 18 | 0.41432 | 0.33932 | MYO1B | 0.33932 | 0.32725 | 0.37647 |
| ENSG00000145545 | 5 | 6633456 | 6669675 | 173 | 20 | 0.41425 | 0.33935 | SRD5A1 | 0.33935 | 0.31088 | 0.36317 |
| ENSG00000019102 | 11 | 124617368 | 124622134 | 17 | 5 | 0.41406 | 0.33942 | VSIG2 | 0.33942 | 0.38542 | 0.44201 |
| ENSG00000110011 | 11 | 63997750 | 64001758 | 12 | 6 | 0.41403 | 0.33943 | DNAJC4 | 0.33943 | 0.47759 | 0.43077 |
| ENSG00000204393 | 20 | 31175281 | 31196695 | 68 | 15 | 0.41401 | 0.33943 | RP11-410N8.4 | 0.33943 | 0.39698 | 0.58025 |
| ENSG00000165929 | 14 | 92246095 | 92333880 | 193 | 19 | 0.41387 | 0.33948 | TC2N | 0.33948 | 0.33754 | 0.3146 |
| ENSG00000160973 | 8 | 145698795 | 145701718 | 3 | 2 | 0.41375 | 0.33953 | FOXH1 | 0.33953 | 0.32477 | 0.43614 |
| ENSG00000113761 | 5 | 176449697 | 176508190 | 137 | 15 | 0.4137 | 0.33955 | ZNF346 | 0.33955 | 0.36157 | 0.3086 |
| ENSG00000221996 | 11 | 4388493 | 4389616 | 4 | 2 | 0.41333 | 0.33968 | OR52B4 | 0.33968 | 0.2946 | 0.44947 |
| ENSG00000074276 | 5 | 175969512 | 176022975 | 220 | 23 | 0.41228 | 0.34007 | CDHR2 | 0.34007 | 0.36068 | 0.40986 |
| ENSG00000128242 | 22 | 30950622 | 30970574 | 74 | 26 | 0.41227 | 0.34007 | GAL3ST1 | 0.34007 | 0.41794 | 0.49589 |
| ENSG00000218357 | 22 | 47857048 | 47882860 | 91 | 11 | 0.4121 | 0.34013 | LL22NC03-75H12.2 | 0.34013 | 0.1869 | 0.16191 |
| ENSG00000186448 | 3 | 44626380 | 44689963 | 109 | 15 | 0.41208 | 0.34014 | ZNF197 | 0.34014 | 0.35847 | 0.43072 |
| ENSG00000183826 | 6 | 38136227 | 38607924 | 1346 | 68 | 0.41194 | 0.34019 | BTBD9 | 0.34019 | 0.35624 | 0.36513 |
| ENSG00000132437 | 7 | 50526134 | 50633154 | 588 | 26 | 0.41184 | 0.34023 | DDC | 0.34023 | 0.32694 | 0.30345 |
| ENSG00000084090 | 2 | 96850597 | 96874563 | 58 | 14 | 0.41135 | 0.34041 | STARD7 | 0.34041 | 0.2318 | 0.12638 |
| ENSG00000186795 | 10 | 118957000 | 118969810 | 75 | 6 | 0.41132 | 0.34042 | KCNK18 | 0.34042 | 0.36082 | 0.30897 |
| ENSG00000152954 | 6 | 24126350 | 24155128 | 84 | 26 | 0.41119 | 0.34047 | NRSN1 | 0.34047 | 0.54099 | 0.42288 |
| ENSG00000239961 | 19 | 54844456 | 54850421 | 34 | 11 | 0.411 | 0.34054 | LILRA4 | 0.34054 | 0.20563 | 0.29049 |
| ENSG00000090372 | 19 | 47222764 | 47250251 | 54 | 11 | 0.41075 | 0.34063 | STRN4 | 0.34063 | 0.26703 | 0.25845 |
| ENSG00000196517 | 1 | 44457172 | 44497139 | 94 | 21 | 0.41011 | 0.34086 | SLC6A9 | 0.34086 | 0.39254 | 0.41304 |
| ENSG00000175899 | 12 | 9220260 | 9268825 | 149 | 10 | 0.40997 | 0.34092 | A2M | 0.34092 | 0.36874 | 0.39339 |
| ENSG00000160326 | 9 | 136336217 | 136344259 | 25 | 9 | 0.40993 | 0.34093 | SLC2A6 | 0.34093 | 0.65611 | 0.59695 |
| ENSG00000198860 | 1 | 184020811 | 184043346 | 36 | 10 | 0.40983 | 0.34096 | TSEN15 | 0.34096 | 0.5061 | 0.64991 |
| ENSG00000111716 | 12 | 21788276 | 21910791 | 220 | 13 | 0.40957 | 0.34106 | LDHB | 0.34106 | 0.34243 | 0.38091 |
| ENSG00000188729 | 3 | 190917030 | 190983404 | 264 | 27 | 0.40943 | 0.34111 | OSTN | 0.34111 | 0.22556 | 0.40865 |
| ENSG00000137672 | 11 | 101322295 | 101743293 | 1652 | 51 | 0.40925 | 0.34118 | TRPC6 | 0.34118 | 0.34258 | 0.35269 |
| ENSG00000162267 | 3 | 52828784 | 52843025 | 32 | 10 | 0.40863 | 0.34141 | ITIH3 | 0.34141 | 0.49682 | 0.55503 |
| ENSG00000123983 | 2 | 223725652 | 223809357 | 266 | 16 | 0.4082 | 0.34156 | ACSL3 | 0.34156 | 0.39917 | 0.47574 |
| ENSG00000143365 | 1 | 151778547 | 151804348 | 67 | 15 | 0.4081 | 0.3416 | RORC | 0.3416 | 0.11717 | 0.13762 |
| ENSG00000183036 | 21 | 41239243 | 41301322 | 332 | 54 | 0.40804 | 0.34162 | PCP4 | 0.34162 | 0.37081 | 0.4096 |
| ENSG00000151611 | 4 | 146539415 | 146581187 | 93 | 17 | 0.40767 | 0.34176 | MMAA | 0.34176 | 0.37415 | 0.39698 |
| ENSG00000255307 | 11 | 6190560 | 6191638 | 2 | 1 | 0.40761 | 0.34178 | OR52B2 | 0.34178 | 0.8873 | 0.97673 |
| ENSG00000117020 | 1 | 243651535 | 244014381 | 635 | 26 | 0.4073 | 0.34189 | AKT3 | 0.34189 | 0.37315 | 0.36219 |
| ENSG00000148219 | 9 | 119187504 | 120177348 | 3500 | 167 | 0.40729 | 0.3419 | ASTN2 | 0.3419 | 0.33257 | 0.34003 |
| ENSG00000164062 | 3 | 49711435 | 49721396 | 9 | 3 | 0.40719 | 0.34193 | APEH | 0.34193 | 0.44696 | 0.19227 |
| ENSG00000145781 | 5 | 115420688 | 115748459 | 1871 | 34 | 0.40704 | 0.34199 | COMMD10 | 0.34199 | 0.34148 | 0.32181 |
| ENSG00000122335 | 6 | 158530536 | 158589312 | 166 | 21 | 0.40703 | 0.34199 | SERAC1 | 0.34199 | 0.39772 | 0.41374 |
| ENSG00000270136 | 1 | 19923473 | 19984549 | 223 | 36 | 0.40701 | 0.342 | MINOS1-NBL1 | 0.342 | 0.46762 | 0.5441 |
| ENSG00000268024 | 17 | 32581900 | 32582306 | 1 | 1 | 0.40701 | 0.342 | AC005549.3 | 0.342 | 0.37444 | 0.43931 |
| ENSG00000173230 | 3 | 121382046 | 121468602 | 168 | 18 | 0.40685 | 0.34206 | GOLGB1 | 0.34206 | 0.2957 | 0.32242 |
| ENSG00000228032 | 14 | 104710541 | 104712473 | 4 | 2 | 0.40673 | 0.3421 | C14orf144 | 0.3421 | 0.14762 | 0.16285 |

| | | | | | | | | | | | |
|---|---|---|---|---|---|---|---|---|---|---|---|
| ENSG00000178922 | 1 | 43916824 | 43919660 | 7 | 4 | 0.40669 | 0.34212 | HYI | 0.34212 | 0.61789 | 0.63117 |
| ENSG00000174038 | 9 | 35041092 | 35045988 | 7 | 3 | 0.40662 | 0.34214 | C9orf131 | 0.34214 | 0.67717 | 0.69835 |
| ENSG00000172667 | 3 | 178735011 | 178790067 | 131 | 14 | 0.40654 | 0.34217 | ZMAT3 | 0.34217 | 0.37686 | 0.34071 |
| ENSG00000108556 | 17 | 4801069 | 4806369 | 27 | 8 | 0.40646 | 0.3422 | CHRNE | 0.3422 | 0.43798 | 0.24522 |
| ENSG00000126106 | 1 | 45100910 | 45140227 | 100 | 8 | 0.40642 | 0.34222 | TMEM53 | 0.34222 | 0.36667 | 0.48357 |
| ENSG00000226976 | 6 | 37012607 | 37013177 | 2 | 1 | 0.40623 | 0.34229 | COX6A1P2 | 0.34229 | 0.53047 | 0.56991 |
| ENSG00000172380 | 1 | 68167149 | 68299150 | 338 | 30 | 0.40619 | 0.3423 | GNG12 | 0.3423 | 0.38927 | 0.40574 |
| ENSG00000151962 | 4 | 155702365 | 155749965 | 116 | 12 | 0.40619 | 0.3423 | RBM46 | 0.3423 | 0.36959 | 0.54231 |
| ENSG00000164107 | 4 | 174446120 | 174451380 | 9 | 4 | 0.40608 | 0.34234 | HAND2 | 0.34234 | 0.80084 | 0.79836 |
| ENSG00000101574 | 18 | 2537524 | 2571508 | 139 | 14 | 0.40558 | 0.34253 | METTL4 | 0.34253 | 0.37724 | 0.4932 |
| ENSG00000004142 | 17 | 26673659 | 26684545 | 15 | 5 | 0.40553 | 0.34255 | POLDIP2 | 0.34255 | 0.55375 | 0.54521 |
| ENSG00000124444 | 19 | 44100544 | 44105309 | 3 | 2 | 0.40547 | 0.34257 | ZNF576 | 0.34257 | 0.55871 | 0.58485 |
| ENSG00000168348 | 14 | 36003248 | 36006260 | 4 | 2 | 0.40534 | 0.34261 | INSM2 | 0.34261 | 0.32694 | 0.16894 |
| ENSG00000126353 | 17 | 38710021 | 38721724 | 26 | 10 | 0.40497 | 0.34275 | CCR7 | 0.34275 | 0.27147 | 0.35007 |
| ENSG00000146386 | 6 | 139349819 | 139364439 | 63 | 13 | 0.40483 | 0.3428 | ABRACL | 0.3428 | 0.074773 | 0.022605 |
| ENSG00000131899 | 17 | 18128901 | 18148189 | 47 | 7 | 0.40463 | 0.34287 | LLGL1 | 0.34287 | 0.31096 | 0.21173 |
| ENSG00000183607 | 2 | 69172364 | 69180102 | 20 | 6 | 0.40458 | 0.34289 | GKN2 | 0.34289 | 0.12436 | 0.34463 |
| ENSG00000122679 | 7 | 45197390 | 45225901 | 225 | 11 | 0.40442 | 0.34295 | RAMP3 | 0.34295 | 0.23031 | 0.18731 |
| ENSG00000164331 | 5 | 72848160 | 72861511 | 46 | 6 | 0.40423 | 0.34302 | ANKRA2 | 0.34302 | 0.32898 | 0.38887 |
| ENSG00000149452 | 11 | 62756626 | 62783311 | 52 | 20 | 0.40391 | 0.34314 | SLC22A8 | 0.34314 | 0.49242 | 0.70381 |
| ENSG00000228672 | 5 | 138727635 | 138730885 | 3 | 1 | 0.40373 | 0.3432 | PROB1 | 0.3432 | 0.57316 | 0.72646 |
| ENSG00000108771 | 17 | 40253422 | 40264751 | 38 | 7 | 0.4037 | 0.34322 | DHX58 | 0.34322 | 0.30183 | 0.43784 |
| ENSG00000137948 | 1 | 92414928 | 92479983 | 242 | 17 | 0.40339 | 0.34333 | BRDT | 0.34333 | 0.30855 | 0.26414 |
| ENSG00000251283 | 4 | 157555646 | 157563606 | 30 | 8 | 0.40332 | 0.34335 | RP11-171N4.2 | 0.34335 | 0.35441 | 0.32511 |
| ENSG00000204442 | 13 | 107820883 | 108519083 | 2872 | 246 | 0.40299 | 0.34348 | FAM155A | 0.34348 | 0.35222 | 0.29386 |
| ENSG00000129988 | 20 | 36974759 | 37005665 | 165 | 24 | 0.40296 | 0.34349 | LBP | 0.34349 | 0.48499 | 0.53565 |
| ENSG00000167508 | 16 | 88718343 | 88729569 | 47 | 13 | 0.40293 | 0.3435 | MVD | 0.3435 | 0.12676 | 0.27689 |
| ENSG00000123144 | 19 | 12841454 | 12845589 | 1 | 1 | 0.40293 | 0.3435 | C19orf43 | 0.3435 | 0.032116 | 0.001243 |
| ENSG00000240053 | 6 | 31637944 | 31641553 | 15 | 7 | 0.40288 | 0.34352 | LY6G5B | 0.34352 | 0.43485 | 0.47798 |
| ENSG00000255835 | 1 | 226074396 | 226111978 | 131 | 19 | 0.40266 | 0.3436 | RP4-559A3.7 | 0.3436 | 0.28241 | 0.32277 |
| ENSG00000169255 | 3 | 160801671 | 160823172 | 45 | 7 | 0.4023 | 0.34373 | B3GALNT1 | 0.34373 | 0.43979 | 0.4704 |
| ENSG00000223547 | 19 | 12175514 | 12192380 | 48 | 5 | 0.40229 | 0.34374 | ZNF844 | 0.34374 | 0.46321 | 0.35868 |
| ENSG00000107485 | 10 | 8095567 | 8117161 | 94 | 13 | 0.40219 | 0.34377 | GATA3 | 0.34377 | 0.37603 | 0.30903 |
| ENSG00000186480 | 7 | 155089486 | 155101945 | 41 | 10 | 0.4021 | 0.34381 | INSIG1 | 0.34381 | 0.36729 | 0.45107 |
| ENSG00000111110 | 12 | 63037762 | 63328817 | 1096 | 114 | 0.40209 | 0.34381 | PPM1H | 0.34381 | 0.3444 | 0.32804 |
| ENSG00000204498 | 6 | 31514647 | 31526606 | 84 | 21 | 0.40193 | 0.34387 | NFKBIL1 | 0.34387 | 0.1135 | 0.1003 |
| ENSG00000186235 | 2 | 239133193 | 239140318 | 39 | 6 | 0.4016 | 0.34399 | AC016757.3 | 0.34399 | 0.3257 | 0.34842 |
| ENSG00000188976 | 1 | 879584 | 894689 | 50 | 9 | 0.40138 | 0.34407 | NOC2L | 0.34407 | 0.46036 | 0.49562 |
| ENSG00000138439 | 2 | 203499901 | 203634480 | 166 | 29 | 0.40137 | 0.34407 | FAM117B | 0.34407 | 0.38223 | 0.45912 |
| ENSG00000145425 | 4 | 152020725 | 152025804 | 7 | 2 | 0.40133 | 0.34409 | RPS3A | 0.34409 | 0.41708 | 0.33841 |
| ENSG00000067704 | 1 | 220267444 | 220321380 | 134 | 19 | 0.40124 | 0.34412 | IARS2 | 0.34412 | 0.36005 | 0.37185 |
| ENSG00000104419 | 8 | 134249414 | 134314265 | 261 | 34 | 0.40125 | 0.34412 | NDRG1 | 0.34412 | 0.42382 | 0.68688 |
| ENSG00000188223 | 19 | 36236579 | 36245420 | 21 | 5 | 0.40111 | 0.34417 | AC002398.9 | 0.34417 | 0.50038 | 0.54166 |
| ENSG00000263417 | 18 | 68297755 | 68317980 | 78 | 14 | 0.40072 | 0.34431 | GTSCR1 | 0.34431 | 0.25047 | 0.39568 |
| ENSG00000181019 | 16 | 69740899 | 69760854 | 62 | 15 | 0.40065 | 0.34434 | NQO1 | 0.34434 | 0.5116 | 0.57824 |
| ENSG00000131242 | 17 | 29718642 | 29865236 | 492 | 58 | 0.40065 | 0.34434 | RAB11FIP4 | 0.34434 | 0.47245 | 0.56069 |
| ENSG00000182149 | 16 | 71879899 | 71962913 | 232 | 25 | 0.40063 | 0.34435 | IST1 | 0.34435 | 0.29377 | 0.33178 |
| ENSG00000169429 | 4 | 74606223 | 74609433 | 7 | 2 | 0.4006 | 0.34436 | IL8 | 0.34436 | 0.33791 | 0.48303 |
| ENSG00000141086 | 16 | 67961543 | 67966317 | 4 | 2 | 0.40029 | 0.34447 | CTRL | 0.34447 | 0.56414 | 0.6729 |
| ENSG00000106927 | 9 | 116822407 | 116840752 | 68 | 9 | 0.4 | 0.34458 | AMBP | 0.34458 | 0.29574 | 0.2578 |
| ENSG00000134775 | 18 | 33877677 | 34360018 | 1393 | 94 | 0.39985 | 0.34463 | FHOD3 | 0.34463 | 0.35126 | 0.39801 |
| ENSG00000095981 | 6 | 39282474 | 39290744 | 30 | 4 | 0.39981 | 0.34465 | KCNK16 | 0.34465 | 0.17189 | 0.20295 |
| ENSG00000173889 | 3 | 169804520 | 169899537 | 163 | 17 | 0.39967 | 0.3447 | PHC3 | 0.3447 | 0.38987 | 0.25826 |
| ENSG00000086289 | 7 | 37723446 | 37991543 | 1174 | 68 | 0.39965 | 0.34471 | EPDR1 | 0.34471 | 0.37119 | 0.35607 |
| ENSG00000105287 | 19 | 47177532 | 47220384 | 112 | 18 | 0.39964 | 0.34471 | PRKD2 | 0.34471 | 0.37921 | 0.3635 |
| ENSG00000105357 | 19 | 50691443 | 50813802 | 604 | 60 | 0.39939 | 0.3448 | MYH14 | 0.3448 | 0.3849 | 0.38974 |
| ENSG00000239388 | 3 | 57302375 | 57326710 | 52 | 10 | 0.39921 | 0.34487 | ASB14 | 0.34487 | 0.40204 | 0.21219 |
| ENSG00000150753 | 5 | 10250033 | 10266524 | 95 | 5 | 0.39918 | 0.34488 | CCT5 | 0.34488 | 0.41325 | 0.47647 |
| ENSG00000125898 | 20 | 814358 | 838106 | 120 | 26 | 0.39912 | 0.3449 | FAM110A | 0.3449 | 0.3772 | 0.23604 |
| ENSG00000122194 | 6 | 161123270 | 161174347 | 153 | 23 | 0.39859 | 0.3451 | PLG | 0.3451 | 0.37648 | 0.39193 |
| ENSG00000125903 | 20 | 207899 | 210527 | 16 | 3 | 0.39859 | 0.3451 | DEFB129 | 0.3451 | 0.17217 | 0.32896 |
| ENSG00000105290 | 19 | 36358801 | 36370693 | 20 | 2 | 0.39841 | 0.34516 | APLP1 | 0.34516 | 0.37459 | 0.47269 |
| ENSG00000197894 | 4 | 99992132 | 100009952 | 100 | 12 | 0.39746 | 0.34551 | ADH5 | 0.34551 | 0.40819 | 0.43402 |
| ENSG00000083896 | 4 | 69176105 | 69215807 | 80 | 15 | 0.39727 | 0.34559 | YTHDC1 | 0.34559 | 0.36298 | 0.31854 |
| ENSG00000170948 | 19 | 8943074 | 8954016 | 27 | 7 | 0.39704 | 0.34567 | MBD3L1 | 0.34567 | 0.37608 | 0.21187 |
| ENSG00000206418 | 18 | 8609443 | 8639379 | 98 | 10 | 0.39618 | 0.34599 | RAB12 | 0.34599 | 0.57801 | 0.54083 |
| ENSG00000091483 | 1 | 241660903 | 241683061 | 39 | 10 | 0.39597 | 0.34606 | FH | 0.34606 | 0.53566 | 0.75718 |
| ENSG00000143107 | 1 | 109255279 | 109285365 | 81 | 19 | 0.39585 | 0.34611 | FNDC7 | 0.34611 | 0.27655 | 0.16407 |
| ENSG00000121848 | 1 | 145611036 | 145696218 | 156 | 10 | 0.39579 | 0.34613 | RNF115 | 0.34613 | 0.36385 | 0.37827 |
| ENSG00000203970 | 6 | 49976851 | 49989694 | 23 | 6 | 0.39573 | 0.34615 | DEFB110 | 0.34615 | 0.47381 | 0.6155 |
| ENSG00000171448 | 9 | 125677845 | 125693779 | 36 | 10 | 0.39564 | 0.34619 | ZBTB26 | 0.34619 | 0.26981 | 0.28343 |
| ENSG00000182896 | 17 | 7258442 | 7260538 | 6 | 3 | 0.39549 | 0.34624 | TMEM95 | 0.34624 | 0.75465 | 0.91494 |
| ENSG00000169499 | 8 | 38758753 | 38831428 | 222 | 48 | 0.39546 | 0.34625 | PLEKHA2 | 0.34625 | 0.34491 | 0.12534 |

| | | | | | | | | | | | |
|---|---|---|---|---|---|---|---|---|---|---|---|
| ENSG00000131795 | 1 | 145507598 | 145513536 | 6 | 3 | 0.39529 | 0.34632 | RBM8A | 0.34632 | 0.52164 | 0.65849 |
| ENSG00000140854 | 16 | 57769642 | 57791162 | 77 | 16 | 0.39507 | 0.3464 | KATNB1 | 0.3464 | 0.26697 | 0.39256 |
| ENSG00000136682 | 2 | 114195268 | 114253766 | 125 | 8 | 0.39488 | 0.34646 | CBWD2 | 0.34646 | 0.41817 | 0.44428 |
| ENSG00000178935 | 19 | 58315209 | 58326281 | 31 | 6 | 0.39491 | 0.34646 | ZNF552 | 0.34646 | 0.40631 | 0.57462 |
| ENSG00000197111 | 12 | 53835525 | 53874946 | 55 | 12 | 0.39485 | 0.34648 | PCBP2 | 0.34648 | 0.4257 | 0.48447 |
| ENSG00000134440 | 18 | 55267888 | 55289445 | 83 | 13 | 0.39434 | 0.34666 | NARS | 0.34666 | 0.39501 | 0.48546 |
| ENSG00000183605 | 10 | 120900279 | 120925179 | 80 | 10 | 0.39405 | 0.34677 | SFXN4 | 0.34677 | 0.49926 | 0.58283 |
| ENSG00000132671 | 20 | 23016057 | 23017314 | 4 | 1 | 0.39406 | 0.34677 | SSTR4 | 0.34677 | 0.4004 | 0.4535 |
| ENSG00000103811 | 15 | 79213400 | 79241916 | 138 | 17 | 0.39401 | 0.34679 | CTSH | 0.34679 | 0.2717 | 0.26004 |
| ENSG00000140093 | 14 | 94749650 | 94759608 | 25 | 10 | 0.39398 | 0.3468 | SERPINA10 | 0.3468 | 0.58482 | 0.66409 |
| ENSG00000167232 | 19 | 23487793 | 23578362 | 180 | 11 | 0.39386 | 0.34684 | ZNF91 | 0.34684 | 0.28108 | 0.10233 |
| ENSG00000204387 | 6 | 31802385 | 31807541 | 22 | 11 | 0.39367 | 0.34691 | C6orf48 | 0.34691 | 0.32258 | 0.25256 |
| ENSG00000080824 | 14 | 102547075 | 102606036 | 112 | 19 | 0.39362 | 0.34693 | HSP90AA1 | 0.34693 | 0.47213 | 0.51754 |
| ENSG00000197757 | 12 | 54384408 | 54424607 | 90 | 18 | 0.39358 | 0.34694 | HOXC6 | 0.34694 | 0.43275 | 0.45243 |
| ENSG00000163354 | 1 | 154990996 | 155006257 | 27 | 7 | 0.39325 | 0.34707 | DCST2 | 0.34707 | 0.32206 | 0.28577 |
| ENSG00000255374 | 12 | 11243886 | 11244912 | 11 | 2 | 0.39323 | 0.34707 | TAS2R43 | 0.34707 | 0.72601 | 0.55722 |
| ENSG00000116885 | 1 | 36881428 | 36916086 | 172 | 9 | 0.39309 | 0.34713 | OSCP1 | 0.34713 | 0.21813 | 0.25051 |
| ENSG00000121207 | 4 | 155548097 | 155674270 | 256 | 13 | 0.39305 | 0.34714 | LRAT | 0.34714 | 0.4067 | 0.4229 |
| ENSG00000047188 | 5 | 112849380 | 112930982 | 416 | 21 | 0.39302 | 0.34715 | YTHDC2 | 0.34715 | 0.38183 | 0.41163 |
| ENSG00000198677 | 5 | 94799599 | 94890711 | 98 | 16 | 0.393 | 0.34716 | TTC37 | 0.34716 | 0.2052 | 0.1301 |
| ENSG00000178761 | 15 | 75192328 | 75199462 | 10 | 4 | 0.39292 | 0.34719 | FAM219B | 0.34719 | 0.36189 | 0.36327 |
| ENSG00000174871 | 11 | 66045661 | 66052772 | 7 | 3 | 0.39285 | 0.34722 | CNIH2 | 0.34722 | 0.56931 | 0.76935 |
| ENSG00000120159 | 9 | 26840683 | 26892802 | 154 | 23 | 0.39261 | 0.3473 | CAAP1 | 0.3473 | 0.39227 | 0.40168 |
| ENSG00000223417 | 11 | 89644279 | 89654935 | 10 | 4 | 0.3924 | 0.34738 | TRIM49D1 | 0.34738 | 0.11044 | 0.18689 |
| ENSG00000149428 | 11 | 118914900 | 118927913 | 23 | 7 | 0.39212 | 0.34748 | HYOU1 | 0.34748 | 0.44318 | 0.53417 |
| ENSG00000167535 | 12 | 49207577 | 49222724 | 26 | 10 | 0.39207 | 0.3475 | CACNB3 | 0.3475 | 0.23743 | 0.1799 |
| ENSG00000117528 | 1 | 94883933 | 94984222 | 148 | 16 | 0.39192 | 0.34756 | ABCD3 | 0.34756 | 0.36061 | 0.35702 |
| ENSG00000130939 | 1 | 10092890 | 10241297 | 386 | 21 | 0.39169 | 0.34764 | UBE4B | 0.34764 | 0.39604 | 0.44612 |
| ENSG00000106028 | 7 | 141438121 | 141487722 | 153 | 5 | 0.39168 | 0.34765 | SSBP1 | 0.34765 | 0.35834 | 0.38145 |
| ENSG00000061656 | 20 | 34203814 | 34208971 | 8 | 3 | 0.39154 | 0.3477 | SPAG4 | 0.3477 | 0.22695 | 0.25104 |
| ENSG00000103275 | 16 | 1355548 | 1377019 | 159 | 15 | 0.39144 | 0.34774 | UBE2I | 0.34774 | 0.4092 | 0.35758 |
| ENSG00000100068 | 22 | 25747385 | 25801344 | 224 | 33 | 0.39129 | 0.34779 | LRP5L | 0.34779 | 0.43104 | 0.34179 |
| ENSG00000153790 | 7 | 25174316 | 25219975 | 216 | 16 | 0.39107 | 0.34787 | C7orf31 | 0.34787 | 0.38336 | 0.44271 |
| ENSG00000141293 | 17 | 46210802 | 46507637 | 630 | 31 | 0.39078 | 0.34798 | SKAP1 | 0.34798 | 0.33532 | 0.3134 |
| ENSG00000109475 | 4 | 109541722 | 109551568 | 23 | 5 | 0.39051 | 0.34808 | RPL34 | 0.34808 | 0.1726 | 0.053786 |
| ENSG00000126746 | 12 | 6775643 | 6798738 | 26 | 9 | 0.3905 | 0.34808 | ZNF384 | 0.34808 | 0.48552 | 0.52705 |
| ENSG00000125735 | 19 | 6663148 | 6670599 | 31 | 11 | 0.39049 | 0.34809 | TNFSF14 | 0.34809 | 0.15568 | 0.009027 |
| ENSG00000268206 | 17 | 26638667 | 26642323 | 2 | 1 | 0.39033 | 0.34815 | AC061975.10 | 0.34815 | 0.34514 | 0.34992 |
| ENSG00000203832 | 1 | 148250249 | 148347506 | 1 | 1 | 0.39018 | 0.3482 | NBPF20 | 0.3482 | 0.3482 | 0.3482 |
| ENSG00000154719 | 21 | 26957968 | 26979829 | 62 | 7 | 0.38992 | 0.3483 | MRPL39 | 0.3483 | 0.22916 | 0.24032 |
| ENSG00000154813 | 3 | 16299485 | 16306479 | 31 | 6 | 0.38982 | 0.34833 | DPH3 | 0.34833 | 0.36377 | 0.43626 |
| ENSG00000204909 | 5 | 147700766 | 147719412 | 43 | 3 | 0.38979 | 0.34834 | SPINK9 | 0.34834 | 0.3733 | 0.34531 |
| ENSG00000175087 | 1 | 26437664 | 26452034 | 44 | 4 | 0.38972 | 0.34837 | PDIK1L | 0.34837 | 0.35368 | 0.37448 |
| ENSG00000139746 | 13 | 79885962 | 79980612 | 328 | 10 | 0.38973 | 0.34837 | RBM26 | 0.34837 | 0.33448 | 0.3555 |
| ENSG00000028277 | 19 | 42590263 | 42700737 | 136 | 10 | 0.38944 | 0.34848 | POU2F2 | 0.34848 | 0.3734 | 0.41038 |
| ENSG00000157766 | 15 | 89346674 | 89418585 | 290 | 45 | 0.38938 | 0.3485 | ACAN | 0.3485 | 0.35141 | 0.6044 |
| ENSG00000132581 | 17 | 26975374 | 26989207 | 18 | 9 | 0.38895 | 0.34866 | SDF2 | 0.34866 | 0.26621 | 0.24221 |
| ENSG00000028203 | 12 | 95611522 | 95696566 | 327 | 23 | 0.38881 | 0.34871 | VEZT | 0.34871 | 0.32322 | 0.27116 |
| ENSG00000164920 | 8 | 99956631 | 99964332 | 6 | 3 | 0.38878 | 0.34872 | OSR2 | 0.34872 | 0.39292 | 0.49195 |
| ENSG00000188375 | 12 | 31944123 | 31945175 | 7 | 4 | 0.38869 | 0.34875 | H3F3C | 0.34875 | 0.42737 | 0.41511 |
| ENSG00000182389 | 2 | 152689290 | 152955593 | 647 | 32 | 0.38856 | 0.3488 | CACNB4 | 0.3488 | 0.3683 | 0.38981 |
| ENSG00000092036 | 14 | 23415437 | 23426370 | 27 | 5 | 0.38842 | 0.34885 | HAUS4 | 0.34885 | 0.59188 | 0.68967 |
| ENSG00000126561 | 17 | 40439565 | 40463961 | 69 | 9 | 0.38842 | 0.34885 | STAT5A | 0.34885 | 0.30174 | 0.27749 |
| ENSG00000146242 | 6 | 83072923 | 83080545 | 19 | 5 | 0.38839 | 0.34886 | TPBG | 0.34886 | 0.53466 | 0.64463 |
| ENSG00000188672 | 1 | 25688740 | 25756683 | 110 | 7 | 0.38838 | 0.34887 | RHCE | 0.34887 | 0.40011 | 0.42447 |
| ENSG00000184588 | 1 | 66258197 | 66840259 | 1807 | 89 | 0.38834 | 0.34888 | PDE4B | 0.34888 | 0.3299 | 0.31336 |
| ENSG00000160716 | 1 | 154540257 | 154552502 | 24 | 9 | 0.38819 | 0.34894 | CHRNB2 | 0.34894 | 0.59386 | 0.62091 |
| ENSG00000188784 | 1 | 20246502 | 20250110 | 19 | 6 | 0.388 | 0.34901 | PLA2G2E | 0.34901 | 0.096553 | 0.11514 |
| ENSG00000134686 | 1 | 33789224 | 33896653 | 377 | 23 | 0.388 | 0.34901 | PHC2 | 0.34901 | 0.37075 | 0.31389 |
| ENSG00000059804 | 12 | 8071826 | 8088871 | 34 | 7 | 0.388 | 0.34901 | SLC2A3 | 0.34901 | 0.68152 | 0.74511 |
| ENSG00000182315 | 19 | 7056220 | 7058651 | 2 | 1 | 0.38793 | 0.34904 | MBD3L3 | 0.34904 | 0.17818 | 0.19594 |
| ENSG00000170074 | 5 | 177134982 | 177210399 | 147 | 17 | 0.38773 | 0.34911 | FAM153A | 0.34911 | 0.29912 | 0.30655 |
| ENSG00000115998 | 2 | 70377012 | 70475747 | 249 | 20 | 0.38765 | 0.34914 | C2orf42 | 0.34914 | 0.32447 | 0.31849 |
| ENSG00000108262 | 17 | 27900487 | 27921072 | 35 | 10 | 0.38677 | 0.34946 | GIT1 | 0.34946 | 0.3203 | 0.24311 |
| ENSG00000173862 | 7 | 33765593 | 33768068 | 6 | 3 | 0.38663 | 0.34951 | RP11-89N17.1 | 0.34951 | 0.062443 | 0.050933 |
| ENSG00000198598 | 12 | 132312938 | 132336328 | 136 | 27 | 0.38659 | 0.34953 | MMP17 | 0.34953 | 0.21264 | 0.06765 |
| ENSG00000105364 | 19 | 10362577 | 10370721 | 21 | 7 | 0.38647 | 0.34958 | MRPL4 | 0.34958 | 0.25686 | 0.17743 |
| ENSG00000133740 | 8 | 86089460 | 86129387 | 73 | 8 | 0.3864 | 0.3496 | E2F5 | 0.3496 | 0.26985 | 0.30079 |
| ENSG00000072163 | 2 | 128395956 | 128439360 | 181 | 33 | 0.38632 | 0.34963 | LIMS2 | 0.34963 | 0.14977 | 0.069605 |
| ENSG00000063177 | 19 | 49118585 | 49122793 | 8 | 3 | 0.38631 | 0.34963 | RPL18 | 0.34963 | 0.11775 | 0.030765 |
| ENSG00000159592 | 1 | 46092976 | 46153785 | 123 | 9 | 0.38624 | 0.34966 | GPBP1L1 | 0.34966 | 0.37047 | 0.36412 |
| ENSG00000196620 | 4 | 69512348 | 69536346 | 104 | 23 | 0.38604 | 0.34973 | UGT2B15 | 0.34973 | 0.31961 | 0.19264 |
| ENSG00000135903 | 2 | 223064607 | 223163715 | 378 | 39 | 0.38601 | 0.34974 | PAX3 | 0.34974 | 0.33194 | 0.47928 |

| | | | | | | | | | | | |
|---|---|---|---|---|---|---|---|---|---|---|---|
| ENSG00000204482 | 6 | 31553901 | 31556686 | 22 | 13 | 0.3858 | 0.34982 | LST1 | 0.34982 | 0.37274 | 0.056578 |
| ENSG00000181045 | 17 | 78193498 | 78227299 | 214 | 29 | 0.3857 | 0.34986 | SLC26A11 | 0.34986 | 0.27953 | 0.25027 |
| ENSG00000205147 | 19 | 4041127 | 4043154 | 10 | 4 | 0.38563 | 0.34989 | AC016586.1 | 0.34989 | 0.2996 | 0.21325 |
| ENSG00000141456 | 17 | 4574679 | 4607632 | 96 | 11 | 0.38548 | 0.34994 | PELP1 | 0.34994 | 0.3556 | 0.36102 |
| ENSG00000146378 | 6 | 132938161 | 132945414 | 43 | 11 | 0.3851 | 0.35008 | TAAR2 | 0.35008 | 0.29894 | 0.37099 |
| ENSG00000175063 | 20 | 44441215 | 44445596 | 18 | 7 | 0.3851 | 0.35008 | UBE2C | 0.35008 | 0.74855 | 0.57455 |
| ENSG00000123094 | 12 | 26111962 | 26232825 | 547 | 33 | 0.38499 | 0.35012 | RASSF8 | 0.35012 | 0.33496 | 0.3411 |
| ENSG00000165948 | 14 | 94547628 | 94570192 | 125 | 15 | 0.38496 | 0.35013 | IFI27L1 | 0.35013 | 0.18227 | 0.092318 |
| ENSG00000206559 | 3 | 28390637 | 28579613 | 348 | 20 | 0.38489 | 0.35016 | ZCWPW2 | 0.35016 | 0.33169 | 0.32697 |
| ENSG00000104881 | 19 | 45882892 | 45909607 | 68 | 17 | 0.38471 | 0.35023 | PPP1R13L | 0.35023 | 0.48351 | 0.60584 |
| ENSG00000100335 | 22 | 39895437 | 39914137 | 23 | 8 | 0.38449 | 0.35031 | MIEF1 | 0.35031 | 0.38646 | 0.443 |
| ENSG00000214414 | 11 | 89443467 | 89451040 | 5 | 3 | 0.38443 | 0.35033 | TRIM77 | 0.35033 | 0.35689 | 0.53648 |
| ENSG00000143315 | 1 | 159997462 | 160001783 | 8 | 5 | 0.38436 | 0.35035 | PIGM | 0.35035 | 0.47979 | 0.51004 |
| ENSG00000244462 | 20 | 34236847 | 34252878 | 35 | 6 | 0.38437 | 0.35035 | RBM12 | 0.35035 | 0.36322 | 0.30122 |
| ENSG00000222012 | 7 | 157406715 | 157411457 | 15 | 5 | 0.38435 | 0.35036 | AC005481.5 | 0.35036 | 0.64227 | 0.53129 |
| ENSG00000143847 | 1 | 202995626 | 203047868 | 196 | 28 | 0.38393 | 0.35052 | PPFIA4 | 0.35052 | 0.2895 | 0.23033 |
| ENSG00000138678 | 4 | 84457067 | 84527028 | 228 | 18 | 0.38385 | 0.35054 | AGPAT9 | 0.35054 | 0.44675 | 0.48528 |
| ENSG00000260914 | 16 | 69333585 | 69354614 | 46 | 7 | 0.38382 | 0.35055 | RP11-343C2.11 | 0.35055 | 0.30745 | 0.28954 |
| ENSG00000117620 | 1 | 100435345 | 100492535 | 158 | 12 | 0.38378 | 0.35057 | SLC35A3 | 0.35057 | 0.33867 | 0.37048 |
| ENSG00000166856 | 12 | 57388230 | 57390468 | 2 | 1 | 0.38363 | 0.35063 | GPR182 | 0.35063 | 0.52982 | 0.54253 |
| ENSG00000165416 | 13 | 53226844 | 53275044 | 157 | 7 | 0.38358 | 0.35064 | SUGT1 | 0.35064 | 0.34133 | 0.33353 |
| ENSG00000143341 | 1 | 185703683 | 186160085 | 740 | 39 | 0.38341 | 0.35071 | HMCN1 | 0.35071 | 0.39384 | 0.33514 |
| ENSG00000187166 | 12 | 48722763 | 48724062 | 7 | 3 | 0.38323 | 0.35078 | H1FNT | 0.35078 | 0.21362 | 0.46427 |
| ENSG00000140553 | 15 | 91473410 | 91497323 | 84 | 11 | 0.38308 | 0.35083 | UNC45A | 0.35083 | 0.51655 | 0.58827 |
| ENSG00000063127 | 19 | 49792895 | 49828482 | 59 | 9 | 0.38305 | 0.35084 | SLC6A16 | 0.35084 | 0.4063 | 0.46301 |
| ENSG00000188486 | 11 | 118964564 | 118966177 | 4 | 1 | 0.38303 | 0.35085 | H2AFX | 0.35085 | 0.58608 | 0.4365 |
| ENSG00000143748 | 1 | 224415036 | 224518089 | 256 | 27 | 0.3829 | 0.3509 | NVL | 0.3509 | 0.3514 | 0.41431 |
| ENSG00000223510 | 17 | 14138990 | 14140179 | 2 | 1 | 0.38275 | 0.35095 | CDRT15 | 0.35095 | 0.46937 | 0.53758 |
| ENSG00000105193 | 19 | 39923847 | 39926588 | 12 | 3 | 0.3827 | 0.35097 | RPS16 | 0.35097 | 0.38389 | 0.34875 |
| ENSG00000139767 | 12 | 119419300 | 119600856 | 823 | 54 | 0.38255 | 0.35103 | SRRM4 | 0.35103 | 0.27557 | 0.1907 |
| ENSG00000169418 | 1 | 153651113 | 153666468 | 25 | 6 | 0.38251 | 0.35104 | NPR1 | 0.35104 | 0.38067 | 0.45319 |
| ENSG00000163162 | 2 | 101887681 | 101925163 | 75 | 13 | 0.38245 | 0.35106 | RNF149 | 0.35106 | 0.26272 | 0.25706 |
| ENSG00000187164 | 10 | 118643742 | 118886097 | 371 | 24 | 0.38234 | 0.35111 | KIAA1598 | 0.35111 | 0.35777 | 0.38563 |
| ENSG00000196235 | 19 | 39926796 | 39967310 | 148 | 13 | 0.38215 | 0.35118 | SUPT5H | 0.35118 | 0.34388 | 0.39006 |
| ENSG00000140807 | 16 | 50582241 | 50670647 | 153 | 25 | 0.38199 | 0.35124 | NKD1 | 0.35124 | 0.35699 | 0.40592 |
| ENSG00000008735 | 22 | 51039114 | 51052409 | 59 | 12 | 0.38194 | 0.35125 | MAPK8IP2 | 0.35125 | 0.35489 | 0.42305 |
| ENSG00000165895 | 11 | 100558384 | 100862668 | 1520 | 60 | 0.38192 | 0.35126 | ARHGAP42 | 0.35126 | 0.35761 | 0.41561 |
| ENSG00000203877 | 6 | 84562985 | 84567234 | 11 | 2 | 0.38184 | 0.35129 | RIPPLY2 | 0.35129 | 0.42556 | 0.45797 |
| ENSG00000169327 | 14 | 21623026 | 21624222 | 7 | 3 | 0.38169 | 0.35135 | OR5AU1 | 0.35135 | 0.5911 | 0.35657 |
| ENSG00000105197 | 19 | 39971052 | 39984422 | 38 | 7 | 0.38121 | 0.35152 | TIMM50 | 0.35152 | 0.43921 | 0.45875 |
| ENSG00000214376 | 11 | 93551398 | 93583697 | 114 | 17 | 0.38062 | 0.35174 | VSTM5 | 0.35174 | 0.28299 | 0.22646 |
| ENSG00000178385 | 2 | 208693027 | 208890284 | 402 | 39 | 0.38058 | 0.35176 | PLEKHM3 | 0.35176 | 0.32398 | 0.30151 |
| ENSG00000197385 | 3 | 32023263 | 32033120 | 48 | 7 | 0.38042 | 0.35182 | ZNF860 | 0.35182 | 0.42517 | 0.49333 |
| ENSG00000136819 | 9 | 132589569 | 132598142 | 59 | 4 | 0.38024 | 0.35188 | C9orf78 | 0.35188 | 0.405 | 0.57668 |
| ENSG00000100258 | 22 | 50941376 | 50946135 | 24 | 7 | 0.38011 | 0.35193 | LMF2 | 0.35193 | 0.706 | 0.79097 |
| ENSG00000187391 | 7 | 77646393 | 79082890 | 6395 | 286 | 0.37992 | 0.352 | MAGI2 | 0.352 | 0.34451 | 0.35178 |
| ENSG00000171223 | 19 | 12902310 | 12904124 | 2 | 1 | 0.37971 | 0.35208 | JUNB | 0.35208 | 0.46009 | 0.52659 |
| ENSG00000082482 | 1 | 215179118 | 215410436 | 809 | 34 | 0.37951 | 0.35215 | KCNK2 | 0.35215 | 0.38188 | 0.39504 |
| ENSG00000136928 | 9 | 101050391 | 101471479 | 1798 | 138 | 0.37943 | 0.35218 | GABBR2 | 0.35218 | 0.41088 | 0.44141 |
| ENSG00000164118 | 4 | 175204828 | 175254531 | 127 | 23 | 0.37924 | 0.35226 | CEP44 | 0.35226 | 0.36903 | 0.32914 |
| ENSG00000125388 | 4 | 2965335 | 3042474 | 295 | 20 | 0.37917 | 0.35228 | GRK4 | 0.35228 | 0.39343 | 0.42239 |
| ENSG00000168702 | 2 | 140988992 | 142889270 | 8855 | 267 | 0.37911 | 0.3523 | LRP1B | 0.3523 | 0.32943 | 0.30706 |
| ENSG00000198826 | 15 | 32907345 | 32932150 | 69 | 14 | 0.37909 | 0.35231 | ARHGAP11A | 0.35231 | 0.489 | 0.63789 |
| ENSG00000185633 | 12 | 57628686 | 57634498 | 1 | 1 | 0.37885 | 0.3524 | NDUFA4L2 | 0.3524 | 0.10836 | 0.23074 |
| ENSG00000113108 | 5 | 139937853 | 139973337 | 43 | 12 | 0.37845 | 0.35255 | APBB3 | 0.35255 | 0.40743 | 0.43604 |
| ENSG00000181656 | 1 | 101003693 | 101007574 | 10 | 4 | 0.37836 | 0.35258 | GPR88 | 0.35258 | 0.14619 | 0.08053 |
| ENSG00000167554 | 19 | 52839498 | 52871031 | 129 | 16 | 0.37832 | 0.3526 | ZNF610 | 0.3526 | 0.28429 | 0.25261 |
| ENSG00000145088 | 3 | 121554030 | 121605373 | 168 | 11 | 0.37829 | 0.35261 | EAF2 | 0.35261 | 0.35766 | 0.40965 |
| ENSG00000167656 | 8 | 143866296 | 143868008 | 8 | 3 | 0.37812 | 0.35267 | LY6D | 0.35267 | 0.64703 | 0.67972 |
| ENSG00000269686 | 2 | 133874577 | 133876625 | 9 | 3 | 0.37797 | 0.35273 | AC011755.1 | 0.35273 | 0.69257 | 0.83094 |
| ENSG00000232040 | 6 | 28539407 | 28583989 | 131 | 22 | 0.37795 | 0.35274 | SCAND3 | 0.35274 | 0.27513 | 0.1565 |
| ENSG00000178795 | 11 | 76927603 | 77012732 | 333 | 20 | 0.37781 | 0.35279 | GDPD4 | 0.35279 | 0.3858 | 0.30953 |
| ENSG00000183864 | 22 | 41829496 | 41843027 | 30 | 5 | 0.37777 | 0.3528 | TOB2 | 0.3528 | 0.52853 | 0.62375 |
| ENSG00000171443 | 19 | 56111388 | 56114504 | 11 | 3 | 0.37757 | 0.35287 | ZNF524 | 0.35287 | 0.084007 | 0.16084 |
| ENSG00000185304 | 2 | 88055474 | 88285309 | 121 | 28 | 0.37751 | 0.3529 | RGPD2 | 0.3529 | 0.47393 | 0.52964 |
| ENSG00000172215 | 3 | 45982425 | 45989845 | 15 | 5 | 0.37727 | 0.35299 | CXCR6 | 0.35299 | 0.30995 | 0.27894 |
| ENSG00000153246 | 2 | 160788519 | 160919121 | 410 | 38 | 0.37702 | 0.35308 | PLA2R1 | 0.35308 | 0.40324 | 0.49251 |
| ENSG00000105258 | 19 | 36604612 | 36606248 | 5 | 3 | 0.37698 | 0.35309 | POLR2I | 0.35309 | 0.36143 | 0.24333 |
| ENSG00000166816 | 16 | 75145758 | 75150669 | 8 | 3 | 0.37696 | 0.3531 | LDHD | 0.3531 | 0.36083 | 0.60415 |
| ENSG00000213886 | 6 | 29523292 | 29527702 | 28 | 5 | 0.3768 | 0.35316 | UBD | 0.35316 | 0.65448 | 0.77453 |
| ENSG00000130414 | 2 | 240831867 | 240964819 | 713 | 47 | 0.37633 | 0.35334 | NDUFA10 | 0.35334 | 0.35335 | 0.22096 |
| ENSG00000013523 | 14 | 77253588 | 77292589 | 131 | 23 | 0.37632 | 0.35334 | ANGEL1 | 0.35334 | 0.42914 | 0.63015 |
| ENSG00000179256 | 12 | 14957584 | 14967116 | 26 | 8 | 0.37627 | 0.35336 | SMCO3 | 0.35336 | 0.55937 | 0.33838 |

| | | | | | | | | | | | |
|---|---|---|---|---|---|---|---|---|---|---|---|
| ENSG00000198093 | 19 | 52392477 | 52408293 | 45 | 7 | 0.37627 | 0.35336 | ZNF649 | 0.35336 | 0.14102 | 0.15908 |
| ENSG00000018625 | 1 | 160085549 | 160113381 | 104 | 21 | 0.37609 | 0.35342 | ATP1A2 | 0.35342 | 0.44076 | 0.30154 |
| ENSG00000196455 | 3 | 130397779 | 130465673 | 135 | 5 | 0.37594 | 0.35348 | PIK3R4 | 0.35348 | 0.36905 | 0.39744 |
| ENSG00000188779 | 15 | 68112042 | 68126899 | 27 | 8 | 0.37593 | 0.35349 | SKOR1 | 0.35349 | 0.53757 | 0.5914 |
| ENSG00000159459 | 15 | 43235095 | 43398311 | 193 | 22 | 0.37577 | 0.35355 | UBR1 | 0.35355 | 0.36455 | 0.3436 |
| ENSG00000104671 | 8 | 30013813 | 30041156 | 67 | 8 | 0.37565 | 0.35359 | DCTN6 | 0.35359 | 0.37099 | 0.44259 |
| ENSG00000084110 | 12 | 96366440 | 96390143 | 125 | 18 | 0.37547 | 0.35366 | HAL | 0.35366 | 0.61238 | 0.71122 |
| ENSG00000122707 | 9 | 36036430 | 36124448 | 196 | 24 | 0.37529 | 0.35372 | RECK | 0.35372 | 0.47182 | 0.52849 |
| ENSG00000090512 | 3 | 186353758 | 186370930 | 51 | 15 | 0.37526 | 0.35373 | FETUB | 0.35373 | 0.27803 | 0.40567 |
| ENSG00000075884 | 2 | 143848931 | 144525921 | 1884 | 114 | 0.37522 | 0.35375 | ARHGAP15 | 0.35375 | 0.31256 | 0.30061 |
| ENSG00000132793 | 20 | 39969560 | 39989222 | 56 | 13 | 0.37515 | 0.35377 | LPIN3 | 0.35377 | 0.29838 | 0.44191 |
| ENSG00000154328 | 8 | 11627148 | 11644855 | 112 | 19 | 0.37509 | 0.3538 | NEIL2 | 0.3538 | 0.40045 | 0.58749 |
| ENSG00000100206 | 22 | 38914954 | 38966291 | 95 | 20 | 0.37505 | 0.35381 | DMC1 | 0.35381 | 0.35336 | 0.37293 |
| ENSG00000060971 | 3 | 38144620 | 38178733 | 62 | 16 | 0.37502 | 0.35382 | ACAA1 | 0.35382 | 0.35404 | 0.48074 |
| ENSG00000138600 | 15 | 50999506 | 51058005 | 190 | 23 | 0.375 | 0.35383 | SPPL2A | 0.35383 | 0.33805 | 0.39369 |
| ENSG00000101210 | 20 | 62119366 | 62130505 | 66 | 18 | 0.37495 | 0.35385 | EEF1A2 | 0.35385 | 0.75628 | 0.61928 |
| ENSG00000257315 | 1 | 203765437 | 203769686 | 7 | 2 | 0.37479 | 0.35391 | ZBED6 | 0.35391 | 0.44632 | 0.36341 |
| ENSG00000198650 | 16 | 71599563 | 71611033 | 31 | 11 | 0.37469 | 0.35394 | TAT | 0.35394 | 0.46696 | 0.41699 |
| ENSG00000167972 | 16 | 2325882 | 2390747 | 106 | 26 | 0.37455 | 0.354 | ABCA3 | 0.354 | 0.37433 | 0.31853 |
| ENSG00000157119 | 3 | 42727011 | 42734036 | 10 | 5 | 0.37443 | 0.35404 | KLHL40 | 0.35404 | 0.31285 | 0.40592 |
| ENSG00000173846 | 1 | 45265897 | 45271662 | 10 | 3 | 0.37433 | 0.35408 | PLK3 | 0.35408 | 0.25896 | 0.22642 |
| ENSG00000176659 | 20 | 58630980 | 58648008 | 49 | 9 | 0.37428 | 0.3541 | C20orf197 | 0.3541 | 0.23228 | 0.46112 |
| ENSG00000152763 | 1 | 67278568 | 67390570 | 291 | 31 | 0.37417 | 0.35414 | WDR78 | 0.35414 | 0.42065 | 0.43115 |
| ENSG00000133121 | 13 | 33677272 | 33924767 | 823 | 58 | 0.3741 | 0.35416 | STARD13 | 0.35416 | 0.34114 | 0.32007 |
| ENSG00000205544 | 17 | 7306294 | 7307456 | 5 | 3 | 0.37411 | 0.35416 | TMEM256 | 0.35416 | 0.69799 | 0.77958 |
| ENSG00000183496 | 15 | 82334119 | 82338482 | 3 | 2 | 0.37398 | 0.35421 | MEX3B | 0.35421 | 0.20909 | 0.3778 |
| ENSG00000186844 | 1 | 152799949 | 152800573 | 1 | 1 | 0.37374 | 0.3543 | LCE1A | 0.3543 | 0.0058845 | 0.04212 |
| ENSG00000180353 | 3 | 121350246 | 121379774 | 72 | 15 | 0.37375 | 0.3543 | HCLS1 | 0.3543 | 0.37122 | 0.46766 |
| ENSG00000100504 | 14 | 51324609 | 51411454 | 329 | 30 | 0.37357 | 0.35436 | PYGL | 0.35436 | 0.243 | 0.21651 |
| ENSG00000102962 | 16 | 57392684 | 57400102 | 27 | 9 | 0.37354 | 0.35437 | CCL22 | 0.35437 | 0.18453 | 0.27967 |
| ENSG00000183779 | 8 | 37553269 | 37557537 | 4 | 2 | 0.3735 | 0.35439 | ZNF703 | 0.35439 | 0.41498 | 0.50485 |
| ENSG00000108688 | 17 | 32597240 | 32599261 | 3 | 2 | 0.37349 | 0.35439 | CCL7 | 0.35439 | 0.33629 | 0.23475 |
| ENSG00000155890 | 3 | 140396881 | 140419992 | 69 | 10 | 0.37285 | 0.35463 | TRIM42 | 0.35463 | 0.44559 | 0.39942 |
| ENSG00000138942 | 22 | 31556168 | 31603005 | 161 | 19 | 0.37268 | 0.35469 | RNF185 | 0.35469 | 0.40193 | 0.43018 |
| ENSG00000198873 | 10 | 120967101 | 121215131 | 893 | 67 | 0.37263 | 0.35471 | GRK5 | 0.35471 | 0.34049 | 0.38005 |
| ENSG00000164209 | 5 | 110073837 | 110100857 | 70 | 13 | 0.37221 | 0.35487 | SLC25A46 | 0.35487 | 0.31971 | 0.29791 |
| ENSG00000137975 | 1 | 86889769 | 86922241 | 190 | 22 | 0.37206 | 0.35493 | CLCA2 | 0.35493 | 0.50902 | 0.56066 |
| ENSG00000197818 | 20 | 48429250 | 48508779 | 268 | 22 | 0.37185 | 0.355 | SLC9A8 | 0.355 | 0.46041 | 0.59308 |
| ENSG00000099940 | 22 | 21213271 | 21245506 | 101 | 13 | 0.37157 | 0.35511 | SNAP29 | 0.35511 | 0.34768 | 0.31753 |
| ENSG00000165071 | 8 | 133697253 | 133772958 | 283 | 14 | 0.37152 | 0.35513 | TMEM71 | 0.35513 | 0.44299 | 0.4726 |
| ENSG00000124380 | 2 | 70120692 | 70132707 | 30 | 5 | 0.37146 | 0.35515 | SNRNP27 | 0.35515 | 0.33054 | 0.32349 |
| ENSG00000121797 | 3 | 46448654 | 46454488 | 11 | 2 | 0.37141 | 0.35517 | CCRL2 | 0.35517 | 0.48896 | 0.55569 |
| ENSG00000136868 | 9 | 115983808 | 116028674 | 167 | 23 | 0.3712 | 0.35524 | SLC31A1 | 0.35524 | 0.30407 | 0.42011 |
| ENSG00000242550 | 18 | 61564408 | 61603345 | 193 | 7 | 0.37108 | 0.35529 | SERPINB10 | 0.35529 | 0.34014 | 0.38069 |
| ENSG00000134323 | 2 | 16080686 | 16087129 | 9 | 3 | 0.37105 | 0.3553 | MYCN | 0.3553 | 0.47733 | 0.58099 |
| ENSG00000270024 | 8 | 67579831 | 67772707 | 188 | 33 | 0.37093 | 0.35534 | C8orf44-SGK3 | 0.35534 | 0.40367 | 0.4196 |
| ENSG00000122694 | 9 | 36136732 | 36163910 | 86 | 20 | 0.37088 | 0.35536 | GLIPR2 | 0.35536 | 0.45494 | 0.44119 |
| ENSG00000185963 | 9 | 95473645 | 95527094 | 160 | 14 | 0.37085 | 0.35537 | BICD2 | 0.35537 | 0.32333 | 0.30396 |
| ENSG00000048471 | 16 | 12070594 | 12668146 | 3029 | 129 | 0.37085 | 0.35537 | SNX29 | 0.35537 | 0.3212 | 0.33362 |
| ENSG00000178149 | 3 | 49052921 | 49059726 | 11 | 6 | 0.37065 | 0.35545 | DALRD3 | 0.35545 | 0.74062 | 0.54807 |
| ENSG00000153495 | 13 | 111968531 | 111996596 | 143 | 28 | 0.37057 | 0.35548 | TEX29 | 0.35548 | 0.3905 | 0.36408 |
| ENSG00000147601 | 8 | 73921099 | 73960357 | 105 | 12 | 0.37043 | 0.35553 | TERF1 | 0.35553 | 0.39348 | 0.4984 |
| ENSG00000101457 | 20 | 44420576 | 44440066 | 63 | 12 | 0.37041 | 0.35554 | DNTTIP1 | 0.35554 | 0.34408 | 0.44703 |
| ENSG00000107890 | 10 | 27280843 | 27389421 | 411 | 20 | 0.37037 | 0.35555 | ANKRD26 | 0.35555 | 0.36908 | 0.35908 |
| ENSG00000140285 | 15 | 49715293 | 49780972 | 242 | 21 | 0.37039 | 0.35555 | FGF7 | 0.35555 | 0.31558 | 0.39174 |
| ENSG00000253309 | 13 | 51909909 | 51938871 | 68 | 9 | 0.37026 | 0.35559 | SERPINE3 | 0.35559 | 0.33908 | 0.41223 |
| ENSG00000142233 | 19 | 49164664 | 49176338 | 35 | 9 | 0.37027 | 0.35559 | NTN5 | 0.35559 | 0.15268 | 0.27777 |
| ENSG00000160695 | 11 | 118938403 | 118952688 | 41 | 5 | 0.37019 | 0.35562 | VPS11 | 0.35562 | 0.3674 | 0.39597 |
| ENSG00000203880 | 20 | 62887094 | 62926855 | 106 | 13 | 0.37018 | 0.35562 | PCMTD2 | 0.35562 | 0.31839 | 0.34243 |
| ENSG00000205420 | 12 | 52880958 | 52887041 | 37 | 5 | 0.37014 | 0.35564 | KRT6A | 0.35564 | 0.33457 | 0.42601 |
| ENSG00000090470 | 15 | 65409717 | 65426174 | 16 | 6 | 0.37 | 0.35569 | PDCD7 | 0.35569 | 0.35229 | 0.27551 |
| ENSG00000173933 | 11 | 66384097 | 66434153 | 64 | 16 | 0.36986 | 0.35574 | RBM4 | 0.35574 | 0.36586 | 0.26817 |
| ENSG00000184984 | 15 | 34260921 | 34357291 | 371 | 21 | 0.36982 | 0.35576 | CHRM5 | 0.35576 | 0.364 | 0.42012 |
| ENSG00000104331 | 8 | 57870492 | 57906403 | 82 | 9 | 0.36949 | 0.35588 | IMPAD1 | 0.35588 | 0.24516 | 0.13602 |
| ENSG00000172927 | 11 | 69061605 | 69182494 | 208 | 33 | 0.36937 | 0.35592 | MYEOV | 0.35592 | 0.33329 | 0.1894 |
| ENSG00000185527 | 17 | 79617489 | 79630142 | 88 | 10 | 0.36938 | 0.35592 | PDE6G | 0.35592 | 0.48593 | 0.55528 |
| ENSG00000144355 | 2 | 172949468 | 172954405 | 8 | 5 | 0.36925 | 0.35597 | DLX1 | 0.35597 | 0.077872 | 0.054013 |
| ENSG00000138336 | 10 | 70320413 | 70454239 | 584 | 23 | 0.36904 | 0.35605 | TET1 | 0.35605 | 0.3685 | 0.38615 |
| ENSG00000204764 | 5 | 170288874 | 170727019 | 992 | 20 | 0.36892 | 0.35609 | RANBP17 | 0.35609 | 0.31568 | 0.30059 |
| ENSG00000140995 | 16 | 90014333 | 90034468 | 116 | 19 | 0.36874 | 0.35616 | DEF8 | 0.35616 | 0.34001 | 0.23001 |
| ENSG00000176641 | 18 | 59475296 | 59561480 | 351 | 30 | 0.36853 | 0.35624 | RNF152 | 0.35624 | 0.2741 | 0.3052 |
| ENSG00000066294 | 1 | 160510885 | 160549306 | 123 | 17 | 0.3685 | 0.35625 | CD84 | 0.35625 | 0.34539 | 0.27805 |
| ENSG00000110711 | 11 | 67250512 | 67258574 | 11 | 3 | 0.36842 | 0.35628 | AIP | 0.35628 | 0.38583 | 0.651 |

| | | | | | | | | | | | |
|---|---|---|---|---|---|---|---|---|---|---|---|
| ENSG00000076928 | 19 | 42387228 | 42434302 | 87 | 15 | 0.36819 | 0.35637 | ARHGEF1 | 0.35637 | 0.41704 | 0.55477 |
| ENSG00000123091 | 1 | 51701943 | 51739127 | 52 | 9 | 0.36814 | 0.35639 | RNF11 | 0.35639 | 0.34674 | 0.29672 |
| ENSG00000044459 | 9 | 17134980 | 17503921 | 1590 | 39 | 0.36811 | 0.35639 | CNTLN | 0.35639 | 0.41129 | 0.40517 |
| ENSG00000163935 | 3 | 52937588 | 53080766 | 371 | 18 | 0.36811 | 0.3564 | SFMBT1 | 0.3564 | 0.35174 | 0.35289 |
| ENSG00000122548 | 7 | 26572740 | 26578407 | 24 | 5 | 0.36798 | 0.35644 | KIAA0087 | 0.35644 | 0.6579 | 0.56862 |
| ENSG00000157869 | 4 | 13362978 | 13485989 | 436 | 13 | 0.36786 | 0.35649 | RAB28 | 0.35649 | 0.38657 | 0.4341 |
| ENSG00000111752 | 12 | 9066492 | 9094063 | 86 | 14 | 0.36772 | 0.35654 | PHC1 | 0.35654 | 0.39544 | 0.50671 |
| ENSG00000268960 | 18 | 45778672 | 45779208 | 3 | 2 | 0.36771 | 0.35654 | AC091150.1 | 0.35654 | 0.35177 | 0.48773 |
| ENSG00000269026 | 19 | 58193413 | 58264943 | 205 | 25 | 0.36771 | 0.35654 | AC003006.7 | 0.35654 | 0.3112 | 0.26979 |
| ENSG00000198914 | 2 | 105471969 | 105476929 | 10 | 4 | 0.36732 | 0.35669 | POU3F3 | 0.35669 | 0.80059 | 0.89696 |
| ENSG00000182150 | 9 | 98637983 | 98776842 | 405 | 22 | 0.36729 | 0.3567 | ERCC6L2 | 0.3567 | 0.37183 | 0.39117 |
| ENSG00000256162 | 6 | 131148546 | 131158275 | 43 | 8 | 0.36665 | 0.35694 | SMLR1 | 0.35694 | 0.62696 | 0.70218 |
| ENSG00000162843 | 1 | 241815580 | 241965435 | 672 | 61 | 0.36663 | 0.35695 | WDR64 | 0.35695 | 0.31284 | 0.31694 |
| ENSG00000076356 | 1 | 208195587 | 208417665 | 694 | 59 | 0.36649 | 0.357 | PLXNA2 | 0.357 | 0.22563 | 0.21743 |
| ENSG00000131471 | 17 | 41003201 | 41010147 | 4 | 2 | 0.36643 | 0.35702 | AOC3 | 0.35702 | 0.63856 | 0.32499 |
| ENSG00000130489 | 22 | 50961997 | 50964868 | 17 | 5 | 0.3664 | 0.35703 | SCO2 | 0.35703 | 0.48389 | 0.73872 |
| ENSG00000164932 | 8 | 104383743 | 104395225 | 48 | 9 | 0.36638 | 0.35704 | CTHRC1 | 0.35704 | 0.36182 | 0.43234 |
| ENSG00000161996 | 16 | 699311 | 717833 | 78 | 5 | 0.36614 | 0.35713 | WDR90 | 0.35713 | 0.35183 | 0.3536 |
| ENSG00000035720 | 4 | 68424446 | 68473055 | 213 | 28 | 0.36606 | 0.35716 | STAP1 | 0.35716 | 0.42733 | 0.43646 |
| ENSG00000173465 | 11 | 65337901 | 65341413 | 6 | 2 | 0.36606 | 0.35716 | SSSCA1 | 0.35716 | 0.11525 | 0.074939 |
| ENSG00000110696 | 11 | 16634679 | 16778428 | 206 | 22 | 0.36595 | 0.3572 | C11orf58 | 0.3572 | 0.39052 | 0.30687 |
| ENSG00000188986 | 9 | 140149625 | 140167998 | 85 | 8 | 0.36566 | 0.35731 | NELFB | 0.35731 | 0.30459 | 0.40462 |
| ENSG00000101446 | 20 | 44141101 | 44144264 | 9 | 3 | 0.36553 | 0.35736 | SPINT3 | 0.35736 | 0.36125 | 0.26465 |
| ENSG00000119922 | 10 | 91061712 | 91069033 | 16 | 6 | 0.36536 | 0.35742 | IFIT2 | 0.35742 | 0.29771 | 0.3147 |
| ENSG00000108854 | 17 | 62538413 | 62658186 | 239 | 12 | 0.36514 | 0.3575 | SMURF2 | 0.3575 | 0.39017 | 0.42255 |
| ENSG00000167306 | 18 | 47349183 | 47721463 | 1607 | 63 | 0.365 | 0.35756 | MYO5B | 0.35756 | 0.37765 | 0.34181 |
| ENSG00000147316 | 8 | 6264113 | 6501144 | 1448 | 95 | 0.36469 | 0.35767 | MCPH1 | 0.35767 | 0.29807 | 0.41392 |
| ENSG00000270466 | 16 | 30389755 | 30393863 | 6 | 3 | 0.36448 | 0.35775 | SEPT1 | 0.35775 | 0.54625 | 0.67427 |
| ENSG00000255798 | 11 | 102552041 | 102554644 | 4 | 2 | 0.36441 | 0.35778 | RP11-817J15.3 | 0.35778 | 0.0067806 | 0.0012457 |
| ENSG00000168807 | 16 | 69221032 | 69342955 | 287 | 20 | 0.36426 | 0.35783 | SNTB2 | 0.35783 | 0.35173 | 0.34002 |
| ENSG00000186895 | 11 | 69624992 | 69633792 | 50 | 7 | 0.36421 | 0.35785 | FGF3 | 0.35785 | 0.51874 | 0.68759 |
| ENSG00000253719 | 12 | 74931551 | 74935223 | 15 | 5 | 0.36406 | 0.35791 | ATXN7L3B | 0.35791 | 0.55888 | 0.65346 |
| ENSG00000147862 | 9 | 14081842 | 14398982 | 900 | 160 | 0.36393 | 0.35796 | NFIB | 0.35796 | 0.34356 | 0.37126 |
| ENSG00000143512 | 1 | 222695602 | 222721445 | 117 | 8 | 0.36389 | 0.35797 | HHIPL2 | 0.35797 | 0.29482 | 0.36796 |
| ENSG00000067365 | 16 | 8715540 | 8740081 | 72 | 13 | 0.36361 | 0.35807 | METTL22 | 0.35807 | 0.075389 | 0.19076 |
| ENSG00000072756 | 3 | 3168600 | 3192563 | 121 | 17 | 0.36353 | 0.35811 | TRNT1 | 0.35811 | 0.34199 | 0.45898 |
| ENSG00000163001 | 2 | 55746740 | 55773015 | 97 | 20 | 0.36334 | 0.35818 | CCDC104 | 0.35818 | 0.33179 | 0.39272 |
| ENSG00000176055 | 5 | 89754020 | 89770585 | 27 | 7 | 0.36323 | 0.35822 | MBLAC2 | 0.35822 | 0.42795 | 0.47343 |
| ENSG00000132470 | 17 | 73717408 | 73753899 | 89 | 19 | 0.36305 | 0.35829 | ITGB4 | 0.35829 | 0.48595 | 0.6303 |
| ENSG00000212658 | 17 | 39458078 | 39459103 | 5 | 1 | 0.36292 | 0.35833 | KRTAP29-1 | 0.35833 | 0.23116 | 0.27583 |
| ENSG00000164647 | 7 | 89783689 | 89794143 | 22 | 6 | 0.36279 | 0.35838 | STEAP1 | 0.35838 | 0.44422 | 0.40898 |
| ENSG00000204469 | 6 | 31588497 | 31605548 | 78 | 19 | 0.36265 | 0.35843 | PRRC2A | 0.35843 | 0.2257 | 0.21562 |
| ENSG00000145016 | 3 | 197398264 | 197476598 | 190 | 12 | 0.36262 | 0.35844 | KIAA0226 | 0.35844 | 0.38114 | 0.50956 |
| ENSG00000164187 | 5 | 36098514 | 36152063 | 77 | 15 | 0.36235 | 0.35855 | LMBRD2 | 0.35855 | 0.22759 | 0.24649 |
| ENSG00000204300 | 11 | 123753633 | 123756349 | 10 | 3 | 0.36223 | 0.35859 | TMEM225 | 0.35859 | 0.31838 | 0.3485 |
| ENSG00000087884 | 11 | 77532155 | 77629478 | 161 | 14 | 0.36218 | 0.35861 | AAMDC | 0.35861 | 0.37889 | 0.24911 |
| ENSG00000170889 | 19 | 54704610 | 54752862 | 382 | 35 | 0.36197 | 0.35869 | RPS9 | 0.35869 | 0.37302 | 0.33986 |
| ENSG00000272741 | 3 | 195975101 | 196045093 | 214 | 16 | 0.3618 | 0.35875 | RP11-447L10.1 | 0.35875 | 0.19302 | 0.16504 |
| ENSG00000054523 | 1 | 10270863 | 10441661 | 600 | 18 | 0.36157 | 0.35884 | KIF1B | 0.35884 | 0.371 | 0.40894 |
| ENSG00000125482 | 9 | 135251008 | 135282209 | 86 | 13 | 0.3614 | 0.3589 | TTF1 | 0.3589 | 0.24462 | 0.055161 |
| ENSG00000198542 | 13 | 102104966 | 102375456 | 909 | 65 | 0.36124 | 0.35896 | ITGBL1 | 0.35896 | 0.36063 | 0.33046 |
| ENSG00000103671 | 15 | 64679947 | 64747502 | 146 | 23 | 0.36122 | 0.35897 | TRIP4 | 0.35897 | 0.39189 | 0.49821 |
| ENSG00000109572 | 4 | 170533784 | 170644824 | 278 | 17 | 0.36107 | 0.35902 | CLCN3 | 0.35902 | 0.40267 | 0.40467 |
| ENSG00000137959 | 1 | 79085607 | 79111830 | 63 | 14 | 0.3607 | 0.35916 | IFI44L | 0.35916 | 0.34955 | 0.20902 |
| ENSG00000131914 | 1 | 26737269 | 26756213 | 54 | 16 | 0.36033 | 0.3593 | LIN28A | 0.3593 | 0.44576 | 0.53808 |
| ENSG00000186146 | 4 | 9446260 | 9452240 | 28 | 4 | 0.36015 | 0.35937 | DEFB131 | 0.35937 | 0.40529 | 0.53882 |
| ENSG00000132670 | 20 | 2844830 | 3019722 | 344 | 19 | 0.36015 | 0.35937 | PTPRA | 0.35937 | 0.37252 | 0.28107 |
| ENSG00000198300 | 19 | 57321445 | 57352096 | 45 | 12 | 0.3601 | 0.35939 | PEG3 | 0.35939 | 0.39207 | 0.26445 |
| ENSG00000165966 | 12 | 41582250 | 41968392 | 1580 | 47 | 0.36003 | 0.35941 | PDZRN4 | 0.35941 | 0.34763 | 0.3699 |
| ENSG00000196532 | 6 | 26045639 | 26046097 | 3 | 2 | 0.35989 | 0.35946 | HIST1H3C | 0.35946 | 0.43452 | 0.45754 |
| ENSG00000168283 | 10 | 22610140 | 22620413 | 5 | 3 | 0.35956 | 0.35959 | BMI1 | 0.35959 | 0.63755 | 0.75617 |
| ENSG00000168447 | 16 | 23289552 | 23392620 | 345 | 38 | 0.35954 | 0.35959 | SCNN1B | 0.35959 | 0.43323 | 0.55547 |
| ENSG00000205302 | 5 | 122110691 | 122165803 | 263 | 17 | 0.3593 | 0.35968 | SNX2 | 0.35968 | 0.36871 | 0.36179 |
| ENSG00000198053 | 20 | 1875154 | 1920543 | 206 | 26 | 0.35926 | 0.3597 | SIRPA | 0.3597 | 0.23401 | 0.198 |
| ENSG00000176619 | 19 | 2427636 | 2456994 | 142 | 17 | 0.35903 | 0.35979 | LMNB2 | 0.35979 | 0.29314 | 0.308 |
| ENSG00000145192 | 3 | 186330712 | 186339107 | 21 | 6 | 0.35893 | 0.35982 | AHSG | 0.35982 | 0.53545 | 0.53226 |
| ENSG00000256715 | 21 | 14741931 | 14745386 | 25 | 6 | 0.35876 | 0.35989 | AL050302.1 | 0.35989 | 0.42923 | 0.45387 |
| ENSG00000092096 | 14 | 23815515 | 23822121 | 6 | 2 | 0.35861 | 0.35994 | SLC22A17 | 0.35994 | 0.89506 | 0.94605 |
| ENSG00000121749 | 12 | 72233487 | 72320629 | 236 | 20 | 0.35843 | 0.36001 | TBC1D15 | 0.36001 | 0.32847 | 0.33626 |
| ENSG00000180066 | 10 | 134257831 | 134262912 | 17 | 2 | 0.35818 | 0.36011 | C10orf91 | 0.36011 | 0.237 | 0.17619 |
| ENSG00000162104 | 16 | 4003388 | 4166186 | 733 | 89 | 0.35804 | 0.36016 | ADCY9 | 0.36016 | 0.36655 | 0.48601 |
| ENSG00000126778 | 14 | 61110133 | 61124977 | 18 | 8 | 0.35801 | 0.36017 | SIX1 | 0.36017 | 0.24673 | 0.23196 |
| ENSG00000126883 | 9 | 134000948 | 134110057 | 254 | 13 | 0.3576 | 0.36032 | NUP214 | 0.36032 | 0.40653 | 0.36881 |

| | | | | | | | | | | | |
|---|---|---|---|---|---|---|---|---|---|---|---|
| ENSG00000185974 | 13 | 114321470 | 114440708 | 93 | 13 | 0.35758 | 0.36033 | GRK1 | 0.36033 | 0.30753 | 0.31947 |
| ENSG00000240224 | 2 | 234621638 | 234681945 | 232 | 22 | 0.35742 | 0.36039 | UGT1A5 | 0.36039 | 0.3769 | 0.21163 |
| ENSG00000167323 | 11 | 3875757 | 4114439 | 691 | 30 | 0.35739 | 0.3604 | STIM1 | 0.3604 | 0.36891 | 0.39299 |
| ENSG00000167653 | 8 | 143751726 | 143764142 | 75 | 5 | 0.35735 | 0.36042 | PSCA | 0.36042 | 0.58295 | 0.76677 |
| ENSG00000183808 | 8 | 94741584 | 94753245 | 21 | 5 | 0.35724 | 0.36046 | RBM12B | 0.36046 | 0.42019 | 0.51909 |
| ENSG00000171596 | 2 | 232387871 | 232395206 | 40 | 4 | 0.35712 | 0.3605 | NMUR1 | 0.3605 | 0.42653 | 0.50778 |
| ENSG00000188199 | 10 | 81462983 | 81474437 | 8 | 3 | 0.35712 | 0.3605 | NUTM2B | 0.3605 | 0.81314 | 0.6753 |
| ENSG00000206344 | 6 | 31165537 | 31171745 | 81 | 13 | 0.35699 | 0.36055 | HCG27 | 0.36055 | 0.43086 | 0.26575 |
| ENSG00000178038 | 3 | 46710487 | 46735194 | 77 | 13 | 0.35691 | 0.36058 | ALS2CL | 0.36058 | 0.56946 | 0.76113 |
| ENSG00000151276 | 3 | 65339200 | 66024509 | 2690 | 175 | 0.3568 | 0.36062 | MAGI1 | 0.36062 | 0.36508 | 0.45007 |
| ENSG00000108219 | 10 | 82213922 | 82292879 | 190 | 31 | 0.35675 | 0.36064 | TSPAN14 | 0.36064 | 0.41407 | 0.33531 |
| ENSG00000162923 | 1 | 224572845 | 224624735 | 103 | 9 | 0.35661 | 0.36069 | WDR26 | 0.36069 | 0.45257 | 0.48967 |
| ENSG00000172508 | 11 | 67182439 | 67193078 | 21 | 6 | 0.35641 | 0.36077 | CARNS1 | 0.36077 | 0.46044 | 0.52027 |
| ENSG00000186400 | 1 | 158548709 | 158549689 | 5 | 1 | 0.3563 | 0.36081 | OR10X1 | 0.36081 | 0.37782 | 0.52746 |
| ENSG00000251349 | 9 | 103204553 | 103339918 | 370 | 21 | 0.35622 | 0.36084 | MSANTD3-TMEFF1 | 0.36084 | 0.4051 | 0.43467 |
| ENSG00000241697 | 9 | 103204560 | 103339918 | 370 | 21 | 0.35622 | 0.36084 | TMEFF1 | 0.36084 | 0.4051 | 0.43467 |
| ENSG00000273170 | 9 | 38540566 | 38577204 | 165 | 40 | 0.35616 | 0.36086 | ANKRD18A | 0.36086 | 0.11195 | 0.21149 |
| ENSG00000184220 | 3 | 99536678 | 99897447 | 353 | 38 | 0.35608 | 0.36089 | CMSS1 | 0.36089 | 0.23606 | 0.14846 |
| ENSG00000213593 | 11 | 57480072 | 57508445 | 59 | 8 | 0.35609 | 0.36089 | TMX2 | 0.36089 | 0.3829 | 0.37337 |
| ENSG00000099968 | 22 | 18111621 | 18213388 | 439 | 24 | 0.35587 | 0.36097 | BCL2L13 | 0.36097 | 0.34791 | 0.26529 |
| ENSG00000214694 | 2 | 39117021 | 39202590 | 186 | 30 | 0.35562 | 0.36106 | ARHGEF33 | 0.36106 | 0.33623 | 0.43871 |
| ENSG00000138613 | 15 | 63568217 | 63601325 | 99 | 21 | 0.35557 | 0.36108 | APH1B | 0.36108 | 0.4262 | 0.50789 |
| ENSG00000248643 | 11 | 66384097 | 66413940 | 35 | 10 | 0.35512 | 0.36125 | RBM14-RBM4 | 0.36125 | 0.30531 | 0.29653 |
| ENSG00000183044 | 16 | 8768422 | 8878432 | 680 | 89 | 0.35441 | 0.36152 | ABAT | 0.36152 | 0.55976 | 0.52362 |
| ENSG00000185681 | 9 | 124922190 | 124962367 | 148 | 26 | 0.35432 | 0.36155 | MORN5 | 0.36155 | 0.37152 | 0.37735 |
| ENSG00000235942 | 1 | 152815344 | 152816459 | 1 | 1 | 0.35419 | 0.3616 | LCE6A | 0.3616 | 0.033522 | 0.025916 |
| ENSG00000205795 | 2 | 10196907 | 10221071 | 110 | 21 | 0.35412 | 0.36163 | CYS1 | 0.36163 | 0.4112 | 0.32193 |
| ENSG00000267748 | 19 | 38779791 | 38795550 | 58 | 12 | 0.35409 | 0.36163 | CTB-102L5.4 | 0.36163 | 0.32891 | 0.44661 |
| ENSG00000231389 | 6 | 33032346 | 33048552 | 421 | 12 | 0.35379 | 0.36175 | HLA-DPA1 | 0.36175 | 0.47612 | 0.59148 |
| ENSG00000068615 | 2 | 86441116 | 86565206 | 472 | 20 | 0.35373 | 0.36177 | REEP1 | 0.36177 | 0.37455 | 0.31225 |
| ENSG00000159202 | 17 | 46985731 | 47006418 | 69 | 5 | 0.35373 | 0.36177 | UBE2Z | 0.36177 | 0.32972 | 0.32281 |
| ENSG00000173334 | 8 | 126442563 | 126450647 | 28 | 13 | 0.35363 | 0.36181 | TRIB1 | 0.36181 | 0.38124 | 0.42506 |
| ENSG00000136114 | 13 | 52951305 | 52980629 | 86 | 6 | 0.35335 | 0.36191 | THSD1 | 0.36191 | 0.33441 | 0.29396 |
| ENSG00000095209 | 9 | 108456825 | 108538893 | 311 | 10 | 0.35326 | 0.36195 | TMEM38B | 0.36195 | 0.34695 | 0.34458 |
| ENSG00000124789 | 6 | 17615269 | 17706656 | 299 | 26 | 0.35316 | 0.36199 | NUP153 | 0.36199 | 0.41181 | 0.45124 |
| ENSG00000185122 | 8 | 145515280 | 145538385 | 59 | 5 | 0.35265 | 0.36217 | HSF1 | 0.36217 | 0.49785 | 0.56792 |
| ENSG00000170260 | 7 | 148936742 | 148952700 | 51 | 13 | 0.35243 | 0.36226 | ZNF212 | 0.36226 | 0.31201 | 0.48799 |
| ENSG00000141971 | 19 | 17516531 | 17544533 | 157 | 21 | 0.35241 | 0.36227 | MVB12A | 0.36227 | 0.37918 | 0.35149 |
| ENSG00000109762 | 4 | 186125391 | 186291339 | 614 | 33 | 0.35193 | 0.36244 | SNX25 | 0.36244 | 0.36913 | 0.43971 |
| ENSG00000166896 | 12 | 58335324 | 58351052 | 61 | 7 | 0.35189 | 0.36246 | XRCC6BP1 | 0.36246 | 0.38903 | 0.38419 |
| ENSG00000063438 | 5 | 304291 | 438406 | 564 | 40 | 0.35156 | 0.36258 | AHRR | 0.36258 | 0.42512 | 0.41047 |
| ENSG00000117525 | 1 | 94994781 | 95007356 | 20 | 5 | 0.35098 | 0.3628 | F3 | 0.3628 | 0.28186 | 0.46283 |
| ENSG00000204478 | 1 | 13736907 | 13747803 | 2 | 1 | 0.35065 | 0.36293 | PRAMEF20 | 0.36293 | 0.36769 | 0.69797 |
| ENSG00000114166 | 3 | 20081515 | 20195896 | 599 | 33 | 0.35065 | 0.36293 | KAT2B | 0.36293 | 0.41016 | 0.42217 |
| ENSG00000103569 | 15 | 58430368 | 58478110 | 203 | 30 | 0.35058 | 0.36295 | AQP9 | 0.36295 | 0.50838 | 0.56356 |
| ENSG00000189157 | 4 | 77135193 | 77204933 | 385 | 24 | 0.35047 | 0.36299 | FAM47E | 0.36299 | 0.38349 | 0.35663 |
| ENSG00000196323 | 11 | 130096572 | 130184581 | 293 | 12 | 0.35048 | 0.36299 | ZBTB44 | 0.36299 | 0.38192 | 0.38134 |
| ENSG00000187837 | 6 | 26055968 | 26056699 | 5 | 3 | 0.35033 | 0.36304 | HIST1H1C | 0.36304 | 0.38608 | 0.4317 |
| ENSG00000127152 | 14 | 99635624 | 99737861 | 317 | 36 | 0.35013 | 0.36312 | BCL11B | 0.36312 | 0.35233 | 0.31793 |
| ENSG00000203896 | 20 | 62366815 | 62370456 | 8 | 2 | 0.35011 | 0.36313 | LIME1 | 0.36313 | 0.22876 | 0.29266 |
| ENSG00000139190 | 12 | 6571403 | 6580153 | 36 | 8 | 0.35001 | 0.36316 | VAMP1 | 0.36316 | 0.34557 | 0.26193 |
| ENSG00000175911 | 17 | 78977135 | 78978932 | 8 | 2 | 0.34976 | 0.36326 | AC127496.1 | 0.36326 | 0.58871 | 0.61791 |
| ENSG00000178776 | 5 | 147260289 | 147286101 | 83 | 15 | 0.34967 | 0.36329 | C5orf46 | 0.36329 | 0.39407 | 0.14557 |
| ENSG00000167371 | 16 | 29823177 | 29827201 | 5 | 2 | 0.34969 | 0.36329 | PRRT2 | 0.36329 | 0.1773 | 0.20722 |
| ENSG00000110906 | 12 | 109886461 | 109915349 | 142 | 12 | 0.34953 | 0.36334 | KCTD10 | 0.36334 | 0.29526 | 0.26507 |
| ENSG00000143256 | 1 | 161070346 | 161087901 | 36 | 6 | 0.34942 | 0.36339 | PFDN2 | 0.36339 | 0.58323 | 0.75015 |
| ENSG00000115825 | 2 | 37477645 | 37551951 | 357 | 20 | 0.34934 | 0.36342 | PRKD3 | 0.36342 | 0.38317 | 0.37435 |
| ENSG00000140563 | 15 | 94774767 | 95023632 | 1017 | 109 | 0.34912 | 0.3635 | MCTP2 | 0.3635 | 0.37528 | 0.44817 |
| ENSG00000110484 | 11 | 62037627 | 62040628 | 6 | 3 | 0.34908 | 0.36351 | SCGB2A2 | 0.36351 | 0.57443 | 0.49928 |
| ENSG00000165458 | 11 | 71934745 | 71950149 | 30 | 8 | 0.34908 | 0.36351 | INPPL1 | 0.36351 | 0.32 | 0.37681 |
| ENSG00000155034 | 7 | 5470966 | 5553429 | 432 | 51 | 0.34895 | 0.36356 | FBXL18 | 0.36356 | 0.37952 | 0.45709 |
| ENSG00000214113 | 6 | 5102827 | 5261172 | 660 | 43 | 0.34892 | 0.36357 | LYRM4 | 0.36357 | 0.31495 | 0.33468 |
| ENSG00000176148 | 11 | 33060963 | 33127489 | 334 | 21 | 0.34893 | 0.36357 | TCP11L1 | 0.36357 | 0.42683 | 0.4142 |
| ENSG00000197937 | 19 | 53627325 | 53662328 | 162 | 34 | 0.34892 | 0.36358 | ZNF347 | 0.36358 | 0.53464 | 0.30007 |
| ENSG00000131149 | 16 | 85645015 | 85709810 | 238 | 50 | 0.34834 | 0.36379 | GSE1 | 0.36379 | 0.47984 | 0.51805 |
| ENSG00000110852 | 12 | 10005583 | 10022735 | 71 | 12 | 0.34818 | 0.36385 | CLEC2B | 0.36385 | 0.59549 | 0.63932 |
| ENSG00000184357 | 6 | 27834570 | 27835359 | 4 | 2 | 0.3481 | 0.36388 | HIST1H1B | 0.36388 | 0.71852 | 0.88373 |
| ENSG00000112159 | 6 | 90352218 | 90529442 | 473 | 32 | 0.34805 | 0.3639 | MDN1 | 0.3639 | 0.38352 | 0.36828 |
| ENSG00000004846 | 7 | 20654830 | 20816658 | 670 | 46 | 0.34784 | 0.36398 | ABCB5 | 0.36398 | 0.34094 | 0.31564 |
| ENSG00000205133 | 8 | 93895758 | 94029901 | 287 | 30 | 0.34783 | 0.36398 | TRIQK | 0.36398 | 0.32036 | 0.33979 |
| ENSG00000085840 | 1 | 52838501 | 52870131 | 38 | 11 | 0.3477 | 0.36403 | ORC1 | 0.36403 | 0.38632 | 0.36875 |
| ENSG00000165841 | 10 | 96447911 | 96613017 | 542 | 18 | 0.34771 | 0.36403 | CYP2C19 | 0.36403 | 0.37751 | 0.38884 |
| ENSG00000172375 | 11 | 118972908 | 118989252 | 50 | 9 | 0.3477 | 0.36403 | C2CD2L | 0.36403 | 0.2806 | 0.31978 |

| | | | | | | | | | | | |
|---|---|---|---|---|---|---|---|---|---|---|---|
| ENSG00000184949 | 22 | 38974125 | 39052634 | 208 | 16 | 0.34739 | 0.36415 | FAM227A | 0.36415 | 0.38016 | 0.39759 |
| ENSG00000111652 | 12 | 6832907 | 6841041 | 35 | 7 | 0.34729 | 0.36419 | COPS7A | 0.36419 | 0.46318 | 0.53756 |
| ENSG00000205330 | 12 | 55714346 | 55715408 | 5 | 2 | 0.34692 | 0.36432 | OR6C1 | 0.36432 | 0.86422 | 0.91631 |
| ENSG00000143476 | 1 | 212208919 | 212280742 | 302 | 24 | 0.34691 | 0.36433 | DTL | 0.36433 | 0.40248 | 0.40765 |
| ENSG00000138698 | 4 | 99182535 | 99365012 | 366 | 19 | 0.34664 | 0.36443 | RAP1GDS1 | 0.36443 | 0.36673 | 0.37276 |
| ENSG00000064115 | 12 | 27126128 | 27167367 | 133 | 25 | 0.34663 | 0.36444 | TM7SF3 | 0.36444 | 0.37933 | 0.41468 |
| ENSG00000176879 | 11 | 4944531 | 4945637 | 8 | 3 | 0.34659 | 0.36445 | OR51G1 | 0.36445 | 0.37552 | 0.39786 |
| ENSG00000170927 | 6 | 51480098 | 51952423 | 1845 | 63 | 0.34657 | 0.36446 | PKHD1 | 0.36446 | 0.35758 | 0.33895 |
| ENSG00000148481 | 10 | 15820169 | 15902519 | 124 | 19 | 0.34637 | 0.36453 | FAM188A | 0.36453 | 0.38584 | 0.15019 |
| ENSG00000172381 | 11 | 57798402 | 57799403 | 5 | 2 | 0.3462 | 0.36459 | OR6Q1 | 0.36459 | 0.50272 | 0.52892 |
| ENSG00000140749 | 16 | 21652609 | 21663981 | 17 | 8 | 0.34621 | 0.36459 | IGSF6 | 0.36459 | 0.76538 | 0.73838 |
| ENSG00000182634 | 11 | 123908773 | 123909717 | 11 | 3 | 0.34608 | 0.36464 | OR10G7 | 0.36464 | 0.28412 | 0.45634 |
| ENSG00000161642 | 12 | 54762917 | 54785082 | 45 | 10 | 0.34609 | 0.36464 | ZNF385A | 0.36464 | 0.47535 | 0.20234 |
| ENSG00000172888 | 3 | 40566369 | 40616176 | 132 | 21 | 0.34606 | 0.36465 | ZNF621 | 0.36465 | 0.35106 | 0.3721 |
| ENSG00000151475 | 4 | 128651533 | 128695448 | 77 | 10 | 0.34605 | 0.36465 | SLC25A31 | 0.36465 | 0.38928 | 0.36421 |
| ENSG00000171476 | 4 | 57514155 | 57548065 | 153 | 24 | 0.34596 | 0.36469 | HOPX | 0.36469 | 0.32421 | 0.07605 |
| ENSG00000120149 | 5 | 174151536 | 174157896 | 23 | 6 | 0.34589 | 0.36471 | MSX2 | 0.36471 | 0.19352 | 0.093464 |
| ENSG00000156860 | 16 | 30669752 | 30682135 | 10 | 5 | 0.3459 | 0.36471 | FBRS | 0.36471 | 0.70904 | 0.71256 |
| ENSG00000174132 | 5 | 99871009 | 99922445 | 125 | 14 | 0.34579 | 0.36475 | FAM174A | 0.36475 | 0.33289 | 0.37783 |
| ENSG00000173918 | 17 | 77018896 | 77045870 | 101 | 12 | 0.3457 | 0.36479 | C1QTNF1 | 0.36479 | 0.41341 | 0.56661 |
| ENSG00000137501 | 11 | 85405267 | 85522184 | 313 | 20 | 0.34561 | 0.36482 | SYTL2 | 0.36482 | 0.39508 | 0.40068 |
| ENSG00000154822 | 3 | 16844159 | 17132086 | 925 | 40 | 0.34538 | 0.3649 | PLCL2 | 0.3649 | 0.36324 | 0.38801 |
| ENSG00000178093 | 19 | 19623227 | 19626838 | 5 | 2 | 0.3454 | 0.3649 | TSSK6 | 0.3649 | 0.61404 | 0.54103 |
| ENSG00000135905 | 2 | 225629807 | 225907162 | 769 | 47 | 0.34532 | 0.36493 | DOCK10 | 0.36493 | 0.38196 | 0.32744 |
| ENSG00000175416 | 5 | 175819456 | 175843570 | 67 | 17 | 0.34461 | 0.36519 | CLTB | 0.36519 | 0.2805 | 0.28912 |
| ENSG00000075290 | 10 | 102222798 | 102243501 | 22 | 5 | 0.34463 | 0.36519 | WNT8B | 0.36519 | 0.53363 | 0.56767 |
| ENSG00000139187 | 12 | 9102640 | 9163356 | 204 | 15 | 0.34454 | 0.36522 | KLRG1 | 0.36522 | 0.36089 | 0.39867 |
| ENSG00000267335 | 19 | 49526210 | 49540152 | 98 | 13 | 0.34404 | 0.36541 | CTB-60B18.6 | 0.36541 | 0.30706 | 0.20569 |
| ENSG00000186049 | 12 | 53001354 | 53012343 | 31 | 10 | 0.34393 | 0.36545 | KRT73 | 0.36545 | 0.45304 | 0.59992 |
| ENSG00000233863 | 8 | 35649365 | 35653384 | 1 | 1 | 0.3438 | 0.3655 | AC012215.1 | 0.3655 | 0.11006 | 0.19767 |
| ENSG00000054277 | 1 | 241753404 | 241840678 | 324 | 58 | 0.34359 | 0.36558 | OPN3 | 0.36558 | 0.50612 | 0.40763 |
| ENSG00000137558 | 8 | 75736772 | 75767264 | 92 | 14 | 0.3432 | 0.36572 | PI15 | 0.36572 | 0.356 | 0.38191 |
| ENSG00000085719 | 8 | 87497059 | 87573726 | 284 | 25 | 0.34299 | 0.3658 | CPNE3 | 0.3658 | 0.21818 | 0.1438 |
| ENSG00000153832 | 2 | 230787018 | 230877825 | 167 | 27 | 0.34296 | 0.36581 | FBXO36 | 0.36581 | 0.36485 | 0.36689 |
| ENSG00000035141 | 2 | 70523107 | 70529222 | 14 | 3 | 0.34274 | 0.3659 | FAM136A | 0.3659 | 0.54246 | 0.7436 |
| ENSG00000146399 | 6 | 132966123 | 132967142 | 3 | 2 | 0.34258 | 0.36596 | TAAR1 | 0.36596 | 0.31499 | 0.21833 |
| ENSG00000196917 | 12 | 123104824 | 123215390 | 383 | 32 | 0.3423 | 0.36606 | HCAR1 | 0.36606 | 0.38841 | 0.41441 |
| ENSG00000186529 | 19 | 15751707 | 15773635 | 194 | 15 | 0.34216 | 0.36611 | CYP4F3 | 0.36611 | 0.37012 | 0.33237 |
| ENSG00000123388 | 12 | 54366910 | 54371427 | 6 | 2 | 0.34169 | 0.36629 | HOXC11 | 0.36629 | 0.17582 | 0.089734 |
| ENSG00000237247 | 19 | 7030589 | 7033022 | 5 | 2 | 0.34164 | 0.36631 | MBD3L5 | 0.36631 | 0.07368 | 0.29209 |
| ENSG00000036257 | 2 | 225334867 | 225450110 | 311 | 15 | 0.34146 | 0.36638 | CUL3 | 0.36638 | 0.3619 | 0.43733 |
| ENSG00000165886 | 10 | 99258625 | 99330966 | 201 | 25 | 0.34137 | 0.36641 | UBTD1 | 0.36641 | 0.15098 | 0.15141 |
| ENSG00000064655 | 20 | 45523263 | 45817492 | 1080 | 99 | 0.34114 | 0.3665 | EYA2 | 0.3665 | 0.39424 | 0.38248 |
| ENSG00000006059 | 17 | 39502344 | 39507064 | 8 | 3 | 0.34103 | 0.36654 | KRT33A | 0.36654 | 0.21608 | 0.25164 |
| ENSG00000137142 | 9 | 38408991 | 38424444 | 71 | 15 | 0.34083 | 0.36662 | IGFBPL1 | 0.36662 | 0.43741 | 0.45892 |
| ENSG00000121933 | 1 | 112025970 | 112106584 | 284 | 33 | 0.34077 | 0.36664 | ADORA3 | 0.36664 | 0.32172 | 0.19741 |
| ENSG00000198835 | 1 | 228337553 | 228347527 | 14 | 5 | 0.3407 | 0.36667 | GJC2 | 0.36667 | 0.42094 | 0.3534 |
| ENSG00000133256 | 4 | 619373 | 664571 | 179 | 15 | 0.34066 | 0.36668 | PDE6B | 0.36668 | 0.29468 | 0.25554 |
| ENSG00000174111 | 17 | 36508111 | 36556019 | 116 | 13 | 0.34066 | 0.36668 | SOCS7 | 0.36668 | 0.37865 | 0.37403 |
| ENSG00000197697 | 6 | 26183958 | 26184454 | 2 | 1 | 0.3406 | 0.3667 | HIST1H2BE | 0.3667 | 0.4737 | 0.54178 |
| ENSG00000108963 | 17 | 1933404 | 1946724 | 48 | 15 | 0.34052 | 0.36673 | DPH1 | 0.36673 | 0.71771 | 0.77523 |
| ENSG00000136244 | 7 | 22765503 | 22771621 | 17 | 4 | 0.34047 | 0.36675 | IL6 | 0.36675 | 0.44178 | 0.54432 |
| ENSG00000261603 | 3 | 46761073 | 46777919 | 68 | 5 | 0.34044 | 0.36676 | PRSS46 | 0.36676 | 0.55941 | 0.67776 |
| ENSG00000095485 | 10 | 101992055 | 102027437 | 124 | 12 | 0.3403 | 0.36681 | CWF19L1 | 0.36681 | 0.42267 | 0.4234 |
| ENSG00000080823 | 14 | 102690837 | 102771537 | 140 | 22 | 0.34015 | 0.36687 | MOK | 0.36687 | 0.30793 | 0.17094 |
| ENSG00000234857 | 11 | 62457747 | 62494856 | 87 | 11 | 0.34008 | 0.3669 | HNRNPUL2-BSCL2 | 0.3669 | 0.41987 | 0.51733 |
| ENSG00000140299 | 15 | 59951345 | 59981733 | 151 | 6 | 0.33984 | 0.36699 | BNIP2 | 0.36699 | 0.41788 | 0.44937 |
| ENSG00000213523 | 5 | 139916925 | 139937895 | 36 | 9 | 0.33977 | 0.36701 | SRA1 | 0.36701 | 0.4427 | 0.40605 |
| ENSG00000232112 | 3 | 48481667 | 48485616 | 15 | 2 | 0.33953 | 0.3671 | TMA7 | 0.3671 | 0.39138 | 0.4238 |
| ENSG00000237330 | 1 | 1006346 | 1009687 | 8 | 3 | 0.33945 | 0.36713 | RNF223 | 0.36713 | 0.51667 | 0.51294 |
| ENSG00000163116 | 4 | 98105244 | 99064391 | 3110 | 39 | 0.33947 | 0.36713 | STPG2 | 0.36713 | 0.36134 | 0.36576 |
| ENSG00000172324 | 11 | 59189416 | 59190449 | 2 | 1 | 0.33945 | 0.36713 | OR5A2 | 0.36713 | 0.099046 | 0.051392 |
| ENSG00000154928 | 3 | 134316643 | 134979309 | 2275 | 152 | 0.33926 | 0.36721 | EPHB1 | 0.36721 | 0.33536 | 0.32769 |
| ENSG00000140961 | 16 | 83981887 | 83999937 | 157 | 22 | 0.3392 | 0.36723 | OSGIN1 | 0.36723 | 0.51933 | 0.68912 |
| ENSG00000078177 | 4 | 40058446 | 40159872 | 281 | 23 | 0.33918 | 0.36724 | N4BP2 | 0.36724 | 0.39015 | 0.34223 |
| ENSG00000148842 | 10 | 104678050 | 104849978 | 522 | 22 | 0.33918 | 0.36724 | CNNM2 | 0.36724 | 0.38031 | 0.40172 |
| ENSG00000177462 | 1 | 248084320 | 248085258 | 17 | 5 | 0.3389 | 0.36734 | OR2T8 | 0.36734 | 0.63543 | 0.46567 |
| ENSG00000261115 | 7 | 140774032 | 141180180 | 1101 | 89 | 0.33857 | 0.36747 | TMEM178B | 0.36747 | 0.29982 | 0.16417 |
| ENSG00000140287 | 15 | 50534144 | 50558223 | 69 | 15 | 0.338 | 0.36768 | HDC | 0.36768 | 0.39405 | 0.36738 |
| ENSG00000205209 | 19 | 35084234 | 35166604 | 443 | 28 | 0.33797 | 0.36769 | SCGB2B2 | 0.36769 | 0.37162 | 0.4787 |
| ENSG00000134463 | 10 | 11784365 | 11806069 | 76 | 10 | 0.33792 | 0.36771 | ECHDC3 | 0.36771 | 0.36383 | 0.29154 |
| ENSG00000105793 | 7 | 89964537 | 90020769 | 233 | 20 | 0.33791 | 0.36772 | GTPBP10 | 0.36772 | 0.37922 | 0.37603 |
| ENSG00000169609 | 15 | 83657193 | 83680393 | 63 | 12 | 0.33776 | 0.36777 | C15orf40 | 0.36777 | 0.275 | 0.3295 |

| | | | | | | | | | | | |
|---|---|---|---|---|---|---|---|---|---|---|---|
| ENSG00000130707 | 9 | 133320316 | 133376661 | 167 | 30 | 0.33755 | 0.36785 | ASS1 | 0.36785 | 0.065611 | 0.17386 |
| ENSG00000186407 | 17 | 72606026 | 72619897 | 51 | 11 | 0.3375 | 0.36787 | CD300E | 0.36787 | 0.45114 | 0.54794 |
| ENSG00000186818 | 19 | 55155340 | 55181810 | 163 | 26 | 0.33711 | 0.36802 | LILRB4 | 0.36802 | 0.35874 | 0.44704 |
| ENSG00000228835 | 18 | 30349758 | 30354376 | 8 | 3 | 0.33688 | 0.36811 | AC012123.1 | 0.36811 | 0.11228 | 0.16358 |
| ENSG00000117475 | 1 | 169337208 | 169365778 | 103 | 14 | 0.33683 | 0.36812 | BLZF1 | 0.36812 | 0.38025 | 0.38867 |
| ENSG00000164690 | 7 | 155592680 | 155604967 | 28 | 12 | 0.33679 | 0.36814 | SHH | 0.36814 | 0.78244 | 0.89281 |
| ENSG00000184350 | 11 | 3248928 | 3253616 | 25 | 9 | 0.33665 | 0.36819 | MRGPRE | 0.36819 | 0.11188 | 0.13269 |
| ENSG00000151338 | 14 | 37667118 | 38021566 | 810 | 19 | 0.33666 | 0.36819 | MIPOL1 | 0.36819 | 0.40257 | 0.43389 |
| ENSG00000070501 | 8 | 42195972 | 42229326 | 82 | 11 | 0.33652 | 0.36824 | POLB | 0.36824 | 0.3181 | 0.29908 |
| ENSG00000101474 | 20 | 24943561 | 24973615 | 101 | 11 | 0.33629 | 0.36833 | APMAP | 0.36833 | 0.32342 | 0.25285 |
| ENSG00000185127 | 6 | 170102233 | 170106401 | 7 | 3 | 0.33622 | 0.36835 | C6orf120 | 0.36835 | 0.036142 | 0.033625 |
| ENSG00000170323 | 8 | 82390654 | 82395498 | 19 | 3 | 0.33617 | 0.36837 | FABP4 | 0.36837 | 0.65148 | 0.81253 |
| ENSG00000185215 | 14 | 103589779 | 103603776 | 39 | 8 | 0.33598 | 0.36844 | TNFAIP2 | 0.36844 | 0.43346 | 0.79135 |
| ENSG00000151665 | 2 | 46808076 | 46844258 | 99 | 15 | 0.33595 | 0.36845 | PIGF | 0.36845 | 0.51137 | 0.48747 |
| ENSG00000147854 | 9 | 6413151 | 6507054 | 304 | 18 | 0.33572 | 0.36854 | UHRF2 | 0.36854 | 0.36883 | 0.39787 |
| ENSG00000267913 | 21 | 45587818 | 45593580 | 12 | 5 | 0.3357 | 0.36855 | AP001055.1 | 0.36855 | 0.46337 | 0.15477 |
| ENSG00000067057 | 10 | 3108525 | 3179904 | 652 | 72 | 0.33552 | 0.36862 | PFKP | 0.36862 | 0.45897 | 0.55404 |
| ENSG00000204161 | 10 | 50362773 | 50396630 | 150 | 28 | 0.33498 | 0.36882 | C10orf128 | 0.36882 | 0.37099 | 0.27386 |
| ENSG00000206562 | 3 | 15422782 | 15482073 | 148 | 16 | 0.33495 | 0.36883 | METTL6 | 0.36883 | 0.39806 | 0.49509 |
| ENSG00000233701 | 3 | 138760944 | 138763734 | 12 | 2 | 0.33484 | 0.36887 | PRR23C | 0.36887 | 0.27718 | 0.24566 |
| ENSG00000088247 | 19 | 6413359 | 6424805 | 45 | 6 | 0.33475 | 0.36891 | KHSRP | 0.36891 | 0.37706 | 0.23789 |
| ENSG00000240204 | 7 | 129142320 | 129152773 | 36 | 11 | 0.33458 | 0.36897 | SMKR1 | 0.36897 | 0.62705 | 0.22643 |
| ENSG00000166226 | 12 | 69979114 | 69995350 | 71 | 11 | 0.3345 | 0.369 | CCT2 | 0.369 | 0.46614 | 0.39197 |
| ENSG00000132681 | 1 | 160121360 | 160156767 | 162 | 34 | 0.33448 | 0.36901 | ATP1A4 | 0.36901 | 0.62212 | 0.66986 |
| ENSG00000137968 | 1 | 75667816 | 76076801 | 1224 | 41 | 0.33436 | 0.36905 | SLC44A5 | 0.36905 | 0.42542 | 0.48624 |
| ENSG00000204388 | 6 | 31795512 | 31798031 | 10 | 6 | 0.33435 | 0.36906 | HSPA1B | 0.36906 | 0.42557 | 0.27285 |
| ENSG00000149212 | 11 | 94898704 | 94965705 | 121 | 16 | 0.33419 | 0.36912 | SESN3 | 0.36912 | 0.36684 | 0.41571 |
| ENSG00000070404 | 19 | 676392 | 683385 | 16 | 4 | 0.33388 | 0.36924 | FSTL3 | 0.36924 | 0.75313 | 0.72021 |
| ENSG00000136918 | 9 | 127615755 | 127620160 | 20 | 4 | 0.33353 | 0.36937 | WDR38 | 0.36937 | 0.41655 | 0.6075 |
| ENSG00000130702 | 20 | 60883011 | 60942368 | 307 | 26 | 0.33349 | 0.36938 | LAMA5 | 0.36938 | 0.37565 | 0.28914 |
| ENSG00000135363 | 11 | 33880122 | 33913836 | 128 | 24 | 0.33337 | 0.36943 | LMO2 | 0.36943 | 0.36602 | 0.43354 |
| ENSG00000138709 | 4 | 128982423 | 129144086 | 428 | 13 | 0.33331 | 0.36945 | LARP1B | 0.36945 | 0.37341 | 0.39874 |
| ENSG00000167580 | 12 | 50344524 | 50352664 | 28 | 6 | 0.33297 | 0.36958 | AQP2 | 0.36958 | 0.5099 | 0.81798 |
| ENSG00000167332 | 11 | 4701401 | 4719084 | 85 | 17 | 0.33293 | 0.36959 | OR51E2 | 0.36959 | 0.25709 | 0.38458 |
| ENSG00000128965 | 15 | 41245160 | 41248710 | 13 | 6 | 0.33292 | 0.3696 | CHAC1 | 0.3696 | 0.63459 | 0.5973 |
| ENSG00000112276 | 6 | 105544697 | 105585049 | 145 | 14 | 0.3327 | 0.36968 | BVES | 0.36968 | 0.38451 | 0.36779 |
| ENSG00000132780 | 1 | 46049518 | 46084566 | 37 | 10 | 0.33264 | 0.3697 | NASP | 0.3697 | 0.36631 | 0.33778 |
| ENSG00000176293 | 19 | 58570607 | 58597677 | 102 | 13 | 0.33227 | 0.36984 | ZNF135 | 0.36984 | 0.48138 | 0.74011 |
| ENSG00000179133 | 10 | 23556124 | 23633774 | 149 | 18 | 0.33205 | 0.36993 | C10orf67 | 0.36993 | 0.17974 | 0.1648 |
| ENSG00000072080 | 2 | 234959323 | 234985778 | 122 | 13 | 0.33197 | 0.36996 | SPP2 | 0.36996 | 0.1654 | 0.13838 |
| ENSG00000172209 | 7 | 107110463 | 107116098 | 7 | 3 | 0.33152 | 0.37013 | GPR22 | 0.37013 | 0.63185 | 0.77312 |
| ENSG00000138379 | 2 | 190920423 | 190927455 | 17 | 3 | 0.33148 | 0.37014 | MSTN | 0.37014 | 0.35595 | 0.4587 |
| ENSG00000136758 | 10 | 27399383 | 27444195 | 197 | 16 | 0.33109 | 0.37029 | YME1L1 | 0.37029 | 0.36257 | 0.29676 |
| ENSG00000157540 | 21 | 38738092 | 38889753 | 434 | 23 | 0.33091 | 0.37035 | DYRK1A | 0.37035 | 0.37053 | 0.41823 |
| ENSG00000256797 | 12 | 10034088 | 10048432 | 57 | 13 | 0.3307 | 0.37044 | KLRF2 | 0.37044 | 0.56589 | 0.61939 |
| ENSG00000148843 | 10 | 105156405 | 105206049 | 95 | 18 | 0.33041 | 0.37054 | PDCD11 | 0.37054 | 0.34418 | 0.52123 |
| ENSG00000084207 | 11 | 67351066 | 67354131 | 13 | 3 | 0.33015 | 0.37064 | GSTP1 | 0.37064 | 0.3388 | 0.31693 |
| ENSG00000037749 | 5 | 153418466 | 153600038 | 564 | 35 | 0.33013 | 0.37065 | MFAP3 | 0.37065 | 0.39235 | 0.37334 |
| ENSG00000143420 | 1 | 150573327 | 150602088 | 66 | 13 | 0.33006 | 0.37068 | ENSA | 0.37068 | 0.43797 | 0.59015 |
| ENSG00000163319 | 4 | 84377085 | 84390888 | 41 | 6 | 0.32987 | 0.37075 | MRPS18C | 0.37075 | 0.39258 | 0.48448 |
| ENSG00000177842 | 3 | 40547483 | 40560227 | 23 | 6 | 0.32978 | 0.37078 | ZNF620 | 0.37078 | 0.33835 | 0.53045 |
| ENSG00000104679 | 8 | 23127633 | 23153792 | 126 | 22 | 0.32944 | 0.37091 | R3HCC1 | 0.37091 | 0.38521 | 0.43339 |
| ENSG00000165970 | 11 | 20620946 | 20680831 | 300 | 31 | 0.3294 | 0.37093 | SLC6A5 | 0.37093 | 0.48658 | 0.52385 |
| ENSG00000241973 | 22 | 21061979 | 21213705 | 515 | 15 | 0.3292 | 0.371 | PI4KA | 0.371 | 0.37908 | 0.33457 |
| ENSG00000136875 | 9 | 116037623 | 116055185 | 65 | 13 | 0.32915 | 0.37102 | PRPF4 | 0.37102 | 0.34167 | 0.44359 |
| ENSG00000180979 | 15 | 42834720 | 42841000 | 12 | 5 | 0.32885 | 0.37113 | LRRC57 | 0.37113 | 0.7121 | 0.74257 |
| ENSG00000101311 | 20 | 6055492 | 6104191 | 177 | 27 | 0.32886 | 0.37113 | FERMT1 | 0.37113 | 0.44917 | 0.47677 |
| ENSG00000170262 | 21 | 33664124 | 33687095 | 54 | 16 | 0.3287 | 0.37119 | MRAP | 0.37119 | 0.6395 | 0.75371 |
| ENSG00000136450 | 17 | 56080721 | 56084707 | 2 | 1 | 0.32836 | 0.37132 | SRSF1 | 0.37132 | 0.13722 | 0.17617 |
| ENSG00000198075 | 2 | 108994367 | 109004513 | 32 | 8 | 0.32823 | 0.37137 | SULT1C4 | 0.37137 | 0.30572 | 0.17226 |
| ENSG00000188659 | 15 | 82555151 | 82577271 | 65 | 16 | 0.32823 | 0.37137 | FAM154B | 0.37137 | 0.30137 | 0.4334 |
| ENSG00000139263 | 12 | 59265931 | 59314303 | 156 | 19 | 0.32779 | 0.37153 | LRIG3 | 0.37153 | 0.22268 | 0.19385 |
| ENSG00000128536 | 7 | 105517242 | 105676877 | 670 | 85 | 0.32774 | 0.37155 | CDHR3 | 0.37155 | 0.40013 | 0.41084 |
| ENSG00000109099 | 17 | 15133095 | 15168643 | 100 | 14 | 0.32772 | 0.37156 | PMP22 | 0.37156 | 0.45965 | 0.51998 |
| ENSG00000167721 | 17 | 2225797 | 2240801 | 27 | 8 | 0.32756 | 0.37162 | TSR1 | 0.37162 | 0.61191 | 0.74613 |
| ENSG00000100242 | 22 | 39130730 | 39190148 | 125 | 27 | 0.32746 | 0.37166 | SUN2 | 0.37166 | 0.41039 | 0.41985 |
| ENSG00000123689 | 1 | 209848765 | 209849733 | 4 | 2 | 0.32734 | 0.37171 | G0S2 | 0.37171 | 0.65364 | 0.81701 |
| ENSG00000146859 | 7 | 134832824 | 134850967 | 51 | 4 | 0.32715 | 0.37178 | TMEM140 | 0.37178 | 0.39346 | 0.48618 |
| ENSG00000253598 | 8 | 82605842 | 82608409 | 8 | 3 | 0.32699 | 0.37184 | SLC10A5 | 0.37184 | 0.42024 | 0.47667 |
| ENSG00000047457 | 3 | 148880197 | 148939842 | 179 | 35 | 0.32684 | 0.37189 | CP | 0.37189 | 0.39942 | 0.20749 |
| ENSG00000269533 | 19 | 57901252 | 57931362 | 69 | 14 | 0.32682 | 0.3719 | AC003002.6 | 0.3719 | 0.32823 | 0.38082 |
| ENSG00000197838 | 19 | 41594377 | 41602100 | 33 | 9 | 0.3267 | 0.37195 | CYP2A13 | 0.37195 | 0.1984 | 0.26713 |
| ENSG00000163071 | 4 | 52917497 | 52963458 | 105 | 12 | 0.32648 | 0.37203 | SPATA18 | 0.37203 | 0.39666 | 0.43303 |

| | | | | | | | | | | | |
|---|---|---|---|---|---|---|---|---|---|---|---|
| ENSG00000102900 | 16 | 56764017 | 56878797 | 296 | 27 | 0.32645 | 0.37204 | NUP93 | 0.37204 | 0.35864 | 0.33663 |
| ENSG00000048991 | 2 | 136289025 | 136482840 | 350 | 21 | 0.32625 | 0.37212 | R3HDM1 | 0.37212 | 0.36726 | 0.42329 |
| ENSG00000143774 | 1 | 228327663 | 228336685 | 14 | 5 | 0.32595 | 0.37223 | GUK1 | 0.37223 | 0.39447 | 0.49235 |
| ENSG00000116459 | 1 | 111991486 | 112005395 | 31 | 7 | 0.32593 | 0.37224 | ATP5F1 | 0.37224 | 0.21066 | 0.21254 |
| ENSG00000170633 | 12 | 121837844 | 121868389 | 104 | 7 | 0.3258 | 0.37229 | RNF34 | 0.37229 | 0.22748 | 0.23298 |
| ENSG00000197535 | 15 | 52599480 | 52821247 | 733 | 24 | 0.32577 | 0.3723 | MYO5A | 0.3723 | 0.38836 | 0.43065 |
| ENSG00000168016 | 3 | 36868311 | 36986548 | 305 | 22 | 0.32561 | 0.37236 | TRANK1 | 0.37236 | 0.46747 | 0.45171 |
| ENSG00000237763 | 1 | 104197912 | 104207173 | 4 | 2 | 0.32535 | 0.37246 | AMY1A | 0.37246 | 0.22665 | 0.054013 |
| ENSG00000111266 | 12 | 12628829 | 12715317 | 284 | 23 | 0.32535 | 0.37246 | DUSP16 | 0.37246 | 0.23631 | 0.21463 |
| ENSG00000070915 | 16 | 56899119 | 56949762 | 271 | 28 | 0.32512 | 0.37254 | SLC12A3 | 0.37254 | 0.40349 | 0.48088 |
| ENSG00000060140 | 12 | 10771538 | 10826917 | 189 | 13 | 0.32491 | 0.37262 | STYK1 | 0.37262 | 0.552 | 0.67538 |
| ENSG00000105186 | 19 | 33087913 | 33167503 | 492 | 23 | 0.32465 | 0.37272 | ANKRD27 | 0.37272 | 0.33157 | 0.33259 |
| ENSG00000176593 | 19 | 58513429 | 58522600 | 37 | 7 | 0.3246 | 0.37274 | CTD-2368P22.1 | 0.37274 | 0.51522 | 0.47187 |
| ENSG00000077984 | 20 | 24929866 | 24940564 | 65 | 9 | 0.32449 | 0.37278 | CST7 | 0.37278 | 0.18435 | 0.2675 |
| ENSG00000126456 | 19 | 50162826 | 50169132 | 13 | 3 | 0.32421 | 0.37289 | IRF3 | 0.37289 | 0.44526 | 0.46337 |
| ENSG00000183914 | 17 | 7620672 | 7737062 | 316 | 32 | 0.32396 | 0.37298 | DNAH2 | 0.37298 | 0.38447 | 0.43706 |
| ENSG00000196361 | 19 | 11562141 | 11591861 | 93 | 14 | 0.32386 | 0.37302 | ELAVL3 | 0.37302 | 0.31639 | 0.17855 |
| ENSG00000186812 | 18 | 32820994 | 32847097 | 51 | 13 | 0.32372 | 0.37308 | ZNF397 | 0.37308 | 0.47024 | 0.29801 |
| ENSG00000080910 | 1 | 196788898 | 196928356 | 522 | 22 | 0.32358 | 0.37313 | CFHR2 | 0.37313 | 0.40331 | 0.52489 |
| ENSG00000145808 | 5 | 128795958 | 129074376 | 852 | 32 | 0.32356 | 0.37314 | ADAMTS19 | 0.37314 | 0.36378 | 0.35103 |
| ENSG00000165209 | 9 | 125871779 | 126030855 | 239 | 33 | 0.32354 | 0.37314 | STRBP | 0.37314 | 0.36188 | 0.38422 |
| ENSG00000205336 | 16 | 57644564 | 57698944 | 268 | 51 | 0.32354 | 0.37314 | GPR56 | 0.37314 | 0.46647 | 0.44593 |
| ENSG00000234409 | 22 | 20135950 | 20138399 | 14 | 4 | 0.32355 | 0.37314 | AC006547.14 | 0.37314 | 0.23071 | 0.31866 |
| ENSG00000111481 | 12 | 54694986 | 54745633 | 75 | 10 | 0.32353 | 0.37315 | COPZ1 | 0.37315 | 0.4119 | 0.52828 |
| ENSG00000273154 | 20 | 62340216 | 62370456 | 113 | 9 | 0.32348 | 0.37317 | RP4-583P15.15 | 0.37317 | 0.32344 | 0.29367 |
| ENSG00000235194 | 14 | 23764852 | 23772057 | 8 | 5 | 0.32325 | 0.37325 | PPP1R3E | 0.37325 | 0.6819 | 0.77259 |
| ENSG00000197714 | 19 | 57791419 | 57804937 | 52 | 8 | 0.32279 | 0.37343 | ZNF460 | 0.37343 | 0.36482 | 0.35123 |
| ENSG00000066185 | 1 | 42896000 | 42921938 | 100 | 10 | 0.32275 | 0.37344 | ZMYND12 | 0.37344 | 0.37613 | 0.34588 |
| ENSG00000068024 | 2 | 239969864 | 240323348 | 1362 | 93 | 0.32273 | 0.37345 | HDAC4 | 0.37345 | 0.41912 | 0.48932 |
| ENSG00000270672 | 7 | 142374104 | 142375550 | 6 | 1 | 0.32227 | 0.37362 | MTRNR2L6 | 0.37362 | 0.64652 | 0.67465 |
| ENSG00000104783 | 19 | 44270685 | 44285409 | 55 | 15 | 0.32208 | 0.3737 | KCNN4 | 0.3737 | 0.67251 | 0.799 |
| ENSG00000129518 | 14 | 34985135 | 35008916 | 90 | 11 | 0.322 | 0.37373 | EAPP | 0.37373 | 0.38465 | 0.28994 |
| ENSG00000198429 | 19 | 11998599 | 12025144 | 73 | 11 | 0.32198 | 0.37374 | ZNF69 | 0.37374 | 0.41244 | 0.25959 |
| ENSG00000103707 | 15 | 65294845 | 65321977 | 73 | 11 | 0.3218 | 0.3738 | MTFMT | 0.3738 | 0.42279 | 0.3273 |
| ENSG00000101544 | 18 | 77866915 | 77905406 | 145 | 14 | 0.32159 | 0.37388 | ADNP2 | 0.37388 | 0.39913 | 0.39321 |
| ENSG00000080345 | 2 | 152266397 | 152364527 | 235 | 25 | 0.32157 | 0.37389 | RIF1 | 0.37389 | 0.39037 | 0.40921 |
| ENSG00000273173 | 15 | 25200133 | 25223729 | 50 | 9 | 0.32154 | 0.3739 | SNURF | 0.3739 | 0.38882 | 0.37933 |
| ENSG00000268936 | 11 | 134855246 | 134856693 | 8 | 5 | 0.32146 | 0.37393 | AP003062.1 | 0.37393 | 0.60312 | 0.30113 |
| ENSG00000196366 | 9 | 139377947 | 139380518 | 5 | 3 | 0.32124 | 0.37401 | C9orf163 | 0.37401 | 0.41681 | 0.60959 |
| ENSG00000048405 | 7 | 126986844 | 127071978 | 329 | 21 | 0.32124 | 0.37402 | ZNF800 | 0.37402 | 0.34548 | 0.39446 |
| ENSG00000131373 | 3 | 15602211 | 15643338 | 101 | 20 | 0.32115 | 0.37405 | HACL1 | 0.37405 | 0.28411 | 0.31853 |
| ENSG00000009765 | 6 | 150690028 | 150727105 | 232 | 23 | 0.32111 | 0.37406 | IYD | 0.37406 | 0.35177 | 0.45288 |
| ENSG00000064607 | 19 | 19101697 | 19144832 | 96 | 11 | 0.32112 | 0.37406 | SUGP2 | 0.37406 | 0.44249 | 0.47356 |
| ENSG00000171757 | 3 | 169511216 | 169530774 | 51 | 5 | 0.32095 | 0.37412 | LRRC34 | 0.37412 | 0.56586 | 0.60715 |
| ENSG00000205174 | 7 | 108524032 | 108524644 | 1 | 1 | 0.32075 | 0.3742 | C7orf66 | 0.3742 | 0.76678 | 0.84382 |
| ENSG00000172037 | 3 | 49158547 | 49170551 | 13 | 8 | 0.3205 | 0.37429 | LAMB2 | 0.37429 | 0.75809 | 0.73048 |
| ENSG00000007372 | 11 | 31806340 | 31839509 | 66 | 11 | 0.32033 | 0.37436 | PAX6 | 0.37436 | 0.30424 | 0.36468 |
| ENSG00000126243 | 19 | 36426260 | 36436669 | 21 | 5 | 0.32023 | 0.3744 | LRFN3 | 0.3744 | 0.34302 | 0.49701 |
| ENSG00000073792 | 3 | 185361527 | 185542844 | 348 | 35 | 0.32021 | 0.37441 | IGF2BP2 | 0.37441 | 0.44653 | 0.43974 |
| ENSG00000204711 | 9 | 72435709 | 72521148 | 254 | 23 | 0.31957 | 0.37465 | C9orf135 | 0.37465 | 0.43644 | 0.53572 |
| ENSG00000134248 | 1 | 110943871 | 110950564 | 20 | 5 | 0.31942 | 0.3747 | LAMTOR5 | 0.3747 | 0.53514 | 0.71264 |
| ENSG00000165650 | 10 | 119040000 | 119134978 | 283 | 13 | 0.31919 | 0.37479 | PDZD8 | 0.37479 | 0.39351 | 0.34893 |
| ENSG00000164176 | 5 | 83236373 | 83680611 | 1097 | 87 | 0.319 | 0.37486 | EDIL3 | 0.37486 | 0.37339 | 0.36383 |
| ENSG00000163530 | 3 | 109012635 | 109035364 | 65 | 17 | 0.31862 | 0.37501 | DPPA2 | 0.37501 | 0.14403 | 0.11079 |
| ENSG00000104823 | 19 | 39306062 | 39322645 | 52 | 10 | 0.31816 | 0.37518 | ECH1 | 0.37518 | 0.51562 | 0.73749 |
| ENSG00000100353 | 22 | 36906897 | 36925483 | 116 | 14 | 0.31816 | 0.37518 | EIF3D | 0.37518 | 0.23312 | 0.065283 |
| ENSG00000197880 | 1 | 23907985 | 23967058 | 131 | 29 | 0.31807 | 0.37522 | MDS2 | 0.37522 | 0.22821 | 0.15019 |
| ENSG00000087301 | 14 | 52897308 | 53019240 | 371 | 20 | 0.31801 | 0.37524 | TXNDC16 | 0.37524 | 0.36314 | 0.37341 |
| ENSG00000182521 | 14 | 55880259 | 55923444 | 249 | 16 | 0.31798 | 0.37525 | TBPL2 | 0.37525 | 0.23751 | 0.17767 |
| ENSG00000005007 | 19 | 18942747 | 18979045 | 70 | 15 | 0.31768 | 0.37537 | UPF1 | 0.37537 | 0.35071 | 0.48046 |
| ENSG00000105426 | 19 | 5158506 | 5340814 | 780 | 92 | 0.31749 | 0.37544 | PTPRS | 0.37544 | 0.42663 | 0.56083 |
| ENSG00000118900 | 16 | 4896666 | 4932361 | 147 | 19 | 0.31736 | 0.37549 | UBN1 | 0.37549 | 0.41648 | 0.45705 |
| ENSG00000268550 | 19 | 14834747 | 14835121 | 2 | 1 | 0.31729 | 0.37551 | AC090427.1 | 0.37551 | 0.49432 | 0.37558 |
| ENSG00000172578 | 3 | 183205319 | 183273477 | 181 | 20 | 0.31724 | 0.37553 | KLHL6 | 0.37553 | 0.33511 | 0.092074 |
| ENSG00000170571 | 5 | 49692026 | 49739082 | 101 | 8 | 0.31702 | 0.37562 | EMB | 0.37562 | 0.38699 | 0.37905 |
| ENSG00000132130 | 17 | 35294084 | 35301917 | 23 | 8 | 0.317 | 0.37562 | LHX1 | 0.37562 | 0.19264 | 0.34442 |
| ENSG00000178904 | 19 | 32896449 | 32976801 | 225 | 27 | 0.31679 | 0.3757 | DPY19L3 | 0.3757 | 0.38819 | 0.42309 |
| ENSG00000112651 | 6 | 43021767 | 43027544 | 17 | 4 | 0.31662 | 0.37577 | MRPL2 | 0.37577 | 0.41639 | 0.43668 |
| ENSG00000160862 | 7 | 99564343 | 99573780 | 16 | 5 | 0.31644 | 0.37583 | AZGP1 | 0.37583 | 0.60779 | 0.44022 |
| ENSG00000206579 | 8 | 56014949 | 56454613 | 1528 | 78 | 0.31644 | 0.37583 | XKR4 | 0.37583 | 0.30968 | 0.20172 |
| ENSG00000171291 | 19 | 11959541 | 11994565 | 120 | 18 | 0.31643 | 0.37584 | ZNF439 | 0.37584 | 0.32321 | 0.35342 |
| ENSG00000187642 | 1 | 910579 | 917497 | 25 | 6 | 0.31612 | 0.37596 | C1orf170 | 0.37596 | 0.32985 | 0.44337 |
| ENSG00000169621 | 2 | 68694693 | 68858004 | 566 | 15 | 0.31582 | 0.37607 | APLF | 0.37607 | 0.38361 | 0.33583 |

| | | | | | | | | | | | |
|---|---|---|---|---|---|---|---|---|---|---|---|
| ENSG00000096872 | 9 | 26947037 | 27062928 | 362 | 27 | 0.31575 | 0.3761 | IFT74 | 0.3761 | 0.37833 | 0.38486 |
| ENSG00000259900 | 16 | 69369179 | 69385633 | 39 | 10 | 0.31574 | 0.3761 | RP11-343C2.7 | 0.3761 | 0.29611 | 0.3052 |
| ENSG00000144029 | 2 | 95752952 | 95815179 | 53 | 11 | 0.31568 | 0.37612 | MRPS5 | 0.37612 | 0.25492 | 0.27366 |
| ENSG00000174339 | 5 | 180166042 | 180167099 | 4 | 2 | 0.3153 | 0.37627 | OR2Y1 | 0.37627 | 0.56344 | 0.54348 |
| ENSG00000119946 | 10 | 101088856 | 101154087 | 173 | 30 | 0.31518 | 0.37631 | CNNM1 | 0.37631 | 0.35524 | 0.30888 |
| ENSG00000104343 | 8 | 74692332 | 74791145 | 168 | 24 | 0.31473 | 0.37648 | UBE2W | 0.37648 | 0.39424 | 0.44348 |
| ENSG00000157890 | 15 | 66187417 | 66546085 | 1283 | 125 | 0.31408 | 0.37673 | MEGF11 | 0.37673 | 0.38628 | 0.38647 |
| ENSG00000213983 | 14 | 24028774 | 24037279 | 9 | 4 | 0.31396 | 0.37678 | AP1G2 | 0.37678 | 0.67555 | 0.66774 |
| ENSG00000196449 | 1 | 38268616 | 38273857 | 9 | 3 | 0.31393 | 0.37679 | YRDC | 0.37679 | 0.4252 | 0.50702 |
| ENSG00000134049 | 18 | 44681413 | 44702745 | 73 | 9 | 0.3139 | 0.3768 | IER3IP1 | 0.3768 | 0.082593 | 0.083522 |
| ENSG00000180440 | 13 | 37248049 | 37271976 | 61 | 12 | 0.31383 | 0.37683 | SERTM1 | 0.37683 | 0.083088 | 0.022942 |
| ENSG00000269529 | 12 | 50690489 | 50691718 | 6 | 2 | 0.31374 | 0.37686 | AC140061.12 | 0.37686 | 0.33294 | 0.26244 |
| ENSG00000188959 | 9 | 112952328 | 112970469 | 86 | 18 | 0.31343 | 0.37698 | C9orf152 | 0.37698 | 0.46737 | 0.59947 |
| ENSG00000179846 | 19 | 45653008 | 45663408 | 27 | 10 | 0.31337 | 0.377 | NKPD1 | 0.377 | 0.49824 | 0.54485 |
| ENSG00000149305 | 11 | 113775399 | 113817287 | 147 | 21 | 0.31282 | 0.37721 | HTR3B | 0.37721 | 0.43056 | 0.50834 |
| ENSG00000099814 | 14 | 105331617 | 105363107 | 102 | 13 | 0.31269 | 0.37726 | CEP170B | 0.37726 | 0.43218 | 0.37897 |
| ENSG00000164093 | 4 | 111538579 | 111563279 | 50 | 11 | 0.31258 | 0.3773 | PITX2 | 0.3773 | 0.38518 | 0.30365 |
| ENSG00000124134 | 20 | 43720951 | 43729753 | 16 | 5 | 0.3122 | 0.37745 | KCNS1 | 0.37745 | 0.57559 | 0.60853 |
| ENSG00000183476 | 15 | 78370150 | 78397251 | 64 | 19 | 0.31209 | 0.37749 | SH2D7 | 0.37749 | 0.47306 | 0.41679 |
| ENSG00000114529 | 3 | 111805182 | 111849851 | 137 | 9 | 0.31198 | 0.37753 | C3orf52 | 0.37753 | 0.44973 | 0.29987 |
| ENSG00000111843 | 6 | 10723148 | 10731362 | 24 | 3 | 0.31193 | 0.37755 | TMEM14C | 0.37755 | 0.60145 | 0.63141 |
| ENSG00000156219 | 4 | 76932337 | 77033955 | 710 | 25 | 0.31178 | 0.3776 | ART3 | 0.3776 | 0.34355 | 0.32099 |
| ENSG00000180433 | 1 | 158724510 | 158725681 | 2 | 1 | 0.31173 | 0.37762 | OR6K6 | 0.37762 | 0.76647 | 0.74222 |
| ENSG00000213218 | 17 | 61949372 | 61951126 | 2 | 1 | 0.31169 | 0.37764 | CSH2 | 0.37764 | 0.69666 | 0.64783 |
| ENSG00000204439 | 6 | 31626075 | 31628549 | 5 | 3 | 0.31166 | 0.37765 | C6orf47 | 0.37765 | 0.45026 | 0.31164 |
| ENSG00000068831 | 11 | 64494383 | 64512928 | 32 | 6 | 0.31161 | 0.37767 | RASGRP2 | 0.37767 | 0.63653 | 0.49265 |
| ENSG00000006210 | 16 | 57406370 | 57418960 | 39 | 11 | 0.31123 | 0.37781 | CX3CL1 | 0.37781 | 0.32709 | 0.26425 |
| ENSG00000078114 | 10 | 21068902 | 21463116 | 1134 | 72 | 0.3112 | 0.37782 | NEBL | 0.37782 | 0.37346 | 0.46386 |
| ENSG00000221932 | 11 | 124789146 | 124790573 | 3 | 2 | 0.31113 | 0.37785 | HEPN1 | 0.37785 | 0.3766 | 0.29988 |
| ENSG00000221852 | 17 | 39182278 | 39183454 | 5 | 3 | 0.31094 | 0.37792 | KRTAP1-5 | 0.37792 | 0.42061 | 0.44373 |
| ENSG00000116337 | 1 | 110158726 | 110174673 | 38 | 10 | 0.31092 | 0.37793 | AMPD2 | 0.37793 | 0.23575 | 0.35852 |
| ENSG00000008311 | 7 | 121715701 | 121784334 | 146 | 19 | 0.31093 | 0.37793 | AASS | 0.37793 | 0.50679 | 0.41897 |
| ENSG00000084636 | 1 | 32117848 | 32169920 | 111 | 19 | 0.31082 | 0.37797 | COL16A1 | 0.37797 | 0.068517 | 0.089765 |
| ENSG00000198785 | 9 | 104331635 | 104500862 | 743 | 29 | 0.31052 | 0.37808 | GRIN3A | 0.37808 | 0.37689 | 0.39625 |
| ENSG00000164978 | 9 | 34329504 | 34343709 | 43 | 9 | 0.31033 | 0.37816 | NUDT2 | 0.37816 | 0.40631 | 0.39747 |
| ENSG00000140379 | 15 | 80253231 | 80263788 | 23 | 3 | 0.31013 | 0.37823 | BCL2A1 | 0.37823 | 0.43915 | 0.42938 |
| ENSG00000091986 | 3 | 112323407 | 112368377 | 141 | 25 | 0.3101 | 0.37824 | CCDC80 | 0.37824 | 0.5137 | 0.64533 |
| ENSG00000164294 | 5 | 54455946 | 54462899 | 16 | 4 | 0.31005 | 0.37826 | GPX8 | 0.37826 | 0.41969 | 0.34894 |
| ENSG00000148156 | 9 | 111616871 | 111619239 | 11 | 4 | 0.30948 | 0.37848 | ACTL7B | 0.37848 | 0.4346 | 0.30851 |
| ENSG00000113013 | 5 | 137890571 | 137911133 | 49 | 9 | 0.30943 | 0.3785 | HSPA9 | 0.3785 | 0.38473 | 0.38631 |
| ENSG00000233404 | 2 | 102508955 | 102511150 | 3 | 2 | 0.30938 | 0.37851 | FLJ20373 | 0.37851 | 0.31177 | 0.27514 |
| ENSG00000101040 | 20 | 45837859 | 45985567 | 294 | 31 | 0.30926 | 0.37856 | ZMYND8 | 0.37856 | 0.30284 | 0.56784 |
| ENSG00000016490 | 1 | 86934051 | 86965972 | 145 | 31 | 0.30898 | 0.37867 | CLCA1 | 0.37867 | 0.47734 | 0.52404 |
| ENSG00000172289 | 11 | 59480296 | 59481337 | 6 | 3 | 0.30891 | 0.3787 | OR10V1 | 0.3787 | 0.19064 | 0.22467 |
| ENSG00000106537 | 7 | 16793160 | 16824161 | 132 | 17 | 0.3088 | 0.37874 | TSPAN13 | 0.37874 | 0.19477 | 0.14672 |
| ENSG00000154134 | 11 | 124735282 | 124751366 | 44 | 16 | 0.30875 | 0.37876 | ROBO3 | 0.37876 | 0.4731 | 0.54466 |
| ENSG00000121446 | 1 | 182419256 | 182529734 | 397 | 18 | 0.30859 | 0.37882 | RGSL1 | 0.37882 | 0.3802 | 0.36693 |
| ENSG00000197632 | 18 | 61538926 | 61571124 | 129 | 4 | 0.30815 | 0.37898 | SERPINB2 | 0.37898 | 0.37386 | 0.38805 |
| ENSG00000104365 | 8 | 42128820 | 42189973 | 144 | 14 | 0.30796 | 0.37906 | IKBKB | 0.37906 | 0.42789 | 0.49066 |
| ENSG00000136371 | 15 | 80125927 | 80189721 | 214 | 27 | 0.30796 | 0.37906 | MTHFS | 0.37906 | 0.41116 | 0.36302 |
| ENSG00000164729 | 17 | 33519539 | 33521412 | 4 | 2 | 0.30771 | 0.37915 | SLC35G3 | 0.37915 | 0.80709 | 0.77415 |
| ENSG00000197903 | 6 | 27106073 | 27114619 | 15 | 7 | 0.30761 | 0.37919 | HIST1H2BK | 0.37919 | 0.41209 | 0.41951 |
| ENSG00000156414 | 14 | 104394799 | 104519004 | 495 | 21 | 0.30754 | 0.37922 | TDRD9 | 0.37922 | 0.39096 | 0.39655 |
| ENSG00000148411 | 9 | 138898383 | 138987131 | 403 | 62 | 0.30727 | 0.37932 | NACC2 | 0.37932 | 0.27346 | 0.21042 |
| ENSG00000197114 | 20 | 62338817 | 62367494 | 106 | 9 | 0.30715 | 0.37936 | ZGPAT | 0.37936 | 0.36155 | 0.31829 |
| ENSG00000105366 | 19 | 51954101 | 51961710 | 16 | 3 | 0.30696 | 0.37944 | SIGLEC8 | 0.37944 | 0.42079 | 0.68151 |
| ENSG00000243440 | 21 | 15964251 | 16031142 | 195 | 19 | 0.30649 | 0.37961 | AF165138.7 | 0.37961 | 0.23371 | 0.31367 |
| ENSG00000262628 | 17 | 2965963 | 2966901 | 4 | 2 | 0.30648 | 0.37962 | OR1D5 | 0.37962 | 0.0016436 | 0.008387 |
| ENSG00000183741 | 22 | 39257455 | 39268319 | 23 | 9 | 0.30641 | 0.37965 | CBX6 | 0.37965 | 0.39981 | 0.66349 |
| ENSG00000183801 | 11 | 7506619 | 7532608 | 130 | 17 | 0.30625 | 0.37971 | OLFML1 | 0.37971 | 0.35127 | 0.27198 |
| ENSG00000116883 | 1 | 36789335 | 36794822 | 14 | 3 | 0.30515 | 0.38013 | RP11-268J15.5 | 0.38013 | 0.34488 | 0.34228 |
| ENSG00000007944 | 6 | 16129356 | 16148479 | 45 | 11 | 0.30489 | 0.38023 | MYLIP | 0.38023 | 0.30809 | 0.47177 |
| ENSG00000079435 | 19 | 42905659 | 42931578 | 16 | 7 | 0.30482 | 0.38025 | LIPE | 0.38025 | 0.60063 | 0.42547 |
| ENSG00000156875 | 1 | 100503653 | 100548933 | 122 | 16 | 0.30432 | 0.38044 | HIAT1 | 0.38044 | 0.34718 | 0.31485 |
| ENSG00000187961 | 1 | 895967 | 901095 | 30 | 8 | 0.30426 | 0.38046 | KLHL17 | 0.38046 | 0.27784 | 0.37974 |
| ENSG00000169715 | 16 | 56659387 | 56661024 | 8 | 4 | 0.30405 | 0.38055 | MT1E | 0.38055 | 0.50617 | 0.47685 |
| ENSG00000268472 | 11 | 112130988 | 112132117 | 4 | 2 | 0.30401 | 0.38056 | AP002884.2 | 0.38056 | 0.57582 | 0.61454 |
| ENSG00000230000 | 7 | 62809239 | 62812151 | 7 | 2 | 0.30371 | 0.38067 | AC006455.1 | 0.38067 | 0.46203 | 0.33599 |
| ENSG00000140527 | 15 | 90234028 | 90286869 | 205 | 14 | 0.30367 | 0.38069 | WDR93 | 0.38069 | 0.31238 | 0.26211 |
| ENSG00000105669 | 19 | 19010323 | 19030206 | 62 | 13 | 0.30356 | 0.38073 | COPE | 0.38073 | 0.46291 | 0.42811 |
| ENSG00000205269 | 6 | 11537938 | 11583757 | 177 | 20 | 0.3035 | 0.38075 | TMEM170B | 0.38075 | 0.35347 | 0.38608 |
| ENSG00000133710 | 5 | 147405246 | 147516852 | 556 | 18 | 0.30344 | 0.38078 | SPINK5 | 0.38078 | 0.34906 | 0.29513 |
| ENSG00000140992 | 16 | 2587965 | 2653189 | 25 | 14 | 0.30333 | 0.38082 | PDPK1 | 0.38082 | 0.4887 | 0.61157 |

| | | | | | | | | | | | |
|---|---|---|---|---|---|---|---|---|---|---|---|
| ENSG00000112312 | 6 | 24775159 | 24786327 | 28 | 12 | 0.30327 | 0.38084 | GMNN | 0.38084 | 0.19995 | 0.3661 |
| ENSG00000154146 | 11 | 124609742 | 124617106 | 15 | 4 | 0.30328 | 0.38084 | NRGN | 0.38084 | 0.46467 | 0.48252 |
| ENSG00000169696 | 17 | 79934683 | 79975282 | 101 | 21 | 0.30268 | 0.38107 | ASPSCR1 | 0.38107 | 0.31286 | 0.27044 |
| ENSG00000105819 | 7 | 102937869 | 102969958 | 55 | 15 | 0.30263 | 0.38109 | PMPCB | 0.38109 | 0.26837 | 0.38467 |
| ENSG00000116132 | 1 | 170631869 | 170708560 | 160 | 24 | 0.30253 | 0.38113 | PRRX1 | 0.38113 | 0.48196 | 0.56951 |
| ENSG00000145309 | 4 | 71200683 | 71202833 | 3 | 2 | 0.30245 | 0.38115 | CABS1 | 0.38115 | 0.67154 | 0.71429 |
| ENSG00000187510 | 12 | 93115281 | 93166231 | 124 | 16 | 0.30236 | 0.38119 | PLEKHG7 | 0.38119 | 0.29042 | 0.29135 |
| ENSG00000186977 | 21 | 31873975 | 31874435 | 2 | 1 | 0.30209 | 0.38129 | KRTAP19-5 | 0.38129 | 0.50861 | 0.53894 |
| ENSG00000197353 | 8 | 143831568 | 143833952 | 10 | 2 | 0.30206 | 0.3813 | LYPD2 | 0.3813 | 0.8321 | 0.7019 |
| ENSG00000105829 | 7 | 93592074 | 93633694 | 75 | 15 | 0.30196 | 0.38134 | BET1 | 0.38134 | 0.47162 | 0.48564 |
| ENSG00000124222 | 20 | 57226328 | 57254582 | 77 | 14 | 0.30194 | 0.38135 | STX16 | 0.38135 | 0.26305 | 0.31652 |
| ENSG00000159210 | 17 | 47006678 | 47022479 | 71 | 8 | 0.30179 | 0.3814 | SNF8 | 0.3814 | 0.3674 | 0.41206 |
| ENSG00000185477 | 4 | 90157537 | 90229161 | 223 | 33 | 0.30178 | 0.38141 | GPRIN3 | 0.38141 | 0.47956 | 0.4995 |
| ENSG00000145020 | 3 | 49454211 | 49460186 | 12 | 3 | 0.30158 | 0.38149 | AMT | 0.38149 | 0.5955 | 0.6205 |
| ENSG00000168118 | 1 | 229406822 | 229441641 | 85 | 10 | 0.30148 | 0.38153 | RAB4A | 0.38153 | 0.41853 | 0.55334 |
| ENSG00000105865 | 7 | 107203929 | 107218906 | 22 | 5 | 0.3008 | 0.38178 | DUS4L | 0.38178 | 0.44737 | 0.53801 |
| ENSG00000148300 | 9 | 136271186 | 136283164 | 57 | 5 | 0.30082 | 0.38178 | REXO4 | 0.38178 | 0.41396 | 0.47695 |
| ENSG00000108786 | 17 | 40701232 | 40707231 | 25 | 5 | 0.30079 | 0.38179 | HSD17B1 | 0.38179 | 0.37318 | 0.43268 |
| ENSG00000127529 | 19 | 15052301 | 15053260 | 3 | 1 | 0.30063 | 0.38185 | OR7C2 | 0.38185 | 0.75871 | 0.74555 |
| ENSG00000186470 | 6 | 26365387 | 26378546 | 101 | 9 | 0.30055 | 0.38188 | BTN3A2 | 0.38188 | 0.35162 | 0.26462 |
| ENSG00000171222 | 20 | 34541539 | 34547394 | 14 | 3 | 0.30042 | 0.38193 | SCAND1 | 0.38193 | 0.38074 | 0.36373 |
| ENSG00000155380 | 1 | 113454469 | 113499635 | 73 | 13 | 0.30035 | 0.38195 | SLC16A1 | 0.38195 | 0.42427 | 0.38763 |
| ENSG00000124641 | 6 | 41873092 | 41888877 | 26 | 9 | 0.29991 | 0.38212 | MED20 | 0.38212 | 0.63207 | 0.6382 |
| ENSG00000215788 | 1 | 6521211 | 6526255 | 17 | 6 | 0.29989 | 0.38213 | TNFRSF25 | 0.38213 | 0.57963 | 0.47791 |
| ENSG00000109084 | 17 | 26646121 | 26655707 | 11 | 4 | 0.29979 | 0.38217 | TMEM97 | 0.38217 | 0.32624 | 0.3305 |
| ENSG00000241794 | 1 | 153028589 | 153030013 | 2 | 1 | 0.29973 | 0.38219 | SPRR2A | 0.38219 | 0.42427 | 0.50593 |
| ENSG00000189334 | 1 | 153586731 | 153589462 | 8 | 2 | 0.29969 | 0.38221 | S100A14 | 0.38221 | 0.44613 | 0.43928 |
| ENSG00000242498 | 15 | 90443159 | 90456188 | 42 | 12 | 0.29963 | 0.38223 | C15orf38 | 0.38223 | 0.43618 | 0.51066 |
| ENSG00000262814 | 17 | 79670387 | 79674556 | 23 | 6 | 0.29962 | 0.38223 | MRPL12 | 0.38223 | 0.21542 | 0.29651 |
| ENSG00000267795 | 16 | 4838398 | 4846492 | 20 | 7 | 0.29945 | 0.3823 | SMIM22 | 0.3823 | 0.43672 | 0.46657 |
| ENSG00000103160 | 16 | 84155886 | 84178797 | 71 | 13 | 0.29937 | 0.38233 | HSDL1 | 0.38233 | 0.24204 | 0.11464 |
| ENSG00000004139 | 17 | 26691378 | 26728065 | 94 | 11 | 0.29932 | 0.38235 | SARM1 | 0.38235 | 0.38355 | 0.40029 |
| ENSG00000120784 | 19 | 38104650 | 38183238 | 202 | 25 | 0.29929 | 0.38236 | ZFP30 | 0.38236 | 0.37348 | 0.37259 |
| ENSG00000089327 | 19 | 35645633 | 35660786 | 69 | 14 | 0.29921 | 0.38239 | FXYD5 | 0.38239 | 0.42284 | 0.65442 |
| ENSG00000123171 | 13 | 52436117 | 52440366 | 11 | 3 | 0.29903 | 0.38246 | CCDC70 | 0.38246 | 0.114 | 0.14699 |
| ENSG00000023330 | 3 | 52232102 | 52248343 | 19 | 3 | 0.29881 | 0.38254 | ALAS1 | 0.38254 | 0.52803 | 0.59226 |
| ENSG00000273291 | 3 | 42850938 | 43097363 | 609 | 34 | 0.29857 | 0.38263 | KRBOX1 | 0.38263 | 0.39918 | 0.43464 |
| ENSG00000197701 | 4 | 53179 | 88099 | 35 | 6 | 0.29859 | 0.38263 | ZNF595 | 0.38263 | 0.38791 | 0.33454 |
| ENSG00000140057 | 14 | 96858448 | 96955764 | 304 | 36 | 0.29857 | 0.38263 | AK7 | 0.38263 | 0.2884 | 0.1054 |
| ENSG00000120868 | 12 | 99038919 | 99129204 | 183 | 27 | 0.29815 | 0.38279 | APAF1 | 0.38279 | 0.25338 | 0.222 |
| ENSG00000144837 | 3 | 119316689 | 119348658 | 145 | 14 | 0.29763 | 0.38299 | PLA1A | 0.38299 | 0.33555 | 0.12643 |
| ENSG00000115464 | 2 | 61414598 | 61697904 | 816 | 28 | 0.29746 | 0.38306 | USP34 | 0.38306 | 0.38788 | 0.4039 |
| ENSG00000188011 | 2 | 242811752 | 242815975 | 20 | 5 | 0.29745 | 0.38306 | CXXC11 | 0.38306 | 0.34966 | 0.21136 |
| ENSG00000255394 | 8 | 11618896 | 11620309 | 8 | 3 | 0.29739 | 0.38308 | C8orf49 | 0.38308 | 0.65007 | 0.49823 |
| ENSG00000156253 | 21 | 30376705 | 30391699 | 40 | 8 | 0.29731 | 0.38311 | RWDD2B | 0.38311 | 0.33842 | 0.38558 |
| ENSG00000268413 | 17 | 42335297 | 42355825 | 59 | 20 | 0.29708 | 0.3832 | AC003043.1 | 0.3832 | 0.40204 | 0.79021 |
| ENSG00000165006 | 9 | 34179003 | 34252521 | 217 | 20 | 0.29705 | 0.38322 | UBAP1 | 0.38322 | 0.34927 | 0.28578 |
| ENSG00000149295 | 11 | 113280318 | 113346413 | 191 | 26 | 0.29703 | 0.38322 | DRD2 | 0.38322 | 0.43623 | 0.36945 |
| ENSG00000197724 | 9 | 96338689 | 96441869 | 560 | 30 | 0.29696 | 0.38325 | PHF2 | 0.38325 | 0.40326 | 0.4271 |
| ENSG00000197361 | 15 | 63889552 | 63894627 | 12 | 6 | 0.29659 | 0.38339 | FBXL22 | 0.38339 | 0.61322 | 0.59388 |
| ENSG00000105497 | 19 | 52074551 | 52092991 | 82 | 18 | 0.29654 | 0.38341 | ZNF175 | 0.38341 | 0.097376 | 0.065527 |
| ENSG00000157911 | 1 | 2336236 | 2345236 | 28 | 9 | 0.29602 | 0.38361 | PEX10 | 0.38361 | 0.49636 | 0.79644 |
| ENSG00000186141 | 1 | 145592605 | 145611025 | 26 | 4 | 0.29578 | 0.3837 | POLR3C | 0.3837 | 0.4338 | 0.43128 |
| ENSG00000197296 | 20 | 42931478 | 42939809 | 16 | 5 | 0.29577 | 0.3837 | FITM2 | 0.3837 | 0.47658 | 0.53467 |
| ENSG00000188859 | 1 | 166026674 | 166136206 | 346 | 21 | 0.29551 | 0.3838 | FAM78B | 0.3838 | 0.41258 | 0.42349 |
| ENSG00000138193 | 10 | 95753746 | 96092580 | 737 | 62 | 0.29506 | 0.38398 | PLCE1 | 0.38398 | 0.41107 | 0.39129 |
| ENSG00000088782 | 20 | 138111 | 139804 | 16 | 1 | 0.29503 | 0.38398 | DEFB127 | 0.38398 | 0.35198 | 0.30873 |
| ENSG00000228083 | 9 | 21239201 | 21239978 | 1 | 1 | 0.29499 | 0.384 | IFNA14 | 0.384 | 0.38267 | 0.48203 |
| ENSG00000205155 | 19 | 36236015 | 36237911 | 4 | 2 | 0.29473 | 0.3841 | PSENEN | 0.3841 | 0.39955 | 0.51709 |
| ENSG00000185888 | 1 | 228003394 | 228034171 | 118 | 20 | 0.29464 | 0.38414 | PRSS38 | 0.38414 | 0.34028 | 0.31963 |
| ENSG00000161533 | 17 | 73937588 | 73975515 | 124 | 26 | 0.29449 | 0.38419 | ACOX1 | 0.38419 | 0.35289 | 0.2592 |
| ENSG00000132164 | 3 | 10857885 | 10982419 | 403 | 42 | 0.29444 | 0.38421 | SLC6A11 | 0.38421 | 0.44282 | 0.38094 |
| ENSG00000268702 | 14 | 23790839 | 23791484 | 3 | 2 | 0.29439 | 0.38423 | AL049829.1 | 0.38423 | 0.81581 | 0.88807 |
| ENSG00000153982 | 17 | 57297828 | 57353328 | 134 | 13 | 0.29435 | 0.38424 | GDPD1 | 0.38424 | 0.37325 | 0.51126 |
| ENSG00000104885 | 19 | 2164148 | 2232577 | 233 | 34 | 0.29417 | 0.38431 | DOT1L | 0.38431 | 0.37393 | 0.32987 |
| ENSG00000169902 | 7 | 65670186 | 65885530 | 522 | 18 | 0.29376 | 0.38447 | TPST1 | 0.38447 | 0.3867 | 0.34271 |
| ENSG00000133105 | 13 | 32313674 | 32377009 | 205 | 19 | 0.29352 | 0.38456 | RXFP2 | 0.38456 | 0.46672 | 0.59643 |
| ENSG00000137648 | 11 | 117947753 | 117992605 | 262 | 24 | 0.29339 | 0.38461 | TMPRSS4 | 0.38461 | 0.41339 | 0.36716 |
| ENSG00000174521 | 19 | 40721965 | 40724306 | 5 | 2 | 0.29329 | 0.38465 | TTC9B | 0.38465 | 0.17468 | 0.23718 |
| ENSG00000176834 | 12 | 118501398 | 118573831 | 329 | 48 | 0.29325 | 0.38467 | VSIG10 | 0.38467 | 0.54322 | 0.61648 |
| ENSG00000124391 | 16 | 88704999 | 88706881 | 4 | 2 | 0.29321 | 0.38468 | IL17C | 0.38468 | 0.22599 | 0.26915 |
| ENSG00000052802 | 4 | 166248775 | 166264312 | 75 | 16 | 0.29316 | 0.3847 | MSMO1 | 0.3847 | 0.35227 | 0.18845 |
| ENSG00000180479 | 19 | 38045684 | 38085673 | 107 | 13 | 0.29292 | 0.38479 | ZNF571 | 0.38479 | 0.4047 | 0.3873 |

| | | | | | | | | | | | |
|---|---|---|---|---|---|---|---|---|---|---|---|
| ENSG00000182827 | 1 | 226332380 | 226374431 | 107 | 15 | 0.29285 | 0.38482 | ACBD3 | 0.38482 | 0.47524 | 0.58515 |
| ENSG00000136870 | 9 | 104161155 | 104172942 | 49 | 7 | 0.29279 | 0.38484 | ZNF189 | 0.38484 | 0.48887 | 0.51602 |
| ENSG00000122718 | 9 | 35957105 | 35958151 | 4 | 1 | 0.29264 | 0.3849 | OR2S2 | 0.3849 | 0.47278 | 0.47339 |
| ENSG00000170324 | 10 | 49364601 | 49482941 | 216 | 23 | 0.29229 | 0.38503 | FRMPD2 | 0.38503 | 0.44357 | 0.46004 |
| ENSG00000105254 | 19 | 36605191 | 36616849 | 33 | 8 | 0.29221 | 0.38506 | TBCB | 0.38506 | 0.32666 | 0.25276 |
| ENSG00000120647 | 12 | 498439 | 551811 | 201 | 36 | 0.29188 | 0.38519 | CCDC77 | 0.38519 | 0.35547 | 0.43969 |
| ENSG00000136231 | 7 | 23349828 | 23510086 | 419 | 45 | 0.29183 | 0.38521 | IGF2BP3 | 0.38521 | 0.38117 | 0.42087 |
| ENSG00000087085 | 7 | 100487615 | 100494594 | 15 | 6 | 0.29143 | 0.38536 | ACHE | 0.38536 | 0.32121 | 0.38324 |
| ENSG00000180269 | 16 | 20042807 | 20085239 | 152 | 31 | 0.2914 | 0.38537 | GPR139 | 0.38537 | 0.40331 | 0.41694 |
| ENSG00000157343 | 6 | 35704809 | 35716856 | 33 | 7 | 0.29125 | 0.38543 | ARMC12 | 0.38543 | 0.51937 | 0.70293 |
| ENSG00000214782 | 11 | 60496789 | 60511685 | 25 | 4 | 0.2908 | 0.3856 | MS4A18 | 0.3856 | 0.4718 | 0.47059 |
| ENSG00000070061 | 9 | 111629797 | 111696396 | 321 | 26 | 0.2902 | 0.38583 | IKBKAP | 0.38583 | 0.45498 | 0.44904 |
| ENSG00000125831 | 20 | 23431040 | 23433494 | 5 | 3 | 0.29016 | 0.38585 | CST11 | 0.38585 | 0.70318 | 0.74762 |
| ENSG00000072609 | 12 | 133398773 | 133532890 | 485 | 32 | 0.29002 | 0.3859 | CHFR | 0.3859 | 0.38562 | 0.43241 |
| ENSG00000134077 | 3 | 9404526 | 9428475 | 35 | 16 | 0.28985 | 0.38596 | THUMPD3 | 0.38596 | 0.48506 | 0.20307 |
| ENSG00000258864 | 5 | 112162910 | 112203279 | 149 | 18 | 0.28978 | 0.38599 | CTC-554D6.1 | 0.38599 | 0.38736 | 0.42844 |
| ENSG00000021776 | 15 | 35147732 | 35262040 | 323 | 20 | 0.28939 | 0.38614 | AQR | 0.38614 | 0.39398 | 0.42953 |
| ENSG00000117519 | 1 | 95362507 | 95392834 | 141 | 30 | 0.28936 | 0.38615 | CNN3 | 0.38615 | 0.46843 | 0.19237 |
| ENSG00000146109 | 6 | 26597180 | 26600278 | 12 | 4 | 0.28937 | 0.38615 | ABT1 | 0.38615 | 0.54732 | 0.67084 |
| ENSG00000142621 | 1 | 15573768 | 15726779 | 537 | 56 | 0.28935 | 0.38616 | FHAD1 | 0.38616 | 0.41803 | 0.39795 |
| ENSG00000187905 | 22 | 21400249 | 21418457 | 87 | 24 | 0.28916 | 0.38623 | AC002472.13 | 0.38623 | 0.68893 | 0.88495 |
| ENSG00000103449 | 16 | 51169886 | 51185278 | 23 | 11 | 0.28902 | 0.38628 | SALL1 | 0.38628 | 0.63756 | 0.66017 |
| ENSG00000171595 | 17 | 72270386 | 72311023 | 152 | 26 | 0.28878 | 0.38638 | DNAI2 | 0.38638 | 0.48089 | 0.74683 |
| ENSG00000149925 | 16 | 30064411 | 30081778 | 16 | 6 | 0.28865 | 0.38643 | ALDOA | 0.38643 | 0.64506 | 0.76757 |
| ENSG00000177054 | 11 | 19138646 | 19197969 | 154 | 16 | 0.28857 | 0.38646 | ZDHHC13 | 0.38646 | 0.4359 | 0.28167 |
| ENSG00000110934 | 12 | 51674822 | 51718452 | 182 | 13 | 0.28855 | 0.38646 | BIN2 | 0.38646 | 0.44474 | 0.4804 |
| ENSG00000269215 | 5 | 39105358 | 39107350 | 2 | 1 | 0.2882 | 0.3866 | AC008964.1 | 0.3866 | 0.51212 | 0.48905 |
| ENSG00000143013 | 1 | 87794151 | 87814606 | 57 | 18 | 0.28815 | 0.38661 | LMO4 | 0.38661 | 0.35957 | 0.30454 |
| ENSG00000197991 | 13 | 61983818 | 62002220 | 69 | 12 | 0.28813 | 0.38662 | PCDH20 | 0.38662 | 0.4634 | 0.49998 |
| ENSG00000179097 | 3 | 88039846 | 88042919 | 5 | 3 | 0.28808 | 0.38664 | HTR1F | 0.38664 | 0.72655 | 0.66466 |
| ENSG00000170786 | 8 | 57212569 | 57233335 | 46 | 10 | 0.28793 | 0.3867 | SDR16C5 | 0.3867 | 0.65201 | 0.7322 |
| ENSG00000141127 | 17 | 18743398 | 18834581 | 360 | 16 | 0.28787 | 0.38672 | PRPSAP2 | 0.38672 | 0.36636 | 0.36171 |
| ENSG00000013588 | 12 | 13043716 | 13070871 | 98 | 18 | 0.28784 | 0.38673 | GPRC5A | 0.38673 | 0.28311 | 0.4308 |
| ENSG00000213859 | 17 | 7255208 | 7258258 | 5 | 3 | 0.28785 | 0.38673 | KCTD11 | 0.38673 | 0.61199 | 0.92768 |
| ENSG00000259332 | 15 | 80137537 | 80216096 | 338 | 25 | 0.28779 | 0.38675 | ST20-MTHFS | 0.38675 | 0.42111 | 0.42348 |
| ENSG00000240303 | 3 | 132276982 | 132379567 | 196 | 23 | 0.28761 | 0.38682 | ACAD11 | 0.38682 | 0.45142 | 0.48517 |
| ENSG00000170011 | 3 | 39850405 | 40301812 | 1679 | 65 | 0.28758 | 0.38684 | MYRIP | 0.38684 | 0.39225 | 0.41616 |
| ENSG00000213782 | 12 | 12966250 | 12982915 | 60 | 15 | 0.28749 | 0.38687 | DDX47 | 0.38687 | 0.2464 | 0.51177 |
| ENSG00000265690 | 16 | 67197288 | 67199718 | 2 | 1 | 0.28736 | 0.38692 | RP11-5A19.5 | 0.38692 | 0.91013 | 0.91496 |
| ENSG00000133639 | 12 | 92536286 | 92539673 | 10 | 5 | 0.28727 | 0.38695 | BTG1 | 0.38695 | 0.1691 | 0.1049 |
| ENSG00000204334 | 2 | 171640296 | 171655481 | 58 | 8 | 0.28663 | 0.3872 | ERICH2 | 0.3872 | 0.32607 | 0.47058 |
| ENSG00000184939 | 16 | 68563993 | 68609975 | 226 | 9 | 0.28661 | 0.3872 | ZFP90 | 0.3872 | 0.36935 | 0.35415 |
| ENSG00000128283 | 22 | 37956454 | 37965412 | 26 | 7 | 0.28658 | 0.38722 | CDC42EP1 | 0.38722 | 0.25332 | 0.30152 |
| ENSG00000147697 | 8 | 130760442 | 130799134 | 132 | 21 | 0.28645 | 0.38727 | GSDMC | 0.38727 | 0.17368 | 0.067944 |
| ENSG00000065989 | 19 | 10527449 | 10580305 | 105 | 15 | 0.28631 | 0.38732 | PDE4A | 0.38732 | 0.45579 | 0.57175 |
| ENSG00000130876 | 19 | 33699570 | 33716756 | 59 | 24 | 0.28615 | 0.38738 | SLC7A10 | 0.38738 | 0.58053 | 0.7459 |
| ENSG00000140280 | 15 | 52015208 | 52043782 | 104 | 11 | 0.28596 | 0.38745 | LYSMD2 | 0.38745 | 0.39462 | 0.38038 |
| ENSG00000204033 | 10 | 85980254 | 85985345 | 17 | 4 | 0.28587 | 0.38749 | LRIT2 | 0.38749 | 0.34074 | 0.15849 |
| ENSG00000143416 | 1 | 151336778 | 151345209 | 24 | 13 | 0.28567 | 0.38757 | SELENBP1 | 0.38757 | 0.75329 | 0.80793 |
| ENSG00000163378 | 3 | 69024365 | 69063112 | 115 | 12 | 0.28552 | 0.38762 | EOGT | 0.38762 | 0.45358 | 0.51616 |
| ENSG00000167700 | 8 | 145734457 | 145736596 | 1 | 1 | 0.28532 | 0.3877 | MFSD3 | 0.3877 | 0.57724 | 0.54477 |
| ENSG00000149534 | 11 | 59855734 | 59863444 | 18 | 3 | 0.28532 | 0.3877 | MS4A2 | 0.3877 | 0.098141 | 0.13253 |
| ENSG00000187210 | 9 | 79034752 | 79122332 | 329 | 29 | 0.28508 | 0.38779 | GCNT1 | 0.38779 | 0.39867 | 0.39117 |
| ENSG00000133985 | 14 | 71108504 | 71142077 | 94 | 24 | 0.28508 | 0.38779 | TTC9 | 0.38779 | 0.70461 | 0.6576 |
| ENSG00000183760 | 19 | 39574553 | 39602133 | 136 | 14 | 0.28503 | 0.38781 | PAPL | 0.38781 | 0.25506 | 0.20288 |
| ENSG00000178852 | 17 | 45400656 | 45518678 | 430 | 18 | 0.28465 | 0.38796 | EFCAB13 | 0.38796 | 0.40554 | 0.41681 |
| ENSG00000179213 | 19 | 51749603 | 51772584 | 82 | 12 | 0.28461 | 0.38797 | SIGLECL1 | 0.38797 | 0.35793 | 0.38066 |
| ENSG00000167941 | 17 | 41831099 | 41836156 | 4 | 2 | 0.2846 | 0.38798 | SOST | 0.38798 | 0.3521 | 0.17098 |
| ENSG00000121270 | 16 | 48200821 | 48281479 | 279 | 15 | 0.28445 | 0.38803 | ABCC11 | 0.38803 | 0.3787 | 0.38595 |
| ENSG00000007202 | 17 | 26941458 | 26972472 | 31 | 13 | 0.28445 | 0.38803 | KIAA0100 | 0.38803 | 0.30152 | 0.19931 |
| ENSG00000136883 | 9 | 116848406 | 116861507 | 54 | 12 | 0.28419 | 0.38813 | KIF12 | 0.38813 | 0.54082 | 0.55786 |
| ENSG00000164163 | 4 | 146019084 | 146050331 | 28 | 11 | 0.28391 | 0.38824 | ABCE1 | 0.38824 | 0.45543 | 0.49697 |
| ENSG00000138684 | 4 | 123533783 | 123542224 | 19 | 6 | 0.28369 | 0.38832 | IL21 | 0.38832 | 0.33291 | 0.26197 |
| ENSG00000178764 | 8 | 123793633 | 123986750 | 496 | 88 | 0.28345 | 0.38842 | ZHX2 | 0.38842 | 0.45698 | 0.5323 |
| ENSG00000250317 | 4 | 25863452 | 25931435 | 225 | 30 | 0.28263 | 0.38873 | SMIM20 | 0.38873 | 0.40693 | 0.45293 |
| ENSG00000175806 | 8 | 9911778 | 10286401 | 1605 | 85 | 0.28259 | 0.38874 | MSRA | 0.38874 | 0.40676 | 0.42523 |
| ENSG00000119737 | 2 | 54080050 | 54087126 | 22 | 9 | 0.28237 | 0.38883 | GPR75 | 0.38883 | 0.37109 | 0.18658 |
| ENSG00000185442 | 15 | 93160673 | 93353114 | 985 | 57 | 0.28228 | 0.38886 | FAM174B | 0.38886 | 0.36271 | 0.37771 |
| ENSG00000011114 | 14 | 93703896 | 93799438 | 336 | 19 | 0.28189 | 0.38901 | BTBD7 | 0.38901 | 0.40952 | 0.47878 |
| ENSG00000020129 | 1 | 36023074 | 36032875 | 13 | 4 | 0.28173 | 0.38908 | NCDN | 0.38908 | 0.46861 | 0.37605 |
| ENSG00000141540 | 17 | 72209653 | 72258155 | 186 | 21 | 0.28164 | 0.38911 | TTYH2 | 0.38911 | 0.2389 | 0.38724 |
| ENSG00000177875 | 12 | 48577366 | 48579709 | 8 | 4 | 0.28161 | 0.38912 | C12orf68 | 0.38912 | 0.47416 | 0.52916 |
| ENSG00000166825 | 15 | 90328120 | 90358633 | 130 | 16 | 0.28156 | 0.38914 | ANPEP | 0.38914 | 0.3489 | 0.37077 |

| | | | | | | | | | | | |
|---|---|---|---|---|---|---|---|---|---|---|---|
| ENSG00000174938 | 16 | 29882480 | 29910868 | 42 | 8 | 0.28133 | 0.38923 | SEZ6L2 | 0.38923 | 0.56032 | 0.632 |
| ENSG00000178586 | 2 | 240984494 | 240985489 | 4 | 2 | 0.28128 | 0.38925 | OR6B3 | 0.38925 | 0.0823 | 0.057494 |
| ENSG00000009307 | 1 | 115259534 | 115301297 | 91 | 13 | 0.28122 | 0.38927 | CSDE1 | 0.38927 | 0.40599 | 0.47173 |
| ENSG00000138468 | 3 | 101043049 | 101232085 | 603 | 12 | 0.28113 | 0.38931 | SENP7 | 0.38931 | 0.39902 | 0.40567 |
| ENSG00000142748 | 1 | 27695603 | 27701315 | 14 | 7 | 0.28094 | 0.38938 | FCN3 | 0.38938 | 0.18911 | 0.14435 |
| ENSG00000120158 | 9 | 4792869 | 4885917 | 283 | 36 | 0.28058 | 0.38951 | RCL1 | 0.38951 | 0.50649 | 0.3984 |
| ENSG00000168710 | 1 | 110527308 | 110566357 | 46 | 13 | 0.28058 | 0.38952 | AHCYL1 | 0.38952 | 0.38474 | 0.17521 |
| ENSG00000002746 | 7 | 43152198 | 43605600 | 1606 | 117 | 0.28042 | 0.38958 | HECW1 | 0.38958 | 0.36855 | 0.33307 |
| ENSG00000126368 | 17 | 38249040 | 38256978 | 17 | 6 | 0.28036 | 0.3896 | NR1D1 | 0.3896 | 0.57746 | 0.48414 |
| ENSG00000265352 | 18 | 69399851 | 69455364 | 319 | 17 | 0.28026 | 0.38964 | RP11-723G8.2 | 0.38964 | 0.411 | 0.4888 |
| ENSG00000163661 | 3 | 157154578 | 157161417 | 9 | 4 | 0.28023 | 0.38965 | PTX3 | 0.38965 | 0.46208 | 0.14551 |
| ENSG00000147364 | 8 | 356428 | 421225 | 314 | 31 | 0.27993 | 0.38976 | FBXO25 | 0.38976 | 0.37484 | 0.38885 |
| ENSG00000177706 | 7 | 192969 | 300711 | 432 | 47 | 0.27977 | 0.38983 | FAM20C | 0.38983 | 0.3469 | 0.34059 |
| ENSG00000120832 | 12 | 107371069 | 107380944 | 24 | 8 | 0.27972 | 0.38984 | MTERFD3 | 0.38984 | 0.71764 | 0.59662 |
| ENSG00000114383 | 3 | 50357458 | 50365682 | 7 | 4 | 0.27947 | 0.38994 | TUSC2 | 0.38994 | 0.33354 | 0.31229 |
| ENSG00000268466 | 14 | 58754751 | 58755865 | 2 | 1 | 0.27935 | 0.38999 | AL132989.1 | 0.38999 | 0.22833 | 0.2103 |
| ENSG00000142556 | 19 | 52516018 | 52533493 | 48 | 9 | 0.27913 | 0.39007 | ZNF614 | 0.39007 | 0.33183 | 0.15499 |
| ENSG00000156735 | 8 | 38034051 | 38070819 | 71 | 4 | 0.27905 | 0.3901 | BAG4 | 0.3901 | 0.42805 | 0.46969 |
| ENSG00000107643 | 10 | 49514698 | 49647403 | 334 | 22 | 0.27906 | 0.3901 | MAPK8 | 0.3901 | 0.39554 | 0.36353 |
| ENSG00000100100 | 22 | 31677579 | 31688520 | 34 | 7 | 0.27905 | 0.3901 | PIK3IP1 | 0.3901 | 0.3365 | 0.38438 |
| ENSG00000156502 | 10 | 70939988 | 70968855 | 87 | 12 | 0.27899 | 0.39013 | SUPV3L1 | 0.39013 | 0.50226 | 0.51414 |
| ENSG00000267157 | 19 | 5784016 | 5785630 | 8 | 3 | 0.27891 | 0.39016 | CTB-54O9.9 | 0.39016 | 0.3491 | 0.44836 |
| ENSG00000182816 | 21 | 31743709 | 31744557 | 4 | 2 | 0.27885 | 0.39018 | KRTAP13-2 | 0.39018 | 0.1974 | 0.27371 |
| ENSG00000179477 | 17 | 7975954 | 7991021 | 62 | 18 | 0.27875 | 0.39022 | ALOX12B | 0.39022 | 0.23325 | 0.17337 |
| ENSG00000133794 | 11 | 13298199 | 13408813 | 306 | 40 | 0.27852 | 0.3903 | ARNTL | 0.3903 | 0.40884 | 0.42732 |
| ENSG00000154240 | 17 | 63631656 | 64188202 | 2457 | 35 | 0.27836 | 0.39037 | CEP112 | 0.39037 | 0.40435 | 0.4335 |
| ENSG00000214435 | 10 | 104629273 | 104661656 | 139 | 16 | 0.27817 | 0.39044 | AS3MT | 0.39044 | 0.43951 | 0.50933 |
| ENSG00000119537 | 18 | 60994959 | 61034743 | 101 | 14 | 0.27798 | 0.39051 | KDSR | 0.39051 | 0.31208 | 0.35311 |
| ENSG00000180011 | 18 | 72907063 | 72921303 | 38 | 8 | 0.27793 | 0.39053 | ZADH2 | 0.39053 | 0.39696 | 0.36874 |
| ENSG00000160679 | 1 | 153606525 | 153618782 | 33 | 5 | 0.27789 | 0.39055 | CHTOP | 0.39055 | 0.4142 | 0.41851 |
| ENSG00000221972 | 3 | 133646992 | 133648656 | 1 | 1 | 0.27776 | 0.3906 | C3orf36 | 0.3906 | 0.67037 | 0.58171 |
| ENSG00000196549 | 3 | 154741913 | 154901497 | 436 | 55 | 0.27774 | 0.3906 | MME | 0.3906 | 0.46825 | 0.4723 |
| ENSG00000134363 | 5 | 52776239 | 52782964 | 15 | 8 | 0.27773 | 0.39061 | FST | 0.39061 | 0.8549 | 0.6344 |
| ENSG00000100505 | 14 | 51441980 | 51562779 | 529 | 44 | 0.2776 | 0.39066 | TRIM9 | 0.39066 | 0.37536 | 0.23832 |
| ENSG00000118971 | 12 | 4382938 | 4414516 | 75 | 21 | 0.2775 | 0.3907 | CCND2 | 0.3907 | 0.46427 | 0.33283 |
| ENSG00000158615 | 1 | 204372515 | 204380919 | 15 | 4 | 0.27731 | 0.39077 | PPP1R15B | 0.39077 | 0.016245 | 0.0011013 |
| ENSG00000130304 | 19 | 17579578 | 17616977 | 139 | 18 | 0.27729 | 0.39078 | SLC27A1 | 0.39078 | 0.51665 | 0.5726 |
| ENSG00000269871 | 17 | 6915928 | 6917755 | 1 | 1 | 0.27723 | 0.3908 | AC040977.1 | 0.3908 | 0.61696 | 0.78945 |
| ENSG00000183624 | 3 | 128997671 | 129025029 | 63 | 4 | 0.27707 | 0.39086 | HMCES | 0.39086 | 0.38039 | 0.47151 |
| ENSG00000132423 | 6 | 99817276 | 99842080 | 100 | 14 | 0.27707 | 0.39086 | COQ3 | 0.39086 | 0.32434 | 0.33409 |
| ENSG00000131747 | 17 | 38544768 | 38574202 | 57 | 6 | 0.27709 | 0.39086 | TOP2A | 0.39086 | 0.27782 | 0.34212 |
| ENSG00000135245 | 7 | 128095903 | 128098472 | 9 | 4 | 0.2769 | 0.39093 | HILPDA | 0.39093 | 0.69065 | 0.7107 |
| ENSG00000003436 | 2 | 188328957 | 188430487 | 306 | 32 | 0.27678 | 0.39097 | TFPI | 0.39097 | 0.44332 | 0.43652 |
| ENSG00000204687 | 6 | 29454474 | 29455738 | 3 | 1 | 0.27679 | 0.39097 | MAS1L | 0.39097 | 0.63397 | 0.76068 |
| ENSG00000141622 | 18 | 43906772 | 44043103 | 428 | 69 | 0.27671 | 0.391 | RNF165 | 0.391 | 0.36297 | 0.37953 |
| ENSG00000214193 | 1 | 36771988 | 36790484 | 29 | 8 | 0.27657 | 0.39106 | SH3D21 | 0.39106 | 0.32009 | 0.28871 |
| ENSG00000166130 | 12 | 99007183 | 99038891 | 100 | 20 | 0.2765 | 0.39108 | IKBIP | 0.39108 | 0.36437 | 0.3538 |
| ENSG00000186230 | 19 | 57946697 | 57956853 | 20 | 8 | 0.27634 | 0.39114 | ZNF749 | 0.39114 | 0.68305 | 0.60333 |
| ENSG00000180043 | 19 | 55866276 | 55874628 | 43 | 10 | 0.27619 | 0.3912 | FAM71E2 | 0.3912 | 0.4278 | 0.44416 |
| ENSG00000164049 | 3 | 48413709 | 48442666 | 64 | 2 | 0.27612 | 0.39123 | FBXW12 | 0.39123 | 0.35754 | 0.3005 |
| ENSG00000091106 | 2 | 32449522 | 32490923 | 114 | 14 | 0.27601 | 0.39127 | NLRC4 | 0.39127 | 0.40829 | 0.42045 |
| ENSG00000153976 | 17 | 13399006 | 13505244 | 461 | 48 | 0.27599 | 0.39128 | HS3ST3A1 | 0.39128 | 0.37358 | 0.42149 |
| ENSG00000213297 | 19 | 12242932 | 12267546 | 48 | 6 | 0.2759 | 0.39131 | ZNF625-ZNF20 | 0.39131 | 0.2682 | 0.20393 |
| ENSG00000140386 | 15 | 76640526 | 77197785 | 1138 | 23 | 0.27576 | 0.39137 | SCAPER | 0.39137 | 0.37906 | 0.36708 |
| ENSG00000116478 | 1 | 32757687 | 32799236 | 32 | 20 | 0.27572 | 0.39138 | HDAC1 | 0.39138 | 0.63636 | 0.73106 |
| ENSG00000056661 | 17 | 36890150 | 36906070 | 50 | 13 | 0.27552 | 0.39146 | PCGF2 | 0.39146 | 0.4396 | 0.36396 |
| ENSG00000180998 | 14 | 53019866 | 53104431 | 236 | 28 | 0.27544 | 0.39149 | GPR137C | 0.39149 | 0.43861 | 0.43899 |
| ENSG00000090554 | 19 | 49977464 | 49989488 | 48 | 5 | 0.27539 | 0.39151 | FLT3LG | 0.39151 | 0.41052 | 0.17586 |
| ENSG00000139610 | 12 | 51722227 | 51740463 | 70 | 11 | 0.27522 | 0.39157 | CELA1 | 0.39157 | 0.39727 | 0.36313 |
| ENSG00000228727 | 6 | 31730576 | 31732628 | 6 | 4 | 0.27512 | 0.39161 | SAPCD1 | 0.39161 | 0.63294 | 0.68384 |
| ENSG00000119953 | 10 | 112050488 | 112064709 | 30 | 8 | 0.27467 | 0.39178 | SMNDC1 | 0.39178 | 0.45392 | 0.59214 |
| ENSG00000184814 | 3 | 138737873 | 138739768 | 1 | 1 | 0.27463 | 0.3918 | PRR23B | 0.3918 | 0.067534 | 0.12905 |
| ENSG00000147592 | 8 | 71547553 | 71581409 | 129 | 17 | 0.27463 | 0.3918 | LACTB2 | 0.3918 | 0.39121 | 0.33989 |
| ENSG00000026751 | 1 | 160709037 | 160724611 | 33 | 6 | 0.2746 | 0.39181 | SLAMF7 | 0.39181 | 0.40925 | 0.34631 |
| ENSG00000150337 | 1 | 149754227 | 149764074 | 9 | 2 | 0.27456 | 0.39183 | FCGR1A | 0.39183 | 0.43736 | 0.4095 |
| ENSG00000122873 | 10 | 60028818 | 60049346 | 79 | 8 | 0.27456 | 0.39183 | CISD1 | 0.39183 | 0.32698 | 0.20452 |
| ENSG00000006652 | 7 | 112063023 | 112121072 | 182 | 27 | 0.27446 | 0.39187 | IFRD1 | 0.39187 | 0.32922 | 0.25732 |
| ENSG00000172244 | 5 | 43486803 | 43515247 | 76 | 11 | 0.27424 | 0.39195 | C5orf34 | 0.39195 | 0.41505 | 0.4652 |
| ENSG00000197414 | 15 | 22736246 | 22746002 | 2 | 1 | 0.27423 | 0.39195 | GOLGA6L1 | 0.39195 | 0.4001 | 0.39405 |
| ENSG00000090539 | 3 | 184097861 | 184108524 | 26 | 9 | 0.27419 | 0.39197 | CHRD | 0.39197 | 0.057275 | 0.093509 |
| ENSG00000148019 | 9 | 80850978 | 80894606 | 84 | 12 | 0.27401 | 0.39204 | CEP78 | 0.39204 | 0.46055 | 0.60714 |
| ENSG00000136839 | 9 | 107379529 | 107380485 | 5 | 2 | 0.27381 | 0.39211 | OR13C9 | 0.39211 | 0.41893 | 0.33065 |
| ENSG00000112164 | 6 | 39016574 | 39055519 | 199 | 26 | 0.2738 | 0.39212 | GLP1R | 0.39212 | 0.42553 | 0.57576 |

| | | | | | | | | | | | |
|---|---|---|---|---|---|---|---|---|---|---|---|
| ENSG00000160117 | 19 | 17392454 | 17398455 | 37 | 5 | 0.27378 | 0.39213 | ANKLE1 | 0.39213 | 0.47382 | 0.57022 |
| ENSG00000213699 | 2 | 26987152 | 27004099 | 30 | 4 | 0.27348 | 0.39224 | SLC35F6 | 0.39224 | 0.59215 | 0.53461 |
| ENSG00000133884 | 11 | 65101225 | 65120720 | 42 | 11 | 0.27338 | 0.39228 | DPF2 | 0.39228 | 0.56433 | 0.71767 |
| ENSG00000170522 | 4 | 110967002 | 111120355 | 626 | 63 | 0.2731 | 0.39239 | ELOVL6 | 0.39239 | 0.34664 | 0.46313 |
| ENSG00000133606 | 7 | 140152840 | 140179369 | 101 | 7 | 0.2731 | 0.39239 | MKRN1 | 0.39239 | 0.39983 | 0.34733 |
| ENSG00000096433 | 6 | 33588142 | 33664351 | 393 | 42 | 0.27298 | 0.39243 | ITPR3 | 0.39243 | 0.41472 | 0.4751 |
| ENSG00000147570 | 8 | 66933795 | 67012751 | 255 | 35 | 0.27274 | 0.39253 | DNAJC5B | 0.39253 | 0.24388 | 0.16833 |
| ENSG00000187553 | 10 | 94821021 | 94828454 | 20 | 6 | 0.27244 | 0.39264 | CYP26C1 | 0.39264 | 0.50848 | 0.59283 |
| ENSG00000133574 | 7 | 150264365 | 150271041 | 19 | 4 | 0.27238 | 0.39267 | GIMAP4 | 0.39267 | 0.46123 | 0.58495 |
| ENSG00000102445 | 13 | 46916139 | 47012325 | 244 | 18 | 0.27237 | 0.39267 | KIAA0226L | 0.39267 | 0.44889 | 0.45605 |
| ENSG00000034510 | 2 | 85132749 | 85133795 | 3 | 1 | 0.27227 | 0.39271 | TMSB10 | 0.39271 | 0.62914 | 0.56513 |
| ENSG00000118197 | 1 | 200593024 | 200639097 | 176 | 22 | 0.27217 | 0.39275 | DDX59 | 0.39275 | 0.54222 | 0.42268 |
| ENSG00000156687 | 8 | 35092975 | 35654068 | 1272 | 51 | 0.27193 | 0.39284 | UNC5D | 0.39284 | 0.40272 | 0.42432 |
| ENSG00000214510 | 5 | 147647743 | 147665817 | 37 | 6 | 0.27114 | 0.39314 | SPINK13 | 0.39314 | 0.26304 | 0.18431 |
| ENSG00000143850 | 1 | 204187979 | 204346793 | 503 | 35 | 0.27093 | 0.39322 | PLEKHA6 | 0.39322 | 0.41247 | 0.38572 |
| ENSG00000188322 | 16 | 28303840 | 28335170 | 56 | 15 | 0.27092 | 0.39323 | SBK1 | 0.39323 | 0.27311 | 0.13564 |
| ENSG00000170365 | 4 | 146402346 | 146479231 | 137 | 16 | 0.27087 | 0.39324 | SMAD1 | 0.39324 | 0.33695 | 0.30134 |
| ENSG00000122417 | 1 | 86814344 | 86862023 | 167 | 18 | 0.27082 | 0.39326 | ODF2L | 0.39326 | 0.41494 | 0.40255 |
| ENSG00000048545 | 6 | 42123144 | 42147794 | 139 | 23 | 0.27081 | 0.39327 | GUCA1A | 0.39327 | 0.29558 | 0.47776 |
| ENSG00000164190 | 5 | 36876861 | 37066515 | 285 | 23 | 0.27075 | 0.39329 | NIPBL | 0.39329 | 0.39896 | 0.36447 |
| ENSG00000142082 | 11 | 215458 | 236931 | 154 | 14 | 0.27048 | 0.3934 | SIRT3 | 0.3934 | 0.43608 | 0.31082 |
| ENSG00000146006 | 5 | 138204612 | 138211057 | 10 | 3 | 0.27024 | 0.39349 | LRRTM2 | 0.39349 | 0.38557 | 0.35677 |
| ENSG00000116044 | 2 | 178092323 | 178257425 | 498 | 33 | 0.26997 | 0.39359 | NFE2L2 | 0.39359 | 0.40016 | 0.434 |
| ENSG00000188643 | 1 | 153579362 | 153585621 | 12 | 3 | 0.26991 | 0.39361 | S100A16 | 0.39361 | 0.40096 | 0.45178 |
| ENSG00000124812 | 6 | 49801970 | 49844809 | 81 | 15 | 0.26923 | 0.39388 | CRISP1 | 0.39388 | 0.30584 | 0.37408 |
| ENSG00000217702 | 2 | 74361599 | 74362521 | 5 | 3 | 0.26918 | 0.3939 | MGC10955 | 0.3939 | 0.71148 | 0.62333 |
| ENSG00000164048 | 3 | 48282590 | 48340743 | 106 | 7 | 0.26912 | 0.39392 | ZNF589 | 0.39392 | 0.38648 | 0.36265 |
| ENSG00000103855 | 15 | 73976307 | 74006859 | 97 | 13 | 0.26906 | 0.39394 | CD276 | 0.39394 | 0.45805 | 0.47853 |
| ENSG00000166595 | 16 | 66965959 | 66968326 | 3 | 2 | 0.26896 | 0.39398 | FAM96B | 0.39398 | 0.54672 | 0.33057 |
| ENSG00000253787 | 5 | 67485704 | 67494235 | 31 | 8 | 0.26832 | 0.39423 | RP11-404L6.2 | 0.39423 | 0.72566 | 0.67734 |
| ENSG00000143033 | 1 | 93544792 | 93604638 | 105 | 18 | 0.26824 | 0.39426 | MTF2 | 0.39426 | 0.4646 | 0.31649 |
| ENSG00000167671 | 19 | 4444996 | 4457819 | 41 | 8 | 0.26817 | 0.39429 | UBXN6 | 0.39429 | 0.52618 | 0.38404 |
| ENSG00000146122 | 6 | 39760142 | 39872648 | 416 | 50 | 0.26808 | 0.39432 | DAAM2 | 0.39432 | 0.33245 | 0.32584 |
| ENSG00000272333 | 19 | 36208921 | 36229779 | 47 | 12 | 0.26778 | 0.39443 | KMT2B | 0.39443 | 0.4854 | 0.53126 |
| ENSG00000184602 | 16 | 11762270 | 11773015 | 49 | 8 | 0.26764 | 0.39449 | SNN | 0.39449 | 0.46164 | 0.69855 |
| ENSG00000260371 | 16 | 69368994 | 69390209 | 45 | 12 | 0.2675 | 0.39454 | RP11-343C2.9 | 0.39454 | 0.30136 | 0.29771 |
| ENSG00000186198 | 15 | 65337708 | 65345734 | 15 | 4 | 0.26749 | 0.39455 | SLC51B | 0.39455 | 0.32011 | 0.23031 |
| ENSG00000133488 | 22 | 30884900 | 30901698 | 83 | 12 | 0.26744 | 0.39456 | SEC14L4 | 0.39456 | 0.41378 | 0.51254 |
| ENSG00000151348 | 11 | 44117099 | 44266979 | 273 | 25 | 0.26738 | 0.39459 | EXT2 | 0.39459 | 0.40301 | 0.45383 |
| ENSG00000100604 | 14 | 93389425 | 93401638 | 49 | 10 | 0.26733 | 0.39461 | CHGA | 0.39461 | 0.56546 | 0.35333 |
| ENSG00000170113 | 15 | 23043277 | 23100005 | 264 | 27 | 0.26726 | 0.39464 | NIPA1 | 0.39464 | 0.28526 | 0.19698 |
| ENSG00000237787 | 3 | 153202284 | 153220486 | 83 | 10 | 0.26712 | 0.39469 | C3orf79 | 0.39469 | 0.47447 | 0.48519 |
| ENSG00000006576 | 7 | 77428122 | 77586818 | 400 | 23 | 0.26707 | 0.39471 | PHTF2 | 0.39471 | 0.45263 | 0.5047 |
| ENSG00000171798 | 10 | 134973951 | 135039916 | 379 | 27 | 0.26697 | 0.39475 | KNDC1 | 0.39475 | 0.49895 | 0.43362 |
| ENSG00000088827 | 20 | 3667617 | 3687775 | 76 | 13 | 0.26685 | 0.39479 | SIGLEC1 | 0.39479 | 0.22865 | 0.16166 |
| ENSG00000127366 | 7 | 141490017 | 141491166 | 4 | 2 | 0.26674 | 0.39483 | TAS2R5 | 0.39483 | 0.35591 | 0.40519 |
| ENSG00000119912 | 10 | 94211441 | 94333833 | 353 | 28 | 0.26594 | 0.39514 | IDE | 0.39514 | 0.4528 | 0.42783 |
| ENSG00000105443 | 19 | 48972289 | 48985571 | 17 | 6 | 0.26586 | 0.39517 | CYTH2 | 0.39517 | 0.16238 | 0.15184 |
| ENSG00000168806 | 15 | 43619974 | 43622803 | 4 | 2 | 0.26575 | 0.39522 | LCMT2 | 0.39522 | 0.55817 | 0.59891 |
| ENSG00000119777 | 2 | 27255778 | 27264563 | 7 | 3 | 0.26559 | 0.39528 | TMEM214 | 0.39528 | 0.44353 | 0.6266 |
| ENSG00000092345 | 3 | 16628299 | 16711813 | 446 | 61 | 0.26558 | 0.39528 | DAZL | 0.39528 | 0.31775 | 0.30362 |
| ENSG00000169903 | 3 | 149191761 | 149221068 | 237 | 22 | 0.26555 | 0.39529 | TM4SF4 | 0.39529 | 0.49208 | 0.62962 |
| ENSG00000205318 | 6 | 10633993 | 10647501 | 17 | 7 | 0.26547 | 0.39532 | GCNT6 | 0.39532 | 0.39952 | 0.62934 |
| ENSG00000070961 | 12 | 89981828 | 90103077 | 233 | 26 | 0.26543 | 0.39534 | ATP2B1 | 0.39534 | 0.39407 | 0.33844 |
| ENSG00000161243 | 19 | 39481354 | 39523425 | 231 | 18 | 0.26539 | 0.39536 | FBXO27 | 0.39536 | 0.3188 | 0.41718 |
| ENSG00000130656 | 16 | 202686 | 204502 | 6 | 3 | 0.26527 | 0.3954 | HBZ | 0.3954 | 0.35865 | 0.43132 |
| ENSG00000268126 | 5 | 147698550 | 147701891 | 14 | 3 | 0.26511 | 0.39546 | AC091948.1 | 0.39546 | 0.35386 | 0.35803 |
| ENSG00000111897 | 6 | 122764499 | 122792967 | 85 | 13 | 0.2651 | 0.39546 | SERINC1 | 0.39546 | 0.43243 | 0.42511 |
| ENSG00000147813 | 8 | 144656955 | 144660819 | 22 | 6 | 0.26505 | 0.39548 | NAPRT1 | 0.39548 | 0.47681 | 0.57932 |
| ENSG00000198408 | 10 | 103544200 | 103578696 | 50 | 10 | 0.26499 | 0.39551 | MGEA5 | 0.39551 | 0.46711 | 0.51467 |
| ENSG00000115461 | 2 | 217536828 | 217560248 | 61 | 8 | 0.26481 | 0.39558 | IGFBP5 | 0.39558 | 0.49283 | 0.70414 |
| ENSG00000196604 | 2 | 130831108 | 130886795 | 84 | 5 | 0.26476 | 0.3956 | POTEF | 0.3956 | 0.41474 | 0.38777 |
| ENSG00000198729 | 6 | 150464212 | 150571493 | 489 | 66 | 0.26475 | 0.3956 | PPP1R14C | 0.3956 | 0.36968 | 0.36752 |
| ENSG00000121858 | 3 | 172223298 | 172241297 | 89 | 13 | 0.26469 | 0.39562 | TNFSF10 | 0.39562 | 0.40315 | 0.55659 |
| ENSG00000212935 | 21 | 45977673 | 45978643 | 8 | 3 | 0.26458 | 0.39567 | KRTAP10-3 | 0.39567 | 0.12708 | 0.32703 |
| ENSG00000137834 | 15 | 66994566 | 67074338 | 204 | 31 | 0.26443 | 0.39572 | SMAD6 | 0.39572 | 0.33345 | 0.46258 |
| ENSG00000166603 | 18 | 58038564 | 58040001 | 3 | 2 | 0.26422 | 0.39581 | MC4R | 0.39581 | 0.010223 | 0.01188 |
| ENSG00000177370 | 17 | 900357 | 906911 | 22 | 7 | 0.26384 | 0.39595 | TIMM22 | 0.39595 | 0.40018 | 0.64562 |
| ENSG00000101577 | 18 | 2916992 | 3013313 | 342 | 28 | 0.26374 | 0.39599 | LPIN2 | 0.39599 | 0.24826 | 0.20292 |
| ENSG00000179294 | 17 | 36827961 | 36831187 | 3 | 2 | 0.26355 | 0.39606 | C17orf96 | 0.39606 | 0.30063 | 0.23128 |
| ENSG00000010295 | 12 | 6647541 | 6665239 | 57 | 8 | 0.26348 | 0.39609 | IFFO1 | 0.39609 | 0.41876 | 0.51811 |
| ENSG00000181481 | 17 | 29295803 | 29326929 | 34 | 6 | 0.26314 | 0.39622 | RNF135 | 0.39622 | 0.35516 | 0.36039 |
| ENSG00000182944 | 22 | 29663998 | 29696515 | 76 | 19 | 0.26288 | 0.39632 | EWSR1 | 0.39632 | 0.44143 | 0.61008 |

| | | | | | | | | | | | |
|---|---|---|---|---|---|---|---|---|---|---|---|
| ENSG00000181038 | 17 | 74722912 | 74730018 | 6 | 3 | 0.26279 | 0.39636 | METTL23 | 0.39636 | 0.91156 | 0.79215 |
| ENSG00000137962 | 1 | 94614544 | 94740624 | 245 | 34 | 0.26265 | 0.39641 | ARHGAP29 | 0.39641 | 0.46425 | 0.39768 |
| ENSG00000010319 | 3 | 52467069 | 52479101 | 18 | 8 | 0.26253 | 0.39645 | SEMA3G | 0.39645 | 0.41227 | 0.44145 |
| ENSG00000180336 | 17 | 42733762 | 42767676 | 46 | 15 | 0.26246 | 0.39648 | C17orf104 | 0.39648 | 0.51577 | 0.54003 |
| ENSG00000255012 | 11 | 56380031 | 56380978 | 2 | 1 | 0.26179 | 0.39674 | OR5M1 | 0.39674 | 0.1234 | 0.028046 |
| ENSG00000169515 | 19 | 46913629 | 46916841 | 9 | 5 | 0.2618 | 0.39674 | CCDC8 | 0.39674 | 0.52502 | 0.24802 |
| ENSG00000163898 | 3 | 185224050 | 185270401 | 141 | 15 | 0.26159 | 0.39682 | LIPH | 0.39682 | 0.35184 | 0.39096 |
| ENSG00000148248 | 9 | 136228325 | 136242970 | 41 | 7 | 0.26122 | 0.39696 | SURF4 | 0.39696 | 0.41443 | 0.35035 |
| ENSG00000105974 | 7 | 116164839 | 116201233 | 117 | 16 | 0.26094 | 0.39707 | CAV1 | 0.39707 | 0.36921 | 0.35854 |
| ENSG00000186907 | 11 | 57228022 | 57245007 | 21 | 8 | 0.26092 | 0.39708 | RTN4RL2 | 0.39708 | 0.37128 | 0.36526 |
| ENSG00000136247 | 7 | 6617065 | 6629005 | 33 | 5 | 0.26089 | 0.39709 | ZDHHC4 | 0.39709 | 0.45079 | 0.45206 |
| ENSG00000131374 | 3 | 17198654 | 18486309 | 3152 | 53 | 0.26085 | 0.3971 | TBC1D5 | 0.3971 | 0.40491 | 0.39834 |
| ENSG00000135093 | 12 | 109460894 | 109525831 | 90 | 16 | 0.26083 | 0.39711 | USP30 | 0.39711 | 0.38454 | 0.45859 |
| ENSG00000189139 | 14 | 44973545 | 44976482 | 8 | 3 | 0.26073 | 0.39715 | FSCB | 0.39715 | 0.14396 | 0.14165 |
| ENSG00000167139 | 15 | 74165949 | 74181555 | 48 | 11 | 0.26056 | 0.39722 | TBC1D21 | 0.39722 | 0.496 | 0.53384 |
| ENSG00000076003 | 2 | 136597196 | 136633996 | 79 | 10 | 0.26053 | 0.39723 | MCM6 | 0.39723 | 0.41126 | 0.47413 |
| ENSG00000163666 | 3 | 57231944 | 57260549 | 51 | 9 | 0.2601 | 0.39739 | HESX1 | 0.39739 | 0.45571 | 0.51875 |
| ENSG00000203756 | 6 | 130152389 | 130182692 | 216 | 21 | 0.26012 | 0.39739 | TMEM244 | 0.39739 | 0.45185 | 0.32873 |
| ENSG00000165244 | 9 | 99148223 | 99180611 | 105 | 9 | 0.26004 | 0.39742 | ZNF367 | 0.39742 | 0.43137 | 0.43246 |
| ENSG00000178607 | 17 | 62116502 | 62208179 | 165 | 19 | 0.26 | 0.39743 | ERN1 | 0.39743 | 0.45068 | 0.54069 |
| ENSG00000060558 | 19 | 3136191 | 3163767 | 221 | 32 | 0.25998 | 0.39744 | GNA15 | 0.39744 | 0.43399 | 0.29126 |
| ENSG00000189325 | 6 | 36283534 | 36304662 | 63 | 22 | 0.25996 | 0.39745 | C6orf222 | 0.39745 | 0.29426 | 0.57777 |
| ENSG00000165695 | 9 | 135600965 | 135754164 | 494 | 38 | 0.25994 | 0.39746 | AK8 | 0.39746 | 0.40532 | 0.36426 |
| ENSG00000108010 | 10 | 131934663 | 131982785 | 258 | 21 | 0.25951 | 0.39762 | GLRX3 | 0.39762 | 0.43929 | 0.047137 |
| ENSG00000205089 | 5 | 132083137 | 132089856 | 7 | 2 | 0.25947 | 0.39764 | CCNI2 | 0.39764 | 0.58994 | 0.60594 |
| ENSG00000101104 | 20 | 43538703 | 43587676 | 108 | 19 | 0.2592 | 0.39774 | PABPC1L | 0.39774 | 0.57867 | 0.66536 |
| ENSG00000254318 | 8 | 94146324 | 94179079 | 88 | 17 | 0.25895 | 0.39784 | C8orf87 | 0.39784 | 0.20844 | 0.30973 |
| ENSG00000125820 | 20 | 21491648 | 21494664 | 9 | 4 | 0.25891 | 0.39785 | NKX2-2 | 0.39785 | 0.012849 | 0.075048 |
| ENSG00000148291 | 9 | 136223428 | 136228045 | 15 | 4 | 0.25889 | 0.39786 | SURF2 | 0.39786 | 0.5195 | 0.43533 |
| ENSG00000145428 | 4 | 154631277 | 154681387 | 230 | 16 | 0.25886 | 0.39787 | RNF175 | 0.39787 | 0.44441 | 0.45954 |
| ENSG00000172345 | 15 | 81601394 | 81616524 | 51 | 13 | 0.25884 | 0.39788 | STARD5 | 0.39788 | 0.025627 | 0.03466 |
| ENSG00000120029 | 10 | 103605356 | 103815950 | 338 | 28 | 0.25873 | 0.39792 | C10orf76 | 0.39792 | 0.4134 | 0.39806 |
| ENSG00000155304 | 21 | 15743436 | 15755805 | 56 | 11 | 0.25858 | 0.39798 | HSPA13 | 0.39798 | 0.29124 | 0.35434 |
| ENSG00000184113 | 22 | 19510547 | 19515068 | 19 | 7 | 0.25856 | 0.39799 | CLDN5 | 0.39799 | 0.47555 | 0.55312 |
| ENSG00000163633 | 4 | 87797358 | 87857354 | 190 | 20 | 0.25853 | 0.398 | C4orf36 | 0.398 | 0.42749 | 0.41976 |
| ENSG00000105697 | 19 | 35771619 | 35776046 | 14 | 5 | 0.25842 | 0.39804 | HAMP | 0.39804 | 0.17269 | 0.14874 |
| ENSG00000127334 | 12 | 68042118 | 68059186 | 32 | 6 | 0.25835 | 0.39807 | DYRK2 | 0.39807 | 0.43944 | 0.55697 |
| ENSG00000268805 | 21 | 46511596 | 46515493 | 18 | 6 | 0.25833 | 0.39808 | PRED57 | 0.39808 | 0.53593 | 0.5823 |
| ENSG00000137831 | 15 | 70946893 | 71055932 | 184 | 15 | 0.25828 | 0.39809 | UACA | 0.39809 | 0.51257 | 0.70679 |
| ENSG00000188060 | 1 | 28918712 | 28921955 | 9 | 5 | 0.25824 | 0.39811 | RAB42 | 0.39811 | 0.52167 | 0.51719 |
| ENSG00000167676 | 19 | 4502204 | 4517716 | 97 | 21 | 0.25825 | 0.39811 | PLIN4 | 0.39811 | 0.30547 | 0.30795 |
| ENSG00000104320 | 8 | 90945564 | 91015456 | 203 | 13 | 0.25822 | 0.39812 | NBN | 0.39812 | 0.38729 | 0.44752 |
| ENSG00000182255 | 11 | 30031288 | 30038570 | 12 | 5 | 0.25818 | 0.39813 | KCNA4 | 0.39813 | 0.50228 | 0.46857 |
| ENSG00000143228 | 1 | 163236366 | 163325554 | 445 | 35 | 0.25809 | 0.39817 | NUF2 | 0.39817 | 0.47391 | 0.48506 |
| ENSG00000083937 | 3 | 87276421 | 87304698 | 85 | 17 | 0.25786 | 0.39826 | CHMP2B | 0.39826 | 0.32003 | 0.32334 |
| ENSG00000179066 | 19 | 35596873 | 35598141 | 12 | 5 | 0.25775 | 0.3983 | AC020907.1 | 0.3983 | 0.20335 | 0.50621 |
| ENSG00000062582 | 7 | 43906157 | 43909492 | 5 | 2 | 0.25767 | 0.39833 | MRPS24 | 0.39833 | 0.29933 | 0.32943 |
| ENSG00000196550 | 1 | 206136916 | 206155151 | 11 | 6 | 0.25761 | 0.39835 | FAM72A | 0.39835 | 0.34828 | 0.34236 |
| ENSG00000236279 | 7 | 139208602 | 139229730 | 35 | 6 | 0.2576 | 0.39836 | CLEC2L | 0.39836 | 0.57664 | 0.41507 |
| ENSG00000006740 | 17 | 12692856 | 12894960 | 938 | 63 | 0.25744 | 0.39842 | ARHGAP44 | 0.39842 | 0.30944 | 0.31795 |
| ENSG00000147471 | 8 | 37620111 | 37637283 | 10 | 6 | 0.25743 | 0.39843 | PROSC | 0.39843 | 0.69734 | 0.94233 |
| ENSG00000008282 | 7 | 105730949 | 105753022 | 64 | 16 | 0.25739 | 0.39844 | SYPL1 | 0.39844 | 0.63701 | 0.61341 |
| ENSG00000196139 | 10 | 5077546 | 5149878 | 457 | 44 | 0.25736 | 0.39845 | AKR1C3 | 0.39845 | 0.39893 | 0.57755 |
| ENSG00000161036 | 7 | 102105376 | 102113615 | 59 | 9 | 0.25733 | 0.39846 | LRWD1 | 0.39846 | 0.47522 | 0.4752 |
| ENSG00000156675 | 8 | 37716136 | 37756985 | 43 | 14 | 0.25726 | 0.39849 | RAB11FIP1 | 0.39849 | 0.073957 | 0.11534 |
| ENSG00000268313 | 1 | 205682497 | 205684153 | 4 | 2 | 0.25722 | 0.39851 | AC119673.1 | 0.39851 | 0.60234 | 0.69781 |
| ENSG00000133687 | 12 | 29653773 | 29937692 | 1237 | 84 | 0.25666 | 0.39872 | TMTC1 | 0.39872 | 0.2504 | 0.21331 |
| ENSG00000072274 | 3 | 195754054 | 195809060 | 280 | 35 | 0.25658 | 0.39875 | TFRC | 0.39875 | 0.36782 | 0.51888 |
| ENSG00000115844 | 2 | 172964167 | 172967628 | 7 | 5 | 0.25647 | 0.39879 | DLX2 | 0.39879 | 0.022956 | 0.025555 |
| ENSG00000085274 | 3 | 169490619 | 169507504 | 24 | 5 | 0.25645 | 0.3988 | MYNN | 0.3988 | 0.38665 | 0.38226 |
| ENSG00000088832 | 20 | 1349622 | 1373806 | 64 | 12 | 0.25635 | 0.39884 | FKBP1A | 0.39884 | 0.60197 | 0.48727 |
| ENSG00000198846 | 8 | 59717977 | 60031767 | 943 | 85 | 0.25631 | 0.39886 | TOX | 0.39886 | 0.37412 | 0.53764 |
| ENSG00000227450 | 17 | 36905613 | 36906969 | 4 | 2 | 0.25623 | 0.39889 | CTB-58E17.5 | 0.39889 | 0.59441 | 0.35294 |
| ENSG00000155850 | 5 | 149340300 | 149373018 | 54 | 8 | 0.25606 | 0.39895 | SLC26A2 | 0.39895 | 0.5096 | 0.5241 |
| ENSG00000165802 | 9 | 140342022 | 140353786 | 18 | 9 | 0.25595 | 0.399 | NSMF | 0.399 | 0.90051 | 0.89153 |
| ENSG00000214670 | 14 | 75158879 | 75179800 | 35 | 5 | 0.25586 | 0.39903 | AC007956.1 | 0.39903 | 0.42729 | 0.50318 |
| ENSG00000103479 | 16 | 53467889 | 53525561 | 144 | 11 | 0.25574 | 0.39908 | RBL2 | 0.39908 | 0.41941 | 0.39766 |
| ENSG00000169288 | 4 | 78783674 | 78873944 | 187 | 21 | 0.25546 | 0.39919 | MRPL1 | 0.39919 | 0.44873 | 0.39351 |
| ENSG00000103512 | 16 | 14927538 | 14990017 | 58 | 18 | 0.25545 | 0.39919 | NOMO1 | 0.39919 | 0.27311 | 0.38337 |
| ENSG00000157106 | 16 | 18816175 | 18937776 | 391 | 16 | 0.25538 | 0.39922 | SMG1 | 0.39922 | 0.4151 | 0.40546 |
| ENSG00000121053 | 17 | 56270098 | 56282535 | 23 | 8 | 0.25523 | 0.39927 | EPX | 0.39927 | 0.71885 | 0.58695 |
| ENSG00000205683 | 14 | 73086004 | 73360809 | 950 | 77 | 0.25498 | 0.39937 | DPF3 | 0.39937 | 0.33144 | 0.37073 |
| ENSG00000111206 | 12 | 2966847 | 2986206 | 41 | 11 | 0.25478 | 0.39945 | FOXM1 | 0.39945 | 0.50636 | 0.42124 |

| | | | | | | | | | | | |
|---|---|---|---|---|---|---|---|---|---|---|---|
| ENSG00000213145 | 14 | 105952654 | 105955284 | 10 | 4 | 0.2546 | 0.39952 | CRIP1 | 0.39952 | 0.27612 | 0.34525 |
| ENSG00000105723 | 19 | 42734338 | 42746777 | 13 | 3 | 0.25458 | 0.39952 | GSK3A | 0.39952 | 0.53587 | 0.60598 |
| ENSG00000099377 | 16 | 30996519 | 31000473 | 8 | 3 | 0.25425 | 0.39965 | HSD3B7 | 0.39965 | 0.71166 | 0.76292 |
| ENSG00000087095 | 17 | 26368763 | 26523407 | 96 | 36 | 0.25405 | 0.39973 | NLK | 0.39973 | 0.40604 | 0.36126 |
| ENSG00000136104 | 13 | 51483814 | 51544592 | 130 | 23 | 0.25397 | 0.39976 | RNASEH2B | 0.39976 | 0.21213 | 0.10537 |
| ENSG00000071073 | 2 | 99235569 | 99347589 | 262 | 23 | 0.25391 | 0.39978 | MGAT4A | 0.39978 | 0.35308 | 0.30973 |
| ENSG00000134809 | 11 | 57295936 | 57298276 | 3 | 1 | 0.25371 | 0.39986 | TIMM10 | 0.39986 | 0.30128 | 0.50487 |
| ENSG00000106397 | 7 | 100849258 | 100861701 | 28 | 8 | 0.2536 | 0.3999 | PLOD3 | 0.3999 | 0.51376 | 0.62689 |
| ENSG00000198865 | 5 | 42756903 | 42802462 | 152 | 16 | 0.25355 | 0.39992 | CCDC152 | 0.39992 | 0.4 | 0.41781 |
| ENSG00000188549 | 15 | 40623653 | 40633168 | 26 | 7 | 0.25349 | 0.39994 | C15orf52 | 0.39994 | 0.61369 | 0.71248 |
| ENSG00000175325 | 5 | 177419236 | 177423243 | 17 | 6 | 0.25348 | 0.39995 | PROP1 | 0.39995 | 0.62325 | 0.37034 |
| ENSG00000182674 | 8 | 73449626 | 73850584 | 1253 | 72 | 0.25345 | 0.39996 | KCNB2 | 0.39996 | 0.29074 | 0.26229 |
| ENSG00000154478 | 10 | 125425871 | 125454123 | 108 | 21 | 0.25329 | 0.40002 | GPR26 | 0.40002 | 0.19652 | 0.28805 |
| ENSG00000179152 | 3 | 44379611 | 44450943 | 192 | 18 | 0.25315 | 0.40008 | TCAIM | 0.40008 | 0.41411 | 0.42122 |
| ENSG00000167842 | 17 | 5389605 | 5394134 | 24 | 7 | 0.25303 | 0.40012 | MIS12 | 0.40012 | 0.40144 | 0.30715 |
| ENSG00000162086 | 16 | 3355406 | 3368852 | 31 | 5 | 0.25267 | 0.40026 | ZNF75A | 0.40026 | 0.54938 | 0.55029 |
| ENSG00000255713 | 17 | 56247017 | 56247940 | 4 | 1 | 0.25253 | 0.40032 | OR4D2 | 0.40032 | 0.31648 | 0.45865 |
| ENSG00000164053 | 3 | 48488114 | 48507115 | 29 | 6 | 0.25241 | 0.40036 | ATRIP | 0.40036 | 0.39817 | 0.41122 |
| ENSG00000117305 | 1 | 24128375 | 24165110 | 55 | 4 | 0.25231 | 0.4004 | HMGCL | 0.4004 | 0.45499 | 0.56663 |
| ENSG00000079785 | 2 | 15731302 | 15771235 | 156 | 13 | 0.25232 | 0.4004 | DDX1 | 0.4004 | 0.40276 | 0.46916 |
| ENSG00000204513 | 1 | 12884468 | 12891264 | 45 | 9 | 0.25205 | 0.4005 | PRAMEF11 | 0.4005 | 0.11839 | 0.40639 |
| ENSG00000212998 | 8 | 94752349 | 94753001 | 1 | 1 | 0.25205 | 0.4005 | RBM12B-AS1 | 0.4005 | 0.35252 | 0.48098 |
| ENSG00000243317 | 7 | 135347244 | 135378166 | 108 | 16 | 0.25191 | 0.40056 | C7orf73 | 0.40056 | 0.36617 | 0.3087 |
| ENSG00000164794 | 8 | 110975874 | 110988076 | 22 | 7 | 0.25189 | 0.40056 | KCNV1 | 0.40056 | 0.58983 | 0.63858 |
| ENSG00000187550 | 19 | 56041100 | 56048456 | 42 | 9 | 0.25153 | 0.4007 | SBK2 | 0.4007 | 0.59872 | 0.20839 |
| ENSG00000143624 | 1 | 153700543 | 153746555 | 99 | 18 | 0.25149 | 0.40072 | INTS3 | 0.40072 | 0.47105 | 0.379 |
| ENSG00000113758 | 5 | 176883609 | 176901402 | 32 | 7 | 0.25111 | 0.40086 | DBN1 | 0.40086 | 0.56546 | 0.46356 |
| ENSG00000121578 | 3 | 118930579 | 118959950 | 131 | 18 | 0.2511 | 0.40087 | B4GALT4 | 0.40087 | 0.32841 | 0.18852 |
| ENSG00000050767 | 5 | 177664619 | 178017556 | 1676 | 143 | 0.25109 | 0.40087 | COL23A1 | 0.40087 | 0.27391 | 0.29244 |
| ENSG00000164080 | 3 | 51575160 | 51702683 | 133 | 32 | 0.25105 | 0.40089 | RAD54L2 | 0.40089 | 0.37348 | 0.43095 |
| ENSG00000122862 | 10 | 70847862 | 70864567 | 57 | 10 | 0.25061 | 0.40106 | SRGN | 0.40106 | 0.59565 | 0.63479 |
| ENSG00000213123 | 3 | 196018090 | 196045170 | 83 | 9 | 0.25014 | 0.40124 | TCTEX1D2 | 0.40124 | 0.19092 | 0.14827 |
| ENSG00000186073 | 15 | 36871812 | 37102449 | 677 | 50 | 0.25012 | 0.40125 | C15orf41 | 0.40125 | 0.40358 | 0.43776 |
| ENSG00000198719 | 6 | 170591294 | 170599561 | 17 | 7 | 0.25002 | 0.40129 | DLL1 | 0.40129 | 0.10979 | 0.10192 |
| ENSG00000273049 | 12 | 54379629 | 54428672 | 110 | 21 | 0.24993 | 0.40132 | RP11-834C11.12 | 0.40132 | 0.3951 | 0.44086 |
| ENSG00000203666 | 1 | 245133007 | 245290466 | 333 | 36 | 0.24971 | 0.40141 | EFCAB2 | 0.40141 | 0.35508 | 0.21105 |
| ENSG00000203872 | 6 | 88054567 | 88075181 | 76 | 7 | 0.24958 | 0.40146 | C6orf163 | 0.40146 | 0.30698 | 0.31665 |
| ENSG00000124193 | 20 | 42086568 | 42092245 | 14 | 5 | 0.24922 | 0.4016 | SRSF6 | 0.4016 | 0.32301 | 0.29355 |
| ENSG00000116288 | 1 | 8014351 | 8045565 | 89 | 15 | 0.24918 | 0.40161 | PARK7 | 0.40161 | 0.3755 | 0.3764 |
| ENSG00000162761 | 1 | 165171104 | 165325952 | 595 | 56 | 0.249 | 0.40168 | LMX1A | 0.40168 | 0.40267 | 0.46503 |
| ENSG00000166704 | 19 | 58488421 | 58514717 | 90 | 9 | 0.24879 | 0.40176 | ZNF606 | 0.40176 | 0.43749 | 0.49919 |
| ENSG00000147576 | 8 | 67342420 | 67383836 | 165 | 17 | 0.24873 | 0.40179 | ADHFE1 | 0.40179 | 0.25251 | 0.31677 |
| ENSG00000151640 | 10 | 134000404 | 134019280 | 87 | 15 | 0.24842 | 0.40191 | DPYSL4 | 0.40191 | 0.52444 | 0.57752 |
| ENSG00000213719 | 6 | 31698358 | 31707540 | 17 | 9 | 0.24836 | 0.40193 | CLIC1 | 0.40193 | 0.63753 | 0.70424 |
| ENSG00000166664 | 15 | 30653443 | 30686052 | 5 | 2 | 0.24828 | 0.40196 | CHRFAM7A | 0.40196 | 0.45905 | 0.3185 |
| ENSG00000105993 | 7 | 157128075 | 157210133 | 378 | 27 | 0.24826 | 0.40197 | DNAJB6 | 0.40197 | 0.31123 | 0.46927 |
| ENSG00000242866 | 15 | 43891596 | 44010458 | 188 | 15 | 0.24806 | 0.40204 | STRC | 0.40204 | 0.35286 | 0.33062 |
| ENSG00000239665 | 10 | 13652047 | 13710299 | 247 | 23 | 0.2477 | 0.40219 | RP11-295P9.3 | 0.40219 | 0.50232 | 0.54776 |
| ENSG00000183784 | 9 | 213108 | 215893 | 19 | 7 | 0.24759 | 0.40223 | C9orf66 | 0.40223 | 0.58036 | 0.53656 |
| ENSG00000113742 | 5 | 173315283 | 173388979 | 200 | 19 | 0.24754 | 0.40224 | CPEB4 | 0.40224 | 0.4242 | 0.40449 |
| ENSG00000267680 | 19 | 44598492 | 44612479 | 56 | 6 | 0.24743 | 0.40229 | ZNF224 | 0.40229 | 0.26522 | 0.32161 |
| ENSG00000100600 | 14 | 93170152 | 93215047 | 107 | 23 | 0.24739 | 0.4023 | LGMN | 0.4023 | 0.53142 | 0.41881 |
| ENSG00000205502 | 15 | 62455734 | 62457482 | 7 | 4 | 0.24739 | 0.4023 | C2CD4B | 0.4023 | 0.33461 | 0.60448 |
| ENSG00000196975 | 2 | 69871557 | 70053596 | 484 | 32 | 0.24734 | 0.40232 | ANXA4 | 0.40232 | 0.45478 | 0.50442 |
| ENSG00000128595 | 7 | 128379346 | 128411861 | 84 | 8 | 0.24726 | 0.40236 | CALU | 0.40236 | 0.53747 | 0.71446 |
| ENSG00000148218 | 9 | 116148597 | 116163613 | 47 | 15 | 0.24714 | 0.4024 | ALAD | 0.4024 | 0.26918 | 0.16449 |
| ENSG00000171428 | 8 | 18027986 | 18081198 | 312 | 33 | 0.2471 | 0.40241 | NAT1 | 0.40241 | 0.39244 | 0.41177 |
| ENSG00000136827 | 9 | 132575223 | 132586413 | 36 | 8 | 0.24698 | 0.40246 | TOR1A | 0.40246 | 0.41503 | 0.54838 |
| ENSG00000130751 | 19 | 47523077 | 47549033 | 104 | 33 | 0.24672 | 0.40256 | NPAS1 | 0.40256 | 0.59027 | 0.69918 |
| ENSG00000205307 | 7 | 100169855 | 100171270 | 2 | 1 | 0.24636 | 0.4027 | SAP25 | 0.4027 | 0.18968 | 0.15125 |
| ENSG00000127954 | 7 | 87905744 | 87936206 | 60 | 9 | 0.2462 | 0.40276 | STEAP4 | 0.40276 | 0.40505 | 0.39626 |
| ENSG00000100077 | 22 | 25960816 | 26125261 | 395 | 19 | 0.24612 | 0.4028 | ADRBK2 | 0.4028 | 0.38265 | 0.35368 |
| ENSG00000133106 | 13 | 43460524 | 43566407 | 412 | 52 | 0.24605 | 0.40282 | EPSTI1 | 0.40282 | 0.46965 | 0.61273 |
| ENSG00000156103 | 8 | 89044237 | 89340254 | 835 | 59 | 0.24569 | 0.40296 | MMP16 | 0.40296 | 0.32569 | 0.26772 |
| ENSG00000188493 | 19 | 41246761 | 41257458 | 44 | 10 | 0.24545 | 0.40305 | C19orf54 | 0.40305 | 0.36403 | 0.27585 |
| ENSG00000177932 | 5 | 178487416 | 178510538 | 55 | 13 | 0.24502 | 0.40322 | ZNF354C | 0.40322 | 0.58624 | 0.64019 |
| ENSG00000164976 | 9 | 34366668 | 34376851 | 26 | 7 | 0.24494 | 0.40325 | KIAA1161 | 0.40325 | 0.47798 | 0.44363 |
| ENSG00000141698 | 17 | 39981335 | 39992523 | 47 | 4 | 0.24473 | 0.40333 | NT5C3B | 0.40333 | 0.43142 | 0.44499 |
| ENSG00000092108 | 14 | 31091318 | 31205018 | 322 | 21 | 0.24462 | 0.40337 | SCFD1 | 0.40337 | 0.42906 | 0.45887 |
| ENSG00000205693 | 12 | 27915671 | 27924209 | 40 | 4 | 0.24456 | 0.4034 | MANSC4 | 0.4034 | 0.27764 | 0.28177 |
| ENSG00000166573 | 18 | 74962505 | 74980858 | 106 | 22 | 0.24454 | 0.40341 | GALR1 | 0.40341 | 0.12397 | 0.40556 |
| ENSG00000249160 | 5 | 10505108 | 10522196 | 80 | 11 | 0.24428 | 0.40351 | RP11-1C1.5 | 0.40351 | 0.44591 | 0.61656 |
| ENSG00000181619 | 14 | 59895740 | 59932060 | 141 | 14 | 0.24413 | 0.40356 | GPR135 | 0.40356 | 0.41722 | 0.32749 |

| | | | | | | | | | | | |
|---|---|---|---|---|---|---|---|---|---|---|---|
| ENSG00000178809 | 7 | 75024337 | 75040279 | 9 | 5 | 0.24381 | 0.40369 | TRIM73 | 0.40369 | 0.63937 | 0.58699 |
| ENSG00000167751 | 19 | 51364824 | 51383823 | 126 | 20 | 0.24368 | 0.40374 | KLK2 | 0.40374 | 0.67647 | 0.78283 |
| ENSG00000176532 | 7 | 29603427 | 29606911 | 15 | 7 | 0.24363 | 0.40376 | PRR15 | 0.40376 | 0.6268 | 0.70636 |
| ENSG00000116667 | 1 | 184356192 | 184598154 | 616 | 36 | 0.24358 | 0.40378 | C1orf21 | 0.40378 | 0.37173 | 0.40189 |
| ENSG00000130714 | 9 | 134378289 | 134399193 | 100 | 8 | 0.24295 | 0.40402 | POMT1 | 0.40402 | 0.27217 | 0.22434 |
| ENSG00000166352 | 11 | 36616051 | 36694823 | 235 | 18 | 0.2429 | 0.40404 | C11orf74 | 0.40404 | 0.3664 | 0.28852 |
| ENSG00000149313 | 11 | 105946228 | 105969437 | 44 | 9 | 0.24272 | 0.40411 | AASDHPPT | 0.40411 | 0.29171 | 0.3375 |
| ENSG00000163993 | 4 | 6694796 | 6698897 | 17 | 4 | 0.24267 | 0.40413 | S100P | 0.40413 | 0.43596 | 0.3625 |
| ENSG00000047410 | 1 | 186280784 | 186344825 | 142 | 14 | 0.24213 | 0.40434 | TPR | 0.40434 | 0.40375 | 0.40895 |
| ENSG00000254206 | 16 | 29392675 | 29415350 | 49 | 12 | 0.2421 | 0.40435 | NPIPB11 | 0.40435 | 0.34433 | 0.43454 |
| ENSG00000257341 | 14 | 105953536 | 105958604 | 8 | 4 | 0.24202 | 0.40438 | CRIP1 | 0.40438 | 0.27612 | 0.34525 |
| ENSG00000075407 | 10 | 38383264 | 38414472 | 147 | 9 | 0.24189 | 0.40443 | ZNF37A | 0.40443 | 0.45394 | 0.50095 |
| ENSG00000203797 | 6 | 110712974 | 110736765 | 110 | 21 | 0.24148 | 0.40459 | DDO | 0.40459 | 0.37264 | 0.58908 |
| ENSG00000115368 | 2 | 190306159 | 190340291 | 205 | 7 | 0.24118 | 0.40471 | WDR75 | 0.40471 | 0.41628 | 0.47817 |
| ENSG00000258708 | 14 | 37641093 | 37643016 | 5 | 3 | 0.241 | 0.40478 | SLC25A21-AS1 | 0.40478 | 0.73723 | 0.7039 |
| ENSG00000170802 | 2 | 48541776 | 48606433 | 256 | 18 | 0.24093 | 0.4048 | FOXN2 | 0.4048 | 0.37255 | 0.31914 |
| ENSG00000196415 | 19 | 840963 | 848175 | 46 | 8 | 0.24095 | 0.4048 | PRTN3 | 0.4048 | 0.50237 | 0.56772 |
| ENSG00000212993 | 8 | 128426535 | 128432311 | 39 | 7 | 0.24076 | 0.40487 | POU5F1B | 0.40487 | 0.79484 | 0.89481 |
| ENSG00000135476 | 12 | 53662083 | 53687427 | 56 | 8 | 0.24071 | 0.40489 | ESPL1 | 0.40489 | 0.55752 | 0.60973 |
| ENSG00000272305 | 3 | 53003135 | 53133469 | 392 | 19 | 0.24056 | 0.40495 | RP11-894J14.5 | 0.40495 | 0.41997 | 0.39619 |
| ENSG00000255837 | 12 | 11149094 | 11150474 | 12 | 2 | 0.2403 | 0.40505 | TAS2R20 | 0.40505 | 0.40355 | 0.43524 |
| ENSG00000151553 | 10 | 116581503 | 116659591 | 264 | 16 | 0.24027 | 0.40506 | FAM160B1 | 0.40506 | 0.34184 | 0.31657 |
| ENSG00000176904 | 11 | 4880886 | 4881794 | 2 | 1 | 0.24008 | 0.40513 | OR51H1P | 0.40513 | 0.69762 | 0.72789 |
| ENSG00000184224 | 11 | 67370351 | 67374177 | 9 | 3 | 0.2401 | 0.40513 | C11orf72 | 0.40513 | 0.60402 | 0.51934 |
| ENSG00000168874 | 2 | 85978467 | 86015189 | 88 | 15 | 0.24 | 0.40516 | ATOH8 | 0.40516 | 0.3533 | 0.27589 |
| ENSG00000152457 | 10 | 14939358 | 14996431 | 161 | 31 | 0.2398 | 0.40524 | DCLRE1C | 0.40524 | 0.53406 | 0.62521 |
| ENSG00000115415 | 2 | 191829084 | 191885686 | 163 | 25 | 0.23949 | 0.40536 | STAT1 | 0.40536 | 0.34954 | 0.30355 |
| ENSG00000196159 | 4 | 126237554 | 126414087 | 524 | 39 | 0.23946 | 0.40537 | FAT4 | 0.40537 | 0.43672 | 0.47176 |
| ENSG00000064999 | 6 | 34857042 | 35059179 | 559 | 37 | 0.23942 | 0.40539 | ANKS1A | 0.40539 | 0.40962 | 0.41024 |
| ENSG00000078098 | 2 | 163027194 | 163101661 | 136 | 10 | 0.23938 | 0.40541 | FAP | 0.40541 | 0.39893 | 0.43701 |
| ENSG00000171435 | 12 | 117890817 | 118406788 | 2130 | 191 | 0.23935 | 0.40542 | KSR2 | 0.40542 | 0.41548 | 0.35909 |
| ENSG00000108424 | 17 | 45726842 | 45762871 | 62 | 11 | 0.2392 | 0.40547 | KPNB1 | 0.40547 | 0.38428 | 0.39287 |
| ENSG00000177674 | 1 | 11796141 | 11814859 | 66 | 11 | 0.23887 | 0.4056 | AGTRAP | 0.4056 | 0.69887 | 0.83181 |
| ENSG00000160194 | 21 | 44299754 | 44333414 | 196 | 18 | 0.23882 | 0.40562 | NDUFV3 | 0.40562 | 0.40355 | 0.40892 |
| ENSG00000123353 | 12 | 56211703 | 56215663 | 3 | 2 | 0.23864 | 0.40569 | ORMDL2 | 0.40569 | 0.97351 | 0.98855 |
| ENSG00000267059 | 19 | 1578338 | 1605444 | 49 | 13 | 0.23854 | 0.40573 | UQCR11 | 0.40573 | 0.42987 | 0.50656 |
| ENSG00000198142 | 2 | 110371911 | 110376563 | 11 | 3 | 0.2382 | 0.40586 | SOWAHC | 0.40586 | 0.64301 | 0.67083 |
| ENSG00000118492 | 6 | 146920101 | 147136598 | 642 | 55 | 0.23819 | 0.40587 | ADGB | 0.40587 | 0.43872 | 0.41863 |
| ENSG00000160360 | 9 | 139221932 | 139254057 | 169 | 27 | 0.23804 | 0.40592 | GPSM1 | 0.40592 | 0.53329 | 0.50726 |
| ENSG00000091127 | 7 | 105080108 | 105162714 | 345 | 39 | 0.23791 | 0.40598 | PUS7 | 0.40598 | 0.57668 | 0.64984 |
| ENSG00000260234 | 3 | 150600784 | 150659530 | 236 | 49 | 0.23747 | 0.40615 | RP11-166N6.3 | 0.40615 | 0.35843 | 0.40561 |
| ENSG00000127837 | 2 | 219128850 | 219134980 | 12 | 4 | 0.23742 | 0.40617 | AAMP | 0.40617 | 0.44762 | 0.47212 |
| ENSG00000111540 | 12 | 56367697 | 56388490 | 42 | 6 | 0.23731 | 0.40621 | RAB5B | 0.40621 | 0.50044 | 0.62193 |
| ENSG00000242441 | 2 | 48844937 | 48960287 | 477 | 46 | 0.23711 | 0.40629 | GTF2A1L | 0.40629 | 0.22636 | 0.21845 |
| ENSG00000242485 | 1 | 1337288 | 1342693 | 18 | 7 | 0.237 | 0.40633 | MRPL20 | 0.40633 | 0.63888 | 0.66843 |
| ENSG00000196569 | 6 | 129204342 | 129837714 | 1978 | 83 | 0.23694 | 0.40635 | LAMA2 | 0.40635 | 0.40522 | 0.41563 |
| ENSG00000100288 | 22 | 51017378 | 51039884 | 121 | 22 | 0.23693 | 0.40636 | CHKB | 0.40636 | 0.51412 | 0.34315 |
| ENSG00000142892 | 1 | 77554675 | 77685115 | 412 | 23 | 0.2365 | 0.40652 | PIGK | 0.40652 | 0.43083 | 0.5086 |
| ENSG00000140451 | 15 | 65107831 | 65117867 | 53 | 9 | 0.23643 | 0.40655 | PIF1 | 0.40655 | 0.48742 | 0.52107 |
| ENSG00000114270 | 3 | 48601506 | 48632700 | 22 | 12 | 0.23639 | 0.40656 | COL7A1 | 0.40656 | 0.59565 | 0.61322 |
| ENSG00000113905 | 3 | 186378005 | 186396029 | 86 | 12 | 0.23626 | 0.40662 | HRG | 0.40662 | 0.37004 | 0.41409 |
| ENSG00000164953 | 8 | 94767072 | 94831462 | 87 | 18 | 0.23598 | 0.40672 | TMEM67 | 0.40672 | 0.38992 | 0.39552 |
| ENSG00000170486 | 12 | 52979373 | 52995322 | 86 | 13 | 0.23593 | 0.40674 | KRT72 | 0.40674 | 0.46834 | 0.52592 |
| ENSG00000151135 | 12 | 107349497 | 107372556 | 60 | 11 | 0.23577 | 0.4068 | C12orf23 | 0.4068 | 0.30769 | 0.31836 |
| ENSG00000119411 | 9 | 116111821 | 116133513 | 142 | 18 | 0.2356 | 0.40687 | BSPRY | 0.40687 | 0.41938 | 0.50263 |
| ENSG00000119919 | 10 | 101292690 | 101296278 | 10 | 3 | 0.23551 | 0.4069 | NKX2-3 | 0.4069 | 0.4639 | 0.53012 |
| ENSG00000176658 | 17 | 30819540 | 31204195 | 955 | 68 | 0.23551 | 0.4069 | MYO1D | 0.4069 | 0.40459 | 0.4242 |
| ENSG00000198739 | 10 | 68685764 | 68859588 | 530 | 40 | 0.2355 | 0.40691 | LRRTM3 | 0.40691 | 0.46208 | 0.52148 |
| ENSG00000164627 | 6 | 39297766 | 39693181 | 1286 | 61 | 0.23543 | 0.40694 | KIF6 | 0.40694 | 0.36292 | 0.41398 |
| ENSG00000259529 | 14 | 24620427 | 24636611 | 38 | 11 | 0.23531 | 0.40698 | RP11-468E2.4 | 0.40698 | 0.58952 | 0.74186 |
| ENSG00000203814 | 1 | 149754245 | 149783928 | 52 | 7 | 0.23494 | 0.40713 | HIST2H2BF | 0.40713 | 0.44916 | 0.4222 |
| ENSG00000176978 | 9 | 140004994 | 140009629 | 20 | 6 | 0.2349 | 0.40714 | DPP7 | 0.40714 | 0.47372 | 0.66845 |
| ENSG00000137807 | 15 | 69706585 | 69740764 | 43 | 9 | 0.23473 | 0.40721 | KIF23 | 0.40721 | 0.42915 | 0.42387 |
| ENSG00000267824 | 16 | 20499024 | 20500981 | 8 | 4 | 0.23448 | 0.4073 | AC137056.1 | 0.4073 | 0.33751 | 0.52984 |
| ENSG00000089060 | 12 | 113736564 | 113797298 | 198 | 19 | 0.23441 | 0.40733 | SLC8B1 | 0.40733 | 0.3718 | 0.29429 |
| ENSG00000197263 | 11 | 124189134 | 124190184 | 3 | 1 | 0.23392 | 0.40752 | OR8D2 | 0.40752 | 0.58462 | 0.68757 |
| ENSG00000178184 | 18 | 77915115 | 78005429 | 282 | 21 | 0.23381 | 0.40757 | PARD6G | 0.40757 | 0.44769 | 0.45539 |
| ENSG00000085265 | 9 | 137801431 | 137809809 | 29 | 4 | 0.23374 | 0.40759 | FCN1 | 0.40759 | 0.55807 | 0.33068 |
| ENSG00000160392 | 19 | 40825443 | 40854434 | 76 | 12 | 0.23363 | 0.40764 | C19orf47 | 0.40764 | 0.39589 | 0.25579 |
| ENSG00000267963 | 2 | 241053907 | 241064304 | 63 | 8 | 0.23347 | 0.4077 | AC013469.1 | 0.4077 | 0.56387 | 0.58385 |
| ENSG00000166863 | 12 | 57403784 | 57422667 | 33 | 6 | 0.23345 | 0.40771 | TAC3 | 0.40771 | 0.47379 | 0.40761 |
| ENSG00000141458 | 18 | 21086148 | 21166862 | 169 | 16 | 0.2332 | 0.4078 | NPC1 | 0.4078 | 0.25474 | 0.28475 |
| ENSG00000125810 | 20 | 23059986 | 23066977 | 13 | 5 | 0.23298 | 0.40789 | CD93 | 0.40789 | 0.1825 | 0.033366 |

| | | | | | | | | | | | |
|---|---|---|---|---|---|---|---|---|---|---|---|
| ENSG00000175832 | 17 | 41605212 | 41656988 | 131 | 13 | 0.23294 | 0.4079 | ETV4 | 0.4079 | 0.45269 | 0.4688 |
| ENSG00000143119 | 1 | 111415775 | 111442550 | 101 | 13 | 0.23249 | 0.40808 | CD53 | 0.40808 | 0.41978 | 0.25888 |
| ENSG00000160208 | 21 | 45079429 | 45115958 | 64 | 6 | 0.23217 | 0.4082 | RRP1B | 0.4082 | 0.41923 | 0.5073 |
| ENSG00000158435 | 2 | 101869264 | 101886778 | 36 | 10 | 0.23215 | 0.40821 | CNOT11 | 0.40821 | 0.38053 | 0.40338 |
| ENSG00000241690 | 20 | 2795614 | 2798712 | 12 | 7 | 0.23212 | 0.40822 | TMEM239 | 0.40822 | 0.36242 | 0.24016 |
| ENSG00000163933 | 3 | 53122499 | 53164478 | 123 | 10 | 0.23209 | 0.40823 | RFT1 | 0.40823 | 0.37654 | 0.45206 |
| ENSG00000174718 | 12 | 32112304 | 32146039 | 175 | 25 | 0.23202 | 0.40826 | KIAA1551 | 0.40826 | 0.42712 | 0.37516 |
| ENSG00000174827 | 1 | 145726918 | 145764074 | 24 | 9 | 0.23181 | 0.40834 | PDZK1 | 0.40834 | 0.62867 | 0.47859 |
| ENSG00000171817 | 19 | 38042308 | 38104998 | 151 | 17 | 0.2315 | 0.40846 | ZNF540 | 0.40846 | 0.38813 | 0.37686 |
| ENSG00000182393 | 19 | 39786965 | 39789313 | 4 | 2 | 0.23143 | 0.40849 | IFNL1 | 0.40849 | 0.15176 | 0.10622 |
| ENSG00000221946 | 19 | 35634154 | 35645204 | 28 | 10 | 0.23131 | 0.40854 | FXYD7 | 0.40854 | 0.49543 | 0.67468 |
| ENSG00000130294 | 2 | 241653181 | 241759725 | 328 | 48 | 0.23098 | 0.40866 | KIF1A | 0.40866 | 0.43365 | 0.47806 |
| ENSG00000110975 | 12 | 33527173 | 33592754 | 258 | 19 | 0.23098 | 0.40867 | SYT10 | 0.40867 | 0.41727 | 0.4535 |
| ENSG00000183921 | 16 | 22177618 | 22203075 | 47 | 10 | 0.23084 | 0.40872 | SDR42E2 | 0.40872 | 0.34296 | 0.30434 |
| ENSG00000118785 | 4 | 88896819 | 88904562 | 29 | 10 | 0.23074 | 0.40876 | SPP1 | 0.40876 | 0.11853 | 0.084855 |
| ENSG00000205106 | 11 | 45792983 | 45793909 | 5 | 3 | 0.23044 | 0.40888 | CTD-2210P24.4 | 0.40888 | 0.76104 | 0.76374 |
| ENSG00000140526 | 15 | 89630690 | 89745591 | 274 | 26 | 0.23031 | 0.40893 | ABHD2 | 0.40893 | 0.43707 | 0.4964 |
| ENSG00000214787 | 11 | 59968726 | 60010561 | 144 | 14 | 0.23022 | 0.40896 | MS4A4E | 0.40896 | 0.45294 | 0.51443 |
| ENSG00000139287 | 12 | 72332626 | 72580398 | 745 | 38 | 0.23015 | 0.40899 | TPH2 | 0.40899 | 0.42435 | 0.47151 |
| ENSG00000068489 | 17 | 57232860 | 57282066 | 103 | 15 | 0.23002 | 0.40904 | PRR11 | 0.40904 | 0.39696 | 0.27098 |
| ENSG00000134769 | 18 | 32073254 | 32471808 | 1018 | 29 | 0.22983 | 0.40911 | DTNA | 0.40911 | 0.42359 | 0.41811 |
| ENSG00000178623 | 2 | 241544848 | 241570676 | 119 | 36 | 0.22974 | 0.40915 | GPR35 | 0.40915 | 0.35489 | 0.27758 |
| ENSG00000134001 | 14 | 67826714 | 67853233 | 100 | 15 | 0.22947 | 0.40925 | EIF2S1 | 0.40925 | 0.39054 | 0.50525 |
| ENSG00000184378 | 3 | 169484709 | 169487683 | 9 | 2 | 0.22939 | 0.40928 | ACTRT3 | 0.40928 | 0.40548 | 0.32804 |
| ENSG00000160991 | 7 | 102073553 | 102097268 | 81 | 18 | 0.2294 | 0.40928 | ORAI2 | 0.40928 | 0.37528 | 0.48111 |
| ENSG00000128815 | 10 | 49892921 | 50191001 | 1160 | 79 | 0.2291 | 0.4094 | WDFY4 | 0.4094 | 0.4423 | 0.50827 |
| ENSG00000168476 | 8 | 21995533 | 21999464 | 11 | 3 | 0.22848 | 0.40964 | REEP4 | 0.40964 | 0.11891 | 0.1609 |
| ENSG00000175215 | 12 | 58213710 | 58240522 | 42 | 9 | 0.22829 | 0.40971 | CTDSP2 | 0.40971 | 0.51531 | 0.55914 |
| ENSG00000100344 | 22 | 44319619 | 44360368 | 188 | 19 | 0.2281 | 0.40978 | PNPLA3 | 0.40978 | 0.2771 | 0.22103 |
| ENSG00000233024 | 16 | 18451943 | 18473188 | 1 | 1 | 0.2278 | 0.4099 | NPIPA7 | 0.4099 | 0.073767 | 0.28034 |
| ENSG00000163636 | 3 | 63996225 | 64009658 | 46 | 12 | 0.22775 | 0.40992 | PSMD6 | 0.40992 | 0.33587 | 0.39702 |
| ENSG00000124215 | 20 | 58533471 | 58609066 | 228 | 36 | 0.2276 | 0.40998 | CDH26 | 0.40998 | 0.4338 | 0.2326 |
| ENSG00000101347 | 20 | 35518632 | 35580246 | 113 | 16 | 0.22751 | 0.41001 | SAMHD1 | 0.41001 | 0.35494 | 0.17145 |
| ENSG00000168310 | 4 | 185308867 | 185395734 | 355 | 69 | 0.22736 | 0.41007 | IRF2 | 0.41007 | 0.4546 | 0.23301 |
| ENSG00000168386 | 3 | 99548985 | 99833357 | 294 | 34 | 0.22717 | 0.41014 | FILIP1L | 0.41014 | 0.3982 | 0.37773 |
| ENSG00000213828 | 2 | 240083493 | 240084053 | 3 | 1 | 0.2271 | 0.41017 | AC017028.1 | 0.41017 | 0.48579 | 0.48165 |
| ENSG00000198633 | 19 | 52932440 | 52955568 | 193 | 9 | 0.22712 | 0.41017 | ZNF534 | 0.41017 | 0.25193 | 0.26107 |
| ENSG00000157077 | 1 | 52608046 | 52812358 | 365 | 23 | 0.22709 | 0.41018 | ZFYVE9 | 0.41018 | 0.38741 | 0.40194 |
| ENSG00000187860 | 22 | 30752624 | 30774647 | 82 | 8 | 0.22692 | 0.41024 | CCDC157 | 0.41024 | 0.46073 | 0.62632 |
| ENSG00000111639 | 12 | 6601150 | 6603007 | 15 | 2 | 0.22679 | 0.41029 | MRPL51 | 0.41029 | 0.40536 | 0.41319 |
| ENSG00000215397 | 20 | 642240 | 656823 | 41 | 12 | 0.22672 | 0.41032 | SCRT2 | 0.41032 | 0.41524 | 0.40577 |
| ENSG00000181072 | 7 | 136553416 | 136705002 | 518 | 32 | 0.22665 | 0.41035 | CHRM2 | 0.41035 | 0.44049 | 0.47754 |
| ENSG00000204428 | 6 | 31644461 | 31651817 | 19 | 7 | 0.2266 | 0.41037 | LY6G5C | 0.41037 | 0.38549 | 0.51793 |
| ENSG00000170748 | 11 | 7110165 | 7112379 | 6 | 2 | 0.22654 | 0.41039 | RBMXL2 | 0.41039 | 0.33683 | 0.30124 |
| ENSG00000214357 | 5 | 172068269 | 172118543 | 154 | 24 | 0.22651 | 0.4104 | NEURL1B | 0.4104 | 0.41725 | 0.44255 |
| ENSG00000013573 | 12 | 31226779 | 31257725 | 198 | 8 | 0.22631 | 0.41048 | DDX11 | 0.41048 | 0.42662 | 0.44592 |
| ENSG00000188786 | 1 | 38275239 | 38325292 | 99 | 19 | 0.22604 | 0.41059 | MTF1 | 0.41059 | 0.39385 | 0.44225 |
| ENSG00000182591 | 21 | 32252966 | 32253874 | 2 | 1 | 0.226 | 0.4106 | KRTAP11-1 | 0.4106 | 0.39185 | 0.69724 |
| ENSG00000088298 | 20 | 33703167 | 33865928 | 449 | 27 | 0.22596 | 0.41062 | EDEM2 | 0.41062 | 0.34681 | 0.3317 |
| ENSG00000091039 | 12 | 76745577 | 76953589 | 434 | 18 | 0.22593 | 0.41063 | OSBPL8 | 0.41063 | 0.39277 | 0.40144 |
| ENSG00000185033 | 15 | 90703836 | 90772911 | 309 | 20 | 0.22552 | 0.41079 | SEMA4B | 0.41079 | 0.42489 | 0.51318 |
| ENSG00000134539 | 12 | 10378657 | 10469850 | 337 | 19 | 0.22526 | 0.41089 | KLRD1 | 0.41089 | 0.4458 | 0.60278 |
| ENSG00000111046 | 12 | 81101277 | 81103253 | 5 | 2 | 0.22504 | 0.41097 | MYF6 | 0.41097 | 0.3383 | 0.35013 |
| ENSG00000115875 | 2 | 38970741 | 38978636 | 12 | 5 | 0.22491 | 0.41102 | SRSF7 | 0.41102 | 0.7289 | 0.74258 |
| ENSG00000129455 | 19 | 51499263 | 51505967 | 44 | 6 | 0.22477 | 0.41108 | KLK8 | 0.41108 | 0.41578 | 0.54148 |
| ENSG00000111358 | 12 | 124118375 | 124147153 | 89 | 11 | 0.22437 | 0.41124 | GTF2H3 | 0.41124 | 0.56756 | 0.55729 |
| ENSG00000148541 | 10 | 61005890 | 61122939 | 291 | 27 | 0.22412 | 0.41133 | FAM13C | 0.41133 | 0.434 | 0.55492 |
| ENSG00000083842 | 19 | 58790317 | 58807254 | 64 | 9 | 0.22413 | 0.41133 | ZNF8 | 0.41133 | 0.54675 | 0.34082 |
| ENSG00000273439 | 19 | 58790318 | 58807254 | 64 | 9 | 0.22413 | 0.41133 | ZNF8 | 0.41133 | 0.54675 | 0.34082 |
| ENSG00000133055 | 1 | 203136939 | 203144941 | 21 | 7 | 0.22345 | 0.41159 | MYBPH | 0.41159 | 0.50219 | 0.63683 |
| ENSG00000134240 | 1 | 120290619 | 120311528 | 58 | 12 | 0.22325 | 0.41167 | HMGCS2 | 0.41167 | 0.47934 | 0.44505 |
| ENSG00000138801 | 4 | 108511433 | 108641608 | 423 | 25 | 0.22322 | 0.41168 | PAPSS1 | 0.41168 | 0.34056 | 0.27198 |
| ENSG00000148926 | 11 | 10326227 | 10328944 | 2 | 1 | 0.22309 | 0.41173 | ADM | 0.41173 | 0.67118 | 0.71137 |
| ENSG00000244355 | 6 | 31683133 | 31685581 | 7 | 3 | 0.223 | 0.41177 | LY6G6D | 0.41177 | 0.73656 | 0.6748 |
| ENSG00000164073 | 4 | 128838960 | 128887150 | 80 | 10 | 0.22294 | 0.41179 | MFSD8 | 0.41179 | 0.31746 | 0.27431 |
| ENSG00000128872 | 15 | 52043758 | 52108565 | 249 | 16 | 0.22281 | 0.41184 | TMOD2 | 0.41184 | 0.41368 | 0.41127 |
| ENSG00000162433 | 1 | 65613232 | 65697828 | 250 | 22 | 0.22269 | 0.41189 | AK4 | 0.41189 | 0.47562 | 0.57822 |
| ENSG00000021574 | 2 | 32288680 | 32382706 | 322 | 21 | 0.22247 | 0.41198 | SPAST | 0.41198 | 0.34075 | 0.3231 |
| ENSG00000107862 | 10 | 104005289 | 104142656 | 267 | 26 | 0.22242 | 0.41199 | GBF1 | 0.41199 | 0.38816 | 0.36112 |
| ENSG00000163067 | 2 | 95831177 | 95850065 | 23 | 8 | 0.2224 | 0.412 | ZNF2 | 0.412 | 0.21649 | 0.26869 |
| ENSG00000135940 | 2 | 98262503 | 98264846 | 1 | 1 | 0.2224 | 0.412 | COX5B | 0.412 | 0.73061 | 0.90282 |
| ENSG00000143179 | 1 | 165796768 | 165880855 | 187 | 29 | 0.22235 | 0.41202 | UCK2 | 0.41202 | 0.41355 | 0.5587 |
| ENSG00000150051 | 10 | 27961804 | 28034989 | 205 | 21 | 0.22233 | 0.41203 | MKX | 0.41203 | 0.2876 | 0.12754 |

| | | | | | | | | | | | |
|---|---|---|---|---|---|---|---|---|---|---|---|
| ENSG00000198844 | 17 | 8213559 | 8225829 | 46 | 12 | 0.22228 | 0.41205 | ARHGEF15 | 0.41205 | 0.54221 | 0.55016 |
| ENSG00000095777 | 10 | 26223196 | 26501456 | 1343 | 24 | 0.22212 | 0.41211 | MYO3A | 0.41211 | 0.39697 | 0.35115 |
| ENSG00000175895 | 8 | 96146032 | 96168912 | 72 | 12 | 0.22201 | 0.41215 | PLEKHF2 | 0.41215 | 0.47188 | 0.3474 |
| ENSG00000181023 | 11 | 5757681 | 5758768 | 5 | 3 | 0.22198 | 0.41217 | OR56B1 | 0.41217 | 0.5784 | 0.71604 |
| ENSG00000179958 | 16 | 30434940 | 30441396 | 4 | 2 | 0.2219 | 0.41219 | DCTPP1 | 0.41219 | 0.74222 | 0.64756 |
| ENSG00000184922 | 17 | 43298811 | 43324687 | 59 | 10 | 0.22145 | 0.41237 | FMNL1 | 0.41237 | 0.4597 | 0.53513 |
| ENSG00000127507 | 19 | 14843205 | 14889353 | 285 | 26 | 0.22132 | 0.41242 | EMR2 | 0.41242 | 0.52812 | 0.55949 |
| ENSG00000142910 | 1 | 32042116 | 32053288 | 26 | 12 | 0.22122 | 0.41246 | TINAGL1 | 0.41246 | 0.61677 | 0.62883 |
| ENSG00000196758 | 2 | 240499995 | 240507788 | 45 | 10 | 0.22119 | 0.41247 | AC079612.1 | 0.41247 | 0.47179 | 0.47682 |
| ENSG00000155511 | 5 | 152869175 | 153193429 | 1311 | 72 | 0.22106 | 0.41252 | GRIA1 | 0.41252 | 0.43449 | 0.50547 |
| ENSG00000078747 | 20 | 32951041 | 33099198 | 277 | 11 | 0.22103 | 0.41254 | ITCH | 0.41254 | 0.4142 | 0.41797 |
| ENSG00000165233 | 9 | 95858500 | 95875565 | 60 | 11 | 0.221 | 0.41255 | C9orf89 | 0.41255 | 0.4579 | 0.33258 |
| ENSG00000182168 | 4 | 96083655 | 96470357 | 1526 | 119 | 0.22098 | 0.41256 | UNC5C | 0.41256 | 0.40196 | 0.36897 |
| ENSG00000105428 | 19 | 5455426 | 5456867 | 9 | 4 | 0.22097 | 0.41256 | ZNRF4 | 0.41256 | 0.77524 | 0.83035 |
| ENSG00000173914 | 11 | 66432469 | 66445392 | 19 | 9 | 0.22084 | 0.41261 | RBM4B | 0.41261 | 0.31005 | 0.27148 |
| ENSG00000013725 | 11 | 60739115 | 60787849 | 240 | 42 | 0.22053 | 0.41273 | CD6 | 0.41273 | 0.3992 | 0.41394 |
| ENSG00000153310 | 8 | 130851839 | 131029375 | 408 | 39 | 0.2203 | 0.41282 | FAM49B | 0.41282 | 0.46728 | 0.27665 |
| ENSG00000123453 | 9 | 136528682 | 136605077 | 322 | 38 | 0.22019 | 0.41286 | SARDH | 0.41286 | 0.52584 | 0.52395 |
| ENSG00000125869 | 20 | 9495005 | 9511171 | 78 | 16 | 0.22015 | 0.41288 | LAMP5 | 0.41288 | 0.60745 | 0.87093 |
| ENSG00000156011 | 8 | 18384811 | 18942240 | 3276 | 170 | 0.21981 | 0.41301 | PSD3 | 0.41301 | 0.36586 | 0.36899 |
| ENSG00000156650 | 10 | 76585340 | 76792380 | 349 | 23 | 0.21981 | 0.41301 | KAT6B | 0.41301 | 0.38425 | 0.36452 |
| ENSG00000112742 | 6 | 80713604 | 80752244 | 170 | 13 | 0.21963 | 0.41308 | TTK | 0.41308 | 0.4503 | 0.47923 |
| ENSG00000160953 | 19 | 1285890 | 1378430 | 326 | 49 | 0.21957 | 0.4131 | MUM1 | 0.4131 | 0.48986 | 0.59475 |
| ENSG00000171236 | 19 | 4536421 | 4540486 | 8 | 4 | 0.21947 | 0.41314 | LRG1 | 0.41314 | 0.76131 | 0.57237 |
| ENSG00000205730 | 16 | 19125254 | 19132946 | 20 | 4 | 0.21939 | 0.41317 | ITPRIPL2 | 0.41317 | 0.63235 | 0.74824 |
| ENSG00000124508 | 6 | 26383324 | 26395102 | 29 | 8 | 0.21937 | 0.41318 | BTN2A2 | 0.41318 | 0.37791 | 0.46336 |
| ENSG00000270188 | 1 | 238107024 | 238108575 | 11 | 2 | 0.21916 | 0.41326 | MTRNR2L11 | 0.41326 | 0.6961 | 0.47373 |
| ENSG00000132676 | 1 | 155657751 | 155708801 | 73 | 12 | 0.21906 | 0.4133 | DAP3 | 0.4133 | 0.4377 | 0.45423 |
| ENSG00000172262 | 5 | 43065278 | 43192123 | 515 | 27 | 0.21898 | 0.41333 | ZNF131 | 0.41333 | 0.44601 | 0.40991 |
| ENSG00000156398 | 10 | 104474295 | 104503249 | 68 | 14 | 0.21879 | 0.41341 | SFXN2 | 0.41341 | 0.43731 | 0.31894 |
| ENSG00000009335 | 7 | 156931607 | 157062066 | 487 | 21 | 0.21863 | 0.41347 | UBE3C | 0.41347 | 0.36968 | 0.28982 |
| ENSG00000167757 | 19 | 51525472 | 51531295 | 14 | 7 | 0.21849 | 0.41352 | KLK11 | 0.41352 | 0.6608 | 0.67625 |
| ENSG00000204899 | 13 | 73282495 | 73301825 | 74 | 9 | 0.2184 | 0.41356 | MZT1 | 0.41356 | 0.40654 | 0.43336 |
| ENSG00000166762 | 15 | 43920701 | 43960316 | 97 | 10 | 0.21837 | 0.41357 | CATSPER2 | 0.41357 | 0.45576 | 0.45677 |
| ENSG00000196407 | 1 | 151819739 | 151826173 | 12 | 2 | 0.21824 | 0.41362 | THEM5 | 0.41362 | 0.15663 | 0.17997 |
| ENSG00000122965 | 12 | 114254543 | 114404176 | 707 | 37 | 0.21812 | 0.41367 | RBM19 | 0.41367 | 0.41919 | 0.46527 |
| ENSG00000115616 | 2 | 103236166 | 103327777 | 231 | 39 | 0.21799 | 0.41372 | SLC9A2 | 0.41372 | 0.50834 | 0.67981 |
| ENSG00000206075 | 18 | 61143994 | 61172318 | 91 | 13 | 0.21742 | 0.41394 | SERPINB5 | 0.41394 | 0.44065 | 0.74917 |
| ENSG00000107651 | 10 | 121652223 | 121702014 | 204 | 14 | 0.21737 | 0.41396 | SEC23IP | 0.41396 | 0.45845 | 0.49834 |
| ENSG00000104818 | 19 | 49535169 | 49536495 | 2 | 1 | 0.21737 | 0.41396 | CGB2 | 0.41396 | 0.27903 | 0.21405 |
| ENSG00000149131 | 11 | 57364860 | 57382326 | 42 | 6 | 0.21729 | 0.41399 | SERPING1 | 0.41399 | 0.6734 | 0.75235 |
| ENSG00000171097 | 9 | 131595221 | 131644773 | 115 | 11 | 0.21725 | 0.41401 | CCBL1 | 0.41401 | 0.4188 | 0.35522 |
| ENSG00000177494 | 3 | 111311747 | 111314290 | 4 | 2 | 0.21685 | 0.41416 | ZBED2 | 0.41416 | 0.33996 | 0.44468 |
| ENSG00000152580 | 3 | 151143172 | 151176497 | 97 | 14 | 0.21686 | 0.41416 | IGSF10 | 0.41416 | 0.50973 | 0.49882 |
| ENSG00000170477 | 12 | 53200333 | 53208335 | 21 | 6 | 0.21678 | 0.41419 | KRT4 | 0.41419 | 0.27564 | 0.41359 |
| ENSG00000118690 | 6 | 109169619 | 109295186 | 398 | 31 | 0.21656 | 0.41428 | ARMC2 | 0.41428 | 0.40066 | 0.46904 |
| ENSG00000092140 | 14 | 31028329 | 31089269 | 153 | 23 | 0.21655 | 0.41428 | G2E3 | 0.41428 | 0.39819 | 0.40548 |
| ENSG00000170396 | 2 | 185463093 | 185804219 | 893 | 30 | 0.21649 | 0.4143 | ZNF804A | 0.4143 | 0.38032 | 0.32865 |
| ENSG00000115183 | 2 | 159825146 | 160089170 | 970 | 58 | 0.21633 | 0.41437 | TANC1 | 0.41437 | 0.40967 | 0.3891 |
| ENSG00000211452 | 1 | 54356912 | 54376759 | 93 | 12 | 0.21617 | 0.41443 | DIO1 | 0.41443 | 0.39448 | 0.50259 |
| ENSG00000188162 | 11 | 17568920 | 17668697 | 273 | 24 | 0.21612 | 0.41445 | OTOG | 0.41445 | 0.39274 | 0.39177 |
| ENSG00000185158 | 17 | 30334891 | 30380523 | 66 | 7 | 0.21602 | 0.41448 | LRRC37B | 0.41448 | 0.41342 | 0.40054 |
| ENSG00000168517 | 17 | 43238067 | 43247407 | 15 | 6 | 0.21604 | 0.41448 | HEXIM2 | 0.41448 | 0.12176 | 0.096716 |
| ENSG00000187556 | 19 | 13972877 | 13991571 | 32 | 8 | 0.21605 | 0.41448 | NANOS3 | 0.41448 | 0.43271 | 0.40851 |
| ENSG00000270099 | 10 | 97620305 | 97684601 | 208 | 27 | 0.21594 | 0.41452 | RP11-248J23.7 | 0.41452 | 0.41725 | 0.47092 |
| ENSG00000081803 | 7 | 121958481 | 122526813 | 1721 | 65 | 0.21584 | 0.41455 | CADPS2 | 0.41455 | 0.41637 | 0.42232 |
| ENSG00000267796 | 19 | 36239262 | 36245420 | 15 | 4 | 0.21566 | 0.41463 | LIN37 | 0.41463 | 0.52338 | 0.53667 |
| ENSG00000176024 | 19 | 52430400 | 52452012 | 99 | 11 | 0.21545 | 0.41471 | ZNF613 | 0.41471 | 0.15126 | 0.086578 |
| ENSG00000256525 | 17 | 62473902 | 62493154 | 50 | 4 | 0.21529 | 0.41477 | POLG2 | 0.41477 | 0.48654 | 0.4954 |
| ENSG00000213967 | 19 | 24097678 | 24127961 | 114 | 17 | 0.21529 | 0.41477 | ZNF726 | 0.41477 | 0.27709 | 0.16687 |
| ENSG00000169871 | 7 | 100728720 | 100735017 | 15 | 5 | 0.21523 | 0.41479 | TRIM56 | 0.41479 | 0.18489 | 0.0084382 |
| ENSG00000132840 | 5 | 78365540 | 78385289 | 47 | 8 | 0.21513 | 0.41483 | BHMT2 | 0.41483 | 0.46866 | 0.61576 |
| ENSG00000234769 | 16 | 64043 | 69452 | 2 | 1 | 0.2151 | 0.41484 | WASH4P | 0.41484 | 0.72998 | 0.42206 |
| ENSG00000166265 | 21 | 27838528 | 27945603 | 347 | 61 | 0.21499 | 0.41489 | CYYR1 | 0.41489 | 0.45572 | 0.37372 |
| ENSG00000116691 | 1 | 12079523 | 12092102 | 25 | 6 | 0.21495 | 0.4149 | MIIP | 0.4149 | 0.57835 | 0.3365 |
| ENSG00000203815 | 1 | 149675978 | 149676609 | 1 | 1 | 0.21496 | 0.4149 | FAM231D | 0.4149 | 0.50166 | 0.54263 |
| ENSG00000184166 | 17 | 2995352 | 2996290 | 3 | 1 | 0.21496 | 0.4149 | OR1D2 | 0.4149 | 0.34308 | 0.29872 |
| ENSG00000134152 | 15 | 34432875 | 34502297 | 298 | 14 | 0.21487 | 0.41494 | KATNBL1 | 0.41494 | 0.42886 | 0.4285 |
| ENSG00000181773 | 1 | 27719148 | 27722318 | 2 | 1 | 0.21474 | 0.41499 | GPR3 | 0.41499 | 0.25124 | 0.26354 |
| ENSG00000010322 | 3 | 52489134 | 52527087 | 89 | 8 | 0.21473 | 0.41499 | NISCH | 0.41499 | 0.402 | 0.37034 |
| ENSG00000087338 | 2 | 70056774 | 70108528 | 108 | 11 | 0.21462 | 0.41503 | GMCL1 | 0.41503 | 0.42612 | 0.39227 |
| ENSG00000198768 | 20 | 57034157 | 57090187 | 180 | 37 | 0.2146 | 0.41504 | APCDD1L | 0.41504 | 0.27974 | 0.20202 |
| ENSG00000157954 | 7 | 5229819 | 5273457 | 156 | 24 | 0.21451 | 0.41507 | WIPI2 | 0.41507 | 0.40884 | 0.58837 |

| | | | | | | | | | | | |
|---|---|---|---|---|---|---|---|---|---|---|---|
| ENSG00000197721 | 1 | 207818458 | 207911761 | 446 | 23 | 0.21413 | 0.41522 | CR1L | 0.41522 | 0.42375 | 0.42918 |
| ENSG00000145879 | 5 | 147691982 | 147695485 | 10 | 2 | 0.21404 | 0.41526 | SPINK7 | 0.41526 | 0.37482 | 0.36249 |
| ENSG00000179299 | 4 | 40751914 | 40812002 | 153 | 17 | 0.21387 | 0.41532 | NSUN7 | 0.41532 | 0.4757 | 0.47836 |
| ENSG00000086696 | 16 | 82068609 | 82132139 | 147 | 25 | 0.21388 | 0.41532 | HSD17B2 | 0.41532 | 0.49153 | 0.77363 |
| ENSG00000020633 | 1 | 25226002 | 25291612 | 148 | 31 | 0.21377 | 0.41536 | RUNX3 | 0.41536 | 0.44966 | 0.4358 |
| ENSG00000100101 | 22 | 38077680 | 38170137 | 280 | 23 | 0.21371 | 0.41539 | NOL12 | 0.41539 | 0.38033 | 0.32211 |
| ENSG00000185697 | 8 | 67474410 | 67526482 | 70 | 7 | 0.21365 | 0.41541 | MYBL1 | 0.41541 | 0.40203 | 0.42764 |
| ENSG00000269554 | 1 | 2121237 | 2123179 | 4 | 2 | 0.21356 | 0.41544 | AL590822.2 | 0.41544 | 0.37777 | 0.31632 |
| ENSG00000147684 | 8 | 125551344 | 125580751 | 137 | 22 | 0.21328 | 0.41555 | NDUFB9 | 0.41555 | 0.45691 | 0.59087 |
| ENSG00000172007 | 4 | 140374386 | 140397763 | 59 | 6 | 0.21323 | 0.41557 | RAB33B | 0.41557 | 0.57525 | 0.62486 |
| ENSG00000117594 | 1 | 209859510 | 209908295 | 121 | 16 | 0.21322 | 0.41558 | HSD11B1 | 0.41558 | 0.45211 | 0.50001 |
| ENSG00000178965 | 1 | 75033795 | 75139422 | 208 | 23 | 0.21281 | 0.41574 | C1orf173 | 0.41574 | 0.46136 | 0.63151 |
| ENSG00000177383 | 3 | 184428156 | 184429836 | 3 | 1 | 0.21273 | 0.41577 | MAGEF1 | 0.41577 | 0.36793 | 0.42886 |
| ENSG00000167625 | 19 | 42724423 | 42732353 | 6 | 2 | 0.2127 | 0.41578 | ZNF526 | 0.41578 | 0.49331 | 0.54115 |
| ENSG00000157500 | 3 | 57261765 | 57307496 | 91 | 13 | 0.21257 | 0.41583 | APPL1 | 0.41583 | 0.34659 | 0.36418 |
| ENSG00000166558 | 16 | 84043272 | 84076241 | 353 | 52 | 0.21258 | 0.41583 | SLC38A8 | 0.41583 | 0.38353 | 0.44911 |
| ENSG00000125633 | 2 | 118673054 | 118771709 | 365 | 24 | 0.21189 | 0.4161 | CCDC93 | 0.4161 | 0.46109 | 0.52715 |
| ENSG00000137491 | 11 | 74811608 | 74917594 | 395 | 30 | 0.21188 | 0.4161 | SLCO2B1 | 0.4161 | 0.41077 | 0.41219 |
| ENSG00000185090 | 1 | 38259474 | 38266809 | 7 | 3 | 0.21163 | 0.4162 | MANEAL | 0.4162 | 0.4837 | 0.46659 |
| ENSG00000100629 | 14 | 80943330 | 81425861 | 1655 | 65 | 0.21108 | 0.41641 | CEP128 | 0.41641 | 0.42481 | 0.43677 |
| ENSG00000198183 | 20 | 31823801 | 31831115 | 27 | 3 | 0.21097 | 0.41645 | BPIFA1 | 0.41645 | 0.46807 | 0.47941 |
| ENSG00000150459 | 13 | 21714653 | 21723221 | 21 | 7 | 0.21086 | 0.4165 | SAP18 | 0.4165 | 0.65505 | 0.83174 |
| ENSG00000206172 | 16 | 226679 | 227521 | 1 | 1 | 0.21086 | 0.4165 | HBA1 | 0.4165 | 0.20789 | 0.14905 |
| ENSG00000183722 | 13 | 39917029 | 40177665 | 987 | 78 | 0.21072 | 0.41655 | LHFP | 0.41655 | 0.48865 | 0.4771 |
| ENSG00000143553 | 1 | 153631130 | 153634306 | 8 | 3 | 0.21065 | 0.41658 | SNAPIN | 0.41658 | 0.45331 | 0.46523 |
| ENSG00000107551 | 10 | 45454855 | 45491339 | 99 | 16 | 0.21063 | 0.41659 | RASSF4 | 0.41659 | 0.5826 | 0.67938 |
| ENSG00000198327 | 6 | 26240561 | 26240976 | 4 | 2 | 0.2106 | 0.4166 | HIST1H4F | 0.4166 | 0.41921 | 0.40214 |
| ENSG00000153071 | 5 | 39371780 | 39462402 | 357 | 38 | 0.21059 | 0.41661 | DAB2 | 0.41661 | 0.32195 | 0.22895 |
| ENSG00000170860 | 3 | 14219858 | 14242619 | 51 | 9 | 0.21043 | 0.41667 | LSM3 | 0.41667 | 0.29702 | 0.38186 |
| ENSG00000142609 | 1 | 1853396 | 1935276 | 460 | 32 | 0.21013 | 0.41678 | C1orf222 | 0.41678 | 0.45524 | 0.47819 |
| ENSG00000099250 | 10 | 33466420 | 33625190 | 665 | 56 | 0.21 | 0.41683 | NRP1 | 0.41683 | 0.35114 | 0.37718 |
| ENSG00000167191 | 16 | 19868616 | 19897489 | 89 | 14 | 0.20993 | 0.41686 | GPRC5B | 0.41686 | 0.49977 | 0.49609 |
| ENSG00000104067 | 15 | 29991571 | 30261068 | 767 | 52 | 0.20992 | 0.41687 | TJP1 | 0.41687 | 0.39263 | 0.40158 |
| ENSG00000135454 | 12 | 58017193 | 58027138 | 26 | 6 | 0.20977 | 0.41692 | B4GALNT1 | 0.41692 | 0.20337 | 0.31642 |
| ENSG00000137269 | 6 | 53659295 | 53788919 | 410 | 21 | 0.20966 | 0.41697 | LRRC1 | 0.41697 | 0.43418 | 0.46844 |
| ENSG00000159263 | 21 | 38071433 | 38122218 | 181 | 46 | 0.20965 | 0.41697 | SIM2 | 0.41697 | 0.62342 | 0.76795 |
| ENSG00000130377 | 19 | 6135258 | 6193112 | 294 | 15 | 0.20948 | 0.41704 | ACSBG2 | 0.41704 | 0.54871 | 0.50968 |
| ENSG00000117013 | 1 | 41249684 | 41306124 | 204 | 43 | 0.20936 | 0.41708 | KCNQ4 | 0.41708 | 0.45103 | 0.51164 |
| ENSG00000213625 | 1 | 65886270 | 65901690 | 59 | 15 | 0.20915 | 0.41717 | LEPROT | 0.41717 | 0.45668 | 0.3499 |
| ENSG00000145331 | 4 | 100467866 | 100485189 | 79 | 12 | 0.20912 | 0.41718 | TRMT10A | 0.41718 | 0.35811 | 0.25485 |
| ENSG00000237651 | 2 | 61372243 | 61391964 | 89 | 7 | 0.20885 | 0.41728 | C2orf74 | 0.41728 | 0.40901 | 0.41858 |
| ENSG00000144847 | 3 | 118619404 | 118864915 | 815 | 45 | 0.20882 | 0.41729 | IGSF11 | 0.41729 | 0.39041 | 0.38205 |
| ENSG00000188211 | 11 | 17373273 | 17398888 | 83 | 14 | 0.20864 | 0.41737 | NCR3LG1 | 0.41737 | 0.4544 | 0.43979 |
| ENSG00000115705 | 2 | 1377995 | 1547483 | 914 | 49 | 0.2086 | 0.41738 | TPO | 0.41738 | 0.48063 | 0.55095 |
| ENSG00000176273 | 10 | 95653730 | 95715819 | 117 | 8 | 0.20859 | 0.41738 | SLC35G1 | 0.41738 | 0.45062 | 0.46375 |
| ENSG00000043462 | 5 | 169673241 | 169725231 | 140 | 44 | 0.2083 | 0.4175 | LCP2 | 0.4175 | 0.58384 | 0.73221 |
| ENSG00000114446 | 3 | 107879659 | 107941417 | 401 | 7 | 0.20815 | 0.41756 | IFT57 | 0.41756 | 0.44801 | 0.51892 |
| ENSG00000156795 | 8 | 124428965 | 124479470 | 339 | 9 | 0.20807 | 0.41759 | WDYHV1 | 0.41759 | 0.3913 | 0.39231 |
| ENSG00000151572 | 12 | 101111304 | 101522419 | 1377 | 88 | 0.2079 | 0.41765 | ANO4 | 0.41765 | 0.39706 | 0.29252 |
| ENSG00000162909 | 1 | 223889295 | 223963720 | 274 | 14 | 0.20777 | 0.4177 | CAPN2 | 0.4177 | 0.44371 | 0.47844 |
| ENSG00000131771 | 17 | 37782993 | 37792879 | 25 | 6 | 0.20778 | 0.4177 | PPP1R1B | 0.4177 | 0.22536 | 0.36031 |
| ENSG00000154864 | 18 | 10666480 | 11148587 | 2089 | 146 | 0.20766 | 0.41775 | PIEZO2 | 0.41775 | 0.44865 | 0.48258 |
| ENSG00000151779 | 2 | 15307032 | 15701454 | 1068 | 27 | 0.20763 | 0.41776 | NBAS | 0.41776 | 0.44061 | 0.45695 |
| ENSG00000166924 | 7 | 100081550 | 100092422 | 25 | 7 | 0.20739 | 0.41785 | NYAP1 | 0.41785 | 0.46067 | 0.5257 |
| ENSG00000167461 | 19 | 16222439 | 16245044 | 82 | 9 | 0.20721 | 0.41792 | RAB8A | 0.41792 | 0.42787 | 0.19969 |
| ENSG00000268662 | 1 | 186364123 | 186365908 | 2 | 1 | 0.20715 | 0.41795 | AL596220.1 | 0.41795 | 0.38186 | 0.33802 |
| ENSG00000125912 | 19 | 3185561 | 3209573 | 103 | 9 | 0.20713 | 0.41796 | NCLN | 0.41796 | 0.52199 | 0.32532 |
| ENSG00000232125 | 2 | 207516345 | 207583120 | 311 | 31 | 0.20708 | 0.41797 | DYTN | 0.41797 | 0.50109 | 0.61836 |
| ENSG00000107745 | 10 | 74127098 | 74385899 | 548 | 12 | 0.20701 | 0.418 | MICU1 | 0.418 | 0.44501 | 0.50754 |
| ENSG00000143493 | 1 | 212113741 | 212208884 | 407 | 22 | 0.20684 | 0.41807 | INTS7 | 0.41807 | 0.4362 | 0.40133 |
| ENSG00000186340 | 6 | 169615875 | 169654139 | 267 | 36 | 0.20658 | 0.41817 | THBS2 | 0.41817 | 0.34392 | 0.28661 |
| ENSG00000087365 | 11 | 65818200 | 65836779 | 27 | 6 | 0.20647 | 0.41821 | SF3B2 | 0.41821 | 0.35112 | 0.38974 |
| ENSG00000137309 | 6 | 34204650 | 34214008 | 18 | 5 | 0.20631 | 0.41827 | HMGA1 | 0.41827 | 0.51324 | 0.51574 |
| ENSG00000075213 | 7 | 83585093 | 84122040 | 2270 | 79 | 0.20626 | 0.41829 | SEMA3A | 0.41829 | 0.43088 | 0.45179 |
| ENSG00000057935 | 2 | 42721709 | 42984087 | 739 | 25 | 0.20624 | 0.4183 | MTA3 | 0.4183 | 0.39883 | 0.27083 |
| ENSG00000219016 | 22 | 48934712 | 48943199 | 40 | 8 | 0.2061 | 0.41836 | CTA-299D3.8 | 0.41836 | 0.66856 | 0.64828 |
| ENSG00000101152 | 20 | 62526518 | 62567384 | 105 | 14 | 0.20607 | 0.41837 | DNAJC5 | 0.41837 | 0.36917 | 0.35618 |
| ENSG00000143198 | 1 | 165600098 | 165631033 | 165 | 25 | 0.2059 | 0.41843 | MGST3 | 0.41843 | 0.26022 | 0.18809 |
| ENSG00000124802 | 6 | 8073593 | 8102811 | 104 | 15 | 0.20571 | 0.41851 | EEF1E1 | 0.41851 | 0.53981 | 0.54702 |
| ENSG00000164287 | 5 | 54408799 | 54469005 | 144 | 15 | 0.20564 | 0.41854 | CDC20B | 0.41854 | 0.42694 | 0.35421 |
| ENSG00000196290 | 2 | 201754050 | 201768655 | 20 | 5 | 0.20525 | 0.41869 | NIF3L1 | 0.41869 | 0.23689 | 0.39039 |
| ENSG00000089157 | 12 | 120634489 | 120639038 | 10 | 4 | 0.20525 | 0.41869 | RPLP0 | 0.41869 | 0.54741 | 0.55256 |
| ENSG00000129749 | 11 | 3686817 | 3692614 | 24 | 5 | 0.20522 | 0.4187 | CHRNA10 | 0.4187 | 0.83226 | 0.94498 |

| | | | | | | | | | | | |
|---|---|---|---|---|---|---|---|---|---|---|---|
| ENSG00000144583 | 2 | 217122588 | 217236750 | 319 | 39 | 0.20516 | 0.41872 | MARCHF4 | 0.41872 | 0.41937 | 0.4765 |
| ENSG00000188997 | 11 | 77882295 | 77899868 | 66 | 10 | 0.20493 | 0.41881 | KCTD21 | 0.41881 | 0.53251 | 0.70046 |
| ENSG00000130479 | 19 | 17830051 | 17845325 | 69 | 9 | 0.20491 | 0.41882 | MAP1S | 0.41882 | 0.24338 | 0.21509 |
| ENSG00000145863 | 5 | 160974069 | 161129599 | 391 | 25 | 0.20478 | 0.41887 | GABRA6 | 0.41887 | 0.43372 | 0.39269 |
| ENSG00000159216 | 21 | 36160098 | 37376965 | 4147 | 211 | 0.20464 | 0.41893 | RUNX1 | 0.41893 | 0.44734 | 0.47007 |
| ENSG00000122482 | 1 | 91380859 | 91487829 | 226 | 18 | 0.20454 | 0.41896 | ZNF644 | 0.41896 | 0.43317 | 0.45672 |
| ENSG00000188612 | 17 | 73163408 | 73179078 | 41 | 11 | 0.2043 | 0.41906 | SUMO2 | 0.41906 | 0.54138 | 0.57224 |
| ENSG00000120697 | 13 | 37523912 | 37574398 | 110 | 13 | 0.2039 | 0.41921 | ALG5 | 0.41921 | 0.3876 | 0.38977 |
| ENSG00000182551 | 2 | 3501132 | 3523507 | 139 | 10 | 0.20359 | 0.41934 | ADI1 | 0.41934 | 0.50795 | 0.66438 |
| ENSG00000154642 | 21 | 19161284 | 19191703 | 61 | 10 | 0.20358 | 0.41934 | C21orf91 | 0.41934 | 0.58292 | 0.66266 |
| ENSG00000118156 | 19 | 48023942 | 48059113 | 56 | 12 | 0.20352 | 0.41936 | ZNF541 | 0.41936 | 0.59963 | 0.77646 |
| ENSG00000152382 | 1 | 166825747 | 166845564 | 75 | 9 | 0.20332 | 0.41944 | TADA1 | 0.41944 | 0.38036 | 0.36924 |
| ENSG00000179218 | 19 | 13049392 | 13055303 | 8 | 4 | 0.2029 | 0.41961 | CALR | 0.41961 | 0.54831 | 0.72083 |
| ENSG00000155366 | 1 | 113243728 | 113250056 | 18 | 6 | 0.2027 | 0.41968 | RHOC | 0.41968 | 0.10715 | 0.20019 |
| ENSG00000171566 | 4 | 155456158 | 155471587 | 21 | 9 | 0.20261 | 0.41972 | PLRG1 | 0.41972 | 0.44476 | 0.60027 |
| ENSG00000021488 | 19 | 33321415 | 33360672 | 140 | 20 | 0.20242 | 0.41979 | SLC7A9 | 0.41979 | 0.2994 | 0.25352 |
| ENSG00000119685 | 14 | 76099968 | 76421421 | 731 | 27 | 0.20237 | 0.41981 | TTLL5 | 0.41981 | 0.46418 | 0.51037 |
| ENSG00000244623 | 7 | 99473610 | 99474680 | 3 | 1 | 0.20215 | 0.4199 | OR2AE1 | 0.4199 | 0.52169 | 0.6924 |
| ENSG00000236637 | 9 | 21186617 | 21187598 | 5 | 2 | 0.20198 | 0.41997 | IFNA4 | 0.41997 | 0.29535 | 0.30892 |
| ENSG00000269408 | 9 | 138150512 | 138151275 | 14 | 3 | 0.20189 | 0.42 | AL390778.1 | 0.42 | 0.20272 | 0.45033 |
| ENSG00000174595 | 7 | 130417401 | 130418888 | 6 | 2 | 0.2018 | 0.42004 | KLF14 | 0.42004 | 0.32509 | 0.31921 |
| ENSG00000198853 | 9 | 35490124 | 35561895 | 138 | 22 | 0.20158 | 0.42012 | RUSC2 | 0.42012 | 0.43105 | 0.40169 |
| ENSG00000089057 | 20 | 4833002 | 4990939 | 493 | 61 | 0.20115 | 0.42029 | SLC23A2 | 0.42029 | 0.55763 | 0.57073 |
| ENSG00000113889 | 3 | 186435065 | 186461743 | 179 | 17 | 0.20106 | 0.42033 | KNG1 | 0.42033 | 0.41879 | 0.49373 |
| ENSG00000165512 | 10 | 45495923 | 45500774 | 19 | 5 | 0.20096 | 0.42036 | ZNF22 | 0.42036 | 0.53718 | 0.43819 |
| ENSG00000139675 | 13 | 53191605 | 53217919 | 93 | 6 | 0.20097 | 0.42036 | HNRNPA1L2 | 0.42036 | 0.38092 | 0.39971 |
| ENSG00000146425 | 6 | 159057506 | 159065771 | 19 | 5 | 0.20092 | 0.42038 | DYNLT1 | 0.42038 | 0.56748 | 0.65169 |
| ENSG00000148832 | 10 | 135192695 | 135205198 | 58 | 12 | 0.20092 | 0.42038 | PAOX | 0.42038 | 0.68306 | 0.85284 |
| ENSG00000183773 | 22 | 21319396 | 21335649 | 59 | 15 | 0.20083 | 0.42042 | AIFM3 | 0.42042 | 0.14249 | 0.22162 |
| ENSG00000078018 | 2 | 210288782 | 210598842 | 863 | 28 | 0.20068 | 0.42047 | MAP2 | 0.42047 | 0.42711 | 0.44549 |
| ENSG00000008324 | 3 | 42623332 | 42636606 | 38 | 7 | 0.2007 | 0.42047 | SS18L2 | 0.42047 | 0.37678 | 0.35686 |
| ENSG00000023608 | 14 | 62229075 | 62263146 | 162 | 15 | 0.20067 | 0.42048 | SNAPC1 | 0.42048 | 0.45757 | 0.47674 |
| ENSG00000100139 | 22 | 38301664 | 38338829 | 64 | 13 | 0.20045 | 0.42056 | MICALL1 | 0.42056 | 0.32795 | 0.35345 |
| ENSG00000175093 | 3 | 140770244 | 140867453 | 230 | 24 | 0.2004 | 0.42058 | SPSB4 | 0.42058 | 0.44468 | 0.48584 |
| ENSG00000177425 | 12 | 79968759 | 80084877 | 164 | 17 | 0.20004 | 0.42072 | PAWR | 0.42072 | 0.438 | 0.43038 |
| ENSG00000196937 | 7 | 120988905 | 121036418 | 122 | 8 | 0.19995 | 0.42076 | FAM3C | 0.42076 | 0.44562 | 0.43877 |
| ENSG00000111885 | 6 | 119498374 | 119670926 | 791 | 26 | 0.19992 | 0.42077 | MAN1A1 | 0.42077 | 0.41667 | 0.40993 |
| ENSG00000150394 | 16 | 61681146 | 62070939 | 1070 | 61 | 0.19989 | 0.42078 | CDH8 | 0.42078 | 0.41999 | 0.40947 |
| ENSG00000178722 | 5 | 60933535 | 61047590 | 397 | 34 | 0.19957 | 0.42091 | C5orf64 | 0.42091 | 0.40028 | 0.42343 |
| ENSG00000162931 | 1 | 228595641 | 228604562 | 10 | 3 | 0.19947 | 0.42095 | TRIM17 | 0.42095 | 0.21547 | 0.19363 |
| ENSG00000081870 | 1 | 54382247 | 54411602 | 71 | 6 | 0.19928 | 0.42102 | HSPB11 | 0.42102 | 0.30057 | 0.34243 |
| ENSG00000122958 | 10 | 70883268 | 70932617 | 162 | 14 | 0.19917 | 0.42106 | VPS26A | 0.42106 | 0.45215 | 0.43774 |
| ENSG00000112245 | 6 | 64231666 | 64293492 | 115 | 15 | 0.19906 | 0.42111 | PTP4A1 | 0.42111 | 0.42583 | 0.4406 |
| ENSG00000176601 | 2 | 135722061 | 135805038 | 299 | 22 | 0.199 | 0.42113 | MAP3K19 | 0.42113 | 0.46899 | 0.51845 |
| ENSG00000120705 | 5 | 137841784 | 137878989 | 87 | 17 | 0.19882 | 0.4212 | ETF1 | 0.4212 | 0.4048 | 0.48598 |
| ENSG00000029725 | 17 | 5185558 | 5289129 | 348 | 26 | 0.1984 | 0.42137 | RABEP1 | 0.42137 | 0.30677 | 0.26679 |
| ENSG00000149564 | 11 | 124622026 | 124632186 | 18 | 7 | 0.19829 | 0.42141 | ESAM | 0.42141 | 0.34699 | 0.36689 |
| ENSG00000198933 | 17 | 45771447 | 45789416 | 47 | 12 | 0.19827 | 0.42142 | TBKBP1 | 0.42142 | 0.4076 | 0.31243 |
| ENSG00000065978 | 1 | 43148098 | 43168020 | 48 | 12 | 0.19822 | 0.42144 | YBX1 | 0.42144 | 0.48891 | 0.48061 |
| ENSG00000143549 | 1 | 154127784 | 154167124 | 54 | 10 | 0.19819 | 0.42145 | TPM3 | 0.42145 | 0.39368 | 0.47967 |
| ENSG00000168631 | 6 | 30908749 | 30921998 | 41 | 13 | 0.19809 | 0.42149 | DPCR1 | 0.42149 | 0.35527 | 0.38019 |
| ENSG00000116580 | 1 | 155719508 | 155829191 | 211 | 16 | 0.19801 | 0.42152 | GON4L | 0.42152 | 0.42769 | 0.45585 |
| ENSG00000185055 | 7 | 105205567 | 105241322 | 122 | 23 | 0.19764 | 0.42166 | EFCAB10 | 0.42166 | 0.45819 | 0.5789 |
| ENSG00000105971 | 7 | 115927434 | 116148595 | 734 | 40 | 0.19727 | 0.42181 | CAV2 | 0.42181 | 0.47033 | 0.53812 |
| ENSG00000166866 | 12 | 57422301 | 57444982 | 57 | 8 | 0.19727 | 0.42181 | MYO1A | 0.42181 | 0.43686 | 0.44478 |
| ENSG00000111245 | 12 | 111348623 | 111358526 | 38 | 6 | 0.197 | 0.42191 | MYL2 | 0.42191 | 0.45716 | 0.48146 |
| ENSG00000117281 | 1 | 145695798 | 145715614 | 25 | 3 | 0.19677 | 0.42201 | CD160 | 0.42201 | 0.49402 | 0.48713 |
| ENSG00000122375 | 10 | 88414314 | 88426605 | 57 | 15 | 0.19674 | 0.42201 | OPN4 | 0.42201 | 0.55773 | 0.49383 |
| ENSG00000166676 | 16 | 10854776 | 10912651 | 166 | 39 | 0.1967 | 0.42203 | TVP23A | 0.42203 | 0.59258 | 0.78813 |
| ENSG00000160551 | 17 | 27717482 | 27878922 | 395 | 19 | 0.19671 | 0.42203 | TAOK1 | 0.42203 | 0.39775 | 0.41343 |
| ENSG00000204695 | 6 | 29274403 | 29275519 | 4 | 2 | 0.1965 | 0.42211 | OR14J1 | 0.42211 | 0.82472 | 0.85256 |
| ENSG00000138115 | 10 | 96796530 | 96829254 | 172 | 19 | 0.1961 | 0.42227 | CYP2C8 | 0.42227 | 0.37956 | 0.34706 |
| ENSG00000177374 | 17 | 1957448 | 1962981 | 14 | 7 | 0.19608 | 0.42227 | HIC1 | 0.42227 | 0.67813 | 0.50273 |
| ENSG00000253903 | 8 | 26943694 | 26944766 | 6 | 3 | 0.19606 | 0.42228 | RP11-521M14.2 | 0.42228 | 0.27483 | 0.26287 |
| ENSG00000165204 | 9 | 125562370 | 125563395 | 3 | 2 | 0.19601 | 0.4223 | OR1K1 | 0.4223 | 0.43968 | 0.69742 |
| ENSG00000143816 | 1 | 228106357 | 228135599 | 103 | 13 | 0.19572 | 0.42242 | WNT9A | 0.42242 | 0.39011 | 0.36446 |
| ENSG00000124233 | 20 | 43835638 | 43838413 | 6 | 2 | 0.19567 | 0.42243 | SEMG1 | 0.42243 | 0.50277 | 0.54009 |
| ENSG00000072778 | 17 | 7120444 | 7128592 | 14 | 4 | 0.19546 | 0.42252 | ACADVL | 0.42252 | 0.43511 | 0.46063 |
| ENSG00000171987 | 11 | 4592653 | 4599050 | 66 | 13 | 0.19523 | 0.42261 | C11orf40 | 0.42261 | 0.4254 | 0.69159 |
| ENSG00000142657 | 1 | 10458649 | 10480201 | 61 | 9 | 0.19516 | 0.42263 | PGD | 0.42263 | 0.26377 | 0.18835 |
| ENSG00000134339 | 11 | 18260770 | 18270190 | 43 | 14 | 0.1951 | 0.42266 | SAA2 | 0.42266 | 0.49559 | 0.70736 |
| ENSG00000119969 | 10 | 96305547 | 96373662 | 177 | 14 | 0.19481 | 0.42277 | HELLS | 0.42277 | 0.42843 | 0.43059 |
| ENSG00000154358 | 1 | 228395831 | 228566577 | 367 | 34 | 0.19455 | 0.42287 | OBSCN | 0.42287 | 0.42198 | 0.3967 |

| | | | | | | | | | | | |
|---|---|---|---|---|---|---|---|---|---|---|---|
| ENSG00000122188 | 1 | 203734304 | 203745361 | 39 | 5 | 0.19433 | 0.42296 | LAX1 | 0.42296 | 0.52959 | 0.60564 |
| ENSG00000182752 | 9 | 118916083 | 119164601 | 530 | 81 | 0.19386 | 0.42314 | PAPPA | 0.42314 | 0.45375 | 0.47506 |
| ENSG00000253426 | 8 | 9009252 | 9025646 | 27 | 4 | 0.19373 | 0.4232 | RP11-10A14.4 | 0.4232 | 0.38847 | 0.49669 |
| ENSG00000008853 | 8 | 22844930 | 22877712 | 88 | 19 | 0.1934 | 0.42332 | RHOBTB2 | 0.42332 | 0.6685 | 0.718 |
| ENSG00000145700 | 5 | 74364100 | 74532703 | 543 | 17 | 0.19316 | 0.42342 | ANKRD31 | 0.42342 | 0.41288 | 0.34917 |
| ENSG00000100106 | 22 | 38093011 | 38172563 | 244 | 26 | 0.19307 | 0.42345 | TRIOBP | 0.42345 | 0.43331 | 0.39943 |
| ENSG00000188483 | 9 | 131937835 | 131940540 | 3 | 1 | 0.19304 | 0.42347 | IER5L | 0.42347 | 0.51162 | 0.62148 |
| ENSG00000173065 | 17 | 27082996 | 27182250 | 193 | 9 | 0.19293 | 0.42351 | FAM222B | 0.42351 | 0.40773 | 0.35641 |
| ENSG00000160193 | 21 | 44263204 | 44299678 | 244 | 23 | 0.19278 | 0.42356 | WDR4 | 0.42356 | 0.43743 | 0.43846 |
| ENSG00000117477 | 1 | 169364108 | 169429907 | 204 | 23 | 0.19275 | 0.42358 | CCDC181 | 0.42358 | 0.44595 | 0.43078 |
| ENSG00000126461 | 19 | 50145382 | 50161899 | 26 | 9 | 0.19274 | 0.42358 | SCAF1 | 0.42358 | 0.21676 | 0.17619 |
| ENSG00000117616 | 1 | 25568728 | 25664704 | 118 | 21 | 0.19245 | 0.42369 | C1orf63 | 0.42369 | 0.43134 | 0.46937 |
| ENSG00000168438 | 6 | 110501344 | 110575478 | 219 | 25 | 0.1924 | 0.42371 | CDC40 | 0.42371 | 0.48081 | 0.46696 |
| ENSG00000257242 | 12 | 92378756 | 92536690 | 510 | 66 | 0.19232 | 0.42375 | C12orf79 | 0.42375 | 0.46207 | 0.45837 |
| ENSG00000265817 | 8 | 95390605 | 95449180 | 204 | 19 | 0.19228 | 0.42376 | FSBP | 0.42376 | 0.44847 | 0.48636 |
| ENSG00000253459 | 14 | 50064193 | 50065882 | 8 | 3 | 0.19222 | 0.42379 | AL139099.1 | 0.42379 | 0.62712 | 0.7328 |
| ENSG00000229183 | 11 | 60989688 | 61018929 | 25 | 11 | 0.19203 | 0.42386 | PGA4 | 0.42386 | 0.5357 | 0.74227 |
| ENSG00000080546 | 6 | 109307640 | 109416022 | 258 | 11 | 0.19184 | 0.42393 | SESN1 | 0.42393 | 0.42349 | 0.39425 |
| ENSG00000165752 | 10 | 134020996 | 134145351 | 635 | 55 | 0.19165 | 0.42401 | STK32C | 0.42401 | 0.27327 | 0.21455 |
| ENSG00000158669 | 8 | 41434706 | 41482520 | 156 | 14 | 0.19148 | 0.42407 | AGPAT6 | 0.42407 | 0.51696 | 0.65858 |
| ENSG00000172889 | 9 | 139553308 | 139567130 | 41 | 8 | 0.19135 | 0.42413 | EGFL7 | 0.42413 | 0.35488 | 0.31628 |
| ENSG00000135439 | 12 | 58118980 | 58135940 | 26 | 8 | 0.19109 | 0.42423 | AGAP2 | 0.42423 | 0.28529 | 0.4265 |
| ENSG00000168925 | 16 | 75252898 | 75258822 | 41 | 8 | 0.19109 | 0.42423 | CTRB1 | 0.42423 | 0.69689 | 0.66863 |
| ENSG00000112139 | 6 | 37600284 | 37667082 | 331 | 37 | 0.19069 | 0.42438 | MDGA1 | 0.42438 | 0.424 | 0.42771 |
| ENSG00000132591 | 17 | 27181956 | 27188085 | 14 | 1 | 0.19064 | 0.4244 | ERAL1 | 0.4244 | 0.24206 | 0.23354 |
| ENSG00000179588 | 16 | 88519725 | 88603424 | 439 | 42 | 0.1906 | 0.42442 | ZFPM1 | 0.42442 | 0.51834 | 0.46842 |
| ENSG00000135549 | 6 | 122793076 | 123047518 | 763 | 41 | 0.1903 | 0.42454 | PKIB | 0.42454 | 0.3989 | 0.26495 |
| ENSG00000181552 | 14 | 21236586 | 21239107 | 22 | 6 | 0.1901 | 0.42462 | EDDM3B | 0.42462 | 0.74133 | 0.89106 |
| ENSG00000187260 | 7 | 151072995 | 151107813 | 144 | 34 | 0.19004 | 0.42464 | WDR86 | 0.42464 | 0.48739 | 0.63389 |
| ENSG00000171150 | 2 | 46926091 | 46990268 | 239 | 27 | 0.18977 | 0.42474 | SOCS5 | 0.42474 | 0.48084 | 0.62453 |
| ENSG00000155918 | 6 | 150339470 | 150346668 | 41 | 6 | 0.18972 | 0.42476 | RAET1L | 0.42476 | 0.17566 | 0.25152 |
| ENSG00000002933 | 7 | 150497491 | 150502208 | 36 | 7 | 0.18973 | 0.42476 | TMEM176A | 0.42476 | 0.68727 | 0.53253 |
| ENSG00000167383 | 19 | 44921685 | 44952766 | 163 | 21 | 0.18968 | 0.42478 | ZNF229 | 0.42478 | 0.41885 | 0.49687 |
| ENSG00000184351 | 21 | 31852018 | 31852663 | 1 | 1 | 0.18963 | 0.4248 | KRTAP19-1 | 0.4248 | 0.47355 | 0.45396 |
| ENSG00000058085 | 1 | 183155373 | 183214035 | 240 | 19 | 0.18959 | 0.42482 | LAMC2 | 0.42482 | 0.44405 | 0.48093 |
| ENSG00000177888 | 1 | 197122810 | 197169672 | 105 | 14 | 0.18952 | 0.42484 | ZBTB41 | 0.42484 | 0.38311 | 0.47788 |
| ENSG00000153443 | 16 | 4658884 | 4665028 | 14 | 4 | 0.18951 | 0.42485 | UBALD1 | 0.42485 | 0.3465 | 0.35476 |
| ENSG00000183726 | 1 | 25664408 | 25688852 | 35 | 9 | 0.18934 | 0.42491 | TMEM50A | 0.42491 | 0.31708 | 0.34843 |
| ENSG00000196946 | 12 | 8290733 | 8341133 | 157 | 10 | 0.18931 | 0.42493 | ZNF705A | 0.42493 | 0.4821 | 0.55285 |
| ENSG00000118689 | 6 | 108881038 | 109005977 | 197 | 24 | 0.18893 | 0.42507 | FOXO3 | 0.42507 | 0.50652 | 0.53943 |
| ENSG00000135655 | 12 | 62654119 | 62811211 | 444 | 25 | 0.18887 | 0.4251 | USP15 | 0.4251 | 0.37287 | 0.31201 |
| ENSG00000106367 | 7 | 100797678 | 100804877 | 34 | 5 | 0.18875 | 0.42514 | AP1S1 | 0.42514 | 0.75218 | 0.69575 |
| ENSG00000182700 | 5 | 139505521 | 139508391 | 1 | 1 | 0.18861 | 0.4252 | IGIP | 0.4252 | 0.6867 | 0.55775 |
| ENSG00000203286 | 9 | 3452305 | 3469181 | 36 | 7 | 0.18854 | 0.42523 | AL365202.1 | 0.42523 | 0.45067 | 0.46626 |
| ENSG00000133302 | 5 | 93954052 | 94075141 | 414 | 22 | 0.18813 | 0.42539 | ANKRD32 | 0.42539 | 0.41694 | 0.461 |
| ENSG00000240038 | 1 | 104096437 | 104122156 | 40 | 15 | 0.18784 | 0.4255 | AMY2B | 0.4255 | 0.30588 | 0.26 |
| ENSG00000197084 | 1 | 152777311 | 152779107 | 15 | 4 | 0.18784 | 0.4255 | LCE1C | 0.4255 | 0.35818 | 0.14967 |
| ENSG00000116985 | 1 | 40222854 | 40254533 | 148 | 21 | 0.18764 | 0.42558 | BMP8B | 0.42558 | 0.34717 | 0.34557 |
| ENSG00000005187 | 16 | 20621565 | 20808903 | 551 | 22 | 0.18763 | 0.42558 | ACSM3 | 0.42558 | 0.43335 | 0.47448 |
| ENSG00000204472 | 6 | 31582961 | 31584798 | 8 | 4 | 0.18757 | 0.42561 | AIF1 | 0.42561 | 0.22573 | 0.20915 |
| ENSG00000175691 | 19 | 2933216 | 2944969 | 57 | 17 | 0.18736 | 0.42569 | ZNF77 | 0.42569 | 0.39233 | 0.50792 |
| ENSG00000137070 | 9 | 34650699 | 34661889 | 23 | 6 | 0.1873 | 0.42571 | IL11RA | 0.42571 | 0.2238 | 0.1898 |
| ENSG00000137080 | 9 | 21165636 | 21166659 | 4 | 2 | 0.18727 | 0.42572 | IFNA21 | 0.42572 | 0.26962 | 0.3132 |
| ENSG00000101331 | 20 | 30598245 | 30619984 | 46 | 12 | 0.18726 | 0.42573 | CCM2L | 0.42573 | 0.4569 | 0.5695 |
| ENSG00000100314 | 22 | 30116073 | 30127828 | 35 | 8 | 0.18695 | 0.42585 | CABP7 | 0.42585 | 0.65739 | 0.6995 |
| ENSG00000157680 | 7 | 137065783 | 137531838 | 1180 | 82 | 0.18693 | 0.42586 | DGKI | 0.42586 | 0.42895 | 0.56113 |
| ENSG00000184601 | 14 | 105046021 | 105056852 | 31 | 6 | 0.18687 | 0.42588 | C14orf180 | 0.42588 | 0.64075 | 0.76931 |
| ENSG00000083814 | 19 | 58231119 | 58238992 | 22 | 7 | 0.18686 | 0.42589 | ZNF671 | 0.42589 | 0.35132 | 0.40441 |
| ENSG00000198176 | 13 | 114239013 | 114295785 | 125 | 13 | 0.18673 | 0.42594 | TFDP1 | 0.42594 | 0.44899 | 0.46983 |
| ENSG00000168237 | 3 | 52321105 | 52329272 | 16 | 4 | 0.18625 | 0.42612 | GLYCTK | 0.42612 | 0.45941 | 0.42744 |
| ENSG00000198169 | 8 | 145946298 | 145981802 | 78 | 21 | 0.18613 | 0.42617 | ZNF251 | 0.42617 | 0.30508 | 0.22057 |
| ENSG00000111602 | 12 | 56810903 | 56843187 | 100 | 8 | 0.18613 | 0.42617 | TIMELESS | 0.42617 | 0.46432 | 0.52457 |
| ENSG00000139318 | 12 | 89741009 | 89747048 | 29 | 6 | 0.18602 | 0.42622 | DUSP6 | 0.42622 | 0.5197 | 0.48506 |
| ENSG00000165548 | 14 | 77582911 | 77725838 | 694 | 53 | 0.186 | 0.42622 | TMEM63C | 0.42622 | 0.49169 | 0.57161 |
| ENSG00000164252 | 5 | 76325076 | 76361059 | 103 | 18 | 0.1858 | 0.4263 | AGGF1 | 0.4263 | 0.51013 | 0.6713 |
| ENSG00000172828 | 16 | 66995140 | 67009051 | 20 | 5 | 0.18565 | 0.42636 | CES3 | 0.42636 | 0.64109 | 0.61267 |
| ENSG00000185087 | 15 | 98980391 | 99057611 | 297 | 20 | 0.18559 | 0.42638 | FAM169B | 0.42638 | 0.39767 | 0.59709 |
| ENSG00000065320 | 17 | 8924859 | 9147317 | 1070 | 100 | 0.18545 | 0.42644 | NTN1 | 0.42644 | 0.40322 | 0.39962 |
| ENSG00000129625 | 5 | 112212084 | 112258236 | 222 | 19 | 0.18498 | 0.42662 | REEP5 | 0.42662 | 0.28697 | 0.30232 |
| ENSG00000126251 | 19 | 35861831 | 35863855 | 6 | 3 | 0.18495 | 0.42664 | GPR42 | 0.42664 | 0.77839 | 0.79353 |
| ENSG00000250038 | 4 | 28363901 | 28404486 | 178 | 19 | 0.18491 | 0.42665 | RP11-180C1.1 | 0.42665 | 0.42751 | 0.37943 |
| ENSG00000196944 | 1 | 248524883 | 248525929 | 15 | 2 | 0.18486 | 0.42667 | OR2T4 | 0.42667 | 0.7253 | 0.66333 |
| ENSG00000182230 | 5 | 175487692 | 175559261 | 132 | 25 | 0.18461 | 0.42677 | FAM153B | 0.42677 | 0.31422 | 0.37586 |

| | | | | | | | | | | | |
|---|---|---|---|---|---|---|---|---|---|---|---|
| ENSG00000111011 | 12 | 122989190 | 123011547 | 42 | 5 | 0.18461 | 0.42677 | RSRC2 | 0.42677 | 0.53055 | 0.59965 |
| ENSG00000100884 | 14 | 24540046 | 24547309 | 13 | 6 | 0.18451 | 0.42681 | CPNE6 | 0.42681 | 0.39284 | 0.28229 |
| ENSG00000198746 | 1 | 27216979 | 27226957 | 17 | 6 | 0.18438 | 0.42686 | GPATCH3 | 0.42686 | 0.62934 | 0.69262 |
| ENSG00000204610 | 6 | 30130993 | 30140473 | 68 | 9 | 0.18437 | 0.42686 | TRIM15 | 0.42686 | 0.43829 | 0.59463 |
| ENSG00000185737 | 10 | 83635070 | 84746935 | 3627 | 187 | 0.18434 | 0.42687 | NRG3 | 0.42687 | 0.44998 | 0.45618 |
| ENSG00000130772 | 1 | 28655513 | 28662476 | 10 | 5 | 0.18428 | 0.4269 | MED18 | 0.4269 | 0.8282 | 0.87823 |
| ENSG00000185627 | 11 | 236546 | 252983 | 113 | 15 | 0.18384 | 0.42707 | PSMD13 | 0.42707 | 0.2882 | 0.33982 |
| ENSG00000172399 | 4 | 120056939 | 120108944 | 201 | 19 | 0.18379 | 0.42709 | MYOZ2 | 0.42709 | 0.54175 | 0.53794 |
| ENSG00000031698 | 1 | 109756540 | 109780791 | 54 | 14 | 0.18358 | 0.42717 | SARS | 0.42717 | 0.61179 | 0.79113 |
| ENSG00000179111 | 17 | 8023908 | 8027410 | 14 | 6 | 0.18349 | 0.42721 | HES7 | 0.42721 | 0.40697 | 0.6418 |
| ENSG00000108587 | 17 | 28804380 | 28854610 | 82 | 10 | 0.1834 | 0.42724 | GOSR1 | 0.42724 | 0.45631 | 0.43215 |
| ENSG00000171530 | 5 | 76986991 | 77164604 | 752 | 40 | 0.18337 | 0.42725 | TBCA | 0.42725 | 0.49185 | 0.48053 |
| ENSG00000186803 | 9 | 21206180 | 21207142 | 3 | 1 | 0.18334 | 0.42727 | IFNA10 | 0.42727 | 0.44045 | 0.37137 |
| ENSG00000173545 | 5 | 16451628 | 16465901 | 37 | 9 | 0.18329 | 0.42729 | ZNF622 | 0.42729 | 0.52254 | 0.59374 |
| ENSG00000178498 | 12 | 57998405 | 58003587 | 9 | 5 | 0.18321 | 0.42732 | DTX3 | 0.42732 | 0.16209 | 0.11663 |
| ENSG00000100304 | 22 | 43562628 | 43583139 | 117 | 21 | 0.18285 | 0.42746 | TTLL12 | 0.42746 | 0.067393 | 0.070098 |
| ENSG00000146090 | 5 | 179527795 | 179636153 | 582 | 79 | 0.18249 | 0.4276 | RASGEF1C | 0.4276 | 0.43413 | 0.54125 |
| ENSG00000197586 | 20 | 25176329 | 25207365 | 81 | 19 | 0.1825 | 0.4276 | ENTPD6 | 0.4276 | 0.33725 | 0.38044 |
| ENSG00000112130 | 6 | 37321748 | 37362514 | 60 | 16 | 0.18234 | 0.42766 | RNF8 | 0.42766 | 0.32108 | 0.10602 |
| ENSG00000160789 | 1 | 156052364 | 156109880 | 158 | 16 | 0.1823 | 0.42767 | LMNA | 0.42767 | 0.46535 | 0.35851 |
| ENSG00000163873 | 1 | 37261128 | 37499730 | 562 | 61 | 0.18144 | 0.42801 | GRIK3 | 0.42801 | 0.41372 | 0.44038 |
| ENSG00000133878 | 8 | 33448856 | 33457624 | 23 | 5 | 0.18119 | 0.42811 | DUSP26 | 0.42811 | 0.67453 | 0.691 |
| ENSG00000144567 | 2 | 220040947 | 220050201 | 20 | 7 | 0.18099 | 0.42819 | FAM134A | 0.42819 | 0.096477 | 0.21494 |
| ENSG00000106049 | 7 | 27565061 | 27702614 | 559 | 17 | 0.18053 | 0.42837 | HIBADH | 0.42837 | 0.43383 | 0.42035 |
| ENSG00000164300 | 5 | 79407050 | 79551898 | 578 | 49 | 0.1805 | 0.42838 | SERINC5 | 0.42838 | 0.46589 | 0.44476 |
| ENSG00000268858 | 20 | 62492879 | 62493217 | 1 | 1 | 0.18045 | 0.4284 | C20ORF135 | 0.4284 | 0.64431 | 0.51376 |
| ENSG00000204435 | 6 | 31633013 | 31638120 | 20 | 8 | 0.18021 | 0.4285 | CSNK2B | 0.4285 | 0.52855 | 0.37158 |
| ENSG00000140199 | 15 | 34525460 | 34630261 | 512 | 22 | 0.18014 | 0.42852 | SLC12A6 | 0.42852 | 0.47538 | 0.48171 |
| ENSG00000255863 | 12 | 49315767 | 49333446 | 51 | 6 | 0.18009 | 0.42854 | AC073610.5 | 0.42854 | 0.52235 | 0.52446 |
| ENSG00000114405 | 3 | 62304648 | 62321888 | 21 | 9 | 0.17989 | 0.42862 | C3orf14 | 0.42862 | 0.62034 | 0.80249 |
| ENSG00000099817 | 19 | 1086594 | 1095598 | 51 | 18 | 0.17989 | 0.42862 | POLR2E | 0.42862 | 0.83196 | 0.8964 |
| ENSG00000105401 | 19 | 10501810 | 10530797 | 72 | 10 | 0.17989 | 0.42862 | CDC37 | 0.42862 | 0.38257 | 0.37948 |
| ENSG00000049656 | 5 | 1317859 | 1345214 | 115 | 18 | 0.17979 | 0.42866 | CLPTM1L | 0.42866 | 0.46597 | 0.27662 |
| ENSG00000029153 | 12 | 27485787 | 27576241 | 334 | 41 | 0.17948 | 0.42878 | ARNTL2 | 0.42878 | 0.42207 | 0.37264 |
| ENSG00000181322 | 3 | 137980279 | 138048728 | 69 | 23 | 0.17933 | 0.42884 | NME9 | 0.42884 | 0.50562 | 0.52048 |
| ENSG00000214188 | 7 | 116593953 | 116738860 | 195 | 38 | 0.17916 | 0.4289 | ST7-OT4 | 0.4289 | 0.41001 | 0.39131 |
| ENSG00000182871 | 21 | 46825052 | 46933634 | 690 | 43 | 0.1791 | 0.42893 | COL18A1 | 0.42893 | 0.48355 | 0.61407 |
| ENSG00000171970 | 19 | 2900896 | 2918474 | 147 | 17 | 0.17905 | 0.42895 | ZNF57 | 0.42895 | 0.38678 | 0.39685 |
| ENSG00000116001 | 2 | 70436576 | 70475792 | 93 | 8 | 0.17881 | 0.42904 | TIA1 | 0.42904 | 0.4572 | 0.45504 |
| ENSG00000184307 | 3 | 113666748 | 113684248 | 40 | 11 | 0.17872 | 0.42908 | ZDHHC23 | 0.42908 | 0.4206 | 0.38513 |
| ENSG00000142252 | 19 | 45582453 | 45594782 | 83 | 5 | 0.17863 | 0.42912 | GEMIN7 | 0.42912 | 0.70478 | 0.77926 |
| ENSG00000267884 | 1 | 166026679 | 166028709 | 6 | 2 | 0.1786 | 0.42913 | AL626787.1 | 0.42913 | 0.44703 | 0.50902 |
| ENSG00000269766 | 11 | 35150091 | 35150215 | 1 | 1 | 0.17841 | 0.4292 | AL356215.1 | 0.4292 | 0.56804 | 0.65342 |
| ENSG00000169203 | 16 | 29495010 | 29517320 | 12 | 2 | 0.17828 | 0.42925 | RP11-231C14.4 | 0.42925 | 0.34105 | 0.3514 |
| ENSG00000170892 | 19 | 54693789 | 54697585 | 10 | 3 | 0.17814 | 0.42931 | TSEN34 | 0.42931 | 0.47799 | 0.28414 |
| ENSG00000151575 | 15 | 56536207 | 56738195 | 673 | 25 | 0.17753 | 0.42954 | TEX9 | 0.42954 | 0.41999 | 0.43125 |
| ENSG00000140522 | 15 | 89753100 | 89764982 | 39 | 8 | 0.17741 | 0.42959 | RLBP1 | 0.42959 | 0.50126 | 0.5391 |
| ENSG00000168389 | 1 | 40420802 | 40435638 | 25 | 11 | 0.17715 | 0.42969 | MFSD2A | 0.42969 | 0.11279 | 0.16526 |
| ENSG00000182541 | 22 | 31608225 | 31676066 | 157 | 23 | 0.17712 | 0.42971 | LIMK2 | 0.42971 | 0.44019 | 0.40176 |
| ENSG00000115267 | 2 | 163123589 | 163175213 | 123 | 6 | 0.17706 | 0.42973 | IFIH1 | 0.42973 | 0.43458 | 0.4108 |
| ENSG00000257949 | 17 | 73975301 | 73996667 | 70 | 14 | 0.17701 | 0.42975 | TEN1 | 0.42975 | 0.52547 | 0.57464 |
| ENSG00000154429 | 1 | 229456758 | 229479041 | 74 | 13 | 0.17698 | 0.42976 | CCSAP | 0.42976 | 0.402 | 0.33776 |
| ENSG00000096060 | 6 | 35541362 | 35696360 | 358 | 27 | 0.17687 | 0.42981 | FKBP5 | 0.42981 | 0.39662 | 0.43845 |
| ENSG00000168101 | 16 | 4743695 | 4745860 | 3 | 1 | 0.17687 | 0.42981 | NUDT16L1 | 0.42981 | 0.46232 | 0.60182 |
| ENSG00000214140 | 17 | 74523668 | 74549660 | 69 | 10 | 0.17641 | 0.42999 | PRCD | 0.42999 | 0.25571 | 0.39512 |
| ENSG00000197102 | 14 | 102430865 | 102517129 | 201 | 21 | 0.17629 | 0.43003 | DYNC1H1 | 0.43003 | 0.43505 | 0.46985 |
| ENSG00000171643 | 5 | 76145826 | 76217475 | 213 | 24 | 0.17628 | 0.43004 | S100Z | 0.43004 | 0.32968 | 0.32741 |
| ENSG00000197993 | 7 | 142638201 | 142659768 | 56 | 9 | 0.17628 | 0.43004 | KEL | 0.43004 | 0.72018 | 0.70774 |
| ENSG00000003509 | 2 | 37458774 | 37480546 | 99 | 16 | 0.17625 | 0.43005 | NDUFAF7 | 0.43005 | 0.44859 | 0.37667 |
| ENSG00000171208 | 16 | 47111614 | 47177908 | 67 | 9 | 0.17597 | 0.43016 | NETO2 | 0.43016 | 0.47947 | 0.54347 |
| ENSG00000172456 | 1 | 59762310 | 60233347 | 1287 | 86 | 0.17592 | 0.43018 | FGGY | 0.43018 | 0.47602 | 0.48327 |
| ENSG00000137171 | 6 | 43008515 | 43042837 | 93 | 13 | 0.17592 | 0.43018 | KLC4 | 0.43018 | 0.44113 | 0.56989 |
| ENSG00000110697 | 11 | 67259239 | 67273734 | 12 | 7 | 0.17564 | 0.43029 | PITPNM1 | 0.43029 | 0.47838 | 0.57516 |
| ENSG00000166192 | 15 | 72406599 | 72433311 | 44 | 7 | 0.17559 | 0.43031 | SENP8 | 0.43031 | 0.47096 | 0.47406 |
| ENSG00000145337 | 4 | 89442136 | 89444964 | 2 | 1 | 0.17545 | 0.43036 | PYURF | 0.43036 | 0.31431 | 0.60735 |
| ENSG00000149600 | 20 | 31290493 | 31331803 | 191 | 12 | 0.17514 | 0.43048 | COMMD7 | 0.43048 | 0.41831 | 0.33397 |
| ENSG00000196648 | 15 | 83098710 | 83108085 | 1 | 1 | 0.1751 | 0.4305 | GOLGA6L20 | 0.4305 | 0.4305 | 0.23317 |
| ENSG00000273395 | 1 | 48226979 | 48227290 | 1 | 1 | 0.17485 | 0.4306 | FLJ00388 | 0.4306 | 0.70103 | 0.6816 |
| ENSG00000198331 | 11 | 125753509 | 125770543 | 59 | 8 | 0.17479 | 0.43062 | HYLS1 | 0.43062 | 0.62999 | 0.67512 |
| ENSG00000120457 | 11 | 128761251 | 128790930 | 140 | 22 | 0.17474 | 0.43064 | KCNJ5 | 0.43064 | 0.45169 | 0.69423 |
| ENSG00000189171 | 1 | 153591263 | 153606568 | 35 | 4 | 0.17455 | 0.43072 | S100A13 | 0.43072 | 0.41714 | 0.42641 |
| ENSG00000075234 | 22 | 46663858 | 46689905 | 99 | 9 | 0.17454 | 0.43072 | TTC38 | 0.43072 | 0.44145 | 0.52695 |
| ENSG00000198752 | 14 | 103398716 | 103523799 | 363 | 33 | 0.17452 | 0.43073 | CDC42BPB | 0.43073 | 0.33091 | 0.40067 |

| | | | | | | | | | | | |
|---|---|---|---|---|---|---|---|---|---|---|---|
| ENSG00000183888 | 1 | 16330731 | 16335302 | 16 | 5 | 0.17446 | 0.43075 | C1orf64 | 0.43075 | 0.62623 | 0.74249 |
| ENSG00000206107 | 21 | 31709331 | 31710012 | 2 | 1 | 0.1744 | 0.43077 | KRTAP27-1 | 0.43077 | 0.64101 | 0.48841 |
| ENSG00000172175 | 18 | 56338618 | 56417371 | 272 | 19 | 0.17395 | 0.43095 | MALT1 | 0.43095 | 0.44061 | 0.38691 |
| ENSG00000152467 | 19 | 58545400 | 58565999 | 48 | 8 | 0.17378 | 0.43102 | ZSCAN1 | 0.43102 | 0.6069 | 0.53478 |
| ENSG00000160223 | 21 | 45642874 | 45660849 | 83 | 16 | 0.17378 | 0.43102 | ICOSLG | 0.43102 | 0.6378 | 0.73571 |
| ENSG00000111269 | 12 | 12764761 | 12798042 | 87 | 13 | 0.17372 | 0.43104 | CREBL2 | 0.43104 | 0.50078 | 0.56585 |
| ENSG00000146376 | 6 | 129897277 | 130031370 | 495 | 51 | 0.17359 | 0.43109 | ARHGAP18 | 0.43109 | 0.56722 | 0.55695 |
| ENSG00000114209 | 3 | 167401086 | 167452727 | 86 | 5 | 0.17358 | 0.4311 | PDCD10 | 0.4311 | 0.44693 | 0.48639 |
| ENSG00000182352 | 17 | 72580818 | 72590348 | 36 | 7 | 0.17345 | 0.43115 | C17orf77 | 0.43115 | 0.36607 | 0.40271 |
| ENSG00000198959 | 20 | 36756863 | 36794980 | 99 | 15 | 0.17327 | 0.43122 | TGM2 | 0.43122 | 0.17543 | 0.24702 |
| ENSG00000158019 | 2 | 28112808 | 28561768 | 1150 | 53 | 0.17325 | 0.43123 | BRE | 0.43123 | 0.43668 | 0.54365 |
| ENSG00000143970 | 2 | 25956622 | 26101385 | 327 | 14 | 0.17301 | 0.43132 | ASXL2 | 0.43132 | 0.42705 | 0.44158 |
| ENSG00000120341 | 1 | 177893091 | 177953438 | 240 | 40 | 0.17298 | 0.43133 | SEC16B | 0.43133 | 0.3232 | 0.34381 |
| ENSG00000124491 | 6 | 6144318 | 6321246 | 793 | 108 | 0.17243 | 0.43155 | F13A1 | 0.43155 | 0.43309 | 0.51963 |
| ENSG00000107130 | 9 | 132934857 | 132999583 | 282 | 33 | 0.17244 | 0.43155 | NCS1 | 0.43155 | 0.44592 | 0.37982 |
| ENSG00000170502 | 4 | 88343734 | 88380606 | 106 | 15 | 0.17231 | 0.4316 | NUDT9 | 0.4316 | 0.50602 | 0.47365 |
| ENSG00000183049 | 10 | 12391481 | 12877545 | 2138 | 194 | 0.17217 | 0.43165 | CAMK1D | 0.43165 | 0.37754 | 0.29489 |
| ENSG00000169490 | 8 | 38846327 | 38854343 | 14 | 3 | 0.17216 | 0.43166 | TM2D2 | 0.43166 | 0.5969 | 0.72984 |
| ENSG00000268301 | 12 | 11187087 | 11187275 | 2 | 1 | 0.17173 | 0.43182 | AC018630.1 | 0.43182 | 0.41334 | 0.39629 |
| ENSG00000186212 | 4 | 77816082 | 77819002 | 8 | 3 | 0.17153 | 0.4319 | SOWAHB | 0.4319 | 0.33595 | 0.32817 |
| ENSG00000205464 | 5 | 81575281 | 81682796 | 191 | 24 | 0.17152 | 0.43191 | ATP6AP1L | 0.43191 | 0.39199 | 0.45423 |
| ENSG00000259060 | 14 | 21051051 | 21077954 | 119 | 29 | 0.17144 | 0.43194 | RNASE11 | 0.43194 | 0.22565 | 0.27646 |
| ENSG00000173464 | 14 | 21051054 | 21078043 | 119 | 29 | 0.17144 | 0.43194 | RNASE11 | 0.43194 | 0.22565 | 0.27646 |
| ENSG00000139160 | 12 | 31800094 | 31826048 | 94 | 23 | 0.17141 | 0.43195 | METTL20 | 0.43195 | 0.35179 | 0.46565 |
| ENSG00000162367 | 1 | 47681962 | 47697892 | 38 | 8 | 0.17067 | 0.43224 | TAL1 | 0.43224 | 0.27169 | 0.20497 |
| ENSG00000114547 | 3 | 125687987 | 125702297 | 36 | 12 | 0.17059 | 0.43227 | ROPN1B | 0.43227 | 0.18933 | 0.21465 |
| ENSG00000253148 | 1 | 192286122 | 192336415 | 138 | 17 | 0.17058 | 0.43228 | RGS21 | 0.43228 | 0.44368 | 0.45866 |
| ENSG00000079974 | 22 | 51205929 | 51222091 | 37 | 6 | 0.17048 | 0.43232 | RABL2B | 0.43232 | 0.23714 | 0.19974 |
| ENSG00000123454 | 9 | 136501482 | 136524466 | 143 | 28 | 0.17037 | 0.43236 | DBH | 0.43236 | 0.48958 | 0.32661 |
| ENSG00000186998 | 22 | 29601840 | 29655586 | 220 | 31 | 0.16988 | 0.43255 | EMID1 | 0.43255 | 0.39802 | 0.41429 |
| ENSG00000154639 | 21 | 18884700 | 18965897 | 285 | 21 | 0.16969 | 0.43263 | CXADR | 0.43263 | 0.47105 | 0.56483 |
| ENSG00000214367 | 4 | 2229191 | 2243891 | 40 | 6 | 0.16958 | 0.43267 | HAUS3 | 0.43267 | 0.45839 | 0.47372 |
| ENSG00000105989 | 7 | 116916685 | 116963343 | 96 | 21 | 0.16932 | 0.43277 | WNT2 | 0.43277 | 0.37105 | 0.28306 |
| ENSG00000128731 | 15 | 28356186 | 28567298 | 285 | 15 | 0.16929 | 0.43279 | HERC2 | 0.43279 | 0.39458 | 0.42259 |
| ENSG00000157064 | 1 | 183217372 | 183387737 | 449 | 30 | 0.16906 | 0.43288 | NMNAT2 | 0.43288 | 0.44689 | 0.45662 |
| ENSG00000134871 | 13 | 110958159 | 111165374 | 1179 | 119 | 0.16897 | 0.43291 | COL4A2 | 0.43291 | 0.52154 | 0.67831 |
| ENSG00000128245 | 22 | 32340447 | 32353590 | 51 | 8 | 0.16895 | 0.43292 | YWHAH | 0.43292 | 0.61636 | 0.5476 |
| ENSG00000174945 | 7 | 2719156 | 2815134 | 492 | 51 | 0.16882 | 0.43297 | AMZ1 | 0.43297 | 0.38663 | 0.40718 |
| ENSG00000113838 | 3 | 186263862 | 186288332 | 64 | 10 | 0.16867 | 0.43303 | TBCCD1 | 0.43303 | 0.42776 | 0.45416 |
| ENSG00000124140 | 20 | 44650356 | 44688784 | 150 | 23 | 0.16867 | 0.43303 | SLC12A5 | 0.43303 | 0.27518 | 0.16439 |
| ENSG00000197345 | 11 | 68658744 | 68671303 | 33 | 7 | 0.1685 | 0.43309 | MRPL21 | 0.43309 | 0.3823 | 0.39941 |
| ENSG00000142684 | 1 | 26496362 | 26498551 | 10 | 3 | 0.16844 | 0.43312 | ZNF593 | 0.43312 | 0.39488 | 0.30923 |
| ENSG00000095787 | 10 | 28821422 | 28912041 | 217 | 20 | 0.16832 | 0.43317 | WAC | 0.43317 | 0.466 | 0.56667 |
| ENSG00000180488 | 1 | 78245309 | 78344106 | 201 | 34 | 0.16813 | 0.43324 | FAM73A | 0.43324 | 0.46359 | 0.47701 |
| ENSG00000154263 | 17 | 67143355 | 67240987 | 270 | 15 | 0.168 | 0.43329 | ABCA10 | 0.43329 | 0.4902 | 0.51585 |
| ENSG00000162241 | 11 | 65142663 | 65151172 | 13 | 5 | 0.16789 | 0.43333 | SLC25A45 | 0.43333 | 0.79288 | 0.84195 |
| ENSG00000187223 | 1 | 152635872 | 152637135 | 2 | 1 | 0.16783 | 0.43336 | LCE2D | 0.43336 | 0.22703 | 0.25562 |
| ENSG00000130164 | 19 | 11200038 | 11244492 | 202 | 26 | 0.16769 | 0.43341 | LDLR | 0.43341 | 0.55585 | 0.46106 |
| ENSG00000182771 | 10 | 87359312 | 88126250 | 2548 | 136 | 0.16765 | 0.43343 | GRID1 | 0.43343 | 0.45238 | 0.45754 |
| ENSG00000142694 | 1 | 36787632 | 36789755 | 3 | 1 | 0.16749 | 0.43349 | EVA1B | 0.43349 | 0.38095 | 0.33502 |
| ENSG00000087237 | 16 | 56995762 | 57017757 | 109 | 22 | 0.16746 | 0.4335 | CETP | 0.4335 | 0.47465 | 0.3431 |
| ENSG00000064835 | 3 | 87308554 | 87325737 | 38 | 9 | 0.16741 | 0.43352 | POU1F1 | 0.43352 | 0.32246 | 0.43282 |
| ENSG00000175294 | 11 | 65784223 | 65793988 | 22 | 4 | 0.16742 | 0.43352 | CATSPER1 | 0.43352 | 0.4393 | 0.48246 |
| ENSG00000112081 | 6 | 36562145 | 36573377 | 55 | 6 | 0.16735 | 0.43355 | SRSF3 | 0.43355 | 0.27364 | 0.29658 |
| ENSG00000147408 | 8 | 19261672 | 19615540 | 1596 | 89 | 0.16732 | 0.43356 | CSGALNACT1 | 0.43356 | 0.40627 | 0.33658 |
| ENSG00000105737 | 19 | 42502473 | 42573650 | 61 | 16 | 0.16719 | 0.43361 | GRIK5 | 0.43361 | 0.49298 | 0.50561 |
| ENSG00000170638 | 22 | 50624344 | 50638027 | 55 | 11 | 0.16696 | 0.4337 | TRABD | 0.4337 | 0.18855 | 0.16143 |
| ENSG00000242366 | 2 | 234668894 | 234681945 | 35 | 10 | 0.16683 | 0.43375 | UGT1A1 | 0.43375 | 0.52137 | 0.24187 |
| ENSG00000137225 | 6 | 44126548 | 44152139 | 132 | 25 | 0.16655 | 0.43386 | CAPN11 | 0.43386 | 0.4385 | 0.62749 |
| ENSG00000188992 | 21 | 15481134 | 15583166 | 479 | 25 | 0.16644 | 0.4339 | LIPI | 0.4339 | 0.49838 | 0.30581 |
| ENSG00000118307 | 12 | 25261354 | 25348096 | 344 | 20 | 0.16629 | 0.43396 | CASC1 | 0.43396 | 0.4436 | 0.50597 |
| ENSG00000176973 | 11 | 65339820 | 65341669 | 2 | 1 | 0.16627 | 0.43397 | FAM89B | 0.43397 | 0.11955 | 0.069316 |
| ENSG00000180357 | 15 | 64752941 | 64978264 | 511 | 24 | 0.16611 | 0.43403 | ZNF609 | 0.43403 | 0.43235 | 0.4373 |
| ENSG00000163982 | 4 | 4190530 | 4228616 | 213 | 25 | 0.16586 | 0.43413 | OTOP1 | 0.43413 | 0.47694 | 0.66265 |
| ENSG00000204256 | 6 | 32936437 | 32949282 | 127 | 28 | 0.16564 | 0.43422 | BRD2 | 0.43422 | 0.61626 | 0.42598 |
| ENSG00000174500 | 3 | 111839688 | 111852152 | 30 | 7 | 0.16562 | 0.43423 | GCSAM | 0.43423 | 0.17851 | 0.18232 |
| ENSG00000033011 | 16 | 5083703 | 5137380 | 196 | 42 | 0.16528 | 0.43436 | ALG1 | 0.43436 | 0.50718 | 0.31015 |
| ENSG00000120685 | 13 | 39584003 | 39612252 | 62 | 12 | 0.16517 | 0.4344 | PROSER1 | 0.4344 | 0.30773 | 0.23023 |
| ENSG00000268279 | 3 | 14176717 | 14189548 | 42 | 9 | 0.1651 | 0.43443 | RP11-434D12.1 | 0.43443 | 0.63268 | 0.7031 |
| ENSG00000145721 | 5 | 96427574 | 96478576 | 91 | 13 | 0.16453 | 0.43466 | LIX1 | 0.43466 | 0.49517 | 0.54523 |
| ENSG00000186160 | 1 | 47533160 | 47583991 | 235 | 10 | 0.16446 | 0.43469 | CYP4Z1 | 0.43469 | 0.43625 | 0.3903 |
| ENSG00000101460 | 20 | 33134658 | 33148149 | 28 | 6 | 0.1644 | 0.43471 | MAP1LC3A | 0.43471 | 0.356 | 0.41886 |
| ENSG00000186468 | 5 | 81569177 | 81574396 | 18 | 6 | 0.16436 | 0.43472 | RPS23 | 0.43472 | 0.48343 | 0.61528 |

| | | | | | | | | | | | |
|---|---|---|---|---|---|---|---|---|---|---|---|
| ENSG00000177291 | 10 | 35894338 | 35897863 | 15 | 5 | 0.16432 | 0.43474 | GJD4 | 0.43474 | 0.056234 | 0.076291 |
| ENSG00000136603 | 3 | 170075466 | 170114623 | 140 | 15 | 0.16398 | 0.43487 | SKIL | 0.43487 | 0.54634 | 0.60489 |
| ENSG00000164054 | 3 | 48509197 | 48542259 | 53 | 12 | 0.16396 | 0.43488 | SHISA5 | 0.43488 | 0.47958 | 0.46653 |
| ENSG00000187678 | 5 | 141689992 | 141706020 | 36 | 9 | 0.16357 | 0.43504 | SPRY4 | 0.43504 | 0.5424 | 0.56484 |
| ENSG00000115866 | 2 | 136664247 | 136743670 | 102 | 6 | 0.16335 | 0.43512 | DARS | 0.43512 | 0.45182 | 0.51033 |
| ENSG00000164099 | 4 | 119201193 | 119274158 | 212 | 30 | 0.16327 | 0.43515 | PRSS12 | 0.43515 | 0.40244 | 0.3625 |
| ENSG00000084734 | 2 | 27719709 | 27746554 | 57 | 11 | 0.1632 | 0.43518 | GCKR | 0.43518 | 0.41218 | 0.52147 |
| ENSG00000119699 | 14 | 76424442 | 76449334 | 59 | 10 | 0.16307 | 0.43523 | TGFB3 | 0.43523 | 0.40174 | 0.35354 |
| ENSG00000150510 | 13 | 51796503 | 51858377 | 196 | 37 | 0.16305 | 0.43524 | FAM124A | 0.43524 | 0.51618 | 0.65267 |
| ENSG00000063660 | 2 | 241375088 | 241407493 | 169 | 36 | 0.16271 | 0.43537 | GPC1 | 0.43537 | 0.65291 | 0.73905 |
| ENSG00000166912 | 15 | 31231144 | 31283810 | 179 | 17 | 0.16263 | 0.4354 | MTMR10 | 0.4354 | 0.49247 | 0.60484 |
| ENSG00000169174 | 1 | 55505221 | 55530525 | 133 | 26 | 0.16208 | 0.43562 | PCSK9 | 0.43562 | 0.46542 | 0.62597 |
| ENSG00000165029 | 9 | 107543283 | 107690518 | 674 | 80 | 0.16165 | 0.43579 | ABCA1 | 0.43579 | 0.60469 | 0.67137 |
| ENSG00000104756 | 8 | 25285366 | 25315992 | 88 | 5 | 0.16161 | 0.43581 | KCTD9 | 0.43581 | 0.47117 | 0.47573 |
| ENSG00000159445 | 1 | 151846060 | 151882284 | 95 | 9 | 0.16143 | 0.43588 | THEM4 | 0.43588 | 0.42851 | 0.50037 |
| ENSG00000113249 | 5 | 156456424 | 156486130 | 162 | 21 | 0.16134 | 0.43591 | HAVCR1 | 0.43591 | 0.50263 | 0.53998 |
| ENSG00000152578 | 11 | 105480721 | 105852819 | 913 | 42 | 0.16113 | 0.43599 | GRIA4 | 0.43599 | 0.46221 | 0.51562 |
| ENSG00000172748 | 8 | 182137 | 197342 | 48 | 10 | 0.16113 | 0.436 | ZNF596 | 0.436 | 0.6462 | 0.65487 |
| ENSG00000180720 | 11 | 46406640 | 46408107 | 1 | 1 | 0.16112 | 0.436 | CHRM4 | 0.436 | 0.31709 | 0.47205 |
| ENSG00000177202 | 19 | 49110000 | 49110970 | 1 | 1 | 0.16086 | 0.4361 | SPACA4 | 0.4361 | 0.056324 | 0.017385 |
| ENSG00000078674 | 8 | 17780349 | 17885478 | 556 | 21 | 0.16049 | 0.43625 | PCM1 | 0.43625 | 0.4262 | 0.37209 |
| ENSG00000151690 | 2 | 191273081 | 191373931 | 306 | 18 | 0.16005 | 0.43642 | MFSD6 | 0.43642 | 0.39244 | 0.35346 |
| ENSG00000184730 | 16 | 28505970 | 28510291 | 11 | 3 | 0.15999 | 0.43644 | APOBR | 0.43644 | 0.82223 | 0.81833 |
| ENSG00000187474 | 19 | 52298416 | 52329442 | 130 | 19 | 0.15992 | 0.43647 | FPR3 | 0.43647 | 0.52137 | 0.67053 |
| ENSG00000197056 | 1 | 35525387 | 35581460 | 88 | 10 | 0.15938 | 0.43669 | ZMYM1 | 0.43669 | 0.42417 | 0.42093 |
| ENSG00000153714 | 9 | 12775020 | 12822130 | 175 | 40 | 0.15936 | 0.43669 | LURAP1L | 0.43669 | 0.48472 | 0.57403 |
| ENSG00000163216 | 1 | 153012201 | 153014407 | 14 | 4 | 0.15935 | 0.4367 | SPRR2D | 0.4367 | 0.64444 | 0.60503 |
| ENSG00000215454 | 21 | 45993606 | 45994987 | 7 | 4 | 0.15931 | 0.43671 | KRTAP10-4 | 0.43671 | 0.54252 | 0.25787 |
| ENSG00000171680 | 1 | 6526152 | 6580121 | 161 | 26 | 0.15922 | 0.43675 | PLEKHG5 | 0.43675 | 0.24261 | 0.16953 |
| ENSG00000153107 | 2 | 112523848 | 112642267 | 390 | 22 | 0.15922 | 0.43675 | ANAPC1 | 0.43675 | 0.43257 | 0.43328 |
| ENSG00000160796 | 3 | 47021173 | 47051193 | 48 | 6 | 0.15906 | 0.43681 | NBEAL2 | 0.43681 | 0.48005 | 0.54698 |
| ENSG00000121281 | 16 | 50280048 | 50352046 | 196 | 32 | 0.15884 | 0.4369 | ADCY7 | 0.4369 | 0.52819 | 0.60761 |
| ENSG00000175265 | 15 | 34671269 | 34880704 | 879 | 30 | 0.15874 | 0.43694 | GOLGA8A | 0.43694 | 0.40841 | 0.38092 |
| ENSG00000108352 | 17 | 38333263 | 38351908 | 40 | 8 | 0.15851 | 0.43703 | RAPGEFL1 | 0.43703 | 0.3731 | 0.41459 |
| ENSG00000147596 | 8 | 70963886 | 70983928 | 66 | 14 | 0.15838 | 0.43708 | PRDM14 | 0.43708 | 0.57238 | 0.51324 |
| ENSG00000198271 | 17 | 39305178 | 39306054 | 6 | 3 | 0.15818 | 0.43716 | KRTAP4-5 | 0.43716 | 0.40776 | 0.22945 |
| ENSG00000125845 | 20 | 6748311 | 6760927 | 32 | 10 | 0.15797 | 0.43724 | BMP2 | 0.43724 | 0.43174 | 0.68339 |
| ENSG00000267970 | 7 | 48887362 | 48891221 | 23 | 6 | 0.15795 | 0.43725 | AC004899.1 | 0.43725 | 0.25321 | 0.2343 |
| ENSG00000144048 | 2 | 73989311 | 74007284 | 19 | 6 | 0.15784 | 0.43729 | DUSP11 | 0.43729 | 0.50963 | 0.64427 |
| ENSG00000150783 | 11 | 112038095 | 112043282 | 15 | 5 | 0.15772 | 0.43734 | TEX12 | 0.43734 | 0.64719 | 0.60845 |
| ENSG00000269741 | 19 | 51499274 | 51512837 | 69 | 11 | 0.15753 | 0.43741 | KLK9 | 0.43741 | 0.47794 | 0.51422 |
| ENSG00000182319 | 8 | 8175258 | 8244008 | 254 | 32 | 0.15746 | 0.43744 | SGK223 | 0.43744 | 0.53005 | 0.53059 |
| ENSG00000197020 | 19 | 21905568 | 21950430 | 217 | 15 | 0.15726 | 0.43752 | ZNF100 | 0.43752 | 0.50654 | 0.61646 |
| ENSG00000267849 | 6 | 44400472 | 44400720 | 2 | 1 | 0.15707 | 0.43759 | AL133262.1 | 0.43759 | 0.75777 | 0.79664 |
| ENSG00000136866 | 9 | 115800660 | 115819039 | 47 | 14 | 0.15708 | 0.43759 | ZFP37 | 0.43759 | 0.51916 | 0.53936 |
| ENSG00000234829 | 9 | 21227242 | 21228221 | 6 | 1 | 0.15707 | 0.4376 | IFNA17 | 0.4376 | 0.40011 | 0.44015 |
| ENSG00000119771 | 2 | 23608088 | 23931481 | 1285 | 110 | 0.15679 | 0.4377 | KLHL29 | 0.4377 | 0.41063 | 0.49493 |
| ENSG00000196581 | 1 | 4714792 | 4852594 | 765 | 86 | 0.1567 | 0.43774 | AJAP1 | 0.43774 | 0.45231 | 0.40151 |
| ENSG00000151693 | 2 | 9346894 | 9545812 | 514 | 28 | 0.15665 | 0.43776 | ASAP2 | 0.43776 | 0.30668 | 0.21842 |
| ENSG00000154079 | 6 | 71276620 | 71299272 | 99 | 7 | 0.15655 | 0.4378 | C6orf57 | 0.4378 | 0.51106 | 0.56005 |
| ENSG00000164050 | 3 | 48445261 | 48471594 | 28 | 4 | 0.1563 | 0.4379 | PLXNB1 | 0.4379 | 0.41272 | 0.3903 |
| ENSG00000110871 | 12 | 120941077 | 120972237 | 122 | 16 | 0.15564 | 0.43816 | COQ5 | 0.43816 | 0.46731 | 0.4862 |
| ENSG00000027869 | 1 | 156776035 | 156786654 | 20 | 4 | 0.15558 | 0.43818 | SH2D2A | 0.43818 | 0.39811 | 0.48035 |
| ENSG00000104112 | 15 | 51973550 | 52013223 | 111 | 19 | 0.15558 | 0.43818 | SCG3 | 0.43818 | 0.33475 | 0.38094 |
| ENSG00000010165 | 1 | 171750788 | 171783163 | 116 | 8 | 0.15542 | 0.43824 | METTL13 | 0.43824 | 0.40894 | 0.44375 |
| ENSG00000239306 | 11 | 66384053 | 66394818 | 12 | 5 | 0.15544 | 0.43824 | RBM14 | 0.43824 | 0.35767 | 0.28236 |
| ENSG00000132274 | 11 | 5710919 | 5758319 | 332 | 28 | 0.15482 | 0.43848 | TRIM22 | 0.43848 | 0.57549 | 0.73596 |
| ENSG00000152726 | 10 | 47894023 | 47949412 | 1 | 1 | 0.15477 | 0.4385 | FAM21B | 0.4385 | 0.50581 | 0.2326 |
| ENSG00000119943 | 10 | 100143322 | 100174941 | 160 | 10 | 0.15446 | 0.43862 | PYROXD2 | 0.43862 | 0.39459 | 0.42217 |
| ENSG00000150760 | 10 | 128593978 | 129250781 | 2230 | 149 | 0.15438 | 0.43866 | DOCK1 | 0.43866 | 0.3761 | 0.41194 |
| ENSG00000203747 | 1 | 161511549 | 161600917 | 405 | 40 | 0.15429 | 0.43869 | FCGR3A | 0.43869 | 0.4598 | 0.49333 |
| ENSG00000163113 | 1 | 149909705 | 149982625 | 101 | 12 | 0.15427 | 0.4387 | OTUD7B | 0.4387 | 0.43531 | 0.5437 |
| ENSG00000166971 | 16 | 53524952 | 53538323 | 22 | 5 | 0.15419 | 0.43873 | AKTIP | 0.43873 | 0.332 | 0.46716 |
| ENSG00000205981 | 3 | 180701497 | 180707562 | 11 | 5 | 0.15403 | 0.43879 | DNAJC19 | 0.43879 | 0.46224 | 0.3109 |
| ENSG00000197914 | 6 | 27798952 | 27799305 | 1 | 1 | 0.15401 | 0.4388 | HIST1H4K | 0.4388 | 0.5969 | 0.71617 |
| ENSG00000167965 | 16 | 2254249 | 2259417 | 13 | 4 | 0.154 | 0.4388 | MLST8 | 0.4388 | 0.71029 | 0.8253 |
| ENSG00000163781 | 3 | 133317019 | 133380737 | 215 | 19 | 0.15387 | 0.43886 | TOPBP1 | 0.43886 | 0.38896 | 0.40223 |
| ENSG00000182600 | 2 | 233721980 | 233743418 | 65 | 13 | 0.15381 | 0.43888 | C2orf82 | 0.43888 | 0.54046 | 0.56701 |
| ENSG00000181220 | 7 | 149169885 | 149194908 | 30 | 4 | 0.15353 | 0.43899 | ZNF746 | 0.43899 | 0.3676 | 0.22991 |
| ENSG00000149658 | 20 | 61826781 | 61847586 | 51 | 7 | 0.15347 | 0.43902 | YTHDF1 | 0.43902 | 0.55801 | 0.67495 |
| ENSG00000116750 | 1 | 192981380 | 193029237 | 50 | 11 | 0.1533 | 0.43908 | UCHL5 | 0.43908 | 0.43174 | 0.38372 |
| ENSG00000164822 | 8 | 6782215 | 6783598 | 9 | 5 | 0.15327 | 0.43909 | DEFA6 | 0.43909 | 0.19998 | 0.21175 |
| ENSG00000206203 | 22 | 19118321 | 19120134 | 5 | 1 | 0.15324 | 0.4391 | TSSK2 | 0.4391 | 0.73417 | 0.81988 |

| | | | | | | | | | | | |
|---|---|---|---|---|---|---|---|---|---|---|---|
| ENSG00000186184 | 13 | 28194903 | 28241548 | 160 | 28 | 0.15293 | 0.43923 | POLR1D | 0.43923 | 0.099835 | 0.071178 |
| ENSG00000229086 | 21 | 34537776 | 34542541 | 17 | 7 | 0.15253 | 0.43938 | C21orf54 | 0.43938 | 0.723 | 0.73456 |
| ENSG00000136270 | 7 | 45139699 | 45151646 | 44 | 10 | 0.15252 | 0.43939 | TBRG4 | 0.43939 | 0.76845 | 0.75917 |
| ENSG00000010539 | 16 | 3272325 | 3286221 | 38 | 6 | 0.15213 | 0.43954 | ZNF200 | 0.43954 | 0.49746 | 0.38323 |
| ENSG00000221968 | 11 | 61640991 | 61659523 | 38 | 5 | 0.15197 | 0.43961 | FADS3 | 0.43961 | 0.20254 | 0.045257 |
| ENSG00000183092 | 14 | 101003486 | 101053750 | 145 | 20 | 0.15175 | 0.43969 | BEGAIN | 0.43969 | 0.43677 | 0.49823 |
| ENSG00000196748 | 6 | 35744371 | 35747329 | 11 | 5 | 0.15165 | 0.43973 | CLPSL2 | 0.43973 | 0.52642 | 0.53538 |
| ENSG00000164024 | 4 | 99916771 | 99983964 | 194 | 14 | 0.15164 | 0.43974 | METAP1 | 0.43974 | 0.44316 | 0.44735 |
| ENSG00000181284 | 17 | 7338762 | 7340998 | 4 | 1 | 0.15147 | 0.4398 | TMEM102 | 0.4398 | 0.20024 | 0.40159 |
| ENSG00000188895 | 17 | 38278551 | 38293042 | 33 | 5 | 0.15147 | 0.4398 | MSL1 | 0.4398 | 0.34545 | 0.37811 |
| ENSG00000185163 | 12 | 132621139 | 132628880 | 26 | 8 | 0.15144 | 0.43982 | DDX51 | 0.43982 | 0.50602 | 0.61431 |
| ENSG00000198952 | 1 | 156219015 | 156252620 | 96 | 11 | 0.15131 | 0.43986 | SMG5 | 0.43986 | 0.41268 | 0.54833 |
| ENSG00000165084 | 8 | 69242957 | 69731257 | 1631 | 89 | 0.15131 | 0.43986 | C8orf34 | 0.43986 | 0.43486 | 0.46286 |
| ENSG00000114841 | 3 | 52350335 | 52434507 | 122 | 12 | 0.15098 | 0.43999 | DNAH1 | 0.43999 | 0.44758 | 0.44181 |
| ENSG00000124107 | 20 | 43880880 | 43883205 | 12 | 4 | 0.15097 | 0.44 | SLPI | 0.44 | 0.6832 | 0.66277 |
| ENSG00000180138 | 13 | 37677398 | 37679803 | 9 | 2 | 0.1509 | 0.44003 | CSNK1A1L | 0.44003 | 0.55972 | 0.62976 |
| ENSG00000140937 | 16 | 64977656 | 65160015 | 561 | 36 | 0.15087 | 0.44004 | CDH11 | 0.44004 | 0.47829 | 0.44741 |
| ENSG00000130700 | 20 | 61038553 | 61051026 | 50 | 11 | 0.15082 | 0.44006 | GATA5 | 0.44006 | 0.45965 | 0.53833 |
| ENSG00000065833 | 6 | 83920108 | 84140797 | 593 | 19 | 0.15075 | 0.44009 | ME1 | 0.44009 | 0.31389 | 0.29914 |
| ENSG00000104643 | 8 | 11141925 | 11185646 | 130 | 10 | 0.15046 | 0.4402 | MTMR9 | 0.4402 | 0.43981 | 0.42695 |
| ENSG00000105854 | 7 | 95034175 | 95064510 | 97 | 13 | 0.15044 | 0.44021 | PON2 | 0.44021 | 0.54046 | 0.59538 |
| ENSG00000187116 | 19 | 54818353 | 54824409 | 14 | 3 | 0.15033 | 0.44025 | LILRA5 | 0.44025 | 0.60757 | 0.53384 |
| ENSG00000079277 | 1 | 47023090 | 47082515 | 146 | 15 | 0.15024 | 0.44029 | MKNK1 | 0.44029 | 0.46261 | 0.50692 |
| ENSG00000117335 | 1 | 207925402 | 207968858 | 124 | 16 | 0.15002 | 0.44037 | CD46 | 0.44037 | 0.37693 | 0.3555 |
| ENSG00000138758 | 4 | 77870856 | 77961537 | 124 | 28 | 0.14996 | 0.4404 | SEPTIN11 | 0.4404 | 0.51861 | 0.55268 |
| ENSG00000118515 | 6 | 134490384 | 134639250 | 526 | 64 | 0.14981 | 0.44046 | SGK1 | 0.44046 | 0.5091 | 0.607 |
| ENSG00000136943 | 9 | 99794940 | 99801925 | 4 | 2 | 0.14979 | 0.44047 | CTSV | 0.44047 | 0.052414 | 0.040743 |
| ENSG00000115009 | 2 | 228678558 | 228682272 | 5 | 3 | 0.14968 | 0.44051 | CCL20 | 0.44051 | 0.60103 | 0.70278 |
| ENSG00000197816 | 9 | 100000765 | 100140806 | 470 | 17 | 0.14946 | 0.44059 | CCDC180 | 0.44059 | 0.45803 | 0.41217 |
| ENSG00000182010 | 10 | 63942794 | 64028466 | 271 | 15 | 0.14928 | 0.44067 | RTKN2 | 0.44067 | 0.45308 | 0.44621 |
| ENSG00000173226 | 3 | 121488610 | 121553926 | 238 | 15 | 0.14909 | 0.44074 | IQCB1 | 0.44074 | 0.46398 | 0.42575 |
| ENSG00000189182 | 12 | 53083410 | 53097247 | 40 | 9 | 0.14908 | 0.44074 | KRT77 | 0.44074 | 0.65548 | 0.86596 |
| ENSG00000163257 | 4 | 17802278 | 17812381 | 19 | 3 | 0.14907 | 0.44075 | DCAF16 | 0.44075 | 0.67191 | 0.7579 |
| ENSG00000124783 | 6 | 7268539 | 7347679 | 597 | 27 | 0.14903 | 0.44076 | SSR1 | 0.44076 | 0.42045 | 0.48681 |
| ENSG00000091428 | 2 | 173600002 | 173917621 | 1054 | 71 | 0.14893 | 0.44081 | RAPGEF4 | 0.44081 | 0.46127 | 0.42052 |
| ENSG00000204140 | 6 | 35748794 | 35761452 | 86 | 17 | 0.14874 | 0.44088 | CLPSL1 | 0.44088 | 0.39603 | 0.54266 |
| ENSG00000171227 | 2 | 120187477 | 120196096 | 41 | 8 | 0.14861 | 0.44093 | TMEM37 | 0.44093 | 0.16832 | 0.18856 |
| ENSG00000196074 | 20 | 58438618 | 58508710 | 193 | 7 | 0.14828 | 0.44106 | SYCP2 | 0.44106 | 0.45129 | 0.40738 |
| ENSG00000105369 | 19 | 42381190 | 42385439 | 6 | 3 | 0.14819 | 0.4411 | CD79A | 0.4411 | 0.66888 | 0.63095 |
| ENSG00000086598 | 12 | 124069078 | 124083116 | 53 | 8 | 0.14772 | 0.44128 | TMED2 | 0.44128 | 0.38087 | 0.45409 |
| ENSG00000177489 | 1 | 247751662 | 247752615 | 6 | 4 | 0.14749 | 0.44137 | OR2G2 | 0.44137 | 0.8674 | 0.8986 |
| ENSG00000163581 | 3 | 170714137 | 170744539 | 89 | 7 | 0.14742 | 0.4414 | SLC2A2 | 0.4414 | 0.39639 | 0.38708 |
| ENSG00000187244 | 19 | 45312328 | 45324673 | 47 | 9 | 0.14734 | 0.44143 | BCAM | 0.44143 | 0.56282 | 0.53408 |
| ENSG00000268217 | 5 | 72894203 | 72896845 | 12 | 4 | 0.14704 | 0.44155 | AC008387.1 | 0.44155 | 0.33696 | 0.2925 |
| ENSG00000011243 | 19 | 15490859 | 15529952 | 79 | 7 | 0.14675 | 0.44166 | AKAP8L | 0.44166 | 0.41131 | 0.40751 |
| ENSG00000106384 | 7 | 100838288 | 100844302 | 15 | 4 | 0.14669 | 0.44169 | MOGAT3 | 0.44169 | 0.53479 | 0.68398 |
| ENSG00000135447 | 12 | 54969171 | 54982443 | 40 | 9 | 0.14669 | 0.44169 | PPP1R1A | 0.44169 | 0.4264 | 0.61362 |
| ENSG00000186207 | 1 | 152483320 | 152484653 | 4 | 2 | 0.14667 | 0.4417 | LCE5A | 0.4417 | 0.35478 | 0.34886 |
| ENSG00000163659 | 3 | 156391024 | 156424559 | 83 | 15 | 0.1466 | 0.44172 | TIPARP | 0.44172 | 0.41273 | 0.41135 |
| ENSG00000139233 | 12 | 66516842 | 66524548 | 20 | 4 | 0.14657 | 0.44174 | LLPH | 0.44174 | 0.459 | 0.47173 |
| ENSG00000205078 | 16 | 77233294 | 77247112 | 93 | 13 | 0.14655 | 0.44174 | SYCE1L | 0.44174 | 0.50971 | 0.40694 |
| ENSG00000134216 | 1 | 111833484 | 111863188 | 230 | 23 | 0.14654 | 0.44175 | CHIA | 0.44175 | 0.44879 | 0.45501 |
| ENSG00000164736 | 8 | 55370495 | 55373448 | 7 | 2 | 0.14649 | 0.44177 | SOX17 | 0.44177 | 0.57916 | 0.73609 |
| ENSG00000227729 | 14 | 104406763 | 104408645 | 7 | 2 | 0.14633 | 0.44183 | RD3L | 0.44183 | 0.63948 | 0.6598 |
| ENSG00000137757 | 11 | 104864962 | 104893895 | 101 | 15 | 0.14584 | 0.44202 | CASP5 | 0.44202 | 0.19695 | 0.042008 |
| ENSG00000138293 | 10 | 51565108 | 51590734 | 33 | 8 | 0.14554 | 0.44214 | NCOA4 | 0.44214 | 0.52083 | 0.54041 |
| ENSG00000215621 | 17 | 79780293 | 79784533 | 11 | 2 | 0.14541 | 0.44219 | AC174470.1 | 0.44219 | 0.64405 | 0.61296 |
| ENSG00000185088 | 15 | 63418071 | 63450220 | 146 | 17 | 0.14532 | 0.44223 | RPS27L | 0.44223 | 0.55034 | 0.52487 |
| ENSG00000255737 | 12 | 58120054 | 58122139 | 1 | 1 | 0.14514 | 0.4423 | AGAP2-AS1 | 0.4423 | 0.42919 | 0.37699 |
| ENSG00000235169 | 1 | 3689352 | 3692546 | 17 | 6 | 0.14512 | 0.44231 | SMIM1 | 0.44231 | 0.75563 | 0.63675 |
| ENSG00000176723 | 16 | 31445569 | 31454346 | 14 | 6 | 0.14509 | 0.44232 | ZNF843 | 0.44232 | 0.5534 | 0.59758 |
| ENSG00000183549 | 16 | 20420856 | 20452658 | 242 | 16 | 0.14493 | 0.44238 | ACSM5 | 0.44238 | 0.52099 | 0.46312 |
| ENSG00000149932 | 16 | 29952206 | 29984373 | 56 | 6 | 0.1448 | 0.44243 | TMEM219 | 0.44243 | 0.51484 | 0.55175 |
| ENSG00000244474 | 2 | 234627424 | 234681945 | 194 | 19 | 0.14479 | 0.44244 | UGT1A4 | 0.44244 | 0.37343 | 0.227 |
| ENSG00000169851 | 4 | 30722037 | 31148422 | 1310 | 63 | 0.14432 | 0.44262 | PCDH7 | 0.44262 | 0.44905 | 0.49276 |
| ENSG00000115307 | 2 | 74753772 | 74757066 | 4 | 2 | 0.14411 | 0.44271 | AUP1 | 0.44271 | 0.64815 | 0.27849 |
| ENSG00000141314 | 17 | 30593195 | 30651680 | 191 | 31 | 0.14385 | 0.44281 | RHBDL3 | 0.44281 | 0.43307 | 0.39198 |
| ENSG00000250506 | 17 | 73996987 | 74002080 | 13 | 6 | 0.14369 | 0.44287 | CDK3 | 0.44287 | 0.35887 | 0.47418 |
| ENSG00000127780 | 17 | 3336164 | 3337135 | 2 | 1 | 0.14366 | 0.44288 | OR1E2 | 0.44288 | 0.62945 | 0.57335 |
| ENSG00000176984 | 11 | 120039685 | 120042387 | 12 | 4 | 0.1436 | 0.44291 | AP000679.2 | 0.44291 | 0.84197 | 0.8229 |
| ENSG00000120662 | 13 | 41790505 | 41837742 | 153 | 12 | 0.14361 | 0.44291 | MTRF1 | 0.44291 | 0.44068 | 0.39809 |
| ENSG00000165023 | 9 | 93372114 | 93405386 | 131 | 23 | 0.14351 | 0.44294 | DIRAS2 | 0.44294 | 0.20186 | 0.16664 |
| ENSG00000130544 | 19 | 7069455 | 7087979 | 97 | 18 | 0.14348 | 0.44296 | ZNF557 | 0.44296 | 0.42059 | 0.23166 |

| | | | | | | | | | | | | |
|---|---|---|---|---|---|---|---|---|---|---|---|---|
| ENSG00000136045 | 12 | 108079509 | 108106944 | 82 | 12 | 0.1432 | 0.44307 | PWP1 | | 0.44307 | 0.41214 | 0.38003 |
| ENSG00000163207 | 1 | 152881021 | 152884362 | 18 | 2 | 0.14316 | 0.44308 | IVL | | 0.44308 | 0.4135 | 0.58497 |
| ENSG00000105609 | 19 | 54754263 | 54761164 | 48 | 13 | 0.14297 | 0.44316 | LILRB5 | | 0.44316 | 0.38269 | 0.53855 |
| ENSG00000065054 | 16 | 2075357 | 2089027 | 47 | 12 | 0.14288 | 0.44319 | SLC9A3R2 | | 0.44319 | 0.64779 | 0.82048 |
| ENSG00000183513 | 2 | 99215773 | 99224978 | 18 | 2 | 0.14284 | 0.44321 | COA5 | | 0.44321 | 0.49373 | 0.34836 |
| ENSG00000196118 | 16 | 30768744 | 30774031 | 11 | 7 | 0.14264 | 0.44329 | C16orf93 | | 0.44329 | 0.67406 | 0.75358 |
| ENSG00000112425 | 6 | 145822719 | 146057160 | 766 | 17 | 0.14257 | 0.44332 | EPM2A | | 0.44332 | 0.46387 | 0.4285 |
| ENSG00000091844 | 6 | 153325594 | 153452384 | 490 | 23 | 0.14256 | 0.44332 | RGS17 | | 0.44332 | 0.56477 | 0.67962 |
| ENSG00000083223 | 9 | 88902648 | 88969369 | 137 | 12 | 0.14256 | 0.44332 | ZCCHC6 | | 0.44332 | 0.45773 | 0.54199 |
| ENSG00000182450 | 11 | 64058774 | 64067503 | 11 | 5 | 0.14256 | 0.44332 | KCNK4 | | 0.44332 | 0.23867 | 0.24382 |
| ENSG00000081087 | 6 | 108362613 | 108487058 | 410 | 59 | 0.14228 | 0.44343 | OSTM1 | | 0.44343 | 0.41227 | 0.35851 |
| ENSG00000258223 | 7 | 141951963 | 141957878 | 30 | 4 | 0.14211 | 0.4435 | PRSS58 | | 0.4435 | 0.47405 | 0.52732 |
| ENSG00000180739 | 19 | 10623623 | 10628607 | 18 | 9 | 0.14209 | 0.44351 | S1PR5 | | 0.44351 | 0.80916 | 0.92364 |
| ENSG00000198003 | 19 | 11531272 | 11546603 | 44 | 11 | 0.14196 | 0.44356 | CCDC151 | | 0.44356 | 0.3939 | 0.66112 |
| ENSG00000254462 | 11 | 57480077 | 57559058 | 143 | 15 | 0.14191 | 0.44358 | TMX2-CTNND1 | | 0.44358 | 0.41954 | 0.40441 |
| ENSG00000184203 | 3 | 195241221 | 195270209 | 96 | 13 | 0.14182 | 0.44361 | PPP1R2 | | 0.44361 | 0.4661 | 0.55668 |
| ENSG00000102967 | 16 | 72042487 | 72058954 | 47 | 11 | 0.14182 | 0.44361 | DHODH | | 0.44361 | 0.32477 | 0.51102 |
| ENSG00000134365 | 1 | 196819371 | 196888102 | 222 | 20 | 0.14175 | 0.44364 | CFHR4 | | 0.44364 | 0.41167 | 0.39268 |
| ENSG00000118245 | 2 | 217724181 | 217724787 | 1 | 1 | 0.14159 | 0.4437 | TNP1 | | 0.4437 | 0.94857 | 0.90165 |
| ENSG00000162745 | 1 | 161952982 | 161993644 | 156 | 29 | 0.1414 | 0.44378 | OLFML2B | | 0.44378 | 0.30242 | 0.39791 |
| ENSG00000008513 | 8 | 134467091 | 134584183 | 515 | 78 | 0.14132 | 0.44381 | ST3GAL1 | | 0.44381 | 0.37767 | 0.35489 |
| ENSG00000154096 | 11 | 119288090 | 119295695 | 20 | 3 | 0.14127 | 0.44383 | THY1 | | 0.44383 | 0.48093 | 0.46317 |
| ENSG00000268387 | 1 | 161190160 | 161192296 | 11 | 4 | 0.14117 | 0.44387 | AL590714.1 | | 0.44387 | 0.7503 | 0.84483 |
| ENSG00000172366 | 16 | 691813 | 698474 | 6 | 3 | 0.14103 | 0.44392 | FAM195A | | 0.44392 | 0.40256 | 0.35716 |
| ENSG00000204839 | 8 | 144648357 | 144655141 | 26 | 7 | 0.141 | 0.44394 | MROH6 | | 0.44394 | 0.347 | 0.61917 |
| ENSG00000160588 | 11 | 118097409 | 118123065 | 108 | 10 | 0.1409 | 0.44397 | MPZL3 | | 0.44397 | 0.4168 | 0.43758 |
| ENSG00000125207 | 12 | 130822432 | 130857182 | 125 | 20 | 0.14069 | 0.44406 | PIWIL1 | | 0.44406 | 0.59925 | 0.74685 |
| ENSG00000197658 | 11 | 62847412 | 62911693 | 304 | 10 | 0.14059 | 0.4441 | SLC22A24 | | 0.4441 | 0.43541 | 0.39753 |
| ENSG00000121871 | 3 | 164904508 | 164914897 | 29 | 7 | 0.14054 | 0.44412 | SLITRK3 | | 0.44412 | 0.53704 | 0.48927 |
| ENSG00000170788 | 10 | 82095861 | 82116511 | 50 | 13 | 0.13995 | 0.44435 | DYDC1 | | 0.44435 | 0.26541 | 0.21015 |
| ENSG00000168148 | 1 | 228612546 | 228613026 | 1 | 1 | 0.13982 | 0.4444 | HIST3H3 | | 0.4444 | 0.12996 | 0.24945 |
| ENSG00000114248 | 3 | 169556967 | 169587718 | 99 | 13 | 0.13959 | 0.44449 | LRRC31 | | 0.44449 | 0.52725 | 0.40258 |
| ENSG00000196934 | 22 | 21737663 | 21743446 | 1 | 1 | 0.13957 | 0.4445 | RIMBP3B | | 0.4445 | 0.4445 | 0.39809 |
| ENSG00000197409 | 6 | 26197068 | 26199521 | 14 | 4 | 0.13954 | 0.44451 | HIST1H3D | | 0.44451 | 0.50993 | 0.52794 |
| ENSG00000156097 | 1 | 110082494 | 110091028 | 12 | 4 | 0.13951 | 0.44452 | GPR61 | | 0.44452 | 0.63015 | 0.81544 |
| ENSG00000271605 | 17 | 62461569 | 62464760 | 3 | 2 | 0.13927 | 0.44462 | MILR1 | | 0.44462 | 0.42323 | 0.41262 |
| ENSG00000196963 | 5 | 140560980 | 140565793 | 36 | 9 | 0.13918 | 0.44465 | PCDHB16 | | 0.44465 | 0.4074 | 0.44228 |
| ENSG00000197479 | 5 | 140579183 | 140582618 | 13 | 5 | 0.13891 | 0.44476 | PCDHB11 | | 0.44476 | 0.3991 | 0.33595 |
| ENSG00000103023 | 16 | 58313901 | 58328951 | 91 | 6 | 0.13889 | 0.44477 | PRSS54 | | 0.44477 | 0.40387 | 0.29466 |
| ENSG00000155100 | 8 | 92082424 | 92099323 | 37 | 8 | 0.13883 | 0.44479 | OTUD6B | | 0.44479 | 0.53208 | 0.58018 |
| ENSG00000176533 | 19 | 2511217 | 2702707 | 1087 | 116 | 0.13872 | 0.44483 | GNG7 | | 0.44483 | 0.43533 | 0.27964 |
| ENSG00000122566 | 7 | 26229547 | 26241149 | 31 | 6 | 0.13856 | 0.4449 | HNRNPA2B1 | | 0.4449 | 0.18902 | 0.18094 |
| ENSG00000145934 | 5 | 166711804 | 167691162 | 2763 | 237 | 0.13849 | 0.44493 | TENM2 | | 0.44493 | 0.41865 | 0.37931 |
| ENSG00000163349 | 1 | 114471814 | 114520426 | 80 | 16 | 0.13841 | 0.44496 | HIPK1 | | 0.44496 | 0.64862 | 0.77843 |
| ENSG00000137161 | 6 | 42896938 | 42907025 | 28 | 7 | 0.13841 | 0.44496 | CNPY3 | | 0.44496 | 0.41329 | 0.26701 |
| ENSG00000204677 | 5 | 177433406 | 177482560 | 139 | 23 | 0.13831 | 0.445 | FAM153C | | 0.445 | 0.44099 | 0.61977 |
| ENSG00000100226 | 22 | 39101728 | 39134304 | 66 | 13 | 0.13798 | 0.44513 | GTPBP1 | | 0.44513 | 0.49319 | 0.35451 |
| ENSG00000138069 | 2 | 65297835 | 65357240 | 238 | 24 | 0.13778 | 0.44521 | RAB1A | | 0.44521 | 0.48415 | 0.55978 |
| ENSG00000058262 | 3 | 127770484 | 127790526 | 48 | 3 | 0.13773 | 0.44523 | SEC61A1 | | 0.44523 | 0.51483 | 0.51231 |
| ENSG00000122733 | 9 | 34957484 | 34982541 | 52 | 13 | 0.13748 | 0.44533 | KIAA1045 | | 0.44533 | 0.66045 | 0.58657 |
| ENSG00000110400 | 11 | 119494120 | 119599794 | 284 | 34 | 0.13737 | 0.44537 | PVRL1 | | 0.44537 | 0.51505 | 0.44974 |
| ENSG00000163295 | 2 | 233320833 | 233324742 | 12 | 3 | 0.13733 | 0.44539 | ALPI | | 0.44539 | 0.78982 | 0.74181 |
| ENSG00000197168 | 13 | 52611093 | 52703214 | 258 | 18 | 0.13719 | 0.44544 | NEK5 | | 0.44544 | 0.41495 | 0.42165 |
| ENSG00000124102 | 20 | 43803517 | 43805185 | 15 | 3 | 0.13716 | 0.44545 | PI3 | | 0.44545 | 0.33515 | 0.31943 |
| ENSG00000188994 | 6 | 87862551 | 87973914 | 359 | 15 | 0.13697 | 0.44553 | ZNF292 | | 0.44553 | 0.45308 | 0.45041 |
| ENSG00000155975 | 8 | 17104080 | 17159936 | 219 | 30 | 0.13691 | 0.44555 | VPS37A | | 0.44555 | 0.52877 | 0.59308 |
| ENSG00000119421 | 9 | 124894745 | 124922098 | 116 | 15 | 0.13676 | 0.44561 | NDUFA8 | | 0.44561 | 0.43695 | 0.53904 |
| ENSG00000133731 | 8 | 82570196 | 82598928 | 91 | 18 | 0.1367 | 0.44563 | IMPA1 | | 0.44563 | 0.48931 | 0.5538 |
| ENSG00000174697 | 7 | 127881337 | 127897681 | 46 | 11 | 0.13663 | 0.44566 | LEP | | 0.44566 | 0.39027 | 0.65772 |
| ENSG00000106278 | 7 | 121513143 | 121702090 | 509 | 34 | 0.13652 | 0.44571 | PTPRZ1 | | 0.44571 | 0.41596 | 0.4253 |
| ENSG00000186188 | 10 | 95326422 | 95364237 | 132 | 25 | 0.13627 | 0.44581 | FFAR4 | | 0.44581 | 0.43323 | 0.36104 |
| ENSG00000268218 | 16 | 89334512 | 89339452 | 22 | 8 | 0.13619 | 0.44584 | AC137932.1 | | 0.44584 | 0.77945 | 0.90084 |
| ENSG00000219073 | 1 | 22303514 | 22325135 | 229 | 28 | 0.13595 | 0.44593 | CELA3B | | 0.44593 | 0.47478 | 0.5554 |
| ENSG00000196859 | 17 | 39114669 | 39123144 | 34 | 6 | 0.13578 | 0.446 | KRT39 | | 0.446 | 0.57927 | 0.55173 |
| ENSG00000248333 | 1 | 1570603 | 1590473 | 39 | 7 | 0.13561 | 0.44607 | CDK11B | | 0.44607 | 0.63206 | 0.68235 |
| ENSG00000198216 | 1 | 181382238 | 181777219 | 1173 | 86 | 0.13555 | 0.44609 | CACNA1E | | 0.44609 | 0.44154 | 0.4093 |
| ENSG00000106648 | 7 | 151653464 | 151717019 | 295 | 29 | 0.13543 | 0.44614 | GALNTL5 | | 0.44614 | 0.2949 | 0.29825 |
| ENSG00000112137 | 6 | 12717893 | 13288645 | 1692 | 79 | 0.13536 | 0.44616 | PHACTR1 | | 0.44616 | 0.46467 | 0.43558 |
| ENSG00000111799 | 6 | 75794042 | 75915767 | 222 | 35 | 0.13528 | 0.44619 | COL12A1 | | 0.44619 | 0.42611 | 0.41952 |
| ENSG00000173338 | 11 | 65360326 | 65363467 | 4 | 2 | 0.13521 | 0.44622 | KCNK7 | | 0.44622 | 0.10619 | 0.10651 |
| ENSG00000130590 | 20 | 62605466 | 62611361 | 19 | 3 | 0.13511 | 0.44626 | SAMD10 | | 0.44626 | 0.40276 | 0.39537 |
| ENSG00000117090 | 1 | 160577890 | 160617085 | 136 | 23 | 0.13504 | 0.44629 | SLAMF1 | | 0.44629 | 0.36773 | 0.23364 |
| ENSG00000142484 | 17 | 4675179 | 4686506 | 40 | 10 | 0.13498 | 0.44631 | TM4SF5 | | 0.44631 | 0.43032 | 0.38203 |

| | | | | | | | | | | | |
|---|---|---|---|---|---|---|---|---|---|---|---|
| ENSG00000186440 | 1 | 158532441 | 158533394 | 5 | 3 | 0.13479 | 0.44639 | OR6P1 | 0.44639 | 0.43284 | 0.53055 |
| ENSG00000162482 | 1 | 19609052 | 19615744 | 30 | 5 | 0.13473 | 0.44641 | AKR7A3 | 0.44641 | 0.54837 | 0.56527 |
| ENSG00000115840 | 2 | 172640880 | 172864766 | 687 | 32 | 0.13466 | 0.44644 | SLC25A12 | 0.44644 | 0.36665 | 0.31443 |
| ENSG00000135541 | 6 | 135604670 | 135818914 | 384 | 30 | 0.13455 | 0.44648 | AHI1 | 0.44648 | 0.49822 | 0.50898 |
| ENSG00000177707 | 3 | 110788918 | 110994410 | 520 | 45 | 0.13439 | 0.44655 | PVRL3 | 0.44655 | 0.4094 | 0.36479 |
| ENSG00000178026 | 22 | 24981588 | 24989175 | 15 | 5 | 0.1343 | 0.44658 | FAM211B | 0.44658 | 0.55158 | 0.62485 |
| ENSG00000112246 | 6 | 100832891 | 100912805 | 215 | 32 | 0.13419 | 0.44663 | SIM1 | 0.44663 | 0.39263 | 0.46227 |
| ENSG00000143514 | 1 | 223967601 | 224033674 | 168 | 15 | 0.13408 | 0.44667 | TP53BP2 | 0.44667 | 0.47075 | 0.47768 |
| ENSG00000268470 | 17 | 76494911 | 76499138 | 32 | 8 | 0.13407 | 0.44667 | DNAH17-AS1 | 0.44667 | 0.51727 | 0.81026 |
| ENSG00000186971 | 21 | 31802572 | 31803216 | 5 | 2 | 0.13405 | 0.44668 | KRTAP13-4 | 0.44668 | 0.4898 | 0.52868 |
| ENSG00000170899 | 6 | 52842751 | 52860176 | 113 | 16 | 0.13392 | 0.44673 | GSTA4 | 0.44673 | 0.49323 | 0.6047 |
| ENSG00000196422 | 9 | 138370925 | 138380739 | 34 | 6 | 0.13368 | 0.44683 | PPP1R26 | 0.44683 | 0.2448 | 0.21143 |
| ENSG00000196511 | 7 | 144149034 | 144533488 | 1227 | 67 | 0.13362 | 0.44685 | TPK1 | 0.44685 | 0.46173 | 0.48628 |
| ENSG00000175048 | 6 | 157802165 | 158099178 | 1272 | 83 | 0.13352 | 0.44689 | ZDHHC14 | 0.44689 | 0.54562 | 0.4755 |
| ENSG00000117399 | 1 | 43824626 | 43828874 | 6 | 3 | 0.13339 | 0.44694 | CDC20 | 0.44694 | 0.4833 | 0.49618 |
| ENSG00000148660 | 10 | 75572259 | 75634343 | 117 | 16 | 0.13333 | 0.44697 | CAMK2G | 0.44697 | 0.43642 | 0.51973 |
| ENSG00000050820 | 16 | 75262928 | 75301951 | 195 | 19 | 0.13317 | 0.44703 | BCAR1 | 0.44703 | 0.50242 | 0.63487 |
| ENSG00000037042 | 17 | 40811323 | 40819024 | 16 | 4 | 0.1331 | 0.44706 | TUBG2 | 0.44706 | 0.31908 | 0.33172 |
| ENSG00000100592 | 14 | 59655364 | 59838123 | 639 | 35 | 0.13298 | 0.4471 | DAAM1 | 0.4471 | 0.44102 | 0.43172 |
| ENSG00000132613 | 16 | 70695107 | 70719969 | 77 | 16 | 0.13298 | 0.44711 | MTSS1L | 0.44711 | 0.50802 | 0.37689 |
| ENSG00000196437 | 19 | 37902057 | 37960177 | 117 | 16 | 0.13286 | 0.44715 | ZNF569 | 0.44715 | 0.44259 | 0.48942 |
| ENSG00000176200 | 11 | 59271049 | 59271984 | 1 | 1 | 0.13274 | 0.4472 | OR4D11 | 0.4472 | 0.47711 | 0.45666 |
| ENSG00000124564 | 6 | 25833294 | 25882514 | 166 | 19 | 0.13264 | 0.44724 | SLC17A3 | 0.44724 | 0.45432 | 0.45042 |
| ENSG00000205858 | 7 | 16566505 | 16621193 | 225 | 35 | 0.13245 | 0.44731 | LRRC72 | 0.44731 | 0.52978 | 0.38535 |
| ENSG00000179988 | 10 | 124713897 | 124757029 | 149 | 16 | 0.13226 | 0.44739 | PSTK | 0.44739 | 0.41403 | 0.27452 |
| ENSG00000091831 | 6 | 151977826 | 152450754 | 1760 | 94 | 0.13224 | 0.4474 | ESR1 | 0.4474 | 0.45312 | 0.47268 |
| ENSG00000159374 | 2 | 74785010 | 74875465 | 119 | 16 | 0.13199 | 0.44749 | M1AP | 0.44749 | 0.39141 | 0.43676 |
| ENSG00000169248 | 4 | 76954835 | 76962568 | 59 | 4 | 0.13196 | 0.44751 | CXCL11 | 0.44751 | 0.40526 | 0.41631 |
| ENSG00000092445 | 15 | 41849873 | 41871536 | 46 | 11 | 0.13169 | 0.44762 | TYRO3 | 0.44762 | 0.63887 | 0.66076 |
| ENSG00000157782 | 12 | 121078355 | 121105127 | 134 | 25 | 0.13144 | 0.44771 | CABP1 | 0.44771 | 0.43977 | 0.54464 |
| ENSG00000185728 | 8 | 64081112 | 64125346 | 82 | 17 | 0.13139 | 0.44773 | YTHDF3 | 0.44773 | 0.42366 | 0.46933 |
| ENSG00000164548 | 7 | 23544399 | 23571660 | 62 | 13 | 0.1313 | 0.44777 | TRA2A | 0.44777 | 0.53484 | 0.5966 |
| ENSG00000197275 | 8 | 95384188 | 95487337 | 418 | 20 | 0.13124 | 0.44779 | RAD54B | 0.44779 | 0.49422 | 0.56704 |
| ENSG00000187862 | 1 | 156549519 | 156556562 | 17 | 4 | 0.13112 | 0.44784 | TTC24 | 0.44784 | 0.61079 | 0.70211 |
| ENSG00000197587 | 1 | 46972668 | 46979898 | 27 | 8 | 0.13094 | 0.44791 | DMBX1 | 0.44791 | 0.39574 | 0.53015 |
| ENSG00000198822 | 7 | 86273230 | 86494200 | 480 | 45 | 0.13095 | 0.44791 | GRM3 | 0.44791 | 0.43871 | 0.42263 |
| ENSG00000161992 | 16 | 610422 | 615528 | 25 | 9 | 0.13031 | 0.44816 | C16orf11 | 0.44816 | 0.51001 | 0.62031 |
| ENSG00000163631 | 4 | 74262831 | 74287129 | 47 | 6 | 0.13017 | 0.44822 | ALB | 0.44822 | 0.45189 | 0.42647 |
| ENSG00000189367 | 6 | 127761488 | 127780536 | 32 | 4 | 0.13015 | 0.44822 | KIAA0408 | 0.44822 | 0.47807 | 0.51205 |
| ENSG00000004866 | 7 | 116593292 | 116870157 | 467 | 54 | 0.12994 | 0.44831 | ST7 | 0.44831 | 0.46687 | 0.53486 |
| ENSG00000269358 | 1 | 224215760 | 224216371 | 3 | 1 | 0.12988 | 0.44833 | AC138393.1 | 0.44833 | 0.66117 | 0.66409 |
| ENSG00000271079 | 7 | 143268894 | 143271480 | 3 | 2 | 0.12984 | 0.44835 | CTAGE15 | 0.44835 | 0.89619 | 0.89781 |
| ENSG00000132970 | 13 | 27131840 | 27263085 | 494 | 29 | 0.12977 | 0.44838 | WASF3 | 0.44838 | 0.36011 | 0.22819 |
| ENSG00000122483 | 1 | 93645476 | 93744287 | 235 | 16 | 0.12968 | 0.44841 | CCDC18 | 0.44841 | 0.45637 | 0.46284 |
| ENSG00000112706 | 6 | 76630832 | 76782395 | 333 | 39 | 0.12968 | 0.44841 | IMPG1 | 0.44841 | 0.42316 | 0.59148 |
| ENSG00000125966 | 20 | 33814457 | 33864801 | 80 | 15 | 0.12966 | 0.44842 | MMP24 | 0.44842 | 0.19489 | 0.15438 |
| ENSG00000183597 | 22 | 20004537 | 20053449 | 187 | 28 | 0.12948 | 0.44849 | TANGO2 | 0.44849 | 0.28477 | 0.13339 |
| ENSG00000168273 | 3 | 52568029 | 52613253 | 117 | 11 | 0.12935 | 0.44854 | SMIM4 | 0.44854 | 0.41723 | 0.37985 |
| ENSG00000146373 | 6 | 125283691 | 125413779 | 331 | 44 | 0.12923 | 0.44859 | RNF217 | 0.44859 | 0.36374 | 0.2749 |
| ENSG00000150540 | 2 | 138721590 | 138773930 | 112 | 8 | 0.12918 | 0.44861 | HNMT | 0.44861 | 0.4834 | 0.37474 |
| ENSG00000124743 | 6 | 53512699 | 53530506 | 79 | 10 | 0.12915 | 0.44862 | KLHL31 | 0.44862 | 0.81908 | 0.92618 |
| ENSG00000196943 | 14 | 24769068 | 24778330 | 28 | 7 | 0.12893 | 0.44871 | NOP9 | 0.44871 | 0.55616 | 0.67267 |
| ENSG00000132485 | 1 | 71528974 | 71546980 | 36 | 9 | 0.12889 | 0.44872 | ZRANB2 | 0.44872 | 0.42069 | 0.31609 |
| ENSG00000071189 | 7 | 17830385 | 17980124 | 386 | 19 | 0.1289 | 0.44872 | SNX13 | 0.44872 | 0.39951 | 0.40302 |
| ENSG00000089154 | 12 | 120565007 | 120632513 | 122 | 10 | 0.12872 | 0.44879 | GCN1L1 | 0.44879 | 0.51718 | 0.64974 |
| ENSG00000205595 | 4 | 75480629 | 75490486 | 8 | 4 | 0.12864 | 0.44882 | AREGB | 0.44882 | 0.87029 | 0.86255 |
| ENSG00000243489 | 21 | 46066331 | 46067564 | 4 | 2 | 0.12862 | 0.44883 | KRTAP10-11 | 0.44883 | 0.848 | 0.83457 |
| ENSG00000139197 | 12 | 7341281 | 7371170 | 161 | 14 | 0.12851 | 0.44887 | PEX5 | 0.44887 | 0.4493 | 0.61539 |
| ENSG00000242220 | 21 | 33948862 | 33959683 | 44 | 7 | 0.12829 | 0.44896 | TCP10L | 0.44896 | 0.49042 | 0.43823 |
| ENSG00000100726 | 16 | 1543345 | 1560458 | 81 | 17 | 0.12813 | 0.44902 | TELO2 | 0.44902 | 0.6834 | 0.87554 |
| ENSG00000189320 | 7 | 135413096 | 135433594 | 78 | 16 | 0.12799 | 0.44908 | FAM180A | 0.44908 | 0.36711 | 0.34647 |
| ENSG00000185101 | 11 | 417933 | 442011 | 87 | 13 | 0.12795 | 0.44909 | ANO9 | 0.44909 | 0.31935 | 0.053915 |
| ENSG00000167257 | 11 | 117103341 | 117157161 | 163 | 20 | 0.12773 | 0.44918 | RNF214 | 0.44918 | 0.45394 | 0.54037 |
| ENSG00000221882 | 17 | 3181193 | 3182268 | 4 | 2 | 0.12771 | 0.44919 | OR3A2 | 0.44919 | 0.36674 | 0.30064 |
| ENSG00000241127 | 7 | 39605975 | 39649919 | 197 | 25 | 0.12754 | 0.44925 | YAE1D1 | 0.44925 | 0.45314 | 0.53702 |
| ENSG00000167779 | 12 | 53491220 | 53496129 | 9 | 4 | 0.12746 | 0.44929 | IGFBP6 | 0.44929 | 0.11534 | 0.046599 |
| ENSG00000166592 | 16 | 66955582 | 66959547 | 6 | 2 | 0.12731 | 0.44935 | RRAD | 0.44935 | 0.26711 | 0.3616 |
| ENSG00000220891 | 22 | 22901750 | 22909007 | 53 | 5 | 0.12711 | 0.44943 | LL22NC03-63E9.3 | 0.44943 | 0.57063 | 0.56532 |
| ENSG00000262664 | 17 | 1945327 | 1946724 | 7 | 3 | 0.12701 | 0.44947 | OVCA2 | 0.44947 | 0.59877 | 0.66685 |
| ENSG00000100554 | 14 | 67761088 | 67826982 | 310 | 14 | 0.12684 | 0.44953 | ATP6V1D | 0.44953 | 0.45977 | 0.46288 |
| ENSG00000090266 | 7 | 140390577 | 140422590 | 72 | 8 | 0.12675 | 0.44957 | NDUFB2 | 0.44957 | 0.45436 | 0.31143 |
| ENSG00000131446 | 5 | 180217541 | 180242652 | 171 | 9 | 0.12652 | 0.44966 | MGAT1 | 0.44966 | 0.45185 | 0.49499 |
| ENSG00000198610 | 10 | 5237425 | 5260912 | 144 | 11 | 0.12631 | 0.44974 | AKR1C4 | 0.44974 | 0.37017 | 0.24401 |

| | | | | | | | | | | | |
|---|---|---|---|---|---|---|---|---|---|---|---|
| ENSG00000159433 | 15 | 42867857 | 43013179 | 247 | 26 | 0.12563 | 0.45001 | STARD9 | 0.45001 | 0.47473 | 0.53891 |
| ENSG00000171044 | 8 | 10753555 | 11058875 | 1166 | 37 | 0.12558 | 0.45003 | XKR6 | 0.45003 | 0.45222 | 0.47742 |
| ENSG00000164047 | 3 | 48264837 | 48266981 | 2 | 1 | 0.12539 | 0.45011 | CAMP | 0.45011 | 0.77364 | 0.72811 |
| ENSG00000132305 | 2 | 86371055 | 86422893 | 185 | 9 | 0.12536 | 0.45012 | IMMT | 0.45012 | 0.47369 | 0.46995 |
| ENSG00000214397 | 15 | 100255908 | 100258029 | 8 | 3 | 0.12535 | 0.45012 | DKFZP779J2370 | 0.45012 | 0.37062 | 0.2073 |
| ENSG00000132953 | 13 | 21351469 | 21477187 | 368 | 18 | 0.12523 | 0.45017 | XPO4 | 0.45017 | 0.422 | 0.39478 |
| ENSG00000185721 | 22 | 31795509 | 31924726 | 400 | 29 | 0.12522 | 0.45018 | DRG1 | 0.45018 | 0.49951 | 0.5684 |
| ENSG00000122696 | 9 | 37879400 | 37904350 | 38 | 12 | 0.12513 | 0.45021 | SLC25A51 | 0.45021 | 0.30283 | 0.36309 |
| ENSG00000172594 | 6 | 123110315 | 123130865 | 71 | 15 | 0.12502 | 0.45025 | SMPDL3A | 0.45025 | 0.73055 | 0.85296 |
| ENSG00000091536 | 17 | 18012020 | 18083116 | 165 | 22 | 0.12497 | 0.45027 | MYO15A | 0.45027 | 0.40054 | 0.40431 |
| ENSG00000131503 | 5 | 139781399 | 139929163 | 179 | 26 | 0.12491 | 0.4503 | ANKHD1 | 0.4503 | 0.43897 | 0.44401 |
| ENSG00000254996 | 5 | 139781493 | 139929154 | 179 | 26 | 0.12491 | 0.4503 | ANKHD1-EIF4EBP3 | 0.4503 | 0.43897 | 0.44401 |
| ENSG00000170054 | 14 | 94929054 | 94946026 | 90 | 21 | 0.12462 | 0.45041 | SERPINA9 | 0.45041 | 0.73134 | 0.34616 |
| ENSG00000166783 | 16 | 15688243 | 15737023 | 104 | 23 | 0.12444 | 0.45049 | KIAA0430 | 0.45049 | 0.50133 | 0.48363 |
| ENSG00000106628 | 7 | 44154286 | 44163957 | 24 | 8 | 0.12437 | 0.45051 | POLD2 | 0.45051 | 0.5315 | 0.41682 |
| ENSG00000110721 | 11 | 67820326 | 67888911 | 115 | 14 | 0.12434 | 0.45052 | CHKA | 0.45052 | 0.42874 | 0.44387 |
| ENSG00000186665 | 17 | 65987217 | 65989765 | 3 | 1 | 0.12435 | 0.45052 | C17orf58 | 0.45052 | 0.48828 | 0.44927 |
| ENSG00000127252 | 3 | 192958914 | 192995527 | 94 | 10 | 0.1243 | 0.45054 | HRASLS | 0.45054 | 0.62277 | 0.63434 |
| ENSG00000172009 | 19 | 2785458 | 2813599 | 70 | 8 | 0.12396 | 0.45067 | THOP1 | 0.45067 | 0.58054 | 0.85367 |
| ENSG00000263264 | 19 | 7413848 | 7448271 | 190 | 19 | 0.12381 | 0.45073 | CTB-133G6.1 | 0.45073 | 0.49218 | 0.48281 |
| ENSG00000141720 | 17 | 36921942 | 36956379 | 95 | 19 | 0.12377 | 0.45075 | PIP4K2B | 0.45075 | 0.38429 | 0.52242 |
| ENSG00000185420 | 1 | 245912642 | 246670614 | 4141 | 110 | 0.12375 | 0.45076 | SMYD3 | 0.45076 | 0.42718 | 0.42697 |
| ENSG00000169862 | 5 | 10971952 | 11904155 | 3078 | 210 | 0.12372 | 0.45077 | CTNND2 | 0.45077 | 0.48409 | 0.40672 |
| ENSG00000154124 | 5 | 14664773 | 14699820 | 65 | 11 | 0.12365 | 0.4508 | FAM105B | 0.4508 | 0.41383 | 0.37331 |
| ENSG00000219200 | 17 | 6915736 | 6917851 | 2 | 1 | 0.12322 | 0.45097 | RNASEK | 0.45097 | 0.61696 | 0.78742 |
| ENSG00000092199 | 14 | 21677295 | 21737653 | 164 | 17 | 0.1227 | 0.45117 | HNRNPC | 0.45117 | 0.51602 | 0.58915 |
| ENSG00000142025 | 19 | 42348806 | 42356401 | 14 | 4 | 0.12267 | 0.45119 | DMRTC2 | 0.45119 | 0.70432 | 0.78234 |
| ENSG00000267110 | 19 | 55666125 | 55672039 | 36 | 6 | 0.12201 | 0.45144 | CTD-2587H24.4 | 0.45144 | 0.69521 | 0.86396 |
| ENSG00000138617 | 15 | 65526798 | 65592956 | 216 | 19 | 0.12183 | 0.45152 | PARP16 | 0.45152 | 0.40007 | 0.32623 |
| ENSG00000175267 | 16 | 22103859 | 22168287 | 112 | 26 | 0.12165 | 0.45159 | VWA3A | 0.45159 | 0.48217 | 0.51933 |
| ENSG00000158006 | 1 | 26286258 | 26324648 | 95 | 7 | 0.12148 | 0.45166 | PAFAH2 | 0.45166 | 0.46749 | 0.43117 |
| ENSG00000144161 | 2 | 112969102 | 113012713 | 66 | 9 | 0.12139 | 0.45169 | ZC3H8 | 0.45169 | 0.55359 | 0.51 |
| ENSG00000146385 | 6 | 132873832 | 132874860 | 1 | 1 | 0.12137 | 0.4517 | TAAR8 | 0.4517 | 0.54416 | 0.28075 |
| ENSG00000185345 | 6 | 161768452 | 163148803 | 6085 | 264 | 0.12124 | 0.45175 | PARK2 | 0.45175 | 0.4874 | 0.55792 |
| ENSG00000136807 | 9 | 130547958 | 130553066 | 14 | 4 | 0.12114 | 0.45179 | CDK9 | 0.45179 | 0.38753 | 0.34529 |
| ENSG00000137992 | 1 | 100652475 | 100715390 | 155 | 13 | 0.12106 | 0.45182 | DBT | 0.45182 | 0.32697 | 0.23353 |
| ENSG00000197822 | 5 | 68788119 | 68853931 | 71 | 11 | 0.12036 | 0.4521 | OCLN | 0.4521 | 0.49016 | 0.4994 |
| ENSG00000184988 | 17 | 41363854 | 41372061 | 16 | 5 | 0.12019 | 0.45216 | TMEM106A | 0.45216 | 0.6958 | 0.69933 |
| ENSG00000136273 | 7 | 47735328 | 48019178 | 1443 | 78 | 0.11985 | 0.4523 | HUS1 | 0.4523 | 0.4945 | 0.50729 |
| ENSG00000171053 | 11 | 125616188 | 125619762 | 26 | 8 | 0.11984 | 0.4523 | PATE1 | 0.4523 | 0.81356 | 0.8746 |
| ENSG00000139173 | 12 | 44229770 | 44783545 | 1672 | 39 | 0.11974 | 0.45234 | TMEM117 | 0.45234 | 0.4259 | 0.39816 |
| ENSG00000080845 | 20 | 34894258 | 35157040 | 315 | 38 | 0.11937 | 0.45249 | DLGAP4 | 0.45249 | 0.50524 | 0.5104 |
| ENSG00000155858 | 5 | 157170703 | 157187717 | 46 | 7 | 0.11916 | 0.45257 | LSM11 | 0.45257 | 0.41535 | 0.65358 |
| ENSG00000138964 | 22 | 44568836 | 44615413 | 231 | 32 | 0.11915 | 0.45258 | PARVG | 0.45258 | 0.55731 | 0.61408 |
| ENSG00000159840 | 7 | 143078173 | 143088204 | 26 | 11 | 0.11877 | 0.45273 | ZYX | 0.45273 | 0.4507 | 0.57364 |
| ENSG00000268171 | 4 | 57276682 | 57277131 | 1 | 1 | 0.11859 | 0.4528 | AC068620.1 | 0.4528 | 0.85442 | 0.89843 |
| ENSG00000171989 | 15 | 59499042 | 59500705 | 13 | 3 | 0.11858 | 0.4528 | LDHAL6B | 0.4528 | 0.33968 | 0.36943 |
| ENSG00000164221 | 5 | 114602885 | 114632528 | 55 | 10 | 0.11857 | 0.45281 | CCDC112 | 0.45281 | 0.57067 | 0.70695 |
| ENSG00000162396 | 1 | 55222571 | 55230187 | 27 | 7 | 0.11852 | 0.45283 | PARS2 | 0.45283 | 0.64017 | 0.88662 |
| ENSG00000251012 | 3 | 118866222 | 118906654 | 178 | 28 | 0.11844 | 0.45286 | RP11-484M3.5 | 0.45286 | 0.39094 | 0.32345 |
| ENSG00000179029 | 17 | 8076555 | 8079717 | 21 | 7 | 0.11799 | 0.45304 | TMEM107 | 0.45304 | 0.3994 | 0.41589 |
| ENSG00000176171 | 10 | 133781578 | 133795435 | 40 | 9 | 0.11793 | 0.45306 | BNIP3 | 0.45306 | 0.45658 | 0.28641 |
| ENSG00000198690 | 15 | 31196055 | 31235311 | 122 | 17 | 0.11792 | 0.45307 | FAN1 | 0.45307 | 0.42991 | 0.50899 |
| ENSG00000270316 | 10 | 104614029 | 104661656 | 182 | 19 | 0.11779 | 0.45312 | C10orf32-ASMT | 0.45312 | 0.46523 | 0.48744 |
| ENSG00000158516 | 7 | 129906667 | 129929638 | 70 | 10 | 0.11765 | 0.45317 | CPA2 | 0.45317 | 0.55016 | 0.53018 |
| ENSG00000108774 | 17 | 40276994 | 40307035 | 70 | 8 | 0.11767 | 0.45317 | RAB5C | 0.45317 | 0.32842 | 0.29671 |
| ENSG00000178951 | 19 | 4044362 | 4066943 | 52 | 10 | 0.11746 | 0.45325 | ZBTB7A | 0.45325 | 0.17099 | 0.17997 |
| ENSG00000126581 | 17 | 40962152 | 40985367 | 9 | 6 | 0.11734 | 0.45329 | BECN1 | 0.45329 | 0.13777 | 0.26628 |
| ENSG00000124171 | 20 | 49348081 | 49373332 | 68 | 14 | 0.11731 | 0.45331 | PARD6B | 0.45331 | 0.43238 | 0.58412 |
| ENSG00000151033 | 10 | 30900718 | 30918691 | 110 | 17 | 0.11728 | 0.45332 | LYZL2 | 0.45332 | 0.47823 | 0.19023 |
| ENSG00000112357 | 6 | 137143717 | 137235075 | 246 | 23 | 0.11725 | 0.45333 | PEX7 | 0.45333 | 0.45309 | 0.48093 |
| ENSG00000100599 | 14 | 92980118 | 93155339 | 707 | 88 | 0.1171 | 0.45339 | RIN3 | 0.45339 | 0.49224 | 0.40641 |
| ENSG00000176231 | 19 | 16059818 | 16060768 | 6 | 2 | 0.11702 | 0.45342 | OR10H4 | 0.45342 | 0.55035 | 0.68592 |
| ENSG00000170379 | 7 | 143318043 | 143427502 | 25 | 8 | 0.11684 | 0.45349 | FAM115C | 0.45349 | 0.53979 | 0.51046 |
| ENSG00000153113 | 5 | 95860971 | 96115299 | 1029 | 75 | 0.11682 | 0.4535 | CAST | 0.4535 | 0.44524 | 0.3892 |
| ENSG00000257591 | 19 | 12251032 | 12267546 | 31 | 4 | 0.11644 | 0.45365 | ZNF625 | 0.45365 | 0.37319 | 0.2182 |
| ENSG00000157349 | 16 | 70323566 | 70369186 | 58 | 27 | 0.11608 | 0.45379 | DDX19B | 0.45379 | 0.4388 | 0.56912 |
| ENSG00000268091 | 12 | 131514221 | 131514769 | 8 | 2 | 0.11607 | 0.4538 | AC078925.1 | 0.4538 | 0.68548 | 0.60419 |
| ENSG00000115020 | 2 | 209130991 | 209223475 | 323 | 13 | 0.11574 | 0.45393 | PIKFYVE | 0.45393 | 0.48127 | 0.49354 |
| ENSG00000197079 | 17 | 39632941 | 39637392 | 17 | 4 | 0.11566 | 0.45396 | KRT35 | 0.45396 | 0.72247 | 0.78664 |
| ENSG00000066926 | 18 | 55215515 | 55254004 | 159 | 28 | 0.1156 | 0.45399 | FECH | 0.45399 | 0.51907 | 0.60629 |
| ENSG00000023697 | 12 | 16064106 | 16190220 | 240 | 33 | 0.11547 | 0.45404 | DERA | 0.45404 | 0.43623 | 0.44357 |
| ENSG00000048140 | 5 | 176074388 | 176086058 | 43 | 7 | 0.1154 | 0.45407 | TSPAN17 | 0.45407 | 0.68347 | 0.69972 |

| | | | | | | | | | | | |
|---|---|---|---|---|---|---|---|---|---|---|---|
| ENSG00000196236 | 22 | 41253081 | 41363838 | 111 | 19 | 0.11529 | 0.45411 | XPNPEP3 | 0.45411 | 0.39764 | 0.41418 |
| ENSG00000107796 | 10 | 90694831 | 90751147 | 164 | 34 | 0.11527 | 0.45412 | ACTA2 | 0.45412 | 0.37399 | 0.31994 |
| ENSG00000203772 | 10 | 135234170 | 135382916 | 694 | 46 | 0.11515 | 0.45416 | SPRN | 0.45416 | 0.43409 | 0.40064 |
| ENSG00000108784 | 17 | 40688190 | 40696467 | 13 | 6 | 0.11494 | 0.45425 | NAGLU | 0.45425 | 0.67831 | 0.55886 |
| ENSG00000196693 | 10 | 43084555 | 43133992 | 134 | 14 | 0.11492 | 0.45426 | ZNF33B | 0.45426 | 0.4563 | 0.47039 |
| ENSG00000112782 | 6 | 45868045 | 46048132 | 633 | 62 | 0.11471 | 0.45434 | CLIC5 | 0.45434 | 0.26675 | 0.31019 |
| ENSG00000166889 | 11 | 59404189 | 59436453 | 59 | 10 | 0.11431 | 0.4545 | PATL1 | 0.4545 | 0.45968 | 0.42879 |
| ENSG00000264187 | 17 | 17105963 | 17140499 | 140 | 20 | 0.1143 | 0.4545 | RP11-45M22.4 | 0.4545 | 0.52307 | 0.51277 |
| ENSG00000124935 | 11 | 62009682 | 62012280 | 9 | 4 | 0.11425 | 0.45452 | SCGB1D2 | 0.45452 | 0.33963 | 0.36599 |
| ENSG00000159882 | 19 | 44507100 | 44518078 | 25 | 4 | 0.11421 | 0.45454 | ZNF230 | 0.45454 | 0.43145 | 0.70639 |
| ENSG00000120160 | 9 | 27284656 | 27297137 | 62 | 10 | 0.11405 | 0.4546 | EQTN | 0.4546 | 0.53697 | 0.60024 |
| ENSG00000179476 | 14 | 45366498 | 45376460 | 20 | 3 | 0.11403 | 0.45461 | C14orf28 | 0.45461 | 0.31443 | 0.45558 |
| ENSG00000130589 | 20 | 62189439 | 62205592 | 84 | 17 | 0.11403 | 0.45461 | HELZ2 | 0.45461 | 0.2429 | 0.34172 |
| ENSG00000184825 | 6 | 27114861 | 27115317 | 3 | 2 | 0.11397 | 0.45463 | HIST1H2AH | 0.45463 | 0.30517 | 0.46055 |
| ENSG00000168781 | 15 | 43825660 | 43882430 | 61 | 10 | 0.11388 | 0.45467 | PPIP5K1 | 0.45467 | 0.44551 | 0.38555 |
| ENSG00000172661 | 10 | 46222648 | 46288409 | 92 | 7 | 0.11382 | 0.45469 | FAM21C | 0.45469 | 0.47634 | 0.48818 |
| ENSG00000116005 | 2 | 70484518 | 70508323 | 58 | 5 | 0.11326 | 0.45491 | PCYOX1 | 0.45491 | 0.4695 | 0.46082 |
| ENSG00000175387 | 18 | 45357922 | 45457515 | 239 | 15 | 0.11323 | 0.45492 | SMAD2 | 0.45492 | 0.42436 | 0.40249 |
| ENSG00000198646 | 20 | 33284722 | 33413452 | 251 | 20 | 0.11299 | 0.45502 | NCOA6 | 0.45502 | 0.3873 | 0.35153 |
| ENSG00000162174 | 11 | 62104920 | 62160882 | 161 | 15 | 0.11288 | 0.45506 | ASRGL1 | 0.45506 | 0.52463 | 0.48466 |
| ENSG00000158710 | 1 | 159887897 | 159895522 | 21 | 7 | 0.11278 | 0.4551 | TAGLN2 | 0.4551 | 0.55262 | 0.31108 |
| ENSG00000179933 | 14 | 23563974 | 23569665 | 25 | 6 | 0.11235 | 0.45527 | C14orf119 | 0.45527 | 0.33532 | 0.16103 |
| ENSG00000183303 | 11 | 7817452 | 7818520 | 14 | 2 | 0.11227 | 0.4553 | OR5P2 | 0.4553 | 0.4239 | 0.40824 |
| ENSG00000140873 | 16 | 77281710 | 77469011 | 1163 | 57 | 0.11223 | 0.45532 | ADAMTS18 | 0.45532 | 0.45071 | 0.43917 |
| ENSG00000149182 | 11 | 47185848 | 47198676 | 31 | 12 | 0.11217 | 0.45534 | ARFGAP2 | 0.45534 | 0.72083 | 0.68818 |
| ENSG00000153130 | 4 | 141178440 | 141306880 | 378 | 39 | 0.11206 | 0.45539 | SCOC | 0.45539 | 0.34021 | 0.36191 |
| ENSG00000163735 | 4 | 74861359 | 74864496 | 5 | 3 | 0.11197 | 0.45542 | CXCL5 | 0.45542 | 0.20342 | 0.33193 |
| ENSG00000168961 | 17 | 25956824 | 25976586 | 69 | 7 | 0.11153 | 0.4556 | LGALS9 | 0.4556 | 0.36359 | 0.38061 |
| ENSG00000173898 | 11 | 66452719 | 66496697 | 42 | 17 | 0.1115 | 0.45561 | SPTBN2 | 0.45561 | 0.37897 | 0.41917 |
| ENSG00000048740 | 10 | 11047259 | 11378674 | 1292 | 101 | 0.11143 | 0.45564 | CELF2 | 0.45564 | 0.32392 | 0.16969 |
| ENSG00000006695 | 17 | 13972813 | 14111994 | 452 | 43 | 0.11131 | 0.45569 | COX10 | 0.45569 | 0.49059 | 0.43519 |
| ENSG00000100028 | 22 | 24951471 | 25005947 | 143 | 22 | 0.11081 | 0.45588 | SNRPD3 | 0.45588 | 0.45405 | 0.35868 |
| ENSG00000124151 | 20 | 46130601 | 46285621 | 439 | 20 | 0.11077 | 0.4559 | NCOA3 | 0.4559 | 0.43933 | 0.44077 |
| ENSG00000137960 | 1 | 78445226 | 78604133 | 474 | 40 | 0.11053 | 0.45599 | GIPC2 | 0.45599 | 0.47772 | 0.54627 |
| ENSG00000075624 | 7 | 5566782 | 5603415 | 160 | 34 | 0.11055 | 0.45599 | ACTB | 0.45599 | 0.50409 | 0.61745 |
| ENSG00000160570 | 19 | 42702750 | 42724292 | 24 | 4 | 0.11028 | 0.4561 | DEDD2 | 0.4561 | 0.49176 | 0.4117 |
| ENSG00000157368 | 16 | 70613798 | 70694585 | 394 | 28 | 0.11001 | 0.4562 | IL34 | 0.4562 | 0.41449 | 0.37023 |
| ENSG00000173113 | 11 | 64083932 | 64085556 | 3 | 1 | 0.10998 | 0.45621 | TRMT112 | 0.45621 | 0.40012 | 0.19628 |
| ENSG00000175820 | 13 | 103381801 | 103389159 | 8 | 4 | 0.11 | 0.45621 | CCDC168 | 0.45621 | 0.54448 | 0.57434 |
| ENSG00000172809 | 17 | 72199721 | 72206794 | 20 | 7 | 0.10995 | 0.45622 | RPL38 | 0.45622 | 0.29929 | 0.22388 |
| ENSG00000120738 | 5 | 137801179 | 137805004 | 5 | 3 | 0.10965 | 0.45634 | EGR1 | 0.45634 | 0.7189 | 0.71696 |
| ENSG00000267908 | 19 | 56751773 | 56758863 | 56 | 16 | 0.10951 | 0.4564 | ZSCAN5D | 0.4564 | 0.95493 | 0.96021 |
| ENSG00000196072 | 10 | 102033713 | 102046469 | 47 | 9 | 0.10947 | 0.45641 | BLOC1S2 | 0.45641 | 0.43385 | 0.28491 |
| ENSG00000183960 | 3 | 19189946 | 19577138 | 657 | 54 | 0.10938 | 0.45645 | KCNH8 | 0.45645 | 0.42793 | 0.3738 |
| ENSG00000108509 | 17 | 4871287 | 4890960 | 60 | 13 | 0.10924 | 0.4565 | CAMTA2 | 0.4565 | 0.2944 | 0.24081 |
| ENSG00000180209 | 16 | 30382255 | 30389312 | 5 | 3 | 0.10905 | 0.45658 | MYLPF | 0.45658 | 0.52342 | 0.67824 |
| ENSG00000106330 | 7 | 100209725 | 100213007 | 8 | 1 | 0.10888 | 0.45665 | MOSPD3 | 0.45665 | 0.40055 | 0.39701 |
| ENSG00000162959 | 2 | 32090129 | 32236299 | 346 | 21 | 0.10874 | 0.4567 | MEMO1 | 0.4567 | 0.45225 | 0.45411 |
| ENSG00000182170 | 11 | 3239174 | 3240043 | 1 | 1 | 0.10875 | 0.4567 | MRGPRG | 0.4567 | 0.02879 | 0.13841 |
| ENSG00000122971 | 12 | 121163538 | 121177811 | 63 | 11 | 0.10832 | 0.45687 | ACADS | 0.45687 | 0.5847 | 0.63912 |
| ENSG00000008196 | 6 | 50786436 | 50815326 | 74 | 12 | 0.10827 | 0.45689 | TFAP2B | 0.45689 | 0.48749 | 0.63077 |
| ENSG00000177606 | 1 | 59246465 | 59249785 | 4 | 2 | 0.10819 | 0.45692 | JUN | 0.45692 | 0.31601 | 0.47626 |
| ENSG00000072310 | 17 | 17713713 | 17740325 | 49 | 6 | 0.10803 | 0.45699 | SREBF1 | 0.45699 | 0.42761 | 0.50461 |
| ENSG00000128510 | 7 | 129932974 | 129964020 | 106 | 15 | 0.108 | 0.457 | CPA4 | 0.457 | 0.43269 | 0.41081 |
| ENSG00000267901 | 14 | 71683031 | 71683462 | 1 | 1 | 0.10799 | 0.457 | AC004817.1 | 0.457 | 0.79689 | 0.88786 |
| ENSG00000180871 | 2 | 218990012 | 219001976 | 25 | 4 | 0.10798 | 0.45701 | CXCR2 | 0.45701 | 0.32358 | 0.19426 |
| ENSG00000204653 | 19 | 51014857 | 51017947 | 13 | 5 | 0.10786 | 0.45705 | ASPDH | 0.45705 | 0.41152 | 0.71472 |
| ENSG00000159496 | 22 | 24030323 | 24041363 | 24 | 6 | 0.10781 | 0.45707 | RGL4 | 0.45707 | 0.34875 | 0.41583 |
| ENSG00000135220 | 4 | 69794181 | 69817509 | 93 | 8 | 0.10779 | 0.45708 | UGT2A3 | 0.45708 | 0.57136 | 0.65587 |
| ENSG00000171862 | 10 | 89622870 | 89731687 | 156 | 34 | 0.1075 | 0.4572 | PTEN | 0.4572 | 0.5324 | 0.64922 |
| ENSG00000140557 | 15 | 92937058 | 93011958 | 251 | 29 | 0.10737 | 0.45725 | ST8SIA2 | 0.45725 | 0.37056 | 0.55241 |
| ENSG00000096996 | 19 | 18169805 | 18209754 | 170 | 20 | 0.10718 | 0.45732 | IL12RB1 | 0.45732 | 0.54965 | 0.61748 |
| ENSG00000070193 | 5 | 44303646 | 44389808 | 188 | 14 | 0.1071 | 0.45735 | FGF10 | 0.45735 | 0.49268 | 0.50355 |
| ENSG00000160207 | 21 | 44949072 | 45079374 | 314 | 15 | 0.10712 | 0.45735 | HSF2BP | 0.45735 | 0.43016 | 0.44048 |
| ENSG00000130720 | 9 | 133777825 | 133814673 | 173 | 24 | 0.10702 | 0.45739 | FIBCD1 | 0.45739 | 0.36612 | 0.32579 |
| ENSG00000161981 | 16 | 103010 | 107669 | 12 | 3 | 0.10699 | 0.4574 | SNRNP25 | 0.4574 | 0.31374 | 0.48612 |
| ENSG00000104231 | 8 | 82613569 | 82645138 | 125 | 13 | 0.10689 | 0.45744 | ZFAND1 | 0.45744 | 0.50405 | 0.56972 |
| ENSG00000170954 | 19 | 53611132 | 53636330 | 176 | 29 | 0.10685 | 0.45745 | ZNF415 | 0.45745 | 0.27621 | 0.24348 |
| ENSG00000172403 | 4 | 119809996 | 119982402 | 660 | 57 | 0.10672 | 0.45751 | SYNPO2 | 0.45751 | 0.46652 | 0.32395 |
| ENSG00000100983 | 20 | 33516236 | 33543620 | 42 | 7 | 0.1067 | 0.45751 | GSS | 0.45751 | 0.41607 | 0.43174 |
| ENSG00000162390 | 1 | 55007930 | 55104865 | 435 | 38 | 0.10662 | 0.45754 | ACOT11 | 0.45754 | 0.44369 | 0.37911 |
| ENSG00000160310 | 21 | 48055079 | 48085036 | 113 | 14 | 0.10661 | 0.45755 | PRMT2 | 0.45755 | 0.33653 | 0.35244 |
| ENSG00000176406 | 8 | 104512976 | 105268322 | 2780 | 31 | 0.10656 | 0.45757 | RIMS2 | 0.45757 | 0.42705 | 0.34227 |

| | | | | | | | | | | | |
|---|---|---|---|---|---|---|---|---|---|---|---|
| ENSG00000185513 | 20 | 42136320 | 42179590 | 127 | 16 | 0.10653 | 0.45758 | L3MBTL1 | 0.45758 | 0.56317 | 0.67917 |
| ENSG00000070031 | 11 | 626431 | 627143 | 1 | 1 | 0.10648 | 0.4576 | SCT | 0.4576 | 0.002698 | 0.0035685 |
| ENSG00000156381 | 14 | 102973179 | 102976136 | 10 | 3 | 0.10641 | 0.45763 | ANKRD9 | 0.45763 | 0.54479 | 0.6441 |
| ENSG00000141569 | 17 | 73876416 | 73893084 | 61 | 8 | 0.10632 | 0.45766 | TRIM65 | 0.45766 | 0.38403 | 0.2157 |
| ENSG00000162961 | 2 | 32092878 | 32264881 | 432 | 23 | 0.1063 | 0.45767 | DPY30 | 0.45767 | 0.45971 | 0.44266 |
| ENSG00000153132 | 4 | 141309609 | 141349122 | 114 | 11 | 0.10627 | 0.45768 | CLGN | 0.45768 | 0.48126 | 0.46908 |
| ENSG00000137077 | 9 | 34709002 | 34710121 | 1 | 1 | 0.10623 | 0.4577 | CCL21 | 0.4577 | 0.013041 | 0.0061598 |
| ENSG00000213339 | 19 | 10812106 | 10824113 | 32 | 8 | 0.10619 | 0.45771 | QTRT1 | 0.45771 | 0.73149 | 0.82519 |
| ENSG00000163521 | 2 | 220101328 | 220110200 | 22 | 8 | 0.10611 | 0.45775 | GLB1L | 0.45775 | 0.49441 | 0.58393 |
| ENSG00000091592 | 17 | 5402747 | 5522744 | 639 | 26 | 0.10606 | 0.45777 | NLRP1 | 0.45777 | 0.49314 | 0.51081 |
| ENSG00000131389 | 3 | 14444076 | 14530857 | 287 | 45 | 0.10599 | 0.4578 | SLC6A6 | 0.4578 | 0.46633 | 0.46189 |
| ENSG00000011590 | 19 | 36195429 | 36207940 | 36 | 10 | 0.10597 | 0.4578 | ZBTB32 | 0.4578 | 0.65007 | 0.67084 |
| ENSG00000118873 | 1 | 220321635 | 220445796 | 275 | 20 | 0.10589 | 0.45783 | RAB3GAP2 | 0.45783 | 0.46293 | 0.47711 |
| ENSG00000144445 | 2 | 210886147 | 211036107 | 153 | 28 | 0.10572 | 0.4579 | KANSL1L | 0.4579 | 0.51743 | 0.52076 |
| ENSG00000176422 | 12 | 56862301 | 56864763 | 5 | 2 | 0.10556 | 0.45796 | SPRYD4 | 0.45796 | 0.5078 | 0.65105 |
| ENSG00000153827 | 2 | 230628554 | 230787955 | 258 | 22 | 0.10545 | 0.45801 | TRIP12 | 0.45801 | 0.42559 | 0.32014 |
| ENSG00000183258 | 5 | 176938578 | 176944470 | 8 | 4 | 0.1052 | 0.45811 | DDX41 | 0.45811 | 0.25628 | 0.20357 |
| ENSG00000170801 | 4 | 8271492 | 8308838 | 156 | 20 | 0.10495 | 0.45821 | HTRA3 | 0.45821 | 0.47099 | 0.22013 |
| ENSG00000181666 | 19 | 37803739 | 37860267 | 140 | 18 | 0.10487 | 0.45824 | HKR1 | 0.45824 | 0.48484 | 0.48186 |
| ENSG00000135821 | 1 | 182350839 | 182361341 | 34 | 5 | 0.1048 | 0.45827 | GLUL | 0.45827 | 0.35915 | 0.36154 |
| ENSG00000179240 | 11 | 76092357 | 76125663 | 132 | 19 | 0.10477 | 0.45828 | RP11-111M22.2 | 0.45828 | 0.38692 | 0.49311 |
| ENSG00000092051 | 14 | 24037244 | 24048009 | 10 | 3 | 0.10477 | 0.45828 | JPH4 | 0.45828 | 0.74244 | 0.74937 |
| ENSG00000157734 | 15 | 64443914 | 64449680 | 16 | 7 | 0.10458 | 0.45836 | SNX22 | 0.45836 | 0.50694 | 0.46769 |
| ENSG00000160741 | 1 | 153920145 | 153931101 | 9 | 2 | 0.1045 | 0.45839 | CRTC2 | 0.45839 | 0.1765 | 0.24222 |
| ENSG00000087077 | 7 | 100464760 | 100471076 | 15 | 8 | 0.10412 | 0.45854 | TRIP6 | 0.45854 | 0.45641 | 0.42018 |
| ENSG00000133030 | 17 | 16945859 | 17120993 | 434 | 34 | 0.10371 | 0.4587 | MPRIP | 0.4587 | 0.47813 | 0.56902 |
| ENSG00000134532 | 12 | 23682440 | 24103966 | 1348 | 130 | 0.10349 | 0.45879 | SOX5 | 0.45879 | 0.49149 | 0.44212 |
| ENSG00000184508 | 15 | 91474148 | 91475799 | 3 | 1 | 0.10338 | 0.45883 | HDDC3 | 0.45883 | 0.32874 | 0.53627 |
| ENSG00000186010 | 19 | 19626545 | 19644285 | 42 | 6 | 0.10251 | 0.45918 | NDUFA13 | 0.45918 | 0.53105 | 0.53225 |
| ENSG00000136463 | 17 | 61678231 | 61685725 | 7 | 4 | 0.10219 | 0.4593 | TACO1 | 0.4593 | 0.90616 | 0.8292 |
| ENSG00000124203 | 20 | 57766075 | 57834168 | 178 | 33 | 0.10202 | 0.45937 | ZNF831 | 0.45937 | 0.39738 | 0.22841 |
| ENSG00000106635 | 7 | 72950686 | 72972332 | 41 | 5 | 0.10198 | 0.45939 | BCL7B | 0.45939 | 0.39617 | 0.37945 |
| ENSG00000145730 | 5 | 102089685 | 102366809 | 734 | 29 | 0.10183 | 0.45944 | PAM | 0.45944 | 0.4382 | 0.37155 |
| ENSG00000131711 | 5 | 71403061 | 71505395 | 245 | 52 | 0.10178 | 0.45947 | MAP1B | 0.45947 | 0.49248 | 0.58326 |
| ENSG00000167981 | 16 | 3486104 | 3493542 | 47 | 8 | 0.10149 | 0.45958 | ZNF597 | 0.45958 | 0.46894 | 0.62725 |
| ENSG00000198685 | 3 | 128290843 | 128294929 | 20 | 7 | 0.10132 | 0.45965 | C3orf27 | 0.45965 | 0.84008 | 0.93191 |
| ENSG00000125454 | 17 | 73269073 | 73285591 | 66 | 9 | 0.10115 | 0.45971 | SLC25A19 | 0.45971 | 0.45511 | 0.48248 |
| ENSG00000173145 | 10 | 96075004 | 96122716 | 142 | 15 | 0.10111 | 0.45973 | NOC3L | 0.45973 | 0.3405 | 0.38356 |
| ENSG00000072182 | 2 | 220378892 | 220403494 | 64 | 13 | 0.10098 | 0.45978 | ASIC4 | 0.45978 | 0.42658 | 0.56703 |
| ENSG00000157335 | 16 | 70207225 | 70221264 | 12 | 7 | 0.10099 | 0.45978 | CLEC18C | 0.45978 | 0.3783 | 0.56023 |
| ENSG00000227152 | 1 | 248604508 | 248605434 | 2 | 1 | 0.1008 | 0.45985 | OR2T7 | 0.45985 | 0.38105 | 0.39903 |
| ENSG00000240972 | 22 | 24236191 | 24237414 | 4 | 1 | 0.10049 | 0.45998 | MIF | 0.45998 | 0.089387 | 0.10356 |
| ENSG00000126247 | 19 | 36630477 | 36641255 | 38 | 17 | 0.10034 | 0.46004 | CAPNS1 | 0.46004 | 0.24173 | 0.36096 |
| ENSG00000123472 | 1 | 47098409 | 47139539 | 94 | 7 | 0.10021 | 0.46009 | ATPAF1 | 0.46009 | 0.47743 | 0.5174 |
| ENSG00000178568 | 2 | 212240446 | 213403565 | 5139 | 206 | 0.099494 | 0.46037 | ERBB4 | 0.46037 | 0.43609 | 0.3787 |
| ENSG00000089220 | 12 | 118573663 | 118583389 | 31 | 9 | 0.099373 | 0.46042 | PEBP1 | 0.46042 | 0.60394 | 0.52349 |
| ENSG00000165406 | 10 | 45950035 | 46090354 | 510 | 11 | 0.099349 | 0.46043 | MARCHF8 | 0.46043 | 0.4725 | 0.48682 |
| ENSG00000164091 | 3 | 52288437 | 52322036 | 64 | 9 | 0.099026 | 0.46056 | WDR82 | 0.46056 | 0.45148 | 0.45273 |
| ENSG00000125571 | 2 | 113670548 | 113676459 | 51 | 4 | 0.099002 | 0.46057 | IL37 | 0.46057 | 0.57036 | 0.67824 |
| ENSG00000160216 | 21 | 45285067 | 45406417 | 306 | 32 | 0.098957 | 0.46059 | AGPAT3 | 0.46059 | 0.43201 | 0.29563 |
| ENSG00000105072 | 19 | 16607122 | 16632163 | 76 | 9 | 0.098914 | 0.4606 | C19orf44 | 0.4606 | 0.34906 | 0.31359 |
| ENSG00000204120 | 2 | 233562009 | 233725285 | 501 | 25 | 0.098671 | 0.4607 | GIGYF2 | 0.4607 | 0.43561 | 0.42239 |
| ENSG00000139547 | 12 | 57345219 | 57353158 | 16 | 5 | 0.098635 | 0.46071 | RDH16 | 0.46071 | 0.82925 | 0.79982 |
| ENSG00000168260 | 14 | 50550369 | 50559361 | 23 | 5 | 0.098612 | 0.46072 | C14orf183 | 0.46072 | 0.21685 | 0.28522 |
| ENSG00000160401 | 9 | 130469268 | 130478281 | 9 | 4 | 0.098525 | 0.46076 | C9orf117 | 0.46076 | 0.70202 | 0.59957 |
| ENSG00000114850 | 3 | 156257929 | 156272973 | 59 | 8 | 0.097551 | 0.46114 | SSR3 | 0.46114 | 0.65891 | 0.47963 |
| ENSG00000269058 | 19 | 16589868 | 16607003 | 79 | 7 | 0.097499 | 0.46117 | CALR3 | 0.46117 | 0.39661 | 0.35596 |
| ENSG00000129422 | 8 | 17501304 | 17658426 | 1016 | 73 | 0.097375 | 0.46121 | MTUS1 | 0.46121 | 0.31574 | 0.17285 |
| ENSG00000133101 | 13 | 37005967 | 37017019 | 26 | 5 | 0.097242 | 0.46127 | CCNA1 | 0.46127 | 0.41031 | 0.5999 |
| ENSG00000115271 | 2 | 163175350 | 163228105 | 57 | 7 | 0.097043 | 0.46135 | GCA | 0.46135 | 0.51752 | 0.51206 |
| ENSG00000165457 | 11 | 71927645 | 71932994 | 9 | 4 | 0.096971 | 0.46137 | FOLR2 | 0.46137 | 0.63625 | 0.56789 |
| ENSG00000204323 | 17 | 73629514 | 73637484 | 13 | 7 | 0.096829 | 0.46143 | SMIM5 | 0.46143 | 0.50492 | 0.54977 |
| ENSG00000205832 | 16 | 4606491 | 4650715 | 112 | 16 | 0.096655 | 0.4615 | C16orf96 | 0.4615 | 0.44744 | 0.55313 |
| ENSG00000164828 | 7 | 855528 | 936072 | 290 | 26 | 0.096405 | 0.4616 | SUN1 | 0.4616 | 0.31535 | 0.13625 |
| ENSG00000204006 | 1 | 51567906 | 51613752 | 69 | 9 | 0.096373 | 0.46161 | C1orf185 | 0.46161 | 0.44219 | 0.46934 |
| ENSG00000122203 | 5 | 175773064 | 175788971 | 36 | 8 | 0.096214 | 0.46168 | KIAA1191 | 0.46168 | 0.49262 | 0.57332 |
| ENSG00000155561 | 7 | 135242667 | 135333505 | 456 | 18 | 0.096098 | 0.46172 | NUP205 | 0.46172 | 0.46179 | 0.42288 |
| ENSG00000137710 | 11 | 110045605 | 110167447 | 268 | 14 | 0.096093 | 0.46172 | RDX | 0.46172 | 0.49132 | 0.48002 |
| ENSG00000070182 | 14 | 65213002 | 65346601 | 566 | 43 | 0.096073 | 0.46173 | SPTB | 0.46173 | 0.4629 | 0.40209 |
| ENSG00000158055 | 1 | 24645812 | 24690972 | 111 | 25 | 0.095764 | 0.46185 | GRHL3 | 0.46185 | 0.59596 | 0.32187 |
| ENSG00000158985 | 5 | 130581186 | 130734140 | 275 | 16 | 0.095683 | 0.46189 | CDC42SE2 | 0.46189 | 0.42252 | 0.43069 |
| ENSG00000101665 | 18 | 46446223 | 46477081 | 115 | 25 | 0.095588 | 0.46192 | SMAD7 | 0.46192 | 0.36352 | 0.27039 |
| ENSG00000130038 | 12 | 3715799 | 3873985 | 730 | 72 | 0.095505 | 0.46196 | EFCAB4B | 0.46196 | 0.53588 | 0.63905 |

| | | | | | | | | | | | |
|---|---|---|---|---|---|---|---|---|---|---|---|
| ENSG00000065618 | 10 | 105791044 | 105845760 | 108 | 18 | 0.095361 | 0.46201 | COL17A1 | 0.46201 | 0.60909 | 0.56777 |
| ENSG00000152292 | 2 | 85645844 | 85664152 | 72 | 16 | 0.095267 | 0.46205 | SH2D6 | 0.46205 | 0.31019 | 0.24202 |
| ENSG00000149573 | 11 | 118124118 | 118135251 | 46 | 5 | 0.095176 | 0.46209 | MPZL2 | 0.46209 | 0.4968 | 0.28237 |
| ENSG00000204671 | 12 | 122656577 | 122658746 | 6 | 2 | 0.095041 | 0.46214 | IL31 | 0.46214 | 0.23325 | 0.17941 |
| ENSG00000148187 | 9 | 125026882 | 125085743 | 57 | 13 | 0.095002 | 0.46216 | MRRF | 0.46216 | 0.64765 | 0.70256 |
| ENSG00000170819 | 3 | 133118839 | 133194066 | 300 | 33 | 0.094766 | 0.46225 | BFSP2 | 0.46225 | 0.58032 | 0.70159 |
| ENSG00000196218 | 19 | 38924339 | 39078204 | 620 | 45 | 0.094581 | 0.46232 | RYR1 | 0.46232 | 0.438 | 0.41156 |
| ENSG00000137842 | 15 | 43415477 | 43477344 | 77 | 16 | 0.094467 | 0.46237 | TMEM62 | 0.46237 | 0.53798 | 0.45341 |
| ENSG00000198791 | 8 | 17086737 | 17104387 | 48 | 10 | 0.094306 | 0.46243 | CNOT7 | 0.46243 | 0.39426 | 0.29197 |
| ENSG00000113924 | 3 | 120347020 | 120401418 | 221 | 36 | 0.094246 | 0.46246 | HGD | 0.46246 | 0.34505 | 0.27085 |
| ENSG00000203908 | 6 | 74072400 | 74073894 | 5 | 2 | 0.094163 | 0.46249 | KHDC3L | 0.46249 | 0.75205 | 0.70631 |
| ENSG00000178691 | 17 | 30264037 | 30328064 | 201 | 8 | 0.094103 | 0.46251 | SUZ12 | 0.46251 | 0.45942 | 0.49677 |
| ENSG00000114026 | 3 | 9791628 | 9829903 | 82 | 14 | 0.093964 | 0.46257 | OGG1 | 0.46257 | 0.37447 | 0.19921 |
| ENSG00000120675 | 13 | 43597339 | 43683045 | 259 | 13 | 0.093869 | 0.46261 | DNAJC15 | 0.46261 | 0.48187 | 0.53304 |
| ENSG00000130561 | 2 | 234216462 | 234255701 | 160 | 20 | 0.093808 | 0.46263 | SAG | 0.46263 | 0.51124 | 0.5497 |
| ENSG00000256870 | 12 | 101549271 | 101604185 | 161 | 18 | 0.09345 | 0.46277 | SLC5A8 | 0.46277 | 0.58819 | 0.70684 |
| ENSG00000091972 | 3 | 112051194 | 112081659 | 101 | 15 | 0.093379 | 0.4628 | CD200 | 0.4628 | 0.59098 | 0.69821 |
| ENSG00000135535 | 6 | 109687717 | 109703762 | 56 | 8 | 0.093391 | 0.4628 | CD164 | 0.4628 | 0.53355 | 0.57322 |
| ENSG00000197921 | 1 | 2460184 | 2461684 | 5 | 3 | 0.093351 | 0.46281 | HES5 | 0.46281 | 0.15051 | 0.22297 |
| ENSG00000171960 | 1 | 43124096 | 43142429 | 35 | 8 | 0.093134 | 0.4629 | PPIH | 0.4629 | 0.45687 | 0.46201 |
| ENSG00000204264 | 6 | 32808494 | 32812480 | 19 | 5 | 0.093086 | 0.46292 | PSMB8 | 0.46292 | 0.77294 | 0.25371 |
| ENSG00000143164 | 1 | 167905021 | 168045081 | 244 | 18 | 0.092988 | 0.46296 | DCAF6 | 0.46296 | 0.48132 | 0.47623 |
| ENSG00000197530 | 1 | 1550795 | 1565990 | 38 | 7 | 0.09286 | 0.46301 | MIB2 | 0.46301 | 0.56655 | 0.58464 |
| ENSG00000014216 | 11 | 64948037 | 64979477 | 77 | 13 | 0.092834 | 0.46302 | CAPN1 | 0.46302 | 0.48375 | 0.47375 |
| ENSG00000240184 | 5 | 140855580 | 140892542 | 107 | 20 | 0.092714 | 0.46307 | PCDHGC3 | 0.46307 | 0.4443 | 0.44105 |
| ENSG00000116661 | 1 | 11708424 | 11715842 | 32 | 16 | 0.09255 | 0.46313 | FBXO2 | 0.46313 | 0.44017 | 0.47016 |
| ENSG00000196539 | 1 | 248636627 | 248637634 | 4 | 2 | 0.092535 | 0.46314 | OR2T3 | 0.46314 | 0.38631 | 0.48814 |
| ENSG00000173706 | 3 | 124684554 | 124774802 | 316 | 27 | 0.0925 | 0.46315 | HEG1 | 0.46315 | 0.49504 | 0.52268 |
| ENSG00000003393 | 2 | 202565277 | 202645912 | 317 | 21 | 0.092446 | 0.46317 | ALS2 | 0.46317 | 0.48552 | 0.48942 |
| ENSG00000143536 | 1 | 152381719 | 152386739 | 8 | 3 | 0.092412 | 0.46319 | CRNN | 0.46319 | 0.70779 | 0.75901 |
| ENSG00000204618 | 6 | 30038043 | 30043664 | 48 | 7 | 0.09232 | 0.46322 | RNF39 | 0.46322 | 0.66905 | 0.68179 |
| ENSG00000167513 | 16 | 88869621 | 88875666 | 27 | 4 | 0.092292 | 0.46323 | CDT1 | 0.46323 | 0.46092 | 0.60912 |
| ENSG00000232774 | 14 | 62037258 | 62125414 | 250 | 39 | 0.092182 | 0.46328 | RP11-47I22.3 | 0.46328 | 0.4491 | 0.47881 |
| ENSG00000070785 | 1 | 45316450 | 45452282 | 417 | 14 | 0.092117 | 0.4633 | EIF2B3 | 0.4633 | 0.48017 | 0.52362 |
| ENSG00000184347 | 5 | 168088745 | 168728133 | 2562 | 194 | 0.092109 | 0.46331 | SLIT3 | 0.46331 | 0.45368 | 0.48836 |
| ENSG00000166796 | 11 | 18433854 | 18473605 | 213 | 13 | 0.091985 | 0.46336 | LDHC | 0.46336 | 0.62788 | 0.64986 |
| ENSG00000107960 | 10 | 105642300 | 105677963 | 117 | 12 | 0.091935 | 0.46337 | OBFC1 | 0.46337 | 0.4641 | 0.51639 |
| ENSG00000172172 | 8 | 121393000 | 121457642 | 117 | 14 | 0.091894 | 0.46339 | MRPL13 | 0.46339 | 0.47944 | 0.51609 |
| ENSG00000116604 | 1 | 156433519 | 156470620 | 80 | 13 | 0.091837 | 0.46341 | MEF2D | 0.46341 | 0.36858 | 0.38383 |
| ENSG00000204604 | 19 | 53341261 | 53360902 | 144 | 16 | 0.091847 | 0.46341 | ZNF468 | 0.46341 | 0.3991 | 0.41876 |
| ENSG00000068971 | 11 | 64685025 | 64701945 | 31 | 13 | 0.091805 | 0.46343 | PPP2R5B | 0.46343 | 0.50052 | 0.47444 |
| ENSG00000164603 | 7 | 112459202 | 112579971 | 236 | 30 | 0.091456 | 0.46357 | C7orf60 | 0.46357 | 0.49126 | 0.52972 |
| ENSG00000010292 | 12 | 6602522 | 6641121 | 163 | 18 | 0.091395 | 0.46359 | NCAPD2 | 0.46359 | 0.42154 | 0.41127 |
| ENSG00000005955 | 17 | 34900737 | 34946278 | 106 | 13 | 0.091287 | 0.46363 | GGNBP2 | 0.46363 | 0.41217 | 0.36869 |
| ENSG00000139352 | 12 | 103351464 | 103354294 | 5 | 3 | 0.091195 | 0.46367 | ASCL1 | 0.46367 | 0.90523 | 0.67512 |
| ENSG00000183137 | 6 | 109416313 | 109485135 | 102 | 12 | 0.091092 | 0.46371 | CEP57L1 | 0.46371 | 0.49523 | 0.4268 |
| ENSG00000151151 | 10 | 59951278 | 60027694 | 335 | 10 | 0.0911 | 0.46371 | IPMK | 0.46371 | 0.43248 | 0.39232 |
| ENSG00000073756 | 1 | 186640923 | 186649559 | 14 | 6 | 0.089925 | 0.46417 | PTGS2 | 0.46417 | 0.83085 | 0.8627 |
| ENSG00000124713 | 6 | 42928496 | 42931618 | 11 | 3 | 0.089598 | 0.4643 | GNMT | 0.4643 | 0.51708 | 0.54598 |
| ENSG00000112234 | 6 | 99316420 | 99395849 | 158 | 18 | 0.089567 | 0.46432 | FBXL4 | 0.46432 | 0.48926 | 0.46886 |
| ENSG00000112981 | 5 | 137450866 | 137475132 | 53 | 13 | 0.089405 | 0.46438 | NME5 | 0.46438 | 0.51091 | 0.48489 |
| ENSG00000136802 | 9 | 131644391 | 131680318 | 79 | 11 | 0.089281 | 0.46443 | LRRC8A | 0.46443 | 0.54841 | 0.54067 |
| ENSG00000167995 | 11 | 61717293 | 61732987 | 60 | 11 | 0.089053 | 0.46452 | BEST1 | 0.46452 | 0.50261 | 0.45712 |
| ENSG00000250151 | 3 | 9834766 | 9877085 | 147 | 17 | 0.089013 | 0.46454 | ARPC4-TTLL3 | 0.46454 | 0.52293 | 0.54786 |
| ENSG00000214518 | 17 | 39210750 | 39211482 | 3 | 2 | 0.088673 | 0.46467 | KRTAP2-2 | 0.46467 | 0.63697 | 0.76641 |
| ENSG00000197536 | 5 | 131746328 | 131811736 | 198 | 19 | 0.088587 | 0.4647 | C5orf56 | 0.4647 | 0.48069 | 0.56502 |
| ENSG00000198492 | 1 | 29063133 | 29096287 | 77 | 9 | 0.08844 | 0.46476 | YTHDF2 | 0.46476 | 0.38113 | 0.37008 |
| ENSG00000066135 | 1 | 44115829 | 44171186 | 98 | 15 | 0.08818 | 0.46487 | KDM4A | 0.46487 | 0.4843 | 0.48758 |
| ENSG00000106631 | 7 | 44178463 | 44180931 | 10 | 4 | 0.087933 | 0.46496 | MYL7 | 0.46496 | 0.63303 | 0.6716 |
| ENSG00000119608 | 14 | 75319736 | 75330537 | 29 | 8 | 0.08789 | 0.46498 | PROX2 | 0.46498 | 0.61657 | 0.68522 |
| ENSG00000184185 | 17 | 21279509 | 21323179 | 64 | 18 | 0.087904 | 0.46498 | KCNJ12 | 0.46498 | 0.68476 | 0.625 |
| ENSG00000176087 | 5 | 139944041 | 139948688 | 7 | 4 | 0.087442 | 0.46516 | SLC35A4 | 0.46516 | 0.3348 | 0.3476 |
| ENSG00000170370 | 10 | 119301955 | 119309056 | 22 | 6 | 0.087118 | 0.46529 | EMX2 | 0.46529 | 0.46098 | 0.64888 |
| ENSG00000221914 | 8 | 26149007 | 26230196 | 206 | 33 | 0.086912 | 0.46537 | PPP2R2A | 0.46537 | 0.56464 | 0.60735 |
| ENSG00000205279 | 5 | 126984736 | 126994322 | 25 | 10 | 0.086812 | 0.46541 | CTXN3 | 0.46541 | 0.50862 | 0.45157 |
| ENSG00000100580 | 14 | 77801364 | 77843452 | 124 | 16 | 0.086763 | 0.46543 | TMED8 | 0.46543 | 0.57497 | 0.70762 |
| ENSG00000149294 | 11 | 112831997 | 113149158 | 1091 | 59 | 0.0867 | 0.46545 | NCAM1 | 0.46545 | 0.50497 | 0.51788 |
| ENSG00000125726 | 19 | 6583194 | 6604114 | 110 | 19 | 0.086307 | 0.46561 | CD70 | 0.46561 | 0.24538 | 0.19242 |
| ENSG00000144771 | 3 | 54952264 | 55001115 | 149 | 21 | 0.086235 | 0.46564 | LRTM1 | 0.46564 | 0.44732 | 0.53566 |
| ENSG00000144736 | 3 | 72798428 | 72911065 | 193 | 12 | 0.086206 | 0.46565 | SHQ1 | 0.46565 | 0.478 | 0.48532 |
| ENSG00000175894 | 21 | 45917775 | 46131495 | 1086 | 63 | 0.086081 | 0.4657 | TSPEAR | 0.4657 | 0.47127 | 0.49106 |
| ENSG00000109208 | 4 | 71226478 | 71232875 | 35 | 8 | 0.086067 | 0.46571 | SMR3A | 0.46571 | 0.52611 | 0.69587 |
| ENSG00000256713 | 11 | 61008514 | 61018920 | 22 | 10 | 0.085912 | 0.46577 | PGA5 | 0.46577 | 0.55181 | 0.60929 |

| | | | | | | | | | | | |
|---|---|---|---|---|---|---|---|---|---|---|---|
| ENSG00000258941 | 14 | 39703106 | 39820397 | 309 | 21 | 0.085618 | 0.46588 | RP11-407N17.3 | 0.46588 | 0.55441 | 0.64261 |
| ENSG00000104321 | 8 | 72932152 | 72987852 | 257 | 18 | 0.085554 | 0.46591 | TRPA1 | 0.46591 | 0.43376 | 0.49711 |
| ENSG00000221949 | 12 | 62995531 | 62997214 | 8 | 3 | 0.085521 | 0.46592 | C12orf61 | 0.46592 | 0.89657 | 0.89916 |
| ENSG00000162415 | 1 | 45482071 | 45771881 | 684 | 26 | 0.085309 | 0.46601 | ZSWIM5 | 0.46601 | 0.47101 | 0.49672 |
| ENSG00000121851 | 1 | 145456236 | 145470388 | 9 | 6 | 0.08527 | 0.46602 | POLR3GL | 0.46602 | 0.21893 | 0.34871 |
| ENSG00000119414 | 9 | 127908852 | 127952218 | 118 | 12 | 0.085255 | 0.46603 | PPP6C | 0.46603 | 0.43687 | 0.40165 |
| ENSG00000101489 | 18 | 34823010 | 35146000 | 906 | 105 | 0.085085 | 0.4661 | CELF4 | 0.4661 | 0.4494 | 0.44567 |
| ENSG00000132661 | 20 | 23331373 | 23335414 | 16 | 5 | 0.085082 | 0.4661 | NXT1 | 0.4661 | 0.5368 | 0.46552 |
| ENSG00000140548 | 15 | 90544624 | 90625438 | 228 | 31 | 0.084995 | 0.46613 | ZNF710 | 0.46613 | 0.35722 | 0.54107 |
| ENSG00000130508 | 2 | 1635659 | 1748624 | 459 | 56 | 0.084918 | 0.46616 | PXDN | 0.46616 | 0.52613 | 0.65232 |
| ENSG00000155330 | 16 | 46830519 | 46865323 | 31 | 7 | 0.084701 | 0.46625 | C16orf87 | 0.46625 | 0.53582 | 0.57281 |
| ENSG00000104904 | 19 | 2269485 | 2273487 | 21 | 6 | 0.084548 | 0.46631 | OAZ1 | 0.46631 | 0.24989 | 0.54124 |
| ENSG00000133816 | 11 | 12115543 | 12285334 | 742 | 83 | 0.084015 | 0.46652 | MICAL2 | 0.46652 | 0.58089 | 0.65888 |
| ENSG00000141252 | 17 | 411908 | 624957 | 830 | 40 | 0.083935 | 0.46655 | VPS53 | 0.46655 | 0.47591 | 0.47189 |
| ENSG00000230778 | 15 | 40573645 | 40574787 | 1 | 1 | 0.083819 | 0.4666 | ANKRD63 | 0.4666 | 0.68045 | 0.7546 |
| ENSG00000121005 | 8 | 75896750 | 75946793 | 201 | 23 | 0.083578 | 0.4667 | CRISPLD1 | 0.4667 | 0.56162 | 0.4905 |
| ENSG00000165916 | 11 | 47440320 | 47447993 | 23 | 6 | 0.083548 | 0.46671 | PSMC3 | 0.46671 | 0.70157 | 0.64095 |
| ENSG00000145741 | 5 | 72794233 | 72801460 | 31 | 4 | 0.083369 | 0.46678 | BTF3 | 0.46678 | 0.5023 | 0.63196 |
| ENSG00000064313 | 8 | 120743015 | 120845103 | 331 | 20 | 0.083379 | 0.46678 | TAF2 | 0.46678 | 0.44539 | 0.48962 |
| ENSG00000215549 | 13 | 26442061 | 26455095 | 8 | 4 | 0.083222 | 0.46684 | AL138815.1 | 0.46684 | 0.432 | 0.49879 |
| ENSG00000169245 | 4 | 76942273 | 76944650 | 12 | 2 | 0.082913 | 0.46696 | CXCL10 | 0.46696 | 0.41984 | 0.39656 |
| ENSG00000125870 | 20 | 16710606 | 16722421 | 39 | 10 | 0.082789 | 0.46701 | SNRPB2 | 0.46701 | 0.53399 | 0.84014 |
| ENSG00000163714 | 3 | 142683339 | 142779567 | 260 | 20 | 0.082632 | 0.46707 | U2SURP | 0.46707 | 0.54997 | 0.6829 |
| ENSG00000204283 | 17 | 75875109 | 75878659 | 11 | 3 | 0.082497 | 0.46713 | FLJ45079 | 0.46713 | 0.66017 | 0.64618 |
| ENSG00000138246 | 3 | 132136370 | 132257876 | 297 | 24 | 0.082431 | 0.46715 | DNAJC13 | 0.46715 | 0.44981 | 0.46343 |
| ENSG00000008294 | 17 | 49039535 | 49198226 | 247 | 21 | 0.082052 | 0.4673 | SPAG9 | 0.4673 | 0.49497 | 0.54365 |
| ENSG00000064989 | 2 | 188207856 | 188313187 | 285 | 18 | 0.081983 | 0.46733 | CALCRL | 0.46733 | 0.44486 | 0.42553 |
| ENSG00000163312 | 4 | 84328496 | 84377009 | 107 | 14 | 0.081845 | 0.46738 | HELQ | 0.46738 | 0.50394 | 0.57152 |
| ENSG00000174963 | 3 | 147103833 | 147124647 | 36 | 9 | 0.081805 | 0.4674 | ZIC4 | 0.4674 | 0.52864 | 0.69156 |
| ENSG00000131808 | 11 | 30252563 | 30256808 | 12 | 2 | 0.081583 | 0.46749 | FSHB | 0.46749 | 0.40673 | 0.43321 |
| ENSG00000071655 | 19 | 1576639 | 1592882 | 37 | 9 | 0.081578 | 0.46749 | MBD3 | 0.46749 | 0.44705 | 0.53388 |
| ENSG00000095066 | 19 | 12873817 | 12983554 | 310 | 24 | 0.081571 | 0.46749 | HOOK2 | 0.46749 | 0.50635 | 0.45058 |
| ENSG00000108239 | 10 | 96162261 | 96295687 | 273 | 15 | 0.081473 | 0.46753 | TBC1D12 | 0.46753 | 0.49028 | 0.49925 |
| ENSG00000162739 | 1 | 160454820 | 160493052 | 134 | 20 | 0.081345 | 0.46758 | SLAMF6 | 0.46758 | 0.55308 | 0.61755 |
| ENSG00000130935 | 17 | 65713949 | 65740647 | 93 | 8 | 0.081133 | 0.46767 | NOL11 | 0.46767 | 0.4242 | 0.4852 |
| ENSG00000125971 | 20 | 33104214 | 33128762 | 30 | 7 | 0.08108 | 0.46769 | DYNLRB1 | 0.46769 | 0.29363 | 0.35092 |
| ENSG00000144199 | 2 | 97749320 | 97760619 | 28 | 5 | 0.080926 | 0.46775 | FAHD2B | 0.46775 | 0.52592 | 0.49897 |
| ENSG00000151229 | 12 | 40148823 | 40499891 | 1058 | 61 | 0.080926 | 0.46775 | SLC2A13 | 0.46775 | 0.50709 | 0.53069 |
| ENSG00000128322 | 22 | 23915312 | 23922495 | 52 | 15 | 0.080937 | 0.46775 | IGLL1 | 0.46775 | 0.4152 | 0.10818 |
| ENSG00000114395 | 3 | 50388126 | 50395891 | 8 | 4 | 0.080886 | 0.46777 | CYB561D2 | 0.46777 | 0.48758 | 0.39409 |
| ENSG00000226288 | 11 | 4608021 | 4609135 | 3 | 2 | 0.080606 | 0.46788 | OR52I2 | 0.46788 | 0.70282 | 0.44313 |
| ENSG00000228120 | 21 | 44579176 | 44582387 | 11 | 3 | 0.080611 | 0.46788 | AP001631.10 | 0.46788 | 0.55365 | 0.56513 |
| ENSG00000166741 | 11 | 114128509 | 114184007 | 141 | 23 | 0.080382 | 0.46797 | NNMT | 0.46797 | 0.32279 | 0.095487 |
| ENSG00000243135 | 2 | 234637754 | 234681945 | 156 | 18 | 0.080213 | 0.46803 | UGT1A3 | 0.46803 | 0.45756 | 0.25743 |
| ENSG00000203485 | 14 | 105155943 | 105185942 | 143 | 23 | 0.080228 | 0.46803 | INF2 | 0.46803 | 0.56804 | 0.78134 |
| ENSG00000172239 | 5 | 43526369 | 43557860 | 84 | 10 | 0.080126 | 0.46807 | PAIP1 | 0.46807 | 0.44761 | 0.3441 |
| ENSG00000163958 | 3 | 195924320 | 195938308 | 78 | 16 | 0.080048 | 0.4681 | ZDHHC19 | 0.4681 | 0.48324 | 0.40448 |
| ENSG00000162444 | 1 | 10057264 | 10076078 | 50 | 6 | 0.079955 | 0.46814 | RBP7 | 0.46814 | 0.34654 | 0.2891 |
| ENSG00000172717 | 14 | 67656110 | 67695267 | 145 | 17 | 0.079689 | 0.46824 | FAM71D | 0.46824 | 0.49581 | 0.49572 |
| ENSG00000196843 | 2 | 97202480 | 97218375 | 25 | 8 | 0.079586 | 0.46828 | ARID5A | 0.46828 | 0.56217 | 0.51151 |
| ENSG00000268863 | 16 | 30996808 | 30997533 | 2 | 1 | 0.079488 | 0.46832 | AC135048.1 | 0.46832 | 0.6908 | 0.75483 |
| ENSG00000151117 | 11 | 18714669 | 18726332 | 26 | 9 | 0.079409 | 0.46835 | TMEM86A | 0.46835 | 0.52524 | 0.56672 |
| ENSG00000064201 | 11 | 2323227 | 2339430 | 81 | 13 | 0.079098 | 0.46848 | TSPAN32 | 0.46848 | 0.37064 | 0.44359 |
| ENSG00000127129 | 1 | 41944446 | 41950344 | 20 | 7 | 0.078726 | 0.46863 | EDN2 | 0.46863 | 0.2679 | 0.59656 |
| ENSG00000165113 | 9 | 86354336 | 86444431 | 248 | 13 | 0.078644 | 0.46866 | GKAP1 | 0.46866 | 0.4628 | 0.48667 |
| ENSG00000257218 | 12 | 120884241 | 120899389 | 58 | 11 | 0.078647 | 0.46866 | GATC | 0.46866 | 0.38027 | 0.47919 |
| ENSG00000117122 | 1 | 17300997 | 17307330 | 18 | 4 | 0.078387 | 0.46876 | MFAP2 | 0.46876 | 0.5569 | 0.7096 |
| ENSG00000173641 | 1 | 16340523 | 16346089 | 46 | 5 | 0.078173 | 0.46885 | HSPB7 | 0.46885 | 0.5106 | 0.76301 |
| ENSG00000174837 | 19 | 6887577 | 6940470 | 360 | 23 | 0.077809 | 0.46899 | EMR1 | 0.46899 | 0.56352 | 0.67224 |
| ENSG00000198477 | 22 | 22838767 | 22863505 | 98 | 8 | 0.077788 | 0.469 | ZNF280B | 0.469 | 0.56416 | 0.57301 |
| ENSG00000105676 | 19 | 19144384 | 19170563 | 72 | 9 | 0.077536 | 0.4691 | ARMC6 | 0.4691 | 0.43784 | 0.46725 |
| ENSG00000103313 | 16 | 3292028 | 3306627 | 68 | 10 | 0.07738 | 0.46916 | MEFV | 0.46916 | 0.53014 | 0.57474 |
| ENSG00000128802 | 10 | 48413092 | 48416853 | 7 | 3 | 0.077091 | 0.46928 | GDF2 | 0.46928 | 0.085256 | 0.023621 |
| ENSG00000113648 | 5 | 134669590 | 134735604 | 120 | 16 | 0.077039 | 0.4693 | H2AFY | 0.4693 | 0.35875 | 0.32176 |
| ENSG00000101546 | 18 | 77794358 | 77806397 | 56 | 9 | 0.07702 | 0.4693 | RBFA | 0.4693 | 0.36597 | 0.38429 |
| ENSG00000259495 | 15 | 80637195 | 80695917 | 374 | 23 | 0.076945 | 0.46933 | RP11-210M15.2 | 0.46933 | 0.50953 | 0.54021 |
| ENSG00000149150 | 11 | 57252007 | 57283259 | 61 | 10 | 0.076774 | 0.4694 | SLC43A1 | 0.4694 | 0.41022 | 0.40937 |
| ENSG00000073910 | 13 | 32605437 | 32870794 | 892 | 74 | 0.076345 | 0.46957 | FRY | 0.46957 | 0.50444 | 0.49532 |
| ENSG00000183979 | 17 | 79858621 | 79860781 | 3 | 2 | 0.076211 | 0.46963 | NPB | 0.46963 | 0.40442 | 0.37358 |
| ENSG00000160959 | 8 | 145743376 | 145750557 | 14 | 5 | 0.076096 | 0.46967 | LRRC14 | 0.46967 | 0.6131 | 0.56566 |
| ENSG00000198054 | 21 | 39493545 | 39560110 | 173 | 21 | 0.076056 | 0.46969 | DSCR8 | 0.46969 | 0.49536 | 0.61175 |
| ENSG00000149930 | 16 | 29984962 | 30003582 | 26 | 4 | 0.07603 | 0.4697 | TAOK2 | 0.4697 | 0.44383 | 0.53607 |
| ENSG00000166483 | 11 | 9595228 | 9615004 | 53 | 13 | 0.076004 | 0.46971 | WEE1 | 0.46971 | 0.42429 | 0.53037 |

| | | | | | | | | | | | |
|---|---|---|---|---|---|---|---|---|---|---|---|
| ENSG00000127184 | 5 | 85913721 | 85916779 | 4 | 2 | 0.075825 | 0.46978 | COX7C | 0.46978 | 0.64192 | 0.70565 |
| ENSG00000102710 | 13 | 37583449 | 37633850 | 84 | 9 | 0.075519 | 0.4699 | SUPT20H | 0.4699 | 0.42072 | 0.44539 |
| ENSG00000175073 | 8 | 67540722 | 67579452 | 56 | 9 | 0.07542 | 0.46994 | VCPIP1 | 0.46994 | 0.48586 | 0.42283 |
| ENSG00000139974 | 14 | 61447832 | 61550451 | 167 | 21 | 0.075388 | 0.46995 | SLC38A6 | 0.46995 | 0.45723 | 0.39894 |
| ENSG00000232560 | 21 | 18811208 | 18821503 | 40 | 6 | 0.07529 | 0.46999 | C21orf37 | 0.46999 | 0.5093 | 0.80489 |
| ENSG00000140326 | 15 | 43015757 | 43029324 | 34 | 5 | 0.074848 | 0.47017 | CDAN1 | 0.47017 | 0.44271 | 0.4315 |
| ENSG00000169045 | 5 | 179041179 | 179061785 | 97 | 17 | 0.074823 | 0.47018 | HNRNPH1 | 0.47018 | 0.43722 | 0.32956 |
| ENSG00000115226 | 2 | 27714750 | 27718112 | 6 | 2 | 0.074546 | 0.47029 | FNDC4 | 0.47029 | 0.56452 | 0.4801 |
| ENSG00000159307 | 22 | 43593289 | 43739394 | 721 | 100 | 0.074438 | 0.47033 | SCUBE1 | 0.47033 | 0.28707 | 0.2717 |
| ENSG00000221937 | 7 | 142919130 | 142920162 | 1 | 1 | 0.074264 | 0.4704 | TAS2R40 | 0.4704 | 0.85311 | 0.92804 |
| ENSG00000198390 | 21 | 31768392 | 31769136 | 1 | 1 | 0.074264 | 0.4704 | KRTAP13-1 | 0.4704 | 0.53853 | 0.58552 |
| ENSG00000147586 | 8 | 80830952 | 80942524 | 197 | 21 | 0.074243 | 0.47041 | MRPS28 | 0.47041 | 0.46818 | 0.46914 |
| ENSG00000262543 | 6 | 138699042 | 138704212 | 30 | 3 | 0.074062 | 0.47048 | RP3-422G23.4 | 0.47048 | 0.55518 | 0.61393 |
| ENSG00000158470 | 20 | 48249482 | 48330415 | 213 | 17 | 0.073962 | 0.47052 | B4GALT5 | 0.47052 | 0.41912 | 0.20431 |
| ENSG00000196591 | 6 | 114254192 | 114332472 | 194 | 26 | 0.073564 | 0.47068 | HDAC2 | 0.47068 | 0.61983 | 0.79656 |
| ENSG00000122557 | 7 | 35672269 | 35735181 | 166 | 16 | 0.073534 | 0.47069 | HERPUD2 | 0.47069 | 0.47483 | 0.3488 |
| ENSG00000213341 | 10 | 101948055 | 101989376 | 108 | 13 | 0.073361 | 0.47076 | CHUK | 0.47076 | 0.44307 | 0.44183 |
| ENSG00000111049 | 12 | 81110691 | 81113452 | 5 | 2 | 0.073023 | 0.47089 | MYF5 | 0.47089 | 0.47776 | 0.40784 |
| ENSG00000127364 | 7 | 141478242 | 141479235 | 3 | 1 | 0.072805 | 0.47098 | TAS2R4 | 0.47098 | 0.37573 | 0.36257 |
| ENSG00000204524 | 19 | 57751973 | 57766503 | 82 | 9 | 0.072736 | 0.47101 | ZNF805 | 0.47101 | 0.50863 | 0.55266 |
| ENSG00000174279 | 2 | 176942200 | 176948641 | 9 | 3 | 0.072647 | 0.47104 | EVX2 | 0.47104 | 0.48109 | 0.16209 |
| ENSG00000106479 | 7 | 149535456 | 149564568 | 64 | 14 | 0.072268 | 0.47119 | ZNF862 | 0.47119 | 0.38117 | 0.47765 |
| ENSG00000141867 | 19 | 15347647 | 15443356 | 109 | 14 | 0.072035 | 0.47129 | BRD4 | 0.47129 | 0.49446 | 0.38744 |
| ENSG00000214978 | 17 | 9694544 | 9725388 | 235 | 34 | 0.071937 | 0.47133 | RP11-477N12.3 | 0.47133 | 0.56318 | 0.68224 |
| ENSG00000269071 | 5 | 138852108 | 138852506 | 1 | 1 | 0.071751 | 0.4714 | AC138517.1 | 0.4714 | 0.17713 | 0.21898 |
| ENSG00000108797 | 17 | 40834631 | 40851832 | 31 | 7 | 0.071618 | 0.47145 | CNTNAP1 | 0.47145 | 0.52153 | 0.53846 |
| ENSG00000122025 | 13 | 28577411 | 28674729 | 401 | 25 | 0.071545 | 0.47148 | FLT3 | 0.47148 | 0.30464 | 0.11556 |
| ENSG00000163590 | 3 | 160473390 | 160796695 | 940 | 31 | 0.071383 | 0.47155 | PPM1L | 0.47155 | 0.48097 | 0.49327 |
| ENSG00000161905 | 17 | 4534197 | 4545589 | 52 | 16 | 0.071275 | 0.47159 | ALOX15 | 0.47159 | 0.17 | 0.34797 |
| ENSG00000196839 | 20 | 43248163 | 43280874 | 102 | 24 | 0.071095 | 0.47166 | ADA | 0.47166 | 0.2855 | 0.20379 |
| ENSG00000008226 | 3 | 38080696 | 38165516 | 295 | 19 | 0.070987 | 0.4717 | DLEC1 | 0.4717 | 0.47161 | 0.51952 |
| ENSG00000165272 | 9 | 33441152 | 33447609 | 23 | 10 | 0.070929 | 0.47173 | AQP3 | 0.47173 | 0.77768 | 0.54866 |
| ENSG00000072415 | 14 | 67707826 | 67802536 | 411 | 16 | 0.070899 | 0.47174 | MPP5 | 0.47174 | 0.4549 | 0.45538 |
| ENSG00000177688 | 6 | 149721495 | 149722177 | 2 | 1 | 0.070854 | 0.47176 | SUMO4 | 0.47176 | 0.6307 | 0.65494 |
| ENSG00000152620 | 5 | 36192694 | 36242381 | 125 | 14 | 0.070659 | 0.47183 | NADK2 | 0.47183 | 0.51903 | 0.50391 |
| ENSG00000133937 | 14 | 95234553 | 95236562 | 5 | 2 | 0.070667 | 0.47183 | GSC | 0.47183 | 0.40749 | 0.61276 |
| ENSG00000180251 | 2 | 103089762 | 103150431 | 273 | 14 | 0.070646 | 0.47184 | SLC9A4 | 0.47184 | 0.40748 | 0.37168 |
| ENSG00000117614 | 1 | 25549170 | 25558993 | 7 | 5 | 0.070597 | 0.47186 | SYF2 | 0.47186 | 0.81858 | 0.927 |
| ENSG00000060749 | 11 | 32914724 | 33014862 | 176 | 17 | 0.070245 | 0.472 | QSER1 | 0.472 | 0.47768 | 0.42366 |
| ENSG00000183798 | 20 | 39988606 | 39995467 | 23 | 5 | 0.070126 | 0.47205 | EMILIN3 | 0.47205 | 0.5987 | 0.52507 |
| ENSG00000068654 | 2 | 86247339 | 86333278 | 240 | 23 | 0.06978 | 0.47218 | POLR1A | 0.47218 | 0.5092 | 0.55857 |
| ENSG00000129197 | 17 | 5322961 | 5336196 | 75 | 6 | 0.069801 | 0.47218 | RPAIN | 0.47218 | 0.28066 | 0.22808 |
| ENSG00000001631 | 7 | 91828283 | 91875480 | 60 | 11 | 0.069746 | 0.4722 | KRIT1 | 0.4722 | 0.56017 | 0.53984 |
| ENSG00000198561 | 11 | 57520715 | 57587018 | 107 | 15 | 0.069482 | 0.4723 | CTNND1 | 0.4723 | 0.47067 | 0.38617 |
| ENSG00000125611 | 2 | 113341817 | 113346852 | 8 | 3 | 0.06939 | 0.47234 | CHCHD5 | 0.47234 | 0.49844 | 0.2457 |
| ENSG00000164236 | 5 | 10564442 | 10650308 | 348 | 33 | 0.069398 | 0.47234 | ANKRD33B | 0.47234 | 0.54786 | 0.65652 |
| ENSG00000163687 | 3 | 58177984 | 58200424 | 92 | 14 | 0.0692 | 0.47242 | DNASE1L3 | 0.47242 | 0.35275 | 0.29624 |
| ENSG00000074935 | 6 | 112391980 | 112408732 | 30 | 5 | 0.06918 | 0.47242 | TUBE1 | 0.47242 | 0.51255 | 0.59398 |
| ENSG00000078246 | 12 | 2986389 | 3050306 | 268 | 6 | 0.069188 | 0.47242 | TULP3 | 0.47242 | 0.36123 | 0.34519 |
| ENSG00000138641 | 4 | 89442199 | 89629693 | 540 | 32 | 0.069106 | 0.47245 | HERC3 | 0.47245 | 0.46047 | 0.47533 |
| ENSG00000099783 | 19 | 8509651 | 8553998 | 166 | 19 | 0.06879 | 0.47258 | HNRNPM | 0.47258 | 0.46304 | 0.69772 |
| ENSG00000143369 | 1 | 150480538 | 150486265 | 14 | 5 | 0.068419 | 0.47273 | ECM1 | 0.47273 | 0.58722 | 0.82028 |
| ENSG00000080822 | 3 | 98216756 | 98241910 | 63 | 13 | 0.068408 | 0.47273 | CLDND1 | 0.47273 | 0.70331 | 0.84712 |
| ENSG00000221989 | 7 | 143806607 | 143807657 | 4 | 2 | 0.068232 | 0.4728 | OR2A2 | 0.4728 | 0.447 | 0.4693 |
| ENSG00000214732 | 6 | 42123279 | 42131156 | 63 | 11 | 0.068045 | 0.47287 | RP1-139D8.6 | 0.47287 | 0.5523 | 0.42282 |
| ENSG00000268925 | 10 | 135341026 | 135342661 | 7 | 4 | 0.068023 | 0.47288 | AL161645.2 | 0.47288 | 0.16473 | 0.3162 |
| ENSG00000184471 | 16 | 1140005 | 1146244 | 37 | 8 | 0.067413 | 0.47313 | C1QTNF8 | 0.47313 | 0.23983 | 0.39665 |
| ENSG00000269624 | 1 | 63073437 | 63075249 | 3 | 1 | 0.067235 | 0.4732 | AL138847.1 | 0.4732 | 0.58032 | 0.51787 |
| ENSG00000101307 | 20 | 1544167 | 1600707 | 102 | 16 | 0.067205 | 0.47321 | SIRPB1 | 0.47321 | 0.62474 | 0.53113 |
| ENSG00000187135 | 19 | 30017406 | 30055386 | 192 | 25 | 0.067172 | 0.47322 | VSTM2B | 0.47322 | 0.63934 | 0.6436 |
| ENSG00000105647 | 19 | 18263928 | 18281350 | 46 | 9 | 0.066911 | 0.47333 | PIK3R2 | 0.47333 | 0.46418 | 0.3149 |
| ENSG00000173302 | 2 | 131486643 | 131487909 | 4 | 2 | 0.066364 | 0.47354 | GPR148 | 0.47354 | 0.42334 | 0.23817 |
| ENSG00000269099 | 13 | 25591541 | 25591675 | 1 | 1 | 0.065972 | 0.4737 | LSP1 | 0.4737 | 0.35504 | 0.42086 |
| ENSG00000182070 | 11 | 5172239 | 5207612 | 209 | 19 | 0.065921 | 0.47372 | OR52A1 | 0.47372 | 0.55352 | 0.74415 |
| ENSG00000119682 | 14 | 75120140 | 75179818 | 80 | 8 | 0.065797 | 0.47377 | AREL1 | 0.47377 | 0.49761 | 0.5992 |
| ENSG00000109684 | 4 | 10488019 | 10686489 | 804 | 38 | 0.065542 | 0.47387 | CLNK | 0.47387 | 0.52783 | 0.52614 |
| ENSG00000106236 | 7 | 98246609 | 98259180 | 33 | 8 | 0.065501 | 0.47389 | NPTX2 | 0.47389 | 0.47434 | 0.49596 |
| ENSG00000139970 | 14 | 60062694 | 60337684 | 633 | 27 | 0.065343 | 0.47395 | RTN1 | 0.47395 | 0.4943 | 0.56277 |
| ENSG00000141198 | 17 | 52976748 | 53039310 | 239 | 21 | 0.065261 | 0.47398 | TOM1L1 | 0.47398 | 0.58274 | 0.6045 |
| ENSG00000214102 | 7 | 141408153 | 141431071 | 35 | 3 | 0.065198 | 0.47401 | WEE2 | 0.47401 | 0.51333 | 0.45805 |
| ENSG00000171533 | 11 | 75297963 | 75380165 | 164 | 17 | 0.065098 | 0.47405 | MAP6 | 0.47405 | 0.56123 | 0.43235 |
| ENSG00000144749 | 3 | 66429221 | 66551687 | 342 | 39 | 0.064941 | 0.47411 | LRIG1 | 0.47411 | 0.48681 | 0.40496 |
| ENSG00000135698 | 16 | 82181403 | 82203831 | 233 | 13 | 0.064771 | 0.47418 | MPHOSPH6 | 0.47418 | 0.38988 | 0.4784 |

| | | | | | | | | | | | |
|---|---|---|---|---|---|---|---|---|---|---|---|
| ENSG00000036828 | 3 | 121902530 | 122005342 | 469 | 34 | 0.064743 | 0.47419 | CASR | 0.47419 | 0.48753 | 0.5413 |
| ENSG00000115756 | 2 | 10443015 | 10567743 | 623 | 76 | 0.064688 | 0.47421 | HPCAL1 | 0.47421 | 0.53407 | 0.55573 |
| ENSG00000164442 | 6 | 139693393 | 139695757 | 11 | 5 | 0.064629 | 0.47423 | CITED2 | 0.47423 | 0.73855 | 0.73315 |
| ENSG00000173705 | 3 | 33191537 | 33260707 | 272 | 21 | 0.064623 | 0.47424 | SUSD5 | 0.47424 | 0.2224 | 0.20171 |
| ENSG00000172269 | 11 | 118967213 | 118979041 | 34 | 7 | 0.064508 | 0.47428 | DPAGT1 | 0.47428 | 0.48247 | 0.31643 |
| ENSG00000178397 | 7 | 6369040 | 6388612 | 108 | 23 | 0.064429 | 0.47431 | FAM220A | 0.47431 | 0.51208 | 0.62516 |
| ENSG00000213366 | 1 | 110210644 | 110252171 | 173 | 32 | 0.064319 | 0.47436 | GSTM2 | 0.47436 | 0.62207 | 0.67939 |
| ENSG00000160161 | 19 | 19649057 | 19657468 | 19 | 3 | 0.064256 | 0.47438 | CILP2 | 0.47438 | 0.49038 | 0.51698 |
| ENSG00000073050 | 19 | 44047192 | 44084625 | 138 | 23 | 0.064234 | 0.47439 | XRCC1 | 0.47439 | 0.562 | 0.47941 |
| ENSG00000136908 | 9 | 130697378 | 130700763 | 7 | 1 | 0.064121 | 0.47444 | DPM2 | 0.47444 | 0.54002 | 0.57137 |
| ENSG00000185652 | 12 | 5541278 | 5630702 | 275 | 40 | 0.064097 | 0.47445 | NTF3 | 0.47445 | 0.13 | 0.15388 |
| ENSG00000127920 | 7 | 93551011 | 93557922 | 28 | 12 | 0.063994 | 0.47449 | GNG11 | 0.47449 | 0.75575 | 0.79386 |
| ENSG00000179909 | 19 | 58208735 | 58220579 | 37 | 8 | 0.063902 | 0.47452 | ZNF154 | 0.47452 | 0.2329 | 0.25481 |
| ENSG00000139291 | 12 | 72079867 | 72097836 | 29 | 7 | 0.063743 | 0.47459 | TMEM19 | 0.47459 | 0.64615 | 0.77527 |
| ENSG00000231213 | 3 | 146294342 | 146324003 | 182 | 16 | 0.063703 | 0.4746 | PLSCR5 | 0.4746 | 0.36816 | 0.42997 |
| ENSG00000196344 | 4 | 100333418 | 100356894 | 93 | 21 | 0.063694 | 0.47461 | ADH7 | 0.47461 | 0.68988 | 0.70466 |
| ENSG00000255221 | 11 | 104963196 | 104972158 | 29 | 6 | 0.063659 | 0.47462 | CARD17 | 0.47462 | 0.67373 | 0.79229 |
| ENSG00000146166 | 6 | 63985856 | 64029882 | 55 | 15 | 0.063512 | 0.47468 | LGSN | 0.47468 | 0.51354 | 0.64018 |
| ENSG00000105329 | 19 | 41807492 | 41859816 | 113 | 15 | 0.063523 | 0.47468 | TGFB1 | 0.47468 | 0.39424 | 0.31645 |
| ENSG00000212127 | 12 | 11090005 | 11324172 | 1539 | 19 | 0.063256 | 0.47478 | TAS2R14 | 0.47478 | 0.46714 | 0.46648 |
| ENSG00000145782 | 5 | 115163893 | 115177555 | 25 | 8 | 0.063186 | 0.47481 | ATG12 | 0.47481 | 0.47696 | 0.65704 |
| ENSG00000073598 | 17 | 33448598 | 33457751 | 22 | 10 | 0.063169 | 0.47482 | FNDC8 | 0.47482 | 0.35317 | 0.50007 |
| ENSG00000141639 | 18 | 48086448 | 48258194 | 457 | 59 | 0.063033 | 0.47487 | MAPK4 | 0.47487 | 0.35155 | 0.47889 |
| ENSG00000171243 | 7 | 16501106 | 16570205 | 335 | 29 | 0.063001 | 0.47488 | SOSTDC1 | 0.47488 | 0.46921 | 0.53967 |
| ENSG00000104889 | 19 | 12917394 | 12924452 | 37 | 5 | 0.062854 | 0.47494 | RNASEH2A | 0.47494 | 0.49725 | 0.33531 |
| ENSG00000196418 | 1 | 247285277 | 247335318 | 209 | 22 | 0.062811 | 0.47496 | ZNF124 | 0.47496 | 0.56053 | 0.41368 |
| ENSG00000086232 | 7 | 6061881 | 6098861 | 169 | 20 | 0.062735 | 0.47499 | EIF2AK1 | 0.47499 | 0.53721 | 0.58769 |
| ENSG00000186105 | 5 | 61874562 | 61877275 | 1 | 1 | 0.062204 | 0.4752 | LRRC70 | 0.4752 | 0.46011 | 0.6318 |
| ENSG00000244025 | 21 | 31863782 | 31864275 | 1 | 1 | 0.062204 | 0.4752 | KRTAP19-3 | 0.4752 | 0.4397 | 0.50989 |
| ENSG00000125780 | 20 | 2276647 | 2321724 | 172 | 25 | 0.062148 | 0.47522 | TGM3 | 0.47522 | 0.38074 | 0.66607 |
| ENSG00000165300 | 13 | 88324870 | 88331871 | 8 | 3 | 0.061954 | 0.4753 | SLITRK5 | 0.4753 | 0.31324 | 0.3437 |
| ENSG00000161040 | 7 | 102453308 | 102715286 | 487 | 29 | 0.061909 | 0.47532 | FBXL13 | 0.47532 | 0.49009 | 0.45584 |
| ENSG00000244291 | 7 | 156431057 | 156433348 | 6 | 3 | 0.060819 | 0.47575 | C7orf13 | 0.47575 | 0.044923 | 0.033147 |
| ENSG00000186973 | 1 | 43610824 | 43622067 | 51 | 9 | 0.060792 | 0.47576 | FAM183A | 0.47576 | 0.56208 | 0.55715 |
| ENSG00000132000 | 19 | 14042000 | 14064204 | 40 | 13 | 0.060801 | 0.47576 | PODNL1 | 0.47576 | 0.66073 | 0.75962 |
| ENSG00000197851 | 7 | 96110938 | 96132835 | 52 | 11 | 0.060696 | 0.4758 | C7orf76 | 0.4758 | 0.42616 | 0.47583 |
| ENSG00000197766 | 19 | 859453 | 863453 | 16 | 4 | 0.060687 | 0.4758 | CFD | 0.4758 | 0.41201 | 0.61454 |
| ENSG00000205754 | 12 | 20968673 | 21245679 | 1534 | 27 | 0.060594 | 0.47584 | SLCO1B7 | 0.47584 | 0.49684 | 0.49743 |
| ENSG00000093217 | 3 | 38388251 | 38462839 | 310 | 21 | 0.060534 | 0.47587 | XYLB | 0.47587 | 0.50241 | 0.49171 |
| ENSG00000204928 | 5 | 145239296 | 145252531 | 34 | 11 | 0.060514 | 0.47587 | GRXCR2 | 0.47587 | 0.096848 | 0.12433 |
| ENSG00000112655 | 6 | 43044006 | 43129457 | 240 | 36 | 0.06024 | 0.47598 | PTK7 | 0.47598 | 0.58702 | 0.61241 |
| ENSG00000108296 | 17 | 36956687 | 36981734 | 87 | 14 | 0.060209 | 0.47599 | CWC25 | 0.47599 | 0.647 | 0.706 |
| ENSG00000131238 | 1 | 40538379 | 40563375 | 106 | 12 | 0.060181 | 0.47601 | PPT1 | 0.47601 | 0.56928 | 0.55777 |
| ENSG00000111218 | 12 | 3490515 | 3703139 | 608 | 88 | 0.060157 | 0.47602 | PRMT8 | 0.47602 | 0.53039 | 0.5219 |
| ENSG00000070831 | 1 | 22379120 | 22419437 | 127 | 17 | 0.059749 | 0.47618 | CDC42 | 0.47618 | 0.55397 | 0.54988 |
| ENSG00000060709 | 12 | 130880682 | 131200826 | 1660 | 163 | 0.059152 | 0.47642 | RIMBP2 | 0.47642 | 0.49678 | 0.36056 |
| ENSG00000178217 | 10 | 82297658 | 82406316 | 496 | 48 | 0.059123 | 0.47643 | SH2D4B | 0.47643 | 0.46693 | 0.52551 |
| ENSG00000169635 | 22 | 21771693 | 21805752 | 13 | 4 | 0.058947 | 0.4765 | HIC2 | 0.4765 | 0.55205 | 0.74429 |
| ENSG00000115361 | 2 | 211052663 | 211090215 | 45 | 8 | 0.058826 | 0.47655 | ACADL | 0.47655 | 0.54715 | 0.63827 |
| ENSG00000138642 | 4 | 89299891 | 89364263 | 110 | 34 | 0.058503 | 0.47667 | HERC6 | 0.47667 | 0.66512 | 0.86277 |
| ENSG00000186191 | 20 | 31667450 | 31699557 | 158 | 18 | 0.058405 | 0.47671 | BPIFB4 | 0.47671 | 0.60079 | 0.69962 |
| ENSG00000166068 | 15 | 38544527 | 38649450 | 380 | 9 | 0.058228 | 0.47678 | SPRED1 | 0.47678 | 0.4705 | 0.42768 |
| ENSG00000101204 | 20 | 61975420 | 62009753 | 235 | 42 | 0.058219 | 0.47679 | CHRNA4 | 0.47679 | 0.26761 | 0.093838 |
| ENSG00000163606 | 3 | 112640056 | 112693969 | 265 | 22 | 0.058182 | 0.4768 | CD200R1 | 0.4768 | 0.24091 | 0.27951 |
| ENSG00000187010 | 1 | 25598884 | 25656936 | 76 | 14 | 0.05812 | 0.47683 | RHD | 0.47683 | 0.42755 | 0.39401 |
| ENSG00000197776 | 14 | 50159823 | 50219870 | 88 | 18 | 0.058004 | 0.47687 | KLHDC1 | 0.47687 | 0.56209 | 0.64853 |
| ENSG00000135847 | 1 | 180244515 | 180472089 | 698 | 23 | 0.057957 | 0.47689 | ACBD6 | 0.47689 | 0.48776 | 0.47629 |
| ENSG00000074211 | 4 | 6322305 | 6565327 | 1101 | 117 | 0.057777 | 0.47696 | PPP2R2C | 0.47696 | 0.48806 | 0.56546 |
| ENSG00000269781 | 7 | 6522922 | 6524873 | 5 | 3 | 0.057508 | 0.47707 | FLJ20306 | 0.47707 | 0.51998 | 0.47711 |
| ENSG00000118363 | 11 | 74660292 | 74690076 | 69 | 12 | 0.057436 | 0.4771 | SPCS2 | 0.4771 | 0.40042 | 0.38464 |
| ENSG00000188177 | 2 | 113033171 | 113097640 | 91 | 17 | 0.057394 | 0.47712 | ZC3H6 | 0.47712 | 0.42451 | 0.45925 |
| ENSG00000221843 | 2 | 27799389 | 27805588 | 12 | 4 | 0.057222 | 0.47718 | C2orf16 | 0.47718 | 0.45299 | 0.47916 |
| ENSG00000143382 | 1 | 150521884 | 150533413 | 37 | 13 | 0.057128 | 0.47722 | ADAMTSL4 | 0.47722 | 0.68405 | 0.72823 |
| ENSG00000180376 | 3 | 56591189 | 56655846 | 310 | 22 | 0.057134 | 0.47722 | CCDC66 | 0.47722 | 0.49402 | 0.53451 |
| ENSG00000176842 | 16 | 54964774 | 54968397 | 7 | 2 | 0.057129 | 0.47722 | IRX5 | 0.47722 | 0.37751 | 0.38011 |
| ENSG00000144406 | 2 | 210636717 | 210864024 | 497 | 31 | 0.057116 | 0.47723 | UNC80 | 0.47723 | 0.5353 | 0.54551 |
| ENSG00000125247 | 13 | 101256181 | 101327347 | 246 | 19 | 0.057072 | 0.47724 | TMTC4 | 0.47724 | 0.44288 | 0.41764 |
| ENSG00000125378 | 14 | 54416454 | 54425479 | 32 | 9 | 0.056995 | 0.47727 | BMP4 | 0.47727 | 0.2251 | 0.2012 |
| ENSG00000114904 | 3 | 52744800 | 52804965 | 156 | 11 | 0.056818 | 0.47734 | NEK4 | 0.47734 | 0.45456 | 0.54583 |
| ENSG00000183346 | 10 | 63422719 | 63526524 | 444 | 27 | 0.056775 | 0.47736 | C10orf107 | 0.47736 | 0.47593 | 0.41981 |
| ENSG00000008197 | 6 | 50681541 | 50740701 | 111 | 11 | 0.056667 | 0.47741 | TFAP2D | 0.47741 | 0.50233 | 0.50333 |
| ENSG00000055211 | 6 | 149887430 | 149912884 | 34 | 7 | 0.056437 | 0.4775 | GINM1 | 0.4775 | 0.45475 | 0.53583 |
| ENSG00000204257 | 6 | 32916390 | 32936871 | 119 | 24 | 0.056386 | 0.47752 | HLA-DMA | 0.47752 | 0.5931 | 0.62541 |

| | | | | | | | | | | | |
|---|---|---|---|---|---|---|---|---|---|---|---|
| ENSG00000113658 | 5 | 135468534 | 135524435 | 161 | 6 | 0.056336 | 0.47754 | SMAD5 | 0.47754 | 0.47657 | 0.50169 |
| ENSG00000261594 | 11 | 74951950 | 74954742 | 6 | 2 | 0.056259 | 0.47757 | TPBGL | 0.47757 | 0.31102 | 0.24617 |
| ENSG00000143977 | 2 | 70508494 | 70520903 | 25 | 3 | 0.055894 | 0.47771 | SNRPG | 0.47771 | 0.52472 | 0.55259 |
| ENSG00000112337 | 6 | 25912982 | 25930946 | 70 | 10 | 0.055798 | 0.47775 | SLC17A2 | 0.47775 | 0.48285 | 0.48864 |
| ENSG00000205208 | 4 | 159587831 | 159593407 | 5 | 3 | 0.055705 | 0.47779 | C4orf46 | 0.47779 | 0.50266 | 0.74155 |
| ENSG00000173805 | 17 | 39873994 | 39890896 | 104 | 11 | 0.055376 | 0.47792 | HAP1 | 0.47792 | 0.59895 | 0.37074 |
| ENSG00000125818 | 20 | 1093906 | 1160596 | 143 | 30 | 0.055298 | 0.47795 | PSMF1 | 0.47795 | 0.5484 | 0.41366 |
| ENSG00000175202 | 15 | 72968124 | 72978490 | 23 | 8 | 0.055133 | 0.47802 | HIGD2B | 0.47802 | 0.70417 | 0.76331 |
| ENSG00000221953 | 1 | 247273462 | 247275719 | 18 | 6 | 0.054951 | 0.47809 | C1orf229 | 0.47809 | 0.66801 | 0.58443 |
| ENSG00000204923 | 2 | 68686551 | 68694390 | 9 | 2 | 0.054876 | 0.47812 | FBXO48 | 0.47812 | 0.32827 | 0.27763 |
| ENSG00000165188 | 9 | 116059373 | 116065656 | 21 | 5 | 0.054854 | 0.47813 | RNF183 | 0.47813 | 0.51614 | 0.44449 |
| ENSG00000170128 | 1 | 200842083 | 200843306 | 2 | 1 | 0.054826 | 0.47814 | GPR25 | 0.47814 | 0.9289 | 0.96439 |
| ENSG00000162517 | 1 | 32095463 | 32110497 | 26 | 4 | 0.054588 | 0.47823 | PEF1 | 0.47823 | 0.58288 | 0.57758 |
| ENSG00000136144 | 13 | 50106082 | 50159719 | 339 | 17 | 0.054597 | 0.47823 | RCBTB1 | 0.47823 | 0.52027 | 0.47088 |
| ENSG00000150967 | 12 | 123405498 | 123466196 | 71 | 18 | 0.054533 | 0.47826 | ABCB9 | 0.47826 | 0.5127 | 0.40949 |
| ENSG00000099338 | 19 | 38826415 | 38861589 | 120 | 17 | 0.054472 | 0.47828 | CATSPERG | 0.47828 | 0.50057 | 0.49063 |
| ENSG00000105677 | 19 | 36036497 | 36038428 | 5 | 3 | 0.054404 | 0.47831 | TMEM147 | 0.47831 | 0.36524 | 0.26859 |
| ENSG00000016864 | 3 | 52728505 | 52740048 | 19 | 5 | 0.054066 | 0.47844 | GLT8D1 | 0.47844 | 0.56194 | 0.5305 |
| ENSG00000166788 | 11 | 18091482 | 18127638 | 146 | 10 | 0.053907 | 0.4785 | SAAL1 | 0.4785 | 0.45673 | 0.42823 |
| ENSG00000186666 | 12 | 50231573 | 50236912 | 12 | 3 | 0.053713 | 0.47858 | BCDIN3D | 0.47858 | 0.27931 | 0.29772 |
| ENSG00000101746 | 18 | 31431064 | 31804916 | 899 | 49 | 0.053687 | 0.47859 | NOL4 | 0.47859 | 0.49908 | 0.55969 |
| ENSG00000034971 | 1 | 171604557 | 171621823 | 89 | 13 | 0.053609 | 0.47862 | MYOC | 0.47862 | 0.43418 | 0.5058 |
| ENSG00000152253 | 2 | 169690642 | 169769881 | 278 | 24 | 0.053424 | 0.4787 | SPC25 | 0.4787 | 0.50397 | 0.49397 |
| ENSG00000087206 | 5 | 176332006 | 176449634 | 327 | 8 | 0.052832 | 0.47893 | UIMC1 | 0.47893 | 0.47911 | 0.487 |
| ENSG00000168263 | 9 | 2717502 | 2730037 | 35 | 9 | 0.052831 | 0.47893 | KCNV2 | 0.47893 | 0.66532 | 0.77106 |
| ENSG00000105953 | 7 | 44646171 | 44748665 | 282 | 25 | 0.052767 | 0.47896 | OGDH | 0.47896 | 0.4729 | 0.43067 |
| ENSG00000257446 | 19 | 12154620 | 12167127 | 19 | 5 | 0.052759 | 0.47896 | ZNF878 | 0.47896 | 0.59818 | 0.51142 |
| ENSG00000111445 | 12 | 118451393 | 118470935 | 59 | 15 | 0.052741 | 0.47897 | RFC5 | 0.47897 | 0.75333 | 0.83377 |
| ENSG00000134551 | 12 | 11081835 | 11084940 | 7 | 2 | 0.052522 | 0.47906 | PRH2 | 0.47906 | 0.7904 | 0.86658 |
| ENSG00000116586 | 1 | 156024543 | 156028301 | 8 | 4 | 0.052214 | 0.47918 | LAMTOR2 | 0.47918 | 0.6217 | 0.85466 |
| ENSG00000106013 | 7 | 117854727 | 117882785 | 105 | 12 | 0.05218 | 0.47919 | ANKRD7 | 0.47919 | 0.42633 | 0.47419 |
| ENSG00000184408 | 7 | 119913722 | 120390385 | 1088 | 45 | 0.051904 | 0.4793 | KCND2 | 0.4793 | 0.46657 | 0.46963 |
| ENSG00000121552 | 3 | 122044091 | 122060819 | 68 | 13 | 0.051813 | 0.47934 | CSTA | 0.47934 | 0.5852 | 0.55664 |
| ENSG00000120800 | 12 | 101673887 | 101780394 | 330 | 26 | 0.051447 | 0.47948 | UTP20 | 0.47948 | 0.50777 | 0.59606 |
| ENSG00000163637 | 3 | 64079543 | 64431152 | 1283 | 98 | 0.051339 | 0.47953 | PRICKLE2 | 0.47953 | 0.51754 | 0.53913 |
| ENSG00000133069 | 1 | 205197304 | 205242471 | 150 | 17 | 0.0513 | 0.47954 | TMCC2 | 0.47954 | 0.5497 | 0.52189 |
| ENSG00000012779 | 10 | 45869661 | 45941561 | 251 | 28 | 0.051302 | 0.47954 | ALOX5 | 0.47954 | 0.49423 | 0.46346 |
| ENSG00000198393 | 12 | 133562951 | 133589154 | 1 | 1 | 0.051158 | 0.4796 | ZNF26 | 0.4796 | 0.34482 | 0.63249 |
| ENSG00000149489 | 11 | 62379194 | 62382592 | 8 | 2 | 0.051144 | 0.47961 | ROM1 | 0.47961 | 0.59581 | 0.4566 |
| ENSG00000112763 | 6 | 26458150 | 26476849 | 70 | 17 | 0.05094 | 0.47969 | BTN2A1 | 0.47969 | 0.47777 | 0.58903 |
| ENSG00000103121 | 16 | 81009698 | 81053875 | 147 | 16 | 0.05072 | 0.47977 | CMC2 | 0.47977 | 0.64898 | 0.75381 |
| ENSG00000136982 | 8 | 120846216 | 120868250 | 106 | 16 | 0.050645 | 0.4798 | DSCC1 | 0.4798 | 0.49197 | 0.37884 |
| ENSG00000206503 | 6 | 29909037 | 29913661 | 303 | 26 | 0.050609 | 0.47982 | HLA-A | 0.47982 | 0.62782 | 0.64172 |
| ENSG00000182544 | 12 | 53645035 | 53648189 | 10 | 3 | 0.050487 | 0.47987 | MFSD5 | 0.47987 | 0.55849 | 0.48168 |
| ENSG00000147642 | 8 | 110586207 | 110704020 | 340 | 23 | 0.050453 | 0.47988 | SYBU | 0.47988 | 0.37089 | 0.25196 |
| ENSG00000122877 | 10 | 64571756 | 64679660 | 337 | 27 | 0.050442 | 0.47989 | EGR2 | 0.47989 | 0.55872 | 0.61243 |
| ENSG00000154814 | 3 | 16306706 | 16391806 | 347 | 24 | 0.049981 | 0.48007 | OXNAD1 | 0.48007 | 0.46214 | 0.4852 |
| ENSG00000006468 | 7 | 13930853 | 14031050 | 455 | 53 | 0.049909 | 0.4801 | ETV1 | 0.4801 | 0.42162 | 0.39948 |
| ENSG00000054793 | 20 | 50213053 | 50385173 | 890 | 34 | 0.049818 | 0.48013 | ATP9A | 0.48013 | 0.45716 | 0.36573 |
| ENSG00000132964 | 13 | 26828276 | 26979375 | 390 | 19 | 0.049779 | 0.48015 | CDK8 | 0.48015 | 0.38143 | 0.34281 |
| ENSG00000125520 | 20 | 62371214 | 62374858 | 7 | 2 | 0.049468 | 0.48027 | SLC2A4RG | 0.48027 | 0.24837 | 0.24958 |
| ENSG00000116014 | 19 | 917287 | 921015 | 7 | 3 | 0.049425 | 0.48029 | KISS1R | 0.48029 | 0.7693 | 0.9055 |
| ENSG00000205221 | 2 | 36923833 | 37041935 | 508 | 63 | 0.049397 | 0.4803 | VIT | 0.4803 | 0.36253 | 0.34768 |
| ENSG00000215045 | 7 | 6537093 | 6591067 | 163 | 34 | 0.049368 | 0.48031 | GRID2IP | 0.48031 | 0.49062 | 0.51127 |
| ENSG00000164318 | 5 | 38258511 | 38465123 | 676 | 62 | 0.049337 | 0.48033 | EGFLAM | 0.48033 | 0.49066 | 0.5321 |
| ENSG00000064225 | 3 | 98451080 | 98540045 | 222 | 33 | 0.048782 | 0.48055 | ST3GAL6 | 0.48055 | 0.4897 | 0.52227 |
| ENSG00000016602 | 1 | 87012761 | 87046437 | 93 | 18 | 0.04876 | 0.48056 | CLCA4 | 0.48056 | 0.52921 | 0.61567 |
| ENSG00000162755 | 1 | 161068151 | 161070136 | 5 | 3 | 0.048349 | 0.48072 | KLHDC9 | 0.48072 | 0.56514 | 0.77853 |
| ENSG00000161544 | 17 | 74523438 | 74547257 | 58 | 9 | 0.048324 | 0.48073 | CYGB | 0.48073 | 0.26235 | 0.39502 |
| ENSG00000136205 | 7 | 47314752 | 47622156 | 954 | 92 | 0.048097 | 0.48082 | TNS3 | 0.48082 | 0.51868 | 0.60192 |
| ENSG00000161542 | 17 | 74305567 | 74380602 | 316 | 35 | 0.048093 | 0.48082 | PRPSAP1 | 0.48082 | 0.40776 | 0.51244 |
| ENSG00000146411 | 6 | 134309835 | 134373774 | 169 | 28 | 0.048068 | 0.48083 | SLC2A12 | 0.48083 | 0.62533 | 0.78002 |
| ENSG00000185361 | 19 | 4639530 | 4655580 | 62 | 14 | 0.047514 | 0.48105 | TNFAIP8L1 | 0.48105 | 0.6003 | 0.73191 |
| ENSG00000154845 | 18 | 9546789 | 9615238 | 238 | 27 | 0.04747 | 0.48107 | PPP4R1 | 0.48107 | 0.33708 | 0.3562 |
| ENSG00000102882 | 16 | 30125426 | 30134827 | 15 | 6 | 0.047367 | 0.48111 | MAPK3 | 0.48111 | 0.73904 | 0.78289 |
| ENSG00000257046 | 12 | 20968608 | 21243179 | 1517 | 27 | 0.04732 | 0.48113 | LST3 | 0.48113 | 0.49617 | 0.50037 |
| ENSG00000171199 | 4 | 71263599 | 71275911 | 40 | 9 | 0.046999 | 0.48126 | PROL1 | 0.48126 | 0.29422 | 0.47135 |
| ENSG00000148737 | 10 | 114710009 | 114927437 | 583 | 70 | 0.046883 | 0.4813 | TCF7L2 | 0.4813 | 0.322 | 0.23931 |
| ENSG00000183791 | 18 | 44554573 | 44556449 | 1 | 1 | 0.046891 | 0.4813 | TCEB3C | 0.4813 | 0.15609 | 0.093256 |
| ENSG00000092009 | 14 | 24974559 | 24977471 | 9 | 3 | 0.046688 | 0.48138 | CMA1 | 0.48138 | 0.78045 | 0.86778 |
| ENSG00000151655 | 10 | 7745232 | 7791483 | 218 | 33 | 0.046549 | 0.48144 | ITIH2 | 0.48144 | 0.39995 | 0.48564 |
| ENSG00000162521 | 1 | 33116743 | 33151812 | 115 | 6 | 0.046486 | 0.48146 | RBBP4 | 0.48146 | 0.47397 | 0.49725 |
| ENSG00000170144 | 2 | 178077291 | 178088686 | 13 | 3 | 0.046434 | 0.48148 | HNRNPA3 | 0.48148 | 0.46452 | 0.50958 |

| | | | | | | | | | | | |
|---|---|---|---|---|---|---|---|---|---|---|---|
| ENSG00000183856 | 1 | 156495197 | 156542396 | 108 | 14 | 0.046232 | 0.48156 | IQGAP3 | 0.48156 | 0.5376 | 0.50654 |
| ENSG00000100211 | 22 | 39052641 | 39069859 | 77 | 6 | 0.046215 | 0.48157 | CBY1 | 0.48157 | 0.44192 | 0.42033 |
| ENSG00000254959 | 7 | 30791753 | 30931696 | 583 | 36 | 0.046188 | 0.48158 | INMT-FAM188B | 0.48158 | 0.42901 | 0.3801 |
| ENSG00000144559 | 3 | 11831916 | 11888393 | 133 | 18 | 0.046161 | 0.48159 | TAMM41 | 0.48159 | 0.50837 | 0.73076 |
| ENSG00000157350 | 16 | 70413338 | 70473140 | 78 | 26 | 0.046136 | 0.4816 | ST3GAL2 | 0.4816 | 0.49031 | 0.51115 |
| ENSG00000139835 | 13 | 113978505 | 114018446 | 166 | 28 | 0.045957 | 0.48167 | GRTP1 | 0.48167 | 0.64609 | 0.66929 |
| ENSG00000171916 | 17 | 18380051 | 18398259 | 15 | 7 | 0.045898 | 0.4817 | LGALS9C | 0.4817 | 0.61847 | 0.5381 |
| ENSG00000268155 | 9 | 5077163 | 5084580 | 23 | 5 | 0.045856 | 0.48171 | AL161450.1 | 0.48171 | 0.3842 | 0.4343 |
| ENSG00000188906 | 12 | 40590546 | 40763087 | 676 | 37 | 0.045826 | 0.48172 | LRRK2 | 0.48172 | 0.49085 | 0.51625 |
| ENSG00000125656 | 19 | 6361463 | 6368919 | 34 | 5 | 0.045801 | 0.48173 | CLPP | 0.48173 | 0.31414 | 0.29458 |
| ENSG00000268083 | 19 | 39308111 | 39330818 | 81 | 14 | 0.04572 | 0.48177 | AC104534.3 | 0.48177 | 0.43622 | 0.71019 |
| ENSG00000177463 | 3 | 14989091 | 15095107 | 217 | 28 | 0.045632 | 0.4818 | NR2C2 | 0.4818 | 0.47471 | 0.39851 |
| ENSG00000171806 | 1 | 169761670 | 169764107 | 8 | 4 | 0.045596 | 0.48182 | METTL18 | 0.48182 | 0.56663 | 0.5005 |
| ENSG00000178695 | 13 | 77454312 | 77460540 | 25 | 12 | 0.045519 | 0.48185 | KCTD12 | 0.48185 | 0.66845 | 0.74783 |
| ENSG00000125686 | 17 | 37560538 | 37607539 | 101 | 10 | 0.045406 | 0.48189 | MED1 | 0.48189 | 0.5218 | 0.53757 |
| ENSG00000171812 | 1 | 36560837 | 36590821 | 82 | 13 | 0.045299 | 0.48193 | COL8A2 | 0.48193 | 0.6019 | 0.71652 |
| ENSG00000257411 | 12 | 56495115 | 56503073 | 12 | 5 | 0.045228 | 0.48196 | RP11-603J24.9 | 0.48196 | 0.65135 | 0.82926 |
| ENSG00000135090 | 12 | 118587606 | 118810750 | 548 | 24 | 0.045084 | 0.48202 | TAOK3 | 0.48202 | 0.48184 | 0.51069 |
| ENSG00000150316 | 11 | 94695787 | 94706776 | 37 | 10 | 0.044756 | 0.48215 | CWC15 | 0.48215 | 0.5849 | 0.65916 |
| ENSG00000164117 | 4 | 175157809 | 175205415 | 117 | 23 | 0.044743 | 0.48216 | FBXO8 | 0.48216 | 0.47237 | 0.4685 |
| ENSG00000197465 | 4 | 144792020 | 144826716 | 113 | 14 | 0.044669 | 0.48219 | GYPE | 0.48219 | 0.49549 | 0.45116 |
| ENSG00000204613 | 6 | 30119722 | 30128711 | 75 | 9 | 0.044618 | 0.48221 | TRIM10 | 0.48221 | 0.65161 | 0.66163 |
| ENSG00000114854 | 3 | 52485118 | 52488086 | 2 | 1 | 0.044298 | 0.48233 | TNNC1 | 0.48233 | 0.25412 | 0.39359 |
| ENSG00000117906 | 15 | 77223960 | 77242601 | 38 | 14 | 0.044261 | 0.48235 | RCN2 | 0.48235 | 0.64694 | 0.62182 |
| ENSG00000077063 | 7 | 117350705 | 117514193 | 406 | 44 | 0.044151 | 0.48239 | CTTNBP2 | 0.48239 | 0.52455 | 0.6129 |
| ENSG00000167258 | 17 | 37617764 | 37721160 | 249 | 15 | 0.043935 | 0.48248 | CDK12 | 0.48248 | 0.48801 | 0.50257 |
| ENSG00000164197 | 5 | 63461671 | 63668696 | 359 | 19 | 0.043898 | 0.48249 | RNF180 | 0.48249 | 0.49711 | 0.51792 |
| ENSG00000160298 | 21 | 47720095 | 47743789 | 88 | 18 | 0.043742 | 0.48256 | C21orf58 | 0.48256 | 0.55932 | 0.67743 |
| ENSG00000153317 | 8 | 131064353 | 131455906 | 1491 | 40 | 0.04362 | 0.4826 | ASAP1 | 0.4826 | 0.49426 | 0.46677 |
| ENSG00000149646 | 20 | 34556512 | 34618622 | 102 | 10 | 0.043614 | 0.48261 | CNBD2 | 0.48261 | 0.43317 | 0.41007 |
| ENSG00000235631 | 7 | 122341718 | 122343021 | 2 | 1 | 0.043504 | 0.48265 | RNF148 | 0.48265 | 0.44706 | 0.56922 |
| ENSG00000242852 | 19 | 12571998 | 12624668 | 225 | 12 | 0.043463 | 0.48267 | ZNF709 | 0.48267 | 0.30367 | 0.23926 |
| ENSG00000083544 | 13 | 60970591 | 61148012 | 295 | 23 | 0.043386 | 0.4827 | TDRD3 | 0.4827 | 0.51375 | 0.57799 |
| ENSG00000128805 | 10 | 49654077 | 49864310 | 855 | 68 | 0.043113 | 0.48281 | ARHGAP22 | 0.48281 | 0.46804 | 0.56396 |
| ENSG00000185187 | 11 | 405716 | 417455 | 37 | 9 | 0.04296 | 0.48287 | SIGIRR | 0.48287 | 0.38295 | 0.36898 |
| ENSG00000135506 | 12 | 58087738 | 58115340 | 61 | 10 | 0.042746 | 0.48295 | OS9 | 0.48295 | 0.52809 | 0.46992 |
| ENSG00000168000 | 11 | 62457747 | 62477317 | 53 | 9 | 0.042622 | 0.483 | BSCL2 | 0.483 | 0.37361 | 0.50497 |
| ENSG00000205439 | 21 | 46077849 | 46078258 | 2 | 1 | 0.042588 | 0.48302 | KRTAP12-3 | 0.48302 | 0.70645 | 0.86284 |
| ENSG00000186335 | 5 | 150694539 | 150727151 | 139 | 15 | 0.042464 | 0.48306 | SLC36A2 | 0.48306 | 0.72869 | 0.87879 |
| ENSG00000147889 | 9 | 21967751 | 21995300 | 52 | 12 | 0.042441 | 0.48307 | CDKN2A | 0.48307 | 0.55621 | 0.70339 |
| ENSG00000114784 | 3 | 40351175 | 40353915 | 13 | 3 | 0.042419 | 0.48308 | EIF1B | 0.48308 | 0.3912 | 0.42029 |
| ENSG00000174527 | 12 | 109785708 | 109893328 | 585 | 25 | 0.042427 | 0.48308 | MYO1H | 0.48308 | 0.5013 | 0.55088 |
| ENSG00000182329 | 2 | 202937978 | 203062585 | 338 | 31 | 0.042403 | 0.48309 | AC079354.1 | 0.48309 | 0.5421 | 0.61018 |
| ENSG00000205413 | 7 | 92728829 | 92747336 | 82 | 14 | 0.042398 | 0.48309 | SAMD9 | 0.48309 | 0.56425 | 0.69239 |
| ENSG00000188171 | 19 | 20802867 | 20844402 | 320 | 14 | 0.042394 | 0.48309 | ZNF626 | 0.48309 | 0.53631 | 0.48027 |
| ENSG00000138944 | 22 | 44639547 | 44708731 | 345 | 54 | 0.042401 | 0.48309 | KIAA1644 | 0.48309 | 0.39205 | 0.61145 |
| ENSG00000149968 | 11 | 102706532 | 102714534 | 13 | 3 | 0.04234 | 0.48311 | MMP3 | 0.48311 | 0.48484 | 0.46694 |
| ENSG00000174007 | 3 | 196433148 | 196439164 | 38 | 6 | 0.042197 | 0.48317 | CEP19 | 0.48317 | 0.19105 | 0.1992 |
| ENSG00000184276 | 11 | 71544246 | 71548756 | 20 | 4 | 0.042056 | 0.48323 | DEFB108B | 0.48323 | 0.50799 | 0.59471 |
| ENSG00000166881 | 12 | 57449426 | 57481846 | 73 | 11 | 0.041867 | 0.4833 | TMEM194A | 0.4833 | 0.50024 | 0.4519 |
| ENSG00000170191 | 20 | 25593571 | 25604811 | 33 | 6 | 0.041741 | 0.48335 | NANP | 0.48335 | 0.54384 | 0.59796 |
| ENSG00000124795 | 6 | 18224099 | 18265054 | 97 | 19 | 0.041604 | 0.48341 | DEK | 0.48341 | 0.52964 | 0.57519 |
| ENSG00000163660 | 3 | 156864297 | 156878549 | 29 | 4 | 0.041367 | 0.4835 | CCNL1 | 0.4835 | 0.54772 | 0.5506 |
| ENSG00000108773 | 17 | 40265126 | 40273376 | 25 | 6 | 0.041042 | 0.48363 | KAT2A | 0.48363 | 0.4033 | 0.3949 |
| ENSG00000215305 | 20 | 2821349 | 2847378 | 100 | 10 | 0.040922 | 0.48368 | VPS16 | 0.48368 | 0.42704 | 0.46525 |
| ENSG00000170345 | 14 | 75745477 | 75748933 | 9 | 3 | 0.040648 | 0.48379 | FOS | 0.48379 | 0.21008 | 0.23237 |
| ENSG00000109736 | 4 | 2932288 | 2936586 | 15 | 9 | 0.040629 | 0.4838 | MFSD10 | 0.4838 | 0.61171 | 0.68298 |
| ENSG00000172640 | 12 | 48596081 | 48597170 | 12 | 2 | 0.040424 | 0.48388 | OR10AD1 | 0.48388 | 0.44404 | 0.46542 |
| ENSG00000167670 | 19 | 4402659 | 4445015 | 123 | 14 | 0.040353 | 0.48391 | CHAF1A | 0.48391 | 0.4669 | 0.50999 |
| ENSG00000132405 | 4 | 6910969 | 7034845 | 555 | 24 | 0.040208 | 0.48396 | TBC1D14 | 0.48396 | 0.47987 | 0.55104 |
| ENSG00000253379 | 8 | 72315675 | 72504260 | 453 | 47 | 0.040113 | 0.484 | RP11-1102P16.1 | 0.484 | 0.43489 | 0.49929 |
| ENSG00000047621 | 12 | 4596894 | 4647674 | 56 | 5 | 0.040087 | 0.48401 | C12orf4 | 0.48401 | 0.63035 | 0.74999 |
| ENSG00000081277 | 1 | 201252580 | 201302121 | 228 | 21 | 0.039699 | 0.48417 | PKP1 | 0.48417 | 0.54762 | 0.58737 |
| ENSG00000023287 | 8 | 53535016 | 53658403 | 185 | 30 | 0.039559 | 0.48422 | RB1CC1 | 0.48422 | 0.5001 | 0.48942 |
| ENSG00000268046 | 18 | 8336437 | 8337038 | 4 | 2 | 0.039552 | 0.48422 | AP001094.1 | 0.48422 | 0.61814 | 0.71272 |
| ENSG00000197980 | 3 | 156543270 | 156763918 | 697 | 33 | 0.03955 | 0.48423 | LEKR1 | 0.48423 | 0.47866 | 0.44932 |
| ENSG00000188624 | 19 | 46623328 | 46627931 | 15 | 4 | 0.039472 | 0.48426 | IGFL3 | 0.48426 | 0.57982 | 0.61294 |
| ENSG00000182218 | 14 | 100111447 | 100146906 | 150 | 18 | 0.039437 | 0.48427 | HHIPL1 | 0.48427 | 0.4728 | 0.22841 |
| ENSG00000134970 | 5 | 114949205 | 114968689 | 38 | 7 | 0.039234 | 0.48435 | TMED7 | 0.48435 | 0.36722 | 0.42962 |
| ENSG00000170270 | 14 | 93669239 | 93673439 | 3 | 1 | 0.039041 | 0.48443 | C14orf142 | 0.48443 | 0.6795 | 0.69038 |
| ENSG00000130202 | 19 | 45349432 | 45392485 | 213 | 30 | 0.03902 | 0.48444 | PVRL2 | 0.48444 | 0.39417 | 0.37749 |
| ENSG00000170779 | 14 | 105475910 | 105487485 | 70 | 6 | 0.038939 | 0.48447 | CDCA4 | 0.48447 | 0.559 | 0.60735 |
| ENSG00000063978 | 4 | 2463947 | 2627047 | 476 | 41 | 0.038843 | 0.48451 | RNF4 | 0.48451 | 0.48073 | 0.51559 |

| | | | | | | | | | | | |
|---|---|---|---|---|---|---|---|---|---|---|---|
| ENSG00000188958 | 3 | 190984957 | 191048325 | 227 | 24 | 0.038738 | 0.48455 | UTS2B | 0.48455 | 0.35241 | 0.50271 |
| ENSG00000204960 | 7 | 155149680 | 155160629 | 82 | 10 | 0.038747 | 0.48455 | BLACE | 0.48455 | 0.48459 | 0.39817 |
| ENSG00000135333 | 6 | 93949738 | 94129265 | 481 | 59 | 0.03871 | 0.48456 | EPHA7 | 0.48456 | 0.48157 | 0.39442 |
| ENSG00000260861 | 20 | 1520790 | 1600655 | 185 | 27 | 0.03871 | 0.48456 | RP4-576H24.4 | 0.48456 | 0.51797 | 0.57986 |
| ENSG00000170412 | 17 | 72420990 | 72447792 | 97 | 21 | 0.038641 | 0.48459 | GPRC5C | 0.48459 | 0.24587 | 0.56543 |
| ENSG00000164089 | 4 | 109663196 | 109684210 | 87 | 9 | 0.038582 | 0.48461 | ETNPPL | 0.48461 | 0.4595 | 0.31487 |
| ENSG00000205592 | 12 | 40787197 | 40964632 | 1102 | 30 | 0.038367 | 0.4847 | MUC19 | 0.4847 | 0.51485 | 0.47665 |
| ENSG00000254636 | 10 | 124214169 | 124216868 | 17 | 4 | 0.038179 | 0.48477 | ARMS2 | 0.48477 | 0.67807 | 0.69103 |
| ENSG00000175489 | 19 | 18501954 | 18508427 | 27 | 7 | 0.038179 | 0.48477 | LRRC25 | 0.48477 | 0.75673 | 0.89995 |
| ENSG00000131094 | 17 | 43037061 | 43045439 | 34 | 9 | 0.038089 | 0.48481 | C1QL1 | 0.48481 | 0.43585 | 0.60781 |
| ENSG00000130255 | 19 | 5674958 | 5691887 | 34 | 8 | 0.038086 | 0.48481 | RPL36 | 0.48481 | 0.44621 | 0.56266 |
| ENSG00000182912 | 21 | 45937099 | 45945835 | 17 | 5 | 0.038048 | 0.48482 | C21orf90 | 0.48482 | 0.43372 | 0.49247 |
| ENSG00000106624 | 7 | 44143960 | 44154161 | 29 | 8 | 0.038013 | 0.48484 | AEBP1 | 0.48484 | 0.2018 | 0.40259 |
| ENSG00000150637 | 18 | 67498394 | 67629039 | 455 | 47 | 0.037786 | 0.48493 | CD226 | 0.48493 | 0.50447 | 0.51418 |
| ENSG00000145391 | 4 | 140417095 | 140527853 | 363 | 53 | 0.037753 | 0.48494 | SETD7 | 0.48494 | 0.59717 | 0.24529 |
| ENSG00000176890 | 18 | 657604 | 673578 | 84 | 7 | 0.037335 | 0.48511 | TYMS | 0.48511 | 0.4154 | 0.097618 |
| ENSG00000138311 | 10 | 64133951 | 64431771 | 1057 | 76 | 0.0373 | 0.48512 | ZNF365 | 0.48512 | 0.5501 | 0.61028 |
| ENSG00000188817 | 3 | 63638344 | 63664696 | 132 | 15 | 0.03706 | 0.48522 | SNTN | 0.48522 | 0.42002 | 0.30862 |
| ENSG00000042304 | 2 | 228474806 | 228498036 | 209 | 21 | 0.036701 | 0.48536 | C2orf83 | 0.48536 | 0.14695 | 0.20976 |
| ENSG00000221983 | 19 | 18682540 | 18688360 | 15 | 3 | 0.035976 | 0.48565 | UBA52 | 0.48565 | 0.64222 | 0.71875 |
| ENSG00000155849 | 7 | 36893961 | 37488852 | 2108 | 129 | 0.035947 | 0.48566 | ELMO1 | 0.48566 | 0.54594 | 0.53375 |
| ENSG00000144597 | 3 | 15468862 | 15484120 | 31 | 11 | 0.035532 | 0.48583 | EAF1 | 0.48583 | 0.37395 | 0.45955 |
| ENSG00000185149 | 4 | 156129781 | 156138230 | 21 | 6 | 0.035224 | 0.48595 | NPY2R | 0.48595 | 0.77126 | 0.79325 |
| ENSG00000175879 | 2 | 176994422 | 176997423 | 6 | 3 | 0.035164 | 0.48597 | HOXD8 | 0.48597 | 0.31006 | 0.35123 |
| ENSG00000171509 | 4 | 159236463 | 159574524 | 1178 | 22 | 0.035184 | 0.48597 | RXFP1 | 0.48597 | 0.4761 | 0.46572 |
| ENSG00000137845 | 15 | 58887403 | 59042177 | 378 | 21 | 0.035172 | 0.48597 | ADAM10 | 0.48597 | 0.48836 | 0.47691 |
| ENSG00000109163 | 4 | 68605046 | 68620078 | 61 | 9 | 0.035136 | 0.48599 | GNRHR | 0.48599 | 0.65961 | 0.74016 |
| ENSG00000175779 | 15 | 38988799 | 38992239 | 22 | 5 | 0.035076 | 0.48601 | C15orf53 | 0.48601 | 0.67228 | 0.64368 |
| ENSG00000141298 | 17 | 27952956 | 28257294 | 513 | 18 | 0.034912 | 0.48607 | SSH2 | 0.48607 | 0.48255 | 0.46858 |
| ENSG00000213563 | 8 | 145751117 | 145754516 | 3 | 1 | 0.034781 | 0.48613 | C8orf82 | 0.48613 | 0.54636 | 0.63148 |
| ENSG00000101079 | 20 | 35280169 | 35374481 | 118 | 39 | 0.034605 | 0.4862 | NDRG3 | 0.4862 | 0.49235 | 0.54761 |
| ENSG00000164086 | 3 | 52082935 | 52090566 | 7 | 4 | 0.034492 | 0.48624 | DUSP7 | 0.48624 | 0.3785 | 0.24875 |
| ENSG00000166295 | 10 | 73975787 | 73995618 | 55 | 8 | 0.034502 | 0.48624 | ANAPC16 | 0.48624 | 0.37651 | 0.47516 |
| ENSG00000114857 | 3 | 42642106 | 42690227 | 133 | 11 | 0.034211 | 0.48635 | NKTR | 0.48635 | 0.5067 | 0.47748 |
| ENSG00000255482 | 11 | 102337986 | 102369532 | 129 | 18 | 0.034129 | 0.48639 | RP11-315O6.2 | 0.48639 | 0.57977 | 0.58314 |
| ENSG00000074660 | 17 | 1537152 | 1549041 | 32 | 8 | 0.034105 | 0.4864 | SCARF1 | 0.4864 | 0.5609 | 0.47846 |
| ENSG00000103037 | 16 | 58549383 | 58554431 | 20 | 5 | 0.034081 | 0.48641 | SETD6 | 0.48641 | 0.27549 | 0.34602 |
| ENSG00000170909 | 19 | 54597933 | 54606000 | 45 | 14 | 0.034021 | 0.48643 | OSCAR | 0.48643 | 0.72321 | 0.85088 |
| ENSG00000198077 | 19 | 41381344 | 41388657 | 95 | 11 | 0.033842 | 0.4865 | CYP2A7 | 0.4865 | 0.62908 | 0.62448 |
| ENSG00000102858 | 16 | 4666494 | 4740975 | 385 | 29 | 0.033765 | 0.48653 | MGRN1 | 0.48653 | 0.47368 | 0.55715 |
| ENSG00000148120 | 9 | 97488983 | 97849441 | 885 | 29 | 0.033666 | 0.48657 | C9orf3 | 0.48657 | 0.48597 | 0.49298 |
| ENSG00000110195 | 11 | 71900602 | 71907345 | 20 | 6 | 0.03356 | 0.48661 | FOLR1 | 0.48661 | 0.58068 | 0.6626 |
| ENSG00000086015 | 1 | 46252659 | 46501796 | 429 | 18 | 0.033168 | 0.48677 | MAST2 | 0.48677 | 0.4877 | 0.50836 |
| ENSG00000183675 | 10 | 48736874 | 48827966 | 23 | 6 | 0.033137 | 0.48678 | PTPN20B | 0.48678 | 0.49285 | 0.61144 |
| ENSG00000197938 | 3 | 98001732 | 98002676 | 3 | 1 | 0.032528 | 0.48703 | OR5H2 | 0.48703 | 0.8702 | 0.75629 |
| ENSG00000157214 | 7 | 89796904 | 89867451 | 284 | 16 | 0.032436 | 0.48706 | STEAP2 | 0.48706 | 0.48011 | 0.52016 |
| ENSG00000125730 | 19 | 6677715 | 6730573 | 253 | 27 | 0.032235 | 0.48714 | C3 | 0.48714 | 0.50573 | 0.26632 |
| ENSG00000117543 | 1 | 101455179 | 101491644 | 53 | 6 | 0.032208 | 0.48715 | DPH5 | 0.48715 | 0.49672 | 0.53247 |
| ENSG00000100221 | 22 | 39081548 | 39097561 | 24 | 8 | 0.032168 | 0.48717 | JOSD1 | 0.48717 | 0.38996 | 0.42217 |
| ENSG00000261086 | 15 | 83209620 | 83231124 | 54 | 13 | 0.032123 | 0.48719 | RP11-379H8.1 | 0.48719 | 0.35401 | 0.25151 |
| ENSG00000130881 | 19 | 33668509 | 33698694 | 109 | 19 | 0.032107 | 0.48719 | LRP3 | 0.48719 | 0.37999 | 0.59127 |
| ENSG00000172663 | 11 | 67231824 | 67236743 | 5 | 3 | 0.031863 | 0.48729 | TMEM134 | 0.48729 | 0.7005 | 0.61128 |
| ENSG00000196260 | 6 | 30899130 | 30899952 | 10 | 4 | 0.031735 | 0.48734 | SFTA2 | 0.48734 | 0.23684 | 0.30996 |
| ENSG00000100938 | 14 | 24701628 | 24708448 | 9 | 5 | 0.031539 | 0.48742 | GMPR2 | 0.48742 | 0.88057 | 0.92414 |
| ENSG00000065615 | 6 | 84569362 | 84677143 | 244 | 14 | 0.031429 | 0.48746 | CYB5R4 | 0.48746 | 0.51775 | 0.5372 |
| ENSG00000105245 | 19 | 41172596 | 41196877 | 88 | 17 | 0.031202 | 0.48755 | NUMBL | 0.48755 | 0.54624 | 0.3838 |
| ENSG00000148488 | 10 | 17360382 | 17496329 | 834 | 40 | 0.030753 | 0.48773 | ST8SIA6 | 0.48773 | 0.3773 | 0.39399 |
| ENSG00000258674 | 19 | 19627036 | 19646885 | 43 | 6 | 0.030694 | 0.48776 | CTC-260F20.3 | 0.48776 | 0.52513 | 0.54652 |
| ENSG00000203791 | 10 | 126436718 | 126480439 | 101 | 15 | 0.030671 | 0.48777 | METTL10 | 0.48777 | 0.46591 | 0.48863 |
| ENSG00000165863 | 10 | 118423207 | 118429775 | 16 | 5 | 0.030517 | 0.48783 | C10orf82 | 0.48783 | 0.51474 | 0.6268 |
| ENSG00000160404 | 9 | 130493803 | 130497604 | 13 | 4 | 0.0304 | 0.48787 | TOR2A | 0.48787 | 0.4274 | 0.51241 |
| ENSG00000188100 | 10 | 88780063 | 88784487 | 22 | 6 | 0.030374 | 0.48788 | FAM25A | 0.48788 | 0.7686 | 0.8012 |
| ENSG00000186919 | 17 | 74068042 | 74079747 | 14 | 6 | 0.030274 | 0.48792 | ZACN | 0.48792 | 0.73265 | 0.48617 |
| ENSG00000133247 | 19 | 55851221 | 55859488 | 36 | 10 | 0.030194 | 0.48796 | SUV420H2 | 0.48796 | 0.45549 | 0.50476 |
| ENSG00000183307 | 22 | 17597189 | 17602257 | 21 | 6 | 0.030154 | 0.48797 | CECR6 | 0.48797 | 0.43197 | 0.21511 |
| ENSG00000137764 | 15 | 67835047 | 68099461 | 635 | 29 | 0.029885 | 0.48808 | MAP2K5 | 0.48808 | 0.45288 | 0.42906 |
| ENSG00000169926 | 15 | 31619058 | 31727868 | 326 | 36 | 0.029831 | 0.4881 | KLF13 | 0.4881 | 0.54619 | 0.70782 |
| ENSG00000161558 | 19 | 48835613 | 48867494 | 132 | 16 | 0.029714 | 0.48815 | TMEM143 | 0.48815 | 0.50912 | 0.60231 |
| ENSG00000196655 | 11 | 118889142 | 118896164 | 34 | 5 | 0.029297 | 0.48831 | TRAPPC4 | 0.48831 | 0.6406 | 0.68223 |
| ENSG00000163468 | 1 | 156278759 | 156337664 | 151 | 12 | 0.028884 | 0.48848 | CCT3 | 0.48848 | 0.54493 | 0.58066 |
| ENSG00000103266 | 16 | 730224 | 732870 | 5 | 2 | 0.02887 | 0.48848 | STUB1 | 0.48848 | 0.26463 | 0.35019 |
| ENSG00000127995 | 7 | 94138531 | 94186331 | 98 | 12 | 0.028864 | 0.48849 | CASD1 | 0.48849 | 0.51492 | 0.56242 |
| ENSG00000157601 | 21 | 42792231 | 42831141 | 233 | 19 | 0.028769 | 0.48852 | MX1 | 0.48852 | 0.41695 | 0.47565 |

| | | | | | | | | | | | |
|---|---|---|---|---|---|---|---|---|---|---|---|
| ENSG00000104290 | 8 | 28351729 | 28431775 | 176 | 7 | 0.028683 | 0.48856 | FZD3 | 0.48856 | 0.57462 | 0.60218 |
| ENSG00000120332 | 1 | 175036994 | 175117202 | 379 | 29 | 0.028547 | 0.48861 | TNN | 0.48861 | 0.62808 | 0.76614 |
| ENSG00000107874 | 10 | 104183002 | 104192418 | 18 | 5 | 0.028554 | 0.48861 | CUEDC2 | 0.48861 | 0.25459 | 0.25942 |
| ENSG00000174482 | 9 | 27948076 | 28670283 | 3023 | 194 | 0.028536 | 0.48862 | LINGO2 | 0.48862 | 0.52198 | 0.54083 |
| ENSG00000139192 | 12 | 6560856 | 6575683 | 63 | 9 | 0.028419 | 0.48866 | TAPBPL | 0.48866 | 0.21871 | 0.21653 |
| ENSG00000137154 | 9 | 19375713 | 19380252 | 13 | 3 | 0.028392 | 0.48867 | RPS6 | 0.48867 | 0.53896 | 0.46249 |
| ENSG00000133275 | 19 | 1941188 | 1981337 | 194 | 18 | 0.02809 | 0.4888 | CSNK1G2 | 0.4888 | 0.52184 | 0.5042 |
| ENSG00000112983 | 5 | 137475455 | 137514675 | 69 | 13 | 0.028008 | 0.48883 | BRD8 | 0.48883 | 0.52526 | 0.52994 |
| ENSG00000172817 | 8 | 65500320 | 65711318 | 421 | 26 | 0.027833 | 0.4889 | CYP7B1 | 0.4889 | 0.4947 | 0.5049 |
| ENSG00000205363 | 15 | 74032141 | 74045088 | 32 | 12 | 0.027785 | 0.48892 | C15orf59 | 0.48892 | 0.63172 | 0.56214 |
| ENSG00000130005 | 19 | 1397091 | 1401569 | 12 | 5 | 0.027713 | 0.48895 | GAMT | 0.48895 | 0.86777 | 0.88508 |
| ENSG00000163644 | 4 | 89178772 | 89205921 | 44 | 11 | 0.027632 | 0.48898 | PPM1K | 0.48898 | 0.83275 | 0.70848 |
| ENSG00000005020 | 7 | 26706681 | 27034858 | 987 | 23 | 0.027575 | 0.489 | SKAP2 | 0.489 | 0.49608 | 0.50432 |
| ENSG00000205531 | 11 | 2965667 | 3013607 | 204 | 20 | 0.027519 | 0.48902 | NAP1L4 | 0.48902 | 0.43531 | 0.46891 |
| ENSG00000267261 | 17 | 40271692 | 40306934 | 85 | 8 | 0.027408 | 0.48907 | CTD-2132N18.3 | 0.48907 | 0.31517 | 0.30014 |
| ENSG00000122304 | 16 | 11369496 | 11370337 | 3 | 2 | 0.027301 | 0.48911 | PRM2 | 0.48911 | 0.58026 | 0.43489 |
| ENSG00000152592 | 4 | 88571459 | 88585513 | 41 | 8 | 0.027011 | 0.48923 | DMP1 | 0.48923 | 0.44272 | 0.50265 |
| ENSG00000149929 | 16 | 30003645 | 30007757 | 3 | 2 | 0.026765 | 0.48932 | HIRIP3 | 0.48932 | 0.53191 | 0.56565 |
| ENSG00000171606 | 19 | 58694396 | 58724928 | 72 | 16 | 0.026414 | 0.48946 | ZNF274 | 0.48946 | 0.30589 | 0.41257 |
| ENSG00000061492 | 5 | 137419581 | 137428054 | 23 | 7 | 0.026021 | 0.48962 | WNT8A | 0.48962 | 0.5209 | 0.57202 |
| ENSG00000155115 | 6 | 111279763 | 111289093 | 82 | 5 | 0.026033 | 0.48962 | GTF3C6 | 0.48962 | 0.54287 | 0.55763 |
| ENSG00000139921 | 14 | 51706880 | 51722759 | 53 | 12 | 0.02578 | 0.48972 | TMX1 | 0.48972 | 0.38375 | 0.43741 |
| ENSG00000221819 | 16 | 90095316 | 90096309 | 9 | 3 | 0.025738 | 0.48973 | C16orf3 | 0.48973 | 0.56593 | 0.71648 |
| ENSG00000065883 | 7 | 39989636 | 40136733 | 378 | 23 | 0.025718 | 0.48974 | CDK13 | 0.48974 | 0.4828 | 0.50705 |
| ENSG00000166167 | 10 | 103113820 | 103317078 | 522 | 16 | 0.025611 | 0.48978 | BTRC | 0.48978 | 0.50023 | 0.52107 |
| ENSG00000151773 | 13 | 44398045 | 44453827 | 264 | 20 | 0.025616 | 0.48978 | CCDC122 | 0.48978 | 0.47854 | 0.39572 |
| ENSG00000079739 | 1 | 64058947 | 64125916 | 207 | 37 | 0.025517 | 0.48982 | PGM1 | 0.48982 | 0.36793 | 0.26777 |
| ENSG00000112214 | 6 | 97010424 | 97064512 | 167 | 13 | 0.025492 | 0.48983 | FHL5 | 0.48983 | 0.47041 | 0.45198 |
| ENSG00000107560 | 10 | 119764427 | 119806114 | 92 | 15 | 0.025432 | 0.48986 | RAB11FIP2 | 0.48986 | 0.4389 | 0.42996 |
| ENSG00000115594 | 2 | 102681004 | 102796334 | 386 | 33 | 0.025364 | 0.48988 | IL1R1 | 0.48988 | 0.52465 | 0.61342 |
| ENSG00000131732 | 5 | 80597409 | 80609116 | 36 | 8 | 0.025277 | 0.48992 | ZCCHC9 | 0.48992 | 0.45616 | 0.37251 |
| ENSG00000185920 | 9 | 98205262 | 98279339 | 195 | 20 | 0.025092 | 0.48999 | PTCH1 | 0.48999 | 0.48454 | 0.47645 |
| ENSG00000136487 | 17 | 61957578 | 61959295 | 5 | 2 | 0.024902 | 0.49007 | GH2 | 0.49007 | 0.48949 | 0.63484 |
| ENSG00000260537 | 16 | 70333097 | 70400163 | 105 | 31 | 0.024816 | 0.4901 | RP11-529K1.3 | 0.4901 | 0.54167 | 0.52759 |
| ENSG00000105227 | 19 | 40899675 | 40919273 | 45 | 12 | 0.024787 | 0.49011 | PRX | 0.49011 | 0.45478 | 0.32328 |
| ENSG00000167766 | 19 | 53097313 | 53193749 | 672 | 44 | 0.024802 | 0.49011 | ZNF83 | 0.49011 | 0.66802 | 0.83605 |
| ENSG00000145979 | 6 | 13266774 | 13328815 | 353 | 19 | 0.024759 | 0.49012 | TBC1D7 | 0.49012 | 0.51138 | 0.43728 |
| ENSG00000205301 | 4 | 141363416 | 141419531 | 212 | 18 | 0.024562 | 0.4902 | RP11-542P2.1 | 0.4902 | 0.52604 | 0.55328 |
| ENSG00000134760 | 18 | 28898052 | 28936992 | 177 | 11 | 0.02457 | 0.4902 | DSG1 | 0.4902 | 0.49169 | 0.38922 |
| ENSG00000117697 | 1 | 212899495 | 212965124 | 365 | 16 | 0.02451 | 0.49022 | NSL1 | 0.49022 | 0.44769 | 0.43516 |
| ENSG00000269028 | 3 | 96335981 | 96337000 | 3 | 1 | 0.024389 | 0.49027 | MTRNR2L12 | 0.49027 | 0.44091 | 0.47872 |
| ENSG00000134470 | 10 | 5990855 | 6020150 | 158 | 29 | 0.024099 | 0.49039 | IL15RA | 0.49039 | 0.61652 | 0.53995 |
| ENSG00000111707 | 12 | 118814185 | 118855840 | 135 | 12 | 0.024087 | 0.49039 | SUDS3 | 0.49039 | 0.4621 | 0.48394 |
| ENSG00000248751 | 22 | 30681227 | 30695471 | 33 | 12 | 0.023946 | 0.49045 | RP1-130H16.18 | 0.49045 | 0.70416 | 0.69701 |
| ENSG00000157191 | 1 | 16767167 | 16786573 | 51 | 13 | 0.023643 | 0.49057 | NECAP2 | 0.49057 | 0.46234 | 0.38057 |
| ENSG00000151876 | 5 | 41925356 | 41941845 | 30 | 9 | 0.023575 | 0.4906 | FBXO4 | 0.4906 | 0.62319 | 0.77708 |
| ENSG00000186117 | 11 | 55578854 | 55579952 | 7 | 2 | 0.023457 | 0.49064 | OR5L1 | 0.49064 | 0.43532 | 0.42494 |
| ENSG00000183735 | 12 | 64845660 | 64895888 | 113 | 16 | 0.023063 | 0.4908 | TBK1 | 0.4908 | 0.42883 | 0.45381 |
| ENSG00000116017 | 19 | 925781 | 975939 | 272 | 34 | 0.023072 | 0.4908 | ARID3A | 0.4908 | 0.52058 | 0.31173 |
| ENSG00000134962 | 4 | 39408473 | 39453156 | 186 | 35 | 0.022717 | 0.49094 | KLB | 0.49094 | 0.29971 | 0.2656 |
| ENSG00000168556 | 4 | 184426147 | 184432249 | 17 | 7 | 0.02266 | 0.49096 | ING2 | 0.49096 | 0.42189 | 0.33282 |
| ENSG00000130638 | 22 | 46067678 | 46241187 | 448 | 29 | 0.022496 | 0.49103 | ATXN10 | 0.49103 | 0.46521 | 0.43672 |
| ENSG00000162576 | 1 | 1288069 | 1297157 | 35 | 6 | 0.022389 | 0.49107 | MXRA8 | 0.49107 | 0.13282 | 0.21391 |
| ENSG00000002549 | 4 | 17578815 | 17609595 | 92 | 5 | 0.022297 | 0.49111 | LAP3 | 0.49111 | 0.45411 | 0.53415 |
| ENSG00000197951 | 19 | 57106632 | 57137849 | 140 | 18 | 0.022216 | 0.49114 | ZNF71 | 0.49114 | 0.32958 | 0.43035 |
| ENSG00000122687 | 7 | 2273866 | 2281840 | 23 | 7 | 0.02211 | 0.49118 | FTSJ2 | 0.49118 | 0.36254 | 0.43051 |
| ENSG00000007402 | 3 | 50400233 | 50541675 | 307 | 20 | 0.021294 | 0.49151 | CACNA2D2 | 0.49151 | 0.48828 | 0.52896 |
| ENSG00000117632 | 1 | 26210672 | 26233482 | 37 | 7 | 0.021224 | 0.49153 | STMN1 | 0.49153 | 0.38126 | 0.41891 |
| ENSG00000203943 | 1 | 84764049 | 84855640 | 211 | 28 | 0.021115 | 0.49158 | SAMD13 | 0.49158 | 0.46408 | 0.42105 |
| ENSG00000142871 | 1 | 86046444 | 86049645 | 6 | 3 | 0.021034 | 0.49161 | CYR61 | 0.49161 | 0.63087 | 0.63033 |
| ENSG00000197712 | 4 | 38869298 | 38947360 | 362 | 25 | 0.020895 | 0.49166 | FAM114A1 | 0.49166 | 0.50879 | 0.49891 |
| ENSG00000232056 | 2 | 10987703 | 10995081 | 46 | 10 | 0.02088 | 0.49167 | AC092687.4 | 0.49167 | 0.3761 | 0.52097 |
| ENSG00000101181 | 20 | 60758085 | 60778624 | 137 | 25 | 0.020547 | 0.4918 | MTG2 | 0.4918 | 0.56897 | 0.62375 |
| ENSG00000133265 | 19 | 55773599 | 55791749 | 40 | 10 | 0.02052 | 0.49181 | HSPBP1 | 0.49181 | 0.67353 | 0.50152 |
| ENSG00000100916 | 14 | 36295524 | 36401531 | 220 | 26 | 0.020502 | 0.49182 | BRMS1L | 0.49182 | 0.47341 | 0.46686 |
| ENSG00000021461 | 7 | 99425636 | 99463718 | 101 | 10 | 0.020411 | 0.49186 | CYP3A43 | 0.49186 | 0.48835 | 0.61891 |
| ENSG00000152284 | 2 | 85360533 | 85537511 | 732 | 64 | 0.020311 | 0.4919 | TCF7L1 | 0.4919 | 0.44539 | 0.52273 |
| ENSG00000163359 | 2 | 238232646 | 238323018 | 252 | 33 | 0.020185 | 0.49195 | COL6A3 | 0.49195 | 0.46288 | 0.499 |
| ENSG00000111052 | 12 | 81186299 | 81331704 | 440 | 30 | 0.019853 | 0.49208 | LIN7A | 0.49208 | 0.52144 | 0.62973 |
| ENSG00000174796 | 4 | 76439156 | 76475683 | 68 | 12 | 0.019801 | 0.4921 | THAP6 | 0.4921 | 0.65889 | 0.6476 |
| ENSG00000272573 | 3 | 52867130 | 52869235 | 8 | 2 | 0.019668 | 0.49215 | MUSTN1 | 0.49215 | 0.69096 | 0.55043 |
| ENSG00000177105 | 11 | 3848208 | 3862213 | 51 | 8 | 0.019568 | 0.49219 | RHOG | 0.49219 | 0.525 | 0.78037 |
| ENSG00000116962 | 1 | 236139130 | 236228462 | 314 | 29 | 0.01946 | 0.49224 | NID1 | 0.49224 | 0.5387 | 0.60061 |

| | | | | | | | | | | | |
|---|---|---|---|---|---|---|---|---|---|---|---|
| ENSG00000178974 | 14 | 55738021 | 55828636 | 356 | 16 | 0.018973 | 0.49243 | FBXO34 | 0.49243 | 0.50785 | 0.54179 |
| ENSG00000169031 | 2 | 228029281 | 228179508 | 950 | 69 | 0.018843 | 0.49248 | COL4A3 | 0.49248 | 0.45279 | 0.51052 |
| ENSG00000251357 | 22 | 24204375 | 24237410 | 142 | 23 | 0.018834 | 0.49249 | AP000350.10 | 0.49249 | 0.24399 | 0.17115 |
| ENSG00000198838 | 15 | 33603163 | 34158303 | 2708 | 205 | 0.018694 | 0.49254 | RYR3 | 0.49254 | 0.54356 | 0.53366 |
| ENSG00000197454 | 1 | 248185250 | 248186188 | 4 | 2 | 0.018514 | 0.49261 | OR2L5 | 0.49261 | 0.17787 | 0.20159 |
| ENSG00000159685 | 3 | 126423063 | 126679249 | 611 | 31 | 0.018533 | 0.49261 | CHCHD6 | 0.49261 | 0.55655 | 0.55267 |
| ENSG00000272896 | 9 | 139685831 | 139695047 | 37 | 7 | 0.018421 | 0.49265 | RP11-216L13.17 | 0.49265 | 0.79484 | 0.83087 |
| ENSG00000113387 | 5 | 32531739 | 32604185 | 197 | 25 | 0.018202 | 0.49274 | SUB1 | 0.49274 | 0.32893 | 0.38944 |
| ENSG00000214300 | 7 | 99905325 | 99919819 | 16 | 4 | 0.018151 | 0.49276 | SPDYE3 | 0.49276 | 0.52089 | 0.46413 |
| ENSG00000100813 | 14 | 23527773 | 23564823 | 98 | 14 | 0.018074 | 0.49279 | ACIN1 | 0.49279 | 0.55498 | 0.48064 |
| ENSG00000116690 | 1 | 186265405 | 186283694 | 37 | 6 | 0.018042 | 0.4928 | PRG4 | 0.4928 | 0.45633 | 0.2341 |
| ENSG00000111700 | 12 | 20963636 | 21243040 | 1552 | 30 | 0.017908 | 0.49286 | SLCO1B3 | 0.49286 | 0.49539 | 0.50359 |
| ENSG00000141526 | 17 | 80186273 | 80219005 | 77 | 25 | 0.017804 | 0.4929 | SLC16A3 | 0.4929 | 0.6435 | 0.70713 |
| ENSG00000135540 | 6 | 138743180 | 139013708 | 862 | 68 | 0.01766 | 0.49296 | NHSL1 | 0.49296 | 0.5444 | 0.64595 |
| ENSG00000031081 | 3 | 119013220 | 119139561 | 491 | 61 | 0.017606 | 0.49298 | ARHGAP31 | 0.49298 | 0.32213 | 0.25096 |
| ENSG00000188487 | 11 | 15133970 | 15268754 | 474 | 53 | 0.017514 | 0.49301 | INSC | 0.49301 | 0.4754 | 0.45007 |
| ENSG00000204957 | 19 | 42746927 | 42749125 | 8 | 2 | 0.017429 | 0.49305 | AC006486.1 | 0.49305 | 0.60043 | 0.63131 |
| ENSG00000138207 | 10 | 95351444 | 95361501 | 40 | 6 | 0.016996 | 0.49322 | RBP4 | 0.49322 | 0.52369 | 0.45206 |
| ENSG00000223638 | 19 | 56270380 | 56274541 | 17 | 7 | 0.016813 | 0.49329 | RFPL4A | 0.49329 | 0.37503 | 0.35205 |
| ENSG00000197487 | 19 | 56687389 | 56697144 | 59 | 16 | 0.016726 | 0.49333 | GALP | 0.49333 | 0.71917 | 0.6897 |
| ENSG00000117650 | 1 | 211836114 | 211848960 | 36 | 10 | 0.016622 | 0.49337 | NEK2 | 0.49337 | 0.56793 | 0.34474 |
| ENSG00000164930 | 8 | 104310661 | 104345094 | 124 | 15 | 0.016413 | 0.49345 | FZD6 | 0.49345 | 0.49658 | 0.43895 |
| ENSG00000235268 | 11 | 94758422 | 94760760 | 7 | 3 | 0.016248 | 0.49352 | KDM4E | 0.49352 | 0.60012 | 0.64701 |
| ENSG00000104863 | 19 | 49617581 | 49621717 | 21 | 3 | 0.016205 | 0.49354 | LIN7B | 0.49354 | 0.58736 | 0.82495 |
| ENSG00000175224 | 11 | 46638826 | 46696368 | 74 | 13 | 0.016045 | 0.4936 | ATG13 | 0.4936 | 0.58084 | 0.60849 |
| ENSG00000135426 | 12 | 55341802 | 55378530 | 195 | 10 | 0.015904 | 0.49366 | TESPA1 | 0.49366 | 0.45075 | 0.59238 |
| ENSG00000258881 | 2 | 71166448 | 71222466 | 194 | 42 | 0.015867 | 0.49367 | AC007040.11 | 0.49367 | 0.63494 | 0.76788 |
| ENSG00000205212 | 17 | 20739760 | 20799453 | 159 | 21 | 0.015819 | 0.49369 | CCDC144NL | 0.49369 | 0.54967 | 0.61908 |
| ENSG00000085377 | 6 | 105725440 | 105850959 | 412 | 32 | 0.015711 | 0.49373 | PREP | 0.49373 | 0.5188 | 0.51305 |
| ENSG00000103145 | 16 | 3072621 | 3074287 | 1 | 1 | 0.015542 | 0.4938 | HCFC1R1 | 0.4938 | 0.68001 | 0.75125 |
| ENSG00000178919 | 9 | 100615536 | 100618986 | 17 | 3 | 0.015186 | 0.49394 | FOXE1 | 0.49394 | 0.43326 | 0.47226 |
| ENSG00000138376 | 2 | 215590370 | 215674428 | 509 | 31 | 0.01488 | 0.49406 | BARD1 | 0.49406 | 0.51385 | 0.4841 |
| ENSG00000185686 | 22 | 22890123 | 22901768 | 55 | 8 | 0.014535 | 0.4942 | PRAME | 0.4942 | 0.51225 | 0.61516 |
| ENSG00000183621 | 10 | 31109136 | 31320866 | 797 | 28 | 0.014406 | 0.49425 | ZNF438 | 0.49425 | 0.54111 | 0.55362 |
| ENSG00000260170 | 15 | 45879549 | 45968512 | 287 | 27 | 0.014399 | 0.49426 | RP11-96O20.4 | 0.49426 | 0.36822 | 0.5038 |
| ENSG00000125462 | 1 | 156374042 | 156400493 | 32 | 10 | 0.014354 | 0.49427 | C1orf61 | 0.49427 | 0.26578 | 0.34489 |
| ENSG00000122912 | 10 | 70237756 | 70287231 | 280 | 16 | 0.01434 | 0.49428 | SLC25A16 | 0.49428 | 0.48347 | 0.49231 |
| ENSG00000134030 | 18 | 46065417 | 46389588 | 1177 | 134 | 0.014339 | 0.49428 | CTIF | 0.49428 | 0.47292 | 0.31733 |
| ENSG00000073754 | 1 | 157800704 | 157868046 | 233 | 20 | 0.014135 | 0.49436 | CD5L | 0.49436 | 0.54593 | 0.67779 |
| ENSG00000049541 | 7 | 73645829 | 73668774 | 52 | 13 | 0.014122 | 0.49437 | RFC2 | 0.49437 | 0.47328 | 0.63877 |
| ENSG00000117016 | 1 | 41086351 | 41131329 | 144 | 21 | 0.014076 | 0.49438 | RIMS3 | 0.49438 | 0.48957 | 0.38066 |
| ENSG00000250067 | 19 | 19627036 | 19648390 | 45 | 6 | 0.013879 | 0.49446 | YJEFN3 | 0.49446 | 0.54146 | 0.53826 |
| ENSG00000094916 | 12 | 54624724 | 54673886 | 54 | 10 | 0.013761 | 0.49451 | CBX5 | 0.49451 | 0.47401 | 0.28625 |
| ENSG00000160961 | 19 | 14800613 | 14844558 | 193 | 24 | 0.013399 | 0.49465 | ZNF333 | 0.49465 | 0.45735 | 0.36008 |
| ENSG00000227051 | 14 | 96505661 | 96560417 | 261 | 21 | 0.013176 | 0.49474 | C14orf132 | 0.49474 | 0.62296 | 0.60348 |
| ENSG00000140829 | 16 | 72127461 | 72146811 | 60 | 9 | 0.013157 | 0.49475 | DHX38 | 0.49475 | 0.61552 | 0.72721 |
| ENSG00000143384 | 1 | 150547032 | 150552066 | 8 | 5 | 0.012936 | 0.49484 | MCL1 | 0.49484 | 0.48578 | 0.47563 |
| ENSG00000138772 | 4 | 79472673 | 79531597 | 155 | 20 | 0.012822 | 0.49488 | ANXA3 | 0.49488 | 0.56209 | 0.61142 |
| ENSG00000175329 | 22 | 35462129 | 35483380 | 118 | 10 | 0.012734 | 0.49492 | ISX | 0.49492 | 0.69236 | 0.75531 |
| ENSG00000211445 | 5 | 150400124 | 150408554 | 32 | 7 | 0.012361 | 0.49507 | GPX3 | 0.49507 | 0.7738 | 0.87765 |
| ENSG00000268133 | 19 | 57874934 | 57909952 | 99 | 17 | 0.012202 | 0.49513 | AC003002.4 | 0.49513 | 0.51251 | 0.50228 |
| ENSG00000129152 | 11 | 17741115 | 17743678 | 6 | 4 | 0.012135 | 0.49516 | MYOD1 | 0.49516 | 0.65183 | 0.709 |
| ENSG00000171503 | 4 | 159593277 | 159630775 | 46 | 17 | 0.012109 | 0.49517 | ETFDH | 0.49517 | 0.49629 | 0.57842 |
| ENSG00000198339 | 6 | 27107076 | 27108418 | 1 | 1 | 0.012032 | 0.4952 | HIST1H4I | 0.4952 | 0.42372 | 0.38804 |
| ENSG00000110955 | 12 | 57031959 | 57039798 | 16 | 5 | 0.011961 | 0.49523 | ATP5B | 0.49523 | 0.5818 | 0.69829 |
| ENSG00000212916 | 1 | 232940643 | 232946092 | 11 | 3 | 0.011669 | 0.49535 | MAP10 | 0.49535 | 0.18139 | 0.18045 |
| ENSG00000121957 | 1 | 109417972 | 109477167 | 184 | 21 | 0.011524 | 0.4954 | GPSM2 | 0.4954 | 0.44067 | 0.34453 |
| ENSG00000256073 | 21 | 33765439 | 33766254 | 3 | 2 | 0.011476 | 0.49542 | C21orf119 | 0.49542 | 0.23767 | 0.065033 |
| ENSG00000042832 | 8 | 133879203 | 134147147 | 1003 | 59 | 0.01115 | 0.49555 | TG | 0.49555 | 0.4962 | 0.51065 |
| ENSG00000108669 | 17 | 76670130 | 76778379 | 272 | 16 | 0.011142 | 0.49555 | CYTH1 | 0.49555 | 0.52771 | 0.61979 |
| ENSG00000248469 | 5 | 175476681 | 175489024 | 19 | 6 | 0.011038 | 0.4956 | RP11-826N14.2 | 0.4956 | 0.58912 | 0.5127 |
| ENSG00000164754 | 8 | 117858174 | 117887105 | 102 | 11 | 0.011034 | 0.4956 | RAD21 | 0.4956 | 0.52013 | 0.47651 |
| ENSG00000091181 | 3 | 3111233 | 3168297 | 345 | 69 | 0.011003 | 0.49561 | IL5RA | 0.49561 | 0.48264 | 0.54998 |
| ENSG00000180953 | 15 | 80191182 | 80216044 | 171 | 13 | 0.010931 | 0.49564 | ST20 | 0.49564 | 0.45409 | 0.44136 |
| ENSG00000169704 | 3 | 128779610 | 128781249 | 1 | 1 | 0.010779 | 0.4957 | GP9 | 0.4957 | 0.74536 | 0.83726 |
| ENSG00000197497 | 19 | 53662466 | 53696639 | 162 | 21 | 0.01071 | 0.49573 | ZNF665 | 0.49573 | 0.53832 | 0.63652 |
| ENSG00000146830 | 7 | 100277130 | 100287071 | 27 | 10 | 0.010619 | 0.49576 | GIGYF1 | 0.49576 | 0.49896 | 0.48765 |
| ENSG00000139044 | 12 | 569530 | 672675 | 585 | 87 | 0.010624 | 0.49576 | B4GALNT3 | 0.49576 | 0.45677 | 0.50231 |
| ENSG00000266202 | 17 | 26125832 | 26220391 | 213 | 21 | 0.01058 | 0.49578 | RP1-66C13.4 | 0.49578 | 0.41368 | 0.38369 |
| ENSG00000159658 | 1 | 47124366 | 47184824 | 140 | 9 | 0.010551 | 0.49579 | EFCAB14 | 0.49579 | 0.43118 | 0.3857 |
| ENSG00000258817 | 11 | 49973943 | 49974971 | 5 | 3 | 0.010434 | 0.49584 | OR4C13 | 0.49584 | 0.45248 | 0.42604 |
| ENSG00000148396 | 9 | 139334549 | 139372141 | 119 | 22 | 0.01034 | 0.49587 | SEC16A | 0.49587 | 0.50463 | 0.62482 |
| ENSG00000094963 | 1 | 171154347 | 171181822 | 179 | 17 | 0.010128 | 0.49596 | FMO2 | 0.49596 | 0.43115 | 0.39869 |

| | | | | | | | | | | | |
|---|---|---|---|---|---|---|---|---|---|---|---|
| ENSG00000065911 | 2 | 74425689 | 74444692 | 42 | 7 | 0.010026 | 0.496 | MTHFD2 | 0.496 | 0.47824 | 0.46299 |
| ENSG00000103174 | 16 | 5074845 | 5084142 | 39 | 10 | 0.0099283 | 0.49604 | NAGPA | 0.49604 | 0.5718 | 0.63966 |
| ENSG00000103034 | 16 | 58496750 | 58547532 | 165 | 46 | 0.0095381 | 0.49619 | NDRG4 | 0.49619 | 0.41787 | 0.55424 |
| ENSG00000111361 | 12 | 124104953 | 124118313 | 48 | 8 | 0.0095212 | 0.4962 | EIF2B1 | 0.4962 | 0.39242 | 0.38388 |
| ENSG00000048544 | 6 | 42174539 | 42185603 | 57 | 5 | 0.0094739 | 0.49622 | MRPS10 | 0.49622 | 0.33243 | 0.32679 |
| ENSG00000013503 | 12 | 106751436 | 106903976 | 419 | 22 | 0.0093787 | 0.49626 | POLR3B | 0.49626 | 0.52434 | 0.56748 |
| ENSG00000170310 | 17 | 9153788 | 9479908 | 1216 | 126 | 0.0092733 | 0.4963 | STX8 | 0.4963 | 0.55243 | 0.5154 |
| ENSG00000196616 | 4 | 100226121 | 100242558 | 49 | 11 | 0.0090317 | 0.4964 | ADH1B | 0.4964 | 0.40489 | 0.4724 |
| ENSG00000143184 | 1 | 168545711 | 168551315 | 31 | 8 | 0.0089791 | 0.49642 | XCL1 | 0.49642 | 0.72786 | 0.8238 |
| ENSG00000187600 | 2 | 46656329 | 46711564 | 220 | 17 | 0.0089719 | 0.49642 | TMEM247 | 0.49642 | 0.51229 | 0.48599 |
| ENSG00000144843 | 3 | 119298115 | 119308792 | 25 | 9 | 0.0087323 | 0.49652 | ADPRH | 0.49652 | 0.044398 | 0.089155 |
| ENSG00000126882 | 9 | 134133463 | 134151934 | 51 | 12 | 0.0085963 | 0.49657 | FAM78A | 0.49657 | 0.37815 | 0.58287 |
| ENSG00000159461 | 16 | 56395364 | 56459450 | 209 | 15 | 0.0085921 | 0.49657 | AMFR | 0.49657 | 0.54675 | 0.52271 |
| ENSG00000224730 | 19 | 55147173 | 55150343 | 37 | 7 | 0.0086051 | 0.49657 | AC009892.10 | 0.49657 | 0.24575 | 0.33288 |
| ENSG00000196230 | 6 | 30687978 | 30693203 | 29 | 8 | 0.008519 | 0.4966 | TUBB | 0.4966 | 0.38547 | 0.42372 |
| ENSG00000073670 | 17 | 42836399 | 42859214 | 65 | 9 | 0.0083645 | 0.49666 | ADAM11 | 0.49666 | 0.49092 | 0.38217 |
| ENSG00000173083 | 4 | 84213614 | 84256306 | 134 | 13 | 0.0079758 | 0.49682 | HPSE | 0.49682 | 0.41754 | 0.29503 |
| ENSG00000185808 | 21 | 38431470 | 38445470 | 58 | 14 | 0.0079598 | 0.49682 | PIGP | 0.49682 | 0.55384 | 0.52374 |
| ENSG00000163900 | 3 | 185194284 | 185216845 | 12 | 6 | 0.0079479 | 0.49683 | TMEM41A | 0.49683 | 0.54328 | 0.67226 |
| ENSG00000093134 | 6 | 133043926 | 133055904 | 46 | 7 | 0.0075871 | 0.49697 | VNN3 | 0.49697 | 0.68443 | 0.84191 |
| ENSG00000178562 | 2 | 204571198 | 204603635 | 63 | 15 | 0.0075066 | 0.49701 | CD28 | 0.49701 | 0.41007 | 0.41098 |
| ENSG00000106511 | 7 | 15650837 | 15726437 | 292 | 27 | 0.0074805 | 0.49702 | MEOX2 | 0.49702 | 0.31457 | 0.1226 |
| ENSG00000131037 | 19 | 55583388 | 55599291 | 85 | 12 | 0.0074372 | 0.49703 | EPS8L1 | 0.49703 | 0.4025 | 0.81454 |
| ENSG00000231031 | 2 | 15830906 | 15884463 | 245 | 29 | 0.007413 | 0.49704 | AC008271.1 | 0.49704 | 0.52949 | 0.55236 |
| ENSG00000118260 | 2 | 208394461 | 208468155 | 140 | 17 | 0.0073149 | 0.49708 | CREB1 | 0.49708 | 0.4784 | 0.48913 |
| ENSG00000184650 | 17 | 8243182 | 8249364 | 39 | 5 | 0.0072964 | 0.49709 | ODF4 | 0.49709 | 0.48307 | 0.43422 |
| ENSG00000185650 | 14 | 69254377 | 69263190 | 20 | 6 | 0.0068944 | 0.49725 | ZFP36L1 | 0.49725 | 0.62401 | 0.41013 |
| ENSG00000108924 | 17 | 53342373 | 53402426 | 164 | 32 | 0.0066964 | 0.49733 | HLF | 0.49733 | 0.61264 | 0.47569 |
| ENSG00000105607 | 19 | 13001840 | 13025021 | 41 | 10 | 0.0066335 | 0.49735 | GCDH | 0.49735 | 0.74373 | 0.74193 |
| ENSG00000143641 | 1 | 230193536 | 230417870 | 917 | 69 | 0.0063462 | 0.49747 | GALNT2 | 0.49747 | 0.5341 | 0.53675 |
| ENSG00000122756 | 9 | 34551430 | 34590121 | 131 | 20 | 0.0061059 | 0.49756 | CNTFR | 0.49756 | 0.5495 | 0.55189 |
| ENSG00000184155 | 1 | 159504793 | 159505842 | 2 | 1 | 0.0059295 | 0.49763 | OR10J5 | 0.49763 | 0.53347 | 0.34639 |
| ENSG00000171204 | 11 | 85339629 | 85347580 | 15 | 6 | 0.0059033 | 0.49764 | TMEM126B | 0.49764 | 0.61913 | 0.73179 |
| ENSG00000128563 | 7 | 102004319 | 102067123 | 172 | 14 | 0.0058776 | 0.49766 | PRKRIP1 | 0.49766 | 0.49474 | 0.4638 |
| ENSG00000183785 | 22 | 18593097 | 18629321 | 135 | 26 | 0.0058633 | 0.49766 | TUBA8 | 0.49766 | 0.73361 | 0.84943 |
| ENSG00000164303 | 4 | 185009859 | 185142383 | 572 | 48 | 0.0058206 | 0.49768 | ENPP6 | 0.49768 | 0.53586 | 0.61065 |
| ENSG00000019169 | 2 | 119699742 | 119752236 | 207 | 26 | 0.0056566 | 0.49774 | MARCO | 0.49774 | 0.29263 | 0.224 |
| ENSG00000143297 | 1 | 157483167 | 157522310 | 162 | 12 | 0.0056051 | 0.49776 | FCRL5 | 0.49776 | 0.6325 | 0.36912 |
| ENSG00000196968 | 10 | 75532049 | 75540009 | 17 | 4 | 0.0056215 | 0.49776 | FUT11 | 0.49776 | 0.5451 | 0.64358 |
| ENSG00000142039 | 19 | 41816094 | 41830788 | 25 | 2 | 0.0055431 | 0.49779 | CCDC97 | 0.49779 | 0.55262 | 0.51931 |
| ENSG00000174137 | 4 | 1619642 | 1686029 | 185 | 36 | 0.0054781 | 0.49781 | FAM53A | 0.49781 | 0.5089 | 0.50726 |
| ENSG00000100201 | 22 | 38879445 | 38903665 | 48 | 13 | 0.0053561 | 0.49786 | DDX17 | 0.49786 | 0.47634 | 0.62363 |
| ENSG00000079102 | 8 | 92967203 | 93115514 | 220 | 10 | 0.0053137 | 0.49788 | RUNX1T1 | 0.49788 | 0.49859 | 0.52361 |
| ENSG00000136003 | 12 | 108956358 | 108963160 | 22 | 5 | 0.005267 | 0.4979 | ISCU | 0.4979 | 0.671 | 0.67652 |
| ENSG00000169221 | 16 | 30368423 | 30381585 | 17 | 7 | 0.0052232 | 0.49792 | TBC1D10B | 0.49792 | 0.44445 | 0.56102 |
| ENSG00000035928 | 4 | 39289076 | 39367995 | 130 | 10 | 0.0047841 | 0.49809 | RFC1 | 0.49809 | 0.4988 | 0.52071 |
| ENSG00000113302 | 5 | 158741791 | 158757895 | 39 | 5 | 0.0046442 | 0.49815 | IL12B | 0.49815 | 0.6041 | 0.78236 |
| ENSG00000212122 | 5 | 112768251 | 112770728 | 18 | 6 | 0.0045897 | 0.49817 | TSSK1B | 0.49817 | 0.45736 | 0.4629 |
| ENSG00000268173 | 19 | 18263968 | 18288927 | 66 | 12 | 0.0045893 | 0.49817 | PIK3R2 | 0.49817 | 0.46418 | 0.3149 |
| ENSG00000188732 | 7 | 23719749 | 23742868 | 89 | 15 | 0.0045014 | 0.4982 | FAM221A | 0.4982 | 0.64467 | 0.67787 |
| ENSG00000093144 | 6 | 127609855 | 127664754 | 134 | 17 | 0.0043508 | 0.49826 | ECHDC1 | 0.49826 | 0.4785 | 0.42345 |
| ENSG00000268553 | 5 | 172036245 | 172036436 | 1 | 1 | 0.0040106 | 0.4984 | AC027309.1 | 0.4984 | 0.52741 | 0.79644 |
| ENSG00000141295 | 17 | 45915058 | 45918699 | 11 | 6 | 0.0039933 | 0.49841 | SCRN2 | 0.49841 | 0.68842 | 0.70294 |
| ENSG00000105605 | 19 | 54412589 | 54447195 | 113 | 26 | 0.0037364 | 0.49851 | CACNG7 | 0.49851 | 0.25752 | 0.051005 |
| ENSG00000140396 | 8 | 71021997 | 71316040 | 527 | 21 | 0.0037215 | 0.49852 | NCOA2 | 0.49852 | 0.48354 | 0.47855 |
| ENSG00000139278 | 12 | 75874460 | 75897633 | 71 | 9 | 0.0036012 | 0.49856 | GLIPR1 | 0.49856 | 0.40158 | 0.36901 |
| ENSG00000206150 | 14 | 21500979 | 21502944 | 16 | 6 | 0.0033365 | 0.49867 | RNASE13 | 0.49867 | 0.69218 | 0.86566 |
| ENSG00000157538 | 21 | 38595721 | 38640262 | 107 | 23 | 0.003295 | 0.49869 | DSCR3 | 0.49869 | 0.57617 | 0.63503 |
| ENSG00000162510 | 1 | 31184124 | 31196434 | 42 | 4 | 0.00324 | 0.49871 | MATN1 | 0.49871 | 0.79373 | 0.84935 |
| ENSG00000126001 | 20 | 34042985 | 34099804 | 121 | 9 | 0.0031549 | 0.49874 | CEP250 | 0.49874 | 0.42625 | 0.28329 |
| ENSG00000132604 | 16 | 69389464 | 69442474 | 108 | 18 | 0.0031287 | 0.49875 | TERF2 | 0.49875 | 0.57564 | 0.54989 |
| ENSG00000240403 | 19 | 55361898 | 55378662 | 116 | 13 | 0.0030343 | 0.49879 | KIR3DL2 | 0.49879 | 0.43666 | 0.45914 |
| ENSG00000183733 | 2 | 71004442 | 71017775 | 65 | 7 | 0.0030156 | 0.4988 | FIGLA | 0.4988 | 0.56881 | 0.52384 |
| ENSG00000182512 | 14 | 95999840 | 96011061 | 52 | 14 | 0.0030123 | 0.4988 | GLRX5 | 0.4988 | 0.4768 | 0.52013 |
| ENSG00000171481 | 9 | 125437315 | 125438432 | 2 | 1 | 0.0027649 | 0.4989 | OR1L3 | 0.4989 | 0.76002 | 0.79319 |
| ENSG00000127578 | 16 | 679239 | 684116 | 21 | 4 | 0.0027192 | 0.49892 | WFIKKN1 | 0.49892 | 0.36524 | 0.33226 |
| ENSG00000268361 | 19 | 44058844 | 44097188 | 111 | 19 | 0.0025611 | 0.49898 | L34079.2 | 0.49898 | 0.55081 | 0.58149 |
| ENSG00000141034 | 17 | 17942606 | 17971718 | 66 | 6 | 0.0024936 | 0.49901 | GID4 | 0.49901 | 0.60338 | 0.62871 |
| ENSG00000149488 | 20 | 2517253 | 2622430 | 478 | 33 | 0.0024524 | 0.49902 | TMC2 | 0.49902 | 0.47542 | 0.45096 |
| ENSG00000134072 | 3 | 9799026 | 9811676 | 21 | 7 | 0.0024101 | 0.49904 | CAMK1 | 0.49904 | 0.36577 | 0.30692 |
| ENSG00000128045 | 4 | 53728457 | 53733000 | 9 | 2 | 0.0023273 | 0.49907 | RASL11B | 0.49907 | 0.83891 | 0.7984 |
| ENSG00000100290 | 22 | 43506754 | 43525718 | 65 | 11 | 0.0023094 | 0.49908 | BIK | 0.49908 | 0.23304 | 0.25824 |
| ENSG00000164691 | 6 | 159455500 | 159466184 | 44 | 6 | 0.0020528 | 0.49918 | TAGAP | 0.49918 | 0.5604 | 0.54213 |

| | | | | | | | | | | | |
|---|---|---|---|---|---|---|---|---|---|---|---|
| ENSG00000153046 | 6 | 4706393 | 4955785 | 878 | 78 | 0.0016718 | 0.49933 | CDYL | 0.49933 | 0.5585 | 0.53988 |
| ENSG00000135213 | 7 | 75046066 | 75115548 | 196 | 27 | 0.0015103 | 0.4994 | POM121C | 0.4994 | 0.52657 | 0.49953 |
| ENSG00000112562 | 6 | 168841831 | 169073984 | 1185 | 98 | 0.0012914 | 0.49948 | SMOC2 | 0.49948 | 0.47743 | 0.41577 |
| ENSG00000164466 | 5 | 174904065 | 174956745 | 224 | 19 | 0.0012363 | 0.49951 | SFXN1 | 0.49951 | 0.60818 | 0.64763 |
| ENSG00000126545 | 4 | 70796799 | 70812289 | 42 | 7 | 0.0010465 | 0.49958 | CSN1S1 | 0.49958 | 0.57756 | 0.6811 |
| ENSG00000100307 | 22 | 39516172 | 39548679 | 122 | 16 | 0.0010255 | 0.49959 | CBX7 | 0.49959 | 0.41961 | 0.25036 |
| ENSG00000125731 | 19 | 6752171 | 6767599 | 43 | 5 | 0.00098348 | 0.49961 | SH2D3A | 0.49961 | 0.49632 | 0.44465 |
| ENSG00000259075 | 12 | 89913185 | 89920039 | 4 | 2 | 0.00094625 | 0.49962 | POC1B-GALNT4 | 0.49962 | 0.59995 | 0.50726 |
| ENSG00000257594 | 12 | 89913185 | 89920039 | 4 | 2 | 0.00094625 | 0.49962 | GALNT4 | 0.49962 | 0.59995 | 0.50726 |
| ENSG00000118246 | 2 | 207630081 | 207657233 | 78 | 13 | 0.00091281 | 0.49964 | FASTKD2 | 0.49964 | 0.56467 | 0.55829 |
| ENSG00000152944 | 12 | 27175479 | 27219276 | 174 | 21 | 0.00087825 | 0.49965 | MED21 | 0.49965 | 0.48789 | 0.50338 |
| ENSG00000154783 | 3 | 14860469 | 14975895 | 573 | 34 | 0.00065959 | 0.49974 | FGD5 | 0.49974 | 0.52662 | 0.52352 |
| ENSG00000108465 | 17 | 46045176 | 46059140 | 35 | 6 | 0.00060896 | 0.49976 | CDK5RAP3 | 0.49976 | 0.58518 | 0.51273 |
| ENSG00000100225 | 22 | 32870663 | 32894818 | 96 | 10 | 0.00058076 | 0.49977 | FBXO7 | 0.49977 | 0.55692 | 0.45225 |
| ENSG00000109738 | 4 | 157997209 | 158093242 | 292 | 25 | 0.00040506 | 0.49984 | GLRB | 0.49984 | 0.54163 | 0.67947 |
| ENSG00000171804 | 19 | 38375463 | 38397317 | 44 | 12 | 0.00014541 | 0.49994 | WDR87 | 0.49994 | 0.54084 | 0.53293 |
| ENSG00000164880 | 7 | 1509913 | 1545489 | 138 | 6 | 0.000003343 | 0.5 | INTS1 | 0.5 | 0.49797 | 0.5956 |
| ENSG00000213512 | 1 | 89597434 | 89641723 | 82 | 8 | -0.00015972 | 0.50006 | GBP7 | 0.50006 | 0.55665 | 0.5528 |
| ENSG00000173769 | 3 | 44283378 | 44373590 | 228 | 16 | -0.00016844 | 0.50007 | TOPAZ1 | 0.50007 | 0.46551 | 0.42849 |
| ENSG00000151500 | 11 | 134118173 | 134123264 | 8 | 4 | -0.00028046 | 0.50011 | THYN1 | 0.50011 | 0.80122 | 0.8894 |
| ENSG00000129009 | 15 | 74466012 | 74469213 | 9 | 2 | -0.00027375 | 0.50011 | ISLR | 0.50011 | 0.53992 | 0.56163 |
| ENSG00000137033 | 9 | 6215805 | 6257983 | 133 | 19 | -0.0003231 | 0.50013 | IL33 | 0.50013 | 0.28253 | 0.26251 |
| ENSG00000232258 | 16 | 8619502 | 8622304 | 16 | 5 | -0.00054483 | 0.50022 | TMEM114 | 0.50022 | 0.37724 | 0.26061 |
| ENSG00000137776 | 15 | 59171244 | 59225852 | 96 | 13 | -0.00060854 | 0.50024 | SLTM | 0.50024 | 0.50943 | 0.649 |
| ENSG00000148357 | 9 | 133046882 | 133309510 | 489 | 87 | -0.0007931 | 0.50032 | HMCN2 | 0.50032 | 0.2877 | 0.29633 |
| ENSG00000265303 | 17 | 57274927 | 57292587 | 40 | 7 | -0.0010108 | 0.5004 | CTD-2510F5.6 | 0.5004 | 0.42425 | 0.43465 |
| ENSG00000169710 | 17 | 80036214 | 80056208 | 57 | 14 | -0.0010004 | 0.5004 | FASN | 0.5004 | 0.588 | 0.72124 |
| ENSG00000139496 | 13 | 25875662 | 25923938 | 66 | 18 | -0.0011108 | 0.50044 | NUPL1 | 0.50044 | 0.53668 | 0.6299 |
| ENSG00000156973 | 2 | 232597135 | 232650982 | 157 | 11 | -0.001241 | 0.5005 | PDE6D | 0.5005 | 0.4606 | 0.49732 |
| ENSG00000091436 | 2 | 173940163 | 174132738 | 778 | 41 | -0.0014067 | 0.50056 | MLTK | 0.50056 | 0.55341 | 0.51488 |
| ENSG00000120885 | 8 | 27454434 | 27472548 | 41 | 7 | -0.0014292 | 0.50057 | CLU | 0.50057 | 0.30396 | 0.20197 |
| ENSG00000117411 | 1 | 44444615 | 44456840 | 22 | 7 | -0.0014631 | 0.50058 | B4GALT2 | 0.50058 | 0.47555 | 0.35349 |
| ENSG00000197451 | 5 | 177631508 | 177638164 | 19 | 7 | -0.0017323 | 0.50069 | HNRNPAB | 0.50069 | 0.69994 | 0.29067 |
| ENSG00000196184 | 1 | 159409512 | 159410600 | 5 | 3 | -0.0017798 | 0.50071 | OR10J1 | 0.50071 | 0.57957 | 0.59271 |
| ENSG00000043514 | 1 | 40306723 | 40349183 | 144 | 17 | -0.0018154 | 0.50072 | TRIT1 | 0.50072 | 0.54472 | 0.54107 |
| ENSG00000204564 | 6 | 30614816 | 30620987 | 15 | 8 | -0.0020312 | 0.50081 | C6orf136 | 0.50081 | 0.16336 | 0.3186 |
| ENSG00000136448 | 17 | 43128978 | 43186384 | 249 | 26 | -0.0022922 | 0.50091 | NMT1 | 0.50091 | 0.43732 | 0.29404 |
| ENSG00000123360 | 12 | 54943134 | 54973023 | 87 | 23 | -0.0023636 | 0.50094 | PDE1B | 0.50094 | 0.55467 | 0.76397 |
| ENSG00000119403 | 9 | 123617977 | 123639606 | 39 | 8 | -0.0024236 | 0.50097 | PHF19 | 0.50097 | 0.23222 | 0.26182 |
| ENSG00000155393 | 16 | 50099852 | 50140298 | 142 | 13 | -0.00286 | 0.50114 | HEATR3 | 0.50114 | 0.52434 | 0.50534 |
| ENSG00000197604 | 10 | 72194585 | 72196314 | 3 | 2 | -0.0031922 | 0.50127 | AC022532.1 | 0.50127 | 0.61191 | 0.68006 |
| ENSG00000104518 | 8 | 144635377 | 144645232 | 45 | 12 | -0.0033317 | 0.50133 | GSDMD | 0.50133 | 0.47749 | 0.57936 |
| ENSG00000173141 | 13 | 21750784 | 21753223 | 5 | 3 | -0.0033646 | 0.50134 | MRP63 | 0.50134 | 0.89963 | 0.92756 |
| ENSG00000183475 | 15 | 101142739 | 101191910 | 146 | 22 | -0.0033685 | 0.50134 | ASB7 | 0.50134 | 0.61428 | 0.26123 |
| ENSG00000126814 | 14 | 61438169 | 61448076 | 12 | 4 | -0.0033846 | 0.50135 | TRMT5 | 0.50135 | 0.63923 | 0.53587 |
| ENSG00000179813 | 13 | 43355686 | 43365683 | 23 | 7 | -0.003517 | 0.5014 | FAM216B | 0.5014 | 0.48424 | 0.51152 |
| ENSG00000196098 | 3 | 98072698 | 98073663 | 2 | 1 | -0.0036865 | 0.50147 | OR5K4 | 0.50147 | 0.83593 | 0.89 |
| ENSG00000121940 | 1 | 109472130 | 109506111 | 143 | 25 | -0.0037394 | 0.50149 | CLCC1 | 0.50149 | 0.4781 | 0.46582 |
| ENSG00000146576 | 7 | 6629648 | 6648357 | 66 | 8 | -0.0038444 | 0.50153 | C7orf26 | 0.50153 | 0.49568 | 0.47432 |
| ENSG00000001167 | 6 | 41040684 | 41067715 | 52 | 8 | -0.0038747 | 0.50155 | NFYA | 0.50155 | 0.66488 | 0.72132 |
| ENSG00000121900 | 1 | 33360195 | 33367039 | 10 | 3 | -0.0042367 | 0.50169 | TMEM54 | 0.50169 | 0.59016 | 0.50853 |
| ENSG00000101438 | 20 | 37353105 | 37358015 | 4 | 2 | -0.0042707 | 0.5017 | SLC32A1 | 0.5017 | 0.67455 | 0.50982 |
| ENSG00000164904 | 5 | 125877533 | 125931110 | 200 | 31 | -0.004322 | 0.50172 | ALDH7A1 | 0.50172 | 0.55935 | 0.71466 |
| ENSG00000256463 | 18 | 76740275 | 76762677 | 56 | 13 | -0.0045983 | 0.50183 | SALL3 | 0.50183 | 0.61095 | 0.58435 |
| ENSG00000163645 | 3 | 150377672 | 150421758 | 137 | 28 | -0.0047141 | 0.50188 | FAM194A | 0.50188 | 0.18682 | 0.21023 |
| ENSG00000142700 | 1 | 50883222 | 50889172 | 6 | 4 | -0.0048004 | 0.50192 | DMRTA2 | 0.50192 | 0.57955 | 0.317 |
| ENSG00000131051 | 20 | 34291531 | 34330234 | 133 | 8 | -0.00486 | 0.50194 | RBM39 | 0.50194 | 0.4496 | 0.40567 |
| ENSG00000153208 | 2 | 112656056 | 112787138 | 512 | 17 | -0.0049021 | 0.50196 | MERTK | 0.50196 | 0.54504 | 0.59073 |
| ENSG00000172340 | 3 | 67410884 | 67705038 | 1319 | 64 | -0.0049163 | 0.50196 | SUCLG2 | 0.50196 | 0.49437 | 0.49553 |
| ENSG00000232629 | 6 | 32723875 | 32731311 | 67 | 5 | -0.0050723 | 0.50202 | HLA-DQB2 | 0.50202 | 0.36866 | 0.036249 |
| ENSG00000100731 | 14 | 71374122 | 71582099 | 696 | 29 | -0.0051924 | 0.50207 | PCNX | 0.50207 | 0.48852 | 0.50485 |
| ENSG00000249693 | 4 | 57396775 | 57469673 | 293 | 33 | -0.0052192 | 0.50208 | THEGL | 0.50208 | 0.55592 | 0.62117 |
| ENSG00000152219 | 11 | 30344598 | 30359774 | 50 | 9 | -0.0053996 | 0.50215 | ARL14EP | 0.50215 | 0.62231 | 0.55363 |
| ENSG00000122786 | 7 | 134429003 | 134655479 | 668 | 52 | -0.0054265 | 0.50216 | CALD1 | 0.50216 | 0.5383 | 0.4876 |
| ENSG00000188694 | 21 | 31653629 | 31655276 | 5 | 2 | -0.0055918 | 0.50223 | KRTAP24-1 | 0.50223 | 0.29038 | 0.30517 |
| ENSG00000162520 | 1 | 33145507 | 33169197 | 73 | 6 | -0.0062998 | 0.50251 | SYNC | 0.50251 | 0.58339 | 0.50748 |
| ENSG00000235531 | 8 | 72740402 | 73030628 | 1292 | 71 | -0.0064629 | 0.50258 | RP11-383H13.1 | 0.50258 | 0.53875 | 0.56992 |
| ENSG00000167874 | 17 | 7758383 | 7759417 | 2 | 1 | -0.0066784 | 0.50266 | TMEM88 | 0.50266 | 0.59558 | 0.70475 |
| ENSG00000144524 | 2 | 232646381 | 232673963 | 71 | 11 | -0.0067342 | 0.50269 | COPS7B | 0.50269 | 0.4914 | 0.31898 |
| ENSG00000165899 | 12 | 80603233 | 80772870 | 424 | 32 | -0.0068069 | 0.50272 | OTOGL | 0.50272 | 0.51386 | 0.56577 |
| ENSG00000215545 | 20 | 29891015 | 29896388 | 13 | 5 | -0.007107 | 0.50284 | DEFB116 | 0.50284 | 0.60765 | 0.66167 |
| ENSG00000163032 | 2 | 17720393 | 17838285 | 463 | 20 | -0.0072257 | 0.50288 | VSNL1 | 0.50288 | 0.45665 | 0.45087 |
| ENSG00000122591 | 7 | 22980878 | 23053749 | 237 | 22 | -0.0073487 | 0.50293 | FAM126A | 0.50293 | 0.4745 | 0.31537 |

| | | | | | | | | | | | |
|---|---|---|---|---|---|---|---|---|---|---|---|
| ENSG00000143621 | 1 | 153634512 | 153643524 | 13 | 3 | -0.0074061 | 0.50295 | ILF2 | 0.50295 | 0.50655 | 0.43945 |
| ENSG00000140450 | 15 | 98462784 | 98517068 | 285 | 37 | -0.0074766 | 0.50298 | ARRDC4 | 0.50298 | 0.54224 | 0.59648 |
| ENSG00000167523 | 16 | 89724210 | 89737680 | 69 | 16 | -0.0076537 | 0.50305 | SPATA33 | 0.50305 | 0.6599 | 0.61936 |
| ENSG00000143768 | 1 | 226124298 | 226129189 | 8 | 3 | -0.0077771 | 0.5031 | LEFTY2 | 0.5031 | 0.55304 | 0.43839 |
| ENSG00000182093 | 21 | 40752170 | 40800454 | 124 | 12 | -0.0077628 | 0.5031 | WRB | 0.5031 | 0.46356 | 0.54157 |
| ENSG00000124092 | 20 | 56071035 | 56100708 | 117 | 27 | -0.0080162 | 0.5032 | CTCFL | 0.5032 | 0.6789 | 0.69367 |
| ENSG00000115468 | 2 | 233470767 | 233547491 | 281 | 30 | -0.0080403 | 0.50321 | EFHD1 | 0.50321 | 0.52072 | 0.42389 |
| ENSG00000259784 | 16 | 2563966 | 2577956 | 11 | 7 | -0.0080884 | 0.50323 | ATP6C | 0.50323 | 0.044922 | 0.070887 |
| ENSG00000213022 | 19 | 51503806 | 51512890 | 37 | 10 | -0.0081353 | 0.50325 | KLK9 | 0.50325 | 0.47794 | 0.51422 |
| ENSG00000114737 | 3 | 50643921 | 50649262 | 10 | 4 | -0.0082452 | 0.50329 | CISH | 0.50329 | 0.61548 | 0.79787 |
| ENSG00000171160 | 10 | 99374310 | 99393344 | 51 | 8 | -0.0082389 | 0.50329 | MORN4 | 0.50329 | 0.55266 | 0.56605 |
| ENSG00000261787 | 8 | 67858736 | 67874825 | 8 | 5 | -0.0083052 | 0.50331 | TCF24 | 0.50331 | 0.74575 | 0.77154 |
| ENSG00000110060 | 11 | 125763381 | 125773116 | 40 | 8 | -0.0084356 | 0.50337 | PUS3 | 0.50337 | 0.54785 | 0.64522 |
| ENSG00000079112 | 8 | 95139399 | 95229531 | 266 | 27 | -0.0085909 | 0.50343 | CDH17 | 0.50343 | 0.48516 | 0.569 |
| ENSG00000007171 | 17 | 26083792 | 26127525 | 130 | 21 | -0.0086359 | 0.50345 | NOS2 | 0.50345 | 0.56726 | 0.5915 |
| ENSG00000110719 | 11 | 67806483 | 67818362 | 27 | 9 | -0.0087233 | 0.50348 | TCIRG1 | 0.50348 | 0.31543 | 0.33833 |
| ENSG00000110799 | 12 | 6058040 | 6233936 | 834 | 69 | -0.0088576 | 0.50353 | VWF | 0.50353 | 0.55297 | 0.50261 |
| ENSG00000144908 | 3 | 125822412 | 125916837 | 452 | 40 | -0.008876 | 0.50354 | ALDH1L1 | 0.50354 | 0.52674 | 0.58783 |
| ENSG00000109832 | 11 | 125773271 | 125793158 | 80 | 14 | -0.008904 | 0.50355 | DDX25 | 0.50355 | 0.50785 | 0.24966 |
| ENSG00000128272 | 22 | 39915700 | 39918691 | 2 | 1 | -0.008987 | 0.50359 | ATF4 | 0.50359 | 0.38461 | 0.35175 |
| ENSG00000082641 | 17 | 46125691 | 46138849 | 29 | 7 | -0.0096793 | 0.50386 | NFE2L1 | 0.50386 | 0.64522 | 0.68738 |
| ENSG00000243678 | 17 | 49230920 | 49249108 | 72 | 16 | -0.0097511 | 0.50389 | NME1-NME2 | 0.50389 | 0.59625 | 0.59262 |
| ENSG00000011052 | 17 | 49230951 | 49249105 | 72 | 16 | -0.0097511 | 0.50389 | NME2 | 0.50389 | 0.59625 | 0.59262 |
| ENSG00000204851 | 19 | 46990163 | 46999755 | 33 | 9 | -0.010033 | 0.504 | PNMAL2 | 0.504 | 0.53139 | 0.44021 |
| ENSG00000157224 | 7 | 90013035 | 90142716 | 434 | 37 | -0.010162 | 0.50405 | CLDN12 | 0.50405 | 0.50794 | 0.50688 |
| ENSG00000186118 | 1 | 47134527 | 47139266 | 8 | 3 | -0.010179 | 0.50406 | TEX38 | 0.50406 | 0.47252 | 0.43271 |
| ENSG00000132196 | 1 | 162760492 | 162782607 | 56 | 17 | -0.010309 | 0.50411 | HSD17B7 | 0.50411 | 0.54135 | 0.59002 |
| ENSG00000144229 | 2 | 137523115 | 138435287 | 3932 | 118 | -0.010378 | 0.50414 | THSD7B | 0.50414 | 0.46445 | 0.43242 |
| ENSG00000259471 | 15 | 66874528 | 66978132 | 533 | 49 | -0.010521 | 0.5042 | RP11-321F6.1 | 0.5042 | 0.57989 | 0.42751 |
| ENSG00000071967 | 2 | 172378757 | 172414643 | 143 | 23 | -0.010796 | 0.50431 | CYBRD1 | 0.50431 | 0.37134 | 0.43889 |
| ENSG00000100557 | 14 | 57936019 | 57960585 | 43 | 9 | -0.010884 | 0.50434 | C14orf105 | 0.50434 | 0.70155 | 0.79693 |
| ENSG00000197857 | 19 | 12335501 | 12405702 | 182 | 13 | -0.010953 | 0.50437 | ZNF44 | 0.50437 | 0.45695 | 0.40501 |
| ENSG00000188523 | 9 | 135285430 | 135448704 | 551 | 69 | -0.011102 | 0.50443 | C9orf171 | 0.50443 | 0.38029 | 0.43519 |
| ENSG00000153140 | 5 | 89688078 | 89705603 | 8 | 4 | -0.011147 | 0.50445 | CETN3 | 0.50445 | 0.48607 | 0.61296 |
| ENSG00000236981 | 11 | 123893720 | 123894655 | 10 | 3 | -0.011187 | 0.50446 | OR10G9 | 0.50446 | 0.58383 | 0.46255 |
| ENSG00000162366 | 1 | 47649265 | 47656716 | 17 | 6 | -0.011228 | 0.50448 | PDZK1IP1 | 0.50448 | 0.4762 | 0.15363 |
| ENSG00000198250 | 10 | 47657581 | 47701443 | 254 | 22 | -0.011412 | 0.50455 | ANTXRL | 0.50455 | 0.65998 | 0.8459 |
| ENSG00000174407 | 20 | 61147660 | 61167971 | 118 | 17 | -0.011405 | 0.50455 | C20orf166 | 0.50455 | 0.53762 | 0.54247 |
| ENSG00000147853 | 9 | 4711155 | 4742043 | 219 | 24 | -0.011708 | 0.50467 | AK3 | 0.50467 | 0.63376 | 0.58318 |
| ENSG00000148572 | 10 | 64893050 | 64914783 | 52 | 13 | -0.011898 | 0.50475 | NRBF2 | 0.50475 | 0.45364 | 0.56638 |
| ENSG00000268351 | 11 | 69587254 | 69587718 | 1 | 1 | -0.012032 | 0.5048 | AP001888.1 | 0.5048 | 0.74655 | 0.83101 |
| ENSG00000185813 | 17 | 79858841 | 79869340 | 9 | 5 | -0.012112 | 0.50483 | PCYT2 | 0.50483 | 0.51594 | 0.23102 |
| ENSG00000000457 | 1 | 169818772 | 169863408 | 172 | 16 | -0.012169 | 0.50485 | SCYL3 | 0.50485 | 0.49418 | 0.50365 |
| ENSG00000125347 | 5 | 131817301 | 131826490 | 47 | 3 | -0.012176 | 0.50486 | IRF1 | 0.50486 | 0.52653 | 0.38793 |
| ENSG00000101222 | 20 | 3758151 | 3762095 | 7 | 3 | -0.012202 | 0.50487 | SPEF1 | 0.50487 | 0.57198 | 0.72671 |
| ENSG00000101292 | 20 | 5282317 | 5297378 | 164 | 7 | -0.012223 | 0.50488 | PROKR2 | 0.50488 | 0.42019 | 0.46679 |
| ENSG00000213214 | 7 | 143883176 | 143991230 | 58 | 21 | -0.012285 | 0.5049 | ARHGEF35 | 0.5049 | 0.62824 | 0.59066 |
| ENSG00000132549 | 8 | 100025494 | 100889808 | 1820 | 24 | -0.012891 | 0.50514 | VPS13B | 0.50514 | 0.50881 | 0.50304 |
| ENSG00000268424 | 5 | 101569696 | 101571279 | 6 | 3 | -0.012937 | 0.50516 | AC008948.1 | 0.50516 | 0.72137 | 0.90715 |
| ENSG00000196832 | 14 | 20665495 | 20666605 | 4 | 1 | -0.012952 | 0.50517 | OR11G2 | 0.50517 | 0.52599 | 0.51387 |
| ENSG00000169612 | 15 | 83654959 | 83659809 | 8 | 4 | -0.013303 | 0.50531 | FAM103A1 | 0.50531 | 0.35953 | 0.44301 |
| ENSG00000126246 | 19 | 36230058 | 36233354 | 15 | 5 | -0.013377 | 0.50534 | IGFLR1 | 0.50534 | 0.38629 | 0.53573 |
| ENSG00000110092 | 11 | 69455855 | 69469242 | 31 | 9 | -0.013419 | 0.50535 | CCND1 | 0.50535 | 0.66098 | 0.5963 |
| ENSG00000143622 | 1 | 155867599 | 155881195 | 26 | 4 | -0.013458 | 0.50537 | RIT1 | 0.50537 | 0.5115 | 0.50941 |
| ENSG00000155465 | 14 | 23242431 | 23299029 | 337 | 40 | -0.013451 | 0.50537 | SLC7A7 | 0.50537 | 0.47318 | 0.18961 |
| ENSG00000119865 | 2 | 68511303 | 68547183 | 96 | 23 | -0.013551 | 0.50541 | CNRIP1 | 0.50541 | 0.38257 | 0.4712 |
| ENSG00000153162 | 6 | 7727030 | 7881655 | 600 | 62 | -0.013573 | 0.50541 | BMP6 | 0.50541 | 0.51493 | 0.58516 |
| ENSG00000189167 | 13 | 32877837 | 32889481 | 35 | 8 | -0.013572 | 0.50541 | ZAR1L | 0.50541 | 0.68114 | 0.50606 |
| ENSG00000165698 | 9 | 135753414 | 135765588 | 22 | 7 | -0.01373 | 0.50548 | C9orf9 | 0.50548 | 0.34128 | 0.15202 |
| ENSG00000169180 | 16 | 28109300 | 28223241 | 146 | 9 | -0.013967 | 0.50557 | XPO6 | 0.50557 | 0.5268 | 0.6141 |
| ENSG00000109794 | 4 | 187025573 | 187093821 | 261 | 33 | -0.014054 | 0.50561 | FAM149A | 0.50561 | 0.61087 | 0.78934 |
| ENSG00000177599 | 19 | 11908482 | 11920437 | 31 | 6 | -0.014319 | 0.50571 | ZNF491 | 0.50571 | 0.60894 | 0.44747 |
| ENSG00000103222 | 16 | 16043434 | 16236931 | 867 | 63 | -0.01437 | 0.50573 | ABCC1 | 0.50573 | 0.47428 | 0.46111 |
| ENSG00000239839 | 8 | 6873391 | 6875823 | 7 | 3 | -0.014544 | 0.5058 | DEFA3 | 0.5058 | 0.4891 | 0.62612 |
| ENSG00000163866 | 1 | 35178338 | 35325417 | 610 | 72 | -0.014559 | 0.50581 | SMIM12 | 0.50581 | 0.59177 | 0.79884 |
| ENSG00000094880 | 5 | 137523339 | 137549032 | 50 | 11 | -0.014672 | 0.50585 | CDC23 | 0.50585 | 0.56146 | 0.55272 |
| ENSG00000115474 | 2 | 233631174 | 233641278 | 31 | 7 | -0.014977 | 0.50597 | KCNJ13 | 0.50597 | 0.48642 | 0.4334 |
| ENSG00000143207 | 1 | 175913967 | 176176629 | 818 | 28 | -0.01517 | 0.50605 | RFWD2 | 0.50605 | 0.51344 | 0.54717 |
| ENSG00000082175 | 11 | 100900355 | 101001255 | 324 | 21 | -0.015348 | 0.50612 | PGR | 0.50612 | 0.60125 | 0.67405 |
| ENSG00000142949 | 1 | 43990858 | 44089343 | 285 | 17 | -0.015551 | 0.5062 | PTPRF | 0.5062 | 0.50771 | 0.52435 |
| ENSG00000169507 | 2 | 165752696 | 165812035 | 209 | 22 | -0.015573 | 0.50621 | SLC38A11 | 0.50621 | 0.43813 | 0.49946 |
| ENSG00000164983 | 8 | 125324231 | 125384933 | 196 | 22 | -0.015819 | 0.50631 | TMEM65 | 0.50631 | 0.49416 | 0.40228 |
| ENSG00000123933 | 4 | 2249159 | 2264021 | 43 | 6 | -0.016068 | 0.50641 | MXD4 | 0.50641 | 0.46353 | 0.46928 |

| | | | | | | | | | | | |
|---|---|---|---|---|---|---|---|---|---|---|---|
| ENSG00000184708 | 22 | 31832963 | 31892094 | 199 | 31 | -0.016202 | 0.50646 | EIF4ENIF1 | 0.50646 | 0.45199 | 0.45462 |
| ENSG00000175318 | 15 | 72452148 | 72490126 | 54 | 10 | -0.016264 | 0.50649 | GRAMD2 | 0.50649 | 0.51781 | 0.51377 |
| ENSG00000108469 | 17 | 73622925 | 73663269 | 74 | 13 | -0.016279 | 0.50649 | RECQL5 | 0.50649 | 0.48402 | 0.62962 |
| ENSG00000136111 | 13 | 75858808 | 76056250 | 605 | 70 | -0.016352 | 0.50652 | TBC1D4 | 0.50652 | 0.42989 | 0.34501 |
| ENSG00000091651 | 16 | 46723555 | 46732306 | 7 | 3 | -0.016333 | 0.50652 | ORC6 | 0.50652 | 0.67182 | 0.68281 |
| ENSG00000183850 | 19 | 23258012 | 23330021 | 199 | 18 | -0.016384 | 0.50654 | ZNF730 | 0.50654 | 0.46116 | 0.47769 |
| ENSG00000135018 | 9 | 86274878 | 86323118 | 164 | 16 | -0.016475 | 0.50657 | UBQLN1 | 0.50657 | 0.51741 | 0.56359 |
| ENSG00000180525 | 10 | 695888 | 711109 | 11 | 7 | -0.016462 | 0.50657 | PRR26 | 0.50657 | 0.61812 | 0.74579 |
| ENSG00000103994 | 15 | 42705021 | 42783321 | 174 | 19 | -0.016506 | 0.50658 | ZNF106 | 0.50658 | 0.52833 | 0.45472 |
| ENSG00000197763 | 3 | 126290622 | 126373998 | 235 | 26 | -0.016537 | 0.5066 | TXNRD3 | 0.5066 | 0.47677 | 0.54541 |
| ENSG00000163510 | 2 | 180809603 | 180871840 | 194 | 16 | -0.016559 | 0.50661 | CWC22 | 0.50661 | 0.50959 | 0.52372 |
| ENSG00000165507 | 10 | 45466429 | 45474258 | 29 | 5 | -0.016616 | 0.50663 | C10orf10 | 0.50663 | 0.44554 | 0.65224 |
| ENSG00000088812 | 20 | 3451687 | 3631769 | 580 | 24 | -0.016669 | 0.50665 | ATRN | 0.50665 | 0.54236 | 0.48635 |
| ENSG00000143845 | 1 | 204100190 | 204121307 | 60 | 19 | -0.016691 | 0.50666 | ETNK2 | 0.50666 | 0.72162 | 0.84816 |
| ENSG00000185899 | 7 | 143140546 | 143141502 | 3 | 1 | -0.0168 | 0.5067 | TAS2R60 | 0.5067 | 0.87555 | 0.83319 |
| ENSG00000137440 | 4 | 15937192 | 15940363 | 10 | 4 | -0.016937 | 0.50676 | FGFBP1 | 0.50676 | 0.56969 | 0.38638 |
| ENSG00000223601 | 10 | 22497743 | 22498950 | 5 | 2 | -0.017225 | 0.50687 | EBLN1 | 0.50687 | 0.64661 | 0.72263 |
| ENSG00000128596 | 7 | 128430811 | 128462186 | 70 | 7 | -0.017255 | 0.50688 | CCDC136 | 0.50688 | 0.51827 | 0.59442 |
| ENSG00000128714 | 2 | 176957619 | 176960666 | 4 | 2 | -0.017423 | 0.50695 | HOXD13 | 0.50695 | 0.21081 | 0.30362 |
| ENSG00000125952 | 14 | 65472892 | 65569413 | 241 | 31 | -0.017516 | 0.50699 | MAX | 0.50699 | 0.49786 | 0.4777 |
| ENSG00000158552 | 2 | 220059959 | 220074373 | 33 | 14 | -0.017667 | 0.50705 | ZFAND2B | 0.50705 | 0.13871 | 0.2969 |
| ENSG00000127220 | 19 | 17402940 | 17421045 | 78 | 13 | -0.017736 | 0.50708 | ABHD8 | 0.50708 | 0.31388 | 0.34079 |
| ENSG00000184661 | 8 | 25316513 | 25365436 | 146 | 10 | -0.017869 | 0.50713 | CDCA2 | 0.50713 | 0.50315 | 0.49971 |
| ENSG00000175426 | 5 | 95726119 | 95769847 | 87 | 12 | -0.017912 | 0.50715 | PCSK1 | 0.50715 | 0.57402 | 0.47599 |
| ENSG00000011275 | 7 | 5659678 | 5821370 | 647 | 27 | -0.017941 | 0.50716 | RNF216 | 0.50716 | 0.49502 | 0.43732 |
| ENSG00000138119 | 10 | 95066186 | 95242074 | 872 | 60 | -0.018014 | 0.50719 | MYOF | 0.50719 | 0.43265 | 0.52411 |
| ENSG00000121903 | 1 | 33938246 | 33962107 | 62 | 6 | -0.018065 | 0.50721 | ZSCAN20 | 0.50721 | 0.5876 | 0.66702 |
| ENSG00000149503 | 11 | 61891445 | 61920635 | 143 | 12 | -0.018067 | 0.50721 | INCENP | 0.50721 | 0.57715 | 0.38092 |
| ENSG00000139223 | 12 | 48866448 | 48866843 | 2 | 1 | -0.018118 | 0.50723 | ANP32D | 0.50723 | 0.78449 | 0.8214 |
| ENSG00000136243 | 7 | 23221446 | 23240630 | 59 | 4 | -0.018299 | 0.5073 | NUPL2 | 0.5073 | 0.41 | 0.46441 |
| ENSG00000062716 | 17 | 57784553 | 57919616 | 275 | 18 | -0.01839 | 0.50734 | VMP1 | 0.50734 | 0.58418 | 0.61079 |
| ENSG00000172602 | 12 | 49250928 | 49259681 | 21 | 7 | -0.018602 | 0.50742 | RND1 | 0.50742 | 0.10373 | 0.074443 |
| ENSG00000128585 | 7 | 130794855 | 131181395 | 1176 | 100 | -0.018774 | 0.50749 | MKLN1 | 0.50749 | 0.52751 | 0.54221 |
| ENSG00000183309 | 8 | 144718183 | 144735900 | 70 | 12 | -0.018827 | 0.50751 | ZNF623 | 0.50751 | 0.69942 | 0.81419 |
| ENSG00000250091 | 12 | 124410971 | 124419531 | 14 | 3 | -0.019019 | 0.50759 | DNAH10OS | 0.50759 | 0.65406 | 0.64157 |
| ENSG00000151287 | 13 | 103418340 | 103426161 | 20 | 7 | -0.019089 | 0.50761 | TEX30 | 0.50761 | 0.51916 | 0.61133 |
| ENSG00000124635 | 6 | 27093676 | 27100541 | 19 | 8 | -0.019242 | 0.50768 | HIST1H2BJ | 0.50768 | 0.38633 | 0.27655 |
| ENSG00000149499 | 11 | 62369690 | 62380237 | 20 | 4 | -0.019247 | 0.50768 | EML3 | 0.50768 | 0.48294 | 0.40121 |
| ENSG00000166147 | 15 | 48700503 | 48938046 | 530 | 18 | -0.019365 | 0.50773 | FBN1 | 0.50773 | 0.48522 | 0.47798 |
| ENSG00000063180 | 19 | 49141199 | 49149569 | 19 | 4 | -0.019422 | 0.50775 | CA11 | 0.50775 | 0.16922 | 0.13209 |
| ENSG00000203852 | 1 | 149824181 | 149825836 | 1 | 1 | -0.019553 | 0.5078 | HIST2H3A | 0.5078 | 0.67549 | 0.64683 |
| ENSG00000216490 | 19 | 18283972 | 18288927 | 11 | 2 | -0.019549 | 0.5078 | IFI30 | 0.5078 | 0.33954 | 0.37613 |
| ENSG00000179918 | 16 | 30454952 | 30457502 | 3 | 2 | -0.01964 | 0.50783 | SEPHS2 | 0.50783 | 0.63681 | 0.58845 |
| ENSG00000117054 | 1 | 76190036 | 76253260 | 308 | 18 | -0.019685 | 0.50785 | ACADM | 0.50785 | 0.51934 | 0.55948 |
| ENSG00000141232 | 17 | 48939584 | 48945339 | 7 | 3 | -0.019668 | 0.50785 | TOB1 | 0.50785 | 0.56177 | 0.39666 |
| ENSG00000136143 | 13 | 48510622 | 48612125 | 489 | 21 | -0.019755 | 0.50788 | SUCLA2 | 0.50788 | 0.53018 | 0.55 |
| ENSG00000268192 | 11 | 119027590 | 119027930 | 1 | 1 | -0.019804 | 0.5079 | AP002956.1 | 0.5079 | 0.57711 | 0.47105 |
| ENSG00000121905 | 1 | 33351595 | 33364042 | 20 | 7 | -0.019973 | 0.50797 | HPCA | 0.50797 | 0.5971 | 0.52883 |
| ENSG00000142619 | 1 | 17575593 | 17610728 | 188 | 29 | -0.020013 | 0.50798 | PADI3 | 0.50798 | 0.56062 | 0.43215 |
| ENSG00000111537 | 12 | 68548548 | 68553527 | 5 | 2 | -0.020742 | 0.50827 | IFNG | 0.50827 | 0.069013 | 0.0096155 |
| ENSG00000102910 | 16 | 48278207 | 48397033 | 166 | 13 | -0.020993 | 0.50837 | LONP2 | 0.50837 | 0.48235 | 0.43496 |
| ENSG00000078140 | 4 | 39699664 | 39784412 | 304 | 16 | -0.021054 | 0.5084 | UBE2K | 0.5084 | 0.54335 | 0.49826 |
| ENSG00000198010 | 8 | 1449532 | 1656642 | 1104 | 54 | -0.02106 | 0.5084 | DLGAP2 | 0.5084 | 0.40747 | 0.39271 |
| ENSG00000112852 | 5 | 140474227 | 140476962 | 8 | 2 | -0.021092 | 0.50841 | PCDHB2 | 0.50841 | 0.70212 | 0.75625 |
| ENSG00000137492 | 11 | 76061000 | 76092015 | 78 | 17 | -0.021111 | 0.50842 | PRKRIR | 0.50842 | 0.57383 | 0.50868 |
| ENSG00000182901 | 1 | 240931554 | 241520530 | 2306 | 168 | -0.021254 | 0.50848 | RGS7 | 0.50848 | 0.61476 | 0.51852 |
| ENSG00000160593 | 11 | 118064455 | 118095809 | 95 | 9 | -0.021462 | 0.50856 | AMICA1 | 0.50856 | 0.57019 | 0.66126 |
| ENSG00000100714 | 14 | 64854749 | 64926722 | 223 | 31 | -0.021853 | 0.50872 | MTHFD1 | 0.50872 | 0.52618 | 0.45597 |
| ENSG00000112146 | 6 | 52916789 | 52965671 | 97 | 15 | -0.021998 | 0.50878 | FBXO9 | 0.50878 | 0.46785 | 0.38566 |
| ENSG00000132323 | 2 | 239079042 | 239112370 | 164 | 18 | -0.022697 | 0.50905 | ILKAP | 0.50905 | 0.51891 | 0.45453 |
| ENSG00000103404 | 16 | 23072727 | 23160591 | 260 | 13 | -0.022702 | 0.50906 | USP31 | 0.50906 | 0.50664 | 0.45685 |
| ENSG00000167614 | 19 | 54926393 | 54948080 | 100 | 22 | -0.022793 | 0.50909 | TTYH1 | 0.50909 | 0.59735 | 0.62702 |
| ENSG00000168137 | 3 | 9439299 | 9520924 | 183 | 17 | -0.022844 | 0.50911 | SETD5 | 0.50911 | 0.51128 | 0.45605 |
| ENSG00000184005 | 1 | 76540404 | 77100286 | 1973 | 150 | -0.022931 | 0.50915 | ST6GALNAC3 | 0.50915 | 0.26697 | 0.2231 |
| ENSG00000157703 | 7 | 138279030 | 138386097 | 531 | 72 | -0.022946 | 0.50915 | SVOPL | 0.50915 | 0.58523 | 0.41615 |
| ENSG00000135919 | 2 | 224839829 | 224904036 | 297 | 30 | -0.022954 | 0.50916 | SERPINE2 | 0.50916 | 0.57337 | 0.6472 |
| ENSG00000110318 | 11 | 101785746 | 101871789 | 171 | 14 | -0.023215 | 0.50926 | KIAA1377 | 0.50926 | 0.51299 | 0.62549 |
| ENSG00000092020 | 14 | 35554673 | 35591723 | 167 | 21 | -0.023274 | 0.50928 | PPP2R3C | 0.50928 | 0.53161 | 0.62761 |
| ENSG00000089692 | 12 | 6881678 | 6887621 | 11 | 6 | -0.023316 | 0.5093 | LAG3 | 0.5093 | 0.5963 | 0.85531 |
| ENSG00000184076 | 22 | 30163358 | 30166402 | 9 | 3 | -0.023339 | 0.50931 | UQCR10 | 0.50931 | 0.71492 | 0.75348 |
| ENSG00000101096 | 20 | 50003494 | 50179370 | 601 | 78 | -0.02337 | 0.50932 | NFATC2 | 0.50932 | 0.32976 | 0.26784 |
| ENSG00000107738 | 10 | 73507316 | 73533255 | 117 | 30 | -0.023915 | 0.50954 | C10orf54 | 0.50954 | 0.68866 | 0.7901 |
| ENSG00000175595 | 16 | 14014014 | 14046202 | 90 | 9 | -0.024016 | 0.50958 | ERCC4 | 0.50958 | 0.47268 | 0.34551 |

| | | | | | | | | | | | |
|---|---|---|---|---|---|---|---|---|---|---|---|
| ENSG00000137216 | 6 | 44094651 | 44123256 | 131 | 20 | -0.024046 | 0.50959 | TMEM63B | 0.50959 | 0.5852 | 0.57471 |
| ENSG00000169067 | 5 | 56775846 | 56778636 | 4 | 2 | -0.024227 | 0.50966 | ACTBL2 | 0.50966 | 0.38339 | 0.098741 |
| ENSG00000106125 | 7 | 30811033 | 30932002 | 495 | 35 | -0.02445 | 0.50975 | FAM188B | 0.50975 | 0.44496 | 0.38625 |
| ENSG00000243896 | 7 | 143955700 | 143956815 | 3 | 1 | -0.024503 | 0.50977 | OR2A7 | 0.50977 | 0.21365 | 0.27519 |
| ENSG00000165511 | 10 | 45493146 | 45496336 | 5 | 2 | -0.024542 | 0.50979 | C10orf25 | 0.50979 | 0.35948 | 0.41872 |
| ENSG00000104946 | 19 | 50380682 | 50392005 | 25 | 6 | -0.024611 | 0.50982 | TBC1D17 | 0.50982 | 0.34306 | 0.43041 |
| ENSG00000196700 | 20 | 62588055 | 62680113 | 252 | 18 | -0.024717 | 0.50986 | ZNF512B | 0.50986 | 0.46327 | 0.46445 |
| ENSG00000151632 | 10 | 5029967 | 5060223 | 226 | 28 | -0.024923 | 0.50994 | AKR1C2 | 0.50994 | 0.60297 | 0.6911 |
| ENSG00000132664 | 20 | 18447771 | 18465287 | 58 | 11 | -0.025155 | 0.51003 | POLR3F | 0.51003 | 0.67601 | 0.81173 |
| ENSG00000110200 | 11 | 71817424 | 71823826 | 12 | 3 | -0.025203 | 0.51005 | ANAPC15 | 0.51005 | 0.69542 | 0.80552 |
| ENSG00000178971 | 17 | 8130191 | 8151362 | 73 | 14 | -0.025292 | 0.51009 | CTC1 | 0.51009 | 0.56818 | 0.55879 |
| ENSG00000181744 | 3 | 143690640 | 143767561 | 266 | 17 | -0.02535 | 0.51011 | C3orf58 | 0.51011 | 0.41132 | 0.41004 |
| ENSG00000116981 | 1 | 40124793 | 40137710 | 26 | 9 | -0.025379 | 0.51012 | NT5C1A | 0.51012 | 0.3366 | 0.26885 |
| ENSG00000162687 | 1 | 196194909 | 196578355 | 782 | 22 | -0.025567 | 0.5102 | KCNT2 | 0.5102 | 0.50693 | 0.48608 |
| ENSG00000185864 | 16 | 21845890 | 21892148 | 3 | 2 | -0.025632 | 0.51022 | NPIPB4 | 0.51022 | 0.31454 | 0.53648 |
| ENSG00000141012 | 16 | 88880142 | 88923378 | 331 | 24 | -0.025715 | 0.51026 | GALNS | 0.51026 | 0.39491 | 0.48858 |
| ENSG00000185670 | 11 | 62515791 | 62521660 | 10 | 4 | -0.02588 | 0.51032 | ZBTB3 | 0.51032 | 0.57107 | 0.50345 |
| ENSG00000119042 | 2 | 200134223 | 200335989 | 298 | 20 | -0.02636 | 0.51051 | SATB2 | 0.51051 | 0.53468 | 0.63425 |
| ENSG00000100285 | 22 | 29876219 | 29887379 | 39 | 10 | -0.026434 | 0.51054 | NEFH | 0.51054 | 0.38225 | 0.24363 |
| ENSG00000184389 | 1 | 33772367 | 33786699 | 50 | 10 | -0.026578 | 0.5106 | A3GALT2 | 0.5106 | 0.4066 | 0.30086 |
| ENSG00000268000 | 19 | 50003781 | 50004614 | 2 | 1 | -0.026591 | 0.51061 | hsa-mir-150 | 0.51061 | 0.60372 | 0.55077 |
| ENSG00000122033 | 13 | 28009776 | 28024739 | 83 | 11 | -0.026885 | 0.51072 | MTIF3 | 0.51072 | 0.50005 | 0.47891 |
| ENSG00000136367 | 14 | 23990066 | 24025401 | 60 | 17 | -0.027027 | 0.51078 | ZFHX2 | 0.51078 | 0.77113 | 0.71536 |
| ENSG00000170782 | 11 | 6897856 | 6898850 | 3 | 2 | -0.027158 | 0.51083 | OR10A4 | 0.51083 | 0.8431 | 0.24192 |
| ENSG00000053770 | 14 | 57735627 | 57756797 | 21 | 10 | -0.027234 | 0.51086 | AP5M1 | 0.51086 | 0.5407 | 0.47789 |
| ENSG00000168502 | 18 | 8705659 | 8832776 | 502 | 64 | -0.027263 | 0.51088 | SOGA2 | 0.51088 | 0.57122 | 0.63093 |
| ENSG00000203857 | 1 | 120049821 | 120057681 | 9 | 3 | -0.027351 | 0.51091 | HSD3B1 | 0.51091 | 0.70709 | 0.76836 |
| ENSG00000129282 | 17 | 34958001 | 34965407 | 25 | 6 | -0.027511 | 0.51097 | MRM1 | 0.51097 | 0.46901 | 0.53685 |
| ENSG00000163263 | 1 | 154171848 | 154178809 | 5 | 3 | -0.027777 | 0.51108 | C1orf189 | 0.51108 | 0.41292 | 0.38282 |
| ENSG00000121988 | 2 | 135894486 | 136288806 | 792 | 15 | -0.027787 | 0.51108 | ZRANB3 | 0.51108 | 0.51699 | 0.50355 |
| ENSG00000245888 | 16 | 27279526 | 27301813 | 53 | 11 | -0.027784 | 0.51108 | CTD-3203P2.2 | 0.51108 | 0.34543 | 0.35033 |
| ENSG00000115107 | 2 | 119981384 | 120023228 | 188 | 27 | -0.027987 | 0.51116 | STEAP3 | 0.51116 | 0.55969 | 0.66799 |
| ENSG00000152433 | 19 | 57874845 | 57890933 | 48 | 11 | -0.028059 | 0.51119 | ZNF547 | 0.51119 | 0.52565 | 0.54332 |
| ENSG00000198099 | 4 | 100044808 | 100078949 | 239 | 10 | -0.028233 | 0.51126 | ADH4 | 0.51126 | 0.55229 | 0.57188 |
| ENSG00000162771 | 1 | 212797789 | 212800120 | 10 | 5 | -0.028286 | 0.51128 | FAM71A | 0.51128 | 0.42864 | 0.35094 |
| ENSG00000228144 | 12 | 66517697 | 66563765 | 149 | 16 | -0.028307 | 0.51129 | TMBIM4 | 0.51129 | 0.54844 | 0.56053 |
| ENSG00000173575 | 15 | 93426526 | 93571237 | 409 | 34 | -0.028574 | 0.5114 | CHD2 | 0.5114 | 0.54782 | 0.70088 |
| ENSG00000165799 | 14 | 21510385 | 21512393 | 6 | 3 | -0.028787 | 0.51148 | RNASE7 | 0.51148 | 0.60147 | 0.93489 |
| ENSG00000128185 | 22 | 20301799 | 20307603 | 15 | 3 | -0.028772 | 0.51148 | DGCR6L | 0.51148 | 0.66677 | 0.62615 |
| ENSG00000081320 | 2 | 196998290 | 197041227 | 168 | 12 | -0.028814 | 0.51149 | STK17B | 0.51149 | 0.50173 | 0.49485 |
| ENSG00000164120 | 4 | 175411328 | 175444305 | 132 | 25 | -0.028806 | 0.51149 | HPGD | 0.51149 | 0.3404 | 0.09718 |
| ENSG00000103047 | 16 | 68877507 | 69119083 | 595 | 30 | -0.029177 | 0.51164 | TANGO6 | 0.51164 | 0.50238 | 0.41652 |
| ENSG00000136297 | 7 | 4945620 | 4998844 | 238 | 27 | -0.029289 | 0.51168 | MMD2 | 0.51168 | 0.69429 | 0.77391 |
| ENSG00000185324 | 16 | 89747145 | 89762772 | 76 | 13 | -0.029317 | 0.51169 | CDK10 | 0.51169 | 0.62285 | 0.57 |
| ENSG00000173744 | 2 | 228336868 | 228425930 | 363 | 24 | -0.029436 | 0.51174 | AGFG1 | 0.51174 | 0.52852 | 0.48956 |
| ENSG00000135976 | 2 | 97779233 | 97930258 | 280 | 16 | -0.029534 | 0.51178 | ANKRD36 | 0.51178 | 0.50591 | 0.52173 |
| ENSG00000156873 | 16 | 30759591 | 30772490 | 20 | 10 | -0.029775 | 0.51188 | PHKG2 | 0.51188 | 0.66523 | 0.80323 |
| ENSG00000004777 | 19 | 36265434 | 36279724 | 42 | 7 | -0.029777 | 0.51188 | ARHGAP33 | 0.51188 | 0.57656 | 0.67715 |
| ENSG00000062524 | 15 | 41795836 | 41806085 | 32 | 8 | -0.029865 | 0.51191 | LTK | 0.51191 | 0.62078 | 0.76326 |
| ENSG00000172061 | 3 | 194075976 | 194090472 | 116 | 9 | -0.029884 | 0.51192 | LRRC15 | 0.51192 | 0.69063 | 0.6651 |
| ENSG00000197119 | 14 | 100757448 | 100772884 | 40 | 8 | -0.02991 | 0.51193 | SLC25A29 | 0.51193 | 0.51972 | 0.33469 |
| ENSG00000150457 | 13 | 21547171 | 21635686 | 334 | 24 | -0.03002 | 0.51197 | LATS2 | 0.51197 | 0.55241 | 0.6493 |
| ENSG00000044090 | 6 | 43005355 | 43021683 | 35 | 10 | -0.030046 | 0.51198 | CUL7 | 0.51198 | 0.47559 | 0.4839 |
| ENSG00000157330 | 1 | 12806163 | 12823847 | 65 | 11 | -0.030265 | 0.51207 | C1orf158 | 0.51207 | 0.26085 | 0.26439 |
| ENSG00000196419 | 22 | 42017123 | 42060044 | 93 | 12 | -0.030703 | 0.51225 | XRCC6 | 0.51225 | 0.61065 | 0.65657 |
| ENSG00000196099 | 11 | 123676043 | 123677095 | 3 | 2 | -0.030782 | 0.51228 | OR6M1 | 0.51228 | 0.024757 | 0.11128 |
| ENSG00000184478 | 11 | 5968570 | 5969591 | 2 | 1 | -0.031001 | 0.51237 | OR56A3 | 0.51237 | 0.69729 | 0.53394 |
| ENSG00000174370 | 11 | 128769460 | 128775930 | 27 | 9 | -0.031011 | 0.51237 | C11orf45 | 0.51237 | 0.57215 | 0.53998 |
| ENSG00000259243 | 15 | 83011379 | 83018198 | 3 | 2 | -0.031153 | 0.51243 | GOLGA6L19 | 0.51243 | 0.80679 | 0.43605 |
| ENSG00000182704 | 11 | 76493295 | 76509198 | 45 | 7 | -0.031179 | 0.51244 | TSKU | 0.51244 | 0.63284 | 0.76838 |
| ENSG00000204049 | 10 | 79626633 | 79629368 | 5 | 2 | -0.031453 | 0.51255 | AL391421.1 | 0.51255 | 0.56188 | 0.66115 |
| ENSG00000196860 | 14 | 58862634 | 58875419 | 12 | 7 | -0.031496 | 0.51256 | TOMM20L | 0.51256 | 0.48774 | 0.3518 |
| ENSG00000168802 | 16 | 69151913 | 69166487 | 33 | 11 | -0.031586 | 0.5126 | CHTF8 | 0.5126 | 0.34012 | 0.26359 |
| ENSG00000189306 | 22 | 42905974 | 42915808 | 52 | 12 | -0.031596 | 0.5126 | RRP7A | 0.5126 | 0.37162 | 0.3972 |
| ENSG00000187624 | 17 | 260118 | 273510 | 52 | 10 | -0.031681 | 0.51264 | C17orf97 | 0.51264 | 0.42711 | 0.64469 |
| ENSG00000156966 | 2 | 232260254 | 232265875 | 15 | 6 | -0.031828 | 0.5127 | B3GNT7 | 0.5127 | 0.67399 | 0.76806 |
| ENSG00000163749 | 4 | 77234154 | 77343021 | 229 | 23 | -0.031915 | 0.51273 | CCDC158 | 0.51273 | 0.47868 | 0.50866 |
| ENSG00000183098 | 13 | 93879095 | 95059655 | 3890 | 191 | -0.031987 | 0.51276 | GPC6 | 0.51276 | 0.51126 | 0.58211 |
| ENSG00000141150 | 17 | 34058668 | 34070540 | 28 | 8 | -0.03206 | 0.51279 | RASL10B | 0.51279 | 0.72312 | 0.76656 |
| ENSG00000136158 | 13 | 80910111 | 80915086 | 6 | 2 | -0.03225 | 0.51286 | SPRY2 | 0.51286 | 0.034234 | 0.072055 |
| ENSG00000198682 | 10 | 89419370 | 89507462 | 293 | 29 | -0.032345 | 0.5129 | PAPSS2 | 0.5129 | 0.6371 | 0.69071 |
| ENSG00000167100 | 17 | 48187404 | 48207246 | 53 | 17 | -0.032639 | 0.51302 | SAMD14 | 0.51302 | 0.4902 | 0.54289 |
| ENSG00000123989 | 2 | 220403669 | 220408509 | 12 | 3 | -0.032688 | 0.51304 | CHPF | 0.51304 | 0.64196 | 0.6649 |

| | | | | | | | | | | | |
|---|---|---|---|---|---|---|---|---|---|---|---|
| ENSG00000213931 | 11 | 5289582 | 5526847 | 1873 | 53 | -0.032886 | 0.51312 | HBE1 | 0.51312 | 0.52844 | 0.56112 |
| ENSG00000044524 | 3 | 89156674 | 89531284 | 851 | 32 | -0.033119 | 0.51321 | EPHA3 | 0.51321 | 0.4987 | 0.48198 |
| ENSG00000100991 | 20 | 33590207 | 33680674 | 319 | 15 | -0.033296 | 0.51328 | TRPC4AP | 0.51328 | 0.51605 | 0.55114 |
| ENSG00000105792 | 7 | 89874488 | 89940377 | 205 | 23 | -0.033374 | 0.51331 | C7orf63 | 0.51331 | 0.521 | 0.5013 |
| ENSG00000163823 | 3 | 46243200 | 46249887 | 13 | 2 | -0.033735 | 0.51346 | CCR1 | 0.51346 | 0.58975 | 0.572 |
| ENSG00000100124 | 22 | 38226862 | 38245334 | 20 | 7 | -0.033785 | 0.51348 | ANKRD54 | 0.51348 | 0.25349 | 0.20098 |
| ENSG00000159398 | 16 | 55880066 | 55989943 | 366 | 38 | -0.034036 | 0.51358 | CES5A | 0.51358 | 0.56846 | 0.50511 |
| ENSG00000147573 | 8 | 67039131 | 67087720 | 108 | 12 | -0.034195 | 0.51364 | TRIM55 | 0.51364 | 0.32813 | 0.18233 |
| ENSG00000141655 | 18 | 59992520 | 60058516 | 283 | 39 | -0.034433 | 0.51373 | TNFRSF11A | 0.51373 | 0.5442 | 0.57013 |
| ENSG00000113621 | 5 | 134209493 | 134237215 | 37 | 10 | -0.034662 | 0.51383 | TXNDC15 | 0.51383 | 0.50973 | 0.51437 |
| ENSG00000181991 | 15 | 89010684 | 89022222 | 11 | 5 | -0.034921 | 0.51393 | MRPS11 | 0.51393 | 0.42964 | 0.50296 |
| ENSG00000109180 | 4 | 48807229 | 48863834 | 79 | 12 | -0.034955 | 0.51394 | OCIAD1 | 0.51394 | 0.53897 | 0.48223 |
| ENSG00000173542 | 4 | 71768043 | 71888166 | 230 | 23 | -0.034969 | 0.51395 | MOB1B | 0.51395 | 0.48207 | 0.46594 |
| ENSG00000131981 | 14 | 55590828 | 55612126 | 71 | 9 | -0.03519 | 0.51404 | LGALS3 | 0.51404 | 0.58009 | 0.55868 |
| ENSG00000167094 | 9 | 130478345 | 130493879 | 34 | 4 | -0.035284 | 0.51407 | TTC16 | 0.51407 | 0.44997 | 0.53536 |
| ENSG00000101216 | 20 | 62218955 | 62258394 | 207 | 22 | -0.035264 | 0.51407 | GMEB2 | 0.51407 | 0.52916 | 0.25474 |
| ENSG00000173273 | 8 | 9413424 | 9639856 | 773 | 14 | -0.035351 | 0.5141 | TNKS | 0.5141 | 0.52237 | 0.53409 |
| ENSG00000144120 | 2 | 120436743 | 120444083 | 44 | 7 | -0.035387 | 0.51411 | TMEM177 | 0.51411 | 0.6598 | 0.68969 |
| ENSG00000163738 | 4 | 74979891 | 75168816 | 343 | 26 | -0.035474 | 0.51415 | MTHFD2L | 0.51415 | 0.56318 | 0.59685 |
| ENSG00000127720 | 12 | 82752276 | 82873015 | 313 | 11 | -0.035519 | 0.51417 | METTL25 | 0.51417 | 0.53306 | 0.53947 |
| ENSG00000273047 | 20 | 62369623 | 62371751 | 2 | 1 | -0.036193 | 0.51444 | RP4-583P15.14 | 0.51444 | 0.2209 | 0.26536 |
| ENSG00000184983 | 22 | 42481529 | 42486959 | 12 | 3 | -0.03669 | 0.51463 | NDUFA6 | 0.51463 | 0.75776 | 0.78124 |
| ENSG00000064932 | 19 | 1107636 | 1174282 | 283 | 33 | -0.036867 | 0.5147 | SBNO2 | 0.5147 | 0.63464 | 0.75216 |
| ENSG00000243509 | 20 | 62328021 | 62330037 | 10 | 2 | -0.036892 | 0.51471 | TNFRSF6B | 0.51471 | 0.26431 | 0.20183 |
| ENSG00000173402 | 3 | 49506146 | 49573048 | 149 | 16 | -0.0369 | 0.51472 | DAG1 | 0.51472 | 0.54999 | 0.56011 |
| ENSG00000188242 | 5 | 466239 | 473213 | 40 | 8 | -0.037186 | 0.51483 | CTD-2228K2.5 | 0.51483 | 0.46686 | 0.24526 |
| ENSG00000187514 | 2 | 232571605 | 232578251 | 33 | 8 | -0.037259 | 0.51486 | PTMA | 0.51486 | 0.52309 | 0.43671 |
| ENSG00000185758 | 4 | 184242917 | 184243579 | 2 | 1 | -0.03727 | 0.51487 | CLDN24 | 0.51487 | 0.84845 | 0.83515 |
| ENSG00000143994 | 2 | 27346682 | 27353680 | 13 | 4 | -0.037366 | 0.5149 | ABHD1 | 0.5149 | 0.83016 | 0.77392 |
| ENSG00000086475 | 10 | 13359424 | 13390297 | 137 | 15 | -0.037467 | 0.51494 | SEPHS1 | 0.51494 | 0.42969 | 0.55969 |
| ENSG00000162643 | 1 | 85464830 | 85598821 | 463 | 42 | -0.037675 | 0.51503 | WDR63 | 0.51503 | 0.44315 | 0.17321 |
| ENSG00000055483 | 17 | 76783463 | 76837523 | 136 | 18 | -0.037864 | 0.5151 | USP36 | 0.5151 | 0.50197 | 0.49738 |
| ENSG00000159596 | 1 | 46152886 | 46160115 | 11 | 4 | -0.03791 | 0.51512 | TMEM69 | 0.51512 | 0.3974 | 0.32371 |
| ENSG00000178921 | 17 | 8150936 | 8173809 | 85 | 16 | -0.037993 | 0.51515 | PFAS | 0.51515 | 0.50793 | 0.43545 |
| ENSG00000206561 | 3 | 15491640 | 15563258 | 203 | 25 | -0.038155 | 0.51522 | COLQ | 0.51522 | 0.53908 | 0.56361 |
| ENSG00000120051 | 10 | 106113522 | 106214848 | 219 | 33 | -0.038372 | 0.5153 | CCDC147 | 0.5153 | 0.53726 | 0.46516 |
| ENSG00000136826 | 9 | 110247133 | 110252763 | 16 | 2 | -0.038466 | 0.51534 | KLF4 | 0.51534 | 0.49717 | 0.14248 |
| ENSG00000155957 | 12 | 66517709 | 66563852 | 150 | 16 | -0.038541 | 0.51537 | TMBIM4 | 0.51537 | 0.54844 | 0.56053 |
| ENSG00000089902 | 14 | 103058998 | 103196913 | 408 | 22 | -0.038642 | 0.51541 | RCOR1 | 0.51541 | 0.4541 | 0.43115 |
| ENSG00000206053 | 16 | 1728257 | 1752281 | 44 | 9 | -0.038682 | 0.51543 | HN1L | 0.51543 | 0.61642 | 0.72591 |
| ENSG00000180708 | 1 | 158389718 | 158390656 | 2 | 1 | -0.038715 | 0.51544 | OR10K2 | 0.51544 | 0.21425 | 0.26419 |
| ENSG00000185610 | 12 | 45408455 | 45444882 | 139 | 16 | -0.038955 | 0.51554 | DBX2 | 0.51554 | 0.57662 | 0.61069 |
| ENSG00000196734 | 1 | 152784447 | 152785585 | 1 | 1 | -0.039364 | 0.5157 | LCE1B | 0.5157 | 0.24892 | 0.11363 |
| ENSG00000156256 | 21 | 30396950 | 30426809 | 46 | 13 | -0.03937 | 0.5157 | USP16 | 0.5157 | 0.55289 | 0.54759 |
| ENSG00000096093 | 6 | 52285106 | 52387892 | 375 | 36 | -0.039696 | 0.51583 | EFHC1 | 0.51583 | 0.57379 | 0.57475 |
| ENSG00000011465 | 12 | 91539025 | 91576900 | 60 | 10 | -0.039972 | 0.51594 | DCN | 0.51594 | 0.32011 | 0.18675 |
| ENSG00000139734 | 13 | 60239717 | 60738121 | 1430 | 35 | -0.040063 | 0.51598 | DIAPH3 | 0.51598 | 0.52238 | 0.51073 |
| ENSG00000087245 | 16 | 55423612 | 55540603 | 658 | 52 | -0.040345 | 0.51609 | MMP2 | 0.51609 | 0.58917 | 0.61927 |
| ENSG00000164270 | 5 | 147830595 | 148056798 | 717 | 42 | -0.040408 | 0.51612 | HTR4 | 0.51612 | 0.55911 | 0.61796 |
| ENSG00000165572 | 13 | 41701705 | 41706882 | 2 | 1 | -0.040444 | 0.51613 | KBTBD6 | 0.51613 | 0.68514 | 0.69066 |
| ENSG00000109107 | 17 | 26900133 | 26904282 | 4 | 2 | -0.040554 | 0.51617 | ALDOC | 0.51617 | 0.43577 | 0.074204 |
| ENSG00000167889 | 17 | 74864538 | 74946475 | 542 | 50 | -0.040651 | 0.51621 | MGAT5B | 0.51621 | 0.57613 | 0.55071 |
| ENSG00000176108 | 17 | 78965398 | 78983317 | 92 | 13 | -0.04068 | 0.51622 | CHMP6 | 0.51622 | 0.57085 | 0.49512 |
| ENSG00000185761 | 19 | 1505017 | 1513603 | 23 | 11 | -0.040669 | 0.51622 | ADAMTSL5 | 0.51622 | 0.62462 | 0.82616 |
| ENSG00000120733 | 5 | 137688285 | 137772717 | 205 | 31 | -0.04069 | 0.51623 | KDM3B | 0.51623 | 0.53683 | 0.62376 |
| ENSG00000100219 | 22 | 29190543 | 29196585 | 14 | 5 | -0.040687 | 0.51623 | XBP1 | 0.51623 | 0.57772 | 0.6279 |
| ENSG00000167778 | 12 | 53458388 | 53473204 | 30 | 9 | -0.041109 | 0.5164 | SPRYD3 | 0.5164 | 0.50272 | 0.38694 |
| ENSG00000135631 | 2 | 73300510 | 73383849 | 294 | 22 | -0.04122 | 0.51644 | RAB11FIP5 | 0.51644 | 0.5715 | 0.51086 |
| ENSG00000198915 | 10 | 43689983 | 43762367 | 304 | 24 | -0.041225 | 0.51644 | RASGEF1A | 0.51644 | 0.47097 | 0.48595 |
| ENSG00000214415 | 7 | 80087987 | 80141336 | 149 | 19 | -0.041266 | 0.51646 | GNAT3 | 0.51646 | 0.58916 | 0.69553 |
| ENSG00000107954 | 10 | 105253736 | 105352309 | 310 | 33 | -0.041663 | 0.51662 | NEURL1 | 0.51662 | 0.4897 | 0.47018 |
| ENSG00000154768 | 17 | 34087916 | 34092098 | 13 | 5 | -0.041724 | 0.51664 | C17orf50 | 0.51664 | 0.65906 | 0.73855 |
| ENSG00000110455 | 11 | 44087475 | 44105772 | 68 | 14 | -0.041736 | 0.51665 | ACCS | 0.51665 | 0.37078 | 0.33549 |
| ENSG00000108846 | 17 | 48712138 | 48769613 | 171 | 39 | -0.041821 | 0.51668 | ABCC3 | 0.51668 | 0.47339 | 0.34657 |
| ENSG00000072195 | 2 | 220299568 | 220363009 | 124 | 20 | -0.041878 | 0.5167 | SPEG | 0.5167 | 0.48337 | 0.49436 |
| ENSG00000114120 | 3 | 140660672 | 140698775 | 48 | 7 | -0.041885 | 0.5167 | SLC25A36 | 0.5167 | 0.54823 | 0.56843 |
| ENSG00000163064 | 2 | 119599747 | 119605254 | 8 | 4 | -0.04189 | 0.51671 | EN1 | 0.51671 | 0.42262 | 0.42097 |
| ENSG00000137770 | 15 | 44719432 | 44821236 | 210 | 14 | -0.041995 | 0.51675 | CTDSPL2 | 0.51675 | 0.50914 | 0.52147 |
| ENSG00000183576 | 14 | 99864083 | 99947216 | 125 | 10 | -0.04213 | 0.5168 | SETD3 | 0.5168 | 0.56828 | 0.52347 |
| ENSG00000137103 | 9 | 35814448 | 35854844 | 52 | 12 | -0.042186 | 0.51682 | TMEM8B | 0.51682 | 0.59072 | 0.73228 |
| ENSG00000183273 | 12 | 119772517 | 119978852 | 639 | 63 | -0.042321 | 0.51688 | CCDC60 | 0.51688 | 0.51642 | 0.5085 |
| ENSG00000158022 | 1 | 26377795 | 26394927 | 45 | 7 | -0.042367 | 0.5169 | TRIM63 | 0.5169 | 0.54592 | 0.62875 |
| ENSG00000144824 | 3 | 111451344 | 111695364 | 1049 | 54 | -0.042391 | 0.51691 | PHLDB2 | 0.51691 | 0.52509 | 0.55399 |

| | | | | | | | | | | | |
|---|---|---|---|---|---|---|---|---|---|---|---|
| ENSG00000075292 | 2 | 71503691 | 71662199 | 591 | 26 | -0.042522 | 0.51696 | ZNF638 | 0.51696 | 0.52486 | 0.55148 |
| ENSG00000185479 | 12 | 52840435 | 52845910 | 22 | 9 | -0.042753 | 0.51705 | KRT6B | 0.51705 | 0.22018 | 0.29612 |
| ENSG00000170190 | 17 | 73083822 | 73102257 | 107 | 16 | -0.042883 | 0.5171 | SLC16A5 | 0.5171 | 0.55031 | 0.62289 |
| ENSG00000099917 | 22 | 20850200 | 20941919 | 301 | 20 | -0.043033 | 0.51716 | MED15 | 0.51716 | 0.53285 | 0.3817 |
| ENSG00000012963 | 14 | 93673401 | 93695561 | 91 | 15 | -0.043194 | 0.51723 | UBR7 | 0.51723 | 0.54916 | 0.53544 |
| ENSG00000147799 | 8 | 145754563 | 145911194 | 320 | 24 | -0.04324 | 0.51724 | ARHGAP39 | 0.51724 | 0.54923 | 0.57705 |
| ENSG00000177409 | 7 | 92759368 | 92777682 | 39 | 11 | -0.04333 | 0.51728 | SAMD9L | 0.51728 | 0.56127 | 0.45834 |
| ENSG00000172795 | 5 | 112312399 | 112356667 | 126 | 19 | -0.043374 | 0.5173 | DCP2 | 0.5173 | 0.57629 | 0.47211 |
| ENSG00000109181 | 4 | 69681711 | 69696914 | 74 | 8 | -0.043413 | 0.51731 | UGT2B10 | 0.51731 | 0.55474 | 0.4888 |
| ENSG00000136122 | 13 | 73302061 | 73330336 | 82 | 17 | -0.043738 | 0.51744 | BORA | 0.51744 | 0.55614 | 0.48581 |
| ENSG00000214929 | 9 | 84603687 | 84610171 | 13 | 6 | -0.043901 | 0.51751 | SPATA31D1 | 0.51751 | 0.42849 | 0.5234 |
| ENSG00000115138 | 2 | 25383722 | 25391772 | 22 | 11 | -0.043981 | 0.51754 | POMC | 0.51754 | 0.79342 | 0.80034 |
| ENSG00000176261 | 1 | 33065773 | 33116504 | 151 | 10 | -0.044085 | 0.51758 | ZBTB8OS | 0.51758 | 0.55296 | 0.57378 |
| ENSG00000172156 | 17 | 32612687 | 32615353 | 11 | 6 | -0.044304 | 0.51767 | CCL11 | 0.51767 | 0.10969 | 0.2126 |
| ENSG00000166948 | 20 | 2361554 | 2413399 | 212 | 28 | -0.044429 | 0.51772 | TGM6 | 0.51772 | 0.44637 | 0.61649 |
| ENSG00000168582 | 2 | 209025464 | 209028300 | 20 | 5 | -0.044498 | 0.51775 | CRYGA | 0.51775 | 0.71908 | 0.73182 |
| ENSG00000254732 | 11 | 57509635 | 57560715 | 84 | 14 | -0.044511 | 0.51775 | RP11-691N7.6 | 0.51775 | 0.44659 | 0.45278 |
| ENSG00000132356 | 5 | 40759481 | 40798476 | 93 | 13 | -0.044649 | 0.51781 | PRKAA1 | 0.51781 | 0.42523 | 0.38897 |
| ENSG00000174373 | 14 | 36007558 | 36278510 | 451 | 19 | -0.04468 | 0.51782 | RALGAPA1 | 0.51782 | 0.50079 | 0.4889 |
| ENSG00000176142 | 3 | 119148347 | 119187677 | 112 | 14 | -0.04492 | 0.51791 | TMEM39A | 0.51791 | 0.55975 | 0.45449 |
| ENSG00000105472 | 19 | 51226586 | 51228974 | 5 | 2 | -0.044926 | 0.51792 | CLEC11A | 0.51792 | 0.84953 | 0.91652 |
| ENSG00000172296 | 20 | 12989627 | 13147411 | 804 | 47 | -0.044936 | 0.51792 | SPTLC3 | 0.51792 | 0.53266 | 0.56464 |
| ENSG00000144655 | 3 | 39183346 | 39196053 | 43 | 11 | -0.044951 | 0.51793 | CSRNP1 | 0.51793 | 0.58644 | 0.51446 |
| ENSG00000137513 | 11 | 78147007 | 78285919 | 403 | 8 | -0.044983 | 0.51794 | NARS2 | 0.51794 | 0.54702 | 0.538 |
| ENSG00000160867 | 5 | 176513887 | 176525145 | 37 | 7 | -0.045041 | 0.51796 | FGFR4 | 0.51796 | 0.19752 | 0.20534 |
| ENSG00000144619 | 3 | 2140497 | 3099645 | 4940 | 324 | -0.045127 | 0.518 | CNTN4 | 0.518 | 0.4896 | 0.50246 |
| ENSG00000114742 | 3 | 39093489 | 39138155 | 154 | 20 | -0.045241 | 0.51804 | WDR48 | 0.51804 | 0.57893 | 0.58834 |
| ENSG00000213533 | 3 | 52870235 | 52931612 | 141 | 12 | -0.04529 | 0.51806 | TMEM110 | 0.51806 | 0.58054 | 0.56004 |
| ENSG00000152767 | 13 | 98794816 | 99102027 | 1417 | 144 | -0.045306 | 0.51807 | FARP1 | 0.51807 | 0.57026 | 0.57219 |
| ENSG00000115008 | 2 | 113531492 | 113542167 | 33 | 4 | -0.045367 | 0.51809 | IL1A | 0.51809 | 0.66832 | 0.73024 |
| ENSG00000184221 | 21 | 34442450 | 34444726 | 5 | 2 | -0.045528 | 0.51816 | OLIG1 | 0.51816 | 0.28768 | 0.25917 |
| ENSG00000166016 | 11 | 34172535 | 34379555 | 787 | 106 | -0.045558 | 0.51817 | ABTB2 | 0.51817 | 0.5897 | 0.50129 |
| ENSG00000006459 | 7 | 139784546 | 139876835 | 284 | 16 | -0.045626 | 0.5182 | KDM7A | 0.5182 | 0.49277 | 0.52219 |
| ENSG00000162298 | 11 | 64889252 | 64902004 | 22 | 9 | -0.045747 | 0.51824 | SYVN1 | 0.51824 | 0.64296 | 0.68207 |
| ENSG00000163374 | 1 | 155629237 | 155658791 | 22 | 13 | -0.046078 | 0.51838 | YY1AP1 | 0.51838 | 0.68854 | 0.46805 |
| ENSG00000015171 | 10 | 180405 | 300577 | 230 | 22 | -0.046359 | 0.51849 | ZMYND11 | 0.51849 | 0.50779 | 0.46435 |
| ENSG00000120055 | 10 | 104209594 | 104211300 | 1 | 1 | -0.046389 | 0.5185 | C10orf95 | 0.5185 | 0.55036 | 0.71835 |
| ENSG00000214954 | 8 | 92114060 | 92231464 | 404 | 16 | -0.04641 | 0.51851 | LRRC69 | 0.51851 | 0.55155 | 0.54002 |
| ENSG00000138032 | 2 | 44395108 | 44471523 | 206 | 21 | -0.046431 | 0.51852 | PPM1B | 0.51852 | 0.55941 | 0.68428 |
| ENSG00000272325 | 6 | 34247456 | 34360451 | 325 | 13 | -0.046483 | 0.51854 | NUDT3 | 0.51854 | 0.56057 | 0.57602 |
| ENSG00000114978 | 2 | 74379655 | 74406025 | 70 | 6 | -0.046836 | 0.51868 | MOB1A | 0.51868 | 0.56854 | 0.57128 |
| ENSG00000272822 | 12 | 49297286 | 49351148 | 193 | 11 | -0.046905 | 0.51871 | RP11-302B13.5 | 0.51871 | 0.51195 | 0.47335 |
| ENSG00000243543 | 20 | 44162835 | 44168134 | 12 | 4 | -0.047015 | 0.51875 | WFDC6 | 0.51875 | 0.12342 | 0.55204 |
| ENSG00000154655 | 18 | 5954705 | 6415236 | 1406 | 99 | -0.047056 | 0.51877 | L3MBTL4 | 0.51877 | 0.54132 | 0.54352 |
| ENSG00000258728 | 9 | 34646642 | 34657110 | 20 | 6 | -0.047221 | 0.51883 | GALT | 0.51883 | 0.3815 | 0.19455 |
| ENSG00000177990 | 12 | 63952693 | 64062719 | 514 | 23 | -0.047226 | 0.51883 | DPY19L2 | 0.51883 | 0.48613 | 0.47651 |
| ENSG00000161929 | 17 | 5112256 | 5138155 | 93 | 14 | -0.047257 | 0.51885 | SCIMP | 0.51885 | 0.41388 | 0.309 |
| ENSG00000111424 | 12 | 48235320 | 48336831 | 380 | 50 | -0.047372 | 0.51889 | VDR | 0.51889 | 0.52343 | 0.4186 |
| ENSG00000025156 | 6 | 122720691 | 122754264 | 86 | 11 | -0.047398 | 0.5189 | HSF2 | 0.5189 | 0.55838 | 0.67741 |
| ENSG00000185823 | 15 | 24920541 | 24928593 | 27 | 8 | -0.047396 | 0.5189 | NPAP1 | 0.5189 | 0.80607 | 0.91449 |
| ENSG00000152049 | 2 | 223916532 | 224063117 | 742 | 34 | -0.047545 | 0.51896 | KCNE4 | 0.51896 | 0.60744 | 0.68039 |
| ENSG00000136002 | 2 | 131594489 | 131804836 | 661 | 47 | -0.047687 | 0.51902 | ARHGEF4 | 0.51902 | 0.54418 | 0.59383 |
| ENSG00000128298 | 22 | 38480896 | 38506677 | 60 | 14 | -0.047775 | 0.51905 | BAIAP2L2 | 0.51905 | 0.73598 | 0.70546 |
| ENSG00000213780 | 6 | 30875961 | 30881883 | 24 | 5 | -0.047883 | 0.5191 | GTF2H4 | 0.5191 | 0.36905 | 0.35017 |
| ENSG00000113971 | 3 | 132276986 | 132441303 | 332 | 27 | -0.048001 | 0.51914 | NPHP3 | 0.51914 | 0.58031 | 0.63264 |
| ENSG00000126775 | 14 | 55833110 | 55878576 | 185 | 17 | -0.048082 | 0.51917 | ATG14 | 0.51917 | 0.50006 | 0.49967 |
| ENSG00000048649 | 11 | 77371041 | 77532063 | 223 | 25 | -0.048154 | 0.5192 | RSF1 | 0.5192 | 0.50529 | 0.4967 |
| ENSG00000174175 | 1 | 169558087 | 169599431 | 145 | 27 | -0.048405 | 0.5193 | SELP | 0.5193 | 0.51205 | 0.46475 |
| ENSG00000182810 | 16 | 68055179 | 68057770 | 2 | 1 | -0.048452 | 0.51932 | DDX28 | 0.51932 | 0.88249 | 0.89375 |
| ENSG00000149577 | 11 | 117049449 | 117068160 | 68 | 11 | -0.048658 | 0.5194 | SIDT2 | 0.5194 | 0.67246 | 0.64228 |
| ENSG00000196482 | 1 | 216676588 | 217311097 | 2490 | 168 | -0.048711 | 0.51943 | ESRRG | 0.51943 | 0.51602 | 0.65085 |
| ENSG00000136156 | 13 | 48807294 | 48837063 | 38 | 7 | -0.048768 | 0.51945 | ITM2B | 0.51945 | 0.42619 | 0.35303 |
| ENSG00000187522 | 10 | 14880163 | 14913740 | 57 | 11 | -0.049053 | 0.51956 | HSPA14 | 0.51956 | 0.39752 | 0.28865 |
| ENSG00000101493 | 18 | 74069644 | 74207146 | 529 | 43 | -0.049313 | 0.51967 | ZNF516 | 0.51967 | 0.68364 | 0.81972 |
| ENSG00000186026 | 19 | 44576297 | 44593766 | 51 | 7 | -0.049405 | 0.5197 | ZNF284 | 0.5197 | 0.44634 | 0.38278 |
| ENSG00000156535 | 6 | 74405508 | 74538040 | 488 | 34 | -0.049453 | 0.51972 | CD109 | 0.51972 | 0.49093 | 0.30508 |
| ENSG00000141433 | 18 | 904944 | 912173 | 37 | 17 | -0.049554 | 0.51976 | ADCYAP1 | 0.51976 | 0.84279 | 0.90488 |
| ENSG00000128408 | 22 | 45809572 | 45828376 | 76 | 7 | -0.049549 | 0.51976 | RIBC2 | 0.51976 | 0.52057 | 0.581 |
| ENSG00000022840 | 12 | 120971283 | 121015397 | 174 | 21 | -0.049627 | 0.51979 | RNF10 | 0.51979 | 0.49183 | 0.47985 |
| ENSG00000134070 | 3 | 10206549 | 10285427 | 294 | 35 | -0.04975 | 0.51984 | IRAK2 | 0.51984 | 0.48752 | 0.46838 |
| ENSG00000170500 | 2 | 100889753 | 100939195 | 211 | 12 | -0.049965 | 0.51992 | LONRF2 | 0.51992 | 0.55403 | 0.54846 |
| ENSG00000122729 | 9 | 32384618 | 32454767 | 206 | 34 | -0.050042 | 0.51996 | ACO1 | 0.51996 | 0.57587 | 0.51971 |
| ENSG00000005810 | 13 | 77618792 | 77901185 | 614 | 26 | -0.050053 | 0.51996 | MYCBP2 | 0.51996 | 0.5259 | 0.60814 |

| | | | | | | | | | | | |
|---|---|---|---|---|---|---|---|---|---|---|---|
| ENSG00000151322 | 14 | 33404139 | 34273382 | 3165 | 315 | -0.050139 | 0.51999 | NPAS3 | 0.51999 | 0.4644 | 0.491 |
| ENSG00000105323 | 19 | 41768401 | 41813503 | 76 | 7 | -0.050119 | 0.51999 | HNRNPUL1 | 0.51999 | 0.5017 | 0.40307 |
| ENSG00000164038 | 4 | 103941025 | 104006986 | 98 | 21 | -0.050268 | 0.52005 | SLC9B2 | 0.52005 | 0.42507 | 0.38963 |
| ENSG00000121316 | 12 | 14656595 | 14721283 | 176 | 26 | -0.050362 | 0.52008 | PLBD1 | 0.52008 | 0.53771 | 0.5443 |
| ENSG00000197106 | 1 | 110693108 | 110744824 | 128 | 18 | -0.050613 | 0.52018 | SLC6A17 | 0.52018 | 0.49189 | 0.52893 |
| ENSG00000164045 | 3 | 48198636 | 48229892 | 45 | 8 | -0.05065 | 0.5202 | CDC25A | 0.5202 | 0.68482 | 0.7116 |
| ENSG00000091317 | 3 | 32522804 | 32544900 | 64 | 14 | -0.050769 | 0.52025 | CMTM6 | 0.52025 | 0.46819 | 0.52262 |
| ENSG00000221938 | 7 | 143826151 | 143827162 | 3 | 1 | -0.050797 | 0.52026 | OR2A14 | 0.52026 | 0.4504 | 0.46403 |
| ENSG00000263812 | 18 | 74240612 | 74322925 | 488 | 78 | -0.050813 | 0.52026 | LINC00908 | 0.52026 | 0.59902 | 0.49809 |
| ENSG00000167792 | 11 | 67374323 | 67380006 | 5 | 3 | -0.051067 | 0.52036 | NDUFV1 | 0.52036 | 0.62249 | 0.56126 |
| ENSG00000105711 | 19 | 35521588 | 35531352 | 18 | 7 | -0.051274 | 0.52045 | SCN1B | 0.52045 | 0.654 | 0.69343 |
| ENSG00000256060 | 19 | 57874879 | 57876721 | 3 | 2 | -0.051343 | 0.52047 | TRAPPC2P1 | 0.52047 | 0.56917 | 0.56296 |
| ENSG00000148468 | 10 | 15253642 | 15413061 | 682 | 56 | -0.051458 | 0.52052 | FAM171A1 | 0.52052 | 0.68383 | 0.70196 |
| ENSG00000153201 | 2 | 109335937 | 109402267 | 151 | 9 | -0.051587 | 0.52057 | RANBP2 | 0.52057 | 0.5424 | 0.59672 |
| ENSG00000147650 | 8 | 105501459 | 105601252 | 197 | 21 | -0.051579 | 0.52057 | LRP12 | 0.52057 | 0.4918 | 0.49721 |
| ENSG00000071205 | 4 | 148653214 | 148993931 | 818 | 53 | -0.051721 | 0.52062 | ARHGAP10 | 0.52062 | 0.4377 | 0.39922 |
| ENSG00000221970 | 7 | 144015218 | 144016150 | 3 | 1 | -0.051869 | 0.52068 | OR2A1 | 0.52068 | 0.72224 | 0.69509 |
| ENSG00000137496 | 11 | 71709587 | 71716761 | 15 | 3 | -0.051977 | 0.52073 | IL18BP | 0.52073 | 0.55073 | 0.61796 |
| ENSG00000185933 | 10 | 105213144 | 105218645 | 16 | 7 | -0.052378 | 0.52089 | CALHM1 | 0.52089 | 0.67254 | 0.54505 |
| ENSG00000083168 | 8 | 41786997 | 41909508 | 214 | 19 | -0.05252 | 0.52094 | KAT6A | 0.52094 | 0.52876 | 0.49537 |
| ENSG00000103740 | 15 | 78459810 | 78538030 | 276 | 45 | -0.05258 | 0.52097 | ACSBG1 | 0.52097 | 0.43739 | 0.34708 |
| ENSG00000263711 | 18 | 70818476 | 70931715 | 351 | 34 | -0.052585 | 0.52097 | RP11-169F17.1 | 0.52097 | 0.54514 | 0.66444 |
| ENSG00000110756 | 11 | 18300223 | 18343745 | 146 | 15 | -0.052672 | 0.521 | HPS5 | 0.521 | 0.55542 | 0.57455 |
| ENSG00000182472 | 19 | 39220827 | 39260544 | 214 | 20 | -0.052847 | 0.52107 | CAPN12 | 0.52107 | 0.59665 | 0.63607 |
| ENSG00000162924 | 2 | 61108656 | 61158745 | 93 | 23 | -0.053304 | 0.52125 | REL | 0.52125 | 0.54253 | 0.59768 |
| ENSG00000167578 | 19 | 41284121 | 41302847 | 81 | 16 | -0.053317 | 0.52126 | RAB4B | 0.52126 | 0.55654 | 0.56017 |
| ENSG00000147883 | 9 | 22002902 | 22009362 | 10 | 4 | -0.053388 | 0.52129 | CDKN2B | 0.52129 | 0.56037 | 0.69498 |
| ENSG00000132510 | 17 | 7743222 | 7758114 | 32 | 12 | -0.053382 | 0.52129 | KDM6B | 0.52129 | 0.73579 | 0.67183 |
| ENSG00000154760 | 17 | 33762115 | 33775856 | 58 | 13 | -0.053627 | 0.52138 | SLFN13 | 0.52138 | 0.48548 | 0.39193 |
| ENSG00000148513 | 10 | 37414785 | 37673039 | 536 | 32 | -0.053658 | 0.5214 | ANKRD30A | 0.5214 | 0.52136 | 0.56243 |
| ENSG00000166689 | 11 | 16799842 | 17035990 | 937 | 55 | -0.053791 | 0.52145 | PLEKHA7 | 0.52145 | 0.56966 | 0.58257 |
| ENSG00000204701 | 6 | 29079668 | 29080603 | 4 | 2 | -0.053844 | 0.52147 | OR2J3 | 0.52147 | 0.83079 | 0.86493 |
| ENSG00000131015 | 6 | 150263136 | 150270371 | 28 | 7 | -0.053903 | 0.52149 | ULBP2 | 0.52149 | 0.61018 | 0.72276 |
| ENSG00000204752 | 19 | 7011514 | 7013338 | 20 | 3 | -0.054187 | 0.52161 | AC025278.1 | 0.52161 | 0.50765 | 0.39977 |
| ENSG00000198518 | 6 | 26204858 | 26206266 | 14 | 4 | -0.054271 | 0.52164 | HIST1H4E | 0.52164 | 0.51923 | 0.51255 |
| ENSG00000171148 | 3 | 9821544 | 9834695 | 35 | 7 | -0.054359 | 0.52168 | TADA3 | 0.52168 | 0.50447 | 0.49089 |
| ENSG00000186166 | 11 | 118868852 | 118886501 | 63 | 9 | -0.054551 | 0.52175 | CCDC84 | 0.52175 | 0.573 | 0.75459 |
| ENSG00000198730 | 11 | 10772534 | 10801290 | 64 | 14 | -0.05456 | 0.52176 | CTR9 | 0.52176 | 0.3404 | 0.16976 |
| ENSG00000171608 | 1 | 9711790 | 9789172 | 174 | 35 | -0.054726 | 0.52182 | PIK3CD | 0.52182 | 0.3715 | 0.46511 |
| ENSG00000214871 | 7 | 23210760 | 23234503 | 65 | 4 | -0.054849 | 0.52187 | AC005082.1 | 0.52187 | 0.50707 | 0.36992 |
| ENSG00000142599 | 1 | 8412457 | 8877702 | 900 | 27 | -0.054941 | 0.52191 | RERE | 0.52191 | 0.53486 | 0.55796 |
| ENSG00000175471 | 5 | 94039446 | 94620279 | 1737 | 131 | -0.055012 | 0.52194 | MCTP1 | 0.52194 | 0.4927 | 0.45687 |
| ENSG00000173638 | 21 | 46913486 | 46964325 | 236 | 25 | -0.055181 | 0.522 | SLC19A1 | 0.522 | 0.53835 | 0.53078 |
| ENSG00000050628 | 1 | 71318036 | 71513491 | 681 | 49 | -0.055254 | 0.52203 | PTGER3 | 0.52203 | 0.50311 | 0.50029 |
| ENSG00000117010 | 1 | 40997233 | 41013841 | 68 | 11 | -0.055354 | 0.52207 | ZNF684 | 0.52207 | 0.52263 | 0.54007 |
| ENSG00000150712 | 5 | 32227100 | 32313115 | 282 | 15 | -0.055445 | 0.52211 | MTMR12 | 0.52211 | 0.48741 | 0.46819 |
| ENSG00000107404 | 1 | 1270656 | 1284730 | 49 | 10 | -0.05551 | 0.52213 | DVL1 | 0.52213 | 0.44547 | 0.21149 |
| ENSG00000011376 | 3 | 45429998 | 45590913 | 374 | 27 | -0.055567 | 0.52216 | LARS2 | 0.52216 | 0.41862 | 0.21894 |
| ENSG00000139537 | 12 | 49297893 | 49325623 | 90 | 7 | -0.055584 | 0.52216 | CCDC65 | 0.52216 | 0.48055 | 0.44033 |
| ENSG00000204394 | 6 | 31745295 | 31763730 | 41 | 17 | -0.055632 | 0.52218 | VARS | 0.52218 | 0.6083 | 0.52672 |
| ENSG00000161057 | 7 | 102984701 | 103009842 | 43 | 8 | -0.05583 | 0.52226 | PSMC2 | 0.52226 | 0.39667 | 0.27209 |
| ENSG00000135070 | 9 | 88879461 | 88897676 | 40 | 7 | -0.055935 | 0.5223 | ISCA1 | 0.5223 | 0.49584 | 0.63211 |
| ENSG00000107566 | 10 | 101909851 | 101948091 | 94 | 10 | -0.056037 | 0.52234 | ERLIN1 | 0.52234 | 0.49048 | 0.55955 |
| ENSG00000185873 | 4 | 69092371 | 69111438 | 31 | 5 | -0.0565 | 0.52253 | TMPRSS11B | 0.52253 | 0.38463 | 0.36289 |
| ENSG00000160688 | 1 | 154955814 | 154965587 | 18 | 3 | -0.056766 | 0.52263 | FLAD1 | 0.52263 | 0.64891 | 0.72533 |
| ENSG00000184385 | 21 | 43522244 | 43528644 | 30 | 4 | -0.05688 | 0.52268 | C21orf128 | 0.52268 | 0.77952 | 0.9058 |
| ENSG00000163412 | 3 | 71724567 | 71803924 | 292 | 32 | -0.056935 | 0.5227 | EIF4E3 | 0.5227 | 0.55269 | 0.59558 |
| ENSG00000120063 | 17 | 63006833 | 63052957 | 36 | 9 | -0.056943 | 0.5227 | GNA13 | 0.5227 | 0.41712 | 0.25675 |
| ENSG00000176919 | 9 | 139839698 | 139841426 | 12 | 3 | -0.056968 | 0.52271 | C8G | 0.52271 | 0.17968 | 0.087195 |
| ENSG00000103168 | 16 | 84211458 | 84220669 | 51 | 11 | -0.057315 | 0.52285 | TAF1C | 0.52285 | 0.18482 | 0.15039 |
| ENSG00000182196 | 12 | 123464607 | 123467456 | 1 | 1 | -0.057433 | 0.5229 | ARL6IP4 | 0.5229 | 0.70308 | 0.63924 |
| ENSG00000270800 | 6 | 34256547 | 34393825 | 377 | 22 | -0.057461 | 0.52291 | RPS10-NUDT3 | 0.52291 | 0.519 | 0.59895 |
| ENSG00000174429 | 8 | 107771711 | 107782473 | 28 | 7 | -0.05766 | 0.52299 | ABRA | 0.52299 | 0.71235 | 0.79994 |
| ENSG00000257860 | 12 | 103545620 | 103562087 | 63 | 13 | -0.057893 | 0.52308 | RP11-552I14.1 | 0.52308 | 0.60175 | 0.69848 |
| ENSG00000184160 | 4 | 3768075 | 3770251 | 8 | 3 | -0.058138 | 0.52318 | ADRA2C | 0.52318 | 0.087661 | 0.06392 |
| ENSG00000197563 | 18 | 59710800 | 59854351 | 471 | 15 | -0.058326 | 0.52326 | PIGN | 0.52326 | 0.50482 | 0.50251 |
| ENSG00000119509 | 9 | 102861538 | 103063282 | 539 | 15 | -0.058382 | 0.52328 | INVS | 0.52328 | 0.53005 | 0.53358 |
| ENSG00000176714 | 2 | 27848506 | 27851879 | 4 | 2 | -0.058761 | 0.52343 | CCDC121 | 0.52343 | 0.65678 | 0.63109 |
| ENSG00000163528 | 3 | 14153580 | 14166370 | 26 | 7 | -0.058842 | 0.52346 | CHCHD4 | 0.52346 | 0.58649 | 0.46042 |
| ENSG00000177673 | 2 | 232457575 | 232458994 | 3 | 2 | -0.059145 | 0.52358 | C2orf57 | 0.52358 | 0.62182 | 0.61476 |
| ENSG00000132155 | 3 | 12625100 | 12705725 | 286 | 17 | -0.059295 | 0.52364 | RAF1 | 0.52364 | 0.5581 | 0.46922 |
| ENSG00000168875 | 3 | 137483579 | 137484396 | 2 | 1 | -0.059295 | 0.52364 | SOX14 | 0.52364 | 0.65267 | 0.70311 |
| ENSG00000080802 | 7 | 135046547 | 135194875 | 300 | 25 | -0.059296 | 0.52364 | CNOT4 | 0.52364 | 0.4494 | 0.4828 |

| | | | | | | | | | | | |
|---|---|---|---|---|---|---|---|---|---|---|---|
| ENSG00000166484 | 17 | 19281034 | 19286857 | 1 | 1 | -0.059442 | 0.5237 | MAPK7 | 0.5237 | 0.92137 | 0.89275 |
| ENSG00000063515 | 22 | 19136089 | 19137796 | 5 | 1 | -0.059469 | 0.52371 | GSC2 | 0.52371 | 0.55367 | 0.053363 |
| ENSG00000181350 | 17 | 16344891 | 16395467 | 197 | 21 | -0.059723 | 0.52381 | FAM211A | 0.52381 | 0.42388 | 0.34738 |
| ENSG00000170540 | 16 | 18802991 | 18813000 | 26 | 6 | -0.059837 | 0.52386 | ARL6IP1 | 0.52386 | 0.6456 | 0.55766 |
| ENSG00000072818 | 17 | 7239848 | 7254797 | 31 | 11 | -0.059872 | 0.52387 | ACAP1 | 0.52387 | 0.79183 | 0.94585 |
| ENSG00000145919 | 5 | 173034517 | 173043663 | 32 | 6 | -0.060022 | 0.52393 | BOD1 | 0.52393 | 0.61339 | 0.24945 |
| ENSG00000131759 | 17 | 38465444 | 38513094 | 72 | 13 | -0.060043 | 0.52394 | RARA | 0.52394 | 0.57723 | 0.51443 |
| ENSG00000124786 | 6 | 8413301 | 8435794 | 89 | 7 | -0.06011 | 0.52397 | SLC35B3 | 0.52397 | 0.69675 | 0.76326 |
| ENSG00000101246 | 20 | 62329996 | 62339377 | 18 | 3 | -0.060264 | 0.52403 | ARFRP1 | 0.52403 | 0.38845 | 0.25081 |
| ENSG00000151413 | 14 | 31959162 | 32330430 | 1459 | 31 | -0.060318 | 0.52405 | NUBPL | 0.52405 | 0.54244 | 0.59799 |
| ENSG00000196372 | 10 | 5680830 | 5708558 | 178 | 42 | -0.060527 | 0.52413 | ASB13 | 0.52413 | 0.36254 | 0.293 |
| ENSG00000123352 | 12 | 49760367 | 49921205 | 406 | 26 | -0.060638 | 0.52418 | SPATS2 | 0.52418 | 0.50945 | 0.55675 |
| ENSG00000070729 | 16 | 57917503 | 58005020 | 374 | 63 | -0.060639 | 0.52418 | CNGB1 | 0.52418 | 0.36673 | 0.27419 |
| ENSG00000105619 | 19 | 54610320 | 54619055 | 32 | 10 | -0.060718 | 0.52421 | TFPT | 0.52421 | 0.62093 | 0.68264 |
| ENSG00000134508 | 18 | 20714528 | 20840431 | 310 | 39 | -0.06076 | 0.52422 | CABLES1 | 0.52422 | 0.56807 | 0.66122 |
| ENSG00000168216 | 6 | 70385694 | 70507003 | 362 | 14 | -0.060833 | 0.52425 | LMBRD1 | 0.52425 | 0.50715 | 0.50012 |
| ENSG00000116127 | 2 | 73612886 | 73837920 | 460 | 16 | -0.061055 | 0.52434 | ALMS1 | 0.52434 | 0.54486 | 0.54327 |
| ENSG00000260916 | 15 | 55632230 | 55700708 | 332 | 20 | -0.06107 | 0.52435 | CCPG1 | 0.52435 | 0.59243 | 0.67265 |
| ENSG00000125458 | 17 | 73126320 | 73127890 | 4 | 2 | -0.061296 | 0.52444 | NT5C | 0.52444 | 0.5773 | 0.61992 |
| ENSG00000128294 | 22 | 26921458 | 26992681 | 326 | 34 | -0.06134 | 0.52446 | TPST2 | 0.52446 | 0.5562 | 0.52727 |
| ENSG00000025293 | 20 | 34359896 | 34538303 | 509 | 14 | -0.06155 | 0.52454 | PHF20 | 0.52454 | 0.51195 | 0.51933 |
| ENSG00000113212 | 5 | 140552243 | 140555957 | 19 | 5 | -0.061803 | 0.52464 | PCDHB7 | 0.52464 | 0.41867 | 0.47786 |
| ENSG00000143771 | 1 | 224544552 | 224567161 | 44 | 5 | -0.061892 | 0.52468 | CNIH4 | 0.52468 | 0.44568 | 0.39993 |
| ENSG00000088899 | 20 | 3143263 | 3154192 | 37 | 16 | -0.0619 | 0.52468 | LZTS3 | 0.52468 | 0.56296 | 0.23811 |
| ENSG00000100522 | 14 | 53241912 | 53258386 | 20 | 4 | -0.062152 | 0.52478 | GNPNAT1 | 0.52478 | 0.5593 | 0.65024 |
| ENSG00000162813 | 1 | 220230824 | 220263804 | 92 | 19 | -0.062173 | 0.52479 | BPNT1 | 0.52479 | 0.58695 | 0.53415 |
| ENSG00000100297 | 22 | 35796056 | 35821423 | 87 | 18 | -0.06221 | 0.5248 | MCM5 | 0.5248 | 0.58267 | 0.13096 |
| ENSG00000100473 | 14 | 31343720 | 31364271 | 66 | 20 | -0.062288 | 0.52483 | COCH | 0.52483 | 0.70397 | 0.58498 |
| ENSG00000091879 | 8 | 6357172 | 6420930 | 491 | 44 | -0.062415 | 0.52488 | ANGPT2 | 0.52488 | 0.53407 | 0.54702 |
| ENSG00000156508 | 6 | 74225473 | 74233520 | 25 | 6 | -0.06249 | 0.52491 | EEF1A1 | 0.52491 | 0.48358 | 0.40921 |
| ENSG00000103152 | 16 | 127006 | 135852 | 33 | 7 | -0.062717 | 0.525 | MPG | 0.525 | 0.71185 | 0.72694 |
| ENSG00000226124 | 2 | 200625267 | 200715896 | 211 | 33 | -0.062783 | 0.52503 | FTCDNL1 | 0.52503 | 0.5774 | 0.5493 |
| ENSG00000171174 | 2 | 28004231 | 28113965 | 230 | 16 | -0.063102 | 0.52516 | RBKS | 0.52516 | 0.55768 | 0.55959 |
| ENSG00000144834 | 3 | 111717511 | 111732734 | 87 | 19 | -0.063247 | 0.52521 | TAGLN3 | 0.52521 | 0.66281 | 0.69413 |
| ENSG00000163512 | 3 | 28356494 | 28390618 | 50 | 9 | -0.063254 | 0.52522 | AZI2 | 0.52522 | 0.58335 | 0.58475 |
| ENSG00000082556 | 8 | 54138284 | 54164257 | 151 | 13 | -0.063342 | 0.52525 | OPRK1 | 0.52525 | 0.45371 | 0.3185 |
| ENSG00000165684 | 9 | 139270029 | 139293249 | 108 | 16 | -0.063503 | 0.52532 | SNAPC4 | 0.52532 | 0.55435 | 0.53514 |
| ENSG00000185339 | 22 | 31002825 | 31023265 | 160 | 23 | -0.063513 | 0.52532 | TCN2 | 0.52532 | 0.58816 | 0.66745 |
| ENSG00000177984 | 9 | 139654086 | 139660707 | 27 | 4 | -0.063709 | 0.5254 | LCN15 | 0.5254 | 0.57007 | 0.43091 |
| ENSG00000131584 | 1 | 1227756 | 1244989 | 74 | 13 | -0.06374 | 0.52541 | ACAP3 | 0.52541 | 0.57983 | 0.75454 |
| ENSG00000143319 | 1 | 156691683 | 156698591 | 13 | 6 | -0.063883 | 0.52547 | ISG20L2 | 0.52547 | 0.90691 | 0.6454 |
| ENSG00000136542 | 2 | 158114110 | 158170723 | 168 | 15 | -0.064288 | 0.52563 | GALNT5 | 0.52563 | 0.46605 | 0.43933 |
| ENSG00000187288 | 3 | 9908398 | 9921938 | 38 | 7 | -0.064321 | 0.52564 | CIDEC | 0.52564 | 0.69928 | 0.6377 |
| ENSG00000105738 | 19 | 38397868 | 38699012 | 998 | 49 | -0.06492 | 0.52588 | SIPA1L3 | 0.52588 | 0.47974 | 0.52617 |
| ENSG00000204941 | 19 | 43670408 | 43690688 | 132 | 13 | -0.065129 | 0.52596 | PSG5 | 0.52596 | 0.59583 | 0.66677 |
| ENSG00000248592 | 3 | 52867137 | 52931578 | 152 | 12 | -0.065176 | 0.52598 | TMEM110-MUSTN1 | 0.52598 | 0.57479 | 0.53757 |
| ENSG00000087087 | 7 | 100472733 | 100486285 | 30 | 6 | -0.065459 | 0.5261 | SRRT | 0.5261 | 0.48952 | 0.37668 |
| ENSG00000068697 | 2 | 20232411 | 20251789 | 59 | 12 | -0.065802 | 0.52623 | LAPTM4A | 0.52623 | 0.35711 | 0.37896 |
| ENSG00000197594 | 6 | 132129156 | 132216295 | 311 | 33 | -0.065968 | 0.5263 | ENPP1 | 0.5263 | 0.58231 | 0.56141 |
| ENSG00000183840 | 2 | 133174147 | 133404132 | 792 | 58 | -0.066032 | 0.52632 | GPR39 | 0.52632 | 0.60569 | 0.70268 |
| ENSG00000233412 | 3 | 97887544 | 97888485 | 7 | 2 | -0.066125 | 0.52636 | OR5H15 | 0.52636 | 0.5032 | 0.66167 |
| ENSG00000145349 | 4 | 114372188 | 114683083 | 919 | 58 | -0.06613 | 0.52636 | CAMK2D | 0.52636 | 0.55537 | 0.62843 |
| ENSG00000162714 | 1 | 247460714 | 247495148 | 114 | 14 | -0.066216 | 0.5264 | ZNF496 | 0.5264 | 0.53611 | 0.47449 |
| ENSG00000242221 | 19 | 43568363 | 43587197 | 126 | 13 | -0.066314 | 0.52644 | PSG2 | 0.52644 | 0.56038 | 0.57512 |
| ENSG00000168743 | 4 | 106815932 | 106925184 | 458 | 28 | -0.066419 | 0.52648 | NPNT | 0.52648 | 0.5273 | 0.57396 |
| ENSG00000089351 | 19 | 35485688 | 35517375 | 106 | 11 | -0.066709 | 0.52659 | GRAMD1A | 0.52659 | 0.55334 | 0.61621 |
| ENSG00000145936 | 5 | 169801675 | 169816681 | 88 | 14 | -0.066767 | 0.52662 | KCNMB1 | 0.52662 | 0.60721 | 0.4467 |
| ENSG00000175854 | 9 | 131037658 | 131051269 | 25 | 4 | -0.066946 | 0.52669 | SWI5 | 0.52669 | 0.66986 | 0.75257 |
| ENSG00000141401 | 18 | 11981024 | 12030876 | 268 | 27 | -0.067063 | 0.52673 | IMPA2 | 0.52673 | 0.64154 | 0.76743 |
| ENSG00000172554 | 2 | 946554 | 1371385 | 2344 | 72 | -0.06708 | 0.52674 | SNTG2 | 0.52674 | 0.52611 | 0.53501 |
| ENSG00000086589 | 5 | 150070356 | 150080669 | 11 | 4 | -0.067473 | 0.5269 | RBM22 | 0.5269 | 0.56206 | 0.63179 |
| ENSG00000111790 | 12 | 27091316 | 27119583 | 79 | 16 | -0.067658 | 0.52697 | FGFR1OP2 | 0.52697 | 0.32301 | 0.33023 |
| ENSG00000101213 | 20 | 62159778 | 62168723 | 36 | 9 | -0.067736 | 0.527 | PTK6 | 0.527 | 0.59345 | 0.54138 |
| ENSG00000213930 | 9 | 34638130 | 34651032 | 24 | 6 | -0.068213 | 0.52719 | GALT | 0.52719 | 0.3815 | 0.19455 |
| ENSG00000124003 | 2 | 223536505 | 223574649 | 121 | 22 | -0.068374 | 0.52726 | MOGAT1 | 0.52726 | 0.68956 | 0.80458 |
| ENSG00000164088 | 3 | 52279841 | 52284613 | 3 | 1 | -0.068478 | 0.5273 | PPM1M | 0.5273 | 0.41304 | 0.48894 |
| ENSG00000146350 | 6 | 121400640 | 121655891 | 1163 | 27 | -0.06849 | 0.5273 | TBC1D32 | 0.5273 | 0.52072 | 0.5442 |
| ENSG00000069998 | 22 | 17618401 | 17646177 | 121 | 23 | -0.068513 | 0.52731 | CECR5 | 0.52731 | 0.38166 | 0.40506 |
| ENSG00000105221 | 19 | 40736224 | 40791443 | 118 | 15 | -0.068838 | 0.52744 | AKT2 | 0.52744 | 0.2805 | 0.23204 |
| ENSG00000132692 | 1 | 156611182 | 156629324 | 43 | 14 | -0.069219 | 0.52759 | BCAN | 0.52759 | 0.28249 | 0.33276 |
| ENSG00000204946 | 7 | 148959262 | 148994393 | 129 | 19 | -0.069237 | 0.5276 | ZNF783 | 0.5276 | 0.47702 | 0.46182 |
| ENSG00000167920 | 17 | 38975358 | 38992522 | 64 | 15 | -0.069281 | 0.52762 | TMEM99 | 0.52762 | 0.71954 | 0.7833 |
| ENSG00000167286 | 11 | 118209669 | 118213459 | 8 | 3 | -0.069517 | 0.52771 | CD3D | 0.52771 | 0.60252 | 0.58108 |

| | | | | | | | | | | | |
|---|---|---|---|---|---|---|---|---|---|---|---|
| ENSG00000077232 | 2 | 183580999 | 183659191 | 232 | 19 | -0.069529 | 0.52772 | DNAJC10 | 0.52772 | 0.51123 | 0.52197 |
| ENSG00000203867 | 10 | 112404155 | 112599227 | 806 | 62 | -0.069789 | 0.52782 | RBM20 | 0.52782 | 0.55081 | 0.60607 |
| ENSG00000125648 | 19 | 6436090 | 6465214 | 131 | 20 | -0.069891 | 0.52786 | SLC25A23 | 0.52786 | 0.43067 | 0.49501 |
| ENSG00000027001 | 13 | 24304328 | 24463558 | 561 | 23 | -0.069923 | 0.52787 | MIPEP | 0.52787 | 0.56117 | 0.57127 |
| ENSG00000160808 | 3 | 46899362 | 46923659 | 51 | 12 | -0.069942 | 0.52788 | MYL3 | 0.52788 | 0.75678 | 0.88993 |
| ENSG00000065675 | 10 | 6469105 | 6622263 | 509 | 61 | -0.069988 | 0.5279 | PRKCQ | 0.5279 | 0.5476 | 0.67082 |
| ENSG00000186280 | 11 | 94706845 | 94732682 | 77 | 16 | -0.070166 | 0.52797 | KDM4D | 0.52797 | 0.58427 | 0.68589 |
| ENSG00000186577 | 6 | 34214157 | 34217247 | 8 | 3 | -0.070186 | 0.52798 | C6orf1 | 0.52798 | 0.61824 | 0.56741 |
| ENSG00000176994 | 17 | 18218624 | 18226517 | 25 | 6 | -0.070464 | 0.52809 | SMCR8 | 0.52809 | 0.68262 | 0.69712 |
| ENSG00000143569 | 1 | 154192655 | 154243986 | 85 | 10 | -0.070523 | 0.52811 | UBAP2L | 0.52811 | 0.52424 | 0.51864 |
| ENSG00000116641 | 1 | 62920399 | 63153969 | 387 | 11 | -0.070638 | 0.52816 | DOCK7 | 0.52816 | 0.5327 | 0.55602 |
| ENSG00000018408 | 3 | 149235022 | 149454501 | 886 | 76 | -0.070732 | 0.52819 | WWTR1 | 0.52819 | 0.57392 | 0.6502 |
| ENSG00000172551 | 12 | 55224303 | 55252177 | 139 | 9 | -0.070792 | 0.52822 | MUCL1 | 0.52822 | 0.69279 | 0.79195 |
| ENSG00000173585 | 3 | 45927996 | 45944667 | 43 | 12 | -0.070821 | 0.52823 | CCR9 | 0.52823 | 0.52186 | 0.53081 |
| ENSG00000151806 | 4 | 44680444 | 44702943 | 56 | 7 | -0.070834 | 0.52824 | GUF1 | 0.52824 | 0.5876 | 0.60769 |
| ENSG00000235711 | 15 | 79575146 | 79590580 | 30 | 8 | -0.070901 | 0.52826 | ANKRD34C | 0.52826 | 0.70595 | 0.72638 |
| ENSG00000142632 | 1 | 16524349 | 16539104 | 71 | 7 | -0.07095 | 0.52828 | ARHGEF19 | 0.52828 | 0.53193 | 0.48184 |
| ENSG00000162769 | 1 | 213031597 | 213072705 | 275 | 12 | -0.071073 | 0.52833 | FLVCR1 | 0.52833 | 0.384 | 0.37818 |
| ENSG00000172409 | 11 | 57416465 | 57429340 | 18 | 4 | -0.07146 | 0.52848 | CLP1 | 0.52848 | 0.6298 | 0.50295 |
| ENSG00000136238 | 7 | 6414154 | 6443608 | 186 | 26 | -0.071486 | 0.52849 | RAC1 | 0.52849 | 0.57693 | 0.54913 |
| ENSG00000167565 | 19 | 40946751 | 40950612 | 21 | 4 | -0.071562 | 0.52852 | SERTAD3 | 0.52852 | 0.36756 | 0.27479 |
| ENSG00000100346 | 22 | 39966758 | 40085742 | 282 | 31 | -0.071695 | 0.52858 | CACNA1I | 0.52858 | 0.52762 | 0.58083 |
| ENSG00000171824 | 1 | 11126675 | 11159938 | 117 | 8 | -0.071774 | 0.52861 | EXOSC10 | 0.52861 | 0.5476 | 0.5258 |
| ENSG00000228716 | 5 | 79922047 | 79950802 | 170 | 7 | -0.071823 | 0.52863 | DHFR | 0.52863 | 0.56995 | 0.56883 |
| ENSG00000188690 | 10 | 127477146 | 127511817 | 77 | 5 | -0.072207 | 0.52878 | UROS | 0.52878 | 0.52775 | 0.49618 |
| ENSG00000139505 | 13 | 25802307 | 25862147 | 125 | 11 | -0.0722 | 0.52878 | MTMR6 | 0.52878 | 0.5883 | 0.63582 |
| ENSG00000135828 | 1 | 182542769 | 182558391 | 41 | 15 | -0.072259 | 0.5288 | RNASEL | 0.5288 | 0.56077 | 0.49487 |
| ENSG00000105679 | 19 | 36024314 | 36036218 | 37 | 9 | -0.07227 | 0.52881 | GAPDHS | 0.52881 | 0.29695 | 0.32406 |
| ENSG00000184292 | 1 | 59041099 | 59043166 | 11 | 3 | -0.072344 | 0.52884 | TACSTD2 | 0.52884 | 0.56416 | 0.63228 |
| ENSG00000164746 | 7 | 48075108 | 48100901 | 111 | 12 | -0.07239 | 0.52885 | C7orf57 | 0.52885 | 0.66309 | 0.68789 |
| ENSG00000009954 | 7 | 72854728 | 72936608 | 140 | 16 | -0.072417 | 0.52887 | BAZ1B | 0.52887 | 0.51421 | 0.56548 |
| ENSG00000107968 | 10 | 30722866 | 30750762 | 70 | 11 | -0.072706 | 0.52898 | MAP3K8 | 0.52898 | 0.64013 | 0.60463 |
| ENSG00000143314 | 1 | 156707095 | 156711382 | 12 | 6 | -0.072765 | 0.529 | MRPL24 | 0.529 | 0.68027 | 0.84588 |
| ENSG00000095370 | 9 | 130500596 | 130541020 | 112 | 16 | -0.07278 | 0.52901 | SH2D3C | 0.52901 | 0.47825 | 0.47152 |
| ENSG00000141219 | 17 | 71228372 | 71245091 | 103 | 5 | -0.073082 | 0.52913 | C17orf80 | 0.52913 | 0.62974 | 0.66098 |
| ENSG00000122085 | 2 | 242011584 | 242041747 | 72 | 10 | -0.073326 | 0.52923 | MTERFD2 | 0.52923 | 0.48174 | 0.51125 |
| ENSG00000174059 | 1 | 208057594 | 208084747 | 101 | 18 | -0.073351 | 0.52924 | CD34 | 0.52924 | 0.49909 | 0.51547 |
| ENSG00000159200 | 21 | 35885440 | 35987441 | 527 | 28 | -0.073392 | 0.52925 | RCAN1 | 0.52925 | 0.54981 | 0.58825 |
| ENSG00000140332 | 15 | 70340129 | 70390515 | 158 | 28 | -0.073692 | 0.52937 | TLE3 | 0.52937 | 0.68855 | 0.85425 |
| ENSG00000145708 | 5 | 76248538 | 76276983 | 75 | 13 | -0.073819 | 0.52942 | CRHBP | 0.52942 | 0.40283 | 0.5168 |
| ENSG00000083817 | 19 | 58082934 | 58090295 | 13 | 6 | -0.073998 | 0.52949 | ZNF416 | 0.52949 | 0.48422 | 0.55165 |
| ENSG00000241595 | 17 | 39405939 | 39406904 | 11 | 3 | -0.074306 | 0.52962 | KRTAP9-4 | 0.52962 | 0.65019 | 0.63826 |
| ENSG00000112290 | 6 | 110421022 | 110501207 | 252 | 13 | -0.074328 | 0.52963 | WASF1 | 0.52963 | 0.55379 | 0.55317 |
| ENSG00000171564 | 4 | 155484108 | 155492238 | 24 | 4 | -0.074398 | 0.52965 | FGB | 0.52965 | 0.49067 | 0.5703 |
| ENSG00000181847 | 3 | 113995760 | 114029135 | 91 | 23 | -0.074521 | 0.5297 | TIGIT | 0.5297 | 0.53087 | 0.54093 |
| ENSG00000268953 | 19 | 31640362 | 31640757 | 2 | 1 | -0.074571 | 0.52972 | AC020952.1 | 0.52972 | 0.32059 | 0.52985 |
| ENSG00000170044 | 3 | 101818088 | 102198685 | 909 | 51 | -0.074601 | 0.52973 | ZPLD1 | 0.52973 | 0.5091 | 0.33453 |
| ENSG00000163808 | 3 | 44803209 | 44914868 | 202 | 22 | -0.074607 | 0.52974 | KIF15 | 0.52974 | 0.38645 | 0.31464 |
| ENSG00000197498 | 6 | 111303218 | 111349466 | 313 | 14 | -0.074625 | 0.52974 | RPF2 | 0.52974 | 0.43436 | 0.37188 |
| ENSG00000198342 | 19 | 12460185 | 12476719 | 16 | 6 | -0.074706 | 0.52978 | ZNF442 | 0.52978 | 0.59112 | 0.57449 |
| ENSG00000133874 | 8 | 33405273 | 33424643 | 62 | 8 | -0.074782 | 0.52981 | RNF122 | 0.52981 | 0.56605 | 0.599 |
| ENSG00000255432 | 11 | 62417166 | 62432650 | 34 | 8 | -0.074811 | 0.52982 | RP11-831H9.11 | 0.52982 | 0.51911 | 0.57423 |
| ENSG00000105996 | 7 | 27139721 | 27142430 | 5 | 2 | -0.074888 | 0.52985 | HOXA2 | 0.52985 | 0.89718 | 0.93458 |
| ENSG00000183470 | 7 | 132333553 | 132413528 | 230 | 29 | -0.074887 | 0.52985 | AC009365.3 | 0.52985 | 0.55861 | 0.58587 |
| ENSG00000171219 | 11 | 64590859 | 64612041 | 57 | 10 | -0.074885 | 0.52985 | CDC42BPG | 0.52985 | 0.81654 | 0.8994 |
| ENSG00000047579 | 6 | 15523032 | 15663289 | 375 | 26 | -0.074922 | 0.52986 | DTNBP1 | 0.52986 | 0.52905 | 0.49587 |
| ENSG00000109079 | 17 | 26662628 | 26674035 | 20 | 7 | -0.075041 | 0.52991 | TNFAIP1 | 0.52991 | 0.3901 | 0.47554 |
| ENSG00000175567 | 11 | 73685712 | 73694352 | 21 | 4 | -0.07534 | 0.53003 | UCP2 | 0.53003 | 0.17038 | 0.55396 |
| ENSG00000188559 | 20 | 20370196 | 20693131 | 575 | 25 | -0.075419 | 0.53006 | RALGAPA2 | 0.53006 | 0.55583 | 0.52922 |
| ENSG00000015133 | 14 | 91737667 | 91884188 | 513 | 62 | -0.075697 | 0.53017 | CCDC88C | 0.53017 | 0.54251 | 0.65273 |
| ENSG00000117500 | 1 | 93615299 | 93646285 | 69 | 11 | -0.0759 | 0.53025 | TMED5 | 0.53025 | 0.60787 | 0.60534 |
| ENSG00000133561 | 7 | 150322463 | 150329473 | 18 | 4 | -0.075923 | 0.53026 | GIMAP6 | 0.53026 | 0.63637 | 0.86361 |
| ENSG00000109790 | 4 | 39046659 | 39128477 | 320 | 9 | -0.076027 | 0.5303 | KLHL5 | 0.5303 | 0.54552 | 0.53721 |
| ENSG00000146530 | 7 | 12370511 | 12443567 | 327 | 22 | -0.0762 | 0.53037 | VWDE | 0.53037 | 0.45863 | 0.44067 |
| ENSG00000204060 | 1 | 41827594 | 41849262 | 70 | 15 | -0.076238 | 0.53039 | FOXO6 | 0.53039 | 0.34515 | 0.34188 |
| ENSG00000174804 | 11 | 86656721 | 86666433 | 22 | 6 | -0.076386 | 0.53044 | FZD4 | 0.53044 | 0.61614 | 0.80498 |
| ENSG00000162645 | 1 | 89571815 | 89616139 | 99 | 7 | -0.076444 | 0.53047 | GBP2 | 0.53047 | 0.5081 | 0.52904 |
| ENSG00000104886 | 19 | 2230002 | 2237703 | 17 | 5 | -0.076527 | 0.5305 | PLEKHJ1 | 0.5305 | 0.7449 | 0.82173 |
| ENSG00000132879 | 1 | 11714432 | 11723384 | 32 | 10 | -0.076725 | 0.53058 | FBXO44 | 0.53058 | 0.39899 | 0.4537 |
| ENSG00000176563 | 17 | 40950810 | 40963605 | 8 | 4 | -0.076729 | 0.53058 | CNTD1 | 0.53058 | 0.42277 | 0.461 |
| ENSG00000107537 | 10 | 13319796 | 13344412 | 138 | 19 | -0.076807 | 0.53061 | PHYH | 0.53061 | 0.57417 | 0.66304 |
| ENSG00000176256 | 1 | 34326076 | 34330392 | 12 | 6 | -0.076818 | 0.53062 | HMGB4 | 0.53062 | 0.5646 | 0.6942 |
| ENSG00000075303 | 7 | 87462883 | 87505672 | 99 | 12 | -0.077282 | 0.5308 | SLC25A40 | 0.5308 | 0.49916 | 0.55984 |

| | | | | | | | | | | | |
|---|---|---|---|---|---|---|---|---|---|---|---|
| ENSG00000254737 | 11 | 123886282 | 123887217 | 9 | 3 | -0.077356 | 0.53083 | OR10G4 | 0.53083 | 0.55202 | 0.50395 |
| ENSG00000206483 | 3 | 126290625 | 126327398 | 102 | 17 | -0.077393 | 0.53084 | TXNRD3NB | 0.53084 | 0.50751 | 0.65866 |
| ENSG00000198326 | 20 | 2795614 | 2800930 | 17 | 8 | -0.077612 | 0.53093 | TMEM239 | 0.53093 | 0.36242 | 0.24016 |
| ENSG00000157379 | 14 | 24759804 | 24769039 | 30 | 6 | -0.077738 | 0.53098 | DHRS1 | 0.53098 | 0.61509 | 0.66824 |
| ENSG00000164485 | 6 | 137464968 | 137494785 | 95 | 17 | -0.078124 | 0.53114 | IL22RA2 | 0.53114 | 0.33494 | 0.44705 |
| ENSG00000160305 | 21 | 47878812 | 47989926 | 495 | 33 | -0.078235 | 0.53118 | DIP2A | 0.53118 | 0.54115 | 0.52109 |
| ENSG00000166477 | 15 | 52230222 | 52264003 | 106 | 10 | -0.078284 | 0.5312 | LEO1 | 0.5312 | 0.57079 | 0.62336 |
| ENSG00000156172 | 8 | 96257147 | 96281429 | 88 | 16 | -0.078338 | 0.53122 | C8orf37 | 0.53122 | 0.43636 | 0.37997 |
| ENSG00000130477 | 19 | 17712137 | 17799401 | 443 | 78 | -0.078366 | 0.53123 | UNC13A | 0.53123 | 0.36511 | 0.20076 |
| ENSG00000123136 | 19 | 14519631 | 14530192 | 18 | 9 | -0.078698 | 0.53136 | DDX39A | 0.53136 | 0.88392 | 0.9101 |
| ENSG00000158517 | 7 | 74188309 | 74203659 | 16 | 5 | -0.078758 | 0.53139 | NCF1 | 0.53139 | 0.60745 | 0.70707 |
| ENSG00000108953 | 17 | 1247566 | 1303672 | 281 | 25 | -0.078952 | 0.53146 | YWHAE | 0.53146 | 0.64188 | 0.4965 |
| ENSG00000170469 | 5 | 138732252 | 138739777 | 14 | 2 | -0.079044 | 0.5315 | SPATA24 | 0.5315 | 0.65979 | 0.71743 |
| ENSG00000175262 | 1 | 11006528 | 11042094 | 139 | 18 | -0.079086 | 0.53152 | C1orf127 | 0.53152 | 0.71225 | 0.76652 |
| ENSG00000150527 | 14 | 39734488 | 39856156 | 355 | 21 | -0.079288 | 0.5316 | CTAGE5 | 0.5316 | 0.58671 | 0.6381 |
| ENSG00000185262 | 17 | 74261283 | 74267375 | 25 | 7 | -0.0793 | 0.5316 | UBALD2 | 0.5316 | 0.67797 | 0.78621 |
| ENSG00000117984 | 11 | 1773982 | 1785222 | 38 | 7 | -0.079327 | 0.53161 | CTSD | 0.53161 | 0.076534 | 0.023205 |
| ENSG00000178537 | 3 | 48894369 | 48936426 | 59 | 10 | -0.079332 | 0.53162 | SLC25A20 | 0.53162 | 0.57961 | 0.58384 |
| ENSG00000171102 | 9 | 136080664 | 136084630 | 22 | 9 | -0.079417 | 0.53165 | OBP2B | 0.53165 | 0.35249 | 0.33635 |
| ENSG00000012660 | 6 | 53132196 | 53213947 | 268 | 11 | -0.079528 | 0.53169 | ELOVL5 | 0.53169 | 0.51013 | 0.42788 |
| ENSG00000212643 | 5 | 112227313 | 112228791 | 7 | 2 | -0.079935 | 0.53186 | ZRSR1 | 0.53186 | 0.56357 | 0.45938 |
| ENSG00000186074 | 17 | 72690452 | 72709117 | 41 | 11 | -0.080102 | 0.53192 | CD300LF | 0.53192 | 0.64914 | 0.57089 |
| ENSG00000213424 | 17 | 38810917 | 38821433 | 18 | 5 | -0.080129 | 0.53193 | KRT222 | 0.53193 | 0.5908 | 0.57718 |
| ENSG00000204518 | 1 | 12704566 | 12727097 | 61 | 7 | -0.080182 | 0.53195 | AADACL4 | 0.53195 | 0.67734 | 0.70396 |
| ENSG00000166961 | 11 | 60524426 | 60544205 | 53 | 20 | -0.080174 | 0.53195 | MS4A15 | 0.53195 | 0.57383 | 0.54353 |
| ENSG00000123684 | 1 | 211916799 | 212004114 | 352 | 24 | -0.080343 | 0.53202 | LPGAT1 | 0.53202 | 0.47631 | 0.39185 |
| ENSG00000096968 | 9 | 4985033 | 5128183 | 430 | 18 | -0.080395 | 0.53204 | JAK2 | 0.53204 | 0.55361 | 0.59333 |
| ENSG00000052344 | 16 | 31142756 | 31147083 | 6 | 4 | -0.080531 | 0.53209 | PRSS8 | 0.53209 | 0.37325 | 0.51338 |
| ENSG00000105131 | 19 | 15337730 | 15344246 | 4 | 2 | -0.080543 | 0.5321 | EPHX3 | 0.5321 | 0.39892 | 0.4064 |
| ENSG00000176783 | 5 | 178977559 | 179037027 | 216 | 26 | -0.080701 | 0.53216 | RUFY1 | 0.53216 | 0.50847 | 0.53128 |
| ENSG00000157107 | 5 | 72251808 | 72386349 | 274 | 16 | -0.080781 | 0.53219 | FCHO2 | 0.53219 | 0.53769 | 0.52893 |
| ENSG00000185869 | 19 | 37379026 | 37407193 | 68 | 8 | -0.080768 | 0.53219 | ZNF829 | 0.53219 | 0.37018 | 0.35843 |
| ENSG00000179950 | 8 | 144898514 | 144912029 | 29 | 8 | -0.080862 | 0.53222 | PUF60 | 0.53222 | 0.55475 | 0.6462 |
| ENSG00000164535 | 7 | 6448757 | 6523821 | 347 | 36 | -0.080978 | 0.53227 | DAGLB | 0.53227 | 0.53568 | 0.53304 |
| ENSG00000167799 | 11 | 67395409 | 67397401 | 4 | 2 | -0.08103 | 0.53229 | NUDT8 | 0.53229 | 0.55839 | 0.41442 |
| ENSG00000108576 | 17 | 28521337 | 28563020 | 79 | 15 | -0.081188 | 0.53235 | SLC6A4 | 0.53235 | 0.57957 | 0.56947 |
| ENSG00000125813 | 20 | 21686297 | 21696620 | 13 | 5 | -0.081354 | 0.53242 | PAX1 | 0.53242 | 0.18197 | 0.079791 |
| ENSG00000263563 | 17 | 21729601 | 21731762 | 8 | 3 | -0.081499 | 0.53248 | UBBP4 | 0.53248 | 0.45103 | 0.51343 |
| ENSG00000155093 | 7 | 157331750 | 158380480 | 4924 | 282 | -0.081662 | 0.53254 | PTPRN2 | 0.53254 | 0.53644 | 0.57856 |
| ENSG00000198799 | 1 | 113615831 | 113674882 | 141 | 19 | -0.081725 | 0.53257 | LRIG2 | 0.53257 | 0.48933 | 0.52966 |
| ENSG00000160613 | 11 | 117075053 | 117103241 | 114 | 19 | -0.081849 | 0.53262 | PCSK7 | 0.53262 | 0.56477 | 0.62881 |
| ENSG00000168116 | 6 | 56911347 | 56920023 | 6 | 2 | -0.081924 | 0.53265 | KIAA1586 | 0.53265 | 0.52203 | 0.60189 |
| ENSG00000102471 | 13 | 80055287 | 80130210 | 145 | 15 | -0.08202 | 0.53268 | NDFIP2 | 0.53268 | 0.53512 | 0.50967 |
| ENSG00000100276 | 22 | 29708922 | 29715668 | 7 | 4 | -0.082027 | 0.53269 | RASL10A | 0.53269 | 0.33428 | 0.35465 |
| ENSG00000091073 | 7 | 76090993 | 76135312 | 152 | 15 | -0.082065 | 0.5327 | DTX2 | 0.5327 | 0.55336 | 0.55872 |
| ENSG00000100280 | 22 | 29723669 | 29819168 | 286 | 25 | -0.082107 | 0.53272 | AP1B1 | 0.53272 | 0.52751 | 0.43779 |
| ENSG00000120656 | 1 | 28915835 | 28969597 | 84 | 12 | -0.082131 | 0.53273 | TAF12 | 0.53273 | 0.5403 | 0.55314 |
| ENSG00000086827 | 11 | 113603909 | 113644533 | 148 | 19 | -0.082126 | 0.53273 | ZW10 | 0.53273 | 0.56574 | 0.58619 |
| ENSG00000170374 | 12 | 53720362 | 53739099 | 36 | 7 | -0.082357 | 0.53282 | SP7 | 0.53282 | 0.57849 | 0.64215 |
| ENSG00000043355 | 13 | 100634026 | 100639018 | 10 | 4 | -0.08255 | 0.5329 | ZIC2 | 0.5329 | 0.7602 | 0.7511 |
| ENSG00000268865 | 12 | 25055243 | 25057615 | 11 | 4 | -0.082629 | 0.53293 | AC026310.1 | 0.53293 | 0.65626 | 0.75712 |
| ENSG00000116117 | 2 | 205410516 | 206484886 | 3114 | 149 | -0.082939 | 0.53305 | PARD3B | 0.53305 | 0.5474 | 0.55208 |
| ENSG00000179695 | 12 | 55845998 | 55846936 | 1 | 1 | -0.083065 | 0.5331 | OR6C2 | 0.5331 | 0.67579 | 0.73954 |
| ENSG00000067182 | 12 | 6437923 | 6451280 | 27 | 7 | -0.083152 | 0.53313 | TNFRSF1A | 0.53313 | 0.74468 | 0.59709 |
| ENSG00000205358 | 16 | 56703726 | 56705041 | 7 | 3 | -0.083161 | 0.53314 | MT1H | 0.53314 | 0.63704 | 0.60555 |
| ENSG00000085449 | 2 | 224720433 | 224810104 | 305 | 14 | -0.083407 | 0.53324 | WDFY1 | 0.53324 | 0.56253 | 0.65695 |
| ENSG00000085871 | 4 | 140586922 | 140661899 | 272 | 35 | -0.083504 | 0.53327 | MGST2 | 0.53327 | 0.70753 | 0.87437 |
| ENSG00000213077 | 17 | 18427880 | 18430160 | 2 | 1 | -0.083521 | 0.53328 | FAM106A | 0.53328 | 0.31861 | 0.42072 |
| ENSG00000213853 | 16 | 10622279 | 10674555 | 266 | 50 | -0.083757 | 0.53338 | EMP2 | 0.53338 | 0.52404 | 0.43642 |
| ENSG00000204449 | 11 | 89764274 | 89806558 | 86 | 12 | -0.083832 | 0.53341 | TRIM49C | 0.53341 | 0.53183 | 0.59913 |
| ENSG00000166225 | 12 | 69864129 | 69973562 | 303 | 19 | -0.083864 | 0.53342 | FRS2 | 0.53342 | 0.53576 | 0.53983 |
| ENSG00000267360 | 19 | 37583921 | 37701414 | 305 | 21 | -0.083943 | 0.53345 | CTC-454I21.3 | 0.53345 | 0.51139 | 0.40357 |
| ENSG00000123999 | 2 | 220433884 | 220440435 | 17 | 6 | -0.08402 | 0.53348 | INHA | 0.53348 | 0.69239 | 0.6423 |
| ENSG00000172331 | 7 | 134331560 | 134364565 | 141 | 8 | -0.084099 | 0.53351 | BPGM | 0.53351 | 0.53851 | 0.49256 |
| ENSG00000176490 | 19 | 2714565 | 2721416 | 17 | 3 | -0.084087 | 0.53351 | DIRAS1 | 0.53351 | 0.72509 | 0.73587 |
| ENSG00000135870 | 1 | 173900352 | 173991435 | 233 | 15 | -0.084109 | 0.53352 | RC3H1 | 0.53352 | 0.55674 | 0.56894 |
| ENSG00000054967 | 11 | 73087309 | 73108519 | 47 | 9 | -0.084142 | 0.53353 | RELT | 0.53353 | 0.54499 | 0.66883 |
| ENSG00000260836 | 15 | 83208783 | 83224682 | 36 | 11 | -0.084399 | 0.53363 | RP11-152F13.10 | 0.53363 | 0.39716 | 0.27695 |
| ENSG00000103326 | 16 | 577717 | 604636 | 95 | 25 | -0.084637 | 0.53373 | CAPN15 | 0.53373 | 0.4538 | 0.48098 |
| ENSG00000100302 | 22 | 35936915 | 35950048 | 30 | 6 | -0.085091 | 0.53391 | RASD2 | 0.53391 | 0.64799 | 0.29831 |
| ENSG00000145569 | 5 | 14581884 | 14615116 | 50 | 10 | -0.085254 | 0.53397 | FAM105A | 0.53397 | 0.5691 | 0.51833 |
| ENSG00000163145 | 4 | 15341442 | 15447790 | 338 | 20 | -0.08528 | 0.53398 | C1QTNF7 | 0.53398 | 0.53483 | 0.484 |
| ENSG00000266208 | 17 | 38516907 | 38520945 | 12 | 3 | -0.085367 | 0.53402 | CTD-2267D19.3 | 0.53402 | 0.56159 | 0.46671 |

| | | | | | | | | | | | |
|---|---|---|---|---|---|---|---|---|---|---|---|
| ENSG00000159217 | 17 | 47074774 | 47133012 | 147 | 21 | -0.08554 | 0.53408 | IGF2BP1 | 0.53408 | 0.48728 | 0.42225 |
| ENSG00000095917 | 16 | 1306060 | 1308532 | 17 | 5 | -0.085795 | 0.53419 | TPSD1 | 0.53419 | 0.5707 | 0.71525 |
| ENSG00000125285 | 13 | 95361886 | 95364389 | 4 | 2 | -0.085836 | 0.5342 | SOX21 | 0.5342 | 0.40175 | 0.20081 |
| ENSG00000178386 | 19 | 44555520 | 44572144 | 34 | 7 | -0.085892 | 0.53422 | ZNF223 | 0.53422 | 0.53206 | 0.45837 |
| ENSG00000124920 | 11 | 61520114 | 61555990 | 71 | 15 | -0.085999 | 0.53427 | MYRF | 0.53427 | 0.65194 | 0.62853 |
| ENSG00000169184 | 22 | 28144265 | 28197486 | 149 | 53 | -0.086111 | 0.53431 | MN1 | 0.53431 | 0.58469 | 0.23049 |
| ENSG00000203780 | 10 | 127585108 | 127698161 | 379 | 20 | -0.086374 | 0.53442 | FANK1 | 0.53442 | 0.52078 | 0.48543 |
| ENSG00000197461 | 7 | 536895 | 559933 | 111 | 19 | -0.086586 | 0.5345 | PDGFA | 0.5345 | 0.71698 | 0.69086 |
| ENSG00000140876 | 16 | 77756411 | 77776157 | 149 | 19 | -0.086609 | 0.53451 | NUDT7 | 0.53451 | 0.66646 | 0.73 |
| ENSG00000171827 | 19 | 37958487 | 37976260 | 27 | 6 | -0.086696 | 0.53454 | ZNF570 | 0.53454 | 0.45027 | 0.46093 |
| ENSG00000142405 | 19 | 54296857 | 54327648 | 170 | 29 | -0.086688 | 0.53454 | NLRP12 | 0.53454 | 0.6388 | 0.83711 |
| ENSG00000188677 | 22 | 44395091 | 44568829 | 791 | 112 | -0.086804 | 0.53459 | PARVB | 0.53459 | 0.41497 | 0.50215 |
| ENSG00000146072 | 6 | 47199268 | 47277641 | 214 | 38 | -0.087021 | 0.53467 | TNFRSF21 | 0.53467 | 0.45983 | 0.28434 |
| ENSG00000092201 | 14 | 21819631 | 21852425 | 70 | 8 | -0.087061 | 0.53469 | SUPT16H | 0.53469 | 0.38588 | 0.18753 |
| ENSG00000158816 | 1 | 20617412 | 20681387 | 274 | 25 | -0.087118 | 0.53471 | VWA5B1 | 0.53471 | 0.52711 | 0.59554 |
| ENSG00000220702 | 22 | 44761431 | 44762164 | 5 | 3 | -0.087193 | 0.53474 | RP1-32I10.10 | 0.53474 | 0.95834 | 0.93434 |
| ENSG00000099840 | 19 | 2096380 | 2099592 | 9 | 5 | -0.087452 | 0.53484 | IZUMO4 | 0.53484 | 0.16971 | 0.20371 |
| ENSG00000107949 | 10 | 127512115 | 127542264 | 56 | 10 | -0.087464 | 0.53485 | BCCIP | 0.53485 | 0.56894 | 0.595 |
| ENSG00000143294 | 1 | 156720402 | 156770607 | 157 | 21 | -0.087611 | 0.53491 | PRCC | 0.53491 | 0.4848 | 0.52587 |
| ENSG00000167680 | 19 | 4542600 | 4559820 | 69 | 18 | -0.087699 | 0.53494 | SEMA6B | 0.53494 | 0.43186 | 0.67392 |
| ENSG00000267857 | 21 | 46520728 | 46534873 | 45 | 9 | -0.087796 | 0.53498 | PRED58 | 0.53498 | 0.52209 | 0.59213 |
| ENSG00000076043 | 11 | 114310108 | 114321001 | 33 | 10 | -0.088256 | 0.53516 | REXO2 | 0.53516 | 0.20597 | 0.37547 |
| ENSG00000134538 | 12 | 21284136 | 21392180 | 502 | 28 | -0.08851 | 0.53526 | SLCO1B1 | 0.53526 | 0.49063 | 0.42564 |
| ENSG00000198286 | 7 | 2945775 | 3083579 | 703 | 90 | -0.088589 | 0.5353 | CARD11 | 0.5353 | 0.2774 | 0.099315 |
| ENSG00000215504 | 11 | 85563600 | 85565986 | 16 | 3 | -0.088591 | 0.5353 | AP000974.1 | 0.5353 | 0.21859 | 0.24271 |
| ENSG00000204403 | 11 | 104756445 | 104769397 | 10 | 6 | -0.088652 | 0.53532 | CASP12 | 0.53532 | 0.80645 | 0.69899 |
| ENSG00000172689 | 11 | 60552821 | 60568778 | 66 | 11 | -0.088668 | 0.53533 | MS4A10 | 0.53533 | 0.62833 | 0.69315 |
| ENSG00000259803 | 16 | 89262406 | 89268072 | 24 | 7 | -0.088725 | 0.53535 | SLC22A31 | 0.53535 | 0.6075 | 0.87507 |
| ENSG00000113719 | 5 | 172261278 | 172379688 | 433 | 64 | -0.089048 | 0.53548 | ERGIC1 | 0.53548 | 0.52793 | 0.32444 |
| ENSG00000188425 | 19 | 46416475 | 46418036 | 3 | 1 | -0.089104 | 0.5355 | NANOS2 | 0.5355 | 0.29585 | 0.076417 |
| ENSG00000125841 | 20 | 327426 | 340304 | 36 | 13 | -0.089103 | 0.5355 | NRSN2 | 0.5355 | 0.78389 | 0.94145 |
| ENSG00000175344 | 15 | 32322691 | 32464722 | 309 | 49 | -0.089207 | 0.53554 | CHRNA7 | 0.53554 | 0.45914 | 0.47216 |
| ENSG00000117479 | 1 | 169433147 | 169455241 | 52 | 15 | -0.089568 | 0.53568 | SLC19A2 | 0.53568 | 0.70961 | 0.78944 |
| ENSG00000114268 | 3 | 48555117 | 48599448 | 39 | 19 | -0.089594 | 0.5357 | PFKFB4 | 0.5357 | 0.35192 | 0.53861 |
| ENSG00000198839 | 7 | 111846643 | 111983151 | 370 | 24 | -0.089855 | 0.5358 | ZNF277 | 0.5358 | 0.51361 | 0.50171 |
| ENSG00000204022 | 10 | 90346510 | 90366733 | 22 | 7 | -0.08993 | 0.53583 | LIPJ | 0.53583 | 0.46822 | 0.3749 |
| ENSG00000121022 | 8 | 67955314 | 67996018 | 22 | 12 | -0.090002 | 0.53586 | COPS5 | 0.53586 | 0.37881 | 0.48611 |
| ENSG00000173517 | 15 | 77400471 | 77712486 | 555 | 23 | -0.090131 | 0.53591 | PEAK1 | 0.53591 | 0.5103 | 0.49308 |
| ENSG00000107719 | 10 | 72238577 | 72328205 | 440 | 48 | -0.090251 | 0.53596 | PALD1 | 0.53596 | 0.44232 | 0.44903 |
| ENSG00000178752 | 2 | 239067623 | 239077541 | 44 | 13 | -0.090345 | 0.53599 | FAM132B | 0.53599 | 0.54351 | 0.57542 |
| ENSG00000188511 | 22 | 49808176 | 50051190 | 1145 | 87 | -0.09037 | 0.536 | C22orf34 | 0.536 | 0.53405 | 0.45992 |
| ENSG00000145491 | 5 | 10441636 | 10472141 | 102 | 10 | -0.090395 | 0.53601 | ROPN1L | 0.53601 | 0.61239 | 0.77585 |
| ENSG00000180900 | 8 | 144873090 | 144897549 | 73 | 16 | -0.090444 | 0.53603 | SCRIB | 0.53603 | 0.61234 | 0.77698 |
| ENSG00000179902 | 1 | 109648573 | 109656479 | 36 | 7 | -0.090475 | 0.53605 | C1orf194 | 0.53605 | 0.57215 | 0.41615 |
| ENSG00000060138 | 12 | 10851683 | 10875911 | 70 | 11 | -0.090696 | 0.53613 | YBX3 | 0.53613 | 0.62938 | 0.69581 |
| ENSG00000048028 | 11 | 113668596 | 113746292 | 212 | 24 | -0.090804 | 0.53618 | USP28 | 0.53618 | 0.51351 | 0.53524 |
| ENSG00000116497 | 1 | 33282368 | 33324476 | 87 | 14 | -0.090892 | 0.53621 | S100PBP | 0.53621 | 0.56153 | 0.56537 |
| ENSG00000164823 | 8 | 90914087 | 90940116 | 66 | 12 | -0.091033 | 0.53627 | OSGIN2 | 0.53627 | 0.34577 | 0.31549 |
| ENSG00000169972 | 1 | 1243947 | 1247057 | 5 | 3 | -0.091059 | 0.53628 | PUSL1 | 0.53628 | 0.50682 | 0.49823 |
| ENSG00000163737 | 4 | 74846794 | 74847841 | 2 | 1 | -0.091165 | 0.53632 | PF4 | 0.53632 | 0.3425 | 0.36107 |
| ENSG00000197771 | 10 | 121588972 | 121652068 | 258 | 11 | -0.091614 | 0.5365 | MCMBP | 0.5365 | 0.5726 | 0.51133 |
| ENSG00000092068 | 14 | 23594504 | 23652883 | 174 | 38 | -0.091791 | 0.53657 | SLC7A8 | 0.53657 | 0.21575 | 0.18551 |
| ENSG00000134262 | 1 | 114437370 | 114447823 | 20 | 7 | -0.09192 | 0.53662 | AP4B1 | 0.53662 | 0.6446 | 0.5023 |
| ENSG00000268537 | 3 | 46927699 | 46930171 | 3 | 2 | -0.09195 | 0.53663 | AC109583.1 | 0.53663 | 0.62858 | 0.68467 |
| ENSG00000174173 | 3 | 101280706 | 101285290 | 13 | 4 | -0.092035 | 0.53666 | TRMT10C | 0.53666 | 0.46599 | 0.5412 |
| ENSG00000197965 | 1 | 167690429 | 167761156 | 234 | 30 | -0.092184 | 0.53672 | MPZL1 | 0.53672 | 0.50363 | 0.39868 |
| ENSG00000111912 | 6 | 126102307 | 126252266 | 316 | 28 | -0.092176 | 0.53672 | NCOA7 | 0.53672 | 0.45218 | 0.34375 |
| ENSG00000240247 | 8 | 6854288 | 6875778 | 36 | 10 | -0.092248 | 0.53675 | DEFA1B | 0.53675 | 0.52495 | 0.60434 |
| ENSG00000115942 | 2 | 201773696 | 201828403 | 91 | 13 | -0.092393 | 0.53681 | ORC2 | 0.53681 | 0.52376 | 0.50857 |
| ENSG00000105655 | 19 | 18545198 | 18549111 | 8 | 2 | -0.092617 | 0.5369 | ISYNA1 | 0.5369 | 0.81782 | 0.83051 |
| ENSG00000136878 | 9 | 132596977 | 132644107 | 396 | 13 | -0.092794 | 0.53697 | USP20 | 0.53697 | 0.43521 | 0.4196 |
| ENSG00000215182 | 11 | 1151580 | 1222364 | 93 | 11 | -0.092979 | 0.53704 | MUC5AC | 0.53704 | 0.44147 | 0.095346 |
| ENSG00000167325 | 11 | 4115937 | 4160106 | 149 | 15 | -0.09309 | 0.53708 | RRM1 | 0.53708 | 0.60894 | 0.55775 |
| ENSG00000134748 | 1 | 52870236 | 52886511 | 23 | 7 | -0.093165 | 0.53711 | PRPF38A | 0.53711 | 0.52671 | 0.51503 |
| ENSG00000158122 | 9 | 99401859 | 99417585 | 16 | 10 | -0.093235 | 0.53714 | AAED1 | 0.53714 | 0.35832 | 0.52566 |
| ENSG00000254706 | 1 | 162335197 | 162351905 | 60 | 20 | -0.093469 | 0.53723 | RP11-565P22.6 | 0.53723 | 0.31156 | 0.16344 |
| ENSG00000137502 | 11 | 82684175 | 82782965 | 243 | 29 | -0.093553 | 0.53727 | RAB30 | 0.53727 | 0.54772 | 0.53575 |
| ENSG00000109851 | 11 | 20177701 | 20182159 | 9 | 4 | -0.093709 | 0.53733 | DBX1 | 0.53733 | 0.48018 | 0.14156 |
| ENSG00000100227 | 22 | 42979727 | 43010968 | 68 | 13 | -0.093712 | 0.53733 | POLDIP3 | 0.53733 | 0.42995 | 0.35536 |
| ENSG00000180613 | 4 | 54965690 | 54968672 | 8 | 2 | -0.093896 | 0.5374 | GSX2 | 0.5374 | 0.61022 | 0.62966 |
| ENSG00000179869 | 7 | 48211055 | 48687092 | 2044 | 70 | -0.093978 | 0.53744 | ABCA13 | 0.53744 | 0.54351 | 0.55711 |
| ENSG00000164199 | 5 | 89825161 | 90460038 | 1781 | 67 | -0.09427 | 0.53755 | GPR98 | 0.53755 | 0.5533 | 0.52763 |
| ENSG00000184619 | 17 | 8271955 | 8280029 | 21 | 5 | -0.094443 | 0.53762 | KRBA2 | 0.53762 | 0.2822 | 0.39911 |

| | | | | | | | | | | | |
|---|---|---|---|---|---|---|---|---|---|---|---|
| ENSG00000186081 | 12 | 52908359 | 52914471 | 27 | 2 | -0.094889 | 0.5378 | KRT5 | 0.5378 | 0.67421 | 0.63476 |
| ENSG00000196182 | 1 | 36805225 | 36851497 | 70 | 10 | -0.095017 | 0.53785 | STK40 | 0.53785 | 0.46418 | 0.30815 |
| ENSG00000188846 | 3 | 40498783 | 40503861 | 19 | 7 | -0.095023 | 0.53785 | RPL14 | 0.53785 | 0.50205 | 0.59827 |
| ENSG00000124678 | 6 | 35085848 | 35116387 | 119 | 13 | -0.095018 | 0.53785 | TCP11 | 0.53785 | 0.61837 | 0.58974 |
| ENSG00000255472 | 14 | 21500977 | 21502054 | 11 | 3 | -0.095062 | 0.53787 | RP11-998D10.1 | 0.53787 | 0.69218 | 0.85122 |
| ENSG00000269267 | 5 | 60411538 | 60411762 | 1 | 1 | -0.095145 | 0.5379 | AC008498.1 | 0.5379 | 0.084733 | 0.099266 |
| ENSG00000168412 | 4 | 187454809 | 187476721 | 100 | 20 | -0.095465 | 0.53803 | MTNR1A | 0.53803 | 0.58487 | 0.72987 |
| ENSG00000167280 | 17 | 77071021 | 77084681 | 79 | 15 | -0.095526 | 0.53805 | ENGASE | 0.53805 | 0.70686 | 0.64737 |
| ENSG00000102683 | 13 | 23755091 | 23899304 | 1188 | 69 | -0.095581 | 0.53807 | SGCG | 0.53807 | 0.55471 | 0.62131 |
| ENSG00000204130 | 10 | 70100864 | 70167051 | 191 | 13 | -0.095588 | 0.53808 | RUFY2 | 0.53808 | 0.58317 | 0.61812 |
| ENSG00000120696 | 13 | 41763969 | 41768702 | 6 | 2 | -0.095599 | 0.53808 | KBTBD7 | 0.53808 | 0.73691 | 0.71814 |
| ENSG00000151893 | 10 | 120433679 | 120514761 | 205 | 27 | -0.095614 | 0.53809 | CACUL1 | 0.53809 | 0.42736 | 0.40134 |
| ENSG00000213722 | 6 | 31694815 | 31698394 | 9 | 4 | -0.095828 | 0.53817 | DDAH2 | 0.53817 | 0.62755 | 0.67109 |
| ENSG00000154133 | 11 | 124753587 | 124768396 | 46 | 14 | -0.095821 | 0.53817 | ROBO4 | 0.53817 | 0.58547 | 0.56765 |
| ENSG00000168329 | 3 | 39304985 | 39323226 | 64 | 19 | -0.095875 | 0.53819 | CX3CR1 | 0.53819 | 0.67976 | 0.85372 |
| ENSG00000153283 | 3 | 111011566 | 111384597 | 1253 | 60 | -0.096234 | 0.53833 | CD96 | 0.53833 | 0.55313 | 0.48302 |
| ENSG00000130731 | 16 | 684429 | 686358 | 5 | 3 | -0.096242 | 0.53834 | C16orf13 | 0.53834 | 0.44646 | 0.31367 |
| ENSG00000185821 | 12 | 55820038 | 55820976 | 1 | 1 | -0.096404 | 0.5384 | OR6C76 | 0.5384 | 0.69526 | 0.7193 |
| ENSG00000172590 | 14 | 23299088 | 23304246 | 27 | 9 | -0.096468 | 0.53843 | MRPL52 | 0.53843 | 0.70412 | 0.67959 |
| ENSG00000080371 | 12 | 72148654 | 72184699 | 31 | 11 | -0.096508 | 0.53844 | RAB21 | 0.53844 | 0.49861 | 0.54805 |
| ENSG00000119121 | 9 | 77337411 | 77503010 | 369 | 39 | -0.096541 | 0.53845 | TRPM6 | 0.53845 | 0.54087 | 0.64508 |
| ENSG00000105173 | 19 | 30302805 | 30315215 | 36 | 11 | -0.096523 | 0.53845 | CCNE1 | 0.53845 | 0.42416 | 0.4159 |
| ENSG00000104218 | 8 | 67974661 | 68108498 | 90 | 33 | -0.096582 | 0.53847 | CSPP1 | 0.53847 | 0.56939 | 0.56215 |
| ENSG00000089847 | 19 | 4183351 | 4224811 | 223 | 31 | -0.096605 | 0.53848 | ANKRD24 | 0.53848 | 0.54006 | 0.63962 |
| ENSG00000077935 | 22 | 45739944 | 45809500 | 213 | 14 | -0.096902 | 0.5386 | SMC1B | 0.5386 | 0.52959 | 0.56812 |
| ENSG00000139055 | 12 | 15066969 | 15092016 | 98 | 6 | -0.097043 | 0.53865 | ERP27 | 0.53865 | 0.33138 | 0.24487 |
| ENSG00000154767 | 3 | 14186647 | 14220283 | 72 | 15 | -0.097101 | 0.53868 | XPC | 0.53868 | 0.5095 | 0.53879 |
| ENSG00000130783 | 12 | 123258874 | 123312075 | 141 | 18 | -0.097107 | 0.53868 | CCDC62 | 0.53868 | 0.55072 | 0.58585 |
| ENSG00000154065 | 18 | 21178890 | 21242849 | 112 | 24 | -0.097096 | 0.53868 | ANKRD29 | 0.53868 | 0.29987 | 0.38305 |
| ENSG00000156831 | 8 | 126103921 | 126379362 | 719 | 53 | -0.09716 | 0.5387 | NSMCE2 | 0.5387 | 0.58192 | 0.57573 |
| ENSG00000251369 | 19 | 58046625 | 58071231 | 29 | 11 | -0.097315 | 0.53876 | ZNF550 | 0.53876 | 0.49341 | 0.50452 |
| ENSG00000105388 | 19 | 42212504 | 42233718 | 55 | 11 | -0.097428 | 0.53881 | CEACAM5 | 0.53881 | 0.49695 | 0.63965 |
| ENSG00000138030 | 2 | 27309615 | 27323640 | 22 | 5 | -0.09752 | 0.53884 | KHK | 0.53884 | 0.60159 | 0.72178 |
| ENSG00000162992 | 2 | 182537815 | 182545603 | 10 | 3 | -0.097517 | 0.53884 | NEUROD1 | 0.53884 | 0.32635 | 0.22581 |
| ENSG00000145283 | 4 | 87744621 | 87770416 | 55 | 13 | -0.097539 | 0.53885 | SLC10A6 | 0.53885 | 0.46918 | 0.54027 |
| ENSG00000101558 | 18 | 9913999 | 9960018 | 167 | 26 | -0.097844 | 0.53897 | VAPA | 0.53897 | 0.48161 | 0.46172 |
| ENSG00000165028 | 9 | 107526438 | 107539738 | 34 | 10 | -0.097989 | 0.53903 | NIPSNAP3B | 0.53903 | 0.77756 | 0.7736 |
| ENSG00000135801 | 1 | 229728858 | 229761794 | 98 | 12 | -0.098309 | 0.53916 | TAF5L | 0.53916 | 0.59522 | 0.60754 |
| ENSG00000116147 | 1 | 175284330 | 175712906 | 1379 | 115 | -0.098403 | 0.53919 | TNR | 0.53919 | 0.56854 | 0.64462 |
| ENSG00000163389 | 3 | 119187785 | 119213555 | 82 | 15 | -0.098632 | 0.53928 | POGLUT1 | 0.53928 | 0.5589 | 0.56976 |
| ENSG00000182652 | 14 | 20215587 | 20216528 | 1 | 1 | -0.09867 | 0.5393 | OR4Q3 | 0.5393 | 0.9231 | 0.92927 |
| ENSG00000124126 | 20 | 47240790 | 47444420 | 652 | 40 | -0.098855 | 0.53937 | PREX1 | 0.53937 | 0.5411 | 0.52395 |
| ENSG00000214021 | 3 | 9849770 | 9896822 | 165 | 21 | -0.098905 | 0.53939 | TTLL3 | 0.53939 | 0.5924 | 0.62693 |
| ENSG00000140968 | 16 | 85932409 | 85956215 | 86 | 13 | -0.098947 | 0.53941 | IRF8 | 0.53941 | 0.28736 | 0.49153 |
| ENSG00000189337 | 1 | 14925200 | 15444539 | 2286 | 219 | -0.098985 | 0.53943 | KAZN | 0.53943 | 0.5386 | 0.60034 |
| ENSG00000127074 | 1 | 192605275 | 192629390 | 82 | 8 | -0.099022 | 0.53944 | RGS13 | 0.53944 | 0.51708 | 0.47855 |
| ENSG00000159788 | 4 | 3294755 | 3441640 | 373 | 41 | -0.099065 | 0.53946 | RGS12 | 0.53946 | 0.54138 | 0.60615 |
| ENSG00000085831 | 1 | 51752930 | 51810788 | 128 | 26 | -0.099245 | 0.53953 | TTC39A | 0.53953 | 0.59835 | 0.6754 |
| ENSG00000113569 | 5 | 37288239 | 37371283 | 181 | 17 | -0.099453 | 0.53961 | NUP155 | 0.53961 | 0.53436 | 0.49603 |
| ENSG00000163406 | 3 | 121612936 | 121662949 | 210 | 6 | -0.099653 | 0.53969 | SLC15A2 | 0.53969 | 0.50081 | 0.44243 |
| ENSG00000079215 | 5 | 36606457 | 36688436 | 225 | 51 | -0.099761 | 0.53973 | SLC1A3 | 0.53973 | 0.58543 | 0.65944 |
| ENSG00000261857 | 19 | 41277553 | 41283395 | 15 | 6 | -0.099856 | 0.53977 | MIA | 0.53977 | 0.5774 | 0.59651 |
| ENSG00000125046 | 3 | 8661086 | 8786726 | 663 | 110 | -0.10025 | 0.53993 | SSUH2 | 0.53993 | 0.64457 | 0.73846 |
| ENSG00000162616 | 1 | 78444859 | 78483648 | 118 | 23 | -0.10033 | 0.53996 | DNAJB4 | 0.53996 | 0.58946 | 0.62509 |
| ENSG00000196363 | 9 | 137000487 | 137025093 | 190 | 14 | -0.10039 | 0.53998 | WDR5 | 0.53998 | 0.43203 | 0.51346 |
| ENSG00000118046 | 19 | 1189406 | 1228428 | 114 | 28 | -0.10068 | 0.5401 | STK11 | 0.5401 | 0.62485 | 0.3697 |
| ENSG00000140987 | 16 | 3432085 | 3451065 | 37 | 6 | -0.10072 | 0.54011 | ZSCAN32 | 0.54011 | 0.71407 | 0.77244 |
| ENSG00000185024 | 14 | 105675623 | 105781926 | 304 | 17 | -0.10101 | 0.54023 | BRF1 | 0.54023 | 0.54294 | 0.56257 |
| ENSG00000239672 | 17 | 49230897 | 49239789 | 39 | 7 | -0.10113 | 0.54028 | NME1 | 0.54028 | 0.56615 | 0.56218 |
| ENSG00000139737 | 13 | 78272023 | 78338377 | 151 | 25 | -0.10139 | 0.54038 | SLAIN1 | 0.54038 | 0.59858 | 0.71936 |
| ENSG00000173376 | 4 | 121956768 | 121994176 | 106 | 19 | -0.10171 | 0.54051 | NDNF | 0.54051 | 0.55101 | 0.59457 |
| ENSG00000140443 | 15 | 99192200 | 99507759 | 1071 | 110 | -0.10189 | 0.54058 | IGF1R | 0.54058 | 0.53534 | 0.33151 |
| ENSG00000115806 | 2 | 171784974 | 171823639 | 106 | 12 | -0.10201 | 0.54063 | GORASP2 | 0.54063 | 0.58051 | 0.59241 |
| ENSG00000033327 | 11 | 77926343 | 78129394 | 453 | 9 | -0.10211 | 0.54067 | GAB2 | 0.54067 | 0.53126 | 0.54933 |
| ENSG00000011260 | 17 | 49337889 | 49375297 | 122 | 14 | -0.1022 | 0.5407 | UTP18 | 0.5407 | 0.50415 | 0.52116 |
| ENSG00000163467 | 1 | 156307105 | 156316786 | 20 | 6 | -0.10236 | 0.54077 | TSACC | 0.54077 | 0.44778 | 0.50817 |
| ENSG00000136381 | 15 | 78729773 | 78793798 | 170 | 17 | -0.10248 | 0.54081 | IREB2 | 0.54081 | 0.52271 | 0.49617 |
| ENSG00000105404 | 19 | 42460833 | 42463542 | 4 | 2 | -0.10265 | 0.54088 | RABAC1 | 0.54088 | 0.44582 | 0.44952 |
| ENSG00000155636 | 2 | 178977151 | 179003738 | 54 | 14 | -0.10273 | 0.54091 | RBM45 | 0.54091 | 0.52362 | 0.72252 |
| ENSG00000085788 | 8 | 38082736 | 38133076 | 60 | 9 | -0.10284 | 0.54095 | DDHD2 | 0.54095 | 0.51852 | 0.51051 |
| ENSG00000147604 | 8 | 74202506 | 74208024 | 21 | 9 | -0.10288 | 0.54097 | RPL7 | 0.54097 | 0.77981 | 0.81919 |
| ENSG00000186431 | 19 | 55385704 | 55401838 | 152 | 21 | -0.103 | 0.54102 | FCAR | 0.54102 | 0.63018 | 0.72504 |
| ENSG00000179604 | 17 | 71279763 | 71308314 | 111 | 22 | -0.10304 | 0.54103 | CDC42EP4 | 0.54103 | 0.54795 | 0.47893 |

| | | | | | | | | | | | |
|---|---|---|---|---|---|---|---|---|---|---|---|
| ENSG00000149136 | 11 | 57093459 | 57103351 | 23 | 5 | -0.10335 | 0.54116 | SSRP1 | 0.54116 | 0.72459 | 0.86992 |
| ENSG00000189159 | 17 | 73131343 | 73164376 | 77 | 11 | -0.10342 | 0.54119 | HN1 | 0.54119 | 0.52437 | 0.55767 |
| ENSG00000137575 | 8 | 59465483 | 59495419 | 59 | 4 | -0.10363 | 0.54127 | SDCBP | 0.54127 | 0.5986 | 0.69004 |
| ENSG00000134287 | 12 | 49329506 | 49351334 | 85 | 8 | -0.10362 | 0.54127 | ARF3 | 0.54127 | 0.52334 | 0.52578 |
| ENSG00000069020 | 5 | 65892176 | 66465423 | 1816 | 98 | -0.10371 | 0.5413 | MAST4 | 0.5413 | 0.43637 | 0.27164 |
| ENSG00000171722 | 1 | 162342993 | 162346660 | 13 | 5 | -0.10381 | 0.54134 | C1orf111 | 0.54134 | 0.3916 | 0.22932 |
| ENSG00000183389 | 11 | 6023216 | 6024424 | 3 | 1 | -0.10406 | 0.54144 | OR56A4 | 0.54144 | 0.76345 | 0.70641 |
| ENSG00000268818 | 22 | 36023040 | 36031181 | 36 | 9 | -0.1041 | 0.54145 | AL049747.1 | 0.54145 | 0.4054 | 0.29165 |
| ENSG00000100147 | 22 | 42196683 | 42222303 | 67 | 11 | -0.10423 | 0.5415 | CCDC134 | 0.5415 | 0.48651 | 0.5667 |
| ENSG00000163909 | 1 | 40089825 | 40105617 | 45 | 17 | -0.10439 | 0.54157 | HEYL | 0.54157 | 0.70971 | 0.70265 |
| ENSG00000153485 | 14 | 93651296 | 93653434 | 4 | 2 | -0.10447 | 0.5416 | TMEM251 | 0.5416 | 0.8179 | 0.51459 |
| ENSG00000115446 | 2 | 99225042 | 99234978 | 21 | 4 | -0.10465 | 0.54167 | UNC50 | 0.54167 | 0.35572 | 0.29278 |
| ENSG00000186453 | 2 | 24397938 | 24423718 | 108 | 14 | -0.10466 | 0.54168 | FAM228A | 0.54168 | 0.52418 | 0.48723 |
| ENSG00000140455 | 15 | 63796793 | 63886839 | 251 | 24 | -0.10482 | 0.54174 | USP3 | 0.54174 | 0.57577 | 0.5284 |
| ENSG00000103657 | 15 | 63900817 | 64126141 | 673 | 22 | -0.10487 | 0.54176 | HERC1 | 0.54176 | 0.56222 | 0.57771 |
| ENSG00000268950 | 1 | 31902192 | 31902614 | 2 | 1 | -0.10502 | 0.54182 | AC114494.1 | 0.54182 | 0.66815 | 0.69427 |
| ENSG00000116584 | 1 | 155916630 | 155976861 | 108 | 16 | -0.10511 | 0.54185 | ARHGEF2 | 0.54185 | 0.55209 | 0.57703 |
| ENSG00000112116 | 6 | 52101479 | 52109335 | 30 | 9 | -0.10528 | 0.54192 | IL17F | 0.54192 | 0.76637 | 0.7767 |
| ENSG00000114638 | 3 | 118892364 | 118924000 | 110 | 23 | -0.10532 | 0.54194 | UPK1B | 0.54194 | 0.56355 | 0.57591 |
| ENSG00000172954 | 2 | 30670092 | 30867091 | 789 | 22 | -0.10563 | 0.54206 | LCLAT1 | 0.54206 | 0.58737 | 0.60812 |
| ENSG00000158458 | 5 | 139226364 | 139422884 | 433 | 45 | -0.10566 | 0.54208 | NRG2 | 0.54208 | 0.52977 | 0.49873 |
| ENSG00000221866 | 7 | 131808091 | 132333447 | 1841 | 146 | -0.10572 | 0.5421 | PLXNA4 | 0.5421 | 0.52367 | 0.49414 |
| ENSG00000177710 | 8 | 11188397 | 11189717 | 11 | 3 | -0.10577 | 0.54212 | SLC35G5 | 0.54212 | 0.46564 | 0.41631 |
| ENSG00000165115 | 9 | 86451613 | 86536342 | 211 | 15 | -0.10581 | 0.54213 | KIF27 | 0.54213 | 0.49206 | 0.54498 |
| ENSG00000006530 | 7 | 141250989 | 141355044 | 175 | 13 | -0.10591 | 0.54217 | AGK | 0.54217 | 0.58263 | 0.6398 |
| ENSG00000028528 | 15 | 64386322 | 64438289 | 110 | 14 | -0.10593 | 0.54218 | SNX1 | 0.54218 | 0.52198 | 0.57212 |
| ENSG00000165630 | 10 | 13628927 | 13672868 | 115 | 18 | -0.10598 | 0.5422 | PRPF18 | 0.5422 | 0.46368 | 0.39418 |
| ENSG00000162975 | 2 | 11052063 | 11054350 | 4 | 2 | -0.10608 | 0.54224 | KCNF1 | 0.54224 | 0.26438 | 0.6194 |
| ENSG00000180116 | 12 | 40019969 | 40302102 | 891 | 29 | -0.10609 | 0.54224 | C12orf40 | 0.54224 | 0.50616 | 0.4739 |
| ENSG00000148671 | 10 | 88727949 | 88743691 | 61 | 10 | -0.1061 | 0.54225 | ADIRF | 0.54225 | 0.37261 | 0.29724 |
| ENSG00000154153 | 5 | 16473147 | 16617167 | 529 | 41 | -0.10635 | 0.54235 | FAM134B | 0.54235 | 0.57453 | 0.61511 |
| ENSG00000095627 | 10 | 115939029 | 115992063 | 146 | 14 | -0.10635 | 0.54235 | TDRD1 | 0.54235 | 0.60439 | 0.61687 |
| ENSG00000140323 | 15 | 40650436 | 40663257 | 33 | 9 | -0.10646 | 0.54239 | DISP2 | 0.54239 | 0.60226 | 0.69147 |
| ENSG00000170255 | 11 | 18955360 | 18961054 | 25 | 5 | -0.10662 | 0.54246 | MRGPRX1 | 0.54246 | 0.45362 | 0.044196 |
| ENSG00000105053 | 19 | 50479724 | 50529203 | 163 | 17 | -0.10665 | 0.54247 | VRK3 | 0.54247 | 0.41542 | 0.38582 |
| ENSG00000144476 | 2 | 237476430 | 237491001 | 37 | 9 | -0.10673 | 0.5425 | ACKR3 | 0.5425 | 0.54547 | 0.53553 |
| ENSG00000090520 | 3 | 186285192 | 186315061 | 107 | 7 | -0.10689 | 0.54256 | DNAJB11 | 0.54256 | 0.56718 | 0.54299 |
| ENSG00000179615 | 12 | 55968199 | 55969128 | 1 | 1 | -0.10699 | 0.5426 | OR2AP1 | 0.5426 | 0.88042 | 0.86352 |
| ENSG00000138448 | 2 | 187454792 | 187545628 | 190 | 25 | -0.10724 | 0.5427 | ITGAV | 0.5427 | 0.53568 | 0.6066 |
| ENSG00000197405 | 19 | 47793280 | 47825323 | 73 | 15 | -0.10744 | 0.54278 | C5AR1 | 0.54278 | 0.71587 | 0.35101 |
| ENSG00000110002 | 11 | 123986069 | 124018428 | 95 | 18 | -0.10753 | 0.54282 | VWA5A | 0.54282 | 0.38991 | 0.56531 |
| ENSG00000155368 | 2 | 120124497 | 120130126 | 28 | 6 | -0.10755 | 0.54283 | DBI | 0.54283 | 0.66549 | 0.77464 |
| ENSG00000110048 | 11 | 59341871 | 59383617 | 102 | 11 | -0.10783 | 0.54293 | OSBP | 0.54293 | 0.53161 | 0.49693 |
| ENSG00000153487 | 13 | 111365083 | 111373421 | 18 | 4 | -0.10805 | 0.54302 | ING1 | 0.54302 | 0.6875 | 0.72376 |
| ENSG00000198160 | 1 | 67390578 | 67454302 | 170 | 16 | -0.1081 | 0.54304 | MIER1 | 0.54304 | 0.58726 | 0.65357 |
| ENSG00000186908 | 12 | 77157368 | 77247476 | 378 | 16 | -0.10811 | 0.54305 | ZDHHC17 | 0.54305 | 0.56357 | 0.58929 |
| ENSG00000254598 | 11 | 11373489 | 11374904 | 2 | 1 | -0.10817 | 0.54307 | CSNK2A3 | 0.54307 | 0.57185 | 0.38024 |
| ENSG00000132297 | 8 | 133073733 | 133123406 | 209 | 33 | -0.10845 | 0.54318 | HHLA1 | 0.54318 | 0.51233 | 0.4084 |
| ENSG00000186889 | 2 | 62727356 | 62739029 | 53 | 10 | -0.10861 | 0.54324 | TMEM17 | 0.54324 | 0.48759 | 0.49275 |
| ENSG00000149923 | 16 | 30087299 | 30096698 | 15 | 4 | -0.10864 | 0.54326 | PPP4C | 0.54326 | 0.69323 | 0.68641 |
| ENSG00000180881 | 12 | 75669759 | 75784708 | 224 | 24 | -0.10876 | 0.5433 | CAPS2 | 0.5433 | 0.35373 | 0.25285 |
| ENSG00000198876 | 9 | 34086385 | 34127397 | 158 | 19 | -0.10881 | 0.54332 | DCAF12 | 0.54332 | 0.55161 | 0.58754 |
| ENSG00000100523 | 14 | 53510686 | 53620000 | 213 | 15 | -0.10888 | 0.54335 | DDHD1 | 0.54335 | 0.52715 | 0.53394 |
| ENSG00000138653 | 4 | 115748919 | 116035032 | 889 | 41 | -0.10919 | 0.54348 | NDST4 | 0.54348 | 0.57139 | 0.53417 |
| ENSG00000104964 | 19 | 3052908 | 3063105 | 41 | 13 | -0.10922 | 0.54349 | AES | 0.54349 | 0.94242 | 0.9292 |
| ENSG00000159692 | 4 | 1205236 | 1243741 | 148 | 18 | -0.10925 | 0.5435 | CTBP1 | 0.5435 | 0.47594 | 0.36192 |
| ENSG00000109381 | 4 | 139949266 | 140098372 | 278 | 27 | -0.10931 | 0.54352 | ELF2 | 0.54352 | 0.50665 | 0.6066 |
| ENSG00000267706 | 19 | 55863669 | 55870884 | 27 | 10 | -0.10929 | 0.54352 | CTD-2105E13.6 | 0.54352 | 0.48042 | 0.42833 |
| ENSG00000125459 | 1 | 155579979 | 155718153 | 143 | 26 | -0.10947 | 0.54359 | MSTO1 | 0.54359 | 0.56172 | 0.56784 |
| ENSG00000171953 | 17 | 17880723 | 17942523 | 108 | 11 | -0.10959 | 0.54363 | ATPAF2 | 0.54363 | 0.5613 | 0.5821 |
| ENSG00000099985 | 22 | 30658818 | 30662829 | 8 | 4 | -0.10964 | 0.54365 | OSM | 0.54365 | 0.54648 | 0.28068 |
| ENSG00000137098 | 9 | 35808042 | 35812269 | 5 | 2 | -0.10966 | 0.54366 | SPAG8 | 0.54366 | 0.82618 | 0.75332 |
| ENSG00000169181 | 16 | 27798850 | 28074830 | 1028 | 104 | -0.10984 | 0.54373 | GSG1L | 0.54373 | 0.57314 | 0.59688 |
| ENSG00000119280 | 1 | 230972865 | 231005335 | 129 | 24 | -0.11 | 0.54379 | C1orf198 | 0.54379 | 0.3932 | 0.12325 |
| ENSG00000135409 | 12 | 53817639 | 53825318 | 13 | 1 | -0.1105 | 0.54399 | AMHR2 | 0.54399 | 0.64098 | 0.73648 |
| ENSG00000165914 | 14 | 91006932 | 91282823 | 1050 | 75 | -0.11048 | 0.54399 | TTC7B | 0.54399 | 0.59077 | 0.64539 |
| ENSG00000132464 | 4 | 71494461 | 71552533 | 109 | 18 | -0.11055 | 0.54401 | ENAM | 0.54401 | 0.50907 | 0.52921 |
| ENSG00000166289 | 19 | 30155963 | 30166364 | 28 | 6 | -0.11068 | 0.54406 | PLEKHF1 | 0.54406 | 0.69429 | 0.68749 |
| ENSG00000168067 | 11 | 64556290 | 64570713 | 15 | 6 | -0.11117 | 0.54426 | MAP4K2 | 0.54426 | 0.84906 | 0.8412 |
| ENSG00000123374 | 12 | 56360553 | 56366568 | 8 | 4 | -0.11133 | 0.54432 | CDK2 | 0.54432 | 0.59914 | 0.56131 |
| ENSG00000078814 | 20 | 33563206 | 33590240 | 88 | 12 | -0.11132 | 0.54432 | MYH7B | 0.54432 | 0.56237 | 0.55562 |
| ENSG00000134138 | 15 | 37181406 | 37393504 | 488 | 84 | -0.11142 | 0.54436 | MEIS2 | 0.54436 | 0.75767 | 0.81596 |
| ENSG00000161594 | 17 | 39991937 | 40004636 | 50 | 5 | -0.11151 | 0.54439 | KLHL10 | 0.54439 | 0.53498 | 0.4961 |

| | | | | | | | | | | | |
|---|---|---|---|---|---|---|---|---|---|---|---|
| ENSG00000117408 | 1 | 44412611 | 44433694 | 34 | 6 | -0.11201 | 0.54459 | IPO13 | 0.54459 | 0.59487 | 0.63085 |
| ENSG00000124587 | 6 | 42931608 | 42946958 | 67 | 10 | -0.11205 | 0.54461 | PEX6 | 0.54461 | 0.56865 | 0.69085 |
| ENSG00000112414 | 6 | 142622991 | 142767403 | 268 | 24 | -0.11206 | 0.54461 | GPR126 | 0.54461 | 0.55954 | 0.59115 |
| ENSG00000099942 | 22 | 21271714 | 21308037 | 129 | 12 | -0.11214 | 0.54464 | CRKL | 0.54464 | 0.45715 | 0.45668 |
| ENSG00000172123 | 17 | 33738079 | 33760302 | 76 | 11 | -0.11232 | 0.54472 | SLFN12 | 0.54472 | 0.57912 | 0.70888 |
| ENSG00000046604 | 18 | 29078006 | 29128971 | 147 | 16 | -0.11233 | 0.54472 | DSG2 | 0.54472 | 0.57208 | 0.80455 |
| ENSG00000013374 | 7 | 151038785 | 151075535 | 160 | 30 | -0.1124 | 0.54475 | NUB1 | 0.54475 | 0.45786 | 0.52763 |
| ENSG00000144283 | 2 | 159313476 | 159539391 | 563 | 26 | -0.11299 | 0.54498 | PKP4 | 0.54498 | 0.54394 | 0.55418 |
| ENSG00000171302 | 17 | 76987799 | 77005949 | 88 | 17 | -0.1132 | 0.54506 | CANT1 | 0.54506 | 0.61893 | 0.42494 |
| ENSG00000128708 | 2 | 172778958 | 172848599 | 203 | 26 | -0.11328 | 0.54509 | HAT1 | 0.54509 | 0.51978 | 0.43065 |
| ENSG00000030419 | 2 | 213864429 | 214017151 | 462 | 31 | -0.11348 | 0.54517 | IKZF2 | 0.54517 | 0.54039 | 0.5359 |
| ENSG00000100528 | 14 | 54893654 | 54908149 | 35 | 5 | -0.11351 | 0.54519 | CNIH1 | 0.54519 | 0.60502 | 0.73733 |
| ENSG00000211456 | 3 | 45730548 | 45786916 | 229 | 11 | -0.11355 | 0.5452 | SACM1L | 0.5452 | 0.42959 | 0.25477 |
| ENSG00000225663 | 17 | 79780287 | 79791178 | 22 | 2 | -0.11372 | 0.54527 | FAM195B | 0.54527 | 0.60329 | 0.62826 |
| ENSG00000140511 | 15 | 89420519 | 89438857 | 84 | 11 | -0.11374 | 0.54528 | HAPLN3 | 0.54528 | 0.49392 | 0.47713 |
| ENSG00000121716 | 7 | 99933737 | 99965356 | 77 | 6 | -0.11394 | 0.54536 | PILRB | 0.54536 | 0.4554 | 0.46094 |
| ENSG00000144840 | 3 | 120405528 | 120461840 | 115 | 17 | -0.11419 | 0.54546 | RABL3 | 0.54546 | 0.60253 | 0.68894 |
| ENSG00000183474 | 5 | 68856035 | 68890550 | 14 | 6 | -0.11436 | 0.54552 | GTF2H2C | 0.54552 | 0.61387 | 0.67549 |
| ENSG00000164405 | 5 | 132202252 | 132203723 | 4 | 2 | -0.11437 | 0.54553 | UQCRQ | 0.54553 | 0.83793 | 0.83146 |
| ENSG00000204278 | 17 | 76227391 | 76237068 | 43 | 10 | -0.11459 | 0.54562 | TMEM235 | 0.54562 | 0.59199 | 0.56608 |
| ENSG00000108439 | 17 | 46018872 | 46025654 | 17 | 7 | -0.11462 | 0.54563 | PNPO | 0.54563 | 0.75806 | 0.84125 |
| ENSG00000267988 | 7 | 45902684 | 45906045 | 14 | 5 | -0.11473 | 0.54567 | AC096582.1 | 0.54567 | 0.42992 | 0.56193 |
| ENSG00000112200 | 6 | 56951642 | 57035105 | 117 | 11 | -0.11499 | 0.54577 | ZNF451 | 0.54577 | 0.56216 | 0.52272 |
| ENSG00000203778 | 6 | 112408802 | 112423993 | 51 | 2 | -0.11498 | 0.54577 | FAM229B | 0.54577 | 0.57518 | 0.58045 |
| ENSG00000172377 | 11 | 57885972 | 57886916 | 1 | 1 | -0.11531 | 0.5459 | OR9I1 | 0.5459 | 0.052275 | 0.05655 |
| ENSG00000143643 | 1 | 231041989 | 231114621 | 332 | 23 | -0.11555 | 0.54599 | TTC13 | 0.54599 | 0.4295 | 0.38009 |
| ENSG00000183090 | 4 | 144498455 | 144621828 | 392 | 24 | -0.11557 | 0.546 | FREM3 | 0.546 | 0.60216 | 0.64134 |
| ENSG00000224982 | 12 | 120031264 | 120079363 | 169 | 13 | -0.11563 | 0.54603 | TMEM233 | 0.54603 | 0.60252 | 0.65663 |
| ENSG00000240747 | 3 | 42850975 | 42984284 | 314 | 27 | -0.11573 | 0.54607 | KRBOX1 | 0.54607 | 0.39918 | 0.43464 |
| ENSG00000161021 | 5 | 179159851 | 179223512 | 209 | 19 | -0.11575 | 0.54607 | MAML1 | 0.54607 | 0.57939 | 0.59054 |
| ENSG00000187664 | 19 | 19366450 | 19373605 | 18 | 3 | -0.11581 | 0.5461 | HAPLN4 | 0.5461 | 0.73112 | 0.8433 |
| ENSG00000172548 | 5 | 156887027 | 156901725 | 53 | 10 | -0.11588 | 0.54613 | NIPAL4 | 0.54613 | 0.45085 | 0.37707 |
| ENSG00000196247 | 7 | 64126511 | 64171404 | 136 | 16 | -0.11597 | 0.54616 | ZNF107 | 0.54616 | 0.49287 | 0.55776 |
| ENSG00000172940 | 3 | 38307303 | 38319806 | 18 | 3 | -0.11634 | 0.54631 | SLC22A13 | 0.54631 | 0.84194 | 0.43528 |
| ENSG00000116209 | 1 | 54497347 | 54519177 | 54 | 11 | -0.11643 | 0.54635 | TMEM59 | 0.54635 | 0.50564 | 0.27031 |
| ENSG00000125753 | 19 | 46009837 | 46030241 | 70 | 13 | -0.11645 | 0.54635 | VASP | 0.54635 | 0.53574 | 0.51331 |
| ENSG00000090857 | 16 | 70147529 | 70195203 | 131 | 22 | -0.11673 | 0.54646 | PDPR | 0.54646 | 0.60144 | 0.55064 |
| ENSG00000152439 | 19 | 58011283 | 58029772 | 143 | 5 | -0.11679 | 0.54649 | ZNF773 | 0.54649 | 0.35338 | 0.37589 |
| ENSG00000143669 | 1 | 235824341 | 236046940 | 352 | 19 | -0.11692 | 0.54654 | LYST | 0.54654 | 0.56828 | 0.57437 |
| ENSG00000080618 | 13 | 46627321 | 46679211 | 184 | 11 | -0.11722 | 0.54666 | CPB2 | 0.54666 | 0.51066 | 0.47979 |
| ENSG00000115145 | 2 | 152973315 | 153032506 | 190 | 16 | -0.11728 | 0.54668 | STAM2 | 0.54668 | 0.61969 | 0.64536 |
| ENSG00000186642 | 11 | 72287185 | 72385635 | 298 | 29 | -0.11727 | 0.54668 | PDE2A | 0.54668 | 0.54085 | 0.54295 |
| ENSG00000106483 | 7 | 37945543 | 38065297 | 554 | 50 | -0.11747 | 0.54675 | SFRP4 | 0.54675 | 0.49285 | 0.43729 |
| ENSG00000152700 | 5 | 133936834 | 133984961 | 57 | 18 | -0.11759 | 0.5468 | SAR1B | 0.5468 | 0.40605 | 0.34533 |
| ENSG00000174405 | 13 | 108859787 | 108870716 | 18 | 5 | -0.11788 | 0.54692 | LIG4 | 0.54692 | 0.75017 | 0.57431 |
| ENSG00000160710 | 1 | 154554538 | 154600475 | 108 | 11 | -0.11805 | 0.54699 | ADAR | 0.54699 | 0.52493 | 0.532 |
| ENSG00000147614 | 8 | 86999552 | 87166457 | 532 | 27 | -0.1186 | 0.5472 | ATP6V0D2 | 0.5472 | 0.41214 | 0.22759 |
| ENSG00000145715 | 5 | 86563705 | 86687748 | 160 | 20 | -0.11871 | 0.54725 | RASA1 | 0.54725 | 0.54391 | 0.54101 |
| ENSG00000137274 | 6 | 3118608 | 3153812 | 118 | 13 | -0.11871 | 0.54725 | BPHL | 0.54725 | 0.47635 | 0.52878 |
| ENSG00000138380 | 2 | 203776937 | 203851786 | 176 | 12 | -0.11883 | 0.54729 | CARF | 0.54729 | 0.54917 | 0.55109 |
| ENSG00000159579 | 16 | 57220049 | 57274387 | 131 | 18 | -0.11893 | 0.54733 | RSPRY1 | 0.54733 | 0.55455 | 0.61705 |
| ENSG00000139218 | 12 | 46312914 | 46385903 | 113 | 19 | -0.11919 | 0.54744 | SCAF11 | 0.54744 | 0.49372 | 0.51674 |
| ENSG00000151136 | 12 | 107712190 | 108053419 | 981 | 75 | -0.11929 | 0.54748 | BTBD11 | 0.54748 | 0.51211 | 0.45655 |
| ENSG00000259132 | 14 | 23415479 | 23451467 | 70 | 9 | -0.1193 | 0.54748 | RP11-298I3.5 | 0.54748 | 0.66721 | 0.72765 |
| ENSG00000176204 | 2 | 76974845 | 77820445 | 3575 | 120 | -0.12001 | 0.54776 | LRRTM4 | 0.54776 | 0.59261 | 0.65143 |
| ENSG00000244411 | 11 | 71238313 | 71239210 | 2 | 1 | -0.12003 | 0.54777 | KRTAP5-7 | 0.54777 | 0.89985 | 0.86994 |
| ENSG00000159224 | 17 | 47035916 | 47045958 | 54 | 7 | -0.12034 | 0.54789 | GIP | 0.54789 | 0.52297 | 0.45782 |
| ENSG00000107020 | 9 | 5357973 | 5437878 | 313 | 25 | -0.12038 | 0.54791 | PLGRKT | 0.54791 | 0.62188 | 0.63885 |
| ENSG00000069018 | 5 | 135548999 | 135732730 | 516 | 27 | -0.1205 | 0.54796 | TRPC7 | 0.54796 | 0.54888 | 0.54552 |
| ENSG00000130803 | 19 | 9251056 | 9274100 | 116 | 11 | -0.12069 | 0.54803 | ZNF317 | 0.54803 | 0.76809 | 0.7193 |
| ENSG00000154803 | 17 | 17115526 | 17140502 | 102 | 16 | -0.12083 | 0.54809 | FLCN | 0.54809 | 0.51295 | 0.47988 |
| ENSG00000183155 | 1 | 202848085 | 202858263 | 38 | 7 | -0.12114 | 0.54821 | RABIF | 0.54821 | 0.6305 | 0.67521 |
| ENSG00000141556 | 17 | 80709940 | 80900724 | 801 | 30 | -0.12137 | 0.5483 | TBCD | 0.5483 | 0.56052 | 0.52482 |
| ENSG00000169740 | 10 | 44139307 | 44144304 | 18 | 5 | -0.12142 | 0.54832 | ZNF32 | 0.54832 | 0.46064 | 0.55745 |
| ENSG00000104894 | 19 | 49838428 | 49846592 | 19 | 4 | -0.12145 | 0.54833 | CD37 | 0.54833 | 0.54499 | 0.56207 |
| ENSG00000181778 | 9 | 71151496 | 71155783 | 9 | 3 | -0.12147 | 0.54834 | TMEM252 | 0.54834 | 0.77805 | 0.66342 |
| ENSG00000169592 | 16 | 30006615 | 30017114 | 28 | 5 | -0.12146 | 0.54834 | INO80E | 0.54834 | 0.66041 | 0.63975 |
| ENSG00000253537 | 5 | 140762467 | 140892546 | 402 | 31 | -0.12192 | 0.54852 | PCDHGA7 | 0.54852 | 0.58888 | 0.52674 |
| ENSG00000147481 | 8 | 50822349 | 51706678 | 3167 | 63 | -0.12199 | 0.54855 | SNTG1 | 0.54855 | 0.55025 | 0.54658 |
| ENSG00000273303 | 3 | 44607051 | 44635665 | 81 | 12 | -0.12223 | 0.54864 | RP11-944L7.5 | 0.54864 | 0.5652 | 0.54544 |
| ENSG00000226894 | 4 | 69054242 | 69083798 | 51 | 6 | -0.12233 | 0.54868 | TMPRSS11BNL | 0.54868 | 0.44726 | 0.46467 |
| ENSG00000145864 | 5 | 160715436 | 160976050 | 596 | 35 | -0.12258 | 0.54878 | GABRB2 | 0.54878 | 0.51682 | 0.59207 |
| ENSG00000141959 | 21 | 45719934 | 45747259 | 54 | 12 | -0.12262 | 0.5488 | PFKL | 0.5488 | 0.45884 | 0.50082 |

| | | | | | | | | | | | |
|---|---|---|---|---|---|---|---|---|---|---|---|
| ENSG00000162526 | 1 | 32817122 | 32829913 | 9 | 6 | -0.12266 | 0.54881 | TSSK3 | 0.54881 | 0.57797 | 0.80635 |
| ENSG00000010318 | 3 | 52444673 | 52457657 | 16 | 7 | -0.12268 | 0.54882 | PHF7 | 0.54882 | 0.73384 | 0.49963 |
| ENSG00000163596 | 2 | 203640690 | 203736708 | 228 | 17 | -0.12279 | 0.54886 | ICA1L | 0.54886 | 0.52271 | 0.53283 |
| ENSG00000153002 | 3 | 148508889 | 148577974 | 271 | 41 | -0.12292 | 0.54891 | CPB1 | 0.54891 | 0.61597 | 0.58799 |
| ENSG00000111199 | 12 | 110220890 | 110271212 | 178 | 27 | -0.12301 | 0.54895 | TRPV4 | 0.54895 | 0.57319 | 0.69416 |
| ENSG00000179833 | 2 | 64858755 | 64978139 | 267 | 21 | -0.12315 | 0.54901 | SERTAD2 | 0.54901 | 0.54957 | 0.58782 |
| ENSG00000115514 | 2 | 99935445 | 99957165 | 52 | 9 | -0.1232 | 0.54903 | TXNDC9 | 0.54903 | 0.5603 | 0.51088 |
| ENSG00000188334 | 19 | 48471303 | 48495427 | 147 | 19 | -0.12322 | 0.54903 | BSPH1 | 0.54903 | 0.55258 | 0.34336 |
| ENSG00000176678 | 16 | 86609974 | 86615303 | 18 | 8 | -0.12355 | 0.54917 | FOXL1 | 0.54917 | 0.52723 | 0.75279 |
| ENSG00000101255 | 20 | 361261 | 378203 | 81 | 31 | -0.1237 | 0.54922 | TRIB3 | 0.54922 | 0.85678 | 0.92032 |
| ENSG00000162913 | 1 | 228391207 | 228401365 | 24 | 5 | -0.12379 | 0.54926 | C1orf145 | 0.54926 | 0.42188 | 0.35531 |
| ENSG00000171056 | 8 | 10581278 | 10697357 | 480 | 25 | -0.12397 | 0.54933 | SOX7 | 0.54933 | 0.47957 | 0.20554 |
| ENSG00000179115 | 19 | 13033293 | 13044851 | 18 | 6 | -0.12435 | 0.54948 | FARSA | 0.54948 | 0.69112 | 0.64342 |
| ENSG00000221916 | 19 | 49621655 | 49622397 | 2 | 1 | -0.12472 | 0.54963 | C19orf73 | 0.54963 | 0.66051 | 0.79812 |
| ENSG00000156521 | 10 | 71897737 | 71906432 | 45 | 9 | -0.12479 | 0.54965 | TYSND1 | 0.54965 | 0.29048 | 0.26625 |
| ENSG00000257040 | 14 | 72456191 | 72458326 | 12 | 3 | -0.12478 | 0.54965 | AC005477.1 | 0.54965 | 0.9065 | 0.89946 |
| ENSG00000215547 | 20 | 29845467 | 29847435 | 5 | 2 | -0.12478 | 0.54965 | DEFB115 | 0.54965 | 0.62755 | 0.62504 |
| ENSG00000185818 | 4 | 2061239 | 2070816 | 12 | 2 | -0.12479 | 0.54966 | NAT8L | 0.54966 | 0.77979 | 0.79804 |
| ENSG00000139350 | 12 | 97301001 | 97347129 | 100 | 9 | -0.12481 | 0.54966 | NEDD1 | 0.54966 | 0.5643 | 0.58641 |
| ENSG00000178826 | 7 | 142977050 | 142985141 | 6 | 4 | -0.12501 | 0.54974 | TMEM139 | 0.54974 | 0.88848 | 0.91744 |
| ENSG00000165490 | 11 | 82611017 | 82669319 | 161 | 18 | -0.12556 | 0.54996 | C11orf82 | 0.54996 | 0.52463 | 0.51021 |
| ENSG00000144214 | 2 | 99900701 | 99921205 | 43 | 9 | -0.12564 | 0.54999 | LYG1 | 0.54999 | 0.49861 | 0.52488 |
| ENSG00000108306 | 17 | 37415384 | 37558776 | 386 | 14 | -0.12613 | 0.55019 | FBXL20 | 0.55019 | 0.55952 | 0.5567 |
| ENSG00000115850 | 2 | 136545410 | 136594750 | 137 | 14 | -0.12616 | 0.5502 | LCT | 0.5502 | 0.55264 | 0.51981 |
| ENSG00000176083 | 1 | 26688125 | 26701013 | 53 | 9 | -0.12618 | 0.55021 | ZNF683 | 0.55021 | 0.66653 | 0.60091 |
| ENSG00000178175 | 5 | 71738479 | 71803554 | 156 | 18 | -0.12622 | 0.55022 | ZNF366 | 0.55022 | 0.60475 | 0.72666 |
| ENSG00000115386 | 2 | 79347488 | 79350545 | 12 | 5 | -0.12631 | 0.55026 | REG1A | 0.55026 | 0.49311 | 0.57575 |
| ENSG00000159593 | 16 | 66836778 | 66907159 | 134 | 14 | -0.12637 | 0.55028 | NAE1 | 0.55028 | 0.5791 | 0.60461 |
| ENSG00000110108 | 11 | 60681346 | 60690915 | 15 | 6 | -0.12667 | 0.5504 | TMEM109 | 0.5504 | 0.60656 | 0.65152 |
| ENSG00000125484 | 9 | 135545422 | 135570342 | 96 | 9 | -0.12707 | 0.55056 | GTF3C4 | 0.55056 | 0.61288 | 0.60411 |
| ENSG00000149575 | 11 | 118032666 | 118047388 | 50 | 14 | -0.12725 | 0.55063 | SCN2B | 0.55063 | 0.75527 | 0.74692 |
| ENSG00000120217 | 9 | 5450503 | 5470566 | 64 | 10 | -0.12735 | 0.55067 | CD274 | 0.55067 | 0.70987 | 0.76895 |
| ENSG00000118898 | 16 | 4932508 | 5010742 | 411 | 49 | -0.12739 | 0.55068 | PPL | 0.55068 | 0.62928 | 0.65575 |
| ENSG00000186732 | 22 | 43807202 | 43903728 | 445 | 54 | -0.12744 | 0.5507 | MPPED1 | 0.5507 | 0.50156 | 0.5396 |
| ENSG00000168288 | 2 | 150426148 | 150444330 | 51 | 8 | -0.12764 | 0.55078 | MMADHC | 0.55078 | 0.51774 | 0.37209 |
| ENSG00000212734 | 17 | 6554971 | 6556791 | 5 | 2 | -0.12764 | 0.55078 | C17orf100 | 0.55078 | 0.51296 | 0.53794 |
| ENSG00000197044 | 19 | 11877815 | 11894893 | 100 | 7 | -0.12797 | 0.55091 | ZNF441 | 0.55091 | 0.59989 | 0.74876 |
| ENSG00000127412 | 7 | 142605267 | 142630905 | 85 | 10 | -0.12819 | 0.551 | TRPV5 | 0.551 | 0.57217 | 0.60564 |
| ENSG00000204293 | 11 | 124252291 | 124253258 | 2 | 1 | -0.12826 | 0.55103 | OR8B2 | 0.55103 | 0.6312 | 0.73343 |
| ENSG00000073060 | 12 | 125261402 | 125367214 | 385 | 69 | -0.12839 | 0.55108 | SCARB1 | 0.55108 | 0.57308 | 0.62371 |
| ENSG00000268193 | 19 | 19017822 | 19045206 | 80 | 17 | -0.12838 | 0.55108 | AC002985.3 | 0.55108 | 0.55014 | 0.4052 |
| ENSG00000122952 | 10 | 58116989 | 58121036 | 10 | 4 | -0.12901 | 0.55132 | ZWINT | 0.55132 | 0.60255 | 0.54909 |
| ENSG00000176399 | 9 | 22446840 | 22455739 | 25 | 6 | -0.12915 | 0.55138 | DMRTA1 | 0.55138 | 0.73447 | 0.59291 |
| ENSG00000167701 | 8 | 145728356 | 145732557 | 7 | 3 | -0.12921 | 0.5514 | GPT | 0.5514 | 0.50162 | 0.44641 |
| ENSG00000105671 | 19 | 19030484 | 19039436 | 27 | 8 | -0.12943 | 0.55149 | DDX49 | 0.55149 | 0.59994 | 0.42144 |
| ENSG00000127452 | 19 | 9920943 | 9938492 | 28 | 15 | -0.12949 | 0.55152 | FBXL12 | 0.55152 | 0.76303 | 0.87909 |
| ENSG00000262484 | 19 | 39399620 | 39402798 | 9 | 3 | -0.12961 | 0.55156 | CCER2 | 0.55156 | 0.11297 | 0.067899 |
| ENSG00000178982 | 19 | 39109735 | 39127595 | 23 | 6 | -0.12963 | 0.55157 | EIF3K | 0.55157 | 0.55081 | 0.51087 |
| ENSG00000167491 | 19 | 19496635 | 19619740 | 218 | 17 | -0.12983 | 0.55165 | GATAD2A | 0.55165 | 0.54918 | 0.58811 |
| ENSG00000081665 | 19 | 19896648 | 19932560 | 121 | 16 | -0.12991 | 0.55168 | ZNF506 | 0.55168 | 0.63147 | 0.6654 |
| ENSG00000137473 | 4 | 147627790 | 147867034 | 482 | 32 | -0.12996 | 0.5517 | TTC29 | 0.5517 | 0.51114 | 0.48557 |
| ENSG00000164877 | 7 | 1468101 | 1499138 | 139 | 17 | -0.12996 | 0.5517 | MICALL2 | 0.5517 | 0.65089 | 0.70823 |
| ENSG00000147548 | 8 | 38127215 | 38239790 | 181 | 15 | -0.13017 | 0.55178 | WHSC1L1 | 0.55178 | 0.53702 | 0.49896 |
| ENSG00000111644 | 12 | 6747241 | 6756626 | 16 | 5 | -0.13018 | 0.55179 | ACRBP | 0.55179 | 0.53743 | 0.59921 |
| ENSG00000122497 | 1 | 148003642 | 148025863 | 1 | 1 | -0.13021 | 0.5518 | NBPF14 | 0.5518 | 0.60482 | 0.60482 |
| ENSG00000132313 | 2 | 86426478 | 86440917 | 42 | 6 | -0.1302 | 0.5518 | MRPL35 | 0.5518 | 0.31837 | 0.35942 |
| ENSG00000233276 | 3 | 49394609 | 49396033 | 4 | 2 | -0.13023 | 0.55181 | GPX1 | 0.55181 | 0.6349 | 0.72975 |
| ENSG00000104413 | 8 | 95653302 | 95719694 | 282 | 18 | -0.13023 | 0.55181 | ESRP1 | 0.55181 | 0.53802 | 0.55757 |
| ENSG00000164512 | 5 | 55395507 | 55529186 | 514 | 49 | -0.13028 | 0.55183 | ANKRD55 | 0.55183 | 0.56172 | 0.53421 |
| ENSG00000147955 | 9 | 34634719 | 34637806 | 7 | 3 | -0.13038 | 0.55187 | SIGMAR1 | 0.55187 | 0.38203 | 0.38667 |
| ENSG00000111215 | 12 | 10977559 | 11324212 | 1883 | 22 | -0.13045 | 0.55189 | PRR4 | 0.55189 | 0.54031 | 0.54362 |
| ENSG00000151881 | 5 | 43444354 | 43483995 | 122 | 11 | -0.13057 | 0.55194 | C5orf28 | 0.55194 | 0.5383 | 0.4816 |
| ENSG00000141985 | 19 | 4360367 | 4400544 | 91 | 16 | -0.13062 | 0.55196 | SH3GL1 | 0.55196 | 0.48341 | 0.6782 |
| ENSG00000056097 | 5 | 32354456 | 32444867 | 291 | 20 | -0.13072 | 0.552 | ZFR | 0.552 | 0.56054 | 0.64074 |
| ENSG00000142789 | 1 | 22328149 | 22339032 | 102 | 18 | -0.13082 | 0.55204 | CELA3A | 0.55204 | 0.41433 | 0.47515 |
| ENSG00000103375 | 16 | 25227052 | 25240261 | 69 | 7 | -0.13098 | 0.5521 | AQP8 | 0.5521 | 0.28214 | 0.23204 |
| ENSG00000244187 | 9 | 139685807 | 139687709 | 8 | 3 | -0.1313 | 0.55223 | TMEM141 | 0.55223 | 0.7567 | 0.82217 |
| ENSG00000142511 | 19 | 51273721 | 51274989 | 1 | 1 | -0.13147 | 0.5523 | GPR32 | 0.5523 | 0.54988 | 0.74073 |
| ENSG00000111907 | 6 | 125440195 | 125585553 | 491 | 37 | -0.13163 | 0.55236 | TPD52L1 | 0.55236 | 0.54277 | 0.59155 |
| ENSG00000088986 | 12 | 120907653 | 120936296 | 103 | 15 | -0.13175 | 0.55241 | DYNLL1 | 0.55241 | 0.52868 | 0.51655 |
| ENSG00000164002 | 1 | 40974413 | 40982228 | 39 | 8 | -0.13182 | 0.55244 | EXO5 | 0.55244 | 0.57958 | 0.61916 |
| ENSG00000154415 | 7 | 113516832 | 113715975 | 453 | 15 | -0.13186 | 0.55245 | PPP1R3A | 0.55245 | 0.55513 | 0.53372 |
| ENSG00000053747 | 18 | 21269407 | 21535030 | 518 | 20 | -0.13199 | 0.5525 | LAMA3 | 0.5525 | 0.59576 | 0.59031 |

| | | | | | | | | | | | |
|---|---|---|---|---|---|---|---|---|---|---|---|
| ENSG00000140092 | 14 | 92335756 | 92414331 | 311 | 37 | -0.13219 | 0.55258 | FBLN5 | 0.55258 | 0.53423 | 0.40316 |
| ENSG00000125827 | 20 | 7957995 | 8000476 | 122 | 13 | -0.13225 | 0.55261 | TMX4 | 0.55261 | 0.31779 | 0.075444 |
| ENSG00000133460 | 22 | 24198890 | 24228496 | 120 | 20 | -0.13251 | 0.55271 | SLC2A11 | 0.55271 | 0.52984 | 0.26375 |
| ENSG00000165626 | 10 | 13480484 | 13570974 | 300 | 22 | -0.13257 | 0.55273 | BEND7 | 0.55273 | 0.57199 | 0.61518 |
| ENSG00000179751 | 19 | 39693471 | 39694906 | 1 | 1 | -0.13299 | 0.5529 | SYCN | 0.5529 | 0.85514 | 0.71554 |
| ENSG00000165219 | 9 | 128024073 | 128129486 | 167 | 11 | -0.13309 | 0.55294 | GAPVD1 | 0.55294 | 0.56612 | 0.58332 |
| ENSG00000116729 | 1 | 68564142 | 68698803 | 611 | 67 | -0.13318 | 0.55297 | WLS | 0.55297 | 0.62001 | 0.56641 |
| ENSG00000197213 | 19 | 56700939 | 56709289 | 45 | 11 | -0.13358 | 0.55313 | ZSCAN5B | 0.55313 | 0.45885 | 0.70914 |
| ENSG00000129667 | 17 | 74466973 | 74497872 | 132 | 20 | -0.13373 | 0.55319 | RHBDF2 | 0.55319 | 0.6482 | 0.39659 |
| ENSG00000128989 | 15 | 52839242 | 52862080 | 85 | 9 | -0.13386 | 0.55324 | ARPP19 | 0.55324 | 0.55058 | 0.62204 |
| ENSG00000116701 | 1 | 183524698 | 183560011 | 100 | 20 | -0.13401 | 0.5533 | NCF2 | 0.5533 | 0.38476 | 0.5135 |
| ENSG00000183706 | 15 | 22382382 | 22383507 | 1 | 1 | -0.134 | 0.5533 | OR4N4 | 0.5533 | 0.092928 | 0.058423 |
| ENSG00000196470 | 16 | 48390275 | 48482313 | 173 | 24 | -0.13403 | 0.55331 | SIAH1 | 0.55331 | 0.54123 | 0.53048 |
| ENSG00000120616 | 10 | 32556679 | 32667726 | 346 | 20 | -0.13413 | 0.55335 | EPC1 | 0.55335 | 0.61612 | 0.73793 |
| ENSG00000116191 | 1 | 178694300 | 178889238 | 418 | 21 | -0.13427 | 0.55341 | RALGPS2 | 0.55341 | 0.55476 | 0.57161 |
| ENSG00000150556 | 2 | 149894621 | 150071776 | 646 | 40 | -0.13453 | 0.55351 | LYPD6B | 0.55351 | 0.52847 | 0.53037 |
| ENSG00000176371 | 15 | 85144217 | 85171027 | 53 | 12 | -0.13487 | 0.55364 | ZSCAN2 | 0.55364 | 0.65161 | 0.6839 |
| ENSG00000181104 | 5 | 76011868 | 76031606 | 21 | 10 | -0.13488 | 0.55365 | F2R | 0.55365 | 0.19595 | 0.25463 |
| ENSG00000140030 | 14 | 88471468 | 88478419 | 21 | 5 | -0.13499 | 0.55369 | GPR65 | 0.55369 | 0.54192 | 0.48671 |
| ENSG00000149792 | 11 | 64889655 | 64894843 | 8 | 5 | -0.13513 | 0.55375 | MRPL49 | 0.55375 | 0.65837 | 0.71379 |
| ENSG00000117153 | 1 | 202860228 | 202897764 | 106 | 9 | -0.1353 | 0.55381 | KLHL12 | 0.55381 | 0.58962 | 0.69259 |
| ENSG00000146904 | 7 | 143087382 | 143105985 | 51 | 15 | -0.13537 | 0.55384 | EPHA1 | 0.55384 | 0.58429 | 0.76957 |
| ENSG00000204640 | 2 | 101086944 | 101099742 | 53 | 4 | -0.13545 | 0.55387 | NMS | 0.55387 | 0.48004 | 0.46596 |
| ENSG00000038358 | 16 | 67906926 | 67918406 | 12 | 3 | -0.13572 | 0.55398 | EDC4 | 0.55398 | 0.76665 | 0.79045 |
| ENSG00000181192 | 10 | 12110971 | 12165224 | 206 | 21 | -0.13604 | 0.55411 | DHTKD1 | 0.55411 | 0.70861 | 0.83116 |
| ENSG00000174721 | 10 | 93666346 | 93669240 | 8 | 3 | -0.13623 | 0.55418 | FGFBP3 | 0.55418 | 0.43253 | 0.60707 |
| ENSG00000170417 | 2 | 103353367 | 103460352 | 239 | 19 | -0.13625 | 0.55419 | TMEM182 | 0.55419 | 0.49772 | 0.47218 |
| ENSG00000144681 | 3 | 36421836 | 36589499 | 784 | 44 | -0.13629 | 0.5542 | STAC | 0.5542 | 0.43376 | 0.40689 |
| ENSG00000141968 | 19 | 6772725 | 6857377 | 401 | 62 | -0.1367 | 0.55437 | VAV1 | 0.55437 | 0.67667 | 0.53807 |
| ENSG00000182177 | 2 | 237102095 | 237173052 | 342 | 19 | -0.13713 | 0.55454 | ASB18 | 0.55454 | 0.54893 | 0.57014 |
| ENSG00000149177 | 11 | 48002113 | 48189670 | 425 | 29 | -0.13716 | 0.55455 | PTPRJ | 0.55455 | 0.53447 | 0.30712 |
| ENSG00000134533 | 12 | 15260717 | 15501609 | 557 | 31 | -0.1372 | 0.55456 | RERG | 0.55456 | 0.55808 | 0.43479 |
| ENSG00000097046 | 1 | 91966408 | 91991321 | 66 | 13 | -0.13734 | 0.55462 | CDC7 | 0.55462 | 0.47554 | 0.47864 |
| ENSG00000132718 | 1 | 155829300 | 155854990 | 43 | 9 | -0.13741 | 0.55465 | SYT11 | 0.55465 | 0.57686 | 0.54518 |
| ENSG00000156968 | 16 | 15489611 | 15507125 | 56 | 13 | -0.13746 | 0.55467 | MPV17L | 0.55467 | 0.2985 | 0.38188 |
| ENSG00000162607 | 1 | 62901968 | 62917475 | 35 | 3 | -0.13762 | 0.55473 | USP1 | 0.55473 | 0.58106 | 0.50989 |
| ENSG00000158050 | 2 | 96808905 | 96811179 | 2 | 1 | -0.13775 | 0.55478 | DUSP2 | 0.55478 | 0.11435 | 0.18263 |
| ENSG00000007062 | 4 | 15964699 | 16086001 | 475 | 72 | -0.1379 | 0.55484 | PROM1 | 0.55484 | 0.65907 | 0.74676 |
| ENSG00000099889 | 22 | 19957419 | 20004331 | 205 | 25 | -0.1382 | 0.55496 | ARVCF | 0.55496 | 0.54232 | 0.49829 |
| ENSG00000268080 | 8 | 94241867 | 94242388 | 1 | 1 | -0.1383 | 0.555 | AC016885.1 | 0.555 | 0.41031 | 0.37122 |
| ENSG00000166716 | 15 | 85291866 | 85349659 | 146 | 22 | -0.13865 | 0.55514 | ZNF592 | 0.55514 | 0.62986 | 0.56143 |
| ENSG00000026025 | 10 | 17270258 | 17279592 | 18 | 6 | -0.13888 | 0.55523 | VIM | 0.55523 | 0.55643 | 0.82112 |
| ENSG00000048828 | 9 | 96214004 | 96328397 | 366 | 23 | -0.13893 | 0.55525 | FAM120A | 0.55525 | 0.47979 | 0.41692 |
| ENSG00000066933 | 15 | 72114632 | 72410918 | 606 | 21 | -0.13896 | 0.55526 | MYO9A | 0.55526 | 0.56507 | 0.55775 |
| ENSG00000164061 | 3 | 49591922 | 49708978 | 211 | 21 | -0.13913 | 0.55532 | BSN | 0.55532 | 0.55729 | 0.56492 |
| ENSG00000243943 | 2 | 27805897 | 27858041 | 127 | 13 | -0.13913 | 0.55533 | ZNF512 | 0.55533 | 0.5834 | 0.57895 |
| ENSG00000128340 | 22 | 37621301 | 37640488 | 81 | 16 | -0.13928 | 0.55538 | RAC2 | 0.55538 | 0.60431 | 0.71423 |
| ENSG00000198055 | 5 | 176830205 | 176869902 | 64 | 24 | -0.13935 | 0.55541 | GRK6 | 0.55541 | 0.59638 | 0.37858 |
| ENSG00000247077 | 12 | 133287405 | 133299228 | 62 | 15 | -0.13933 | 0.55541 | PGAM5 | 0.55541 | 0.62906 | 0.81668 |
| ENSG00000174946 | 3 | 150915619 | 150920988 | 13 | 4 | -0.13954 | 0.55549 | GPR171 | 0.55549 | 0.76127 | 0.87612 |
| ENSG00000166266 | 11 | 107879459 | 107978503 | 248 | 24 | -0.13957 | 0.5555 | CUL5 | 0.5555 | 0.6011 | 0.59935 |
| ENSG00000138463 | 3 | 122513642 | 122599986 | 224 | 28 | -0.13966 | 0.55553 | DIRC2 | 0.55553 | 0.41601 | 0.42084 |
| ENSG00000156639 | 6 | 37787275 | 38122400 | 1034 | 25 | -0.13966 | 0.55554 | ZFAND3 | 0.55554 | 0.54692 | 0.57299 |
| ENSG00000187416 | 7 | 103969104 | 104549001 | 2096 | 112 | -0.13972 | 0.55556 | LHFPL3 | 0.55556 | 0.55064 | 0.55909 |
| ENSG00000113328 | 5 | 162864575 | 162873157 | 12 | 4 | -0.13988 | 0.55562 | CCNG1 | 0.55562 | 0.78923 | 0.83963 |
| ENSG00000116205 | 1 | 54519260 | 54578192 | 135 | 15 | -0.14001 | 0.55567 | TCEANC2 | 0.55567 | 0.46149 | 0.31231 |
| ENSG00000116539 | 1 | 155305059 | 155532598 | 150 | 47 | -0.14046 | 0.55585 | ASH1L | 0.55585 | 0.52431 | 0.49179 |
| ENSG00000205883 | 8 | 11839830 | 11842099 | 10 | 3 | -0.14046 | 0.55585 | DEFB135 | 0.55585 | 0.73074 | 0.82177 |
| ENSG00000196542 | 3 | 161062580 | 161090668 | 127 | 18 | -0.1405 | 0.55587 | SPTSSB | 0.55587 | 0.68968 | 0.76791 |
| ENSG00000269858 | 19 | 41304901 | 41314338 | 33 | 10 | -0.1405 | 0.55587 | EGLN2 | 0.55587 | 0.50264 | 0.60154 |
| ENSG00000049449 | 11 | 31833939 | 32127301 | 840 | 73 | -0.14056 | 0.55589 | RCN1 | 0.55589 | 0.54428 | 0.5627 |
| ENSG00000115841 | 2 | 38150330 | 38294285 | 535 | 25 | -0.14078 | 0.55598 | RMDN2 | 0.55598 | 0.53993 | 0.50295 |
| ENSG00000118946 | 13 | 58205944 | 58303445 | 256 | 30 | -0.14079 | 0.55598 | PCDH17 | 0.55598 | 0.597 | 0.58728 |
| ENSG00000189079 | 12 | 46123448 | 46301823 | 190 | 25 | -0.14089 | 0.55602 | ARID2 | 0.55602 | 0.53443 | 0.43761 |
| ENSG00000186103 | 3 | 121289551 | 121305481 | 67 | 7 | -0.14093 | 0.55604 | ARGFX | 0.55604 | 0.58338 | 0.67746 |
| ENSG00000173366 | 3 | 52255097 | 52265206 | 12 | 6 | -0.14105 | 0.55609 | TLR9 | 0.55609 | 0.5759 | 0.49632 |
| ENSG00000149289 | 11 | 109964087 | 110042566 | 228 | 17 | -0.14106 | 0.55609 | ZC3H12C | 0.55609 | 0.49664 | 0.47845 |
| ENSG00000116679 | 1 | 185265520 | 185286461 | 16 | 6 | -0.1411 | 0.5561 | IVNS1ABP | 0.5561 | 0.79327 | 0.81937 |
| ENSG00000184731 | 2 | 38814 | 46870 | 14 | 6 | -0.14117 | 0.55613 | FAM110C | 0.55613 | 0.57517 | 0.642 |
| ENSG00000070388 | 19 | 639879 | 643703 | 24 | 6 | -0.14121 | 0.55615 | FGF22 | 0.55615 | 0.60833 | 0.80589 |
| ENSG00000263002 | 19 | 44645710 | 44663156 | 32 | 8 | -0.14121 | 0.55615 | ZNF234 | 0.55615 | 0.58049 | 0.53495 |
| ENSG00000158457 | 7 | 128784712 | 128808671 | 83 | 9 | -0.14135 | 0.5562 | TSPAN33 | 0.5562 | 0.55259 | 0.5266 |
| ENSG00000084731 | 2 | 26149471 | 26205618 | 208 | 31 | -0.14152 | 0.55627 | KIF3C | 0.55627 | 0.59366 | 0.33968 |

| | | | | | | | | | | | |
|---|---|---|---|---|---|---|---|---|---|---|---|
| ENSG00000214841 | 7 | 34685709 | 34699772 | 68 | 12 | -0.14158 | 0.55629 | AC005493.1 | 0.55629 | 0.41063 | 0.35409 |
| ENSG00000079263 | 2 | 231067826 | 231223762 | 480 | 34 | -0.14179 | 0.55638 | SP140 | 0.55638 | 0.2929 | 0.27489 |
| ENSG00000160145 | 3 | 123798870 | 124445172 | 2336 | 99 | -0.14192 | 0.55643 | KALRN | 0.55643 | 0.54845 | 0.50565 |
| ENSG00000139364 | 12 | 125671382 | 126146917 | 1730 | 161 | -0.14194 | 0.55644 | TMEM132B | 0.55644 | 0.57396 | 0.56841 |
| ENSG00000131115 | 19 | 44711700 | 44741421 | 54 | 10 | -0.14206 | 0.55648 | ZNF227 | 0.55648 | 0.66314 | 0.68247 |
| ENSG00000170852 | 7 | 32907784 | 32933743 | 76 | 9 | -0.14217 | 0.55653 | KBTBD2 | 0.55653 | 0.61885 | 0.59662 |
| ENSG00000136816 | 9 | 132565432 | 132573560 | 23 | 5 | -0.14233 | 0.55659 | TOR1B | 0.55659 | 0.51956 | 0.61503 |
| ENSG00000215356 | 8 | 7783859 | 7812271 | 1 | 1 | -0.14235 | 0.5566 | ZNF705B | 0.5566 | 0.5566 | 0.72897 |
| ENSG00000100614 | 14 | 60712470 | 60765805 | 121 | 10 | -0.14254 | 0.55667 | PPM1A | 0.55667 | 0.53886 | 0.4961 |
| ENSG00000164211 | 5 | 110831731 | 110848288 | 25 | 7 | -0.14255 | 0.55668 | STARD4 | 0.55668 | 0.58694 | 0.7587 |
| ENSG00000258890 | 17 | 62502706 | 62538579 | 57 | 7 | -0.14299 | 0.55685 | CEP95 | 0.55685 | 0.59278 | 0.53361 |
| ENSG00000138231 | 3 | 137879854 | 137893791 | 19 | 3 | -0.14322 | 0.55694 | DBR1 | 0.55694 | 0.43477 | 0.41528 |
| ENSG00000203836 | 1 | 147574428 | 147634886 | 16 | 5 | -0.14332 | 0.55698 | NBPF24 | 0.55698 | 0.6468 | 0.5241 |
| ENSG00000186687 | 5 | 130506503 | 130541119 | 106 | 11 | -0.14345 | 0.55703 | LYRM7 | 0.55703 | 0.43816 | 0.37994 |
| ENSG00000166828 | 16 | 23194036 | 23228204 | 139 | 15 | -0.14351 | 0.55706 | SCNN1G | 0.55706 | 0.56793 | 0.55855 |
| ENSG00000176396 | 19 | 40028890 | 40030870 | 7 | 3 | -0.14354 | 0.55707 | EID2 | 0.55707 | 0.58915 | 0.56195 |
| ENSG00000125744 | 19 | 45988547 | 46000319 | 38 | 7 | -0.14363 | 0.5571 | RTN2 | 0.5571 | 0.84131 | 0.87622 |
| ENSG00000197745 | 11 | 62063754 | 62066536 | 13 | 5 | -0.14374 | 0.55715 | SCGB1D4 | 0.55715 | 0.50596 | 0.45993 |
| ENSG00000213402 | 11 | 67202981 | 67205538 | 4 | 2 | -0.14393 | 0.55722 | PTPRCAP | 0.55722 | 0.65294 | 0.65273 |
| ENSG00000165434 | 11 | 74041363 | 74109518 | 190 | 17 | -0.14399 | 0.55724 | PGM2L1 | 0.55724 | 0.6027 | 0.66674 |
| ENSG00000269964 | 6 | 78400375 | 78636691 | 957 | 25 | -0.14409 | 0.55728 | MEI4 | 0.55728 | 0.56354 | 0.58069 |
| ENSG00000240682 | 3 | 128846258 | 128880136 | 54 | 15 | -0.14416 | 0.55731 | ISY1 | 0.55731 | 0.55922 | 0.6377 |
| ENSG00000116698 | 1 | 183441351 | 183567381 | 272 | 30 | -0.14424 | 0.55734 | SMG7 | 0.55734 | 0.5503 | 0.69439 |
| ENSG00000176046 | 16 | 28548606 | 28550495 | 6 | 2 | -0.14432 | 0.55738 | NUPR1 | 0.55738 | 0.72965 | 0.76989 |
| ENSG00000188888 | 17 | 36481413 | 36499730 | 37 | 10 | -0.14455 | 0.55747 | GPR179 | 0.55747 | 0.66026 | 0.59783 |
| ENSG00000122678 | 7 | 44111846 | 44122139 | 24 | 5 | -0.14466 | 0.55751 | POLM | 0.55751 | 0.58222 | 0.4399 |
| ENSG00000168872 | 16 | 70380732 | 70407286 | 46 | 12 | -0.14468 | 0.55752 | DDX19A | 0.55752 | 0.50847 | 0.58879 |
| ENSG00000139549 | 12 | 49483204 | 49488602 | 8 | 4 | -0.14482 | 0.55757 | DHH | 0.55757 | 0.45218 | 0.43481 |
| ENSG00000181790 | 8 | 143530791 | 143626370 | 276 | 44 | -0.14523 | 0.55774 | BAI1 | 0.55774 | 0.31729 | 0.1744 |
| ENSG00000131050 | 20 | 31749574 | 31769218 | 38 | 4 | -0.14524 | 0.55774 | BPIFA2 | 0.55774 | 0.61819 | 0.68131 |
| ENSG00000078124 | 11 | 76571911 | 76737841 | 570 | 11 | -0.14539 | 0.5578 | ACER3 | 0.5578 | 0.58376 | 0.62234 |
| ENSG00000133398 | 5 | 6371994 | 6378707 | 19 | 5 | -0.14548 | 0.55783 | MED10 | 0.55783 | 0.32062 | 0.28368 |
| ENSG00000157020 | 3 | 10334815 | 10362862 | 90 | 15 | -0.14573 | 0.55793 | SEC13 | 0.55793 | 0.575 | 0.66621 |
| ENSG00000081923 | 18 | 55313658 | 55470333 | 655 | 66 | -0.1459 | 0.558 | ATP8B1 | 0.558 | 0.57471 | 0.59516 |
| ENSG00000105246 | 19 | 4229495 | 4237528 | 43 | 5 | -0.14591 | 0.558 | EBI3 | 0.558 | 0.60819 | 0.70062 |
| ENSG00000248789 | 5 | 51303865 | 51376821 | 290 | 12 | -0.14601 | 0.55804 | CTD-2203A3.1 | 0.55804 | 0.56816 | 0.53422 |
| ENSG00000151729 | 4 | 186064395 | 186071536 | 18 | 7 | -0.14611 | 0.55808 | SLC25A4 | 0.55808 | 0.81502 | 0.696 |
| ENSG00000128610 | 7 | 121941448 | 121950745 | 31 | 11 | -0.1461 | 0.55808 | FEZF1 | 0.55808 | 0.53452 | 0.37864 |
| ENSG00000142552 | 19 | 50030875 | 50050219 | 122 | 3 | -0.14634 | 0.55817 | RCN3 | 0.55817 | 0.57802 | 0.60165 |
| ENSG00000082074 | 5 | 39105338 | 39274630 | 572 | 35 | -0.14638 | 0.55819 | FYB | 0.55819 | 0.54285 | 0.5175 |
| ENSG00000171570 | 19 | 41284147 | 41314103 | 119 | 21 | -0.14639 | 0.55819 | RAB4B-EGLN2 | 0.55819 | 0.55176 | 0.6225 |
| ENSG00000110107 | 11 | 60658202 | 60674060 | 34 | 7 | -0.14645 | 0.55822 | PRPF19 | 0.55822 | 0.59808 | 0.64549 |
| ENSG00000160111 | 19 | 17003758 | 17137625 | 533 | 67 | -0.14647 | 0.55822 | CPAMD8 | 0.55822 | 0.60895 | 0.76943 |
| ENSG00000147872 | 9 | 19108373 | 19149288 | 152 | 28 | -0.14652 | 0.55825 | PLIN2 | 0.55825 | 0.46351 | 0.32881 |
| ENSG00000164893 | 8 | 87226281 | 87333375 | 309 | 22 | -0.14665 | 0.5583 | SLC7A13 | 0.5583 | 0.34819 | 0.25846 |
| ENSG00000165629 | 10 | 7830092 | 7849778 | 73 | 11 | -0.14674 | 0.55833 | ATP5C1 | 0.55833 | 0.38907 | 0.41472 |
| ENSG00000162695 | 1 | 101361632 | 101447309 | 158 | 15 | -0.14681 | 0.55836 | SLC30A7 | 0.55836 | 0.53384 | 0.52366 |
| ENSG00000123096 | 12 | 26274924 | 26452223 | 609 | 62 | -0.14692 | 0.5584 | SSPN | 0.5584 | 0.41657 | 0.35098 |
| ENSG00000156269 | 4 | 80146848 | 80247204 | 162 | 21 | -0.14699 | 0.55843 | NAA11 | 0.55843 | 0.48405 | 0.55963 |
| ENSG00000197852 | 1 | 112223252 | 112298446 | 328 | 35 | -0.14701 | 0.55844 | FAM212B | 0.55844 | 0.62139 | 0.64563 |
| ENSG00000114902 | 3 | 52738971 | 52742182 | 7 | 2 | -0.14706 | 0.55846 | SPCS1 | 0.55846 | 0.45916 | 0.49911 |
| ENSG00000186458 | 20 | 238377 | 241737 | 17 | 2 | -0.1471 | 0.55847 | DEFB132 | 0.55847 | 0.6721 | 0.19635 |
| ENSG00000188010 | 2 | 39103103 | 39156213 | 108 | 15 | -0.14718 | 0.5585 | MORN2 | 0.5585 | 0.61875 | 0.59598 |
| ENSG00000132424 | 6 | 99845927 | 99873207 | 68 | 10 | -0.14718 | 0.5585 | PNISR | 0.5585 | 0.39858 | 0.39439 |
| ENSG00000125375 | 14 | 50779044 | 50802276 | 85 | 6 | -0.14726 | 0.55854 | ATP5S | 0.55854 | 0.59517 | 0.58874 |
| ENSG00000164326 | 5 | 71014990 | 71016875 | 4 | 2 | -0.1479 | 0.55879 | CARTPT | 0.55879 | 0.45506 | 0.65282 |
| ENSG00000127527 | 19 | 16466050 | 16582896 | 285 | 26 | -0.14797 | 0.55882 | EPS15L1 | 0.55882 | 0.56958 | 0.54316 |
| ENSG00000164764 | 8 | 73976775 | 74036323 | 206 | 21 | -0.14825 | 0.55893 | SBSPON | 0.55893 | 0.54256 | 0.59949 |
| ENSG00000178372 | 10 | 5540658 | 5541533 | 3 | 1 | -0.14828 | 0.55894 | CALML5 | 0.55894 | 0.62285 | 0.42526 |
| ENSG00000185787 | 15 | 79102829 | 79190475 | 437 | 22 | -0.14841 | 0.55899 | MORF4L1 | 0.55899 | 0.53797 | 0.58353 |
| ENSG00000138640 | 4 | 89647106 | 90032549 | 1264 | 45 | -0.1486 | 0.55907 | FAM13A | 0.55907 | 0.56991 | 0.54302 |
| ENSG00000116133 | 1 | 55315306 | 55352891 | 152 | 19 | -0.14907 | 0.55925 | DHCR24 | 0.55925 | 0.53201 | 0.69909 |
| ENSG00000169252 | 5 | 148206156 | 148208196 | 11 | 4 | -0.14909 | 0.55926 | ADRB2 | 0.55926 | 0.58241 | 0.80138 |
| ENSG00000172464 | 11 | 56408942 | 56409947 | 2 | 1 | -0.14919 | 0.5593 | OR5AP2 | 0.5593 | 0.091832 | 0.031839 |
| ENSG00000149532 | 11 | 61170121 | 61197503 | 34 | 9 | -0.14924 | 0.55932 | CPSF7 | 0.55932 | 0.65112 | 0.72251 |
| ENSG00000204661 | 5 | 179068545 | 179072047 | 13 | 6 | -0.14953 | 0.55943 | C5orf60 | 0.55943 | 0.3466 | 0.42796 |
| ENSG00000158169 | 9 | 97861336 | 98079991 | 263 | 20 | -0.14959 | 0.55946 | FANCC | 0.55946 | 0.55255 | 0.54347 |
| ENSG00000148288 | 9 | 136028340 | 136039332 | 43 | 10 | -0.14963 | 0.55947 | GBGT1 | 0.55947 | 0.43972 | 0.33444 |
| ENSG00000241241 | 17 | 39257754 | 39258461 | 4 | 2 | -0.14968 | 0.55949 | KRTAP4-16P | 0.55949 | 0.38098 | 0.38729 |
| ENSG00000259207 | 17 | 45331212 | 45421658 | 330 | 30 | -0.14967 | 0.55949 | ITGB3 | 0.55949 | 0.59963 | 0.65778 |
| ENSG00000196544 | 17 | 8091652 | 8093564 | 2 | 1 | -0.14988 | 0.55957 | C17orf59 | 0.55957 | 0.5089 | 0.55056 |
| ENSG00000129515 | 14 | 35030300 | 35099389 | 223 | 20 | -0.15002 | 0.55963 | SNX6 | 0.55963 | 0.58188 | 0.59053 |
| ENSG00000141434 | 18 | 29765032 | 29800367 | 124 | 17 | -0.15011 | 0.55966 | MEP1B | 0.55966 | 0.64762 | 0.69357 |

| | | | | | | | | | | | |
|---|---|---|---|---|---|---|---|---|---|---|---|
| ENSG00000182698 | 2 | 220192131 | 220197899 | 30 | 3 | -0.15026 | 0.55972 | RESP18 | 0.55972 | 0.62962 | 0.67226 |
| ENSG00000139990 | 14 | 69517598 | 69619867 | 203 | 24 | -0.15032 | 0.55974 | DCAF5 | 0.55974 | 0.5429 | 0.51269 |
| ENSG00000137075 | 9 | 36336393 | 36487545 | 307 | 29 | -0.15045 | 0.5598 | RNF38 | 0.5598 | 0.56389 | 0.54221 |
| ENSG00000153404 | 5 | 140373 | 190085 | 226 | 21 | -0.15063 | 0.55987 | PLEKHG4B | 0.55987 | 0.61302 | 0.65073 |
| ENSG00000254093 | 8 | 10622473 | 10697394 | 322 | 19 | -0.15079 | 0.55993 | PINX1 | 0.55993 | 0.58661 | 0.59311 |
| ENSG00000138442 | 2 | 203739505 | 203879521 | 353 | 15 | -0.15098 | 0.56001 | WDR12 | 0.56001 | 0.56822 | 0.57644 |
| ENSG00000079950 | 6 | 132767006 | 132834337 | 161 | 25 | -0.15132 | 0.56014 | STX7 | 0.56014 | 0.47136 | 0.36631 |
| ENSG00000139797 | 13 | 98828039 | 98829519 | 3 | 1 | -0.15152 | 0.56022 | RNF113B | 0.56022 | 0.45103 | 0.39289 |
| ENSG00000130349 | 6 | 107349407 | 107372546 | 113 | 14 | -0.15156 | 0.56023 | C6orf203 | 0.56023 | 0.47833 | 0.42184 |
| ENSG00000213658 | 16 | 28996147 | 29002104 | 11 | 5 | -0.15161 | 0.56025 | LAT | 0.56025 | 0.4796 | 0.5999 |
| ENSG00000165525 | 14 | 50249997 | 50319921 | 95 | 21 | -0.15177 | 0.56031 | NEMF | 0.56031 | 0.56159 | 0.59364 |
| ENSG00000142534 | 19 | 49999622 | 50002946 | 9 | 2 | -0.15188 | 0.56036 | RPS11 | 0.56036 | 0.5579 | 0.54419 |
| ENSG00000166004 | 11 | 93394805 | 93463522 | 163 | 15 | -0.152 | 0.56041 | KIAA1731 | 0.56041 | 0.62084 | 0.58116 |
| ENSG00000101447 | 20 | 37554955 | 37581703 | 57 | 8 | -0.15201 | 0.56041 | FAM83D | 0.56041 | 0.63527 | 0.69827 |
| ENSG00000197641 | 18 | 61254223 | 61271873 | 27 | 7 | -0.15209 | 0.56044 | SERPINB13 | 0.56044 | 0.71612 | 0.56527 |
| ENSG00000074047 | 2 | 121493199 | 121750229 | 786 | 84 | -0.15221 | 0.56049 | GLI2 | 0.56049 | 0.51179 | 0.30409 |
| ENSG00000090863 | 16 | 74485856 | 74641012 | 572 | 26 | -0.15226 | 0.56051 | GLG1 | 0.56051 | 0.52012 | 0.46222 |
| ENSG00000189362 | 2 | 191369068 | 191399448 | 74 | 12 | -0.15238 | 0.56055 | TMEM194B | 0.56055 | 0.47862 | 0.54161 |
| ENSG00000259753 | 17 | 45331263 | 45421658 | 329 | 29 | -0.1526 | 0.56064 | ITGB3 | 0.56064 | 0.59963 | 0.65778 |
| ENSG00000158764 | 1 | 160914815 | 160924589 | 23 | 5 | -0.15265 | 0.56066 | ITLN2 | 0.56066 | 0.79062 | 0.85962 |
| ENSG00000118579 | 4 | 17616254 | 17635728 | 78 | 6 | -0.15265 | 0.56066 | MED28 | 0.56066 | 0.5304 | 0.50519 |
| ENSG00000163607 | 3 | 112709765 | 112733907 | 76 | 17 | -0.1527 | 0.56068 | GTPBP8 | 0.56068 | 0.44834 | 0.35004 |
| ENSG00000004455 | 1 | 33473585 | 33546597 | 142 | 12 | -0.15296 | 0.56078 | AK2 | 0.56078 | 0.51372 | 0.46333 |
| ENSG00000183562 | 11 | 3011093 | 3012563 | 7 | 1 | -0.15294 | 0.56078 | AC131971.1 | 0.56078 | 0.41901 | 0.4945 |
| ENSG00000178694 | 3 | 93781760 | 93847389 | 67 | 19 | -0.15297 | 0.56079 | NSUN3 | 0.56079 | 0.46688 | 0.50995 |
| ENSG00000081014 | 15 | 51200869 | 51298097 | 287 | 16 | -0.1531 | 0.56084 | AP4E1 | 0.56084 | 0.54759 | 0.5519 |
| ENSG00000164708 | 7 | 44102326 | 44105186 | 8 | 2 | -0.15319 | 0.56088 | PGAM2 | 0.56088 | 0.56213 | 0.62492 |
| ENSG00000168004 | 11 | 63228876 | 63258666 | 64 | 11 | -0.15339 | 0.56096 | HRASLS5 | 0.56096 | 0.27003 | 0.13859 |
| ENSG00000123444 | 11 | 47593749 | 47600567 | 8 | 5 | -0.15345 | 0.56098 | KBTBD4 | 0.56098 | 0.61116 | 0.63419 |
| ENSG00000170091 | 5 | 173472607 | 173670504 | 783 | 88 | -0.15374 | 0.56109 | NSG2 | 0.56109 | 0.5931 | 0.64932 |
| ENSG00000131669 | 9 | 95883771 | 95896570 | 61 | 13 | -0.15376 | 0.5611 | NINJ1 | 0.5611 | 0.49219 | 0.57627 |
| ENSG00000161270 | 19 | 36316866 | 36360189 | 113 | 21 | -0.15402 | 0.5612 | NPHS1 | 0.5612 | 0.62182 | 0.67632 |
| ENSG00000072786 | 5 | 171469077 | 171615390 | 651 | 55 | -0.15403 | 0.56121 | STK10 | 0.56121 | 0.68326 | 0.50146 |
| ENSG00000164344 | 4 | 187130133 | 187179625 | 221 | 21 | -0.15412 | 0.56124 | KLKB1 | 0.56124 | 0.61317 | 0.64776 |
| ENSG00000226763 | 19 | 44100769 | 44118649 | 35 | 7 | -0.15414 | 0.56125 | SRRM5 | 0.56125 | 0.57366 | 0.66873 |
| ENSG00000178084 | 3 | 183770835 | 183778459 | 30 | 7 | -0.15434 | 0.56133 | HTR3C | 0.56133 | 0.78819 | 0.6737 |
| ENSG00000111642 | 12 | 6679249 | 6716642 | 99 | 9 | -0.15441 | 0.56136 | CHD4 | 0.56136 | 0.55572 | 0.59828 |
| ENSG00000184945 | 2 | 241631262 | 241637900 | 14 | 6 | -0.15461 | 0.56144 | AQP12A | 0.56144 | 0.51577 | 0.51997 |
| ENSG00000048342 | 4 | 15471489 | 15603180 | 331 | 33 | -0.15465 | 0.56145 | CC2D2A | 0.56145 | 0.55897 | 0.59504 |
| ENSG00000254647 | 11 | 2181009 | 2182571 | 5 | 2 | -0.15499 | 0.56158 | INS | 0.56158 | 0.020591 | 0.26865 |
| ENSG00000120451 | 11 | 130745331 | 130786404 | 263 | 12 | -0.15501 | 0.56159 | SNX19 | 0.56159 | 0.50987 | 0.5116 |
| ENSG00000258838 | 10 | 50723247 | 50747584 | 76 | 15 | -0.15515 | 0.56165 | ERCC6-PGBD3 | 0.56165 | 0.55862 | 0.53813 |
| ENSG00000254470 | 11 | 65543364 | 65548273 | 8 | 3 | -0.15566 | 0.56185 | AP5B1 | 0.56185 | 0.72482 | 0.78618 |
| ENSG00000147533 | 8 | 41347915 | 41368499 | 78 | 14 | -0.1557 | 0.56186 | GOLGA7 | 0.56186 | 0.57887 | 0.54004 |
| ENSG00000184575 | 12 | 64798130 | 64844907 | 77 | 13 | -0.15578 | 0.5619 | XPOT | 0.5619 | 0.61967 | 0.48219 |
| ENSG00000164841 | 8 | 109619079 | 109799844 | 605 | 37 | -0.15587 | 0.56193 | TMEM74 | 0.56193 | 0.59604 | 0.59134 |
| ENSG00000137693 | 11 | 101981192 | 102104154 | 325 | 17 | -0.15599 | 0.56198 | YAP1 | 0.56198 | 0.53598 | 0.48697 |
| ENSG00000198925 | 2 | 220074494 | 220094439 | 43 | 8 | -0.15607 | 0.56201 | ATG9A | 0.56201 | 0.53851 | 0.36108 |
| ENSG00000077080 | 7 | 100240720 | 100254084 | 23 | 9 | -0.15609 | 0.56202 | ACTL6B | 0.56202 | 0.51762 | 0.55959 |
| ENSG00000173715 | 11 | 66511922 | 66610987 | 116 | 22 | -0.15613 | 0.56203 | C11orf80 | 0.56203 | 0.58523 | 0.60579 |
| ENSG00000186185 | 17 | 43002077 | 43025082 | 77 | 19 | -0.1562 | 0.56206 | KIF18B | 0.56206 | 0.52181 | 0.57725 |
| ENSG00000066777 | 8 | 68085747 | 68255912 | 153 | 25 | -0.1566 | 0.56222 | ARFGEF1 | 0.56222 | 0.6076 | 0.61186 |
| ENSG00000136811 | 9 | 131217465 | 131263571 | 83 | 8 | -0.15678 | 0.56229 | ODF2 | 0.56229 | 0.51694 | 0.5808 |
| ENSG00000185651 | 22 | 21903736 | 21978323 | 134 | 6 | -0.15681 | 0.5623 | UBE2L3 | 0.5623 | 0.54404 | 0.60466 |
| ENSG00000099875 | 19 | 2037470 | 2051243 | 38 | 11 | -0.15693 | 0.56235 | MKNK2 | 0.56235 | 0.51943 | 0.18639 |
| ENSG00000184566 | 11 | 33902189 | 33903338 | 4 | 2 | -0.15706 | 0.5624 | AC132216.1 | 0.5624 | 0.53218 | 0.26645 |
| ENSG00000154678 | 7 | 31790793 | 32338941 | 2296 | 130 | -0.15709 | 0.56241 | PDE1C | 0.56241 | 0.53103 | 0.35528 |
| ENSG00000117133 | 1 | 84944942 | 84963473 | 85 | 16 | -0.15727 | 0.56248 | RPF1 | 0.56248 | 0.62398 | 0.47587 |
| ENSG00000257093 | 7 | 141356528 | 141401953 | 106 | 13 | -0.15727 | 0.56248 | KIAA1147 | 0.56248 | 0.58764 | 0.6017 |
| ENSG00000253305 | 5 | 140787770 | 140892546 | 321 | 27 | -0.15791 | 0.56274 | PCDHGB6 | 0.56274 | 0.56012 | 0.48597 |
| ENSG00000105552 | 19 | 49298319 | 49314286 | 47 | 10 | -0.15794 | 0.56275 | BCAT2 | 0.56275 | 0.57717 | 0.75061 |
| ENSG00000100142 | 22 | 38348614 | 38437922 | 96 | 9 | -0.15797 | 0.56276 | POLR2F | 0.56276 | 0.57654 | 0.30063 |
| ENSG00000214842 | 2 | 17691851 | 17699706 | 34 | 4 | -0.15804 | 0.56279 | RAD51AP2 | 0.56279 | 0.60938 | 0.57684 |
| ENSG00000145723 | 5 | 102421704 | 102455855 | 122 | 7 | -0.15813 | 0.56282 | GIN1 | 0.56282 | 0.654 | 0.69259 |
| ENSG00000067225 | 15 | 72491370 | 72524164 | 62 | 12 | -0.15816 | 0.56284 | PKM | 0.56284 | 0.65694 | 0.68023 |
| ENSG00000125775 | 20 | 1290619 | 1309883 | 82 | 24 | -0.15818 | 0.56284 | SDCBP2 | 0.56284 | 0.28155 | 0.16711 |
| ENSG00000126500 | 11 | 63870660 | 63886645 | 65 | 13 | -0.15819 | 0.56285 | FLRT1 | 0.56285 | 0.62656 | 0.67601 |
| ENSG00000179715 | 12 | 47473386 | 47630443 | 569 | 45 | -0.15826 | 0.56287 | PCED1B | 0.56287 | 0.62803 | 0.39919 |
| ENSG00000212724 | 17 | 39215495 | 39216344 | 3 | 1 | -0.1583 | 0.56289 | KRTAP2-3 | 0.56289 | 0.66418 | 0.77271 |
| ENSG00000135517 | 12 | 56843286 | 56862950 | 49 | 14 | -0.15842 | 0.56294 | MIP | 0.56294 | 0.49955 | 0.51329 |
| ENSG00000108344 | 17 | 38137050 | 38154213 | 76 | 9 | -0.15842 | 0.56294 | PSMD3 | 0.56294 | 0.69187 | 0.76245 |
| ENSG00000260286 | 6 | 24797549 | 24799117 | 4 | 2 | -0.1586 | 0.56301 | C6orf229 | 0.56301 | 0.51373 | 0.48086 |
| ENSG00000253767 | 5 | 140772381 | 140892546 | 372 | 30 | -0.15867 | 0.56304 | PCDHGA8 | 0.56304 | 0.56631 | 0.52345 |

| | | | | | | | | | | | |
|---|---|---|---|---|---|---|---|---|---|---|---|
| ENSG00000272658 | 14 | 24778161 | 24781252 | 3 | 1 | -0.1587 | 0.56305 | LTB4R2 | 0.56305 | 0.54878 | 0.65034 |
| ENSG00000170681 | 9 | 103340361 | 103350188 | 42 | 9 | -0.1589 | 0.56313 | MURC | 0.56313 | 0.51274 | 0.49067 |
| ENSG00000122420 | 1 | 78769568 | 79005434 | 715 | 56 | -0.15897 | 0.56315 | PTGFR | 0.56315 | 0.51482 | 0.37855 |
| ENSG00000253953 | 5 | 140767452 | 140892546 | 385 | 32 | -0.15897 | 0.56315 | PCDHGB4 | 0.56315 | 0.55417 | 0.50389 |
| ENSG00000123191 | 13 | 52506809 | 52585630 | 229 | 22 | -0.15905 | 0.56318 | ATP7B | 0.56318 | 0.59378 | 0.59103 |
| ENSG00000157999 | 7 | 6071007 | 6076017 | 11 | 4 | -0.15932 | 0.56329 | ANKRD61 | 0.56329 | 0.4252 | 0.59324 |
| ENSG00000141519 | 17 | 78010435 | 78074412 | 341 | 30 | -0.15935 | 0.5633 | CCDC40 | 0.5633 | 0.32069 | 0.14901 |
| ENSG00000141744 | 17 | 37824234 | 37826728 | 4 | 2 | -0.1594 | 0.56332 | PNMT | 0.56332 | 0.91613 | 0.81556 |
| ENSG00000125885 | 20 | 5931298 | 5975852 | 160 | 20 | -0.15949 | 0.56336 | MCM8 | 0.56336 | 0.58799 | 0.19627 |
| ENSG00000185250 | 6 | 109711418 | 109762374 | 154 | 14 | -0.15951 | 0.56337 | PPIL6 | 0.56337 | 0.57258 | 0.59127 |
| ENSG00000116703 | 1 | 186412698 | 186430254 | 30 | 4 | -0.15961 | 0.56341 | PDC | 0.56341 | 0.55555 | 0.6174 |
| ENSG00000156304 | 21 | 33043346 | 33104388 | 157 | 20 | -0.15967 | 0.56343 | SCAF4 | 0.56343 | 0.53461 | 0.51167 |
| ENSG00000032389 | 2 | 3192696 | 3381653 | 908 | 32 | -0.15977 | 0.56347 | TSSC1 | 0.56347 | 0.61517 | 0.63504 |
| ENSG00000124228 | 20 | 47835884 | 47860614 | 98 | 11 | -0.15988 | 0.56351 | DDX27 | 0.56351 | 0.58474 | 0.5916 |
| ENSG00000100652 | 14 | 70242134 | 70264006 | 53 | 7 | -0.15989 | 0.56352 | SLC10A1 | 0.56352 | 0.69587 | 0.71678 |
| ENSG00000062650 | 10 | 88195013 | 88281572 | 279 | 22 | -0.16019 | 0.56363 | WAPAL | 0.56363 | 0.60357 | 0.6271 |
| ENSG00000135175 | 7 | 97613996 | 97620481 | 41 | 5 | -0.16026 | 0.56366 | OCM2 | 0.56366 | 0.81642 | 0.94804 |
| ENSG00000152061 | 1 | 174128548 | 174964445 | 2182 | 24 | -0.16029 | 0.56367 | RABGAP1L | 0.56367 | 0.5675 | 0.55136 |
| ENSG00000002919 | 17 | 46180719 | 46200436 | 68 | 9 | -0.16028 | 0.56367 | SNX11 | 0.56367 | 0.61606 | 0.64053 |
| ENSG00000072694 | 1 | 161551101 | 161648444 | 524 | 30 | -0.16033 | 0.56369 | FCGR2B | 0.56369 | 0.56239 | 0.54975 |
| ENSG00000159079 | 21 | 33964389 | 33985176 | 75 | 9 | -0.16049 | 0.56375 | C21orf59 | 0.56375 | 0.58645 | 0.50398 |
| ENSG00000135365 | 11 | 45950871 | 46142985 | 394 | 19 | -0.1605 | 0.56376 | PHF21A | 0.56376 | 0.57167 | 0.54436 |
| ENSG00000135116 | 12 | 117293949 | 117319246 | 42 | 11 | -0.16061 | 0.5638 | HRK | 0.5638 | 0.7031 | 0.63208 |
| ENSG00000137393 | 6 | 18368779 | 18469105 | 585 | 34 | -0.16073 | 0.56385 | RNF144B | 0.56385 | 0.51931 | 0.43989 |
| ENSG00000166848 | 16 | 75681684 | 75795770 | 390 | 22 | -0.16074 | 0.56385 | TERF2IP | 0.56385 | 0.62961 | 0.65846 |
| ENSG00000087263 | 16 | 56485402 | 56513012 | 69 | 10 | -0.16081 | 0.56388 | OGFOD1 | 0.56388 | 0.62546 | 0.62067 |
| ENSG00000184564 | 13 | 86366925 | 86373623 | 29 | 4 | -0.16105 | 0.56397 | SLITRK6 | 0.56397 | 0.41859 | 0.52513 |
| ENSG00000139656 | 13 | 44717679 | 44735393 | 38 | 8 | -0.16123 | 0.56404 | SMIM2 | 0.56404 | 0.67732 | 0.77596 |
| ENSG00000001036 | 6 | 143815948 | 143832827 | 41 | 5 | -0.16138 | 0.5641 | FUCA2 | 0.5641 | 0.6027 | 0.78135 |
| ENSG00000137812 | 15 | 40886218 | 40956540 | 127 | 13 | -0.1618 | 0.56427 | CASC5 | 0.56427 | 0.63742 | 0.71824 |
| ENSG00000171681 | 12 | 14518610 | 14651697 | 403 | 26 | -0.16202 | 0.56435 | ATF7IP | 0.56435 | 0.59486 | 0.61598 |
| ENSG00000130475 | 19 | 17858527 | 17899377 | 190 | 27 | -0.16209 | 0.56438 | FCHO1 | 0.56438 | 0.6043 | 0.664 |
| ENSG00000102699 | 13 | 24995064 | 25086948 | 647 | 15 | -0.16221 | 0.56443 | PARP4 | 0.56443 | 0.62788 | 0.67323 |
| ENSG00000172840 | 16 | 66912492 | 66929657 | 48 | 10 | -0.16231 | 0.56447 | PDP2 | 0.56447 | 0.5087 | 0.37359 |
| ENSG00000214513 | 2 | 73429386 | 73439641 | 56 | 8 | -0.16236 | 0.56449 | NOTO | 0.56449 | 0.68712 | 0.63153 |
| ENSG00000139832 | 13 | 111175417 | 111214080 | 231 | 25 | -0.16253 | 0.56456 | RAB20 | 0.56456 | 0.61947 | 0.74182 |
| ENSG00000230062 | 6 | 46714654 | 46727243 | 27 | 8 | -0.16259 | 0.56458 | ANKRD66 | 0.56458 | 0.56006 | 0.72113 |
| ENSG00000219545 | 7 | 7680342 | 8043689 | 1842 | 143 | -0.16263 | 0.5646 | RPA3-AS1 | 0.5646 | 0.53628 | 0.43982 |
| ENSG00000125787 | 20 | 3024268 | 3026393 | 7 | 3 | -0.16285 | 0.56468 | GNRH2 | 0.56468 | 0.067996 | 0.11691 |
| ENSG00000163235 | 2 | 70674412 | 70781325 | 408 | 45 | -0.16294 | 0.56472 | TGFA | 0.56472 | 0.64493 | 0.73504 |
| ENSG00000125122 | 16 | 67241042 | 67260951 | 28 | 6 | -0.16303 | 0.56475 | LRRC29 | 0.56475 | 0.62486 | 0.65565 |
| ENSG00000169495 | 8 | 38831683 | 38846181 | 33 | 6 | -0.16337 | 0.56489 | HTRA4 | 0.56489 | 0.53339 | 0.67727 |
| ENSG00000133863 | 8 | 30689060 | 30748122 | 159 | 15 | -0.16378 | 0.56505 | TEX15 | 0.56505 | 0.58272 | 0.61353 |
| ENSG00000130254 | 19 | 5587010 | 5624057 | 106 | 16 | -0.1638 | 0.56506 | SAFB2 | 0.56506 | 0.73593 | 0.82613 |
| ENSG00000101425 | 20 | 36888551 | 36965907 | 248 | 59 | -0.16391 | 0.5651 | BPI | 0.5651 | 0.70384 | 0.60125 |
| ENSG00000221826 | 19 | 43225790 | 43244721 | 69 | 9 | -0.16397 | 0.56512 | PSG3 | 0.56512 | 0.52023 | 0.54103 |
| ENSG00000234127 | 6 | 30152232 | 30181204 | 147 | 20 | -0.16398 | 0.56513 | TRIM26 | 0.56513 | 0.56725 | 0.52426 |
| ENSG00000204361 | 11 | 114549108 | 114579362 | 122 | 20 | -0.16401 | 0.56514 | NXPE2 | 0.56514 | 0.40172 | 0.3565 |
| ENSG00000186583 | 8 | 145086582 | 145101933 | 37 | 10 | -0.16418 | 0.5652 | SPATC1 | 0.5652 | 0.21293 | 0.2614 |
| ENSG00000008382 | 19 | 4343524 | 4360083 | 64 | 14 | -0.16435 | 0.56527 | MPND | 0.56527 | 0.7349 | 0.78518 |
| ENSG00000143612 | 1 | 154179182 | 154193104 | 25 | 6 | -0.16443 | 0.5653 | C1orf43 | 0.5653 | 0.47582 | 0.4678 |
| ENSG00000114648 | 3 | 47324407 | 47388306 | 73 | 13 | -0.16446 | 0.56532 | KLHL18 | 0.56532 | 0.49002 | 0.57326 |
| ENSG00000184502 | 17 | 39868578 | 39872221 | 13 | 4 | -0.16447 | 0.56532 | GAST | 0.56532 | 0.40757 | 0.37351 |
| ENSG00000215644 | 17 | 79762008 | 79771889 | 20 | 4 | -0.16454 | 0.56535 | GCGR | 0.56535 | 0.61602 | 0.63538 |
| ENSG00000250722 | 5 | 42799982 | 42887494 | 289 | 27 | -0.16483 | 0.56546 | SEPP1 | 0.56546 | 0.56142 | 0.55697 |
| ENSG00000182718 | 15 | 60639333 | 60695082 | 181 | 31 | -0.16499 | 0.56552 | ANXA2 | 0.56552 | 0.55346 | 0.58376 |
| ENSG00000170442 | 12 | 52643084 | 52702947 | 284 | 20 | -0.16503 | 0.56554 | KRT86 | 0.56554 | 0.68011 | 0.70305 |
| ENSG00000213759 | 4 | 70065669 | 70080449 | 65 | 11 | -0.16517 | 0.56559 | UGT2B11 | 0.56559 | 0.54934 | 0.57797 |
| ENSG00000171855 | 9 | 21077104 | 21077943 | 1 | 1 | -0.16518 | 0.5656 | IFNB1 | 0.5656 | 0.73386 | 0.75557 |
| ENSG00000196565 | 11 | 5274420 | 5667019 | 2701 | 95 | -0.16518 | 0.5656 | HBG2 | 0.5656 | 0.67011 | 0.69255 |
| ENSG00000129451 | 19 | 51515995 | 51523431 | 58 | 6 | -0.16546 | 0.56571 | KLK10 | 0.56571 | 0.57244 | 0.52392 |
| ENSG00000175221 | 19 | 867962 | 893218 | 130 | 18 | -0.16565 | 0.56578 | MED16 | 0.56578 | 0.63939 | 0.66878 |
| ENSG00000155097 | 8 | 104033291 | 104085279 | 135 | 18 | -0.16585 | 0.56586 | ATP6V1C1 | 0.56586 | 0.47948 | 0.37829 |
| ENSG00000188827 | 16 | 3631182 | 3661599 | 98 | 5 | -0.16586 | 0.56587 | SLX4 | 0.56587 | 0.55543 | 0.54482 |
| ENSG00000197301 | 12 | 66245120 | 66275947 | 73 | 15 | -0.16595 | 0.5659 | RP11-366L20.2 | 0.5659 | 0.67843 | 0.67566 |
| ENSG00000102595 | 13 | 96453834 | 96705736 | 622 | 23 | -0.16594 | 0.5659 | UGGT2 | 0.5659 | 0.58708 | 0.61704 |
| ENSG00000188257 | 1 | 20301925 | 20306932 | 16 | 7 | -0.166 | 0.56592 | PLA2G2A | 0.56592 | 0.7551 | 0.72576 |
| ENSG00000166402 | 11 | 8040791 | 8127659 | 291 | 49 | -0.16611 | 0.56596 | TUB | 0.56596 | 0.61942 | 0.67408 |
| ENSG00000090932 | 19 | 39989535 | 39999121 | 28 | 4 | -0.16649 | 0.56611 | DLL3 | 0.56611 | 0.66135 | 0.5229 |
| ENSG00000250361 | 4 | 144917257 | 145061844 | 485 | 40 | -0.16656 | 0.56614 | GYPB | 0.56614 | 0.45401 | 0.44726 |
| ENSG00000052126 | 12 | 19282648 | 19529334 | 762 | 37 | -0.16688 | 0.56627 | PLEKHA5 | 0.56627 | 0.56756 | 0.56308 |
| ENSG00000206127 | 15 | 32737307 | 32747835 | 1 | 1 | -0.16696 | 0.5663 | GOLGA8O | 0.5663 | 0.5663 | 0.15008 |
| ENSG00000187033 | 3 | 169629360 | 169656963 | 88 | 14 | -0.1673 | 0.56643 | SAMD7 | 0.56643 | 0.5859 | 0.40792 |

| | | | | | | | | | | | |
|---|---|---|---|---|---|---|---|---|---|---|---|
| ENSG00000140545 | 15 | 89441916 | 89456642 | 69 | 14 | -0.16729 | 0.56643 | MFGE8 | 0.56643 | 0.74839 | 0.8278 |
| ENSG00000004776 | 19 | 36245469 | 36248980 | 5 | 3 | -0.16739 | 0.56647 | HSPB6 | 0.56647 | 0.54685 | 0.51618 |
| ENSG00000115306 | 2 | 54683422 | 54896812 | 744 | 30 | -0.16745 | 0.56649 | SPTBN1 | 0.56649 | 0.59234 | 0.617 |
| ENSG00000117877 | 19 | 45909467 | 45914024 | 24 | 6 | -0.16744 | 0.56649 | CD3EAP | 0.56649 | 0.45663 | 0.53749 |
| ENSG00000258677 | 8 | 74600926 | 74791089 | 438 | 28 | -0.16749 | 0.56651 | RP11-463D19.2 | 0.56651 | 0.58575 | 0.56122 |
| ENSG00000177666 | 11 | 818902 | 825573 | 16 | 5 | -0.16752 | 0.56652 | PNPLA2 | 0.56652 | 0.59556 | 0.71342 |
| ENSG00000139988 | 14 | 68168603 | 68201169 | 129 | 16 | -0.16761 | 0.56655 | RDH12 | 0.56655 | 0.58618 | 0.55453 |
| ENSG00000077152 | 1 | 202300785 | 202311108 | 15 | 4 | -0.16779 | 0.56663 | UBE2T | 0.56663 | 0.73704 | 0.71639 |
| ENSG00000166035 | 15 | 58702768 | 58861151 | 669 | 92 | -0.16798 | 0.5667 | LIPC | 0.5667 | 0.55464 | 0.44884 |
| ENSG00000165521 | 14 | 89078775 | 89259096 | 394 | 21 | -0.168 | 0.56671 | EML5 | 0.56671 | 0.57646 | 0.5845 |
| ENSG00000182606 | 3 | 42055294 | 42267381 | 711 | 45 | -0.16806 | 0.56673 | TRAK1 | 0.56673 | 0.5978 | 0.6472 |
| ENSG00000161791 | 12 | 50031724 | 50101948 | 122 | 11 | -0.16813 | 0.56676 | FMNL3 | 0.56676 | 0.61228 | 0.64569 |
| ENSG00000166263 | 17 | 53046088 | 53241646 | 557 | 19 | -0.16813 | 0.56676 | STXBP4 | 0.56676 | 0.58583 | 0.55794 |
| ENSG00000145908 | 5 | 150273954 | 150284545 | 33 | 2 | -0.16819 | 0.56678 | ZNF300 | 0.56678 | 0.84377 | 0.87871 |
| ENSG00000012983 | 14 | 50885219 | 51027844 | 283 | 23 | -0.16821 | 0.56679 | MAP4K5 | 0.56679 | 0.54396 | 0.58574 |
| ENSG00000126860 | 17 | 29644578 | 29648902 | 13 | 2 | -0.16845 | 0.56688 | EVI2A | 0.56688 | 0.68122 | 0.68179 |
| ENSG00000175970 | 12 | 121148238 | 121161443 | 62 | 13 | -0.16849 | 0.5669 | UNC119B | 0.5669 | 0.55735 | 0.61752 |
| ENSG00000090273 | 1 | 27226729 | 27273353 | 91 | 13 | -0.16858 | 0.56694 | NUDC | 0.56694 | 0.6068 | 0.67366 |
| ENSG00000099866 | 19 | 489176 | 505347 | 95 | 20 | -0.16865 | 0.56696 | MADCAM1 | 0.56696 | 0.76701 | 0.68359 |
| ENSG00000139304 | 12 | 80799774 | 81072802 | 933 | 48 | -0.16892 | 0.56707 | PTPRQ | 0.56707 | 0.5798 | 0.58666 |
| ENSG00000137275 | 6 | 3064225 | 3115421 | 168 | 27 | -0.16897 | 0.56709 | RIPK1 | 0.56709 | 0.58975 | 0.77533 |
| ENSG00000214517 | 11 | 73882144 | 73965748 | 203 | 13 | -0.16907 | 0.56713 | PPME1 | 0.56713 | 0.63129 | 0.69714 |
| ENSG00000188676 | 8 | 39792133 | 39873910 | 495 | 40 | -0.16949 | 0.5673 | IDO2 | 0.5673 | 0.51 | 0.45343 |
| ENSG00000185860 | 1 | 162794248 | 162838605 | 233 | 33 | -0.16957 | 0.56733 | C1orf110 | 0.56733 | 0.63715 | 0.6904 |
| ENSG00000159189 | 1 | 22970123 | 22974603 | 26 | 5 | -0.16976 | 0.5674 | C1QC | 0.5674 | 0.55182 | 0.49381 |
| ENSG00000101445 | 20 | 37434348 | 37551667 | 307 | 42 | -0.16976 | 0.5674 | PPP1R16B | 0.5674 | 0.54138 | 0.42044 |
| ENSG00000167759 | 19 | 51559463 | 51568371 | 31 | 8 | -0.16987 | 0.56744 | KLK13 | 0.56744 | 0.75924 | 0.55494 |
| ENSG00000172794 | 17 | 72666717 | 72743474 | 179 | 35 | -0.16994 | 0.56747 | RAB37 | 0.56747 | 0.65219 | 0.82936 |
| ENSG00000112293 | 6 | 24424793 | 24495433 | 560 | 35 | -0.17001 | 0.5675 | GPLD1 | 0.5675 | 0.54143 | 0.56193 |
| ENSG00000160957 | 8 | 145736667 | 145743229 | 17 | 4 | -0.1702 | 0.56757 | RECQL4 | 0.56757 | 0.5553 | 0.57592 |
| ENSG00000139200 | 12 | 6802958 | 6809966 | 6 | 3 | -0.17028 | 0.5676 | PIANP | 0.5676 | 0.33499 | 0.27262 |
| ENSG00000117713 | 1 | 27022524 | 27108595 | 139 | 9 | -0.1703 | 0.56761 | ARID1A | 0.56761 | 0.57503 | 0.55147 |
| ENSG00000033030 | 12 | 122957417 | 122985518 | 50 | 7 | -0.17034 | 0.56763 | ZCCHC8 | 0.56763 | 0.55185 | 0.55432 |
| ENSG00000222007 | 2 | 236682499 | 236692031 | 40 | 3 | -0.17044 | 0.56767 | AC064874.1 | 0.56767 | 0.6124 | 0.59631 |
| ENSG00000101189 | 20 | 61427805 | 61431945 | 16 | 6 | -0.17044 | 0.56767 | MRGBP | 0.56767 | 0.59325 | 0.69729 |
| ENSG00000169884 | 12 | 49359123 | 49365546 | 14 | 6 | -0.17047 | 0.56768 | WNT10B | 0.56768 | 0.53439 | 0.5354 |
| ENSG00000115884 | 2 | 20400558 | 20425194 | 47 | 16 | -0.17056 | 0.56772 | SDC1 | 0.56772 | 0.73602 | 0.80663 |
| ENSG00000166926 | 11 | 60102304 | 60164069 | 178 | 17 | -0.17058 | 0.56772 | MS4A6E | 0.56772 | 0.26912 | 0.23894 |
| ENSG00000108753 | 17 | 36046435 | 36105237 | 170 | 47 | -0.1707 | 0.56777 | HNF1B | 0.56777 | 0.63236 | 0.47792 |
| ENSG00000107341 | 9 | 33817565 | 33920402 | 354 | 20 | -0.17149 | 0.56808 | UBE2R2 | 0.56808 | 0.58476 | 0.61655 |
| ENSG00000145087 | 3 | 120626919 | 121143608 | 1903 | 31 | -0.17156 | 0.56811 | STXBP5L | 0.56811 | 0.58229 | 0.5875 |
| ENSG00000164164 | 4 | 146031990 | 146101313 | 82 | 21 | -0.17163 | 0.56813 | OTUD4 | 0.56813 | 0.49889 | 0.50102 |
| ENSG00000136932 | 9 | 100666771 | 100684852 | 47 | 7 | -0.17161 | 0.56813 | C9orf156 | 0.56813 | 0.63619 | 0.6669 |
| ENSG00000070748 | 10 | 50817141 | 50901925 | 235 | 39 | -0.17167 | 0.56815 | CHAT | 0.56815 | 0.54418 | 0.64212 |
| ENSG00000104870 | 19 | 50010073 | 50029590 | 65 | 11 | -0.17166 | 0.56815 | FCGRT | 0.56815 | 0.64412 | 0.62264 |
| ENSG00000167236 | 17 | 34340096 | 34345005 | 24 | 5 | -0.17186 | 0.56823 | CCL23 | 0.56823 | 0.72699 | 0.5859 |
| ENSG00000122034 | 13 | 27998681 | 28009958 | 47 | 8 | -0.17192 | 0.56825 | GTF3A | 0.56825 | 0.54312 | 0.55949 |
| ENSG00000128534 | 7 | 117824086 | 117832878 | 59 | 5 | -0.17203 | 0.56829 | NAA38 | 0.56829 | 0.59803 | 0.59215 |
| ENSG00000013810 | 4 | 1723227 | 1746898 | 164 | 16 | -0.17209 | 0.56832 | TACC3 | 0.56832 | 0.54829 | 0.43841 |
| ENSG00000197057 | 4 | 36283244 | 36347378 | 244 | 42 | -0.17223 | 0.56837 | DTHD1 | 0.56837 | 0.50603 | 0.55382 |
| ENSG00000033627 | 17 | 40610862 | 40674629 | 90 | 17 | -0.17248 | 0.56847 | ATP6V0A1 | 0.56847 | 0.57408 | 0.69292 |
| ENSG00000188816 | 10 | 124907638 | 124910188 | 5 | 1 | -0.17254 | 0.56849 | HMX2 | 0.56849 | 0.515 | 0.46921 |
| ENSG00000213901 | 2 | 219940051 | 220035549 | 188 | 18 | -0.17336 | 0.56882 | SLC23A3 | 0.56882 | 0.58174 | 0.45376 |
| ENSG00000183971 | 16 | 2059927 | 2070756 | 42 | 10 | -0.17358 | 0.5689 | NPW | 0.5689 | 0.8133 | 0.86081 |
| ENSG00000126804 | 14 | 64970430 | 65000408 | 101 | 10 | -0.17364 | 0.56893 | ZBTB1 | 0.56893 | 0.66669 | 0.77461 |
| ENSG00000259030 | 1 | 74663919 | 75009666 | 800 | 58 | -0.17371 | 0.56895 | FPGT-TNNI3K | 0.56895 | 0.60431 | 0.63212 |
| ENSG00000116783 | 1 | 74663947 | 75010112 | 800 | 58 | -0.17371 | 0.56895 | TNNI3K | 0.56895 | 0.60234 | 0.63212 |
| ENSG00000076248 | 12 | 109535379 | 109548797 | 28 | 10 | -0.17372 | 0.56896 | UNG | 0.56896 | 0.63389 | 0.77759 |
| ENSG00000100413 | 22 | 41921808 | 41940610 | 49 | 7 | -0.17381 | 0.56899 | POLR3H | 0.56899 | 0.54407 | 0.61583 |
| ENSG00000215428 | 9 | 133274029 | 133282639 | 36 | 11 | -0.17388 | 0.56902 | AL354898.1 | 0.56902 | 0.59644 | 0.59665 |
| ENSG00000083457 | 17 | 3617922 | 3704537 | 372 | 48 | -0.17391 | 0.56903 | ITGAE | 0.56903 | 0.64204 | 0.67621 |
| ENSG00000036549 | 1 | 78028101 | 78149104 | 235 | 26 | -0.17415 | 0.56913 | ZZZ3 | 0.56913 | 0.54358 | 0.56066 |
| ENSG00000082269 | 6 | 71122644 | 71270877 | 487 | 17 | -0.17422 | 0.56915 | FAM135A | 0.56915 | 0.57682 | 0.55959 |
| ENSG00000165304 | 9 | 36572859 | 36677678 | 218 | 14 | -0.17445 | 0.56925 | MELK | 0.56925 | 0.56719 | 0.54639 |
| ENSG00000119630 | 14 | 75408537 | 75422487 | 21 | 10 | -0.17448 | 0.56925 | PGF | 0.56925 | 0.26195 | 0.24435 |
| ENSG00000143797 | 2 | 8992820 | 9143942 | 259 | 15 | -0.17448 | 0.56926 | MBOAT2 | 0.56926 | 0.57961 | 0.49174 |
| ENSG00000134899 | 13 | 103497194 | 103528345 | 85 | 12 | -0.17491 | 0.56942 | ERCC5 | 0.56942 | 0.39689 | 0.22716 |
| ENSG00000131779 | 1 | 145516252 | 145523730 | 3 | 1 | -0.17493 | 0.56943 | PEX11B | 0.56943 | 0.5873 | 0.68945 |
| ENSG00000243667 | 2 | 68350068 | 68384692 | 60 | 9 | -0.17496 | 0.56944 | WDR92 | 0.56944 | 0.64057 | 0.71851 |
| ENSG00000227124 | 3 | 75758794 | 75834734 | 185 | 14 | -0.17495 | 0.56944 | ZNF717 | 0.56944 | 0.59666 | 0.57096 |
| ENSG00000110717 | 11 | 67798084 | 67804111 | 16 | 6 | -0.17515 | 0.56952 | NDUFS8 | 0.56952 | 0.33551 | 0.31807 |
| ENSG00000176937 | 11 | 4824663 | 4825847 | 5 | 2 | -0.17519 | 0.56954 | OR52R1 | 0.56954 | 0.60797 | 0.719 |
| ENSG00000124383 | 2 | 71357444 | 71377231 | 75 | 7 | -0.17555 | 0.56967 | MPHOSPH10 | 0.56967 | 0.59266 | 0.45947 |

| | | | | | | | | | | | |
|---|---|---|---|---|---|---|---|---|---|---|---|
| ENSG00000076924 | 19 | 7684411 | 7694451 | 25 | 3 | -0.17567 | 0.56972 | XAB2 | 0.56972 | 0.427 | 0.43502 |
| ENSG00000095932 | 19 | 3474405 | 3480540 | 20 | 7 | -0.17585 | 0.5698 | C19orf77 | 0.5698 | 0.34719 | 0.34645 |
| ENSG00000175183 | 12 | 77252495 | 77272840 | 89 | 16 | -0.17602 | 0.56986 | CSRP2 | 0.56986 | 0.52951 | 0.50869 |
| ENSG00000116678 | 1 | 65886248 | 66107242 | 775 | 29 | -0.17607 | 0.56988 | LEPR | 0.56988 | 0.53068 | 0.44752 |
| ENSG00000221933 | 7 | 143771275 | 143772288 | 3 | 1 | -0.17617 | 0.56992 | OR2A25 | 0.56992 | 0.53012 | 0.45784 |
| ENSG00000170099 | 14 | 94770585 | 94789731 | 68 | 12 | -0.17626 | 0.56995 | SERPINA6 | 0.56995 | 0.73802 | 0.78795 |
| ENSG00000104695 | 8 | 30631973 | 30671830 | 129 | 14 | -0.17639 | 0.57001 | PPP2CB | 0.57001 | 0.6421 | 0.74825 |
| ENSG00000271810 | 1 | 113245236 | 113254055 | 26 | 8 | -0.1773 | 0.57036 | RP11-426L16.10 | 0.57036 | 0.15032 | 0.1772 |
| ENSG00000205643 | 22 | 46639908 | 46646576 | 31 | 5 | -0.17752 | 0.57045 | CDPF1 | 0.57045 | 0.56856 | 0.52905 |
| ENSG00000166947 | 15 | 43398423 | 43513481 | 136 | 24 | -0.17755 | 0.57046 | EPB42 | 0.57046 | 0.43944 | 0.37417 |
| ENSG00000124249 | 20 | 43374421 | 43379675 | 16 | 5 | -0.17756 | 0.57047 | KCNK15 | 0.57047 | 0.72557 | 0.88198 |
| ENSG00000172179 | 6 | 22287480 | 22297730 | 27 | 8 | -0.17775 | 0.57054 | PRL | 0.57054 | 0.21613 | 0.062473 |
| ENSG00000137261 | 6 | 24544332 | 24646383 | 394 | 43 | -0.17788 | 0.57059 | KIAA0319 | 0.57059 | 0.52553 | 0.53969 |
| ENSG00000007541 | 16 | 616995 | 634136 | 77 | 5 | -0.17814 | 0.57069 | PIGQ | 0.57069 | 0.56252 | 0.50295 |
| ENSG00000113739 | 5 | 172741716 | 172756506 | 31 | 8 | -0.17819 | 0.57071 | STC2 | 0.57071 | 0.56529 | 0.62928 |
| ENSG00000171345 | 17 | 39679869 | 39684560 | 8 | 1 | -0.17837 | 0.57078 | KRT19 | 0.57078 | 0.61583 | 0.82308 |
| ENSG00000071242 | 6 | 166822852 | 167319939 | 2198 | 173 | -0.17841 | 0.5708 | RPS6KA2 | 0.5708 | 0.62752 | 0.49403 |
| ENSG00000021645 | 14 | 78708734 | 80330762 | 4650 | 243 | -0.17858 | 0.57087 | NRXN3 | 0.57087 | 0.49151 | 0.52408 |
| ENSG00000163743 | 4 | 76404247 | 76439974 | 103 | 12 | -0.17872 | 0.57092 | RCHY1 | 0.57092 | 0.55033 | 0.57585 |
| ENSG00000108278 | 17 | 34842473 | 34855154 | 23 | 5 | -0.17871 | 0.57092 | ZNHIT3 | 0.57092 | 0.45901 | 0.49836 |
| ENSG00000215114 | 8 | 59323823 | 59364060 | 111 | 17 | -0.17895 | 0.57101 | UBXN2B | 0.57101 | 0.57054 | 0.73178 |
| ENSG00000164332 | 5 | 158690089 | 158713044 | 45 | 10 | -0.17903 | 0.57104 | UBLCP1 | 0.57104 | 0.60266 | 0.60293 |
| ENSG00000129151 | 11 | 27062272 | 27149356 | 328 | 24 | -0.17933 | 0.57116 | BBOX1 | 0.57116 | 0.55517 | 0.51993 |
| ENSG00000033178 | 4 | 68478370 | 68566897 | 349 | 14 | -0.17936 | 0.57117 | UBA6 | 0.57117 | 0.60266 | 0.65554 |
| ENSG00000183206 | 18 | 14507338 | 14543584 | 117 | 17 | -0.17937 | 0.57118 | POTEC | 0.57118 | 0.60615 | 0.59953 |
| ENSG00000135932 | 2 | 231577560 | 231685790 | 203 | 20 | -0.17941 | 0.57119 | CAB39 | 0.57119 | 0.50808 | 0.45633 |
| ENSG00000100379 | 22 | 37447779 | 37459430 | 19 | 7 | -0.17946 | 0.57121 | KCTD17 | 0.57121 | 0.015746 | 0.017913 |
| ENSG00000106993 | 9 | 4679559 | 4708398 | 93 | 9 | -0.17967 | 0.57129 | CDC37L1 | 0.57129 | 0.36056 | 0.39528 |
| ENSG00000092098 | 14 | 24615892 | 24629870 | 35 | 13 | -0.17966 | 0.57129 | RNF31 | 0.57129 | 0.67559 | 0.754 |
| ENSG00000198400 | 1 | 156785432 | 156851642 | 183 | 31 | -0.17997 | 0.57141 | NTRK1 | 0.57141 | 0.49462 | 0.28111 |
| ENSG00000144061 | 2 | 110879888 | 110962643 | 178 | 12 | -0.18001 | 0.57143 | NPHP1 | 0.57143 | 0.59789 | 0.52069 |
| ENSG00000083838 | 19 | 58985384 | 58992597 | 13 | 5 | -0.18021 | 0.57151 | ZNF446 | 0.57151 | 0.65741 | 0.78147 |
| ENSG00000249967 | 10 | 99344131 | 99433667 | 247 | 29 | -0.18027 | 0.57153 | PI4K2A | 0.57153 | 0.6189 | 0.6996 |
| ENSG00000152413 | 5 | 78668459 | 78810040 | 375 | 25 | -0.18054 | 0.57164 | HOMER1 | 0.57164 | 0.48499 | 0.37447 |
| ENSG00000072803 | 5 | 171288553 | 171433877 | 257 | 17 | -0.18061 | 0.57166 | FBXW11 | 0.57166 | 0.59932 | 0.70541 |
| ENSG00000105810 | 7 | 92234235 | 92465908 | 322 | 34 | -0.18061 | 0.57166 | CDK6 | 0.57166 | 0.63889 | 0.7259 |
| ENSG00000166436 | 11 | 8633584 | 8693413 | 160 | 18 | -0.18067 | 0.57169 | TRIM66 | 0.57169 | 0.51834 | 0.30966 |
| ENSG00000116954 | 1 | 39303870 | 39325495 | 65 | 14 | -0.18077 | 0.57173 | RRAGC | 0.57173 | 0.57486 | 0.27912 |
| ENSG00000147535 | 8 | 38120648 | 38126761 | 9 | 3 | -0.18096 | 0.5718 | PPAPDC1B | 0.5718 | 0.50748 | 0.54583 |
| ENSG00000198833 | 6 | 90036344 | 90062567 | 60 | 20 | -0.18114 | 0.57187 | UBE2J1 | 0.57187 | 0.64427 | 0.57737 |
| ENSG00000090615 | 12 | 133345495 | 133405444 | 309 | 27 | -0.18113 | 0.57187 | GOLGA3 | 0.57187 | 0.58849 | 0.54522 |
| ENSG00000150776 | 11 | 111944810 | 111955874 | 15 | 6 | -0.18115 | 0.57188 | C11orf57 | 0.57188 | 0.69669 | 0.76687 |
| ENSG00000258973 | 14 | 24801516 | 24806588 | 19 | 5 | -0.18115 | 0.57188 | RP11-934B9.3 | 0.57188 | 0.33966 | 0.10034 |
| ENSG00000143507 | 1 | 221874766 | 221915518 | 101 | 22 | -0.18138 | 0.57196 | DUSP10 | 0.57196 | 0.55713 | 0.64857 |
| ENSG00000268628 | 20 | 19738296 | 19738679 | 2 | 1 | -0.18147 | 0.572 | AL121761.2 | 0.572 | 0.82498 | 0.80987 |
| ENSG00000188280 | 22 | 20692438 | 20716938 | 49 | 5 | -0.18146 | 0.572 | FAM230A | 0.572 | 0.62559 | 0.76438 |
| ENSG00000179862 | 1 | 41326729 | 41328018 | 7 | 1 | -0.18162 | 0.57206 | CITED4 | 0.57206 | 0.41712 | 0.38679 |
| ENSG00000183814 | 1 | 226418850 | 226497570 | 155 | 21 | -0.18177 | 0.57212 | LIN9 | 0.57212 | 0.61828 | 0.59745 |
| ENSG00000138083 | 2 | 45168902 | 45173216 | 19 | 6 | -0.18181 | 0.57213 | SIX3 | 0.57213 | 0.40973 | 0.32411 |
| ENSG00000182575 | 17 | 47653220 | 47661189 | 13 | 4 | -0.1818 | 0.57213 | NXPH3 | 0.57213 | 0.88327 | 0.7598 |
| ENSG00000152404 | 11 | 107197071 | 107328572 | 575 | 25 | -0.18192 | 0.57218 | CWF19L2 | 0.57218 | 0.56476 | 0.505 |
| ENSG00000146147 | 6 | 53794780 | 54131078 | 856 | 54 | -0.18216 | 0.57227 | MLIP | 0.57227 | 0.59173 | 0.62169 |
| ENSG00000196196 | 9 | 35906189 | 35907138 | 6 | 4 | -0.18218 | 0.57228 | HRCT1 | 0.57228 | 0.87751 | 0.90065 |
| ENSG00000072864 | 16 | 15737124 | 15820210 | 236 | 29 | -0.18229 | 0.57232 | NDE1 | 0.57232 | 0.53928 | 0.60254 |
| ENSG00000184697 | 16 | 3064713 | 3070072 | 17 | 4 | -0.18245 | 0.57239 | CLDN6 | 0.57239 | 0.91624 | 0.53204 |
| ENSG00000184208 | 22 | 42084943 | 42094140 | 16 | 3 | -0.18247 | 0.57239 | C22orf46 | 0.57239 | 0.75033 | 0.74768 |
| ENSG00000159212 | 21 | 36041688 | 36090525 | 115 | 14 | -0.18267 | 0.57247 | CLIC6 | 0.57247 | 0.64259 | 0.65019 |
| ENSG00000175482 | 11 | 67118248 | 67124443 | 9 | 3 | -0.18276 | 0.57251 | POLD4 | 0.57251 | 0.80634 | 0.94191 |
| ENSG00000198483 | 1 | 145549230 | 145568526 | 43 | 7 | -0.18292 | 0.57257 | ANKRD35 | 0.57257 | 0.59382 | 0.64928 |
| ENSG00000204444 | 6 | 31620193 | 31625987 | 15 | 6 | -0.183 | 0.5726 | APOM | 0.5726 | 0.33743 | 0.31656 |
| ENSG00000180332 | 13 | 45766988 | 45775175 | 14 | 2 | -0.18306 | 0.57262 | KCTD4 | 0.57262 | 0.66733 | 0.69426 |
| ENSG00000242110 | 5 | 33986283 | 34008220 | 67 | 10 | -0.18308 | 0.57263 | AMACR | 0.57263 | 0.62626 | 0.79647 |
| ENSG00000087053 | 11 | 95566046 | 95658479 | 304 | 23 | -0.18314 | 0.57266 | MTMR2 | 0.57266 | 0.6103 | 0.63419 |
| ENSG00000186918 | 8 | 28203102 | 28260218 | 73 | 13 | -0.18341 | 0.57276 | ZNF395 | 0.57276 | 0.68469 | 0.72546 |
| ENSG00000259399 | 20 | 35202956 | 35240787 | 41 | 16 | -0.18347 | 0.57278 | TGIF2-C20orf24 | 0.57278 | 0.56014 | 0.51486 |
| ENSG00000173627 | 1 | 183615541 | 183622451 | 42 | 3 | -0.18354 | 0.57281 | APOBEC4 | 0.57281 | 0.6013 | 0.61898 |
| ENSG00000182272 | 11 | 369796 | 382116 | 50 | 9 | -0.18354 | 0.57281 | B4GALNT4 | 0.57281 | 0.28982 | 0.15374 |
| ENSG00000125409 | 17 | 15207128 | 15244958 | 119 | 15 | -0.18356 | 0.57282 | TEKT3 | 0.57282 | 0.75432 | 0.8886 |
| ENSG00000112041 | 6 | 35465651 | 35480715 | 35 | 9 | -0.18371 | 0.57288 | TULP1 | 0.57288 | 0.57763 | 0.34239 |
| ENSG00000134900 | 13 | 103249353 | 103331521 | 134 | 23 | -0.18402 | 0.573 | TPP2 | 0.573 | 0.61068 | 0.5147 |
| ENSG00000166260 | 17 | 53029263 | 53046146 | 80 | 9 | -0.18436 | 0.57313 | COX11 | 0.57313 | 0.54414 | 0.53003 |
| ENSG00000204438 | 6 | 31629006 | 31634060 | 34 | 13 | -0.18455 | 0.57321 | GPANK1 | 0.57321 | 0.41699 | 0.30599 |
| ENSG00000168803 | 15 | 43622872 | 43646096 | 67 | 8 | -0.1848 | 0.57331 | ADAL | 0.57331 | 0.6316 | 0.62026 |

| | | | | | | | | | | | |
|---|---|---|---|---|---|---|---|---|---|---|---|
| ENSG00000163428 | 3 | 120043356 | 120068186 | 88 | 13 | -0.18488 | 0.57334 | LRRC58 | 0.57334 | 0.54494 | 0.53665 |
| ENSG00000144747 | 3 | 69068978 | 69101484 | 60 | 9 | -0.1849 | 0.57335 | TMF1 | 0.57335 | 0.54593 | 0.63246 |
| ENSG00000185955 | 7 | 100054238 | 100061894 | 9 | 3 | -0.18492 | 0.57335 | C7orf61 | 0.57335 | 0.64268 | 0.62242 |
| ENSG00000106302 | 7 | 123469037 | 123517532 | 101 | 18 | -0.18498 | 0.57338 | HYAL4 | 0.57338 | 0.63432 | 0.61243 |
| ENSG00000263155 | 15 | 57884139 | 57977562 | 424 | 33 | -0.18505 | 0.57341 | MYZAP | 0.57341 | 0.54725 | 0.55407 |
| ENSG00000092841 | 12 | 56551945 | 56557280 | 6 | 3 | -0.18508 | 0.57342 | MYL6 | 0.57342 | 0.81533 | 0.71512 |
| ENSG00000152022 | 1 | 145477085 | 145501669 | 22 | 7 | -0.18532 | 0.57351 | LIX1L | 0.57351 | 0.39071 | 0.46104 |
| ENSG00000253846 | 5 | 140792743 | 140892546 | 312 | 27 | -0.18538 | 0.57353 | PCDHGA10 | 0.57353 | 0.58848 | 0.47921 |
| ENSG00000118513 | 6 | 135502453 | 135540311 | 73 | 11 | -0.18537 | 0.57353 | MYB | 0.57353 | 0.72834 | 0.5128 |
| ENSG00000111261 | 12 | 12482198 | 12503475 | 110 | 13 | -0.18536 | 0.57353 | MANSC1 | 0.57353 | 0.72198 | 0.65887 |
| ENSG00000141068 | 17 | 25783670 | 25953461 | 393 | 39 | -0.18536 | 0.57353 | KSR1 | 0.57353 | 0.55761 | 0.65188 |
| ENSG00000161526 | 17 | 73663196 | 73704142 | 99 | 18 | -0.18545 | 0.57356 | SAP30BP | 0.57356 | 0.52503 | 0.58644 |
| ENSG00000197930 | 14 | 53106634 | 53162618 | 160 | 18 | -0.18547 | 0.57357 | ERO1L | 0.57357 | 0.53803 | 0.51806 |
| ENSG00000173335 | 20 | 23583047 | 23586513 | 1 | 1 | -0.18555 | 0.5736 | CST9 | 0.5736 | 0.67428 | 0.93455 |
| ENSG00000178913 | 5 | 140698057 | 140700330 | 3 | 2 | -0.1859 | 0.57374 | TAF7 | 0.57374 | 0.41812 | 0.263 |
| ENSG00000266964 | 19 | 35629712 | 35634013 | 11 | 2 | -0.18594 | 0.57375 | FXYD1 | 0.57375 | 0.65623 | 0.71673 |
| ENSG00000091947 | 17 | 42088556 | 42101314 | 35 | 9 | -0.18599 | 0.57377 | TMEM101 | 0.57377 | 0.64945 | 0.70652 |
| ENSG00000160200 | 21 | 44473301 | 44497053 | 94 | 18 | -0.18598 | 0.57377 | CBS | 0.57377 | 0.69509 | 0.82471 |
| ENSG00000145384 | 4 | 120238405 | 120243545 | 47 | 4 | -0.18609 | 0.57381 | FABP2 | 0.57381 | 0.60294 | 0.59979 |
| ENSG00000114650 | 3 | 47455203 | 47518616 | 81 | 13 | -0.18617 | 0.57385 | SCAP | 0.57385 | 0.59402 | 0.58353 |
| ENSG00000125850 | 20 | 17937623 | 18039832 | 477 | 36 | -0.18629 | 0.57389 | OVOL2 | 0.57389 | 0.62581 | 0.64799 |
| ENSG00000164619 | 7 | 33944523 | 34195484 | 610 | 57 | -0.18639 | 0.57393 | BMPER | 0.57393 | 0.52539 | 0.38216 |
| ENSG00000077420 | 10 | 26727132 | 26856732 | 516 | 53 | -0.18646 | 0.57396 | APBB1IP | 0.57396 | 0.64605 | 0.70873 |
| ENSG00000080573 | 19 | 10070237 | 10121147 | 397 | 42 | -0.18688 | 0.57412 | COL5A3 | 0.57412 | 0.60009 | 0.49565 |
| ENSG00000138386 | 2 | 191511472 | 191557492 | 124 | 17 | -0.18703 | 0.57418 | NAB1 | 0.57418 | 0.61556 | 0.63049 |
| ENSG00000132849 | 1 | 62208149 | 62629592 | 1586 | 103 | -0.18714 | 0.57423 | INADL | 0.57423 | 0.5924 | 0.56139 |
| ENSG00000213689 | 3 | 48506445 | 48509044 | 3 | 2 | -0.18718 | 0.57424 | TREX1 | 0.57424 | 0.42036 | 0.42255 |
| ENSG00000181195 | 8 | 57349233 | 57359293 | 42 | 7 | -0.1872 | 0.57425 | PENK | 0.57425 | 0.46054 | 0.42587 |
| ENSG00000119326 | 9 | 111704851 | 111775809 | 242 | 25 | -0.18721 | 0.57425 | CTNNAL1 | 0.57425 | 0.56996 | 0.4882 |
| ENSG00000162702 | 1 | 200374068 | 200379184 | 3 | 2 | -0.18724 | 0.57427 | ZNF281 | 0.57427 | 0.45115 | 0.383 |
| ENSG00000016391 | 3 | 53846362 | 53880417 | 122 | 23 | -0.18744 | 0.57434 | CHDH | 0.57434 | 0.47437 | 0.62688 |
| ENSG00000130313 | 19 | 17622438 | 17632097 | 32 | 7 | -0.18756 | 0.57439 | PGLS | 0.57439 | 0.54092 | 0.30857 |
| ENSG00000131042 | 19 | 54777675 | 54785039 | 58 | 14 | -0.18763 | 0.57442 | LILRB2 | 0.57442 | 0.51898 | 0.43178 |
| ENSG00000100170 | 22 | 32439019 | 32509016 | 244 | 16 | -0.18771 | 0.57445 | SLC5A1 | 0.57445 | 0.59886 | 0.60061 |
| ENSG00000138796 | 4 | 108910870 | 108956331 | 131 | 9 | -0.18807 | 0.57459 | HADH | 0.57459 | 0.59111 | 0.56572 |
| ENSG00000205362 | 16 | 56672578 | 56673999 | 8 | 2 | -0.18808 | 0.57459 | MT1A | 0.57459 | 0.3007 | 0.44266 |
| ENSG00000075218 | 22 | 46692638 | 46726707 | 106 | 8 | -0.18817 | 0.57463 | GTSE1 | 0.57463 | 0.52346 | 0.54706 |
| ENSG00000135373 | 11 | 34642640 | 34682604 | 128 | 33 | -0.1882 | 0.57464 | EHF | 0.57464 | 0.52717 | 0.29151 |
| ENSG00000206531 | 3 | 112534556 | 112565703 | 140 | 20 | -0.18837 | 0.57471 | CD200R1L | 0.57471 | 0.59327 | 0.59129 |
| ENSG00000249437 | 5 | 70264310 | 70320941 | 16 | 6 | -0.18841 | 0.57472 | NAIP | 0.57472 | 0.69899 | 0.80172 |
| ENSG00000070669 | 7 | 97481430 | 97501854 | 79 | 15 | -0.18844 | 0.57473 | ASNS | 0.57473 | 0.66857 | 0.75186 |
| ENSG00000119650 | 14 | 76368479 | 76550928 | 462 | 32 | -0.18848 | 0.57475 | IFT43 | 0.57475 | 0.5996 | 0.63889 |
| ENSG00000156574 | 10 | 72192071 | 72207707 | 44 | 10 | -0.18879 | 0.57487 | NODAL | 0.57487 | 0.65291 | 0.71049 |
| ENSG00000169214 | 1 | 247875045 | 247876105 | 7 | 3 | -0.1889 | 0.57492 | OR6F1 | 0.57492 | 0.93963 | 0.93566 |
| ENSG00000185475 | 11 | 62554887 | 62557877 | 5 | 2 | -0.18893 | 0.57493 | TMEM179B | 0.57493 | 0.85578 | 0.5975 |
| ENSG00000128463 | 15 | 34517200 | 34522357 | 24 | 7 | -0.18893 | 0.57493 | EMC4 | 0.57493 | 0.73572 | 0.72112 |
| ENSG00000102970 | 16 | 57438679 | 57449974 | 49 | 12 | -0.18905 | 0.57497 | CCL17 | 0.57497 | 0.047754 | 0.018534 |
| ENSG00000186766 | 10 | 129535499 | 129539450 | 19 | 4 | -0.18906 | 0.57498 | FOXI2 | 0.57498 | 0.51743 | 0.64758 |
| ENSG00000100376 | 22 | 45704849 | 45737836 | 122 | 13 | -0.18927 | 0.57506 | FAM118A | 0.57506 | 0.61842 | 0.46118 |
| ENSG00000118454 | 1 | 70726271 | 70820417 | 184 | 24 | -0.18935 | 0.57509 | ANKRD13C | 0.57509 | 0.59285 | 0.64503 |
| ENSG00000162692 | 1 | 101185298 | 101204601 | 35 | 16 | -0.1894 | 0.57511 | VCAM1 | 0.57511 | 0.24369 | 0.114 |
| ENSG00000221883 | 3 | 48955221 | 48956818 | 2 | 1 | -0.18958 | 0.57518 | ARIH2OS | 0.57518 | 0.61781 | 0.59298 |
| ENSG00000243251 | 10 | 50723151 | 50747086 | 74 | 15 | -0.18961 | 0.57519 | PGBD3 | 0.57519 | 0.557 | 0.5304 |
| ENSG00000110427 | 11 | 33563618 | 33695648 | 328 | 33 | -0.18999 | 0.57534 | KIAA1549L | 0.57534 | 0.44942 | 0.5178 |
| ENSG00000131408 | 19 | 50832910 | 50886239 | 134 | 14 | -0.19007 | 0.57537 | NR1H2 | 0.57537 | 0.70998 | 0.83699 |
| ENSG00000085276 | 3 | 168801287 | 169381406 | 2085 | 137 | -0.19015 | 0.5754 | MECOM | 0.5754 | 0.56165 | 0.52718 |
| ENSG00000255994 | 14 | 70037483 | 70038032 | 1 | 1 | -0.19014 | 0.5754 | CCDC177 | 0.5754 | 0.78288 | 0.83395 |
| ENSG00000137801 | 15 | 39873280 | 39891667 | 27 | 4 | -0.1902 | 0.57542 | THBS1 | 0.57542 | 0.80181 | 0.79831 |
| ENSG00000119616 | 14 | 75179847 | 75203394 | 35 | 6 | -0.19059 | 0.57558 | FCF1 | 0.57558 | 0.57056 | 0.52301 |
| ENSG00000113712 | 5 | 148871760 | 148931007 | 105 | 16 | -0.19064 | 0.57559 | CSNK1A1 | 0.57559 | 0.57722 | 0.6634 |
| ENSG00000241404 | 6 | 32132360 | 32136058 | 15 | 6 | -0.19087 | 0.57569 | EGFL8 | 0.57569 | 0.023094 | 0.04735 |
| ENSG00000166813 | 15 | 90152020 | 90198682 | 132 | 13 | -0.19095 | 0.57572 | KIF7 | 0.57572 | 0.59503 | 0.55991 |
| ENSG00000105383 | 19 | 51728320 | 51747115 | 38 | 12 | -0.19109 | 0.57577 | CD33 | 0.57577 | 0.10291 | 0.28145 |
| ENSG00000131778 | 1 | 146714291 | 146767443 | 257 | 21 | -0.19128 | 0.57585 | CHD1L | 0.57585 | 0.54928 | 0.5491 |
| ENSG00000184489 | 8 | 142402093 | 142441620 | 122 | 17 | -0.19135 | 0.57587 | PTP4A3 | 0.57587 | 0.55606 | 0.49254 |
| ENSG00000162231 | 11 | 62559595 | 62573774 | 14 | 6 | -0.19154 | 0.57595 | NXF1 | 0.57595 | 0.74708 | 0.68324 |
| ENSG00000127084 | 9 | 95709733 | 95798518 | 297 | 26 | -0.19169 | 0.57601 | FGD3 | 0.57601 | 0.64262 | 0.63341 |
| ENSG00000029534 | 8 | 41510739 | 41754280 | 829 | 71 | -0.19203 | 0.57614 | ANK1 | 0.57614 | 0.56672 | 0.59285 |
| ENSG00000170315 | 17 | 16284112 | 16286059 | 2 | 1 | -0.19216 | 0.57619 | UBB | 0.57619 | 0.088099 | 0.11932 |
| ENSG00000083642 | 13 | 33160564 | 33352157 | 472 | 13 | -0.19234 | 0.57626 | PDS5B | 0.57626 | 0.57761 | 0.5837 |
| ENSG00000149403 | 11 | 120382468 | 120859613 | 1531 | 144 | -0.19239 | 0.57628 | GRIK4 | 0.57628 | 0.51888 | 0.38933 |
| ENSG00000165312 | 10 | 23728198 | 23731308 | 4 | 2 | -0.1924 | 0.57629 | OTUD1 | 0.57629 | 0.8712 | 0.88588 |
| ENSG00000233608 | 2 | 239756673 | 239795893 | 158 | 44 | -0.19245 | 0.5763 | TWIST2 | 0.5763 | 0.47592 | 0.49021 |

| | | | | | | | | | | | |
|---|---|---|---|---|---|---|---|---|---|---|---|
| ENSG00000100056 | 22 | 19117792 | 19132197 | 39 | 10 | -0.19264 | 0.57638 | DGCR14 | 0.57638 | 0.76446 | 0.13536 |
| ENSG00000140479 | 15 | 101840818 | 102065405 | 1184 | 146 | -0.19297 | 0.57651 | PCSK6 | 0.57651 | 0.49625 | 0.5627 |
| ENSG00000135049 | 9 | 88161455 | 88356944 | 729 | 20 | -0.193 | 0.57652 | AGTPBP1 | 0.57652 | 0.58667 | 0.62864 |
| ENSG00000255633 | 6 | 62284008 | 62284534 | 8 | 3 | -0.19301 | 0.57653 | MTRNR2L9 | 0.57653 | 0.27914 | 0.23635 |
| ENSG00000227507 | 6 | 31548302 | 31550299 | 1 | 1 | -0.1932 | 0.5766 | LTB | 0.5766 | 0.12886 | 0.081067 |
| ENSG00000128918 | 15 | 58245622 | 58790065 | 2227 | 144 | -0.19324 | 0.57661 | ALDH1A2 | 0.57661 | 0.57081 | 0.58361 |
| ENSG00000255663 | 11 | 114271404 | 114314654 | 154 | 18 | -0.1934 | 0.57668 | RP11-212D19.4 | 0.57668 | 0.55336 | 0.46903 |
| ENSG00000168530 | 2 | 211154874 | 211179914 | 65 | 7 | -0.19343 | 0.57669 | MYL1 | 0.57669 | 0.48622 | 0.38256 |
| ENSG00000178222 | 4 | 1050038 | 1107350 | 354 | 13 | -0.19344 | 0.57669 | RNF212 | 0.57669 | 0.63886 | 0.71834 |
| ENSG00000163531 | 1 | 204797779 | 204991950 | 611 | 58 | -0.19363 | 0.57677 | NFASC | 0.57677 | 0.50002 | 0.31066 |
| ENSG00000198225 | 6 | 63921351 | 63922929 | 11 | 3 | -0.19364 | 0.57677 | FKBP1C | 0.57677 | 0.67087 | 0.73683 |
| ENSG00000175155 | 17 | 57409050 | 57479090 | 182 | 23 | -0.19377 | 0.57682 | YPEL2 | 0.57682 | 0.64729 | 0.82647 |
| ENSG00000181392 | 19 | 36494209 | 36499695 | 9 | 4 | -0.19377 | 0.57682 | SYNE4 | 0.57682 | 0.73726 | 0.6356 |
| ENSG00000169314 | 22 | 24105208 | 24108048 | 6 | 3 | -0.19382 | 0.57684 | C22orf15 | 0.57684 | 0.89132 | 0.9076 |
| ENSG00000064703 | 1 | 112297867 | 112310638 | 58 | 8 | -0.19387 | 0.57686 | DDX20 | 0.57686 | 0.50948 | 0.34087 |
| ENSG00000077238 | 16 | 27324989 | 27376099 | 193 | 24 | -0.19387 | 0.57686 | IL4R | 0.57686 | 0.43087 | 0.4486 |
| ENSG00000172150 | 17 | 3100813 | 3101742 | 5 | 2 | -0.1939 | 0.57687 | OR1A2 | 0.57687 | 0.75086 | 0.829 |
| ENSG00000176953 | 16 | 28962128 | 28978418 | 30 | 8 | -0.19408 | 0.57694 | NFATC2IP | 0.57694 | 0.49065 | 0.52641 |
| ENSG00000136738 | 10 | 17686124 | 17757913 | 344 | 18 | -0.19411 | 0.57696 | STAM | 0.57696 | 0.63428 | 0.71839 |
| ENSG00000272869 | 5 | 112196885 | 112228776 | 158 | 15 | -0.19414 | 0.57697 | CTC-487M23.8 | 0.57697 | 0.55744 | 0.46437 |
| ENSG00000267120 | 19 | 36230153 | 36236333 | 27 | 9 | -0.1942 | 0.57699 | AD000671.6 | 0.57699 | 0.36371 | 0.52854 |
| ENSG00000269881 | 16 | 299546 | 319942 | 74 | 10 | -0.19438 | 0.57706 | ITFG3 | 0.57706 | 0.39761 | 0.33681 |
| ENSG00000110077 | 11 | 59939081 | 59952139 | 29 | 5 | -0.19444 | 0.57708 | MS4A6A | 0.57708 | 0.54264 | 0.57297 |
| ENSG00000171617 | 5 | 73923234 | 73937249 | 38 | 7 | -0.19462 | 0.57716 | ENC1 | 0.57716 | 0.55269 | 0.80823 |
| ENSG00000142945 | 1 | 45205490 | 45233439 | 75 | 12 | -0.19474 | 0.5772 | KIF2C | 0.5772 | 0.53164 | 0.58697 |
| ENSG00000147324 | 8 | 8640864 | 8751155 | 512 | 38 | -0.19479 | 0.57722 | MFHAS1 | 0.57722 | 0.63078 | 0.55356 |
| ENSG00000124568 | 6 | 25783125 | 25832287 | 162 | 16 | -0.19489 | 0.57726 | SLC17A1 | 0.57726 | 0.46901 | 0.44848 |
| ENSG00000228570 | 10 | 81601114 | 81610632 | 14 | 6 | -0.19488 | 0.57726 | NUTM2E | 0.57726 | 0.42771 | 0.51819 |
| ENSG00000125843 | 20 | 3801178 | 3805949 | 8 | 2 | -0.19506 | 0.57733 | AP5S1 | 0.57733 | 0.72401 | 0.75422 |
| ENSG00000162929 | 2 | 61293006 | 61391960 | 277 | 21 | -0.1951 | 0.57734 | KIAA1841 | 0.57734 | 0.5924 | 0.57882 |
| ENSG00000039560 | 5 | 34656342 | 34832732 | 649 | 69 | -0.19524 | 0.5774 | RAI14 | 0.5774 | 0.61469 | 0.52291 |
| ENSG00000034239 | 8 | 49623348 | 49647870 | 61 | 17 | -0.19535 | 0.57744 | EFCAB1 | 0.57744 | 0.71027 | 0.73152 |
| ENSG00000104356 | 8 | 99129525 | 99172062 | 170 | 22 | -0.19563 | 0.57755 | POP1 | 0.57755 | 0.54807 | 0.53753 |
| ENSG00000185056 | 5 | 173400782 | 173433143 | 167 | 20 | -0.19572 | 0.57758 | C5orf47 | 0.57758 | 0.5187 | 0.54404 |
| ENSG00000156049 | 9 | 80037995 | 80263223 | 760 | 89 | -0.19594 | 0.57767 | GNA14 | 0.57767 | 0.52327 | 0.54599 |
| ENSG00000130755 | 19 | 39818993 | 39833012 | 42 | 9 | -0.19599 | 0.57769 | GMFG | 0.57769 | 0.54233 | 0.52444 |
| ENSG00000114204 | 3 | 167159577 | 167196792 | 129 | 13 | -0.1961 | 0.57773 | SERPINI2 | 0.57773 | 0.37624 | 0.2983 |
| ENSG00000136717 | 2 | 127805603 | 127864931 | 276 | 27 | -0.1964 | 0.57785 | BIN1 | 0.57785 | 0.54583 | 0.52261 |
| ENSG00000107014 | 9 | 5299868 | 5304969 | 14 | 4 | -0.19648 | 0.57788 | RLN2 | 0.57788 | 0.82005 | 0.50924 |
| ENSG00000243449 | 4 | 2043689 | 2045697 | 4 | 2 | -0.19661 | 0.57793 | C4orf48 | 0.57793 | 0.70319 | 0.78825 |
| ENSG00000197410 | 4 | 155153399 | 155412930 | 888 | 52 | -0.19661 | 0.57793 | DCHS2 | 0.57793 | 0.64311 | 0.69314 |
| ENSG00000116863 | 1 | 36554476 | 36559533 | 6 | 4 | -0.19662 | 0.57794 | ADPRHL2 | 0.57794 | 0.73495 | 0.63264 |
| ENSG00000123384 | 12 | 57522276 | 57607134 | 131 | 22 | -0.19684 | 0.57802 | LRP1 | 0.57802 | 0.50151 | 0.4136 |
| ENSG00000118564 | 4 | 15606162 | 15683302 | 274 | 17 | -0.1971 | 0.57813 | FBXL5 | 0.57813 | 0.5744 | 0.60141 |
| ENSG00000147912 | 9 | 37510889 | 37588871 | 242 | 33 | -0.19721 | 0.57817 | FBXO10 | 0.57817 | 0.521 | 0.39312 |
| ENSG00000144339 | 2 | 192813769 | 193060435 | 586 | 63 | -0.19725 | 0.57818 | TMEFF2 | 0.57818 | 0.51296 | 0.52735 |
| ENSG00000059122 | 16 | 2961938 | 3001209 | 261 | 27 | -0.19724 | 0.57818 | FLYWCH1 | 0.57818 | 0.74291 | 0.75023 |
| ENSG00000161999 | 16 | 731671 | 734529 | 4 | 2 | -0.19737 | 0.57823 | JMJD8 | 0.57823 | 0.27971 | 0.35391 |
| ENSG00000110497 | 11 | 46417964 | 46615675 | 235 | 24 | -0.19759 | 0.57832 | AMBRA1 | 0.57832 | 0.57103 | 0.55742 |
| ENSG00000156411 | 14 | 104378625 | 104394606 | 80 | 10 | -0.19777 | 0.57839 | C14orf2 | 0.57839 | 0.61717 | 0.6856 |
| ENSG00000152760 | 1 | 67218142 | 67244470 | 89 | 11 | -0.19814 | 0.57853 | TCTEX1D1 | 0.57853 | 0.53783 | 0.57258 |
| ENSG00000204392 | 6 | 31765173 | 31774761 | 37 | 14 | -0.19812 | 0.57853 | LSM2 | 0.57853 | 0.33398 | 0.43031 |
| ENSG00000175398 | 12 | 56030644 | 56031638 | 3 | 1 | -0.19813 | 0.57853 | OR10P1 | 0.57853 | 0.24 | 0.39199 |
| ENSG00000070269 | 14 | 56955072 | 57117324 | 689 | 56 | -0.19817 | 0.57854 | TMEM260 | 0.57854 | 0.56314 | 0.71745 |
| ENSG00000143919 | 2 | 44589089 | 44999731 | 1172 | 38 | -0.19829 | 0.57859 | CAMKMT | 0.57859 | 0.58532 | 0.59616 |
| ENSG00000145819 | 5 | 142149949 | 142608576 | 1282 | 83 | -0.19841 | 0.57864 | ARHGAP26 | 0.57864 | 0.56248 | 0.3583 |
| ENSG00000128928 | 15 | 40697686 | 40728146 | 101 | 11 | -0.19853 | 0.57869 | IVD | 0.57869 | 0.60062 | 0.66818 |
| ENSG00000151623 | 4 | 148999913 | 149365850 | 1006 | 73 | -0.19864 | 0.57873 | NR3C2 | 0.57873 | 0.50464 | 0.51637 |
| ENSG00000171451 | 18 | 65173819 | 65184217 | 22 | 8 | -0.19876 | 0.57877 | DSEL | 0.57877 | 0.73703 | 0.56424 |
| ENSG00000167103 | 9 | 130683158 | 130693076 | 18 | 5 | -0.19884 | 0.57881 | PIP5KL1 | 0.57881 | 0.55779 | 0.57945 |
| ENSG00000187098 | 3 | 69788586 | 70017488 | 671 | 24 | -0.19926 | 0.57897 | MITF | 0.57897 | 0.58893 | 0.62887 |
| ENSG00000118194 | 1 | 201328136 | 201346890 | 89 | 13 | -0.19942 | 0.57903 | TNNT2 | 0.57903 | 0.64359 | 0.64677 |
| ENSG00000134759 | 18 | 33709407 | 33757909 | 148 | 14 | -0.19945 | 0.57904 | ELP2 | 0.57904 | 0.47105 | 0.49308 |
| ENSG00000243696 | 3 | 52847298 | 52869745 | 57 | 6 | -0.19971 | 0.57914 | RP5-966M1.6 | 0.57914 | 0.48447 | 0.60242 |
| ENSG00000268467 | 11 | 107462471 | 107463949 | 5 | 2 | -0.19974 | 0.57916 | AP000889.3 | 0.57916 | 0.64382 | 0.51751 |
| ENSG00000099822 | 19 | 589893 | 617159 | 218 | 51 | -0.19977 | 0.57917 | HCN2 | 0.57917 | 0.90221 | 0.88903 |
| ENSG00000135926 | 2 | 219138915 | 219157309 | 55 | 4 | -0.20027 | 0.57937 | TMBIM1 | 0.57937 | 0.64829 | 0.58289 |
| ENSG00000105520 | 19 | 11466062 | 11476374 | 16 | 6 | -0.20032 | 0.57938 | DKFZP761J1410 | 0.57938 | 0.62664 | 0.55778 |
| ENSG00000255245 | 11 | 117690878 | 117747382 | 167 | 43 | -0.20034 | 0.57939 | FXYD6-FXYD2 | 0.57939 | 0.7659 | 0.73212 |
| ENSG00000257065 | 6 | 144128229 | 144165754 | 127 | 15 | -0.20057 | 0.57948 | RP3-468K18.5 | 0.57948 | 0.69703 | 0.76534 |
| ENSG00000122735 | 9 | 34457412 | 34520982 | 122 | 19 | -0.20057 | 0.57948 | DNAI1 | 0.57948 | 0.58165 | 0.53221 |
| ENSG00000149054 | 11 | 6947635 | 7005863 | 236 | 22 | -0.20078 | 0.57956 | ZNF215 | 0.57956 | 0.5772 | 0.46177 |
| ENSG00000166268 | 12 | 70219084 | 70352877 | 623 | 44 | -0.20088 | 0.5796 | MYRFL | 0.5796 | 0.52327 | 0.52609 |

| | | | | | | | | | | | |
|---|---|---|---|---|---|---|---|---|---|---|---|
| ENSG00000215568 | 22 | 17442826 | 17489112 | 186 | 15 | -0.2009 | 0.57961 | GAB4 | 0.57961 | 0.481 | 0.4513 |
| ENSG00000108654 | 17 | 62495734 | 62504317 | 18 | 4 | -0.20139 | 0.5798 | DDX5 | 0.5798 | 0.63145 | 0.49963 |
| ENSG00000151778 | 13 | 44947801 | 44971850 | 39 | 9 | -0.20141 | 0.57981 | SERP2 | 0.57981 | 0.47161 | 0.43716 |
| ENSG00000182795 | 1 | 207191866 | 207206101 | 12 | 7 | -0.20158 | 0.57988 | C1orf116 | 0.57988 | 0.67256 | 0.63742 |
| ENSG00000179454 | 14 | 45393522 | 45511525 | 248 | 12 | -0.20184 | 0.57998 | KLHL28 | 0.57998 | 0.50361 | 0.53456 |
| ENSG00000111786 | 12 | 120899471 | 120907596 | 34 | 10 | -0.20188 | 0.57999 | SRSF9 | 0.57999 | 0.6088 | 0.4546 |
| ENSG00000168955 | 2 | 228226872 | 228246711 | 98 | 20 | -0.20189 | 0.58 | TM4SF20 | 0.58 | 0.54112 | 0.48816 |
| ENSG00000143324 | 1 | 180601140 | 180859387 | 554 | 28 | -0.20234 | 0.58018 | XPR1 | 0.58018 | 0.61741 | 0.58133 |
| ENSG00000183560 | 11 | 94038759 | 94040887 | 6 | 1 | -0.20255 | 0.58026 | FOLR4 | 0.58026 | 0.79795 | 0.81934 |
| ENSG00000176884 | 9 | 140032842 | 140063207 | 65 | 13 | -0.20297 | 0.58042 | GRIN1 | 0.58042 | 0.73149 | 0.73553 |
| ENSG00000136425 | 15 | 78396948 | 78423886 | 78 | 15 | -0.20299 | 0.58043 | CIB2 | 0.58043 | 0.61738 | 0.71141 |
| ENSG00000186417 | 15 | 51633826 | 51700210 | 332 | 31 | -0.20304 | 0.58045 | GLDN | 0.58045 | 0.61441 | 0.58588 |
| ENSG00000115970 | 2 | 43393800 | 43823185 | 1281 | 38 | -0.20308 | 0.58046 | THADA | 0.58046 | 0.58389 | 0.56475 |
| ENSG00000205186 | 8 | 82370576 | 82373814 | 10 | 5 | -0.20312 | 0.58048 | FABP9 | 0.58048 | 0.94342 | 0.88372 |
| ENSG00000125457 | 17 | 73262309 | 73267308 | 6 | 2 | -0.20317 | 0.5805 | MIF4GD | 0.5805 | 0.6479 | 0.54926 |
| ENSG00000130203 | 19 | 45409011 | 45412650 | 6 | 3 | -0.2033 | 0.58055 | APOE | 0.58055 | 0.26566 | 0.29514 |
| ENSG00000146112 | 6 | 30644166 | 30655672 | 33 | 8 | -0.20334 | 0.58056 | PPP1R18 | 0.58056 | 0.60692 | 0.65438 |
| ENSG00000121381 | 12 | 10961693 | 10962767 | 1 | 1 | -0.20343 | 0.5806 | TAS2R9 | 0.5806 | 0.67515 | 0.62023 |
| ENSG00000118434 | 6 | 88757507 | 88776550 | 30 | 5 | -0.20349 | 0.58062 | SPACA1 | 0.58062 | 0.70662 | 0.77335 |
| ENSG00000114315 | 3 | 193853934 | 193856521 | 3 | 1 | -0.20372 | 0.58071 | HES1 | 0.58071 | 0.39874 | 0.49711 |
| ENSG00000139618 | 13 | 32889611 | 32973805 | 221 | 25 | -0.20402 | 0.58083 | BRCA2 | 0.58083 | 0.54072 | 0.63025 |
| ENSG00000214265 | 15 | 25200181 | 25245423 | 121 | 14 | -0.20402 | 0.58083 | SNURF | 0.58083 | 0.38882 | 0.37933 |
| ENSG00000105668 | 19 | 36157715 | 36169367 | 50 | 8 | -0.20402 | 0.58083 | UPK1A | 0.58083 | 0.7212 | 0.74688 |
| ENSG00000248235 | 8 | 22446787 | 22459597 | 30 | 5 | -0.20446 | 0.581 | AC037459.4 | 0.581 | 0.7256 | 0.83216 |
| ENSG00000168661 | 19 | 35417807 | 35436074 | 46 | 8 | -0.20462 | 0.58107 | ZNF30 | 0.58107 | 0.54922 | 0.62152 |
| ENSG00000204616 | 6 | 30070674 | 30080883 | 80 | 20 | -0.20493 | 0.58119 | TRIM31 | 0.58119 | 0.56013 | 0.73002 |
| ENSG00000130958 | 9 | 99082988 | 99145992 | 263 | 14 | -0.20497 | 0.5812 | SLC35D2 | 0.5812 | 0.54764 | 0.54938 |
| ENSG00000153707 | 9 | 8314246 | 10612723 | 11171 | 502 | -0.20503 | 0.58123 | PTPRD | 0.58123 | 0.43808 | 0.39874 |
| ENSG00000165659 | 13 | 72012098 | 72441330 | 872 | 87 | -0.20537 | 0.58136 | DACH1 | 0.58136 | 0.57874 | 0.53614 |
| ENSG00000126259 | 19 | 36346763 | 36358011 | 12 | 3 | -0.20546 | 0.58139 | KIRREL2 | 0.58139 | 0.38437 | 0.42229 |
| ENSG00000164749 | 8 | 76320149 | 76479078 | 664 | 38 | -0.20566 | 0.58147 | HNF4G | 0.58147 | 0.63042 | 0.64218 |
| ENSG00000160746 | 3 | 43396351 | 43733086 | 747 | 40 | -0.20575 | 0.58151 | ANO10 | 0.58151 | 0.55599 | 0.55863 |
| ENSG00000113100 | 5 | 26880709 | 27121257 | 721 | 41 | -0.20588 | 0.58156 | CDH9 | 0.58156 | 0.61553 | 0.65411 |
| ENSG00000113615 | 5 | 133984479 | 134063513 | 107 | 29 | -0.20618 | 0.58167 | SEC24A | 0.58167 | 0.53735 | 0.51197 |
| ENSG00000112667 | 6 | 43193367 | 43197222 | 11 | 4 | -0.20625 | 0.5817 | DNPH1 | 0.5817 | 0.57041 | 0.54376 |
| ENSG00000198081 | 18 | 5289018 | 5297052 | 28 | 5 | -0.20628 | 0.58172 | ZBTB14 | 0.58172 | 0.3408 | 0.46218 |
| ENSG00000136695 | 2 | 113816215 | 113822959 | 50 | 3 | -0.20646 | 0.58179 | IL36RN | 0.58179 | 0.71867 | 0.61486 |
| ENSG00000205808 | 9 | 4662298 | 4665256 | 9 | 5 | -0.20669 | 0.58188 | PPAPDC2 | 0.58188 | 0.2164 | 0.31489 |
| ENSG00000153347 | 5 | 94727048 | 94786158 | 220 | 30 | -0.20674 | 0.58189 | FAM81B | 0.58189 | 0.41854 | 0.32982 |
| ENSG00000251258 | 6 | 112668532 | 112672498 | 15 | 3 | -0.20681 | 0.58192 | RFPL4B | 0.58192 | 0.33685 | 0.45716 |
| ENSG00000081154 | 3 | 101292939 | 101313281 | 64 | 11 | -0.20691 | 0.58196 | PCNP | 0.58196 | 0.6676 | 0.62471 |
| ENSG00000180190 | 8 | 439803 | 495781 | 236 | 37 | -0.20696 | 0.58198 | TDRP | 0.58198 | 0.60205 | 0.63917 |
| ENSG00000113845 | 3 | 119217379 | 119243937 | 121 | 18 | -0.20709 | 0.58203 | TIMMDC1 | 0.58203 | 0.63227 | 0.70686 |
| ENSG00000141384 | 18 | 23805900 | 23971649 | 550 | 27 | -0.20714 | 0.58205 | TAF4B | 0.58205 | 0.57969 | 0.5583 |
| ENSG00000157093 | 3 | 42438570 | 42452092 | 31 | 8 | -0.2074 | 0.58215 | LYZL4 | 0.58215 | 0.55721 | 0.51506 |
| ENSG00000115896 | 2 | 198669426 | 199437305 | 1699 | 79 | -0.20752 | 0.5822 | PLCL1 | 0.5822 | 0.58596 | 0.62829 |
| ENSG00000196326 | 10 | 5196335 | 5244449 | 290 | 18 | -0.20771 | 0.58227 | AKR1CL1 | 0.58227 | 0.55533 | 0.53428 |
| ENSG00000185559 | 14 | 101192042 | 101201539 | 29 | 13 | -0.20771 | 0.58227 | DLK1 | 0.58227 | 0.73774 | 0.29267 |
| ENSG00000139725 | 12 | 122215664 | 122241309 | 84 | 12 | -0.20775 | 0.58229 | RHOF | 0.58229 | 0.74175 | 0.82617 |
| ENSG00000081237 | 1 | 198607801 | 198726545 | 378 | 30 | -0.20778 | 0.5823 | PTPRC | 0.5823 | 0.5952 | 0.68466 |
| ENSG00000154734 | 21 | 28208066 | 28217728 | 37 | 7 | -0.20803 | 0.5824 | ADAMTS1 | 0.5824 | 0.74038 | 0.62023 |
| ENSG00000111860 | 6 | 118781935 | 119031238 | 695 | 24 | -0.20809 | 0.58242 | CEP85L | 0.58242 | 0.5802 | 0.62611 |
| ENSG00000175110 | 3 | 138724648 | 139076065 | 1057 | 56 | -0.20814 | 0.58244 | MRPS22 | 0.58244 | 0.5649 | 0.55681 |
| ENSG00000118482 | 6 | 64345725 | 64489229 | 253 | 21 | -0.20835 | 0.58252 | PHF3 | 0.58252 | 0.45509 | 0.38444 |
| ENSG00000167615 | 19 | 54960065 | 54973217 | 67 | 11 | -0.20834 | 0.58252 | LENG8 | 0.58252 | 0.72271 | 0.65817 |
| ENSG00000152601 | 3 | 151961617 | 152183569 | 442 | 29 | -0.20849 | 0.58258 | MBNL1 | 0.58258 | 0.62984 | 0.70322 |
| ENSG00000182050 | 12 | 86372516 | 87232681 | 2274 | 51 | -0.20852 | 0.58259 | MGAT4C | 0.58259 | 0.57147 | 0.56411 |
| ENSG00000077800 | 7 | 72742167 | 72772634 | 75 | 7 | -0.20855 | 0.5826 | FKBP6 | 0.5826 | 0.60996 | 0.51571 |
| ENSG00000138068 | 2 | 37394963 | 37423741 | 263 | 14 | -0.20865 | 0.58264 | SULT6B1 | 0.58264 | 0.58383 | 0.57296 |
| ENSG00000115159 | 2 | 157291802 | 157470247 | 368 | 21 | -0.20869 | 0.58265 | GPD2 | 0.58265 | 0.5671 | 0.46085 |
| ENSG00000144231 | 2 | 128603840 | 128615731 | 46 | 8 | -0.20889 | 0.58273 | POLR2D | 0.58273 | 0.67693 | 0.59537 |
| ENSG00000168917 | 3 | 136537489 | 136574734 | 126 | 6 | -0.20901 | 0.58278 | SLC35G2 | 0.58278 | 0.61869 | 0.67494 |
| ENSG00000177669 | 8 | 29989340 | 30002202 | 47 | 6 | -0.20903 | 0.58279 | MBOAT4 | 0.58279 | 0.64939 | 0.71836 |
| ENSG00000101161 | 20 | 62612488 | 62664453 | 132 | 13 | -0.20928 | 0.58288 | PRPF6 | 0.58288 | 0.57195 | 0.51848 |
| ENSG00000175591 | 11 | 72929343 | 72947397 | 43 | 14 | -0.20943 | 0.58294 | P2RY2 | 0.58294 | 0.76033 | 0.68094 |
| ENSG00000160299 | 21 | 47744036 | 47865682 | 657 | 29 | -0.20949 | 0.58297 | PCNT | 0.58297 | 0.58297 | 0.57913 |
| ENSG00000153086 | 2 | 135596117 | 135659604 | 157 | 11 | -0.20953 | 0.58298 | ACMSD | 0.58298 | 0.59004 | 0.74249 |
| ENSG00000237102 | 16 | 67244251 | 67260930 | 20 | 6 | -0.20956 | 0.58299 | AC040160.1 | 0.58299 | 0.61906 | 0.63625 |
| ENSG00000149781 | 11 | 63974150 | 63991354 | 37 | 10 | -0.20958 | 0.583 | FERMT3 | 0.583 | 0.72482 | 0.57512 |
| ENSG00000133477 | 22 | 40390953 | 40439533 | 80 | 13 | -0.20956 | 0.583 | FAM83F | 0.583 | 0.63645 | 0.58299 |
| ENSG00000178773 | 16 | 89642176 | 89663654 | 127 | 26 | -0.20973 | 0.58306 | CPNE7 | 0.58306 | 0.58781 | 0.74802 |
| ENSG00000137414 | 6 | 17600586 | 17611950 | 26 | 6 | -0.20983 | 0.5831 | FAM8A1 | 0.5831 | 0.59906 | 0.7284 |
| ENSG00000168062 | 11 | 64755415 | 64764517 | 20 | 7 | -0.20983 | 0.5831 | BATF2 | 0.5831 | 0.66425 | 0.75189 |

| | | | | | | | | | | | |
|---|---|---|---|---|---|---|---|---|---|---|---|
| ENSG00000186469 | 14 | 52292913 | 52446060 | 734 | 75 | -0.20996 | 0.58315 | GNG2 | 0.58315 | 0.55344 | 0.57578 |
| ENSG00000105784 | 7 | 87256864 | 87461611 | 373 | 18 | -0.2102 | 0.58324 | RUNDC3B | 0.58324 | 0.60726 | 0.62497 |
| ENSG00000134815 | 19 | 47852538 | 47885961 | 131 | 17 | -0.21028 | 0.58328 | DHX34 | 0.58328 | 0.65206 | 0.74185 |
| ENSG00000141579 | 17 | 80787311 | 80798454 | 39 | 7 | -0.21034 | 0.5833 | ZNF750 | 0.5833 | 0.65426 | 0.63875 |
| ENSG00000011105 | 12 | 3186521 | 3395730 | 1135 | 99 | -0.21038 | 0.58331 | TSPAN9 | 0.58331 | 0.50342 | 0.49188 |
| ENSG00000204246 | 9 | 107298030 | 107299137 | 3 | 1 | -0.21053 | 0.58337 | OR13C3 | 0.58337 | 0.59513 | 0.70483 |
| ENSG00000161849 | 12 | 52771596 | 52779436 | 62 | 4 | -0.2106 | 0.5834 | KRT84 | 0.5834 | 0.72629 | 0.76341 |
| ENSG00000173110 | 1 | 161494036 | 161496681 | 11 | 5 | -0.21079 | 0.58348 | HSPA6 | 0.58348 | 0.44153 | 0.71474 |
| ENSG00000050555 | 9 | 133884469 | 133969860 | 388 | 45 | -0.21132 | 0.58368 | LAMC3 | 0.58368 | 0.58171 | 0.59377 |
| ENSG00000167311 | 11 | 3659733 | 3663546 | 25 | 7 | -0.21153 | 0.58376 | ART5 | 0.58376 | 0.95996 | 0.96698 |
| ENSG00000187824 | 17 | 10602332 | 10633633 | 135 | 3 | -0.21167 | 0.58382 | TMEM220 | 0.58382 | 0.60146 | 0.65092 |
| ENSG00000164778 | 7 | 155250824 | 155257526 | 36 | 10 | -0.21174 | 0.58384 | EN2 | 0.58384 | 0.78152 | 0.77338 |
| ENSG00000119392 | 9 | 131266979 | 131304567 | 65 | 9 | -0.21177 | 0.58386 | GLE1 | 0.58386 | 0.58163 | 0.50068 |
| ENSG00000164591 | 5 | 150040436 | 150058927 | 55 | 3 | -0.21228 | 0.58405 | MYOZ3 | 0.58405 | 0.52136 | 0.54307 |
| ENSG00000154654 | 21 | 22370633 | 22915650 | 2122 | 103 | -0.21229 | 0.58406 | NCAM2 | 0.58406 | 0.60695 | 0.60351 |
| ENSG00000262209 | 5 | 140749831 | 140892546 | 446 | 36 | -0.21233 | 0.58407 | PCDHGB3 | 0.58407 | 0.59754 | 0.50737 |
| ENSG00000113492 | 5 | 34998206 | 35048198 | 218 | 24 | -0.21258 | 0.58417 | AGXT2 | 0.58417 | 0.62689 | 0.6547 |
| ENSG00000196724 | 19 | 58433252 | 58446761 | 38 | 8 | -0.21262 | 0.58419 | ZNF418 | 0.58419 | 0.5236 | 0.55539 |
| ENSG00000187764 | 9 | 91975702 | 92113045 | 606 | 49 | -0.21273 | 0.58423 | SEMA4D | 0.58423 | 0.56028 | 0.66559 |
| ENSG00000143368 | 1 | 149895209 | 149900236 | 13 | 2 | -0.21295 | 0.58432 | SF3B4 | 0.58432 | 0.74734 | 0.69313 |
| ENSG00000106392 | 7 | 7196565 | 7288282 | 431 | 26 | -0.21298 | 0.58433 | C1GALT1 | 0.58433 | 0.64588 | 0.62654 |
| ENSG00000171877 | 15 | 44162962 | 44487450 | 544 | 21 | -0.2132 | 0.58441 | FRMD5 | 0.58441 | 0.57742 | 0.58256 |
| ENSG00000168334 | 3 | 39224701 | 39234087 | 17 | 6 | -0.21322 | 0.58442 | XIRP1 | 0.58442 | 0.59826 | 0.43491 |
| ENSG00000131059 | 20 | 31805116 | 31815564 | 12 | 6 | -0.21329 | 0.58445 | BPIFA3 | 0.58445 | 0.51179 | 0.53898 |
| ENSG00000134744 | 1 | 52873954 | 53019159 | 299 | 20 | -0.21336 | 0.58448 | ZCCHC11 | 0.58448 | 0.59089 | 0.58306 |
| ENSG00000171136 | 19 | 14138960 | 14141854 | 10 | 3 | -0.21338 | 0.58448 | RLN3 | 0.58448 | 0.72205 | 0.78273 |
| ENSG00000197782 | 19 | 40575059 | 40596845 | 77 | 7 | -0.21337 | 0.58448 | ZNF780A | 0.58448 | 0.61989 | 0.6571 |
| ENSG00000146151 | 6 | 55299167 | 55444012 | 749 | 29 | -0.21358 | 0.58456 | HMGCLL1 | 0.58456 | 0.61696 | 0.65665 |
| ENSG00000033800 | 15 | 68346517 | 68483096 | 215 | 24 | -0.21365 | 0.58459 | PIAS1 | 0.58459 | 0.62726 | 0.67173 |
| ENSG00000178234 | 7 | 151722759 | 151819425 | 159 | 9 | -0.2137 | 0.58461 | GALNT11 | 0.58461 | 0.63109 | 0.73998 |
| ENSG00000127533 | 19 | 16999671 | 17003417 | 9 | 3 | -0.2137 | 0.58461 | F2RL3 | 0.58461 | 0.62649 | 0.72156 |
| ENSG00000163348 | 1 | 154929502 | 154936329 | 7 | 4 | -0.21372 | 0.58462 | PYGO2 | 0.58462 | 0.77348 | 0.90452 |
| ENSG00000177731 | 17 | 18148150 | 18162230 | 24 | 7 | -0.21372 | 0.58462 | FLII | 0.58462 | 0.60681 | 0.42069 |
| ENSG00000049283 | 17 | 48609904 | 48621111 | 41 | 12 | -0.21389 | 0.58468 | EPN3 | 0.58468 | 0.83708 | 0.89849 |
| ENSG00000169992 | 17 | 7308193 | 7323179 | 41 | 17 | -0.214 | 0.58473 | NLGN2 | 0.58473 | 0.46506 | 0.63714 |
| ENSG00000096717 | 10 | 69644427 | 69678147 | 115 | 14 | -0.21412 | 0.58478 | SIRT1 | 0.58478 | 0.60257 | 0.64151 |
| ENSG00000232237 | 1 | 201083081 | 201096312 | 41 | 9 | -0.21417 | 0.58479 | ASCL5 | 0.58479 | 0.42367 | 0.36015 |
| ENSG00000104689 | 8 | 23047965 | 23082639 | 93 | 18 | -0.21416 | 0.58479 | TNFRSF10A | 0.58479 | 0.55746 | 0.34337 |
| ENSG00000138622 | 15 | 73612200 | 73661605 | 155 | 29 | -0.21467 | 0.58499 | HCN4 | 0.58499 | 0.65492 | 0.67993 |
| ENSG00000185128 | 17 | 36283971 | 36294915 | 1 | 1 | -0.2147 | 0.585 | TBC1D3F | 0.585 | 0.97735 | 0.99063 |
| ENSG00000166797 | 15 | 64364758 | 64386217 | 69 | 9 | -0.21485 | 0.58506 | FAM96A | 0.58506 | 0.67339 | 0.6908 |
| ENSG00000186193 | 9 | 139956581 | 139965040 | 31 | 6 | -0.21489 | 0.58507 | SAPCD2 | 0.58507 | 0.63242 | 0.72293 |
| ENSG00000138363 | 2 | 216176540 | 216214487 | 124 | 13 | -0.21494 | 0.58509 | ATIC | 0.58509 | 0.59225 | 0.59292 |
| ENSG00000120658 | 13 | 43787654 | 44361044 | 1705 | 82 | -0.21493 | 0.58509 | ENOX1 | 0.58509 | 0.60379 | 0.66939 |
| ENSG00000116761 | 1 | 70876901 | 70905534 | 83 | 19 | -0.21499 | 0.58511 | CTH | 0.58511 | 0.6689 | 0.70797 |
| ENSG00000080608 | 9 | 2720469 | 2844241 | 422 | 48 | -0.21505 | 0.58514 | KIAA0020 | 0.58514 | 0.63285 | 0.70297 |
| ENSG00000248801 | 8 | 69215703 | 69243726 | 59 | 11 | -0.21515 | 0.58517 | RP11-664D7.4 | 0.58517 | 0.73923 | 0.82921 |
| ENSG00000253731 | 5 | 140753651 | 140892546 | 438 | 36 | -0.21525 | 0.58521 | PCDHGA6 | 0.58521 | 0.59529 | 0.51937 |
| ENSG00000153406 | 16 | 4511681 | 4545764 | 145 | 10 | -0.21535 | 0.58525 | NMRAL1 | 0.58525 | 0.66774 | 0.69375 |
| ENSG00000169474 | 1 | 152956557 | 152958289 | 7 | 1 | -0.21537 | 0.58526 | SPRR1A | 0.58526 | 0.5807 | 0.61465 |
| ENSG00000168066 | 11 | 64532078 | 64546258 | 20 | 5 | -0.2154 | 0.58527 | SF1 | 0.58527 | 0.69325 | 0.6446 |
| ENSG00000100341 | 22 | 44275558 | 44287893 | 69 | 11 | -0.21538 | 0.58527 | PNPLA5 | 0.58527 | 0.55722 | 0.52943 |
| ENSG00000081853 | 5 | 140718539 | 140892546 | 515 | 31 | -0.21553 | 0.58532 | PCDHGA2 | 0.58532 | 0.58272 | 0.51779 |
| ENSG00000163399 | 1 | 116915290 | 116952883 | 76 | 7 | -0.21564 | 0.58536 | ATP1A1 | 0.58536 | 0.63262 | 0.65009 |
| ENSG00000186976 | 22 | 43924624 | 44208217 | 879 | 38 | -0.21571 | 0.58539 | EFCAB6 | 0.58539 | 0.56624 | 0.5909 |
| ENSG00000258724 | 8 | 10582909 | 10697357 | 473 | 25 | -0.21575 | 0.58541 | SOX7 | 0.58541 | 0.47957 | 0.20554 |
| ENSG00000117419 | 1 | 44686742 | 44820932 | 231 | 25 | -0.21583 | 0.58544 | ERI3 | 0.58544 | 0.5536 | 0.58287 |
| ENSG00000136697 | 2 | 113825547 | 113833427 | 47 | 9 | -0.21596 | 0.58549 | IL1F10 | 0.58549 | 0.49491 | 0.6064 |
| ENSG00000182601 | 16 | 25703347 | 26149009 | 1836 | 161 | -0.21596 | 0.58549 | HS3ST4 | 0.58549 | 0.65736 | 0.742 |
| ENSG00000228567 | 19 | 53769929 | 53770972 | 1 | 1 | -0.21598 | 0.5855 | VN1R4 | 0.5855 | 0.18498 | 0.23389 |
| ENSG00000144962 | 3 | 172607148 | 172859058 | 776 | 58 | -0.21613 | 0.58555 | SPATA16 | 0.58555 | 0.63117 | 0.67532 |
| ENSG00000175445 | 8 | 19759228 | 19824769 | 328 | 24 | -0.21613 | 0.58556 | LPL | 0.58556 | 0.65541 | 0.75514 |
| ENSG00000137642 | 11 | 121322912 | 121504402 | 432 | 60 | -0.2162 | 0.58558 | SORL1 | 0.58558 | 0.4943 | 0.4163 |
| ENSG00000144130 | 2 | 113479063 | 113500456 | 56 | 7 | -0.21659 | 0.58573 | NT5DC4 | 0.58573 | 0.85057 | 0.95008 |
| ENSG00000126218 | 13 | 113777128 | 113803843 | 64 | 23 | -0.21663 | 0.58575 | F10 | 0.58575 | 0.79083 | 0.868 |
| ENSG00000171161 | 1 | 249132409 | 249143716 | 16 | 5 | -0.2167 | 0.58578 | ZNF672 | 0.58578 | 0.83979 | 0.83811 |
| ENSG00000184574 | 12 | 6728001 | 6745613 | 40 | 14 | -0.21676 | 0.5858 | LPAR5 | 0.5858 | 0.56943 | 0.6267 |
| ENSG00000089169 | 12 | 113008184 | 113336686 | 712 | 68 | -0.2169 | 0.58586 | RPH3A | 0.58586 | 0.56046 | 0.23298 |
| ENSG00000242114 | 22 | 30821518 | 30825045 | 9 | 4 | -0.21707 | 0.58592 | MTFP1 | 0.58592 | 0.73646 | 0.92282 |
| ENSG00000112679 | 6 | 291630 | 351355 | 74 | 18 | -0.21718 | 0.58597 | DUSP22 | 0.58597 | 0.7445 | 0.75041 |
| ENSG00000124882 | 4 | 75230860 | 75254468 | 86 | 8 | -0.21725 | 0.58599 | EREG | 0.58599 | 0.7273 | 0.7556 |
| ENSG00000213741 | 14 | 50043390 | 50065408 | 65 | 8 | -0.2175 | 0.58609 | RPS29 | 0.58609 | 0.61526 | 0.66953 |
| ENSG00000249931 | 15 | 32684983 | 32695396 | 2 | 1 | -0.21755 | 0.58611 | GOLGA8K | 0.58611 | 0.54133 | 0.073142 |

| | | | | | | | | | | | |
|---|---|---|---|---|---|---|---|---|---|---|---|
| ENSG00000104827 | 19 | 49526126 | 49527590 | 14 | 2 | -0.21778 | 0.5862 | CGB | 0.5862 | 0.37964 | 0.23306 |
| ENSG00000112494 | 6 | 167684657 | 167729507 | 301 | 31 | -0.21783 | 0.58622 | UNC93A | 0.58622 | 0.60868 | 0.56489 |
| ENSG00000185662 | 5 | 171212876 | 171221602 | 39 | 9 | -0.21794 | 0.58626 | SMIM23 | 0.58626 | 0.42689 | 0.43335 |
| ENSG00000175287 | 9 | 131683174 | 131704320 | 55 | 4 | -0.21797 | 0.58627 | PHYHD1 | 0.58627 | 0.60154 | 0.61585 |
| ENSG00000135423 | 12 | 56864736 | 56882198 | 27 | 6 | -0.21796 | 0.58627 | GLS2 | 0.58627 | 0.70696 | 0.68615 |
| ENSG00000215271 | 14 | 23741666 | 23768656 | 56 | 14 | -0.21809 | 0.58632 | HOMEZ | 0.58632 | 0.61563 | 0.65133 |
| ENSG00000139445 | 12 | 109715784 | 109747025 | 67 | 11 | -0.21817 | 0.58635 | FOXN4 | 0.58635 | 0.64915 | 0.71649 |
| ENSG00000052723 | 1 | 115312100 | 115323306 | 11 | 5 | -0.21824 | 0.58638 | SIKE1 | 0.58638 | 0.29028 | 0.38716 |
| ENSG00000270617 | 7 | 43906157 | 43946188 | 122 | 18 | -0.21831 | 0.58641 | URGCP-MRPS24 | 0.58641 | 0.60503 | 0.56982 |
| ENSG00000268461 | 19 | 20047769 | 20048639 | 3 | 2 | -0.21832 | 0.58641 | AC007204.1 | 0.58641 | 0.82079 | 0.5526 |
| ENSG00000146005 | 5 | 139175406 | 139224051 | 75 | 9 | -0.21851 | 0.58649 | PSD2 | 0.58649 | 0.51752 | 0.22745 |
| ENSG00000254221 | 5 | 140729828 | 140892546 | 492 | 38 | -0.21863 | 0.58653 | PCDHGB1 | 0.58653 | 0.59754 | 0.50543 |
| ENSG00000001461 | 1 | 24742284 | 24799466 | 178 | 17 | -0.21864 | 0.58654 | NIPAL3 | 0.58654 | 0.67971 | 0.67825 |
| ENSG00000235162 | 12 | 105629068 | 105789875 | 645 | 39 | -0.21868 | 0.58655 | C12orf75 | 0.58655 | 0.59509 | 0.65073 |
| ENSG00000168928 | 16 | 75237994 | 75241083 | 10 | 5 | -0.21892 | 0.58664 | CTRB2 | 0.58664 | 0.72689 | 0.58735 |
| ENSG00000111325 | 12 | 123459127 | 123464590 | 7 | 2 | -0.21893 | 0.58665 | OGFOD2 | 0.58665 | 0.71853 | 0.6145 |
| ENSG00000187175 | 21 | 46101491 | 46102078 | 5 | 2 | -0.21935 | 0.58681 | KRTAP12-1 | 0.58681 | 0.80459 | 0.56959 |
| ENSG00000204178 | 1 | 25757388 | 25826700 | 125 | 5 | -0.21952 | 0.58688 | TMEM57 | 0.58688 | 0.55441 | 0.51333 |
| ENSG00000044115 | 5 | 137946656 | 138270723 | 666 | 31 | -0.21954 | 0.58689 | CTNNA1 | 0.58689 | 0.58875 | 0.5988 |
| ENSG00000156509 | 8 | 101145588 | 101158028 | 31 | 5 | -0.21958 | 0.5869 | FBXO43 | 0.5869 | 0.53772 | 0.61123 |
| ENSG00000106852 | 9 | 124964856 | 124991905 | 33 | 12 | -0.21961 | 0.58691 | LHX6 | 0.58691 | 0.66924 | 0.60386 |
| ENSG00000153767 | 3 | 120461484 | 120501916 | 88 | 17 | -0.21983 | 0.587 | GTF2E1 | 0.587 | 0.46642 | 0.4231 |
| ENSG00000253910 | 5 | 140739703 | 140892546 | 467 | 35 | -0.22005 | 0.58708 | PCDHGB2 | 0.58708 | 0.58319 | 0.52088 |
| ENSG00000059377 | 7 | 139476850 | 139720125 | 637 | 91 | -0.22013 | 0.58711 | TBXAS1 | 0.58711 | 0.48627 | 0.47557 |
| ENSG00000135144 | 12 | 113494514 | 113535833 | 195 | 37 | -0.22011 | 0.58711 | DTX1 | 0.58711 | 0.62278 | 0.53852 |
| ENSG00000144724 | 3 | 61547243 | 62283288 | 2669 | 182 | -0.22058 | 0.58729 | PTPRG | 0.58729 | 0.643 | 0.65178 |
| ENSG00000130957 | 9 | 97321002 | 97356075 | 223 | 20 | -0.22061 | 0.5873 | FBP2 | 0.5873 | 0.64907 | 0.60059 |
| ENSG00000167685 | 19 | 56643968 | 56672262 | 111 | 17 | -0.22066 | 0.58732 | ZNF444 | 0.58732 | 0.7583 | 0.79766 |
| ENSG00000197893 | 10 | 115348475 | 115423886 | 318 | 28 | -0.22079 | 0.58737 | NRAP | 0.58737 | 0.61138 | 0.74478 |
| ENSG00000033170 | 14 | 65877310 | 66210839 | 956 | 33 | -0.22082 | 0.58738 | FUT8 | 0.58738 | 0.58982 | 0.56183 |
| ENSG00000178645 | 10 | 50887697 | 50918307 | 120 | 12 | -0.22095 | 0.58744 | C10orf53 | 0.58744 | 0.69708 | 0.75025 |
| ENSG00000172350 | 11 | 119019722 | 119033360 | 26 | 5 | -0.22099 | 0.58745 | ABCG4 | 0.58745 | 0.59682 | 0.43103 |
| ENSG00000115661 | 2 | 220110177 | 220115059 | 7 | 3 | -0.22116 | 0.58751 | STK16 | 0.58751 | 0.545 | 0.58279 |
| ENSG00000160908 | 7 | 99084142 | 99097947 | 26 | 8 | -0.22122 | 0.58754 | ZNF394 | 0.58754 | 0.69948 | 0.80197 |
| ENSG00000261147 | 15 | 90792788 | 90815440 | 82 | 7 | -0.22133 | 0.58758 | RP11-697E2.6 | 0.58758 | 0.54576 | 0.4501 |
| ENSG00000183150 | 12 | 12813825 | 12849141 | 98 | 19 | -0.22137 | 0.5876 | GPR19 | 0.5876 | 0.49836 | 0.45626 |
| ENSG00000070366 | 17 | 1963133 | 2207065 | 786 | 26 | -0.22144 | 0.58762 | SMG6 | 0.58762 | 0.60223 | 0.61677 |
| ENSG00000189164 | 19 | 37861963 | 37883968 | 34 | 7 | -0.22151 | 0.58765 | ZNF527 | 0.58765 | 0.41573 | 0.44973 |
| ENSG00000177663 | 22 | 17565844 | 17596583 | 119 | 32 | -0.22163 | 0.5877 | IL17RA | 0.5877 | 0.51172 | 0.31498 |
| ENSG00000249961 | 16 | 66788879 | 66835523 | 106 | 8 | -0.22182 | 0.58777 | CCDC79 | 0.58777 | 0.62653 | 0.66568 |
| ENSG00000132182 | 3 | 13357737 | 13461809 | 358 | 17 | -0.22206 | 0.58787 | NUP210 | 0.58787 | 0.64132 | 0.73164 |
| ENSG00000066468 | 10 | 123237848 | 123357972 | 248 | 47 | -0.22211 | 0.58789 | FGFR2 | 0.58789 | 0.3845 | 0.45186 |
| ENSG00000167768 | 12 | 53068520 | 53074191 | 16 | 6 | -0.22239 | 0.58799 | KRT1 | 0.58799 | 0.72957 | 0.76971 |
| ENSG00000027075 | 14 | 61654277 | 62017694 | 1246 | 103 | -0.22237 | 0.58799 | PRKCH | 0.58799 | 0.59217 | 0.56305 |
| ENSG00000198721 | 6 | 4115923 | 4135831 | 71 | 9 | -0.223 | 0.58823 | ECI2 | 0.58823 | 0.68323 | 0.72209 |
| ENSG00000167880 | 17 | 74000583 | 74023533 | 75 | 15 | -0.22324 | 0.58833 | EVPL | 0.58833 | 0.45941 | 0.46203 |
| ENSG00000088727 | 3 | 47269516 | 47324941 | 93 | 12 | -0.22329 | 0.58835 | KIF9 | 0.58835 | 0.59943 | 0.55516 |
| ENSG00000231256 | 17 | 41857803 | 41864980 | 23 | 2 | -0.22329 | 0.58835 | C17orf105 | 0.58835 | 0.41922 | 0.28293 |
| ENSG00000140836 | 16 | 72816784 | 73093597 | 857 | 108 | -0.22344 | 0.5884 | ZFHX3 | 0.5884 | 0.61188 | 0.57945 |
| ENSG00000188282 | 2 | 218899683 | 218955304 | 167 | 17 | -0.22356 | 0.58845 | RUFY4 | 0.58845 | 0.52147 | 0.53521 |
| ENSG00000215372 | 8 | 7213039 | 7243080 | 5 | 3 | -0.22357 | 0.58845 | ZNF705G | 0.58845 | 0.36446 | 0.060195 |
| ENSG00000198951 | 22 | 42454358 | 42466846 | 27 | 6 | -0.22369 | 0.5885 | NAGA | 0.5885 | 0.76074 | 0.80076 |
| ENSG00000158955 | 17 | 44910567 | 44964096 | 142 | 18 | -0.22409 | 0.58866 | WNT9B | 0.58866 | 0.69749 | 0.7291 |
| ENSG00000058453 | 1 | 17066768 | 17299474 | 426 | 30 | -0.22415 | 0.58868 | CROCC | 0.58868 | 0.61301 | 0.62048 |
| ENSG00000204956 | 5 | 140710252 | 140892546 | 530 | 34 | -0.22442 | 0.58878 | PCDHGA1 | 0.58878 | 0.57461 | 0.50145 |
| ENSG00000115657 | 2 | 220074490 | 220083712 | 11 | 5 | -0.22456 | 0.58884 | ABCB6 | 0.58884 | 0.55407 | 0.31579 |
| ENSG00000013293 | 3 | 170177372 | 170303863 | 342 | 38 | -0.22467 | 0.58888 | SLC7A14 | 0.58888 | 0.69101 | 0.75338 |
| ENSG00000118985 | 5 | 95220802 | 95297775 | 289 | 11 | -0.22492 | 0.58898 | ELL2 | 0.58898 | 0.58934 | 0.54814 |
| ENSG00000139651 | 12 | 53574484 | 53584596 | 31 | 7 | -0.22519 | 0.58908 | ZNF740 | 0.58908 | 0.51414 | 0.46594 |
| ENSG00000211448 | 14 | 80663873 | 80854100 | 411 | 39 | -0.2256 | 0.58924 | DIO2 | 0.58924 | 0.68681 | 0.75893 |
| ENSG00000064300 | 17 | 47572655 | 47592379 | 66 | 22 | -0.22568 | 0.58928 | NGFR | 0.58928 | 0.74996 | 0.57724 |
| ENSG00000140632 | 16 | 4853204 | 4897343 | 162 | 27 | -0.22574 | 0.5893 | GLYR1 | 0.5893 | 0.56369 | 0.46414 |
| ENSG00000163516 | 2 | 220094479 | 220101391 | 20 | 6 | -0.22623 | 0.58949 | ANKZF1 | 0.58949 | 0.53984 | 0.52923 |
| ENSG00000228049 | 7 | 102277474 | 102312076 | 6 | 3 | -0.22654 | 0.58961 | POLR2J2 | 0.58961 | 0.46903 | 0.79127 |
| ENSG00000267645 | 7 | 102277496 | 102312088 | 6 | 3 | -0.22654 | 0.58961 | POLR2J2 | 0.58961 | 0.46903 | 0.79127 |
| ENSG00000157654 | 9 | 112542589 | 112934792 | 1459 | 131 | -0.22654 | 0.58961 | PALM2-AKAP2 | 0.58961 | 0.48877 | 0.5119 |
| ENSG00000170162 | 6 | 117586721 | 117594728 | 23 | 4 | -0.22661 | 0.58963 | VGLL2 | 0.58963 | 0.49706 | 0.28241 |
| ENSG00000155760 | 2 | 202899310 | 202903160 | 8 | 4 | -0.22664 | 0.58965 | FZD7 | 0.58965 | 0.58228 | 0.65705 |
| ENSG00000102837 | 13 | 53602894 | 53626192 | 78 | 19 | -0.22672 | 0.58968 | OLFM4 | 0.58968 | 0.80246 | 0.83123 |
| ENSG00000162727 | 1 | 248308450 | 248309388 | 6 | 2 | -0.22677 | 0.5897 | OR2M5 | 0.5897 | 0.23996 | 0.21376 |
| ENSG00000133958 | 14 | 93799565 | 94174222 | 963 | 29 | -0.227 | 0.58979 | UNC79 | 0.58979 | 0.60445 | 0.6083 |
| ENSG00000142102 | 11 | 289135 | 296107 | 26 | 8 | -0.22738 | 0.58994 | ATHL1 | 0.58994 | 0.67371 | 0.51268 |
| ENSG00000163322 | 4 | 84382092 | 84444501 | 221 | 12 | -0.22745 | 0.58996 | FAM175A | 0.58996 | 0.56505 | 0.5645 |

| | | | | | | | | | | | |
|---|---|---|---|---|---|---|---|---|---|---|---|
| ENSG00000172765 | 3 | 129366635 | 129612419 | 238 | 23 | -0.2276 | 0.59002 | TMCC1 | 0.59002 | 0.5829 | 0.56239 |
| ENSG00000153037 | 5 | 112196919 | 112205485 | 45 | 9 | -0.22761 | 0.59002 | SRP19 | 0.59002 | 0.52534 | 0.52367 |
| ENSG00000088881 | 20 | 2673524 | 2740753 | 198 | 28 | -0.22769 | 0.59006 | EBF4 | 0.59006 | 0.56336 | 0.43549 |
| ENSG00000139714 | 12 | 122089024 | 122110537 | 58 | 12 | -0.22781 | 0.5901 | MORN3 | 0.5901 | 0.41406 | 0.40966 |
| ENSG00000205328 | 12 | 55794214 | 55795289 | 2 | 1 | -0.22786 | 0.59012 | OR6C65 | 0.59012 | 0.78965 | 0.82257 |
| ENSG00000189266 | 1 | 24285599 | 24289952 | 9 | 2 | -0.22819 | 0.59025 | PNRC2 | 0.59025 | 0.72476 | 0.70815 |
| ENSG00000273171 | 17 | 26691310 | 26695033 | 13 | 4 | -0.22824 | 0.59027 | CTB-96E2.2 | 0.59027 | 0.62334 | 0.49882 |
| ENSG00000203965 | 1 | 63989043 | 64038364 | 94 | 11 | -0.2283 | 0.59029 | EFCAB7 | 0.59029 | 0.64706 | 0.6152 |
| ENSG00000136942 | 9 | 127620159 | 127624260 | 8 | 3 | -0.22836 | 0.59032 | RPL35 | 0.59032 | 0.40286 | 0.62416 |
| ENSG00000076604 | 17 | 27071002 | 27077974 | 11 | 3 | -0.22841 | 0.59033 | TRAF4 | 0.59033 | 0.32415 | 0.28295 |
| ENSG00000142207 | 21 | 33683329 | 33765335 | 158 | 13 | -0.22886 | 0.59051 | URB1 | 0.59051 | 0.56346 | 0.34555 |
| ENSG00000214872 | 11 | 57308979 | 57318108 | 26 | 5 | -0.22902 | 0.59057 | SMTNL1 | 0.59057 | 0.63252 | 0.64427 |
| ENSG00000176887 | 2 | 5832799 | 5841516 | 25 | 8 | -0.22904 | 0.59058 | SOX11 | 0.59058 | 0.63962 | 0.33111 |
| ENSG00000178531 | 19 | 7989372 | 7991051 | 7 | 4 | -0.22905 | 0.59058 | CTXN1 | 0.59058 | 0.95296 | 0.96027 |
| ENSG00000006071 | 11 | 17414432 | 17498449 | 325 | 34 | -0.22913 | 0.59062 | ABCC8 | 0.59062 | 0.57278 | 0.63694 |
| ENSG00000267881 | 19 | 42212531 | 42265190 | 134 | 15 | -0.22915 | 0.59062 | CEA | 0.59062 | 0.58038 | 0.69841 |
| ENSG00000126934 | 19 | 4090319 | 4124126 | 96 | 20 | -0.22919 | 0.59064 | MAP2K2 | 0.59064 | 0.78957 | 0.54967 |
| ENSG00000145824 | 5 | 134906373 | 134914969 | 14 | 5 | -0.2293 | 0.59068 | CXCL14 | 0.59068 | 0.38168 | 0.40953 |
| ENSG00000124209 | 20 | 56884752 | 56942563 | 210 | 23 | -0.2293 | 0.59068 | RAB22A | 0.59068 | 0.51533 | 0.20323 |
| ENSG00000103489 | 16 | 17195626 | 17564738 | 1369 | 113 | -0.22931 | 0.59069 | XYLT1 | 0.59069 | 0.60534 | 0.61407 |
| ENSG00000113407 | 5 | 33440802 | 33469644 | 93 | 14 | -0.22934 | 0.5907 | TARS | 0.5907 | 0.69583 | 0.75314 |
| ENSG00000165410 | 14 | 35179593 | 35184029 | 13 | 4 | -0.22943 | 0.59073 | CFL2 | 0.59073 | 0.47773 | 0.70335 |
| ENSG00000164751 | 8 | 77892494 | 77913280 | 54 | 11 | -0.22951 | 0.59076 | PEX2 | 0.59076 | 0.60303 | 0.58386 |
| ENSG00000029363 | 6 | 136578001 | 136610989 | 33 | 11 | -0.22953 | 0.59077 | BCLAF1 | 0.59077 | 0.57386 | 0.69507 |
| ENSG00000069275 | 1 | 205681947 | 205719404 | 80 | 19 | -0.2297 | 0.59084 | NUCKS1 | 0.59084 | 0.58668 | 0.67507 |
| ENSG00000100122 | 22 | 26995242 | 27014052 | 69 | 11 | -0.22982 | 0.59088 | CRYBB1 | 0.59088 | 0.55389 | 0.51921 |
| ENSG00000011405 | 11 | 17099277 | 17229530 | 232 | 30 | -0.22986 | 0.5909 | PIK3C2A | 0.5909 | 0.50354 | 0.50625 |
| ENSG00000187257 | 7 | 77325760 | 77412339 | 278 | 16 | -0.22991 | 0.59092 | RSBN1L | 0.59092 | 0.58479 | 0.57569 |
| ENSG00000141161 | 17 | 33474836 | 33516364 | 134 | 20 | -0.23002 | 0.59096 | UNC45B | 0.59096 | 0.58735 | 0.68152 |
| ENSG00000145555 | 5 | 16665395 | 16936372 | 1148 | 124 | -0.23006 | 0.59098 | MYO10 | 0.59098 | 0.58646 | 0.74489 |
| ENSG00000227345 | 10 | 51026325 | 51130715 | 172 | 13 | -0.23038 | 0.5911 | PARG | 0.5911 | 0.62056 | 0.69226 |
| ENSG00000187840 | 8 | 37887859 | 37917883 | 39 | 8 | -0.2304 | 0.59111 | EIF4EBP1 | 0.59111 | 0.42916 | 0.48818 |
| ENSG00000183186 | 19 | 405438 | 409139 | 21 | 8 | -0.23053 | 0.59116 | C2CD4C | 0.59116 | 0.86374 | 0.92839 |
| ENSG00000135525 | 6 | 136663875 | 136871957 | 376 | 16 | -0.23058 | 0.59118 | MAP7 | 0.59118 | 0.67009 | 0.62788 |
| ENSG00000196387 | 12 | 133656424 | 133684130 | 24 | 9 | -0.23077 | 0.59125 | ZNF140 | 0.59125 | 0.78124 | 0.83891 |
| ENSG00000196531 | 12 | 57106212 | 57125412 | 37 | 7 | -0.2308 | 0.59126 | NACA | 0.59126 | 0.68124 | 0.66112 |
| ENSG00000111725 | 12 | 120105558 | 120119435 | 23 | 4 | -0.23089 | 0.5913 | PRKAB1 | 0.5913 | 0.78803 | 0.82501 |
| ENSG00000136859 | 9 | 129849611 | 129885162 | 60 | 11 | -0.23108 | 0.59137 | ANGPTL2 | 0.59137 | 0.60318 | 0.65932 |
| ENSG00000019549 | 8 | 49830249 | 49834299 | 3 | 2 | -0.23116 | 0.59141 | SNAI2 | 0.59141 | 0.89015 | 0.78692 |
| ENSG00000125450 | 17 | 73201754 | 73231853 | 65 | 9 | -0.23125 | 0.59144 | NUP85 | 0.59144 | 0.63172 | 0.62057 |
| ENSG00000198723 | 19 | 7557862 | 7573336 | 85 | 16 | -0.23132 | 0.59147 | C19orf45 | 0.59147 | 0.37636 | 0.30136 |
| ENSG00000109321 | 4 | 75310851 | 75320726 | 5 | 3 | -0.23139 | 0.5915 | AREG | 0.5915 | 0.91405 | 0.92935 |
| ENSG00000182831 | 16 | 9185505 | 9215497 | 102 | 15 | -0.23147 | 0.59152 | C16orf72 | 0.59152 | 0.73448 | 0.73505 |
| ENSG00000181626 | 18 | 12093852 | 12129748 | 130 | 9 | -0.23152 | 0.59155 | ANKRD62 | 0.59155 | 0.60575 | 0.61262 |
| ENSG00000115163 | 2 | 26987157 | 27023935 | 98 | 9 | -0.23156 | 0.59156 | CENPA | 0.59156 | 0.54936 | 0.47829 |
| ENSG00000241978 | 9 | 112542769 | 112934792 | 1458 | 131 | -0.23156 | 0.59156 | AKAP2 | 0.59156 | 0.48574 | 0.50775 |
| ENSG00000168032 | 3 | 40428647 | 40470110 | 125 | 9 | -0.23159 | 0.59157 | ENTPD3 | 0.59157 | 0.59083 | 0.5764 |
| ENSG00000142197 | 21 | 37529080 | 37666572 | 679 | 53 | -0.23196 | 0.59171 | DOPEY2 | 0.59171 | 0.59218 | 0.64628 |
| ENSG00000131943 | 19 | 30191721 | 30206364 | 57 | 13 | -0.23197 | 0.59172 | C19orf12 | 0.59172 | 0.80699 | 0.90165 |
| ENSG00000183421 | 21 | 43159529 | 43187266 | 112 | 18 | -0.23224 | 0.59182 | RIPK4 | 0.59182 | 0.58905 | 0.84814 |
| ENSG00000112761 | 6 | 112375275 | 112392171 | 31 | 7 | -0.23229 | 0.59185 | WISP3 | 0.59185 | 0.5506 | 0.60061 |
| ENSG00000144401 | 2 | 208445355 | 208490652 | 94 | 8 | -0.23232 | 0.59186 | METTL21A | 0.59186 | 0.27493 | 0.15702 |
| ENSG00000104388 | 8 | 61429416 | 61536186 | 318 | 15 | -0.23242 | 0.59189 | RAB2A | 0.59189 | 0.57669 | 0.58542 |
| ENSG00000075856 | 12 | 108916357 | 108955176 | 150 | 13 | -0.23243 | 0.5919 | SART3 | 0.5919 | 0.6395 | 0.83881 |
| ENSG00000167881 | 17 | 74035184 | 74068734 | 76 | 7 | -0.23259 | 0.59196 | SRP68 | 0.59196 | 0.51509 | 0.64416 |
| ENSG00000142528 | 19 | 50528971 | 50556671 | 54 | 12 | -0.23258 | 0.59196 | ZNF473 | 0.59196 | 0.70994 | 0.68508 |
| ENSG00000130684 | 20 | 25654851 | 25677477 | 54 | 7 | -0.23285 | 0.59206 | ZNF337 | 0.59206 | 0.65272 | 0.67824 |
| ENSG00000070718 | 8 | 42010464 | 42029191 | 38 | 7 | -0.23292 | 0.59209 | AP3M2 | 0.59209 | 0.5559 | 0.56135 |
| ENSG00000126264 | 19 | 36393382 | 36395279 | 6 | 2 | -0.23365 | 0.59237 | HCST | 0.59237 | 0.77516 | 0.52157 |
| ENSG00000108294 | 17 | 36908989 | 36920484 | 35 | 12 | -0.23367 | 0.59238 | PSMB3 | 0.59238 | 0.48362 | 0.36941 |
| ENSG00000132031 | 2 | 20191872 | 20212455 | 48 | 11 | -0.23379 | 0.59243 | MATN3 | 0.59243 | 0.51254 | 0.45289 |
| ENSG00000163520 | 3 | 13573824 | 13679922 | 481 | 52 | -0.23386 | 0.59245 | FBLN2 | 0.59245 | 0.70345 | 0.70044 |
| ENSG00000184719 | 10 | 90033621 | 90344287 | 766 | 42 | -0.23423 | 0.5926 | RNLS | 0.5926 | 0.60364 | 0.62418 |
| ENSG00000143575 | 1 | 154244987 | 154248351 | 3 | 2 | -0.23427 | 0.59261 | HAX1 | 0.59261 | 0.54596 | 0.49363 |
| ENSG00000142544 | 19 | 51600863 | 51611627 | 37 | 8 | -0.23429 | 0.59262 | CTU1 | 0.59262 | 0.59063 | 0.50005 |
| ENSG00000198829 | 3 | 151591431 | 151602405 | 37 | 10 | -0.23434 | 0.59264 | SUCNR1 | 0.59264 | 0.73057 | 0.74873 |
| ENSG00000157510 | 5 | 148651434 | 148721365 | 159 | 41 | -0.23456 | 0.59272 | AFAP1L1 | 0.59272 | 0.5349 | 0.43018 |
| ENSG00000120709 | 5 | 137667624 | 137685416 | 39 | 12 | -0.23464 | 0.59276 | FAM53C | 0.59276 | 0.45523 | 0.41492 |
| ENSG00000152749 | 13 | 95254157 | 95286899 | 80 | 12 | -0.23475 | 0.5928 | GPR180 | 0.5928 | 0.6374 | 0.64433 |
| ENSG00000179627 | 14 | 105266933 | 105271049 | 12 | 4 | -0.23492 | 0.59286 | ZBTB42 | 0.59286 | 0.60587 | 0.66493 |
| ENSG00000062725 | 17 | 58520520 | 58603580 | 62 | 15 | -0.23495 | 0.59288 | APPBP2 | 0.59288 | 0.6917 | 0.69943 |
| ENSG00000004468 | 4 | 15779898 | 15854853 | 262 | 17 | -0.23531 | 0.59302 | CD38 | 0.59302 | 0.52787 | 0.49302 |
| ENSG00000142539 | 19 | 50919056 | 50931666 | 49 | 15 | -0.23537 | 0.59304 | CTD-2545M3.6 | 0.59304 | 0.81027 | 0.93699 |

| | | | | | | | | | | | |
|---|---|---|---|---|---|---|---|---|---|---|---|
| ENSG00000131068 | 20 | 29956421 | 29961726 | 7 | 3 | -0.23536 | 0.59304 | DEFB118 | 0.59304 | 0.50782 | 0.51922 |
| ENSG00000139636 | 12 | 49490919 | 49504683 | 13 | 3 | -0.23543 | 0.59306 | LMBR1L | 0.59306 | 0.51765 | 0.42594 |
| ENSG00000173156 | 11 | 66824289 | 66839484 | 33 | 6 | -0.23547 | 0.59308 | RHOD | 0.59308 | 0.70769 | 0.84706 |
| ENSG00000197846 | 6 | 26199748 | 26200942 | 10 | 4 | -0.23551 | 0.59309 | HIST1H2BF | 0.59309 | 0.51398 | 0.51257 |
| ENSG00000084112 | 12 | 109176466 | 109251366 | 193 | 27 | -0.23557 | 0.59312 | SSH1 | 0.59312 | 0.55966 | 0.51604 |
| ENSG00000100445 | 14 | 24908972 | 24912111 | 19 | 2 | -0.23558 | 0.59312 | SDR39U1 | 0.59312 | 0.57424 | 0.52183 |
| ENSG00000066322 | 1 | 43829068 | 43833696 | 4 | 2 | -0.23587 | 0.59323 | ELOVL1 | 0.59323 | 0.39089 | 0.45091 |
| ENSG00000176239 | 11 | 5372738 | 5373676 | 19 | 3 | -0.23593 | 0.59326 | OR51B6 | 0.59326 | 0.17935 | 0.25072 |
| ENSG00000101363 | 20 | 35918041 | 35945663 | 34 | 12 | -0.23598 | 0.59328 | MANBAL | 0.59328 | 0.58438 | 0.31616 |
| ENSG00000173157 | 12 | 43747669 | 43945724 | 723 | 33 | -0.23602 | 0.59329 | ADAMTS20 | 0.59329 | 0.6226 | 0.62244 |
| ENSG00000179755 | 16 | 34256507 | 34257278 | 1 | 1 | -0.2363 | 0.5934 | CTD-2144E22.5 | 0.5934 | 0.78988 | 0.73005 |
| ENSG00000133678 | 10 | 81838402 | 81852313 | 59 | 10 | -0.23637 | 0.59343 | TMEM254 | 0.59343 | 0.70554 | 0.73355 |
| ENSG00000164885 | 7 | 150750899 | 150755617 | 11 | 4 | -0.23643 | 0.59345 | CDK5 | 0.59345 | 0.64493 | 0.76813 |
| ENSG00000167969 | 16 | 2289396 | 2302301 | 21 | 8 | -0.23643 | 0.59345 | ECI1 | 0.59345 | 0.54568 | 0.71697 |
| ENSG00000072121 | 14 | 68194091 | 68283307 | 241 | 28 | -0.23646 | 0.59346 | ZFYVE26 | 0.59346 | 0.60291 | 0.67653 |
| ENSG00000173153 | 11 | 64073044 | 64084215 | 13 | 5 | -0.23649 | 0.59348 | ESRRA | 0.59348 | 0.38995 | 0.1827 |
| ENSG00000137252 | 6 | 55039050 | 55147418 | 420 | 26 | -0.23658 | 0.59351 | HCRTR2 | 0.59351 | 0.59766 | 0.56523 |
| ENSG00000269514 | 12 | 48592170 | 48595814 | 26 | 4 | -0.23659 | 0.59351 | DKFZP779L1853 | 0.59351 | 0.45279 | 0.4681 |
| ENSG00000213903 | 14 | 24780656 | 24787242 | 12 | 5 | -0.23679 | 0.59359 | LTB4R | 0.59359 | 0.61781 | 0.63402 |
| ENSG00000268432 | 5 | 157160003 | 157161727 | 5 | 1 | -0.23692 | 0.59364 | AC026407.1 | 0.59364 | 0.69807 | 0.78168 |
| ENSG00000154978 | 7 | 55503749 | 55640681 | 747 | 14 | -0.23705 | 0.59369 | VOPP1 | 0.59369 | 0.60865 | 0.63567 |
| ENSG00000225830 | 10 | 50663414 | 50747584 | 196 | 25 | -0.23704 | 0.59369 | ERCC6 | 0.59369 | 0.55754 | 0.5297 |
| ENSG00000255767 | 2 | 3579574 | 3595547 | 49 | 11 | -0.23706 | 0.5937 | RP13-512J5.1 | 0.5937 | 0.60845 | 0.37923 |
| ENSG00000120211 | 9 | 5231419 | 5235304 | 7 | 2 | -0.23712 | 0.59372 | INSL4 | 0.59372 | 0.57047 | 0.51003 |
| ENSG00000128791 | 18 | 9334765 | 9402418 | 244 | 20 | -0.23729 | 0.59379 | TWSG1 | 0.59379 | 0.59331 | 0.59426 |
| ENSG00000226180 | 16 | 87727144 | 87729753 | 12 | 3 | -0.23737 | 0.59381 | AC010536.1 | 0.59381 | 0.97012 | 0.79785 |
| ENSG00000213471 | 15 | 90792762 | 90808199 | 74 | 6 | -0.23737 | 0.59382 | TTLL13 | 0.59382 | 0.52294 | 0.47771 |
| ENSG00000174738 | 3 | 23986751 | 24022109 | 113 | 16 | -0.23751 | 0.59387 | NR1D2 | 0.59387 | 0.59738 | 0.635 |
| ENSG00000183891 | 2 | 20096404 | 20101747 | 10 | 4 | -0.23787 | 0.59401 | TTC32 | 0.59401 | 0.695 | 0.85602 |
| ENSG00000149634 | 20 | 44515128 | 44516274 | 4 | 2 | -0.23794 | 0.59404 | SPATA25 | 0.59404 | 0.85045 | 0.89453 |
| ENSG00000184909 | 1 | 178511950 | 178517579 | 27 | 4 | -0.23796 | 0.59405 | C1ORF220 | 0.59405 | 0.75822 | 0.74893 |
| ENSG00000109424 | 4 | 141480588 | 141489959 | 22 | 4 | -0.23808 | 0.59409 | UCP1 | 0.59409 | 0.77997 | 0.75074 |
| ENSG00000077157 | 1 | 202317827 | 202561834 | 368 | 15 | -0.23842 | 0.59422 | PPP1R12B | 0.59422 | 0.61542 | 0.63764 |
| ENSG00000090382 | 12 | 69742121 | 69748014 | 22 | 6 | -0.2385 | 0.59425 | LYZ | 0.59425 | 0.58725 | 0.67903 |
| ENSG00000173451 | 12 | 72056789 | 72074419 | 30 | 7 | -0.23849 | 0.59425 | THAP2 | 0.59425 | 0.67453 | 0.67198 |
| ENSG00000115255 | 19 | 1491165 | 1497926 | 13 | 3 | -0.23853 | 0.59427 | REEP6 | 0.59427 | 0.53687 | 0.39112 |
| ENSG00000163281 | 4 | 44684217 | 44728612 | 130 | 10 | -0.23874 | 0.59435 | GNPDA2 | 0.59435 | 0.59685 | 0.62503 |
| ENSG00000153802 | 4 | 68686594 | 68749750 | 174 | 23 | -0.23875 | 0.59435 | TMPRSS11D | 0.59435 | 0.48515 | 0.5949 |
| ENSG00000105255 | 19 | 4304597 | 4323840 | 69 | 15 | -0.23879 | 0.59437 | FSD1 | 0.59437 | 0.37655 | 0.25437 |
| ENSG00000165566 | 13 | 25735822 | 25746426 | 32 | 6 | -0.23905 | 0.59447 | AMER2 | 0.59447 | 0.43545 | 0.45308 |
| ENSG00000063015 | 17 | 27281919 | 27333458 | 80 | 13 | -0.23922 | 0.59453 | SEZ6 | 0.59453 | 0.59253 | 0.65882 |
| ENSG00000197121 | 2 | 197697728 | 197792520 | 176 | 10 | -0.23928 | 0.59455 | PGAP1 | 0.59455 | 0.42572 | 0.14401 |
| ENSG00000141452 | 18 | 21083473 | 21111746 | 61 | 9 | -0.23974 | 0.59473 | C18orf8 | 0.59473 | 0.40422 | 0.4614 |
| ENSG00000155249 | 14 | 20403767 | 20404842 | 7 | 2 | -0.23986 | 0.59478 | OR4K1 | 0.59478 | 0.67255 | 0.56794 |
| ENSG00000173077 | 9 | 117904097 | 118164923 | 1096 | 63 | -0.24001 | 0.59484 | 44166 | 0.59484 | 0.6385 | 0.67034 |
| ENSG00000198901 | 15 | 91509270 | 91538859 | 113 | 12 | -0.24022 | 0.59492 | PRC1 | 0.59492 | 0.66803 | 0.71375 |
| ENSG00000224689 | 19 | 9800600 | 9811452 | 38 | 4 | -0.24046 | 0.59501 | ZNF812 | 0.59501 | 0.61258 | 0.62139 |
| ENSG00000123240 | 10 | 13141449 | 13180291 | 146 | 26 | -0.24066 | 0.59509 | OPTN | 0.59509 | 0.54615 | 0.068869 |
| ENSG00000188610 | 1 | 120837756 | 120855681 | 9 | 3 | -0.24076 | 0.59513 | FAM72B | 0.59513 | 0.5822 | 0.61199 |
| ENSG00000126351 | 17 | 38214543 | 38250120 | 73 | 16 | -0.24082 | 0.59515 | THRA | 0.59515 | 0.68381 | 0.66495 |
| ENSG00000140848 | 16 | 57126449 | 57181878 | 221 | 43 | -0.24087 | 0.59517 | CPNE2 | 0.59517 | 0.43741 | 0.33461 |
| ENSG00000226974 | 13 | 64320934 | 64321323 | 1 | 1 | -0.24094 | 0.5952 | AL445989.1 | 0.5952 | 0.53245 | 0.39182 |
| ENSG00000116016 | 2 | 46520806 | 46613836 | 294 | 35 | -0.24129 | 0.59534 | EPAS1 | 0.59534 | 0.68504 | 0.73432 |
| ENSG00000180389 | 13 | 28519343 | 28519727 | 3 | 2 | -0.24132 | 0.59535 | ATP5EP2 | 0.59535 | 0.79778 | 0.48173 |
| ENSG00000068305 | 15 | 100017370 | 100256671 | 683 | 50 | -0.24145 | 0.5954 | MEF2A | 0.5954 | 0.33255 | 0.36456 |
| ENSG00000258659 | 11 | 5640994 | 5665628 | 97 | 14 | -0.24164 | 0.59547 | TRIM34 | 0.59547 | 0.78707 | 0.90995 |
| ENSG00000142166 | 21 | 34696734 | 34732168 | 91 | 14 | -0.24169 | 0.59549 | IFNAR1 | 0.59549 | 0.74804 | 0.69098 |
| ENSG00000176566 | 8 | 88882973 | 88886296 | 10 | 6 | -0.24172 | 0.5955 | DCAF4L2 | 0.5955 | 0.4535 | 0.57052 |
| ENSG00000067221 | 15 | 74275547 | 74286963 | 30 | 7 | -0.24188 | 0.59556 | STOML1 | 0.59556 | 0.57975 | 0.4547 |
| ENSG00000272790 | 11 | 112041253 | 112071394 | 102 | 17 | -0.24189 | 0.59557 | AP002884.3 | 0.59557 | 0.6275 | 0.6541 |
| ENSG00000178096 | 1 | 149859440 | 149872351 | 36 | 3 | -0.24201 | 0.59561 | BOLA1 | 0.59561 | 0.63717 | 0.68625 |
| ENSG00000143851 | 1 | 202116141 | 202130716 | 49 | 19 | -0.24253 | 0.59582 | PTPN7 | 0.59582 | 0.71001 | 0.30392 |
| ENSG00000152518 | 2 | 43449541 | 43453748 | 10 | 4 | -0.24264 | 0.59586 | ZFP36L2 | 0.59586 | 0.31102 | 0.21207 |
| ENSG00000164256 | 5 | 23507264 | 23528706 | 46 | 14 | -0.24264 | 0.59586 | PRDM9 | 0.59586 | 0.69157 | 0.7602 |
| ENSG00000204612 | 9 | 79634571 | 79635869 | 1 | 1 | -0.24275 | 0.5959 | FOXB2 | 0.5959 | 0.98248 | 0.96098 |
| ENSG00000246366 | 8 | 71520812 | 71575514 | 191 | 19 | -0.24303 | 0.59601 | RP11-382J12.1 | 0.59601 | 0.4916 | 0.42656 |
| ENSG00000125871 | 20 | 17949556 | 17971765 | 85 | 11 | -0.24304 | 0.59601 | MGME1 | 0.59601 | 0.62653 | 0.56839 |
| ENSG00000260259 | 16 | 89749028 | 89752977 | 24 | 6 | -0.24314 | 0.59605 | RP11-368I7.4 | 0.59605 | 0.58467 | 0.61534 |
| ENSG00000118162 | 19 | 47978401 | 47987525 | 13 | 6 | -0.24346 | 0.59617 | KPTN | 0.59617 | 0.86729 | 0.90769 |
| ENSG00000130226 | 7 | 153584182 | 154685995 | 4594 | 357 | -0.24358 | 0.59622 | DPP6 | 0.59622 | 0.59534 | 0.63908 |
| ENSG00000137806 | 15 | 41679551 | 41694717 | 45 | 9 | -0.24366 | 0.59625 | NDUFAF1 | 0.59625 | 0.63535 | 0.66089 |
| ENSG00000154274 | 4 | 37455563 | 37625117 | 849 | 75 | -0.24371 | 0.59627 | C4orf19 | 0.59627 | 0.55951 | 0.6259 |
| ENSG00000156858 | 16 | 30662038 | 30667761 | 3 | 2 | -0.24392 | 0.59635 | PRR14 | 0.59635 | 0.50741 | 0.6636 |

| | | | | | | | | | | | |
|---|---|---|---|---|---|---|---|---|---|---|---|
| ENSG00000183269 | 11 | 5877904 | 5878964 | 3 | 1 | -0.244 | 0.59639 | OR52E8 | 0.59639 | 0.64432 | 0.74114 |
| ENSG00000152102 | 2 | 131805449 | 131851033 | 52 | 6 | -0.2442 | 0.59646 | FAM168B | 0.59646 | 0.62798 | 0.63594 |
| ENSG00000159256 | 21 | 37692487 | 37758446 | 245 | 15 | -0.24419 | 0.59646 | MORC3 | 0.59646 | 0.64719 | 0.68481 |
| ENSG00000198556 | 7 | 99070464 | 99101273 | 48 | 12 | -0.24441 | 0.59654 | ZNF789 | 0.59654 | 0.75635 | 0.77604 |
| ENSG00000040933 | 2 | 99061317 | 99210853 | 266 | 16 | -0.24452 | 0.59659 | INPP4A | 0.59659 | 0.64503 | 0.59845 |
| ENSG00000128915 | 15 | 60711808 | 60771359 | 143 | 14 | -0.24453 | 0.59659 | NARG2 | 0.59659 | 0.63293 | 0.68349 |
| ENSG00000162076 | 16 | 2933187 | 2949383 | 102 | 12 | -0.24475 | 0.59668 | FLYWCH2 | 0.59668 | 0.79788 | 0.8783 |
| ENSG00000145390 | 4 | 120133742 | 120216672 | 292 | 19 | -0.24479 | 0.59669 | USP53 | 0.59669 | 0.58532 | 0.58299 |
| ENSG00000196776 | 3 | 107762145 | 107809872 | 108 | 19 | -0.24483 | 0.59671 | CD47 | 0.59671 | 0.71627 | 0.17365 |
| ENSG00000108839 | 17 | 6899384 | 6914055 | 54 | 7 | -0.24502 | 0.59678 | ALOX12 | 0.59678 | 0.66723 | 0.76282 |
| ENSG00000146414 | 6 | 146185381 | 146285559 | 190 | 11 | -0.24504 | 0.59679 | SHPRH | 0.59679 | 0.56686 | 0.56354 |
| ENSG00000144230 | 2 | 128403439 | 128410213 | 27 | 9 | -0.2454 | 0.59693 | GPR17 | 0.59693 | 0.13939 | 0.078447 |
| ENSG00000173237 | 11 | 66742748 | 66744475 | 2 | 1 | -0.24557 | 0.59699 | C11orf86 | 0.59699 | 0.24137 | 0.64378 |
| ENSG00000105672 | 19 | 36132647 | 36135773 | 5 | 2 | -0.24555 | 0.59699 | ETV2 | 0.59699 | 0.91119 | 0.8282 |
| ENSG00000083099 | 6 | 90277863 | 90348472 | 180 | 26 | -0.2456 | 0.597 | LYRM2 | 0.597 | 0.61564 | 0.52472 |
| ENSG00000140319 | 15 | 40327940 | 40331389 | 13 | 5 | -0.24562 | 0.59701 | SRP14 | 0.59701 | 0.38365 | 0.37027 |
| ENSG00000253485 | 5 | 140743898 | 140892546 | 457 | 37 | -0.24582 | 0.59709 | PCDHGA5 | 0.59709 | 0.60239 | 0.50549 |
| ENSG00000127663 | 19 | 4969125 | 5153606 | 647 | 39 | -0.24582 | 0.59709 | KDM4B | 0.59709 | 0.65236 | 0.68354 |
| ENSG00000197959 | 1 | 171810621 | 172387606 | 1606 | 71 | -0.24588 | 0.59711 | DNM3 | 0.59711 | 0.59693 | 0.58687 |
| ENSG00000177764 | 20 | 277737 | 280965 | 8 | 3 | -0.24591 | 0.59712 | ZCCHC3 | 0.59712 | 0.5464 | 0.32454 |
| ENSG00000159650 | 3 | 126200124 | 126236616 | 175 | 36 | -0.246 | 0.59716 | UROC1 | 0.59716 | 0.65534 | 0.82766 |
| ENSG00000137270 | 6 | 52991762 | 53013627 | 87 | 17 | -0.24601 | 0.59716 | GCM1 | 0.59716 | 0.48137 | 0.46652 |
| ENSG00000166405 | 11 | 8127597 | 8190602 | 242 | 22 | -0.24624 | 0.59725 | RIC3 | 0.59725 | 0.56888 | 0.60881 |
| ENSG00000163535 | 2 | 201374731 | 201448505 | 188 | 23 | -0.24631 | 0.59728 | SGOL2 | 0.59728 | 0.65764 | 0.7261 |
| ENSG00000235705 | 10 | 81266126 | 81270678 | 23 | 7 | -0.24634 | 0.59729 | AL133481.1 | 0.59729 | 0.44586 | 0.49119 |
| ENSG00000055955 | 3 | 52846991 | 52865495 | 40 | 5 | -0.24636 | 0.5973 | ITIH4 | 0.5973 | 0.5381 | 0.59147 |
| ENSG00000111731 | 12 | 22601517 | 22697480 | 184 | 7 | -0.24653 | 0.59736 | C2CD5 | 0.59736 | 0.60608 | 0.54884 |
| ENSG00000138180 | 10 | 95256389 | 95288849 | 153 | 19 | -0.24663 | 0.5974 | CEP55 | 0.5974 | 0.45316 | 0.26065 |
| ENSG00000068885 | 3 | 159974774 | 160117668 | 245 | 28 | -0.24687 | 0.5975 | IFT80 | 0.5975 | 0.62408 | 0.67703 |
| ENSG00000267022 | 19 | 44529506 | 44591471 | 150 | 12 | -0.24687 | 0.5975 | ZNF223 | 0.5975 | 0.53206 | 0.45837 |
| ENSG00000172650 | 10 | 75434033 | 75457639 | 20 | 6 | -0.2469 | 0.59751 | AGAP5 | 0.59751 | 0.77734 | 0.86819 |
| ENSG00000261934 | 5 | 140782520 | 140892546 | 341 | 30 | -0.24701 | 0.59755 | PCDHGA9 | 0.59755 | 0.55532 | 0.48732 |
| ENSG00000133422 | 22 | 31321117 | 31364284 | 66 | 9 | -0.24705 | 0.59757 | MORC2 | 0.59757 | 0.52487 | 0.5833 |
| ENSG00000170606 | 5 | 132387654 | 132442141 | 172 | 14 | -0.24711 | 0.59759 | HSPA4 | 0.59759 | 0.63207 | 0.49146 |
| ENSG00000148384 | 9 | 139323071 | 139334274 | 60 | 11 | -0.24712 | 0.59759 | INPP5E | 0.59759 | 0.74344 | 0.73888 |
| ENSG00000127249 | 3 | 193119866 | 193310900 | 557 | 46 | -0.24718 | 0.59762 | ATP13A4 | 0.59762 | 0.61829 | 0.59742 |
| ENSG00000130402 | 19 | 39138289 | 39222223 | 282 | 19 | -0.24733 | 0.59767 | ACTN4 | 0.59767 | 0.56306 | 0.52415 |
| ENSG00000116473 | 1 | 112084840 | 112259313 | 651 | 46 | -0.24738 | 0.59769 | RAP1A | 0.59769 | 0.59131 | 0.49378 |
| ENSG00000066455 | 14 | 93260576 | 93306308 | 199 | 14 | -0.24759 | 0.59777 | GOLGA5 | 0.59777 | 0.63895 | 0.72397 |
| ENSG00000244486 | 22 | 20778874 | 20792146 | 52 | 10 | -0.24758 | 0.59777 | SCARF2 | 0.59777 | 0.72578 | 0.73432 |
| ENSG00000181555 | 3 | 47057919 | 47205457 | 253 | 14 | -0.24773 | 0.59783 | SETD2 | 0.59783 | 0.59305 | 0.56943 |
| ENSG00000116741 | 1 | 192778169 | 192781403 | 6 | 2 | -0.24803 | 0.59794 | RGS2 | 0.59794 | 0.64726 | 0.34538 |
| ENSG00000157851 | 2 | 27070615 | 27173219 | 197 | 13 | -0.24842 | 0.59809 | DPYSL5 | 0.59809 | 0.49015 | 0.46597 |
| ENSG00000138668 | 4 | 83273651 | 83295656 | 63 | 7 | -0.24866 | 0.59819 | HNRNPD | 0.59819 | 0.70745 | 0.86186 |
| ENSG00000077254 | 1 | 78161672 | 78225537 | 116 | 21 | -0.24871 | 0.59821 | USP33 | 0.59821 | 0.42399 | 0.43617 |
| ENSG00000166862 | 22 | 36959968 | 37099603 | 453 | 58 | -0.24888 | 0.59827 | CACNG2 | 0.59827 | 0.23592 | 0.024215 |
| ENSG00000182224 | 17 | 7761064 | 7765600 | 9 | 4 | -0.24898 | 0.59831 | CYB5D1 | 0.59831 | 0.63055 | 0.71475 |
| ENSG00000187736 | 2 | 219940039 | 220025587 | 177 | 17 | -0.24906 | 0.59834 | NHEJ1 | 0.59834 | 0.59816 | 0.62951 |
| ENSG00000170820 | 2 | 49189296 | 49381676 | 894 | 38 | -0.2491 | 0.59836 | FSHR | 0.59836 | 0.6043 | 0.57867 |
| ENSG00000185252 | 22 | 20748405 | 20762745 | 56 | 5 | -0.24923 | 0.59841 | ZNF74 | 0.59841 | 0.65338 | 0.81925 |
| ENSG00000073921 | 11 | 85668727 | 85780924 | 403 | 23 | -0.24933 | 0.59845 | PICALM | 0.59845 | 0.57664 | 0.5061 |
| ENSG00000130312 | 19 | 17403418 | 17417652 | 61 | 11 | -0.24942 | 0.59848 | MRPL34 | 0.59848 | 0.37996 | 0.33951 |
| ENSG00000269404 | 19 | 50922195 | 50934570 | 45 | 13 | -0.2494 | 0.59848 | SPIB | 0.59848 | 0.8221 | 0.94996 |
| ENSG00000129158 | 11 | 17809595 | 18034709 | 481 | 47 | -0.24948 | 0.59851 | SERGEF | 0.59851 | 0.61768 | 0.67501 |
| ENSG00000132975 | 13 | 27329341 | 27334922 | 17 | 5 | -0.24955 | 0.59853 | GPR12 | 0.59853 | 0.56523 | 0.56831 |
| ENSG00000165973 | 11 | 20691117 | 21597227 | 3451 | 267 | -0.24956 | 0.59854 | NELL1 | 0.59854 | 0.64734 | 0.62206 |
| ENSG00000073711 | 3 | 135684515 | 135866733 | 351 | 23 | -0.24961 | 0.59856 | PPP2R3A | 0.59856 | 0.59485 | 0.60773 |
| ENSG00000100218 | 22 | 23401593 | 23487208 | 199 | 17 | -0.24965 | 0.59857 | RTDR1 | 0.59857 | 0.49284 | 0.41648 |
| ENSG00000171561 | 11 | 74799758 | 74800799 | 1 | 1 | -0.24973 | 0.5986 | OR2AT4 | 0.5986 | 0.28751 | 0.53891 |
| ENSG00000178031 | 9 | 18473892 | 18910948 | 2150 | 154 | -0.24992 | 0.59868 | ADAMTSL1 | 0.59868 | 0.61133 | 0.61951 |
| ENSG00000154493 | 10 | 128113566 | 128359079 | 865 | 90 | -0.24994 | 0.59868 | C10orf90 | 0.59868 | 0.634 | 0.64315 |
| ENSG00000212126 | 12 | 11138512 | 11139511 | 3 | 2 | -0.25004 | 0.59872 | TAS2R50 | 0.59872 | 0.39831 | 0.46463 |
| ENSG00000205744 | 19 | 6467218 | 6482568 | 60 | 13 | -0.25018 | 0.59878 | DENND1C | 0.59878 | 0.70316 | 0.55314 |
| ENSG00000099795 | 19 | 14676890 | 14682874 | 15 | 5 | -0.25018 | 0.59878 | NDUFB7 | 0.59878 | 0.61388 | 0.4536 |
| ENSG00000112367 | 6 | 110012499 | 110146631 | 374 | 33 | -0.25041 | 0.59886 | FIG4 | 0.59886 | 0.61721 | 0.68916 |
| ENSG00000186452 | 12 | 51236703 | 51281667 | 183 | 14 | -0.25039 | 0.59886 | TMPRSS12 | 0.59886 | 0.61855 | 0.71825 |
| ENSG00000140740 | 16 | 21963981 | 21994981 | 37 | 10 | -0.25039 | 0.59886 | UQCRC2 | 0.59886 | 0.51603 | 0.49097 |
| ENSG00000114473 | 3 | 197615946 | 197687013 | 251 | 21 | -0.25043 | 0.59887 | IQCG | 0.59887 | 0.64032 | 0.69058 |
| ENSG00000243147 | 2 | 27994584 | 28210954 | 420 | 19 | -0.25047 | 0.59889 | MRPL33 | 0.59889 | 0.6151 | 0.58975 |
| ENSG00000247596 | 3 | 52262626 | 52273276 | 29 | 9 | -0.25049 | 0.59889 | TWF2 | 0.59889 | 0.5475 | 0.4867 |
| ENSG00000108039 | 10 | 111624524 | 111683311 | 115 | 16 | -0.25063 | 0.59895 | XPNPEP1 | 0.59895 | 0.64368 | 0.80707 |
| ENSG00000205707 | 12 | 25348150 | 25362579 | 44 | 6 | -0.25062 | 0.59895 | LYRM5 | 0.59895 | 0.63962 | 0.62531 |
| ENSG00000076067 | 12 | 56915713 | 56984745 | 223 | 23 | -0.2507 | 0.59898 | RBMS2 | 0.59898 | 0.56831 | 0.6147 |

| | | | | | | | | | | | |
|---|---|---|---|---|---|---|---|---|---|---|---|
| ENSG00000069122 | 6 | 46820249 | 46922680 | 365 | 42 | -0.25075 | 0.599 | GPR116 | 0.599 | 0.7084 | 0.86216 |
| ENSG00000125505 | 19 | 54677107 | 54693733 | 62 | 19 | -0.25075 | 0.599 | MBOAT7 | 0.599 | 0.30424 | 0.26187 |
| ENSG00000145687 | 5 | 80708840 | 81047616 | 843 | 38 | -0.25104 | 0.59911 | SSBP2 | 0.59911 | 0.61044 | 0.64123 |
| ENSG00000100417 | 22 | 41972898 | 41985894 | 24 | 5 | -0.25112 | 0.59914 | PMM1 | 0.59914 | 0.57442 | 0.65458 |
| ENSG00000171790 | 1 | 41481269 | 41488909 | 33 | 9 | -0.25122 | 0.59918 | SLFNL1 | 0.59918 | 0.58792 | 0.53598 |
| ENSG00000169122 | 8 | 58907068 | 59116838 | 656 | 54 | -0.25167 | 0.59935 | FAM110B | 0.59935 | 0.64441 | 0.72567 |
| ENSG00000073584 | 17 | 38781214 | 38804760 | 42 | 10 | -0.25194 | 0.59946 | SMARCE1 | 0.59946 | 0.56723 | 0.56863 |
| ENSG00000075711 | 3 | 196769431 | 197026171 | 737 | 27 | -0.25203 | 0.59949 | DLG1 | 0.59949 | 0.57355 | 0.53985 |
| ENSG00000110660 | 11 | 107661717 | 107799019 | 678 | 35 | -0.25217 | 0.59954 | SLC35F2 | 0.59954 | 0.57874 | 0.49195 |
| ENSG00000158869 | 1 | 161185024 | 161190489 | 20 | 8 | -0.25219 | 0.59955 | FCER1G | 0.59955 | 0.77133 | 0.85492 |
| ENSG00000134882 | 13 | 99853028 | 100038688 | 542 | 29 | -0.2525 | 0.59967 | UBAC2 | 0.59967 | 0.64514 | 0.66712 |
| ENSG00000112208 | 6 | 57037124 | 57054631 | 24 | 4 | -0.25293 | 0.59984 | BAG2 | 0.59984 | 0.54106 | 0.42252 |
| ENSG00000267596 | 17 | 34323476 | 34329084 | 27 | 7 | -0.25295 | 0.59985 | CCL15 | 0.59985 | 0.39694 | 0.49765 |
| ENSG00000124767 | 6 | 38643701 | 38670917 | 90 | 17 | -0.25299 | 0.59986 | GLO1 | 0.59986 | 0.55161 | 0.48493 |
| ENSG00000163939 | 3 | 52579368 | 52719933 | 395 | 12 | -0.25302 | 0.59987 | PBRM1 | 0.59987 | 0.57036 | 0.51933 |
| ENSG00000162069 | 16 | 3077683 | 3086927 | 33 | 6 | -0.253 | 0.59987 | CCDC64B | 0.59987 | 0.67021 | 0.88036 |
| ENSG00000100228 | 22 | 23487513 | 23506537 | 65 | 5 | -0.25312 | 0.59991 | RAB36 | 0.59991 | 0.61495 | 0.7288 |
| ENSG00000204248 | 6 | 33130458 | 33160276 | 110 | 18 | -0.25344 | 0.60003 | COL11A2 | 0.60003 | 0.55279 | 0.62539 |
| ENSG00000180304 | 15 | 64979772 | 64995480 | 28 | 7 | -0.25345 | 0.60004 | OAZ2 | 0.60004 | 0.54496 | 0.62504 |
| ENSG00000160201 | 21 | 44513066 | 44527697 | 35 | 5 | -0.25358 | 0.60009 | U2AF1 | 0.60009 | 0.79226 | 0.82671 |
| ENSG00000268730 | 15 | 99859300 | 99859828 | 2 | 1 | -0.25361 | 0.6001 | AC022819.2 | 0.6001 | 0.66621 | 0.79536 |
| ENSG00000077009 | 19 | 3933101 | 3942414 | 44 | 14 | -0.25365 | 0.60012 | NMRK2 | 0.60012 | 0.58946 | 0.52744 |
| ENSG00000184682 | 11 | 1910375 | 1912084 | 5 | 2 | -0.25383 | 0.60019 | C11orf89 | 0.60019 | 0.31087 | 0.51879 |
| ENSG00000166143 | 15 | 41107650 | 41120907 | 39 | 8 | -0.25406 | 0.60028 | PPP1R14D | 0.60028 | 0.70105 | 0.81312 |
| ENSG00000214447 | 17 | 42977135 | 42982758 | 16 | 3 | -0.25408 | 0.60028 | FAM187A | 0.60028 | 0.69971 | 0.75079 |
| ENSG00000001629 | 7 | 91875548 | 92030698 | 212 | 20 | -0.2541 | 0.60029 | ANKIB1 | 0.60029 | 0.59694 | 0.63968 |
| ENSG00000143147 | 1 | 168053997 | 168106821 | 125 | 13 | -0.25478 | 0.60055 | GPR161 | 0.60055 | 0.68863 | 0.68996 |
| ENSG00000198198 | 1 | 43855553 | 43918321 | 101 | 11 | -0.25479 | 0.60056 | SZT2 | 0.60056 | 0.59976 | 0.56056 |
| ENSG00000100823 | 14 | 20923350 | 20925927 | 4 | 2 | -0.2548 | 0.60056 | APEX1 | 0.60056 | 0.22781 | 0.37288 |
| ENSG00000131126 | 19 | 43905648 | 43922763 | 58 | 7 | -0.25493 | 0.60061 | TEX101 | 0.60061 | 0.70963 | 0.82871 |
| ENSG00000104901 | 19 | 49865040 | 49878373 | 50 | 4 | -0.25512 | 0.60069 | DKKL1 | 0.60069 | 0.61274 | 0.68862 |
| ENSG00000106128 | 7 | 30978284 | 31032869 | 213 | 37 | -0.25518 | 0.60071 | GHRHR | 0.60071 | 0.63359 | 0.53637 |
| ENSG00000011451 | 19 | 15532319 | 15560762 | 25 | 9 | -0.25523 | 0.60073 | WIZ | 0.60073 | 0.72139 | 0.79331 |
| ENSG00000261247 | 15 | 30427352 | 30437770 | 4 | 2 | -0.25558 | 0.60086 | GOLGA8T | 0.60086 | 0.68278 | 0.76684 |
| ENSG00000178567 | 3 | 37027357 | 37034795 | 10 | 3 | -0.2558 | 0.60095 | EPM2AIP1 | 0.60095 | 0.72746 | 0.64874 |
| ENSG00000269343 | 19 | 58331099 | 58370018 | 118 | 16 | -0.2568 | 0.60133 | ZNF587B | 0.60133 | 0.5836 | 0.61818 |
| ENSG00000067798 | 12 | 78224685 | 78606790 | 1518 | 81 | -0.25681 | 0.60134 | NAV3 | 0.60134 | 0.65822 | 0.6856 |
| ENSG00000027697 | 6 | 137518621 | 137540586 | 40 | 9 | -0.25685 | 0.60135 | IFNGR1 | 0.60135 | 0.55771 | 0.20854 |
| ENSG00000174233 | 12 | 49159975 | 49182820 | 41 | 5 | -0.25695 | 0.60139 | ADCY6 | 0.60139 | 0.71806 | 0.73755 |
| ENSG00000178772 | 3 | 194060494 | 194072057 | 80 | 9 | -0.25703 | 0.60142 | CPN2 | 0.60142 | 0.50316 | 0.67759 |
| ENSG00000119720 | 14 | 90742580 | 90798481 | 137 | 17 | -0.25719 | 0.60149 | NRDE2 | 0.60149 | 0.55922 | 0.417 |
| ENSG00000081059 | 5 | 133450402 | 133487556 | 88 | 11 | -0.25768 | 0.60167 | TCF7 | 0.60167 | 0.5892 | 0.63865 |
| ENSG00000155816 | 1 | 240177648 | 240638489 | 1959 | 106 | -0.258 | 0.60179 | FMN2 | 0.60179 | 0.54042 | 0.56298 |
| ENSG00000119041 | 2 | 197627756 | 197664449 | 52 | 13 | -0.258 | 0.6018 | GTF3C3 | 0.6018 | 0.76194 | 0.76849 |
| ENSG00000172367 | 11 | 119056166 | 119060932 | 3 | 2 | -0.25801 | 0.6018 | PDZD3 | 0.6018 | 0.73898 | 0.76331 |
| ENSG00000170276 | 11 | 111782966 | 111789574 | 12 | 4 | -0.25805 | 0.60181 | HSPB2 | 0.60181 | 0.78768 | 0.88312 |
| ENSG00000189280 | 1 | 35220648 | 35224113 | 12 | 5 | -0.25808 | 0.60183 | GJB5 | 0.60183 | 0.93968 | 0.74253 |
| ENSG00000269795 | 1 | 243388058 | 243395911 | 25 | 6 | -0.25817 | 0.60186 | AC092782.1 | 0.60186 | 0.52779 | 0.555 |
| ENSG00000166153 | 12 | 100597447 | 100660857 | 166 | 23 | -0.25838 | 0.60194 | DEPDC4 | 0.60194 | 0.61733 | 0.61751 |
| ENSG00000040531 | 17 | 3539762 | 3564836 | 144 | 15 | -0.25837 | 0.60194 | CTNS | 0.60194 | 0.65562 | 0.60938 |
| ENSG00000254122 | 5 | 140797427 | 140892546 | 303 | 27 | -0.2584 | 0.60195 | PCDHGB7 | 0.60195 | 0.55567 | 0.48605 |
| ENSG00000138686 | 4 | 122745595 | 122791652 | 104 | 15 | -0.25868 | 0.60206 | BBS7 | 0.60206 | 0.626 | 0.69048 |
| ENSG00000104044 | 15 | 28000021 | 28344504 | 1102 | 49 | -0.2588 | 0.6021 | OCA2 | 0.6021 | 0.60467 | 0.67373 |
| ENSG00000165449 | 10 | 61410523 | 61495760 | 281 | 34 | -0.25896 | 0.60217 | SLC16A9 | 0.60217 | 0.59932 | 0.56826 |
| ENSG00000251201 | 5 | 114914339 | 114961858 | 91 | 11 | -0.25899 | 0.60218 | TMED7-TICAM2 | 0.60218 | 0.59418 | 0.55876 |
| ENSG00000243414 | 5 | 114914339 | 114961876 | 91 | 11 | -0.25899 | 0.60218 | TICAM2 | 0.60218 | 0.59418 | 0.55876 |
| ENSG00000163382 | 1 | 156561554 | 156564091 | 13 | 1 | -0.25933 | 0.60231 | APOA1BP | 0.60231 | 0.6276 | 0.7133 |
| ENSG00000196365 | 19 | 5691845 | 5720583 | 127 | 19 | -0.25934 | 0.60231 | LONP1 | 0.60231 | 0.52351 | 0.63791 |
| ENSG00000069966 | 15 | 52413117 | 52483566 | 238 | 22 | -0.2596 | 0.60241 | GNB5 | 0.60241 | 0.61986 | 0.63165 |
| ENSG00000239732 | 3 | 52255096 | 52273183 | 38 | 11 | -0.25962 | 0.60242 | TLR9 | 0.60242 | 0.5759 | 0.49632 |
| ENSG00000147689 | 8 | 124191200 | 124222314 | 74 | 15 | -0.2597 | 0.60245 | FAM83A | 0.60245 | 0.59857 | 0.48704 |
| ENSG00000171962 | 17 | 17876127 | 17920203 | 91 | 10 | -0.25977 | 0.60248 | LRRC48 | 0.60248 | 0.61778 | 0.62257 |
| ENSG00000182463 | 20 | 51588946 | 52111869 | 1668 | 181 | -0.25984 | 0.60251 | TSHZ2 | 0.60251 | 0.60612 | 0.66629 |
| ENSG00000176390 | 17 | 29096406 | 29151794 | 126 | 12 | -0.26002 | 0.60257 | CRLF3 | 0.60257 | 0.57269 | 0.49298 |
| ENSG00000148143 | 9 | 109625378 | 109775915 | 307 | 43 | -0.26004 | 0.60258 | ZNF462 | 0.60258 | 0.64329 | 0.7156 |
| ENSG00000132005 | 19 | 14072350 | 14117851 | 40 | 13 | -0.2602 | 0.60264 | RFX1 | 0.60264 | 0.56526 | 0.7285 |
| ENSG00000139178 | 12 | 7242183 | 7261869 | 61 | 14 | -0.26021 | 0.60265 | C1RL | 0.60265 | 0.67065 | 0.51598 |
| ENSG00000204421 | 6 | 31686425 | 31689622 | 8 | 4 | -0.2604 | 0.60272 | LY6G6C | 0.60272 | 0.63147 | 0.68679 |
| ENSG00000183087 | 13 | 114523522 | 114567046 | 271 | 27 | -0.26044 | 0.60274 | GAS6 | 0.60274 | 0.59989 | 0.75601 |
| ENSG00000120254 | 6 | 151186685 | 151423023 | 1164 | 90 | -0.26061 | 0.6028 | MTHFD1L | 0.6028 | 0.59286 | 0.60488 |
| ENSG00000164659 | 7 | 86506222 | 86689015 | 480 | 16 | -0.26066 | 0.60282 | KIAA1324L | 0.60282 | 0.63489 | 0.53002 |
| ENSG00000030066 | 11 | 47799639 | 47870107 | 181 | 18 | -0.26071 | 0.60284 | NUP160 | 0.60284 | 0.61044 | 0.60537 |
| ENSG00000134419 | 16 | 18792617 | 18801705 | 8 | 3 | -0.26081 | 0.60288 | RPS15A | 0.60288 | 0.64824 | 0.59375 |

| | | | | | | | | | | | |
|---|---|---|---|---|---|---|---|---|---|---|---|
| ENSG00000015532 | 17 | 48423453 | 48440499 | 72 | 9 | -0.26086 | 0.6029 | XYLT2 | 0.6029 | 0.53571 | 0.62875 |
| ENSG00000188056 | 6 | 41196062 | 41206620 | 95 | 9 | -0.2611 | 0.60299 | TREML4 | 0.60299 | 0.44858 | 0.45653 |
| ENSG00000148297 | 9 | 136205160 | 136214986 | 28 | 10 | -0.26122 | 0.60304 | MED22 | 0.60304 | 0.53098 | 0.46201 |
| ENSG00000164100 | 4 | 118954773 | 119179803 | 842 | 27 | -0.26167 | 0.60321 | NDST3 | 0.60321 | 0.61952 | 0.65294 |
| ENSG00000262576 | 5 | 140734768 | 140892546 | 479 | 38 | -0.26173 | 0.60324 | PCDHGA4 | 0.60324 | 0.58384 | 0.52124 |
| ENSG00000150093 | 10 | 33189247 | 33294720 | 355 | 31 | -0.26174 | 0.60324 | ITGB1 | 0.60324 | 0.62565 | 0.66301 |
| ENSG00000162426 | 1 | 8377886 | 8404227 | 97 | 16 | -0.26183 | 0.60327 | SLC45A1 | 0.60327 | 0.57092 | 0.1745 |
| ENSG00000119718 | 14 | 75469614 | 75476292 | 11 | 2 | -0.26185 | 0.60328 | EIF2B2 | 0.60328 | 0.75214 | 0.70822 |
| ENSG00000130748 | 19 | 47549165 | 47551888 | 7 | 2 | -0.26195 | 0.60332 | TMEM160 | 0.60332 | 0.55899 | 0.47702 |
| ENSG00000100412 | 22 | 41865129 | 41924993 | 91 | 17 | -0.26199 | 0.60334 | ACO2 | 0.60334 | 0.60565 | 0.60262 |
| ENSG00000183579 | 22 | 29279580 | 29453475 | 379 | 34 | -0.2621 | 0.60338 | ZNRF3 | 0.60338 | 0.62854 | 0.69968 |
| ENSG00000112237 | 6 | 99990256 | 100016849 | 102 | 15 | -0.2623 | 0.60346 | CCNC | 0.60346 | 0.68224 | 0.68046 |
| ENSG00000126391 | 11 | 65154070 | 65180996 | 40 | 12 | -0.26231 | 0.60346 | FRMD8 | 0.60346 | 0.60939 | 0.81244 |
| ENSG00000138138 | 10 | 89511269 | 89601100 | 200 | 14 | -0.26243 | 0.6035 | ATAD1 | 0.6035 | 0.62528 | 0.58477 |
| ENSG00000167106 | 9 | 130702858 | 130742792 | 103 | 10 | -0.26244 | 0.60351 | FAM102A | 0.60351 | 0.58378 | 0.64245 |
| ENSG00000163209 | 1 | 152974223 | 152976332 | 23 | 2 | -0.2626 | 0.60357 | SPRR3 | 0.60357 | 0.67857 | 0.65322 |
| ENSG00000204193 | 9 | 113065801 | 113100127 | 150 | 35 | -0.26286 | 0.60367 | TXNDC8 | 0.60367 | 0.70445 | 0.76057 |
| ENSG00000198879 | 10 | 7200586 | 7453450 | 1037 | 70 | -0.26285 | 0.60367 | SFMBT2 | 0.60367 | 0.63596 | 0.63099 |
| ENSG00000151790 | 4 | 156775890 | 156841558 | 336 | 20 | -0.26288 | 0.60368 | TDO2 | 0.60368 | 0.60821 | 0.68449 |
| ENSG00000083828 | 19 | 58281023 | 58331307 | 168 | 20 | -0.26292 | 0.6037 | ZNF586 | 0.6037 | 0.49962 | 0.51662 |
| ENSG00000186075 | 17 | 38024417 | 38034149 | 38 | 3 | -0.26296 | 0.60371 | ZPBP2 | 0.60371 | 0.71598 | 0.78422 |
| ENSG00000172197 | 6 | 20100935 | 20212670 | 405 | 63 | -0.26305 | 0.60374 | MBOAT1 | 0.60374 | 0.5778 | 0.67867 |
| ENSG00000215845 | 1 | 161007421 | 161008780 | 2 | 1 | -0.26308 | 0.60375 | TSTD1 | 0.60375 | 0.90834 | 0.79406 |
| ENSG00000167930 | 16 | 284545 | 319942 | 128 | 19 | -0.26319 | 0.6038 | ITFG3 | 0.6038 | 0.39761 | 0.33681 |
| ENSG00000166426 | 15 | 78632666 | 78640572 | 22 | 8 | -0.26324 | 0.60382 | CRABP1 | 0.60382 | 0.6968 | 0.51769 |
| ENSG00000128274 | 22 | 43088127 | 43117304 | 190 | 26 | -0.26334 | 0.60385 | A4GALT | 0.60385 | 0.68686 | 0.73313 |
| ENSG00000006118 | 11 | 60691935 | 60704631 | 52 | 6 | -0.26343 | 0.60389 | TMEM132A | 0.60389 | 0.64576 | 0.6817 |
| ENSG00000102786 | 13 | 51928213 | 52028400 | 211 | 23 | -0.26349 | 0.60391 | INTS6 | 0.60391 | 0.55802 | 0.49007 |
| ENSG00000138107 | 10 | 104238986 | 104262482 | 83 | 5 | -0.26371 | 0.604 | ACTR1A | 0.604 | 0.68322 | 0.74421 |
| ENSG00000169783 | 15 | 77905369 | 78113242 | 735 | 46 | -0.2637 | 0.604 | LINGO1 | 0.604 | 0.61164 | 0.68932 |
| ENSG00000086548 | 19 | 42254885 | 42276113 | 57 | 10 | -0.26372 | 0.604 | CEACAM6 | 0.604 | 0.65769 | 0.66426 |
| ENSG00000125823 | 20 | 23420322 | 23425567 | 16 | 5 | -0.26405 | 0.60413 | CSTL1 | 0.60413 | 0.67123 | 0.76225 |
| ENSG00000112378 | 6 | 138409642 | 138428648 | 102 | 21 | -0.26427 | 0.60421 | PERP | 0.60421 | 0.74426 | 0.72865 |
| ENSG00000158286 | 1 | 6265535 | 6281359 | 46 | 7 | -0.26434 | 0.60424 | RNF207 | 0.60424 | 0.5864 | 0.37054 |
| ENSG00000272195 | 1 | 245132652 | 245134390 | 7 | 2 | -0.26519 | 0.60457 | RP11-156E8.1 | 0.60457 | 0.26215 | 0.053408 |
| ENSG00000117425 | 1 | 45285516 | 45308735 | 69 | 12 | -0.26536 | 0.60463 | PTCH2 | 0.60463 | 0.36775 | 0.3862 |
| ENSG00000152359 | 5 | 74969949 | 75013313 | 228 | 14 | -0.26539 | 0.60465 | POC5 | 0.60465 | 0.61696 | 0.59186 |
| ENSG00000138297 | 10 | 51592080 | 51623365 | 17 | 6 | -0.26542 | 0.60466 | TIMM23 | 0.60466 | 0.51845 | 0.5074 |
| ENSG00000162704 | 1 | 183592401 | 183604892 | 85 | 3 | -0.26547 | 0.60468 | ARPC5 | 0.60468 | 0.65359 | 0.62698 |
| ENSG00000196155 | 16 | 67311413 | 67323402 | 17 | 6 | -0.26566 | 0.60475 | PLEKHG4 | 0.60475 | 0.80801 | 0.86326 |
| ENSG00000152475 | 19 | 58878985 | 58892427 | 52 | 9 | -0.26575 | 0.60478 | ZNF837 | 0.60478 | 0.54082 | 0.64792 |
| ENSG00000141873 | 19 | 2732202 | 2740150 | 17 | 5 | -0.26579 | 0.6048 | SLC39A3 | 0.6048 | 0.81467 | 0.85755 |
| ENSG00000179038 | 11 | 66963391 | 66964638 | 1 | 1 | -0.26605 | 0.6049 | AP001885.1 | 0.6049 | 0.81943 | 0.82074 |
| ENSG00000164411 | 6 | 87992696 | 88038996 | 196 | 14 | -0.26673 | 0.60516 | GJB7 | 0.60516 | 0.56554 | 0.5734 |
| ENSG00000170523 | 12 | 52708085 | 52715182 | 58 | 6 | -0.2668 | 0.60519 | KRT83 | 0.60519 | 0.70394 | 0.78332 |
| ENSG00000256394 | 4 | 156750881 | 156787425 | 161 | 20 | -0.26708 | 0.60529 | ASIC5 | 0.60529 | 0.60018 | 0.77748 |
| ENSG00000160838 | 1 | 156890442 | 156902886 | 42 | 5 | -0.26721 | 0.60535 | LRRC71 | 0.60535 | 0.56164 | 0.39813 |
| ENSG00000089876 | 10 | 127524906 | 127585005 | 92 | 15 | -0.26746 | 0.60544 | DHX32 | 0.60544 | 0.62708 | 0.67824 |
| ENSG00000160349 | 9 | 138413284 | 138418378 | 22 | 7 | -0.26758 | 0.60549 | LCN1 | 0.60549 | 0.93319 | 0.46765 |
| ENSG00000124216 | 20 | 48599536 | 48605423 | 31 | 3 | -0.26764 | 0.60551 | SNAI1 | 0.60551 | 0.76681 | 0.66233 |
| ENSG00000171360 | 17 | 39592698 | 39597173 | 31 | 6 | -0.26768 | 0.60553 | KRT38 | 0.60553 | 0.50286 | 0.52989 |
| ENSG00000165915 | 11 | 47428683 | 47438047 | 23 | 5 | -0.26778 | 0.60557 | SLC39A13 | 0.60557 | 0.45811 | 0.60678 |
| ENSG00000169900 | 16 | 31227283 | 31228680 | 1 | 1 | -0.26787 | 0.6056 | PYDC1 | 0.6056 | 0.80714 | 0.72233 |
| ENSG00000138780 | 4 | 106629935 | 106768885 | 379 | 13 | -0.2679 | 0.60561 | GSTCD | 0.60561 | 0.60465 | 0.59834 |
| ENSG00000126787 | 14 | 55614830 | 55658396 | 156 | 20 | -0.26792 | 0.60562 | DLGAP5 | 0.60562 | 0.52966 | 0.52574 |
| ENSG00000102794 | 13 | 77522632 | 77532777 | 19 | 7 | -0.26795 | 0.60563 | IRG1 | 0.60563 | 0.80591 | 0.7456 |
| ENSG00000198750 | 7 | 74807499 | 74867509 | 10 | 4 | -0.26803 | 0.60566 | GATSL2 | 0.60566 | 0.66034 | 0.70713 |
| ENSG00000148290 | 9 | 136218610 | 136223552 | 16 | 6 | -0.26822 | 0.60573 | SURF1 | 0.60573 | 0.56458 | 0.47655 |
| ENSG00000189052 | 19 | 49547141 | 49548568 | 6 | 3 | -0.2683 | 0.60577 | CGB5 | 0.60577 | 0.12253 | 0.33873 |
| ENSG00000116977 | 1 | 236681300 | 236716281 | 307 | 12 | -0.26837 | 0.60579 | LGALS8 | 0.60579 | 0.68071 | 0.85186 |
| ENSG00000273452 | 17 | 12626199 | 12661542 | 111 | 23 | -0.26838 | 0.6058 | AC005358.1 | 0.6058 | 0.28575 | 0.023439 |
| ENSG00000196497 | 14 | 24649425 | 24658170 | 20 | 7 | -0.26842 | 0.60581 | IPO4 | 0.60581 | 0.65595 | 0.62572 |
| ENSG00000213337 | 2 | 97502348 | 97523832 | 28 | 4 | -0.2685 | 0.60584 | ANKRD39 | 0.60584 | 0.59207 | 0.51674 |
| ENSG00000196104 | 4 | 167654535 | 168155947 | 1936 | 68 | -0.2685 | 0.60584 | SPOCK3 | 0.60584 | 0.6079 | 0.68696 |
| ENSG00000084754 | 2 | 26413504 | 26467594 | 39 | 10 | -0.26868 | 0.60591 | HADHA | 0.60591 | 0.5433 | 0.55622 |
| ENSG00000120327 | 5 | 140602931 | 140605858 | 10 | 6 | -0.269 | 0.60603 | PCDHB14 | 0.60603 | 0.085579 | 0.13238 |
| ENSG00000215186 | 15 | 72947079 | 72958735 | 16 | 10 | -0.26899 | 0.60603 | GOLGA6B | 0.60603 | 0.69066 | 0.72853 |
| ENSG00000146707 | 7 | 76239303 | 76256578 | 123 | 10 | -0.26901 | 0.60604 | POMZP3 | 0.60604 | 0.52108 | 0.51288 |
| ENSG00000108591 | 17 | 17991200 | 18011285 | 40 | 7 | -0.26902 | 0.60604 | DRG2 | 0.60604 | 0.74117 | 0.753 |
| ENSG00000087502 | 12 | 29490285 | 29534122 | 102 | 13 | -0.26919 | 0.60611 | ERGIC2 | 0.60611 | 0.60836 | 0.49836 |
| ENSG00000214253 | 7 | 100882739 | 100895597 | 58 | 5 | -0.26934 | 0.60617 | FIS1 | 0.60617 | 0.77728 | 0.79826 |
| ENSG00000180483 | 20 | 29964967 | 29978406 | 48 | 11 | -0.26934 | 0.60617 | DEFB119 | 0.60617 | 0.57191 | 0.59311 |
| ENSG00000183844 | 21 | 42676139 | 42729358 | 192 | 29 | -0.26934 | 0.60617 | FAM3B | 0.60617 | 0.6023 | 0.58529 |

| | | | | | | | | | | | |
|---|---|---|---|---|---|---|---|---|---|---|---|
| ENSG00000155754 | 2 | 202352148 | 202483901 | 504 | 20 | -0.26949 | 0.60622 | ALS2CR11 | 0.60622 | 0.59351 | 0.59757 |
| ENSG00000198373 | 16 | 69796209 | 69975644 | 558 | 30 | -0.26947 | 0.60622 | WWP2 | 0.60622 | 0.59764 | 0.60212 |
| ENSG00000120210 | 9 | 5131979 | 5185668 | 178 | 6 | -0.26973 | 0.60632 | INSL6 | 0.60632 | 0.59395 | 0.59413 |
| ENSG00000269711 | 19 | 7743387 | 7747740 | 15 | 6 | -0.26988 | 0.60637 | CTD-3214H19.16 | 0.60637 | 0.90913 | 0.84481 |
| ENSG00000145888 | 5 | 151202074 | 151304403 | 224 | 15 | -0.27027 | 0.60652 | GLRA1 | 0.60652 | 0.68163 | 0.70234 |
| ENSG00000100911 | 14 | 24612574 | 24616779 | 8 | 5 | -0.27038 | 0.60657 | PSME2 | 0.60657 | 0.91872 | 0.77901 |
| ENSG00000268750 | 19 | 58331087 | 58370309 | 119 | 16 | -0.27049 | 0.60661 | CTD-2583A14.10 | 0.60661 | 0.58745 | 0.61118 |
| ENSG00000253873 | 5 | 140800762 | 140891835 | 295 | 26 | -0.27069 | 0.60669 | PCDHGA11 | 0.60669 | 0.56691 | 0.47566 |
| ENSG00000123975 | 9 | 91926113 | 91931618 | 19 | 8 | -0.27072 | 0.6067 | CKS2 | 0.6067 | 0.86748 | 0.814 |
| ENSG00000121764 | 1 | 32083287 | 32098119 | 37 | 5 | -0.27082 | 0.60673 | HCRTR1 | 0.60673 | 0.50306 | 0.52598 |
| ENSG00000080819 | 3 | 98239976 | 98312567 | 208 | 35 | -0.27087 | 0.60676 | CPOX | 0.60676 | 0.69918 | 0.74864 |
| ENSG00000173432 | 11 | 18287721 | 18291524 | 33 | 9 | -0.2709 | 0.60676 | SAA1 | 0.60676 | 0.74864 | 0.58854 |
| ENSG00000187908 | 10 | 124320181 | 124403252 | 347 | 25 | -0.27115 | 0.60686 | DMBT1 | 0.60686 | 0.62917 | 0.67172 |
| ENSG00000181616 | 11 | 5565719 | 5566779 | 7 | 2 | -0.27128 | 0.60691 | OR52H1 | 0.60691 | 0.80143 | 0.52589 |
| ENSG00000141499 | 17 | 7589389 | 7606820 | 45 | 8 | -0.27137 | 0.60695 | WRAP53 | 0.60695 | 0.6691 | 0.58319 |
| ENSG00000168385 | 2 | 242254515 | 242293442 | 132 | 21 | -0.2715 | 0.607 | 44076 | 0.607 | 0.62329 | 0.62005 |
| ENSG00000185264 | 22 | 37387163 | 37403882 | 89 | 13 | -0.27185 | 0.60713 | TEX33 | 0.60713 | 0.67557 | 0.57698 |
| ENSG00000131773 | 8 | 136469700 | 136668965 | 498 | 27 | -0.27236 | 0.60733 | KHDRBS3 | 0.60733 | 0.61893 | 0.60197 |
| ENSG00000181908 | 11 | 64216546 | 64219126 | 4 | 2 | -0.27243 | 0.60735 | AP003774.4 | 0.60735 | 0.48507 | 0.36981 |
| ENSG00000144366 | 2 | 189156396 | 189460653 | 728 | 29 | -0.27244 | 0.60736 | GULP1 | 0.60736 | 0.60742 | 0.6558 |
| ENSG00000169118 | 15 | 64457716 | 64648442 | 443 | 23 | -0.2725 | 0.60738 | CSNK1G1 | 0.60738 | 0.62354 | 0.61505 |
| ENSG00000127054 | 1 | 1246965 | 1260071 | 25 | 7 | -0.27286 | 0.60752 | CPSF3L | 0.60752 | 0.49554 | 0.52162 |
| ENSG00000104859 | 19 | 45542298 | 45574214 | 70 | 11 | -0.27294 | 0.60755 | CLASRP | 0.60755 | 0.58175 | 0.60054 |
| ENSG00000143412 | 1 | 150954493 | 150968110 | 35 | 11 | -0.27306 | 0.6076 | ANXA9 | 0.6076 | 0.72935 | 0.80215 |
| ENSG00000163930 | 3 | 52435029 | 52444366 | 6 | 2 | -0.27311 | 0.60762 | BAP1 | 0.60762 | 0.56827 | 0.60094 |
| ENSG00000058729 | 5 | 96496571 | 96518964 | 94 | 16 | -0.27318 | 0.60764 | RIOK2 | 0.60764 | 0.72817 | 0.76425 |
| ENSG00000197713 | 2 | 210867289 | 210886300 | 25 | 8 | -0.27376 | 0.60787 | RPE | 0.60787 | 0.5466 | 0.65103 |
| ENSG00000136986 | 8 | 124025404 | 124054663 | 86 | 14 | -0.27389 | 0.60792 | DERL1 | 0.60792 | 0.64016 | 0.64565 |
| ENSG00000086730 | 7 | 73613982 | 73644161 | 42 | 17 | -0.27398 | 0.60795 | LAT2 | 0.60795 | 0.35555 | 0.46344 |
| ENSG00000187715 | 3 | 127634075 | 127706514 | 199 | 23 | -0.27408 | 0.60799 | KBTBD12 | 0.60799 | 0.59402 | 0.51123 |
| ENSG00000254245 | 5 | 140723601 | 140892546 | 503 | 33 | -0.27425 | 0.60805 | PCDHGA3 | 0.60805 | 0.59048 | 0.51019 |
| ENSG00000130158 | 19 | 11309971 | 11373157 | 160 | 23 | -0.27433 | 0.60808 | DOCK6 | 0.60808 | 0.64976 | 0.35687 |
| ENSG00000115112 | 2 | 121974163 | 122042783 | 214 | 32 | -0.27436 | 0.6081 | TFCP2L1 | 0.6081 | 0.54958 | 0.62442 |
| ENSG00000116032 | 19 | 1000418 | 1009731 | 89 | 18 | -0.2744 | 0.60811 | GRIN3B | 0.60811 | 0.1635 | 0.32647 |
| ENSG00000168303 | 7 | 40165622 | 40174258 | 10 | 3 | -0.27441 | 0.60812 | MPLKIP | 0.60812 | 0.57606 | 0.67213 |
| ENSG00000178734 | 13 | 76445187 | 76457947 | 83 | 10 | -0.27443 | 0.60812 | C13orf45 | 0.60812 | 0.66323 | 0.67092 |
| ENSG00000102554 | 13 | 73629114 | 73651676 | 63 | 15 | -0.27449 | 0.60815 | KLF5 | 0.60815 | 0.78448 | 0.74291 |
| ENSG00000140521 | 15 | 89859534 | 89878092 | 55 | 8 | -0.27454 | 0.60816 | POLG | 0.60816 | 0.67348 | 0.62471 |
| ENSG00000134779 | 18 | 34359987 | 34409158 | 117 | 7 | -0.27473 | 0.60824 | TPGS2 | 0.60824 | 0.55612 | 0.5323 |
| ENSG00000113073 | 5 | 139739787 | 139754728 | 21 | 8 | -0.27477 | 0.60825 | SLC4A9 | 0.60825 | 0.73654 | 0.21599 |
| ENSG00000083720 | 5 | 41730167 | 41870621 | 219 | 14 | -0.27504 | 0.60836 | OXCT1 | 0.60836 | 0.64326 | 0.66537 |
| ENSG00000118965 | 2 | 20110021 | 20189892 | 169 | 20 | -0.27514 | 0.6084 | WDR35 | 0.6084 | 0.58246 | 0.64265 |
| ENSG00000228368 | 7 | 116605900 | 116607526 | 4 | 2 | -0.27515 | 0.6084 | AC106873.4 | 0.6084 | 0.44432 | 0.3881 |
| ENSG00000186684 | 2 | 127941696 | 127977654 | 98 | 18 | -0.27537 | 0.60848 | CYP27C1 | 0.60848 | 0.66368 | 0.77967 |
| ENSG00000197885 | 3 | 23933151 | 23988082 | 243 | 19 | -0.27535 | 0.60848 | NKIRAS1 | 0.60848 | 0.59958 | 0.6277 |
| ENSG00000170920 | 19 | 9236688 | 9237626 | 4 | 1 | -0.27576 | 0.60863 | OR7G3 | 0.60863 | 0.68923 | 0.63982 |
| ENSG00000167034 | 8 | 23536206 | 23540440 | 12 | 4 | -0.27585 | 0.60867 | NKX3-1 | 0.60867 | 0.14821 | 0.59087 |
| ENSG00000188340 | 1 | 158746472 | 158747425 | 2 | 1 | -0.27594 | 0.6087 | OR6N2 | 0.6087 | 0.43474 | 0.57559 |
| ENSG00000025772 | 20 | 43570771 | 43589127 | 22 | 5 | -0.27597 | 0.60872 | TOMM34 | 0.60872 | 0.61083 | 0.60784 |
| ENSG00000177479 | 3 | 48956254 | 49023815 | 95 | 14 | -0.27617 | 0.60879 | ARIH2 | 0.60879 | 0.54413 | 0.55278 |
| ENSG00000055917 | 2 | 20448452 | 20551995 | 261 | 17 | -0.27635 | 0.60886 | PUM2 | 0.60886 | 0.63428 | 0.63824 |
| ENSG00000261130 | 16 | 15489611 | 15609205 | 260 | 31 | -0.27644 | 0.6089 | RP11-1021N1.1 | 0.6089 | 0.57398 | 0.62485 |
| ENSG00000088356 | 20 | 30532758 | 30539895 | 8 | 2 | -0.27649 | 0.60891 | PDRG1 | 0.60891 | 0.73482 | 0.70309 |
| ENSG00000115649 | 2 | 220036619 | 220042828 | 10 | 4 | -0.27651 | 0.60892 | CNPPD1 | 0.60892 | 0.15489 | 0.20887 |
| ENSG00000170881 | 8 | 125486979 | 125500155 | 35 | 10 | -0.27677 | 0.60902 | RNF139 | 0.60902 | 0.35807 | 0.34193 |
| ENSG00000072444 | 10 | 48001603 | 48055018 | 28 | 5 | -0.27723 | 0.6092 | ASAH2C | 0.6092 | 0.4614 | 0.35928 |
| ENSG00000168070 | 11 | 64704989 | 64739557 | 86 | 11 | -0.27726 | 0.60921 | C11orf85 | 0.60921 | 0.65334 | 0.58297 |
| ENSG00000203864 | 1 | 117236734 | 117249225 | 14 | 7 | -0.27743 | 0.60927 | C1orf137 | 0.60927 | 0.28528 | 0.16998 |
| ENSG00000150045 | 12 | 9980077 | 9997606 | 46 | 7 | -0.2775 | 0.6093 | KLRF1 | 0.6093 | 0.63971 | 0.48075 |
| ENSG00000177981 | 12 | 48541571 | 48574996 | 84 | 12 | -0.27749 | 0.6093 | ASB8 | 0.6093 | 0.57227 | 0.61109 |
| ENSG00000130560 | 9 | 138824815 | 138853226 | 113 | 13 | -0.27758 | 0.60933 | UBAC1 | 0.60933 | 0.69253 | 0.78052 |
| ENSG00000253159 | 5 | 140810185 | 140892546 | 266 | 26 | -0.27763 | 0.60935 | PCDHGA12 | 0.60935 | 0.58722 | 0.47309 |
| ENSG00000174498 | 15 | 65619465 | 65670378 | 150 | 25 | -0.27788 | 0.60945 | IGDCC3 | 0.60945 | 0.67683 | 0.75179 |
| ENSG00000118702 | 20 | 35879489 | 35890238 | 26 | 6 | -0.27789 | 0.60945 | GHRH | 0.60945 | 0.17801 | 0.21493 |
| ENSG00000155085 | 6 | 109814059 | 110012420 | 811 | 17 | -0.27792 | 0.60946 | AK9 | 0.60946 | 0.63204 | 0.62556 |
| ENSG00000106327 | 7 | 100218039 | 100240402 | 54 | 9 | -0.27808 | 0.60953 | TFR2 | 0.60953 | 0.52873 | 0.47117 |
| ENSG00000205329 | 12 | 55725485 | 55726420 | 1 | 1 | -0.27828 | 0.6096 | OR6C3 | 0.6096 | 0.8651 | 0.88807 |
| ENSG00000221845 | 7 | 47694842 | 47701246 | 9 | 3 | -0.2783 | 0.60961 | C7orf65 | 0.60961 | 0.68464 | 0.68975 |
| ENSG00000206536 | 3 | 98109510 | 98110475 | 2 | 1 | -0.27835 | 0.60963 | OR5K3 | 0.60963 | 0.69041 | 0.81418 |
| ENSG00000101605 | 18 | 3066805 | 3220106 | 729 | 77 | -0.27836 | 0.60963 | MYOM1 | 0.60963 | 0.59756 | 0.6779 |
| ENSG00000230510 | 19 | 46984053 | 47104457 | 479 | 67 | -0.27866 | 0.60975 | PPP5D1 | 0.60975 | 0.59815 | 0.59561 |
| ENSG00000150787 | 11 | 112097088 | 112140678 | 140 | 7 | -0.27868 | 0.60976 | PTS | 0.60976 | 0.60576 | 0.66865 |
| ENSG00000260342 | 16 | 18799385 | 18812841 | 34 | 6 | -0.2787 | 0.60976 | RP11-1035H13.3 | 0.60976 | 0.6413 | 0.54903 |

| | | | | | | | | | | | |
|---|---|---|---|---|---|---|---|---|---|---|---|
| ENSG00000181378 | 2 | 219867568 | 219906249 | 67 | 11 | -0.27871 | 0.60977 | CCDC108 | 0.60977 | 0.66056 | 0.65899 |
| ENSG00000204889 | 17 | 39133968 | 39143387 | 91 | 6 | -0.27875 | 0.60978 | KRT40 | 0.60978 | 0.6683 | 0.60139 |
| ENSG00000091129 | 7 | 107788068 | 108097161 | 1174 | 54 | -0.27904 | 0.60989 | NRCAM | 0.60989 | 0.5663 | 0.58381 |
| ENSG00000080031 | 19 | 55692616 | 55720874 | 154 | 32 | -0.27902 | 0.60989 | PTPRH | 0.60989 | 0.86998 | 0.91853 |
| ENSG00000170017 | 3 | 105085753 | 105295744 | 754 | 35 | -0.27905 | 0.6099 | ALCAM | 0.6099 | 0.38505 | 0.36733 |
| ENSG00000136197 | 7 | 42948325 | 42951904 | 11 | 2 | -0.27909 | 0.60991 | C7orf25 | 0.60991 | 0.74194 | 0.71996 |
| ENSG00000261949 | 19 | 49927006 | 49932075 | 9 | 5 | -0.27913 | 0.60993 | GFY | 0.60993 | 0.16678 | 0.090244 |
| ENSG00000177025 | 19 | 58469805 | 58485902 | 77 | 8 | -0.27945 | 0.61005 | C19orf18 | 0.61005 | 0.46461 | 0.47303 |
| ENSG00000187730 | 1 | 1950780 | 1962192 | 29 | 6 | -0.27962 | 0.61011 | GABRD | 0.61011 | 0.56149 | 0.56073 |
| ENSG00000179088 | 12 | 103631369 | 103889749 | 703 | 34 | -0.27966 | 0.61013 | C12orf42 | 0.61013 | 0.54323 | 0.58959 |
| ENSG00000156958 | 15 | 49447853 | 49660066 | 719 | 21 | -0.27967 | 0.61013 | GALK2 | 0.61013 | 0.61138 | 0.61455 |
| ENSG00000169951 | 16 | 30565085 | 30569819 | 6 | 4 | -0.27975 | 0.61017 | ZNF764 | 0.61017 | 0.55877 | 0.85234 |
| ENSG00000159733 | 4 | 2271309 | 2420390 | 731 | 51 | -0.27997 | 0.61025 | ZFYVE28 | 0.61025 | 0.65271 | 0.68839 |
| ENSG00000171316 | 8 | 61591337 | 61779465 | 440 | 24 | -0.28 | 0.61026 | CHD7 | 0.61026 | 0.62587 | 0.67239 |
| ENSG00000170871 | 4 | 6783102 | 6885897 | 168 | 26 | -0.28009 | 0.6103 | KIAA0232 | 0.6103 | 0.68817 | 0.73443 |
| ENSG00000256812 | 16 | 55600584 | 55601599 | 1 | 1 | -0.2801 | 0.6103 | CAPNS2 | 0.6103 | 0.74158 | 0.73787 |
| ENSG00000120314 | 5 | 140044261 | 140053709 | 21 | 6 | -0.28013 | 0.61031 | WDR55 | 0.61031 | 0.69701 | 0.7631 |
| ENSG00000170558 | 18 | 25530930 | 25757410 | 588 | 42 | -0.28015 | 0.61032 | CDH2 | 0.61032 | 0.62293 | 0.67741 |
| ENSG00000086189 | 5 | 61683081 | 61699766 | 52 | 7 | -0.28021 | 0.61034 | DIMT1 | 0.61034 | 0.63636 | 0.6604 |
| ENSG00000137815 | 15 | 41700606 | 41775761 | 187 | 19 | -0.28023 | 0.61035 | RTF1 | 0.61035 | 0.64302 | 0.68347 |
| ENSG00000084628 | 1 | 31652592 | 31712401 | 281 | 14 | -0.28044 | 0.61043 | NKAIN1 | 0.61043 | 0.62956 | 0.47352 |
| ENSG00000180098 | 1 | 28879597 | 28905051 | 42 | 11 | -0.28083 | 0.61058 | TRNAU1AP | 0.61058 | 0.71947 | 0.67149 |
| ENSG00000115598 | 2 | 102803433 | 102856462 | 141 | 18 | -0.28086 | 0.61059 | IL1RL2 | 0.61059 | 0.62899 | 0.62489 |
| ENSG00000100650 | 14 | 70193617 | 70238722 | 129 | 18 | -0.28102 | 0.61065 | SRSF5 | 0.61065 | 0.66022 | 0.69577 |
| ENSG00000083307 | 8 | 102504660 | 102681954 | 602 | 50 | -0.28112 | 0.61069 | GRHL2 | 0.61069 | 0.49804 | 0.5521 |
| ENSG00000084073 | 1 | 40723779 | 40759856 | 59 | 12 | -0.2812 | 0.61072 | ZMPSTE24 | 0.61072 | 0.71778 | 0.33354 |
| ENSG00000248993 | 6 | 32905141 | 32920899 | 105 | 11 | -0.2812 | 0.61072 | XXbac-BPG181M17.5 | 0.61072 | 0.55449 | 0.55006 |
| ENSG00000259522 | 14 | 24649601 | 24659622 | 21 | 7 | -0.28122 | 0.61073 | RP11-468E2.2 | 0.61073 | 0.64023 | 0.6453 |
| ENSG00000129480 | 14 | 31915242 | 31926716 | 28 | 5 | -0.28124 | 0.61074 | DTD2 | 0.61074 | 0.74888 | 0.81716 |
| ENSG00000158545 | 16 | 88636789 | 88698374 | 253 | 27 | -0.28127 | 0.61075 | ZC3H18 | 0.61075 | 0.63171 | 0.43654 |
| ENSG00000166343 | 10 | 75183339 | 75193319 | 7 | 3 | -0.28139 | 0.61079 | MSS51 | 0.61079 | 0.76847 | 0.8172 |
| ENSG00000132109 | 11 | 4406127 | 4414926 | 42 | 10 | -0.28148 | 0.61083 | TRIM21 | 0.61083 | 0.45333 | 0.47123 |
| ENSG00000160408 | 9 | 130647600 | 130667687 | 57 | 16 | -0.28159 | 0.61087 | ST6GALNAC6 | 0.61087 | 0.65637 | 0.82135 |
| ENSG00000104129 | 15 | 41060067 | 41099675 | 82 | 12 | -0.28172 | 0.61092 | DNAJC17 | 0.61092 | 0.58349 | 0.64471 |
| ENSG00000186150 | 1 | 110655062 | 110656569 | 2 | 1 | -0.28214 | 0.61108 | UBL4B | 0.61108 | 0.80685 | 0.71734 |
| ENSG00000253797 | 13 | 52598827 | 52607736 | 12 | 3 | -0.28219 | 0.6111 | UTP14C | 0.6111 | 0.58969 | 0.54365 |
| ENSG00000174227 | 4 | 492989 | 533985 | 170 | 11 | -0.28224 | 0.61112 | PIGG | 0.61112 | 0.46136 | 0.43463 |
| ENSG00000106086 | 7 | 30067020 | 30170096 | 310 | 13 | -0.28247 | 0.61121 | PLEKHA8 | 0.61121 | 0.64761 | 0.72618 |
| ENSG00000256966 | 9 | 37512544 | 37592466 | 252 | 27 | -0.28259 | 0.61125 | RP11-613M10.8 | 0.61125 | 0.487 | 0.38023 |
| ENSG00000106689 | 9 | 126763949 | 126795580 | 97 | 12 | -0.28258 | 0.61125 | LHX2 | 0.61125 | 0.6792 | 0.25981 |
| ENSG00000141576 | 17 | 74138534 | 74236454 | 280 | 28 | -0.28273 | 0.61131 | RNF157 | 0.61131 | 0.60668 | 0.57466 |
| ENSG00000120318 | 5 | 141032968 | 141061788 | 97 | 22 | -0.28278 | 0.61133 | ARAP3 | 0.61133 | 0.5291 | 0.34095 |
| ENSG00000074800 | 1 | 8921061 | 8939308 | 33 | 5 | -0.28297 | 0.6114 | ENO1 | 0.6114 | 0.49259 | 0.45405 |
| ENSG00000115234 | 2 | 27593389 | 27599995 | 11 | 3 | -0.28299 | 0.61141 | SNX17 | 0.61141 | 0.75339 | 0.83059 |
| ENSG00000112079 | 6 | 36461669 | 36515247 | 157 | 17 | -0.28309 | 0.61145 | STK38 | 0.61145 | 0.63009 | 0.64143 |
| ENSG00000187742 | 9 | 91933421 | 91974557 | 93 | 19 | -0.28313 | 0.61146 | SECISBP2 | 0.61146 | 0.64682 | 0.67601 |
| ENSG00000138430 | 2 | 174937175 | 175113426 | 604 | 21 | -0.28341 | 0.61157 | OLA1 | 0.61157 | 0.60064 | 0.60602 |
| ENSG00000197261 | 6 | 49518113 | 49529620 | 61 | 7 | -0.28375 | 0.6117 | C6orf141 | 0.6117 | 0.46887 | 0.43736 |
| ENSG00000100565 | 14 | 77292715 | 77336645 | 185 | 40 | -0.28379 | 0.61171 | C14orf166B | 0.61171 | 0.3302 | 0.17328 |
| ENSG00000137200 | 6 | 37400995 | 37450603 | 80 | 16 | -0.28407 | 0.61182 | CMTR1 | 0.61182 | 0.5986 | 0.53954 |
| ENSG00000076201 | 3 | 47422501 | 47454931 | 37 | 5 | -0.28418 | 0.61186 | PTPN23 | 0.61186 | 0.60892 | 0.62322 |
| ENSG00000005238 | 9 | 35104109 | 35116338 | 20 | 8 | -0.28435 | 0.61193 | FAM214B | 0.61193 | 0.65468 | 0.68565 |
| ENSG00000177994 | 2 | 54557171 | 54610879 | 160 | 25 | -0.28454 | 0.612 | C2orf73 | 0.612 | 0.67095 | 0.68451 |
| ENSG00000047617 | 12 | 5641035 | 6055398 | 1598 | 119 | -0.28464 | 0.61204 | ANO2 | 0.61204 | 0.57427 | 0.58749 |
| ENSG00000196535 | 17 | 27400528 | 27507430 | 234 | 23 | -0.28465 | 0.61204 | MYO18A | 0.61204 | 0.60611 | 0.61098 |
| ENSG00000108443 | 17 | 57970447 | 58027925 | 123 | 14 | -0.28468 | 0.61206 | RPS6KB1 | 0.61206 | 0.59647 | 0.6565 |
| ENSG00000259066 | 14 | 93651318 | 93684932 | 109 | 20 | -0.28481 | 0.61211 | RP11-371E8.4 | 0.61211 | 0.72424 | 0.53175 |
| ENSG00000137731 | 11 | 117671559 | 117699413 | 121 | 25 | -0.28491 | 0.61214 | FXYD2 | 0.61214 | 0.40616 | 0.52829 |
| ENSG00000259133 | 14 | 57115683 | 57197253 | 366 | 37 | -0.2849 | 0.61214 | RP11-1085N6.3 | 0.61214 | 0.4904 | 0.32704 |
| ENSG00000114115 | 3 | 139236276 | 139258671 | 67 | 15 | -0.28503 | 0.61219 | RBP1 | 0.61219 | 0.42754 | 0.51354 |
| ENSG00000006047 | 17 | 7191571 | 7197934 | 17 | 3 | -0.28516 | 0.61224 | YBX2 | 0.61224 | 0.94061 | 0.97627 |
| ENSG00000089685 | 17 | 76210267 | 76221717 | 85 | 11 | -0.28535 | 0.61231 | BIRC5 | 0.61231 | 0.63865 | 0.78872 |
| ENSG00000187605 | 2 | 74229840 | 74335303 | 235 | 16 | -0.28548 | 0.61236 | TET3 | 0.61236 | 0.61168 | 0.70051 |
| ENSG00000141013 | 16 | 90086037 | 90111383 | 152 | 19 | -0.28558 | 0.6124 | GAS8 | 0.6124 | 0.67616 | 0.62695 |
| ENSG00000108511 | 17 | 46671639 | 46682354 | 23 | 9 | -0.2856 | 0.61241 | HOXB6 | 0.61241 | 0.7785 | 0.87658 |
| ENSG00000134504 | 18 | 24034874 | 24237365 | 700 | 92 | -0.28562 | 0.61242 | KCTD1 | 0.61242 | 0.65278 | 0.81537 |
| ENSG00000110171 | 11 | 6469843 | 6495689 | 59 | 13 | -0.28579 | 0.61248 | TRIM3 | 0.61248 | 0.8372 | 0.93962 |
| ENSG00000138785 | 4 | 106603784 | 106817143 | 554 | 15 | -0.28587 | 0.61251 | INTS12 | 0.61251 | 0.62261 | 0.62662 |
| ENSG00000214114 | 1 | 39328636 | 39347289 | 48 | 8 | -0.28591 | 0.61252 | MYCBP | 0.61252 | 0.65119 | 0.66457 |
| ENSG00000134186 | 1 | 109234945 | 109244425 | 8 | 5 | -0.28598 | 0.61255 | PRPF38B | 0.61255 | 0.82305 | 0.2866 |
| ENSG00000213190 | 1 | 151030234 | 151040970 | 20 | 6 | -0.28627 | 0.61267 | MLLT11 | 0.61267 | 0.77525 | 0.87944 |
| ENSG00000121989 | 2 | 148602086 | 148688393 | 138 | 16 | -0.28633 | 0.61269 | ACVR2A | 0.61269 | 0.63244 | 0.61272 |
| ENSG00000124232 | 20 | 43935491 | 43945803 | 30 | 7 | -0.28642 | 0.61272 | RBPJL | 0.61272 | 0.1898 | 0.29412 |

| | | | | | | | | | | | |
|---|---|---|---|---|---|---|---|---|---|---|---|
| ENSG00000110786 | 11 | 18749475 | 18814268 | 209 | 39 | -0.28649 | 0.61275 | PTPN5 | 0.61275 | 0.50372 | 0.42378 |
| ENSG00000170484 | 12 | 52959566 | 52967609 | 53 | 14 | -0.28674 | 0.61284 | KRT74 | 0.61284 | 0.72454 | 0.36057 |
| ENSG00000168065 | 11 | 64323098 | 64340347 | 21 | 3 | -0.28679 | 0.61286 | SLC22A11 | 0.61286 | 0.64708 | 0.41193 |
| ENSG00000100410 | 22 | 41855721 | 41864729 | 23 | 3 | -0.28693 | 0.61292 | PHF5A | 0.61292 | 0.58814 | 0.50518 |
| ENSG00000181009 | 11 | 5798864 | 5799897 | 2 | 1 | -0.287 | 0.61294 | OR52N5 | 0.61294 | 0.97529 | 0.94958 |
| ENSG00000116990 | 1 | 40361098 | 40367928 | 20 | 10 | -0.28717 | 0.61301 | MYCL | 0.61301 | 0.58904 | 0.63503 |
| ENSG00000213953 | 2 | 187361840 | 187365393 | 12 | 4 | -0.28729 | 0.61305 | AC018867.1 | 0.61305 | 0.79322 | 0.79918 |
| ENSG00000150977 | 12 | 123899936 | 123921264 | 80 | 19 | -0.28794 | 0.6133 | RILPL2 | 0.6133 | 0.68391 | 0.71085 |
| ENSG00000163072 | 2 | 169643049 | 169722024 | 260 | 26 | -0.28797 | 0.61332 | NOSTRIN | 0.61332 | 0.7295 | 0.72876 |
| ENSG00000197181 | 8 | 22132810 | 22215076 | 267 | 14 | -0.28818 | 0.6134 | PIWIL2 | 0.6134 | 0.62342 | 0.63773 |
| ENSG00000222018 | 21 | 35772615 | 35773370 | 2 | 1 | -0.2883 | 0.61344 | C21orf140 | 0.61344 | 0.80299 | 0.82326 |
| ENSG00000166510 | 18 | 52568740 | 52626739 | 155 | 19 | -0.28839 | 0.61347 | CCDC68 | 0.61347 | 0.4641 | 0.42968 |
| ENSG00000197753 | 6 | 35773070 | 35801651 | 98 | 10 | -0.28849 | 0.61351 | LHFPL5 | 0.61351 | 0.54652 | 0.51536 |
| ENSG00000182952 | 6 | 26538633 | 26546482 | 23 | 5 | -0.28856 | 0.61354 | HMGN4 | 0.61354 | 0.52617 | 0.54146 |
| ENSG00000187193 | 16 | 56716336 | 56718108 | 11 | 4 | -0.28892 | 0.61368 | MT1X | 0.61368 | 0.7193 | 0.50494 |
| ENSG00000125878 | 20 | 584441 | 591042 | 17 | 7 | -0.28891 | 0.61368 | TCF15 | 0.61368 | 0.82779 | 0.89263 |
| ENSG00000112304 | 6 | 24667263 | 24705293 | 168 | 22 | -0.28909 | 0.61375 | ACOT13 | 0.61375 | 0.61825 | 0.55718 |
| ENSG00000186283 | 1 | 179050512 | 179067158 | 62 | 16 | -0.28928 | 0.61382 | TOR3A | 0.61382 | 0.66352 | 0.47885 |
| ENSG00000132911 | 5 | 151771093 | 151812929 | 165 | 8 | -0.28928 | 0.61382 | NMUR2 | 0.61382 | 0.60713 | 0.68354 |
| ENSG00000163009 | 2 | 10281509 | 10351851 | 351 | 37 | -0.28973 | 0.61399 | C2orf48 | 0.61399 | 0.63138 | 0.76322 |
| ENSG00000105617 | 19 | 54658899 | 54663620 | 9 | 6 | -0.28985 | 0.61403 | LENG1 | 0.61403 | 0.13482 | 0.18427 |
| ENSG00000120896 | 8 | 22402499 | 22433301 | 81 | 24 | -0.28994 | 0.61407 | SORBS3 | 0.61407 | 0.64143 | 0.45755 |
| ENSG00000175324 | 8 | 38020839 | 38034248 | 34 | 7 | -0.29033 | 0.61422 | LSM1 | 0.61422 | 0.52161 | 0.52776 |
| ENSG00000183770 | 3 | 138663066 | 138665982 | 6 | 2 | -0.29054 | 0.6143 | FOXL2 | 0.6143 | 0.50487 | 0.54793 |
| ENSG00000169246 | 16 | 21413558 | 21459888 | 3 | 2 | -0.29059 | 0.61432 | NPIPB3 | 0.61432 | 0.89018 | 0.84302 |
| ENSG00000164188 | 5 | 36248536 | 36302216 | 180 | 21 | -0.29064 | 0.61434 | RANBP3L | 0.61434 | 0.6063 | 0.67045 |
| ENSG00000175573 | 11 | 65684279 | 65686588 | 5 | 1 | -0.29065 | 0.61434 | C11orf68 | 0.61434 | 0.86418 | 0.89103 |
| ENSG00000106829 | 9 | 82186688 | 82341658 | 353 | 28 | -0.29069 | 0.61436 | TLE4 | 0.61436 | 0.59601 | 0.60742 |
| ENSG00000170145 | 11 | 111473115 | 111601577 | 325 | 14 | -0.29076 | 0.61438 | SIK2 | 0.61438 | 0.60743 | 0.62921 |
| ENSG00000044574 | 9 | 127997132 | 128003609 | 7 | 3 | -0.29081 | 0.6144 | HSPA5 | 0.6144 | 0.28316 | 0.38728 |
| ENSG00000198924 | 10 | 115594488 | 115614142 | 17 | 5 | -0.29105 | 0.61449 | DCLRE1A | 0.61449 | 0.60737 | 0.3361 |
| ENSG00000179163 | 1 | 24171567 | 24194784 | 61 | 7 | -0.29137 | 0.61462 | FUCA1 | 0.61462 | 0.46194 | 0.58548 |
| ENSG00000120526 | 8 | 110253148 | 110346614 | 274 | 13 | -0.29147 | 0.61465 | NUDCD1 | 0.61465 | 0.6401 | 0.72852 |
| ENSG00000256269 | 11 | 118955576 | 118964259 | 18 | 3 | -0.29148 | 0.61466 | HMBS | 0.61466 | 0.60118 | 0.41272 |
| ENSG00000171163 | 1 | 249144205 | 249153343 | 11 | 2 | -0.29154 | 0.61468 | ZNF692 | 0.61468 | 0.6494 | 0.77136 |
| ENSG00000165617 | 14 | 59100685 | 59115039 | 28 | 8 | -0.29155 | 0.61469 | DACT1 | 0.61469 | 0.70258 | 0.64088 |
| ENSG00000223865 | 6 | 33043703 | 33054978 | 350 | 19 | -0.29177 | 0.61477 | HLA-DPB1 | 0.61477 | 0.58454 | 0.58144 |
| ENSG00000171773 | 19 | 17566234 | 17571763 | 27 | 5 | -0.29197 | 0.61484 | NXNL1 | 0.61484 | 0.74627 | 0.67932 |
| ENSG00000163565 | 1 | 158969758 | 159024945 | 279 | 19 | -0.292 | 0.61485 | IFI16 | 0.61485 | 0.63175 | 0.67934 |
| ENSG00000151849 | 13 | 25457171 | 25497018 | 140 | 21 | -0.29208 | 0.61489 | CENPJ | 0.61489 | 0.75449 | 0.19165 |
| ENSG00000185940 | 11 | 1651033 | 1652160 | 17 | 5 | -0.29222 | 0.61494 | KRTAP5-5 | 0.61494 | 0.91778 | 0.75185 |
| ENSG00000145041 | 3 | 51433298 | 51534010 | 163 | 16 | -0.29224 | 0.61495 | VPRBP | 0.61495 | 0.65834 | 0.6618 |
| ENSG00000133808 | 11 | 12297627 | 12380691 | 453 | 34 | -0.29226 | 0.61496 | MICALCL | 0.61496 | 0.68101 | 0.75616 |
| ENSG00000198580 | 7 | 6713376 | 6715991 | 12 | 5 | -0.29236 | 0.61499 | AC073343.1 | 0.61499 | 0.39162 | 0.29558 |
| ENSG00000102924 | 16 | 49311828 | 49315742 | 12 | 5 | -0.29234 | 0.61499 | CBLN1 | 0.61499 | 0.63927 | 0.22776 |
| ENSG00000072422 | 10 | 62629196 | 62761198 | 314 | 33 | -0.29237 | 0.615 | RHOBTB1 | 0.615 | 0.74028 | 0.7637 |
| ENSG00000158793 | 1 | 161087876 | 161095235 | 12 | 5 | -0.2924 | 0.61501 | NIT1 | 0.61501 | 0.49855 | 0.62621 |
| ENSG00000188931 | 1 | 161334521 | 161337664 | 18 | 1 | -0.29243 | 0.61502 | C1orf192 | 0.61502 | 0.66642 | 0.6711 |
| ENSG00000164087 | 3 | 52109269 | 52188706 | 119 | 18 | -0.29259 | 0.61508 | POC1A | 0.61508 | 0.66713 | 0.66795 |
| ENSG00000122390 | 16 | 3493611 | 3536963 | 233 | 22 | -0.29261 | 0.61509 | NAA60 | 0.61509 | 0.51044 | 0.58108 |
| ENSG00000112539 | 6 | 165693153 | 165723096 | 107 | 14 | -0.29267 | 0.61511 | C6orf118 | 0.61511 | 0.6899 | 0.67531 |
| ENSG00000173611 | 9 | 127704887 | 127905785 | 459 | 27 | -0.2927 | 0.61513 | SCAI | 0.61513 | 0.632 | 0.61924 |
| ENSG00000174483 | 11 | 66278077 | 66301098 | 53 | 7 | -0.29281 | 0.61517 | BBS1 | 0.61517 | 0.68386 | 0.73515 |
| ENSG00000148965 | 11 | 18252896 | 18258440 | 17 | 5 | -0.2929 | 0.6152 | SAA4 | 0.6152 | 0.61985 | 0.64566 |
| ENSG00000172782 | 17 | 72873428 | 72889708 | 53 | 12 | -0.29327 | 0.61534 | FADS6 | 0.61534 | 0.83859 | 0.76862 |
| ENSG00000141096 | 16 | 68009566 | 68014732 | 8 | 4 | -0.29329 | 0.61535 | DPEP3 | 0.61535 | 0.88559 | 0.90202 |
| ENSG00000113594 | 5 | 38475065 | 38608456 | 247 | 25 | -0.29343 | 0.6154 | LIFR | 0.6154 | 0.57126 | 0.71951 |
| ENSG00000197323 | 1 | 114935399 | 115053781 | 358 | 22 | -0.2935 | 0.61543 | TRIM33 | 0.61543 | 0.61103 | 0.65907 |
| ENSG00000188647 | 9 | 72324438 | 72374875 | 88 | 14 | -0.29366 | 0.61549 | PTAR1 | 0.61549 | 0.71553 | 0.75654 |
| ENSG00000187231 | 2 | 179966419 | 180129517 | 506 | 24 | -0.29421 | 0.6157 | SESTD1 | 0.6157 | 0.65916 | 0.64853 |
| ENSG00000149548 | 11 | 124824017 | 124911385 | 316 | 19 | -0.29428 | 0.61573 | CCDC15 | 0.61573 | 0.50481 | 0.4719 |
| ENSG00000156206 | 15 | 81299374 | 81441516 | 493 | 39 | -0.29438 | 0.61577 | C15orf26 | 0.61577 | 0.65828 | 0.67742 |
| ENSG00000158315 | 1 | 39351479 | 39407471 | 215 | 32 | -0.29463 | 0.61586 | RHBDL2 | 0.61586 | 0.66378 | 0.64879 |
| ENSG00000168234 | 18 | 21572737 | 21715574 | 448 | 47 | -0.29469 | 0.61589 | TTC39C | 0.61589 | 0.62183 | 0.57046 |
| ENSG00000188674 | 2 | 209030067 | 209054797 | 129 | 22 | -0.29472 | 0.6159 | C2orf80 | 0.6159 | 0.7113 | 0.62981 |
| ENSG00000135902 | 2 | 233390703 | 233401377 | 33 | 7 | -0.29482 | 0.61594 | CHRND | 0.61594 | 0.25676 | 0.16734 |
| ENSG00000243335 | 7 | 66093868 | 66276446 | 742 | 23 | -0.29484 | 0.61594 | KCTD7 | 0.61594 | 0.63282 | 0.62819 |
| ENSG00000116663 | 1 | 11724181 | 11734411 | 46 | 9 | -0.29488 | 0.61596 | FBXO6 | 0.61596 | 0.57072 | 0.37728 |
| ENSG00000135472 | 12 | 50260679 | 50298000 | 160 | 30 | -0.29488 | 0.61596 | FAIM2 | 0.61596 | 0.61694 | 0.59207 |
| ENSG00000198105 | 10 | 38091751 | 38147034 | 217 | 13 | -0.29492 | 0.61597 | ZNF248 | 0.61597 | 0.61476 | 0.58462 |
| ENSG00000177275 | 1 | 248097071 | 248098057 | 1 | 1 | -0.29499 | 0.616 | OR2AJ1 | 0.616 | 0.47233 | 0.62671 |
| ENSG00000120156 | 9 | 27109139 | 27230173 | 787 | 63 | -0.2951 | 0.61604 | TEK | 0.61604 | 0.60791 | 0.65077 |
| ENSG00000211455 | 12 | 27396901 | 27478892 | 259 | 30 | -0.29512 | 0.61605 | STK38L | 0.61605 | 0.59694 | 0.55388 |

| | | | | | | | | | | | |
|---|---|---|---|---|---|---|---|---|---|---|---|
| ENSG00000147647 | 8 | 105342552 | 105479281 | 467 | 56 | -0.29521 | 0.61608 | DPYS | 0.61608 | 0.46973 | 0.31032 |
| ENSG00000163600 | 2 | 204801471 | 204826300 | 76 | 17 | -0.29531 | 0.61612 | ICOS | 0.61612 | 0.61831 | 0.64479 |
| ENSG00000175606 | 8 | 74884672 | 74895018 | 45 | 6 | -0.29547 | 0.61618 | TMEM70 | 0.61618 | 0.7051 | 0.75747 |
| ENSG00000131748 | 17 | 37793318 | 37819737 | 75 | 9 | -0.29552 | 0.6162 | STARD3 | 0.6162 | 0.68878 | 0.63346 |
| ENSG00000155926 | 8 | 134048973 | 134115298 | 328 | 25 | -0.29562 | 0.61624 | SLA | 0.61624 | 0.71293 | 0.6726 |
| ENSG00000165283 | 9 | 35099888 | 35103154 | 6 | 2 | -0.29565 | 0.61625 | STOML2 | 0.61625 | 0.76099 | 0.72739 |
| ENSG00000119812 | 2 | 33808725 | 33824449 | 49 | 11 | -0.29607 | 0.61641 | FAM98A | 0.61641 | 0.66773 | 0.7051 |
| ENSG00000169432 | 2 | 167051695 | 167232503 | 693 | 47 | -0.29625 | 0.61648 | SCN9A | 0.61648 | 0.60917 | 0.55287 |
| ENSG00000040633 | 17 | 7138347 | 7143041 | 5 | 3 | -0.29637 | 0.61653 | PHF23 | 0.61653 | 0.82883 | 0.75817 |
| ENSG00000196345 | 3 | 44596685 | 44624975 | 97 | 13 | -0.2964 | 0.61654 | ZKSCAN7 | 0.61654 | 0.5823 | 0.5739 |
| ENSG00000141447 | 18 | 21742008 | 21977844 | 666 | 75 | -0.2965 | 0.61658 | OSBPL1A | 0.61658 | 0.61934 | 0.68339 |
| ENSG00000157315 | 16 | 69377151 | 69385712 | 20 | 8 | -0.29662 | 0.61662 | TMED6 | 0.61662 | 0.34558 | 0.31505 |
| ENSG00000169218 | 1 | 38076951 | 38100595 | 76 | 15 | -0.29669 | 0.61665 | RSPO1 | 0.61665 | 0.65432 | 0.75848 |
| ENSG00000173210 | 5 | 148521046 | 148640105 | 271 | 28 | -0.29675 | 0.61667 | ABLIM3 | 0.61667 | 0.68129 | 0.77695 |
| ENSG00000196800 | 5 | 147549296 | 147554961 | 24 | 4 | -0.29716 | 0.61683 | SPINK14 | 0.61683 | 0.42843 | 0.40853 |
| ENSG00000057252 | 1 | 179262925 | 179327815 | 339 | 22 | -0.29725 | 0.61686 | SOAT1 | 0.61686 | 0.68377 | 0.63044 |
| ENSG00000172530 | 16 | 87982850 | 88110924 | 760 | 23 | -0.29728 | 0.61688 | BANP | 0.61688 | 0.63933 | 0.69223 |
| ENSG00000128654 | 2 | 177134123 | 177202753 | 109 | 14 | -0.29735 | 0.6169 | MTX2 | 0.6169 | 0.64428 | 0.61722 |
| ENSG00000051382 | 3 | 138372860 | 138553780 | 181 | 31 | -0.29745 | 0.61694 | PIK3CB | 0.61694 | 0.6343 | 0.63334 |
| ENSG00000149380 | 11 | 73946846 | 74022702 | 242 | 22 | -0.29766 | 0.61702 | P4HA3 | 0.61702 | 0.66067 | 0.63313 |
| ENSG00000181333 | 11 | 93754527 | 93846917 | 222 | 21 | -0.29782 | 0.61708 | HEPHL1 | 0.61708 | 0.59978 | 0.6189 |
| ENSG00000233438 | 2 | 27760253 | 27790011 | 92 | 11 | -0.29817 | 0.61721 | AC109829.1 | 0.61721 | 0.53482 | 0.51372 |
| ENSG00000088386 | 13 | 99336055 | 99404908 | 249 | 52 | -0.29819 | 0.61722 | SLC15A1 | 0.61722 | 0.67737 | 0.73666 |
| ENSG00000187266 | 19 | 11488236 | 11495018 | 4 | 2 | -0.29836 | 0.61728 | EPOR | 0.61728 | 0.52067 | 0.68564 |
| ENSG00000163126 | 2 | 97490263 | 97523671 | 52 | 6 | -0.29839 | 0.6173 | ANKRD23 | 0.6173 | 0.59173 | 0.51697 |
| ENSG00000196900 | 5 | 125967414 | 125971979 | 18 | 3 | -0.29862 | 0.61739 | C5orf48 | 0.61739 | 0.27852 | 0.25077 |
| ENSG00000183323 | 5 | 68576002 | 68628636 | 166 | 15 | -0.2987 | 0.61742 | CCDC125 | 0.61742 | 0.68043 | 0.6945 |
| ENSG00000128973 | 15 | 68499330 | 68549549 | 110 | 7 | -0.29883 | 0.61746 | CLN6 | 0.61746 | 0.51078 | 0.49222 |
| ENSG00000187796 | 9 | 139256355 | 139268133 | 45 | 12 | -0.29899 | 0.61753 | CARD9 | 0.61753 | 0.35426 | 0.37335 |
| ENSG00000178665 | 7 | 55955169 | 56009918 | 195 | 27 | -0.29903 | 0.61754 | ZNF713 | 0.61754 | 0.68346 | 0.73953 |
| ENSG00000184924 | 2 | 25012603 | 25016251 | 13 | 5 | -0.29917 | 0.6176 | PTRHD1 | 0.6176 | 0.70434 | 0.65958 |
| ENSG00000179023 | 1 | 18807424 | 18812478 | 27 | 6 | -0.29927 | 0.61763 | KLHDC7A | 0.61763 | 0.76445 | 0.87048 |
| ENSG00000172939 | 3 | 38206580 | 38296979 | 123 | 30 | -0.2993 | 0.61764 | OXSR1 | 0.61764 | 0.66414 | 0.639 |
| ENSG00000142541 | 19 | 49990811 | 49995565 | 14 | 3 | -0.29932 | 0.61765 | RPL13A | 0.61765 | 0.4792 | 0.4199 |
| ENSG00000165813 | 10 | 115880621 | 115933979 | 127 | 13 | -0.29958 | 0.61775 | C10orf118 | 0.61775 | 0.65486 | 0.66837 |
| ENSG00000143155 | 1 | 168148171 | 168169950 | 65 | 7 | -0.29968 | 0.61779 | TIPRL | 0.61779 | 0.56807 | 0.35106 |
| ENSG00000153044 | 5 | 68485375 | 68506184 | 55 | 5 | -0.29972 | 0.6178 | CENPH | 0.6178 | 0.66116 | 0.70055 |
| ENSG00000165496 | 14 | 47120222 | 47121028 | 2 | 1 | -0.29986 | 0.61786 | RPL10L | 0.61786 | 0.52349 | 0.53375 |
| ENSG00000148803 | 10 | 135168658 | 135171529 | 9 | 2 | -0.29991 | 0.61788 | FUOM | 0.61788 | 0.56741 | 0.65345 |
| ENSG00000185379 | 17 | 33426811 | 33448541 | 38 | 13 | -0.29995 | 0.61789 | RAD51D | 0.61789 | 0.54645 | 0.50872 |
| ENSG00000183793 | 16 | 15457516 | 15474904 | 39 | 12 | -0.3 | 0.61791 | NPIPA5 | 0.61791 | 0.45776 | 0.35213 |
| ENSG00000105613 | 19 | 12944765 | 12985765 | 103 | 10 | -0.30002 | 0.61792 | MAST1 | 0.61792 | 0.5111 | 0.50776 |
| ENSG00000025770 | 22 | 50946645 | 50961901 | 70 | 15 | -0.3003 | 0.61803 | NCAPH2 | 0.61803 | 0.56114 | 0.86164 |
| ENSG00000268614 | 19 | 7598481 | 7604945 | 10 | 5 | -0.30051 | 0.61811 | CTD-2207O23.10 | 0.61811 | 0.7432 | 0.82036 |
| ENSG00000182107 | 14 | 61744088 | 61748558 | 12 | 6 | -0.3006 | 0.61814 | TMEM30B | 0.61814 | 0.63898 | 0.66336 |
| ENSG00000116704 | 1 | 67465015 | 67519782 | 181 | 17 | -0.3008 | 0.61822 | SLC35D1 | 0.61822 | 0.54794 | 0.55391 |
| ENSG00000133019 | 1 | 239549865 | 240078750 | 1580 | 115 | -0.30092 | 0.61826 | CHRM3 | 0.61826 | 0.68839 | 0.72664 |
| ENSG00000148700 | 10 | 111756126 | 111895323 | 265 | 21 | -0.30097 | 0.61828 | ADD3 | 0.61828 | 0.63418 | 0.69726 |
| ENSG00000214087 | 17 | 79648204 | 79650954 | 13 | 3 | -0.30103 | 0.6183 | ARL16 | 0.6183 | 0.31321 | 0.28994 |
| ENSG00000069956 | 15 | 52244303 | 52358462 | 339 | 28 | -0.30119 | 0.61836 | MAPK6 | 0.61836 | 0.62074 | 0.6576 |
| ENSG00000144852 | 3 | 119499331 | 119537332 | 152 | 25 | -0.30145 | 0.61847 | NR1I2 | 0.61847 | 0.56719 | 0.43146 |
| ENSG00000173762 | 17 | 80272744 | 80275478 | 11 | 5 | -0.3015 | 0.61848 | CD7 | 0.61848 | 0.51 | 0.6279 |
| ENSG00000143954 | 2 | 79252812 | 79255631 | 23 | 6 | -0.30167 | 0.61855 | REG3G | 0.61855 | 0.51978 | 0.70657 |
| ENSG00000103353 | 16 | 23568392 | 23585710 | 58 | 9 | -0.30172 | 0.61857 | UBFD1 | 0.61857 | 0.73927 | 0.82092 |
| ENSG00000141337 | 17 | 66255323 | 66418872 | 518 | 68 | -0.30184 | 0.61861 | ARSG | 0.61861 | 0.56523 | 0.56569 |
| ENSG00000010803 | 1 | 41492872 | 41707826 | 682 | 27 | -0.30188 | 0.61863 | SCMH1 | 0.61863 | 0.61441 | 0.6296 |
| ENSG00000148942 | 11 | 26688566 | 26744974 | 211 | 37 | -0.30192 | 0.61864 | SLC5A12 | 0.61864 | 0.6457 | 0.69422 |
| ENSG00000166024 | 10 | 99894387 | 100004654 | 234 | 20 | -0.30197 | 0.61866 | R3HCC1L | 0.61866 | 0.59928 | 0.55885 |
| ENSG00000170142 | 3 | 23847394 | 23932807 | 220 | 28 | -0.3021 | 0.61871 | UBE2E1 | 0.61871 | 0.58456 | 0.5217 |
| ENSG00000174032 | 13 | 45967451 | 45992590 | 70 | 11 | -0.30219 | 0.61874 | SLC25A30 | 0.61874 | 0.51034 | 0.53822 |
| ENSG00000132481 | 17 | 73870242 | 73874656 | 13 | 3 | -0.30222 | 0.61876 | TRIM47 | 0.61876 | 0.47101 | 0.29088 |
| ENSG00000142178 | 21 | 44834395 | 44847008 | 58 | 8 | -0.30233 | 0.6188 | SIK1 | 0.6188 | 0.60672 | 0.67696 |
| ENSG00000114054 | 3 | 135969148 | 136056738 | 162 | 20 | -0.30244 | 0.61884 | PCCB | 0.61884 | 0.62818 | 0.66847 |
| ENSG00000166262 | 15 | 49619159 | 49913128 | 1075 | 27 | -0.30245 | 0.61884 | FAM227B | 0.61884 | 0.60735 | 0.60107 |
| ENSG00000117154 | 1 | 18434240 | 18704977 | 1134 | 137 | -0.30251 | 0.61887 | IGSF21 | 0.61887 | 0.63586 | 0.62962 |
| ENSG00000166454 | 16 | 81069452 | 81080963 | 22 | 5 | -0.3028 | 0.61898 | ATMIN | 0.61898 | 0.76221 | 0.76532 |
| ENSG00000188760 | 2 | 220408385 | 220415317 | 14 | 4 | -0.30292 | 0.61903 | TMEM198 | 0.61903 | 0.6903 | 0.64028 |
| ENSG00000131944 | 19 | 33463115 | 33469128 | 26 | 7 | -0.30311 | 0.6191 | C19orf40 | 0.6191 | 0.70921 | 0.59707 |
| ENSG00000131233 | 1 | 39330175 | 39347289 | 47 | 8 | -0.30335 | 0.61919 | GJA9 | 0.61919 | 0.63815 | 0.66079 |
| ENSG00000109046 | 17 | 25621102 | 25640657 | 56 | 8 | -0.30347 | 0.61924 | WSB1 | 0.61924 | 0.4942 | 0.46561 |
| ENSG00000257008 | 17 | 72363546 | 72368761 | 16 | 6 | -0.30361 | 0.61929 | GPR142 | 0.61929 | 0.94556 | 0.88864 |
| ENSG00000268310 | 17 | 76210462 | 76220740 | 79 | 11 | -0.30375 | 0.61934 | AC087645.1 | 0.61934 | 0.64143 | 0.80256 |
| ENSG00000064195 | 17 | 48067369 | 48072588 | 25 | 7 | -0.30392 | 0.61941 | DLX3 | 0.61941 | 0.78741 | 0.75619 |

| | | | | | | | | | | | |
|---|---|---|---|---|---|---|---|---|---|---|---|
| ENSG00000137875 | 15 | 52401460 | 52404972 | 13 | 3 | -0.30422 | 0.61952 | BCL2L10 | 0.61952 | 0.52249 | 0.5327 |
| ENSG00000155970 | 8 | 16884747 | 16980153 | 358 | 30 | -0.30451 | 0.61963 | MICU3 | 0.61963 | 0.63706 | 0.63916 |
| ENSG00000114626 | 3 | 127391778 | 127399768 | 12 | 5 | -0.30453 | 0.61964 | ABTB1 | 0.61964 | 0.82546 | 0.80376 |
| ENSG00000171865 | 2 | 3592383 | 3606206 | 46 | 12 | -0.30464 | 0.61968 | RNASEH1 | 0.61968 | 0.34715 | 0.21263 |
| ENSG00000063244 | 19 | 56165512 | 56186081 | 61 | 12 | -0.30556 | 0.62003 | U2AF2 | 0.62003 | 0.36146 | 0.26534 |
| ENSG00000108001 | 10 | 131633547 | 131762105 | 358 | 86 | -0.30567 | 0.62007 | EBF3 | 0.62007 | 0.65058 | 0.58042 |
| ENSG00000102890 | 16 | 67233014 | 67237932 | 13 | 3 | -0.3057 | 0.62008 | ELMO3 | 0.62008 | 0.62864 | 0.71838 |
| ENSG00000074219 | 19 | 49843852 | 49865714 | 59 | 7 | -0.30577 | 0.62011 | TEAD2 | 0.62011 | 0.599 | 0.58081 |
| ENSG00000160447 | 9 | 131464802 | 131483197 | 18 | 10 | -0.30587 | 0.62015 | PKN3 | 0.62015 | 0.49195 | 0.066612 |
| ENSG00000181396 | 17 | 80347099 | 80376513 | 108 | 20 | -0.30593 | 0.62017 | OGFOD3 | 0.62017 | 0.48836 | 0.64626 |
| ENSG00000178257 | 16 | 11367144 | 11367452 | 1 | 1 | -0.30601 | 0.6202 | PRM3 | 0.6202 | 0.57668 | 0.4348 |
| ENSG00000212807 | 7 | 143929004 | 143929936 | 1 | 1 | -0.30627 | 0.6203 | OR2A42 | 0.6203 | 0.52098 | 0.31006 |
| ENSG00000135678 | 12 | 69235977 | 69365350 | 450 | 59 | -0.30626 | 0.6203 | CPM | 0.6203 | 0.63638 | 0.66959 |
| ENSG00000157578 | 21 | 40777770 | 40817731 | 119 | 10 | -0.30633 | 0.62032 | LCA5L | 0.62032 | 0.6832 | 0.65326 |
| ENSG00000236624 | 1 | 45959538 | 45965751 | 13 | 4 | -0.3064 | 0.62035 | CCDC163P | 0.62035 | 0.68389 | 0.57387 |
| ENSG00000164244 | 5 | 126853301 | 126890781 | 125 | 10 | -0.30641 | 0.62035 | PRRC1 | 0.62035 | 0.57988 | 0.55909 |
| ENSG00000121417 | 19 | 58141761 | 58154147 | 90 | 5 | -0.30647 | 0.62038 | ZNF211 | 0.62038 | 0.68098 | 0.63379 |
| ENSG00000198366 | 6 | 26020718 | 26021186 | 1 | 1 | -0.30653 | 0.6204 | HIST1H3A | 0.6204 | 0.51792 | 0.34862 |
| ENSG00000204427 | 6 | 31654726 | 31671221 | 43 | 11 | -0.30691 | 0.62054 | ABHD16A | 0.62054 | 0.59273 | 0.59859 |
| ENSG00000183723 | 16 | 66648653 | 66730610 | 226 | 15 | -0.30709 | 0.62061 | CMTM4 | 0.62061 | 0.59386 | 0.62038 |
| ENSG00000137265 | 6 | 391739 | 411447 | 84 | 19 | -0.3072 | 0.62065 | IRF4 | 0.62065 | 0.60538 | 0.79843 |
| ENSG00000183828 | 14 | 105639275 | 105647660 | 7 | 3 | -0.30726 | 0.62068 | NUDT14 | 0.62068 | 0.80648 | 0.76606 |
| ENSG00000177301 | 1 | 111136202 | 111174096 | 112 | 13 | -0.30741 | 0.62074 | KCNA2 | 0.62074 | 0.6058 | 0.58322 |
| ENSG00000143248 | 1 | 163080911 | 163291577 | 936 | 54 | -0.3076 | 0.62081 | RGS5 | 0.62081 | 0.63865 | 0.65116 |
| ENSG00000113273 | 5 | 78073032 | 78281910 | 890 | 43 | -0.3077 | 0.62085 | ARSB | 0.62085 | 0.51448 | 0.54236 |
| ENSG00000154645 | 21 | 19273580 | 19639690 | 1403 | 123 | -0.30776 | 0.62087 | CHODL | 0.62087 | 0.67747 | 0.72946 |
| ENSG00000135222 | 4 | 70820974 | 70826725 | 13 | 4 | -0.30779 | 0.62088 | CSN2 | 0.62088 | 0.65855 | 0.65689 |
| ENSG00000120913 | 8 | 22435792 | 22455538 | 63 | 14 | -0.30793 | 0.62093 | PDLIM2 | 0.62093 | 0.80253 | 0.81357 |
| ENSG00000120694 | 13 | 31710762 | 31736525 | 73 | 10 | -0.30794 | 0.62093 | HSPH1 | 0.62093 | 0.7081 | 0.76628 |
| ENSG00000119669 | 14 | 77490888 | 77495034 | 9 | 4 | -0.30808 | 0.62099 | IRF2BPL | 0.62099 | 0.53067 | 0.47983 |
| ENSG00000106078 | 7 | 51083909 | 51384515 | 696 | 42 | -0.30811 | 0.621 | COBL | 0.621 | 0.63303 | 0.56598 |
| ENSG00000113851 | 3 | 3190676 | 3221394 | 136 | 20 | -0.3084 | 0.62111 | CRBN | 0.62111 | 0.44849 | 0.53726 |
| ENSG00000168916 | 5 | 123972608 | 124084500 | 366 | 44 | -0.30844 | 0.62113 | ZNF608 | 0.62113 | 0.6344 | 0.46785 |
| ENSG00000180287 | 1 | 242246288 | 242687998 | 2031 | 112 | -0.30848 | 0.62114 | PLD5 | 0.62114 | 0.65106 | 0.69121 |
| ENSG00000172831 | 16 | 66968347 | 66978999 | 12 | 4 | -0.30847 | 0.62114 | CES2 | 0.62114 | 0.58515 | 0.46015 |
| ENSG00000182611 | 6 | 27782112 | 27782607 | 4 | 1 | -0.30857 | 0.62118 | HIST1H2AJ | 0.62118 | 0.64197 | 0.71499 |
| ENSG00000089012 | 20 | 1609798 | 1638425 | 119 | 17 | -0.30888 | 0.62129 | SIRPG | 0.62129 | 0.69944 | 0.77301 |
| ENSG00000158483 | 11 | 71498556 | 71512282 | 93 | 9 | -0.30904 | 0.62135 | FAM86C1 | 0.62135 | 0.59548 | 0.62438 |
| ENSG00000165516 | 14 | 50234326 | 50249909 | 23 | 6 | -0.30909 | 0.62137 | KLHDC2 | 0.62137 | 0.7455 | 0.61208 |
| ENSG00000261796 | 3 | 128806418 | 128879879 | 103 | 17 | -0.3091 | 0.62138 | ISY1-RAB43 | 0.62138 | 0.64155 | 0.67109 |
| ENSG00000165983 | 10 | 16478942 | 16555736 | 319 | 19 | -0.30917 | 0.6214 | PTER | 0.6214 | 0.66182 | 0.48469 |
| ENSG00000145216 | 4 | 54243810 | 55161439 | 3164 | 150 | -0.30922 | 0.62142 | FIP1L1 | 0.62142 | 0.62137 | 0.61742 |
| ENSG00000205442 | 9 | 24542950 | 24545944 | 7 | 2 | -0.30928 | 0.62144 | IZUMO3 | 0.62144 | 0.62066 | 0.68402 |
| ENSG00000204866 | 19 | 46646363 | 46664556 | 50 | 9 | -0.30926 | 0.62144 | IGFL2 | 0.62144 | 0.60864 | 0.67719 |
| ENSG00000182156 | 17 | 77704681 | 77716021 | 52 | 12 | -0.30933 | 0.62147 | ENPP7 | 0.62147 | 0.72697 | 0.69725 |
| ENSG00000166340 | 11 | 6634000 | 6640692 | 13 | 5 | -0.3094 | 0.62149 | TPP1 | 0.62149 | 0.087846 | 0.07732 |
| ENSG00000168748 | 16 | 66878282 | 66888056 | 15 | 4 | -0.30968 | 0.6216 | CA7 | 0.6216 | 0.43105 | 0.45664 |
| ENSG00000124541 | 6 | 42989383 | 43001894 | 40 | 8 | -0.30971 | 0.62161 | RRP36 | 0.62161 | 0.68807 | 0.66978 |
| ENSG00000177076 | 9 | 19408925 | 19452018 | 142 | 12 | -0.30975 | 0.62162 | ACER2 | 0.62162 | 0.61686 | 0.70661 |
| ENSG00000140612 | 15 | 85212775 | 85259947 | 119 | 22 | -0.30982 | 0.62165 | SEC11A | 0.62165 | 0.64359 | 0.64254 |
| ENSG00000168746 | 20 | 43080624 | 43093984 | 34 | 10 | -0.30988 | 0.62168 | C20orf62 | 0.62168 | 0.42319 | 0.24788 |
| ENSG00000175518 | 11 | 5535623 | 5537935 | 9 | 2 | -0.3101 | 0.62176 | UBQLNL | 0.62176 | 0.69315 | 0.65553 |
| ENSG00000140691 | 16 | 31469401 | 31478487 | 16 | 7 | -0.31024 | 0.62181 | ARMC5 | 0.62181 | 0.78733 | 0.7912 |
| ENSG00000145147 | 4 | 20254883 | 20622184 | 1124 | 105 | -0.31049 | 0.62191 | SLIT2 | 0.62191 | 0.5333 | 0.33209 |
| ENSG00000052795 | 4 | 159690290 | 159829201 | 249 | 19 | -0.31049 | 0.62191 | FNIP2 | 0.62191 | 0.60187 | 0.61792 |
| ENSG00000168612 | 20 | 44509866 | 44513905 | 13 | 6 | -0.31077 | 0.62201 | ZSWIM1 | 0.62201 | 0.84991 | 0.89523 |
| ENSG00000084072 | 1 | 40157854 | 40229586 | 239 | 44 | -0.31091 | 0.62207 | PPIE | 0.62207 | 0.49286 | 0.43551 |
| ENSG00000177575 | 12 | 7623409 | 7656489 | 87 | 18 | -0.31092 | 0.62207 | CD163 | 0.62207 | 0.54521 | 0.7174 |
| ENSG00000204634 | 2 | 101624079 | 101869328 | 679 | 40 | -0.31116 | 0.62216 | TBC1D8 | 0.62216 | 0.63587 | 0.63656 |
| ENSG00000267942 | 12 | 66291518 | 66317967 | 44 | 11 | -0.31128 | 0.62221 | AC090673.2 | 0.62221 | 0.59105 | 0.59919 |
| ENSG00000204396 | 6 | 31733367 | 31745108 | 40 | 17 | -0.31143 | 0.62226 | VWA7 | 0.62226 | 0.62714 | 0.60088 |
| ENSG00000269279 | 1 | 116915890 | 116926718 | 28 | 7 | -0.31148 | 0.62228 | AL136376.1 | 0.62228 | 0.66688 | 0.64441 |
| ENSG00000268643 | 19 | 42738735 | 42759284 | 32 | 4 | -0.31157 | 0.62232 | AC006486.9 | 0.62232 | 0.64845 | 0.5636 |
| ENSG00000155792 | 8 | 120885957 | 121063152 | 884 | 31 | -0.31213 | 0.62253 | DEPTOR | 0.62253 | 0.6154 | 0.63963 |
| ENSG00000146013 | 5 | 137588068 | 137610360 | 57 | 13 | -0.31217 | 0.62254 | GFRA3 | 0.62254 | 0.49246 | 0.52178 |
| ENSG00000127337 | 12 | 69753483 | 69784576 | 161 | 14 | -0.31221 | 0.62256 | YEATS4 | 0.62256 | 0.62472 | 0.60245 |
| ENSG00000140153 | 14 | 102605840 | 102691184 | 173 | 28 | -0.3122 | 0.62256 | WDR20 | 0.62256 | 0.5592 | 0.51167 |
| ENSG00000187758 | 4 | 100197524 | 100212185 | 37 | 9 | -0.31231 | 0.6226 | ADH1A | 0.6226 | 0.37418 | 0.39714 |
| ENSG00000188738 | 2 | 186603355 | 186698017 | 247 | 11 | -0.31238 | 0.62262 | FSIP2 | 0.62262 | 0.64711 | 0.63406 |
| ENSG00000259202 | 15 | 67435072 | 67439277 | 10 | 4 | -0.31239 | 0.62263 | RP11-342M21.2 | 0.62263 | 0.57265 | 0.20604 |
| ENSG00000185352 | 13 | 96743093 | 97485671 | 1976 | 71 | -0.31243 | 0.62264 | HS6ST3 | 0.62264 | 0.62981 | 0.65394 |
| ENSG00000143473 | 1 | 210856555 | 211307457 | 1656 | 88 | -0.31265 | 0.62273 | KCNH1 | 0.62273 | 0.61618 | 0.63273 |
| ENSG00000188785 | 19 | 57901218 | 57912786 | 26 | 9 | -0.31285 | 0.6228 | ZNF548 | 0.6228 | 0.45975 | 0.3737 |

| | | | | | | | | | | | |
|---|---|---|---|---|---|---|---|---|---|---|---|
| ENSG00000177303 | 17 | 73496342 | 73511664 | 55 | 9 | -0.31299 | 0.62286 | CASKIN2 | 0.62286 | 0.69683 | 0.77097 |
| ENSG00000166391 | 11 | 75428864 | 75444003 | 43 | 10 | -0.31306 | 0.62288 | MOGAT2 | 0.62288 | 0.87508 | 0.9172 |
| ENSG00000106331 | 7 | 127250346 | 127255982 | 20 | 5 | -0.31307 | 0.62289 | PAX4 | 0.62289 | 0.734 | 0.71672 |
| ENSG00000070444 | 17 | 2287354 | 2304412 | 53 | 9 | -0.31322 | 0.62294 | MNT | 0.62294 | 0.55099 | 0.56493 |
| ENSG00000162222 | 11 | 62495541 | 62507765 | 28 | 5 | -0.31332 | 0.62298 | TTC9C | 0.62298 | 0.48005 | 0.39783 |
| ENSG00000117707 | 1 | 214156524 | 214214595 | 142 | 25 | -0.31344 | 0.62303 | PROX1 | 0.62303 | 0.62179 | 0.6343 |
| ENSG00000180902 | 2 | 242673994 | 242708231 | 177 | 24 | -0.31359 | 0.62308 | D2HGDH | 0.62308 | 0.78629 | 0.86406 |
| ENSG00000164056 | 4 | 124317950 | 124324910 | 22 | 6 | -0.31358 | 0.62308 | SPRY1 | 0.62308 | 0.46358 | 0.42557 |
| ENSG00000265590 | 21 | 33951132 | 33984591 | 119 | 16 | -0.31363 | 0.6231 | AP000275.65 | 0.6231 | 0.58118 | 0.47422 |
| ENSG00000177000 | 1 | 11845780 | 11866977 | 97 | 13 | -0.31386 | 0.62318 | MTHFR | 0.62318 | 0.56814 | 0.64271 |
| ENSG00000105248 | 19 | 4247076 | 4269087 | 97 | 6 | -0.31415 | 0.62329 | CCDC94 | 0.62329 | 0.48338 | 0.35627 |
| ENSG00000212659 | 17 | 39421591 | 39422893 | 14 | 2 | -0.3149 | 0.62358 | KRTAP9-6 | 0.62358 | 0.7358 | 0.62369 |
| ENSG00000166455 | 16 | 81087102 | 81110872 | 78 | 8 | -0.31494 | 0.6236 | C16orf46 | 0.6236 | 0.6051 | 0.67171 |
| ENSG00000166886 | 12 | 57482677 | 57489259 | 19 | 10 | -0.31509 | 0.62365 | NAB2 | 0.62365 | 0.47693 | 0.43158 |
| ENSG00000164934 | 8 | 104426942 | 104455681 | 83 | 9 | -0.3151 | 0.62366 | DCAF13 | 0.62366 | 0.65631 | 0.62628 |
| ENSG00000124731 | 6 | 41235664 | 41254457 | 70 | 16 | -0.31544 | 0.62378 | TREM1 | 0.62378 | 0.47508 | 0.55249 |
| ENSG00000137574 | 8 | 56685701 | 56738007 | 190 | 14 | -0.31548 | 0.6238 | TGS1 | 0.6238 | 0.62263 | 0.705 |
| ENSG00000256349 | 11 | 66276550 | 66301084 | 57 | 7 | -0.31549 | 0.6238 | CTD-3074O7.11 | 0.6238 | 0.68525 | 0.73515 |
| ENSG00000163875 | 1 | 37958176 | 37980375 | 96 | 18 | -0.31556 | 0.62383 | MEAF6 | 0.62383 | 0.55981 | 0.55391 |
| ENSG00000120798 | 12 | 95414005 | 95467479 | 157 | 24 | -0.31557 | 0.62384 | NR2C1 | 0.62384 | 0.612 | 0.64247 |
| ENSG00000171540 | 5 | 76924538 | 76935513 | 25 | 4 | -0.31564 | 0.62386 | OTP | 0.62386 | 0.61602 | 0.41949 |
| ENSG00000087448 | 12 | 27932953 | 27955973 | 53 | 12 | -0.31578 | 0.62392 | KLHL42 | 0.62392 | 0.48186 | 0.49041 |
| ENSG00000135968 | 2 | 109065017 | 109125871 | 287 | 16 | -0.31586 | 0.62394 | GCC2 | 0.62394 | 0.66228 | 0.65112 |
| ENSG00000184635 | 19 | 20011722 | 20046384 | 127 | 17 | -0.31586 | 0.62394 | ZNF93 | 0.62394 | 0.64586 | 0.59963 |
| ENSG00000162989 | 2 | 155554811 | 155714863 | 577 | 54 | -0.31589 | 0.62396 | KCNJ3 | 0.62396 | 0.63868 | 0.69164 |
| ENSG00000272162 | 6 | 10748027 | 10930656 | 539 | 49 | -0.3163 | 0.62411 | RP11-637O19.3 | 0.62411 | 0.66797 | 0.68584 |
| ENSG00000185009 | 10 | 75881524 | 75910821 | 73 | 9 | -0.31629 | 0.62411 | AP3M1 | 0.62411 | 0.62103 | 0.59281 |
| ENSG00000087274 | 4 | 2845584 | 2931803 | 188 | 23 | -0.31643 | 0.62416 | ADD1 | 0.62416 | 0.62809 | 0.57576 |
| ENSG00000198471 | 3 | 187416047 | 187420345 | 22 | 6 | -0.31653 | 0.6242 | RTP2 | 0.6242 | 0.62993 | 0.60752 |
| ENSG00000060566 | 19 | 4153598 | 4173051 | 76 | 16 | -0.31659 | 0.62422 | CREB3L3 | 0.62422 | 0.63469 | 0.74261 |
| ENSG00000113810 | 3 | 160117062 | 160152750 | 84 | 15 | -0.31673 | 0.62428 | SMC4 | 0.62428 | 0.6911 | 0.65443 |
| ENSG00000163946 | 3 | 56654161 | 56717265 | 298 | 18 | -0.31685 | 0.62432 | FAM208A | 0.62432 | 0.64356 | 0.60611 |
| ENSG00000205352 | 12 | 53835389 | 53840429 | 9 | 4 | -0.31695 | 0.62436 | PRR13 | 0.62436 | 0.57605 | 0.65603 |
| ENSG00000133065 | 1 | 205758221 | 205782876 | 63 | 14 | -0.31699 | 0.62438 | SLC41A1 | 0.62438 | 0.79866 | 0.84854 |
| ENSG00000145040 | 3 | 48599160 | 48601206 | 3 | 2 | -0.3174 | 0.62453 | UCN2 | 0.62453 | 0.40695 | 0.62757 |
| ENSG00000135842 | 1 | 184759858 | 184943682 | 435 | 30 | -0.31783 | 0.62469 | FAM129A | 0.62469 | 0.7191 | 0.79218 |
| ENSG00000197416 | 8 | 82437216 | 82443613 | 35 | 7 | -0.31792 | 0.62473 | FABP12 | 0.62473 | 0.60258 | 0.63839 |
| ENSG00000105649 | 19 | 18307594 | 18314884 | 18 | 5 | -0.31828 | 0.62486 | RAB3A | 0.62486 | 0.32938 | 0.16646 |
| ENSG00000115295 | 2 | 29320571 | 29412509 | 228 | 27 | -0.31871 | 0.62503 | CLIP4 | 0.62503 | 0.38155 | 0.37246 |
| ENSG00000162654 | 1 | 89646831 | 89664615 | 29 | 4 | -0.31903 | 0.62515 | GBP4 | 0.62515 | 0.56825 | 0.53825 |
| ENSG00000111850 | 6 | 88032301 | 88052043 | 134 | 12 | -0.31908 | 0.62517 | SMIM8 | 0.62517 | 0.58853 | 0.56341 |
| ENSG00000167987 | 11 | 60897728 | 60929089 | 138 | 8 | -0.31948 | 0.62532 | VPS37C | 0.62532 | 0.58639 | 0.59189 |
| ENSG00000125844 | 20 | 17594323 | 17662940 | 155 | 27 | -0.31973 | 0.62541 | RRBP1 | 0.62541 | 0.49969 | 0.35952 |
| ENSG00000187735 | 8 | 54879112 | 54935089 | 129 | 10 | -0.31998 | 0.62551 | TCEA1 | 0.62551 | 0.64465 | 0.61472 |
| ENSG00000107679 | 10 | 124134212 | 124191867 | 176 | 19 | -0.31998 | 0.62551 | PLEKHA1 | 0.62551 | 0.65256 | 0.70236 |
| ENSG00000228696 | 17 | 44352150 | 44439130 | 227 | 8 | -0.32054 | 0.62572 | ARL17B | 0.62572 | 0.64299 | 0.65745 |
| ENSG00000183230 | 10 | 67672276 | 69455927 | 7751 | 197 | -0.3206 | 0.62574 | CTNNA3 | 0.62574 | 0.61412 | 0.6143 |
| ENSG00000183250 | 21 | 46352729 | 46359828 | 34 | 6 | -0.32059 | 0.62574 | C21orf67 | 0.62574 | 0.69321 | 0.85557 |
| ENSG00000117228 | 1 | 89518002 | 89531043 | 47 | 9 | -0.32098 | 0.62589 | GBP1 | 0.62589 | 0.67106 | 0.69537 |
| ENSG00000135749 | 1 | 233119181 | 233431459 | 1233 | 69 | -0.32115 | 0.62595 | PCNXL2 | 0.62595 | 0.5705 | 0.59687 |
| ENSG00000271043 | 5 | 79945819 | 79946855 | 3 | 1 | -0.32126 | 0.62599 | MTRNR2L2 | 0.62599 | 0.56583 | 0.53957 |
| ENSG00000163904 | 3 | 185300284 | 185351339 | 91 | 16 | -0.32164 | 0.62614 | SENP2 | 0.62614 | 0.58623 | 0.56945 |
| ENSG00000188393 | 12 | 10051272 | 10084980 | 130 | 31 | -0.3217 | 0.62616 | CLEC2A | 0.62616 | 0.52389 | 0.40379 |
| ENSG00000132300 | 2 | 86333305 | 86369280 | 101 | 11 | -0.32179 | 0.62619 | PTCD3 | 0.62619 | 0.60476 | 0.58062 |
| ENSG00000115488 | 2 | 233897382 | 233899767 | 7 | 2 | -0.32183 | 0.62621 | NEU2 | 0.62621 | 0.83509 | 0.84842 |
| ENSG00000155252 | 10 | 99344131 | 99436191 | 255 | 29 | -0.32207 | 0.6263 | PI4K2A | 0.6263 | 0.6189 | 0.6996 |
| ENSG00000136492 | 17 | 59758627 | 59940882 | 350 | 22 | -0.32207 | 0.6263 | BRIP1 | 0.6263 | 0.64284 | 0.68268 |
| ENSG00000065717 | 19 | 2997636 | 3047633 | 271 | 44 | -0.32224 | 0.62636 | TLE2 | 0.62636 | 0.73092 | 0.87417 |
| ENSG00000172031 | 1 | 92495539 | 92529093 | 85 | 13 | -0.32224 | 0.62637 | EPHX4 | 0.62637 | 0.36007 | 0.28192 |
| ENSG00000151079 | 12 | 4918342 | 4960277 | 243 | 35 | -0.32225 | 0.62637 | KCNA6 | 0.62637 | 0.63959 | 0.69893 |
| ENSG00000100399 | 22 | 41625517 | 41636938 | 24 | 6 | -0.3224 | 0.62643 | CHADL | 0.62643 | 0.70854 | 0.73404 |
| ENSG00000185668 | 1 | 38509523 | 38512450 | 2 | 1 | -0.32254 | 0.62648 | POU3F1 | 0.62648 | 0.66651 | 0.49162 |
| ENSG00000129933 | 19 | 19431490 | 19469563 | 89 | 10 | -0.32272 | 0.62655 | MAU2 | 0.62655 | 0.60699 | 0.62059 |
| ENSG00000244045 | 17 | 26684604 | 26708716 | 58 | 8 | -0.3228 | 0.62658 | TMEM199 | 0.62658 | 0.49467 | 0.38466 |
| ENSG00000135052 | 9 | 88641061 | 88715088 | 315 | 25 | -0.32297 | 0.62664 | GOLM1 | 0.62664 | 0.68505 | 0.72128 |
| ENSG00000148925 | 11 | 13409548 | 13484844 | 225 | 17 | -0.32312 | 0.6267 | BTBD10 | 0.6267 | 0.62293 | 0.56845 |
| ENSG00000268975 | 19 | 41277553 | 41292590 | 47 | 12 | -0.3232 | 0.62673 | MIA-RAB4B | 0.62673 | 0.54472 | 0.55986 |
| ENSG00000087470 | 12 | 32832134 | 32898486 | 327 | 7 | -0.32327 | 0.62675 | DNM1L | 0.62675 | 0.66374 | 0.75289 |
| ENSG00000138400 | 2 | 207602487 | 207630271 | 144 | 11 | -0.32334 | 0.62678 | MDH1B | 0.62678 | 0.60326 | 0.60895 |
| ENSG00000156076 | 12 | 65444406 | 65515346 | 148 | 17 | -0.32333 | 0.62678 | WIF1 | 0.62678 | 0.68464 | 0.54212 |
| ENSG00000089225 | 12 | 114791736 | 114846247 | 197 | 36 | -0.32335 | 0.62678 | TBX5 | 0.62678 | 0.66813 | 0.46355 |
| ENSG00000188379 | 9 | 21384254 | 21385396 | 1 | 1 | -0.32365 | 0.6269 | IFNA2 | 0.6269 | 0.38643 | 0.40176 |
| ENSG00000087088 | 19 | 49458072 | 49465055 | 29 | 7 | -0.32367 | 0.6269 | BAX | 0.6269 | 0.7167 | 0.82736 |

| | | | | | | | | | | | |
|---|---|---|---|---|---|---|---|---|---|---|---|
| ENSG00000167183 | 17 | 46029333 | 46035244 | 22 | 6 | -0.32368 | 0.62691 | PRR15L | 0.62691 | 0.6761 | 0.71906 |
| ENSG00000159176 | 1 | 201452658 | 201478584 | 119 | 16 | -0.32384 | 0.62697 | CSRP1 | 0.62697 | 0.40836 | 0.16001 |
| ENSG00000258064 | 12 | 72067984 | 72092748 | 42 | 9 | -0.32383 | 0.62697 | RP11-293I14.2 | 0.62697 | 0.62604 | 0.73591 |
| ENSG00000047346 | 15 | 52873514 | 53002014 | 297 | 25 | -0.3239 | 0.62699 | FAM214A | 0.62699 | 0.71705 | 0.70861 |
| ENSG00000119986 | 10 | 99437181 | 99447080 | 39 | 7 | -0.32392 | 0.627 | AVPI1 | 0.627 | 0.68307 | 0.66973 |
| ENSG00000267883 | 21 | 47183565 | 47190005 | 12 | 4 | -0.32402 | 0.62704 | PRED60 | 0.62704 | 0.78615 | 0.8214 |
| ENSG00000124780 | 6 | 39266777 | 39282329 | 48 | 16 | -0.3241 | 0.62707 | KCNK17 | 0.62707 | 0.50121 | 0.39532 |
| ENSG00000204422 | 6 | 31654739 | 31681849 | 75 | 18 | -0.32426 | 0.62713 | XXbac-BPG32J3.20 | 0.62713 | 0.61988 | 0.58963 |
| ENSG00000183454 | 16 | 9852376 | 10276611 | 1969 | 56 | -0.32432 | 0.62715 | GRIN2A | 0.62715 | 0.64758 | 0.66052 |
| ENSG00000148386 | 9 | 138555168 | 138558268 | 11 | 5 | -0.32477 | 0.62732 | LCN9 | 0.62732 | 0.7984 | 0.80992 |
| ENSG00000168763 | 2 | 97481982 | 97499648 | 29 | 5 | -0.3248 | 0.62733 | CNNM3 | 0.62733 | 0.57576 | 0.60427 |
| ENSG00000055163 | 5 | 156693089 | 156822606 | 460 | 27 | -0.32509 | 0.62744 | CYFIP2 | 0.62744 | 0.51974 | 0.32306 |
| ENSG00000173480 | 19 | 58411664 | 58427978 | 55 | 9 | -0.32542 | 0.62757 | ZNF417 | 0.62757 | 0.62164 | 0.59379 |
| ENSG00000111845 | 6 | 10694928 | 10710015 | 20 | 10 | -0.32587 | 0.62774 | PAK1IP1 | 0.62774 | 0.7992 | 0.73793 |
| ENSG00000174606 | 1 | 213165524 | 213189168 | 40 | 8 | -0.32617 | 0.62785 | ANGEL2 | 0.62785 | 0.65967 | 0.63599 |
| ENSG00000108878 | 17 | 65040706 | 65052909 | 31 | 7 | -0.32631 | 0.6279 | CACNG1 | 0.6279 | 0.79681 | 0.89117 |
| ENSG00000105397 | 19 | 10461209 | 10491352 | 101 | 17 | -0.32636 | 0.62792 | TYK2 | 0.62792 | 0.66687 | 0.68232 |
| ENSG00000137101 | 9 | 35609530 | 35646807 | 92 | 19 | -0.32637 | 0.62793 | CD72 | 0.62793 | 0.57851 | 0.3165 |
| ENSG00000039319 | 5 | 79703832 | 79775169 | 190 | 11 | -0.32644 | 0.62795 | ZFYVE16 | 0.62795 | 0.61398 | 0.61967 |
| ENSG00000131626 | 11 | 70116806 | 70230509 | 305 | 9 | -0.32654 | 0.62799 | PPFIA1 | 0.62799 | 0.62407 | 0.62424 |
| ENSG00000204673 | 19 | 50372295 | 50381716 | 16 | 3 | -0.32664 | 0.62803 | AKT1S1 | 0.62803 | 0.48487 | 0.42697 |
| ENSG00000141524 | 17 | 76106539 | 76128488 | 98 | 31 | -0.32669 | 0.62805 | TMC6 | 0.62805 | 0.68019 | 0.40485 |
| ENSG00000159723 | 16 | 67516474 | 67517716 | 3 | 1 | -0.32672 | 0.62806 | AGRP | 0.62806 | 0.65049 | 0.60747 |
| ENSG00000181029 | 19 | 7745729 | 7747744 | 6 | 3 | -0.32686 | 0.62811 | TRAPPC5 | 0.62811 | 0.90089 | 0.85135 |
| ENSG00000185950 | 13 | 110406184 | 110438915 | 77 | 14 | -0.32688 | 0.62812 | IRS2 | 0.62812 | 0.62553 | 0.77836 |
| ENSG00000253857 | 8 | 56074043 | 56077287 | 5 | 2 | -0.32698 | 0.62816 | RP11-386G21.2 | 0.62816 | 0.85661 | 0.91551 |
| ENSG00000035664 | 15 | 64199235 | 64364232 | 610 | 33 | -0.32718 | 0.62823 | DAPK2 | 0.62823 | 0.65241 | 0.71279 |
| ENSG00000136643 | 1 | 213224589 | 213448116 | 357 | 21 | -0.32734 | 0.62829 | RPS6KC1 | 0.62829 | 0.61853 | 0.54837 |
| ENSG00000179750 | 22 | 39378352 | 39388809 | 31 | 8 | -0.32737 | 0.6283 | APOBEC3B | 0.6283 | 0.76638 | 0.6707 |
| ENSG00000184647 | 8 | 10383056 | 10411676 | 214 | 22 | -0.32738 | 0.62831 | PRSS55 | 0.62831 | 0.76461 | 0.80643 |
| ENSG00000168811 | 3 | 159706537 | 159713806 | 19 | 6 | -0.32741 | 0.62832 | IL12A | 0.62832 | 0.76803 | 0.26151 |
| ENSG00000145833 | 5 | 134094469 | 134190823 | 136 | 29 | -0.32749 | 0.62835 | DDX46 | 0.62835 | 0.60445 | 0.53878 |
| ENSG00000178997 | 15 | 41474923 | 41522941 | 252 | 12 | -0.32774 | 0.62844 | EXD1 | 0.62844 | 0.68771 | 0.71409 |
| ENSG00000146834 | 7 | 100026413 | 100031741 | 16 | 4 | -0.32832 | 0.62867 | MEPCE | 0.62867 | 0.75946 | 0.64081 |
| ENSG00000171992 | 5 | 149980642 | 150038782 | 125 | 23 | -0.32837 | 0.62868 | SYNPO | 0.62868 | 0.67857 | 0.71441 |
| ENSG00000129534 | 14 | 45672393 | 45722743 | 97 | 11 | -0.32836 | 0.62868 | MIS18BP1 | 0.62868 | 0.58257 | 0.67602 |
| ENSG00000120837 | 12 | 104510855 | 104532067 | 46 | 10 | -0.32868 | 0.6288 | NFYB | 0.6288 | 0.64463 | 0.70834 |
| ENSG00000197467 | 10 | 71561644 | 71724031 | 824 | 117 | -0.32906 | 0.62894 | COL13A1 | 0.62894 | 0.51219 | 0.50514 |
| ENSG00000272916 | 10 | 75556272 | 75571409 | 21 | 5 | -0.32914 | 0.62897 | RP11-574K11.31 | 0.62897 | 0.75576 | 0.5803 |
| ENSG00000149435 | 20 | 23965690 | 23969416 | 23 | 8 | -0.32912 | 0.62897 | GGTLC1 | 0.62897 | 0.56836 | 0.5557 |
| ENSG00000205086 | 2 | 42162508 | 42181406 | 49 | 14 | -0.3292 | 0.629 | C2orf91 | 0.629 | 0.80779 | 0.84593 |
| ENSG00000185155 | 1 | 226411319 | 226414755 | 2 | 1 | -0.32924 | 0.62901 | MIXL1 | 0.62901 | 0.57272 | 0.70878 |
| ENSG00000001460 | 1 | 24683489 | 24743424 | 190 | 19 | -0.32943 | 0.62908 | STPG1 | 0.62908 | 0.61528 | 0.60361 |
| ENSG00000119655 | 14 | 74942895 | 74960880 | 53 | 6 | -0.32964 | 0.62916 | NPC2 | 0.62916 | 0.66422 | 0.69506 |
| ENSG00000099995 | 22 | 30727977 | 30752936 | 82 | 12 | -0.32964 | 0.62916 | SF3A1 | 0.62916 | 0.57368 | 0.54288 |
| ENSG00000181092 | 3 | 186560463 | 186576252 | 64 | 18 | -0.32971 | 0.62919 | ADIPOQ | 0.62919 | 0.68654 | 0.70723 |
| ENSG00000130305 | 7 | 72716514 | 72722864 | 4 | 2 | -0.32974 | 0.6292 | NSUN5 | 0.6292 | 0.29352 | 0.52668 |
| ENSG00000257207 | 2 | 111160511 | 111230652 | 10 | 6 | -0.3298 | 0.62922 | LIMS3 | 0.62922 | 0.6298 | 0.55767 |
| ENSG00000222038 | 2 | 131369054 | 131415664 | 32 | 15 | -0.33016 | 0.62936 | POTEJ | 0.62936 | 0.65 | 0.52154 |
| ENSG00000174473 | 4 | 172733405 | 173962710 | 3536 | 120 | -0.33028 | 0.6294 | GALNTL6 | 0.6294 | 0.63696 | 0.66536 |
| ENSG00000142627 | 1 | 16450832 | 16482582 | 52 | 13 | -0.33028 | 0.62941 | EPHA2 | 0.62941 | 0.6536 | 0.73474 |
| ENSG00000101115 | 20 | 50400581 | 50419059 | 89 | 19 | -0.33042 | 0.62946 | SALL4 | 0.62946 | 0.35174 | 0.44019 |
| ENSG00000166507 | 10 | 75561669 | 75571589 | 15 | 5 | -0.33066 | 0.62955 | NDST2 | 0.62955 | 0.75205 | 0.58635 |
| ENSG00000175727 | 12 | 122516628 | 122631894 | 228 | 8 | -0.33075 | 0.62958 | MLXIP | 0.62958 | 0.56256 | 0.46363 |
| ENSG00000177694 | 3 | 174156363 | 175523428 | 5203 | 252 | -0.3308 | 0.6296 | NAALADL2 | 0.6296 | 0.64012 | 0.66144 |
| ENSG00000164104 | 4 | 174252846 | 174256276 | 4 | 2 | -0.33083 | 0.62962 | HMGB2 | 0.62962 | 0.67969 | 0.68979 |
| ENSG00000177548 | 16 | 28915742 | 28947847 | 49 | 12 | -0.33121 | 0.62976 | RABEP2 | 0.62976 | 0.6772 | 0.69088 |
| ENSG00000143153 | 1 | 169074935 | 169101960 | 139 | 24 | -0.33132 | 0.6298 | ATP1B1 | 0.6298 | 0.65395 | 0.64651 |
| ENSG00000164651 | 7 | 20823906 | 20826505 | 1 | 1 | -0.33132 | 0.6298 | SP8 | 0.6298 | 0.37189 | 0.34326 |
| ENSG00000221855 | 7 | 143174966 | 143175889 | 2 | 1 | -0.33146 | 0.62985 | TAS2R41 | 0.62985 | 0.51946 | 0.41016 |
| ENSG00000188848 | 4 | 42112955 | 42154895 | 124 | 15 | -0.33156 | 0.62989 | BEND4 | 0.62989 | 0.71206 | 0.66896 |
| ENSG00000162227 | 11 | 62538775 | 62554814 | 36 | 7 | -0.33157 | 0.62989 | TAF6L | 0.62989 | 0.52828 | 0.62191 |
| ENSG00000219492 | 4 | 9385743 | 9390709 | 40 | 8 | -0.33163 | 0.62991 | RP11-1396O13.13 | 0.62991 | 0.64974 | 0.67664 |
| ENSG00000141560 | 17 | 80674559 | 80688204 | 50 | 12 | -0.33175 | 0.62996 | FN3KRP | 0.62996 | 0.47068 | 0.42501 |
| ENSG00000134697 | 1 | 38032417 | 38061536 | 77 | 14 | -0.33182 | 0.62999 | GNL2 | 0.62999 | 0.66585 | 0.7285 |
| ENSG00000185246 | 14 | 45553302 | 45585485 | 55 | 6 | -0.33182 | 0.62999 | PRPF39 | 0.62999 | 0.65184 | 0.64445 |
| ENSG00000113811 | 3 | 53918437 | 53926015 | 26 | 6 | -0.33191 | 0.63002 | SELK | 0.63002 | 0.63906 | 0.68669 |
| ENSG00000108179 | 10 | 81107225 | 81115093 | 22 | 6 | -0.33196 | 0.63004 | PPIF | 0.63004 | 0.49964 | 0.27716 |
| ENSG00000163539 | 3 | 33537737 | 33759848 | 475 | 32 | -0.33208 | 0.63008 | CLASP2 | 0.63008 | 0.65394 | 0.67915 |
| ENSG00000149782 | 11 | 64018995 | 64036622 | 31 | 12 | -0.3321 | 0.63009 | PLCB3 | 0.63009 | 0.27594 | 0.29803 |
| ENSG00000135763 | 1 | 229761981 | 229795946 | 75 | 10 | -0.3322 | 0.63013 | URB2 | 0.63013 | 0.71136 | 0.66508 |
| ENSG00000160179 | 21 | 43619799 | 43717354 | 429 | 63 | -0.33236 | 0.63019 | ABCG1 | 0.63019 | 0.82456 | 0.82059 |
| ENSG00000136485 | 17 | 61627822 | 61671639 | 57 | 12 | -0.33241 | 0.63021 | DCAF7 | 0.63021 | 0.76821 | 0.77057 |

| | | | | | | | | | | | |
|---|---|---|---|---|---|---|---|---|---|---|---|
| ENSG00000124356 | 2 | 74056086 | 74100786 | 135 | 20 | -0.33252 | 0.63025 | STAMBP | 0.63025 | 0.40118 | 0.52083 |
| ENSG00000087116 | 5 | 178537852 | 178772431 | 1083 | 132 | -0.33254 | 0.63026 | ADAMTS2 | 0.63026 | 0.63717 | 0.62082 |
| ENSG00000108582 | 17 | 28705923 | 28797007 | 131 | 10 | -0.33258 | 0.63027 | CPD | 0.63027 | 0.62584 | 0.47942 |
| ENSG00000230453 | 9 | 33524392 | 33573007 | 187 | 27 | -0.33264 | 0.6303 | ANKRD18B | 0.6303 | 0.54636 | 0.50004 |
| ENSG00000120669 | 13 | 36742345 | 36871979 | 348 | 32 | -0.33284 | 0.63037 | SOHLH2 | 0.63037 | 0.51265 | 0.43787 |
| ENSG00000250709 | 13 | 36742931 | 36871979 | 348 | 32 | -0.33284 | 0.63037 | CCDC169-SOHLH2 | 0.63037 | 0.52147 | 0.44595 |
| ENSG00000096384 | 6 | 44214824 | 44221620 | 26 | 9 | -0.33317 | 0.6305 | HSP90AB1 | 0.6305 | 0.77059 | 0.66223 |
| ENSG00000060642 | 1 | 27113963 | 27124889 | 16 | 5 | -0.33321 | 0.63051 | PIGV | 0.63051 | 0.62451 | 0.64821 |
| ENSG00000187871 | 6 | 55192267 | 55267291 | 672 | 24 | -0.33331 | 0.63055 | GFRAL | 0.63055 | 0.52506 | 0.55637 |
| ENSG00000250733 | 8 | 140943416 | 140946190 | 8 | 4 | -0.3337 | 0.6307 | C8orf17 | 0.6307 | 0.25809 | 0.26019 |
| ENSG00000175467 | 11 | 65729160 | 65747299 | 57 | 8 | -0.33383 | 0.63075 | SART1 | 0.63075 | 0.67364 | 0.68998 |
| ENSG00000073849 | 3 | 186648274 | 186796341 | 682 | 85 | -0.3339 | 0.63077 | ST6GAL1 | 0.63077 | 0.64159 | 0.66953 |
| ENSG00000198108 | 5 | 129240165 | 129522327 | 579 | 23 | -0.3339 | 0.63077 | CHSY3 | 0.63077 | 0.65142 | 0.6588 |
| ENSG00000120688 | 13 | 41635410 | 41658137 | 28 | 12 | -0.33402 | 0.63082 | WBP4 | 0.63082 | 0.74719 | 0.87813 |
| ENSG00000175315 | 11 | 65779312 | 65780976 | 3 | 2 | -0.3341 | 0.63085 | CST6 | 0.63085 | 0.5233 | 0.56572 |
| ENSG00000120586 | 10 | 18098352 | 18200091 | 35 | 11 | -0.33417 | 0.63087 | MRC1 | 0.63087 | 0.68131 | 0.78105 |
| ENSG00000160799 | 3 | 46963216 | 47023500 | 89 | 10 | -0.3342 | 0.63089 | CCDC12 | 0.63089 | 0.61871 | 0.65407 |
| ENSG00000204775 | 8 | 145317003 | 145331153 | 3 | 1 | -0.33433 | 0.63093 | KM-PA-2 | 0.63093 | 0.8135 | 0.8135 |
| ENSG00000259316 | 15 | 64472547 | 64665980 | 481 | 24 | -0.33437 | 0.63095 | CTD-2116N17.1 | 0.63095 | 0.6324 | 0.61784 |
| ENSG00000141030 | 17 | 17150141 | 17184607 | 131 | 13 | -0.3345 | 0.631 | COPS3 | 0.631 | 0.69397 | 0.70221 |
| ENSG00000105204 | 19 | 40315990 | 40324841 | 17 | 4 | -0.3345 | 0.631 | DYRK1B | 0.631 | 0.70681 | 0.75529 |
| ENSG00000144031 | 2 | 71205510 | 71212626 | 24 | 8 | -0.33461 | 0.63104 | ANKRD53 | 0.63104 | 0.3057 | 0.47783 |
| ENSG00000181690 | 8 | 57073463 | 57123883 | 111 | 14 | -0.33489 | 0.63114 | PLAG1 | 0.63114 | 0.63067 | 0.68194 |
| ENSG00000143194 | 1 | 166958346 | 166991451 | 97 | 15 | -0.33493 | 0.63116 | MAEL | 0.63116 | 0.70878 | 0.72705 |
| ENSG00000134042 | 18 | 48324574 | 48351772 | 149 | 29 | -0.33498 | 0.63118 | MRO | 0.63118 | 0.48712 | 0.53975 |
| ENSG00000149021 | 11 | 62172575 | 62190667 | 68 | 15 | -0.335 | 0.63119 | SCGB1A1 | 0.63119 | 0.74 | 0.63883 |
| ENSG00000176383 | 12 | 122688090 | 122693499 | 14 | 6 | -0.33501 | 0.63119 | B3GNT4 | 0.63119 | 0.88193 | 0.67417 |
| ENSG00000181929 | 12 | 49396057 | 49412980 | 19 | 3 | -0.33512 | 0.63123 | PRKAG1 | 0.63123 | 0.61541 | 0.40485 |
| ENSG00000066629 | 14 | 100204030 | 100408397 | 652 | 47 | -0.33518 | 0.63125 | EML1 | 0.63125 | 0.58906 | 0.54144 |
| ENSG00000158156 | 1 | 28285973 | 28294607 | 16 | 8 | -0.33522 | 0.63127 | XKR8 | 0.63127 | 0.6081 | 0.76454 |
| ENSG00000204548 | 20 | 29992648 | 30000641 | 20 | 4 | -0.33525 | 0.63128 | DEFB121 | 0.63128 | 0.71567 | 0.62996 |
| ENSG00000138346 | 10 | 70173821 | 70231879 | 272 | 16 | -0.33528 | 0.63129 | DNA2 | 0.63129 | 0.70398 | 0.70836 |
| ENSG00000005075 | 7 | 102113565 | 102119354 | 35 | 8 | -0.33529 | 0.6313 | POLR2J | 0.6313 | 0.49682 | 0.53826 |
| ENSG00000148450 | 10 | 23384435 | 23410942 | 79 | 13 | -0.33546 | 0.63136 | MSRB2 | 0.63136 | 0.70004 | 0.72021 |
| ENSG00000134460 | 10 | 6052652 | 6104288 | 259 | 32 | -0.3356 | 0.63141 | IL2RA | 0.63141 | 0.66646 | 0.7116 |
| ENSG00000197355 | 9 | 139971953 | 139978991 | 10 | 4 | -0.33562 | 0.63142 | UAP1L1 | 0.63142 | 0.71733 | 0.68847 |
| ENSG00000171649 | 19 | 58089824 | 58105145 | 29 | 4 | -0.33571 | 0.63145 | ZIK1 | 0.63145 | 0.60537 | 0.62715 |
| ENSG00000129968 | 19 | 1876809 | 1885546 | 23 | 3 | -0.33573 | 0.63146 | ABHD17A | 0.63146 | 0.6885 | 0.27831 |
| ENSG00000143167 | 1 | 167022073 | 167059868 | 124 | 23 | -0.33576 | 0.63147 | GPA33 | 0.63147 | 0.6334 | 0.7497 |
| ENSG00000187627 | 2 | 87135076 | 87241104 | 59 | 12 | -0.33579 | 0.63149 | RGPD1 | 0.63149 | 0.65809 | 0.48929 |
| ENSG00000157551 | 21 | 39529128 | 39679279 | 592 | 25 | -0.3358 | 0.63149 | KCNJ15 | 0.63149 | 0.66269 | 0.63796 |
| ENSG00000157005 | 3 | 187386694 | 187388187 | 6 | 2 | -0.33603 | 0.63158 | SST | 0.63158 | 0.52269 | 0.44651 |
| ENSG00000100605 | 14 | 93403259 | 93582665 | 571 | 32 | -0.33608 | 0.6316 | ITPK1 | 0.6316 | 0.66387 | 0.78 |
| ENSG00000079691 | 6 | 25279306 | 25620758 | 1529 | 80 | -0.33631 | 0.63168 | LRRC16A | 0.63168 | 0.64512 | 0.69304 |
| ENSG00000196166 | 8 | 38368352 | 38386180 | 73 | 8 | -0.33661 | 0.6318 | C8orf86 | 0.6318 | 0.55337 | 0.65094 |
| ENSG00000154710 | 7 | 66147151 | 66276451 | 516 | 20 | -0.33689 | 0.6319 | RABGEF1 | 0.6319 | 0.64426 | 0.64069 |
| ENSG00000172575 | 15 | 38780304 | 38857776 | 220 | 27 | -0.33696 | 0.63193 | RASGRP1 | 0.63193 | 0.6582 | 0.72241 |
| ENSG00000189120 | 17 | 45922279 | 45933240 | 30 | 6 | -0.33699 | 0.63194 | SP6 | 0.63194 | 0.62717 | 0.72395 |
| ENSG00000053918 | 11 | 2465914 | 2870339 | 1352 | 162 | -0.33717 | 0.63201 | KCNQ1 | 0.63201 | 0.61302 | 0.74992 |
| ENSG00000163331 | 2 | 159651829 | 159719293 | 330 | 24 | -0.33729 | 0.63205 | DAPL1 | 0.63205 | 0.67745 | 0.5624 |
| ENSG00000111863 | 6 | 11712287 | 11807279 | 396 | 40 | -0.33732 | 0.63206 | ADTRP | 0.63206 | 0.66322 | 0.83744 |
| ENSG00000154122 | 5 | 14704910 | 14871887 | 532 | 46 | -0.33733 | 0.63207 | ANKH | 0.63207 | 0.66211 | 0.67409 |
| ENSG00000115738 | 2 | 8818975 | 8824583 | 13 | 5 | -0.33741 | 0.6321 | ID2 | 0.6321 | 0.90754 | 0.84199 |
| ENSG00000197768 | 9 | 140145713 | 140147934 | 7 | 3 | -0.33744 | 0.63211 | C9orf173 | 0.63211 | 0.40785 | 0.38802 |
| ENSG00000076513 | 12 | 110436991 | 110477568 | 99 | 20 | -0.3376 | 0.63217 | ANKRD13A | 0.63217 | 0.71635 | 0.76059 |
| ENSG00000135723 | 16 | 67263290 | 67281561 | 25 | 8 | -0.33768 | 0.6322 | FHOD1 | 0.6322 | 0.64853 | 0.67394 |
| ENSG00000197646 | 9 | 5510545 | 5571282 | 169 | 31 | -0.33771 | 0.63221 | PDCD1LG2 | 0.63221 | 0.62113 | 0.68728 |
| ENSG00000025039 | 6 | 90074355 | 90121989 | 124 | 24 | -0.33783 | 0.63226 | RRAGD | 0.63226 | 0.51587 | 0.49333 |
| ENSG00000135437 | 12 | 56114151 | 56118489 | 8 | 3 | -0.33789 | 0.63228 | RDH5 | 0.63228 | 0.90981 | 0.95672 |
| ENSG00000122711 | 9 | 33218363 | 33248565 | 114 | 10 | -0.33857 | 0.63253 | SPINK4 | 0.63253 | 0.66779 | 0.79227 |
| ENSG00000165671 | 5 | 176560026 | 176727216 | 472 | 23 | -0.3388 | 0.63262 | NSD1 | 0.63262 | 0.66713 | 0.68586 |
| ENSG00000084764 | 2 | 27193480 | 27250064 | 77 | 8 | -0.33931 | 0.63281 | MAPRE3 | 0.63281 | 0.54056 | 0.53444 |
| ENSG00000135913 | 2 | 219314974 | 219433084 | 268 | 11 | -0.3393 | 0.63281 | USP37 | 0.63281 | 0.63273 | 0.65218 |
| ENSG00000119760 | 2 | 27873679 | 27886676 | 13 | 3 | -0.33938 | 0.63284 | SUPT7L | 0.63284 | 0.74057 | 0.83852 |
| ENSG00000104613 | 8 | 19674651 | 19709594 | 116 | 19 | -0.33966 | 0.63294 | INTS10 | 0.63294 | 0.64505 | 0.51556 |
| ENSG00000062370 | 19 | 44830708 | 44871377 | 168 | 19 | -0.33999 | 0.63307 | ZNF112 | 0.63307 | 0.72428 | 0.75274 |
| ENSG00000213096 | 19 | 24216276 | 24312654 | 384 | 21 | -0.34048 | 0.63325 | ZNF254 | 0.63325 | 0.6558 | 0.62968 |
| ENSG00000066056 | 1 | 43766664 | 43788779 | 36 | 5 | -0.34053 | 0.63327 | TIE1 | 0.63327 | 0.5896 | 0.60969 |
| ENSG00000016402 | 6 | 137321108 | 137366298 | 113 | 24 | -0.34073 | 0.63334 | IL20RA | 0.63334 | 0.6985 | 0.30109 |
| ENSG00000161265 | 19 | 36233365 | 36236346 | 12 | 6 | -0.34082 | 0.63338 | U2AF1L4 | 0.63338 | 0.40155 | 0.53275 |
| ENSG00000066557 | 1 | 70610488 | 70671303 | 104 | 14 | -0.34102 | 0.63345 | LRRC40 | 0.63345 | 0.63056 | 0.63619 |
| ENSG00000132478 | 17 | 73780681 | 73821886 | 116 | 17 | -0.34107 | 0.63347 | UNK | 0.63347 | 0.74013 | 0.711 |
| ENSG00000067955 | 16 | 67063019 | 67134961 | 117 | 13 | -0.34109 | 0.63348 | CBFB | 0.63348 | 0.64326 | 0.7029 |

| | | | | | | | | | | | |
|---|---|---|---|---|---|---|---|---|---|---|---|
| ENSG00000165271 | 9 | 33461439 | 33473928 | 32 | 9 | -0.34113 | 0.6335 | NOL6 | 0.6335 | 0.67511 | 0.75193 |
| ENSG00000133661 | 10 | 81697496 | 81742370 | 292 | 13 | -0.34115 | 0.63351 | SFTPD | 0.63351 | 0.70389 | 0.78546 |
| ENSG00000132855 | 1 | 63063158 | 63071830 | 9 | 5 | -0.34123 | 0.63354 | ANGPTL3 | 0.63354 | 0.46861 | 0.57702 |
| ENSG00000124532 | 6 | 24403153 | 24425810 | 144 | 12 | -0.34125 | 0.63354 | MRS2 | 0.63354 | 0.59337 | 0.52846 |
| ENSG00000198535 | 15 | 62359176 | 62363116 | 9 | 3 | -0.34136 | 0.63358 | C2CD4A | 0.63358 | 0.67019 | 0.76957 |
| ENSG00000134184 | 1 | 110230436 | 110251661 | 95 | 18 | -0.34138 | 0.63359 | GSTM1 | 0.63359 | 0.5652 | 0.70218 |
| ENSG00000137601 | 4 | 170314426 | 170533780 | 402 | 27 | -0.34151 | 0.63364 | NEK1 | 0.63364 | 0.66379 | 0.6452 |
| ENSG00000225996 | 1 | 150521897 | 150524367 | 4 | 2 | -0.3416 | 0.63367 | AL356356.1 | 0.63367 | 0.68629 | 0.76009 |
| ENSG00000153294 | 6 | 47653600 | 47689757 | 170 | 9 | -0.34159 | 0.63367 | GPR115 | 0.63367 | 0.5767 | 0.63569 |
| ENSG00000158104 | 12 | 122277433 | 122301502 | 69 | 12 | -0.34167 | 0.6337 | HPD | 0.6337 | 0.68462 | 0.81995 |
| ENSG00000134489 | 18 | 22040593 | 22059921 | 65 | 18 | -0.34166 | 0.6337 | HRH4 | 0.6337 | 0.72324 | 0.77687 |
| ENSG00000149300 | 11 | 111788756 | 111797596 | 20 | 4 | -0.34193 | 0.6338 | C11orf52 | 0.6338 | 0.73084 | 0.88555 |
| ENSG00000197872 | 2 | 16730727 | 16847599 | 349 | 58 | -0.34213 | 0.63387 | FAM49A | 0.63387 | 0.64703 | 0.62531 |
| ENSG00000141564 | 17 | 78518619 | 78940171 | 2038 | 72 | -0.34236 | 0.63396 | RPTOR | 0.63396 | 0.66663 | 0.68957 |
| ENSG00000179886 | 8 | 144680074 | 144682485 | 6 | 3 | -0.34259 | 0.63404 | TIGD5 | 0.63404 | 0.79335 | 0.78801 |
| ENSG00000170214 | 5 | 159343790 | 159399551 | 180 | 23 | -0.34264 | 0.63407 | ADRA1B | 0.63407 | 0.57167 | 0.57372 |
| ENSG00000149657 | 20 | 60697517 | 60710434 | 26 | 5 | -0.34285 | 0.63414 | LSM14B | 0.63414 | 0.72522 | 0.76663 |
| ENSG00000248710 | 3 | 159945241 | 160167617 | 436 | 35 | -0.34346 | 0.63438 | RP11-432B6.3 | 0.63438 | 0.71742 | 0.75253 |
| ENSG00000139597 | 13 | 32974861 | 33002315 | 69 | 16 | -0.34354 | 0.6344 | N4BP2L1 | 0.6344 | 0.64889 | 0.53587 |
| ENSG00000158092 | 3 | 136581050 | 136668665 | 257 | 9 | -0.34402 | 0.63458 | NCK1 | 0.63458 | 0.64959 | 0.68004 |
| ENSG00000148444 | 10 | 22604903 | 22609235 | 2 | 1 | -0.344 | 0.63458 | COMMD3 | 0.63458 | 0.69396 | 0.61938 |
| ENSG00000100393 | 22 | 41487790 | 41576081 | 176 | 16 | -0.34424 | 0.63467 | EP300 | 0.63467 | 0.63379 | 0.6226 |
| ENSG00000186143 | 2 | 27359723 | 27362278 | 7 | 4 | -0.34429 | 0.63469 | C2orf53 | 0.63469 | 0.53287 | 0.6839 |
| ENSG00000170166 | 2 | 177015950 | 177017954 | 4 | 2 | -0.3446 | 0.6348 | HOXD4 | 0.6348 | 0.29354 | 0.21435 |
| ENSG00000174151 | 1 | 110036674 | 110045554 | 18 | 3 | -0.34463 | 0.63481 | CYB561D1 | 0.63481 | 0.81092 | 0.86675 |
| ENSG00000116685 | 1 | 11979648 | 11986485 | 23 | 7 | -0.34476 | 0.63486 | KIAA2013 | 0.63486 | 0.47527 | 0.29125 |
| ENSG00000169344 | 16 | 20344374 | 20367623 | 65 | 17 | -0.34489 | 0.63491 | UMOD | 0.63491 | 0.078303 | 0.040859 |
| ENSG00000188554 | 17 | 41322498 | 41363708 | 91 | 4 | -0.34499 | 0.63495 | NBR1 | 0.63495 | 0.62409 | 0.68073 |
| ENSG00000171262 | 15 | 38746328 | 38779911 | 151 | 17 | -0.34526 | 0.63505 | FAM98B | 0.63505 | 0.6235 | 0.45974 |
| ENSG00000025423 | 12 | 57145945 | 57181574 | 72 | 14 | -0.34528 | 0.63506 | HSD17B6 | 0.63506 | 0.59499 | 0.66337 |
| ENSG00000172428 | 2 | 241065980 | 241076224 | 73 | 10 | -0.34547 | 0.63513 | MYEOV2 | 0.63513 | 0.75205 | 0.71475 |
| ENSG00000104368 | 8 | 42032236 | 42065242 | 72 | 13 | -0.34565 | 0.6352 | PLAT | 0.6352 | 0.62911 | 0.72169 |
| ENSG00000152128 | 2 | 135213330 | 135476570 | 833 | 46 | -0.34574 | 0.63523 | TMEM163 | 0.63523 | 0.55705 | 0.5239 |
| ENSG00000182885 | 16 | 57702099 | 57723975 | 93 | 19 | -0.34576 | 0.63524 | GPR97 | 0.63524 | 0.64815 | 0.6921 |
| ENSG00000134684 | 1 | 33240840 | 33283754 | 91 | 11 | -0.34582 | 0.63526 | YARS | 0.63526 | 0.67791 | 0.72479 |
| ENSG00000196277 | 3 | 6811688 | 7783215 | 3781 | 225 | -0.34581 | 0.63526 | GRM7 | 0.63526 | 0.65162 | 0.68126 |
| ENSG00000159128 | 21 | 34775202 | 34851655 | 276 | 17 | -0.34586 | 0.63528 | IFNGR2 | 0.63528 | 0.70356 | 0.75565 |
| ENSG00000154007 | 1 | 76384558 | 76398116 | 66 | 13 | -0.3459 | 0.63529 | ASB17 | 0.63529 | 0.68396 | 0.68405 |
| ENSG00000268344 | 13 | 26452422 | 26452679 | 1 | 1 | -0.34592 | 0.6353 | AL138815.2 | 0.6353 | 0.56828 | 0.43831 |
| ENSG00000108523 | 17 | 4843303 | 4848517 | 13 | 6 | -0.34606 | 0.63535 | RNF167 | 0.63535 | 0.23752 | 0.14949 |
| ENSG00000196526 | 4 | 7760441 | 7941653 | 1066 | 43 | -0.34621 | 0.63541 | AFAP1 | 0.63541 | 0.59252 | 0.62656 |
| ENSG00000170035 | 2 | 181831975 | 181941312 | 477 | 17 | -0.34626 | 0.63543 | UBE2E3 | 0.63543 | 0.6468 | 0.66697 |
| ENSG00000151617 | 4 | 148402069 | 148466106 | 127 | 16 | -0.34633 | 0.63545 | EDNRA | 0.63545 | 0.74641 | 0.8525 |
| ENSG00000167797 | 11 | 67273968 | 67276120 | 3 | 1 | -0.34632 | 0.63545 | CDK2AP2 | 0.63545 | 0.55785 | 0.51129 |
| ENSG00000161896 | 6 | 33689444 | 33714762 | 122 | 20 | -0.34638 | 0.63547 | IP6K3 | 0.63547 | 0.62729 | 0.7265 |
| ENSG00000173947 | 1 | 111888910 | 111895635 | 31 | 5 | -0.34647 | 0.63551 | PIFO | 0.63551 | 0.13268 | 0.091993 |
| ENSG00000104812 | 19 | 49471382 | 49496567 | 50 | 9 | -0.34665 | 0.63557 | GYS1 | 0.63557 | 0.32873 | 0.36807 |
| ENSG00000213057 | 1 | 178511887 | 178518024 | 32 | 5 | -0.34713 | 0.63575 | C1orf220 | 0.63575 | 0.75822 | 0.74893 |
| ENSG00000108828 | 17 | 41166622 | 41177140 | 15 | 4 | -0.34715 | 0.63576 | VAT1 | 0.63576 | 0.82132 | 0.89506 |
| ENSG00000113441 | 5 | 96271098 | 96373219 | 349 | 17 | -0.34719 | 0.63578 | LNPEP | 0.63578 | 0.69733 | 0.73912 |
| ENSG00000072840 | 4 | 5712924 | 5830772 | 790 | 72 | -0.34725 | 0.6358 | EVC | 0.6358 | 0.7001 | 0.74658 |
| ENSG00000117139 | 1 | 202696526 | 202778598 | 249 | 23 | -0.34729 | 0.63581 | KDM5B | 0.63581 | 0.66258 | 0.74506 |
| ENSG00000164961 | 8 | 126036502 | 126104082 | 302 | 12 | -0.34733 | 0.63583 | KIAA0196 | 0.63583 | 0.66007 | 0.73643 |
| ENSG00000175646 | 16 | 11374693 | 11375207 | 2 | 1 | -0.34732 | 0.63583 | PRM1 | 0.63583 | 0.59712 | 0.53805 |
| ENSG00000180592 | 10 | 21802407 | 21814611 | 15 | 6 | -0.34736 | 0.63584 | SKIDA1 | 0.63584 | 0.63193 | 0.63832 |
| ENSG00000183688 | 17 | 289769 | 295730 | 28 | 9 | -0.34738 | 0.63585 | FAM101B | 0.63585 | 0.64606 | 0.55314 |
| ENSG00000165682 | 12 | 10138241 | 10166023 | 168 | 20 | -0.34748 | 0.63588 | CLEC1B | 0.63588 | 0.52192 | 0.28995 |
| ENSG00000187608 | 1 | 948803 | 949920 | 7 | 2 | -0.34752 | 0.6359 | ISG15 | 0.6359 | 0.65785 | 0.81998 |
| ENSG00000153234 | 2 | 157180944 | 157198860 | 25 | 7 | -0.34795 | 0.63606 | NR4A2 | 0.63606 | 0.59232 | 0.57006 |
| ENSG00000234465 | 19 | 44080952 | 44088116 | 27 | 6 | -0.34794 | 0.63606 | PINLYP | 0.63606 | 0.4472 | 0.5789 |
| ENSG00000103540 | 16 | 19535133 | 19564730 | 118 | 13 | -0.34799 | 0.63608 | CCP110 | 0.63608 | 0.5022 | 0.45656 |
| ENSG00000130517 | 19 | 18451397 | 18480760 | 121 | 24 | -0.34806 | 0.6361 | PGPEP1 | 0.6361 | 0.75598 | 0.83659 |
| ENSG00000107105 | 9 | 23690102 | 23826335 | 432 | 41 | -0.34807 | 0.63611 | ELAVL2 | 0.63611 | 0.69621 | 0.48375 |
| ENSG00000140297 | 15 | 59887074 | 59932438 | 220 | 31 | -0.34807 | 0.63611 | GCNT3 | 0.63611 | 0.69023 | 0.55157 |
| ENSG00000139613 | 12 | 56556767 | 56583351 | 23 | 8 | -0.3481 | 0.63612 | SMARCC2 | 0.63612 | 0.76627 | 0.77389 |
| ENSG00000185085 | 11 | 62414320 | 62420774 | 4 | 2 | -0.34842 | 0.63624 | INTS5 | 0.63624 | 0.32821 | 0.55571 |
| ENSG00000164123 | 4 | 159814286 | 159959912 | 506 | 20 | -0.34857 | 0.63629 | C4orf45 | 0.63629 | 0.69433 | 0.72564 |
| ENSG00000164949 | 8 | 95261481 | 95274578 | 36 | 7 | -0.3486 | 0.6363 | GEM | 0.6363 | 0.67224 | 0.64751 |
| ENSG00000165091 | 9 | 75136717 | 75451267 | 935 | 37 | -0.34861 | 0.63631 | TMC1 | 0.63631 | 0.58698 | 0.55947 |
| ENSG00000183783 | 4 | 44175926 | 44450824 | 791 | 50 | -0.34871 | 0.63634 | KCTD8 | 0.63634 | 0.61756 | 0.59659 |
| ENSG00000182957 | 13 | 24553944 | 24881212 | 1632 | 176 | -0.34868 | 0.63634 | SPATA13 | 0.63634 | 0.68035 | 0.68945 |
| ENSG00000197915 | 1 | 152184558 | 152196669 | 77 | 9 | -0.34884 | 0.63639 | HRNR | 0.63639 | 0.52753 | 0.5438 |
| ENSG00000143224 | 1 | 161136200 | 161147803 | 21 | 7 | -0.34909 | 0.63649 | PPOX | 0.63649 | 0.88187 | 0.88304 |

| | | | | | | | | | | | |
|---|---|---|---|---|---|---|---|---|---|---|---|
| ENSG00000160013 | 19 | 47123725 | 47128375 | 7 | 2 | -0.34928 | 0.63656 | PTGIR | 0.63656 | 0.85123 | 0.83041 |
| ENSG00000177238 | 16 | 31225342 | 31236510 | 8 | 3 | -0.34931 | 0.63657 | TRIM72 | 0.63657 | 0.7415 | 0.72698 |
| ENSG00000105171 | 19 | 30094924 | 30108144 | 64 | 13 | -0.34934 | 0.63658 | POP4 | 0.63658 | 0.57776 | 0.45109 |
| ENSG00000075790 | 7 | 107220422 | 107269615 | 81 | 11 | -0.34975 | 0.63674 | BCAP29 | 0.63674 | 0.57056 | 0.54679 |
| ENSG00000110448 | 11 | 60869867 | 60895324 | 67 | 13 | -0.34982 | 0.63676 | CD5 | 0.63676 | 0.68203 | 0.57181 |
| ENSG00000055118 | 7 | 150642049 | 150675403 | 94 | 16 | -0.35013 | 0.63688 | KCNH2 | 0.63688 | 0.6782 | 0.74075 |
| ENSG00000196990 | 9 | 136444079 | 136451319 | 1 | 1 | -0.35018 | 0.6369 | FAM163B | 0.6369 | 0.97332 | 0.88343 |
| ENSG00000124766 | 6 | 21593972 | 21598847 | 8 | 4 | -0.35047 | 0.63701 | SOX4 | 0.63701 | 0.90613 | 0.75769 |
| ENSG00000075539 | 4 | 48499378 | 48782339 | 442 | 24 | -0.35064 | 0.63707 | FRYL | 0.63707 | 0.63334 | 0.63929 |
| ENSG00000004838 | 3 | 50378541 | 50384283 | 6 | 3 | -0.35066 | 0.63708 | ZMYND10 | 0.63708 | 0.32628 | 0.36502 |
| ENSG00000059573 | 10 | 97365696 | 97416463 | 201 | 24 | -0.35072 | 0.6371 | ALDH18A1 | 0.6371 | 0.69165 | 0.75566 |
| ENSG00000179284 | 19 | 13075973 | 13085567 | 27 | 11 | -0.35073 | 0.6371 | DAND5 | 0.6371 | 0.63445 | 0.52361 |
| ENSG00000109956 | 11 | 134248398 | 134281812 | 159 | 17 | -0.35105 | 0.63722 | B3GAT1 | 0.63722 | 0.66022 | 0.55517 |
| ENSG00000203876 | 10 | 111701029 | 111713649 | 21 | 2 | -0.35123 | 0.63729 | RP11-451M19.3 | 0.63729 | 0.71617 | 0.7489 |
| ENSG00000077585 | 1 | 236305832 | 236385165 | 308 | 25 | -0.3513 | 0.63732 | GPR137B | 0.63732 | 0.70353 | 0.78904 |
| ENSG00000184232 | 11 | 120081475 | 120101041 | 97 | 17 | -0.35134 | 0.63733 | OAF | 0.63733 | 0.60309 | 0.62512 |
| ENSG00000134285 | 12 | 49315301 | 49320257 | 11 | 3 | -0.35142 | 0.63736 | FKBP11 | 0.63736 | 0.5113 | 0.51163 |
| ENSG00000145861 | 5 | 159774758 | 159797648 | 76 | 12 | -0.35172 | 0.63747 | C1QTNF2 | 0.63747 | 0.60331 | 0.68293 |
| ENSG00000140832 | 16 | 71660064 | 71676017 | 35 | 8 | -0.35182 | 0.63751 | MARVELD3 | 0.63751 | 0.70202 | 0.72957 |
| ENSG00000133216 | 1 | 23037332 | 23241818 | 540 | 62 | -0.35191 | 0.63755 | EPHB2 | 0.63755 | 0.62773 | 0.70218 |
| ENSG00000105610 | 19 | 12995237 | 12997995 | 4 | 2 | -0.35193 | 0.63755 | KLF1 | 0.63755 | 0.74431 | 0.6747 |
| ENSG00000215749 | 15 | 82930438 | 82939159 | 9 | 3 | -0.3522 | 0.63765 | GOLGA6L18 | 0.63765 | 0.77214 | 0.70256 |
| ENSG00000182253 | 15 | 99638420 | 99675798 | 211 | 24 | -0.35228 | 0.63769 | SYNM | 0.63769 | 0.56812 | 0.54818 |
| ENSG00000236032 | 3 | 97868170 | 97869249 | 5 | 1 | -0.35238 | 0.63772 | OR5H14 | 0.63772 | 0.57882 | 0.54308 |
| ENSG00000158220 | 3 | 138153428 | 138200528 | 89 | 18 | -0.35238 | 0.63772 | ESYT3 | 0.63772 | 0.59884 | 0.57428 |
| ENSG00000136938 | 9 | 100745643 | 100778225 | 57 | 11 | -0.35239 | 0.63773 | ANP32B | 0.63773 | 0.68151 | 0.707 |
| ENSG00000172500 | 11 | 65651212 | 65656010 | 11 | 2 | -0.35249 | 0.63776 | FIBP | 0.63776 | 0.85596 | 0.90528 |
| ENSG00000169918 | 15 | 31775329 | 32162992 | 1241 | 41 | -0.35263 | 0.63782 | OTUD7A | 0.63782 | 0.67206 | 0.68579 |
| ENSG00000165417 | 14 | 81641796 | 81687721 | 115 | 19 | -0.35269 | 0.63784 | GTF2A1 | 0.63784 | 0.63147 | 0.56901 |
| ENSG00000089006 | 20 | 17922241 | 17949623 | 115 | 11 | -0.35271 | 0.63785 | SNX5 | 0.63785 | 0.61504 | 0.73256 |
| ENSG00000153250 | 2 | 161128662 | 161350305 | 567 | 33 | -0.3529 | 0.63792 | RBMS1 | 0.63792 | 0.63598 | 0.68867 |
| ENSG00000196935 | 12 | 64238073 | 64541613 | 694 | 83 | -0.35294 | 0.63793 | SRGAP1 | 0.63793 | 0.63307 | 0.58358 |
| ENSG00000178460 | 8 | 67782984 | 67834283 | 30 | 15 | -0.35294 | 0.63794 | MCMDC2 | 0.63794 | 0.82315 | 0.67188 |
| ENSG00000174792 | 4 | 76481258 | 76491095 | 36 | 9 | -0.35327 | 0.63806 | C4orf26 | 0.63806 | 0.65303 | 0.55646 |
| ENSG00000152092 | 1 | 176826438 | 177134109 | 828 | 60 | -0.35343 | 0.63812 | ASTN1 | 0.63812 | 0.6569 | 0.67265 |
| ENSG00000264058 | 17 | 38785049 | 38821393 | 55 | 10 | -0.35357 | 0.63817 | KRT222 | 0.63817 | 0.5908 | 0.57718 |
| ENSG00000189366 | 3 | 125648118 | 125655882 | 52 | 6 | -0.35364 | 0.6382 | ALG1L | 0.6382 | 0.64084 | 0.59685 |
| ENSG00000082497 | 1 | 210406144 | 210419976 | 18 | 7 | -0.35423 | 0.63842 | SERTAD4 | 0.63842 | 0.80932 | 0.58096 |
| ENSG00000196653 | 3 | 44754135 | 44765323 | 44 | 4 | -0.35423 | 0.63842 | ZNF502 | 0.63842 | 0.67655 | 0.53519 |
| ENSG00000185640 | 12 | 53215194 | 53228079 | 26 | 9 | -0.35436 | 0.63846 | KRT79 | 0.63846 | 0.51232 | 0.48339 |
| ENSG00000089123 | 20 | 13246709 | 13619587 | 1309 | 61 | -0.3549 | 0.63867 | TASP1 | 0.63867 | 0.67952 | 0.68144 |
| ENSG00000184254 | 15 | 101417919 | 101456831 | 89 | 17 | -0.355 | 0.6387 | ALDH1A3 | 0.6387 | 0.39043 | 0.46415 |
| ENSG00000257743 | 7 | 141811549 | 141922124 | 369 | 45 | -0.35537 | 0.63884 | RP11-1220K2.2 | 0.63884 | 0.58123 | 0.4062 |
| ENSG00000198523 | 6 | 118869461 | 118881893 | 23 | 5 | -0.35567 | 0.63896 | PLN | 0.63896 | 0.66 | 0.56828 |
| ENSG00000203697 | 1 | 223711349 | 223853436 | 332 | 34 | -0.35573 | 0.63898 | CAPN8 | 0.63898 | 0.68819 | 0.64325 |
| ENSG00000180776 | 13 | 21950263 | 22033509 | 264 | 17 | -0.35591 | 0.63905 | ZDHHC20 | 0.63905 | 0.62882 | 0.67904 |
| ENSG00000047365 | 4 | 35949843 | 36246131 | 856 | 65 | -0.35625 | 0.63917 | ARAP2 | 0.63917 | 0.65335 | 0.66403 |
| ENSG00000164933 | 8 | 104410863 | 104427417 | 63 | 8 | -0.35637 | 0.63922 | SLC25A32 | 0.63922 | 0.60987 | 0.59017 |
| ENSG00000168528 | 1 | 31882412 | 31907525 | 80 | 10 | -0.35646 | 0.63925 | SERINC2 | 0.63925 | 0.70935 | 0.75482 |
| ENSG00000108799 | 17 | 40852293 | 40897071 | 53 | 7 | -0.35645 | 0.63925 | EZH1 | 0.63925 | 0.6842 | 0.63262 |
| ENSG00000101265 | 20 | 4760669 | 4804291 | 244 | 37 | -0.35676 | 0.63937 | RASSF2 | 0.63937 | 0.45645 | 0.62928 |
| ENSG00000136279 | 7 | 44084239 | 44109055 | 84 | 10 | -0.35688 | 0.63941 | DBNL | 0.63941 | 0.64079 | 0.71804 |
| ENSG00000172572 | 12 | 20522179 | 20837315 | 1218 | 90 | -0.35721 | 0.63953 | PDE3A | 0.63953 | 0.7068 | 0.73358 |
| ENSG00000181541 | 4 | 151503077 | 151505843 | 3 | 2 | -0.35723 | 0.63954 | MAB21L2 | 0.63954 | 0.20712 | 0.13046 |
| ENSG00000111653 | 12 | 6759446 | 6772314 | 23 | 7 | -0.35743 | 0.63961 | ING4 | 0.63961 | 0.64477 | 0.56764 |
| ENSG00000175334 | 11 | 65769550 | 65771620 | 11 | 3 | -0.35753 | 0.63965 | BANF1 | 0.63965 | 0.64662 | 0.66243 |
| ENSG00000155052 | 2 | 124782864 | 125672864 | 3153 | 131 | -0.35754 | 0.63966 | CNTNAP5 | 0.63966 | 0.63956 | 0.67759 |
| ENSG00000238243 | 1 | 248031277 | 248060449 | 165 | 25 | -0.35766 | 0.6397 | OR2W3 | 0.6397 | 0.71366 | 0.76543 |
| ENSG00000108468 | 17 | 46147414 | 46178883 | 100 | 12 | -0.35778 | 0.63975 | CBX1 | 0.63975 | 0.62963 | 0.62298 |
| ENSG00000175544 | 11 | 67219877 | 67226699 | 16 | 6 | -0.35782 | 0.63976 | CABP4 | 0.63976 | 0.76954 | 0.70894 |
| ENSG00000100029 | 22 | 30972612 | 31003070 | 173 | 33 | -0.3582 | 0.6399 | PES1 | 0.6399 | 0.64027 | 0.6674 |
| ENSG00000160097 | 1 | 33327869 | 33338083 | 16 | 5 | -0.35821 | 0.63991 | FNDC5 | 0.63991 | 0.50645 | 0.52086 |
| ENSG00000108622 | 17 | 62079954 | 62097994 | 26 | 12 | -0.35851 | 0.64002 | ICAM2 | 0.64002 | 0.44638 | 0.45836 |
| ENSG00000151615 | 4 | 147560045 | 147563626 | 10 | 4 | -0.35853 | 0.64003 | POU4F2 | 0.64003 | 0.82993 | 0.87702 |
| ENSG00000134758 | 18 | 29671818 | 29711524 | 107 | 15 | -0.35861 | 0.64006 | RNF138 | 0.64006 | 0.68545 | 0.76337 |
| ENSG00000171201 | 4 | 71235810 | 71255961 | 116 | 11 | -0.35863 | 0.64007 | SMR3B | 0.64007 | 0.50777 | 0.60389 |
| ENSG00000156689 | 11 | 58601542 | 58671688 | 374 | 11 | -0.35927 | 0.6403 | GLYATL2 | 0.6403 | 0.41343 | 0.22549 |
| ENSG00000103035 | 16 | 74330673 | 74340186 | 17 | 9 | -0.35932 | 0.64032 | PSMD7 | 0.64032 | 0.50754 | 0.49215 |
| ENSG00000013583 | 12 | 13127798 | 13153207 | 73 | 11 | -0.35933 | 0.64033 | HEBP1 | 0.64033 | 0.70762 | 0.76265 |
| ENSG00000164946 | 9 | 14734664 | 14910993 | 859 | 83 | -0.35944 | 0.64037 | FREM1 | 0.64037 | 0.69707 | 0.77882 |
| ENSG00000108578 | 17 | 28575218 | 28619074 | 88 | 15 | -0.3595 | 0.64039 | BLMH | 0.64039 | 0.56949 | 0.5163 |
| ENSG00000178636 | 4 | 120113928 | 120133799 | 72 | 13 | -0.35965 | 0.64045 | RP11-455G16.1 | 0.64045 | 0.73186 | 0.72616 |
| ENSG00000021762 | 11 | 3108346 | 3187969 | 485 | 65 | -0.35978 | 0.64049 | OSBPL5 | 0.64049 | 0.64901 | 0.73128 |

| | | | | | | | | | | | |
|---|---|---|---|---|---|---|---|---|---|---|---|
| ENSG00000155666 | 16 | 27214807 | 27233089 | 33 | 8 | -0.35992 | 0.64054 | KDM8 | 0.64054 | 0.58424 | 0.66068 |
| ENSG00000180432 | 3 | 42897497 | 42917633 | 54 | 10 | -0.36 | 0.64058 | CYP8B1 | 0.64058 | 0.6857 | 0.57131 |
| ENSG00000167131 | 17 | 42976510 | 42982758 | 19 | 4 | -0.36023 | 0.64066 | CCDC103 | 0.64066 | 0.71134 | 0.75079 |
| ENSG00000182676 | 17 | 79791368 | 79792926 | 3 | 1 | -0.3604 | 0.64073 | PPP1R27 | 0.64073 | 0.57038 | 0.60747 |
| ENSG00000109220 | 4 | 54875956 | 54930857 | 92 | 9 | -0.36056 | 0.64078 | CHIC2 | 0.64078 | 0.68438 | 0.69238 |
| ENSG00000177885 | 17 | 73314157 | 73401790 | 281 | 13 | -0.36056 | 0.64079 | GRB2 | 0.64079 | 0.64257 | 0.61677 |
| ENSG00000108557 | 17 | 17584787 | 17714767 | 236 | 32 | -0.36061 | 0.6408 | RAI1 | 0.6408 | 0.64803 | 0.7041 |
| ENSG00000185624 | 17 | 79801035 | 79818570 | 40 | 3 | -0.3606 | 0.6408 | P4HB | 0.6408 | 0.672 | 0.6183 |
| ENSG00000124201 | 20 | 47854483 | 47894963 | 176 | 14 | -0.3608 | 0.64087 | ZNFX1 | 0.64087 | 0.61184 | 0.56134 |
| ENSG00000073605 | 17 | 38060848 | 38076107 | 64 | 5 | -0.36105 | 0.64097 | GSDMB | 0.64097 | 0.65432 | 0.7576 |
| ENSG00000182307 | 8 | 146277764 | 146281416 | 7 | 4 | -0.36116 | 0.64101 | C8orf33 | 0.64101 | 0.24911 | 0.23037 |
| ENSG00000134333 | 11 | 18415935 | 18429972 | 61 | 5 | -0.36119 | 0.64102 | LDHA | 0.64102 | 0.81254 | 0.73285 |
| ENSG00000006327 | 16 | 3068446 | 3072384 | 11 | 3 | -0.3612 | 0.64102 | TNFRSF12A | 0.64102 | 0.86933 | 0.6063 |
| ENSG00000168679 | 1 | 110905470 | 110933704 | 78 | 15 | -0.36127 | 0.64105 | SLC16A4 | 0.64105 | 0.55579 | 0.57235 |
| ENSG00000137727 | 11 | 110447766 | 110583912 | 402 | 25 | -0.36129 | 0.64106 | ARHGAP20 | 0.64106 | 0.66823 | 0.74404 |
| ENSG00000080493 | 4 | 72053003 | 72437804 | 847 | 47 | -0.36135 | 0.64108 | SLC4A4 | 0.64108 | 0.68004 | 0.72726 |
| ENSG00000186891 | 1 | 1138888 | 1142071 | 11 | 2 | -0.36147 | 0.64113 | TNFRSF18 | 0.64113 | 0.45768 | 0.34971 |
| ENSG00000181418 | 12 | 49388932 | 49393092 | 7 | 4 | -0.36165 | 0.64119 | DDN | 0.64119 | 0.45246 | 0.442 |
| ENSG00000135924 | 2 | 220143989 | 220151622 | 20 | 5 | -0.36165 | 0.6412 | DNAJB2 | 0.6412 | 0.52601 | 0.40949 |
| ENSG00000257355 | 19 | 12128703 | 12163782 | 41 | 21 | -0.3617 | 0.64121 | CTD-2006C1.10 | 0.64121 | 0.76048 | 0.49944 |
| ENSG00000141753 | 17 | 38599702 | 38613983 | 39 | 9 | -0.36201 | 0.64133 | IGFBP4 | 0.64133 | 0.62769 | 0.59828 |
| ENSG00000008118 | 1 | 209757062 | 209787283 | 112 | 19 | -0.36222 | 0.6414 | CAMK1G | 0.6414 | 0.57576 | 0.77984 |
| ENSG00000139182 | 12 | 7282294 | 7311541 | 63 | 10 | -0.36271 | 0.64159 | CLSTN3 | 0.64159 | 0.52064 | 0.53998 |
| ENSG00000108684 | 17 | 31340105 | 32501983 | 4140 | 301 | -0.36274 | 0.6416 | ASIC2 | 0.6416 | 0.62613 | 0.65529 |
| ENSG00000184492 | 2 | 114256661 | 114258728 | 13 | 5 | -0.3628 | 0.64162 | FOXD4L1 | 0.64162 | 0.67046 | 0.6504 |
| ENSG00000140525 | 15 | 89787180 | 89860492 | 171 | 11 | -0.36297 | 0.64169 | FANCI | 0.64169 | 0.63373 | 0.65306 |
| ENSG00000165105 | 9 | 85594500 | 85678092 | 194 | 14 | -0.36349 | 0.64188 | RASEF | 0.64188 | 0.65837 | 0.65107 |
| ENSG00000226490 | 8 | 142524738 | 142528837 | 21 | 6 | -0.36357 | 0.64191 | AC138647.1 | 0.64191 | 0.48315 | 0.45348 |
| ENSG00000213002 | 8 | 71485681 | 71486671 | 4 | 2 | -0.36365 | 0.64194 | AC120194.1 | 0.64194 | 0.82507 | 0.81312 |
| ENSG00000151923 | 10 | 121334199 | 121356541 | 38 | 9 | -0.36375 | 0.64198 | TIAL1 | 0.64198 | 0.60737 | 0.73786 |
| ENSG00000122224 | 1 | 160765864 | 160798051 | 103 | 10 | -0.36377 | 0.64199 | LY9 | 0.64199 | 0.61379 | 0.62769 |
| ENSG00000163818 | 3 | 45864808 | 45957534 | 215 | 33 | -0.36394 | 0.64205 | LZTFL1 | 0.64205 | 0.65936 | 0.65471 |
| ENSG00000166949 | 15 | 67356101 | 67487533 | 456 | 64 | -0.36404 | 0.64209 | SMAD3 | 0.64209 | 0.70805 | 0.52886 |
| ENSG00000161798 | 12 | 50355653 | 50359464 | 11 | 8 | -0.36416 | 0.64213 | AQP5 | 0.64213 | 0.77194 | 0.82045 |
| ENSG00000258466 | 14 | 68126537 | 68157084 | 88 | 15 | -0.36428 | 0.64218 | RP11-1012A1.4 | 0.64218 | 0.69111 | 0.66046 |
| ENSG00000117586 | 1 | 173152873 | 173176452 | 69 | 10 | -0.36469 | 0.64233 | TNFSF4 | 0.64233 | 0.66275 | 0.67126 |
| ENSG00000143543 | 1 | 153946745 | 153950164 | 2 | 1 | -0.36478 | 0.64236 | JTB | 0.64236 | 0.27294 | 0.20874 |
| ENSG00000168306 | 3 | 58490863 | 58523046 | 74 | 19 | -0.36495 | 0.64242 | ACOX2 | 0.64242 | 0.72114 | 0.69122 |
| ENSG00000119979 | 10 | 120863598 | 120897496 | 109 | 5 | -0.3651 | 0.64248 | FAM45A | 0.64248 | 0.6575 | 0.69679 |
| ENSG00000163630 | 3 | 63213991 | 63602597 | 1683 | 111 | -0.36525 | 0.64254 | SYNPR | 0.64254 | 0.8313 | 0.8283 |
| ENSG00000143756 | 1 | 224301789 | 224349749 | 161 | 14 | -0.36607 | 0.64284 | FBXO28 | 0.64284 | 0.65239 | 0.67409 |
| ENSG00000129473 | 14 | 23767999 | 23780968 | 27 | 13 | -0.36614 | 0.64287 | BCL2L2 | 0.64287 | 0.70256 | 0.84131 |
| ENSG00000157837 | 12 | 121200313 | 121342174 | 573 | 24 | -0.3662 | 0.64289 | SPPL3 | 0.64289 | 0.64094 | 0.66521 |
| ENSG00000166960 | 18 | 30517366 | 31021065 | 1136 | 31 | -0.36632 | 0.64294 | CCDC178 | 0.64294 | 0.66659 | 0.6557 |
| ENSG00000181374 | 17 | 32683471 | 32685629 | 2 | 1 | -0.3666 | 0.64304 | CCL13 | 0.64304 | 0.78957 | 0.61449 |
| ENSG00000165806 | 10 | 115438942 | 115490662 | 186 | 21 | -0.36664 | 0.64305 | CASP7 | 0.64305 | 0.72364 | 0.80064 |
| ENSG00000105722 | 19 | 42751724 | 42759309 | 12 | 2 | -0.3667 | 0.64308 | ERF | 0.64308 | 0.69666 | 0.56705 |
| ENSG00000167646 | 19 | 55670031 | 55678090 | 45 | 8 | -0.36675 | 0.6431 | DNAAF3 | 0.6431 | 0.79049 | 0.80324 |
| ENSG00000269883 | 14 | 77607026 | 77609339 | 6 | 2 | -0.36695 | 0.64317 | AC007375.1 | 0.64317 | 0.78099 | 0.83528 |
| ENSG00000122859 | 10 | 71331454 | 71332994 | 6 | 2 | -0.36718 | 0.64326 | NEUROG3 | 0.64326 | 0.30204 | 0.27218 |
| ENSG00000176635 | 22 | 30476163 | 30573064 | 238 | 26 | -0.36744 | 0.64335 | HORMAD2 | 0.64335 | 0.67414 | 0.62695 |
| ENSG00000253710 | 13 | 52586534 | 52603800 | 31 | 7 | -0.36811 | 0.64361 | ALG11 | 0.64361 | 0.59662 | 0.53544 |
| ENSG00000119242 | 12 | 124403207 | 124457378 | 131 | 10 | -0.36815 | 0.64362 | CCDC92 | 0.64362 | 0.6785 | 0.66273 |
| ENSG00000099625 | 19 | 1228286 | 1238026 | 22 | 7 | -0.36818 | 0.64363 | C19orf26 | 0.64363 | 0.37478 | 0.12785 |
| ENSG00000181074 | 11 | 5775923 | 5776959 | 6 | 2 | -0.36828 | 0.64367 | OR52N4 | 0.64367 | 0.81478 | 0.85347 |
| ENSG00000167747 | 19 | 51300961 | 51308186 | 44 | 9 | -0.36865 | 0.6438 | C19orf48 | 0.6438 | 0.61868 | 0.8381 |
| ENSG00000148341 | 9 | 131769315 | 131790582 | 34 | 8 | -0.36868 | 0.64382 | SH3GLB2 | 0.64382 | 0.87187 | 0.88755 |
| ENSG00000008300 | 3 | 48673902 | 48700348 | 21 | 5 | -0.36883 | 0.64387 | CELSR3 | 0.64387 | 0.64294 | 0.7859 |
| ENSG00000119844 | 2 | 64751465 | 64820139 | 165 | 10 | -0.3692 | 0.64401 | AFTPH | 0.64401 | 0.68418 | 0.70938 |
| ENSG00000142875 | 1 | 84543745 | 84704181 | 297 | 19 | -0.36961 | 0.64417 | PRKACB | 0.64417 | 0.62827 | 0.65467 |
| ENSG00000170509 | 4 | 88224941 | 88244058 | 67 | 15 | -0.36963 | 0.64417 | HSD17B13 | 0.64417 | 0.70505 | 0.61784 |
| ENSG00000163879 | 1 | 38022520 | 38032458 | 31 | 8 | -0.36964 | 0.64418 | DNALI1 | 0.64418 | 0.64537 | 0.64539 |
| ENSG00000163491 | 3 | 27151576 | 27410951 | 935 | 25 | -0.36982 | 0.64424 | NEK10 | 0.64424 | 0.66002 | 0.65954 |
| ENSG00000066379 | 6 | 30026676 | 30032686 | 44 | 7 | -0.36989 | 0.64427 | ZNRD1 | 0.64427 | 0.58972 | 0.59922 |
| ENSG00000156026 | 10 | 74451889 | 74647452 | 236 | 16 | -0.37004 | 0.64432 | MCU | 0.64432 | 0.64316 | 0.62054 |
| ENSG00000140598 | 15 | 82422571 | 82555104 | 522 | 31 | -0.37006 | 0.64433 | EFTUD1 | 0.64433 | 0.64575 | 0.65565 |
| ENSG00000196466 | 19 | 12500830 | 12512085 | 16 | 4 | -0.3701 | 0.64435 | ZNF799 | 0.64435 | 0.31211 | 0.14731 |
| ENSG00000183580 | 5 | 15500305 | 15939900 | 1038 | 56 | -0.37013 | 0.64436 | FBXL7 | 0.64436 | 0.66581 | 0.71708 |
| ENSG00000075826 | 10 | 102246399 | 102289628 | 31 | 11 | -0.37043 | 0.64447 | SEC31B | 0.64447 | 0.27286 | 0.31966 |
| ENSG00000072210 | 17 | 19551449 | 19580911 | 83 | 12 | -0.37042 | 0.64447 | ALDH3A2 | 0.64447 | 0.67685 | 0.67653 |
| ENSG00000131845 | 19 | 57862675 | 57871266 | 32 | 11 | -0.37047 | 0.64448 | ZNF304 | 0.64448 | 0.653 | 0.5293 |
| ENSG00000197506 | 9 | 86890372 | 86955672 | 250 | 35 | -0.37101 | 0.64469 | SLC28A3 | 0.64469 | 0.11438 | 0.14183 |
| ENSG00000093000 | 22 | 45559722 | 45583896 | 113 | 11 | -0.37106 | 0.6447 | NUP50 | 0.6447 | 0.72792 | 0.80352 |

| | | | | | | | | | | | |
|---|---|---|---|---|---|---|---|---|---|---|---|
| ENSG00000168952 | 14 | 25278862 | 25519503 | 752 | 68 | -0.3712 | 0.64476 | STXBP6 | 0.64476 | 0.6942 | 0.76482 |
| ENSG00000021300 | 11 | 73357223 | 73373864 | 65 | 14 | -0.37124 | 0.64477 | PLEKHB1 | 0.64477 | 0.78273 | 0.67455 |
| ENSG00000105270 | 19 | 36505562 | 36524245 | 32 | 11 | -0.37159 | 0.6449 | CLIP3 | 0.6449 | 0.28787 | 0.26071 |
| ENSG00000070614 | 5 | 149865381 | 149937773 | 175 | 22 | -0.37172 | 0.64495 | NDST1 | 0.64495 | 0.69059 | 0.82148 |
| ENSG00000150201 | 10 | 43867090 | 43871783 | 11 | 5 | -0.37174 | 0.64496 | FXYD4 | 0.64496 | 0.67417 | 0.6415 |
| ENSG00000197162 | 16 | 30585061 | 30597092 | 17 | 8 | -0.37187 | 0.645 | ZNF785 | 0.645 | 0.71284 | 0.79563 |
| ENSG00000115525 | 2 | 86066267 | 86116137 | 175 | 16 | -0.37195 | 0.64503 | ST3GAL5 | 0.64503 | 0.66104 | 0.72339 |
| ENSG00000256407 | 1 | 54606897 | 54665746 | 216 | 37 | -0.3723 | 0.64517 | RP11-446E24.4 | 0.64517 | 0.33914 | 0.2658 |
| ENSG00000224838 | 5 | 59726565 | 59726939 | 1 | 1 | -0.37239 | 0.6452 | FKSG52 | 0.6452 | 0.28213 | 0.32496 |
| ENSG00000214167 | 17 | 65027509 | 65028198 | 1 | 1 | -0.37239 | 0.6452 | AC005544.1 | 0.6452 | 0.91604 | 0.84367 |
| ENSG00000100918 | 14 | 24641062 | 24649463 | 19 | 5 | -0.37261 | 0.64528 | REC8 | 0.64528 | 0.50543 | 0.55455 |
| ENSG00000139083 | 12 | 11802788 | 12048336 | 797 | 99 | -0.37269 | 0.64531 | ETV6 | 0.64531 | 0.62134 | 0.50775 |
| ENSG00000187811 | 11 | 71292956 | 71318487 | 184 | 15 | -0.37282 | 0.64536 | AP000867.1 | 0.64536 | 0.49475 | 0.46566 |
| ENSG00000075223 | 7 | 80371854 | 80551675 | 445 | 58 | -0.37287 | 0.64538 | SEMA3C | 0.64538 | 0.73338 | 0.7825 |
| ENSG00000148057 | 9 | 86237964 | 86259045 | 63 | 12 | -0.3731 | 0.64546 | IDNK | 0.64546 | 0.67263 | 0.73318 |
| ENSG00000172775 | 16 | 57186378 | 57220028 | 70 | 14 | -0.3731 | 0.64546 | FAM192A | 0.64546 | 0.56173 | 0.51552 |
| ENSG00000143079 | 1 | 112938803 | 113006078 | 195 | 8 | -0.3733 | 0.64554 | CTTNBP2NL | 0.64554 | 0.65477 | 0.59399 |
| ENSG00000204632 | 6 | 29794744 | 29798902 | 77 | 7 | -0.37347 | 0.6456 | HLA-G | 0.6456 | 0.82411 | 0.76183 |
| ENSG00000158716 | 1 | 159750722 | 159752333 | 6 | 2 | -0.37366 | 0.64567 | DUSP23 | 0.64567 | 0.92922 | 0.49627 |
| ENSG00000220575 | 7 | 154858777 | 154863267 | 19 | 5 | -0.37375 | 0.6457 | HTR5A-AS1 | 0.6457 | 0.54647 | 0.63987 |
| ENSG00000166803 | 15 | 64657193 | 64679886 | 65 | 11 | -0.37374 | 0.6457 | KIAA0101 | 0.6457 | 0.6758 | 0.64551 |
| ENSG00000144711 | 3 | 12938719 | 13114617 | 587 | 80 | -0.37382 | 0.64573 | IQSEC1 | 0.64573 | 0.68724 | 0.76237 |
| ENSG00000160255 | 21 | 46305868 | 46351904 | 253 | 49 | -0.37397 | 0.64579 | ITGB2 | 0.64579 | 0.78887 | 0.85871 |
| ENSG00000128609 | 7 | 123177051 | 123198309 | 44 | 7 | -0.37405 | 0.64582 | NDUFA5 | 0.64582 | 0.82485 | 0.92517 |
| ENSG00000154380 | 1 | 225674537 | 225840844 | 444 | 18 | -0.37409 | 0.64583 | ENAH | 0.64583 | 0.64002 | 0.65096 |
| ENSG00000182333 | 10 | 90424198 | 90438571 | 47 | 12 | -0.37421 | 0.64588 | LIPF | 0.64588 | 0.60636 | 0.72689 |
| ENSG00000159713 | 16 | 67423712 | 67427438 | 9 | 3 | -0.37433 | 0.64592 | TPPP3 | 0.64592 | 0.74737 | 0.71213 |
| ENSG00000129646 | 17 | 74270130 | 74303761 | 142 | 15 | -0.37434 | 0.64592 | QRICH2 | 0.64592 | 0.42384 | 0.44417 |
| ENSG00000104131 | 15 | 44829255 | 44855227 | 46 | 5 | -0.3744 | 0.64595 | EIF3J | 0.64595 | 0.6881 | 0.73408 |
| ENSG00000138172 | 10 | 105206543 | 105212660 | 9 | 5 | -0.37456 | 0.64601 | CALHM2 | 0.64601 | 0.51785 | 0.61623 |
| ENSG00000168061 | 11 | 64808373 | 64812300 | 15 | 4 | -0.37456 | 0.64601 | SAC3D1 | 0.64601 | 0.74113 | 0.84277 |
| ENSG00000145965 | 6 | 3982909 | 3984372 | 5 | 2 | -0.37466 | 0.64604 | C6ORF50 | 0.64604 | 0.82982 | 0.87291 |
| ENSG00000135637 | 2 | 74699113 | 74710535 | 16 | 9 | -0.37481 | 0.6461 | CCDC142 | 0.6461 | 0.46735 | 0.40456 |
| ENSG00000168427 | 2 | 239047363 | 239061588 | 48 | 13 | -0.37497 | 0.64616 | KLHL30 | 0.64616 | 0.58954 | 0.30271 |
| ENSG00000152464 | 10 | 15139179 | 15181817 | 203 | 23 | -0.37508 | 0.6462 | RPP38 | 0.6462 | 0.7318 | 0.83298 |
| ENSG00000166165 | 14 | 103985996 | 103989448 | 5 | 2 | -0.37526 | 0.64627 | CKB | 0.64627 | 0.73042 | 0.74532 |
| ENSG00000146858 | 7 | 138710452 | 138720775 | 57 | 4 | -0.37538 | 0.64631 | ZC3HAV1L | 0.64631 | 0.52531 | 0.32414 |
| ENSG00000137878 | 15 | 57884106 | 58006943 | 521 | 36 | -0.37543 | 0.64633 | GCOM1 | 0.64633 | 0.61253 | 0.63406 |
| ENSG00000156127 | 14 | 75988768 | 76013358 | 68 | 12 | -0.37548 | 0.64635 | BATF | 0.64635 | 0.44015 | 0.70248 |
| ENSG00000169762 | 4 | 16162128 | 16229033 | 246 | 26 | -0.37584 | 0.64648 | TAPT1 | 0.64648 | 0.6894 | 0.81694 |
| ENSG00000110917 | 12 | 121124672 | 121139667 | 64 | 18 | -0.37583 | 0.64648 | MLEC | 0.64648 | 0.66519 | 0.57309 |
| ENSG00000168418 | 16 | 84255823 | 84273356 | 95 | 20 | -0.37589 | 0.6465 | KCNG4 | 0.6465 | 0.36073 | 0.3046 |
| ENSG00000188051 | 19 | 17546318 | 17559376 | 54 | 11 | -0.37597 | 0.64653 | TMEM221 | 0.64653 | 0.78479 | 0.58972 |
| ENSG00000197651 | 12 | 91299399 | 91348953 | 104 | 19 | -0.37612 | 0.64658 | CCER1 | 0.64658 | 0.66592 | 0.52367 |
| ENSG00000196796 | 16 | 29049923 | 29064047 | 9 | 5 | -0.37629 | 0.64665 | CTB-134H23.2 | 0.64665 | 0.91614 | 0.75586 |
| ENSG00000237440 | 19 | 20718631 | 20748615 | 174 | 20 | -0.37643 | 0.6467 | ZNF737 | 0.6467 | 0.65704 | 0.61364 |
| ENSG00000108852 | 17 | 41952725 | 41987068 | 103 | 9 | -0.37648 | 0.64672 | MPP2 | 0.64672 | 0.6339 | 0.53952 |
| ENSG00000105641 | 19 | 17982782 | 18005983 | 83 | 10 | -0.37662 | 0.64677 | SLC5A5 | 0.64677 | 0.4962 | 0.4782 |
| ENSG00000127564 | 16 | 3018025 | 3030540 | 17 | 5 | -0.37669 | 0.6468 | PKMYT1 | 0.6468 | 0.50341 | 0.38311 |
| ENSG00000170515 | 12 | 56498103 | 56507691 | 15 | 7 | -0.37673 | 0.64681 | PA2G4 | 0.64681 | 0.6856 | 0.83274 |
| ENSG00000249773 | 7 | 55954970 | 56022932 | 243 | 29 | -0.37687 | 0.64686 | MRPS17 | 0.64686 | 0.70795 | 0.72389 |
| ENSG00000198585 | 3 | 131100515 | 131107674 | 10 | 6 | -0.37696 | 0.6469 | NUDT16 | 0.6469 | 0.67323 | 0.86538 |
| ENSG00000188822 | 1 | 24197016 | 24285549 | 286 | 16 | -0.37731 | 0.64703 | CNR2 | 0.64703 | 0.64593 | 0.6802 |
| ENSG00000164867 | 7 | 150688083 | 150711676 | 45 | 11 | -0.37735 | 0.64704 | NOS3 | 0.64704 | 0.78839 | 0.83925 |
| ENSG00000074071 | 16 | 1821891 | 1823156 | 6 | 3 | -0.37735 | 0.64704 | MRPS34 | 0.64704 | 0.80747 | 0.85947 |
| ENSG00000236609 | 7 | 6655248 | 6663921 | 22 | 6 | -0.3774 | 0.64706 | ZNF853 | 0.64706 | 0.48315 | 0.33681 |
| ENSG00000269476 | 19 | 58388133 | 58427959 | 118 | 18 | -0.37746 | 0.64708 | CTD-2583A14.9 | 0.64708 | 0.67426 | 0.69398 |
| ENSG00000179832 | 8 | 145202919 | 145316843 | 141 | 19 | -0.37755 | 0.64712 | MROH1 | 0.64712 | 0.6529 | 0.51127 |
| ENSG00000174695 | 5 | 82348665 | 82373682 | 49 | 9 | -0.3777 | 0.64717 | TMEM167A | 0.64717 | 0.63341 | 0.6 |
| ENSG00000062282 | 11 | 75470557 | 75512579 | 50 | 6 | -0.37781 | 0.64721 | DGAT2 | 0.64721 | 0.73596 | 0.81941 |
| ENSG00000142583 | 1 | 9095166 | 9148537 | 270 | 27 | -0.37794 | 0.64726 | SLC2A5 | 0.64726 | 0.76882 | 0.85167 |
| ENSG00000115194 | 2 | 27476552 | 27498685 | 35 | 8 | -0.378 | 0.64729 | SLC30A3 | 0.64729 | 0.78545 | 0.79721 |
| ENSG00000126861 | 17 | 29599031 | 29624557 | 49 | 6 | -0.37801 | 0.64729 | OMG | 0.64729 | 0.64842 | 0.64342 |
| ENSG00000179094 | 17 | 8043790 | 8059824 | 42 | 12 | -0.37842 | 0.64744 | PER1 | 0.64744 | 0.5583 | 0.44454 |
| ENSG00000143520 | 1 | 152321211 | 152332482 | 26 | 5 | -0.37882 | 0.64759 | FLG2 | 0.64759 | 0.62029 | 0.70535 |
| ENSG00000109689 | 4 | 26859300 | 27027003 | 342 | 27 | -0.37914 | 0.64771 | STIM2 | 0.64771 | 0.61761 | 0.60346 |
| ENSG00000142856 | 1 | 63906441 | 64059392 | 308 | 16 | -0.37917 | 0.64772 | ITGB3BP | 0.64772 | 0.63964 | 0.66198 |
| ENSG00000011021 | 1 | 11866207 | 11903201 | 223 | 16 | -0.3792 | 0.64773 | CLCN6 | 0.64773 | 0.68371 | 0.71667 |
| ENSG00000113318 | 5 | 79950467 | 80172279 | 780 | 27 | -0.37938 | 0.6478 | MSH3 | 0.6478 | 0.64116 | 0.6444 |
| ENSG00000104450 | 8 | 101170134 | 101271506 | 229 | 17 | -0.37954 | 0.64785 | SPAG1 | 0.64785 | 0.63205 | 0.6424 |
| ENSG00000178209 | 8 | 144989321 | 145050902 | 177 | 10 | -0.37955 | 0.64786 | PLEC | 0.64786 | 0.63992 | 0.65718 |
| ENSG00000183734 | 11 | 2289725 | 2292182 | 7 | 2 | -0.37955 | 0.64786 | ASCL2 | 0.64786 | 0.91785 | 0.77505 |
| ENSG00000186765 | 17 | 79495422 | 79504156 | 58 | 11 | -0.37972 | 0.64792 | FSCN2 | 0.64792 | 0.50229 | 0.4121 |

| | | | | | | | | | | | |
|---|---|---|---|---|---|---|---|---|---|---|---|
| ENSG00000069849 | 3 | 141594966 | 141645356 | 163 | 21 | -0.37998 | 0.64802 | ATP1B3 | 0.64802 | 0.68777 | 0.71304 |
| ENSG00000143125 | 1 | 110993822 | 110999976 | 21 | 6 | -0.38008 | 0.64806 | PROK1 | 0.64806 | 0.8607 | 0.94597 |
| ENSG00000119328 | 9 | 111696461 | 111713024 | 53 | 10 | -0.38019 | 0.6481 | FAM206A | 0.6481 | 0.68182 | 0.65004 |
| ENSG00000160214 | 21 | 45209394 | 45225174 | 88 | 10 | -0.38032 | 0.64815 | RRP1 | 0.64815 | 0.63173 | 0.62048 |
| ENSG00000174586 | 19 | 58865723 | 58874214 | 17 | 5 | -0.38043 | 0.64819 | ZNF497 | 0.64819 | 0.62703 | 0.77356 |
| ENSG00000075089 | 12 | 100592900 | 100635643 | 120 | 22 | -0.38086 | 0.64835 | ACTR6 | 0.64835 | 0.6535 | 0.71768 |
| ENSG00000083067 | 9 | 73143979 | 74061820 | 3217 | 131 | -0.38108 | 0.64843 | TRPM3 | 0.64843 | 0.65719 | 0.65491 |
| ENSG00000204084 | 1 | 38326369 | 38412729 | 233 | 30 | -0.38128 | 0.6485 | INPP5B | 0.6485 | 0.69558 | 0.64547 |
| ENSG00000255098 | 8 | 11870545 | 11873043 | 37 | 6 | -0.38134 | 0.64852 | RP11-481A20.11 | 0.64852 | 0.47181 | 0.4018 |
| ENSG00000213918 | 16 | 3661729 | 3730144 | 192 | 19 | -0.38145 | 0.64856 | DNASE1 | 0.64856 | 0.66641 | 0.63753 |
| ENSG00000143793 | 1 | 228288427 | 228293112 | 5 | 3 | -0.38147 | 0.64857 | C1orf35 | 0.64857 | 0.4622 | 0.31883 |
| ENSG00000204021 | 10 | 90484301 | 90512543 | 178 | 10 | -0.38165 | 0.64864 | LIPK | 0.64864 | 0.60705 | 0.65251 |
| ENSG00000118997 | 2 | 196602427 | 196933536 | 779 | 26 | -0.38189 | 0.64873 | DNAH7 | 0.64873 | 0.65326 | 0.64833 |
| ENSG00000160883 | 5 | 176307870 | 176326333 | 47 | 17 | -0.38215 | 0.64883 | HK3 | 0.64883 | 0.7442 | 0.86639 |
| ENSG00000255622 | 5 | 140535577 | 140538639 | 10 | 4 | -0.38223 | 0.64885 | PCDHB17 | 0.64885 | 0.57691 | 0.57851 |
| ENSG00000187838 | 17 | 7293046 | 7307416 | 45 | 13 | -0.38221 | 0.64885 | TMEM256-PLSCR3 | 0.64885 | 0.889 | 0.86329 |
| ENSG00000262481 | 17 | 7293053 | 7307416 | 45 | 13 | -0.38221 | 0.64885 | C17orf61-PLSCR3 | 0.64885 | 0.889 | 0.86329 |
| ENSG00000198336 | 17 | 45277812 | 45301045 | 44 | 10 | -0.3823 | 0.64888 | MYL4 | 0.64888 | 0.60134 | 0.87323 |
| ENSG00000184374 | 8 | 120007691 | 120118821 | 624 | 28 | -0.3826 | 0.64899 | COLEC10 | 0.64899 | 0.62793 | 0.62738 |
| ENSG00000127804 | 17 | 2308856 | 2415185 | 301 | 38 | -0.38262 | 0.649 | METTL16 | 0.649 | 0.65147 | 0.64361 |
| ENSG00000166377 | 18 | 76829285 | 77138283 | 1057 | 29 | -0.38268 | 0.64902 | ATP9B | 0.64902 | 0.63383 | 0.57686 |
| ENSG00000159322 | 15 | 73043710 | 73078187 | 80 | 13 | -0.38274 | 0.64904 | ADPGK | 0.64904 | 0.6848 | 0.73463 |
| ENSG00000259003 | 14 | 23025534 | 23027939 | 14 | 6 | -0.38275 | 0.64905 | AE000662.92 | 0.64905 | 0.94663 | 0.94927 |
| ENSG00000171459 | 9 | 125512019 | 125513062 | 8 | 2 | -0.38282 | 0.64907 | OR1L6 | 0.64907 | 0.79484 | 0.81955 |
| ENSG00000120729 | 5 | 137203480 | 137223540 | 20 | 7 | -0.38295 | 0.64912 | MYOT | 0.64912 | 0.84574 | 0.83295 |
| ENSG00000187650 | 19 | 5904869 | 5910864 | 34 | 7 | -0.38303 | 0.64915 | VMAC | 0.64915 | 0.7698 | 0.84073 |
| ENSG00000105556 | 19 | 305575 | 344798 | 238 | 18 | -0.38346 | 0.64931 | MIER2 | 0.64931 | 0.7838 | 0.84337 |
| ENSG00000138316 | 10 | 72432559 | 72522197 | 486 | 57 | -0.38352 | 0.64933 | ADAMTS14 | 0.64933 | 0.58206 | 0.64341 |
| ENSG00000171466 | 19 | 9759330 | 9785776 | 58 | 6 | -0.38358 | 0.64935 | ZNF562 | 0.64935 | 0.61105 | 0.64488 |
| ENSG00000151474 | 10 | 13685706 | 14504141 | 3802 | 263 | -0.38364 | 0.64938 | FRMD4A | 0.64938 | 0.67915 | 0.75799 |
| ENSG00000169488 | 14 | 20443603 | 20444818 | 14 | 3 | -0.38369 | 0.6494 | OR4K15 | 0.6494 | 0.72918 | 0.67552 |
| ENSG00000241123 | 21 | 45999332 | 46000481 | 7 | 2 | -0.38378 | 0.64943 | KRTAP10-5 | 0.64943 | 0.61284 | 0.27547 |
| ENSG00000132541 | 8 | 99114572 | 99129469 | 74 | 14 | -0.38381 | 0.64944 | HRSP12 | 0.64944 | 0.70496 | 0.69355 |
| ENSG00000123901 | 11 | 94110477 | 94134585 | 119 | 19 | -0.38392 | 0.64948 | GPR83 | 0.64948 | 0.69228 | 0.69183 |
| ENSG00000107077 | 9 | 6720863 | 7175648 | 2400 | 115 | -0.38397 | 0.6495 | KDM4C | 0.6495 | 0.67457 | 0.68066 |
| ENSG00000244607 | 3 | 42734155 | 42814745 | 195 | 19 | -0.38398 | 0.64951 | CCDC13 | 0.64951 | 0.66939 | 0.65838 |
| ENSG00000243073 | 1 | 12939045 | 12946025 | 49 | 7 | -0.38441 | 0.64966 | PRAMEF4 | 0.64966 | 0.67931 | 0.61646 |
| ENSG00000174325 | 2 | 189598882 | 189654831 | 189 | 9 | -0.38443 | 0.64967 | DIRC1 | 0.64967 | 0.67958 | 0.67513 |
| ENSG00000102854 | 16 | 810762 | 818865 | 54 | 11 | -0.38465 | 0.64975 | MSLN | 0.64975 | 0.62895 | 0.64764 |
| ENSG00000184524 | 11 | 787104 | 790123 | 8 | 3 | -0.38478 | 0.6498 | CEND1 | 0.6498 | 0.37555 | 0.46237 |
| ENSG00000111012 | 12 | 58156117 | 58162769 | 6 | 2 | -0.38485 | 0.64982 | CYP27B1 | 0.64982 | 0.76704 | 0.60438 |
| ENSG00000110328 | 11 | 11292423 | 11643552 | 1696 | 173 | -0.38489 | 0.64984 | GALNT18 | 0.64984 | 0.63153 | 0.59641 |
| ENSG00000176749 | 17 | 30813637 | 30818274 | 4 | 2 | -0.38501 | 0.64988 | CDK5R1 | 0.64988 | 0.089655 | 0.20554 |
| ENSG00000270106 | 1 | 231664399 | 231954990 | 766 | 61 | -0.38525 | 0.64998 | TSNAX-DISC1 | 0.64998 | 0.46326 | 0.49203 |
| ENSG00000164393 | 6 | 47624172 | 47665533 | 225 | 7 | -0.3854 | 0.65003 | GPR111 | 0.65003 | 0.65187 | 0.57406 |
| ENSG00000141179 | 17 | 53828340 | 53920191 | 293 | 13 | -0.38568 | 0.65013 | PCTP | 0.65013 | 0.69522 | 0.80907 |
| ENSG00000086200 | 5 | 61699799 | 61924409 | 501 | 24 | -0.38571 | 0.65015 | IPO11 | 0.65015 | 0.65754 | 0.65773 |
| ENSG00000185133 | 22 | 31518717 | 31530682 | 28 | 8 | -0.3859 | 0.65021 | INPP5J | 0.65021 | 0.28919 | 0.26061 |
| ENSG00000172671 | 10 | 46111043 | 46168228 | 204 | 11 | -0.38598 | 0.65025 | ZFAND4 | 0.65025 | 0.60944 | 0.64609 |
| ENSG00000180626 | 17 | 5082831 | 5095178 | 33 | 11 | -0.38599 | 0.65025 | ZNF594 | 0.65025 | 0.70414 | 0.73282 |
| ENSG00000105700 | 19 | 18668572 | 18680197 | 39 | 5 | -0.38607 | 0.65028 | KXD1 | 0.65028 | 0.64765 | 0.73512 |
| ENSG00000230989 | 16 | 83841448 | 83853342 | 43 | 11 | -0.38624 | 0.65034 | HSBP1 | 0.65034 | 0.78343 | 0.47582 |
| ENSG00000196131 | 19 | 53761545 | 53765090 | 17 | 4 | -0.38625 | 0.65034 | VN1R2 | 0.65034 | 0.27967 | 0.22672 |
| ENSG00000129195 | 17 | 6347735 | 6354789 | 35 | 11 | -0.3864 | 0.6504 | FAM64A | 0.6504 | 0.8949 | 0.67882 |
| ENSG00000074201 | 11 | 77225981 | 77348850 | 178 | 27 | -0.38651 | 0.65044 | CLNS1A | 0.65044 | 0.64432 | 0.68398 |
| ENSG00000251606 | 5 | 96424779 | 96519354 | 280 | 39 | -0.3868 | 0.65055 | CTD-2215E18.1 | 0.65055 | 0.66899 | 0.59561 |
| ENSG00000062598 | 20 | 44994688 | 45061704 | 86 | 10 | -0.38685 | 0.65057 | ELMO2 | 0.65057 | 0.67887 | 0.70114 |
| ENSG00000100055 | 22 | 37678068 | 37711382 | 85 | 24 | -0.38696 | 0.65061 | CYTH4 | 0.65061 | 0.53157 | 0.79016 |
| ENSG00000021826 | 2 | 211342406 | 211543831 | 578 | 36 | -0.38714 | 0.65067 | CPS1 | 0.65067 | 0.65758 | 0.66143 |
| ENSG00000176720 | 2 | 242498136 | 242513546 | 63 | 18 | -0.38721 | 0.6507 | BOK | 0.6507 | 0.70288 | 0.66361 |
| ENSG00000000971 | 1 | 196621008 | 196716634 | 273 | 16 | -0.38723 | 0.65071 | CFH | 0.65071 | 0.67128 | 0.73316 |
| ENSG00000116747 | 1 | 193028552 | 193060907 | 41 | 10 | -0.38737 | 0.65076 | TROVE2 | 0.65076 | 0.4928 | 0.36835 |
| ENSG00000172915 | 13 | 35516424 | 36247159 | 2233 | 68 | -0.38776 | 0.6509 | NBEA | 0.6509 | 0.65734 | 0.62741 |
| ENSG00000165118 | 9 | 86553226 | 86571901 | 49 | 12 | -0.38793 | 0.65097 | C9orf64 | 0.65097 | 0.70771 | 0.68176 |
| ENSG00000162836 | 1 | 147101453 | 147142618 | 128 | 16 | -0.38799 | 0.65099 | ACP6 | 0.65099 | 0.66428 | 0.63759 |
| ENSG00000101098 | 20 | 43380449 | 43438979 | 113 | 16 | -0.38862 | 0.65122 | RIMS4 | 0.65122 | 0.69011 | 0.73441 |
| ENSG00000196209 | 20 | 1451386 | 1472233 | 87 | 12 | -0.38873 | 0.65126 | SIRPB2 | 0.65126 | 0.74131 | 0.80689 |
| ENSG00000125895 | 20 | 1161205 | 1166059 | 14 | 9 | -0.38877 | 0.65128 | TMEM74B | 0.65128 | 0.44506 | 0.25857 |
| ENSG00000103502 | 16 | 29869678 | 29875057 | 5 | 3 | -0.38919 | 0.65143 | CDIPT | 0.65143 | 0.64032 | 0.57823 |
| ENSG00000113083 | 5 | 121398890 | 121413980 | 28 | 5 | -0.3892 | 0.65144 | LOX | 0.65144 | 0.39003 | 0.66499 |
| ENSG00000140534 | 15 | 90118713 | 90174287 | 157 | 14 | -0.3894 | 0.65151 | TICRR | 0.65151 | 0.591 | 0.5405 |
| ENSG00000152454 | 19 | 58452201 | 58459043 | 12 | 4 | -0.38942 | 0.65152 | ZNF256 | 0.65152 | 0.37872 | 0.55273 |
| ENSG00000182584 | 20 | 32254304 | 32256331 | 3 | 1 | -0.38958 | 0.65158 | ACTL10 | 0.65158 | 0.91854 | 0.92569 |

| | | | | | | | | | | | |
|---|---|---|---|---|---|---|---|---|---|---|---|
| ENSG00000008710 | 16 | 2138711 | 2185899 | 149 | 19 | -0.38985 | 0.65168 | PKD1 | 0.65168 | 0.62094 | 0.60859 |
| ENSG00000179148 | 17 | 7999218 | 8022365 | 91 | 26 | -0.38991 | 0.6517 | ALOXE3 | 0.6517 | 0.39761 | 0.52476 |
| ENSG00000143226 | 1 | 161475220 | 161493803 | 78 | 14 | -0.39043 | 0.65189 | FCGR2A | 0.65189 | 0.71419 | 0.77043 |
| ENSG00000104983 | 19 | 46498339 | 46524576 | 136 | 23 | -0.39055 | 0.65194 | CCDC61 | 0.65194 | 0.46859 | 0.43483 |
| ENSG00000169967 | 2 | 128056306 | 128146041 | 273 | 19 | -0.39061 | 0.65196 | MAP3K2 | 0.65196 | 0.65667 | 0.67211 |
| ENSG00000100987 | 20 | 25051521 | 25062996 | 45 | 4 | -0.39071 | 0.65199 | VSX1 | 0.65199 | 0.73262 | 0.77545 |
| ENSG00000198718 | 14 | 45431411 | 45543634 | 223 | 14 | -0.39079 | 0.65202 | FAM179B | 0.65202 | 0.64038 | 0.61617 |
| ENSG00000140795 | 16 | 46740891 | 46824319 | 90 | 15 | -0.39077 | 0.65202 | MYLK3 | 0.65202 | 0.67128 | 0.64291 |
| ENSG00000111727 | 12 | 104458235 | 104498639 | 110 | 17 | -0.39119 | 0.65217 | HCFC2 | 0.65217 | 0.59812 | 0.50544 |
| ENSG00000144802 | 3 | 101546835 | 101579866 | 46 | 13 | -0.3912 | 0.65218 | NFKBIZ | 0.65218 | 0.56424 | 0.52897 |
| ENSG00000034677 | 8 | 101269288 | 101348446 | 166 | 9 | -0.3913 | 0.65221 | RNF19A | 0.65221 | 0.74178 | 0.83826 |
| ENSG00000148690 | 10 | 95427640 | 95462329 | 110 | 12 | -0.39161 | 0.65233 | FRA10AC1 | 0.65233 | 0.61655 | 0.61302 |
| ENSG00000103089 | 16 | 74746853 | 74808729 | 236 | 39 | -0.39165 | 0.65234 | FA2H | 0.65234 | 0.76115 | 0.71523 |
| ENSG00000242574 | 6 | 32902406 | 32908847 | 46 | 7 | -0.39173 | 0.65237 | HLA-DMB | 0.65237 | 0.49703 | 0.55679 |
| ENSG00000131437 | 5 | 132028320 | 132073330 | 98 | 7 | -0.39203 | 0.65248 | KIF3A | 0.65248 | 0.70891 | 0.75353 |
| ENSG00000038427 | 5 | 82767284 | 82878122 | 300 | 40 | -0.39215 | 0.65253 | VCAN | 0.65253 | 0.43484 | 0.45605 |
| ENSG00000168439 | 11 | 63952744 | 63972015 | 55 | 13 | -0.39216 | 0.65253 | STIP1 | 0.65253 | 0.69221 | 0.65272 |
| ENSG00000013016 | 2 | 31456880 | 31492313 | 129 | 16 | -0.39239 | 0.65262 | EHD3 | 0.65262 | 0.52463 | 0.20693 |
| ENSG00000026950 | 6 | 26402465 | 26415444 | 57 | 6 | -0.39296 | 0.65283 | BTN3A1 | 0.65283 | 0.6172 | 0.6063 |
| ENSG00000164916 | 7 | 4683388 | 4811074 | 389 | 62 | -0.39346 | 0.65301 | FOXK1 | 0.65301 | 0.80601 | 0.62491 |
| ENSG00000272781 | 14 | 47311134 | 48143999 | 2775 | 88 | -0.39349 | 0.65302 | MDGA2 | 0.65302 | 0.67768 | 0.68896 |
| ENSG00000158417 | 2 | 99953816 | 100017789 | 163 | 19 | -0.39353 | 0.65304 | EIF5B | 0.65304 | 0.64783 | 0.63385 |
| ENSG00000134046 | 18 | 51679079 | 51751158 | 127 | 20 | -0.39367 | 0.65309 | MBD2 | 0.65309 | 0.61804 | 0.70913 |
| ENSG00000107372 | 9 | 74966341 | 74980163 | 21 | 7 | -0.39405 | 0.65323 | ZFAND5 | 0.65323 | 0.59661 | 0.58783 |
| ENSG00000269422 | 16 | 88894366 | 88894545 | 2 | 1 | -0.39418 | 0.65328 | AC092384.1 | 0.65328 | 0.72872 | 0.53799 |
| ENSG00000123838 | 1 | 207277607 | 207318317 | 132 | 19 | -0.39423 | 0.65329 | C4BPA | 0.65329 | 0.68142 | 0.67124 |
| ENSG00000142089 | 11 | 319669 | 327537 | 53 | 12 | -0.3944 | 0.65336 | IFITM3 | 0.65336 | 0.47394 | 0.51925 |
| ENSG00000103642 | 15 | 63413999 | 63434260 | 70 | 8 | -0.39442 | 0.65336 | LACTB | 0.65336 | 0.62617 | 0.52474 |
| ENSG00000081307 | 3 | 132373290 | 132396941 | 31 | 8 | -0.39469 | 0.65346 | UBA5 | 0.65346 | 0.51458 | 0.47503 |
| ENSG00000147475 | 8 | 37594117 | 37616619 | 9 | 6 | -0.39474 | 0.65348 | ERLIN2 | 0.65348 | 0.59178 | 0.62609 |
| ENSG00000160113 | 19 | 17342692 | 17356749 | 46 | 16 | -0.3948 | 0.65351 | NR2F6 | 0.65351 | 0.8961 | 0.90321 |
| ENSG00000104808 | 19 | 49436939 | 49448226 | 60 | 5 | -0.39481 | 0.65351 | DHDH | 0.65351 | 0.88363 | 0.89209 |
| ENSG00000177542 | 11 | 790475 | 798316 | 14 | 7 | -0.39483 | 0.65352 | SLC25A22 | 0.65352 | 0.63849 | 0.52009 |
| ENSG00000267848 | 20 | 62585007 | 62585495 | 1 | 1 | -0.39506 | 0.6536 | AL118506.1 | 0.6536 | 0.075333 | 0.15235 |
| ENSG00000121039 | 8 | 74206847 | 74237516 | 108 | 18 | -0.39509 | 0.65361 | RDH10 | 0.65361 | 0.68721 | 0.75113 |
| ENSG00000115257 | 19 | 1481427 | 1490751 | 31 | 9 | -0.39512 | 0.65362 | PCSK4 | 0.65362 | 0.2459 | 0.15629 |
| ENSG00000081148 | 3 | 100941390 | 101039404 | 206 | 14 | -0.39522 | 0.65366 | IMPG2 | 0.65366 | 0.68047 | 0.67065 |
| ENSG00000188385 | 10 | 133918175 | 133998313 | 511 | 47 | -0.39532 | 0.6537 | JAKMIP3 | 0.6537 | 0.3705 | 0.37126 |
| ENSG00000213762 | 19 | 58125601 | 58134724 | 22 | 5 | -0.39552 | 0.65377 | ZNF134 | 0.65377 | 0.79053 | 0.73859 |
| ENSG00000105852 | 7 | 94989256 | 95025680 | 71 | 14 | -0.39573 | 0.65385 | PON3 | 0.65385 | 0.54137 | 0.51837 |
| ENSG00000008394 | 12 | 16500076 | 16762193 | 609 | 52 | -0.39576 | 0.65386 | MGST1 | 0.65386 | 0.64842 | 0.67353 |
| ENSG00000100985 | 20 | 44637547 | 44645200 | 24 | 6 | -0.39577 | 0.65386 | MMP9 | 0.65386 | 0.51308 | 0.45901 |
| ENSG00000135480 | 12 | 52626304 | 52645970 | 63 | 11 | -0.39603 | 0.65396 | KRT7 | 0.65396 | 0.59789 | 0.77515 |
| ENSG00000174446 | 15 | 66782473 | 66790151 | 17 | 7 | -0.39619 | 0.65402 | SNAPC5 | 0.65402 | 0.27979 | 0.17443 |
| ENSG00000061676 | 2 | 183773843 | 183903586 | 358 | 25 | -0.39697 | 0.65431 | NCKAP1 | 0.65431 | 0.65195 | 0.66475 |
| ENSG00000215910 | 1 | 11821844 | 11849642 | 169 | 13 | -0.39709 | 0.65435 | C1orf167 | 0.65435 | 0.6724 | 0.63977 |
| ENSG00000224132 | 2 | 238165734 | 238166319 | 1 | 1 | -0.39723 | 0.6544 | AC112715.2 | 0.6544 | 0.52743 | 0.67926 |
| ENSG00000204217 | 2 | 203241659 | 203432474 | 402 | 23 | -0.39751 | 0.65451 | BMPR2 | 0.65451 | 0.65638 | 0.65515 |
| ENSG00000125508 | 20 | 62172163 | 62178857 | 27 | 7 | -0.39764 | 0.65455 | SRMS | 0.65455 | 0.1919 | 0.4263 |
| ENSG00000140104 | 14 | 105452112 | 105476819 | 105 | 14 | -0.3978 | 0.65461 | C14orf79 | 0.65461 | 0.61253 | 0.59415 |
| ENSG00000116194 | 1 | 178818840 | 178840187 | 42 | 9 | -0.39794 | 0.65466 | ANGPTL1 | 0.65466 | 0.58929 | 0.57204 |
| ENSG00000215193 | 22 | 18560689 | 18613905 | 214 | 34 | -0.39806 | 0.65471 | PEX26 | 0.65471 | 0.6245 | 0.88436 |
| ENSG00000073578 | 5 | 218356 | 256815 | 359 | 5 | -0.39827 | 0.65479 | SDHA | 0.65479 | 0.69908 | 0.75588 |
| ENSG00000165794 | 14 | 21467414 | 21470030 | 8 | 3 | -0.39835 | 0.65481 | SLC39A2 | 0.65481 | 0.0025943 | 0.00015066 |
| ENSG00000126856 | 16 | 90122974 | 90158480 | 128 | 17 | -0.39849 | 0.65487 | PRDM7 | 0.65487 | 0.57468 | 0.66007 |
| ENSG00000099246 | 10 | 27793197 | 27831143 | 66 | 11 | -0.39862 | 0.65491 | RAB18 | 0.65491 | 0.66897 | 0.54286 |
| ENSG00000144278 | 2 | 154728426 | 155310361 | 2145 | 56 | -0.39886 | 0.655 | GALNT13 | 0.655 | 0.6659 | 0.69343 |
| ENSG00000113532 | 5 | 100142639 | 100238970 | 243 | 16 | -0.3989 | 0.65502 | ST8SIA4 | 0.65502 | 0.62716 | 0.60521 |
| ENSG00000184345 | 3 | 51895645 | 51897440 | 5 | 3 | -0.39904 | 0.65507 | IQCF2 | 0.65507 | 0.58876 | 0.66117 |
| ENSG00000165156 | 8 | 124260696 | 124287781 | 19 | 8 | -0.39919 | 0.65512 | ZHX1 | 0.65512 | 0.60419 | 0.50469 |
| ENSG00000174799 | 4 | 56815037 | 56899529 | 211 | 25 | -0.3994 | 0.6552 | CEP135 | 0.6552 | 0.6197 | 0.31831 |
| ENSG00000132963 | 13 | 29233241 | 29253062 | 51 | 7 | -0.39963 | 0.65528 | POMP | 0.65528 | 0.78547 | 0.88953 |
| ENSG00000167654 | 19 | 3879862 | 3928077 | 172 | 40 | -0.39963 | 0.65528 | ATCAY | 0.65528 | 0.86959 | 0.85735 |
| ENSG00000117410 | 1 | 44440159 | 44443967 | 8 | 4 | -0.39981 | 0.65535 | ATP6V0B | 0.65535 | 0.44114 | 0.40936 |
| ENSG00000162599 | 1 | 61330931 | 61928465 | 1815 | 179 | -0.39981 | 0.65535 | NFIA | 0.65535 | 0.68869 | 0.65029 |
| ENSG00000109072 | 17 | 26691290 | 26700110 | 24 | 5 | -0.39985 | 0.65537 | VTN | 0.65537 | 0.63611 | 0.41089 |
| ENSG00000164900 | 7 | 150845676 | 150871832 | 59 | 12 | -0.39995 | 0.6554 | GBX1 | 0.6554 | 0.62994 | 0.84046 |
| ENSG00000155744 | 2 | 201838441 | 201936394 | 217 | 20 | -0.40061 | 0.65565 | FAM126B | 0.65565 | 0.6564 | 0.65711 |
| ENSG00000139915 | 14 | 47308826 | 48144157 | 2780 | 89 | -0.40081 | 0.65572 | MDGA2 | 0.65572 | 0.67768 | 0.68896 |
| ENSG00000184809 | 21 | 40969074 | 40984748 | 47 | 14 | -0.40101 | 0.65579 | C21orf88 | 0.65579 | 0.81771 | 0.9043 |
| ENSG00000135775 | 1 | 230778235 | 230829728 | 200 | 32 | -0.40119 | 0.65586 | COG2 | 0.65586 | 0.573 | 0.47627 |
| ENSG00000124227 | 20 | 56793551 | 56803709 | 25 | 7 | -0.40119 | 0.65586 | ANKRD60 | 0.65586 | 0.45263 | 0.46026 |
| ENSG00000175906 | 17 | 41476327 | 41478505 | 6 | 2 | -0.40121 | 0.65587 | ARL4D | 0.65587 | 0.72046 | 0.86466 |

| | | | | | | | | | | | |
|---|---|---|---|---|---|---|---|---|---|---|---|
| ENSG00000174748 | 3 | 23958036 | 23965183 | 15 | 6 | -0.40142 | 0.65595 | RPL15 | 0.65595 | 0.63647 | 0.61562 |
| ENSG00000254901 | 19 | 19287712 | 19303400 | 67 | 10 | -0.40152 | 0.65598 | MEF2BNB | 0.65598 | 0.73206 | 0.80595 |
| ENSG00000176945 | 3 | 195447753 | 195467994 | 141 | 22 | -0.40169 | 0.65604 | MUC20 | 0.65604 | 0.70835 | 0.67318 |
| ENSG00000076053 | 11 | 114270752 | 114284925 | 38 | 6 | -0.40177 | 0.65607 | RBM7 | 0.65607 | 0.70612 | 0.55892 |
| ENSG00000163040 | 2 | 132285248 | 132291239 | 22 | 7 | -0.40274 | 0.65643 | CCDC74A | 0.65643 | 0.73971 | 0.67011 |
| ENSG00000162194 | 11 | 62430287 | 62439727 | 21 | 4 | -0.40281 | 0.65645 | C11orf48 | 0.65645 | 0.67012 | 0.54396 |
| ENSG00000186474 | 19 | 51532348 | 51538486 | 16 | 4 | -0.40285 | 0.65647 | KLK12 | 0.65647 | 0.43206 | 0.66362 |
| ENSG00000170961 | 8 | 122624356 | 122653630 | 90 | 11 | -0.4029 | 0.65649 | HAS2 | 0.65649 | 0.78486 | 0.81579 |
| ENSG00000267906 | 7 | 16828866 | 16829246 | 1 | 1 | -0.40293 | 0.6565 | AC073333.1 | 0.6565 | 0.025162 | 0.060189 |
| ENSG00000123106 | 12 | 28286182 | 28732883 | 1470 | 22 | -0.40296 | 0.65651 | CCDC91 | 0.65651 | 0.67418 | 0.67769 |
| ENSG00000131558 | 7 | 132937829 | 133751342 | 1666 | 37 | -0.4031 | 0.65656 | EXOC4 | 0.65656 | 0.67764 | 0.69428 |
| ENSG00000080298 | 9 | 3218297 | 3526004 | 924 | 48 | -0.40312 | 0.65657 | RFX3 | 0.65657 | 0.65186 | 0.66027 |
| ENSG00000135124 | 12 | 121647660 | 121671909 | 105 | 13 | -0.40312 | 0.65657 | P2RX4 | 0.65657 | 0.60885 | 0.50858 |
| ENSG00000185115 | 15 | 29560353 | 29562033 | 3 | 2 | -0.40313 | 0.65657 | NDNL2 | 0.65657 | 0.69593 | 0.70322 |
| ENSG00000127824 | 2 | 220114433 | 220142892 | 77 | 10 | -0.40337 | 0.65666 | TUBA4A | 0.65666 | 0.65995 | 0.50767 |
| ENSG00000110375 | 11 | 118795873 | 118829269 | 118 | 24 | -0.40342 | 0.65668 | UPK2 | 0.65668 | 0.65907 | 0.75538 |
| ENSG00000256771 | 19 | 19976695 | 20005483 | 119 | 14 | -0.40343 | 0.65668 | ZNF253 | 0.65668 | 0.69479 | 0.73252 |
| ENSG00000144674 | 3 | 37284668 | 37408242 | 283 | 15 | -0.40354 | 0.65672 | GOLGA4 | 0.65672 | 0.64119 | 0.64734 |
| ENSG00000113048 | 5 | 71515236 | 71616473 | 178 | 20 | -0.40363 | 0.65676 | MRPS27 | 0.65676 | 0.62053 | 0.64593 |
| ENSG00000179626 | 12 | 55944983 | 55946028 | 1 | 1 | -0.40375 | 0.6568 | OR6C4 | 0.6568 | 0.60597 | 0.73321 |
| ENSG00000173093 | 12 | 111284573 | 111345339 | 228 | 28 | -0.40378 | 0.65681 | CCDC63 | 0.65681 | 0.65368 | 0.7073 |
| ENSG00000183682 | 1 | 39957318 | 39991607 | 105 | 21 | -0.40379 | 0.65682 | BMP8A | 0.65682 | 0.80262 | 0.84658 |
| ENSG00000171813 | 10 | 134210672 | 134231367 | 104 | 22 | -0.40394 | 0.65687 | PWWP2B | 0.65687 | 0.59413 | 0.47714 |
| ENSG00000152217 | 18 | 42260138 | 42648475 | 859 | 93 | -0.40401 | 0.6569 | SETBP1 | 0.6569 | 0.67598 | 0.64547 |
| ENSG00000184117 | 22 | 29950797 | 29977326 | 95 | 10 | -0.40405 | 0.65691 | NIPSNAP1 | 0.65691 | 0.65078 | 0.63331 |
| ENSG00000153914 | 5 | 65435799 | 65479443 | 71 | 13 | -0.40416 | 0.65695 | SREK1 | 0.65695 | 0.7097 | 0.77049 |
| ENSG00000100890 | 14 | 35591052 | 35743271 | 604 | 26 | -0.40416 | 0.65695 | KIAA0391 | 0.65695 | 0.66018 | 0.65486 |
| ENSG00000121690 | 11 | 33037410 | 33055128 | 74 | 7 | -0.40428 | 0.657 | DEPDC7 | 0.657 | 0.62663 | 0.62235 |
| ENSG00000008838 | 17 | 38175350 | 38217468 | 107 | 13 | -0.40447 | 0.65707 | MED24 | 0.65707 | 0.67914 | 0.67626 |
| ENSG00000170903 | 11 | 105866350 | 105893130 | 57 | 11 | -0.40454 | 0.65709 | MSANTD4 | 0.65709 | 0.6937 | 0.70455 |
| ENSG00000159873 | 22 | 29168662 | 29185283 | 84 | 13 | -0.40461 | 0.65712 | CCDC117 | 0.65712 | 0.54122 | 0.68241 |
| ENSG00000176248 | 9 | 140069236 | 140082989 | 29 | 3 | -0.40478 | 0.65718 | ANAPC2 | 0.65718 | 0.79312 | 0.70133 |
| ENSG00000140688 | 16 | 31500792 | 31520630 | 22 | 8 | -0.40485 | 0.6572 | C16orf58 | 0.6572 | 0.74423 | 0.82842 |
| ENSG00000180210 | 11 | 46740730 | 46761056 | 63 | 9 | -0.40504 | 0.65728 | F2 | 0.65728 | 0.65991 | 0.70031 |
| ENSG00000171496 | 9 | 125329778 | 125330838 | 6 | 2 | -0.40507 | 0.65729 | OR1L8 | 0.65729 | 0.73795 | 0.83736 |
| ENSG00000185278 | 1 | 173837220 | 173872687 | 117 | 14 | -0.40546 | 0.65743 | ZBTB37 | 0.65743 | 0.74569 | 0.76114 |
| ENSG00000215131 | 16 | 3543484 | 3545480 | 6 | 2 | -0.40546 | 0.65743 | C16orf90 | 0.65743 | 0.35731 | 0.52408 |
| ENSG00000105639 | 19 | 17935589 | 17958880 | 85 | 25 | -0.40553 | 0.65745 | JAK3 | 0.65745 | 0.52958 | 0.49253 |
| ENSG00000111371 | 12 | 46576846 | 46663800 | 213 | 41 | -0.40556 | 0.65747 | SLC38A1 | 0.65747 | 0.70535 | 0.75955 |
| ENSG00000161677 | 19 | 51009255 | 51014610 | 18 | 7 | -0.40563 | 0.65749 | JOSD2 | 0.65749 | 0.68141 | 0.65401 |
| ENSG00000175662 | 17 | 17746828 | 17875736 | 228 | 10 | -0.40608 | 0.65766 | TOM1L2 | 0.65766 | 0.63496 | 0.63836 |
| ENSG00000258744 | 14 | 24970800 | 24977894 | 27 | 7 | -0.40618 | 0.65769 | RP11-80A15.1 | 0.65769 | 0.76491 | 0.88127 |
| ENSG00000117748 | 1 | 28218035 | 28241257 | 70 | 6 | -0.40646 | 0.6578 | RPA2 | 0.6578 | 0.74167 | 0.80877 |
| ENSG00000151948 | 12 | 129337972 | 129469509 | 633 | 72 | -0.40656 | 0.65783 | GLT1D1 | 0.65783 | 0.35677 | 0.32747 |
| ENSG00000267106 | 19 | 9731967 | 9756572 | 42 | 7 | -0.40656 | 0.65783 | C19orf82 | 0.65783 | 0.70494 | 0.68999 |
| ENSG00000182103 | 11 | 82443053 | 82444906 | 7 | 2 | -0.40699 | 0.65799 | FAM181B | 0.65799 | 0.49315 | 0.48409 |
| ENSG00000144827 | 3 | 111697857 | 111712210 | 73 | 9 | -0.40703 | 0.65801 | ABHD10 | 0.65801 | 0.5171 | 0.57297 |
| ENSG00000268358 | 5 | 111755280 | 111756310 | 8 | 4 | -0.40703 | 0.65801 | EPB41L4A-AS2 | 0.65801 | 0.8558 | 0.82595 |
| ENSG00000269307 | 19 | 17378227 | 17393775 | 70 | 12 | -0.40729 | 0.6581 | CTD-2278I10.6 | 0.6581 | 0.58859 | 0.65238 |
| ENSG00000166794 | 15 | 64448011 | 64455404 | 20 | 6 | -0.40736 | 0.65813 | PPIB | 0.65813 | 0.48882 | 0.48594 |
| ENSG00000100632 | 14 | 69846848 | 69865344 | 63 | 14 | -0.40743 | 0.65815 | ERH | 0.65815 | 0.75694 | 0.80314 |
| ENSG00000182173 | 17 | 73512141 | 73520820 | 63 | 10 | -0.40748 | 0.65817 | TSEN54 | 0.65817 | 0.63843 | 0.82141 |
| ENSG00000145832 | 5 | 135170338 | 135224326 | 213 | 30 | -0.40762 | 0.65822 | SLC25A48 | 0.65822 | 0.7295 | 0.83694 |
| ENSG00000170160 | 17 | 16592851 | 16707767 | 132 | 16 | -0.40779 | 0.65829 | CCDC144A | 0.65829 | 0.65983 | 0.7312 |
| ENSG00000083520 | 13 | 73329540 | 73356234 | 48 | 13 | -0.40785 | 0.65831 | DIS3 | 0.65831 | 0.5456 | 0.54681 |
| ENSG00000189269 | 22 | 23950639 | 23974508 | 76 | 13 | -0.40799 | 0.65836 | C22orf43 | 0.65836 | 0.61927 | 0.53106 |
| ENSG00000187801 | 1 | 40915774 | 40929390 | 63 | 20 | -0.40803 | 0.65838 | ZFP69B | 0.65838 | 0.40674 | 0.71002 |
| ENSG00000197157 | 7 | 127292234 | 127732661 | 791 | 33 | -0.40828 | 0.65847 | SND1 | 0.65847 | 0.66006 | 0.66681 |
| ENSG00000180884 | 19 | 35447258 | 35454953 | 23 | 7 | -0.40838 | 0.6585 | ZNF792 | 0.6585 | 0.4639 | 0.54381 |
| ENSG00000069329 | 16 | 46690054 | 46723430 | 24 | 5 | -0.40841 | 0.65851 | VPS35 | 0.65851 | 0.67286 | 0.77256 |
| ENSG00000127511 | 19 | 16940211 | 16991164 | 129 | 21 | -0.40861 | 0.65859 | SIN3B | 0.65859 | 0.71399 | 0.78904 |
| ENSG00000267314 | 19 | 5904852 | 5914718 | 48 | 10 | -0.40877 | 0.65865 | AC104532.2 | 0.65865 | 0.76996 | 0.85456 |
| ENSG00000165424 | 10 | 81142081 | 81205383 | 273 | 43 | -0.40881 | 0.65866 | ZCCHC24 | 0.65866 | 0.67974 | 0.75783 |
| ENSG00000126603 | 16 | 4364762 | 4389598 | 59 | 16 | -0.40895 | 0.65871 | GLIS2 | 0.65871 | 0.5829 | 0.43514 |
| ENSG00000188883 | 7 | 139137436 | 139168458 | 141 | 16 | -0.40918 | 0.6588 | KLRG2 | 0.6588 | 0.68845 | 0.72247 |
| ENSG00000269753 | 1 | 2258581 | 2259042 | 4 | 2 | -0.40952 | 0.65892 | AL589739.1 | 0.65892 | 0.78045 | 0.86961 |
| ENSG00000259164 | 14 | 77564654 | 77691805 | 519 | 46 | -0.40951 | 0.65892 | RP11-463C8.4 | 0.65892 | 0.53304 | 0.55917 |
| ENSG00000268656 | 2 | 27998851 | 28009186 | 18 | 8 | -0.40966 | 0.65897 | AC110084.1 | 0.65897 | 0.61939 | 0.41296 |
| ENSG00000187569 | 12 | 7864050 | 7870155 | 27 | 8 | -0.40968 | 0.65898 | DPPA3 | 0.65898 | 0.34137 | 0.2115 |
| ENSG00000250510 | 12 | 6930711 | 6939136 | 28 | 14 | -0.40974 | 0.659 | GPR162 | 0.659 | 0.70815 | 0.54073 |
| ENSG00000124103 | 20 | 55092243 | 55101198 | 21 | 6 | -0.40977 | 0.65901 | FAM209A | 0.65901 | 0.74853 | 0.85499 |
| ENSG00000154001 | 14 | 63838075 | 64010092 | 550 | 30 | -0.40978 | 0.65902 | PPP2R5E | 0.65902 | 0.70029 | 0.774 |
| ENSG00000134321 | 2 | 7005937 | 7038370 | 62 | 12 | -0.40981 | 0.65903 | RSAD2 | 0.65903 | 0.73506 | 0.79474 |

| | | | | | | | | | | | |
|---|---|---|---|---|---|---|---|---|---|---|---|
| ENSG00000100519 | 14 | 53173890 | 53195305 | 32 | 7 | -0.41028 | 0.6592 | PSMC6 | 0.6592 | 0.67665 | 0.61435 |
| ENSG00000162591 | 1 | 3406484 | 3528059 | 589 | 89 | -0.41063 | 0.65933 | MEGF6 | 0.65933 | 0.64349 | 0.68864 |
| ENSG00000164638 | 7 | 5314000 | 5346501 | 201 | 30 | -0.41077 | 0.65938 | SLC29A4 | 0.65938 | 0.66144 | 0.71602 |
| ENSG00000152254 | 2 | 169757750 | 169766505 | 19 | 6 | -0.41092 | 0.65943 | G6PC2 | 0.65943 | 0.38907 | 0.28503 |
| ENSG00000162006 | 16 | 819428 | 833370 | 93 | 18 | -0.41098 | 0.65946 | MSLNL | 0.65946 | 0.73384 | 0.75491 |
| ENSG00000107807 | 10 | 102889257 | 102897545 | 17 | 6 | -0.41122 | 0.65954 | TLX1 | 0.65954 | 0.16848 | 0.633 |
| ENSG00000126091 | 1 | 44171495 | 44396831 | 655 | 27 | -0.41129 | 0.65957 | ST3GAL3 | 0.65957 | 0.67703 | 0.64982 |
| ENSG00000109471 | 4 | 123372625 | 123377880 | 6 | 3 | -0.41154 | 0.65966 | IL2 | 0.65966 | 0.25926 | 0.13182 |
| ENSG00000116199 | 1 | 178994939 | 179045697 | 197 | 20 | -0.41167 | 0.65971 | FAM20B | 0.65971 | 0.64798 | 0.69185 |
| ENSG00000113119 | 5 | 140019012 | 140024993 | 12 | 4 | -0.41166 | 0.65971 | TMCO6 | 0.65971 | 0.59871 | 0.52587 |
| ENSG00000077235 | 16 | 27470876 | 27561234 | 137 | 16 | -0.41171 | 0.65972 | GTF3C1 | 0.65972 | 0.70073 | 0.73056 |
| ENSG00000189332 | 11 | 18230685 | 18236109 | 10 | 3 | -0.41173 | 0.65973 | RP11-113D6.10 | 0.65973 | 0.69098 | 0.56143 |
| ENSG00000136935 | 9 | 127640646 | 127710771 | 153 | 16 | -0.41206 | 0.65985 | GOLGA1 | 0.65985 | 0.63849 | 0.55771 |
| ENSG00000129167 | 11 | 18039111 | 18063973 | 51 | 6 | -0.4123 | 0.65994 | TPH1 | 0.65994 | 0.73451 | 0.65866 |
| ENSG00000144460 | 2 | 226265364 | 226518734 | 549 | 39 | -0.41247 | 0.66 | NYAP2 | 0.66 | 0.696 | 0.70335 |
| ENSG00000178287 | 8 | 7705398 | 7726389 | 1 | 1 | -0.41246 | 0.66 | SPAG11A | 0.66 | 0.66 | 0.81871 |
| ENSG00000125449 | 17 | 73106047 | 73126360 | 57 | 13 | -0.41257 | 0.66004 | ARMC7 | 0.66004 | 0.59364 | 0.59336 |
| ENSG00000228278 | 9 | 117092149 | 117095532 | 10 | 3 | -0.41283 | 0.66014 | ORM2 | 0.66014 | 0.19986 | 0.087182 |
| ENSG00000167105 | 17 | 48348767 | 48358844 | 38 | 14 | -0.41291 | 0.66017 | TMEM92 | 0.66017 | 0.60698 | 0.5511 |
| ENSG00000092847 | 1 | 36335409 | 36395211 | 84 | 14 | -0.41321 | 0.66027 | AGO1 | 0.66027 | 0.63738 | 0.54896 |
| ENSG00000148948 | 11 | 40135753 | 41481323 | 4479 | 197 | -0.41331 | 0.66031 | LRRC4C | 0.66031 | 0.61205 | 0.62466 |
| ENSG00000178279 | 16 | 11361605 | 11363390 | 6 | 2 | -0.41338 | 0.66033 | TNP2 | 0.66033 | 0.48979 | 0.44709 |
| ENSG00000161813 | 12 | 50786166 | 50873787 | 201 | 15 | -0.41362 | 0.66042 | LARP4 | 0.66042 | 0.6668 | 0.65176 |
| ENSG00000160202 | 21 | 44589118 | 44592915 | 11 | 5 | -0.41369 | 0.66045 | CRYAA | 0.66045 | 0.67081 | 0.57005 |
| ENSG00000070778 | 14 | 88932122 | 89021077 | 240 | 14 | -0.41395 | 0.66054 | PTPN21 | 0.66054 | 0.68553 | 0.71664 |
| ENSG00000167720 | 17 | 2206677 | 2228554 | 49 | 12 | -0.41411 | 0.6606 | SRR | 0.6606 | 0.52153 | 0.56463 |
| ENSG00000157796 | 4 | 39184024 | 39287430 | 249 | 13 | -0.41474 | 0.66083 | WDR19 | 0.66083 | 0.64664 | 0.60773 |
| ENSG00000168569 | 11 | 62539101 | 62559493 | 44 | 9 | -0.41489 | 0.66089 | TMEM223 | 0.66089 | 0.53548 | 0.6418 |
| ENSG00000135587 | 6 | 109761966 | 109765122 | 9 | 4 | -0.41491 | 0.6609 | SMPD2 | 0.6609 | 0.70181 | 0.65423 |
| ENSG00000198369 | 2 | 65537985 | 65659771 | 387 | 28 | -0.41495 | 0.66091 | SPRED2 | 0.66091 | 0.66317 | 0.69936 |
| ENSG00000141577 | 17 | 79163393 | 79196799 | 153 | 24 | -0.41528 | 0.66103 | AZI1 | 0.66103 | 0.46618 | 0.24636 |
| ENSG00000166831 | 15 | 65032091 | 65067786 | 105 | 15 | -0.41558 | 0.66114 | RBPMS2 | 0.66114 | 0.65873 | 0.68723 |
| ENSG00000105514 | 19 | 11432722 | 11456946 | 78 | 9 | -0.41574 | 0.6612 | RAB3D | 0.6612 | 0.72017 | 0.87144 |
| ENSG00000159915 | 19 | 44754318 | 44779470 | 54 | 8 | -0.41577 | 0.66121 | ZNF233 | 0.66121 | 0.72704 | 0.71203 |
| ENSG00000171033 | 8 | 79428374 | 79517502 | 224 | 21 | -0.41598 | 0.66129 | PKIA | 0.66129 | 0.68186 | 0.66659 |
| ENSG00000132768 | 1 | 44435672 | 44439041 | 3 | 1 | -0.41606 | 0.66132 | DPH2 | 0.66132 | 0.45041 | 0.53231 |
| ENSG00000197013 | 19 | 21679484 | 21739072 | 230 | 13 | -0.41627 | 0.66139 | ZNF429 | 0.66139 | 0.66748 | 0.63907 |
| ENSG00000153029 | 1 | 181003067 | 181031074 | 93 | 17 | -0.41641 | 0.66145 | MR1 | 0.66145 | 0.61334 | 0.51923 |
| ENSG00000172752 | 3 | 130064359 | 130203688 | 571 | 31 | -0.41648 | 0.66147 | COL6A5 | 0.66147 | 0.6754 | 0.66596 |
| ENSG00000197568 | 1 | 70820488 | 70851022 | 63 | 10 | -0.41656 | 0.6615 | HHLA3 | 0.6615 | 0.75841 | 0.76109 |
| ENSG00000164265 | 5 | 147250245 | 147261754 | 79 | 13 | -0.4166 | 0.66151 | SCGB3A2 | 0.66151 | 0.56259 | 0.52576 |
| ENSG00000130348 | 6 | 107077453 | 107116292 | 184 | 22 | -0.4166 | 0.66151 | QRSL1 | 0.66151 | 0.68934 | 0.77775 |
| ENSG00000100372 | 22 | 41165634 | 41215403 | 75 | 7 | -0.41694 | 0.66164 | SLC25A17 | 0.66164 | 0.5486 | 0.44511 |
| ENSG00000121351 | 12 | 21507893 | 21532912 | 120 | 14 | -0.41707 | 0.66169 | IAPP | 0.66169 | 0.54053 | 0.35213 |
| ENSG00000267168 | 17 | 74729060 | 74734604 | 5 | 3 | -0.4173 | 0.66177 | RP11-318A15.7 | 0.66177 | 0.5223 | 0.62568 |
| ENSG00000185022 | 22 | 38597889 | 38612518 | 23 | 6 | -0.41738 | 0.6618 | MAFF | 0.6618 | 0.25535 | 0.18484 |
| ENSG00000075240 | 22 | 46971909 | 47075688 | 510 | 34 | -0.41746 | 0.66183 | GRAMD4 | 0.66183 | 0.63264 | 0.72877 |
| ENSG00000166800 | 11 | 18477371 | 18501147 | 86 | 17 | -0.41802 | 0.66203 | LDHAL6A | 0.66203 | 0.42521 | 0.30366 |
| ENSG00000186951 | 22 | 46546424 | 46639653 | 277 | 33 | -0.41803 | 0.66204 | PPARA | 0.66204 | 0.71611 | 0.73224 |
| ENSG00000007306 | 19 | 42177235 | 42210895 | 134 | 13 | -0.41808 | 0.66206 | CEACAM7 | 0.66206 | 0.66589 | 0.74792 |
| ENSG00000052841 | 11 | 43380482 | 43516483 | 360 | 33 | -0.41821 | 0.6621 | TTC17 | 0.6621 | 0.65612 | 0.66838 |
| ENSG00000132698 | 1 | 156030951 | 156040295 | 8 | 5 | -0.41831 | 0.66214 | RAB25 | 0.66214 | 0.54735 | 0.81824 |
| ENSG00000253206 | 8 | 50448945 | 50466973 | 31 | 3 | -0.41844 | 0.66219 | RP11-738G5.2 | 0.66219 | 0.77634 | 0.74136 |
| ENSG00000123600 | 2 | 172173919 | 172291312 | 248 | 29 | -0.41847 | 0.6622 | METTL8 | 0.6622 | 0.57935 | 0.54659 |
| ENSG00000035681 | 8 | 59496063 | 59572403 | 215 | 21 | -0.41903 | 0.6624 | NSMAF | 0.6624 | 0.68603 | 0.70084 |
| ENSG00000231861 | 3 | 98216448 | 98217496 | 2 | 1 | -0.41906 | 0.66242 | OR5K2 | 0.66242 | 0.85348 | 0.86108 |
| ENSG00000100395 | 22 | 41601209 | 41627275 | 103 | 6 | -0.41913 | 0.66244 | L3MBTL2 | 0.66244 | 0.64091 | 0.70437 |
| ENSG00000057149 | 18 | 61322431 | 61329197 | 19 | 8 | -0.41927 | 0.66249 | SERPINB3 | 0.66249 | 0.58937 | 0.66006 |
| ENSG00000091136 | 7 | 107564244 | 107643700 | 219 | 39 | -0.41937 | 0.66253 | LAMB1 | 0.66253 | 0.63033 | 0.58215 |
| ENSG00000120693 | 13 | 37418968 | 37494902 | 227 | 32 | -0.41947 | 0.66256 | SMAD9 | 0.66256 | 0.54629 | 0.47936 |
| ENSG00000137868 | 15 | 74471807 | 74504608 | 92 | 14 | -0.4195 | 0.66257 | STRA6 | 0.66257 | 0.63122 | 0.50529 |
| ENSG00000113916 | 3 | 187439165 | 187463515 | 79 | 16 | -0.41955 | 0.66259 | BCL6 | 0.66259 | 0.75079 | 0.75942 |
| ENSG00000154162 | 5 | 21750782 | 22853731 | 3284 | 113 | -0.41957 | 0.6626 | CDH12 | 0.6626 | 0.68751 | 0.70592 |
| ENSG00000258539 | 10 | 126305649 | 126480296 | 383 | 30 | -0.41961 | 0.66262 | RP11-12J10.3 | 0.66262 | 0.69744 | 0.74739 |
| ENSG00000185238 | 11 | 20409076 | 20530840 | 384 | 24 | -0.41963 | 0.66262 | PRMT3 | 0.66262 | 0.6501 | 0.41669 |
| ENSG00000152332 | 1 | 162467041 | 162499419 | 108 | 6 | -0.41966 | 0.66263 | UHMK1 | 0.66263 | 0.71583 | 0.5656 |
| ENSG00000144191 | 2 | 98962618 | 99015064 | 214 | 21 | -0.4197 | 0.66265 | CNGA3 | 0.66265 | 0.72136 | 0.74976 |
| ENSG00000119596 | 14 | 75230069 | 75322244 | 186 | 15 | -0.4198 | 0.66268 | YLPM1 | 0.66268 | 0.64401 | 0.64988 |
| ENSG00000160229 | 19 | 20959110 | 20991922 | 312 | 10 | -0.41991 | 0.66272 | ZNF66 | 0.66272 | 0.58644 | 0.62488 |
| ENSG00000181499 | 11 | 123813492 | 123814580 | 4 | 1 | -0.42006 | 0.66278 | OR6T1 | 0.66278 | 0.38879 | 0.34908 |
| ENSG00000186714 | 11 | 32623792 | 32816204 | 712 | 25 | -0.42033 | 0.66288 | CCDC73 | 0.66288 | 0.63444 | 0.63568 |
| ENSG00000139579 | 12 | 56615799 | 56623638 | 13 | 6 | -0.42048 | 0.66293 | NABP2 | 0.66293 | 0.69661 | 0.5379 |
| ENSG00000109618 | 4 | 25121627 | 25162204 | 85 | 16 | -0.42051 | 0.66294 | SEPSECS | 0.66294 | 0.72676 | 0.82279 |

| | | | | | | | | | | | |
|---|---|---|---|---|---|---|---|---|---|---|---|
| ENSG00000241935 | 10 | 99344080 | 99372559 | 87 | 20 | -0.4205 | 0.66294 | HOGA1 | 0.66294 | 0.63653 | 0.66484 |
| ENSG00000100575 | 14 | 58875212 | 58894332 | 20 | 4 | -0.42069 | 0.66301 | TIMM9 | 0.66301 | 0.68994 | 0.62382 |
| ENSG00000123349 | 12 | 53689075 | 53693234 | 16 | 5 | -0.42078 | 0.66304 | PFDN5 | 0.66304 | 0.67924 | 0.66743 |
| ENSG00000139131 | 12 | 32880424 | 32908836 | 95 | 7 | -0.42081 | 0.66305 | YARS2 | 0.66305 | 0.70382 | 0.81147 |
| ENSG00000105866 | 7 | 21467652 | 21554440 | 377 | 32 | -0.42097 | 0.66311 | SP4 | 0.66311 | 0.64561 | 0.62931 |
| ENSG00000124721 | 6 | 38683117 | 38998301 | 1386 | 60 | -0.42106 | 0.66314 | DNAH8 | 0.66314 | 0.63419 | 0.65255 |
| ENSG00000112309 | 6 | 71566382 | 71666741 | 335 | 46 | -0.42117 | 0.66318 | B3GAT2 | 0.66318 | 0.64556 | 0.61573 |
| ENSG00000103528 | 16 | 19179293 | 19279652 | 321 | 50 | -0.4213 | 0.66323 | SYT17 | 0.66323 | 0.73588 | 0.7359 |
| ENSG00000212670 | 1 | 54569968 | 54571750 | 5 | 3 | -0.42147 | 0.66329 | AL161915.1 | 0.66329 | 0.63921 | 0.35356 |
| ENSG00000188725 | 5 | 60453536 | 60458301 | 9 | 5 | -0.42151 | 0.66331 | SMIM15 | 0.66331 | 0.20763 | 0.11769 |
| ENSG00000167702 | 8 | 145691426 | 145699585 | 9 | 5 | -0.42169 | 0.66337 | KIFC2 | 0.66337 | 0.44436 | 0.24391 |
| ENSG00000137824 | 15 | 41028082 | 41048049 | 43 | 6 | -0.42183 | 0.66343 | RMDN3 | 0.66343 | 0.64939 | 0.62261 |
| ENSG00000079385 | 19 | 43011456 | 43065386 | 53 | 8 | -0.42185 | 0.66343 | CEACAM1 | 0.66343 | 0.6117 | 0.50154 |
| ENSG00000165660 | 10 | 126490354 | 126525239 | 72 | 13 | -0.42205 | 0.6635 | FAM175B | 0.6635 | 0.58264 | 0.54009 |
| ENSG00000163534 | 1 | 157764193 | 157789895 | 79 | 9 | -0.42225 | 0.66358 | FCRL1 | 0.66358 | 0.65372 | 0.61784 |
| ENSG00000143702 | 1 | 243287730 | 243418650 | 295 | 22 | -0.42226 | 0.66358 | CEP170 | 0.66358 | 0.64195 | 0.61065 |
| ENSG00000155833 | 9 | 105757593 | 105780770 | 117 | 11 | -0.42225 | 0.66358 | CYLC2 | 0.66358 | 0.57842 | 0.44641 |
| ENSG00000188372 | 7 | 76026835 | 76071388 | 238 | 30 | -0.42256 | 0.66369 | ZP3 | 0.66369 | 0.57155 | 0.63451 |
| ENSG00000170906 | 19 | 54606036 | 54612564 | 32 | 6 | -0.42292 | 0.66382 | NDUFA3 | 0.66382 | 0.65086 | 0.82818 |
| ENSG00000125249 | 13 | 98086476 | 98121382 | 68 | 11 | -0.42354 | 0.66405 | RAP2A | 0.66405 | 0.73559 | 0.62183 |
| ENSG00000108932 | 17 | 66263167 | 66287408 | 78 | 15 | -0.42357 | 0.66406 | SLC16A6 | 0.66406 | 0.71467 | 0.75479 |
| ENSG00000166448 | 7 | 98444111 | 98468394 | 61 | 10 | -0.4237 | 0.66411 | TMEM130 | 0.66411 | 0.40731 | 0.13454 |
| ENSG00000213889 | 19 | 45992035 | 46005768 | 51 | 9 | -0.42375 | 0.66413 | PPM1N | 0.66413 | 0.79983 | 0.83805 |
| ENSG00000171815 | 5 | 140430979 | 140433512 | 3 | 2 | -0.42383 | 0.66415 | PCDHB1 | 0.66415 | 0.83614 | 0.89641 |
| ENSG00000186960 | 14 | 29241910 | 29282493 | 78 | 23 | -0.42384 | 0.66416 | C14orf23 | 0.66416 | 0.60617 | 0.47936 |
| ENSG00000127022 | 5 | 179105629 | 179157926 | 181 | 20 | -0.42415 | 0.66427 | CANX | 0.66427 | 0.59199 | 0.66143 |
| ENSG00000131067 | 20 | 33432523 | 33460663 | 67 | 10 | -0.42418 | 0.66428 | GGT7 | 0.66428 | 0.54603 | 0.47208 |
| ENSG00000130035 | 12 | 4829507 | 4960277 | 495 | 54 | -0.42427 | 0.66432 | GALNT8 | 0.66432 | 0.71342 | 0.75485 |
| ENSG00000120071 | 17 | 44107282 | 44302733 | 1255 | 6 | -0.42441 | 0.66437 | KANSL1 | 0.66437 | 0.66732 | 0.65845 |
| ENSG00000163499 | 2 | 219854911 | 219858143 | 1 | 1 | -0.4245 | 0.6644 | CRYBA2 | 0.6644 | 0.027076 | 0.044144 |
| ENSG00000101443 | 20 | 44098346 | 44110172 | 49 | 9 | -0.42479 | 0.6645 | WFDC2 | 0.6645 | 0.44387 | 0.41448 |
| ENSG00000237515 | 16 | 12995477 | 13334272 | 1608 | 103 | -0.42492 | 0.66455 | SHISA9 | 0.66455 | 0.65443 | 0.63859 |
| ENSG00000183281 | 2 | 87229682 | 87248975 | 6 | 2 | -0.42525 | 0.66467 | PLGLB1 | 0.66467 | 0.62345 | 0.59522 |
| ENSG00000142513 | 19 | 51293672 | 51298481 | 24 | 5 | -0.42532 | 0.6647 | ACPT | 0.6647 | 0.70168 | 0.82888 |
| ENSG00000160181 | 21 | 43766466 | 43771237 | 21 | 6 | -0.42566 | 0.66482 | TFF2 | 0.66482 | 0.11344 | 0.15178 |
| ENSG00000267318 | 17 | 58018269 | 58050462 | 65 | 12 | -0.42581 | 0.66488 | RP11-178C3.1 | 0.66488 | 0.6808 | 0.73214 |
| ENSG00000230301 | 3 | 97983088 | 97984200 | 12 | 3 | -0.42584 | 0.66489 | OR5H6 | 0.66489 | 0.52901 | 0.69693 |
| ENSG00000204887 | 17 | 39185949 | 39186356 | 6 | 2 | -0.42585 | 0.66489 | KRTAP1-4 | 0.66489 | 0.55872 | 0.4225 |
| ENSG00000212899 | 17 | 39149686 | 39150385 | 7 | 3 | -0.42616 | 0.665 | KRTAP3-3 | 0.665 | 0.51029 | 0.56035 |
| ENSG00000128591 | 7 | 128470431 | 128499328 | 53 | 11 | -0.42655 | 0.66515 | FLNC | 0.66515 | 0.69258 | 0.77819 |
| ENSG00000109911 | 11 | 31531297 | 31805546 | 444 | 16 | -0.42657 | 0.66516 | ELP4 | 0.66516 | 0.66446 | 0.68294 |
| ENSG00000102891 | 16 | 56598961 | 56602869 | 22 | 9 | -0.42673 | 0.66521 | MT4 | 0.66521 | 0.67801 | 0.65559 |
| ENSG00000105948 | 7 | 138818490 | 138876732 | 154 | 8 | -0.42692 | 0.66528 | TTC26 | 0.66528 | 0.73394 | 0.7949 |
| ENSG00000101076 | 20 | 42984340 | 43061485 | 266 | 39 | -0.42691 | 0.66528 | HNF4A | 0.66528 | 0.60904 | 0.59521 |
| ENSG00000134253 | 1 | 117653682 | 117665209 | 38 | 8 | -0.42729 | 0.66542 | TRIM45 | 0.66542 | 0.78825 | 0.82968 |
| ENSG00000154227 | 15 | 100940600 | 101085200 | 784 | 28 | -0.42748 | 0.66548 | CERS3 | 0.66548 | 0.66808 | 0.76639 |
| ENSG00000178201 | 19 | 57966542 | 57967854 | 2 | 1 | -0.4275 | 0.66549 | VN1R1 | 0.66549 | 0.69136 | 0.69488 |
| ENSG00000135211 | 7 | 77423045 | 77427897 | 15 | 6 | -0.42758 | 0.66552 | TMEM60 | 0.66552 | 0.51213 | 0.5 |
| ENSG00000180822 | 6 | 3231637 | 3303607 | 433 | 20 | -0.4277 | 0.66557 | PSMG4 | 0.66557 | 0.69832 | 0.69362 |
| ENSG00000197956 | 1 | 153507075 | 153508720 | 3 | 2 | -0.4279 | 0.66564 | S100A6 | 0.66564 | 0.88585 | 0.88921 |
| ENSG00000139323 | 12 | 89813495 | 89919801 | 162 | 18 | -0.42796 | 0.66566 | POC1B | 0.66566 | 0.62433 | 0.54566 |
| ENSG00000258790 | 14 | 35591755 | 35786680 | 767 | 24 | -0.42808 | 0.6657 | KIAA0391 | 0.6657 | 0.66018 | 0.65486 |
| ENSG00000110844 | 12 | 49962001 | 50038449 | 121 | 16 | -0.42814 | 0.66572 | PRPF40B | 0.66572 | 0.64096 | 0.65662 |
| ENSG00000167283 | 11 | 118271869 | 118302211 | 69 | 12 | -0.42824 | 0.66576 | ATP5L | 0.66576 | 0.72274 | 0.71172 |
| ENSG00000110172 | 11 | 89934328 | 89956532 | 89 | 9 | -0.42831 | 0.66579 | CHORDC1 | 0.66579 | 0.71352 | 0.74692 |
| ENSG00000118276 | 18 | 29202210 | 29265799 | 135 | 18 | -0.42831 | 0.66579 | B4GALT6 | 0.66579 | 0.74549 | 0.79118 |
| ENSG00000159166 | 1 | 201342372 | 201368736 | 120 | 22 | -0.42839 | 0.66582 | LAD1 | 0.66582 | 0.61 | 0.58872 |
| ENSG00000126602 | 16 | 3701640 | 3767598 | 234 | 38 | -0.42851 | 0.66586 | TRAP1 | 0.66586 | 0.69122 | 0.63933 |
| ENSG00000165138 | 9 | 101493611 | 101559247 | 139 | 17 | -0.42856 | 0.66588 | ANKS6 | 0.66588 | 0.66899 | 0.73041 |
| ENSG00000127980 | 7 | 92116334 | 92157845 | 73 | 11 | -0.4288 | 0.66597 | PEX1 | 0.66597 | 0.73116 | 0.70385 |
| ENSG00000120686 | 13 | 38923986 | 38937140 | 28 | 4 | -0.42885 | 0.66598 | UFM1 | 0.66598 | 0.60299 | 0.6139 |
| ENSG00000180828 | 8 | 65492814 | 65496181 | 7 | 3 | -0.42894 | 0.66602 | BHLHE22 | 0.66602 | 0.56738 | 0.57132 |
| ENSG00000185129 | 5 | 139487362 | 139496321 | 6 | 3 | -0.42913 | 0.66609 | PURA | 0.66609 | 0.61848 | 0.7648 |
| ENSG00000115966 | 2 | 175936978 | 176033110 | 278 | 14 | -0.42918 | 0.6661 | ATF2 | 0.6661 | 0.67752 | 0.72089 |
| ENSG00000115109 | 2 | 120770581 | 120936695 | 365 | 14 | -0.42934 | 0.66616 | EPB41L5 | 0.66616 | 0.68879 | 0.67726 |
| ENSG00000184029 | 21 | 39323728 | 39493454 | 507 | 54 | -0.42947 | 0.66621 | DSCR4 | 0.66621 | 0.6718 | 0.69344 |
| ENSG00000079387 | 12 | 48436681 | 48500091 | 198 | 19 | -0.42949 | 0.66622 | SENP1 | 0.66622 | 0.69203 | 0.50923 |
| ENSG00000172789 | 12 | 54426637 | 54429142 | 6 | 2 | -0.42958 | 0.66625 | HOXC5 | 0.66625 | 0.77271 | 0.87277 |
| ENSG00000129465 | 14 | 24805227 | 24809251 | 16 | 2 | -0.4296 | 0.66626 | RIPK3 | 0.66626 | 0.28296 | 0.091549 |
| ENSG00000123843 | 1 | 207262187 | 207273338 | 21 | 6 | -0.42964 | 0.66627 | C4BPB | 0.66627 | 0.37012 | 0.37809 |
| ENSG00000113272 | 5 | 157158205 | 157168456 | 47 | 9 | -0.42978 | 0.66632 | THG1L | 0.66632 | 0.70664 | 0.72512 |
| ENSG00000179965 | 16 | 30418618 | 30440920 | 28 | 12 | -0.42988 | 0.66636 | ZNF771 | 0.66636 | 0.71991 | 0.73236 |
| ENSG00000179051 | 1 | 17733256 | 17766220 | 87 | 12 | -0.42995 | 0.66638 | RCC2 | 0.66638 | 0.58882 | 0.661 |

| | | | | | | | | | | | | |
|---|---|---|---|---|---|---|---|---|---|---|---|---|
| ENSG00000256061 | 15 | 55702723 | 55800432 | 399 | 26 | -0.42993 | 0.66638 | DYX1C1 | | 0.66638 | 0.68648 | 0.6237 |
| ENSG00000179242 | 20 | 59827482 | 60515673 | 3470 | 274 | -0.43001 | 0.66641 | CDH4 | | 0.66641 | 0.70003 | 0.71733 |
| ENSG00000095321 | 9 | 131857089 | 131873468 | 32 | 4 | -0.43004 | 0.66642 | CRAT | | 0.66642 | 0.65347 | 0.71396 |
| ENSG00000215712 | 6 | 157710418 | 157744633 | 126 | 21 | -0.43036 | 0.66653 | TMEM242 | | 0.66653 | 0.65463 | 0.55551 |
| ENSG00000092929 | 17 | 73823306 | 73840798 | 79 | 12 | -0.43045 | 0.66657 | UNC13D | | 0.66657 | 0.50948 | 0.40118 |
| ENSG00000204659 | 5 | 179105584 | 179107975 | 4 | 2 | -0.43063 | 0.66663 | CBY3 | | 0.66663 | 0.83839 | 0.80125 |
| ENSG00000120690 | 13 | 41506056 | 41635576 | 316 | 21 | -0.43064 | 0.66663 | ELF1 | | 0.66663 | 0.68035 | 0.69791 |
| ENSG00000205060 | 7 | 133974084 | 134001803 | 88 | 4 | -0.43067 | 0.66665 | SLC35B4 | | 0.66665 | 0.71179 | 0.64105 |
| ENSG00000067369 | 15 | 43699407 | 43802926 | 248 | 14 | -0.43112 | 0.66681 | TP53BP1 | | 0.66681 | 0.67739 | 0.68157 |
| ENSG00000120049 | 10 | 103585731 | 103603677 | 32 | 11 | -0.43129 | 0.66687 | KCNIP2 | | 0.66687 | 0.66341 | 0.54834 |
| ENSG00000204978 | 19 | 41949063 | 41950670 | 5 | 2 | -0.4313 | 0.66687 | C19orf69 | | 0.66687 | 0.16907 | 0.22895 |
| ENSG00000170419 | 7 | 54610018 | 54638773 | 106 | 17 | -0.43135 | 0.66689 | VSTM2A | | 0.66689 | 0.77868 | 0.79925 |
| ENSG00000130643 | 10 | 135138927 | 135150475 | 35 | 12 | -0.43134 | 0.66689 | CALY | | 0.66689 | 0.64473 | 0.68452 |
| ENSG00000184454 | 1 | 24882602 | 24935819 | 274 | 27 | -0.4314 | 0.66691 | NCMAP | | 0.66691 | 0.67587 | 0.68192 |
| ENSG00000204644 | 6 | 29640169 | 29648887 | 63 | 11 | -0.43139 | 0.66691 | ZFP57 | | 0.66691 | 0.84561 | 0.88346 |
| ENSG00000213822 | 19 | 51979838 | 51994206 | 83 | 9 | -0.43149 | 0.66694 | CEACAM18 | | 0.66694 | 0.51966 | 0.13579 |
| ENSG00000088035 | 1 | 63833261 | 63904233 | 170 | 21 | -0.43162 | 0.66699 | ALG6 | | 0.66699 | 0.66736 | 0.80963 |
| ENSG00000171931 | 17 | 18647326 | 18682662 | 128 | 20 | -0.43174 | 0.66703 | FBXW10 | | 0.66703 | 0.65198 | 0.5376 |
| ENSG00000133067 | 1 | 202163029 | 202288909 | 362 | 36 | -0.43183 | 0.66707 | LGR6 | | 0.66707 | 0.69059 | 0.65201 |
| ENSG00000198890 | 1 | 107599267 | 107601916 | 3 | 2 | -0.43187 | 0.66708 | PRMT6 | | 0.66708 | 0.58827 | 0.70483 |
| ENSG00000036565 | 8 | 20002366 | 20040717 | 170 | 23 | -0.43209 | 0.66716 | SLC18A1 | | 0.66716 | 0.68748 | 0.66281 |
| ENSG00000144741 | 3 | 66119285 | 66438540 | 800 | 51 | -0.43211 | 0.66717 | SLC25A26 | | 0.66717 | 0.63397 | 0.55857 |
| ENSG00000184402 | 20 | 60718822 | 60757540 | 209 | 15 | -0.43212 | 0.66717 | SS18L1 | | 0.66717 | 0.69633 | 0.71416 |
| ENSG00000048052 | 7 | 18126572 | 19042039 | 2785 | 175 | -0.43223 | 0.66721 | HDAC9 | | 0.66721 | 0.63913 | 0.60159 |
| ENSG00000162998 | 2 | 183698002 | 183731890 | 115 | 15 | -0.43255 | 0.66733 | FRZB | | 0.66733 | 0.32225 | 0.36733 |
| ENSG00000100234 | 22 | 33197687 | 33259030 | 137 | 28 | -0.43257 | 0.66733 | TIMP3 | | 0.66733 | 0.70849 | 0.49581 |
| ENSG00000171612 | 1 | 9599541 | 9645237 | 80 | 12 | -0.43264 | 0.66736 | SLC25A33 | | 0.66736 | 0.73985 | 0.73455 |
| ENSG00000198056 | 12 | 57125380 | 57146157 | 39 | 6 | -0.4328 | 0.66742 | PRIM1 | | 0.66742 | 0.65169 | 0.67797 |
| ENSG00000267757 | 19 | 46144752 | 46146098 | 4 | 2 | -0.43294 | 0.66747 | C19orf83 | | 0.66747 | 0.00023494 | 3.7961E-06 |
| ENSG00000180263 | 12 | 95470525 | 95611258 | 567 | 30 | -0.43306 | 0.66751 | FGD6 | | 0.66751 | 0.64192 | 0.62074 |
| ENSG00000123609 | 2 | 152126979 | 152146571 | 88 | 11 | -0.4331 | 0.66753 | NMI | | 0.66753 | 0.74237 | 0.73952 |
| ENSG00000049323 | 2 | 33172039 | 33624576 | 2158 | 90 | -0.43315 | 0.66755 | LTBP1 | | 0.66755 | 0.67113 | 0.70118 |
| ENSG00000176155 | 17 | 80059336 | 80170706 | 625 | 11 | -0.43318 | 0.66756 | CCDC57 | | 0.66756 | 0.67202 | 0.69181 |
| ENSG00000121671 | 11 | 45868669 | 45904798 | 83 | 15 | -0.43331 | 0.6676 | CRY2 | | 0.6676 | 0.74585 | 0.76093 |
| ENSG00000271321 | 7 | 143452182 | 143454789 | 5 | 3 | -0.43338 | 0.66763 | CTAGE6 | | 0.66763 | 0.57557 | 0.55361 |
| ENSG00000182334 | 11 | 7846584 | 7847519 | 4 | 2 | -0.43352 | 0.66768 | OR5P3 | | 0.66768 | 0.66996 | 0.43733 |
| ENSG00000162614 | 1 | 78354198 | 78409580 | 134 | 21 | -0.43356 | 0.66769 | NEXN | | 0.66769 | 0.6086 | 0.5585 |
| ENSG00000108423 | 17 | 57936851 | 57970304 | 96 | 13 | -0.43355 | 0.66769 | TUBD1 | | 0.66769 | 0.64241 | 0.62396 |
| ENSG00000145416 | 4 | 164445450 | 165305202 | 3447 | 129 | -0.43387 | 0.66781 | MARCHF1 | | 0.66781 | 0.18133 | 0.30778 |
| ENSG00000177192 | 12 | 132413745 | 132428406 | 38 | 9 | -0.43388 | 0.66781 | PUS1 | | 0.66781 | 0.39578 | 0.28588 |
| ENSG00000067208 | 1 | 92974253 | 93257961 | 839 | 17 | -0.43393 | 0.66783 | EVI5 | | 0.66783 | 0.65638 | 0.66012 |
| ENSG00000138669 | 4 | 82009837 | 82136218 | 405 | 22 | -0.43413 | 0.6679 | PRKG2 | | 0.6679 | 0.71121 | 0.77172 |
| ENSG00000159423 | 1 | 19197926 | 19229275 | 154 | 23 | -0.43442 | 0.66801 | ALDH4A1 | | 0.66801 | 0.78251 | 0.95415 |
| ENSG00000110931 | 12 | 121675497 | 121736111 | 204 | 37 | -0.43443 | 0.66801 | CAMKK2 | | 0.66801 | 0.70437 | 0.78881 |
| ENSG00000143811 | 1 | 226107578 | 226111978 | 8 | 4 | -0.43446 | 0.66802 | PYCR2 | | 0.66802 | 0.36413 | 0.45787 |
| ENSG00000177465 | 14 | 74058410 | 74063200 | 19 | 8 | -0.43446 | 0.66802 | ACOT4 | | 0.66802 | 0.37854 | 0.43521 |
| ENSG00000141854 | 19 | 14183348 | 14185874 | 7 | 2 | -0.43446 | 0.66802 | hsa-mir-1199 | | 0.66802 | 0.73211 | 0.86351 |
| ENSG00000162441 | 1 | 9982173 | 10003465 | 59 | 11 | -0.43452 | 0.66805 | LZIC | | 0.66805 | 0.51435 | 0.36976 |
| ENSG00000196712 | 17 | 29421945 | 29709134 | 487 | 12 | -0.43461 | 0.66808 | NF1 | | 0.66808 | 0.67294 | 0.68379 |
| ENSG00000267937 | 21 | 38888740 | 38888973 | 1 | 1 | -0.43467 | 0.6681 | AP001421.1 | | 0.6681 | 0.78887 | 0.82669 |
| ENSG00000198060 | 10 | 94050920 | 94113721 | 107 | 14 | -0.43471 | 0.66811 | | 43895 | 0.66811 | 0.70408 | 0.72325 |
| ENSG00000142185 | 21 | 45770046 | 45862964 | 220 | 27 | -0.43477 | 0.66814 | TRPM2 | | 0.66814 | 0.6966 | 0.69309 |
| ENSG00000120725 | 5 | 138282409 | 138629246 | 549 | 23 | -0.43497 | 0.66821 | SIL1 | | 0.66821 | 0.66066 | 0.62919 |
| ENSG00000058600 | 16 | 22308730 | 22346424 | 41 | 19 | -0.43501 | 0.66822 | POLR3E | | 0.66822 | 0.76549 | 0.80236 |
| ENSG00000206549 | 3 | 46753605 | 46854064 | 271 | 13 | -0.43525 | 0.66831 | PRSS50 | | 0.66831 | 0.70483 | 0.73248 |
| ENSG00000166479 | 18 | 66340925 | 66382535 | 123 | 9 | -0.43544 | 0.66838 | TMX3 | | 0.66838 | 0.63196 | 0.61712 |
| ENSG00000156299 | 21 | 32490734 | 32932290 | 1592 | 124 | -0.43555 | 0.66842 | TIAM1 | | 0.66842 | 0.65719 | 0.62735 |
| ENSG00000109323 | 4 | 103552660 | 103682151 | 262 | 17 | -0.43561 | 0.66844 | MANBA | | 0.66844 | 0.67409 | 0.7086 |
| ENSG00000167202 | 15 | 78276378 | 78370066 | 375 | 27 | -0.43613 | 0.66863 | TBC1D2B | | 0.66863 | 0.62829 | 0.62272 |
| ENSG00000089916 | 14 | 76618259 | 76720685 | 431 | 28 | -0.43616 | 0.66864 | GPATCH2L | | 0.66864 | 0.66946 | 0.63314 |
| ENSG00000174957 | 11 | 55944094 | 55945032 | 1 | 1 | -0.43633 | 0.6687 | OR5J2 | | 0.6687 | 0.030539 | 0.032729 |
| ENSG00000178685 | 8 | 145051321 | 145086940 | 84 | 8 | -0.4365 | 0.66876 | PARP10 | | 0.66876 | 0.7506 | 0.55324 |
| ENSG00000166220 | 10 | 72530995 | 72545157 | 86 | 24 | -0.43689 | 0.6689 | TBATA | | 0.6689 | 0.77843 | 0.62623 |
| ENSG00000125977 | 20 | 32676104 | 32700138 | 31 | 7 | -0.43702 | 0.66895 | EIF2S2 | | 0.66895 | 0.62774 | 0.62106 |
| ENSG00000225940 | 5 | 55807394 | 55902059 | 487 | 62 | -0.43707 | 0.66897 | AC022431.2 | | 0.66897 | 0.59103 | 0.54445 |
| ENSG00000104998 | 19 | 14142560 | 14163743 | 71 | 9 | -0.43752 | 0.66913 | IL27RA | | 0.66913 | 0.7808 | 0.8348 |
| ENSG00000214708 | 17 | 30468245 | 30470154 | 6 | 2 | -0.43763 | 0.66917 | AC090616.2 | | 0.66917 | 0.72993 | 0.6641 |
| ENSG00000166963 | 15 | 43803156 | 43823818 | 25 | 6 | -0.43777 | 0.66922 | MAP1A | | 0.66922 | 0.60687 | 0.54747 |
| ENSG00000057608 | 10 | 5807186 | 5884095 | 267 | 31 | -0.43791 | 0.66928 | GDI2 | | 0.66928 | 0.30771 | 0.1353 |
| ENSG00000177791 | 10 | 75391412 | 75401515 | 21 | 5 | -0.43799 | 0.6693 | MYOZ1 | | 0.6693 | 0.82942 | 0.89283 |
| ENSG00000109674 | 4 | 178230990 | 178284097 | 207 | 38 | -0.43805 | 0.66933 | NEIL3 | | 0.66933 | 0.54761 | 0.4243 |
| ENSG00000257987 | 12 | 49121218 | 49159569 | 99 | 16 | -0.43809 | 0.66934 | LINC00935 | | 0.66934 | 0.67867 | 0.7496 |
| ENSG00000146918 | 7 | 158424003 | 158497520 | 248 | 17 | -0.4383 | 0.66942 | NCAPG2 | | 0.66942 | 0.74839 | 0.7568 |

| | | | | | | | | | | | |
|---|---|---|---|---|---|---|---|---|---|---|---|
| ENSG00000214960 | 7 | 16130817 | 16460947 | 1167 | 45 | -0.43853 | 0.6695 | ISPD | 0.6695 | 0.66232 | 0.66397 |
| ENSG00000159720 | 16 | 67471917 | 67515140 | 60 | 11 | -0.43853 | 0.6695 | ATP6V0D1 | 0.6695 | 0.69339 | 0.6674 |
| ENSG00000129991 | 19 | 55663137 | 55669141 | 24 | 7 | -0.4386 | 0.66953 | TNNI3 | 0.66953 | 0.68366 | 0.89009 |
| ENSG00000143502 | 1 | 223394161 | 223537544 | 368 | 25 | -0.43868 | 0.66955 | SUSD4 | 0.66955 | 0.66429 | 0.60129 |
| ENSG00000171914 | 15 | 62682725 | 63136830 | 1448 | 130 | -0.43867 | 0.66955 | TLN2 | 0.66955 | 0.62493 | 0.62219 |
| ENSG00000254445 | 11 | 111783460 | 111797595 | 30 | 6 | -0.43902 | 0.66968 | HSPB2-C11orf52 | 0.66968 | 0.80088 | 0.89315 |
| ENSG00000159588 | 1 | 46085716 | 46089729 | 3 | 2 | -0.43926 | 0.66976 | CCDC17 | 0.66976 | 0.4201 | 0.38448 |
| ENSG00000178662 | 2 | 166326157 | 166545917 | 519 | 34 | -0.43935 | 0.66979 | CSRNP3 | 0.66979 | 0.7067 | 0.72039 |
| ENSG00000174891 | 3 | 157823644 | 158263519 | 1519 | 20 | -0.4395 | 0.66985 | RSRC1 | 0.66985 | 0.64061 | 0.64266 |
| ENSG00000176974 | 17 | 18231187 | 18266856 | 99 | 11 | -0.43957 | 0.66988 | SHMT1 | 0.66988 | 0.69708 | 0.78984 |
| ENSG00000203859 | 1 | 119957554 | 119965658 | 26 | 5 | -0.43961 | 0.66989 | HSD3B2 | 0.66989 | 0.81472 | 0.8908 |
| ENSG00000178243 | 9 | 138235095 | 138238408 | 13 | 3 | -0.43973 | 0.66993 | C9orf62 | 0.66993 | 0.7435 | 0.64434 |
| ENSG00000183153 | 17 | 38517235 | 38520067 | 8 | 3 | -0.43997 | 0.67002 | GJD3 | 0.67002 | 0.55704 | 0.45967 |
| ENSG00000115365 | 2 | 211295973 | 211342376 | 127 | 11 | -0.44006 | 0.67005 | LANCL1 | 0.67005 | 0.71076 | 0.73114 |
| ENSG00000177047 | 9 | 21140631 | 21142144 | 2 | 1 | -0.44019 | 0.6701 | IFNW1 | 0.6701 | 0.44266 | 0.48227 |
| ENSG00000090861 | 16 | 70286198 | 70323446 | 43 | 17 | -0.44024 | 0.67012 | AARS | 0.67012 | 0.68198 | 0.62932 |
| ENSG00000119314 | 9 | 114980715 | 115095947 | 407 | 13 | -0.44042 | 0.67019 | PTBP3 | 0.67019 | 0.66532 | 0.68774 |
| ENSG00000172653 | 17 | 34181955 | 34195895 | 32 | 6 | -0.44045 | 0.67019 | C17orf66 | 0.67019 | 0.71042 | 0.78486 |
| ENSG00000127603 | 1 | 39546988 | 39952849 | 988 | 36 | -0.44046 | 0.6702 | MACF1 | 0.6702 | 0.68048 | 0.67685 |
| ENSG00000077522 | 1 | 236849754 | 236927931 | 324 | 35 | -0.44046 | 0.6702 | ACTN2 | 0.6702 | 0.75486 | 0.81131 |
| ENSG00000100325 | 22 | 30184597 | 30234271 | 168 | 19 | -0.44048 | 0.6702 | ASCC2 | 0.6702 | 0.69737 | 0.66812 |
| ENSG00000168077 | 8 | 27491385 | 27534293 | 184 | 15 | -0.44064 | 0.67026 | SCARA3 | 0.67026 | 0.6664 | 0.8 |
| ENSG00000153064 | 4 | 102332443 | 102995969 | 2093 | 90 | -0.44096 | 0.67038 | BANK1 | 0.67038 | 0.68411 | 0.69357 |
| ENSG00000131116 | 19 | 44111371 | 44124026 | 20 | 5 | -0.44106 | 0.67041 | ZNF428 | 0.67041 | 0.58036 | 0.73674 |
| ENSG00000150873 | 2 | 11273179 | 11286916 | 66 | 8 | -0.44136 | 0.67052 | C2orf50 | 0.67052 | 0.82409 | 0.84428 |
| ENSG00000104081 | 15 | 40380091 | 40401093 | 39 | 11 | -0.44135 | 0.67052 | BMF | 0.67052 | 0.06581 | 0.073235 |
| ENSG00000153006 | 5 | 64013971 | 64064512 | 98 | 11 | -0.44145 | 0.67056 | SREK1IP1 | 0.67056 | 0.71111 | 0.71642 |
| ENSG00000148082 | 9 | 91628060 | 91793682 | 546 | 38 | -0.44145 | 0.67056 | SHC3 | 0.67056 | 0.71181 | 0.74396 |
| ENSG00000175322 | 18 | 14057456 | 14132489 | 189 | 23 | -0.44154 | 0.67059 | ZNF519 | 0.67059 | 0.62821 | 0.6167 |
| ENSG00000198722 | 9 | 35161999 | 35405335 | 433 | 28 | -0.44161 | 0.67061 | UNC13B | 0.67061 | 0.67557 | 0.70803 |
| ENSG00000164241 | 5 | 126378250 | 126409184 | 78 | 8 | -0.44174 | 0.67066 | C5orf63 | 0.67066 | 0.6493 | 0.52501 |
| ENSG00000129696 | 8 | 33330904 | 33371119 | 165 | 13 | -0.44175 | 0.67067 | TTI2 | 0.67067 | 0.66051 | 0.70852 |
| ENSG00000148337 | 9 | 130928343 | 130966662 | 105 | 11 | -0.4419 | 0.67072 | CIZ1 | 0.67072 | 0.7509 | 0.83227 |
| ENSG00000005156 | 17 | 33307513 | 33332083 | 22 | 6 | -0.44189 | 0.67072 | LIG3 | 0.67072 | 0.63947 | 0.50447 |
| ENSG00000268154 | 1 | 161035655 | 161035943 | 2 | 1 | -0.44218 | 0.67082 | AL591806.1 | 0.67082 | 0.93128 | 0.90043 |
| ENSG00000118298 | 1 | 150229554 | 150237478 | 10 | 6 | -0.44222 | 0.67084 | CA14 | 0.67084 | 0.78352 | 0.91284 |
| ENSG00000183401 | 19 | 11455360 | 11465620 | 16 | 5 | -0.4423 | 0.67086 | CCDC159 | 0.67086 | 0.56893 | 0.67994 |
| ENSG00000136051 | 12 | 105501102 | 105562912 | 193 | 21 | -0.44231 | 0.67087 | KIAA1033 | 0.67087 | 0.68111 | 0.70973 |
| ENSG00000142961 | 1 | 47073387 | 47082563 | 24 | 3 | -0.44242 | 0.67091 | MOB3C | 0.67091 | 0.70983 | 0.55975 |
| ENSG00000086570 | 5 | 150883654 | 150948505 | 247 | 42 | -0.44244 | 0.67092 | FAT2 | 0.67092 | 0.78982 | 0.8113 |
| ENSG00000196350 | 19 | 22469210 | 22499978 | 69 | 12 | -0.44247 | 0.67092 | ZNF729 | 0.67092 | 0.66712 | 0.72905 |
| ENSG00000205177 | 11 | 33719807 | 33722347 | 16 | 7 | -0.44262 | 0.67098 | C11orf91 | 0.67098 | 0.85284 | 0.51309 |
| ENSG00000148483 | 10 | 17794251 | 17842867 | 43 | 12 | -0.44264 | 0.67099 | TMEM236 | 0.67099 | 0.55936 | 0.71337 |
| ENSG00000116128 | 1 | 147013182 | 147098017 | 298 | 48 | -0.44271 | 0.67101 | BCL9 | 0.67101 | 0.76634 | 0.75267 |
| ENSG00000105048 | 19 | 55644162 | 55660722 | 65 | 17 | -0.4428 | 0.67105 | TNNT1 | 0.67105 | 0.81698 | 0.90307 |
| ENSG00000035687 | 1 | 244571796 | 244615436 | 136 | 22 | -0.44284 | 0.67106 | ADSS | 0.67106 | 0.66047 | 0.61008 |
| ENSG00000072042 | 14 | 68143518 | 68162531 | 45 | 12 | -0.44296 | 0.6711 | RDH11 | 0.6711 | 0.62472 | 0.65904 |
| ENSG00000189050 | 17 | 58029601 | 58042122 | 29 | 8 | -0.44299 | 0.67111 | RNFT1 | 0.67111 | 0.65105 | 0.70636 |
| ENSG00000041802 | 3 | 194361517 | 194393206 | 184 | 13 | -0.44308 | 0.67115 | LSG1 | 0.67115 | 0.74956 | 0.7965 |
| ENSG00000155897 | 8 | 131792547 | 132054672 | 943 | 65 | -0.44309 | 0.67115 | ADCY8 | 0.67115 | 0.66267 | 0.68189 |
| ENSG00000110446 | 11 | 60704556 | 60720002 | 49 | 12 | -0.44315 | 0.67117 | SLC15A3 | 0.67117 | 0.67268 | 0.66045 |
| ENSG00000076826 | 19 | 7660788 | 7683190 | 64 | 9 | -0.44368 | 0.67136 | CAMSAP3 | 0.67136 | 0.68356 | 0.58772 |
| ENSG00000203734 | 6 | 139117063 | 139225207 | 404 | 47 | -0.44371 | 0.67137 | ECT2L | 0.67137 | 0.60824 | 0.60501 |
| ENSG00000251184 | 9 | 131703757 | 131719300 | 17 | 3 | -0.44412 | 0.67152 | RP11-101E3.5 | 0.67152 | 0.67588 | 0.6475 |
| ENSG00000178177 | 4 | 17842822 | 18023499 | 441 | 15 | -0.44443 | 0.67163 | LCORL | 0.67163 | 0.70627 | 0.70055 |
| ENSG00000189152 | 17 | 19030782 | 19062489 | 5 | 3 | -0.4446 | 0.6717 | GRAPL | 0.6717 | 0.42795 | 0.66685 |
| ENSG00000115935 | 2 | 175424300 | 175547644 | 214 | 31 | -0.4449 | 0.6718 | WIPF1 | 0.6718 | 0.67364 | 0.58764 |
| ENSG00000145022 | 3 | 49449639 | 49453908 | 9 | 4 | -0.44497 | 0.67183 | TCTA | 0.67183 | 0.61514 | 0.59491 |
| ENSG00000149346 | 20 | 10415951 | 10617477 | 647 | 30 | -0.44503 | 0.67185 | SLX4IP | 0.67185 | 0.72348 | 0.75398 |
| ENSG00000167130 | 9 | 131843379 | 131852717 | 13 | 4 | -0.44529 | 0.67195 | DOLPP1 | 0.67195 | 0.73863 | 0.74029 |
| ENSG00000170175 | 17 | 7348380 | 7361026 | 39 | 7 | -0.4453 | 0.67195 | CHRNB1 | 0.67195 | 0.51799 | 0.55517 |
| ENSG00000115523 | 2 | 85912298 | 85925977 | 72 | 10 | -0.44532 | 0.67196 | GNLY | 0.67196 | 0.66168 | 0.68564 |
| ENSG00000266614 | 18 | 73971121 | 73972069 | 5 | 1 | -0.4455 | 0.67202 | RP11-94B19.4 | 0.67202 | 0.73947 | 0.73181 |
| ENSG00000077312 | 19 | 41256542 | 41271294 | 36 | 11 | -0.44555 | 0.67204 | SNRPA | 0.67204 | 0.47563 | 0.44504 |
| ENSG00000186318 | 11 | 117156402 | 117186975 | 67 | 19 | -0.44561 | 0.67206 | BACE1 | 0.67206 | 0.6209 | 0.37748 |
| ENSG00000127831 | 2 | 219283815 | 219318018 | 57 | 8 | -0.44564 | 0.67207 | VIL1 | 0.67207 | 0.6649 | 0.59764 |
| ENSG00000138081 | 2 | 48016455 | 48132932 | 341 | 19 | -0.44588 | 0.67216 | FBXO11 | 0.67216 | 0.67373 | 0.5965 |
| ENSG00000102738 | 13 | 41303432 | 41345309 | 37 | 7 | -0.44593 | 0.67218 | MRPS31 | 0.67218 | 0.71176 | 0.83265 |
| ENSG00000197580 | 11 | 112046190 | 112095422 | 150 | 21 | -0.44598 | 0.67219 | BCO2 | 0.67219 | 0.65211 | 0.67476 |
| ENSG00000148339 | 9 | 130830480 | 130871524 | 155 | 4 | -0.44635 | 0.67233 | SLC25A25 | 0.67233 | 0.64691 | 0.70774 |
| ENSG00000166148 | 12 | 63539014 | 63544722 | 17 | 6 | -0.44644 | 0.67236 | AVPR1A | 0.67236 | 0.49547 | 0.54681 |
| ENSG00000131351 | 19 | 17160539 | 17186435 | 112 | 16 | -0.44681 | 0.67249 | HAUS8 | 0.67249 | 0.76989 | 0.85554 |
| ENSG00000147454 | 8 | 23386318 | 23432976 | 312 | 22 | -0.44727 | 0.67266 | SLC25A37 | 0.67266 | 0.67619 | 0.66213 |

| | | | | | | | | | | | |
|---|---|---|---|---|---|---|---|---|---|---|---|
| ENSG00000174013 | 3 | 196295482 | 196315930 | 67 | 12 | -0.4476 | 0.67278 | FBXO45 | 0.67278 | 0.54046 | 0.095906 |
| ENSG00000087510 | 20 | 55204358 | 55214339 | 18 | 7 | -0.44762 | 0.67279 | TFAP2C | 0.67279 | 0.63567 | 0.68098 |
| ENSG00000035499 | 5 | 59892739 | 59996017 | 242 | 23 | -0.44769 | 0.67281 | DEPDC1B | 0.67281 | 0.6657 | 0.62123 |
| ENSG00000160886 | 8 | 143781529 | 143786545 | 19 | 3 | -0.44776 | 0.67284 | LY6K | 0.67284 | 0.66969 | 0.55749 |
| ENSG00000162512 | 1 | 31342314 | 31381608 | 119 | 29 | -0.44797 | 0.67291 | SDC3 | 0.67291 | 0.73895 | 0.84809 |
| ENSG00000134057 | 5 | 68462837 | 68474072 | 40 | 5 | -0.44796 | 0.67291 | CCNB1 | 0.67291 | 0.61127 | 0.66852 |
| ENSG00000137185 | 6 | 28192664 | 28201260 | 27 | 7 | -0.44827 | 0.67302 | ZSCAN9 | 0.67302 | 0.74618 | 0.83879 |
| ENSG00000090674 | 19 | 7587512 | 7598895 | 23 | 8 | -0.4484 | 0.67307 | MCOLN1 | 0.67307 | 0.84876 | 0.88724 |
| ENSG00000165323 | 11 | 92085262 | 92629618 | 1357 | 57 | -0.44849 | 0.6731 | FAT3 | 0.6731 | 0.68388 | 0.71745 |
| ENSG00000262304 | 17 | 3468738 | 3539543 | 323 | 38 | -0.44856 | 0.67312 | SHPK | 0.67312 | 0.44399 | 0.62358 |
| ENSG00000214063 | 11 | 842808 | 867116 | 58 | 13 | -0.44872 | 0.67318 | TSPAN4 | 0.67318 | 0.76095 | 0.73329 |
| ENSG00000126262 | 19 | 35934809 | 35942669 | 30 | 10 | -0.44875 | 0.67319 | FFAR2 | 0.67319 | 0.63266 | 0.78872 |
| ENSG00000113657 | 5 | 146770374 | 146889619 | 154 | 36 | -0.44882 | 0.67322 | DPYSL3 | 0.67322 | 0.43922 | 0.32155 |
| ENSG00000187833 | 2 | 74011316 | 74044274 | 16 | 4 | -0.44886 | 0.67324 | C2orf78 | 0.67324 | 0.68012 | 0.86771 |
| ENSG00000196266 | 1 | 159283460 | 159284449 | 2 | 1 | -0.4489 | 0.67325 | OR10J3 | 0.67325 | 0.20357 | 0.2155 |
| ENSG00000084070 | 1 | 40810522 | 40888998 | 225 | 36 | -0.44943 | 0.67344 | SMAP2 | 0.67344 | 0.65533 | 0.66221 |
| ENSG00000173213 | 18 | 47225 | 73545 | 60 | 5 | -0.44967 | 0.67352 | RP11-683L23.1 | 0.67352 | 0.69425 | 0.6203 |
| ENSG00000147687 | 8 | 125500726 | 125551699 | 125 | 18 | -0.44978 | 0.67357 | TATDN1 | 0.67357 | 0.68037 | 0.5135 |
| ENSG00000178252 | 3 | 49044495 | 49053386 | 6 | 2 | -0.44983 | 0.67359 | WDR6 | 0.67359 | 0.58724 | 0.51543 |
| ENSG00000161939 | 17 | 6915954 | 6920839 | 10 | 2 | -0.44985 | 0.67359 | RNASEK-C17orf49 | 0.67359 | 0.6477 | 0.7956 |
| ENSG00000189099 | 4 | 152198323 | 152212605 | 47 | 7 | -0.45052 | 0.67383 | PRSS48 | 0.67383 | 0.6178 | 0.69282 |
| ENSG00000149948 | 12 | 66217911 | 66360075 | 286 | 36 | -0.45054 | 0.67384 | HMGA2 | 0.67384 | 0.66302 | 0.62806 |
| ENSG00000171988 | 10 | 64926981 | 65225722 | 707 | 31 | -0.45063 | 0.67387 | JMJD1C | 0.67387 | 0.66184 | 0.66424 |
| ENSG00000092850 | 1 | 36549676 | 36553876 | 4 | 2 | -0.4507 | 0.6739 | TEKT2 | 0.6739 | 0.89459 | 0.75667 |
| ENSG00000132958 | 13 | 19997017 | 20110903 | 325 | 17 | -0.45098 | 0.674 | TPTE2 | 0.674 | 0.64422 | 0.53274 |
| ENSG00000178718 | 15 | 75246757 | 75249805 | 2 | 1 | -0.45099 | 0.674 | RPP25 | 0.674 | 0.37996 | 0.33792 |
| ENSG00000103148 | 16 | 134273 | 188859 | 187 | 25 | -0.45104 | 0.67402 | NPRL3 | 0.67402 | 0.74957 | 0.7133 |
| ENSG00000174903 | 11 | 66036004 | 66044963 | 20 | 6 | -0.45122 | 0.67408 | RAB1B | 0.67408 | 0.66938 | 0.61931 |
| ENSG00000138722 | 4 | 90800683 | 90875780 | 232 | 25 | -0.45133 | 0.67412 | MMRN1 | 0.67412 | 0.57583 | 0.45637 |
| ENSG00000149735 | 11 | 64701943 | 64703360 | 3 | 2 | -0.45147 | 0.67418 | GPHA2 | 0.67418 | 0.65161 | 0.55447 |
| ENSG00000188176 | 17 | 4487294 | 4511614 | 127 | 30 | -0.45149 | 0.67418 | SMTNL2 | 0.67418 | 0.85896 | 0.83912 |
| ENSG00000166140 | 15 | 41099284 | 41106767 | 28 | 5 | -0.45154 | 0.6742 | ZFYVE19 | 0.6742 | 0.61985 | 0.71965 |
| ENSG00000173572 | 19 | 56403065 | 56443702 | 191 | 24 | -0.45156 | 0.67421 | NLRP13 | 0.67421 | 0.72655 | 0.69746 |
| ENSG00000144468 | 2 | 227700297 | 227863931 | 371 | 15 | -0.45184 | 0.67431 | RHBDD1 | 0.67431 | 0.68268 | 0.69485 |
| ENSG00000178802 | 15 | 75182346 | 75191798 | 13 | 3 | -0.45192 | 0.67434 | MPI | 0.67434 | 0.49952 | 0.35047 |
| ENSG00000179165 | 6 | 36358328 | 36410666 | 202 | 18 | -0.45198 | 0.67436 | PXT1 | 0.67436 | 0.65981 | 0.73762 |
| ENSG00000134853 | 4 | 55095264 | 55164414 | 170 | 14 | -0.45208 | 0.6744 | PDGFRA | 0.6744 | 0.60936 | 0.45152 |
| ENSG00000105122 | 19 | 15562435 | 15575382 | 25 | 13 | -0.45224 | 0.67445 | RASAL3 | 0.67445 | 0.84139 | 0.9058 |
| ENSG00000257017 | 16 | 72088491 | 72094954 | 23 | 7 | -0.45237 | 0.6745 | HP | 0.6745 | 0.74015 | 0.66512 |
| ENSG00000168522 | 8 | 42889337 | 42940931 | 87 | 11 | -0.45239 | 0.67451 | FNTA | 0.67451 | 0.72672 | 0.74312 |
| ENSG00000140265 | 15 | 43650370 | 43663223 | 33 | 5 | -0.45268 | 0.67461 | ZSCAN29 | 0.67461 | 0.67236 | 0.68172 |
| ENSG00000164220 | 5 | 75911307 | 75919259 | 33 | 8 | -0.45272 | 0.67462 | F2RL2 | 0.67462 | 0.65072 | 0.4949 |
| ENSG00000148200 | 9 | 127279554 | 127533589 | 225 | 22 | -0.45289 | 0.67469 | NR6A1 | 0.67469 | 0.69198 | 0.6884 |
| ENSG00000123119 | 8 | 91803778 | 91971636 | 454 | 18 | -0.45295 | 0.67471 | NECAB1 | 0.67471 | 0.70881 | 0.71334 |
| ENSG00000131653 | 16 | 2205699 | 2228130 | 42 | 14 | -0.45306 | 0.67475 | TRAF7 | 0.67475 | 0.63 | 0.64505 |
| ENSG00000145423 | 4 | 154701744 | 154710272 | 17 | 5 | -0.4531 | 0.67476 | SFRP2 | 0.67476 | 0.52524 | 0.50962 |
| ENSG00000111647 | 12 | 100422233 | 100536626 | 359 | 25 | -0.45313 | 0.67477 | UHRF1BP1L | 0.67477 | 0.65787 | 0.69683 |
| ENSG00000101057 | 20 | 42295754 | 42345136 | 157 | 14 | -0.45312 | 0.67477 | MYBL2 | 0.67477 | 0.70276 | 0.69696 |
| ENSG00000124205 | 20 | 57875482 | 57901047 | 68 | 17 | -0.45312 | 0.67477 | EDN3 | 0.67477 | 0.85465 | 0.95641 |
| ENSG00000165443 | 10 | 60936350 | 61007534 | 127 | 9 | -0.45341 | 0.67487 | PHYHIPL | 0.67487 | 0.66239 | 0.69614 |
| ENSG00000178502 | 17 | 40009797 | 40021684 | 45 | 7 | -0.45341 | 0.67487 | KLHL11 | 0.67487 | 0.66991 | 0.64086 |
| ENSG00000146842 | 7 | 129804555 | 129847610 | 93 | 16 | -0.45346 | 0.67489 | TMEM209 | 0.67489 | 0.71806 | 0.87495 |
| ENSG00000164362 | 5 | 1253262 | 1295184 | 110 | 33 | -0.45379 | 0.67501 | TERT | 0.67501 | 0.29448 | 0.33841 |
| ENSG00000131620 | 11 | 69924408 | 70035634 | 532 | 56 | -0.45393 | 0.67506 | ANO1 | 0.67506 | 0.63688 | 0.5392 |
| ENSG00000169469 | 1 | 153003678 | 153005376 | 19 | 1 | -0.45416 | 0.67514 | SPRR1B | 0.67514 | 0.64166 | 0.65153 |
| ENSG00000119383 | 9 | 131873229 | 131911225 | 121 | 6 | -0.45419 | 0.67515 | PPP2R4 | 0.67515 | 0.67865 | 0.66648 |
| ENSG00000129467 | 14 | 24787555 | 24804299 | 66 | 13 | -0.45427 | 0.67518 | ADCY4 | 0.67518 | 0.48008 | 0.13295 |
| ENSG00000154016 | 17 | 18923986 | 18950950 | 14 | 7 | -0.4543 | 0.67519 | GRAP | 0.67519 | 0.31173 | 0.6396 |
| ENSG00000160072 | 1 | 1407143 | 1433228 | 132 | 11 | -0.45439 | 0.67522 | ATAD3B | 0.67522 | 0.78472 | 0.83401 |
| ENSG00000152952 | 3 | 145787227 | 145881440 | 296 | 10 | -0.45442 | 0.67524 | PLOD2 | 0.67524 | 0.65226 | 0.66806 |
| ENSG00000116903 | 1 | 231468480 | 231473598 | 14 | 4 | -0.45449 | 0.67526 | EXOC8 | 0.67526 | 0.48899 | 0.36641 |
| ENSG00000230873 | 6 | 17102489 | 17131603 | 92 | 9 | -0.45459 | 0.6753 | STMND1 | 0.6753 | 0.76516 | 0.78758 |
| ENSG00000235750 | 1 | 175126123 | 175162135 | 120 | 14 | -0.4547 | 0.67534 | KIAA0040 | 0.67534 | 0.69206 | 0.52732 |
| ENSG00000144224 | 2 | 136499189 | 136542625 | 97 | 9 | -0.45484 | 0.67539 | UBXN4 | 0.67539 | 0.65995 | 0.61296 |
| ENSG00000011143 | 17 | 56282803 | 56296966 | 17 | 6 | -0.45516 | 0.6755 | MKS1 | 0.6755 | 0.40428 | 0.54346 |
| ENSG00000083535 | 13 | 73356197 | 73590591 | 596 | 31 | -0.45539 | 0.67559 | PIBF1 | 0.67559 | 0.67875 | 0.67983 |
| ENSG00000119689 | 14 | 75348594 | 75370448 | 71 | 10 | -0.45544 | 0.6756 | DLST | 0.6756 | 0.58624 | 0.51126 |
| ENSG00000139971 | 14 | 58466453 | 58764857 | 850 | 53 | -0.4556 | 0.67566 | C14orf37 | 0.67566 | 0.65133 | 0.67337 |
| ENSG00000101639 | 18 | 12991361 | 13125051 | 442 | 23 | -0.45559 | 0.67566 | CEP192 | 0.67566 | 0.68801 | 0.67133 |
| ENSG00000107104 | 9 | 470291 | 746105 | 1511 | 48 | -0.45569 | 0.67569 | KANK1 | 0.67569 | 0.73091 | 0.64199 |
| ENSG00000165898 | 14 | 74960423 | 74963809 | 10 | 3 | -0.45588 | 0.67576 | ISCA2 | 0.67576 | 0.66406 | 0.75515 |
| ENSG00000132821 | 20 | 36531499 | 36573752 | 131 | 19 | -0.4561 | 0.67584 | VSTM2L | 0.67584 | 0.686 | 0.77318 |
| ENSG00000116299 | 1 | 109656301 | 109749401 | 402 | 44 | -0.45649 | 0.67598 | KIAA1324 | 0.67598 | 0.70879 | 0.65405 |

| | | | | | | | | | | | |
|---|---|---|---|---|---|---|---|---|---|---|---|
| ENSG00000088726 | 3 | 12775024 | 12810956 | 129 | 22 | -0.45648 | 0.67598 | TMEM40 | 0.67598 | 0.6985 | 0.79238 |
| ENSG00000132321 | 2 | 237232794 | 237416185 | 492 | 30 | -0.45656 | 0.676 | IQCA1 | 0.676 | 0.70187 | 0.75587 |
| ENSG00000102931 | 16 | 57279010 | 57287516 | 13 | 3 | -0.45663 | 0.67603 | ARL2BP | 0.67603 | 0.70186 | 0.6441 |
| ENSG00000187140 | 1 | 63788730 | 63790797 | 3 | 2 | -0.45697 | 0.67615 | FOXD3 | 0.67615 | 0.40887 | 0.63472 |
| ENSG00000143751 | 1 | 226170403 | 226187032 | 23 | 9 | -0.45696 | 0.67615 | SDE2 | 0.67615 | 0.43744 | 0.26222 |
| ENSG00000055957 | 3 | 52811603 | 52826078 | 45 | 5 | -0.4571 | 0.6762 | ITIH1 | 0.6762 | 0.61258 | 0.49744 |
| ENSG00000187461 | 6 | 13469513 | 13486415 | 38 | 11 | -0.45712 | 0.67621 | AL583828.1 | 0.67621 | 0.88917 | 0.88419 |
| ENSG00000104154 | 15 | 45771809 | 45815005 | 47 | 8 | -0.45773 | 0.67643 | SLC30A4 | 0.67643 | 0.55067 | 0.64992 |
| ENSG00000101624 | 18 | 12661832 | 12702776 | 147 | 14 | -0.45773 | 0.67643 | CEP76 | 0.67643 | 0.72917 | 0.75063 |
| ENSG00000153563 | 2 | 87011729 | 87035519 | 71 | 8 | -0.45781 | 0.67646 | CD8A | 0.67646 | 0.82062 | 0.90082 |
| ENSG00000187672 | 3 | 55542336 | 56502391 | 3243 | 117 | -0.45784 | 0.67647 | ERC2 | 0.67647 | 0.6789 | 0.69018 |
| ENSG00000167333 | 11 | 4619902 | 4629489 | 37 | 7 | -0.45789 | 0.67648 | TRIM68 | 0.67648 | 0.68777 | 0.60758 |
| ENSG00000108515 | 17 | 4851387 | 4860426 | 14 | 6 | -0.45794 | 0.6765 | ENO3 | 0.6765 | 0.19528 | 0.25037 |
| ENSG00000081052 | 2 | 227867427 | 228028829 | 563 | 40 | -0.45804 | 0.67654 | COL4A4 | 0.67654 | 0.68599 | 0.68964 |
| ENSG00000057019 | 3 | 98514785 | 98620533 | 276 | 19 | -0.45808 | 0.67655 | DCBLD2 | 0.67655 | 0.62721 | 0.64573 |
| ENSG00000100294 | 22 | 43528212 | 43539400 | 29 | 8 | -0.45806 | 0.67655 | MCAT | 0.67655 | 0.51237 | 0.20251 |
| ENSG00000170222 | 17 | 10600911 | 10614550 | 37 | 3 | -0.45816 | 0.67658 | ADPRM | 0.67658 | 0.60822 | 0.63005 |
| ENSG00000005812 | 13 | 77566740 | 77601330 | 133 | 12 | -0.45848 | 0.6767 | FBXL3 | 0.6767 | 0.7058 | 0.66029 |
| ENSG00000272968 | 7 | 5023349 | 5112852 | 432 | 23 | -0.45869 | 0.67677 | RBAK-RBAKDN | 0.67677 | 0.67231 | 0.70196 |
| ENSG00000112659 | 6 | 43149913 | 43192325 | 107 | 16 | -0.45931 | 0.677 | CUL9 | 0.677 | 0.67276 | 0.66875 |
| ENSG00000106348 | 7 | 128032331 | 128050306 | 66 | 15 | -0.45935 | 0.67701 | IMPDH1 | 0.67701 | 0.66391 | 0.45173 |
| ENSG00000124091 | 20 | 55066548 | 55100981 | 94 | 9 | -0.45949 | 0.67706 | GCNT7 | 0.67706 | 0.73105 | 0.84807 |
| ENSG00000241399 | 2 | 160625364 | 160654753 | 137 | 17 | -0.45982 | 0.67718 | CD302 | 0.67718 | 0.69639 | 0.69389 |
| ENSG00000136999 | 8 | 120428546 | 120436593 | 42 | 3 | -0.45993 | 0.67722 | NOV | 0.67722 | 0.6583 | 0.679 |
| ENSG00000145029 | 3 | 49460379 | 49466759 | 9 | 4 | -0.46007 | 0.67727 | NICN1 | 0.67727 | 0.51693 | 0.656 |
| ENSG00000183011 | 17 | 7760003 | 7788556 | 79 | 14 | -0.4603 | 0.67735 | LSMD1 | 0.67735 | 0.72061 | 0.74517 |
| ENSG00000137936 | 1 | 94027347 | 94312706 | 1022 | 35 | -0.46047 | 0.67741 | BCAR3 | 0.67741 | 0.65303 | 0.64602 |
| ENSG00000181355 | 6 | 9596343 | 10211841 | 2155 | 119 | -0.46057 | 0.67745 | OFCC1 | 0.67745 | 0.70912 | 0.72993 |
| ENSG00000198862 | 21 | 30300466 | 30365270 | 163 | 24 | -0.46069 | 0.67749 | LTN1 | 0.67749 | 0.65621 | 0.66058 |
| ENSG00000118491 | 6 | 144185573 | 144259483 | 196 | 29 | -0.46103 | 0.67761 | ZC2HC1B | 0.67761 | 0.71297 | 0.78527 |
| ENSG00000261609 | 16 | 81348557 | 81424489 | 438 | 32 | -0.46121 | 0.67768 | GAN | 0.67768 | 0.74257 | 0.77914 |
| ENSG00000149357 | 11 | 71796941 | 71814433 | 43 | 5 | -0.46148 | 0.67777 | LAMTOR1 | 0.67777 | 0.70337 | 0.74283 |
| ENSG00000172167 | 8 | 121457640 | 121554373 | 263 | 10 | -0.46151 | 0.67778 | MTBP | 0.67778 | 0.68564 | 0.72696 |
| ENSG00000186354 | 9 | 91605778 | 91611055 | 18 | 7 | -0.46158 | 0.67781 | C9orf47 | 0.67781 | 0.63816 | 0.72451 |
| ENSG00000172497 | 5 | 80625824 | 80689998 | 230 | 40 | -0.46173 | 0.67786 | ACOT12 | 0.67786 | 0.6671 | 0.69464 |
| ENSG00000266302 | 17 | 16593575 | 16707767 | 126 | 15 | -0.46178 | 0.67788 | RP11-219A15.1 | 0.67788 | 0.65928 | 0.726 |
| ENSG00000161013 | 5 | 179224597 | 179233952 | 25 | 6 | -0.46192 | 0.67793 | MGAT4B | 0.67793 | 0.86 | 0.82714 |
| ENSG00000076344 | 16 | 318300 | 325980 | 30 | 8 | -0.46198 | 0.67795 | RGS11 | 0.67795 | 0.28601 | 0.43229 |
| ENSG00000085721 | 16 | 15153879 | 15188174 | 109 | 18 | -0.46204 | 0.67798 | RRN3 | 0.67798 | 0.61322 | 0.37661 |
| ENSG00000002016 | 12 | 1021243 | 1099219 | 300 | 37 | -0.46208 | 0.67799 | RAD52 | 0.67799 | 0.73743 | 0.76497 |
| ENSG00000111802 | 6 | 24650205 | 24667261 | 86 | 15 | -0.46218 | 0.67802 | TDP2 | 0.67802 | 0.7651 | 0.84005 |
| ENSG00000143127 | 1 | 145524891 | 145543868 | 26 | 8 | -0.46261 | 0.67818 | ITGA10 | 0.67818 | 0.74927 | 0.66126 |
| ENSG00000167536 | 17 | 27224799 | 27230089 | 6 | 2 | -0.46263 | 0.67818 | DHRS13 | 0.67818 | 0.17361 | 0.21667 |
| ENSG00000214226 | 17 | 54869274 | 54916134 | 146 | 13 | -0.46279 | 0.67824 | C17orf67 | 0.67824 | 0.7222 | 0.73563 |
| ENSG00000078487 | 7 | 99998449 | 100026615 | 52 | 5 | -0.46288 | 0.67827 | ZCWPW1 | 0.67827 | 0.67015 | 0.60484 |
| ENSG00000112280 | 6 | 70924764 | 71012786 | 435 | 36 | -0.4629 | 0.67828 | COL9A1 | 0.67828 | 0.70128 | 0.77675 |
| ENSG00000179562 | 7 | 127220672 | 127233665 | 42 | 11 | -0.46329 | 0.67842 | GCC1 | 0.67842 | 0.67501 | 0.74102 |
| ENSG00000185100 | 14 | 105190523 | 105213662 | 72 | 11 | -0.46393 | 0.67865 | ADSSL1 | 0.67865 | 0.70967 | 0.75622 |
| ENSG00000127124 | 1 | 41972036 | 42501596 | 1581 | 67 | -0.46408 | 0.6787 | HIVEP3 | 0.6787 | 0.69611 | 0.66593 |
| ENSG00000106609 | 7 | 66386212 | 66423538 | 113 | 14 | -0.46411 | 0.67872 | TMEM248 | 0.67872 | 0.69572 | 0.64745 |
| ENSG00000196967 | 19 | 37597636 | 37663643 | 194 | 13 | -0.46421 | 0.67875 | ZNF585A | 0.67875 | 0.69926 | 0.66306 |
| ENSG00000162341 | 11 | 68816365 | 68858072 | 215 | 15 | -0.46442 | 0.67883 | TPCN2 | 0.67883 | 0.65756 | 0.68886 |
| ENSG00000114544 | 3 | 125725198 | 125820404 | 417 | 19 | -0.4645 | 0.67886 | SLC41A3 | 0.67886 | 0.63841 | 0.54537 |
| ENSG00000170088 | 4 | 165995574 | 166129701 | 591 | 34 | -0.46462 | 0.6789 | TMEM192 | 0.6789 | 0.7286 | 0.75993 |
| ENSG00000163541 | 2 | 84650647 | 84687169 | 112 | 10 | -0.46471 | 0.67893 | SUCLG1 | 0.67893 | 0.68178 | 0.68247 |
| ENSG00000139914 | 14 | 24600484 | 24602058 | 2 | 1 | -0.46472 | 0.67893 | FITM1 | 0.67893 | 0.85847 | 0.86816 |
| ENSG00000066136 | 1 | 41157320 | 41237275 | 245 | 22 | -0.46474 | 0.67894 | NFYC | 0.67894 | 0.61338 | 0.5896 |
| ENSG00000164600 | 7 | 31377075 | 31380508 | 7 | 3 | -0.46495 | 0.67902 | NEUROD6 | 0.67902 | 0.56056 | 0.38447 |
| ENSG00000071537 | 14 | 81937893 | 82000205 | 96 | 23 | -0.46507 | 0.67906 | SEL1L | 0.67906 | 0.81579 | 0.73929 |
| ENSG00000179636 | 14 | 21492268 | 21504435 | 63 | 17 | -0.4652 | 0.67911 | TPPP2 | 0.67911 | 0.80002 | 0.89417 |
| ENSG00000269165 | 10 | 52094298 | 52095876 | 4 | 2 | -0.46574 | 0.6793 | AC069547.1 | 0.6793 | 0.36343 | 0.54017 |
| ENSG00000233539 | 7 | 46713383 | 46799449 | 371 | 32 | -0.46578 | 0.67931 | AC011294.3 | 0.67931 | 0.8095 | 0.81513 |
| ENSG00000105726 | 19 | 19756007 | 19774502 | 53 | 14 | -0.46596 | 0.67938 | ATP13A1 | 0.67938 | 0.76701 | 0.72392 |
| ENSG00000269449 | 7 | 95171457 | 95173854 | 8 | 5 | -0.46607 | 0.67942 | AC002451.1 | 0.67942 | 0.60417 | 0.52525 |
| ENSG00000256655 | 8 | 128959126 | 128960591 | 2 | 1 | -0.4662 | 0.67947 | TMEM75 | 0.67947 | 0.13816 | 0.15059 |
| ENSG00000164323 | 4 | 186080819 | 186130658 | 209 | 12 | -0.46627 | 0.67949 | KIAA1430 | 0.67949 | 0.70658 | 0.74806 |
| ENSG00000087076 | 19 | 49316274 | 49339935 | 104 | 20 | -0.46631 | 0.6795 | HSD17B14 | 0.6795 | 0.73044 | 0.80045 |
| ENSG00000105894 | 7 | 136912088 | 137028611 | 372 | 48 | -0.46634 | 0.67951 | PTN | 0.67951 | 0.68347 | 0.74979 |
| ENSG00000143862 | 1 | 202102532 | 202113869 | 23 | 6 | -0.46637 | 0.67952 | ARL8A | 0.67952 | 0.27562 | 0.28834 |
| ENSG00000146587 | 7 | 5085452 | 5109119 | 99 | 16 | -0.46637 | 0.67952 | RBAK | 0.67952 | 0.64692 | 0.52877 |
| ENSG00000138002 | 2 | 27667238 | 27712656 | 84 | 8 | -0.46647 | 0.67956 | IFT172 | 0.67956 | 0.66174 | 0.62243 |
| ENSG00000090581 | 16 | 1401924 | 1413352 | 35 | 12 | -0.46647 | 0.67956 | GNPTG | 0.67956 | 0.83209 | 0.84871 |
| ENSG00000198830 | 1 | 26798941 | 26802463 | 5 | 2 | -0.4665 | 0.67957 | HMGN2 | 0.67957 | 0.66875 | 0.71506 |

| | | | | | | | | | | | |
|---|---|---|---|---|---|---|---|---|---|---|---|
| ENSG00000103932 | 15 | 41809374 | 41836467 | 97 | 14 | -0.46655 | 0.67959 | RPAP1 | 0.67959 | 0.66651 | 0.69378 |
| ENSG00000160691 | 1 | 154934774 | 154946871 | 16 | 4 | -0.46663 | 0.67962 | SHC1 | 0.67962 | 0.71956 | 0.83801 |
| ENSG00000268076 | 12 | 25381515 | 25390840 | 37 | 11 | -0.467 | 0.67975 | AC087239.1 | 0.67975 | 0.70455 | 0.7577 |
| ENSG00000175105 | 3 | 88188254 | 88193815 | 16 | 3 | -0.46709 | 0.67978 | ZNF654 | 0.67978 | 0.7376 | 0.74115 |
| ENSG00000185972 | 9 | 36169389 | 36171329 | 6 | 3 | -0.46707 | 0.67978 | CCIN | 0.67978 | 0.38498 | 0.32452 |
| ENSG00000162885 | 1 | 235613238 | 235667781 | 157 | 15 | -0.46711 | 0.67979 | B3GALNT2 | 0.67979 | 0.66129 | 0.55999 |
| ENSG00000125447 | 17 | 73232694 | 73258444 | 64 | 9 | -0.46714 | 0.6798 | GGA3 | 0.6798 | 0.67499 | 0.62331 |
| ENSG00000137713 | 11 | 111597632 | 111637151 | 89 | 15 | -0.46727 | 0.67985 | PPP2R1B | 0.67985 | 0.67424 | 0.66175 |
| ENSG00000213445 | 11 | 65405568 | 65418401 | 23 | 7 | -0.46735 | 0.67987 | SIPA1 | 0.67987 | 0.81662 | 0.82905 |
| ENSG00000172469 | 6 | 96025419 | 96057333 | 125 | 12 | -0.46794 | 0.68009 | MANEA | 0.68009 | 0.65709 | 0.70236 |
| ENSG00000067560 | 3 | 49396578 | 49450431 | 139 | 15 | -0.46822 | 0.68019 | RHOA | 0.68019 | 0.64824 | 0.65286 |
| ENSG00000197403 | 1 | 158735466 | 158736492 | 9 | 3 | -0.46827 | 0.6802 | OR6N1 | 0.6802 | 0.81209 | 0.61082 |
| ENSG00000146802 | 7 | 112402437 | 112430647 | 52 | 9 | -0.4683 | 0.68022 | TMEM168 | 0.68022 | 0.7016 | 0.64345 |
| ENSG00000135912 | 2 | 219575568 | 219620139 | 100 | 6 | -0.46838 | 0.68024 | TTLL4 | 0.68024 | 0.66369 | 0.65865 |
| ENSG00000144868 | 3 | 132757235 | 133116636 | 1329 | 48 | -0.46921 | 0.68054 | TMEM108 | 0.68054 | 0.69643 | 0.78038 |
| ENSG00000240871 | 17 | 39240459 | 39241396 | 11 | 3 | -0.46925 | 0.68056 | KRTAP4-7 | 0.68056 | 0.62085 | 0.55741 |
| ENSG00000113812 | 3 | 53901093 | 53916229 | 52 | 9 | -0.46929 | 0.68057 | ACTR8 | 0.68057 | 0.65176 | 0.52613 |
| ENSG00000145675 | 5 | 67511548 | 67597649 | 291 | 41 | -0.46932 | 0.68058 | PIK3R1 | 0.68058 | 0.62479 | 0.26639 |
| ENSG00000135953 | 2 | 103332299 | 103353347 | 47 | 8 | -0.46945 | 0.68063 | MFSD9 | 0.68063 | 0.58874 | 0.74435 |
| ENSG00000130487 | 22 | 50986462 | 50989451 | 15 | 5 | -0.46964 | 0.68069 | KLHDC7B | 0.68069 | 0.29717 | 0.48762 |
| ENSG00000128519 | 7 | 122634759 | 122635754 | 2 | 1 | -0.46965 | 0.6807 | TAS2R16 | 0.6807 | 0.41451 | 0.31906 |
| ENSG00000129295 | 8 | 133584320 | 133687838 | 188 | 20 | -0.46984 | 0.68076 | LRRC6 | 0.68076 | 0.63751 | 0.64279 |
| ENSG00000130943 | 22 | 46651560 | 46659219 | 32 | 4 | -0.4699 | 0.68079 | PKDREJ | 0.68079 | 0.55075 | 0.48713 |
| ENSG00000266958 | 19 | 45690646 | 45720191 | 179 | 33 | -0.47 | 0.68082 | AC006126.3 | 0.68082 | 0.85637 | 0.89448 |
| ENSG00000084676 | 2 | 24714783 | 24993571 | 774 | 29 | -0.47022 | 0.6809 | NCOA1 | 0.6809 | 0.67856 | 0.65995 |
| ENSG00000198919 | 3 | 108308529 | 108413693 | 184 | 20 | -0.47023 | 0.68091 | DZIP3 | 0.68091 | 0.66416 | 0.67972 |
| ENSG00000198382 | 11 | 75526212 | 75854239 | 655 | 20 | -0.47027 | 0.68092 | UVRAG | 0.68092 | 0.74048 | 0.7689 |
| ENSG00000188038 | 16 | 67918708 | 67922758 | 4 | 2 | -0.47031 | 0.68093 | NRN1L | 0.68093 | 0.73023 | 0.78136 |
| ENSG00000167992 | 11 | 61025762 | 61062896 | 34 | 12 | -0.47047 | 0.68099 | VWCE | 0.68099 | 0.61628 | 0.59242 |
| ENSG00000268275 | 6 | 41653964 | 41654127 | 1 | 1 | -0.4705 | 0.681 | AL035588.1 | 0.681 | 0.49156 | 0.86126 |
| ENSG00000138395 | 2 | 202655184 | 202760273 | 332 | 43 | -0.47055 | 0.68102 | CDK15 | 0.68102 | 0.72914 | 0.71275 |
| ENSG00000255330 | 6 | 127759551 | 127840146 | 131 | 21 | -0.47063 | 0.68105 | SOGA3 | 0.68105 | 0.685 | 0.78014 |
| ENSG00000214338 | 6 | 127759551 | 127840500 | 131 | 21 | -0.47063 | 0.68105 | SOGA3 | 0.68105 | 0.685 | 0.78014 |
| ENSG00000151461 | 10 | 11962021 | 12085169 | 482 | 23 | -0.47074 | 0.68109 | UPF2 | 0.68109 | 0.67099 | 0.8271 |
| ENSG00000162889 | 1 | 206858289 | 206907628 | 140 | 23 | -0.47094 | 0.68116 | MAPKAPK2 | 0.68116 | 0.59774 | 0.58343 |
| ENSG00000125124 | 16 | 56500748 | 56554195 | 120 | 15 | -0.47097 | 0.68117 | BBS2 | 0.68117 | 0.6338 | 0.66189 |
| ENSG00000084463 | 12 | 14939410 | 14956474 | 36 | 9 | -0.47106 | 0.6812 | WBP11 | 0.6812 | 0.41318 | 0.37334 |
| ENSG00000179709 | 19 | 56459198 | 56499995 | 191 | 28 | -0.47115 | 0.68123 | NLRP8 | 0.68123 | 0.65266 | 0.46293 |
| ENSG00000166887 | 15 | 42450899 | 42500514 | 117 | 12 | -0.47133 | 0.6813 | VPS39 | 0.6813 | 0.71583 | 0.64257 |
| ENSG00000167800 | 11 | 67398774 | 67407031 | 38 | 7 | -0.47137 | 0.68131 | TBX10 | 0.68131 | 0.60302 | 0.2853 |
| ENSG00000198265 | 17 | 65066554 | 65242105 | 375 | 19 | -0.47143 | 0.68133 | HELZ | 0.68133 | 0.68149 | 0.69916 |
| ENSG00000109118 | 17 | 27232268 | 27278789 | 50 | 10 | -0.47155 | 0.68138 | PHF12 | 0.68138 | 0.6213 | 0.48714 |
| ENSG00000253976 | 8 | 56047763 | 56054918 | 27 | 7 | -0.47171 | 0.68143 | RP11-386G21.1 | 0.68143 | 0.76029 | 0.81313 |
| ENSG00000050748 | 5 | 179660143 | 179719099 | 182 | 25 | -0.47173 | 0.68144 | MAPK9 | 0.68144 | 0.66179 | 0.5583 |
| ENSG00000056487 | 22 | 45277042 | 45405880 | 498 | 50 | -0.47186 | 0.68149 | PHF21B | 0.68149 | 0.46085 | 0.65873 |
| ENSG00000100031 | 22 | 24979718 | 25024972 | 136 | 13 | -0.47254 | 0.68173 | GGT1 | 0.68173 | 0.65475 | 0.60125 |
| ENSG00000101150 | 20 | 62496596 | 62522898 | 70 | 12 | -0.47284 | 0.68184 | TPD52L2 | 0.68184 | 0.66323 | 0.49469 |
| ENSG00000167977 | 16 | 2732476 | 2759031 | 80 | 14 | -0.47318 | 0.68196 | KCTD5 | 0.68196 | 0.81593 | 0.86537 |
| ENSG00000141101 | 16 | 69775770 | 69788843 | 32 | 8 | -0.47319 | 0.68196 | NOB1 | 0.68196 | 0.72615 | 0.60308 |
| ENSG00000018699 | 2 | 32853099 | 33046118 | 850 | 42 | -0.47321 | 0.68197 | TTC27 | 0.68197 | 0.71113 | 0.73906 |
| ENSG00000171763 | 15 | 45694529 | 45713617 | 34 | 6 | -0.47329 | 0.682 | SPATA5L1 | 0.682 | 0.88038 | 0.90826 |
| ENSG00000173826 | 17 | 61600695 | 61626338 | 46 | 12 | -0.47339 | 0.68203 | KCNH6 | 0.68203 | 0.76848 | 0.7938 |
| ENSG00000164430 | 6 | 74123238 | 74161999 | 104 | 30 | -0.47357 | 0.68209 | MB21D1 | 0.68209 | 0.73171 | 0.6344 |
| ENSG00000124370 | 2 | 71336814 | 71357369 | 88 | 9 | -0.47357 | 0.6821 | MCEE | 0.6821 | 0.7022 | 0.75146 |
| ENSG00000128606 | 7 | 102553438 | 102585396 | 55 | 9 | -0.47372 | 0.68215 | LRRC17 | 0.68215 | 0.66017 | 0.63769 |
| ENSG00000138092 | 2 | 25016005 | 25045245 | 115 | 18 | -0.47376 | 0.68216 | CENPO | 0.68216 | 0.73818 | 0.74103 |
| ENSG00000147724 | 8 | 139142266 | 139509065 | 1454 | 120 | -0.47375 | 0.68216 | FAM135B | 0.68216 | 0.65145 | 0.62139 |
| ENSG00000151012 | 4 | 139085251 | 139163503 | 233 | 25 | -0.47382 | 0.68219 | SLC7A11 | 0.68219 | 0.72783 | 0.71757 |
| ENSG00000152556 | 12 | 48498922 | 48540187 | 117 | 16 | -0.47383 | 0.68219 | PFKM | 0.68219 | 0.63758 | 0.69631 |
| ENSG00000125538 | 2 | 113587328 | 113594480 | 16 | 4 | -0.47395 | 0.68223 | IL1B | 0.68223 | 0.65336 | 0.5979 |
| ENSG00000181026 | 15 | 89164527 | 89175513 | 60 | 10 | -0.47415 | 0.6823 | AEN | 0.6823 | 0.77973 | 0.74421 |
| ENSG00000142784 | 1 | 27561007 | 27635110 | 130 | 19 | -0.47428 | 0.68235 | WDTC1 | 0.68235 | 0.64687 | 0.65101 |
| ENSG00000139344 | 12 | 96337071 | 96362370 | 131 | 12 | -0.47436 | 0.68238 | AMDHD1 | 0.68238 | 0.6878 | 0.66411 |
| ENSG00000103966 | 15 | 42190950 | 42264776 | 244 | 32 | -0.4744 | 0.68239 | EHD4 | 0.68239 | 0.61421 | 0.41527 |
| ENSG00000128253 | 22 | 32586422 | 32600718 | 60 | 11 | -0.47462 | 0.68247 | RFPL2 | 0.68247 | 0.59257 | 0.48734 |
| ENSG00000198252 | 14 | 53196898 | 53241716 | 107 | 12 | -0.47467 | 0.68249 | STYX | 0.68249 | 0.67026 | 0.66691 |
| ENSG00000161860 | 19 | 13009600 | 13030090 | 35 | 10 | -0.47478 | 0.68253 | SYCE2 | 0.68253 | 0.63518 | 0.73579 |
| ENSG00000003400 | 2 | 202047604 | 202094129 | 92 | 11 | -0.47482 | 0.68254 | CASP10 | 0.68254 | 0.67467 | 0.80601 |
| ENSG00000261652 | 15 | 55700746 | 55710962 | 57 | 10 | -0.47498 | 0.6826 | C15orf65 | 0.6826 | 0.61698 | 0.57912 |
| ENSG00000158008 | 1 | 26346375 | 26362955 | 29 | 8 | -0.47512 | 0.68265 | EXTL1 | 0.68265 | 0.73284 | 0.72354 |
| ENSG00000198796 | 18 | 56148479 | 56296189 | 602 | 64 | -0.47539 | 0.68275 | ALPK2 | 0.68275 | 0.69153 | 0.59758 |
| ENSG00000165816 | 10 | 115999018 | 116051272 | 149 | 23 | -0.47548 | 0.68278 | VWA2 | 0.68278 | 0.6887 | 0.68423 |
| ENSG00000174876 | 1 | 104230037 | 104239302 | 1 | 1 | -0.47554 | 0.6828 | AMY1B | 0.6828 | 0.040026 | 0.18213 |

| | | | | | | | | | | | |
|---|---|---|---|---|---|---|---|---|---|---|---|
| ENSG00000110090 | 11 | 68522088 | 68611878 | 252 | 25 | -0.47561 | 0.68282 | CPT1A | 0.68282 | 0.62392 | 0.53027 |
| ENSG00000073803 | 3 | 185000729 | 185206885 | 616 | 28 | -0.47567 | 0.68285 | MAP3K13 | 0.68285 | 0.67821 | 0.70362 |
| ENSG00000117601 | 1 | 173872947 | 173886516 | 34 | 7 | -0.47604 | 0.68298 | SERPINC1 | 0.68298 | 0.70402 | 0.59775 |
| ENSG00000114573 | 3 | 113465866 | 113530903 | 112 | 5 | -0.47613 | 0.68301 | ATP6V1A | 0.68301 | 0.62352 | 0.6183 |
| ENSG00000158717 | 16 | 88762903 | 88772829 | 58 | 17 | -0.47622 | 0.68304 | RNF166 | 0.68304 | 0.82398 | 0.79444 |
| ENSG00000198074 | 7 | 134212344 | 134226160 | 61 | 11 | -0.47629 | 0.68306 | AKR1B10 | 0.68306 | 0.56661 | 0.41218 |
| ENSG00000187848 | 12 | 133195366 | 133198972 | 21 | 7 | -0.47633 | 0.68308 | P2RX2 | 0.68308 | 0.75101 | 0.78898 |
| ENSG00000130711 | 9 | 133539981 | 133558368 | 65 | 15 | -0.47643 | 0.68311 | PRDM12 | 0.68311 | 0.69248 | 0.81002 |
| ENSG00000198517 | 7 | 1570350 | 1582679 | 38 | 7 | -0.47662 | 0.68318 | MAFK | 0.68318 | 0.73774 | 0.75113 |
| ENSG00000259305 | 8 | 124238431 | 124279627 | 40 | 15 | -0.47678 | 0.68324 | ZHX1-C8ORF76 | 0.68324 | 0.63659 | 0.6938 |
| ENSG00000134644 | 1 | 31404353 | 31538838 | 337 | 21 | -0.47698 | 0.68331 | PUM1 | 0.68331 | 0.69467 | 0.68007 |
| ENSG00000137522 | 11 | 71639747 | 71708643 | 230 | 12 | -0.47707 | 0.68334 | RNF121 | 0.68334 | 0.66115 | 0.6618 |
| ENSG00000158488 | 1 | 158323254 | 158327343 | 7 | 3 | -0.47711 | 0.68336 | CD1E | 0.68336 | 0.5704 | 0.66393 |
| ENSG00000100490 | 14 | 50796310 | 50883179 | 280 | 28 | -0.47721 | 0.68339 | CDKL1 | 0.68339 | 0.74984 | 0.74888 |
| ENSG00000167528 | 12 | 48733791 | 48745197 | 26 | 6 | -0.4778 | 0.6836 | ZNF641 | 0.6836 | 0.56861 | 0.48221 |
| ENSG00000163393 | 1 | 116519119 | 116612675 | 269 | 25 | -0.47812 | 0.68372 | SLC22A15 | 0.68372 | 0.5898 | 0.59072 |
| ENSG00000179919 | 12 | 55614809 | 55615759 | 1 | 1 | -0.47835 | 0.6838 | OR10A7 | 0.6838 | 0.91732 | 0.91765 |
| ENSG00000198042 | 8 | 33342268 | 33358778 | 62 | 8 | -0.47851 | 0.68386 | MAK16 | 0.68386 | 0.68216 | 0.67037 |
| ENSG00000174943 | 16 | 29916333 | 29938356 | 39 | 6 | -0.47856 | 0.68388 | KCTD13 | 0.68388 | 0.61209 | 0.65822 |
| ENSG00000132153 | 3 | 47844399 | 47891685 | 49 | 6 | -0.4786 | 0.68389 | DHX30 | 0.68389 | 0.80063 | 0.85268 |
| ENSG00000183196 | 16 | 75510949 | 75529282 | 64 | 20 | -0.47874 | 0.68394 | CHST6 | 0.68394 | 0.68362 | 0.87843 |
| ENSG00000112183 | 6 | 17281577 | 17294106 | 26 | 9 | -0.47884 | 0.68397 | RBM24 | 0.68397 | 0.56842 | 0.47968 |
| ENSG00000127947 | 7 | 77166592 | 77269388 | 339 | 19 | -0.47882 | 0.68397 | PTPN12 | 0.68397 | 0.66863 | 0.61653 |
| ENSG00000136040 | 12 | 94542499 | 94701451 | 667 | 69 | -0.47884 | 0.68398 | PLXNC1 | 0.68398 | 0.74718 | 0.80605 |
| ENSG00000143515 | 1 | 154298029 | 154323783 | 72 | 11 | -0.47893 | 0.684 | ATP8B2 | 0.684 | 0.55099 | 0.34504 |
| ENSG00000166111 | 12 | 109304658 | 109459045 | 366 | 33 | -0.47909 | 0.68406 | SVOP | 0.68406 | 0.7244 | 0.77611 |
| ENSG00000100503 | 14 | 51186481 | 51297839 | 346 | 45 | -0.47932 | 0.68414 | NIN | 0.68414 | 0.76358 | 0.65688 |
| ENSG00000135111 | 12 | 115108059 | 115121969 | 37 | 12 | -0.47941 | 0.68418 | TBX3 | 0.68418 | 0.27957 | 0.31583 |
| ENSG00000150782 | 11 | 112013974 | 112034840 | 61 | 13 | -0.4797 | 0.68428 | IL18 | 0.68428 | 0.56842 | 0.58921 |
| ENSG00000148634 | 10 | 69681665 | 69835105 | 363 | 23 | -0.47973 | 0.68429 | HERC4 | 0.68429 | 0.68271 | 0.68312 |
| ENSG00000126467 | 19 | 50243010 | 50266587 | 66 | 14 | -0.47974 | 0.68429 | TSKS | 0.68429 | 0.70673 | 0.82473 |
| ENSG00000166747 | 16 | 71762913 | 71843104 | 238 | 24 | -0.48001 | 0.68439 | AP1G1 | 0.68439 | 0.68698 | 0.62162 |
| ENSG00000273294 | 5 | 33987279 | 34124633 | 325 | 29 | -0.48005 | 0.6844 | RP11-1084J3.4 | 0.6844 | 0.66865 | 0.71977 |
| ENSG00000162881 | 2 | 42989642 | 42991401 | 7 | 2 | -0.48006 | 0.68441 | OXER1 | 0.68441 | 0.48293 | 0.22229 |
| ENSG00000175066 | 3 | 141882414 | 141944449 | 162 | 27 | -0.48014 | 0.68444 | GK5 | 0.68444 | 0.7558 | 0.7649 |
| ENSG00000149506 | 11 | 60635035 | 60643166 | 16 | 5 | -0.48038 | 0.68452 | ZP1 | 0.68452 | 0.5736 | 0.76582 |
| ENSG00000177830 | 11 | 867357 | 915058 | 123 | 14 | -0.48057 | 0.68459 | CHID1 | 0.68459 | 0.67094 | 0.6979 |
| ENSG00000161920 | 17 | 4634723 | 4636905 | 5 | 2 | -0.48061 | 0.6846 | MED11 | 0.6846 | 0.8028 | 0.71426 |
| ENSG00000163646 | 3 | 150643950 | 150690786 | 199 | 27 | -0.48075 | 0.68465 | CLRN1 | 0.68465 | 0.73315 | 0.76662 |
| ENSG00000110075 | 11 | 68228186 | 68382802 | 398 | 19 | -0.48085 | 0.68469 | PPP6R3 | 0.68469 | 0.60155 | 0.44217 |
| ENSG00000214688 | 10 | 73471458 | 73497581 | 64 | 13 | -0.48101 | 0.68474 | C10orf105 | 0.68474 | 0.83353 | 0.81835 |
| ENSG00000181789 | 3 | 128968449 | 128996614 | 52 | 6 | -0.48106 | 0.68476 | COPG1 | 0.68476 | 0.65677 | 0.60413 |
| ENSG00000100926 | 14 | 24658349 | 24682679 | 90 | 16 | -0.48124 | 0.68483 | TM9SF1 | 0.68483 | 0.73547 | 0.72503 |
| ENSG00000254692 | 14 | 24658363 | 24682679 | 90 | 16 | -0.48124 | 0.68483 | TM9SF1 | 0.68483 | 0.73547 | 0.72503 |
| ENSG00000267618 | 17 | 33338368 | 33448512 | 143 | 24 | -0.48131 | 0.68485 | RAD51L3-RFFL | 0.68485 | 0.65748 | 0.6022 |
| ENSG00000163923 | 3 | 186838736 | 186898696 | 251 | 35 | -0.48142 | 0.68489 | RPL39L | 0.68489 | 0.64811 | 0.69845 |
| ENSG00000140905 | 16 | 81115566 | 81130008 | 90 | 4 | -0.48149 | 0.68492 | GCSH | 0.68492 | 0.67554 | 0.82737 |
| ENSG00000133393 | 16 | 15959577 | 15982482 | 201 | 8 | -0.48155 | 0.68494 | FOPNL | 0.68494 | 0.61665 | 0.56806 |
| ENSG00000101198 | 20 | 61872136 | 61904046 | 156 | 30 | -0.48171 | 0.68499 | NKAIN4 | 0.68499 | 0.39269 | 0.16024 |
| ENSG00000196405 | 14 | 100437786 | 100610573 | 551 | 32 | -0.48176 | 0.68501 | EVL | 0.68501 | 0.63945 | 0.45974 |
| ENSG00000173401 | 12 | 75728419 | 75764340 | 76 | 9 | -0.48178 | 0.68502 | GLIPR1L1 | 0.68502 | 0.61414 | 0.54368 |
| ENSG00000145623 | 5 | 38845960 | 38945698 | 351 | 39 | -0.48212 | 0.68514 | OSMR | 0.68514 | 0.61019 | 0.55608 |
| ENSG00000131849 | 19 | 58944181 | 58951589 | 9 | 3 | -0.48214 | 0.68515 | ZNF132 | 0.68515 | 0.72465 | 0.76161 |
| ENSG00000229544 | 10 | 126135592 | 126138753 | 13 | 2 | -0.48232 | 0.68521 | NKX1-2 | 0.68521 | 0.30461 | 0.4605 |
| ENSG00000100462 | 14 | 23389720 | 23398794 | 23 | 4 | -0.48235 | 0.68522 | PRMT5 | 0.68522 | 0.79246 | 0.72924 |
| ENSG00000112685 | 6 | 485133 | 693117 | 866 | 50 | -0.48253 | 0.68529 | EXOC2 | 0.68529 | 0.70153 | 0.64099 |
| ENSG00000179455 | 15 | 23810454 | 23873064 | 171 | 19 | -0.48259 | 0.68531 | MKRN3 | 0.68531 | 0.53845 | 0.63506 |
| ENSG00000126107 | 1 | 45468212 | 45477001 | 22 | 4 | -0.48264 | 0.68532 | HECTD3 | 0.68532 | 0.58953 | 0.56383 |
| ENSG00000164185 | 5 | 121465208 | 121515312 | 91 | 25 | -0.48266 | 0.68533 | ZNF474 | 0.68533 | 0.73066 | 0.87108 |
| ENSG00000168907 | 15 | 42433332 | 42448839 | 61 | 14 | -0.48266 | 0.68533 | PLA2G4F | 0.68533 | 0.39288 | 0.60602 |
| ENSG00000183020 | 11 | 924894 | 1012239 | 350 | 14 | -0.48314 | 0.6855 | AP2A2 | 0.6855 | 0.62355 | 0.48882 |
| ENSG00000163218 | 1 | 153302596 | 153321316 | 87 | 7 | -0.48332 | 0.68556 | PGLYRP4 | 0.68556 | 0.68125 | 0.57624 |
| ENSG00000173207 | 1 | 154947129 | 154951725 | 2 | 1 | -0.48349 | 0.68562 | CKS1B | 0.68562 | 0.44668 | 0.61149 |
| ENSG00000124159 | 20 | 43922085 | 43937169 | 61 | 9 | -0.4836 | 0.68567 | MATN4 | 0.68567 | 0.75236 | 0.31939 |
| ENSG00000171819 | 1 | 11249398 | 11256038 | 17 | 3 | -0.48379 | 0.68573 | ANGPTL7 | 0.68573 | 0.70299 | 0.74152 |
| ENSG00000198825 | 10 | 121485609 | 121588652 | 296 | 16 | -0.48411 | 0.68585 | INPP5F | 0.68585 | 0.71132 | 0.6989 |
| ENSG00000161249 | 19 | 35988122 | 36004560 | 74 | 18 | -0.48433 | 0.68593 | DMKN | 0.68593 | 0.48804 | 0.37225 |
| ENSG00000105771 | 19 | 44235301 | 44259142 | 32 | 9 | -0.48435 | 0.68593 | SMG9 | 0.68593 | 0.67614 | 0.87728 |
| ENSG00000144369 | 2 | 187558698 | 187630685 | 147 | 20 | -0.48437 | 0.68594 | FAM171B | 0.68594 | 0.64639 | 0.68055 |
| ENSG00000109686 | 4 | 152023903 | 152246784 | 715 | 28 | -0.48442 | 0.68596 | SH3D19 | 0.68596 | 0.69672 | 0.67903 |
| ENSG00000108272 | 17 | 34948228 | 34957235 | 19 | 4 | -0.48443 | 0.68596 | DHRS11 | 0.68596 | 0.48669 | 0.48154 |
| ENSG00000171944 | 11 | 5152922 | 5153872 | 1 | 1 | -0.48483 | 0.6861 | OR52A5 | 0.6861 | 0.69051 | 0.37209 |
| ENSG00000069424 | 1 | 6051526 | 6161253 | 415 | 62 | -0.48491 | 0.68613 | KCNAB2 | 0.68613 | 0.54953 | 0.54383 |

| | | | | | | | | | | | |
|---|---|---|---|---|---|---|---|---|---|---|---|
| ENSG00000164902 | 5 | 125935960 | 125962944 | 107 | 9 | -0.48517 | 0.68622 | PHAX | 0.68622 | 0.57028 | 0.35825 |
| ENSG00000100483 | 14 | 50575350 | 50583318 | 45 | 8 | -0.48516 | 0.68622 | VCPKMT | 0.68622 | 0.62357 | 0.5731 |
| ENSG00000169398 | 8 | 141667999 | 142012315 | 683 | 16 | -0.4852 | 0.68623 | PTK2 | 0.68623 | 0.67377 | 0.63069 |
| ENSG00000135040 | 9 | 88556061 | 88637213 | 125 | 26 | -0.48574 | 0.68642 | NAA35 | 0.68642 | 0.74002 | 0.7657 |
| ENSG00000117461 | 1 | 46505812 | 46642160 | 234 | 23 | -0.48587 | 0.68647 | PIK3R3 | 0.68647 | 0.66304 | 0.51287 |
| ENSG00000087589 | 20 | 54987168 | 55034396 | 150 | 24 | -0.48613 | 0.68656 | CASS4 | 0.68656 | 0.84635 | 0.91934 |
| ENSG00000113356 | 5 | 89767565 | 89810370 | 113 | 18 | -0.48618 | 0.68658 | POLR3G | 0.68658 | 0.69442 | 0.60049 |
| ENSG00000154518 | 2 | 176040986 | 176049335 | 36 | 6 | -0.48633 | 0.68663 | ATP5G3 | 0.68663 | 0.7643 | 0.75146 |
| ENSG00000129103 | 7 | 56131695 | 56148363 | 56 | 13 | -0.48652 | 0.6867 | SUMF2 | 0.6867 | 0.69302 | 0.76012 |
| ENSG00000197935 | 6 | 28962562 | 28973093 | 29 | 8 | -0.48658 | 0.68672 | ZNF311 | 0.68672 | 0.67482 | 0.81374 |
| ENSG00000117114 | 1 | 81771845 | 82458120 | 2633 | 171 | -0.487 | 0.68687 | LPHN2 | 0.68687 | 0.70546 | 0.76677 |
| ENSG00000214706 | 3 | 50325163 | 50330349 | 4 | 2 | -0.48708 | 0.6869 | IFRD2 | 0.6869 | 0.22096 | 0.24349 |
| ENSG00000188001 | 3 | 188665003 | 189043093 | 929 | 74 | -0.48712 | 0.68691 | TPRG1 | 0.68691 | 0.67309 | 0.44366 |
| ENSG00000196578 | 3 | 97806017 | 97806946 | 6 | 3 | -0.48719 | 0.68694 | OR5AC2 | 0.68694 | 0.56143 | 0.49402 |
| ENSG00000141933 | 19 | 507299 | 519654 | 65 | 18 | -0.48754 | 0.68706 | TPGS1 | 0.68706 | 0.35859 | 0.5172 |
| ENSG00000157450 | 15 | 59157374 | 59389618 | 584 | 33 | -0.48763 | 0.68709 | RNF111 | 0.68709 | 0.70106 | 0.74954 |
| ENSG00000164889 | 7 | 150754297 | 150773614 | 56 | 12 | -0.48788 | 0.68718 | SLC4A2 | 0.68718 | 0.65104 | 0.87856 |
| ENSG00000051108 | 16 | 56965960 | 56977798 | 40 | 8 | -0.48791 | 0.68719 | HERPUD1 | 0.68719 | 0.69838 | 0.79146 |
| ENSG00000171695 | 20 | 62714733 | 62715712 | 3 | 1 | -0.48792 | 0.6872 | C20orf201 | 0.6872 | 0.86769 | 0.8977 |
| ENSG00000079819 | 6 | 131160487 | 131384462 | 830 | 32 | -0.48803 | 0.68724 | EPB41L2 | 0.68724 | 0.68181 | 0.72625 |
| ENSG00000129493 | 14 | 31760994 | 31889788 | 228 | 22 | -0.48827 | 0.68732 | HEATR5A | 0.68732 | 0.71678 | 0.78654 |
| ENSG00000177613 | 10 | 53455247 | 53459355 | 6 | 2 | -0.48853 | 0.68741 | CSTF2T | 0.68741 | 0.73437 | 0.92979 |
| ENSG00000180992 | 6 | 44081194 | 44095194 | 53 | 10 | -0.48859 | 0.68743 | MRPL14 | 0.68743 | 0.50169 | 0.47937 |
| ENSG00000135824 | 1 | 182615239 | 182653711 | 105 | 14 | -0.48866 | 0.68746 | RGS8 | 0.68746 | 0.57896 | 0.45317 |
| ENSG00000139168 | 12 | 42705880 | 42719920 | 32 | 6 | -0.4887 | 0.68747 | ZCRB1 | 0.68747 | 0.69025 | 0.70984 |
| ENSG00000146731 | 7 | 56119323 | 56131682 | 41 | 11 | -0.48877 | 0.6875 | CCT6A | 0.6875 | 0.72458 | 0.79466 |
| ENSG00000140044 | 14 | 75894419 | 75940814 | 143 | 26 | -0.48879 | 0.68751 | JDP2 | 0.68751 | 0.76136 | 0.60757 |
| ENSG00000067191 | 17 | 37329709 | 37353956 | 69 | 23 | -0.48894 | 0.68756 | CACNB1 | 0.68756 | 0.71736 | 0.69142 |
| ENSG00000215475 | 13 | 46354405 | 46425871 | 310 | 32 | -0.48931 | 0.68769 | SIAH3 | 0.68769 | 0.68491 | 0.59694 |
| ENSG00000164122 | 4 | 177134824 | 177198722 | 306 | 22 | -0.48952 | 0.68776 | ASB5 | 0.68776 | 0.66148 | 0.74775 |
| ENSG00000159708 | 16 | 67360701 | 67419106 | 123 | 13 | -0.48958 | 0.68778 | LRRC36 | 0.68778 | 0.68447 | 0.7165 |
| ENSG00000140406 | 15 | 81293295 | 81296342 | 4 | 2 | -0.48973 | 0.68784 | MESDC1 | 0.68784 | 0.72795 | 0.69221 |
| ENSG00000158525 | 7 | 129984630 | 130008571 | 104 | 15 | -0.49006 | 0.68795 | CPA5 | 0.68795 | 0.56014 | 0.64391 |
| ENSG00000100583 | 14 | 77843032 | 77857840 | 54 | 7 | -0.49024 | 0.68802 | SAMD15 | 0.68802 | 0.6983 | 0.66728 |
| ENSG00000109929 | 11 | 121163162 | 121179403 | 20 | 5 | -0.49063 | 0.68816 | SC5D | 0.68816 | 0.7359 | 0.54694 |
| ENSG00000144285 | 2 | 166845670 | 166984523 | 446 | 27 | -0.49079 | 0.68821 | SCN1A | 0.68821 | 0.72077 | 0.77954 |
| ENSG00000152642 | 3 | 32147181 | 32210205 | 301 | 19 | -0.49079 | 0.68821 | GPD1L | 0.68821 | 0.7736 | 0.80187 |
| ENSG00000167910 | 8 | 59402737 | 59412795 | 29 | 7 | -0.4908 | 0.68822 | CYP7A1 | 0.68822 | 0.79758 | 0.88674 |
| ENSG00000164379 | 6 | 1312675 | 1314992 | 10 | 4 | -0.49086 | 0.68824 | FOXQ1 | 0.68824 | 0.81561 | 0.5114 |
| ENSG00000172830 | 11 | 67070919 | 67080078 | 15 | 7 | -0.4909 | 0.68825 | SSH3 | 0.68825 | 0.85881 | 0.85665 |
| ENSG00000269185 | 7 | 108540389 | 108540655 | 1 | 1 | -0.49104 | 0.6883 | FLJ00325 | 0.6883 | 0.74077 | 0.82085 |
| ENSG00000157445 | 3 | 54156574 | 55108584 | 3257 | 205 | -0.49119 | 0.68835 | CACNA2D3 | 0.68835 | 0.63777 | 0.66035 |
| ENSG00000039139 | 5 | 13690440 | 13944652 | 1054 | 54 | -0.49147 | 0.68845 | DNAH5 | 0.68845 | 0.72337 | 0.73442 |
| ENSG00000170873 | 8 | 125563031 | 125740730 | 727 | 84 | -0.49153 | 0.68847 | MTSS1 | 0.68847 | 0.58168 | 0.47349 |
| ENSG00000146909 | 7 | 156742417 | 156765876 | 163 | 13 | -0.49156 | 0.68849 | NOM1 | 0.68849 | 0.76775 | 0.75284 |
| ENSG00000153975 | 6 | 116956781 | 116989957 | 142 | 15 | -0.49171 | 0.68854 | ZUFSP | 0.68854 | 0.67837 | 0.67152 |
| ENSG00000057294 | 12 | 32943679 | 33049774 | 403 | 31 | -0.49181 | 0.68857 | PKP2 | 0.68857 | 0.72681 | 0.6985 |
| ENSG00000134294 | 12 | 46751972 | 46766650 | 29 | 5 | -0.49198 | 0.68863 | SLC38A2 | 0.68863 | 0.51463 | 0.59384 |
| ENSG00000131873 | 15 | 101715928 | 101792137 | 291 | 23 | -0.4921 | 0.68867 | CHSY1 | 0.68867 | 0.68398 | 0.73728 |
| ENSG00000125398 | 17 | 70117161 | 70122561 | 10 | 3 | -0.49216 | 0.6887 | SOX9 | 0.6887 | 0.80606 | 0.59942 |
| ENSG00000157933 | 1 | 2160134 | 2241558 | 271 | 45 | -0.49236 | 0.68877 | SKI | 0.68877 | 0.73823 | 0.71249 |
| ENSG00000198133 | 14 | 67913801 | 68000456 | 368 | 53 | -0.49254 | 0.68883 | TMEM229B | 0.68883 | 0.71629 | 0.74223 |
| ENSG00000198546 | 10 | 135121979 | 135166033 | 144 | 23 | -0.49256 | 0.68884 | ZNF511 | 0.68884 | 0.66202 | 0.42774 |
| ENSG00000269035 | 19 | 17547420 | 17571613 | 94 | 13 | -0.49296 | 0.68898 | CTD-2521M24.10 | 0.68898 | 0.79874 | 0.57289 |
| ENSG00000166171 | 10 | 103330317 | 103369425 | 80 | 10 | -0.49324 | 0.68908 | DPCD | 0.68908 | 0.64013 | 0.58025 |
| ENSG00000088387 | 13 | 99445741 | 99738879 | 1110 | 34 | -0.49333 | 0.68911 | DOCK9 | 0.68911 | 0.72792 | 0.77159 |
| ENSG00000228804 | 3 | 187420101 | 187451637 | 124 | 19 | -0.49337 | 0.68912 | RP11-211G3.3 | 0.68912 | 0.68469 | 0.68696 |
| ENSG00000114113 | 3 | 139171726 | 139199589 | 77 | 10 | -0.49366 | 0.68923 | RBP2 | 0.68923 | 0.68805 | 0.59308 |
| ENSG00000007174 | 17 | 11501748 | 11873065 | 1217 | 99 | -0.49373 | 0.68925 | DNAH9 | 0.68925 | 0.63367 | 0.54544 |
| ENSG00000268163 | 19 | 57949338 | 57988904 | 182 | 16 | -0.49387 | 0.6893 | AC004076.9 | 0.6893 | 0.66688 | 0.44239 |
| ENSG00000244588 | 20 | 1206700 | 1277065 | 231 | 31 | -0.49414 | 0.6894 | RAD21L1 | 0.6894 | 0.53796 | 0.5033 |
| ENSG00000042493 | 2 | 85621871 | 85645555 | 120 | 22 | -0.49424 | 0.68943 | CAPG | 0.68943 | 0.71517 | 0.59003 |
| ENSG00000167110 | 9 | 131018108 | 131038274 | 44 | 8 | -0.49429 | 0.68945 | GOLGA2 | 0.68945 | 0.64119 | 0.69237 |
| ENSG00000123700 | 17 | 68164814 | 68176189 | 13 | 5 | -0.49465 | 0.68958 | KCNJ2 | 0.68958 | 0.7892 | 0.86659 |
| ENSG00000132563 | 5 | 137774706 | 137782658 | 25 | 6 | -0.49475 | 0.68961 | REEP2 | 0.68961 | 0.695 | 0.65991 |
| ENSG00000026103 | 10 | 90750414 | 90775542 | 120 | 14 | -0.49476 | 0.68962 | FAS | 0.68962 | 0.7954 | 0.83896 |
| ENSG00000135404 | 12 | 56119107 | 56123491 | 6 | 3 | -0.49483 | 0.68964 | CD63 | 0.68964 | 0.87334 | 0.97746 |
| ENSG00000065665 | 10 | 12171636 | 12211960 | 151 | 11 | -0.49495 | 0.68968 | SEC61A2 | 0.68968 | 0.6636 | 0.71078 |
| ENSG00000188015 | 1 | 153519805 | 153521848 | 20 | 2 | -0.49509 | 0.68973 | S100A3 | 0.68973 | 0.8486 | 0.89185 |
| ENSG00000143061 | 1 | 117117031 | 117210375 | 144 | 27 | -0.49511 | 0.68974 | IGSF3 | 0.68974 | 0.56188 | 0.46504 |
| ENSG00000110619 | 11 | 3022152 | 3078843 | 197 | 19 | -0.49529 | 0.6898 | CARS | 0.6898 | 0.66296 | 0.61774 |
| ENSG00000177700 | 11 | 837356 | 842545 | 36 | 3 | -0.49534 | 0.68982 | POLR2L | 0.68982 | 0.62687 | 0.71764 |
| ENSG00000254469 | 11 | 71576555 | 71639700 | 351 | 8 | -0.49547 | 0.68987 | RP11-849H4.2 | 0.68987 | 0.68434 | 0.6798 |

| | | | | | | | | | | | |
|---|---|---|---|---|---|---|---|---|---|---|---|
| ENSG00000103197 | 16 | 2097466 | 2138716 | 91 | 28 | -0.49556 | 0.6899 | TSC2 | 0.6899 | 0.77978 | 0.82317 |
| ENSG00000005421 | 7 | 94926988 | 95025673 | 244 | 42 | -0.49572 | 0.68995 | PON1 | 0.68995 | 0.67435 | 0.58886 |
| ENSG00000006128 | 7 | 97361220 | 97369784 | 29 | 4 | -0.4958 | 0.68998 | TAC1 | 0.68998 | 0.78534 | 0.84737 |
| ENSG00000240654 | 13 | 24881304 | 24896673 | 104 | 15 | -0.49587 | 0.69001 | C1QTNF9 | 0.69001 | 0.69667 | 0.69082 |
| ENSG00000150630 | 4 | 177604689 | 177713881 | 386 | 18 | -0.49603 | 0.69006 | VEGFC | 0.69006 | 0.67088 | 0.5991 |
| ENSG00000164684 | 8 | 81540686 | 81787016 | 415 | 19 | -0.49603 | 0.69006 | ZNF704 | 0.69006 | 0.69593 | 0.72141 |
| ENSG00000054267 | 1 | 235294949 | 235491534 | 580 | 17 | -0.49625 | 0.69014 | ARID4B | 0.69014 | 0.71682 | 0.66619 |
| ENSG00000188229 | 9 | 140135665 | 140138159 | 6 | 4 | -0.49628 | 0.69015 | TUBB4B | 0.69015 | 0.60985 | 0.46095 |
| ENSG00000121486 | 1 | 185087220 | 185126204 | 38 | 8 | -0.4963 | 0.69016 | TRMT1L | 0.69016 | 0.76614 | 0.83771 |
| ENSG00000163568 | 1 | 159032274 | 159116886 | 187 | 27 | -0.49641 | 0.6902 | AIM2 | 0.6902 | 0.63988 | 0.54006 |
| ENSG00000226757 | 4 | 122685740 | 122687962 | 9 | 3 | -0.49644 | 0.69021 | AC079341.1 | 0.69021 | 0.83318 | 0.82209 |
| ENSG00000004059 | 7 | 127228399 | 127231759 | 10 | 3 | -0.49651 | 0.69023 | ARF5 | 0.69023 | 0.69746 | 0.73575 |
| ENSG00000177684 | 6 | 49927962 | 49931818 | 12 | 3 | -0.49653 | 0.69024 | DEFB114 | 0.69024 | 0.62361 | 0.72127 |
| ENSG00000167077 | 22 | 42095503 | 42195460 | 306 | 15 | -0.49653 | 0.69024 | MEI1 | 0.69024 | 0.69071 | 0.67616 |
| ENSG00000163877 | 1 | 38000050 | 38019903 | 54 | 12 | -0.49659 | 0.69026 | SNIP1 | 0.69026 | 0.66682 | 0.62015 |
| ENSG00000011523 | 2 | 65283500 | 65314138 | 97 | 18 | -0.49678 | 0.69033 | CEP68 | 0.69033 | 0.72247 | 0.5933 |
| ENSG00000113319 | 5 | 80256491 | 80525975 | 980 | 96 | -0.49701 | 0.69041 | RASGRF2 | 0.69041 | 0.65277 | 0.67592 |
| ENSG00000140830 | 16 | 72078188 | 72128330 | 162 | 20 | -0.49701 | 0.69041 | TXNL4B | 0.69041 | 0.63304 | 0.65857 |
| ENSG00000184012 | 21 | 42836478 | 42903043 | 348 | 36 | -0.49704 | 0.69042 | TMPRSS2 | 0.69042 | 0.7326 | 0.82338 |
| ENSG00000101407 | 20 | 36611409 | 36661870 | 74 | 14 | -0.49713 | 0.69045 | TTI1 | 0.69045 | 0.65668 | 0.75218 |
| ENSG00000222001 | 2 | 233877324 | 233880595 | 9 | 4 | -0.49721 | 0.69048 | AC106876.2 | 0.69048 | 0.86896 | 0.93457 |
| ENSG00000122778 | 7 | 138516126 | 138666064 | 495 | 48 | -0.49721 | 0.69048 | KIAA1549 | 0.69048 | 0.71556 | 0.7744 |
| ENSG00000205863 | 13 | 24465238 | 24476794 | 57 | 6 | -0.49722 | 0.69048 | C1QTNF9B | 0.69048 | 0.6825 | 0.68211 |
| ENSG00000110876 | 12 | 109015686 | 109027735 | 45 | 15 | -0.49723 | 0.69049 | SELPLG | 0.69049 | 0.27194 | 0.3498 |
| ENSG00000189129 | 10 | 81891438 | 81905115 | 63 | 14 | -0.49728 | 0.6905 | PLAC9 | 0.6905 | 0.49316 | 0.43976 |
| ENSG00000205409 | 11 | 5862158 | 5863182 | 10 | 2 | -0.49743 | 0.69056 | OR52E6 | 0.69056 | 0.87766 | 0.89473 |
| ENSG00000111701 | 12 | 7801996 | 7818499 | 103 | 19 | -0.49746 | 0.69057 | APOBEC1 | 0.69057 | 0.57676 | 0.56066 |
| ENSG00000228768 | 17 | 29898161 | 29902678 | 6 | 2 | -0.49775 | 0.69067 | AC003101.1 | 0.69067 | 0.89445 | 0.83247 |
| ENSG00000197723 | 17 | 40274756 | 40275371 | 1 | 1 | -0.49812 | 0.6908 | HSPB9 | 0.6908 | 0.48851 | 0.44933 |
| ENSG00000183166 | 7 | 71244476 | 71912136 | 2359 | 71 | -0.49822 | 0.69083 | CALN1 | 0.69083 | 0.68678 | 0.71875 |
| ENSG00000147465 | 8 | 38001167 | 38008783 | 9 | 4 | -0.4983 | 0.69086 | STAR | 0.69086 | 0.81539 | 0.79004 |
| ENSG00000197555 | 14 | 71787166 | 72207946 | 706 | 39 | -0.49832 | 0.69087 | SIPA1L1 | 0.69087 | 0.71934 | 0.77256 |
| ENSG00000090924 | 19 | 39903225 | 39919054 | 27 | 9 | -0.49843 | 0.69091 | PLEKHG2 | 0.69091 | 0.55628 | 0.55075 |
| ENSG00000170465 | 12 | 52862300 | 52867569 | 30 | 5 | -0.49852 | 0.69094 | KRT6C | 0.69094 | 0.36459 | 0.35075 |
| ENSG00000104447 | 8 | 116420724 | 116821899 | 638 | 27 | -0.4986 | 0.69097 | TRPS1 | 0.69097 | 0.6927 | 0.63769 |
| ENSG00000168876 | 11 | 94226795 | 94232749 | 20 | 7 | -0.49859 | 0.69097 | ANKRD49 | 0.69097 | 0.69863 | 0.71493 |
| ENSG00000241962 | 2 | 99757948 | 99939204 | 456 | 25 | -0.49918 | 0.69118 | C2orf15 | 0.69118 | 0.70336 | 0.69758 |
| ENSG00000107036 | 9 | 5629025 | 5776557 | 477 | 26 | -0.4992 | 0.69118 | KIAA1432 | 0.69118 | 0.71383 | 0.69805 |
| ENSG00000237765 | 4 | 15683285 | 15707188 | 72 | 11 | -0.49924 | 0.69119 | FAM200B | 0.69119 | 0.64304 | 0.34477 |
| ENSG00000113525 | 5 | 131877136 | 131892530 | 16 | 3 | -0.49922 | 0.69119 | IL5 | 0.69119 | 0.73854 | 0.79258 |
| ENSG00000237896 | 7 | 80804824 | 80828431 | 117 | 16 | -0.49928 | 0.69121 | AC005008.2 | 0.69121 | 0.62345 | 0.61271 |
| ENSG00000163870 | 3 | 127291912 | 127317094 | 31 | 14 | -0.49931 | 0.69122 | TPRA1 | 0.69122 | 0.82362 | 0.53771 |
| ENSG00000100568 | 14 | 68113792 | 68141548 | 83 | 13 | -0.4993 | 0.69122 | VTI1B | 0.69122 | 0.68702 | 0.75589 |
| ENSG00000138594 | 15 | 52121825 | 52239492 | 419 | 16 | -0.4995 | 0.69129 | TMOD3 | 0.69129 | 0.65222 | 0.62904 |
| ENSG00000053900 | 4 | 25378835 | 25420120 | 169 | 13 | -0.49966 | 0.69134 | ANAPC4 | 0.69134 | 0.68779 | 0.70778 |
| ENSG00000198887 | 9 | 72873937 | 72969804 | 274 | 15 | -0.49965 | 0.69134 | SMC5 | 0.69134 | 0.73131 | 0.738 |
| ENSG00000140320 | 15 | 40731920 | 40760441 | 38 | 10 | -0.49964 | 0.69134 | BAHD1 | 0.69134 | 0.83554 | 0.75354 |
| ENSG00000142449 | 19 | 8130286 | 8214730 | 515 | 52 | -0.49992 | 0.69143 | FBN3 | 0.69143 | 0.79938 | 0.87636 |
| ENSG00000112759 | 6 | 44187242 | 44201888 | 27 | 10 | -0.50004 | 0.69148 | SLC29A1 | 0.69148 | 0.65803 | 0.74497 |
| ENSG00000107829 | 10 | 103370423 | 103455052 | 138 | 17 | -0.50004 | 0.69148 | FBXW4 | 0.69148 | 0.65723 | 0.6519 |
| ENSG00000128789 | 18 | 12658737 | 12725739 | 231 | 18 | -0.50008 | 0.69149 | PSMG2 | 0.69149 | 0.72888 | 0.80671 |
| ENSG00000105393 | 19 | 17378159 | 17392058 | 61 | 11 | -0.50028 | 0.69156 | BABAM1 | 0.69156 | 0.60167 | 0.67722 |
| ENSG00000163449 | 2 | 216946589 | 216967506 | 50 | 8 | -0.50032 | 0.69157 | TMEM169 | 0.69157 | 0.81314 | 0.64436 |
| ENSG00000198590 | 3 | 37427760 | 37476988 | 96 | 16 | -0.50047 | 0.69163 | C3orf35 | 0.69163 | 0.73626 | 0.77325 |
| ENSG00000165588 | 14 | 57267425 | 57277197 | 7 | 4 | -0.50071 | 0.69171 | OTX2 | 0.69171 | 0.66697 | 0.51632 |
| ENSG00000117971 | 15 | 78916461 | 79020096 | 294 | 24 | -0.50076 | 0.69173 | CHRNB4 | 0.69173 | 0.58761 | 0.57676 |
| ENSG00000085415 | 18 | 12947132 | 12987535 | 150 | 20 | -0.50079 | 0.69174 | SEH1L | 0.69174 | 0.74643 | 0.82683 |
| ENSG00000180198 | 1 | 28832455 | 28865812 | 104 | 12 | -0.50114 | 0.69186 | RCC1 | 0.69186 | 0.67764 | 0.65526 |
| ENSG00000180872 | 6 | 50011288 | 50016364 | 18 | 6 | -0.50126 | 0.69191 | DEFB112 | 0.69191 | 0.87465 | 0.68735 |
| ENSG00000124191 | 20 | 42543504 | 42698256 | 523 | 67 | -0.50134 | 0.69193 | TOX2 | 0.69193 | 0.80782 | 0.80362 |
| ENSG00000161204 | 3 | 183903811 | 183911800 | 30 | 8 | -0.50146 | 0.69198 | ABCF3 | 0.69198 | 0.7989 | 0.89217 |
| ENSG00000104660 | 8 | 29952914 | 30034724 | 231 | 21 | -0.50156 | 0.69201 | LEPROTL1 | 0.69201 | 0.69775 | 0.67921 |
| ENSG00000028116 | 2 | 58134786 | 58387055 | 574 | 29 | -0.50178 | 0.69209 | VRK2 | 0.69209 | 0.69337 | 0.64707 |
| ENSG00000184517 | 16 | 75182390 | 75206134 | 130 | 13 | -0.50178 | 0.69209 | ZFP1 | 0.69209 | 0.78821 | 0.75043 |
| ENSG00000186047 | 13 | 51285144 | 51418075 | 454 | 36 | -0.5018 | 0.6921 | DLEU7 | 0.6921 | 0.72704 | 0.76326 |
| ENSG00000169223 | 5 | 176758563 | 176778853 | 41 | 8 | -0.50235 | 0.69229 | LMAN2 | 0.69229 | 0.69562 | 0.61263 |
| ENSG00000104375 | 8 | 99413631 | 99955055 | 923 | 42 | -0.50246 | 0.69233 | STK3 | 0.69233 | 0.69506 | 0.70009 |
| ENSG00000137210 | 6 | 10747992 | 10852986 | 260 | 29 | -0.50259 | 0.69237 | TMEM14B | 0.69237 | 0.71172 | 0.71338 |
| ENSG00000181894 | 19 | 58637619 | 58666477 | 102 | 19 | -0.50259 | 0.69237 | ZNF329 | 0.69237 | 0.81519 | 0.75011 |
| ENSG00000197324 | 14 | 23340822 | 23350789 | 11 | 5 | -0.50291 | 0.69249 | LRP10 | 0.69249 | 0.51769 | 0.56084 |
| ENSG00000105849 | 7 | 19735085 | 19748710 | 43 | 8 | -0.50295 | 0.6925 | TWISTNB | 0.6925 | 0.67012 | 0.62643 |
| ENSG00000105372 | 19 | 42363988 | 42376994 | 30 | 3 | -0.50297 | 0.69251 | RPS19 | 0.69251 | 0.66429 | 0.71829 |
| ENSG00000136531 | 2 | 166095912 | 166248818 | 472 | 36 | -0.50304 | 0.69253 | SCN2A | 0.69253 | 0.65855 | 0.65984 |

| | | | | | | | | | | | |
|---|---|---|---|---|---|---|---|---|---|---|---|
| ENSG00000121807 | 3 | 46395225 | 46402419 | 20 | 5 | -0.50325 | 0.69261 | CCR2 | 0.69261 | 0.66604 | 0.60716 |
| ENSG00000158792 | 16 | 89762751 | 89768113 | 19 | 7 | -0.50332 | 0.69263 | SPATA2L | 0.69263 | 0.64543 | 0.64371 |
| ENSG00000186790 | 1 | 47881744 | 47883723 | 4 | 2 | -0.50357 | 0.69272 | FOXE3 | 0.69272 | 0.33818 | 0.23799 |
| ENSG00000092010 | 14 | 24605367 | 24608176 | 7 | 2 | -0.50363 | 0.69274 | PSME1 | 0.69274 | 0.8473 | 0.79293 |
| ENSG00000170027 | 7 | 75956116 | 75988348 | 84 | 12 | -0.50365 | 0.69275 | YWHAG | 0.69275 | 0.67512 | 0.70418 |
| ENSG00000180869 | 1 | 85093913 | 85100703 | 14 | 6 | -0.50369 | 0.69276 | C1orf180 | 0.69276 | 0.19736 | 0.11026 |
| ENSG00000157985 | 2 | 236402733 | 237040444 | 1817 | 136 | -0.50384 | 0.69281 | AGAP1 | 0.69281 | 0.69383 | 0.65934 |
| ENSG00000175121 | 20 | 43738093 | 43743813 | 23 | 6 | -0.50386 | 0.69282 | WFDC5 | 0.69282 | 0.51593 | 0.39841 |
| ENSG00000107736 | 10 | 73156691 | 73575702 | 1480 | 177 | -0.50392 | 0.69284 | CDH23 | 0.69284 | 0.77613 | 0.78699 |
| ENSG00000187713 | 9 | 140098534 | 140100090 | 2 | 1 | -0.50409 | 0.6929 | TMEM203 | 0.6929 | 0.34559 | 0.56874 |
| ENSG00000180448 | 19 | 1065922 | 1086627 | 95 | 20 | -0.50437 | 0.693 | HMHA1 | 0.693 | 0.8085 | 0.93348 |
| ENSG00000173041 | 7 | 63980262 | 64023484 | 217 | 16 | -0.5045 | 0.69305 | ZNF680 | 0.69305 | 0.70523 | 0.72346 |
| ENSG00000125485 | 9 | 135468384 | 135545788 | 185 | 24 | -0.50453 | 0.69305 | DDX31 | 0.69305 | 0.65762 | 0.62414 |
| ENSG00000131475 | 17 | 40925454 | 40931617 | 3 | 2 | -0.5048 | 0.69315 | VPS25 | 0.69315 | 0.6757 | 0.72907 |
| ENSG00000167791 | 11 | 67286383 | 67290899 | 5 | 3 | -0.50511 | 0.69326 | CABP2 | 0.69326 | 0.55528 | 0.49834 |
| ENSG00000090905 | 16 | 24741016 | 24838953 | 259 | 21 | -0.5051 | 0.69326 | TNRC6A | 0.69326 | 0.66594 | 0.66214 |
| ENSG00000135249 | 7 | 105172532 | 105208124 | 100 | 12 | -0.50526 | 0.69331 | RINT1 | 0.69331 | 0.67564 | 0.5894 |
| ENSG00000268791 | 4 | 6693782 | 6694189 | 3 | 2 | -0.50533 | 0.69334 | AC093323.1 | 0.69334 | 0.40426 | 0.30314 |
| ENSG00000176912 | 18 | 641320 | 658340 | 62 | 12 | -0.50534 | 0.69334 | C18orf56 | 0.69334 | 0.70437 | 0.56955 |
| ENSG00000118473 | 1 | 66999066 | 67213982 | 853 | 26 | -0.50542 | 0.69337 | SGIP1 | 0.69337 | 0.6979 | 0.73929 |
| ENSG00000214814 | 8 | 124864227 | 125132302 | 734 | 49 | -0.50564 | 0.69344 | FER1L6 | 0.69344 | 0.6745 | 0.48943 |
| ENSG00000251139 | 4 | 185734773 | 185742443 | 21 | 9 | -0.50565 | 0.69345 | RP11-701P16.2 | 0.69345 | 0.69228 | 0.35754 |
| ENSG00000113211 | 5 | 140529683 | 140532868 | 21 | 8 | -0.5057 | 0.69347 | PCDHB6 | 0.69347 | 0.62242 | 0.67199 |
| ENSG00000187778 | 12 | 49950327 | 49961936 | 23 | 6 | -0.50581 | 0.6935 | MCRS1 | 0.6935 | 0.69112 | 0.68256 |
| ENSG00000160062 | 1 | 33005028 | 33071540 | 188 | 10 | -0.50589 | 0.69353 | ZBTB8A | 0.69353 | 0.70477 | 0.71482 |
| ENSG00000162496 | 1 | 12627939 | 12677737 | 214 | 41 | -0.50607 | 0.6936 | DHRS3 | 0.6936 | 0.78125 | 0.70802 |
| ENSG00000010810 | 6 | 111981535 | 112194655 | 686 | 38 | -0.50609 | 0.6936 | FYN | 0.6936 | 0.64526 | 0.60383 |
| ENSG00000102934 | 16 | 57290004 | 57318599 | 111 | 16 | -0.50608 | 0.6936 | PLLP | 0.6936 | 0.56787 | 0.63696 |
| ENSG00000010379 | 12 | 329789 | 372039 | 211 | 35 | -0.50615 | 0.69362 | SLC6A13 | 0.69362 | 0.71981 | 0.79771 |
| ENSG00000100296 | 22 | 29901868 | 29951205 | 159 | 17 | -0.50626 | 0.69366 | THOC5 | 0.69366 | 0.54577 | 0.51998 |
| ENSG00000137573 | 8 | 70378859 | 70573150 | 609 | 80 | -0.50642 | 0.69372 | SULF1 | 0.69372 | 0.77164 | 0.88142 |
| ENSG00000170631 | 8 | 146155744 | 146176274 | 78 | 10 | -0.50652 | 0.69376 | ZNF16 | 0.69376 | 0.71452 | 0.72492 |
| ENSG00000137817 | 15 | 72533522 | 72565340 | 84 | 13 | -0.50666 | 0.6938 | PARP6 | 0.6938 | 0.70223 | 0.71129 |
| ENSG00000188089 | 15 | 42273780 | 42343388 | 374 | 37 | -0.50667 | 0.69381 | PLA2G4E | 0.69381 | 0.78845 | 0.84565 |
| ENSG00000106526 | 7 | 149941005 | 150020814 | 240 | 20 | -0.50705 | 0.69394 | ACTR3C | 0.69394 | 0.7335 | 0.76441 |
| ENSG00000213949 | 5 | 52083730 | 52255040 | 702 | 70 | -0.50733 | 0.69404 | ITGA1 | 0.69404 | 0.75413 | 0.79107 |
| ENSG00000165832 | 10 | 116697952 | 116737430 | 114 | 16 | -0.50749 | 0.69409 | TRUB1 | 0.69409 | 0.7061 | 0.75529 |
| ENSG00000036530 | 14 | 100150641 | 100193638 | 135 | 18 | -0.50758 | 0.69413 | CYP46A1 | 0.69413 | 0.61521 | 0.44502 |
| ENSG00000114948 | 2 | 207308263 | 207485851 | 392 | 39 | -0.50762 | 0.69414 | ADAM23 | 0.69414 | 0.67845 | 0.64493 |
| ENSG00000268852 | 17 | 80214131 | 80216080 | 5 | 3 | -0.50775 | 0.69419 | AC132872.2 | 0.69419 | 0.45505 | 0.40542 |
| ENSG00000135205 | 7 | 76751751 | 76958850 | 774 | 33 | -0.50789 | 0.69423 | CCDC146 | 0.69423 | 0.73076 | 0.72301 |
| ENSG00000180901 | 17 | 73028670 | 73061984 | 145 | 21 | -0.50788 | 0.69423 | KCTD2 | 0.69423 | 0.65175 | 0.71165 |
| ENSG00000138303 | 10 | 73856278 | 73976892 | 230 | 21 | -0.50806 | 0.69429 | ASCC1 | 0.69429 | 0.74404 | 0.61198 |
| ENSG00000119004 | 2 | 204103663 | 204163009 | 140 | 15 | -0.50807 | 0.6943 | CYP20A1 | 0.6943 | 0.70936 | 0.7607 |
| ENSG00000167996 | 11 | 61727190 | 61735132 | 19 | 8 | -0.50808 | 0.6943 | FTH1 | 0.6943 | 0.43252 | 0.60491 |
| ENSG00000170209 | 11 | 113258513 | 113271140 | 42 | 8 | -0.50808 | 0.6943 | ANKK1 | 0.6943 | 0.74649 | 0.81741 |
| ENSG00000189127 | 5 | 79852574 | 79866307 | 44 | 10 | -0.50824 | 0.69436 | ANKRD34B | 0.69436 | 0.86584 | 0.90709 |
| ENSG00000066813 | 16 | 20547548 | 20587749 | 192 | 19 | -0.50825 | 0.69436 | ACSM2B | 0.69436 | 0.71755 | 0.8027 |
| ENSG00000198130 | 2 | 191054461 | 191208919 | 519 | 13 | -0.50828 | 0.69437 | HIBCH | 0.69437 | 0.69911 | 0.67544 |
| ENSG00000212933 | 21 | 46074130 | 46074576 | 2 | 1 | -0.50859 | 0.69448 | KRTAP12-4 | 0.69448 | 0.71015 | 0.85896 |
| ENSG00000115677 | 2 | 242166679 | 242256476 | 274 | 25 | -0.50868 | 0.69451 | HDLBP | 0.69451 | 0.69847 | 0.61445 |
| ENSG00000163915 | 3 | 185430316 | 185447575 | 23 | 8 | -0.50868 | 0.69451 | C3orf65 | 0.69451 | 0.68299 | 0.3676 |
| ENSG00000142273 | 19 | 45281126 | 45303891 | 74 | 14 | -0.50938 | 0.69476 | CBLC | 0.69476 | 0.71918 | 0.77135 |
| ENSG00000253304 | 1 | 29445940 | 29450447 | 13 | 4 | -0.50959 | 0.69483 | TMEM200B | 0.69483 | 0.77322 | 0.86231 |
| ENSG00000162817 | 1 | 220863187 | 220872499 | 37 | 8 | -0.50959 | 0.69483 | C1orf115 | 0.69483 | 0.80968 | 0.80845 |
| ENSG00000178732 | 3 | 194115550 | 194119995 | 17 | 3 | -0.50961 | 0.69484 | GP5 | 0.69484 | 0.58747 | 0.61421 |
| ENSG00000188747 | 9 | 140317802 | 140328858 | 37 | 5 | -0.5098 | 0.69491 | NOXA1 | 0.69491 | 0.7581 | 0.72778 |
| ENSG00000126803 | 14 | 65002623 | 65009955 | 27 | 6 | -0.5099 | 0.69494 | HSPA2 | 0.69494 | 0.65823 | 0.70681 |
| ENSG00000177212 | 1 | 248436073 | 248437138 | 1 | 1 | -0.51007 | 0.695 | OR2T33 | 0.695 | 0.65793 | 0.6529 |
| ENSG00000205745 | 7 | 36118694 | 36125346 | 33 | 11 | -0.51016 | 0.69503 | PP13004 | 0.69503 | 0.54268 | 0.54545 |
| ENSG00000110344 | 11 | 118230300 | 118269926 | 86 | 14 | -0.51022 | 0.69505 | UBE4A | 0.69505 | 0.68039 | 0.64384 |
| ENSG00000121864 | 3 | 179040779 | 179053323 | 28 | 8 | -0.51033 | 0.69509 | ZNF639 | 0.69509 | 0.8867 | 0.91459 |
| ENSG00000073712 | 14 | 53323986 | 53419153 | 245 | 26 | -0.51102 | 0.69533 | FERMT2 | 0.69533 | 0.68102 | 0.72423 |
| ENSG00000197208 | 5 | 131630136 | 131679899 | 124 | 13 | -0.51121 | 0.6954 | SLC22A4 | 0.6954 | 0.71753 | 0.76746 |
| ENSG00000168398 | 14 | 96671016 | 96710666 | 194 | 38 | -0.51136 | 0.69545 | BDKRB2 | 0.69545 | 0.57351 | 0.47596 |
| ENSG00000120533 | 8 | 110346553 | 110358182 | 28 | 8 | -0.51137 | 0.69546 | ENY2 | 0.69546 | 0.86164 | 0.87255 |
| ENSG00000120915 | 8 | 27348296 | 27403081 | 77 | 16 | -0.5115 | 0.6955 | EPHX2 | 0.6955 | 0.56771 | 0.57661 |
| ENSG00000168461 | 18 | 9708162 | 9862548 | 726 | 109 | -0.51162 | 0.69554 | RAB31 | 0.69554 | 0.80532 | 0.45671 |
| ENSG00000060982 | 12 | 24964295 | 25102393 | 620 | 50 | -0.51174 | 0.69558 | BCAT1 | 0.69558 | 0.82922 | 0.88684 |
| ENSG00000168314 | 3 | 39508689 | 39570970 | 331 | 10 | -0.51177 | 0.69559 | MOBP | 0.69559 | 0.71176 | 0.76654 |
| ENSG00000206535 | 3 | 100120037 | 100175163 | 98 | 21 | -0.51185 | 0.69562 | LNP1 | 0.69562 | 0.6354 | 0.56319 |
| ENSG00000173253 | 9 | 1050354 | 1057552 | 38 | 10 | -0.51194 | 0.69565 | DMRT2 | 0.69565 | 0.67746 | 0.84219 |
| ENSG00000054965 | 11 | 73111532 | 73309234 | 457 | 24 | -0.51194 | 0.69565 | FAM168A | 0.69565 | 0.68633 | 0.64398 |

| | | | | | | | | | | | |
|---|---|---|---|---|---|---|---|---|---|---|---|
| ENSG00000186827 | 1 | 1146706 | 1149518 | 9 | 3 | -0.51223 | 0.69576 | TNFRSF4 | 0.69576 | 0.88455 | 0.38254 |
| ENSG00000185046 | 12 | 99120235 | 100378432 | 3616 | 187 | -0.51231 | 0.69578 | ANKS1B | 0.69578 | 0.70167 | 0.67955 |
| ENSG00000115526 | 2 | 101008327 | 101034118 | 80 | 13 | -0.51237 | 0.6958 | CHST10 | 0.6958 | 0.70798 | 0.75685 |
| ENSG00000204713 | 6 | 28870779 | 28891766 | 77 | 14 | -0.51239 | 0.69581 | TRIM27 | 0.69581 | 0.59408 | 0.6053 |
| ENSG00000267140 | 18 | 33023835 | 33077904 | 126 | 20 | -0.51242 | 0.69582 | RP11-322E11.6 | 0.69582 | 0.6871 | 0.54468 |
| ENSG00000141858 | 19 | 14198652 | 14201848 | 3 | 2 | -0.51252 | 0.69586 | SAMD1 | 0.69586 | 0.90111 | 0.84276 |
| ENSG00000204703 | 6 | 29053985 | 29055090 | 2 | 1 | -0.51262 | 0.69589 | OR2B3 | 0.69589 | 0.93889 | 0.90645 |
| ENSG00000104973 | 19 | 50321539 | 50342073 | 52 | 9 | -0.51264 | 0.6959 | MED25 | 0.6959 | 0.80619 | 0.84619 |
| ENSG00000182013 | 19 | 46969748 | 46974820 | 34 | 5 | -0.51321 | 0.6961 | PNMAL1 | 0.6961 | 0.37282 | 0.37976 |
| ENSG00000180801 | 4 | 114821440 | 114900883 | 281 | 25 | -0.5136 | 0.69624 | ARSJ | 0.69624 | 0.71256 | 0.66692 |
| ENSG00000266076 | 17 | 63534220 | 63637151 | 307 | 45 | -0.51373 | 0.69628 | CTD-2535L24.2 | 0.69628 | 0.78832 | 0.85612 |
| ENSG00000138741 | 4 | 122800182 | 122872909 | 149 | 22 | -0.51378 | 0.6963 | TRPC3 | 0.6963 | 0.56317 | 0.27317 |
| ENSG00000197136 | 11 | 65383244 | 65404910 | 36 | 13 | -0.5138 | 0.6963 | PCNXL3 | 0.6963 | 0.62168 | 0.46888 |
| ENSG00000180096 | 16 | 30389454 | 30407312 | 32 | 17 | -0.51379 | 0.6963 | SEPTIN1 | 0.6963 | 0.54625 | 0.67427 |
| ENSG00000184178 | 4 | 53739149 | 54232242 | 1423 | 63 | -0.51381 | 0.69631 | SCFD2 | 0.69631 | 0.69851 | 0.73265 |
| ENSG00000126249 | 19 | 34895289 | 34917073 | 64 | 13 | -0.51397 | 0.69636 | PDCD2L | 0.69636 | 0.74791 | 0.74062 |
| ENSG00000130997 | 4 | 2073645 | 2243848 | 473 | 15 | -0.51398 | 0.69637 | POLN | 0.69637 | 0.68297 | 0.71611 |
| ENSG00000137700 | 11 | 118894824 | 118901616 | 29 | 10 | -0.51401 | 0.69638 | SLC37A4 | 0.69638 | 0.7471 | 0.6608 |
| ENSG00000069248 | 1 | 229577045 | 229644103 | 191 | 11 | -0.51403 | 0.69639 | NUP133 | 0.69639 | 0.6618 | 0.51929 |
| ENSG00000103264 | 16 | 87360593 | 87425748 | 243 | 37 | -0.5141 | 0.69641 | FBXO31 | 0.69641 | 0.77625 | 0.85946 |
| ENSG00000085552 | 1 | 159896829 | 159915386 | 40 | 10 | -0.51421 | 0.69645 | IGSF9 | 0.69645 | 0.38306 | 0.51631 |
| ENSG00000004700 | 12 | 21621845 | 21654603 | 147 | 25 | -0.51435 | 0.6965 | RECQL | 0.6965 | 0.80717 | 0.86174 |
| ENSG00000268085 | 8 | 23412287 | 23420651 | 48 | 9 | -0.51462 | 0.69659 | AC051642.1 | 0.69659 | 0.7252 | 0.74792 |
| ENSG00000173040 | 4 | 5544499 | 5711275 | 700 | 61 | -0.51495 | 0.69671 | EVC2 | 0.69671 | 0.74759 | 0.79217 |
| ENSG00000092871 | 17 | 33333009 | 33416338 | 100 | 17 | -0.51506 | 0.69675 | RFFL | 0.69675 | 0.64076 | 0.68913 |
| ENSG00000006194 | 16 | 3313743 | 3351401 | 62 | 9 | -0.51518 | 0.69679 | ZNF263 | 0.69679 | 0.71732 | 0.63656 |
| ENSG00000161570 | 17 | 34198495 | 34207797 | 21 | 5 | -0.51522 | 0.6968 | CCL5 | 0.6968 | 0.71946 | 0.7229 |
| ENSG00000152939 | 5 | 68710939 | 68740157 | 58 | 5 | -0.51531 | 0.69683 | MARVELD2 | 0.69683 | 0.72862 | 0.77047 |
| ENSG00000163751 | 3 | 148583043 | 148614983 | 107 | 14 | -0.51534 | 0.69684 | CPA3 | 0.69684 | 0.78296 | 0.77502 |
| ENSG00000198911 | 22 | 42229109 | 42303312 | 216 | 29 | -0.51544 | 0.69688 | SREBF2 | 0.69688 | 0.69029 | 0.68754 |
| ENSG00000160256 | 21 | 46359925 | 46396904 | 150 | 21 | -0.51559 | 0.69693 | FAM207A | 0.69693 | 0.66564 | 0.80664 |
| ENSG00000112319 | 6 | 133561736 | 133853258 | 1003 | 62 | -0.51582 | 0.69701 | EYA4 | 0.69701 | 0.71376 | 0.72997 |
| ENSG00000135740 | 16 | 67271586 | 67306093 | 50 | 12 | -0.51581 | 0.69701 | SLC9A5 | 0.69701 | 0.76471 | 0.74207 |
| ENSG00000131788 | 1 | 145575233 | 145586546 | 7 | 4 | -0.51597 | 0.69706 | PIAS3 | 0.69706 | 0.5535 | 0.54827 |
| ENSG00000187630 | 14 | 24439148 | 24475617 | 209 | 24 | -0.51598 | 0.69706 | DHRS4L2 | 0.69706 | 0.74967 | 0.82312 |
| ENSG00000197124 | 19 | 20107867 | 20150315 | 192 | 17 | -0.51613 | 0.69712 | ZNF682 | 0.69712 | 0.69583 | 0.53622 |
| ENSG00000163701 | 3 | 9944296 | 9958086 | 46 | 4 | -0.51621 | 0.69715 | IL17RE | 0.69715 | 0.59519 | 0.30315 |
| ENSG00000112053 | 6 | 35911291 | 35992645 | 220 | 32 | -0.51634 | 0.69719 | SLC26A8 | 0.69719 | 0.72249 | 0.83003 |
| ENSG00000157778 | 7 | 1606966 | 1610641 | 15 | 3 | -0.51639 | 0.69721 | PSMG3 | 0.69721 | 0.72992 | 0.91964 |
| ENSG00000137078 | 9 | 35649292 | 35650937 | 2 | 1 | -0.51638 | 0.69721 | SIT1 | 0.69721 | 0.10203 | 0.026478 |
| ENSG00000085231 | 5 | 68646811 | 68665840 | 29 | 6 | -0.51641 | 0.69722 | TAF9 | 0.69722 | 0.69766 | 0.63863 |
| ENSG00000080603 | 16 | 30709530 | 30755602 | 39 | 12 | -0.51668 | 0.69731 | SRCAP | 0.69731 | 0.65399 | 0.69162 |
| ENSG00000118217 | 1 | 161736084 | 161933860 | 618 | 19 | -0.51682 | 0.69736 | ATF6 | 0.69736 | 0.61257 | 0.52672 |
| ENSG00000006634 | 7 | 87505531 | 87538856 | 79 | 12 | -0.51686 | 0.69737 | DBF4 | 0.69737 | 0.57835 | 0.52798 |
| ENSG00000079335 | 1 | 100810584 | 100985833 | 327 | 22 | -0.51695 | 0.69741 | CDC14A | 0.69741 | 0.73666 | 0.71504 |
| ENSG00000197170 | 17 | 65334032 | 65362743 | 73 | 13 | -0.51711 | 0.69746 | PSMD12 | 0.69746 | 0.76448 | 0.77359 |
| ENSG00000180919 | 11 | 6128914 | 6130065 | 1 | 1 | -0.51722 | 0.6975 | OR56B4 | 0.6975 | 0.63195 | 0.68369 |
| ENSG00000111641 | 12 | 6666029 | 6677857 | 21 | 8 | -0.51725 | 0.69751 | NOP2 | 0.69751 | 0.57629 | 0.5044 |
| ENSG00000169894 | 7 | 100547187 | 100611118 | 67 | 15 | -0.51767 | 0.69766 | MUC3A | 0.69766 | 0.42635 | 0.5218 |
| ENSG00000179262 | 19 | 13056669 | 13064456 | 17 | 8 | -0.51781 | 0.69771 | RAD23A | 0.69771 | 0.61612 | 0.69273 |
| ENSG00000138074 | 2 | 27422455 | 27435826 | 24 | 8 | -0.51798 | 0.69776 | SLC5A6 | 0.69776 | 0.76962 | 0.73872 |
| ENSG00000068796 | 5 | 61601989 | 61833076 | 604 | 20 | -0.51799 | 0.69777 | KIF2A | 0.69777 | 0.7329 | 0.72224 |
| ENSG00000107854 | 10 | 93558069 | 93625033 | 166 | 18 | -0.518 | 0.69777 | TNKS2 | 0.69777 | 0.69786 | 0.70049 |
| ENSG00000169877 | 16 | 31539185 | 31540124 | 5 | 2 | -0.5181 | 0.69781 | AHSP | 0.69781 | 0.79868 | 0.78729 |
| ENSG00000205710 | 17 | 4802713 | 4806227 | 10 | 4 | -0.5181 | 0.69781 | C17orf107 | 0.69781 | 0.4542 | 0.24088 |
| ENSG00000152977 | 3 | 147111209 | 147228080 | 306 | 23 | -0.51813 | 0.69782 | ZIC1 | 0.69782 | 0.69511 | 0.73719 |
| ENSG00000180957 | 22 | 28247657 | 28316122 | 97 | 18 | -0.51834 | 0.69789 | PITPNB | 0.69789 | 0.79263 | 0.83895 |
| ENSG00000014257 | 3 | 132036211 | 132087142 | 258 | 20 | -0.51863 | 0.69799 | ACPP | 0.69799 | 0.68059 | 0.68743 |
| ENSG00000148225 | 9 | 116075502 | 116102578 | 76 | 14 | -0.51868 | 0.69801 | WDR31 | 0.69801 | 0.72878 | 0.59169 |
| ENSG00000091164 | 18 | 54264439 | 54318831 | 154 | 16 | -0.51872 | 0.69802 | TXNL1 | 0.69802 | 0.72393 | 0.77014 |
| ENSG00000047662 | 4 | 17630929 | 17783135 | 467 | 15 | -0.5188 | 0.69805 | FAM184B | 0.69805 | 0.72148 | 0.69759 |
| ENSG00000162746 | 1 | 161691353 | 161697933 | 37 | 10 | -0.51894 | 0.6981 | FCRLB | 0.6981 | 0.57827 | 0.46564 |
| ENSG00000119723 | 14 | 74416629 | 74430373 | 64 | 11 | -0.51895 | 0.6981 | COQ6 | 0.6981 | 0.76033 | 0.80759 |
| ENSG00000131165 | 16 | 89710839 | 89724253 | 105 | 19 | -0.51923 | 0.6982 | CHMP1A | 0.6982 | 0.62962 | 0.67548 |
| ENSG00000165637 | 10 | 76969912 | 76991206 | 43 | 11 | -0.51925 | 0.69821 | VDAC2 | 0.69821 | 0.51418 | 0.55376 |
| ENSG00000105202 | 19 | 40325098 | 40337054 | 25 | 7 | -0.5197 | 0.69836 | FBL | 0.69836 | 0.83615 | 0.93227 |
| ENSG00000119729 | 2 | 46768945 | 46810260 | 112 | 20 | -0.51974 | 0.69838 | RHOQ | 0.69838 | 0.54871 | 0.45197 |
| ENSG00000185862 | 17 | 29630784 | 29641130 | 20 | 4 | -0.51998 | 0.69846 | EVI2B | 0.69846 | 0.70039 | 0.69502 |
| ENSG00000130177 | 13 | 115000362 | 115038198 | 106 | 11 | -0.51999 | 0.69847 | CDC16 | 0.69847 | 0.69931 | 0.82272 |
| ENSG00000186834 | 17 | 43224684 | 43229468 | 9 | 4 | -0.52006 | 0.69849 | HEXIM1 | 0.69849 | 0.43724 | 0.14447 |
| ENSG00000273167 | 13 | 24553839 | 24895169 | 1732 | 181 | -0.52053 | 0.69865 | RP11-307N16.6 | 0.69865 | 0.67389 | 0.68604 |
| ENSG00000152133 | 2 | 37311594 | 37326387 | 48 | 8 | -0.52057 | 0.69867 | GPATCH11 | 0.69867 | 0.72634 | 0.76123 |
| ENSG00000249471 | 19 | 58962971 | 58969200 | 13 | 5 | -0.52061 | 0.69868 | ZNF324B | 0.69868 | 0.73564 | 0.69196 |

| | | | | | | | | | | | |
|---|---|---|---|---|---|---|---|---|---|---|---|
| ENSG00000185924 | 17 | 1837971 | 1928639 | 455 | 51 | -0.52087 | 0.69877 | RTN4RL1 | 0.69877 | 0.65439 | 0.71415 |
| ENSG00000163285 | 4 | 46037786 | 46126098 | 317 | 18 | -0.5209 | 0.69878 | GABRG1 | 0.69878 | 0.66277 | 0.68553 |
| ENSG00000204410 | 6 | 31707725 | 31732622 | 77 | 19 | -0.52089 | 0.69878 | MSH5 | 0.69878 | 0.63223 | 0.67194 |
| ENSG00000198842 | 1 | 167063282 | 167098402 | 155 | 21 | -0.52126 | 0.69891 | DUSP27 | 0.69891 | 0.56048 | 0.55682 |
| ENSG00000142512 | 19 | 51913275 | 51921057 | 36 | 9 | -0.52131 | 0.69892 | SIGLEC10 | 0.69892 | 0.63658 | 0.82489 |
| ENSG00000121769 | 1 | 31838472 | 31849697 | 23 | 7 | -0.52143 | 0.69897 | FABP3 | 0.69897 | 0.80273 | 0.77529 |
| ENSG00000105220 | 19 | 34850385 | 34893061 | 114 | 20 | -0.52159 | 0.69902 | GPI | 0.69902 | 0.71554 | 0.77612 |
| ENSG00000184321 | 11 | 5423827 | 5424777 | 13 | 2 | -0.52168 | 0.69905 | OR51J1 | 0.69905 | 0.81011 | 0.62435 |
| ENSG00000226887 | 4 | 53588785 | 53617807 | 60 | 5 | -0.5219 | 0.69913 | ERVMER34-1 | 0.69913 | 0.72665 | 0.83817 |
| ENSG00000154511 | 1 | 93307724 | 93427057 | 245 | 12 | -0.52198 | 0.69916 | FAM69A | 0.69916 | 0.68386 | 0.65323 |
| ENSG00000009844 | 6 | 142468367 | 142545826 | 91 | 21 | -0.52204 | 0.69918 | VTA1 | 0.69918 | 0.64874 | 0.64579 |
| ENSG00000117643 | 1 | 25943959 | 26112698 | 389 | 34 | -0.52213 | 0.69921 | MAN1C1 | 0.69921 | 0.71132 | 0.72076 |
| ENSG00000168918 | 2 | 233924677 | 234116549 | 728 | 60 | -0.52215 | 0.69922 | INPP5D | 0.69922 | 0.75567 | 0.85885 |
| ENSG00000162490 | 1 | 11751786 | 11785914 | 138 | 29 | -0.52236 | 0.69929 | DRAXIN | 0.69929 | 0.61908 | 0.64635 |
| ENSG00000180817 | 10 | 71962586 | 71993667 | 146 | 20 | -0.52244 | 0.69932 | PPA1 | 0.69932 | 0.80431 | 0.77867 |
| ENSG00000152932 | 5 | 57878048 | 58155213 | 1041 | 57 | -0.52259 | 0.69937 | RAB3C | 0.69937 | 0.68273 | 0.64969 |
| ENSG00000254979 | 11 | 57154838 | 57191532 | 71 | 12 | -0.52283 | 0.69945 | RP11-872D17.8 | 0.69945 | 0.62165 | 0.66411 |
| ENSG00000151176 | 12 | 113796371 | 113827203 | 61 | 6 | -0.52284 | 0.69946 | PLBD2 | 0.69946 | 0.624 | 0.63221 |
| ENSG00000228157 | 17 | 18996287 | 19000150 | 1 | 1 | -0.52296 | 0.6995 | AC007952.5 | 0.6995 | 0.9759 | 0.75907 |
| ENSG00000143469 | 1 | 210111538 | 210337636 | 664 | 29 | -0.52312 | 0.69955 | SYT14 | 0.69955 | 0.69947 | 0.69507 |
| ENSG00000049883 | 5 | 71616194 | 71656052 | 50 | 10 | -0.52339 | 0.69965 | PTCD2 | 0.69965 | 0.55161 | 0.54085 |
| ENSG00000087299 | 14 | 50704281 | 50779266 | 207 | 17 | -0.5234 | 0.69965 | L2HGDH | 0.69965 | 0.73266 | 0.76078 |
| ENSG00000179178 | 1 | 43735665 | 43739673 | 9 | 5 | -0.52354 | 0.6997 | TMEM125 | 0.6997 | 0.66236 | 0.72213 |
| ENSG00000229859 | 11 | 60970779 | 60980570 | 17 | 8 | -0.5236 | 0.69972 | PGA3 | 0.69972 | 0.71314 | 0.54007 |
| ENSG00000112855 | 5 | 140071011 | 140078889 | 14 | 4 | -0.52365 | 0.69974 | HARS2 | 0.69974 | 0.7938 | 0.78724 |
| ENSG00000127325 | 12 | 70037140 | 70093256 | 290 | 18 | -0.52368 | 0.69975 | BEST3 | 0.69975 | 0.67415 | 0.61385 |
| ENSG00000068137 | 17 | 40819932 | 40829048 | 13 | 6 | -0.52408 | 0.69989 | PLEKHH3 | 0.69989 | 0.40195 | 0.34645 |
| ENSG00000196792 | 14 | 31363005 | 31495607 | 410 | 23 | -0.52421 | 0.69993 | STRN3 | 0.69993 | 0.6631 | 0.6918 |
| ENSG00000160014 | 19 | 47104331 | 47114050 | 22 | 7 | -0.52426 | 0.69995 | CALM3 | 0.69995 | 0.73169 | 0.81519 |
| ENSG00000126062 | 3 | 50392180 | 50397041 | 3 | 2 | -0.52429 | 0.69996 | TMEM115 | 0.69996 | 0.40352 | 0.38412 |
| ENSG00000125815 | 20 | 23471738 | 23476655 | 20 | 6 | -0.5244 | 0.7 | CST8 | 0.7 | 0.70407 | 0.74003 |
| ENSG00000172057 | 17 | 38077294 | 38083854 | 11 | 5 | -0.52456 | 0.70006 | ORMDL3 | 0.70006 | 0.58711 | 0.79538 |
| ENSG00000176236 | 10 | 15137384 | 15139318 | 7 | 3 | -0.52482 | 0.70015 | C10orf111 | 0.70015 | 0.88388 | 0.89207 |
| ENSG00000269095 | 19 | 17346749 | 17366257 | 85 | 19 | -0.52484 | 0.70015 | AC010646.3 | 0.70015 | 0.86146 | 0.92014 |
| ENSG00000173239 | 10 | 90562487 | 90580303 | 65 | 19 | -0.52485 | 0.70016 | LIPM | 0.70016 | 0.62511 | 0.55045 |
| ENSG00000198824 | 13 | 115079988 | 115092796 | 34 | 6 | -0.52495 | 0.70019 | CHAMP1 | 0.70019 | 0.67854 | 0.73565 |
| ENSG00000111981 | 6 | 150285143 | 150294846 | 55 | 6 | -0.525 | 0.70021 | ULBP1 | 0.70021 | 0.75412 | 0.83808 |
| ENSG00000269113 | 1 | 48226200 | 48462567 | 649 | 34 | -0.52534 | 0.70033 | TRABD2B | 0.70033 | 0.71394 | 0.7878 |
| ENSG00000117450 | 1 | 45976708 | 45988719 | 43 | 6 | -0.52551 | 0.70038 | PRDX1 | 0.70038 | 0.68462 | 0.60912 |
| ENSG00000115694 | 2 | 242432089 | 242449145 | 64 | 14 | -0.52553 | 0.70039 | STK25 | 0.70039 | 0.7688 | 0.78133 |
| ENSG00000112701 | 6 | 76311225 | 76427997 | 353 | 21 | -0.52557 | 0.70041 | SENP6 | 0.70041 | 0.71969 | 0.74625 |
| ENSG00000163501 | 2 | 219919142 | 219925189 | 15 | 3 | -0.52586 | 0.70051 | IHH | 0.70051 | 0.83659 | 0.84259 |
| ENSG00000180934 | 11 | 6047901 | 6048971 | 2 | 1 | -0.52591 | 0.70052 | OR56A1 | 0.70052 | 0.65665 | 0.74538 |
| ENSG00000183067 | 21 | 41117334 | 41174023 | 347 | 19 | -0.52598 | 0.70055 | IGSF5 | 0.70055 | 0.66651 | 0.66928 |
| ENSG00000165887 | 10 | 99332198 | 99343641 | 33 | 8 | -0.52629 | 0.70066 | ANKRD2 | 0.70066 | 0.56893 | 0.7652 |
| ENSG00000185684 | 12 | 132568828 | 132613029 | 110 | 10 | -0.52635 | 0.70068 | EP400NL | 0.70068 | 0.68586 | 0.63969 |
| ENSG00000125970 | 20 | 32581452 | 32696114 | 179 | 16 | -0.52635 | 0.70068 | RALY | 0.70068 | 0.69233 | 0.70739 |
| ENSG00000178591 | 20 | 67891 | 77735 | 16 | 8 | -0.52641 | 0.7007 | DEFB125 | 0.7007 | 0.8946 | 0.78211 |
| ENSG00000078618 | 1 | 52254863 | 52344477 | 199 | 26 | -0.52654 | 0.70074 | NRD1 | 0.70074 | 0.78791 | 0.75932 |
| ENSG00000254709 | 22 | 23229960 | 23238287 | 7 | 4 | -0.52657 | 0.70075 | IGLL5 | 0.70075 | 0.55182 | 0.58776 |
| ENSG00000184154 | 11 | 71791382 | 71821828 | 75 | 6 | -0.5268 | 0.70084 | LRTOMT | 0.70084 | 0.7032 | 0.77122 |
| ENSG00000113966 | 3 | 97483365 | 97519953 | 101 | 13 | -0.52697 | 0.70089 | ARL6 | 0.70089 | 0.67619 | 0.68365 |
| ENSG00000168564 | 4 | 184365744 | 184369351 | 9 | 4 | -0.52706 | 0.70092 | CDKN2AIP | 0.70092 | 0.70055 | 0.70747 |
| ENSG00000269897 | 10 | 22605317 | 22618471 | 9 | 4 | -0.5272 | 0.70097 | COMMD3-BMI1 | 0.70097 | 0.66153 | 0.66654 |
| ENSG00000132874 | 18 | 42792960 | 43263072 | 1710 | 108 | -0.52745 | 0.70106 | SLC14A2 | 0.70106 | 0.77088 | 0.75508 |
| ENSG00000273259 | 14 | 95058395 | 95090983 | 170 | 27 | -0.52749 | 0.70107 | RP11-986E7.7 | 0.70107 | 0.62418 | 0.48865 |
| ENSG00000182223 | 4 | 48492269 | 48496406 | 14 | 4 | -0.52792 | 0.70122 | ZAR1 | 0.70122 | 0.71216 | 0.66643 |
| ENSG00000262621 | 16 | 3415099 | 3536960 | 492 | 35 | -0.52804 | 0.70126 | NAA60 | 0.70126 | 0.51044 | 0.58108 |
| ENSG00000160539 | 9 | 134165081 | 134184649 | 75 | 6 | -0.52822 | 0.70133 | PPAPDC3 | 0.70133 | 0.45317 | 0.45484 |
| ENSG00000101224 | 20 | 3767578 | 3786762 | 60 | 10 | -0.52822 | 0.70133 | CDC25B | 0.70133 | 0.7123 | 0.79811 |
| ENSG00000050730 | 4 | 122052563 | 122148621 | 306 | 18 | -0.52838 | 0.70138 | TNIP3 | 0.70138 | 0.73577 | 0.77097 |
| ENSG00000220032 | 18 | 74207477 | 74210045 | 5 | 3 | -0.52837 | 0.70138 | RP11-17M16.1 | 0.70138 | 0.9387 | 0.87774 |
| ENSG00000188800 | 1 | 40711619 | 40717363 | 10 | 6 | -0.52841 | 0.70139 | TMCO2 | 0.70139 | 0.58728 | 0.45298 |
| ENSG00000184908 | 1 | 16370272 | 16383803 | 140 | 15 | -0.52853 | 0.70143 | CLCNKB | 0.70143 | 0.87182 | 0.8996 |
| ENSG00000146955 | 7 | 140103843 | 140126050 | 121 | 26 | -0.52856 | 0.70144 | RAB19 | 0.70144 | 0.79766 | 0.76031 |
| ENSG00000204709 | 6 | 28911654 | 28912314 | 4 | 2 | -0.52862 | 0.70146 | C6orf100 | 0.70146 | 0.64507 | 0.66435 |
| ENSG00000267060 | 17 | 41120105 | 41132545 | 27 | 6 | -0.52859 | 0.70146 | PTGES3L | 0.70146 | 0.75466 | 0.85827 |
| ENSG00000197969 | 9 | 79792269 | 80036457 | 666 | 28 | -0.52863 | 0.70147 | VPS13A | 0.70147 | 0.67479 | 0.69036 |
| ENSG00000103260 | 16 | 765115 | 769655 | 22 | 4 | -0.52927 | 0.70169 | METRN | 0.70169 | 0.44561 | 0.29626 |
| ENSG00000104611 | 8 | 19171128 | 19253729 | 281 | 33 | -0.52983 | 0.70188 | SH2D4A | 0.70188 | 0.84024 | 0.45539 |
| ENSG00000187546 | 7 | 15239943 | 15601640 | 1334 | 78 | -0.53014 | 0.70199 | AGMO | 0.70199 | 0.61969 | 0.46366 |
| ENSG00000181035 | 19 | 19174808 | 19223697 | 103 | 13 | -0.53026 | 0.70204 | SLC25A42 | 0.70204 | 0.6906 | 0.68539 |
| ENSG00000118292 | 1 | 150240600 | 150253327 | 18 | 6 | -0.53045 | 0.7021 | C1orf54 | 0.7021 | 0.84845 | 0.89832 |

| | | | | | | | | | | | |
|---|---|---|---|---|---|---|---|---|---|---|---|
| ENSG00000196872 | 2 | 99410309 | 99552722 | 411 | 23 | -0.53045 | 0.7021 | KIAA1211L | 0.7021 | 0.67933 | 0.66586 |
| ENSG00000154258 | 17 | 66970629 | 67057205 | 205 | 25 | -0.53086 | 0.70224 | ABCA9 | 0.70224 | 0.7928 | 0.70743 |
| ENSG00000204950 | 11 | 61276272 | 61278482 | 5 | 1 | -0.53114 | 0.70234 | LRRC10B | 0.70234 | 0.44593 | 0.44264 |
| ENSG00000163618 | 3 | 62384022 | 62861054 | 2107 | 109 | -0.53142 | 0.70244 | CADPS | 0.70244 | 0.68586 | 0.61938 |
| ENSG00000162892 | 1 | 207070788 | 207077484 | 17 | 4 | -0.53164 | 0.70251 | IL24 | 0.70251 | 0.67519 | 0.72284 |
| ENSG00000121904 | 1 | 33979609 | 34631443 | 2087 | 158 | -0.53173 | 0.70254 | CSMD2 | 0.70254 | 0.69425 | 0.71235 |
| ENSG00000166181 | 11 | 43333513 | 43366079 | 69 | 17 | -0.53226 | 0.70273 | API5 | 0.70273 | 0.63579 | 0.48142 |
| ENSG00000117400 | 1 | 43803478 | 43818443 | 15 | 4 | -0.53258 | 0.70284 | MPL | 0.70284 | 0.62961 | 0.57476 |
| ENSG00000197375 | 5 | 131705444 | 131731306 | 96 | 10 | -0.53267 | 0.70287 | SLC22A5 | 0.70287 | 0.71345 | 0.7034 |
| ENSG00000233822 | 6 | 27806323 | 27823487 | 36 | 7 | -0.53284 | 0.70293 | HIST1H2BN | 0.70293 | 0.71758 | 0.75301 |
| ENSG00000198298 | 10 | 44101855 | 44113351 | 47 | 9 | -0.53284 | 0.70293 | ZNF485 | 0.70293 | 0.8089 | 0.75486 |
| ENSG00000177889 | 12 | 93799449 | 93836038 | 61 | 16 | -0.53287 | 0.70294 | UBE2N | 0.70294 | 0.77386 | 0.8791 |
| ENSG00000197128 | 19 | 57978031 | 57988938 | 45 | 5 | -0.53288 | 0.70294 | ZNF772 | 0.70294 | 0.61545 | 0.43055 |
| ENSG00000156162 | 8 | 95731931 | 95806064 | 269 | 20 | -0.53297 | 0.70297 | DPY19L4 | 0.70297 | 0.63161 | 0.44947 |
| ENSG00000152782 | 10 | 91342745 | 91405215 | 268 | 27 | -0.53307 | 0.70301 | PANK1 | 0.70301 | 0.73987 | 0.80098 |
| ENSG00000196961 | 19 | 50270225 | 50310370 | 135 | 20 | -0.53307 | 0.70301 | AP2A1 | 0.70301 | 0.77116 | 0.83547 |
| ENSG00000118369 | 11 | 77899858 | 77925757 | 65 | 10 | -0.53318 | 0.70304 | USP35 | 0.70304 | 0.55686 | 0.60936 |
| ENSG00000164663 | 6 | 41757634 | 41863099 | 241 | 25 | -0.53333 | 0.7031 | USP49 | 0.7031 | 0.72852 | 0.69936 |
| ENSG00000163938 | 3 | 52715172 | 52728508 | 28 | 3 | -0.53343 | 0.70313 | GNL3 | 0.70313 | 0.59933 | 0.58381 |
| ENSG00000055609 | 7 | 151832010 | 152133090 | 435 | 14 | -0.53353 | 0.70317 | KMT2C | 0.70317 | 0.72494 | 0.74792 |
| ENSG00000131477 | 17 | 40910465 | 40915059 | 11 | 3 | -0.53356 | 0.70318 | RAMP2 | 0.70318 | 0.78349 | 0.73422 |
| ENSG00000011258 | 17 | 49254786 | 49337524 | 221 | 17 | -0.53371 | 0.70323 | MBTD1 | 0.70323 | 0.66358 | 0.68175 |
| ENSG00000075336 | 18 | 71815746 | 71826197 | 52 | 4 | -0.53385 | 0.70328 | TIMM21 | 0.70328 | 0.66939 | 0.61921 |
| ENSG00000027644 | 1 | 156809855 | 156828810 | 27 | 10 | -0.53437 | 0.70346 | INSRR | 0.70346 | 0.44221 | 0.57495 |
| ENSG00000244067 | 6 | 52614897 | 52628367 | 50 | 6 | -0.53447 | 0.70349 | GSTA2 | 0.70349 | 0.72406 | 0.71562 |
| ENSG00000204611 | 19 | 52616344 | 52643175 | 96 | 15 | -0.53494 | 0.70365 | ZNF616 | 0.70365 | 0.57752 | 0.61481 |
| ENSG00000100196 | 22 | 38864067 | 38879452 | 48 | 16 | -0.5351 | 0.70371 | KDELR3 | 0.70371 | 0.80117 | 0.8037 |
| ENSG00000022277 | 20 | 55043647 | 55093943 | 129 | 11 | -0.53516 | 0.70373 | RTFDC1 | 0.70373 | 0.76417 | 0.84816 |
| ENSG00000182154 | 9 | 140445651 | 140447007 | 2 | 1 | -0.53532 | 0.70379 | MRPL41 | 0.70379 | 0.4368 | 0.40392 |
| ENSG00000087269 | 4 | 2939660 | 2965112 | 86 | 12 | -0.53558 | 0.70387 | NOP14 | 0.70387 | 0.72987 | 0.68323 |
| ENSG00000267976 | 21 | 37858165 | 37860058 | 15 | 7 | -0.53557 | 0.70387 | AP000695.1 | 0.70387 | 0.89889 | 0.98038 |
| ENSG00000186868 | 17 | 43971748 | 44105700 | 855 | 6 | -0.53573 | 0.70393 | MAPT | 0.70393 | 0.70756 | 0.7001 |
| ENSG00000132669 | 20 | 19867165 | 19983101 | 463 | 59 | -0.53581 | 0.70396 | RIN2 | 0.70396 | 0.83142 | 0.77292 |
| ENSG00000114771 | 3 | 151531825 | 151546276 | 94 | 9 | -0.53607 | 0.70404 | AADAC | 0.70404 | 0.67243 | 0.79993 |
| ENSG00000198033 | 13 | 19747910 | 19755992 | 71 | 8 | -0.53624 | 0.7041 | TUBA3C | 0.7041 | 0.80091 | 0.90574 |
| ENSG00000175756 | 1 | 1309110 | 1310875 | 11 | 4 | -0.53634 | 0.70414 | AURKAIP1 | 0.70414 | 0.12353 | 0.1644 |
| ENSG00000240720 | 7 | 91741473 | 91808845 | 132 | 8 | -0.53638 | 0.70415 | LRRD1 | 0.70415 | 0.66596 | 0.68271 |
| ENSG00000109805 | 4 | 17812525 | 17846485 | 77 | 9 | -0.5364 | 0.70416 | NCAPG | 0.70416 | 0.58015 | 0.65144 |
| ENSG00000173908 | 17 | 38948455 | 38956211 | 23 | 7 | -0.53662 | 0.70423 | KRT28 | 0.70423 | 0.82032 | 0.81802 |
| ENSG00000160818 | 1 | 156564279 | 156571288 | 41 | 4 | -0.53699 | 0.70436 | GPATCH4 | 0.70436 | 0.67891 | 0.71634 |
| ENSG00000108702 | 17 | 32687347 | 32690250 | 10 | 3 | -0.53763 | 0.70458 | CCL1 | 0.70458 | 0.69748 | 0.40099 |
| ENSG00000205517 | 19 | 11495017 | 11530018 | 108 | 12 | -0.53786 | 0.70466 | RGL3 | 0.70466 | 0.6511 | 0.66308 |
| ENSG00000054116 | 1 | 36602173 | 36615098 | 28 | 4 | -0.53788 | 0.70467 | TRAPPC3 | 0.70467 | 0.80587 | 0.6703 |
| ENSG00000149483 | 11 | 61129473 | 61136981 | 5 | 3 | -0.53811 | 0.70475 | TMEM138 | 0.70475 | 0.72017 | 0.74307 |
| ENSG00000268626 | 17 | 72346906 | 72348271 | 6 | 3 | -0.53814 | 0.70476 | AC103809.2 | 0.70476 | 0.74055 | 0.85735 |
| ENSG00000122254 | 16 | 22825498 | 22927659 | 349 | 51 | -0.53832 | 0.70482 | HS3ST2 | 0.70482 | 0.80323 | 0.85597 |
| ENSG00000187581 | 14 | 93813537 | 93814702 | 9 | 3 | -0.5385 | 0.70488 | COX8C | 0.70488 | 0.75216 | 0.66131 |
| ENSG00000058804 | 1 | 54231133 | 54304533 | 155 | 14 | -0.53883 | 0.705 | NDC1 | 0.705 | 0.73689 | 0.78218 |
| ENSG00000025708 | 22 | 50964181 | 50968485 | 16 | 8 | -0.53893 | 0.70503 | TYMP | 0.70503 | 0.50865 | 0.5999 |
| ENSG00000118007 | 3 | 136055077 | 136471220 | 796 | 26 | -0.5391 | 0.70509 | STAG1 | 0.70509 | 0.69522 | 0.69089 |
| ENSG00000164532 | 7 | 35242042 | 35293758 | 149 | 8 | -0.53909 | 0.70509 | TBX20 | 0.70509 | 0.76449 | 0.86698 |
| ENSG00000159885 | 19 | 44529494 | 44537265 | 23 | 8 | -0.53921 | 0.70513 | ZNF222 | 0.70513 | 0.64804 | 0.5045 |
| ENSG00000205020 | 17 | 34538310 | 34540280 | 2 | 1 | -0.53937 | 0.70518 | CCL4L1 | 0.70518 | 0.71458 | 0.67075 |
| ENSG00000167900 | 17 | 76170160 | 76183314 | 67 | 14 | -0.53973 | 0.70531 | TK1 | 0.70531 | 0.78495 | 0.87094 |
| ENSG00000222022 | 2 | 238330063 | 238333919 | 9 | 3 | -0.53985 | 0.70535 | AC112721.1 | 0.70535 | 0.71486 | 0.52785 |
| ENSG00000187867 | 19 | 14164177 | 14169971 | 14 | 8 | -0.53984 | 0.70535 | PALM3 | 0.70535 | 0.81441 | 0.7901 |
| ENSG00000108379 | 17 | 44839872 | 44910520 | 154 | 29 | -0.53993 | 0.70538 | WNT3 | 0.70538 | 0.76551 | 0.72913 |
| ENSG00000217128 | 5 | 130977407 | 131132710 | 262 | 13 | -0.53997 | 0.70539 | FNIP1 | 0.70539 | 0.72972 | 0.73618 |
| ENSG00000172346 | 22 | 41956767 | 41973745 | 44 | 5 | -0.54003 | 0.70541 | CSDC2 | 0.70541 | 0.69659 | 0.62341 |
| ENSG00000128422 | 17 | 39775689 | 39781094 | 8 | 5 | -0.54012 | 0.70544 | KRT17 | 0.70544 | 0.50046 | 0.45562 |
| ENSG00000158887 | 1 | 161274525 | 161279762 | 16 | 6 | -0.54016 | 0.70546 | MPZ | 0.70546 | 0.74579 | 0.86183 |
| ENSG00000250641 | 6 | 31674681 | 31685695 | 37 | 15 | -0.54031 | 0.70551 | MEGT1 | 0.70551 | 0.64695 | 0.6433 |
| ENSG00000066697 | 9 | 103189438 | 103213511 | 80 | 7 | -0.54031 | 0.70551 | MSANTD3 | 0.70551 | 0.67445 | 0.70067 |
| ENSG00000215277 | 14 | 23654525 | 23742686 | 426 | 36 | -0.54054 | 0.70559 | C14orf164 | 0.70559 | 0.74649 | 0.81552 |
| ENSG00000145642 | 5 | 63986135 | 64013917 | 48 | 9 | -0.54068 | 0.70564 | FAM159B | 0.70564 | 0.66778 | 0.58851 |
| ENSG00000139508 | 13 | 29274201 | 29293107 | 40 | 7 | -0.54068 | 0.70564 | SLC46A3 | 0.70564 | 0.76276 | 0.84364 |
| ENSG00000170423 | 12 | 53231588 | 53242876 | 45 | 7 | -0.54075 | 0.70566 | KRT78 | 0.70566 | 0.58746 | 0.43016 |
| ENSG00000115263 | 2 | 162999392 | 163008914 | 12 | 2 | -0.54077 | 0.70567 | GCG | 0.70567 | 0.88517 | 0.86289 |
| ENSG00000119535 | 1 | 36931644 | 36948879 | 45 | 11 | -0.54102 | 0.70575 | CSF3R | 0.70575 | 0.60445 | 0.53122 |
| ENSG00000119632 | 14 | 94594116 | 94596590 | 6 | 4 | -0.54103 | 0.70576 | IFI27L2 | 0.70576 | 0.95751 | 0.93785 |
| ENSG00000145242 | 4 | 66185281 | 66536213 | 1498 | 72 | -0.54127 | 0.70584 | EPHA5 | 0.70584 | 0.70496 | 0.67641 |
| ENSG00000135521 | 6 | 144164481 | 144184949 | 47 | 8 | -0.54136 | 0.70587 | LTV1 | 0.70587 | 0.62214 | 0.56385 |
| ENSG00000111186 | 12 | 1639057 | 1756409 | 529 | 108 | -0.54139 | 0.70588 | WNT5B | 0.70588 | 0.82312 | 0.82221 |

| | | | | | | | | | | | |
|---|---|---|---|---|---|---|---|---|---|---|---|
| ENSG00000163159 | 1 | 151142463 | 151167797 | 25 | 9 | -0.5415 | 0.70592 | VPS72 | 0.70592 | 0.89661 | 0.88762 |
| ENSG00000172046 | 3 | 49145479 | 49158371 | 13 | 5 | -0.54155 | 0.70594 | USP19 | 0.70594 | 0.62635 | 0.64797 |
| ENSG00000213865 | 8 | 67579850 | 67597799 | 17 | 5 | -0.54187 | 0.70604 | C8orf44 | 0.70604 | 0.54068 | 0.54478 |
| ENSG00000006607 | 2 | 242295658 | 242434256 | 456 | 34 | -0.542 | 0.70609 | FARP2 | 0.70609 | 0.69058 | 0.72863 |
| ENSG00000136044 | 12 | 105567074 | 105630016 | 241 | 23 | -0.54203 | 0.7061 | APPL2 | 0.7061 | 0.73174 | 0.73343 |
| ENSG00000184544 | 17 | 9674751 | 9694614 | 79 | 11 | -0.54211 | 0.70613 | DHRS7C | 0.70613 | 0.66534 | 0.77047 |
| ENSG00000067334 | 1 | 94333373 | 94345474 | 65 | 10 | -0.54219 | 0.70615 | DNTTIP2 | 0.70615 | 0.43561 | 0.30662 |
| ENSG00000114956 | 2 | 74153953 | 74186088 | 102 | 10 | -0.54227 | 0.70618 | DGUOK | 0.70618 | 0.76453 | 0.90551 |
| ENSG00000179387 | 4 | 141445312 | 141474924 | 54 | 9 | -0.54226 | 0.70618 | ELMOD2 | 0.70618 | 0.68366 | 0.7204 |
| ENSG00000161381 | 17 | 37219556 | 37310647 | 383 | 34 | -0.54226 | 0.70618 | PLXDC1 | 0.70618 | 0.69986 | 0.64359 |
| ENSG00000150893 | 13 | 39261266 | 39460074 | 531 | 63 | -0.54245 | 0.70625 | FREM2 | 0.70625 | 0.7976 | 0.86695 |
| ENSG00000157429 | 16 | 71498453 | 71598992 | 188 | 27 | -0.54245 | 0.70625 | ZNF19 | 0.70625 | 0.67282 | 0.68788 |
| ENSG00000189375 | 17 | 18538319 | 18565263 | 87 | 14 | -0.54267 | 0.70632 | TBC1D28 | 0.70632 | 0.80551 | 0.91673 |
| ENSG00000113141 | 5 | 140026643 | 140042064 | 32 | 5 | -0.54289 | 0.7064 | IK | 0.7064 | 0.68271 | 0.57263 |
| ENSG00000175520 | 11 | 5528530 | 5531215 | 6 | 3 | -0.54292 | 0.70641 | UBQLN3 | 0.70641 | 0.60391 | 0.34049 |
| ENSG00000162236 | 11 | 62574369 | 62599560 | 59 | 6 | -0.54292 | 0.70641 | STX5 | 0.70641 | 0.72034 | 0.76354 |
| ENSG00000118257 | 2 | 206546714 | 206662857 | 315 | 58 | -0.54296 | 0.70642 | NRP2 | 0.70642 | 0.77024 | 0.83324 |
| ENSG00000182853 | 17 | 4688580 | 4689728 | 4 | 2 | -0.54305 | 0.70645 | VMO1 | 0.70645 | 0.49736 | 0.26705 |
| ENSG00000203760 | 6 | 126661320 | 126670021 | 11 | 4 | -0.54327 | 0.70653 | CENPW | 0.70653 | 0.596 | 0.67495 |
| ENSG00000138356 | 2 | 201450591 | 201541787 | 348 | 36 | -0.54338 | 0.70656 | AOX1 | 0.70656 | 0.71692 | 0.69089 |
| ENSG00000115593 | 2 | 88367299 | 88412906 | 173 | 26 | -0.54361 | 0.70664 | SMYD1 | 0.70664 | 0.80675 | 0.89869 |
| ENSG00000254726 | 1 | 156041804 | 156051789 | 18 | 8 | -0.54381 | 0.70671 | MEX3A | 0.70671 | 0.80267 | 0.61893 |
| ENSG00000166166 | 14 | 103995521 | 104003410 | 21 | 4 | -0.54387 | 0.70673 | TRMT61A | 0.70673 | 0.76168 | 0.77591 |
| ENSG00000205334 | 2 | 27928653 | 27938599 | 17 | 7 | -0.544 | 0.70678 | AC074091.13 | 0.70678 | 0.63253 | 0.54781 |
| ENSG00000204176 | 10 | 46955444 | 46971400 | 1 | 1 | -0.54406 | 0.7068 | SYT15 | 0.7068 | 0.77466 | 0.91865 |
| ENSG00000162378 | 1 | 53192126 | 53293014 | 398 | 22 | -0.54415 | 0.70683 | ZYG11B | 0.70683 | 0.67273 | 0.67464 |
| ENSG00000122643 | 7 | 33053742 | 33102409 | 208 | 12 | -0.54417 | 0.70684 | NT5C3A | 0.70684 | 0.74076 | 0.72655 |
| ENSG00000197566 | 17 | 16524051 | 16557170 | 110 | 10 | -0.54432 | 0.70689 | ZNF624 | 0.70689 | 0.73076 | 0.73088 |
| ENSG00000120471 | 11 | 128804626 | 128813040 | 31 | 11 | -0.54467 | 0.70701 | TP53AIP1 | 0.70701 | 0.90801 | 0.63012 |
| ENSG00000205359 | 5 | 101707486 | 101834720 | 545 | 17 | -0.54469 | 0.70702 | SLCO6A1 | 0.70702 | 0.69447 | 0.69099 |
| ENSG00000119929 | 10 | 101462315 | 101515891 | 99 | 17 | -0.54472 | 0.70703 | CUTC | 0.70703 | 0.68172 | 0.73878 |
| ENSG00000001561 | 6 | 46097730 | 46114436 | 59 | 7 | -0.54477 | 0.70705 | ENPP4 | 0.70705 | 0.72837 | 0.74631 |
| ENSG00000188191 | 7 | 588834 | 767287 | 785 | 63 | -0.54478 | 0.70705 | PRKAR1B | 0.70705 | 0.74061 | 0.77304 |
| ENSG00000149743 | 11 | 63991271 | 63993726 | 6 | 3 | -0.54489 | 0.70708 | TRPT1 | 0.70708 | 0.6341 | 0.47437 |
| ENSG00000091704 | 7 | 130020180 | 130027955 | 22 | 5 | -0.54504 | 0.70714 | CPA1 | 0.70714 | 0.7288 | 0.72965 |
| ENSG00000174080 | 11 | 66330934 | 66336312 | 16 | 4 | -0.54504 | 0.70714 | CTSF | 0.70714 | 0.46081 | 0.50616 |
| ENSG00000120075 | 17 | 46668619 | 46671323 | 8 | 4 | -0.5453 | 0.70723 | HOXB5 | 0.70723 | 0.73917 | 0.84884 |
| ENSG00000100442 | 14 | 45584803 | 45604522 | 25 | 6 | -0.54549 | 0.70729 | FKBP3 | 0.70729 | 0.671 | 0.81085 |
| ENSG00000164327 | 5 | 38938021 | 39074510 | 358 | 15 | -0.54572 | 0.70737 | RICTOR | 0.70737 | 0.71237 | 0.78386 |
| ENSG00000229117 | 12 | 56510370 | 56511727 | 3 | 2 | -0.54577 | 0.70739 | RPL41 | 0.70739 | 0.8209 | 0.82741 |
| ENSG00000107819 | 10 | 102790991 | 102800998 | 21 | 10 | -0.54587 | 0.70742 | SFXN3 | 0.70742 | 0.33472 | 0.46345 |
| ENSG00000161807 | 19 | 9225274 | 9226439 | 9 | 3 | -0.54587 | 0.70742 | OR7G1 | 0.70742 | 0.42775 | 0.63501 |
| ENSG00000168995 | 19 | 51645556 | 51656783 | 16 | 5 | -0.54596 | 0.70745 | SIGLEC7 | 0.70745 | 0.37253 | 0.61554 |
| ENSG00000100422 | 22 | 47080308 | 47134158 | 253 | 38 | -0.54632 | 0.70758 | CERK | 0.70758 | 0.61336 | 0.62424 |
| ENSG00000269367 | 12 | 42624050 | 42628075 | 7 | 4 | -0.54657 | 0.70766 | AC020629.1 | 0.70766 | 0.91693 | 0.89697 |
| ENSG00000020577 | 14 | 55033815 | 55260033 | 702 | 119 | -0.54659 | 0.70767 | SAMD4A | 0.70767 | 0.80552 | 0.82975 |
| ENSG00000168785 | 4 | 99391518 | 99579780 | 662 | 61 | -0.54678 | 0.70773 | TSPAN5 | 0.70773 | 0.7296 | 0.75942 |
| ENSG00000128513 | 7 | 124462440 | 124570037 | 348 | 12 | -0.54687 | 0.70777 | POT1 | 0.70777 | 0.70567 | 0.7084 |
| ENSG00000248712 | 11 | 119060347 | 119067479 | 14 | 5 | -0.54703 | 0.70782 | CCDC153 | 0.70782 | 0.71485 | 0.80027 |
| ENSG00000248905 | 15 | 33057747 | 33486897 | 2277 | 133 | -0.54715 | 0.70786 | FMN1 | 0.70786 | 0.69652 | 0.68137 |
| ENSG00000179456 | 1 | 244214585 | 244220778 | 7 | 4 | -0.5472 | 0.70788 | ZBTB18 | 0.70788 | 0.78786 | 0.97114 |
| ENSG00000154582 | 8 | 74851404 | 74884522 | 119 | 17 | -0.54727 | 0.7079 | TCEB1 | 0.7079 | 0.74297 | 0.84437 |
| ENSG00000173894 | 17 | 77751931 | 77761782 | 24 | 11 | -0.5477 | 0.70805 | CBX2 | 0.70805 | 0.695 | 0.62515 |
| ENSG00000143753 | 1 | 224363458 | 224381143 | 55 | 11 | -0.54775 | 0.70807 | DEGS1 | 0.70807 | 0.74338 | 0.74131 |
| ENSG00000243137 | 19 | 43696854 | 43711451 | 147 | 14 | -0.54783 | 0.7081 | PSG4 | 0.7081 | 0.73141 | 0.72871 |
| ENSG00000018189 | 4 | 71569921 | 71673032 | 382 | 14 | -0.54789 | 0.70812 | RUFY3 | 0.70812 | 0.68584 | 0.73103 |
| ENSG00000092529 | 15 | 42640301 | 42704516 | 184 | 16 | -0.54797 | 0.70814 | CAPN3 | 0.70814 | 0.67613 | 0.65687 |
| ENSG00000115211 | 2 | 27587219 | 27593353 | 18 | 8 | -0.54804 | 0.70817 | EIF2B4 | 0.70817 | 0.77444 | 0.84666 |
| ENSG00000229644 | 10 | 36810649 | 36813162 | 8 | 3 | -0.54845 | 0.70831 | NAMPTL | 0.70831 | 0.38143 | 0.48665 |
| ENSG00000109576 | 4 | 170981373 | 171012850 | 92 | 15 | -0.54852 | 0.70833 | AADAT | 0.70833 | 0.74213 | 0.71716 |
| ENSG00000172845 | 2 | 174771187 | 174830430 | 209 | 25 | -0.54857 | 0.70835 | SP3 | 0.70835 | 0.67988 | 0.58335 |
| ENSG00000130340 | 6 | 158244296 | 158366109 | 459 | 23 | -0.5491 | 0.70853 | SNX9 | 0.70853 | 0.71509 | 0.73633 |
| ENSG00000186038 | 3 | 183814852 | 183824783 | 31 | 6 | -0.54916 | 0.70855 | HTR3E | 0.70855 | 0.82912 | 0.83563 |
| ENSG00000005844 | 16 | 30483979 | 30534506 | 111 | 24 | -0.54934 | 0.70861 | ITGAL | 0.70861 | 0.71875 | 0.69862 |
| ENSG00000120942 | 1 | 11333263 | 11356106 | 41 | 5 | -0.54935 | 0.70862 | UBIAD1 | 0.70862 | 0.82948 | 0.79087 |
| ENSG00000125144 | 16 | 56700643 | 56701977 | 9 | 4 | -0.54942 | 0.70864 | MT1G | 0.70864 | 0.51282 | 0.56651 |
| ENSG00000146054 | 5 | 180620924 | 180632293 | 69 | 11 | -0.54944 | 0.70865 | TRIM7 | 0.70865 | 0.66616 | 0.6396 |
| ENSG00000178789 | 17 | 72517313 | 72527613 | 33 | 6 | -0.54949 | 0.70867 | CD300LB | 0.70867 | 0.84782 | 0.89076 |
| ENSG00000174898 | 19 | 5720688 | 5778745 | 235 | 46 | -0.54975 | 0.70876 | CATSPERD | 0.70876 | 0.72288 | 0.68842 |
| ENSG00000166275 | 10 | 104613980 | 104624718 | 32 | 8 | -0.54979 | 0.70877 | C10orf32 | 0.70877 | 0.47341 | 0.41865 |
| ENSG00000120952 | 1 | 12916941 | 12921764 | 74 | 13 | -0.54994 | 0.70882 | PRAMEF2 | 0.70882 | 0.36653 | 0.29009 |
| ENSG00000183160 | 12 | 108983622 | 108992096 | 25 | 8 | -0.55027 | 0.70893 | TMEM119 | 0.70893 | 0.70151 | 0.69105 |
| ENSG00000186314 | 5 | 144851362 | 145214932 | 1104 | 41 | -0.55028 | 0.70894 | PRELID2 | 0.70894 | 0.6794 | 0.57497 |

| | | | | | | | | | | | |
|---|---|---|---|---|---|---|---|---|---|---|---|
| ENSG00000161835 | 12 | 52400724 | 52409673 | 16 | 8 | -0.55051 | 0.70901 | GRASP | 0.70901 | 0.54285 | 0.21846 |
| ENSG00000156218 | 15 | 84322838 | 84708594 | 1150 | 45 | -0.55064 | 0.70906 | ADAMTSL3 | 0.70906 | 0.71819 | 0.69843 |
| ENSG00000232268 | 11 | 4615037 | 4616243 | 6 | 2 | -0.55073 | 0.70909 | OR52I1 | 0.70909 | 0.6994 | 0.60165 |
| ENSG00000160124 | 3 | 122078438 | 122102078 | 74 | 14 | -0.55082 | 0.70912 | CCDC58 | 0.70912 | 0.74207 | 0.79117 |
| ENSG00000164825 | 8 | 6728097 | 6735544 | 83 | 5 | -0.55088 | 0.70914 | DEFB1 | 0.70914 | 0.71649 | 0.91048 |
| ENSG00000166446 | 16 | 80631803 | 80838226 | 844 | 62 | -0.55119 | 0.70925 | CDYL2 | 0.70925 | 0.52875 | 0.3617 |
| ENSG00000205765 | 5 | 41904290 | 41921738 | 26 | 7 | -0.55129 | 0.70928 | C5orf51 | 0.70928 | 0.71961 | 0.69189 |
| ENSG00000156136 | 4 | 71858255 | 71896631 | 83 | 13 | -0.55165 | 0.70941 | DCK | 0.70941 | 0.66576 | 0.47914 |
| ENSG00000129460 | 14 | 23938897 | 23979071 | 120 | 13 | -0.55171 | 0.70943 | NGDN | 0.70943 | 0.83489 | 0.8414 |
| ENSG00000135776 | 1 | 229652329 | 229694442 | 79 | 9 | -0.55184 | 0.70947 | ABCB10 | 0.70947 | 0.71757 | 0.71802 |
| ENSG00000071626 | 19 | 1407568 | 1435719 | 54 | 11 | -0.55186 | 0.70948 | DAZAP1 | 0.70948 | 0.73441 | 0.7356 |
| ENSG00000135482 | 12 | 56511943 | 56516278 | 4 | 2 | -0.55208 | 0.70955 | ZC3H10 | 0.70955 | 0.8264 | 0.78649 |
| ENSG00000182533 | 3 | 8775486 | 8883492 | 589 | 57 | -0.55212 | 0.70957 | CAV3 | 0.70957 | 0.73112 | 0.59229 |
| ENSG00000124787 | 6 | 4994966 | 5004297 | 53 | 8 | -0.55213 | 0.70957 | RPP40 | 0.70957 | 0.75563 | 0.65213 |
| ENSG00000171049 | 19 | 52255279 | 52273779 | 82 | 12 | -0.55226 | 0.70962 | FPR2 | 0.70962 | 0.71617 | 0.80283 |
| ENSG00000253506 | 17 | 59667794 | 59668563 | 4 | 1 | -0.55248 | 0.70969 | NACA2 | 0.70969 | 0.052252 | 0.076038 |
| ENSG00000137337 | 6 | 30667584 | 30685666 | 119 | 11 | -0.5525 | 0.7097 | MDC1 | 0.7097 | 0.63364 | 0.4936 |
| ENSG00000169313 | 3 | 151055168 | 151102600 | 179 | 26 | -0.55277 | 0.70979 | P2RY12 | 0.70979 | 0.50457 | 0.46765 |
| ENSG00000248672 | 2 | 160628362 | 160761221 | 506 | 39 | -0.5528 | 0.7098 | LY75-CD302 | 0.7098 | 0.71155 | 0.70578 |
| ENSG00000054219 | 2 | 160628362 | 160761260 | 506 | 39 | -0.5528 | 0.7098 | LY75 | 0.7098 | 0.71155 | 0.70578 |
| ENSG00000109920 | 11 | 47738072 | 47788995 | 141 | 19 | -0.55296 | 0.70986 | FNBP4 | 0.70986 | 0.72414 | 0.7176 |
| ENSG00000187773 | 18 | 72102963 | 72125179 | 89 | 22 | -0.55299 | 0.70986 | FAM69C | 0.70986 | 0.70616 | 0.76382 |
| ENSG00000109756 | 4 | 160025330 | 160281321 | 610 | 36 | -0.55309 | 0.7099 | RAPGEF2 | 0.7099 | 0.71387 | 0.71193 |
| ENSG00000099341 | 19 | 38865176 | 38874464 | 29 | 6 | -0.55319 | 0.70993 | PSMD8 | 0.70993 | 0.83914 | 0.72933 |
| ENSG00000185883 | 16 | 2563871 | 2570219 | 6 | 4 | -0.55321 | 0.70994 | ATP6V0C | 0.70994 | 0.020994 | 0.066328 |
| ENSG00000134480 | 5 | 86687311 | 86708836 | 30 | 6 | -0.55336 | 0.70999 | CCNH | 0.70999 | 0.68307 | 0.70999 |
| ENSG00000132763 | 1 | 45965725 | 45976739 | 25 | 5 | -0.55356 | 0.71006 | MMACHC | 0.71006 | 0.69746 | 0.65445 |
| ENSG00000178199 | 6 | 149768794 | 149806197 | 194 | 11 | -0.55364 | 0.71009 | ZC3H12D | 0.71009 | 0.74942 | 0.72949 |
| ENSG00000183628 | 22 | 18893541 | 18901751 | 48 | 11 | -0.55371 | 0.71011 | DGCR6 | 0.71011 | 0.7187 | 0.79975 |
| ENSG00000074266 | 11 | 85955586 | 85989855 | 111 | 16 | -0.55378 | 0.71014 | EED | 0.71014 | 0.72001 | 0.69332 |
| ENSG00000144426 | 2 | 203879602 | 204091101 | 382 | 26 | -0.55389 | 0.71017 | NBEAL1 | 0.71017 | 0.70453 | 0.69201 |
| ENSG00000134283 | 12 | 42632249 | 42853517 | 706 | 23 | -0.55393 | 0.71019 | PPHLN1 | 0.71019 | 0.70968 | 0.7237 |
| ENSG00000061337 | 8 | 20103676 | 20161474 | 217 | 43 | -0.55401 | 0.71021 | LZTS1 | 0.71021 | 0.53181 | 0.48522 |
| ENSG00000186628 | 15 | 83424114 | 83474822 | 146 | 23 | -0.55401 | 0.71021 | FSD2 | 0.71021 | 0.68443 | 0.67897 |
| ENSG00000250021 | 15 | 90377540 | 90456114 | 234 | 19 | -0.55418 | 0.71027 | C15orf38-AP3S2 | 0.71027 | 0.66268 | 0.68029 |
| ENSG00000150756 | 5 | 10226442 | 10250009 | 143 | 24 | -0.55437 | 0.71034 | FAM173B | 0.71034 | 0.59633 | 0.54022 |
| ENSG00000134109 | 3 | 5229331 | 5261642 | 60 | 18 | -0.55467 | 0.71044 | EDEM1 | 0.71044 | 0.64932 | 0.79616 |
| ENSG00000124818 | 6 | 47749718 | 47800516 | 179 | 16 | -0.55471 | 0.71045 | OPN5 | 0.71045 | 0.73839 | 0.69334 |
| ENSG00000165215 | 7 | 73183328 | 73184600 | 2 | 1 | -0.55483 | 0.71049 | CLDN3 | 0.71049 | 0.40514 | 0.41364 |
| ENSG00000179292 | 11 | 66059341 | 66064135 | 10 | 6 | -0.555 | 0.71055 | TMEM151A | 0.71055 | 0.76716 | 0.88647 |
| ENSG00000137825 | 15 | 41785591 | 41795747 | 32 | 9 | -0.5553 | 0.71066 | ITPKA | 0.71066 | 0.73671 | 0.78786 |
| ENSG00000164008 | 1 | 43232940 | 43263968 | 61 | 12 | -0.55536 | 0.71067 | C1orf50 | 0.71067 | 0.78886 | 0.79805 |
| ENSG00000268657 | 14 | 92038788 | 92041383 | 11 | 4 | -0.5556 | 0.71076 | AL133373.1 | 0.71076 | 0.56812 | 0.55139 |
| ENSG00000186924 | 21 | 31973414 | 31973612 | 2 | 1 | -0.55591 | 0.71086 | KRTAP22-1 | 0.71086 | 0.81368 | 0.72896 |
| ENSG00000146926 | 7 | 150872785 | 150884919 | 39 | 7 | -0.55596 | 0.71088 | ASB10 | 0.71088 | 0.66824 | 0.80641 |
| ENSG00000198001 | 12 | 44152747 | 44183346 | 90 | 11 | -0.55614 | 0.71094 | IRAK4 | 0.71094 | 0.77102 | 0.81885 |
| ENSG00000102452 | 13 | 101706130 | 102068843 | 1577 | 87 | -0.55615 | 0.71094 | NALCN | 0.71094 | 0.68876 | 0.66917 |
| ENSG00000264278 | 18 | 74506688 | 74534251 | 99 | 18 | -0.5562 | 0.71096 | RP11-162A12.2 | 0.71096 | 0.68802 | 0.73448 |
| ENSG00000174990 | 16 | 87921625 | 87970135 | 298 | 47 | -0.55636 | 0.71102 | CA5A | 0.71102 | 0.75916 | 0.7842 |
| ENSG00000072041 | 12 | 85253492 | 85307394 | 91 | 14 | -0.55673 | 0.71115 | SLC6A15 | 0.71115 | 0.72487 | 0.55651 |
| ENSG00000054690 | 14 | 68000018 | 68056329 | 186 | 37 | -0.55677 | 0.71116 | PLEKHH1 | 0.71116 | 0.67153 | 0.65839 |
| ENSG00000085563 | 7 | 87133175 | 87342611 | 517 | 39 | -0.55691 | 0.71121 | ABCB1 | 0.71121 | 0.75127 | 0.80433 |
| ENSG00000143498 | 1 | 222731244 | 222763275 | 81 | 11 | -0.55706 | 0.71126 | TAF1A | 0.71126 | 0.58367 | 0.62491 |
| ENSG00000206384 | 3 | 130279178 | 130396999 | 245 | 37 | -0.55707 | 0.71126 | COL6A6 | 0.71126 | 0.62948 | 0.49636 |
| ENSG00000115808 | 2 | 37070783 | 37193615 | 383 | 22 | -0.55732 | 0.71134 | STRN | 0.71134 | 0.71697 | 0.71115 |
| ENSG00000167595 | 19 | 36249044 | 36261930 | 43 | 12 | -0.55731 | 0.71134 | C19orf55 | 0.71134 | 0.60267 | 0.56072 |
| ENSG00000254673 | 8 | 42873567 | 42932477 | 96 | 10 | -0.55762 | 0.71145 | RP11-598P20.5 | 0.71145 | 0.72071 | 0.74144 |
| ENSG00000187866 | 9 | 71394964 | 71398609 | 10 | 4 | -0.55783 | 0.71152 | FAM122A | 0.71152 | 0.69336 | 0.8004 |
| ENSG00000155256 | 10 | 99496878 | 99520664 | 88 | 9 | -0.55787 | 0.71153 | ZFYVE27 | 0.71153 | 0.6216 | 0.46615 |
| ENSG00000135413 | 12 | 55024595 | 55028679 | 2 | 1 | -0.55788 | 0.71154 | LACRT | 0.71154 | 0.81931 | 0.75332 |
| ENSG00000090621 | 1 | 40026488 | 40042462 | 40 | 10 | -0.55808 | 0.7116 | PABPC4 | 0.7116 | 0.77377 | 0.80826 |
| ENSG00000135945 | 2 | 100016938 | 100106497 | 221 | 18 | -0.55809 | 0.71161 | REV1 | 0.71161 | 0.72539 | 0.75354 |
| ENSG00000130413 | 11 | 8413418 | 8615836 | 747 | 28 | -0.55822 | 0.71165 | STK33 | 0.71165 | 0.70524 | 0.69714 |
| ENSG00000134827 | 11 | 59620273 | 59634048 | 14 | 7 | -0.55829 | 0.71168 | TCN1 | 0.71168 | 0.94442 | 0.96521 |
| ENSG00000167315 | 18 | 47309869 | 47340330 | 84 | 13 | -0.55834 | 0.71169 | ACAA2 | 0.71169 | 0.72969 | 0.73595 |
| ENSG00000185674 | 2 | 99858709 | 99871745 | 25 | 9 | -0.55866 | 0.7118 | LYG2 | 0.7118 | 0.55132 | 0.58551 |
| ENSG00000120370 | 1 | 170501270 | 170522587 | 36 | 10 | -0.55869 | 0.71181 | GORAB | 0.71181 | 0.59473 | 0.70476 |
| ENSG00000197790 | 11 | 4566421 | 4567374 | 5 | 3 | -0.55886 | 0.71187 | OR52M1 | 0.71187 | 0.89354 | 0.75193 |
| ENSG00000135272 | 7 | 114562209 | 114659256 | 180 | 40 | -0.55914 | 0.71197 | MDFIC | 0.71197 | 0.48145 | 0.23428 |
| ENSG00000213694 | 9 | 91606362 | 91619925 | 39 | 15 | -0.5595 | 0.71209 | S1PR3 | 0.71209 | 0.67115 | 0.76658 |
| ENSG00000120519 | 4 | 147175127 | 147443123 | 634 | 28 | -0.55956 | 0.71211 | SLC10A7 | 0.71211 | 0.73748 | 0.79999 |
| ENSG00000092969 | 1 | 218519577 | 218617961 | 213 | 43 | -0.55993 | 0.71224 | TGFB2 | 0.71224 | 0.71761 | 0.7179 |
| ENSG00000169605 | 2 | 69201705 | 69208106 | 31 | 5 | -0.55997 | 0.71225 | GKN1 | 0.71225 | 0.74228 | 0.67163 |

| | | | | | | | | | | | |
|---|---|---|---|---|---|---|---|---|---|---|---|
| ENSG00000117151 | 1 | 85015289 | 85040163 | 79 | 12 | -0.56006 | 0.71228 | CTBS | 0.71228 | 0.73376 | 0.8052 |
| ENSG00000112837 | 6 | 85397069 | 85474237 | 215 | 22 | -0.56006 | 0.71228 | TBX18 | 0.71228 | 0.63995 | 0.67411 |
| ENSG00000141574 | 17 | 80278900 | 80291950 | 72 | 6 | -0.56016 | 0.71231 | SECTM1 | 0.71231 | 0.6277 | 0.68059 |
| ENSG00000121741 | 13 | 20532810 | 20665968 | 331 | 19 | -0.5602 | 0.71233 | ZMYM2 | 0.71233 | 0.66627 | 0.45875 |
| ENSG00000204983 | 7 | 142457319 | 142460923 | 5 | 1 | -0.56028 | 0.71235 | PRSS1 | 0.71235 | 0.6559 | 0.67954 |
| ENSG00000024526 | 1 | 68939835 | 68962904 | 73 | 10 | -0.56035 | 0.71238 | DEPDC1 | 0.71238 | 0.68585 | 0.6091 |
| ENSG00000163833 | 3 | 121311966 | 121349139 | 145 | 28 | -0.56042 | 0.7124 | FBXO40 | 0.7124 | 0.71413 | 0.69575 |
| ENSG00000129559 | 14 | 24686058 | 24701660 | 36 | 13 | -0.56042 | 0.7124 | NEDD8 | 0.7124 | 0.84198 | 0.80961 |
| ENSG00000038219 | 4 | 13570362 | 13629347 | 116 | 19 | -0.56082 | 0.71254 | BOD1L1 | 0.71254 | 0.73558 | 0.81066 |
| ENSG00000137699 | 11 | 119981983 | 120056237 | 182 | 26 | -0.56122 | 0.71268 | TRIM29 | 0.71268 | 0.61994 | 0.67204 |
| ENSG00000154781 | 3 | 14693271 | 14714166 | 88 | 11 | -0.56131 | 0.71271 | CCDC174 | 0.71271 | 0.67296 | 0.52432 |
| ENSG00000158555 | 11 | 75145685 | 75236948 | 247 | 27 | -0.56135 | 0.71272 | GDPD5 | 0.71272 | 0.49603 | 0.594 |
| ENSG00000074181 | 19 | 15270444 | 15311792 | 170 | 14 | -0.56168 | 0.71283 | NOTCH3 | 0.71283 | 0.70737 | 0.63659 |
| ENSG00000162604 | 1 | 62146718 | 62191095 | 144 | 17 | -0.56171 | 0.71284 | TM2D1 | 0.71284 | 0.73209 | 0.48116 |
| ENSG00000106404 | 7 | 100875373 | 100882101 | 16 | 5 | -0.56171 | 0.71284 | CLDN15 | 0.71284 | 0.65577 | 0.76988 |
| ENSG00000167641 | 19 | 38741877 | 38747231 | 19 | 4 | -0.56188 | 0.7129 | PPP1R14A | 0.7129 | 0.8323 | 0.92554 |
| ENSG00000198445 | 22 | 17071667 | 17073700 | 3 | 1 | -0.56192 | 0.71291 | CCT8L2 | 0.71291 | 0.38485 | 0.2187 |
| ENSG00000166923 | 15 | 33010175 | 33026870 | 67 | 10 | -0.56202 | 0.71295 | GREM1 | 0.71295 | 0.44449 | 0.38366 |
| ENSG00000174021 | 1 | 84964008 | 84972248 | 32 | 9 | -0.56206 | 0.71296 | GNG5 | 0.71296 | 0.60128 | 0.58456 |
| ENSG00000268162 | 12 | 57390745 | 57391335 | 2 | 1 | -0.56204 | 0.71296 | HBCBP | 0.71296 | 0.44407 | 0.56206 |
| ENSG00000196547 | 15 | 91445448 | 91465814 | 71 | 13 | -0.56218 | 0.713 | MAN2A2 | 0.713 | 0.72257 | 0.46725 |
| ENSG00000164398 | 5 | 131142683 | 131347936 | 372 | 16 | -0.56228 | 0.71304 | ACSL6 | 0.71304 | 0.75247 | 0.76397 |
| ENSG00000111832 | 6 | 116892530 | 116918838 | 73 | 11 | -0.56232 | 0.71305 | RWDD1 | 0.71305 | 0.78228 | 0.82494 |
| ENSG00000138771 | 4 | 77356253 | 77704406 | 1191 | 66 | -0.56247 | 0.7131 | SHROOM3 | 0.7131 | 0.73623 | 0.75443 |
| ENSG00000169717 | 1 | 2938046 | 2939465 | 5 | 2 | -0.56256 | 0.71313 | ACTRT2 | 0.71313 | 0.88222 | 0.90262 |
| ENSG00000108840 | 17 | 42154114 | 42201070 | 95 | 12 | -0.56283 | 0.71323 | HDAC5 | 0.71323 | 0.79605 | 0.86693 |
| ENSG00000165828 | 10 | 135160650 | 135166187 | 25 | 6 | -0.56292 | 0.71326 | PRAP1 | 0.71326 | 0.63774 | 0.66071 |
| ENSG00000115457 | 2 | 217497551 | 217529159 | 69 | 19 | -0.56303 | 0.71329 | IGFBP2 | 0.71329 | 0.63576 | 0.62047 |
| ENSG00000116649 | 1 | 11114641 | 11120081 | 10 | 3 | -0.56306 | 0.7133 | SRM | 0.7133 | 0.3983 | 0.32805 |
| ENSG00000268098 | 21 | 37402368 | 37417818 | 58 | 10 | -0.56315 | 0.71333 | AP000688.1 | 0.71333 | 0.80021 | 0.79657 |
| ENSG00000167771 | 11 | 63678693 | 63684316 | 8 | 2 | -0.56318 | 0.71334 | RCOR2 | 0.71334 | 0.88496 | 0.88945 |
| ENSG00000161654 | 17 | 42112003 | 42144987 | 65 | 14 | -0.56322 | 0.71336 | LSM12 | 0.71336 | 0.68941 | 0.64919 |
| ENSG00000185863 | 9 | 140065310 | 140066861 | 3 | 2 | -0.5635 | 0.71345 | TMEM210 | 0.71345 | 0.79153 | 0.80643 |
| ENSG00000128710 | 2 | 176973518 | 176984670 | 29 | 9 | -0.56366 | 0.71351 | HOXD10 | 0.71351 | 0.28204 | 0.29085 |
| ENSG00000061918 | 4 | 156680144 | 156728743 | 165 | 32 | -0.56373 | 0.71353 | GUCY1B3 | 0.71353 | 0.73751 | 0.7572 |
| ENSG00000076662 | 19 | 10444452 | 10450499 | 29 | 4 | -0.56382 | 0.71356 | ICAM3 | 0.71356 | 0.70311 | 0.46315 |
| ENSG00000117152 | 1 | 163038565 | 163046592 | 18 | 7 | -0.56405 | 0.71364 | RGS4 | 0.71364 | 0.67453 | 0.75214 |
| ENSG00000163507 | 3 | 108268716 | 108308491 | 109 | 14 | -0.56407 | 0.71365 | KIAA1524 | 0.71365 | 0.74822 | 0.76153 |
| ENSG00000110675 | 11 | 107461817 | 107537505 | 217 | 35 | -0.56407 | 0.71365 | ELMOD1 | 0.71365 | 0.68904 | 0.66865 |
| ENSG00000130347 | 6 | 107018903 | 107077373 | 272 | 16 | -0.56415 | 0.71367 | RTN4IP1 | 0.71367 | 0.70156 | 0.72084 |
| ENSG00000092200 | 14 | 21756098 | 21819460 | 220 | 29 | -0.56422 | 0.7137 | RPGRIP1 | 0.7137 | 0.70054 | 0.74709 |
| ENSG00000125498 | 19 | 55281263 | 55295774 | 14 | 5 | -0.56438 | 0.71375 | KIR2DL1 | 0.71375 | 0.41701 | 0.27633 |
| ENSG00000215915 | 1 | 1385069 | 1405538 | 108 | 13 | -0.56448 | 0.71379 | ATAD3C | 0.71379 | 0.78046 | 0.82936 |
| ENSG00000162494 | 1 | 13801445 | 13840543 | 204 | 43 | -0.56464 | 0.71384 | LRRC38 | 0.71384 | 0.72034 | 0.72001 |
| ENSG00000204922 | 11 | 62437745 | 62441159 | 10 | 2 | -0.56464 | 0.71384 | C11orf83 | 0.71384 | 0.6541 | 0.63089 |
| ENSG00000248487 | 3 | 52005442 | 52015212 | 13 | 6 | -0.56469 | 0.71386 | ABHD14A | 0.71386 | 0.87981 | 0.76171 |
| ENSG00000171501 | 9 | 125315391 | 125316493 | 13 | 4 | -0.56477 | 0.71388 | OR1N2 | 0.71388 | 0.67878 | 0.73691 |
| ENSG00000258315 | 17 | 6917814 | 6920844 | 9 | 2 | -0.56482 | 0.7139 | C17orf49 | 0.7139 | 0.70139 | 0.79031 |
| ENSG00000105479 | 19 | 48799714 | 48825151 | 107 | 13 | -0.56485 | 0.71391 | CCDC114 | 0.71391 | 0.65359 | 0.68714 |
| ENSG00000122299 | 16 | 11844442 | 11891123 | 154 | 23 | -0.56501 | 0.71397 | ZC3H7A | 0.71397 | 0.82138 | 0.91249 |
| ENSG00000184887 | 14 | 105714827 | 105717430 | 7 | 2 | -0.56509 | 0.71399 | BTBD6 | 0.71399 | 0.5082 | 0.50992 |
| ENSG00000144043 | 2 | 71213068 | 71222075 | 32 | 7 | -0.56527 | 0.71406 | TEX261 | 0.71406 | 0.80291 | 0.5368 |
| ENSG00000160446 | 9 | 131483148 | 131486406 | 6 | 4 | -0.5655 | 0.71413 | ZDHHC12 | 0.71413 | 0.3014 | 0.24858 |
| ENSG00000240021 | 1 | 178482212 | 178517734 | 108 | 11 | -0.56557 | 0.71416 | TEX35 | 0.71416 | 0.74751 | 0.6443 |
| ENSG00000160180 | 21 | 43731777 | 43735761 | 9 | 4 | -0.56585 | 0.71425 | TFF3 | 0.71425 | 0.62894 | 0.79815 |
| ENSG00000169258 | 5 | 176022803 | 176037134 | 20 | 9 | -0.56629 | 0.7144 | GPRIN1 | 0.7144 | 0.56796 | 0.51478 |
| ENSG00000028137 | 1 | 12227060 | 12269285 | 166 | 20 | -0.56633 | 0.71442 | TNFRSF1B | 0.71442 | 0.4413 | 0.22814 |
| ENSG00000225921 | 6 | 13615559 | 13632971 | 28 | 7 | -0.56633 | 0.71442 | NOL7 | 0.71442 | 0.83883 | 0.87652 |
| ENSG00000172115 | 7 | 25159710 | 25164980 | 9 | 3 | -0.56648 | 0.71446 | CYCS | 0.71446 | 0.5351 | 0.55328 |
| ENSG00000157833 | 2 | 26395960 | 26412532 | 31 | 11 | -0.5666 | 0.71451 | GAREML | 0.71451 | 0.6826 | 0.62127 |
| ENSG00000173588 | 12 | 94700225 | 94853764 | 349 | 29 | -0.56664 | 0.71452 | CCDC41 | 0.71452 | 0.71333 | 0.66468 |
| ENSG00000182327 | 17 | 4692254 | 4693685 | 3 | 2 | -0.56722 | 0.71472 | GLTPD2 | 0.71472 | 0.36541 | 0.23247 |
| ENSG00000269375 | 14 | 101359265 | 101360646 | 6 | 4 | -0.56729 | 0.71474 | AL117190.3 | 0.71474 | 0.28743 | 0.16097 |
| ENSG00000110700 | 11 | 17095936 | 17099334 | 8 | 3 | -0.56732 | 0.71475 | RPS13 | 0.71475 | 0.77933 | 0.73993 |
| ENSG00000164007 | 1 | 43198764 | 43205925 | 18 | 6 | -0.56744 | 0.71479 | CLDN19 | 0.71479 | 0.63582 | 0.71949 |
| ENSG00000170390 | 4 | 150999426 | 151178609 | 655 | 41 | -0.56745 | 0.7148 | DCLK2 | 0.7148 | 0.68732 | 0.64982 |
| ENSG00000136720 | 2 | 128994290 | 129076151 | 324 | 45 | -0.56749 | 0.71481 | HS6ST1 | 0.71481 | 0.70249 | 0.77674 |
| ENSG00000158850 | 1 | 161141100 | 161147287 | 9 | 3 | -0.56768 | 0.71487 | B4GALT3 | 0.71487 | 0.84703 | 0.90497 |
| ENSG00000168291 | 3 | 58413357 | 58419584 | 30 | 7 | -0.56775 | 0.7149 | PDHB | 0.7149 | 0.74687 | 0.7705 |
| ENSG00000153093 | 2 | 111490150 | 111875799 | 1243 | 75 | -0.56778 | 0.71491 | ACOXL | 0.71491 | 0.72682 | 0.74315 |
| ENSG00000107521 | 10 | 100175955 | 100206684 | 132 | 19 | -0.56791 | 0.71495 | HPS1 | 0.71495 | 0.76145 | 0.76065 |
| ENSG00000186510 | 1 | 16345370 | 16360545 | 97 | 13 | -0.56804 | 0.715 | CLCNKA | 0.715 | 0.72064 | 0.79927 |
| ENSG00000167207 | 16 | 50727514 | 50766988 | 77 | 10 | -0.56842 | 0.71513 | NOD2 | 0.71513 | 0.67034 | 0.70384 |

| | | | | | | | | | | | |
|---|---|---|---|---|---|---|---|---|---|---|---|
| ENSG00000177302 | 17 | 18174742 | 18218321 | 89 | 14 | -0.56859 | 0.71518 | TOP3A | 0.71518 | 0.68573 | 0.68729 |
| ENSG00000196335 | 7 | 23749786 | 23872132 | 476 | 28 | -0.56874 | 0.71523 | STK31 | 0.71523 | 0.73349 | 0.7126 |
| ENSG00000197892 | 8 | 28924796 | 29120641 | 481 | 44 | -0.56875 | 0.71524 | KIF13B | 0.71524 | 0.65006 | 0.6402 |
| ENSG00000260001 | 19 | 7981030 | 7983982 | 4 | 2 | -0.56876 | 0.71524 | TGFBR3L | 0.71524 | 0.93633 | 0.97931 |
| ENSG00000110025 | 11 | 64794880 | 64808044 | 58 | 8 | -0.56888 | 0.71528 | SNX15 | 0.71528 | 0.80212 | 0.87888 |
| ENSG00000148730 | 10 | 72164135 | 72188374 | 54 | 11 | -0.56891 | 0.71529 | EIF4EBP2 | 0.71529 | 0.52503 | 0.60033 |
| ENSG00000162407 | 1 | 56960419 | 57110974 | 618 | 44 | -0.56906 | 0.71534 | PPAP2B | 0.71534 | 0.75252 | 0.81179 |
| ENSG00000197635 | 2 | 162848751 | 162931052 | 229 | 30 | -0.56907 | 0.71535 | DPP4 | 0.71535 | 0.76534 | 0.75951 |
| ENSG00000181495 | 5 | 32788945 | 32791819 | 6 | 2 | -0.56937 | 0.71545 | AC026703.1 | 0.71545 | 0.77985 | 0.84047 |
| ENSG00000174628 | 16 | 19727778 | 19868907 | 305 | 15 | -0.56938 | 0.71545 | IQCK | 0.71545 | 0.71889 | 0.69783 |
| ENSG00000172000 | 19 | 2867333 | 2878515 | 61 | 14 | -0.56948 | 0.71549 | ZNF556 | 0.71549 | 0.79023 | 0.82734 |
| ENSG00000161850 | 12 | 52787715 | 52800139 | 38 | 10 | -0.56952 | 0.7155 | KRT82 | 0.7155 | 0.71989 | 0.70538 |
| ENSG00000185404 | 2 | 231191899 | 231268447 | 367 | 27 | -0.56973 | 0.71557 | SP140L | 0.71557 | 0.76851 | 0.79781 |
| ENSG00000130713 | 9 | 133569108 | 133580248 | 23 | 10 | -0.56987 | 0.71562 | EXOSC2 | 0.71562 | 0.49264 | 0.57199 |
| ENSG00000182346 | 13 | 106118216 | 106143383 | 146 | 15 | -0.57006 | 0.71568 | DAOA | 0.71568 | 0.6541 | 0.42672 |
| ENSG00000081189 | 5 | 88013975 | 88199922 | 381 | 31 | -0.57014 | 0.71571 | MEF2C | 0.71571 | 0.74176 | 0.75212 |
| ENSG00000185829 | 17 | 44594068 | 44657088 | 43 | 8 | -0.5706 | 0.71586 | ARL17A | 0.71586 | 0.7048 | 0.67631 |
| ENSG00000205882 | 8 | 11850692 | 11853821 | 16 | 6 | -0.57065 | 0.71588 | DEFB134 | 0.71588 | 0.48222 | 0.70168 |
| ENSG00000196352 | 1 | 207494853 | 207534311 | 86 | 8 | -0.57071 | 0.7159 | CD55 | 0.7159 | 0.71701 | 0.74837 |
| ENSG00000099364 | 16 | 30934376 | 30960104 | 34 | 5 | -0.57072 | 0.71591 | FBXL19 | 0.71591 | 0.75613 | 0.81022 |
| ENSG00000135250 | 7 | 104751151 | 105039755 | 599 | 22 | -0.57089 | 0.71596 | SRPK2 | 0.71596 | 0.72188 | 0.734 |
| ENSG00000154917 | 3 | 133543083 | 133614680 | 227 | 50 | -0.57134 | 0.71611 | RAB6B | 0.71611 | 0.74068 | 0.81697 |
| ENSG00000129204 | 17 | 5019733 | 5078329 | 146 | 18 | -0.57132 | 0.71611 | USP6 | 0.71611 | 0.7586 | 0.75314 |
| ENSG00000157014 | 3 | 10289707 | 10322902 | 110 | 15 | -0.57143 | 0.71614 | TATDN2 | 0.71614 | 0.73442 | 0.69955 |
| ENSG00000255152 | 6 | 31707797 | 31732628 | 76 | 19 | -0.57143 | 0.71615 | MSH5-SAPCD1 | 0.71615 | 0.63223 | 0.67194 |
| ENSG00000106034 | 7 | 120628731 | 120937498 | 856 | 39 | -0.57183 | 0.71628 | CPED1 | 0.71628 | 0.72634 | 0.783 |
| ENSG00000176922 | 11 | 4869427 | 4870514 | 9 | 3 | -0.572 | 0.71634 | OR51S1 | 0.71634 | 0.7108 | 0.76544 |
| ENSG00000268797 | 19 | 41307202 | 41404187 | 568 | 44 | -0.57204 | 0.71635 | CTC-490E21.12 | 0.71635 | 0.72706 | 0.70697 |
| ENSG00000095203 | 9 | 111934255 | 112083244 | 424 | 65 | -0.57207 | 0.71636 | EPB41L4B | 0.71636 | 0.72672 | 0.83185 |
| ENSG00000100330 | 22 | 30279144 | 30426855 | 439 | 30 | -0.57206 | 0.71636 | MTMR3 | 0.71636 | 0.69087 | 0.66574 |
| ENSG00000172824 | 16 | 67022492 | 67043661 | 30 | 11 | -0.57217 | 0.7164 | CES4A | 0.7164 | 0.77363 | 0.74558 |
| ENSG00000176909 | 19 | 49215999 | 49222978 | 28 | 6 | -0.57225 | 0.71642 | MAMSTR | 0.71642 | 0.70466 | 0.80914 |
| ENSG00000171792 | 12 | 2985424 | 2998626 | 42 | 3 | -0.57252 | 0.71651 | RHNO1 | 0.71651 | 0.60513 | 0.52565 |
| ENSG00000106038 | 7 | 27282164 | 27290112 | 13 | 5 | -0.57268 | 0.71657 | EVX1 | 0.71657 | 0.72771 | 0.74918 |
| ENSG00000133703 | 12 | 25357723 | 25403870 | 167 | 16 | -0.57269 | 0.71657 | KRAS | 0.71657 | 0.70923 | 0.76738 |
| ENSG00000154589 | 8 | 74903587 | 74941322 | 130 | 17 | -0.57278 | 0.7166 | LY96 | 0.7166 | 0.74313 | 0.81249 |
| ENSG00000004660 | 17 | 3763609 | 3798185 | 137 | 40 | -0.57367 | 0.71691 | CAMKK1 | 0.71691 | 0.79812 | 0.76585 |
| ENSG00000178127 | 18 | 9102628 | 9134343 | 78 | 14 | -0.57381 | 0.71695 | NDUFV2 | 0.71695 | 0.55685 | 0.53475 |
| ENSG00000154743 | 3 | 12525931 | 12581122 | 258 | 13 | -0.57383 | 0.71696 | TSEN2 | 0.71696 | 0.5171 | 0.37827 |
| ENSG00000092208 | 14 | 39583427 | 39606177 | 94 | 18 | -0.57384 | 0.71696 | GEMIN2 | 0.71696 | 0.71143 | 0.73502 |
| ENSG00000132185 | 1 | 161676762 | 161684142 | 33 | 7 | -0.57406 | 0.71703 | FCRLA | 0.71703 | 0.80574 | 0.70032 |
| ENSG00000198788 | 11 | 1074875 | 1104419 | 201 | 22 | -0.57408 | 0.71704 | MUC2 | 0.71704 | 0.52136 | 0.07112 |
| ENSG00000205971 | 7 | 1732446 | 1733961 | 5 | 2 | -0.57417 | 0.71707 | AC074389.6 | 0.71707 | 0.51484 | 0.77476 |
| ENSG00000197599 | 16 | 1484384 | 1494557 | 80 | 16 | -0.57431 | 0.71712 | CCDC154 | 0.71712 | 0.76717 | 0.88674 |
| ENSG00000225697 | 3 | 48663156 | 48672926 | 5 | 3 | -0.57439 | 0.71715 | SLC26A6 | 0.71715 | 0.76359 | 0.82483 |
| ENSG00000119820 | 2 | 32502979 | 32541663 | 98 | 19 | -0.57448 | 0.71718 | YIPF4 | 0.71718 | 0.7325 | 0.67997 |
| ENSG00000166292 | 17 | 53796988 | 53809482 | 12 | 5 | -0.57447 | 0.71718 | TMEM100 | 0.71718 | 0.79067 | 0.94196 |
| ENSG00000119013 | 2 | 201936156 | 201950473 | 42 | 7 | -0.57468 | 0.71725 | NDUFB3 | 0.71725 | 0.70206 | 0.66585 |
| ENSG00000225119 | 10 | 38717074 | 38736820 | 39 | 8 | -0.57473 | 0.71726 | LINC00999 | 0.71726 | 0.5743 | 0.61876 |
| ENSG00000136152 | 13 | 46039060 | 46110765 | 273 | 14 | -0.57475 | 0.71727 | COG3 | 0.71727 | 0.74462 | 0.74462 |
| ENSG00000158828 | 1 | 20959948 | 20978004 | 97 | 13 | -0.57488 | 0.71731 | PINK1 | 0.71731 | 0.77451 | 0.74945 |
| ENSG00000170561 | 5 | 2745959 | 2752969 | 11 | 5 | -0.57509 | 0.71739 | IRX2 | 0.71739 | 0.51523 | 0.62145 |
| ENSG00000176986 | 10 | 75504120 | 75531919 | 50 | 9 | -0.57578 | 0.71762 | SEC24C | 0.71762 | 0.69837 | 0.77929 |
| ENSG00000127530 | 19 | 14909958 | 14911023 | 6 | 2 | -0.57594 | 0.71767 | OR7C1 | 0.71767 | 0.79009 | 0.91917 |
| ENSG00000163082 | 2 | 223289236 | 223425667 | 360 | 57 | -0.5762 | 0.71776 | SGPP2 | 0.71776 | 0.72756 | 0.75569 |
| ENSG00000113205 | 5 | 140480234 | 140483406 | 13 | 3 | -0.57659 | 0.71789 | PCDHB3 | 0.71789 | 0.75781 | 0.7463 |
| ENSG00000164985 | 9 | 15464064 | 15511017 | 128 | 18 | -0.57657 | 0.71789 | PSIP1 | 0.71789 | 0.70721 | 0.68465 |
| ENSG00000143252 | 1 | 161284047 | 161332984 | 288 | 8 | -0.57674 | 0.71794 | SDHC | 0.71794 | 0.69923 | 0.72753 |
| ENSG00000176920 | 19 | 49199228 | 49209207 | 45 | 5 | -0.57681 | 0.71797 | FUT2 | 0.71797 | 0.7594 | 0.78877 |
| ENSG00000159640 | 17 | 61554422 | 61599205 | 65 | 16 | -0.57703 | 0.71804 | ACE | 0.71804 | 0.47514 | 0.64982 |
| ENSG00000167840 | 17 | 5008836 | 5026411 | 61 | 17 | -0.57716 | 0.71808 | ZNF232 | 0.71808 | 0.70787 | 0.80108 |
| ENSG00000164162 | 4 | 145888264 | 146019693 | 154 | 24 | -0.57744 | 0.71818 | ANAPC10 | 0.71818 | 0.69118 | 0.7139 |
| ENSG00000108298 | 17 | 37356536 | 37360980 | 6 | 2 | -0.57747 | 0.71819 | RPL19 | 0.71819 | 0.57192 | 0.41454 |
| ENSG00000128000 | 19 | 40534167 | 40562116 | 49 | 9 | -0.57748 | 0.71819 | ZNF780B | 0.71819 | 0.82749 | 0.76729 |
| ENSG00000255501 | 11 | 105008448 | 105010453 | 3 | 2 | -0.57754 | 0.71821 | CARD18 | 0.71821 | 0.68863 | 0.76905 |
| ENSG00000160325 | 9 | 136325089 | 136335970 | 34 | 10 | -0.57777 | 0.71829 | CACFD1 | 0.71829 | 0.51931 | 0.72607 |
| ENSG00000265118 | 17 | 29632231 | 29645849 | 23 | 3 | -0.57777 | 0.71829 | CTD-2370N5.3 | 0.71829 | 0.71292 | 0.68445 |
| ENSG00000187957 | 2 | 230222345 | 230579274 | 1464 | 124 | -0.57808 | 0.71839 | DNER | 0.71839 | 0.67909 | 0.6709 |
| ENSG00000132475 | 17 | 73772515 | 73781974 | 25 | 7 | -0.57811 | 0.71841 | H3F3B | 0.71841 | 0.79448 | 0.62969 |
| ENSG00000132846 | 5 | 76367897 | 76383148 | 58 | 13 | -0.57835 | 0.71849 | ZBED3 | 0.71849 | 0.60685 | 0.5812 |
| ENSG00000055044 | 2 | 203130439 | 203168389 | 90 | 10 | -0.57851 | 0.71854 | NOP58 | 0.71854 | 0.75093 | 0.74804 |
| ENSG00000122852 | 10 | 81370695 | 81375196 | 33 | 6 | -0.57853 | 0.71855 | SFTPA1 | 0.71855 | 0.70205 | 0.69326 |
| ENSG00000188133 | 9 | 32783497 | 32787397 | 9 | 4 | -0.57888 | 0.71867 | TMEM215 | 0.71867 | 0.77639 | 0.8452 |

| | | | | | | | | | | | |
|---|---|---|---|---|---|---|---|---|---|---|---|
| ENSG00000206073 | 18 | 61304493 | 61311532 | 4 | 2 | -0.57904 | 0.71872 | SERPINB4 | 0.71872 | 0.5519 | 0.44335 |
| ENSG00000124422 | 17 | 20902910 | 20947073 | 193 | 14 | -0.57925 | 0.71879 | USP22 | 0.71879 | 0.69628 | 0.61852 |
| ENSG00000160801 | 3 | 46919236 | 46945287 | 39 | 11 | -0.57945 | 0.71886 | PTH1R | 0.71886 | 0.67681 | 0.85035 |
| ENSG00000133835 | 5 | 118788138 | 118972894 | 567 | 30 | -0.57952 | 0.71888 | HSD17B4 | 0.71888 | 0.73051 | 0.75423 |
| ENSG00000185000 | 8 | 145539954 | 145550573 | 18 | 5 | -0.57968 | 0.71893 | DGAT1 | 0.71893 | 0.58806 | 0.40312 |
| ENSG00000111181 | 12 | 299243 | 323736 | 128 | 26 | -0.57981 | 0.71898 | SLC6A12 | 0.71898 | 0.52401 | 0.56578 |
| ENSG00000138152 | 10 | 124030821 | 124097677 | 258 | 20 | -0.57988 | 0.719 | BTBD16 | 0.719 | 0.7869 | 0.81963 |
| ENSG00000123636 | 2 | 160175490 | 160473203 | 957 | 56 | -0.58009 | 0.71907 | BAZ2B | 0.71907 | 0.71613 | 0.66119 |
| ENSG00000154035 | 17 | 21142183 | 21156722 | 45 | 11 | -0.58016 | 0.7191 | C17orf103 | 0.7191 | 0.71288 | 0.53364 |
| ENSG00000204571 | 11 | 71292901 | 71314399 | 169 | 14 | -0.58024 | 0.71912 | KRTAP5-11 | 0.71912 | 0.61576 | 0.51372 |
| ENSG00000204414 | 17 | 61986957 | 61996198 | 28 | 6 | -0.58032 | 0.71915 | CSHL1 | 0.71915 | 0.48755 | 0.36694 |
| ENSG00000137814 | 15 | 42841008 | 42862192 | 51 | 9 | -0.58052 | 0.71922 | HAUS2 | 0.71922 | 0.68108 | 0.74317 |
| ENSG00000136052 | 12 | 105196331 | 105352522 | 552 | 24 | -0.58077 | 0.7193 | SLC41A2 | 0.7193 | 0.71908 | 0.72301 |
| ENSG00000167985 | 11 | 61197514 | 61215001 | 29 | 4 | -0.58085 | 0.71933 | SDHAF2 | 0.71933 | 0.78419 | 0.75538 |
| ENSG00000054179 | 9 | 139942550 | 139948497 | 32 | 14 | -0.58102 | 0.71939 | ENTPD2 | 0.71939 | 0.6344 | 0.79627 |
| ENSG00000165097 | 6 | 18155560 | 18224084 | 305 | 30 | -0.58109 | 0.71941 | KDM1B | 0.71941 | 0.77965 | 0.83122 |
| ENSG00000214756 | 11 | 62432781 | 62435968 | 4 | 2 | -0.58154 | 0.71956 | METTL12 | 0.71956 | 0.66499 | 0.64301 |
| ENSG00000138061 | 2 | 38294116 | 38337044 | 182 | 23 | -0.58165 | 0.7196 | CYP1B1 | 0.7196 | 0.73572 | 0.79621 |
| ENSG00000168060 | 11 | 64812290 | 64826021 | 20 | 7 | -0.58166 | 0.7196 | NAALADL1 | 0.7196 | 0.77666 | 0.84786 |
| ENSG00000105374 | 19 | 51874860 | 51875969 | 1 | 1 | -0.58165 | 0.7196 | NKG7 | 0.7196 | 0.47224 | 0.5499 |
| ENSG00000144648 | 3 | 42846244 | 42929466 | 230 | 24 | -0.58199 | 0.71971 | ACKR2 | 0.71971 | 0.69194 | 0.68043 |
| ENSG00000168096 | 16 | 4746513 | 4784379 | 148 | 17 | -0.58203 | 0.71973 | ANKS3 | 0.71973 | 0.74979 | 0.73253 |
| ENSG00000167377 | 16 | 71481500 | 71496998 | 32 | 8 | -0.58206 | 0.71974 | ZNF23 | 0.71974 | 0.71621 | 0.79017 |
| ENSG00000172086 | 2 | 88326724 | 88355248 | 66 | 10 | -0.5823 | 0.71982 | KRCC1 | 0.71982 | 0.74608 | 0.67198 |
| ENSG00000128266 | 22 | 23412540 | 23467224 | 123 | 13 | -0.58229 | 0.71982 | GNAZ | 0.71982 | 0.59855 | 0.46659 |
| ENSG00000133392 | 16 | 15797029 | 15950890 | 702 | 66 | -0.58242 | 0.71986 | MYH11 | 0.71986 | 0.67693 | 0.78597 |
| ENSG00000075785 | 3 | 128444965 | 128533639 | 268 | 16 | -0.58244 | 0.71987 | RAB7A | 0.71987 | 0.74564 | 0.85682 |
| ENSG00000170915 | 6 | 52226219 | 52272575 | 190 | 38 | -0.58263 | 0.71993 | PAQR8 | 0.71993 | 0.76727 | 0.81808 |
| ENSG00000189058 | 3 | 195295573 | 195311076 | 82 | 19 | -0.58271 | 0.71996 | APOD | 0.71996 | 0.87151 | 0.86831 |
| ENSG00000064419 | 7 | 128594948 | 128695198 | 264 | 20 | -0.58355 | 0.72024 | TNPO3 | 0.72024 | 0.69705 | 0.65185 |
| ENSG00000144134 | 2 | 114384806 | 114400973 | 64 | 11 | -0.58365 | 0.72027 | RABL2A | 0.72027 | 0.73587 | 0.67024 |
| ENSG00000160185 | 21 | 43824008 | 43867791 | 220 | 26 | -0.58366 | 0.72028 | UBASH3A | 0.72028 | 0.73833 | 0.87454 |
| ENSG00000196628 | 18 | 52889562 | 53332018 | 807 | 53 | -0.58384 | 0.72034 | TCF4 | 0.72034 | 0.72261 | 0.75049 |
| ENSG00000231824 | 18 | 5145284 | 5197502 | 203 | 36 | -0.58408 | 0.72042 | C18orf42 | 0.72042 | 0.88481 | 0.92932 |
| ENSG00000128313 | 22 | 36113919 | 36125530 | 61 | 7 | -0.58424 | 0.72047 | APOL5 | 0.72047 | 0.65568 | 0.48058 |
| ENSG00000086065 | 9 | 33264940 | 33281977 | 22 | 5 | -0.58436 | 0.72051 | CHMP5 | 0.72051 | 0.78735 | 0.79853 |
| ENSG00000180938 | 8 | 125985540 | 125991631 | 9 | 2 | -0.58442 | 0.72053 | ZNF572 | 0.72053 | 0.62173 | 0.67435 |
| ENSG00000168589 | 16 | 80574631 | 80584657 | 52 | 7 | -0.58442 | 0.72053 | DYNLRB2 | 0.72053 | 0.68592 | 0.65487 |
| ENSG00000119139 | 9 | 71736209 | 71870124 | 573 | 46 | -0.58449 | 0.72055 | TJP2 | 0.72055 | 0.71498 | 0.73331 |
| ENSG00000153561 | 2 | 86947296 | 87005164 | 120 | 12 | -0.58469 | 0.72062 | RMND5A | 0.72062 | 0.72129 | 0.74572 |
| ENSG00000198848 | 16 | 55836763 | 55867249 | 116 | 22 | -0.58478 | 0.72065 | CES1 | 0.72065 | 0.83904 | 0.25836 |
| ENSG00000131044 | 20 | 30458505 | 30532764 | 193 | 31 | -0.58487 | 0.72068 | TTLL9 | 0.72068 | 0.69313 | 0.60722 |
| ENSG00000121067 | 17 | 47676246 | 47755596 | 118 | 28 | -0.5855 | 0.7209 | SPOP | 0.7209 | 0.72309 | 0.56234 |
| ENSG00000015413 | 16 | 89679716 | 89704839 | 138 | 22 | -0.5856 | 0.72093 | DPEP1 | 0.72093 | 0.70161 | 0.68098 |
| ENSG00000130956 | 9 | 99212483 | 99253618 | 90 | 9 | -0.58599 | 0.72106 | HABP4 | 0.72106 | 0.7053 | 0.70083 |
| ENSG00000112167 | 6 | 39071840 | 39082965 | 34 | 13 | -0.58612 | 0.7211 | SAYSD1 | 0.7211 | 0.59991 | 0.70413 |
| ENSG00000152910 | 16 | 76311176 | 76593135 | 1347 | 56 | -0.58627 | 0.72115 | CNTNAP4 | 0.72115 | 0.70948 | 0.637 |
| ENSG00000213920 | 14 | 24683143 | 24685276 | 7 | 4 | -0.58638 | 0.72119 | MDP1 | 0.72119 | 0.81617 | 0.73611 |
| ENSG00000139220 | 12 | 81652045 | 82153332 | 1258 | 60 | -0.58641 | 0.7212 | PPFIA2 | 0.7212 | 0.75159 | 0.74568 |
| ENSG00000212900 | 17 | 39155447 | 39156138 | 9 | 2 | -0.58687 | 0.72136 | KRTAP3-2 | 0.72136 | 0.43608 | 0.49068 |
| ENSG00000251692 | 16 | 1535887 | 1538982 | 28 | 7 | -0.58705 | 0.72141 | PTX4 | 0.72141 | 0.94171 | 0.89416 |
| ENSG00000172818 | 11 | 65554493 | 65564690 | 29 | 7 | -0.5873 | 0.7215 | OVOL1 | 0.7215 | 0.76275 | 0.87305 |
| ENSG00000127472 | 1 | 20354672 | 20417683 | 240 | 28 | -0.58757 | 0.72159 | PLA2G5 | 0.72159 | 0.75004 | 0.77342 |
| ENSG00000102921 | 16 | 48572637 | 48654059 | 182 | 7 | -0.58769 | 0.72163 | N4BP1 | 0.72163 | 0.68922 | 0.67395 |
| ENSG00000136895 | 9 | 129986544 | 130155939 | 337 | 27 | -0.58779 | 0.72166 | GARNL3 | 0.72166 | 0.7288 | 0.69079 |
| ENSG00000177143 | 18 | 580367 | 581524 | 4 | 2 | -0.5878 | 0.72167 | CETN1 | 0.72167 | 0.86093 | 0.70637 |
| ENSG00000168958 | 2 | 228189867 | 228222550 | 128 | 14 | -0.58792 | 0.72171 | MFF | 0.72171 | 0.72561 | 0.69905 |
| ENSG00000161203 | 3 | 183892477 | 183901879 | 24 | 8 | -0.58804 | 0.72175 | AP2M1 | 0.72175 | 0.74343 | 0.80416 |
| ENSG00000050393 | 6 | 13786789 | 13814800 | 63 | 17 | -0.58814 | 0.72178 | MCUR1 | 0.72178 | 0.69805 | 0.416 |
| ENSG00000059145 | 16 | 1413206 | 1464752 | 161 | 19 | -0.58822 | 0.72181 | UNKL | 0.72181 | 0.72876 | 0.79983 |
| ENSG00000096063 | 6 | 35800743 | 35889119 | 257 | 10 | -0.58828 | 0.72183 | SRPK1 | 0.72183 | 0.71119 | 0.69711 |
| ENSG00000257921 | 12 | 58166811 | 58180829 | 33 | 3 | -0.58864 | 0.72195 | RP11-571M6.15 | 0.72195 | 0.72569 | 0.66959 |
| ENSG00000186001 | 3 | 197518097 | 197615307 | 348 | 38 | -0.58869 | 0.72197 | LRCH3 | 0.72197 | 0.73824 | 0.76515 |
| ENSG00000249679 | 4 | 186392670 | 186393417 | 4 | 2 | -0.58869 | 0.72197 | RP11-279O9.4 | 0.72197 | 0.64535 | 0.60206 |
| ENSG00000137497 | 11 | 71713910 | 71791739 | 191 | 10 | -0.58884 | 0.72202 | NUMA1 | 0.72202 | 0.71112 | 0.69999 |
| ENSG00000249751 | 5 | 138784245 | 138842328 | 10 | 5 | -0.58897 | 0.72206 | ECSCR | 0.72206 | 0.7434 | 0.50042 |
| ENSG00000168026 | 3 | 39149152 | 39180394 | 135 | 17 | -0.58964 | 0.72229 | TTC21A | 0.72229 | 0.7359 | 0.7102 |
| ENSG00000260007 | 15 | 68486139 | 68522056 | 76 | 6 | -0.58966 | 0.72229 | RP11-315D16.2 | 0.72229 | 0.71997 | 0.68556 |
| ENSG00000243444 | 9 | 112403068 | 112713755 | 1395 | 93 | -0.58968 | 0.7223 | PALM2 | 0.7223 | 0.72247 | 0.7265 |
| ENSG00000185198 | 19 | 685521 | 695460 | 80 | 12 | -0.58985 | 0.72235 | PRSS57 | 0.72235 | 0.59547 | 0.40241 |
| ENSG00000269510 | 9 | 35042390 | 35042824 | 1 | 1 | -0.58999 | 0.7224 | FLJ00273 | 0.7224 | 0.72971 | 0.68248 |
| ENSG00000006744 | 17 | 12895708 | 12921504 | 175 | 8 | -0.59002 | 0.72241 | ELAC2 | 0.72241 | 0.68837 | 0.62174 |
| ENSG00000138735 | 4 | 120415550 | 120550146 | 484 | 19 | -0.5901 | 0.72244 | PDE5A | 0.72244 | 0.70691 | 0.71565 |

| | | | | | | | | | | | |
|---|---|---|---|---|---|---|---|---|---|---|---|
| ENSG00000169251 | 3 | 160822484 | 160971320 | 416 | 24 | -0.59013 | 0.72245 | NMD3 | 0.72245 | 0.70198 | 0.68928 |
| ENSG00000111057 | 12 | 53342655 | 53346685 | 6 | 3 | -0.59014 | 0.72245 | KRT18 | 0.72245 | 0.78858 | 0.60413 |
| ENSG00000205669 | 14 | 74077649 | 74086592 | 36 | 9 | -0.59012 | 0.72245 | ACOT6 | 0.72245 | 0.67065 | 0.57558 |
| ENSG00000177239 | 9 | 139981379 | 140003635 | 72 | 13 | -0.5903 | 0.72251 | MAN1B1 | 0.72251 | 0.61847 | 0.68198 |
| ENSG00000167604 | 19 | 36378555 | 36393205 | 38 | 6 | -0.59045 | 0.72256 | NFKBID | 0.72256 | 0.59182 | 0.52065 |
| ENSG00000268735 | 13 | 24322243 | 24323901 | 12 | 5 | -0.59053 | 0.72258 | AL139080.1 | 0.72258 | 0.57108 | 0.52797 |
| ENSG00000080986 | 18 | 2571510 | 2616634 | 146 | 18 | -0.59072 | 0.72265 | NDC80 | 0.72265 | 0.66984 | 0.36428 |
| ENSG00000185716 | 16 | 22018959 | 22098855 | 107 | 20 | -0.59086 | 0.72269 | C16orf52 | 0.72269 | 0.74038 | 0.71575 |
| ENSG00000173163 | 2 | 62115859 | 62374382 | 375 | 36 | -0.59101 | 0.72274 | COMMD1 | 0.72274 | 0.67837 | 0.6014 |
| ENSG00000127914 | 7 | 91570181 | 91739987 | 396 | 9 | -0.59122 | 0.72281 | AKAP9 | 0.72281 | 0.70625 | 0.71574 |
| ENSG00000112078 | 6 | 36410544 | 36458920 | 131 | 16 | -0.59134 | 0.72285 | KCTD20 | 0.72285 | 0.73461 | 0.71545 |
| ENSG00000130307 | 19 | 17359985 | 17393595 | 146 | 23 | -0.59133 | 0.72285 | USHBP1 | 0.72285 | 0.65278 | 0.77661 |
| ENSG00000186970 | 21 | 31812597 | 31813070 | 3 | 2 | -0.59154 | 0.72292 | KRTAP15-1 | 0.72292 | 0.46683 | 0.48397 |
| ENSG00000116774 | 1 | 114522063 | 114524876 | 6 | 4 | -0.59193 | 0.72305 | OLFML3 | 0.72305 | 0.78469 | 0.84081 |
| ENSG00000180999 | 1 | 172389828 | 172437971 | 152 | 19 | -0.59192 | 0.72305 | C1orf105 | 0.72305 | 0.74488 | 0.715 |
| ENSG00000032444 | 19 | 7598890 | 7626650 | 147 | 18 | -0.59214 | 0.72312 | PNPLA6 | 0.72312 | 0.76688 | 0.8446 |
| ENSG00000171217 | 6 | 155585147 | 155597682 | 77 | 11 | -0.59221 | 0.72314 | CLDN20 | 0.72314 | 0.73479 | 0.75602 |
| ENSG00000162039 | 16 | 1883984 | 1934295 | 437 | 6 | -0.59244 | 0.72322 | MEIOB | 0.72322 | 0.76439 | 0.79695 |
| ENSG00000213578 | 15 | 75118888 | 75124141 | 7 | 3 | -0.5925 | 0.72324 | CPLX3 | 0.72324 | 0.21203 | 0.10045 |
| ENSG00000071994 | 6 | 170884383 | 170893780 | 32 | 6 | -0.59253 | 0.72325 | PDCD2 | 0.72325 | 0.76158 | 0.74319 |
| ENSG00000124214 | 20 | 47729878 | 47804904 | 211 | 21 | -0.59274 | 0.72332 | STAU1 | 0.72332 | 0.75029 | 0.73776 |
| ENSG00000109332 | 4 | 103715540 | 103790053 | 210 | 12 | -0.59281 | 0.72334 | UBE2D3 | 0.72334 | 0.74235 | 0.74893 |
| ENSG00000179195 | 12 | 124456392 | 124499986 | 95 | 10 | -0.59288 | 0.72337 | ZNF664 | 0.72337 | 0.7401 | 0.77095 |
| ENSG00000121966 | 2 | 136871919 | 136875735 | 5 | 2 | -0.593 | 0.72341 | CXCR4 | 0.72341 | 0.81455 | 0.64515 |
| ENSG00000177432 | 4 | 89617066 | 89619386 | 8 | 4 | -0.59307 | 0.72343 | NAP1L5 | 0.72343 | 0.47348 | 0.37057 |
| ENSG00000183431 | 1 | 38422647 | 38456593 | 114 | 12 | -0.59328 | 0.7235 | SF3A3 | 0.7235 | 0.75008 | 0.83042 |
| ENSG00000183625 | 3 | 46205096 | 46308197 | 313 | 16 | -0.59327 | 0.7235 | CCR3 | 0.7235 | 0.72538 | 0.76648 |
| ENSG00000102678 | 13 | 22245522 | 22278637 | 62 | 9 | -0.59328 | 0.7235 | FGF9 | 0.7235 | 0.76653 | 0.65746 |
| ENSG00000164975 | 9 | 15422702 | 15465951 | 101 | 15 | -0.59334 | 0.72352 | SNAPC3 | 0.72352 | 0.67173 | 0.70582 |
| ENSG00000114126 | 3 | 141663277 | 141868386 | 857 | 20 | -0.59337 | 0.72353 | TFDP2 | 0.72353 | 0.73171 | 0.73764 |
| ENSG00000089063 | 20 | 5080486 | 5093749 | 31 | 7 | -0.59339 | 0.72354 | TMEM230 | 0.72354 | 0.78952 | 0.73807 |
| ENSG00000269042 | 11 | 108130255 | 108136480 | 7 | 2 | -0.59343 | 0.72355 | AP001925.1 | 0.72355 | 0.73508 | 0.78935 |
| ENSG00000090920 | 19 | 40353963 | 40440533 | 363 | 38 | -0.59346 | 0.72356 | FCGBP | 0.72356 | 0.49157 | 0.56772 |
| ENSG00000272410 | 3 | 10291056 | 10327480 | 119 | 17 | -0.59352 | 0.72358 | RP11-438J1.1 | 0.72358 | 0.70833 | 0.74462 |
| ENSG00000234545 | 7 | 92190107 | 92219708 | 73 | 11 | -0.59358 | 0.7236 | FAM133B | 0.7236 | 0.76722 | 0.73331 |
| ENSG00000106443 | 7 | 11013499 | 11209250 | 544 | 21 | -0.59364 | 0.72362 | PHF14 | 0.72362 | 0.74112 | 0.78823 |
| ENSG00000130787 | 12 | 123319000 | 123347507 | 66 | 12 | -0.59368 | 0.72364 | HIP1R | 0.72364 | 0.72224 | 0.60061 |
| ENSG00000171793 | 1 | 41445007 | 41478235 | 107 | 22 | -0.59375 | 0.72366 | CTPS1 | 0.72366 | 0.77753 | 0.79531 |
| ENSG00000104327 | 8 | 91070836 | 91107703 | 96 | 12 | -0.59385 | 0.7237 | CALB1 | 0.7237 | 0.66411 | 0.71333 |
| ENSG00000137822 | 15 | 43661419 | 43699293 | 77 | 9 | -0.5939 | 0.72371 | TUBGCP4 | 0.72371 | 0.72362 | 0.72801 |
| ENSG00000198420 | 7 | 143548468 | 143599291 | 55 | 11 | -0.59445 | 0.72389 | FAM115A | 0.72389 | 0.75875 | 0.66018 |
| ENSG00000126653 | 17 | 28442539 | 28513493 | 132 | 9 | -0.59464 | 0.72396 | NSRP1 | 0.72396 | 0.74553 | 0.77906 |
| ENSG00000163788 | 3 | 43328004 | 43466256 | 222 | 32 | -0.59473 | 0.72399 | SNRK | 0.72399 | 0.71342 | 0.74239 |
| ENSG00000143786 | 1 | 224622362 | 224928251 | 908 | 47 | -0.59478 | 0.724 | CNIH3 | 0.724 | 0.70773 | 0.67583 |
| ENSG00000178700 | 3 | 93766680 | 93782233 | 14 | 6 | -0.5949 | 0.72404 | DHFRL1 | 0.72404 | 0.75959 | 0.71129 |
| ENSG00000163885 | 3 | 126113782 | 126155399 | 197 | 33 | -0.59494 | 0.72406 | CCDC37 | 0.72406 | 0.87539 | 0.95219 |
| ENSG00000205476 | 14 | 99977603 | 100070363 | 429 | 27 | -0.59499 | 0.72407 | CCDC85C | 0.72407 | 0.57728 | 0.46841 |
| ENSG00000196367 | 7 | 98475556 | 98610866 | 321 | 19 | -0.59503 | 0.72409 | TRRAP | 0.72409 | 0.75988 | 0.77563 |
| ENSG00000135677 | 12 | 65107225 | 65153227 | 91 | 9 | -0.59515 | 0.72413 | GNS | 0.72413 | 0.7231 | 0.74438 |
| ENSG00000157881 | 1 | 2439972 | 2458039 | 52 | 9 | -0.59526 | 0.72417 | PANK4 | 0.72417 | 0.55648 | 0.26835 |
| ENSG00000169905 | 1 | 179809102 | 179846938 | 68 | 11 | -0.59535 | 0.7242 | TOR1AIP2 | 0.7242 | 0.6525 | 0.40712 |
| ENSG00000213281 | 1 | 115247090 | 115259515 | 24 | 7 | -0.59545 | 0.72423 | NRAS | 0.72423 | 0.6148 | 0.67159 |
| ENSG00000174547 | 11 | 66202546 | 66234209 | 36 | 9 | -0.59558 | 0.72427 | MRPL11 | 0.72427 | 0.79702 | 0.8151 |
| ENSG00000187792 | 22 | 24083769 | 24093279 | 26 | 7 | -0.59597 | 0.7244 | ZNF70 | 0.7244 | 0.86455 | 0.89978 |
| ENSG00000256618 | 17 | 22022437 | 22023991 | 6 | 2 | -0.59605 | 0.72443 | MTRNR2L1 | 0.72443 | 0.76221 | 0.75084 |
| ENSG00000185838 | 22 | 19770747 | 19842462 | 184 | 29 | -0.59604 | 0.72443 | GNB1L | 0.72443 | 0.51753 | 0.29559 |
| ENSG00000154920 | 17 | 48450581 | 48458844 | 27 | 4 | -0.59612 | 0.72445 | EME1 | 0.72445 | 0.74385 | 0.7712 |
| ENSG00000178301 | 11 | 77300436 | 77321400 | 25 | 9 | -0.59628 | 0.72451 | AQP11 | 0.72451 | 0.67169 | 0.7231 |
| ENSG00000161547 | 17 | 74730197 | 74733456 | 2 | 1 | -0.59649 | 0.72458 | SRSF2 | 0.72458 | 0.5223 | 0.60534 |
| ENSG00000099365 | 16 | 31000577 | 31021949 | 53 | 12 | -0.59652 | 0.72459 | STX1B | 0.72459 | 0.74954 | 0.78292 |
| ENSG00000141367 | 17 | 57697219 | 57773671 | 182 | 14 | -0.59654 | 0.72459 | CLTC | 0.72459 | 0.76654 | 0.77202 |
| ENSG00000188636 | 22 | 44888452 | 44894178 | 29 | 3 | -0.59656 | 0.7246 | LDOC1L | 0.7246 | 0.57463 | 0.65636 |
| ENSG00000161572 | 17 | 34261522 | 34270703 | 18 | 5 | -0.59665 | 0.72463 | LYZL6 | 0.72463 | 0.82216 | 0.55202 |
| ENSG00000162894 | 1 | 207077731 | 207096592 | 35 | 8 | -0.59681 | 0.72468 | FAIM3 | 0.72468 | 0.75751 | 0.773 |
| ENSG00000132139 | 17 | 34071530 | 34079897 | 30 | 7 | -0.59684 | 0.72469 | GAS2L2 | 0.72469 | 0.6532 | 0.72821 |
| ENSG00000139330 | 12 | 91444268 | 91451760 | 51 | 12 | -0.59685 | 0.7247 | KERA | 0.7247 | 0.91542 | 0.88933 |
| ENSG00000268111 | 20 | 44095910 | 44107452 | 45 | 8 | -0.59692 | 0.72472 | AL031663.1 | 0.72472 | 0.51899 | 0.42917 |
| ENSG00000173391 | 12 | 10310902 | 10324737 | 59 | 9 | -0.59703 | 0.72476 | OLR1 | 0.72476 | 0.54954 | 0.74894 |
| ENSG00000258928 | 14 | 52382863 | 52385289 | 8 | 4 | -0.59705 | 0.72476 | RP11-463J10.2 | 0.72476 | 0.83386 | 0.42085 |
| ENSG00000203995 | 1 | 53308183 | 53360670 | 182 | 14 | -0.59721 | 0.72482 | ZYG11A | 0.72482 | 0.74309 | 0.78049 |
| ENSG00000101132 | 20 | 52824386 | 52844591 | 50 | 10 | -0.5975 | 0.72491 | PFDN4 | 0.72491 | 0.52746 | 0.53914 |
| ENSG00000134107 | 3 | 5020801 | 5027008 | 23 | 5 | -0.5977 | 0.72498 | BHLHE40 | 0.72498 | 0.64118 | 0.89065 |
| ENSG00000166750 | 17 | 33570055 | 33600674 | 118 | 19 | -0.59769 | 0.72498 | SLFN5 | 0.72498 | 0.80165 | 0.82237 |

| | | | | | | | | | | | |
|---|---|---|---|---|---|---|---|---|---|---|---|
| ENSG00000155099 | 8 | 92006024 | 92053292 | 103 | 14 | -0.59792 | 0.72505 | TMEM55A | 0.72505 | 0.75787 | 0.77274 |
| ENSG00000124469 | 19 | 43084393 | 43099207 | 17 | 9 | -0.59793 | 0.72506 | CEACAM8 | 0.72506 | 0.56615 | 0.57687 |
| ENSG00000080709 | 5 | 113696642 | 113832337 | 360 | 40 | -0.59803 | 0.72509 | KCNN2 | 0.72509 | 0.7728 | 0.79947 |
| ENSG00000165689 | 9 | 139296377 | 139305061 | 73 | 13 | -0.59802 | 0.72509 | SDCCAG3 | 0.72509 | 0.77265 | 0.71054 |
| ENSG00000120664 | 13 | 36920568 | 36943738 | 81 | 12 | -0.59804 | 0.72509 | SPG20OS | 0.72509 | 0.80319 | 0.79001 |
| ENSG00000159167 | 8 | 23699428 | 23712320 | 33 | 6 | -0.59814 | 0.72513 | STC1 | 0.72513 | 0.64985 | 0.71496 |
| ENSG00000184009 | 17 | 79476997 | 79490873 | 68 | 14 | -0.59833 | 0.72519 | ACTG1 | 0.72519 | 0.75977 | 0.4927 |
| ENSG00000214700 | 12 | 27233991 | 27235447 | 10 | 4 | -0.59854 | 0.72526 | C12orf71 | 0.72526 | 0.5574 | 0.59035 |
| ENSG00000151364 | 11 | 77726761 | 77757237 | 116 | 13 | -0.5986 | 0.72528 | KCTD14 | 0.72528 | 0.6483 | 0.50409 |
| ENSG00000149100 | 11 | 32605344 | 32627808 | 77 | 13 | -0.59867 | 0.7253 | EIF3M | 0.7253 | 0.78679 | 0.56288 |
| ENSG00000163517 | 3 | 13521224 | 13547916 | 94 | 27 | -0.59882 | 0.72535 | HDAC11 | 0.72535 | 0.87396 | 0.86305 |
| ENSG00000164776 | 7 | 56148440 | 56160689 | 57 | 12 | -0.59912 | 0.72545 | PHKG1 | 0.72545 | 0.77946 | 0.75352 |
| ENSG00000167904 | 8 | 56608983 | 56685966 | 242 | 24 | -0.59925 | 0.7255 | TMEM68 | 0.7255 | 0.8061 | 0.82344 |
| ENSG00000162676 | 1 | 92940319 | 92952433 | 31 | 8 | -0.59961 | 0.72562 | GFI1 | 0.72562 | 0.65924 | 0.63078 |
| ENSG00000020922 | 11 | 94152895 | 94227074 | 219 | 22 | -0.59961 | 0.72562 | MRE11A | 0.72562 | 0.74657 | 0.78188 |
| ENSG00000116141 | 1 | 220701568 | 220837803 | 312 | 16 | -0.59987 | 0.7257 | MARK1 | 0.7257 | 0.72485 | 0.73787 |
| ENSG00000136011 | 12 | 103981051 | 104160505 | 758 | 90 | -0.59993 | 0.72572 | STAB2 | 0.72572 | 0.82746 | 0.69759 |
| ENSG00000123908 | 8 | 141541264 | 141645718 | 360 | 44 | -0.59995 | 0.72573 | AGO2 | 0.72573 | 0.69388 | 0.77745 |
| ENSG00000186943 | 9 | 107331449 | 107332411 | 4 | 2 | -0.6002 | 0.72581 | OR13C8 | 0.72581 | 0.37015 | 0.40819 |
| ENSG00000151208 | 10 | 79550549 | 79686378 | 439 | 20 | -0.60021 | 0.72582 | DLG5 | 0.72582 | 0.74754 | 0.70195 |
| ENSG00000117899 | 15 | 81239667 | 81282219 | 129 | 11 | -0.60028 | 0.72584 | MESDC2 | 0.72584 | 0.72699 | 0.74381 |
| ENSG00000187980 | 1 | 20487746 | 20503917 | 55 | 12 | -0.60054 | 0.72593 | PLA2G2C | 0.72593 | 0.79165 | 0.96292 |
| ENSG00000147613 | 8 | 87060602 | 87100850 | 106 | 15 | -0.60072 | 0.72599 | PSKH2 | 0.72599 | 0.71266 | 0.69147 |
| ENSG00000168488 | 16 | 28834356 | 28848558 | 34 | 4 | -0.60087 | 0.72604 | ATXN2L | 0.72604 | 0.80353 | 0.785 |
| ENSG00000264194 | 17 | 10048831 | 10050277 | 7 | 3 | -0.60093 | 0.72606 | AC000003.2 | 0.72606 | 0.99113 | 0.99673 |
| ENSG00000160563 | 9 | 134735494 | 134955295 | 400 | 35 | -0.60113 | 0.72612 | MED27 | 0.72612 | 0.75432 | 0.81273 |
| ENSG00000181444 | 7 | 149461271 | 149470568 | 30 | 7 | -0.60158 | 0.72627 | ZNF467 | 0.72627 | 0.57643 | 0.30258 |
| ENSG00000160050 | 1 | 32665987 | 32670988 | 4 | 2 | -0.60163 | 0.72629 | CCDC28B | 0.72629 | 0.0044296 | 0.0042207 |
| ENSG00000155640 | 10 | 98741041 | 98745582 | 4 | 2 | -0.60163 | 0.72629 | C10orf12 | 0.72629 | 0.84409 | 0.87412 |
| ENSG00000081913 | 18 | 60382672 | 60647666 | 531 | 26 | -0.60174 | 0.72633 | PHLPP1 | 0.72633 | 0.73701 | 0.65946 |
| ENSG00000102580 | 13 | 96329393 | 96447243 | 190 | 31 | -0.6018 | 0.72635 | DNAJC3 | 0.72635 | 0.77232 | 0.81587 |
| ENSG00000154553 | 4 | 186422851 | 186456766 | 86 | 9 | -0.6019 | 0.72638 | PDLIM3 | 0.72638 | 0.79705 | 0.83341 |
| ENSG00000105875 | 7 | 134868590 | 134896316 | 91 | 6 | -0.60194 | 0.72639 | WDR91 | 0.72639 | 0.71763 | 0.68474 |
| ENSG00000152409 | 5 | 78532012 | 78623038 | 191 | 14 | -0.60199 | 0.72641 | JMY | 0.72641 | 0.71645 | 0.74998 |
| ENSG00000130545 | 19 | 6463788 | 6467232 | 10 | 3 | -0.60205 | 0.72643 | CRB3 | 0.72643 | 0.89859 | 0.64165 |
| ENSG00000198218 | 3 | 49067140 | 49131796 | 99 | 18 | -0.60235 | 0.72653 | QRICH1 | 0.72653 | 0.77894 | 0.75582 |
| ENSG00000182747 | 6 | 137243402 | 137246777 | 2 | 1 | -0.60246 | 0.72657 | SLC35D3 | 0.72657 | 0.64452 | 0.66892 |
| ENSG00000116871 | 1 | 36621180 | 36646450 | 40 | 8 | -0.60267 | 0.72664 | MAP7D1 | 0.72664 | 0.72545 | 0.84374 |
| ENSG00000100744 | 14 | 96829814 | 96853625 | 42 | 8 | -0.60281 | 0.72668 | GSKIP | 0.72668 | 0.84592 | 0.89973 |
| ENSG00000100897 | 14 | 24583404 | 24594451 | 21 | 4 | -0.60302 | 0.72675 | DCAF11 | 0.72675 | 0.71242 | 0.69715 |
| ENSG00000137714 | 11 | 110300607 | 110335605 | 124 | 10 | -0.60311 | 0.72678 | FDX1 | 0.72678 | 0.81074 | 0.83705 |
| ENSG00000146221 | 6 | 44246480 | 44265458 | 88 | 11 | -0.60325 | 0.72683 | TCTE1 | 0.72683 | 0.75858 | 0.80221 |
| ENSG00000070018 | 12 | 12268959 | 12419946 | 413 | 25 | -0.60351 | 0.72692 | LRP6 | 0.72692 | 0.75315 | 0.86853 |
| ENSG00000205497 | 11 | 4967355 | 4968356 | 7 | 2 | -0.60355 | 0.72693 | OR51A4 | 0.72693 | 0.48521 | 0.41551 |
| ENSG00000100403 | 22 | 41697526 | 41756151 | 109 | 12 | -0.60473 | 0.72732 | ZC3H7B | 0.72732 | 0.77114 | 0.79843 |
| ENSG00000127329 | 12 | 70910630 | 71031220 | 544 | 60 | -0.60483 | 0.72735 | PTPRB | 0.72735 | 0.66724 | 0.70025 |
| ENSG00000171475 | 17 | 38375556 | 38440388 | 141 | 13 | -0.60481 | 0.72735 | WIPF2 | 0.72735 | 0.68878 | 0.65575 |
| ENSG00000105549 | 19 | 361750 | 376013 | 125 | 37 | -0.60485 | 0.72736 | THEG | 0.72736 | 0.88914 | 0.95821 |
| ENSG00000110422 | 11 | 33278218 | 33378569 | 250 | 23 | -0.60497 | 0.7274 | HIPK3 | 0.7274 | 0.76421 | 0.7431 |
| ENSG00000112175 | 6 | 55618443 | 55740362 | 279 | 36 | -0.6051 | 0.72744 | BMP5 | 0.72744 | 0.69162 | 0.65544 |
| ENSG00000133773 | 12 | 82617460 | 82752584 | 373 | 25 | -0.60522 | 0.72748 | CCDC59 | 0.72748 | 0.72589 | 0.72483 |
| ENSG00000106615 | 7 | 151163098 | 151217206 | 116 | 12 | -0.60533 | 0.72752 | RHEB | 0.72752 | 0.69799 | 0.6222 |
| ENSG00000188342 | 13 | 45694650 | 45858237 | 368 | 14 | -0.60535 | 0.72753 | GTF2F2 | 0.72753 | 0.74234 | 0.74895 |
| ENSG00000178202 | 11 | 108342832 | 108369159 | 55 | 9 | -0.60552 | 0.72758 | KDELC2 | 0.72758 | 0.65753 | 0.64323 |
| ENSG00000213995 | 13 | 111267881 | 111292340 | 141 | 12 | -0.6055 | 0.72758 | CARKD | 0.72758 | 0.76305 | 0.89249 |
| ENSG00000131864 | 19 | 57630506 | 57643294 | 35 | 5 | -0.60559 | 0.72761 | USP29 | 0.72761 | 0.68896 | 0.78494 |
| ENSG00000168610 | 17 | 40465342 | 40540586 | 177 | 17 | -0.60582 | 0.72768 | STAT3 | 0.72768 | 0.70854 | 0.70171 |
| ENSG00000131375 | 3 | 15247659 | 15294425 | 52 | 17 | -0.60592 | 0.72772 | CAPN7 | 0.72772 | 0.79868 | 0.72715 |
| ENSG00000255524 | 16 | 28648975 | 28670003 | 31 | 7 | -0.60603 | 0.72775 | NPIPB8 | 0.72775 | 0.73284 | 0.75814 |
| ENSG00000110237 | 11 | 73019334 | 73080136 | 167 | 13 | -0.60616 | 0.7278 | ARHGEF17 | 0.7278 | 0.73064 | 0.65591 |
| ENSG00000001630 | 7 | 91741465 | 91772266 | 68 | 4 | -0.6062 | 0.72781 | CYP51A1 | 0.72781 | 0.63786 | 0.67526 |
| ENSG00000006831 | 12 | 1797740 | 1897844 | 351 | 21 | -0.6062 | 0.72781 | ADIPOR2 | 0.72781 | 0.76181 | 0.73299 |
| ENSG00000118418 | 6 | 79910962 | 79944406 | 88 | 12 | -0.60645 | 0.72789 | HMGN3 | 0.72789 | 0.76065 | 0.84377 |
| ENSG00000166211 | 12 | 101869199 | 101880775 | 48 | 13 | -0.60647 | 0.7279 | SPIC | 0.7279 | 0.70008 | 0.75329 |
| ENSG00000108651 | 17 | 30187923 | 30228784 | 116 | 13 | -0.60654 | 0.72792 | UTP6 | 0.72792 | 0.70581 | 0.77937 |
| ENSG00000273217 | 5 | 130761584 | 131132676 | 653 | 17 | -0.60698 | 0.72807 | CTC-432M15.3 | 0.72807 | 0.71163 | 0.69092 |
| ENSG00000153560 | 3 | 33429828 | 33482863 | 168 | 17 | -0.60706 | 0.72809 | UBP1 | 0.72809 | 0.70445 | 0.68251 |
| ENSG00000145248 | 4 | 48485360 | 48491213 | 14 | 3 | -0.60705 | 0.72809 | SLC10A4 | 0.72809 | 0.7337 | 0.73465 |
| ENSG00000005700 | 6 | 82879700 | 82957471 | 244 | 13 | -0.60704 | 0.72809 | IBTK | 0.72809 | 0.77782 | 0.86325 |
| ENSG00000142444 | 19 | 11039409 | 11044211 | 10 | 5 | -0.60708 | 0.7281 | C19orf52 | 0.7281 | 0.80829 | 0.74761 |
| ENSG00000171843 | 9 | 20341663 | 20622542 | 570 | 70 | -0.60736 | 0.72819 | MLLT3 | 0.72819 | 0.63987 | 0.56648 |
| ENSG00000131018 | 6 | 152442819 | 152958936 | 2024 | 136 | -0.60742 | 0.72821 | SYNE1 | 0.72821 | 0.71931 | 0.6911 |
| ENSG00000160870 | 7 | 99302660 | 99332819 | 74 | 5 | -0.60743 | 0.72822 | CYP3A7 | 0.72822 | 0.73999 | 0.76441 |

| | | | | | | | | | | | |
|---|---|---|---|---|---|---|---|---|---|---|---|
| ENSG00000138279 | 10 | 75134859 | 75173834 | 91 | 4 | -0.60747 | 0.72823 | ANXA7 | 0.72823 | 0.74968 | 0.76538 |
| ENSG00000089335 | 19 | 35168544 | 35177302 | 38 | 7 | -0.60751 | 0.72824 | ZNF302 | 0.72824 | 0.63226 | 0.65492 |
| ENSG00000151327 | 14 | 35514113 | 35582336 | 308 | 22 | -0.60762 | 0.72828 | FAM177A1 | 0.72828 | 0.72443 | 0.7407 |
| ENSG00000142188 | 21 | 34804792 | 34852318 | 201 | 13 | -0.60771 | 0.72831 | TMEM50B | 0.72831 | 0.71344 | 0.74291 |
| ENSG00000124733 | 6 | 42979832 | 42981706 | 5 | 3 | -0.60793 | 0.72838 | MEA1 | 0.72838 | 0.81031 | 0.7812 |
| ENSG00000082126 | 2 | 202509593 | 202563417 | 165 | 37 | -0.60798 | 0.7284 | MPP4 | 0.7284 | 0.61935 | 0.61698 |
| ENSG00000174939 | 16 | 29911696 | 29931185 | 32 | 9 | -0.60805 | 0.72842 | ASPHD1 | 0.72842 | 0.60136 | 0.63872 |
| ENSG00000255292 | 11 | 111957627 | 112064528 | 289 | 34 | -0.60806 | 0.72843 | SDHD | 0.72843 | 0.73129 | 0.77644 |
| ENSG00000214941 | 17 | 15879874 | 15903031 | 75 | 6 | -0.60807 | 0.72843 | ZSWIM7 | 0.72843 | 0.85626 | 0.87372 |
| ENSG00000174600 | 12 | 108681821 | 108733118 | 194 | 37 | -0.60813 | 0.72845 | CMKLR1 | 0.72845 | 0.68075 | 0.57577 |
| ENSG00000159625 | 16 | 57728705 | 57765717 | 134 | 19 | -0.60815 | 0.72846 | CCDC135 | 0.72846 | 0.55196 | 0.64956 |
| ENSG00000156671 | 10 | 76859344 | 76941881 | 193 | 22 | -0.60825 | 0.72849 | SAMD8 | 0.72849 | 0.76928 | 0.72656 |
| ENSG00000244617 | 2 | 70187226 | 70189397 | 4 | 2 | -0.60849 | 0.72857 | ASPRV1 | 0.72857 | 0.4864 | 0.33787 |
| ENSG00000114656 | 3 | 128628709 | 128721533 | 255 | 22 | -0.60866 | 0.72863 | KIAA1257 | 0.72863 | 0.75557 | 0.77182 |
| ENSG00000197683 | 21 | 31691452 | 31692607 | 5 | 2 | -0.60869 | 0.72863 | KRTAP26-1 | 0.72863 | 0.5732 | 0.53224 |
| ENSG00000081985 | 1 | 67773047 | 67862583 | 316 | 22 | -0.60876 | 0.72866 | IL12RB2 | 0.72866 | 0.73574 | 0.80662 |
| ENSG00000262633 | 17 | 45000499 | 45124520 | 485 | 25 | -0.60876 | 0.72866 | RP11-156P1.2 | 0.72866 | 0.7408 | 0.72339 |
| ENSG00000106266 | 7 | 2291405 | 2393953 | 386 | 39 | -0.60882 | 0.72868 | SNX8 | 0.72868 | 0.65523 | 0.75006 |
| ENSG00000136193 | 7 | 29959719 | 30029905 | 161 | 22 | -0.60924 | 0.72882 | SCRN1 | 0.72882 | 0.66697 | 0.63609 |
| ENSG00000154265 | 17 | 67240452 | 67323385 | 191 | 18 | -0.60984 | 0.72902 | ABCA5 | 0.72902 | 0.72209 | 0.79288 |
| ENSG00000155329 | 5 | 132332677 | 132362296 | 100 | 6 | -0.60991 | 0.72904 | ZCCHC10 | 0.72904 | 0.71936 | 0.83011 |
| ENSG00000162572 | 1 | 1215816 | 1227409 | 63 | 23 | -0.61002 | 0.72907 | SCNN1D | 0.72907 | 0.8457 | 0.83316 |
| ENSG00000171195 | 4 | 71296209 | 71348714 | 209 | 18 | -0.61021 | 0.72914 | MUC7 | 0.72914 | 0.70819 | 0.69652 |
| ENSG00000256018 | 6 | 26271146 | 26271612 | 1 | 1 | -0.6104 | 0.7292 | HIST1H3G | 0.7292 | 0.167 | 0.2708 |
| ENSG00000197381 | 21 | 46493768 | 46646478 | 461 | 28 | -0.61047 | 0.72922 | ADARB1 | 0.72922 | 0.75298 | 0.84295 |
| ENSG00000159714 | 16 | 67428322 | 67450736 | 44 | 10 | -0.61049 | 0.72923 | ZDHHC1 | 0.72923 | 0.70847 | 0.74699 |
| ENSG00000135387 | 11 | 34073230 | 34122703 | 163 | 5 | -0.61073 | 0.72931 | CAPRIN1 | 0.72931 | 0.71979 | 0.75653 |
| ENSG00000163220 | 1 | 153330330 | 153333503 | 3 | 2 | -0.61125 | 0.72948 | S100A9 | 0.72948 | 0.64879 | 0.44113 |
| ENSG00000269067 | 19 | 23158270 | 23185978 | 85 | 6 | -0.61126 | 0.72949 | ZNF728 | 0.72949 | 0.75023 | 0.76625 |
| ENSG00000267954 | 22 | 24124467 | 24126145 | 4 | 2 | -0.6113 | 0.7295 | AP000349.1 | 0.7295 | 0.83239 | 0.93268 |
| ENSG00000177879 | 5 | 115177178 | 115249778 | 148 | 21 | -0.61153 | 0.72957 | AP3S1 | 0.72957 | 0.7339 | 0.87383 |
| ENSG00000152242 | 18 | 43753519 | 43846949 | 186 | 12 | -0.61154 | 0.72958 | C18orf25 | 0.72958 | 0.72019 | 0.61345 |
| ENSG00000172613 | 11 | 67159176 | 67165881 | 16 | 7 | -0.61156 | 0.72959 | RAD9A | 0.72959 | 0.47103 | 0.63718 |
| ENSG00000144063 | 2 | 110841447 | 110874143 | 59 | 15 | -0.61167 | 0.72962 | MALL | 0.72962 | 0.79554 | 0.78612 |
| ENSG00000143303 | 1 | 156698234 | 156706752 | 18 | 6 | -0.61182 | 0.72967 | RRNAD1 | 0.72967 | 0.70079 | 0.82467 |
| ENSG00000185942 | 8 | 63161150 | 63912211 | 2627 | 66 | -0.61181 | 0.72967 | NKAIN3 | 0.72967 | 0.73099 | 0.73346 |
| ENSG00000181982 | 4 | 24807739 | 24981826 | 604 | 40 | -0.61188 | 0.72969 | CCDC149 | 0.72969 | 0.77009 | 0.8062 |
| ENSG00000106025 | 7 | 120427376 | 120498456 | 128 | 15 | -0.61208 | 0.72976 | TSPAN12 | 0.72976 | 0.75025 | 0.84405 |
| ENSG00000115604 | 2 | 102927989 | 103015218 | 399 | 21 | -0.61218 | 0.72979 | IL18R1 | 0.72979 | 0.7185 | 0.64983 |
| ENSG00000175395 | 10 | 38238500 | 38265561 | 81 | 12 | -0.61233 | 0.72984 | ZNF25 | 0.72984 | 0.72234 | 0.66424 |
| ENSG00000121653 | 11 | 45907202 | 45928016 | 36 | 13 | -0.61249 | 0.72989 | MAPK8IP1 | 0.72989 | 0.50571 | 0.49933 |
| ENSG00000269383 | 2 | 100986705 | 100987007 | 1 | 1 | -0.61251 | 0.7299 | AC012493.2 | 0.7299 | 0.51812 | 0.53587 |
| ENSG00000135547 | 6 | 126068810 | 126082415 | 33 | 6 | -0.61261 | 0.72993 | HEY2 | 0.72993 | 0.72195 | 0.77 |
| ENSG00000165506 | 14 | 50091892 | 50101948 | 36 | 5 | -0.61277 | 0.72999 | DNAAF2 | 0.72999 | 0.84444 | 0.83137 |
| ENSG00000203710 | 1 | 207669492 | 207813992 | 279 | 19 | -0.61281 | 0.73 | CR1 | 0.73 | 0.74184 | 0.73785 |
| ENSG00000167272 | 12 | 121016567 | 121019201 | 13 | 5 | -0.61321 | 0.73013 | POP5 | 0.73013 | 0.49134 | 0.39282 |
| ENSG00000183292 | 2 | 131328402 | 131341734 | 2 | 1 | -0.61323 | 0.73014 | AC140481.2 | 0.73014 | 0.68441 | 0.68441 |
| ENSG00000091140 | 7 | 107531415 | 107572175 | 84 | 7 | -0.61357 | 0.73025 | DLD | 0.73025 | 0.72336 | 0.6934 |
| ENSG00000185294 | 17 | 43922256 | 43924438 | 17 | 2 | -0.61361 | 0.73026 | SPPL2C | 0.73026 | 0.64709 | 0.65714 |
| ENSG00000268179 | 1 | 861264 | 866445 | 33 | 10 | -0.61392 | 0.73037 | AL645608.1 | 0.73037 | 0.7996 | 0.73926 |
| ENSG00000106991 | 9 | 130577291 | 130617035 | 99 | 23 | -0.61398 | 0.73039 | ENG | 0.73039 | 0.66574 | 0.68985 |
| ENSG00000105088 | 19 | 9964394 | 10047228 | 305 | 26 | -0.614 | 0.73039 | OLFM2 | 0.73039 | 0.54913 | 0.60709 |
| ENSG00000110492 | 11 | 46402306 | 46405375 | 3 | 1 | -0.61406 | 0.73041 | MDK | 0.73041 | 0.69428 | 0.45623 |
| ENSG00000166569 | 18 | 56942388 | 56985881 | 169 | 23 | -0.61408 | 0.73042 | CPLX4 | 0.73042 | 0.74965 | 0.72459 |
| ENSG00000136147 | 13 | 50069746 | 50103123 | 142 | 15 | -0.61428 | 0.73048 | PHF11 | 0.73048 | 0.73283 | 0.78007 |
| ENSG00000167526 | 16 | 89627065 | 89630950 | 18 | 2 | -0.61435 | 0.73051 | RPL13 | 0.73051 | 0.70817 | 0.88086 |
| ENSG00000176597 | 3 | 182971032 | 183016292 | 64 | 17 | -0.61441 | 0.73053 | B3GNT5 | 0.73053 | 0.78128 | 0.78342 |
| ENSG00000175764 | 9 | 124584207 | 124855885 | 1096 | 56 | -0.61456 | 0.73058 | TTLL11 | 0.73058 | 0.68339 | 0.59229 |
| ENSG00000126858 | 17 | 30469473 | 30580393 | 184 | 16 | -0.61458 | 0.73058 | RHOT1 | 0.73058 | 0.73446 | 0.7767 |
| ENSG00000163697 | 4 | 40812044 | 41218731 | 1624 | 82 | -0.61461 | 0.73059 | APBB2 | 0.73059 | 0.76496 | 0.76877 |
| ENSG00000101441 | 20 | 23666277 | 23669677 | 14 | 4 | -0.61463 | 0.7306 | CST4 | 0.7306 | 0.57026 | 0.63724 |
| ENSG00000167074 | 22 | 41763337 | 41795330 | 72 | 7 | -0.61466 | 0.73061 | TEF | 0.73061 | 0.76554 | 0.79265 |
| ENSG00000180264 | 9 | 127213423 | 127239379 | 76 | 9 | -0.61468 | 0.73062 | GPR144 | 0.73062 | 0.81074 | 0.86841 |
| ENSG00000159792 | 16 | 67927175 | 67963581 | 48 | 10 | -0.61474 | 0.73064 | PSKH1 | 0.73064 | 0.7099 | 0.7104 |
| ENSG00000116035 | 2 | 71127720 | 71160576 | 122 | 22 | -0.61496 | 0.73071 | VAX2 | 0.73071 | 0.82268 | 0.88563 |
| ENSG00000174705 | 5 | 171752185 | 171881527 | 514 | 23 | -0.61523 | 0.7308 | SH3PXD2B | 0.7308 | 0.60177 | 0.48097 |
| ENSG00000136813 | 9 | 114122972 | 114247025 | 346 | 23 | -0.61549 | 0.73088 | KIAA0368 | 0.73088 | 0.70716 | 0.63378 |
| ENSG00000095380 | 9 | 100819021 | 100845357 | 54 | 8 | -0.61552 | 0.73089 | NANS | 0.73089 | 0.66062 | 0.70961 |
| ENSG00000163694 | 4 | 40425272 | 40632892 | 807 | 111 | -0.61568 | 0.73095 | RBM47 | 0.73095 | 0.73289 | 0.67552 |
| ENSG00000156239 | 21 | 30244513 | 30257693 | 75 | 8 | -0.61588 | 0.73101 | N6AMT1 | 0.73101 | 0.71093 | 0.7927 |
| ENSG00000238083 | 17 | 44588877 | 44633016 | 32 | 7 | -0.61609 | 0.73108 | LRRC37A2 | 0.73108 | 0.78021 | 0.67161 |
| ENSG00000159111 | 17 | 45900638 | 45908900 | 47 | 8 | -0.61619 | 0.73111 | MRPL10 | 0.73111 | 0.72917 | 0.73831 |
| ENSG00000196296 | 16 | 28889726 | 28915830 | 49 | 8 | -0.61622 | 0.73113 | ATP2A1 | 0.73113 | 0.7091 | 0.72326 |

| | | | | | | | | | | | |
|---|---|---|---|---|---|---|---|---|---|---|---|
| ENSG00000187323 | 18 | 49866542 | 51057784 | 5048 | 121 | -0.61625 | 0.73113 | DCC | 0.73113 | 0.75111 | 0.74293 |
| ENSG00000101413 | 20 | 36661948 | 36720768 | 132 | 11 | -0.61627 | 0.73114 | RPRD1B | 0.73114 | 0.72135 | 0.74304 |
| ENSG00000259009 | 19 | 48337701 | 48364769 | 198 | 16 | -0.61647 | 0.73121 | TPRX2P | 0.73121 | 0.85365 | 0.89762 |
| ENSG00000222033 | 2 | 171568961 | 171571077 | 5 | 3 | -0.61658 | 0.73124 | LINC01124 | 0.73124 | 0.14285 | 0.31282 |
| ENSG00000006025 | 17 | 45884738 | 45899200 | 52 | 9 | -0.61683 | 0.73133 | OSBPL7 | 0.73133 | 0.77017 | 0.45307 |
| ENSG00000188076 | 11 | 193080 | 194573 | 5 | 3 | -0.61686 | 0.73134 | SCGB1C1 | 0.73134 | 0.37247 | 0.21901 |
| ENSG00000073734 | 2 | 169779448 | 169887832 | 331 | 26 | -0.61693 | 0.73136 | ABCB11 | 0.73136 | 0.71256 | 0.70172 |
| ENSG00000177034 | 5 | 79275584 | 79287082 | 31 | 5 | -0.61723 | 0.73146 | MTX3 | 0.73146 | 0.73138 | 0.75186 |
| ENSG00000145868 | 5 | 147763498 | 147822399 | 104 | 10 | -0.61726 | 0.73147 | FBXO38 | 0.73147 | 0.66318 | 0.60861 |
| ENSG00000124302 | 19 | 34112861 | 34264414 | 594 | 70 | -0.61736 | 0.7315 | CHST8 | 0.7315 | 0.77196 | 0.78932 |
| ENSG00000172216 | 20 | 48807376 | 48809212 | 3 | 2 | -0.6175 | 0.73155 | CEBPB | 0.73155 | 0.81633 | 0.92196 |
| ENSG00000221910 | 7 | 143632259 | 143633372 | 7 | 2 | -0.61771 | 0.73162 | OR2F2 | 0.73162 | 0.7323 | 0.57233 |
| ENSG00000184500 | 3 | 93591881 | 93692910 | 105 | 22 | -0.61776 | 0.73163 | PROS1 | 0.73163 | 0.75547 | 0.74678 |
| ENSG00000165124 | 9 | 113127531 | 113342160 | 1031 | 63 | -0.61796 | 0.7317 | SVEP1 | 0.7317 | 0.75784 | 0.80039 |
| ENSG00000114993 | 2 | 74652963 | 74669549 | 8 | 5 | -0.61802 | 0.73172 | RTKN | 0.73172 | 0.14866 | 0.36542 |
| ENSG00000167618 | 19 | 55009100 | 55021897 | 98 | 26 | -0.61804 | 0.73173 | LAIR2 | 0.73173 | 0.17758 | 0.28772 |
| ENSG00000153666 | 15 | 23255242 | 23268823 | 4 | 2 | -0.6181 | 0.73174 | GOLGA8I | 0.73174 | 0.74064 | 0.63658 |
| ENSG00000143373 | 1 | 151254094 | 151264656 | 9 | 3 | -0.61815 | 0.73176 | ZNF687 | 0.73176 | 0.70557 | 0.86272 |
| ENSG00000162148 | 11 | 61248592 | 61258403 | 17 | 5 | -0.61813 | 0.73176 | PPP1R32 | 0.73176 | 0.72482 | 0.69858 |
| ENSG00000100385 | 22 | 37521878 | 37571094 | 161 | 38 | -0.61815 | 0.73176 | IL2RB | 0.73176 | 0.83098 | 0.68357 |
| ENSG00000065613 | 10 | 105726959 | 105788991 | 208 | 13 | -0.6182 | 0.73178 | SLK | 0.73178 | 0.7527 | 0.78107 |
| ENSG00000170837 | 3 | 71803201 | 71805647 | 2 | 1 | -0.61844 | 0.73186 | GPR27 | 0.73186 | 0.80768 | 0.75621 |
| ENSG00000166946 | 15 | 43477316 | 43487396 | 10 | 4 | -0.61859 | 0.73191 | CCNDBP1 | 0.73191 | 0.5453 | 0.55949 |
| ENSG00000143418 | 1 | 150933059 | 150947479 | 36 | 7 | -0.61873 | 0.73195 | CERS2 | 0.73195 | 0.72181 | 0.77676 |
| ENSG00000139842 | 13 | 113862552 | 113919399 | 143 | 7 | -0.61884 | 0.73199 | CUL4A | 0.73199 | 0.69674 | 0.62634 |
| ENSG00000244754 | 13 | 33006554 | 33112970 | 193 | 21 | -0.61895 | 0.73203 | N4BP2L2 | 0.73203 | 0.74955 | 0.77192 |
| ENSG00000177595 | 11 | 799179 | 809753 | 42 | 7 | -0.61898 | 0.73204 | PIDD | 0.73204 | 0.79915 | 0.5571 |
| ENSG00000102908 | 16 | 69598997 | 69738569 | 378 | 20 | -0.61927 | 0.73213 | NFAT5 | 0.73213 | 0.7651 | 0.71065 |
| ENSG00000169155 | 9 | 129567285 | 129600489 | 59 | 18 | -0.61939 | 0.73217 | ZBTB43 | 0.73217 | 0.69718 | 0.83964 |
| ENSG00000058063 | 3 | 182511288 | 182639423 | 229 | 12 | -0.61962 | 0.73225 | ATP11B | 0.73225 | 0.77132 | 0.74014 |
| ENSG00000255526 | 14 | 24683161 | 24701558 | 43 | 14 | -0.61967 | 0.73226 | NEDD8-MDP1 | 0.73226 | 0.79761 | 0.82422 |
| ENSG00000119950 | 10 | 111967363 | 112047123 | 238 | 22 | -0.61984 | 0.73232 | MXI1 | 0.73232 | 0.73322 | 0.75132 |
| ENSG00000127586 | 16 | 838046 | 850737 | 98 | 18 | -0.61993 | 0.73235 | CHTF18 | 0.73235 | 0.81076 | 0.77569 |
| ENSG00000241852 | 8 | 22457114 | 22461663 | 11 | 4 | -0.62 | 0.73237 | C8orf58 | 0.73237 | 0.69639 | 0.79341 |
| ENSG00000133110 | 13 | 38136720 | 38172981 | 110 | 17 | -0.62013 | 0.73241 | POSTN | 0.73241 | 0.70436 | 0.43703 |
| ENSG00000170231 | 5 | 159614374 | 159665742 | 312 | 26 | -0.6203 | 0.73247 | FABP6 | 0.73247 | 0.73783 | 0.72938 |
| ENSG00000123427 | 12 | 58165275 | 58176324 | 23 | 3 | -0.62039 | 0.7325 | METTL21B | 0.7325 | 0.73871 | 0.62823 |
| ENSG00000104381 | 8 | 75233365 | 75401107 | 488 | 34 | -0.62062 | 0.73258 | GDAP1 | 0.73258 | 0.69698 | 0.73064 |
| ENSG00000196371 | 11 | 94277017 | 94283063 | 17 | 6 | -0.62064 | 0.73258 | FUT4 | 0.73258 | 0.73352 | 0.8644 |
| ENSG00000100902 | 14 | 35747839 | 35786699 | 149 | 18 | -0.62084 | 0.73265 | PSMA6 | 0.73265 | 0.65881 | 0.66892 |
| ENSG00000185049 | 4 | 1984441 | 2043630 | 145 | 19 | -0.62088 | 0.73266 | NELFA | 0.73266 | 0.71284 | 0.74979 |
| ENSG00000006016 | 19 | 18683030 | 18718551 | 118 | 15 | -0.62103 | 0.73271 | CRLF1 | 0.73271 | 0.74617 | 0.76974 |
| ENSG00000159905 | 19 | 44455375 | 44471861 | 57 | 11 | -0.62158 | 0.73289 | ZNF221 | 0.73289 | 0.69281 | 0.81079 |
| ENSG00000108433 | 17 | 45000483 | 45105003 | 436 | 23 | -0.62176 | 0.73295 | GOSR2 | 0.73295 | 0.73152 | 0.73197 |
| ENSG00000137720 | 11 | 111749659 | 111756699 | 13 | 2 | -0.62183 | 0.73297 | C11orf1 | 0.73297 | 0.88588 | 0.91865 |
| ENSG00000153157 | 6 | 10748027 | 10979553 | 766 | 60 | -0.62184 | 0.73298 | SYCP2L | 0.73298 | 0.7485 | 0.76261 |
| ENSG00000158987 | 5 | 130759614 | 130970929 | 386 | 15 | -0.62206 | 0.73305 | RAPGEF6 | 0.73305 | 0.70778 | 0.66948 |
| ENSG00000178952 | 16 | 28853732 | 28857729 | 7 | 2 | -0.62222 | 0.7331 | TUFM | 0.7331 | 0.81044 | 0.78752 |
| ENSG00000109065 | 17 | 72766686 | 72772506 | 21 | 3 | -0.62236 | 0.73315 | NAT9 | 0.73315 | 0.78406 | 0.78771 |
| ENSG00000204683 | 10 | 21414692 | 21435488 | 37 | 9 | -0.62245 | 0.73318 | C10orf113 | 0.73318 | 0.7229 | 0.67225 |
| ENSG00000163956 | 4 | 3508103 | 3534286 | 174 | 23 | -0.62254 | 0.73321 | LRPAP1 | 0.73321 | 0.78229 | 0.77351 |
| ENSG00000198046 | 19 | 56950694 | 56989434 | 85 | 11 | -0.62265 | 0.73324 | ZNF667 | 0.73324 | 0.81087 | 0.84583 |
| ENSG00000185900 | 8 | 42948658 | 42978577 | 44 | 6 | -0.62288 | 0.73332 | POMK | 0.73332 | 0.7641 | 0.78443 |
| ENSG00000161653 | 17 | 42081914 | 42086431 | 13 | 4 | -0.62291 | 0.73333 | NAGS | 0.73333 | 0.82621 | 0.71701 |
| ENSG00000111344 | 12 | 113536624 | 113574044 | 177 | 15 | -0.62298 | 0.73335 | RASAL1 | 0.73335 | 0.62643 | 0.58896 |
| ENSG00000124657 | 6 | 27925019 | 27925960 | 4 | 2 | -0.62317 | 0.73341 | OR2B6 | 0.73341 | 0.96734 | 0.96803 |
| ENSG00000144810 | 3 | 99357319 | 99518070 | 408 | 32 | -0.62343 | 0.7335 | COL8A1 | 0.7335 | 0.71557 | 0.69359 |
| ENSG00000156042 | 10 | 75013517 | 75118617 | 363 | 5 | -0.62358 | 0.73355 | TTC18 | 0.73355 | 0.7377 | 0.74973 |
| ENSG00000151150 | 10 | 61786056 | 62493248 | 2612 | 91 | -0.62386 | 0.73364 | ANK3 | 0.73364 | 0.75842 | 0.76777 |
| ENSG00000107263 | 9 | 134452157 | 134615461 | 512 | 29 | -0.62432 | 0.73379 | RAPGEF1 | 0.73379 | 0.71808 | 0.64199 |
| ENSG00000258875 | 14 | 91709103 | 91717426 | 33 | 4 | -0.62435 | 0.7338 | CTD-2547L24.3 | 0.7338 | 0.80037 | 0.51746 |
| ENSG00000197077 | 22 | 25348697 | 25593415 | 787 | 62 | -0.62434 | 0.7338 | KIAA1671 | 0.7338 | 0.74963 | 0.74146 |
| ENSG00000187720 | 15 | 71389291 | 72075722 | 2506 | 179 | -0.62439 | 0.73381 | THSD4 | 0.73381 | 0.7927 | 0.81659 |
| ENSG00000141522 | 17 | 79825597 | 79829282 | 7 | 2 | -0.62442 | 0.73382 | ARHGDIA | 0.73382 | 0.67742 | 0.63936 |
| ENSG00000196772 | 1 | 247978102 | 247979031 | 3 | 2 | -0.62455 | 0.73387 | OR14A16 | 0.73387 | 0.48655 | 0.69309 |
| ENSG00000099958 | 22 | 24176690 | 24181315 | 22 | 5 | -0.62471 | 0.73392 | DERL3 | 0.73392 | 0.69547 | 0.81008 |
| ENSG00000185418 | 15 | 102193801 | 102264807 | 180 | 22 | -0.62478 | 0.73394 | TARSL2 | 0.73394 | 0.75151 | 0.76678 |
| ENSG00000103510 | 16 | 31127075 | 31142714 | 32 | 5 | -0.6248 | 0.73395 | KAT8 | 0.73395 | 0.64277 | 0.56262 |
| ENSG00000104728 | 8 | 1772142 | 1906807 | 888 | 108 | -0.62484 | 0.73396 | ARHGEF10 | 0.73396 | 0.81367 | 0.85391 |
| ENSG00000140750 | 16 | 24930706 | 25026987 | 210 | 17 | -0.62488 | 0.73398 | ARHGAP17 | 0.73398 | 0.75557 | 0.80373 |
| ENSG00000171224 | 10 | 71390007 | 71393352 | 26 | 6 | -0.62491 | 0.73399 | C10orf35 | 0.73399 | 0.7248 | 0.71998 |
| ENSG00000139910 | 14 | 26912299 | 27066960 | 328 | 15 | -0.62502 | 0.73402 | NOVA1 | 0.73402 | 0.76446 | 0.78913 |
| ENSG00000126261 | 19 | 34919257 | 34960853 | 109 | 19 | -0.62515 | 0.73406 | UBA2 | 0.73406 | 0.70972 | 0.77895 |

| | | | | | | | | | | | | |
|---|---|---|---|---|---|---|---|---|---|---|---|---|
| ENSG00000212997 | 8 | 95558771 | 95560630 | 11 | 3 | -0.62554 | 0.73419 | AC023632.1 | | 0.73419 | 0.85165 | 0.86629 |
| ENSG00000184640 | 17 | 75276651 | 75496678 | 787 | 78 | -0.62575 | 0.73426 | | 44083 | 0.73426 | 0.76368 | 0.77394 |
| ENSG00000187123 | 2 | 150186499 | 150330662 | 325 | 31 | -0.6258 | 0.73428 | LYPD6 | | 0.73428 | 0.71833 | 0.5978 |
| ENSG00000184986 | 14 | 105992940 | 105996539 | 5 | 2 | -0.62588 | 0.7343 | TMEM121 | | 0.7343 | 0.49069 | 0.10416 |
| ENSG00000167360 | 11 | 5443341 | 5444436 | 13 | 2 | -0.62597 | 0.73433 | OR51Q1 | | 0.73433 | 0.93494 | 0.94351 |
| ENSG00000273045 | 2 | 99757948 | 99767950 | 40 | 6 | -0.62614 | 0.73439 | C2ORF15 | | 0.73439 | 0.70336 | 0.69758 |
| ENSG00000122507 | 7 | 33168856 | 33645680 | 1702 | 57 | -0.62613 | 0.73439 | BBS9 | | 0.73439 | 0.74256 | 0.72074 |
| ENSG00000141580 | 17 | 80572438 | 80606429 | 139 | 24 | -0.62616 | 0.73439 | WDR45B | | 0.73439 | 0.76784 | 0.69452 |
| ENSG00000167850 | 17 | 72537247 | 72542282 | 23 | 6 | -0.62635 | 0.73446 | CD300C | | 0.73446 | 0.74547 | 0.59741 |
| ENSG00000120215 | 9 | 5890802 | 5910606 | 56 | 14 | -0.62669 | 0.73457 | MLANA | | 0.73457 | 0.73537 | 0.87099 |
| ENSG00000169435 | 4 | 74437267 | 74486348 | 154 | 12 | -0.62679 | 0.7346 | RASSF6 | | 0.7346 | 0.79991 | 0.88183 |
| ENSG00000117091 | 1 | 160648536 | 160681641 | 164 | 24 | -0.62684 | 0.73462 | CD48 | | 0.73462 | 0.56729 | 0.46347 |
| ENSG00000253831 | 1 | 157061836 | 157069600 | 36 | 10 | -0.62694 | 0.73465 | ETV3L | | 0.73465 | 0.61121 | 0.83876 |
| ENSG00000186393 | 17 | 38922490 | 38928414 | 15 | 5 | -0.62713 | 0.73471 | KRT26 | | 0.73471 | 0.77394 | 0.84154 |
| ENSG00000163814 | 3 | 45123770 | 45187914 | 262 | 65 | -0.62732 | 0.73478 | CDCP1 | | 0.73478 | 0.61511 | 0.76642 |
| ENSG00000134389 | 1 | 196946667 | 196978804 | 67 | 11 | -0.62738 | 0.73479 | CFHR5 | | 0.73479 | 0.66505 | 0.49491 |
| ENSG00000203724 | 1 | 197871777 | 197876497 | 4 | 2 | -0.62738 | 0.73479 | C1orf53 | | 0.73479 | 0.18901 | 0.13213 |
| ENSG00000138185 | 10 | 97471536 | 97637023 | 553 | 21 | -0.62735 | 0.73479 | ENTPD1 | | 0.73479 | 0.76506 | 0.74215 |
| ENSG00000184361 | 17 | 43331760 | 43339479 | 25 | 5 | -0.62742 | 0.73481 | SPATA32 | | 0.73481 | 0.64144 | 0.59977 |
| ENSG00000166925 | 7 | 100060982 | 100076902 | 22 | 4 | -0.62747 | 0.73482 | TSC22D4 | | 0.73482 | 0.58296 | 0.54443 |
| ENSG00000135604 | 6 | 144471663 | 144509507 | 78 | 12 | -0.6276 | 0.73487 | STX11 | | 0.73487 | 0.86102 | 0.93303 |
| ENSG00000010030 | 6 | 36322419 | 36356164 | 163 | 37 | -0.62797 | 0.73499 | ETV7 | | 0.73499 | 0.68234 | 0.60859 |
| ENSG00000167011 | 7 | 100813774 | 100823557 | 51 | 10 | -0.62808 | 0.73503 | NAT16 | | 0.73503 | 0.77116 | 0.71644 |
| ENSG00000136536 | 2 | 160569000 | 160627538 | 273 | 17 | -0.62821 | 0.73507 | | 43897 | 0.73507 | 0.72254 | 0.7299 |
| ENSG00000170855 | 12 | 120881764 | 120884215 | 4 | 1 | -0.62828 | 0.73509 | TRIAP1 | | 0.73509 | 0.25427 | 0.43929 |
| ENSG00000169385 | 14 | 21423611 | 21424595 | 1 | 1 | -0.62831 | 0.7351 | RNASE2 | | 0.7351 | 0.60039 | 0.067319 |
| ENSG00000129007 | 15 | 68483043 | 68498417 | 29 | 6 | -0.62845 | 0.73515 | CALML4 | | 0.73515 | 0.77568 | 0.73414 |
| ENSG00000117407 | 1 | 44398992 | 44402913 | 10 | 4 | -0.62863 | 0.73521 | ARTN | | 0.73521 | 0.70158 | 0.71068 |
| ENSG00000165476 | 10 | 65281123 | 65384883 | 273 | 19 | -0.62892 | 0.7353 | REEP3 | | 0.7353 | 0.75052 | 0.7676 |
| ENSG00000189169 | 21 | 46117087 | 46117959 | 2 | 1 | -0.62891 | 0.7353 | KRTAP10-12 | | 0.7353 | 0.13302 | 0.35001 |
| ENSG00000140993 | 16 | 3348832 | 3355729 | 6 | 2 | -0.62926 | 0.73541 | TIGD7 | | 0.73541 | 0.55961 | 0.6789 |
| ENSG00000143409 | 1 | 150969025 | 150980851 | 23 | 7 | -0.62971 | 0.73556 | FAM63A | | 0.73556 | 0.73488 | 0.87192 |
| ENSG00000125445 | 17 | 73257755 | 73262454 | 16 | 2 | -0.62972 | 0.73556 | MRPS7 | | 0.73556 | 0.6313 | 0.58747 |
| ENSG00000130513 | 19 | 18485541 | 18499986 | 65 | 13 | -0.62994 | 0.73563 | GDF15 | | 0.73563 | 0.7744 | 0.8316 |
| ENSG00000136631 | 1 | 150039369 | 150117505 | 129 | 7 | -0.63011 | 0.73569 | VPS45 | | 0.73569 | 0.72624 | 0.74423 |
| ENSG00000100593 | 14 | 77940740 | 77965210 | 85 | 14 | -0.63047 | 0.73581 | ISM2 | | 0.73581 | 0.79257 | 0.81366 |
| ENSG00000074590 | 12 | 106457118 | 106533811 | 258 | 36 | -0.63053 | 0.73582 | NUAK1 | | 0.73582 | 0.84311 | 0.84348 |
| ENSG00000228146 | 16 | 3194220 | 3199964 | 16 | 2 | -0.6305 | 0.73582 | CASP16 | | 0.73582 | 0.42888 | 0.51006 |
| ENSG00000141748 | 17 | 37313147 | 37323737 | 61 | 13 | -0.63059 | 0.73585 | ARL5C | | 0.73585 | 0.73967 | 0.76127 |
| ENSG00000089818 | 12 | 8234807 | 8250367 | 31 | 10 | -0.63083 | 0.73592 | NECAP1 | | 0.73592 | 0.779 | 0.8928 |
| ENSG00000164175 | 5 | 33944721 | 33984835 | 87 | 14 | -0.63098 | 0.73597 | SLC45A2 | | 0.73597 | 0.77451 | 0.75296 |
| ENSG00000100811 | 14 | 100704635 | 100749129 | 120 | 13 | -0.63107 | 0.736 | YY1 | | 0.736 | 0.74835 | 0.73119 |
| ENSG00000162398 | 1 | 55271736 | 55307925 | 165 | 12 | -0.63132 | 0.73608 | C1orf177 | | 0.73608 | 0.69634 | 0.74772 |
| ENSG00000110218 | 11 | 93862094 | 93915138 | 165 | 18 | -0.63139 | 0.73611 | PANX1 | | 0.73611 | 0.63247 | 0.63377 |
| ENSG00000133250 | 19 | 8575462 | 8579048 | 17 | 2 | -0.63139 | 0.73611 | ZNF414 | | 0.73611 | 0.76293 | 0.80118 |
| ENSG00000166482 | 17 | 19286755 | 19290553 | 4 | 2 | -0.63146 | 0.73613 | MFAP4 | | 0.73613 | 0.87257 | 0.88208 |
| ENSG00000143257 | 1 | 161199456 | 161208092 | 33 | 10 | -0.63153 | 0.73615 | NR1I3 | | 0.73615 | 0.77405 | 0.86646 |
| ENSG00000165269 | 9 | 33384765 | 33402643 | 65 | 17 | -0.6316 | 0.73618 | AQP7 | | 0.73618 | 0.8182 | 0.65171 |
| ENSG00000204544 | 6 | 30951495 | 30957680 | 63 | 13 | -0.63181 | 0.73624 | MUC21 | | 0.73624 | 0.54478 | 0.38453 |
| ENSG00000174165 | 11 | 66288108 | 66313709 | 62 | 8 | -0.63186 | 0.73626 | ZDHHC24 | | 0.73626 | 0.71611 | 0.69673 |
| ENSG00000165672 | 10 | 120927215 | 120938345 | 16 | 6 | -0.6319 | 0.73627 | PRDX3 | | 0.73627 | 0.3475 | 0.40108 |
| ENSG00000197953 | 3 | 151451704 | 151479127 | 114 | 14 | -0.63209 | 0.73634 | AADACL2 | | 0.73634 | 0.79342 | 0.80159 |
| ENSG00000132646 | 20 | 5095599 | 5107272 | 27 | 3 | -0.63211 | 0.73634 | PCNA | | 0.73634 | 0.68857 | 0.70865 |
| ENSG00000176381 | 6 | 166719168 | 166721936 | 8 | 2 | -0.63221 | 0.73637 | PRR18 | | 0.73637 | 0.78423 | 0.70742 |
| ENSG00000110723 | 11 | 108376158 | 108464465 | 266 | 29 | -0.63219 | 0.73637 | EXPH5 | | 0.73637 | 0.76614 | 0.79925 |
| ENSG00000196642 | 9 | 139702374 | 139735639 | 55 | 7 | -0.63227 | 0.73639 | RABL6 | | 0.73639 | 0.67788 | 0.65025 |
| ENSG00000196154 | 1 | 153516089 | 153522612 | 32 | 2 | -0.63233 | 0.73641 | S100A4 | | 0.73641 | 0.85695 | 0.89753 |
| ENSG00000074842 | 19 | 4641386 | 4670382 | 137 | 17 | -0.63245 | 0.73645 | C19orf10 | | 0.73645 | 0.70375 | 0.75416 |
| ENSG00000115216 | 2 | 27650657 | 27665126 | 25 | 3 | -0.63249 | 0.73647 | NRBP1 | | 0.73647 | 0.6803 | 0.7337 |
| ENSG00000107614 | 10 | 17184253 | 17244053 | 260 | 10 | -0.63253 | 0.73648 | TRDMT1 | | 0.73648 | 0.76663 | 0.8037 |
| ENSG00000160714 | 1 | 154521053 | 154531504 | 15 | 3 | -0.63265 | 0.73652 | UBE2Q1 | | 0.73652 | 0.753 | 0.79073 |
| ENSG00000182759 | 8 | 144501352 | 144512576 | 69 | 15 | -0.63281 | 0.73657 | MAFA | | 0.73657 | 0.57087 | 0.65733 |
| ENSG00000213221 | 9 | 139253932 | 139258241 | 22 | 7 | -0.63286 | 0.73659 | DNLZ | | 0.73659 | 0.33168 | 0.3218 |
| ENSG00000121297 | 19 | 31765851 | 31840453 | 171 | 38 | -0.63304 | 0.73665 | TSHZ3 | | 0.73665 | 0.68432 | 0.4806 |
| ENSG00000177335 | 8 | 144120626 | 144141359 | 87 | 6 | -0.63325 | 0.73672 | C8orf31 | | 0.73672 | 0.7666 | 0.78865 |
| ENSG00000109320 | 4 | 103422486 | 103538459 | 237 | 25 | -0.63391 | 0.73693 | NFKB1 | | 0.73693 | 0.73794 | 0.72014 |
| ENSG00000141378 | 17 | 57751997 | 57784987 | 65 | 10 | -0.63397 | 0.73695 | PTRH2 | | 0.73695 | 0.6955 | 0.69175 |
| ENSG00000141738 | 17 | 37894180 | 37903544 | 12 | 3 | -0.63406 | 0.73698 | GRB7 | | 0.73698 | 0.79459 | 0.91247 |
| ENSG00000177558 | 19 | 35715703 | 35719632 | 18 | 6 | -0.63423 | 0.73704 | FAM187B | | 0.73704 | 0.7449 | 0.734 |
| ENSG00000106636 | 7 | 44240567 | 44253893 | 29 | 6 | -0.63458 | 0.73715 | YKT6 | | 0.73715 | 0.73877 | 0.66286 |
| ENSG00000137040 | 9 | 6011043 | 6015618 | 6 | 2 | -0.63458 | 0.73715 | RANBP6 | | 0.73715 | 0.5228 | 0.65672 |
| ENSG00000261701 | 16 | 72088522 | 72111145 | 70 | 14 | -0.63472 | 0.73719 | HPR | | 0.73719 | 0.68518 | 0.66223 |
| ENSG00000106123 | 7 | 142552792 | 142568847 | 45 | 8 | -0.63474 | 0.7372 | EPHB6 | | 0.7372 | 0.68025 | 0.44522 |

| | | | | | | | | | | | |
|---|---|---|---|---|---|---|---|---|---|---|---|
| ENSG00000160323 | 9 | 136279478 | 136324508 | 165 | 21 | -0.63483 | 0.73723 | ADAMTS13 | 0.73723 | 0.63683 | 0.55085 |
| ENSG00000146243 | 6 | 79577189 | 79656157 | 179 | 19 | -0.63487 | 0.73724 | IRAK1BP1 | 0.73724 | 0.71968 | 0.72469 |
| ENSG00000178163 | 4 | 10441498 | 10459034 | 53 | 17 | -0.63499 | 0.73728 | ZNF518B | 0.73728 | 0.74894 | 0.77262 |
| ENSG00000179532 | 11 | 6518490 | 6614988 | 305 | 39 | -0.63502 | 0.73729 | DNHD1 | 0.73729 | 0.6776 | 0.49645 |
| ENSG00000101349 | 20 | 9518036 | 9819689 | 1137 | 90 | -0.635 | 0.73729 | PAK7 | 0.73729 | 0.73238 | 0.73676 |
| ENSG00000151304 | 5 | 121297656 | 121411265 | 239 | 14 | -0.63503 | 0.7373 | SRFBP1 | 0.7373 | 0.6743 | 0.71494 |
| ENSG00000135299 | 6 | 90142889 | 90343553 | 409 | 40 | -0.63504 | 0.7373 | ANKRD6 | 0.7373 | 0.75778 | 0.6954 |
| ENSG00000172732 | 11 | 65624597 | 65635124 | 28 | 4 | -0.63506 | 0.73731 | MUS81 | 0.73731 | 0.80248 | 0.85891 |
| ENSG00000133321 | 11 | 63304281 | 63313934 | 9 | 5 | -0.6352 | 0.73735 | RARRES3 | 0.73735 | 0.98463 | 0.98868 |
| ENSG00000157219 | 7 | 154862034 | 154877459 | 69 | 11 | -0.63553 | 0.73746 | HTR5A | 0.73746 | 0.65287 | 0.67739 |
| ENSG00000029364 | 14 | 69864732 | 69929105 | 167 | 21 | -0.63614 | 0.73766 | SLC39A9 | 0.73766 | 0.71425 | 0.78668 |
| ENSG00000163161 | 2 | 128014866 | 128051752 | 101 | 19 | -0.63642 | 0.73775 | ERCC3 | 0.73775 | 0.75636 | 0.76694 |
| ENSG00000197901 | 11 | 62703857 | 62752455 | 130 | 16 | -0.63654 | 0.73779 | SLC22A6 | 0.73779 | 0.72192 | 0.72365 |
| ENSG00000103274 | 16 | 10837643 | 10863208 | 77 | 13 | -0.63669 | 0.73784 | NUBP1 | 0.73784 | 0.76111 | 0.80315 |
| ENSG00000158158 | 2 | 97426639 | 97477628 | 88 | 11 | -0.6372 | 0.738 | CNNM4 | 0.738 | 0.67826 | 0.67695 |
| ENSG00000131876 | 15 | 101821715 | 101835487 | 47 | 5 | -0.63728 | 0.73803 | SNRPA1 | 0.73803 | 0.71671 | 0.68585 |
| ENSG00000105011 | 19 | 14230321 | 14247768 | 38 | 4 | -0.6374 | 0.73807 | ASF1B | 0.73807 | 0.73482 | 0.77424 |
| ENSG00000114491 | 3 | 124449213 | 124464040 | 73 | 12 | -0.63753 | 0.73811 | UMPS | 0.73811 | 0.72293 | 0.84077 |
| ENSG00000085662 | 7 | 134127102 | 134144036 | 44 | 12 | -0.63756 | 0.73812 | AKR1B1 | 0.73812 | 0.83562 | 0.66501 |
| ENSG00000167005 | 16 | 56463045 | 56486111 | 37 | 8 | -0.6376 | 0.73813 | NUDT21 | 0.73813 | 0.65978 | 0.62518 |
| ENSG00000106799 | 9 | 101866320 | 101916474 | 103 | 12 | -0.63768 | 0.73816 | TGFBR1 | 0.73816 | 0.74156 | 0.67718 |
| ENSG00000160471 | 19 | 55860674 | 55866182 | 11 | 7 | -0.63794 | 0.73824 | COX6B2 | 0.73824 | 0.50197 | 0.47902 |
| ENSG00000204642 | 6 | 29690552 | 29706305 | 181 | 19 | -0.63851 | 0.73843 | HLA-F | 0.73843 | 0.75854 | 0.83674 |
| ENSG00000167741 | 17 | 4460222 | 4464113 | 15 | 7 | -0.63883 | 0.73853 | GGT6 | 0.73853 | 0.74235 | 0.3464 |
| ENSG00000214097 | 3 | 196233748 | 196242237 | 28 | 6 | -0.63886 | 0.73854 | SMCO1 | 0.73854 | 0.86557 | 0.94319 |
| ENSG00000164172 | 5 | 52391509 | 52405893 | 63 | 7 | -0.63892 | 0.73856 | MOCS2 | 0.73856 | 0.82251 | 0.90409 |
| ENSG00000268365 | 1 | 240629183 | 240643504 | 64 | 8 | -0.63893 | 0.73857 | AL646016.1 | 0.73857 | 0.63381 | 0.73582 |
| ENSG00000203883 | 20 | 62679076 | 62680994 | 1 | 1 | -0.63904 | 0.7386 | SOX18 | 0.7386 | 0.70714 | 0.80885 |
| ENSG00000101109 | 20 | 43595115 | 43708600 | 256 | 13 | -0.63911 | 0.73862 | STK4 | 0.73862 | 0.74089 | 0.76707 |
| ENSG00000160318 | 19 | 51870352 | 51872257 | 1 | 1 | -0.63996 | 0.7389 | CLDND2 | 0.7389 | 0.16871 | 0.52227 |
| ENSG00000196557 | 16 | 1203241 | 1271771 | 303 | 55 | -0.64013 | 0.73896 | CACNA1H | 0.73896 | 0.85171 | 0.88784 |
| ENSG00000213977 | 17 | 3566196 | 3571976 | 17 | 5 | -0.64014 | 0.73896 | TAX1BP3 | 0.73896 | 0.77962 | 0.7581 |
| ENSG00000176753 | 15 | 40542865 | 40545170 | 7 | 1 | -0.64054 | 0.73909 | C15orf56 | 0.73909 | 0.64397 | 0.36835 |
| ENSG00000170325 | 11 | 129769601 | 129872730 | 328 | 36 | -0.64057 | 0.7391 | PRDM10 | 0.7391 | 0.66352 | 0.49633 |
| ENSG00000156232 | 15 | 83478380 | 83503611 | 83 | 18 | -0.64057 | 0.7391 | WHAMM | 0.7391 | 0.75357 | 0.74374 |
| ENSG00000268800 | 5 | 162866447 | 162873821 | 13 | 4 | -0.64065 | 0.73913 | AC112205.1 | 0.73913 | 0.79407 | 0.86039 |
| ENSG00000161395 | 17 | 37827375 | 37853050 | 82 | 6 | -0.64068 | 0.73913 | PGAP3 | 0.73913 | 0.82766 | 0.88858 |
| ENSG00000106328 | 7 | 127231463 | 127242198 | 36 | 9 | -0.64076 | 0.73916 | FSCN3 | 0.73916 | 0.70355 | 0.74982 |
| ENSG00000158485 | 1 | 158297741 | 158301321 | 11 | 5 | -0.64089 | 0.7392 | CD1B | 0.7392 | 0.8811 | 0.8492 |
| ENSG00000018869 | 19 | 56887413 | 56904914 | 63 | 21 | -0.64101 | 0.73924 | ZNF582 | 0.73924 | 0.84188 | 0.65221 |
| ENSG00000141946 | 19 | 57645464 | 57656570 | 58 | 8 | -0.64105 | 0.73925 | ZIM3 | 0.73925 | 0.84379 | 0.80406 |
| ENSG00000130940 | 1 | 10696661 | 10856707 | 525 | 80 | -0.64105 | 0.73926 | CASZ1 | 0.73926 | 0.74577 | 0.8105 |
| ENSG00000168131 | 6 | 27878963 | 27880174 | 3 | 1 | -0.64111 | 0.73927 | OR2B2 | 0.73927 | 0.9391 | 0.97437 |
| ENSG00000184330 | 1 | 153389000 | 153395701 | 17 | 3 | -0.64144 | 0.73938 | S100A7A | 0.73938 | 0.57623 | 0.45568 |
| ENSG00000100558 | 14 | 67853700 | 67878917 | 96 | 15 | -0.64144 | 0.73938 | PLEK2 | 0.73938 | 0.74087 | 0.68514 |
| ENSG00000121406 | 19 | 58038693 | 58068910 | 32 | 11 | -0.64146 | 0.73939 | ZNF549 | 0.73939 | 0.43738 | 0.47899 |
| ENSG00000171502 | 1 | 86194916 | 86622626 | 1820 | 61 | -0.64149 | 0.7394 | COL24A1 | 0.7394 | 0.76509 | 0.78757 |
| ENSG00000180914 | 3 | 8792094 | 8811314 | 92 | 24 | -0.64154 | 0.73941 | OXTR | 0.73941 | 0.86401 | 0.90276 |
| ENSG00000221874 | 19 | 53431728 | 53466076 | 170 | 27 | -0.64158 | 0.73943 | ZNF321P | 0.73943 | 0.85854 | 0.92138 |
| ENSG00000159917 | 19 | 44732882 | 44809199 | 161 | 16 | -0.64178 | 0.73949 | ZNF235 | 0.73949 | 0.71808 | 0.70524 |
| ENSG00000268744 | 19 | 12490003 | 12512088 | 46 | 7 | -0.64191 | 0.73953 | CTD-3105H18.14 | 0.73953 | 0.46824 | 0.16279 |
| ENSG00000266258 | 18 | 68002675 | 68019695 | 67 | 13 | -0.64224 | 0.73964 | RP11-41O4.1 | 0.73964 | 0.85688 | 0.84976 |
| ENSG00000204514 | 19 | 58360099 | 58400442 | 137 | 15 | -0.64232 | 0.73967 | ZNF814 | 0.73967 | 0.70738 | 0.70274 |
| ENSG00000248511 | 4 | 93218402 | 93225329 | 7 | 2 | -0.64257 | 0.73975 | RP11-9B6.1 | 0.73975 | 0.85316 | 0.86995 |
| ENSG00000184302 | 14 | 60975669 | 60979568 | 5 | 3 | -0.64259 | 0.73975 | SIX6 | 0.73975 | 0.77242 | 0.79132 |
| ENSG00000163686 | 3 | 58223233 | 58281420 | 263 | 17 | -0.64262 | 0.73977 | ABHD6 | 0.73977 | 0.64418 | 0.64393 |
| ENSG00000163762 | 3 | 149036285 | 149052201 | 50 | 15 | -0.64309 | 0.73992 | TM4SF18 | 0.73992 | 0.78282 | 0.69249 |
| ENSG00000166183 | 14 | 104552016 | 104579098 | 101 | 18 | -0.64313 | 0.73993 | ASPG | 0.73993 | 0.58363 | 0.51826 |
| ENSG00000149591 | 11 | 117070037 | 117075498 | 21 | 6 | -0.64322 | 0.73996 | TAGLN | 0.73996 | 0.60416 | 0.57898 |
| ENSG00000172270 | 19 | 571297 | 583493 | 109 | 14 | -0.64338 | 0.74001 | BSG | 0.74001 | 0.78439 | 0.56575 |
| ENSG00000180628 | 10 | 92979908 | 93044088 | 170 | 8 | -0.64346 | 0.74004 | PCGF5 | 0.74004 | 0.63039 | 0.34412 |
| ENSG00000151332 | 14 | 36767770 | 36789882 | 42 | 5 | -0.64349 | 0.74005 | MBIP | 0.74005 | 0.77241 | 0.71597 |
| ENSG00000198758 | 1 | 110292702 | 110306649 | 47 | 14 | -0.64353 | 0.74006 | EPS8L3 | 0.74006 | 0.83664 | 0.89092 |
| ENSG00000104140 | 15 | 41164412 | 41166487 | 2 | 1 | -0.64358 | 0.74008 | RHOV | 0.74008 | 0.54018 | 0.72381 |
| ENSG00000101188 | 20 | 61340189 | 61394123 | 279 | 30 | -0.64359 | 0.74008 | NTSR1 | 0.74008 | 0.75513 | 0.66353 |
| ENSG00000150594 | 10 | 112836790 | 112840658 | 7 | 4 | -0.64372 | 0.74012 | ADRA2A | 0.74012 | 0.90845 | 0.9324 |
| ENSG00000132950 | 13 | 20397622 | 20437776 | 154 | 9 | -0.64399 | 0.74021 | ZMYM5 | 0.74021 | 0.7512 | 0.77247 |
| ENSG00000163793 | 2 | 27498289 | 27504367 | 14 | 6 | -0.64437 | 0.74033 | DNAJC5G | 0.74033 | 0.76758 | 0.7897 |
| ENSG00000168672 | 8 | 127564687 | 127570638 | 6 | 3 | -0.64436 | 0.74033 | FAM84B | 0.74033 | 0.88565 | 0.75537 |
| ENSG00000258740 | 14 | 75705319 | 75735986 | 62 | 16 | -0.64458 | 0.7404 | RP11-293M10.1 | 0.7404 | 0.4273 | 0.2102 |
| ENSG00000198104 | 1 | 248550910 | 248551836 | 6 | 2 | -0.6446 | 0.74041 | OR2T6 | 0.74041 | 0.87755 | 0.7909 |
| ENSG00000187049 | 11 | 61159159 | 61166335 | 13 | 4 | -0.6451 | 0.74057 | TMEM216 | 0.74057 | 0.62976 | 0.63625 |
| ENSG00000125337 | 6 | 168396921 | 168445769 | 266 | 33 | -0.64532 | 0.74064 | KIF25 | 0.74064 | 0.69877 | 0.77994 |

| | | | | | | | | | | | | |
|---|---|---|---|---|---|---|---|---|---|---|---|---|
| ENSG00000117598 | 1 | 99355806 | 99470588 | 505 | 46 | -0.64557 | 0.74072 | LPPR5 | | 0.74072 | 0.68941 | 0.57561 |
| ENSG00000103507 | 16 | 31117428 | 31124110 | 9 | 3 | -0.64563 | 0.74074 | BCKDK | | 0.74074 | 0.66054 | 0.77484 |
| ENSG00000129595 | 5 | 111478138 | 111755013 | 973 | 52 | -0.64567 | 0.74075 | EPB41L4A | | 0.74075 | 0.73666 | 0.73753 |
| ENSG00000137726 | 11 | 117707693 | 117748201 | 107 | 26 | -0.64569 | 0.74076 | FXYD6 | | 0.74076 | 0.72721 | 0.79253 |
| ENSG00000106012 | 7 | 2598632 | 2654368 | 334 | 26 | -0.64574 | 0.74078 | IQCE | | 0.74078 | 0.6084 | 0.50784 |
| ENSG00000181924 | 11 | 73583712 | 73588033 | 12 | 6 | -0.64579 | 0.74079 | COA4 | | 0.74079 | 0.87458 | 0.91122 |
| ENSG00000203727 | 6 | 147830063 | 148058683 | 1187 | 75 | -0.64583 | 0.7408 | SAMD5 | | 0.7408 | 0.74413 | 0.77977 |
| ENSG00000179388 | 8 | 22545172 | 22550815 | 8 | 4 | -0.64588 | 0.74082 | EGR3 | | 0.74082 | 0.94698 | 0.76943 |
| ENSG00000236311 | 10 | 102849078 | 102890883 | 148 | 19 | -0.64593 | 0.74084 | TLX1NB | | 0.74084 | 0.767 | 0.75823 |
| ENSG00000167703 | 17 | 1472561 | 1532180 | 298 | 61 | -0.64598 | 0.74085 | SLC43A2 | | 0.74085 | 0.51268 | 0.30862 |
| ENSG00000137955 | 1 | 76251879 | 76260764 | 29 | 9 | -0.64625 | 0.74094 | RABGGTB | | 0.74094 | 0.78816 | 0.77658 |
| ENSG00000170085 | 5 | 175665365 | 175772994 | 325 | 19 | -0.64638 | 0.74098 | SIMC1 | | 0.74098 | 0.71483 | 0.65598 |
| ENSG00000115241 | 2 | 27604061 | 27632554 | 56 | 6 | -0.64659 | 0.74105 | PPM1G | | 0.74105 | 0.7598 | 0.74516 |
| ENSG00000136848 | 9 | 124329336 | 124547809 | 525 | 70 | -0.64657 | 0.74105 | DAB2IP | | 0.74105 | 0.62636 | 0.45621 |
| ENSG00000159251 | 15 | 35080297 | 35088340 | 26 | 14 | -0.64658 | 0.74105 | ACTC1 | | 0.74105 | 0.011909 | 0.017271 |
| ENSG00000104884 | 19 | 45853095 | 45874176 | 82 | 13 | -0.64689 | 0.74115 | ERCC2 | | 0.74115 | 0.70807 | 0.78266 |
| ENSG00000144455 | 3 | 3742498 | 4508965 | 3434 | 126 | -0.64698 | 0.74118 | SUMF1 | | 0.74118 | 0.7832 | 0.79215 |
| ENSG00000089177 | 20 | 16252749 | 16554078 | 1233 | 58 | -0.6471 | 0.74121 | KIF16B | | 0.74121 | 0.74185 | 0.62779 |
| ENSG00000176018 | 5 | 89811428 | 89825401 | 36 | 5 | -0.64729 | 0.74128 | LYSMD3 | | 0.74128 | 0.6373 | 0.69318 |
| ENSG00000076650 | 19 | 33571786 | 33621448 | 179 | 13 | -0.64732 | 0.74129 | GPATCH1 | | 0.74129 | 0.78216 | 0.84006 |
| ENSG00000187954 | 8 | 145674965 | 145691060 | 21 | 10 | -0.64746 | 0.74133 | CYHR1 | | 0.74133 | 0.14157 | 0.085614 |
| ENSG00000161973 | 17 | 8633252 | 8648537 | 61 | 8 | -0.64752 | 0.74135 | CCDC42 | | 0.74135 | 0.83502 | 0.9103 |
| ENSG00000104852 | 19 | 49588676 | 49611862 | 61 | 18 | -0.64801 | 0.74151 | SNRNP70 | | 0.74151 | 0.62981 | 0.75839 |
| ENSG00000179021 | 3 | 88198893 | 88217879 | 53 | 4 | -0.64863 | 0.74171 | C3orf38 | | 0.74171 | 0.73738 | 0.76744 |
| ENSG00000132702 | 1 | 156589086 | 156595517 | 10 | 2 | -0.64868 | 0.74173 | HAPLN2 | | 0.74173 | 0.81007 | 0.75799 |
| ENSG00000165120 | 7 | 129847700 | 129856683 | 16 | 6 | -0.64869 | 0.74173 | SSMEM1 | | 0.74173 | 0.82405 | 0.83883 |
| ENSG00000151704 | 11 | 128706210 | 128737268 | 86 | 15 | -0.64871 | 0.74174 | KCNJ1 | | 0.74174 | 0.88708 | 0.72924 |
| ENSG00000189114 | 19 | 45682003 | 45685059 | 6 | 2 | -0.64871 | 0.74174 | BLOC1S3 | | 0.74174 | 0.83506 | 0.72941 |
| ENSG00000228474 | 2 | 27293340 | 27294641 | 1 | 1 | -0.6489 | 0.7418 | OST4 | | 0.7418 | 0.71402 | 0.67512 |
| ENSG00000145494 | 5 | 1801514 | 1816719 | 65 | 9 | -0.64891 | 0.7418 | NDUFS6 | | 0.7418 | 0.76325 | 0.72399 |
| ENSG00000171133 | 9 | 114089763 | 114090800 | 2 | 1 | -0.64896 | 0.74182 | OR2K2 | | 0.74182 | 0.7295 | 0.48058 |
| ENSG00000160460 | 19 | 40972148 | 41082370 | 322 | 46 | -0.64932 | 0.74193 | SPTBN4 | | 0.74193 | 0.64397 | 0.54468 |
| ENSG00000115993 | 2 | 202241930 | 202316302 | 181 | 15 | -0.64937 | 0.74195 | TRAK2 | | 0.74195 | 0.79475 | 0.83561 |
| ENSG00000112964 | 5 | 42423879 | 42721979 | 637 | 38 | -0.64956 | 0.74201 | GHR | | 0.74201 | 0.73088 | 0.76318 |
| ENSG00000099785 | 19 | 8478154 | 8503901 | 105 | 22 | -0.64955 | 0.74201 | | 43892 | 0.74201 | 0.72233 | 0.43796 |
| ENSG00000114631 | 3 | 127348024 | 127391652 | 44 | 14 | -0.6497 | 0.74206 | PODXL2 | | 0.74206 | 0.7971 | 0.78885 |
| ENSG00000023839 | 10 | 101542489 | 101611949 | 235 | 21 | -0.64974 | 0.74207 | ABCC2 | | 0.74207 | 0.76148 | 0.7222 |
| ENSG00000119906 | 10 | 102672326 | 102724893 | 106 | 16 | -0.64976 | 0.74208 | FAM178A | | 0.74208 | 0.78653 | 0.82341 |
| ENSG00000149273 | 11 | 75110530 | 75133324 | 47 | 11 | -0.64979 | 0.74209 | RPS3 | | 0.74209 | 0.80986 | 0.77975 |
| ENSG00000133466 | 22 | 37576207 | 37595425 | 86 | 11 | -0.64985 | 0.74211 | C1QTNF6 | | 0.74211 | 0.74679 | 0.75989 |
| ENSG00000111450 | 12 | 131274145 | 131323811 | 183 | 11 | -0.64995 | 0.74214 | STX2 | | 0.74214 | 0.847 | 0.90236 |
| ENSG00000228300 | 19 | 1275026 | 1279248 | 17 | 7 | -0.65038 | 0.74228 | C19orf24 | | 0.74228 | 0.6496 | 0.0928 |
| ENSG00000164989 | 9 | 15552895 | 16061661 | 1848 | 20 | -0.65085 | 0.74243 | CCDC171 | | 0.74243 | 0.76185 | 0.78817 |
| ENSG00000172159 | 9 | 85857905 | 86153461 | 1319 | 87 | -0.65085 | 0.74243 | FRMD3 | | 0.74243 | 0.76916 | 0.80975 |
| ENSG00000134851 | 4 | 56262124 | 56319564 | 209 | 19 | -0.65094 | 0.74246 | TMEM165 | | 0.74246 | 0.79441 | 0.79424 |
| ENSG00000164136 | 4 | 142557752 | 142655140 | 174 | 20 | -0.65102 | 0.74248 | IL15 | | 0.74248 | 0.61427 | 0.55644 |
| ENSG00000104147 | 15 | 41601466 | 41624819 | 76 | 9 | -0.6512 | 0.74254 | OIP5 | | 0.74254 | 0.74298 | 0.76805 |
| ENSG00000205609 | 16 | 28390900 | 28415200 | 7 | 3 | -0.65173 | 0.74271 | EIF3CL | | 0.74271 | 0.64481 | 0.60211 |
| ENSG00000134852 | 4 | 56294070 | 56413305 | 521 | 14 | -0.65181 | 0.74274 | CLOCK | | 0.74274 | 0.73734 | 0.7662 |
| ENSG00000135972 | 2 | 105654441 | 105716418 | 275 | 8 | -0.65207 | 0.74282 | MRPS9 | | 0.74282 | 0.72844 | 0.72809 |
| ENSG00000105705 | 19 | 19386827 | 19431653 | 101 | 11 | -0.6521 | 0.74283 | SUGP1 | | 0.74283 | 0.73324 | 0.70948 |
| ENSG00000134443 | 18 | 56887400 | 56898006 | 47 | 4 | -0.65213 | 0.74284 | GRP | | 0.74284 | 0.8781 | 0.76651 |
| ENSG00000264324 | 2 | 74438731 | 74590504 | 237 | 40 | -0.65245 | 0.74294 | RP11-287D1.3 | | 0.74294 | 0.72848 | 0.69481 |
| ENSG00000165131 | 7 | 142636440 | 142637955 | 5 | 1 | -0.65256 | 0.74298 | C7orf34 | | 0.74298 | 0.51629 | 0.59252 |
| ENSG00000127377 | 7 | 151125921 | 151137899 | 30 | 9 | -0.65277 | 0.74305 | CRYGN | | 0.74305 | 0.83831 | 0.94194 |
| ENSG00000243627 | 21 | 35791013 | 35796264 | 11 | 4 | -0.65311 | 0.74316 | AP000322.53 | | 0.74316 | 0.68721 | 0.75654 |
| ENSG00000167395 | 16 | 31085743 | 31095517 | 15 | 4 | -0.65315 | 0.74317 | ZNF646 | | 0.74317 | 0.83506 | 0.85176 |
| ENSG00000123297 | 12 | 58176372 | 58201854 | 41 | 7 | -0.65327 | 0.74321 | TSFM | | 0.74321 | 0.74906 | 0.69822 |
| ENSG00000113209 | 5 | 140514800 | 140517703 | 6 | 3 | -0.65343 | 0.74326 | PCDHB5 | | 0.74326 | 0.83847 | 0.80569 |
| ENSG00000112299 | 6 | 133002729 | 133035188 | 177 | 15 | -0.65341 | 0.74326 | VNN1 | | 0.74326 | 0.63439 | 0.58104 |
| ENSG00000165923 | 11 | 47681143 | 47736941 | 130 | 15 | -0.65351 | 0.74329 | AGBL2 | | 0.74329 | 0.71826 | 0.72423 |
| ENSG00000151418 | 1 | 198492352 | 198510075 | 37 | 10 | -0.65357 | 0.74331 | ATP6V1G3 | | 0.74331 | 0.80243 | 0.81705 |
| ENSG00000181585 | 3 | 46742823 | 46752376 | 29 | 8 | -0.65379 | 0.74338 | TMIE | | 0.74338 | 0.75981 | 0.55935 |
| ENSG00000182670 | 21 | 38445526 | 38575413 | 493 | 18 | -0.65379 | 0.74338 | TTC3 | | 0.74338 | 0.73139 | 0.73225 |
| ENSG00000088538 | 3 | 50712672 | 51421629 | 1350 | 23 | -0.65383 | 0.74339 | DOCK3 | | 0.74339 | 0.74609 | 0.75883 |
| ENSG00000181634 | 9 | 117546915 | 117568406 | 34 | 11 | -0.65414 | 0.74349 | TNFSF15 | | 0.74349 | 0.79173 | 0.76346 |
| ENSG00000162373 | 1 | 49193195 | 49242641 | 89 | 17 | -0.65436 | 0.74356 | BEND5 | | 0.74356 | 0.70691 | 0.40332 |
| ENSG00000114541 | 3 | 69219141 | 69591734 | 1402 | 94 | -0.65439 | 0.74357 | FRMD4B | | 0.74357 | 0.76707 | 0.77479 |
| ENSG00000117533 | 1 | 171669300 | 171711387 | 123 | 11 | -0.65446 | 0.74359 | VAMP4 | | 0.74359 | 0.77041 | 0.79221 |
| ENSG00000258247 | 12 | 95770984 | 95774672 | 38 | 6 | -0.65444 | 0.74359 | RP11-167N24.6 | | 0.74359 | 0.87524 | 0.90398 |
| ENSG00000230185 | 9 | 115196096 | 115249484 | 220 | 11 | -0.65485 | 0.74372 | C9orf147 | | 0.74372 | 0.78832 | 0.83039 |
| ENSG00000130775 | 1 | 28199055 | 28213196 | 54 | 10 | -0.655 | 0.74377 | THEMIS2 | | 0.74377 | 0.69063 | 0.69753 |
| ENSG00000136274 | 7 | 45120037 | 45128513 | 32 | 7 | -0.65506 | 0.74378 | NACAD | | 0.74378 | 0.81406 | 0.78478 |

| | | | | | | | | | | | |
|---|---|---|---|---|---|---|---|---|---|---|---|
| ENSG00000198538 | 19 | 53300662 | 53360853 | 467 | 39 | -0.65518 | 0.74383 | ZNF28 | 0.74383 | 0.64092 | 0.62009 |
| ENSG00000143442 | 1 | 151375200 | 151431941 | 118 | 11 | -0.65536 | 0.74388 | POGZ | 0.74388 | 0.7392 | 0.75468 |
| ENSG00000204397 | 11 | 104912053 | 104972158 | 191 | 17 | -0.65542 | 0.7439 | CARD16 | 0.7439 | 0.81033 | 0.84879 |
| ENSG00000197746 | 10 | 73576055 | 73611126 | 147 | 22 | -0.65558 | 0.74395 | PSAP | 0.74395 | 0.84643 | 0.88791 |
| ENSG00000169504 | 1 | 25071848 | 25170815 | 135 | 14 | -0.6557 | 0.74399 | CLIC4 | 0.74399 | 0.72896 | 0.64862 |
| ENSG00000198793 | 1 | 11166592 | 11322564 | 398 | 12 | -0.65573 | 0.744 | MTOR | 0.744 | 0.73062 | 0.732 |
| ENSG00000051128 | 19 | 19040010 | 19052070 | 32 | 10 | -0.65586 | 0.74404 | HOMER3 | 0.74404 | 0.46306 | 0.41771 |
| ENSG00000164597 | 7 | 106842000 | 107204959 | 948 | 22 | -0.65593 | 0.74406 | COG5 | 0.74406 | 0.76374 | 0.76157 |
| ENSG00000099910 | 22 | 20783528 | 20850170 | 182 | 17 | -0.65592 | 0.74406 | KLHL22 | 0.74406 | 0.81157 | 0.80792 |
| ENSG00000240065 | 6 | 32811913 | 32827362 | 134 | 17 | -0.65597 | 0.74408 | PSMB9 | 0.74408 | 0.62847 | 0.32044 |
| ENSG00000103415 | 16 | 4524691 | 4560348 | 117 | 9 | -0.65604 | 0.7441 | HMOX2 | 0.7441 | 0.65546 | 0.69682 |
| ENSG00000225683 | 6 | 163586446 | 163612840 | 88 | 15 | -0.65609 | 0.74412 | AL078585.1 | 0.74412 | 0.89155 | 0.96612 |
| ENSG00000103226 | 16 | 16326352 | 16388668 | 82 | 8 | -0.65612 | 0.74413 | NOMO3 | 0.74413 | 0.75432 | 0.77228 |
| ENSG00000152990 | 4 | 22346694 | 22517677 | 645 | 47 | -0.65621 | 0.74415 | GPR125 | 0.74415 | 0.67053 | 0.86972 |
| ENSG00000136169 | 13 | 50018429 | 50069138 | 180 | 18 | -0.65638 | 0.74421 | SETDB2 | 0.74421 | 0.76492 | 0.80152 |
| ENSG00000136891 | 9 | 103064359 | 103115221 | 143 | 13 | -0.6566 | 0.74428 | TEX10 | 0.74428 | 0.73455 | 0.6853 |
| ENSG00000103091 | 16 | 74907468 | 75034071 | 488 | 54 | -0.65673 | 0.74432 | WDR59 | 0.74432 | 0.71382 | 0.63279 |
| ENSG00000078070 | 3 | 182733006 | 182833863 | 248 | 19 | -0.6569 | 0.74438 | MCCC1 | 0.74438 | 0.75958 | 0.7647 |
| ENSG00000171604 | 5 | 139026884 | 139063467 | 40 | 11 | -0.65697 | 0.7444 | CXXC5 | 0.7444 | 0.68065 | 0.67296 |
| ENSG00000103018 | 16 | 69458428 | 69500169 | 165 | 20 | -0.65711 | 0.74444 | CYB5B | 0.74444 | 0.75413 | 0.66965 |
| ENSG00000240694 | 8 | 26362202 | 26371608 | 20 | 7 | -0.65714 | 0.74445 | PNMA2 | 0.74445 | 0.84862 | 0.87036 |
| ENSG00000128311 | 22 | 37406900 | 37415681 | 48 | 9 | -0.65742 | 0.74454 | TST | 0.74454 | 0.54179 | 0.28466 |
| ENSG00000164500 | 7 | 50135632 | 50199426 | 241 | 18 | -0.65746 | 0.74456 | C7orf72 | 0.74456 | 0.78323 | 0.80408 |
| ENSG00000051009 | 11 | 6232565 | 6255941 | 57 | 10 | -0.65756 | 0.74459 | FAM160A2 | 0.74459 | 0.86038 | 0.94301 |
| ENSG00000149823 | 11 | 64856796 | 64879332 | 40 | 11 | -0.65755 | 0.74459 | VPS51 | 0.74459 | 0.77861 | 0.8151 |
| ENSG00000256053 | 14 | 104029299 | 104073860 | 140 | 15 | -0.65778 | 0.74466 | APOPT1 | 0.74466 | 0.76211 | 0.82968 |
| ENSG00000164128 | 4 | 164245113 | 164265984 | 70 | 14 | -0.65789 | 0.7447 | NPY1R | 0.7447 | 0.7911 | 0.91469 |
| ENSG00000118729 | 1 | 116242628 | 116311402 | 373 | 22 | -0.65805 | 0.74475 | CASQ2 | 0.74475 | 0.8057 | 0.78359 |
| ENSG00000174579 | 3 | 135867764 | 135916083 | 76 | 15 | -0.65812 | 0.74477 | MSL2 | 0.74477 | 0.74521 | 0.80079 |
| ENSG00000184954 | 12 | 55862984 | 55863922 | 3 | 2 | -0.65814 | 0.74477 | OR6C70 | 0.74477 | 0.81394 | 0.82092 |
| ENSG00000118193 | 1 | 200520628 | 200589862 | 226 | 24 | -0.65844 | 0.74487 | KIF14 | 0.74487 | 0.7305 | 0.76319 |
| ENSG00000169372 | 12 | 94071151 | 94288616 | 643 | 58 | -0.65847 | 0.74488 | CRADD | 0.74488 | 0.73018 | 0.64581 |
| ENSG00000068001 | 3 | 50355221 | 50360337 | 3 | 2 | -0.65892 | 0.74503 | HYAL2 | 0.74503 | 0.36695 | 0.30524 |
| ENSG00000196341 | 11 | 124179708 | 124180733 | 5 | 2 | -0.65904 | 0.74506 | OR8D1 | 0.74506 | 0.50145 | 0.6283 |
| ENSG00000135686 | 16 | 84682131 | 84701292 | 100 | 17 | -0.65937 | 0.74517 | KLHL36 | 0.74517 | 0.65614 | 0.63001 |
| ENSG00000197614 | 12 | 8789942 | 8815484 | 94 | 7 | -0.65948 | 0.74521 | MFAP5 | 0.74521 | 0.71428 | 0.75791 |
| ENSG00000038274 | 5 | 162930120 | 162946342 | 50 | 11 | -0.65962 | 0.74525 | MAT2B | 0.74525 | 0.56435 | 0.65873 |
| ENSG00000106714 | 9 | 39072764 | 39288312 | 419 | 18 | -0.65963 | 0.74526 | CNTNAP3 | 0.74526 | 0.74789 | 0.73554 |
| ENSG00000172780 | 3 | 128806412 | 128841644 | 42 | 9 | -0.65978 | 0.7453 | RAB43 | 0.7453 | 0.67928 | 0.6832 |
| ENSG00000175115 | 11 | 65837834 | 66012218 | 357 | 20 | -0.65986 | 0.74533 | PACS1 | 0.74533 | 0.71816 | 0.69928 |
| ENSG00000129221 | 17 | 6297013 | 6338519 | 266 | 40 | -0.65997 | 0.74536 | AIPL1 | 0.74536 | 0.89398 | 0.92145 |
| ENSG00000166105 | 11 | 134144139 | 134189458 | 165 | 28 | -0.66031 | 0.74547 | GLB1L3 | 0.74547 | 0.79261 | 0.90244 |
| ENSG00000257482 | 7 | 63505821 | 63538927 | 176 | 11 | -0.66047 | 0.74552 | ZNF727 | 0.74552 | 0.73977 | 0.71371 |
| ENSG00000181830 | 11 | 45825623 | 45834566 | 32 | 10 | -0.66061 | 0.74557 | SLC35C1 | 0.74557 | 0.8908 | 0.84696 |
| ENSG00000272047 | 6 | 158589384 | 158620376 | 82 | 11 | -0.66079 | 0.74563 | GTF2H5 | 0.74563 | 0.70738 | 0.78499 |
| ENSG00000261611 | 16 | 71481503 | 71523214 | 76 | 14 | -0.66087 | 0.74565 | AC010547.9 | 0.74565 | 0.72371 | 0.77917 |
| ENSG00000067606 | 1 | 1981909 | 2116834 | 499 | 39 | -0.66097 | 0.74568 | PRKCZ | 0.74568 | 0.71647 | 0.71945 |
| ENSG00000215346 | 8 | 10983980 | 10987745 | 13 | 4 | -0.66119 | 0.74575 | AF131215.5 | 0.74575 | 0.86604 | 0.8086 |
| ENSG00000137818 | 15 | 69745123 | 69748255 | 6 | 3 | -0.66124 | 0.74577 | RPLP1 | 0.74577 | 0.47328 | 0.40198 |
| ENSG00000133424 | 22 | 33558212 | 34318829 | 3118 | 204 | -0.66129 | 0.74579 | LARGE | 0.74579 | 0.76381 | 0.80784 |
| ENSG00000170689 | 17 | 46698518 | 46703839 | 15 | 4 | -0.66148 | 0.74585 | HOXB9 | 0.74585 | 0.85499 | 0.88763 |
| ENSG00000143556 | 1 | 153430220 | 153433177 | 8 | 3 | -0.66154 | 0.74587 | S100A7 | 0.74587 | 0.87051 | 0.88459 |
| ENSG00000105516 | 19 | 49133287 | 49140695 | 14 | 4 | -0.66164 | 0.7459 | DBP | 0.7459 | 0.41332 | 0.11416 |
| ENSG00000120337 | 1 | 173009100 | 173020103 | 40 | 8 | -0.66173 | 0.74593 | TNFSF18 | 0.74593 | 0.72737 | 0.63921 |
| ENSG00000258792 | 14 | 90095265 | 90109015 | 42 | 9 | -0.66196 | 0.746 | RP11-944C7.1 | 0.746 | 0.6217 | 0.58315 |
| ENSG00000065809 | 10 | 14560556 | 14816896 | 1151 | 99 | -0.66206 | 0.74603 | FAM107B | 0.74603 | 0.7012 | 0.60817 |
| ENSG00000234237 | 1 | 180941695 | 180949868 | 14 | 2 | -0.66207 | 0.74604 | AL162431.1 | 0.74604 | 0.85553 | 0.87742 |
| ENSG00000189180 | 10 | 38299578 | 38354016 | 171 | 17 | -0.66212 | 0.74605 | ZNF33A | 0.74605 | 0.70326 | 0.68994 |
| ENSG00000139697 | 12 | 123773656 | 123849390 | 311 | 15 | -0.66224 | 0.74609 | SBNO1 | 0.74609 | 0.79013 | 0.81912 |
| ENSG00000157557 | 21 | 40177231 | 40196879 | 49 | 10 | -0.66226 | 0.7461 | ETS2 | 0.7461 | 0.69807 | 0.50149 |
| ENSG00000097033 | 1 | 87170259 | 87213867 | 114 | 18 | -0.66231 | 0.74611 | SH3GLB1 | 0.74611 | 0.72933 | 0.75814 |
| ENSG00000053501 | 19 | 17326155 | 17330638 | 10 | 5 | -0.6628 | 0.74627 | USE1 | 0.74627 | 0.92285 | 0.91177 |
| ENSG00000177201 | 1 | 248457918 | 248458880 | 8 | 3 | -0.66292 | 0.74631 | OR2T12 | 0.74631 | 0.33045 | 0.33972 |
| ENSG00000107864 | 10 | 93806449 | 94050844 | 412 | 22 | -0.66323 | 0.74641 | CPEB3 | 0.74641 | 0.73707 | 0.76513 |
| ENSG00000181649 | 11 | 2949503 | 2950685 | 3 | 2 | -0.6638 | 0.74659 | PHLDA2 | 0.74659 | 0.38376 | 0.72624 |
| ENSG00000014914 | 1 | 149900543 | 149908791 | 17 | 3 | -0.664 | 0.74665 | MTMR11 | 0.74665 | 0.71986 | 0.64206 |
| ENSG00000108528 | 17 | 4840425 | 4843546 | 4 | 2 | -0.66408 | 0.74668 | SLC25A11 | 0.74668 | 0.21591 | 0.1544 |
| ENSG00000132600 | 16 | 68344877 | 68392466 | 118 | 19 | -0.6642 | 0.74672 | PRMT7 | 0.74672 | 0.79897 | 0.82253 |
| ENSG00000154646 | 21 | 19641433 | 19858197 | 805 | 89 | -0.66436 | 0.74677 | TMPRSS15 | 0.74677 | 0.80413 | 0.84094 |
| ENSG00000149654 | 20 | 44802372 | 44937137 | 338 | 50 | -0.66441 | 0.74679 | CDH22 | 0.74679 | 0.58064 | 0.51312 |
| ENSG00000063854 | 16 | 1845621 | 1877195 | 350 | 14 | -0.66486 | 0.74693 | HAGH | 0.74693 | 0.82649 | 0.84124 |
| ENSG00000117226 | 1 | 89472349 | 89488577 | 45 | 8 | -0.66505 | 0.74699 | GBP3 | 0.74699 | 0.81395 | 0.84683 |
| ENSG00000132463 | 4 | 71681499 | 71705662 | 99 | 10 | -0.66507 | 0.747 | GRSF1 | 0.747 | 0.67405 | 0.68702 |

| | | | | | | | | | | | |
|---|---|---|---|---|---|---|---|---|---|---|---|
| ENSG00000109184 | 4 | 52709166 | 52783003 | 98 | 9 | -0.66527 | 0.74706 | DCUN1D4 | 0.74706 | 0.75918 | 0.75896 |
| ENSG00000094914 | 12 | 53701240 | 53718648 | 28 | 4 | -0.66531 | 0.74707 | AAAS | 0.74707 | 0.73944 | 0.70565 |
| ENSG00000188343 | 8 | 94710789 | 94743755 | 52 | 14 | -0.66533 | 0.74708 | FAM92A1 | 0.74708 | 0.59623 | 0.59574 |
| ENSG00000113552 | 5 | 141371314 | 141392606 | 105 | 8 | -0.6654 | 0.7471 | GNPDA1 | 0.7471 | 0.75801 | 0.64692 |
| ENSG00000163884 | 3 | 126061478 | 126076285 | 24 | 7 | -0.66543 | 0.74711 | KLF15 | 0.74711 | 0.85426 | 0.88658 |
| ENSG00000114745 | 3 | 39138150 | 39149854 | 38 | 10 | -0.66577 | 0.74722 | GORASP1 | 0.74722 | 0.73841 | 0.70113 |
| ENSG00000176787 | 11 | 5079880 | 5080857 | 7 | 4 | -0.6658 | 0.74723 | OR52E2 | 0.74723 | 0.71838 | 0.68173 |
| ENSG00000108309 | 17 | 42385781 | 42396039 | 24 | 10 | -0.66584 | 0.74724 | RUNDC3A | 0.74724 | 0.89784 | 0.90161 |
| ENSG00000165102 | 8 | 42995556 | 43057998 | 72 | 15 | -0.66586 | 0.74725 | HGSNAT | 0.74725 | 0.80763 | 0.79649 |
| ENSG00000008277 | 7 | 87563458 | 87832204 | 578 | 26 | -0.66595 | 0.74728 | ADAM22 | 0.74728 | 0.73777 | 0.71885 |
| ENSG00000189376 | 8 | 124232196 | 124253638 | 19 | 5 | -0.66605 | 0.74731 | C8orf76 | 0.74731 | 0.66704 | 0.75352 |
| ENSG00000147676 | 8 | 120177273 | 120257913 | 297 | 31 | -0.66615 | 0.74734 | MAL2 | 0.74734 | 0.7692 | 0.67798 |
| ENSG00000073111 | 3 | 127317066 | 127341276 | 19 | 8 | -0.66618 | 0.74735 | MCM2 | 0.74735 | 0.75966 | 0.71671 |
| ENSG00000167945 | 16 | 855443 | 863861 | 45 | 14 | -0.66641 | 0.74743 | PRR25 | 0.74743 | 0.65986 | 0.8287 |
| ENSG00000221837 | 21 | 46047040 | 46048294 | 19 | 4 | -0.66653 | 0.74746 | KRTAP10-9 | 0.74746 | 0.60965 | 0.46142 |
| ENSG00000158373 | 6 | 26158349 | 26171577 | 21 | 6 | -0.66679 | 0.74755 | HIST1H2BD | 0.74755 | 0.70457 | 0.59706 |
| ENSG00000089250 | 12 | 117645947 | 117889975 | 803 | 64 | -0.66682 | 0.74756 | NOS1 | 0.74756 | 0.74433 | 0.75362 |
| ENSG00000092148 | 14 | 31569318 | 31677010 | 172 | 18 | -0.66688 | 0.74758 | HECTD1 | 0.74758 | 0.73799 | 0.81606 |
| ENSG00000100941 | 14 | 39644387 | 39652422 | 24 | 7 | -0.66699 | 0.74761 | PNN | 0.74761 | 0.89981 | 0.8331 |
| ENSG00000197006 | 16 | 21608539 | 21668794 | 93 | 22 | -0.66706 | 0.74763 | METTL9 | 0.74763 | 0.69894 | 0.48031 |
| ENSG00000107262 | 9 | 33247818 | 33264761 | 30 | 5 | -0.66707 | 0.74764 | BAG1 | 0.74764 | 0.77604 | 0.7686 |
| ENSG00000235878 | 21 | 47612391 | 47613673 | 13 | 5 | -0.66707 | 0.74764 | AP001468.1 | 0.74764 | 0.81631 | 0.27518 |
| ENSG00000174106 | 12 | 65563351 | 65642107 | 94 | 14 | -0.66748 | 0.74777 | LEMD3 | 0.74777 | 0.71741 | 0.52738 |
| ENSG00000254553 | 1 | 32930658 | 33066393 | 329 | 14 | -0.66778 | 0.74786 | RP1-27O5.3 | 0.74786 | 0.76566 | 0.83443 |
| ENSG00000167098 | 20 | 31571579 | 31592239 | 119 | 10 | -0.66791 | 0.74791 | SUN5 | 0.74791 | 0.7252 | 0.52342 |
| ENSG00000130429 | 7 | 98971872 | 98992424 | 57 | 9 | -0.66795 | 0.74792 | ARPC1B | 0.74792 | 0.802 | 0.83162 |
| ENSG00000183260 | 20 | 62492566 | 62494341 | 6 | 2 | -0.66848 | 0.74809 | ABHD16B | 0.74809 | 0.67167 | 0.53874 |
| ENSG00000174885 | 11 | 278365 | 285359 | 24 | 10 | -0.66858 | 0.74812 | NLRP6 | 0.74812 | 0.63436 | 0.46359 |
| ENSG00000227782 | 17 | 15943505 | 15943994 | 1 | 1 | -0.66884 | 0.7482 | AC002553.1 | 0.7482 | 0.78414 | 0.78067 |
| ENSG00000164508 | 6 | 25726291 | 25726790 | 6 | 2 | -0.6692 | 0.74832 | HIST1H2AA | 0.74832 | 0.70651 | 0.71218 |
| ENSG00000163584 | 3 | 170582664 | 170588272 | 17 | 4 | -0.6694 | 0.74838 | RPL22L1 | 0.74838 | 0.89987 | 0.84502 |
| ENSG00000204219 | 1 | 23707554 | 23751233 | 139 | 22 | -0.66954 | 0.74842 | TCEA3 | 0.74842 | 0.56286 | 0.3094 |
| ENSG00000006125 | 17 | 33905065 | 34053436 | 479 | 26 | -0.66955 | 0.74843 | AP2B1 | 0.74843 | 0.77558 | 0.76986 |
| ENSG00000100416 | 22 | 46726772 | 46753237 | 71 | 12 | -0.66959 | 0.74844 | TRMU | 0.74844 | 0.73258 | 0.63496 |
| ENSG00000253293 | 7 | 27210210 | 27219880 | 13 | 4 | -0.66963 | 0.74845 | HOXA10 | 0.74845 | 0.5536 | 0.52778 |
| ENSG00000112578 | 6 | 41888926 | 41900784 | 25 | 8 | -0.66964 | 0.74846 | BYSL | 0.74846 | 0.57954 | 0.49992 |
| ENSG00000160703 | 11 | 119037277 | 119054725 | 20 | 7 | -0.66972 | 0.74848 | NLRX1 | 0.74848 | 0.7316 | 0.68224 |
| ENSG00000082153 | 2 | 201675317 | 201688569 | 23 | 5 | -0.66986 | 0.74853 | BZW1 | 0.74853 | 0.62964 | 0.56373 |
| ENSG00000162409 | 1 | 57110995 | 57181008 | 288 | 15 | -0.66994 | 0.74855 | PRKAA2 | 0.74855 | 0.64244 | 0.55674 |
| ENSG00000117481 | 1 | 46805849 | 46830824 | 187 | 3 | -0.66997 | 0.74856 | NSUN4 | 0.74856 | 0.76131 | 0.72407 |
| ENSG00000180974 | 11 | 5905501 | 5906527 | 4 | 1 | -0.66999 | 0.74857 | OR52E4 | 0.74857 | 0.67322 | 0.65271 |
| ENSG00000159231 | 21 | 37507210 | 37518864 | 65 | 9 | -0.67064 | 0.74877 | CBR3 | 0.74877 | 0.78347 | 0.8327 |
| ENSG00000105707 | 19 | 35531410 | 35557475 | 102 | 15 | -0.67069 | 0.74879 | HPN | 0.74879 | 0.68968 | 0.7712 |
| ENSG00000113558 | 5 | 133484633 | 133512729 | 62 | 9 | -0.67087 | 0.74885 | SKP1 | 0.74885 | 0.75905 | 0.79853 |
| ENSG00000221858 | 7 | 143792141 | 143793186 | 2 | 1 | -0.67111 | 0.74892 | OR2A12 | 0.74892 | 0.55771 | 0.4956 |
| ENSG00000187531 | 17 | 79869815 | 79879199 | 4 | 2 | -0.67124 | 0.74897 | SIRT7 | 0.74897 | 0.29005 | 0.075806 |
| ENSG00000135436 | 12 | 49976668 | 49999422 | 39 | 9 | -0.67151 | 0.74905 | FAM186B | 0.74905 | 0.72242 | 0.69166 |
| ENSG00000160307 | 21 | 48018875 | 48025121 | 30 | 7 | -0.67164 | 0.74909 | S100B | 0.74909 | 0.36497 | 0.40845 |
| ENSG00000135248 | 7 | 128349115 | 128371797 | 113 | 23 | -0.67182 | 0.74915 | FAM71F1 | 0.74915 | 0.82293 | 0.84825 |
| ENSG00000111961 | 6 | 148593440 | 148873186 | 1227 | 121 | -0.67195 | 0.74919 | SASH1 | 0.74919 | 0.72453 | 0.72198 |
| ENSG00000158747 | 1 | 19967048 | 19984945 | 84 | 25 | -0.67197 | 0.7492 | NBL1 | 0.7492 | 0.71499 | 0.57444 |
| ENSG00000175182 | 3 | 184053714 | 184064063 | 26 | 10 | -0.67197 | 0.7492 | FAM131A | 0.7492 | 0.67493 | 0.21798 |
| ENSG00000091513 | 3 | 133464800 | 133497850 | 124 | 17 | -0.67207 | 0.74923 | TF | 0.74923 | 0.79681 | 0.79624 |
| ENSG00000157800 | 7 | 139993493 | 140104233 | 460 | 51 | -0.67234 | 0.74932 | SLC37A3 | 0.74932 | 0.66256 | 0.55238 |
| ENSG00000174282 | 17 | 7362685 | 7387582 | 80 | 12 | -0.67245 | 0.74935 | ZBTB4 | 0.74935 | 0.81758 | 0.78564 |
| ENSG00000164756 | 8 | 117962512 | 118188953 | 492 | 33 | -0.67248 | 0.74936 | SLC30A8 | 0.74936 | 0.80288 | 0.81917 |
| ENSG00000114013 | 3 | 121774213 | 121839983 | 127 | 25 | -0.67261 | 0.7494 | CD86 | 0.7494 | 0.7888 | 0.87047 |
| ENSG00000135114 | 12 | 121458095 | 121477045 | 57 | 15 | -0.673 | 0.74953 | OASL | 0.74953 | 0.70152 | 0.78203 |
| ENSG00000141200 | 17 | 51900239 | 51902573 | 14 | 5 | -0.67306 | 0.74954 | KIF2B | 0.74954 | 0.62511 | 0.69138 |
| ENSG00000113721 | 5 | 149493400 | 149535435 | 132 | 24 | -0.67334 | 0.74963 | PDGFRB | 0.74963 | 0.3088 | 0.2112 |
| ENSG00000077327 | 10 | 22634399 | 22743153 | 167 | 12 | -0.6734 | 0.74965 | SPAG6 | 0.74965 | 0.74594 | 0.71704 |
| ENSG00000186364 | 1 | 145586115 | 145589439 | 10 | 5 | -0.67348 | 0.74968 | NUDT17 | 0.74968 | 0.50986 | 0.52694 |
| ENSG00000164161 | 4 | 145567173 | 145666423 | 208 | 16 | -0.67352 | 0.74969 | HHIP | 0.74969 | 0.75111 | 0.78251 |
| ENSG00000120088 | 17 | 43699267 | 43913194 | 1169 | 10 | -0.67368 | 0.74974 | CRHR1 | 0.74974 | 0.76402 | 0.73476 |
| ENSG00000188771 | 11 | 112118876 | 112131583 | 41 | 5 | -0.67389 | 0.74981 | PLET1 | 0.74981 | 0.58137 | 0.61304 |
| ENSG00000116748 | 1 | 115215719 | 115238239 | 79 | 21 | -0.67398 | 0.74984 | AMPD1 | 0.74984 | 0.80421 | 0.80543 |
| ENSG00000168259 | 17 | 40128451 | 40173394 | 75 | 14 | -0.67399 | 0.74984 | DNAJC7 | 0.74984 | 0.82997 | 0.86068 |
| ENSG00000144895 | 3 | 150264465 | 150302029 | 119 | 18 | -0.67411 | 0.74988 | EIF2A | 0.74988 | 0.80306 | 0.8421 |
| ENSG00000115339 | 2 | 166604101 | 166651192 | 73 | 10 | -0.67442 | 0.74998 | GALNT3 | 0.74998 | 0.88192 | 0.85418 |
| ENSG00000051180 | 15 | 40986972 | 41024354 | 93 | 9 | -0.6745 | 0.75 | RAD51 | 0.75 | 0.75622 | 0.77474 |
| ENSG00000168453 | 8 | 21971928 | 21990897 | 64 | 13 | -0.67468 | 0.75006 | HR | 0.75006 | 0.44892 | 0.14624 |
| ENSG00000042317 | 14 | 88851268 | 88936694 | 129 | 18 | -0.6747 | 0.75007 | SPATA7 | 0.75007 | 0.74421 | 0.72465 |
| ENSG00000057593 | 13 | 113760105 | 113774995 | 29 | 8 | -0.67494 | 0.75014 | F7 | 0.75014 | 0.82138 | 0.69134 |

| | | | | | | | | | | | |
|---|---|---|---|---|---|---|---|---|---|---|---|
| ENSG00000269407 | 20 | 23330015 | 23334066 | 14 | 4 | -0.67509 | 0.75019 | AL096677.1 | 0.75019 | 0.47189 | 0.47052 |
| ENSG00000138670 | 4 | 82347547 | 82965397 | 2474 | 96 | -0.67515 | 0.75021 | RASGEF1B | 0.75021 | 0.74907 | 0.72904 |
| ENSG00000168209 | 10 | 74033678 | 74035794 | 4 | 2 | -0.67532 | 0.75026 | DDIT4 | 0.75026 | 0.74017 | 0.61506 |
| ENSG00000251322 | 22 | 51112843 | 51171726 | 165 | 40 | -0.67536 | 0.75028 | SHANK3 | 0.75028 | 0.73586 | 0.60739 |
| ENSG00000198466 | 19 | 58361225 | 58376480 | 68 | 10 | -0.67541 | 0.75029 | ZNF587 | 0.75029 | 0.73806 | 0.70718 |
| ENSG00000069345 | 16 | 46989299 | 47007699 | 26 | 9 | -0.67575 | 0.7504 | DNAJA2 | 0.7504 | 0.66546 | 0.69742 |
| ENSG00000049540 | 7 | 73442119 | 73484237 | 98 | 24 | -0.67615 | 0.75053 | ELN | 0.75053 | 0.77853 | 0.92754 |
| ENSG00000131236 | 1 | 40505905 | 40538321 | 135 | 10 | -0.67634 | 0.75059 | CAP1 | 0.75059 | 0.76629 | 0.71694 |
| ENSG00000108883 | 17 | 42927311 | 42977030 | 169 | 19 | -0.67635 | 0.75059 | EFTUD2 | 0.75059 | 0.76761 | 0.79736 |
| ENSG00000104499 | 8 | 143915663 | 143997922 | 453 | 19 | -0.67645 | 0.75062 | GML | 0.75062 | 0.75977 | 0.82441 |
| ENSG00000255112 | 18 | 11851395 | 11854448 | 6 | 4 | -0.67662 | 0.75068 | CHMP1B | 0.75068 | 0.95056 | 0.93038 |
| ENSG00000106948 | 9 | 117096436 | 117156685 | 138 | 21 | -0.67665 | 0.75069 | AKNA | 0.75069 | 0.58468 | 0.3287 |
| ENSG00000168092 | 11 | 117014983 | 117047610 | 138 | 18 | -0.67666 | 0.75069 | PAFAH1B2 | 0.75069 | 0.66773 | 0.66692 |
| ENSG00000184845 | 5 | 174867042 | 174871211 | 8 | 3 | -0.67673 | 0.75071 | DRD1 | 0.75071 | 0.22848 | 0.17163 |
| ENSG00000064393 | 7 | 139246316 | 139477577 | 516 | 52 | -0.67685 | 0.75075 | HIPK2 | 0.75075 | 0.72479 | 0.6831 |
| ENSG00000162009 | 16 | 1122756 | 1131454 | 59 | 14 | -0.67684 | 0.75075 | SSTR5 | 0.75075 | 0.73986 | 0.50244 |
| ENSG00000124596 | 6 | 41001366 | 41065526 | 159 | 14 | -0.6772 | 0.75086 | OARD1 | 0.75086 | 0.78778 | 0.80899 |
| ENSG00000187187 | 19 | 40490041 | 40523514 | 95 | 9 | -0.67723 | 0.75087 | ZNF546 | 0.75087 | 0.79708 | 0.82647 |
| ENSG00000154114 | 11 | 120894781 | 120961484 | 131 | 13 | -0.67735 | 0.75091 | TBCEL | 0.75091 | 0.75695 | 0.70595 |
| ENSG00000146215 | 6 | 43267448 | 43276535 | 15 | 4 | -0.67738 | 0.75092 | CRIP3 | 0.75092 | 0.75811 | 0.41604 |
| ENSG00000153094 | 2 | 111876955 | 111926024 | 119 | 17 | -0.67747 | 0.75095 | BCL2L11 | 0.75095 | 0.5828 | 0.42568 |
| ENSG00000079246 | 2 | 216972187 | 217071026 | 272 | 23 | -0.6775 | 0.75096 | XRCC5 | 0.75096 | 0.70203 | 0.66808 |
| ENSG00000196878 | 1 | 209788215 | 209825811 | 160 | 26 | -0.67763 | 0.751 | LAMB3 | 0.751 | 0.76087 | 0.81762 |
| ENSG00000132694 | 1 | 156904632 | 157015162 | 283 | 17 | -0.67786 | 0.75107 | ARHGEF11 | 0.75107 | 0.74336 | 0.78574 |
| ENSG00000188687 | 2 | 74443369 | 74570541 | 201 | 39 | -0.67799 | 0.75111 | SLC4A5 | 0.75111 | 0.75389 | 0.73451 |
| ENSG00000104722 | 8 | 24770525 | 24776607 | 5 | 3 | -0.67805 | 0.75113 | NEFM | 0.75113 | 0.42777 | 0.35411 |
| ENSG00000108826 | 17 | 48445218 | 48450575 | 19 | 5 | -0.67818 | 0.75117 | MRPL27 | 0.75117 | 0.71442 | 0.67212 |
| ENSG00000255823 | 11 | 10529434 | 10530723 | 6 | 3 | -0.67819 | 0.75118 | MTRNR2L8 | 0.75118 | 0.28487 | 0.32293 |
| ENSG00000168769 | 4 | 106067032 | 106200973 | 320 | 30 | -0.67835 | 0.75123 | TET2 | 0.75123 | 0.76778 | 0.76326 |
| ENSG00000120937 | 1 | 11917521 | 11918988 | 4 | 2 | -0.67872 | 0.75134 | NPPB | 0.75134 | 0.91751 | 0.94689 |
| ENSG00000127948 | 7 | 75528518 | 75616173 | 169 | 35 | -0.67888 | 0.75139 | POR | 0.75139 | 0.77452 | 0.69003 |
| ENSG00000135720 | 16 | 66754796 | 66785701 | 70 | 5 | -0.67886 | 0.75139 | DYNC1LI2 | 0.75139 | 0.70312 | 0.66227 |
| ENSG00000156886 | 16 | 31404633 | 31437826 | 62 | 12 | -0.67912 | 0.75147 | ITGAD | 0.75147 | 0.75488 | 0.80837 |
| ENSG00000078579 | 8 | 16849678 | 16859690 | 60 | 13 | -0.67916 | 0.75148 | FGF20 | 0.75148 | 0.5856 | 0.4316 |
| ENSG00000145332 | 4 | 88081255 | 88161466 | 210 | 26 | -0.67919 | 0.75149 | KLHL8 | 0.75149 | 0.79098 | 0.81716 |
| ENSG00000143457 | 1 | 150618701 | 150669630 | 145 | 8 | -0.67973 | 0.75166 | GOLPH3L | 0.75166 | 0.78152 | 0.77705 |
| ENSG00000177868 | 1 | 43272723 | 43282954 | 33 | 9 | -0.68001 | 0.75175 | CCDC23 | 0.75175 | 0.725 | 0.81316 |
| ENSG00000116039 | 2 | 71163012 | 71192536 | 113 | 24 | -0.68013 | 0.75179 | ATP6V1B1 | 0.75179 | 0.43679 | 0.55914 |
| ENSG00000186115 | 19 | 15988833 | 16008930 | 100 | 11 | -0.68013 | 0.75179 | CYP4F2 | 0.75179 | 0.67288 | 0.55555 |
| ENSG00000134874 | 13 | 96230457 | 96296957 | 113 | 21 | -0.68018 | 0.75181 | DZIP1 | 0.75181 | 0.73899 | 0.78456 |
| ENSG00000018510 | 2 | 178257372 | 178408564 | 390 | 16 | -0.68034 | 0.75185 | AGPS | 0.75185 | 0.73184 | 0.71663 |
| ENSG00000175213 | 11 | 46722368 | 46727462 | 8 | 2 | -0.68045 | 0.75189 | ZNF408 | 0.75189 | 0.62507 | 0.69324 |
| ENSG00000204700 | 6 | 29141311 | 29142351 | 8 | 3 | -0.68078 | 0.75199 | OR2J2 | 0.75199 | 0.7142 | 0.76036 |
| ENSG00000167754 | 19 | 51446559 | 51456349 | 56 | 16 | -0.68085 | 0.75202 | KLK5 | 0.75202 | 0.63195 | 0.79301 |
| ENSG00000146047 | 6 | 25727137 | 25727573 | 4 | 1 | -0.68091 | 0.75204 | HIST1H2BA | 0.75204 | 0.70651 | 0.71236 |
| ENSG00000196228 | 2 | 108863651 | 108881807 | 51 | 11 | -0.68126 | 0.75215 | SULT1C3 | 0.75215 | 0.50951 | 0.36299 |
| ENSG00000007545 | 16 | 1662326 | 1727909 | 138 | 15 | -0.68138 | 0.75219 | CRAMP1L | 0.75219 | 0.79054 | 0.82208 |
| ENSG00000188290 | 1 | 934342 | 935552 | 4 | 2 | -0.68163 | 0.75226 | HES4 | 0.75226 | 0.64628 | 0.62822 |
| ENSG00000196453 | 7 | 149128454 | 149158214 | 65 | 9 | -0.68164 | 0.75227 | ZNF777 | 0.75227 | 0.8279 | 0.67383 |
| ENSG00000138109 | 10 | 96698415 | 96749147 | 213 | 19 | -0.68169 | 0.75228 | CYP2C9 | 0.75228 | 0.76109 | 0.75486 |
| ENSG00000256223 | 12 | 133707161 | 133736051 | 80 | 10 | -0.68169 | 0.75228 | ZNF10 | 0.75228 | 0.71286 | 0.7773 |
| ENSG00000108590 | 17 | 6546635 | 6554954 | 46 | 8 | -0.68176 | 0.7523 | MED31 | 0.7523 | 0.61453 | 0.54425 |
| ENSG00000243660 | 10 | 43932282 | 43991517 | 235 | 23 | -0.68196 | 0.75237 | ZNF487 | 0.75237 | 0.77197 | 0.74073 |
| ENSG00000150907 | 13 | 41129804 | 41240734 | 224 | 15 | -0.68208 | 0.7524 | FOXO1 | 0.7524 | 0.79396 | 0.8083 |
| ENSG00000173250 | 5 | 145892666 | 145895753 | 3 | 1 | -0.6821 | 0.75241 | GPR151 | 0.75241 | 0.79541 | 0.78749 |
| ENSG00000177426 | 18 | 3411606 | 3458409 | 144 | 40 | -0.68237 | 0.7525 | TGIF1 | 0.7525 | 0.65734 | 0.71866 |
| ENSG00000152936 | 12 | 25562241 | 25801513 | 1051 | 62 | -0.68244 | 0.75252 | IFLTD1 | 0.75252 | 0.80352 | 0.63738 |
| ENSG00000070950 | 3 | 8817088 | 9005457 | 891 | 34 | -0.68276 | 0.75262 | RAD18 | 0.75262 | 0.77143 | 0.80498 |
| ENSG00000180061 | 19 | 55824169 | 55845416 | 85 | 10 | -0.68276 | 0.75262 | TMEM150B | 0.75262 | 0.48761 | 0.37506 |
| ENSG00000096080 | 6 | 43639040 | 43655528 | 48 | 8 | -0.68299 | 0.75269 | MRPS18A | 0.75269 | 0.75325 | 0.81578 |
| ENSG00000109272 | 4 | 74718906 | 74719872 | 5 | 1 | -0.68312 | 0.75273 | PF4V1 | 0.75273 | 0.61233 | 0.86993 |
| ENSG00000187689 | 4 | 71384257 | 71398459 | 87 | 8 | -0.68316 | 0.75275 | AMTN | 0.75275 | 0.76226 | 0.81133 |
| ENSG00000150764 | 11 | 111797868 | 111893308 | 135 | 18 | -0.68326 | 0.75278 | DIXDC1 | 0.75278 | 0.76516 | 0.83646 |
| ENSG00000143443 | 1 | 151020216 | 151024462 | 3 | 2 | -0.68328 | 0.75279 | C1orf56 | 0.75279 | 0.92541 | 0.92201 |
| ENSG00000119285 | 1 | 236712305 | 236767804 | 403 | 16 | -0.6833 | 0.75279 | HEATR1 | 0.75279 | 0.72719 | 0.73563 |
| ENSG00000119927 | 10 | 113909624 | 113975135 | 152 | 20 | -0.68332 | 0.7528 | GPAM | 0.7528 | 0.83486 | 0.94461 |
| ENSG00000121892 | 4 | 39824483 | 39979576 | 436 | 28 | -0.68336 | 0.75281 | PDS5A | 0.75281 | 0.77449 | 0.74823 |
| ENSG00000251655 | 12 | 11504757 | 11548500 | 178 | 18 | -0.68378 | 0.75294 | PRB1 | 0.75294 | 0.73885 | 0.70036 |
| ENSG00000130520 | 19 | 18417040 | 18434084 | 50 | 11 | -0.6838 | 0.75295 | LSM4 | 0.75295 | 0.67543 | 0.83363 |
| ENSG00000141385 | 18 | 12328943 | 12377313 | 130 | 13 | -0.68382 | 0.75296 | AFG3L2 | 0.75296 | 0.80607 | 0.90214 |
| ENSG00000058799 | 1 | 54317392 | 54356407 | 148 | 14 | -0.68388 | 0.75297 | YIPF1 | 0.75297 | 0.70747 | 0.64432 |
| ENSG00000169247 | 5 | 148303202 | 148442726 | 467 | 40 | -0.6839 | 0.75298 | SH3TC2 | 0.75298 | 0.7019 | 0.47732 |
| ENSG00000181896 | 19 | 19779605 | 19794318 | 40 | 10 | -0.68416 | 0.75306 | ZNF101 | 0.75306 | 0.41772 | 0.42124 |

| | | | | | | | | | | | |
|---|---|---|---|---|---|---|---|---|---|---|---|
| ENSG00000182963 | 17 | 42875816 | 42908184 | 67 | 13 | -0.6843 | 0.75311 | GJC1 | 0.75311 | 0.62174 | 0.67085 |
| ENSG00000134193 | 1 | 120336641 | 120354283 | 57 | 10 | -0.68438 | 0.75313 | REG4 | 0.75313 | 0.72306 | 0.72441 |
| ENSG00000124602 | 6 | 40994772 | 41006928 | 20 | 4 | -0.68444 | 0.75315 | UNC5CL | 0.75315 | 0.8398 | 0.91297 |
| ENSG00000116560 | 1 | 35641979 | 35658749 | 29 | 2 | -0.68472 | 0.75324 | SFPQ | 0.75324 | 0.69189 | 0.73341 |
| ENSG00000100342 | 22 | 36649056 | 36663576 | 81 | 12 | -0.68502 | 0.75333 | APOL1 | 0.75333 | 0.86179 | 0.91072 |
| ENSG00000180638 | 17 | 19581601 | 19622292 | 112 | 20 | -0.68504 | 0.75334 | SLC47A2 | 0.75334 | 0.83427 | 0.83734 |
| ENSG00000260903 | 20 | 30555805 | 30591190 | 89 | 24 | -0.68509 | 0.75335 | XKR7 | 0.75335 | 0.72643 | 0.72031 |
| ENSG00000157017 | 3 | 10327359 | 10334631 | 15 | 7 | -0.68513 | 0.75337 | GHRL | 0.75337 | 0.68802 | 0.73749 |
| ENSG00000131370 | 3 | 15296360 | 15382875 | 319 | 47 | -0.68525 | 0.75341 | SH3BP5 | 0.75341 | 0.79965 | 0.83302 |
| ENSG00000114988 | 2 | 97371666 | 97405801 | 45 | 7 | -0.68536 | 0.75344 | LMAN2L | 0.75344 | 0.62645 | 0.68604 |
| ENSG00000183977 | 3 | 20021453 | 20053822 | 143 | 18 | -0.68554 | 0.7535 | PP2D1 | 0.7535 | 0.74142 | 0.73775 |
| ENSG00000258472 | 17 | 26782770 | 26941215 | 275 | 33 | -0.68567 | 0.75354 | RP11-192H23.4 | 0.75354 | 0.75347 | 0.76129 |
| ENSG00000165501 | 14 | 50065415 | 50081390 | 62 | 6 | -0.68575 | 0.75356 | LRR1 | 0.75356 | 0.74576 | 0.76622 |
| ENSG00000114686 | 3 | 131181056 | 131221827 | 96 | 15 | -0.68592 | 0.75362 | MRPL3 | 0.75362 | 0.73796 | 0.75426 |
| ENSG00000105519 | 19 | 5911718 | 5915888 | 14 | 6 | -0.68592 | 0.75362 | CAPS | 0.75362 | 0.78769 | 0.87766 |
| ENSG00000166913 | 20 | 43514317 | 43537173 | 45 | 5 | -0.686 | 0.75364 | YWHAB | 0.75364 | 0.5026 | 0.45734 |
| ENSG00000133657 | 3 | 194123401 | 194219093 | 253 | 21 | -0.68603 | 0.75365 | ATP13A3 | 0.75365 | 0.5095 | 0.10808 |
| ENSG00000133858 | 12 | 72003252 | 72061505 | 120 | 11 | -0.68605 | 0.75366 | ZFC3H1 | 0.75366 | 0.73796 | 0.75015 |
| ENSG00000158773 | 1 | 161009041 | 161015767 | 26 | 6 | -0.68641 | 0.75377 | USF1 | 0.75377 | 0.83911 | 0.82356 |
| ENSG00000137944 | 1 | 89401456 | 89458636 | 129 | 11 | -0.68644 | 0.75378 | CCBL2 | 0.75378 | 0.85256 | 0.76997 |
| ENSG00000128218 | 22 | 24094930 | 24096655 | 7 | 2 | -0.68642 | 0.75378 | VPREB3 | 0.75378 | 0.84467 | 0.90063 |
| ENSG00000182141 | 19 | 21473962 | 21512227 | 135 | 13 | -0.68657 | 0.75382 | ZNF708 | 0.75382 | 0.81424 | 0.75822 |
| ENSG00000101280 | 20 | 853296 | 896977 | 239 | 42 | -0.68672 | 0.75387 | ANGPT4 | 0.75387 | 0.74633 | 0.83781 |
| ENSG00000186265 | 3 | 112182815 | 112218408 | 55 | 8 | -0.68689 | 0.75392 | BTLA | 0.75392 | 0.707 | 0.74164 |
| ENSG00000004897 | 17 | 45195069 | 45266788 | 189 | 26 | -0.68689 | 0.75392 | CDC27 | 0.75392 | 0.70712 | 0.75322 |
| ENSG00000172164 | 8 | 121547985 | 121825513 | 765 | 49 | -0.68695 | 0.75394 | SNTB1 | 0.75394 | 0.74055 | 0.71006 |
| ENSG00000168003 | 11 | 62623518 | 62656352 | 49 | 13 | -0.68697 | 0.75395 | SLC3A2 | 0.75395 | 0.76163 | 0.85481 |
| ENSG00000163626 | 4 | 73921797 | 73935472 | 17 | 7 | -0.68699 | 0.75396 | COX18 | 0.75396 | 0.68362 | 0.70976 |
| ENSG00000131508 | 5 | 138906016 | 139008018 | 239 | 21 | -0.68724 | 0.75403 | UBE2D2 | 0.75403 | 0.76078 | 0.73157 |
| ENSG00000213213 | 9 | 139690802 | 139702193 | 35 | 8 | -0.68758 | 0.75414 | CCDC183 | 0.75414 | 0.76378 | 0.83239 |
| ENSG00000169860 | 3 | 152552736 | 152555841 | 6 | 2 | -0.68759 | 0.75415 | P2RY1 | 0.75415 | 0.76917 | 0.85037 |
| ENSG00000183520 | 1 | 38474930 | 38490496 | 38 | 15 | -0.68804 | 0.75429 | UTP11L | 0.75429 | 0.79941 | 0.81887 |
| ENSG00000139433 | 12 | 110288748 | 110318293 | 100 | 14 | -0.68831 | 0.75437 | GLTP | 0.75437 | 0.6988 | 0.74466 |
| ENSG00000144395 | 2 | 197504278 | 197628214 | 275 | 15 | -0.68842 | 0.75441 | CCDC150 | 0.75441 | 0.73251 | 0.62968 |
| ENSG00000204420 | 6 | 31686371 | 31694491 | 18 | 6 | -0.68842 | 0.75441 | C6orf25 | 0.75441 | 0.58937 | 0.68551 |
| ENSG00000165733 | 10 | 43278249 | 43330385 | 184 | 17 | -0.68847 | 0.75442 | BMS1 | 0.75442 | 0.70605 | 0.75418 |
| ENSG00000129351 | 19 | 10764937 | 10803093 | 78 | 10 | -0.68858 | 0.75446 | ILF3 | 0.75446 | 0.80029 | 0.82502 |
| ENSG00000099992 | 22 | 30687979 | 30723035 | 81 | 20 | -0.68867 | 0.75448 | TBC1D10A | 0.75448 | 0.66923 | 0.68407 |
| ENSG00000104164 | 15 | 45879321 | 45908197 | 46 | 12 | -0.68885 | 0.75454 | BLOC1S6 | 0.75454 | 0.64411 | 0.68098 |
| ENSG00000113430 | 5 | 1877541 | 1887350 | 50 | 9 | -0.68905 | 0.7546 | IRX4 | 0.7546 | 0.94765 | 0.93648 |
| ENSG00000184530 | 6 | 127840600 | 127912962 | 332 | 17 | -0.68912 | 0.75463 | C6orf58 | 0.75463 | 0.76398 | 0.75076 |
| ENSG00000186575 | 22 | 29999545 | 30094587 | 228 | 25 | -0.68925 | 0.75467 | NF2 | 0.75467 | 0.71557 | 0.67288 |
| ENSG00000186867 | 4 | 122250467 | 122302214 | 259 | 23 | -0.68945 | 0.75473 | QRFPR | 0.75473 | 0.81515 | 0.8392 |
| ENSG00000162511 | 1 | 31205316 | 31230667 | 71 | 22 | -0.68975 | 0.75482 | LAPTM5 | 0.75482 | 0.77101 | 0.64022 |
| ENSG00000006712 | 19 | 39876492 | 39881835 | 10 | 3 | -0.6898 | 0.75484 | PAF1 | 0.75484 | 0.87506 | 0.90335 |
| ENSG00000184678 | 1 | 149856010 | 149858232 | 4 | 2 | -0.68983 | 0.75485 | HIST2H2BE | 0.75485 | 0.58574 | 0.63621 |
| ENSG00000197050 | 19 | 37498759 | 37621216 | 368 | 12 | -0.68991 | 0.75487 | ZNF420 | 0.75487 | 0.79061 | 0.77771 |
| ENSG00000196639 | 3 | 11178779 | 11305243 | 429 | 57 | -0.69016 | 0.75495 | HRH1 | 0.75495 | 0.75556 | 0.66268 |
| ENSG00000131400 | 19 | 50861734 | 50869087 | 15 | 3 | -0.69014 | 0.75495 | NAPSA | 0.75495 | 0.46349 | 0.58898 |
| ENSG00000128923 | 15 | 59063391 | 59154099 | 231 | 23 | -0.69019 | 0.75496 | FAM63B | 0.75496 | 0.76563 | 0.77004 |
| ENSG00000241128 | 1 | 247886401 | 247887345 | 4 | 2 | -0.69058 | 0.75509 | OR14A2 | 0.75509 | 0.80778 | 0.93988 |
| ENSG00000126524 | 7 | 66452664 | 66460588 | 38 | 15 | -0.69076 | 0.75514 | SBDS | 0.75514 | 0.81279 | 0.67458 |
| ENSG00000090612 | 12 | 133707570 | 133783698 | 219 | 17 | -0.69082 | 0.75516 | ZNF268 | 0.75516 | 0.78015 | 0.82002 |
| ENSG00000127838 | 2 | 219135115 | 219211516 | 225 | 15 | -0.69084 | 0.75517 | PNKD | 0.75517 | 0.7647 | 0.76193 |
| ENSG00000189045 | 5 | 74907284 | 74967671 | 182 | 22 | -0.69114 | 0.75526 | ANKDD1B | 0.75526 | 0.75203 | 0.70554 |
| ENSG00000165801 | 14 | 21538429 | 21558399 | 61 | 10 | -0.69114 | 0.75526 | ARHGEF40 | 0.75526 | 0.91851 | 0.96684 |
| ENSG00000051341 | 3 | 121150278 | 121264853 | 290 | 16 | -0.69131 | 0.75532 | POLQ | 0.75532 | 0.78302 | 0.75566 |
| ENSG00000161202 | 3 | 183873176 | 183891398 | 41 | 9 | -0.69136 | 0.75533 | DVL3 | 0.75533 | 0.75259 | 0.73509 |
| ENSG00000159164 | 1 | 149874870 | 149889434 | 24 | 7 | -0.69152 | 0.75538 | SV2A | 0.75538 | 0.71169 | 0.69833 |
| ENSG00000136159 | 13 | 48611703 | 48621358 | 21 | 7 | -0.69155 | 0.75539 | NUDT15 | 0.75539 | 0.7976 | 0.68521 |
| ENSG00000185414 | 2 | 99797542 | 99814089 | 48 | 8 | -0.69193 | 0.75551 | MRPL30 | 0.75551 | 0.78707 | 0.7911 |
| ENSG00000183172 | 22 | 42475695 | 42480288 | 4 | 1 | -0.69195 | 0.75551 | SMDT1 | 0.75551 | 0.79187 | 0.76824 |
| ENSG00000206262 | 3 | 138666076 | 138672293 | 25 | 3 | -0.69198 | 0.75552 | C3orf72 | 0.75552 | 0.47973 | 0.52402 |
| ENSG00000115165 | 2 | 158271131 | 158345473 | 182 | 25 | -0.69232 | 0.75563 | CYTIP | 0.75563 | 0.77636 | 0.74788 |
| ENSG00000234284 | 5 | 178450753 | 178462065 | 47 | 5 | -0.69243 | 0.75567 | ZNF879 | 0.75567 | 0.72408 | 0.6538 |
| ENSG00000100036 | 22 | 31031639 | 31065003 | 167 | 32 | -0.69255 | 0.7557 | SLC35E4 | 0.7557 | 0.79254 | 0.7767 |
| ENSG00000145451 | 4 | 175558065 | 175750465 | 868 | 61 | -0.69269 | 0.75575 | GLRA3 | 0.75575 | 0.77161 | 0.75057 |
| ENSG00000160856 | 1 | 157644111 | 157670647 | 56 | 9 | -0.69301 | 0.75585 | FCRL3 | 0.75585 | 0.70938 | 0.76205 |
| ENSG00000267001 | 19 | 2717767 | 2740046 | 80 | 15 | -0.69319 | 0.7559 | AC006538.4 | 0.7559 | 0.80458 | 0.84348 |
| ENSG00000133028 | 17 | 10583654 | 10601692 | 29 | 4 | -0.69321 | 0.75591 | SCO1 | 0.75591 | 0.74816 | 0.61518 |
| ENSG00000166959 | 11 | 60467047 | 60483284 | 20 | 6 | -0.69322 | 0.75592 | MS4A8 | 0.75592 | 0.58743 | 0.58398 |
| ENSG00000196862 | 2 | 108443388 | 108507297 | 108 | 11 | -0.69332 | 0.75595 | RGPD4 | 0.75595 | 0.74586 | 0.73928 |
| ENSG00000160882 | 8 | 143953772 | 143961262 | 54 | 7 | -0.69361 | 0.75604 | CYP11B1 | 0.75604 | 0.70952 | 0.68619 |

| | | | | | | | | | | | | |
|---|---|---|---|---|---|---|---|---|---|---|---|---|
| ENSG00000164402 | 5 | 132086509 | 132142933 | 114 | 9 | -0.69377 | 0.75609 | | 44082 | 0.75609 | 0.80402 | 0.74654 |
| ENSG00000104960 | 19 | 50353992 | 50364001 | 37 | 10 | -0.69395 | 0.75614 | PTOV1 | | 0.75614 | 0.61654 | 0.66289 |
| ENSG00000174928 | 3 | 155480401 | 155524140 | 139 | 18 | -0.69403 | 0.75617 | C3orf33 | | 0.75617 | 0.613 | 0.36721 |
| ENSG00000160917 | 7 | 99036545 | 99054994 | 49 | 7 | -0.6941 | 0.75619 | CPSF4 | | 0.75619 | 0.81318 | 0.82366 |
| ENSG00000169548 | 22 | 22868060 | 22874613 | 59 | 8 | -0.69446 | 0.7563 | ZNF280A | | 0.7563 | 0.34577 | 0.45407 |
| ENSG00000115459 | 2 | 85581517 | 85618875 | 115 | 20 | -0.69447 | 0.75631 | ELMOD3 | | 0.75631 | 0.61244 | 0.42067 |
| ENSG00000105926 | 7 | 24612887 | 24733812 | 456 | 12 | -0.69447 | 0.75631 | MPP6 | | 0.75631 | 0.74069 | 0.81129 |
| ENSG00000239704 | 17 | 15339332 | 15406922 | 228 | 30 | -0.6947 | 0.75638 | CDRT4 | | 0.75638 | 0.82258 | 0.87735 |
| ENSG00000165810 | 5 | 180467225 | 180488523 | 88 | 13 | -0.69475 | 0.75639 | BTNL9 | | 0.75639 | 0.77442 | 0.90365 |
| ENSG00000177752 | 4 | 44624086 | 44680573 | 260 | 15 | -0.69492 | 0.75645 | YIPF7 | | 0.75645 | 0.72286 | 0.62388 |
| ENSG00000184445 | 12 | 123011793 | 123110943 | 169 | 20 | -0.69496 | 0.75646 | KNTC1 | | 0.75646 | 0.74063 | 0.73368 |
| ENSG00000205846 | 12 | 8608522 | 8630926 | 106 | 4 | -0.69535 | 0.75658 | CLEC6A | | 0.75658 | 0.77611 | 0.79172 |
| ENSG00000157593 | 6 | 44221833 | 44225291 | 8 | 4 | -0.69538 | 0.75659 | SLC35B2 | | 0.75659 | 0.82776 | 0.70745 |
| ENSG00000034533 | 3 | 130732719 | 130746493 | 29 | 3 | -0.69542 | 0.75661 | ASTE1 | | 0.75661 | 0.77693 | 0.87372 |
| ENSG00000004975 | 17 | 7128660 | 7137864 | 14 | 4 | -0.69547 | 0.75662 | DVL2 | | 0.75662 | 0.66432 | 0.65838 |
| ENSG00000256100 | 11 | 63742092 | 63755818 | 24 | 3 | -0.69565 | 0.75668 | AP000721.4 | | 0.75668 | 0.84039 | 0.87561 |
| ENSG00000188580 | 6 | 124125286 | 125146803 | 3272 | 215 | -0.69571 | 0.75669 | NKAIN2 | | 0.75669 | 0.75146 | 0.76999 |
| ENSG00000174996 | 11 | 66024765 | 66035331 | 17 | 5 | -0.69568 | 0.75669 | KLC2 | | 0.75669 | 0.58957 | 0.56618 |
| ENSG00000161904 | 6 | 33738979 | 33756913 | 88 | 11 | -0.69573 | 0.7567 | LEMD2 | | 0.7567 | 0.80319 | 0.70751 |
| ENSG00000241468 | 7 | 99046098 | 99063954 | 40 | 6 | -0.69572 | 0.7567 | ATP5J2 | | 0.7567 | 0.77207 | 0.78507 |
| ENSG00000170727 | 8 | 145486055 | 145515082 | 66 | 9 | -0.69571 | 0.7567 | BOP1 | | 0.7567 | 0.58728 | 0.54851 |
| ENSG00000205850 | 13 | 24895264 | 24895566 | 1 | 1 | -0.69573 | 0.7567 | AL359736.1 | | 0.7567 | 0.7494 | 0.72539 |
| ENSG00000102805 | 13 | 77564795 | 77576652 | 48 | 8 | -0.69572 | 0.7567 | CLN5 | | 0.7567 | 0.71504 | 0.69424 |
| ENSG00000258873 | 19 | 57665389 | 57678811 | 69 | 17 | -0.69579 | 0.75672 | DUXA | | 0.75672 | 0.66587 | 0.82968 |
| ENSG00000167210 | 18 | 44056935 | 44236996 | 619 | 65 | -0.69634 | 0.75689 | LOXHD1 | | 0.75689 | 0.77564 | 0.73625 |
| ENSG00000179954 | 19 | 55999771 | 56030465 | 176 | 29 | -0.69659 | 0.75697 | SSC5D | | 0.75697 | 0.66153 | 0.49847 |
| ENSG00000161509 | 17 | 72838162 | 72857627 | 52 | 5 | -0.69662 | 0.75698 | GRIN2C | | 0.75698 | 0.85385 | 0.86889 |
| ENSG00000114520 | 3 | 125165495 | 125239041 | 288 | 17 | -0.69668 | 0.757 | SNX4 | | 0.757 | 0.78157 | 0.84239 |
| ENSG00000169756 | 2 | 109150857 | 109303702 | 534 | 21 | -0.69673 | 0.75701 | LIMS1 | | 0.75701 | 0.7437 | 0.7563 |
| ENSG00000108406 | 17 | 57642886 | 57685706 | 90 | 13 | -0.69675 | 0.75702 | DHX40 | | 0.75702 | 0.78056 | 0.82428 |
| ENSG00000167916 | 17 | 38854243 | 38860002 | 7 | 3 | -0.69689 | 0.75706 | KRT24 | | 0.75706 | 0.75101 | 0.79177 |
| ENSG00000205268 | 8 | 66629745 | 66754557 | 411 | 27 | -0.69707 | 0.75712 | PDE7A | | 0.75712 | 0.76701 | 0.77405 |
| ENSG00000233436 | 11 | 57510986 | 57519253 | 7 | 4 | -0.69708 | 0.75713 | BTBD18 | | 0.75713 | 0.46983 | 0.4102 |
| ENSG00000173621 | 11 | 66624118 | 66627946 | 2 | 1 | -0.69709 | 0.75713 | LRFN4 | | 0.75713 | 0.59752 | 0.61661 |
| ENSG00000104866 | 19 | 45594654 | 45651335 | 187 | 16 | -0.69716 | 0.75715 | PPP1R37 | | 0.75715 | 0.66691 | 0.68495 |
| ENSG00000170037 | 17 | 7835419 | 7853236 | 50 | 3 | -0.69729 | 0.75719 | CNTROB | | 0.75719 | 0.87747 | 0.88129 |
| ENSG00000104921 | 19 | 7753644 | 7767032 | 177 | 12 | -0.69729 | 0.75719 | FCER2 | | 0.75719 | 0.73347 | 0.72856 |
| ENSG00000167046 | 20 | 61272071 | 61274656 | 6 | 2 | -0.6976 | 0.75729 | RP11-93B14.6 | | 0.75729 | 0.78592 | 0.58852 |
| ENSG00000075035 | 12 | 108523248 | 108644314 | 371 | 30 | -0.69788 | 0.75737 | WSCD2 | | 0.75737 | 0.7938 | 0.88426 |
| ENSG00000173482 | 18 | 7566780 | 8406859 | 2785 | 166 | -0.69802 | 0.75742 | PTPRM | | 0.75742 | 0.7563 | 0.75551 |
| ENSG00000135929 | 2 | 219646472 | 219680016 | 80 | 5 | -0.69838 | 0.75753 | CYP27A1 | | 0.75753 | 0.75516 | 0.71638 |
| ENSG00000185988 | 19 | 1524073 | 1535455 | 68 | 10 | -0.69854 | 0.75758 | PLK5 | | 0.75758 | 0.7755 | 0.7827 |
| ENSG00000174807 | 11 | 66081958 | 66084515 | 3 | 1 | -0.69862 | 0.75761 | CD248 | | 0.75761 | 0.88046 | 0.90258 |
| ENSG00000185052 | 20 | 19193290 | 19703581 | 1850 | 113 | -0.69869 | 0.75763 | SLC24A3 | | 0.75763 | 0.69907 | 0.73796 |
| ENSG00000126088 | 1 | 45477819 | 45481247 | 5 | 2 | -0.69877 | 0.75765 | UROD | | 0.75765 | 0.55543 | 0.59675 |
| ENSG00000106608 | 7 | 43915493 | 43966010 | 171 | 20 | -0.69944 | 0.75786 | URGCP | | 0.75786 | 0.73605 | 0.69276 |
| ENSG00000221955 | 3 | 124801480 | 124998021 | 786 | 79 | -0.69948 | 0.75787 | SLC12A8 | | 0.75787 | 0.70078 | 0.72393 |
| ENSG00000179044 | 16 | 67218269 | 67224107 | 11 | 4 | -0.69948 | 0.75788 | EXOC3L1 | | 0.75788 | 0.83315 | 0.73544 |
| ENSG00000153147 | 4 | 144434616 | 144478639 | 89 | 9 | -0.69955 | 0.7579 | SMARCA5 | | 0.7579 | 0.77714 | 0.82065 |
| ENSG00000112249 | 6 | 100956070 | 101329248 | 1123 | 25 | -0.69957 | 0.7579 | ASCC3 | | 0.7579 | 0.74848 | 0.75753 |
| ENSG00000124224 | 20 | 56806188 | 56884495 | 235 | 19 | -0.69978 | 0.75797 | PPP4R1L | | 0.75797 | 0.7502 | 0.73429 |
| ENSG00000172901 | 5 | 115298151 | 115363316 | 305 | 34 | -0.69993 | 0.75801 | AQPEP | | 0.75801 | 0.71681 | 0.90854 |
| ENSG00000129474 | 14 | 23440383 | 23451851 | 20 | 4 | -0.70039 | 0.75816 | AJUBA | | 0.75816 | 0.76167 | 0.65962 |
| ENSG00000167363 | 17 | 80693451 | 80709073 | 47 | 10 | -0.70045 | 0.75818 | FN3K | | 0.75818 | 0.84435 | 0.71814 |
| ENSG00000101310 | 20 | 18488137 | 18542059 | 221 | 16 | -0.70051 | 0.75819 | SEC23B | | 0.75819 | 0.801 | 0.80527 |
| ENSG00000101197 | 20 | 61867235 | 61871859 | 13 | 5 | -0.70058 | 0.75822 | BIRC7 | | 0.75822 | 0.78473 | 0.76693 |
| ENSG00000236669 | 7 | 157318477 | 157320019 | 6 | 3 | -0.70062 | 0.75823 | AC006372.1 | | 0.75823 | 0.49507 | 0.3862 |
| ENSG00000109472 | 4 | 166282346 | 166419472 | 678 | 31 | -0.70083 | 0.75829 | CPE | | 0.75829 | 0.76066 | 0.76092 |
| ENSG00000269402 | 17 | 30506379 | 30506885 | 2 | 1 | -0.70092 | 0.75832 | AC116407.2 | | 0.75832 | 0.69229 | 0.68261 |
| ENSG00000084693 | 2 | 27265232 | 27293490 | 53 | 12 | -0.70103 | 0.75836 | AGBL5 | | 0.75836 | 0.70109 | 0.60883 |
| ENSG00000198558 | 6 | 27840926 | 27841289 | 1 | 1 | -0.70117 | 0.7584 | HIST1H4L | | 0.7584 | 0.79959 | 0.91325 |
| ENSG00000173068 | 9 | 16409501 | 16870841 | 1726 | 92 | -0.70145 | 0.75849 | BNC2 | | 0.75849 | 0.79392 | 0.79349 |
| ENSG00000119698 | 14 | 94612465 | 94746072 | 255 | 23 | -0.7015 | 0.7585 | PPP4R4 | | 0.7585 | 0.77976 | 0.79211 |
| ENSG00000106400 | 7 | 100860949 | 100867471 | 22 | 5 | -0.70163 | 0.75854 | ZNHIT1 | | 0.75854 | 0.65349 | 0.59449 |
| ENSG00000155827 | 9 | 104296133 | 104325622 | 63 | 15 | -0.70182 | 0.7586 | RNF20 | | 0.7586 | 0.89743 | 0.88612 |
| ENSG00000184497 | 13 | 114462216 | 114514926 | 253 | 17 | -0.70204 | 0.75867 | TMEM255B | | 0.75867 | 0.64469 | 0.49015 |
| ENSG00000110047 | 11 | 64619114 | 64655768 | 65 | 10 | -0.70238 | 0.75878 | EHD1 | | 0.75878 | 0.70826 | 0.75141 |
| ENSG00000037280 | 5 | 180028506 | 180076624 | 233 | 45 | -0.70242 | 0.75879 | FLT4 | | 0.75879 | 0.78781 | 0.74327 |
| ENSG00000221823 | 2 | 68405989 | 68483369 | 163 | 10 | -0.70252 | 0.75882 | PPP3R1 | | 0.75882 | 0.75675 | 0.7616 |
| ENSG00000075945 | 1 | 169890467 | 170054349 | 564 | 12 | -0.7026 | 0.75885 | KIFAP3 | | 0.75885 | 0.72945 | 0.7046 |
| ENSG00000134762 | 18 | 28569974 | 28622781 | 196 | 22 | -0.70272 | 0.75889 | DSC3 | | 0.75889 | 0.57923 | 0.56766 |
| ENSG00000100162 | 22 | 42334725 | 42343168 | 31 | 5 | -0.7029 | 0.75894 | CENPM | | 0.75894 | 0.86331 | 0.87346 |
| ENSG00000125637 | 2 | 113914902 | 113966973 | 199 | 37 | -0.70298 | 0.75896 | PSD4 | | 0.75896 | 0.8297 | 0.83365 |

| | | | | | | | | | | | |
|---|---|---|---|---|---|---|---|---|---|---|---|
| ENSG00000108018 | 10 | 108333421 | 108924292 | 1758 | 128 | -0.70309 | 0.759 | SORCS1 | 0.759 | 0.75566 | 0.74401 |
| ENSG00000172071 | 2 | 88856259 | 88927094 | 97 | 16 | -0.7032 | 0.75903 | EIF2AK3 | 0.75903 | 0.79802 | 0.8589 |
| ENSG00000095319 | 9 | 131709978 | 131769375 | 81 | 10 | -0.70357 | 0.75915 | NUP188 | 0.75915 | 0.74896 | 0.70593 |
| ENSG00000088888 | 20 | 3827487 | 3849280 | 75 | 25 | -0.70379 | 0.75922 | MAVS | 0.75922 | 0.84246 | 0.90968 |
| ENSG00000140395 | 15 | 78570177 | 78592136 | 61 | 13 | -0.70391 | 0.75926 | WDR61 | 0.75926 | 0.65279 | 0.28417 |
| ENSG00000116237 | 1 | 6281253 | 6296032 | 37 | 6 | -0.70394 | 0.75927 | ICMT | 0.75927 | 0.70528 | 0.76921 |
| ENSG00000163154 | 1 | 151129105 | 151132225 | 4 | 2 | -0.70416 | 0.75933 | TNFAIP8L2 | 0.75933 | 0.94091 | 0.91809 |
| ENSG00000147896 | 9 | 27524312 | 27526496 | 15 | 6 | -0.70415 | 0.75933 | IFNK | 0.75933 | 0.67564 | 0.75168 |
| ENSG00000135519 | 12 | 49932940 | 49952091 | 45 | 7 | -0.70419 | 0.75934 | KCNH3 | 0.75934 | 0.63174 | 0.61291 |
| ENSG00000131910 | 1 | 27237980 | 27240457 | 5 | 2 | -0.70422 | 0.75935 | NR0B2 | 0.75935 | 0.57721 | 0.55951 |
| ENSG00000095585 | 10 | 97951458 | 98031344 | 310 | 53 | -0.70436 | 0.75939 | BLNK | 0.75939 | 0.80802 | 0.86132 |
| ENSG00000204138 | 1 | 28696114 | 28826881 | 500 | 14 | -0.7045 | 0.75944 | PHACTR4 | 0.75944 | 0.75133 | 0.76287 |
| ENSG00000118094 | 11 | 118528026 | 118550399 | 56 | 5 | -0.70452 | 0.75945 | TREH | 0.75945 | 0.74376 | 0.75063 |
| ENSG00000126267 | 19 | 36139125 | 36149763 | 31 | 7 | -0.70453 | 0.75945 | COX6B1 | 0.75945 | 0.73242 | 0.82224 |
| ENSG00000107959 | 10 | 3179920 | 3215003 | 188 | 22 | -0.70458 | 0.75946 | PITRM1 | 0.75946 | 0.78716 | 0.59668 |
| ENSG00000176623 | 8 | 87480486 | 87526586 | 162 | 14 | -0.7046 | 0.75947 | RMDN1 | 0.75947 | 0.78302 | 0.77076 |
| ENSG00000165972 | 12 | 96260826 | 96336752 | 364 | 33 | -0.70461 | 0.75947 | CCDC38 | 0.75947 | 0.80258 | 0.8144 |
| ENSG00000133316 | 11 | 62599814 | 62609281 | 45 | 7 | -0.70469 | 0.7595 | WDR74 | 0.7595 | 0.75396 | 0.75853 |
| ENSG00000217930 | 16 | 4381550 | 4405608 | 60 | 13 | -0.70524 | 0.75967 | PAM16 | 0.75967 | 0.74922 | 0.78346 |
| ENSG00000072832 | 4 | 5749811 | 5894785 | 818 | 91 | -0.70529 | 0.75968 | CRMP1 | 0.75968 | 0.76543 | 0.79287 |
| ENSG00000107863 | 10 | 24872538 | 25012597 | 252 | 21 | -0.7055 | 0.75975 | ARHGAP21 | 0.75975 | 0.75725 | 0.5482 |
| ENSG00000103599 | 15 | 67547138 | 67794598 | 482 | 29 | -0.70558 | 0.75978 | IQCH | 0.75978 | 0.73223 | 0.69746 |
| ENSG00000184608 | 8 | 11225911 | 11296167 | 363 | 30 | -0.70575 | 0.75983 | C8orf12 | 0.75983 | 0.39547 | 0.36765 |
| ENSG00000174099 | 12 | 65672423 | 65882024 | 377 | 31 | -0.70586 | 0.75986 | MSRB3 | 0.75986 | 0.76332 | 0.72809 |
| ENSG00000173406 | 1 | 57460451 | 59012406 | 5270 | 292 | -0.70597 | 0.7599 | DAB1 | 0.7599 | 0.7945 | 0.73537 |
| ENSG00000137494 | 11 | 82904781 | 82971736 | 170 | 20 | -0.70651 | 0.76006 | ANKRD42 | 0.76006 | 0.77293 | 0.65744 |
| ENSG00000113161 | 5 | 74632154 | 74657929 | 57 | 10 | -0.70682 | 0.76016 | HMGCR | 0.76016 | 0.80754 | 0.63435 |
| ENSG00000119446 | 9 | 124999903 | 125027118 | 96 | 12 | -0.70685 | 0.76017 | RBM18 | 0.76017 | 0.79353 | 0.73692 |
| ENSG00000177108 | 14 | 77597613 | 77609077 | 34 | 12 | -0.70684 | 0.76017 | ZDHHC22 | 0.76017 | 0.85408 | 0.87621 |
| ENSG00000187695 | 3 | 128628717 | 128690173 | 192 | 24 | -0.70691 | 0.76019 | RP11-723O4.6 | 0.76019 | 0.76153 | 0.76731 |
| ENSG00000110887 | 12 | 109252708 | 109294819 | 163 | 28 | -0.70712 | 0.76025 | DAO | 0.76025 | 0.76565 | 0.76894 |
| ENSG00000152942 | 5 | 68665120 | 68710628 | 118 | 11 | -0.70717 | 0.76027 | RAD17 | 0.76027 | 0.74894 | 0.72906 |
| ENSG00000189319 | 10 | 126307861 | 126432838 | 272 | 27 | -0.70746 | 0.76036 | FAM53B | 0.76036 | 0.7384 | 0.76172 |
| ENSG00000107789 | 10 | 89264632 | 89313217 | 116 | 19 | -0.70758 | 0.7604 | MINPP1 | 0.7604 | 0.74784 | 0.73781 |
| ENSG00000128228 | 22 | 21996550 | 21998587 | 7 | 3 | -0.70776 | 0.76045 | SDF2L1 | 0.76045 | 0.79636 | 0.78576 |
| ENSG00000162244 | 3 | 52027616 | 52029958 | 4 | 2 | -0.70781 | 0.76047 | RPL29 | 0.76047 | 0.8723 | 0.84634 |
| ENSG00000122218 | 1 | 160259063 | 160313190 | 89 | 15 | -0.70784 | 0.76048 | COPA | 0.76048 | 0.78147 | 0.79407 |
| ENSG00000147459 | 8 | 25042238 | 25275598 | 786 | 63 | -0.70791 | 0.7605 | DOCK5 | 0.7605 | 0.71182 | 0.62876 |
| ENSG00000196704 | 17 | 66243715 | 66253297 | 33 | 10 | -0.70793 | 0.7605 | AMZ2 | 0.7605 | 0.75021 | 0.75915 |
| ENSG00000114107 | 3 | 138213186 | 138313380 | 163 | 17 | -0.70795 | 0.76051 | CEP70 | 0.76051 | 0.75584 | 0.72162 |
| ENSG00000145692 | 5 | 78407602 | 78428108 | 47 | 11 | -0.708 | 0.76053 | BHMT | 0.76053 | 0.70103 | 0.63789 |
| ENSG00000143889 | 2 | 38789120 | 38830728 | 115 | 16 | -0.70817 | 0.76058 | HNRNPLL | 0.76058 | 0.74414 | 0.62477 |
| ENSG00000135100 | 12 | 121416346 | 121440315 | 119 | 20 | -0.7082 | 0.76059 | HNF1A | 0.76059 | 0.74099 | 0.73746 |
| ENSG00000158796 | 1 | 161090764 | 161102478 | 16 | 6 | -0.70834 | 0.76063 | DEDD | 0.76063 | 0.60247 | 0.59877 |
| ENSG00000197622 | 1 | 151023447 | 151042801 | 38 | 11 | -0.70857 | 0.76071 | CDC42SE1 | 0.76071 | 0.8497 | 0.91346 |
| ENSG00000095464 | 10 | 95372345 | 95425767 | 205 | 35 | -0.70858 | 0.76071 | PDE6C | 0.76071 | 0.82337 | 0.8376 |
| ENSG00000197696 | 15 | 85198360 | 85201794 | 10 | 3 | -0.70858 | 0.76071 | NMB | 0.76071 | 0.67259 | 0.74066 |
| ENSG00000186174 | 11 | 118764584 | 118796317 | 58 | 17 | -0.70879 | 0.76077 | BCL9L | 0.76077 | 0.70498 | 0.71002 |
| ENSG00000173679 | 9 | 125423845 | 125424927 | 1 | 1 | -0.70888 | 0.7608 | OR1L1 | 0.7608 | 0.68579 | 0.81742 |
| ENSG00000196663 | 14 | 102829300 | 102968818 | 527 | 22 | -0.70886 | 0.7608 | TECPR2 | 0.7608 | 0.7262 | 0.67654 |
| ENSG00000082805 | 12 | 1099675 | 1605099 | 2105 | 47 | -0.70897 | 0.76083 | ERC1 | 0.76083 | 0.78583 | 0.81387 |
| ENSG00000134326 | 2 | 6980701 | 7006766 | 32 | 15 | -0.70902 | 0.76084 | CMPK2 | 0.76084 | 0.85288 | 0.76308 |
| ENSG00000105618 | 19 | 54618837 | 54635140 | 61 | 16 | -0.7093 | 0.76093 | PRPF31 | 0.76093 | 0.51668 | 0.53264 |
| ENSG00000034693 | 6 | 143771944 | 143811147 | 76 | 8 | -0.70936 | 0.76095 | PEX3 | 0.76095 | 0.8063 | 0.83914 |
| ENSG00000136160 | 13 | 78469616 | 78493903 | 59 | 12 | -0.7094 | 0.76096 | EDNRB | 0.76096 | 0.78914 | 0.85311 |
| ENSG00000179841 | 14 | 64932217 | 64941221 | 12 | 5 | -0.70953 | 0.761 | AKAP5 | 0.761 | 0.71711 | 0.76621 |
| ENSG00000204852 | 12 | 111051832 | 111087235 | 73 | 9 | -0.70978 | 0.76108 | TCTN1 | 0.76108 | 0.71192 | 0.74156 |
| ENSG00000108604 | 17 | 61909444 | 61920425 | 22 | 3 | -0.70979 | 0.76108 | SMARCD2 | 0.76108 | 0.81599 | 0.82549 |
| ENSG00000110395 | 11 | 119076752 | 119178859 | 177 | 16 | -0.71008 | 0.76117 | CBL | 0.76117 | 0.7881 | 0.8369 |
| ENSG00000064489 | 19 | 19256376 | 19302967 | 215 | 18 | -0.71007 | 0.76117 | MEF2BNB-MEF2B | 0.76117 | 0.72622 | 0.80305 |
| ENSG00000104723 | 8 | 15274724 | 15624158 | 1921 | 72 | -0.71009 | 0.76118 | TUSC3 | 0.76118 | 0.68819 | 0.69793 |
| ENSG00000166743 | 16 | 20634559 | 20710212 | 221 | 22 | -0.7102 | 0.76121 | ACSM1 | 0.76121 | 0.71582 | 0.65373 |
| ENSG00000149926 | 16 | 30035748 | 30064299 | 69 | 10 | -0.71035 | 0.76126 | FAM57B | 0.76126 | 0.76754 | 0.74217 |
| ENSG00000268423 | 19 | 47050313 | 47104118 | 171 | 29 | -0.71063 | 0.76134 | AC011551.3 | 0.76134 | 0.79621 | 0.73893 |
| ENSG00000064547 | 19 | 19734477 | 19739739 | 5 | 2 | -0.71071 | 0.76137 | LPAR2 | 0.76137 | 0.75778 | 0.89856 |
| ENSG00000185591 | 12 | 53773960 | 53810230 | 105 | 8 | -0.71078 | 0.76139 | SP1 | 0.76139 | 0.75258 | 0.69843 |
| ENSG00000138078 | 2 | 44543420 | 44589001 | 163 | 16 | -0.71089 | 0.76142 | PREPL | 0.76142 | 0.75716 | 0.73825 |
| ENSG00000167978 | 16 | 2802330 | 2822539 | 35 | 8 | -0.71096 | 0.76145 | SRRM2 | 0.76145 | 0.90244 | 0.84908 |
| ENSG00000164649 | 7 | 21940518 | 21985702 | 170 | 26 | -0.71111 | 0.76149 | CDCA7L | 0.76149 | 0.80973 | 0.86295 |
| ENSG00000103111 | 16 | 77224732 | 77236302 | 54 | 9 | -0.71143 | 0.76159 | MON1B | 0.76159 | 0.63141 | 0.45165 |
| ENSG00000068079 | 17 | 41158742 | 41166473 | 18 | 5 | -0.71153 | 0.76162 | IFI35 | 0.76162 | 0.84705 | 0.89433 |
| ENSG00000258436 | 14 | 21058352 | 21058982 | 1 | 1 | -0.71178 | 0.7617 | RNASE12 | 0.7617 | 0.15418 | 0.24087 |
| ENSG00000181449 | 3 | 181429714 | 181432221 | 3 | 2 | -0.71216 | 0.76182 | SOX2 | 0.76182 | 0.19832 | 0.070649 |

| | | | | | | | | | | | |
|---|---|---|---|---|---|---|---|---|---|---|---|
| ENSG00000170876 | 3 | 14166440 | 14185179 | 46 | 13 | -0.71233 | 0.76187 | TMEM43 | 0.76187 | 0.59921 | 0.59087 |
| ENSG00000053372 | 1 | 19578033 | 19586622 | 59 | 9 | -0.71247 | 0.76191 | MRTO4 | 0.76191 | 0.68318 | 0.58217 |
| ENSG00000164506 | 6 | 147525561 | 147706866 | 345 | 18 | -0.7125 | 0.76192 | STXBP5 | 0.76192 | 0.76899 | 0.80587 |
| ENSG00000156853 | 16 | 30613879 | 30635333 | 17 | 8 | -0.71258 | 0.76195 | ZNF689 | 0.76195 | 0.78653 | 0.75948 |
| ENSG00000168701 | 16 | 67261006 | 67263181 | 1 | 1 | -0.71275 | 0.762 | TMEM208 | 0.762 | 0.71706 | 0.61126 |
| ENSG00000273398 | 2 | 68358370 | 68488362 | 247 | 13 | -0.71305 | 0.76209 | RP11-474G23.1 | 0.76209 | 0.75826 | 0.73215 |
| ENSG00000196169 | 17 | 72322349 | 72351959 | 121 | 22 | -0.71322 | 0.76215 | KIF19 | 0.76215 | 0.824 | 0.85262 |
| ENSG00000243130 | 19 | 43511808 | 43530664 | 101 | 7 | -0.71329 | 0.76217 | PSG11 | 0.76217 | 0.72046 | 0.75418 |
| ENSG00000170790 | 11 | 6890924 | 6891976 | 8 | 2 | -0.71332 | 0.76218 | OR10A2 | 0.76218 | 0.41441 | 0.17633 |
| ENSG00000100478 | 14 | 31494312 | 31562818 | 233 | 35 | -0.71333 | 0.76218 | AP4S1 | 0.76218 | 0.83382 | 0.8615 |
| ENSG00000092531 | 15 | 42783431 | 42837547 | 105 | 12 | -0.7134 | 0.7622 | SNAP23 | 0.7622 | 0.74333 | 0.76429 |
| ENSG00000198185 | 20 | 45129701 | 45142198 | 26 | 6 | -0.7134 | 0.7622 | ZNF334 | 0.7622 | 0.6163 | 0.70058 |
| ENSG00000137193 | 6 | 37137979 | 37143202 | 9 | 4 | -0.71354 | 0.76225 | PIM1 | 0.76225 | 0.85428 | 0.28552 |
| ENSG00000103351 | 16 | 3550924 | 3589048 | 96 | 17 | -0.71366 | 0.76228 | CLUAP1 | 0.76228 | 0.66837 | 0.52441 |
| ENSG00000176101 | 9 | 140083099 | 140084822 | 4 | 1 | -0.71371 | 0.7623 | SSNA1 | 0.7623 | 0.79918 | 0.64048 |
| ENSG00000156466 | 8 | 97154562 | 97173020 | 38 | 10 | -0.71374 | 0.76231 | GDF6 | 0.76231 | 0.91841 | 0.93486 |
| ENSG00000197183 | 20 | 31030862 | 31172876 | 294 | 23 | -0.71398 | 0.76238 | C20orf112 | 0.76238 | 0.75641 | 0.68193 |
| ENSG00000160321 | 19 | 22115760 | 22193751 | 331 | 13 | -0.71419 | 0.76245 | ZNF208 | 0.76245 | 0.7709 | 0.8016 |
| ENSG00000175376 | 11 | 65764016 | 65769647 | 22 | 4 | -0.71451 | 0.76254 | EIF1AD | 0.76254 | 0.64726 | 0.63655 |
| ENSG00000179761 | 17 | 27277531 | 27384234 | 226 | 42 | -0.7145 | 0.76254 | PIPOX | 0.76254 | 0.81304 | 0.83946 |
| ENSG00000239264 | 6 | 7881483 | 8026646 | 716 | 66 | -0.71474 | 0.76262 | TXNDC5 | 0.76262 | 0.72699 | 0.758 |
| ENSG00000153391 | 18 | 33032194 | 33077955 | 108 | 19 | -0.71491 | 0.76267 | INO80C | 0.76267 | 0.69644 | 0.56715 |
| ENSG00000187066 | 11 | 64852451 | 64856847 | 7 | 3 | -0.71495 | 0.76268 | TMEM262 | 0.76268 | 0.82802 | 0.87285 |
| ENSG00000111077 | 12 | 53440753 | 53458156 | 43 | 11 | -0.71545 | 0.76283 | TENC1 | 0.76283 | 0.50531 | 0.27141 |
| ENSG00000171467 | 6 | 43274872 | 43337216 | 116 | 16 | -0.71585 | 0.76296 | ZNF318 | 0.76296 | 0.77769 | 0.71596 |
| ENSG00000172955 | 4 | 100123795 | 100140694 | 38 | 8 | -0.7159 | 0.76297 | ADH6 | 0.76297 | 0.65905 | 0.58735 |
| ENSG00000133943 | 14 | 91526677 | 91691976 | 540 | 54 | -0.71593 | 0.76298 | C14orf159 | 0.76298 | 0.8001 | 0.90421 |
| ENSG00000087494 | 12 | 28111017 | 28125638 | 49 | 10 | -0.7167 | 0.76322 | PTHLH | 0.76322 | 0.8471 | 0.89559 |
| ENSG00000165555 | 14 | 77860364 | 77889860 | 90 | 14 | -0.71671 | 0.76322 | NOXRED1 | 0.76322 | 0.77135 | 0.78644 |
| ENSG00000123737 | 4 | 122722472 | 122738176 | 45 | 9 | -0.71673 | 0.76323 | EXOSC9 | 0.76323 | 0.77757 | 0.75439 |
| ENSG00000002745 | 7 | 120965421 | 120981158 | 52 | 11 | -0.7168 | 0.76325 | WNT16 | 0.76325 | 0.62352 | 0.57503 |
| ENSG00000126822 | 14 | 65170820 | 65213610 | 105 | 13 | -0.71686 | 0.76327 | PLEKHG3 | 0.76327 | 0.77938 | 0.68063 |
| ENSG00000043093 | 3 | 182655862 | 182703741 | 67 | 11 | -0.71706 | 0.76333 | DCUN1D1 | 0.76333 | 0.76993 | 0.78547 |
| ENSG00000177427 | 17 | 18163848 | 18169866 | 13 | 5 | -0.71734 | 0.76342 | MIEF2 | 0.76342 | 0.70526 | 0.58933 |
| ENSG00000105717 | 19 | 19672522 | 19729725 | 87 | 14 | -0.71734 | 0.76342 | PBX4 | 0.76342 | 0.69839 | 0.73816 |
| ENSG00000168495 | 8 | 22102617 | 22112113 | 28 | 6 | -0.71737 | 0.76343 | POLR3D | 0.76343 | 0.87485 | 0.87126 |
| ENSG00000162040 | 16 | 1961464 | 1968441 | 36 | 4 | -0.71743 | 0.76345 | HS3ST6 | 0.76345 | 0.87394 | 0.8332 |
| ENSG00000258465 | 1 | 160187254 | 160254920 | 105 | 11 | -0.71792 | 0.7636 | DCAF8 | 0.7636 | 0.80544 | 0.85843 |
| ENSG00000108825 | 17 | 41102543 | 41132545 | 55 | 9 | -0.71809 | 0.76365 | PTGES3L-AARSD1 | 0.76365 | 0.75795 | 0.80167 |
| ENSG00000171574 | 19 | 58912871 | 58929694 | 46 | 6 | -0.71842 | 0.76375 | ZNF584 | 0.76375 | 0.71895 | 0.72305 |
| ENSG00000100418 | 22 | 41994032 | 42017100 | 42 | 9 | -0.71859 | 0.7638 | DESI1 | 0.7638 | 0.58143 | 0.59159 |
| ENSG00000139890 | 14 | 23352374 | 23356895 | 10 | 6 | -0.71861 | 0.76381 | REM2 | 0.76381 | 0.1919 | 0.59307 |
| ENSG00000116754 | 1 | 70671365 | 70718735 | 66 | 8 | -0.71863 | 0.76382 | SRSF11 | 0.76382 | 0.72142 | 0.66814 |
| ENSG00000125503 | 19 | 55602281 | 55628927 | 71 | 18 | -0.71879 | 0.76387 | PPP1R12C | 0.76387 | 0.82444 | 0.61969 |
| ENSG00000082929 | 4 | 5526296 | 5529528 | 9 | 3 | -0.71892 | 0.7639 | C4orf6 | 0.7639 | 0.85053 | 0.83484 |
| ENSG00000144451 | 2 | 214149113 | 215275225 | 3770 | 84 | -0.71904 | 0.76394 | SPAG16 | 0.76394 | 0.76197 | 0.77878 |
| ENSG00000256762 | 17 | 44076616 | 44077060 | 1 | 1 | -0.71923 | 0.764 | STH | 0.764 | 0.72423 | 0.72374 |
| ENSG00000258484 | 15 | 69110560 | 69239150 | 248 | 28 | -0.71931 | 0.76403 | SPESP1 | 0.76403 | 0.80231 | 0.86669 |
| ENSG00000100138 | 22 | 42069934 | 42086508 | 45 | 5 | -0.71935 | 0.76404 | NHP2L1 | 0.76404 | 0.74808 | 0.71079 |
| ENSG00000066855 | 8 | 66556969 | 66683496 | 426 | 26 | -0.71937 | 0.76405 | MTFR1 | 0.76405 | 0.76373 | 0.77735 |
| ENSG00000121594 | 3 | 119243140 | 119278449 | 131 | 26 | -0.71955 | 0.7641 | CD80 | 0.7641 | 0.16488 | 0.13564 |
| ENSG00000116030 | 2 | 203070903 | 203103331 | 86 | 13 | -0.7196 | 0.76411 | SUMO1 | 0.76411 | 0.76732 | 0.76348 |
| ENSG00000256825 | 12 | 133707570 | 133778955 | 205 | 17 | -0.71966 | 0.76413 | CTD-2140B24.4 | 0.76413 | 0.78896 | 0.83522 |
| ENSG00000116721 | 1 | 12851546 | 12856777 | 42 | 11 | -0.71969 | 0.76414 | PRAMEF1 | 0.76414 | 0.62693 | 0.47957 |
| ENSG00000213999 | 19 | 19256377 | 19303400 | 217 | 18 | -0.71978 | 0.76417 | MEF2B | 0.76417 | 0.72622 | 0.80305 |
| ENSG00000158161 | 1 | 28296855 | 28415207 | 297 | 20 | -0.72012 | 0.76428 | EYA3 | 0.76428 | 0.79774 | 0.85899 |
| ENSG00000150773 | 11 | 111934734 | 111944998 | 16 | 4 | -0.72035 | 0.76435 | PIH1D2 | 0.76435 | 0.72515 | 0.84513 |
| ENSG00000126453 | 19 | 50168823 | 50177173 | 11 | 3 | -0.72055 | 0.76441 | BCL2L12 | 0.76441 | 0.64159 | 0.76085 |
| ENSG00000144034 | 2 | 73956231 | 73964527 | 13 | 2 | -0.72086 | 0.7645 | TPRKB | 0.7645 | 0.71318 | 0.70913 |
| ENSG00000187446 | 15 | 41523037 | 41574043 | 153 | 16 | -0.72088 | 0.76451 | CHP1 | 0.76451 | 0.7716 | 0.66806 |
| ENSG00000197362 | 7 | 148766735 | 148787874 | 63 | 11 | -0.72101 | 0.76455 | ZNF786 | 0.76455 | 0.83394 | 0.86891 |
| ENSG00000162105 | 11 | 70313961 | 70963623 | 2124 | 144 | -0.72101 | 0.76455 | SHANK2 | 0.76455 | 0.77559 | 0.79926 |
| ENSG00000161048 | 7 | 102740223 | 102790007 | 85 | 14 | -0.72106 | 0.76456 | NAPEPLD | 0.76456 | 0.67859 | 0.58295 |
| ENSG00000256950 | 12 | 122326715 | 122359376 | 96 | 12 | -0.72105 | 0.76456 | RP11-87C12.2 | 0.76456 | 0.78905 | 0.8431 |
| ENSG00000099622 | 19 | 1259384 | 1274879 | 25 | 5 | -0.72148 | 0.76469 | CIRBP | 0.76469 | 0.24235 | 0.061442 |
| ENSG00000111262 | 12 | 5019071 | 5040527 | 62 | 12 | -0.72171 | 0.76476 | KCNA1 | 0.76476 | 0.84834 | 0.82378 |
| ENSG00000160685 | 1 | 154975127 | 154990998 | 23 | 9 | -0.72172 | 0.76477 | ZBTB7B | 0.76477 | 0.64282 | 0.46565 |
| ENSG00000206013 | 11 | 298200 | 299526 | 5 | 3 | -0.72182 | 0.7648 | IFITM5 | 0.7648 | 0.44872 | 0.43822 |
| ENSG00000114302 | 3 | 48782030 | 48885279 | 122 | 21 | -0.72186 | 0.76481 | PRKAR2A | 0.76481 | 0.75536 | 0.77303 |
| ENSG00000006015 | 19 | 18699495 | 18703146 | 13 | 3 | -0.722 | 0.76485 | C19orf60 | 0.76485 | 0.76552 | 0.74147 |
| ENSG00000180035 | 16 | 30389427 | 30411429 | 35 | 18 | -0.72216 | 0.7649 | ZNF48 | 0.7649 | 0.66751 | 0.72858 |
| ENSG00000012048 | 17 | 41196312 | 41277500 | 208 | 7 | -0.72216 | 0.7649 | BRCA1 | 0.7649 | 0.77933 | 0.75477 |
| ENSG00000117751 | 1 | 28157289 | 28178187 | 33 | 7 | -0.72232 | 0.76495 | PPP1R8 | 0.76495 | 0.70026 | 0.64622 |

| | | | | | | | | | | | |
|---|---|---|---|---|---|---|---|---|---|---|---|
| ENSG00000213316 | 5 | 179220981 | 179223648 | 7 | 3 | -0.72237 | 0.76497 | LTC4S | 0.76497 | 0.80274 | 0.73625 |
| ENSG00000115947 | 2 | 148687968 | 148779147 | 174 | 14 | -0.72244 | 0.76499 | ORC4 | 0.76499 | 0.76905 | 0.74948 |
| ENSG00000160813 | 7 | 100032905 | 100034188 | 1 | 1 | -0.72248 | 0.765 | PPP1R35 | 0.765 | 0.65549 | 0.64303 |
| ENSG00000260238 | 1 | 156182773 | 156213123 | 111 | 11 | -0.72253 | 0.76502 | PMF1-BGLAP | 0.76502 | 0.76601 | 0.75303 |
| ENSG00000160783 | 1 | 156182784 | 156212874 | 111 | 11 | -0.72253 | 0.76502 | PMF1 | 0.76502 | 0.75791 | 0.75303 |
| ENSG00000180767 | 3 | 126243126 | 126262134 | 97 | 15 | -0.72257 | 0.76503 | CHST13 | 0.76503 | 0.78482 | 0.75114 |
| ENSG00000169019 | 4 | 47452885 | 47465736 | 25 | 5 | -0.72291 | 0.76513 | COMMD8 | 0.76513 | 0.73989 | 0.69867 |
| ENSG00000269089 | 11 | 61735453 | 61736755 | 3 | 1 | -0.72291 | 0.76513 | AP003733.1 | 0.76513 | 0.46232 | 0.67197 |
| ENSG00000177084 | 12 | 133200348 | 133263951 | 236 | 14 | -0.72311 | 0.76519 | POLE | 0.76519 | 0.80462 | 0.81892 |
| ENSG00000127616 | 19 | 11071598 | 11176071 | 304 | 25 | -0.72335 | 0.76527 | SMARCA4 | 0.76527 | 0.72423 | 0.83562 |
| ENSG00000037474 | 5 | 6599352 | 6633404 | 181 | 18 | -0.72343 | 0.76529 | NSUN2 | 0.76529 | 0.61645 | 0.57407 |
| ENSG00000117758 | 1 | 28099694 | 28150963 | 76 | 8 | -0.72351 | 0.76532 | STX12 | 0.76532 | 0.78475 | 0.81614 |
| ENSG00000184344 | 12 | 7842378 | 7848372 | 39 | 12 | -0.72365 | 0.76536 | GDF3 | 0.76536 | 0.53157 | 0.5339 |
| ENSG00000119523 | 9 | 101978708 | 101984238 | 10 | 2 | -0.72368 | 0.76537 | ALG2 | 0.76537 | 0.64332 | 0.7789 |
| ENSG00000113790 | 3 | 184908412 | 184999778 | 188 | 18 | -0.72379 | 0.7654 | EHHADH | 0.7654 | 0.77553 | 0.735 |
| ENSG00000155287 | 10 | 101370282 | 101380366 | 24 | 4 | -0.7238 | 0.7654 | SLC25A28 | 0.7654 | 0.59316 | 0.62849 |
| ENSG00000167196 | 15 | 76196200 | 76227609 | 57 | 6 | -0.72383 | 0.76542 | FBXO22 | 0.76542 | 0.76372 | 0.82629 |
| ENSG00000102606 | 13 | 111766906 | 111958084 | 515 | 49 | -0.72403 | 0.76548 | ARHGEF7 | 0.76548 | 0.84032 | 0.82785 |
| ENSG00000204424 | 6 | 31674640 | 31685581 | 36 | 14 | -0.72472 | 0.76569 | LY6G6F | 0.76569 | 0.64695 | 0.6433 |
| ENSG00000100591 | 14 | 77924213 | 77935817 | 64 | 8 | -0.72471 | 0.76569 | AHSA1 | 0.76569 | 0.78499 | 0.83697 |
| ENSG00000137124 | 9 | 38392661 | 38398658 | 29 | 11 | -0.72486 | 0.76573 | ALDH1B1 | 0.76573 | 0.45124 | 0.3979 |
| ENSG00000170248 | 3 | 33839844 | 33911194 | 289 | 12 | -0.72494 | 0.76576 | PDCD6IP | 0.76576 | 0.8122 | 0.83503 |
| ENSG00000187242 | 17 | 39017555 | 39023462 | 13 | 6 | -0.72531 | 0.76587 | KRT12 | 0.76587 | 0.65986 | 0.65503 |
| ENSG00000111837 | 6 | 10762956 | 10838764 | 183 | 23 | -0.7259 | 0.76605 | MAK | 0.76605 | 0.69044 | 0.70063 |
| ENSG00000103591 | 15 | 67493371 | 67547533 | 113 | 10 | -0.72621 | 0.76615 | AAGAB | 0.76615 | 0.75073 | 0.71387 |
| ENSG00000204516 | 6 | 31462658 | 31478901 | 181 | 20 | -0.72642 | 0.76621 | MICB | 0.76621 | 0.57716 | 0.4445 |
| ENSG00000124574 | 6 | 43395104 | 43418168 | 57 | 14 | -0.72649 | 0.76623 | ABCC10 | 0.76623 | 0.7207 | 0.74706 |
| ENSG00000168778 | 12 | 124155660 | 124192948 | 206 | 10 | -0.72651 | 0.76624 | TCTN2 | 0.76624 | 0.79401 | 0.81213 |
| ENSG00000112462 | 6 | 29341200 | 29343068 | 10 | 4 | -0.7267 | 0.7663 | OR12D3 | 0.7663 | 0.77302 | 0.84535 |
| ENSG00000185739 | 16 | 4239375 | 4292081 | 250 | 39 | -0.72689 | 0.76635 | SRL | 0.76635 | 0.86797 | 0.81358 |
| ENSG00000170807 | 7 | 123295861 | 123304344 | 21 | 6 | -0.72695 | 0.76637 | LMOD2 | 0.76637 | 0.81934 | 0.74112 |
| ENSG00000109881 | 11 | 27352374 | 27385415 | 103 | 18 | -0.72706 | 0.7664 | CCDC34 | 0.7664 | 0.86354 | 0.90366 |
| ENSG00000123243 | 10 | 7601232 | 7708961 | 567 | 60 | -0.72715 | 0.76643 | ITIH5 | 0.76643 | 0.74859 | 0.77251 |
| ENSG00000163344 | 1 | 154897210 | 154909467 | 27 | 5 | -0.72717 | 0.76644 | PMVK | 0.76644 | 0.87042 | 0.87649 |
| ENSG00000178233 | 6 | 44238203 | 44275243 | 155 | 17 | -0.72765 | 0.76659 | TMEM151B | 0.76659 | 0.79066 | 0.80484 |
| ENSG00000172867 | 12 | 53038342 | 53045948 | 12 | 6 | -0.72784 | 0.76664 | KRT2 | 0.76664 | 0.91149 | 0.92801 |
| ENSG00000153902 | 19 | 35615417 | 35633355 | 55 | 6 | -0.72805 | 0.76671 | LGI4 | 0.76671 | 0.78743 | 0.65301 |
| ENSG00000143545 | 1 | 153954127 | 153958834 | 8 | 5 | -0.7284 | 0.76681 | RAB13 | 0.76681 | 0.30433 | 0.15895 |
| ENSG00000101752 | 18 | 19284918 | 19450918 | 311 | 21 | -0.7288 | 0.76694 | MIB1 | 0.76694 | 0.81407 | 0.80553 |
| ENSG00000255093 | 11 | 111322064 | 111328876 | 22 | 6 | -0.72886 | 0.76695 | RP11-794P6.2 | 0.76695 | 0.52743 | 0.66125 |
| ENSG00000136146 | 13 | 48627459 | 48669267 | 121 | 18 | -0.72937 | 0.76711 | MED4 | 0.76711 | 0.79722 | 0.82238 |
| ENSG00000179314 | 17 | 5675554 | 6027747 | 1625 | 179 | -0.72935 | 0.76711 | WSCD1 | 0.76711 | 0.77823 | 0.75233 |
| ENSG00000109182 | 4 | 48988264 | 49064098 | 143 | 12 | -0.72943 | 0.76713 | CWH43 | 0.76713 | 0.78761 | 0.81059 |
| ENSG00000222020 | 2 | 240323130 | 240324058 | 4 | 2 | -0.73021 | 0.76737 | AC062017.1 | 0.76737 | 0.60199 | 0.68828 |
| ENSG00000115760 | 2 | 32582096 | 32843966 | 644 | 28 | -0.73052 | 0.76747 | BIRC6 | 0.76747 | 0.79339 | 0.78684 |
| ENSG00000176401 | 19 | 40021630 | 40023494 | 8 | 3 | -0.73065 | 0.7675 | EID2B | 0.7675 | 0.65304 | 0.53547 |
| ENSG00000114738 | 3 | 50648951 | 50686720 | 70 | 12 | -0.732 | 0.76792 | MAPKAPK3 | 0.76792 | 0.77048 | 0.79793 |
| ENSG00000115042 | 2 | 96068469 | 96082364 | 11 | 8 | -0.73238 | 0.76803 | FAHD2A | 0.76803 | 0.8662 | 0.6722 |
| ENSG00000107140 | 9 | 35605367 | 35610038 | 9 | 5 | -0.73243 | 0.76805 | TESK1 | 0.76805 | 0.84271 | 0.70973 |
| ENSG00000082146 | 2 | 202252581 | 202345569 | 201 | 17 | -0.73253 | 0.76808 | STRADB | 0.76808 | 0.76388 | 0.82298 |
| ENSG00000224361 | 2 | 23729913 | 23747214 | 69 | 16 | -0.73322 | 0.76829 | AC011239.1 | 0.76829 | 0.75352 | 0.65999 |
| ENSG00000125534 | 20 | 62152077 | 62153559 | 7 | 3 | -0.7335 | 0.76837 | PPDPF | 0.76837 | 0.81545 | 0.83144 |
| ENSG00000151239 | 12 | 44187526 | 44200178 | 27 | 7 | -0.73398 | 0.76852 | TWF1 | 0.76852 | 0.87103 | 0.84791 |
| ENSG00000203691 | 1 | 225600404 | 225602040 | 4 | 2 | -0.73402 | 0.76853 | AC092811.1 | 0.76853 | 0.1954 | 0.38298 |
| ENSG00000176769 | 10 | 132890654 | 133109984 | 1202 | 131 | -0.73409 | 0.76855 | TCERG1L | 0.76855 | 0.64203 | 0.65021 |
| ENSG00000172059 | 2 | 10182976 | 10194963 | 42 | 9 | -0.73445 | 0.76866 | KLF11 | 0.76866 | 0.76288 | 0.59931 |
| ENSG00000149609 | 20 | 32250090 | 32253425 | 1 | 1 | -0.73457 | 0.7687 | C20orf144 | 0.7687 | 0.88056 | 0.9603 |
| ENSG00000158186 | 3 | 138066539 | 138124375 | 149 | 14 | -0.73472 | 0.76874 | MRAS | 0.76874 | 0.7472 | 0.75827 |
| ENSG00000112796 | 6 | 46126924 | 46138708 | 48 | 7 | -0.73483 | 0.76878 | ENPP5 | 0.76878 | 0.82621 | 0.57723 |
| ENSG00000167434 | 17 | 58227297 | 58248260 | 29 | 8 | -0.73493 | 0.76881 | CA4 | 0.76881 | 0.84198 | 0.84563 |
| ENSG00000163170 | 2 | 74362525 | 74375121 | 42 | 6 | -0.73512 | 0.76887 | BOLA3 | 0.76887 | 0.70677 | 0.62435 |
| ENSG00000088205 | 2 | 118572226 | 118589955 | 59 | 13 | -0.7354 | 0.76895 | DDX18 | 0.76895 | 0.77911 | 0.82472 |
| ENSG00000158715 | 1 | 205626979 | 205649587 | 63 | 18 | -0.73542 | 0.76896 | SLC45A3 | 0.76896 | 0.84457 | 0.90383 |
| ENSG00000174669 | 11 | 66129992 | 66139685 | 23 | 4 | -0.73546 | 0.76897 | SLC29A2 | 0.76897 | 0.9147 | 0.91724 |
| ENSG00000172638 | 11 | 65633912 | 65641063 | 16 | 3 | -0.7357 | 0.76904 | EFEMP2 | 0.76904 | 0.8323 | 0.87679 |
| ENSG00000205356 | 7 | 97843936 | 97881563 | 110 | 12 | -0.73574 | 0.76906 | TECPR1 | 0.76906 | 0.68428 | 0.59777 |
| ENSG00000163214 | 2 | 39024871 | 39103075 | 183 | 15 | -0.73587 | 0.76909 | DHX57 | 0.76909 | 0.7875 | 0.76653 |
| ENSG00000165078 | 8 | 68334360 | 68658620 | 997 | 74 | -0.73587 | 0.7691 | CPA6 | 0.7691 | 0.70845 | 0.63793 |
| ENSG00000198482 | 19 | 53030905 | 53067717 | 167 | 18 | -0.73599 | 0.76913 | ZNF808 | 0.76913 | 0.78547 | 0.86629 |
| ENSG00000136514 | 3 | 187086120 | 187089864 | 21 | 8 | -0.73609 | 0.76916 | RTP4 | 0.76916 | 0.69697 | 0.73985 |
| ENSG00000268688 | 2 | 37068327 | 37068530 | 1 | 1 | -0.73621 | 0.7692 | AC007382.1 | 0.7692 | 0.57534 | 0.52149 |
| ENSG00000128944 | 15 | 40674922 | 40686447 | 35 | 5 | -0.73623 | 0.76921 | KNSTRN | 0.76921 | 0.63116 | 0.58325 |
| ENSG00000114021 | 3 | 100053545 | 100075710 | 67 | 12 | -0.73632 | 0.76923 | NIT2 | 0.76923 | 0.90289 | 0.93677 |

| | | | | | | | | | | | |
|---|---|---|---|---|---|---|---|---|---|---|---|
| ENSG00000132716 | 1 | 160185505 | 160254920 | 106 | 11 | -0.7364 | 0.76926 | DCAF8 | 0.76926 | 0.80544 | 0.85843 |
| ENSG00000196793 | 10 | 44051792 | 44070066 | 52 | 6 | -0.73641 | 0.76926 | ZNF239 | 0.76926 | 0.72059 | 0.70771 |
| ENSG00000157895 | 12 | 121440225 | 121454305 | 70 | 15 | -0.73654 | 0.7693 | C12orf43 | 0.7693 | 0.79675 | 0.79245 |
| ENSG00000158941 | 8 | 22462145 | 22479027 | 46 | 4 | -0.73683 | 0.76939 | CCAR2 | 0.76939 | 0.81312 | 0.81813 |
| ENSG00000103496 | 16 | 31044210 | 31054296 | 19 | 4 | -0.73693 | 0.76942 | STX4 | 0.76942 | 0.77705 | 0.82884 |
| ENSG00000180245 | 4 | 110749150 | 110765760 | 57 | 6 | -0.73721 | 0.7695 | RRH | 0.7695 | 0.78983 | 0.87017 |
| ENSG00000187535 | 16 | 1560428 | 1662111 | 358 | 22 | -0.73722 | 0.76951 | IFT140 | 0.76951 | 0.73143 | 0.80144 |
| ENSG00000114487 | 3 | 108677086 | 108836989 | 591 | 17 | -0.73726 | 0.76952 | MORC1 | 0.76952 | 0.78656 | 0.75483 |
| ENSG00000198018 | 10 | 101419263 | 101465997 | 110 | 12 | -0.73728 | 0.76952 | ENTPD7 | 0.76952 | 0.81191 | 0.81435 |
| ENSG00000170734 | 6 | 43543887 | 43586701 | 80 | 14 | -0.73729 | 0.76953 | POLH | 0.76953 | 0.78944 | 0.7956 |
| ENSG00000114853 | 3 | 42695176 | 42709072 | 32 | 7 | -0.7374 | 0.76956 | ZBTB47 | 0.76956 | 0.656 | 0.54089 |
| ENSG00000126231 | 13 | 113812968 | 113826694 | 43 | 8 | -0.73764 | 0.76963 | PROZ | 0.76963 | 0.74958 | 0.55183 |
| ENSG00000198435 | 9 | 140194083 | 140196703 | 4 | 2 | -0.73767 | 0.76964 | NRARP | 0.76964 | 0.84178 | 0.55473 |
| ENSG00000084710 | 2 | 25264999 | 25378243 | 322 | 29 | -0.7377 | 0.76965 | EFR3B | 0.76965 | 0.76776 | 0.83636 |
| ENSG00000178363 | 10 | 5566924 | 5568225 | 5 | 2 | -0.73806 | 0.76976 | CALML3 | 0.76976 | 0.821 | 0.9244 |
| ENSG00000197647 | 19 | 12125547 | 12146556 | 20 | 13 | -0.73804 | 0.76976 | ZNF433 | 0.76976 | 0.90703 | 0.41087 |
| ENSG00000133818 | 11 | 14299472 | 14386052 | 146 | 27 | -0.73813 | 0.76978 | RRAS2 | 0.76978 | 0.75567 | 0.74278 |
| ENSG00000049130 | 12 | 88886570 | 88974628 | 204 | 11 | -0.73811 | 0.76978 | KITLG | 0.76978 | 0.77563 | 0.8142 |
| ENSG00000100647 | 14 | 70078313 | 70181859 | 206 | 31 | -0.73821 | 0.76981 | KIAA0247 | 0.76981 | 0.75906 | 0.76424 |
| ENSG00000161082 | 19 | 3224701 | 3297074 | 338 | 35 | -0.73822 | 0.76981 | CELF5 | 0.76981 | 0.7457 | 0.86073 |
| ENSG00000138073 | 2 | 27353624 | 27357543 | 9 | 3 | -0.7383 | 0.76983 | PREB | 0.76983 | 0.66224 | 0.66897 |
| ENSG00000173261 | 5 | 145463949 | 145483932 | 60 | 5 | -0.73869 | 0.76995 | PLAC8L1 | 0.76995 | 0.76256 | 0.76736 |
| ENSG00000131152 | 16 | 87336482 | 87367879 | 153 | 25 | -0.73923 | 0.77012 | RP11-178L8.4 | 0.77012 | 0.81853 | 0.82451 |
| ENSG00000085365 | 5 | 77656339 | 77776562 | 394 | 15 | -0.73947 | 0.77019 | SCAMP1 | 0.77019 | 0.7898 | 0.80303 |
| ENSG00000183647 | 19 | 58111253 | 58124090 | 27 | 8 | -0.7396 | 0.77023 | ZNF530 | 0.77023 | 0.77233 | 0.81565 |
| ENSG00000163440 | 4 | 56422692 | 56458379 | 161 | 14 | -0.73966 | 0.77025 | PDCL2 | 0.77025 | 0.62464 | 0.4859 |
| ENSG00000071082 | 2 | 101618177 | 101640494 | 64 | 9 | -0.74007 | 0.77037 | RPL31 | 0.77037 | 0.82617 | 0.80469 |
| ENSG00000090975 | 12 | 123468027 | 123634562 | 234 | 30 | -0.7404 | 0.77047 | PITPNM2 | 0.77047 | 0.77893 | 0.81855 |
| ENSG00000125691 | 17 | 37004118 | 37010096 | 16 | 7 | -0.74044 | 0.77048 | RPL23 | 0.77048 | 0.90518 | 0.89737 |
| ENSG00000261732 | 16 | 1682260 | 1736716 | 110 | 12 | -0.74051 | 0.7705 | LA16c-431H6.6 | 0.7705 | 0.67677 | 0.73186 |
| ENSG00000143437 | 1 | 150782181 | 150849244 | 129 | 15 | -0.74058 | 0.77052 | ARNT | 0.77052 | 0.79402 | 0.8134 |
| ENSG00000071246 | 14 | 77228532 | 77249354 | 73 | 19 | -0.74073 | 0.77057 | VASH1 | 0.77057 | 0.61372 | 0.29768 |
| ENSG00000131148 | 16 | 85805364 | 85833214 | 86 | 10 | -0.74075 | 0.77058 | EMC8 | 0.77058 | 0.77284 | 0.85377 |
| ENSG00000119547 | 18 | 55102917 | 55158529 | 183 | 26 | -0.74077 | 0.77058 | ONECUT2 | 0.77058 | 0.83288 | 0.91804 |
| ENSG00000110801 | 12 | 122326637 | 122356203 | 81 | 11 | -0.74094 | 0.77063 | PSMD9 | 0.77063 | 0.80113 | 0.8321 |
| ENSG00000132693 | 1 | 159682079 | 159684379 | 8 | 4 | -0.7411 | 0.77069 | CRP | 0.77069 | 0.65657 | 0.64433 |
| ENSG00000106299 | 7 | 123321989 | 123389121 | 178 | 15 | -0.7412 | 0.77071 | WASL | 0.77071 | 0.74388 | 0.70593 |
| ENSG00000117834 | 1 | 48688357 | 48714316 | 54 | 12 | -0.74129 | 0.77074 | SLC5A9 | 0.77074 | 0.77948 | 0.76736 |
| ENSG00000163832 | 3 | 47537130 | 47555251 | 39 | 6 | -0.74136 | 0.77076 | ELP6 | 0.77076 | 0.7715 | 0.64904 |
| ENSG00000168172 | 8 | 42752075 | 42885682 | 295 | 11 | -0.74177 | 0.77089 | HOOK3 | 0.77089 | 0.76166 | 0.78372 |
| ENSG00000167617 | 19 | 54976210 | 54984411 | 31 | 6 | -0.74211 | 0.77099 | CDC42EP5 | 0.77099 | 0.65607 | 0.26655 |
| ENSG00000149922 | 16 | 30097114 | 30103208 | 11 | 5 | -0.74215 | 0.771 | TBX6 | 0.771 | 0.73642 | 0.71896 |
| ENSG00000165474 | 13 | 20761609 | 20767037 | 18 | 7 | -0.74217 | 0.77101 | GJB2 | 0.77101 | 0.65924 | 0.47526 |
| ENSG00000114654 | 3 | 128720472 | 128759585 | 61 | 14 | -0.74241 | 0.77108 | EFCC1 | 0.77108 | 0.79757 | 0.73522 |
| ENSG00000140274 | 15 | 45406519 | 45410619 | 9 | 2 | -0.74253 | 0.77112 | DUOXA2 | 0.77112 | 0.88779 | 0.88082 |
| ENSG00000118137 | 11 | 116706467 | 116708666 | 11 | 5 | -0.7426 | 0.77114 | APOA1 | 0.77114 | 0.7953 | 0.87696 |
| ENSG00000102780 | 13 | 42614176 | 42830714 | 894 | 41 | -0.74284 | 0.77121 | DGKH | 0.77121 | 0.72278 | 0.70738 |
| ENSG00000139219 | 12 | 48366748 | 48398269 | 124 | 17 | -0.74334 | 0.77136 | COL2A1 | 0.77136 | 0.63051 | 0.39173 |
| ENSG00000060491 | 20 | 61436187 | 61445352 | 51 | 10 | -0.74334 | 0.77136 | OGFR | 0.77136 | 0.65377 | 0.281 |
| ENSG00000127081 | 9 | 95607874 | 95640304 | 108 | 15 | -0.74362 | 0.77145 | ZNF484 | 0.77145 | 0.75417 | 0.72403 |
| ENSG00000166257 | 11 | 123499895 | 123525952 | 103 | 19 | -0.74365 | 0.77145 | SCN3B | 0.77145 | 0.84419 | 0.82901 |
| ENSG00000197561 | 19 | 851014 | 856242 | 10 | 6 | -0.74381 | 0.7715 | ELANE | 0.7715 | 0.37207 | 0.63527 |
| ENSG00000247121 | 5 | 96149731 | 96271513 | 705 | 34 | -0.74425 | 0.77164 | CTD-2260A17.2 | 0.77164 | 0.72562 | 0.62952 |
| ENSG00000232850 | 9 | 130890612 | 130891481 | 2 | 1 | -0.74436 | 0.77167 | AL590708.2 | 0.77167 | 0.90635 | 0.84428 |
| ENSG00000183762 | 22 | 29469066 | 29564321 | 389 | 36 | -0.74435 | 0.77167 | KREMEN1 | 0.77167 | 0.77929 | 0.74241 |
| ENSG00000149196 | 11 | 86013253 | 86056969 | 203 | 15 | -0.74454 | 0.77172 | C11orf73 | 0.77172 | 0.71362 | 0.56414 |
| ENSG00000184752 | 12 | 95290831 | 95397546 | 672 | 45 | -0.74457 | 0.77174 | NDUFA12 | 0.77174 | 0.7753 | 0.75612 |
| ENSG00000102878 | 16 | 67197288 | 67203848 | 5 | 3 | -0.74482 | 0.77181 | HSF4 | 0.77181 | 0.89439 | 0.92122 |
| ENSG00000166965 | 15 | 91498100 | 91506349 | 37 | 8 | -0.74488 | 0.77183 | RCCD1 | 0.77183 | 0.68086 | 0.55795 |
| ENSG00000168818 | 4 | 4417469 | 4544073 | 352 | 19 | -0.74524 | 0.77194 | STX18 | 0.77194 | 0.80491 | 0.81507 |
| ENSG00000139209 | 12 | 47158546 | 47226191 | 261 | 32 | -0.74527 | 0.77195 | SLC38A4 | 0.77195 | 0.76986 | 0.75473 |
| ENSG00000079557 | 4 | 74347400 | 74369691 | 73 | 12 | -0.74532 | 0.77196 | AFM | 0.77196 | 0.74066 | 0.82378 |
| ENSG00000175336 | 12 | 56754353 | 56756607 | 4 | 2 | -0.74557 | 0.77204 | APOF | 0.77204 | 0.22909 | 0.26405 |
| ENSG00000162613 | 1 | 78409740 | 78444794 | 84 | 14 | -0.74566 | 0.77206 | FUBP1 | 0.77206 | 0.66337 | 0.6133 |
| ENSG00000187024 | 9 | 130455257 | 130487152 | 36 | 10 | -0.74568 | 0.77207 | PTRH1 | 0.77207 | 0.68273 | 0.53584 |
| ENSG00000139620 | 12 | 49047184 | 49076021 | 82 | 11 | -0.7457 | 0.77207 | KANSL2 | 0.77207 | 0.73371 | 0.72364 |
| ENSG00000114744 | 3 | 149456257 | 149470286 | 33 | 6 | -0.74594 | 0.77215 | COMMD2 | 0.77215 | 0.87299 | 0.89864 |
| ENSG00000069431 | 12 | 21950335 | 22094336 | 377 | 37 | -0.74598 | 0.77216 | ABCC9 | 0.77216 | 0.79463 | 0.86038 |
| ENSG00000104529 | 8 | 144661867 | 144681711 | 87 | 10 | -0.74604 | 0.77218 | EEF1D | 0.77218 | 0.64754 | 0.65367 |
| ENSG00000162300 | 11 | 64851695 | 64855870 | 3 | 1 | -0.74613 | 0.77221 | ZFPL1 | 0.77221 | 0.84581 | 0.87674 |
| ENSG00000177082 | 15 | 85185999 | 85197574 | 31 | 7 | -0.74614 | 0.77221 | WDR73 | 0.77221 | 0.69663 | 0.72311 |
| ENSG00000111530 | 12 | 67663061 | 67713731 | 183 | 17 | -0.74624 | 0.77224 | CAND1 | 0.77224 | 0.7459 | 0.78913 |
| ENSG00000104848 | 19 | 49570675 | 49576198 | 28 | 5 | -0.74632 | 0.77226 | KCNA7 | 0.77226 | 0.69774 | 0.51946 |

| | | | | | | | | | | | |
|---|---|---|---|---|---|---|---|---|---|---|---|
| ENSG00000179873 | 19 | 56296770 | 56348166 | 358 | 47 | -0.74639 | 0.77228 | NLRP11 | 0.77228 | 0.67939 | 0.70913 |
| ENSG00000183751 | 16 | 2022038 | 2032934 | 66 | 8 | -0.74641 | 0.77229 | TBL3 | 0.77229 | 0.93428 | 0.96411 |
| ENSG00000096401 | 6 | 44355262 | 44418163 | 195 | 27 | -0.74661 | 0.77235 | CDC5L | 0.77235 | 0.79646 | 0.91293 |
| ENSG00000003249 | 16 | 90071273 | 90086536 | 83 | 19 | -0.74765 | 0.77267 | DBNDD1 | 0.77267 | 0.53365 | 0.28941 |
| ENSG00000103550 | 16 | 19714902 | 19729557 | 37 | 4 | -0.74771 | 0.77268 | KNOP1 | 0.77268 | 0.72125 | 0.64887 |
| ENSG00000137273 | 6 | 1390069 | 1395832 | 25 | 10 | -0.74781 | 0.77271 | FOXF2 | 0.77271 | 0.848 | 0.79076 |
| ENSG00000135596 | 6 | 109765265 | 109787171 | 65 | 13 | -0.74814 | 0.77281 | MICAL1 | 0.77281 | 0.84743 | 0.74586 |
| ENSG00000168135 | 22 | 38822332 | 38851205 | 101 | 21 | -0.74827 | 0.77285 | KCNJ4 | 0.77285 | 0.7401 | 0.75081 |
| ENSG00000184881 | 11 | 5344541 | 5345582 | 10 | 2 | -0.74842 | 0.7729 | OR51B2 | 0.7729 | 0.81879 | 0.43562 |
| ENSG00000101343 | 20 | 20015012 | 20036690 | 86 | 10 | -0.74847 | 0.77291 | CRNKL1 | 0.77291 | 0.84418 | 0.84843 |
| ENSG00000167815 | 19 | 12907634 | 12912694 | 18 | 4 | -0.74863 | 0.77296 | PRDX2 | 0.77296 | 0.56717 | 0.37403 |
| ENSG00000148719 | 10 | 74092588 | 74114988 | 71 | 4 | -0.74874 | 0.77299 | DNAJB12 | 0.77299 | 0.78231 | 0.71039 |
| ENSG00000183255 | 21 | 46269500 | 46293752 | 123 | 19 | -0.74881 | 0.77301 | PTTG1IP | 0.77301 | 0.82477 | 0.77396 |
| ENSG00000087903 | 19 | 5993175 | 6199583 | 692 | 43 | -0.74883 | 0.77302 | RFX2 | 0.77302 | 0.72979 | 0.72125 |
| ENSG00000071677 | 2 | 238475217 | 238475818 | 1 | 1 | -0.74909 | 0.7731 | PRLH | 0.7731 | 0.60958 | 0.50945 |
| ENSG00000124575 | 6 | 26234440 | 26235216 | 4 | 2 | -0.74949 | 0.77322 | HIST1H1D | 0.77322 | 0.44277 | 0.48191 |
| ENSG00000124275 | 5 | 7851299 | 7906138 | 289 | 27 | -0.74975 | 0.7733 | MTRR | 0.7733 | 0.83979 | 0.78795 |
| ENSG00000104517 | 8 | 103265240 | 103425069 | 551 | 30 | -0.74974 | 0.7733 | UBR5 | 0.7733 | 0.7946 | 0.80191 |
| ENSG00000129270 | 17 | 34083268 | 34122711 | 105 | 13 | -0.74975 | 0.7733 | MMP28 | 0.7733 | 0.73752 | 0.76391 |
| ENSG00000106244 | 7 | 98989671 | 99006452 | 26 | 5 | -0.7498 | 0.77331 | PDAP1 | 0.77331 | 0.82583 | 0.84139 |
| ENSG00000103742 | 15 | 65673802 | 65715410 | 119 | 26 | -0.74988 | 0.77334 | IGDCC4 | 0.77334 | 0.74524 | 0.65699 |
| ENSG00000166920 | 15 | 45722727 | 45740959 | 29 | 9 | -0.74991 | 0.77335 | C15orf48 | 0.77335 | 0.86571 | 0.84452 |
| ENSG00000075239 | 11 | 107992243 | 108018503 | 61 | 11 | -0.74997 | 0.77336 | ACAT1 | 0.77336 | 0.86857 | 0.85322 |
| ENSG00000114646 | 3 | 47603729 | 47622282 | 31 | 5 | -0.75005 | 0.77339 | CSPG5 | 0.77339 | 0.69066 | 0.82561 |
| ENSG00000152465 | 10 | 15144583 | 15210692 | 304 | 29 | -0.7504 | 0.77349 | NMT2 | 0.77349 | 0.81531 | 0.8384 |
| ENSG00000168787 | 6 | 29364416 | 29365448 | 9 | 2 | -0.75047 | 0.77352 | OR12D2 | 0.77352 | 0.7705 | 0.90206 |
| ENSG00000130766 | 1 | 28586038 | 28609002 | 55 | 11 | -0.75054 | 0.77354 | SESN2 | 0.77354 | 0.59617 | 0.31638 |
| ENSG00000101405 | 20 | 3052266 | 3053163 | 4 | 2 | -0.75055 | 0.77354 | OXT | 0.77354 | 0.096345 | 0.029543 |
| ENSG00000168081 | 8 | 28174503 | 28200872 | 114 | 8 | -0.75075 | 0.7736 | PNOC | 0.7736 | 0.67966 | 0.80345 |
| ENSG00000165912 | 11 | 47199076 | 47207994 | 22 | 6 | -0.75085 | 0.77363 | PACSIN3 | 0.77363 | 0.76703 | 0.73053 |
| ENSG00000239789 | 7 | 56019512 | 56024192 | 15 | 5 | -0.75094 | 0.77365 | MRPS17 | 0.77365 | 0.70795 | 0.72389 |
| ENSG00000117148 | 1 | 18081808 | 18153558 | 204 | 26 | -0.75116 | 0.77372 | ACTL8 | 0.77372 | 0.81294 | 0.85954 |
| ENSG00000268223 | 5 | 115387163 | 115395815 | 76 | 4 | -0.75144 | 0.7738 | ARL14EPL | 0.7738 | 0.92384 | 0.84051 |
| ENSG00000078401 | 6 | 12290596 | 12297427 | 26 | 8 | -0.75177 | 0.7739 | EDN1 | 0.7739 | 0.81228 | 0.7582 |
| ENSG00000100023 | 22 | 22006559 | 22054304 | 206 | 24 | -0.75178 | 0.77391 | PPIL2 | 0.77391 | 0.84466 | 0.81937 |
| ENSG00000168830 | 6 | 87647024 | 87726349 | 178 | 19 | -0.75205 | 0.77399 | HTR1E | 0.77399 | 0.8461 | 0.90717 |
| ENSG00000137204 | 6 | 43263432 | 43273276 | 17 | 6 | -0.75235 | 0.77408 | SLC22A7 | 0.77408 | 0.63873 | 0.28791 |
| ENSG00000170312 | 10 | 62538089 | 62554610 | 66 | 8 | -0.75254 | 0.77414 | CDK1 | 0.77414 | 0.83542 | 0.88429 |
| ENSG00000151233 | 12 | 42475647 | 42538681 | 191 | 22 | -0.753 | 0.77428 | GXYLT1 | 0.77428 | 0.79915 | 0.8158 |
| ENSG00000170482 | 5 | 138702885 | 138720242 | 22 | 5 | -0.75308 | 0.7743 | SLC23A1 | 0.7743 | 0.54266 | 0.34729 |
| ENSG00000186439 | 6 | 123537483 | 123958238 | 1439 | 69 | -0.75308 | 0.7743 | TRDN | 0.7743 | 0.76646 | 0.70603 |
| ENSG00000173473 | 3 | 47626762 | 47823596 | 284 | 12 | -0.7532 | 0.77434 | SMARCC1 | 0.77434 | 0.77979 | 0.80549 |
| ENSG00000111237 | 12 | 110928902 | 110939922 | 22 | 5 | -0.75332 | 0.77437 | VPS29 | 0.77437 | 0.77427 | 0.84054 |
| ENSG00000131746 | 17 | 38632080 | 38657849 | 94 | 20 | -0.75341 | 0.7744 | TNS4 | 0.7744 | 0.78005 | 0.84171 |
| ENSG00000173599 | 11 | 66615704 | 66725847 | 230 | 27 | -0.75363 | 0.77446 | PC | 0.77446 | 0.79681 | 0.74695 |
| ENSG00000168676 | 16 | 67323331 | 67360666 | 57 | 10 | -0.75379 | 0.77451 | KCTD19 | 0.77451 | 0.70857 | 0.76116 |
| ENSG00000143105 | 1 | 111059839 | 111061797 | 6 | 3 | -0.75384 | 0.77453 | KCNA10 | 0.77453 | 0.83292 | 0.94055 |
| ENSG00000169813 | 10 | 43881065 | 43904614 | 94 | 15 | -0.75409 | 0.7746 | HNRNPF | 0.7746 | 0.76209 | 0.76002 |
| ENSG00000187701 | 1 | 248813232 | 248814185 | 6 | 2 | -0.75416 | 0.77462 | OR2T27 | 0.77462 | 0.6264 | 0.35972 |
| ENSG00000099769 | 16 | 1840414 | 1844972 | 28 | 5 | -0.75427 | 0.77466 | IGFALS | 0.77466 | 0.75629 | 0.78367 |
| ENSG00000103199 | 16 | 4798240 | 4817625 | 92 | 12 | -0.75441 | 0.7747 | ZNF500 | 0.7747 | 0.73123 | 0.66357 |
| ENSG00000152595 | 4 | 88742563 | 88767969 | 105 | 17 | -0.75478 | 0.77481 | MEPE | 0.77481 | 0.75819 | 0.6461 |
| ENSG00000133020 | 17 | 10293639 | 10325267 | 52 | 8 | -0.75477 | 0.77481 | MYH8 | 0.77481 | 0.81446 | 0.796 |
| ENSG00000174343 | 4 | 40337346 | 40357234 | 129 | 12 | -0.75531 | 0.77497 | CHRNA9 | 0.77497 | 0.71793 | 0.79395 |
| ENSG00000187051 | 22 | 39925098 | 39928860 | 3 | 1 | -0.75548 | 0.77502 | RPS19BP1 | 0.77502 | 0.44399 | 0.41314 |
| ENSG00000139514 | 13 | 30083547 | 30169825 | 288 | 23 | -0.75565 | 0.77507 | SLC7A1 | 0.77507 | 0.77999 | 0.85433 |
| ENSG00000081377 | 9 | 99252523 | 99382112 | 151 | 23 | -0.75608 | 0.7752 | CDC14B | 0.7752 | 0.78841 | 0.7827 |
| ENSG00000268211 | 9 | 131062367 | 131064212 | 8 | 2 | -0.75608 | 0.7752 | AL359091.2 | 0.7752 | 0.7736 | 0.75655 |
| ENSG00000142864 | 1 | 67873493 | 67896098 | 37 | 8 | -0.75631 | 0.77527 | SERBP1 | 0.77527 | 0.87571 | 0.85191 |
| ENSG00000107447 | 10 | 98064085 | 98098321 | 100 | 12 | -0.75631 | 0.77527 | DNTT | 0.77527 | 0.64676 | 0.62885 |
| ENSG00000107938 | 10 | 127408084 | 127452712 | 59 | 10 | -0.75639 | 0.77529 | EDRF1 | 0.77529 | 0.62126 | 0.68178 |
| ENSG00000112818 | 6 | 46761127 | 46807515 | 146 | 14 | -0.75663 | 0.77536 | MEP1A | 0.77536 | 0.80438 | 0.77811 |
| ENSG00000171421 | 5 | 1798500 | 1801480 | 26 | 5 | -0.75672 | 0.77539 | MRPL36 | 0.77539 | 0.71307 | 0.71079 |
| ENSG00000115233 | 2 | 162164549 | 162268228 | 144 | 17 | -0.75737 | 0.77559 | PSMD14 | 0.77559 | 0.7807 | 0.77056 |
| ENSG00000189377 | 19 | 42932440 | 42947200 | 24 | 3 | -0.75759 | 0.77565 | CXCL17 | 0.77565 | 0.70922 | 0.79225 |
| ENSG00000166206 | 15 | 26788693 | 27184686 | 1532 | 110 | -0.75777 | 0.77571 | GABRB3 | 0.77571 | 0.78156 | 0.80113 |
| ENSG00000155367 | 1 | 113252616 | 113258099 | 15 | 8 | -0.75785 | 0.77573 | PPM1J | 0.77573 | 0.72468 | 0.054525 |
| ENSG00000175602 | 11 | 65657875 | 65659105 | 2 | 1 | -0.75796 | 0.77576 | CCDC85B | 0.77576 | 0.90214 | 0.89267 |
| ENSG00000159314 | 17 | 43471275 | 43511787 | 86 | 4 | -0.75802 | 0.77578 | ARHGAP27 | 0.77578 | 0.7904 | 0.83841 |
| ENSG00000183145 | 21 | 38378450 | 38391959 | 58 | 10 | -0.7582 | 0.77583 | RIPPLY3 | 0.77583 | 0.65574 | 0.77728 |
| ENSG00000206047 | 8 | 6835171 | 6837602 | 2 | 1 | -0.75827 | 0.77586 | DEFA1 | 0.77586 | 0.063978 | 0.044141 |
| ENSG00000115363 | 2 | 75696428 | 75796848 | 313 | 36 | -0.75848 | 0.77592 | EVA1A | 0.77592 | 0.79888 | 0.75641 |
| ENSG00000114480 | 3 | 81538850 | 81811312 | 506 | 31 | -0.75861 | 0.77596 | GBE1 | 0.77596 | 0.81915 | 0.80776 |

| | | | | | | | | | | | |
|---|---|---|---|---|---|---|---|---|---|---|---|
| ENSG00000131634 | 16 | 1578689 | 1605581 | 105 | 14 | -0.7587 | 0.77598 | TMEM204 | 0.77598 | 0.76091 | 0.69073 |
| ENSG00000186866 | 21 | 46683843 | 46707813 | 77 | 8 | -0.75868 | 0.77598 | POFUT2 | 0.77598 | 0.7865 | 0.79 |
| ENSG00000175455 | 3 | 123616152 | 123680564 | 179 | 20 | -0.75877 | 0.776 | CCDC14 | 0.776 | 0.82908 | 0.85569 |
| ENSG00000176845 | 17 | 81037567 | 81052864 | 70 | 22 | -0.75878 | 0.77601 | METRNL | 0.77601 | 0.55436 | 0.28269 |
| ENSG00000255104 | 17 | 15603006 | 15648092 | 167 | 26 | -0.75894 | 0.77605 | ZNF286A | 0.77605 | 0.7551 | 0.72273 |
| ENSG00000175564 | 11 | 73711326 | 73720480 | 31 | 6 | -0.75906 | 0.77609 | UCP3 | 0.77609 | 0.74144 | 0.44954 |
| ENSG00000169718 | 17 | 80015382 | 80023763 | 15 | 7 | -0.7591 | 0.7761 | DUS1L | 0.7761 | 0.816 | 0.76321 |
| ENSG00000186847 | 17 | 39738531 | 39743173 | 9 | 1 | -0.75919 | 0.77613 | KRT14 | 0.77613 | 0.83953 | 0.82723 |
| ENSG00000167612 | 12 | 52281744 | 52285448 | 18 | 7 | -0.75928 | 0.77616 | ANKRD33 | 0.77616 | 0.60995 | 0.6239 |
| ENSG00000064545 | 19 | 19229978 | 19249322 | 56 | 11 | -0.75928 | 0.77616 | TMEM161A | 0.77616 | 0.65056 | 0.80659 |
| ENSG00000169727 | 17 | 80008567 | 80015346 | 17 | 5 | -0.75936 | 0.77618 | GPS1 | 0.77618 | 0.88148 | 0.74127 |
| ENSG00000128683 | 2 | 171669723 | 171717661 | 180 | 24 | -0.75943 | 0.7762 | GAD1 | 0.7762 | 0.70308 | 0.79667 |
| ENSG00000096746 | 10 | 70090931 | 70102948 | 34 | 7 | -0.75958 | 0.77625 | HNRNPH3 | 0.77625 | 0.7972 | 0.66977 |
| ENSG00000138079 | 2 | 44502599 | 44548633 | 185 | 19 | -0.75991 | 0.77635 | SLC3A1 | 0.77635 | 0.83107 | 0.84562 |
| ENSG00000166669 | 16 | 10420291 | 10577495 | 652 | 40 | -0.75994 | 0.77636 | ATF7IP2 | 0.77636 | 0.80861 | 0.78887 |
| ENSG00000131095 | 17 | 42982376 | 42994305 | 51 | 13 | -0.75996 | 0.77636 | GFAP | 0.77636 | 0.73002 | 0.71216 |
| ENSG00000180257 | 19 | 53430388 | 53466164 | 180 | 27 | -0.76031 | 0.77646 | ZNF816 | 0.77646 | 0.85048 | 0.92797 |
| ENSG00000105499 | 19 | 48551100 | 48614074 | 392 | 50 | -0.76076 | 0.7766 | PLA2G4C | 0.7766 | 0.6386 | 0.539 |
| ENSG00000164830 | 8 | 107282473 | 107764922 | 1545 | 65 | -0.76097 | 0.77666 | OXR1 | 0.77666 | 0.77206 | 0.78774 |
| ENSG00000198453 | 19 | 37407231 | 37489602 | 437 | 16 | -0.76095 | 0.77666 | ZNF568 | 0.77666 | 0.78025 | 0.74558 |
| ENSG00000180475 | 11 | 57995354 | 57996390 | 4 | 2 | -0.7612 | 0.77673 | OR10Q1 | 0.77673 | 0.32856 | 0.16348 |
| ENSG00000177556 | 5 | 151121877 | 151152093 | 45 | 11 | -0.76145 | 0.77681 | ATOX1 | 0.77681 | 0.78647 | 0.75195 |
| ENSG00000204814 | 9 | 44867571 | 44870528 | 20 | 3 | -0.76146 | 0.77681 | RP11-160N1.10 | 0.77681 | 0.80935 | 0.80223 |
| ENSG00000116329 | 1 | 29138654 | 29190208 | 201 | 23 | -0.76164 | 0.77686 | OPRD1 | 0.77686 | 0.88399 | 0.91929 |
| ENSG00000140279 | 15 | 45384848 | 45406542 | 45 | 5 | -0.76184 | 0.77692 | DUOX2 | 0.77692 | 0.8761 | 0.91956 |
| ENSG00000186017 | 19 | 36936021 | 36980804 | 138 | 14 | -0.76188 | 0.77693 | ZNF566 | 0.77693 | 0.81926 | 0.80539 |
| ENSG00000137073 | 9 | 33921691 | 34048947 | 510 | 20 | -0.762 | 0.77697 | UBAP2 | 0.77697 | 0.7675 | 0.73636 |
| ENSG00000113749 | 5 | 175085033 | 175113245 | 55 | 14 | -0.76233 | 0.77707 | HRH2 | 0.77707 | 0.76243 | 0.85036 |
| ENSG00000175536 | 11 | 74202757 | 74204778 | 7 | 2 | -0.76239 | 0.77709 | LIPT2 | 0.77709 | 0.75849 | 0.70739 |
| ENSG00000100351 | 22 | 40297086 | 40369725 | 87 | 21 | -0.76245 | 0.7771 | GRAP2 | 0.7771 | 0.82815 | 0.90443 |
| ENSG00000168936 | 4 | 1717679 | 1723085 | 26 | 6 | -0.7629 | 0.77724 | TMEM129 | 0.77724 | 0.60005 | 0.51755 |
| ENSG00000171723 | 14 | 66974125 | 67648520 | 2579 | 20 | -0.76318 | 0.77732 | GPHN | 0.77732 | 0.76188 | 0.77308 |
| ENSG00000103227 | 16 | 903634 | 1031318 | 851 | 38 | -0.76316 | 0.77732 | LMF1 | 0.77732 | 0.72994 | 0.79156 |
| ENSG00000005059 | 4 | 110481361 | 110609874 | 397 | 23 | -0.7632 | 0.77733 | CCDC109B | 0.77733 | 0.7038 | 0.59551 |
| ENSG00000109445 | 4 | 142142041 | 142155851 | 28 | 4 | -0.76322 | 0.77733 | ZNF330 | 0.77733 | 0.85487 | 0.89724 |
| ENSG00000144821 | 3 | 108099216 | 108248169 | 440 | 39 | -0.76323 | 0.77734 | MYH15 | 0.77734 | 0.78884 | 0.83182 |
| ENSG00000099860 | 19 | 2476120 | 2478257 | 7 | 3 | -0.76345 | 0.7774 | GADD45B | 0.7774 | 0.81432 | 0.18288 |
| ENSG00000187951 | 15 | 30916697 | 31065196 | 366 | 16 | -0.76371 | 0.77748 | ARHGAP11B | 0.77748 | 0.75909 | 0.75223 |
| ENSG00000187555 | 16 | 8985951 | 9058371 | 230 | 24 | -0.76375 | 0.77749 | USP7 | 0.77749 | 0.83821 | 0.82947 |
| ENSG00000196171 | 1 | 158669369 | 158670486 | 1 | 1 | -0.76378 | 0.7775 | OR6K2 | 0.7775 | 0.70206 | 0.67941 |
| ENSG00000130856 | 18 | 74534563 | 74682683 | 406 | 25 | -0.7638 | 0.77751 | ZNF236 | 0.77751 | 0.79358 | 0.77865 |
| ENSG00000100316 | 22 | 39708887 | 39716394 | 34 | 9 | -0.76382 | 0.77751 | RPL3 | 0.77751 | 0.58957 | 0.62846 |
| ENSG00000122012 | 5 | 75378997 | 75649764 | 1036 | 70 | -0.76406 | 0.77758 | SV2C | 0.77758 | 0.80483 | 0.82916 |
| ENSG00000234444 | 7 | 63767837 | 63815238 | 264 | 22 | -0.76406 | 0.77758 | ZNF736 | 0.77758 | 0.75657 | 0.75605 |
| ENSG00000156345 | 9 | 90581356 | 90589668 | 18 | 6 | -0.76404 | 0.77758 | CDK20 | 0.77758 | 0.47924 | 0.38994 |
| ENSG00000170545 | 12 | 51638342 | 51664674 | 83 | 11 | -0.76409 | 0.77759 | SMAGP | 0.77759 | 0.88308 | 0.87573 |
| ENSG00000158769 | 1 | 160965001 | 160991138 | 94 | 9 | -0.76433 | 0.77766 | F11R | 0.77766 | 0.74505 | 0.78935 |
| ENSG00000171867 | 20 | 4666882 | 4682236 | 49 | 10 | -0.7643 | 0.77766 | PRNP | 0.77766 | 0.8707 | 0.90529 |
| ENSG00000205436 | 14 | 103566481 | 103576896 | 41 | 7 | -0.76487 | 0.77783 | EXOC3L4 | 0.77783 | 0.81909 | 0.83798 |
| ENSG00000181143 | 19 | 8959520 | 9092018 | 562 | 43 | -0.76501 | 0.77787 | MUC16 | 0.77787 | 0.78218 | 0.79871 |
| ENSG00000111247 | 12 | 4647950 | 4669214 | 39 | 7 | -0.7655 | 0.77801 | RAD51AP1 | 0.77801 | 0.76337 | 0.83624 |
| ENSG00000099282 | 10 | 71211229 | 71267425 | 277 | 18 | -0.7656 | 0.77804 | TSPAN15 | 0.77804 | 0.79857 | 0.76779 |
| ENSG00000131495 | 5 | 140018325 | 140027370 | 23 | 6 | -0.76596 | 0.77815 | NDUFA2 | 0.77815 | 0.58637 | 0.54038 |
| ENSG00000171282 | 17 | 79373540 | 79433357 | 207 | 46 | -0.76597 | 0.77815 | RP11-1055B8.7 | 0.77815 | 0.89184 | 0.88753 |
| ENSG00000112306 | 6 | 133135580 | 133138703 | 26 | 9 | -0.7663 | 0.77825 | RPS12 | 0.77825 | 0.91754 | 0.92574 |
| ENSG00000166347 | 18 | 71920530 | 71959251 | 197 | 19 | -0.76642 | 0.77829 | CYB5A | 0.77829 | 0.79994 | 0.68709 |
| ENSG00000224997 | 14 | 104177607 | 104179149 | 3 | 2 | -0.76666 | 0.77836 | AL049840.1 | 0.77836 | 0.87058 | 0.88841 |
| ENSG00000141664 | 18 | 60190240 | 60254942 | 203 | 26 | -0.76673 | 0.77838 | ZCCHC2 | 0.77838 | 0.79 | 0.78042 |
| ENSG00000182938 | 17 | 72931814 | 72946087 | 58 | 15 | -0.76684 | 0.77841 | OTOP3 | 0.77841 | 0.57546 | 0.33425 |
| ENSG00000121310 | 1 | 53361656 | 53392884 | 43 | 11 | -0.76702 | 0.77846 | ECHDC2 | 0.77846 | 0.8182 | 0.78806 |
| ENSG00000162738 | 1 | 160370376 | 160398468 | 73 | 14 | -0.76707 | 0.77848 | VANGL2 | 0.77848 | 0.7758 | 0.80276 |
| ENSG00000171202 | 11 | 85359011 | 85367591 | 17 | 4 | -0.7672 | 0.77852 | TMEM126A | 0.77852 | 0.87462 | 0.83618 |
| ENSG00000006282 | 17 | 48620419 | 48633213 | 55 | 9 | -0.76723 | 0.77853 | SPATA20 | 0.77853 | 0.82971 | 0.9025 |
| ENSG00000165868 | 10 | 118430703 | 118502085 | 287 | 21 | -0.76795 | 0.77874 | HSPA12A | 0.77874 | 0.75405 | 0.7058 |
| ENSG00000173011 | 4 | 7043626 | 7059679 | 64 | 13 | -0.76814 | 0.7788 | TADA2B | 0.7788 | 0.8179 | 0.81875 |
| ENSG00000142156 | 21 | 47401651 | 47424964 | 124 | 22 | -0.76829 | 0.77884 | COL6A1 | 0.77884 | 0.79394 | 0.86824 |
| ENSG00000135372 | 11 | 34127149 | 34169217 | 94 | 6 | -0.76832 | 0.77885 | NAT10 | 0.77885 | 0.75485 | 0.75419 |
| ENSG00000072858 | 3 | 113251143 | 113348425 | 392 | 39 | -0.76836 | 0.77886 | SIDT1 | 0.77886 | 0.79128 | 0.76538 |
| ENSG00000090104 | 1 | 192544857 | 192549161 | 13 | 5 | -0.76864 | 0.77895 | RGS1 | 0.77895 | 0.16746 | 0.098005 |
| ENSG00000272297 | 4 | 187347700 | 187476464 | 822 | 74 | -0.76878 | 0.77899 | RP11-215A19.2 | 0.77899 | 0.79469 | 0.8232 |
| ENSG00000186260 | 16 | 14165178 | 14360630 | 416 | 21 | -0.76883 | 0.779 | MKL2 | 0.779 | 0.7787 | 0.7777 |
| ENSG00000138136 | 10 | 102986733 | 102989551 | 2 | 1 | -0.76886 | 0.77901 | LBX1 | 0.77901 | 0.67027 | 0.85299 |
| ENSG00000161010 | 5 | 179261436 | 179289173 | 107 | 15 | -0.76907 | 0.77907 | C5orf45 | 0.77907 | 0.79728 | 0.76886 |

| | | | | | | | | | | | |
|---|---|---|---|---|---|---|---|---|---|---|---|
| ENSG00000176040 | 3 | 111753690 | 111800116 | 153 | 21 | -0.76934 | 0.77915 | TMPRSS7 | 0.77915 | 0.8519 | 0.81202 |
| ENSG00000186432 | 3 | 160212783 | 160283376 | 161 | 18 | -0.76939 | 0.77917 | KPNA4 | 0.77917 | 0.78388 | 0.77855 |
| ENSG00000213996 | 19 | 19375173 | 19384200 | 35 | 8 | -0.7694 | 0.77917 | TM6SF2 | 0.77917 | 0.72548 | 0.85383 |
| ENSG00000171747 | 19 | 39292311 | 39304004 | 45 | 13 | -0.76938 | 0.77917 | LGALS4 | 0.77917 | 0.83994 | 0.80876 |
| ENSG00000167895 | 17 | 76126851 | 76139049 | 69 | 15 | -0.76945 | 0.77919 | TMC8 | 0.77919 | 0.81485 | 0.86965 |
| ENSG00000182783 | 1 | 248721784 | 248722797 | 6 | 1 | -0.76962 | 0.77924 | OR2T29 | 0.77924 | 0.8733 | 0.54069 |
| ENSG00000113522 | 5 | 131891711 | 131980313 | 127 | 7 | -0.76966 | 0.77925 | RAD50 | 0.77925 | 0.73429 | 0.70038 |
| ENSG00000197977 | 6 | 10980992 | 11044547 | 180 | 19 | -0.77 | 0.77935 | ELOVL2 | 0.77935 | 0.71027 | 0.70681 |
| ENSG00000122140 | 9 | 138391830 | 138396519 | 19 | 6 | -0.77004 | 0.77936 | MRPS2 | 0.77936 | 0.72957 | 0.74787 |
| ENSG00000068745 | 3 | 48725436 | 48777786 | 101 | 14 | -0.77013 | 0.77939 | IP6K2 | 0.77939 | 0.78496 | 0.81646 |
| ENSG00000009709 | 1 | 18957500 | 19075360 | 471 | 76 | -0.77021 | 0.77941 | PAX7 | 0.77941 | 0.84207 | 0.82094 |
| ENSG00000136240 | 7 | 6485584 | 6523873 | 160 | 31 | -0.77039 | 0.77947 | KDELR2 | 0.77947 | 0.62604 | 0.55154 |
| ENSG00000173728 | 1 | 244515937 | 244552965 | 186 | 23 | -0.7706 | 0.77953 | C1orf100 | 0.77953 | 0.72194 | 0.83142 |
| ENSG00000064651 | 5 | 127419458 | 127525380 | 204 | 17 | -0.77068 | 0.77955 | SLC12A2 | 0.77955 | 0.76891 | 0.74243 |
| ENSG00000143452 | 1 | 150670536 | 150693364 | 77 | 5 | -0.77072 | 0.77956 | HORMAD1 | 0.77956 | 0.78082 | 0.77274 |
| ENSG00000258643 | 14 | 23776044 | 23794578 | 39 | 12 | -0.77072 | 0.77956 | BCL2L2-PABPN1 | 0.77956 | 0.80667 | 0.88043 |
| ENSG00000104957 | 19 | 13842574 | 13874110 | 110 | 14 | -0.77086 | 0.7796 | CCDC130 | 0.7796 | 0.55299 | 0.39752 |
| ENSG00000152147 | 2 | 38978676 | 39012142 | 96 | 13 | -0.77137 | 0.77976 | GEMIN6 | 0.77976 | 0.75133 | 0.77238 |
| ENSG00000110958 | 12 | 57057127 | 57082159 | 74 | 13 | -0.77143 | 0.77977 | PTGES3 | 0.77977 | 0.70084 | 0.669 |
| ENSG00000181007 | 19 | 36874022 | 36909558 | 107 | 10 | -0.77143 | 0.77977 | ZFP82 | 0.77977 | 0.79668 | 0.81076 |
| ENSG00000265818 | 6 | 8015959 | 8102763 | 364 | 28 | -0.77148 | 0.77979 | EEF1E1-BLOC1S5 | 0.77979 | 0.7849 | 0.76645 |
| ENSG00000012223 | 3 | 46477136 | 46526724 | 201 | 25 | -0.77168 | 0.77985 | LTF | 0.77985 | 0.75964 | 0.705 |
| ENSG00000160345 | 9 | 138387027 | 138393580 | 23 | 4 | -0.77195 | 0.77993 | C9orf116 | 0.77993 | 0.58411 | 0.65434 |
| ENSG00000173389 | 3 | 51928892 | 51937351 | 26 | 4 | -0.7721 | 0.77997 | IQCF1 | 0.77997 | 0.89918 | 0.84238 |
| ENSG00000196267 | 19 | 52657117 | 52674896 | 53 | 12 | -0.77208 | 0.77997 | ZNF836 | 0.77997 | 0.91623 | 0.85872 |
| ENSG00000100485 | 14 | 50583847 | 50698276 | 451 | 26 | -0.77326 | 0.78032 | SOS2 | 0.78032 | 0.75614 | 0.74819 |
| ENSG00000178538 | 8 | 61099906 | 61193971 | 354 | 28 | -0.77337 | 0.78035 | CA8 | 0.78035 | 0.71861 | 0.61797 |
| ENSG00000197428 | 11 | 4660945 | 4662068 | 5 | 2 | -0.77347 | 0.78038 | OR51D1 | 0.78038 | 0.22474 | 0.2919 |
| ENSG00000260869 | 16 | 30545548 | 30569600 | 38 | 13 | -0.77353 | 0.78039 | AC002310.13 | 0.78039 | 0.89807 | 0.9348 |
| ENSG00000154856 | 18 | 10454625 | 10489945 | 138 | 20 | -0.7735 | 0.78039 | APCDD1 | 0.78039 | 0.85361 | 0.85715 |
| ENSG00000167863 | 17 | 73034958 | 73043080 | 47 | 11 | -0.77374 | 0.78046 | ATP5H | 0.78046 | 0.7721 | 0.71235 |
| ENSG00000101782 | 18 | 21032787 | 21066567 | 99 | 17 | -0.77376 | 0.78046 | RIOK3 | 0.78046 | 0.66017 | 0.68857 |
| ENSG00000186522 | 2 | 110300559 | 110371783 | 226 | 25 | -0.77406 | 0.78055 | 44084 | 0.78055 | 0.78187 | 0.80854 |
| ENSG00000081791 | 5 | 141303373 | 141321612 | 44 | 5 | -0.7743 | 0.78062 | KIAA0141 | 0.78062 | 0.86136 | 0.9125 |
| ENSG00000074527 | 12 | 96051583 | 96184930 | 625 | 48 | -0.77439 | 0.78065 | NTN4 | 0.78065 | 0.73232 | 0.865 |
| ENSG00000185518 | 15 | 91643180 | 91844539 | 732 | 56 | -0.77441 | 0.78066 | SV2B | 0.78066 | 0.73919 | 0.80822 |
| ENSG00000065325 | 17 | 9725523 | 9795419 | 329 | 28 | -0.77447 | 0.78067 | GLP2R | 0.78067 | 0.7128 | 0.78018 |
| ENSG00000075886 | 2 | 132233666 | 132240507 | 27 | 2 | -0.77457 | 0.7807 | TUBA3D | 0.7807 | 0.84416 | 0.84068 |
| ENSG00000127588 | 16 | 848041 | 850733 | 19 | 6 | -0.77469 | 0.78074 | GNG13 | 0.78074 | 0.79104 | 0.74071 |
| ENSG00000198019 | 1 | 120926979 | 120935937 | 9 | 4 | -0.77474 | 0.78075 | FCGR1B | 0.78075 | 0.8777 | 0.74895 |
| ENSG00000184056 | 15 | 91541646 | 91565833 | 81 | 11 | -0.7749 | 0.7808 | VPS33B | 0.7808 | 0.74724 | 0.096064 |
| ENSG00000196739 | 9 | 116917840 | 117074791 | 608 | 101 | -0.77504 | 0.78084 | COL27A1 | 0.78084 | 0.60563 | 0.53675 |
| ENSG00000105438 | 19 | 48885827 | 48894810 | 32 | 8 | -0.7752 | 0.78089 | KDELR1 | 0.78089 | 0.59821 | 0.42374 |
| ENSG00000120645 | 12 | 175931 | 287626 | 428 | 65 | -0.77525 | 0.78091 | IQSEC3 | 0.78091 | 0.64204 | 0.64328 |
| ENSG00000176515 | 6 | 1080164 | 1105181 | 103 | 18 | -0.77541 | 0.78095 | AL033381.1 | 0.78095 | 0.90624 | 0.96163 |
| ENSG00000162430 | 1 | 26126667 | 26144713 | 57 | 5 | -0.77545 | 0.78096 | SEPN1 | 0.78096 | 0.69705 | 0.77603 |
| ENSG00000168158 | 16 | 3405889 | 3406924 | 7 | 3 | -0.77563 | 0.78101 | OR2C1 | 0.78101 | 0.73311 | 0.61139 |
| ENSG00000123411 | 12 | 56401443 | 56432219 | 42 | 7 | -0.77563 | 0.78102 | IKZF4 | 0.78102 | 0.79769 | 0.72416 |
| ENSG00000185437 | 21 | 40817781 | 40887433 | 308 | 20 | -0.77564 | 0.78102 | SH3BGR | 0.78102 | 0.7762 | 0.77598 |
| ENSG00000185019 | 20 | 3088219 | 3140842 | 163 | 19 | -0.77574 | 0.78105 | UBOX5 | 0.78105 | 0.71561 | 0.68209 |
| ENSG00000057757 | 1 | 24104895 | 24114722 | 9 | 3 | -0.77577 | 0.78106 | PITHD1 | 0.78106 | 0.91676 | 0.72261 |
| ENSG00000166311 | 11 | 6411655 | 6416228 | 10 | 3 | -0.77599 | 0.78112 | SMPD1 | 0.78112 | 0.39416 | 0.51908 |
| ENSG00000182459 | 17 | 80317123 | 80321652 | 24 | 5 | -0.77605 | 0.78114 | TEX19 | 0.78114 | 0.69273 | 0.6057 |
| ENSG00000236939 | 8 | 104145191 | 104153703 | 26 | 8 | -0.77634 | 0.78123 | C8orf56 | 0.78123 | 0.7647 | 0.71305 |
| ENSG00000065328 | 10 | 13203554 | 13253104 | 185 | 37 | -0.77651 | 0.78127 | MCM10 | 0.78127 | 0.8161 | 0.92382 |
| ENSG00000146383 | 6 | 132891461 | 132892498 | 3 | 1 | -0.77657 | 0.78129 | TAAR6 | 0.78129 | 0.15106 | 0.24254 |
| ENSG00000185615 | 16 | 333152 | 337215 | 22 | 9 | -0.77657 | 0.78129 | PDIA2 | 0.78129 | 0.17738 | 0.21399 |
| ENSG00000099994 | 22 | 24577227 | 24585078 | 28 | 10 | -0.77664 | 0.78131 | SUSD2 | 0.78131 | 0.8539 | 0.90369 |
| ENSG00000164733 | 8 | 11700033 | 11726957 | 159 | 19 | -0.77687 | 0.78138 | CTSB | 0.78138 | 0.84346 | 0.82982 |
| ENSG00000178966 | 9 | 86595626 | 86618985 | 46 | 9 | -0.77716 | 0.78147 | RMI1 | 0.78147 | 0.81 | 0.81793 |
| ENSG00000138028 | 2 | 27321757 | 27341995 | 51 | 10 | -0.77749 | 0.78156 | CGREF1 | 0.78156 | 0.70353 | 0.69651 |
| ENSG00000155903 | 3 | 141205889 | 141334184 | 212 | 19 | -0.77761 | 0.7816 | RASA2 | 0.7816 | 0.81452 | 0.74937 |
| ENSG00000172232 | 19 | 825097 | 832017 | 31 | 7 | -0.77759 | 0.7816 | AZU1 | 0.7816 | 0.56935 | 0.46443 |
| ENSG00000144909 | 3 | 125247702 | 125313934 | 280 | 17 | -0.77765 | 0.78161 | OSBPL11 | 0.78161 | 0.75547 | 0.79938 |
| ENSG00000121454 | 1 | 180199421 | 180249380 | 237 | 44 | -0.77789 | 0.78168 | LHX4 | 0.78168 | 0.65488 | 0.65218 |
| ENSG00000110057 | 11 | 67758575 | 67772452 | 31 | 6 | -0.77801 | 0.78172 | UNC93B1 | 0.78172 | 0.70897 | 0.63244 |
| ENSG00000154188 | 8 | 108261721 | 108510283 | 959 | 62 | -0.77805 | 0.78173 | ANGPT1 | 0.78173 | 0.81322 | 0.84281 |
| ENSG00000166532 | 12 | 8834196 | 8935691 | 154 | 27 | -0.77804 | 0.78173 | RIMKLB | 0.78173 | 0.77122 | 0.85763 |
| ENSG00000065548 | 2 | 187350883 | 187374090 | 53 | 9 | -0.77823 | 0.78178 | ZC3H15 | 0.78178 | 0.79279 | 0.77152 |
| ENSG00000138075 | 2 | 44039611 | 44066004 | 104 | 16 | -0.77867 | 0.78191 | ABCG5 | 0.78191 | 0.69283 | 0.61992 |
| ENSG00000169570 | 5 | 118173017 | 118324240 | 339 | 35 | -0.77867 | 0.78191 | DTWD2 | 0.78191 | 0.79413 | 0.83076 |
| ENSG00000171124 | 19 | 5842899 | 5851485 | 32 | 9 | -0.77892 | 0.78199 | FUT3 | 0.78199 | 0.51499 | 0.8498 |
| ENSG00000107443 | 10 | 97803151 | 97820627 | 51 | 15 | -0.77934 | 0.78211 | CCNJ | 0.78211 | 0.75573 | 0.82355 |

| | | | | | | | | | | | |
|---|---|---|---|---|---|---|---|---|---|---|---|
| ENSG00000079101 | 18 | 596988 | 650334 | 255 | 52 | -0.77934 | 0.78211 | CLUL1 | 0.78211 | 0.83966 | 0.83612 |
| ENSG00000108588 | 17 | 61822610 | 61853711 | 84 | 8 | -0.77983 | 0.78225 | CCDC47 | 0.78225 | 0.83349 | 0.87639 |
| ENSG00000236294 | 7 | 113091127 | 113091457 | 1 | 1 | -0.77998 | 0.7823 | TSRM | 0.7823 | 0.62723 | 0.57006 |
| ENSG00000156006 | 8 | 18248755 | 18258728 | 51 | 9 | -0.78 | 0.7823 | NAT2 | 0.7823 | 0.89934 | 0.90181 |
| ENSG00000175198 | 13 | 100741269 | 101182686 | 998 | 48 | -0.78001 | 0.78231 | PCCA | 0.78231 | 0.70695 | 0.66916 |
| ENSG00000090238 | 16 | 30103635 | 30108236 | 7 | 4 | -0.78012 | 0.78234 | YPEL3 | 0.78234 | 0.84308 | 0.75036 |
| ENSG00000162670 | 1 | 190066792 | 190446759 | 847 | 36 | -0.78023 | 0.78237 | BRINP3 | 0.78237 | 0.7596 | 0.71146 |
| ENSG00000206527 | 3 | 123209667 | 123304032 | 293 | 18 | -0.78035 | 0.78241 | PTPLB | 0.78241 | 0.76958 | 0.67601 |
| ENSG00000184898 | 2 | 152104454 | 152118393 | 42 | 7 | -0.78043 | 0.78243 | RBM43 | 0.78243 | 0.82799 | 0.75767 |
| ENSG00000261190 | 16 | 52039399 | 52112386 | 217 | 31 | -0.7805 | 0.78245 | C16orf97 | 0.78245 | 0.66042 | 0.63431 |
| ENSG00000005249 | 7 | 106685094 | 106802256 | 316 | 23 | -0.78057 | 0.78247 | PRKAR2B | 0.78247 | 0.80668 | 0.81477 |
| ENSG00000026297 | 6 | 167342992 | 167370679 | 113 | 25 | -0.78064 | 0.78249 | RNASET2 | 0.78249 | 0.73489 | 0.74538 |
| ENSG00000197045 | 14 | 54941202 | 54955914 | 22 | 6 | -0.7811 | 0.78263 | GMFB | 0.78263 | 0.74723 | 0.76913 |
| ENSG00000259224 | 17 | 7384721 | 7386383 | 6 | 2 | -0.78151 | 0.78275 | SLC35G6 | 0.78275 | 0.85578 | 0.82186 |
| ENSG00000084774 | 2 | 27440258 | 27466811 | 53 | 11 | -0.78162 | 0.78278 | CAD | 0.78278 | 0.81425 | 0.7647 |
| ENSG00000064490 | 19 | 19303008 | 19312678 | 20 | 6 | -0.78187 | 0.78285 | RFXANK | 0.78285 | 0.78623 | 0.83355 |
| ENSG00000144152 | 2 | 112895962 | 112945791 | 156 | 28 | -0.78206 | 0.78291 | FBLN7 | 0.78291 | 0.83396 | 0.88842 |
| ENSG00000095906 | 16 | 1832902 | 1839192 | 34 | 9 | -0.78221 | 0.78295 | NUBP2 | 0.78295 | 0.79222 | 0.77034 |
| ENSG00000171469 | 19 | 9715356 | 9732075 | 33 | 5 | -0.7822 | 0.78295 | ZNF561 | 0.78295 | 0.71832 | 0.51082 |
| ENSG00000078967 | 7 | 43966037 | 43995735 | 111 | 18 | -0.78257 | 0.78306 | UBE2D4 | 0.78306 | 0.86284 | 0.89776 |
| ENSG00000204370 | 11 | 111957497 | 111990353 | 66 | 12 | -0.78263 | 0.78308 | SDHD | 0.78308 | 0.73129 | 0.77644 |
| ENSG00000132677 | 1 | 156339003 | 156355011 | 60 | 12 | -0.7827 | 0.7831 | RHBG | 0.7831 | 0.62174 | 0.54439 |
| ENSG00000180644 | 10 | 72357104 | 72362531 | 17 | 7 | -0.78305 | 0.7832 | PRF1 | 0.7832 | 0.72562 | 0.72341 |
| ENSG00000104524 | 8 | 144686083 | 144691943 | 15 | 6 | -0.78346 | 0.78332 | PYCRL | 0.78332 | 0.8512 | 0.86651 |
| ENSG00000204880 | 17 | 39253233 | 39254393 | 21 | 4 | -0.78367 | 0.78338 | KRTAP4-8 | 0.78338 | 0.5017 | 0.42162 |
| ENSG00000188596 | 12 | 96883349 | 97269333 | 1321 | 42 | -0.78369 | 0.78339 | C12orf55 | 0.78339 | 0.78852 | 0.80988 |
| ENSG00000215251 | 20 | 3127165 | 3140543 | 36 | 8 | -0.78394 | 0.78346 | FASTKD5 | 0.78346 | 0.61985 | 0.67141 |
| ENSG00000048392 | 8 | 103216730 | 103251346 | 134 | 16 | -0.78404 | 0.78349 | RRM2B | 0.78349 | 0.82395 | 0.77394 |
| ENSG00000169064 | 3 | 166958075 | 167099134 | 415 | 29 | -0.7841 | 0.78351 | ZBBX | 0.78351 | 0.78123 | 0.72903 |
| ENSG00000100722 | 14 | 89029253 | 89079853 | 105 | 17 | -0.78433 | 0.78358 | ZC3H14 | 0.78358 | 0.80884 | 0.76342 |
| ENSG00000123095 | 12 | 26272959 | 26278060 | 9 | 4 | -0.78445 | 0.78361 | BHLHE41 | 0.78361 | 0.34836 | 0.29345 |
| ENSG00000150526 | 14 | 39699435 | 39722859 | 70 | 13 | -0.78457 | 0.78365 | MIA2 | 0.78365 | 0.75824 | 0.81526 |
| ENSG00000157823 | 15 | 90373831 | 90437574 | 172 | 16 | -0.78477 | 0.78371 | AP3S2 | 0.78371 | 0.76263 | 0.74005 |
| ENSG00000109171 | 4 | 48343339 | 48428229 | 289 | 15 | -0.78484 | 0.78372 | SLAIN2 | 0.78372 | 0.81841 | 0.86298 |
| ENSG00000162623 | 1 | 75198836 | 75232362 | 143 | 18 | -0.78485 | 0.78373 | TYW3 | 0.78373 | 0.74008 | 0.65138 |
| ENSG00000196844 | 11 | 125646008 | 125648714 | 12 | 3 | -0.78498 | 0.78377 | PATE2 | 0.78377 | 0.93023 | 0.85972 |
| ENSG00000119714 | 14 | 91698876 | 91720269 | 80 | 9 | -0.78504 | 0.78378 | GPR68 | 0.78378 | 0.76119 | 0.55341 |
| ENSG00000179142 | 8 | 143991975 | 143999259 | 77 | 9 | -0.78506 | 0.78379 | CYP11B2 | 0.78379 | 0.79711 | 0.79302 |
| ENSG00000220008 | 19 | 2289774 | 2308156 | 79 | 17 | -0.78519 | 0.78383 | LINGO3 | 0.78383 | 0.55175 | 0.27823 |
| ENSG00000141294 | 17 | 45908993 | 45915079 | 23 | 8 | -0.78528 | 0.78385 | LRRC46 | 0.78385 | 0.72808 | 0.7114 |
| ENSG00000197272 | 16 | 28510683 | 28523372 | 37 | 7 | -0.78555 | 0.78394 | IL27 | 0.78394 | 0.7735 | 0.83784 |
| ENSG00000173818 | 17 | 78388965 | 78411886 | 96 | 22 | -0.78595 | 0.78405 | ENDOV | 0.78405 | 0.86831 | 0.91343 |
| ENSG00000176853 | 8 | 124780696 | 124827692 | 116 | 13 | -0.78602 | 0.78407 | FAM91A1 | 0.78407 | 0.79368 | 0.8338 |
| ENSG00000197889 | 10 | 15001438 | 15030049 | 222 | 16 | -0.78623 | 0.78413 | MEIG1 | 0.78413 | 0.71716 | 0.55232 |
| ENSG00000176302 | 11 | 118842417 | 118852001 | 28 | 6 | -0.78683 | 0.78431 | FOXR1 | 0.78431 | 0.8206 | 0.76585 |
| ENSG00000065308 | 6 | 52362200 | 52441713 | 396 | 36 | -0.78699 | 0.78435 | TRAM2 | 0.78435 | 0.72874 | 0.70411 |
| ENSG00000111581 | 12 | 69080514 | 69136785 | 129 | 18 | -0.78697 | 0.78435 | NUP107 | 0.78435 | 0.80161 | 0.79611 |
| ENSG00000188269 | 19 | 14903302 | 14946188 | 180 | 31 | -0.78708 | 0.78438 | OR7A5 | 0.78438 | 0.57921 | 0.39882 |
| ENSG00000179673 | 17 | 45055523 | 45056614 | 2 | 1 | -0.78711 | 0.78439 | RPRML | 0.78439 | 0.80843 | 0.79305 |
| ENSG00000164308 | 5 | 96211643 | 96255420 | 284 | 11 | -0.78731 | 0.78445 | ERAP2 | 0.78445 | 0.82251 | 0.88044 |
| ENSG00000171724 | 16 | 77822427 | 78014004 | 938 | 118 | -0.78733 | 0.78445 | VAT1L | 0.78445 | 0.50263 | 0.52388 |
| ENSG00000162664 | 1 | 90460671 | 90501090 | 65 | 15 | -0.78734 | 0.78446 | ZNF326 | 0.78446 | 0.74917 | 0.64315 |
| ENSG00000144580 | 2 | 219433303 | 219461803 | 64 | 6 | -0.78746 | 0.78449 | RQCD1 | 0.78449 | 0.7442 | 0.70675 |
| ENSG00000213672 | 3 | 48701364 | 48723797 | 39 | 8 | -0.78747 | 0.7845 | NCKIPSD | 0.7845 | 0.73832 | 0.74288 |
| ENSG00000178750 | 3 | 93733213 | 93747454 | 17 | 5 | -0.78782 | 0.7846 | STX19 | 0.7846 | 0.75464 | 0.79358 |
| ENSG00000124784 | 6 | 7389729 | 7418270 | 82 | 11 | -0.78786 | 0.78461 | RIOK1 | 0.78461 | 0.84523 | 0.87685 |
| ENSG00000138434 | 2 | 182756560 | 182795465 | 74 | 8 | -0.78857 | 0.78482 | SSFA2 | 0.78482 | 0.7932 | 0.83385 |
| ENSG00000269920 | 10 | 97733786 | 97792432 | 212 | 31 | -0.78887 | 0.7849 | RP11-690P14.4 | 0.7849 | 0.70038 | 0.5372 |
| ENSG00000188649 | 10 | 97733786 | 97792441 | 212 | 31 | -0.78887 | 0.7849 | CC2D2B | 0.7849 | 0.70038 | 0.5372 |
| ENSG00000156453 | 5 | 141232938 | 141258811 | 49 | 19 | -0.78901 | 0.78495 | PCDH1 | 0.78495 | 0.9315 | 0.83133 |
| ENSG00000076242 | 3 | 37034823 | 37107380 | 178 | 15 | -0.78917 | 0.78499 | MLH1 | 0.78499 | 0.76756 | 0.77716 |
| ENSG00000179468 | 7 | 142723245 | 142724282 | 1 | 1 | -0.78953 | 0.7851 | OR9A2 | 0.7851 | 0.21897 | 0.30733 |
| ENSG00000229729 | 3 | 88108424 | 88188123 | 204 | 5 | -0.78959 | 0.78512 | RP11-159G9.5 | 0.78512 | 0.76107 | 0.79666 |
| ENSG00000102531 | 13 | 49550048 | 49783915 | 335 | 21 | -0.78967 | 0.78514 | FNDC3A | 0.78514 | 0.78825 | 0.74594 |
| ENSG00000130921 | 12 | 123717463 | 123742506 | 101 | 10 | -0.78992 | 0.78521 | C12orf65 | 0.78521 | 0.73958 | 0.7942 |
| ENSG00000129317 | 12 | 44122410 | 44152620 | 107 | 11 | -0.78996 | 0.78522 | PUS7L | 0.78522 | 0.79989 | 0.76982 |
| ENSG00000163527 | 3 | 31574130 | 31679112 | 253 | 18 | -0.79003 | 0.78525 | STT3B | 0.78525 | 0.79873 | 0.73735 |
| ENSG00000186881 | 9 | 107266455 | 107267547 | 9 | 3 | -0.79004 | 0.78525 | OR13F1 | 0.78525 | 0.80846 | 0.81181 |
| ENSG00000124019 | 2 | 225243415 | 225266802 | 57 | 11 | -0.79012 | 0.78527 | FAM124B | 0.78527 | 0.77355 | 0.7946 |
| ENSG00000169105 | 15 | 40763160 | 40765353 | 1 | 1 | -0.79022 | 0.7853 | CHST14 | 0.7853 | 0.76171 | 0.71233 |
| ENSG00000128607 | 7 | 129710350 | 129775560 | 232 | 27 | -0.79066 | 0.78543 | KLHDC10 | 0.78543 | 0.77458 | 0.85877 |
| ENSG00000163320 | 3 | 88101094 | 88199035 | 255 | 6 | -0.79086 | 0.78549 | CGGBP1 | 0.78549 | 0.77123 | 0.79483 |
| ENSG00000109390 | 4 | 140188034 | 140223705 | 82 | 7 | -0.79178 | 0.78575 | NDUFC1 | 0.78575 | 0.85439 | 0.88952 |

| | | | | | | | | | | | | |
|---|---|---|---|---|---|---|---|---|---|---|---|---|
| ENSG00000172301 | 17 | 30178883 | 30186356 | 16 | 6 | -0.79183 | 0.78577 | COPRS | | 0.78577 | 0.85302 | 0.83947 |
| ENSG00000146247 | 6 | 79645584 | 79787953 | 327 | 21 | -0.79197 | 0.78581 | PHIP | | 0.78581 | 0.75239 | 0.74945 |
| ENSG00000155714 | 16 | 21995186 | 22012433 | 19 | 8 | -0.79198 | 0.78581 | PDZD9 | | 0.78581 | 0.68124 | 0.70415 |
| ENSG00000103056 | 16 | 68392231 | 68482591 | 183 | 25 | -0.79195 | 0.78581 | SMPD3 | | 0.78581 | 0.80988 | 0.83978 |
| ENSG00000173120 | 11 | 66886740 | 67025558 | 178 | 31 | -0.792 | 0.78582 | KDM2A | | 0.78582 | 0.79403 | 0.7693 |
| ENSG00000132639 | 20 | 10199478 | 10288066 | 243 | 36 | -0.79201 | 0.78582 | SNAP25 | | 0.78582 | 0.87584 | 0.8546 |
| ENSG00000112118 | 6 | 52128807 | 52149679 | 77 | 15 | -0.79211 | 0.78585 | MCM3 | | 0.78585 | 0.76477 | 0.75821 |
| ENSG00000106080 | 7 | 30050203 | 30066300 | 36 | 7 | -0.79214 | 0.78586 | FKBP14 | | 0.78586 | 0.76732 | 0.72433 |
| ENSG00000177363 | 11 | 62453874 | 62457371 | 4 | 2 | -0.79233 | 0.78592 | LRRN4CL | | 0.78592 | 0.52053 | 0.58396 |
| ENSG00000182362 | 21 | 47706251 | 47717665 | 116 | 11 | -0.79267 | 0.78602 | YBEY | | 0.78602 | 0.72017 | 0.69597 |
| ENSG00000100167 | 22 | 42372276 | 42394225 | 56 | 13 | -0.79293 | 0.78609 | | 44077 | 0.78609 | 0.80304 | 0.82749 |
| ENSG00000225190 | 17 | 43513266 | 43568115 | 147 | 7 | -0.79309 | 0.78614 | PLEKHM1 | | 0.78614 | 0.77583 | 0.78857 |
| ENSG00000114200 | 3 | 165490692 | 165555260 | 219 | 24 | -0.79314 | 0.78615 | BCHE | | 0.78615 | 0.85149 | 0.90391 |
| ENSG00000167346 | 11 | 4726157 | 5013659 | 1485 | 56 | -0.79318 | 0.78616 | MMP26 | | 0.78616 | 0.8049 | 0.80545 |
| ENSG00000166780 | 16 | 15528152 | 15718885 | 340 | 41 | -0.79355 | 0.78627 | C16orf45 | | 0.78627 | 0.76659 | 0.70359 |
| ENSG00000185664 | 12 | 56347889 | 56367101 | 23 | 9 | -0.79378 | 0.78634 | PMEL | | 0.78634 | 0.68054 | 0.62195 |
| ENSG00000162598 | 1 | 60452941 | 60539442 | 227 | 38 | -0.7938 | 0.78635 | C1orf87 | | 0.78635 | 0.83166 | 0.77834 |
| ENSG00000100678 | 14 | 70510934 | 70655787 | 664 | 41 | -0.79392 | 0.78638 | SLC8A3 | | 0.78638 | 0.75189 | 0.63806 |
| ENSG00000131409 | 19 | 51020149 | 51071302 | 171 | 43 | -0.79393 | 0.78638 | LRRC4B | | 0.78638 | 0.78232 | 0.81874 |
| ENSG00000115252 | 2 | 183004763 | 183387919 | 1490 | 59 | -0.79422 | 0.78646 | PDE1A | | 0.78646 | 0.73614 | 0.74786 |
| ENSG00000268714 | 5 | 115358088 | 115394625 | 226 | 26 | -0.79428 | 0.78648 | CTD-2287O16.3 | | 0.78648 | 0.75756 | 0.74194 |
| ENSG00000164129 | 4 | 164265091 | 164273086 | 26 | 5 | -0.79449 | 0.78654 | NPY5R | | 0.78654 | 0.93085 | 0.94933 |
| ENSG00000164400 | 5 | 131409483 | 131411859 | 5 | 3 | -0.79449 | 0.78654 | CSF2 | | 0.78654 | 0.34163 | 0.32221 |
| ENSG00000131871 | 15 | 101811022 | 101817705 | 14 | 6 | -0.79488 | 0.78666 | VIMP | | 0.78666 | 0.72845 | 0.67118 |
| ENSG00000222047 | 10 | 75669727 | 75682535 | 41 | 9 | -0.795 | 0.78669 | C10orf55 | | 0.78669 | 0.71553 | 0.5487 |
| ENSG00000105825 | 7 | 93514709 | 93520303 | 23 | 4 | -0.79524 | 0.78676 | TFPI2 | | 0.78676 | 0.79259 | 0.85354 |
| ENSG00000175782 | 12 | 69139886 | 69187508 | 128 | 25 | -0.79541 | 0.78681 | SLC35E3 | | 0.78681 | 0.76729 | 0.5885 |
| ENSG00000073331 | 4 | 113206665 | 113363776 | 515 | 63 | -0.7955 | 0.78684 | ALPK1 | | 0.78684 | 0.39253 | 0.36515 |
| ENSG00000146360 | 6 | 110299514 | 110301921 | 3 | 1 | -0.79551 | 0.78684 | GPR6 | | 0.78684 | 0.79612 | 0.67397 |
| ENSG00000107249 | 9 | 3824127 | 4348392 | 2578 | 193 | -0.7956 | 0.78687 | GLIS3 | | 0.78687 | 0.78242 | 0.78876 |
| ENSG00000100105 | 22 | 31721790 | 31742218 | 42 | 12 | -0.79567 | 0.78689 | PATZ1 | | 0.78689 | 0.51573 | 0.63153 |
| ENSG00000179008 | 14 | 60863187 | 60982261 | 209 | 16 | -0.7957 | 0.7869 | C14orf39 | | 0.7869 | 0.77577 | 0.77921 |
| ENSG00000165309 | 10 | 23216953 | 23326518 | 328 | 26 | -0.79582 | 0.78693 | ARMC3 | | 0.78693 | 0.78159 | 0.81576 |
| ENSG00000259517 | 16 | 54279456 | 54304658 | 51 | 8 | -0.796 | 0.78698 | RP11-324D17.1 | | 0.78698 | 0.8582 | 0.84908 |
| ENSG00000176281 | 14 | 20388741 | 20389818 | 2 | 1 | -0.79601 | 0.78699 | OR4K5 | | 0.78699 | 0.44025 | 0.46382 |
| ENSG00000100479 | 14 | 50110273 | 50155140 | 122 | 7 | -0.79604 | 0.787 | POLE2 | | 0.787 | 0.81588 | 0.83511 |
| ENSG00000007129 | 19 | 42055886 | 42093197 | 105 | 13 | -0.79626 | 0.78706 | CEACAM21 | | 0.78706 | 0.86831 | 0.92604 |
| ENSG00000171408 | 6 | 136172834 | 136516712 | 755 | 60 | -0.79679 | 0.78721 | PDE7B | | 0.78721 | 0.83181 | 0.86438 |
| ENSG00000166670 | 11 | 102641234 | 102651359 | 39 | 9 | -0.79709 | 0.7873 | MMP10 | | 0.7873 | 0.29836 | 0.11645 |
| ENSG00000184209 | 12 | 123942188 | 123957701 | 51 | 9 | -0.79715 | 0.78732 | SNRNP35 | | 0.78732 | 0.71874 | 0.71968 |
| ENSG00000102796 | 13 | 52342129 | 52378293 | 82 | 19 | -0.79717 | 0.78732 | DHRS12 | | 0.78732 | 0.78934 | 0.8116 |
| ENSG00000072201 | 4 | 54325468 | 54567572 | 935 | 61 | -0.79718 | 0.78733 | LNX1 | | 0.78733 | 0.81868 | 0.85316 |
| ENSG00000101384 | 20 | 10618332 | 10654694 | 123 | 27 | -0.79718 | 0.78733 | JAG1 | | 0.78733 | 0.78434 | 0.76221 |
| ENSG00000186493 | 5 | 2752245 | 2755508 | 8 | 3 | -0.79735 | 0.78738 | C5orf38 | | 0.78738 | 0.5911 | 0.55849 |
| ENSG00000159674 | 4 | 1160720 | 1202750 | 130 | 27 | -0.79745 | 0.7874 | SPON2 | | 0.7874 | 0.81462 | 0.88172 |
| ENSG00000091010 | 5 | 145718587 | 145720083 | 1 | 1 | -0.79743 | 0.7874 | POU4F3 | | 0.7874 | 0.73403 | 0.727 |
| ENSG00000124571 | 6 | 43490072 | 43543812 | 108 | 14 | -0.79792 | 0.78754 | XPO5 | | 0.78754 | 0.8268 | 0.8319 |
| ENSG00000157240 | 7 | 90893783 | 90898123 | 6 | 2 | -0.79805 | 0.78758 | FZD1 | | 0.78758 | 0.94765 | 0.89775 |
| ENSG00000083782 | 12 | 91357456 | 91398803 | 132 | 13 | -0.79819 | 0.78762 | EPYC | | 0.78762 | 0.76944 | 0.77344 |
| ENSG00000132122 | 1 | 48761044 | 48937845 | 385 | 35 | -0.79823 | 0.78763 | SPATA6 | | 0.78763 | 0.78336 | 0.76871 |
| ENSG00000132872 | 18 | 40847843 | 40857615 | 11 | 4 | -0.79843 | 0.78769 | SYT4 | | 0.78769 | 0.63083 | 0.47869 |
| ENSG00000165055 | 7 | 128116783 | 128146656 | 133 | 24 | -0.7985 | 0.78771 | METTL2B | | 0.78771 | 0.71225 | 0.73524 |
| ENSG00000154127 | 11 | 122526383 | 122685181 | 612 | 87 | -0.7985 | 0.78771 | UBASH3B | | 0.78771 | 0.71408 | 0.58943 |
| ENSG00000270149 | 1 | 160967747 | 161008752 | 139 | 17 | -0.79854 | 0.78772 | RP11-544M22.13 | | 0.78772 | 0.81175 | 0.84087 |
| ENSG00000179580 | 16 | 2016824 | 2018976 | 11 | 2 | -0.79855 | 0.78772 | RNF151 | | 0.78772 | 0.88567 | 0.93885 |
| ENSG00000078081 | 3 | 182840001 | 182881627 | 184 | 23 | -0.7988 | 0.7878 | LAMP3 | | 0.7878 | 0.86085 | 0.69799 |
| ENSG00000165805 | 12 | 88373816 | 88427814 | 143 | 13 | -0.7988 | 0.7878 | C12orf50 | | 0.7878 | 0.71307 | 0.66327 |
| ENSG00000169752 | 15 | 76228310 | 76352136 | 273 | 7 | -0.79885 | 0.78781 | NRG4 | | 0.78781 | 0.77776 | 0.76802 |
| ENSG00000167476 | 19 | 2252251 | 2269758 | 72 | 16 | -0.79894 | 0.78784 | JSRP1 | | 0.78784 | 0.53924 | 0.56718 |
| ENSG00000142168 | 21 | 33031935 | 33041244 | 23 | 7 | -0.79894 | 0.78784 | SOD1 | | 0.78784 | 0.7192 | 0.66527 |
| ENSG00000158865 | 16 | 24857162 | 24922949 | 227 | 29 | -0.79905 | 0.78787 | SLC5A11 | | 0.78787 | 0.80991 | 0.82123 |
| ENSG00000267216 | 19 | 58790332 | 58826563 | 126 | 26 | -0.79941 | 0.78797 | AC010642.1 | | 0.78797 | 0.65497 | 0.46901 |
| ENSG00000181562 | 14 | 21214051 | 21216539 | 16 | 3 | -0.79961 | 0.78803 | EDDM3A | | 0.78803 | 0.93943 | 0.92627 |
| ENSG00000132972 | 13 | 25338290 | 25454059 | 363 | 19 | -0.80004 | 0.78816 | RNF17 | | 0.78816 | 0.77152 | 0.78869 |
| ENSG00000125812 | 20 | 23342787 | 23353700 | 31 | 6 | -0.80051 | 0.78829 | GZF1 | | 0.78829 | 0.66946 | 0.65452 |
| ENSG00000051620 | 6 | 138724668 | 138743334 | 52 | 11 | -0.80055 | 0.7883 | HEBP2 | | 0.7883 | 0.68098 | 0.56684 |
| ENSG00000169006 | 2 | 11798304 | 11810290 | 40 | 10 | -0.80058 | 0.78831 | NTSR2 | | 0.78831 | 0.872 | 0.90368 |
| ENSG00000123685 | 1 | 212859760 | 212873327 | 54 | 15 | -0.80092 | 0.78841 | BATF3 | | 0.78841 | 0.6714 | 0.68259 |
| ENSG00000131435 | 5 | 131593364 | 131609147 | 52 | 8 | -0.80108 | 0.78846 | PDLIM4 | | 0.78846 | 0.84762 | 0.84466 |
| ENSG00000168140 | 16 | 4421849 | 4433529 | 29 | 7 | -0.80116 | 0.78848 | VASN | | 0.78848 | 0.90195 | 0.91734 |
| ENSG00000174374 | 7 | 74441226 | 74490064 | 30 | 8 | -0.80158 | 0.7886 | WBSCR16 | | 0.7886 | 0.7907 | 0.78275 |
| ENSG00000162522 | 1 | 33207486 | 33240571 | 64 | 16 | -0.80209 | 0.78875 | KIAA1522 | | 0.78875 | 0.64142 | 0.53309 |
| ENSG00000133246 | 19 | 8554940 | 8567996 | 37 | 8 | -0.80229 | 0.78881 | PRAM1 | | 0.78881 | 0.81339 | 0.78823 |

| | | | | | | | | | | | |
|---|---|---|---|---|---|---|---|---|---|---|---|
| ENSG00000184156 | 8 | 133133108 | 133493200 | 1247 | 105 | -0.80242 | 0.78885 | KCNQ3 | 0.78885 | 0.77099 | 0.76807 |
| ENSG00000142149 | 21 | 33245628 | 33416946 | 695 | 67 | -0.80242 | 0.78885 | HUNK | 0.78885 | 0.88224 | 0.8878 |
| ENSG00000198944 | 5 | 132149033 | 132152488 | 6 | 2 | -0.80248 | 0.78886 | SOWAHA | 0.78886 | 0.9282 | 0.8507 |
| ENSG00000242265 | 7 | 94285637 | 94299007 | 20 | 7 | -0.80278 | 0.78895 | PEG10 | 0.78895 | 0.8511 | 0.86709 |
| ENSG00000130201 | 19 | 45715879 | 45737469 | 116 | 8 | -0.80278 | 0.78895 | EXOC3L2 | 0.78895 | 0.724 | 0.86369 |
| ENSG00000143631 | 1 | 152274651 | 152297679 | 116 | 10 | -0.80306 | 0.78903 | FLG | 0.78903 | 0.7558 | 0.70119 |
| ENSG00000118322 | 5 | 159990127 | 160279221 | 1025 | 64 | -0.80306 | 0.78903 | ATP10B | 0.78903 | 0.78541 | 0.7635 |
| ENSG00000161405 | 17 | 37921198 | 38020441 | 269 | 7 | -0.8031 | 0.78904 | IKZF3 | 0.78904 | 0.79643 | 0.83211 |
| ENSG00000106460 | 7 | 12250867 | 12282993 | 194 | 9 | -0.80314 | 0.78905 | TMEM106B | 0.78905 | 0.9018 | 0.94826 |
| ENSG00000154721 | 21 | 27011584 | 27089874 | 255 | 25 | -0.80313 | 0.78905 | JAM2 | 0.78905 | 0.72096 | 0.61737 |
| ENSG00000182400 | 14 | 39617015 | 39639736 | 83 | 15 | -0.80321 | 0.78907 | TRAPPC6B | 0.78907 | 0.80027 | 0.7842 |
| ENSG00000171462 | 6 | 43418090 | 43424370 | 18 | 6 | -0.80366 | 0.7892 | DLK2 | 0.7892 | 0.93033 | 0.95116 |
| ENSG00000144227 | 2 | 139428342 | 139537918 | 393 | 49 | -0.8038 | 0.78924 | NXPH2 | 0.78924 | 0.76384 | 0.68073 |
| ENSG00000007255 | 19 | 45666186 | 45681495 | 41 | 8 | -0.80378 | 0.78924 | TRAPPC6A | 0.78924 | 0.65223 | 0.74226 |
| ENSG00000119541 | 18 | 61056423 | 61089716 | 84 | 9 | -0.80385 | 0.78926 | VPS4B | 0.78926 | 0.76618 | 0.66847 |
| ENSG00000242715 | 13 | 36801182 | 36871977 | 213 | 12 | -0.80409 | 0.78933 | CCDC169 | 0.78933 | 0.80534 | 0.75933 |
| ENSG00000198909 | 17 | 61699775 | 61773663 | 113 | 7 | -0.80454 | 0.78946 | MAP3K3 | 0.78946 | 0.83047 | 0.8721 |
| ENSG00000176681 | 17 | 44370099 | 44415160 | 30 | 9 | -0.80457 | 0.78947 | LRRC37A | 0.78947 | 0.64655 | 0.63511 |
| ENSG00000006555 | 1 | 55245385 | 55266940 | 69 | 13 | -0.80467 | 0.78949 | TTC22 | 0.78949 | 0.9236 | 0.87934 |
| ENSG00000154174 | 3 | 100082275 | 100120242 | 103 | 16 | -0.80464 | 0.78949 | TOMM70A | 0.78949 | 0.81016 | 0.79534 |
| ENSG00000138738 | 4 | 121606074 | 121844025 | 968 | 39 | -0.80466 | 0.78949 | PRDM5 | 0.78949 | 0.79009 | 0.81075 |
| ENSG00000065361 | 12 | 56473641 | 56497289 | 46 | 13 | -0.80471 | 0.78951 | ERBB3 | 0.78951 | 0.77489 | 0.82174 |
| ENSG00000100577 | 14 | 77787227 | 77797940 | 43 | 12 | -0.80476 | 0.78952 | GSTZ1 | 0.78952 | 0.74914 | 0.75964 |
| ENSG00000169855 | 3 | 78646390 | 79816965 | 3150 | 112 | -0.805 | 0.78959 | ROBO1 | 0.78959 | 0.79402 | 0.77241 |
| ENSG00000171189 | 21 | 30909254 | 31312351 | 1276 | 63 | -0.80525 | 0.78966 | GRIK1 | 0.78966 | 0.77689 | 0.78338 |
| ENSG00000124449 | 19 | 44220162 | 44224173 | 8 | 2 | -0.80529 | 0.78967 | IRGC | 0.78967 | 0.94882 | 0.96527 |
| ENSG00000180929 | 3 | 51989330 | 51991509 | 5 | 2 | -0.80546 | 0.78972 | GPR62 | 0.78972 | 0.57617 | 0.77873 |
| ENSG00000196136 | 14 | 95078714 | 95090392 | 37 | 13 | -0.80575 | 0.78981 | SERPINA3 | 0.78981 | 0.48975 | 0.58904 |
| ENSG00000142751 | 1 | 27202624 | 27216788 | 23 | 5 | -0.80604 | 0.78989 | GPN2 | 0.78989 | 0.73086 | 0.74781 |
| ENSG00000162928 | 2 | 61244360 | 61279125 | 69 | 15 | -0.80614 | 0.78992 | PEX13 | 0.78992 | 0.83468 | 0.82187 |
| ENSG00000008323 | 12 | 6419602 | 6437672 | 55 | 8 | -0.80618 | 0.78993 | PLEKHG6 | 0.78993 | 0.38563 | 0.31619 |
| ENSG00000155189 | 8 | 6565878 | 6617184 | 204 | 23 | -0.8067 | 0.79008 | AGPAT5 | 0.79008 | 0.87747 | 0.92383 |
| ENSG00000188766 | 19 | 38879061 | 38886881 | 21 | 3 | -0.8067 | 0.79008 | SPRED3 | 0.79008 | 0.79606 | 0.75738 |
| ENSG00000107281 | 9 | 139933922 | 139940655 | 13 | 5 | -0.80679 | 0.79011 | NPDC1 | 0.79011 | 0.80725 | 0.79155 |
| ENSG00000170667 | 7 | 102122892 | 102158228 | 13 | 6 | -0.80685 | 0.79012 | RASA4B | 0.79012 | 0.80471 | 0.49881 |
| ENSG00000214128 | 7 | 138482695 | 138522846 | 118 | 13 | -0.80732 | 0.79026 | TMEM213 | 0.79026 | 0.66648 | 0.63767 |
| ENSG00000130733 | 19 | 11033444 | 11039357 | 6 | 4 | -0.80748 | 0.7903 | YIPF2 | 0.7903 | 0.84873 | 0.85349 |
| ENSG00000187147 | 1 | 44870866 | 45117396 | 771 | 86 | -0.80786 | 0.79042 | RNF220 | 0.79042 | 0.76523 | 0.8114 |
| ENSG00000259040 | 6 | 7881755 | 8064597 | 887 | 59 | -0.808 | 0.79046 | BLOC1S5-TXNDC5 | 0.79046 | 0.77168 | 0.77937 |
| ENSG00000196993 | 16 | 28763108 | 28784128 | 18 | 8 | -0.80838 | 0.79056 | NPIPB9 | 0.79056 | 0.86631 | 0.83125 |
| ENSG00000198700 | 1 | 201798269 | 201853422 | 120 | 19 | -0.8086 | 0.79063 | IPO9 | 0.79063 | 0.84105 | 0.86946 |
| ENSG00000138071 | 2 | 65454887 | 65498387 | 134 | 15 | -0.80901 | 0.79075 | ACTR2 | 0.79075 | 0.83592 | 0.84805 |
| ENSG00000087460 | 20 | 57414773 | 57486247 | 218 | 44 | -0.80912 | 0.79078 | GNAS | 0.79078 | 0.83743 | 0.8433 |
| ENSG00000095539 | 10 | 102729275 | 102745628 | 32 | 7 | -0.80939 | 0.79085 | SEMA4G | 0.79085 | 0.83778 | 0.86636 |
| ENSG00000187398 | 11 | 24518516 | 25104150 | 2764 | 106 | -0.80948 | 0.79088 | LUZP2 | 0.79088 | 0.81035 | 0.78489 |
| ENSG00000182685 | 16 | 2259254 | 2261951 | 15 | 4 | -0.80956 | 0.7909 | BRICD5 | 0.7909 | 0.7793 | 0.83046 |
| ENSG00000076984 | 19 | 7968728 | 7979363 | 46 | 15 | -0.80972 | 0.79095 | MAP2K7 | 0.79095 | 0.9412 | 0.99075 |
| ENSG00000138435 | 2 | 175612320 | 175629200 | 51 | 7 | -0.80977 | 0.79096 | CHRNA1 | 0.79096 | 0.83003 | 0.96799 |
| ENSG00000156234 | 4 | 78432907 | 78532988 | 181 | 33 | -0.80976 | 0.79096 | CXCL13 | 0.79096 | 0.86403 | 0.87197 |
| ENSG00000188735 | 12 | 122150658 | 122220907 | 236 | 23 | -0.80999 | 0.79103 | TMEM120B | 0.79103 | 0.7607 | 0.74824 |
| ENSG00000003987 | 8 | 17155539 | 17271037 | 662 | 48 | -0.8101 | 0.79106 | MTMR7 | 0.79106 | 0.75358 | 0.68871 |
| ENSG00000124688 | 6 | 43597277 | 43608689 | 25 | 8 | -0.81063 | 0.79121 | MAD2L1BP | 0.79121 | 0.86031 | 0.85827 |
| ENSG00000135778 | 1 | 233086351 | 233119628 | 86 | 10 | -0.81102 | 0.79132 | NTPCR | 0.79132 | 0.80324 | 0.71479 |
| ENSG00000131979 | 14 | 55308726 | 55369570 | 242 | 22 | -0.81117 | 0.79137 | GCH1 | 0.79137 | 0.83525 | 0.82103 |
| ENSG00000166562 | 18 | 56806709 | 56826068 | 92 | 13 | -0.81128 | 0.7914 | SEC11C | 0.7914 | 0.77303 | 0.86355 |
| ENSG00000100448 | 14 | 25042728 | 25045466 | 7 | 3 | -0.8114 | 0.79143 | CTSG | 0.79143 | 0.93067 | 0.9404 |
| ENSG00000169291 | 1 | 154442248 | 154474589 | 52 | 9 | -0.81175 | 0.79153 | SHE | 0.79153 | 0.83732 | 0.54778 |
| ENSG00000184486 | 6 | 99282580 | 99286660 | 8 | 3 | -0.81222 | 0.79167 | POU3F2 | 0.79167 | 0.78909 | 0.71804 |
| ENSG00000185189 | 8 | 144915764 | 144924200 | 11 | 5 | -0.81235 | 0.7917 | NRBP2 | 0.7917 | 0.81611 | 0.80607 |
| ENSG00000185290 | 7 | 56182374 | 56184093 | 5 | 2 | -0.81237 | 0.79171 | NUPR1L | 0.79171 | 0.70072 | 0.75789 |
| ENSG00000135226 | 4 | 70146217 | 70160768 | 57 | 5 | -0.8125 | 0.79175 | UGT2B28 | 0.79175 | 0.79253 | 0.77683 |
| ENSG00000138190 | 10 | 94590935 | 94819250 | 719 | 31 | -0.81273 | 0.79181 | EXOC6 | 0.79181 | 0.76589 | 0.7678 |
| ENSG00000105352 | 19 | 42125344 | 42133442 | 50 | 11 | -0.81276 | 0.79182 | CEACAM4 | 0.79182 | 0.74904 | 0.87649 |
| ENSG00000139292 | 12 | 71833550 | 71980090 | 678 | 33 | -0.81282 | 0.79184 | LGR5 | 0.79184 | 0.78777 | 0.75495 |
| ENSG00000168397 | 2 | 242576628 | 242613272 | 184 | 27 | -0.81322 | 0.79195 | ATG4B | 0.79195 | 0.80522 | 0.68567 |
| ENSG00000225968 | 7 | 1727755 | 1787590 | 185 | 24 | -0.8133 | 0.79198 | ELFN1 | 0.79198 | 0.68547 | 0.51833 |
| ENSG00000178188 | 16 | 28857921 | 28885526 | 79 | 6 | -0.81396 | 0.79216 | SH2B1 | 0.79216 | 0.79622 | 0.75669 |
| ENSG00000176393 | 1 | 201951500 | 201975275 | 136 | 18 | -0.81405 | 0.79219 | RNPEP | 0.79219 | 0.79891 | 0.68373 |
| ENSG00000134780 | 11 | 61447905 | 61514473 | 143 | 21 | -0.81404 | 0.79219 | DAGLA | 0.79219 | 0.84347 | 0.85408 |
| ENSG00000177989 | 22 | 50968139 | 50971009 | 5 | 2 | -0.81406 | 0.7922 | ODF3B | 0.7922 | 0.59749 | 0.49866 |
| ENSG00000104892 | 19 | 45836692 | 45854778 | 71 | 14 | -0.81445 | 0.79231 | KLC3 | 0.79231 | 0.88955 | 0.90763 |
| ENSG00000101082 | 20 | 35240721 | 35274619 | 40 | 19 | -0.81464 | 0.79236 | SLA2 | 0.79236 | 0.76664 | 0.6591 |
| ENSG00000131016 | 6 | 151561134 | 151679692 | 603 | 61 | -0.81532 | 0.79255 | AKAP12 | 0.79255 | 0.78457 | 0.75638 |

| | | | | | | | | | | | |
|---|---|---|---|---|---|---|---|---|---|---|---|
| ENSG00000160932 | 8 | 144099399 | 144105249 | 18 | 6 | -0.81537 | 0.79257 | LY6E | 0.79257 | 0.7057 | 0.8231 |
| ENSG00000141646 | 18 | 48494410 | 48611415 | 160 | 15 | -0.81546 | 0.79259 | SMAD4 | 0.79259 | 0.81456 | 0.83682 |
| ENSG00000154175 | 3 | 100468000 | 100712359 | 723 | 58 | -0.81565 | 0.79265 | ABI3BP | 0.79265 | 0.76824 | 0.73264 |
| ENSG00000065060 | 6 | 34759857 | 34850915 | 318 | 20 | -0.81565 | 0.79265 | UHRF1BP1 | 0.79265 | 0.80192 | 0.71726 |
| ENSG00000183423 | 4 | 110769358 | 110793471 | 63 | 8 | -0.81601 | 0.79275 | LRIT3 | 0.79275 | 0.84865 | 0.88671 |
| ENSG00000166173 | 15 | 71123863 | 71146498 | 46 | 9 | -0.81606 | 0.79277 | LARP6 | 0.79277 | 0.89334 | 0.9488 |
| ENSG00000136931 | 9 | 127243516 | 127269709 | 55 | 7 | -0.81635 | 0.79285 | NR5A1 | 0.79285 | 0.75658 | 0.83298 |
| ENSG00000101144 | 20 | 55743804 | 55841685 | 390 | 30 | -0.8164 | 0.79286 | BMP7 | 0.79286 | 0.79908 | 0.82337 |
| ENSG00000086506 | 16 | 230452 | 231180 | 3 | 1 | -0.81642 | 0.79287 | HBQ1 | 0.79287 | 0.17238 | 0.14198 |
| ENSG00000256646 | 7 | 42948872 | 42971773 | 48 | 4 | -0.81679 | 0.79297 | PSMA2 | 0.79297 | 0.76318 | 0.79939 |
| ENSG00000099953 | 22 | 24110413 | 24126503 | 43 | 12 | -0.81678 | 0.79297 | MMP11 | 0.79297 | 0.89396 | 0.94933 |
| ENSG00000152463 | 10 | 15074226 | 15115851 | 155 | 27 | -0.81696 | 0.79303 | OLAH | 0.79303 | 0.81532 | 0.68948 |
| ENSG00000101452 | 20 | 37590942 | 37668366 | 147 | 14 | -0.81711 | 0.79307 | DHX35 | 0.79307 | 0.75442 | 0.64734 |
| ENSG00000112303 | 6 | 133065009 | 133084598 | 90 | 20 | -0.8174 | 0.79315 | VNN2 | 0.79315 | 0.83728 | 0.88229 |
| ENSG00000183032 | 14 | 37147636 | 37642071 | 1738 | 113 | -0.81743 | 0.79316 | SLC25A21 | 0.79316 | 0.81439 | 0.82479 |
| ENSG00000183495 | 12 | 132434508 | 132565005 | 229 | 21 | -0.81789 | 0.79329 | EP400 | 0.79329 | 0.82193 | 0.75785 |
| ENSG00000114779 | 3 | 52002526 | 52017425 | 17 | 8 | -0.818 | 0.79332 | ABHD14B | 0.79332 | 0.87244 | 0.7841 |
| ENSG00000221880 | 17 | 39190142 | 39191107 | 4 | 1 | -0.81812 | 0.79335 | KRTAP1-3 | 0.79335 | 0.67881 | 0.49986 |
| ENSG00000117322 | 1 | 207627575 | 207663240 | 95 | 14 | -0.81813 | 0.79336 | CR2 | 0.79336 | 0.86924 | 0.90935 |
| ENSG00000188037 | 7 | 143013219 | 143049176 | 125 | 23 | -0.81831 | 0.79341 | CLCN1 | 0.79341 | 0.73275 | 0.78174 |
| ENSG00000166189 | 10 | 103825147 | 103827792 | 3 | 2 | -0.81851 | 0.79347 | HPS6 | 0.79347 | 0.14354 | 0.30078 |
| ENSG00000150995 | 3 | 4535032 | 4889524 | 1337 | 155 | -0.81859 | 0.79349 | ITPR1 | 0.79349 | 0.85824 | 0.91825 |
| ENSG00000185619 | 4 | 699537 | 764428 | 327 | 31 | -0.81878 | 0.79355 | PCGF3 | 0.79355 | 0.79591 | 0.78908 |
| ENSG00000121895 | 4 | 38968365 | 39034542 | 360 | 34 | -0.81882 | 0.79356 | TMEM156 | 0.79356 | 0.76792 | 0.76643 |
| ENSG00000186190 | 20 | 31643230 | 31661434 | 70 | 13 | -0.81902 | 0.79361 | BPIFB3 | 0.79361 | 0.81227 | 0.78541 |
| ENSG00000131096 | 17 | 42030106 | 42081837 | 175 | 26 | -0.81937 | 0.79371 | PYY | 0.79371 | 0.8048 | 0.86351 |
| ENSG00000176387 | 16 | 67464555 | 67471456 | 12 | 4 | -0.8196 | 0.79378 | HSD11B2 | 0.79378 | 0.76989 | 0.71346 |
| ENSG00000100564 | 14 | 68048672 | 68067004 | 71 | 12 | -0.81982 | 0.79384 | PIGH | 0.79384 | 0.6706 | 0.78801 |
| ENSG00000108961 | 17 | 8191815 | 8193410 | 5 | 2 | -0.81987 | 0.79386 | RANGRF | 0.79386 | 0.37099 | 0.53051 |
| ENSG00000111711 | 12 | 21654715 | 21671342 | 93 | 16 | -0.81998 | 0.79389 | GOLT1B | 0.79389 | 0.78663 | 0.48138 |
| ENSG00000080189 | 20 | 44978167 | 44993043 | 17 | 5 | -0.82001 | 0.7939 | SLC35C2 | 0.7939 | 0.9041 | 0.93895 |
| ENSG00000205659 | 14 | 74551499 | 74667936 | 478 | 20 | -0.82034 | 0.79399 | LIN52 | 0.79399 | 0.79584 | 0.81049 |
| ENSG00000240857 | 2 | 18735989 | 18741946 | 14 | 8 | -0.82052 | 0.79404 | RDH14 | 0.79404 | 0.75916 | 0.51506 |
| ENSG00000010818 | 6 | 143072604 | 143266338 | 535 | 47 | -0.82061 | 0.79406 | HIVEP2 | 0.79406 | 0.7665 | 0.6304 |
| ENSG00000129538 | 14 | 21269387 | 21271437 | 3 | 1 | -0.82073 | 0.7941 | RNASE1 | 0.7941 | 0.95445 | 0.9111 |
| ENSG00000177627 | 12 | 48876286 | 48890295 | 78 | 11 | -0.82079 | 0.79412 | C12orf54 | 0.79412 | 0.86316 | 0.87227 |
| ENSG00000180008 | 14 | 55493948 | 55516206 | 45 | 6 | -0.82112 | 0.79421 | SOCS4 | 0.79421 | 0.84268 | 0.88397 |
| ENSG00000165059 | 9 | 71627469 | 71629039 | 4 | 2 | -0.82125 | 0.79425 | PRKACG | 0.79425 | 0.60239 | 0.83173 |
| ENSG00000185338 | 16 | 11348262 | 11350036 | 5 | 3 | -0.82124 | 0.79425 | SOCS1 | 0.79425 | 0.3281 | 0.40011 |
| ENSG00000153574 | 2 | 88991162 | 89050452 | 153 | 10 | -0.82141 | 0.79429 | RPIA | 0.79429 | 0.81785 | 0.81702 |
| ENSG00000179057 | 11 | 18725852 | 18747777 | 91 | 14 | -0.82141 | 0.79429 | IGSF22 | 0.79429 | 0.80081 | 0.72767 |
| ENSG00000168913 | 9 | 34521038 | 34523039 | 5 | 3 | -0.82157 | 0.79434 | ENHO | 0.79434 | 0.4958 | 0.71261 |
| ENSG00000205884 | 8 | 11831446 | 11832108 | 5 | 2 | -0.82183 | 0.79441 | DEFB136 | 0.79441 | 0.87769 | 0.90382 |
| ENSG00000184743 | 11 | 63391559 | 63439393 | 83 | 8 | -0.82184 | 0.79442 | ATL3 | 0.79442 | 0.83539 | 0.84781 |
| ENSG00000134917 | 11 | 130274820 | 130298888 | 86 | 24 | -0.82203 | 0.79447 | ADAMTS8 | 0.79447 | 0.69541 | 0.37327 |
| ENSG00000185760 | 6 | 73331520 | 73908574 | 1755 | 122 | -0.82209 | 0.79449 | KCNQ5 | 0.79449 | 0.78077 | 0.74662 |
| ENSG00000187545 | 1 | 12952727 | 12958101 | 17 | 4 | -0.82217 | 0.79451 | PRAMEF10 | 0.79451 | 0.66153 | 0.63252 |
| ENSG00000089775 | 14 | 64915824 | 64971931 | 153 | 12 | -0.82238 | 0.79457 | ZBTB25 | 0.79457 | 0.75189 | 0.69278 |
| ENSG00000134056 | 5 | 68513587 | 68525956 | 44 | 6 | -0.82267 | 0.79465 | MRPS36 | 0.79465 | 0.79566 | 0.78168 |
| ENSG00000136834 | 9 | 125239220 | 125240235 | 6 | 2 | -0.82308 | 0.79477 | OR1J1 | 0.79477 | 0.93535 | 0.96658 |
| ENSG00000188529 | 1 | 24291294 | 24307417 | 12 | 4 | -0.82337 | 0.79485 | SRSF10 | 0.79485 | 0.70237 | 0.7234 |
| ENSG00000181631 | 3 | 151044100 | 151047336 | 12 | 4 | -0.82345 | 0.79487 | P2RY13 | 0.79487 | 0.94802 | 0.94422 |
| ENSG00000175352 | 11 | 9002123 | 9025596 | 74 | 10 | -0.82355 | 0.7949 | NRIP3 | 0.7949 | 0.67871 | 0.71243 |
| ENSG00000187753 | 9 | 88835180 | 88874572 | 103 | 16 | -0.82382 | 0.79498 | C9orf153 | 0.79498 | 0.68322 | 0.54846 |
| ENSG00000104442 | 8 | 66514694 | 66546442 | 157 | 11 | -0.82393 | 0.79501 | ARMC1 | 0.79501 | 0.83723 | 0.85186 |
| ENSG00000134873 | 13 | 96085858 | 96232013 | 621 | 36 | -0.82391 | 0.79501 | CLDN10 | 0.79501 | 0.80378 | 0.82204 |
| ENSG00000244038 | 1 | 20978270 | 20988000 | 39 | 8 | -0.82439 | 0.79514 | DDOST | 0.79514 | 0.77691 | 0.68767 |
| ENSG00000168765 | 1 | 110198703 | 110208118 | 24 | 7 | -0.82439 | 0.79514 | GSTM4 | 0.79514 | 0.57654 | 0.31023 |
| ENSG00000073737 | 2 | 169921299 | 169952677 | 36 | 12 | -0.82443 | 0.79515 | DHRS9 | 0.79515 | 0.71916 | 0.62919 |
| ENSG00000159450 | 1 | 152078793 | 152086556 | 10 | 5 | -0.82462 | 0.79521 | TCHH | 0.79521 | 0.70848 | 0.68008 |
| ENSG00000164855 | 7 | 1581871 | 1600457 | 72 | 8 | -0.82464 | 0.79521 | TMEM184A | 0.79521 | 0.85253 | 0.84982 |
| ENSG00000146729 | 7 | 56019486 | 56067874 | 181 | 25 | -0.82464 | 0.79521 | GBAS | 0.79521 | 0.81436 | 0.80987 |
| ENSG00000167414 | 19 | 47137333 | 47137942 | 3 | 2 | -0.82481 | 0.79526 | GNG8 | 0.79526 | 0.92167 | 0.81553 |
| ENSG00000183066 | 22 | 42394729 | 42454460 | 127 | 12 | -0.82488 | 0.79528 | WBP2NL | 0.79528 | 0.76413 | 0.80704 |
| ENSG00000100796 | 14 | 91923955 | 91976898 | 193 | 14 | -0.82492 | 0.79529 | SMEK1 | 0.79529 | 0.83403 | 0.84744 |
| ENSG00000183570 | 21 | 47063608 | 47362368 | 874 | 26 | -0.82504 | 0.79533 | PCBP3 | 0.79533 | 0.76521 | 0.77892 |
| ENSG00000122512 | 7 | 6012870 | 6048756 | 170 | 30 | -0.82536 | 0.79542 | PMS2 | 0.79542 | 0.80133 | 0.78911 |
| ENSG00000148943 | 11 | 27516123 | 27528320 | 28 | 7 | -0.82541 | 0.79543 | LIN7C | 0.79543 | 0.8001 | 0.69122 |
| ENSG00000129910 | 16 | 89238175 | 89261900 | 106 | 16 | -0.82552 | 0.79546 | CDH15 | 0.79546 | 0.80201 | 0.92485 |
| ENSG00000243725 | 1 | 55181495 | 55208330 | 207 | 7 | -0.82592 | 0.79557 | TTC4 | 0.79557 | 0.83516 | 0.86353 |
| ENSG00000177692 | 21 | 34860497 | 34864027 | 9 | 3 | -0.82624 | 0.79567 | DNAJC28 | 0.79567 | 0.92083 | 0.90995 |
| ENSG00000165949 | 14 | 94571182 | 94583033 | 51 | 15 | -0.82638 | 0.79571 | IFI27 | 0.79571 | 0.58265 | 0.58228 |
| ENSG00000163719 | 3 | 9691117 | 9744077 | 98 | 16 | -0.82703 | 0.79589 | MTMR14 | 0.79589 | 0.7687 | 0.56126 |

| | | | | | | | | | | | | |
|---|---|---|---|---|---|---|---|---|---|---|---|---|
| ENSG00000176894 | 12 | 133264192 | 133297276 | 148 | 20 | -0.82713 | 0.79592 | PXMP2 | | 0.79592 | 0.76434 | 0.79254 |
| ENSG00000183654 | 5 | 16067248 | 16180871 | 219 | 29 | -0.82718 | 0.79593 | | 43901 | 0.79593 | 0.74178 | 0.70861 |
| ENSG00000165714 | 12 | 12510013 | 12619840 | 457 | 41 | -0.82716 | 0.79593 | LOH12CR1 | | 0.79593 | 0.73196 | 0.71285 |
| ENSG00000141469 | 18 | 43304092 | 43332485 | 128 | 8 | -0.82718 | 0.79593 | SLC14A1 | | 0.79593 | 0.68755 | 0.66189 |
| ENSG00000173124 | 10 | 96953957 | 96988685 | 133 | 10 | -0.82734 | 0.79598 | C10orf129 | | 0.79598 | 0.82109 | 0.85434 |
| ENSG00000108592 | 17 | 61896793 | 61907372 | 25 | 4 | -0.82747 | 0.79602 | FTSJ3 | | 0.79602 | 0.82259 | 0.85797 |
| ENSG00000089050 | 20 | 18467184 | 18477887 | 36 | 11 | -0.82752 | 0.79603 | RBBP9 | | 0.79603 | 0.78813 | 0.71113 |
| ENSG00000138411 | 2 | 197059094 | 197458416 | 1257 | 65 | -0.82768 | 0.79607 | HECW2 | | 0.79607 | 0.78371 | 0.80036 |
| ENSG00000184432 | 3 | 139074442 | 139108574 | 99 | 10 | -0.82784 | 0.79612 | COPB2 | | 0.79612 | 0.8058 | 0.78506 |
| ENSG00000235645 | 10 | 71444655 | 71452208 | 40 | 7 | -0.82798 | 0.79616 | RP11-242G20.1 | | 0.79616 | 0.73787 | 0.62531 |
| ENSG00000154727 | 21 | 27106881 | 27144771 | 66 | 10 | -0.82814 | 0.7962 | GABPA | | 0.7962 | 0.78689 | 0.83308 |
| ENSG00000163608 | 3 | 112721287 | 112738708 | 49 | 14 | -0.82823 | 0.79623 | C3orf17 | | 0.79623 | 0.41382 | 0.39633 |
| ENSG00000167749 | 19 | 51409608 | 51413994 | 22 | 7 | -0.82843 | 0.79629 | KLK4 | | 0.79629 | 0.74063 | 0.79762 |
| ENSG00000146555 | 7 | 3341080 | 4308632 | 4083 | 119 | -0.82855 | 0.79632 | SDK1 | | 0.79632 | 0.80338 | 0.83984 |
| ENSG00000178055 | 3 | 46871032 | 46875585 | 7 | 2 | -0.82861 | 0.79634 | PRSS42 | | 0.79634 | 0.86688 | 0.92884 |
| ENSG00000186591 | 7 | 129470572 | 129592789 | 505 | 15 | -0.82881 | 0.79639 | UBE2H | | 0.79639 | 0.77056 | 0.80766 |
| ENSG00000167767 | 12 | 52562780 | 52585784 | 70 | 23 | -0.82885 | 0.7964 | KRT80 | | 0.7964 | 0.67094 | 0.60497 |
| ENSG00000122068 | 3 | 197464050 | 197514467 | 122 | 17 | -0.82893 | 0.79643 | FYTTD1 | | 0.79643 | 0.76666 | 0.73221 |
| ENSG00000120860 | 12 | 102406705 | 102455927 | 120 | 17 | -0.82892 | 0.79643 | CCDC53 | | 0.79643 | 0.78579 | 0.78512 |
| ENSG00000108381 | 17 | 3375668 | 3406713 | 117 | 25 | -0.82907 | 0.79647 | ASPA | | 0.79647 | 0.85013 | 0.92916 |
| ENSG00000110583 | 11 | 63706431 | 63724800 | 46 | 6 | -0.82953 | 0.7966 | NAA40 | | 0.7966 | 0.80586 | 0.84529 |
| ENSG00000186197 | 1 | 236511562 | 236648214 | 761 | 58 | -0.82967 | 0.79664 | EDARADD | | 0.79664 | 0.83599 | 0.92759 |
| ENSG00000144668 | 3 | 37493606 | 37865005 | 1115 | 90 | -0.82994 | 0.79671 | ITGA9 | | 0.79671 | 0.78758 | 0.78481 |
| ENSG00000145050 | 3 | 51422478 | 51426828 | 10 | 3 | -0.83059 | 0.7969 | MANF | | 0.7969 | 0.6344 | 0.75544 |
| ENSG00000144290 | 2 | 162280843 | 162841792 | 1384 | 27 | -0.83064 | 0.79691 | SLC4A10 | | 0.79691 | 0.81699 | 0.80689 |
| ENSG00000105750 | 19 | 21106028 | 21133503 | 178 | 9 | -0.83109 | 0.79704 | ZNF85 | | 0.79704 | 0.79795 | 0.80141 |
| ENSG00000213203 | 7 | 150413645 | 150421372 | 16 | 6 | -0.83116 | 0.79706 | GIMAP1 | | 0.79706 | 0.84165 | 0.86639 |
| ENSG00000146070 | 6 | 46671938 | 46703430 | 58 | 13 | -0.83194 | 0.79728 | PLA2G7 | | 0.79728 | 0.86646 | 0.7529 |
| ENSG00000179528 | 2 | 74724644 | 74730443 | 3 | 2 | -0.83232 | 0.79739 | LBX2 | | 0.79739 | 0.11896 | 0.55577 |
| ENSG00000164037 | 4 | 103806205 | 103940896 | 348 | 9 | -0.83242 | 0.79741 | SLC9B1 | | 0.79741 | 0.80593 | 0.80708 |
| ENSG00000109066 | 17 | 72772622 | 72835918 | 181 | 19 | -0.83254 | 0.79745 | TMEM104 | | 0.79745 | 0.85153 | 0.8798 |
| ENSG00000185069 | 12 | 53161939 | 53171129 | 33 | 8 | -0.83257 | 0.79746 | KRT76 | | 0.79746 | 0.89204 | 0.85415 |
| ENSG00000186723 | 19 | 15917817 | 15918936 | 5 | 2 | -0.83278 | 0.79752 | OR10H1 | | 0.79752 | 0.39595 | 0.32532 |
| ENSG00000170542 | 6 | 2887500 | 2903514 | 61 | 21 | -0.83288 | 0.79754 | SERPINB9 | | 0.79754 | 0.73992 | 0.86309 |
| ENSG00000101400 | 20 | 31995761 | 32031698 | 63 | 9 | -0.83288 | 0.79754 | SNTA1 | | 0.79754 | 0.83667 | 0.85529 |
| ENSG00000146433 | 6 | 158957468 | 159056460 | 423 | 28 | -0.83305 | 0.79759 | TMEM181 | | 0.79759 | 0.8153 | 0.83051 |
| ENSG00000143842 | 1 | 204042243 | 204096863 | 172 | 41 | -0.83331 | 0.79766 | SOX13 | | 0.79766 | 0.83655 | 0.90689 |
| ENSG00000113460 | 5 | 34915481 | 34926101 | 21 | 7 | -0.83339 | 0.79769 | BRIX1 | | 0.79769 | 0.75993 | 0.82819 |
| ENSG00000187097 | 14 | 74424713 | 74486102 | 306 | 21 | -0.83342 | 0.7977 | ENTPD5 | | 0.7977 | 0.80199 | 0.83123 |
| ENSG00000066248 | 2 | 233743396 | 233877982 | 642 | 50 | -0.83362 | 0.79775 | NGEF | | 0.79775 | 0.80426 | 0.8485 |
| ENSG00000136828 | 9 | 129677053 | 129985445 | 653 | 23 | -0.83372 | 0.79778 | RALGPS1 | | 0.79778 | 0.8337 | 0.84372 |
| ENSG00000189056 | 7 | 103112231 | 103629963 | 2306 | 110 | -0.83393 | 0.79784 | RELN | | 0.79784 | 0.81636 | 0.83475 |
| ENSG00000188315 | 3 | 49306035 | 49315342 | 15 | 4 | -0.834 | 0.79786 | C3orf62 | | 0.79786 | 0.85187 | 0.92526 |
| ENSG00000173227 | 11 | 66774249 | 66818334 | 62 | 23 | -0.83408 | 0.79788 | SYT12 | | 0.79788 | 0.78566 | 0.79045 |
| ENSG00000141682 | 18 | 57567180 | 57571538 | 10 | 5 | -0.83436 | 0.79796 | PMAIP1 | | 0.79796 | 0.87531 | 0.79172 |
| ENSG00000161133 | 22 | 20704868 | 20745048 | 127 | 23 | -0.83444 | 0.79798 | USP41 | | 0.79798 | 0.83779 | 0.73657 |
| ENSG00000185305 | 5 | 53179775 | 53606412 | 1562 | 91 | -0.83465 | 0.79804 | ARL15 | | 0.79804 | 0.8007 | 0.79021 |
| ENSG00000149557 | 11 | 125315646 | 125366213 | 141 | 22 | -0.83469 | 0.79805 | FEZ1 | | 0.79805 | 0.80719 | 0.81368 |
| ENSG00000108733 | 17 | 33901814 | 33905882 | 12 | 5 | -0.8348 | 0.79808 | PEX12 | | 0.79808 | 0.81766 | 0.83852 |
| ENSG00000116668 | 1 | 185126212 | 185260897 | 178 | 15 | -0.83491 | 0.79812 | SWT1 | | 0.79812 | 0.80782 | 0.79597 |
| ENSG00000113805 | 3 | 74311719 | 74570291 | 631 | 81 | -0.83503 | 0.79815 | CNTN3 | | 0.79815 | 0.7977 | 0.75118 |
| ENSG00000221813 | 7 | 143701022 | 143702068 | 2 | 1 | -0.83504 | 0.79815 | OR6B1 | | 0.79815 | 0.84816 | 0.79619 |
| ENSG00000161682 | 17 | 42430583 | 42441243 | 17 | 9 | -0.83503 | 0.79815 | FAM171A2 | | 0.79815 | 0.98513 | 0.90713 |
| ENSG00000138756 | 4 | 79697496 | 79837526 | 276 | 11 | -0.83511 | 0.79817 | BMP2K | | 0.79817 | 0.79867 | 0.83243 |
| ENSG00000174428 | 7 | 74508364 | 74565623 | 42 | 16 | -0.83518 | 0.79819 | GTF2IRD2B | | 0.79819 | 0.71739 | 0.61761 |
| ENSG00000125995 | 20 | 34287194 | 34288906 | 2 | 1 | -0.83587 | 0.79839 | ROMO1 | | 0.79839 | 0.2133 | 0.33387 |
| ENSG00000182552 | 4 | 184560788 | 184580378 | 145 | 19 | -0.83592 | 0.7984 | RWDD4 | | 0.7984 | 0.7231 | 0.69055 |
| ENSG00000158106 | 8 | 144451057 | 144466390 | 55 | 10 | -0.83597 | 0.79841 | RHPN1 | | 0.79841 | 0.91409 | 0.88264 |
| ENSG00000101463 | 20 | 24449835 | 24647252 | 522 | 51 | -0.83619 | 0.79848 | SYNDIG1 | | 0.79848 | 0.42151 | 0.45764 |
| ENSG00000112977 | 5 | 10679342 | 10761384 | 314 | 15 | -0.83634 | 0.79852 | DAP | | 0.79852 | 0.85482 | 0.88577 |
| ENSG00000130559 | 9 | 138700333 | 138799074 | 156 | 21 | -0.83647 | 0.79856 | CAMSAP1 | | 0.79856 | 0.807 | 0.82374 |
| ENSG00000187122 | 10 | 98757795 | 98945677 | 585 | 69 | -0.83649 | 0.79856 | SLIT1 | | 0.79856 | 0.82371 | 0.50141 |
| ENSG00000106018 | 7 | 158820866 | 158937649 | 546 | 29 | -0.83652 | 0.79857 | VIPR2 | | 0.79857 | 0.82735 | 0.72249 |
| ENSG00000161031 | 19 | 15579456 | 15609767 | 113 | 21 | -0.8366 | 0.79859 | PGLYRP2 | | 0.79859 | 0.77157 | 0.81442 |
| ENSG00000115355 | 2 | 55514978 | 55647057 | 490 | 22 | -0.8367 | 0.79862 | CCDC88A | | 0.79862 | 0.82589 | 0.82446 |
| ENSG00000187987 | 6 | 28399707 | 28411279 | 37 | 7 | -0.83671 | 0.79862 | ZSCAN23 | | 0.79862 | 0.88449 | 0.80553 |
| ENSG00000080854 | 11 | 133778459 | 133826880 | 146 | 28 | -0.8368 | 0.79865 | IGSF9B | | 0.79865 | 0.85562 | 0.85718 |
| ENSG00000158901 | 20 | 44179792 | 44207965 | 187 | 4 | -0.83691 | 0.79868 | WFDC8 | | 0.79868 | 0.74621 | 0.66831 |
| ENSG00000215298 | 8 | 23430157 | 23432974 | 14 | 3 | -0.83717 | 0.79875 | FP15737 | | 0.79875 | 0.82987 | 0.82857 |
| ENSG00000182979 | 14 | 105886159 | 105937066 | 65 | 26 | -0.83721 | 0.79876 | MTA1 | | 0.79876 | 0.58294 | 0.55191 |
| ENSG00000181061 | 3 | 42798669 | 42846023 | 120 | 19 | -0.83782 | 0.79894 | HIGD1A | | 0.79894 | 0.79157 | 0.82256 |
| ENSG00000117640 | 1 | 26145131 | 26159432 | 37 | 3 | -0.83816 | 0.79903 | MTFR1L | | 0.79903 | 0.83415 | 0.76211 |
| ENSG00000269205 | 16 | 2031633 | 2032918 | 8 | 2 | -0.83815 | 0.79903 | AC005606.1 | | 0.79903 | 0.88537 | 0.94898 |

| | | | | | | | | | | | |
|---|---|---|---|---|---|---|---|---|---|---|---|
| ENSG00000197343 | 7 | 99156029 | 99174076 | 35 | 10 | -0.83855 | 0.79914 | ZNF655 | 0.79914 | 0.86708 | 0.8408 |
| ENSG00000083857 | 4 | 187508937 | 187647876 | 538 | 65 | -0.83864 | 0.79916 | FAT1 | 0.79916 | 0.80359 | 0.85892 |
| ENSG00000150681 | 1 | 192127587 | 192154945 | 49 | 12 | -0.83901 | 0.79927 | RGS18 | 0.79927 | 0.83997 | 0.83279 |
| ENSG00000103356 | 16 | 23533335 | 23569052 | 136 | 13 | -0.83907 | 0.79928 | EARS2 | 0.79928 | 0.79382 | 0.8154 |
| ENSG00000253320 | 8 | 103876528 | 103990104 | 386 | 35 | -0.83926 | 0.79934 | KB-1507C5.2 | 0.79934 | 0.8206 | 0.7303 |
| ENSG00000170627 | 12 | 54849734 | 54867386 | 59 | 6 | -0.83931 | 0.79935 | GTSF1 | 0.79935 | 0.79744 | 0.74607 |
| ENSG00000100146 | 22 | 38366693 | 38383429 | 10 | 3 | -0.83994 | 0.79953 | SOX10 | 0.79953 | 0.53291 | 0.57557 |
| ENSG00000182175 | 15 | 93586636 | 93632433 | 240 | 41 | -0.84026 | 0.79962 | RGMA | 0.79962 | 0.84955 | 0.818 |
| ENSG00000113649 | 5 | 145826874 | 145891524 | 139 | 18 | -0.84051 | 0.79969 | TCERG1 | 0.79969 | 0.78367 | 0.7334 |
| ENSG00000184611 | 2 | 163227917 | 163695240 | 1113 | 65 | -0.84059 | 0.79971 | KCNH7 | 0.79971 | 0.79126 | 0.77828 |
| ENSG00000175746 | 15 | 39542885 | 39547046 | 17 | 5 | -0.84061 | 0.79972 | C15orf54 | 0.79972 | 0.854 | 0.75611 |
| ENSG00000166897 | 22 | 37764000 | 37823505 | 254 | 26 | -0.84061 | 0.79972 | ELFN2 | 0.79972 | 0.68777 | 0.86526 |
| ENSG00000144893 | 3 | 150803484 | 151154860 | 1038 | 46 | -0.84078 | 0.79977 | MED12L | 0.79977 | 0.80958 | 0.80456 |
| ENSG00000171848 | 2 | 10262455 | 10271545 | 19 | 5 | -0.84115 | 0.79987 | RRM2 | 0.79987 | 0.64383 | 0.49752 |
| ENSG00000014164 | 8 | 144519825 | 144623623 | 340 | 34 | -0.8412 | 0.79988 | ZC3H3 | 0.79988 | 0.78735 | 0.74158 |
| ENSG00000140988 | 16 | 2012053 | 2014861 | 27 | 5 | -0.8412 | 0.79988 | RPS2 | 0.79988 | 0.88162 | 0.94171 |
| ENSG00000170004 | 17 | 7788124 | 7816078 | 47 | 14 | -0.8412 | 0.79988 | CHD3 | 0.79988 | 0.5975 | 0.77617 |
| ENSG00000162896 | 1 | 207101863 | 207119811 | 40 | 7 | -0.84162 | 0.8 | PIGR | 0.8 | 0.79692 | 0.83953 |
| ENSG00000255529 | 15 | 57884231 | 58074960 | 782 | 68 | -0.84162 | 0.8 | POLR2M | 0.8 | 0.77231 | 0.70546 |
| ENSG00000108405 | 17 | 3799886 | 3819794 | 69 | 14 | -0.84164 | 0.80001 | P2RX1 | 0.80001 | 0.82685 | 0.91473 |
| ENSG00000250741 | 2 | 18736811 | 18770830 | 81 | 15 | -0.84183 | 0.80006 | NT5C1B-RDH14 | 0.80006 | 0.80512 | 0.67326 |
| ENSG00000185013 | 2 | 18737050 | 18770838 | 81 | 15 | -0.84183 | 0.80006 | NT5C1B | 0.80006 | 0.80512 | 0.69629 |
| ENSG00000255054 | 1 | 26137961 | 26150235 | 40 | 4 | -0.84209 | 0.80013 | RP1-317E23.6 | 0.80013 | 0.7978 | 0.75244 |
| ENSG00000170577 | 2 | 45232300 | 45236569 | 11 | 5 | -0.84211 | 0.80014 | SIX2 | 0.80014 | 0.86505 | 0.84429 |
| ENSG00000165076 | 7 | 141536086 | 141541287 | 30 | 3 | -0.84216 | 0.80015 | PRSS37 | 0.80015 | 0.77909 | 0.76572 |
| ENSG00000180574 | 12 | 10658201 | 10675734 | 76 | 7 | -0.84218 | 0.80016 | EIF2S3L | 0.80016 | 0.77793 | 0.8326 |
| ENSG00000088992 | 12 | 117476728 | 117537284 | 229 | 23 | -0.84227 | 0.80018 | TESC | 0.80018 | 0.7941 | 0.81518 |
| ENSG00000168546 | 8 | 21547915 | 21669869 | 318 | 58 | -0.84241 | 0.80022 | GFRA2 | 0.80022 | 0.48504 | 0.34906 |
| ENSG00000176253 | 14 | 20502003 | 20502919 | 2 | 1 | -0.84241 | 0.80022 | OR4K13 | 0.80022 | 0.88598 | 0.87108 |
| ENSG00000120992 | 8 | 54958938 | 55014577 | 143 | 11 | -0.84247 | 0.80024 | LYPLA1 | 0.80024 | 0.79365 | 0.74293 |
| ENSG00000269709 | 1 | 48569306 | 48569878 | 2 | 1 | -0.84263 | 0.80028 | AL109659.1 | 0.80028 | 0.58665 | 0.67218 |
| ENSG00000182372 | 8 | 1703944 | 1734738 | 123 | 12 | -0.84278 | 0.80033 | CLN8 | 0.80033 | 0.84034 | 0.86717 |
| ENSG00000112297 | 6 | 106959730 | 107018326 | 336 | 24 | -0.84291 | 0.80036 | AIM1 | 0.80036 | 0.80343 | 0.84102 |
| ENSG00000129355 | 19 | 10677138 | 10679735 | 7 | 2 | -0.84317 | 0.80043 | CDKN2D | 0.80043 | 0.92663 | 0.95965 |
| ENSG00000145907 | 5 | 151150606 | 151192346 | 81 | 12 | -0.84362 | 0.80056 | G3BP1 | 0.80056 | 0.80071 | 0.77228 |
| ENSG00000019485 | 11 | 45115564 | 45247734 | 322 | 28 | -0.84364 | 0.80056 | PRDM11 | 0.80056 | 0.82742 | 0.80447 |
| ENSG00000165125 | 7 | 142568956 | 142583507 | 58 | 4 | -0.84374 | 0.80059 | TRPV6 | 0.80059 | 0.84185 | 0.70376 |
| ENSG00000172404 | 22 | 41255553 | 41258130 | 3 | 2 | -0.84384 | 0.80062 | DNAJB7 | 0.80062 | 0.42382 | 0.47253 |
| ENSG00000197448 | 7 | 142941186 | 142967947 | 14 | 8 | -0.84413 | 0.8007 | GSTK1 | 0.8007 | 0.89698 | 0.86692 |
| ENSG00000166037 | 11 | 95523129 | 95565857 | 176 | 15 | -0.84416 | 0.80071 | CEP57 | 0.80071 | 0.82742 | 0.79118 |
| ENSG00000166323 | 11 | 108179246 | 108338258 | 383 | 14 | -0.84416 | 0.80071 | C11orf65 | 0.80071 | 0.7808 | 0.81303 |
| ENSG00000165917 | 11 | 47459308 | 47470730 | 42 | 7 | -0.84441 | 0.80078 | RAPSN | 0.80078 | 0.79365 | 0.7622 |
| ENSG00000223609 | 11 | 5253908 | 5256600 | 4 | 2 | -0.84451 | 0.80081 | HBD | 0.80081 | 0.65039 | 0.91265 |
| ENSG00000221887 | 18 | 61616535 | 61649008 | 135 | 17 | -0.84467 | 0.80085 | HMSD | 0.80085 | 0.72542 | 0.64549 |
| ENSG00000156990 | 3 | 9879533 | 9886286 | 11 | 5 | -0.84478 | 0.80088 | RPUSD3 | 0.80088 | 0.49416 | 0.62446 |
| ENSG00000221818 | 8 | 25699246 | 25902913 | 702 | 84 | -0.84477 | 0.80088 | EBF2 | 0.80088 | 0.84926 | 0.87609 |
| ENSG00000114757 | 3 | 179512746 | 179754841 | 763 | 44 | -0.8448 | 0.80089 | PEX5L | 0.80089 | 0.82571 | 0.84573 |
| ENSG00000258292 | 12 | 96196875 | 96217085 | 74 | 13 | -0.8449 | 0.80092 | RP11-536G4.1 | 0.80092 | 0.85678 | 0.89797 |
| ENSG00000015592 | 8 | 27092840 | 27115937 | 111 | 14 | -0.84495 | 0.80093 | STMN4 | 0.80093 | 0.88169 | 0.92851 |
| ENSG00000159495 | 15 | 43568478 | 43594453 | 47 | 6 | -0.84497 | 0.80093 | TGM7 | 0.80093 | 0.79442 | 0.76459 |
| ENSG00000147432 | 8 | 42552519 | 42592550 | 241 | 7 | -0.84503 | 0.80095 | CHRNB3 | 0.80095 | 0.83622 | 0.8857 |
| ENSG00000077498 | 11 | 88910620 | 89028927 | 373 | 19 | -0.84513 | 0.80098 | TYR | 0.80098 | 0.7688 | 0.71921 |
| ENSG00000215203 | 4 | 42895284 | 43032675 | 478 | 38 | -0.84527 | 0.80102 | GRXCR1 | 0.80102 | 0.78662 | 0.70565 |
| ENSG00000135636 | 2 | 71680852 | 71913898 | 926 | 95 | -0.84566 | 0.80113 | DYSF | 0.80113 | 0.81564 | 0.73969 |
| ENSG00000112592 | 6 | 170863390 | 170881958 | 53 | 7 | -0.84608 | 0.80125 | TBP | 0.80125 | 0.79667 | 0.78015 |
| ENSG00000174453 | 2 | 215275789 | 215443683 | 484 | 30 | -0.84643 | 0.80134 | VWC2L | 0.80134 | 0.8115 | 0.82827 |
| ENSG00000169733 | 17 | 80005778 | 80009707 | 20 | 3 | -0.84646 | 0.80135 | RFNG | 0.80135 | 0.91858 | 0.69665 |
| ENSG00000100084 | 22 | 19318221 | 19435224 | 290 | 18 | -0.8465 | 0.80136 | HIRA | 0.80136 | 0.78846 | 0.74377 |
| ENSG00000272762 | 3 | 51994317 | 52008032 | 14 | 4 | -0.84662 | 0.8014 | RP11-155D18.14 | 0.8014 | 0.83941 | 0.80078 |
| ENSG00000085382 | 6 | 105175968 | 105307794 | 390 | 17 | -0.84668 | 0.80141 | HACE1 | 0.80141 | 0.8017 | 0.79643 |
| ENSG00000110367 | 11 | 118620034 | 118661858 | 117 | 14 | -0.84711 | 0.80153 | DDX6 | 0.80153 | 0.82788 | 0.81125 |
| ENSG00000149311 | 11 | 108093211 | 108239829 | 236 | 12 | -0.84734 | 0.8016 | ATM | 0.8016 | 0.82091 | 0.82955 |
| ENSG00000268059 | 6 | 13615728 | 13621126 | 6 | 2 | -0.84743 | 0.80162 | AL441883.1 | 0.80162 | 0.75151 | 0.86032 |
| ENSG00000130768 | 1 | 28261504 | 28285668 | 81 | 11 | -0.84754 | 0.80165 | SMPDL3B | 0.80165 | 0.79049 | 0.81451 |
| ENSG00000182742 | 17 | 46652875 | 46657473 | 14 | 4 | -0.8476 | 0.80167 | HOXB4 | 0.80167 | 0.82909 | 0.82953 |
| ENSG00000198711 | 1 | 54703710 | 54704736 | 5 | 3 | -0.84775 | 0.80171 | SSBP3-AS1 | 0.80171 | 0.71624 | 0.79726 |
| ENSG00000138031 | 2 | 25042038 | 25142708 | 434 | 27 | -0.84773 | 0.80171 | ADCY3 | 0.80171 | 0.83908 | 0.87114 |
| ENSG00000188611 | 10 | 51942538 | 52008370 | 91 | 13 | -0.84816 | 0.80182 | ASAH2 | 0.80182 | 0.75663 | 0.39259 |
| ENSG00000198121 | 9 | 113635543 | 113800981 | 609 | 40 | -0.84823 | 0.80184 | LPAR1 | 0.80184 | 0.79891 | 0.83932 |
| ENSG00000198522 | 2 | 27851114 | 27874375 | 51 | 8 | -0.84855 | 0.80193 | GPN1 | 0.80193 | 0.72227 | 0.66484 |
| ENSG00000110243 | 11 | 116660083 | 116663136 | 11 | 4 | -0.84853 | 0.80193 | APOA5 | 0.80193 | 0.88665 | 0.90165 |
| ENSG00000047578 | 16 | 27561454 | 27791690 | 344 | 17 | -0.84864 | 0.80196 | KIAA0556 | 0.80196 | 0.82322 | 0.84757 |
| ENSG00000169379 | 3 | 93698983 | 93774512 | 80 | 17 | -0.84868 | 0.80197 | ARL13B | 0.80197 | 0.8268 | 0.8105 |

| | | | | | | | | | | | |
|---|---|---|---|---|---|---|---|---|---|---|---|
| ENSG00000149798 | 11 | 65082289 | 65089900 | 13 | 6 | -0.84884 | 0.80202 | CDC42EP2 | 0.80202 | 0.54609 | 0.4633 |
| ENSG00000166359 | 19 | 33622996 | 33666701 | 182 | 23 | -0.84885 | 0.80202 | WDR88 | 0.80202 | 0.7149 | 0.74123 |
| ENSG00000214736 | 6 | 41755400 | 41757636 | 6 | 3 | -0.84888 | 0.80203 | TOMM6 | 0.80203 | 0.79313 | 0.72396 |
| ENSG00000055950 | 10 | 102729215 | 102747272 | 37 | 8 | -0.84899 | 0.80206 | MRPL43 | 0.80206 | 0.85486 | 0.85147 |
| ENSG00000159346 | 1 | 202909951 | 202927700 | 68 | 14 | -0.84926 | 0.80213 | ADIPOR1 | 0.80213 | 0.85139 | 0.84739 |
| ENSG00000180425 | 11 | 114262165 | 114271139 | 31 | 7 | -0.84941 | 0.80217 | C11orf71 | 0.80217 | 0.70611 | 0.51934 |
| ENSG00000013392 | 6 | 83903098 | 83908651 | 7 | 3 | -0.84954 | 0.80221 | RWDD2A | 0.80221 | 0.0048274 | 0.0034603 |
| ENSG00000089486 | 16 | 4560676 | 4588829 | 77 | 9 | -0.8499 | 0.80231 | CDIP1 | 0.80231 | 0.81637 | 0.76925 |
| ENSG00000116120 | 2 | 223435255 | 223521056 | 191 | 20 | -0.8502 | 0.80239 | FARSB | 0.80239 | 0.80605 | 0.82523 |
| ENSG00000140284 | 15 | 50474393 | 50528592 | 262 | 21 | -0.85023 | 0.8024 | SLC27A2 | 0.8024 | 0.75176 | 0.75948 |
| ENSG00000124659 | 6 | 42712219 | 42714558 | 3 | 1 | -0.85057 | 0.8025 | TBCC | 0.8025 | 0.5649 | 0.40505 |
| ENSG00000196814 | 9 | 129089128 | 129269320 | 512 | 35 | -0.8506 | 0.8025 | MVB12B | 0.8025 | 0.82039 | 0.85632 |
| ENSG00000162775 | 1 | 110881128 | 110889299 | 13 | 4 | -0.85095 | 0.8026 | RBM15 | 0.8026 | 0.47459 | 0.44594 |
| ENSG00000177646 | 3 | 128598439 | 128634910 | 117 | 15 | -0.85097 | 0.80261 | ACAD9 | 0.80261 | 0.81727 | 0.80837 |
| ENSG00000157992 | 2 | 27665233 | 27669348 | 5 | 1 | -0.85119 | 0.80267 | KRTCAP3 | 0.80267 | 0.7358 | 0.69621 |
| ENSG00000178878 | 12 | 12878851 | 12982909 | 403 | 75 | -0.85121 | 0.80267 | APOLD1 | 0.80267 | 0.71195 | 0.75969 |
| ENSG00000013275 | 19 | 40476912 | 40487348 | 33 | 6 | -0.85128 | 0.80269 | PSMC4 | 0.80269 | 0.7784 | 0.83378 |
| ENSG00000121410 | 19 | 58856544 | 58864865 | 15 | 2 | -0.85136 | 0.80272 | A1BG | 0.80272 | 0.79166 | 0.76526 |
| ENSG00000134243 | 1 | 109852192 | 109940573 | 159 | 13 | -0.85144 | 0.80274 | SORT1 | 0.80274 | 0.80513 | 0.85293 |
| ENSG00000159337 | 15 | 42359207 | 42386752 | 134 | 9 | -0.85148 | 0.80275 | PLA2G4D | 0.80275 | 0.84039 | 0.9343 |
| ENSG00000124702 | 6 | 42981951 | 42989036 | 9 | 3 | -0.8519 | 0.80286 | KLHDC3 | 0.80286 | 0.72531 | 0.7568 |
| ENSG00000177732 | 20 | 306207 | 310865 | 20 | 4 | -0.85198 | 0.80289 | SOX12 | 0.80289 | 0.87633 | 0.82669 |
| ENSG00000256500 | 14 | 104029299 | 104152261 | 362 | 21 | -0.85226 | 0.80296 | RP11-73M18.2 | 0.80296 | 0.80866 | 0.81351 |
| ENSG00000153066 | 16 | 11772936 | 11836734 | 201 | 21 | -0.85238 | 0.803 | TXNDC11 | 0.803 | 0.71812 | 0.76427 |
| ENSG00000171444 | 5 | 112357796 | 112824527 | 2135 | 63 | -0.85247 | 0.80302 | MCC | 0.80302 | 0.80596 | 0.8156 |
| ENSG00000174951 | 19 | 49251268 | 49258647 | 22 | 5 | -0.85322 | 0.80323 | FUT1 | 0.80323 | 0.92973 | 0.95723 |
| ENSG00000143401 | 1 | 150190717 | 150208504 | 37 | 7 | -0.85332 | 0.80326 | ANP32E | 0.80326 | 0.80352 | 0.8267 |
| ENSG00000099203 | 19 | 10943114 | 10946994 | 3 | 2 | -0.85344 | 0.80329 | TMED1 | 0.80329 | 0.87817 | 0.89536 |
| ENSG00000133706 | 5 | 145492601 | 145562223 | 284 | 5 | -0.85346 | 0.8033 | LARS | 0.8033 | 0.79383 | 0.79198 |
| ENSG00000140463 | 15 | 72978527 | 73030817 | 142 | 21 | -0.85369 | 0.80336 | BBS4 | 0.80336 | 0.75926 | 0.73765 |
| ENSG00000117280 | 1 | 205737114 | 205744588 | 22 | 8 | -0.85377 | 0.80338 | RAB7L1 | 0.80338 | 0.74536 | 0.68022 |
| ENSG00000104969 | 19 | 2754712 | 2783369 | 101 | 9 | -0.85394 | 0.80343 | SGTA | 0.80343 | 0.81257 | 0.76425 |
| ENSG00000130244 | 19 | 38893775 | 38899728 | 30 | 4 | -0.85398 | 0.80344 | FAM98C | 0.80344 | 0.68143 | 0.70182 |
| ENSG00000176624 | 18 | 48700920 | 48744674 | 124 | 11 | -0.85411 | 0.80348 | MEX3C | 0.80348 | 0.81447 | 0.76526 |
| ENSG00000136478 | 17 | 62224587 | 62340661 | 341 | 18 | -0.85419 | 0.8035 | TEX2 | 0.8035 | 0.76213 | 0.72367 |
| ENSG00000003137 | 2 | 72356367 | 72375167 | 40 | 10 | -0.85491 | 0.8037 | CYP26B1 | 0.8037 | 0.40986 | 0.49485 |
| ENSG00000204704 | 6 | 29011990 | 29013017 | 3 | 1 | -0.8549 | 0.8037 | OR2W1 | 0.8037 | 0.80418 | 0.81551 |
| ENSG00000213921 | 19 | 40267235 | 40276775 | 17 | 5 | -0.85493 | 0.80371 | LEUTX | 0.80371 | 0.71299 | 0.71061 |
| ENSG00000156515 | 10 | 71029740 | 71161638 | 542 | 72 | -0.85553 | 0.80387 | HK1 | 0.80387 | 0.73554 | 0.7393 |
| ENSG00000173702 | 3 | 124624289 | 124672663 | 214 | 42 | -0.85577 | 0.80394 | MUC13 | 0.80394 | 0.8687 | 0.85756 |
| ENSG00000119138 | 9 | 72999503 | 73029540 | 58 | 17 | -0.85585 | 0.80396 | KLF9 | 0.80396 | 0.83142 | 0.89635 |
| ENSG00000163249 | 2 | 208576264 | 208626563 | 159 | 10 | -0.8561 | 0.80403 | CCNYL1 | 0.80403 | 0.86215 | 0.90758 |
| ENSG00000115568 | 2 | 219502639 | 219524378 | 59 | 7 | -0.85623 | 0.80406 | ZNF142 | 0.80406 | 0.86515 | 0.86374 |
| ENSG00000135407 | 12 | 58191159 | 58212487 | 30 | 10 | -0.85622 | 0.80406 | AVIL | 0.80406 | 0.65693 | 0.66546 |
| ENSG00000164741 | 8 | 12940870 | 13373167 | 2635 | 89 | -0.85624 | 0.80407 | DLC1 | 0.80407 | 0.82771 | 0.83409 |
| ENSG00000102935 | 16 | 49521435 | 49891830 | 1265 | 121 | -0.85628 | 0.80408 | ZNF423 | 0.80408 | 0.81332 | 0.81647 |
| ENSG00000085733 | 11 | 70244510 | 70282690 | 90 | 10 | -0.85633 | 0.80409 | CTTN | 0.80409 | 0.72616 | 0.77805 |
| ENSG00000184995 | 9 | 21480841 | 21482312 | 6 | 2 | -0.85639 | 0.80411 | IFNE | 0.80411 | 0.61895 | 0.68345 |
| ENSG00000108342 | 17 | 38171614 | 38174066 | 12 | 3 | -0.85644 | 0.80412 | CSF3 | 0.80412 | 0.63334 | 0.61344 |
| ENSG00000255639 | 12 | 4766800 | 4853747 | 329 | 24 | -0.85661 | 0.80417 | RP11-234B24.6 | 0.80417 | 0.84198 | 0.78449 |
| ENSG00000231192 | 3 | 97851542 | 97852483 | 3 | 1 | -0.85747 | 0.80441 | OR5H1 | 0.80441 | 0.58888 | 0.65063 |
| ENSG00000181222 | 17 | 7387685 | 7417933 | 148 | 11 | -0.85812 | 0.80459 | POLR2A | 0.80459 | 0.73503 | 0.66219 |
| ENSG00000104953 | 19 | 2977444 | 2995177 | 87 | 14 | -0.85822 | 0.80461 | TLE6 | 0.80461 | 0.85861 | 0.88412 |
| ENSG00000134222 | 1 | 109822178 | 109825808 | 9 | 5 | -0.85835 | 0.80465 | PSRC1 | 0.80465 | 0.93489 | 0.94341 |
| ENSG00000171823 | 12 | 1675159 | 1703331 | 87 | 29 | -0.85857 | 0.80471 | FBXL14 | 0.80471 | 0.96091 | 0.91716 |
| ENSG00000126070 | 1 | 36396319 | 36538101 | 191 | 21 | -0.85896 | 0.80482 | AGO3 | 0.80482 | 0.80301 | 0.82372 |
| ENSG00000055332 | 2 | 37326353 | 37384208 | 219 | 21 | -0.85913 | 0.80487 | EIF2AK2 | 0.80487 | 0.8254 | 0.81743 |
| ENSG00000259363 | 15 | 100347228 | 100417353 | 513 | 19 | -0.85922 | 0.80489 | CTD-2054N24.2 | 0.80489 | 0.61912 | 0.56594 |
| ENSG00000206043 | 18 | 71983048 | 72026422 | 115 | 10 | -0.85945 | 0.80495 | C18orf63 | 0.80495 | 0.84523 | 0.85895 |
| ENSG00000160191 | 21 | 44073746 | 44195619 | 639 | 50 | -0.8595 | 0.80497 | PDE9A | 0.80497 | 0.8219 | 0.84977 |
| ENSG00000180988 | 11 | 5841544 | 5842578 | 11 | 2 | -0.85955 | 0.80498 | OR52N2 | 0.80498 | 0.91674 | 0.94682 |
| ENSG00000105856 | 7 | 106809406 | 106842974 | 72 | 7 | -0.85978 | 0.80504 | HBP1 | 0.80504 | 0.77871 | 0.81311 |
| ENSG00000183379 | 14 | 74872596 | 74892805 | 74 | 12 | -0.86041 | 0.80522 | SYNDIG1L | 0.80522 | 0.78466 | 0.82684 |
| ENSG00000248485 | 1 | 161228517 | 161255240 | 128 | 15 | -0.86061 | 0.80527 | PCP4L1 | 0.80527 | 0.83267 | 0.87132 |
| ENSG00000254986 | 11 | 66247484 | 66277130 | 64 | 5 | -0.86083 | 0.80533 | DPP3 | 0.80533 | 0.78359 | 0.77179 |
| ENSG00000150076 | 10 | 32832297 | 33171802 | 1346 | 26 | -0.86104 | 0.80539 | C10ORF68 | 0.80539 | 0.79561 | 0.78971 |
| ENSG00000154640 | 21 | 18965971 | 18985265 | 24 | 6 | -0.86112 | 0.80541 | BTG3 | 0.80541 | 0.79301 | 0.65682 |
| ENSG00000164530 | 6 | 36922209 | 36932613 | 39 | 9 | -0.86134 | 0.80547 | PI16 | 0.80547 | 0.95664 | 0.86003 |
| ENSG00000174145 | 4 | 37245842 | 37451087 | 724 | 61 | -0.86148 | 0.80551 | KIAA1239 | 0.80551 | 0.82532 | 0.79257 |
| ENSG00000114023 | 3 | 122103023 | 122131181 | 95 | 14 | -0.86181 | 0.8056 | FAM162A | 0.8056 | 0.86097 | 0.82241 |
| ENSG00000126233 | 8 | 143822362 | 143823829 | 2 | 1 | -0.86197 | 0.80565 | SLURP1 | 0.80565 | 0.75873 | 0.77489 |
| ENSG00000163554 | 1 | 158580496 | 158656488 | 215 | 24 | -0.86208 | 0.80568 | SPTA1 | 0.80568 | 0.8046 | 0.81275 |
| ENSG00000124693 | 6 | 26031817 | 26032288 | 1 | 1 | -0.86216 | 0.8057 | HIST1H3B | 0.8057 | 0.33191 | 0.40199 |

| | | | | | | | | | | | | |
|---|---|---|---|---|---|---|---|---|---|---|---|---|
| ENSG00000037897 | 12 | 58162254 | 58166576 | 6 | 2 | -0.86231 | 0.80574 | METTL1 | | 0.80574 | 0.76252 | 0.62973 |
| ENSG00000166123 | 16 | 46918290 | 46965209 | 41 | 19 | -0.86236 | 0.80576 | GPT2 | | 0.80576 | 0.74042 | 0.79236 |
| ENSG00000108829 | 17 | 48452420 | 48474914 | 67 | 8 | -0.86241 | 0.80577 | LRRC59 | | 0.80577 | 0.80493 | 0.85066 |
| ENSG00000006074 | 17 | 34391640 | 34399392 | 12 | 5 | -0.86246 | 0.80578 | CCL18 | | 0.80578 | 0.54516 | 0.54541 |
| ENSG00000182180 | 10 | 75006510 | 75012451 | 19 | 2 | -0.86249 | 0.80579 | MRPS16 | | 0.80579 | 0.75206 | 0.71194 |
| ENSG00000141258 | 17 | 2240792 | 2284352 | 182 | 25 | -0.86312 | 0.80597 | SGSM2 | | 0.80597 | 0.79356 | 0.79401 |
| ENSG00000177697 | 11 | 832843 | 839831 | 56 | 4 | -0.86317 | 0.80598 | CD151 | | 0.80598 | 0.60155 | 0.71457 |
| ENSG00000091009 | 5 | 145583113 | 145718814 | 241 | 22 | -0.86359 | 0.80609 | RBM27 | | 0.80609 | 0.8028 | 0.80887 |
| ENSG00000269064 | 1 | 179443716 | 179457805 | 158 | 10 | -0.86365 | 0.80611 | AL160286.1 | | 0.80611 | 0.84442 | 0.91152 |
| ENSG00000006453 | 7 | 97920963 | 98030380 | 329 | 12 | -0.86366 | 0.80611 | BAIAP2L1 | | 0.80611 | 0.78848 | 0.75887 |
| ENSG00000249115 | 19 | 36103646 | 36116251 | 23 | 8 | -0.86364 | 0.80611 | HAUS5 | | 0.80611 | 0.79276 | 0.82706 |
| ENSG00000185104 | 1 | 50905150 | 51425935 | 838 | 28 | -0.86401 | 0.80621 | FAF1 | | 0.80621 | 0.8159 | 0.82111 |
| ENSG00000131188 | 5 | 176873446 | 176883283 | 13 | 5 | -0.86423 | 0.80627 | PRR7 | | 0.80627 | 0.42026 | 0.49326 |
| ENSG00000257138 | 7 | 141672431 | 141673573 | 3 | 1 | -0.86434 | 0.8063 | TAS2R38 | | 0.8063 | 0.71084 | 0.11301 |
| ENSG00000173418 | 20 | 19997760 | 20014299 | 36 | 10 | -0.86469 | 0.80639 | NAA20 | | 0.80639 | 0.84507 | 0.86866 |
| ENSG00000160584 | 11 | 116714118 | 116969153 | 1100 | 28 | -0.86544 | 0.8066 | SIK3 | | 0.8066 | 0.8161 | 0.84266 |
| ENSG00000010610 | 12 | 6896024 | 6929974 | 162 | 26 | -0.86576 | 0.80669 | CD4 | | 0.80669 | 0.82357 | 0.79149 |
| ENSG00000106462 | 7 | 148504475 | 148581413 | 174 | 21 | -0.86594 | 0.80674 | EZH2 | | 0.80674 | 0.80343 | 0.87386 |
| ENSG00000215009 | 12 | 7456880 | 7481320 | 54 | 11 | -0.86594 | 0.80674 | ACSM4 | | 0.80674 | 0.86098 | 0.90426 |
| ENSG00000163395 | 1 | 201159953 | 201198080 | 175 | 32 | -0.86638 | 0.80686 | IGFN1 | | 0.80686 | 0.79638 | 0.82143 |
| ENSG00000248835 | 1 | 54635333 | 54637997 | 12 | 4 | -0.86643 | 0.80687 | AL357673.1 | | 0.80687 | 0.79259 | 0.87337 |
| ENSG00000141002 | 16 | 89940000 | 89977792 | 191 | 13 | -0.86645 | 0.80688 | TCF25 | | 0.80688 | 0.86994 | 0.91612 |
| ENSG00000166927 | 11 | 60145955 | 60163424 | 40 | 8 | -0.86659 | 0.80692 | MS4A7 | | 0.80692 | 0.78871 | 0.82332 |
| ENSG00000160472 | 19 | 55888204 | 55889612 | 5 | 3 | -0.86682 | 0.80698 | TMEM190 | | 0.80698 | 0.45943 | 0.71372 |
| ENSG00000109606 | 4 | 24519064 | 24586173 | 130 | 20 | -0.86698 | 0.80702 | DHX15 | | 0.80702 | 0.77375 | 0.93276 |
| ENSG00000254873 | 11 | 118267810 | 118272610 | 7 | 3 | -0.86705 | 0.80704 | RP11-770J1.5 | | 0.80704 | 0.7126 | 0.73473 |
| ENSG00000174989 | 12 | 117348761 | 117468953 | 263 | 14 | -0.86745 | 0.80715 | FBXW8 | | 0.80715 | 0.76384 | 0.81624 |
| ENSG00000155096 | 8 | 103838585 | 103906092 | 236 | 23 | -0.8675 | 0.80717 | AZIN1 | | 0.80717 | 0.80275 | 0.83938 |
| ENSG00000123080 | 1 | 51426417 | 51440305 | 18 | 6 | -0.86779 | 0.80725 | CDKN2C | | 0.80725 | 0.81944 | 0.55947 |
| ENSG00000104897 | 19 | 2236520 | 2248678 | 54 | 14 | -0.86812 | 0.80733 | SF3A2 | | 0.80733 | 0.8762 | 0.8034 |
| ENSG00000168301 | 3 | 58477841 | 58488087 | 25 | 9 | -0.86859 | 0.80746 | KCTD6 | | 0.80746 | 0.63193 | 0.64913 |
| ENSG00000182318 | 19 | 58838385 | 58853698 | 51 | 3 | -0.86876 | 0.80751 | ZSCAN22 | | 0.80751 | 0.75484 | 0.81604 |
| ENSG00000101367 | 20 | 31407699 | 31438211 | 98 | 19 | -0.86925 | 0.80765 | MAPRE1 | | 0.80765 | 0.77482 | 0.70405 |
| ENSG00000108759 | 17 | 39616063 | 39623681 | 48 | 6 | -0.86939 | 0.80768 | KRT32 | | 0.80768 | 0.74864 | 0.74811 |
| ENSG00000166164 | 16 | 50347398 | 50402845 | 159 | 18 | -0.86942 | 0.80769 | BRD7 | | 0.80769 | 0.85923 | 0.81696 |
| ENSG00000139266 | 12 | 58148881 | 58154190 | 3 | 2 | -0.86953 | 0.80772 | | 43899 | 0.80772 | 0.3655 | 0.56706 |
| ENSG00000161921 | 17 | 4636821 | 4643217 | 33 | 5 | -0.86965 | 0.80775 | CXCL16 | | 0.80775 | 0.8861 | 0.73412 |
| ENSG00000157326 | 14 | 24422795 | 24438488 | 85 | 11 | -0.86975 | 0.80778 | DHRS4 | | 0.80778 | 0.77612 | 0.8521 |
| ENSG00000115073 | 2 | 98272431 | 98280570 | 10 | 2 | -0.86997 | 0.80784 | ACTR1B | | 0.80784 | 0.77868 | 0.83358 |
| ENSG00000110442 | 11 | 36295051 | 36310999 | 102 | 13 | -0.87008 | 0.80787 | COMMD9 | | 0.80787 | 0.80699 | 0.8682 |
| ENSG00000116176 | 16 | 1271651 | 1275257 | 31 | 9 | -0.8702 | 0.8079 | TPSG1 | | 0.8079 | 0.8361 | 0.78643 |
| ENSG00000164924 | 8 | 101928753 | 101965616 | 108 | 14 | -0.87024 | 0.80791 | YWHAZ | | 0.80791 | 0.81363 | 0.82652 |
| ENSG00000143337 | 1 | 179851177 | 179894135 | 126 | 13 | -0.87028 | 0.80792 | TOR1AIP1 | | 0.80792 | 0.78889 | 0.8468 |
| ENSG00000112531 | 6 | 163835032 | 163999628 | 425 | 21 | -0.87028 | 0.80793 | QKI | | 0.80793 | 0.85431 | 0.90204 |
| ENSG00000185669 | 16 | 88744090 | 88752901 | 23 | 7 | -0.87053 | 0.80799 | SNAI3 | | 0.80799 | 0.71401 | 0.60044 |
| ENSG00000106686 | 9 | 4553386 | 4666674 | 613 | 55 | -0.87066 | 0.80803 | SPATA6L | | 0.80803 | 0.64553 | 0.48177 |
| ENSG00000168993 | 4 | 778745 | 819986 | 206 | 30 | -0.87084 | 0.80808 | CPLX1 | | 0.80808 | 0.85815 | 0.76607 |
| ENSG00000100836 | 14 | 23790498 | 23795394 | 12 | 6 | -0.87083 | 0.80808 | PABPN1 | | 0.80808 | 0.87679 | 0.88383 |
| ENSG00000158321 | 7 | 69063905 | 70258054 | 2885 | 82 | -0.87087 | 0.80809 | AUTS2 | | 0.80809 | 0.83296 | 0.83598 |
| ENSG00000166033 | 10 | 124221041 | 124274424 | 215 | 29 | -0.87137 | 0.80822 | HTRA1 | | 0.80822 | 0.80927 | 0.8611 |
| ENSG00000164418 | 6 | 101846664 | 102517958 | 2513 | 127 | -0.8715 | 0.80826 | GRIK2 | | 0.80826 | 0.82242 | 0.8527 |
| ENSG00000109534 | 4 | 110736666 | 110745893 | 24 | 5 | -0.8716 | 0.80829 | GAR1 | | 0.80829 | 0.86561 | 0.91202 |
| ENSG00000133104 | 13 | 36875775 | 36944317 | 230 | 15 | -0.87169 | 0.80831 | SPG20 | | 0.80831 | 0.84082 | 0.80208 |
| ENSG00000130762 | 1 | 3370990 | 3397677 | 107 | 13 | -0.87193 | 0.80838 | ARHGEF16 | | 0.80838 | 0.76209 | 0.57712 |
| ENSG00000198920 | 17 | 6481468 | 6544247 | 210 | 19 | -0.87194 | 0.80838 | KIAA0753 | | 0.80838 | 0.78692 | 0.70838 |
| ENSG00000139133 | 12 | 34175216 | 34182629 | 24 | 5 | -0.87209 | 0.80842 | ALG10 | | 0.80842 | 0.83997 | 0.83634 |
| ENSG00000135341 | 6 | 91223292 | 91296764 | 226 | 19 | -0.87227 | 0.80847 | MAP3K7 | | 0.80847 | 0.7737 | 0.85704 |
| ENSG00000108448 | 17 | 18601311 | 18639431 | 109 | 20 | -0.87233 | 0.80848 | TRIM16L | | 0.80848 | 0.86671 | 0.95923 |
| ENSG00000154252 | 2 | 242716240 | 242743623 | 88 | 14 | -0.87238 | 0.8085 | GAL3ST2 | | 0.8085 | 0.77339 | 0.54937 |
| ENSG00000164342 | 4 | 186990306 | 187009223 | 57 | 14 | -0.87258 | 0.80856 | TLR3 | | 0.80856 | 0.89375 | 0.84232 |
| ENSG00000146674 | 7 | 45951949 | 45961473 | 23 | 6 | -0.87263 | 0.80857 | IGFBP3 | | 0.80857 | 0.87427 | 0.59238 |
| ENSG00000188536 | 16 | 222846 | 223709 | 1 | 1 | -0.87275 | 0.8086 | HBA2 | | 0.8086 | 0.31657 | 0.17281 |
| ENSG00000099991 | 22 | 24407642 | 24574596 | 296 | 22 | -0.87277 | 0.80861 | CABIN1 | | 0.80861 | 0.76061 | 0.76626 |
| ENSG00000197756 | 2 | 217362912 | 217443903 | 279 | 35 | -0.87293 | 0.80865 | RPL37A | | 0.80865 | 0.83403 | 0.87494 |
| ENSG00000184613 | 12 | 44902058 | 45315631 | 1163 | 57 | -0.87295 | 0.80865 | NELL2 | | 0.80865 | 0.79963 | 0.79274 |
| ENSG00000144645 | 3 | 31699382 | 32119072 | 1612 | 133 | -0.87295 | 0.80866 | OSBPL10 | | 0.80866 | 0.78332 | 0.85149 |
| ENSG00000254413 | 22 | 51007298 | 51021394 | 66 | 11 | -0.87302 | 0.80867 | CHKB-CPT1B | | 0.80867 | 0.60269 | 0.41874 |
| ENSG00000104055 | 15 | 43524793 | 43559055 | 60 | 9 | -0.8731 | 0.8087 | TGM5 | | 0.8087 | 0.74793 | 0.7892 |
| ENSG00000143595 | 1 | 154293566 | 154297801 | 14 | 4 | -0.87317 | 0.80871 | AQP10 | | 0.80871 | 0.75789 | 0.69694 |
| ENSG00000188428 | 6 | 8013800 | 8064647 | 230 | 19 | -0.87326 | 0.80874 | BLOC1S5 | | 0.80874 | 0.85174 | 0.85079 |
| ENSG00000166118 | 11 | 133710526 | 133715433 | 30 | 5 | -0.87331 | 0.80875 | SPATA19 | | 0.80875 | 0.72444 | 0.72278 |
| ENSG00000139977 | 14 | 57857262 | 57882635 | 34 | 6 | -0.87335 | 0.80876 | NAA30 | | 0.80876 | 0.94768 | 0.97359 |
| ENSG00000162620 | 1 | 74491699 | 74663871 | 283 | 17 | -0.87354 | 0.80882 | LRRIQ3 | | 0.80882 | 0.80346 | 0.80713 |

| | | | | | | | | | | | | |
|---|---|---|---|---|---|---|---|---|---|---|---|---|
| ENSG00000177675 | 12 | 7499281 | 7632493 | 375 | 39 | -0.87364 | 0.80884 | CD163L1 | | 0.80884 | 0.87334 | 0.87004 |
| ENSG00000242259 | 22 | 19338891 | 19435755 | 231 | 21 | -0.87361 | 0.80884 | C22orf39 | | 0.80884 | 0.78058 | 0.69904 |
| ENSG00000092439 | 15 | 50844670 | 50979012 | 554 | 19 | -0.87371 | 0.80886 | TRPM7 | | 0.80886 | 0.81093 | 0.77404 |
| ENSG00000221869 | 8 | 48649471 | 48651648 | 2 | 1 | -0.87392 | 0.80892 | CEBPD | | 0.80892 | 0.25063 | 0.091523 |
| ENSG00000111801 | 6 | 26440700 | 26453643 | 36 | 10 | -0.87396 | 0.80893 | BTN3A3 | | 0.80893 | 0.81294 | 0.72295 |
| ENSG00000198569 | 9 | 140125209 | 140131006 | 18 | 8 | -0.87403 | 0.80895 | SLC34A3 | | 0.80895 | 0.57267 | 0.41731 |
| ENSG00000150593 | 10 | 112631565 | 112659764 | 68 | 15 | -0.87422 | 0.809 | PDCD4 | | 0.809 | 0.82931 | 0.77768 |
| ENSG00000169193 | 7 | 23636998 | 23684327 | 91 | 17 | -0.87437 | 0.80904 | CCDC126 | | 0.80904 | 0.78945 | 0.8658 |
| ENSG00000241878 | 22 | 32014477 | 32058418 | 179 | 17 | -0.87437 | 0.80904 | PISD | | 0.80904 | 0.80301 | 0.84503 |
| ENSG00000115275 | 2 | 74688184 | 74692537 | 4 | 2 | -0.87458 | 0.8091 | MOGS | | 0.8091 | 0.49675 | 0.38679 |
| ENSG00000187616 | 9 | 136379708 | 136393734 | 32 | 12 | -0.8747 | 0.80913 | TMEM8C | | 0.80913 | 0.73656 | 0.90815 |
| ENSG00000264424 | 17 | 10346607 | 10372876 | 39 | 7 | -0.87483 | 0.80917 | MYH4 | | 0.80917 | 0.87473 | 0.87898 |
| ENSG00000151365 | 11 | 77774907 | 77779397 | 27 | 3 | -0.87489 | 0.80918 | THRSP | | 0.80918 | 0.91555 | 0.90843 |
| ENSG00000004939 | 17 | 42325753 | 42345509 | 54 | 18 | -0.87508 | 0.80923 | SLC4A1 | | 0.80923 | 0.64027 | 0.68441 |
| ENSG00000197024 | 7 | 148823508 | 148880116 | 211 | 14 | -0.87514 | 0.80925 | ZNF398 | | 0.80925 | 0.83062 | 0.87794 |
| ENSG00000171700 | 20 | 62704534 | 62711323 | 12 | 5 | -0.87563 | 0.80938 | RGS19 | | 0.80938 | 0.84869 | 0.90905 |
| ENSG00000139436 | 12 | 110367607 | 110434194 | 168 | 22 | -0.87593 | 0.80947 | GIT2 | | 0.80947 | 0.80995 | 0.76276 |
| ENSG00000138821 | 4 | 103172198 | 103352415 | 641 | 38 | -0.87618 | 0.80953 | SLC39A8 | | 0.80953 | 0.78584 | 0.75938 |
| ENSG00000167769 | 19 | 6306153 | 6333640 | 129 | 21 | -0.87616 | 0.80953 | ACER1 | | 0.80953 | 0.88234 | 0.7932 |
| ENSG00000126215 | 14 | 104163946 | 104181841 | 41 | 10 | -0.8764 | 0.80959 | XRCC3 | | 0.80959 | 0.79192 | 0.87845 |
| ENSG00000126709 | 1 | 27992572 | 27998729 | 19 | 5 | -0.87656 | 0.80964 | IFI6 | | 0.80964 | 0.84397 | 0.86103 |
| ENSG00000076108 | 12 | 56989380 | 57030600 | 59 | 13 | -0.87668 | 0.80967 | BAZ2A | | 0.80967 | 0.78034 | 0.663 |
| ENSG00000125863 | 20 | 10381657 | 10414870 | 137 | 9 | -0.87682 | 0.80971 | MKKS | | 0.80971 | 0.84846 | 0.9251 |
| ENSG00000115556 | 2 | 219472488 | 219501907 | 59 | 8 | -0.87695 | 0.80974 | PLCD4 | | 0.80974 | 0.78756 | 0.77431 |
| ENSG00000115761 | 2 | 10710892 | 10830101 | 651 | 25 | -0.87709 | 0.80978 | NOL10 | | 0.80978 | 0.81313 | 0.85391 |
| ENSG00000250588 | 3 | 158680024 | 159615155 | 2309 | 106 | -0.87716 | 0.8098 | IQCJ-SCHIP1 | | 0.8098 | 0.82308 | 0.8173 |
| ENSG00000177150 | 18 | 13663346 | 13726662 | 248 | 28 | -0.87737 | 0.80986 | FAM210A | | 0.80986 | 0.82526 | 0.86813 |
| ENSG00000168487 | 8 | 22022249 | 22069839 | 180 | 31 | -0.8775 | 0.80989 | BMP1 | | 0.80989 | 0.76826 | 0.61634 |
| ENSG00000173838 | 17 | 60778675 | 60885705 | 330 | 22 | -0.87748 | 0.80989 | | 43900 | 0.80989 | 0.72202 | 0.55567 |
| ENSG00000181826 | 4 | 37592422 | 37687998 | 424 | 52 | -0.8776 | 0.80992 | RELL1 | | 0.80992 | 0.85382 | 0.92115 |
| ENSG00000163817 | 3 | 45796942 | 45838027 | 99 | 31 | -0.87795 | 0.81001 | SLC6A20 | | 0.81001 | 0.81005 | 0.79456 |
| ENSG00000186517 | 1 | 161016736 | 161039760 | 66 | 14 | -0.87799 | 0.81002 | ARHGAP30 | | 0.81002 | 0.87985 | 0.91755 |
| ENSG00000158480 | 20 | 48519928 | 48532080 | 37 | 8 | -0.87799 | 0.81002 | SPATA2 | | 0.81002 | 0.82361 | 0.77102 |
| ENSG00000138182 | 10 | 91461367 | 91534700 | 296 | 11 | -0.87839 | 0.81014 | KIF20B | | 0.81014 | 0.80349 | 0.81031 |
| ENSG00000135127 | 12 | 120427673 | 120532298 | 254 | 17 | -0.87859 | 0.81019 | CCDC64 | | 0.81019 | 0.81642 | 0.83648 |
| ENSG00000124006 | 2 | 220415451 | 220436581 | 55 | 10 | -0.87893 | 0.81028 | OBSL1 | | 0.81028 | 0.7492 | 0.61806 |
| ENSG00000102893 | 16 | 47495034 | 47735434 | 229 | 20 | -0.87901 | 0.8103 | PHKB | | 0.8103 | 0.82193 | 0.81571 |
| ENSG00000104804 | 19 | 49384222 | 49401990 | 84 | 11 | -0.87911 | 0.81033 | TULP2 | | 0.81033 | 0.89717 | 0.93667 |
| ENSG00000170242 | 11 | 11862970 | 11980870 | 274 | 20 | -0.87917 | 0.81034 | USP47 | | 0.81034 | 0.80006 | 0.75815 |
| ENSG00000186153 | 16 | 78133310 | 79246564 | 7906 | 390 | -0.87914 | 0.81034 | WWOX | | 0.81034 | 0.8006 | 0.80546 |
| ENSG00000266714 | 17 | 73584139 | 73622929 | 223 | 30 | -0.87935 | 0.81039 | MYO15B | | 0.81039 | 0.88268 | 0.85987 |
| ENSG00000151743 | 12 | 31824071 | 31882108 | 164 | 29 | -0.8794 | 0.81041 | AMN1 | | 0.81041 | 0.75152 | 0.64945 |
| ENSG00000115652 | 2 | 106709759 | 106810795 | 209 | 38 | -0.87975 | 0.8105 | UXS1 | | 0.8105 | 0.47677 | 0.40937 |
| ENSG00000109854 | 11 | 20385231 | 20405329 | 67 | 12 | -0.87976 | 0.8105 | HTATIP2 | | 0.8105 | 0.75614 | 0.81209 |
| ENSG00000138100 | 2 | 27505260 | 27530307 | 47 | 10 | -0.87982 | 0.81052 | TRIM54 | | 0.81052 | 0.85646 | 0.82553 |
| ENSG00000141098 | 16 | 67708434 | 67753324 | 76 | 12 | -0.87998 | 0.81056 | GFOD2 | | 0.81056 | 0.86387 | 0.88497 |
| ENSG00000189030 | 1 | 156268415 | 156269428 | 2 | 1 | -0.88029 | 0.81065 | VHLL | | 0.81065 | 0.44057 | 0.4637 |
| ENSG00000188032 | 19 | 14192431 | 14196687 | 8 | 3 | -0.88037 | 0.81067 | C19orf67 | | 0.81067 | 0.84822 | 0.81643 |
| ENSG00000102547 | 13 | 49882786 | 50018262 | 453 | 36 | -0.88047 | 0.8107 | CAB39L | | 0.8107 | 0.85753 | 0.88946 |
| ENSG00000167378 | 19 | 44088521 | 44100287 | 19 | 9 | -0.88064 | 0.81074 | IRGQ | | 0.81074 | 0.66381 | 0.48074 |
| ENSG00000145495 | 5 | 10353815 | 10440500 | 135 | 14 | -0.88091 | 0.81082 | | 43896 | 0.81082 | 0.81106 | 0.74648 |
| ENSG00000154059 | 18 | 22006580 | 22033499 | 83 | 12 | -0.88094 | 0.81083 | IMPACT | | 0.81083 | 0.84754 | 0.82398 |
| ENSG00000214022 | 7 | 150065278 | 150071133 | 18 | 3 | -0.8812 | 0.8109 | REPIN1 | | 0.8109 | 0.80188 | 0.82778 |
| ENSG00000213614 | 15 | 72635775 | 72668817 | 65 | 8 | -0.88141 | 0.81095 | HEXA | | 0.81095 | 0.90163 | 0.8972 |
| ENSG00000213186 | 3 | 160150233 | 160203561 | 121 | 18 | -0.88145 | 0.81096 | TRIM59 | | 0.81096 | 0.75007 | 0.77394 |
| ENSG00000215527 | 18 | 12658042 | 12663470 | 24 | 4 | -0.88166 | 0.81102 | AP005482.1 | | 0.81102 | 0.76589 | 0.7949 |
| ENSG00000120708 | 5 | 135364584 | 135399507 | 89 | 9 | -0.88178 | 0.81105 | TGFBI | | 0.81105 | 0.80118 | 0.85943 |
| ENSG00000182264 | 19 | 49244109 | 49250166 | 27 | 2 | -0.88203 | 0.81112 | IZUMO1 | | 0.81112 | 0.84089 | 0.93208 |
| ENSG00000128891 | 15 | 40820882 | 40857256 | 109 | 10 | -0.88205 | 0.81113 | C15orf57 | | 0.81113 | 0.83055 | 0.8151 |
| ENSG00000123124 | 8 | 87354967 | 87490649 | 471 | 25 | -0.88214 | 0.81115 | WWP1 | | 0.81115 | 0.81383 | 0.8325 |
| ENSG00000156110 | 10 | 75910960 | 76469061 | 1141 | 27 | -0.88216 | 0.81115 | ADK | | 0.81115 | 0.79466 | 0.7662 |
| ENSG00000000419 | 20 | 49551404 | 49575092 | 32 | 9 | -0.88212 | 0.81115 | DPM1 | | 0.81115 | 0.8299 | 0.87261 |
| ENSG00000117501 | 1 | 170904612 | 171033906 | 523 | 25 | -0.88222 | 0.81117 | MROH9 | | 0.81117 | 0.83082 | 0.84154 |
| ENSG00000270033 | 17 | 60349848 | 60360276 | 62 | 11 | -0.88221 | 0.81117 | RP11-51L5.7 | | 0.81117 | 0.79373 | 0.75235 |
| ENSG00000186925 | 21 | 31913854 | 31914183 | 2 | 1 | -0.88231 | 0.8112 | KRTAP19-6 | | 0.8112 | 0.77227 | 0.75977 |
| ENSG00000150938 | 2 | 36583069 | 36778278 | 616 | 54 | -0.88255 | 0.81126 | CRIM1 | | 0.81126 | 0.85354 | 0.89165 |
| ENSG00000138449 | 2 | 190425305 | 190448484 | 61 | 11 | -0.88256 | 0.81126 | SLC40A1 | | 0.81126 | 0.65828 | 0.63538 |
| ENSG00000162976 | 2 | 11295324 | 11319000 | 114 | 20 | -0.8826 | 0.81127 | PQLC3 | | 0.81127 | 0.87246 | 0.8976 |
| ENSG00000173535 | 8 | 22941868 | 22974950 | 137 | 21 | -0.88305 | 0.8114 | TNFRSF10C | | 0.8114 | 0.72246 | 0.75519 |
| ENSG00000129159 | 11 | 17756359 | 17804602 | 143 | 24 | -0.88307 | 0.8114 | KCNC1 | | 0.8114 | 0.88451 | 0.90932 |
| ENSG00000104237 | 8 | 55528627 | 55543394 | 21 | 4 | -0.88315 | 0.81142 | RP1 | | 0.81142 | 0.85653 | 0.89641 |
| ENSG00000133789 | 11 | 9685624 | 9774538 | 383 | 16 | -0.88316 | 0.81142 | SWAP70 | | 0.81142 | 0.82191 | 0.85971 |
| ENSG00000139826 | 13 | 108870727 | 108886603 | 48 | 7 | -0.88315 | 0.81142 | ABHD13 | | 0.81142 | 0.7388 | 0.60973 |

| | | | | | | | | | | | | |
|---|---|---|---|---|---|---|---|---|---|---|---|---|
| ENSG00000163239 | 1 | 154474695 | 154520623 | 187 | 11 | -0.88338 | 0.81148 | TDRD10 | | 0.81148 | 0.82098 | 0.81322 |
| ENSG00000258986 | 14 | 104941015 | 105071984 | 561 | 39 | -0.88369 | 0.81157 | TMEM179 | | 0.81157 | 0.86031 | 0.88387 |
| ENSG00000167112 | 9 | 131071382 | 131085021 | 24 | 8 | -0.88381 | 0.8116 | TRUB2 | | 0.8116 | 0.85233 | 0.84422 |
| ENSG00000180386 | 17 | 39431911 | 39432949 | 5 | 1 | -0.88387 | 0.81162 | KRTAP9-7 | | 0.81162 | 0.74698 | 0.47139 |
| ENSG00000029639 | 6 | 155578643 | 155635627 | 283 | 21 | -0.88395 | 0.81164 | TFB1M | | 0.81164 | 0.77696 | 0.75037 |
| ENSG00000147649 | 8 | 98656407 | 98740998 | 201 | 19 | -0.88404 | 0.81166 | MTDH | | 0.81166 | 0.84095 | 0.89884 |
| ENSG00000151006 | 16 | 31094746 | 31100949 | 14 | 4 | -0.88436 | 0.81175 | PRSS53 | | 0.81175 | 0.87228 | 0.82678 |
| ENSG00000170180 | 4 | 145030457 | 145061904 | 159 | 13 | -0.88473 | 0.81185 | GYPA | | 0.81185 | 0.77689 | 0.65728 |
| ENSG00000105518 | 19 | 11453452 | 11457194 | 1 | 1 | -0.88492 | 0.8119 | TMEM205 | | 0.8119 | 0.5308 | 0.72368 |
| ENSG00000121068 | 17 | 59477257 | 59486827 | 36 | 5 | -0.88528 | 0.812 | TBX2 | | 0.812 | 0.77245 | 0.73293 |
| ENSG00000267496 | 17 | 41994576 | 41995326 | 7 | 1 | -0.88532 | 0.81201 | FAM215A | | 0.81201 | 0.74536 | 0.82706 |
| ENSG00000106305 | 7 | 6048876 | 6063465 | 88 | 18 | -0.88559 | 0.81208 | AIMP2 | | 0.81208 | 0.78844 | 0.70003 |
| ENSG00000177151 | 1 | 248801588 | 248802559 | 1 | 1 | -0.88566 | 0.8121 | OR2T35 | | 0.8121 | 0.85464 | 0.51823 |
| ENSG00000147421 | 8 | 28747911 | 28922281 | 255 | 19 | -0.8857 | 0.81211 | HMBOX1 | | 0.81211 | 0.80593 | 0.82635 |
| ENSG00000174109 | 16 | 1469745 | 1479345 | 26 | 7 | -0.88598 | 0.81219 | C16orf91 | | 0.81219 | 0.71766 | 0.76931 |
| ENSG00000172938 | 11 | 68747490 | 68748455 | 2 | 1 | -0.88603 | 0.8122 | MRGPRD | | 0.8122 | 0.28896 | 0.25799 |
| ENSG00000205835 | 3 | 190570666 | 190610218 | 148 | 27 | -0.88628 | 0.81227 | GMNC | | 0.81227 | 0.75333 | 0.81743 |
| ENSG00000165487 | 13 | 22066836 | 22178353 | 379 | 21 | -0.88629 | 0.81227 | MICU2 | | 0.81227 | 0.75724 | 0.6533 |
| ENSG00000151929 | 10 | 121410882 | 121437331 | 100 | 15 | -0.88632 | 0.81228 | BAG3 | | 0.81228 | 0.88941 | 0.93575 |
| ENSG00000188833 | 9 | 140328816 | 140336268 | 15 | 6 | -0.88642 | 0.81231 | ENTPD8 | | 0.81231 | 0.79446 | 0.8063 |
| ENSG00000197233 | 9 | 125273081 | 125274022 | 3 | 2 | -0.88654 | 0.81234 | OR1J2 | | 0.81234 | 0.95015 | 0.86255 |
| ENSG00000183473 | 22 | 37600278 | 37608362 | 21 | 6 | -0.88671 | 0.81238 | SSTR3 | | 0.81238 | 0.73691 | 0.67919 |
| ENSG00000122986 | 12 | 111065646 | 111142755 | 176 | 15 | -0.88692 | 0.81244 | HVCN1 | | 0.81244 | 0.81215 | 0.82344 |
| ENSG00000109846 | 11 | 111779289 | 111794446 | 30 | 7 | -0.88696 | 0.81245 | CRYAB | | 0.81245 | 0.85516 | 0.88561 |
| ENSG00000136696 | 2 | 113779668 | 113810444 | 116 | 14 | -0.88749 | 0.81259 | IL36B | | 0.81259 | 0.78697 | 0.74663 |
| ENSG00000126214 | 14 | 104028233 | 104167888 | 409 | 23 | -0.88759 | 0.81262 | KLC1 | | 0.81262 | 0.80341 | 0.84609 |
| ENSG00000156273 | 21 | 30566392 | 31003071 | 1017 | 63 | -0.88766 | 0.81264 | BACH1 | | 0.81264 | 0.83901 | 0.88916 |
| ENSG00000141026 | 17 | 17380300 | 17396540 | 39 | 8 | -0.888 | 0.81273 | MED9 | | 0.81273 | 0.81393 | 0.72071 |
| ENSG00000186806 | 19 | 51834790 | 51845378 | 27 | 6 | -0.88816 | 0.81277 | VSIG10L | | 0.81277 | 0.24597 | 0.10543 |
| ENSG00000105855 | 7 | 20370325 | 20455377 | 382 | 23 | -0.88832 | 0.81282 | ITGB8 | | 0.81282 | 0.73364 | 0.71492 |
| ENSG00000163171 | 2 | 37869032 | 37965611 | 465 | 50 | -0.88838 | 0.81283 | CDC42EP3 | | 0.81283 | 0.78026 | 0.66313 |
| ENSG00000135622 | 2 | 74881355 | 74909186 | 34 | 8 | -0.88848 | 0.81286 | SEMA4F | | 0.81286 | 0.80188 | 0.80927 |
| ENSG00000183696 | 7 | 48128225 | 48148330 | 46 | 11 | -0.88852 | 0.81287 | UPP1 | | 0.81287 | 0.60468 | 0.60828 |
| ENSG00000176900 | 11 | 4903049 | 4904113 | 1 | 1 | -0.88901 | 0.813 | OR51T1 | | 0.813 | 0.53699 | 0.42736 |
| ENSG00000068976 | 11 | 64513861 | 64527769 | 31 | 4 | -0.88911 | 0.81303 | PYGM | | 0.81303 | 0.56374 | 0.6561 |
| ENSG00000120742 | 3 | 150259781 | 150321015 | 224 | 25 | -0.88931 | 0.81308 | SERP1 | | 0.81308 | 0.836 | 0.86876 |
| ENSG00000161911 | 6 | 41117080 | 41122075 | 5 | 1 | -0.88939 | 0.8131 | TREML1 | | 0.8131 | 0.68331 | 0.61923 |
| ENSG00000125779 | 20 | 3869486 | 3907605 | 111 | 15 | -0.88944 | 0.81312 | PANK2 | | 0.81312 | 0.91919 | 0.94835 |
| ENSG00000169062 | 13 | 115047059 | 115071283 | 85 | 19 | -0.88961 | 0.81316 | UPF3A | | 0.81316 | 0.78857 | 0.82131 |
| ENSG00000158748 | 1 | 19991780 | 20006055 | 50 | 10 | -0.88974 | 0.8132 | HTR6 | | 0.8132 | 0.85472 | 0.86142 |
| ENSG00000268182 | 19 | 57154513 | 57167134 | 61 | 9 | -0.89003 | 0.81328 | SMIM17 | | 0.81328 | 0.37546 | 0.39663 |
| ENSG00000102886 | 16 | 30116131 | 30125177 | 25 | 10 | -0.89016 | 0.81331 | GDPD3 | | 0.81331 | 0.7359 | 0.75639 |
| ENSG00000167637 | 19 | 44331444 | 44353307 | 118 | 9 | -0.89015 | 0.81331 | ZNF283 | | 0.81331 | 0.8058 | 0.87962 |
| ENSG00000038002 | 4 | 178351924 | 178363657 | 30 | 8 | -0.89049 | 0.8134 | AGA | | 0.8134 | 0.6637 | 0.61498 |
| ENSG00000137251 | 6 | 54172657 | 54254950 | 264 | 25 | -0.89064 | 0.81344 | TINAG | | 0.81344 | 0.82791 | 0.82656 |
| ENSG00000136457 | 17 | 48541857 | 48546327 | 11 | 4 | -0.89068 | 0.81345 | CHAD | | 0.81345 | 0.86543 | 0.89335 |
| ENSG00000087008 | 4 | 8368009 | 8442450 | 353 | 28 | -0.89074 | 0.81347 | ACOX3 | | 0.81347 | 0.8203 | 0.86339 |
| ENSG00000188981 | 4 | 3246096 | 3273465 | 100 | 11 | -0.89088 | 0.8135 | MSANTD1 | | 0.8135 | 0.84525 | 0.87535 |
| ENSG00000214645 | 9 | 110539471 | 110540419 | 1 | 1 | -0.89087 | 0.8135 | AL162389.1 | | 0.8135 | 0.79452 | 0.60724 |
| ENSG00000163131 | 1 | 150702672 | 150738433 | 85 | 9 | -0.8912 | 0.81359 | CTSS | | 0.81359 | 0.8375 | 0.8336 |
| ENSG00000151687 | 2 | 190539016 | 190625919 | 203 | 22 | -0.89138 | 0.81364 | ANKAR | | 0.81364 | 0.80378 | 0.85036 |
| ENSG00000145287 | 4 | 84011201 | 84058228 | 108 | 18 | -0.89144 | 0.81365 | PLAC8 | | 0.81365 | 0.70365 | 0.64277 |
| ENSG00000187821 | 4 | 185939995 | 185941958 | 6 | 2 | -0.89163 | 0.8137 | HELT | | 0.8137 | 0.47913 | 0.6845 |
| ENSG00000273025 | 15 | 72559087 | 72612287 | 132 | 18 | -0.89215 | 0.81384 | CELF6 | | 0.81384 | 0.81322 | 0.85255 |
| ENSG00000034713 | 16 | 75600249 | 75611779 | 29 | 7 | -0.8923 | 0.81388 | GABARAPL2 | | 0.81388 | 0.8988 | 0.92224 |
| ENSG00000243710 | 1 | 43637820 | 43720029 | 378 | 31 | -0.89249 | 0.81393 | WDR65 | | 0.81393 | 0.82707 | 0.83238 |
| ENSG00000174156 | 6 | 52761437 | 52774483 | 44 | 8 | -0.89315 | 0.81411 | GSTA3 | | 0.81411 | 0.82495 | 0.86133 |
| ENSG00000100439 | 14 | 23067146 | 23081265 | 60 | 13 | -0.89325 | 0.81414 | ABHD4 | | 0.81414 | 0.816 | 0.069018 |
| ENSG00000182854 | 15 | 102358366 | 102359350 | 1 | 1 | -0.89348 | 0.8142 | OR4F15 | | 0.8142 | 0.50244 | 0.54054 |
| ENSG00000175137 | 1 | 249104648 | 249120832 | 29 | 4 | -0.89371 | 0.81426 | SH3BP5L | | 0.81426 | 0.8084 | 0.83167 |
| ENSG00000136059 | 3 | 38029550 | 38048679 | 61 | 7 | -0.89369 | 0.81426 | VILL | | 0.81426 | 0.83014 | 0.88693 |
| ENSG00000111276 | 12 | 12867992 | 12875305 | 17 | 7 | -0.89371 | 0.81426 | CDKN1B | | 0.81426 | 0.64769 | 0.59732 |
| ENSG00000117036 | 1 | 157090983 | 157108266 | 21 | 8 | -0.89393 | 0.81432 | ETV3 | | 0.81432 | 0.89084 | 0.9178 |
| ENSG00000204930 | 9 | 35816388 | 35828744 | 17 | 7 | -0.89411 | 0.81437 | FAM221B | | 0.81437 | 0.58988 | 0.68747 |
| ENSG00000164818 | 7 | 766338 | 829190 | 303 | 21 | -0.89415 | 0.81438 | HEATR2 | | 0.81438 | 0.87181 | 0.85291 |
| ENSG00000166169 | 10 | 103338639 | 103348027 | 16 | 2 | -0.89442 | 0.81445 | POLL | | 0.81445 | 0.7409 | 0.63765 |
| ENSG00000165392 | 8 | 30891317 | 31031285 | 368 | 26 | -0.89453 | 0.81448 | WRN | | 0.81448 | 0.80498 | 0.8165 |
| ENSG00000117791 | 1 | 220921567 | 220958150 | 119 | 11 | -0.89456 | 0.81449 | | 43892 | 0.81449 | 0.72233 | 0.43796 |
| ENSG00000170677 | 18 | 67956137 | 67997436 | 172 | 25 | -0.89459 | 0.8145 | SOCS6 | | 0.8145 | 0.82552 | 0.8626 |
| ENSG00000105656 | 19 | 18553473 | 18632937 | 210 | 13 | -0.89472 | 0.81453 | ELL | | 0.81453 | 0.83733 | 0.84594 |
| ENSG00000144045 | 2 | 74745258 | 74753463 | 6 | 4 | -0.89478 | 0.81455 | DQX1 | | 0.81455 | 0.62715 | 0.32354 |
| ENSG00000165282 | 9 | 35088685 | 35096591 | 8 | 4 | -0.8949 | 0.81458 | PIGO | | 0.81458 | 0.8229 | 0.82029 |
| ENSG00000197445 | 16 | 73160536 | 73178346 | 39 | 14 | -0.89587 | 0.81484 | C16orf47 | | 0.81484 | 0.84824 | 0.89558 |

| | | | | | | | | | | | |
|---|---|---|---|---|---|---|---|---|---|---|---|
| ENSG00000268040 | 21 | 47347679 | 47352477 | 15 | 4 | -0.896 | 0.81487 | PRED62 | 0.81487 | 0.58754 | 0.58747 |
| ENSG00000176925 | 11 | 4842551 | 4843686 | 3 | 2 | -0.89611 | 0.8149 | OR51F2 | 0.8149 | 0.68405 | 0.70977 |
| ENSG00000117480 | 1 | 46859937 | 46879520 | 39 | 9 | -0.89648 | 0.815 | FAAH | 0.815 | 0.84074 | 0.7629 |
| ENSG00000259658 | 15 | 102277302 | 102285913 | 20 | 5 | -0.89648 | 0.815 | RP11-89K11.1 | 0.815 | 0.6167 | 0.60388 |
| ENSG00000166398 | 19 | 34745442 | 34846491 | 299 | 21 | -0.89682 | 0.81509 | KIAA0355 | 0.81509 | 0.80243 | 0.75506 |
| ENSG00000164744 | 7 | 48026745 | 48068716 | 102 | 8 | -0.89708 | 0.81516 | SUN3 | 0.81516 | 0.84005 | 0.76371 |
| ENSG00000023902 | 1 | 150121373 | 150136916 | 39 | 7 | -0.89712 | 0.81517 | PLEKHO1 | 0.81517 | 0.82488 | 0.84795 |
| ENSG00000123607 | 2 | 166713985 | 166810353 | 238 | 24 | -0.89719 | 0.81519 | TTC21B | 0.81519 | 0.8245 | 0.78958 |
| ENSG00000107438 | 10 | 96997329 | 97050781 | 186 | 27 | -0.8972 | 0.81519 | PDLIM1 | 0.81519 | 0.86649 | 0.86646 |
| ENSG00000091664 | 11 | 22359643 | 22401049 | 151 | 36 | -0.89731 | 0.81522 | SLC17A6 | 0.81522 | 0.74285 | 0.75727 |
| ENSG00000133193 | 17 | 71203492 | 71232892 | 144 | 9 | -0.89791 | 0.81538 | FAM104A | 0.81538 | 0.72059 | 0.77189 |
| ENSG00000129292 | 8 | 133787618 | 133861052 | 129 | 13 | -0.89793 | 0.81539 | PHF20L1 | 0.81539 | 0.80787 | 0.76095 |
| ENSG00000136939 | 9 | 125486269 | 125487204 | 2 | 1 | -0.89809 | 0.81543 | OR1L4 | 0.81543 | 0.7993 | 0.79697 |
| ENSG00000136444 | 17 | 48556161 | 48563336 | 12 | 3 | -0.89813 | 0.81544 | RSAD1 | 0.81544 | 0.90542 | 0.88483 |
| ENSG00000129514 | 14 | 38059189 | 38069245 | 26 | 8 | -0.89825 | 0.81547 | FOXA1 | 0.81547 | 0.61524 | 0.73392 |
| ENSG00000087253 | 16 | 55542910 | 55620582 | 258 | 15 | -0.89832 | 0.81549 | LPCAT2 | 0.81549 | 0.84472 | 0.83308 |
| ENSG00000168703 | 20 | 43752066 | 43753106 | 4 | 2 | -0.89884 | 0.81563 | WFDC12 | 0.81563 | 0.36793 | 0.38768 |
| ENSG00000100319 | 22 | 30126945 | 30163000 | 130 | 13 | -0.89909 | 0.8157 | ZMAT5 | 0.8157 | 0.76096 | 0.73077 |
| ENSG00000136535 | 2 | 162272605 | 162282381 | 10 | 4 | -0.89921 | 0.81573 | TBR1 | 0.81573 | 0.41869 | 0.41427 |
| ENSG00000100908 | 14 | 24608174 | 24610797 | 4 | 2 | -0.89938 | 0.81578 | EMC9 | 0.81578 | 0.87376 | 0.80069 |
| ENSG00000158292 | 1 | 6307406 | 6321035 | 37 | 3 | -0.89945 | 0.81579 | GPR153 | 0.81579 | 0.68795 | 0.3582 |
| ENSG00000007080 | 19 | 18043825 | 18054800 | 34 | 10 | -0.89949 | 0.8158 | CCDC124 | 0.8158 | 0.63251 | 0.72678 |
| ENSG00000153303 | 6 | 168456425 | 168482237 | 135 | 20 | -0.89952 | 0.81581 | FRMD1 | 0.81581 | 0.75121 | 0.77957 |
| ENSG00000135077 | 5 | 156512843 | 156569880 | 237 | 25 | -0.89969 | 0.81586 | HAVCR2 | 0.81586 | 0.842 | 0.84741 |
| ENSG00000144744 | 3 | 69103881 | 69129559 | 45 | 10 | -0.90008 | 0.81596 | UBA3 | 0.81596 | 0.6274 | 0.39159 |
| ENSG00000087586 | 20 | 54944445 | 54967393 | 55 | 19 | -0.90011 | 0.81597 | AURKA | 0.81597 | 0.96991 | 0.98515 |
| ENSG00000125814 | 20 | 23355159 | 23402125 | 202 | 15 | -0.90032 | 0.81602 | NAPB | 0.81602 | 0.80485 | 0.76912 |
| ENSG00000159363 | 1 | 17312453 | 17338423 | 58 | 12 | -0.90037 | 0.81604 | ATP13A2 | 0.81604 | 0.8163 | 0.84972 |
| ENSG00000093167 | 3 | 37094117 | 37225180 | 301 | 13 | -0.9004 | 0.81605 | LRRFIP2 | 0.81605 | 0.78828 | 0.81359 |
| ENSG00000145284 | 4 | 83550692 | 83720010 | 935 | 61 | -0.9006 | 0.8161 | SCD5 | 0.8161 | 0.79224 | 0.76142 |
| ENSG00000162032 | 16 | 1826713 | 1843701 | 67 | 13 | -0.90063 | 0.81611 | SPSB3 | 0.81611 | 0.7625 | 0.80801 |
| ENSG00000099385 | 16 | 30844947 | 30906281 | 100 | 13 | -0.90082 | 0.81616 | BCL7C | 0.81616 | 0.81456 | 0.81669 |
| ENSG00000124529 | 6 | 26027124 | 26027480 | 3 | 1 | -0.90102 | 0.81621 | HIST1H4B | 0.81621 | 0.38416 | 0.38112 |
| ENSG00000139531 | 12 | 56390964 | 56400425 | 16 | 8 | -0.90123 | 0.81627 | SUOX | 0.81627 | 0.62744 | 0.59486 |
| ENSG00000155026 | 7 | 5965181 | 6010314 | 81 | 30 | -0.90129 | 0.81628 | RSPH10B | 0.81628 | 0.67628 | 0.40057 |
| ENSG00000242950 | 7 | 92097694 | 92107300 | 24 | 8 | -0.90142 | 0.81632 | ERVW-1 | 0.81632 | 0.7001 | 0.73713 |
| ENSG00000172425 | 11 | 118398187 | 118401912 | 1 | 1 | -0.90173 | 0.8164 | TTC36 | 0.8164 | 0.16279 | 0.15242 |
| ENSG00000008018 | 6 | 170844205 | 170862429 | 65 | 8 | -0.90193 | 0.81645 | PSMB1 | 0.81645 | 0.77571 | 0.75634 |
| ENSG00000198807 | 14 | 37126773 | 37148920 | 87 | 15 | -0.90208 | 0.81649 | PAX9 | 0.81649 | 0.88149 | 0.91589 |
| ENSG00000168229 | 14 | 52734431 | 52743442 | 42 | 10 | -0.90214 | 0.81651 | PTGDR | 0.81651 | 0.45018 | 0.51153 |
| ENSG00000105696 | 19 | 18718240 | 18731849 | 61 | 13 | -0.90218 | 0.81652 | TMEM59L | 0.81652 | 0.67297 | 0.48808 |
| ENSG00000105402 | 19 | 47990894 | 48018497 | 43 | 8 | -0.90259 | 0.81663 | NAPA | 0.81663 | 0.84414 | 0.82127 |
| ENSG00000203546 | 14 | 31803518 | 31926647 | 257 | 20 | -0.90268 | 0.81665 | RP11-176H8.1 | 0.81665 | 0.82559 | 0.83903 |
| ENSG00000011422 | 19 | 44150247 | 44174699 | 124 | 24 | -0.90273 | 0.81667 | PLAUR | 0.81667 | 0.75899 | 0.85877 |
| ENSG00000178401 | 12 | 49740700 | 49751309 | 34 | 3 | -0.90284 | 0.81669 | DNAJC22 | 0.81669 | 0.63255 | 0.47718 |
| ENSG00000135486 | 12 | 54673977 | 54680872 | 7 | 3 | -0.90291 | 0.81671 | HNRNPA1 | 0.81671 | 0.51428 | 0.57578 |
| ENSG00000141582 | 17 | 77806955 | 77813228 | 13 | 5 | -0.90292 | 0.81671 | CBX4 | 0.81671 | 0.60524 | 0.51542 |
| ENSG00000010270 | 7 | 38217824 | 38270272 | 180 | 21 | -0.90297 | 0.81673 | STARD3NL | 0.81673 | 0.76756 | 0.88622 |
| ENSG00000155530 | 7 | 133812052 | 133949343 | 433 | 18 | -0.9031 | 0.81676 | LRGUK | 0.81676 | 0.81723 | 0.8437 |
| ENSG00000160994 | 19 | 15121556 | 15134081 | 104 | 20 | -0.90348 | 0.81686 | CCDC105 | 0.81686 | 0.84997 | 0.96881 |
| ENSG00000106803 | 9 | 101984346 | 101992897 | 35 | 6 | -0.90362 | 0.8169 | SEC61B | 0.8169 | 0.75419 | 0.74955 |
| ENSG00000167851 | 17 | 72462555 | 72480933 | 87 | 16 | -0.90387 | 0.81697 | CD300A | 0.81697 | 0.83726 | 0.79509 |
| ENSG00000138795 | 4 | 108968701 | 109090112 | 312 | 24 | -0.90392 | 0.81698 | LEF1 | 0.81698 | 0.81273 | 0.83529 |
| ENSG00000172660 | 17 | 34136459 | 34191619 | 157 | 14 | -0.90408 | 0.81702 | TAF15 | 0.81702 | 0.78119 | 0.80498 |
| ENSG00000188993 | 4 | 52859868 | 52883786 | 28 | 8 | -0.90414 | 0.81704 | LRRC66 | 0.81704 | 0.85424 | 0.79943 |
| ENSG00000145725 | 5 | 102455853 | 102548500 | 229 | 13 | -0.90431 | 0.81709 | PPIP5K2 | 0.81709 | 0.79295 | 0.76938 |
| ENSG00000129315 | 12 | 49082247 | 49110681 | 78 | 12 | -0.9049 | 0.81724 | CCNT1 | 0.81724 | 0.73321 | 0.71451 |
| ENSG00000131097 | 17 | 42923721 | 42927848 | 13 | 5 | -0.90488 | 0.81724 | HIGD1B | 0.81724 | 0.78396 | 0.80133 |
| ENSG00000111729 | 12 | 8276228 | 8291203 | 47 | 9 | -0.90495 | 0.81725 | CLEC4A | 0.81725 | 0.87388 | 0.85123 |
| ENSG00000130779 | 12 | 122755979 | 122907179 | 259 | 26 | -0.90493 | 0.81725 | CLIP1 | 0.81725 | 0.82816 | 0.83638 |
| ENSG00000040199 | 16 | 71671738 | 71758604 | 222 | 21 | -0.905 | 0.81727 | PHLPP2 | 0.81727 | 0.80142 | 0.81929 |
| ENSG00000139637 | 12 | 53693470 | 53700961 | 13 | 4 | -0.90504 | 0.81728 | C12orf10 | 0.81728 | 0.7515 | 0.70197 |
| ENSG00000196850 | 12 | 110969120 | 111021125 | 131 | 12 | -0.90509 | 0.81729 | PPTC7 | 0.81729 | 0.83524 | 0.86468 |
| ENSG00000172426 | 6 | 43612783 | 43640336 | 105 | 13 | -0.90513 | 0.8173 | RSPH9 | 0.8173 | 0.83034 | 0.82424 |
| ENSG00000123415 | 12 | 54558529 | 54582778 | 66 | 9 | -0.90513 | 0.8173 | SMUG1 | 0.8173 | 0.82863 | 0.8126 |
| ENSG00000255537 | 11 | 125365110 | 125369424 | 9 | 4 | -0.9058 | 0.81748 | AP000708.1 | 0.81748 | 0.58663 | 0.78111 |
| ENSG00000181031 | 17 | 62293 | 236045 | 1265 | 159 | -0.90602 | 0.81754 | RPH3AL | 0.81754 | 0.86728 | 0.89529 |
| ENSG00000100532 | 14 | 54976530 | 55005567 | 46 | 11 | -0.90634 | 0.81762 | CGRRF1 | 0.81762 | 0.84613 | 0.84504 |
| ENSG00000136770 | 10 | 22045466 | 22292698 | 465 | 26 | -0.90667 | 0.81771 | DNAJC1 | 0.81771 | 0.79177 | 0.7883 |
| ENSG00000115221 | 2 | 160956177 | 161128399 | 536 | 41 | -0.9068 | 0.81774 | ITGB6 | 0.81774 | 0.82279 | 0.8471 |
| ENSG00000184164 | 22 | 50311815 | 50321188 | 104 | 6 | -0.90697 | 0.81779 | CRELD2 | 0.81779 | 0.76766 | 0.81418 |
| ENSG00000163684 | 3 | 58291974 | 58310422 | 90 | 11 | -0.90763 | 0.81796 | RPP14 | 0.81796 | 0.76797 | 0.79903 |
| ENSG00000205002 | 8 | 117950438 | 117956726 | 12 | 3 | -0.90768 | 0.81798 | AARD | 0.81798 | 0.86028 | 0.8667 |

| | | | | | | | | | | | |
|---|---|---|---|---|---|---|---|---|---|---|---|
| ENSG00000171401 | 17 | 39657233 | 39661957 | 14 | 5 | -0.90782 | 0.81801 | KRT13 | 0.81801 | 0.78391 | 0.78264 |
| ENSG00000043591 | 10 | 115803806 | 115806667 | 4 | 2 | -0.90809 | 0.81808 | ADRB1 | 0.81808 | 0.88111 | 0.81261 |
| ENSG00000183780 | 1 | 234040679 | 234460262 | 1868 | 129 | -0.9087 | 0.81825 | SLC35F3 | 0.81825 | 0.77275 | 0.72459 |
| ENSG00000143390 | 1 | 151313116 | 151319833 | 6 | 3 | -0.90892 | 0.8183 | RFX5 | 0.8183 | 0.89229 | 0.85385 |
| ENSG00000138085 | 2 | 27434895 | 27440046 | 11 | 3 | -0.90915 | 0.81836 | ATRAID | 0.81836 | 0.81814 | 0.74672 |
| ENSG00000116918 | 1 | 231664399 | 231702270 | 63 | 14 | -0.90919 | 0.81838 | TSNAX | 0.81838 | 0.82123 | 0.87418 |
| ENSG00000120659 | 13 | 43136872 | 43182149 | 132 | 16 | -0.90924 | 0.81839 | TNFSF11 | 0.81839 | 0.77844 | 0.78557 |
| ENSG00000175634 | 11 | 67195931 | 67202872 | 14 | 3 | -0.9093 | 0.81841 | RPS6KB2 | 0.81841 | 0.58496 | 0.49819 |
| ENSG00000268402 | 6 | 143759575 | 143764295 | 14 | 3 | -0.90947 | 0.81845 | AL031320.1 | 0.81845 | 0.90108 | 0.8773 |
| ENSG00000175003 | 6 | 160542821 | 160579750 | 116 | 25 | -0.90955 | 0.81847 | SLC22A1 | 0.81847 | 0.83505 | 0.86455 |
| ENSG00000103876 | 15 | 80444832 | 80479288 | 156 | 21 | -0.90975 | 0.81852 | FAH | 0.81852 | 0.75397 | 0.71011 |
| ENSG00000163347 | 3 | 190023490 | 190040264 | 83 | 20 | -0.90978 | 0.81853 | CLDN1 | 0.81853 | 0.76546 | 0.54622 |
| ENSG00000169169 | 19 | 50194155 | 50216988 | 76 | 13 | -0.90981 | 0.81854 | CPT1C | 0.81854 | 0.918 | 0.91921 |
| ENSG00000107562 | 10 | 44793038 | 44881941 | 409 | 44 | -0.91034 | 0.81868 | CXCL12 | 0.81868 | 0.85064 | 0.5549 |
| ENSG00000162552 | 1 | 22443798 | 22470462 | 92 | 16 | -0.9104 | 0.81869 | WNT4 | 0.81869 | 0.78967 | 0.73963 |
| ENSG00000139343 | 12 | 96252706 | 96297606 | 197 | 27 | -0.91039 | 0.81869 | SNRPF | 0.81869 | 0.73506 | 0.75486 |
| ENSG00000186825 | 2 | 132552534 | 132559234 | 24 | 4 | -0.91045 | 0.81871 | C2orf27B | 0.81871 | 0.55462 | 0.49038 |
| ENSG00000229474 | 15 | 44957930 | 45003514 | 45 | 19 | -0.91046 | 0.81871 | PATL2 | 0.81871 | 0.65397 | 0.52298 |
| ENSG00000083290 | 17 | 19674142 | 19771249 | 202 | 12 | -0.91051 | 0.81872 | ULK2 | 0.81872 | 0.83655 | 0.81888 |
| ENSG00000172935 | 11 | 68771863 | 68780877 | 19 | 7 | -0.91084 | 0.81881 | MRGPRF | 0.81881 | 0.73445 | 0.54049 |
| ENSG00000179914 | 1 | 160846329 | 160854960 | 34 | 2 | -0.91091 | 0.81883 | ITLN1 | 0.81883 | 0.80811 | 0.80177 |
| ENSG00000235098 | 1 | 1353800 | 1357149 | 8 | 2 | -0.91104 | 0.81886 | ANKRD65 | 0.81886 | 0.92663 | 0.83344 |
| ENSG00000269855 | 19 | 58907457 | 58908407 | 2 | 1 | -0.91104 | 0.81886 | AC012313.1 | 0.81886 | 0.51325 | 0.70075 |
| ENSG00000104714 | 8 | 564746 | 688106 | 671 | 43 | -0.91131 | 0.81893 | ERICH1 | 0.81893 | 0.82522 | 0.82719 |
| ENSG00000116396 | 1 | 110753965 | 110825722 | 235 | 31 | -0.91145 | 0.81897 | KCNC4 | 0.81897 | 0.80428 | 0.84083 |
| ENSG00000166348 | 10 | 75257296 | 75385711 | 265 | 13 | -0.91197 | 0.81911 | USP54 | 0.81911 | 0.81864 | 0.82366 |
| ENSG00000109189 | 4 | 53457138 | 53525502 | 101 | 10 | -0.91204 | 0.81913 | USP46 | 0.81913 | 0.77636 | 0.72846 |
| ENSG00000168395 | 2 | 242641450 | 242668893 | 149 | 19 | -0.9121 | 0.81914 | ING5 | 0.81914 | 0.7341 | 0.77217 |
| ENSG00000136237 | 7 | 22157856 | 22396763 | 784 | 52 | -0.91211 | 0.81914 | RAPGEF5 | 0.81914 | 0.82703 | 0.86555 |
| ENSG00000120440 | 6 | 167738574 | 167772991 | 122 | 22 | -0.91251 | 0.81925 | TTLL2 | 0.81925 | 0.76746 | 0.82028 |
| ENSG00000161940 | 17 | 6926339 | 6933609 | 16 | 5 | -0.9125 | 0.81925 | BCL6B | 0.81925 | 0.87693 | 0.80243 |
| ENSG00000110245 | 11 | 116700422 | 116703788 | 26 | 8 | -0.9126 | 0.81927 | APOC3 | 0.81927 | 0.80824 | 0.8974 |
| ENSG00000164404 | 5 | 132196873 | 132202576 | 15 | 4 | -0.91264 | 0.81929 | GDF9 | 0.81929 | 0.75094 | 0.82652 |
| ENSG00000040275 | 5 | 169010638 | 169031782 | 76 | 8 | -0.91276 | 0.81932 | SPDL1 | 0.81932 | 0.75243 | 0.79018 |
| ENSG00000214029 | 12 | 133694740 | 133707059 | 47 | 8 | -0.91296 | 0.81937 | ZNF891 | 0.81937 | 0.85414 | 0.80602 |
| ENSG00000137860 | 15 | 45544428 | 45568149 | 74 | 4 | -0.91303 | 0.81939 | SLC28A2 | 0.81939 | 0.71119 | 0.6509 |
| ENSG00000170613 | 5 | 156588857 | 156593275 | 9 | 3 | -0.91331 | 0.81946 | FAM71B | 0.81946 | 0.79619 | 0.81095 |
| ENSG00000112062 | 6 | 35995488 | 36079013 | 228 | 20 | -0.91331 | 0.81946 | MAPK14 | 0.81946 | 0.81987 | 0.7723 |
| ENSG00000122728 | 9 | 32629452 | 32635667 | 7 | 4 | -0.91332 | 0.81946 | TAF1L | 0.81946 | 0.90984 | 0.94529 |
| ENSG00000256453 | 5 | 140050379 | 140053171 | 3 | 2 | -0.91335 | 0.81947 | DND1 | 0.81947 | 0.74124 | 0.7663 |
| ENSG00000127561 | 16 | 2039661 | 2044276 | 11 | 3 | -0.91405 | 0.81965 | SYNGR3 | 0.81965 | 0.90406 | 0.93246 |
| ENSG00000248385 | 19 | 54573149 | 54584634 | 64 | 10 | -0.91412 | 0.81967 | TARM1 | 0.81967 | 0.88626 | 0.91717 |
| ENSG00000176244 | 10 | 15119522 | 15130775 | 66 | 15 | -0.91424 | 0.8197 | ACBD7 | 0.8197 | 0.89935 | 0.90806 |
| ENSG00000212719 | 17 | 21428051 | 21477722 | 41 | 8 | -0.91425 | 0.81971 | C17orf51 | 0.81971 | 0.85853 | 0.83983 |
| ENSG00000124074 | 16 | 67696848 | 67701168 | 4 | 2 | -0.9149 | 0.81988 | ENKD1 | 0.81988 | 0.97292 | 0.8626 |
| ENSG00000216560 | 4 | 3578596 | 3592438 | 16 | 2 | -0.91541 | 0.82001 | LINC00955 | 0.82001 | 0.66761 | 0.56495 |
| ENSG00000126337 | 17 | 39642388 | 39648798 | 21 | 8 | -0.91546 | 0.82003 | KRT36 | 0.82003 | 0.63205 | 0.81523 |
| ENSG00000102904 | 16 | 67840668 | 67866051 | 40 | 8 | -0.91555 | 0.82005 | TSNAXIP1 | 0.82005 | 0.84227 | 0.88384 |
| ENSG00000128276 | 22 | 32750872 | 32757148 | 41 | 14 | -0.91566 | 0.82008 | RFPL3 | 0.82008 | 0.89541 | 0.90422 |
| ENSG00000187676 | 13 | 31774073 | 31906413 | 406 | 31 | -0.91586 | 0.82013 | B3GALTL | 0.82013 | 0.83308 | 0.83541 |
| ENSG00000099381 | 16 | 30968615 | 30996437 | 50 | 8 | -0.9159 | 0.82014 | SETD1A | 0.82014 | 0.78164 | 0.78514 |
| ENSG00000152377 | 5 | 136310987 | 136934068 | 1946 | 101 | -0.91595 | 0.82015 | SPOCK1 | 0.82015 | 0.80901 | 0.8102 |
| ENSG00000171603 | 1 | 9789084 | 9884584 | 179 | 22 | -0.91659 | 0.82032 | CLSTN1 | 0.82032 | 0.78134 | 0.75846 |
| ENSG00000107831 | 10 | 103529899 | 103535854 | 3 | 2 | -0.91661 | 0.82033 | FGF8 | 0.82033 | 0.64995 | 0.64822 |
| ENSG00000255974 | 19 | 41349443 | 41356352 | 45 | 12 | -0.91665 | 0.82034 | CYP2A6 | 0.82034 | 0.61825 | 0.79271 |
| ENSG00000134256 | 1 | 117544382 | 117579167 | 102 | 23 | -0.91669 | 0.82035 | CD101 | 0.82035 | 0.83174 | 0.93923 |
| ENSG00000121879 | 3 | 178865902 | 178957881 | 252 | 19 | -0.91669 | 0.82035 | PIK3CA | 0.82035 | 0.79225 | 0.65395 |
| ENSG00000135632 | 2 | 73441350 | 73454365 | 28 | 8 | -0.91673 | 0.82036 | SMYD5 | 0.82036 | 0.69722 | 0.63337 |
| ENSG00000165338 | 10 | 93170096 | 93274586 | 207 | 12 | -0.91675 | 0.82036 | HECTD2 | 0.82036 | 0.81647 | 0.81392 |
| ENSG00000156413 | 19 | 5830621 | 5839742 | 42 | 9 | -0.91675 | 0.82036 | FUT6 | 0.82036 | 0.82958 | 0.71056 |
| ENSG00000167123 | 9 | 131174030 | 131199626 | 74 | 5 | -0.9177 | 0.82061 | CERCAM | 0.82061 | 0.79291 | 0.77554 |
| ENSG00000182108 | 16 | 11022748 | 11036317 | 18 | 7 | -0.91781 | 0.82064 | DEXI | 0.82064 | 0.71684 | 0.48596 |
| ENSG00000141570 | 17 | 77765931 | 77775482 | 18 | 6 | -0.91787 | 0.82066 | CBX8 | 0.82066 | 0.59103 | 0.60362 |
| ENSG00000114439 | 3 | 107241783 | 107530171 | 653 | 37 | -0.91793 | 0.82067 | BBX | 0.82067 | 0.83853 | 0.83066 |
| ENSG00000107362 | 9 | 74477368 | 74525847 | 50 | 9 | -0.91812 | 0.82072 | ABHD17B | 0.82072 | 0.86768 | 0.87244 |
| ENSG00000143321 | 1 | 156711899 | 156736717 | 73 | 17 | -0.91837 | 0.82079 | HDGF | 0.82079 | 0.72684 | 0.6293 |
| ENSG00000067167 | 8 | 71485677 | 71520622 | 101 | 15 | -0.91882 | 0.82091 | TRAM1 | 0.82091 | 0.85854 | 0.81753 |
| ENSG00000166133 | 15 | 40861499 | 40866659 | 11 | 5 | -0.91885 | 0.82091 | RPUSD2 | 0.82091 | 0.84265 | 0.8001 |
| ENSG00000175311 | 16 | 21244986 | 21263750 | 45 | 15 | -0.91886 | 0.82092 | ANKS4B | 0.82092 | 0.63167 | 0.35931 |
| ENSG00000120907 | 8 | 26605667 | 26724790 | 548 | 55 | -0.91906 | 0.82097 | ADRA1A | 0.82097 | 0.84158 | 0.88734 |
| ENSG00000143398 | 1 | 151170425 | 151222012 | 91 | 11 | -0.91918 | 0.821 | PIP5K1A | 0.821 | 0.81825 | 0.85839 |
| ENSG00000124496 | 6 | 42192669 | 42419789 | 612 | 46 | -0.9193 | 0.82103 | TRERF1 | 0.82103 | 0.75374 | 0.80896 |
| ENSG00000156050 | 14 | 74398204 | 74417117 | 84 | 10 | -0.91933 | 0.82104 | FAM161B | 0.82104 | 0.81589 | 0.83913 |

| | | | | | | | | | | | |
|---|---|---|---|---|---|---|---|---|---|---|---|
| ENSG00000106070 | 7 | 50657760 | 50861159 | 747 | 34 | -0.91937 | 0.82105 | GRB10 | 0.82105 | 0.79803 | 0.77403 |
| ENSG00000176697 | 11 | 27676440 | 27743605 | 139 | 20 | -0.91943 | 0.82106 | BDNF | 0.82106 | 0.79073 | 0.8833 |
| ENSG00000106591 | 7 | 42971799 | 42988557 | 54 | 8 | -0.91946 | 0.82107 | MRPL32 | 0.82107 | 0.86325 | 0.83748 |
| ENSG00000116183 | 1 | 176432307 | 176814735 | 917 | 39 | -0.91948 | 0.82108 | PAPPA2 | 0.82108 | 0.82538 | 0.83027 |
| ENSG00000081760 | 12 | 125549925 | 125627873 | 256 | 30 | -0.91961 | 0.82111 | AACS | 0.82111 | 0.85395 | 0.81096 |
| ENSG00000143126 | 1 | 109792641 | 109818372 | 87 | 16 | -0.91982 | 0.82117 | CELSR2 | 0.82117 | 0.86681 | 0.86267 |
| ENSG00000175485 | 11 | 6220376 | 6221489 | 7 | 3 | -0.92001 | 0.82122 | OR52W1 | 0.82122 | 0.97042 | 0.97425 |
| ENSG00000203733 | 6 | 142454227 | 142456288 | 3 | 2 | -0.92016 | 0.82126 | GJE1 | 0.82126 | 0.93241 | 0.89476 |
| ENSG00000261514 | 17 | 42015731 | 42016327 | 1 | 1 | -0.92071 | 0.8214 | RP11-527L4.2 | 0.8214 | 0.87505 | 0.86775 |
| ENSG00000101470 | 20 | 44451853 | 44462384 | 35 | 8 | -0.92105 | 0.82149 | TNNC2 | 0.82149 | 0.86423 | 0.855 |
| ENSG00000131187 | 5 | 176829141 | 176836577 | 16 | 7 | -0.9213 | 0.82155 | F12 | 0.82155 | 0.67727 | 0.36571 |
| ENSG00000116791 | 1 | 75171170 | 75199092 | 92 | 18 | -0.92254 | 0.82188 | CRYZ | 0.82188 | 0.88504 | 0.92673 |
| ENSG00000160294 | 21 | 47655047 | 47706211 | 181 | 21 | -0.92271 | 0.82192 | MCM3AP | 0.82192 | 0.80197 | 0.79089 |
| ENSG00000237136 | 4 | 146601356 | 146692184 | 322 | 35 | -0.92331 | 0.82208 | C4orf51 | 0.82208 | 0.69698 | 0.67967 |
| ENSG00000109927 | 11 | 120971882 | 121062202 | 336 | 25 | -0.92331 | 0.82208 | TECTA | 0.82208 | 0.81772 | 0.85947 |
| ENSG00000118707 | 20 | 35201891 | 35222353 | 20 | 10 | -0.92347 | 0.82212 | TGIF2 | 0.82212 | 0.36339 | 0.4345 |
| ENSG00000026508 | 11 | 35160417 | 35253949 | 369 | 63 | -0.9236 | 0.82215 | CD44 | 0.82215 | 0.51573 | 0.44144 |
| ENSG00000203907 | 6 | 74078278 | 74104856 | 54 | 15 | -0.92363 | 0.82216 | OOEP | 0.82216 | 0.85419 | 0.83669 |
| ENSG00000144848 | 3 | 112251356 | 112280893 | 119 | 10 | -0.92391 | 0.82223 | ATG3 | 0.82223 | 0.83108 | 0.8733 |
| ENSG00000131480 | 17 | 40996617 | 41002724 | 4 | 2 | -0.92394 | 0.82224 | AOC2 | 0.82224 | 0.18657 | 0.39131 |
| ENSG00000167394 | 16 | 31072164 | 31085641 | 29 | 5 | -0.92437 | 0.82235 | ZNF668 | 0.82235 | 0.83226 | 0.81858 |
| ENSG00000141446 | 18 | 19109242 | 19180845 | 97 | 12 | -0.92435 | 0.82235 | ESCO1 | 0.82235 | 0.86449 | 0.87703 |
| ENSG00000135617 | 2 | 73455134 | 73460366 | 17 | 5 | -0.92471 | 0.82244 | PRADC1 | 0.82244 | 0.57532 | 0.57254 |
| ENSG00000136490 | 17 | 61773262 | 61778532 | 9 | 4 | -0.92481 | 0.82247 | LIMD2 | 0.82247 | 0.879 | 0.91274 |
| ENSG00000129028 | 15 | 71173681 | 71185124 | 15 | 7 | -0.92524 | 0.82258 | THAP10 | 0.82258 | 0.81227 | 0.8502 |
| ENSG00000177508 | 16 | 54317216 | 54320675 | 4 | 2 | -0.92536 | 0.82261 | IRX3 | 0.82261 | 0.73827 | 0.62395 |
| ENSG00000180185 | 16 | 1876968 | 1890208 | 143 | 4 | -0.92548 | 0.82264 | FAHD1 | 0.82264 | 0.81253 | 0.79495 |
| ENSG00000095794 | 10 | 35415719 | 35501886 | 245 | 12 | -0.92573 | 0.82271 | CREM | 0.82271 | 0.82661 | 0.78213 |
| ENSG00000160285 | 21 | 47608055 | 47648738 | 163 | 20 | -0.9258 | 0.82272 | LSS | 0.82272 | 0.67691 | 0.48667 |
| ENSG00000140945 | 16 | 82660408 | 83830204 | 7186 | 357 | -0.92609 | 0.8228 | CDH13 | 0.8228 | 0.86951 | 0.8732 |
| ENSG00000176014 | 18 | 12307668 | 12344319 | 128 | 10 | -0.92625 | 0.82284 | TUBB6 | 0.82284 | 0.77424 | 0.8389 |
| ENSG00000058056 | 3 | 179370543 | 179507189 | 432 | 46 | -0.92632 | 0.82286 | USP13 | 0.82286 | 0.82639 | 0.91368 |
| ENSG00000112658 | 6 | 43139037 | 43149243 | 16 | 4 | -0.92637 | 0.82287 | SRF | 0.82287 | 0.80533 | 0.66423 |
| ENSG00000106952 | 9 | 117656003 | 117692697 | 135 | 11 | -0.92645 | 0.82289 | TNFSF8 | 0.82289 | 0.85162 | 0.92272 |
| ENSG00000183054 | 2 | 111271389 | 111334762 | 1 | 1 | -0.92647 | 0.8229 | RGPD6 | 0.8229 | 0.8229 | 0.69669 |
| ENSG00000139668 | 13 | 52158644 | 52336171 | 262 | 28 | -0.92653 | 0.82292 | WDFY2 | 0.82292 | 0.82852 | 0.81906 |
| ENSG00000078795 | 5 | 137223657 | 137278436 | 115 | 17 | -0.9266 | 0.82293 | PKD2L2 | 0.82293 | 0.83502 | 0.8419 |
| ENSG00000259371 | 14 | 24590560 | 24601038 | 21 | 2 | -0.92665 | 0.82295 | RP11-468E2.6 | 0.82295 | 0.80757 | 0.79719 |
| ENSG00000151445 | 14 | 77893018 | 77924295 | 109 | 16 | -0.92677 | 0.82298 | VIPAS39 | 0.82298 | 0.81085 | 0.80365 |
| ENSG00000166851 | 16 | 23688977 | 23701688 | 32 | 8 | -0.92694 | 0.82302 | PLK1 | 0.82302 | 0.83344 | 0.86829 |
| ENSG00000163069 | 4 | 52886872 | 52904648 | 39 | 7 | -0.92713 | 0.82307 | SGCB | 0.82307 | 0.81373 | 0.81907 |
| ENSG00000114933 | 2 | 206858445 | 206951027 | 249 | 25 | -0.92721 | 0.82309 | INO80D | 0.82309 | 0.72499 | 0.64009 |
| ENSG00000140471 | 15 | 101099574 | 101143435 | 267 | 17 | -0.92729 | 0.82311 | LINS | 0.82311 | 0.78347 | 0.83343 |
| ENSG00000122861 | 10 | 75668935 | 75677255 | 27 | 7 | -0.92748 | 0.82316 | PLAU | 0.82316 | 0.72077 | 0.58908 |
| ENSG00000142530 | 19 | 50970042 | 50980010 | 49 | 8 | -0.92762 | 0.8232 | FAM71E1 | 0.8232 | 0.89189 | 0.95122 |
| ENSG00000180697 | 3 | 126245842 | 126277808 | 167 | 20 | -0.92769 | 0.82321 | C3orf22 | 0.82321 | 0.74098 | 0.71017 |
| ENSG00000184672 | 8 | 85095022 | 85834079 | 2123 | 54 | -0.92816 | 0.82334 | RALYL | 0.82334 | 0.82012 | 0.84165 |
| ENSG00000182585 | 4 | 75174190 | 75181024 | 10 | 2 | -0.92823 | 0.82336 | EPGN | 0.82336 | 0.81413 | 0.74436 |
| ENSG00000263203 | 16 | 69152757 | 69154331 | 4 | 2 | -0.92828 | 0.82337 | CHTF8 | 0.82337 | 0.34012 | 0.26359 |
| ENSG00000143379 | 1 | 150898739 | 150937213 | 75 | 11 | -0.92842 | 0.8234 | SETDB1 | 0.8234 | 0.79707 | 0.8118 |
| ENSG00000107338 | 9 | 37919131 | 38069210 | 482 | 25 | -0.92843 | 0.82341 | SHB | 0.82341 | 0.86806 | 0.87217 |
| ENSG00000162927 | 2 | 61167357 | 61245394 | 156 | 28 | -0.92846 | 0.82342 | PUS10 | 0.82342 | 0.81701 | 0.76672 |
| ENSG00000100461 | 14 | 23369854 | 23388393 | 61 | 7 | -0.92861 | 0.82346 | RBM23 | 0.82346 | 0.62863 | 0.68664 |
| ENSG00000149187 | 11 | 47487496 | 47587121 | 188 | 33 | -0.92878 | 0.8235 | CELF1 | 0.8235 | 0.8428 | 0.84665 |
| ENSG00000081248 | 1 | 201008642 | 201081694 | 304 | 48 | -0.92888 | 0.82352 | CACNA1S | 0.82352 | 0.79082 | 0.73835 |
| ENSG00000204052 | 6 | 43474707 | 43478424 | 9 | 2 | -0.92927 | 0.82362 | LRRC73 | 0.82362 | 0.89934 | 0.87714 |
| ENSG00000171621 | 1 | 9352939 | 9429591 | 345 | 68 | -0.92974 | 0.82375 | SPSB1 | 0.82375 | 0.84925 | 0.86795 |
| ENSG00000177800 | 1 | 229385383 | 229387557 | 4 | 2 | -0.92979 | 0.82376 | TMEM78 | 0.82376 | 0.81526 | 0.86211 |
| ENSG00000158805 | 16 | 89786808 | 89807311 | 153 | 21 | -0.92993 | 0.8238 | ZNF276 | 0.8238 | 0.83301 | 0.86513 |
| ENSG00000135898 | 2 | 231772033 | 231825781 | 155 | 34 | -0.93026 | 0.82388 | GPR55 | 0.82388 | 0.91477 | 0.93967 |
| ENSG00000106780 | 9 | 123363091 | 123476748 | 153 | 14 | -0.93028 | 0.82389 | MEGF9 | 0.82389 | 0.80186 | 0.74881 |
| ENSG00000143353 | 1 | 219347186 | 219386207 | 78 | 14 | -0.93042 | 0.82392 | LYPLAL1 | 0.82392 | 0.85452 | 0.85845 |
| ENSG00000187726 | 11 | 73661364 | 73681411 | 93 | 17 | -0.93042 | 0.82392 | DNAJB13 | 0.82392 | 0.66771 | 0.49702 |
| ENSG00000103423 | 16 | 4475806 | 4506776 | 82 | 11 | -0.93049 | 0.82394 | DNAJA3 | 0.82394 | 0.78962 | 0.74864 |
| ENSG00000106245 | 7 | 99006264 | 99017239 | 25 | 6 | -0.93057 | 0.82396 | BUD31 | 0.82396 | 0.87351 | 0.86776 |
| ENSG00000232838 | 20 | 18118517 | 18123813 | 24 | 8 | -0.93059 | 0.82397 | PET117 | 0.82397 | 0.57621 | 0.56017 |
| ENSG00000162722 | 1 | 248020501 | 248041507 | 109 | 16 | -0.93062 | 0.82398 | TRIM58 | 0.82398 | 0.84621 | 0.84288 |
| ENSG00000269190 | 19 | 39432044 | 39466550 | 138 | 20 | -0.93087 | 0.82404 | FBXO17 | 0.82404 | 0.53823 | 0.35319 |
| ENSG00000115085 | 2 | 98330023 | 98356325 | 55 | 9 | -0.93096 | 0.82406 | ZAP70 | 0.82406 | 0.83791 | 0.86278 |
| ENSG00000149972 | 11 | 98891683 | 100229616 | 6480 | 233 | -0.93108 | 0.82409 | CNTN5 | 0.82409 | 0.81677 | 0.82085 |
| ENSG00000169714 | 3 | 128888327 | 128902765 | 17 | 7 | -0.93118 | 0.82412 | CNBP | 0.82412 | 0.8014 | 0.71874 |
| ENSG00000169020 | 4 | 666225 | 668127 | 8 | 4 | -0.93146 | 0.82419 | ATP5I | 0.82419 | 0.30021 | 0.3171 |
| ENSG00000214860 | 17 | 18280976 | 18292961 | 58 | 4 | -0.93162 | 0.82423 | EVPLL | 0.82423 | 0.87438 | 0.88892 |

| | | | | | | | | | | | |
|---|---|---|---|---|---|---|---|---|---|---|---|
| ENSG00000121377 | 12 | 10954131 | 10955226 | 3 | 1 | -0.93179 | 0.82428 | TAS2R7 | 0.82428 | 0.63378 | 0.57455 |
| ENSG00000169484 | 14 | 20482419 | 20483352 | 2 | 1 | -0.93189 | 0.8243 | OR4K14 | 0.8243 | 0.86355 | 0.90784 |
| ENSG00000174780 | 4 | 57333081 | 57369839 | 137 | 14 | -0.9325 | 0.82446 | SRP72 | 0.82446 | 0.85987 | 0.88609 |
| ENSG00000119457 | 9 | 115641200 | 115653193 | 39 | 14 | -0.93264 | 0.8245 | SLC46A2 | 0.8245 | 0.87065 | 0.84453 |
| ENSG00000168297 | 3 | 58318607 | 58411748 | 458 | 21 | -0.93277 | 0.82453 | PXK | 0.82453 | 0.83635 | 0.8458 |
| ENSG00000214216 | 3 | 158680717 | 158984096 | 723 | 42 | -0.93279 | 0.82454 | IQCJ | 0.82454 | 0.8271 | 0.8463 |
| ENSG00000172869 | 5 | 118373467 | 118584833 | 372 | 37 | -0.93292 | 0.82457 | DMXL1 | 0.82457 | 0.77292 | 0.79973 |
| ENSG00000159899 | 9 | 35792151 | 35809729 | 33 | 5 | -0.93292 | 0.82457 | NPR2 | 0.82457 | 0.77279 | 0.61142 |
| ENSG00000196199 | 13 | 20207788 | 20247599 | 116 | 13 | -0.93291 | 0.82457 | MPHOSPH8 | 0.82457 | 0.85164 | 0.8737 |
| ENSG00000205678 | 4 | 65140975 | 65275186 | 580 | 19 | -0.93313 | 0.82462 | TECRL | 0.82462 | 0.84171 | 0.85018 |
| ENSG00000120262 | 6 | 151815165 | 151942328 | 609 | 42 | -0.93343 | 0.8247 | CCDC170 | 0.8247 | 0.85313 | 0.81784 |
| ENSG00000104687 | 8 | 30535583 | 30585443 | 152 | 17 | -0.93362 | 0.82475 | GSR | 0.82475 | 0.88441 | 0.81574 |
| ENSG00000184007 | 1 | 32372022 | 32410457 | 56 | 10 | -0.93379 | 0.82479 | PTP4A2 | 0.82479 | 0.82289 | 0.70018 |
| ENSG00000187733 | 1 | 104293028 | 104301312 | 2 | 1 | -0.93407 | 0.82487 | AMY1C | 0.82487 | 0.36446 | 0.43511 |
| ENSG00000184792 | 22 | 31089769 | 31303811 | 714 | 55 | -0.93409 | 0.82487 | OSBP2 | 0.82487 | 0.84063 | 0.82526 |
| ENSG00000138111 | 10 | 104221149 | 104236802 | 69 | 6 | -0.93416 | 0.82489 | TMEM180 | 0.82489 | 0.77954 | 0.71646 |
| ENSG00000187566 | 6 | 18120718 | 18122851 | 6 | 2 | -0.9342 | 0.8249 | NHLRC1 | 0.8249 | 0.90709 | 0.96745 |
| ENSG00000127423 | 1 | 26158414 | 26185903 | 80 | 3 | -0.93476 | 0.82505 | AUNIP | 0.82505 | 0.77759 | 0.76725 |
| ENSG00000089195 | 20 | 5917881 | 5931182 | 67 | 11 | -0.93513 | 0.82514 | TRMT6 | 0.82514 | 0.60099 | 0.42677 |
| ENSG00000157470 | 15 | 59664892 | 59815748 | 578 | 44 | -0.9356 | 0.82526 | FAM81A | 0.82526 | 0.7857 | 0.71308 |
| ENSG00000181754 | 1 | 110046797 | 110052360 | 8 | 3 | -0.93573 | 0.82529 | AMIGO1 | 0.82529 | 0.7521 | 0.80375 |
| ENSG00000164898 | 7 | 139024203 | 139031065 | 23 | 3 | -0.93576 | 0.8253 | C7orf55 | 0.8253 | 0.8291 | 0.8773 |
| ENSG00000269302 | 4 | 187111913 | 187112626 | 3 | 1 | -0.9362 | 0.82541 | AC110771.1 | 0.82541 | 0.87525 | 0.92655 |
| ENSG00000156486 | 8 | 99439250 | 99445076 | 7 | 3 | -0.93632 | 0.82544 | KCNS2 | 0.82544 | 0.89216 | 0.90131 |
| ENSG00000076321 | 1 | 173684080 | 173755840 | 115 | 13 | -0.93632 | 0.82545 | KLHL20 | 0.82545 | 0.82266 | 0.85148 |
| ENSG00000181588 | 19 | 1554668 | 1568057 | 23 | 11 | -0.9364 | 0.82547 | MEX3D | 0.82547 | 0.5749 | 0.71483 |
| ENSG00000127990 | 7 | 94214542 | 94285521 | 106 | 10 | -0.93697 | 0.82561 | SGCE | 0.82561 | 0.82654 | 0.81485 |
| ENSG00000164399 | 5 | 131396222 | 131398897 | 3 | 1 | -0.93709 | 0.82564 | IL3 | 0.82564 | 0.34094 | 0.36837 |
| ENSG00000150990 | 12 | 125431371 | 125473668 | 317 | 37 | -0.9374 | 0.82572 | DHX37 | 0.82572 | 0.91126 | 0.91711 |
| ENSG00000117215 | 1 | 20438432 | 20446050 | 24 | 10 | -0.93742 | 0.82573 | PLA2G2D | 0.82573 | 0.68247 | 0.78027 |
| ENSG00000269360 | 15 | 93749295 | 93751277 | 8 | 4 | -0.93754 | 0.82576 | AC112693.2 | 0.82576 | 0.15601 | 0.26419 |
| ENSG00000166246 | 16 | 4784273 | 4799397 | 75 | 8 | -0.93789 | 0.82585 | C16orf71 | 0.82585 | 0.84744 | 0.81339 |
| ENSG00000267545 | 19 | 45683080 | 45705702 | 111 | 21 | -0.93788 | 0.82585 | AC005779.2 | 0.82585 | 0.72047 | 0.86259 |
| ENSG00000129596 | 5 | 115140430 | 115152651 | 20 | 7 | -0.93801 | 0.82588 | CDO1 | 0.82588 | 0.9624 | 0.86417 |
| ENSG00000197774 | 16 | 1823208 | 1831709 | 31 | 11 | -0.93814 | 0.82591 | EME2 | 0.82591 | 0.80555 | 0.81313 |
| ENSG00000205189 | 8 | 81397854 | 81438500 | 115 | 17 | -0.93881 | 0.82608 | ZBTB10 | 0.82608 | 0.79411 | 0.85007 |
| ENSG00000005882 | 17 | 48172101 | 48189516 | 64 | 13 | -0.93877 | 0.82608 | PDK2 | 0.82608 | 0.87057 | 0.83855 |
| ENSG00000168256 | 17 | 40163400 | 40177659 | 20 | 6 | -0.93952 | 0.82627 | NKIRAS2 | 0.82627 | 0.80205 | 0.83322 |
| ENSG00000131473 | 17 | 40023161 | 40086795 | 153 | 15 | -0.93957 | 0.82628 | ACLY | 0.82628 | 0.7842 | 0.76925 |
| ENSG00000082258 | 2 | 135675805 | 135716912 | 109 | 11 | -0.93972 | 0.82632 | CCNT2 | 0.82632 | 0.84652 | 0.709 |
| ENSG00000173991 | 17 | 37820440 | 37822808 | 5 | 2 | -0.93971 | 0.82632 | TCAP | 0.82632 | 0.89695 | 0.79658 |
| ENSG00000251695 | 12 | 69186125 | 69187743 | 4 | 2 | -0.93996 | 0.82638 | AC124890.1 | 0.82638 | 0.37651 | 0.44704 |
| ENSG00000129636 | 16 | 47188298 | 47498060 | 366 | 23 | -0.93994 | 0.82638 | ITFG1 | 0.82638 | 0.82359 | 0.78104 |
| ENSG00000168646 | 17 | 63524681 | 63557765 | 89 | 21 | -0.94001 | 0.82639 | AXIN2 | 0.82639 | 0.87836 | 0.72532 |
| ENSG00000106144 | 7 | 142985308 | 143004789 | 20 | 7 | -0.94027 | 0.82646 | CASP2 | 0.82646 | 0.83985 | 0.93922 |
| ENSG00000183648 | 14 | 92582466 | 92588261 | 49 | 5 | -0.94041 | 0.8265 | NDUFB1 | 0.8265 | 0.51946 | 0.46866 |
| ENSG00000181577 | 6 | 43968317 | 43973695 | 29 | 7 | -0.94068 | 0.82657 | C6orf223 | 0.82657 | 0.90707 | 0.98895 |
| ENSG00000216937 | 10 | 32735068 | 32863492 | 536 | 23 | -0.94101 | 0.82665 | CCDC7 | 0.82665 | 0.8507 | 0.81098 |
| ENSG00000170743 | 11 | 7260009 | 7490273 | 955 | 84 | -0.9412 | 0.8267 | SYT9 | 0.8267 | 0.63358 | 0.35619 |
| ENSG00000179241 | 11 | 35965531 | 36253686 | 1137 | 115 | -0.94121 | 0.8267 | LDLRAD3 | 0.8267 | 0.8289 | 0.86075 |
| ENSG00000173262 | 12 | 7965108 | 8043744 | 432 | 46 | -0.94134 | 0.82674 | SLC2A14 | 0.82674 | 0.85897 | 0.88991 |
| ENSG00000163888 | 3 | 183977001 | 183979251 | 4 | 2 | -0.94139 | 0.82675 | CAMK2N2 | 0.82675 | 0.83563 | 0.93432 |
| ENSG00000227268 | 10 | 89618918 | 89623194 | 1 | 1 | -0.9416 | 0.8268 | KLLN | 0.8268 | 0.82991 | 0.77646 |
| ENSG00000141510 | 17 | 7565097 | 7590856 | 102 | 21 | -0.94187 | 0.82687 | TP53 | 0.82687 | 0.6599 | 0.086922 |
| ENSG00000069188 | 17 | 71330523 | 71640228 | 1217 | 141 | -0.94192 | 0.82688 | SDK2 | 0.82688 | 0.82747 | 0.88815 |
| ENSG00000214681 | 3 | 51907737 | 51909600 | 3 | 1 | -0.94258 | 0.82705 | IQCF5 | 0.82705 | 0.65397 | 0.72496 |
| ENSG00000105983 | 7 | 156461646 | 156685924 | 917 | 31 | -0.94266 | 0.82707 | LMBR1 | 0.82707 | 0.81724 | 0.79283 |
| ENSG00000177352 | 3 | 49199968 | 49203754 | 5 | 2 | -0.94358 | 0.82731 | CCDC71 | 0.82731 | 0.93233 | 0.86837 |
| ENSG00000221995 | 17 | 27400537 | 27418537 | 46 | 6 | -0.94366 | 0.82733 | TIAF1 | 0.82733 | 0.83363 | 0.84992 |
| ENSG00000127993 | 7 | 92158087 | 92167319 | 20 | 5 | -0.9438 | 0.82737 | RBM48 | 0.82737 | 0.61901 | 0.64352 |
| ENSG00000164919 | 8 | 100885428 | 100906290 | 67 | 8 | -0.94393 | 0.8274 | COX6C | 0.8274 | 0.8177 | 0.83106 |
| ENSG00000168280 | 2 | 149632819 | 149883273 | 254 | 15 | -0.94406 | 0.82743 | KIF5C | 0.82743 | 0.80155 | 0.81004 |
| ENSG00000184916 | 14 | 105607318 | 105635161 | 63 | 17 | -0.9441 | 0.82744 | JAG2 | 0.82744 | 0.79525 | 0.81341 |
| ENSG00000102870 | 16 | 30789778 | 30798523 | 2 | 1 | -0.94416 | 0.82746 | ZNF629 | 0.82746 | 0.79414 | 0.76865 |
| ENSG00000140948 | 16 | 87439852 | 87525651 | 380 | 19 | -0.94448 | 0.82754 | ZCCHC14 | 0.82754 | 0.79704 | 0.79038 |
| ENSG00000142661 | 1 | 24382525 | 24438665 | 210 | 23 | -0.94518 | 0.82772 | MYOM3 | 0.82772 | 0.94278 | 0.94378 |
| ENSG00000127863 | 13 | 24144509 | 24250232 | 427 | 41 | -0.94529 | 0.82774 | TNFRSF19 | 0.82774 | 0.79549 | 0.63182 |
| ENSG00000125788 | 20 | 123010 | 126392 | 7 | 3 | -0.94528 | 0.82774 | DEFB126 | 0.82774 | 0.51044 | 0.47625 |
| ENSG00000013561 | 5 | 141337893 | 141369856 | 103 | 10 | -0.94536 | 0.82776 | RNF14 | 0.82776 | 0.80158 | 0.8332 |
| ENSG00000079999 | 19 | 10596796 | 10614417 | 51 | 9 | -0.94549 | 0.8278 | KEAP1 | 0.8278 | 0.78275 | 0.87445 |
| ENSG00000198892 | 1 | 201857808 | 201861434 | 7 | 3 | -0.94553 | 0.82781 | SHISA4 | 0.82781 | 0.92996 | 0.90688 |
| ENSG00000136275 | 7 | 47834889 | 47859445 | 100 | 16 | -0.94586 | 0.82789 | C7orf69 | 0.82789 | 0.77034 | 0.69155 |
| ENSG00000075388 | 11 | 69587797 | 69590171 | 10 | 4 | -0.9463 | 0.828 | FGF4 | 0.828 | 0.75925 | 0.83817 |

| | | | | | | | | | | | | |
|---|---|---|---|---|---|---|---|---|---|---|---|---|
| ENSG00000139567 | 12 | 52300692 | 52317145 | 51 | 12 | -0.94636 | 0.82802 | ACVRL1 | | 0.82802 | 0.85262 | 0.88673 |
| ENSG00000267873 | 12 | 52709128 | 52709790 | 2 | 1 | -0.94645 | 0.82804 | AC121757.1 | | 0.82804 | 0.69115 | 0.7567 |
| ENSG00000166200 | 15 | 49398268 | 49447858 | 152 | 19 | -0.94708 | 0.8282 | COPS2 | | 0.8282 | 0.83237 | 0.81066 |
| ENSG00000170684 | 19 | 45574758 | 45579846 | 6 | 3 | -0.94713 | 0.82821 | ZNF296 | | 0.82821 | 0.48637 | 0.73506 |
| ENSG00000132704 | 1 | 157715523 | 157746922 | 70 | 13 | -0.94727 | 0.82825 | FCRL2 | | 0.82825 | 0.79699 | 0.72993 |
| ENSG00000174886 | 19 | 5891287 | 5904017 | 57 | 14 | -0.94739 | 0.82828 | NDUFA11 | | 0.82828 | 0.79664 | 0.86624 |
| ENSG00000178078 | 19 | 4324040 | 4342783 | 86 | 14 | -0.94752 | 0.82831 | STAP2 | | 0.82831 | 0.82384 | 0.5921 |
| ENSG00000169194 | 5 | 131991955 | 131996802 | 9 | 2 | -0.94759 | 0.82833 | IL13 | | 0.82833 | 0.41471 | 0.75313 |
| ENSG00000155275 | 4 | 8437867 | 8495258 | 259 | 22 | -0.94776 | 0.82837 | TRMT44 | | 0.82837 | 0.83316 | 0.8385 |
| ENSG00000091157 | 18 | 54318574 | 54698828 | 1052 | 50 | -0.94786 | 0.8284 | WDR7 | | 0.8284 | 0.83085 | 0.79681 |
| ENSG00000184207 | 16 | 2261998 | 2264808 | 12 | 4 | -0.94796 | 0.82843 | PGP | | 0.82843 | 0.78489 | 0.83377 |
| ENSG00000110713 | 11 | 3692313 | 3819022 | 435 | 28 | -0.94846 | 0.82855 | NUP98 | | 0.82855 | 0.87046 | 0.95521 |
| ENSG00000151532 | 10 | 114206756 | 114578503 | 968 | 101 | -0.94876 | 0.82863 | VTI1A | | 0.82863 | 0.83866 | 0.71841 |
| ENSG00000136280 | 7 | 45039074 | 45116068 | 302 | 17 | -0.94898 | 0.82868 | CCM2 | | 0.82868 | 0.81252 | 0.8524 |
| ENSG00000147852 | 9 | 2621834 | 2660053 | 155 | 29 | -0.94902 | 0.82869 | VLDLR | | 0.82869 | 0.8707 | 0.92641 |
| ENSG00000106004 | 7 | 27180671 | 27183287 | 9 | 4 | -0.94911 | 0.82872 | HOXA5 | | 0.82872 | 0.9364 | 0.97328 |
| ENSG00000163655 | 3 | 155588325 | 155661815 | 243 | 15 | -0.94916 | 0.82873 | GMPS | | 0.82873 | 0.82873 | 0.79093 |
| ENSG00000167104 | 20 | 31619454 | 31631853 | 37 | 6 | -0.9492 | 0.82874 | BPIFB6 | | 0.82874 | 0.58569 | 0.53303 |
| ENSG00000187997 | 17 | 76142434 | 76162258 | 113 | 25 | -0.94929 | 0.82876 | C17orf99 | | 0.82876 | 0.77729 | 0.87975 |
| ENSG00000128254 | 22 | 32329507 | 32341504 | 51 | 14 | -0.94972 | 0.82887 | C22orf24 | | 0.82887 | 0.4406 | 0.43662 |
| ENSG00000157087 | 3 | 10365707 | 10749716 | 1489 | 124 | -0.94999 | 0.82894 | ATP2B2 | | 0.82894 | 0.83719 | 0.76693 |
| ENSG00000177992 | 9 | 90497741 | 90503814 | 22 | 6 | -0.95007 | 0.82896 | SPATA31E1 | | 0.82896 | 0.85843 | 0.87958 |
| ENSG00000082781 | 3 | 124480795 | 124620265 | 531 | 30 | -0.95032 | 0.82903 | ITGB5 | | 0.82903 | 0.85362 | 0.86221 |
| ENSG00000155621 | 9 | 74526426 | 74600970 | 113 | 21 | -0.95034 | 0.82903 | C9orf85 | | 0.82903 | 0.79686 | 0.86002 |
| ENSG00000146278 | 6 | 89790470 | 89794879 | 15 | 4 | -0.95062 | 0.8291 | PNRC1 | | 0.8291 | 0.85512 | 0.90071 |
| ENSG00000009830 | 14 | 77741299 | 77787227 | 164 | 28 | -0.95068 | 0.82912 | POMT2 | | 0.82912 | 0.86265 | 0.80972 |
| ENSG00000248919 | 7 | 99017372 | 99063820 | 116 | 13 | -0.95075 | 0.82913 | ATP5J2-PTCD1 | | 0.82913 | 0.84412 | 0.80682 |
| ENSG00000120093 | 17 | 46626232 | 46682274 | 123 | 24 | -0.95082 | 0.82915 | HOXB3 | | 0.82915 | 0.86261 | 0.91895 |
| ENSG00000158201 | 18 | 19230858 | 19284766 | 101 | 11 | -0.95111 | 0.82922 | ABHD3 | | 0.82922 | 0.83338 | 0.8388 |
| ENSG00000137841 | 15 | 40570377 | 40600136 | 81 | 18 | -0.95135 | 0.82929 | PLCB2 | | 0.82929 | 0.67733 | 0.80065 |
| ENSG00000167772 | 19 | 8428173 | 8439257 | 30 | 9 | -0.95144 | 0.82931 | ANGPTL4 | | 0.82931 | 0.84987 | 0.46927 |
| ENSG00000122545 | 7 | 35840542 | 35944917 | 225 | 17 | -0.95162 | 0.82935 | | 44081 | 0.82935 | 0.84372 | 0.82253 |
| ENSG00000204920 | 19 | 44472014 | 44502477 | 132 | 13 | -0.95194 | 0.82944 | ZNF155 | | 0.82944 | 0.80712 | 0.76759 |
| ENSG00000163602 | 3 | 72420976 | 72496069 | 107 | 27 | -0.95212 | 0.82948 | RYBP | | 0.82948 | 0.84877 | 0.95514 |
| ENSG00000119686 | 14 | 76044960 | 76129557 | 244 | 42 | -0.9522 | 0.8295 | FLVCR2 | | 0.8295 | 0.88573 | 0.83887 |
| ENSG00000187634 | 1 | 860260 | 879955 | 90 | 21 | -0.95236 | 0.82954 | SAMD11 | | 0.82954 | 0.74752 | 0.68684 |
| ENSG00000177234 | 10 | 122357721 | 122359629 | 9 | 4 | -0.95244 | 0.82956 | C10orf85 | | 0.82956 | 0.93892 | 0.87511 |
| ENSG00000198586 | 2 | 171847333 | 172087824 | 736 | 26 | -0.95246 | 0.82957 | TLK1 | | 0.82957 | 0.83308 | 0.81283 |
| ENSG00000162377 | 1 | 53152508 | 53164038 | 31 | 6 | -0.95251 | 0.82958 | COA7 | | 0.82958 | 0.63542 | 0.61158 |
| ENSG00000174851 | 11 | 66052051 | 66056641 | 5 | 2 | -0.95251 | 0.82958 | YIF1A | | 0.82958 | 0.56594 | 0.85934 |
| ENSG00000204614 | 6 | 30103885 | 30116512 | 132 | 11 | -0.95257 | 0.82959 | TRIM40 | | 0.82959 | 0.79302 | 0.69377 |
| ENSG00000258405 | 19 | 52956829 | 53015407 | 402 | 23 | -0.95266 | 0.82962 | ZNF578 | | 0.82962 | 0.81534 | 0.75864 |
| ENSG00000159214 | 1 | 44457031 | 44462200 | 10 | 3 | -0.9528 | 0.82965 | CCDC24 | | 0.82965 | 0.25281 | 0.35879 |
| ENSG00000091732 | 7 | 129658126 | 129691291 | 121 | 11 | -0.95294 | 0.82969 | ZC3HC1 | | 0.82969 | 0.83333 | 0.66978 |
| ENSG00000214194 | 7 | 112756773 | 112758668 | 7 | 3 | -0.95299 | 0.8297 | LINC00998 | | 0.8297 | 0.89816 | 0.90915 |
| ENSG00000182083 | 2 | 240968841 | 240969906 | 2 | 1 | -0.95309 | 0.82973 | OR6B2 | | 0.82973 | 0.090348 | 0.10034 |
| ENSG00000214248 | 19 | 7963689 | 7968427 | 17 | 5 | -0.95322 | 0.82976 | AC010336.1 | | 0.82976 | 0.94874 | 0.98902 |
| ENSG00000160888 | 19 | 13261229 | 13265722 | 11 | 6 | -0.95359 | 0.82986 | IER2 | | 0.82986 | 0.47406 | 0.45194 |
| ENSG00000137752 | 11 | 104896170 | 104972158 | 256 | 16 | -0.95382 | 0.82991 | CASP1 | | 0.82991 | 0.81602 | 0.83501 |
| ENSG00000102901 | 16 | 67862060 | 67881714 | 26 | 5 | -0.9541 | 0.82998 | CENPT | | 0.82998 | 0.83785 | 0.8355 |
| ENSG00000272636 | 17 | 5810 | 31427 | 82 | 11 | -0.95407 | 0.82998 | DOC2B | | 0.82998 | 0.85899 | 0.88746 |
| ENSG00000151917 | 6 | 56819773 | 56892140 | 171 | 11 | -0.95419 | 0.83001 | BEND6 | | 0.83001 | 0.83953 | 0.84207 |
| ENSG00000174326 | 17 | 6944949 | 6947242 | 4 | 2 | -0.95421 | 0.83001 | SLC16A11 | | 0.83001 | 0.79942 | 0.81601 |
| ENSG00000090061 | 14 | 99947506 | 100001381 | 93 | 13 | -0.9544 | 0.83006 | CCNK | | 0.83006 | 0.90068 | 0.91503 |
| ENSG00000273155 | 2 | 99771461 | 99811761 | 119 | 12 | -0.95452 | 0.83009 | MRPL30 | | 0.83009 | 0.78707 | 0.7911 |
| ENSG00000105889 | 7 | 22459063 | 22672544 | 829 | 66 | -0.95464 | 0.83012 | STEAP1B | | 0.83012 | 0.76488 | 0.73884 |
| ENSG00000146007 | 5 | 140078265 | 140086248 | 15 | 2 | -0.95527 | 0.83028 | ZMAT2 | | 0.83028 | 0.82063 | 0.82848 |
| ENSG00000226479 | 2 | 120978854 | 120980984 | 3 | 2 | -0.95537 | 0.8303 | TMEM185B | | 0.8303 | 0.68832 | 0.5759 |
| ENSG00000170537 | 16 | 18995256 | 19075264 | 281 | 18 | -0.95538 | 0.83031 | TMC7 | | 0.83031 | 0.81683 | 0.83202 |
| ENSG00000118507 | 6 | 131456806 | 131604675 | 384 | 33 | -0.95555 | 0.83035 | AKAP7 | | 0.83035 | 0.84436 | 0.88624 |
| ENSG00000205670 | 21 | 35747779 | 35780164 | 168 | 18 | -0.95558 | 0.83036 | SMIM11 | | 0.83036 | 0.84932 | 0.8965 |
| ENSG00000183542 | 12 | 10559983 | 10562356 | 8 | 2 | -0.95574 | 0.8304 | KLRC4 | | 0.8304 | 0.94343 | 0.94123 |
| ENSG00000172543 | 11 | 65647280 | 65651212 | 9 | 4 | -0.95587 | 0.83043 | CTSW | | 0.83043 | 0.76637 | 0.89028 |
| ENSG00000198855 | 12 | 108908962 | 108919614 | 23 | 8 | -0.95618 | 0.83051 | FICD | | 0.83051 | 0.83451 | 0.84915 |
| ENSG00000131013 | 6 | 149825869 | 149867174 | 61 | 9 | -0.95677 | 0.83066 | PPIL4 | | 0.83066 | 0.85688 | 0.83048 |
| ENSG00000100596 | 14 | 77972340 | 78083116 | 558 | 43 | -0.95679 | 0.83066 | SPTLC2 | | 0.83066 | 0.87436 | 0.9186 |
| ENSG00000129128 | 4 | 177241115 | 177253396 | 57 | 9 | -0.95682 | 0.83067 | SPCS3 | | 0.83067 | 0.77091 | 0.65993 |
| ENSG00000085832 | 1 | 51819935 | 51985000 | 613 | 22 | -0.95749 | 0.83084 | EPS15 | | 0.83084 | 0.82399 | 0.85234 |
| ENSG00000187013 | 17 | 59489112 | 59490641 | 3 | 1 | -0.95748 | 0.83084 | C17orf82 | | 0.83084 | 0.66643 | 0.71914 |
| ENSG00000204160 | 1 | 27153201 | 27184093 | 53 | 8 | -0.95751 | 0.83085 | ZDHHC18 | | 0.83085 | 0.79491 | 0.78442 |
| ENSG00000197223 | 2 | 68268262 | 68338080 | 114 | 22 | -0.95758 | 0.83086 | C1D | | 0.83086 | 0.81455 | 0.51045 |
| ENSG00000110074 | 11 | 126138950 | 126148026 | 20 | 8 | -0.95779 | 0.83092 | FOXRED1 | | 0.83092 | 0.95902 | 0.9721 |
| ENSG00000128833 | 15 | 52484519 | 52587995 | 343 | 35 | -0.95804 | 0.83098 | MYO5C | | 0.83098 | 0.82494 | 0.8894 |

| | | | | | | | | | | | |
|---|---|---|---|---|---|---|---|---|---|---|---|
| ENSG00000204186 | 2 | 207139387 | 207179148 | 82 | 11 | -0.95843 | 0.83108 | ZDBF2 | 0.83108 | 0.78151 | 0.76724 |
| ENSG00000204619 | 6 | 30034486 | 30038110 | 19 | 5 | -0.95853 | 0.8311 | PPP1R11 | 0.8311 | 0.60447 | 0.6549 |
| ENSG00000125740 | 19 | 45971253 | 45978437 | 25 | 6 | -0.95851 | 0.8311 | FOSB | 0.8311 | 0.92191 | 0.87534 |
| ENSG00000067836 | 16 | 4846969 | 4852951 | 13 | 4 | -0.95873 | 0.83115 | ROGDI | 0.83115 | 0.62556 | 0.37344 |
| ENSG00000269417 | 2 | 85122209 | 85122345 | 2 | 1 | -0.95874 | 0.83116 | AC022210.1 | 0.83116 | 0.83783 | 0.69762 |
| ENSG00000179981 | 18 | 72922710 | 73001905 | 268 | 35 | -0.95875 | 0.83116 | TSHZ1 | 0.83116 | 0.72638 | 0.63327 |
| ENSG00000269237 | 19 | 21587961 | 21674068 | 321 | 15 | -0.95888 | 0.83119 | CTD-2561J22.3 | 0.83119 | 0.81551 | 0.81766 |
| ENSG00000112379 | 6 | 138483058 | 138665800 | 584 | 33 | -0.95905 | 0.83123 | KIAA1244 | 0.83123 | 0.81728 | 0.84892 |
| ENSG00000236882 | 5 | 95187936 | 95195837 | 43 | 12 | -0.95912 | 0.83125 | C5orf27 | 0.83125 | 0.64133 | 0.737 |
| ENSG00000214655 | 10 | 75545340 | 75561551 | 12 | 4 | -0.95921 | 0.83127 | ZSWIM8 | 0.83127 | 0.61475 | 0.6191 |
| ENSG00000205809 | 12 | 10579453 | 10594899 | 74 | 10 | -0.95936 | 0.83131 | KLRC2 | 0.83131 | 0.94934 | 0.96387 |
| ENSG00000157827 | 2 | 153191751 | 153506348 | 1166 | 65 | -0.95939 | 0.83132 | FMNL2 | 0.83132 | 0.83925 | 0.87421 |
| ENSG00000139437 | 12 | 110338069 | 110421646 | 221 | 23 | -0.95939 | 0.83132 | TCHP | 0.83132 | 0.82263 | 0.7902 |
| ENSG00000175697 | 3 | 119883492 | 120003941 | 399 | 27 | -0.95947 | 0.83134 | GPR156 | 0.83134 | 0.86663 | 0.86755 |
| ENSG00000212999 | 8 | 93895865 | 93898013 | 4 | 2 | -0.9595 | 0.83135 | AC117834.1 | 0.83135 | 0.37832 | 0.47502 |
| ENSG00000129691 | 8 | 37962760 | 38001594 | 53 | 20 | -0.95955 | 0.83136 | ASH2L | 0.83136 | 0.81183 | 0.83128 |
| ENSG00000213888 | 22 | 31742875 | 31744670 | 7 | 4 | -0.95982 | 0.83143 | AC005003.1 | 0.83143 | 0.41679 | 0.65218 |
| ENSG00000107625 | 10 | 70661034 | 70706603 | 171 | 23 | -0.96014 | 0.83151 | DDX50 | 0.83151 | 0.77738 | 0.81123 |
| ENSG00000224226 | 17 | 34493061 | 34503996 | 2 | 1 | -0.96018 | 0.83152 | TBC1D3B | 0.83152 | 0.071944 | 0.14656 |
| ENSG00000215529 | 20 | 31446729 | 31549006 | 373 | 35 | -0.96018 | 0.83152 | EFCAB8 | 0.83152 | 0.82687 | 0.84542 |
| ENSG00000116198 | 1 | 3728645 | 3773778 | 338 | 19 | -0.96032 | 0.83155 | CEP104 | 0.83155 | 0.84888 | 0.4679 |
| ENSG00000206538 | 3 | 86987119 | 87040269 | 125 | 12 | -0.96047 | 0.83159 | VGLL3 | 0.83159 | 0.84778 | 0.82008 |
| ENSG00000164112 | 4 | 122680088 | 122686582 | 24 | 6 | -0.96047 | 0.83159 | TMEM155 | 0.83159 | 0.80194 | 0.81642 |
| ENSG00000145198 | 3 | 183948217 | 183960117 | 23 | 5 | -0.96052 | 0.8316 | VWA5B2 | 0.8316 | 0.72943 | 0.86533 |
| ENSG00000261717 | 16 | 75446582 | 75498604 | 282 | 16 | -0.96052 | 0.8316 | RP11-77K12.1 | 0.8316 | 0.83412 | 0.82966 |
| ENSG00000100453 | 14 | 25100160 | 25103473 | 18 | 3 | -0.96081 | 0.83168 | GZMB | 0.83168 | 0.3565 | 0.32547 |
| ENSG00000137876 | 15 | 55473004 | 55489265 | 43 | 9 | -0.96118 | 0.83177 | RSL24D1 | 0.83177 | 0.8648 | 0.91981 |
| ENSG00000130045 | 9 | 91150016 | 91199189 | 120 | 15 | -0.96182 | 0.83193 | NXNL2 | 0.83193 | 0.74735 | 0.27045 |
| ENSG00000072134 | 17 | 19118928 | 19240028 | 207 | 13 | -0.96238 | 0.83207 | EPN2 | 0.83207 | 0.84456 | 0.85867 |
| ENSG00000185630 | 1 | 164524821 | 164868533 | 1067 | 119 | -0.96257 | 0.83212 | PBX1 | 0.83212 | 0.83858 | 0.84393 |
| ENSG00000178999 | 17 | 8108056 | 8113918 | 19 | 8 | -0.96261 | 0.83213 | AURKB | 0.83213 | 0.64352 | 0.55819 |
| ENSG00000124782 | 6 | 7107830 | 7252213 | 360 | 43 | -0.96268 | 0.83215 | RREB1 | 0.83215 | 0.85113 | 0.79449 |
| ENSG00000196123 | 16 | 67209505 | 67217943 | 13 | 5 | -0.96321 | 0.83228 | KIAA0895L | 0.83228 | 0.90658 | 0.81995 |
| ENSG00000135048 | 9 | 74298282 | 74431606 | 508 | 33 | -0.96336 | 0.83232 | TMEM2 | 0.83232 | 0.88657 | 0.92045 |
| ENSG00000007516 | 16 | 1383602 | 1399439 | 69 | 19 | -0.96341 | 0.83233 | BAIAP3 | 0.83233 | 0.88323 | 0.67442 |
| ENSG00000119711 | 14 | 74523553 | 74551196 | 112 | 11 | -0.96424 | 0.83254 | ALDH6A1 | 0.83254 | 0.83656 | 0.82348 |
| ENSG00000162897 | 1 | 207131310 | 207143970 | 40 | 5 | -0.96501 | 0.83273 | FCAMR | 0.83273 | 0.8475 | 0.90656 |
| ENSG00000122223 | 1 | 160799950 | 160832692 | 108 | 9 | -0.96542 | 0.83283 | CD244 | 0.83283 | 0.93044 | 0.92752 |
| ENSG00000106588 | 7 | 42956460 | 42971822 | 31 | 4 | -0.96554 | 0.83286 | PSMA2 | 0.83286 | 0.76318 | 0.79939 |
| ENSG00000132465 | 4 | 71521258 | 71547534 | 33 | 7 | -0.96607 | 0.833 | IGJ | 0.833 | 0.79637 | 0.61605 |
| ENSG00000115648 | 2 | 238394071 | 238463961 | 388 | 18 | -0.96634 | 0.83306 | MLPH | 0.83306 | 0.83405 | 0.79643 |
| ENSG00000088451 | 13 | 95226308 | 95248511 | 65 | 6 | -0.96637 | 0.83307 | TGDS | 0.83307 | 0.81937 | 0.79318 |
| ENSG00000205038 | 8 | 110374706 | 110542559 | 571 | 20 | -0.9664 | 0.83308 | PKHD1L1 | 0.83308 | 0.82944 | 0.80371 |
| ENSG00000111834 | 6 | 116937642 | 116954148 | 58 | 12 | -0.96644 | 0.83309 | RSPH4A | 0.83309 | 0.75852 | 0.7389 |
| ENSG00000121064 | 17 | 55055466 | 55084129 | 64 | 10 | -0.96673 | 0.83316 | SCPEP1 | 0.83316 | 0.8641 | 0.75581 |
| ENSG00000102743 | 13 | 41363548 | 41384247 | 65 | 8 | -0.96689 | 0.8332 | SLC25A15 | 0.8332 | 0.83749 | 0.85049 |
| ENSG00000074755 | 17 | 3907739 | 4046314 | 535 | 18 | -0.96691 | 0.8332 | ZZEF1 | 0.8332 | 0.85252 | 0.86614 |
| ENSG00000085741 | 11 | 75897369 | 75921780 | 81 | 16 | -0.96711 | 0.83326 | WNT11 | 0.83326 | 0.78439 | 0.81479 |
| ENSG00000139644 | 12 | 50101508 | 50158717 | 190 | 14 | -0.9677 | 0.8334 | TMBIM6 | 0.8334 | 0.81876 | 0.76807 |
| ENSG00000168724 | 5 | 34929698 | 34959069 | 61 | 13 | -0.9678 | 0.83343 | DNAJC21 | 0.83343 | 0.79656 | 0.83439 |
| ENSG00000170445 | 5 | 140052758 | 140071609 | 45 | 8 | -0.96782 | 0.83343 | HARS | 0.83343 | 0.78205 | 0.77592 |
| ENSG00000198168 | 11 | 22835345 | 22851845 | 31 | 6 | -0.96807 | 0.83349 | SVIP | 0.83349 | 0.86698 | 0.86812 |
| ENSG00000062822 | 19 | 50887461 | 50921273 | 92 | 10 | -0.96829 | 0.83355 | POLD1 | 0.83355 | 0.8535 | 0.90814 |
| ENSG00000058091 | 7 | 90095738 | 90839905 | 2637 | 85 | -0.96833 | 0.83356 | CDK14 | 0.83356 | 0.83546 | 0.84384 |
| ENSG00000106410 | 7 | 144094333 | 144107320 | 43 | 6 | -0.96833 | 0.83356 | NOBOX | 0.83356 | 0.75282 | 0.74858 |
| ENSG00000114735 | 3 | 50606583 | 50622366 | 22 | 6 | -0.96853 | 0.83361 | HEMK1 | 0.83361 | 0.89887 | 0.89457 |
| ENSG00000139626 | 12 | 53585102 | 53601091 | 39 | 10 | -0.9688 | 0.83368 | ITGB7 | 0.83368 | 0.57344 | 0.3288 |
| ENSG00000128165 | 22 | 50919985 | 50924869 | 17 | 9 | -0.96903 | 0.83373 | ADM2 | 0.83373 | 0.80946 | 0.55105 |
| ENSG00000106785 | 9 | 100831557 | 100881494 | 145 | 27 | -0.96936 | 0.83382 | TRIM14 | 0.83382 | 0.83985 | 0.83108 |
| ENSG00000148950 | 11 | 31453948 | 31531192 | 94 | 8 | -0.96944 | 0.83384 | IMMP1L | 0.83384 | 0.82564 | 0.85501 |
| ENSG00000198315 | 6 | 28109688 | 28127250 | 53 | 7 | -0.96955 | 0.83386 | ZKSCAN8 | 0.83386 | 0.85325 | 0.82562 |
| ENSG00000188626 | 15 | 28947100 | 28983530 | 79 | 4 | -0.96952 | 0.83386 | GOLGA8M | 0.83386 | 0.81096 | 0.78295 |
| ENSG00000087191 | 17 | 61904512 | 61909379 | 10 | 3 | -0.96962 | 0.83388 | PSMC5 | 0.83388 | 0.81115 | 0.84364 |
| ENSG00000143554 | 1 | 153746830 | 153752633 | 11 | 6 | -0.96978 | 0.83392 | SLC27A3 | 0.83392 | 0.46978 | 0.23668 |
| ENSG00000063322 | 19 | 39881943 | 39891277 | 30 | 5 | -0.97028 | 0.83405 | MED29 | 0.83405 | 0.85619 | 0.93009 |
| ENSG00000137804 | 15 | 41624892 | 41673248 | 118 | 14 | -0.97073 | 0.83416 | NUSAP1 | 0.83416 | 0.79715 | 0.76136 |
| ENSG00000267508 | 19 | 44886459 | 44905774 | 137 | 11 | -0.97083 | 0.83418 | ZNF285 | 0.83418 | 0.84864 | 0.92311 |
| ENSG00000268280 | 5 | 55224468 | 55224569 | 1 | 1 | -0.9709 | 0.8342 | AC008914.1 | 0.8342 | 0.51786 | 0.22146 |
| ENSG00000114786 | 3 | 52009066 | 52023199 | 22 | 11 | -0.97115 | 0.83426 | ABHD14A-ACY1 | 0.83426 | 0.89963 | 0.87976 |
| ENSG00000243989 | 3 | 52009066 | 52023213 | 22 | 11 | -0.97115 | 0.83426 | ACY1 | 0.83426 | 0.89963 | 0.87976 |
| ENSG00000163586 | 2 | 88422510 | 88427635 | 10 | 4 | -0.97131 | 0.8343 | FABP1 | 0.8343 | 0.89791 | 0.92463 |
| ENSG00000007264 | 19 | 3777971 | 3802127 | 89 | 22 | -0.97135 | 0.83431 | MATK | 0.83431 | 0.73111 | 0.40214 |
| ENSG00000196754 | 1 | 153533584 | 153540366 | 13 | 4 | -0.97151 | 0.83435 | S100A2 | 0.83435 | 0.90085 | 0.83263 |

| | | | | | | | | | | | |
|---|---|---|---|---|---|---|---|---|---|---|---|
| ENSG00000253250 | 8 | 91970865 | 91997485 | 67 | 12 | -0.97165 | 0.83439 | C8orf88 | 0.83439 | 0.68212 | 0.70699 |
| ENSG00000111144 | 12 | 96394606 | 96437298 | 143 | 17 | -0.97177 | 0.83442 | LTA4H | 0.83442 | 0.85133 | 0.79395 |
| ENSG00000145780 | 5 | 114856608 | 114880591 | 38 | 9 | -0.97186 | 0.83444 | FEM1C | 0.83444 | 0.74166 | 0.84122 |
| ENSG00000123739 | 4 | 110631145 | 110651233 | 33 | 8 | -0.97231 | 0.83455 | PLA2G12A | 0.83455 | 0.80112 | 0.86484 |
| ENSG00000173456 | 11 | 119205237 | 119208023 | 9 | 4 | -0.97233 | 0.83456 | RNF26 | 0.83456 | 0.084992 | 0.0067395 |
| ENSG00000090263 | 7 | 140705854 | 140715028 | 24 | 5 | -0.9724 | 0.83457 | MRPS33 | 0.83457 | 0.835 | 0.83905 |
| ENSG00000085978 | 2 | 234118697 | 234204320 | 277 | 33 | -0.97263 | 0.83463 | ATG16L1 | 0.83463 | 0.85449 | 0.86019 |
| ENSG00000072062 | 19 | 14202500 | 14228896 | 37 | 9 | -0.97291 | 0.8347 | PRKACA | 0.8347 | 0.80783 | 0.85774 |
| ENSG00000141642 | 18 | 48494361 | 48514491 | 29 | 8 | -0.97303 | 0.83473 | ELAC1 | 0.83473 | 0.92153 | 0.94807 |
| ENSG00000100253 | 22 | 50925213 | 50929077 | 31 | 12 | -0.97314 | 0.83476 | MIOX | 0.83476 | 0.95517 | 0.67835 |
| ENSG00000163815 | 3 | 45043040 | 45077563 | 123 | 11 | -0.9732 | 0.83477 | CLEC3B | 0.83477 | 0.83918 | 0.86799 |
| ENSG00000163811 | 2 | 29117509 | 29171088 | 206 | 16 | -0.97346 | 0.83484 | WDR43 | 0.83484 | 0.85137 | 0.84797 |
| ENSG00000164124 | 4 | 159122756 | 159176563 | 88 | 17 | -0.9735 | 0.83485 | TMEM144 | 0.83485 | 0.86134 | 0.50646 |
| ENSG00000151065 | 12 | 2055220 | 2113701 | 245 | 13 | -0.97394 | 0.83496 | DCP1B | 0.83496 | 0.77907 | 0.85388 |
| ENSG00000254004 | 19 | 37001597 | 37019562 | 49 | 10 | -0.97405 | 0.83498 | ZNF260 | 0.83498 | 0.76207 | 0.73815 |
| ENSG00000118160 | 19 | 47931274 | 47975425 | 89 | 23 | -0.97412 | 0.835 | SLC8A2 | 0.835 | 0.81346 | 0.77037 |
| ENSG00000113456 | 5 | 34905369 | 34919094 | 24 | 6 | -0.9742 | 0.83502 | RAD1 | 0.83502 | 0.78023 | 0.76544 |
| ENSG00000109680 | 4 | 26578059 | 26756973 | 226 | 20 | -0.97456 | 0.83511 | TBC1D19 | 0.83511 | 0.83185 | 0.85957 |
| ENSG00000170043 | 17 | 7833663 | 7835441 | 5 | 3 | -0.97454 | 0.83511 | TRAPPC1 | 0.83511 | 0.80829 | 0.7707 |
| ENSG00000172457 | 11 | 56510303 | 56511287 | 3 | 1 | -0.9747 | 0.83515 | OR9G4 | 0.83515 | 0.12792 | 0.06724 |
| ENSG00000140350 | 15 | 69070874 | 69113236 | 109 | 20 | -0.9749 | 0.83519 | ANP32A | 0.83519 | 0.75124 | 0.80363 |
| ENSG00000168658 | 2 | 98703579 | 98929762 | 704 | 41 | -0.97521 | 0.83527 | VWA3B | 0.83527 | 0.83918 | 0.80778 |
| ENSG00000133895 | 11 | 64570982 | 64578766 | 18 | 7 | -0.97527 | 0.83529 | MEN1 | 0.83529 | 0.96962 | 0.895 |
| ENSG00000175920 | 4 | 3465033 | 3503200 | 365 | 36 | -0.97538 | 0.83531 | DOK7 | 0.83531 | 0.83826 | 0.78122 |
| ENSG00000164106 | 4 | 174305852 | 174327531 | 45 | 9 | -0.97546 | 0.83533 | SCRG1 | 0.83533 | 0.82861 | 0.81828 |
| ENSG00000141084 | 16 | 67757005 | 67840555 | 126 | 18 | -0.97565 | 0.83538 | RANBP10 | 0.83538 | 0.84453 | 0.84567 |
| ENSG00000136213 | 7 | 2443223 | 2474242 | 75 | 13 | -0.97568 | 0.83539 | CHST12 | 0.83539 | 0.84124 | 0.88367 |
| ENSG00000142173 | 21 | 47518011 | 47552763 | 202 | 26 | -0.97579 | 0.83541 | COL6A2 | 0.83541 | 0.7164 | 0.57151 |
| ENSG00000171703 | 20 | 62681189 | 62703700 | 62 | 15 | -0.97601 | 0.83547 | TCEA2 | 0.83547 | 0.883 | 0.89451 |
| ENSG00000132466 | 4 | 73939093 | 74124515 | 169 | 30 | -0.97611 | 0.8355 | ANKRD17 | 0.8355 | 0.82461 | 0.81401 |
| ENSG00000177459 | 8 | 99076539 | 99105838 | 105 | 28 | -0.97636 | 0.83556 | C8orf47 | 0.83556 | 0.90763 | 0.92398 |
| ENSG00000122970 | 12 | 110562140 | 110656602 | 216 | 19 | -0.97663 | 0.83562 | IFT81 | 0.83562 | 0.84068 | 0.85618 |
| ENSG00000106246 | 7 | 99014362 | 99063787 | 126 | 14 | -0.97672 | 0.83565 | PTCD1 | 0.83565 | 0.84892 | 0.81582 |
| ENSG00000127314 | 12 | 69004619 | 69054372 | 123 | 18 | -0.97679 | 0.83566 | RAP1B | 0.83566 | 0.80155 | 0.75205 |
| ENSG00000197123 | 7 | 63688852 | 63727309 | 194 | 22 | -0.97691 | 0.83569 | ZNF679 | 0.83569 | 0.78266 | 0.80179 |
| ENSG00000141349 | 17 | 42148103 | 42153709 | 11 | 5 | -0.97726 | 0.83578 | G6PC3 | 0.83578 | 0.75271 | 0.76484 |
| ENSG00000119878 | 2 | 46843555 | 46852881 | 31 | 7 | -0.97735 | 0.8358 | CRIPT | 0.8358 | 0.58704 | 0.28639 |
| ENSG00000170624 | 5 | 155297354 | 156194799 | 2945 | 105 | -0.97734 | 0.8358 | SGCD | 0.8358 | 0.85812 | 0.85634 |
| ENSG00000168394 | 6 | 32812986 | 32821755 | 93 | 11 | -0.97757 | 0.83586 | TAP1 | 0.83586 | 0.63222 | 0.32631 |
| ENSG00000187479 | 11 | 43946892 | 43965888 | 89 | 25 | -0.97776 | 0.8359 | C11orf96 | 0.8359 | 0.7144 | 0.88222 |
| ENSG00000234511 | 5 | 169659451 | 169679579 | 30 | 8 | -0.97788 | 0.83593 | C5orf58 | 0.83593 | 0.84007 | 0.61555 |
| ENSG00000158089 | 2 | 31133333 | 31378068 | 994 | 71 | -0.97805 | 0.83597 | GALNT14 | 0.83597 | 0.82339 | 0.76283 |
| ENSG00000136694 | 2 | 113763038 | 113765621 | 6 | 2 | -0.97836 | 0.83605 | IL36A | 0.83605 | 0.84782 | 0.85908 |
| ENSG00000260230 | 9 | 111892573 | 111929571 | 154 | 23 | -0.97834 | 0.83605 | FRRS1L | 0.83605 | 0.79966 | 0.63955 |
| ENSG00000226174 | 14 | 105864916 | 105916443 | 50 | 20 | -0.97865 | 0.83612 | TEX22 | 0.83612 | 0.8281 | 0.49084 |
| ENSG00000162073 | 16 | 3019246 | 3023490 | 6 | 3 | -0.97867 | 0.83613 | PAQR4 | 0.83613 | 0.64911 | 0.35848 |
| ENSG00000178860 | 8 | 72753784 | 72756703 | 10 | 4 | -0.9787 | 0.83614 | MSC | 0.83614 | 0.82673 | 0.94146 |
| ENSG00000111231 | 12 | 110890289 | 110907073 | 45 | 8 | -0.97877 | 0.83615 | GPN3 | 0.83615 | 0.83325 | 0.85186 |
| ENSG00000006451 | 7 | 39663082 | 39747723 | 214 | 28 | -0.97879 | 0.83616 | RALA | 0.83616 | 0.82751 | 0.71687 |
| ENSG00000166822 | 16 | 75476952 | 75499395 | 126 | 15 | -0.9788 | 0.83616 | TMEM170A | 0.83616 | 0.82912 | 0.82138 |
| ENSG00000169071 | 9 | 94325373 | 94712444 | 1753 | 93 | -0.97909 | 0.83623 | ROR2 | 0.83623 | 0.82439 | 0.84501 |
| ENSG00000166582 | 17 | 16245848 | 16256970 | 23 | 4 | -0.9791 | 0.83624 | CENPV | 0.83624 | 0.93762 | 0.93763 |
| ENSG00000197150 | 7 | 150725510 | 150744869 | 60 | 12 | -0.97971 | 0.83638 | ABCB8 | 0.83638 | 0.85658 | 0.82449 |
| ENSG00000126821 | 14 | 64150932 | 64194757 | 84 | 8 | -0.98016 | 0.8365 | SGPP1 | 0.8365 | 0.8065 | 0.81942 |
| ENSG00000187091 | 3 | 38048987 | 38071253 | 81 | 16 | -0.98045 | 0.83657 | PLCD1 | 0.83657 | 0.90288 | 0.7928 |
| ENSG00000136960 | 8 | 120569326 | 120685693 | 309 | 32 | -0.98044 | 0.83657 | ENPP2 | 0.83657 | 0.87136 | 0.87612 |
| ENSG00000143839 | 1 | 204123944 | 204135465 | 45 | 9 | -0.9805 | 0.83658 | REN | 0.83658 | 0.79102 | 0.81951 |
| ENSG00000122787 | 7 | 137687070 | 137802732 | 316 | 30 | -0.98059 | 0.8366 | AKR1D1 | 0.8366 | 0.84661 | 0.73241 |
| ENSG00000166326 | 11 | 35684353 | 35829775 | 220 | 34 | -0.98058 | 0.8366 | TRIM44 | 0.8366 | 0.84486 | 0.85505 |
| ENSG00000153558 | 3 | 33318517 | 33445154 | 351 | 23 | -0.98072 | 0.83664 | FBXL2 | 0.83664 | 0.82616 | 0.80058 |
| ENSG00000114346 | 3 | 172468472 | 172539264 | 153 | 15 | -0.98084 | 0.83666 | ECT2 | 0.83666 | 0.79972 | 0.7074 |
| ENSG00000168813 | 19 | 32836500 | 32878573 | 97 | 17 | -0.98141 | 0.8368 | ZNF507 | 0.8368 | 0.85467 | 0.83236 |
| ENSG00000218819 | 2 | 21346789 | 21366144 | 30 | 10 | -0.98144 | 0.83681 | TDRD15 | 0.83681 | 0.86464 | 0.68304 |
| ENSG00000123992 | 2 | 220238268 | 220264744 | 84 | 11 | -0.98166 | 0.83687 | DNPEP | 0.83687 | 0.83537 | 0.86737 |
| ENSG00000122592 | 7 | 27193335 | 27197555 | 11 | 5 | -0.98171 | 0.83688 | HOXA7 | 0.83688 | 0.92241 | 0.94899 |
| ENSG00000170476 | 5 | 138723169 | 138725770 | 1 | 1 | -0.9818 | 0.8369 | MZB1 | 0.8369 | 0.61275 | 0.65584 |
| ENSG00000112530 | 6 | 163148164 | 163736524 | 2130 | 136 | -0.98184 | 0.83691 | PACRG | 0.83691 | 0.86018 | 0.88828 |
| ENSG00000172137 | 16 | 71392616 | 71424341 | 119 | 16 | -0.98187 | 0.83692 | CALB2 | 0.83692 | 0.8552 | 0.83665 |
| ENSG00000116171 | 1 | 53392901 | 53517375 | 533 | 28 | -0.98213 | 0.83698 | SCP2 | 0.83698 | 0.85339 | 0.8504 |
| ENSG00000173852 | 7 | 34968488 | 35077883 | 447 | 20 | -0.98233 | 0.83703 | DPY19L1 | 0.83703 | 0.86784 | 0.90561 |
| ENSG00000128482 | 17 | 19314438 | 19320589 | 10 | 2 | -0.98254 | 0.83708 | RNF112 | 0.83708 | 0.72084 | 0.86185 |
| ENSG00000149292 | 11 | 113185251 | 113254266 | 158 | 21 | -0.98268 | 0.83712 | TTC12 | 0.83712 | 0.87595 | 0.89962 |
| ENSG00000113520 | 5 | 132009678 | 132018368 | 27 | 4 | -0.98277 | 0.83714 | IL4 | 0.83714 | 0.76811 | 0.77822 |

| | | | | | | | | | | | |
|---|---|---|---|---|---|---|---|---|---|---|---|
| ENSG00000121361 | 12 | 21917889 | 21928515 | 6 | 4 | -0.98333 | 0.83728 | KCNJ8 | 0.83728 | 0.89186 | 0.84054 |
| ENSG00000160058 | 1 | 32830704 | 32860332 | 28 | 16 | -0.98348 | 0.83732 | BSDC1 | 0.83732 | 0.91464 | 0.92272 |
| ENSG00000112486 | 6 | 167525295 | 167553184 | 106 | 19 | -0.98359 | 0.83734 | CCR6 | 0.83734 | 0.81019 | 0.6228 |
| ENSG00000169896 | 16 | 31271311 | 31344213 | 214 | 16 | -0.98358 | 0.83734 | ITGAM | 0.83734 | 0.84321 | 0.81139 |
| ENSG00000101182 | 20 | 60711791 | 60718496 | 17 | 2 | -0.98369 | 0.83737 | PSMA7 | 0.83737 | 0.8358 | 0.77159 |
| ENSG00000147873 | 9 | 21304613 | 21305312 | 1 | 1 | -0.98423 | 0.8375 | IFNA5 | 0.8375 | 0.36255 | 0.44625 |
| ENSG00000119636 | 14 | 74486056 | 74549566 | 272 | 18 | -0.98473 | 0.83762 | CCDC176 | 0.83762 | 0.82652 | 0.8297 |
| ENSG00000163795 | 2 | 27600098 | 27603657 | 3 | 2 | -0.98498 | 0.83768 | ZNF513 | 0.83768 | 0.79057 | 0.77083 |
| ENSG00000120278 | 6 | 150920999 | 151164799 | 1097 | 71 | -0.98496 | 0.83768 | PLEKHG1 | 0.83768 | 0.87525 | 0.86361 |
| ENSG00000130528 | 19 | 49654455 | 49658681 | 18 | 7 | -0.98535 | 0.83777 | HRC | 0.83777 | 0.45079 | 0.39249 |
| ENSG00000074410 | 15 | 63613577 | 63674360 | 157 | 26 | -0.98542 | 0.83779 | CA12 | 0.83779 | 0.88291 | 0.87281 |
| ENSG00000162777 | 1 | 111729796 | 111747157 | 51 | 18 | -0.98546 | 0.8378 | DENND2D | 0.8378 | 0.90702 | 0.90821 |
| ENSG00000227059 | 12 | 133794898 | 133812681 | 47 | 12 | -0.98566 | 0.83785 | ANHX | 0.83785 | 0.92366 | 0.91191 |
| ENSG00000186854 | 2 | 85048774 | 85134132 | 341 | 21 | -0.9857 | 0.83786 | TRABD2A | 0.83786 | 0.81876 | 0.77618 |
| ENSG00000255103 | 1 | 39876151 | 39882154 | 13 | 5 | -0.98573 | 0.83787 | KIAA0754 | 0.83787 | 0.73188 | 0.61575 |
| ENSG00000115687 | 2 | 242045514 | 242089679 | 193 | 27 | -0.98588 | 0.8379 | PASK | 0.8379 | 0.80483 | 0.8642 |
| ENSG00000225485 | 17 | 36584662 | 36668628 | 303 | 37 | -0.98585 | 0.8379 | ARHGAP23 | 0.8379 | 0.72813 | 0.28495 |
| ENSG00000131153 | 16 | 85709804 | 85723679 | 58 | 9 | -0.98652 | 0.83806 | GINS2 | 0.83806 | 0.84385 | 0.84152 |
| ENSG00000183283 | 12 | 51632076 | 51665146 | 100 | 13 | -0.98668 | 0.8381 | DAZAP2 | 0.8381 | 0.90982 | 0.88791 |
| ENSG00000011638 | 16 | 21169698 | 21191937 | 56 | 10 | -0.98732 | 0.83826 | TMEM159 | 0.83826 | 0.80777 | 0.79083 |
| ENSG00000100150 | 22 | 32149944 | 32303012 | 459 | 23 | -0.98755 | 0.83831 | DEPDC5 | 0.83831 | 0.8102 | 0.75649 |
| ENSG00000136715 | 2 | 128698791 | 128785694 | 210 | 22 | -0.98756 | 0.83832 | SAP130 | 0.83832 | 0.87495 | 0.73272 |
| ENSG00000128694 | 2 | 190611386 | 190627953 | 33 | 9 | -0.98784 | 0.83839 | OSGEPL1 | 0.83839 | 0.80823 | 0.84168 |
| ENSG00000196233 | 10 | 98592017 | 98740800 | 232 | 15 | -0.9879 | 0.8384 | LCOR | 0.8384 | 0.82402 | 0.84977 |
| ENSG00000184937 | 11 | 32409321 | 32457176 | 180 | 17 | -0.98858 | 0.83857 | WT1 | 0.83857 | 0.89275 | 0.90718 |
| ENSG00000163623 | 4 | 85413140 | 85419603 | 12 | 5 | -0.98873 | 0.8386 | NKX6-1 | 0.8386 | 0.38905 | 0.10472 |
| ENSG00000135900 | 2 | 224822121 | 224832431 | 21 | 4 | -0.98943 | 0.83877 | MRPL44 | 0.83877 | 0.84127 | 0.8325 |
| ENSG00000124788 | 6 | 16299343 | 16761722 | 1612 | 178 | -0.98941 | 0.83877 | ATXN1 | 0.83877 | 0.85144 | 0.86339 |
| ENSG00000224867 | 1 | 220603286 | 220608023 | 18 | 6 | -0.98969 | 0.83884 | AC096644.1 | 0.83884 | 0.8515 | 0.90795 |
| ENSG00000141448 | 18 | 19749404 | 19782491 | 52 | 13 | -0.98974 | 0.83885 | GATA6 | 0.83885 | 0.88103 | 0.93316 |
| ENSG00000138286 | 10 | 74927924 | 75004262 | 210 | 5 | -0.98994 | 0.8389 | FAM149B1 | 0.8389 | 0.83385 | 0.80979 |
| ENSG00000169925 | 9 | 136895427 | 136933657 | 147 | 19 | -0.98998 | 0.83891 | BRD3 | 0.83891 | 0.87073 | 0.73312 |
| ENSG00000148426 | 10 | 11865338 | 11914276 | 178 | 29 | -0.99032 | 0.83899 | PROSER2 | 0.83899 | 0.91099 | 0.83991 |
| ENSG00000055208 | 6 | 149539777 | 149732749 | 685 | 42 | -0.99039 | 0.83901 | TAB2 | 0.83901 | 0.8566 | 0.8439 |
| ENSG00000011295 | 17 | 15902694 | 15948329 | 133 | 11 | -0.99046 | 0.83903 | TTC19 | 0.83903 | 0.82935 | 0.8114 |
| ENSG00000161681 | 19 | 51165084 | 51222707 | 171 | 40 | -0.99089 | 0.83913 | SHANK1 | 0.83913 | 0.92883 | 0.98973 |
| ENSG00000103064 | 16 | 68298433 | 68335722 | 71 | 12 | -0.99105 | 0.83917 | SLC7A6 | 0.83917 | 0.87683 | 0.89277 |
| ENSG00000198521 | 19 | 21987752 | 22034927 | 160 | 18 | -0.99113 | 0.83919 | ZNF43 | 0.83919 | 0.83933 | 0.86199 |
| ENSG00000136929 | 9 | 100689073 | 100707138 | 32 | 4 | -0.9916 | 0.8393 | HEMGN | 0.8393 | 0.7881 | 0.74359 |
| ENSG00000132837 | 5 | 78293438 | 78531861 | 724 | 38 | -0.99171 | 0.83933 | DMGDH | 0.83933 | 0.83877 | 0.81171 |
| ENSG00000077458 | 11 | 95502106 | 95523573 | 80 | 11 | -0.99179 | 0.83935 | FAM76B | 0.83935 | 0.80395 | 0.74989 |
| ENSG00000198083 | 17 | 39411636 | 39412616 | 2 | 1 | -0.99193 | 0.83938 | KRTAP9-9 | 0.83938 | 0.64166 | 0.66756 |
| ENSG00000170956 | 19 | 42300369 | 42315591 | 58 | 4 | -0.99197 | 0.83939 | CEACAM3 | 0.83939 | 0.81261 | 0.81921 |
| ENSG00000177476 | 1 | 247768856 | 247769838 | 4 | 2 | -0.9923 | 0.83947 | OR2G3 | 0.83947 | 0.91886 | 0.79837 |
| ENSG00000100320 | 22 | 36134783 | 36424473 | 414 | 25 | -0.99231 | 0.83948 | RBFOX2 | 0.83948 | 0.86976 | 0.85544 |
| ENSG00000143742 | 1 | 225965531 | 225978168 | 38 | 9 | -0.99243 | 0.83951 | SRP9 | 0.83951 | 0.46388 | 0.47878 |
| ENSG00000105701 | 19 | 18642561 | 18654887 | 14 | 9 | -0.99292 | 0.83963 | FKBP8 | 0.83963 | 0.93638 | 0.89852 |
| ENSG00000143878 | 2 | 20646835 | 20649200 | 6 | 4 | -0.99298 | 0.83964 | RHOB | 0.83964 | 0.60926 | 0.65181 |
| ENSG00000118271 | 18 | 29171689 | 29178974 | 10 | 2 | -0.99325 | 0.83971 | TTR | 0.83971 | 0.93665 | 0.96892 |
| ENSG00000166685 | 17 | 71189129 | 71204646 | 58 | 8 | -0.99349 | 0.83977 | COG1 | 0.83977 | 0.89631 | 0.83847 |
| ENSG00000186298 | 12 | 111157485 | 111180744 | 62 | 11 | -0.99365 | 0.8398 | PPP1CC | 0.8398 | 0.81313 | 0.81617 |
| ENSG00000138798 | 4 | 110834040 | 110933422 | 235 | 18 | -0.9938 | 0.83984 | EGF | 0.83984 | 0.84447 | 0.90272 |
| ENSG00000181652 | 7 | 150709297 | 150721586 | 29 | 12 | -0.994 | 0.83989 | ATG9B | 0.83989 | 0.8202 | 0.89226 |
| ENSG00000273003 | 11 | 64781654 | 64808042 | 99 | 14 | -0.99416 | 0.83993 | RP11-399J13.3 | 0.83993 | 0.88347 | 0.88638 |
| ENSG00000165119 | 9 | 86582998 | 86595569 | 25 | 7 | -0.9942 | 0.83994 | HNRNPK | 0.83994 | 0.82936 | 0.76786 |
| ENSG00000184040 | 10 | 18041218 | 18089855 | 21 | 9 | -0.99425 | 0.83995 | TMEM236 | 0.83995 | 0.55936 | 0.71337 |
| ENSG00000249915 | 5 | 271736 | 353971 | 382 | 21 | -0.99433 | 0.83997 | PDCD6 | 0.83997 | 0.85747 | 0.84658 |
| ENSG00000103355 | 16 | 2833954 | 2837949 | 12 | 4 | -0.99433 | 0.83997 | PRSS33 | 0.83997 | 0.95432 | 0.94468 |
| ENSG00000171446 | 17 | 38933060 | 38938786 | 29 | 7 | -0.99441 | 0.83999 | KRT27 | 0.83999 | 0.84126 | 0.84788 |
| ENSG00000196374 | 6 | 27782822 | 27783267 | 1 | 1 | -0.99446 | 0.84 | HIST1H2BM | 0.84 | 0.66976 | 0.70868 |
| ENSG00000198732 | 14 | 70320848 | 70499083 | 759 | 66 | -0.99451 | 0.84001 | SMOC1 | 0.84001 | 0.85402 | 0.88242 |
| ENSG00000170279 | 7 | 148287657 | 148312952 | 89 | 18 | -0.99458 | 0.84003 | C7orf33 | 0.84003 | 0.8927 | 0.93054 |
| ENSG00000148204 | 9 | 126118449 | 126142603 | 101 | 20 | -0.9948 | 0.84008 | CRB2 | 0.84008 | 0.91252 | 0.85818 |
| ENSG00000005102 | 17 | 41717756 | 41739322 | 80 | 12 | -0.99481 | 0.84009 | MEOX1 | 0.84009 | 0.90441 | 0.88577 |
| ENSG00000213551 | 10 | 74943120 | 75008620 | 175 | 5 | -0.99491 | 0.84011 | DNAJC9 | 0.84011 | 0.82808 | 0.78486 |
| ENSG00000204149 | 10 | 51748078 | 51770259 | 36 | 6 | -0.99502 | 0.84014 | AGAP6 | 0.84014 | 0.75075 | 0.80651 |
| ENSG00000115590 | 2 | 102608306 | 102645006 | 153 | 11 | -0.99525 | 0.84019 | IL1R2 | 0.84019 | 0.75258 | 0.69145 |
| ENSG00000173726 | 1 | 235272651 | 235292251 | 47 | 7 | -0.99568 | 0.8403 | TOMM20 | 0.8403 | 0.863 | 0.87102 |
| ENSG00000135312 | 6 | 78171948 | 78173490 | 5 | 2 | -0.99582 | 0.84033 | HTR1B | 0.84033 | 0.49508 | 0.47536 |
| ENSG00000149761 | 11 | 63993413 | 63998158 | 16 | 6 | -0.99587 | 0.84034 | NUDT22 | 0.84034 | 0.53236 | 0.4566 |
| ENSG00000007216 | 17 | 26800311 | 26824799 | 44 | 17 | -0.99615 | 0.84041 | SLC13A2 | 0.84041 | 0.70315 | 0.8066 |
| ENSG00000198795 | 18 | 22641890 | 22932154 | 885 | 84 | -0.99615 | 0.84041 | ZNF521 | 0.84041 | 0.85722 | 0.80388 |
| ENSG00000179674 | 3 | 160394948 | 160396233 | 4 | 2 | -0.99641 | 0.84047 | ARL14 | 0.84047 | 0.62387 | 0.68052 |

| | | | | | | | | | | | |
|---|---|---|---|---|---|---|---|---|---|---|---|
| ENSG00000160226 | 21 | 45748827 | 45759285 | 32 | 10 | -0.99652 | 0.8405 | C21orf2 | 0.8405 | 0.62819 | 0.56526 |
| ENSG00000157259 | 7 | 92076767 | 92088150 | 19 | 7 | -0.99678 | 0.84056 | GATAD1 | 0.84056 | 0.67013 | 0.72528 |
| ENSG00000105767 | 19 | 44126520 | 44143991 | 37 | 10 | -0.99675 | 0.84056 | CADM4 | 0.84056 | 0.59268 | 0.82495 |
| ENSG00000196504 | 2 | 153508107 | 153574511 | 183 | 18 | -0.99699 | 0.84061 | PRPF40A | 0.84061 | 0.88376 | 0.91763 |
| ENSG00000170616 | 8 | 145554228 | 145559943 | 8 | 5 | -0.99698 | 0.84061 | SCRT1 | 0.84061 | 0.46099 | 0.22834 |
| ENSG00000198554 | 14 | 55405668 | 55493823 | 237 | 18 | -0.99698 | 0.84061 | WDHD1 | 0.84061 | 0.85783 | 0.85888 |
| ENSG00000188452 | 2 | 182401403 | 182545392 | 468 | 33 | -0.99699 | 0.84062 | CERKL | 0.84062 | 0.75289 | 0.56265 |
| ENSG00000146021 | 5 | 136953189 | 137071779 | 247 | 29 | -0.99827 | 0.84093 | KLHL3 | 0.84093 | 0.87837 | 0.84796 |
| ENSG00000164713 | 7 | 97881691 | 97937162 | 167 | 11 | -0.9984 | 0.84096 | BRI3 | 0.84096 | 0.81018 | 0.78499 |
| ENSG00000231738 | 12 | 85408094 | 85430055 | 34 | 8 | -0.99846 | 0.84097 | TSPAN19 | 0.84097 | 0.86633 | 0.81425 |
| ENSG00000115946 | 2 | 68384976 | 68403370 | 38 | 5 | -0.99855 | 0.84099 | PNO1 | 0.84099 | 0.79114 | 0.77008 |
| ENSG00000151503 | 11 | 134020014 | 134095348 | 199 | 28 | -0.99878 | 0.84105 | NCAPD3 | 0.84105 | 0.87638 | 0.89104 |
| ENSG00000173272 | 2 | 132222473 | 132250316 | 194 | 11 | -0.99882 | 0.84106 | MZT2A | 0.84106 | 0.85699 | 0.82429 |
| ENSG00000161573 | 17 | 34303529 | 34308532 | 22 | 6 | -0.99892 | 0.84108 | CCL16 | 0.84108 | 0.34598 | 0.34678 |
| ENSG00000135951 | 2 | 99613724 | 99771427 | 401 | 22 | -0.99894 | 0.84109 | TSGA10 | 0.84109 | 0.84251 | 0.85807 |
| ENSG00000181915 | 10 | 64564516 | 64568238 | 6 | 2 | -0.99922 | 0.84116 | ADO | 0.84116 | 0.54755 | 0.70982 |
| ENSG00000160633 | 19 | 5623046 | 5668489 | 95 | 13 | -0.99946 | 0.84121 | SAFB | 0.84121 | 0.69089 | 0.57919 |
| ENSG00000196652 | 7 | 99102274 | 99132323 | 51 | 11 | -0.99974 | 0.84128 | ZKSCAN5 | 0.84128 | 0.82434 | 0.79914 |
| ENSG00000123395 | 12 | 52463030 | 52471278 | 31 | 5 | -0.99983 | 0.8413 | C12orf44 | 0.8413 | 0.62002 | 0.64716 |
| ENSG00000166069 | 15 | 38214140 | 38259925 | 77 | 16 | -0.99987 | 0.84131 | TMCO5A | 0.84131 | 0.86667 | 0.90909 |
| ENSG00000107611 | 10 | 16865963 | 17171830 | 1417 | 89 | -1.0002 | 0.84139 | CUBN | 0.84139 | 0.86841 | 0.91124 |
| ENSG00000136574 | 8 | 11534468 | 11617511 | 290 | 32 | -1.0005 | 0.84146 | GATA4 | 0.84146 | 0.8094 | 0.50814 |
| ENSG00000197134 | 19 | 22235254 | 22274282 | 204 | 8 | -1.0006 | 0.8415 | ZNF257 | 0.8415 | 0.83795 | 0.81518 |
| ENSG00000020181 | 8 | 37641709 | 37702414 | 160 | 37 | -1.0007 | 0.84151 | GPR124 | 0.84151 | 0.70597 | 0.72248 |
| ENSG00000095564 | 10 | 93683526 | 93790082 | 246 | 20 | -1.0008 | 0.84153 | BTAF1 | 0.84153 | 0.85526 | 0.80561 |
| ENSG00000175826 | 17 | 7146910 | 7155810 | 10 | 3 | -1.0013 | 0.84165 | CTDNEP1 | 0.84165 | 0.88123 | 0.8555 |
| ENSG00000243955 | 6 | 52656462 | 52668708 | 34 | 6 | -1.0014 | 0.84167 | GSTA1 | 0.84167 | 0.62757 | 0.61218 |
| ENSG00000100628 | 14 | 94400499 | 94443137 | 167 | 27 | -1.0017 | 0.84176 | ASB2 | 0.84176 | 0.88272 | 0.93632 |
| ENSG00000163479 | 1 | 155978839 | 155990750 | 23 | 7 | -1.0018 | 0.84178 | SSR2 | 0.84178 | 0.83761 | 0.85693 |
| ENSG00000057468 | 1 | 76262567 | 76378923 | 538 | 23 | -1.0018 | 0.84179 | MSH4 | 0.84179 | 0.82467 | 0.82056 |
| ENSG00000205560 | 22 | 51007290 | 51017899 | 54 | 9 | -1.002 | 0.84183 | CPT1B | 0.84183 | 0.63987 | 0.43188 |
| ENSG00000266953 | 19 | 34887220 | 34900269 | 31 | 10 | -1.0027 | 0.84201 | RP11-618P17.4 | 0.84201 | 0.82588 | 0.72697 |
| ENSG00000260362 | 16 | 9760742 | 9770700 | 77 | 7 | -1.0031 | 0.8421 | RP11-297M9.1 | 0.8421 | 0.85904 | 0.93253 |
| ENSG00000250799 | 19 | 36290890 | 36304201 | 30 | 13 | -1.0036 | 0.84222 | PRODH2 | 0.84222 | 0.84985 | 0.84302 |
| ENSG00000141027 | 17 | 15932471 | 16121499 | 451 | 17 | -1.0037 | 0.84224 | NCOR1 | 0.84224 | 0.85848 | 0.85445 |
| ENSG00000132388 | 17 | 4172554 | 4269923 | 179 | 12 | -1.0039 | 0.84229 | UBE2G1 | 0.84229 | 0.86052 | 0.8622 |
| ENSG00000188086 | 3 | 46783582 | 46854048 | 129 | 11 | -1.0044 | 0.8424 | PRSS45 | 0.8424 | 0.76361 | 0.72732 |
| ENSG00000113282 | 5 | 157212751 | 157286183 | 232 | 16 | -1.0045 | 0.84243 | CLINT1 | 0.84243 | 0.86489 | 0.86858 |
| ENSG00000196562 | 20 | 46285092 | 46415360 | 582 | 92 | -1.0045 | 0.84243 | SULF2 | 0.84243 | 0.87194 | 0.89372 |
| ENSG00000140067 | 14 | 94385240 | 94395954 | 38 | 9 | -1.0046 | 0.84246 | FAM181A | 0.84246 | 0.74609 | 0.8104 |
| ENSG00000242616 | 9 | 114423615 | 114432526 | 40 | 9 | -1.0048 | 0.8425 | GNG10 | 0.8425 | 0.93088 | 0.93765 |
| ENSG00000168286 | 16 | 67876213 | 67878097 | 2 | 1 | -1.0052 | 0.8426 | THAP11 | 0.8426 | 0.8026 | 0.8462 |
| ENSG00000205864 | 11 | 1718425 | 1718985 | 3 | 1 | -1.0053 | 0.84263 | KRTAP5-6 | 0.84263 | 0.031447 | 0.013172 |
| ENSG00000153015 | 5 | 64064757 | 64314590 | 617 | 24 | -1.0054 | 0.84265 | CWC27 | 0.84265 | 0.82962 | 0.83221 |
| ENSG00000112499 | 6 | 160592093 | 160698670 | 388 | 24 | -1.0055 | 0.84266 | SLC22A2 | 0.84266 | 0.91847 | 0.95364 |
| ENSG00000135862 | 1 | 182992595 | 183114727 | 556 | 11 | -1.0056 | 0.8427 | LAMC1 | 0.8427 | 0.83864 | 0.84326 |
| ENSG00000163288 | 4 | 46995740 | 47428461 | 1264 | 75 | -1.0057 | 0.84272 | GABRB1 | 0.84272 | 0.82854 | 0.79329 |
| ENSG00000006747 | 7 | 12610203 | 12693228 | 343 | 46 | -1.0058 | 0.84275 | SCIN | 0.84275 | 0.75976 | 0.64353 |
| ENSG00000088543 | 3 | 50595462 | 50608458 | 19 | 6 | -1.0059 | 0.84278 | C3orf18 | 0.84278 | 0.86995 | 0.85737 |
| ENSG00000153774 | 16 | 75327596 | 75467383 | 709 | 23 | -1.0061 | 0.84283 | CFDP1 | 0.84283 | 0.83403 | 0.84684 |
| ENSG00000107758 | 10 | 75196186 | 75255782 | 65 | 12 | -1.0062 | 0.84284 | PPP3CB | 0.84284 | 0.84243 | 0.82432 |
| ENSG00000103978 | 15 | 42502730 | 42565861 | 112 | 16 | -1.0066 | 0.84293 | TMEM87A | 0.84293 | 0.84172 | 0.80585 |
| ENSG00000184779 | 15 | 82821158 | 82824972 | 1 | 1 | -1.0069 | 0.843 | RPS17 | 0.843 | 0.55173 | 0.69932 |
| ENSG00000205922 | 19 | 1752372 | 1780987 | 104 | 29 | -1.0073 | 0.84311 | ONECUT3 | 0.84311 | 0.9277 | 0.76761 |
| ENSG00000133640 | 12 | 85430092 | 85657002 | 385 | 19 | -1.0074 | 0.84312 | LRRIQ1 | 0.84312 | 0.83415 | 0.85757 |
| ENSG00000167065 | 22 | 31048038 | 31063877 | 84 | 19 | -1.0074 | 0.84312 | DUSP18 | 0.84312 | 0.758 | 0.80514 |
| ENSG00000108849 | 17 | 42018172 | 42019836 | 3 | 1 | -1.0075 | 0.84314 | PPY | 0.84314 | 0.86587 | 0.8947 |
| ENSG00000100802 | 14 | 23456110 | 23479375 | 39 | 9 | -1.0075 | 0.84316 | C14orf93 | 0.84316 | 0.94299 | 0.93884 |
| ENSG00000086102 | 9 | 33290509 | 33371155 | 120 | 19 | -1.0081 | 0.84329 | NFX1 | 0.84329 | 0.83098 | 0.84287 |
| ENSG00000129993 | 16 | 88941266 | 89043612 | 536 | 41 | -1.0081 | 0.8433 | CBFA2T3 | 0.8433 | 0.83758 | 0.72759 |
| ENSG00000175354 | 18 | 12785477 | 12929642 | 504 | 40 | -1.0082 | 0.84332 | PTPN2 | 0.84332 | 0.85423 | 0.92239 |
| ENSG00000114030 | 3 | 122140796 | 122233792 | 274 | 18 | -1.0087 | 0.84345 | KPNA1 | 0.84345 | 0.84969 | 0.87038 |
| ENSG00000185332 | 17 | 79285074 | 79304474 | 91 | 11 | -1.0087 | 0.84345 | TMEM105 | 0.84345 | 0.83079 | 0.91571 |
| ENSG00000108094 | 10 | 35297479 | 35379570 | 243 | 12 | -1.0092 | 0.84357 | CUL2 | 0.84357 | 0.85255 | 0.86997 |
| ENSG00000254505 | 14 | 24678789 | 24683075 | 12 | 7 | -1.0092 | 0.84357 | CHMP4A | 0.84357 | 0.71057 | 0.74908 |
| ENSG00000267802 | 14 | 52436014 | 52436247 | 1 | 1 | -1.0094 | 0.8436 | AL358333.1 | 0.8436 | 0.75153 | 0.70168 |
| ENSG00000100033 | 22 | 18900294 | 18924066 | 139 | 22 | -1.0095 | 0.84364 | PRODH | 0.84364 | 0.84477 | 0.69587 |
| ENSG00000090659 | 19 | 7804879 | 7812464 | 28 | 7 | -1.01 | 0.84375 | CD209 | 0.84375 | 0.36322 | 0.16043 |
| ENSG00000101193 | 20 | 61569471 | 61576996 | 27 | 9 | -1.01 | 0.84375 | GID8 | 0.84375 | 0.85831 | 0.15378 |
| ENSG00000164114 | 4 | 156263810 | 156298122 | 154 | 14 | -1.0102 | 0.8438 | MAP9 | 0.8438 | 0.86117 | 0.90291 |
| ENSG00000176209 | 8 | 42396298 | 42409603 | 47 | 6 | -1.0105 | 0.84386 | SMIM19 | 0.84386 | 0.8728 | 0.84611 |
| ENSG00000012504 | 12 | 100867486 | 100958191 | 188 | 25 | -1.0105 | 0.84387 | NR1H4 | 0.84387 | 0.81029 | 0.31192 |
| ENSG00000177469 | 17 | 40554470 | 40575535 | 83 | 12 | -1.0105 | 0.84387 | PTRF | 0.84387 | 0.89832 | 0.89677 |

| | | | | | | | | | | | |
|---|---|---|---|---|---|---|---|---|---|---|---|
| ENSG00000197386 | 4 | 3076408 | 3245676 | 430 | 25 | -1.0108 | 0.84394 | HTT | 0.84394 | 0.86734 | 0.85278 |
| ENSG00000112419 | 6 | 143857982 | 144152322 | 1011 | 61 | -1.0108 | 0.84395 | PHACTR2 | 0.84395 | 0.81043 | 0.79653 |
| ENSG00000183643 | 15 | 93014884 | 93044347 | 99 | 21 | -1.0108 | 0.84395 | C15orf32 | 0.84395 | 0.62432 | 0.44378 |
| ENSG00000163486 | 1 | 206516200 | 206637783 | 272 | 18 | -1.0109 | 0.84397 | SRGAP2 | 0.84397 | 0.84931 | 0.70103 |
| ENSG00000114388 | 3 | 50384761 | 50388522 | 1 | 1 | -1.011 | 0.844 | NPRL2 | 0.844 | 0.4463 | 0.33631 |
| ENSG00000258839 | 16 | 89978527 | 89987385 | 63 | 11 | -1.0113 | 0.84407 | MC1R | 0.84407 | 0.93616 | 0.92422 |
| ENSG00000221963 | 22 | 36044442 | 36064456 | 84 | 11 | -1.0118 | 0.84417 | APOL6 | 0.84417 | 0.7543 | 0.79619 |
| ENSG00000105063 | 19 | 55741148 | 55770363 | 94 | 16 | -1.0126 | 0.84437 | PPP6R1 | 0.84437 | 0.92237 | 0.94658 |
| ENSG00000146833 | 7 | 99474581 | 99517223 | 123 | 10 | -1.0126 | 0.84438 | TRIM4 | 0.84438 | 0.79763 | 0.73116 |
| ENSG00000070476 | 3 | 126156444 | 126194762 | 150 | 18 | -1.0131 | 0.8445 | ZXDC | 0.8445 | 0.83878 | 0.78923 |
| ENSG00000131398 | 19 | 50815194 | 50836772 | 71 | 18 | -1.0132 | 0.84451 | KCNC3 | 0.84451 | 0.88666 | 0.78342 |
| ENSG00000088367 | 20 | 34679426 | 34820721 | 178 | 21 | -1.0135 | 0.84459 | EPB41L1 | 0.84459 | 0.84651 | 0.79851 |
| ENSG00000158023 | 12 | 122355768 | 122441833 | 236 | 25 | -1.0136 | 0.8446 | WDR66 | 0.8446 | 0.83205 | 0.89096 |
| ENSG00000075413 | 14 | 103851729 | 103970168 | 359 | 13 | -1.0141 | 0.84473 | MARK3 | 0.84473 | 0.82023 | 0.79994 |
| ENSG00000134453 | 10 | 6130950 | 6159420 | 112 | 23 | -1.0142 | 0.84476 | RBM17 | 0.84476 | 0.8868 | 0.90931 |
| ENSG00000169607 | 2 | 113493930 | 113522254 | 61 | 8 | -1.0143 | 0.84478 | CKAP2L | 0.84478 | 0.8389 | 0.86438 |
| ENSG00000171180 | 1 | 248402231 | 248403166 | 1 | 1 | -1.0144 | 0.8448 | OR2M4 | 0.8448 | 0.76541 | 0.8742 |
| ENSG00000165238 | 9 | 95947198 | 96082854 | 448 | 36 | -1.0147 | 0.84487 | WNK2 | 0.84487 | 0.86203 | 0.88886 |
| ENSG00000149179 | 11 | 46958240 | 47185936 | 407 | 24 | -1.0148 | 0.84491 | C11orf49 | 0.84491 | 0.80983 | 0.836 |
| ENSG00000124067 | 16 | 67977377 | 68003504 | 33 | 9 | -1.0155 | 0.84506 | SLC12A4 | 0.84506 | 0.84393 | 0.80384 |
| ENSG00000122641 | 7 | 41724712 | 41742706 | 17 | 6 | -1.016 | 0.84518 | INHBA | 0.84518 | 0.93836 | 0.9682 |
| ENSG00000159842 | 17 | 906758 | 1132315 | 1258 | 145 | -1.016 | 0.84518 | ABR | 0.84518 | 0.88028 | 0.90023 |
| ENSG00000215217 | 5 | 7830491 | 7851603 | 87 | 13 | -1.016 | 0.84519 | C5orf49 | 0.84519 | 0.90227 | 0.87202 |
| ENSG00000108830 | 17 | 41177258 | 41184057 | 12 | 3 | -1.0161 | 0.84521 | RND2 | 0.84521 | 0.76549 | 0.86266 |
| ENSG00000125703 | 1 | 63249806 | 63331184 | 173 | 11 | -1.0163 | 0.84527 | ATG4C | 0.84527 | 0.84108 | 0.86156 |
| ENSG00000125967 | 20 | 32244893 | 32262269 | 15 | 5 | -1.0163 | 0.84527 | NECAB3 | 0.84527 | 0.93397 | 0.90651 |
| ENSG00000078237 | 12 | 4430371 | 4462338 | 110 | 16 | -1.0168 | 0.84537 | C12orf5 | 0.84537 | 0.83224 | 0.50429 |
| ENSG00000150281 | 16 | 30907928 | 30914881 | 11 | 5 | -1.0171 | 0.84544 | CTF1 | 0.84544 | 0.94255 | 0.85481 |
| ENSG00000161609 | 19 | 49891475 | 49921251 | 99 | 9 | -1.0171 | 0.84546 | CCDC155 | 0.84546 | 0.80949 | 0.66263 |
| ENSG00000111880 | 6 | 89319985 | 89673348 | 834 | 23 | -1.0173 | 0.8455 | RNGTT | 0.8455 | 0.82672 | 0.8899 |
| ENSG00000204979 | 11 | 60282886 | 60310194 | 102 | 12 | -1.0173 | 0.8455 | MS4A13 | 0.8455 | 0.78129 | 0.83945 |
| ENSG00000079393 | 10 | 76854192 | 76868979 | 49 | 12 | -1.0178 | 0.84562 | DUSP13 | 0.84562 | 0.86616 | 0.73294 |
| ENSG00000205927 | 21 | 34398153 | 34401504 | 9 | 4 | -1.0179 | 0.84564 | OLIG2 | 0.84564 | 0.72902 | 0.74837 |
| ENSG00000115602 | 2 | 102927962 | 102968497 | 202 | 17 | -1.0182 | 0.8457 | IL1RL1 | 0.8457 | 0.84688 | 0.81072 |
| ENSG00000163328 | 2 | 175296966 | 175351822 | 132 | 16 | -1.0182 | 0.84572 | GPR155 | 0.84572 | 0.76722 | 0.58146 |
| ENSG00000118402 | 6 | 80624529 | 80657297 | 126 | 12 | -1.0186 | 0.84579 | ELOVL4 | 0.84579 | 0.89849 | 0.85292 |
| ENSG00000075914 | 3 | 45016733 | 45077558 | 199 | 16 | -1.0188 | 0.84585 | EXOSC7 | 0.84585 | 0.84017 | 0.86688 |
| ENSG00000133063 | 1 | 203181955 | 203242769 | 195 | 30 | -1.019 | 0.84589 | CHIT1 | 0.84589 | 0.72653 | 0.75041 |
| ENSG00000164611 | 5 | 159848829 | 159855748 | 16 | 5 | -1.0191 | 0.84591 | PTTG1 | 0.84591 | 0.897 | 0.95238 |
| ENSG00000172728 | 8 | 33228342 | 33330940 | 395 | 22 | -1.0191 | 0.84593 | FUT10 | 0.84593 | 0.82784 | 0.74493 |
| ENSG00000106976 | 9 | 130965658 | 131017527 | 135 | 21 | -1.0193 | 0.84596 | DNM1 | 0.84596 | 0.86805 | 0.86526 |
| ENSG00000064601 | 20 | 44518783 | 44527459 | 28 | 6 | -1.0192 | 0.84596 | CTSA | 0.84596 | 0.86898 | 0.88462 |
| ENSG00000172936 | 3 | 38179969 | 38184513 | 5 | 3 | -1.0194 | 0.84598 | MYD88 | 0.84598 | 0.42118 | 0.55027 |
| ENSG00000137857 | 15 | 45422131 | 45457774 | 104 | 16 | -1.0194 | 0.84599 | DUOX1 | 0.84599 | 0.90583 | 0.94034 |
| ENSG00000174177 | 16 | 88772871 | 88781794 | 77 | 15 | -1.0196 | 0.84603 | CTU2 | 0.84603 | 0.82596 | 0.77542 |
| ENSG00000122406 | 1 | 93297582 | 93307481 | 12 | 5 | -1.0196 | 0.84605 | RPL5 | 0.84605 | 0.58617 | 0.55027 |
| ENSG00000130522 | 19 | 18390563 | 18392432 | 4 | 2 | -1.0202 | 0.84618 | JUND | 0.84618 | 0.75951 | 0.53021 |
| ENSG00000182447 | 3 | 161214596 | 161221730 | 30 | 3 | -1.0202 | 0.84619 | OTOL1 | 0.84619 | 0.84313 | 0.88486 |
| ENSG00000105398 | 19 | 48373723 | 48389654 | 77 | 11 | -1.0202 | 0.84619 | SULT2A1 | 0.84619 | 0.80658 | 0.83286 |
| ENSG00000134905 | 13 | 111293759 | 111365950 | 361 | 18 | -1.0205 | 0.84626 | CARS2 | 0.84626 | 0.85569 | 0.87706 |
| ENSG00000175216 | 11 | 46764598 | 46867847 | 184 | 21 | -1.0206 | 0.84628 | CKAP5 | 0.84628 | 0.8449 | 0.83019 |
| ENSG00000196353 | 3 | 131252399 | 132004254 | 3465 | 100 | -1.0209 | 0.84634 | CPNE4 | 0.84634 | 0.84359 | 0.84534 |
| ENSG00000172183 | 15 | 89179384 | 89199714 | 101 | 19 | -1.0211 | 0.84639 | ISG20 | 0.84639 | 0.89531 | 0.84343 |
| ENSG00000167037 | 22 | 25202236 | 25323545 | 633 | 73 | -1.0211 | 0.8464 | SGSM1 | 0.8464 | 0.83898 | 0.85239 |
| ENSG00000104522 | 8 | 144694788 | 144700218 | 19 | 4 | -1.0221 | 0.84664 | TSTA3 | 0.84664 | 0.92241 | 0.89322 |
| ENSG00000165995 | 10 | 18429606 | 18830798 | 1869 | 122 | -1.0222 | 0.84665 | CACNB2 | 0.84665 | 0.84025 | 0.80011 |
| ENSG00000166575 | 11 | 86748886 | 87034800 | 1828 | 40 | -1.0224 | 0.8467 | TMEM135 | 0.8467 | 0.85498 | 0.85247 |
| ENSG00000204856 | 12 | 110906169 | 110928190 | 65 | 10 | -1.0228 | 0.84679 | FAM216A | 0.84679 | 0.83337 | 0.82876 |
| ENSG00000132003 | 19 | 13906274 | 13943044 | 94 | 9 | -1.0231 | 0.84687 | ZSWIM4 | 0.84687 | 0.78771 | 0.52926 |
| ENSG00000264813 | 17 | 61562184 | 61599209 | 52 | 12 | -1.0232 | 0.84688 | ACE | 0.84688 | 0.47514 | 0.64982 |
| ENSG00000176907 | 8 | 40010989 | 40012821 | 3 | 2 | -1.0233 | 0.84691 | C8orf4 | 0.84691 | 0.89956 | 0.80955 |
| ENSG00000198898 | 7 | 116451124 | 116562103 | 220 | 25 | -1.0234 | 0.84693 | CAPZA2 | 0.84693 | 0.81983 | 0.81989 |
| ENSG00000125879 | 20 | 16729003 | 16750707 | 126 | 23 | -1.0238 | 0.84703 | OTOR | 0.84703 | 0.78061 | 0.74316 |
| ENSG00000187145 | 1 | 150266289 | 150281414 | 65 | 9 | -1.024 | 0.84708 | MRPS21 | 0.84708 | 0.89431 | 0.88537 |
| ENSG00000198520 | 1 | 45140364 | 45191263 | 117 | 35 | -1.0241 | 0.84712 | C1orf228 | 0.84712 | 0.71256 | 0.66557 |
| ENSG00000005801 | 11 | 3360491 | 3400448 | 164 | 13 | -1.0242 | 0.84712 | ZNF195 | 0.84712 | 0.86346 | 0.84538 |
| ENSG00000162144 | 11 | 61116217 | 61129771 | 4 | 2 | -1.0243 | 0.84715 | CYB561A3 | 0.84715 | 0.76262 | 0.48163 |
| ENSG00000171051 | 19 | 52248425 | 52307363 | 290 | 38 | -1.0243 | 0.84715 | FPR1 | 0.84715 | 0.78462 | 0.70609 |
| ENSG00000151498 | 11 | 134123389 | 134135749 | 39 | 9 | -1.0245 | 0.84719 | ACAD8 | 0.84719 | 0.81011 | 0.86484 |
| ENSG00000124875 | 4 | 74702214 | 74714781 | 32 | 12 | -1.0245 | 0.84721 | CXCL6 | 0.84721 | 0.72325 | 0.82452 |
| ENSG00000100426 | 22 | 50247490 | 50283726 | 178 | 12 | -1.0247 | 0.84724 | ZBED4 | 0.84724 | 0.83914 | 0.85026 |
| ENSG00000203972 | 6 | 49467671 | 49494760 | 95 | 9 | -1.0247 | 0.84725 | GLYATL3 | 0.84725 | 0.85593 | 0.70433 |
| ENSG00000116830 | 1 | 117602925 | 117650075 | 126 | 26 | -1.025 | 0.84731 | TTF2 | 0.84731 | 0.88283 | 0.91915 |

| | | | | | | | | | | | |
|---|---|---|---|---|---|---|---|---|---|---|---|
| ENSG00000138496 | 3 | 122246771 | 122283424 | 110 | 15 | -1.025 | 0.84732 | PARP9 | 0.84732 | 0.77134 | 0.292 |
| ENSG00000033867 | 3 | 27414214 | 27525911 | 447 | 25 | -1.0251 | 0.84733 | SLC4A7 | 0.84733 | 0.84294 | 0.82563 |
| ENSG00000120925 | 8 | 42704780 | 42752433 | 74 | 12 | -1.0251 | 0.84735 | RNF170 | 0.84735 | 0.84275 | 0.83062 |
| ENSG00000108510 | 17 | 60019966 | 60142643 | 202 | 22 | -1.0254 | 0.8474 | MED13 | 0.8474 | 0.83343 | 0.86724 |
| ENSG00000105954 | 7 | 25264189 | 25268105 | 25 | 7 | -1.0254 | 0.84742 | NPVF | 0.84742 | 0.78553 | 0.65258 |
| ENSG00000221829 | 9 | 35073832 | 35080013 | 6 | 3 | -1.0255 | 0.84744 | FANCG | 0.84744 | 0.883 | 0.83249 |
| ENSG00000081181 | 14 | 68086515 | 68118437 | 88 | 12 | -1.0258 | 0.84751 | ARG2 | 0.84751 | 0.84601 | 0.75838 |
| ENSG00000182379 | 12 | 57610578 | 57620232 | 21 | 6 | -1.0259 | 0.84753 | NXPH4 | 0.84753 | 0.46104 | 0.44727 |
| ENSG00000231924 | 19 | 43370616 | 43383974 | 82 | 7 | -1.026 | 0.84754 | PSG1 | 0.84754 | 0.85807 | 0.84719 |
| ENSG00000010704 | 6 | 26087509 | 26098571 | 32 | 11 | -1.0261 | 0.84759 | HFE | 0.84759 | 0.76001 | 0.64593 |
| ENSG00000176915 | 12 | 133302254 | 133338474 | 187 | 29 | -1.0262 | 0.8476 | ANKLE2 | 0.8476 | 0.77418 | 0.81209 |
| ENSG00000135443 | 12 | 52753790 | 52761265 | 27 | 9 | -1.0264 | 0.84765 | KRT85 | 0.84765 | 0.86259 | 0.84013 |
| ENSG00000144036 | 2 | 72403113 | 73053170 | 1159 | 26 | -1.0266 | 0.8477 | EXOC6B | 0.8477 | 0.84017 | 0.81765 |
| ENSG00000189298 | 6 | 28317691 | 28336947 | 79 | 15 | -1.0266 | 0.8477 | ZKSCAN3 | 0.8477 | 0.89244 | 0.92056 |
| ENSG00000121390 | 13 | 20248896 | 20357142 | 273 | 19 | -1.0274 | 0.84787 | PSPC1 | 0.84787 | 0.83743 | 0.82686 |
| ENSG00000204347 | 17 | 72352555 | 72358085 | 23 | 5 | -1.0277 | 0.84794 | BTBD17 | 0.84794 | 0.84696 | 0.91341 |
| ENSG00000175600 | 7 | 40174575 | 40900362 | 1528 | 79 | -1.0277 | 0.84796 | SUGCT | 0.84796 | 0.85807 | 0.86512 |
| ENSG00000176971 | 11 | 27015628 | 27018630 | 5 | 3 | -1.0278 | 0.84799 | FIBIN | 0.84799 | 0.71408 | 0.74577 |
| ENSG00000188739 | 1 | 235294498 | 235324772 | 78 | 10 | -1.0279 | 0.84801 | RBM34 | 0.84801 | 0.88291 | 0.8753 |
| ENSG00000017797 | 18 | 9475007 | 9538114 | 241 | 27 | -1.028 | 0.84803 | RALBP1 | 0.84803 | 0.78443 | 0.73685 |
| ENSG00000171132 | 2 | 45878484 | 46415129 | 2240 | 202 | -1.0283 | 0.8481 | PRKCE | 0.8481 | 0.85319 | 0.85147 |
| ENSG00000005486 | 7 | 75471920 | 75518244 | 124 | 21 | -1.0286 | 0.84816 | RHBDD2 | 0.84816 | 0.88958 | 0.9572 |
| ENSG00000138814 | 4 | 101944566 | 102269435 | 838 | 45 | -1.0287 | 0.84819 | PPP3CA | 0.84819 | 0.86093 | 0.86366 |
| ENSG00000250305 | 8 | 12803151 | 12889012 | 598 | 36 | -1.0293 | 0.84832 | KIAA1456 | 0.84832 | 0.81544 | 0.85757 |
| ENSG00000072736 | 16 | 68118654 | 68263162 | 299 | 12 | -1.0295 | 0.84838 | NFATC3 | 0.84838 | 0.85011 | 0.86116 |
| ENSG00000148482 | 10 | 18240768 | 18332221 | 398 | 45 | -1.0298 | 0.84844 | SLC39A12 | 0.84844 | 0.8239 | 0.78908 |
| ENSG00000167748 | 19 | 51322404 | 51327043 | 21 | 5 | -1.0299 | 0.84848 | KLK1 | 0.84848 | 0.85413 | 0.82305 |
| ENSG00000143183 | 1 | 165696032 | 165796992 | 264 | 31 | -1.03 | 0.8485 | TMCO1 | 0.8485 | 0.82475 | 0.69326 |
| ENSG00000179403 | 1 | 1370241 | 1378262 | 21 | 5 | -1.0301 | 0.84853 | VWA1 | 0.84853 | 0.80237 | 0.89332 |
| ENSG00000155729 | 2 | 201353675 | 201384507 | 92 | 21 | -1.0303 | 0.84857 | KCTD18 | 0.84857 | 0.72139 | 0.78004 |
| ENSG00000196979 | 9 | 132902889 | 132906498 | 18 | 4 | -1.0304 | 0.84858 | AL360004.1 | 0.84858 | 0.90491 | 0.35558 |
| ENSG00000125257 | 13 | 95672083 | 95953687 | 1386 | 98 | -1.0306 | 0.84863 | ABCC4 | 0.84863 | 0.86766 | 0.82799 |
| ENSG00000112033 | 6 | 35310335 | 35395968 | 262 | 25 | -1.0309 | 0.8487 | PPARD | 0.8487 | 0.83085 | 0.77934 |
| ENSG00000120068 | 17 | 46688739 | 46692478 | 15 | 7 | -1.0309 | 0.8487 | HOXB8 | 0.8487 | 0.81947 | 0.86149 |
| ENSG00000124570 | 6 | 2948393 | 2972090 | 113 | 16 | -1.031 | 0.84872 | SERPINB6 | 0.84872 | 0.86804 | 0.82052 |
| ENSG00000268519 | 12 | 77949297 | 77950568 | 5 | 1 | -1.031 | 0.84874 | AC073528.1 | 0.84874 | 0.80407 | 0.8571 |
| ENSG00000162736 | 1 | 160313062 | 160328742 | 33 | 13 | -1.0315 | 0.84884 | NCSTN | 0.84884 | 0.79061 | 0.85521 |
| ENSG00000164853 | 7 | 1272543 | 1276954 | 5 | 2 | -1.0315 | 0.84884 | UNCX | 0.84884 | 0.79907 | 0.81652 |
| ENSG00000213619 | 11 | 47586888 | 47606114 | 24 | 10 | -1.0323 | 0.84904 | NDUFS3 | 0.84904 | 0.54475 | 0.73586 |
| ENSG00000104549 | 8 | 126010739 | 126034525 | 106 | 11 | -1.0325 | 0.84908 | SQLE | 0.84908 | 0.87636 | 0.85291 |
| ENSG00000177398 | 21 | 43483068 | 43563563 | 494 | 63 | -1.0326 | 0.8491 | UMODL1 | 0.8491 | 0.89069 | 0.92981 |
| ENSG00000197616 | 14 | 23851199 | 23877486 | 80 | 21 | -1.033 | 0.84919 | MYH6 | 0.84919 | 0.77943 | 0.76318 |
| ENSG00000259112 | 11 | 77728017 | 77790911 | 272 | 15 | -1.0332 | 0.84925 | NDUFC2-KCTD14 | 0.84925 | 0.79277 | 0.67503 |
| ENSG00000186965 | 21 | 31859362 | 31859755 | 2 | 1 | -1.0332 | 0.84925 | KRTAP19-2 | 0.84925 | 0.42954 | 0.46883 |
| ENSG00000224531 | 6 | 11094266 | 11138964 | 191 | 17 | -1.0334 | 0.84928 | SMIM13 | 0.84928 | 0.90037 | 0.96383 |
| ENSG00000197019 | 19 | 40927499 | 40931932 | 9 | 3 | -1.0333 | 0.84928 | SERTAD1 | 0.84928 | 0.33558 | 0.37438 |
| ENSG00000185432 | 12 | 51317255 | 51326300 | 22 | 8 | -1.034 | 0.84943 | METTL7A | 0.84943 | 0.57618 | 0.65945 |
| ENSG00000159261 | 21 | 37832919 | 37948867 | 537 | 41 | -1.0341 | 0.84947 | CLDN14 | 0.84947 | 0.86854 | 0.9265 |
| ENSG00000117868 | 7 | 158523686 | 158622944 | 507 | 18 | -1.0343 | 0.8495 | ESYT2 | 0.8495 | 0.85746 | 0.87401 |
| ENSG00000237693 | 5 | 150226085 | 150280298 | 230 | 7 | -1.0346 | 0.84957 | IRGM | 0.84957 | 0.83629 | 0.82352 |
| ENSG00000124226 | 20 | 48552948 | 48570429 | 51 | 12 | -1.0346 | 0.84957 | RNF114 | 0.84957 | 0.76191 | 0.8123 |
| ENSG00000105141 | 19 | 15160195 | 15169104 | 45 | 7 | -1.0351 | 0.84969 | CASP14 | 0.84969 | 0.89465 | 0.71447 |
| ENSG00000215262 | 8 | 36641842 | 36793646 | 304 | 17 | -1.0352 | 0.8497 | KCNU1 | 0.8497 | 0.89439 | 0.89111 |
| ENSG00000214940 | 16 | 18411799 | 18441131 | 1 | 1 | -1.0351 | 0.8497 | NPIPA8 | 0.8497 | 0.8497 | 0.8497 |
| ENSG00000169397 | 14 | 21359558 | 21360507 | 3 | 2 | -1.0355 | 0.84977 | RNASE3 | 0.84977 | 0.78874 | 0.54663 |
| ENSG00000140678 | 16 | 31366455 | 31394318 | 66 | 19 | -1.0355 | 0.84978 | ITGAX | 0.84978 | 0.79069 | 0.84636 |
| ENSG00000114316 | 3 | 49315264 | 49378145 | 106 | 17 | -1.0355 | 0.84979 | USP4 | 0.84979 | 0.87314 | 0.86165 |
| ENSG00000196458 | 12 | 133498019 | 133532892 | 45 | 9 | -1.0359 | 0.84988 | ZNF605 | 0.84988 | 0.74505 | 0.62326 |
| ENSG00000175785 | 14 | 94184644 | 94254827 | 250 | 59 | -1.036 | 0.8499 | PRIMA1 | 0.8499 | 0.81871 | 0.79129 |
| ENSG00000172053 | 3 | 49133365 | 49142553 | 11 | 4 | -1.0362 | 0.84994 | QARS | 0.84994 | 0.56618 | 0.69973 |
| ENSG00000166317 | 10 | 75404639 | 75423561 | 36 | 5 | -1.0363 | 0.84998 | SYNPO2L | 0.84998 | 0.82394 | 0.89614 |
| ENSG00000074695 | 18 | 56995055 | 57027194 | 94 | 11 | -1.0364 | 0.85 | LMAN1 | 0.85 | 0.89173 | 0.87994 |
| ENSG00000231068 | 21 | 32090843 | 32091095 | 1 | 1 | -1.0364 | 0.85 | KRTAP21-3 | 0.85 | 0.75593 | 0.77636 |
| ENSG00000131686 | 1 | 9005926 | 9035151 | 134 | 30 | -1.0366 | 0.85003 | CA6 | 0.85003 | 0.94832 | 0.94735 |
| ENSG00000135298 | 6 | 69345259 | 70099403 | 2379 | 74 | -1.0366 | 0.85004 | BAI3 | 0.85004 | 0.85753 | 0.86926 |
| ENSG00000167380 | 19 | 44669226 | 44682534 | 15 | 5 | -1.0366 | 0.85004 | ZNF226 | 0.85004 | 0.74995 | 0.77842 |
| ENSG00000214050 | 8 | 28205804 | 28347835 | 314 | 50 | -1.0367 | 0.85006 | FBXO16 | 0.85006 | 0.8715 | 0.9149 |
| ENSG00000153291 | 6 | 46620678 | 46645930 | 42 | 8 | -1.0369 | 0.85011 | SLC25A27 | 0.85011 | 0.83687 | 0.94256 |
| ENSG00000107731 | 10 | 72972327 | 73062621 | 378 | 48 | -1.0371 | 0.85015 | UNC5B | 0.85015 | 0.8767 | 0.95108 |
| ENSG00000134058 | 5 | 68530668 | 68573250 | 148 | 15 | -1.0371 | 0.85016 | CDK7 | 0.85016 | 0.85349 | 0.81026 |
| ENSG00000124467 | 19 | 43256838 | 43359843 | 554 | 23 | -1.0371 | 0.85016 | PSG8 | 0.85016 | 0.85299 | 0.83817 |
| ENSG00000205853 | 22 | 32755893 | 32778039 | 140 | 19 | -1.0373 | 0.8502 | RFPL3S | 0.8502 | 0.84836 | 0.93364 |
| ENSG00000196533 | 1 | 206238872 | 206288647 | 95 | 12 | -1.0375 | 0.85026 | C1orf186 | 0.85026 | 0.49703 | 0.25273 |

| | | | | | | | | | | | |
|---|---|---|---|---|---|---|---|---|---|---|---|
| ENSG00000107882 | 10 | 104263744 | 104393292 | 358 | 17 | -1.0375 | 0.85026 | SUFU | 0.85026 | 0.81846 | 0.83055 |
| ENSG00000177200 | 16 | 53088945 | 53363062 | 787 | 22 | -1.0378 | 0.85032 | CHD9 | 0.85032 | 0.85422 | 0.80341 |
| ENSG00000144182 | 2 | 99771418 | 99779620 | 30 | 6 | -1.0388 | 0.85055 | LIPT1 | 0.85055 | 0.82341 | 0.82807 |
| ENSG00000184992 | 12 | 125478246 | 125515777 | 241 | 25 | -1.039 | 0.8506 | BRI3BP | 0.8506 | 0.79304 | 0.82092 |
| ENSG00000129450 | 19 | 51628165 | 51639908 | 24 | 10 | -1.039 | 0.8506 | SIGLEC9 | 0.8506 | 0.44495 | 0.6996 |
| ENSG00000031003 | 5 | 137273649 | 137387650 | 306 | 18 | -1.0393 | 0.85067 | FAM13B | 0.85067 | 0.84125 | 0.82652 |
| ENSG00000121853 | 3 | 172162923 | 172166246 | 9 | 3 | -1.0394 | 0.85068 | GHSR | 0.85068 | 0.67627 | 0.41291 |
| ENSG00000124507 | 6 | 34433916 | 34503006 | 191 | 33 | -1.0398 | 0.85078 | PACSIN1 | 0.85078 | 0.87157 | 0.93364 |
| ENSG00000005108 | 7 | 11409984 | 11871824 | 1966 | 131 | -1.0398 | 0.85079 | THSD7A | 0.85079 | 0.8859 | 0.88082 |
| ENSG00000146700 | 7 | 76018651 | 76039012 | 99 | 12 | -1.0401 | 0.85086 | SRCRB4D | 0.85086 | 0.8328 | 0.64564 |
| ENSG00000216588 | 19 | 45116940 | 45140081 | 72 | 16 | -1.0402 | 0.85088 | IGSF23 | 0.85088 | 0.73465 | 0.77597 |
| ENSG00000103549 | 16 | 30773066 | 30787628 | 17 | 7 | -1.0403 | 0.8509 | RNF40 | 0.8509 | 0.78013 | 0.74182 |
| ENSG00000145014 | 3 | 194308402 | 194354418 | 301 | 24 | -1.0408 | 0.85101 | TMEM44 | 0.85101 | 0.90714 | 0.91283 |
| ENSG00000197702 | 11 | 12398732 | 12552348 | 532 | 77 | -1.0409 | 0.85105 | PARVA | 0.85105 | 0.85075 | 0.51263 |
| ENSG00000154930 | 20 | 24986868 | 25039616 | 135 | 27 | -1.0409 | 0.85105 | ACSS1 | 0.85105 | 0.66564 | 0.74062 |
| ENSG00000019144 | 11 | 118477155 | 118528741 | 100 | 15 | -1.041 | 0.85107 | PHLDB1 | 0.85107 | 0.76283 | 0.64356 |
| ENSG00000105612 | 19 | 12986025 | 12992282 | 10 | 3 | -1.0413 | 0.85112 | DNASE2 | 0.85112 | 0.78694 | 0.67872 |
| ENSG00000164219 | 5 | 114546527 | 114598569 | 137 | 20 | -1.0417 | 0.85123 | PGGT1B | 0.85123 | 0.82409 | 0.72382 |
| ENSG00000185608 | 22 | 19419425 | 19423598 | 15 | 4 | -1.0418 | 0.85124 | MRPL40 | 0.85124 | 0.80167 | 0.69957 |
| ENSG00000182446 | 17 | 79523913 | 79615495 | 561 | 19 | -1.042 | 0.8513 | NPLOC4 | 0.8513 | 0.83182 | 0.68175 |
| ENSG00000204897 | 17 | 38904273 | 38911584 | 18 | 6 | -1.0424 | 0.85138 | KRT25 | 0.85138 | 0.8305 | 0.86574 |
| ENSG00000205629 | 16 | 25123050 | 25189552 | 145 | 10 | -1.0425 | 0.8514 | LCMT1 | 0.8514 | 0.83764 | 0.84927 |
| ENSG00000014919 | 10 | 101471601 | 101491857 | 39 | 6 | -1.0426 | 0.85144 | COX15 | 0.85144 | 0.79225 | 0.73849 |
| ENSG00000197165 | 16 | 28603264 | 28608430 | 29 | 7 | -1.0431 | 0.85155 | SULT1A2 | 0.85155 | 0.91983 | 0.9265 |
| ENSG00000143772 | 1 | 226819391 | 226927024 | 226 | 36 | -1.0432 | 0.85156 | ITPKB | 0.85156 | 0.81134 | 0.84123 |
| ENSG00000150672 | 11 | 83166055 | 85338966 | 6733 | 200 | -1.0431 | 0.85156 | DLG2 | 0.85156 | 0.8653 | 0.86263 |
| ENSG00000176340 | 11 | 63742079 | 63744015 | 2 | 1 | -1.0433 | 0.8516 | COX8A | 0.8516 | 0.81291 | 0.84909 |
| ENSG00000101049 | 20 | 42187608 | 42216877 | 99 | 25 | -1.0434 | 0.85161 | SGK2 | 0.85161 | 0.85416 | 0.86957 |
| ENSG00000055732 | 1 | 85483765 | 85514182 | 71 | 15 | -1.0436 | 0.85168 | MCOLN3 | 0.85168 | 0.7869 | 0.71379 |
| ENSG00000132952 | 13 | 31191830 | 31233686 | 36 | 16 | -1.0437 | 0.8517 | USPL1 | 0.8517 | 0.84817 | 0.81299 |
| ENSG00000067141 | 15 | 73344051 | 73597547 | 649 | 28 | -1.0438 | 0.8517 | NEO1 | 0.8517 | 0.86097 | 0.88274 |
| ENSG00000116731 | 1 | 14026693 | 14151574 | 417 | 31 | -1.0439 | 0.85174 | PRDM2 | 0.85174 | 0.84764 | 0.70059 |
| ENSG00000196584 | 7 | 152341864 | 152373250 | 125 | 20 | -1.0439 | 0.85174 | XRCC2 | 0.85174 | 0.87479 | 0.92803 |
| ENSG00000149043 | 11 | 1848709 | 1858751 | 41 | 11 | -1.0439 | 0.85174 | SYT8 | 0.85174 | 0.65827 | 0.75302 |
| ENSG00000156959 | 3 | 9543481 | 9595486 | 241 | 23 | -1.0441 | 0.85177 | LHFPL4 | 0.85177 | 0.84632 | 0.85933 |
| ENSG00000171403 | 17 | 39722096 | 39728310 | 13 | 3 | -1.0442 | 0.85181 | KRT9 | 0.85181 | 0.90435 | 0.89578 |
| ENSG00000083844 | 19 | 57702868 | 57724724 | 54 | 13 | -1.0442 | 0.85181 | ZNF264 | 0.85181 | 0.88627 | 0.78053 |
| ENSG00000152672 | 2 | 71035775 | 71047732 | 43 | 12 | -1.0443 | 0.85182 | CLEC4F | 0.85182 | 0.49455 | 0.67732 |
| ENSG00000214239 | 6 | 159047471 | 159049322 | 17 | 4 | -1.0443 | 0.85183 | AL591025.1 | 0.85183 | 0.67938 | 0.66807 |
| ENSG00000188655 | 14 | 21024252 | 21029090 | 15 | 5 | -1.0449 | 0.85196 | RNASE9 | 0.85196 | 0.18047 | 0.062671 |
| ENSG00000267819 | 12 | 131780067 | 131782763 | 6 | 2 | -1.0456 | 0.85212 | AC092850.1 | 0.85212 | 0.6314 | 0.53193 |
| ENSG00000180543 | 8 | 98285717 | 98290176 | 13 | 5 | -1.0456 | 0.85213 | TSPYL5 | 0.85213 | 0.73401 | 0.68656 |
| ENSG00000205084 | 16 | 75572015 | 75590184 | 83 | 9 | -1.0457 | 0.85215 | TMEM231 | 0.85215 | 0.89706 | 0.91702 |
| ENSG00000158014 | 1 | 26363743 | 26372624 | 20 | 3 | -1.0466 | 0.85235 | SLC30A2 | 0.85235 | 0.8265 | 0.65851 |
| ENSG00000146592 | 7 | 28338940 | 28865511 | 1841 | 167 | -1.0466 | 0.85235 | CREB5 | 0.85235 | 0.86074 | 0.87012 |
| ENSG00000214013 | 15 | 42565431 | 42645864 | 275 | 15 | -1.0468 | 0.8524 | GANC | 0.8524 | 0.84936 | 0.85243 |
| ENSG00000109991 | 11 | 57105848 | 57138073 | 106 | 13 | -1.0469 | 0.85243 | P2RX3 | 0.85243 | 0.8386 | 0.86485 |
| ENSG00000134258 | 1 | 117686209 | 117753556 | 224 | 31 | -1.0469 | 0.85244 | VTCN1 | 0.85244 | 0.52245 | 0.43747 |
| ENSG00000157353 | 16 | 70488324 | 70514177 | 29 | 15 | -1.0471 | 0.85246 | FUK | 0.85246 | 0.51002 | 0.30308 |
| ENSG00000168904 | 15 | 99791567 | 99930934 | 499 | 33 | -1.0471 | 0.85248 | LRRC28 | 0.85248 | 0.71441 | 0.59589 |
| ENSG00000141562 | 17 | 80416056 | 80448413 | 96 | 16 | -1.0474 | 0.85254 | NARF | 0.85254 | 0.89414 | 0.83358 |
| ENSG00000140941 | 16 | 87417601 | 87438385 | 86 | 18 | -1.0475 | 0.85257 | MAP1LC3B | 0.85257 | 0.74248 | 0.84763 |
| ENSG00000164626 | 6 | 39156749 | 39197226 | 132 | 29 | -1.0479 | 0.85265 | KCNK5 | 0.85265 | 0.7072 | 0.3315 |
| ENSG00000166321 | 10 | 74870217 | 74891586 | 60 | 3 | -1.048 | 0.85268 | NUDT13 | 0.85268 | 0.84189 | 0.83933 |
| ENSG00000115504 | 2 | 62900986 | 63273622 | 484 | 40 | -1.0481 | 0.8527 | EHBP1 | 0.8527 | 0.84105 | 0.8221 |
| ENSG00000103995 | 15 | 49005125 | 49103343 | 191 | 23 | -1.0481 | 0.8527 | CEP152 | 0.8527 | 0.88001 | 0.85441 |
| ENSG00000075188 | 12 | 102467967 | 102513902 | 118 | 17 | -1.0486 | 0.85281 | NUP37 | 0.85281 | 0.83451 | 0.86081 |
| ENSG00000145476 | 4 | 187112674 | 187134610 | 108 | 14 | -1.0486 | 0.85283 | CYP4V2 | 0.85283 | 0.88827 | 0.9089 |
| ENSG00000109917 | 11 | 116648436 | 116658766 | 30 | 9 | -1.049 | 0.85292 | ZNF259 | 0.85292 | 0.88426 | 0.90928 |
| ENSG00000182613 | 5 | 180581943 | 180582890 | 3 | 2 | -1.0493 | 0.85299 | OR2V2 | 0.85299 | 0.47366 | 0.56391 |
| ENSG00000085760 | 2 | 55463731 | 55496483 | 120 | 17 | -1.0495 | 0.85302 | MTIF2 | 0.85302 | 0.88423 | 0.92334 |
| ENSG00000153933 | 17 | 54911460 | 54946036 | 101 | 7 | -1.0497 | 0.85307 | DGKE | 0.85307 | 0.80802 | 0.838 |
| ENSG00000038945 | 8 | 15965387 | 16424999 | 1872 | 67 | -1.0499 | 0.85311 | MSR1 | 0.85311 | 0.88613 | 0.88975 |
| ENSG00000178307 | 17 | 21100593 | 21117937 | 43 | 10 | -1.0501 | 0.85317 | TMEM11 | 0.85317 | 0.92069 | 0.92733 |
| ENSG00000168175 | 14 | 55518349 | 55536910 | 37 | 9 | -1.0502 | 0.85319 | MAPK1IP1L | 0.85319 | 0.72342 | 0.72283 |
| ENSG00000108602 | 17 | 19641297 | 19652256 | 19 | 6 | -1.0502 | 0.85319 | ALDH3A1 | 0.85319 | 0.90366 | 0.95052 |
| ENSG00000117395 | 1 | 43629846 | 43736607 | 464 | 36 | -1.0505 | 0.85326 | EBNA1BP2 | 0.85326 | 0.83199 | 0.8277 |
| ENSG00000236334 | 1 | 143767144 | 143767881 | 1 | 1 | -1.0507 | 0.8533 | PPIAL4G | 0.8533 | 0.53384 | 0.41399 |
| ENSG00000051825 | 12 | 123636867 | 123728561 | 332 | 14 | -1.051 | 0.85338 | MPHOSPH9 | 0.85338 | 0.8602 | 0.87446 |
| ENSG00000136936 | 9 | 100437191 | 100459639 | 25 | 10 | -1.0516 | 0.8535 | XPA | 0.8535 | 0.88328 | 0.85153 |
| ENSG00000197279 | 6 | 28048753 | 28057341 | 23 | 5 | -1.0518 | 0.85356 | ZNF165 | 0.85356 | 0.83922 | 0.83952 |
| ENSG00000156475 | 5 | 145967936 | 146464347 | 1341 | 64 | -1.0525 | 0.85371 | PPP2R2B | 0.85371 | 0.87727 | 0.89131 |
| ENSG00000163075 | 2 | 120302008 | 120419827 | 211 | 22 | -1.0529 | 0.85381 | PCDP1 | 0.85381 | 0.80651 | 0.77655 |

| | | | | | | | | | | | |
|---|---|---|---|---|---|---|---|---|---|---|---|
| ENSG00000163947 | 3 | 56761446 | 57113357 | 1221 | 108 | -1.053 | 0.85384 | ARHGEF3 | 0.85384 | 0.88537 | 0.88627 |
| ENSG00000113391 | 5 | 92953775 | 93447404 | 792 | 24 | -1.0533 | 0.85389 | FAM172A | 0.85389 | 0.82191 | 0.81383 |
| ENSG00000134352 | 5 | 55230923 | 55290821 | 125 | 17 | -1.0533 | 0.85391 | IL6ST | 0.85391 | 0.88622 | 0.69618 |
| ENSG00000124157 | 20 | 43849941 | 43853099 | 10 | 2 | -1.0535 | 0.85395 | SEMG2 | 0.85395 | 0.62468 | 0.63863 |
| ENSG00000171119 | 19 | 5823813 | 5828335 | 13 | 5 | -1.0537 | 0.854 | NRTN | 0.854 | 0.92911 | 0.83164 |
| ENSG00000137656 | 11 | 116618886 | 116643704 | 105 | 20 | -1.0542 | 0.85409 | BUD13 | 0.85409 | 0.87342 | 0.86795 |
| ENSG00000167634 | 19 | 55434877 | 55477680 | 237 | 48 | -1.0544 | 0.85416 | NLRP7 | 0.85416 | 0.93927 | 0.98485 |
| ENSG00000178927 | 17 | 80400465 | 80408705 | 33 | 10 | -1.0545 | 0.85417 | C17orf62 | 0.85417 | 0.89341 | 0.88685 |
| ENSG00000099326 | 19 | 59073298 | 59084942 | 23 | 5 | -1.0547 | 0.85421 | MZF1 | 0.85421 | 0.90653 | 0.93187 |
| ENSG00000179104 | 12 | 83080659 | 83528649 | 1468 | 97 | -1.0549 | 0.85427 | TMTC2 | 0.85427 | 0.76235 | 0.80332 |
| ENSG00000170848 | 19 | 43406231 | 43423715 | 85 | 9 | -1.0549 | 0.85427 | PSG6 | 0.85427 | 0.85315 | 0.86351 |
| ENSG00000144481 | 2 | 234826043 | 234928166 | 366 | 58 | -1.0551 | 0.85432 | TRPM8 | 0.85432 | 0.65288 | 0.64748 |
| ENSG00000139155 | 12 | 20848289 | 20906320 | 277 | 33 | -1.0552 | 0.85433 | SLCO1C1 | 0.85433 | 0.84118 | 0.81087 |
| ENSG00000103363 | 16 | 2821415 | 2827298 | 24 | 3 | -1.0559 | 0.85449 | TCEB2 | 0.85449 | 0.9621 | 0.94427 |
| ENSG00000268265 | 21 | 34947431 | 34947645 | 1 | 1 | -1.0559 | 0.8545 | AP000304.1 | 0.8545 | 0.26821 | 0.32568 |
| ENSG00000141255 | 17 | 3343313 | 3417146 | 350 | 33 | -1.056 | 0.85452 | SPATA22 | 0.85452 | 0.89916 | 0.93642 |
| ENSG00000172062 | 5 | 70220768 | 70249769 | 5 | 3 | -1.0561 | 0.85454 | SMN1 | 0.85454 | 0.89183 | 0.93901 |
| ENSG00000175745 | 5 | 92919043 | 92930321 | 6 | 4 | -1.0562 | 0.85457 | NR2F1 | 0.85457 | 0.4025 | 0.61476 |
| ENSG00000104142 | 15 | 41186628 | 41196173 | 30 | 5 | -1.0568 | 0.8547 | VPS18 | 0.8547 | 0.68793 | 0.68629 |
| ENSG00000160183 | 21 | 43791999 | 43816955 | 138 | 34 | -1.0575 | 0.85485 | TMPRSS3 | 0.85485 | 0.85098 | 0.35106 |
| ENSG00000133313 | 18 | 72163051 | 72188366 | 115 | 18 | -1.0576 | 0.85488 | CNDP2 | 0.85488 | 0.91216 | 0.90186 |
| ENSG00000047849 | 3 | 47892182 | 48130769 | 333 | 18 | -1.0582 | 0.85502 | MAP4 | 0.85502 | 0.82302 | 0.80537 |
| ENSG00000159208 | 1 | 150254953 | 150259505 | 11 | 3 | -1.0585 | 0.85508 | C1orf51 | 0.85508 | 0.90531 | 0.89175 |
| ENSG00000235978 | 3 | 4855978 | 4928977 | 301 | 30 | -1.0588 | 0.85515 | AC018816.3 | 0.85515 | 0.84794 | 0.56067 |
| ENSG00000105486 | 19 | 48618702 | 48673860 | 268 | 15 | -1.0593 | 0.85526 | LIG1 | 0.85526 | 0.85972 | 0.9152 |
| ENSG00000133116 | 13 | 33590207 | 33640282 | 208 | 16 | -1.0594 | 0.85529 | KL | 0.85529 | 0.85091 | 0.89212 |
| ENSG00000100626 | 14 | 69725994 | 69821183 | 389 | 45 | -1.0595 | 0.85531 | GALNT16 | 0.85531 | 0.90364 | 0.87344 |
| ENSG00000198657 | 11 | 124293807 | 124294789 | 5 | 2 | -1.0604 | 0.85553 | OR8B4 | 0.85553 | 0.73906 | 0.57626 |
| ENSG00000165030 | 9 | 94171327 | 94186144 | 31 | 9 | -1.0606 | 0.85555 | NFIL3 | 0.85555 | 0.72456 | 0.79778 |
| ENSG00000134146 | 15 | 35509546 | 35838394 | 843 | 39 | -1.0606 | 0.85555 | DPH6 | 0.85555 | 0.86203 | 0.87912 |
| ENSG00000184277 | 15 | 102161847 | 102192594 | 72 | 18 | -1.0605 | 0.85555 | TM2D3 | 0.85555 | 0.7056 | 0.42199 |
| ENSG00000136810 | 9 | 113006091 | 113018920 | 51 | 18 | -1.0606 | 0.85557 | TXN | 0.85557 | 0.93797 | 0.9724 |
| ENSG00000043143 | 5 | 133860003 | 133918918 | 196 | 33 | -1.0607 | 0.85559 | JADE2 | 0.85559 | 0.78124 | 0.69537 |
| ENSG00000148334 | 9 | 130882972 | 130890741 | 12 | 6 | -1.0609 | 0.85563 | PTGES2 | 0.85563 | 0.82837 | 0.82244 |
| ENSG00000072571 | 5 | 162887209 | 162918947 | 79 | 12 | -1.061 | 0.85564 | HMMR | 0.85564 | 0.8514 | 0.83248 |
| ENSG00000125695 | 17 | 61780205 | 61829457 | 132 | 9 | -1.061 | 0.85566 | RP11-51F16.8 | 0.85566 | 0.88261 | 0.87711 |
| ENSG00000168575 | 8 | 42273993 | 42397069 | 250 | 34 | -1.0615 | 0.85576 | SLC20A2 | 0.85576 | 0.86336 | 0.86759 |
| ENSG00000182899 | 3 | 197676858 | 197683481 | 17 | 3 | -1.0622 | 0.85594 | RPL35A | 0.85594 | 0.73303 | 0.80893 |
| ENSG00000013297 | 3 | 170136653 | 170578169 | 1416 | 81 | -1.0624 | 0.85597 | CLDN11 | 0.85597 | 0.86378 | 0.87579 |
| ENSG00000164105 | 4 | 174291120 | 174298683 | 9 | 2 | -1.0624 | 0.85597 | SAP30 | 0.85597 | 0.72552 | 0.72825 |
| ENSG00000137413 | 6 | 42018251 | 42055199 | 171 | 32 | -1.0624 | 0.85598 | TAF8 | 0.85598 | 0.92825 | 0.98741 |
| ENSG00000105538 | 19 | 49223844 | 49243978 | 47 | 7 | -1.0625 | 0.85598 | RASIP1 | 0.85598 | 0.82106 | 0.87746 |
| ENSG00000172986 | 3 | 72937224 | 73047289 | 422 | 30 | -1.0625 | 0.856 | GXYLT2 | 0.856 | 0.7404 | 0.62977 |
| ENSG00000256235 | 5 | 150157508 | 150176296 | 69 | 7 | -1.0625 | 0.856 | SMIM3 | 0.856 | 0.87023 | 0.88213 |
| ENSG00000165409 | 14 | 81421333 | 81612646 | 872 | 54 | -1.0625 | 0.85601 | TSHR | 0.85601 | 0.80175 | 0.66146 |
| ENSG00000005483 | 7 | 104654626 | 104754808 | 164 | 12 | -1.0627 | 0.85604 | KMT2E | 0.85604 | 0.86019 | 0.83108 |
| ENSG00000122882 | 10 | 74889913 | 74928813 | 124 | 3 | -1.0628 | 0.85607 | ECD | 0.85607 | 0.84825 | 0.84604 |
| ENSG00000073350 | 17 | 73521161 | 73571289 | 271 | 25 | -1.0629 | 0.85608 | LLGL2 | 0.85608 | 0.86546 | 0.91585 |
| ENSG00000269636 | 5 | 150157867 | 150176297 | 68 | 7 | -1.0629 | 0.85609 | AC010441.1 | 0.85609 | 0.86963 | 0.88213 |
| ENSG00000146963 | 7 | 139025105 | 139108198 | 332 | 7 | -1.063 | 0.8561 | C7orf55-LUC7L2 | 0.8561 | 0.8605 | 0.87923 |
| ENSG00000053438 | 20 | 36149617 | 36152092 | 1 | 1 | -1.0634 | 0.8562 | NNAT | 0.8562 | 0.18579 | 0.20398 |
| ENSG00000143374 | 1 | 150459887 | 150480078 | 75 | 10 | -1.0638 | 0.85629 | TARS2 | 0.85629 | 0.81939 | 0.82017 |
| ENSG00000150628 | 4 | 177105789 | 177116822 | 39 | 11 | -1.0639 | 0.85632 | SPATA4 | 0.85632 | 0.76064 | 0.66809 |
| ENSG00000115091 | 2 | 114647537 | 114720173 | 292 | 17 | -1.064 | 0.85633 | ACTR3 | 0.85633 | 0.86587 | 0.71873 |
| ENSG00000137177 | 6 | 17759414 | 17987854 | 820 | 32 | -1.0641 | 0.85636 | KIF13A | 0.85636 | 0.86463 | 0.88787 |
| ENSG00000137404 | 6 | 30655824 | 30659197 | 13 | 5 | -1.0642 | 0.85638 | NRM | 0.85638 | 0.67277 | 0.69603 |
| ENSG00000156265 | 21 | 30449792 | 30548210 | 362 | 20 | -1.0642 | 0.85638 | MAP3K7CL | 0.85638 | 0.8699 | 0.89311 |
| ENSG00000178928 | 19 | 48304500 | 48322308 | 86 | 18 | -1.0647 | 0.85649 | TPRX1 | 0.85649 | 0.90932 | 0.9764 |
| ENSG00000224153 | 3 | 184119325 | 184157436 | 206 | 31 | -1.0647 | 0.8565 | RP11-433C9.2 | 0.8565 | 0.87082 | 0.57026 |
| ENSG00000163291 | 4 | 79808281 | 79860592 | 170 | 8 | -1.0649 | 0.85654 | PAQR3 | 0.85654 | 0.83622 | 0.81485 |
| ENSG00000154839 | 18 | 47901365 | 47920543 | 72 | 7 | -1.0651 | 0.85658 | SKA1 | 0.85658 | 0.86232 | 0.80607 |
| ENSG00000014123 | 6 | 96969471 | 97003152 | 55 | 10 | -1.0652 | 0.8566 | UFL1 | 0.8566 | 0.80149 | 0.73839 |
| ENSG00000115977 | 2 | 69688532 | 69901481 | 441 | 32 | -1.0653 | 0.85662 | AAK1 | 0.85662 | 0.82802 | 0.71962 |
| ENSG00000175348 | 11 | 8968841 | 8986558 | 62 | 8 | -1.0655 | 0.85668 | TMEM9B | 0.85668 | 0.93126 | 0.92657 |
| ENSG00000159761 | 16 | 67700719 | 67702661 | 1 | 1 | -1.0661 | 0.8568 | C16orf86 | 0.8568 | 0.95746 | 0.92427 |
| ENSG00000151502 | 11 | 134094539 | 134117686 | 47 | 12 | -1.0663 | 0.85686 | VPS26B | 0.85686 | 0.88017 | 0.9113 |
| ENSG00000179134 | 19 | 39833036 | 39876350 | 112 | 13 | -1.0664 | 0.85689 | SAMD4B | 0.85689 | 0.84481 | 0.7819 |
| ENSG00000141429 | 18 | 33161081 | 33291798 | 449 | 18 | -1.067 | 0.85702 | GALNT1 | 0.85702 | 0.8551 | 0.83353 |
| ENSG00000146733 | 7 | 56078744 | 56119297 | 132 | 19 | -1.0671 | 0.85704 | PSPH | 0.85704 | 0.84192 | 0.84766 |
| ENSG00000100365 | 22 | 37257030 | 37274057 | 79 | 15 | -1.0672 | 0.85705 | NCF4 | 0.85705 | 0.74312 | 0.78976 |
| ENSG00000221986 | 1 | 109834987 | 109849663 | 40 | 10 | -1.0674 | 0.8571 | MYBPHL | 0.8571 | 0.90012 | 0.94687 |
| ENSG00000127955 | 7 | 79763271 | 79848718 | 238 | 34 | -1.0676 | 0.85716 | GNAI1 | 0.85716 | 0.82427 | 0.91775 |
| ENSG00000117245 | 1 | 20990507 | 21044510 | 261 | 44 | -1.0677 | 0.85717 | KIF17 | 0.85717 | 0.91695 | 0.9239 |

| | | | | | | | | | | | |
|---|---|---|---|---|---|---|---|---|---|---|---|
| ENSG00000146910 | 7 | 155266901 | 155326557 | 224 | 56 | -1.0677 | 0.85717 | CNPY1 | 0.85717 | 0.8997 | 0.94427 |
| ENSG00000153165 | 2 | 107021446 | 107084832 | 120 | 27 | -1.0678 | 0.8572 | RGPD3 | 0.8572 | 0.90502 | 0.86529 |
| ENSG00000204848 | 9 | 39884975 | 39891210 | 1 | 1 | -1.0678 | 0.8572 | SPATA31A2 | 0.8572 | 0.8572 | 0.8572 |
| ENSG00000161610 | 17 | 40336078 | 40337470 | 1 | 1 | -1.0678 | 0.8572 | HCRT | 0.8572 | 0.39589 | 0.36904 |
| ENSG00000143195 | 1 | 166882443 | 166944719 | 116 | 21 | -1.0685 | 0.85735 | ILDR2 | 0.85735 | 0.87162 | 0.84061 |
| ENSG00000254087 | 8 | 56792372 | 56923940 | 518 | 32 | -1.0686 | 0.85738 | LYN | 0.85738 | 0.84474 | 0.86185 |
| ENSG00000230610 | 1 | 38292232 | 38293886 | 2 | 1 | -1.069 | 0.85746 | AL929472.1 | 0.85746 | 0.39576 | 0.41047 |
| ENSG00000142182 | 21 | 45666222 | 45682099 | 73 | 21 | -1.069 | 0.85746 | DNMT3L | 0.85746 | 0.75341 | 0.6192 |
| ENSG00000164944 | 8 | 95499921 | 95565757 | 298 | 18 | -1.0692 | 0.85752 | KIAA1429 | 0.85752 | 0.8563 | 0.82613 |
| ENSG00000256591 | 11 | 61196692 | 61253294 | 93 | 9 | -1.0699 | 0.85766 | RP11-286N22.8 | 0.85766 | 0.84432 | 0.72808 |
| ENSG00000170340 | 2 | 62423248 | 62451866 | 55 | 11 | -1.0701 | 0.85771 | B3GNT2 | 0.85771 | 0.90024 | 0.92285 |
| ENSG00000137266 | 6 | 3269196 | 3457256 | 863 | 77 | -1.0703 | 0.85775 | SLC22A23 | 0.85775 | 0.84203 | 0.85305 |
| ENSG00000119772 | 2 | 25455845 | 25565459 | 220 | 45 | -1.0705 | 0.85779 | DNMT3A | 0.85779 | 0.92535 | 0.9374 |
| ENSG00000112212 | 6 | 41010293 | 41012076 | 3 | 1 | -1.0706 | 0.85781 | TSPO2 | 0.85781 | 0.83745 | 0.88341 |
| ENSG00000256632 | 12 | 133264635 | 133291498 | 106 | 14 | -1.0705 | 0.85781 | RP13-672B3.2 | 0.85781 | 0.77535 | 0.77911 |
| ENSG00000145860 | 5 | 158584417 | 158637061 | 118 | 21 | -1.071 | 0.85791 | RNF145 | 0.85791 | 0.83199 | 0.78385 |
| ENSG00000204352 | 9 | 96080481 | 96108696 | 131 | 21 | -1.071 | 0.85792 | C9orf129 | 0.85792 | 0.81874 | 0.83622 |
| ENSG00000110063 | 11 | 126173647 | 126215644 | 93 | 18 | -1.0711 | 0.85793 | DCPS | 0.85793 | 0.93645 | 0.95271 |
| ENSG00000028839 | 6 | 134273308 | 134311570 | 101 | 11 | -1.0711 | 0.85794 | TBPL1 | 0.85794 | 0.89196 | 0.79726 |
| ENSG00000165782 | 14 | 20925878 | 20929771 | 4 | 2 | -1.0712 | 0.85795 | TMEM55B | 0.85795 | 0.16299 | 0.41627 |
| ENSG00000137166 | 6 | 41514164 | 41570122 | 158 | 24 | -1.0717 | 0.85807 | FOXP4 | 0.85807 | 0.9184 | 0.91393 |
| ENSG00000083454 | 17 | 3575493 | 3599698 | 90 | 16 | -1.0718 | 0.85808 | P2RX5 | 0.85808 | 0.94012 | 0.9501 |
| ENSG00000272772 | 5 | 133502861 | 133561762 | 181 | 11 | -1.0721 | 0.85817 | CTD-2410N18.5 | 0.85817 | 0.86374 | 0.87507 |
| ENSG00000153292 | 6 | 46965440 | 47010099 | 109 | 23 | -1.0725 | 0.85824 | GPR110 | 0.85824 | 0.72729 | 0.83131 |
| ENSG00000106665 | 7 | 73703805 | 73820273 | 365 | 29 | -1.0726 | 0.85828 | CLIP2 | 0.85828 | 0.87514 | 0.92803 |
| ENSG00000143387 | 1 | 150768684 | 150780799 | 13 | 4 | -1.0734 | 0.85845 | CTSK | 0.85845 | 0.87155 | 0.8724 |
| ENSG00000131966 | 14 | 58666798 | 58701750 | 93 | 14 | -1.0734 | 0.85846 | ACTR10 | 0.85846 | 0.84937 | 0.70318 |
| ENSG00000136286 | 7 | 45002261 | 45018697 | 46 | 8 | -1.0735 | 0.85848 | MYO1G | 0.85848 | 0.90415 | 0.83751 |
| ENSG00000109519 | 4 | 7060633 | 7069924 | 48 | 8 | -1.0738 | 0.85855 | GRPEL1 | 0.85855 | 0.81143 | 0.79193 |
| ENSG00000153012 | 4 | 25000469 | 25032501 | 159 | 25 | -1.0739 | 0.85856 | LGI2 | 0.85856 | 0.87505 | 0.86379 |
| ENSG00000157554 | 21 | 39751949 | 40033704 | 1178 | 100 | -1.074 | 0.8586 | ERG | 0.8586 | 0.87806 | 0.80582 |
| ENSG00000128524 | 7 | 128502880 | 128505898 | 14 | 5 | -1.0741 | 0.85861 | ATP6V1F | 0.85861 | 0.9364 | 0.91999 |
| ENSG00000197249 | 14 | 94843084 | 94857030 | 49 | 16 | -1.0742 | 0.85864 | SERPINA1 | 0.85864 | 0.88825 | 0.95765 |
| ENSG00000113575 | 5 | 133530025 | 133561833 | 116 | 7 | -1.0744 | 0.85868 | PPP2CA | 0.85868 | 0.85225 | 0.87952 |
| ENSG00000163362 | 1 | 200860176 | 200884863 | 81 | 17 | -1.0746 | 0.85871 | C1orf106 | 0.85871 | 0.9136 | 0.93383 |
| ENSG00000196456 | 7 | 150065879 | 150109558 | 117 | 15 | -1.075 | 0.8588 | ZNF775 | 0.8588 | 0.87587 | 0.89608 |
| ENSG00000174564 | 3 | 136665072 | 136729927 | 122 | 10 | -1.0751 | 0.85884 | IL20RB | 0.85884 | 0.84257 | 0.83177 |
| ENSG00000109061 | 17 | 10395624 | 10421860 | 42 | 9 | -1.0755 | 0.85893 | MYH1 | 0.85893 | 0.88268 | 0.91511 |
| ENSG00000174437 | 12 | 110718561 | 110788898 | 188 | 15 | -1.0757 | 0.85898 | ATP2A2 | 0.85898 | 0.85069 | 0.8584 |
| ENSG00000145354 | 4 | 103790135 | 103810399 | 40 | 4 | -1.0761 | 0.85905 | CISD2 | 0.85905 | 0.84327 | 0.88708 |
| ENSG00000243452 | 1 | 148555979 | 148596267 | 6 | 1 | -1.0761 | 0.85906 | NBPF15 | 0.85906 | 0.87046 | 0.87522 |
| ENSG00000113361 | 5 | 31193857 | 31329253 | 293 | 36 | -1.0761 | 0.85906 | CDH6 | 0.85906 | 0.88774 | 0.85371 |
| ENSG00000149809 | 11 | 64879317 | 64883856 | 9 | 2 | -1.0761 | 0.85906 | TM7SF2 | 0.85906 | 0.67295 | 0.70553 |
| ENSG00000203737 | 1 | 174417212 | 174418683 | 1 | 1 | -1.0763 | 0.8591 | GPR52 | 0.8591 | 0.6792 | 0.63487 |
| ENSG00000163251 | 2 | 208627310 | 208634287 | 14 | 6 | -1.0764 | 0.85912 | FZD5 | 0.85912 | 0.8815 | 0.83352 |
| ENSG00000162493 | 1 | 13909960 | 13944452 | 140 | 34 | -1.0768 | 0.85921 | PDPN | 0.85921 | 0.73608 | 0.63807 |
| ENSG00000187889 | 1 | 57184477 | 57285369 | 369 | 56 | -1.0769 | 0.85924 | C1orf168 | 0.85924 | 0.68389 | 0.48494 |
| ENSG00000176927 | 17 | 28256218 | 28435470 | 287 | 14 | -1.0769 | 0.85924 | EFCAB5 | 0.85924 | 0.86163 | 0.84952 |
| ENSG00000169679 | 2 | 111395275 | 111435691 | 44 | 9 | -1.077 | 0.85925 | BUB1 | 0.85925 | 0.82879 | 0.74152 |
| ENSG00000197140 | 8 | 38964509 | 39142430 | 448 | 16 | -1.077 | 0.85925 | ADAM32 | 0.85925 | 0.87169 | 0.86413 |
| ENSG00000266997 | 18 | 47322163 | 47363993 | 133 | 18 | -1.0773 | 0.85932 | RP11-886H22.1 | 0.85932 | 0.84621 | 0.79632 |
| ENSG00000132406 | 4 | 4237269 | 4249950 | 123 | 6 | -1.0782 | 0.85952 | TMEM128 | 0.85952 | 0.88666 | 0.91591 |
| ENSG00000021355 | 6 | 2832566 | 2842240 | 38 | 8 | -1.0784 | 0.85957 | SERPINB1 | 0.85957 | 0.86631 | 0.91646 |
| ENSG00000137434 | 6 | 10671651 | 10695030 | 25 | 12 | -1.0785 | 0.85959 | C6orf52 | 0.85959 | 0.86275 | 0.53785 |
| ENSG00000110148 | 11 | 6280966 | 6293357 | 78 | 13 | -1.0788 | 0.85966 | CCKBR | 0.85966 | 0.93265 | 0.85312 |
| ENSG00000135823 | 1 | 180941861 | 180992047 | 112 | 13 | -1.0788 | 0.85967 | STX6 | 0.85967 | 0.90867 | 0.92426 |
| ENSG00000179397 | 1 | 244617679 | 244804479 | 404 | 45 | -1.0793 | 0.85978 | C1orf101 | 0.85978 | 0.84632 | 0.9011 |
| ENSG00000038210 | 4 | 25162263 | 25280714 | 422 | 32 | -1.0793 | 0.85978 | PI4K2B | 0.85978 | 0.85743 | 0.88393 |
| ENSG00000148702 | 10 | 115310596 | 115349361 | 206 | 35 | -1.0795 | 0.85983 | HABP2 | 0.85983 | 0.9012 | 0.91785 |
| ENSG00000134343 | 11 | 26210829 | 26684835 | 2609 | 85 | -1.0797 | 0.85985 | ANO3 | 0.85985 | 0.86296 | 0.88586 |
| ENSG00000154548 | 6 | 89805678 | 89827800 | 87 | 9 | -1.0799 | 0.8599 | SRSF12 | 0.8599 | 0.8825 | 0.89889 |
| ENSG00000180318 | 12 | 85673885 | 85695562 | 42 | 7 | -1.0799 | 0.8599 | ALX1 | 0.8599 | 0.79398 | 0.7839 |
| ENSG00000102898 | 16 | 67880635 | 67906470 | 49 | 11 | -1.0808 | 0.86011 | NUTF2 | 0.86011 | 0.81974 | 0.80084 |
| ENSG00000065526 | 1 | 16174359 | 16266955 | 194 | 23 | -1.081 | 0.86015 | SPEN | 0.86015 | 0.81632 | 0.69658 |
| ENSG00000185480 | 12 | 102513956 | 102591298 | 184 | 19 | -1.0812 | 0.8602 | PARPBP | 0.8602 | 0.87681 | 0.87396 |
| ENSG00000119048 | 5 | 133706870 | 133727683 | 42 | 5 | -1.0817 | 0.8603 | UBE2B | 0.8603 | 0.84916 | 0.84805 |
| ENSG00000187105 | 14 | 73945189 | 74025651 | 247 | 48 | -1.0818 | 0.86032 | HEATR4 | 0.86032 | 0.80475 | 0.74981 |
| ENSG00000213799 | 19 | 53837002 | 53858122 | 132 | 26 | -1.0824 | 0.86047 | ZNF845 | 0.86047 | 0.96548 | 0.96952 |
| ENSG00000109670 | 4 | 153242410 | 153457253 | 499 | 13 | -1.0825 | 0.86048 | FBXW7 | 0.86048 | 0.86858 | 0.91664 |
| ENSG00000249481 | 6 | 44310397 | 44344904 | 103 | 12 | -1.0825 | 0.86048 | SPATS1 | 0.86048 | 0.87437 | 0.87363 |
| ENSG00000067900 | 18 | 18526867 | 18691812 | 154 | 14 | -1.0825 | 0.86048 | ROCK1 | 0.86048 | 0.8347 | 0.81036 |
| ENSG00000177455 | 16 | 28943260 | 28950667 | 8 | 4 | -1.0828 | 0.86056 | CD19 | 0.86056 | 0.57083 | 0.62856 |
| ENSG00000163848 | 3 | 124944405 | 125094198 | 371 | 13 | -1.0832 | 0.86064 | ZNF148 | 0.86064 | 0.8759 | 0.87766 |

| | | | | | | | | | | | |
|---|---|---|---|---|---|---|---|---|---|---|---|
| ENSG00000141736 | 17 | 37844167 | 37886679 | 65 | 12 | -1.0833 | 0.86067 | ERBB2 | 0.86067 | 0.82226 | 0.87471 |
| ENSG00000228198 | 1 | 248366332 | 248367392 | 2 | 1 | -1.0835 | 0.86071 | OR2M3 | 0.86071 | 0.79796 | 0.86479 |
| ENSG00000011332 | 19 | 38701646 | 38720354 | 77 | 15 | -1.0835 | 0.86072 | DPF1 | 0.86072 | 0.9257 | 0.94086 |
| ENSG00000175106 | 17 | 15341205 | 15466909 | 421 | 31 | -1.0838 | 0.86078 | TVP23C | 0.86078 | 0.87947 | 0.91734 |
| ENSG00000183484 | 14 | 105515728 | 105531782 | 28 | 4 | -1.0844 | 0.86091 | GPR132 | 0.86091 | 0.7684 | 0.79583 |
| ENSG00000111058 | 12 | 81331594 | 81650533 | 1254 | 35 | -1.0845 | 0.86093 | ACSS3 | 0.86093 | 0.89569 | 0.89711 |
| ENSG00000100883 | 14 | 35451163 | 35498773 | 236 | 20 | -1.0845 | 0.86094 | SRP54 | 0.86094 | 0.84494 | 0.8563 |
| ENSG00000135702 | 16 | 75562433 | 75569145 | 23 | 5 | -1.0846 | 0.86094 | CHST5 | 0.86094 | 0.89614 | 0.9027 |
| ENSG00000139174 | 12 | 42852140 | 42984157 | 506 | 49 | -1.0846 | 0.86095 | PRICKLE1 | 0.86095 | 0.84682 | 0.87401 |
| ENSG00000132286 | 11 | 6502677 | 6505909 | 8 | 2 | -1.0848 | 0.861 | TIMM10B | 0.861 | 0.86534 | 0.87736 |
| ENSG00000003402 | 2 | 201980827 | 202041410 | 172 | 18 | -1.0849 | 0.86102 | CFLAR | 0.86102 | 0.83307 | 0.80847 |
| ENSG00000100348 | 22 | 36863083 | 36878077 | 86 | 12 | -1.085 | 0.86104 | TXN2 | 0.86104 | 0.91516 | 0.90446 |
| ENSG00000116151 | 1 | 2252692 | 2323146 | 354 | 41 | -1.0855 | 0.86116 | MORN1 | 0.86116 | 0.8611 | 0.87698 |
| ENSG00000205250 | 16 | 67226072 | 67232821 | 10 | 3 | -1.0859 | 0.86123 | E2F4 | 0.86123 | 0.76031 | 0.72721 |
| ENSG00000184979 | 22 | 18632666 | 18660164 | 130 | 14 | -1.0858 | 0.86123 | USP18 | 0.86123 | 0.88817 | 0.7896 |
| ENSG00000170959 | 11 | 30851916 | 31391357 | 873 | 26 | -1.0864 | 0.86135 | DCDC1 | 0.86135 | 0.85975 | 0.86108 |
| ENSG00000124535 | 6 | 2765648 | 2787186 | 68 | 12 | -1.0865 | 0.86138 | WRNIP1 | 0.86138 | 0.8713 | 0.9183 |
| ENSG00000116350 | 1 | 29474255 | 29508499 | 77 | 16 | -1.0866 | 0.8614 | SRSF4 | 0.8614 | 0.90941 | 0.88708 |
| ENSG00000116717 | 1 | 68150744 | 68154021 | 10 | 4 | -1.0867 | 0.86141 | GADD45A | 0.86141 | 0.56548 | 0.54602 |
| ENSG00000223474 | 1 | 26145212 | 26147288 | 7 | 3 | -1.087 | 0.86148 | AL020996.1 | 0.86148 | 0.8071 | 0.75337 |
| ENSG00000166478 | 11 | 9481866 | 9550071 | 177 | 19 | -1.087 | 0.86148 | ZNF143 | 0.86148 | 0.89926 | 0.90761 |
| ENSG00000148935 | 11 | 22647188 | 22834601 | 587 | 37 | -1.0874 | 0.86157 | GAS2 | 0.86157 | 0.8614 | 0.84774 |
| ENSG00000135269 | 7 | 115850547 | 115898837 | 195 | 14 | -1.0876 | 0.86162 | TES | 0.86162 | 0.89487 | 0.92039 |
| ENSG00000167562 | 19 | 53059075 | 53090427 | 161 | 30 | -1.0877 | 0.86163 | ZNF701 | 0.86163 | 0.87138 | 0.9042 |
| ENSG00000057657 | 6 | 106534195 | 106557814 | 54 | 15 | -1.0877 | 0.86164 | PRDM1 | 0.86164 | 0.94575 | 0.96859 |
| ENSG00000178104 | 1 | 144836157 | 145076186 | 28 | 8 | -1.088 | 0.86169 | PDE4DIP | 0.86169 | 0.86169 | 0.75463 |
| ENSG00000102543 | 13 | 49822047 | 49867618 | 100 | 7 | -1.0881 | 0.86172 | CDADC1 | 0.86172 | 0.88166 | 0.89284 |
| ENSG00000165807 | 14 | 65016620 | 65056098 | 144 | 20 | -1.0882 | 0.86174 | PPP1R36 | 0.86174 | 0.82635 | 0.76375 |
| ENSG00000088179 | 2 | 120517207 | 120741394 | 489 | 22 | -1.0882 | 0.86175 | PTPN4 | 0.86175 | 0.86618 | 0.84206 |
| ENSG00000185303 | 10 | 81315608 | 81320153 | 43 | 12 | -1.0882 | 0.86175 | SFTPA2 | 0.86175 | 0.80544 | 0.83571 |
| ENSG00000138794 | 4 | 110609785 | 110624739 | 35 | 9 | -1.0884 | 0.86179 | CASP6 | 0.86179 | 0.87586 | 0.87308 |
| ENSG00000157637 | 17 | 79218800 | 79269347 | 196 | 30 | -1.0885 | 0.86181 | SLC38A10 | 0.86181 | 0.7981 | 0.70642 |
| ENSG00000183318 | 17 | 8656424 | 8661877 | 14 | 4 | -1.0887 | 0.86185 | SPDYE4 | 0.86185 | 0.85107 | 0.75901 |
| ENSG00000127554 | 16 | 2034208 | 2037750 | 20 | 3 | -1.0892 | 0.86196 | GFER | 0.86196 | 0.86202 | 0.95233 |
| ENSG00000070985 | 11 | 2425745 | 2444275 | 91 | 17 | -1.0894 | 0.862 | TRPM5 | 0.862 | 0.76005 | 0.77639 |
| ENSG00000004864 | 7 | 95749532 | 95951459 | 310 | 14 | -1.0897 | 0.86208 | SLC25A13 | 0.86208 | 0.84712 | 0.82908 |
| ENSG00000088808 | 14 | 104200089 | 104313927 | 301 | 11 | -1.0897 | 0.86208 | PPP1R13B | 0.86208 | 0.86415 | 0.88838 |
| ENSG00000143869 | 2 | 20866424 | 20873418 | 26 | 4 | -1.0898 | 0.86211 | GDF7 | 0.86211 | 0.83136 | 0.54681 |
| ENSG00000188112 | 6 | 42068856 | 42110357 | 204 | 37 | -1.0901 | 0.86218 | C6orf132 | 0.86218 | 0.86628 | 0.91445 |
| ENSG00000198643 | 3 | 58619673 | 58652575 | 134 | 17 | -1.0904 | 0.86224 | FAM3D | 0.86224 | 0.89914 | 0.9266 |
| ENSG00000143032 | 1 | 91177096 | 91182794 | 12 | 7 | -1.0905 | 0.86225 | BARHL2 | 0.86225 | 0.89917 | 0.81587 |
| ENSG00000151689 | 2 | 191208196 | 191236391 | 125 | 10 | -1.0905 | 0.86226 | INPP1 | 0.86226 | 0.82266 | 0.68988 |
| ENSG00000255346 | 15 | 69222864 | 69355083 | 230 | 23 | -1.0907 | 0.86229 | NOX5 | 0.86229 | 0.85807 | 0.84727 |
| ENSG00000257950 | 17 | 3566357 | 3599488 | 106 | 19 | -1.0912 | 0.8624 | P2RX5-TAX1BP3 | 0.8624 | 0.91246 | 0.91804 |
| ENSG00000154975 | 17 | 49707674 | 50237377 | 1437 | 85 | -1.0913 | 0.86242 | CA10 | 0.86242 | 0.8623 | 0.86088 |
| ENSG00000132965 | 13 | 31309645 | 31338556 | 111 | 19 | -1.0913 | 0.86244 | ALOX5AP | 0.86244 | 0.88203 | 0.97602 |
| ENSG00000151025 | 10 | 25463991 | 25891155 | 2074 | 60 | -1.0916 | 0.86249 | GPR158 | 0.86249 | 0.85768 | 0.84088 |
| ENSG00000185008 | 3 | 75955846 | 77699115 | 6303 | 243 | -1.0917 | 0.86253 | ROBO2 | 0.86253 | 0.88456 | 0.90666 |
| ENSG00000197177 | 10 | 134884433 | 134945179 | 329 | 42 | -1.0919 | 0.86255 | GPR123 | 0.86255 | 0.87359 | 0.81509 |
| ENSG00000105467 | 19 | 48867652 | 48879633 | 27 | 9 | -1.0921 | 0.86261 | SYNGR4 | 0.86261 | 0.77358 | 0.68919 |
| ENSG00000171714 | 11 | 22214722 | 22304903 | 175 | 14 | -1.0928 | 0.86275 | ANO5 | 0.86275 | 0.88841 | 0.85688 |
| ENSG00000177380 | 19 | 49622646 | 49654283 | 112 | 16 | -1.0935 | 0.86292 | PPFIA3 | 0.86292 | 0.75752 | 0.53147 |
| ENSG00000170549 | 5 | 3596168 | 3601517 | 17 | 4 | -1.0936 | 0.86294 | IRX1 | 0.86294 | 0.51321 | 0.64004 |
| ENSG00000145386 | 4 | 122737599 | 122745087 | 19 | 6 | -1.0937 | 0.86296 | CCNA2 | 0.86296 | 0.73145 | 0.73859 |
| ENSG00000179922 | 19 | 56132108 | 56135967 | 6 | 2 | -1.0939 | 0.863 | ZNF784 | 0.863 | 0.44214 | 0.12859 |
| ENSG00000092607 | 1 | 119425669 | 119532179 | 323 | 28 | -1.0944 | 0.86311 | TBX15 | 0.86311 | 0.89589 | 0.91373 |
| ENSG00000205281 | 15 | 82635130 | 82641706 | 3 | 1 | -1.0945 | 0.86312 | GOLGA6L10 | 0.86312 | 0.54149 | 0.5493 |
| ENSG00000184774 | 1 | 202789394 | 202796353 | 26 | 3 | -1.0945 | 0.86314 | RP11-480I12.4 | 0.86314 | 0.90685 | 0.90994 |
| ENSG00000137221 | 6 | 43445261 | 43474294 | 69 | 9 | -1.0945 | 0.86314 | TJAP1 | 0.86314 | 0.90401 | 0.94108 |
| ENSG00000198843 | 3 | 150320662 | 150348222 | 69 | 14 | -1.0947 | 0.86318 | SELT | 0.86318 | 0.85125 | 0.88813 |
| ENSG00000124196 | 20 | 42354804 | 42355638 | 1 | 1 | -1.0948 | 0.8632 | GTSF1L | 0.8632 | 0.24799 | 0.59357 |
| ENSG00000185436 | 1 | 24480647 | 24514449 | 163 | 23 | -1.0949 | 0.86322 | IFNLR1 | 0.86322 | 0.88012 | 0.96771 |
| ENSG00000173530 | 8 | 22993101 | 23021543 | 114 | 22 | -1.095 | 0.86325 | TNFRSF10D | 0.86325 | 0.76305 | 0.76355 |
| ENSG00000077097 | 3 | 25639475 | 25706398 | 137 | 11 | -1.0952 | 0.86328 | TOP2B | 0.86328 | 0.8102 | 0.8545 |
| ENSG00000154025 | 17 | 18853658 | 18925908 | 174 | 40 | -1.0952 | 0.86329 | SLC5A10 | 0.86329 | 0.8715 | 0.83957 |
| ENSG00000152380 | 5 | 79783788 | 79838382 | 157 | 26 | -1.0954 | 0.86332 | FAM151B | 0.86332 | 0.82801 | 0.84545 |
| ENSG00000076641 | 8 | 81880045 | 82024303 | 360 | 45 | -1.0954 | 0.86333 | PAG1 | 0.86333 | 0.86074 | 0.85647 |
| ENSG00000151612 | 4 | 146678779 | 146859787 | 428 | 45 | -1.0956 | 0.86336 | ZNF827 | 0.86336 | 0.89771 | 0.34053 |
| ENSG00000185917 | 21 | 37406839 | 37451687 | 183 | 24 | -1.0956 | 0.86337 | SETD4 | 0.86337 | 0.88253 | 0.87581 |
| ENSG00000172478 | 2 | 241825465 | 241836306 | 94 | 7 | -1.0962 | 0.86351 | C2orf54 | 0.86351 | 0.81219 | 0.81104 |
| ENSG00000113645 | 5 | 167718656 | 167899308 | 786 | 89 | -1.0962 | 0.86351 | WWC1 | 0.86351 | 0.85247 | 0.8049 |
| ENSG00000142002 | 19 | 4675236 | 4724685 | 114 | 25 | -1.0965 | 0.86358 | DPP9 | 0.86358 | 0.91797 | 0.93244 |
| ENSG00000008869 | 2 | 37195526 | 37311485 | 398 | 30 | -1.0966 | 0.86359 | HEATR5B | 0.86359 | 0.84913 | 0.867 |

| | | | | | | | | | | | |
|---|---|---|---|---|---|---|---|---|---|---|---|
| ENSG00000259024 | 17 | 15339338 | 15466875 | 425 | 32 | -1.0966 | 0.8636 | TVP23C-CDRT4 | 0.8636 | 0.88723 | 0.91735 |
| ENSG00000186088 | 7 | 76940068 | 77045717 | 378 | 28 | -1.0967 | 0.86362 | GSAP | 0.86362 | 0.80111 | 0.85918 |
| ENSG00000113296 | 5 | 79287134 | 79379110 | 256 | 15 | -1.097 | 0.86368 | THBS4 | 0.86368 | 0.88008 | 0.84851 |
| ENSG00000165502 | 14 | 50085237 | 50087403 | 9 | 2 | -1.0975 | 0.86378 | RPL36AL | 0.86378 | 0.85134 | 0.7817 |
| ENSG00000236699 | 4 | 106473777 | 106629250 | 464 | 40 | -1.0978 | 0.86384 | ARHGEF38 | 0.86384 | 0.87462 | 0.80671 |
| ENSG00000120137 | 5 | 167975500 | 168006605 | 48 | 10 | -1.0978 | 0.86386 | PANK3 | 0.86386 | 0.69322 | 0.57825 |
| ENSG00000267740 | 19 | 5866182 | 5903798 | 224 | 27 | -1.0978 | 0.86386 | AC024592.12 | 0.86386 | 0.8475 | 0.80244 |
| ENSG00000112773 | 6 | 82201156 | 82462491 | 879 | 44 | -1.0981 | 0.86392 | FAM46A | 0.86392 | 0.84206 | 0.80787 |
| ENSG00000124279 | 5 | 7859272 | 7869150 | 35 | 9 | -1.0982 | 0.86394 | FASTKD3 | 0.86394 | 0.93249 | 0.87009 |
| ENSG00000111319 | 12 | 6456009 | 6486896 | 130 | 34 | -1.0989 | 0.86409 | SCNN1A | 0.86409 | 0.94744 | 0.96642 |
| ENSG00000112640 | 6 | 42952237 | 42980080 | 93 | 14 | -1.0991 | 0.86413 | PPP2R5D | 0.86413 | 0.79641 | 0.74769 |
| ENSG00000101220 | 20 | 3734155 | 3749034 | 69 | 22 | -1.0992 | 0.86416 | C20orf27 | 0.86416 | 0.63646 | 0.6142 |
| ENSG00000170923 | 19 | 9212945 | 9213982 | 1 | 1 | -1.0994 | 0.8642 | OR7G2 | 0.8642 | 0.33318 | 0.48999 |
| ENSG00000077616 | 11 | 89864683 | 89926062 | 328 | 17 | -1.0995 | 0.86423 | NAALAD2 | 0.86423 | 0.87851 | 0.87481 |
| ENSG00000167968 | 16 | 2285817 | 2288712 | 6 | 4 | -1.0995 | 0.86423 | DNASE1L2 | 0.86423 | 0.88476 | 0.78463 |
| ENSG00000175170 | 20 | 25744102 | 25848861 | 146 | 16 | -1.1 | 0.86433 | FAM182B | 0.86433 | 0.86363 | 0.88081 |
| ENSG00000122644 | 7 | 12726481 | 12730559 | 17 | 7 | -1.1 | 0.86434 | ARL4A | 0.86434 | 0.70729 | 0.44096 |
| ENSG00000151131 | 12 | 105380088 | 105443515 | 274 | 16 | -1.1001 | 0.86436 | C12orf45 | 0.86436 | 0.90808 | 0.91168 |
| ENSG00000100578 | 14 | 58894103 | 59015216 | 152 | 34 | -1.1003 | 0.8644 | KIAA0586 | 0.8644 | 0.86339 | 0.88552 |
| ENSG00000177873 | 3 | 40518604 | 40531727 | 27 | 7 | -1.1004 | 0.86443 | ZNF619 | 0.86443 | 0.61214 | 0.58197 |
| ENSG00000143458 | 1 | 151043054 | 151098018 | 154 | 20 | -1.1007 | 0.86449 | GABPB2 | 0.86449 | 0.85167 | 0.87568 |
| ENSG00000213465 | 11 | 64781585 | 64789656 | 22 | 6 | -1.1013 | 0.86461 | ARL2 | 0.86461 | 0.92764 | 0.87202 |
| ENSG00000154723 | 21 | 27088815 | 27107984 | 35 | 10 | -1.1013 | 0.86461 | ATP5J | 0.86461 | 0.88037 | 0.86609 |
| ENSG00000131652 | 16 | 3074028 | 3077756 | 10 | 4 | -1.1014 | 0.86465 | THOC6 | 0.86465 | 0.67671 | 0.77544 |
| ENSG00000188807 | 1 | 9648932 | 9674935 | 43 | 7 | -1.1021 | 0.86478 | TMEM201 | 0.86478 | 0.883 | 0.71294 |
| ENSG00000204520 | 6 | 31371356 | 31383092 | 222 | 27 | -1.1022 | 0.8648 | MICA | 0.8648 | 0.86568 | 0.81657 |
| ENSG00000153179 | 12 | 65004293 | 65091347 | 308 | 20 | -1.1023 | 0.86483 | RASSF3 | 0.86483 | 0.86422 | 0.88664 |
| ENSG00000197241 | 1 | 9063359 | 9086404 | 84 | 21 | -1.1025 | 0.86488 | SLC2A7 | 0.86488 | 0.89394 | 0.91111 |
| ENSG00000171873 | 20 | 4201329 | 4229721 | 112 | 23 | -1.1027 | 0.86492 | ADRA1D | 0.86492 | 0.8389 | 0.87786 |
| ENSG00000104435 | 8 | 80523049 | 80578410 | 190 | 25 | -1.1027 | 0.86493 | STMN2 | 0.86493 | 0.83104 | 0.85083 |
| ENSG00000092964 | 8 | 26371791 | 26515694 | 706 | 37 | -1.1028 | 0.86494 | DPYSL2 | 0.86494 | 0.88415 | 0.90959 |
| ENSG00000259431 | 14 | 24025216 | 24029480 | 3 | 1 | -1.1028 | 0.86494 | THTPA | 0.86494 | 0.56087 | 0.65724 |
| ENSG00000140488 | 15 | 72577068 | 72612470 | 77 | 9 | -1.1029 | 0.86496 | CELF6 | 0.86496 | 0.81322 | 0.85255 |
| ENSG00000119638 | 14 | 75548822 | 75594047 | 83 | 6 | -1.1029 | 0.86497 | NEK9 | 0.86497 | 0.86839 | 0.86866 |
| ENSG00000121774 | 1 | 32479430 | 32526451 | 32 | 18 | -1.1035 | 0.86509 | KHDRBS1 | 0.86509 | 0.78079 | 0.31206 |
| ENSG00000130383 | 19 | 5865837 | 5903798 | 227 | 27 | -1.1035 | 0.86509 | FUT5 | 0.86509 | 0.85728 | 0.82618 |
| ENSG00000180089 | 19 | 55738007 | 55741647 | 11 | 4 | -1.1035 | 0.8651 | TMEM86B | 0.8651 | 0.94771 | 0.97828 |
| ENSG00000100644 | 14 | 62162231 | 62214976 | 118 | 19 | -1.1037 | 0.86513 | HIF1A | 0.86513 | 0.88073 | 0.88816 |
| ENSG00000197728 | 12 | 56435637 | 56438116 | 5 | 3 | -1.1043 | 0.86527 | RPS26 | 0.86527 | 0.80985 | 0.81973 |
| ENSG00000180777 | 18 | 14728271 | 14852737 | 395 | 21 | -1.1044 | 0.86528 | ANKRD30B | 0.86528 | 0.87915 | 0.88614 |
| ENSG00000138381 | 2 | 190526111 | 190535557 | 28 | 7 | -1.1049 | 0.86539 | ASNSD1 | 0.86539 | 0.8964 | 0.90792 |
| ENSG00000145736 | 5 | 70330784 | 70363516 | 14 | 5 | -1.1049 | 0.86539 | GTF2H2 | 0.86539 | 0.87803 | 0.68983 |
| ENSG00000119977 | 10 | 97423158 | 97453900 | 94 | 18 | -1.1049 | 0.86539 | TCTN3 | 0.86539 | 0.86199 | 0.86006 |
| ENSG00000108551 | 17 | 17397751 | 17399709 | 7 | 3 | -1.1049 | 0.8654 | RASD1 | 0.8654 | 0.7924 | 0.7631 |
| ENSG00000101247 | 20 | 13765596 | 13799067 | 114 | 17 | -1.105 | 0.86543 | NDUFAF5 | 0.86543 | 0.87181 | 0.90588 |
| ENSG00000103066 | 16 | 68279207 | 68294961 | 38 | 7 | -1.1052 | 0.86546 | PLA2G15 | 0.86546 | 0.92885 | 0.8663 |
| ENSG00000163812 | 3 | 44956749 | 45017677 | 119 | 16 | -1.1053 | 0.86549 | ZDHHC3 | 0.86549 | 0.81993 | 0.65378 |
| ENSG00000090376 | 12 | 66582659 | 66648402 | 160 | 26 | -1.1055 | 0.86552 | IRAK3 | 0.86552 | 0.93214 | 0.90147 |
| ENSG00000167261 | 16 | 68021297 | 68034489 | 22 | 6 | -1.1055 | 0.86553 | DPEP2 | 0.86553 | 0.93729 | 0.92632 |
| ENSG00000156194 | 4 | 76781020 | 76823724 | 136 | 21 | -1.1056 | 0.86556 | PPEF2 | 0.86556 | 0.62589 | 0.37869 |
| ENSG00000258311 | 12 | 56109820 | 56118487 | 16 | 6 | -1.1058 | 0.8656 | RP11-644F5.10 | 0.8656 | 0.94339 | 0.94897 |
| ENSG00000250210 | 8 | 142091 | 150563 | 2 | 1 | -1.1059 | 0.86561 | RP11-585F1.10 | 0.86561 | 0.28292 | 0.39125 |
| ENSG00000132357 | 5 | 40841286 | 40860275 | 41 | 8 | -1.1063 | 0.86569 | CARD6 | 0.86569 | 0.8021 | 0.70487 |
| ENSG00000186832 | 17 | 39766030 | 39772151 | 9 | 3 | -1.1062 | 0.86569 | KRT16 | 0.86569 | 0.84078 | 0.63543 |
| ENSG00000165732 | 10 | 70715884 | 70744829 | 74 | 16 | -1.1064 | 0.86573 | DDX21 | 0.86573 | 0.85677 | 0.89574 |
| ENSG00000012061 | 19 | 45910591 | 45982086 | 266 | 30 | -1.107 | 0.86585 | ERCC1 | 0.86585 | 0.81269 | 0.8184 |
| ENSG00000180347 | 7 | 31553704 | 31698334 | 505 | 31 | -1.1071 | 0.86587 | CCDC129 | 0.86587 | 0.89736 | 0.91483 |
| ENSG00000167014 | 15 | 45248900 | 45271427 | 95 | 8 | -1.1074 | 0.86595 | C15orf43 | 0.86595 | 0.81331 | 0.61248 |
| ENSG00000181781 | 19 | 463346 | 474983 | 66 | 10 | -1.1075 | 0.86595 | ODF3L2 | 0.86595 | 0.95148 | 0.97977 |
| ENSG00000124523 | 6 | 13574816 | 13614790 | 80 | 13 | -1.1083 | 0.86613 | SIRT5 | 0.86613 | 0.83228 | 0.85273 |
| ENSG00000165480 | 13 | 21727734 | 21750741 | 119 | 18 | -1.1088 | 0.86625 | SKA3 | 0.86625 | 0.87825 | 0.87427 |
| ENSG00000136449 | 17 | 48585745 | 48608862 | 87 | 13 | -1.1088 | 0.86625 | MYCBPAP | 0.86625 | 0.80639 | 0.90688 |
| ENSG00000154237 | 15 | 101459420 | 101610317 | 582 | 44 | -1.1089 | 0.86627 | LRRK1 | 0.86627 | 0.80791 | 0.89888 |
| ENSG00000241685 | 7 | 98923521 | 98985787 | 198 | 20 | -1.1094 | 0.86637 | ARPC1A | 0.86637 | 0.87498 | 0.88083 |
| ENSG00000078403 | 10 | 21823094 | 22032559 | 223 | 31 | -1.1094 | 0.86637 | MLLT10 | 0.86637 | 0.84442 | 0.82301 |
| ENSG00000105143 | 19 | 15060846 | 15133802 | 600 | 47 | -1.1094 | 0.86637 | SLC1A6 | 0.86637 | 0.8695 | 0.91616 |
| ENSG00000169217 | 16 | 30362087 | 30366682 | 6 | 3 | -1.1095 | 0.86639 | CD2BP2 | 0.86639 | 0.49557 | 0.45153 |
| ENSG00000115053 | 2 | 232318242 | 232348352 | 114 | 15 | -1.1096 | 0.86642 | NCL | 0.86642 | 0.64568 | 0.54907 |
| ENSG00000006704 | 7 | 73868120 | 74016931 | 398 | 42 | -1.1101 | 0.86652 | GTF2IRD1 | 0.86652 | 0.79358 | 0.88361 |
| ENSG00000221864 | 21 | 46086106 | 46086844 | 9 | 3 | -1.1104 | 0.86658 | KRTAP12-2 | 0.86658 | 0.81272 | 0.82076 |
| ENSG00000139985 | 14 | 70918874 | 70926622 | 36 | 7 | -1.1104 | 0.86659 | ADAM21 | 0.86659 | 0.91959 | 0.92641 |
| ENSG00000183130 | 1 | 248789420 | 248790491 | 3 | 1 | -1.1105 | 0.8666 | OR2T11 | 0.8666 | 0.66597 | 0.28644 |
| ENSG00000103343 | 16 | 3451235 | 3459370 | 23 | 6 | -1.1105 | 0.8666 | ZNF174 | 0.8666 | 0.74539 | 0.62373 |

| | | | | | | | | | | | |
|---|---|---|---|---|---|---|---|---|---|---|---|
| ENSG00000174327 | 17 | 6939394 | 6943440 | 10 | 3 | -1.1109 | 0.8667 | SLC16A13 | 0.8667 | 0.86442 | 0.82624 |
| ENSG00000139718 | 12 | 122242086 | 122270562 | 65 | 12 | -1.1115 | 0.86681 | SETD1B | 0.86681 | 0.8349 | 0.76198 |
| ENSG00000138326 | 10 | 79793518 | 79816570 | 50 | 10 | -1.1116 | 0.86684 | RPS24 | 0.86684 | 0.87405 | 0.82756 |
| ENSG00000011600 | 19 | 36395303 | 36399197 | 23 | 6 | -1.1119 | 0.86691 | TYROBP | 0.86691 | 0.75984 | 0.48072 |
| ENSG00000134830 | 19 | 47835404 | 47846606 | 46 | 5 | -1.1124 | 0.86701 | C5AR2 | 0.86701 | 0.86152 | 0.80619 |
| ENSG00000117009 | 1 | 241695434 | 241758944 | 266 | 44 | -1.1128 | 0.8671 | KMO | 0.8671 | 0.84878 | 0.77697 |
| ENSG00000133983 | 14 | 70791798 | 70826448 | 114 | 16 | -1.1129 | 0.86712 | COX16 | 0.86712 | 0.83921 | 0.87692 |
| ENSG00000116221 | 1 | 54649714 | 54691137 | 111 | 24 | -1.113 | 0.86714 | MRPL37 | 0.86714 | 0.90046 | 0.94371 |
| ENSG00000171159 | 9 | 130922539 | 130926207 | 6 | 2 | -1.1133 | 0.86721 | C9orf16 | 0.86721 | 0.77161 | 0.71256 |
| ENSG00000182858 | 22 | 50293877 | 50312106 | 118 | 10 | -1.1134 | 0.86723 | ALG12 | 0.86723 | 0.8244 | 0.8191 |
| ENSG00000165526 | 11 | 126071993 | 126081587 | 20 | 6 | -1.1135 | 0.86725 | RPUSD4 | 0.86725 | 0.82557 | 0.88387 |
| ENSG00000064933 | 2 | 190649107 | 190742355 | 222 | 22 | -1.1139 | 0.86734 | PMS1 | 0.86734 | 0.90318 | 0.91686 |
| ENSG00000176715 | 16 | 89154783 | 89222254 | 426 | 25 | -1.1142 | 0.8674 | ACSF3 | 0.8674 | 0.92121 | 0.95528 |
| ENSG00000185274 | 7 | 70597155 | 71178585 | 2360 | 144 | -1.1145 | 0.86746 | WBSCR17 | 0.86746 | 0.88189 | 0.90136 |
| ENSG00000164109 | 4 | 120976763 | 120988229 | 39 | 8 | -1.1146 | 0.86749 | MAD2L1 | 0.86749 | 0.85463 | 0.73188 |
| ENSG00000010017 | 6 | 13621730 | 13711796 | 145 | 22 | -1.1146 | 0.86749 | RANBP9 | 0.86749 | 0.87541 | 0.88598 |
| ENSG00000112559 | 6 | 41604620 | 41621984 | 91 | 18 | -1.1147 | 0.86752 | MDFI | 0.86752 | 0.92625 | 0.97016 |
| ENSG00000162592 | 1 | 3668962 | 3688209 | 87 | 18 | -1.1149 | 0.86755 | CCDC27 | 0.86755 | 0.91568 | 0.81192 |
| ENSG00000151164 | 12 | 110939460 | 110969891 | 94 | 12 | -1.115 | 0.86756 | RAD9B | 0.86756 | 0.85835 | 0.82643 |
| ENSG00000101435 | 20 | 23545369 | 23549386 | 23 | 2 | -1.115 | 0.86758 | CST9L | 0.86758 | 0.89045 | 0.90594 |
| ENSG00000138459 | 3 | 112280556 | 112304424 | 110 | 12 | -1.1151 | 0.86759 | SLC35A5 | 0.86759 | 0.84705 | 0.84765 |
| ENSG00000116406 | 1 | 184659365 | 184724047 | 144 | 16 | -1.1157 | 0.86773 | EDEM3 | 0.86773 | 0.87693 | 0.89265 |
| ENSG00000029559 | 4 | 88720733 | 88733074 | 46 | 11 | -1.1157 | 0.86773 | IBSP | 0.86773 | 0.85709 | 0.80632 |
| ENSG00000160785 | 1 | 156163880 | 156182587 | 34 | 6 | -1.1158 | 0.86774 | SLC25A44 | 0.86774 | 0.81344 | 0.75604 |
| ENSG00000197046 | 18 | 43405477 | 43424045 | 45 | 10 | -1.1159 | 0.86776 | SIGLEC15 | 0.86776 | 0.8122 | 0.53128 |
| ENSG00000196420 | 1 | 153509623 | 153514241 | 7 | 2 | -1.1162 | 0.86783 | S100A5 | 0.86783 | 0.84453 | 0.88293 |
| ENSG00000103249 | 16 | 1494935 | 1525581 | 226 | 23 | -1.1163 | 0.86786 | CLCN7 | 0.86786 | 0.9225 | 0.95361 |
| ENSG00000065135 | 1 | 110091233 | 110136975 | 92 | 18 | -1.1167 | 0.86793 | GNAI3 | 0.86793 | 0.81021 | 0.75472 |
| ENSG00000108312 | 17 | 42282401 | 42298994 | 34 | 7 | -1.1167 | 0.86794 | UBTF | 0.86794 | 0.87061 | 0.86776 |
| ENSG00000165632 | 10 | 7860467 | 8058590 | 733 | 40 | -1.1169 | 0.86797 | TAF3 | 0.86797 | 0.87509 | 0.85174 |
| ENSG00000134901 | 13 | 103436631 | 103451357 | 44 | 6 | -1.1168 | 0.86797 | KDELC1 | 0.86797 | 0.66978 | 0.54989 |
| ENSG00000155313 | 21 | 17102344 | 17252377 | 347 | 12 | -1.1168 | 0.86797 | USP25 | 0.86797 | 0.88267 | 0.89379 |
| ENSG00000119471 | 9 | 115142217 | 115234690 | 354 | 17 | -1.1171 | 0.86802 | HSDL2 | 0.86802 | 0.86071 | 0.86071 |
| ENSG00000166342 | 18 | 70409549 | 70535381 | 572 | 43 | -1.1171 | 0.86803 | NETO1 | 0.86803 | 0.84683 | 0.85871 |
| ENSG00000172687 | 19 | 21541732 | 21562104 | 114 | 6 | -1.1172 | 0.86804 | ZNF738 | 0.86804 | 0.88278 | 0.88317 |
| ENSG00000101391 | 20 | 31946645 | 31989367 | 71 | 10 | -1.1173 | 0.86806 | CDK5RAP1 | 0.86806 | 0.88261 | 0.92822 |
| ENSG00000168038 | 3 | 41288090 | 42003922 | 2795 | 43 | -1.1174 | 0.8681 | ULK4 | 0.8681 | 0.85455 | 0.84456 |
| ENSG00000236027 | 11 | 125658006 | 125661495 | 10 | 4 | -1.118 | 0.86822 | PATE3 | 0.86822 | 0.69947 | 0.83619 |
| ENSG00000159763 | 7 | 142829170 | 142836839 | 19 | 5 | -1.1186 | 0.86834 | PIP | 0.86834 | 0.92154 | 0.86186 |
| ENSG00000104472 | 8 | 141521397 | 141527236 | 17 | 6 | -1.1187 | 0.86836 | CHRAC1 | 0.86836 | 0.71213 | 0.46292 |
| ENSG00000109062 | 17 | 72744791 | 72765492 | 34 | 9 | -1.1199 | 0.86863 | SLC9A3R1 | 0.86863 | 0.82057 | 0.78124 |
| ENSG00000176402 | 7 | 99520892 | 99527243 | 23 | 6 | -1.1201 | 0.86867 | GJC3 | 0.86867 | 0.75994 | 0.69653 |
| ENSG00000128590 | 7 | 108210012 | 108215294 | 10 | 3 | -1.1202 | 0.86868 | DNAJB9 | 0.86868 | 0.81376 | 0.77259 |
| ENSG00000120334 | 1 | 173768688 | 173793858 | 34 | 6 | -1.1204 | 0.86873 | CENPL | 0.86873 | 0.93936 | 0.93597 |
| ENSG00000158786 | 1 | 20465819 | 20476879 | 28 | 13 | -1.1208 | 0.8688 | PLA2G2F | 0.8688 | 0.90738 | 0.91115 |
| ENSG00000012124 | 19 | 35810164 | 35838258 | 122 | 35 | -1.1209 | 0.86883 | CD22 | 0.86883 | 0.83779 | 0.6616 |
| ENSG00000100949 | 14 | 24734744 | 24740945 | 13 | 3 | -1.1209 | 0.86884 | RABGGTA | 0.86884 | 0.84126 | 0.86439 |
| ENSG00000183653 | 21 | 30968360 | 31003067 | 141 | 21 | -1.1212 | 0.8689 | GRIK1-AS2 | 0.8689 | 0.9191 | 0.83129 |
| ENSG00000261427 | 16 | 19125324 | 19184900 | 172 | 32 | -1.1215 | 0.86896 | CTD-2349B8.1 | 0.86896 | 0.88548 | 0.89752 |
| ENSG00000117691 | 1 | 212606229 | 212619714 | 53 | 9 | -1.1216 | 0.86897 | NENF | 0.86897 | 0.87878 | 0.8026 |
| ENSG00000197775 | 14 | 24407940 | 24409196 | 6 | 2 | -1.1218 | 0.86903 | DHRS4-AS1 | 0.86903 | 0.78248 | 0.80755 |
| ENSG00000197576 | 7 | 27168126 | 27170418 | 10 | 2 | -1.1219 | 0.86905 | HOXA4 | 0.86905 | 0.9711 | 0.96385 |
| ENSG00000255046 | 8 | 11655437 | 11660077 | 34 | 9 | -1.1219 | 0.86905 | RP11-297N6.4 | 0.86905 | 0.72691 | 0.60487 |
| ENSG00000183530 | 22 | 32072242 | 32146126 | 208 | 22 | -1.1224 | 0.86915 | PRR14L | 0.86915 | 0.86746 | 0.89286 |
| ENSG00000198353 | 12 | 54410715 | 54449813 | 90 | 14 | -1.1224 | 0.86916 | HOXC4 | 0.86916 | 0.8847 | 0.84269 |
| ENSG00000197329 | 2 | 64319786 | 64371588 | 155 | 19 | -1.1228 | 0.86925 | PELI1 | 0.86925 | 0.88956 | 0.91971 |
| ENSG00000075461 | 17 | 64961026 | 65029514 | 244 | 43 | -1.123 | 0.86929 | CACNG4 | 0.86929 | 0.86996 | 0.75989 |
| ENSG00000152492 | 3 | 191046866 | 191116459 | 362 | 25 | -1.1231 | 0.8693 | CCDC50 | 0.8693 | 0.83039 | 0.86626 |
| ENSG00000183709 | 19 | 39759154 | 39760732 | 7 | 3 | -1.1234 | 0.86936 | IFNL2 | 0.86936 | 0.4008 | 0.12824 |
| ENSG00000173653 | 11 | 66610306 | 66614017 | 6 | 4 | -1.1235 | 0.86939 | RCE1 | 0.86939 | 0.80281 | 0.45139 |
| ENSG00000197769 | 1 | 242158792 | 242162375 | 11 | 2 | -1.1236 | 0.86941 | MAP1LC3C | 0.86941 | 0.91958 | 0.94811 |
| ENSG00000173013 | 4 | 7042579 | 7044728 | 3 | 2 | -1.1236 | 0.86941 | CCDC96 | 0.86941 | 0.69861 | 0.78717 |
| ENSG00000144504 | 2 | 241418839 | 241508626 | 365 | 47 | -1.1239 | 0.86948 | ANKMY1 | 0.86948 | 0.81868 | 0.69458 |
| ENSG00000272647 | 7 | 99156291 | 99205433 | 100 | 14 | -1.1242 | 0.86955 | GS1-259H13.10 | 0.86955 | 0.86088 | 0.87152 |
| ENSG00000162669 | 1 | 91726323 | 91870426 | 413 | 26 | -1.1245 | 0.86959 | HFM1 | 0.86959 | 0.86408 | 0.86251 |
| ENSG00000166847 | 16 | 23652713 | 23681195 | 72 | 11 | -1.1248 | 0.86966 | DCTN5 | 0.86966 | 0.90105 | 0.92375 |
| ENSG00000059769 | 9 | 114393632 | 114432508 | 171 | 22 | -1.1248 | 0.86967 | DNAJC25 | 0.86967 | 0.88215 | 0.80521 |
| ENSG00000244115 | 9 | 114393666 | 114432509 | 171 | 22 | -1.1248 | 0.86967 | DNAJC25-GNG10 | 0.86967 | 0.88215 | 0.81005 |
| ENSG00000111229 | 12 | 110872630 | 110888227 | 53 | 10 | -1.1249 | 0.86969 | ARPC3 | 0.86969 | 0.86515 | 0.83399 |
| ENSG00000105808 | 7 | 102220093 | 102257204 | 1 | 1 | -1.125 | 0.8697 | RASA4 | 0.8697 | 0.8697 | 0.8697 |
| ENSG00000205236 | 7 | 102222970 | 102283316 | 1 | 1 | -1.125 | 0.8697 | RP11-514P8.6 | 0.8697 | 0.8697 | 0.3689 |
| ENSG00000163961 | 3 | 196195654 | 196230639 | 140 | 15 | -1.125 | 0.86971 | RNF168 | 0.86971 | 0.88862 | 0.93918 |
| ENSG00000167862 | 17 | 73008765 | 73017356 | 46 | 9 | -1.125 | 0.86971 | ICT1 | 0.86971 | 0.83959 | 0.79599 |

| | | | | | | | | | | | |
|---|---|---|---|---|---|---|---|---|---|---|---|
| ENSG00000167460 | 19 | 16177831 | 16213813 | 105 | 19 | -1.1251 | 0.86973 | TPM4 | 0.86973 | 0.84477 | 0.80933 |
| ENSG00000269955 | 7 | 139026106 | 139107345 | 324 | 7 | -1.1253 | 0.86978 | LUC7L2 | 0.86978 | 0.85973 | 0.87854 |
| ENSG00000122966 | 12 | 120123595 | 120315095 | 573 | 34 | -1.1254 | 0.86978 | CIT | 0.86978 | 0.85414 | 0.87188 |
| ENSG00000267385 | 19 | 4528439 | 4540079 | 44 | 8 | -1.1255 | 0.86981 | CTB-50L17.14 | 0.86981 | 0.77326 | 0.56634 |
| ENSG00000177511 | 18 | 55018044 | 55038962 | 56 | 8 | -1.1258 | 0.86987 | ST8SIA3 | 0.86987 | 0.93153 | 0.89688 |
| ENSG00000007520 | 16 | 1399241 | 1401912 | 7 | 3 | -1.1258 | 0.86988 | TSR3 | 0.86988 | 0.8501 | 0.86965 |
| ENSG00000135837 | 1 | 179923873 | 180084015 | 496 | 22 | -1.1261 | 0.86994 | CEP350 | 0.86994 | 0.87317 | 0.87478 |
| ENSG00000170275 | 3 | 33155471 | 33189265 | 89 | 15 | -1.1263 | 0.86997 | CRTAP | 0.86997 | 0.85605 | 0.63067 |
| ENSG00000151067 | 12 | 2079952 | 2802108 | 2337 | 208 | -1.1264 | 0.87001 | CACNA1C | 0.87001 | 0.86031 | 0.86992 |
| ENSG00000146063 | 5 | 180649499 | 180662809 | 44 | 10 | -1.1266 | 0.87005 | TRIM41 | 0.87005 | 0.32994 | 0.33657 |
| ENSG00000008405 | 12 | 107385142 | 107487607 | 275 | 16 | -1.1267 | 0.87006 | CRY1 | 0.87006 | 0.86117 | 0.88178 |
| ENSG00000141449 | 18 | 18822203 | 19105378 | 346 | 27 | -1.1267 | 0.87008 | GREB1L | 0.87008 | 0.88326 | 0.88293 |
| ENSG00000268927 | 16 | 70695570 | 70699739 | 13 | 8 | -1.1269 | 0.87011 | FLJ00418 | 0.87011 | 0.7634 | 0.47979 |
| ENSG00000204947 | 7 | 148799876 | 148823438 | 64 | 8 | -1.127 | 0.87012 | ZNF425 | 0.87012 | 0.84144 | 0.86093 |
| ENSG00000123338 | 12 | 54891495 | 54937726 | 130 | 34 | -1.1269 | 0.87012 | NCKAP1L | 0.87012 | 0.87739 | 0.82745 |
| ENSG00000070413 | 22 | 19023795 | 19109967 | 452 | 19 | -1.1276 | 0.87025 | DGCR2 | 0.87025 | 0.883 | 0.87395 |
| ENSG00000111196 | 12 | 10758612 | 10766222 | 32 | 5 | -1.1276 | 0.87026 | MAGOHB | 0.87026 | 0.93755 | 0.82021 |
| ENSG00000103365 | 16 | 23474863 | 23533316 | 159 | 15 | -1.1276 | 0.87026 | GGA2 | 0.87026 | 0.87752 | 0.87313 |
| ENSG00000143340 | 1 | 179712298 | 179785333 | 263 | 29 | -1.1278 | 0.8703 | FAM163A | 0.8703 | 0.5147 | 0.53413 |
| ENSG00000168393 | 2 | 242615157 | 242626406 | 45 | 12 | -1.1284 | 0.87043 | DTYMK | 0.87043 | 0.73141 | 0.74147 |
| ENSG00000197870 | 12 | 11418857 | 11422739 | 14 | 4 | -1.1287 | 0.87048 | PRB3 | 0.87048 | 0.66117 | 0.21471 |
| ENSG00000116809 | 1 | 16268364 | 16302627 | 74 | 17 | -1.1288 | 0.8705 | ZBTB17 | 0.8705 | 0.86935 | 0.87892 |
| ENSG00000151062 | 12 | 1901123 | 2028002 | 564 | 57 | -1.1288 | 0.87051 | CACNA2D4 | 0.87051 | 0.92257 | 0.91732 |
| ENSG00000267699 | 18 | 48494389 | 48584514 | 125 | 12 | -1.1289 | 0.87053 | RP11-729L2.2 | 0.87053 | 0.85435 | 0.8885 |
| ENSG00000186334 | 5 | 150656323 | 150683327 | 142 | 13 | -1.1289 | 0.87054 | SLC36A3 | 0.87054 | 0.81742 | 0.78742 |
| ENSG00000124225 | 20 | 56223448 | 56286592 | 189 | 40 | -1.1293 | 0.87062 | PMEPA1 | 0.87062 | 0.63445 | 0.05824 |
| ENSG00000148606 | 10 | 79734907 | 79789303 | 90 | 13 | -1.13 | 0.87076 | POLR3A | 0.87076 | 0.89263 | 0.88712 |
| ENSG00000105991 | 7 | 27132612 | 27135615 | 6 | 4 | -1.13 | 0.87077 | HOXA1 | 0.87077 | 0.82409 | 0.83057 |
| ENSG00000100934 | 14 | 39501123 | 39578850 | 235 | 28 | -1.1301 | 0.87079 | SEC23A | 0.87079 | 0.87285 | 0.86145 |
| ENSG00000149927 | 16 | 30016830 | 30034591 | 42 | 6 | -1.1304 | 0.87084 | DOC2A | 0.87084 | 0.7314 | 0.7275 |
| ENSG00000128335 | 22 | 36622256 | 36636000 | 81 | 17 | -1.1306 | 0.87089 | APOL2 | 0.87089 | 0.89817 | 0.95831 |
| ENSG00000221909 | 7 | 99143931 | 99156159 | 10 | 3 | -1.1309 | 0.87095 | FAM200A | 0.87095 | 0.79471 | 0.85779 |
| ENSG00000068097 | 17 | 58120555 | 58156292 | 42 | 4 | -1.1309 | 0.87096 | HEATR6 | 0.87096 | 0.85885 | 0.78444 |
| ENSG00000236320 | 17 | 33875144 | 33885117 | 37 | 9 | -1.1317 | 0.87111 | SLFN14 | 0.87111 | 0.80562 | 0.84134 |
| ENSG00000108700 | 17 | 32646055 | 32648421 | 5 | 1 | -1.1323 | 0.87124 | CCL8 | 0.87124 | 0.86415 | 0.6612 |
| ENSG00000188306 | 3 | 169539710 | 169555563 | 52 | 9 | -1.1328 | 0.87136 | LRRIQ4 | 0.87136 | 0.8777 | 0.76338 |
| ENSG00000189067 | 16 | 11641853 | 11730237 | 418 | 43 | -1.1333 | 0.87146 | LITAF | 0.87146 | 0.8852 | 0.85411 |
| ENSG00000174791 | 11 | 66097713 | 66104311 | 20 | 5 | -1.1334 | 0.87147 | RIN1 | 0.87147 | 0.88134 | 0.9512 |
| ENSG00000206474 | 6 | 29407083 | 29408731 | 16 | 4 | -1.1336 | 0.87152 | OR10C1 | 0.87152 | 0.88077 | 0.95812 |
| ENSG00000165655 | 10 | 77157588 | 77161664 | 8 | 4 | -1.1342 | 0.87164 | ZNF503 | 0.87164 | 0.61255 | 0.7317 |
| ENSG00000196116 | 9 | 100174232 | 100258407 | 194 | 18 | -1.1343 | 0.87166 | TDRD7 | 0.87166 | 0.90687 | 0.71343 |
| ENSG00000121350 | 12 | 21590549 | 21623300 | 276 | 10 | -1.1346 | 0.87173 | PYROXD1 | 0.87173 | 0.86419 | 0.81745 |
| ENSG00000134569 | 11 | 46878419 | 46940193 | 140 | 17 | -1.1347 | 0.87175 | LRP4 | 0.87175 | 0.89133 | 0.85593 |
| ENSG00000117682 | 1 | 26758773 | 26797785 | 76 | 17 | -1.1348 | 0.87177 | DHDDS | 0.87177 | 0.84898 | 0.70554 |
| ENSG00000264116 | 18 | 73118967 | 73130995 | 43 | 9 | -1.1356 | 0.87195 | RP11-321M21.3 | 0.87195 | 0.91852 | 0.97472 |
| ENSG00000248458 | 1 | 67131547 | 67142710 | 37 | 8 | -1.1357 | 0.87196 | AL139147.1 | 0.87196 | 0.89796 | 0.79006 |
| ENSG00000181163 | 5 | 170814120 | 170838141 | 86 | 7 | -1.1361 | 0.87204 | NPM1 | 0.87204 | 0.83397 | 0.85389 |
| ENSG00000167522 | 16 | 89334038 | 89556969 | 572 | 31 | -1.1361 | 0.87204 | ANKRD11 | 0.87204 | 0.9106 | 0.93804 |
| ENSG00000177535 | 1 | 247614255 | 247615308 | 7 | 3 | -1.1362 | 0.87206 | OR2B11 | 0.87206 | 0.807 | 0.87278 |
| ENSG00000162849 | 1 | 245318287 | 245872733 | 2604 | 213 | -1.1362 | 0.87207 | KIF26B | 0.87207 | 0.91097 | 0.8852 |
| ENSG00000170632 | 7 | 102715328 | 102740205 | 37 | 13 | -1.1364 | 0.87211 | ARMC10 | 0.87211 | 0.73422 | 0.55473 |
| ENSG00000072682 | 5 | 131527531 | 131631008 | 281 | 19 | -1.1369 | 0.87221 | P4HA2 | 0.87221 | 0.91193 | 0.88404 |
| ENSG00000148400 | 9 | 139388896 | 139440314 | 171 | 39 | -1.137 | 0.87224 | NOTCH1 | 0.87224 | 0.80073 | 0.84865 |
| ENSG00000133318 | 11 | 63448918 | 63527363 | 125 | 13 | -1.1372 | 0.87227 | RTN3 | 0.87227 | 0.86406 | 0.84621 |
| ENSG00000164011 | 1 | 43312280 | 43318148 | 2 | 1 | -1.1372 | 0.87228 | ZNF691 | 0.87228 | 0.78284 | 0.82031 |
| ENSG00000165066 | 8 | 41502697 | 41508855 | 22 | 7 | -1.1376 | 0.87235 | NKX6-3 | 0.87235 | 0.89113 | 0.88204 |
| ENSG00000011009 | 1 | 24117460 | 24122029 | 4 | 1 | -1.1377 | 0.87237 | LYPLA2 | 0.87237 | 0.8574 | 0.68465 |
| ENSG00000165821 | 14 | 21989232 | 22005350 | 48 | 12 | -1.1383 | 0.87251 | SALL2 | 0.87251 | 0.91294 | 0.75552 |
| ENSG00000198863 | 17 | 41132582 | 41145707 | 15 | 5 | -1.1383 | 0.87251 | RUNDC1 | 0.87251 | 0.87269 | 0.84722 |
| ENSG00000138764 | 4 | 78078304 | 78354542 | 1759 | 29 | -1.1385 | 0.87255 | CCNG2 | 0.87255 | 0.87604 | 0.89994 |
| ENSG00000111728 | 12 | 22216707 | 22589975 | 1396 | 66 | -1.1385 | 0.87255 | ST8SIA1 | 0.87255 | 0.88092 | 0.87933 |
| ENSG00000117069 | 1 | 77333126 | 77531396 | 562 | 36 | -1.1388 | 0.87261 | ST6GALNAC5 | 0.87261 | 0.8677 | 0.91856 |
| ENSG00000155066 | 2 | 95940201 | 95957056 | 22 | 9 | -1.139 | 0.87265 | PROM2 | 0.87265 | 0.4211 | 0.27129 |
| ENSG00000160505 | 19 | 56347944 | 56393220 | 255 | 42 | -1.139 | 0.87266 | NLRP4 | 0.87266 | 0.87053 | 0.98112 |
| ENSG00000134398 | 16 | 23701647 | 23724821 | 52 | 16 | -1.14 | 0.87286 | ERN2 | 0.87286 | 0.87536 | 0.90521 |
| ENSG00000165553 | 14 | 77731826 | 77737655 | 11 | 4 | -1.1401 | 0.87288 | NGB | 0.87288 | 0.75621 | 0.79786 |
| ENSG00000137218 | 6 | 41737914 | 41754280 | 43 | 12 | -1.1413 | 0.87313 | FRS3 | 0.87313 | 0.81735 | 0.58394 |
| ENSG00000072071 | 19 | 14260750 | 14316999 | 143 | 26 | -1.1414 | 0.87315 | LPHN1 | 0.87315 | 0.85562 | 0.85073 |
| ENSG00000186838 | 19 | 40005753 | 40011326 | 18 | 8 | -1.1415 | 0.87317 | SELV | 0.87317 | 0.63315 | 0.64135 |
| ENSG00000182973 | 3 | 32726637 | 32815367 | 462 | 23 | -1.1417 | 0.8732 | CNOT10 | 0.8732 | 0.88774 | 0.92075 |
| ENSG00000143006 | 1 | 53925072 | 53933161 | 30 | 7 | -1.1418 | 0.87324 | DMRTB1 | 0.87324 | 0.78738 | 0.79831 |
| ENSG00000183386 | 1 | 38462442 | 38471278 | 21 | 10 | -1.142 | 0.87328 | FHL3 | 0.87328 | 0.71515 | 0.83476 |
| ENSG00000134516 | 5 | 169064251 | 169510386 | 1620 | 118 | -1.142 | 0.87328 | DOCK2 | 0.87328 | 0.87525 | 0.76595 |

| | | | | | | | | | | | |
|---|---|---|---|---|---|---|---|---|---|---|---|
| ENSG00000196502 | 16 | 28616903 | 28634946 | 99 | 9 | -1.142 | 0.87328 | SULT1A1 | 0.87328 | 0.89607 | 0.90411 |
| ENSG00000143947 | 2 | 55459039 | 55462989 | 14 | 6 | -1.1422 | 0.87332 | RPS27A | 0.87332 | 0.95266 | 0.93523 |
| ENSG00000128829 | 15 | 40226347 | 40327797 | 372 | 31 | -1.1427 | 0.87341 | EIF2AK4 | 0.87341 | 0.83286 | 0.83014 |
| ENSG00000065357 | 12 | 56321103 | 56347811 | 20 | 5 | -1.1428 | 0.87344 | DGKA | 0.87344 | 0.8853 | 0.88446 |
| ENSG00000172243 | 12 | 10269376 | 10282857 | 86 | 8 | -1.1432 | 0.87352 | CLEC7A | 0.87352 | 0.81301 | 0.48146 |
| ENSG00000100207 | 22 | 42556019 | 42739622 | 431 | 51 | -1.1433 | 0.87354 | TCF20 | 0.87354 | 0.84403 | 0.85919 |
| ENSG00000099330 | 19 | 17337013 | 17340028 | 28 | 6 | -1.1436 | 0.8736 | OCEL1 | 0.8736 | 0.90556 | 0.91546 |
| ENSG00000182111 | 7 | 57509883 | 57533252 | 124 | 6 | -1.1437 | 0.87363 | ZNF716 | 0.87363 | 0.84534 | 0.83206 |
| ENSG00000204020 | 10 | 90521163 | 90537999 | 88 | 16 | -1.1443 | 0.87375 | LIPN | 0.87375 | 0.91242 | 0.8668 |
| ENSG00000164082 | 3 | 51741086 | 51752629 | 9 | 5 | -1.1446 | 0.87381 | GRM2 | 0.87381 | 0.50306 | 0.45242 |
| ENSG00000188279 | 10 | 49199649 | 49207818 | 30 | 7 | -1.1446 | 0.87381 | FAM25C | 0.87381 | 0.88736 | 0.88324 |
| ENSG00000187790 | 14 | 45605143 | 45670093 | 127 | 12 | -1.1447 | 0.87383 | FANCM | 0.87383 | 0.84458 | 0.83508 |
| ENSG00000168014 | 11 | 73723763 | 73882255 | 430 | 24 | -1.1448 | 0.87386 | C2CD3 | 0.87386 | 0.88391 | 0.83675 |
| ENSG00000049239 | 1 | 9294834 | 9331396 | 128 | 22 | -1.1449 | 0.87387 | H6PD | 0.87387 | 0.89212 | 0.92715 |
| ENSG00000169738 | 17 | 79993012 | 79995608 | 15 | 1 | -1.1449 | 0.87388 | DCXR | 0.87388 | 0.61888 | 0.69783 |
| ENSG00000117385 | 1 | 43212006 | 43232755 | 57 | 13 | -1.1456 | 0.87402 | LEPRE1 | 0.87402 | 0.82562 | 0.75704 |
| ENSG00000089639 | 19 | 19740285 | 19754476 | 27 | 10 | -1.1456 | 0.87402 | GMIP | 0.87402 | 0.89602 | 0.87836 |
| ENSG00000124243 | 20 | 49411431 | 49493714 | 312 | 33 | -1.146 | 0.87411 | BCAS4 | 0.87411 | 0.91214 | 0.90886 |
| ENSG00000050327 | 7 | 144052381 | 144077725 | 20 | 6 | -1.1462 | 0.87414 | ARHGEF5 | 0.87414 | 0.7288 | 0.66359 |
| ENSG00000161800 | 12 | 50370706 | 50426919 | 126 | 10 | -1.1463 | 0.87416 | RACGAP1 | 0.87416 | 0.90761 | 0.85655 |
| ENSG00000255408 | 5 | 140180783 | 140391929 | 546 | 21 | -1.1464 | 0.87419 | PCDHA3 | 0.87419 | 0.87125 | 0.89707 |
| ENSG00000143870 | 2 | 10923517 | 10978103 | 441 | 21 | -1.1465 | 0.8742 | PDIA6 | 0.8742 | 0.81052 | 0.81758 |
| ENSG00000198374 | 6 | 27833034 | 27833606 | 1 | 1 | -1.1465 | 0.8742 | HIST1H2AL | 0.8742 | 0.7335 | 0.85694 |
| ENSG00000217455 | 7 | 3179868 | 3214287 | 218 | 32 | -1.1469 | 0.87429 | AC091801.1 | 0.87429 | 0.74139 | 0.43924 |
| ENSG00000174899 | 3 | 157261035 | 157395538 | 490 | 38 | -1.1476 | 0.87443 | C3orf55 | 0.87443 | 0.86406 | 0.84165 |
| ENSG00000117266 | 1 | 205473723 | 205501921 | 141 | 36 | -1.1476 | 0.87444 | CDK18 | 0.87444 | 0.94393 | 0.95254 |
| ENSG00000188069 | 11 | 4790209 | 4791168 | 19 | 3 | -1.1478 | 0.87447 | OR51F1 | 0.87447 | 0.88514 | 0.89664 |
| ENSG00000105976 | 7 | 116312444 | 116438440 | 228 | 39 | -1.1481 | 0.87453 | MET | 0.87453 | 0.91797 | 0.92392 |
| ENSG00000186297 | 15 | 27111510 | 27194354 | 265 | 36 | -1.1481 | 0.87454 | GABRA5 | 0.87454 | 0.68798 | 0.59864 |
| ENSG00000125826 | 20 | 388142 | 411610 | 101 | 19 | -1.1482 | 0.87456 | RBCK1 | 0.87456 | 0.86095 | 0.58447 |
| ENSG00000161179 | 22 | 21982378 | 21984353 | 4 | 2 | -1.1482 | 0.87456 | YDJC | 0.87456 | 0.42894 | 0.7042 |
| ENSG00000136261 | 7 | 16685756 | 16746148 | 215 | 31 | -1.1483 | 0.87458 | BZW2 | 0.87458 | 0.91953 | 0.7081 |
| ENSG00000255359 | 11 | 22868477 | 22881972 | 48 | 5 | -1.1483 | 0.87458 | CCDC179 | 0.87458 | 0.71808 | 0.68293 |
| ENSG00000122884 | 10 | 74766975 | 74856732 | 200 | 7 | -1.1486 | 0.87465 | P4HA1 | 0.87465 | 0.86813 | 0.86011 |
| ENSG00000197520 | 1 | 222910549 | 222924147 | 22 | 6 | -1.1492 | 0.87475 | FAM177B | 0.87475 | 0.82652 | 0.86254 |
| ENSG00000158411 | 2 | 99777890 | 99797521 | 49 | 7 | -1.1493 | 0.87479 | MITD1 | 0.87479 | 0.85598 | 0.79986 |
| ENSG00000169676 | 4 | 9783258 | 9785632 | 5 | 2 | -1.1496 | 0.87485 | DRD5 | 0.87485 | 0.6632 | 0.66518 |
| ENSG00000171206 | 10 | 104404253 | 104418164 | 26 | 9 | -1.1498 | 0.87489 | TRIM8 | 0.87489 | 0.85718 | 0.79764 |
| ENSG00000143106 | 1 | 109941653 | 109969062 | 54 | 9 | -1.1499 | 0.87491 | PSMA5 | 0.87491 | 0.86857 | 0.88514 |
| ENSG00000183354 | 9 | 5881596 | 6007901 | 372 | 24 | -1.1501 | 0.87496 | KIAA2026 | 0.87496 | 0.87435 | 0.90635 |
| ENSG00000089091 | 20 | 18364011 | 18447925 | 281 | 22 | -1.1503 | 0.87499 | DZANK1 | 0.87499 | 0.83083 | 0.85414 |
| ENSG00000177096 | 22 | 42470255 | 42475445 | 12 | 3 | -1.1505 | 0.87502 | FAM109B | 0.87502 | 0.78449 | 0.76412 |
| ENSG00000183086 | 7 | 74379083 | 74438803 | 23 | 10 | -1.1509 | 0.87511 | GATSL1 | 0.87511 | 0.91968 | 0.96403 |
| ENSG00000100401 | 22 | 41641615 | 41682255 | 113 | 14 | -1.1509 | 0.87512 | RANGAP1 | 0.87512 | 0.8818 | 0.84107 |
| ENSG00000213516 | 1 | 89445139 | 89458643 | 22 | 10 | -1.1515 | 0.87523 | RBMXL1 | 0.87523 | 0.95621 | 0.92797 |
| ENSG00000139641 | 12 | 56512034 | 56538455 | 41 | 15 | -1.1515 | 0.87523 | ESYT1 | 0.87523 | 0.7979 | 0.80522 |
| ENSG00000173218 | 1 | 116184574 | 116240845 | 218 | 36 | -1.1521 | 0.87536 | VANGL1 | 0.87536 | 0.75482 | 0.70302 |
| ENSG00000267909 | 14 | 70038216 | 70040339 | 6 | 2 | -1.1524 | 0.87543 | CCDC177 | 0.87543 | 0.78288 | 0.83395 |
| ENSG00000106006 | 7 | 27185015 | 27190222 | 20 | 5 | -1.1531 | 0.87556 | HOXA6 | 0.87556 | 0.91179 | 0.97282 |
| ENSG00000204572 | 11 | 71276609 | 71279980 | 35 | 5 | -1.1531 | 0.87556 | KRTAP5-10 | 0.87556 | 0.89292 | 0.96149 |
| ENSG00000204969 | 5 | 140174444 | 140391929 | 557 | 22 | -1.1534 | 0.87563 | PCDHA2 | 0.87563 | 0.8762 | 0.89015 |
| ENSG00000166025 | 11 | 94439597 | 94609918 | 430 | 37 | -1.1535 | 0.87565 | AMOTL1 | 0.87565 | 0.87096 | 0.84988 |
| ENSG00000140043 | 14 | 74318547 | 74353530 | 180 | 13 | -1.1536 | 0.87566 | PTGR2 | 0.87566 | 0.86749 | 0.88715 |
| ENSG00000165388 | 10 | 48355024 | 48373866 | 39 | 12 | -1.1539 | 0.87572 | ZNF488 | 0.87572 | 0.76007 | 0.52776 |
| ENSG00000161664 | 17 | 42247815 | 42256451 | 21 | 4 | -1.154 | 0.87575 | ASB16 | 0.87575 | 0.93951 | 0.89864 |
| ENSG00000185261 | 5 | 93488671 | 93954309 | 783 | 78 | -1.1543 | 0.8758 | KIAA0825 | 0.8758 | 0.86182 | 0.85094 |
| ENSG00000165685 | 12 | 10323141 | 10344400 | 96 | 19 | -1.1543 | 0.87581 | TMEM52B | 0.87581 | 0.94508 | 0.85871 |
| ENSG00000124459 | 19 | 44416781 | 44439430 | 75 | 8 | -1.1543 | 0.87581 | ZNF45 | 0.87581 | 0.8767 | 0.82683 |
| ENSG00000088833 | 20 | 1422807 | 1454487 | 82 | 19 | -1.1543 | 0.87581 | NSFL1C | 0.87581 | 0.85448 | 0.90724 |
| ENSG00000196503 | 4 | 57371509 | 57390614 | 73 | 14 | -1.1545 | 0.87585 | ARL9 | 0.87585 | 0.84884 | 0.68408 |
| ENSG00000170473 | 12 | 56295197 | 56326402 | 49 | 7 | -1.1548 | 0.87592 | WIBG | 0.87592 | 0.87123 | 0.91129 |
| ENSG00000214413 | 10 | 112658488 | 112679032 | 57 | 14 | -1.1549 | 0.87594 | BBIP1 | 0.87594 | 0.86179 | 0.85675 |
| ENSG00000157111 | 5 | 72416119 | 72427644 | 39 | 13 | -1.155 | 0.87595 | TMEM171 | 0.87595 | 0.93595 | 0.93823 |
| ENSG00000163902 | 3 | 128338817 | 128399918 | 224 | 22 | -1.1551 | 0.87598 | RPN1 | 0.87598 | 0.9029 | 0.90422 |
| ENSG00000189410 | 1 | 21046225 | 21059330 | 37 | 9 | -1.1561 | 0.87618 | SH2D5 | 0.87618 | 0.91096 | 0.96619 |
| ENSG00000171209 | 4 | 71108305 | 71117145 | 76 | 5 | -1.1563 | 0.87623 | CSN3 | 0.87623 | 0.85131 | 0.66858 |
| ENSG00000003147 | 7 | 8152814 | 8302317 | 688 | 80 | -1.1569 | 0.87634 | ICA1 | 0.87634 | 0.87923 | 0.90551 |
| ENSG00000148158 | 9 | 115513118 | 115643951 | 407 | 21 | -1.1569 | 0.87634 | SNX30 | 0.87634 | 0.88829 | 0.92115 |
| ENSG00000064687 | 19 | 1040102 | 1065571 | 132 | 23 | -1.1569 | 0.87635 | ABCA7 | 0.87635 | 0.94046 | 0.94087 |
| ENSG00000187533 | 4 | 70999333 | 71042516 | 202 | 26 | -1.157 | 0.87636 | C4orf40 | 0.87636 | 0.93766 | 0.9126 |
| ENSG00000169189 | 16 | 27236312 | 27280115 | 121 | 13 | -1.1574 | 0.87646 | NSMCE1 | 0.87646 | 0.81506 | 0.75571 |
| ENSG00000168615 | 8 | 38854388 | 38962663 | 246 | 22 | -1.1577 | 0.87651 | ADAM9 | 0.87651 | 0.84689 | 0.86412 |
| ENSG00000214854 | 22 | 39656865 | 39657063 | 2 | 1 | -1.1581 | 0.87659 | AL031590.1 | 0.87659 | 0.51635 | 0.59004 |

| | | | | | | | | | | | |
|---|---|---|---|---|---|---|---|---|---|---|---|
| ENSG00000176246 | 14 | 20528204 | 20529142 | 10 | 2 | -1.1583 | 0.87662 | OR4L1 | 0.87662 | 0.74183 | 0.76293 |
| ENSG00000117592 | 1 | 173446405 | 173457946 | 16 | 5 | -1.1586 | 0.87669 | PRDX6 | 0.87669 | 0.92293 | 0.5254 |
| ENSG00000108061 | 10 | 112679301 | 112773425 | 220 | 26 | -1.1587 | 0.87671 | SHOC2 | 0.87671 | 0.88224 | 0.88149 |
| ENSG00000099810 | 9 | 21802542 | 21931646 | 349 | 33 | -1.1588 | 0.87672 | MTAP | 0.87672 | 0.89079 | 0.8103 |
| ENSG00000268208 | 14 | 78227711 | 78228033 | 3 | 1 | -1.1588 | 0.87674 | AC008372.1 | 0.87674 | 0.95635 | 0.95801 |
| ENSG00000155428 | 7 | 72430016 | 72439997 | 8 | 4 | -1.1595 | 0.87688 | TRIM74 | 0.87688 | 0.89754 | 0.30102 |
| ENSG00000166233 | 15 | 72766667 | 72879692 | 193 | 20 | -1.1596 | 0.87689 | ARIH1 | 0.87689 | 0.87683 | 0.89554 |
| ENSG00000136522 | 3 | 179306073 | 179322442 | 36 | 5 | -1.1605 | 0.87707 | MRPL47 | 0.87707 | 0.93298 | 0.96206 |
| ENSG00000075643 | 18 | 33767482 | 33852120 | 328 | 32 | -1.1605 | 0.87708 | MOCOS | 0.87708 | 0.87278 | 0.81473 |
| ENSG00000175143 | 1 | 248569296 | 248570405 | 1 | 1 | -1.1606 | 0.8771 | OR2T1 | 0.8771 | 0.65726 | 0.68377 |
| ENSG00000204542 | 6 | 31079000 | 31080336 | 16 | 5 | -1.1609 | 0.87717 | C6orf15 | 0.87717 | 0.55998 | 0.18591 |
| ENSG00000100197 | 22 | 42522501 | 42526908 | 32 | 7 | -1.1611 | 0.8772 | CYP2D6 | 0.8772 | 0.83118 | 0.80871 |
| ENSG00000166411 | 15 | 78423840 | 78464291 | 109 | 17 | -1.1611 | 0.87721 | IDH3A | 0.87721 | 0.83405 | 0.85829 |
| ENSG00000177728 | 17 | 73437240 | 73496171 | 201 | 19 | -1.1612 | 0.87722 | KIAA0195 | 0.87722 | 0.87686 | 0.87745 |
| ENSG00000089053 | 12 | 121746048 | 121837699 | 342 | 37 | -1.1613 | 0.87724 | ANAPC5 | 0.87724 | 0.84875 | 0.81608 |
| ENSG00000117569 | 1 | 97187221 | 97289294 | 268 | 20 | -1.162 | 0.87737 | PTBP2 | 0.87737 | 0.86787 | 0.88957 |
| ENSG00000163689 | 3 | 58703092 | 59035810 | 583 | 33 | -1.1621 | 0.8774 | C3orf67 | 0.8774 | 0.87286 | 0.88203 |
| ENSG00000152223 | 18 | 43427574 | 43547240 | 264 | 25 | -1.1623 | 0.87745 | EPG5 | 0.87745 | 0.90968 | 0.86237 |
| ENSG00000057663 | 6 | 106632351 | 106773666 | 450 | 26 | -1.1625 | 0.87748 | ATG5 | 0.87748 | 0.87645 | 0.88908 |
| ENSG00000160439 | 19 | 55550476 | 55582659 | 237 | 32 | -1.1629 | 0.87757 | RDH13 | 0.87757 | 0.80395 | 0.83494 |
| ENSG00000184599 | 1 | 113263041 | 113269857 | 20 | 5 | -1.1631 | 0.87761 | FAM19A3 | 0.87761 | 0.66141 | 0.096398 |
| ENSG00000163913 | 3 | 129158968 | 129239198 | 267 | 12 | -1.1631 | 0.87761 | IFT122 | 0.87761 | 0.88641 | 0.92852 |
| ENSG00000248369 | 4 | 133356957 | 133368390 | 23 | 4 | -1.1636 | 0.8777 | RP11-62N21.1 | 0.8777 | 0.87048 | 0.85214 |
| ENSG00000019186 | 20 | 52769988 | 52790512 | 120 | 23 | -1.1643 | 0.87786 | CYP24A1 | 0.87786 | 0.86044 | 0.86158 |
| ENSG00000049245 | 1 | 7831329 | 7841492 | 21 | 6 | -1.1647 | 0.87792 | VAMP3 | 0.87792 | 0.93159 | 0.95113 |
| ENSG00000178473 | 10 | 5406972 | 5416169 | 64 | 13 | -1.1658 | 0.87814 | UCN3 | 0.87814 | 0.88879 | 0.20755 |
| ENSG00000166869 | 16 | 23765948 | 23770272 | 8 | 4 | -1.1658 | 0.87816 | CHP2 | 0.87816 | 0.55585 | 0.76349 |
| ENSG00000164292 | 5 | 95049226 | 95160087 | 247 | 40 | -1.1659 | 0.87817 | RHOBTB3 | 0.87817 | 0.7534 | 0.79383 |
| ENSG00000129990 | 19 | 55682567 | 55691809 | 66 | 8 | -1.1661 | 0.87821 | SYT5 | 0.87821 | 0.81381 | 0.78918 |
| ENSG00000158486 | 16 | 20944433 | 21170762 | 617 | 44 | -1.1666 | 0.87831 | DNAH3 | 0.87831 | 0.86798 | 0.82622 |
| ENSG00000137094 | 9 | 34989638 | 34998897 | 10 | 6 | -1.1668 | 0.87836 | DNAJB5 | 0.87836 | 0.77257 | 0.6443 |
| ENSG00000136634 | 1 | 206940947 | 206945839 | 18 | 5 | -1.1671 | 0.87841 | IL10 | 0.87841 | 0.81797 | 0.82089 |
| ENSG00000148672 | 10 | 88810243 | 88854623 | 131 | 18 | -1.1671 | 0.87841 | GLUD1 | 0.87841 | 0.8845 | 0.88868 |
| ENSG00000100336 | 22 | 36585172 | 36600886 | 133 | 15 | -1.1675 | 0.87849 | APOL4 | 0.87849 | 0.86667 | 0.63033 |
| ENSG00000215883 | 1 | 54638009 | 54665709 | 71 | 16 | -1.1681 | 0.87861 | CYB5RL | 0.87861 | 0.91354 | 0.91469 |
| ENSG00000140986 | 16 | 1993975 | 2007607 | 76 | 24 | -1.1685 | 0.8787 | RPL3L | 0.8787 | 0.8516 | 0.92687 |
| ENSG00000213892 | 19 | 45202421 | 45213986 | 37 | 10 | -1.1686 | 0.87871 | CEACAM16 | 0.87871 | 0.80658 | 0.65089 |
| ENSG00000196329 | 7 | 150419341 | 150447121 | 54 | 9 | -1.1688 | 0.87877 | GIMAP5 | 0.87877 | 0.89884 | 0.89623 |
| ENSG00000163558 | 3 | 169940153 | 170023769 | 362 | 13 | -1.1694 | 0.87887 | PRKCI | 0.87887 | 0.87598 | 0.82353 |
| ENSG00000090097 | 3 | 51991470 | 52002426 | 15 | 4 | -1.1696 | 0.87893 | PCBP4 | 0.87893 | 0.699 | 0.81028 |
| ENSG00000213047 | 1 | 197473878 | 197744826 | 409 | 20 | -1.1697 | 0.87894 | DENND1B | 0.87894 | 0.86916 | 0.87332 |
| ENSG00000156052 | 9 | 80331003 | 80646374 | 634 | 27 | -1.1698 | 0.87895 | GNAQ | 0.87895 | 0.88817 | 0.89579 |
| ENSG00000090054 | 9 | 94794281 | 94877666 | 332 | 23 | -1.1702 | 0.87904 | SPTLC1 | 0.87904 | 0.89087 | 0.89555 |
| ENSG00000135577 | 6 | 142379467 | 142409936 | 66 | 10 | -1.1705 | 0.87911 | NMBR | 0.87911 | 0.8537 | 0.85483 |
| ENSG00000189339 | 1 | 1592939 | 1624167 | 87 | 15 | -1.1711 | 0.87922 | SLC35E2B | 0.87922 | 0.80523 | 0.69886 |
| ENSG00000135697 | 16 | 81272053 | 81324747 | 229 | 32 | -1.1711 | 0.87922 | BCMO1 | 0.87922 | 0.92591 | 0.97792 |
| ENSG00000172757 | 11 | 65590493 | 65629497 | 89 | 17 | -1.1712 | 0.87923 | CFL1 | 0.87923 | 0.90731 | 0.91991 |
| ENSG00000204963 | 5 | 140213969 | 140391929 | 454 | 23 | -1.1716 | 0.87931 | PCDHA7 | 0.87931 | 0.89917 | 0.91914 |
| ENSG00000183077 | 17 | 76183398 | 76203782 | 144 | 29 | -1.1723 | 0.87946 | AFMID | 0.87946 | 0.89369 | 0.84965 |
| ENSG00000175707 | 1 | 27276053 | 27286897 | 18 | 6 | -1.1723 | 0.87947 | C1orf172 | 0.87947 | 0.79701 | 0.76033 |
| ENSG00000083799 | 16 | 50775961 | 50835846 | 112 | 14 | -1.1724 | 0.87949 | CYLD | 0.87949 | 0.78448 | 0.77556 |
| ENSG00000185842 | 1 | 225083964 | 225586996 | 1808 | 31 | -1.1728 | 0.87957 | DNAH14 | 0.87957 | 0.87629 | 0.85881 |
| ENSG00000196141 | 2 | 201170604 | 201346986 | 470 | 47 | -1.1729 | 0.87958 | SPATS2L | 0.87958 | 0.88085 | 0.90787 |
| ENSG00000116752 | 1 | 115110178 | 115124260 | 46 | 15 | -1.1731 | 0.87963 | BCAS2 | 0.87963 | 0.94258 | 0.94609 |
| ENSG00000136854 | 9 | 130374544 | 130457460 | 106 | 19 | -1.1732 | 0.87964 | STXBP1 | 0.87964 | 0.91497 | 0.89882 |
| ENSG00000075673 | 13 | 25254549 | 25285921 | 206 | 27 | -1.1735 | 0.8797 | ATP12A | 0.8797 | 0.90796 | 0.96668 |
| ENSG00000205085 | 7 | 128312342 | 128326929 | 75 | 9 | -1.1737 | 0.87975 | FAM71F2 | 0.87975 | 0.8902 | 0.9135 |
| ENSG00000103154 | 16 | 84002237 | 84036381 | 308 | 24 | -1.1739 | 0.87979 | NECAB2 | 0.87979 | 0.9206 | 0.81399 |
| ENSG00000254685 | 1 | 74663896 | 74699770 | 118 | 8 | -1.174 | 0.87981 | FPGT | 0.87981 | 0.83783 | 0.83339 |
| ENSG00000177683 | 7 | 108194987 | 108210194 | 44 | 6 | -1.1741 | 0.87983 | THAP5 | 0.87983 | 0.88274 | 0.82595 |
| ENSG00000140675 | 16 | 31494323 | 31502181 | 7 | 3 | -1.1743 | 0.87986 | SLC5A2 | 0.87986 | 0.90558 | 0.81741 |
| ENSG00000198734 | 1 | 169483404 | 169555826 | 392 | 33 | -1.1745 | 0.8799 | F5 | 0.8799 | 0.88201 | 0.85071 |
| ENSG00000158683 | 7 | 47814250 | 47988088 | 793 | 51 | -1.1747 | 0.87994 | PKD1L1 | 0.87994 | 0.82564 | 0.39628 |
| ENSG00000132522 | 17 | 7214643 | 7218883 | 11 | 5 | -1.1747 | 0.87995 | GPS2 | 0.87995 | 0.9342 | 0.95138 |
| ENSG00000239332 | 2 | 47043836 | 47086146 | 179 | 39 | -1.1749 | 0.87998 | LINC01119 | 0.87998 | 0.83106 | 0.68096 |
| ENSG00000105185 | 19 | 33071974 | 33078358 | 18 | 7 | -1.1751 | 0.88003 | PDCD5 | 0.88003 | 0.83236 | 0.81634 |
| ENSG00000008516 | 16 | 3096682 | 3110727 | 69 | 13 | -1.1756 | 0.88012 | MMP25 | 0.88012 | 0.78471 | 0.65647 |
| ENSG00000124562 | 6 | 34725183 | 34741571 | 67 | 9 | -1.1756 | 0.88013 | SNRPC | 0.88013 | 0.88313 | 0.89341 |
| ENSG00000145703 | 5 | 75699074 | 76003957 | 1195 | 99 | -1.1757 | 0.88014 | IQGAP2 | 0.88014 | 0.81815 | 0.82349 |
| ENSG00000104765 | 8 | 26240414 | 26363152 | 566 | 59 | -1.1758 | 0.88017 | BNIP3L | 0.88017 | 0.89108 | 0.88186 |
| ENSG00000132640 | 20 | 11871371 | 11907257 | 97 | 9 | -1.1762 | 0.88024 | BTBD3 | 0.88024 | 0.89495 | 0.91197 |
| ENSG00000007384 | 16 | 108058 | 126354 | 59 | 12 | -1.1763 | 0.88025 | RHBDF1 | 0.88025 | 0.66743 | 0.54417 |
| ENSG00000117600 | 1 | 99729509 | 99775146 | 131 | 20 | -1.1763 | 0.88026 | LPPR4 | 0.88026 | 0.92069 | 0.95027 |

| | | | | | | | | | | | |
|---|---|---|---|---|---|---|---|---|---|---|---|
| ENSG00000066739 | 14 | 96747595 | 96830207 | 206 | 17 | -1.1763 | 0.88026 | ATG2B | 0.88026 | 0.87259 | 0.72648 |
| ENSG00000121335 | 12 | 11544476 | 11653975 | 486 | 29 | -1.1764 | 0.88028 | PRB2 | 0.88028 | 0.92149 | 0.95251 |
| ENSG00000101126 | 20 | 49505585 | 49547958 | 104 | 12 | -1.1765 | 0.8803 | ADNP | 0.8803 | 0.91481 | 0.9064 |
| ENSG00000116874 | 1 | 119573839 | 119683294 | 327 | 24 | -1.1766 | 0.88033 | WARS2 | 0.88033 | 0.89817 | 0.91736 |
| ENSG00000164967 | 9 | 34610483 | 34612101 | 2 | 1 | -1.1769 | 0.88039 | RPP25L | 0.88039 | 0.33921 | 0.33195 |
| ENSG00000089693 | 12 | 6857170 | 6876641 | 78 | 13 | -1.1772 | 0.88044 | MLF2 | 0.88044 | 0.83872 | 0.81347 |
| ENSG00000137747 | 11 | 117771358 | 117800174 | 144 | 23 | -1.1774 | 0.88049 | TMPRSS13 | 0.88049 | 0.8704 | 0.86332 |
| ENSG00000134376 | 1 | 197170592 | 197447585 | 501 | 46 | -1.1787 | 0.88074 | CRB1 | 0.88074 | 0.86935 | 0.86821 |
| ENSG00000089356 | 19 | 35606732 | 35615228 | 45 | 10 | -1.1787 | 0.88074 | FXYD3 | 0.88074 | 0.81203 | 0.50256 |
| ENSG00000111704 | 12 | 7940390 | 7948655 | 71 | 7 | -1.1789 | 0.88078 | NANOG | 0.88078 | 0.84541 | 0.80126 |
| ENSG00000144233 | 2 | 128619204 | 128643496 | 46 | 6 | -1.179 | 0.88079 | AMMECR1L | 0.88079 | 0.78215 | 0.79218 |
| ENSG00000196220 | 3 | 9022275 | 9404737 | 1334 | 150 | -1.179 | 0.8808 | SRGAP3 | 0.8808 | 0.8741 | 0.87109 |
| ENSG00000132912 | 5 | 150088002 | 150138671 | 124 | 5 | -1.179 | 0.88081 | DCTN4 | 0.88081 | 0.86458 | 0.83225 |
| ENSG00000196730 | 9 | 90112143 | 90323548 | 891 | 81 | -1.1792 | 0.88084 | DAPK1 | 0.88084 | 0.87464 | 0.89434 |
| ENSG00000185585 | 9 | 127539437 | 127577164 | 76 | 20 | -1.1792 | 0.88084 | OLFML2A | 0.88084 | 0.76631 | 0.73078 |
| ENSG00000111796 | 12 | 9747147 | 9760482 | 70 | 13 | -1.1795 | 0.8809 | KLRB1 | 0.8809 | 0.85801 | 0.8093 |
| ENSG00000260027 | 17 | 46684594 | 46710934 | 85 | 20 | -1.1797 | 0.88094 | HOXB7 | 0.88094 | 0.86061 | 0.91346 |
| ENSG00000180549 | 9 | 139924626 | 139927462 | 9 | 3 | -1.1798 | 0.88096 | FUT7 | 0.88096 | 0.88825 | 0.53328 |
| ENSG00000215462 | 13 | 50007495 | 50020554 | 41 | 8 | -1.18 | 0.881 | AL136218.1 | 0.881 | 0.93942 | 0.93582 |
| ENSG00000104918 | 19 | 7733930 | 7735334 | 7 | 3 | -1.18 | 0.881 | RETN | 0.881 | 0.9154 | 0.94276 |
| ENSG00000136732 | 2 | 127413509 | 127454246 | 206 | 34 | -1.1806 | 0.88113 | GYPC | 0.88113 | 0.77524 | 0.86473 |
| ENSG00000167642 | 19 | 38734675 | 38783254 | 193 | 28 | -1.1806 | 0.88113 | SPINT2 | 0.88113 | 0.88218 | 0.74331 |
| ENSG00000203757 | 1 | 158686958 | 158687953 | 4 | 2 | -1.1812 | 0.88124 | OR6K3 | 0.88124 | 0.83586 | 0.74479 |
| ENSG00000168309 | 3 | 58549844 | 58613337 | 256 | 32 | -1.1813 | 0.88126 | FAM107A | 0.88126 | 0.91366 | 0.8961 |
| ENSG00000187607 | 17 | 15602891 | 15640874 | 148 | 24 | -1.1814 | 0.88127 | ZNF286A | 0.88127 | 0.7551 | 0.72273 |
| ENSG00000196981 | 3 | 122130715 | 122134882 | 6 | 3 | -1.1819 | 0.88138 | WDR5B | 0.88138 | 0.84893 | 0.8442 |
| ENSG00000005243 | 17 | 46103533 | 46115392 | 26 | 10 | -1.182 | 0.88139 | COPZ2 | 0.88139 | 0.74141 | 0.68101 |
| ENSG00000175582 | 11 | 73386683 | 73472182 | 314 | 25 | -1.1825 | 0.8815 | RAB6A | 0.8815 | 0.90368 | 0.90117 |
| ENSG00000119522 | 9 | 126141933 | 126692431 | 1552 | 57 | -1.1826 | 0.88151 | DENND1A | 0.88151 | 0.88714 | 0.91156 |
| ENSG00000116031 | 2 | 71057347 | 71062952 | 31 | 8 | -1.1826 | 0.88152 | CD207 | 0.88152 | 0.52142 | 0.65475 |
| ENSG00000143633 | 1 | 231359509 | 231376933 | 45 | 11 | -1.1828 | 0.88156 | C1orf131 | 0.88156 | 0.50845 | 0.48796 |
| ENSG00000185798 | 3 | 196281056 | 196295545 | 41 | 4 | -1.1834 | 0.88167 | WDR53 | 0.88167 | 0.86444 | 0.88982 |
| ENSG00000269515 | 10 | 44873687 | 44878739 | 25 | 7 | -1.184 | 0.8818 | AL137026.1 | 0.8818 | 0.88406 | 0.91531 |
| ENSG00000149090 | 11 | 35453370 | 35551848 | 368 | 42 | -1.184 | 0.8818 | PAMR1 | 0.8818 | 0.88073 | 0.85267 |
| ENSG00000158864 | 1 | 161166894 | 161184185 | 30 | 13 | -1.1846 | 0.88191 | NDUFS2 | 0.88191 | 0.8326 | 0.89383 |
| ENSG00000158882 | 1 | 161195793 | 161200408 | 4 | 2 | -1.1849 | 0.88196 | TOMM40L | 0.88196 | 0.73129 | 0.83907 |
| ENSG00000197912 | 16 | 89557325 | 89624176 | 261 | 11 | -1.1852 | 0.88202 | SPG7 | 0.88202 | 0.88802 | 0.89183 |
| ENSG00000164587 | 5 | 149822753 | 149829319 | 35 | 3 | -1.1852 | 0.88203 | RPS14 | 0.88203 | 0.87008 | 0.85462 |
| ENSG00000069943 | 15 | 55611158 | 55647846 | 147 | 20 | -1.1854 | 0.88207 | PIGB | 0.88207 | 0.81283 | 0.80601 |
| ENSG00000187783 | 10 | 45406648 | 45432450 | 77 | 11 | -1.1855 | 0.88208 | TMEM72 | 0.88208 | 0.85705 | 0.82561 |
| ENSG00000129235 | 17 | 6544078 | 6547861 | 16 | 4 | -1.1857 | 0.88212 | TXNDC17 | 0.88212 | 0.77486 | 0.6773 |
| ENSG00000189134 | 6 | 28227098 | 28228736 | 6 | 3 | -1.1858 | 0.88215 | NKAPL | 0.88215 | 0.9562 | 0.92465 |
| ENSG00000183668 | 19 | 43715943 | 43773682 | 388 | 23 | -1.186 | 0.8822 | PSG9 | 0.8822 | 0.87517 | 0.89639 |
| ENSG00000204967 | 5 | 140186659 | 140391929 | 519 | 20 | -1.1861 | 0.88221 | PCDHA4 | 0.88221 | 0.9027 | 0.91155 |
| ENSG00000129173 | 11 | 19245610 | 19263167 | 39 | 16 | -1.1865 | 0.88228 | E2F8 | 0.88228 | 0.71313 | 0.49094 |
| ENSG00000121898 | 10 | 125465723 | 125699783 | 1160 | 110 | -1.1873 | 0.88244 | CPXM2 | 0.88244 | 0.83177 | 0.81369 |
| ENSG00000146281 | 6 | 89855769 | 89875284 | 61 | 7 | -1.1876 | 0.8825 | PM20D2 | 0.8825 | 0.88176 | 0.63008 |
| ENSG00000174417 | 8 | 110098850 | 110131813 | 97 | 15 | -1.1878 | 0.88254 | TRHR | 0.88254 | 0.85658 | 0.86975 |
| ENSG00000117859 | 1 | 52042851 | 52254889 | 446 | 33 | -1.1879 | 0.88257 | OSBPL9 | 0.88257 | 0.9042 | 0.92027 |
| ENSG00000233041 | 15 | 40643234 | 40648635 | 19 | 6 | -1.1881 | 0.88259 | PHGR1 | 0.88259 | 0.80818 | 0.64279 |
| ENSG00000070087 | 3 | 149682691 | 149768575 | 269 | 23 | -1.1883 | 0.88265 | PFN2 | 0.88265 | 0.87662 | 0.86606 |
| ENSG00000171453 | 6 | 43477440 | 43497323 | 36 | 6 | -1.1885 | 0.88268 | POLR1C | 0.88268 | 0.86076 | 0.85111 |
| ENSG00000168824 | 4 | 4349867 | 4420785 | 334 | 45 | -1.1888 | 0.88274 | NSG1 | 0.88274 | 0.84668 | 0.51086 |
| ENSG00000176022 | 1 | 1167629 | 1170421 | 9 | 3 | -1.1892 | 0.88281 | B3GALT6 | 0.88281 | 0.93725 | 0.9534 |
| ENSG00000115268 | 19 | 1438358 | 1440583 | 1 | 1 | -1.1896 | 0.8829 | RPS15 | 0.8829 | 0.73762 | 0.39296 |
| ENSG00000258588 | 11 | 5617955 | 5665628 | 261 | 36 | -1.1897 | 0.88293 | TRIM6-TRIM34 | 0.88293 | 0.90188 | 0.60216 |
| ENSG00000114279 | 3 | 191857184 | 192485553 | 2060 | 157 | -1.1898 | 0.88294 | FGF12 | 0.88294 | 0.87247 | 0.89552 |
| ENSG00000170464 | 5 | 138744279 | 138780180 | 46 | 5 | -1.1901 | 0.883 | DNAJC18 | 0.883 | 0.85996 | 0.79488 |
| ENSG00000080224 | 3 | 96533425 | 97471304 | 1519 | 81 | -1.1902 | 0.88301 | EPHA6 | 0.88301 | 0.88011 | 0.89546 |
| ENSG00000150656 | 18 | 72201675 | 72254448 | 174 | 47 | -1.1901 | 0.88301 | CNDP1 | 0.88301 | 0.93177 | 0.95047 |
| ENSG00000184571 | 22 | 25115001 | 25170687 | 250 | 28 | -1.1903 | 0.88303 | PIWIL3 | 0.88303 | 0.90357 | 0.86923 |
| ENSG00000136897 | 9 | 104149915 | 104160896 | 28 | 8 | -1.1905 | 0.88307 | MRPL50 | 0.88307 | 0.6966 | 0.65901 |
| ENSG00000257184 | 7 | 27203154 | 27219632 | 31 | 10 | -1.1906 | 0.8831 | RP1-170O19.20 | 0.8831 | 0.70054 | 0.58649 |
| ENSG00000136378 | 15 | 79051545 | 79103773 | 248 | 18 | -1.1907 | 0.88312 | ADAMTS7 | 0.88312 | 0.91488 | 0.86585 |
| ENSG00000172568 | 5 | 156768608 | 156772729 | 17 | 5 | -1.1911 | 0.8832 | FNDC9 | 0.8832 | 0.72185 | 0.68713 |
| ENSG00000124701 | 6 | 41021043 | 41032250 | 34 | 6 | -1.1912 | 0.8832 | APOBEC2 | 0.8832 | 0.846 | 0.81206 |
| ENSG00000269047 | 16 | 1031252 | 1031663 | 1 | 1 | -1.1916 | 0.8833 | AC009041.2 | 0.8833 | 0.11624 | 0.44128 |
| ENSG00000166928 | 11 | 60146003 | 60185161 | 121 | 12 | -1.1923 | 0.88342 | MS4A14 | 0.88342 | 0.88839 | 0.89254 |
| ENSG00000135845 | 1 | 172339329 | 172413230 | 155 | 28 | -1.1924 | 0.88344 | PIGC | 0.88344 | 0.84563 | 0.80797 |
| ENSG00000104626 | 8 | 8859657 | 8974256 | 420 | 32 | -1.1929 | 0.88354 | ERI1 | 0.88354 | 0.91092 | 0.90139 |
| ENSG00000109501 | 4 | 6271576 | 6304992 | 207 | 14 | -1.1931 | 0.88358 | WFS1 | 0.88358 | 0.87407 | 0.81942 |
| ENSG00000089280 | 16 | 31191431 | 31203127 | 13 | 6 | -1.1931 | 0.88359 | FUS | 0.88359 | 0.72906 | 0.71489 |
| ENSG00000111328 | 12 | 123745528 | 123756881 | 32 | 7 | -1.1932 | 0.88361 | CDK2AP1 | 0.88361 | 0.83535 | 0.825 |

| | | | | | | | | | | | |
|---|---|---|---|---|---|---|---|---|---|---|---|
| ENSG00000130816 | 19 | 10244021 | 10341962 | 288 | 26 | -1.1934 | 0.88364 | DNMT1 | 0.88364 | 0.86303 | 0.76045 |
| ENSG00000135185 | 7 | 86825478 | 86849903 | 93 | 10 | -1.1939 | 0.88375 | TMEM243 | 0.88375 | 0.90424 | 0.94488 |
| ENSG00000155158 | 9 | 15163620 | 15307358 | 555 | 53 | -1.1943 | 0.88382 | TTC39B | 0.88382 | 0.90195 | 0.86897 |
| ENSG00000042781 | 1 | 215796236 | 216596738 | 2524 | 154 | -1.1945 | 0.88387 | USH2A | 0.88387 | 0.8778 | 0.877 |
| ENSG00000176194 | 18 | 12254318 | 12277594 | 94 | 20 | -1.1947 | 0.88389 | CIDEA | 0.88389 | 0.88738 | 0.90979 |
| ENSG00000105392 | 19 | 48322703 | 48346587 | 207 | 21 | -1.1947 | 0.8839 | CRX | 0.8839 | 0.86933 | 0.90724 |
| ENSG00000140006 | 14 | 64063757 | 64108579 | 133 | 11 | -1.1949 | 0.88394 | WDR89 | 0.88394 | 0.8763 | 0.91011 |
| ENSG00000262180 | 1 | 186369704 | 186370587 | 2 | 1 | -1.1951 | 0.88397 | OCLM | 0.88397 | 0.35124 | 0.32331 |
| ENSG00000132554 | 8 | 100973164 | 101143496 | 324 | 29 | -1.1952 | 0.88399 | RGS22 | 0.88399 | 0.89117 | 0.87609 |
| ENSG00000166329 | 17 | 55821842 | 55822673 | 4 | 1 | -1.1953 | 0.88402 | CCDC182 | 0.88402 | 0.89184 | 0.8358 |
| ENSG00000124608 | 6 | 44267391 | 44281063 | 40 | 9 | -1.1963 | 0.8842 | AARS2 | 0.8842 | 0.85889 | 0.81385 |
| ENSG00000107902 | 10 | 126150403 | 126306457 | 999 | 64 | -1.1965 | 0.88424 | LHPP | 0.88424 | 0.92081 | 0.88536 |
| ENSG00000108100 | 10 | 35535953 | 35860852 | 614 | 27 | -1.1969 | 0.88432 | CCNY | 0.88432 | 0.91012 | 0.82499 |
| ENSG00000136379 | 15 | 80972025 | 81047962 | 273 | 24 | -1.1969 | 0.88432 | ABHD17C | 0.88432 | 0.87682 | 0.94883 |
| ENSG00000144535 | 2 | 232825955 | 233209060 | 1097 | 31 | -1.197 | 0.88434 | DIS3L2 | 0.88434 | 0.85933 | 0.72766 |
| ENSG00000119684 | 14 | 75480467 | 75518235 | 65 | 5 | -1.1971 | 0.88436 | MLH3 | 0.88436 | 0.83858 | 0.85607 |
| ENSG00000166170 | 14 | 104022881 | 104029168 | 12 | 3 | -1.1974 | 0.88443 | BAG5 | 0.88443 | 0.8667 | 0.84715 |
| ENSG00000132801 | 20 | 44486256 | 44507761 | 105 | 10 | -1.1977 | 0.88449 | ZSWIM3 | 0.88449 | 0.85361 | 0.90539 |
| ENSG00000144867 | 3 | 133502877 | 133544616 | 107 | 21 | -1.1979 | 0.88452 | SRPRB | 0.88452 | 0.889 | 0.88295 |
| ENSG00000158246 | 1 | 27331511 | 27339327 | 11 | 2 | -1.198 | 0.88455 | FAM46B | 0.88455 | 0.78681 | 0.83309 |
| ENSG00000134259 | 1 | 115828539 | 115880857 | 170 | 37 | -1.198 | 0.88455 | NGF | 0.88455 | 0.86504 | 0.8994 |
| ENSG00000171320 | 8 | 27629466 | 27670157 | 179 | 10 | -1.198 | 0.88455 | ESCO2 | 0.88455 | 0.90584 | 0.93713 |
| ENSG00000116833 | 1 | 199996730 | 200146552 | 618 | 65 | -1.1982 | 0.88457 | NR5A2 | 0.88457 | 0.82952 | 0.64256 |
| ENSG00000215014 | 1 | 1510355 | 1511373 | 6 | 2 | -1.1983 | 0.8846 | AL645728.1 | 0.8846 | 0.91364 | 0.93466 |
| ENSG00000169087 | 3 | 122458846 | 122512671 | 249 | 23 | -1.1987 | 0.88468 | HSPBAP1 | 0.88468 | 0.8697 | 0.84439 |
| ENSG00000106178 | 7 | 75440983 | 75452674 | 75 | 16 | -1.1988 | 0.8847 | CCL24 | 0.8847 | 0.4355 | 0.36893 |
| ENSG00000170832 | 17 | 58256455 | 58499831 | 278 | 11 | -1.1994 | 0.88482 | USP32 | 0.88482 | 0.87973 | 0.9208 |
| ENSG00000184787 | 21 | 46188955 | 46221934 | 142 | 21 | -1.1996 | 0.88485 | UBE2G2 | 0.88485 | 0.82794 | 0.85077 |
| ENSG00000176454 | 15 | 34651106 | 34659479 | 31 | 11 | -1.1997 | 0.88486 | LPCAT4 | 0.88486 | 0.41933 | 0.46476 |
| ENSG00000145826 | 5 | 135257968 | 135290723 | 155 | 20 | -1.2002 | 0.88497 | LECT2 | 0.88497 | 0.80905 | 0.88876 |
| ENSG00000204767 | 5 | 169291268 | 169407744 | 341 | 55 | -1.2006 | 0.88505 | FAM196B | 0.88505 | 0.89335 | 0.84483 |
| ENSG00000108255 | 17 | 27573881 | 27581512 | 11 | 4 | -1.2011 | 0.88515 | CRYBA1 | 0.88515 | 0.84743 | 0.88456 |
| ENSG00000165475 | 13 | 20977806 | 21099996 | 542 | 52 | -1.2014 | 0.8852 | CRYL1 | 0.8852 | 0.87483 | 0.80976 |
| ENSG00000170100 | 16 | 89284118 | 89295363 | 76 | 7 | -1.2015 | 0.88522 | ZNF778 | 0.88522 | 0.89991 | 0.91088 |
| ENSG00000162383 | 1 | 53552855 | 53608289 | 334 | 37 | -1.2017 | 0.88526 | SLC1A7 | 0.88526 | 0.85693 | 0.89042 |
| ENSG00000188167 | 3 | 33131913 | 33138293 | 6 | 4 | -1.2018 | 0.88527 | TMPPE | 0.88527 | 0.6691 | 0.72006 |
| ENSG00000213937 | 16 | 3062457 | 3064506 | 3 | 1 | -1.2019 | 0.8853 | CLDN9 | 0.8853 | 0.88804 | 0.52088 |
| ENSG00000173212 | 1 | 116654376 | 116677861 | 88 | 15 | -1.2021 | 0.88534 | MAB21L3 | 0.88534 | 0.78249 | 0.80684 |
| ENSG00000169660 | 17 | 80376194 | 80400521 | 68 | 14 | -1.2022 | 0.88536 | HEXDC | 0.88536 | 0.86887 | 0.84278 |
| ENSG00000170439 | 12 | 56075330 | 56078395 | 13 | 5 | -1.2024 | 0.88539 | METTL7B | 0.88539 | 0.87708 | 0.90509 |
| ENSG00000181610 | 17 | 55916842 | 55927417 | 41 | 13 | -1.2023 | 0.88539 | MRPS23 | 0.88539 | 0.89384 | 0.7581 |
| ENSG00000143434 | 1 | 151104161 | 151119104 | 47 | 10 | -1.2025 | 0.88541 | SEMA6C | 0.88541 | 0.89097 | 0.90287 |
| ENSG00000255561 | 11 | 111744780 | 111751967 | 19 | 6 | -1.2028 | 0.88546 | FDXACB1 | 0.88546 | 0.89072 | 0.91474 |
| ENSG00000170236 | 15 | 50792759 | 50838905 | 197 | 20 | -1.2029 | 0.8855 | USP50 | 0.8855 | 0.86985 | 0.88615 |
| ENSG00000167397 | 16 | 31102163 | 31107301 | 9 | 3 | -1.2033 | 0.88557 | VKORC1 | 0.88557 | 0.83579 | 0.78777 |
| ENSG00000180354 | 7 | 30174426 | 30202378 | 115 | 20 | -1.2036 | 0.88562 | MTURN | 0.88562 | 0.91543 | 0.89489 |
| ENSG00000197153 | 6 | 27858093 | 27860884 | 8 | 3 | -1.2036 | 0.88563 | HIST1H3J | 0.88563 | 0.97134 | 0.94726 |
| ENSG00000170266 | 3 | 33038100 | 33138722 | 346 | 29 | -1.204 | 0.8857 | GLB1 | 0.8857 | 0.88639 | 0.93184 |
| ENSG00000131941 | 19 | 33469499 | 33555794 | 436 | 38 | -1.204 | 0.88571 | RHPN2 | 0.88571 | 0.88353 | 0.86359 |
| ENSG00000170153 | 4 | 141780961 | 142134031 | 1378 | 82 | -1.2041 | 0.88573 | RNF150 | 0.88573 | 0.90196 | 0.92399 |
| ENSG00000149308 | 11 | 108027942 | 108093369 | 116 | 12 | -1.2042 | 0.88575 | NPAT | 0.88575 | 0.88026 | 0.89043 |
| ENSG00000268130 | 13 | 113755563 | 113756608 | 3 | 2 | -1.2045 | 0.88579 | AL137002.1 | 0.88579 | 0.46604 | 0.7598 |
| ENSG00000198604 | 14 | 35221937 | 35344853 | 479 | 30 | -1.2048 | 0.88586 | BAZ1A | 0.88586 | 0.82767 | 0.90262 |
| ENSG00000132341 | 12 | 131356424 | 131362223 | 19 | 3 | -1.2048 | 0.88587 | RAN | 0.88587 | 0.7633 | 0.82321 |
| ENSG00000166451 | 16 | 81040103 | 81066719 | 62 | 14 | -1.2051 | 0.88592 | CENPN | 0.88592 | 0.78228 | 0.69208 |
| ENSG00000173548 | 15 | 75940247 | 75954642 | 22 | 7 | -1.2058 | 0.88605 | SNX33 | 0.88605 | 0.77302 | 0.26606 |
| ENSG00000262302 | 17 | 7150148 | 7165408 | 34 | 5 | -1.2059 | 0.88606 | RP1-4G17.5 | 0.88606 | 0.87094 | 0.89153 |
| ENSG00000145362 | 4 | 113739265 | 114304896 | 1873 | 133 | -1.2062 | 0.88613 | ANK2 | 0.88613 | 0.88146 | 0.89296 |
| ENSG00000182534 | 17 | 74668633 | 74707098 | 150 | 14 | -1.2062 | 0.88613 | MXRA7 | 0.88613 | 0.87935 | 0.86654 |
| ENSG00000104325 | 8 | 91013633 | 91064320 | 131 | 13 | -1.2063 | 0.88614 | DECR1 | 0.88614 | 0.81453 | 0.7877 |
| ENSG00000266173 | 17 | 61780192 | 61819330 | 105 | 9 | -1.2064 | 0.88617 | STRADA | 0.88617 | 0.87096 | 0.86324 |
| ENSG00000183640 | 21 | 32185016 | 32185570 | 1 | 1 | -1.2071 | 0.8863 | KRTAP8-1 | 0.8863 | 0.9028 | 0.90577 |
| ENSG00000196821 | 6 | 34555065 | 34664636 | 301 | 19 | -1.2071 | 0.88631 | C6orf106 | 0.88631 | 0.87794 | 0.88852 |
| ENSG00000166974 | 18 | 32556892 | 32723434 | 424 | 37 | -1.2073 | 0.88633 | MAPRE2 | 0.88633 | 0.87349 | 0.86442 |
| ENSG00000139946 | 14 | 56584532 | 56768244 | 371 | 31 | -1.2073 | 0.88634 | PELI2 | 0.88634 | 0.93647 | 0.9539 |
| ENSG00000157765 | 4 | 25656923 | 25680370 | 60 | 19 | -1.2075 | 0.88638 | SLC34A2 | 0.88638 | 0.66868 | 0.58651 |
| ENSG00000196132 | 20 | 62783144 | 62873604 | 346 | 51 | -1.2078 | 0.88644 | MYT1 | 0.88644 | 0.85536 | 0.86352 |
| ENSG00000075151 | 1 | 21132963 | 21503377 | 916 | 28 | -1.208 | 0.88647 | EIF4G3 | 0.88647 | 0.88645 | 0.89372 |
| ENSG00000204710 | 11 | 64937625 | 64940715 | 6 | 3 | -1.2081 | 0.88649 | SPDYC | 0.88649 | 0.44512 | 0.56716 |
| ENSG00000006283 | 17 | 48638429 | 48704835 | 162 | 24 | -1.2089 | 0.88664 | CACNA1G | 0.88664 | 0.90965 | 0.90539 |
| ENSG00000109193 | 4 | 70676498 | 70725870 | 135 | 19 | -1.2091 | 0.88669 | SULT1E1 | 0.88669 | 0.88335 | 0.92342 |
| ENSG00000167914 | 17 | 38119226 | 38134019 | 69 | 10 | -1.2093 | 0.88672 | GSDMA | 0.88672 | 0.8279 | 0.83562 |
| ENSG00000129562 | 14 | 23033805 | 23058175 | 184 | 11 | -1.2094 | 0.88675 | DAD1 | 0.88675 | 0.93375 | 0.96703 |

| | | | | | | | | | | | |
|---|---|---|---|---|---|---|---|---|---|---|---|
| ENSG00000145945 | 6 | 3849620 | 3851551 | 3 | 2 | -1.2096 | 0.88679 | FAM50B | 0.88679 | 0.44327 | 0.66185 |
| ENSG00000012232 | 8 | 28457986 | 28613116 | 387 | 35 | -1.21 | 0.88685 | EXTL3 | 0.88685 | 0.87387 | 0.87019 |
| ENSG00000268882 | 10 | 135192720 | 135193102 | 1 | 1 | -1.2107 | 0.887 | AL360181.1 | 0.887 | 0.56598 | 0.73833 |
| ENSG00000130287 | 19 | 19322782 | 19363042 | 117 | 15 | -1.2108 | 0.88701 | NCAN | 0.88701 | 0.88339 | 0.92296 |
| ENSG00000144713 | 3 | 12875984 | 12883087 | 34 | 5 | -1.2109 | 0.88703 | RPL32 | 0.88703 | 0.95206 | 0.91183 |
| ENSG00000129946 | 19 | 416583 | 460996 | 255 | 38 | -1.2112 | 0.88708 | SHC2 | 0.88708 | 0.93972 | 0.97937 |
| ENSG00000153956 | 7 | 81575760 | 82073114 | 1807 | 113 | -1.2115 | 0.88714 | CACNA2D1 | 0.88714 | 0.88658 | 0.86888 |
| ENSG00000179603 | 7 | 126078652 | 126893348 | 2452 | 69 | -1.2118 | 0.88721 | GRM8 | 0.88721 | 0.87722 | 0.86452 |
| ENSG00000165644 | 10 | 76993727 | 76995788 | 2 | 1 | -1.212 | 0.88725 | COMTD1 | 0.88725 | 0.39212 | 0.5222 |
| ENSG00000197408 | 19 | 41497204 | 41524303 | 158 | 19 | -1.2122 | 0.88727 | CYP2B6 | 0.88727 | 0.93089 | 0.91917 |
| ENSG00000159063 | 11 | 77811982 | 77850706 | 275 | 11 | -1.2122 | 0.88728 | ALG8 | 0.88728 | 0.89084 | 0.88592 |
| ENSG00000188582 | 3 | 142668006 | 142682178 | 44 | 13 | -1.2126 | 0.88736 | PAQR9 | 0.88736 | 0.8365 | 0.58901 |
| ENSG00000103335 | 16 | 88781751 | 88851619 | 462 | 56 | -1.2127 | 0.88737 | PIEZO1 | 0.88737 | 0.89732 | 0.89739 |
| ENSG00000121766 | 1 | 31769842 | 31837783 | 184 | 12 | -1.2127 | 0.88738 | ZCCHC17 | 0.88738 | 0.87576 | 0.90754 |
| ENSG00000244476 | 6 | 11102722 | 11111965 | 40 | 8 | -1.213 | 0.88744 | ERVFRD-1 | 0.88744 | 0.84711 | 0.91411 |
| ENSG00000163577 | 3 | 170606204 | 170626482 | 47 | 7 | -1.2132 | 0.88748 | EIF5A2 | 0.88748 | 0.82726 | 0.80027 |
| ENSG00000214595 | 2 | 54950636 | 55199157 | 957 | 73 | -1.2134 | 0.88751 | EML6 | 0.88751 | 0.86457 | 0.84919 |
| ENSG00000125798 | 20 | 22561643 | 22566093 | 8 | 1 | -1.214 | 0.88762 | FOXA2 | 0.88762 | 0.70646 | 0.81175 |
| ENSG00000138375 | 2 | 217277137 | 217347776 | 291 | 21 | -1.2147 | 0.88777 | SMARCAL1 | 0.88777 | 0.87528 | 0.93142 |
| ENSG00000145982 | 6 | 5261277 | 5771813 | 2014 | 163 | -1.215 | 0.88782 | FARS2 | 0.88782 | 0.86094 | 0.83918 |
| ENSG00000053108 | 5 | 132532147 | 132948255 | 1288 | 140 | -1.215 | 0.88783 | FSTL4 | 0.88783 | 0.83055 | 0.8543 |
| ENSG00000101473 | 20 | 44470360 | 44486045 | 57 | 16 | -1.2158 | 0.88797 | ACOT8 | 0.88797 | 0.91134 | 0.92947 |
| ENSG00000174100 | 17 | 36452989 | 36479101 | 48 | 8 | -1.2158 | 0.88798 | MRPL45 | 0.88798 | 0.91737 | 0.85692 |
| ENSG00000101236 | 20 | 3907956 | 3996229 | 265 | 15 | -1.2161 | 0.88802 | RNF24 | 0.88802 | 0.87551 | 0.88983 |
| ENSG00000125618 | 2 | 113973574 | 114036527 | 205 | 25 | -1.217 | 0.8882 | PAX8 | 0.8882 | 0.88638 | 0.83707 |
| ENSG00000267173 | 19 | 44833848 | 44905774 | 367 | 32 | -1.2171 | 0.88821 | CTC-512J12.6 | 0.88821 | 0.89222 | 0.86728 |
| ENSG00000130726 | 19 | 59055458 | 59062089 | 6 | 3 | -1.2171 | 0.88822 | TRIM28 | 0.88822 | 0.9481 | 0.95374 |
| ENSG00000186501 | 1 | 27648651 | 27662891 | 29 | 7 | -1.2172 | 0.88824 | TMEM222 | 0.88824 | 0.76608 | 0.45197 |
| ENSG00000269591 | 13 | 20268547 | 20269240 | 2 | 1 | -1.2179 | 0.88836 | AL354808.2 | 0.88836 | 0.81825 | 0.82476 |
| ENSG00000255275 | 1 | 19175767 | 19247615 | 403 | 51 | -1.2181 | 0.88841 | RP13-279N23.2 | 0.88841 | 0.93988 | 0.97374 |
| ENSG00000204970 | 5 | 140165876 | 140391929 | 577 | 23 | -1.2185 | 0.88849 | PCDHA1 | 0.88849 | 0.89004 | 0.91035 |
| ENSG00000165280 | 9 | 35056061 | 35073246 | 36 | 7 | -1.2188 | 0.88853 | VCP | 0.88853 | 0.82398 | 0.81493 |
| ENSG00000231887 | 12 | 11033610 | 11036854 | 12 | 2 | -1.2191 | 0.8886 | PRH1 | 0.8886 | 0.82805 | 0.81538 |
| ENSG00000128645 | 2 | 177053307 | 177055688 | 4 | 2 | -1.2192 | 0.88861 | HOXD1 | 0.88861 | 0.62582 | 0.51365 |
| ENSG00000176225 | 18 | 67671029 | 67873181 | 610 | 21 | -1.2192 | 0.88861 | RTTN | 0.88861 | 0.90386 | 0.91002 |
| ENSG00000226397 | 12 | 25146358 | 25150453 | 17 | 9 | -1.2192 | 0.88862 | C12orf77 | 0.88862 | 0.97113 | 0.9787 |
| ENSG00000099219 | 9 | 5765076 | 5833117 | 193 | 17 | -1.2196 | 0.88869 | ERMP1 | 0.88869 | 0.86218 | 0.88359 |
| ENSG00000112394 | 6 | 111408781 | 111552397 | 295 | 19 | -1.22 | 0.88877 | SLC16A10 | 0.88877 | 0.90346 | 0.89221 |
| ENSG00000070159 | 9 | 112137746 | 112260590 | 399 | 36 | -1.2205 | 0.88886 | PTPN3 | 0.88886 | 0.86323 | 0.85221 |
| ENSG00000166199 | 11 | 43902361 | 43941816 | 175 | 21 | -1.2206 | 0.88888 | ALKBH3 | 0.88888 | 0.88783 | 0.91932 |
| ENSG00000269559 | 4 | 75023327 | 75024085 | 1 | 1 | -1.2207 | 0.8889 | AC093677.1 | 0.8889 | 0.6881 | 0.78268 |
| ENSG00000115353 | 2 | 75273590 | 75426826 | 494 | 53 | -1.2211 | 0.88897 | TACR1 | 0.88897 | 0.90276 | 0.87233 |
| ENSG00000165355 | 14 | 39866873 | 39901704 | 100 | 11 | -1.2214 | 0.88904 | FBXO33 | 0.88904 | 0.91554 | 0.89767 |
| ENSG00000170291 | 17 | 7154735 | 7163259 | 21 | 4 | -1.2214 | 0.88904 | ELP5 | 0.88904 | 0.85423 | 0.90727 |
| ENSG00000135597 | 6 | 139224630 | 139309398 | 352 | 17 | -1.2217 | 0.88908 | REPS1 | 0.88908 | 0.90141 | 0.84531 |
| ENSG00000164742 | 7 | 45613739 | 45762715 | 507 | 47 | -1.2222 | 0.88919 | ADCY1 | 0.88919 | 0.90184 | 0.89692 |
| ENSG00000033122 | 1 | 70034081 | 70617628 | 1894 | 73 | -1.2223 | 0.88921 | LRRC7 | 0.88921 | 0.87078 | 0.86925 |
| ENSG00000128656 | 2 | 175664091 | 175870097 | 337 | 21 | -1.2226 | 0.88925 | CHN1 | 0.88925 | 0.89955 | 0.91385 |
| ENSG00000113303 | 5 | 180326077 | 180377906 | 236 | 16 | -1.2226 | 0.88927 | BTNL8 | 0.88927 | 0.86568 | 0.83534 |
| ENSG00000124762 | 6 | 36644305 | 36655116 | 44 | 10 | -1.2228 | 0.88929 | CDKN1A | 0.88929 | 0.86888 | 0.83271 |
| ENSG00000186652 | 11 | 57154267 | 57158130 | 3 | 2 | -1.2237 | 0.88948 | PRG2 | 0.88948 | 0.73437 | 0.81198 |
| ENSG00000138834 | 16 | 1756184 | 1820318 | 193 | 27 | -1.2239 | 0.88951 | MAPK8IP3 | 0.88951 | 0.88157 | 0.87964 |
| ENSG00000111737 | 12 | 120532899 | 120555306 | 82 | 17 | -1.2247 | 0.88965 | RAB35 | 0.88965 | 0.93008 | 0.90074 |
| ENSG00000118655 | 1 | 114447763 | 114456708 | 20 | 6 | -1.2259 | 0.88988 | DCLRE1B | 0.88988 | 0.76182 | 0.5446 |
| ENSG00000036054 | 3 | 99979844 | 100044095 | 135 | 13 | -1.2261 | 0.88992 | TBC1D23 | 0.88992 | 0.91283 | 0.90141 |
| ENSG00000099999 | 22 | 30773835 | 30817760 | 147 | 23 | -1.2263 | 0.88996 | RNF215 | 0.88996 | 0.79449 | 0.73174 |
| ENSG00000092295 | 14 | 24718320 | 24733638 | 41 | 15 | -1.2266 | 0.89001 | TGM1 | 0.89001 | 0.95828 | 0.91969 |
| ENSG00000171560 | 4 | 155504278 | 155511918 | 12 | 6 | -1.2267 | 0.89003 | FGA | 0.89003 | 0.74581 | 0.61679 |
| ENSG00000011007 | 1 | 24069645 | 24088549 | 24 | 11 | -1.2268 | 0.89005 | TCEB3 | 0.89005 | 0.69911 | 0.67086 |
| ENSG00000023445 | 11 | 102188215 | 102210134 | 31 | 6 | -1.2272 | 0.89013 | BIRC3 | 0.89013 | 0.79187 | 0.73664 |
| ENSG00000181963 | 11 | 4470525 | 4471591 | 1 | 1 | -1.2276 | 0.8902 | OR52K2 | 0.8902 | 0.22806 | 0.52481 |
| ENSG00000196465 | 12 | 56546040 | 56553431 | 11 | 5 | -1.2276 | 0.8902 | MYL6B | 0.8902 | 0.70535 | 0.77108 |
| ENSG00000162595 | 1 | 68511645 | 68517314 | 13 | 5 | -1.2278 | 0.89023 | DIRAS3 | 0.89023 | 0.87576 | 0.76341 |
| ENSG00000256043 | 4 | 156845270 | 156875069 | 152 | 25 | -1.2281 | 0.89029 | CTSO | 0.89029 | 0.86426 | 0.70513 |
| ENSG00000184923 | 10 | 88985205 | 88996665 | 18 | 6 | -1.2284 | 0.89036 | NUTM2A | 0.89036 | 0.92937 | 0.96675 |
| ENSG00000117360 | 1 | 150293925 | 150325671 | 113 | 14 | -1.2285 | 0.89038 | PRPF3 | 0.89038 | 0.85288 | 0.86958 |
| ENSG00000134531 | 12 | 13349650 | 13369708 | 52 | 14 | -1.2289 | 0.89044 | EMP1 | 0.89044 | 0.96194 | 0.97404 |
| ENSG00000111726 | 12 | 22199108 | 22218608 | 50 | 9 | -1.229 | 0.89046 | CMAS | 0.89046 | 0.70225 | 0.82684 |
| ENSG00000106263 | 7 | 2393721 | 2420380 | 115 | 14 | -1.2292 | 0.89051 | EIF3B | 0.89051 | 0.93656 | 0.92841 |
| ENSG00000115286 | 19 | 1383526 | 1395583 | 45 | 8 | -1.2296 | 0.89058 | NDUFS7 | 0.89058 | 0.87139 | 0.79914 |
| ENSG00000114450 | 3 | 179116990 | 179169378 | 129 | 18 | -1.2308 | 0.89079 | GNB4 | 0.89079 | 0.86908 | 0.89943 |
| ENSG00000116984 | 1 | 236958610 | 237067281 | 311 | 15 | -1.2309 | 0.89082 | MTR | 0.89082 | 0.87678 | 0.86255 |
| ENSG00000150687 | 11 | 86502101 | 86663952 | 728 | 64 | -1.2314 | 0.89091 | PRSS23 | 0.89091 | 0.9173 | 0.94379 |

| | | | | | | | | | | | | |
|---|---|---|---|---|---|---|---|---|---|---|---|---|
| ENSG00000119965 | 10 | 124690419 | 124713919 | 39 | 9 | -1.2315 | 0.89094 | C10orf88 | | 0.89094 | 0.83547 | 0.82894 |
| ENSG00000116785 | 1 | 196743925 | 196763203 | 109 | 3 | -1.2319 | 0.89101 | CFHR3 | | 0.89101 | 0.8595 | 0.82905 |
| ENSG00000187815 | 1 | 40942887 | 40962015 | 72 | 15 | -1.232 | 0.89103 | ZFP69 | | 0.89103 | 0.72717 | 0.70727 |
| ENSG00000173557 | 2 | 26785450 | 26802400 | 49 | 9 | -1.2327 | 0.89116 | C2orf70 | | 0.89116 | 0.94846 | 0.83515 |
| ENSG00000124593 | 6 | 41748087 | 41757879 | 33 | 8 | -1.2334 | 0.8913 | PRICKLE4 | | 0.8913 | 0.78189 | 0.73402 |
| ENSG00000177614 | 1 | 230457392 | 230561475 | 455 | 45 | -1.234 | 0.8914 | PGBD5 | | 0.8914 | 0.83578 | 0.78233 |
| ENSG00000141371 | 17 | 58469790 | 58508782 | 59 | 4 | -1.2342 | 0.89143 | C17orf64 | | 0.89143 | 0.93059 | 0.93705 |
| ENSG00000070882 | 7 | 24836158 | 25021253 | 606 | 68 | -1.2344 | 0.89147 | OSBPL3 | | 0.89147 | 0.91206 | 0.83455 |
| ENSG00000065150 | 13 | 98605912 | 98676551 | 156 | 19 | -1.2345 | 0.89148 | IPO5 | | 0.89148 | 0.86981 | 0.90803 |
| ENSG00000008988 | 8 | 56979854 | 56987069 | 31 | 5 | -1.2345 | 0.8915 | RPS20 | | 0.8915 | 0.9385 | 0.94668 |
| ENSG00000196605 | 19 | 9862669 | 9903856 | 85 | 11 | -1.2353 | 0.89164 | ZNF846 | | 0.89164 | 0.84119 | 0.83026 |
| ENSG00000175535 | 10 | 118305443 | 118327367 | 37 | 7 | -1.2354 | 0.89165 | PNLIP | | 0.89165 | 0.8547 | 0.89988 |
| ENSG00000085117 | 11 | 44585977 | 44641913 | 249 | 44 | -1.2355 | 0.89168 | CD82 | | 0.89168 | 0.91619 | 0.86429 |
| ENSG00000066027 | 1 | 212458879 | 212535200 | 216 | 24 | -1.2356 | 0.8917 | PPP2R5A | | 0.8917 | 0.87814 | 0.89886 |
| ENSG00000185028 | 5 | 191626 | 195468 | 13 | 4 | -1.2356 | 0.8917 | LRRC14B | | 0.8917 | 0.96112 | 0.94127 |
| ENSG00000164304 | 6 | 7326889 | 7389976 | 254 | 21 | -1.237 | 0.89195 | CAGE1 | | 0.89195 | 0.85393 | 0.7559 |
| ENSG00000154997 | 7 | 55861237 | 55930482 | 271 | 24 | -1.237 | 0.89196 | | 44088 | 0.89196 | 0.89094 | 0.87203 |
| ENSG00000188157 | 1 | 955503 | 991496 | 150 | 19 | -1.2371 | 0.89197 | AGRN | | 0.89197 | 0.83968 | 0.8354 |
| ENSG00000172432 | 6 | 43573053 | 43596899 | 45 | 10 | -1.2371 | 0.89197 | GTPBP2 | | 0.89197 | 0.76456 | 0.82441 |
| ENSG00000135424 | 12 | 56078352 | 56109827 | 74 | 8 | -1.2371 | 0.89197 | ITGA7 | | 0.89197 | 0.89863 | 0.9301 |
| ENSG00000197444 | 10 | 50942689 | 50970425 | 103 | 13 | -1.2374 | 0.89204 | OGDHL | | 0.89204 | 0.88434 | 0.88684 |
| ENSG00000183908 | 11 | 56949221 | 56959191 | 32 | 7 | -1.2375 | 0.89205 | LRRC55 | | 0.89205 | 0.84615 | 0.84223 |
| ENSG00000167113 | 9 | 131084815 | 131096351 | 31 | 7 | -1.2375 | 0.89206 | COQ4 | | 0.89206 | 0.91345 | 0.90541 |
| ENSG00000260092 | 16 | 75562430 | 75579326 | 78 | 9 | -1.2376 | 0.89206 | RP11-77K12.7 | | 0.89206 | 0.87815 | 0.91093 |
| ENSG00000125245 | 13 | 99906968 | 99913998 | 15 | 6 | -1.2382 | 0.89218 | GPR18 | | 0.89218 | 0.71906 | 0.51241 |
| ENSG00000188107 | 6 | 64429876 | 66417118 | 7033 | 128 | -1.2388 | 0.89229 | EYS | | 0.89229 | 0.89638 | 0.90078 |
| ENSG00000163125 | 1 | 150335567 | 150449042 | 439 | 17 | -1.2397 | 0.89245 | RPRD2 | | 0.89245 | 0.88896 | 0.88006 |
| ENSG00000180530 | 21 | 16333556 | 16437321 | 234 | 31 | -1.2399 | 0.8925 | NRIP1 | | 0.8925 | 0.92909 | 0.87447 |
| ENSG00000165140 | 9 | 97365415 | 97402531 | 161 | 19 | -1.24 | 0.89252 | FBP1 | | 0.89252 | 0.87397 | 0.80261 |
| ENSG00000166090 | 14 | 23842018 | 23845612 | 12 | 4 | -1.2407 | 0.89264 | IL25 | | 0.89264 | 0.92505 | 0.9199 |
| ENSG00000114423 | 3 | 105374305 | 105588396 | 664 | 28 | -1.2411 | 0.89271 | CBLB | | 0.89271 | 0.87653 | 0.85107 |
| ENSG00000126254 | 19 | 36119932 | 36128588 | 18 | 6 | -1.2411 | 0.89271 | RBM42 | | 0.89271 | 0.87017 | 0.85039 |
| ENSG00000132518 | 17 | 7905912 | 7923657 | 42 | 11 | -1.2411 | 0.89272 | GUCY2D | | 0.89272 | 0.87863 | 0.81441 |
| ENSG00000239382 | 19 | 36500022 | 36505141 | 2 | 1 | -1.2419 | 0.89286 | ALKBH6 | | 0.89286 | 0.68264 | 0.53107 |
| ENSG00000184709 | 9 | 140063210 | 140064503 | 4 | 2 | -1.2419 | 0.89287 | LRRC26 | | 0.89287 | 0.7436 | 0.76913 |
| ENSG00000117115 | 1 | 17393256 | 17445948 | 292 | 17 | -1.242 | 0.89288 | PADI2 | | 0.89288 | 0.89878 | 0.93541 |
| ENSG00000158473 | 1 | 158149737 | 158154686 | 10 | 3 | -1.2423 | 0.89294 | CD1D | | 0.89294 | 0.86381 | 0.91521 |
| ENSG00000205495 | 11 | 5067756 | 5068691 | 7 | 2 | -1.2425 | 0.89298 | OR52J3 | | 0.89298 | 0.70971 | 0.71754 |
| ENSG00000185875 | 10 | 25305587 | 25315593 | 35 | 6 | -1.2428 | 0.89303 | THNSL1 | | 0.89303 | 0.94389 | 0.96257 |
| ENSG00000141441 | 18 | 29704840 | 30050447 | 1079 | 53 | -1.243 | 0.89306 | GAREM | | 0.89306 | 0.90064 | 0.89947 |
| ENSG00000168282 | 14 | 50087489 | 50090198 | 2 | 1 | -1.2431 | 0.89308 | MGAT2 | | 0.89308 | 0.8711 | 0.79024 |
| ENSG00000004779 | 16 | 23592323 | 23607677 | 66 | 8 | -1.2431 | 0.89308 | NDUFAB1 | | 0.89308 | 0.90289 | 0.93034 |
| ENSG00000124440 | 19 | 46800303 | 46846690 | 204 | 36 | -1.2431 | 0.89309 | HIF3A | | 0.89309 | 0.72348 | 0.54349 |
| ENSG00000168005 | 11 | 63580860 | 63595190 | 33 | 6 | -1.2434 | 0.89313 | C11orf84 | | 0.89313 | 0.83202 | 0.95072 |
| ENSG00000167740 | 17 | 4046462 | 4090605 | 150 | 17 | -1.2436 | 0.89317 | CYB5D2 | | 0.89317 | 0.8768 | 0.82479 |
| ENSG00000186716 | 22 | 23521891 | 23660224 | 458 | 39 | -1.2439 | 0.89322 | BCR | | 0.89322 | 0.89375 | 0.93109 |
| ENSG00000172058 | 5 | 70196492 | 70214357 | 3 | 1 | -1.2444 | 0.89333 | SERF1A | | 0.89333 | 0.89076 | 0.61076 |
| ENSG00000107815 | 10 | 102747124 | 102754158 | 16 | 3 | -1.2448 | 0.8934 | C10orf2 | | 0.8934 | 0.86407 | 0.8613 |
| ENSG00000090686 | 1 | 22004791 | 22110099 | 404 | 32 | -1.2449 | 0.89342 | USP48 | | 0.89342 | 0.89347 | 0.8729 |
| ENSG00000187775 | 17 | 76419778 | 76573476 | 978 | 90 | -1.2452 | 0.89347 | DNAH17 | | 0.89347 | 0.90448 | 0.89621 |
| ENSG00000101052 | 20 | 42219571 | 42275936 | 176 | 13 | -1.2455 | 0.89353 | IFT52 | | 0.89353 | 0.90582 | 0.88467 |
| ENSG00000204962 | 5 | 140220907 | 140391929 | 438 | 21 | -1.2461 | 0.89364 | PCDHA8 | | 0.89364 | 0.92432 | 0.92946 |
| ENSG00000071909 | 2 | 171034655 | 171511681 | 1835 | 172 | -1.2465 | 0.8937 | MYO3B | | 0.8937 | 0.88656 | 0.93528 |
| ENSG00000105778 | 7 | 32535038 | 33078516 | 2314 | 42 | -1.2465 | 0.89371 | AVL9 | | 0.89371 | 0.89349 | 0.89959 |
| ENSG00000100027 | 22 | 22051833 | 22090123 | 126 | 29 | -1.2466 | 0.89373 | YPEL1 | | 0.89373 | 0.88214 | 0.90424 |
| ENSG00000173193 | 3 | 122399465 | 122449687 | 185 | 23 | -1.2471 | 0.89382 | PARP14 | | 0.89382 | 0.88478 | 0.90074 |
| ENSG00000104980 | 19 | 7991603 | 8008805 | 82 | 9 | -1.2475 | 0.89388 | TIMM44 | | 0.89388 | 0.9486 | 0.96833 |
| ENSG00000167182 | 17 | 45973516 | 46006323 | 63 | 13 | -1.2475 | 0.89389 | SP2 | | 0.89389 | 0.89684 | 0.84955 |
| ENSG00000161634 | 12 | 55038375 | 55042277 | 12 | 5 | -1.2477 | 0.89392 | DCD | | 0.89392 | 0.73122 | 0.82404 |
| ENSG00000118407 | 6 | 76001575 | 76203454 | 358 | 22 | -1.248 | 0.89398 | FILIP1 | | 0.89398 | 0.90333 | 0.90166 |
| ENSG00000111224 | 12 | 3900213 | 3982608 | 279 | 19 | -1.2486 | 0.89409 | PARP11 | | 0.89409 | 0.92035 | 0.9462 |
| ENSG00000015153 | 12 | 42550906 | 42632151 | 177 | 20 | -1.2491 | 0.89418 | YAF2 | | 0.89418 | 0.88959 | 0.90842 |
| ENSG00000183439 | 4 | 165875598 | 165898820 | 58 | 8 | -1.2491 | 0.89419 | TRIM61 | | 0.89419 | 0.85321 | 0.66788 |
| ENSG00000205300 | 20 | 4050799 | 4055812 | 40 | 13 | -1.2491 | 0.89419 | RP11-352D3.2 | | 0.89419 | 0.81412 | 0.66731 |
| ENSG00000137203 | 6 | 10393419 | 10419892 | 58 | 13 | -1.2494 | 0.89424 | TFAP2A | | 0.89424 | 0.95651 | 0.85472 |
| ENSG00000039537 | 5 | 41142336 | 41261540 | 415 | 28 | -1.2502 | 0.89438 | C6 | | 0.89438 | 0.89494 | 0.9133 |
| ENSG00000112787 | 12 | 133066137 | 133161774 | 511 | 33 | -1.2503 | 0.89441 | FBRSL1 | | 0.89441 | 0.88776 | 0.93011 |
| ENSG00000173917 | 17 | 46618256 | 46623441 | 13 | 8 | -1.2503 | 0.89441 | HOXB2 | | 0.89441 | 0.95126 | 0.94225 |
| ENSG00000143882 | 2 | 10861775 | 10925236 | 243 | 36 | -1.2505 | 0.89444 | ATP6V1C2 | | 0.89444 | 0.9475 | 0.95494 |
| ENSG00000134215 | 1 | 108113782 | 108507766 | 1489 | 98 | -1.2506 | 0.89447 | VAV3 | | 0.89447 | 0.87303 | 0.8537 |
| ENSG00000229937 | 7 | 18066405 | 18067486 | 2 | 1 | -1.2506 | 0.89447 | PRPS1L1 | | 0.89447 | 0.0053685 | 0.0052675 |
| ENSG00000133315 | 11 | 63766030 | 63933578 | 456 | 60 | -1.251 | 0.89454 | MACROD1 | | 0.89454 | 0.88052 | 0.89056 |
| ENSG00000072364 | 5 | 132211071 | 132299326 | 245 | 16 | -1.2515 | 0.89462 | AFF4 | | 0.89462 | 0.9018 | 0.90334 |

| | | | | | | | | | | | |
|---|---|---|---|---|---|---|---|---|---|---|---|
| ENSG00000100612 | 14 | 60610838 | 60636574 | 86 | 4 | -1.252 | 0.89471 | DHRS7 | 0.89471 | 0.91206 | 0.92903 |
| ENSG00000108256 | 17 | 27582854 | 27621136 | 73 | 10 | -1.2524 | 0.89479 | NUFIP2 | 0.89479 | 0.88937 | 0.89039 |
| ENSG00000074964 | 1 | 17866330 | 18024369 | 331 | 66 | -1.2526 | 0.89482 | ARHGEF10L | 0.89482 | 0.90095 | 0.90074 |
| ENSG00000119640 | 14 | 75519924 | 75536186 | 30 | 3 | -1.2527 | 0.89484 | ACYP1 | 0.89484 | 0.91568 | 0.89925 |
| ENSG00000170584 | 5 | 162873532 | 162887146 | 28 | 7 | -1.2528 | 0.89485 | NUDCD2 | 0.89485 | 0.80883 | 0.83272 |
| ENSG00000180861 | 12 | 13524023 | 13540101 | 60 | 7 | -1.2529 | 0.89488 | C12orf36 | 0.89488 | 0.82602 | 0.91018 |
| ENSG00000217075 | 2 | 36758948 | 36779411 | 72 | 13 | -1.2531 | 0.89492 | AC007401.2 | 0.89492 | 0.92582 | 0.9219 |
| ENSG00000170049 | 17 | 7825177 | 7833121 | 18 | 6 | -1.2531 | 0.89492 | KCNAB3 | 0.89492 | 0.72192 | 0.7507 |
| ENSG00000165609 | 10 | 12207324 | 12238143 | 84 | 18 | -1.2535 | 0.89498 | NUDT5 | 0.89498 | 0.93847 | 0.91179 |
| ENSG00000124664 | 6 | 34505579 | 34524110 | 51 | 13 | -1.2538 | 0.89504 | SPDEF | 0.89504 | 0.80821 | 0.82884 |
| ENSG00000171396 | 17 | 39315910 | 39316983 | 1 | 1 | -1.2541 | 0.8951 | KRTAP4-4 | 0.8951 | 0.67171 | 0.24113 |
| ENSG00000157456 | 15 | 59397277 | 59417244 | 46 | 10 | -1.2542 | 0.89512 | CCNB2 | 0.89512 | 0.85329 | 0.8471 |
| ENSG00000103024 | 16 | 1820287 | 1821731 | 1 | 1 | -1.2547 | 0.8952 | NME3 | 0.8952 | 0.82002 | 0.87015 |
| ENSG00000182261 | 11 | 7980971 | 7986973 | 22 | 7 | -1.2548 | 0.89522 | NLRP10 | 0.89522 | 0.79569 | 0.6519 |
| ENSG00000071243 | 7 | 120590803 | 120617270 | 51 | 6 | -1.2549 | 0.89524 | ING3 | 0.89524 | 0.93064 | 0.9345 |
| ENSG00000237353 | 11 | 125703211 | 125709964 | 34 | 7 | -1.255 | 0.89525 | PATE4 | 0.89525 | 0.83215 | 0.85436 |
| ENSG00000173905 | 3 | 167726465 | 167813763 | 277 | 23 | -1.2557 | 0.89539 | GOLIM4 | 0.89539 | 0.90447 | 0.91156 |
| ENSG00000108106 | 19 | 55912652 | 55919145 | 22 | 9 | -1.2567 | 0.89557 | UBE2S | 0.89557 | 0.91638 | 0.6225 |
| ENSG00000081842 | 5 | 140207563 | 140391929 | 469 | 23 | -1.2575 | 0.89571 | PCDHA6 | 0.89571 | 0.92392 | 0.88736 |
| ENSG00000154914 | 17 | 9548015 | 9633008 | 296 | 33 | -1.2581 | 0.89582 | USP43 | 0.89582 | 0.9106 | 0.94702 |
| ENSG00000269650 | 20 | 34756461 | 34758180 | 7 | 2 | -1.2581 | 0.89582 | AL121895.1 | 0.89582 | 0.95367 | 0.84755 |
| ENSG00000140743 | 16 | 22357257 | 22448486 | 82 | 23 | -1.2582 | 0.89584 | CDR2 | 0.89584 | 0.89022 | 0.83794 |
| ENSG00000204965 | 5 | 140201222 | 140391929 | 476 | 23 | -1.2594 | 0.89606 | PCDHA5 | 0.89606 | 0.87399 | 0.91654 |
| ENSG00000159259 | 21 | 37757676 | 37791313 | 135 | 14 | -1.2597 | 0.89612 | CHAF1B | 0.89612 | 0.83206 | 0.91563 |
| ENSG00000107816 | 10 | 102756375 | 102767593 | 32 | 7 | -1.2606 | 0.89627 | LZTS2 | 0.89627 | 0.87241 | 0.7236 |
| ENSG00000163914 | 3 | 129247483 | 129254012 | 17 | 5 | -1.2612 | 0.89639 | RHO | 0.89639 | 0.94565 | 0.93115 |
| ENSG00000137463 | 4 | 140187317 | 140201492 | 32 | 5 | -1.2613 | 0.89639 | MGARP | 0.89639 | 0.86019 | 0.84524 |
| ENSG00000213463 | 14 | 70838148 | 70883778 | 155 | 12 | -1.2614 | 0.89642 | SYNJ2BP | 0.89642 | 0.89348 | 0.93386 |
| ENSG00000175390 | 11 | 7991798 | 8023409 | 137 | 28 | -1.2615 | 0.89644 | EIF3F | 0.89644 | 0.9276 | 0.90982 |
| ENSG00000157884 | 2 | 26804070 | 26864236 | 165 | 24 | -1.2616 | 0.89645 | CIB4 | 0.89645 | 0.92833 | 0.87012 |
| ENSG00000119787 | 2 | 38522022 | 38604427 | 270 | 20 | -1.2617 | 0.89647 | ATL2 | 0.89647 | 0.89343 | 0.86494 |
| ENSG00000204702 | 6 | 29068386 | 29069658 | 7 | 2 | -1.2617 | 0.89648 | OR2J1 | 0.89648 | 0.88708 | 0.89711 |
| ENSG00000183873 | 3 | 38589548 | 38691164 | 345 | 51 | -1.2619 | 0.8965 | SCN5A | 0.8965 | 0.89888 | 0.85689 |
| ENSG00000155307 | 21 | 15857549 | 15955723 | 416 | 42 | -1.2621 | 0.89654 | SAMSN1 | 0.89654 | 0.77027 | 0.78172 |
| ENSG00000065154 | 10 | 126085872 | 126107545 | 89 | 11 | -1.2622 | 0.89656 | OAT | 0.89656 | 0.91245 | 0.49617 |
| ENSG00000010278 | 12 | 6308881 | 6347425 | 70 | 11 | -1.2624 | 0.89659 | CD9 | 0.89659 | 0.86469 | 0.79497 |
| ENSG00000129071 | 3 | 129149787 | 129158878 | 24 | 5 | -1.2625 | 0.89661 | MBD4 | 0.89661 | 0.91004 | 0.96908 |
| ENSG00000214456 | 19 | 4522543 | 4535236 | 54 | 11 | -1.2624 | 0.89661 | PLIN5 | 0.89661 | 0.79236 | 0.64646 |
| ENSG00000179399 | 13 | 92050929 | 93519490 | 4948 | 212 | -1.2628 | 0.89667 | GPC5 | 0.89667 | 0.89076 | 0.94993 |
| ENSG00000108506 | 17 | 59942731 | 60005377 | 120 | 15 | -1.2629 | 0.89669 | INTS2 | 0.89669 | 0.89192 | 0.89826 |
| ENSG00000148516 | 10 | 31607424 | 31818742 | 264 | 18 | -1.2631 | 0.89673 | ZEB1 | 0.89673 | 0.87644 | 0.89154 |
| ENSG00000169299 | 4 | 37828255 | 37864558 | 218 | 19 | -1.2635 | 0.8968 | PGM2 | 0.8968 | 0.8457 | 0.86839 |
| ENSG00000133612 | 7 | 150782918 | 150841523 | 170 | 25 | -1.264 | 0.89688 | AGAP3 | 0.89688 | 0.87635 | 0.88449 |
| ENSG00000225885 | 8 | 119294378 | 119412844 | 424 | 30 | -1.2641 | 0.89691 | AC023590.1 | 0.89691 | 0.89703 | 0.87857 |
| ENSG00000178015 | 5 | 94955782 | 94957846 | 3 | 2 | -1.2644 | 0.89696 | GPR150 | 0.89696 | 0.49072 | 0.079076 |
| ENSG00000229615 | 2 | 87089023 | 87092827 | 8 | 3 | -1.2645 | 0.89697 | AC111200.1 | 0.89697 | 0.51656 | 0.44316 |
| ENSG00000196408 | 16 | 2028918 | 2034193 | 33 | 6 | -1.2645 | 0.89698 | NOXO1 | 0.89698 | 0.9093 | 0.95354 |
| ENSG00000112874 | 5 | 102884556 | 102898494 | 46 | 7 | -1.2649 | 0.89705 | NUDT12 | 0.89705 | 0.97814 | 0.97686 |
| ENSG00000150551 | 2 | 133402426 | 133429152 | 72 | 4 | -1.265 | 0.89706 | LYPD1 | 0.89706 | 0.90735 | 0.8598 |
| ENSG00000074582 | 2 | 219523487 | 219528166 | 10 | 3 | -1.2651 | 0.89708 | BCS1L | 0.89708 | 0.87743 | 0.89527 |
| ENSG00000122376 | 10 | 88853918 | 88951225 | 314 | 20 | -1.2651 | 0.89708 | FAM35A | 0.89708 | 0.89772 | 0.90619 |
| ENSG00000156587 | 11 | 57319129 | 57335757 | 56 | 10 | -1.2655 | 0.89715 | UBE2L6 | 0.89715 | 0.80081 | 0.80361 |
| ENSG00000078808 | 1 | 1152288 | 1167411 | 117 | 6 | -1.2658 | 0.8972 | SDF4 | 0.8972 | 0.92418 | 0.71714 |
| ENSG00000128699 | 2 | 190635049 | 190649097 | 33 | 6 | -1.2659 | 0.89723 | ORMDL1 | 0.89723 | 0.79639 | 0.79297 |
| ENSG00000160087 | 1 | 1189289 | 1209265 | 61 | 10 | -1.266 | 0.89725 | UBE2J2 | 0.89725 | 0.97624 | 0.95917 |
| ENSG00000171505 | 9 | 125288637 | 125289572 | 6 | 3 | -1.2666 | 0.89736 | OR1N1 | 0.89736 | 0.60718 | 0.74113 |
| ENSG00000258653 | 14 | 74318674 | 74398981 | 385 | 17 | -1.267 | 0.89742 | RP5-1021I20.4 | 0.89742 | 0.89372 | 0.86812 |
| ENSG00000165837 | 13 | 46108657 | 46189874 | 250 | 33 | -1.2672 | 0.89747 | FAM194B | 0.89747 | 0.85586 | 0.86155 |
| ENSG00000129003 | 15 | 62144588 | 62352672 | 601 | 24 | -1.2673 | 0.89748 | VPS13C | 0.89748 | 0.89248 | 0.89857 |
| ENSG00000197322 | 17 | 32901142 | 32906388 | 15 | 5 | -1.2674 | 0.89749 | C17orf102 | 0.89749 | 0.94768 | 0.86367 |
| ENSG00000140990 | 16 | 2009509 | 2011976 | 25 | 5 | -1.2689 | 0.89776 | NDUFB10 | 0.89776 | 0.87299 | 0.91264 |
| ENSG00000066084 | 12 | 50898768 | 51142450 | 540 | 20 | -1.2694 | 0.89785 | DIP2B | 0.89785 | 0.90412 | 0.8961 |
| ENSG00000181852 | 12 | 56598285 | 56615717 | 26 | 13 | -1.2696 | 0.89788 | RNF41 | 0.89788 | 0.91923 | 0.69952 |
| ENSG00000106617 | 7 | 151253197 | 151574210 | 1295 | 150 | -1.2697 | 0.89791 | PRKAG2 | 0.89791 | 0.91328 | 0.9247 |
| ENSG00000161011 | 5 | 179233388 | 179265078 | 157 | 28 | -1.2702 | 0.898 | SQSTM1 | 0.898 | 0.90505 | 0.90661 |
| ENSG00000135362 | 11 | 36317838 | 36486754 | 655 | 67 | -1.2703 | 0.89802 | PRR5L | 0.89802 | 0.85939 | 0.82353 |
| ENSG00000136573 | 8 | 11351510 | 11422113 | 276 | 30 | -1.2705 | 0.89804 | BLK | 0.89804 | 0.8609 | 0.61398 |
| ENSG00000248483 | 5 | 93070508 | 93077343 | 7 | 2 | -1.2706 | 0.89807 | POU5F2 | 0.89807 | 0.94694 | 0.93969 |
| ENSG00000109047 | 17 | 9799637 | 9808938 | 68 | 9 | -1.2708 | 0.89811 | RCVRN | 0.89811 | 0.6561 | 0.72682 |
| ENSG00000175505 | 11 | 67131639 | 67141648 | 24 | 4 | -1.2709 | 0.89812 | CLCF1 | 0.89812 | 0.8967 | 0.87205 |
| ENSG00000143995 | 2 | 66660584 | 66801001 | 308 | 46 | -1.2714 | 0.8982 | MEIS1 | 0.8982 | 0.91307 | 0.88009 |
| ENSG00000128284 | 22 | 36536372 | 36562225 | 112 | 19 | -1.272 | 0.89831 | APOL3 | 0.89831 | 0.46456 | 0.52132 |
| ENSG00000168228 | 4 | 25314407 | 25372005 | 185 | 18 | -1.2723 | 0.89837 | ZCCHC4 | 0.89837 | 0.87384 | 0.87815 |

| | | | | | | | | | | | |
|---|---|---|---|---|---|---|---|---|---|---|---|
| ENSG00000005471 | 7 | 87031013 | 87109751 | 227 | 19 | -1.2726 | 0.89842 | ABCB4 | 0.89842 | 0.92804 | 0.93029 |
| ENSG00000142453 | 19 | 10982189 | 11033453 | 94 | 14 | -1.2727 | 0.89843 | CARM1 | 0.89843 | 0.89379 | 0.86288 |
| ENSG00000076555 | 12 | 109554400 | 109706031 | 560 | 45 | -1.2729 | 0.89847 | ACACB | 0.89847 | 0.90164 | 0.89625 |
| ENSG00000100842 | 14 | 23825611 | 23834961 | 26 | 6 | -1.2733 | 0.89855 | EFS | 0.89855 | 0.81622 | 0.95146 |
| ENSG00000108231 | 10 | 95517566 | 95557916 | 117 | 20 | -1.2736 | 0.89859 | LGI1 | 0.89859 | 0.30089 | 0.11632 |
| ENSG00000137198 | 6 | 16238811 | 16295780 | 200 | 30 | -1.2736 | 0.8986 | GMPR | 0.8986 | 0.90201 | 0.93963 |
| ENSG00000024422 | 19 | 48216600 | 48246391 | 130 | 21 | -1.2737 | 0.89861 | EHD2 | 0.89861 | 0.93336 | 0.90315 |
| ENSG00000152127 | 2 | 134877554 | 135212192 | 1125 | 91 | -1.2742 | 0.89869 | MGAT5 | 0.89869 | 0.75851 | 0.68792 |
| ENSG00000137745 | 11 | 102813724 | 102826463 | 27 | 5 | -1.2741 | 0.89869 | MMP13 | 0.89869 | 0.97839 | 0.95454 |
| ENSG00000164645 | 7 | 88423420 | 88425035 | 14 | 7 | -1.2743 | 0.89872 | C7orf62 | 0.89872 | 0.3742 | 0.21935 |
| ENSG00000172803 | 11 | 65601112 | 65624367 | 45 | 13 | -1.2743 | 0.89872 | SNX32 | 0.89872 | 0.87491 | 0.89668 |
| ENSG00000196778 | 11 | 4510109 | 4511138 | 3 | 1 | -1.2743 | 0.89873 | OR52K1 | 0.89873 | 0.864 | 0.81871 |
| ENSG00000205403 | 4 | 110661852 | 110723335 | 184 | 18 | -1.2747 | 0.89879 | CFI | 0.89879 | 0.93237 | 0.95504 |
| ENSG00000164434 | 6 | 123100620 | 123105219 | 12 | 3 | -1.275 | 0.89885 | FABP7 | 0.89885 | 0.95186 | 0.79192 |
| ENSG00000204681 | 6 | 29523406 | 29601753 | 314 | 35 | -1.2751 | 0.89886 | GABBR1 | 0.89886 | 0.91239 | 0.91747 |
| ENSG00000146670 | 11 | 64833772 | 64851636 | 50 | 13 | -1.2752 | 0.89888 | CDCA5 | 0.89888 | 0.87977 | 0.85386 |
| ENSG00000170921 | 17 | 61086917 | 61505060 | 779 | 27 | -1.2753 | 0.8989 | TANC2 | 0.8989 | 0.90677 | 0.91836 |
| ENSG00000125864 | 20 | 17474550 | 17549865 | 313 | 51 | -1.2755 | 0.89894 | BFSP1 | 0.89894 | 0.9175 | 0.79352 |
| ENSG00000163806 | 2 | 29005383 | 29073477 | 159 | 16 | -1.2757 | 0.89897 | SPDYA | 0.89897 | 0.91914 | 0.92653 |
| ENSG00000205076 | 19 | 39261611 | 39264132 | 4 | 2 | -1.2758 | 0.89898 | LGALS7 | 0.89898 | 0.88857 | 0.83065 |
| ENSG00000040341 | 8 | 74332604 | 74659943 | 1189 | 48 | -1.2758 | 0.89899 | STAU2 | 0.89899 | 0.91064 | 0.91905 |
| ENSG00000151718 | 4 | 184020446 | 184241930 | 494 | 47 | -1.2759 | 0.899 | WWC2 | 0.899 | 0.89991 | 0.9209 |
| ENSG00000149541 | 11 | 62382768 | 62389647 | 8 | 3 | -1.2761 | 0.89903 | B3GAT3 | 0.89903 | 0.66057 | 0.45328 |
| ENSG00000130147 | 2 | 235860617 | 235964358 | 351 | 42 | -1.2763 | 0.89908 | SH3BP4 | 0.89908 | 0.89831 | 0.88145 |
| ENSG00000170950 | 6 | 49753366 | 49755053 | 1 | 1 | -1.2764 | 0.8991 | PGK2 | 0.8991 | 0.43755 | 0.33813 |
| ENSG00000198948 | 4 | 170907748 | 170954182 | 131 | 20 | -1.2769 | 0.89917 | MFAP3L | 0.89917 | 0.87215 | 0.83661 |
| ENSG00000204961 | 5 | 140227048 | 140391929 | 425 | 21 | -1.2769 | 0.89918 | PCDHA9 | 0.89918 | 0.91985 | 0.91748 |
| ENSG00000134318 | 2 | 11319887 | 11488456 | 594 | 20 | -1.277 | 0.89919 | ROCK2 | 0.89919 | 0.8852 | 0.90331 |
| ENSG00000175766 | 5 | 176057683 | 176073642 | 22 | 8 | -1.2771 | 0.89922 | EIF4E1B | 0.89922 | 0.83646 | 0.75236 |
| ENSG00000105550 | 19 | 49258816 | 49261587 | 13 | 5 | -1.2771 | 0.89922 | FGF21 | 0.89922 | 0.94906 | 0.96315 |
| ENSG00000107295 | 9 | 17579080 | 17797127 | 1038 | 56 | -1.2771 | 0.89923 | SH3GL2 | 0.89923 | 0.89323 | 0.83069 |
| ENSG00000017427 | 12 | 102789645 | 102874423 | 161 | 24 | -1.2773 | 0.89925 | IGF1 | 0.89925 | 0.92023 | 0.94916 |
| ENSG00000134490 | 18 | 20777108 | 21017925 | 620 | 51 | -1.2774 | 0.89928 | TMEM241 | 0.89928 | 0.91187 | 0.91214 |
| ENSG00000162399 | 1 | 55464606 | 55476556 | 37 | 9 | -1.2776 | 0.8993 | BSND | 0.8993 | 0.76228 | 0.77642 |
| ENSG00000118523 | 6 | 132269316 | 132272513 | 6 | 4 | -1.278 | 0.89938 | CTGF | 0.89938 | 0.036823 | 0.066391 |
| ENSG00000268965 | 17 | 76422409 | 76422834 | 3 | 2 | -1.2782 | 0.8994 | AC061992.1 | 0.8994 | 0.7608 | 0.58654 |
| ENSG00000171234 | 4 | 69917081 | 69978705 | 404 | 11 | -1.2782 | 0.89941 | UGT2B7 | 0.89941 | 0.90418 | 0.90328 |
| ENSG00000241233 | 11 | 71249071 | 71250253 | 2 | 1 | -1.2783 | 0.89943 | KRTAP5-8 | 0.89943 | 0.92116 | 0.90276 |
| ENSG00000163297 | 4 | 80822303 | 81046608 | 653 | 41 | -1.2785 | 0.89947 | ANTXR2 | 0.89947 | 0.89939 | 0.79092 |
| ENSG00000112984 | 5 | 137514408 | 137523404 | 12 | 6 | -1.2786 | 0.89947 | KIF20A | 0.89947 | 0.57286 | 0.52704 |
| ENSG00000180909 | 11 | 6173006 | 6173818 | 1 | 1 | -1.2787 | 0.8995 | OR52B1P | 0.8995 | 0.87619 | 0.91867 |
| ENSG00000156738 | 11 | 60223225 | 60238233 | 41 | 8 | -1.2791 | 0.89956 | MS4A1 | 0.89956 | 0.95814 | 0.98109 |
| ENSG00000064042 | 4 | 41361624 | 41702061 | 988 | 76 | -1.2791 | 0.89958 | LIMCH1 | 0.89958 | 0.79411 | 0.5866 |
| ENSG00000198670 | 6 | 160952515 | 161087407 | 306 | 24 | -1.2798 | 0.89968 | LPA | 0.89968 | 0.90097 | 0.90481 |
| ENSG00000061987 | 12 | 62860597 | 62991363 | 398 | 25 | -1.2797 | 0.89968 | MON2 | 0.89968 | 0.9029 | 0.93667 |
| ENSG00000106258 | 7 | 99245817 | 99277621 | 48 | 12 | -1.2798 | 0.89969 | CYP3A5 | 0.89969 | 0.88554 | 0.83915 |
| ENSG00000086619 | 1 | 236378855 | 236445319 | 435 | 13 | -1.2799 | 0.89971 | ERO1LB | 0.89971 | 0.92484 | 0.91647 |
| ENSG00000110203 | 11 | 71825915 | 71850936 | 120 | 14 | -1.28 | 0.89973 | FOLR3 | 0.89973 | 0.89088 | 0.89945 |
| ENSG00000187595 | 17 | 40177594 | 40217383 | 87 | 11 | -1.2802 | 0.89977 | ZNF385C | 0.89977 | 0.87037 | 0.86672 |
| ENSG00000178053 | 3 | 158288952 | 158325041 | 148 | 14 | -1.2806 | 0.89984 | MLF1 | 0.89984 | 0.84899 | 0.84295 |
| ENSG00000164134 | 4 | 140222609 | 140341187 | 285 | 17 | -1.281 | 0.8999 | NAA15 | 0.8999 | 0.87595 | 0.85408 |
| ENSG00000129757 | 11 | 2904443 | 2907111 | 5 | 3 | -1.2814 | 0.89998 | CDKN1C | 0.89998 | 0.1762 | 0.56189 |
| ENSG00000150722 | 2 | 182818968 | 182996125 | 463 | 50 | -1.2817 | 0.90002 | PPP1R1C | 0.90002 | 0.91955 | 0.94613 |
| ENSG00000197376 | 12 | 48901137 | 48923680 | 122 | 14 | -1.2816 | 0.90002 | OR8S1 | 0.90002 | 0.91626 | 0.90407 |
| ENSG00000179583 | 16 | 10971055 | 11026079 | 183 | 38 | -1.282 | 0.90007 | CIITA | 0.90007 | 0.92012 | 0.72184 |
| ENSG00000128059 | 4 | 57259528 | 57301781 | 143 | 16 | -1.2828 | 0.90022 | PPAT | 0.90022 | 0.9175 | 0.93484 |
| ENSG00000221954 | 11 | 50003009 | 50004071 | 2 | 1 | -1.2828 | 0.90022 | OR4C12 | 0.90022 | 0.47932 | 0.46361 |
| ENSG00000073969 | 17 | 44668035 | 44834830 | 121 | 13 | -1.2831 | 0.90028 | NSF | 0.90028 | 0.91742 | 0.89368 |
| ENSG00000185621 | 3 | 197687071 | 197770591 | 294 | 19 | -1.2843 | 0.90048 | LMLN | 0.90048 | 0.87896 | 0.80127 |
| ENSG00000079689 | 6 | 25652464 | 25702011 | 180 | 28 | -1.2843 | 0.90048 | SCGN | 0.90048 | 0.91269 | 0.89016 |
| ENSG00000203690 | 6 | 167770521 | 167797998 | 200 | 30 | -1.2845 | 0.90052 | TCP10 | 0.90052 | 0.8954 | 0.90528 |
| ENSG00000149115 | 11 | 57067112 | 57092426 | 75 | 12 | -1.285 | 0.90061 | TNKS1BP1 | 0.90061 | 0.86461 | 0.8815 |
| ENSG00000113555 | 5 | 141323150 | 141349304 | 61 | 13 | -1.2852 | 0.90064 | PCDH12 | 0.90064 | 0.88305 | 0.90584 |
| ENSG00000165072 | 9 | 72658497 | 72841886 | 652 | 20 | -1.2853 | 0.90066 | MAMDC2 | 0.90066 | 0.88006 | 0.82241 |
| ENSG00000148346 | 9 | 130911350 | 130915734 | 7 | 4 | -1.2857 | 0.90073 | LCN2 | 0.90073 | 0.77587 | 0.84959 |
| ENSG00000197037 | 7 | 99214569 | 99230030 | 19 | 4 | -1.2858 | 0.90074 | ZSCAN25 | 0.90074 | 0.74787 | 0.84507 |
| ENSG00000180758 | 1 | 9160364 | 9189229 | 102 | 14 | -1.2862 | 0.90081 | GPR157 | 0.90081 | 0.92715 | 0.91041 |
| ENSG00000203795 | 10 | 124670217 | 124672627 | 5 | 2 | -1.2865 | 0.90087 | FAM24A | 0.90087 | 0.90699 | 0.93754 |
| ENSG00000170605 | 12 | 55523465 | 55524586 | 5 | 1 | -1.2867 | 0.90089 | OR9K2 | 0.90089 | 0.91253 | 0.93698 |
| ENSG00000117118 | 1 | 17345217 | 17380665 | 132 | 12 | -1.2867 | 0.90091 | SDHB | 0.90091 | 0.93716 | 0.95076 |
| ENSG00000227184 | 8 | 144939497 | 144952632 | 42 | 16 | -1.2868 | 0.90092 | EPPK1 | 0.90092 | 0.91518 | 0.92124 |
| ENSG00000118495 | 6 | 144261437 | 144385735 | 363 | 45 | -1.2872 | 0.90099 | PLAGL1 | 0.90099 | 0.93113 | 0.94212 |
| ENSG00000158691 | 6 | 28346732 | 28367511 | 74 | 10 | -1.2874 | 0.90101 | ZSCAN12 | 0.90101 | 0.88096 | 0.85178 |

| | | | | | | | | | | | |
|---|---|---|---|---|---|---|---|---|---|---|---|
| ENSG00000102539 | 13 | 49794474 | 49796513 | 2 | 1 | -1.2874 | 0.90102 | MLNR | 0.90102 | 0.69427 | 0.78357 |
| ENSG00000125821 | 20 | 18568537 | 18744561 | 798 | 24 | -1.2875 | 0.90104 | DTD1 | 0.90104 | 0.92129 | 0.91963 |
| ENSG00000172006 | 19 | 2819872 | 2835771 | 80 | 13 | -1.2875 | 0.90105 | ZNF554 | 0.90105 | 0.87695 | 0.88121 |
| ENSG00000196109 | 19 | 22361893 | 22379753 | 54 | 5 | -1.2878 | 0.90109 | ZNF676 | 0.90109 | 0.85511 | 0.81063 |
| ENSG00000109861 | 11 | 88026760 | 88070955 | 180 | 21 | -1.288 | 0.90113 | CTSC | 0.90113 | 0.86015 | 0.80568 |
| ENSG00000116711 | 1 | 186798085 | 186958113 | 534 | 45 | -1.2881 | 0.90115 | PLA2G4A | 0.90115 | 0.91903 | 0.94389 |
| ENSG00000214212 | 19 | 10947251 | 10980466 | 63 | 14 | -1.2881 | 0.90115 | C19orf38 | 0.90115 | 0.89633 | 0.88952 |
| ENSG00000167264 | 16 | 68021649 | 68113223 | 166 | 16 | -1.2882 | 0.90117 | DUS2 | 0.90117 | 0.89579 | 0.902 |
| ENSG00000129347 | 19 | 10663761 | 10676713 | 64 | 10 | -1.2884 | 0.9012 | KRI1 | 0.9012 | 0.95755 | 0.97071 |
| ENSG00000159348 | 1 | 202930997 | 202936408 | 12 | 3 | -1.2887 | 0.90124 | CYB5R1 | 0.90124 | 0.9021 | 0.88253 |
| ENSG00000130511 | 19 | 18529674 | 18545372 | 32 | 7 | -1.2891 | 0.90131 | SSBP4 | 0.90131 | 0.88039 | 0.87259 |
| ENSG00000147536 | 8 | 41386725 | 41402565 | 50 | 12 | -1.2892 | 0.90133 | GINS4 | 0.90133 | 0.83194 | 0.84623 |
| ENSG00000197584 | 3 | 177990720 | 178562217 | 1916 | 99 | -1.2897 | 0.90142 | KCNMB2 | 0.90142 | 0.84182 | 0.82426 |
| ENSG00000081818 | 5 | 140501581 | 140505201 | 10 | 3 | -1.2898 | 0.90145 | PCDHB4 | 0.90145 | 0.84461 | 0.82623 |
| ENSG00000107833 | 10 | 103541082 | 103543170 | 3 | 1 | -1.2903 | 0.90153 | NPM3 | 0.90153 | 0.84906 | 0.53108 |
| ENSG00000134160 | 15 | 31293264 | 31453476 | 638 | 69 | -1.2904 | 0.90155 | TRPM1 | 0.90155 | 0.9262 | 0.93602 |
| ENSG00000106436 | 7 | 101256605 | 101272576 | 60 | 14 | -1.2906 | 0.90157 | MYL10 | 0.90157 | 0.65609 | 0.36477 |
| ENSG00000136352 | 14 | 36985602 | 36990354 | 10 | 4 | -1.2906 | 0.90157 | NKX2-1 | 0.90157 | 0.74021 | 0.81446 |
| ENSG00000159131 | 21 | 34876238 | 34915797 | 79 | 18 | -1.2909 | 0.90163 | GART | 0.90163 | 0.88247 | 0.83905 |
| ENSG00000213639 | 2 | 28974506 | 29025806 | 143 | 14 | -1.291 | 0.90165 | PPP1CB | 0.90165 | 0.85867 | 0.84564 |
| ENSG00000175985 | 14 | 69951409 | 69995215 | 130 | 23 | -1.291 | 0.90165 | PLEKHD1 | 0.90165 | 0.92952 | 0.9425 |
| ENSG00000104805 | 19 | 49403307 | 49426629 | 105 | 23 | -1.2914 | 0.90172 | NUCB1 | 0.90172 | 0.90167 | 0.95735 |
| ENSG00000184110 | 16 | 28699879 | 28747051 | 25 | 11 | -1.2919 | 0.9018 | EIF3C | 0.9018 | 0.92602 | 0.88204 |
| ENSG00000171055 | 2 | 36778570 | 36873230 | 382 | 29 | -1.2924 | 0.9019 | FEZ2 | 0.9019 | 0.89762 | 0.81653 |
| ENSG00000269430 | 21 | 45879814 | 45882078 | 6 | 2 | -1.2926 | 0.90192 | LRRC3DN | 0.90192 | 0.33574 | 0.55968 |
| ENSG00000171310 | 12 | 104849073 | 105155792 | 1182 | 127 | -1.2928 | 0.90196 | CHST11 | 0.90196 | 0.93442 | 0.93526 |
| ENSG00000185344 | 12 | 124196865 | 124246302 | 175 | 10 | -1.2928 | 0.90196 | ATP6V0A2 | 0.90196 | 0.91779 | 0.89624 |
| ENSG00000174456 | 12 | 110465872 | 110511491 | 142 | 23 | -1.2928 | 0.90197 | C12orf76 | 0.90197 | 0.90212 | 0.86592 |
| ENSG00000163346 | 1 | 154916552 | 154928599 | 16 | 8 | -1.2929 | 0.90198 | PBXIP1 | 0.90198 | 0.89697 | 0.87892 |
| ENSG00000174371 | 1 | 242011269 | 242058450 | 274 | 21 | -1.2932 | 0.90203 | EXO1 | 0.90203 | 0.89757 | 0.8939 |
| ENSG00000100852 | 14 | 32545320 | 32628934 | 74 | 32 | -1.2935 | 0.90208 | ARHGAP5 | 0.90208 | 0.87141 | 0.53638 |
| ENSG00000082397 | 18 | 5392383 | 5630699 | 817 | 71 | -1.2938 | 0.90212 | EPB41L3 | 0.90212 | 0.88166 | 0.66813 |
| ENSG00000157168 | 8 | 31496902 | 32622548 | 4249 | 157 | -1.2939 | 0.90214 | NRG1 | 0.90214 | 0.90393 | 0.90243 |
| ENSG00000126457 | 19 | 50179043 | 50192286 | 25 | 15 | -1.2942 | 0.9022 | PRMT1 | 0.9022 | 0.9244 | 0.83226 |
| ENSG00000143363 | 1 | 150980896 | 151008189 | 99 | 26 | -1.2945 | 0.90225 | PRUNE | 0.90225 | 0.93352 | 0.93393 |
| ENSG00000180875 | 1 | 240652873 | 240775449 | 487 | 84 | -1.2945 | 0.90225 | GREM2 | 0.90225 | 0.87258 | 0.87188 |
| ENSG00000249034 | 5 | 140240341 | 140243224 | 11 | 4 | -1.2953 | 0.9024 | AC005609.1 | 0.9024 | 0.76641 | 0.7413 |
| ENSG00000237524 | 2 | 127656472 | 127659673 | 17 | 3 | -1.2956 | 0.90245 | AC114783.1 | 0.90245 | 0.85114 | 0.86109 |
| ENSG00000090530 | 3 | 189674517 | 189840226 | 770 | 32 | -1.2957 | 0.90247 | LEPREL1 | 0.90247 | 0.90513 | 0.92161 |
| ENSG00000101084 | 20 | 35234137 | 35240960 | 13 | 7 | -1.296 | 0.90251 | C20orf24 | 0.90251 | 0.52593 | 0.64994 |
| ENSG00000151240 | 10 | 320130 | 735683 | 1130 | 61 | -1.2961 | 0.90253 | DIP2C | 0.90253 | 0.86519 | 0.88709 |
| ENSG00000142065 | 19 | 36827162 | 36870101 | 119 | 12 | -1.2967 | 0.90262 | ZFP14 | 0.90262 | 0.91362 | 0.89145 |
| ENSG00000089159 | 12 | 120648250 | 120703574 | 87 | 13 | -1.2971 | 0.9027 | PXN | 0.9027 | 0.85468 | 0.79626 |
| ENSG00000112851 | 5 | 65222303 | 65378377 | 232 | 20 | -1.2975 | 0.90278 | ERBB2IP | 0.90278 | 0.91693 | 0.861 |
| ENSG00000110536 | 11 | 47586982 | 47595013 | 10 | 5 | -1.2978 | 0.90283 | PTPMT1 | 0.90283 | 0.63997 | 0.72658 |
| ENSG00000255439 | 16 | 31094760 | 31106277 | 24 | 6 | -1.298 | 0.90286 | RP11-196G11.1 | 0.90286 | 0.81612 | 0.80948 |
| ENSG00000065600 | 1 | 212537273 | 212588243 | 157 | 24 | -1.2981 | 0.90287 | TMEM206 | 0.90287 | 0.88765 | 0.90031 |
| ENSG00000168143 | 6 | 54711569 | 54806820 | 263 | 24 | -1.2981 | 0.90288 | FAM83B | 0.90288 | 0.88168 | 0.86679 |
| ENSG00000188710 | 9 | 133768815 | 133769225 | 3 | 2 | -1.2984 | 0.90292 | QRFP | 0.90292 | 0.80543 | 0.81618 |
| ENSG00000167971 | 16 | 2227184 | 2246526 | 24 | 6 | -1.2985 | 0.90294 | CASKIN1 | 0.90294 | 0.69625 | 0.79957 |
| ENSG00000235109 | 6 | 28292470 | 28324048 | 129 | 20 | -1.2992 | 0.90306 | ZSCAN31 | 0.90306 | 0.88358 | 0.91708 |
| ENSG00000117143 | 1 | 162531323 | 162569627 | 68 | 12 | -1.2997 | 0.90315 | UAP1 | 0.90315 | 0.90366 | 0.92188 |
| ENSG00000198440 | 19 | 56909335 | 56947404 | 89 | 12 | -1.2999 | 0.90318 | ZNF583 | 0.90318 | 0.85915 | 0.83762 |
| ENSG00000228208 | 1 | 218683438 | 218699320 | 57 | 3 | -1.3 | 0.9032 | C1orf143 | 0.9032 | 0.90161 | 0.90011 |
| ENSG00000249590 | 22 | 30805244 | 30824977 | 52 | 13 | -1.3001 | 0.90322 | RP4-539M6.19 | 0.90322 | 0.88203 | 0.92242 |
| ENSG00000241598 | 11 | 1642188 | 1643368 | 11 | 6 | -1.3002 | 0.90324 | KRTAP5-4 | 0.90324 | 0.95376 | 0.76684 |
| ENSG00000133872 | 8 | 29920528 | 29940723 | 40 | 7 | -1.3006 | 0.9033 | TMEM66 | 0.9033 | 0.89595 | 0.86586 |
| ENSG00000167962 | 16 | 2047655 | 2059824 | 66 | 11 | -1.3009 | 0.90335 | ZNF598 | 0.90335 | 0.85501 | 0.9012 |
| ENSG00000112029 | 6 | 153291664 | 153304714 | 49 | 8 | -1.3011 | 0.90339 | FBXO5 | 0.90339 | 0.94766 | 0.89629 |
| ENSG00000153233 | 12 | 71031853 | 71314623 | 956 | 86 | -1.3012 | 0.9034 | PTPRR | 0.9034 | 0.89504 | 0.8738 |
| ENSG00000232748 | 16 | 31072813 | 31073451 | 1 | 1 | -1.3012 | 0.9034 | ZNF668 | 0.9034 | 0.83226 | 0.81858 |
| ENSG00000107242 | 9 | 71320575 | 71624092 | 1007 | 44 | -1.3025 | 0.90363 | PIP5K1B | 0.90363 | 0.91636 | 0.94138 |
| ENSG00000145734 | 5 | 70751442 | 70863649 | 294 | 11 | -1.3034 | 0.90378 | BDP1 | 0.90378 | 0.90566 | 0.90958 |
| ENSG00000235034 | 19 | 51152702 | 51162567 | 37 | 12 | -1.3036 | 0.90381 | C19orf81 | 0.90381 | 0.96064 | 0.965 |
| ENSG00000168621 | 5 | 37812779 | 37839788 | 91 | 19 | -1.304 | 0.90389 | GDNF | 0.90389 | 0.88537 | 0.84485 |
| ENSG00000134265 | 18 | 10525902 | 10552758 | 72 | 6 | -1.3044 | 0.90394 | NAPG | 0.90394 | 0.89686 | 0.8985 |
| ENSG00000164631 | 7 | 6728064 | 6746554 | 64 | 14 | -1.3046 | 0.90399 | ZNF12 | 0.90399 | 0.90795 | 0.76473 |
| ENSG00000197747 | 1 | 151955391 | 151966866 | 22 | 5 | -1.3047 | 0.904 | S100A10 | 0.904 | 0.70569 | 0.65379 |
| ENSG00000137709 | 11 | 120107349 | 120190653 | 235 | 34 | -1.3051 | 0.90407 | POU2F3 | 0.90407 | 0.85719 | 0.77372 |
| ENSG00000196284 | 6 | 44777054 | 45345690 | 1996 | 30 | -1.3052 | 0.90409 | SUPT3H | 0.90409 | 0.8982 | 0.90004 |
| ENSG00000141741 | 17 | 37884749 | 37887040 | 3 | 1 | -1.3052 | 0.90409 | MIEN1 | 0.90409 | 0.82617 | 0.90577 |
| ENSG00000255168 | 1 | 145209145 | 145319796 | 1 | 1 | -1.3053 | 0.9041 | RP11-458D21.5 | 0.9041 | 0.30883 | 0.30883 |
| ENSG00000143393 | 1 | 151264273 | 151300191 | 37 | 7 | -1.3054 | 0.90412 | PI4KB | 0.90412 | 0.87287 | 0.86319 |

| | | | | | | | | | | | |
|---|---|---|---|---|---|---|---|---|---|---|---|
| ENSG00000204694 | 6 | 29393281 | 29424848 | 158 | 14 | -1.3057 | 0.90417 | OR11A1 | 0.90417 | 0.95913 | 0.94186 |
| ENSG00000145740 | 5 | 68389473 | 68426896 | 113 | 11 | -1.3061 | 0.90425 | SLC30A5 | 0.90425 | 0.88541 | 0.86766 |
| ENSG00000138892 | 22 | 50453552 | 50495364 | 214 | 41 | -1.3061 | 0.90425 | TTLL8 | 0.90425 | 0.39238 | 0.37405 |
| ENSG00000125522 | 20 | 62737173 | 62738524 | 5 | 2 | -1.3063 | 0.90428 | NPBWR2 | 0.90428 | 0.79546 | 0.88525 |
| ENSG00000136206 | 7 | 44040488 | 44049721 | 24 | 9 | -1.3068 | 0.90436 | SPDYE1 | 0.90436 | 0.9774 | 0.97517 |
| ENSG00000096395 | 6 | 33762450 | 33771788 | 39 | 8 | -1.3072 | 0.90442 | MLN | 0.90442 | 0.76111 | 0.88271 |
| ENSG00000082996 | 3 | 149530495 | 149679926 | 351 | 24 | -1.3072 | 0.90443 | RNF13 | 0.90443 | 0.92855 | 0.93485 |
| ENSG00000153789 | 16 | 85131965 | 85146114 | 101 | 18 | -1.3073 | 0.90445 | FAM92B | 0.90445 | 0.84083 | 0.85241 |
| ENSG00000166046 | 12 | 106695707 | 106740793 | 129 | 16 | -1.3079 | 0.90455 | TCP11L2 | 0.90455 | 0.91475 | 0.82882 |
| ENSG00000100038 | 22 | 22311397 | 22337213 | 74 | 9 | -1.308 | 0.90456 | TOP3B | 0.90456 | 0.95648 | 0.96723 |
| ENSG00000170456 | 12 | 31535157 | 31744031 | 904 | 27 | -1.3082 | 0.90461 | DENND5B | 0.90461 | 0.89831 | 0.91306 |
| ENSG00000130193 | 8 | 143808621 | 143818345 | 31 | 9 | -1.309 | 0.90474 | THEM6 | 0.90474 | 0.8035 | 0.80966 |
| ENSG00000180921 | 8 | 144806103 | 144815971 | 34 | 6 | -1.3098 | 0.90487 | FAM83H | 0.90487 | 0.9252 | 0.95448 |
| ENSG00000101670 | 18 | 47087069 | 47119272 | 90 | 21 | -1.3098 | 0.90487 | LIPG | 0.90487 | 0.90142 | 0.91157 |
| ENSG00000099956 | 22 | 24129150 | 24176703 | 341 | 22 | -1.31 | 0.9049 | SMARCB1 | 0.9049 | 0.89935 | 0.88022 |
| ENSG00000172493 | 4 | 87856154 | 88062206 | 651 | 36 | -1.3102 | 0.90494 | AFF1 | 0.90494 | 0.91391 | 0.92053 |
| ENSG00000168434 | 16 | 23399814 | 23464501 | 137 | 13 | -1.3104 | 0.90497 | COG7 | 0.90497 | 0.92081 | 0.93726 |
| ENSG00000105146 | 19 | 57742377 | 57746916 | 18 | 4 | -1.3105 | 0.90498 | AURKC | 0.90498 | 0.76938 | 0.64989 |
| ENSG00000101191 | 20 | 61509090 | 61569304 | 233 | 28 | -1.311 | 0.90508 | DIDO1 | 0.90508 | 0.90511 | 0.73684 |
| ENSG00000140367 | 15 | 76135622 | 76193419 | 89 | 6 | -1.3112 | 0.90511 | UBE2Q2 | 0.90511 | 0.92538 | 0.90525 |
| ENSG00000269808 | 19 | 55738587 | 55741647 | 9 | 3 | -1.3114 | 0.90513 | AC010327.2 | 0.90513 | 0.94771 | 0.97802 |
| ENSG00000132254 | 11 | 6496910 | 6502666 | 4 | 2 | -1.3115 | 0.90516 | ARFIP2 | 0.90516 | 0.83049 | 0.84258 |
| ENSG00000078399 | 7 | 27202054 | 27210117 | 20 | 9 | -1.3119 | 0.90522 | HOXA9 | 0.90522 | 0.94431 | 0.82017 |
| ENSG00000264472 | 18 | 66567018 | 66569492 | 8 | 4 | -1.3125 | 0.90532 | RP11-861L17.3 | 0.90532 | 0.94824 | 0.93228 |
| ENSG00000249158 | 5 | 140248689 | 140391929 | 363 | 18 | -1.3131 | 0.90543 | PCDHA11 | 0.90543 | 0.92576 | 0.92372 |
| ENSG00000168490 | 8 | 22077222 | 22089854 | 45 | 15 | -1.3133 | 0.90545 | PHYHIP | 0.90545 | 0.93278 | 0.87952 |
| ENSG00000147869 | 9 | 14719722 | 14722715 | 16 | 5 | -1.3132 | 0.90545 | CER1 | 0.90545 | 0.96928 | 0.18051 |
| ENSG00000115137 | 2 | 25166505 | 25194963 | 63 | 13 | -1.3134 | 0.90548 | DNAJC27 | 0.90548 | 0.93988 | 0.93338 |
| ENSG00000198089 | 22 | 31884674 | 32014572 | 550 | 22 | -1.3136 | 0.9055 | SFI1 | 0.9055 | 0.87926 | 0.87228 |
| ENSG00000137288 | 6 | 33662070 | 33679504 | 82 | 9 | -1.3142 | 0.90561 | UQCC2 | 0.90561 | 0.88913 | 0.90805 |
| ENSG00000115423 | 2 | 84743579 | 85046713 | 871 | 23 | -1.3142 | 0.90562 | DNAH6 | 0.90562 | 0.90159 | 0.88562 |
| ENSG00000146232 | 6 | 44225903 | 44233500 | 19 | 6 | -1.3142 | 0.90562 | NFKBIE | 0.90562 | 0.69199 | 0.67805 |
| ENSG00000163191 | 1 | 152004982 | 152020383 | 24 | 4 | -1.3146 | 0.90567 | S100A11 | 0.90567 | 0.90828 | 0.77732 |
| ENSG00000145685 | 5 | 77781038 | 78065844 | 820 | 71 | -1.3146 | 0.90567 | LHFPL2 | 0.90567 | 0.81751 | 0.87472 |
| ENSG00000164609 | 5 | 159828648 | 159848718 | 70 | 10 | -1.3154 | 0.90581 | SLU7 | 0.90581 | 0.92961 | 0.89894 |
| ENSG00000128050 | 4 | 57301907 | 57327534 | 81 | 10 | -1.3157 | 0.90586 | PAICS | 0.90586 | 0.93506 | 0.92736 |
| ENSG00000248099 | 19 | 17927321 | 17932383 | 12 | 4 | -1.3157 | 0.90586 | INSL3 | 0.90586 | 0.90753 | 0.93348 |
| ENSG00000136783 | 9 | 107509969 | 107522403 | 36 | 9 | -1.3159 | 0.90589 | NIPSNAP3A | 0.90589 | 0.94324 | 0.8852 |
| ENSG00000249141 | 6 | 167271582 | 167369612 | 362 | 40 | -1.3162 | 0.90595 | RP11-514O12.4 | 0.90595 | 0.90908 | 0.94097 |
| ENSG00000215041 | 17 | 7218947 | 7232712 | 32 | 9 | -1.3163 | 0.90596 | NEURL4 | 0.90596 | 0.93701 | 0.92975 |
| ENSG00000266967 | 17 | 41102543 | 41116515 | 18 | 5 | -1.3169 | 0.90606 | AARSD1 | 0.90606 | 0.73019 | 0.7112 |
| ENSG00000189007 | 6 | 143743969 | 143771810 | 103 | 4 | -1.3177 | 0.90619 | ADAT2 | 0.90619 | 0.88614 | 0.91346 |
| ENSG00000167770 | 11 | 63753325 | 63769283 | 28 | 5 | -1.3176 | 0.90619 | OTUB1 | 0.90619 | 0.88615 | 0.92571 |
| ENSG00000111605 | 12 | 69633317 | 69668138 | 84 | 13 | -1.3181 | 0.90627 | CPSF6 | 0.90627 | 0.88827 | 0.87759 |
| ENSG00000196683 | 7 | 22852251 | 22862470 | 73 | 10 | -1.3182 | 0.90629 | TOMM7 | 0.90629 | 0.85266 | 0.87636 |
| ENSG00000134690 | 1 | 38158090 | 38175391 | 59 | 10 | -1.3183 | 0.9063 | CDCA8 | 0.9063 | 0.92777 | 0.97262 |
| ENSG00000171517 | 1 | 85277285 | 85358896 | 304 | 48 | -1.3186 | 0.90635 | LPAR3 | 0.90635 | 0.91258 | 0.9298 |
| ENSG00000146410 | 6 | 136552162 | 136571473 | 21 | 13 | -1.3187 | 0.90636 | MTFR2 | 0.90636 | 0.92228 | 0.94587 |
| ENSG00000158445 | 20 | 47980414 | 48099184 | 467 | 47 | -1.3192 | 0.90645 | KCNB1 | 0.90645 | 0.88373 | 0.86294 |
| ENSG00000178882 | 12 | 124457788 | 124800570 | 1194 | 100 | -1.3198 | 0.90655 | FAM101A | 0.90655 | 0.91754 | 0.9257 |
| ENSG00000145293 | 4 | 83351715 | 83382244 | 112 | 24 | -1.32 | 0.90658 | ENOPH1 | 0.90658 | 0.9454 | 0.9657 |
| ENSG00000155868 | 5 | 156564423 | 156586030 | 87 | 12 | -1.3203 | 0.90663 | MED7 | 0.90663 | 0.91643 | 0.90366 |
| ENSG00000157227 | 14 | 23305766 | 23318236 | 36 | 13 | -1.3218 | 0.90688 | MMP14 | 0.90688 | 0.76908 | 0.70598 |
| ENSG00000125510 | 20 | 62711526 | 62731996 | 74 | 22 | -1.322 | 0.90691 | OPRL1 | 0.90691 | 0.87477 | 0.90688 |
| ENSG00000088053 | 19 | 55525073 | 55549632 | 143 | 16 | -1.3225 | 0.907 | GP6 | 0.907 | 0.92407 | 0.96767 |
| ENSG00000130338 | 6 | 158733692 | 158932860 | 834 | 31 | -1.3237 | 0.9072 | TULP4 | 0.9072 | 0.90727 | 0.91608 |
| ENSG00000179837 | 3 | 51428731 | 51435330 | 8 | 2 | -1.324 | 0.90725 | RBM15B | 0.90725 | 0.75474 | 0.75944 |
| ENSG00000107581 | 10 | 120794356 | 120840316 | 89 | 12 | -1.3241 | 0.90727 | EIF3A | 0.90727 | 0.91959 | 0.92467 |
| ENSG00000167107 | 17 | 48503519 | 48552206 | 127 | 16 | -1.3242 | 0.90728 | ACSF2 | 0.90728 | 0.95307 | 0.97091 |
| ENSG00000169604 | 2 | 69240310 | 69476459 | 679 | 82 | -1.3242 | 0.90729 | ANTXR1 | 0.90729 | 0.88349 | 0.91027 |
| ENSG00000163435 | 1 | 201977073 | 201986316 | 29 | 9 | -1.3247 | 0.90737 | ELF3 | 0.90737 | 0.94176 | 0.86661 |
| ENSG00000007376 | 16 | 834974 | 838397 | 12 | 5 | -1.325 | 0.90742 | RPUSD1 | 0.90742 | 0.71274 | 0.80173 |
| ENSG00000132170 | 3 | 12328867 | 12475855 | 332 | 33 | -1.3251 | 0.90743 | PPARG | 0.90743 | 0.93176 | 0.91024 |
| ENSG00000145901 | 5 | 150409506 | 150473138 | 237 | 36 | -1.3253 | 0.90747 | TNIP1 | 0.90747 | 0.88925 | 0.95678 |
| ENSG00000167967 | 16 | 2273567 | 2285743 | 18 | 5 | -1.3257 | 0.90754 | E4F1 | 0.90754 | 0.90689 | 0.74696 |
| ENSG00000132256 | 11 | 5684425 | 5959849 | 1784 | 56 | -1.326 | 0.90758 | TRIM5 | 0.90758 | 0.91035 | 0.9215 |
| ENSG00000173452 | 7 | 19758933 | 19813221 | 224 | 20 | -1.3264 | 0.90765 | TMEM196 | 0.90765 | 0.89692 | 0.90078 |
| ENSG00000182095 | 7 | 5346421 | 5465045 | 488 | 72 | -1.3265 | 0.90767 | TNRC18 | 0.90767 | 0.87642 | 0.71841 |
| ENSG00000121274 | 16 | 50186829 | 50269221 | 208 | 20 | -1.3269 | 0.90774 | PAPD5 | 0.90774 | 0.87694 | 0.82687 |
| ENSG00000147439 | 8 | 22477931 | 22526661 | 128 | 15 | -1.3275 | 0.90782 | BIN3 | 0.90782 | 0.88825 | 0.88534 |
| ENSG00000004799 | 7 | 95212811 | 95225803 | 48 | 8 | -1.3281 | 0.90793 | PDK4 | 0.90793 | 0.74848 | 0.77695 |
| ENSG00000268941 | 20 | 57209828 | 57210772 | 5 | 2 | -1.3285 | 0.90799 | MGC4294 | 0.90799 | 0.8685 | 0.82731 |
| ENSG00000006837 | 5 | 133541305 | 133706738 | 373 | 13 | -1.3287 | 0.90803 | CDKL3 | 0.90803 | 0.89246 | 0.89604 |

| | | | | | | | | | | | |
|---|---|---|---|---|---|---|---|---|---|---|---|
| ENSG00000060339 | 10 | 70480769 | 70552134 | 182 | 20 | -1.3289 | 0.90806 | CCAR1 | 0.90806 | 0.90542 | 0.85471 |
| ENSG00000165757 | 10 | 30301729 | 30404423 | 284 | 37 | -1.329 | 0.90808 | KIAA1462 | 0.90808 | 0.93066 | 0.93963 |
| ENSG00000133997 | 14 | 71047974 | 71067384 | 70 | 10 | -1.3292 | 0.90811 | MED6 | 0.90811 | 0.82649 | 0.87511 |
| ENSG00000014641 | 2 | 63815743 | 63834331 | 35 | 6 | -1.3293 | 0.90812 | MDH1 | 0.90812 | 0.83406 | 0.75253 |
| ENSG00000158234 | 3 | 138327448 | 138352218 | 29 | 10 | -1.3295 | 0.90815 | FAIM | 0.90815 | 0.81524 | 0.86819 |
| ENSG00000123268 | 12 | 51157493 | 51214905 | 162 | 18 | -1.3299 | 0.90823 | ATF1 | 0.90823 | 0.93045 | 0.91801 |
| ENSG00000167925 | 17 | 40340817 | 40346531 | 5 | 3 | -1.3304 | 0.9083 | GHDC | 0.9083 | 0.53848 | 0.28354 |
| ENSG00000127445 | 19 | 9945933 | 9960358 | 30 | 8 | -1.3314 | 0.90848 | PIN1 | 0.90848 | 0.84565 | 0.89944 |
| ENSG00000166159 | 12 | 1929433 | 1945918 | 76 | 26 | -1.3319 | 0.90856 | LRTM2 | 0.90856 | 0.9338 | 0.85328 |
| ENSG00000212673 | 1 | 32379174 | 32380745 | 1 | 1 | -1.3322 | 0.9086 | AL136115.1 | 0.9086 | 0.85547 | 0.75741 |
| ENSG00000075275 | 22 | 46756731 | 46933067 | 980 | 96 | -1.333 | 0.90874 | CELSR1 | 0.90874 | 0.86172 | 0.9272 |
| ENSG00000204381 | 11 | 111411005 | 111432264 | 35 | 7 | -1.3332 | 0.90877 | LAYN | 0.90877 | 0.78659 | 0.6521 |
| ENSG00000125319 | 17 | 42219274 | 42239844 | 41 | 7 | -1.3337 | 0.90885 | C17orf53 | 0.90885 | 0.94166 | 0.91397 |
| ENSG00000143494 | 1 | 213123862 | 213165379 | 149 | 24 | -1.3338 | 0.90887 | VASH2 | 0.90887 | 0.93616 | 0.87272 |
| ENSG00000143028 | 1 | 110009180 | 110024759 | 51 | 5 | -1.3344 | 0.90896 | SYPL2 | 0.90896 | 0.87963 | 0.87105 |
| ENSG00000185156 | 17 | 8700436 | 8702667 | 6 | 3 | -1.3348 | 0.90902 | MFSD6L | 0.90902 | 0.85394 | 0.42387 |
| ENSG00000158477 | 1 | 158223927 | 158228059 | 20 | 7 | -1.3355 | 0.90915 | CD1A | 0.90915 | 0.98311 | 0.97384 |
| ENSG00000107147 | 9 | 138594031 | 138684992 | 423 | 49 | -1.3362 | 0.90925 | KCNT1 | 0.90925 | 0.88557 | 0.84482 |
| ENSG00000139163 | 12 | 22778009 | 22843599 | 132 | 14 | -1.3362 | 0.90926 | ETNK1 | 0.90926 | 0.94583 | 0.9586 |
| ENSG00000110104 | 11 | 60609544 | 60618554 | 39 | 12 | -1.3366 | 0.90932 | CCDC86 | 0.90932 | 0.93268 | 0.91604 |
| ENSG00000165623 | 10 | 13263767 | 13276334 | 85 | 19 | -1.3369 | 0.90937 | UCMA | 0.90937 | 0.85236 | 0.64428 |
| ENSG00000175206 | 1 | 11905766 | 11908402 | 20 | 5 | -1.3376 | 0.90948 | NPPA | 0.90948 | 0.84744 | 0.87751 |
| ENSG00000251664 | 5 | 140255058 | 140391929 | 351 | 19 | -1.3376 | 0.90948 | PCDHA12 | 0.90948 | 0.92211 | 0.93276 |
| ENSG00000110169 | 11 | 6452279 | 6463847 | 23 | 5 | -1.3376 | 0.90949 | HPX | 0.90949 | 0.91424 | 0.9364 |
| ENSG00000107164 | 9 | 133454352 | 133513739 | 169 | 25 | -1.338 | 0.90956 | FUBP3 | 0.90956 | 0.92421 | 0.97119 |
| ENSG00000143858 | 1 | 202559724 | 202679545 | 323 | 42 | -1.3385 | 0.90963 | SYT2 | 0.90963 | 0.92173 | 0.91415 |
| ENSG00000243729 | 6 | 29323007 | 29399744 | 363 | 25 | -1.3385 | 0.90964 | OR5V1 | 0.90964 | 0.92451 | 0.93411 |
| ENSG00000171484 | 9 | 125390858 | 125391852 | 9 | 3 | -1.3389 | 0.90969 | OR1B1 | 0.90969 | 0.9803 | 0.97553 |
| ENSG00000174516 | 11 | 66234216 | 66244808 | 20 | 4 | -1.3389 | 0.90969 | PELI3 | 0.90969 | 0.88045 | 0.85341 |
| ENSG00000078549 | 7 | 31092076 | 31151089 | 177 | 40 | -1.3391 | 0.90973 | ADCYAP1R1 | 0.90973 | 0.93126 | 0.93041 |
| ENSG00000106565 | 7 | 150488373 | 150498448 | 48 | 11 | -1.3394 | 0.90978 | TMEM176B | 0.90978 | 0.82886 | 0.62594 |
| ENSG00000144560 | 3 | 11597544 | 11762220 | 696 | 59 | -1.3398 | 0.90984 | VGLL4 | 0.90984 | 0.86684 | 0.91026 |
| ENSG00000142676 | 1 | 24018269 | 24022915 | 13 | 4 | -1.3399 | 0.90986 | RPL11 | 0.90986 | 0.64968 | 0.56718 |
| ENSG00000010361 | 19 | 50310126 | 50320633 | 29 | 9 | -1.3407 | 0.90999 | FUZ | 0.90999 | 0.88605 | 0.82525 |
| ENSG00000196932 | 10 | 63166401 | 63213208 | 141 | 22 | -1.3411 | 0.91005 | TMEM26 | 0.91005 | 0.90395 | 0.92473 |
| ENSG00000183103 | 19 | 42341148 | 42348760 | 16 | 2 | -1.3421 | 0.91021 | LYPD4 | 0.91021 | 0.80073 | 0.7949 |
| ENSG00000114867 | 3 | 184032283 | 184053146 | 34 | 11 | -1.3422 | 0.91023 | EIF4G1 | 0.91023 | 0.90256 | 0.78647 |
| ENSG00000109586 | 4 | 174089904 | 174245118 | 298 | 31 | -1.3424 | 0.91026 | GALNT7 | 0.91026 | 0.90566 | 0.88794 |
| ENSG00000106304 | 7 | 123565286 | 123611468 | 131 | 20 | -1.3427 | 0.91031 | SPAM1 | 0.91031 | 0.93201 | 0.953 |
| ENSG00000112561 | 6 | 41651716 | 41703997 | 175 | 30 | -1.3427 | 0.91032 | TFEB | 0.91032 | 0.82586 | 0.86562 |
| ENSG00000150768 | 11 | 111895538 | 111935114 | 76 | 10 | -1.3431 | 0.91038 | DLAT | 0.91038 | 0.8857 | 0.85063 |
| ENSG00000100012 | 22 | 30843946 | 30868036 | 133 | 12 | -1.3434 | 0.91043 | SEC14L3 | 0.91043 | 0.897 | 0.78731 |
| ENSG00000170485 | 2 | 101436614 | 101613291 | 596 | 64 | -1.3436 | 0.91046 | NPAS2 | 0.91046 | 0.9166 | 0.94652 |
| ENSG00000139117 | 12 | 39040624 | 39301232 | 933 | 33 | -1.3437 | 0.91047 | CPNE8 | 0.91047 | 0.93025 | 0.92703 |
| ENSG00000145335 | 4 | 90645250 | 90759466 | 493 | 33 | -1.3438 | 0.91049 | SNCA | 0.91049 | 0.85652 | 0.84032 |
| ENSG00000158825 | 1 | 20915441 | 20945401 | 147 | 22 | -1.3439 | 0.9105 | CDA | 0.9105 | 0.91852 | 0.92383 |
| ENSG00000129354 | 19 | 10683347 | 10697991 | 85 | 9 | -1.3447 | 0.91063 | AP1M2 | 0.91063 | 0.94178 | 0.97156 |
| ENSG00000187626 | 6 | 28212401 | 28227011 | 43 | 12 | -1.345 | 0.91068 | ZKSCAN4 | 0.91068 | 0.91071 | 0.8973 |
| ENSG00000129116 | 4 | 169418217 | 169849608 | 1588 | 113 | -1.3458 | 0.91082 | PALLD | 0.91082 | 0.91249 | 0.92467 |
| ENSG00000120903 | 8 | 27317279 | 27337400 | 93 | 18 | -1.346 | 0.91084 | CHRNA2 | 0.91084 | 0.93433 | 0.9538 |
| ENSG00000165688 | 9 | 139305110 | 139318213 | 54 | 11 | -1.3465 | 0.91093 | PMPCA | 0.91093 | 0.80291 | 0.78058 |
| ENSG00000141568 | 17 | 80477589 | 80602538 | 530 | 55 | -1.3478 | 0.91113 | FOXK2 | 0.91113 | 0.88587 | 0.80689 |
| ENSG00000172073 | 2 | 88824169 | 88829102 | 14 | 3 | -1.3478 | 0.91115 | TEX37 | 0.91115 | 0.90477 | 0.92973 |
| ENSG00000170946 | 11 | 31391387 | 31453396 | 113 | 14 | -1.3495 | 0.91141 | DNAJC24 | 0.91141 | 0.8882 | 0.87535 |
| ENSG00000147588 | 8 | 82352561 | 82359758 | 16 | 4 | -1.35 | 0.91149 | PMP2 | 0.91149 | 0.95981 | 0.9453 |
| ENSG00000009780 | 1 | 28052490 | 28089633 | 82 | 5 | -1.35 | 0.9115 | FAM76A | 0.9115 | 0.88982 | 0.92073 |
| ENSG00000110628 | 11 | 2920951 | 2946476 | 129 | 30 | -1.3506 | 0.91158 | SLC22A18 | 0.91158 | 0.7617 | 0.59922 |
| ENSG00000116096 | 2 | 73114489 | 73119287 | 2 | 1 | -1.3507 | 0.91161 | SPR | 0.91161 | 0.87103 | 0.6387 |
| ENSG00000215067 | 17 | 6779954 | 6915668 | 426 | 57 | -1.3514 | 0.91172 | AC027763.2 | 0.91172 | 0.92357 | 0.95145 |
| ENSG00000196188 | 1 | 206317459 | 206332104 | 24 | 3 | -1.3516 | 0.91175 | CTSE | 0.91175 | 0.84787 | 0.59276 |
| ENSG00000079459 | 8 | 11653082 | 11696818 | 303 | 23 | -1.3517 | 0.91177 | FDFT1 | 0.91177 | 0.87052 | 0.77258 |
| ENSG00000203326 | 19 | 53868946 | 53895470 | 170 | 26 | -1.3519 | 0.91179 | ZNF525 | 0.91179 | 0.96248 | 0.15563 |
| ENSG00000204174 | 10 | 47083534 | 47088320 | 1 | 1 | -1.3519 | 0.9118 | NPY4R | 0.9118 | 0.9118 | 0.9118 |
| ENSG00000175220 | 11 | 46698630 | 46722165 | 40 | 14 | -1.3528 | 0.91194 | ARHGAP1 | 0.91194 | 0.72475 | 0.6506 |
| ENSG00000147485 | 8 | 52232138 | 52722005 | 1959 | 63 | -1.353 | 0.91197 | PXDNL | 0.91197 | 0.88962 | 0.86619 |
| ENSG00000128383 | 22 | 39348746 | 39359188 | 30 | 11 | -1.353 | 0.91197 | APOBEC3A | 0.91197 | 0.90531 | 0.92188 |
| ENSG00000229972 | 3 | 51851620 | 51864876 | 31 | 4 | -1.3531 | 0.91198 | IQCF3 | 0.91198 | 0.82923 | 0.79205 |
| ENSG00000185130 | 6 | 27775257 | 27775709 | 2 | 1 | -1.3531 | 0.91199 | HIST1H2BL | 0.91199 | 0.71684 | 0.71709 |
| ENSG00000179270 | 2 | 29283842 | 29297127 | 66 | 10 | -1.3536 | 0.91206 | C2orf71 | 0.91206 | 0.88112 | 0.92028 |
| ENSG00000118855 | 3 | 158449987 | 158547508 | 295 | 25 | -1.3535 | 0.91206 | MFSD1 | 0.91206 | 0.86141 | 0.87371 |
| ENSG00000005001 | 16 | 2902728 | 2908171 | 16 | 5 | -1.3537 | 0.91209 | PRSS22 | 0.91209 | 0.90886 | 0.50135 |
| ENSG00000176476 | 16 | 28565236 | 28603111 | 125 | 16 | -1.3538 | 0.91209 | CCDC101 | 0.91209 | 0.88514 | 0.8825 |
| ENSG00000106560 | 7 | 150382785 | 150390729 | 16 | 6 | -1.3538 | 0.9121 | GIMAP2 | 0.9121 | 0.9485 | 0.92295 |

| | | | | | | | | | | | |
|---|---|---|---|---|---|---|---|---|---|---|---|
| ENSG00000169744 | 4 | 16503164 | 16900432 | 1313 | 135 | -1.355 | 0.9123 | LDB2 | 0.9123 | 0.91388 | 0.93208 |
| ENSG00000144566 | 3 | 19988571 | 20026667 | 172 | 20 | -1.3553 | 0.91234 | RAB5A | 0.91234 | 0.91097 | 0.87713 |
| ENSG00000205220 | 16 | 67968405 | 67970990 | 2 | 1 | -1.3556 | 0.91238 | PSMB10 | 0.91238 | 0.56756 | 0.72554 |
| ENSG00000143222 | 1 | 161122566 | 161128646 | 10 | 4 | -1.3558 | 0.91243 | UFC1 | 0.91243 | 0.92385 | 0.7993 |
| ENSG00000132842 | 5 | 77296349 | 77590579 | 753 | 25 | -1.356 | 0.91245 | AP3B1 | 0.91245 | 0.91631 | 0.93314 |
| ENSG00000196275 | 7 | 74210483 | 74267847 | 73 | 17 | -1.3579 | 0.91274 | GTF2IRD2 | 0.91274 | 0.84808 | 0.84496 |
| ENSG00000205649 | 4 | 70894130 | 70902255 | 27 | 6 | -1.3587 | 0.91288 | HTN3 | 0.91288 | 0.93459 | 0.95016 |
| ENSG00000182087 | 19 | 1009647 | 1021117 | 97 | 18 | -1.3587 | 0.91288 | TMEM259 | 0.91288 | 0.77984 | 0.41369 |
| ENSG00000254440 | 6 | 138537129 | 138539627 | 12 | 2 | -1.3589 | 0.91291 | PBOV1 | 0.91291 | 0.9094 | 0.78788 |
| ENSG00000204991 | 16 | 89884587 | 89937727 | 271 | 21 | -1.3592 | 0.91295 | SPIRE2 | 0.91295 | 0.91148 | 0.91753 |
| ENSG00000099957 | 22 | 21364097 | 21383119 | 84 | 21 | -1.3594 | 0.91299 | P2RX6 | 0.91299 | 0.79327 | 0.6406 |
| ENSG00000130653 | 9 | 140354404 | 140444986 | 201 | 49 | -1.3598 | 0.91305 | PNPLA7 | 0.91305 | 0.8588 | 0.88535 |
| ENSG00000145839 | 5 | 135227935 | 135231516 | 9 | 3 | -1.3607 | 0.9132 | IL9 | 0.9132 | 0.95911 | 0.90411 |
| ENSG00000213714 | 20 | 55108302 | 55111576 | 12 | 4 | -1.3611 | 0.91325 | FAM209B | 0.91325 | 0.90174 | 0.93467 |
| ENSG00000166091 | 14 | 23846017 | 23848981 | 8 | 4 | -1.3611 | 0.91326 | CMTM5 | 0.91326 | 0.91012 | 0.9342 |
| ENSG00000163611 | 3 | 113161565 | 113234034 | 395 | 21 | -1.3618 | 0.91337 | SPICE1 | 0.91337 | 0.92619 | 0.90311 |
| ENSG00000273266 | 12 | 54447661 | 54449814 | 8 | 4 | -1.3619 | 0.91338 | HOXC4 | 0.91338 | 0.8847 | 0.84269 |
| ENSG00000268759 | 15 | 32774822 | 32780287 | 1 | 1 | -1.362 | 0.9134 | AC135983.2 | 0.9134 | 0.9134 | 0.90308 |
| ENSG00000206530 | 3 | 113005777 | 113160457 | 671 | 26 | -1.3621 | 0.91342 | WDR52 | 0.91342 | 0.92272 | 0.93215 |
| ENSG00000124493 | 6 | 33989628 | 34123399 | 670 | 57 | -1.3625 | 0.91348 | GRM4 | 0.91348 | 0.95808 | 0.95987 |
| ENSG00000123545 | 6 | 97337189 | 97345757 | 33 | 9 | -1.3625 | 0.91348 | NDUFAF4 | 0.91348 | 0.8814 | 0.9201 |
| ENSG00000197591 | 1 | 248004199 | 248005254 | 8 | 1 | -1.3633 | 0.91361 | OR11L1 | 0.91361 | 0.89668 | 0.89356 |
| ENSG00000204479 | 1 | 13716092 | 13719089 | 3 | 1 | -1.3634 | 0.91363 | PRAMEF17 | 0.91363 | 0.70179 | 0.53587 |
| ENSG00000196586 | 6 | 76458909 | 76629254 | 396 | 24 | -1.3637 | 0.91366 | MYO6 | 0.91366 | 0.88929 | 0.83249 |
| ENSG00000131931 | 8 | 42691817 | 42698468 | 11 | 3 | -1.3636 | 0.91366 | THAP1 | 0.91366 | 0.86284 | 0.84522 |
| ENSG00000196781 | 9 | 84198598 | 84304220 | 420 | 84 | -1.3637 | 0.91366 | TLE1 | 0.91366 | 0.84284 | 0.78096 |
| ENSG00000215455 | 21 | 45958864 | 45960078 | 12 | 2 | -1.3639 | 0.91369 | KRTAP10-1 | 0.91369 | 0.46062 | 0.19301 |
| ENSG00000181867 | 5 | 121187650 | 121188519 | 1 | 1 | -1.3639 | 0.9137 | FTMT | 0.9137 | 0.44016 | 0.38471 |
| ENSG00000138606 | 15 | 45459412 | 45493373 | 60 | 10 | -1.3639 | 0.9137 | SHF | 0.9137 | 0.86905 | 0.67215 |
| ENSG00000175984 | 1 | 115125469 | 115213043 | 263 | 23 | -1.364 | 0.91372 | DENND2C | 0.91372 | 0.88887 | 0.89773 |
| ENSG00000115762 | 2 | 131862420 | 132111282 | 435 | 16 | -1.3641 | 0.91373 | PLEKHB2 | 0.91373 | 0.91561 | 0.89654 |
| ENSG00000244694 | 6 | 47845764 | 48036425 | 713 | 48 | -1.3644 | 0.91378 | PTCHD4 | 0.91378 | 0.91375 | 0.92536 |
| ENSG00000161513 | 17 | 72858619 | 72869156 | 34 | 3 | -1.3655 | 0.91396 | FDXR | 0.91396 | 0.85816 | 0.875 |
| ENSG00000163017 | 2 | 74119441 | 74146992 | 143 | 15 | -1.3659 | 0.91402 | ACTG2 | 0.91402 | 0.9212 | 0.56021 |
| ENSG00000196812 | 6 | 28092338 | 28097860 | 16 | 4 | -1.3661 | 0.91405 | ZSCAN16 | 0.91405 | 0.74577 | 0.81417 |
| ENSG00000004399 | 3 | 129274018 | 129325661 | 155 | 19 | -1.3663 | 0.91407 | PLXND1 | 0.91407 | 0.94047 | 0.94854 |
| ENSG00000152495 | 5 | 110559351 | 110830584 | 844 | 42 | -1.3665 | 0.91411 | CAMK4 | 0.91411 | 0.92557 | 0.90995 |
| ENSG00000269866 | 16 | 70258261 | 70258641 | 1 | 1 | -1.3671 | 0.9142 | FKSG63 | 0.9142 | 0.46327 | 0.43905 |
| ENSG00000198894 | 14 | 77564440 | 77583630 | 51 | 9 | -1.3673 | 0.91423 | CIPC | 0.91423 | 0.88211 | 0.87939 |
| ENSG00000136925 | 9 | 100362362 | 100395962 | 36 | 12 | -1.3675 | 0.91426 | TSTD2 | 0.91426 | 0.90296 | 0.85217 |
| ENSG00000119725 | 14 | 74353320 | 74399214 | 210 | 17 | -1.368 | 0.91434 | ZNF410 | 0.91434 | 0.89998 | 0.88397 |
| ENSG00000182572 | 6 | 27839623 | 27840099 | 2 | 1 | -1.3686 | 0.91445 | HIST1H3I | 0.91445 | 0.79872 | 0.91663 |
| ENSG00000151465 | 10 | 12237964 | 12292588 | 171 | 21 | -1.3687 | 0.91446 | CDC123 | 0.91446 | 0.9232 | 0.92709 |
| ENSG00000272214 | 12 | 121407641 | 121410095 | 4 | 2 | -1.369 | 0.9145 | AC079602.1 | 0.9145 | 0.5604 | 0.62147 |
| ENSG00000133317 | 11 | 63273556 | 63284246 | 32 | 10 | -1.3699 | 0.91464 | LGALS12 | 0.91464 | 0.86809 | 0.88821 |
| ENSG00000143185 | 1 | 168510003 | 168513235 | 29 | 9 | -1.3704 | 0.91471 | XCL2 | 0.91471 | 0.76463 | 0.80676 |
| ENSG00000007047 | 19 | 45582546 | 45808541 | 865 | 61 | -1.3704 | 0.91471 | MARK4 | 0.91471 | 0.93334 | 0.94546 |
| ENSG00000184381 | 22 | 38507502 | 38601697 | 353 | 18 | -1.3703 | 0.91471 | PLA2G6 | 0.91471 | 0.91189 | 0.82356 |
| ENSG00000183559 | 10 | 124457225 | 124459338 | 7 | 3 | -1.3705 | 0.91473 | C10orf120 | 0.91473 | 0.64693 | 0.52678 |
| ENSG00000149571 | 11 | 126293254 | 126873355 | 2262 | 164 | -1.3706 | 0.91476 | KIRREL3 | 0.91476 | 0.89085 | 0.81982 |
| ENSG00000100603 | 14 | 78183942 | 78227550 | 206 | 21 | -1.3708 | 0.91478 | SNW1 | 0.91478 | 0.92753 | 0.94319 |
| ENSG00000112365 | 6 | 109783797 | 109804440 | 57 | 13 | -1.3709 | 0.91479 | ZBTB24 | 0.91479 | 0.90757 | 0.79718 |
| ENSG00000162735 | 1 | 160246602 | 160256138 | 10 | 4 | -1.3718 | 0.91494 | PEX19 | 0.91494 | 0.77554 | 0.71983 |
| ENSG00000214921 | 17 | 42325758 | 42327236 | 3 | 2 | -1.372 | 0.91496 | AC003102.1 | 0.91496 | 0.83407 | 0.79913 |
| ENSG00000100053 | 22 | 25595817 | 25603330 | 30 | 12 | -1.372 | 0.91497 | CRYBB3 | 0.91497 | 0.8774 | 0.50466 |
| ENSG00000197798 | 11 | 126081309 | 126132881 | 115 | 21 | -1.3723 | 0.91502 | FAM118B | 0.91502 | 0.93339 | 0.9348 |
| ENSG00000176058 | 9 | 140086069 | 140098645 | 14 | 2 | -1.3724 | 0.91503 | TPRN | 0.91503 | 0.56537 | 0.61792 |
| ENSG00000137145 | 9 | 19230433 | 19374139 | 502 | 29 | -1.3726 | 0.91507 | DENND4C | 0.91507 | 0.90338 | 0.87305 |
| ENSG00000166145 | 15 | 41136216 | 41150405 | 48 | 7 | -1.373 | 0.91512 | SPINT1 | 0.91512 | 0.8431 | 0.79611 |
| ENSG00000163286 | 2 | 233271553 | 233275424 | 21 | 8 | -1.373 | 0.91513 | ALPPL2 | 0.91513 | 0.59644 | 0.54939 |
| ENSG00000171951 | 2 | 224461658 | 224467221 | 9 | 4 | -1.3736 | 0.91521 | SCG2 | 0.91521 | 0.87816 | 0.91621 |
| ENSG00000104731 | 16 | 87730091 | 87799598 | 401 | 28 | -1.3744 | 0.91534 | KLHDC4 | 0.91534 | 0.92056 | 0.92287 |
| ENSG00000104324 | 8 | 97657455 | 98161882 | 924 | 56 | -1.3745 | 0.91536 | CPQ | 0.91536 | 0.92505 | 0.90126 |
| ENSG00000134324 | 2 | 11817721 | 11967535 | 548 | 49 | -1.3746 | 0.91538 | LPIN1 | 0.91538 | 0.87425 | 0.85069 |
| ENSG00000172680 | 8 | 57025501 | 57026541 | 2 | 1 | -1.3755 | 0.91551 | MOS | 0.91551 | 0.97163 | 0.95451 |
| ENSG00000172985 | 2 | 109745804 | 110262207 | 1399 | 79 | -1.3756 | 0.91552 | SH3RF3 | 0.91552 | 0.91629 | 0.92665 |
| ENSG00000104970 | 19 | 55043909 | 55057053 | 57 | 21 | -1.3756 | 0.91553 | KIR3DX1 | 0.91553 | 0.87714 | 0.74033 |
| ENSG00000026652 | 6 | 161551011 | 161695093 | 521 | 49 | -1.3763 | 0.91564 | AGPAT4 | 0.91564 | 0.86737 | 0.86345 |
| ENSG00000225950 | 19 | 49559050 | 49568333 | 15 | 7 | -1.3765 | 0.91567 | NTF4 | 0.91567 | 0.50338 | 0.35831 |
| ENSG00000176956 | 8 | 144239331 | 144242128 | 7 | 2 | -1.3766 | 0.91568 | LY6H | 0.91568 | 0.85426 | 0.87909 |
| ENSG00000186654 | 22 | 45064593 | 45133561 | 437 | 63 | -1.3766 | 0.91568 | PRR5 | 0.91568 | 0.89111 | 0.82044 |
| ENSG00000152684 | 5 | 52083774 | 52099880 | 42 | 9 | -1.3774 | 0.9158 | PELO | 0.9158 | 0.93372 | 0.97461 |
| ENSG00000100345 | 22 | 36677327 | 36784063 | 359 | 40 | -1.3776 | 0.91584 | MYH9 | 0.91584 | 0.92518 | 0.96296 |

| | | | | | | | | | | | |
|---|---|---|---|---|---|---|---|---|---|---|---|
| ENSG00000104369 | 8 | 75146935 | 75233563 | 203 | 31 | -1.3791 | 0.91606 | JPH1 | 0.91606 | 0.91032 | 0.85382 |
| ENSG00000163485 | 1 | 203059782 | 203136533 | 202 | 31 | -1.3791 | 0.91607 | ADORA1 | 0.91607 | 0.8924 | 0.75702 |
| ENSG00000165181 | 9 | 114448453 | 114557288 | 473 | 21 | -1.3797 | 0.91615 | C9orf84 | 0.91615 | 0.90154 | 0.92261 |
| ENSG00000111696 | 12 | 104164231 | 104234975 | 383 | 23 | -1.3805 | 0.91629 | NT5DC3 | 0.91629 | 0.69685 | 0.2052 |
| ENSG00000105483 | 19 | 48684027 | 48759203 | 353 | 31 | -1.3808 | 0.91633 | CARD8 | 0.91633 | 0.94841 | 0.97818 |
| ENSG00000135046 | 9 | 75766673 | 75785309 | 47 | 13 | -1.3815 | 0.91643 | ANXA1 | 0.91643 | 0.94257 | 0.91629 |
| ENSG00000151892 | 10 | 117816444 | 118032979 | 763 | 63 | -1.382 | 0.91652 | GFRA1 | 0.91652 | 0.82917 | 0.78195 |
| ENSG00000160190 | 21 | 43916118 | 44001550 | 413 | 27 | -1.3822 | 0.91654 | SLC37A1 | 0.91654 | 0.9229 | 0.95494 |
| ENSG00000115310 | 2 | 55199325 | 55339757 | 506 | 32 | -1.3824 | 0.91657 | RTN4 | 0.91657 | 0.90248 | 0.86501 |
| ENSG00000100425 | 22 | 50166931 | 50221160 | 165 | 18 | -1.383 | 0.91666 | BRD1 | 0.91666 | 0.91807 | 0.92292 |
| ENSG00000162819 | 1 | 222885895 | 222908538 | 35 | 10 | -1.3836 | 0.91676 | BROX | 0.91676 | 0.89376 | 0.9081 |
| ENSG00000157426 | 4 | 57204456 | 57253666 | 205 | 20 | -1.3839 | 0.9168 | AASDH | 0.9168 | 0.89598 | 0.85914 |
| ENSG00000188613 | 10 | 120789228 | 120793854 | 7 | 3 | -1.3848 | 0.91695 | NANOS1 | 0.91695 | 0.94106 | 0.98453 |
| ENSG00000160188 | 21 | 43892596 | 43916464 | 80 | 21 | -1.3852 | 0.917 | RSPH1 | 0.917 | 0.91817 | 0.92412 |
| ENSG00000026559 | 20 | 49620193 | 49639666 | 58 | 10 | -1.3855 | 0.91705 | KCNG1 | 0.91705 | 0.97447 | 0.98832 |
| ENSG00000181904 | 5 | 134181370 | 134195427 | 17 | 9 | -1.3867 | 0.91724 | C5orf24 | 0.91724 | 0.63962 | 0.59772 |
| ENSG00000126773 | 14 | 60558629 | 60635851 | 239 | 9 | -1.3882 | 0.91746 | PCNXL4 | 0.91746 | 0.92778 | 0.93852 |
| ENSG00000064886 | 1 | 111743393 | 111786062 | 175 | 23 | -1.3884 | 0.9175 | CHI3L2 | 0.9175 | 0.94879 | 0.83729 |
| ENSG00000153898 | 1 | 85391268 | 85462796 | 311 | 21 | -1.3888 | 0.91755 | MCOLN2 | 0.91755 | 0.9401 | 0.95417 |
| ENSG00000101421 | 20 | 32399110 | 32442172 | 50 | 8 | -1.3893 | 0.91763 | CHMP4B | 0.91763 | 0.86442 | 0.94691 |
| ENSG00000125629 | 2 | 118846028 | 118868573 | 65 | 21 | -1.3898 | 0.91771 | INSIG2 | 0.91771 | 0.87929 | 0.84776 |
| ENSG00000171471 | 12 | 116997186 | 117014425 | 71 | 18 | -1.3909 | 0.91788 | MAP1LC3B2 | 0.91788 | 0.90901 | 0.83075 |
| ENSG00000163141 | 1 | 151009046 | 151020076 | 32 | 10 | -1.3914 | 0.91794 | BNIPL | 0.91794 | 0.96427 | 0.93583 |
| ENSG00000115685 | 2 | 242088991 | 242123067 | 86 | 16 | -1.3916 | 0.91798 | PPP1R7 | 0.91798 | 0.92087 | 0.86379 |
| ENSG00000171954 | 19 | 15619304 | 15663128 | 175 | 37 | -1.3916 | 0.91798 | CYP4F22 | 0.91798 | 0.93458 | 0.87626 |
| ENSG00000105278 | 19 | 3804022 | 3869030 | 277 | 27 | -1.3919 | 0.91802 | ZFR2 | 0.91802 | 0.93385 | 0.94827 |
| ENSG00000069206 | 8 | 24298443 | 24384483 | 296 | 22 | -1.3921 | 0.91806 | ADAM7 | 0.91806 | 0.89854 | 0.81613 |
| ENSG00000197681 | 17 | 36337711 | 36358166 | 10 | 5 | -1.3921 | 0.91806 | TBC1D3 | 0.91806 | 0.80578 | 0.93427 |
| ENSG00000197548 | 3 | 11313995 | 11599139 | 790 | 23 | -1.3924 | 0.9181 | ATG7 | 0.9181 | 0.90457 | 0.91226 |
| ENSG00000250120 | 5 | 140235595 | 140391929 | 397 | 22 | -1.3929 | 0.91817 | PCDHA10 | 0.91817 | 0.90964 | 0.92558 |
| ENSG00000137500 | 11 | 82970139 | 82997450 | 87 | 11 | -1.3929 | 0.91818 | CCDC90B | 0.91818 | 0.91001 | 0.89191 |
| ENSG00000137207 | 6 | 43479565 | 43484728 | 11 | 4 | -1.3936 | 0.91829 | YIPF3 | 0.91829 | 0.86725 | 0.86002 |
| ENSG00000140543 | 15 | 89054790 | 89089906 | 102 | 11 | -1.3937 | 0.91829 | DET1 | 0.91829 | 0.91075 | 0.88252 |
| ENSG00000168827 | 3 | 158362067 | 158410364 | 199 | 10 | -1.3945 | 0.91842 | GFM1 | 0.91842 | 0.93047 | 0.93476 |
| ENSG00000170425 | 17 | 15848231 | 15879060 | 80 | 17 | -1.3952 | 0.91852 | ADORA2B | 0.91852 | 0.89785 | 0.90823 |
| ENSG00000165716 | 9 | 139607022 | 139618502 | 47 | 12 | -1.3952 | 0.91853 | FAM69B | 0.91853 | 0.6479 | 0.20862 |
| ENSG00000136944 | 9 | 129376722 | 129463311 | 278 | 39 | -1.3953 | 0.91854 | LMX1B | 0.91854 | 0.94708 | 0.92887 |
| ENSG00000146374 | 6 | 127439749 | 127518910 | 165 | 12 | -1.3956 | 0.91859 | RSPO3 | 0.91859 | 0.91628 | 0.90356 |
| ENSG00000197061 | 6 | 26104104 | 26104518 | 1 | 1 | -1.3957 | 0.9186 | HIST1H4C | 0.9186 | 0.73303 | 0.62061 |
| ENSG00000133112 | 13 | 45911008 | 45915505 | 11 | 3 | -1.396 | 0.91864 | TPT1 | 0.91864 | 0.85912 | 0.89634 |
| ENSG00000198954 | 10 | 70748487 | 70776738 | 106 | 16 | -1.397 | 0.91879 | KIAA1279 | 0.91879 | 0.94931 | 0.96107 |
| ENSG00000248383 | 5 | 140306302 | 140391929 | 246 | 20 | -1.3972 | 0.91882 | PCDHAC1 | 0.91882 | 0.92569 | 0.95685 |
| ENSG00000119147 | 2 | 106679702 | 106694615 | 47 | 15 | -1.3973 | 0.91884 | C2orf40 | 0.91884 | 0.67095 | 0.73367 |
| ENSG00000156113 | 10 | 78629359 | 79398353 | 2472 | 174 | -1.3976 | 0.91889 | KCNMA1 | 0.91889 | 0.93822 | 0.94704 |
| ENSG00000206026 | 18 | 73121431 | 73139658 | 76 | 10 | -1.3982 | 0.91897 | SMIM21 | 0.91897 | 0.92062 | 0.97777 |
| ENSG00000198601 | 1 | 248343288 | 248344331 | 3 | 1 | -1.3988 | 0.91906 | OR2M2 | 0.91906 | 0.75555 | 0.6996 |
| ENSG00000123219 | 5 | 64813593 | 64858998 | 185 | 14 | -1.4 | 0.91924 | CENPK | 0.91924 | 0.94259 | 0.94203 |
| ENSG00000107165 | 9 | 12685439 | 12710290 | 78 | 9 | -1.4004 | 0.9193 | TYRP1 | 0.9193 | 0.92553 | 0.91513 |
| ENSG00000125149 | 16 | 67143861 | 67182442 | 50 | 12 | -1.4006 | 0.91934 | C16orf70 | 0.91934 | 0.90551 | 0.88238 |
| ENSG00000166073 | 15 | 40091233 | 40213093 | 302 | 33 | -1.4009 | 0.91938 | GPR176 | 0.91938 | 0.9128 | 0.92685 |
| ENSG00000117174 | 1 | 86115106 | 86174116 | 152 | 22 | -1.4011 | 0.91941 | ZNHIT6 | 0.91941 | 0.88823 | 0.91327 |
| ENSG00000242802 | 7 | 4815253 | 4833943 | 129 | 12 | -1.4016 | 0.91948 | AP5Z1 | 0.91948 | 0.87974 | 0.8391 |
| ENSG00000234965 | 22 | 42307297 | 42310570 | 13 | 2 | -1.4018 | 0.91952 | SHISA8 | 0.91952 | 0.781 | 0.8488 |
| ENSG00000183134 | 11 | 60618413 | 60623444 | 27 | 8 | -1.4021 | 0.91956 | PTGDR2 | 0.91956 | 0.82666 | 0.91106 |
| ENSG00000129187 | 4 | 183811213 | 183839089 | 160 | 21 | -1.4023 | 0.91958 | DCTD | 0.91958 | 0.94209 | 0.42158 |
| ENSG00000136010 | 12 | 105413568 | 105478355 | 315 | 25 | -1.4029 | 0.91968 | ALDH1L2 | 0.91968 | 0.91258 | 0.91223 |
| ENSG00000094755 | 5 | 170190354 | 170241051 | 233 | 35 | -1.403 | 0.91969 | GABRP | 0.91969 | 0.81424 | 0.78148 |
| ENSG00000185236 | 19 | 8454865 | 8469318 | 67 | 14 | -1.4038 | 0.9198 | RAB11B | 0.9198 | 0.93015 | 0.93869 |
| ENSG00000187772 | 6 | 105404923 | 105531207 | 201 | 28 | -1.4039 | 0.91982 | LIN28B | 0.91982 | 0.90373 | 0.869 |
| ENSG00000163798 | 2 | 27886338 | 27917840 | 41 | 14 | -1.4047 | 0.91994 | SLC4A1AP | 0.91994 | 0.87259 | 0.81116 |
| ENSG00000119913 | 10 | 114043493 | 114064793 | 57 | 17 | -1.4054 | 0.92005 | TECTB | 0.92005 | 0.9296 | 0.94398 |
| ENSG00000145358 | 4 | 101107027 | 101111939 | 35 | 9 | -1.4057 | 0.9201 | DDIT4L | 0.9201 | 0.93502 | 0.8714 |
| ENSG00000188603 | 16 | 28477983 | 28506896 | 44 | 7 | -1.4067 | 0.92025 | CLN3 | 0.92025 | 0.79692 | 0.83016 |
| ENSG00000148735 | 10 | 115511213 | 115543188 | 114 | 12 | -1.4073 | 0.92032 | PLEKHS1 | 0.92032 | 0.92506 | 0.85533 |
| ENSG00000087074 | 19 | 49375649 | 49379314 | 21 | 4 | -1.4072 | 0.92032 | PPP1R15A | 0.92032 | 0.97018 | 0.94824 |
| ENSG00000269197 | 8 | 70850403 | 70850635 | 1 | 1 | -1.4084 | 0.9205 | AC090574.1 | 0.9205 | 0.049976 | 0.027429 |
| ENSG00000134909 | 11 | 128834955 | 129149219 | 831 | 28 | -1.4089 | 0.92057 | ARHGAP32 | 0.92057 | 0.92242 | 0.91869 |
| ENSG00000108107 | 19 | 55896713 | 55914612 | 118 | 21 | -1.409 | 0.92058 | RPL28 | 0.92058 | 0.90456 | 0.72996 |
| ENSG00000072518 | 11 | 63606400 | 63678491 | 154 | 13 | -1.4093 | 0.92062 | MARK2 | 0.92062 | 0.90626 | 0.93998 |
| ENSG00000106546 | 7 | 17338246 | 17385776 | 61 | 18 | -1.4093 | 0.92063 | AHR | 0.92063 | 0.89276 | 0.92749 |
| ENSG00000137441 | 4 | 15961866 | 15970932 | 35 | 10 | -1.4095 | 0.92065 | FGFBP2 | 0.92065 | 0.80967 | 0.50279 |
| ENSG00000168243 | 1 | 235710987 | 235814054 | 438 | 42 | -1.41 | 0.92074 | GNG4 | 0.92074 | 0.89311 | 0.90253 |
| ENSG00000082196 | 5 | 34019553 | 34043937 | 75 | 8 | -1.4101 | 0.92075 | C1QTNF3 | 0.92075 | 0.88459 | 0.85396 |

| | | | | | | | | | | | |
|---|---|---|---|---|---|---|---|---|---|---|---|
| ENSG00000164695 | 8 | 82644669 | 82671750 | 76 | 17 | -1.4102 | 0.92076 | CHMP4C | 0.92076 | 0.84424 | 0.79445 |
| ENSG00000071203 | 11 | 60260251 | 60274903 | 75 | 3 | -1.4105 | 0.9208 | MS4A12 | 0.9208 | 0.9444 | 0.97062 |
| ENSG00000125846 | 20 | 18269121 | 18297640 | 100 | 18 | -1.4105 | 0.92081 | ZNF133 | 0.92081 | 0.92749 | 0.94813 |
| ENSG00000180878 | 11 | 6226796 | 6232362 | 10 | 4 | -1.4106 | 0.92082 | C11orf42 | 0.92082 | 0.91364 | 0.95764 |
| ENSG00000188505 | 19 | 39687601 | 39692524 | 12 | 2 | -1.4112 | 0.9209 | NCCRP1 | 0.9209 | 0.85706 | 0.79717 |
| ENSG00000109265 | 4 | 57036361 | 57194791 | 668 | 83 | -1.4112 | 0.92091 | KIAA1211 | 0.92091 | 0.92811 | 0.93096 |
| ENSG00000172673 | 6 | 128029217 | 128239776 | 445 | 34 | -1.4115 | 0.92095 | THEMIS | 0.92095 | 0.90787 | 0.92405 |
| ENSG00000173950 | 3 | 194789008 | 194991896 | 857 | 48 | -1.4122 | 0.92105 | XXYLT1 | 0.92105 | 0.89849 | 0.85296 |
| ENSG00000137285 | 6 | 3224495 | 3231964 | 13 | 5 | -1.4129 | 0.92116 | TUBB2B | 0.92116 | 0.85249 | 0.65742 |
| ENSG00000165185 | 9 | 115249127 | 115431677 | 446 | 44 | -1.4132 | 0.9212 | KIAA1958 | 0.9212 | 0.93469 | 0.94587 |
| ENSG00000119703 | 14 | 75530873 | 75545126 | 30 | 4 | -1.4139 | 0.9213 | ZC2HC1C | 0.9213 | 0.89519 | 0.89964 |
| ENSG00000004809 | 6 | 110745890 | 110797844 | 237 | 36 | -1.4141 | 0.92133 | SLC22A16 | 0.92133 | 0.89319 | 0.85264 |
| ENSG00000176177 | 22 | 40139049 | 40289868 | 219 | 19 | -1.4147 | 0.92142 | ENTHD1 | 0.92142 | 0.90211 | 0.91113 |
| ENSG00000225399 | 3 | 49297518 | 49298744 | 4 | 2 | -1.4148 | 0.92143 | RP11-3B7.1 | 0.92143 | 0.93822 | 0.94594 |
| ENSG00000166401 | 18 | 61637159 | 61672278 | 106 | 28 | -1.4147 | 0.92143 | SERPINB8 | 0.92143 | 0.85634 | 0.82324 |
| ENSG00000169856 | 15 | 53049637 | 53083273 | 96 | 23 | -1.4153 | 0.92151 | ONECUT1 | 0.92151 | 0.85037 | 0.83635 |
| ENSG00000112699 | 6 | 1624041 | 2245926 | 1931 | 119 | -1.4158 | 0.92159 | GMDS | 0.92159 | 0.88916 | 0.81672 |
| ENSG00000049192 | 5 | 64444563 | 64777747 | 963 | 43 | -1.4165 | 0.92168 | ADAMTS6 | 0.92168 | 0.92176 | 0.92238 |
| ENSG00000268421 | 10 | 131308884 | 131309525 | 4 | 2 | -1.4165 | 0.92168 | AL355531.2 | 0.92168 | 0.27647 | 0.18435 |
| ENSG00000156261 | 21 | 30428126 | 30446118 | 37 | 12 | -1.4178 | 0.92187 | CCT8 | 0.92187 | 0.92276 | 0.85264 |
| ENSG00000168754 | 2 | 97541620 | 97684175 | 389 | 27 | -1.4179 | 0.92189 | FAM178B | 0.92189 | 0.91554 | 0.90723 |
| ENSG00000181085 | 8 | 144798429 | 144804628 | 24 | 10 | -1.418 | 0.9219 | MAPK15 | 0.9219 | 0.96323 | 0.96799 |
| ENSG00000213066 | 6 | 167412670 | 167466201 | 335 | 10 | -1.4184 | 0.92196 | FGFR1OP | 0.92196 | 0.90669 | 0.87427 |
| ENSG00000171587 | 21 | 41382926 | 42219065 | 3842 | 228 | -1.4184 | 0.92196 | DSCAM | 0.92196 | 0.92196 | 0.92833 |
| ENSG00000130592 | 11 | 1874200 | 1913497 | 215 | 35 | -1.4189 | 0.92204 | LSP1 | 0.92204 | 0.35504 | 0.42086 |
| ENSG00000184990 | 14 | 105219437 | 105234831 | 44 | 8 | -1.4194 | 0.92212 | SIVA1 | 0.92212 | 0.8048 | 0.86831 |
| ENSG00000106341 | 7 | 31726329 | 31748069 | 73 | 15 | -1.4196 | 0.92214 | PPP1R17 | 0.92214 | 0.9187 | 0.92352 |
| ENSG00000171786 | 1 | 160336857 | 160342638 | 18 | 6 | -1.4198 | 0.92217 | NHLH1 | 0.92217 | 0.86166 | 0.81503 |
| ENSG00000226104 | 19 | 10751715 | 10752093 | 3 | 2 | -1.42 | 0.92219 | AC011475.1 | 0.92219 | 0.93982 | 0.88301 |
| ENSG00000176009 | 11 | 8959119 | 8964580 | 30 | 4 | -1.4204 | 0.92225 | ASCL3 | 0.92225 | 0.95538 | 0.94905 |
| ENSG00000205339 | 11 | 9406169 | 9469673 | 143 | 15 | -1.4212 | 0.92237 | IPO7 | 0.92237 | 0.92912 | 0.83546 |
| ENSG00000128039 | 4 | 56212276 | 56239263 | 94 | 16 | -1.4217 | 0.92244 | SRD5A3 | 0.92244 | 0.93063 | 0.91228 |
| ENSG00000120899 | 8 | 27168999 | 27316903 | 477 | 26 | -1.4218 | 0.92245 | PTK2B | 0.92245 | 0.95005 | 0.95474 |
| ENSG00000066405 | 3 | 137717577 | 137752494 | 78 | 11 | -1.422 | 0.92249 | CLDN18 | 0.92249 | 0.89884 | 0.80469 |
| ENSG00000148153 | 9 | 115446206 | 115480516 | 85 | 12 | -1.4222 | 0.92251 | INIP | 0.92251 | 0.97783 | 0.98542 |
| ENSG00000100979 | 20 | 44527399 | 44540794 | 37 | 14 | -1.4227 | 0.92259 | PLTP | 0.92259 | 0.89731 | 0.58024 |
| ENSG00000170498 | 1 | 204159469 | 204165614 | 29 | 5 | -1.4229 | 0.92262 | KISS1 | 0.92262 | 0.56322 | 0.60998 |
| ENSG00000128626 | 19 | 39421188 | 39423802 | 4 | 2 | -1.4231 | 0.92264 | MRPS12 | 0.92264 | 0.19768 | 0.22055 |
| ENSG00000108599 | 17 | 19807615 | 19881656 | 145 | 12 | -1.4234 | 0.92269 | AKAP10 | 0.92269 | 0.91688 | 0.89349 |
| ENSG00000183831 | 1 | 173578700 | 173639001 | 65 | 16 | -1.4236 | 0.92272 | ANKRD45 | 0.92272 | 0.90677 | 0.802 |
| ENSG00000214562 | 10 | 89117425 | 89130452 | 21 | 8 | -1.4246 | 0.92287 | NUTM2D | 0.92287 | 0.96403 | 0.97759 |
| ENSG00000256040 | 9 | 119160439 | 119162885 | 8 | 4 | -1.425 | 0.92291 | PAPPA-AS1 | 0.92291 | 0.57075 | 0.42395 |
| ENSG00000008083 | 6 | 15246527 | 15522252 | 833 | 54 | -1.4253 | 0.92296 | JARID2 | 0.92296 | 0.93529 | 0.90874 |
| ENSG00000082212 | 18 | 48405419 | 48474691 | 158 | 15 | -1.4257 | 0.92303 | ME2 | 0.92303 | 0.92221 | 0.92979 |
| ENSG00000259384 | 17 | 61994560 | 61996179 | 4 | 2 | -1.4261 | 0.92308 | GH1 | 0.92308 | 0.41165 | 0.38485 |
| ENSG00000065057 | 16 | 2089816 | 2097867 | 8 | 3 | -1.4267 | 0.92317 | NTHL1 | 0.92317 | 0.7095 | 0.64064 |
| ENSG00000268674 | 1 | 16865561 | 16866070 | 1 | 1 | -1.4269 | 0.9232 | FAM231B | 0.9232 | 0.84982 | 0.90476 |
| ENSG00000100209 | 22 | 29138019 | 29153503 | 48 | 11 | -1.4273 | 0.92326 | HSCB | 0.92326 | 0.62039 | 0.43889 |
| ENSG00000131951 | 14 | 60386431 | 60530277 | 331 | 14 | -1.4279 | 0.92333 | LRRC9 | 0.92333 | 0.93441 | 0.93302 |
| ENSG00000099256 | 10 | 25137536 | 25241533 | 387 | 45 | -1.4282 | 0.92338 | PRTFDC1 | 0.92338 | 0.94851 | 0.97697 |
| ENSG00000187959 | 17 | 71244588 | 71258491 | 42 | 11 | -1.4282 | 0.92338 | CPSF4L | 0.92338 | 0.51239 | 0.54301 |
| ENSG00000132024 | 19 | 14017014 | 14041692 | 46 | 16 | -1.4284 | 0.92341 | CC2D1A | 0.92341 | 0.33146 | 0.28522 |
| ENSG00000170348 | 14 | 75598173 | 75643334 | 89 | 7 | -1.429 | 0.9235 | TMED10 | 0.9235 | 0.90104 | 0.9131 |
| ENSG00000196268 | 19 | 21579921 | 21610375 | 136 | 12 | -1.429 | 0.9235 | ZNF493 | 0.9235 | 0.92631 | 0.92192 |
| ENSG00000116922 | 1 | 38147242 | 38157914 | 48 | 5 | -1.4291 | 0.92351 | C1orf109 | 0.92351 | 0.92709 | 0.95783 |
| ENSG00000196091 | 12 | 101962131 | 102079796 | 547 | 48 | -1.4293 | 0.92354 | MYBPC1 | 0.92354 | 0.73773 | 0.75013 |
| ENSG00000186300 | 19 | 2841433 | 2860482 | 67 | 10 | -1.4294 | 0.92355 | ZNF555 | 0.92355 | 0.82804 | 0.90358 |
| ENSG00000092978 | 1 | 217600334 | 217804424 | 659 | 56 | -1.4295 | 0.92357 | GPATCH2 | 0.92357 | 0.8899 | 0.83081 |
| ENSG00000203661 | 1 | 248651885 | 248652898 | 4 | 1 | -1.4301 | 0.92365 | OR2T5 | 0.92365 | 0.92406 | 0.52043 |
| ENSG00000184163 | 1 | 1177826 | 1182102 | 14 | 3 | -1.4305 | 0.92371 | FAM132A | 0.92371 | 0.9537 | 0.96498 |
| ENSG00000173930 | 5 | 101569690 | 101632253 | 277 | 24 | -1.4318 | 0.9239 | SLCO4C1 | 0.9239 | 0.92113 | 0.91873 |
| ENSG00000197557 | 2 | 178477720 | 178483694 | 37 | 7 | -1.4323 | 0.92398 | TTC30A | 0.92398 | 0.92464 | 0.91156 |
| ENSG00000157741 | 7 | 138915102 | 138992982 | 134 | 17 | -1.4327 | 0.92403 | UBN2 | 0.92403 | 0.9414 | 0.958 |
| ENSG00000156427 | 5 | 170846660 | 170884627 | 93 | 17 | -1.433 | 0.92408 | FGF18 | 0.92408 | 0.97206 | 0.96281 |
| ENSG00000198931 | 16 | 88875747 | 88878352 | 8 | 4 | -1.4337 | 0.92417 | APRT | 0.92417 | 0.51028 | 0.63241 |
| ENSG00000244687 | 20 | 48697661 | 48732496 | 88 | 12 | -1.4343 | 0.92425 | UBE2V1 | 0.92425 | 0.95503 | 0.94519 |
| ENSG00000186132 | 2 | 120059801 | 120124404 | 273 | 25 | -1.4348 | 0.92432 | C2orf76 | 0.92432 | 0.92252 | 0.92346 |
| ENSG00000138483 | 3 | 107096188 | 107097481 | 2 | 1 | -1.436 | 0.9245 | CCDC54 | 0.9245 | 0.92012 | 0.89504 |
| ENSG00000135392 | 12 | 56214744 | 56224608 | 22 | 13 | -1.4367 | 0.9246 | DNAJC14 | 0.9246 | 0.97804 | 0.99282 |
| ENSG00000187741 | 16 | 89803957 | 89883065 | 597 | 20 | -1.4379 | 0.92477 | FANCA | 0.92477 | 0.91989 | 0.90175 |
| ENSG00000110429 | 11 | 33762485 | 33796089 | 78 | 14 | -1.4382 | 0.92481 | FBXO3 | 0.92481 | 0.96852 | 0.95498 |
| ENSG00000250486 | 4 | 165878100 | 165880273 | 6 | 2 | -1.4398 | 0.92503 | FAM218A | 0.92503 | 0.78898 | 0.59519 |
| ENSG00000152208 | 4 | 93225550 | 94695707 | 4206 | 128 | -1.4398 | 0.92504 | GRID2 | 0.92504 | 0.88978 | 0.82743 |

| | | | | | | | | | | | |
|---|---|---|---|---|---|---|---|---|---|---|---|
| ENSG00000133812 | 11 | 9800214 | 10315754 | 1809 | 36 | -1.44 | 0.92506 | SBF2 | 0.92506 | 0.93601 | 0.93526 |
| ENSG00000142515 | 19 | 51358171 | 51364020 | 37 | 10 | -1.44 | 0.92506 | KLK3 | 0.92506 | 0.73839 | 0.73998 |
| ENSG00000160868 | 7 | 99354604 | 99381888 | 28 | 9 | -1.441 | 0.9252 | CYP3A4 | 0.9252 | 0.86204 | 0.85855 |
| ENSG00000162877 | 1 | 205797150 | 205819260 | 42 | 14 | -1.4428 | 0.92547 | PM20D1 | 0.92547 | 0.93611 | 0.87229 |
| ENSG00000196510 | 12 | 110810705 | 110841535 | 114 | 15 | -1.4428 | 0.92547 | ANAPC7 | 0.92547 | 0.8881 | 0.8807 |
| ENSG00000181617 | 4 | 71091788 | 71100969 | 17 | 5 | -1.443 | 0.92549 | FDCSP | 0.92549 | 0.86539 | 0.90694 |
| ENSG00000204913 | 17 | 38097727 | 38101000 | 9 | 4 | -1.443 | 0.92549 | LRRC3C | 0.92549 | 0.87033 | 0.78764 |
| ENSG00000204655 | 6 | 29624758 | 29640149 | 103 | 16 | -1.4436 | 0.92558 | MOG | 0.92558 | 0.86667 | 0.8556 |
| ENSG00000169282 | 3 | 155755490 | 156256545 | 1592 | 73 | -1.4443 | 0.92567 | KCNAB1 | 0.92567 | 0.93262 | 0.94214 |
| ENSG00000173597 | 4 | 70586880 | 70653679 | 184 | 23 | -1.4462 | 0.92594 | SULT1B1 | 0.92594 | 0.93701 | 0.95126 |
| ENSG00000180346 | 4 | 90033968 | 90036050 | 4 | 1 | -1.4481 | 0.92621 | TIGD2 | 0.92621 | 0.3938 | 0.37978 |
| ENSG00000158428 | 2 | 219221579 | 219232822 | 41 | 10 | -1.4487 | 0.92629 | C2orf62 | 0.92629 | 0.90414 | 0.75972 |
| ENSG00000143344 | 1 | 183605220 | 183897665 | 986 | 46 | -1.4509 | 0.92659 | RGL1 | 0.92659 | 0.89944 | 0.84867 |
| ENSG00000171109 | 3 | 179065480 | 179112719 | 108 | 15 | -1.4509 | 0.9266 | MFN1 | 0.9266 | 0.93387 | 0.92578 |
| ENSG00000146416 | 6 | 143381633 | 143661441 | 596 | 58 | -1.451 | 0.92661 | AIG1 | 0.92661 | 0.93802 | 0.94018 |
| ENSG00000147434 | 8 | 42607763 | 42651535 | 121 | 12 | -1.4516 | 0.92669 | CHRNA6 | 0.92669 | 0.93209 | 0.95134 |
| ENSG00000160007 | 19 | 47421933 | 47508334 | 96 | 18 | -1.4519 | 0.92673 | ARHGAP35 | 0.92673 | 0.89174 | 0.85362 |
| ENSG00000116062 | 2 | 47922669 | 48037240 | 343 | 32 | -1.4523 | 0.92679 | MSH6 | 0.92679 | 0.93959 | 0.92518 |
| ENSG00000103426 | 16 | 4390252 | 4470495 | 200 | 25 | -1.4528 | 0.92686 | CORO7-PAM16 | 0.92686 | 0.91289 | 0.90642 |
| ENSG00000102871 | 16 | 67188083 | 67194201 | 6 | 3 | -1.4529 | 0.92687 | TRADD | 0.92687 | 0.86256 | 0.92987 |
| ENSG00000121075 | 17 | 59529765 | 59562471 | 82 | 19 | -1.4533 | 0.92693 | TBX4 | 0.92693 | 0.90805 | 0.89474 |
| ENSG00000123146 | 19 | 14491313 | 14519537 | 93 | 29 | -1.4533 | 0.92693 | CD97 | 0.92693 | 0.73535 | 0.7695 |
| ENSG00000166523 | 12 | 8685901 | 8693559 | 16 | 3 | -1.4533 | 0.92694 | CLEC4E | 0.92694 | 0.98094 | 0.97756 |
| ENSG00000145949 | 6 | 2663863 | 2751200 | 475 | 38 | -1.4535 | 0.92696 | MYLK4 | 0.92696 | 0.93689 | 0.95565 |
| ENSG00000175166 | 3 | 184016497 | 184026842 | 22 | 7 | -1.4541 | 0.92704 | PSMD2 | 0.92704 | 0.90686 | 0.9684 |
| ENSG00000197943 | 16 | 81772702 | 81991899 | 1497 | 124 | -1.4542 | 0.92706 | PLCG2 | 0.92706 | 0.90232 | 0.93105 |
| ENSG00000065882 | 4 | 37892708 | 38140796 | 948 | 79 | -1.4544 | 0.92708 | TBC1D1 | 0.92708 | 0.9308 | 0.91117 |
| ENSG00000265264 | 11 | 6502715 | 6530208 | 84 | 9 | -1.4547 | 0.92713 | TIMM10B | 0.92713 | 0.86534 | 0.87736 |
| ENSG00000168078 | 8 | 27667137 | 27695612 | 182 | 5 | -1.4564 | 0.92736 | PBK | 0.92736 | 0.92444 | 0.92094 |
| ENSG00000262874 | 19 | 51891543 | 51893828 | 17 | 1 | -1.4566 | 0.92739 | C19orf84 | 0.92739 | 0.90205 | 0.84758 |
| ENSG00000188026 | 12 | 123955925 | 124018265 | 223 | 25 | -1.4576 | 0.92753 | RILPL1 | 0.92753 | 0.88889 | 0.84709 |
| ENSG00000262246 | 16 | 4404543 | 4475706 | 183 | 22 | -1.4576 | 0.92753 | CORO7 | 0.92753 | 0.91648 | 0.91439 |
| ENSG00000184058 | 22 | 19744226 | 19771116 | 111 | 17 | -1.4577 | 0.92753 | TBX1 | 0.92753 | 0.93794 | 0.95321 |
| ENSG00000255038 | 11 | 65834748 | 65837090 | 3 | 2 | -1.458 | 0.92759 | RP11-1167A19.2 | 0.92759 | 0.52915 | 0.43647 |
| ENSG00000222028 | 14 | 23511376 | 23513269 | 6 | 3 | -1.4587 | 0.92768 | PSMB11 | 0.92768 | 0.98339 | 0.97286 |
| ENSG00000125414 | 17 | 10424465 | 10453274 | 76 | 9 | -1.4589 | 0.9277 | MYH2 | 0.9277 | 0.92855 | 0.91714 |
| ENSG00000269259 | 10 | 102882536 | 102883624 | 3 | 2 | -1.4591 | 0.92774 | HUG1 | 0.92774 | 0.72932 | 0.52684 |
| ENSG00000175283 | 9 | 131707809 | 131709898 | 1 | 1 | -1.4596 | 0.9278 | DOLK | 0.9278 | 0.64356 | 0.65858 |
| ENSG00000171241 | 16 | 46614466 | 46655538 | 37 | 19 | -1.4599 | 0.92785 | SHCBP1 | 0.92785 | 0.91972 | 0.94236 |
| ENSG00000155111 | 6 | 110931181 | 111137161 | 538 | 24 | -1.46 | 0.92786 | CDK19 | 0.92786 | 0.92631 | 0.92678 |
| ENSG00000167693 | 17 | 702553 | 883010 | 1250 | 144 | -1.4607 | 0.92795 | NXN | 0.92795 | 0.89809 | 0.90602 |
| ENSG00000255819 | 12 | 10524952 | 10562745 | 125 | 12 | -1.4617 | 0.92809 | KLRC4-KLRK1 | 0.92809 | 0.9385 | 0.97208 |
| ENSG00000154305 | 1 | 222791428 | 222841354 | 139 | 14 | -1.4623 | 0.92817 | MIA3 | 0.92817 | 0.95608 | 0.96582 |
| ENSG00000131142 | 19 | 8117651 | 8127534 | 57 | 14 | -1.4629 | 0.92825 | CCL25 | 0.92825 | 0.96999 | 0.96203 |
| ENSG00000162600 | 1 | 58881056 | 59012474 | 606 | 43 | -1.4636 | 0.92835 | OMA1 | 0.92835 | 0.91449 | 0.90946 |
| ENSG00000261832 | 16 | 28477974 | 28503333 | 37 | 6 | -1.4636 | 0.92835 | CLN3 | 0.92835 | 0.79692 | 0.83016 |
| ENSG00000079257 | 3 | 158363611 | 158390482 | 109 | 9 | -1.4652 | 0.92856 | LXN | 0.92856 | 0.91706 | 0.92918 |
| ENSG00000173786 | 17 | 40118759 | 40129749 | 24 | 9 | -1.4659 | 0.92867 | CNP | 0.92867 | 0.83511 | 0.84614 |
| ENSG00000165959 | 14 | 95648277 | 95786243 | 584 | 48 | -1.4661 | 0.92869 | CLMN | 0.92869 | 0.9249 | 0.93032 |
| ENSG00000151005 | 4 | 164392257 | 164395047 | 2 | 1 | -1.467 | 0.92881 | TKTL2 | 0.92881 | 0.60492 | 0.52816 |
| ENSG00000141497 | 17 | 4643319 | 4649411 | 21 | 9 | -1.468 | 0.92895 | ZMYND15 | 0.92895 | 0.93342 | 0.7439 |
| ENSG00000060688 | 1 | 31732417 | 31769662 | 128 | 9 | -1.4687 | 0.92905 | SNRNP40 | 0.92905 | 0.88606 | 0.89671 |
| ENSG00000159055 | 21 | 33640530 | 33651380 | 15 | 4 | -1.4689 | 0.92906 | MIS18A | 0.92906 | 0.96797 | 0.81103 |
| ENSG00000176871 | 12 | 118470712 | 118500235 | 85 | 10 | -1.4695 | 0.92915 | WSB2 | 0.92915 | 0.72681 | 0.4436 |
| ENSG00000205018 | 16 | 89006197 | 89017932 | 30 | 8 | -1.4697 | 0.92917 | RP11-830F9.6 | 0.92917 | 0.87111 | 0.867 |
| ENSG00000157216 | 1 | 54691105 | 54879152 | 532 | 72 | -1.4697 | 0.92918 | SSBP3 | 0.92918 | 0.85018 | 0.6791 |
| ENSG00000189350 | 2 | 29179477 | 29284239 | 459 | 36 | -1.4697 | 0.92918 | FAM179A | 0.92918 | 0.95888 | 0.97809 |
| ENSG00000151090 | 3 | 24158651 | 24536773 | 1266 | 127 | -1.4701 | 0.92923 | THRB | 0.92923 | 0.91542 | 0.89582 |
| ENSG00000180730 | 13 | 26618735 | 26625169 | 16 | 8 | -1.4708 | 0.92933 | SHISA2 | 0.92933 | 0.32292 | 0.60644 |
| ENSG00000184900 | 21 | 46225532 | 46238694 | 55 | 13 | -1.4716 | 0.92943 | SUMO3 | 0.92943 | 0.92142 | 0.94569 |
| ENSG00000048540 | 12 | 16701307 | 16763528 | 86 | 24 | -1.4718 | 0.92946 | LMO3 | 0.92946 | 0.96054 | 0.97949 |
| ENSG00000173372 | 1 | 22962999 | 22966101 | 6 | 3 | -1.4721 | 0.92951 | C1QA | 0.92951 | 0.68791 | 0.49548 |
| ENSG00000177917 | 2 | 153574407 | 153617767 | 154 | 20 | -1.4721 | 0.92951 | ARL6IP6 | 0.92951 | 0.92381 | 0.9117 |
| ENSG00000181215 | 4 | 5958845 | 5991545 | 225 | 31 | -1.4726 | 0.92957 | C4orf50 | 0.92957 | 0.79493 | 0.87394 |
| ENSG00000222005 | 2 | 47043807 | 47049796 | 29 | 7 | -1.4726 | 0.92958 | LINC01118 | 0.92958 | 0.72718 | 0.63074 |
| ENSG00000168539 | 11 | 62676151 | 62689279 | 33 | 9 | -1.4733 | 0.92966 | CHRM1 | 0.92966 | 0.90836 | 0.87537 |
| ENSG00000078142 | 18 | 39535171 | 39667794 | 255 | 16 | -1.4734 | 0.92968 | PIK3C3 | 0.92968 | 0.93497 | 0.91284 |
| ENSG00000172977 | 11 | 65479467 | 65487075 | 23 | 6 | -1.4736 | 0.92971 | KAT5 | 0.92971 | 0.86167 | 0.83647 |
| ENSG00000085063 | 11 | 33719807 | 33757991 | 119 | 23 | -1.4738 | 0.92974 | CD59 | 0.92974 | 0.95085 | 0.65591 |
| ENSG00000198231 | 17 | 61850963 | 61896677 | 111 | 8 | -1.4739 | 0.92974 | DDX42 | 0.92974 | 0.88702 | 0.85545 |
| ENSG00000123643 | 5 | 150816607 | 150871942 | 213 | 24 | -1.475 | 0.92989 | SLC36A1 | 0.92989 | 0.91982 | 0.92669 |
| ENSG00000171794 | 10 | 135043778 | 135045062 | 7 | 3 | -1.4756 | 0.92997 | UTF1 | 0.92997 | 0.23513 | 0.15543 |
| ENSG00000158560 | 7 | 95401866 | 95739634 | 1052 | 76 | -1.4759 | 0.93002 | DYNC1I1 | 0.93002 | 0.92502 | 0.8955 |

| | | | | | | | | | | | |
|---|---|---|---|---|---|---|---|---|---|---|---|
| ENSG00000169764 | 2 | 64068074 | 64118696 | 144 | 23 | -1.4761 | 0.93005 | UGP2 | 0.93005 | 0.91564 | 0.9259 |
| ENSG00000181001 | 11 | 5809084 | 5810046 | 11 | 4 | -1.4766 | 0.93011 | OR52N1 | 0.93011 | 0.95136 | 0.93909 |
| ENSG00000177602 | 17 | 3627211 | 3630067 | 11 | 3 | -1.4767 | 0.93012 | GSG2 | 0.93012 | 0.9246 | 0.90442 |
| ENSG00000074370 | 17 | 3827169 | 3867736 | 179 | 20 | -1.4771 | 0.93017 | ATP2A3 | 0.93017 | 0.96394 | 0.96782 |
| ENSG00000239389 | 5 | 140261793 | 140391929 | 343 | 20 | -1.4771 | 0.93018 | PCDHA13 | 0.93018 | 0.91497 | 0.92463 |
| ENSG00000175548 | 12 | 38710380 | 38717784 | 26 | 6 | -1.4781 | 0.93031 | ALG10B | 0.93031 | 0.73006 | 0.77846 |
| ENSG00000167088 | 18 | 19192228 | 19210417 | 29 | 5 | -1.4791 | 0.93044 | SNRPD1 | 0.93044 | 0.92003 | 0.87297 |
| ENSG00000203963 | 1 | 67557848 | 67697536 | 499 | 37 | -1.4798 | 0.93054 | C1orf141 | 0.93054 | 0.93736 | 0.92593 |
| ENSG00000165891 | 12 | 77415027 | 77459360 | 125 | 33 | -1.4799 | 0.93056 | E2F7 | 0.93056 | 0.83907 | 0.90839 |
| ENSG00000112576 | 6 | 41902671 | 42018095 | 468 | 57 | -1.4804 | 0.93062 | CCND3 | 0.93062 | 0.94517 | 0.95473 |
| ENSG00000220256 | 2 | 240684554 | 240724577 | 269 | 22 | -1.481 | 0.9307 | AC093802.1 | 0.9307 | 0.9579 | 0.8589 |
| ENSG00000150667 | 15 | 39892232 | 40075031 | 514 | 24 | -1.4818 | 0.9308 | FSIP1 | 0.9308 | 0.92933 | 0.93379 |
| ENSG00000164451 | 6 | 116850174 | 116880031 | 72 | 10 | -1.4827 | 0.93092 | FAM26D | 0.93092 | 0.93766 | 0.93957 |
| ENSG00000187068 | 3 | 184795838 | 184870802 | 245 | 25 | -1.4841 | 0.93111 | C3orf70 | 0.93111 | 0.95083 | 0.92513 |
| ENSG00000254536 | 10 | 135204338 | 135233999 | 141 | 19 | -1.4843 | 0.93113 | RP11-108K14.8 | 0.93113 | 0.91124 | 0.9058 |
| ENSG00000108091 | 10 | 61548521 | 61666414 | 524 | 22 | -1.4846 | 0.93118 | CCDC6 | 0.93118 | 0.92612 | 0.88375 |
| ENSG00000269175 | 1 | 101455761 | 101457653 | 1 | 1 | -1.4848 | 0.9312 | AC093157.1 | 0.9312 | 0.50601 | 0.54865 |
| ENSG00000196208 | 2 | 11674242 | 11782914 | 402 | 47 | -1.4853 | 0.93126 | GREB1 | 0.93126 | 0.92394 | 0.81489 |
| ENSG00000176294 | 14 | 20187174 | 20296531 | 283 | 37 | -1.4855 | 0.93129 | OR4N2 | 0.93129 | 0.94637 | 0.95936 |
| ENSG00000170293 | 3 | 32280171 | 32411817 | 581 | 81 | -1.4856 | 0.9313 | CMTM8 | 0.9313 | 0.9248 | 0.97413 |
| ENSG00000268175 | 3 | 149689066 | 149690535 | 4 | 2 | -1.4861 | 0.93137 | AC117395.1 | 0.93137 | 0.85679 | 0.90273 |
| ENSG00000246223 | 14 | 98391947 | 98444461 | 211 | 26 | -1.4862 | 0.93139 | C14orf64 | 0.93139 | 0.9557 | 0.96645 |
| ENSG00000101654 | 18 | 13726659 | 13764557 | 117 | 16 | -1.4866 | 0.93144 | RNMT | 0.93144 | 0.86374 | 0.8012 |
| ENSG00000171853 | 2 | 3383446 | 3488865 | 407 | 38 | -1.4871 | 0.9315 | TRAPPC12 | 0.9315 | 0.93337 | 0.93761 |
| ENSG00000154451 | 1 | 89724633 | 89738544 | 21 | 5 | -1.4871 | 0.93151 | GBP5 | 0.93151 | 0.94937 | 0.95672 |
| ENSG00000198680 | 9 | 25676396 | 25678856 | 18 | 5 | -1.4876 | 0.93157 | TUSC1 | 0.93157 | 0.93297 | 0.9641 |
| ENSG00000204446 | 9 | 89763559 | 89774471 | 46 | 8 | -1.4879 | 0.93161 | C9orf170 | 0.93161 | 0.89024 | 0.8975 |
| ENSG00000010256 | 3 | 48636435 | 48648409 | 9 | 5 | -1.4883 | 0.93167 | UQCRC1 | 0.93167 | 0.57839 | 0.83816 |
| ENSG00000263588 | 18 | 19668021 | 19678741 | 61 | 10 | -1.489 | 0.93176 | RP11-595B24.2 | 0.93176 | 0.93765 | 0.9414 |
| ENSG00000229415 | 14 | 36942493 | 36983034 | 80 | 14 | -1.4894 | 0.9318 | SFTA3 | 0.9318 | 0.9844 | 0.97817 |
| ENSG00000258529 | 11 | 111657010 | 111750149 | 199 | 20 | -1.4899 | 0.93187 | ALG9 | 0.93187 | 0.93206 | 0.93263 |
| ENSG00000272442 | 6 | 44240931 | 44346694 | 350 | 40 | -1.4903 | 0.93193 | RP11-444E17.6 | 0.93193 | 0.93077 | 0.94155 |
| ENSG00000113597 | 5 | 64920543 | 64962060 | 161 | 12 | -1.4906 | 0.93196 | TRAPPC13 | 0.93196 | 0.94241 | 0.94819 |
| ENSG00000136754 | 10 | 27035522 | 27150016 | 442 | 22 | -1.4906 | 0.93197 | ABI1 | 0.93197 | 0.91194 | 0.85675 |
| ENSG00000163046 | 2 | 132905164 | 133015542 | 155 | 20 | -1.4912 | 0.93205 | ANKRD30BL | 0.93205 | 0.91352 | 0.91982 |
| ENSG00000255468 | 11 | 66115421 | 66132275 | 44 | 7 | -1.4917 | 0.93211 | RP11-867G23.8 | 0.93211 | 0.92861 | 0.91858 |
| ENSG00000167118 | 9 | 131133598 | 131153015 | 37 | 7 | -1.4919 | 0.93214 | URM1 | 0.93214 | 0.94966 | 0.9601 |
| ENSG00000135916 | 2 | 231729354 | 231743963 | 63 | 13 | -1.4922 | 0.93218 | ITM2C | 0.93218 | 0.95497 | 0.83035 |
| ENSG00000187151 | 11 | 101761405 | 101787253 | 42 | 8 | -1.4935 | 0.93234 | ANGPTL5 | 0.93234 | 0.74802 | 0.61701 |
| ENSG00000105732 | 19 | 42572629 | 42585701 | 14 | 6 | -1.4941 | 0.93243 | ZNF574 | 0.93243 | 0.78695 | 0.68266 |
| ENSG00000147419 | 8 | 27590835 | 27630170 | 166 | 13 | -1.4945 | 0.93248 | CCDC25 | 0.93248 | 0.93886 | 0.95244 |
| ENSG00000109919 | 11 | 47638867 | 47664175 | 38 | 12 | -1.4946 | 0.93249 | MTCH2 | 0.93249 | 0.8911 | 0.75085 |
| ENSG00000116353 | 1 | 29519385 | 29557454 | 86 | 16 | -1.495 | 0.93254 | MECR | 0.93254 | 0.94901 | 0.95452 |
| ENSG00000137504 | 11 | 85370752 | 85393951 | 46 | 11 | -1.4955 | 0.9326 | CREBZF | 0.9326 | 0.94464 | 0.92224 |
| ENSG00000197859 | 9 | 136397286 | 136440641 | 173 | 23 | -1.4969 | 0.93279 | ADAMTSL2 | 0.93279 | 0.95271 | 0.99146 |
| ENSG00000086848 | 11 | 111652919 | 111742305 | 183 | 19 | -1.4979 | 0.93292 | ALG9 | 0.93292 | 0.93206 | 0.93263 |
| ENSG00000141956 | 21 | 43218385 | 43299591 | 445 | 37 | -1.4981 | 0.93295 | PRDM15 | 0.93295 | 0.97483 | 0.9823 |
| ENSG00000119705 | 14 | 78174414 | 78227447 | 269 | 19 | -1.4982 | 0.93296 | SLIRP | 0.93296 | 0.93119 | 0.9507 |
| ENSG00000132434 | 7 | 55433141 | 55501435 | 219 | 14 | -1.4991 | 0.93307 | LANCL2 | 0.93307 | 0.87451 | 0.55406 |
| ENSG00000156282 | 21 | 31538241 | 31538971 | 1 | 1 | -1.4993 | 0.9331 | CLDN17 | 0.9331 | 0.52055 | 0.50898 |
| ENSG00000159228 | 21 | 37442239 | 37445464 | 14 | 5 | -1.5005 | 0.93326 | CBR1 | 0.93326 | 0.89764 | 0.93365 |
| ENSG00000268926 | 9 | 98534605 | 98537013 | 10 | 4 | -1.5016 | 0.9334 | DKFZP434H0512 | 0.9334 | 0.87057 | 0.79355 |
| ENSG00000176222 | 19 | 44376515 | 44405537 | 32 | 10 | -1.5018 | 0.93343 | ZNF404 | 0.93343 | 0.93749 | 0.86376 |
| ENSG00000113140 | 5 | 151040657 | 151066726 | 70 | 15 | -1.5022 | 0.93347 | SPARC | 0.93347 | 0.92731 | 0.97457 |
| ENSG00000135218 | 7 | 79998891 | 80308593 | 1008 | 51 | -1.5029 | 0.93357 | CD36 | 0.93357 | 0.94414 | 0.94777 |
| ENSG00000163421 | 3 | 71820807 | 71834357 | 31 | 10 | -1.503 | 0.93358 | PROK2 | 0.93358 | 0.94882 | 0.91404 |
| ENSG00000166452 | 11 | 8932686 | 8941631 | 37 | 6 | -1.5031 | 0.9336 | AKIP1 | 0.9336 | 0.88434 | 0.91585 |
| ENSG00000167114 | 9 | 131102925 | 131123749 | 53 | 10 | -1.5039 | 0.9337 | SLC27A4 | 0.9337 | 0.95012 | 0.95406 |
| ENSG00000174348 | 1 | 53527854 | 53551174 | 121 | 14 | -1.5043 | 0.93374 | PODN | 0.93374 | 0.94638 | 0.96763 |
| ENSG00000174640 | 3 | 133651540 | 133771028 | 416 | 63 | -1.5061 | 0.93399 | SLCO2A1 | 0.93399 | 0.92131 | 0.92904 |
| ENSG00000109205 | 4 | 71062213 | 71070293 | 25 | 5 | -1.5062 | 0.93399 | ODAM | 0.93399 | 0.90563 | 0.92199 |
| ENSG00000253368 | 1 | 27320198 | 27327389 | 8 | 3 | -1.5067 | 0.93406 | TRNP1 | 0.93406 | 0.88232 | 0.90774 |
| ENSG00000178828 | 1 | 20140523 | 20141771 | 6 | 4 | -1.5071 | 0.93411 | RNF186 | 0.93411 | 0.30327 | 0.18375 |
| ENSG00000165092 | 9 | 75515578 | 75695358 | 369 | 32 | -1.5075 | 0.93416 | ALDH1A1 | 0.93416 | 0.88746 | 0.83254 |
| ENSG00000162782 | 1 | 179560748 | 179660407 | 377 | 20 | -1.5082 | 0.93425 | TDRD5 | 0.93425 | 0.92706 | 0.94468 |
| ENSG00000148343 | 9 | 131798900 | 131834361 | 43 | 14 | -1.5089 | 0.93434 | FAM73B | 0.93434 | 0.92683 | 0.94338 |
| ENSG00000206199 | 3 | 149478892 | 149686172 | 563 | 27 | -1.5091 | 0.93437 | ANKUB1 | 0.93437 | 0.95031 | 0.96799 |
| ENSG00000248405 | 22 | 45098113 | 45258586 | 1277 | 126 | -1.5092 | 0.93438 | PRR5-ARHGAP8 | 0.93438 | 0.96198 | 0.97459 |
| ENSG00000197364 | 1 | 153409534 | 153412425 | 10 | 4 | -1.5102 | 0.9345 | S100A7L2 | 0.9345 | 0.79524 | 0.83332 |
| ENSG00000142224 | 1 | 206972215 | 207016324 | 148 | 24 | -1.5104 | 0.93453 | IL19 | 0.93453 | 0.93328 | 0.90138 |
| ENSG00000249581 | 4 | 17516788 | 17528727 | 38 | 8 | -1.5108 | 0.93458 | CLRN2 | 0.93458 | 0.92671 | 0.90797 |
| ENSG00000213809 | 12 | 10524952 | 10544473 | 66 | 7 | -1.511 | 0.93461 | KLRK1 | 0.93461 | 0.91021 | 0.96556 |
| ENSG00000268485 | 2 | 242167319 | 242168850 | 3 | 2 | -1.5118 | 0.93471 | AC104841.2 | 0.93471 | 0.69834 | 0.44559 |

| | | | | | | | | | | | |
|---|---|---|---|---|---|---|---|---|---|---|---|
| ENSG00000239779 | 2 | 74685456 | 74688018 | 1 | 1 | -1.5125 | 0.9348 | WBP1 | 0.9348 | 0.66883 | 0.69935 |
| ENSG00000245848 | 19 | 33790840 | 33793470 | 6 | 2 | -1.5126 | 0.93481 | CEBPA | 0.93481 | 0.78506 | 0.80047 |
| ENSG00000182985 | 11 | 115039938 | 115375675 | 745 | 64 | -1.5131 | 0.93487 | CADM1 | 0.93487 | 0.95744 | 0.95064 |
| ENSG00000185736 | 10 | 1228073 | 1779670 | 2772 | 209 | -1.5133 | 0.93489 | ADARB2 | 0.93489 | 0.9338 | 0.92796 |
| ENSG00000157927 | 7 | 4836686 | 4923350 | 438 | 32 | -1.5135 | 0.93492 | RADIL | 0.93492 | 0.94489 | 0.96441 |
| ENSG00000137821 | 15 | 71145578 | 71342414 | 263 | 21 | -1.5141 | 0.93501 | LRRC49 | 0.93501 | 0.93457 | 0.94619 |
| ENSG00000100526 | 14 | 54863567 | 54886936 | 47 | 8 | -1.5154 | 0.93517 | CDKN3 | 0.93517 | 0.8842 | 0.82163 |
| ENSG00000109625 | 4 | 8594387 | 8621488 | 232 | 32 | -1.5159 | 0.93523 | CPZ | 0.93523 | 0.91308 | 0.94608 |
| ENSG00000124207 | 20 | 47662849 | 47713489 | 153 | 14 | -1.516 | 0.93524 | CSE1L | 0.93524 | 0.93595 | 0.93479 |
| ENSG00000162779 | 1 | 179334855 | 179523870 | 1325 | 22 | -1.5169 | 0.93535 | AXDND1 | 0.93535 | 0.93916 | 0.94683 |
| ENSG00000241322 | 17 | 15468798 | 15522826 | 245 | 28 | -1.5173 | 0.93541 | CDRT1 | 0.93541 | 0.90428 | 0.93986 |
| ENSG00000249428 | 4 | 2451700 | 2464668 | 46 | 10 | -1.5175 | 0.93543 | RP11-503N18.3 | 0.93543 | 0.92726 | 0.90809 |
| ENSG00000151952 | 12 | 129556270 | 130388211 | 3641 | 313 | -1.5181 | 0.9355 | TMEM132D | 0.9355 | 0.91392 | 0.84955 |
| ENSG00000161652 | 19 | 50655805 | 50666452 | 16 | 7 | -1.5181 | 0.9355 | IZUMO2 | 0.9355 | 0.79101 | 0.67889 |
| ENSG00000115963 | 2 | 151324709 | 151395525 | 171 | 26 | -1.5182 | 0.93552 | RND3 | 0.93552 | 0.95027 | 0.94905 |
| ENSG00000106153 | 7 | 56169262 | 56174269 | 16 | 5 | -1.5188 | 0.93559 | CHCHD2 | 0.93559 | 0.77609 | 0.73635 |
| ENSG00000005436 | 2 | 75879126 | 75938115 | 180 | 16 | -1.5198 | 0.93572 | GCFC2 | 0.93572 | 0.96153 | 0.91543 |
| ENSG00000099821 | 19 | 617223 | 633597 | 108 | 18 | -1.5198 | 0.93572 | POLRMT | 0.93572 | 0.88846 | 0.75452 |
| ENSG00000121236 | 11 | 5617339 | 5634188 | 115 | 28 | -1.5207 | 0.93583 | TRIM6 | 0.93583 | 0.90865 | 0.51574 |
| ENSG00000267952 | 19 | 7571938 | 7584508 | 45 | 10 | -1.522 | 0.93599 | CTD-2207O23.12 | 0.93599 | 0.89891 | 0.48912 |
| ENSG00000174944 | 3 | 150929905 | 150996255 | 172 | 18 | -1.5224 | 0.93605 | P2RY14 | 0.93605 | 0.93562 | 0.96084 |
| ENSG00000172482 | 2 | 241807896 | 241819919 | 59 | 12 | -1.5225 | 0.93606 | AGXT | 0.93606 | 0.83503 | 0.8381 |
| ENSG00000103429 | 16 | 14726672 | 14763093 | 54 | 16 | -1.5226 | 0.93607 | BFAR | 0.93607 | 0.86543 | 0.70839 |
| ENSG00000163482 | 2 | 219536749 | 219567439 | 41 | 7 | -1.5228 | 0.9361 | STK36 | 0.9361 | 0.94659 | 0.8878 |
| ENSG00000139800 | 13 | 100615218 | 100624163 | 22 | 8 | -1.5235 | 0.93619 | ZIC5 | 0.93619 | 0.92909 | 0.94586 |
| ENSG00000248874 | 5 | 23951457 | 24178372 | 963 | 27 | -1.5237 | 0.93621 | C5orf17 | 0.93621 | 0.939 | 0.94826 |
| ENSG00000147894 | 9 | 27546544 | 27573864 | 127 | 18 | -1.524 | 0.93625 | C9orf72 | 0.93625 | 0.86941 | 0.87492 |
| ENSG00000148484 | 10 | 16632610 | 16859527 | 901 | 73 | -1.5242 | 0.93627 | RSU1 | 0.93627 | 0.94981 | 0.95202 |
| ENSG00000177468 | 6 | 137813336 | 137815531 | 2 | 1 | -1.5249 | 0.93636 | OLIG3 | 0.93636 | 0.23401 | 0.20874 |
| ENSG00000138759 | 4 | 78978724 | 79465423 | 2062 | 52 | -1.5251 | 0.93638 | FRAS1 | 0.93638 | 0.94176 | 0.94193 |
| ENSG00000134757 | 18 | 29027758 | 29058665 | 138 | 17 | -1.5258 | 0.93647 | DSG3 | 0.93647 | 0.87445 | 0.84303 |
| ENSG00000172922 | 11 | 65482367 | 65488418 | 15 | 4 | -1.5261 | 0.93651 | RNASEH2C | 0.93651 | 0.8461 | 0.8245 |
| ENSG00000055130 | 7 | 148395006 | 148498128 | 238 | 21 | -1.5265 | 0.93655 | CUL1 | 0.93655 | 0.94992 | 0.94612 |
| ENSG00000198015 | 12 | 93861264 | 93897545 | 153 | 14 | -1.5265 | 0.93655 | MRPL42 | 0.93655 | 0.93794 | 0.92703 |
| ENSG00000213231 | 14 | 96152754 | 96158980 | 36 | 12 | -1.5264 | 0.93655 | TCL1B | 0.93655 | 0.92901 | 0.80291 |
| ENSG00000163431 | 1 | 201865580 | 201915715 | 146 | 26 | -1.527 | 0.93661 | LMOD1 | 0.93661 | 0.94106 | 0.50989 |
| ENSG00000079805 | 19 | 10828755 | 10944164 | 357 | 29 | -1.5273 | 0.93666 | DNM2 | 0.93666 | 0.93111 | 0.93621 |
| ENSG00000165985 | 10 | 16555742 | 16564004 | 28 | 8 | -1.5277 | 0.9367 | C1QL3 | 0.9367 | 0.35791 | 0.395 |
| ENSG00000180539 | 9 | 139921916 | 139931234 | 26 | 5 | -1.529 | 0.93687 | C9orf139 | 0.93687 | 0.85822 | 0.39061 |
| ENSG00000112186 | 6 | 17393447 | 17558023 | 492 | 47 | -1.5296 | 0.93694 | CAP2 | 0.93694 | 0.95116 | 0.96616 |
| ENSG00000120833 | 12 | 93963590 | 93977263 | 21 | 7 | -1.5299 | 0.93698 | SOCS2 | 0.93698 | 0.26789 | 0.16498 |
| ENSG00000159352 | 1 | 151227179 | 151239955 | 25 | 5 | -1.53 | 0.93699 | PSMD4 | 0.93699 | 0.86186 | 0.87061 |
| ENSG00000177182 | 8 | 61969717 | 62414204 | 1608 | 70 | -1.5313 | 0.93715 | CLVS1 | 0.93715 | 0.91535 | 0.87822 |
| ENSG00000213347 | 5 | 176728462 | 176739758 | 26 | 9 | -1.5315 | 0.93717 | MXD3 | 0.93717 | 0.90955 | 0.86638 |
| ENSG00000269676 | 12 | 133049334 | 133050726 | 20 | 8 | -1.5319 | 0.93723 | MUC8 | 0.93723 | 0.76984 | 0.38224 |
| ENSG00000241484 | 22 | 45098355 | 45258665 | 1280 | 130 | -1.5321 | 0.93725 | ARHGAP8 | 0.93725 | 0.96147 | 0.97507 |
| ENSG00000203837 | 10 | 118187379 | 118237469 | 177 | 21 | -1.5322 | 0.93726 | PNLIPRP3 | 0.93726 | 0.9682 | 0.93028 |
| ENSG00000131844 | 5 | 70883115 | 70954531 | 195 | 18 | -1.5323 | 0.93728 | MCCC2 | 0.93728 | 0.95652 | 0.9673 |
| ENSG00000107186 | 9 | 13105703 | 13279589 | 279 | 30 | -1.533 | 0.93737 | MPDZ | 0.93737 | 0.88246 | 0.85167 |
| ENSG00000174514 | 1 | 205538013 | 205572046 | 150 | 24 | -1.5334 | 0.93741 | MFSD4 | 0.93741 | 0.93522 | 0.8945 |
| ENSG00000101138 | 20 | 54967427 | 54979518 | 31 | 8 | -1.5337 | 0.93744 | CSTF1 | 0.93744 | 0.96918 | 0.96999 |
| ENSG00000136688 | 2 | 113730780 | 113743242 | 19 | 5 | -1.5343 | 0.93752 | IL36G | 0.93752 | 0.8587 | 0.89134 |
| ENSG00000164040 | 4 | 129190397 | 129209984 | 33 | 10 | -1.5347 | 0.93757 | PGRMC2 | 0.93757 | 0.93207 | 0.95528 |
| ENSG00000204595 | 19 | 54135310 | 54140263 | 16 | 5 | -1.5347 | 0.93758 | DPRX | 0.93758 | 0.76022 | 0.44859 |
| ENSG00000178719 | 8 | 145064226 | 145067583 | 5 | 2 | -1.5354 | 0.93765 | GRINA | 0.93765 | 0.62154 | 0.70715 |
| ENSG00000083093 | 16 | 23614488 | 23652631 | 115 | 9 | -1.5357 | 0.93769 | PALB2 | 0.93769 | 0.96119 | 0.96895 |
| ENSG00000172146 | 17 | 3118915 | 3119844 | 7 | 3 | -1.5359 | 0.93771 | OR1A1 | 0.93771 | 0.7669 | 0.78828 |
| ENSG00000108474 | 17 | 16120505 | 16252115 | 344 | 16 | -1.5362 | 0.93776 | PIGL | 0.93776 | 0.93818 | 0.9388 |
| ENSG00000112031 | 6 | 153308497 | 153323820 | 54 | 9 | -1.5364 | 0.93778 | MTRF1L | 0.93778 | 0.95451 | 0.9383 |
| ENSG00000072657 | 12 | 72481046 | 73059422 | 1528 | 68 | -1.537 | 0.93785 | TRHDE | 0.93785 | 0.94561 | 0.94364 |
| ENSG00000121413 | 19 | 58595205 | 58629794 | 198 | 16 | -1.537 | 0.93785 | ZSCAN18 | 0.93785 | 0.89822 | 0.88382 |
| ENSG00000119318 | 9 | 110045418 | 110094475 | 205 | 23 | -1.5374 | 0.9379 | RAD23B | 0.9379 | 0.94042 | 0.9494 |
| ENSG00000181856 | 17 | 7184986 | 7191576 | 11 | 5 | -1.5374 | 0.9379 | SLC2A4 | 0.9379 | 0.9268 | 0.97205 |
| ENSG00000261915 | 17 | 7215980 | 7222493 | 18 | 6 | -1.5376 | 0.93792 | RP11-542C16.2 | 0.93792 | 0.94593 | 0.95039 |
| ENSG00000183019 | 19 | 7741514 | 7744712 | 11 | 6 | -1.5378 | 0.93795 | C19orf59 | 0.93795 | 0.92127 | 0.87034 |
| ENSG00000165795 | 14 | 21484922 | 21539031 | 187 | 56 | -1.5381 | 0.93799 | NDRG2 | 0.93799 | 0.95422 | 0.77536 |
| ENSG00000159197 | 21 | 35736323 | 35743688 | 27 | 7 | -1.5382 | 0.938 | KCNE2 | 0.938 | 0.95713 | 0.87624 |
| ENSG00000148824 | 10 | 135207598 | 135234811 | 133 | 17 | -1.5383 | 0.93801 | MTG1 | 0.93801 | 0.93779 | 0.92661 |
| ENSG00000173825 | 11 | 65122238 | 65125084 | 5 | 3 | -1.5394 | 0.93815 | TIGD3 | 0.93815 | 0.78031 | 0.71559 |
| ENSG00000160075 | 1 | 1477053 | 1510249 | 123 | 11 | -1.5399 | 0.9382 | SSU72 | 0.9382 | 0.96674 | 0.97025 |
| ENSG00000172020 | 3 | 115342171 | 115440337 | 266 | 50 | -1.5406 | 0.93829 | GAP43 | 0.93829 | 0.6299 | 0.45336 |
| ENSG00000158481 | 1 | 158259576 | 158263420 | 3 | 2 | -1.5425 | 0.93852 | CD1C | 0.93852 | 0.97169 | 0.94727 |
| ENSG00000178934 | 19 | 39279851 | 39282389 | 8 | 3 | -1.5424 | 0.93852 | LGALS7B | 0.93852 | 0.92465 | 0.93541 |

| | | | | | | | | | | | |
|---|---|---|---|---|---|---|---|---|---|---|---|
| ENSG00000106683 | 7 | 73497263 | 73536855 | 85 | 16 | -1.5425 | 0.93853 | LIMK1 | 0.93853 | 0.94266 | 0.96747 |
| ENSG00000167186 | 16 | 19078921 | 19091417 | 51 | 10 | -1.5429 | 0.93858 | COQ7 | 0.93858 | 0.89746 | 0.90191 |
| ENSG00000136881 | 9 | 104122699 | 104145801 | 63 | 8 | -1.5435 | 0.93865 | BAAT | 0.93865 | 0.90636 | 0.8996 |
| ENSG00000141668 | 18 | 70203915 | 70305756 | 363 | 31 | -1.544 | 0.93871 | CBLN2 | 0.93871 | 0.95036 | 0.93394 |
| ENSG00000178206 | 2 | 132044330 | 132050127 | 13 | 4 | -1.5455 | 0.93888 | CYP4F31P | 0.93888 | 0.92441 | 0.86135 |
| ENSG00000130167 | 19 | 11406824 | 11437672 | 82 | 11 | -1.5457 | 0.93892 | TSPAN16 | 0.93892 | 0.95861 | 0.96435 |
| ENSG00000105559 | 19 | 49340354 | 49371889 | 102 | 15 | -1.5464 | 0.93899 | PLEKHA4 | 0.93899 | 0.94927 | 0.94342 |
| ENSG00000180787 | 17 | 4981543 | 4999669 | 51 | 10 | -1.5475 | 0.93913 | ZFP3 | 0.93913 | 0.71355 | 0.80881 |
| ENSG00000025796 | 6 | 108188960 | 108279393 | 201 | 13 | -1.5493 | 0.93934 | SEC63 | 0.93934 | 0.90779 | 0.87446 |
| ENSG00000121644 | 1 | 244816237 | 244872335 | 156 | 19 | -1.5496 | 0.93938 | DESI2 | 0.93938 | 0.91502 | 0.7253 |
| ENSG00000119661 | 14 | 74111578 | 74170435 | 199 | 20 | -1.5497 | 0.93939 | DNAL1 | 0.93939 | 0.87856 | 0.90633 |
| ENSG00000180509 | 21 | 35818988 | 35884573 | 233 | 30 | -1.5523 | 0.93971 | KCNE1 | 0.93971 | 0.94784 | 0.94125 |
| ENSG00000250479 | 22 | 24108021 | 24110630 | 15 | 5 | -1.5527 | 0.93975 | CHCHD10 | 0.93975 | 0.88448 | 0.91147 |
| ENSG00000138593 | 15 | 49280673 | 49338760 | 130 | 15 | -1.5534 | 0.93984 | SECISBP2L | 0.93984 | 0.95421 | 0.96317 |
| ENSG00000100220 | 22 | 32783569 | 32808242 | 101 | 19 | -1.5535 | 0.93985 | RTCB | 0.93985 | 0.84302 | 0.75357 |
| ENSG00000136842 | 9 | 100263462 | 100364030 | 300 | 42 | -1.5538 | 0.93989 | TMOD1 | 0.93989 | 0.94618 | 0.95836 |
| ENSG00000151914 | 6 | 56322785 | 56819426 | 1123 | 31 | -1.5541 | 0.93992 | DST | 0.93992 | 0.93272 | 0.94203 |
| ENSG00000175938 | 16 | 30960387 | 30967782 | 5 | 3 | -1.5546 | 0.93998 | ORAI3 | 0.93998 | 0.82215 | 0.79874 |
| ENSG00000121210 | 4 | 154387498 | 154557863 | 549 | 62 | -1.5549 | 0.94001 | KIAA0922 | 0.94001 | 0.95316 | 0.97027 |
| ENSG00000120242 | 9 | 21409146 | 21410184 | 2 | 1 | -1.5549 | 0.94002 | IFNA8 | 0.94002 | 0.71679 | 0.47705 |
| ENSG00000103494 | 16 | 53631595 | 53737850 | 297 | 22 | -1.5562 | 0.94017 | RPGRIP1L | 0.94017 | 0.9035 | 0.86238 |
| ENSG00000249860 | 10 | 57358750 | 57360488 | 7 | 3 | -1.5569 | 0.94025 | MTRNR2L5 | 0.94025 | 0.90903 | 0.85962 |
| ENSG00000149243 | 11 | 75133438 | 75141674 | 7 | 3 | -1.5578 | 0.94036 | KLHL35 | 0.94036 | 0.92356 | 0.85825 |
| ENSG00000069764 | 16 | 14766405 | 14788526 | 21 | 8 | -1.5597 | 0.94058 | PLA2G10 | 0.94058 | 0.93903 | 0.92718 |
| ENSG00000103184 | 16 | 5008318 | 5069159 | 287 | 25 | -1.5604 | 0.94067 | SEC14L5 | 0.94067 | 0.94729 | 0.92078 |
| ENSG00000206102 | 21 | 32410478 | 32410795 | 1 | 1 | -1.5607 | 0.9407 | KRTAP19-8 | 0.9407 | 0.81029 | 0.7998 |
| ENSG00000154473 | 10 | 124913793 | 124924886 | 27 | 4 | -1.5625 | 0.94091 | BUB3 | 0.94091 | 0.79014 | 0.67724 |
| ENSG00000134115 | 3 | 1134260 | 1445901 | 1694 | 120 | -1.5627 | 0.94094 | CNTN6 | 0.94094 | 0.95461 | 0.96066 |
| ENSG00000139908 | 14 | 24674903 | 24677568 | 2 | 1 | -1.5631 | 0.94099 | TSSK4 | 0.94099 | 0.67746 | 0.76412 |
| ENSG00000143217 | 1 | 161040785 | 161059389 | 46 | 12 | -1.5642 | 0.94112 | PVRL4 | 0.94112 | 0.94768 | 0.87977 |
| ENSG00000148154 | 9 | 114659046 | 114697649 | 217 | 31 | -1.5657 | 0.94129 | UGCG | 0.94129 | 0.93809 | 0.95186 |
| ENSG00000165152 | 9 | 104235453 | 104295819 | 219 | 31 | -1.5662 | 0.94135 | TMEM246 | 0.94135 | 0.9396 | 0.97609 |
| ENSG00000169946 | 8 | 106330920 | 106816760 | 1288 | 97 | -1.5665 | 0.94139 | ZFPM2 | 0.94139 | 0.94211 | 0.93767 |
| ENSG00000103241 | 16 | 86544133 | 86548076 | 22 | 8 | -1.5681 | 0.94157 | FOXF1 | 0.94157 | 0.96504 | 0.97949 |
| ENSG00000171103 | 2 | 29072687 | 29093167 | 49 | 9 | -1.5682 | 0.94158 | TRMT61B | 0.94158 | 0.89464 | 0.90818 |
| ENSG00000114767 | 3 | 51967446 | 51975957 | 15 | 5 | -1.5683 | 0.9416 | RRP9 | 0.9416 | 0.67858 | 0.8023 |
| ENSG00000153060 | 16 | 10721358 | 10788802 | 455 | 45 | -1.5685 | 0.94161 | TEKT5 | 0.94161 | 0.88788 | 0.82861 |
| ENSG00000170929 | 19 | 9203855 | 9204889 | 1 | 1 | -1.5692 | 0.9417 | OR1M1 | 0.9417 | 0.75273 | 0.48319 |
| ENSG00000180773 | 11 | 92877341 | 92931130 | 153 | 20 | -1.5698 | 0.94177 | SLC36A4 | 0.94177 | 0.92626 | 0.89743 |
| ENSG00000135241 | 7 | 108110866 | 108210110 | 290 | 18 | -1.5702 | 0.94181 | PNPLA8 | 0.94181 | 0.95616 | 0.89436 |
| ENSG00000159958 | 22 | 42321045 | 42322822 | 8 | 3 | -1.5701 | 0.94181 | TNFRSF13C | 0.94181 | 0.79396 | 0.87193 |
| ENSG00000188293 | 19 | 46733009 | 46734500 | 3 | 2 | -1.572 | 0.94202 | IGFL1 | 0.94202 | 0.90866 | 0.91111 |
| ENSG00000236980 | 3 | 49215065 | 49229291 | 25 | 7 | -1.5723 | 0.94206 | C3orf84 | 0.94206 | 0.94736 | 0.95348 |
| ENSG00000197321 | 10 | 29746267 | 30025710 | 1312 | 91 | -1.5727 | 0.9421 | SVIL | 0.9421 | 0.94047 | 0.9444 |
| ENSG00000146409 | 6 | 133090509 | 133119701 | 113 | 12 | -1.5733 | 0.94217 | SLC18B1 | 0.94217 | 0.94149 | 0.95262 |
| ENSG00000167635 | 19 | 36705504 | 36729676 | 100 | 10 | -1.5733 | 0.94217 | ZNF146 | 0.94217 | 0.94189 | 0.94172 |
| ENSG00000118557 | 16 | 72146056 | 72210777 | 194 | 22 | -1.5737 | 0.94223 | PMFBP1 | 0.94223 | 0.90932 | 0.87722 |
| ENSG00000187021 | 10 | 118349897 | 118368687 | 91 | 20 | -1.5742 | 0.94228 | PNLIPRP1 | 0.94228 | 0.94406 | 0.95985 |
| ENSG00000121388 | 13 | 19756173 | 19761496 | 47 | 3 | -1.5749 | 0.94236 | RP11-408E5.4 | 0.94236 | 0.89234 | 0.88135 |
| ENSG00000215853 | 1 | 152126071 | 152131704 | 8 | 4 | -1.5756 | 0.94244 | RPTN | 0.94244 | 0.81078 | 0.85684 |
| ENSG00000267941 | 7 | 65939543 | 65940709 | 3 | 1 | -1.5762 | 0.94251 | AC008267.1 | 0.94251 | 0.70895 | 0.67183 |
| ENSG00000151366 | 11 | 77779350 | 77791265 | 44 | 7 | -1.5783 | 0.94275 | NDUFC2 | 0.94275 | 0.9264 | 0.90562 |
| ENSG00000100030 | 22 | 22108789 | 22221970 | 246 | 27 | -1.5788 | 0.94281 | MAPK1 | 0.94281 | 0.94504 | 0.95169 |
| ENSG00000118960 | 2 | 20760208 | 20850849 | 427 | 42 | -1.58 | 0.94294 | HS1BP3 | 0.94294 | 0.95711 | 0.98022 |
| ENSG00000086967 | 19 | 50936160 | 50969578 | 138 | 23 | -1.5802 | 0.94297 | MYBPC2 | 0.94297 | 0.92723 | 0.96645 |
| ENSG00000137338 | 6 | 28249314 | 28270326 | 66 | 13 | -1.5813 | 0.94309 | PGBD1 | 0.94309 | 0.93641 | 0.93222 |
| ENSG00000119487 | 9 | 128199672 | 128469513 | 588 | 27 | -1.5816 | 0.94313 | MAPKAP1 | 0.94313 | 0.93614 | 0.93705 |
| ENSG00000145194 | 3 | 183967438 | 184010819 | 110 | 17 | -1.5831 | 0.9433 | ECE2 | 0.9433 | 0.9641 | 0.93767 |
| ENSG00000124098 | 20 | 54933971 | 54943719 | 38 | 7 | -1.5832 | 0.94331 | FAM210B | 0.94331 | 0.95891 | 0.96982 |
| ENSG00000145536 | 5 | 5140443 | 5320417 | 923 | 101 | -1.5835 | 0.94334 | ADAMTS16 | 0.94334 | 0.94652 | 0.95291 |
| ENSG00000197653 | 12 | 124247042 | 124420753 | 440 | 48 | -1.5852 | 0.94353 | DNAH10 | 0.94353 | 0.9492 | 0.94721 |
| ENSG00000204175 | 10 | 46994087 | 47005643 | 11 | 2 | -1.5856 | 0.94359 | GPRIN2 | 0.94359 | 0.94564 | 0.9395 |
| ENSG00000212864 | 9 | 140114707 | 140116033 | 2 | 1 | -1.5861 | 0.94364 | RNF208 | 0.94364 | 0.3698 | 0.48308 |
| ENSG00000149527 | 1 | 2357419 | 2436969 | 326 | 43 | -1.5865 | 0.94369 | PLCH2 | 0.94369 | 0.95959 | 0.91296 |
| ENSG00000264545 | 9 | 21802635 | 22032985 | 591 | 60 | -1.5885 | 0.94392 | RP11-145E5.5 | 0.94392 | 0.94551 | 0.90118 |
| ENSG00000261567 | 16 | 85391069 | 85393055 | 6 | 3 | -1.5891 | 0.94398 | RP11-680G10.1 | 0.94398 | 0.60689 | 0.80758 |
| ENSG00000101695 | 18 | 29598335 | 29653176 | 144 | 22 | -1.5897 | 0.94405 | RNF125 | 0.94405 | 0.96239 | 0.8973 |
| ENSG00000169840 | 13 | 28366780 | 28368905 | 2 | 1 | -1.5909 | 0.94419 | GSX1 | 0.94419 | 0.56385 | 0.57701 |
| ENSG00000198626 | 1 | 237205505 | 237997288 | 3371 | 213 | -1.5914 | 0.94424 | RYR2 | 0.94424 | 0.95409 | 0.94433 |
| ENSG00000131697 | 1 | 5922871 | 6052533 | 456 | 23 | -1.5917 | 0.94428 | NPHP4 | 0.94428 | 0.95518 | 0.95685 |
| ENSG00000176798 | 11 | 5020213 | 5021160 | 6 | 2 | -1.5927 | 0.94439 | OR51L1 | 0.94439 | 0.83798 | 0.80829 |
| ENSG00000040731 | 5 | 24487209 | 24645087 | 723 | 33 | -1.5933 | 0.94445 | CDH10 | 0.94445 | 0.86657 | 0.60145 |
| ENSG00000146469 | 6 | 153071933 | 153080900 | 9 | 4 | -1.5936 | 0.94449 | VIP | 0.94449 | 0.87086 | 0.12466 |

| | | | | | | | | | | | |
|---|---|---|---|---|---|---|---|---|---|---|---|
| ENSG00000163710 | 3 | 142534764 | 142608045 | 224 | 34 | -1.5945 | 0.94458 | PCOLCE2 | 0.94458 | 0.9643 | 0.96836 |
| ENSG00000273274 | 1 | 32930670 | 32962287 | 43 | 12 | -1.5947 | 0.94461 | ZBTB8B | 0.94461 | 0.94418 | 0.88697 |
| ENSG00000074803 | 15 | 48483861 | 48596275 | 173 | 13 | -1.5958 | 0.94473 | SLC12A1 | 0.94473 | 0.92927 | 0.92261 |
| ENSG00000116882 | 1 | 119911402 | 119936753 | 58 | 9 | -1.5959 | 0.94474 | HAO2 | 0.94474 | 0.90067 | 0.90549 |
| ENSG00000065427 | 16 | 75661622 | 75682541 | 85 | 14 | -1.5968 | 0.94485 | KARS | 0.94485 | 0.95108 | 0.94652 |
| ENSG00000168591 | 17 | 42264338 | 42269099 | 11 | 4 | -1.5968 | 0.94485 | TMUB2 | 0.94485 | 0.81901 | 0.89836 |
| ENSG00000258947 | 16 | 89987800 | 90005169 | 83 | 15 | -1.5977 | 0.94494 | TUBB3 | 0.94494 | 0.93948 | 0.81802 |
| ENSG00000163576 | 3 | 19920964 | 19988517 | 259 | 19 | -1.599 | 0.94509 | EFHB | 0.94509 | 0.97343 | 0.97079 |
| ENSG00000064205 | 20 | 43343485 | 43357150 | 28 | 10 | -1.5998 | 0.94517 | WISP2 | 0.94517 | 0.90857 | 0.79496 |
| ENSG00000124104 | 20 | 44462449 | 44471914 | 34 | 10 | -1.6 | 0.9452 | SNX21 | 0.9452 | 0.92655 | 0.87823 |
| ENSG00000256861 | 12 | 122692326 | 122750970 | 92 | 13 | -1.6013 | 0.94535 | RP11-512M8.5 | 0.94535 | 0.93037 | 0.90015 |
| ENSG00000162994 | 2 | 55401927 | 55459699 | 174 | 21 | -1.6018 | 0.9454 | CLHC1 | 0.9454 | 0.95983 | 0.97278 |
| ENSG00000165996 | 10 | 17631958 | 17659376 | 87 | 15 | -1.6018 | 0.9454 | PTPLA | 0.9454 | 0.84742 | 0.8294 |
| ENSG00000107984 | 10 | 54074056 | 54077802 | 10 | 3 | -1.6022 | 0.94544 | DKK1 | 0.94544 | 0.89099 | 0.9177 |
| ENSG00000231145 | 2 | 190642909 | 190643157 | 2 | 1 | -1.6023 | 0.94545 | AC013468.1 | 0.94545 | 0.78294 | 0.81575 |
| ENSG00000139180 | 12 | 4758261 | 4798454 | 150 | 21 | -1.6029 | 0.94552 | NDUFA9 | 0.94552 | 0.91776 | 0.70639 |
| ENSG00000132259 | 11 | 6255995 | 6265659 | 24 | 9 | -1.6042 | 0.94566 | CNGA4 | 0.94566 | 0.90085 | 0.88543 |
| ENSG00000166313 | 11 | 6416355 | 6440644 | 52 | 21 | -1.6062 | 0.94588 | APBB1 | 0.94588 | 0.84205 | 0.75503 |
| ENSG00000176595 | 8 | 1922044 | 1955102 | 209 | 25 | -1.6072 | 0.94599 | KBTBD11 | 0.94599 | 0.91985 | 0.36298 |
| ENSG00000100350 | 22 | 36883237 | 36903148 | 83 | 12 | -1.6087 | 0.94616 | FOXRED2 | 0.94616 | 0.95823 | 0.65753 |
| ENSG00000166930 | 11 | 60197062 | 60222687 | 137 | 17 | -1.61 | 0.9463 | MS4A5 | 0.9463 | 0.96094 | 0.97478 |
| ENSG00000129353 | 19 | 10713133 | 10755235 | 141 | 16 | -1.611 | 0.94641 | SLC44A2 | 0.94641 | 0.95927 | 0.98109 |
| ENSG00000091490 | 4 | 25749055 | 25865382 | 315 | 49 | -1.6113 | 0.94644 | SEL1L3 | 0.94644 | 0.9813 | 0.96813 |
| ENSG00000171847 | 12 | 8373856 | 8380214 | 36 | 9 | -1.6113 | 0.94644 | FAM90A1 | 0.94644 | 0.72551 | 0.11811 |
| ENSG00000112406 | 6 | 139456249 | 139501939 | 131 | 13 | -1.6117 | 0.94649 | HECA | 0.94649 | 0.91767 | 0.87031 |
| ENSG00000068784 | 2 | 45615819 | 45839304 | 766 | 30 | -1.6128 | 0.94661 | SRBD1 | 0.94661 | 0.9369 | 0.89607 |
| ENSG00000106992 | 9 | 130628759 | 130640022 | 27 | 12 | -1.6144 | 0.94678 | AK1 | 0.94678 | 0.9334 | 0.8235 |
| ENSG00000139719 | 12 | 122714111 | 122751068 | 58 | 11 | -1.615 | 0.94684 | VPS33A | 0.94684 | 0.94112 | 0.94203 |
| ENSG00000128487 | 17 | 19912657 | 20222339 | 917 | 21 | -1.615 | 0.94684 | SPECC1 | 0.94684 | 0.95436 | 0.96032 |
| ENSG00000158859 | 1 | 161154098 | 161168846 | 23 | 9 | -1.6167 | 0.94703 | ADAMTS4 | 0.94703 | 0.92555 | 0.88912 |
| ENSG00000185722 | 17 | 4067201 | 4167274 | 218 | 18 | -1.6195 | 0.94732 | ANKFY1 | 0.94732 | 0.95724 | 0.94065 |
| ENSG00000143921 | 2 | 44066103 | 44105605 | 324 | 13 | -1.6201 | 0.94739 | ABCG8 | 0.94739 | 0.95142 | 0.92021 |
| ENSG00000184271 | 12 | 51580719 | 51611477 | 86 | 18 | -1.6209 | 0.94749 | POU6F1 | 0.94749 | 0.91339 | 0.43506 |
| ENSG00000243232 | 5 | 140345820 | 140391936 | 77 | 18 | -1.622 | 0.9476 | PCDHAC2 | 0.9476 | 0.96616 | 0.96465 |
| ENSG00000104899 | 19 | 2249308 | 2252072 | 8 | 3 | -1.6224 | 0.94764 | AMH | 0.94764 | 0.95863 | 0.70565 |
| ENSG00000120805 | 12 | 101786898 | 101801598 | 38 | 12 | -1.6225 | 0.94765 | ARL1 | 0.94765 | 0.87352 | 0.82633 |
| ENSG00000169228 | 5 | 176728199 | 176730745 | 5 | 3 | -1.6231 | 0.94772 | RAB24 | 0.94772 | 0.91708 | 0.85774 |
| ENSG00000175575 | 11 | 73587744 | 73638790 | 209 | 22 | -1.6234 | 0.94775 | PAAF1 | 0.94775 | 0.93674 | 0.91185 |
| ENSG00000258869 | 14 | 98215601 | 98230326 | 67 | 12 | -1.6238 | 0.94779 | RP11-204N11.1 | 0.94779 | 0.89491 | 0.78271 |
| ENSG00000173598 | 12 | 93771659 | 93797024 | 93 | 14 | -1.6248 | 0.9479 | NUDT4 | 0.9479 | 0.94068 | 0.88085 |
| ENSG00000134202 | 1 | 110276554 | 110284384 | 19 | 4 | -1.625 | 0.94792 | GSTM3 | 0.94792 | 0.92786 | 0.93121 |
| ENSG00000196636 | 7 | 96745902 | 96811075 | 164 | 13 | -1.625 | 0.94792 | ACN9 | 0.94792 | 0.94156 | 0.95947 |
| ENSG00000169439 | 8 | 97505579 | 97624000 | 387 | 44 | -1.6261 | 0.94803 | SDC2 | 0.94803 | 0.93951 | 0.94449 |
| ENSG00000176903 | 14 | 74178494 | 74181128 | 4 | 2 | -1.6264 | 0.94807 | PNMA1 | 0.94807 | 0.97869 | 0.98933 |
| ENSG00000188064 | 22 | 46316242 | 46373009 | 306 | 45 | -1.6272 | 0.94816 | WNT7B | 0.94816 | 0.98621 | 0.98313 |
| ENSG00000116218 | 1 | 179519674 | 179545087 | 117 | 16 | -1.6274 | 0.94818 | NPHS2 | 0.94818 | 0.95442 | 0.94947 |
| ENSG00000104228 | 8 | 27142404 | 27168836 | 52 | 10 | -1.6281 | 0.94825 | TRIM35 | 0.94825 | 0.95969 | 0.93284 |
| ENSG00000178035 | 3 | 49061758 | 49066841 | 6 | 4 | -1.6285 | 0.94829 | IMPDH2 | 0.94829 | 0.74374 | 0.74804 |
| ENSG00000255384 | 11 | 118303836 | 118305921 | 3 | 2 | -1.6296 | 0.9484 | RP11-770J1.4 | 0.9484 | 0.75808 | 0.65995 |
| ENSG00000178403 | 4 | 113434672 | 113437328 | 5 | 3 | -1.6297 | 0.94841 | NEUROG2 | 0.94841 | 0.98745 | 0.97932 |
| ENSG00000184811 | 17 | 1182957 | 1204281 | 150 | 23 | -1.6301 | 0.94846 | TUSC5 | 0.94846 | 0.95125 | 0.9491 |
| ENSG00000159377 | 1 | 151372010 | 151374420 | 3 | 1 | -1.6306 | 0.94851 | PSMB4 | 0.94851 | 0.80094 | 0.82589 |
| ENSG00000173626 | 6 | 116816152 | 116866773 | 166 | 14 | -1.6317 | 0.94863 | TRAPPC3L | 0.94863 | 0.9505 | 0.92485 |
| ENSG00000140254 | 15 | 45409569 | 45422136 | 26 | 9 | -1.6319 | 0.94865 | DUOXA1 | 0.94865 | 0.92207 | 0.92591 |
| ENSG00000144857 | 3 | 112929850 | 113006303 | 296 | 39 | -1.6333 | 0.94879 | BOC | 0.94879 | 0.92839 | 0.92464 |
| ENSG00000187003 | 9 | 111624603 | 111626035 | 3 | 2 | -1.6336 | 0.94883 | ACTL7A | 0.94883 | 0.4629 | 0.31804 |
| ENSG00000115364 | 2 | 75873909 | 75917977 | 154 | 18 | -1.6349 | 0.94896 | MRPL19 | 0.94896 | 0.93569 | 0.90048 |
| ENSG00000108641 | 17 | 19240867 | 19281495 | 76 | 9 | -1.635 | 0.94898 | B9D1 | 0.94898 | 0.92638 | 0.89775 |
| ENSG00000128617 | 7 | 128412545 | 128415844 | 7 | 4 | -1.6355 | 0.94903 | OPN1SW | 0.94903 | 0.5535 | 0.63633 |
| ENSG00000183955 | 12 | 123868320 | 123893905 | 74 | 14 | -1.6355 | 0.94903 | SETD8 | 0.94903 | 0.92607 | 0.91407 |
| ENSG00000142208 | 14 | 105235686 | 105262088 | 94 | 23 | -1.6365 | 0.94913 | AKT1 | 0.94913 | 0.95669 | 0.84736 |
| ENSG00000164010 | 1 | 43282795 | 43310660 | 55 | 14 | -1.6369 | 0.94917 | ERMAP | 0.94917 | 0.93821 | 0.82545 |
| ENSG00000117308 | 1 | 24122089 | 24127271 | 2 | 1 | -1.6378 | 0.94927 | GALE | 0.94927 | 0.5402 | 0.58348 |
| ENSG00000198355 | 22 | 50354161 | 50357728 | 14 | 4 | -1.6379 | 0.94928 | PIM3 | 0.94928 | 0.8751 | 0.87908 |
| ENSG00000179059 | 4 | 188916925 | 188926204 | 43 | 12 | -1.6386 | 0.94935 | ZFP42 | 0.94935 | 0.85703 | 0.85013 |
| ENSG00000164675 | 7 | 123092454 | 123175131 | 149 | 16 | -1.6396 | 0.94945 | IQUB | 0.94945 | 0.93981 | 0.91491 |
| ENSG00000172116 | 2 | 87042462 | 87089047 | 157 | 19 | -1.6402 | 0.94952 | CD8B | 0.94952 | 0.94209 | 0.80878 |
| ENSG00000139144 | 12 | 18400548 | 18801348 | 1321 | 56 | -1.6408 | 0.94958 | PIK3C2G | 0.94958 | 0.92346 | 0.86599 |
| ENSG00000204688 | 6 | 29424958 | 29432105 | 18 | 8 | -1.6413 | 0.94963 | OR2H1 | 0.94963 | 0.90774 | 0.85782 |
| ENSG00000111291 | 12 | 13093709 | 13105081 | 38 | 10 | -1.6416 | 0.94966 | GPRC5D | 0.94966 | 0.9328 | 0.66367 |
| ENSG00000115207 | 2 | 27548716 | 27579868 | 35 | 7 | -1.6424 | 0.94975 | GTF3C2 | 0.94975 | 0.91525 | 0.91398 |
| ENSG00000161671 | 19 | 50979657 | 50986608 | 25 | 8 | -1.6424 | 0.94975 | EMC10 | 0.94975 | 0.9203 | 0.91949 |
| ENSG00000117593 | 1 | 173793641 | 173827684 | 51 | 11 | -1.6426 | 0.94976 | DARS2 | 0.94976 | 0.94779 | 0.90294 |

| | | | | | | | | | | | |
|---|---|---|---|---|---|---|---|---|---|---|---|
| ENSG00000162344 | 11 | 69513000 | 69519410 | 9 | 4 | -1.6436 | 0.94987 | FGF19 | 0.94987 | 0.87526 | 0.73405 |
| ENSG00000123307 | 12 | 55413729 | 55423798 | 23 | 6 | -1.6443 | 0.94994 | NEUROD4 | 0.94994 | 0.8893 | 0.82562 |
| ENSG00000134201 | 1 | 110254864 | 110318050 | 174 | 29 | -1.6454 | 0.95006 | GSTM5 | 0.95006 | 0.86871 | 0.84776 |
| ENSG00000181135 | 8 | 144766622 | 144796068 | 102 | 17 | -1.646 | 0.95012 | ZNF707 | 0.95012 | 0.95265 | 0.90274 |
| ENSG00000161267 | 3 | 197236654 | 197300194 | 287 | 28 | -1.6468 | 0.9502 | BDH1 | 0.9502 | 0.98215 | 0.99281 |
| ENSG00000101298 | 20 | 1246960 | 1289972 | 150 | 29 | -1.6468 | 0.95021 | SNPH | 0.95021 | 0.93556 | 0.71392 |
| ENSG00000169427 | 8 | 140613081 | 140715299 | 336 | 47 | -1.6471 | 0.95023 | KCNK9 | 0.95023 | 0.97174 | 0.9553 |
| ENSG00000165934 | 14 | 92588281 | 92630755 | 136 | 13 | -1.6472 | 0.95024 | CPSF2 | 0.95024 | 0.88506 | 0.72595 |
| ENSG00000156931 | 3 | 184529931 | 184770402 | 407 | 27 | -1.6484 | 0.95036 | VPS8 | 0.95036 | 0.94866 | 0.96758 |
| ENSG00000175581 | 11 | 73498361 | 73576178 | 349 | 22 | -1.6509 | 0.95062 | MRPL48 | 0.95062 | 0.96265 | 0.96366 |
| ENSG00000133739 | 8 | 86019382 | 86058311 | 47 | 8 | -1.6519 | 0.95072 | LRRCC1 | 0.95072 | 0.92621 | 0.89221 |
| ENSG00000099331 | 19 | 17186591 | 17325346 | 634 | 61 | -1.6519 | 0.95072 | MYO9B | 0.95072 | 0.94691 | 0.96496 |
| ENSG00000176771 | 2 | 133429374 | 134326034 | 2584 | 187 | -1.652 | 0.95073 | NCKAP5 | 0.95073 | 0.95932 | 0.96468 |
| ENSG00000182732 | 14 | 72399156 | 73030654 | 2610 | 131 | -1.652 | 0.95073 | RGS6 | 0.95073 | 0.94526 | 0.95184 |
| ENSG00000159871 | 19 | 44300081 | 44331358 | 155 | 23 | -1.6521 | 0.95075 | LYPD5 | 0.95075 | 0.94572 | 0.89364 |
| ENSG00000134278 | 18 | 12446511 | 12658133 | 795 | 33 | -1.6531 | 0.95085 | SPIRE1 | 0.95085 | 0.95732 | 0.95496 |
| ENSG00000139880 | 14 | 23516271 | 23526747 | 28 | 10 | -1.6532 | 0.95086 | CDH24 | 0.95086 | 0.93459 | 0.93062 |
| ENSG00000186063 | 1 | 222841355 | 222886552 | 92 | 18 | -1.6534 | 0.95088 | AIDA | 0.95088 | 0.9615 | 0.94124 |
| ENSG00000080007 | 6 | 74104471 | 74127292 | 39 | 10 | -1.6535 | 0.95088 | DDX43 | 0.95088 | 0.90176 | 0.83994 |
| ENSG00000113763 | 5 | 176237478 | 176307897 | 182 | 37 | -1.6554 | 0.95107 | UNC5A | 0.95107 | 0.90432 | 0.84171 |
| ENSG00000109101 | 17 | 26833261 | 26865914 | 73 | 10 | -1.6556 | 0.9511 | FOXN1 | 0.9511 | 0.90091 | 0.81783 |
| ENSG00000087495 | 20 | 58152564 | 58422766 | 1091 | 85 | -1.6557 | 0.95111 | PHACTR3 | 0.95111 | 0.90217 | 0.87 |
| ENSG00000182631 | 5 | 33936491 | 33939023 | 3 | 1 | -1.6558 | 0.95112 | RXFP3 | 0.95112 | 0.60754 | 0.66827 |
| ENSG00000142459 | 19 | 7895119 | 7929862 | 115 | 18 | -1.6564 | 0.95118 | EVI5L | 0.95118 | 0.96573 | 0.98328 |
| ENSG00000185909 | 3 | 49209044 | 49213917 | 10 | 3 | -1.6571 | 0.95125 | KLHDC8B | 0.95125 | 0.92068 | 0.93295 |
| ENSG00000112182 | 6 | 90636248 | 91006627 | 912 | 82 | -1.6576 | 0.9513 | BACH2 | 0.9513 | 0.90198 | 0.88748 |
| ENSG00000255587 | 6 | 36683256 | 36699870 | 58 | 11 | -1.6579 | 0.95133 | RAB44 | 0.95133 | 0.75609 | 0.8135 |
| ENSG00000204918 | 13 | 57721622 | 57724643 | 1 | 1 | -1.6586 | 0.9514 | PRR20B | 0.9514 | 0.78495 | 0.82142 |
| ENSG00000005884 | 17 | 48133332 | 48167845 | 97 | 14 | -1.6587 | 0.95142 | ITGA3 | 0.95142 | 0.92756 | 0.93691 |
| ENSG00000100711 | 14 | 104182067 | 104200005 | 47 | 12 | -1.6589 | 0.95143 | ZFYVE21 | 0.95143 | 0.96161 | 0.93741 |
| ENSG00000180053 | 8 | 23559964 | 23564111 | 14 | 4 | -1.6598 | 0.95153 | NKX2-6 | 0.95153 | 0.70097 | 0.40495 |
| ENSG00000137872 | 15 | 47476298 | 48066420 | 1891 | 63 | -1.6601 | 0.95155 | SEMA6D | 0.95155 | 0.95358 | 0.94542 |
| ENSG00000173166 | 2 | 204259068 | 204400133 | 268 | 21 | -1.6608 | 0.95163 | RAPH1 | 0.95163 | 0.95433 | 0.96279 |
| ENSG00000205030 | 11 | 55594695 | 55595630 | 9 | 2 | -1.6611 | 0.95165 | OR5L2 | 0.95165 | 0.44552 | 0.28507 |
| ENSG00000213121 | 6 | 153552455 | 153668623 | 516 | 33 | -1.6617 | 0.95172 | AL590867.1 | 0.95172 | 0.95822 | 0.9117 |
| ENSG00000159753 | 16 | 67678822 | 67691472 | 19 | 5 | -1.6628 | 0.95182 | RLTPR | 0.95182 | 0.80971 | 0.53384 |
| ENSG00000162851 | 1 | 246703862 | 246729626 | 208 | 24 | -1.6631 | 0.95186 | TFB2M | 0.95186 | 0.97934 | 0.85529 |
| ENSG00000075568 | 2 | 98372799 | 98612388 | 596 | 22 | -1.664 | 0.95194 | TMEM131 | 0.95194 | 0.9461 | 0.94083 |
| ENSG00000134007 | 14 | 70989075 | 71001732 | 31 | 6 | -1.6652 | 0.95207 | ADAM20 | 0.95207 | 0.95349 | 0.95568 |
| ENSG00000219438 | 22 | 48885272 | 49246724 | 2383 | 175 | -1.6654 | 0.95208 | FAM19A5 | 0.95208 | 0.94399 | 0.94566 |
| ENSG00000103522 | 16 | 27413483 | 27462115 | 210 | 41 | -1.6662 | 0.95216 | IL21R | 0.95216 | 0.97422 | 0.97893 |
| ENSG00000159023 | 1 | 29213603 | 29446553 | 561 | 28 | -1.6665 | 0.9522 | EPB41 | 0.9522 | 0.9468 | 0.96394 |
| ENSG00000138271 | 3 | 151011891 | 151034740 | 68 | 14 | -1.6668 | 0.95223 | GPR87 | 0.95223 | 0.95439 | 0.96002 |
| ENSG00000111732 | 12 | 8754762 | 8765467 | 22 | 13 | -1.6678 | 0.95233 | AICDA | 0.95233 | 0.9496 | 0.95096 |
| ENSG00000126550 | 4 | 70916119 | 70924562 | 34 | 8 | -1.6691 | 0.95245 | HTN1 | 0.95245 | 0.95634 | 0.95919 |
| ENSG00000126838 | 12 | 9301436 | 9360966 | 178 | 24 | -1.6694 | 0.95248 | PZP | 0.95248 | 0.95214 | 0.95564 |
| ENSG00000082438 | 2 | 165510134 | 165700189 | 523 | 29 | -1.6696 | 0.9525 | COBLL1 | 0.9525 | 0.94695 | 0.95937 |
| ENSG00000165730 | 10 | 70587298 | 70655188 | 234 | 30 | -1.6706 | 0.9526 | STOX1 | 0.9526 | 0.95893 | 0.96769 |
| ENSG00000213988 | 19 | 20188803 | 20237885 | 143 | 23 | -1.6714 | 0.95268 | ZNF90 | 0.95268 | 0.96062 | 0.95998 |
| ENSG00000089048 | 20 | 13694969 | 13765532 | 250 | 27 | -1.6718 | 0.95272 | ESF1 | 0.95272 | 0.93635 | 0.93076 |
| ENSG00000112218 | 6 | 97242002 | 97285353 | 142 | 17 | -1.6723 | 0.95277 | GPR63 | 0.95277 | 0.94297 | 0.92765 |
| ENSG00000176435 | 14 | 38723308 | 38725574 | 1 | 1 | -1.6726 | 0.9528 | CLEC14A | 0.9528 | 0.29567 | 0.50504 |
| ENSG00000146809 | 7 | 123207064 | 123277951 | 189 | 22 | -1.6735 | 0.95289 | ASB15 | 0.95289 | 0.94791 | 0.95194 |
| ENSG00000013288 | 4 | 6576902 | 6625089 | 169 | 27 | -1.6739 | 0.95292 | MAN2B2 | 0.95292 | 0.78094 | 0.7244 |
| ENSG00000197706 | 12 | 55640982 | 55642086 | 8 | 2 | -1.674 | 0.95293 | OR6C74 | 0.95293 | 0.95857 | 0.9751 |
| ENSG00000164334 | 5 | 118965254 | 118971517 | 33 | 4 | -1.6748 | 0.95301 | FAM170A | 0.95301 | 0.92926 | 0.91234 |
| ENSG00000113595 | 5 | 64885507 | 64921802 | 140 | 13 | -1.6763 | 0.95316 | TRIM23 | 0.95316 | 0.95251 | 0.95675 |
| ENSG00000184047 | 12 | 122692210 | 122712081 | 33 | 7 | -1.6764 | 0.95317 | DIABLO | 0.95317 | 0.85632 | 0.83828 |
| ENSG00000124208 | 20 | 48697661 | 48770174 | 157 | 19 | -1.6766 | 0.95319 | TMEM189-UBE2V1 | 0.95319 | 0.92302 | 0.95848 |
| ENSG00000240849 | 20 | 48697663 | 48770335 | 157 | 19 | -1.6766 | 0.95319 | TMEM189 | 0.95319 | 0.92302 | 0.95848 |
| ENSG00000214643 | 6 | 49913814 | 49917157 | 14 | 6 | -1.6776 | 0.95328 | DEFB133 | 0.95328 | 0.72198 | 0.68239 |
| ENSG00000123342 | 12 | 56229217 | 56236750 | 12 | 6 | -1.6777 | 0.95329 | MMP19 | 0.95329 | 0.98309 | 0.98891 |
| ENSG00000076706 | 11 | 119179241 | 119192231 | 20 | 11 | -1.6789 | 0.95341 | MCAM | 0.95341 | 0.87874 | 0.8946 |
| ENSG00000144712 | 3 | 12837971 | 12913415 | 304 | 28 | -1.6789 | 0.95342 | CAND2 | 0.95342 | 0.95309 | 0.9315 |
| ENSG00000115592 | 2 | 219687106 | 219696809 | 11 | 5 | -1.6792 | 0.95344 | PRKAG3 | 0.95344 | 0.97637 | 0.89971 |
| ENSG00000166086 | 11 | 133938820 | 134021896 | 343 | 18 | -1.6792 | 0.95344 | JAM3 | 0.95344 | 0.92734 | 0.92266 |
| ENSG00000205047 | 16 | 87731749 | 87739290 | 43 | 10 | -1.68 | 0.95353 | FLJ00104 | 0.95353 | 0.97837 | 0.9292 |
| ENSG00000174292 | 17 | 7283853 | 7293093 | 49 | 14 | -1.6803 | 0.95355 | TNK1 | 0.95355 | 0.9227 | 0.92504 |
| ENSG00000187527 | 3 | 192992579 | 193096632 | 399 | 28 | -1.6812 | 0.95364 | ATP13A5 | 0.95364 | 0.94175 | 0.92685 |
| ENSG00000222000 | 2 | 98947852 | 98972468 | 72 | 11 | -1.6814 | 0.95366 | AC092675.3 | 0.95366 | 0.90688 | 0.78975 |
| ENSG00000235931 | 10 | 61715188 | 61720671 | 23 | 4 | -1.6823 | 0.95374 | C10orf40 | 0.95374 | 0.97515 | 0.94905 |
| ENSG00000065457 | 16 | 75630879 | 75657198 | 122 | 19 | -1.6823 | 0.95375 | ADAT1 | 0.95375 | 0.93278 | 0.93392 |
| ENSG00000196689 | 17 | 3468738 | 3500392 | 148 | 25 | -1.6831 | 0.95382 | TRPV1 | 0.95382 | 0.93833 | 0.83073 |

| | | | | | | | | | | | |
|---|---|---|---|---|---|---|---|---|---|---|---|
| ENSG00000130724 | 19 | 59062933 | 59066491 | 7 | 4 | -1.683 | 0.95382 | CHMP2A | 0.95382 | 0.93129 | 0.94799 |
| ENSG00000262526 | 17 | 7143746 | 7147954 | 7 | 3 | -1.6832 | 0.95383 | CTD-2545G14.7 | 0.95383 | 0.85224 | 0.78234 |
| ENSG00000171014 | 11 | 123810250 | 123811344 | 3 | 2 | -1.6841 | 0.95392 | OR4D5 | 0.95392 | 0.4958 | 0.39611 |
| ENSG00000205327 | 12 | 55886147 | 55887100 | 4 | 2 | -1.6842 | 0.95393 | OR6C68 | 0.95393 | 0.82556 | 0.81569 |
| ENSG00000104415 | 8 | 134203282 | 134242587 | 148 | 32 | -1.685 | 0.95401 | WISP1 | 0.95401 | 0.94291 | 0.95943 |
| ENSG00000163481 | 2 | 219528587 | 219537134 | 22 | 3 | -1.6859 | 0.95409 | RNF25 | 0.95409 | 0.93887 | 0.89823 |
| ENSG00000143190 | 1 | 167190066 | 167396582 | 468 | 27 | -1.6885 | 0.95434 | POU2F1 | 0.95434 | 0.96794 | 0.96829 |
| ENSG00000198211 | 16 | 89985573 | 90002500 | 74 | 16 | -1.6898 | 0.95447 | TUBB3 | 0.95447 | 0.93948 | 0.81802 |
| ENSG00000131650 | 16 | 3013945 | 3018384 | 8 | 4 | -1.6903 | 0.95451 | KREMEN2 | 0.95451 | 0.72532 | 0.55339 |
| ENSG00000049246 | 1 | 7844380 | 7905237 | 257 | 22 | -1.6905 | 0.95453 | PER3 | 0.95453 | 0.95805 | 0.98249 |
| ENSG00000149328 | 11 | 134201768 | 134248235 | 168 | 49 | -1.6905 | 0.95453 | GLB1L2 | 0.95453 | 0.92178 | 0.93303 |
| ENSG00000187806 | 15 | 72690614 | 72700370 | 14 | 3 | -1.6907 | 0.95455 | TMEM202 | 0.95455 | 0.91361 | 0.88012 |
| ENSG00000165061 | 8 | 40388109 | 40755352 | 1359 | 89 | -1.6908 | 0.95456 | ZMAT4 | 0.95456 | 0.95232 | 0.96764 |
| ENSG00000069399 | 19 | 45250962 | 45263301 | 43 | 12 | -1.6911 | 0.9546 | BCL3 | 0.9546 | 0.86945 | 0.64008 |
| ENSG00000179055 | 9 | 107456660 | 107457766 | 7 | 2 | -1.6915 | 0.95463 | OR13D1 | 0.95463 | 0.88159 | 0.80845 |
| ENSG00000170265 | 7 | 148892577 | 148923339 | 115 | 18 | -1.6955 | 0.95501 | ZNF282 | 0.95501 | 0.8583 | 0.80742 |
| ENSG00000100003 | 22 | 30792846 | 30821305 | 88 | 16 | -1.6977 | 0.95522 | SEC14L2 | 0.95522 | 0.91433 | 0.87952 |
| ENSG00000133328 | 11 | 63320242 | 63330855 | 13 | 5 | -1.6982 | 0.95527 | HRASLS2 | 0.95527 | 0.99659 | 0.99709 |
| ENSG00000080503 | 9 | 2015342 | 2193624 | 751 | 101 | -1.6989 | 0.95533 | SMARCA2 | 0.95533 | 0.90245 | 0.81324 |
| ENSG00000174917 | 19 | 5678432 | 5680907 | 3 | 1 | -1.7002 | 0.95546 | C19orf70 | 0.95546 | 0.50377 | 0.52992 |
| ENSG00000189433 | 1 | 35225342 | 35229325 | 17 | 9 | -1.7005 | 0.95549 | GJB4 | 0.95549 | 0.84233 | 0.53755 |
| ENSG00000187942 | 1 | 22138758 | 22151714 | 42 | 17 | -1.7015 | 0.95557 | LDLRAD2 | 0.95557 | 0.92937 | 0.967 |
| ENSG00000049247 | 1 | 7903143 | 7913572 | 49 | 15 | -1.702 | 0.95562 | UTS2 | 0.95562 | 0.98547 | 0.98844 |
| ENSG00000132561 | 8 | 98881068 | 99048944 | 570 | 43 | -1.7022 | 0.95564 | MATN2 | 0.95564 | 0.96339 | 0.96393 |
| ENSG00000101412 | 20 | 32263489 | 32274210 | 17 | 9 | -1.7044 | 0.95585 | E2F1 | 0.95585 | 0.86697 | 0.9266 |
| ENSG00000152056 | 2 | 224616403 | 224702744 | 402 | 32 | -1.7059 | 0.95599 | AP1S3 | 0.95599 | 0.96965 | 0.95885 |
| ENSG00000074317 | 5 | 176047085 | 176057530 | 20 | 7 | -1.7066 | 0.95605 | SNCB | 0.95605 | 0.86108 | 0.82818 |
| ENSG00000232196 | 16 | 3421053 | 3422283 | 2 | 1 | -1.7084 | 0.95622 | MTRNR2L4 | 0.95622 | 0.82979 | 0.69942 |
| ENSG00000174720 | 4 | 113558120 | 113578748 | 48 | 14 | -1.7087 | 0.95624 | LARP7 | 0.95624 | 0.95089 | 0.94549 |
| ENSG00000120162 | 9 | 27325207 | 27529779 | 782 | 78 | -1.709 | 0.95627 | MOB3B | 0.95627 | 0.95736 | 0.97677 |
| ENSG00000134627 | 11 | 94277006 | 94354587 | 258 | 24 | -1.7089 | 0.95627 | PIWIL4 | 0.95627 | 0.95723 | 0.98282 |
| ENSG00000170296 | 17 | 7143333 | 7146089 | 6 | 3 | -1.7098 | 0.95635 | GABARAP | 0.95635 | 0.87292 | 0.77772 |
| ENSG00000185112 | 3 | 194406622 | 194409762 | 13 | 3 | -1.7105 | 0.95642 | FAM43A | 0.95642 | 0.80579 | 0.83393 |
| ENSG00000123505 | 6 | 111195973 | 111216916 | 76 | 10 | -1.7108 | 0.95644 | AMD1 | 0.95644 | 0.95442 | 0.96925 |
| ENSG00000115204 | 2 | 27532360 | 27548547 | 20 | 8 | -1.7116 | 0.95651 | MPV17 | 0.95651 | 0.9017 | 0.92737 |
| ENSG00000187944 | 2 | 197669726 | 197675000 | 7 | 3 | -1.7127 | 0.95662 | C2orf66 | 0.95662 | 0.79564 | 0.73882 |
| ENSG00000254997 | 11 | 71259466 | 71260653 | 8 | 3 | -1.7153 | 0.95685 | KRTAP5-9 | 0.95685 | 0.90252 | 0.9258 |
| ENSG00000196357 | 19 | 36673188 | 36737159 | 227 | 18 | -1.717 | 0.95701 | ZNF565 | 0.95701 | 0.94427 | 0.84687 |
| ENSG00000135164 | 7 | 86781677 | 86825653 | 166 | 11 | -1.7182 | 0.95712 | DMTF1 | 0.95712 | 0.96314 | 0.94578 |
| ENSG00000169230 | 5 | 176730775 | 176733960 | 9 | 4 | -1.7186 | 0.95715 | PRELID1 | 0.95715 | 0.94398 | 0.87366 |
| ENSG00000205116 | 1 | 1361508 | 1363167 | 10 | 4 | -1.719 | 0.95719 | TMEM88B | 0.95719 | 0.89738 | 0.88559 |
| ENSG00000103061 | 16 | 68318406 | 68344849 | 42 | 11 | -1.7192 | 0.95721 | SLC7A6OS | 0.95721 | 0.86185 | 0.83292 |
| ENSG00000099960 | 22 | 21383007 | 21387129 | 9 | 4 | -1.7198 | 0.95727 | SLC7A4 | 0.95727 | 0.95956 | 0.89431 |
| ENSG00000145526 | 5 | 19473060 | 20575982 | 3629 | 118 | -1.7205 | 0.95733 | CDH18 | 0.95733 | 0.96738 | 0.96793 |
| ENSG00000175946 | 8 | 124657767 | 124665190 | 48 | 9 | -1.7216 | 0.95743 | KLHL38 | 0.95743 | 0.92574 | 0.95002 |
| ENSG00000162891 | 1 | 207038699 | 207042568 | 8 | 4 | -1.7221 | 0.95748 | IL20 | 0.95748 | 0.49816 | 0.55938 |
| ENSG00000160339 | 9 | 137772654 | 137779924 | 47 | 10 | -1.7225 | 0.95751 | FCN2 | 0.95751 | 0.93517 | 0.97201 |
| ENSG00000265629 | 2 | 14775215 | 14790933 | 20 | 4 | -1.7226 | 0.95752 | AC011897.1 | 0.95752 | 0.92976 | 0.90828 |
| ENSG00000100601 | 14 | 78138747 | 78174363 | 180 | 23 | -1.7227 | 0.95753 | ALKBH1 | 0.95753 | 0.96395 | 0.95595 |
| ENSG00000137106 | 9 | 37422663 | 37436987 | 29 | 8 | -1.7241 | 0.95766 | GRHPR | 0.95766 | 0.8361 | 0.97072 |
| ENSG00000198858 | 19 | 896503 | 913240 | 87 | 24 | -1.7243 | 0.95768 | R3HDM4 | 0.95768 | 0.93673 | 0.8996 |
| ENSG00000088826 | 20 | 4101627 | 4168394 | 352 | 46 | -1.7244 | 0.95768 | SMOX | 0.95768 | 0.97525 | 0.99617 |
| ENSG00000168903 | 5 | 180415845 | 180433727 | 53 | 11 | -1.725 | 0.95774 | BTNL3 | 0.95774 | 0.97944 | 0.9925 |
| ENSG00000126549 | 4 | 70861647 | 70868287 | 17 | 7 | -1.7259 | 0.95781 | STATH | 0.95781 | 0.94947 | 0.92992 |
| ENSG00000148773 | 10 | 129894923 | 129924649 | 116 | 22 | -1.7267 | 0.95789 | MKI67 | 0.95789 | 0.93196 | 0.82987 |
| ENSG00000124198 | 20 | 47538427 | 47653230 | 307 | 17 | -1.7295 | 0.95814 | ARFGEF2 | 0.95814 | 0.96194 | 0.96168 |
| ENSG00000157388 | 3 | 53528683 | 53847760 | 846 | 84 | -1.7304 | 0.95822 | CACNA1D | 0.95822 | 0.93949 | 0.85772 |
| ENSG00000143951 | 2 | 63348518 | 64054977 | 1617 | 35 | -1.7307 | 0.95825 | WDPCP | 0.95825 | 0.96423 | 0.95324 |
| ENSG00000251537 | 17 | 15474805 | 15554967 | 313 | 28 | -1.7315 | 0.95832 | RP11-385D13.1 | 0.95832 | 0.97528 | 0.98856 |
| ENSG00000198674 | 11 | 123864870 | 123865868 | 4 | 2 | -1.7332 | 0.95847 | OR10G6 | 0.95847 | 0.63415 | 0.51252 |
| ENSG00000169224 | 1 | 247670360 | 247740992 | 278 | 37 | -1.7336 | 0.9585 | GCSAML | 0.9585 | 0.94754 | 0.97589 |
| ENSG00000119283 | 1 | 231297858 | 231357302 | 222 | 36 | -1.7345 | 0.95859 | TRIM67 | 0.95859 | 0.96296 | 0.93333 |
| ENSG00000164692 | 7 | 94023873 | 94060544 | 112 | 22 | -1.7359 | 0.95871 | COL1A2 | 0.95871 | 0.98307 | 0.97945 |
| ENSG00000197785 | 1 | 1447531 | 1470067 | 116 | 15 | -1.7366 | 0.95877 | ATAD3A | 0.95877 | 0.97422 | 0.96815 |
| ENSG00000168795 | 9 | 37438111 | 37465396 | 49 | 14 | -1.7366 | 0.95877 | ZBTB5 | 0.95877 | 0.90907 | 0.78118 |
| ENSG00000162981 | 2 | 14772810 | 14790933 | 22 | 4 | -1.7372 | 0.95882 | FAM84A | 0.95882 | 0.92526 | 0.90464 |
| ENSG00000165678 | 10 | 85899196 | 85913001 | 46 | 11 | -1.7372 | 0.95882 | GHITM | 0.95882 | 0.93399 | 0.83907 |
| ENSG00000145990 | 6 | 13358062 | 13487894 | 421 | 39 | -1.7389 | 0.95897 | GFOD1 | 0.95897 | 0.97103 | 0.98826 |
| ENSG00000105967 | 7 | 115575202 | 115799950 | 615 | 34 | -1.7397 | 0.95905 | TFEC | 0.95905 | 0.96512 | 0.96958 |
| ENSG00000171766 | 15 | 45653322 | 45694525 | 89 | 6 | -1.7404 | 0.95911 | GATM | 0.95911 | 0.91617 | 0.94258 |
| ENSG00000109016 | 17 | 21026677 | 21096578 | 104 | 15 | -1.7415 | 0.9592 | DHRS7B | 0.9592 | 0.96297 | 0.95646 |
| ENSG00000104976 | 19 | 7985201 | 7988135 | 6 | 3 | -1.7437 | 0.95939 | SNAPC2 | 0.95939 | 0.96909 | 0.97466 |
| ENSG00000139211 | 12 | 47469490 | 47473734 | 10 | 4 | -1.7443 | 0.95945 | AMIGO2 | 0.95945 | 0.97679 | 0.98547 |

| | | | | | | | | | | | |
|---|---|---|---|---|---|---|---|---|---|---|---|
| ENSG00000232070 | 14 | 21567096 | 21571883 | 15 | 6 | -1.7448 | 0.95949 | TMEM253 | 0.95949 | 0.95401 | 0.80066 |
| ENSG00000172236 | 16 | 1290697 | 1292555 | 15 | 6 | -1.7449 | 0.9595 | TPSAB1 | 0.9595 | 0.90614 | 0.79797 |
| ENSG00000166527 | 12 | 8662071 | 8674962 | 38 | 5 | -1.745 | 0.95951 | CLEC4D | 0.95951 | 0.94912 | 0.98319 |
| ENSG00000046889 | 8 | 68864353 | 69149265 | 1104 | 82 | -1.7452 | 0.95952 | PREX2 | 0.95952 | 0.96 | 0.9562 |
| ENSG00000008283 | 17 | 61509665 | 61523739 | 23 | 6 | -1.7459 | 0.95959 | CYB561 | 0.95959 | 0.85319 | 0.84581 |
| ENSG00000100034 | 22 | 22273793 | 22307209 | 79 | 12 | -1.7468 | 0.95966 | PPM1F | 0.95966 | 0.97292 | 0.96602 |
| ENSG00000118308 | 12 | 25173936 | 25261268 | 371 | 58 | -1.747 | 0.95968 | LRMP | 0.95968 | 0.94619 | 0.91434 |
| ENSG00000106633 | 7 | 44183872 | 44237769 | 158 | 29 | -1.7474 | 0.95971 | GCK | 0.95971 | 0.93533 | 0.88656 |
| ENSG00000185989 | 13 | 114747194 | 114898086 | 860 | 38 | -1.7495 | 0.95989 | RASA3 | 0.95989 | 0.94224 | 0.96533 |
| ENSG00000165804 | 14 | 21558205 | 21572881 | 38 | 12 | -1.7502 | 0.95996 | ZNF219 | 0.95996 | 0.91231 | 0.83425 |
| ENSG00000163739 | 4 | 74735110 | 74736959 | 6 | 3 | -1.7511 | 0.96003 | CXCL1 | 0.96003 | 0.89426 | 0.8312 |
| ENSG00000198576 | 8 | 143692405 | 143696833 | 15 | 3 | -1.7511 | 0.96004 | ARC | 0.96004 | 0.92641 | 0.95591 |
| ENSG00000162543 | 1 | 20512578 | 20522541 | 12 | 4 | -1.7546 | 0.96034 | UBXN10 | 0.96034 | 0.96701 | 0.63337 |
| ENSG00000178033 | 6 | 116832809 | 116845955 | 44 | 7 | -1.7547 | 0.96034 | FAM26E | 0.96034 | 0.94449 | 0.94984 |
| ENSG00000196090 | 20 | 40701392 | 41818610 | 3760 | 252 | -1.7548 | 0.96035 | PTPRT | 0.96035 | 0.97697 | 0.98595 |
| ENSG00000197734 | 14 | 78227173 | 78236085 | 45 | 11 | -1.755 | 0.96037 | C14orf178 | 0.96037 | 0.96409 | 0.95652 |
| ENSG00000141934 | 19 | 281040 | 291393 | 46 | 12 | -1.7552 | 0.96039 | PPAP2C | 0.96039 | 0.98042 | 0.9677 |
| ENSG00000255641 | 12 | 10564911 | 10588600 | 90 | 15 | -1.7559 | 0.96045 | NKG2-E | 0.96045 | 0.93946 | 0.95522 |
| ENSG00000144115 | 2 | 88469835 | 88486146 | 60 | 16 | -1.7566 | 0.9605 | THNSL2 | 0.9605 | 0.94287 | 0.95342 |
| ENSG00000182934 | 11 | 126132814 | 126139039 | 12 | 3 | -1.7572 | 0.96055 | SRPR | 0.96055 | 0.91892 | 0.97382 |
| ENSG00000130540 | 22 | 44220389 | 44258398 | 147 | 15 | -1.758 | 0.96063 | SULT4A1 | 0.96063 | 0.9419 | 0.91197 |
| ENSG00000189430 | 19 | 55417508 | 55427508 | 57 | 11 | -1.7602 | 0.96081 | NCR1 | 0.96081 | 0.9409 | 0.91545 |
| ENSG00000140682 | 16 | 31482906 | 31489281 | 5 | 2 | -1.7616 | 0.96093 | TGFB1I1 | 0.96093 | 0.96667 | 0.8329 |
| ENSG00000180113 | 6 | 46655612 | 46672056 | 21 | 9 | -1.7633 | 0.96107 | TDRD6 | 0.96107 | 0.7687 | 0.76499 |
| ENSG00000211450 | 11 | 57508825 | 57510986 | 3 | 1 | -1.7636 | 0.9611 | C11orf31 | 0.9611 | 0.3865 | 0.44862 |
| ENSG00000152078 | 1 | 95582894 | 95663163 | 283 | 25 | -1.7642 | 0.96115 | TMEM56 | 0.96115 | 0.97325 | 0.96679 |
| ENSG00000181885 | 17 | 7163222 | 7167302 | 11 | 5 | -1.7655 | 0.96126 | CLDN7 | 0.96126 | 0.89363 | 0.90533 |
| ENSG00000110987 | 12 | 122457328 | 122499948 | 139 | 26 | -1.7707 | 0.9617 | BCL7A | 0.9617 | 0.95563 | 0.96595 |
| ENSG00000178150 | 19 | 48675575 | 48790865 | 648 | 46 | -1.7707 | 0.9617 | ZNF114 | 0.9617 | 0.9769 | 0.98298 |
| ENSG00000102977 | 16 | 67691415 | 67694713 | 5 | 2 | -1.7716 | 0.96177 | ACD | 0.96177 | 0.95849 | 0.79402 |
| ENSG00000168497 | 2 | 192699028 | 192711981 | 26 | 9 | -1.7726 | 0.96185 | SDPR | 0.96185 | 0.96157 | 0.88228 |
| ENSG00000064692 | 5 | 121647049 | 121799914 | 272 | 19 | -1.7728 | 0.96187 | SNCAIP | 0.96187 | 0.9599 | 0.96361 |
| ENSG00000055813 | 2 | 56411258 | 56613308 | 715 | 58 | -1.7733 | 0.96191 | CCDC85A | 0.96191 | 0.97517 | 0.9483 |
| ENSG00000212128 | 12 | 11060525 | 11062161 | 2 | 1 | -1.7734 | 0.96192 | TAS2R13 | 0.96192 | 0.88045 | 0.87937 |
| ENSG00000261794 | 15 | 30896329 | 30906764 | 17 | 4 | -1.7735 | 0.96193 | GOLGA8H | 0.96193 | 0.82351 | 0.8225 |
| ENSG00000176571 | 8 | 87878670 | 88627447 | 2590 | 67 | -1.7738 | 0.96195 | CNBD1 | 0.96195 | 0.95985 | 0.96592 |
| ENSG00000132915 | 5 | 149237519 | 149324356 | 280 | 36 | -1.774 | 0.96196 | PDE6A | 0.96196 | 0.87021 | 0.87122 |
| ENSG00000176029 | 11 | 8941623 | 8954553 | 40 | 7 | -1.7752 | 0.96206 | C11orf16 | 0.96206 | 0.94344 | 0.92695 |
| ENSG00000178602 | 2 | 241078446 | 241083979 | 46 | 8 | -1.7766 | 0.96219 | OTOS | 0.96219 | 0.93717 | 0.85077 |
| ENSG00000249459 | 17 | 18561742 | 18585575 | 83 | 11 | -1.7774 | 0.96225 | ZNF286B | 0.96225 | 0.9669 | 0.94029 |
| ENSG00000164406 | 5 | 132208014 | 132210738 | 10 | 3 | -1.7777 | 0.96228 | LEAP2 | 0.96228 | 0.85436 | 0.84334 |
| ENSG00000174562 | 19 | 51328545 | 51340469 | 46 | 16 | -1.7782 | 0.96231 | KLK15 | 0.96231 | 0.858 | 0.93327 |
| ENSG00000164032 | 4 | 100869243 | 100871545 | 7 | 3 | -1.7801 | 0.96247 | H2AFZ | 0.96247 | 0.98586 | 0.98288 |
| ENSG00000187272 | 17 | 39394297 | 39395256 | 24 | 1 | -1.7803 | 0.96249 | KRTAP9-8 | 0.96249 | 0.62053 | 0.57773 |
| ENSG00000142677 | 1 | 24446261 | 24469611 | 75 | 16 | -1.7805 | 0.9625 | IL22RA1 | 0.9625 | 0.9756 | 0.90811 |
| ENSG00000132467 | 4 | 71554196 | 71556267 | 4 | 2 | -1.781 | 0.96254 | UTP3 | 0.96254 | 0.82073 | 0.79451 |
| ENSG00000137409 | 6 | 36935917 | 36954074 | 95 | 13 | -1.7817 | 0.9626 | MTCH1 | 0.9626 | 0.97308 | 0.97735 |
| ENSG00000100784 | 14 | 91336799 | 91526980 | 509 | 28 | -1.7829 | 0.9627 | RPS6KA5 | 0.9627 | 0.97016 | 0.97071 |
| ENSG00000124257 | 20 | 44517264 | 44519926 | 8 | 3 | -1.7829 | 0.9627 | NEURL2 | 0.9627 | 0.82132 | 0.88906 |
| ENSG00000221859 | 21 | 46057273 | 46058370 | 19 | 4 | -1.7853 | 0.9629 | KRTAP10-10 | 0.9629 | 0.85983 | 0.73352 |
| ENSG00000131061 | 20 | 32319463 | 32380075 | 112 | 16 | -1.7855 | 0.96291 | ZNF341 | 0.96291 | 0.98428 | 0.98883 |
| ENSG00000189043 | 7 | 10971578 | 10979883 | 34 | 9 | -1.7895 | 0.96323 | NDUFA4 | 0.96323 | 0.96713 | 0.93424 |
| ENSG00000204669 | 9 | 74666292 | 74687733 | 77 | 15 | -1.7941 | 0.9636 | C9orf57 | 0.9636 | 0.86483 | 0.67888 |
| ENSG00000119715 | 14 | 76776957 | 76968178 | 882 | 80 | -1.7945 | 0.96363 | ESRRB | 0.96363 | 0.9467 | 0.91449 |
| ENSG00000205810 | 12 | 10564911 | 10573194 | 22 | 5 | -1.7947 | 0.96365 | KLRC3 | 0.96365 | 0.96973 | 0.92891 |
| ENSG00000198816 | 19 | 7580178 | 7585912 | 17 | 8 | -1.7947 | 0.96365 | ZNF358 | 0.96365 | 0.88747 | 0.67035 |
| ENSG00000114455 | 3 | 108015376 | 108097132 | 343 | 24 | -1.7967 | 0.96381 | HHLA2 | 0.96381 | 0.97224 | 0.97942 |
| ENSG00000118849 | 3 | 158414681 | 158450485 | 148 | 19 | -1.7975 | 0.96387 | RARRES1 | 0.96387 | 0.97125 | 0.97034 |
| ENSG00000171495 | 5 | 40998119 | 41071444 | 316 | 30 | -1.7976 | 0.96388 | MROH2B | 0.96388 | 0.94795 | 0.89756 |
| ENSG00000232653 | 15 | 32885657 | 32896179 | 7 | 4 | -1.7978 | 0.96389 | GOLGA8N | 0.96389 | 0.90139 | 0.80189 |
| ENSG00000162191 | 11 | 62443970 | 62446567 | 3 | 2 | -1.8019 | 0.96422 | UBXN1 | 0.96422 | 0.76325 | 0.66044 |
| ENSG00000070190 | 4 | 100737990 | 100791311 | 223 | 28 | -1.8035 | 0.96434 | DAPP1 | 0.96434 | 0.97736 | 0.93582 |
| ENSG00000135441 | 12 | 56109820 | 56113871 | 8 | 4 | -1.8044 | 0.96442 | BLOC1S1 | 0.96442 | 0.95521 | 0.93605 |
| ENSG00000186910 | 14 | 94908801 | 94919127 | 53 | 10 | -1.8049 | 0.96445 | SERPINA11 | 0.96445 | 0.97635 | 0.92155 |
| ENSG00000176742 | 11 | 5220965 | 5221930 | 4 | 2 | -1.805 | 0.96446 | OR51V1 | 0.96446 | 0.98009 | 0.95608 |
| ENSG00000176485 | 11 | 63340667 | 63384355 | 89 | 22 | -1.8055 | 0.9645 | PLA2G16 | 0.9645 | 0.98492 | 0.99518 |
| ENSG00000127483 | 1 | 21069154 | 21113816 | 124 | 17 | -1.8061 | 0.96455 | HP1BP3 | 0.96455 | 0.97502 | 0.96698 |
| ENSG00000213185 | 10 | 124608594 | 124639157 | 43 | 15 | -1.8071 | 0.96462 | FAM24B | 0.96462 | 0.95024 | 0.93575 |
| ENSG00000109270 | 4 | 100799493 | 100815647 | 33 | 9 | -1.8074 | 0.96465 | LAMTOR3 | 0.96465 | 0.98591 | 0.9855 |
| ENSG00000116544 | 1 | 35331037 | 35395186 | 130 | 35 | -1.8081 | 0.9647 | DLGAP3 | 0.9647 | 0.96633 | 0.9453 |
| ENSG00000104497 | 8 | 82711816 | 82755101 | 183 | 14 | -1.809 | 0.96478 | SNX16 | 0.96478 | 0.97335 | 0.97476 |
| ENSG00000163792 | 2 | 27371872 | 27376378 | 8 | 3 | -1.8105 | 0.96489 | TCF23 | 0.96489 | 0.51946 | 0.55525 |
| ENSG00000133119 | 13 | 34392186 | 34540695 | 434 | 31 | -1.8119 | 0.965 | RFC3 | 0.965 | 0.96208 | 0.93163 |

| | | | | | | | | | | | |
|---|---|---|---|---|---|---|---|---|---|---|---|
| ENSG00000184313 | 1 | 55107463 | 55175939 | 265 | 36 | -1.8123 | 0.96503 | MROH7 | 0.96503 | 0.98774 | 0.98838 |
| ENSG00000175592 | 11 | 65659520 | 65668044 | 27 | 7 | -1.8126 | 0.96506 | FOSL1 | 0.96506 | 0.92079 | 0.91167 |
| ENSG00000138658 | 4 | 113460492 | 113558151 | 247 | 26 | -1.8155 | 0.96527 | C4orf21 | 0.96527 | 0.97985 | 0.97448 |
| ENSG00000105997 | 7 | 27145803 | 27192200 | 134 | 19 | -1.8163 | 0.96534 | HOXA3 | 0.96534 | 0.95819 | 0.97172 |
| ENSG00000109133 | 4 | 41937137 | 41962589 | 77 | 8 | -1.8167 | 0.96537 | TMEM33 | 0.96537 | 0.9611 | 0.8217 |
| ENSG00000178804 | 3 | 129262057 | 129270310 | 24 | 5 | -1.817 | 0.96539 | H1FOO | 0.96539 | 0.87179 | 0.95211 |
| ENSG00000166415 | 15 | 53805938 | 54055075 | 992 | 48 | -1.8172 | 0.96541 | WDR72 | 0.96541 | 0.9722 | 0.97104 |
| ENSG00000237190 | 5 | 133737778 | 133747589 | 9 | 4 | -1.8185 | 0.9655 | CDKN2AIPNL | 0.9655 | 0.85803 | 0.85434 |
| ENSG00000140650 | 16 | 8882680 | 8943188 | 417 | 27 | -1.8191 | 0.96555 | PMM2 | 0.96555 | 0.98237 | 0.98247 |
| ENSG00000156885 | 16 | 31439052 | 31439967 | 3 | 1 | -1.8195 | 0.96558 | COX6A2 | 0.96558 | 0.30815 | 0.65643 |
| ENSG00000198877 | 11 | 55540914 | 55541858 | 4 | 2 | -1.82 | 0.96562 | OR5D13 | 0.96562 | 0.55723 | 0.60108 |
| ENSG00000187456 | 17 | 34245070 | 34257777 | 36 | 4 | -1.8203 | 0.96564 | RDM1 | 0.96564 | 0.90589 | 0.82767 |
| ENSG00000165943 | 14 | 93648541 | 93651273 | 9 | 5 | -1.8236 | 0.96589 | MOAP1 | 0.96589 | 0.81519 | 0.50695 |
| ENSG00000203783 | 1 | 153190060 | 153191793 | 5 | 1 | -1.8239 | 0.96592 | PRR9 | 0.96592 | 0.95852 | 0.93114 |
| ENSG00000108823 | 17 | 48241575 | 48253292 | 44 | 10 | -1.8258 | 0.96606 | SGCA | 0.96606 | 0.92125 | 0.81324 |
| ENSG00000174744 | 11 | 66104804 | 66112596 | 14 | 4 | -1.8262 | 0.96609 | BRMS1 | 0.96609 | 0.95015 | 0.93436 |
| ENSG00000205213 | 11 | 27387508 | 27494322 | 246 | 35 | -1.8265 | 0.96611 | LGR4 | 0.96611 | 0.96243 | 0.95328 |
| ENSG00000105373 | 19 | 48248779 | 48260315 | 71 | 12 | -1.8268 | 0.96614 | GLTSCR2 | 0.96614 | 0.97038 | 0.96842 |
| ENSG00000188263 | 22 | 50432942 | 50451088 | 81 | 13 | -1.8275 | 0.96619 | IL17REL | 0.96619 | 0.94095 | 0.97942 |
| ENSG00000160683 | 11 | 118754475 | 118768508 | 37 | 13 | -1.8292 | 0.96631 | CXCR5 | 0.96631 | 0.90657 | 0.79926 |
| ENSG00000172967 | 22 | 17264302 | 17302589 | 117 | 21 | -1.8292 | 0.96632 | XKR3 | 0.96632 | 0.96797 | 0.94916 |
| ENSG00000183778 | 21 | 40928369 | 41045064 | 429 | 46 | -1.8303 | 0.9664 | B3GALT5 | 0.9664 | 0.94983 | 0.90515 |
| ENSG00000253251 | 5 | 64920592 | 64926718 | 21 | 5 | -1.832 | 0.96653 | CTC-534A2.2 | 0.96653 | 0.94666 | 0.93985 |
| ENSG00000243716 | 16 | 22490442 | 22547842 | 3 | 2 | -1.8353 | 0.96677 | NPIPB5 | 0.96677 | 0.83518 | 0.6543 |
| ENSG00000185372 | 5 | 180551357 | 180552304 | 1 | 1 | -1.8357 | 0.9668 | OR2V1 | 0.9668 | 0.94425 | 0.83738 |
| ENSG00000213020 | 19 | 53206066 | 53238326 | 188 | 19 | -1.8405 | 0.96715 | ZNF611 | 0.96715 | 0.95438 | 0.91135 |
| ENSG00000160803 | 1 | 156005092 | 156023585 | 29 | 9 | -1.8421 | 0.96727 | UBQLN4 | 0.96727 | 0.87672 | 0.87179 |
| ENSG00000064666 | 19 | 1026298 | 1039068 | 96 | 19 | -1.8429 | 0.96733 | CNN2 | 0.96733 | 0.96295 | 0.97146 |
| ENSG00000123977 | 2 | 228735770 | 228789060 | 272 | 17 | -1.843 | 0.96734 | DAW1 | 0.96734 | 0.95538 | 0.94211 |
| ENSG00000257390 | 12 | 56151059 | 56221330 | 105 | 25 | -1.8453 | 0.9675 | RP11-762I7.5 | 0.9675 | 0.97481 | 0.98119 |
| ENSG00000184677 | 1 | 22778344 | 22857650 | 160 | 18 | -1.8486 | 0.96774 | ZBTB40 | 0.96774 | 0.97773 | 0.89615 |
| ENSG00000064012 | 2 | 202098166 | 202152434 | 182 | 29 | -1.849 | 0.96777 | CASP8 | 0.96777 | 0.97679 | 0.98232 |
| ENSG00000143318 | 1 | 160160285 | 160171676 | 25 | 9 | -1.8521 | 0.968 | CASQ1 | 0.968 | 0.99399 | 0.86863 |
| ENSG00000064270 | 16 | 84402133 | 84497793 | 780 | 84 | -1.8521 | 0.968 | ATP2C2 | 0.968 | 0.97188 | 0.96981 |
| ENSG00000154102 | 16 | 85723690 | 85784735 | 264 | 19 | -1.8531 | 0.96807 | C16orf74 | 0.96807 | 0.96146 | 0.97087 |
| ENSG00000134545 | 12 | 10594863 | 10607284 | 66 | 10 | -1.8539 | 0.96812 | KLRC1 | 0.96812 | 0.9615 | 0.96161 |
| ENSG00000173221 | 5 | 95087023 | 95158709 | 179 | 30 | -1.8546 | 0.96817 | GLRX | 0.96817 | 0.93788 | 0.94511 |
| ENSG00000182909 | 19 | 54972981 | 54974894 | 7 | 3 | -1.8545 | 0.96817 | LENG9 | 0.96817 | 0.66954 | 0.70751 |
| ENSG00000105928 | 7 | 24737972 | 24809244 | 326 | 22 | -1.8549 | 0.9682 | DFNA5 | 0.9682 | 0.93512 | 0.94913 |
| ENSG00000233538 | 2 | 232316906 | 232317864 | 3 | 2 | -1.8564 | 0.9683 | AC017104.2 | 0.9683 | 0.14053 | 0.45616 |
| ENSG00000083807 | 19 | 58990879 | 59023780 | 61 | 14 | -1.857 | 0.96834 | SLC27A5 | 0.96834 | 0.93054 | 0.93374 |
| ENSG00000139112 | 12 | 10365057 | 10375727 | 35 | 13 | -1.8575 | 0.96838 | GABARAPL1 | 0.96838 | 0.85335 | 0.93834 |
| ENSG00000183853 | 1 | 157963063 | 158070052 | 394 | 62 | -1.8583 | 0.96844 | KIRREL | 0.96844 | 0.95192 | 0.97835 |
| ENSG00000122547 | 7 | 36192758 | 36341152 | 523 | 48 | -1.8588 | 0.96847 | EEPD1 | 0.96847 | 0.95513 | 0.81881 |
| ENSG00000156482 | 8 | 99037079 | 99058697 | 73 | 13 | -1.8616 | 0.96867 | RPL30 | 0.96867 | 0.98096 | 0.9758 |
| ENSG00000173039 | 11 | 65421067 | 65430565 | 15 | 5 | -1.8622 | 0.96871 | RELA | 0.96871 | 0.89739 | 0.88513 |
| ENSG00000100364 | 22 | 45586219 | 45636650 | 243 | 29 | -1.8654 | 0.96894 | KIAA0930 | 0.96894 | 0.97085 | 0.97075 |
| ENSG00000140939 | 16 | 67204057 | 67209643 | 5 | 2 | -1.867 | 0.96905 | NOL3 | 0.96905 | 0.86756 | 0.87131 |
| ENSG00000144331 | 2 | 180306709 | 180726232 | 1751 | 109 | -1.8677 | 0.9691 | ZNF385B | 0.9691 | 0.98886 | 0.99088 |
| ENSG00000205323 | 12 | 56146247 | 56211540 | 89 | 19 | -1.8685 | 0.96915 | SARNP | 0.96915 | 0.9772 | 0.97046 |
| ENSG00000138161 | 10 | 124591665 | 124639146 | 67 | 21 | -1.8694 | 0.96921 | CUZD1 | 0.96921 | 0.91552 | 0.90606 |
| ENSG00000086159 | 12 | 50360977 | 50370922 | 17 | 5 | -1.8711 | 0.96933 | AQP6 | 0.96933 | 0.95709 | 0.84408 |
| ENSG00000155749 | 2 | 202152994 | 202222121 | 202 | 22 | -1.8718 | 0.96938 | ALS2CR12 | 0.96938 | 0.96229 | 0.9661 |
| ENSG00000179913 | 19 | 17905637 | 17923891 | 66 | 14 | -1.872 | 0.9694 | B3GNT3 | 0.9694 | 0.95945 | 0.98198 |
| ENSG00000128655 | 2 | 178487980 | 178973066 | 1873 | 46 | -1.8743 | 0.96955 | PDE11A | 0.96955 | 0.97228 | 0.97638 |
| ENSG00000163785 | 3 | 133784147 | 133969689 | 626 | 30 | -1.8749 | 0.9696 | RYK | 0.9696 | 0.96671 | 0.95748 |
| ENSG00000271723 | 1 | 55107459 | 55207981 | 491 | 48 | -1.8759 | 0.96967 | MROH7-TTC4 | 0.96967 | 0.97665 | 0.9739 |
| ENSG00000113389 | 5 | 32689176 | 32791819 | 332 | 35 | -1.8764 | 0.9697 | NPR3 | 0.9697 | 0.96873 | 0.96751 |
| ENSG00000171940 | 20 | 52183604 | 52226446 | 116 | 28 | -1.8769 | 0.96974 | ZNF217 | 0.96974 | 0.95152 | 0.97894 |
| ENSG00000140835 | 16 | 71559136 | 71572649 | 20 | 8 | -1.8801 | 0.96995 | CHST4 | 0.96995 | 0.78775 | 0.69278 |
| ENSG00000181408 | 17 | 80332153 | 80333462 | 2 | 1 | -1.8806 | 0.96998 | UTS2R | 0.96998 | 0.40721 | 0.53478 |
| ENSG00000185519 | 1 | 16384264 | 16400127 | 113 | 15 | -1.8812 | 0.97003 | FAM131C | 0.97003 | 0.92355 | 0.91879 |
| ENSG00000127928 | 7 | 93220885 | 93540577 | 1027 | 58 | -1.8813 | 0.97004 | GNGT1 | 0.97004 | 0.97679 | 0.97529 |
| ENSG00000102981 | 16 | 67694849 | 67696681 | 2 | 1 | -1.8831 | 0.97015 | PARD6A | 0.97015 | 0.95867 | 0.88842 |
| ENSG00000163629 | 4 | 87515468 | 87736324 | 697 | 18 | -1.8831 | 0.97016 | PTPN13 | 0.97016 | 0.97375 | 0.97465 |
| ENSG00000197309 | 11 | 124055923 | 124056952 | 6 | 2 | -1.8837 | 0.9702 | OR10D3 | 0.9702 | 0.62097 | 0.66768 |
| ENSG00000084444 | 12 | 13197218 | 13295455 | 217 | 26 | -1.8863 | 0.97037 | KIAA1467 | 0.97037 | 0.97495 | 0.97203 |
| ENSG00000137364 | 6 | 18128542 | 18155305 | 97 | 14 | -1.8868 | 0.9704 | TPMT | 0.9704 | 0.9326 | 0.94181 |
| ENSG00000103196 | 16 | 84853590 | 84954374 | 540 | 55 | -1.8884 | 0.97052 | CRISPLD2 | 0.97052 | 0.68432 | 0.077458 |
| ENSG00000117602 | 1 | 24829387 | 24867530 | 125 | 16 | -1.889 | 0.97055 | RCAN3 | 0.97055 | 0.96756 | 0.96631 |
| ENSG00000149596 | 20 | 42740335 | 42816218 | 347 | 45 | -1.8916 | 0.97073 | JPH2 | 0.97073 | 0.97192 | 0.85956 |
| ENSG00000005469 | 7 | 86974997 | 87029111 | 146 | 21 | -1.8932 | 0.97083 | CROT | 0.97083 | 0.97058 | 0.98832 |
| ENSG00000151552 | 4 | 17461884 | 17513857 | 296 | 30 | -1.8932 | 0.97084 | QDPR | 0.97084 | 0.95906 | 0.9539 |

| | | | | | | | | | | | |
|---|---|---|---|---|---|---|---|---|---|---|---|
| ENSG00000063761 | 14 | 78266426 | 78401355 | 660 | 42 | -1.9001 | 0.97129 | ADCK1 | 0.97129 | 0.98412 | 0.98836 |
| ENSG00000131469 | 17 | 41150290 | 41154976 | 10 | 2 | -1.9008 | 0.97134 | RPL27 | 0.97134 | 0.90567 | 0.87077 |
| ENSG00000130558 | 9 | 137967268 | 138013025 | 247 | 42 | -1.9017 | 0.97139 | OLFM1 | 0.97139 | 0.75365 | 0.67781 |
| ENSG00000136518 | 3 | 179280668 | 179306196 | 65 | 10 | -1.903 | 0.97148 | ACTL6A | 0.97148 | 0.97222 | 0.96538 |
| ENSG00000183287 | 18 | 57098172 | 57364612 | 1166 | 94 | -1.9032 | 0.97149 | CCBE1 | 0.97149 | 0.97424 | 0.97987 |
| ENSG00000169991 | 1 | 19230775 | 19283180 | 218 | 42 | -1.9085 | 0.97184 | IFFO2 | 0.97184 | 0.97614 | 0.94626 |
| ENSG00000169550 | 11 | 26580580 | 26593780 | 57 | 14 | -1.911 | 0.97199 | MUC15 | 0.97199 | 0.93146 | 0.91104 |
| ENSG00000204169 | 10 | 51464162 | 51486327 | 24 | 6 | -1.912 | 0.97206 | AGAP7 | 0.97206 | 0.92906 | 0.81592 |
| ENSG00000165379 | 14 | 42076773 | 42373752 | 1278 | 39 | -1.9134 | 0.97215 | LRFN5 | 0.97215 | 0.97133 | 0.97022 |
| ENSG00000152795 | 4 | 83343717 | 83351294 | 34 | 8 | -1.9145 | 0.97222 | HNRNPDL | 0.97222 | 0.97667 | 0.98181 |
| ENSG00000164366 | 5 | 196986 | 218330 | 129 | 16 | -1.9157 | 0.9723 | CCDC127 | 0.9723 | 0.91487 | 0.85924 |
| ENSG00000017260 | 3 | 130569439 | 130735556 | 346 | 26 | -1.916 | 0.97232 | ATP2C1 | 0.97232 | 0.95205 | 0.95337 |
| ENSG00000144218 | 2 | 100162323 | 100759201 | 1525 | 56 | -1.9192 | 0.97252 | AFF3 | 0.97252 | 0.9812 | 0.98082 |
| ENSG00000179168 | 19 | 38874905 | 38878722 | 12 | 4 | -1.9244 | 0.97285 | GGN | 0.97285 | 0.80874 | 0.7895 |
| ENSG00000214788 | 11 | 59705928 | 59712807 | 7 | 3 | -1.9246 | 0.97286 | OOSP1 | 0.97286 | 0.96249 | 0.93177 |
| ENSG00000263053 | 17 | 79369275 | 79375658 | 14 | 6 | -1.9254 | 0.97291 | RP11-1055B8.6 | 0.97291 | 0.74561 | 0.7521 |
| ENSG00000066044 | 19 | 8023463 | 8070543 | 113 | 16 | -1.9257 | 0.97293 | ELAVL1 | 0.97293 | 0.97622 | 0.97685 |
| ENSG00000146648 | 7 | 55086714 | 55324313 | 799 | 84 | -1.9263 | 0.97296 | EGFR | 0.97296 | 0.97935 | 0.95734 |
| ENSG00000164031 | 4 | 100817405 | 100867883 | 100 | 17 | -1.9277 | 0.97305 | DNAJB14 | 0.97305 | 0.97859 | 0.98055 |
| ENSG00000164171 | 5 | 52285156 | 52390609 | 454 | 32 | -1.9286 | 0.97311 | ITGA2 | 0.97311 | 0.94464 | 0.88807 |
| ENSG00000255552 | 6 | 31679548 | 31681842 | 3 | 2 | -1.9308 | 0.97325 | LY6G6E | 0.97325 | 0.57002 | 0.63769 |
| ENSG00000136521 | 3 | 179322478 | 179345435 | 46 | 8 | -1.9318 | 0.97331 | NDUFB5 | 0.97331 | 0.94558 | 0.93403 |
| ENSG00000151023 | 10 | 25270908 | 25305089 | 72 | 13 | -1.9318 | 0.97331 | ENKUR | 0.97331 | 0.96249 | 0.95321 |
| ENSG00000213398 | 16 | 67973653 | 67978034 | 3 | 2 | -1.9323 | 0.97334 | LCAT | 0.97334 | 0.69534 | 0.75774 |
| ENSG00000179144 | 7 | 150211918 | 150218161 | 32 | 8 | -1.9344 | 0.97347 | GIMAP7 | 0.97347 | 0.78683 | 0.57939 |
| ENSG00000254535 | 4 | 135117488 | 135122903 | 14 | 3 | -1.9345 | 0.97348 | PABPC4L | 0.97348 | 0.90747 | 0.93721 |
| ENSG00000115758 | 2 | 10580094 | 10588630 | 21 | 6 | -1.9354 | 0.97353 | ODC1 | 0.97353 | 0.84097 | 0.75774 |
| ENSG00000138443 | 2 | 204192942 | 204312446 | 235 | 29 | -1.9372 | 0.97364 | ABI2 | 0.97364 | 0.97173 | 0.95805 |
| ENSG00000221886 | 5 | 159820155 | 159827104 | 10 | 4 | -1.9395 | 0.97378 | C5orf54 | 0.97378 | 0.84647 | 0.86789 |
| ENSG00000197860 | 5 | 64961755 | 65018862 | 123 | 16 | -1.9408 | 0.97386 | SGTB | 0.97386 | 0.95419 | 0.89393 |
| ENSG00000185453 | 19 | 48673949 | 48700877 | 107 | 18 | -1.9437 | 0.97404 | C19orf68 | 0.97404 | 0.98588 | 0.9766 |
| ENSG00000221888 | 1 | 247920676 | 247921982 | 3 | 2 | -1.9511 | 0.97448 | OR1C1 | 0.97448 | 0.89671 | 0.92905 |
| ENSG00000178988 | 4 | 6709428 | 6711607 | 1 | 1 | -1.9532 | 0.9746 | MRFAP1L1 | 0.9746 | 0.46221 | 0.29913 |
| ENSG00000110244 | 11 | 116691419 | 116694022 | 16 | 8 | -1.9536 | 0.97463 | APOA4 | 0.97463 | 0.77751 | 0.86423 |
| ENSG00000271092 | 1 | 95583479 | 95712781 | 456 | 29 | -1.9559 | 0.97476 | TMEM56-RWDD3 | 0.97476 | 0.98045 | 0.97284 |
| ENSG00000188522 | 17 | 18872102 | 18908117 | 76 | 18 | -1.956 | 0.97477 | FAM83G | 0.97477 | 0.86639 | 0.8573 |
| ENSG00000168675 | 18 | 13217497 | 13652754 | 1447 | 161 | -1.9572 | 0.97484 | LDLRAD4 | 0.97484 | 0.97573 | 0.98006 |
| ENSG00000225362 | 15 | 71402583 | 71407833 | 12 | 2 | -1.9577 | 0.97487 | CT62 | 0.97487 | 0.75889 | 0.77074 |
| ENSG00000162852 | 1 | 246729746 | 246831886 | 373 | 34 | -1.9592 | 0.97496 | CNST | 0.97496 | 0.96814 | 0.97721 |
| ENSG00000142279 | 19 | 34971874 | 34997258 | 99 | 24 | -1.9603 | 0.97502 | WTIP | 0.97502 | 0.97797 | 0.85997 |
| ENSG00000070495 | 17 | 74708919 | 74722866 | 21 | 6 | -1.966 | 0.97535 | JMJD6 | 0.97535 | 0.88972 | 0.85934 |
| ENSG00000078699 | 20 | 32077881 | 32237842 | 161 | 28 | -1.9681 | 0.97547 | CBFA2T2 | 0.97547 | 0.97099 | 0.96496 |
| ENSG00000109654 | 4 | 154073494 | 154260472 | 539 | 52 | -1.9706 | 0.97562 | TRIM2 | 0.97562 | 0.97968 | 0.94202 |
| ENSG00000162601 | 1 | 59120411 | 59165764 | 203 | 13 | -1.9717 | 0.97568 | MYSM1 | 0.97568 | 0.98172 | 0.97483 |
| ENSG00000138592 | 15 | 50716577 | 50793280 | 301 | 19 | -1.9733 | 0.97577 | USP8 | 0.97577 | 0.97485 | 0.97951 |
| ENSG00000174093 | 17 | 36337717 | 36421858 | 105 | 24 | -1.9732 | 0.97577 | RP11-1407O15.2 | 0.97577 | 0.96955 | 0.96332 |
| ENSG00000136937 | 9 | 100395908 | 100436030 | 36 | 12 | -1.9739 | 0.9758 | NCBP1 | 0.9758 | 0.94899 | 0.94775 |
| ENSG00000143952 | 2 | 64119280 | 64246206 | 474 | 38 | -1.9747 | 0.97585 | VPS54 | 0.97585 | 0.97278 | 0.96683 |
| ENSG00000148985 | 11 | 3818954 | 3847601 | 117 | 12 | -1.9754 | 0.97589 | PGAP2 | 0.97589 | 0.96262 | 0.90291 |
| ENSG00000111321 | 12 | 6484211 | 6500733 | 61 | 19 | -1.9777 | 0.97602 | LTBR | 0.97602 | 0.96555 | 0.9652 |
| ENSG00000249853 | 6 | 114376750 | 114664209 | 651 | 63 | -1.9804 | 0.97617 | HS3ST5 | 0.97617 | 0.97499 | 0.97664 |
| ENSG00000013306 | 17 | 42396993 | 42402238 | 22 | 7 | -1.9839 | 0.97637 | SLC25A39 | 0.97637 | 0.86485 | 0.95176 |
| ENSG00000175161 | 3 | 85008132 | 86123579 | 3309 | 63 | -1.9841 | 0.97638 | CADM2 | 0.97638 | 0.97779 | 0.97491 |
| ENSG00000171860 | 12 | 8210898 | 8219067 | 13 | 5 | -1.9885 | 0.97662 | C3AR1 | 0.97662 | 0.97553 | 0.96149 |
| ENSG00000125430 | 17 | 14204400 | 14252721 | 216 | 56 | -1.9918 | 0.9768 | HS3ST3B1 | 0.9768 | 0.98224 | 0.97231 |
| ENSG00000214160 | 3 | 183960089 | 183967336 | 17 | 7 | -1.9931 | 0.97687 | ALG3 | 0.97687 | 0.91316 | 0.82479 |
| ENSG00000140478 | 15 | 75575176 | 75586816 | 6 | 4 | -1.9936 | 0.9769 | GOLGA6D | 0.9769 | 0.040172 | 0.0077722 |
| ENSG00000109705 | 4 | 13542454 | 13546674 | 7 | 4 | -1.9968 | 0.97708 | NKX3-2 | 0.97708 | 0.95828 | 0.96696 |
| ENSG00000153214 | 2 | 112812800 | 112876895 | 199 | 30 | -1.9991 | 0.9772 | TMEM87B | 0.9772 | 0.95982 | 0.96991 |
| ENSG00000111404 | 12 | 18233803 | 18473041 | 1050 | 44 | -1.9992 | 0.97721 | RERGL | 0.97721 | 0.97766 | 0.97829 |
| ENSG00000113593 | 5 | 64859063 | 64883376 | 99 | 10 | -1.9998 | 0.97724 | PPWD1 | 0.97724 | 0.96943 | 0.96106 |
| ENSG00000268556 | 2 | 105867759 | 105869601 | 8 | 3 | -2.0012 | 0.97731 | AC012360.1 | 0.97731 | 0.81953 | 0.7641 |
| ENSG00000100450 | 14 | 25075686 | 25078905 | 6 | 4 | -2.0022 | 0.97737 | GZMH | 0.97737 | 0.90996 | 0.92094 |
| ENSG00000135829 | 1 | 182808504 | 182856886 | 116 | 14 | -2.013 | 0.97794 | DHX9 | 0.97794 | 0.97354 | 0.97866 |
| ENSG00000150347 | 10 | 63661059 | 63856703 | 536 | 57 | -2.0132 | 0.97795 | ARID5B | 0.97795 | 0.9775 | 0.98038 |
| ENSG00000167723 | 17 | 3413796 | 3461289 | 288 | 40 | -2.0135 | 0.97797 | TRPV3 | 0.97797 | 0.98739 | 0.9937 |
| ENSG00000134884 | 13 | 107194021 | 107220512 | 31 | 7 | -2.0163 | 0.97812 | ARGLU1 | 0.97812 | 0.91191 | 0.84053 |
| ENSG00000214556 | 17 | 36991341 | 36997708 | 24 | 7 | -2.0176 | 0.97818 | C17orf98 | 0.97818 | 0.9553 | 0.91063 |
| ENSG00000213585 | 5 | 133307606 | 133340824 | 99 | 15 | -2.0178 | 0.97819 | VDAC1 | 0.97819 | 0.95951 | 0.89425 |
| ENSG00000155269 | 4 | 8560452 | 8621486 | 412 | 49 | -2.018 | 0.9782 | GPR78 | 0.9782 | 0.97062 | 0.97314 |
| ENSG00000196498 | 12 | 124808961 | 125052135 | 933 | 116 | -2.0196 | 0.97829 | NCOR2 | 0.97829 | 0.96517 | 0.98687 |
| ENSG00000156030 | 14 | 74181825 | 74256988 | 221 | 38 | -2.0208 | 0.97835 | ELMSAN1 | 0.97835 | 0.98875 | 0.99392 |
| ENSG00000103248 | 16 | 86563782 | 86588841 | 111 | 16 | -2.0225 | 0.97844 | MTHFSD | 0.97844 | 0.9168 | 0.90289 |

| | | | | | | | | | | | |
|---|---|---|---|---|---|---|---|---|---|---|---|
| ENSG00000232382 | 3 | 98188324 | 98189420 | 2 | 1 | -2.0227 | 0.97845 | OR5K1 | 0.97845 | 0.95924 | 0.94515 |
| ENSG00000181234 | 12 | 128751948 | 129192460 | 1715 | 190 | -2.0238 | 0.9785 | TMEM132C | 0.9785 | 0.98988 | 0.98871 |
| ENSG00000255181 | 8 | 144788864 | 144790279 | 2 | 1 | -2.0264 | 0.97864 | CCDC166 | 0.97864 | 0.98506 | 0.9761 |
| ENSG00000152953 | 4 | 5053169 | 5502725 | 2365 | 104 | -2.0321 | 0.97893 | STK32B | 0.97893 | 0.97803 | 0.97641 |
| ENSG00000133805 | 11 | 10329860 | 10529126 | 939 | 77 | -2.0332 | 0.97899 | AMPD3 | 0.97899 | 0.96859 | 0.90426 |
| ENSG00000129744 | 11 | 3666358 | 3685646 | 168 | 24 | -2.0415 | 0.9794 | ART1 | 0.9794 | 0.96455 | 0.97592 |
| ENSG00000083123 | 6 | 80816364 | 81055987 | 800 | 23 | -2.042 | 0.97942 | BCKDHB | 0.97942 | 0.9579 | 0.93469 |
| ENSG00000111305 | 12 | 13236494 | 13256619 | 28 | 13 | -2.0424 | 0.97945 | GSG1 | 0.97945 | 0.98731 | 0.98485 |
| ENSG00000213934 | 11 | 5269313 | 5271122 | 21 | 6 | -2.0457 | 0.97961 | HBG1 | 0.97961 | 0.89497 | 0.75989 |
| ENSG00000213023 | 19 | 51124564 | 51171651 | 174 | 33 | -2.0482 | 0.97973 | SYT3 | 0.97973 | 0.97317 | 0.98339 |
| ENSG00000065371 | 3 | 123687368 | 123711025 | 63 | 12 | -2.0502 | 0.97983 | ROPN1 | 0.97983 | 0.68568 | 0.64078 |
| ENSG00000269558 | 7 | 65112081 | 65113280 | 2 | 1 | -2.052 | 0.97991 | AC104057.1 | 0.97991 | 0.66994 | 0.49572 |
| ENSG00000119574 | 19 | 59024897 | 59050278 | 41 | 11 | -2.0527 | 0.97995 | ZBTB45 | 0.97995 | 0.98741 | 0.98191 |
| ENSG00000117298 | 1 | 21543740 | 21671997 | 396 | 40 | -2.0543 | 0.98003 | ECE1 | 0.98003 | 0.98149 | 0.96705 |
| ENSG00000007237 | 17 | 9813926 | 10101868 | 1373 | 173 | -2.0543 | 0.98003 | GAS7 | 0.98003 | 0.98196 | 0.9342 |
| ENSG00000104059 | 15 | 29412457 | 29862927 | 1742 | 113 | -2.0568 | 0.98015 | FAM189A1 | 0.98015 | 0.98307 | 0.96893 |
| ENSG00000205090 | 1 | 1470554 | 1475833 | 9 | 4 | -2.0626 | 0.98042 | TMEM240 | 0.98042 | 0.97733 | 0.9799 |
| ENSG00000071991 | 18 | 64168320 | 64271375 | 278 | 22 | -2.0633 | 0.98046 | CDH19 | 0.98046 | 0.97838 | 0.9641 |
| ENSG00000031823 | 19 | 5916150 | 5978153 | 116 | 24 | -2.0663 | 0.9806 | RANBP3 | 0.9806 | 0.94827 | 0.91204 |
| ENSG00000106355 | 7 | 32524951 | 32534895 | 52 | 9 | -2.0688 | 0.98072 | LSM5 | 0.98072 | 0.9592 | 0.96825 |
| ENSG00000184227 | 14 | 74003818 | 74010498 | 22 | 7 | -2.0699 | 0.98077 | ACOT1 | 0.98077 | 0.97962 | 0.92959 |
| ENSG00000162734 | 1 | 160175127 | 160185166 | 13 | 7 | -2.0715 | 0.98084 | PEA15 | 0.98084 | 0.97672 | 0.99532 |
| ENSG00000188542 | 2 | 241499471 | 241503431 | 15 | 6 | -2.0766 | 0.98108 | DUSP28 | 0.98108 | 0.82948 | 0.58753 |
| ENSG00000141404 | 18 | 11688955 | 11885684 | 817 | 46 | -2.0779 | 0.98114 | GNAL | 0.98114 | 0.96895 | 0.9306 |
| ENSG00000171017 | 19 | 7953390 | 7966901 | 56 | 10 | -2.0796 | 0.98122 | LRRC8E | 0.98122 | 0.97625 | 0.98968 |
| ENSG00000221962 | 3 | 152057484 | 152058779 | 3 | 1 | -2.0807 | 0.98127 | TMEM14E | 0.98127 | 0.55336 | 0.49875 |
| ENSG00000166368 | 11 | 6912721 | 6913830 | 3 | 2 | -2.0822 | 0.98134 | OR2D2 | 0.98134 | 0.60347 | 0.58261 |
| ENSG00000142798 | 1 | 22148738 | 22263790 | 434 | 43 | -2.0876 | 0.98158 | HSPG2 | 0.98158 | 0.98191 | 0.9831 |
| ENSG00000095587 | 10 | 98124363 | 98273675 | 500 | 48 | -2.0907 | 0.98172 | TLL2 | 0.98172 | 0.98336 | 0.89558 |
| ENSG00000101333 | 20 | 9049410 | 9461889 | 1057 | 83 | -2.092 | 0.98178 | PLCB4 | 0.98178 | 0.98548 | 0.98527 |
| ENSG00000185271 | 14 | 20896970 | 20903801 | 32 | 7 | -2.0921 | 0.98179 | KLHL33 | 0.98179 | 0.89938 | 0.52631 |
| ENSG00000169957 | 16 | 30535325 | 30538142 | 3 | 2 | -2.0923 | 0.98179 | ZNF768 | 0.98179 | 0.99511 | 0.97779 |
| ENSG00000183317 | 1 | 38179552 | 38230805 | 227 | 24 | -2.0936 | 0.98185 | EPHA10 | 0.98185 | 0.95279 | 0.93514 |
| ENSG00000189190 | 19 | 53267448 | 53290044 | 144 | 29 | -2.0953 | 0.98193 | ZNF600 | 0.98193 | 0.97316 | 0.97004 |
| ENSG00000173421 | 3 | 49235861 | 49295537 | 101 | 14 | -2.097 | 0.982 | CCDC36 | 0.982 | 0.97952 | 0.96159 |
| ENSG00000188219 | 2 | 131975882 | 132022965 | 30 | 7 | -2.0977 | 0.98203 | POTEE | 0.98203 | 0.96664 | 0.97014 |
| ENSG00000101417 | 20 | 32294512 | 32308125 | 32 | 7 | -2.1026 | 0.98225 | PXMP4 | 0.98225 | 0.97757 | 0.96726 |
| ENSG00000101596 | 18 | 2655737 | 2805015 | 681 | 24 | -2.1053 | 0.98237 | SMCHD1 | 0.98237 | 0.97948 | 0.70271 |
| ENSG00000157542 | 21 | 38979678 | 39288749 | 1226 | 88 | -2.1062 | 0.98241 | KCNJ6 | 0.98241 | 0.9842 | 0.98784 |
| ENSG00000084674 | 2 | 21224301 | 21266945 | 80 | 21 | -2.109 | 0.98253 | APOB | 0.98253 | 0.97389 | 0.96299 |
| ENSG00000128564 | 7 | 100805790 | 100808874 | 2 | 1 | -2.1098 | 0.98256 | VGF | 0.98256 | 0.68686 | 0.78216 |
| ENSG00000262730 | 17 | 7829249 | 7833940 | 13 | 4 | -2.1103 | 0.98258 | RP11-1099M24.7 | 0.98258 | 0.80122 | 0.77057 |
| ENSG00000135838 | 1 | 182758428 | 182799519 | 119 | 14 | -2.1111 | 0.98262 | NPL | 0.98262 | 0.97779 | 0.96724 |
| ENSG00000100804 | 14 | 23485752 | 23504439 | 28 | 6 | -2.1116 | 0.98264 | PSMB5 | 0.98264 | 0.98859 | 0.99334 |
| ENSG00000141837 | 19 | 13317256 | 13734804 | 1839 | 178 | -2.1128 | 0.98269 | CACNA1A | 0.98269 | 0.989 | 0.99332 |
| ENSG00000178057 | 3 | 49057892 | 49060928 | 3 | 2 | -2.1151 | 0.98279 | NDUFAF3 | 0.98279 | 0.73694 | 0.65973 |
| ENSG00000196242 | 1 | 247693434 | 247697141 | 19 | 6 | -2.1184 | 0.98293 | OR2C3 | 0.98293 | 0.89368 | 0.9258 |
| ENSG00000255062 | 11 | 126164193 | 126174213 | 26 | 8 | -2.1214 | 0.98306 | RP11-712L6.5 | 0.98306 | 0.98622 | 0.97869 |
| ENSG00000154764 | 3 | 13857755 | 13921618 | 290 | 39 | -2.1291 | 0.98338 | WNT7A | 0.98338 | 0.96341 | 0.90498 |
| ENSG00000162650 | 1 | 110026101 | 110035426 | 13 | 4 | -2.1303 | 0.98343 | ATXN7L2 | 0.98343 | 0.81712 | 0.86099 |
| ENSG00000132507 | 17 | 7210318 | 7215774 | 16 | 5 | -2.1318 | 0.98349 | EIF5A | 0.98349 | 0.93577 | 0.95622 |
| ENSG00000069974 | 15 | 55495164 | 55611311 | 470 | 43 | -2.1339 | 0.98357 | RAB27A | 0.98357 | 0.97069 | 0.96896 |
| ENSG00000157060 | 1 | 182869000 | 182922660 | 133 | 17 | -2.1365 | 0.98368 | SHCBP1L | 0.98368 | 0.97341 | 0.79496 |
| ENSG00000146233 | 6 | 46517541 | 46620523 | 331 | 24 | -2.1441 | 0.98399 | CYP39A1 | 0.98399 | 0.9859 | 0.98599 |
| ENSG00000196083 | 3 | 190231840 | 190375843 | 571 | 70 | -2.1469 | 0.9841 | IL1RAP | 0.9841 | 0.98858 | 0.98595 |
| ENSG00000022556 | 19 | 55464498 | 55512510 | 326 | 38 | -2.1472 | 0.98411 | NLRP2 | 0.98411 | 0.96245 | 0.96693 |
| ENSG00000118200 | 1 | 200708686 | 200829832 | 348 | 25 | -2.1482 | 0.98415 | CAMSAP2 | 0.98415 | 0.98909 | 0.9845 |
| ENSG00000153048 | 16 | 8946799 | 8962866 | 111 | 20 | -2.1513 | 0.98428 | CARHSP1 | 0.98428 | 0.97837 | 0.9695 |
| ENSG00000177125 | 9 | 129622944 | 129648157 | 38 | 12 | -2.1557 | 0.98444 | ZBTB34 | 0.98444 | 0.972 | 0.9608 |
| ENSG00000108821 | 17 | 48260650 | 48278993 | 57 | 17 | -2.1572 | 0.9845 | COL1A1 | 0.9845 | 0.98033 | 0.97022 |
| ENSG00000180316 | 6 | 36210980 | 36276372 | 204 | 41 | -2.1622 | 0.9847 | PNPLA1 | 0.9847 | 0.98061 | 0.91231 |
| ENSG00000162594 | 1 | 67632083 | 67725662 | 293 | 30 | -2.1725 | 0.98509 | IL23R | 0.98509 | 0.97536 | 0.9629 |
| ENSG00000141391 | 18 | 12407895 | 12432237 | 106 | 20 | -2.1729 | 0.98511 | SLMO1 | 0.98511 | 0.9886 | 0.99653 |
| ENSG00000111859 | 6 | 11183531 | 11382581 | 763 | 92 | -2.1766 | 0.98525 | NEDD9 | 0.98525 | 0.98699 | 0.99614 |
| ENSG00000132932 | 13 | 25946209 | 26599989 | 2218 | 139 | -2.1783 | 0.98531 | ATP8A2 | 0.98531 | 0.97198 | 0.97184 |
| ENSG00000134996 | 9 | 77703459 | 77762181 | 236 | 33 | -2.1787 | 0.98532 | OSTF1 | 0.98532 | 0.98516 | 0.88752 |
| ENSG00000114670 | 3 | 130745694 | 131069309 | 847 | 33 | -2.1802 | 0.98538 | NEK11 | 0.98538 | 0.98614 | 0.99013 |
| ENSG00000167359 | 11 | 5461800 | 5462744 | 7 | 3 | -2.1884 | 0.98568 | OR51I1 | 0.98568 | 0.91615 | 0.87348 |
| ENSG00000130734 | 19 | 10654571 | 10664094 | 29 | 9 | -2.2007 | 0.98612 | ATG4D | 0.98612 | 0.95105 | 0.96457 |
| ENSG00000204657 | 6 | 29555683 | 29556745 | 6 | 3 | -2.201 | 0.98613 | OR2H2 | 0.98613 | 0.92732 | 0.92514 |
| ENSG00000172932 | 11 | 67056018 | 67069956 | 19 | 9 | -2.208 | 0.98638 | ANKRD13D | 0.98638 | 0.78772 | 0.76755 |
| ENSG00000189184 | 4 | 138440072 | 138453648 | 37 | 8 | -2.2084 | 0.98639 | PCDH18 | 0.98639 | 0.98323 | 0.98815 |
| ENSG00000215421 | 18 | 72265106 | 72777627 | 1071 | 43 | -2.2093 | 0.98642 | ZNF407 | 0.98642 | 0.98864 | 0.98326 |

| | | | | | | | | | | | |
|---|---|---|---|---|---|---|---|---|---|---|---|
| ENSG00000184857 | 16 | 8874241 | 8891505 | 114 | 17 | -2.2101 | 0.98645 | TMEM186 | 0.98645 | 0.97751 | 0.96059 |
| ENSG00000178980 | 19 | 48281829 | 48287943 | 13 | 5 | -2.2139 | 0.98658 | SEPW1 | 0.98658 | 0.99242 | 0.99183 |
| ENSG00000170298 | 17 | 20352708 | 20370852 | 33 | 8 | -2.2157 | 0.98664 | LGALS9B | 0.98664 | 0.9849 | 0.98664 |
| ENSG00000128594 | 7 | 127667124 | 127672160 | 11 | 5 | -2.2176 | 0.98671 | LRRC4 | 0.98671 | 0.9129 | 0.95921 |
| ENSG00000178075 | 3 | 113547029 | 113666021 | 334 | 38 | -2.2181 | 0.98672 | GRAMD1C | 0.98672 | 0.98627 | 0.9722 |
| ENSG00000101439 | 20 | 23608534 | 23619110 | 15 | 9 | -2.2215 | 0.98684 | CST3 | 0.98684 | 0.69752 | 0.46491 |
| ENSG00000167468 | 19 | 1103936 | 1106787 | 11 | 6 | -2.2264 | 0.98701 | GPX4 | 0.98701 | 0.92754 | 0.85395 |
| ENSG00000134812 | 11 | 59596741 | 59612974 | 26 | 9 | -2.2316 | 0.98718 | GIF | 0.98718 | 0.51037 | 0.20533 |
| ENSG00000179002 | 1 | 19166093 | 19186176 | 133 | 24 | -2.2398 | 0.98745 | TAS1R2 | 0.98745 | 0.96843 | 0.94409 |
| ENSG00000030582 | 17 | 42422614 | 42430470 | 16 | 6 | -2.2442 | 0.98759 | GRN | 0.98759 | 0.98156 | 0.98492 |
| ENSG00000268482 | 9 | 35361056 | 35361262 | 1 | 1 | -2.2571 | 0.988 | AL160274.1 | 0.988 | 0.52732 | 0.40365 |
| ENSG00000165202 | 9 | 125376948 | 125378026 | 4 | 2 | -2.2619 | 0.98815 | OR1Q1 | 0.98815 | 0.96852 | 0.94075 |
| ENSG00000213417 | 17 | 39221369 | 39222131 | 1 | 1 | -2.2636 | 0.9882 | KRTAP2-4 | 0.9882 | 0.70718 | 0.73154 |
| ENSG00000166793 | 11 | 57412560 | 57417417 | 5 | 3 | -2.264 | 0.98821 | YPEL4 | 0.98821 | 0.59932 | 0.60327 |
| ENSG00000065970 | 12 | 8185299 | 8208099 | 46 | 10 | -2.2641 | 0.98821 | FOXJ2 | 0.98821 | 0.99773 | 0.98721 |
| ENSG00000163132 | 4 | 4861393 | 4865663 | 23 | 7 | -2.2668 | 0.9883 | MSX1 | 0.9883 | 0.98325 | 0.75602 |
| ENSG00000172382 | 16 | 2762419 | 2770552 | 10 | 5 | -2.2688 | 0.98836 | PRSS27 | 0.98836 | 0.6233 | 0.64437 |
| ENSG00000176809 | 17 | 62850430 | 62915598 | 16 | 9 | -2.2771 | 0.98861 | LRRC37A3 | 0.98861 | 0.98242 | 0.97608 |
| ENSG00000184985 | 4 | 7194265 | 7744554 | 3531 | 297 | -2.2866 | 0.98889 | SORCS2 | 0.98889 | 0.99084 | 0.99582 |
| ENSG00000173431 | 14 | 21525981 | 21526614 | 2 | 1 | -2.2912 | 0.98903 | RNASE8 | 0.98903 | 0.89205 | 0.93705 |
| ENSG00000223874 | 2 | 217735495 | 217736362 | 2 | 1 | -2.2956 | 0.98915 | AC007557.1 | 0.98915 | 0.97765 | 0.96508 |
| ENSG00000163155 | 1 | 151132224 | 151138424 | 8 | 4 | -2.299 | 0.98925 | LYSMD1 | 0.98925 | 0.83438 | 0.9216 |
| ENSG00000143603 | 1 | 154669931 | 154842756 | 529 | 90 | -2.3011 | 0.98931 | KCNN3 | 0.98931 | 0.9944 | 0.99687 |
| ENSG00000134247 | 1 | 117452679 | 117532980 | 266 | 25 | -2.3093 | 0.98954 | PTGFRN | 0.98954 | 0.99047 | 0.98663 |
| ENSG00000112273 | 6 | 22569678 | 22571892 | 13 | 5 | -2.3126 | 0.98963 | HDGFL1 | 0.98963 | 0.37019 | 0.5515 |
| ENSG00000163156 | 1 | 151129140 | 151142773 | 16 | 8 | -2.3205 | 0.98984 | SCNM1 | 0.98984 | 0.90518 | 0.93012 |
| ENSG00000167755 | 19 | 51461887 | 51472929 | 32 | 11 | -2.3317 | 0.99014 | KLK6 | 0.99014 | 0.98908 | 0.78448 |
| ENSG00000221926 | 17 | 15531274 | 15587625 | 158 | 27 | -2.3329 | 0.99017 | TRIM16 | 0.99017 | 0.9711 | 0.97974 |
| ENSG00000172771 | 3 | 129120164 | 129147494 | 139 | 12 | -2.3418 | 0.99041 | EFCAB12 | 0.99041 | 0.98895 | 0.97813 |
| ENSG00000134369 | 1 | 201592411 | 201796102 | 555 | 49 | -2.3434 | 0.99044 | NAV1 | 0.99044 | 0.99533 | 0.98452 |
| ENSG00000177300 | 4 | 184239220 | 184241927 | 3 | 2 | -2.3462 | 0.99052 | CLDN22 | 0.99052 | 0.75543 | 0.82114 |
| ENSG00000122481 | 1 | 95699711 | 95712781 | 47 | 11 | -2.3716 | 0.99115 | RWDD3 | 0.99115 | 0.982 | 0.98842 |
| ENSG00000105370 | 19 | 51883163 | 51891214 | 19 | 5 | -2.3936 | 0.99166 | LIM2 | 0.99166 | 0.96292 | 0.5362 |
| ENSG00000156804 | 8 | 124510129 | 124553446 | 156 | 34 | -2.4028 | 0.99187 | FBXO32 | 0.99187 | 0.98913 | 0.96348 |
| ENSG00000104133 | 15 | 44854894 | 44955876 | 95 | 31 | -2.4261 | 0.99237 | SPG11 | 0.99237 | 0.97758 | 0.95101 |
| ENSG00000229809 | 16 | 30580667 | 30584055 | 1 | 1 | -2.4372 | 0.9926 | ZNF688 | 0.9926 | 0.74669 | 0.71855 |
| ENSG00000260456 | 16 | 87117168 | 87351022 | 1272 | 83 | -2.4388 | 0.99263 | C16orf95 | 0.99263 | 0.99271 | 0.99199 |
| ENSG00000150455 | 11 | 126152960 | 126168740 | 42 | 11 | -2.4574 | 0.993 | TIRAP | 0.993 | 0.98487 | 0.97267 |
| ENSG00000132329 | 2 | 238767536 | 238820756 | 252 | 43 | -2.4715 | 0.99327 | RAMP1 | 0.99327 | 0.99675 | 0.99302 |
| ENSG00000136267 | 7 | 14184674 | 15014402 | 3354 | 166 | -2.4764 | 0.99336 | DGKB | 0.99336 | 0.99402 | 0.99537 |
| ENSG00000179071 | 11 | 85394893 | 85397320 | 5 | 2 | -2.4813 | 0.99345 | CCDC89 | 0.99345 | 0.93816 | 0.91379 |
| ENSG00000103067 | 16 | 68263014 | 68272005 | 5 | 2 | -2.4833 | 0.99349 | ESRP2 | 0.99349 | 0.82615 | 0.89448 |
| ENSG00000100263 | 22 | 29655841 | 29664198 | 11 | 6 | -2.486 | 0.99354 | RHBDD3 | 0.99354 | 0.87812 | 0.58883 |
| ENSG00000189068 | 19 | 54544079 | 54567207 | 159 | 29 | -2.5004 | 0.9938 | VSTM1 | 0.9938 | 0.99368 | 0.79218 |
| ENSG00000158258 | 3 | 139654027 | 140296239 | 2146 | 155 | -2.5074 | 0.99392 | CLSTN2 | 0.99392 | 0.99359 | 0.99618 |
| ENSG00000182621 | 20 | 8112824 | 8949003 | 2920 | 185 | -2.5152 | 0.99405 | PLCB1 | 0.99405 | 0.9923 | 0.98066 |
| ENSG00000104879 | 19 | 45809672 | 45826235 | 79 | 17 | -2.5194 | 0.99412 | CKM | 0.99412 | 0.98932 | 0.9747 |
| ENSG00000135414 | 12 | 56137064 | 56150911 | 16 | 7 | -2.5216 | 0.99416 | GDF11 | 0.99416 | 0.93874 | 0.9367 |
| ENSG00000214919 | 3 | 155459933 | 155461515 | 2 | 1 | -2.5387 | 0.99444 | AC104472.1 | 0.99444 | 0.74255 | 0.70242 |
| ENSG00000197532 | 1 | 158516918 | 158517895 | 2 | 1 | -2.5499 | 0.99461 | OR6Y1 | 0.99461 | 0.56163 | 0.50179 |
| ENSG00000119681 | 14 | 74964873 | 75079306 | 330 | 37 | -2.555 | 0.99469 | LTBP2 | 0.99469 | 0.99457 | 0.98842 |
| ENSG00000268146 | 1 | 244227632 | 244231211 | 9 | 4 | -2.5759 | 0.995 | AL590483.1 | 0.995 | 0.96038 | 0.76706 |
| ENSG00000172023 | 2 | 79312156 | 79315145 | 10 | 5 | -2.6012 | 0.99536 | REG1B | 0.99536 | 0.99221 | 0.8353 |
| ENSG00000174684 | 11 | 66112843 | 66115163 | 1 | 1 | -2.6121 | 0.9955 | B3GNT1 | 0.9955 | 0.96502 | 0.92001 |
| ENSG00000185532 | 10 | 52750945 | 54058110 | 5107 | 244 | -2.6169 | 0.99556 | PRKG1 | 0.99556 | 0.9943 | 0.99485 |
| ENSG00000137090 | 9 | 841690 | 969090 | 819 | 65 | -2.6482 | 0.99595 | DMRT1 | 0.99595 | 0.99117 | 0.98235 |
| ENSG00000143258 | 1 | 161129240 | 161135513 | 7 | 4 | -2.6748 | 0.99626 | USP21 | 0.99626 | 0.88592 | 0.84434 |
| ENSG00000205138 | 19 | 36486090 | 36487220 | 2 | 1 | -2.6946 | 0.99648 | SDHAF1 | 0.99648 | 0.75443 | 0.6879 |
| ENSG00000101282 | 20 | 939095 | 982907 | 227 | 34 | -2.7081 | 0.99662 | RSPO4 | 0.99662 | 0.99933 | 0.98724 |
| ENSG00000183248 | 19 | 7933605 | 7939326 | 12 | 4 | -2.7507 | 0.99703 | CTD-3193O13.9 | 0.99703 | 0.96819 | 0.99256 |
| ENSG00000077092 | 3 | 25215823 | 25639423 | 1910 | 110 | -2.7531 | 0.99705 | RARB | 0.99705 | 0.99761 | 0.99782 |
| ENSG00000261459 | 16 | 30537244 | 30546173 | 6 | 3 | -2.7605 | 0.99711 | ZNF747 | 0.99711 | 0.97499 | 0.97611 |
| ENSG00000269490 | 6 | 33663196 | 33663474 | 2 | 1 | -2.8632 | 0.9979 | SBP1 | 0.9979 | 0.90317 | 0.89615 |
| ENSG00000169955 | 16 | 30537244 | 30546668 | 9 | 5 | -2.9258 | 0.99828 | ZNF747 | 0.99828 | 0.97499 | 0.97611 |
| ENSG00000127774 | 17 | 3572109 | 3572962 | 1 | 1 | -2.9478 | 0.9984 | EMC6 | 0.9984 | 0.84843 | 0.82224 |
| ENSG00000172208 | 11 | 48266608 | 48267622 | 2 | 1 | -3.1204 | 0.9991 | OR4X2 | 0.9991 | 0.060686 | 0.058548 |
| ENSG00000134817 | 11 | 57001051 | 57004709 | 9 | 3 | -3.3123 | 0.99954 | APLNR | 0.99954 | 0.8562 | 0.85531 |
| ENSG00000224420 | 19 | 50191921 | 50193832 | 1 | 1 | -3.4316 | 0.9997 | ADM5 | 0.9997 | 0.90979 | 0.88956 |
