## Supplementary Table 5 for "A higher burden of multiple sclerosis genetic risk confers an earlier onset"

**Supplementary Table 5. Gene list submissions to the DAVID Functional Annotat**

| Column Header | Key |
|---|---|
| **Cohort** | Source of gene list |
| **Threshold** | Inclusion threshold of either Genes or SNPs (given the data |
| **nGenes** | Number of unique genes contained within the specified thr |
| **nGenes (DAVID)** | Number of unique genes identied and retained by the DAVI |

| **Analysis** | **SNP-based test (GS1)** | **Gene-based test (GS2)** |
|---|---|---|
| **Threshold** | **Top 2.5%** | **Top 5%** |
| **nGenes** | **986** | **952** |
| **nGenes (DAVID)** | **950** | **872** |
| | CSMD1 | SSB |
| | MGMT | TRAFD1 |
| | UBR3 | HECTD4 |
| | LRRFIP1 | MMP8 |
| | THSD7B | NAA25 |
| | HMCN2 | UBR3 |
| | ZNF652 | ZDHHC11B |
| | CNTN4 | TACC1 |
| | HLA-C | ERP29 |
| | SCN7A | ARMC4 |
| | AGBL1 | CBLL1 |
| | CASC9 | KLHL23 |
| | DPP10 | C4B |
| | AGR2 | KIR3DL3 |
| | LINC00907 | METTL5 |
| | RBFOX1 | CLRN3 |
| | PHOSPHO2-KLHL23 | TMEM116 |
| | CHCHD6 | SLC26A4 |
| | ABTB2 | METTL17 |
| | TMEM178B | GJB6 |
| | GSE1 | ZNF652 |
| | PLXDC2 | SUPT4H1 |
| | GALNT12 | FAM109A |
| | DPP6 | PTPN18 |
| | CATSPER3 | CATSPER3 |
| | BLACE | TEAD1 |
| | DIRC3 | PTPN11 |
| | CBLL1 | AGR3 |
| | TXNRD3 | LPPR1 |
| | LRRTM4 | RP11-162P23.2 |
| | FAM83E | C1orf204 |
| | FAXC | MMP27 |
| | SYT16 | PPM1E |
| | CA1 | KRTAP9-3 |
| | EML2 | HLA-C |

| | |
|---|---|
| METTL5 | CFB |
| RIMS1 | MEST |
| MAPKBP1 | LRRC56 |
| CNTN2 | PAPOLA |
| PIK3R6 | CFB |
| ESYT2 | RP11-302M6.4 |
| PI16 | PARP15 |
| ROBO2 | AC073657.1 |
| DOPEY1 | PACRGL |
| KCNQ1 | PLA2G4B |
| KERA | JMJD7-PLA2G4B |
| PPIG | LIMD1 |
| TENM2 | CUX2 |
| CNTN1 | HIATL2 |
| ARMC4 | CNTN2 |
| MTTP | OR4D10 |
| HLA-DQA1 | TBXA2R |
| FAM109A | SH2B3 |
| MAST4 | CYP21A2 |
| FLJ40194 | CRLS1 |
| GLT8D2 | PHGDH |
| MTG2 | IPO8 |
| HRH3 | FANCE |
| MMP8 | APOBEC3G |
| CASZ1 | TNPO2 |
| ATP10D | C2 |
| UNC13C | COL25A1 |
| CUX2 | DENND6B |
| SCHIP1 | STK19 |
| XRCC5 | PDGFRL |
| ANK3 | EHMT2 |
| HLA-DQB1 | GGCT |
| XRCC4 | PTDSS2 |
| UTRN | TPPP |
| CACNG2 | CA9 |
| KCNIP4 | C4A |
| ATXN7L1 | AC011530.4 |
| SUSD5 | ASNA1 |
| C4BPA | COPG2 |
| DDAH1 | MGA |
| BARX2 | HS2ST1 |
| LPP | FBXW9 |
| CLYBL | THPO |
| APOBEC3D | AC074212.3 |
| LOC254128 | ARHGEF39 |
| PDXDC1 | SOX15 |
| PDE8A | TNRC6C |

| | |
|---|---|
| MAGI1 | KLF3 |
| NOTCH4 | APOBEC3F |
| CMBL | SYDE2 |
| BPIFC | TNXB |
| C4B | SLC16A14 |
| TRAPPC9 | HLA-DQA1 |
| RPL10A | EZR |
| NOX3 | SELENOF |
| MUC22 | HPS4 |
| GABBR2 | CCDC107 |
| NEDD4 | ZNF510 |
| CPAMD8 | RPL27A |
| FRRS1 | ZNRF2 |
| TEAD3 | BBS5 |
| CNOT6 | RP11-724O16.1 |
| TAX1BP1 | KLHL41 |
| RFTN1 | HLA-DRA |
| DHRS7C | RPL6 |
| AHI1 | GLI1 |
| SERPINB12 | CHD8 |
| RAB27B | TSGA13 |
| PTPRO | MGMT |
| PPFIBP1 | STMN3 |
| PCLO | OR11H4 |
| SLCO3A1 | ZNF648 |
| KLHL25 | TH |
| PAK2 | MCM7 |
| ASTN2 | SLC17A9 |
| RALYL | DRAM1 |
| ZNRF1 | NT5DC1 |
| TEAD1 | MAN2B1 |
| SRPK2 | C12orf49 |
| SLC9A3 | ZDHHC11 |
| C6orf7 | POLR2H |
| EHD4 | CCNF |
| WWOX | PHOSPHO2 |
| RBMS3 | SET |
| SNX29 | UTRN |
| ME3 | TEX14 |
| CSMD3 | NUTM2G |
| PDGFRL | UBE2U |
| EZR | IGFBP7 |
| RTL1 | CNOT2 |
| GLTSCR1L | SIX5 |
| MYO5B | C19orf66 |
| FBN2 | CCDC173 |
| LINC00222 | TMEM30C |

| | |
|---|---|
| EEPD1 | SCLT1 |
| PGM3 | EIF3I |
| IQCK | MED30 |
| LUZP2 | SPTBN5 |
| TCF7L2 | PDP1 |
| PVRL2 | TEKT1 |
| HLA-DRB6 | LRRC8C |
| MYO1C | CTD-2192J16.22 |
| DYNC1I2 | SMARCD1 |
| CRLS1 | TRIM37 |
| GUCY1A2 | POLR2B |
| KLHL23 | GADL1 |
| VAV2 | CSMD3 |
| CLASP1 | AGPAT1 |
| FANCE | OR2Z1 |
| TTC28 | CCL28 |
| COLEC12 | RIC8B |
| PLCH1 | ACSL5 |
| TSGA13 | AL645730.2 |
| TOM1L2 | SRRD |
| ZBTB46 | KCMF1 |
| NRCAM | TMEM18 |
| TEX14 | BSX |
| APH1B | SENP3 |
| SYNDIG1 | PIK3R6 |
| HLA-DRA | VAPB |
| MTUS2 | CUX1 |
| MGA | PARL |
| C6orf10 | MALRD1 |
| NRXN1 | ZNF782 |
| SLC26A4 | SLC35E2 |
| LOC285696 | CNOT6 |
| TNXB | NUFIP1 |
| CCDC11 | OAS1 |
| AP2A2 | MTTP |
| AGR3 | TBC1D22B |
| COL6A5 | TMEM39B |
| NDST3 | SEPTIN4 |
| SLC39A11 | MAPKAPK5 |
| CELF2 | C1QL2 |
| TBC1D24 | SKIV2L |
| SEMA5A | ALDH2 |
| DSTYK | APOBEC3D |
| TP73 | SNX27 |
| SGK223 | ACTR3B |
| NOL10 | SLC9A9 |
| COPG2 | RP5-1052I5.2 |

| | |
|---|---|
| BZRAP1 | NR2C2AP |
| PSD3 | AC011366.3 |
| BBOX1 | AP001652.1 |
| MYO3B | HDAC3 |
| PHKB | COLEC12 |
| C9orf92 | TOP1 |
| KCNG4 | GBA2 |
| UBE3D | SLC8A1 |
| METTL17 | PYHIN1 |
| RAD51C | SP9 |
| SYNPO2 | CNGA1 |
| KIR3DL3 | ICAM4 |
| LOC101928991 | TRA2B |
| ABCG2 | TMEM217 |
| SLC8A1 | CRIPAK |
| CCDC173 | ATF6B |
| SAMD12 | SYNGAP1 |
| IFT140 | ITGB1BP1 |
| CNGA3 | IRX6 |
| UGT1A10 | LRRN4 |
| PTPRE | SLC26A3 |
| ITPR2 | RAD51C |
| MAP3K5 | TSN |
| MEST | FKSG48 |
| NTN1 | RB1 |
| TENM4 | BZRAP1 |
| TRAT1 | IQCC |
| COL5A1 | EEF2 |
| DGKG | TBC1D3C |
| CTNND2 | ZFP91 |
| JMJD7-PLA2G4B | PHRF1 |
| SNRPN | RNF103 |
| GPSM3 | ANGPTL6 |
| SLC26A3 | AC069547.2 |
| SMARCC1 | SMIM14 |
| PALM2 | THUMPD2 |
| KCNC2 | C11orf35 |
| GRHL2 | ATP1B2 |
| XIRP2 | TXLNA |
| DDX60L | SSX2IP |
| PTPRT | ARHGAP9 |
| CHEK2 | NELFE |
| FGD4 | PPIG |
| DCLK3 | KLHL25 |
| AKAP6 | IL10RA |
| PALLD | ZNF335 |
| CACNA1C | ZFP91-CNTF |

| | |
|---|---|
| ZBTB7C | BCL11A |
| RIT2 | MRC2 |
| CUX1 | PLGLB2 |
| PRUNE2 | MTNR1B |
| PAPOLA | OR11H6 |
| TENM3 | NTS |
| GALNT3 | RFX4 |
| CYP2B6 | GJA3 |
| BCL11B | GPC2 |
| FASTKD1 | ATP11A |
| TRIML1 | PTGDS |
| MDFIC | FGR |
| LOC100616530 | CAPZA3 |
| ANXA13 | GNPTAB |
| RNFT2 | STRIP2 |
| RABEP1 | ADAP1 |
| ATP2B4 | BUB1B |
| PPM1E | CLC |
| LOC728730 | LGI3 |
| OSMR | RASSF9 |
| FAM135B | TP53I11 |
| C9orf3 | GDF10 |
| CCDC81 | DMWD |
| CDH13 | LCNL1 |
| PDS5B | INPP5A |
| NKAIN1 | LARP4B |
| ERC1 | ATP6V0E1 |
| APOBEC3F | GAS2L3 |
| DAB1 | TP53INP1 |
| DNAJC5 | TMEM218 |
| ANKDD1A | RCBTB2 |
| OBSCN | FLI1 |
| NTM | HRH3 |
| KPNA3 | ARL4C |
| SPTBN5 | CASP4 |
| FAM19A1 | SLC13A5 |
| ROR1 | WDR83 |
| PHOSPHO2 | OAS3 |
| TGM2 | EIF4A1 |
| MCF2L | PPP5C |
| HECTD4 | AFAP1L2 |
| GTPBP4 | LPP |
| MAML1 | CIAPIN1 |
| ARMC9 | DMPK |
| TUSC3 | AC026740.1 |
| DYNC2H1 | TBC1D3H |
| FHIT | RPL10A |

| | |
|---|---|
| ALDH2 | DDX24 |
| TMEM130 | NGLY1 |
| CNTNAP2 | AP1AR |
| SYDE2 | RP11-81K2.1 |
| NLGN1 | MRPL9 |
| NAA25 | EPO |
| BTNL2 | COX19 |
| C19orf66 | SLC35B1 |
| ST8SIA2 | SLC9A3 |
| PRSS12 | CCDC3 |
| SMIM14 | DOPEY1 |
| LRP1B | FGFR3 |
| PCSK5 | PAH |
| CNGB1 | FBXO46 |
| SEPTIN4 | DCDC2C |
| NOS1AP | C16orf80 |
| SLC24A1 | SLC39A10 |
| PARG | TMEM234 |
| SUSD3 | PDE8A |
| TMEM243 | MTERF |
| CRISPLD2 | RNH1 |
| TMEM116 | GOLGA6A |
| SPATA5L1 | APOC4 |
| DPP8 | APOC4-APOC2 |
| PARL | COL15A1 |
| INPP5A | METAP2 |
| MINK1 | AC021860.1 |
| PTPN18 | MYC |
| GRIN2B | UBL3 |
| AGBL3 | KLHL24 |
| KCNT2 | SSTR1 |
| TGM6 | C5 |
| NELL1 | TDG |
| DLGAP2 | OAS2 |
| CYP21A2 | KCNH5 |
| FARS2 | ALX3 |
| RMND1 | SKA2 |
| IDI2-AS1 | DCT |
| TBL1XR1 | KPNA6 |
| MSI2 | PIGX |
| FNTB | VAMP5 |
| CTNNA3 | MAN1A2 |
| TACC1 | FGL2 |
| FIGNL2 | WDR17 |
| DDRGK1 | POLR2C |
| H2AFJ | SHISA7 |
| DOCK5 | UPP2 |

| | |
|---|---|
| DISP1 | OR5M8 |
| CLRN3 | STRIP1 |
| CHID1 | AL162407.1 |
| FBXO39 | SIGLEC5 |
| MROH2A | SLC17A7 |
| ERP29 | DGCR8 |
| DDHD1 | TFIP11 |
| LINC00959 | BRINP2 |
| COL25A1 | FASLG |
| CECR2 | APOC2 |
| FBXW9 | PPP6R2 |
| C8orf69 | INHBB |
| HS2ST1 | APBA1 |
| OR4D6 | DFNB59 |
| LMO7 | OSTC |
| BACE2 | C17orf112 |
| PDE10A | HSH2D |
| PBX2 | ZDHHC6 |
| NUP88 | CILP |
| CDHR5 | PLEKHM2 |
| SPATA17 | HMGB1 |
| SLC9A9 | HRAS |
| TRAFD1 | SLC44A4 |
| TPD52L2 | ANKRD50 |
| POLR2B | AL589765.1 |
| FIBCD1 | RIIAD1 |
| PACRG | ENOSF1 |
| LOC339862 | PRR16 |
| NEDD4L | AC069368.3 |
| CEP128 | FRG2B |
| MTMR4 | CDHR5 |
| MOB3A | FBXO39 |
| CDH12 | NPM2 |
| SPEF2 | ZNF714 |
| LOC285556 | PCLO |
| NTF3 | IFITM10 |
| CYFIP1 | TRIM52 |
| RBFOX3 | AC138655.1 |
| DENND3 | SPNS2 |
| NCAM1 | OR8I2 |
| KCMF1 | CAPRIN2 |
| PRKD2 | LCE2A |
| MICU1 | SCYL1 |
| TTC37 | KIR2DL4 |
| CD247 | ZNF362 |
| PKP3 | C2orf68 |
| TRIM37 | C17orf78 |

| | |
|---|---|
| TWIST1 | BCL10 |
| KCNC4 | RAB40B |
| KCNJ10 | SIPA1L2 |
| FRMD8 | CSNK1G3 |
| AKT3 | MPEG1 |
| RPA3-AS1 | ZNF791 |
| HGD | SIGLEC5 |
| CRIPAK | QSOX2 |
| ARHGEF39 | HGF |
| WIPF3 | CELF3 |
| FBXL17 | MLYCD |
| NEURL1B | CALM2 |
| LOC285500 | ATXN2 |
| SIRT2 | OR5B2 |
| LEKR1 | GUCY1A2 |
| MUC5AC | OR5M9 |
| SESTD1 | PPRC1 |
| HDGFRP3 | TEP1 |
| NLRP9 | USP10 |
| ATRNL1 | MYCT1 |
| HLA-DRB1 | EMR3 |
| ALDH1L1 | CDCA3 |
| ENPEP | CLASP1 |
| ESYT3 | FGF2 |
| TBC1D32 | KCNIP4 |
| C1orf198 | MAPKBP1 |
| CHRNE | FAM150B |
| RGL2 | CSNK2A2 |
| CPEB3 | CMBL |
| KCNJ3 | OR4D6 |
| SLAMF6 | MPC1 |
| THSD7A | KIAA0947 |
| LOC100506403 | CLMP |
| FAR1 | CTB-129P6.11 |
| AKAP13 | SERINC3 |
| RUFY1 | CHMP3 |
| LINC00466 | TRAT1 |
| PTPLAD1 | TRIP13 |
| OSBPL10 | C4orf32 |
| NUFIP1 | PPT2-EGFL8 |
| GOLGA6L2 | DSC1 |
| FGF14 | PPM1D |
| PLS1 | NIPAL1 |
| SUPT4H1 | FHL2 |
| HLA-DOB | CXCR1 |
| EPHB1 | ZNF430 |
| EBF1 | SPRR2G |

| | |
|---|---|
| ZDHHC11B | FAM159A |
| PICALM | HYDIN |
| ATE1 | NUP35 |
| FOXP2 | PWP2 |
| IGFBP7 | TRAF3IP1 |
| WDR62 | HAO1 |
| CCL28 | DHPS |
| CLGN | MYADM |
| KLHL41 | GPX6 |
| GGA3 | CASP9 |
| LRRC2 | FAM20A |
| SDK1 | RNF103-CHMP3 |
| FYB | GOLGA7B |
| MAP3K11 | SCGB2A1 |
| LOC100132146 | NRROS |
| TINCR | PIK3C2B |
| C1orf52 | CABLES2 |
| EMR3 | BEAN1 |
| BLOC1S5 | CRELD1 |
| TNXA | RSPH6A |
| MAML3 | CLPTM1 |
| TMC2 | IFIT1B |
| LPPR1 | ME3 |
| RRP12 | GLIS1 |
| C10orf112 | FAM213B |
| SYNE3 | ATP10D |
| ELP2 | LOXL1 |
| PLB1 | RP11-761B3.1 |
| ZDHHC19 | KHDC1L |
| GMEB1 | IFIT1 |
| RBPJ | NOLC1 |
| PAX5 | WDR41 |
| TANGO2 | FRK |
| CEP89 | SLC26A9 |
| HCG23 | LCN6 |
| SYTL2 | CPEB2 |
| PRC1 | CIR1 |
| TRPV2 | MYPOP |
| MED30 | HLA-DOB |
| SLC35F1 | USP2 |
| SSX2IP | MMS19 |
| CCDC107 | EPPIN |
| DDX25 | ZNF765 |
| SND1 | EFNA3 |
| FOXP1 | PIGZ |
| CYBA | BMP10 |
| CCDC148 | AC004528.1 |

| | |
|---|---|
| SKA2 | LTA |
| FAM155A | LPAR6 |
| C2orf61 | MUC17 |
| SLC25A12 | BRD9 |
| USP45 | TRIO |
| ELK3 | OR51A2 |
| DCLK1 | ZNF490 |
| GADL1 | ATXN7L1 |
| HLA-DQA2 | SYCP3 |
| BTAF1 | LCN10 |
| SNAPC4 | PGM3 |
| BCL10 | LDB1 |
| TSNARE1 | IFIT5 |
| FGD5 | AC026202.1 |
| ACAD11 | ZNF628 |
| LSAMP | SIRT2 |
| ARID1B | PDX1 |
| USP6NL | FN1 |
| LARS2 | DUSP14 |
| CAPS2 | OR5M3 |
| MEGF11 | RP4-734P14.4 |
| KRTAP9-3 | MPDU1 |
| TIMP2 | SLAMF8 |
| CCDC68 | CTNNA2 |
| CLPX | MOB3A |
| SP100 | SLC25A34 |
| STAG3 | PHLDA1 |
| TNFAIP8 | RP11-429E11.3 |
| CHIA | HMOX1 |
| CEP85L | DOK4 |
| MAGI2 | FAXC |
| MYLK | DNAJC16 |
| LINC00333 | OR8H3 |
| MBTPS1 | C9orf142 |
| NUP35 | HIST1H1E |
| AOAH | SOD2 |
| PAH | ENTPD4 |
| RASSF8 | RPAP3 |
| ZSCAN5A | PLA2G12B |
| TSPAN18 | NCOA5 |
| TOB2 | IL31RA |
| PRLR | MIDN |
| KLHL2 | KAT7 |
| CNTNAP5 | CRH |
| COL22A1 | ZNF354A |
| HIBCH | INPP4B |
| FANCC | CLEC19A |

| | |
|---|---|
| PDGFC | PPFIBP1 |
| ZNF552 | ZBTB22 |
| TP63 | IRF2BP1 |
| TAP2 | GNE |
| ETV5 | RPS7 |
| UQCRB | GSTT2 |
| SMAP1 | MUC6 |
| CD320 | ZBTB2 |
| RIMBP2 | RMND1 |
| RYR2 | DOK2 |
| SYCP2L | CDX1 |
| LOC101059906 | EIF1 |
| LOC100500773 | MSH2 |
| CAMK2G | OR5T3 |
| CALM2 | RP11-144F15.1 |
| SH3BP2 | CALCOCO2 |
| ALMS1 | ESD |
| PPP6R2 | CNTN1 |
| GRIA2 | ROS1 |
| TRIM66 | POU5F1 |
| ZNF343 | GJB3 |
| NBAS | RP11-20I23.1 |
| MYO1E | RGP1 |
| CDH23 | ANKRD28 |
| PRR16 | TNFSF12 |
| VPS54 | TNFSF12-TNFSF13 |
| SMIM3 | DYNC2H1 |
| C16orf72 | UQCRB |
| WDR37 | C2CD4D |
| DHX30 | CTB-186H2.3 |
| ANKMY2 | CHEK2 |
| KLC1 | CACNG6 |
| SLC24A4 | WFDC10A |
| ARHGAP10 | ZNF184 |
| POLD3 | CACNG5 |
| ST5 | EFNA3 |
| RNLS | JAK1 |
| AHCYL2 | CEBPE |
| PCDH15 | PRRG2 |
| MAP4 | NLRP9 |
| B3GALT1 | SCGB1D1 |
| RSU1 | AKIRIN2 |
| NINJ2 | SPATA17 |
| BCAS3 | DSE |
| NFKBIB | ARRDC2 |
| ST18 | ARFGAP1 |
| ASAP1 | GPNMB |

| | |
|---|---|
| LINC00276 | EMC3 |
| GSG1L | GSTT2B |
| ZFAND2A | MCFD2 |
| ATF6B | SLC22A9 |
| PRDM6 | PCIF1 |
| TUBGCP3 | MAN2A1 |
| SQRDL | PSORS1C1 |
| MAPKAPK5 | SMCO4 |
| PSORS1C3 | FGF23 |
| PTPRM | FHIT |
| AUH | CTC-435M10.3 |
| HSPA4L | ATP5SL |
| ITFG1 | AGR2 |
| SLC35E2 | DDAH1 |
| C20orf196 | RPS18 |
| CTTNBP2NL | SLC2A9 |
| KCNH5 | TFCP2 |
| ADCK3 | WFDC10B |
| CUEDC1 | RFESD |
| VEGFA | RP11-650K20.3 |
| PRICKLE1 | KIAA1199 |
| KLF3 | FBXL15 |
| VIPR1 | OR8K5 |
| LOC338667 | EEF1B2 |
| ADAMTSL1 | LRRC23 |
| CAMTA1 | AHNAK2 |
| MOCOS | RNF219 |
| FLJ13197 | SFT2D3 |
| KRR1 | ADAM33 |
| RBL1 | ATP2B4 |
| NR3C1 | BCKDHA |
| LRRC56 | C4orf33 |
| DSCAM | ITPA |
| MKX | OR5M11 |
| TPPP | MNX1 |
| PLCL1 | ZG16B |
| HKDC1 | C13orf35 |
| ZFP91 | SLC26A10 |
| TAPBP | TJP3 |
| QSOX1 | CDK5RAP2 |
| SLC16A14 | AL450307.1 |
| MCU | AC022498.1 |
| USP10 | HNRNPR |
| SLC9C1 | PLCG1 |
| FREM3 | ALDH3B2 |
| PALM2-AKAP2 | ZNF479 |
| ESRRG | HTR2A |

| | |
|---|---|
| PARVA | CCDC33 |
| SGSM3 | CA2 |
| RPAP3 | ADCK5 |
| DNAJB4 | CRYGS |
| COBLL1 | RP11-15E18.4 |
| B3GALT4 | SOX8 |
| PLSCR2 | SPN |
| TCF3 | FER |
| GNE | TMPO |
| SCG5 | NT5E |
| PDE3B | RP11-351M8.1 |
| CA9 | DLGAP1 |
| SIPA1L2 | NOS1AP |
| ADSL | GNB2L1 |
| TMTC1 | ZFP62 |
| RUNX1 | LRRC27 |
| RICTOR | C21orf33 |
| ADAMTS14 | CTBP2 |
| TBC1D16 | PPT2 |
| UPP2 | DCAF4 |
| TNPO2 | SLC45A4 |
| FOXJ3 | LACTBL1 |
| RPL27A | PGLYRP1 |
| COL9A2 | EXOC7 |
| SSB | MAS1 |
| MACC1 | AL355390.1 |
| FAM189A2 | RNF123 |
| JAKMIP2 | FKBP4 |
| LRRC28 | SELL |
| TPP2 | UBE3D |
| ITGBL1 | PHLDA3 |
| JPH3 | SEMA4A |
| FOXN3 | VSIG8 |
| MYRIP | PRKD1 |
| ERBB4 | HIST1H3H |
| PPP1R16B | FAM189A2 |
| ADARB2 | CPO |
| LOC100130480 | KCNC2 |
| BRSK2 | RFX6 |
| CDK17 | ITGA8 |
| LY86-AS1 | C7orf43 |
| NEBL | COL10A1 |
| TACC2 | EML2 |
| CPLX2 | MYL12A |
| SLC44A4 | DCDC2B |
| HIP1 | B3GNT8 |
| METTL21C | SKOR2 |

| | |
|---|---|
| RBM20 | CENPBD1 |
| SEC1P | FOSL2 |
| WDR46 | SLC28A1 |
| SORCS3 | TIFA |
| CHST11 | COPS4 |
| ADRB2 | CTC-786C10.1 |
| CLEC17A | IL6R |
| CCSER1 | TRAPPC10 |
| GCNT2 | NREP |
| CPPED1 | ARRB1 |
| IPO8 | DKFZP434E1119 |
| PARK2 | EPPIN-WFDC6 |
| FOCAD | POMGNT1 |
| SIGIRR | RBL1 |
| DOK6 | C1S |
| PPM1H | LHX9 |
| LTBP3 | TMEM178A |
| AGAP1 | GCNT2 |
| HS3ST4 | NELFCD |
| BIVM | MAL |
| RASGRF2 | PTPRO |
| COLEC11 | FOXI1 |
| KIAA1199 | PFN4 |
| ZNF532 | RP1-228P16.5 |
| CELA3A | FAM205A |
| WDFY4 | LOXL2 |
| FAM81A | IFITM1 |
| MAN2A1 | RNASE4 |
| L3MBTL4 | TTPAL |
| TRIO | SGK494 |
| GPM6A | TTC31 |
| PPT2 | WDR81 |
| ZHX3 | NKX1-1 |
| RYR3 | GALNT15 |
| DGKZ | AP002348.1 |
| NCOA5 | ABCD4 |
| SLC1A2 | ZBTB48 |
| CDH3 | AC023469.1 |
| UBE2U | PKN2 |
| SORBS2 | PBX2 |
| VPS52 | ZNF436 |
| SULT2B1 | OR8H2 |
| CORIN | ZNF532 |
| DLGAP1 | S1PR4 |
| RNF144B | AC087477.1 |
| MYO18B | RP11-403P17.5 |
| GPR133 | GGH |

| | |
|---|---|
| GABRG3 | OR5F1 |
| DIAPH1 | SLC13A1 |
| PLD2 | KCNJ10 |
| TESC | TADA2A |
| DOCK10 | LAMP1 |
| MAN1A2 | ITGA11 |
| RBPMS | GATA2 |
| C7orf10 | LONRF1 |
| TBXA2R | GIPC3 |
| FHDC1 | TUBAL3 |
| PRKCE | CPNE5 |
| EFNA5 | LIN54 |
| RAD51B | RP11-105C20.2 |
| RBBP5 | OR52D1 |
| PTDSS2 | EPHB3 |
| OR5A1 | CLEC5A |
| XPO7 | PRB4 |
| INTS8 | PRSS35 |
| APCDD1L | CDHR4 |
| NT5DC1 | PRR24 |
| TEKT1 | ARHGDIB |
| CKAP4 | AL163636.6 |
| MPP7 | CEP72 |
| NFXL1 | LZTR1 |
| PCDH9 | ACAD10 |
| ZHX2 | KLHDC8A |
| ABCC5 | JAZF1 |
| APBA1 | SH2B2 |
| ASIC2 | RALGAPB |
| CCDC154 | CD320 |
| C1orf21 | KIF26A |
| WDPCP | MBLAC1 |
| COL6A6 | PPP1R9B |
| RORA | SPSB2 |
| SRL | EFNA1 |
| RPS18 | TM4SF19 |
| LINC00871 | CCDC102A |
| GJB6 | USP42 |
| NGLY1 | PDE10A |
| EIF3D | C1orf123 |
| KIF26A | OR5AS1 |
| HS3ST1 | TTLL7 |
| LARP4B | OR5M10 |
| GPR123 | TMEM41B |
| KIAA1644 | DYNC1I2 |
| FBXO15 | DDX23 |
| ATP11A | GPR75-ASB3 |

| | |
|---|---|
| ABCA13 | ZHX3 |
| TMPO | THAP9 |
| MYH14 | OR10AG1 |
| ELOF1 | ASPN |
| ITPK1 | RASSF7 |
| C12orf49 | THBD |
| LOC100996485 | OR6J1 |
| CTNNA2 | FCER1A |
| SLC6A15 | CCDC106 |
| SLC2A9 | CCDC81 |
| CSRNP2 | STON2 |
| KLB | LSAMP |
| RNF43 | HDGFRP3 |
| ANKRD30A | ACP5 |
| IL10RA | RP11-796G6.2 |
| OR4D10 | USP6NL |
| NCOR2 | SEC22A |
| BMPR1B | CLTA |
| C8orf87 | LCE1F |
| DTX3L | ERMN |
| CDK14 | MCM4 |
| TJP3 | CCDC39 |
| ZBP1 | PHF10 |
| MYO5A | SORBS2 |
| TECRL | ATN1 |
| SYNE2 | C20orf141 |
| GABRA2 | WDR1 |
| SEC62 | CBLN3 |
| DCC | ZNF391 |
| IMMP2L | PITX1 |
| LMCD1 | RBPJ |
| NKAIN2 | OR8H1 |
| VAT1L | CXXC1 |
| COL9A3 | FANCD2 |
| SMYD3 | ZNF804B |
| APP | NFXL1 |
| ZNF385D | CLK1 |
| MIR3689B | RP11-763F8.1 |
| INPP4B | SWSAP1 |
| ROBO1 | NADK |
| NFIC | RTL1 |
| CAPRIN2 | BAI2 |
| RPA3 | DFFA |
| TIMM21 | TEAD3 |
| UNC5C | NLRP5 |
| FBLN1 | GRK7 |
| MYOM2 | PPCDC |

| | |
|---|---|
| CIAPIN1 | TMEM173 |
| ZNRD1-AS1 | LCE3E |
| CPNE4 | DRD4 |
| CEP57 | TM4SF19 |
| RAP1A | ECM2 |
| SAMD7 | SLC4A11 |
| COLGALT2 | SCEL |
| RPL31 | IP6K1 |
| COL21A1 | BAIAP2 |
| TNKS2 | C1orf52 |
| ATP10A | LRP8 |
| SATB1 | FAM212A |
| LINC00911 | LRRC45 |
| AATK | PGAM1 |
| C1orf95 | AP4M1 |
| SLC12A5 | AC011298.1 |
| CCDC33 | SAP30L |
| SMAD3 | RGCC |
| MAP3K8 | EXOSC5 |
| GLI1 | ST14 |
| ZBTB22 | LPXN |
| FKBPL | AC068039.1 |
| SOX6 | SHMT2 |
| SHISA6 | AMBN |
| SIRPD | TRMT10B |
| SYTL3 | EPS8L2 |
| SPNS3 | CBWD1 |
| SLC26A5 | ATP6V0A4 |
| MCM6 | UCKL1 |
| SLC39A10 | ZNF681 |
| ZDHHC21 | INHBC |
| ANO3 | GIPR |
| SLC19A1 | ZDHHC16 |
| ATXN7L3 | MFSD12 |
| MACROD2 | OR14I1 |
| LIMK2 | COL3A1 |
| FAM154A | C18orf54 |
| NRXN3 | PSMD1 |
| OGDH | URAD |
| AMPH | KHDRBS2 |
| ATP2C2 | GPX5 |
| ADAM12 | GZMK |
| TNRC6B | FANCD2OS |
| LIMD1 | CAPN13 |
| YWHAQ | PHF1 |
| SH3RF3 | C19orf25 |
| LILRP2 | MC2R |

| | |
|---|---|
| ANKRD10 | RP11-219B4.5 |
| NBEA | GJA4 |
| DNAH11 | SH2D1B |
| KCNG3 | PRSS21 |
| EVC | TBC1D9B |
| ALK | LCN6 |
| BBS5 | OR4A16 |
| EFCAB1 | FBXO41 |
| PLA2G12B | C8B |
| DAB2 | LRRC3B |
| DCDC2C | GLYATL1 |
| MAPRE2 | MAFB |
| ARRB1 | SNRPB |
| HMGB1 | GPR31 |
| TFCP2 | MDM1 |
| CRTAC1 | VKORC1L1 |
| PIEZO2 | ANG |
| MKL2 | ILVBL |
| PITPNC1 | NYNRIN |
| LOC100652824 | LTBP3 |
| CHST3 | AIMP1 |
| ZBTB2 | CARD10 |
| CLVS2 | CD40 |
| ARMC8 | CCDC9 |
| TBX19 | LMO1 |
| XAF1 | TPI1 |
| TNFRSF13B | CLVS2 |
| LOC101929065 | PGPEP1L |
| BAIAP2L1 | TUBA1C |
| AFAP1L2 | COX4I2 |
| GMDS | ERLEC1 |
| HIST4H4 | CGN |
| HERPUD2 | CTDP1 |
| LINGO2 | FMO4 |
| CSTF3 | OR2AG2 |
| AGL | TCF19 |
| INSR | SFRP1 |
| PAK7 | LRCH1 |
| GLCCI1 | CLEC4C |
| MMP16 | ACOT7 |
| ADRBK1 | KLHL14 |
| APBB1IP | ENO2 |
| GLI3 | KBTBD8 |
| PIK3CB | OR10A3 |
| BNC2 | MBD3L4 |
| AHNAK2 | HBEGF |
| GGT5 | AOC1 |

| | |
|---|---|
| ACOT7 | KCNA3 |
| SYNGAP1 | LMAN1L |
| LARGE | PSMA1 |
| STAB1 | AVP |
| JAK1 | LUZPP1 |
| MELK | SLC9C1 |
| PITX1 | TTLL1 |
| PARP15 | UBR4 |
| MAP2 | TTC34 |
| TMEM132B | NUCB2 |
| ZNF782 | ZNF763 |
| SCARB1 | CTC-429P9.4 |
| TNRC6C | CSRNP2 |
| ABRACL | ZNF383 |
| HSD17B4 | ZNF141 |
| C18orf42 | MED23 |
| MTHFR | DNAJB1 |
| ACOT13 | CHPT1 |
| TGFBR3 | CORIN |
| LRCH1 | ZNF763 |
| NOX4 | MRPL54 |
| TXK | Z97053.1 |
| PIGX | RP11-187E13.1 |
| NRROS | OXSM |
| NEB | ARMC8 |
| MREG | PIP5K1C |
| SELENOF | RUSC1 |
| NALCN-AS1 | SHC4 |
| KANK3 | BARX2 |
| GRID1 | MASP1 |
| SHC4 | COPS6 |
| PPIL1 | ERMARD |
| ITGAE | WNT5A |
| MAT1A | OR5I1 |
| DSCAML1 | AL031663.2 |
| DENND6B | ZNF627 |
| ACVR1C | SYT6 |
| FMN2 | RP11-272B17.2 |
| FAM73A | EFNA4 |
| DOCK3 | MBD3L2 |
| SPNS2 | CTD-2192J16.24 |
| PACRGL | C17orf74 |
| TNPO1 | UBE3A |
| RB1 | WFDC11 |
| NPLOC4 | KATNAL2 |
| DLG2 | RAB27B |
| RAI14 | NOX4 |

MGAM
DDX10
JRKL-AS1
INPP5K
EPC2
MSRB3
ATP8B3
LARS
LIAS
RIMKLA
PIK3C2B
MECOM
PDZD2
SLC30A9
MYO7B
KDM2A
MOK
C5
RELN
TUBGCP5
TRERF1
LRRC16A
PCSK2
FAM134B
LRP5
EPB41L3
ZNF821
ATP6V1G2-DDX39B
CCDC60
PCBP3
IQCJ-SCHIP1
FLJ42393
LINC00710
SLC8A1-AS1
UGT1A8
PLA2G4B
UGDH-AS1
CHURC1-FNTB
BZRAP1-AS1
BLOC1S5-TXNDC5
NPHP3-ACAD11
LARS2-AS1
GATS
JMJD7
MESTIT1
LOC100506374
SLC9A9-AS1
CNFN
RIMS1
APIP
LHX3
SP110
HDHD2
MRI1
SCN4A
AC018755.1
GAA
TTLL6
QPRT
CTD-3088G3.8
MREG
EIF4E
PAFAH1B1
FBXO24
NBPF4
THOC7
MPP7
MAMDC4
SF3B14
CDK4
CAT

ZFP91-CNTF
JAKMIP2-AS1
SNURF
LRRTM3
PPT2-EGFL8
MAGI1-AS1
ERICH1-AS1
TBC1D8
HCG25
UGT1A9
EEF1E1-MUTED

**:ion tool**

source)
:eshold
ID Functional Annotation Tool

**Gene- & SNP-based tests (GS3)**
**Union**
**1679**
**1574**
SSB
TRAFD1
HECTD4
MMP8
NAA25
UBR3
ZDHHC11B
TACC1
ERP29
ARMC4
CBLL1
KLHL23
C4B
KIR3DL3
METTL5
CLRN3
TMEM116
SLC26A4
METTL17
GJB6
ZNF652
SUPT4H1
FAM109A
PTPN18
CATSPER3
TEAD1
PTPN11
AGR3
LPPR1
RP11-162P23.2
C1orf204
MMP27
PPM1E
KRTAP9-3
HLA-C

CFB
MEST
LRRC56
PAPOLA
RP11-302M6.4
PARP15
AC073657.1
PACRGL
PLA2G4B
JMJD7-PLA2G4B
LIMD1
CUX2
HIATL2
CNTN2
OR4D10
TBXA2R
SH2B3
CYP21A2
CRLS1
PHGDH
IPO8
FANCE
APOBEC3G
TNPO2
C2
COL25A1
DENND6B
STK19
PDGFRL
EHMT2
GGCT
PTDSS2
TPPP
CA9
C4A
AC011530.4
ASNA1
COPG2
MGA
HS2ST1
FBXW9
THPO
AC074212.3
ARHGEF39
SOX15
TNRC6C
KLF3

APOBEC3F
SYDE2
TNXB
SLC16A14
HLA-DQA1
EZR
SELENOF
HPS4
CCDC107
ZNF510
RPL27A
ZNRF2
BBS5
RP11-724O16.1
KLHL41
HLA-DRA
RPL6
GLI1
CHD8
TSGA13
MGMT
STMN3
OR11H4
ZNF648
TH
MCM7
SLC17A9
DRAM1
NT5DC1
MAN2B1
C12orf49
ZDHHC11
POLR2H
CCNF
PHOSPHO2
SET
UTRN
TEX14
NUTM2G
UBE2U
IGFBP7
CNOT2
SIX5
C19orf66
CCDC173
TMEM30C
SCLT1

EIF3I
MED30
SPTBN5
PDP1
TEKT1
LRRC8C
CTD-2192J16.22
SMARCD1
TRIM37
POLR2B
GADL1
CSMD3
AGPAT1
OR2Z1
CCL28
RIC8B
ACSL5
AL645730.2
SRRD
KCMF1
TMEM18
BSX
SENP3
PIK3R6
VAPB
CUX1
PARL
MALRD1
ZNF782
SLC35E2
CNOT6
NUFIP1
OAS1
MTTP
TBC1D22B
TMEM39B
SEPTIN4
MAPKAPK5
C1QL2
SKIV2L
ALDH2
APOBEC3D
SNX27
ACTR3B
SLC9A9
RP5-1052I5.2
NR2C2AP

AC011366.3
AP001652.1
HDAC3
COLEC12
TOP1
GBA2
SLC8A1
PYHIN1
SP9
CNGA1
ICAM4
TRA2B
TMEM217
CRIPAK
ATF6B
SYNGAP1
ITGB1BP1
IRX6
LRRN4
SLC26A3
RAD51C
TSN
FKSG48
RB1
BZRAP1
IQCC
EEF2
TBC1D3C
ZFP91
PHRF1
RNF103
ANGPTL6
AC069547.2
SMIM14
THUMPD2
C11orf35
ATP1B2
TXLNA
SSX2IP
ARHGAP9
NELFE
PPIG
KLHL25
IL10RA
ZNF335
ZFP91-CNTF
BCL11A

MRC2
PLGLB2
MTNR1B
OR11H6
NTS
RFX4
GJA3
GPC2
ATP11A
PTGDS
FGR
CAPZA3
GNPTAB
STRIP2
ADAP1
BUB1B
CLC
LGI3
RASSF9
TP53I11
GDF10
DMWD
LCNL1
INPP5A
LARP4B
ATP6V0E1
GAS2L3
TP53INP1
TMEM218
RCBTB2
FLI1
HRH3
ARL4C
CASP4
SLC13A5
WDR83
OAS3
EIF4A1
PPP5C
AFAP1L2
LPP
CIAPIN1
DMPK
AC026740.1
TBC1D3H
RPL10A
DDX24

NGLY1
AP1AR
RP11-81K2.1
MRPL9
EPO
COX19
SLC35B1
SLC9A3
CCDC3
DOPEY1
FGFR3
PAH
FBXO46
DCDC2C
C16orf80
SLC39A10
TMEM234
PDE8A
MTERF
RNH1
GOLGA6A
APOC4
APOC4-APOC2
COL15A1
METAP2
AC021860.1
MYC
UBL3
KLHL24
SSTR1
C5
TDG
OAS2
KCNH5
ALX3
SKA2
DCT
KPNA6
PIGX
VAMP5
MAN1A2
FGL2
WDR17
POLR2C
SHISA7
UPP2
OR5M8

STRIP1
AL162407.1
SIGLEC5
SLC17A7
DGCR8
TFIP11
BRINP2
FASLG
APOC2
PPP6R2
INHBB
APBA1
DFNB59
OSTC
C17orf112
HSH2D
ZDHHC6
CILP
PLEKHM2
HMGB1
HRAS
SLC44A4
ANKRD50
AL589765.1
RIIAD1
ENOSF1
PRR16
AC069368.3
FRG2B
CDHR5
FBXO39
NPM2
ZNF714
PCLO
IFITM10
TRIM52
AC138655.1
SPNS2
OR8I2
CAPRIN2
LCE2A
SCYL1
KIR2DL4
ZNF362
C2orf68
C17orf78
BCL10

RAB40B
SIPA1L2
CSNK1G3
MPEG1
ZNF791
QSOX2
HGF
CELF3
MLYCD
CALM2
ATXN2
OR5B2
GUCY1A2
OR5M9
PPRC1
TEP1
USP10
MYCT1
EMR3
CDCA3
CLASP1
FGF2
KCNIP4
MAPKBP1
FAM150B
CSNK2A2
CMBL
OR4D6
MPC1
KIAA0947
CLMP
CTB-129P6.11
SERINC3
CHMP3
TRAT1
TRIP13
C4orf32
PPT2-EGFL8
DSC1
PPM1D
NIPAL1
FHL2
CXCR1
ZNF430
SPRR2G
FAM159A
HYDIN

NUP35
PWP2
TRAF3IP1
HAO1
DHPS
MYADM
GPX6
CASP9
FAM20A
RNF103-CHMP3
GOLGA7B
SCGB2A1
NRROS
PIK3C2B
CABLES2
BEAN1
CRELD1
RSPH6A
CLPTM1
IFIT1B
ME3
GLIS1
FAM213B
ATP10D
LOXL1
RP11-761B3.1
KHDC1L
IFIT1
NOLC1
WDR41
FRK
SLC26A9
LCN6
CPEB2
CIR1
MYPOP
HLA-DOB
USP2
MMS19
EPPIN
ZNF765
EFNA3
PIGZ
BMP10
AC004528.1
LTA
LPAR6

MUC17
BRD9
TRIO
OR51A2
ZNF490
ATXN7L1
SYCP3
LCN10
PGM3
LDB1
IFIT5
AC026202.1
ZNF628
SIRT2
PDX1
FN1
DUSP14
OR5M3
RP4-734P14.4
MPDU1
SLAMF8
CTNNA2
MOB3A
SLC25A34
PHLDA1
RP11-429E11.3
HMOX1
DOK4
FAXC
DNAJC16
OR8H3
C9orf142
HIST1H1E
SOD2
ENTPD4
RPAP3
PLA2G12B
NCOA5
IL31RA
MIDN
KAT7
CRH
ZNF354A
INPP4B
CLEC19A
PPFIBP1
ZBTB22

IRF2BP1
GNE
RPS7
GSTT2
MUC6
ZBTB2
RMND1
DOK2
CDX1
EIF1
MSH2
OR5T3
RP11-144F15.1
CALCOCO2
ESD
CNTN1
ROS1
POU5F1
GJB3
RP11-20I23.1
RGP1
ANKRD28
TNFSF12
TNFSF12-TNFSF13
DYNC2H1
UQCRB
C2CD4D
CTB-186H2.3
CHEK2
CACNG6
WFDC10A
ZNF184
CACNG5
JAK1
CEBPE
PRRG2
NLRP9
SCGB1D1
AKIRIN2
SPATA17
DSE
ARRDC2
ARFGAP1
GPNMB
EMC3
GSTT2B
MCFD2

SLC22A9
PCIF1
MAN2A1
PSORS1C1
SMCO4
FGF23
FHIT
CTC-435M10.3
ATP5SL
AGR2
DDAH1
RPS18
SLC2A9
TFCP2
WFDC10B
RFESD
RP11-650K20.3
KIAA1199
FBXL15
OR8K5
EEF1B2
LRRC23
AHNAK2
RNF219
SFT2D3
ADAM33
ATP2B4
BCKDHA
C4orf33
ITPA
OR5M11
MNX1
ZG16B
C13orf35
SLC26A10
TJP3
CDK5RAP2
AL450307.1
AC022498.1
HNRNPR
PLCG1
ALDH3B2
ZNF479
HTR2A
CCDC33
CA2
ADCK5

CRYGS
RP11-15E18.4
SOX8
SPN
FER
TMPO
NT5E
RP11-351M8.1
DLGAP1
NOS1AP
GNB2L1
ZFP62
LRRC27
C21orf33
CTBP2
PPT2
DCAF4
SLC45A4
LACTBL1
PGLYRP1
EXOC7
MAS1
AL355390.1
RNF123
FKBP4
SELL
UBE3D
PHLDA3
SEMA4A
VSIG8
PRKD1
HIST1H3H
FAM189A2
CPO
KCNC2
RFX6
ITGA8
C7orf43
COL10A1
EML2
MYL12A
DCDC2B
B3GNT8
SKOR2
CENPBD1
FOSL2
SLC28A1

TIFA
COPS4
CTC-786C10.1
IL6R
TRAPPC10
NREP
ARRB1
DKFZP434E1119
EPPIN-WFDC6
POMGNT1
RBL1
C1S
LHX9
TMEM178A
GCNT2
NELFCD
MAL
PTPRO
FOXI1
PFN4
RP1-228P16.5
FAM205A
LOXL2
IFITM1
RNASE4
TTPAL
SGK494
TTC31
WDR81
NKX1-1
GALNT15
AP002348.1
ABCD4
ZBTB48
AC023469.1
PKN2
PBX2
ZNF436
OR8H2
ZNF532
S1PR4
AC087477.1
RP11-403P17.5
GGH
OR5F1
SLC13A1
KCNJ10

TADA2A
LAMP1
ITGA11
GATA2
LONRF1
GIPC3
TUBAL3
CPNE5
LIN54
RP11-105C20.2
OR52D1
EPHB3
CLEC5A
PRB4
PRSS35
CDHR4
PRR24
ARHGDIB
AL163636.6
CEP72
LZTR1
ACAD10
KLHDC8A
JAZF1
SH2B2
RALGAPB
CD320
KIF26A
MBLAC1
PPP1R9B
SPSB2
EFNA1
TM4SF19
CCDC102A
USP42
PDE10A
C1orf123
OR5AS1
TTLL7
OR5M10
TMEM41B
DYNC1I2
DDX23
GPR75-ASB3
ZHX3
THAP9
OR10AG1

ASPN
RASSF7
THBD
OR6J1
FCER1A
CCDC106
CCDC81
STON2
LSAMP
HDGFRP3
ACP5
RP11-796G6.2
USP6NL
SEC22A
CLTA
LCE1F
ERMN
MCM4
CCDC39
PHF10
SORBS2
ATN1
C20orf141
WDR1
CBLN3
ZNF391
PITX1
RBPJ
OR8H1
CXXC1
FANCD2
ZNF804B
NFXL1
CLK1
RP11-763F8.1
SWSAP1
NADK
RTL1
BAI2
DFFA
TEAD3
NLRP5
GRK7
PPCDC
TMEM173
LCE3E
DRD4

ECM2
SLC4A11
SCEL
IP6K1
BAIAP2
C1orf52
LRP8
FAM212A
LRRC45
PGAM1
AP4M1
AC011298.1
SAP30L
RGCC
EXOSC5
ST14
LPXN
AC068039.1
SHMT2
AMBN
TRMT10B
EPS8L2
CBWD1
ATP6V0A4
UCKL1
ZNF681
INHBC
GIPR
ZDHHC16
MFSD12
OR14I1
COL3A1
C18orf54
PSMD1
URAD
KHDRBS2
GPX5
GZMK
FANCD2OS
CAPN13
PHF1
C19orf25
MC2R
RP11-219B4.5
GJA4
SH2D1B
PRSS21

TBC1D9B
OR4A16
FBXO41
C8B
LRRC3B
GLYATL1
MAFB
SNRPB
GPR31
MDM1
VKORC1L1
ANG
ILVBL
NYNRIN
LTBP3
AIMP1
CARD10
CD40
CCDC9
LMO1
TPI1
CLVS2
PGPEP1L
TUBA1C
COX4I2
ERLEC1
CGN
CTDP1
FMO4
OR2AG2
TCF19
SFRP1
LRCH1
CLEC4C
ACOT7
KLHL14
ENO2
KBTBD8
OR10A3
MBD3L4
HBEGF
AOC1
KCNA3
LMAN1L
PSMA1
AVP
LUZPP1

SLC9C1
TTLL1
UBR4
TTC34
NUCB2
ZNF763
CTC-429P9.4
CSRNP2
ZNF383
ZNF141
MED23
DNAJB1
CHPT1
CORIN
MRPL54
Z97053.1
RP11-187E13.1
OXSM
ARMC8
PIP5K1C
RUSC1
SHC4
BARX2
MASP1
COPS6
ERMARD
WNT5A
OR5I1
AL031663.2
ZNF627
SYT6
RP11-272B17.2
EFNA4
MBD3L2
CTD-2192J16.24
C17orf74
UBE3A
WFDC11
KATNAL2
RAB27B
NOX4
CNFN
RIMS1
APIP
LHX3
SP110
HDHD2

MRI1
SCN4A
AC018755.1
GAA
TTLL6
QPRT
CTD-3088G3.8
MREG
EIF4E
PAFAH1B1
FBXO24
NBPF4
THOC7
MPP7
MAMDC4
SF3B14
CDK4
CAT
CSMD1
LRRFIP1
THSD7B
HMCN2
CNTN4
SCN7A
AGBL1
CASC9
DPP10
LINC00907
RBFOX1
PHOSPHO2-KLHL23
CHCHD6
ABTB2
TMEM178B
GSE1
PLXDC2
GALNT12
DPP6
BLACE
DIRC3
TXNRD3
LRRTM4
FAM83E
SYT16
CA1
ESYT2
PI16
ROBO2

KCNQ1
KERA
TENM2
MAST4
FLJ40194
GLT8D2
MTG2
CASZ1
UNC13C
SCHIP1
XRCC5
ANK3
HLA-DQB1
XRCC4
CACNG2
SUSD5
C4BPA
CLYBL
LOC254128
PDXDC1
MAGI1
NOTCH4
BPIFC
TRAPPC9
NOX3
MUC22
GABBR2
NEDD4
CPAMD8
FRRS1
TAX1BP1
RFTN1
DHRS7C
AHI1
SERPINB12
SLCO3A1
PAK2
ASTN2
RALYL
ZNRF1
SRPK2
C6orf7
EHD4
WWOX
RBMS3
SNX29
GLTSCR1L

MYO5B
FBN2
LINC00222
EEPD1
IQCK
LUZP2
TCF7L2
PVRL2
HLA-DRB6
MYO1C
VAV2
TTC28
PLCH1
TOM1L2
ZBTB46
NRCAM
APH1B
SYNDIG1
MTUS2
C6orf10
NRXN1
LOC285696
CCDC11
AP2A2
COL6A5
NDST3
SLC39A11
CELF2
TBC1D24
SEMA5A
DSTYK
TP73
SGK223
NOL10
PSD3
BBOX1
MYO3B
PHKB
C9orf92
KCNG4
SYNPO2
LOC101928991
ABCG2
SAMD12
IFT140
CNGA3
UGT1A10

PTPRE
ITPR2
MAP3K5
NTN1
TENM4
COL5A1
DGKG
CTNND2
SNRPN
GPSM3
SMARCC1
PALM2
GRHL2
XIRP2
DDX60L
PTPRT
FGD4
DCLK3
AKAP6
PALLD
CACNA1C
ZBTB7C
RIT2
PRUNE2
TENM3
GALNT3
CYP2B6
BCL11B
FASTKD1
TRIML1
MDFIC
LOC100616530
ANXA13
RNFT2
RABEP1
LOC728730
OSMR
FAM135B
C9orf3
CDH13
PDS5B
NKAIN1
ERC1
DAB1
DNAJC5
ANKDD1A
OBSCN

NTM
KPNA3
FAM19A1
ROR1
TGM2
MCF2L
GTPBP4
MAML1
ARMC9
TUSC3
TMEM130
CNTNAP2
NLGN1
BTNL2
ST8SIA2
PRSS12
LRP1B
PCSK5
CNGB1
SLC24A1
PARG
SUSD3
TMEM243
CRISPLD2
SPATA5L1
DPP8
MINK1
GRIN2B
AGBL3
KCNT2
TGM6
NELL1
DLGAP2
FARS2
IDI2-AS1
TBL1XR1
MSI2
FNTB
CTNNA3
FIGNL2
DDRGK1
H2AFJ
DOCK5
DISP1
CHID1
MROH2A
DDHD1

LINC00959
CECR2
C8orf69
LMO7
BACE2
NUP88
TPD52L2
FIBCD1
PACRG
LOC339862
NEDD4L
CEP128
MTMR4
CDH12
SPEF2
LOC285556
NTF3
CYFIP1
RBFOX3
DENND3
NCAM1
PRKD2
MICU1
TTC37
CD247
PKP3
TWIST1
KCNC4
FRMD8
AKT3
RPA3-AS1
HGD
WIPF3
FBXL17
NEURL1B
LOC285500
LEKR1
MUC5AC
SESTD1
ATRNL1
HLA-DRB1
ALDH1L1
ENPEP
ESYT3
TBC1D32
C1orf198
CHRNE

RGL2
CPEB3
KCNJ3
SLAMF6
THSD7A
LOC100506403
FAR1
AKAP13
RUFY1
LINC00466
PTPLAD1
OSBPL10
GOLGA6L2
FGF14
PLS1
EPHB1
EBF1
PICALM
ATE1
FOXP2
WDR62
CLGN
GGA3
LRRC2
SDK1
FYB
MAP3K11
LOC100132146
TINCR
BLOC1S5
TNXA
MAML3
TMC2
RRP12
C10orf112
SYNE3
ELP2
PLB1
ZDHHC19
GMEB1
PAX5
TANGO2
CEP89
HCG23
SYTL2
PRC1
TRPV2

SLC35F1
DDX25
SND1
FOXP1
CYBA
CCDC148
FAM155A
C2orf61
SLC25A12
USP45
ELK3
DCLK1
HLA-DQA2
BTAF1
SNAPC4
TSNARE1
FGD5
ACAD11
ARID1B
LARS2
CAPS2
MEGF11
TIMP2
CCDC68
CLPX
SP100
STAG3
TNFAIP8
CHIA
CEP85L
MAGI2
MYLK
LINC00333
MBTPS1
AOAH
RASSF8
ZSCAN5A
TSPAN18
TOB2
PRLR
KLHL2
CNTNAP5
COL22A1
HIBCH
FANCC
PDGFC
ZNF552

TP63
TAP2
ETV5
SMAP1
RIMBP2
RYR2
SYCP2L
LOC101059906
LOC100500773
CAMK2G
SH3BP2
ALMS1
GRIA2
TRIM66
ZNF343
NBAS
MYO1E
CDH23
VPS54
SMIM3
C16orf72
WDR37
DHX30
ANKMY2
KLC1
SLC24A4
ARHGAP10
POLD3
ST5
RNLS
AHCYL2
PCDH15
MAP4
B3GALT1
RSU1
NINJ2
BCAS3
NFKBIB
ST18
ASAP1
LINC00276
GSG1L
ZFAND2A
PRDM6
TUBGCP3
SQRDL
PSORS1C3

PTPRM
AUH
HSPA4L
ITFG1
C20orf196
CTTNBP2NL
ADCK3
CUEDC1
VEGFA
PRICKLE1
VIPR1
LOC338667
ADAMTSL1
CAMTA1
MOCOS
FLJ13197
KRR1
NR3C1
DSCAM
MKX
PLCL1
HKDC1
TAPBP
QSOX1
MCU
FREM3
PALM2-AKAP2
ESRRG
PARVA
SGSM3
DNAJB4
COBLL1
B3GALT4
PLSCR2
TCF3
SCG5
PDE3B
ADSL
TMTC1
RUNX1
RICTOR
ADAMTS14
TBC1D16
FOXJ3
COL9A2
SSB
MACC1

JAKMIP2
LRRC28
TPP2
ITGBL1
JPH3
FOXN3
MYRIP
ERBB4
PPP1R16B
ADARB2
LOC100130480
BRSK2
CDK17
LY86-AS1
NEBL
TACC2
CPLX2
HIP1
METTL21C
RBM20
SEC1P
WDR46
SORCS3
CHST11
ADRB2
CLEC17A
CCSER1
CPPED1
PARK2
FOCAD
SIGIRR
DOK6
PPM1H
AGAP1
HS3ST4
BIVM
RASGRF2
COLEC11
CELA3A
WDFY4
FAM81A
L3MBTL4
GPM6A
RYR3
DGKZ
SLC1A2
CDH3

VPS52
SULT2B1
RNF144B
MYO18B
GPR133
GABRG3
DIAPH1
PLD2
TESC
DOCK10
RBPMS
C7orf10
FHDC1
PRKCE
EFNA5
RAD51B
RBBP5
OR5A1
XPO7
INTS8
APCDD1L
CKAP4
PCDH9
ZHX2
ABCC5
ASIC2
CCDC154
C1orf21
WDPCP
COL6A6
RORA
SRL
LINC00871
EIF3D
HS3ST1
GPR123
KIAA1644
FBXO15
ABCA13
MYH14
ELOF1
ITPK1
LOC100996485
SLC6A15
KLB
RNF43
ANKRD30A

NCOR2
BMPR1B
C8orf87
DTX3L
CDK14
ZBP1
MYO5A
TECRL
SYNE2
GABRA2
SEC62
DCC
IMMP2L
LMCD1
NKAIN2
VAT1L
COL9A3
SMYD3
APP
ZNF385D
MIR3689B
ROBO1
NFIC
RPA3
TIMM21
UNC5C
FBLN1
MYOM2
ZNRD1-AS1
CPNE4
CEP57
RAP1A
SAMD7
COLGALT2
RPL31
COL21A1
TNKS2
ATP10A
SATB1
LINC00911
AATK
C1orf95
SLC12A5
SMAD3
MAP3K8
FKBPL
SOX6

SHISA6
SIRPD
SYTL3
SPNS3
SLC26A5
MCM6
ZDHHC21
ANO3
SLC19A1
ATXN7L3
MACROD2
LIMK2
FAM154A
NRXN3
OGDH
AMPH
ATP2C2
ADAM12
TNRC6B
YWHAQ
SH3RF3
LILRP2
ANKRD10
NBEA
DNAH11
KCNG3
EVC
ALK
EFCAB1
DAB2
MAPRE2
CRTAC1
PIEZO2
MKL2
PITPNC1
LOC100652824
CHST3
TBX19
XAF1
TNFRSF13B
LOC101929065
BAIAP2L1
GMDS
HIST4H4
HERPUD2
LINGO2
CSTF3

AGL
INSR
PAK7
GLCCI1
MMP16
ADRBK1
APBB1IP
GLI3
PIK3CB
BNC2
GGT5
LARGE
STAB1
MELK
MAP2
TMEM132B
SCARB1
ABRACL
HSD17B4
C18orf42
MTHFR
ACOT13
TGFBR3
TXK
NEB
NALCN-AS1
KANK3
GRID1
PPIL1
ITGAE
MAT1A
DSCAML1
ACVR1C
FMN2
FAM73A
DOCK3
TNPO1
NPLOC4
DLG2
RAI14
MGAM
DDX10
JRKL-AS1
INPP5K
EPC2
MSRB3
ATP8B3

LARS
LIAS
RIMKLA
MECOM
PDZD2
SLC30A9
MYO7B
KDM2A
MOK
RELN
TUBGCP5
TRERF1
LRRC16A
PCSK2
FAM134B
LRP5
EPB41L3
ZNF821
ATP6V1G2-DDX39B
CCDC60
PCBP3
IQCJ-SCHIP1
FLJ42393
LINC00710
SLC8A1-AS1
UGT1A8
UGDH-AS1
CHURC1-FNTB
BZRAP1-AS1
BLOC1S5-TXNDC5
NPHP3-ACAD11
LARS2-AS1
GATS
JMJD7
MESTIT1
LOC100506374
SLC9A9-AS1
JAKMIP2-AS1
SNURF
LRRTM3
MAGI1-AS1
ERICH1-AS1
TBC1D8
HCG25
UGT1A9
EEF1E1-MUTED
