## Supplementary Table 6 for "A higher burden of multiple sclerosis genetic risk confers an earlier onset"

**Supplementary Table 6. Pathway Analysis Results for top 2.5% genes associated with AAO from the individual SNP-level (GS1) analyses**

| Column Header | Key |
|---|---|
| Category | Pathway Database |
| Term | Pathway name/Identifier |
| Count | Number of genes in the submitted list in the specified term |
| % | Percentage of identified genes in the submitted list associated with the specified term |
| Pvalue | Significance level associated with the EASE enrichment score for the term |
| Genes | List of genes present in the term |
| List Total | Number of genes from the submitted list present in the category |
| Pop Hits | Number of genes involved in the specified term (category-specific) |
| Pop Total | Number of genes in the human genome background (category-specific) |
| Fold Enrichment | Ratio of the proportion of count to list total and population hits to population total |
| Bonferroni | Bonferroni adjustment of p-value |
| Benjamini | Benjamini adjustment of p-value |
| FDR | False Discovery Rate of p-value (percent form) |

| Category | Term | Count | % | PValue | Genes | List Total | Pop Hits | Pop Total | Fold Enrichment | Bonferroni | Benjamini | FDR (%) |
|---|---|---|---|---|---|---|---|---|---|---|---|---|
| REACTOME_PATHWAY | R-HSA-419037:R-HSA-419037 | 9 | 0.947368421 | 6.67E-04 | NCAM1, COL9A2, COL9A3, COL6A6, COL6A5, CNTN2, ST8SIA2, CACNA1C, COL5A1 | 430 | 42 | 9075 | 4.522425249 | 0.349039705 | 0.349039705 | 0.990486099 |
| REACTOME_PATHWAY | R-HSA-202433:R-HSA-202433 | 8 | 0.842105263 | 0.001272259 | FYB, HLA-DQB1, PAK2, HLA-DRB1, CD247, HLA-DQA2, HLA-DQA1, HLA-DRA | 430 | 36 | 9075 | 4.689922481 | 0.558943837 | 0.335879406 | 1.880131742 |
| KEGG_PATHWAY | hsa04514:Cell adhesion molecules | 17 | 1.789473684 | 0.001737596 | HLA-DQB1, PTPRM, HLA-DRB1, NRXN3, NLGN1, HLA-C, NRXN1, CDH3, HLA-DQA2, HLA-DQA1, NCAM1, NRCAM, CNTN2, CNTN1, CNTNAP2, HLA-DOB, HLA-DRA | 343 | 142 | 6879 | 2.400997824 | 0.364858231 | 0.364858231 | 2.248893226 |
| REACTOME_PATHWAY | R-HSA-202430:R-HSA-202430 | 6 | 0.631578947 | 0.003113091 | HLA-DQB1, HLA-DRB1, CD247, HLA-DQA2, HLA-DQA1, HLA-DRA | 430 | 22 | 9075 | 5.755813953 | 0.865318146 | 0.487410512 | 4.542187678 |
| REACTOME_PATHWAY | R-HSA-1650814:R-HSA-1650814 | 10 | 1.052631579 | 0.004049213 | COL9A2, COL9A3, COLGALT2, COL21A1, ADAMTS14, COL6A6, COL6A5, COL22A1, COL25A1, COL5A1 | 430 | 67 | 9075 | 3.149947935 | 0.926386683 | 0.479118303 | 5.869934356 |
| KEGG_PATHWAY | hsa05332:Graft-versus-host diseas | 7 | 0.736842105 | 0.005115849 | HLA-DQB1, HLA-DRB1, HLA-C, HLA-DQA2, HLA-DOB, HLA-DQA1, HLA-DRA | 343 | 33 | 6879 | 4.254174397 | 0.737804227 | 0.487949443 | 6.488108197 |
| REACTOME_PATHWAY | R-HSA-202427:R-HSA-202427 | 6 | 0.631578947 | 0.005589055 | HLA-DQB1, HLA-DRB1, CD247, HLA-DQA2, HLA-DQA1, HLA-DRA | 430 | 25 | 9075 | 5.065116279 | 0.972781084 | 0.513621728 | 8.016551688 |
| REACTOME_PATHWAY | R-HSA-389948:R-HSA-389948 | 6 | 0.631578947 | 0.006656478 | HLA-DQB1, HLA-DRB1, CD247, HLA-DQA2, HLA-DQA1, HLA-DRA | 430 | 26 | 9075 | 4.870304114 | 0.986355574 | 0.511168469 | 9.477693961 |
| KEGG_PATHWAY | hsa05169:Epstein-Barr virus infecti | 14 | 1.473684211 | 0.007329795 | HLA-DQB1, HLA-DRB1, PIK3CB, NFKBIB, HLA-C, RB1, HLA-DQA2, HLA-DQA1, NEDD4, JAK1, RBPJ, NCOR2, AKT3, HLA-DRA | 343 | 122 | 6879 | 2.301438608 | 0.853410967 | 0.472729071 | 9.17348673 |
| KEGG_PATHWAY | hsa04672:Intestinal immune netwo | 8 | 0.842105263 | 0.007966385 | HLA-DQB1, HLA-DRB1, TNFRSF13B, HLA-DQA2, HLA-DOB, CCL28, HLA-DQA1, HLA-DRA | 343 | 47 | 6879 | 3.413684015 | 0.876009786 | 0.406600943 | 9.932340164 |
| KEGG_PATHWAY | hsa05330:Allograft rejection | 7 | 0.736842105 | 0.009089065 | HLA-DQB1, HLA-DRB1, HLA-C, HLA-DQA2, HLA-DOB, HLA-DQA1, HLA-DRA | 343 | 37 | 6879 | 3.794263651 | 0.907735199 | 0.379120659 | 11.25639149 |
| KEGG_PATHWAY | hsa04024:cAMP signaling pathway | 19 | 2 | 0.009387479 | PLD2, PIK3CB, CAMK2G, PDE3B, GABBR2, CNGB1, VAV2, CNGA3, GLI3, GLI1, ADRB2, ATP2B4, GRIN2B, GRIA2, RYR2, RAP1A, CACNA1C, CALM2, AKT3 | 343 | 198 | 6879 | 1.924507465 | 0.914710577 | 0.336538292 | 11.60529391 |
| REACTOME_PATHWAY | R-HSA-1912408:R-HSA-1912408 | 6 | 0.631578947 | 0.010700249 | TNRC6C, MAML1, NOTCH4, MAML3, TNRC6B, RBPJ | 430 | 29 | 9075 | 4.366479551 | 0.999009534 | 0.627750422 | 14.8189132 |
| KEGG_PATHWAY | hsa04974:Protein digestion and ab | 11 | 1.157894737 | 0.011543067 | CELA3A, SLC8A1, COL9A2, COL9A3, COL21A1, COL6A6, COL6A5, SLC9A3, COL22A1, KCNQ1, COL5A1 | 343 | 88 | 6879 | 2.506924198 | 0.951697102 | 0.351371287 | 14.08820612 |
| KEGG_PATHWAY | hsa04612:Antigen processing and p | 10 | 1.052631579 | 0.012549085 | HLA-DQB1, HLA-DRB1, TAP2, KIR3DL3, HLA-C, HLA-DQA2, HLA-DOB, HLA-DQA1, HLA-DRA, TAPBP | 343 | 76 | 6879 | 2.638867577 | 0.962970328 | 0.337678684 | 15.22479919 |
| REACTOME_PATHWAY | R-HSA-376172:R-HSA-376172 | 4 | 0.421052632 | 0.013028927 | DCC, DSCAML1, NTN1, DSCAM | 430 | 11 | 9075 | 7.674418605 | 0.999782357 | 0.651487714 | 17.75981565 |
| KEGG_PATHWAY | hsa05310:Asthma | 6 | 0.631578947 | 0.015061909 | HLA-DQB1, HLA-DRB1, HLA-DQA2, HLA-DOB, HLA-DQA1, HLA-DRA | 343 | 30 | 6879 | 4.011078717 | 0.980957163 | 0.356039731 | 18.00338887 |
| REACTOME_PATHWAY | R-HSA-186797:R-HSA-186797 | 6 | 0.631578947 | 0.01616397 | COL9A2, COL9A3, COL6A6, COL6A5, PDGFC, COL5A1 | 430 | 32 | 9075 | 3.957122093 | 0.999971861 | 0.687846729 | 21.56964993 |
| KEGG_PATHWAY | hsa04940:Type I diabetes mellitus | 7 | 0.736842105 | 0.01666801 | HLA-DQB1, HLA-DRB1, HLA-C, HLA-DQA2, HLA-DOB, HLA-DQA1, HLA-DRA | 343 | 42 | 6879 | 3.342565598 | 0.987562191 | 0.35512671 | 19.73503684 |
| KEGG_PATHWAY | hsa05150:Staphylococcus aureus ir | 8 | 0.842105263 | 0.016693158 | HLA-DQB1, HLA-DRB1, C4B, C5, HLA-DQA2, HLA-DOB, HLA-DQA1, HLA-DRA | 343 | 54 | 6879 | 2.97116942 | 0.987644936 | 0.329295281 | 19.76188011 |
| REACTOME_PATHWAY | R-HSA-936837:R-HSA-936837 | 8 | 0.842105263 | 0.017238235 | ATP2C2, ATP2B4, CAMK2G, ATP10A, ATP11A, ATP10D, ATP8B3, CALM2 | 430 | 57 | 9075 | 2.962056304 | 0.999986061 | 0.673094238 | 22.83680399 |
| REACTOME_PATHWAY | R-HSA-350054:R-HSA-350054 | 4 | 0.421052632 | 0.021052407 | MAML1, NOTCH4, MAML3, RBPJ | 430 | 13 | 9075 | 6.493738819 | 0.999998856 | 0.711698138 | 27.18320015 |
| REACTOME_PATHWAY | R-HSA-425561:R-HSA-425561 | 4 | 0.421052632 | 0.021052407 | SLC8A1, SLC24A4, SLC24A1, CALM2 | 430 | 13 | 9075 | 6.493738819 | 0.999998856 | 0.711698138 | 27.18320015 |
| REACTOME_PATHWAY | R-HSA-193648:R-HSA-193648 | 7 | 0.736842105 | 0.02241025 | OBSCN, RASGRF2, TRIO, AKAP13, VAV2, MCF2L, FGD4 | 430 | 47 | 9075 | 3.143245918 | 0.999999531 | 0.703134367 | 28.67458207 |
| REACTOME_PATHWAY | R-HSA-877300:R-HSA-877300 | 10 | 1.052631579 | 0.022767686 | NCAM1, HLA-DQB1, SP100, HLA-DRB1, CAMK2G, JAK1, HLA-C, HLA-DQA2, HLA-DQA1, HLA-DRA | 430 | 88 | 9075 | 2.398255814 | 0.99999963 | 0.679906579 | 29.06240522 |
| KEGG_PATHWAY | hsa04070:Phosphatidylinositol sigr | 11 | 1.157894737 | 0.023191176 | INPP5K, PIK3CB, PIK3C2B, DGKG, DGKZ, INPP4B, ITPK1, INPP5A, MTMR4, CALM2, ITPR2 | 343 | 98 | 6879 | 2.251115607 | 0.997810733 | 0.399713972 | 26.42671755 |
| KEGG_PATHWAY | hsa05140:Leishmaniasis | 9 | 0.947368421 | 0.02381573 | HLA-DQB1, CYBA, HLA-DRB1, NFKBIB, JAK1, HLA-DQA2, HLA-DOB, HLA-DQA1, HLA-DRA | 343 | 71 | 6879 | 2.54223299 | 0.998147317 | 0.383645087 | 27.03959843 |
| KEGG_PATHWAY | hsa04330:Notch signaling pathway | 7 | 0.736842105 | 0.030449932 | MAML1, DTX3L, NOTCH4, APH1B, MAML3, RBPJ, NCOR2 | 343 | 48 | 6879 | 2.924744898 | 0.999687512 | 0.43813637 | 33.26508346 |
| KEGG_PATHWAY | hsa04014:Ras signaling pathway | 19 | 2 | 0.03200999 | PLD2, PIK3CB, FGF14, RGL2, JMJD7-PLA2G4B, PAK2, GRIN2B, RASGRF2, VEGFA, PLA2G12B, RAP1A, PDGFC, EFNA5, SYNGAP1, PLA2G4B, INSR, AKT3, CALM2, SHC4 | 343 | 226 | 6879 | 1.686072912 | 0.999794743 | 0.432255118 | 34.65593237 |
| KEGG_PATHWAY | hsa04145:Phagosome | 14 | 1.473684211 | 0.035396226 | HLA-DQB1, HLA-DRB1, HLA-C, COLEC12, COLEC11, HLA-DQA2, HLA-DQA1, CYBA, TAP2, DYNC2H1, SCARB1, HLA-DOB, DYNC1I2, HLA-DRA | 343 | 150 | 6879 | 1.871836735 | 0.999917761 | 0.444489391 | 37.58325717 |
| KEGG_PATHWAY | hsa05321:Inflammatory bowel dise | 8 | 0.842105263 | 0.038520985 | HLA-DQB1, HLA-DRB1, SMAD3, RORA, HLA-DQA2, HLA-DOB, HLA-DQA1, HLA-DRA | 343 | 64 | 6879 | 2.506924198 | 0.999964739 | 0.452888427 | 40.17661747 |
| KEGG_PATHWAY | hsa05320:Autoimmune thyroid dis | 7 | 0.736842105 | 0.042882778 | HLA-DQB1, HLA-DRB1, HLA-C, HLA-DQA2, HLA-DOB, HLA-DQA1, HLA-DRA | 343 | 52 | 6879 | 2.699764521 | 0.999989238 | 0.470343391 | 43.63048762 |
| KEGG_PATHWAY | hsa04713:Circadian entrainment | 10 | 1.052631579 | 0.045794812 | NOS1AP, GRIN2B, GRIA2, CAMK2G, RYR3, GUCY1A2, RYR2, CACNA1C, KCNJ3, CALM2 | 343 | 95 | 6879 | 2.111094062 | 0.999995141 | 0.47477879 | 45.8328258 |
| REACTOME_PATHWAY | R-HSA-5218920:R-HSA-5218920 | 5 | 0.526315789 | 0.04590614 | PAK2, RICTOR, VAV2, AKT3, CALM2 | 430 | 29 | 9075 | 3.638732959 | 1 | 0.884482332 | 50.3725882 |
| KEGG_PATHWAY | hsa04925:Aldosterone synthesis ar | 9 | 0.947368421 | 0.047161191 | PRKD2, CYP21A2, CAMK2G, ATF6B, SCARB1, PRKCE, CACNA1C, CALM2, ITPR2 | 343 | 81 | 6879 | 2.228377065 | 0.999996657 | 0.467642155 | 46.83856549 |
| REACTOME_PATHWAY | R-HSA-5578775:R-HSA-5578775 | 7 | 0.736842105 | 0.047692042 | SLC8A1, ATP2B4, CAMK2G, RYR3, RYR2, CALM2, ITPR2 | 430 | 56 | 9075 | 2.638081395 | 1 | 0.876899884 | 51.73967082 |
| KEGG_PATHWAY | hsa04022:cGMP-PKG signaling patl | 14 | 1.473684211 | 0.05034014 | SLC8A1, PDE3B, CNGB1, PRKCE, ITPR2, ADRB2, ATP2B4, ATF6B, GUCY1A2, CACNA1C, INSR, AKT3, CALM2, MYLK | 343 | 158 | 6879 | 1.777060191 | 0.999998603 | 0.473736438 | 49.11210206 |
| REACTOME_PATHWAY | R-HSA-1296072:R-HSA-1296072 | 6 | 0.631578947 | 0.050789169 | KCNC2, KCNC4, KCNG3, KCNQ1, KCNG4, KCNH5 | 430 | 43 | 9075 | 2.944835046 | 1 | 0.876898623 | 54.02751817 |
| REACTOME_PATHWAY | R-HSA-2644606:R-HSA-2644606 | 7 | 0.736842105 | 0.051274843 | TBL1XR1, MAML1, APH1B, NEURL1B, MAML3, RBPJ, NCOR2 | 430 | 57 | 9075 | 2.591799266 | 1 | 0.863425417 | 54.37696968 |
| REACTOME_PATHWAY | R-HSA-2894862:R-HSA-2894862 | 7 | 0.736842105 | 0.051274843 | TBL1XR1, MAML1, APH1B, NEURL1B, MAML3, RBPJ, NCOR2 | 430 | 57 | 9075 | 2.591799266 | 1 | 0.863425417 | 54.37696968 |
| REACTOME_PATHWAY | R-HSA-373752:R-HSA-373752 | 4 | 0.421052632 | 0.057911463 | DCC, EZR, ROBO1, NTN1 | 430 | 19 | 9075 | 4.443084455 | 1 | 0.881286898 | 58.91050642 |
| REACTOME_PATHWAY | R-HSA-442982:R-HSA-442982 | 4 | 0.421052632 | 0.057911463 | GRIN2B, RASGRF2, CAMK2G, CALM2 | 430 | 19 | 9075 | 4.443084455 | 1 | 0.881286898 | 58.91050642 |
| KEGG_PATHWAY | hsa00562:Inositol phosphate meta | 8 | 0.842105263 | 0.061672246 | INPP5K, PIK3CB, PIK3C2B, PLCH1, INPP4B, ITPK1, INPP5A, MTMR4 | 343 | 71 | 6879 | 2.259762658 | 0.999999939 | 0.5300793 | 56.50621008 |
| KEGG_PATHWAY | hsa04520:Adherens junction | 8 | 0.842105263 | 0.061672246 | PTPRM, LMO7, SMAD3, SSX2IP, INSR, TCF7L2, CTNNA3, CTNNA2 | 343 | 71 | 6879 | 2.259762658 | 0.999999939 | 0.5300793 | 56.50621008 |
| KEGG_PATHWAY | hsa05416:Viral myocarditis | 7 | 0.736842105 | 0.062356111 | HLA-DQB1, HLA-DRB1, HLA-C, HLA-DQA2, HLA-DOB, HLA-DQA1, HLA-DRA | 343 | 57 | 6879 | 2.462943072 | 0.99999995 | 0.518394693 | 56.91897659 |
| REACTOME_PATHWAY | R-HSA-2132295:R-HSA-2132295 | 11 | 1.157894737 | 0.062789079 | HLA-DQB1, AP2A2, HLA-DRB1, KLC1, DYNC2H1, KIF26A, HLA-DQA2, HLA-DOB, HLA-DQA1, DYNC1I2, HLA-DRA | 430 | 122 | 9075 | 1.902878384 | 1 | 0.888592014 | 61.97055192 |
| REACTOME_PATHWAY | R-HSA-3000178:R-HSA-3000178 | 8 | 0.842105263 | 0.06332452 | NCAM1, COL9A2, APP, COL9A3, TNXB, COL6A6, COL6A5, COL5A1 | 430 | 75 | 9075 | 2.251162791 | 1 | 0.877934982 | 62.29319438 |
| REACTOME_PATHWAY | R-HSA-5576892:R-HSA-5576892 | 6 | 0.631578947 | 0.064583591 | FGF14, CAMK2G, CACNG2, SCN7A, CACNA1C, CALM2 | 430 | 46 | 9075 | 2.752780586 | 1 | 0.870520735 | 63.041841 |
| BIOCARTA | h_PDZsPathway:Synaptic Proteins | 4 | 0.421052632 | 0.06754667 | NCAM1, NRXN3, PCLO, DLG2 | 88 | 18 | 1625 | 4.103535354 | 0.999986185 | 0.999986185 | 57.02153133 |
| KEGG_PATHWAY | hsa05322:Systemic lupus erythema | 12 | 1.263157895 | 0.069169234 | HLA-DQB1, HIST4H4, GRIN2B, HLA-DRB1, C4B, C5, SSB, H2AFJ, HLA-DQA2, HLA-DOB, HLA-DQA1, HLA-DRA | 343 | 134 | 6879 | 1.796005396 | 0.999999992 | 0.541362903 | 60.83821792 |
| REACTOME_PATHWAY | R-HSA-416482:R-HSA-416482 | 8 | 0.842105263 | 0.070925171 | OBSCN, RASGRF2, TBXA2R, TRIO, AKAP13, VAV2, MCF2L, FGD4 | 430 | 77 | 9075 | 2.19269103 | 1 | 0.883531382 | 66.60630178 |
| REACTOME_PATHWAY | R-HSA-1810476:R-HSA-1810476 | 4 | 0.421052632 | 0.074209561 | HMGB1, APP, NFKBIB, ZBP1 | 430 | 21 | 9075 | 4.019933555 | 1 | 0.884172595 | 68.3237124 |
| BIOCARTA | h_gcrpathway:Corticosteroids and | 4 | 0.421052632 | 0.077209215 | ADRB2, CORIN, NR3C1, CALM2 | 88 | 19 | 1625 | 3.887559809 | 0.999997391 | 0.998384675 | 62.1011587 |
| BIOCARTA | h_tob1Pathway:Role of Tob in T-ce | 4 | 0.421052632 | 0.077209215 | CD247, SMAD3, TGFBR3, TOB2 | 88 | 19 | 1625 | 3.887559809 | 0.999997391 | 0.998384675 | 62.1011587 |
| REACTOME_PATHWAY | R-HSA-418886:R-HSA-418886 | 3 | 0.315789474 | 0.078036191 | DCC, UNC5C, NTN1 | 430 | 10 | 9075 | 6.331395349 | 1 | 0.886596658 | 70.22062945 |
| REACTOME_PATHWAY | R-HSA-1442490:R-HSA-1442490 | 7 | 0.736842105 | 0.080861046 | COL9A2, COL9A3, COL6A6, COL6A5, MMP8, COL25A1, COL5A1 | 430 | 64 | 9075 | 2.308321221 | 1 | 0.885669066 | 71.55236682 |
| REACTOME_PATHWAY | R-HSA-1660499:R-HSA-1660499 | 5 | 0.526315789 | 0.081368156 | INPP5K, PIK3CB, PIK3C2B, INPP4B, PIK3R6 | 430 | 35 | 9075 | 3.014950166 | 1 | 0.877408362 | 71.78547246 |
| REACTOME_PATHWAY | R-HSA-114508:R-HSA-114508 | 4 | 0.421052632 | 0.083037912 | DGKG, DGKZ, PRKCE, ITPR2 | 430 | 22 | 9075 | 3.837209302 | 1 | 0.873116519 | 72.54048532 |
| REACTOME_PATHWAY | R-HSA-1257604:R-HSA-1257604 | 8 | 0.842105263 | 0.087684908 | TNRC6C, KLB, ERBB4, PIK3CB, RICTOR, TNRC6B, TRAT1, AKT3 | 430 | 81 | 9075 | 2.084409991 | 1 | 0.878449945 | 74.54368045 |
| KEGG_PATHWAY | hsa04510:Focal adhesion | 16 | 1.684210526 | 0.087943108 | TNXB, DIAPH1, PIK3CB, VAV2, COL5A1, PAK2, COL6A6, COL6A5, VEGFA, RAP1A, RELN, PDGFC, MYLK, AKT3, PARVA, SHC4 | 343 | 206 | 6879 | 1.557700473 | 1 | 0.6175022 | 69.99942262 |
| KEGG_PATHWAY | hsa04972:Pancreatic secretion | 9 | 0.947368421 | 0.09033886 | CELA3A, SLC26A3, ATP2B4, PLA2G12B, RAP1A, RYR2, RAB27B, KCNQ1, ITPR2 | 343 | 93 | 6879 | 1.940844541 | 1 | 0.613441816 | 71.01390416 |
| REACTOME_PATHWAY | R-HSA-211728:R-HSA-211728 | 2 | 0.210526316 | 0.092315707 | PAK2, ARHGAP10 | 430 | 2 | 9075 | 21.10465116 | 1 | 0.8832348 | 76.40359099 |

| | | | | | | | | | | | | |
|---|---|---|---|---|---|---|---|---|---|---|---|---|
| KEGG_PATHWAY | hsa05145:Toxoplasmosis | 10 | 1.052631579 | 0.09588132 | HLA-DQB1, HLA-DRB1, IL10RA, NFKBIB, JAK1, HLA-DQA2, HLA-DOB, AKT3, HLA-DQA1, HLA-DRA | 343 | 110 | 6879 | 1.823217599 | 1 | 0.622561721 | 73.24064591 |
| KEGG_PATHWAY | hsa00564:Glycerophospholipid me | 9 | 0.947368421 | 0.099216815 | PLD2, CRLS1, JMJD7-PLA2G4B, PLB1, DGKG, PLA2G12B, DGKZ, PLA2G4B, PTDSS2 | 343 | 95 | 6879 | 1.899984656 | 1 | 0.622431205 | 74.50343371 |
| KEGG_PATHWAY | hsa04015:Rap1 signaling pathway | 16 | 1.684210526 | 0.099477471 | FYB, MAGI2, MAGI1, PIK3CB, FGF14, SIPA1L2, APBB1IP, PRKD2, GRIN2B, VEGFA, RAP1A, PDGFC, EFNA5, INSR, AKT3, CALM2 | 343 | 210 | 6879 | 1.528029988 | 1 | 0.610550418 | 74.59975939 |
