## Supplementary Table 7 for "A higher burden of multiple sclerosis genetic risk confers an earlier onset"

**Supplementary Table 7. Pathway Analysis Results for the top %5 genes associated with AAO from the gene-based tests (GS2)**

| Column Header | Key |
|---|---|
| Category | Pathway Database |
| Term | Pathway name/Identifier |
| Count | Number of genes in the submitted list in the specified term |
| % | Percentage of identified genes in the submitted list associated with the specified term |
| Pvalue | Significance level associated with the EASE enrichment score for the term |
| Genes | List of genes present in the term |
| List Total | Number of genes from the submitted list present in the category |
| Pop Hits | Number of genes involved in the specified term (category-specific) |
| Pop Total | Number of genes in the human genome background (category-specific) |
| Fold Enrichment | Ratio of the proportion of count to list total and population hits to population total |
| Bonferroni | Bonferroni adjustment of p-value |
| Benjamini | Benjamini adjustment of p-value |
| FDR | False Discovery Rate of p-value (percent form) |

| Category | Term | Count | % | PValue | Genes | List Total | Pop Hits | Pop Total | Fold Enrichment | Bonferroni | Benjamini | FDR |
|---|---|---|---|---|---|---|---|---|---|---|---|---|
| BIOCARTA | h_compPathway:Complement Path | 7 | 0.802752294 | 3.23E-04 | MASP1, C4A, C4B, CFB, C5, C1S, C2 | 84 | 20 | 1625 | 6.770833333 | 0.059818465 | 0.059818465 | 0.40091772 |
| REACTOME_PATHWAY | R-HSA-174577:R-HSA-174577 | 5 | 0.573394495 | 3.29E-04 | C4A, C4B, CFB, C5, C2 | 442 | 8 | 9075 | 12.83229638 | 0.200442582 | 0.200442582 | 0.493583697 |
| REACTOME_PATHWAY | R-HSA-167242:R-HSA-167242 | 7 | 0.802752294 | 6.33E-04 | POLR2H, NELFCD, SUPT4H1, NELFE, CTDP1, POLR2C, POLR2B | 442 | 23 | 9075 | 6.248770411 | 0.349637909 | 0.193549697 | 0.947123558 |
| KEGG_PATHWAY | hsa05150:Staphylococcus aureus ir | 10 | 1.146788991 | 9.68E-04 | MASP1, C4A, C4B, CFB, C5, C1S, C2, HLA-DOB, HLA-DQA1, HLA-DRA | 332 | 54 | 6879 | 3.837014726 | 0.220248954 | 0.220248954 | 1.255067547 |
| BIOCARTA | h_lectinPathway:Lectin Induced Co | 5 | 0.573394495 | 0.003179064 | MASP1, C4A, C4B, C5, C2 | 84 | 13 | 1625 | 7.44047619 | 0.455653017 | 0.262201259 | 3.883452961 |
| REACTOME_PATHWAY | R-HSA-167158:R-HSA-167158 | 7 | 0.802752294 | 0.004604257 | POLR2H, NELFCD, SUPT4H1, NELFE, CTDP1, POLR2C, POLR2B | 442 | 33 | 9075 | 4.35520362 | 0.956435385 | 0.648133467 | 6.696409304 |
| REACTOME_PATHWAY | R-HSA-113418:R-HSA-113418 | 7 | 0.802752294 | 0.004604257 | POLR2H, NELFCD, SUPT4H1, NELFE, CTDP1, POLR2C, POLR2B | 442 | 33 | 9075 | 4.35520362 | 0.956435385 | 0.648133467 | 6.696409304 |
| BIOCARTA | h_classicPathway:Classical Comple | 5 | 0.573394495 | 0.005610551 | C4A, C4B, C5, C1S, C2 | 84 | 15 | 1625 | 6.448412698 | 0.658576838 | 0.301074327 | 6.759546744 |
| REACTOME_PATHWAY | R-HSA-167243:R-HSA-167243 | 7 | 0.802752294 | 0.007162245 | POLR2H, NELFCD, SUPT4H1, NELFE, CTDP1, POLR2C, POLR2B | 442 | 36 | 9075 | 3.992269985 | 0.992408026 | 0.704818812 | 10.23344443 |
| REACTOME_PATHWAY | R-HSA-167238:R-HSA-167238 | 7 | 0.802752294 | 0.007162245 | POLR2H, NELFCD, SUPT4H1, NELFE, CTDP1, POLR2C, POLR2B | 442 | 36 | 9075 | 3.992269985 | 0.992408026 | 0.704818812 | 10.23344443 |
| REACTOME_PATHWAY | R-HSA-167287:R-HSA-167287 | 7 | 0.802752294 | 0.009356155 | POLR2H, NELFCD, SUPT4H1, NELFE, CTDP1, POLR2C, POLR2B | 442 | 38 | 9075 | 3.782150512 | 0.998309507 | 0.721000987 | 13.1669385 |
| REACTOME_PATHWAY | R-HSA-167290:R-HSA-167290 | 7 | 0.802752294 | 0.009356155 | POLR2H, NELFCD, SUPT4H1, NELFE, CTDP1, POLR2C, POLR2B | 442 | 38 | 9075 | 3.782150512 | 0.998309507 | 0.721000987 | 13.1669385 |
| REACTOME_PATHWAY | R-HSA-977606:R-HSA-977606 | 6 | 0.688073394 | 0.010304572 | C8B, C4A, C4B, CFB, C5, C2 | 442 | 28 | 9075 | 4.399644473 | 0.99911781 | 0.690310132 | 14.40715976 |
| REACTOME_PATHWAY | R-HSA-5654706:R-HSA-5654706 | 5 | 0.573394495 | 0.014333977 | HRAS, FGFR3, FGF23, FGF2, PTPN11 | 442 | 20 | 9075 | 5.132918552 | 0.999944726 | 0.753515703 | 19.49425221 |
| KEGG_PATHWAY | hsa04610:Complement and coagul | 9 | 1.032110092 | 0.017068899 | C8B, THBD, MASP1, C4A, C4B, CFB, C5, C1S, C2 | 332 | 69 | 6879 | 2.702592981 | 0.988020719 | 0.890550096 | 20.11859178 |
| REACTOME_PATHWAY | R-HSA-5654227:R-HSA-5654227 | 4 | 0.458715596 | 0.022644622 | FGFR3, PLCG1, FGF23, FGF2 | 442 | 13 | 9075 | 6.317438218 | 0.999999824 | 0.856877953 | 29.10797493 |
| REACTOME_PATHWAY | R-HSA-167246:R-HSA-167246 | 7 | 0.802752294 | 0.022949439 | POLR2H, NELFCD, SUPT4H1, NELFE, CTDP1, POLR2C, POLR2B | 442 | 46 | 9075 | 3.124385206 | 0.999999858 | 0.82650116 | 29.4393189 |
| KEGG_PATHWAY | hsa05322:Systemic lupus erythema | 13 | 1.490825688 | 0.027379892 | C8B, C4A, C4B, C5, SNRPB, SSB, C1S, C2, CD40, HLA-DOB, HIST1H3H, HLA-DQA1, HLA-DRA | 332 | 134 | 6879 | 2.010137565 | 0.999203082 | 0.907287585 | 30.38691642 |
| REACTOME_PATHWAY | R-HSA-167200:R-HSA-167200 | 7 | 0.802752294 | 0.027737023 | POLR2H, NELFCD, SUPT4H1, NELFE, CTDP1, POLR2C, POLR2B | 442 | 48 | 9075 | 2.994202489 | 0.999999995 | 0.851913136 | 34.4575575 |
| KEGG_PATHWAY | hsa05330:Allograft rejection | 6 | 0.688073394 | 0.030778102 | FASLG, HLA-C, CD40, HLA-DOB, HLA-DQA1, HLA-DRA | 332 | 37 | 6879 | 3.359980462 | 0.999675834 | 0.865818778 | 33.49435542 |
| REACTOME_PATHWAY | R-HSA-167152:R-HSA-167152 | 7 | 0.802752294 | 0.033141514 | POLR2H, NELFCD, SUPT4H1, NELFE, CTDP1, POLR2C, POLR2B | 442 | 50 | 9075 | 2.874434389 | 1 | 0.875120093 | 39.72131079 |
| REACTOME_PATHWAY | R-HSA-190861:R-HSA-190861 | 6 | 0.688073394 | 0.035471004 | GJB3, TUBAL3, GJB6, GJA4, TUBA1C, GJA3 | 442 | 38 | 9075 | 3.241843296 | 1 | 0.870429323 | 41.86610943 |
| REACTOME_PATHWAY | R-HSA-427601:R-HSA-427601 | 3 | 0.344036697 | 0.042059193 | SLC26A4, SLC26A3, SLC26A9 | 442 | 7 | 9075 | 8.799288946 | 1 | 0.894000309 | 47.5526854 |
| REACTOME_PATHWAY | R-HSA-909733:R-HSA-909733 | 8 | 0.917431193 | 0.043151777 | IFIT1, IFITM1, OAS3, JAK1, OAS1, HLA-C, OAS2, PTPN11 | 442 | 67 | 9075 | 2.451543189 | 1 | 0.882269726 | 48.44396778 |
| KEGG_PATHWAY | hsa04740:Olfactory transduction | 28 | 3.211009174 | 0.044272914 | OR10A3, OR8H3, OR4D10, OR52D1, OR8K5, OR8I2, OR5AS1, OR5F1, OR5I1, OR2AG2, OR5M10, OR5N | 332 | 399 | 6879 | 1.454026633 | 0.999991173 | 0.902464624 | 44.61288261 |
| BIOCARTA | h_eifPathway:Eukaryotic protein tr | 4 | 0.458715596 | 0.044445513 | EIF4E, EIF4A1, EEF2, EIF1 | 84 | 16 | 1625 | 4.836309524 | 0.999830647 | 0.885923016 | 43.1946362 |
| KEGG_PATHWAY | hsa05219:Bladder cancer | 6 | 0.688073394 | 0.045338811 | HRAS, FGFR3, HBEGF, RB1, CDK4, MYC | 332 | 41 | 6879 | 3.03217749 | 0.999993374 | 0.862950661 | 45.41344972 |
| REACTOME_PATHWAY | R-HSA-75955:R-HSA-75955 | 7 | 0.802752294 | 0.045888535 | POLR2H, NELFCD, SUPT4H1, NELFE, CTDP1, POLR2C, POLR2B | 442 | 54 | 9075 | 2.661513323 | 1 | 0.880733104 | 50.61482518 |
| KEGG_PATHWAY | hsa04940:Type I diabetes mellitus | 6 | 0.688073394 | 0.049522996 | FASLG, HLA-C, HLA-DOB, HLA-DQA1, LTA, HLA-DRA | 332 | 42 | 6879 | 2.959982788 | 0.999997857 | 0.84506765 | 48.45390587 |
| REACTOME_PATHWAY | R-HSA-112382:R-HSA-112382 | 7 | 0.802752294 | 0.053262603 | POLR2H, NELFCD, SUPT4H1, NELFE, CTDP1, POLR2C, POLR2B | 442 | 56 | 9075 | 2.566459276 | 1 | 0.901996682 | 56.04700963 |
| KEGG_PATHWAY | hsa05310:Asthma | 5 | 0.573394495 | 0.053787188 | FCER1A, CD40, HLA-DOB, HLA-DQA1, HLA-DRA | 332 | 30 | 6879 | 3.453313253 | 0.999999325 | 0.830706229 | 51.39096876 |
| REACTOME_PATHWAY | R-HSA-5654704:R-HSA-5654704 | 4 | 0.458715596 | 0.054036083 | HRAS, FGFR3, FGF23, FGF2 | 442 | 18 | 9075 | 4.562594268 | 1 | 0.89125683 | 56.58325891 |
| REACTOME_PATHWAY | R-HSA-5676594:R-HSA-5676594 | 4 | 0.458715596 | 0.054036083 | TNFSF12-TNFSF13, TNFSF12, CD40, LTA | 442 | 18 | 9075 | 4.562594268 | 1 | 0.89125683 | 56.58325891 |
| REACTOME_PATHWAY | R-HSA-5654710:R-HSA-5654710 | 4 | 0.458715596 | 0.054036083 | FGFR3, FGF23, FGF2, PTPN11 | 442 | 18 | 9075 | 4.562594268 | 1 | 0.89125683 | 56.58325891 |
| KEGG_PATHWAY | hsa04145:Phagosome | 13 | 1.490825688 | 0.057029246 | ATP6V0E1, MRC2, HLA-C, COLEC12, HLA-DQA1, LAMP1, DYNC2H1, TUBAL3, ATP6V0A4, HLA-DOB, TUI | 332 | 150 | 6879 | 1.795722892 | 0.999999721 | 0.813025407 | 53.51974288 |
| KEGG_PATHWAY | hsa05168:Herpes simplex infection | 15 | 1.720183486 | 0.057226165 | IFIT1B, C5, OAS3, FASLG, HLA-C, OAS1, OAS2, HLA-DQA1, PTPN11, CSNK2A2, IFIT1, JAK1, HLA-DOB, LT | 332 | 183 | 6879 | 1.69835078 | 0.999999735 | 0.780075666 | 53.64622659 |
| REACTOME_PATHWAY | R-HSA-112411:R-HSA-112411 | 3 | 0.344036697 | 0.067627705 | JAK1, IL6R, PTPN11 | 442 | 9 | 9075 | 6.843891403 | 1 | 0.928740447 | 65.06515107 |
| KEGG_PATHWAY | hsa04151:PI3K-Akt signaling pathw | 24 | 2.752293578 | 0.06873036 | HRAS, TNXB, FGFR3, EFNA1, COL3A1, EFNA3, ITGA11, PKN2, FASLG, FGF23, IL6R, HGF, CDK4, EIF4E, C | 332 | 345 | 6879 | 1.441382923 | 0.999999989 | 0.810552039 | 60.50742535 |
| KEGG_PATHWAY | hsa00590:Arachidonic acid metabo | 7 | 0.802752294 | 0.071714904 | JMJD7-PLA2G4B, GPX6, PTGDS, GPX5, PLA2G12B, FAM213B, PLA2G4B | 332 | 61 | 6879 | 2.377691092 | 0.999999995 | 0.796837829 | 62.1272777 |
| KEGG_PATHWAY | hsa05332:Graft-versus-host disease | 5 | 0.573394495 | 0.071885818 | FASLG, HLA-C, HLA-DOB, HLA-DQA1, HLA-DRA | 332 | 33 | 6879 | 3.139375685 | 0.999999995 | 0.771174313 | 62.21815724 |
| BIOCARTA | h_bbcellPathway:Bystander B Cell / | 3 | 0.344036697 | 0.073470478 | FASLG, CD40, HLA-DRA | 84 | 9 | 1625 | 6.448412698 | 0.999999532 | 0.945796941 | 61.29665789 |
| REACTOME_PATHWAY | R-HSA-5655302:R-HSA-5655302 | 5 | 0.573394495 | 0.074096589 | HRAS, PLCG1, FGF23, CUX1, FGF2 | 442 | 33 | 9075 | 3.110859729 | 1 | 0.936149673 | 68.53368393 |
| KEGG_PATHWAY | hsa04014:Ras signaling pathway | 17 | 1.949541284 | 0.077451642 | HRAS, FGFR3, EFNA1, EFNA3, FGF23, FASLG, HGF, PTPN11, JMJD7-PLA2G4B, PLCG1, PLA2G12B, EFNA | 332 | 226 | 6879 | 1.558575008 | 0.999999999 | 0.772332706 | 65.06989716 |
| BIOCARTA | h_bArrestin-srcPathway:Roles of Ã | 4 | 0.458715596 | 0.078208159 | HRAS, FGR, ARRB1, KCNA3 | 84 | 20 | 1625 | 3.869047619 | 0.999999824 | 0.925157889 | 63.68773651 |
| REACTOME_PATHWAY | R-HSA-3299685:R-HSA-3299685 | 5 | 0.573394495 | 0.080908277 | NOX4, GPX6, GPX5, CAT, SOD2 | 442 | 34 | 9075 | 3.019363854 | 1 | 0.942979224 | 71.83679079 |
| REACTOME_PATHWAY | R-HSA-110056:R-HSA-110056 | 3 | 0.344036697 | 0.081887367 | JAK1, IL6R, PTPN11 | 442 | 10 | 9075 | 6.159502262 | 1 | 0.936860257 | 72.28404072 |
| BIOCARTA | h_il6Pathway:IL 6 signaling pathwa | 4 | 0.458715596 | 0.087999337 | HRAS, JAK1, IL6R, PTPN11 | 84 | 21 | 1625 | 3.684807256 | 0.999999977 | 0.919008544 | 68.20469758 |
| REACTOME_PATHWAY | R-HSA-5654712:R-HSA-5654712 | 4 | 0.458715596 | 0.088557518 | HRAS, FGF23, FGF2, PTPN11 | 442 | 22 | 9075 | 3.733031674 | 1 | 0.942839142 | 75.15900901 |
| REACTOME_PATHWAY | R-HSA-5654732:R-HSA-5654732 | 4 | 0.458715596 | 0.088557518 | FGFR3, FGF23, FGF2, PTPN11 | 442 | 22 | 9075 | 3.733031674 | 1 | 0.942839142 | 75.15900901 |

| | | | | | | | | | | | | |
|---|---|---|---|---|---|---|---|---|---|---|---|---|
| REACTOME_PATHWAY | R-HSA-381753:R-HSA-381753 | 28 | 3.211009174 | 0.093820458 | OR10A3, OR8H3, OR4D10, OR52D1, OR8K5, OR8I2, OR5AS1, OR5F1, OR6J1, OR5I1, OR2AG2, OR5M1[illegible] | 442 | 426 | 9075 | 1.349499713 | 1 | 0.945437992 | 77.22829141 |
| REACTOME_PATHWAY | R-HSA-1059683:R-HSA-1059683 | 3 | 0.344036697 | 0.09696414 | JAK1, IL6R, PTPN11 | 442 | 11 | 9075 | 5.599547511 | 1 | 0.944176054 | 78.38636026 |
| REACTOME_PATHWAY | R-HSA-5654719:R-HSA-5654719 | 4 | 0.458715596 | 0.098336404 | HRAS, FGF23, FGF2, PTPN11 | 442 | 23 | 9075 | 3.570725949 | 1 | 0.939881299 | 78.87443371 |
| REACTOME_PATHWAY | R-HSA-429947:R-HSA-429947 | 4 | 0.458715596 | 0.098336404 | EIF4E, EIF4A1, CNOT2, CNOT6 | 442 | 23 | 9075 | 3.570725949 | 1 | 0.939881299 | 78.87443371 |
| REACTOME_PATHWAY | R-HSA-203927:R-HSA-203927 | 4 | 0.458715596 | 0.098336404 | POLR2H, DGCR8, POLR2C, POLR2B | 442 | 23 | 9075 | 3.570725949 | 1 | 0.939881299 | 78.87443371 |
| REACTOME_PATHWAY | R-HSA-5654693:R-HSA-5654693 | 4 | 0.458715596 | 0.098336404 | HRAS, FGF23, FGF2, PTPN11 | 442 | 23 | 9075 | 3.570725949 | 1 | 0.939881299 | 78.87443371 |
