## Supplementary Table 8 for "A higher burden of multiple sclerosis genetic risk confers an earlier onset"

**Supplementary Table 8. Pathway Analysis Results for the union (GS3) of the top genes from the SNP-level and Gene-based analyses (see ST4)**

| Column Header | Key |
|---|---|
| **Category** | Pathway Database |
| **Term** | Pathway name/Identifier |
| **Count** | Number of genes in the submitted list in the specified term |
| **%** | Percentage of identified genes in the submitted list associated with the specified term |
| **Pvalue** | Significance level associated with the EASE enrichment score for the term |
| **Genes** | List of genes present in the term |
| **List Total** | Number of genes from the submitted list present in the category |
| **Pop Hits** | Number of genes involved in the specified term (category-specific) |
| **Pop Total** | Number of genes in the human genome background (category-specific) |
| **Fold Enrichment** | Ratio of the proportion of count to list total and population hits to population total |
| **Bonferroni** | Bonferroni adjustment of p-value |
| **Benjamini** | Benjamini adjustment of p-value |
| **FDR** | False Discovery Rate of p-value (percent form) |

| Category | Term | Count | % | PValue | Genes | List Total | Pop Hits | Pop Total | Fold Enrichment | Bonferroni | Benjamini | FDR |
|---|---|---|---|---|---|---|---|---|---|---|---|---|
| KEGG_PATHWAY | hsa05150:Staphylococcus aureus infection | 13 | 0.82592122 | 0.001317045 | HLA-DQB1, MASP1, C4A, HLA-DRB1, C4B, CFB, C5, C1S, C2, HLA-DQA2, HLA-DOB, HLA-DQA1, HLA-DI | 574 | 54 | 6879 | 2.885114208 | 0.307673593 | 0.307673593 | 1.726551035 |
| REACTOME_PATHWAY | R-HSA-174577:R-HSA-174577 | 5 | 0.317662008 | 0.002399156 | C4A, C4B, CFB, C5, C2 | 745 | 8 | 9075 | 7.613255034 | 0.873277173 | 0.873277173 | 3.653740927 |
| REACTOME_PATHWAY | R-HSA-419037:R-HSA-419037 | 10 | 0.635324015 | 0.006054814 | NCAM1, COL9A2, COL9A3, COL6A6, COL6A5, COL3A1, CNTN2, ST8SIA2, CACNA1C, COL5A1 | 745 | 42 | 9075 | 2.900287632 | 0.994608697 | 0.92657451 | 8.981706017 |
| BIOCARTA | h_compPathway:Complement Pathway | 7 | 0.444726811 | 0.006153355 | MASP1, C4A, C4B, CFB, C5, C1S, C2 | 146 | 20 | 1625 | 3.895547945 | 0.74907712 | 0.74907712 | 7.577363584 |
| REACTOME_PATHWAY | R-HSA-202433:R-HSA-202433 | 9 | 0.571791614 | 0.007543799 | FYB, HLA-DQB1, PAK2, HLA-DRB1, PLCG1, CD247, HLA-DQA2, HLA-DQA1, HLA-DRA | 745 | 36 | 9075 | 3.045302013 | 0.998514883 | 0.885908443 | 11.07178474 |
| REACTOME_PATHWAY | R-HSA-1650814:R-HSA-1650814 | 13 | 0.82592122 | 0.007609629 | COL9A2, COL9A3, COLGALT2, COL6A6, COL21A1, ADAMTS14, COL6A5, COL3A1, COL22A1, COL15A1, | 745 | 67 | 9075 | 2.363517981 | 0.998597232 | 0.806470831 | 11.16314538 |
| REACTOME_PATHWAY | R-HSA-167242:R-HSA-167242 | 7 | 0.444726811 | 0.008945057 | POLR2H, NELFCD, SUPT4H1, NELFE, CTDP1, POLR2C, POLR2B | 745 | 23 | 9075 | 3.70732419 | 0.999559395 | 0.786788846 | 12.99763541 |
| KEGG_PATHWAY | hsa05330:Allograft rejection | 9 | 0.571791614 | 0.00976962 | HLA-DQB1, HLA-DRB1, FASLG, HLA-C, CD40, HLA-DQA2, HLA-DOB, HLA-DQA1, HLA-DRA | 574 | 37 | 6879 | 2.915105 | 0.935373196 | 0.745781975 | 12.16770176 |
| KEGG_PATHWAY | hsa05310:Asthma | 8 | 0.508259212 | 0.010043273 | FCER1A, HLA-DQB1, HLA-DRB1, CD40, HLA-DQA2, HLA-DOB, HLA-DQA1, HLA-DRA | 574 | 30 | 6879 | 3.195818815 | 0.940169469 | 0.608882165 | 12.48792638 |
| KEGG_PATHWAY | hsa04014:Ras signaling pathway | 30 | 1.905972046 | 0.012222755 | HRAS, FGFR3, FGF14, EFNA1, EFNA3, FASLG, RGL2, JMJD7-PLA2G4B, GRIN2B, PAK2, PLA2G12B, PDG | 574 | 226 | 6879 | 1.590839012 | 0.967650634 | 0.575901787 | 15.00006326 |
| KEGG_PATHWAY | hsa04974:Protein digestion and absorption | 15 | 0.952986023 | 0.013358314 | CELA3A, SLC8A1, COL21A1, ATP1B2, SLC9A3, COL3A1, COL22A1, COL15A1, COL5A1, COL9A2, COL9A | 574 | 88 | 6879 | 2.042781913 | 0.976531277 | 0.52783054 | 16.28236858 |
| KEGG_PATHWAY | hsa04514:Cell adhesion molecules (CAMs) | 21 | 1.334180432 | 0.013500959 | HLA-DQB1, PTPRM, HLA-DRB1, NRXN3, SELL, NLGN1, HLA-C, CD40, NRXN1, CDH3, HLA-DQA2, HLA-D | 574 | 142 | 6879 | 1.772329097 | 0.977459151 | 0.468510066 | 16.44217603 |
| REACTOME_PATHWAY | R-HSA-389948:R-HSA-389948 | 7 | 0.444726811 | 0.016534101 | HLA-DQB1, HLA-DRB1, CD247, HLA-DQA2, HLA-DQA1, HLA-DRA, PTPN11 | 745 | 26 | 9075 | 3.279556014 | 0.999999407 | 0.908342678 | 22.76763837 |
| KEGG_PATHWAY | hsa05332:Graft-versus-host disease | 8 | 0.508259212 | 0.016964141 | HLA-DQB1, HLA-DRB1, FASLG, HLA-C, HLA-DQA2, HLA-DOB, HLA-DQA1, HLA-DRA | 574 | 33 | 6879 | 2.905289832 | 0.991550101 | 0.494366392 | 20.2365983 |
| REACTOME_PATHWAY | R-HSA-3928665:R-HSA-3928665 | 10 | 0.635324015 | 0.018943536 | AP2A2, CLTA, EFNA1, EFNA3, APH1B, EFNA5, EFNA4, VAV2, EPHB3, EPHB1 | 745 | 50 | 9075 | 2.436241611 | 0.999999928 | 0.904599865 | 25.64818872 |
| KEGG_PATHWAY | hsa04940:Type I diabetes mellitus | 9 | 0.571791614 | 0.020705812 | HLA-DQB1, HLA-DRB1, FASLG, HLA-C, HLA-DQA2, HLA-DOB, HLA-DQA1, LTA, HLA-DRA | 574 | 42 | 6879 | 2.568068691 | 0.997084075 | 0.517944496 | 24.15671972 |
| BIOCARTA | h_lectinPathway:Lectin Induced Complement Pathway | 5 | 0.317662008 | 0.022969103 | MASP1, C4A, C4B, C5, C2 | 146 | 13 | 1625 | 4.280821918 | 0.994511436 | 0.925915159 | 25.66962815 |
| REACTOME_PATHWAY | R-HSA-977606:R-HSA-977606 | 7 | 0.444726811 | 0.023535655 | C8B, C4A, C4B, CFB, C5, C4BPA, C2 | 745 | 28 | 9075 | 3.045302013 | 0.999999999 | 0.922720924 | 30.86196884 |
| KEGG_PATHWAY | hsa05169:Epstein-Barr virus infection | 18 | 1.143583227 | 0.023858967 | HLA-DQB1, HLA-DRB1, PIK3CB, NFKBIB, HLA-C, RB1, CD40, HLA-DQA2, HLA-DQA1, NEDD4, PSMD1, J | 574 | 122 | 6879 | 1.768178443 | 0.998814213 | 0.526968961 | 27.32115896 |
| KEGG_PATHWAY | hsa04015:Rap1 signaling pathway | 27 | 1.715374841 | 0.025787957 | HRAS, FGFR3, FGF14, EFNA1, EFNA3, APBB1IP, GRIN2B, PFN4, PDGFC, INSR, FGF2, AKT3, FYB, MAGI | 574 | 210 | 6879 | 1.540841215 | 0.999317151 | 0.517572025 | 29.19641165 |
| KEGG_PATHWAY | hsa05322:Systemic lupus erythematosus | 19 | 1.207115629 | 0.028567002 | HLA-DQB1, HIST4H4, C4A, HLA-DRB1, C4B, C5, SSB, H2AFJ, CD40, C1S, HLA-DQA2, HLA-DQA1, C8B, G | 574 | 134 | 6879 | 1.699269333 | 0.999692259 | 0.520549094 | 31.81951192 |
| REACTOME_PATHWAY | R-HSA-202430:R-HSA-202430 | 6 | 0.381194409 | 0.029777702 | HLA-DQB1, HLA-DRB1, CD247, HLA-DQA2, HLA-DQA1, HLA-DRA | 745 | 22 | 9075 | 3.322147651 | 1 | 0.944348719 | 37.40221966 |
| REACTOME_PATHWAY | R-HSA-1257604:R-HSA-1257604 | 13 | 0.82592122 | 0.031562491 | FGFR3, KLB, ERBB4, PIK3CB, FGF23, RICTOR, TRAT1, PTPN11, TNRC6C, HBEGF, TNRC6B, FGF2, AKT3 | 745 | 81 | 9075 | 1.9550087 | 1 | 0.936589996 | 39.16301456 |
| REACTOME_PATHWAY | R-HSA-5576892:R-HSA-5576892 | 9 | 0.571791614 | 0.031627864 | FGF14, CAMK2G, CACNG6, CACNG5, CACNG2, SCN7A, CACNA1C, SCN4A, CALM2 | 745 | 46 | 9075 | 2.383279837 | 1 | 0.918948615 | 39.22662029 |
| BIOCARTA | h_classicPathway:Classical Complement Pathway | 5 | 0.317662008 | 0.038114048 | C4A, C4B, C5, C1S, C2 | 146 | 15 | 1625 | 3.710045662 | 0.999834163 | 0.945059309 | 39.10935301 |
| KEGG_PATHWAY | hsa04672:Intestinal immune network for IgA production | 9 | 0.571791614 | 0.03834821 | HLA-DQB1, HLA-DRB1, TNFRSF13B, CD40, HLA-DQA2, HLA-DOB, CCL28, HLA-DQA1, HLA-DRA | 574 | 47 | 6879 | 2.294869894 | 0.99998172 | 0.597130042 | 40.35420477 |
| BIOCARTA | h_bbcellPathway:Bystander B Cell Activation | 4 | 0.254129606 | 0.039137337 | HLA-DRB1, FASLG, CD40, HLA-DRA | 146 | 9 | 1625 | 4.946727549 | 0.999869342 | 0.893086191 | 39.93116792 |
| KEGG_PATHWAY | hsa04722:Neurotrophin signaling pathway | 17 | 1.080050826 | 0.039849201 | HRAS, NTF3, PIK3CB, NFKBIB, CAMK2G, FASLG, TP73, PTPN11, MAP3K5, PLCG1, SH2B3, RAP1A, SH2I | 574 | 120 | 6879 | 1.697778746 | 0.999988178 | 0.582191401 | 41.57283971 |
| REACTOME_PATHWAY | R-HSA-3000178:R-HSA-3000178 | 12 | 0.762388818 | 0.041072005 | ASPN, NCAM1, COL9A2, APP, COL9A3, TNXB, COL6A6, COL6A5, ITGA8, COL3A1, COL5A1, FN1 | 745 | 75 | 9075 | 1.948993289 | 1 | 0.950493393 | 47.789408 |
| KEGG_PATHWAY | hsa04145:Phagosome | 20 | 1.27064803 | 0.042353488 | HLA-DQB1, ATP6V0E1, HLA-DRB1, MRC2, HLA-C, COLEC12, COLEC11, HLA-DQA2, HLA-DQA1, LAMP1 | 574 | 150 | 6879 | 1.597909408 | 0.999994295 | 0.577869137 | 43.55492642 |
| REACTOME_PATHWAY | R-HSA-418886:R-HSA-418886 | 4 | 0.254129606 | 0.042576153 | DCC, UNC5C, NTN1, PTPN11 | 745 | 10 | 9075 | 4.872483221 | 1 | 0.943769386 | 49.04413091 |
| KEGG_PATHWAY | hsa04330:Notch signaling pathway | 9 | 0.571791614 | 0.042804826 | CIR1, CTBP2, MAML1, DTX3L, NOTCH4, APH1B, MAML3, RBPJ, NCOR2 | 574 | 48 | 6879 | 2.247060105 | 0.999994998 | 0.55679025 | 43.90546991 |
| REACTOME_PATHWAY | R-HSA-186797:R-HSA-186797 | 7 | 0.444726811 | 0.042916121 | COL9A2, COL9A3, COL6A6, COL6A5, COL3A1, PDGFC, COL5A1 | 745 | 32 | 9075 | 2.664639262 | 1 | 0.932425042 | 49.32378789 |
| REACTOME_PATHWAY | R-HSA-167158:R-HSA-167158 | 7 | 0.444726811 | 0.048963019 | POLR2H, NELFCD, SUPT4H1, NELFE, CTDP1, POLR2C, POLR2B | 745 | 33 | 9075 | 2.583892617 | 1 | 0.94376787 | 54.06430063 |
| REACTOME_PATHWAY | R-HSA-113418:R-HSA-113418 | 7 | 0.444726811 | 0.048963019 | POLR2H, NELFCD, SUPT4H1, NELFE, CTDP1, POLR2C, POLR2B | 745 | 33 | 9075 | 2.583892617 | 1 | 0.94376787 | 54.06430063 |
| REACTOME_PATHWAY | R-HSA-202427:R-HSA-202427 | 6 | 0.381194409 | 0.049087347 | HLA-DQB1, HLA-DRB1, CD247, HLA-DQA2, HLA-DQA1, HLA-DRA | 745 | 25 | 9075 | 2.923489933 | 1 | 0.93315658 | 54.15726678 |
| REACTOME_PATHWAY | R-HSA-877300:R-HSA-877300 | 13 | 0.82592122 | 0.054635218 | NCAM1, HLA-DQB1, SP100, HLA-DRB1, CAMK2G, OAS3, JAK1, OAS1, HLA-C, OAS2, HLA-DQA2, HLA-D | 745 | 88 | 9075 | 1.799496644 | 1 | 0.941706828 | 58.13102266 |
| REACTOME_PATHWAY | R-HSA-376172:R-HSA-376172 | 4 | 0.254129606 | 0.055067673 | DCC, DSCAML1, NTN1, DSCAM | 745 | 11 | 9075 | 4.429530201 | 1 | 0.933211694 | 58.42683105 |
| REACTOME_PATHWAY | R-HSA-2682334:R-HSA-2682334 | 6 | 0.381194409 | 0.05683812 | EFNA1, EFNA3, EFNA5, EFNA4, EPHB3, EPHB1 | 745 | 26 | 9075 | 2.811048012 | 1 | 0.929255457 | 59.61758641 |
| KEGG_PATHWAY | hsa04070:Phosphatidylinositol signaling system | 14 | 0.889453621 | 0.061990804 | PIK3CB, PIK3C2B, PIP5K1C, ITPR2, PLCG1, INPP5K, DGKG, DGKZ, INPP4B, ITPK1, IP6K1, INPP5A, MTM | 574 | 98 | 6879 | 1.712045794 | 0.999999982 | 0.672386919 | 57.07451936 |
| REACTOME_PATHWAY | R-HSA-3928663:R-HSA-3928663 | 7 | 0.444726811 | 0.062555153 | EFNA1, EFNA3, VEGFA, EFNA5, MYH14, EFNA4, MYL12A | 745 | 35 | 9075 | 2.436241611 | 1 | 0.937817781 | 63.24848219 |
| KEGG_PATHWAY | hsa04024:cAMP signaling pathway | 24 | 1.524777637 | 0.063408262 | PLD2, ATP1B2, PIK3CB, CAMK2G, PDE3B, GABBR2, CNGB1, VAV2, CNGA3, GLI3, CNGA1, GLI1, ADRB | 574 | 198 | 6879 | 1.452644916 | 0.999999988 | 0.658735915 | 57.92838237 |
| KEGG_PATHWAY | hsa05320:Autoimmune thyroid disease | 9 | 0.571791614 | 0.063983153 | HLA-DQB1, HLA-DRB1, FASLG, HLA-C, CD40, HLA-DQA2, HLA-DOB, HLA-DQA1, HLA-DRA | 574 | 52 | 6879 | 2.074209327 | 0.99999999 | 0.641163334 | 58.26388899 |
| KEGG_PATHWAY | hsa00590:Arachidonic acid metabolism | 10 | 0.635324015 | 0.064004165 | GGT5, JMJD7-PLA2G4B, GPX6, PLB1, PTGDS, CYP2B6, GPX5, PLA2G12B, FAM213B, PLA2G4B | 574 | 61 | 6879 | 1.964642714 | 0.99999999 | 0.621400413 | 58.27626854 |
| KEGG_PATHWAY | hsa04151:PI3K-Akt signaling pathway | 38 | 2.414231258 | 0.06415422 | HRAS, FGFR3, FGF14, EFNA1, OSMR, EFNA3, COL3A1, ITGA11, FASLG, CASP9, COL6A6, COL6A5, ATF6 | 574 | 345 | 6879 | 1.320012119 | 0.999999991 | 0.603448321 | 58.36457719 |
| REACTOME_PATHWAY | R-HSA-442982:R-HSA-442982 | 5 | 0.317662008 | 0.064564262 | HRAS, GRIN2B, RASGRF2, CAMK2G, CALM2 | 745 | 19 | 9075 | 3.205581067 | 1 | 0.934994132 | 64.4502341 |
| KEGG_PATHWAY | hsa00562:Inositol phosphate metabolism | 11 | 0.698856417 | 0.068305099 | TPI1, INPP5K, PLCG1, PIK3CB, PIK3C2B, PLCH1, PIP5K1C, INPP4B, ITPK1, INPP5A, MTMR4 | 574 | 71 | 6879 | 1.85672572 | 0.999999997 | 0.609357579 | 60.73998919 |
| REACTOME_PATHWAY | R-HSA-167243:R-HSA-167243 | 7 | 0.444726811 | 0.070105589 | POLR2H, NELFCD, SUPT4H1, NELFE, CTDP1, POLR2C, POLR2B | 745 | 36 | 9075 | 2.368568233 | 1 | 0.941649821 | 67.57705217 |
| REACTOME_PATHWAY | R-HSA-2514859:R-HSA-2514859 | 7 | 0.444726811 | 0.070105589 | FNTB, METAP2, CHURC1-FNTB, GRK7, CNGB1, CNGA1, CALM2 | 745 | 36 | 9075 | 2.368568233 | 1 | 0.941649821 | 67.57705217 |
| REACTOME_PATHWAY | R-HSA-167238:R-HSA-167238 | 7 | 0.444726811 | 0.070105589 | POLR2H, NELFCD, SUPT4H1, NELFE, CTDP1, POLR2C, POLR2B | 745 | 36 | 9075 | 2.368568233 | 1 | 0.941649821 | 67.57705217 |
| KEGG_PATHWAY | hsa05220:Chronic myeloid leukemia | 11 | 0.698856417 | 0.073718237 | HRAS, CTBP2, PIK3CB, RB1, RUNX1, MECOM, CDK4, MYC, AKT3, SHC4, PTPN11 | 574 | 72 | 6879 | 1.830937863 | 0.999999999 | 0.621346295 | 63.64967294 |
| REACTOME_PATHWAY | R-HSA-5654706:R-HSA-5654706 | 5 | 0.317662008 | 0.075685902 | HRAS, FGFR3, FGF23, FGF2, PTPN11 | 745 | 20 | 9075 | 3.045302013 | 1 | 0.947283156 | 70.46442578 |
| REACTOME_PATHWAY | R-HSA-1442490:R-HSA-1442490 | 10 | 0.635324015 | 0.075856304 | COL9A2, COL9A3, COL6A6, COL6A5, COL3A1, MMP8, COL15A1, COL25A1, COL5A1, COL10A1 | 745 | 64 | 9075 | 1.903313758 | 1 | 0.940798558 | 70.54868829 |
| REACTOME_PATHWAY | R-HSA-69052:R-HSA-69052 | 3 | 0.190597205 | 0.080666664 | MCM7, MCM4, MCM6 | 745 | 6 | 9075 | 6.090604027 | 1 | 0.944604882 | 72.83664856 |
| REACTOME_PATHWAY | R-HSA-1912408:R-HSA-1912408 | 6 | 0.381194409 | 0.084033947 | TNRC6C, MAML1, NOTCH4, MAML3, TNRC6B, RBPJ | 745 | 29 | 9075 | 2.520249942 | 1 | 0.945161302 | 74.33810591 |
| REACTOME_PATHWAY | R-HSA-425561:R-HSA-425561 | 4 | 0.254129606 | 0.084534597 | SLC8A1, SLC24A4, SLC24A1, CALM2 | 745 | 13 | 9075 | 3.748064017 | 1 | 0.93998986 | 74.55459672 |
| REACTOME_PATHWAY | R-HSA-350054:R-HSA-350054 | 4 | 0.254129606 | 0.084534597 | MAML1, NOTCH4, MAML3, RBPJ | 745 | 13 | 9075 | 3.748064017 | 1 | 0.93998986 | 74.55459672 |
| REACTOME_PATHWAY | R-HSA-190372:R-HSA-190372 | 4 | 0.254129606 | 0.084534597 | GALNT3, FGFR3, FGF23, FGF2 | 745 | 13 | 9075 | 3.748064017 | 1 | 0.93998986 | 74.55459672 |
| REACTOME_PATHWAY | R-HSA-5654227:R-HSA-5654227 | 4 | 0.254129606 | 0.084534597 | FGFR3, PLCG1, FGF23, FGF2 | 745 | 13 | 9075 | 3.748064017 | 1 | 0.93998986 | 74.55459672 |

| | | | | | | | | | | | | |
|---|---|---|---|---|---|---|---|---|---|---|---|---|
| REACTOME_PATHWAY | R-HSA-5578775:R-HSA-5578775 | 9 | 0.571791614 | 0.084667826 | SLC8A1, ATP2B4, ATP1B2, CAMK2G, RYR3, RYR2, CALM2, ITPR2, DMPK | 745 | 56 | 9075 | 1.957694151 | 1 | 0.933943116 | 74.61191894 |
| REACTOME_PATHWAY | R-HSA-167287:R-HSA-167287 | 7 | 0.444726811 | 0.086708222 | POLR2H, NELFCD, SUPT4H1, NELFE, CTDP1, POLR2C, POLR2B | 745 | 38 | 9075 | 2.243906747 | 1 | 0.932100066 | 75.47485375 |
| REACTOME_PATHWAY | R-HSA-167290:R-HSA-167290 | 7 | 0.444726811 | 0.086708222 | POLR2H, NELFCD, SUPT4H1, NELFE, CTDP1, POLR2C, POLR2B | 745 | 38 | 9075 | 2.243906747 | 1 | 0.932100066 | 75.47485375 |
| REACTOME_PATHWAY | R-HSA-5218921:R-HSA-5218921 | 5 | 0.317662008 | 0.0877009 | HRAS, PLCG1, VEGFA, CALM2, ITPR2 | 745 | 21 | 9075 | 2.900287632 | 1 | 0.928010736 | 75.88468742 |
| KEGG_PATHWAY | hsa05223:Non-small cell lung cancer | 9 | 0.571791614 | 0.09070558 | FHIT, HRAS, CASP9, PLCG1, PIK3CB, RB1, CDK4, ALK, AKT3 | 574 | 56 | 6879 | 1.926051518 | 1 | 0.684450175 | 71.53718018 |
| REACTOME_PATHWAY | R-HSA-2894862:R-HSA-2894862 | 9 | 0.571791614 | 0.091805093 | TBL1XR1, HDAC3, MAML1, APH1B, NEURL1B, MAML3, RBPJ, MYC, NCOR2 | 745 | 57 | 9075 | 1.92334864 | 1 | 0.930847762 | 77.51208597 |
| REACTOME_PATHWAY | R-HSA-936837:R-HSA-936837 | 9 | 0.571791614 | 0.091805093 | ATP2C2, ATP2B4, ATP1B2, CAMK2G, ATP10A, ATP11A, ATP10D, ATP8B3, CALM2 | 745 | 57 | 9075 | 1.92334864 | 1 | 0.930847762 | 77.51208597 |
| REACTOME_PATHWAY | R-HSA-2644606:R-HSA-2644606 | 9 | 0.571791614 | 0.091805093 | TBL1XR1, HDAC3, MAML1, APH1B, NEURL1B, MAML3, RBPJ, MYC, NCOR2 | 745 | 57 | 9075 | 1.92334864 | 1 | 0.930847762 | 77.51208597 |
| REACTOME_PATHWAY | R-HSA-216083:R-HSA-216083 | 12 | 0.762388818 | 0.091953413 | COL9A2, COL9A3, COL6A6, ICAM4, COL6A5, ITGAE, ITGA8, COL3A1, ITGA11, COL5A1, COL10A1, FN1 | 745 | 86 | 9075 | 1.699703449 | 1 | 0.925156169 | 77.56892801 |
| KEGG_PATHWAY | hsa04612:Antigen processing and presentation | 11 | 0.698856417 | 0.09799326 | HLA-DQB1, HLA-DRB1, TAP2, KIR3DL3, HLA-C, HLA-DQA2, HLA-DOB, HLA-DQA1, KIR2DL4, HLA-DRA, | 574 | 76 | 6879 | 1.734572712 | 1 | 0.698481665 | 74.40856175 |
| KEGG_PATHWAY | hsa05416:Viral myocarditis | 9 | 0.571791614 | 0.098258155 | HLA-DQB1, CASP9, HLA-DRB1, HLA-C, CD40, HLA-DQA2, HLA-DOB, HLA-DQA1, HLA-DRA | 574 | 57 | 6879 | 1.892261141 | 1 | 0.684704615 | 74.50770184 |
