## Supplementary Table 9 for "A higher burden of multiple sclerosis genetic risk confers an earlier onset"

**Supplementary Table 9. Functional Annotation Clustering Results for the union (GS3) of the top genes from the SNP-level and Gene-based analyses (see ST4)**

| Column Header | Key |
|---|---|
| **Annotation Cluster** | Name of cluster, sorted by descending Enrichment score |
| **Enrichment Score** | EASE enrichment score for functional annotation cluster |
| **Category** | Pathway Database |
| **Term** | Pathway name/Identifier |
| **Count** | Number of genes in the submitted list in the specified term |
| **%** | Percentage of identified genes in the submitted list associated with the specified term |
| **PValue** | Significance level associated with the EASE enrichment score for the term |
| **Genes** | List of genes present in the term |
| **List Total** | Number of genes from the submitted list present in the category |
| **Pop Hits** | Number of genes involved in the specified term (category-specific) |
| **Pop Total** | Number of genes in the human genome background (category-specific) |
| **Fold Enrichment** | Ratio of the proportion of count to list total and population hits to population total |
| **Bonferroni** | Bonferroni adjustment of p-value |
| **Benjamini** | Benjamini adjustment of p-value |
| **FDR** | False Discovery Rate of p-value (percent form) |

Annotation Cluster 1 **Enrichment Score: 3.8978262119731335**

| Category | Term | Count | % | PValue | Genes | List Total | Pop Hits | Pop Total | Fold Enrichment | Bonferroni | Benjamini | FDR |
|---|---|---|---|---|---|---|---|---|---|---|---|---|
| GOTERM_CC_DIRECT | GO:0005886~plasma membrane | 383 | 24.33290978 | 5.74E-05 | SLC9A9, XRCC5, HRAS, CHMP3, ATP1B2, EFNA1, OSMR, SLC9A3, EFNA3, UTRN, SYT6, ZNRF2, APP, AT | 1425 | 4121 | 18224 | 1.18857065 | 0.038655922 | 0.038655922 | 0.086284383 |
| UP_KEYWORDS | Membrane | 626 | 39.77128335 | 1.53E-04 | SLC9A9, HRAS, OSMR, VKORC1L1, RUSC1, SLC9A3, VPS52, APP, GRIN2B, TECRL, RAB27B, ROS1, ITFG | 1535 | 7494 | 20581 | 1.120001843 | 0.072781834 | 0.006277414 | 0.220047629 |
| UP_KEYWORDS | Cell membrane | 287 | 18.23379924 | 2.30E-04 | HRAS, ATP1B2, EFNA1, SLC9A3, RUSC1, EFNA3, UTRN, SYT6, MEGF11, ZNRF2, ATP2B4, GRIN2B, S1PI | 1535 | 3175 | 20581 | 1.211982251 | 0.107341223 | 0.008696647 | 0.330474651 |

Annotation Cluster 2 **Enrichment Score: 3.651023631992672**

| Category | Term | Count | % | PValue | Genes | List Total | Pop Hits | Pop Total | Fold Enrichment | Bonferroni | Benjamini | FDR |
|---|---|---|---|---|---|---|---|---|---|---|---|---|
| UP_KEYWORDS | Cell junction | 86 | 5.463786531 | 1.25E-06 | KCNC2, CPEB3, RUSC1, UTRN, COPS4, PIP5K1C, DSTYK, SYT6, GABBR2, FER, ZNRF1, GJA4, ZNRF2, GJ/ | 1535 | 675 | 20581 | 1.708255278 | 6.15E-04 | 1.54E-04 | 0.00179425 |
| UP_KEYWORDS | Synapse | 50 | 3.176620076 | 2.46E-05 | STON2, KCNC2, SYNDIG1, CPEB3, RUSC1, COPS4, UTRN, DSCAML1, SYT6, GABBR2, ZNRF1, GSG1L, RI | 1535 | 357 | 20581 | 1.877845601 | 0.01205396 | 0.002422501 | 0.035346789 |
| GOTERM_CC_DIRECT | GO:0030054~cell junction | 60 | 3.811944091 | 1.02E-04 | KCNC2, XRCC4, CPEB3, RUSC1, UTRN, COPS4, DSTYK, SYT6, GABBR2, FER, ZNRF1, IL31RA, ZNRF2, HN | 1425 | 459 | 18224 | 1.671734893 | 0.067383482 | 0.034279276 | 0.152634718 |
| UP_KEYWORDS | Postsynaptic cell membrane | 24 | 1.524777637 | 0.007359395 | KCNC2, GABRA2, DLGAP1, GABRG3, SYNDIG1, CPEB3, DLGAP2, RUSC1, UTRN, CACNG5, NLGN1, PSD | 1535 | 179 | 20581 | 1.797696213 | 0.973789574 | 0.09881585 | 10.07186658 |
| GOTERM_CC_DIRECT | GO:0045211~postsynaptic membrane | 26 | 1.65184244 | 0.024217148 | KCNC2, GABRA2, GABRG3, DLGAP1, SYNDIG1, DLGAP2, CPEB3, RUSC1, UTRN, NLGN1, PSD3, CACNG | 1425 | 211 | 18224 | 1.575867631 | 0.999999952 | 0.429589046 | 30.84201449 |

Annotation Cluster 3 Enrichment Score: 2.9994206148541176

| Category | Term | Count | % | PValue | Genes | List Total | Pop Hits | Pop Total | Fold Enrichment | Bonferroni | Benjamini | FDR |
|---|---|---|---|---|---|---|---|---|---|---|---|---|
| UP_KEYWORDS | SH3 domain | 32 | 2.033036849 | 3.04E-04 | FRK, FGR, MYO7B, RUSC1, ASAP1, MACC1, AMPH, SORBS2, DLG2, ARHGAP9, FYB, OBSCN, BAIAP2L1 | 1535 | 215 | 20581 | 1.995582153 | 0.139140637 | 0.010644664 | 0.435812758 |
| UP_SEQ_FEATURE | domain:SH3 | 27 | 1.715374841 | 5.81E-04 | FRK, FGR, RUSC1, ASAP1, MACC1, MCF2L, AMPH, DLG2, ARHGAP9, FYB, OBSCN, BAIAP2L1, BAIAP2, | 1499 | 175 | 20063 | 2.064999523 | 0.906127004 | 0.166386637 | 1.075702475 |
| INTERPRO | IPR001452:Src homology-3 domain | 32 | 2.033036849 | 8.55E-04 | FRK, FGR, MYO7B, RUSC1, ASAP1, MCF2L, AMPH, SORBS2, DLG2, ARHGAP9, FYB, OBSCN, BAIAP2L1, | 1452 | 218 | 18559 | 1.876209973 | 0.824595 | 0.293993997 | 1.463443477 |
| SMART | SM00326:SH3 | 30 | 1.905972046 | 0.00666089 | FRK, FGR, MYO7B, RUSC1, ASAP1, MCF2L, AMPH, SORBS2, DLG2, FYB, BAIAP2L1, BAIAP2, MYO1E, RI | 878 | 206 | 10057 | 1.668122609 | 0.937557982 | 0.500116003 | 8.945921022 |

Annotation Cluster 4 Enrichment Score: 2.8455446039146026

| Category | Term | Count | % | PValue | Genes | List Total | Pop Hits | Pop Total | Fold Enrichment | Bonferroni | Benjamini | FDR |
|---|---|---|---|---|---|---|---|---|---|---|---|---|
| INTERPRO | IPR001849:Pleckstrin homology domain | 38 | 2.414231258 | 6.32E-04 | PLEKHM2, ASAP1, AKAP13, AFAP1L2, APBB1IP, MCF2L, ADAP1, PLCL1, FAM109A, PLCH1, SH2B3, SH: | 1452 | 271 | 18559 | 1.792265154 | 0.723708267 | 0.274993379 | 1.083522682 |
| INTERPRO | IPR011993:Pleckstrin homology-like domain | 53 | 3.367217281 | 9.36E-04 | PLEKHM2, MYO7B, NBEA, MCF2L, DAB2, DAB1, FAM109A, AGAP1, DOCK10, AKT3, ARHGAP9, SHC4, | 1452 | 427 | 18559 | 1.586484926 | 0.851006664 | 0.271894228 | 1.599546182 |
| SMART | SM00233:PH | 38 | 2.414231258 | 0.002562552 | PLEKHM2, ASAP1, AKAP13, AFAP1L2, APBB1IP, MCF2L, ADAP1, PLCL1, FAM109A, PLCH1, SH2B3, SH: | 878 | 264 | 10057 | 1.648745427 | 0.655211444 | 0.298785404 | 3.534059519 |
| UP_SEQ_FEATURE | domain:PH | 32 | 2.033036849 | 0.002736631 | PLEKHM2, ASAP1, AKAP13, SLC26A10, APBB1IP, MCF2L, PLCL1, FAM109A, PLCH1, SH2B3, SH2B2, AG | 1499 | 245 | 20063 | 1.748147745 | 0.99998579 | 0.384478841 | 4.97451739 |

Annotation Cluster 5 Enrichment Score: 2.70711641328601

| Category | Term | Count | % | PValue | Genes | List Total | Pop Hits | Pop Total | Fold Enrichment | Bonferroni | Benjamini | FDR |
|---|---|---|---|---|---|---|---|---|---|---|---|---|
| UP_SEQ_FEATURE | domain:Fibronectin type-III 3 | 23 | 1.461245235 | 1.44E-07 | DCC, OBSCN, PTPRM, TNXB, TNXA, OSMR, RIMBP2, SDK1, DSCAML1, PTPRT, PTPRO, IL31RA, NRCAN | 1499 | 84 | 20063 | 3.664736809 | 5.86E-04 | 2.93E-04 | 2.68E-04 |
| UP_SEQ_FEATURE | domain:Fibronectin type-III 4 | 19 | 1.207115629 | 4.75E-07 | DCC, OBSCN, PTPRM, TNXB, TNXA, OSMR, SDK1, DSCAML1, PTPRT, PTPRO, IL31RA, NRCAM, MYOM | 1499 | 63 | 20063 | 4.036521702 | 0.001933088 | 6.45E-04 | 8.85E-04 |
| UP_SEQ_FEATURE | domain:Fibronectin type-III 2 | 27 | 1.715374841 | 4.98E-06 | DCC, OSMR, DSCAML1, EPHB3, EPHB1, IL31RA, NRCAM, MYOM2, ROBO1, ROBO2, INSR, ROS1, DSCA | 1499 | 133 | 20063 | 2.717104636 | 0.020067079 | 0.004046024 | 0.00926652 |
| UP_SEQ_FEATURE | domain:Fibronectin type-III 1 | 27 | 1.715374841 | 5.74E-06 | DCC, OSMR, DSCAML1, EPHB3, EPHB1, IL31RA, NRCAM, MYOM2, ROBO1, ROBO2, INSR, ROS1, DSCA | 1499 | 134 | 20063 | 2.696827736 | 0.023118689 | 0.003890764 | 0.010692209 |
| SMART | SM00060:FN3 | 30 | 1.905972046 | 3.31E-05 | DCC, OSMR, DSCAML1, EPHB3, EPHB1, IL31RA, NRCAM, MYOM2, ROBO1, ROBO2, INSR, ROS1, FN1, | 878 | 150 | 10057 | 2.290888383 | 0.013633442 | 0.013633442 | 0.046373282 |
| INTERPRO | IPR003961:Fibronectin, type III | 32 | 2.033036849 | 3.50E-04 | DCC, OSMR, DSCAML1, ASTN2, EPHB3, EPHB1, IL31RA, NRCAM, MYOM2, ROBO1, IL10RA, ROBO2, IN | 1452 | 207 | 18559 | 1.975911952 | 0.508931419 | 0.299237143 | 0.600519348 |
| UP_SEQ_FEATURE | domain:Fibronectin type-III 5 | 12 | 0.762388818 | 4.51E-04 | DCC, NRCAM, TNXB, MYOM2, TNXA, SDK1, DSCAML1, PTPRO, ROS1, DSCAM, FN1, IL31RA | 1499 | 46 | 20063 | 3.491545088 | 0.840949456 | 0.142051413 | 0.836962336 |
| UP_SEQ_FEATURE | domain:Fibronectin type-III 6 | 9 | 0.571791614 | 0.001972411 | DCC, TNXB, TNXA, SDK1, DSCAML1, PTPRO, ROS1, DSCAM, FN1 | 1499 | 32 | 20063 | 3.764322048 | 0.999678194 | 0.360298424 | 3.609444792 |
| UP_SEQ_FEATURE | domain:Ig-like C2-type 5 | 12 | 0.762388818 | 0.002179772 | NCAM1, MYOM2, ROBO1, CNTN2, SDK1, CNTN1, DSCAML1, ROBO2, CNTN4, PALLD, MYLK, DSCAM | 1499 | 55 | 20063 | 2.920201346 | 0.999861951 | 0.35878791 | 3.981661788 |
| UP_SEQ_FEATURE | domain:Ig-like C2-type 4 | 13 | 0.82592122 | 0.010308277 | DCC, SDK1, DSCAML1, PALLD, NCAM1, MYOM2, ROBO1, CNTN2, CNTN1, CNTN4, ROBO2, MYLK, DSC | 1499 | 76 | 20063 | 2.28941224 | 1 | 0.661130877 | 17.54628962 |
| UP_SEQ_FEATURE | domain:Fibronectin type-III 7 | 6 | 0.381194409 | 0.034910027 | TNXB, TNXA, SDK1, PTPRO, ROS1, FN1 | 1499 | 25 | 20063 | 3.212221481 | 1 | 0.873506966 | 48.39911556 |
| UP_SEQ_FEATURE | domain:Fibronectin type-III 8 | 6 | 0.381194409 | 0.034910027 | TNXB, TNXA, SDK1, PTPRO, ROS1, FN1 | 1499 | 25 | 20063 | 3.212221481 | 1 | 0.873506966 | 48.39911556 |
| UP_SEQ_FEATURE | domain:Fibronectin type-III 9 | 5 | 0.317662008 | 0.040730353 | TNXB, TNXA, SDK1, ROS1, FN1 | 1499 | 18 | 20063 | 3.717848936 | 1 | 0.895465471 | 53.89571119 |
| UP_SEQ_FEATURE | domain:Fibronectin type-III 11 | 4 | 0.254129606 | 0.043558648 | TNXB, TNXA, SDK1, FN1 | 1499 | 11 | 20063 | 4.867002244 | 1 | 0.899352497 | 56.36206388 |
| UP_SEQ_FEATURE | domain:Fibronectin type-III 10 | 4 | 0.254129606 | 0.043558648 | TNXB, TNXA, SDK1, FN1 | 1499 | 11 | 20063 | 4.867002244 | 1 | 0.899352497 | 56.36206388 |
| UP_SEQ_FEATURE | domain:Fibronectin type-III 13 | 4 | 0.254129606 | 0.043558648 | TNXB, TNXA, SDK1, FN1 | 1499 | 11 | 20063 | 4.867002244 | 1 | 0.899352497 | 56.36206388 |
| UP_SEQ_FEATURE | domain:Fibronectin type-III 16 | 3 | 0.190597205 | 0.115475174 | TNXB, TNXA, FN1 | 1499 | 8 | 20063 | 5.019096064 | 1 | 0.986545152 | 89.81978955 |
| UP_SEQ_FEATURE | domain:Fibronectin type-III 15 | 3 | 0.190597205 | 0.141381957 | TNXB, TNXA, FN1 | 1499 | 9 | 20063 | 4.461418724 | 1 | 0.992174796 | 94.14703211 |
| UP_SEQ_FEATURE | domain:Fibronectin type-III 14 | 3 | 0.190597205 | 0.141381957 | TNXB, TNXA, FN1 | 1499 | 9 | 20063 | 4.461418724 | 1 | 0.992174796 | 94.14703211 |
| UP_SEQ_FEATURE | domain:Fibronectin type-III 12 | 3 | 0.190597205 | 0.168352902 | TNXB, TNXA, SDK1 | 1499 | 10 | 20063 | 4.015276851 | 1 | 0.995833277 | 96.76935141 |

Annotation Cluster 6 Enrichment Score: 2.7036236930619433

| Category | Term | Count | % | PValue | Genes | List Total | Pop Hits | Pop Total | Fold Enrichment | Bonferroni | Benjamini | FDR |
|---|---|---|---|---|---|---|---|---|---|---|---|---|
| UP_KEYWORDS | Glycoprotein | 400 | 25.41296061 | 9.00E-05 | SLC9A9, MASP1, OSMR, LTBP3, SLC9A3, HMCN2, APP, GRIN2B, ROS1, ITFG1, CDH23, CELA3A, PTPRN | 1535 | 4551 | 20581 | 1.178450239 | 0.043400644 | 0.004917936 | 0.129264988 |
| UP_SEQ_FEATURE | glycosylation site:N-linked (GlcNAc...) | 374 | 23.76111817 | 1.30E-04 | SLC9A9, MASP1, ATP1B2, EFNA1, OSMR, LTBP3, FAM20A, NELL1, B3GALT4, SLC9A3, EFNA3, LUZP2, I | 1499 | 4234 | 20063 | 1.182265425 | 0.411536242 | 0.064131208 | 0.242105368 |
| UP_KEYWORDS | Membrane | 626 | 39.77128335 | 1.53E-04 | SLC9A9, HRAS, OSMR, VKORC1L1, RUSC1, SLC9A3, VPS52, APP, GRIN2B, TECRL, RAB27B, ROS1, ITFG | 1535 | 7494 | 20581 | 1.120001843 | 0.072781834 | 0.006277414 | 0.220047629 |
| UP_SEQ_FEATURE | transmembrane region | 426 | 27.06480305 | 0.002003705 | SLC9A9, VKORC1L1, OSMR, SLC9A3, APP, GRIN2B, TECRL, ROS1, ITFG1, CDH23, PTPRM, TMEM132B, | 1499 | 5056 | 20063 | 1.127708293 | 0.999716778 | 0.349466227 | 3.665706385 |
| UP_SEQ_FEATURE | topological domain:Extracellular | 244 | 15.50190597 | 0.004128957 | TMEM18, ATP1B2, OSMR, SLC9A3, APP, ATP2B4, GRIN2B, S1PR4, CLEC4C, PI16, ROS1, GRID1, CDH2: | 1499 | 2787 | 20063 | 1.171782743 | 0.999999952 | 0.452207751 | 7.414634023 |
| UP_KEYWORDS | Transmembrane helix | 466 | 29.60609911 | 0.004350845 | SLC9A9, OSMR, VKORC1L1, SLC9A3, APP, GRIN2B, TECRL, ROS1, ITFG1, CDH23, PTPRM, TMEM132B, | 1535 | 5634 | 20581 | 1.108988817 | 0.883474652 | 0.073900026 | 6.074304539 |
| UP_KEYWORDS | Transmembrane | 467 | 29.66963151 | 0.004651018 | SLC9A9, OSMR, VKORC1L1, SLC9A3, APP, GRIN2B, TECRL, ROS1, ITFG1, CDH23, PTPRM, TMEM132B, | 1535 | 5651 | 20581 | 1.108025272 | 0.899570638 | 0.073748999 | 6.480455252 |
| UP_SEQ_FEATURE | topological domain:Cytoplasmic | 293 | 18.61499365 | 0.0095743 | TMEM18, ATP1B2, OSMR, VAPB, B3GALT4, SLC9A3, SYT6, HS2ST1, APP, ATP2B4, GRIN2B, S1PR4, CL | 1499 | 3456 | 20063 | 1.134718477 | 1 | 0.643406229 | 16.40023541 |
| GOTERM_CC_DIRECT | GO:0016021~integral component of membrane | 421 | 26.74714104 | 0.161538689 | SLC9A9, VKORC1L1, OSMR, SLC9A3, APP, TECRL, ROS1, ITFG1, CDH23, PTPRM, TMEM132B, CDHR4, | 1425 | 5163 | 18224 | 1.042818707 | 1 | 0.826955778 | 92.93726984 |

Annotation Cluster 7 Enrichment Score: 2.454567448652333

| Category | Term | Count | % | PValue | Genes | List Total | Pop Hits | Pop Total | Fold Enrichment | Bonferroni | Benjamini | FDR |
|---|---|---|---|---|---|---|---|---|---|---|---|---|
| UP_KEYWORDS | Glycoprotein | 400 | 25.41296061 | 9.00E-05 | SLC9A9, MASP1, OSMR, LTBP3, SLC9A3, HMCN2, APP, GRIN2B, ROS1, ITFG1, CDH23, CELA3A, PTPRN | 1535 | 4551 | 20581 | 1.178450239 | 0.043400644 | 0.004917936 | 0.129264988 |

| | | | | | | | | | | | | |
|---|---|---|---|---|---|---|---|---|---|---|---|---|
| UP_SEQ_FEATURE | glycosylation site:N-linked (GlcNAc...) | 374 | 23.76111817 | 1.30E-04 | SLC9A9, MASP1, ATP1B2, EFNA1, OSMR, LTBP3, FAM20A, NELL1, B3GALT4, SLC9A3, EFNA3, LUZP2, | 1499 | 4234 | 20063 | 1.182265425 | 0.411536242 | 0.064131208 | 0.242105368 |
| UP_SEQ_FEATURE | signal peptide | 290 | 18.42439644 | 0.003092704 | CHIA, MASP1, EFNA1, OSMR, FAM20A, LTBP3, NELL1, EFNA3, LUZP2, MEGF11, ERLEC1, APP, BRINP2 | 1499 | 3346 | 20063 | 1.160022202 | 0.999996682 | 0.396279639 | 5.604280276 |
| UP_KEYWORDS | Disulfide bond | 289 | 18.36086404 | 0.014367859 | CHIA, MASP1, ATP1B2, VKORC1L1, EFNA1, OSMR, FAM20A, LTBP3, NELL1, EFNA3, SYT6, MEGF11, H | 1535 | 3434 | 20581 | 1.128380688 | 0.999203062 | 0.156229384 | 18.7785442 |
| UP_SEQ_FEATURE | disulfide bond | 242 | 15.37484117 | 0.043421868 | CHIA, MASP1, ATP1B2, EFNA1, OSMR, LTBP3, NELL1, EFNA3, MEGF11, APP, MAT1A, CLEC4C, RAB27 | 1499 | 2917 | 20063 | 1.110383954 | 1 | 0.901539651 | 56.24571815 |
| UP_KEYWORDS | Signal | 333 | 21.15628971 | 0.08287004 | CHIA, MASP1, EFNA1, OSMR, LTBP3, FAM20A, NELL1, SLC9A3, EFNA3, LUZP2, MEGF11, WDR81, ERL | 1535 | 4160 | 20581 | 1.073270014 | 1 | 0.390978199 | 71.15559184 |

Annotation Cluster 8 Enrichment Score: 1.9046636302574544

| Category | Term | Count | % | PValue | Genes | List Total | Pop Hits | Pop Total | Fold Enrichment | Bonferroni | Benjamini | FDR |
|---|---|---|---|---|---|---|---|---|---|---|---|---|
| UP_KEYWORDS | Innate immunity | 35 | 2.223634053 | 0.00104109 | HMGB1, IFITM1, FGR, MASP1, C5, PGLYRP1, OAS3, OAS1, APOBEC3G, OAS2, C1S, APOBEC3F, APOBE | 1535 | 261 | 20581 | 1.797983202 | 0.401617357 | 0.029755662 | 1.485870782 |
| UP_KEYWORDS | Immunity | 55 | 3.494282084 | 0.003905934 | CHIA, MASP1, PGLYRP1, APOBEC3G, APOBEC3F, IL31RA, APOBEC3D, MAP3K5, TMEM173, CASP4, M | 1535 | 500 | 20581 | 1.474859935 | 0.854764114 | 0.074272943 | 5.469294836 |
| GOTERM_BP_DIRECT | GO:0045087~innate immune response | 36 | 2.287166455 | 0.475111786 | HMGB1, FRK, FGR, PGLYRP1, APOBEC3G, FER, C1S, APOBEC3F, APOBEC3D, SERINC3, APP, MAP3K5, | 1342 | 430 | 16792 | 1.047572176 | 1 | 0.999899747 | 99.99938349 |

Annotation Cluster 9 Enrichment Score: 1.880181659715394

| Category | Term | Count | % | PValue | Genes | List Total | Pop Hits | Pop Total | Fold Enrichment | Bonferroni | Benjamini | FDR |
|---|---|---|---|---|---|---|---|---|---|---|---|---|
| UP_SEQ_FEATURE | nucleotide phosphate-binding region:ATP | 102 | 6.480304956 | 8.91E-04 | RAD51C, MOK, RAD51B, DSTYK, CLK1, DMPK, CSNK2A2, MAP3K5, DDX23, MAT1A, DDX25, MAP3K8, | 1499 | 994 | 20063 | 1.373434738 | 0.973500535 | 0.214976832 | 1.646043167 |
| UP_KEYWORDS | ATP-binding | 135 | 8.576874206 | 0.001184081 | XRCC5, MOK, RAD51C, RAD51B, DSTYK, CLK1, DMPK, CSNK2A2, MAP3K5, ATP2B4, DDX23, MAT1A, D | 1535 | 1391 | 20581 | 1.301261952 | 0.442391172 | 0.031929018 | 1.688338142 |
| INTERPRO | IPR011009:Protein kinase-like domain | 61 | 3.875476493 | 0.002484327 | MOK, CDK17, DSTYK, CLK1, FER, WDR81, ACVR1C, DMPK, CSNK2A2, MAP3K5, PAK2, MAP3K8, ROS1, | 1452 | 531 | 18559 | 1.468328638 | 0.993650679 | 0.514595388 | 4.194574248 |
| UP_SEQ_FEATURE | domain:Protein kinase | 54 | 3.430749682 | 0.002490722 | MOK, CDK17, DSTYK, CLK1, FER, ACVR1C, DMPK, CSNK2A2, MAP3K5, PAK2, MAP3K8, ROS1, INSR, AI | 1499 | 478 | 20063 | 1.51202894 | 0.999961211 | 0.383492056 | 4.53727649 |
| UP_SEQ_FEATURE | binding site:ATP | 61 | 3.875476493 | 0.002733633 | MOK, CDK17, DSTYK, CLK1, FER, ACVR1C, DMPK, CSNK2A2, MAP3K5, PAK2, MAT1A, MAP3K8, ITPK1 | 1499 | 558 | 20063 | 1.463153452 | 0.999985615 | 0.397572497 | 4.969199144 |
| UP_KEYWORDS | Kinase | 76 | 4.828462516 | 0.003417353 | MOK, CDK17, PHKB, NELL1, PIP5K1C, DSTYK, CLK1, FER, ACVR1C, DMPK, CSNK2A2, MAP3K5, PAK2, | 1535 | 735 | 20581 | 1.386386581 | 0.815044517 | 0.067902949 | 4.800723756 |
| INTERPRO | IPR000719:Protein kinase, catalytic domain | 55 | 3.494282084 | 0.005798532 | MOK, CDK17, DSTYK, CLK1, FER, ACVR1C, DMPK, CSNK2A2, MAP3K5, PAK2, MAP3K8, ROS1, INSR, AI | 1452 | 487 | 18559 | 1.443516272 | 0.999992707 | 0.626826503 | 9.532755467 |
| INTERPRO | IPR017441:Protein kinase, ATP binding site | 44 | 2.795425667 | 0.009556683 | FRK, MOK, CDK17, FGFR3, ERBB4, FGR, CAMK2G, BRSK2, DSTYK, CLK1, FER, EPHB3, EPHB1, ACVR1C, | 1452 | 381 | 18559 | 1.476099578 | 0.999999997 | 0.704986973 | 15.24667239 |
| INTERPRO | IPR008271:Serine/threonine-protein kinase, active site | 36 | 2.287166455 | 0.019196671 | SGK494, MOK, CDK17, MAPKAPK5, CAMK2G, BRSK2, DSTYK, CHEK2, CLK1, ACVR1C, DMPK, CSNK2A2 | 1452 | 312 | 18559 | 1.474809282 | 1 | 0.755382557 | 28.38870804 |
| GOTERM_MF_DIRECT | GO:0005524~ATP binding | 141 | 8.958068615 | 0.019218066 | XRCC5, MOK, RAD51C, RAD51B, DSTYK, CLK1, DMPK, CSNK2A2, MAP3K5, ATP2B4, DDX23, MAT1A, D | 1340 | 1495 | 16881 | 1.188150052 | 1 | 0.875017748 | 27.12454213 |
| UP_KEYWORDS | Nucleotide-binding | 156 | 9.911054638 | 0.025803752 | XRCC5, MOK, RAD51C, FHIT, HRAS, RAD51B, DSTYK, CLK1, CNGB1, DMPK, CSNK2A2, MAP3K5, ATP2I | 1535 | 1788 | 20581 | 1.169809588 | 0.999997472 | 0.231277825 | 31.3203094 |
| UP_SEQ_FEATURE | active site:Proton acceptor | 65 | 4.129606099 | 0.026248327 | MOK, CDK17, DSTYK, SULT2B1, CLK1, FER, ACVR1C, DMPK, CSNK2A2, MAP3K5, PAK2, ANG, MAP3K8 | 1499 | 672 | 20063 | 1.294608112 | 1 | 0.845550682 | 39.05904154 |
| UP_KEYWORDS | Serine/threonine-protein kinase | 41 | 2.604828463 | 0.027926823 | SGK494, MOK, CDK17, MAPKAPK5, CAMK2G, BRSK2, DSTYK, CHEK2, CLK1, ACVR1C, DMPK, CSNK2A2 | 1535 | 393 | 20581 | 1.398779952 | 0.999999138 | 0.231618121 | 33.44034091 |
| GOTERM_BP_DIRECT | GO:0018105~peptidyl-serine phosphorylation | 17 | 1.080050826 | 0.040534778 | MAPKAPK5, PKN2, BRSK2, RICTOR, CLK1, CHEK2, PRKCE, DMPK, PRKD1, MAST4, PRKD2, PAK2, DCLK | 1342 | 125 | 16792 | 1.701722802 | 1 | 0.921414456 | 53.7052685 |
| GOTERM_MF_DIRECT | GO:0004674~protein serine/threonine kinase activity | 39 | 2.477763659 | 0.073809984 | SGK494, MOK, CDK17, FAM20A, MAPKAPK5, CAMK2G, BRSK2, DSTYK, CHEK2, CLK1, ACVR1C, DMPK, | 1340 | 376 | 16881 | 1.306682677 | 1 | 0.909734861 | 71.35723513 |
| GOTERM_MF_DIRECT | GO:0004672~protein kinase activity | 37 | 2.350698856 | 0.086942169 | SGK494, MOK, FASTKD1, CDK17, MAPKAPK5, BRSK2, CHEK2, CLK1, DMPK, CSNK2A2, SGK223, MAP3I | 1340 | 359 | 16881 | 1.298376502 | 1 | 0.9213553 | 77.3071614 |
| GOTERM_BP_DIRECT | GO:0006468~protein phosphorylation | 45 | 2.858958069 | 0.111798207 | SGK494, FRK, MOK, FASTKD1, CDK17, FGR, PHKB, BRSK2, AKAP13, CHEK2, FER, ACVR1C, DMPK, CSN | 1342 | 456 | 16792 | 1.23480273 | 1 | 0.979326343 | 88.99199788 |
| SMART | SM00220:S_TKc | 37 | 2.350698856 | 0.209187455 | SGK494, MOK, CDK17, MAPKAPK5, CAMK2G, BRSK2, DSTYK, CHEK2, CLK1, ACVR1C, DMPK, CSNK2A2 | 878 | 359 | 10057 | 1.180541367 | 1 | 0.922935232 | 96.27857634 |

Annotation Cluster 10 Enrichment Score: 1.8682521248851618

| Category | Term | Count | % | PValue | Genes | List Total | Pop Hits | Pop Total | Fold Enrichment | Bonferroni | Benjamini | FDR |
|---|---|---|---|---|---|---|---|---|---|---|---|---|
| UP_SEQ_FEATURE | domain:EGF-like 3 | 17 | 1.080050826 | 1.28E-04 | TNXB, TNXA, NRXN3, LTBP3, NELL1, LRP1B, ASTN2, NRXN1, MEGF11, TENM4, STAB1, TENM2, NOTC | 1499 | 76 | 20063 | 2.993846775 | 0.406681263 | 0.071861943 | 0.238358262 |
| UP_SEQ_FEATURE | domain:EGF-like 1 | 22 | 1.397712834 | 2.06E-04 | CRELD1, ATRNL1, NRXN3, CNTNAP5, LTBP3, NELL1, LRP1B, ASTN2, NRXN1, MEGF11, TENM4, FBLN1 | 1499 | 120 | 20063 | 2.453780298 | 0.567483071 | 0.088921284 | 0.382419351 |
| UP_SEQ_FEATURE | domain:EGF-like 2 | 18 | 1.143583227 | 2.78E-04 | TNXB, TNXA, ATRNL1, NRXN3, CNTNAP5, ASTN2, NRXN1, MEGF11, TENM4, THBD, STAB1, TENM2, | 1499 | 89 | 20063 | 2.706928214 | 0.677391816 | 0.106966937 | 0.515845795 |
| UP_KEYWORDS | EGF-like domain | 34 | 2.160101652 | 4.09E-04 | MALRD1, CRELD1, MASP1, LTBP3, NELL1, ASTN2, C1S, MEGF11, ITGBL1, HMCN2, ADAM33, CNTNAP | 1535 | 238 | 20581 | 1.915402513 | 0.182456215 | 0.013340274 | 0.585545546 |
| UP_SEQ_FEATURE | domain:EGF-like 8 | 9 | 0.571791614 | 4.38E-04 | TENM4, TNXB, TNXA, STAB1, TENM2, TENM3, LRP1B, RELN, MEGF11 | 1499 | 26 | 20063 | 4.633011751 | 0.831955537 | 0.149676819 | 0.812023442 |
| UP_SEQ_FEATURE | domain:EGF-like 7 | 9 | 0.571791614 | 7.62E-04 | TENM4, TNXB, TNXA, STAB1, TENM2, TENM3, LRP1B, RELN, MEGF11 | 1499 | 28 | 20063 | 4.302082341 | 0.955092992 | 0.198807781 | 1.408593823 |
| UP_SEQ_FEATURE | domain:EGF-like 4 | 13 | 0.82592122 | 0.001196875 | TNXB, TNXA, NELL1, LRP1B, MEGF11, TENM4, THBD, STAB1, TENM2, TENM3, NOTCH4, RELN, LRP5 | 1499 | 59 | 20063 | 2.949073394 | 0.9923862 | 0.262774939 | 2.20520249 |
| UP_SEQ_FEATURE | domain:EGF-like 6 | 11 | 0.698856417 | 0.001240368 | TENM4, TNXB, TNXA, STAB1, TENM2, NOTCH4, TENM3, LRP1B, RELN, FBN2, MEGF11 | 1499 | 44 | 20063 | 3.346064043 | 0.993623609 | 0.25722488 | 2.284462464 |
| UP_SEQ_FEATURE | domain:EGF-like 5 | 10 | 0.635324015 | 0.003303428 | TENM4, THBD, TNXB, TNXA, STAB1, TENM2, TENM3, LRP1B, RELN, MEGF11 | 1499 | 42 | 20063 | 3.18672766 | 0.999998597 | 0.404498479 | 5.975110111 |
| INTERPRO | IPR000742:Epidermal growth factor-like domain | 31 | 1.969504447 | 0.004178651 | MALRD1, CRELD1, LTBP3, NELL1, MEGF11, ITGBL1, HMCN2, ADAM33, CNTNAP2, FBN2, PCSK5, TNXI | 1452 | 231 | 18559 | 1.715290449 | 0.999799998 | 0.57331891 | 6.959625542 |
| UP_SEQ_FEATURE | domain:EGF-like 10 | 6 | 0.381194409 | 0.006673928 | TNXB, TNXA, STAB1, NOTCH4, LRP1B, MEGF11 | 1499 | 17 | 20063 | 4.723855119 | 1 | 0.585137079 | 11.72233433 |
| SMART | SM00181:EGF | 27 | 1.715374841 | 0.012271549 | MALRD1, CRELD1, LTBP3, NELL1, MEGF11, ITGBL1, HMCN2, CNTNAP2, FBN2, PCSK5, TNXB, ATRNL1 | 878 | 188 | 10057 | 1.645052828 | 0.994049014 | 0.641146009 | 15.89848819 |
| UP_SEQ_FEATURE | domain:EGF-like 12 | 5 | 0.317662008 | 0.012814284 | TNXB, TNXA, STAB1, LRP1B, MEGF11 | 1499 | 13 | 20063 | 5.147790835 | 1 | 0.696950976 | 21.34822866 |
| UP_SEQ_FEATURE | domain:EGF-like 11 | 5 | 0.317662008 | 0.016897452 | TNXB, TNXA, STAB1, LRP1B, MEGF11 | 1499 | 14 | 20063 | 4.78009149 | 1 | 0.750483062 | 27.18978829 |
| UP_SEQ_FEATURE | domain:EGF-like 14 | 5 | 0.317662008 | 0.021706961 | TNXB, TNXA, STAB1, LRP1B, MEGF11 | 1499 | 15 | 20063 | 4.461418724 | 1 | 0.808965085 | 33.54388138 |
| UP_SEQ_FEATURE | domain:EGF-like 16 | 4 | 0.254129606 | 0.024801433 | TNXB, TNXA, STAB1, NOTCH4 | 1499 | 9 | 20063 | 5.948558298 | 1 | 0.833799715 | 37.35073803 |
| UP_SEQ_FEATURE | domain:EGF-like 13 | 5 | 0.317662008 | 0.033608344 | TNXB, TNXA, STAB1, LRP1B, MEGF11 | 1499 | 17 | 20063 | 3.936545933 | 1 | 0.874845696 | 47.08771362 |
| INTERPRO | IPR013032:EGF-like, conserved site | 24 | 1.524777637 | 0.038209594 | MALRD1, TNXB, MASP1, CRELD1, ATRNL1, LTBP3, SELL, NELL1, LRP1B, MEGF11, HMCN2, TENM4, FE | 1452 | 199 | 18559 | 1.541509199 | 1 | 0.868906876 | 48.88764236 |
| INTERPRO | IPR026823:Complement Clr-like EGF domain | 6 | 0.381194409 | 0.063218072 | HMCN2, FBLN1, THBD, LTBP3, LRP1B, FBN2 | 1452 | 28 | 18559 | 2.738931523 | 1 | 0.929813379 | 67.53502315 |
| UP_SEQ_FEATURE | domain:EGF-like 9 | 5 | 0.317662008 | 0.099617679 | TNXB, TNXA, STAB1, LRP1B, MEGF11 | 1499 | 24 | 20063 | 2.788386702 | 1 | 0.980888024 | 85.82758991 |
| INTERPRO | IPR018097:EGF-like calcium-binding, conserved site | 13 | 0.82592122 | 0.100659471 | HMCN2, FBLN1, THBD, MASP1, CRELD1, CRTAC1, LTBP3, NELL1, NOTCH4, LRP1B, LRP8, C1S, FBN2 | 1452 | 102 | 18559 | 1.629037703 | 1 | 0.963843964 | 83.92135994 |
| INTERPRO | IPR000152:EGF-type aspartate/asparagine hydroxylation site | 13 | 0.82592122 | 0.118073864 | HMCN2, FBLN1, THBD, CRELD1, NRXN3, LTBP3, NELL1, NOTCH4, LRP1B, LRP8, C1S, NRXN1, FBN2 | 1452 | 105 | 18559 | 1.582493769 | 1 | 0.968368246 | 88.51957324 |
| INTERPRO | IPR001881:EGF-like calcium-binding | 15 | 0.952986023 | 0.120678686 | MASP1, CRELD1, LTBP3, CRTAC1, NELL1, LRP1B, C1S, HMCN2, FBLN1, THBD, STAB1, TENM2, NOTCH | 1452 | 127 | 18559 | 1.509647296 | 1 | 0.969430882 | 89.08991998 |
| INTERPRO | IPR009030:Insulin-like growth factor binding protein, N-terminal | 16 | 1.016518424 | 0.126632726 | ERBB4, CRELD1, CRTAC1, LTBP3, IGFBP7, NELL1, LRP1B, EPHB1, HMCN2, FBLN1, THBD, NOTCH4, LR | 1452 | 139 | 18559 | 1.471272569 | 1 | 0.969379898 | 90.2949824 |
| SMART | SM00179:EGF_CA | 15 | 0.952986023 | 0.215996514 | MASP1, CRELD1, LTBP3, CRTAC1, NELL1, LRP1B, C1S, HMCN2, FBLN1, THBD, STAB1, TENM2, NOTCH | 878 | 127 | 10057 | 1.35288684 | 1 | 0.914828972 | 96.70355505 |
| UP_SEQ_FEATURE | domain:EGF-like 15 | 3 | 0.190597205 | 0.252812379 | TNXB, TNXA, STAB1 | 1499 | 13 | 20063 | 3.088674501 | 1 | 0.999215754 | 99.56015612 |
| UP_SEQ_FEATURE | domain:EGF-like 6; calcium-binding | 4 | 0.254129606 | 0.306643203 | FBLN1, THBD, LTBP3, NELL1 | 1499 | 26 | 20063 | 2.059116334 | 1 | 0.999736313 | 99.89068919 |
| UP_SEQ_FEATURE | domain:EGF-like 5; calcium-binding | 5 | 0.317662008 | 0.314756392 | FBLN1, LTBP3, NELL1, NOTCH4, FBN2 | 1499 | 38 | 20063 | 1.761086338 | 1 | 0.999777961 | 99.91220228 |
| UP_SEQ_FEATURE | domain:EGF-like 2; calcium-binding | 6 | 0.381194409 | 0.437966708 | FBLN1, CRELD1, LTBP3, NELL1, LRP1B, LRP8 | 1499 | 58 | 20063 | 1.384578225 | 1 | 0.999975557 | 99.99780899 |

Annotation Cluster 11 Enrichment Score: 1.792846440615368

| Category | Term | Count | % | PValue | Genes | List Total | Pop Hits | Pop Total | Fold Enrichment | Bonferroni | Benjamini | FDR |
|---|---|---|---|---|---|---|---|---|---|---|---|---|
| UP_KEYWORDS | LIM domain | 13 | 0.82592122 | 0.006623988 | LPXN, LMO1, LIMK2, PRICKLE1, LPP, LHX3, LMCD1, FHL2, LMO7, LIMD1, LHX9, SCEL, NEBL | 1535 | 72 | 20581 | 2.420855954 | 0.962239432 | 0.094517721 | 9.109597307 |
| INTERPRO | IPR001781:Zinc finger, LIM-type | 13 | 0.82592122 | 0.009538358 | LPXN, LMO1, LIMK2, PRICKLE1, LPP, LHX3, LMCD1, FHL2, LMO7, LIMD1, LHX9, SCEL, NEBL | 1452 | 72 | 18559 | 2.307803413 | 0.999999997 | 0.727361698 | 15.2196541 |
| UP_SEQ_FEATURE | domain:LIM zinc-binding 1 | 10 | 0.635324015 | 0.010862514 | LPXN, LMO1, LIMK2, PRICKLE1, LPP, LHX3, LMCD1, FHL2, LIMD1, LHX9 | 1499 | 50 | 20063 | 2.676851234 | 1 | 0.66209877 | 18.40182252 |
| UP_SEQ_FEATURE | domain:LIM zinc-binding 2 | 10 | 0.635324015 | 0.010862514 | LPXN, LMO1, LIMK2, PRICKLE1, LPP, LHX3, LMCD1, FHL2, LIMD1, LHX9 | 1499 | 50 | 20063 | 2.676851234 | 1 | 0.66209877 | 18.40182252 |
| SMART | SM00132:LIM | 13 | 0.82592122 | 0.020963839 | LPXN, LMO1, LIMK2, PRICKLE1, LPP, LHX3, LMCD1, FHL2, LMO7, LIMD1, LHX9, SCEL, NEBL | 878 | 72 | 10057 | 2.068163123 | 0.999848129 | 0.623539553 | 25.70274403 |
| UP_SEQ_FEATURE | domain:LIM zinc-binding 3 | 5 | 0.317662008 | 0.111942946 | LPXN, PRICKLE1, LPP, FHL2, LIMD1 | 1499 | 25 | 20063 | 2.676851234 | 1 | 0.986145531 | 89.03560923 |

Annotation Cluster 12 Enrichment Score: 1.6823971256748071

| Category | Term | Count | % | PValue | Genes | List Total | Pop Hits | Pop Total | Fold Enrichment | Bonferroni | Benjamini | FDR |
|---|---|---|---|---|---|---|---|---|---|---|---|---|
| GOTERM_BP_DIRECT | GO:1902358~sulfate transmembrane transport | 6 | 0.381194409 | 0.001585146 | SLC26A4, SLC26A3, SLC26A5, SLC26A9, SLC13A1, SLC26A10 | 1342 | 12 | 16792 | 6.25633383 | 0.998397256 | 0.723959494 | 2.909420921 |
| GOTERM_BP_DIRECT | GO:0051453~regulation of intracellular pH | 10 | 0.635324015 | 0.001676507 | SLC26A4, SLC9A9, SLC26A3, SLC4A11, SLC26A5, SLC9A3, SLC26A9, SLC9C1, CA2, SLC26A10 | 1342 | 36 | 16792 | 3.475741017 | 0.998894323 | 0.678433053 | 3.074642855 |
| GOTERM_MF_DIRECT | GO:0015106~bicarbonate transmembrane transporter activity | 6 | 0.381194409 | 0.004768165 | SLC26A4, SLC26A3, SLC4A11, SLC26A5, SLC26A9, SLC26A10 | 1340 | 15 | 16881 | 5.039104478 | 0.997859093 | 0.953730063 | 7.497562812 |
| GOTERM_BP_DIRECT | GO:0042391~regulation of membrane potential | 14 | 0.889453621 | 0.006064849 | ASIC2, CNGB1, SLC26A10, CNGA3, RIMS1, CNGA1, SLC26A4, SLC26A3, SLC26A5, NEDD4, SLC26A9, NI | 1342 | 75 | 16792 | 2.335697963 | 1 | 0.786154291 | 10.70471531 |
| GOTERM_MF_DIRECT | GO:0015301~anion:anion antiporter activity | 7 | 0.444726811 | 0.00767379 | SLC26A4, SLC26A3, SLC4A11, SLC26A5, SLC22A9, SLC26A9, SLC26A10 | 1340 | 23 | 16881 | 3.834101233 | 0.999950153 | 0.808159743 | 11.80418412 |
| INTERPRO | IPR002645:STAS domain | 5 | 0.317662008 | 0.007855377 | SLC26A4, SLC26A3, SLC26A5, SLC26A9, SLC26A10 | 1452 | 11 | 18559 | 5.809854746 | 0.999999892 | 0.682024266 | 12.70345322 |
| INTERPRO | IPR001902:Sulphate anion transporter | 5 | 0.317662008 | 0.007855377 | SLC26A4, SLC26A3, SLC26A5, SLC26A9, SLC26A10 | 1452 | 11 | 18559 | 5.809854746 | 0.999999892 | 0.682024266 | 12.70345322 |
| INTERPRO | IPR011547:Sulphate transporter | 5 | 0.317662008 | 0.007855377 | SLC26A4, SLC26A3, SLC26A5, SLC26A9, SLC26A10 | 1452 | 11 | 18559 | 5.809854746 | 0.999999892 | 0.682024266 | 12.70345322 |
| GOTERM_MF_DIRECT | GO:0019531~oxalate transmembrane transporter activity | 5 | 0.317662008 | 0.008265321 | SLC26A4, SLC26A3, SLC26A5, SLC26A9, SLC26A10 | 1340 | 11 | 16881 | 5.726255088 | 0.999976847 | 0.782328353 | 12.65755025 |

| | | | | | | | | | | | | |
|---|---|---|---|---|---|---|---|---|---|---|---|---|
| GOTERM_MF_DIRECT | GO:0015116~sulfate transmembrane transporter activity | 5 | 0.317662008 | 0.008265321 | SLC26A4, SLC26A3, SLC26A5, SLC26A9, SLC26A10 | 1340 | 11 | 16881 | 5.726255088 | 0.999976847 | 0.782328353 | 12.65755025 |
| GOTERM_MF_DIRECT | GO:0008271~secondary active sulfate transmembrane transporter activity | 5 | 0.317662008 | 0.008265321 | SLC26A4, SLC26A3, SLC26A5, SLC26A9, SLC26A10 | 1340 | 11 | 16881 | 5.726255088 | 0.999976847 | 0.782328353 | 12.65755025 |
| GOTERM_BP_DIRECT | GO:0019532~oxalate transport | 5 | 0.317662008 | 0.008465773 | SLC26A4, SLC26A3, SLC26A5, SLC26A9, SLC26A10 | 1342 | 11 | 16792 | 5.687576209 | 1 | 0.83722078 | 14.63500841 |
| UP_SEQ_FEATURE | domain:STAS | 5 | 0.317662008 | 0.012814284 | SLC26A4, SLC26A3, SLC26A5, SLC26A9, SLC26A10 | 1499 | 13 | 20063 | 5.147790835 | 1 | 0.696950976 | 21.34822866 |
| GOTERM_BP_DIRECT | GO:0015701~bicarbonate transport | 9 | 0.571791614 | 0.02166779 | SLC26A4, SLC26A3, SLC4A11, SLC26A5, CA9, SLC26A9, CA2, SLC26A10, CA1 | 1342 | 44 | 16792 | 2.559409294 | 1 | 0.879488939 | 33.4828265 |
| INTERPRO | IPR018045:Sulphate anion transporter, conserved site | 3 | 0.190597205 | 0.098565829 | SLC26A4, SLC26A3, SLC26A5 | 1452 | 7 | 18559 | 5.477863046 | 1 | 0.966768877 | 83.2642154 |
| REACTOME_PATHWAY | R-HSA-427601:R-HSA-427601 | 3 | 0.190597205 | 0.106954574 | SLC26A4, SLC26A3, SLC26A9 | 745 | 7 | 9075 | 5.220517737 | 1 | 0.942800819 | 82.67217365 |
| GOTERM_BP_DIRECT | GO:1902476~chloride transmembrane transport | 10 | 0.635324015 | 0.324355997 | SLC26A4, SLC17A7, SLC26A3, GABRA2, GABRG3, SLC26A5, SLC12A5, SLC26A9, ANO3, SLC26A10 | 1342 | 93 | 16792 | 1.345448136 | 1 | 0.999066693 | 99.93227605 |
| GOTERM_MF_DIRECT | GO:0005254~chloride channel activity | 6 | 0.381194409 | 0.428601648 | SLC26A4, SLC26A3, GABRG3, SLC26A5, SLC26A9, SLC26A10 | 1340 | 54 | 16881 | 1.399751244 | 1 | 0.994969124 | 99.98911896 |
| GOTERM_BP_DIRECT | GO:0006821~chloride transport | 4 | 0.254129606 | 0.629622755 | SLC26A3, GABRA2, GABRG3, SLC26A9 | 1342 | 40 | 16792 | 1.251266766 | 1 | 0.999992094 | 99.99999906 |
| UP_KEYWORDS | Chloride | 6 | 0.381194409 | 0.642486481 | SLC26A4, SLC26A3, GABRA2, GABRG3, SLC26A10, RUNX1 | 1535 | 73 | 20581 | 1.102012405 | 1 | 0.925761065 | 99.99996198 |
| Annotation Cluster 13 | Enrichment Score: 1.5795181084483119 | | | | | | | | | | | |
| Category | Term | Count | % | PValue | Genes | List Total | Pop Hits | Pop Total | Fold Enrichment | Bonferroni | Benjamini | FDR |
| SMART | SM00060:FN3 | 30 | 1.905972046 | 3.31E-05 | DCC, OSMR, DSCAML1, EPHB3, EPHB1, IL31RA, NRCAM, MYOM2, ROBO1, ROBO2, INSR, ROS1, FN1, | 878 | 150 | 10057 | 2.290888383 | 0.013633442 | 0.013633442 | 0.046373282 |
| INTERPRO | IPR013098:Immunoglobulin I-set | 25 | 1.588310038 | 2.34E-04 | DCC, OBSCN, ADAMTSL1, FGFR3, IGFBP7, SDK1, DSCAML1, TRIO, PALLD, HMCN2, LINGO2, NRCAM, I | 1452 | 141 | 18559 | 2.266255397 | 0.379334192 | 0.379334192 | 0.403150572 |
| UP_SEQ_FEATURE | domain:Ig-like C2-type 5 | 12 | 0.762388818 | 0.002179772 | NCAM1, MYOM2, ROBO1, CNTN2, SDK1, CNTN1, DSCAML1, ROBO2, CNTN4, PALLD, MYLK, DSCAM | 1499 | 55 | 20063 | 2.920201346 | 0.999861951 | 0.35878791 | 3.981661788 |
| INTERPRO | IPR003598:Immunoglobulin subtype 2 | 33 | 2.09656925 | 0.003960156 | DCC, FGFR3, ADAMTSL1, DSCAML1, HMCN2, LINGO2, NRCAM, MYOM2, ROBO1, ROBO2, UNC5C, DS | 1452 | 250 | 18559 | 1.687181818 | 0.999687513 | 0.59211582 | 6.607322126 |
| UP_SEQ_FEATURE | domain:Ig-like C2-type 4 | 13 | 0.82592122 | 0.010308277 | DCC, SDK1, DSCAML1, PALLD, NCAM1, MYOM2, ROBO1, CNTN2, CNTN1, CNTN4, ROBO2, MYLK, DS( | 1499 | 76 | 20063 | 2.28941224 | 1 | 0.661130877 | 17.54628962 |
| SMART | SM00408:IGc2 | 33 | 2.09656925 | 0.017530716 | DCC, FGFR3, ADAMTSL1, DSCAML1, HMCN2, LINGO2, NRCAM, MYOM2, ROBO1, ROBO2, UNC5C, DS | 878 | 250 | 10057 | 1.511986333 | 0.999350802 | 0.649550585 | 21.96474259 |
| UP_SEQ_FEATURE | domain:Ig-like C2-type 3 | 17 | 1.080050826 | 0.028994492 | DCC, FGFR3, SDK1, DSCAML1, PALLD, NCAM1, MYOM2, ROBO1, LSAMP, KIR3DL3, CNTN2, CNTN1, R | 1499 | 128 | 20063 | 1.777596523 | 1 | 0.864304339 | 42.18083932 |
| UP_SEQ_FEATURE | domain:Ig-like C2-type 1 | 23 | 1.461245235 | 0.047803256 | DCC, CLMP, ADAMTSL1, FGFR3, ICAM4, SDK1, DSCAML1, PALLD, NCAM1, MYOM2, ROBO1, SIGLEC5, | 1499 | 202 | 20063 | 1.52394996 | 1 | 0.914828551 | 59.83040836 |
| UP_SEQ_FEATURE | domain:Ig-like C2-type 2 | 23 | 1.461245235 | 0.049995237 | DCC, CLMP, ADAMTSL1, FGFR3, ICAM4, SDK1, DSCAML1, PALLD, NCAM1, MYOM2, ROBO1, SIGLEC5, | 1499 | 203 | 20063 | 1.516442817 | 1 | 0.9192852 | 61.51772062 |
| UP_KEYWORDS | Immunoglobulin domain | 46 | 2.92249047 | 0.266974508 | DCC, HLA-DQB1, ADAMTSL1, FGFR3, IGFBP7, DSCAML1, TAPBP, HMCN2, LINGO2, NRCAM, MYOM2, | 1535 | 546 | 20581 | 1.129596354 | 1 | 0.694841938 | 98.84784901 |
| INTERPRO | IPR003599:Immunoglobulin subtype | 39 | 2.477763659 | 0.582487288 | DCC, FGFR3, ADAMTSL1, IGFBP7, DSCAML1, HMCN2, LINGO2, NRCAM, MYOM2, ROBO1, ROBO2, UN | 1452 | 497 | 18559 | 1.002989008 | 1 | 0.999921301 | 99.99997081 |
| INTERPRO | IPR013783:Immunoglobulin-like fold | 70 | 4.447268107 | 0.763712272 | OSMR, IGFBP7, TAPBP, IL31RA, HMCN2, NRCAM, ROBO1, ROBO2, UNC5C, ROS1, INSR, DSCAM, PTPI | 1452 | 951 | 18559 | 0.940817698 | 1 | 0.999998818 | 100 |
| SMART | SM00409:IG | 39 | 2.477763659 | 0.829165935 | DCC, FGFR3, ADAMTSL1, IGFBP7, DSCAML1, HMCN2, LINGO2, NRCAM, MYOM2, ROBO1, ROBO2, UN | 878 | 497 | 10057 | 0.898839506 | 1 | 0.999191005 | 100 |
| INTERPRO | IPR007110:Immunoglobulin-like domain | 51 | 3.240152478 | 0.933250924 | IGFBP7, TAPBP, HMCN2, NRCAM, ROBO1, ROBO2, UNC5C, DSCAM, CLMP, PTPRM, ICAM4, BTNL2, S | 1452 | 771 | 18559 | 0.845480807 | 1 | 1 | 100 |
| Annotation Cluster 14 | Enrichment Score: 1.5328830217604308 | | | | | | | | | | | |
| Category | Term | Count | % | PValue | Genes | List Total | Pop Hits | Pop Total | Fold Enrichment | Bonferroni | Benjamini | FDR |
| INTERPRO | IPR000008:C2 calcium-dependent membrane targeting | 27 | 1.715374841 | 4.62E-04 | C2CD4D, SYT6, ESYT2, RIMS1, ESYT3, PLCL1, JMJD7-PLA2G4B, PLCH1, NEDD4L, CPNE4, PIK3CB, CPNE | 1452 | 165 | 18559 | 2.091547708 | 0.609046266 | 0.268790562 | 0.792275508 |
| SMART | SM00239:C2 | 23 | 1.461245235 | 0.001351571 | CPNE4, CPNE5, PIK3C2B, PKN2, C2CD4D, SYT6, PRKCE, ESYT2, RIMS1, PCLO, ESYT3, PLCL1, JMJD7-PL | 878 | 127 | 10057 | 2.074426488 | 0.429522412 | 0.244700332 | 1.878687129 |
| UP_SEQ_FEATURE | domain:C2 | 14 | 0.889453621 | 0.00750672 | PIK3C2B, PKN2, PRKCE, PLCL1, JMJD7-PLA2G4B, SYDE2, PLCG1, JMJD7, PLCH1, CCDC33, INPP4B, NEI | 1499 | 82 | 20063 | 2.285116907 | 1 | 0.605448756 | 13.09026384 |
| UP_SEQ_FEATURE | domain:C2 2 | 11 | 0.698856417 | 0.011366908 | CPNE4, CPNE5, SYT6, SYT16, SYTL2, SYTL3, UNC13C, ESYT2, RIMS1, PCLO, ESYT3 | 1499 | 59 | 20063 | 2.495369795 | 1 | 0.669989988 | 19.1731107 |
| UP_SEQ_FEATURE | domain:C2 1 | 11 | 0.698856417 | 0.011366908 | CPNE4, CPNE5, SYT6, SYT16, SYTL2, SYTL3, UNC13C, ESYT2, RIMS1, PCLO, ESYT3 | 1499 | 59 | 20063 | 2.495369795 | 1 | 0.669989988 | 19.1731107 |
| GOTERM_MF_DIRECT | GO:0030276~clathrin binding | 10 | 0.635324015 | 0.019988779 | SCLT1, SMAP1, PICALM, TOM1L2, C2CD4D, SYT6, SYT16, SYTL2, SYTL3, HIP1 | 1340 | 52 | 16881 | 2.422646383 | 1 | 0.864309855 | 28.05273132 |
| GOTERM_MF_DIRECT | GO:0005544~calcium-dependent phospholipid binding | 10 | 0.635324015 | 0.037953259 | JMJD7-PLA2G4B, ANXA13, C2CD4D, SYT6, SYT16, SYTL2, SYTL3, ESYT2, PCLO, PLA2G4B | 1340 | 58 | 16881 | 2.172027792 | 1 | 0.89583066 | 46.78937009 |
| GOTERM_CC_DIRECT | GO:0070382~exocytic vesicle | 4 | 0.254129606 | 0.106975873 | ANXA13, SYT6, SYTL2, SYTL3 | 1425 | 15 | 18224 | 3.410339181 | 1 | 0.762933266 | 81.76750106 |
| GOTERM_BP_DIRECT | GO:0048791~calcium ion-regulated exocytosis of neurotransmitter | 6 | 0.381194409 | 0.197476361 | C2CD4D, SYT6, SYT16, SYTL2, SYTL3, RIMS1 | 1342 | 39 | 16792 | 1.925025794 | 1 | 0.994079886 | 98.33356912 |
| GOTERM_BP_DIRECT | GO:0017158~regulation of calcium ion-dependent exocytosis | 5 | 0.317662008 | 0.304824482 | C2CD4D, SYT6, SYT16, SYTL2, SYTL3 | 1342 | 35 | 16792 | 1.787523951 | 1 | 0.998883596 | 99.88489553 |
| GOTERM_BP_DIRECT | GO:0006906~vesicle fusion | 7 | 0.444726811 | 0.331242169 | VAMP5, C2CD4D, SYT6, SYT16, SYTL2, SYTL3, TSNARE1 | 1342 | 59 | 16792 | 1.48455379 | 1 | 0.99909306 | 99.94403229 |
| GOTERM_MF_DIRECT | GO:0019905~syntaxin binding | 8 | 0.508259212 | 0.411629991 | VPS54, VPS52, C2CD4D, SYT6, SYT16, SYTL2, SYTL3, TXLNA | 1340 | 77 | 16881 | 1.308858306 | 1 | 0.994736045 | 99.98246347 |
| Annotation Cluster 15 | Enrichment Score: 1.4617588023179386 | | | | | | | | | | | |
| Category | Term | Count | % | PValue | Genes | List Total | Pop Hits | Pop Total | Fold Enrichment | Bonferroni | Benjamini | FDR |
| UP_KEYWORDS | Hydroxylation | 16 | 1.016518424 | 0.004367341 | MASP1, COL3A1, RPL27A, COL15A1, COL25A1, C1S, COL5A1, TNKS2, COL9A2, AMBN, ADRB2, THBD, | 1535 | 95 | 20581 | 2.258158752 | 0.884422586 | 0.071706574 | 6.0966675 |
| UP_KEYWORDS | Collagen | 16 | 1.016518424 | 0.005868355 | COL21A1, COL3A1, MMP8, COL22A1, COL15A1, COL25A1, COLEC12, COLEC11, COL5A1, RPS18, COL9 | 1535 | 98 | 20581 | 2.189031443 | 0.945065907 | 0.089353673 | 8.110888902 |
| REACTOME_PATHWAY | R-HSA-419037:R-HSA-419037 | 10 | 0.635324015 | 0.006054814 | NCAM1, COL9A2, COL9A3, COL6A6, COL6A5, COL3A1, CNTN2, ST8SIA2, CACNA1C, COL5A1 | 745 | 42 | 9075 | 2.900287632 | 0.994608697 | 0.92657451 | 8.981706017 |
| REACTOME_PATHWAY | R-HSA-1650814:R-HSA-1650814 | 13 | 0.82592122 | 0.007609629 | COL9A2, COL9A3, COLGALT2, COL6A6, COL21A1, ADAMTS14, COL6A5, COL3A1, COL22A1, COL15A1, | 745 | 67 | 9075 | 2.363517981 | 0.998597232 | 0.806470831 | 11.16314538 |
| INTERPRO | IPR008160:Collagen triple helix repeat | 14 | 0.889453621 | 0.010826983 | COL21A1, COL22A1, COL3A1, COL15A1, COL25A1, COLEC12, COLEC11, COL5A1, COL9A2, COL9A3, CC | 1452 | 82 | 18559 | 2.182238124 | 1 | 0.688194966 | 17.09990857 |
| GOTERM_BP_DIRECT | GO:0030574~collagen catabolic process | 12 | 0.762388818 | 0.011825078 | ADAMTS14, COL6A6, COL6A5, MRC2, COL3A1, MMP8, MMP27, COL15A1, COL25A1, MMP16, COL5A | 1342 | 64 | 16792 | 2.346125186 | 1 | 0.843728965 | 19.86021227 |
| GOTERM_CC_DIRECT | GO:0005581~collagen trimer | 15 | 0.952986023 | 0.01184733 | COL21A1, MMP8, COL22A1, COL3A1, COL15A1, COL25A1, COLEC12, COLEC11, COL5A1, RPS18, COL9 | 1425 | 92 | 18224 | 2.085125858 | 0.999721951 | 0.310762108 | 16.41288464 |
| KEGG_PATHWAY | hsa04974:Protein digestion and absorption | 15 | 0.952986023 | 0.013358314 | CELA3A, SLC8A1, COL21A1, ATP1B2, SLC9A3, COL3A1, COL22A1, COL15A1, COL5A1, COL9A2, COL9A: | 574 | 88 | 6879 | 2.042781913 | 0.976531277 | 0.52783054 | 16.28236858 |
| REACTOME_PATHWAY | R-HSA-3000178:R-HSA-3000178 | 12 | 0.762388818 | 0.041072005 | ASPN, NCAM1, COL9A2, APP, COL9A3, TNXB, COL6A6, COL6A5, ITGA8, COL3A1, COL5A1, FN1 | 745 | 75 | 9075 | 1.948993289 | 1 | 0.950493393 | 47.789408 |
| REACTOME_PATHWAY | R-HSA-186797:R-HSA-186797 | 7 | 0.444726811 | 0.042916121 | COL9A2, COL9A3, COL6A6, COL6A5, COL3A1, PDGFC, COL5A1 | 745 | 32 | 9075 | 2.664639262 | 1 | 0.932425042 | 49.32378789 |
| REACTOME_PATHWAY | R-HSA-1442490:R-HSA-1442490 | 10 | 0.635324015 | 0.075856304 | COL9A2, COL9A3, COL6A6, COL6A5, COL3A1, MMP8, COL15A1, COL25A1, COL5A1, COL10A1 | 745 | 64 | 9075 | 1.903313758 | 1 | 0.940798558 | 70.54868829 |
| UP_SEQ_FEATURE | region of interest:Triple-helical region | 5 | 0.317662008 | 0.087964253 | COL6A6, COL6A5, COL3A1, COL5A1, COL10A1 | 1499 | 23 | 20063 | 2.909620907 | 1 | 0.973774641 | 81.99335137 |
| REACTOME_PATHWAY | R-HSA-216083:R-HSA-216083 | 12 | 0.762388818 | 0.091953413 | COL9A2, COL9A3, COL6A6, ICAM4, COL6A5, ITGAE, ITGA8, COL3A1, ITGA11, COL5A1, COL10A1, FN1 | 745 | 86 | 9075 | 1.699703449 | 1 | 0.925156169 | 77.56892801 |
| UP_SEQ_FEATURE | region of interest:Nonhelical region | 3 | 0.190597205 | 0.115475174 | COL6A6, COL6A5, COL5A1 | 1499 | 8 | 20063 | 5.019096064 | 1 | 0.986545152 | 89.81978955 |
| REACTOME_PATHWAY | R-HSA-2022090:R-HSA-2022090 | 6 | 0.381194409 | 0.324345318 | COL6A6, COL6A5, COL3A1, COL15A1, COL5A1, COL10A1 | 745 | 46 | 9075 | 1.588853224 | 1 | 0.986738072 | 99.77015158 |
| GOTERM_CC_DIRECT | GO:0005788~endoplasmic reticulum lumen | 18 | 1.143583227 | 0.331530693 | WNT5A, ADAMTSL1, COL21A1, COL3A1, ERP29, COL22A1, COL15A1, ESD, COL25A1, ERLEC1, COL5A1 | 1425 | 192 | 18224 | 1.198947368 | 1 | 0.913591342 | 99.76625604 |
| KEGG_PATHWAY | hsa04512:ECM-receptor interaction | 9 | 0.571791614 | 0.43971047 | TNXB, COL6A6, COL6A5, ITGA8, COL3A1, ITGA11, RELN, COL5A1, FN1 | 574 | 87 | 6879 | 1.239757299 | 1 | 0.857339979 | 99.95265689 |
| Annotation Cluster 16 | Enrichment Score: 1.4597407040762473 | | | | | | | | | | | |
| Category | Term | Count | % | PValue | Genes | List Total | Pop Hits | Pop Total | Fold Enrichment | Bonferroni | Benjamini | FDR |
| INTERPRO | IPR022398:Peptidase S8, subtilisin, His-active site | 4 | 0.254129606 | 0.02801118 | PCSK2, TPP2, PCSK5, MBTPS1 | 1452 | 9 | 18559 | 5.680746863 | 1 | 0.835669019 | 38.7028787 |
| INTERPRO | IPR023828:Peptidase S8, subtilisin, Ser-active site | 4 | 0.254129606 | 0.02801118 | PCSK2, TPP2, PCSK5, MBTPS1 | 1452 | 9 | 18559 | 5.680746863 | 1 | 0.835669019 | 38.7028787 |
| INTERPRO | IPR015500:Peptidase S8, subtilisin-related | 4 | 0.254129606 | 0.037733621 | PCSK2, TPP2, PCSK5, MBTPS1 | 1452 | 10 | 18559 | 5.112672176 | 1 | 0.87239738 | 48.45013759 |
| INTERPRO | IPR000209:Peptidase S8/S53 domain | 4 | 0.254129606 | 0.048937963 | PCSK2, TPP2, PCSK5, MBTPS1 | 1452 | 11 | 18559 | 4.647883797 | 1 | 0.901677162 | 57.86890772 |
| Annotation Cluster 17 | Enrichment Score: 1.4109899123165597 | | | | | | | | | | | |
| Category | Term | Count | % | PValue | Genes | List Total | Pop Hits | Pop Total | Fold Enrichment | Bonferroni | Benjamini | FDR |
| GOTERM_BP_DIRECT | GO:0007169~transmembrane receptor protein tyrosine kinase signaling pathway | 18 | 1.143583227 | 0.001452636 | FRK, NTF3, FGR, ERBB4, PIK3CB, AHI1, PTPRT, ALK, TRAT1, IL31RA, DOK2, DOK4, ROR1, JAK1, TXK, R | 1342 | 96 | 16792 | 2.346125186 | 0.997254047 | 0.771085515 | 2.669310354 |
| INTERPRO | IPR008266:Tyrosine-protein kinase, active site | 16 | 1.016518424 | 0.006736062 | OBSCN, FRK, FGFR3, FGR, ERBB4, FER, EPHB3, ALK, EPHB1, SGK223, ROR1, JAK1, TXK, ROS1, INSR, A | 1452 | 95 | 18559 | 2.152704074 | 0.99999893 | 0.652677077 | 10.99121131 |
| INTERPRO | IPR001245:Serine-threonine/tyrosine-protein kinase catalytic domain | 20 | 1.27064803 | 0.012839086 | FRK, FGFR3, LIMK2, FGR, ERBB4, DSTYK, FER, EPHB3, ALK, EPHB1, ACVR1C, TEX14, ROR1, JAK1, TXK, | 1452 | 140 | 18559 | 1.825954349 | 1 | 0.681066827 | 19.95744212 |
| INTERPRO | IPR020635:Tyrosine-protein kinase, catalytic domain | 13 | 0.82592122 | 0.032340322 | FRK, FGFR3, ERBB4, FGR, JAK1, TXK, FER, EPHB3, ALK, INSR, ROS1, EPHB1, MAP3K11 | 1452 | 85 | 18559 | 1.954845244 | 1 | 0.851993198 | 43.23988248 |
| GOTERM_BP_DIRECT | GO:0038083~peptidyl-tyrosine autophosphorylation | 8 | 0.508259212 | 0.037149508 | FRK, ERBB4, FGR, JAK1, TXK, FER, INSR, ROS1 | 1342 | 40 | 16792 | 2.502533532 | 1 | 0.916021944 | 50.56886859 |
| UP_KEYWORDS | Tyrosine-protein kinase | 15 | 0.952986023 | 0.038819676 | FRK, FGFR3, FGR, ERBB4, DSTYK, FER, CLK1, EPHB3, ALK, EPHB1, SGK223, JAK1, TXK, ROS1, INSR | 1535 | 112 | 20581 | 1.795689856 | 0.999999997 | 0.277707066 | 43.39250743 |
| GOTERM_BP_DIRECT | GO:0018108~peptidyl-tyrosine phosphorylation | 19 | 1.207115629 | 0.06262344 | FGFR3, FGR, ERBB4, DSTYK, FGF23, FER, CLK1, EPHB3, ALK, EPHB1, SGK223, SCYL1, ROR1, HBEGF, RI | 1342 | 153 | 16792 | 1.55386069 | 1 | 0.959221387 | 69.98969873 |
| SMART | SM00219:TyrKc | 13 | 0.82592122 | 0.064279896 | FRK, FGFR3, ERBB4, FGR, JAK1, TXK, FER, EPHB3, ALK, INSR, ROS1, EPHB1, MAP3K11 | 878 | 85 | 10057 | 1.751855822 | 1 | 0.860464785 | 60.6099512 |
| GOTERM_MF_DIRECT | GO:0004715~non-membrane spanning protein tyrosine kinase activity | 8 | 0.508259212 | 0.068790676 | SGK223, FRK, FGR, JAK1, TXK, FER, CLK1, MELK | 1340 | 46 | 16881 | 2.19091499 | 1 | 0.90464245 | 68.71843328 |
| GOTERM_BP_DIRECT | GO:0046777~protein autophosphorylation | 20 | 1.27064803 | 0.093071636 | FGFR3, FGR, ERBB4, MAPKAPK5, CAMK2G, MINK1, FER, CLK1, CHEK2, EPHB3, ALK, EPHB1, PRKD1, P | 1342 | 172 | 16792 | 1.454961356 | 1 | 0.97867594 | 83.7685833 |
| UP_SEQ_FEATURE | domain:SH2 | 12 | 0.762388818 | 0.10193482 | FRK, FGR, SH2B3, JAK1, SH2B2, TXK, FER, VAV2, SH2D1B, SHC4, SH3BP2, HSH2D | 1499 | 96 | 20063 | 1.673032021 | 1 | 0.982000683 | 86.49152294 |
| GOTERM_MF_DIRECT | GO:0004713~protein tyrosine kinase activity | 16 | 1.016518424 | 0.105268069 | FGFR3, FGR, ERBB4, DSTYK, FGF23, FER, ALK, SCYL1, HBEGF, JAK1, TXK, CUX1, FGF2, ROS1, INSR, MA | 1340 | 133 | 16881 | 1.515520144 | 1 | 0.942780728 | 83.69541874 |

| | | | | | | | | | | | | |
|---|---|---|---|---|---|---|---|---|---|---|---|---|
| GOTERM_CC_DIRECT | GO:0031234~extrinsic component of cytoplasmic side of plasma membrane | 8 | 0.508259212 | 0.280224908 | FRK, FGR, JAK1, TXK, FER, ESYT2, MCF2L, ESYT3 | 1425 | 68 | 18224 | 1.504561404 | 1 | 0.886060292 | 99.28904161 |

Annotation Cluster 18 Enrichment Score: 1.4032170134337703

| Category | Term | Count | % | PValue | Genes | List Total | Pop Hits | Pop Total | Fold Enrichment | Bonferroni | Benjamini | FDR |
|---|---|---|---|---|---|---|---|---|---|---|---|---|
| UP_SEQ_FEATURE | domain:EGF-like 8 | 9 | 0.571791614 | 4.38E-04 | TENM4, TNXB, TNXA, STAB1, TENM2, TENM3, LRP1B, RELN, MEGF11 | 1499 | 26 | 20063 | 4.633011751 | 0.831955537 | 0.149676819 | 0.812023442 |
| UP_SEQ_FEATURE | domain:EGF-like 7 | 9 | 0.571791614 | 7.62E-04 | TENM4, TNXB, TNXA, STAB1, TENM2, TENM3, LRP1B, RELN, MEGF11 | 1499 | 28 | 20063 | 4.302082341 | 0.955092992 | 0.198807781 | 1.408593823 |
| INTERPRO | IPR022385:Rhs repeat-associated core | 3 | 0.190597205 | 0.017372155 | TENM4, TENM2, TENM3 | 1452 | 3 | 18559 | 12.78168044 | 1 | 0.732918975 | 26.05887113 |
| UP_SEQ_FEATURE | domain:Teneurin N-terminal | 3 | 0.190597205 | 0.030197446 | TENM4, TENM2, TENM3 | 1499 | 4 | 20063 | 10.03819213 | 1 | 0.862258025 | 43.5001235 |
| UP_SEQ_FEATURE | repeat:YD 20 | 3 | 0.190597205 | 0.030197446 | TENM4, TENM2, TENM3 | 1499 | 4 | 20063 | 10.03819213 | 1 | 0.862258025 | 43.5001235 |
| UP_SEQ_FEATURE | repeat:YD 21 | 3 | 0.190597205 | 0.030197446 | TENM4, TENM2, TENM3 | 1499 | 4 | 20063 | 10.03819213 | 1 | 0.862258025 | 43.5001235 |
| UP_SEQ_FEATURE | repeat:YD 22 | 3 | 0.190597205 | 0.030197446 | TENM4, TENM2, TENM3 | 1499 | 4 | 20063 | 10.03819213 | 1 | 0.862258025 | 43.5001235 |
| UP_SEQ_FEATURE | repeat:YD 23 | 3 | 0.190597205 | 0.030197446 | TENM4, TENM2, TENM3 | 1499 | 4 | 20063 | 10.03819213 | 1 | 0.862258025 | 43.5001235 |
| UP_SEQ_FEATURE | repeat:YD 17 | 3 | 0.190597205 | 0.030197446 | TENM4, TENM2, TENM3 | 1499 | 4 | 20063 | 10.03819213 | 1 | 0.862258025 | 43.5001235 |
| UP_SEQ_FEATURE | repeat:YD 18 | 3 | 0.190597205 | 0.030197446 | TENM4, TENM2, TENM3 | 1499 | 4 | 20063 | 10.03819213 | 1 | 0.862258025 | 43.5001235 |
| UP_SEQ_FEATURE | repeat:YD 19 | 3 | 0.190597205 | 0.030197446 | TENM4, TENM2, TENM3 | 1499 | 4 | 20063 | 10.03819213 | 1 | 0.862258025 | 43.5001235 |
| UP_SEQ_FEATURE | repeat:YD 2 | 3 | 0.190597205 | 0.030197446 | TENM4, TENM2, TENM3 | 1499 | 4 | 20063 | 10.03819213 | 1 | 0.862258025 | 43.5001235 |
| UP_SEQ_FEATURE | repeat:YD 7 | 3 | 0.190597205 | 0.030197446 | TENM4, TENM2, TENM3 | 1499 | 4 | 20063 | 10.03819213 | 1 | 0.862258025 | 43.5001235 |
| UP_SEQ_FEATURE | repeat:YD 8 | 3 | 0.190597205 | 0.030197446 | TENM4, TENM2, TENM3 | 1499 | 4 | 20063 | 10.03819213 | 1 | 0.862258025 | 43.5001235 |
| UP_SEQ_FEATURE | repeat:YD 9 | 3 | 0.190597205 | 0.030197446 | TENM4, TENM2, TENM3 | 1499 | 4 | 20063 | 10.03819213 | 1 | 0.862258025 | 43.5001235 |
| UP_SEQ_FEATURE | repeat:YD 3 | 3 | 0.190597205 | 0.030197446 | TENM4, TENM2, TENM3 | 1499 | 4 | 20063 | 10.03819213 | 1 | 0.862258025 | 43.5001235 |
| UP_SEQ_FEATURE | repeat:YD 4 | 3 | 0.190597205 | 0.030197446 | TENM4, TENM2, TENM3 | 1499 | 4 | 20063 | 10.03819213 | 1 | 0.862258025 | 43.5001235 |
| UP_SEQ_FEATURE | repeat:YD 5 | 3 | 0.190597205 | 0.030197446 | TENM4, TENM2, TENM3 | 1499 | 4 | 20063 | 10.03819213 | 1 | 0.862258025 | 43.5001235 |
| UP_SEQ_FEATURE | repeat:YD 6 | 3 | 0.190597205 | 0.030197446 | TENM4, TENM2, TENM3 | 1499 | 4 | 20063 | 10.03819213 | 1 | 0.862258025 | 43.5001235 |
| UP_SEQ_FEATURE | repeat:YD 14 | 3 | 0.190597205 | 0.030197446 | TENM4, TENM2, TENM3 | 1499 | 4 | 20063 | 10.03819213 | 1 | 0.862258025 | 43.5001235 |
| UP_SEQ_FEATURE | repeat:YD 13 | 3 | 0.190597205 | 0.030197446 | TENM4, TENM2, TENM3 | 1499 | 4 | 20063 | 10.03819213 | 1 | 0.862258025 | 43.5001235 |
| UP_SEQ_FEATURE | repeat:YD 16 | 3 | 0.190597205 | 0.030197446 | TENM4, TENM2, TENM3 | 1499 | 4 | 20063 | 10.03819213 | 1 | 0.862258025 | 43.5001235 |
| UP_SEQ_FEATURE | repeat:YD 15 | 3 | 0.190597205 | 0.030197446 | TENM4, TENM2, TENM3 | 1499 | 4 | 20063 | 10.03819213 | 1 | 0.862258025 | 43.5001235 |
| UP_SEQ_FEATURE | repeat:YD 10 | 3 | 0.190597205 | 0.030197446 | TENM4, TENM2, TENM3 | 1499 | 4 | 20063 | 10.03819213 | 1 | 0.862258025 | 43.5001235 |
| UP_SEQ_FEATURE | repeat:YD 1 | 3 | 0.190597205 | 0.030197446 | TENM4, TENM2, TENM3 | 1499 | 4 | 20063 | 10.03819213 | 1 | 0.862258025 | 43.5001235 |
| UP_SEQ_FEATURE | repeat:YD 12 | 3 | 0.190597205 | 0.030197446 | TENM4, TENM2, TENM3 | 1499 | 4 | 20063 | 10.03819213 | 1 | 0.862258025 | 43.5001235 |
| UP_SEQ_FEATURE | repeat:YD 11 | 3 | 0.190597205 | 0.030197446 | TENM4, TENM2, TENM3 | 1499 | 4 | 20063 | 10.03819213 | 1 | 0.862258025 | 43.5001235 |
| INTERPRO | IPR009471:Teneurin intracellular, N-terminal | 3 | 0.190597205 | 0.03294801 | TENM4, TENM2, TENM3 | 1452 | 4 | 18559 | 9.586260331 | 1 | 0.849368775 | 43.85082476 |
| INTERPRO | IPR006530:YD repeat | 3 | 0.190597205 | 0.03294801 | TENM4, TENM2, TENM3 | 1452 | 4 | 18559 | 9.586260331 | 1 | 0.849368775 | 43.85082476 |
| COG_ONTOLOGY | Cell envelope biogenesis, outer membrane | 5 | 0.317662008 | 0.073466494 | TENM4, GALNT3, TENM2, TENM3, GALNT12 | 166 | 20 | 2026 | 3.051204819 | 0.912992612 | 0.705029852 | 48.35953234 |
| GOTERM_BP_DIRECT | GO:0097264~self proteolysis | 3 | 0.190597205 | 0.102249192 | TENM4, TENM2, TENM3 | 1342 | 7 | 16792 | 5.362571854 | 1 | 0.979902008 | 86.56787542 |
| INTERPRO | IPR011042:Six-bladed beta-propeller, TolB-like | 7 | 0.444726811 | 0.106457144 | TENM4, TENM2, TENM3, LRP1B, LRP8, ROS1, LRP5 | 1452 | 42 | 18559 | 2.130280073 | 1 | 0.967194194 | 85.61657385 |
| INTERPRO | IPR008969:Carboxypeptidase-like, regulatory domain | 4 | 0.254129606 | 0.12452823 | TENM4, TENM2, TENM3, CILP | 1452 | 16 | 18559 | 3.19542011 | 1 | 0.970194765 | 89.88414493 |
| UP_SEQ_FEATURE | repeat:NHL 5 | 3 | 0.190597205 | 0.252812379 | TENM4, TENM2, TENM3 | 1499 | 13 | 20063 | 3.088674501 | 1 | 0.999215754 | 99.56015612 |
| UP_SEQ_FEATURE | repeat:NHL 2 | 3 | 0.190597205 | 0.281381718 | TENM4, TENM2, TENM3 | 1499 | 14 | 20063 | 2.868054894 | 1 | 0.999562574 | 99.78716607 |
| UP_SEQ_FEATURE | repeat:NHL 3 | 3 | 0.190597205 | 0.281381718 | TENM4, TENM2, TENM3 | 1499 | 14 | 20063 | 2.868054894 | 1 | 0.999562574 | 99.78716607 |
| UP_SEQ_FEATURE | repeat:NHL 1 | 3 | 0.190597205 | 0.281381718 | TENM4, TENM2, TENM3 | 1499 | 14 | 20063 | 2.868054894 | 1 | 0.999562574 | 99.78716607 |
| UP_SEQ_FEATURE | repeat:NHL 4 | 3 | 0.190597205 | 0.281381718 | TENM4, TENM2, TENM3 | 1499 | 14 | 20063 | 2.868054894 | 1 | 0.999562574 | 99.78716607 |
| GOTERM_BP_DIRECT | GO:0007157~heterophilic cell-cell adhesion via plasma membrane cell adhesion molecules | 5 | 0.317662008 | 0.573549763 | TENM4, TENM2, TENM3, NLGN1, NRXN1 | 1342 | 50 | 16792 | 1.251266766 | 1 | 0.999976807 | 99.99998708 |

Annotation Cluster 19 Enrichment Score: 1.3942238696620632

| Category | Term | Count | % | PValue | Genes | List Total | Pop Hits | Pop Total | Fold Enrichment | Bonferroni | Benjamini | FDR |
|---|---|---|---|---|---|---|---|---|---|---|---|---|
| UP_KEYWORDS | Non-syndromic deafness | 14 | 0.889453621 | 0.012051873 | DIAPH1, GIPC3, GJB3, PCDH15, GJB6, HGF, GRHL2, MSRB3, SLC26A4, TBC1D24, SLC26A5, DFNB59, M | 1535 | 87 | 20581 | 2.157579842 | 0.997465268 | 0.145558325 | 15.99216479 |
| UP_KEYWORDS | Deafness | 24 | 1.524777637 | 0.022666023 | FGFR3, DIAPH1, GIPC3, GJB3, ALMS1, PCDH15, SIX5, KCNJ10, HGF, GJB6, LARS2, GRHL2, MSRB3, PTF | 1535 | 198 | 20581 | 1.625190011 | 0.999987663 | 0.217855661 | 28.07178426 |
| GOTERM_BP_DIRECT | GO:0007605~sensory perception of sound | 17 | 1.080050826 | 0.06470853 | ZNF354A, DIAPH1, TH, ASIC2, GJB3, ALMS1, PCDH15, GJB6, SLC26A4, EML2, SLC26A5, DFNB59, WDR | 1342 | 133 | 16792 | 1.599363536 | 1 | 0.961989633 | 71.20808345 |
| UP_KEYWORDS | Hearing | 6 | 0.381194409 | 0.149869449 | SLC26A5, DIAPH1, PCDH15, DFNB59, GJB6, CDH23 | 1535 | 38 | 20581 | 2.11702383 | 1 | 0.536836065 | 90.30435297 |

Annotation Cluster 20 Enrichment Score: 1.3225431830927539

| Category | Term | Count | % | PValue | Genes | List Total | Pop Hits | Pop Total | Fold Enrichment | Bonferroni | Benjamini | FDR |
|---|---|---|---|---|---|---|---|---|---|---|---|---|
| INTERPRO | IPR000998:MAM domain | 5 | 0.317662008 | 0.047036072 | MALRD1, MAMDC4, PTPRM, PTPRT, ALK | 1452 | 18 | 18559 | 3.550466789 | 1 | 0.908380405 | 56.3937071 |
| UP_SEQ_FEATURE | domain:MAM 2 | 3 | 0.190597205 | 0.04785967 | MALRD1, MAMDC4, ALK | 1499 | 5 | 20063 | 8.030553702 | 1 | 0.912489302 | 59.8746983 |
| UP_SEQ_FEATURE | domain:MAM 1 | 3 | 0.190597205 | 0.04785967 | MALRD1, MAMDC4, ALK | 1499 | 5 | 20063 | 8.030553702 | 1 | 0.912489302 | 59.8746983 |

Annotation Cluster 21 Enrichment Score: 1.3176268492067542

| Category | Term | Count | % | PValue | Genes | List Total | Pop Hits | Pop Total | Fold Enrichment | Bonferroni | Benjamini | FDR |
|---|---|---|---|---|---|---|---|---|---|---|---|---|
| UP_SEQ_FEATURE | domain:LDL-receptor class A 2 | 8 | 0.508259212 | 0.002998743 | MALRD1, MAMDC4, CORIN, CD320, ST14, LRP1B, LRP8, LRP5 | 1499 | 27 | 20063 | 3.965705532 | 0.999995129 | 0.393309998 | 5.438482116 |
| UP_SEQ_FEATURE | domain:LDL-receptor class A 3 | 7 | 0.444726811 | 0.003503471 | MALRD1, MAMDC4, CORIN, ST14, LRP1B, LRP8, LRP5 | 1499 | 21 | 20063 | 4.461418724 | 0.999999381 | 0.411061214 | 6.325868296 |
| INTERPRO | IPR023415:Low-density lipoprotein (LDL) receptor class A, conserved site | 9 | 0.571791614 | 0.011050967 | MALRD1, C8B, MAMDC4, CORIN, CD320, ST14, LRP1B, LRP8, LRP5 | 1452 | 40 | 18559 | 2.875878099 | 1 | 0.659153573 | 17.42269325 |
| INTERPRO | IPR002172:Low-density lipoprotein (LDL) receptor class A repeat | 10 | 0.635324015 | 0.011072101 | MALRD1, C8B, MAMDC4, CORIN, CD320, ST14, LRP1B, LRP8, ALK, LRP5 | 1452 | 48 | 18559 | 2.662850092 | 1 | 0.642769959 | 17.45308938 |
| UP_SEQ_FEATURE | domain:LDL-receptor class A 1 | 7 | 0.444726811 | 0.013012927 | MALRD1, CORIN, CD320, ST14, LRP1B, LRP8, LRP5 | 1499 | 27 | 20063 | 3.469992341 | 1 | 0.694420403 | 21.64238885 |
| SMART | SM00192:LDLa | 10 | 0.635324015 | 0.016295154 | MALRD1, C8B, MAMDC4, CORIN, CD320, ST14, LRP1B, LRP8, ALK, LRP5 | 878 | 46 | 10057 | 2.490096068 | 0.998906304 | 0.679016458 | 20.57725257 |
| UP_SEQ_FEATURE | domain:LDL-receptor class A 4 | 5 | 0.317662008 | 0.027270834 | MALRD1, CORIN, ST14, LRP1B, LRP8 | 1499 | 16 | 20063 | 4.182580053 | 1 | 0.851735679 | 40.23959954 |
| UP_SEQ_FEATURE | domain:LDL-receptor class A 7 | 4 | 0.254129606 | 0.043558648 | MALRD1, CORIN, LRP1B, LRP8 | 1499 | 11 | 20063 | 4.867002244 | 1 | 0.899352497 | 56.36206388 |
| UP_SEQ_FEATURE | domain:LDL-receptor class A 6 | 4 | 0.254129606 | 0.043558648 | MALRD1, CORIN, LRP1B, LRP8 | 1499 | 11 | 20063 | 4.867002244 | 1 | 0.899352497 | 56.36206388 |
| UP_SEQ_FEATURE | domain:LDL-receptor class A 5 | 4 | 0.254129606 | 0.05493803 | MALRD1, CORIN, LRP1B, LRP8 | 1499 | 12 | 20063 | 4.461418724 | 1 | 0.93330171 | 65.07970204 |
| INTERPRO | IPR011042:Six-bladed beta-propeller, TolB-like | 7 | 0.444726811 | 0.106457144 | TENM4, TENM2, TENM3, LRP1B, LRP8, ROS1, LRP5 | 1452 | 42 | 18559 | 2.130280073 | 1 | 0.967194194 | 85.61657385 |
| INTERPRO | IPR000033:LDLR class B repeat | 4 | 0.254129606 | 0.10711664 | LRP1B, LRP8, ROS1, LRP5 | 1452 | 15 | 18559 | 3.408448118 | 1 | 0.964558329 | 85.79836453 |
| SMART | SM00135:LY | 4 | 0.254129606 | 0.136924544 | LRP1B, LRP8, ROS1, LRP5 | 878 | 15 | 10057 | 3.054517844 | 1 | 0.895997004 | 87.31686152 |
| UP_SEQ_FEATURE | repeat:LDL-receptor class B 5 | 3 | 0.190597205 | 0.252812379 | LRP1B, LRP8, LRP5 | 1499 | 13 | 20063 | 3.088674501 | 1 | 0.999215754 | 99.56015612 |
| UP_SEQ_FEATURE | repeat:LDL-receptor class B 4 | 3 | 0.190597205 | 0.281381718 | LRP1B, LRP8, LRP5 | 1499 | 14 | 20063 | 2.868054894 | 1 | 0.999562574 | 99.78716607 |
| UP_SEQ_FEATURE | repeat:LDL-receptor class B 3 | 3 | 0.190597205 | 0.281381718 | LRP1B, LRP8, LRP5 | 1499 | 14 | 20063 | 2.868054894 | 1 | 0.999562574 | 99.78716607 |
| UP_SEQ_FEATURE | repeat:LDL-receptor class B 1 | 3 | 0.190597205 | 0.281381718 | LRP1B, LRP8, LRP5 | 1499 | 14 | 20063 | 2.868054894 | 1 | 0.999562574 | 99.78716607 |
| UP_SEQ_FEATURE | repeat:LDL-receptor class B 2 | 3 | 0.190597205 | 0.281381718 | LRP1B, LRP8, LRP5 | 1499 | 14 | 20063 | 2.868054894 | 1 | 0.999562574 | 99.78716607 |

Annotation Cluster 22 Enrichment Score: 1.281664139954155

| Category | Term | Count | % | PValue | Genes | List Total | Pop Hits | Pop Total | Fold Enrichment | Bonferroni | Benjamini | FDR |
|---|---|---|---|---|---|---|---|---|---|---|---|---|
| GOTERM_BP_DIRECT | GO:0035418~protein localization to synapse | 6 | 0.381194409 | 0.0024075 | KLC1, ASIC2, NLGN1, RELN, NRXN1, PCLO | 1342 | 13 | 16792 | 5.775077382 | 0.999943372 | 0.623899993 | 4.387053375 |
| GOTERM_BP_DIRECT | GO:0097120~receptor localization to synapse | 4 | 0.254129606 | 0.029616388 | NLGN1, RELN, NRXN1, DLG2 | 1342 | 9 | 16792 | 5.561185627 | 1 | 0.891131644 | 42.85292149 |
| GOTERM_BP_DIRECT | GO:0097114~NMDA glutamate receptor clustering | 3 | 0.190597205 | 0.054161082 | NLGN1, RELN, NRXN1 | 1342 | 5 | 16792 | 7.507600596 | 1 | 0.950810698 | 64.52569176 |
| GOTERM_BP_DIRECT | GO:2000463~positive regulation of excitatory postsynaptic potential | 5 | 0.317662008 | 0.070133242 | NLGN1, RELN, CUX2, NRXN1, RIMS1 | 1342 | 20 | 16792 | 3.128166915 | 1 | 0.967620814 | 74.16238249 |
| GOTERM_BP_DIRECT | GO:0097119~postsynaptic density protein 95 clustering | 3 | 0.190597205 | 0.077013635 | NLGN1, RELN, NRXN1 | 1342 | 6 | 16792 | 6.25633383 | 1 | 0.971924876 | 77.49799337 |

| | | | | | | | | | | | | |
|---|---|---|---|---|---|---|---|---|---|---|---|---|
| GOTERM_BP_DIRECT | GO:2000310~regulation of N-methyl-D-aspartate selective glutamate receptor activity | 4 | 0.254129606 | 0.112429822 | CRH, NLGN1, RELN, NRXN1 | 1342 | 15 | 16792 | 3.336711376 | 1 | 0.97916148 | 89.13678205 |
| GOTERM_BP_DIRECT | GO:0051968~positive regulation of synaptic transmission, glutamatergic | 3 | 0.190597205 | 0.455189596 | NLGN1, RELN, NRXN1 | 1342 | 19 | 16792 | 1.975684367 | 1 | 0.999853888 | 99.99876676 |

Annotation Cluster 23 Enrichment Score: 1.279994250993593

| Category | Term | Count | % | PValue | Genes | List Total | Pop Hits | Pop Total | Fold Enrichment | Bonferroni | Benjamini | FDR |
|---|---|---|---|---|---|---|---|---|---|---|---|---|
| GOTERM_CC_DIRECT | GO:0016459~myosin complex | 11 | 0.698856417 | 0.004717196 | MYO5A, MYO18B, MYO1C, MYO7B, CGN, CCDC102A, MYO1E, MYO3B, MYH14, MYL12A, MYO5B | 1425 | 50 | 18224 | 2.813529825 | 0.961162975 | 0.255697235 | 6.865695203 |
| GOTERM_MF_DIRECT | GO:0003774~motor activity | 12 | 0.762388818 | 0.014099669 | MYO5A, MYO18B, MYO1C, MYO7B, CGN, KLC1, CCDC102A, MYO1E, MYO3B, DYNC2H1, MYH14, MY( | 1340 | 66 | 16881 | 2.290502035 | 0.999999988 | 0.838963111 | 20.66922667 |
| INTERPRO | IPR000048:IQ motif, EF-hand binding site | 14 | 0.889453621 | 0.017337062 | MYO5A, CAMTA1, OBSCN, MYO1C, MYO7B, MYO1E, MYO3B, MYO18B, RASGRF2, SPATA17, MYH14, | 1452 | 87 | 18559 | 2.05682214 | 1 | 0.745431779 | 26.01336681 |
| UP_SEQ_FEATURE | domain:Myosin head-like | 8 | 0.508259212 | 0.018120624 | MYO5A, MYO18B, MYO1C, MYO7B, MYO1E, MYO3B, MYH14, MYO5B | 1499 | 37 | 20063 | 2.893893226 | 1 | 0.767864757 | 28.85818874 |
| UP_SEQ_FEATURE | domain:IQ 2 | 7 | 0.444726811 | 0.033297191 | MYO5A, CAMTA1, MYO1C, MYO7B, MYO3B, SPATA17, MYO5B | 1499 | 33 | 20063 | 2.839084642 | 1 | 0.880205871 | 46.76960017 |
| UP_SEQ_FEATURE | domain:IQ 1 | 7 | 0.444726811 | 0.033297191 | MYO5A, CAMTA1, MYO1C, MYO7B, MYO3B, SPATA17, MYO5B | 1499 | 33 | 20063 | 2.839084642 | 1 | 0.880205871 | 46.76960017 |
| INTERPRO | IPR001609:Myosin head, motor domain | 8 | 0.508259212 | 0.033653208 | MYO5A, MYO18B, MYO1C, MYO7B, MYO1E, MYO3B, MYH14, MYO5B | 1452 | 40 | 18559 | 2.556336088 | 1 | 0.84769389 | 44.55203464 |
| UP_KEYWORDS | Myosin | 9 | 0.571791614 | 0.033824701 | MYO5A, MYO18B, MYO1C, MYO7B, MYO1E, MYO3B, MYH14, MYL12A, MYO5B | 1535 | 51 | 20581 | 2.366085457 | 0.999999957 | 0.253593513 | 39.01460773 |
| SMART | SM00242:MYSc | 8 | 0.508259212 | 0.055288539 | MYO5A, MYO18B, MYO1C, MYO7B, MYO1E, MYO3B, MYH14, MYO5B | 878 | 40 | 10057 | 2.290888383 | 1 | 0.860117481 | 54.95711434 |
| GOTERM_MF_DIRECT | GO:0000146~microfilament motor activity | 5 | 0.317662008 | 0.068735926 | MYO5A, MYO1C, MYO1E, MYH14, MYO5B | 1340 | 20 | 16881 | 3.149440299 | 1 | 0.910182677 | 68.68843034 |
| UP_SEQ_FEATURE | domain:IQ 3 | 5 | 0.317662008 | 0.077014528 | MYO5A, CAMTA1, MYO7B, SPATA17, MYO5B | 1499 | 22 | 20063 | 3.041876402 | 1 | 0.965442434 | 77.51261931 |
| UP_SEQ_FEATURE | domain:IQ 6 | 3 | 0.190597205 | 0.115475174 | MYO5A, MYO7B, MYO5B | 1499 | 8 | 20063 | 5.019096064 | 1 | 0.986545152 | 89.81978955 |
| UP_KEYWORDS | Motor protein | 15 | 0.952986023 | 0.130849239 | MYO5A, DNAH11, MYO1C, MYO7B, MYO1E, MYO3B, MYL12A, MYO18B, SLC26A5, KLC1, DYNC2H1, I | 1535 | 135 | 20581 | 1.48975751 | 1 | 0.502599778 | 86.6745121 |
| GOTERM_BP_DIRECT | GO:0030048~actin filament-based movement | 4 | 0.254129606 | 0.14961134 | MYO5A, MYO1E, WIPF3, MYH14 | 1342 | 17 | 16792 | 2.944157097 | 1 | 0.98677382 | 95.10160843 |
| SMART | SM00015:IQ | 8 | 0.508259212 | 0.163970142 | MYO5A, OBSCN, MYO1C, MYO7B, MYO3B, SPATA17, MYH14, MYO5B | 878 | 52 | 10057 | 1.762221833 | 1 | 0.909055687 | 91.8841458 |
| UP_SEQ_FEATURE | domain:IQ 5 | 3 | 0.190597205 | 0.196084959 | MYO5A, MYO7B, MYO5B | 1499 | 11 | 20063 | 3.650251683 | 1 | 0.997732105 | 98.28192692 |
| UP_SEQ_FEATURE | domain:IQ 4 | 3 | 0.190597205 | 0.309852953 | MYO5A, MYO7B, MYO5B | 1499 | 15 | 20063 | 2.676851234 | 1 | 0.999751312 | 99.89973676 |

Annotation Cluster 24 Enrichment Score: 1.2727440414350037

| Category | Term | Count | % | PValue | Genes | List Total | Pop Hits | Pop Total | Fold Enrichment | Bonferroni | Benjamini | FDR |
|---|---|---|---|---|---|---|---|---|---|---|---|---|
| UP_SEQ_FEATURE | domain:EGF-like 10 | 6 | 0.381194409 | 0.006673928 | TNXB, TNXA, STAB1, NOTCH4, LRP1B, MEGF11 | 1499 | 17 | 20063 | 4.723855119 | 1 | 0.585137079 | 11.72233433 |
| UP_SEQ_FEATURE | domain:EGF-like 16 | 4 | 0.254129606 | 0.024801433 | TNXB, TNXA, STAB1, NOTCH4 | 1499 | 9 | 20063 | 5.948558298 | 1 | 0.833799715 | 37.35073803 |
| UP_SEQ_FEATURE | domain:EGF-like 19 | 3 | 0.190597205 | 0.115475174 | TNXB, TNXA, NOTCH4 | 1499 | 8 | 20063 | 5.019096064 | 1 | 0.986545152 | 89.81978955 |
| UP_SEQ_FEATURE | domain:EGF-like 18 | 3 | 0.190597205 | 0.115475174 | TNXB, TNXA, NOTCH4 | 1499 | 8 | 20063 | 5.019096064 | 1 | 0.986545152 | 89.81978955 |
| UP_SEQ_FEATURE | domain:EGF-like 17 | 3 | 0.190597205 | 0.196084959 | TNXB, TNXA, NOTCH4 | 1499 | 11 | 20063 | 3.650251683 | 1 | 0.997732105 | 98.28192692 |

Annotation Cluster 25 Enrichment Score: 1.2669161441836099

| Category | Term | Count | % | PValue | Genes | List Total | Pop Hits | Pop Total | Fold Enrichment | Bonferroni | Benjamini | FDR |
|---|---|---|---|---|---|---|---|---|---|---|---|---|
| REACTOME_PATHWAY | R-HSA-3928665:R-HSA-3928665 | 10 | 0.635324015 | 0.018943536 | AP2A2, CLTA, EFNA1, EFNA3, APH1B, EFNA5, EFNA4, VAV2, EPHB3, EPHB1 | 745 | 50 | 9075 | 2.436241611 | 0.999999928 | 0.904599865 | 25.64818872 |
| INTERPRO | IPR001799:Ephrin | 4 | 0.254129606 | 0.019809027 | EFNA1, EFNA3, EFNA5, EFNA4 | 1452 | 8 | 18559 | 6.39084022 | 1 | 0.754217829 | 29.15503772 |
| INTERPRO | IPR019765:Ephrin, conserved site | 4 | 0.254129606 | 0.019809027 | EFNA1, EFNA3, EFNA5, EFNA4 | 1452 | 8 | 18559 | 6.39084022 | 1 | 0.754217829 | 29.15503772 |
| GOTERM_BP_DIRECT | GO:0048013~ephrin receptor signaling pathway | 13 | 0.82592122 | 0.040173141 | AP2A2, HRAS, GRIN2B, EFNA1, EFNA3, APH1B, EFNA5, EFNA4, MYL12A, VAV2, EPHB3, EPHB1, PTPN | 1342 | 86 | 16792 | 1.891449763 | 1 | 0.922631723 | 53.37942724 |
| REACTOME_PATHWAY | R-HSA-2682334:R-HSA-2682334 | 6 | 0.381194409 | 0.05683812 | EFNA1, EFNA3, EFNA5, EFNA4, EPHB3, EPHB1 | 745 | 26 | 9075 | 2.811048012 | 1 | 0.929255457 | 59.61758641 |
| REACTOME_PATHWAY | R-HSA-3928663:R-HSA-3928663 | 7 | 0.444726811 | 0.062555153 | EFNA1, EFNA3, VEGFA, EFNA5, MYH14, EFNA4, MYL12A | 745 | 35 | 9075 | 2.436241611 | 1 | 0.937817781 | 63.24848219 |
| INTERPRO | IPR008972:Cupredoxin | 4 | 0.254129606 | 0.202402908 | EFNA1, EFNA3, EFNA5, EFNA4 | 1452 | 20 | 18559 | 2.556336088 | 1 | 0.987485679 | 97.96754682 |
| GOTERM_MF_DIRECT | GO:0046875~ephrin receptor binding | 4 | 0.254129606 | 0.34074396 | EFNA1, EFNA3, EFNA5, EFNA4 | 1340 | 26 | 16881 | 1.938117107 | 1 | 0.991275515 | 99.88792371 |

Annotation Cluster 26 Enrichment Score: 1.2394591787413343

| Category | Term | Count | % | PValue | Genes | List Total | Pop Hits | Pop Total | Fold Enrichment | Bonferroni | Benjamini | FDR |
|---|---|---|---|---|---|---|---|---|---|---|---|---|
| GOTERM_BP_DIRECT | GO:0090630~activation of GTPase activity | 13 | 0.82592122 | 0.02234305 | WNT5A, TBC1D3C, TBC1D8, NTF3, TBC1D3H, RALGAPB, TBC1D22B, USP6NL, TBC1D16, SYDE2, SGSN | 1342 | 79 | 16792 | 2.059046577 | 1 | 0.875505559 | 34.33214225 |
| UP_SEQ_FEATURE | domain:Rab-GAP TBC | 9 | 0.571791614 | 0.024590691 | TBC1D3C, TBC1D16, TBC1D24, TBC1D8, TBC1D3H, SGSM3, TBC1D22B, USP6NL, TBC1D9B | 1499 | 48 | 20063 | 2.509548032 | 1 | 0.836491714 | 37.09817317 |
| GOTERM_BP_DIRECT | GO:1902017~regulation of cilium assembly | 8 | 0.508259212 | 0.025221778 | TBC1D16, TBC1D24, TBC1D8, SGSM3, TBC1D22B, USP6NL, TBC1D9B, IFT140 | 1342 | 37 | 16792 | 2.705441656 | 1 | 0.87937231 | 37.83911544 |
| INTERPRO | IPR000195:Rab-GTPase-TBC domain | 10 | 0.635324015 | 0.031765642 | TBC1D3C, TBC1D16, TBC1D24, TBC1D8, TBC1D3H, SGSM3, TBC1D22B, TBC1D32, USP6NL, TBC1D9B | 1452 | 57 | 18559 | 2.242400077 | 1 | 0.855021561 | 42.65636723 |
| GOTERM_MF_DIRECT | GO:0017137~Rab GTPase binding | 18 | 1.143583227 | 0.036221427 | TBC1D3C, MYO5A, TBC1D8, TBC1D3H, HPS4, VPS52, RGP1, RIMS1, TBC1D22B, USP6NL, TBC1D16, M | 1340 | 134 | 16881 | 1.692236578 | 1 | 0.895576476 | 45.20576753 |
| GOTERM_BP_DIRECT | GO:0031338~regulation of vesicle fusion | 8 | 0.508259212 | 0.05228818 | TBC1D3C, TBC1D16, TBC1D8, TBC1D3H, SGSM3, TBC1D22B, USP6NL, TBC1D9B | 1342 | 43 | 16792 | 2.327938169 | 1 | 0.947362681 | 63.19526767 |
| SMART | SM00164:TBC | 9 | 0.571791614 | 0.065898112 | TBC1D3C, TBC1D16, TBC1D24, TBC1D8, TBC1D3H, SGSM3, TBC1D22B, USP6NL, TBC1D9B | 878 | 50 | 10057 | 2.061799544 | 1 | 0.829352137 | 61.55450513 |
| GOTERM_CC_DIRECT | GO:0012505~endomembrane system | 14 | 0.889453621 | 0.108010766 | TBC1D3C, TBC1D8, TBC1D3H, SGSM3, WDFY4, ATF6B, JAK1, PIP5K1C, NBEA, TBC1D22B, TSNARE1, N | 1425 | 114 | 18224 | 1.570550939 | 1 | 0.760146294 | 82.08276636 |
| UP_KEYWORDS | GTPase activation | 20 | 1.27064803 | 0.149244449 | ARFGAP1, TBC1D3C, TBC1D8, TBC1D3H, RALGAPB, SIPA1L2, ASAP1, TBC1D22B, ADAP1, USP6NL, TB( | 1535 | 197 | 20581 | 1.361199755 | 1 | 0.538664253 | 90.20140443 |
| GOTERM_MF_DIRECT | GO:0005096~GTPase activator activity | 24 | 1.524777637 | 0.455149856 | ARFGAP1, TBC1D3C, TBC1D8, TBC1D3H, RALGAPB, SIPA1L2, ASAP1, RIC8B, TBC1D22B, ADAP1, USP6 | 1340 | 279 | 16881 | 1.083678382 | 1 | 0.995910822 | 99.99499091 |

Annotation Cluster 27 Enrichment Score: 1.2342177632525675

| Category | Term | Count | % | PValue | Genes | List Total | Pop Hits | Pop Total | Fold Enrichment | Bonferroni | Benjamini | FDR |
|---|---|---|---|---|---|---|---|---|---|---|---|---|
| UP_SEQ_FEATURE | domain:PID | 7 | 0.444726811 | 0.01554069 | DAB2, DAB1, NOS1AP, ITGB1BP1, EPS8L2, APBA1, SHC4 | 1499 | 28 | 20063 | 3.346064043 | 1 | 0.742651487 | 25.29588398 |
| INTERPRO | IPR006020:Phosphotyrosine interaction domain | 7 | 0.444726811 | 0.097331541 | DAB2, DAB1, NOS1AP, ITGB1BP1, EPS8L2, APBA1, SHC4 | 1452 | 41 | 18559 | 2.182238124 | 1 | 0.968925205 | 82.86503344 |
| SMART | SM00462:PTB | 7 | 0.444726811 | 0.13110607 | DAB2, DAB1, NOS1AP, ITGB1BP1, EPS8L2, APBA1, SHC4 | 878 | 40 | 10057 | 2.004527335 | 1 | 0.902982905 | 86.06377579 |

Annotation Cluster 28 Enrichment Score: 1.2251715919275556

| Category | Term | Count | % | PValue | Genes | List Total | Pop Hits | Pop Total | Fold Enrichment | Bonferroni | Benjamini | FDR |
|---|---|---|---|---|---|---|---|---|---|---|---|---|
| UP_SEQ_FEATURE | chain:Cytosolic phospholipase A2 beta | 3 | 0.190597205 | 0.015883223 | JMJD7-PLA2G4B, JMJD7, PLA2G4B | 1499 | 3 | 20063 | 13.38425617 | 1 | 0.742973401 | 25.77837608 |
| UP_SEQ_FEATURE | chain:JmjC domain-containing protein 7 | 3 | 0.190597205 | 0.015883223 | JMJD7-PLA2G4B, JMJD7, PLA2G4B | 1499 | 3 | 20063 | 13.38425617 | 1 | 0.742973401 | 25.77837608 |
| UP_SEQ_FEATURE | domain:PLA2c | 3 | 0.190597205 | 0.115475174 | JMJD7-PLA2G4B, JMJD7, PLA2G4B | 1499 | 8 | 20063 | 5.019096064 | 1 | 0.986545152 | 89.81978955 |
| UP_SEQ_FEATURE | domain:JmjC | 4 | 0.254129606 | 0.431467168 | JMJD7-PLA2G4B, KDM2A, JMJD7, PLA2G4B | 1499 | 32 | 20063 | 1.673032021 | 1 | 0.999972527 | 99.99728592 |

Annotation Cluster 29 Enrichment Score: 1.1992517973379717

| Category | Term | Count | % | PValue | Genes | List Total | Pop Hits | Pop Total | Fold Enrichment | Bonferroni | Benjamini | FDR |
|---|---|---|---|---|---|---|---|---|---|---|---|---|
| UP_KEYWORDS | Calcium transport | 16 | 1.016518424 | 0.005868355 | MICU1, SLC8A1, TRPV2, CACNG6, CACNG5, MCU, CACNG2, ITPR2, ATP2C2, ATP2B4, SLC24A4, CATSF | 1535 | 98 | 20581 | 2.189031443 | 0.945065907 | 0.089353673 | 8.110888902 |
| UP_KEYWORDS | Calcium channel | 10 | 0.635324015 | 0.055308229 | CATSPER3, RYR3, TRPV2, CACNG6, CACNG5, MCU, RYR2, CACNG2, CACNA1C, ITPR2 | 1535 | 66 | 20581 | 2.031487514 | 1 | 0.322661056 | 55.8558576 |
| GOTERM_BP_DIRECT | GO:0070588~calcium ion transmembrane transport | 14 | 0.889453621 | 0.154153054 | SLC8A1, TMC2, TRPV2, CACNG6, CACNG5, CACNG2, ATP2C2, ATP2B4, GPM6A, SLC24A4, CATSPER3, | 1342 | 119 | 16792 | 1.472078548 | 1 | 0.98819559 | 95.56627766 |
| GOTERM_MF_DIRECT | GO:0005262~calcium channel activity | 8 | 0.508259212 | 0.318958785 | SLC24A4, GPM6A, TRPV2, CACNG6, SLC24A1, MCU, RYR2, CACNG2 | 1340 | 70 | 16881 | 1.439744136 | 1 | 0.991347833 | 99.80956737 |

Annotation Cluster 30 Enrichment Score: 1.1934914194389876

| Category | Term | Count | % | PValue | Genes | List Total | Pop Hits | Pop Total | Fold Enrichment | Bonferroni | Benjamini | FDR |
|---|---|---|---|---|---|---|---|---|---|---|---|---|
| UP_SEQ_FEATURE | repeat:WD 11 | 8 | 0.508259212 | 0.005648113 | EML2, ELP2, WDR17, WDR62, TEP1, WDR1, MAPKBP1, PWP2 | 1499 | 30 | 20063 | 3.569134979 | 1 | 0.548651894 | 10.00933774 |
| UP_SEQ_FEATURE | repeat:WD 10 | 8 | 0.508259212 | 0.006834225 | EML2, ELP2, WDR17, WDR62, TEP1, WDR1, MAPKBP1, PWP2 | 1499 | 31 | 20063 | 3.454001592 | 1 | 0.582242751 | 11.98720941 |
| UP_SEQ_FEATURE | repeat:WD 12 | 6 | 0.381194409 | 0.020813723 | ELP2, WDR17, WDR62, TEP1, MAPKBP1, PWP2 | 1499 | 22 | 20063 | 3.650251683 | 1 | 0.80746553 | 32.40493656 |
| UP_SEQ_FEATURE | repeat:WD 9 | 8 | 0.508259212 | 0.043008582 | EML2, ELP2, WDR17, WDR62, TEP1, WDR1, MAPKBP1, PWP2 | 1499 | 44 | 20063 | 2.433501122 | 1 | 0.902251553 | 55.89239107 |
| UP_SEQ_FEATURE | repeat:WD 8 | 9 | 0.571791614 | 0.076652365 | EML2, TBL1XR1, ELP2, WDR17, WDR62, TEP1, WDR1, MAPKBP1, PWP2 | 1499 | 60 | 20063 | 2.007638426 | 1 | 0.966072786 | 77.34775645 |
| UP_SEQ_FEATURE | repeat:WD 14 | 4 | 0.254129606 | 0.081396345 | ELP2, WDR62, TEP1, PWP2 | 1499 | 14 | 20063 | 3.824073192 | 1 | 0.967410363 | 79.41941972 |
| UP_SEQ_FEATURE | repeat:WD 13 | 4 | 0.254129606 | 0.146780167 | ELP2, WDR62, TEP1, PWP2 | 1499 | 18 | 20063 | 2.974279149 | 1 | 0.993080316 | 94.79554201 |
| UP_SEQ_FEATURE | repeat:WD 7 | 15 | 0.952986023 | 0.23740349 | TBL1XR1, ELP2, AHI1, PWP2, FBXW9, EML2, WDR83, WDR37, WDR17, WDR62, TEP1, PAFAH1B1, WI | 1499 | 151 | 20063 | 1.329561871 | 1 | 0.998901856 | 99.35677208 |

| | | | | | | | | | | | | |
|---|---|---|---|---|---|---|---|---|---|---|---|---|
| UP_SEQ_FEATURE | repeat:WD 6 | 18 | 1.143583227 | 0.295475061 | TBL1XR1, ELP2, RBBP5, AHI1, PWP2, WDR83, EML2, FBXW9, WDR37, WDR17, WDR62, TEP1, PAFAH | 1499 | 196 | 20063 | 1.229166383 | 1 | 0.999697997 | 99.8528107 |
| INTERPRO | IPR011047:Quinonprotein alcohol dehydrogenase-like superfamily | 5 | 0.317662008 | 0.523338292 | EML2, WDR17, WDR62, TEP1, PWP2 | 1452 | 48 | 18559 | 1.331425046 | 1 | 0.999818079 | 99.9997139 |

Annotation Cluster 31 Enrichment Score: 1.1780515847589232

| Category | Term | Count | % | PValue | Genes | List Total | Pop Hits | Pop Total | Fold Enrichment | Bonferroni | Benjamini | FDR |
|---|---|---|---|---|---|---|---|---|---|---|---|---|
| KEGG_PATHWAY | hsa05150:Staphylococcus aureus infection | 13 | 0.82592122 | 0.001317045 | HLA-DQB1, MASP1, C4A, HLA-DRB1, C4B, CFB, C5, C1S, C2, HLA-DQA2, HLA-DOB, HLA-DQA1, HLA-DI | 574 | 54 | 6879 | 2.885114208 | 0.307673593 | 0.307673593 | 1.726551035 |
| REACTOME_PATHWAY | R-HSA-174577:R-HSA-174577 | 5 | 0.317662008 | 0.002399156 | C4A, C4B, CFB, C5, C2 | 745 | 8 | 9075 | 7.613255034 | 0.873277173 | 0.873277173 | 3.653740927 |
| BIOCARTA | h_compPathway:Complement Pathway | 7 | 0.444726811 | 0.006153355 | MASP1, C4A, C4B, CFB, C5, C1S, C2 | 146 | 20 | 1625 | 3.895547945 | 0.74907712 | 0.74907712 | 7.577363584 |
| INTERPRO | IPR000020:Anaphylatoxin/fibulin | 4 | 0.254129606 | 0.013136595 | FBLN1, C4A, C4B, C5 | 1452 | 7 | 18559 | 7.303817395 | 1 | 0.673949747 | 20.37199691 |
| INTERPRO | IPR008930:Terpenoid cyclases/protein prenyltransferase alpha-alpha toroid | 6 | 0.381194409 | 0.013355289 | FNTB, C4A, CHURC1-FNTB, CPAMD8, C4B, C5 | 1452 | 19 | 18559 | 4.036320139 | 1 | 0.665094685 | 20.67543948 |
| SMART | SM00104:ANATO | 4 | 0.254129606 | 0.01770198 | FBLN1, C4A, C4B, C5 | 878 | 7 | 10057 | 6.545395379 | 0.999396112 | 0.604068952 | 22.15527991 |
| UP_SEQ_FEATURE | domain:NTR | 6 | 0.381194409 | 0.020813723 | C4A, SFRP1, C4B, C5, TIMP2, NTN1 | 1499 | 22 | 20063 | 3.650251683 | 1 | 0.80746553 | 32.40493656 |
| BIOCARTA | h_lectinPathway:Lectin Induced Complement Pathway | 5 | 0.317662008 | 0.022969103 | MASP1, C4A, C4B, C5, C2 | 146 | 13 | 1625 | 4.280821918 | 0.994511436 | 0.925915159 | 25.66962815 |
| REACTOME_PATHWAY | R-HSA-977606:R-HSA-977606 | 7 | 0.444726811 | 0.023535655 | C8B, C4A, C4B, CFB, C5, C4BPA, C2 | 745 | 28 | 9075 | 3.045302013 | 0.999999999 | 0.922720924 | 30.86196884 |
| UP_KEYWORDS | Complement pathway | 7 | 0.444726811 | 0.024923369 | C8B, C4A, C4B, C5, C4BPA, C1S, C2 | 1535 | 31 | 20581 | 3.027571714 | 0.999996054 | 0.228352753 | 30.422895 |
| GOTERM_BP_DIRECT | GO:0030449~regulation of complement activation | 7 | 0.444726811 | 0.028941218 | C8B, C4A, C4B, CFB, C5, C4BPA, C2 | 1342 | 30 | 16792 | 2.919622454 | 1 | 0.894395922 | 42.10833464 |
| INTERPRO | IPR001134:Netrin domain | 6 | 0.381194409 | 0.029818296 | C4A, SFRP1, C4B, C5, TIMP2, NTN1 | 1452 | 23 | 18559 | 3.334351419 | 1 | 0.84523538 | 40.63679163 |
| UP_SEQ_FEATURE | domain:Anaphylatoxin-like | 3 | 0.190597205 | 0.030197446 | C4A, C4B, C5 | 1499 | 4 | 20063 | 10.03819213 | 1 | 0.862258025 | 43.5001235 |
| BIOCARTA | h_classicPathway:Classical Complement Pathway | 5 | 0.317662008 | 0.038114048 | C4A, C4B, C5, C1S, C2 | 146 | 15 | 1625 | 3.710045662 | 0.999834163 | 0.945059309 | 39.10935301 |
| INTERPRO | IPR018933:Netrin module, non-TIMP type | 5 | 0.317662008 | 0.047036072 | C4A, SFRP1, C4B, C5, NTN1 | 1452 | 18 | 18559 | 3.550466789 | 1 | 0.908380405 | 56.3937071 |
| INTERPRO | IPR008993:Tissue inhibitor of metalloproteinases-like, OB-fold | 6 | 0.381194409 | 0.048072362 | C4A, SFRP1, C4B, C5, TIMP2, NTN1 | 1452 | 26 | 18559 | 2.949618563 | 1 | 0.907993354 | 57.20343137 |
| INTERPRO | IPR018081:Anaphylatoxin | 3 | 0.190597205 | 0.052094013 | C4A, C4B, C5 | 1452 | 5 | 18559 | 7.669008264 | 1 | 0.910917985 | 60.21365103 |
| INTERPRO | IPR001840:Complement C3a/C4a/C5a anaphylatoxin | 3 | 0.190597205 | 0.052094013 | C4A, C4B, C5 | 1452 | 5 | 18559 | 7.669008264 | 1 | 0.910917985 | 60.21365103 |
| SMART | SM00643:C345C | 5 | 0.317662008 | 0.054650294 | C4A, SFRP1, C4B, C5, NTN1 | 878 | 17 | 10057 | 3.368953504 | 1 | 0.880002734 | 54.5285064 |
| INTERPRO | IPR009048:Alpha-macroglobulin, receptor-binding | 4 | 0.254129606 | 0.061563084 | C4A, CPAMD8, C4B, C5 | 1452 | 12 | 18559 | 4.260560147 | 1 | 0.928463219 | 66.53267445 |
| INTERPRO | IPR002890:Alpha-macroglobulin, N-terminal | 4 | 0.254129606 | 0.061563084 | C4A, CPAMD8, C4B, C5 | 1452 | 12 | 18559 | 4.260560147 | 1 | 0.928463219 | 66.53267445 |
| INTERPRO | IPR001599:Alpha-2-macroglobulin | 4 | 0.254129606 | 0.075530428 | C4A, CPAMD8, C4B, C5 | 1452 | 13 | 18559 | 3.932824751 | 1 | 0.950903684 | 74.15178854 |
| INTERPRO | IPR011626:A-macroglobulin complement component | 4 | 0.254129606 | 0.075530428 | C4A, CPAMD8, C4B, C5 | 1452 | 13 | 18559 | 3.932824751 | 1 | 0.950903684 | 74.15178854 |
| INTERPRO | IPR011625:Alpha-2-macroglobulin, N-terminal 2 | 4 | 0.254129606 | 0.075530428 | C4A, CPAMD8, C4B, C5 | 1452 | 13 | 18559 | 3.932824751 | 1 | 0.950903684 | 74.15178854 |
| SMART | SM01361:SM01361 | 4 | 0.254129606 | 0.080222515 | C4A, CPAMD8, C4B, C5 | 878 | 12 | 10057 | 3.818147304 | 1 | 0.870152845 | 69.04491501 |
| SMART | SM01359:SM01359 | 4 | 0.254129606 | 0.09778689 | C4A, CPAMD8, C4B, C5 | 878 | 13 | 10057 | 3.524443666 | 1 | 0.869135048 | 76.37834183 |
| SMART | SM01360:SM01360 | 4 | 0.254129606 | 0.09778689 | C4A, CPAMD8, C4B, C5 | 878 | 13 | 10057 | 3.524443666 | 1 | 0.869135048 | 76.37834183 |
| GOTERM_BP_DIRECT | GO:2000427~positive regulation of apoptotic cell clearance | 3 | 0.190597205 | 0.102249192 | C4A, C4B, C2 | 1342 | 7 | 16792 | 5.362571854 | 1 | 0.979902008 | 86.56787542 |
| KEGG_PATHWAY | hsa04610:Complement and coagulation cascades | 10 | 0.635324015 | 0.117365253 | C8B, THBD, MASP1, C4A, C4B, CFB, C5, C4BPA, C1S, C2 | 574 | 69 | 6879 | 1.736858052 | 1 | 0.711765078 | 80.79151757 |
| INTERPRO | IPR019742:Alpha-2-macroglobulin, conserved site | 3 | 0.190597205 | 0.152477215 | C4A, CPAMD8, C4B | 1452 | 9 | 18559 | 4.260560147 | 1 | 0.977197811 | 94.2155735 |
| INTERPRO | IPR019565:Alpha-2-macroglobulin, thiol-ester bond-forming | 3 | 0.190597205 | 0.21054648 | C4A, CPAMD8, C4B | 1452 | 11 | 18559 | 3.485912847 | 1 | 0.98882506 | 98.29690195 |
| UP_KEYWORDS | Complement alternate pathway | 3 | 0.190597205 | 0.223724927 | C8B, CFB, C5 | 1535 | 12 | 20581 | 3.351954397 | 1 | 0.643645778 | 97.37385033 |
| GOTERM_BP_DIRECT | GO:0006956~complement activation | 10 | 0.635324015 | 0.257876058 | C8B, MASP1, C4A, C4B, CFB, C5, RGCC, COLEC11, C1S, C2 | 1342 | 87 | 16792 | 1.438237662 | 1 | 0.997641615 | 99.61155271 |
| GOTERM_BP_DIRECT | GO:0006957~complement activation, alternative pathway | 3 | 0.190597205 | 0.278666743 | C8B, CFB, C5 | 1342 | 13 | 16792 | 2.887538691 | 1 | 0.998183139 | 99.77109605 |
| KEGG_PATHWAY | hsa05133:Pertussis | 7 | 0.444726811 | 0.602572242 | C4A, C4B, C5, C4BPA, C1S, C2, CALM2 | 574 | 75 | 6879 | 1.118536585 | 1 | 0.905757229 | 99.99949397 |
| GOTERM_MF_DIRECT | GO:0004866~endopeptidase inhibitor activity | 4 | 0.254129606 | 0.624909352 | C4A, CPAMD8, C4B, C5 | 1340 | 40 | 16881 | 1.259776119 | 1 | 0.999226802 | 99.99998863 |
| REACTOME_PATHWAY | R-HSA-166663:R-HSA-166663 | 6 | 0.381194409 | 0.747152262 | MASP1, C4A, C4B, COLEC11, C1S, C2 | 745 | 75 | 9075 | 0.974496644 | 1 | 0.999785292 | 99.99999994 |
| GOTERM_BP_DIRECT | GO:0010951~negative regulation of endopeptidase activity | 9 | 0.571791614 | 0.759821218 | WFDC10A, APP, C4A, CPAMD8, C4B, C5, EPPIN, SERPINB12, TIMP2 | 1342 | 121 | 16792 | 0.930694289 | 1 | 0.999999726 | 100 |
| GOTERM_BP_DIRECT | GO:0006958~complement activation, classical pathway | 7 | 0.444726811 | 0.812196743 | C8B, C4A, C4B, C5, C4BPA, C1S, C2 | 1342 | 99 | 16792 | 0.884734077 | 1 | 0.999999935 | 100 |
| GOTERM_CC_DIRECT | GO:0072562~blood microparticle | 6 | 0.381194409 | 0.993565127 | C4A, C4B, CFB, C4BPA, C1S, FN1 | 1425 | 152 | 18224 | 0.504819945 | 1 | 0.999999782 | 100 |

Annotation Cluster 32 Enrichment Score: 1.1715054145170507

| Category | Term | Count | % | PValue | Genes | List Total | Pop Hits | Pop Total | Fold Enrichment | Bonferroni | Benjamini | FDR |
|---|---|---|---|---|---|---|---|---|---|---|---|---|
| GOTERM_MF_DIRECT | GO:0032395~MHC class II receptor activity | 7 | 0.444726811 | 6.59E-04 | HLA-DQB1, HLA-DRB1, HLA-C, HLA-DQA2, HLA-DOB, HLA-DQA1, HLA-DRA | 1340 | 15 | 16881 | 5.878955224 | 0.571631785 | 0.571631785 | 1.069180313 |
| KEGG_PATHWAY | hsa05150:Staphylococcus aureus infection | 13 | 0.82592122 | 0.001317045 | HLA-DQB1, MASP1, C4A, HLA-DRB1, C4B, CFB, C5, C1S, C2, HLA-DQA2, HLA-DOB, HLA-DQA1, HLA-DI | 574 | 54 | 6879 | 2.885114208 | 0.307673593 | 0.307673593 | 1.726551035 |
| GOTERM_CC_DIRECT | GO:0030666~endocytic vesicle membrane | 14 | 0.889453621 | 0.001572948 | WNT5A, HLA-DQB1, HLA-DRB1, CAMK2G, COLEC12, CACNG2, HLA-DQA2, HLA-DQA1, AP2A2, GRIA2 | 1425 | 66 | 18224 | 2.712769803 | 0.66090193 | 0.143152336 | 2.340195043 |
| INTERPRO | IPR014745:MHC class II, alpha/beta chain, N-terminal | 7 | 0.444726811 | 0.004405062 | HLA-DQB1, HLA-DRB1, HLA-C, HLA-DQA2, HLA-DOB, HLA-DQA1, HLA-DRA | 1452 | 21 | 18559 | 4.260560147 | 0.999874058 | 0.557951371 | 7.323371046 |
| GOTERM_CC_DIRECT | GO:0042613~MHC class II protein complex | 7 | 0.444726811 | 0.005645959 | HLA-DQB1, HLA-DRB1, HLA-C, HLA-DQA2, HLA-DOB, HLA-DQA1, HLA-DRA | 1425 | 22 | 18224 | 4.069154705 | 0.979549854 | 0.242582285 | 8.164527001 |
| UP_KEYWORDS | MHC II | 6 | 0.381194409 | 0.006625563 | HLA-DQB1, HLA-DRB1, HLA-DQA2, HLA-DOB, HLA-DQA1, HLA-DRA | 1535 | 17 | 20581 | 4.732170914 | 0.96226893 | 0.09189052 | 9.111667955 |
| REACTOME_PATHWAY | R-HSA-202433:R-HSA-202433 | 9 | 0.571791614 | 0.007543799 | FYB, HLA-DQB1, PAK2, HLA-DRB1, PLCG1, CD247, HLA-DQA2, HLA-DQA1, HLA-DRA | 745 | 36 | 9075 | 3.045302013 | 0.998514883 | 0.885908443 | 11.07178474 |
| UP_SEQ_FEATURE | domain:Ig-like C1-type | 8 | 0.508259212 | 0.008195533 | HLA-DQB1, HLA-DRB1, HLA-C, HLA-DQA2, HLA-DOB, HLA-DQA1, HLA-DRA, TAPBP | 1499 | 32 | 20063 | 3.346064043 | 1 | 0.605861919 | 14.2065142 |
| GOTERM_BP_DIRECT | GO:0002504~antigen processing and presentation of peptide or polysaccharide antigen via MHC cla | 6 | 0.381194409 | 0.008847005 | HLA-DQB1, HLA-DRB1, HLA-DQA2, HLA-DOB, HLA-DQA1, HLA-DRA | 1342 | 17 | 16792 | 4.416235645 | 1 | 0.835130413 | 15.24381531 |
| KEGG_PATHWAY | hsa05330:Allograft rejection | 9 | 0.571791614 | 0.00976962 | HLA-DQB1, HLA-DRB1, FASLG, HLA-C, CD40, HLA-DQA2, HLA-DOB, HLA-DQA1, HLA-DRA | 574 | 37 | 6879 | 2.915105 | 0.935373196 | 0.745781975 | 12.16770176 |
| GOTERM_BP_DIRECT | GO:0060333~interferon-gamma-mediated signaling pathway | 13 | 0.82592122 | 0.010014208 | NCAM1, HLA-DQB1, SP100, HLA-DRB1, CAMK2G, OAS3, JAK1, OAS1, HLA-C, OAS2, HLA-DQA2, HLA-I | 1342 | 71 | 16792 | 2.291051825 | 1 | 0.817563222 | 17.08233523 |
| KEGG_PATHWAY | hsa05310:Asthma | 8 | 0.508259212 | 0.010043273 | FCER1A, HLA-DQB1, HLA-DRB1, CD40, HLA-DQA2, HLA-DOB, HLA-DQA1, HLA-DRA | 574 | 30 | 6879 | 3.195818815 | 0.940169469 | 0.608882165 | 12.48792638 |
| UP_SEQ_FEATURE | region of interest:Connecting peptide | 8 | 0.508259212 | 0.011500544 | HLA-DQB1, CRYGS, HLA-DRB1, HLA-C, HLA-DQA2, HLA-DOB, HLA-DQA1, HLA-DRA | 1499 | 34 | 20063 | 3.149236746 | 1 | 0.665679333 | 19.37629939 |
| GOTERM_CC_DIRECT | GO:0030669~clathrin-coated endocytic vesicle membrane | 9 | 0.571791614 | 0.012770555 | WNT5A, HLA-DQB1, AP2A2, HLA-DRB1, HBEGF, CLVS2, HLA-DQA2, HLA-DQA1, HLA-DRA | 1425 | 41 | 18224 | 2.807291399 | 0.999853703 | 0.318805587 | 17.57997322 |
| KEGG_PATHWAY | hsa04514:Cell adhesion molecules (CAMs) | 21 | 1.334180432 | 0.013500959 | HLA-DQB1, PTPRM, HLA-DRB1, NRXN3, SELL, NLGN1, HLA-C, CD40, NRXN1, CDH3, HLA-DQA2, HLA-I | 574 | 142 | 6879 | 1.772329097 | 0.977459151 | 0.468510066 | 16.44217603 |
| REACTOME_PATHWAY | R-HSA-389948:R-HSA-389948 | 7 | 0.444726811 | 0.016534101 | HLA-DQB1, HLA-DRB1, CD247, HLA-DQA2, HLA-DQA1, HLA-DRA, PTPN11 | 745 | 26 | 9075 | 3.279556014 | 0.999999407 | 0.908342678 | 22.76763837 |
| KEGG_PATHWAY | hsa05332:Graft-versus-host disease | 8 | 0.508259212 | 0.016964141 | HLA-DQB1, HLA-DRB1, FASLG, HLA-C, HLA-DQA2, HLA-DOB, HLA-DQA1, HLA-DRA | 574 | 33 | 6879 | 2.905289832 | 0.991550101 | 0.494366392 | 20.2365983 |
| GOTERM_MF_DIRECT | GO:0042605~peptide antigen binding | 7 | 0.444726811 | 0.020402689 | HLA-DQB1, HLA-DRB1, MAML1, HLA-C, HLA-DQA1, HLA-DRA, TAPBP | 1340 | 28 | 16881 | 3.149440299 | 1 | 0.849458353 | 28.54662204 |
| KEGG_PATHWAY | hsa04940:Type I diabetes mellitus | 9 | 0.571791614 | 0.020705812 | HLA-DQB1, HLA-DRB1, FASLG, HLA-C, HLA-DQA2, HLA-DOB, HLA-DQA1, LTA, HLA-DRA | 574 | 42 | 6879 | 2.568068691 | 0.997084075 | 0.517944496 | 24.15671972 |
| GOTERM_CC_DIRECT | GO:0071556~integral component of lumenal side of endoplasmic reticulum membrane | 7 | 0.444726811 | 0.022515215 | HLA-DQB1, HLA-DRB1, HLA-C, HLA-DQA2, HLA-DQA1, HLA-DRA, TAPBP | 1425 | 29 | 18224 | 3.086944949 | 0.999999839 | 0.416945088 | 29.00511235 |
| KEGG_PATHWAY | hsa05169:Epstein-Barr virus infection | 18 | 1.143583227 | 0.023858967 | HLA-DQB1, HLA-DRB1, PIK3CB, NFKBIB, HLA-C, RB1, CD40, HLA-DQA2, HLA-DQA1, NEDD4, PSMD1, J | 574 | 122 | 6879 | 1.768178443 | 0.998814213 | 0.526968961 | 27.32115896 |
| GOTERM_CC_DIRECT | GO:0032588~trans-Golgi network membrane | 13 | 0.82592122 | 0.027323357 | HLA-DQB1, CLTA, APP, HLA-DRB1, VPS54, VPS52, RGP1, AP4M1, CLVS2, HLA-DQA2, HLA-DQA1, HLA | 1425 | 83 | 18224 | 2.003060664 | 0.999999995 | 0.438272377 | 34.08068369 |
| KEGG_PATHWAY | hsa05322:Systemic lupus erythematosus | 19 | 1.207115629 | 0.028567002 | HLA-DQB1, HIST4H4, C4A, HLA-DRB1, C4B, C5, SSB, H2AFJ, CD40, C1S, HLA-DQA2, HLA-DQA1, C8B, ( | 574 | 134 | 6879 | 1.699269333 | 0.999692259 | 0.520549094 | 31.81951192 |
| REACTOME_PATHWAY | R-HSA-202430:R-HSA-202430 | 6 | 0.381194409 | 0.029777702 | HLA-DQB1, HLA-DRB1, CD247, HLA-DQA2, HLA-DQA1, HLA-DRA | 745 | 22 | 9075 | 3.322147651 | 1 | 0.944348719 | 37.40221966 |
| KEGG_PATHWAY | hsa04672:Intestinal immune network for IgA production | 9 | 0.571791614 | 0.03834821 | HLA-DQB1, HLA-DRB1, TNFRSF13B, CD40, HLA-DQA2, HLA-DOB, CCL28, HLA-DQA1, HLA-DRA | 574 | 47 | 6879 | 2.294869894 | 0.99998172 | 0.597130042 | 40.35420477 |
| BIOCARTA | h_bbcellPathway:Bystander B Cell Activation | 4 | 0.254129606 | 0.039137337 | HLA-DRB1, FASLG, CD40, HLA-DRA | 146 | 9 | 1625 | 4.946727549 | 0.999869342 | 0.893086191 | 39.93116792 |
| KEGG_PATHWAY | hsa04145:Phagosome | 20 | 1.27064803 | 0.042353488 | HLA-DQB1, ATP6V0E1, HLA-DRB1, MRC2, HLA-C, COLEC12, COLEC11, HLA-DQA2, HLA-DQA1, LAMP1 | 574 | 150 | 6879 | 1.597909408 | 0.999994295 | 0.577869137 | 43.55492642 |
| REACTOME_PATHWAY | R-HSA-202427:R-HSA-202427 | 6 | 0.381194409 | 0.049087347 | HLA-DQB1, HLA-DRB1, CD247, HLA-DQA2, HLA-DQA1, HLA-DRA | 745 | 25 | 9075 | 2.923489933 | 1 | 0.93315658 | 54.15726678 |
| REACTOME_PATHWAY | R-HSA-877300:R-HSA-877300 | 13 | 0.82592122 | 0.054635218 | NCAM1, HLA-DQB1, SP100, HLA-DRB1, CAMK2G, OAS3, JAK1, OAS1, HLA-C, OAS2, HLA-DQA2, HLA-I | 745 | 88 | 9075 | 1.799496644 | 1 | 0.941706828 | 58.13102266 |
| KEGG_PATHWAY | hsa05320:Autoimmune thyroid disease | 9 | 0.571791614 | 0.063983153 | HLA-DQB1, HLA-DRB1, FASLG, HLA-C, CD40, HLA-DQA2, HLA-DOB, HLA-DQA1, HLA-DRA | 574 | 52 | 6879 | 2.074209327 | 0.99999999 | 0.641163334 | 58.26388899 |
| INTERPRO | IPR001003:MHC class II, alpha chain, N-terminal | 3 | 0.190597205 | 0.074157476 | HLA-DQA2, HLA-DQA1, HLA-DRA | 1452 | 6 | 18559 | 6.39084022 | 1 | 0.950899407 | 73.48245133 |
| GOTERM_BP_DIRECT | GO:0050852~T cell receptor signaling pathway | 18 | 1.143583227 | 0.08200477 | HLA-DQB1, FYB, BCL10, HRAS, HLA-DRB1, PIK3CB, CD247, RFTN1, HLA-DQA2, HLA-DQA1, TRAT1, PS | 1342 | 148 | 16792 | 1.521810932 | 1 | 0.972086755 | 79.65802954 |
| SMART | SM00920:SM00920 | 3 | 0.190597205 | 0.089972819 | HLA-DQA2, HLA-DQA1, HLA-DRA | 878 | 6 | 10057 | 5.727220957 | 1 | 0.886243657 | 73.34191906 |
| GOTERM_BP_DIRECT | GO:0031295~T cell costimulation | 11 | 0.698856417 | 0.091215102 | HLA-DQB1, PAK2, HLA-DRB1, CD247, MAP3K8, RICTOR, HLA-DQA2, HLA-DQA1, SPN, HLA-DRA, PTPN | 1342 | 78 | 16792 | 1.764606978 | 1 | 0.978548762 | 83.1389007 |
| INTERPRO | IPR003006:Immunoglobulin/major histocompatibility complex, conserved site | 9 | 0.571791614 | 0.094177414 | HLA-DQB1, NRCAM, OBSCN, HLA-DRB1, HLA-C, HLA-DQA2, HLA-DOB, HLA-DQA1, HLA-DRA | 1452 | 60 | 18559 | 1.917252066 | 1 | 0.966956911 | 81.80382384 |
| KEGG_PATHWAY | hsa04612:Antigen processing and presentation | 11 | 0.698856417 | 0.09799326 | HLA-DQB1, HLA-DRB1, TAP2, KIR3DL3, HLA-C, HLA-DQA2, HLA-DOB, HLA-DQA1, KIR2DL4, HLA-DRA, | 574 | 76 | 6879 | 1.734572712 | 1 | 0.698481665 | 74.40856175 |
| KEGG_PATHWAY | hsa05416:Viral myocarditis | 9 | 0.571791614 | 0.098258155 | HLA-DQB1, CASP9, HLA-DRB1, HLA-C, CD40, HLA-DQA2, HLA-DOB, HLA-DQA1, HLA-DRA | 574 | 57 | 6879 | 1.892261141 | 1 | 0.684704615 | 74.50770184 |

| | | | | | | | | | | | | |
|---|---|---|---|---|---|---|---|---|---|---|---|---|
| BIOCARTA | h_asbcellPathway:Antigen Dependent B Cell Activation | 4 | 0.254129606 | 0.102515867 | HLA-DRB1, FASLG, CD40, HLA-DRA | 146 | 13 | 1625 | 3.424657534 | 1 | 0.992136793 | 74.86228576 |
| INTERPRO | IPR000353:MHC class II, beta chain, N-terminal | 4 | 0.254129606 | 0.10711664 | HLA-DQB1, HLA-DRB1, HLA-C, HLA-DOB | 1452 | 15 | 18559 | 3.408448118 | 1 | 0.964558329 | 85.79836453 |
| UP_SEQ_FEATURE | region of interest:Alpha-2 | 4 | 0.254129606 | 0.112252812 | HLA-C, HLA-DQA2, HLA-DQA1, HLA-DRA | 1499 | 16 | 20063 | 3.346064043 | 1 | 0.985793841 | 89.10662529 |
| UP_SEQ_FEATURE | region of interest:Alpha-1 | 4 | 0.254129606 | 0.112252812 | HLA-C, HLA-DQA2, HLA-DQA1, HLA-DRA | 1499 | 16 | 20063 | 3.346064043 | 1 | 0.985793841 | 89.10662529 |
| GOTERM_BP_DIRECT | GO:0019886~antigen processing and presentation of exogenous peptide antigen via MHC class II | 12 | 0.762388818 | 0.115298742 | HLA-DQB1, AP2A2, CLTA, HLA-DRB1, KLC1, DYNC2H1, KIF26A, HLA-DQA2, HLA-DOB, DYNC1I2, HLA-I | 1342 | 92 | 16792 | 1.632087086 | 1 | 0.980027837 | 89.772035 |
| UP_SEQ_FEATURE | region of interest:Beta-1 | 3 | 0.190597205 | 0.115475174 | HLA-DQB1, HLA-DRB1, HLA-DOB | 1499 | 8 | 20063 | 5.019096064 | 1 | 0.986545152 | 89.81978955 |
| UP_SEQ_FEATURE | region of interest:Beta-2 | 3 | 0.190597205 | 0.115475174 | HLA-DQB1, HLA-DRB1, HLA-DOB | 1499 | 8 | 20063 | 5.019096064 | 1 | 0.986545152 | 89.81978955 |
| KEGG_PATHWAY | hsa05168:Herpes simplex infection | 21 | 1.334180432 | 0.125975075 | HLA-DQB1, IFIT1B, SP100, HLA-DRB1, NFKBIB, C5, OAS3, FASLG, OAS1, HLA-C, OAS2, HLA-DQA2, HL/ | 574 | 183 | 6879 | 1.3752499 | 1 | 0.714128076 | 83.12538498 |
| INTERPRO | IPR011162:MHC classes I/II-like antigen recognition protein | 7 | 0.444726811 | 0.13633501 | HLA-DQB1, HLA-DRB1, HLA-C, HLA-DQA2, HLA-DOB, HLA-DQA1, HLA-DRA | 1452 | 45 | 18559 | 1.988261402 | 1 | 0.972452845 | 91.99396315 |
| SMART | SM00921:SM00921 | 4 | 0.254129606 | 0.136924544 | HLA-DQB1, HLA-DRB1, HLA-C, HLA-DOB | 878 | 15 | 10057 | 3.054517844 | 1 | 0.895997004 | 87.31686152 |
| INTERPRO | IPR003597:Immunoglobulin C1-set | 9 | 0.571791614 | 0.159871009 | HLA-DQB1, HLA-DRB1, BTNL2, HLA-C, HLA-DQA2, HLA-DOB, HLA-DQA1, HLA-DRA, TAPBP | 1452 | 68 | 18559 | 1.691693 | 1 | 0.978748572 | 95.02601696 |
| REACTOME_PATHWAY | R-HSA-2132295:R-HSA-2132295 | 14 | 0.889453621 | 0.19898493 | HLA-DQB1, CLTA, HLA-DRB1, HLA-DQA2, HLA-DQA1, AP2A2, KLC1, DYNC2H1, TUBAL3, KIF26A, HLA- | 745 | 122 | 9075 | 1.397843547 | 1 | 0.968861998 | 96.78759058 |
| GOTERM_CC_DIRECT | GO:0012507~ER to Golgi transport vesicle membrane | 7 | 0.444726811 | 0.218328573 | HLA-DQB1, HLA-DRB1, MCFD2, HLA-C, HLA-DQA2, HLA-DQA1, HLA-DRA | 1425 | 52 | 18224 | 1.721565452 | 1 | 0.860222329 | 97.54083336 |
| KEGG_PATHWAY | hsa05140:Leishmaniasis | 9 | 0.571791614 | 0.236805384 | HLA-DQB1, CYBA, HLA-DRB1, NFKBIB, JAK1, HLA-DQA2, HLA-DOB, HLA-DQA1, HLA-DRA | 574 | 71 | 6879 | 1.519139226 | 1 | 0.819780785 | 97.18797821 |
| BIOCARTA | h_ctla4Pathway:The Co-Stimulatory Signal During T-cell Activation | 4 | 0.254129606 | 0.236849351 | HLA-DRB1, CD247, HLA-DRA, PTPN11 | 146 | 19 | 1625 | 2.343186734 | 1 | 0.997653232 | 96.82774838 |
| INTERPRO | IPR001039:MHC class I, alpha chain, alpha1/alpha2 | 3 | 0.190597205 | 0.240343159 | HLA-DQB1, HLA-DRB1, HLA-C | 1452 | 12 | 18559 | 3.19542011 | 1 | 0.993507746 | 99.12220937 |
| BIOCARTA | h_blymphocytePathway:B Lymphocyte Cell Surface Molecules | 3 | 0.190597205 | 0.256709931 | HLA-DRB1, CD40, HLA-DRA | 146 | 11 | 1625 | 3.035491905 | 1 | 0.997621435 | 97.7344762 |
| BIOCARTA | h_tcraPathway:Lck and Fyn tyrosine kinases in initiation of TCR Activation | 3 | 0.190597205 | 0.256709931 | HLA-DRB1, CD247, HLA-DRA | 146 | 11 | 1625 | 3.035491905 | 1 | 0.997621435 | 97.7344762 |
| KEGG_PATHWAY | hsa05321:Inflammatory bowel disease (IBD) | 8 | 0.508259212 | 0.282384506 | HLA-DQB1, HLA-DRB1, SMAD3, RORA, HLA-DQA2, HLA-DOB, HLA-DQA1, HLA-DRA | 574 | 64 | 6879 | 1.49804007 | 1 | 0.808560127 | 98.75375936 |
| KEGG_PATHWAY | hsa05145:Toxoplasmosis | 12 | 0.762388818 | 0.307968201 | HLA-DQB1, CASP9, HLA-DRB1, IL10RA, NFKBIB, JAK1, CD40, HLA-DQA2, HLA-DOB, AKT3, HLA-DQA1, | 574 | 110 | 6879 | 1.307380424 | 1 | 0.819454721 | 99.22864104 |
| KEGG_PATHWAY | hsa05323:Rheumatoid arthritis | 10 | 0.635324015 | 0.310043235 | HLA-DQB1, ATP6V0E1, HLA-DRB1, VEGFA, ACP5, ATP6V0A4, HLA-DQA2, HLA-DOB, HLA-DQA1, HLA- | 574 | 88 | 6879 | 1.361854609 | 1 | 0.806709205 | 99.25865255 |
| REACTOME_PATHWAY | R-HSA-202424:R-HSA-202424 | 11 | 0.698856417 | 0.314011936 | HLA-DQB1, PSMA1, BCL10, HLA-DRB1, PIK3CB, CD247, PSMD1, HLA-DQA2, TRAT1, HLA-DQA1, HLA-I | 745 | 101 | 9075 | 1.326666224 | 1 | 0.986723104 | 99.70920526 |
| SMART | SM00407:IGc1 | 8 | 0.508259212 | 0.337760779 | HLA-DQB1, HLA-DRB1, BTNL2, HLA-C, HLA-DQA2, HLA-DOB, HLA-DQA1, HLA-DRA | 878 | 65 | 10057 | 1.409777466 | 1 | 0.947600066 | 99.69087539 |
| GOTERM_CC_DIRECT | GO:0030658~transport vesicle membrane | 5 | 0.317662008 | 0.345446659 | HLA-DQB1, HLA-DRB1, HLA-DQA2, HLA-DQA1, HLA-DRA | 1425 | 38 | 18224 | 1.682733149 | 1 | 0.920480856 | 99.82966527 |
| GOTERM_MF_DIRECT | GO:0023026~MHC class II protein complex binding | 3 | 0.190597205 | 0.366112403 | HLA-DRB1, HLA-DOB, HLA-DRA | 1340 | 16 | 16881 | 2.362080224 | 1 | 0.992748943 | 99.94089411 |
| GOTERM_BP_DIRECT | GO:0002474~antigen processing and presentation of peptide antigen via MHC class I | 4 | 0.254129606 | 0.433527357 | HLA-DQB1, TAP2, HLA-C, TAPBP | 1342 | 30 | 16792 | 1.668355688 | 1 | 0.99982233 | 99.99745192 |
| GOTERM_BP_DIRECT | GO:0019882~antigen processing and presentation | 6 | 0.381194409 | 0.450529588 | HLA-DQB1, HLA-DRB1, HLA-C, HLA-DQA2, HLA-DQA1, HLA-DRA | 1342 | 55 | 16792 | 1.36501829 | 1 | 0.999854297 | 99.99855492 |
| KEGG_PATHWAY | hsa05166:HTLV-I infection | 23 | 1.461245235 | 0.456633617 | WNT5A, HLA-DQB1, HRAS, HLA-DRB1, PIK3CB, SMAD3, HLA-C, RB1, CD40, CHEK2, CDK4, HLA-DQA2, | 574 | 254 | 6879 | 1.085194381 | 1 | 0.864954237 | 99.96843283 |
| KEGG_PATHWAY | hsa05164:Influenza A | 16 | 1.016518424 | 0.484555565 | HLA-DQB1, HLA-DRB1, PIK3CB, NFKBIB, OAS3, FASLG, OAS1, OAS2, HLA-DQA2, HLA-DQA1, CASP9, J/ | 574 | 174 | 6879 | 1.102006488 | 1 | 0.874761278 | 99.98427949 |
| KEGG_PATHWAY | hsa05152:Tuberculosis | 16 | 1.016518424 | 0.512177259 | HLA-DQB1, BCL10, HLA-DRB1, CAMK2G, MRC2, HLA-DQA2, HLA-DQA1, LAMP1, CASP9, IL10RA, JAK1 | 574 | 177 | 6879 | 1.083328412 | 1 | 0.875831464 | 99.99240785 |
| BIOCARTA | h_th1th2Pathway:Th1/Th2 Differentiation | 3 | 0.190597205 | 0.544922095 | HLA-DRB1, CD40, HLA-DRA | 146 | 20 | 1625 | 1.669520548 | 1 | 0.998543273 | 99.99568461 |
| BIOCARTA | h_cskPathway:Activation of Csk by cAMP-dependent Protein Kinase Inhibits Signaling through the T | 3 | 0.190597205 | 0.59803996 | HLA-DRB1, CD247, HLA-DRA | 146 | 22 | 1625 | 1.517745953 | 1 | 0.999123698 | 99.99911513 |
| GOTERM_BP_DIRECT | GO:0007018~microtubule-based movement | 7 | 0.444726811 | 0.635884168 | AP2A2, CLTA, SNX29, KLC1, DYNC2H1, KIF26A, DYNC1I2 | 1342 | 81 | 16792 | 1.08134165 | 1 | 0.99999331 | 99.99999932 |
| INTERPRO | IPR011161:MHC class I-like antigen recognition | 3 | 0.190597205 | 0.654832406 | HLA-DQB1, HLA-DRB1, HLA-C | 1452 | 28 | 18559 | 1.369465762 | 1 | 0.999978858 | 99.9999989 |
| UP_KEYWORDS | Lysosome | 18 | 1.143583227 | 0.76361001 | HLA-DQB1, HLA-DRB1, GGH, PPT2, FASLG, ACP5, ZNRF1, HLA-DQA2, HLA-DQA1, ZNRF2, LAMP1, CHI | 1535 | 263 | 20581 | 0.91764531 | 1 | 0.959355187 | 99.9999999 |

Annotation Cluster 33 Enrichment Score: 1.1657600473186358

| Category | Term | Count | % | PValue | Genes | List Total | Pop Hits | Pop Total | Fold Enrichment | Bonferroni | Benjamini | FDR |
|---|---|---|---|---|---|---|---|---|---|---|---|---|
| GOTERM_BP_DIRECT | GO:0022400~regulation of rhodopsin mediated signaling pathway | 7 | 0.444726811 | 0.02477501 | FNTB, METAP2, CHURC1-FNTB, GRK7, CNGB1, CNGA1, CALM2 | 1342 | 29 | 16792 | 3.02029909 | 1 | 0.880014043 | 37.30671907 |
| REACTOME_PATHWAY | R-HSA-2514859:R-HSA-2514859 | 7 | 0.444726811 | 0.070105589 | FNTB, METAP2, CHURC1-FNTB, GRK7, CNGB1, CNGA1, CALM2 | 745 | 36 | 9075 | 2.368568233 | 1 | 0.941649821 | 67.57705217 |
| KEGG_PATHWAY | hsa04744:Phototransduction | 5 | 0.317662008 | 0.18321177 | SLC24A1, GRK7, CNGB1, CNGA1, CALM2 | 574 | 27 | 6879 | 2.219318622 | 1 | 0.756239373 | 93.10520151 |

Annotation Cluster 34 Enrichment Score: 1.1579288265190144

| Category | Term | Count | % | PValue | Genes | List Total | Pop Hits | Pop Total | Fold Enrichment | Bonferroni | Benjamini | FDR |
|---|---|---|---|---|---|---|---|---|---|---|---|---|
| GOTERM_BP_DIRECT | GO:0032793~positive regulation of CREB transcription factor activity | 5 | 0.317662008 | 0.027094445 | PRKD1, PRKD2, VEGFA, RELN, LRP8 | 1342 | 15 | 16792 | 4.17088922 | 1 | 0.88753056 | 40.02450887 |
| GOTERM_BP_DIRECT | GO:0010595~positive regulation of endothelial cell migration | 9 | 0.571791614 | 0.027729439 | WNT5A, PRKD1, PRKD2, ITGB1BP1, VEGFA, SCARB1, BCAS3, FGF2, FOXP1 | 1342 | 46 | 16792 | 2.448130629 | 1 | 0.888529041 | 40.74889044 |
| GOTERM_BP_DIRECT | GO:1901727~positive regulation of histone deacetylase activity | 3 | 0.190597205 | 0.054161082 | PRKD1, PRKD2, VEGFA | 1342 | 5 | 16792 | 7.507600596 | 1 | 0.950810698 | 64.52569176 |
| GOTERM_BP_DIRECT | GO:0038033~positive regulation of endothelial cell chemotaxis by VEGF-activated vascular endothe | 3 | 0.190597205 | 0.054161082 | PRKD1, PRKD2, VEGFA | 1342 | 5 | 16792 | 7.507600596 | 1 | 0.950810698 | 64.52569176 |
| GOTERM_BP_DIRECT | GO:0043536~positive regulation of blood vessel endothelial cell migration | 5 | 0.317662008 | 0.059736606 | PRKD1, PRKD2, PLCG1, VEGFA, FGF2 | 1342 | 19 | 16792 | 3.292807279 | 1 | 0.954273847 | 68.22209258 |
| GOTERM_BP_DIRECT | GO:0048010~vascular endothelial growth factor receptor signaling pathway | 9 | 0.571791614 | 0.214944848 | PRKD1, CYBA, PRKD2, PAK2, PIK3CB, BAIAP2, VEGFA, CYFIP1, VAV2 | 1342 | 72 | 16792 | 1.564083458 | 1 | 0.994899878 | 98.89362632 |
| GOTERM_BP_DIRECT | GO:0035924~cellular response to vascular endothelial growth factor stimulus | 4 | 0.254129606 | 0.277162598 | PRKD1, PRKD2, ITGB1BP1, VEGFA | 1342 | 23 | 16792 | 2.176116115 | 1 | 0.998164437 | 99.76204729 |

Annotation Cluster 35 Enrichment Score: 1.144874211276258

| Category | Term | Count | % | PValue | Genes | List Total | Pop Hits | Pop Total | Fold Enrichment | Bonferroni | Benjamini | FDR |
|---|---|---|---|---|---|---|---|---|---|---|---|---|
| UP_SEQ_FEATURE | region of interest:Triple-helical region 1 (COL1) | 3 | 0.190597205 | 0.068290808 | COL9A2, COL9A3, COL15A1 | 1499 | 6 | 20063 | 6.692128085 | 1 | 0.956349522 | 73.20770389 |
| UP_SEQ_FEATURE | region of interest:Triple-helical region 2 (COL2) | 3 | 0.190597205 | 0.068290808 | COL9A2, COL9A3, COL15A1 | 1499 | 6 | 20063 | 6.692128085 | 1 | 0.956349522 | 73.20770389 |
| UP_SEQ_FEATURE | region of interest:Nonhelical region 3 (NC3) | 3 | 0.190597205 | 0.068290808 | COL9A2, COL9A3, COL15A1 | 1499 | 6 | 20063 | 6.692128085 | 1 | 0.956349522 | 73.20770389 |
| UP_SEQ_FEATURE | region of interest:Nonhelical region 2 (NC2) | 3 | 0.190597205 | 0.068290808 | COL9A2, COL9A3, COL15A1 | 1499 | 6 | 20063 | 6.692128085 | 1 | 0.956349522 | 73.20770389 |
| UP_SEQ_FEATURE | region of interest:Nonhelical region 1 (NC1) | 3 | 0.190597205 | 0.068290808 | COL9A2, COL9A3, COL15A1 | 1499 | 6 | 20063 | 6.692128085 | 1 | 0.956349522 | 73.20770389 |
| UP_SEQ_FEATURE | region of interest:Triple-helical region 3 (COL3) | 3 | 0.190597205 | 0.090979486 | COL9A2, COL9A3, COL15A1 | 1499 | 7 | 20063 | 5.736109787 | 1 | 0.975279715 | 83.07009667 |

Annotation Cluster 36 Enrichment Score: 1.136864519281062

| Category | Term | Count | % | PValue | Genes | List Total | Pop Hits | Pop Total | Fold Enrichment | Bonferroni | Benjamini | FDR |
|---|---|---|---|---|---|---|---|---|---|---|---|---|
| UP_KEYWORDS | Notch signaling pathway | 10 | 0.635324015 | 0.012211682 | APP, ITGB1BP1, MAML1, NOTCH4, APH1B, CNTN1, TP63, NEURL1B, MAML3, RBPJ | 1535 | 51 | 20581 | 2.628983841 | 0.99765957 | 0.143857523 | 16.18725195 |
| GOTERM_BP_DIRECT | GO:0007221~positive regulation of transcription of Notch receptor target | 4 | 0.254129606 | 0.013924741 | MAML1, NOTCH4, MAML3, RBPJ | 1342 | 7 | 16792 | 7.150095806 | 1 | 0.849880395 | 22.97082761 |
| KEGG_PATHWAY | hsa04330:Notch signaling pathway | 9 | 0.571791614 | 0.042804826 | CIR1, CTBP2, MAML1, DTX3L, NOTCH4, APH1B, MAML3, RBPJ, NCOR2 | 574 | 48 | 6879 | 2.247060105 | 0.999994998 | 0.55679025 | 43.90546991 |
| REACTOME_PATHWAY | R-HSA-1912408:R-HSA-1912408 | 6 | 0.381194409 | 0.084033947 | TNRC6C, MAML1, NOTCH4, MAML3, TNRC6B, RBPJ | 745 | 29 | 9075 | 2.520249942 | 1 | 0.945161302 | 74.33810591 |
| REACTOME_PATHWAY | R-HSA-350054:R-HSA-350054 | 4 | 0.254129606 | 0.084534597 | MAML1, NOTCH4, MAML3, RBPJ | 745 | 13 | 9075 | 3.748064017 | 1 | 0.93998986 | 74.55459672 |
| REACTOME_PATHWAY | R-HSA-2644606:R-HSA-2644606 | 9 | 0.571791614 | 0.091805093 | TBL1XR1, HDAC3, MAML1, APH1B, NEURL1B, MAML3, RBPJ, MYC, NCOR2 | 745 | 57 | 9075 | 1.92334864 | 1 | 0.930847762 | 77.51208597 |
| REACTOME_PATHWAY | R-HSA-2894862:R-HSA-2894862 | 9 | 0.571791614 | 0.091805093 | TBL1XR1, HDAC3, MAML1, APH1B, NEURL1B, MAML3, RBPJ, MYC, NCOR2 | 745 | 57 | 9075 | 1.92334864 | 1 | 0.930847762 | 77.51208597 |
| REACTOME_PATHWAY | R-HSA-2122947:R-HSA-2122947 | 7 | 0.444726811 | 0.184281663 | TBL1XR1, HDAC3, MAML1, MAML3, RBPJ, MYC, NCOR2 | 745 | 47 | 9075 | 1.814222476 | 1 | 0.967765171 | 95.74165425 |
| GOTERM_BP_DIRECT | GO:0007219~Notch signaling pathway | 13 | 0.82592122 | 0.206711154 | GMDS, MAML1, APH1B, TP63, APP, ITGB1BP1, NOTCH4, SUSD5, CNTN1, NEURL1B, MAML3, RBPJ, M | 1342 | 115 | 16792 | 1.414475475 | 1 | 0.994430483 | 98.65650734 |
| REACTOME_PATHWAY | R-HSA-2197563:R-HSA-2197563 | 3 | 0.190597205 | 0.25776897 | MAML1, MAML3, RBPJ | 745 | 12 | 9075 | 3.045302013 | 1 | 0.980629858 | 99.01396146 |

Annotation Cluster 37 Enrichment Score: 1.1282026364252316

| Category | Term | Count | % | PValue | Genes | List Total | Pop Hits | Pop Total | Fold Enrichment | Bonferroni | Benjamini | FDR |
|---|---|---|---|---|---|---|---|---|---|---|---|---|
| INTERPRO | IPR004012:RUN | 5 | 0.317662008 | 0.056017614 | SNX29, PLEKHM2, SGSM3, RUSC1, RUFY1 | 1452 | 19 | 18559 | 3.363600116 | 1 | 0.913084975 | 62.95737967 |
| SMART | SM00593:RUN | 5 | 0.317662008 | 0.065594927 | SNX29, PLEKHM2, SGSM3, RUSC1, RUFY1 | 878 | 18 | 10057 | 3.18178942 | 1 | 0.846959329 | 61.37915277 |
| UP_SEQ_FEATURE | domain:RUN | 4 | 0.254129606 | 0.112252812 | PLEKHM2, SGSM3, RUSC1, RUFY1 | 1499 | 16 | 20063 | 3.346064043 | 1 | 0.985793841 | 89.10662529 |

Annotation Cluster 38 Enrichment Score: 1.0980003176077864

| Category | Term | Count | % | PValue | Genes | List Total | Pop Hits | Pop Total | Fold Enrichment | Bonferroni | Benjamini | FDR |
|---|---|---|---|---|---|---|---|---|---|---|---|---|
| UP_KEYWORDS | Antiport | 8 | 0.508259212 | 0.030329973 | SLC9A9, SLC26A3, SLC8A1, SLC24A4, SLC9A3, SLC24A1, SLC9C1, SLC26A10 | 1535 | 41 | 20581 | 2.61615953 | 0.999999746 | 0.241243985 | 35.76651165 |
| REACTOME_PATHWAY | R-HSA-425561:R-HSA-425561 | 4 | 0.254129606 | 0.084534597 | SLC8A1, SLC24A4, SLC24A1, CALM2 | 745 | 13 | 9075 | 3.748064017 | 1 | 0.93998986 | 74.55459672 |
| UP_SEQ_FEATURE | repeat:Alpha-2 | 3 | 0.190597205 | 0.090979486 | SLC8A1, SLC24A4, SLC24A1 | 1499 | 7 | 20063 | 5.736109787 | 1 | 0.975279715 | 83.07009667 |

| | | | | | | | | | | | | |
|---|---|---|---|---|---|---|---|---|---|---|---|---|
| UP_SEQ_FEATURE | repeat:Alpha-1 | 3 | 0.190597205 | 0.090979486 | SLC8A1, SLC24A4, SLC24A1 | 1499 | 7 | 20063 | 5.736109787 | 1 | 0.975279715 | 83.07009667 |
| INTERPRO | IPR004837:Sodium/calcium exchanger membrane region | 3 | 0.190597205 | 0.152477215 | SLC8A1, SLC24A4, SLC24A1 | 1452 | 9 | 18559 | 4.260560147 | 1 | 0.977197811 | 94.2155735 |
| Annotation Cluster 39 | Enrichment Score: 1.0867855026186746 | | | | | | | | | | | |
| Category | Term | Count | % | PValue | Genes | List Total | Pop Hits | Pop Total | Fold Enrichment | Bonferroni | Benjamini | FDR |
| UP_KEYWORDS | Metal-binding | 311 | 19.75857687 | 0.004318894 | PDP1, MASP1, UTRN, ZNF782, SYT6, RORA, ZNRF1, OGDH, CIAPIN1, BBOX1, ZNRF2, ZFP91, MAP3K5, | 1535 | 3640 | 20581 | 1.145558041 | 0.881616475 | 0.075988334 | 6.030975661 |
| UP_KEYWORDS | Zinc | 194 | 12.3252859 | 0.077610127 | UTRN, ZNF782, RORA, ZNRF1, BBOX1, ZNRF2, ZFP91, APP, GRIN2B, RNF219, SAP30L, DDAH1, ZNF39 | 1535 | 2348 | 20581 | 1.107800942 | 1 | 0.377582659 | 68.68472309 |
| GOTERM_MF_DIRECT | GO:0046872~metal ion binding | 174 | 11.05463787 | 0.231352347 | PDP1, ZNF821, ZNF782, PDE3B, OGDH, CIAPIN1, CNOT6, CLYBL, NT5DC1, DMPK, ZNF184, ZFP91, ATI | 1340 | 2069 | 16881 | 1.059454059 | 1 | 0.981330987 | 98.63012676 |
| UP_KEYWORDS | Zinc-finger | 132 | 8.386277001 | 0.579819998 | ZNF821, UTRN, ZNF782, TRIM52, RORA, ZNRF1, NFXL1, ZNRF2, GATA2, ZNF184, ZFP91, RNF219, SAF | 1535 | 1781 | 20581 | 0.993729322 | 1 | 0.89956125 | 99.99961269 |
| Annotation Cluster 40 | Enrichment Score: 1.0606959343150066 | | | | | | | | | | | |
| Category | Term | Count | % | PValue | Genes | List Total | Pop Hits | Pop Total | Fold Enrichment | Bonferroni | Benjamini | FDR |
| UP_SEQ_FEATURE | repeat:ARM 8 | 6 | 0.381194409 | 0.020813723 | ARMC8, PKP3, CTNND2, ARMC4, KPNA6, KPNA3 | 1499 | 22 | 20063 | 3.650251683 | 1 | 0.80746553 | 32.40493656 |
| UP_SEQ_FEATURE | repeat:ARM 7 | 6 | 0.381194409 | 0.029697127 | ARMC8, PKP3, CTNND2, ARMC4, KPNA6, KPNA3 | 1499 | 24 | 20063 | 3.346064043 | 1 | 0.861994879 | 42.95492064 |
| UP_SEQ_FEATURE | repeat:ARM 6 | 6 | 0.381194409 | 0.040653474 | ARMC8, PKP3, CTNND2, ARMC4, KPNA6, KPNA3 | 1499 | 26 | 20063 | 3.088674501 | 1 | 0.898159112 | 53.82686364 |
| UP_SEQ_FEATURE | repeat:ARM 9 | 5 | 0.317662008 | 0.040730353 | ARMC8, CTNND2, ARMC4, KPNA6, KPNA3 | 1499 | 18 | 20063 | 3.717848936 | 1 | 0.895465471 | 53.89571119 |
| UP_SEQ_FEATURE | repeat:ARM 5 | 6 | 0.381194409 | 0.06112943 | ARMC8, PKP3, CTNND2, ARMC4, KPNA6, KPNA3 | 1499 | 29 | 20063 | 2.769156449 | 1 | 0.944239431 | 69.10224933 |
| UP_SEQ_FEATURE | repeat:ARM 4 | 6 | 0.381194409 | 0.077488406 | ARMC8, PKP3, CTNND2, ARMC4, KPNA6, KPNA3 | 1499 | 31 | 20063 | 2.590501194 | 1 | 0.964990454 | 77.72662228 |
| UP_SEQ_FEATURE | repeat:ARM 3 | 6 | 0.381194409 | 0.127402072 | ARMC8, PKP3, CTNND2, ARMC4, KPNA6, KPNA3 | 1499 | 36 | 20063 | 2.230709362 | 1 | 0.990202065 | 92.09361764 |
| UP_SEQ_FEATURE | repeat:ARM 2 | 6 | 0.381194409 | 0.138810246 | ARMC8, PKP3, CTNND2, ARMC4, KPNA6, KPNA3 | 1499 | 37 | 20063 | 2.17041992 | 1 | 0.991709818 | 93.81185873 |
| INTERPRO | IPR000225:Armadillo | 6 | 0.381194409 | 0.243592225 | ARMC8, PKP3, CTNND2, ARMC4, KPNA6, KPNA3 | 1452 | 43 | 18559 | 1.783490294 | 1 | 0.993717651 | 99.18468896 |
| SMART | SM00185:ARM | 6 | 0.381194409 | 0.250388262 | ARMC8, PKP3, CTNND2, ARMC4, KPNA6, KPNA3 | 878 | 39 | 10057 | 1.762221833 | 1 | 0.917233195 | 98.24264431 |
| UP_SEQ_FEATURE | repeat:ARM 1 | 4 | 0.254129606 | 0.411057136 | ARMC8, PKP3, CTNND2, ARMC4 | 1499 | 31 | 20063 | 1.727000796 | 1 | 0.999959883 | 99.99476603 |
| Annotation Cluster 41 | Enrichment Score: 1.0584304788411694 | | | | | | | | | | | |
| Category | Term | Count | % | PValue | Genes | List Total | Pop Hits | Pop Total | Fold Enrichment | Bonferroni | Benjamini | FDR |
| UP_SEQ_FEATURE | domain:RabBD | 4 | 0.254129606 | 0.05493803 | MYRIP, SYTL2, SYTL3, RIMS1 | 1499 | 12 | 20063 | 4.461418724 | 1 | 0.93330171 | 65.07970204 |
| INTERPRO | IPR010911:Rab-binding domain | 4 | 0.254129606 | 0.061563084 | MYRIP, SYTL2, SYTL3, RIMS1 | 1452 | 12 | 18559 | 4.260560147 | 1 | 0.928463219 | 66.53267445 |
| GOTERM_BP_DIRECT | GO:0048791~calcium ion-regulated exocytosis of neurotransmitter | 6 | 0.381194409 | 0.197476361 | C2CD4D, SYT6, SYT16, SYTL2, SYTL3, RIMS1 | 1342 | 39 | 16792 | 1.925025794 | 1 | 0.994079886 | 98.33356912 |
| Annotation Cluster 42 | Enrichment Score: 1.041327968009083 | | | | | | | | | | | |
| Category | Term | Count | % | PValue | Genes | List Total | Pop Hits | Pop Total | Fold Enrichment | Bonferroni | Benjamini | FDR |
| UP_KEYWORDS | SH2 domain | 14 | 0.889453621 | 0.058269489 | FRK, FGR, FER, VAV2, PTPN11, HSH2D, PLCG1, SH2B3, JAK1, TXK, SH2B2, SH2D1B, SHC4, SH3BP2 | 1535 | 108 | 20581 | 1.738050428 | 1 | 0.329660122 | 57.80342814 |
| UP_SEQ_FEATURE | domain:SH2 | 12 | 0.762388818 | 0.10193482 | FRK, FGR, SH2B3, JAK1, SH2B2, TXK, FER, VAV2, SH2D1B, SHC4, SH3BP2, HSH2D | 1499 | 96 | 20063 | 1.673032021 | 1 | 0.982000683 | 86.49152294 |
| INTERPRO | IPR000980:SH2 domain | 14 | 0.889453621 | 0.102969996 | FRK, FGR, FER, VAV2, PTPN11, HSH2D, PLCG1, SH2B3, JAK1, TXK, SH2B2, SH2D1B, SHC4, SH3BP2 | 1452 | 113 | 18559 | 1.583571028 | 1 | 0.964876595 | 84.61833573 |
| SMART | SM00252:SH2 | 14 | 0.889453621 | 0.111741286 | FRK, FGR, FER, VAV2, PTPN11, HSH2D, PLCG1, SH2B3, JAK1, TXK, SH2B2, SH2D1B, SHC4, SH3BP2 | 878 | 103 | 10057 | 1.556914435 | 1 | 0.893028668 | 81.01629135 |
| Annotation Cluster 43 | Enrichment Score: 1.025967611903064 | | | | | | | | | | | |
| Category | Term | Count | % | PValue | Genes | List Total | Pop Hits | Pop Total | Fold Enrichment | Bonferroni | Benjamini | FDR |
| UP_SEQ_FEATURE | repeat:TPR 8 | 9 | 0.571791614 | 0.030705913 | IFIT1B, IFIT1, TTC34, IFIT5, TTC37, TMTC1, TTC28, ACAD11, IFT140 | 1499 | 50 | 20063 | 2.409166111 | 1 | 0.862588343 | 44.04915269 |
| UP_SEQ_FEATURE | repeat:TPR 6 | 11 | 0.698856417 | 0.034392439 | RPAP3, IFIT1B, IFIT1, TTC34, KLC1, IFIT5, TTC37, TMTC1, TTC28, ACAD11, IFT140 | 1499 | 70 | 20063 | 2.103240255 | 1 | 0.877085218 | 47.88138933 |
| UP_SEQ_FEATURE | repeat:TPR 4 | 14 | 0.889453621 | 0.034944322 | IFIT1B, TTC34, TTC37, NAA25, TTC28, RPAP3, INTS8, IFIT1, KLC1, IFIT5, TMTC1, ACAD11, LONRF1, IFT | 1499 | 100 | 20063 | 1.873795864 | 1 | 0.870034453 | 48.43324623 |
| UP_SEQ_FEATURE | repeat:TPR 7 | 10 | 0.635324015 | 0.036284082 | RPAP3, IFIT1B, IFIT1, TTC34, IFIT5, TTC37, TMTC1, TTC28, ACAD11, IFT140 | 1499 | 61 | 20063 | 2.194140356 | 1 | 0.876403637 | 49.75002818 |
| UP_SEQ_FEATURE | repeat:TPR 3 | 18 | 1.143583227 | 0.055550395 | IFIT1B, TTC31, TTC34, FKBP4, TTC37, NAA25, TTC28, RPAP3, INTS8, IFIT1, KLC1, IFIT5, TMTC1, LONR | 1499 | 150 | 20063 | 1.60611074 | 1 | 0.933249286 | 65.49861466 |
| UP_SEQ_FEATURE | repeat:TPR 5 | 11 | 0.698856417 | 0.064261478 | RPAP3, IFIT1B, IFIT1, TTC34, KLC1, IFIT5, TTC37, TMTC1, TTC28, ACAD11, IFT140 | 1499 | 78 | 20063 | 1.887523306 | 1 | 0.950502729 | 70.96608249 |
| UP_SEQ_FEATURE | repeat:TPR 10 | 5 | 0.317662008 | 0.077014528 | IFIT1, TTC37, TMTC1, TTC28, ACAD11 | 1499 | 22 | 20063 | 3.041876402 | 1 | 0.965442434 | 77.51261931 |
| UP_SEQ_FEATURE | repeat:TPR 9 | 6 | 0.381194409 | 0.086464925 | IFIT1, TTC37, TMTC1, TTC28, ACAD11, IFT140 | 1499 | 32 | 20063 | 2.509548032 | 1 | 0.972978814 | 81.43411928 |
| INTERPRO | IPR013026:Tetratricopeptide repeat-containing domain | 16 | 1.016518424 | 0.091634258 | IFIT1B, TTC31, NPHP3-ACAD11, TTC34, FKBP4, TTC37, NAA25, TTC28, RPAP3, IFIT1, KLC1, IFIT5, TMT | 1452 | 132 | 18559 | 1.549294599 | 1 | 0.965624615 | 80.90341031 |
| UP_SEQ_FEATURE | repeat:TPR 2 | 18 | 1.143583227 | 0.105654155 | IFIT1B, TTC31, TTC34, FKBP4, TTC37, NAA25, TTC28, RPAP3, INTS8, IFIT1, KLC1, IFIT5, TMTC1, LONR | 1499 | 164 | 20063 | 1.469003726 | 1 | 0.983990392 | 87.49605943 |
| UP_SEQ_FEATURE | repeat:TPR 1 | 18 | 1.143583227 | 0.105654155 | IFIT1B, TTC31, TTC34, FKBP4, TTC37, NAA25, TTC28, RPAP3, INTS8, IFIT1, KLC1, IFIT5, TMTC1, LONR | 1499 | 164 | 20063 | 1.469003726 | 1 | 0.983990392 | 87.49605943 |
| UP_KEYWORDS | TPR repeat | 17 | 1.080050826 | 0.18080595 | IFIT1B, TTC31, TTC34, FKBP4, TTC37, NAA25, TTC28, RPAP3, INTS8, IFIT1, KLC1, IFIT5, TMTC1, LONR | 1535 | 167 | 20581 | 1.364867659 | 1 | 0.590913322 | 94.30876527 |
| INTERPRO | IPR019734:Tetratricopeptide repeat | 15 | 0.952986023 | 0.190652672 | IFIT1B, TTC31, NPHP3-ACAD11, TTC34, FKBP4, TTC37, TTC28, RPAP3, IFIT1, KLC1, IFIT5, TMTC1, LON | 1452 | 138 | 18559 | 1.389313091 | 1 | 0.985273397 | 97.38521487 |
| SMART | SM00028:TPR | 15 | 0.952986023 | 0.250895332 | IFIT1B, TTC31, NPHP3-ACAD11, TTC34, FKBP4, TTC37, TTC28, RPAP3, IFIT1, KLC1, IFIT5, TMTC1, LON | 878 | 131 | 10057 | 1.311577318 | 1 | 0.913413175 | 98.25924068 |
| UP_SEQ_FEATURE | repeat:TPR 11 | 3 | 0.190597205 | 0.309852953 | TTC37, TTC28, ACAD11 | 1499 | 15 | 20063 | 2.676851234 | 1 | 0.999751312 | 99.89973676 |
| INTERPRO | IPR011990:Tetratricopeptide-like helical | 19 | 1.207115629 | 0.440306514 | IFIT1B, TTC31, CSTF3, NPHP3-ACAD11, TTC34, FKBP4, CCNF, TTC37, NAA25, TTC28, RPAP3, INTS8, IF | 1452 | 218 | 18559 | 1.113999671 | 1 | 0.999507443 | 99.99545089 |
| Annotation Cluster 44 | Enrichment Score: 1.018541165828759 | | | | | | | | | | | |
| Category | Term | Count | % | PValue | Genes | List Total | Pop Hits | Pop Total | Fold Enrichment | Bonferroni | Benjamini | FDR |
| GOTERM_CC_DIRECT | GO:0033017~sarcoplasmic reticulum membrane | 9 | 0.571791614 | 0.003942654 | SYNE2, NOS1AP, CAMK2G, RYR3, RYR2, KLHL41, DHRS7C, ITPR2, DMPK | 1425 | 34 | 18224 | 3.385263158 | 0.93372597 | 0.237685223 | 5.769443319 |
| UP_SEQ_FEATURE | domain:MIR 4 | 3 | 0.190597205 | 0.068290808 | RYR3, RYR2, ITPR2 | 1499 | 6 | 20063 | 6.692128085 | 1 | 0.956349522 | 73.20770389 |
| UP_SEQ_FEATURE | domain:MIR 5 | 3 | 0.190597205 | 0.068290808 | RYR3, RYR2, ITPR2 | 1499 | 6 | 20063 | 6.692128085 | 1 | 0.956349522 | 73.20770389 |
| INTERPRO | IPR014821:Inositol 1,4,5-trisphosphate/ryanodine receptor | 3 | 0.190597205 | 0.074157476 | RYR3, RYR2, ITPR2 | 1452 | 6 | 18559 | 6.39084022 | 1 | 0.950899407 | 73.48245133 |
| INTERPRO | IPR000699:Intracellular calcium-release channel | 3 | 0.190597205 | 0.074157476 | RYR3, RYR2, ITPR2 | 1452 | 6 | 18559 | 6.39084022 | 1 | 0.950899407 | 73.48245133 |
| INTERPRO | IPR015925:Ryanodine receptor-related | 3 | 0.190597205 | 0.074157476 | RYR3, RYR2, ITPR2 | 1452 | 6 | 18559 | 6.39084022 | 1 | 0.950899407 | 73.48245133 |
| INTERPRO | IPR013662:RyR/IP3R Homology associated domain | 3 | 0.190597205 | 0.074157476 | RYR3, RYR2, ITPR2 | 1452 | 6 | 18559 | 6.39084022 | 1 | 0.950899407 | 73.48245133 |
| REACTOME_PATHWAY | R-HSA-5578775:R-HSA-5578775 | 9 | 0.571791614 | 0.084667826 | SLC8A1, ATP2B4, ATP1B2, CAMK2G, RYR3, RYR2, CALM2, ITPR2, DMPK | 745 | 56 | 9075 | 1.957694151 | 1 | 0.933943116 | 74.61191894 |
| GOTERM_BP_DIRECT | GO:0051209~release of sequestered calcium ion into cytosol | 7 | 0.444726811 | 0.105118938 | JPH3, RYR3, RYR2, PRKCE, FGF2, HTR2A, ITPR2 | 1342 | 41 | 16792 | 2.136309113 | 1 | 0.978745442 | 87.34491028 |
| GOTERM_BP_DIRECT | GO:0071313~cellular response to caffeine | 3 | 0.190597205 | 0.129342022 | SLC8A1, RYR3, RYR2 | 1342 | 8 | 16792 | 4.692250373 | 1 | 0.984429287 | 92.4062792 |
| GOTERM_BP_DIRECT | GO:1903779~regulation of cardiac conduction | 8 | 0.508259212 | 0.15678727 | SLC8A1, ATP2B4, CORIN, ATP1B2, RYR3, RYR2, ITPR2, DMPK | 1342 | 56 | 16792 | 1.787523951 | 1 | 0.988808795 | 95.81634027 |
| UP_SEQ_FEATURE | domain:MIR 3 | 3 | 0.190597205 | 0.168352902 | RYR3, RYR2, ITPR2 | 1499 | 10 | 20063 | 4.015276851 | 1 | 0.995833277 | 96.76935141 |
| UP_SEQ_FEATURE | domain:MIR 1 | 3 | 0.190597205 | 0.168352902 | RYR3, RYR2, ITPR2 | 1499 | 10 | 20063 | 4.015276851 | 1 | 0.995833277 | 96.76935141 |
| UP_SEQ_FEATURE | domain:MIR 2 | 3 | 0.190597205 | 0.168352902 | RYR3, RYR2, ITPR2 | 1499 | 10 | 20063 | 4.015276851 | 1 | 0.995833277 | 96.76935141 |
| INTERPRO | IPR016093:MIR motif | 3 | 0.190597205 | 0.181163113 | RYR3, RYR2, ITPR2 | 1452 | 10 | 18559 | 3.834504132 | 1 | 0.984209236 | 96.8037019 |
| SMART | SM00472:MIR | 3 | 0.190597205 | 0.214788783 | RYR3, RYR2, ITPR2 | 878 | 10 | 10057 | 3.436332574 | 1 | 0.918626146 | 96.63162789 |
| GOTERM_BP_DIRECT | GO:0006816~calcium ion transport | 9 | 0.571791614 | 0.259701346 | CATSPER3, CAMK2G, RYR3, CACNG6, SLC24A1, RYR2, GJA4, CDH23, ITPR2 | 1342 | 76 | 16792 | 1.481763276 | 1 | 0.997687404 | 99.62895458 |
| Annotation Cluster 45 | Enrichment Score: 1.013149993396764 | | | | | | | | | | | |
| Category | Term | Count | % | PValue | Genes | List Total | Pop Hits | Pop Total | Fold Enrichment | Bonferroni | Benjamini | FDR |
| UP_SEQ_FEATURE | domain:Collagen-like 3 | 4 | 0.254129606 | 0.05493803 | COL21A1, COL22A1, COL25A1, COLEC12 | 1499 | 12 | 20063 | 4.461418724 | 1 | 0.93330171 | 65.07970204 |
| UP_SEQ_FEATURE | domain:Collagen-like 6 | 3 | 0.190597205 | 0.068290808 | COL21A1, COL22A1, COL25A1 | 1499 | 6 | 20063 | 6.692128085 | 1 | 0.956349522 | 73.20770389 |
| UP_SEQ_FEATURE | domain:Collagen-like 5 | 3 | 0.190597205 | 0.115475174 | COL21A1, COL22A1, COL25A1 | 1499 | 8 | 20063 | 5.019096064 | 1 | 0.986545152 | 89.81978955 |
| UP_SEQ_FEATURE | domain:Collagen-like 4 | 3 | 0.190597205 | 0.115475174 | COL21A1, COL22A1, COL25A1 | 1499 | 8 | 20063 | 5.019096064 | 1 | 0.986545152 | 89.81978955 |
| UP_SEQ_FEATURE | domain:Collagen-like 2 | 4 | 0.254129606 | 0.129105198 | COL21A1, COL22A1, COL25A1, COLEC12 | 1499 | 17 | 20063 | 3.149236746 | 1 | 0.990097609 | 92.37606141 |
| UP_SEQ_FEATURE | domain:Collagen-like 1 | 4 | 0.254129606 | 0.129105198 | COL21A1, COL22A1, COL25A1, COLEC12 | 1499 | 17 | 20063 | 3.149236746 | 1 | 0.990097609 | 92.37606141 |

Annotation Cluster 46 Enrichment Score: 1.0117828940691853

| Category | Term | Count | % | PValue | Genes | List Total | Pop Hits | Pop Total | Fold Enrichment | Bonferroni | Benjamini | FDR |
|---|---|---|---|---|---|---|---|---|---|---|---|---|
| UP_SEQ_FEATURE | domain:CRAL-TRIO | 6 | 0.381194409 | 0.06112943 | PRUNE2, TTPAL, SESTD1, TRIO, CLVS2, MCF2L | 1499 | 29 | 20063 | 2.769156449 | 1 | 0.944239431 | 69.10224933 |
| INTERPRO | IPR001251:CRAL-TRIO domain | 6 | 0.381194409 | 0.080741576 | PRUNE2, TTPAL, SESTD1, TRIO, CLVS2, MCF2L | 1452 | 30 | 18559 | 2.556336088 | 1 | 0.95804179 | 76.55025977 |
| SMART | SM00516:SEC14 | 5 | 0.317662008 | 0.186768549 | PRUNE2, TTPAL, TRIO, CLVS2, MCF2L | 878 | 26 | 10057 | 2.202777291 | 1 | 0.913821974 | 94.49250451 |

Annotation Cluster 47 Enrichment Score: 0.9935635865988873

| Category | Term | Count | % | PValue | Genes | List Total | Pop Hits | Pop Total | Fold Enrichment | Bonferroni | Benjamini | FDR |
|---|---|---|---|---|---|---|---|---|---|---|---|---|
| UP_SEQ_FEATURE | repeat:Spectrin 1 | 7 | 0.444726811 | 0.010787552 | SYNE3, SYNE2, SESTD1, SPTBN5, UTRN, TRIO, AKAP6 | 1499 | 26 | 20063 | 3.603453584 | 1 | 0.668593726 | 18.28660397 |
| UP_SEQ_FEATURE | repeat:Spectrin 2 | 7 | 0.444726811 | 0.010787552 | SYNE3, SYNE2, SESTD1, SPTBN5, UTRN, TRIO, AKAP6 | 1499 | 26 | 20063 | 3.603453584 | 1 | 0.668593726 | 18.28660397 |
| INTERPRO | IPR002017:Spectrin repeat | 7 | 0.444726811 | 0.010988224 | SYNE3, SYNE2, SPTBN5, UTRN, TRIO, AKAP6, MCF2L | 1452 | 25 | 18559 | 3.578870523 | 1 | 0.674921127 | 17.33239456 |
| INTERPRO | IPR018159:Spectrin/alpha-actinin | 7 | 0.444726811 | 0.022573624 | SYNE3, SYNE2, SPTBN5, UTRN, TRIO, AKAP6, MCF2L | 1452 | 29 | 18559 | 3.08523321 | 1 | 0.7873336 | 32.51962137 |
| SMART | SM00150:SPEC | 7 | 0.444726811 | 0.036122622 | SYNE3, SYNE2, SPTBN5, UTRN, TRIO, AKAP6, MCF2L | 878 | 29 | 10057 | 2.764865289 | 0.999999766 | 0.782777814 | 40.30454807 |
| INTERPRO | IPR001715:Calponin homology domain | 11 | 0.698856417 | 0.071276486 | GAS2L3, SYNE2, SPTBN5, LRCH1, UTRN, PLS1, LMO7, SPEF2, MAPRE2, VAV2, PARVA | 1452 | 76 | 18559 | 1.849980064 | 1 | 0.947611529 | 72.02448224 |
| UP_SEQ_FEATURE | repeat:Spectrin 3 | 5 | 0.317662008 | 0.077014528 | SYNE2, SESTD1, SPTBN5, UTRN, TRIO | 1499 | 22 | 20063 | 3.041876402 | 1 | 0.965442434 | 77.51261931 |
| UP_SEQ_FEATURE | domain:CH 1 | 5 | 0.317662008 | 0.138464537 | SYNE2, SPTBN5, UTRN, PLS1, PARVA | 1499 | 27 | 20063 | 2.478565958 | 1 | 0.991915016 | 93.76544136 |
| UP_SEQ_FEATURE | domain:CH 2 | 5 | 0.317662008 | 0.138464537 | SYNE2, SPTBN5, UTRN, PLS1, PARVA | 1499 | 27 | 20063 | 2.478565958 | 1 | 0.991915016 | 93.76544136 |
| UP_SEQ_FEATURE | domain:CH | 6 | 0.381194409 | 0.201812683 | GAS2L3, LRCH1, LMO7, SPEF2, MAPRE2, VAV2 | 1499 | 42 | 20063 | 1.912036596 | 1 | 0.997977455 | 98.49609191 |
| UP_SEQ_FEATURE | repeat:Spectrin 4 | 4 | 0.254129606 | 0.203783314 | SYNE2, SPTBN5, UTRN, TRIO | 1499 | 21 | 20063 | 2.549382128 | 1 | 0.998029241 | 98.56374302 |
| SMART | SM00033:CH | 9 | 0.571791614 | 0.214786914 | GAS2L3, SYNE2, SPTBN5, LRCH1, UTRN, PLS1, LMO7, VAV2, PARVA | 878 | 66 | 10057 | 1.561969352 | 1 | 0.92369384 | 96.63151546 |
| UP_SEQ_FEATURE | repeat:Spectrin 6 | 3 | 0.190597205 | 0.252812379 | SYNE2, SPTBN5, UTRN | 1499 | 13 | 20063 | 3.088674501 | 1 | 0.999215754 | 99.56015612 |
| UP_SEQ_FEATURE | repeat:Spectrin 9 | 3 | 0.190597205 | 0.252812379 | SYNE2, SPTBN5, UTRN | 1499 | 13 | 20063 | 3.088674501 | 1 | 0.999215754 | 99.56015612 |
| UP_SEQ_FEATURE | repeat:Spectrin 7 | 3 | 0.190597205 | 0.252812379 | SYNE2, SPTBN5, UTRN | 1499 | 13 | 20063 | 3.088674501 | 1 | 0.999215754 | 99.56015612 |
| UP_SEQ_FEATURE | repeat:Spectrin 8 | 3 | 0.190597205 | 0.252812379 | SYNE2, SPTBN5, UTRN | 1499 | 13 | 20063 | 3.088674501 | 1 | 0.999215754 | 99.56015612 |
| INTERPRO | IPR001589:Actinin-type, actin-binding, conserved site | 4 | 0.254129606 | 0.266441517 | SYNE2, SPTBN5, UTRN, PLS1 | 1452 | 23 | 18559 | 2.222900946 | 1 | 0.994762562 | 99.51934037 |
| UP_SEQ_FEATURE | repeat:Spectrin 5 | 3 | 0.190597205 | 0.281381718 | SYNE2, SPTBN5, UTRN | 1499 | 14 | 20063 | 2.868054894 | 1 | 0.999562574 | 99.78716607 |
| UP_SEQ_FEATURE | domain:Actin-binding | 3 | 0.190597205 | 0.446401725 | SYNE2, SPTBN5, UTRN | 1499 | 20 | 20063 | 2.007638426 | 1 | 0.999979603 | 99.99834666 |

Annotation Cluster 48 Enrichment Score: 0.9863511046454562

| Category | Term | Count | % | PValue | Genes | List Total | Pop Hits | Pop Total | Fold Enrichment | Bonferroni | Benjamini | FDR |
|---|---|---|---|---|---|---|---|---|---|---|---|---|
| INTERPRO | IPR005821:Ion transport domain | 18 | 1.143583227 | 0.003460672 | KCNC2, KCNC4, TRPV2, KCNA3, CNGA3, CNGA1, ITPR2, CATSPER3, RYR3, RYR2, SLC9C1, SCN7A, CAC | 1452 | 106 | 18559 | 2.170474037 | 0.999133645 | 0.585798168 | 5.797227034 |
| GOTERM_CC_DIRECT | GO:0008076~voltage-gated potassium channel complex | 14 | 0.889453621 | 0.018850118 | KCNC2, KCNC4, KCNA3, KCNJ3, KCNIP4, KCNT2, CNTN2, CNTNAP2, KCNG3, DPP6, KCNQ1, KCNG4, D | 1425 | 88 | 18224 | 2.034577352 | 0.9999979 | 0.383815567 | 24.89361812 |
| UP_KEYWORDS | Potassium | 16 | 1.016518424 | 0.056955474 | BCKDHA, KCNC2, KCNC4, ATP1B2, SLC12A5, KCNA3, KCNJ10, KCNJ3, KCNIP4, SLC24A4, KCNT2, MAT | 1535 | 129 | 20581 | 1.662985127 | 1 | 0.327015409 | 56.94930622 |
| GOTERM_BP_DIRECT | GO:0071805~potassium ion transmembrane transport | 16 | 1.016518424 | 0.058368412 | SLC9A9, KCNC2, KCNC4, SLC9A3, SLC12A5, KCNA3, CNGB1, CNGA3, CNGA1, KCNIP4, KCNT2, SLC9C1, | 1342 | 121 | 16792 | 1.654567624 | 1 | 0.954430464 | 67.3503542 |
| UP_KEYWORDS | Ion channel | 36 | 2.287166455 | 0.062597245 | KCNC2, KCNC4, LRRC8C, TRPV2, KCNA3, KCNJ10, CNGB1, KCNJ3, KCNIP4, GRIN2B, SCN7A, KCNG3, K | 1535 | 359 | 20581 | 1.344516527 | 1 | 0.338918622 | 60.50633309 |
| GOTERM_MF_DIRECT | GO:0005249~voltage-gated potassium channel activity | 9 | 0.571791614 | 0.086068369 | KCNC2, KCNC4, KCNT2, CNGB1, CNGA3, KCNQ1, KCNG4, CNGA1, KCNH5 | 1340 | 58 | 16881 | 1.954825013 | 1 | 0.923616308 | 76.95043823 |
| UP_KEYWORDS | Potassium transport | 14 | 0.889453621 | 0.091073749 | KCNC2, KCNC4, ATP1B2, SLC12A5, KCNA3, KCNJ10, KCNJ3, KCNIP4, KCNT2, SLC24A4, KCNG3, KCNG4 | 1535 | 116 | 20581 | 1.618184882 | 1 | 0.414312214 | 74.6499252 |
| UP_KEYWORDS | Voltage-gated channel | 17 | 1.080050826 | 0.093336643 | KCNC2, KCNC4, CACNG6, CACNG5, KCNA3, KCNJ10, CACNG2, KCNJ3, KCNIP4, CATSPER3, SCN7A, KC | 1535 | 150 | 20581 | 1.51955266 | 1 | 0.418861186 | 75.54202408 |
| GOTERM_BP_DIRECT | GO:0034765~regulation of ion transmembrane transport | 14 | 0.889453621 | 0.104748524 | KCNC4, CACNG6, ASIC2, KCNA3, KCNJ10, KCNJ3, NEDD4, CATSPER3, NEDD4L, SCN7A, KCNG3, KCNG | 1342 | 111 | 16792 | 1.578174299 | 1 | 0.979139755 | 87.24706054 |
| UP_SEQ_FEATURE | short sequence motif:Selectivity filter | 8 | 0.508259212 | 0.130636951 | KCNC4, KCNA3, KCNJ10, KCNG3, KCNQ1, KCNG4, KCNJ3, KCNH5 | 1499 | 57 | 20063 | 1.878492094 | 1 | 0.990301186 | 92.62190514 |
| INTERPRO | IPR027359:Voltage-dependent potassium channel, four helix bundle domain | 8 | 0.508259212 | 0.135496435 | KCNC2, KCNC4, KCNA3, SCN7A, KCNG3, CACNA1C, KCNG4, SCN4A | 1452 | 55 | 18559 | 1.859153519 | 1 | 0.974168847 | 91.8589893 |
| REACTOME_PATHWAY | R-HSA-1296072:R-HSA-1296072 | 7 | 0.444726811 | 0.136621639 | KCNC2, KCNC4, KCNA3, KCNG3, KCNQ1, KCNG4, KCNH5 | 745 | 43 | 9075 | 1.982987358 | 1 | 0.957506262 | 89.73443219 |
| GOTERM_MF_DIRECT | GO:0005251~delayed rectifier potassium channel activity | 6 | 0.381194409 | 0.153094276 | KCNC2, KCNC4, KCNA3, KCNG3, KCNQ1, KCNG4 | 1340 | 36 | 16881 | 2.099626866 | 1 | 0.958803137 | 93.34289311 |
| INTERPRO | IPR003968:Potassium channel, voltage dependent, Kv | 5 | 0.317662008 | 0.156166817 | KCNC2, KCNC4, KCNA3, KCNG3, KCNG4 | 1452 | 27 | 18559 | 2.366977859 | 1 | 0.978452388 | 94.63439394 |
| UP_KEYWORDS | Potassium channel | 9 | 0.571791614 | 0.184788174 | KCNC2, KCNC4, KCNT2, KCNA3, KCNG3, KCNQ1, KCNG4, KCNIP4, KCNH5 | 1535 | 74 | 20581 | 1.630680518 | 1 | 0.596435465 | 94.69370905 |
| UP_SEQ_FEATURE | region of interest:Segment H5 (pore-forming) | 4 | 0.254129606 | 0.348675573 | KCNG3, KCNQ1, KCNG4, KCNH5 | 1499 | 28 | 20063 | 1.912036596 | 1 | 0.999870952 | 99.96588361 |
| INTERPRO | IPR003131:Potassium channel tetramerisation-type BTB domain | 5 | 0.317662008 | 0.572211454 | KCNC2, KCNC4, KCNA3, KCNG3, KCNG4 | 1452 | 51 | 18559 | 1.253105926 | 1 | 0.999911791 | 99.99995562 |
| GOTERM_BP_DIRECT | GO:0006813~potassium ion transport | 6 | 0.381194409 | 0.793938347 | KCNC4, SLC24A4, ATP1B2, KCNA3, KCNJ10, KCNJ3 | 1342 | 82 | 16792 | 0.915561048 | 1 | 0.999999894 | 100 |

Annotation Cluster 49 Enrichment Score: 0.983638424286999

| Category | Term | Count | % | PValue | Genes | List Total | Pop Hits | Pop Total | Fold Enrichment | Bonferroni | Benjamini | FDR |
|---|---|---|---|---|---|---|---|---|---|---|---|---|
| UP_SEQ_FEATURE | domain:TSP N-terminal | 5 | 0.317662008 | 0.087964253 | COL21A1, NELL1, COL22A1, COL15A1, COL5A1 | 1499 | 23 | 20063 | 2.909620907 | 1 | 0.973774641 | 81.99335137 |
| INTERPRO | IPR001791:Laminin G domain | 9 | 0.571791614 | 0.094177414 | COL21A1, NRXN3, CNTNAP5, NELL1, COL22A1, COL15A1, CNTNAP2, NRXN1, COL5A1 | 1452 | 60 | 18559 | 1.917252066 | 1 | 0.966956911 | 81.80382384 |
| SMART | SM00210:TSPN | 5 | 0.317662008 | 0.135154884 | COL21A1, NELL1, COL22A1, COL15A1, COL5A1 | 878 | 23 | 10057 | 2.490096068 | 1 | 0.901499408 | 86.94728128 |

Annotation Cluster 50 Enrichment Score: 0.9765273025152105

| Category | Term | Count | % | PValue | Genes | List Total | Pop Hits | Pop Total | Fold Enrichment | Bonferroni | Benjamini | FDR |
|---|---|---|---|---|---|---|---|---|---|---|---|---|
| UP_KEYWORDS | Sushi | 10 | 0.635324015 | 0.021830106 | MASP1, CFB, SELL, CSMD3, SUSD5, C4BPA, C1S, C2, SUSD3, CSMD1 | 1535 | 56 | 20581 | 2.394253141 | 0.999981196 | 0.214794657 | 27.18254642 |
| INTERPRO | IPR000436:Sushi/SCR/CCP | 10 | 0.635324015 | 0.031765642 | MASP1, CFB, SELL, CSMD3, SUSD5, C4BPA, C1S, C2, SUSD3, CSMD1 | 1452 | 57 | 18559 | 2.242400077 | 1 | 0.855021561 | 42.65636723 |
| UP_SEQ_FEATURE | domain:Sushi 1 | 8 | 0.508259212 | 0.047781845 | MASP1, CFB, SELL, CSMD3, C4BPA, C1S, C2, CSMD1 | 1499 | 45 | 20063 | 2.379423319 | 1 | 0.917316304 | 59.81358719 |
| UP_SEQ_FEATURE | domain:Sushi 2 | 8 | 0.508259212 | 0.047781845 | MASP1, CFB, SELL, CSMD3, C4BPA, C1S, C2, CSMD1 | 1499 | 45 | 20063 | 2.379423319 | 1 | 0.917316304 | 59.81358719 |
| SMART | SM00032:CCP | 9 | 0.571791614 | 0.094527742 | MASP1, CFB, SELL, CSMD3, C4BPA, C1S, C2, SUSD3, CSMD1 | 878 | 54 | 10057 | 1.909073652 | 1 | 0.872602958 | 75.15321205 |
| KEGG_PATHWAY | hsa04610:Complement and coagulation cascades | 10 | 0.635324015 | 0.117365253 | C8B, THBD, MASP1, C4A, C4B, CFB, C5, C4BPA, C1S, C2 | 574 | 69 | 6879 | 1.736858052 | 1 | 0.711765078 | 80.79151757 |
| UP_SEQ_FEATURE | domain:Sushi 3 | 5 | 0.317662008 | 0.213760162 | CFB, CSMD3, C4BPA, C2, CSMD1 | 1499 | 32 | 20063 | 2.091290027 | 1 | 0.998342018 | 98.86429271 |
| SMART | SM00042:CUB | 7 | 0.444726811 | 0.237857607 | MASP1, ATRNL1, ST14, CSMD3, PDGFC, C1S, CSMD1 | 878 | 48 | 10057 | 1.670439446 | 1 | 0.922841082 | 97.78272296 |
| INTERPRO | IPR000859:CUB domain | 7 | 0.444726811 | 0.258216774 | MASP1, ATRNL1, ST14, CSMD3, PDGFC, C1S, CSMD1 | 1452 | 55 | 18559 | 1.626759329 | 1 | 0.994686576 | 99.41755411 |
| UP_SEQ_FEATURE | domain:CUB 1 | 5 | 0.317662008 | 0.280328105 | MASP1, ST14, CSMD3, C1S, CSMD1 | 1499 | 36 | 20063 | 1.858924468 | 1 | 0.999586083 | 99.78128014 |
| UP_SEQ_FEATURE | domain:CUB 2 | 5 | 0.317662008 | 0.280328105 | MASP1, ST14, CSMD3, C1S, CSMD1 | 1499 | 36 | 20063 | 1.858924468 | 1 | 0.999586083 | 99.78128014 |

Annotation Cluster 51 Enrichment Score: 0.9691938271679943

| Category | Term | Count | % | PValue | Genes | List Total | Pop Hits | Pop Total | Fold Enrichment | Bonferroni | Benjamini | FDR |
|---|---|---|---|---|---|---|---|---|---|---|---|---|
| INTERPRO | IPR010919:SAND domain-like | 4 | 0.254129606 | 0.061563084 | SKOR2, SP100, GMEB1, SP110 | 1452 | 12 | 18559 | 4.260560147 | 1 | 0.928463219 | 66.53267445 |
| UP_SEQ_FEATURE | domain:SAND | 3 | 0.190597205 | 0.115475174 | SP100, GMEB1, SP110 | 1499 | 8 | 20063 | 5.019096064 | 1 | 0.986545152 | 89.81978955 |
| INTERPRO | IPR000770:SAND domain | 3 | 0.190597205 | 0.124818096 | SP100, GMEB1, SP110 | 1452 | 8 | 18559 | 4.793130165 | 1 | 0.969089745 | 89.94168933 |
| SMART | SM00258:SAND | 3 | 0.190597205 | 0.149671168 | SP100, GMEB1, SP110 | 878 | 8 | 10057 | 4.295415718 | 1 | 0.909556504 | 89.70530327 |

Annotation Cluster 52 Enrichment Score: 0.9471771161172486

| Category | Term | Count | % | PValue | Genes | List Total | Pop Hits | Pop Total | Fold Enrichment | Bonferroni | Benjamini | FDR |
|---|---|---|---|---|---|---|---|---|---|---|---|---|
| GOTERM_BP_DIRECT | GO:0086064~cell communication by electrical coupling involved in cardiac conduction | 4 | 0.254129606 | 0.079466112 | SLC8A1, ATP1B2, RYR2, CACNA1C | 1342 | 13 | 16792 | 3.850051588 | 1 | 0.973004972 | 78.58514095 |
| GOTERM_BP_DIRECT | GO:0060402~calcium ion transport into cytosol | 4 | 0.254129606 | 0.095362747 | SLC8A1, JPH3, RYR2, CACNA1C | 1342 | 14 | 16792 | 3.575047903 | 1 | 0.979190754 | 84.51500586 |
| GOTERM_BP_DIRECT | GO:0010881~regulation of cardiac muscle contraction by regulation of the release of sequestered c | 4 | 0.254129606 | 0.190067532 | SLC8A1, RYR2, CACNA1C, CALM2 | 1342 | 19 | 16792 | 2.634245823 | 1 | 0.993271247 | 98.02272728 |

Annotation Cluster 53 Enrichment Score: 0.9452507230492354

| Category | Term | Count | % | PValue | Genes | List Total | Pop Hits | Pop Total | Fold Enrichment | Bonferroni | Benjamini | FDR |
|---|---|---|---|---|---|---|---|---|---|---|---|---|
| INTERPRO | IPR006652:Kelch repeat type 1 | 11 | 0.698856417 | 0.026029096 | LZTR1, ATRNL1, KLHDC8A, KLHL25, KLHL14, KBTBD8, KLHL24, KLHL23, KLHL41, KLHL2, PHOSPHO2-K | 1452 | 64 | 18559 | 2.196851326 | 1 | 0.822797973 | 36.51354465 |

| | | | | | | | | | | | | |
|---|---|---|---|---|---|---|---|---|---|---|---|---|
| UP_KEYWORDS | Kelch repeat | 11 | 0.698856417 | 0.040307364 | LZTR1, ATRNL1, KLHDC8A, KLHL25, KLHL14, KBTBD8, KLHL24, KLHL23, KLHL41, KLHL2, PHOSPHO2-K | 1535 | 72 | 20581 | 2.048416576 | 0.999999998 | 0.279022233 | 44.63875807 |
| UP_SEQ_FEATURE | repeat:Kelch 5 | 10 | 0.635324015 | 0.0558001 | LZTR1, ATRNL1, KLHDC8A, KLHL25, KLHL14, KBTBD8, KLHL24, KLHL23, KLHL41, KLHL2 | 1499 | 66 | 20063 | 2.027917602 | 1 | 0.931987053 | 65.66806653 |
| SMART | SM00612:Kelch | 10 | 0.635324015 | 0.062358534 | LZTR1, KLHDC8A, KLHL25, KLHL14, KBTBD8, KLHL24, KLHL23, KLHL41, KLHL2, PHOSPHO2-KLHL23 | 878 | 58 | 10057 | 1.974903778 | 1 | 0.871964979 | 59.46047774 |
| UP_SEQ_FEATURE | repeat:Kelch 4 | 10 | 0.635324015 | 0.075549907 | LZTR1, ATRNL1, KLHDC8A, KLHL25, KLHL14, KBTBD8, KLHL24, KLHL23, KLHL41, KLHL2 | 1499 | 70 | 20063 | 1.912036596 | 1 | 0.965541316 | 76.83882976 |
| UP_SEQ_FEATURE | repeat:Kelch 1 | 10 | 0.635324015 | 0.081079423 | LZTR1, ATRNL1, KLHDC8A, KLHL25, KLHL14, KBTBD8, KLHL24, KLHL23, KLHL41, KLHL2 | 1499 | 71 | 20063 | 1.885106503 | 1 | 0.968061686 | 79.2868101 |
| UP_SEQ_FEATURE | repeat:Kelch 2 | 10 | 0.635324015 | 0.081079423 | LZTR1, ATRNL1, KLHDC8A, KLHL25, KLHL14, KBTBD8, KLHL24, KLHL23, KLHL41, KLHL2 | 1499 | 71 | 20063 | 1.885106503 | 1 | 0.968061686 | 79.2868101 |
| UP_SEQ_FEATURE | repeat:Kelch 3 | 10 | 0.635324015 | 0.081079423 | LZTR1, ATRNL1, KLHDC8A, KLHL25, KLHL14, KBTBD8, KLHL24, KLHL23, KLHL41, KLHL2 | 1499 | 71 | 20063 | 1.885106503 | 1 | 0.968061686 | 79.2868101 |
| UP_SEQ_FEATURE | repeat:Kelch 6 | 8 | 0.508259212 | 0.090405888 | LZTR1, ATRNL1, KLHDC8A, KLHL25, KLHL14, KLHL24, KLHL23, KLHL2 | 1499 | 52 | 20063 | 2.059116334 | 1 | 0.975547056 | 82.87007484 |
| INTERPRO | IPR017096:Kelch-like protein, gigaxonin | 7 | 0.444726811 | 0.106457144 | KLHL25, KBTBD8, KLHL24, KLHL23, KLHL41, KLHL2, PHOSPHO2-KLHL23 | 1452 | 42 | 18559 | 2.130280073 | 1 | 0.967194194 | 85.61657385 |
| INTERPRO | IPR011333:BTB/POZ fold | 21 | 1.334180432 | 0.106643305 | ZBTB22, KCNC2, ZBTB48, KCNC4, ZBTB46, ZBTB7C, LZTR1, ABTB2, KCNA3, KLHL2, RCBTB2, KLHL25, K | 1452 | 190 | 18559 | 1.412712049 | 1 | 0.965717836 | 85.66811016 |
| INTERPRO | IPR011705:BTB/Kelch-associated | 9 | 0.571791614 | 0.10887681 | RCBTB2, KLHL25, KLHL14, KBTBD8, KLHL24, KLHL23, KLHL41, KLHL2, PHOSPHO2-KLHL23 | 1452 | 62 | 18559 | 1.855405225 | 1 | 0.964897833 | 86.273018 |
| INTERPRO | IPR000210:BTB/POZ-like | 20 | 1.27064803 | 0.111279991 | ZBTB22, KCNC2, ZBTB48, KCNC4, ZBTB46, ZBTB7C, LZTR1, ABTB2, KCNA3, KLHL2, RCBTB2, KLHL25, K | 1452 | 180 | 18559 | 1.420186716 | 1 | 0.965941596 | 86.8969803 |
| INTERPRO | IPR015915:Kelch-type beta propeller | 8 | 0.508259212 | 0.117307501 | LZTR1, ATRNL1, KLHDC8A, KLHL25, KLHL14, KBTBD8, KLHL23, KLHL41 | 1452 | 53 | 18559 | 1.929310255 | 1 | 0.969090813 | 88.34649766 |
| GOTERM_CC_DIRECT | GO:0031463~Cul3-RING ubiquitin ligase complex | 9 | 0.571791614 | 0.150343227 | KLHDC8A, KLHL25, KLHL14, KBTBD8, KLHL24, KLHL23, KLHL41, KLHL2, PHOSPHO2-KLHL23 | 1425 | 67 | 18224 | 1.717894737 | 1 | 0.821286042 | 91.37763429 |
| PIR_SUPERFAMILY | PIRSF037037:kelch-like protein, gigaxonin type | 7 | 0.444726811 | 0.172986325 | KLHL25, KBTBD8, KLHL24, KLHL23, KLHL41, KLHL2, PHOSPHO2-KLHL23 | 154 | 42 | 1692 | 1.831168831 | 1 | 0.999997966 | 89.30481131 |
| INTERPRO | IPR015916:Galactose oxidase, beta-propeller | 5 | 0.317662008 | 0.187653892 | KLHDC8A, KLHL24, KLHL23, KLHL2, PHOSPHO2-KLHL23 | 1452 | 29 | 18559 | 2.203738007 | 1 | 0.985407354 | 97.21320178 |
| SMART | SM00225:BTB | 20 | 1.27064803 | 0.194758863 | ZBTB22, KCNC2, ZBTB48, KCNC4, ZBTB46, ZBTB7C, LZTR1, ABTB2, KCNA3, KLHL2, RCBTB2, KLHL25, K | 878 | 176 | 10057 | 1.301641127 | 1 | 0.917674325 | 95.20463638 |
| SMART | SM00875:SM00875 | 8 | 0.508259212 | 0.266520611 | KLHL25, KLHL14, KBTBD8, KLHL24, KLHL23, KLHL41, KLHL2, PHOSPHO2-KLHL23 | 878 | 60 | 10057 | 1.527258922 | 1 | 0.91572631 | 98.70471286 |
| UP_SEQ_FEATURE | domain:BTB | 13 | 0.82592122 | 0.389748568 | ZBTB22, ZBTB48, ZBTB46, ZBTB7C, KLHL25, ABTB2, KLHL14, KBTBD8, KLHL24, ZBTB2, KLHL23, KLHL4 | 1499 | 144 | 20063 | 1.208300904 | 1 | 0.999942573 | 99.98985531 |
| UP_SEQ_FEATURE | domain:BACK | 3 | 0.190597205 | 0.666722975 | KBTBD8, KLHL24, KLHL23 | 1499 | 30 | 20063 | 1.338425617 | 1 | 0.999999924 | 99.99999987 |

Annotation Cluster 54 Enrichment Score: 0.9402144690451104

| Category | Term | Count | % | PValue | Genes | List Total | Pop Hits | Pop Total | Fold Enrichment | Bonferroni | Benjamini | FDR |
|---|---|---|---|---|---|---|---|---|---|---|---|---|
| GOTERM_BP_DIRECT | GO:0006607~NLS-bearing protein import into nucleus | 6 | 0.381194409 | 0.032304474 | FYB, KPNA6, TNPO2, KPNA3, NUP35, TNPO1 | 1342 | 23 | 16792 | 3.264174172 | 1 | 0.907354962 | 45.72847736 |
| GOTERM_MF_DIRECT | GO:0008565~protein transporter activity | 10 | 0.635324015 | 0.115049311 | AP2A2, ITGB1BP1, RUFY1, RAP1A, KPNA6, IPO8, TNPO2, KPNA3, TNPO1, SEC62 | 1340 | 72 | 16881 | 1.749689055 | 1 | 0.945563836 | 86.3708911 |
| COG_ONTOLOGY | Intracellular trafficking and secretion | 7 | 0.444726811 | 0.121118171 | COPG2, ARMC8, SEC22A, KPNA6, TNPO2, KPNA3, TNPO1 | 166 | 42 | 2026 | 2.034136546 | 0.983938891 | 0.747695395 | 67.31172809 |
| GOTERM_MF_DIRECT | GO:0008139~nuclear localization sequence binding | 4 | 0.254129606 | 0.38528857 | KPNA6, TNPO2, KPNA3, TNPO1 | 1340 | 28 | 16881 | 1.799680171 | 1 | 0.993568434 | 99.96418258 |

Annotation Cluster 55 Enrichment Score: 0.9383921999468369

| Category | Term | Count | % | PValue | Genes | List Total | Pop Hits | Pop Total | Fold Enrichment | Bonferroni | Benjamini | FDR |
|---|---|---|---|---|---|---|---|---|---|---|---|---|
| UP_SEQ_FEATURE | metal ion-binding site:Magnesium | 16 | 1.016518424 | 0.029588733 | ATP10A, ATP11A, ATP10D, CNOT6, CLYBL, NT5DC1, PGM3, ATP2C2, ITPA, ATP2B4, MAT1A, ENO2, P | 1499 | 118 | 20063 | 1.814814396 | 1 | 0.865403312 | 42.83614752 |
| INTERPRO | IPR006539:Phospholipid-transporting P-type ATPase, subfamily IV | 4 | 0.254129606 | 0.090748497 | ATP10A, ATP11A, ATP10D, ATP8B3 | 1452 | 14 | 18559 | 3.651908697 | 1 | 0.966451574 | 80.58006846 |
| REACTOME_PATHWAY | R-HSA-936837:R-HSA-936837 | 9 | 0.571791614 | 0.091805093 | ATP2C2, ATP2B4, ATP1B2, CAMK2G, ATP10A, ATP11A, ATP10D, ATP8B3, CALM2 | 745 | 57 | 9075 | 1.92334864 | 1 | 0.930847762 | 77.51208597 |
| GOTERM_MF_DIRECT | GO:0004012~phospholipid-translocating ATPase activity | 4 | 0.254129606 | 0.093864395 | ATP10A, ATP11A, ATP10D, ATP8B3 | 1340 | 14 | 16881 | 3.599360341 | 1 | 0.93258866 | 79.95544857 |
| INTERPRO | IPR023214:HAD-like domain | 11 | 0.698856417 | 0.099498347 | ATP2C2, ATP2B4, ATP10A, PHOSPHO2, ATP11A, HDHD2, ATP10D, ACAD10, CTDP1, ATP8B3, NT5DC1 | 1452 | 81 | 18559 | 1.735783764 | 1 | 0.966076548 | 83.55997502 |
| INTERPRO | IPR023299:P-type ATPase, cytoplasmic domain N | 6 | 0.381194409 | 0.146628746 | ATP2C2, ATP2B4, ATP10A, ATP11A, ATP10D, ATP8B3 | 1452 | 36 | 18559 | 2.130280073 | 1 | 0.976485603 | 93.48804981 |
| INTERPRO | IPR008250:P-type ATPase, A domain | 6 | 0.381194409 | 0.146628746 | ATP2C2, ATP2B4, ATP10A, ATP11A, ATP10D, ATP8B3 | 1452 | 36 | 18559 | 2.130280073 | 1 | 0.976485603 | 93.48804981 |
| INTERPRO | IPR001757:Cation-transporting P-type ATPase | 6 | 0.381194409 | 0.146628746 | ATP2C2, ATP2B4, ATP10A, ATP11A, ATP10D, ATP8B3 | 1452 | 36 | 18559 | 2.130280073 | 1 | 0.976485603 | 93.48804981 |
| INTERPRO | IPR018303:P-type ATPase, phosphorylation site | 6 | 0.381194409 | 0.146628746 | ATP2C2, ATP2B4, ATP10A, ATP11A, ATP10D, ATP8B3 | 1452 | 36 | 18559 | 2.130280073 | 1 | 0.976485603 | 93.48804981 |
| UP_SEQ_FEATURE | active site:4-aspartylphosphate intermediate | 6 | 0.381194409 | 0.175516912 | ATP2C2, ATP2B4, ATP10A, ATP11A, ATP10D, ATP8B3 | 1499 | 40 | 20063 | 2.007638426 | 1 | 0.996356761 | 97.25002043 |
| GOTERM_BP_DIRECT | GO:0045332~phospholipid translocation | 4 | 0.254129606 | 0.254883081 | ATP10A, ATP11A, ATP10D, ATP8B3 | 1342 | 22 | 16792 | 2.275030484 | 1 | 0.997516594 | 99.5813364 |

Annotation Cluster 56 Enrichment Score: 0.9306718200624204

| Category | Term | Count | % | PValue | Genes | List Total | Pop Hits | Pop Total | Fold Enrichment | Bonferroni | Benjamini | FDR |
|---|---|---|---|---|---|---|---|---|---|---|---|---|
| GOTERM_MF_DIRECT | GO:0035615~clathrin adaptor activity | 5 | 0.317662008 | 0.005608559 | AP2A2, DAB2, PICALM, ARRB1, HIP1 | 1340 | 10 | 16881 | 6.298880597 | 0.999277583 | 0.910271745 | 8.763034184 |
| GOTERM_BP_DIRECT | GO:0072583~clathrin-mediated endocytosis | 5 | 0.317662008 | 0.070133242 | AP2A2, CLTA, PICALM, PIP5K1C, HIP1 | 1342 | 20 | 16792 | 3.128166915 | 1 | 0.967620814 | 74.16238249 |
| GOTERM_CC_DIRECT | GO:0030122~AP-2 adaptor complex | 3 | 0.190597205 | 0.270087858 | AP2A2, PICALM, HIP1 | 1425 | 13 | 18224 | 2.951255061 | 1 | 0.882518563 | 99.1225738 |
| UP_KEYWORDS | Coated pit | 6 | 0.381194409 | 0.330707369 | AP2A2, CLTA, DAB2, APP, PICALM, ARRB1 | 1535 | 51 | 20581 | 1.577390305 | 1 | 0.759285913 | 99.68827111 |
| GOTERM_CC_DIRECT | GO:0005905~clathrin-coated pit | 5 | 0.317662008 | 0.632290672 | DAB2, APP, PICALM, ARRB1, CDHR5 | 1425 | 55 | 18224 | 1.16261563 | 1 | 0.982870771 | 99.99997089 |

Annotation Cluster 57 Enrichment Score: 0.9172550433751184

| Category | Term | Count | % | PValue | Genes | List Total | Pop Hits | Pop Total | Fold Enrichment | Bonferroni | Benjamini | FDR |
|---|---|---|---|---|---|---|---|---|---|---|---|---|
| GOTERM_MF_DIRECT | GO:0019706~protein-cysteine S-palmitoyltransferase activity | 7 | 0.444726811 | 0.017150904 | ZDHHC16, ZDHHC21, ZDHHC6, ZDHHC11, ZDHHC11B, GOLGA7B, ZDHHC19 | 1340 | 27 | 16881 | 3.266086235 | 1 | 0.867674481 | 24.57919287 |
| UP_SEQ_FEATURE | zinc finger region:DHHC-type | 6 | 0.381194409 | 0.040653474 | ZDHHC16, ZDHHC21, ZDHHC6, ZDHHC11, ZDHHC11B, ZDHHC19 | 1499 | 26 | 20063 | 3.088674501 | 1 | 0.898159112 | 53.82686364 |
| INTERPRO | IPR001594:Zinc finger, DHHC-type, palmitoyltransferase | 6 | 0.381194409 | 0.055345855 | ZDHHC16, ZDHHC21, ZDHHC6, ZDHHC11, ZDHHC11B, ZDHHC19 | 1452 | 27 | 18559 | 2.840373431 | 1 | 0.91490727 | 62.50063669 |
| UP_KEYWORDS | Acyltransferase | 15 | 0.952986023 | 0.397677575 | KAT7, ZDHHC6, ZDHHC11B, GGCT, ZDHHC19, GLYATL1, OXSM, ATE1, GGT5, ZDHHC16, ZDHHC21, ZD | 1535 | 171 | 20581 | 1.17612435 | 1 | 0.80268188 | 99.93149343 |
| GOTERM_MF_DIRECT | GO:0016409~palmitoyltransferase activity | 3 | 0.190597205 | 0.423794435 | ZDHHC16, ZDHHC21, ZDHHC6 | 1340 | 18 | 16881 | 2.099626866 | 1 | 0.994962 | 99.9875262 |
| GOTERM_BP_DIRECT | GO:0018345~protein palmitoylation | 3 | 0.190597205 | 0.482295951 | ZDHHC16, ZDHHC21, ZDHHC6 | 1342 | 20 | 16792 | 1.876900149 | 1 | 0.999906174 | 99.99952298 |

Annotation Cluster 58 Enrichment Score: 0.8750792476074271

| Category | Term | Count | % | PValue | Genes | List Total | Pop Hits | Pop Total | Fold Enrichment | Bonferroni | Benjamini | FDR |
|---|---|---|---|---|---|---|---|---|---|---|---|---|
| UP_KEYWORDS | Aminopeptidase | 6 | 0.381194409 | 0.0534311 | METAP2, TPP2, DPP8, ENPEP, DPP6, C9ORF3 | 1535 | 28 | 20581 | 2.873103769 | 1 | 0.32452478 | 54.5783346 |
| GOTERM_MF_DIRECT | GO:0004177~aminopeptidase activity | 5 | 0.317662008 | 0.162064275 | METAP2, TPP2, ENPEP, DPP6, C9ORF3 | 1340 | 27 | 16881 | 2.33291874 | 1 | 0.959342525 | 94.40395583 |
| GOTERM_MF_DIRECT | GO:0070006~metalloaminopeptidase activity | 4 | 0.254129606 | 0.273703815 | METAP2, MMP16, ENPEP, C9ORF3 | 1340 | 23 | 16881 | 2.19091499 | 1 | 0.987991546 | 99.45633131 |

Annotation Cluster 59 Enrichment Score: 0.8735877622482812

| Category | Term | Count | % | PValue | Genes | List Total | Pop Hits | Pop Total | Fold Enrichment | Bonferroni | Benjamini | FDR |
|---|---|---|---|---|---|---|---|---|---|---|---|---|
| INTERPRO | IPR001202:WW domain | 9 | 0.571791614 | 0.052286556 | MAGI2, DGCR8, MAGI1, PCIF1, NEDD4, UTRN, NEDD4L, WWOX, ARHGAP9 | 1452 | 53 | 18559 | 2.170474037 | 1 | 0.906949304 | 60.35264319 |
| SMART | SM00456:WW | 8 | 0.508259212 | 0.130998208 | MAGI2, DGCR8, MAGI1, PCIF1, NEDD4, UTRN, NEDD4L, WWOX | 878 | 49 | 10057 | 1.870112965 | 1 | 0.911780257 | 86.03949658 |
| UP_SEQ_FEATURE | domain:WW 1 | 5 | 0.317662008 | 0.167214296 | MAGI2, MAGI1, NEDD4, NEDD4L, WWOX | 1499 | 29 | 20063 | 2.307630374 | 1 | 0.995830461 | 96.68599419 |
| UP_SEQ_FEATURE | domain:WW 2 | 5 | 0.317662008 | 0.167214296 | MAGI2, MAGI1, NEDD4, NEDD4L, WWOX | 1499 | 29 | 20063 | 2.307630374 | 1 | 0.995830461 | 96.68599419 |
| UP_SEQ_FEATURE | domain:WW | 4 | 0.254129606 | 0.223799438 | DGCR8, PCIF1, UTRN, ARHGAP9 | 1499 | 22 | 20063 | 2.433501122 | 1 | 0.998542208 | 99.10597811 |

Annotation Cluster 60 Enrichment Score: 0.871017372933464

| Category | Term | Count | % | PValue | Genes | List Total | Pop Hits | Pop Total | Fold Enrichment | Bonferroni | Benjamini | FDR |
|---|---|---|---|---|---|---|---|---|---|---|---|---|
| KEGG_PATHWAY | hsa04070:Phosphatidylinositol signaling system | 14 | 0.889453621 | 0.061990804 | PIK3CB, PIK3C2B, PIP5K1C, ITPR2, PLCG1, INPP5K, DGKG, DGKZ, INPP4B, ITPK1, IP6K1, INPP5A, MTM | 574 | 98 | 6879 | 1.712045794 | 0.999999982 | 0.672386919 | 57.07451936 |
| KEGG_PATHWAY | hsa00562:Inositol phosphate metabolism | 11 | 0.698856417 | 0.068305099 | TPI1, INPP5K, PLCG1, PIK3CB, PIK3C2B, PLCH1, PIP5K1C, INPP4B, ITPK1, INPP5A, MTMR4 | 574 | 71 | 6879 | 1.85672572 | 0.999999997 | 0.609357579 | 60.73998919 |
| REACTOME_PATHWAY | R-HSA-1660499:R-HSA-1660499 | 6 | 0.381194409 | 0.154917718 | INPP5K, PIK3CB, PIK3C2B, PIP5K1C, INPP4B, PIK3R6 | 745 | 35 | 9075 | 2.088207095 | 1 | 0.96274302 | 92.63389211 |
| GOTERM_BP_DIRECT | GO:0043647~inositol phosphate metabolic process | 7 | 0.444726811 | 0.169568809 | PLCG1, PLCH1, INPP4B, IP6K1, ITPK1, INPP5A, CALM2 | 1342 | 47 | 16792 | 1.8635888 | 1 | 0.99100827 | 96.85156394 |
| GOTERM_BP_DIRECT | GO:0006661~phosphatidylinositol biosynthetic process | 8 | 0.508259212 | 0.178053189 | INPP5K, PIK3CB, PIK3C2B, PIP5K1C, INPP4B, PIK3R6, FGF2, MTMR4 | 1342 | 58 | 16792 | 1.725885195 | 1 | 0.991942736 | 97.39931738 |
| GOTERM_CC_DIRECT | GO:0005942~phosphatidylinositol 3-kinase complex | 3 | 0.190597205 | 0.299998128 | PIK3CB, PIK3C2B, PIK3R6 | 1425 | 14 | 18224 | 2.740451128 | 1 | 0.896567074 | 99.53241285 |

Annotation Cluster 61 Enrichment Score: 0.8669079190708627

| Category | Term | Count | % | PValue | Genes | List Total | Pop Hits | Pop Total | Fold Enrichment | Bonferroni | Benjamini | FDR |
|---|---|---|---|---|---|---|---|---|---|---|---|---|
| UP_SEQ_FEATURE | repeat:WD 1 | 30 | 1.905972046 | 0.042380238 | NBEA, DMWD, WDR81, FBXW9, WDR37, WDPCP, WDR17, EIF3I, PAFAH1B1, BCAS3, MAPKBP1, IFT1 | 1499 | 278 | 20063 | 1.444344191 | 1 | 0.901802906 | 55.35002832 |

| | | | | | | | | | | | | |
|---|---|---|---|---|---|---|---|---|---|---|---|---|
| UP_SEQ_FEATURE | repeat:WD 2 | 30 | 1.905972046 | 0.042380238 | NBEA, DMWD, WDR81, FBXW9, WDR37, WDPCP, WDR17, EIF3I, PAFAH1B1, BCAS3, MAPKBP1, IFT14 | 1499 | 278 | 20063 | 1.444344191 | 1 | 0.901802906 | 55.35002832 |
| UP_KEYWORDS | WD repeat | 30 | 1.905972046 | 0.047051238 | NBEA, DMWD, WDR81, FBXW9, WDR37, WDPCP, WDR17, EIF3I, PAFAH1B1, BCAS3, MAPKBP1, IFT14 | 1535 | 281 | 20581 | 1.4314396 | 1 | 0.302320629 | 49.97462321 |
| UP_SEQ_FEATURE | repeat:WD 8 | 9 | 0.571791614 | 0.076652365 | EML2, TBL1XR1, ELP2, WDR17, WDR62, TEP1, WDR1, MAPKBP1, PWP2 | 1499 | 60 | 20063 | 2.007638426 | 1 | 0.966072786 | 77.34775645 |
| UP_SEQ_FEATURE | repeat:WD 5 | 25 | 1.588310038 | 0.078909878 | NBEA, DMWD, WDR81, FBXW9, WDR37, WDR17, EIF3I, PAFAH1B1, MAPKBP1, IFT140, TBL1XR1, EL | 1499 | 237 | 20063 | 1.411841368 | 1 | 0.966010957 | 78.35705682 |
| INTERPRO | IPR017986:WD40-repeat-containing domain | 32 | 2.033036849 | 0.082341075 | NBEA, DMWD, WDR81, FBXW9, WDR37, WDPCP, WDR17, EIF3I, PAFAH1B1, BCAS3, MAPKBP1, IFT14 | 1452 | 306 | 18559 | 1.336646321 | 1 | 0.958323969 | 77.2433264 |
| UP_SEQ_FEATURE | repeat:WD 3 | 27 | 1.715374841 | 0.105820209 | NBEA, DMWD, WDR81, FBXW9, WDR37, WDR17, EIF3I, PAFAH1B1, BCAS3, MAPKBP1, IFT140, TBL1X | 1499 | 269 | 20063 | 1.343401177 | 1 | 0.983495653 | 87.53921655 |
| UP_SEQ_FEATURE | repeat:WD 4 | 25 | 1.588310038 | 0.143020503 | NBEA, DMWD, WDR81, FBXW9, WDR37, WDR17, EIF3I, PAFAH1B1, MAPKBP1, IFT140, TBL1XR1, EL | 1499 | 255 | 20063 | 1.312181978 | 1 | 0.992350513 | 94.35154558 |
| INTERPRO | IPR001680:WD40 repeat | 27 | 1.715374841 | 0.14889993 | NBEA, DMWD, WDR81, FBXW9, WDR37, WDR17, EIF3I, PAFAH1B1, MAPKBP1, IFT140, TBL1XR1, EL | 1452 | 268 | 18559 | 1.287706612 | 1 | 0.976932776 | 93.78025216 |
| INTERPRO | IPR015943:WD40/YVTN repeat-like-containing domain | 32 | 2.033036849 | 0.168434466 | NBEA, DMWD, WDR81, SEMA5A, FBXW9, WDR37, WDPCP, WDR17, EIF3I, PAFAH1B1, BCAS3, MAPK | 1452 | 331 | 18559 | 1.23569116 | 1 | 0.980727347 | 95.83080024 |
| UP_SEQ_FEATURE | repeat:WD 7 | 15 | 0.952986023 | 0.23740349 | TBL1XR1, ELP2, AHI1, PWP2, FBXW9, EML2, WDR83, WDR37, WDR17, WDR62, TEP1, PAFAH1B1, W | 1499 | 151 | 20063 | 1.329561871 | 1 | 0.998901856 | 99.35677208 |
| UP_SEQ_FEATURE | repeat:WD 6 | 18 | 1.143583227 | 0.295475061 | TBL1XR1, ELP2, RBBP5, AHI1, PWP2, WDR83, EML2, FBXW9, WDR37, WDR17, WDR62, TEP1, PAFAH | 1499 | 196 | 20063 | 1.229166383 | 1 | 0.999697997 | 99.8528107 |
| SMART | SM00320:WD40 | 27 | 1.715374841 | 0.305352649 | NBEA, DMWD, WDR81, FBXW9, WDR37, WDR17, EIF3I, PAFAH1B1, MAPKBP1, IFT140, TBL1XR1, EL | 878 | 267 | 10057 | 1.158314351 | 1 | 0.936020572 | 99.39591398 |
| INTERPRO | IPR020472:G-protein beta WD-40 repeat | 9 | 0.571791614 | 0.443265207 | WDR83, FBXW9, TBL1XR1, WDR37, WDR17, PAFAH1B1, WDR1, WDR41, PWP2 | 1452 | 93 | 18559 | 1.236936817 | 1 | 0.999493279 | 99.99584786 |
| INTERPRO | IPR019775:WD40 repeat, conserved site | 13 | 0.82592122 | 0.653799036 | TBL1XR1, WDFY4, PWP2, FBXW9, WDR83, WDR37, WDR17, TEP1, EIF3I, PAFAH1B1, WDR1, WDR41, | 1452 | 166 | 18559 | 1.000974974 | 1 | 0.999979348 | 99.99999884 |

Annotation Cluster 62 Enrichment Score: 0.8661001960078794

| Category | Term | Count | % | PValue | Genes | List Total | Pop Hits | Pop Total | Fold Enrichment | Bonferroni | Benjamini | FDR |
|---|---|---|---|---|---|---|---|---|---|---|---|---|
| INTERPRO | IPR018952:2'-5'-oligoadenylate synthetase 1, domain 2/C-terminal | 3 | 0.190597205 | 0.03294801 | OAS3, OAS1, OAS2 | 1452 | 4 | 18559 | 9.586260331 | 1 | 0.849368775 | 43.85082476 |
| INTERPRO | IPR026774:2'-5'-oligoadenylate synthase | 3 | 0.190597205 | 0.03294801 | OAS3, OAS1, OAS2 | 1452 | 4 | 18559 | 9.586260331 | 1 | 0.849368775 | 43.85082476 |
| INTERPRO | IPR006117:2-5-oligoadenylate synthetase, conserved site | 3 | 0.190597205 | 0.03294801 | OAS3, OAS1, OAS2 | 1452 | 4 | 18559 | 9.586260331 | 1 | 0.849368775 | 43.85082476 |
| GOTERM_MF_DIRECT | GO:0001730~2'-5'-oligoadenylate synthetase activity | 3 | 0.190597205 | 0.033858033 | OAS3, OAS1, OAS2 | 1340 | 4 | 16881 | 9.448320896 | 1 | 0.902835344 | 42.97318508 |
| GOTERM_BP_DIRECT | GO:0045071~negative regulation of viral genome replication | 8 | 0.508259212 | 0.037149508 | SRPK2, IFIT1, IFITM1, C19ORF66, OAS3, APOBEC3G, OAS1, APOBEC3F | 1342 | 40 | 16792 | 2.502533532 | 1 | 0.916021944 | 50.56886859 |
| INTERPRO | IPR002934:Nucleotidyl transferase domain | 4 | 0.254129606 | 0.037733621 | PAPOLA, OAS3, OAS1, OAS2 | 1452 | 10 | 18559 | 5.112672176 | 1 | 0.87239738 | 48.45013759 |
| INTERPRO | IPR006116:2-5-oligoadenylate synthetase, N-terminal | 3 | 0.190597205 | 0.052094013 | OAS3, OAS1, OAS2 | 1452 | 5 | 18559 | 7.669008264 | 1 | 0.910917985 | 60.21365103 |
| GOTERM_BP_DIRECT | GO:0060337~type I interferon signaling pathway | 9 | 0.571791614 | 0.136085444 | IFIT1, SP100, IFITM1, OAS3, JAK1, OAS1, HLA-C, OAS2, XAF1 | 1342 | 64 | 16792 | 1.75959389 | 1 | 0.984691111 | 93.42937922 |
| REACTOME_PATHWAY | R-HSA-909733:R-HSA-909733 | 9 | 0.571791614 | 0.180750907 | IFIT1, IFITM1, OAS3, JAK1, OAS1, HLA-C, OAS2, XAF1, PTPN11 | 745 | 67 | 9075 | 1.636281679 | 1 | 0.967584355 | 95.44690026 |
| GOTERM_BP_DIRECT | GO:0051607~defense response to virus | 16 | 1.016518424 | 0.338877826 | IFIT1B, AIMP1, IFITM1, EXOSC5, OAS3, APOBEC3G, OAS1, OAS2, CD40, APOBEC3F, APOBEC3D, SERI | 1342 | 165 | 16792 | 1.213349591 | 1 | 0.999186194 | 99.95480218 |
| UP_KEYWORDS | Antiviral defense | 11 | 0.698856417 | 0.364003194 | SERINC3, IFIT1, IFITM1, IFIT5, C19ORF66, OAS3, APOBEC3G, OAS1, OAS2, APOBEC3F, APOBEC3D | 1535 | 116 | 20581 | 1.271430978 | 1 | 0.785342148 | 99.85027974 |
| GOTERM_MF_DIRECT | GO:0016779~nucleotidyltransferase activity | 3 | 0.190597205 | 0.643050074 | OAS3, OAS1, OAS2 | 1340 | 27 | 16881 | 1.399751244 | 1 | 0.999414249 | 99.99999493 |
| KEGG_PATHWAY | hsa05160:Hepatitis C | 10 | 0.635324015 | 0.789096345 | IFIT1B, IFIT1, HRAS, PIK3CB, OAS3, JAK1, OAS1, SCARB1, OAS2, AKT3 | 574 | 133 | 6879 | 0.901076734 | 1 | 0.933721654 | 99.99999988 |
| UP_KEYWORDS | Nucleotidyltransferase | 5 | 0.317662008 | 0.849456394 | POLD3, OAS3, OAS1, OAS2, POLR2B | 1535 | 79 | 20581 | 0.84859605 | 1 | 0.9802036 | 100 |
| GOTERM_BP_DIRECT | GO:0009615~response to virus | 7 | 0.444726811 | 0.881702322 | IFIT1, IFITM1, FGR, OAS3, OAS1, OAS2, DCLK1 | 1342 | 110 | 16792 | 0.796260669 | 1 | 0.999999997 | 100 |

Annotation Cluster 63 Enrichment Score: 0.8531731350342785

| Category | Term | Count | % | PValue | Genes | List Total | Pop Hits | Pop Total | Fold Enrichment | Bonferroni | Benjamini | FDR |
|---|---|---|---|---|---|---|---|---|---|---|---|---|
| INTERPRO | IPR002469:Peptidase S9B, dipeptidylpeptidase IV N-terminal | 3 | 0.190597205 | 0.074157476 | DPP10, DPP8, DPP6 | 1452 | 6 | 18559 | 6.39084022 | 1 | 0.950899407 | 73.48245133 |
| GOTERM_MF_DIRECT | GO:0008236~serine-type peptidase activity | 9 | 0.571791614 | 0.123901561 | IMMP2L, GZMK, ST14, DPP10, RELN, DPP8, DPP6, PRSS12, PRSS21 | 1340 | 63 | 16881 | 1.799680171 | 1 | 0.944043815 | 88.43160277 |
| INTERPRO | IPR001375:Peptidase S9, prolyl oligopeptidase, catalytic domain | 3 | 0.190597205 | 0.152477215 | DPP10, DPP8, DPP6 | 1452 | 9 | 18559 | 4.260560147 | 1 | 0.977197811 | 94.2155735 |
| GOTERM_MF_DIRECT | GO:0008239~dipeptidyl-peptidase activity | 3 | 0.190597205 | 0.275975666 | DPP10, DPP8, DPP6 | 1340 | 13 | 16881 | 2.90717566 | 1 | 0.987941205 | 99.48340702 |

Annotation Cluster 64 Enrichment Score: 0.8502407996108761

| Category | Term | Count | % | PValue | Genes | List Total | Pop Hits | Pop Total | Fold Enrichment | Bonferroni | Benjamini | FDR |
|---|---|---|---|---|---|---|---|---|---|---|---|---|
| SMART | SM00327:VWA | 11 | 0.698856417 | 0.093879577 | XRCC5, COL21A1, COL6A6, CPNE4, CPNE5, CFB, COL6A5, ITGAE, COL22A1, ITGA11, C2 | 878 | 72 | 10057 | 1.749984181 | 1 | 0.883895156 | 74.90263597 |
| INTERPRO | IPR002035:von Willebrand factor, type A | 12 | 0.762388818 | 0.135683065 | XRCC5, HMCN2, COL21A1, COL6A6, CPNE4, CPNE5, CFB, COL6A5, ITGAE, COL22A1, ITGA11, C2 | 1452 | 97 | 18559 | 1.581238817 | 1 | 0.973135297 | 91.88921267 |
| UP_SEQ_FEATURE | domain:VWFA | 8 | 0.508259212 | 0.22089243 | COL21A1, CPNE4, CPNE5, CFB, ITGAE, COL22A1, ITGA11, C2 | 1499 | 66 | 20063 | 1.622334081 | 1 | 0.998459197 | 99.04153403 |

Annotation Cluster 65 Enrichment Score: 0.8487723472276086

| Category | Term | Count | % | PValue | Genes | List Total | Pop Hits | Pop Total | Fold Enrichment | Bonferroni | Benjamini | FDR |
|---|---|---|---|---|---|---|---|---|---|---|---|---|
| UP_SEQ_FEATURE | domain:F-box | 10 | 0.635324015 | 0.092852167 | FBXW9, FBXO39, KDM2A, CCNF, FBXO41, FBXO15, FBXO24, FBXL15, FBXL17, FBXO46 | 1499 | 73 | 20063 | 1.833459749 | 1 | 0.976351712 | 83.70785108 |
| INTERPRO | IPR001810:F-box domain, cyclin-like | 10 | 0.635324015 | 0.122257906 | FBXW9, FBXO39, KDM2A, CCNF, FBXO41, FBXO15, FBXO24, FBXL15, FBXL17, FBXO46 | 1452 | 74 | 18559 | 1.727254114 | 1 | 0.969498576 | 89.42259255 |
| SMART | SM00256:FBOX | 6 | 0.381194409 | 0.250388262 | FBXW9, CCNF, FBXO15, FBXO24, FBXL17, FBXO46 | 878 | 39 | 10057 | 1.762221833 | 1 | 0.917233195 | 98.24264431 |

Annotation Cluster 66 Enrichment Score: 0.8440140189583796

| Category | Term | Count | % | PValue | Genes | List Total | Pop Hits | Pop Total | Fold Enrichment | Bonferroni | Benjamini | FDR |
|---|---|---|---|---|---|---|---|---|---|---|---|---|
| GOTERM_BP_DIRECT | GO:0010332~response to gamma radiation | 6 | 0.381194409 | 0.096887579 | FANCD2, TP63, CHEK2, MYC, TP73, SOD2 | 1342 | 31 | 16792 | 2.421806644 | 1 | 0.977473058 | 84.99365066 |
| GOTERM_BP_DIRECT | GO:0006978~DNA damage response, signal transduction by p53 class mediator resulting in transcri | 4 | 0.254129606 | 0.130552062 | SP100, TP63, CHEK2, TP73 | 1342 | 16 | 16792 | 3.128166915 | 1 | 0.984597812 | 92.60031809 |
| GOTERM_BP_DIRECT | GO:0042771~intrinsic apoptotic signaling pathway in response to DNA damage by p53 class mediat | 5 | 0.317662008 | 0.232223465 | MSH2, TP63, CHEK2, PHLDA3, TP73 | 1342 | 31 | 16792 | 2.018172203 | 1 | 0.996350009 | 99.26883989 |

Annotation Cluster 67 Enrichment Score: 0.8231142172212605

| Category | Term | Count | % | PValue | Genes | List Total | Pop Hits | Pop Total | Fold Enrichment | Bonferroni | Benjamini | FDR |
|---|---|---|---|---|---|---|---|---|---|---|---|---|
| UP_KEYWORDS | Guanine-nucleotide releasing factor | 19 | 1.207115629 | 0.028755629 | DENND6B, ARHGEF39, HPS4, PSD3, TRIO, AKAP13, RGP1, VAV2, DOCK5, RIC8B, DOCK3, MCF2L, RGL | 1535 | 149 | 20581 | 1.709721706 | 0.999999434 | 0.233849598 | 34.25130949 |
| UP_SEQ_FEATURE | domain:PH 1 | 7 | 0.444726811 | 0.060683568 | RASGRF2, PLCG1, TRIO, AFAP1L2, FGD5, ADAP1, FGD4 | 1499 | 38 | 20063 | 2.465520874 | 1 | 0.94483998 | 68.8278947 |
| GOTERM_MF_DIRECT | GO:0005089~Rho guanyl-nucleotide exchange factor activity | 11 | 0.698856417 | 0.082375744 | OBSCN, RASGRF2, ARHGEF39, TRIO, AKAP13, FGD5, VAV2, EPS8L2, MCF2L, FGD4, RGL2 | 1340 | 77 | 16881 | 1.799680171 | 1 | 0.92354259 | 75.38401668 |
| GOTERM_MF_DIRECT | GO:0005085~guanyl-nucleotide exchange factor activity | 15 | 0.952986023 | 0.083912031 | OBSCN, HPS4, TRIO, AKAP13, VAV2, DOCK5, RIC8B, DOCK3, MCF2L, RGL2, RASGRF2, TBXA2R, FGD5, | 1340 | 118 | 16881 | 1.601410321 | 1 | 0.92281595 | 76.0474755 |
| GOTERM_BP_DIRECT | GO:0035023~regulation of Rho protein signal transduction | 11 | 0.698856417 | 0.110396446 | OBSCN, RASGRF2, ARHGEF39, TRIO, AKAP13, FGD5, VAV2, EPS8L2, MCF2L, FGD4, ARHGDIB | 1342 | 81 | 16792 | 1.699251164 | 1 | 0.980202354 | 88.66412471 |
| UP_SEQ_FEATURE | domain:DH | 9 | 0.571791614 | 0.133033314 | OBSCN, RASGRF2, ARHGEF39, AKAP13, SLC26A10, FGD5, VAV2, MCF2L, FGD4 | 1499 | 68 | 20063 | 1.77144567 | 1 | 0.99080442 | 92.99152475 |
| UP_SEQ_FEATURE | domain:PH 2 | 6 | 0.381194409 | 0.138810246 | RASGRF2, TRIO, AFAP1L2, FGD5, ADAP1, FGD4 | 1499 | 37 | 20063 | 2.17041992 | 1 | 0.991709818 | 93.81185873 |
| REACTOME_PATHWAY | R-HSA-193648:R-HSA-193648 | 7 | 0.444726811 | 0.184281663 | OBSCN, RASGRF2, TRIO, AKAP13, VAV2, MCF2L, FGD4 | 745 | 47 | 9075 | 1.814222476 | 1 | 0.967765171 | 95.74165425 |
| INTERPRO | IPR000219:Dbl homology (DH) domain | 9 | 0.571791614 | 0.188908315 | OBSCN, RASGRF2, ARHGEF39, TRIO, AKAP13, FGD5, VAV2, MCF2L, FGD4 | 1452 | 71 | 18559 | 1.620213014 | 1 | 0.985249905 | 97.28641452 |
| SMART | SM00325:RhoGEF | 9 | 0.571791614 | 0.23897857 | OBSCN, RASGRF2, ARHGEF39, TRIO, AKAP13, FGD5, VAV2, MCF2L, FGD4 | 878 | 68 | 10057 | 1.516029077 | 1 | 0.91942214 | 97.8280184 |
| GOTERM_BP_DIRECT | GO:0051056~regulation of small GTPase mediated signal transduction | 13 | 0.82592122 | 0.383339103 | OBSCN, RALGAPB, SIPA1L2, TRIO, AKAP13, VAV2, MCF2L, RASGRF2, SYDE2, ARHGAP9, ARHGAP10, A | 1342 | 134 | 16792 | 1.213915519 | 1 | 0.999570193 | 99.98762961 |
| REACTOME_PATHWAY | R-HSA-416482:R-HSA-416482 | 8 | 0.508259212 | 0.446043138 | OBSCN, RASGRF2, TBXA2R, TRIO, AKAP13, VAV2, MCF2L, FGD4 | 745 | 77 | 9075 | 1.265580058 | 1 | 0.995755348 | 99.98940886 |
| REACTOME_PATHWAY | R-HSA-194840:R-HSA-194840 | 11 | 0.698856417 | 0.569781636 | OBSCN, SYDE2, RASGRF2, TRIO, AKAP13, VAV2, MCF2L, FGD4, ARHGDIB, ARHGAP10, ARHGAP9 | 745 | 124 | 9075 | 1.080591037 | 1 | 0.99886285 | 99.99978928 |

Annotation Cluster 68 Enrichment Score: 0.8143187665578262

| Category | Term | Count | % | PValue | Genes | List Total | Pop Hits | Pop Total | Fold Enrichment | Bonferroni | Benjamini | FDR |
|---|---|---|---|---|---|---|---|---|---|---|---|---|
| GOTERM_BP_DIRECT | GO:0006801~superoxide metabolic process | 5 | 0.317662008 | 0.070133242 | NOX4, CYBA, IMMP2L, NRROS, SOD2 | 1342 | 20 | 16792 | 3.128166915 | 1 | 0.967620814 | 74.16238249 |
| GOTERM_BP_DIRECT | GO:0042554~superoxide anion generation | 4 | 0.254129606 | 0.095362747 | NOX4, NOX3, CYBA, SOD2 | 1342 | 14 | 16792 | 3.575047903 | 1 | 0.979190754 | 84.51500586 |
| REACTOME_PATHWAY | R-HSA-3299685:R-HSA-3299685 | 6 | 0.381194409 | 0.141723066 | NOX4, CYBA, GPX6, GPX5, CAT, SOD2 | 745 | 34 | 9075 | 2.149624951 | 1 | 0.956253548 | 90.63514575 |
| GOTERM_BP_DIRECT | GO:0000302~response to reactive oxygen species | 6 | 0.381194409 | 0.197476361 | NOX4, CYBA, GPX5, TRA2B, CAT, SOD2 | 1342 | 39 | 16792 | 1.925025794 | 1 | 0.994079886 | 98.33356912 |
| GOTERM_BP_DIRECT | GO:0014823~response to activity | 5 | 0.317662008 | 0.453051782 | CYBA, TH, MAS1, CAT, SOD2 | 1342 | 43 | 16792 | 1.454961356 | 1 | 0.999859396 | 99.99867351 |

Annotation Cluster 69 Enrichment Score: 0.7915425040917351

| Category | Term | Count | % | PValue | Genes | List Total | Pop Hits | Pop Total | Fold Enrichment | Bonferroni | Benjamini | FDR |
|---|---|---|---|---|---|---|---|---|---|---|---|---|
| UP_KEYWORDS | cGMP-binding | 3 | 0.190597205 | 0.140976755 | PDE10A, CNGA3, CNGA1 | 1535 | 9 | 20581 | 4.46927253 | 1 | 0.527236179 | 88.74028342 |

| | | | | | | | | | | | | |
|---|---|---|---|---|---|---|---|---|---|---|---|---|
| GOTERM_MF_DIRECT | GO:0030553~cGMP binding | 4 | 0.254129606 | 0.147428081 | PDE10A, CNGB1, CNGA3, CNGA1 | 1340 | 17 | 16881 | 2.964179104 | 1 | 0.955300311 | 92.5782433 |
| UP_KEYWORDS | cGMP | 4 | 0.254129606 | 0.20306953 | PDE3B, PDE10A, CNGA3, CNGA1 | 1535 | 21 | 20581 | 2.553870017 | 1 | 0.618901854 | 96.16974569 |
| Annotation Cluster 70 | Enrichment Score: 0.7862842645078107 | | | | | | | | | | | |
| Category | Term | Count | % | PValue | Genes | List Total | Pop Hits | Pop Total | Fold Enrichment | Bonferroni | Benjamini | FDR |
| INTERPRO | IPR001478:PDZ domain | 18 | 1.143583227 | 0.098007326 | MAGI2, LIMK2, MAGI1, SNX27, SIPA1L2, GIPC3, LMO7, SYNPO2, MPP7, RIMS1, PCLO, MAST4, PPP1R | 1452 | 155 | 18559 | 1.48432418 | 1 | 0.967916819 | 83.08468728 |
| UP_SEQ_FEATURE | domain:PDZ 6 | 3 | 0.190597205 | 0.115475174 | MAGI2, MAGI1, PDZD2 | 1499 | 8 | 20063 | 5.019096064 | 1 | 0.986545152 | 89.81978955 |
| UP_SEQ_FEATURE | domain:PDZ | 13 | 0.82592122 | 0.136976498 | MAGI1, LIMK2, SNX27, SIPA1L2, GIPC3, LMO7, SYNPO2, MPP7, PCLO, RIMS1, PPP1R9B, MAST4, AHN | 1499 | 113 | 20063 | 1.539781683 | 1 | 0.991770733 | 93.56185866 |
| UP_SEQ_FEATURE | domain:PDZ 5 | 3 | 0.190597205 | 0.141381957 | MAGI2, MAGI1, PDZD2 | 1499 | 9 | 20063 | 4.461418724 | 1 | 0.992174796 | 94.14703211 |
| SMART | SM00228:PDZ | 18 | 1.143583227 | 0.1523725 | MAGI2, LIMK2, MAGI1, SNX27, SIPA1L2, GIPC3, LMO7, SYNPO2, MPP7, RIMS1, PCLO, MAST4, PPP1R | 878 | 149 | 10057 | 1.383758084 | 1 | 0.906116485 | 90.15454109 |
| UP_SEQ_FEATURE | domain:PDZ 3 | 5 | 0.317662008 | 0.182317954 | MAGI2, MAGI1, PDZD2, TJP3, DLG2 | 1499 | 30 | 20063 | 2.230709362 | 1 | 0.997015518 | 97.64305732 |
| UP_SEQ_FEATURE | domain:PDZ 1 | 6 | 0.381194409 | 0.201812683 | MAGI2, MAGI1, PDZD2, TJP3, DLG2, APBA1 | 1499 | 42 | 20063 | 1.912036596 | 1 | 0.997977455 | 98.49609191 |
| UP_SEQ_FEATURE | domain:PDZ 2 | 6 | 0.381194409 | 0.201812683 | MAGI2, MAGI1, PDZD2, TJP3, DLG2, APBA1 | 1499 | 42 | 20063 | 1.912036596 | 1 | 0.997977455 | 98.49609191 |
| UP_SEQ_FEATURE | domain:PDZ 4 | 3 | 0.190597205 | 0.338081545 | MAGI2, MAGI1, PDZD2 | 1499 | 16 | 20063 | 2.509548032 | 1 | 0.99984201 | 99.95392837 |
| Annotation Cluster 71 | Enrichment Score: 0.7770419873711857 | | | | | | | | | | | |
| Category | Term | Count | % | PValue | Genes | List Total | Pop Hits | Pop Total | Fold Enrichment | Bonferroni | Benjamini | FDR |
| INTERPRO | IPR003350:Homeodomain protein CUT | 3 | 0.190597205 | 0.098565829 | SATB1, CUX2, CUX1 | 1452 | 7 | 18559 | 5.477863046 | 1 | 0.966768877 | 83.2642154 |
| SMART | SM01109:SM01109 | 3 | 0.190597205 | 0.118881154 | SATB1, CUX2, CUX1 | 878 | 7 | 10057 | 4.909046534 | 1 | 0.898086663 | 83.04758981 |
| INTERPRO | IPR010982:Lambda repressor-like, DNA-binding domain | 4 | 0.254129606 | 0.398138933 | SATB1, POU5F1, CUX2, CUX1 | 1452 | 29 | 18559 | 1.762990406 | 1 | 0.999231959 | 99.98410096 |
| Annotation Cluster 72 | Enrichment Score: 0.7756801967255452 | | | | | | | | | | | |
| Category | Term | Count | % | PValue | Genes | List Total | Pop Hits | Pop Total | Fold Enrichment | Bonferroni | Benjamini | FDR |
| UP_SEQ_FEATURE | domain:Importin N-terminal | 4 | 0.254129606 | 0.081396345 | IPO8, TNPO2, XPO7, TNPO1 | 1499 | 14 | 20063 | 3.824073192 | 1 | 0.967410363 | 79.41941972 |
| INTERPRO | IPR001494:Importin-beta, N-terminal | 4 | 0.254129606 | 0.142873304 | IPO8, TNPO2, XPO7, TNPO1 | 1452 | 17 | 18559 | 3.007454221 | 1 | 0.975007669 | 92.97634335 |
| SMART | SM00913:SM00913 | 4 | 0.254129606 | 0.158188967 | IPO8, TNPO2, XPO7, TNPO1 | 878 | 16 | 10057 | 2.863610478 | 1 | 0.907643213 | 91.06073261 |
| GOTERM_MF_DIRECT | GO:0008536~Ran GTPase binding | 4 | 0.254129606 | 0.429088653 | IPO8, TNPO2, XPO7, TNPO1 | 1340 | 30 | 16881 | 1.679701493 | 1 | 0.99481267 | 99.9892692 |
| Annotation Cluster 73 | Enrichment Score: 0.7751851274036656 | | | | | | | | | | | |
| Category | Term | Count | % | PValue | Genes | List Total | Pop Hits | Pop Total | Fold Enrichment | Bonferroni | Benjamini | FDR |
| UP_SEQ_FEATURE | metal ion-binding site:Calcium 2; via carbonyl oxygen | 7 | 0.444726811 | 0.067391001 | ATP2C2, MASP1, PLCH1, MMP8, MMP27, COLEC12, AOC1 | 1499 | 39 | 20063 | 2.40230239 | 1 | 0.955964041 | 72.72179948 |
| UP_SEQ_FEATURE | metal ion-binding site:Calcium 2 | 8 | 0.508259212 | 0.130636951 | ATP2C2, MASP1, PLCH1, MMP8, MMP27, SYT6, COLEC12, AOC1 | 1499 | 57 | 20063 | 1.878492094 | 1 | 0.990301186 | 92.62190514 |
| UP_SEQ_FEATURE | metal ion-binding site:Calcium 3; via carbonyl oxygen | 4 | 0.254129606 | 0.203783314 | MASP1, PLCH1, MMP8, MMP27 | 1499 | 21 | 20063 | 2.549382128 | 1 | 0.998029241 | 98.56374302 |
| UP_SEQ_FEATURE | metal ion-binding site:Calcium 3 | 5 | 0.317662008 | 0.213760162 | MASP1, PLCH1, MMP27, SYT6, COLEC12 | 1499 | 32 | 20063 | 2.091290027 | 1 | 0.998342018 | 98.86429271 |
| UP_SEQ_FEATURE | metal ion-binding site:Calcium 1; via carbonyl oxygen | 5 | 0.317662008 | 0.213760162 | MASP1, MMP8, SYT6, COLEC12, AOC1 | 1499 | 32 | 20063 | 2.091290027 | 1 | 0.998342018 | 98.86429271 |
| UP_SEQ_FEATURE | metal ion-binding site:Calcium 1 | 6 | 0.381194409 | 0.272395002 | MASP1, MMP8, MMP27, SYT6, COLEC12, AOC1 | 1499 | 47 | 20063 | 1.708628447 | 1 | 0.999508885 | 99.73175042 |
| Annotation Cluster 74 | Enrichment Score: 0.7716367818396228 | | | | | | | | | | | |
| Category | Term | Count | % | PValue | Genes | List Total | Pop Hits | Pop Total | Fold Enrichment | Bonferroni | Benjamini | FDR |
| GOTERM_BP_DIRECT | GO:0045869~negative regulation of single stranded viral RNA replication via double stranded DNA | 4 | 0.254129606 | 0.013924741 | INPP5K, APOBEC3G, APOBEC3F, APOBEC3D | 1342 | 7 | 16792 | 7.150095806 | 1 | 0.849880395 | 22.97082761 |
| GOTERM_BP_DIRECT | GO:0070383~DNA cytosine deamination | 3 | 0.190597205 | 0.102249192 | APOBEC3G, APOBEC3F, APOBEC3D | 1342 | 7 | 16792 | 5.362571854 | 1 | 0.979902008 | 86.56787542 |
| GOTERM_BP_DIRECT | GO:0010529~negative regulation of transposition | 3 | 0.190597205 | 0.129342022 | APOBEC3G, APOBEC3F, APOBEC3D | 1342 | 8 | 16792 | 4.692250373 | 1 | 0.984429287 | 92.4062792 |
| GOTERM_MF_DIRECT | GO:0016814~hydrolase activity, acting on carbon-nitrogen (but not peptide) bonds, in cyclic amidin | 3 | 0.190597205 | 0.215252361 | APOBEC3G, APOBEC3F, APOBEC3D | 1340 | 11 | 16881 | 3.435753053 | 1 | 0.977661265 | 98.07921944 |
| INTERPRO | IPR013158:APOBEC-like, N-terminal | 3 | 0.190597205 | 0.240343159 | APOBEC3G, APOBEC3F, APOBEC3D | 1452 | 12 | 18559 | 3.19542011 | 1 | 0.993507746 | 99.12220937 |
| GOTERM_CC_DIRECT | GO:0000932~cytoplasmic mRNA processing body | 9 | 0.571791614 | 0.263539486 | TOP1, RBPMS, EIF4E, CNOT2, APOBEC3G, LIMD1, TNRC6B, APOBEC3F, APOBEC3D | 1425 | 78 | 18224 | 1.47562753 | 1 | 0.885425091 | 98.99640224 |
| INTERPRO | IPR016192:APOBEC/CMP deaminase, zinc-binding | 3 | 0.190597205 | 0.300235879 | APOBEC3G, APOBEC3F, APOBEC3D | 1452 | 14 | 18559 | 2.738931523 | 1 | 0.996713111 | 99.78670791 |
| INTERPRO | IPR002125:CMP/dCMP deaminase, zinc-binding | 3 | 0.190597205 | 0.38814783 | APOBEC3G, APOBEC3F, APOBEC3D | 1452 | 17 | 18559 | 2.255590666 | 1 | 0.999121106 | 99.97888706 |
| INTERPRO | IPR016193:Cytidine deaminase-like | 3 | 0.190597205 | 0.38814783 | APOBEC3G, APOBEC3F, APOBEC3D | 1452 | 17 | 18559 | 2.255590666 | 1 | 0.999121106 | 99.97888706 |
| Annotation Cluster 75 | Enrichment Score: 0.7690696652264886 | | | | | | | | | | | |
| Category | Term | Count | % | PValue | Genes | List Total | Pop Hits | Pop Total | Fold Enrichment | Bonferroni | Benjamini | FDR |
| GOTERM_BP_DIRECT | GO:0042554~superoxide anion generation | 4 | 0.254129606 | 0.095362747 | NOX4, NOX3, CYBA, SOD2 | 1342 | 14 | 16792 | 3.575047903 | 1 | 0.979190754 | 84.51500586 |
| GOTERM_MF_DIRECT | GO:0016175~superoxide-generating NADPH oxidase activity | 3 | 0.190597205 | 0.215252361 | NOX4, NOX3, CYBA | 1340 | 11 | 16881 | 3.435753053 | 1 | 0.977661265 | 98.07921944 |
| GOTERM_CC_DIRECT | GO:0043020~NADPH oxidase complex | 3 | 0.190597205 | 0.240139902 | NOX4, NOX3, CYBA | 1425 | 12 | 18224 | 3.197192982 | 1 | 0.87708635 | 98.39341651 |
| Annotation Cluster 76 | Enrichment Score: 0.7665875028419051 | | | | | | | | | | | |
| Category | Term | Count | % | PValue | Genes | List Total | Pop Hits | Pop Total | Fold Enrichment | Bonferroni | Benjamini | FDR |
| GOTERM_BP_DIRECT | GO:0035725~sodium ion transmembrane transport | 10 | 0.635324015 | 0.126305668 | SLC17A7, SLC8A1, SLC4A11, SLC24A4, SLC24A1, ASIC2, SLC13A1, SCN7A, TRAPPC10, SCN4A | 1342 | 73 | 16792 | 1.714064063 | 1 | 0.984235011 | 91.89795842 |
| UP_KEYWORDS | Sodium transport | 13 | 0.82592122 | 0.162003652 | SLC9A9, SLC17A7, SLC8A1, SLC13A5, SLC24A4, ATP1B2, SLC9A3, ASIC2, SLC13A1, SLC9C1, SLC6A15, S | 1535 | 117 | 20581 | 1.48975751 | 1 | 0.557065113 | 92.11419414 |
| GOTERM_BP_DIRECT | GO:0006814~sodium ion transport | 10 | 0.635324015 | 0.19653222 | SLC13A5, SLC4A11, ATP1B2, CATSPER3, SLC13A1, SLC6A15, NEDD4L, SCN7A, TRAPPC10, SCN4A | 1342 | 81 | 16792 | 1.544773785 | 1 | 0.994092168 | 98.29670022 |
| UP_KEYWORDS | Sodium | 13 | 0.82592122 | 0.213436448 | SLC9A9, SLC17A7, SLC8A1, SLC13A5, SLC24A4, ATP1B2, SLC9A3, ASIC2, SLC13A1, SLC9C1, SLC6A15, S | 1535 | 124 | 20581 | 1.405658296 | 1 | 0.630137668 | 96.82677876 |
| Annotation Cluster 77 | Enrichment Score: 0.7624651966423669 | | | | | | | | | | | |
| Category | Term | Count | % | PValue | Genes | List Total | Pop Hits | Pop Total | Fold Enrichment | Bonferroni | Benjamini | FDR |
| UP_SEQ_FEATURE | domain:Ras-associating | 6 | 0.381194409 | 0.105957656 | SNX27, RASSF9, RASSF8, RASSF7, APBB1IP, RGL2 | 1499 | 34 | 20063 | 2.36192756 | 1 | 0.982975092 | 87.57483224 |
| INTERPRO | IPR000159:Ras-association | 6 | 0.381194409 | 0.199945394 | SNX27, RASSF9, RASSF8, RASSF7, APBB1IP, RGL2 | 1452 | 40 | 18559 | 1.917252066 | 1 | 0.987786283 | 97.85692652 |
| SMART | SM00314:RA | 5 | 0.317662008 | 0.243534595 | RASSF9, RASSF8, RASSF7, APBB1IP, RGL2 | 878 | 29 | 10057 | 1.974903778 | 1 | 0.919375577 | 98.00341705 |
| Annotation Cluster 78 | Enrichment Score: 0.7612566098411084 | | | | | | | | | | | |
| Category | Term | Count | % | PValue | Genes | List Total | Pop Hits | Pop Total | Fold Enrichment | Bonferroni | Benjamini | FDR |
| UP_SEQ_FEATURE | domain:Fibrinogen C-terminal | 7 | 0.444726811 | 0.033297191 | ANGPTL6, TNXB, TNXA, CNTNAP5, CNTNAP2, FGL2, FIBCD1 | 1499 | 33 | 20063 | 2.839084642 | 1 | 0.880205871 | 46.76960017 |
| INTERPRO | IPR002181:Fibrinogen, alpha/beta/gamma chain, C-terminal globular domain | 6 | 0.381194409 | 0.10057291 | ANGPTL6, TNXB, CNTNAP5, CNTNAP2, FGL2, FIBCD1 | 1452 | 32 | 18559 | 2.396565083 | 1 | 0.96556653 | 83.89467935 |
| INTERPRO | IPR020837:Fibrinogen, conserved site | 4 | 0.254129606 | 0.181920088 | ANGPTL6, TNXB, FGL2, FIBCD1 | 1452 | 19 | 18559 | 2.690880093 | 1 | 0.983843451 | 96.85422473 |
| INTERPRO | IPR014715:Fibrinogen, alpha/beta/gamma chain, C-terminal globular, subdomain 2 | 4 | 0.254129606 | 0.288330568 | ANGPTL6, TNXB, FGL2, FIBCD1 | 1452 | 24 | 18559 | 2.130280073 | 1 | 0.996225253 | 99.71477266 |
| SMART | SM00186:FBG | 4 | 0.254129606 | 0.349436793 | ANGPTL6, TNXB, FGL2, FIBCD1 | 878 | 24 | 10057 | 1.909073652 | 1 | 0.948880606 | 99.75911925 |
| INTERPRO | IPR014716:Fibrinogen, alpha/beta/gamma chain, C-terminal globular, subdomain 1 | 4 | 0.254129606 | 0.440993055 | ANGPTL6, TNXB, FGL2, FIBCD1 | 1452 | 31 | 18559 | 1.649249089 | 1 | 0.99949101 | 99.99554607 |
| Annotation Cluster 79 | Enrichment Score: 0.760066233494109 | | | | | | | | | | | |
| Category | Term | Count | % | PValue | Genes | List Total | Pop Hits | Pop Total | Fold Enrichment | Bonferroni | Benjamini | FDR |
| GOTERM_CC_DIRECT | GO:0005913~cell-cell adherens junction | 33 | 2.09656925 | 0.097538718 | VAPB, ASAP1, LMO7, CDH3, ESYT2, PDXDC1, EZR, PICALM, PAK2, RPL6, SND1, SSX2IP, LRRFIP1, MKL | 1425 | 323 | 18224 | 1.306592798 | 1 | 0.742283069 | 78.64388471 |
| GOTERM_MF_DIRECT | GO:0098641~cadherin binding involved in cell-cell adhesion | 29 | 1.842439644 | 0.163855414 | VAPB, ASAP1, ESYT2, PDXDC1, EZR, PICALM, PAK2, RPL6, SND1, LRRFIP1, MKL2, EHD4, COBLL1, BAI | 1340 | 290 | 16881 | 1.259776119 | 1 | 0.959087207 | 94.59584679 |
| GOTERM_BP_DIRECT | GO:0098609~cell-cell adhesion | 25 | 1.588310038 | 0.328218804 | VAPB, ASAP1, ESYT2, PDXDC1, PICALM, PAK2, RPL6, SND1, LRRFIP1, MKL2, EHD4, COBLL1, BAIAP2L1 | 1342 | 271 | 16792 | 1.154305135 | 1 | 0.999130757 | 99.93913083 |
| Annotation Cluster 80 | Enrichment Score: 0.7592844035663446 | | | | | | | | | | | |
| Category | Term | Count | % | PValue | Genes | List Total | Pop Hits | Pop Total | Fold Enrichment | Bonferroni | Benjamini | FDR |

| | | | | | | | | | | | | |
|---|---|---|---|---|---|---|---|---|---|---|---|---|
| UP_SEQ_FEATURE | repeat:HEAT 6 | 6 | 0.381194409 | 0.069040521 | MMS19, BTAF1, COPG2, CLASP1, TNPO2, TNPO1 | 1499 | 30 | 20063 | 2.676851234 | 1 | 0.956418546 | 73.60628933 |
| UP_SEQ_FEATURE | repeat:HEAT 7 | 5 | 0.317662008 | 0.099617679 | MMS19, BTAF1, CLASP1, TNPO2, TNPO1 | 1499 | 24 | 20063 | 2.788386702 | 1 | 0.980888024 | 85.82758991 |
| UP_SEQ_FEATURE | repeat:HEAT 5 | 6 | 0.381194409 | 0.116444601 | MMS19, BTAF1, COPG2, CLASP1, TNPO2, TNPO1 | 1499 | 35 | 20063 | 2.294443915 | 1 | 0.986063482 | 90.0255427 |
| UP_SEQ_FEATURE | repeat:HEAT 3 | 7 | 0.444726811 | 0.167150556 | MMS19, BTAF1, COPG2, SCYL1, CLASP1, TNPO2, TNPO1 | 1499 | 50 | 20063 | 1.873795864 | 1 | 0.995987067 | 96.68126817 |
| UP_SEQ_FEATURE | repeat:HEAT 4 | 6 | 0.381194409 | 0.215430056 | MMS19, BTAF1, COPG2, CLASP1, TNPO2, TNPO1 | 1499 | 43 | 20063 | 1.867570628 | 1 | 0.998366147 | 98.90837523 |
| UP_SEQ_FEATURE | repeat:HEAT 2 | 7 | 0.444726811 | 0.26372798 | MMS19, BTAF1, COPG2, SCYL1, CLASP1, TNPO2, TNPO1 | 1499 | 58 | 20063 | 1.615341262 | 1 | 0.999402318 | 99.66557931 |
| UP_SEQ_FEATURE | repeat:HEAT 1 | 7 | 0.444726811 | 0.26372798 | MMS19, BTAF1, COPG2, SCYL1, CLASP1, TNPO2, TNPO1 | 1499 | 58 | 20063 | 1.615341262 | 1 | 0.999402318 | 99.66557931 |
| UP_SEQ_FEATURE | repeat:HEAT 8 | 3 | 0.190597205 | 0.420182925 | BTAF1, TNPO2, TNPO1 | 1499 | 19 | 20063 | 2.113303606 | 1 | 0.999963936 | 99.99608664 |

Annotation Cluster 81 Enrichment Score: 0.7524949665868461

| Category | Term | Count | % | PValue | Genes | List Total | Pop Hits | Pop Total | Fold Enrichment | Bonferroni | Benjamini | FDR |
|---|---|---|---|---|---|---|---|---|---|---|---|---|
| GOTERM_BP_DIRECT | GO:0048015~phosphatidylinositol-mediated signaling | 16 | 1.016518424 | 0.021094128 | FGFR3, KLB, ERBB4, PIK3CB, PIK3C2B, FGF23, RICTOR, TRAT1, PTPN11, TNRC6C, EZR, PLCH1, HBEGF, | 1342 | 106 | 16792 | 1.888704552 | 1 | 0.891152191 | 32.75313971 |
| REACTOME_PATHWAY | R-HSA-1257604:R-HSA-1257604 | 13 | 0.82592122 | 0.031562491 | FGFR3, KLB, ERBB4, PIK3CB, FGF23, RICTOR, TRAT1, PTPN11, TNRC6C, HBEGF, TNRC6B, FGF2, AKT3 | 745 | 81 | 9075 | 1.9550087 | 1 | 0.936589996 | 39.16301456 |
| GOTERM_MF_DIRECT | GO:0046934~phosphatidylinositol-4,5-bisphosphate 3-kinase activity | 10 | 0.635324015 | 0.054693371 | FGFR3, KLB, ERBB4, PIK3CB, FGF23, HBEGF, PIK3R6, FGF2, TRAT1, PTPN11 | 1340 | 62 | 16881 | 2.031896967 | 1 | 0.880855654 | 60.03398528 |
| GOTERM_BP_DIRECT | GO:0046854~phosphatidylinositol phosphorylation | 13 | 0.82592122 | 0.070590365 | FGFR3, KLB, ERBB4, PIK3CB, PIK3C2B, FGF23, HBEGF, PIP5K1C, PIK3R6, IP6K1, FGF2, TRAT1, PTPN11 | 1342 | 94 | 16792 | 1.730475315 | 1 | 0.965780134 | 74.39776479 |
| REACTOME_PATHWAY | R-HSA-5654706:R-HSA-5654706 | 5 | 0.317662008 | 0.075685902 | HRAS, FGFR3, FGF23, FGF2, PTPN11 | 745 | 20 | 9075 | 3.045302013 | 1 | 0.947283156 | 70.46442578 |
| REACTOME_PATHWAY | R-HSA-5654227:R-HSA-5654227 | 4 | 0.254129606 | 0.084534597 | FGFR3, PLCG1, FGF23, FGF2 | 745 | 13 | 9075 | 3.748064017 | 1 | 0.93998986 | 74.55459672 |
| REACTOME_PATHWAY | R-HSA-190372:R-HSA-190372 | 4 | 0.254129606 | 0.084534597 | GALNT3, FGFR3, FGF23, FGF2 | 745 | 13 | 9075 | 3.748064017 | 1 | 0.93998986 | 74.55459672 |
| GOTERM_BP_DIRECT | GO:0014066~regulation of phosphatidylinositol 3-kinase signaling | 11 | 0.698856417 | 0.091215102 | PPP1R16B, FGFR3, KLB, ERBB4, PIK3CB, FGF23, HBEGF, PIP5K1C, FGF2, TRAT1, PTPN11 | 1342 | 78 | 16792 | 1.764606978 | 1 | 0.978548762 | 83.1389007 |
| GOTERM_BP_DIRECT | GO:0036092~phosphatidylinositol-3-phosphate biosynthetic process | 8 | 0.508259212 | 0.092769044 | FGFR3, KLB, PIK3CB, PIK3C2B, FGF23, INPP4B, FGF2, PTPN11 | 1342 | 49 | 16792 | 2.042884516 | 1 | 0.979191124 | 83.66749388 |
| REACTOME_PATHWAY | R-HSA-5654712:R-HSA-5654712 | 5 | 0.317662008 | 0.100572722 | HRAS, KLB, FGF23, FGF2, PTPN11 | 745 | 22 | 9075 | 2.768456376 | 1 | 0.936856669 | 80.65070886 |
| REACTOME_PATHWAY | R-HSA-5654719:R-HSA-5654719 | 5 | 0.317662008 | 0.114258653 | HRAS, KLB, FGF23, FGF2, PTPN11 | 745 | 23 | 9075 | 2.648088707 | 1 | 0.94927107 | 84.74275571 |
| REACTOME_PATHWAY | R-HSA-5654228:R-HSA-5654228 | 4 | 0.254129606 | 0.119256117 | KLB, PLCG1, FGF23, FGF2 | 745 | 15 | 9075 | 3.248322148 | 1 | 0.951858561 | 86.02350013 |
| GOTERM_MF_DIRECT | GO:0016303~1-phosphatidylinositol-3-kinase activity | 7 | 0.444726811 | 0.122039443 | FGFR3, KLB, PIK3CB, PIK3C2B, FGF23, FGF2, PTPN11 | 1340 | 43 | 16881 | 2.050798334 | 1 | 0.946946164 | 88.02408102 |
| GOTERM_MF_DIRECT | GO:0005088~Ras guanyl-nucleotide exchange factor activity | 14 | 0.889453621 | 0.123421209 | FGFR3, KLB, ERBB4, SPTBN5, CAMK2G, FGF23, RGL2, NCAM1, GRIN2B, RASGRF2, JAK1, HBEGF, FGF2 | 1340 | 115 | 16881 | 1.533640493 | 1 | 0.946106367 | 88.32774278 |
| REACTOME_PATHWAY | R-HSA-2219530:R-HSA-2219530 | 9 | 0.571791614 | 0.123668686 | FGFR3, KLB, ERBB4, PIK3CB, FGF23, HBEGF, FGF2, TRAT1, PTPN11 | 745 | 61 | 9075 | 1.797227418 | 1 | 0.953502879 | 87.07003772 |
| REACTOME_PATHWAY | R-HSA-5655302:R-HSA-5655302 | 6 | 0.381194409 | 0.129044387 | HRAS, PLCG1, FGF23, LRRFIP1, CUX1, FGF2 | 745 | 33 | 9075 | 2.214765101 | 1 | 0.952484493 | 88.24594866 |
| REACTOME_PATHWAY | R-HSA-5673001:R-HSA-5673001 | 14 | 0.889453621 | 0.153855991 | HRAS, FGFR3, KLB, ERBB4, SPTBN5, CAMK2G, FGF23, NCAM1, GRIN2B, RASGRF2, JAK1, HBEGF, FGF2 | 745 | 116 | 9075 | 1.4701458 | 1 | 0.964609583 | 92.48917283 |
| REACTOME_PATHWAY | R-HSA-109704:R-HSA-109704 | 6 | 0.381194409 | 0.168597094 | FGFR3, KLB, PIK3CB, FGF23, FGF2, PTPN11 | 745 | 36 | 9075 | 2.030201342 | 1 | 0.968317138 | 94.27979077 |
| REACTOME_PATHWAY | R-HSA-5654704:R-HSA-5654704 | 4 | 0.254129606 | 0.179024176 | HRAS, FGFR3, FGF23, FGF2 | 745 | 18 | 9075 | 2.706935123 | 1 | 0.96863751 | 95.29589998 |
| REACTOME_PATHWAY | R-HSA-5654710:R-HSA-5654710 | 4 | 0.254129606 | 0.179024176 | FGFR3, FGF23, FGF2, PTPN11 | 745 | 18 | 9075 | 2.706935123 | 1 | 0.96863751 | 95.29589998 |
| GOTERM_BP_DIRECT | GO:0000165~MAPK cascade | 26 | 1.65184244 | 0.202621395 | HRAS, FGFR3, ERBB4, EFNA1, CAMK2G, MAPKAPK5, IL31RA, MAP3K5, GRIN2B, PSMD1, MYC, FGF2, | 1342 | 262 | 16792 | 1.241715111 | 1 | 0.994427331 | 98.52157339 |
| GOTERM_BP_DIRECT | GO:0008543~fibroblast growth factor receptor signaling pathway | 10 | 0.635324015 | 0.206305242 | GALNT3, POLR2H, FGFR3, KLB, CEP57, FGF23, FGF2, POLR2C, POLR2B, PTPN11 | 1342 | 82 | 16792 | 1.525935081 | 1 | 0.994526039 | 98.64365497 |
| REACTOME_PATHWAY | R-HSA-5654720:R-HSA-5654720 | 4 | 0.254129606 | 0.222508853 | KLB, FGF23, FGF2, PTPN11 | 745 | 20 | 9075 | 2.436241611 | 1 | 0.97605155 | 97.97589271 |
| REACTOME_PATHWAY | R-HSA-2033514:R-HSA-2033514 | 3 | 0.190597205 | 0.25776897 | FGFR3, FGF23, FGF2 | 745 | 12 | 9075 | 3.045302013 | 1 | 0.980629858 | 99.01396146 |
| REACTOME_PATHWAY | R-HSA-1839130:R-HSA-1839130 | 3 | 0.190597205 | 0.25776897 | FGFR3, FGF23, FGF2 | 745 | 12 | 9075 | 3.045302013 | 1 | 0.980629858 | 99.01396146 |
| REACTOME_PATHWAY | R-HSA-5654732:R-HSA-5654732 | 4 | 0.254129606 | 0.267777496 | FGFR3, FGF23, FGF2, PTPN11 | 745 | 22 | 9075 | 2.214765101 | 1 | 0.981694453 | 99.20103115 |
| REACTOME_PATHWAY | R-HSA-5654693:R-HSA-5654693 | 4 | 0.254129606 | 0.290805322 | HRAS, FGF23, FGF2, PTPN11 | 745 | 23 | 9075 | 2.118470966 | 1 | 0.985326045 | 99.51304972 |
| REACTOME_PATHWAY | R-HSA-5654733:R-HSA-5654733 | 4 | 0.254129606 | 0.313961679 | KLB, FGF23, FGF2, PTPN11 | 745 | 24 | 9075 | 2.030201342 | 1 | 0.987465623 | 99.70887495 |
| REACTOME_PATHWAY | R-HSA-5654700:R-HSA-5654700 | 4 | 0.254129606 | 0.337155016 | HRAS, FGF23, FGF2, PTPN11 | 745 | 25 | 9075 | 1.948993289 | 1 | 0.988626944 | 99.82914512 |
| REACTOME_PATHWAY | R-HSA-5654699:R-HSA-5654699 | 4 | 0.254129606 | 0.360300997 | HRAS, FGF23, FGF2, PTPN11 | 745 | 26 | 9075 | 1.874032008 | 1 | 0.990771933 | 99.90150167 |
| REACTOME_PATHWAY | R-HSA-5654219:R-HSA-5654219 | 3 | 0.190597205 | 0.382038787 | PLCG1, FGF23, FGF2 | 745 | 16 | 9075 | 2.28397651 | 1 | 0.993175733 | 99.94235427 |
| REACTOME_PATHWAY | R-HSA-5655253:R-HSA-5655253 | 4 | 0.254129606 | 0.383322497 | HRAS, PLCG1, FGF23, FGF2 | 745 | 27 | 9075 | 1.804623415 | 1 | 0.992910838 | 99.94418221 |
| REACTOME_PATHWAY | R-HSA-5654221:R-HSA-5654221 | 3 | 0.190597205 | 0.440984278 | PLCG1, FGF23, FGF2 | 745 | 18 | 9075 | 2.030201342 | 1 | 0.995645245 | 99.98780669 |
| REACTOME_PATHWAY | R-HSA-5654688:R-HSA-5654688 | 3 | 0.190597205 | 0.523193953 | HRAS, FGF23, FGF2 | 745 | 21 | 9075 | 1.740172579 | 1 | 0.998059206 | 99.99896335 |
| REACTOME_PATHWAY | R-HSA-5654689:R-HSA-5654689 | 3 | 0.190597205 | 0.523193953 | FGF23, FGF2, PTPN11 | 745 | 21 | 9075 | 1.740172579 | 1 | 0.998059206 | 99.99896335 |
| REACTOME_PATHWAY | R-HSA-5654695:R-HSA-5654695 | 3 | 0.190597205 | 0.573300387 | FGF23, FGF2, PTPN11 | 745 | 23 | 9075 | 1.588853224 | 1 | 0.998865833 | 99.99981446 |
| REACTOME_PATHWAY | R-HSA-5654726:R-HSA-5654726 | 3 | 0.190597205 | 0.641057799 | FGF23, FGF2, PTPN11 | 745 | 26 | 9075 | 1.405524006 | 1 | 0.999312355 | 99.99998727 |
| REACTOME_PATHWAY | R-HSA-5654727:R-HSA-5654727 | 3 | 0.190597205 | 0.661676623 | FGF23, FGF2, PTPN11 | 745 | 27 | 9075 | 1.353467562 | 1 | 0.999422165 | 99.99999491 |

Annotation Cluster 82 Enrichment Score: 0.7500753334052698

| Category | Term | Count | % | PValue | Genes | List Total | Pop Hits | Pop Total | Fold Enrichment | Bonferroni | Benjamini | FDR |
|---|---|---|---|---|---|---|---|---|---|---|---|---|
| UP_SEQ_FEATURE | domain:Sushi 1 | 8 | 0.508259212 | 0.047781845 | MASP1, CFB, SELL, CSMD3, C4BPA, C1S, C2, CSMD1 | 1499 | 45 | 20063 | 2.379423319 | 1 | 0.917316304 | 59.81358719 |
| UP_SEQ_FEATURE | domain:Sushi 2 | 8 | 0.508259212 | 0.047781845 | MASP1, CFB, SELL, CSMD3, C4BPA, C1S, C2, CSMD1 | 1499 | 45 | 20063 | 2.379423319 | 1 | 0.917316304 | 59.81358719 |
| SMART | SM00032:CCP | 9 | 0.571791614 | 0.094527742 | MASP1, CFB, SELL, CSMD3, C4BPA, C1S, C2, SUSD3, CSMD1 | 878 | 54 | 10057 | 1.909073652 | 1 | 0.872602958 | 75.15321205 |
| UP_SEQ_FEATURE | domain:Sushi 3 | 5 | 0.317662008 | 0.213760162 | CFB, CSMD3, C4BPA, C2, CSMD1 | 1499 | 32 | 20063 | 2.091290027 | 1 | 0.998342018 | 98.86429271 |
| UP_SEQ_FEATURE | domain:Sushi 8 | 3 | 0.190597205 | 0.224314344 | CSMD3, C4BPA, CSMD1 | 1499 | 12 | 20063 | 3.346064043 | 1 | 0.998506684 | 99.11695651 |
| UP_SEQ_FEATURE | domain:Sushi 7 | 3 | 0.190597205 | 0.224314344 | CSMD3, C4BPA, CSMD1 | 1499 | 12 | 20063 | 3.346064043 | 1 | 0.998506684 | 99.11695651 |
| UP_SEQ_FEATURE | domain:Sushi 6 | 3 | 0.190597205 | 0.252812379 | CSMD3, C4BPA, CSMD1 | 1499 | 13 | 20063 | 3.088674501 | 1 | 0.999215754 | 99.56015612 |
| UP_SEQ_FEATURE | domain:Sushi 5 | 3 | 0.190597205 | 0.496754612 | CSMD3, C4BPA, CSMD1 | 1499 | 22 | 20063 | 1.825125841 | 1 | 0.999992869 | 99.99971995 |
| UP_SEQ_FEATURE | domain:Sushi 4 | 3 | 0.190597205 | 0.609053963 | CSMD3, C4BPA, CSMD1 | 1499 | 27 | 20063 | 1.487139575 | 1 | 0.999999615 | 99.99999746 |

Annotation Cluster 83 Enrichment Score: 0.7348304994735374

| Category | Term | Count | % | PValue | Genes | List Total | Pop Hits | Pop Total | Fold Enrichment | Bonferroni | Benjamini | FDR |
|---|---|---|---|---|---|---|---|---|---|---|---|---|
| UP_SEQ_FEATURE | domain:Ig-like C2-type 6 | 7 | 0.444726811 | 0.054389761 | CNTN2, SDK1, CNTN1, DSCAML1, CNTN4, MYLK, DSCAM | 1499 | 37 | 20063 | 2.532156573 | 1 | 0.933575362 | 64.70055728 |
| UP_SEQ_FEATURE | domain:Ig-like C2-type 9 | 3 | 0.190597205 | 0.168352902 | DSCAML1, MYLK, DSCAM | 1499 | 10 | 20063 | 4.015276851 | 1 | 0.995833277 | 96.76935141 |
| UP_SEQ_FEATURE | domain:Ig-like C2-type 8 | 3 | 0.190597205 | 0.252812379 | DSCAML1, MYLK, DSCAM | 1499 | 13 | 20063 | 3.088674501 | 1 | 0.999215754 | 99.56015612 |
| UP_SEQ_FEATURE | domain:Ig-like C2-type 7 | 3 | 0.190597205 | 0.496754612 | DSCAML1, MYLK, DSCAM | 1499 | 22 | 20063 | 1.825125841 | 1 | 0.999992869 | 99.99971995 |

Annotation Cluster 84 Enrichment Score: 0.7216659071080604

| Category | Term | Count | % | PValue | Genes | List Total | Pop Hits | Pop Total | Fold Enrichment | Bonferroni | Benjamini | FDR |
|---|---|---|---|---|---|---|---|---|---|---|---|---|
| GOTERM_BP_DIRECT | GO:0035176~social behavior | 8 | 0.508259212 | 0.085069458 | HRAS, AVP, NRXN3, DRD4, TH, CNTNAP2, NRXN1, GRID1 | 1342 | 48 | 16792 | 2.08544461 | 1 | 0.97479149 | 80.88549296 |
| INTERPRO | IPR001791:Laminin G domain | 9 | 0.571791614 | 0.094177414 | COL21A1, NRXN3, CNTNAP5, NELL1, COL22A1, COL15A1, CNTNAP2, NRXN1, COL5A1 | 1452 | 60 | 18559 | 1.917252066 | 1 | 0.966956911 | 81.80382384 |
| GOTERM_BP_DIRECT | GO:0071625~vocalization behavior | 4 | 0.254129606 | 0.095362747 | NRXN3, CNTNAP2, MYH14, NRXN1 | 1342 | 14 | 16792 | 3.575047903 | 1 | 0.979190754 | 84.51500586 |
| UP_SEQ_FEATURE | domain:Laminin G-like 4 | 4 | 0.254129606 | 0.096318691 | NRXN3, CNTNAP5, CNTNAP2, NRXN1 | 1499 | 15 | 20063 | 3.569134979 | 1 | 0.978830827 | 84.82886297 |
| GOTERM_BP_DIRECT | GO:0030534~adult behavior | 5 | 0.317662008 | 0.149305576 | SLC1A2, NRXN3, PAX5, CNTNAP2, NRXN1 | 1342 | 26 | 16792 | 2.406282242 | 1 | 0.987393063 | 95.06872418 |
| UP_SEQ_FEATURE | domain:Laminin G-like 3 | 4 | 0.254129606 | 0.184213726 | NRXN3, CNTNAP5, CNTNAP2, NRXN1 | 1499 | 20 | 20063 | 2.676851234 | 1 | 0.997091045 | 97.74275297 |
| UP_SEQ_FEATURE | domain:Laminin G-like | 3 | 0.190597205 | 0.224314344 | NRXN3, NRXN1, COL5A1 | 1499 | 12 | 20063 | 3.346064043 | 1 | 0.998506684 | 99.11695651 |
| INTERPRO | IPR003585:Neurexin/syndecan/glycophorin C | 3 | 0.190597205 | 0.300235879 | NRXN3, CNTNAP2, NRXN1 | 1452 | 14 | 18559 | 2.738931523 | 1 | 0.996713111 | 99.78670791 |
| SMART | SM00282:LamG | 6 | 0.381194409 | 0.302200631 | NRXN3, CNTNAP5, NELL1, CNTNAP2, NRXN1, COL5A1 | 878 | 42 | 10057 | 1.636348845 | 1 | 0.937043509 | 99.35631974 |
| SMART | SM00294:4.1m | 3 | 0.190597205 | 0.348428193 | NRXN3, CNTNAP2, NRXN1 | 878 | 14 | 10057 | 2.454523267 | 1 | 0.950860625 | 99.75382927 |
| UP_SEQ_FEATURE | domain:Laminin G-like 2 | 4 | 0.254129606 | 0.390424019 | NRXN3, CNTNAP5, CNTNAP2, NRXN1 | 1499 | 30 | 20063 | 1.784567489 | 1 | 0.999941097 | 99.99006236 |
| UP_SEQ_FEATURE | domain:Laminin G-like 1 | 4 | 0.254129606 | 0.390424019 | NRXN3, CNTNAP5, CNTNAP2, NRXN1 | 1499 | 30 | 20063 | 1.784567489 | 1 | 0.999941097 | 99.99006236 |

Annotation Cluster 85 Enrichment Score: 0.6921248708308873

| Category | Term | Count | % | PValue | Genes | List Total | Pop Hits | Pop Total | Fold Enrichment | Bonferroni | Benjamini | FDR |
|---|---|---|---|---|---|---|---|---|---|---|---|---|
| UP_SEQ_FEATURE | zinc finger region:C4-type | 10 | 0.635324015 | 0.173954795 | ARFGAP1, SMAP1, FHL2, ASAP1, SUPT4H1, AGAP1, PCLO, POLR2B, CLPX, ADAP1 | 1499 | 84 | 20063 | 1.59336383 | 1 | 0.99630144 | 97.15137009 |
| UP_SEQ_FEATURE | domain:Arf-GAP | 5 | 0.317662008 | 0.182317954 | ARFGAP1, SMAP1, ASAP1, AGAP1, ADAP1 | 1499 | 30 | 20063 | 2.230709362 | 1 | 0.997015518 | 97.64305732 |
| INTERPRO | IPR001164:Arf GTPase activating protein | 5 | 0.317662008 | 0.20410194 | ARFGAP1, SMAP1, ASAP1, AGAP1, ADAP1 | 1452 | 30 | 18559 | 2.130280073 | 1 | 0.987480548 | 98.04085625 |
| SMART | SM00105:ArfGap | 5 | 0.317662008 | 0.263261614 | ARFGAP1, SMAP1, ASAP1, AGAP1, ADAP1 | 878 | 30 | 10057 | 1.909073652 | 1 | 0.916767204 | 98.62163174 |
| Annotation Cluster 86 | Enrichment Score: 0.6920705179849828 | | | | | | | | | | | |
| Category | Term | Count | % | PValue | Genes | List Total | Pop Hits | Pop Total | Fold Enrichment | Bonferroni | Benjamini | FDR |
| GOTERM_MF_DIRECT | GO:0005249~voltage-gated potassium channel activity | 9 | 0.571791614 | 0.086068369 | KCNC2, KCNC4, KCNT2, CNGB1, CNGA3, KCNQ1, KCNG4, CNGA1, KCNH5 | 1340 | 58 | 16881 | 1.954825013 | 1 | 0.923616308 | 76.95043823 |
| GOTERM_MF_DIRECT | GO:0005223~intracellular cGMP activated cation channel activity | 3 | 0.190597205 | 0.101058014 | CNGB1, CNGA3, CNGA1 | 1340 | 7 | 16881 | 5.399040512 | 1 | 0.942403751 | 82.39836695 |
| GOTERM_MF_DIRECT | GO:0030553~cGMP binding | 4 | 0.254129606 | 0.147428081 | PDE10A, CNGB1, CNGA3, CNGA1 | 1340 | 17 | 16881 | 2.964179104 | 1 | 0.955300311 | 92.5782433 |
| GOTERM_MF_DIRECT | GO:0005222~intracellular cAMP activated cation channel activity | 3 | 0.190597205 | 0.156104872 | CNGB1, CNGA3, CNGA1 | 1340 | 9 | 16881 | 4.199253731 | 1 | 0.95771482 | 93.7184473 |
| INTERPRO | IPR014710:RmlC-like jelly roll fold | 7 | 0.444726811 | 0.158203042 | MAGI1, SLC9C1, HGD, CNGB1, CNGA3, CNGA1, KCNH5 | 1452 | 47 | 18559 | 1.903654534 | 1 | 0.978705814 | 94.85312827 |
| INTERPRO | IPR000595:Cyclic nucleotide-binding domain | 5 | 0.317662008 | 0.291317942 | SLC9C1, CNGB1, CNGA3, CNGA1, KCNH5 | 1452 | 35 | 18559 | 1.825954349 | 1 | 0.996314081 | 99.73471085 |
| INTERPRO | IPR018490:Cyclic nucleotide-binding-like | 5 | 0.317662008 | 0.345840694 | SLC9C1, CNGB1, CNGA3, CNGA1, KCNH5 | 1452 | 38 | 18559 | 1.681800058 | 1 | 0.998248604 | 99.93320085 |
| INTERPRO | IPR018488:Cyclic nucleotide-binding, conserved site | 3 | 0.190597205 | 0.359292003 | CNGB1, CNGA3, CNGA1 | 1452 | 16 | 18559 | 2.396565083 | 1 | 0.998586361 | 99.95329916 |
| SMART | SM00100:cNMP | 4 | 0.254129606 | 0.515286341 | CNGB1, CNGA3, CNGA1, KCNH5 | 878 | 31 | 10057 | 1.477992505 | 1 | 0.983706843 | 99.99611283 |
| Annotation Cluster 87 | Enrichment Score: 0.690109744899569 | | | | | | | | | | | |
| Category | Term | Count | % | PValue | Genes | List Total | Pop Hits | Pop Total | Fold Enrichment | Bonferroni | Benjamini | FDR |
| UP_SEQ_FEATURE | domain:LisH | 5 | 0.317662008 | 0.152581901 | TBL1XR1, NOLC1, ARMC9, PAFAH1B1, KATNAL2 | 1499 | 28 | 20063 | 2.390045745 | 1 | 0.994186424 | 95.41645503 |
| INTERPRO | IPR006594:LisH dimerisation motif | 5 | 0.317662008 | 0.171658274 | TBL1XR1, NOLC1, ARMC9, PAFAH1B1, KATNAL2 | 1452 | 28 | 18559 | 2.282442936 | 1 | 0.981504563 | 96.10065501 |
| SMART | SM00667:LisH | 4 | 0.254129606 | 0.324715487 | TBL1XR1, ARMC9, PAFAH1B1, KATNAL2 | 878 | 23 | 10057 | 1.992076855 | 1 | 0.94264858 | 99.59361942 |
| Annotation Cluster 88 | Enrichment Score: 0.6869532496862514 | | | | | | | | | | | |
| Category | Term | Count | % | PValue | Genes | List Total | Pop Hits | Pop Total | Fold Enrichment | Bonferroni | Benjamini | FDR |
| UP_SEQ_FEATURE | domain:DHR-2 | 3 | 0.190597205 | 0.196084959 | DOCK5, DOCK10, DOCK3 | 1499 | 11 | 20063 | 3.650251683 | 1 | 0.997732105 | 98.28192692 |
| UP_SEQ_FEATURE | domain:DHR-1 | 3 | 0.190597205 | 0.196084959 | DOCK5, DOCK10, DOCK3 | 1499 | 11 | 20063 | 3.650251683 | 1 | 0.997732105 | 98.28192692 |
| INTERPRO | IPR027357:DHR-2 domain | 3 | 0.190597205 | 0.21054648 | DOCK5, DOCK10, DOCK3 | 1452 | 11 | 18559 | 3.485912847 | 1 | 0.98882506 | 98.29690195 |
| INTERPRO | IPR027007:DHR-1 domain | 3 | 0.190597205 | 0.21054648 | DOCK5, DOCK10, DOCK3 | 1452 | 11 | 18559 | 3.485912847 | 1 | 0.98882506 | 98.29690195 |
| INTERPRO | IPR026791:Dedicator of cytokinesis | 3 | 0.190597205 | 0.21054648 | DOCK5, DOCK10, DOCK3 | 1452 | 11 | 18559 | 3.485912847 | 1 | 0.98882506 | 98.29690195 |
| INTERPRO | IPR010703:Dedicator of cytokinesis C-terminal | 3 | 0.190597205 | 0.21054648 | DOCK5, DOCK10, DOCK3 | 1452 | 11 | 18559 | 3.485912847 | 1 | 0.98882506 | 98.29690195 |
| Annotation Cluster 89 | Enrichment Score: 0.6740105470799573 | | | | | | | | | | | |
| Category | Term | Count | % | PValue | Genes | List Total | Pop Hits | Pop Total | Fold Enrichment | Bonferroni | Benjamini | FDR |
| UP_SEQ_FEATURE | zinc finger region:Phorbol-ester/DAG-type 1 | 5 | 0.317662008 | 0.077014528 | PRKD1, PRKD2, DGKG, DGKZ, PRKCE | 1499 | 22 | 20063 | 3.041876402 | 1 | 0.965442434 | 77.51261931 |
| UP_SEQ_FEATURE | zinc finger region:Phorbol-ester/DAG-type 2 | 5 | 0.317662008 | 0.077014528 | PRKD1, PRKD2, DGKG, DGKZ, PRKCE | 1499 | 22 | 20063 | 3.041876402 | 1 | 0.965442434 | 77.51261931 |
| GOTERM_MF_DIRECT | GO:0004697~protein kinase C activity | 4 | 0.254129606 | 0.110705756 | PRKD1, PRKD2, PKN2, PRKCE | 1340 | 15 | 16881 | 3.359402985 | 1 | 0.945064187 | 85.23815992 |
| INTERPRO | IPR002219:Protein kinase C-like, phorbol ester/diacylglycerol binding | 8 | 0.508259212 | 0.255985382 | PRKD1, PRKD2, DGKG, DGKZ, AKAP13, PRKCE, UNC13C, VAV2 | 1452 | 66 | 18559 | 1.549294599 | 1 | 0.99464614 | 99.38662286 |
| REACTOME_PATHWAY | R-HSA-114508:R-HSA-114508 | 4 | 0.254129606 | 0.267777496 | DGKG, DGKZ, PRKCE, ITPR2 | 745 | 22 | 9075 | 2.214765101 | 1 | 0.981694453 | 99.20103115 |
| INTERPRO | IPR020454:Diacylglycerol/phorbol-ester binding | 4 | 0.254129606 | 0.310344049 | PRKD1, PRKD2, DGKG, PRKCE | 1452 | 25 | 18559 | 2.045068871 | 1 | 0.99714473 | 99.83399779 |
| SMART | SM00109:C1 | 8 | 0.508259212 | 0.337760779 | PRKD1, PRKD2, DGKG, DGKZ, AKAP13, PRKCE, UNC13C, VAV2 | 878 | 65 | 10057 | 1.409777466 | 1 | 0.947600066 | 99.69087539 |
| UP_SEQ_FEATURE | zinc finger region:Phorbol-ester/DAG-type | 3 | 0.190597205 | 0.859326528 | AKAP13, UNC13C, VAV2 | 1499 | 45 | 20063 | 0.892283745 | 1 | 1 | 100 |
| Annotation Cluster 90 | Enrichment Score: 0.6726291988046023 | | | | | | | | | | | |
| Category | Term | Count | % | PValue | Genes | List Total | Pop Hits | Pop Total | Fold Enrichment | Bonferroni | Benjamini | FDR |
| GOTERM_BP_DIRECT | GO:0007154~cell communication | 7 | 0.444726811 | 0.078708897 | SLC8A1, FREM3, ANG, GJB3, GJB6, GJA4, GJA3 | 1342 | 38 | 16792 | 2.304965095 | 1 | 0.973085911 | 78.25489913 |
| REACTOME_PATHWAY | R-HSA-190861:R-HSA-190861 | 6 | 0.381194409 | 0.197276695 | GJB3, TUBAL3, GJB6, GJA4, TUBA1C, GJA3 | 745 | 38 | 9075 | 1.92334864 | 1 | 0.969791919 | 96.67977579 |
| INTERPRO | IPR017990:Connexin, conserved site | 4 | 0.254129606 | 0.202402908 | GJB3, GJB6, GJA4, GJA3 | 1452 | 20 | 18559 | 2.556336088 | 1 | 0.987485679 | 97.96754682 |
| GOTERM_CC_DIRECT | GO:0005922~connexon complex | 4 | 0.254129606 | 0.223132717 | GJB3, GJB6, GJA4, GJA3 | 1425 | 21 | 18224 | 2.435956558 | 1 | 0.860697902 | 97.75863612 |
| INTERPRO | IPR019570:Gap junction protein, cysteine-rich domain | 4 | 0.254129606 | 0.223384015 | GJB3, GJB6, GJA4, GJA3 | 1452 | 21 | 18559 | 2.434605798 | 1 | 0.991063045 | 98.71595892 |
| INTERPRO | IPR013092:Connexin, N-terminal | 4 | 0.254129606 | 0.223384015 | GJB3, GJB6, GJA4, GJA3 | 1452 | 21 | 18559 | 2.434605798 | 1 | 0.991063045 | 98.71595892 |
| INTERPRO | IPR000500:Connexin | 4 | 0.254129606 | 0.223384015 | GJB3, GJB6, GJA4, GJA3 | 1452 | 21 | 18559 | 2.434605798 | 1 | 0.991063045 | 98.71595892 |
| SMART | SM00037:CNX | 4 | 0.254129606 | 0.250966048 | GJB3, GJB6, GJA4, GJA3 | 878 | 20 | 10057 | 2.290888383 | 1 | 0.909142145 | 98.26154363 |
| SMART | SM01089:SM01089 | 4 | 0.254129606 | 0.275368192 | GJB3, GJB6, GJA4, GJA3 | 878 | 21 | 10057 | 2.18179846 | 1 | 0.919704205 | 98.9074061 |
| UP_KEYWORDS | Gap junction | 4 | 0.254129606 | 0.347650918 | GJB3, GJB6, GJA4, GJA3 | 1535 | 28 | 20581 | 1.915402513 | 1 | 0.770696723 | 99.78435861 |
| Annotation Cluster 91 | Enrichment Score: 0.6700918077441579 | | | | | | | | | | | |
| Category | Term | Count | % | PValue | Genes | List Total | Pop Hits | Pop Total | Fold Enrichment | Bonferroni | Benjamini | FDR |
| REACTOME_PATHWAY | R-HSA-112411:R-HSA-112411 | 3 | 0.190597205 | 0.164663088 | JAK1, IL6R, PTPN11 | 745 | 9 | 9075 | 4.060402685 | 1 | 0.967886345 | 93.8456742 |
| REACTOME_PATHWAY | R-HSA-110056:R-HSA-110056 | 3 | 0.190597205 | 0.195184822 | JAK1, IL6R, PTPN11 | 745 | 10 | 9075 | 3.654362416 | 1 | 0.972434902 | 96.54313707 |
| REACTOME_PATHWAY | R-HSA-1059683:R-HSA-1059683 | 3 | 0.190597205 | 0.226321966 | JAK1, IL6R, PTPN11 | 745 | 11 | 9075 | 3.322147651 | 1 | 0.974724651 | 98.12437066 |
| BIOCARTA | h_il6Pathway:IL 6 signaling pathway | 4 | 0.254129606 | 0.286988018 | HRAS, JAK1, IL6R, PTPN11 | 146 | 21 | 1625 | 2.120026093 | 1 | 0.997057304 | 98.66773664 |
| Annotation Cluster 92 | Enrichment Score: 0.6611911234131698 | | | | | | | | | | | |
| Category | Term | Count | % | PValue | Genes | List Total | Pop Hits | Pop Total | Fold Enrichment | Bonferroni | Benjamini | FDR |
| REACTOME_PATHWAY | R-HSA-5576892:R-HSA-5576892 | 9 | 0.571791614 | 0.031627864 | FGF14, CAMK2G, CACNG6, CACNG5, CACNG2, SCN7A, CACNA1C, SCN4A, CALM2 | 745 | 46 | 9075 | 2.383279837 | 1 | 0.918948615 | 39.22662029 |
| UP_KEYWORDS | Calcium channel | 10 | 0.635324015 | 0.055308229 | CATSPER3, RYR3, TRPV2, CACNG6, CACNG5, MCU, RYR2, CACNG2, CACNA1C, ITPR2 | 1535 | 66 | 20581 | 2.031487514 | 1 | 0.322661056 | 55.8558576 |
| KEGG_PATHWAY | hsa05412:Arrhythmogenic right ventricular cardiomyopathy (ARVC) | 10 | 0.635324015 | 0.102238477 | ITGA8, CACNG6, CACNG5, ITGA11, RYR2, CACNG2, CACNA1C, TCF7L2, CTNNA3, CTNNA2 | 574 | 67 | 6879 | 1.788704561 | 1 | 0.685673228 | 75.95527842 |
| INTERPRO | IPR008368:Voltage-dependent calcium channel, gamma subunit | 3 | 0.190597205 | 0.124818096 | CACNG6, CACNG5, CACNG2 | 1452 | 8 | 18559 | 4.793130165 | 1 | 0.969089745 | 89.94168933 |
| REACTOME_PATHWAY | R-HSA-5576893:R-HSA-5576893 | 5 | 0.317662008 | 0.176129003 | CACNG6, CACNG5, CACNG2, CACNA1C, KCNQ1 | 745 | 27 | 9075 | 2.255779269 | 1 | 0.968920245 | 95.03216328 |
| REACTOME_PATHWAY | R-HSA-5576894:R-HSA-5576894 | 5 | 0.317662008 | 0.176129003 | CACNG6, CACNG5, CACNG2, CACNA1C, KCNIP4 | 745 | 27 | 9075 | 2.255779269 | 1 | 0.968920245 | 95.03216328 |
| GOTERM_MF_DIRECT | GO:0005245~voltage-gated calcium channel activity | 6 | 0.381194409 | 0.208082988 | CATSPER3, TMC2, CACNG6, CACNG5, CACNG2, CACNA1C | 1340 | 40 | 16881 | 1.889664179 | 1 | 0.977578794 | 97.77217822 |
| INTERPRO | IPR004031:PMP-22/EMP/MP20/Claudin | 6 | 0.381194409 | 0.273693941 | CACNG6, CACNG5, CACNG2, GSG1L, TMEM178A, TMEM178B | 1452 | 45 | 18559 | 1.704224059 | 1 | 0.994979774 | 99.59754266 |
| GOTERM_BP_DIRECT | GO:0061337~cardiac conduction | 6 | 0.381194409 | 0.288801495 | CACNG6, CACNG5, CACNG2, CACNA1C, KCNQ1, KCNIP4 | 1342 | 45 | 16792 | 1.668355688 | 1 | 0.998483379 | 99.82409367 |
| KEGG_PATHWAY | hsa04261:Adrenergic signaling in cardiomyocytes | 14 | 0.889453621 | 0.363163157 | ATP1B2, CAMK2G, CACNG6, CACNG5, CACNG2, ADRB2, ATP2B4, ATF6B, RYR2, SCN7A, CACNA1C, KC | 574 | 138 | 6879 | 1.215800636 | 1 | 0.834456429 | 99.74282954 |
| KEGG_PATHWAY | hsa04260:Cardiac muscle contraction | 8 | 0.508259212 | 0.435157537 | ATP1B2, CACNG6, CACNG5, COX4I2, RYR2, CACNG2, CACNA1C, UQCRB | 574 | 75 | 6879 | 1.278327526 | 1 | 0.86358988 | 99.94731276 |
| KEGG_PATHWAY | hsa05410:Hypertrophic cardiomyopathy (HCM) | 7 | 0.444726811 | 0.641285595 | ITGA8, CACNG6, CACNG5, ITGA11, RYR2, CACNG2, CACNA1C | 574 | 78 | 6879 | 1.075515947 | 1 | 0.907787245 | 99.99986939 |
| GOTERM_CC_DIRECT | GO:0005891~voltage-gated calcium channel complex | 3 | 0.190597205 | 0.673537956 | CACNG6, CACNG2, CACNA1C | 1425 | 29 | 18224 | 1.322976407 | 1 | 0.987027859 | 99.99999514 |
| KEGG_PATHWAY | hsa05414:Dilated cardiomyopathy | 7 | 0.444726811 | 0.711573422 | ITGA8, CACNG6, CACNG5, ITGA11, RYR2, CACNG2, CACNA1C | 574 | 84 | 6879 | 0.99869338 | 1 | 0.921967009 | 99.99999268 |
| Annotation Cluster 93 | Enrichment Score: 0.6600590693023942 | | | | | | | | | | | |
| Category | Term | Count | % | PValue | Genes | List Total | Pop Hits | Pop Total | Fold Enrichment | Bonferroni | Benjamini | FDR |
| GOTERM_BP_DIRECT | GO:0015701~bicarbonate transport | 9 | 0.571791614 | 0.02166779 | SLC26A4, SLC26A3, SLC4A11, SLC26A5, CA9, SLC26A9, CA2, SLC26A10, CA1 | 1342 | 44 | 16792 | 2.559409294 | 1 | 0.879488939 | 33.4828265 |
| GOTERM_BP_DIRECT | GO:0006730~one-carbon metabolic process | 7 | 0.444726811 | 0.028941218 | SHMT2, ALDH1L1, CA9, MAT1A, CA2, AHCYL2, CA1 | 1342 | 30 | 16792 | 2.919622454 | 1 | 0.894395922 | 42.10833464 |

| | | | | | | | | | | | | |
|---|---|---|---|---|---|---|---|---|---|---|---|---|
| INTERPRO | IPR018338:Carbonic anhydrase, alpha-class, conserved site | 3 | 0.190597205 | 0.240343159 | CA9, CA2, CA1 | 1452 | 12 | 18559 | 3.19542011 | 1 | 0.993507746 | 99.12220937 |
| REACTOME_PATHWAY | R-HSA-1475029:R-HSA-1475029 | 3 | 0.190597205 | 0.25776897 | CA9, CA2, CA1 | 745 | 12 | 9075 | 3.045302013 | 1 | 0.980629858 | 99.01396146 |
| GOTERM_MF_DIRECT | GO:0004089~carbonate dehydratase activity | 3 | 0.190597205 | 0.306327544 | CA9, CA2, CA1 | 1340 | 14 | 16881 | 2.699520256 | 1 | 0.990937537 | 99.74303163 |
| INTERPRO | IPR023561:Carbonic anhydrase, alpha-class | 3 | 0.190597205 | 0.359292003 | CA9, CA2, CA1 | 1452 | 16 | 18559 | 2.396565083 | 1 | 0.998586361 | 99.95329916 |
| INTERPRO | IPR001148:Alpha carbonic anhydrase | 3 | 0.190597205 | 0.416411215 | CA9, CA2, CA1 | 1452 | 18 | 18559 | 2.130280073 | 1 | 0.999358607 | 99.99065213 |
| KEGG_PATHWAY | hsa00910:Nitrogen metabolism | 3 | 0.190597205 | 0.420016493 | CA9, CA2, CA1 | 574 | 17 | 6879 | 2.114880098 | 1 | 0.86107968 | 99.925265 |
| SMART | SM01057:SM01057 | 3 | 0.190597205 | 0.443781593 | CA9, CA2, CA1 | 878 | 17 | 10057 | 2.021372102 | 1 | 0.977711781 | 99.97323083 |
| UP_SEQ_FEATURE | metal ion-binding site:Zinc; catalytic | 9 | 0.571791614 | 0.755829495 | ADAMTS14, CA9, ADAM33, MMP16, ENPEP, CA2, ADAM12, CA1, C9ORF3 | 1499 | 129 | 20063 | 0.933785314 | 1 | 0.999999998 | 100 |
| Annotation Cluster 94 | Enrichment Score: 0.6572286582333988 | | | | | | | | | | | |
| Category | Term | Count | % | PValue | Genes | List Total | Pop Hits | Pop Total | Fold Enrichment | Bonferroni | Benjamini | FDR |
| UP_KEYWORDS | Antiport | 8 | 0.508259212 | 0.030329973 | SLC9A9, SLC26A3, SLC8A1, SLC24A4, SLC9A3, SLC24A1, SLC9C1, SLC26A10 | 1535 | 41 | 20581 | 2.61615953 | 0.999999746 | 0.241243985 | 35.76651165 |
| INTERPRO | IPR018422:Cation/H+ exchanger, CPA1 family | 3 | 0.190597205 | 0.21054648 | SLC9A9, SLC9A3, SLC9C1 | 1452 | 11 | 18559 | 3.485912847 | 1 | 0.98882506 | 98.29690195 |
| GOTERM_MF_DIRECT | GO:0015386~potassium:proton antiporter activity | 3 | 0.190597205 | 0.215252361 | SLC9A9, SLC9A3, SLC9C1 | 1340 | 11 | 16881 | 3.435753053 | 1 | 0.977661265 | 98.07921944 |
| GOTERM_MF_DIRECT | GO:0015385~sodium:proton antiporter activity | 3 | 0.190597205 | 0.245546669 | SLC9A9, SLC9A3, SLC9C1 | 1340 | 12 | 16881 | 3.149440299 | 1 | 0.983715887 | 98.98915326 |
| GOTERM_BP_DIRECT | GO:0098719~sodium ion import across plasma membrane | 3 | 0.190597205 | 0.309217409 | SLC9A9, SLC9A3, SLC9C1 | 1342 | 14 | 16792 | 2.681285927 | 1 | 0.998943716 | 99.8977055 |
| INTERPRO | IPR006153:Cation/H+ exchanger | 3 | 0.190597205 | 0.329946531 | SLC9A9, SLC9A3, SLC9C1 | 1452 | 15 | 18559 | 2.556336088 | 1 | 0.997808954 | 99.89898659 |
| GOTERM_BP_DIRECT | GO:1902600~hydrogen ion transmembrane transport | 5 | 0.317662008 | 0.728433511 | SLC9A9, SLC9A3, COX4I2, SLC9C1, UQCRB | 1342 | 61 | 16792 | 1.025628497 | 1 | 0.999999304 | 100 |
| Annotation Cluster 95 | Enrichment Score: 0.6544038354621932 | | | | | | | | | | | |
| Category | Term | Count | % | PValue | Genes | List Total | Pop Hits | Pop Total | Fold Enrichment | Bonferroni | Benjamini | FDR |
| KEGG_PATHWAY | hsa05231:Choline metabolism in cancer | 13 | 0.82592122 | 0.132906849 | PLD2, JMJD7-PLA2G4B, HRAS, PLCG1, PIK3CB, SLC44A4, DGKG, DGKZ, PIP5K1C, PDGFC, CHPT1, PLA2 | 574 | 101 | 6879 | 1.542536309 | 1 | 0.722928533 | 84.81078298 |
| KEGG_PATHWAY | hsa04370:VEGF signaling pathway | 8 | 0.508259212 | 0.242926693 | JMJD7-PLA2G4B, HRAS, CASP9, PLCG1, PIK3CB, VEGFA, PLA2G4B, AKT3 | 574 | 61 | 6879 | 1.571714171 | 1 | 0.821903579 | 97.47186175 |
| KEGG_PATHWAY | hsa04664:Fc epsilon RI signaling pathway | 8 | 0.508259212 | 0.337105544 | FCER1A, JMJD7-PLA2G4B, HRAS, PLCG1, PIK3CB, VAV2, PLA2G4B, AKT3 | 574 | 68 | 6879 | 1.409920066 | 1 | 0.819506134 | 99.56310482 |
| Annotation Cluster 96 | Enrichment Score: 0.6437567079733363 | | | | | | | | | | | |
| Category | Term | Count | % | PValue | Genes | List Total | Pop Hits | Pop Total | Fold Enrichment | Bonferroni | Benjamini | FDR |
| UP_SEQ_FEATURE | binding site:PAPS | 3 | 0.190597205 | 0.115475174 | NDST3, HS3ST1, SULT2B1 | 1499 | 8 | 20063 | 5.019096064 | 1 | 0.986545152 | 89.81978955 |
| UP_SEQ_FEATURE | nucleotide phosphate-binding region:PAPS | 5 | 0.317662008 | 0.230008575 | NDST3, HS3ST1, CHST11, SULT2B1, CHST3 | 1499 | 33 | 20063 | 2.027917602 | 1 | 0.998656293 | 99.23015828 |
| REACTOME_PATHWAY | R-HSA-2022928:R-HSA-2022928 | 5 | 0.317662008 | 0.246651598 | GPC2, NDST3, HS3ST4, HS3ST1, HS2ST1 | 745 | 31 | 9075 | 1.964710976 | 1 | 0.979063889 | 98.75849944 |
| INTERPRO | IPR000863:Sulfotransferase domain | 5 | 0.317662008 | 0.273380883 | NDST3, HS3ST4, HS3ST1, SULT2B1, CHST3 | 1452 | 34 | 18559 | 1.879658888 | 1 | 0.995128204 | 99.59193842 |
| GOTERM_MF_DIRECT | GO:0008146~sulfotransferase activity | 5 | 0.317662008 | 0.337389631 | HS3ST4, HS3ST1, CHST11, CHST3, HS2ST1 | 1340 | 37 | 16881 | 1.702400161 | 1 | 0.991865213 | 99.87825426 |
| Annotation Cluster 97 | Enrichment Score: 0.6426690841908343 | | | | | | | | | | | |
| Category | Term | Count | % | PValue | Genes | List Total | Pop Hits | Pop Total | Fold Enrichment | Bonferroni | Benjamini | FDR |
| INTERPRO | IPR008144:Guanylate kinase | 5 | 0.317662008 | 0.087988078 | MAGI2, MAGI1, MPP7, TJP3, DLG2 | 1452 | 22 | 18559 | 2.904927373 | 1 | 0.9647498 | 79.53899787 |
| UP_SEQ_FEATURE | domain:Guanylate kinase-like | 5 | 0.317662008 | 0.099617679 | MAGI2, MAGI1, MPP7, TJP3, DLG2 | 1499 | 24 | 20063 | 2.788386702 | 1 | 0.980888024 | 85.82758991 |
| INTERPRO | IPR020590:Guanylate kinase, conserved site | 4 | 0.254129606 | 0.12452823 | MAGI2, MAGI1, MPP7, DLG2 | 1452 | 16 | 18559 | 3.19542011 | 1 | 0.970194765 | 89.88414493 |
| INTERPRO | IPR008145:Guanylate kinase/L-type calcium channel | 5 | 0.317662008 | 0.141230378 | MAGI2, MAGI1, MPP7, TJP3, DLG2 | 1452 | 26 | 18559 | 2.458015469 | 1 | 0.974946358 | 92.74077646 |
| SMART | SM00072:GuKc | 5 | 0.317662008 | 0.168894626 | MAGI2, MAGI1, MPP7, TJP3, DLG2 | 878 | 25 | 10057 | 2.290888383 | 1 | 0.909209601 | 92.52939011 |
| UP_SEQ_FEATURE | domain:PDZ 3 | 5 | 0.317662008 | 0.182317954 | MAGI2, MAGI1, PDZD2, TJP3, DLG2 | 1499 | 30 | 20063 | 2.230709362 | 1 | 0.997015518 | 97.64305732 |
| UP_SEQ_FEATURE | domain:PDZ 1 | 6 | 0.381194409 | 0.201812683 | MAGI2, MAGI1, PDZD2, TJP3, DLG2, APBA1 | 1499 | 42 | 20063 | 1.912036596 | 1 | 0.997977455 | 98.49609191 |
| UP_SEQ_FEATURE | domain:PDZ 2 | 6 | 0.381194409 | 0.201812683 | MAGI2, MAGI1, PDZD2, TJP3, DLG2, APBA1 | 1499 | 42 | 20063 | 1.912036596 | 1 | 0.997977455 | 98.49609191 |
| GOTERM_CC_DIRECT | GO:0005923~bicellular tight junction | 9 | 0.571791614 | 0.665556106 | CLMP, MAGI2, MAGI1, CGN, ANK3, MPP7, TJP3, CDK4, AOC1 | 1425 | 113 | 18224 | 1.018574755 | 1 | 0.986092818 | 99.99999301 |
| KEGG_PATHWAY | hsa04530:Tight junction | 7 | 0.444726811 | 0.742966974 | MAGI1, CGN, MYH14, MYL12A, PRKCE, TJP3, CDK4 | 574 | 87 | 6879 | 0.964255677 | 1 | 0.924110259 | 99.9999984 |
| UP_KEYWORDS | Tight junction | 5 | 0.317662008 | 0.891635869 | CLMP, MAGI1, CGN, MPP7, TJP3 | 1535 | 86 | 20581 | 0.779524278 | 1 | 0.988151089 | 100 |
| Annotation Cluster 98 | Enrichment Score: 0.6421224730798776 | | | | | | | | | | | |
| Category | Term | Count | % | PValue | Genes | List Total | Pop Hits | Pop Total | Fold Enrichment | Bonferroni | Benjamini | FDR |
| REACTOME_PATHWAY | R-HSA-196757:R-HSA-196757 | 4 | 0.254129606 | 0.138282201 | SHMT2, MTHFR, ALDH1L1, SLC19A1 | 745 | 16 | 9075 | 3.045302013 | 1 | 0.955919877 | 90.03615516 |
| GOTERM_BP_DIRECT | GO:0046655~folic acid metabolic process | 4 | 0.254129606 | 0.169488965 | SHMT2, MTHFR, ALDH1L1, SLC19A1 | 1342 | 18 | 16792 | 2.780592813 | 1 | 0.991507908 | 96.84592511 |
| KEGG_PATHWAY | hsa00670:One carbon pool by folate | 3 | 0.190597205 | 0.505504318 | SHMT2, MTHFR, ALDH1L1 | 574 | 20 | 6879 | 1.797648084 | 1 | 0.873583558 | 99.99091469 |
| Annotation Cluster 99 | Enrichment Score: 0.6346740451691963 | | | | | | | | | | | |
| Category | Term | Count | % | PValue | Genes | List Total | Pop Hits | Pop Total | Fold Enrichment | Bonferroni | Benjamini | FDR |
| GOTERM_BP_DIRECT | GO:0003356~regulation of cilium beat frequency | 3 | 0.190597205 | 0.054161082 | DNAH11, ARMC4, CCDC39 | 1342 | 5 | 16792 | 7.507600596 | 1 | 0.950810698 | 64.52569176 |
| UP_KEYWORDS | Ciliopathy | 12 | 0.762388818 | 0.338017025 | DNAH11, TRAF3IP1, BBS5, EVC, HYDIN, WDPCP, DYNC2H1, ARMC4, AHI1, ALMS1, CCDC39, IFT140 | 1535 | 126 | 20581 | 1.276935009 | 1 | 0.761213131 | 99.73378383 |
| GOTERM_BP_DIRECT | GO:0003341~cilium movement | 4 | 0.254129606 | 0.367179625 | DNAH11, HYDIN, ARMC4, CCDC39 | 1342 | 27 | 16792 | 1.853728542 | 1 | 0.999438729 | 99.97997975 |
| UP_KEYWORDS | Primary ciliary dyskinesia | 4 | 0.254129606 | 0.43032774 | DNAH11, HYDIN, ARMC4, CCDC39 | 1535 | 32 | 20581 | 1.675977199 | 1 | 0.819567068 | 99.9692478 |
| Annotation Cluster 100 | Enrichment Score: 0.629271708602199 | | | | | | | | | | | |
| Category | Term | Count | % | PValue | Genes | List Total | Pop Hits | Pop Total | Fold Enrichment | Bonferroni | Benjamini | FDR |
| INTERPRO | IPR026823:Complement Clr-like EGF domain | 6 | 0.381194409 | 0.063218072 | HMCN2, FBLN1, THBD, LTBP3, LRP1B, FBN2 | 1452 | 28 | 18559 | 2.738931523 | 1 | 0.929813379 | 67.53502315 |
| UP_SEQ_FEATURE | domain:EGF-like 9; calcium-binding | 4 | 0.254129606 | 0.112252812 | FBLN1, LTBP3, NOTCH4, FBN2 | 1499 | 16 | 20063 | 3.346064043 | 1 | 0.985793841 | 89.10662529 |
| UP_SEQ_FEATURE | domain:EGF-like 13; calcium-binding | 3 | 0.190597205 | 0.196084959 | LTBP3, NOTCH4, FBN2 | 1499 | 11 | 20063 | 3.650251683 | 1 | 0.997732105 | 98.28192692 |
| REACTOME_PATHWAY | R-HSA-2129379:R-HSA-2129379 | 6 | 0.381194409 | 0.197276695 | BMP10, FBLN1, LTBP3, ITGA8, FBN2, FN1 | 745 | 38 | 9075 | 1.92334864 | 1 | 0.969791919 | 96.67977579 |
| UP_SEQ_FEATURE | domain:EGF-like 8; calcium-binding | 4 | 0.254129606 | 0.264825721 | FBLN1, LTBP3, NOTCH4, FBN2 | 1499 | 24 | 20063 | 2.230709362 | 1 | 0.999397541 | 99.6747422 |
| UP_SEQ_FEATURE | domain:EGF-like 11; calcium-binding | 3 | 0.190597205 | 0.281381718 | LTBP3, NOTCH4, FBN2 | 1499 | 14 | 20063 | 2.868054894 | 1 | 0.999562574 | 99.78716607 |
| UP_SEQ_FEATURE | domain:EGF-like 12; calcium-binding | 3 | 0.190597205 | 0.309852953 | LTBP3, NOTCH4, FBN2 | 1499 | 15 | 20063 | 2.676851234 | 1 | 0.999751312 | 99.89973676 |
| UP_SEQ_FEATURE | domain:EGF-like 5; calcium-binding | 5 | 0.317662008 | 0.314756392 | FBLN1, LTBP3, NELL1, NOTCH4, FBN2 | 1499 | 38 | 20063 | 1.761086338 | 1 | 0.999777961 | 99.91220228 |
| UP_SEQ_FEATURE | domain:EGF-like 7; calcium-binding | 4 | 0.254129606 | 0.348675573 | FBLN1, LTBP3, NOTCH4, FBN2 | 1499 | 28 | 20063 | 1.912036596 | 1 | 0.999870952 | 99.96588361 |
| UP_SEQ_FEATURE | domain:EGF-like 4; calcium-binding | 3 | 0.190597205 | 0.732683833 | FBLN1, LTBP3, FBN2 | 1499 | 34 | 20063 | 1.18096378 | 1 | 0.999999994 | 100 |
| Annotation Cluster 101 | Enrichment Score: 0.6262762931247365 | | | | | | | | | | | |
| Category | Term | Count | % | PValue | Genes | List Total | Pop Hits | Pop Total | Fold Enrichment | Bonferroni | Benjamini | FDR |
| INTERPRO | IPR015655:Protein phosphatase 2C | 4 | 0.254129606 | 0.142873304 | PDP1, PPM1D, PPM1E, PPM1H | 1452 | 17 | 18559 | 3.007454221 | 1 | 0.975007669 | 92.97634335 |
| INTERPRO | IPR001932:Protein phosphatase 2C (PP2C)-like | 4 | 0.254129606 | 0.202402908 | PDP1, PPM1D, PPM1E, PPM1H | 1452 | 20 | 18559 | 2.556336088 | 1 | 0.987485679 | 97.96754682 |
| INTERPRO | IPR000222:Protein phosphatase 2C, manganese/magnesium aspartate binding site | 3 | 0.190597205 | 0.21054648 | PDP1, PPM1D, PPM1E | 1452 | 11 | 18559 | 3.485912847 | 1 | 0.98882506 | 98.29690195 |
| GOTERM_BP_DIRECT | GO:0035970~peptidyl-threonine dephosphorylation | 3 | 0.190597205 | 0.217492002 | PDP1, PPM1D, PPM1E | 1342 | 11 | 16792 | 3.412545726 | 1 | 0.994970664 | 98.95856193 |
| SMART | SM00332:PP2Cc | 4 | 0.254129606 | 0.250966048 | PDP1, PPM1D, PPM1E, PPM1H | 878 | 20 | 10057 | 2.290888383 | 1 | 0.909142145 | 98.26154363 |
| UP_SEQ_FEATURE | domain:PP2C-like | 3 | 0.190597205 | 0.252812379 | PPM1D, PPM1E, PPM1H | 1499 | 13 | 20063 | 3.088674501 | 1 | 0.999215754 | 99.56015612 |
| GOTERM_MF_DIRECT | GO:0004722~protein serine/threonine phosphatase activity | 6 | 0.381194409 | 0.49169244 | PDP1, PPM1D, PPM1E, PPM1H, MTMR4, PPP5C | 1340 | 58 | 16881 | 1.303216675 | 1 | 0.997091856 | 99.99838517 |
| Annotation Cluster 102 | Enrichment Score: 0.6232986808997257 | | | | | | | | | | | |
| Category | Term | Count | % | PValue | Genes | List Total | Pop Hits | Pop Total | Fold Enrichment | Bonferroni | Benjamini | FDR |

| | | | | | | | | | | | | |
|---|---|---|---|---|---|---|---|---|---|---|---|---|
| UP_SEQ_FEATURE | domain:SH3 1 | 6 | 0.381194409 | 0.162891242 | MYO7B, SORBS2, RIMBP2, SH3RF3, TRIO, VAV2 | 1499 | 39 | 20063 | 2.059116334 | 1 | 0.995503081 | 96.35059573 |
| UP_SEQ_FEATURE | domain:SH3 2 | 6 | 0.381194409 | 0.175516912 | MYO7B, SORBS2, RIMBP2, SH3RF3, TRIO, VAV2 | 1499 | 40 | 20063 | 2.007638426 | 1 | 0.996356761 | 97.25002043 |
| UP_SEQ_FEATURE | domain:SH3 3 | 3 | 0.190597205 | 0.471940505 | SORBS2, RIMBP2, SH3RF3 | 1499 | 21 | 20063 | 1.912036596 | 1 | 0.999988315 | 99.99931382 |

Annotation Cluster 103 Enrichment Score: 0.6163518832473759

| Category | Term | Count | % | PValue | Genes | List Total | Pop Hits | Pop Total | Fold Enrichment | Bonferroni | Benjamini | FDR |
|---|---|---|---|---|---|---|---|---|---|---|---|---|
| INTERPRO | IPR017131:Small ribonucleoprotein associated, SmB/SmN | 3 | 0.190597205 | 0.017372155 | SNRPN, SNRPB, SNURF | 1452 | 3 | 18559 | 12.78168044 | 1 | 0.732918975 | 26.05887113 |
| PIR_SUPERFAMILY | PIRSF037187:small nuclear ribonucleoprotein associated protein, SmB/SmN types | 3 | 0.190597205 | 0.022931929 | SNRPN, SNRPB, SNURF | 154 | 3 | 1692 | 10.98701299 | 0.959297101 | 0.959297101 | 23.89328549 |
| UP_SEQ_FEATURE | region of interest:Repeat-rich region | 3 | 0.190597205 | 0.068290808 | SNRPN, SNRPB, SNURF | 1499 | 6 | 20063 | 6.692128085 | 1 | 0.956349522 | 73.20770389 |
| GOTERM_CC_DIRECT | GO:0005682~U5 snRNP | 4 | 0.254129606 | 0.142694809 | DDX23, SNRPN, SNRPB, SNURF | 1425 | 17 | 18224 | 3.009122807 | 1 | 0.808463793 | 90.13330158 |
| GOTERM_CC_DIRECT | GO:0005687~U4 snRNP | 3 | 0.190597205 | 0.210362584 | SNRPN, SNRPB, SNURF | 1425 | 11 | 18224 | 3.48784689 | 1 | 0.855087693 | 97.13563386 |
| INTERPRO | IPR010920:Like-Sm (LSM) domain | 4 | 0.254129606 | 0.288330568 | ATXN2, SNRPN, SNRPB, SNURF | 1452 | 24 | 18559 | 2.130280073 | 1 | 0.996225253 | 99.71477266 |
| GOTERM_CC_DIRECT | GO:0071004~U2-type prespliceosome | 3 | 0.190597205 | 0.387870196 | SNRPN, SNRPB, SNURF | 1425 | 17 | 18224 | 2.256842105 | 1 | 0.932626049 | 99.93783647 |
| GOTERM_CC_DIRECT | GO:0030532~small nuclear ribonucleoprotein complex | 3 | 0.190597205 | 0.387870196 | SNRPN, SNRPB, SNURF | 1425 | 17 | 18224 | 2.256842105 | 1 | 0.932626049 | 99.93783647 |
| GOTERM_CC_DIRECT | GO:0005685~U1 snRNP | 3 | 0.190597205 | 0.443702274 | SNRPN, SNRPB, SNURF | 1425 | 19 | 18224 | 2.019279778 | 1 | 0.950543354 | 99.9852526 |
| GOTERM_CC_DIRECT | GO:0005686~U2 snRNP | 3 | 0.190597205 | 0.470539298 | SNRPN, SNRPB, SNURF | 1425 | 20 | 18224 | 1.918315789 | 1 | 0.957813619 | 99.99299039 |
| INTERPRO | IPR001163:Ribonucleoprotein LSM domain | 3 | 0.190597205 | 0.496890596 | SNRPN, SNRPB, SNURF | 1452 | 21 | 18559 | 1.825954349 | 1 | 0.99974334 | 99.99927469 |
| GOTERM_CC_DIRECT | GO:0046540~U4/U6 x U5 tri-snRNP complex | 3 | 0.190597205 | 0.546136899 | SNRPN, SNRPB, SNURF | 1425 | 23 | 18224 | 1.668100686 | 1 | 0.971191183 | 99.99930944 |
| SMART | SM00651:Sm | 3 | 0.190597205 | 0.557775484 | SNRPN, SNRPB, SNURF | 878 | 21 | 10057 | 1.636348845 | 1 | 0.987692982 | 99.99892618 |
| GOTERM_CC_DIRECT | GO:0071013~catalytic step 2 spliceosome | 8 | 0.508259212 | 0.585482445 | TFIP11, WDR83, DDX23, SNRPN, PPIL1, SNRPB, HNRNPR, SNURF | 1425 | 92 | 18224 | 1.112067124 | 1 | 0.977205769 | 99.99982348 |
| GOTERM_CC_DIRECT | GO:0005681~spliceosomal complex | 6 | 0.381194409 | 0.867364989 | TFIP11, WDR83, SNRPN, SNRPB, HNRNPR, SNURF | 1425 | 94 | 18224 | 0.816304591 | 1 | 0.998962067 | 100 |

Annotation Cluster 104 Enrichment Score: 0.6113523580145223

| Category | Term | Count | % | PValue | Genes | List Total | Pop Hits | Pop Total | Fold Enrichment | Bonferroni | Benjamini | FDR |
|---|---|---|---|---|---|---|---|---|---|---|---|---|
| GOTERM_BP_DIRECT | GO:0018095~protein polyglutamylation | 3 | 0.190597205 | 0.157835061 | TTLL6, TTLL7, TTLL1 | 1342 | 9 | 16792 | 4.17088922 | 1 | 0.988848565 | 95.91204581 |
| UP_SEQ_FEATURE | domain:TTL | 3 | 0.190597205 | 0.281381718 | TTLL6, TTLL7, TTLL1 | 1499 | 14 | 20063 | 2.868054894 | 1 | 0.999562574 | 99.78716607 |
| INTERPRO | IPR004344:Tubulin-tyrosine ligase | 3 | 0.190597205 | 0.329946531 | TTLL6, TTLL7, TTLL1 | 1452 | 15 | 18559 | 2.556336088 | 1 | 0.997808954 | 99.89898659 |

Annotation Cluster 105 Enrichment Score: 0.6092517923574313

| Category | Term | Count | % | PValue | Genes | List Total | Pop Hits | Pop Total | Fold Enrichment | Bonferroni | Benjamini | FDR |
|---|---|---|---|---|---|---|---|---|---|---|---|---|
| INTERPRO | IPR011011:Zinc finger, FYVE/PHD-type | 18 | 1.143583227 | 0.048779355 | PHRF1, SP100, RUFY1, PHF10, SP110, RIMS1, PCLO, CXXC1, MYRIP, TRIM66, KDM2A, PHF1, TCF19, S | 1452 | 141 | 18559 | 1.631703886 | 1 | 0.906102626 | 57.74770343 |
| INTERPRO | IPR019786:Zinc finger, PHD-type, conserved site | 9 | 0.571791614 | 0.150689403 | TRIM66, PHRF1, SP100, KDM2A, PHF1, TCF19, SP110, NFXL1, CXXC1 | 1452 | 67 | 18559 | 1.716942149 | 1 | 0.977066525 | 94.00173268 |
| UP_SEQ_FEATURE | zinc finger region:PHD-type | 6 | 0.381194409 | 0.347176799 | TRIM66, PHRF1, KDM2A, TCF19, SP110, CXXC1 | 1499 | 52 | 20063 | 1.54433725 | 1 | 0.999870681 | 99.96439184 |
| INTERPRO | IPR001965:Zinc finger, PHD-type | 9 | 0.571791614 | 0.394686338 | TRIM66, PHRF1, SP100, KDM2A, PHF1, PHF10, TCF19, SP110, CXXC1 | 1452 | 89 | 18559 | 1.292529483 | 1 | 0.99920761 | 99.98245446 |
| INTERPRO | IPR019787:Zinc finger, PHD-finger | 8 | 0.508259212 | 0.423140487 | TRIM66, PHRF1, SP100, KDM2A, PHF1, PHF10, SP110, CXXC1 | 1452 | 79 | 18559 | 1.294347386 | 1 | 0.999395263 | 99.99234506 |
| SMART | SM00249:PHD | 9 | 0.571791614 | 0.518660426 | TRIM66, PHRF1, SP100, KDM2A, PHF1, PHF10, TCF19, SP110, CXXC1 | 878 | 89 | 10057 | 1.158314351 | 1 | 0.983436425 | 99.99647554 |

Annotation Cluster 106 Enrichment Score: 0.591633800292143

| Category | Term | Count | % | PValue | Genes | List Total | Pop Hits | Pop Total | Fold Enrichment | Bonferroni | Benjamini | FDR |
|---|---|---|---|---|---|---|---|---|---|---|---|---|
| INTERPRO | IPR016186:C-type lectin-like | 13 | 0.82592122 | 0.112088272 | CLEC19A, CLEC17A, THBD, ATRNL1, STAB1, SELL, MRC2, COL15A1, SUSD5, CLEC4C, COLEC12, COLEC | 1452 | 104 | 18559 | 1.597710055 | 1 | 0.965210624 | 87.10076792 |
| INTERPRO | IPR016187:C-type lectin fold | 13 | 0.82592122 | 0.164872584 | CLEC19A, CLEC17A, THBD, ATRNL1, STAB1, SELL, MRC2, COL15A1, SUSD5, CLEC4C, COLEC12, COLEC | 1452 | 112 | 18559 | 1.483587908 | 1 | 0.979729591 | 95.51223002 |
| INTERPRO | IPR018378:C-type lectin, conserved site | 6 | 0.381194409 | 0.258671838 | CLEC17A, SELL, MRC2, COLEC12, CLEC4C, COLEC11 | 1452 | 44 | 18559 | 1.742956424 | 1 | 0.994502281 | 99.42367904 |
| INTERPRO | IPR001304:C-type lectin | 10 | 0.635324015 | 0.259450731 | CLEC19A, CLEC17A, THBD, ATRNL1, SELL, MRC2, COLEC12, CLEC4C, COLEC11, CLEC5A | 1452 | 89 | 18559 | 1.43614387 | 1 | 0.994357599 | 99.43402199 |
| UP_KEYWORDS | Lectin | 16 | 1.016518424 | 0.293593884 | GALNT3, LMAN1L, ATRNL1, SELL, MRC2, COLEC12, COLEC11, CLC, CLEC19A, ZG16B, CLEC17A, SIGLE | 1535 | 171 | 20581 | 1.25453264 | 1 | 0.713703274 | 99.32293445 |
| UP_SEQ_FEATURE | domain:C-type lectin | 8 | 0.508259212 | 0.414082462 | CLEC17A, THBD, ATRNL1, SELL, COLEC12, CLEC4C, COLEC11, CLEC5A | 1499 | 82 | 20063 | 1.30578109 | 1 | 0.999961867 | 99.99524462 |
| SMART | SM00034:CLECT | 9 | 0.571791614 | 0.478869188 | CLEC17A, THBD, ATRNL1, SELL, MRC2, COLEC12, CLEC4C, COLEC11, CLEC5A | 878 | 86 | 10057 | 1.198720665 | 1 | 0.980158327 | 99.9892648 |

Annotation Cluster 107 Enrichment Score: 0.587850033111461

| Category | Term | Count | % | PValue | Genes | List Total | Pop Hits | Pop Total | Fold Enrichment | Bonferroni | Benjamini | FDR |
|---|---|---|---|---|---|---|---|---|---|---|---|---|
| UP_SEQ_FEATURE | metal ion-binding site:Copper | 4 | 0.254129606 | 0.096318691 | APP, LOXL2, AOC1, LOXL1 | 1499 | 15 | 20063 | 3.569134979 | 1 | 0.978830827 | 84.82886297 |
| UP_KEYWORDS | TPQ | 3 | 0.190597205 | 0.11513475 | LOXL2, AOC1, LOXL1 | 1535 | 8 | 20581 | 5.027931596 | 1 | 0.477132414 | 82.760473 |
| GOTERM_MF_DIRECT | GO:0005507~copper ion binding | 6 | 0.381194409 | 0.460414638 | DCT, ANG, LOXL2, AOC1, LOXL1, METTL17 | 1340 | 56 | 16881 | 1.349760128 | 1 | 0.996105828 | 99.99572438 |
| UP_KEYWORDS | Copper | 4 | 0.254129606 | 0.872034662 | APP, LOXL2, AOC1, LOXL1 | 1535 | 65 | 20581 | 0.825096467 | 1 | 0.98509165 | 100 |

Annotation Cluster 108 Enrichment Score: 0.5834782813527617

| Category | Term | Count | % | PValue | Genes | List Total | Pop Hits | Pop Total | Fold Enrichment | Bonferroni | Benjamini | FDR |
|---|---|---|---|---|---|---|---|---|---|---|---|---|
| UP_SEQ_FEATURE | domain:TSP type-1 2 | 7 | 0.444726811 | 0.116172944 | THSD7A, SEMA5A, C8B, ADAMTSL1, ADAMTS14, UNC5C, THSD7B | 1499 | 45 | 20063 | 2.081995404 | 1 | 0.986418742 | 89.96828602 |
| UP_SEQ_FEATURE | domain:TSP type-1 1 | 7 | 0.444726811 | 0.116172944 | THSD7A, SEMA5A, C8B, ADAMTSL1, ADAMTS14, UNC5C, THSD7B | 1499 | 45 | 20063 | 2.081995404 | 1 | 0.986418742 | 89.96828602 |
| UP_SEQ_FEATURE | domain:TSP type-1 4 | 5 | 0.317662008 | 0.213760162 | THSD7A, SEMA5A, ADAMTSL1, ADAMTS14, THSD7B | 1499 | 32 | 20063 | 2.091290027 | 1 | 0.998342018 | 98.86429271 |
| UP_SEQ_FEATURE | domain:TSP type-1 7 | 3 | 0.190597205 | 0.224314344 | THSD7A, SEMA5A, THSD7B | 1499 | 12 | 20063 | 3.346064043 | 1 | 0.998506684 | 99.11695651 |
| INTERPRO | IPR000884:Thrombospondin, type 1 repeat | 8 | 0.508259212 | 0.243871751 | THSD7A, SEMA5A, C8B, ADAMTSL1, ADAMTS14, CILP, UNC5C, THSD7B | 1452 | 65 | 18559 | 1.5731299 | 1 | 0.993472929 | 99.18986381 |
| UP_SEQ_FEATURE | domain:TSP type-1 3 | 5 | 0.317662008 | 0.297481785 | THSD7A, SEMA5A, ADAMTSL1, ADAMTS14, THSD7B | 1499 | 37 | 20063 | 1.808683266 | 1 | 0.999703932 | 99.86042408 |
| SMART | SM00209:TSP1 | 8 | 0.508259212 | 0.337760779 | THSD7A, SEMA5A, C8B, ADAMTSL1, ADAMTS14, CILP, UNC5C, THSD7B | 878 | 65 | 10057 | 1.409777466 | 1 | 0.947600066 | 99.69087539 |
| UP_SEQ_FEATURE | domain:TSP type-1 6 | 3 | 0.190597205 | 0.393340669 | THSD7A, SEMA5A, THSD7B | 1499 | 18 | 20063 | 2.230709362 | 1 | 0.999938317 | 99.99091136 |
| REACTOME_PATHWAY | R-HSA-5173214:R-HSA-5173214 | 5 | 0.317662008 | 0.398740874 | THSD7A, SEMA5A, ADAMTSL1, ADAMTS14, THSD7B | 745 | 39 | 9075 | 1.56169334 | 1 | 0.994184004 | 99.96229719 |
| UP_SEQ_FEATURE | domain:TSP type-1 5 | 3 | 0.190597205 | 0.588210376 | THSD7A, SEMA5A, THSD7B | 1499 | 26 | 20063 | 1.54433725 | 1 | 0.999999408 | 99.99999331 |

Annotation Cluster 109 Enrichment Score: 0.5773558647178297

| Category | Term | Count | % | PValue | Genes | List Total | Pop Hits | Pop Total | Fold Enrichment | Bonferroni | Benjamini | FDR |
|---|---|---|---|---|---|---|---|---|---|---|---|---|
| GOTERM_MF_DIRECT | GO:0004553~hydrolase activity, hydrolyzing O-glycosyl compounds | 7 | 0.444726811 | 0.028104901 | GBA2, CHID1, CHIA, KLB, GNE, MGAM, GAA | 1340 | 30 | 16881 | 2.939477612 | 1 | 0.898863333 | 37.17637491 |
| UP_KEYWORDS | Glycosidase | 9 | 0.571791614 | 0.266294438 | MAN2A1, GBA2, CHIA, MAN1A2, MGAM, GAA, TDG, MAN2B1, AGL | 1535 | 82 | 20581 | 1.471589735 | 1 | 0.696576754 | 98.83239103 |
| INTERPRO | IPR017853:Glycoside hydrolase, superfamily | 6 | 0.381194409 | 0.416156137 | CHID1, CHIA, KLB, MGAM, GAA, AGL | 1452 | 54 | 18559 | 1.420186716 | 1 | 0.999416139 | 99.99058149 |
| INTERPRO | IPR013781:Glycoside hydrolase, catalytic domain | 4 | 0.254129606 | 0.561005805 | CHID1, CHIA, KLB, AGL | 1452 | 37 | 18559 | 1.381803291 | 1 | 0.999888413 | 99.99993072 |
| GOTERM_BP_DIRECT | GO:0005975~carbohydrate metabolic process | 13 | 0.82592122 | 0.742767446 | GALNT3, PGM3, CHID1, CHIA, KLB, MGAM, GAA, ALDH2, CHST3, PARG, ST8SIA2, MAN2B1, INSR | 1342 | 174 | 16792 | 0.93485448 | 1 | 0.999999549 | 100 |

Annotation Cluster 110 Enrichment Score: 0.5758572385796689

| Category | Term | Count | % | PValue | Genes | List Total | Pop Hits | Pop Total | Fold Enrichment | Bonferroni | Benjamini | FDR |
|---|---|---|---|---|---|---|---|---|---|---|---|---|
| UP_SEQ_FEATURE | repeat:ANK 13 | 4 | 0.254129606 | 0.096318691 | ANKRD28, ANK3, ANKRD50, TNKS2 | 1499 | 15 | 20063 | 3.569134979 | 1 | 0.978830827 | 84.82886297 |
| UP_SEQ_FEATURE | repeat:ANK 15 | 4 | 0.254129606 | 0.096318691 | ANKRD28, ANK3, ANKRD50, TNKS2 | 1499 | 15 | 20063 | 3.569134979 | 1 | 0.978830827 | 84.82886297 |
| UP_SEQ_FEATURE | repeat:ANK 14 | 4 | 0.254129606 | 0.096318691 | ANKRD28, ANK3, ANKRD50, TNKS2 | 1499 | 15 | 20063 | 3.569134979 | 1 | 0.978830827 | 84.82886297 |
| UP_SEQ_FEATURE | repeat:ANK 11 | 5 | 0.317662008 | 0.111942946 | ANKRD28, ANK3, ANKRD50, ANKDD1A, TNKS2 | 1499 | 25 | 20063 | 2.676851234 | 1 | 0.986145531 | 89.03560923 |
| UP_SEQ_FEATURE | repeat:ANK 12 | 4 | 0.254129606 | 0.146780167 | ANKRD28, ANK3, ANKRD50, TNKS2 | 1499 | 18 | 20063 | 2.974279149 | 1 | 0.993080316 | 94.79554201 |
| UP_SEQ_FEATURE | repeat:ANK 10 | 5 | 0.317662008 | 0.167214296 | ANKRD28, ANK3, ANKRD50, ANKDD1A, TNKS2 | 1499 | 29 | 20063 | 2.307630374 | 1 | 0.995830461 | 96.68599419 |
| UP_SEQ_FEATURE | repeat:ANK 18 | 3 | 0.190597205 | 0.224314344 | ANKRD28, ANK3, ANKRD50 | 1499 | 12 | 20063 | 3.346064043 | 1 | 0.998506684 | 99.11695651 |
| UP_SEQ_FEATURE | repeat:ANK 19 | 3 | 0.190597205 | 0.224314344 | ANKRD28, ANK3, ANKRD50 | 1499 | 12 | 20063 | 3.346064043 | 1 | 0.998506684 | 99.11695651 |

| | | | | | | | | | | | | |
|---|---|---|---|---|---|---|---|---|---|---|---|---|
| UP_SEQ_FEATURE | repeat:ANK 17 | 3 | 0.190597205 | 0.224314344 | ANKRD28, ANK3, ANKRD50 | 1499 | 12 | 20063 | 3.346064043 | 1 | 0.998506684 | 99.11695651 |
| UP_SEQ_FEATURE | repeat:ANK 16 | 3 | 0.190597205 | 0.252812379 | ANKRD28, ANK3, ANKRD50 | 1499 | 13 | 20063 | 3.088674501 | 1 | 0.999215754 | 99.56015612 |
| UP_SEQ_FEATURE | repeat:ANK 6 | 10 | 0.635324015 | 0.319644821 | ANKRD28, ANK3, ANKRD50, NFKBIB, TRPV2, ANKRD30A, RAI14, ANKDD1A, EHMT2, TNKS2 | 1499 | 99 | 20063 | 1.351945068 | 1 | 0.999792283 | 99.9231597 |
| UP_SEQ_FEATURE | repeat:ANK 7 | 7 | 0.444726811 | 0.330001336 | ANKRD28, ANK3, ANKRD50, RAI14, ANKDD1A, EHMT2, TNKS2 | 1499 | 63 | 20063 | 1.487139575 | 1 | 0.99983716 | 99.94225038 |
| UP_SEQ_FEATURE | repeat:ANK 1 | 21 | 1.334180432 | 0.365234582 | CAMTA1, NFKBIB, TRPV2, ABTB2, ANKMY2, ANKRD30A, ANKRD10, RAI14, ASAP1, EHMT2, KANK3, T | 1499 | 244 | 20063 | 1.151923687 | 1 | 0.999908795 | 99.97887881 |
| UP_SEQ_FEATURE | repeat:ANK 9 | 5 | 0.317662008 | 0.366918666 | ANKRD28, ANK3, ANKRD50, ANKDD1A, TNKS2 | 1499 | 41 | 20063 | 1.632226362 | 1 | 0.999905193 | 99.97989816 |
| UP_SEQ_FEATURE | repeat:ANK 5 | 13 | 0.82592122 | 0.371251661 | PPP1R16B, ANKRD28, ANK3, ANKRD50, NFKBIB, TRPV2, NOTCH4, ANKRD30A, RAI14, ANKDD1A, EHI | 1499 | 142 | 20063 | 1.225319227 | 1 | 0.999909504 | 99.98231115 |
| UP_SEQ_FEATURE | repeat:ANK 2 | 21 | 1.334180432 | 0.372233004 | CAMTA1, NFKBIB, TRPV2, ABTB2, ANKMY2, ANKRD30A, ANKRD10, RAI14, ASAP1, EHMT2, KANK3, T | 1499 | 245 | 20063 | 1.147221957 | 1 | 0.999908186 | 99.9828182 |
| UP_SEQ_FEATURE | repeat:ANK 3 | 18 | 1.143583227 | 0.377697805 | CAMTA1, NFKBIB, TRPV2, ABTB2, ANKMY2, ANKRD30A, RAI14, ANKRD10, EHMT2, KANK3, TNKS2, T | 1499 | 207 | 20063 | 1.163848363 | 1 | 0.999919247 | 99.98539953 |
| UP_SEQ_FEATURE | repeat:ANK 4 | 15 | 0.952986023 | 0.391674696 | TRPV2, ABTB2, NFKBIB, ANKRD30A, RAI14, ANKRD10, KANK3, EHMT2, TNKS2, PPP1R16B, ANKRD28 | 1499 | 170 | 20063 | 1.18096378 | 1 | 0.999940707 | 99.99043522 |
| UP_KEYWORDS | ANK repeat | 22 | 1.397712834 | 0.41084207 | CAMTA1, GPR75-ASB3, NFKBIB, TRPV2, ABTB2, ANKMY2, ANKRD30A, ANKRD10, RAI14, ASAP1, EHN | 1535 | 264 | 20581 | 1.117318132 | 1 | 0.810111636 | 99.9501344 |
| INTERPRO | IPR002110:Ankyrin repeat | 22 | 1.397712834 | 0.434908333 | CAMTA1, GPR75-ASB3, NFKBIB, TRPV2, ABTB2, ANKMY2, ANKRD30A, ANKRD10, RAI14, ASAP1, EHN | 1452 | 255 | 18559 | 1.102733214 | 1 | 0.99949343 | 99.9946329 |
| UP_SEQ_FEATURE | repeat:ANK 8 | 5 | 0.317662008 | 0.452921694 | ANKRD28, ANK3, ANKRD50, ANKDD1A, TNKS2 | 1499 | 46 | 20063 | 1.454810453 | 1 | 0.999980985 | 99.99867395 |
| INTERPRO | IPR020683:Ankyrin repeat-containing domain | 22 | 1.397712834 | 0.507459444 | CAMTA1, GPR75-ASB3, NFKBIB, TRPV2, ABTB2, ANKMY2, ANKRD30A, ANKRD10, RAI14, ASAP1, EHN | 1452 | 265 | 18559 | 1.06112064 | 1 | 0.999780377 | 99.99949687 |
| SMART | SM00248:ANK | 21 | 1.334180432 | 0.684651205 | GPR75-ASB3, NFKBIB, TRPV2, ABTB2, ANKMY2, ANKRD30A, ANKRD10, RAI14, ASAP1, EHMT2, KANK | 878 | 249 | 10057 | 0.96603727 | 1 | 0.994200067 | 99.99999063 |

Annotation Cluster 111 Enrichment Score: 0.5688210920290462

| Category | Term | Count | % | PValue | Genes | List Total | Pop Hits | Pop Total | Fold Enrichment | Bonferroni | Benjamini | FDR |
|---|---|---|---|---|---|---|---|---|---|---|---|---|
| GOTERM_CC_DIRECT | GO:0016514~SWI/SNF complex | 4 | 0.254129606 | 0.106975873 | SMARCC1, SMARCD1, RB1, ARID1B | 1425 | 15 | 18224 | 3.410339181 | 1 | 0.762933266 | 81.76750106 |
| GOTERM_CC_DIRECT | GO:0071565~nBAF complex | 3 | 0.190597205 | 0.299998128 | SMARCC1, SMARCD1, ARID1B | 1425 | 14 | 18224 | 2.740451128 | 1 | 0.896567074 | 99.53241285 |
| REACTOME_PATHWAY | R-HSA-3214858:R-HSA-3214858 | 7 | 0.444726811 | 0.612537021 | HIST4H4, SMARCC1, SMARCD1, H2AFJ, ARID1B, CDK4, HIST1H3H | 745 | 77 | 9075 | 1.10738255 | 1 | 0.999114428 | 99.99995838 |

Annotation Cluster 112 Enrichment Score: 0.5419268242288833

| Category | Term | Count | % | PValue | Genes | List Total | Pop Hits | Pop Total | Fold Enrichment | Bonferroni | Benjamini | FDR |
|---|---|---|---|---|---|---|---|---|---|---|---|---|
| INTERPRO | IPR002404:Insulin receptor substrate-1, PTB | 3 | 0.190597205 | 0.240343159 | DOK2, DOK4, DOK6 | 1452 | 12 | 18559 | 3.19542011 | 1 | 0.993507746 | 99.12220937 |
| SMART | SM00310:PTBI | 3 | 0.190597205 | 0.24825504 | DOK2, DOK4, DOK6 | 878 | 11 | 10057 | 3.123938704 | 1 | 0.919512377 | 98.17120167 |
| UP_SEQ_FEATURE | domain:IRS-type PTB | 3 | 0.190597205 | 0.252812379 | DOK2, DOK4, DOK6 | 1499 | 13 | 20063 | 3.088674501 | 1 | 0.999215754 | 99.56015612 |
| GOTERM_MF_DIRECT | GO:0005158~insulin receptor binding | 4 | 0.254129606 | 0.45057234 | DOK2, DOK4, DOK6, PTPN11 | 1340 | 31 | 16881 | 1.625517573 | 1 | 0.995917818 | 99.99425876 |

Annotation Cluster 113 Enrichment Score: 0.540821012386633

| Category | Term | Count | % | PValue | Genes | List Total | Pop Hits | Pop Total | Fold Enrichment | Bonferroni | Benjamini | FDR |
|---|---|---|---|---|---|---|---|---|---|---|---|---|
| GOTERM_BP_DIRECT | GO:0018149~peptide cross-linking | 8 | 0.508259212 | 0.100843926 | LCE1F, COL3A1, TGM2, SPRR2G, LCE2A, LCE3E, TGM6, FN1 | 1342 | 50 | 16792 | 2.002026826 | 1 | 0.979455619 | 86.1711107 |
| GOTERM_BP_DIRECT | GO:0030216~keratinocyte differentiation | 9 | 0.571791614 | 0.259701346 | WNT5A, LCE1F, ST14, TP63, SPRR2G, LCE2A, LCE3E, RBPJ, SCEL | 1342 | 76 | 16792 | 1.481763276 | 1 | 0.997687404 | 99.62895458 |
| GOTERM_CC_DIRECT | GO:0001533~cornified envelope | 6 | 0.381194409 | 0.289030902 | LCE1F, CNFN, SPRR2G, LCE2A, LCE3E, SCEL | 1425 | 46 | 18224 | 1.668100686 | 1 | 0.888108118 | 99.40922217 |
| UP_KEYWORDS | Keratinization | 5 | 0.317662008 | 0.330934577 | LCE1F, CNFN, SPRR2G, LCE2A, LCE3E | 1535 | 39 | 20581 | 1.718950973 | 1 | 0.757115337 | 99.68978855 |
| GOTERM_BP_DIRECT | GO:0031424~keratinization | 6 | 0.381194409 | 0.337056282 | LCE1F, CNFN, SPRR2G, LCE2A, LCE3E, CDH3 | 1342 | 48 | 16792 | 1.564083458 | 1 | 0.999172051 | 99.95242734 |
| INTERPRO | IPR026075:Small proline-rich protein/late cornified envelope protein | 3 | 0.190597205 | 0.673846737 | LCE1F, LCE2A, LCE3E | 1452 | 29 | 18559 | 1.322242804 | 1 | 0.999985922 | 99.99999959 |

Annotation Cluster 114 Enrichment Score: 0.5407597781255395

| Category | Term | Count | % | PValue | Genes | List Total | Pop Hits | Pop Total | Fold Enrichment | Bonferroni | Benjamini | FDR |
|---|---|---|---|---|---|---|---|---|---|---|---|---|
| GOTERM_BP_DIRECT | GO:0051552~flavone metabolic process | 3 | 0.190597205 | 0.077013635 | UGT1A10, UGT1A9, UGT1A8 | 1342 | 6 | 16792 | 6.25633383 | 1 | 0.971924876 | 77.49799337 |
| GOTERM_BP_DIRECT | GO:1904224~negative regulation of glucuronosyltransferase activity | 3 | 0.190597205 | 0.129342022 | UGT1A10, UGT1A9, UGT1A8 | 1342 | 8 | 16792 | 4.692250373 | 1 | 0.984429287 | 92.4062792 |
| GOTERM_BP_DIRECT | GO:2001030~negative regulation of cellular glucuronidation | 3 | 0.190597205 | 0.129342022 | UGT1A10, UGT1A9, UGT1A8 | 1342 | 8 | 16792 | 4.692250373 | 1 | 0.984429287 | 92.4062792 |
| UP_SEQ_FEATURE | chain:UDP-glucuronosyltransferase 1-1 | 3 | 0.190597205 | 0.141381957 | UGT1A10, UGT1A9, UGT1A8 | 1499 | 9 | 20063 | 4.461418724 | 1 | 0.992174796 | 94.14703211 |
| UP_SEQ_FEATURE | chain:UDP-glucuronosyltransferase 1-10 | 3 | 0.190597205 | 0.141381957 | UGT1A10, UGT1A9, UGT1A8 | 1499 | 9 | 20063 | 4.461418724 | 1 | 0.992174796 | 94.14703211 |
| UP_SEQ_FEATURE | chain:UDP-glucuronosyltransferase 1-3 | 3 | 0.190597205 | 0.141381957 | UGT1A10, UGT1A9, UGT1A8 | 1499 | 9 | 20063 | 4.461418724 | 1 | 0.992174796 | 94.14703211 |
| UP_SEQ_FEATURE | chain:UDP-glucuronosyltransferase 1-4 | 3 | 0.190597205 | 0.141381957 | UGT1A10, UGT1A9, UGT1A8 | 1499 | 9 | 20063 | 4.461418724 | 1 | 0.992174796 | 94.14703211 |
| UP_SEQ_FEATURE | chain:UDP-glucuronosyltransferase 1-5 | 3 | 0.190597205 | 0.141381957 | UGT1A10, UGT1A9, UGT1A8 | 1499 | 9 | 20063 | 4.461418724 | 1 | 0.992174796 | 94.14703211 |
| UP_SEQ_FEATURE | chain:UDP-glucuronosyltransferase 1-6 | 3 | 0.190597205 | 0.141381957 | UGT1A10, UGT1A9, UGT1A8 | 1499 | 9 | 20063 | 4.461418724 | 1 | 0.992174796 | 94.14703211 |
| UP_SEQ_FEATURE | chain:UDP-glucuronosyltransferase 1-7 | 3 | 0.190597205 | 0.141381957 | UGT1A10, UGT1A9, UGT1A8 | 1499 | 9 | 20063 | 4.461418724 | 1 | 0.992174796 | 94.14703211 |
| UP_SEQ_FEATURE | chain:UDP-glucuronosyltransferase 1-8 | 3 | 0.190597205 | 0.141381957 | UGT1A10, UGT1A9, UGT1A8 | 1499 | 9 | 20063 | 4.461418724 | 1 | 0.992174796 | 94.14703211 |
| UP_SEQ_FEATURE | chain:UDP-glucuronosyltransferase 1-9 | 3 | 0.190597205 | 0.141381957 | UGT1A10, UGT1A9, UGT1A8 | 1499 | 9 | 20063 | 4.461418724 | 1 | 0.992174796 | 94.14703211 |
| GOTERM_BP_DIRECT | GO:0045922~negative regulation of fatty acid metabolic process | 3 | 0.190597205 | 0.157835061 | UGT1A10, UGT1A9, UGT1A8 | 1342 | 9 | 16792 | 4.17088922 | 1 | 0.988848565 | 95.91204581 |
| GOTERM_BP_DIRECT | GO:0052697~xenobiotic glucuronidation | 3 | 0.190597205 | 0.157835061 | UGT1A10, UGT1A9, UGT1A8 | 1342 | 9 | 16792 | 4.17088922 | 1 | 0.988848565 | 95.91204581 |
| KEGG_PATHWAY | hsa00983:Drug metabolism - other enzymes | 6 | 0.381194409 | 0.33655786 | UGT1A10, UGT1A9, ITPA, UGT1A8, UPP2, UCKL1 | 574 | 46 | 6879 | 1.563172247 | 1 | 0.823512924 | 99.55831052 |
| KEGG_PATHWAY | hsa00982:Drug metabolism - cytochrome P450 | 8 | 0.508259212 | 0.337105544 | FMO4, UGT1A10, UGT1A9, GSTT2B, UGT1A8, CYP2B6, ALDH3B2, GSTT2 | 574 | 68 | 6879 | 1.409920066 | 1 | 0.819506134 | 99.56310482 |
| GOTERM_BP_DIRECT | GO:0052695~cellular glucuronidation | 3 | 0.190597205 | 0.369341045 | UGT1A10, UGT1A9, UGT1A8 | 1342 | 16 | 16792 | 2.346125186 | 1 | 0.999452961 | 99.98121486 |
| KEGG_PATHWAY | hsa00053:Ascorbate and aldarate metabolism | 4 | 0.254129606 | 0.393289979 | UGT1A10, UGT1A9, UGT1A8, ALDH2 | 574 | 27 | 6879 | 1.775454897 | 1 | 0.85189427 | 99.86445497 |
| INTERPRO | IPR002213:UDP-glucuronosyl/UDP-glucosyltransferase | 3 | 0.190597205 | 0.522103424 | UGT1A10, UGT1A9, UGT1A8 | 1452 | 22 | 18559 | 1.742956424 | 1 | 0.999830232 | 99.99970086 |
| GOTERM_BP_DIRECT | GO:0052696~flavonoid glucuronidation | 3 | 0.190597205 | 0.533917803 | UGT1A10, UGT1A9, UGT1A8 | 1342 | 22 | 16792 | 1.706272863 | 1 | 0.999959782 | 99.99993247 |
| GOTERM_MF_DIRECT | GO:0001972~retinoic acid binding | 3 | 0.190597205 | 0.554560917 | UGT1A10, UGT1A9, UGT1A8 | 1340 | 23 | 16881 | 1.643186243 | 1 | 0.998615147 | 99.99981242 |
| KEGG_PATHWAY | hsa00980:Metabolism of xenobiotics by cytochrome P450 | 7 | 0.444726811 | 0.589179065 | UGT1A10, UGT1A9, GSTT2B, UGT1A8, CYP2B6, ALDH3B2, GSTT2 | 574 | 74 | 6879 | 1.133651945 | 1 | 0.901687646 | 99.99921586 |
| GOTERM_MF_DIRECT | GO:0015020~glucuronosyltransferase activity | 3 | 0.190597205 | 0.681808794 | UGT1A10, UGT1A9, UGT1A8 | 1340 | 29 | 16881 | 1.303216675 | 1 | 0.999650851 | 99.99999922 |
| KEGG_PATHWAY | hsa00860:Porphyrin and chlorophyll metabolism | 4 | 0.254129606 | 0.691748101 | UGT1A10, UGT1A9, UGT1A8, HMOX1 | 574 | 42 | 6879 | 1.141363863 | 1 | 0.919995736 | 99.99998239 |
| KEGG_PATHWAY | hsa00140:Steroid hormone biosynthesis | 5 | 0.317662008 | 0.723544614 | UGT1A10, UGT1A9, UGT1A8, CYP21A2, SULT2B1 | 574 | 58 | 6879 | 1.033131083 | 1 | 0.920032927 | 99.99999582 |
| KEGG_PATHWAY | hsa00040:Pentose and glucuronate interconversions | 3 | 0.190597205 | 0.774020179 | UGT1A10, UGT1A9, UGT1A8 | 574 | 33 | 6879 | 1.089483687 | 1 | 0.931241659 | 99.99999971 |
| KEGG_PATHWAY | hsa05204:Chemical carcinogenesis | 6 | 0.381194409 | 0.807720874 | UGT1A10, UGT1A9, GSTT2B, UGT1A8, ALDH3B2, GSTT2 | 574 | 80 | 6879 | 0.898824042 | 1 | 0.942573517 | 99.99999997 |
| KEGG_PATHWAY | hsa00830:Retinol metabolism | 4 | 0.254129606 | 0.911280046 | UGT1A10, UGT1A9, UGT1A8, CYP2B6 | 574 | 64 | 6879 | 0.749020035 | 1 | 0.976588296 | 100 |
| GOTERM_BP_DIRECT | GO:0008152~metabolic process | 8 | 0.508259212 | 0.983629909 | UGT1A10, UGT1A9, UGT1A8, MAN1A2, ACAD10, OXSM, ENOSF1, AUH | 1342 | 168 | 16792 | 0.595841317 | 1 | 1 | 100 |

Annotation Cluster 115 Enrichment Score: 0.527521646117892

| Category | Term | Count | % | PValue | Genes | List Total | Pop Hits | Pop Total | Fold Enrichment | Bonferroni | Benjamini | FDR |
|---|---|---|---|---|---|---|---|---|---|---|---|---|
| GOTERM_BP_DIRECT | GO:0009953~dorsal/ventral pattern formation | 7 | 0.444726811 | 0.038591652 | DISP1, DYNC2H1, DSCAML1, BMPR1B, FBXL15, GLI3, GLI1 | 1342 | 32 | 16792 | 2.737146051 | 1 | 0.920691032 | 51.92881104 |
| GOTERM_CC_DIRECT | GO:0097546~ciliary base | 5 | 0.317662008 | 0.155935436 | MOK, TRAF3IP1, ARMC4, GLI3, GLI1 | 1425 | 27 | 18224 | 2.368291098 | 1 | 0.824176922 | 92.19296837 |
| GOTERM_CC_DIRECT | GO:0005930~axoneme | 9 | 0.571791614 | 0.286463388 | TRAF3IP1, WDPCP, DYNC2H1, ARMC4, MAP4, CCDC39, GLI3, IFT140, GLI1 | 1425 | 80 | 18224 | 1.438736842 | 1 | 0.890118762 | 99.3763022 |
| GOTERM_CC_DIRECT | GO:0097542~ciliary tip | 5 | 0.317662008 | 0.453898592 | TRAF3IP1, DYNC2H1, GLI3, IFT140, GLI1 | 1425 | 44 | 18224 | 1.453269537 | 1 | 0.953972965 | 99.98883492 |
| GOTERM_CC_DIRECT | GO:0072372~primary cilium | 7 | 0.444726811 | 0.576859702 | TRAF3IP1, DYNC2H1, AHI1, GLI3, TNPO1, IFT140, GLI1 | 1425 | 78 | 18224 | 1.147710301 | 1 | 0.975670642 | 99.9997594 |
| REACTOME_PATHWAY | R-HSA-5620924:R-HSA-5620924 | 5 | 0.317662008 | 0.656549207 | TRAF3IP1, DYNC2H1, TNPO1, TUBA1C, IFT140 | 745 | 54 | 9075 | 1.127889635 | 1 | 0.999396057 | 99.99999358 |
| REACTOME_PATHWAY | R-HSA-5610787:R-HSA-5610787 | 5 | 0.317662008 | 0.684770189 | DYNC2H1, GLI3, TUBA1C, IFT140, GLI1 | 745 | 56 | 9075 | 1.087607862 | 1 | 0.999488853 | 99.9999983 |

Annotation Cluster 116 Enrichment Score: 0.5198290113964804

| Category | Term | Count | % | PValue | Genes | List Total | Pop Hits | Pop Total | Fold Enrichment | Bonferroni | Benjamini | FDR |
|---|---|---|---|---|---|---|---|---|---|---|---|---|
| UP_SEQ_FEATURE | zinc finger region:FYVE-type | 6 | 0.381194409 | 0.162891242 | MYRIP, RUFY1, FGD5, RIMS1, MTMR4, FGD4 | 1499 | 39 | 20063 | 2.059116334 | 1 | 0.995503081 | 96.35059573 |
| INTERPRO | IPR017455:Zinc finger, FYVE-related | 5 | 0.317662008 | 0.273380883 | RUFY1, FGD5, RIMS1, MTMR4, FGD4 | 1452 | 34 | 18559 | 1.879658888 | 1 | 0.995128204 | 99.59193842 |
| INTERPRO | IPR000306:Zinc finger, FYVE-type | 4 | 0.254129606 | 0.398138933 | RUFY1, FGD5, MTMR4, FGD4 | 1452 | 29 | 18559 | 1.762990406 | 1 | 0.999231959 | 99.98410096 |

| Category | Term | Count | % | PValue | Genes | List Total | Pop Hits | Pop Total | Fold Enrichment | Bonferroni | Benjamini | FDR |
|---|---|---|---|---|---|---|---|---|---|---|---|---|
| SMART | SM00064:FYVE | 4 | 0.254129606 | 0.469876181 | RUFY1, FGD5, MTMR4, FGD4 | 878 | 29 | 10057 | 1.579923023 | 1 | 0.98037612 | 99.98635394 |

Annotation Cluster 117 Enrichment Score: 0.502309147655733

| Category | Term | Count | % | PValue | Genes | List Total | Pop Hits | Pop Total | Fold Enrichment | Bonferroni | Benjamini | FDR |
|---|---|---|---|---|---|---|---|---|---|---|---|---|
| KEGG_PATHWAY | hsa05220:Chronic myeloid leukemia | 11 | 0.698856417 | 0.073718237 | HRAS, CTBP2, PIK3CB, RB1, RUNX1, MECOM, CDK4, MYC, AKT3, SHC4, PTPN11 | 574 | 72 | 6879 | 1.830937863 | 0.999999999 | 0.621346295 | 63.64967294 |
| KEGG_PATHWAY | hsa05223:Non-small cell lung cancer | 9 | 0.571791614 | 0.09070558 | FHIT, HRAS, CASP9, PLCG1, PIK3CB, RB1, CDK4, ALK, AKT3 | 574 | 56 | 6879 | 1.926051518 | 1 | 0.684450175 | 71.53718018 |
| KEGG_PATHWAY | hsa05218:Melanoma | 10 | 0.635324015 | 0.133646665 | HRAS, FGF14, PIK3CB, FGF23, PDGFC, RB1, HGF, CDK4, FGF2, AKT3 | 574 | 71 | 6879 | 1.687932473 | 1 | 0.713728093 | 84.98115595 |
| KEGG_PATHWAY | hsa05213:Endometrial cancer | 8 | 0.508259212 | 0.138234773 | HRAS, CASP9, PIK3CB, MYC, TCF7L2, AKT3, CTNNA3, CTNNA2 | 574 | 52 | 6879 | 1.843741624 | 1 | 0.694537601 | 85.89892266 |
| KEGG_PATHWAY | hsa05214:Glioma | 9 | 0.571791614 | 0.17057648 | HRAS, PLCG1, PIK3CB, CAMK2G, RB1, CDK4, AKT3, CALM2, SHC4 | 574 | 65 | 6879 | 1.659367462 | 1 | 0.746692885 | 91.55451237 |
| KEGG_PATHWAY | hsa04370:VEGF signaling pathway | 8 | 0.508259212 | 0.242926693 | JMJD7-PLA2G4B, HRAS, CASP9, PLCG1, PIK3CB, VEGFA, PLA2G4B, AKT3 | 574 | 61 | 6879 | 1.571714171 | 1 | 0.821903579 | 97.47186175 |
| KEGG_PATHWAY | hsa05210:Colorectal cancer | 8 | 0.508259212 | 0.255888644 | DCC, CASP9, PIK3CB, MSH2, SMAD3, MYC, TCF7L2, AKT3 | 574 | 62 | 6879 | 1.546363943 | 1 | 0.820569824 | 97.98772254 |
| KEGG_PATHWAY | hsa05212:Pancreatic cancer | 8 | 0.508259212 | 0.295878578 | CASP9, PIK3CB, VEGFA, SMAD3, JAK1, RB1, CDK4, AKT3 | 574 | 65 | 6879 | 1.474993299 | 1 | 0.820413004 | 99.03026366 |
| KEGG_PATHWAY | hsa05230:Central carbon metabolism in cancer | 7 | 0.444726811 | 0.444575161 | HRAS, FGFR3, PIK3CB, HKDC1, PGAM1, MYC, AKT3 | 574 | 64 | 6879 | 1.310785061 | 1 | 0.858162467 | 99.95781009 |
| KEGG_PATHWAY | hsa05221:Acute myeloid leukemia | 6 | 0.381194409 | 0.504669684 | HRAS, PIK3CB, RUNX1, MYC, TCF7L2, AKT3 | 574 | 56 | 6879 | 1.284034345 | 1 | 0.881222098 | 99.99070994 |
| KEGG_PATHWAY | hsa05222:Small cell lung cancer | 8 | 0.508259212 | 0.570086853 | FHIT, CASP9, PIK3CB, RB1, CDK4, MYC, AKT3, FN1 | 574 | 85 | 6879 | 1.127936052 | 1 | 0.896134027 | 99.99857076 |
| KEGG_PATHWAY | hsa04919:Thyroid hormone signaling pathway | 10 | 0.635324015 | 0.627871598 | HDAC3, HRAS, MED30, CASP9, PLCG1, ATP1B2, PIK3CB, NOTCH4, MYC, AKT3 | 574 | 115 | 6879 | 1.042114831 | 1 | 0.90912102 | 99.99978783 |
| KEGG_PATHWAY | hsa05216:Thyroid cancer | 3 | 0.190597205 | 0.708784455 | HRAS, MYC, TCF7L2 | 574 | 29 | 6879 | 1.239757299 | 1 | 0.923361548 | 99.99999169 |
| BIOCARTA | h_raccycdPathway:Influence of Ras and Rho proteins on G1 to S Transition | 3 | 0.190597205 | 0.71022717 | HRAS, RB1, CDK4 | 146 | 27 | 1625 | 1.236681887 | 1 | 0.999446308 | 99.99998643 |
| KEGG_PATHWAY | hsa05215:Prostate cancer | 7 | 0.444726811 | 0.752869128 | HRAS, CASP9, PIK3CB, PDGFC, RB1, TCF7L2, AKT3 | 574 | 88 | 6879 | 0.953298226 | 1 | 0.925724733 | 99.99999905 |
| KEGG_PATHWAY | hsa05161:Hepatitis B | 11 | 0.698856417 | 0.778601278 | HRAS, CASP9, PIK3CB, ATF6B, YWHAQ, JAK1, FASLG, RB1, CDK4, MYC, AKT3 | 574 | 145 | 6879 | 0.909155353 | 1 | 0.93256725 | 99.99999978 |

Annotation Cluster 118 Enrichment Score: 0.4964936806301937

| Category | Term | Count | % | PValue | Genes | List Total | Pop Hits | Pop Total | Fold Enrichment | Bonferroni | Benjamini | FDR |
|---|---|---|---|---|---|---|---|---|---|---|---|---|
| KEGG_PATHWAY | hsa00590:Arachidonic acid metabolism | 10 | 0.635324015 | 0.064004165 | GGT5, JMJD7-PLA2G4B, GPX6, PLB1, PTGDS, CYP2B6, GPX5, PLA2G12B, FAM213B, PLA2G4B | 574 | 61 | 6879 | 1.964642714 | 0.99999999 | 0.621400413 | 58.27626854 |
| KEGG_PATHWAY | hsa00565:Ether lipid metabolism | 7 | 0.444726811 | 0.168089656 | PLD2, JMJD7-PLA2G4B, PLB1, PLA2G12B, PAFAH1B1, CHPT1, PLA2G4B | 574 | 45 | 6879 | 1.864227642 | 1 | 0.750348724 | 91.21368741 |
| GOTERM_MF_DIRECT | GO:0004623~phospholipase A2 activity | 5 | 0.317662008 | 0.228574246 | JMJD7-PLA2G4B, PLB1, PLA2G12B, PAFAH1B1, PLA2G4B | 1340 | 31 | 16881 | 2.031896967 | 1 | 0.982063487 | 98.54712272 |
| REACTOME_PATHWAY | R-HSA-1483166:R-HSA-1483166 | 5 | 0.317662008 | 0.265185991 | PLD2, DDHD1, JMJD7-PLA2G4B, PLA2G4B, AGPAT1 | 745 | 32 | 9075 | 1.903313758 | 1 | 0.981959386 | 99.15607088 |
| KEGG_PATHWAY | hsa00564:Glycerophospholipid metabolism | 11 | 0.698856417 | 0.265634372 | PLD2, CRLS1, JMJD7-PLA2G4B, PLB1, DGKG, PLA2G12B, DGKZ, CHPT1, PLA2G4B, AGPAT1, PTDSS2 | 574 | 95 | 6879 | 1.38765817 | 1 | 0.815302238 | 98.30946624 |
| GOTERM_BP_DIRECT | GO:0006654~phosphatidic acid biosynthetic process | 5 | 0.317662008 | 0.304824482 | PLD2, DDHD1, JMJD7-PLA2G4B, PLA2G4B, AGPAT1 | 1342 | 35 | 16792 | 1.787523951 | 1 | 0.998883596 | 99.88489553 |
| GOTERM_MF_DIRECT | GO:0004622~lysophospholipase activity | 4 | 0.254129606 | 0.318352206 | JMJD7-PLA2G4B, PLB1, CLC, PLA2G4B | 1340 | 25 | 16881 | 2.015641791 | 1 | 0.992028299 | 99.80678278 |
| KEGG_PATHWAY | hsa00592:alpha-Linolenic acid metabolism | 4 | 0.254129606 | 0.346446755 | JMJD7-PLA2G4B, PLB1, PLA2G12B, PLA2G4B | 574 | 25 | 6879 | 1.917491289 | 1 | 0.825374079 | 99.63781753 |
| UP_KEYWORDS | Lipid degradation | 10 | 0.635324015 | 0.392909297 | PLD2, PLCL1, DDHD1, JMJD7-PLA2G4B, PLB1, PLCG1, PLCH1, APOC2, PAFAH1B1, PLA2G4B | 1535 | 106 | 20581 | 1.264888452 | 1 | 0.80184725 | 99.92327279 |
| KEGG_PATHWAY | hsa00591:Linoleic acid metabolism | 4 | 0.254129606 | 0.439221562 | JMJD7-PLA2G4B, PLB1, PLA2G12B, PLA2G4B | 574 | 29 | 6879 | 1.653009732 | 1 | 0.860273836 | 99.95210804 |
| GOTERM_BP_DIRECT | GO:0036151~phosphatidylcholine acyl-chain remodeling | 3 | 0.190597205 | 0.646843231 | JMJD7-PLA2G4B, PLB1, PLA2G4B | 1342 | 27 | 16792 | 1.390296407 | 1 | 0.999994812 | 99.99999961 |
| REACTOME_PATHWAY | R-HSA-1482788:R-HSA-1482788 | 3 | 0.190597205 | 0.661676623 | JMJD7-PLA2G4B, PLB1, PLA2G4B | 745 | 27 | 9075 | 1.353467562 | 1 | 0.999422165 | 99.99999491 |
| GOTERM_BP_DIRECT | GO:0006644~phospholipid metabolic process | 4 | 0.254129606 | 0.816499245 | JMJD7-PLA2G4B, PLA2G12B, PLA2G4B, AGPAT1 | 1342 | 54 | 16792 | 0.926864271 | 1 | 0.999999946 | 100 |

Annotation Cluster 119 Enrichment Score: 0.4951785962359787

| Category | Term | Count | % | PValue | Genes | List Total | Pop Hits | Pop Total | Fold Enrichment | Bonferroni | Benjamini | FDR |
|---|---|---|---|---|---|---|---|---|---|---|---|---|
| UP_KEYWORDS | GPI-anchor | 15 | 0.952986023 | 0.142218023 | CPO, EFNA1, EFNA3, NCAM1, CDH13, GPC2, LSAMP, CNTN2, CNTN1, CNTN4, EFNA5, EFNA4, NT5E, I | 1535 | 137 | 20581 | 1.468009225 | 1 | 0.527066368 | 88.97186822 |
| GOTERM_CC_DIRECT | GO:0031225~anchored component of membrane | 12 | 0.762388818 | 0.267848455 | NCAM1, CDH13, GPC2, LSAMP, EFNA3, CNTN2, CNTN1, CNTN4, EFNA4, NT5E, NTM, PRSS21 | 1425 | 113 | 18224 | 1.358099674 | 1 | 0.882561971 | 99.08119484 |
| UP_SEQ_FEATURE | lipid moiety-binding region:GPI-anchor amidated serine | 6 | 0.381194409 | 0.392708318 | EFNA1, CNTN1, CNTN4, EFNA4, NT5E, PRSS21 | 1499 | 55 | 20063 | 1.460100673 | 1 | 0.999939905 | 99.99073333 |
| UP_SEQ_FEATURE | propeptide:Removed in mature form | 17 | 1.080050826 | 0.698830309 | HRAS, EFNA1, EFNA3, UBL3, CDH13, GPC2, LSAMP, CNTN2, CNTN1, GRK7, RAP1A, CNTN4, EFNA5, EI | 1499 | 237 | 20063 | 0.96005213 | 1 | 0.999999974 | 99.99999998 |

Annotation Cluster 120 Enrichment Score: 0.49082359645712575

| Category | Term | Count | % | PValue | Genes | List Total | Pop Hits | Pop Total | Fold Enrichment | Bonferroni | Benjamini | FDR |
|---|---|---|---|---|---|---|---|---|---|---|---|---|
| INTERPRO | IPR001005:SANT/Myb domain | 6 | 0.381194409 | 0.243592225 | TADA2A, SMARCC1, SNAPC4, MYPOP, TRERF1, NCOR2 | 1452 | 43 | 18559 | 1.783490294 | 1 | 0.993717651 | 99.18468896 |
| SMART | SM00717:SANT | 6 | 0.381194409 | 0.3198451 | TADA2A, SMARCC1, SNAPC4, MYPOP, TRERF1, NCOR2 | 878 | 43 | 10057 | 1.59829422 | 1 | 0.942521627 | 99.5505322 |
| INTERPRO | IPR017884:SANT domain | 4 | 0.254129606 | 0.332402354 | TADA2A, SMARCC1, TRERF1, NCOR2 | 1452 | 26 | 18559 | 1.966412376 | 1 | 0.997830839 | 99.90517831 |
| UP_SEQ_FEATURE | domain:SANT | 3 | 0.190597205 | 0.420182925 | TADA2A, SMARCC1, TRERF1 | 1499 | 19 | 20063 | 2.113303606 | 1 | 0.999963936 | 99.99608664 |

Annotation Cluster 121 Enrichment Score: 0.48909222643974

| Category | Term | Count | % | PValue | Genes | List Total | Pop Hits | Pop Total | Fold Enrichment | Bonferroni | Benjamini | FDR |
|---|---|---|---|---|---|---|---|---|---|---|---|---|
| UP_SEQ_FEATURE | repeat:LRR 9 | 15 | 0.952986023 | 0.126601287 | ASPN, NLRP5, KERA, LRRN4, LRRC2, LRRC8C, LRRC28, RNH1, ECM2, LINGO2, SYNE2, LRCH1, LRRTM4 | 1499 | 134 | 20063 | 1.498237631 | 1 | 0.99027503 | 91.95742084 |
| INTERPRO | IPR001611:Leucine-rich repeat | 27 | 1.715374841 | 0.128153727 | ASPN, CEP72, KERA, LRRC8C, RNH1, CNOT6, LINGO2, LRRC56, LRRTM4, LRRTM3, NRROS, LGI3, FBXL | 1452 | 263 | 18559 | 1.312187726 | 1 | 0.969402577 | 90.58206982 |
| UP_SEQ_FEATURE | repeat:LRR 10 | 13 | 0.82592122 | 0.156549444 | LINGO2, ASPN, NLRP5, SYNE2, LRRTM4, LRRN4, KERA, LRRTM3, LRRC8C, LRRC28, RNH1, NRROS, ECI | 1499 | 116 | 20063 | 1.499959743 | 1 | 0.994770123 | 95.79996828 |
| UP_SEQ_FEATURE | repeat:LRR 8 | 15 | 0.952986023 | 0.215461565 | ASPN, NLRP5, KERA, LRRN4, LRRC2, LRRC8C, LRRC28, RNH1, ECM2, LINGO2, SYNE2, LRCH1, LRRTM4 | 1499 | 148 | 20063 | 1.35651245 | 1 | 0.998298885 | 98.90919125 |
| UP_SEQ_FEATURE | repeat:LRR 1 | 29 | 1.842439644 | 0.22037709 | ASPN, CEP72, KERA, LRRC8C, RNH1, CNOT6, LINGO2, LRRTM4, LRRC56, LRRTM3, NRROS, LGI3, HECT | 1499 | 321 | 20063 | 1.209169561 | 1 | 0.998496105 | 99.02966054 |
| UP_SEQ_FEATURE | repeat:LRR 2 | 29 | 1.842439644 | 0.224571036 | ASPN, CEP72, KERA, LRRC8C, RNH1, CNOT6, LINGO2, LRRTM4, LRRC56, LRRTM3, NRROS, LGI3, HECT | 1499 | 322 | 20063 | 1.205414376 | 1 | 0.998457753 | 99.12238171 |
| INTERPRO | IPR003591:Leucine-rich repeat, typical subtype | 18 | 1.143583227 | 0.228579993 | ASPN, CEP72, RSU1, KERA, LRRN4, LRRC2, LRRC8C, LRRC28, LRRC27, ECM2, CNOT6, LINGO2, LRRTM | 1452 | 178 | 18559 | 1.292529483 | 1 | 0.991760398 | 98.85618784 |
| UP_SEQ_FEATURE | repeat:LRR 7 | 16 | 1.016518424 | 0.335340399 | ASPN, RSU1, NLRP5, KERA, LRRN4, LRRC2, LRRC8C, LRRC28, RNH1, ECM2, LINGO2, SYNE2, LRCH1, LI | 1499 | 176 | 20063 | 1.216750561 | 1 | 0.99984257 | 99.95024324 |
| UP_SEQ_FEATURE | repeat:LRR 11 | 10 | 0.635324015 | 0.340925784 | ASPN, LINGO2, NLRP5, SYNE2, KERA, LRRC8C, LRRC28, RNH1, NRROS, ECM2 | 1499 | 101 | 20063 | 1.325173878 | 1 | 0.999849042 | 99.9574782 |
| UP_SEQ_FEATURE | repeat:LRR 12 | 9 | 0.571791614 | 0.357777287 | ASPN, LINGO2, NLRP5, SYNE2, KERA, LRRC8C, RNH1, NRROS, ECM2 | 1499 | 90 | 20063 | 1.338425617 | 1 | 0.999889348 | 99.97374785 |
| UP_KEYWORDS | Leucine-rich repeat | 26 | 1.65184244 | 0.37625675 | ASPN, CEP72, KERA, LRRC8C, RNH1, CNOT6, LINGO2, LRRC56, LRRTM4, LRRTM3, NRROS, LGI3, FBXL | 1535 | 311 | 20581 | 1.120910795 | 1 | 0.792437977 | 99.88679866 |
| SMART | SM00369:LRR_TYP | 18 | 1.143583227 | 0.383947148 | ASPN, CEP72, RSU1, KERA, LRRN4, LRRC2, LRRC8C, LRRC28, LRRC27, ECM2, CNOT6, LINGO2, LRRTM | 878 | 178 | 10057 | 1.158314351 | 1 | 0.96093411 | 99.88783385 |
| UP_SEQ_FEATURE | repeat:LRR 3 | 24 | 1.524777637 | 0.488714984 | ASPN, RSU1, NLRP5, KERA, LRRN4, LRRC8C, LRRC2, NLRP9, LRRC28, RNH1, LRRC27, ECM2, CNOT6, L | 1499 | 302 | 20063 | 1.063649497 | 1 | 0.999991508 | 99.99962382 |
| UP_SEQ_FEATURE | repeat:LRR 13 | 7 | 0.444726811 | 0.492233124 | LINGO2, NLRP5, SYNE2, LRRC8C, RNH1, NRROS, ECM2 | 1499 | 75 | 20063 | 1.249197243 | 1 | 0.999992082 | 99.9996692 |
| UP_SEQ_FEATURE | repeat:LRR 5 | 18 | 1.143583227 | 0.561203279 | ASPN, RSU1, NLRP5, KERA, LRRN4, LRRC2, LRRC8C, LRRC28, RNH1, ECM2, LRRC23, LINGO2, SYNE2, I | 1499 | 231 | 20063 | 1.042929052 | 1 | 0.999998806 | 99.99997817 |
| UP_SEQ_FEATURE | repeat:LRR 6 | 16 | 1.016518424 | 0.601777983 | ASPN, RSU1, NLRP5, KERA, LRRN4, LRRC2, LRRC8C, LRRC28, RNH1, ECM2, LINGO2, SYNE2, LRCH1, LI | 1499 | 209 | 20063 | 1.024632051 | 1 | 0.999999613 | 99.99999642 |
| UP_SEQ_FEATURE | repeat:LRR 4 | 19 | 1.207115629 | 0.6330267 | ASPN, RSU1, NLRP5, KERA, LRRN4, LRRC2, LRRC8C, LRRC28, RNH1, LRRC27, ECM2, LRRC23, LINGO2, | 1499 | 255 | 20063 | 0.997258303 | 1 | 0.999999784 | 99.99999922 |
| UP_SEQ_FEATURE | repeat:LRR 14 | 4 | 0.254129606 | 0.835355947 | LRRC8C, RNH1, NRROS, ECM2 | 1499 | 60 | 20063 | 0.892283745 | 1 | 1 | 100 |

Annotation Cluster 122 Enrichment Score: 0.4813268107462214

| Category | Term | Count | % | PValue | Genes | List Total | Pop Hits | Pop Total | Fold Enrichment | Bonferroni | Benjamini | FDR |
|---|---|---|---|---|---|---|---|---|---|---|---|---|
| INTERPRO | IPR018525:Mini-chromosome maintenance, conserved site | 3 | 0.190597205 | 0.074157476 | MCM7, MCM4, MCM6 | 1452 | 6 | 18559 | 6.39084022 | 1 | 0.950899407 | 73.48245133 |
| REACTOME_PATHWAY | R-HSA-69052:R-HSA-69052 | 3 | 0.190597205 | 0.080666664 | MCM7, MCM4, MCM6 | 745 | 6 | 9075 | 6.090604027 | 1 | 0.944604882 | 72.83664856 |
| UP_SEQ_FEATURE | domain:MCM | 3 | 0.190597205 | 0.115475174 | MCM7, MCM4, MCM6 | 1499 | 8 | 20063 | 5.019096064 | 1 | 0.986545152 | 89.81978955 |
| GOTERM_CC_DIRECT | GO:0042555~MCM complex | 3 | 0.190597205 | 0.152335336 | MCM7, MCM4, MCM6 | 1425 | 9 | 18224 | 4.262923977 | 1 | 0.820989742 | 91.67678383 |
| INTERPRO | IPR001208:Mini-chromosome maintenance, DNA-dependent ATPase | 3 | 0.190597205 | 0.152477215 | MCM7, MCM4, MCM6 | 1452 | 9 | 18559 | 4.260560147 | 1 | 0.977197811 | 94.2155735 |
| SMART | SM00350:MCM | 3 | 0.190597205 | 0.181795767 | MCM7, MCM4, MCM6 | 878 | 9 | 10057 | 3.818147304 | 1 | 0.91362254 | 94.00098142 |
| GOTERM_BP_DIRECT | GO:0006268~DNA unwinding involved in DNA replication | 3 | 0.190597205 | 0.187332308 | MCM7, MCM4, MCM6 | 1342 | 10 | 16792 | 3.753800298 | 1 | 0.993324158 | 97.8946819 |
| REACTOME_PATHWAY | R-HSA-176974:R-HSA-176974 | 3 | 0.190597205 | 0.25776897 | MCM7, MCM4, MCM6 | 745 | 12 | 9075 | 3.045302013 | 1 | 0.980629858 | 99.01396146 |
| GOTERM_MF_DIRECT | GO:0003678~DNA helicase activity | 4 | 0.254129606 | 0.295978459 | CHD8, MCM7, MCM4, MCM6 | 1340 | 24 | 16881 | 2.099626866 | 1 | 0.990464678 | 99.67284689 |
| REACTOME_PATHWAY | R-HSA-68867:R-HSA-68867 | 3 | 0.190597205 | 0.351569214 | MCM7, MCM4, MCM6 | 745 | 15 | 9075 | 2.436241611 | 1 | 0.990504147 | 99.87847443 |
| KEGG_PATHWAY | hsa03030:DNA replication | 5 | 0.317662008 | 0.352268222 | POLD3, MCM7, MCM4, MCM6, RPA3 | 574 | 36 | 6879 | 1.664488966 | 1 | 0.827264895 | 99.6782069 |

| | | | | | | | | | | | | |
|---|---|---|---|---|---|---|---|---|---|---|---|---|
| REACTOME_PATHWAY | R-HSA-69300:R-HSA-69300 | 3 | 0.190597205 | 0.4118776 | MCM7, MCM4, MCM6 | 745 | 17 | 9075 | 2.149624951 | 1 | 0.994414386 | 99.97322637 |
| GOTERM_MF_DIRECT | GO:0004003~ATP-dependent DNA helicase activity | 4 | 0.254129606 | 0.471716789 | XRCC5, MCM7, MCM4, MCM6 | 1340 | 32 | 16881 | 1.574720149 | 1 | 0.996515702 | 99.99697244 |
| BIOCARTA | h_mcmPathway:CDK Regulation of DNA Replication | 3 | 0.190597205 | 0.487157755 | MCM7, MCM4, MCM6 | 146 | 18 | 1625 | 1.855022831 | 1 | 0.99850206 | 99.98015888 |
| REACTOME_PATHWAY | R-HSA-68962:R-HSA-68962 | 4 | 0.254129606 | 0.494350406 | MCM7, MCM4, MCM6, RPA3 | 745 | 32 | 9075 | 1.522651007 | 1 | 0.997324648 | 99.99742427 |
| REACTOME_PATHWAY | R-HSA-176187:R-HSA-176187 | 4 | 0.254129606 | 0.594512654 | MCM7, MCM4, MCM6, RPA3 | 745 | 37 | 9075 | 1.316887357 | 1 | 0.999082287 | 99.99991581 |
| REACTOME_PATHWAY | R-HSA-68949:R-HSA-68949 | 6 | 0.381194409 | 0.702306032 | PSMA1, MCM7, PSMD1, RB1, MCM4, MCM6 | 745 | 71 | 9075 | 1.029397864 | 1 | 0.999580528 | 99.9999993 |
| UP_KEYWORDS | DNA replication | 7 | 0.444726811 | 0.71059327 | POLD3, KAT7, MCM7, NFIC, MCM4, MCM6, RPA3 | 1535 | 94 | 20581 | 0.998454501 | 1 | 0.948562441 | 99.99999818 |
| GOTERM_BP_DIRECT | GO:0000082~G1/S transition of mitotic cell cycle | 8 | 0.508259212 | 0.715382825 | EIF4E, MCM7, CAMK2G, RB1, CDK4, MCM4, MCM6, RPA3 | 1342 | 102 | 16792 | 0.981385699 | 1 | 0.999999037 | 99.99999999 |
| INTERPRO | IPR012340:Nucleic acid-binding, OB-fold | 6 | 0.381194409 | 0.72910402 | POLR2H, MCM7, AIMP1, MCM4, MCM6, RPA3 | 1452 | 77 | 18559 | 0.995975099 | 1 | 0.999996792 | 99.99999998 |
| GOTERM_BP_DIRECT | GO:0006270~DNA replication initiation | 3 | 0.190597205 | 0.736948356 | MCM7, MCM4, MCM6 | 1342 | 32 | 16792 | 1.173062593 | 1 | 0.999999468 | 100 |
| GOTERM_BP_DIRECT | GO:0006260~DNA replication | 9 | 0.571791614 | 0.93469029 | POLD3, TOP1, KAT7, MCM7, SET, NFIC, MCM4, MCM6, RPA3 | 1342 | 155 | 16792 | 0.726541993 | 1 | 1 | 100 |

Annotation Cluster 123 Enrichment Score: 0.4783612608011878

| Category | Term | Count | % | PValue | Genes | List Total | Pop Hits | Pop Total | Fold Enrichment | Bonferroni | Benjamini | FDR |
|---|---|---|---|---|---|---|---|---|---|---|---|---|
| REACTOME_PATHWAY | R-HSA-210993:R-HSA-210993 | 4 | 0.254129606 | 0.179024176 | DOK2, HRAS, PIK3CB, PTPN11 | 745 | 18 | 9075 | 2.706935123 | 1 | 0.96863751 | 95.29589998 |
| KEGG_PATHWAY | hsa05211:Renal cell carcinoma | 8 | 0.508259212 | 0.309510066 | HRAS, PAK2, PIK3CB, VEGFA, RAP1A, HGF, AKT3, PTPN11 | 574 | 66 | 6879 | 1.452644916 | 1 | 0.811110384 | 99.25104605 |
| REACTOME_PATHWAY | R-HSA-186763:R-HSA-186763 | 4 | 0.254129606 | 0.428718955 | HRAS, PLCG1, PIK3CB, PTPN11 | 745 | 29 | 9075 | 1.680166628 | 1 | 0.995251408 | 99.98293202 |
| GOTERM_BP_DIRECT | GO:0050900~leukocyte migration | 11 | 0.698856417 | 0.513802023 | DOK2, HRAS, THBD, AIMP1, PLCG1, ATP1B2, PIK3CB, SELL, SPN, FN1, PTPN11 | 1342 | 122 | 16792 | 1.128191346 | 1 | 0.9999437 | 99.99985174 |

Annotation Cluster 124 Enrichment Score: 0.46983772989602895

| Category | Term | Count | % | PValue | Genes | List Total | Pop Hits | Pop Total | Fold Enrichment | Bonferroni | Benjamini | FDR |
|---|---|---|---|---|---|---|---|---|---|---|---|---|
| UP_SEQ_FEATURE | domain:FH2 | 3 | 0.190597205 | 0.309852953 | FMN2, DIAPH1, FHDC1 | 1499 | 15 | 20063 | 2.676851234 | 1 | 0.999751312 | 99.89973676 |
| INTERPRO | IPR015425:Actin-binding FH2 | 3 | 0.190597205 | 0.329946531 | FMN2, DIAPH1, FHDC1 | 1452 | 15 | 18559 | 2.556336088 | 1 | 0.997808954 | 99.89898659 |
| SMART | SM00498:FH2 | 3 | 0.190597205 | 0.380967216 | FMN2, DIAPH1, FHDC1 | 878 | 15 | 10057 | 2.290888383 | 1 | 0.961720177 | 99.87998128 |

Annotation Cluster 125 Enrichment Score: 0.4666930913746874

| Category | Term | Count | % | PValue | Genes | List Total | Pop Hits | Pop Total | Fold Enrichment | Bonferroni | Benjamini | FDR |
|---|---|---|---|---|---|---|---|---|---|---|---|---|
| INTERPRO | IPR008983:Tumour necrosis factor-like domain | 8 | 0.508259212 | 0.135496435 | CBLN3, FASLG, C1QL2, TNFSF12-TNFSF13, TNFSF12, LTA, COL10A1, CAPRIN2 | 1452 | 55 | 18559 | 1.859153519 | 1 | 0.974168847 | 91.8589893 |
| SMART | SM00207:TNF | 4 | 0.254129606 | 0.158188967 | FASLG, TNFSF12-TNFSF13, TNFSF12, LTA | 878 | 16 | 10057 | 2.863610478 | 1 | 0.907643213 | 91.06073261 |
| INTERPRO | IPR006052:Tumour necrosis factor | 4 | 0.254129606 | 0.16204075 | FASLG, TNFSF12-TNFSF13, TNFSF12, LTA | 1452 | 18 | 18559 | 2.840373431 | 1 | 0.97906801 | 95.24273789 |
| REACTOME_PATHWAY | R-HSA-5676594:R-HSA-5676594 | 4 | 0.254129606 | 0.179024176 | TNFSF12-TNFSF13, TNFSF12, CD40, LTA | 745 | 18 | 9075 | 2.706935123 | 1 | 0.96863751 | 95.29589998 |
| INTERPRO | IPR021184:Tumour necrosis factor, conserved site | 3 | 0.190597205 | 0.240343159 | FASLG, TNFSF12-TNFSF13, LTA | 1452 | 12 | 18559 | 3.19542011 | 1 | 0.993507746 | 99.12220937 |
| GOTERM_BP_DIRECT | GO:0097191~extrinsic apoptotic signaling pathway | 6 | 0.381194409 | 0.24197365 | SMAD3, FASLG, IL6R, TNFSF12-TNFSF13, TNFSF12, WWOX | 1342 | 42 | 16792 | 1.787523951 | 1 | 0.996860818 | 99.42362248 |
| GOTERM_MF_DIRECT | GO:0005164~tumor necrosis factor receptor binding | 4 | 0.254129606 | 0.407310579 | TRIM37, FASLG, TNFSF12, LTA | 1340 | 29 | 16881 | 1.737622234 | 1 | 0.994563544 | 99.98024206 |
| GOTERM_BP_DIRECT | GO:0097190~apoptotic signaling pathway | 7 | 0.444726811 | 0.504892839 | MAP3K5, FASLG, TNFSF12-TNFSF13, TNFSF12, CD40, SPN, HIP1 | 1342 | 71 | 16792 | 1.23364329 | 1 | 0.99993671 | 99.99979213 |
| UP_SEQ_FEATURE | site:Cleavage | 6 | 0.381194409 | 0.606649789 | FASLG, PDGFC, MUC5AC, TNFSF12-TNFSF13, TNFSF12, MUC17 | 1499 | 70 | 20063 | 1.147221957 | 1 | 0.999999599 | 99.99999715 |
| REACTOME_PATHWAY | R-HSA-5668541:R-HSA-5668541 | 6 | 0.381194409 | 0.677914455 | PSMA1, PSMD1, TNFSF12-TNFSF13, TNFSF12, CD40, LTA | 745 | 69 | 9075 | 1.059235483 | 1 | 0.999505482 | 99.99999763 |
| GOTERM_BP_DIRECT | GO:0033209~tumor necrosis factor-mediated signaling pathway | 8 | 0.508259212 | 0.841122043 | PSMA1, TNFRSF13B, PSMD1, FASLG, TNFSF12-TNFSF13, TNFSF12, CD40, LTA | 1342 | 118 | 16792 | 0.848316452 | 1 | 0.999999981 | 100 |
| UP_KEYWORDS | Cytokine | 9 | 0.571791614 | 0.975767193 | BMP10, AIMP1, GDF10, FASLG, TNFSF12-TNFSF13, TNFSF12, CCL28, LTA, THPO | 1535 | 190 | 20581 | 0.635107149 | 1 | 0.998820408 | 100 |

Annotation Cluster 126 Enrichment Score: 0.45951895868265086

| Category | Term | Count | % | PValue | Genes | List Total | Pop Hits | Pop Total | Fold Enrichment | Bonferroni | Benjamini | FDR |
|---|---|---|---|---|---|---|---|---|---|---|---|---|
| GOTERM_BP_DIRECT | GO:0050434~positive regulation of viral transcription | 8 | 0.508259212 | 0.006668158 | POLR2H, MDFIC, NELFCD, SUPT4H1, NELFE, CTDP1, POLR2C, POLR2B | 1342 | 29 | 16792 | 3.451770389 | 1 | 0.797429149 | 11.70812671 |
| REACTOME_PATHWAY | R-HSA-167242:R-HSA-167242 | 7 | 0.444726811 | 0.008945057 | POLR2H, NELFCD, SUPT4H1, NELFE, CTDP1, POLR2C, POLR2B | 745 | 23 | 9075 | 3.70732419 | 0.999559395 | 0.786788846 | 12.99763541 |
| REACTOME_PATHWAY | R-HSA-167158:R-HSA-167158 | 7 | 0.444726811 | 0.048963019 | POLR2H, NELFCD, SUPT4H1, NELFE, CTDP1, POLR2C, POLR2B | 745 | 33 | 9075 | 2.583892617 | 1 | 0.94376787 | 54.06430063 |
| REACTOME_PATHWAY | R-HSA-113418:R-HSA-113418 | 7 | 0.444726811 | 0.048963019 | POLR2H, NELFCD, SUPT4H1, NELFE, CTDP1, POLR2C, POLR2B | 745 | 33 | 9075 | 2.583892617 | 1 | 0.94376787 | 54.06430063 |
| REACTOME_PATHWAY | R-HSA-167238:R-HSA-167238 | 7 | 0.444726811 | 0.070105589 | POLR2H, NELFCD, SUPT4H1, NELFE, CTDP1, POLR2C, POLR2B | 745 | 36 | 9075 | 2.368568233 | 1 | 0.941649821 | 67.57705217 |
| REACTOME_PATHWAY | R-HSA-167243:R-HSA-167243 | 7 | 0.444726811 | 0.070105589 | POLR2H, NELFCD, SUPT4H1, NELFE, CTDP1, POLR2C, POLR2B | 745 | 36 | 9075 | 2.368568233 | 1 | 0.941649821 | 67.57705217 |
| REACTOME_PATHWAY | R-HSA-167287:R-HSA-167287 | 7 | 0.444726811 | 0.086708222 | POLR2H, NELFCD, SUPT4H1, NELFE, CTDP1, POLR2C, POLR2B | 745 | 38 | 9075 | 2.243906747 | 1 | 0.932100066 | 75.47485375 |
| REACTOME_PATHWAY | R-HSA-167290:R-HSA-167290 | 7 | 0.444726811 | 0.086708222 | POLR2H, NELFCD, SUPT4H1, NELFE, CTDP1, POLR2C, POLR2B | 745 | 38 | 9075 | 2.243906747 | 1 | 0.932100066 | 75.47485375 |
| REACTOME_PATHWAY | R-HSA-167246:R-HSA-167246 | 7 | 0.444726811 | 0.171795892 | POLR2H, NELFCD, SUPT4H1, NELFE, CTDP1, POLR2C, POLR2B | 745 | 46 | 9075 | 1.853662095 | 1 | 0.968224804 | 94.61148095 |
| GOTERM_BP_DIRECT | GO:0031047~gene silencing by RNA | 13 | 0.82592122 | 0.174820284 | POLR2H, HIST4H4, NUP88, CNOT2, IPO8, TSN, POLR2C, POLR2B, DGCR8, SND1, NUP35, TNRC6B, HIS | 1342 | 111 | 16792 | 1.465447564 | 1 | 0.991626075 | 97.20219587 |
| GOTERM_MF_DIRECT | GO:0001055~RNA polymerase II activity | 3 | 0.190597205 | 0.185341556 | POLR2H, POLR2C, POLR2B | 1340 | 10 | 16881 | 3.779328358 | 1 | 0.967403101 | 96.46510877 |
| REACTOME_PATHWAY | R-HSA-167200:R-HSA-167200 | 7 | 0.444726811 | 0.197111671 | POLR2H, NELFCD, SUPT4H1, NELFE, CTDP1, POLR2C, POLR2B | 745 | 48 | 9075 | 1.776426174 | 1 | 0.971627683 | 96.66918295 |
| REACTOME_PATHWAY | R-HSA-167152:R-HSA-167152 | 7 | 0.444726811 | 0.223710612 | POLR2H, NELFCD, SUPT4H1, NELFE, CTDP1, POLR2C, POLR2B | 745 | 50 | 9075 | 1.705369128 | 1 | 0.975056644 | 98.02383427 |
| GOTERM_BP_DIRECT | GO:0006368~transcription elongation from RNA polymerase II promoter | 10 | 0.635324015 | 0.247234136 | POLR2H, ELP2, ELOF1, NELFCD, THOC7, SUPT4H1, NELFE, CTDP1, POLR2C, POLR2B | 1342 | 86 | 16792 | 1.454961356 | 1 | 0.997201108 | 99.49368856 |
| REACTOME_PATHWAY | R-HSA-75955:R-HSA-75955 | 7 | 0.444726811 | 0.27997412 | POLR2H, NELFCD, SUPT4H1, NELFE, CTDP1, POLR2C, POLR2B | 745 | 54 | 9075 | 1.579045488 | 1 | 0.984300959 | 99.38413124 |
| REACTOME_PATHWAY | R-HSA-203927:R-HSA-203927 | 4 | 0.254129606 | 0.290805322 | POLR2H, DGCR8, POLR2C, POLR2B | 745 | 23 | 9075 | 2.118470966 | 1 | 0.985326045 | 99.51304972 |
| REACTOME_PATHWAY | R-HSA-112382:R-HSA-112382 | 7 | 0.444726811 | 0.309233672 | POLR2H, NELFCD, SUPT4H1, NELFE, CTDP1, POLR2C, POLR2B | 745 | 56 | 9075 | 1.522651007 | 1 | 0.987200618 | 99.67618249 |
| REACTOME_PATHWAY | R-HSA-452723:R-HSA-452723 | 4 | 0.254129606 | 0.406149504 | POLR2H, POU5F1, POLR2C, POLR2B | 745 | 28 | 9075 | 1.740172579 | 1 | 0.994208576 | 99.96888746 |
| GOTERM_CC_DIRECT | GO:0005665~DNA-directed RNA polymerase II, core complex | 3 | 0.190597205 | 0.416123539 | POLR2H, POLR2C, POLR2B | 1425 | 18 | 18224 | 2.131461988 | 1 | 0.941776784 | 99.96946348 |
| REACTOME_PATHWAY | R-HSA-168325:R-HSA-168325 | 5 | 0.317662008 | 0.473861631 | POLR2H, NUP88, NUP35, POLR2C, POLR2B | 745 | 43 | 9075 | 1.416419541 | 1 | 0.996632729 | 99.99523341 |
| GOTERM_BP_DIRECT | GO:0010467~gene expression | 5 | 0.317662008 | 0.54058622 | NOX4, POLR2H, HNRNPR, POLR2C, POLR2B | 1342 | 48 | 16792 | 1.303402881 | 1 | 0.999960786 | 99.99994836 |
| GOTERM_BP_DIRECT | GO:0006283~transcription-coupled nucleotide-excision repair | 7 | 0.444726811 | 0.546198696 | POLD3, POLR2H, COPS6, COPS4, POLR2C, POLR2B, RPA3 | 1342 | 74 | 16792 | 1.183630725 | 1 | 0.999963138 | 99.99995892 |
| REACTOME_PATHWAY | R-HSA-72203:R-HSA-72203 | 4 | 0.254129606 | 0.594512654 | POLR2H, HNRNPR, POLR2C, POLR2B | 745 | 37 | 9075 | 1.316887357 | 1 | 0.999082287 | 99.99991581 |
| REACTOME_PATHWAY | R-HSA-674695:R-HSA-674695 | 7 | 0.444726811 | 0.62527355 | POLR2H, NELFCD, SUPT4H1, NELFE, CTDP1, POLR2C, POLR2B | 745 | 78 | 9075 | 1.093185338 | 1 | 0.999218062 | 99.99997521 |
| REACTOME_PATHWAY | R-HSA-72165:R-HSA-72165 | 5 | 0.317662008 | 0.626677987 | POLR2H, DDX23, SNRPB, POLR2C, POLR2B | 745 | 52 | 9075 | 1.171270005 | 1 | 0.999191845 | 99.99997661 |
| REACTOME_PATHWAY | R-HSA-5578749:R-HSA-5578749 | 9 | 0.571791614 | 0.648915037 | POLR2H, HIST4H4, NUP88, IPO8, H2AFJ, NUP35, POLR2C, POLR2B, HIST1H3H | 745 | 106 | 9075 | 1.034253514 | 1 | 0.999336896 | 99.99999097 |
| REACTOME_PATHWAY | R-HSA-6781823:R-HSA-6781823 | 5 | 0.317662008 | 0.656549207 | POLR2H, COPS6, COPS4, POLR2C, POLR2B | 745 | 54 | 9075 | 1.127889635 | 1 | 0.999396057 | 99.99999358 |
| REACTOME_PATHWAY | R-HSA-77075:R-HSA-77075 | 3 | 0.190597205 | 0.661676623 | POLR2H, POLR2C, POLR2B | 745 | 27 | 9075 | 1.353467562 | 1 | 0.999422165 | 99.99999491 |
| REACTOME_PATHWAY | R-HSA-167160:R-HSA-167160 | 3 | 0.190597205 | 0.661676623 | POLR2H, POLR2C, POLR2B | 745 | 27 | 9075 | 1.353467562 | 1 | 0.999422165 | 99.99999491 |
| REACTOME_PATHWAY | R-HSA-72086:R-HSA-72086 | 3 | 0.190597205 | 0.700044674 | POLR2H, POLR2C, POLR2B | 745 | 29 | 9075 | 1.260124971 | 1 | 0.999584509 | 99.99999921 |
| REACTOME_PATHWAY | R-HSA-5601884:R-HSA-5601884 | 3 | 0.190597205 | 0.700044674 | POLR2H, POLR2C, POLR2B | 745 | 29 | 9075 | 1.260124971 | 1 | 0.999584509 | 99.99999921 |
| KEGG_PATHWAY | hsa00240:Pyrimidine metabolism | 8 | 0.508259212 | 0.747832645 | POLD3, POLR2H, UPP2, UCKL1, ENTPD4, NT5E, POLR2C, POLR2B | 574 | 101 | 6879 | 0.949253113 | 1 | 0.924199436 | 99.99999876 |
| GOTERM_BP_DIRECT | GO:0006370~7-methylguanosine mRNA capping | 3 | 0.190597205 | 0.752420257 | POLR2H, POLR2C, POLR2B | 1342 | 33 | 16792 | 1.137515242 | 1 | 0.99999965 | 100 |
| KEGG_PATHWAY | hsa03020:RNA polymerase | 3 | 0.190597205 | 0.759021636 | POLR2H, POLR2C, POLR2B | 574 | 32 | 6879 | 1.123530052 | 1 | 0.927873992 | 99.99999932 |
| REACTOME_PATHWAY | R-HSA-6782210:R-HSA-6782210 | 5 | 0.317662008 | 0.791248407 | POLD3, POLR2H, POLR2C, POLR2B, RPA3 | 745 | 65 | 9075 | 0.937016004 | 1 | 0.999895383 | 100 |
| UP_KEYWORDS | DNA-directed RNA polymerase | 3 | 0.190597205 | 0.798375592 | POLR2H, POLR2C, POLR2B | 1535 | 39 | 20581 | 1.031370584 | 1 | 0.969596803 | 99.99999999 |
| REACTOME_PATHWAY | R-HSA-6782135:R-HSA-6782135 | 5 | 0.317662008 | 0.801076949 | POLD3, POLR2H, POLR2C, POLR2B, RPA3 | 745 | 66 | 9075 | 0.922818792 | 1 | 0.999892345 | 100 |
| GOTERM_MF_DIRECT | GO:0003899~DNA-directed RNA polymerase activity | 3 | 0.190597205 | 0.815420863 | POLR2H, POLR2C, POLR2B | 1340 | 38 | 16881 | 0.994560094 | 1 | 0.999967919 | 100 |
| GOTERM_BP_DIRECT | GO:0042795~snRNA transcription from RNA polymerase II promoter | 5 | 0.317662008 | 0.820481035 | POLR2H, INTS8, SNAPC4, POLR2C, POLR2B | 1342 | 70 | 16792 | 0.893761976 | 1 | 0.999999951 | 100 |
| REACTOME_PATHWAY | R-HSA-6781827:R-HSA-6781827 | 3 | 0.190597205 | 0.893717217 | POLR2H, POLR2C, POLR2B | 745 | 45 | 9075 | 0.812080537 | 1 | 0.999988885 | 100 |
| REACTOME_PATHWAY | R-HSA-76042:R-HSA-76042 | 3 | 0.190597205 | 0.900727187 | POLR2H, POLR2C, POLR2B | 745 | 46 | 9075 | 0.794426612 | 1 | 0.99998899 | 100 |
| REACTOME_PATHWAY | R-HSA-167161:R-HSA-167161 | 3 | 0.190597205 | 0.900727187 | POLR2H, POLR2C, POLR2B | 745 | 46 | 9075 | 0.794426612 | 1 | 0.99998899 | 100 |
| REACTOME_PATHWAY | R-HSA-75953:R-HSA-75953 | 3 | 0.190597205 | 0.900727187 | POLR2H, POLR2C, POLR2B | 745 | 46 | 9075 | 0.794426612 | 1 | 0.99998899 | 100 |

| | | | | | | | | | | | | |
|---|---|---|---|---|---|---|---|---|---|---|---|---|
| REACTOME_PATHWAY | R-HSA-167162:R-HSA-167162 | 3 | 0.190597205 | 0.900727187 | POLR2H, POLR2C, POLR2B | 745 | 46 | 9075 | 0.794426612 | 1 | 0.99998899 | 100 |
| REACTOME_PATHWAY | R-HSA-73779:R-HSA-73779 | 3 | 0.190597205 | 0.900727187 | POLR2H, POLR2C, POLR2B | 745 | 46 | 9075 | 0.794426612 | 1 | 0.99998899 | 100 |
| REACTOME_PATHWAY | R-HSA-167172:R-HSA-167172 | 3 | 0.190597205 | 0.900727187 | POLR2H, POLR2C, POLR2B | 745 | 46 | 9075 | 0.794426612 | 1 | 0.99998899 | 100 |
| REACTOME_PATHWAY | R-HSA-73776:R-HSA-73776 | 3 | 0.190597205 | 0.900727187 | POLR2H, POLR2C, POLR2B | 745 | 46 | 9075 | 0.794426612 | 1 | 0.99998899 | 100 |
| Annotation Cluster 127 | Enrichment Score: 0.4590439542675749 | | | | | | | | | | | |
| Category | Term | Count | % | PValue | Genes | List Total | Pop Hits | Pop Total | Fold Enrichment | Bonferroni | Benjamini | FDR |
| REACTOME_PATHWAY | R-HSA-442982:R-HSA-442982 | 5 | 0.317662008 | 0.064564262 | HRAS, GRIN2B, RASGRF2, CAMK2G, CALM2 | 745 | 19 | 9075 | 3.205581067 | 1 | 0.934994132 | 64.4502341 |
| KEGG_PATHWAY | hsa04713:Circadian entrainment | 11 | 0.698856417 | 0.265634372 | NOS1AP, GRIN2B, GRIA2, CAMK2G, RYR3, MTNR1B, GUCY1A2, RYR2, CACNA1C, KCNJ3, CALM2 | 574 | 95 | 6879 | 1.38765817 | 1 | 0.815302238 | 98.30946624 |
| KEGG_PATHWAY | hsa04720:Long-term potentiation | 8 | 0.508259212 | 0.309510066 | HRAS, GRIN2B, GRIA2, CAMK2G, RAP1A, CACNA1C, CALM2, ITPR2 | 574 | 66 | 6879 | 1.452644916 | 1 | 0.811110384 | 99.25104605 |
| REACTOME_PATHWAY | R-HSA-442729:R-HSA-442729 | 3 | 0.190597205 | 0.4118776 | GRIN2B, CAMK2G, CALM2 | 745 | 17 | 9075 | 2.149624951 | 1 | 0.994414386 | 99.97322637 |
| REACTOME_PATHWAY | R-HSA-438066:R-HSA-438066 | 3 | 0.190597205 | 0.4118776 | GRIN2B, GRIA2, CAMK2G | 745 | 17 | 9075 | 2.149624951 | 1 | 0.994414386 | 99.97322637 |
| KEGG_PATHWAY | hsa05031:Amphetamine addiction | 7 | 0.444726811 | 0.474715525 | GRIN2B, GRIA2, CAMK2G, ATF6B, TH, CACNA1C, CALM2 | 574 | 66 | 6879 | 1.271064302 | 1 | 0.873137199 | 99.97981643 |
| KEGG_PATHWAY | hsa04728:Dopaminergic synapse | 11 | 0.698856417 | 0.631856699 | GRIN2B, GRIA2, CAMK2G, DRD4, ATF6B, TH, CACNA1C, KCNJ3, AKT3, CALM2, ITPR2 | 574 | 128 | 6879 | 1.029902548 | 1 | 0.907717951 | 99.99981597 |
| KEGG_PATHWAY | hsa05030:Cocaine addiction | 4 | 0.254129606 | 0.787242896 | GRIN2B, GRIA2, ATF6B, TH | 574 | 49 | 6879 | 0.978311882 | 1 | 0.934962719 | 99.99999987 |
| Annotation Cluster 128 | Enrichment Score: 0.451507632558959 | | | | | | | | | | | |
| Category | Term | Count | % | PValue | Genes | List Total | Pop Hits | Pop Total | Fold Enrichment | Bonferroni | Benjamini | FDR |
| UP_KEYWORDS | Serine protease | 17 | 1.080050826 | 0.060544833 | CELA3A, MASP1, CORIN, CFB, C1S, PCSK2, GZMK, PARL, TPP2, ST14, RELN, DPP8, C2, PCSK5, MBTPS1 | 1535 | 141 | 20581 | 1.616545383 | 1 | 0.333114493 | 59.24523947 |
| UP_SEQ_FEATURE | domain:Peptidase S1 | 12 | 0.762388818 | 0.189976853 | CELA3A, GZMK, MASP1, CORIN, CFB, ST14, C1S, C2, HGF, PRSS35, PRSS12, PRSS21 | 1499 | 109 | 20063 | 1.473496092 | 1 | 0.997418482 | 98.02189334 |
| INTERPRO | IPR001254:Peptidase S1 | 12 | 0.762388818 | 0.3366357 | CELA3A, GZMK, MASP1, CORIN, CFB, ST14, C1S, C2, HGF, PRSS35, PRSS12, PRSS21 | 1452 | 120 | 18559 | 1.278168044 | 1 | 0.99793748 | 99.91502043 |
| INTERPRO | IPR009003:Trypsin-like cysteine/serine peptidase domain | 12 | 0.762388818 | 0.417827738 | CELA3A, GZMK, MASP1, CORIN, CFB, ST14, C1S, C2, HGF, PRSS35, PRSS12, PRSS21 | 1452 | 128 | 18559 | 1.198282541 | 1 | 0.999348224 | 99.9910354 |
| UP_SEQ_FEATURE | active site:Charge relay system | 17 | 1.080050826 | 0.448612486 | CELA3A, MASP1, CORIN, CFB, ESD, C1S, PCSK2, GZMK, TPP2, ST14, DPP8, C2, MBTPS1, PRSS35, PCSK | 1499 | 203 | 20063 | 1.120849039 | 1 | 0.999980098 | 99.99846536 |
| INTERPRO | IPR018114:Peptidase S1, trypsin family, active site | 9 | 0.571791614 | 0.549442631 | CELA3A, GZMK, MASP1, CFB, ST14, C2, PRSS35, PRSS12, PRSS21 | 1452 | 102 | 18559 | 1.127795333 | 1 | 0.999871473 | 99.99989157 |
| GOTERM_MF_DIRECT | GO:0004252~serine-type endopeptidase activity | 21 | 1.334180432 | 0.555032815 | CELA3A, MASP1, C4A, CORIN, C4B, CFB, MMP8, COLEC11, C1S, HGF, PCSK2, IMMP2L, GZMK, PARL, T | 1340 | 255 | 16881 | 1.037462687 | 1 | 0.998568936 | 99.99981564 |
| SMART | SM00020:Tryp_SPc | 11 | 0.698856417 | 0.585862668 | CELA3A, GZMK, MASP1, CORIN, CFB, ST14, C1S, C2, HGF, PRSS12, PRSS21 | 878 | 118 | 10057 | 1.067786958 | 1 | 0.990254673 | 99.99957214 |
| INTERPRO | IPR001314:Peptidase S1A, chymotrypsin-type | 9 | 0.571791614 | 0.666174335 | CELA3A, GZMK, MASP1, CFB, C1S, C2, HGF, PRSS12, PRSS21 | 1452 | 113 | 18559 | 1.018009947 | 1 | 0.999983181 | 99.99999938 |
| Annotation Cluster 129 | Enrichment Score: 0.45115498208065613 | | | | | | | | | | | |
| Category | Term | Count | % | PValue | Genes | List Total | Pop Hits | Pop Total | Fold Enrichment | Bonferroni | Benjamini | FDR |
| GOTERM_CC_DIRECT | GO:0000118~histone deacetylase complex | 7 | 0.444726811 | 0.058221368 | TBL1XR1, HDAC3, CIR1, SAP30L, MECOM, TRERF1, NCOR2 | 1425 | 36 | 18224 | 2.486705653 | 1 | 0.583888683 | 59.43810679 |
| REACTOME_PATHWAY | R-HSA-2122947:R-HSA-2122947 | 7 | 0.444726811 | 0.184281663 | TBL1XR1, HDAC3, MAML1, MAML3, RBPJ, MYC, NCOR2 | 745 | 47 | 9075 | 1.814222476 | 1 | 0.967765171 | 95.74165425 |
| REACTOME_PATHWAY | R-HSA-400206:R-HSA-400206 | 3 | 0.190597205 | 0.496697086 | TBL1XR1, HDAC3, NCOR2 | 745 | 20 | 9075 | 1.827181208 | 1 | 0.997272698 | 99.9976034 |
| REACTOME_PATHWAY | R-HSA-381340:R-HSA-381340 | 7 | 0.444726811 | 0.637766142 | TBL1XR1, HDAC3, MED30, EBF1, MED23, CDK4, NCOR2 | 745 | 79 | 9075 | 1.079347549 | 1 | 0.99930909 | 99.99998534 |
| GOTERM_BP_DIRECT | GO:0044255~cellular lipid metabolic process | 4 | 0.254129606 | 0.646449839 | TBL1XR1, HDAC3, FAM135B, NCOR2 | 1342 | 41 | 16792 | 1.220748064 | 1 | 0.999994926 | 99.99999961 |
| REACTOME_PATHWAY | R-HSA-3214815:R-HSA-3214815 | 6 | 0.381194409 | 0.893753329 | TBL1XR1, HDAC3, HIST4H4, SAP30L, NCOR2, HIST1H3H | 745 | 94 | 9075 | 0.777523918 | 1 | 0.999988134 | 100 |
| Annotation Cluster 130 | Enrichment Score: 0.4503864849165277 | | | | | | | | | | | |
| Category | Term | Count | % | PValue | Genes | List Total | Pop Hits | Pop Total | Fold Enrichment | Bonferroni | Benjamini | FDR |
| INTERPRO | IPR008983:Tumour necrosis factor-like domain | 8 | 0.508259212 | 0.135496435 | CBLN3, FASLG, C1QL2, TNFSF12-TNFSF13, TNFSF12, LTA, COL10A1, CAPRIN2 | 1452 | 55 | 18559 | 1.859153519 | 1 | 0.974168847 | 91.8589893 |
| UP_SEQ_FEATURE | domain:C1q | 4 | 0.254129606 | 0.431467168 | CBLN3, C1QL2, COL10A1, CAPRIN2 | 1499 | 32 | 20063 | 1.673032021 | 1 | 0.999972527 | 99.99728592 |
| INTERPRO | IPR001073:Complement C1q protein | 4 | 0.254129606 | 0.5028588 | CBLN3, C1QL2, COL10A1, CAPRIN2 | 1452 | 34 | 18559 | 1.503727111 | 1 | 0.999766402 | 99.99940947 |
| SMART | SM00110:C1Q | 4 | 0.254129606 | 0.53719552 | CBLN3, C1QL2, COL10A1, CAPRIN2 | 878 | 32 | 10057 | 1.431805239 | 1 | 0.985110005 | 99.99796789 |
| Annotation Cluster 131 | Enrichment Score: 0.44816168253866084 | | | | | | | | | | | |
| Category | Term | Count | % | PValue | Genes | List Total | Pop Hits | Pop Total | Fold Enrichment | Bonferroni | Benjamini | FDR |
| REACTOME_PATHWAY | R-HSA-5674135:R-HSA-5674135 | 6 | 0.381194409 | 0.227483216 | WDR83, HRAS, ARRB1, RAP1A, APBB1IP, FN1 | 745 | 40 | 9075 | 1.827181208 | 1 | 0.973716361 | 98.1675235 |
| REACTOME_PATHWAY | R-HSA-354194:R-HSA-354194 | 3 | 0.190597205 | 0.351569214 | RAP1A, APBB1IP, FN1 | 745 | 15 | 9075 | 2.436241611 | 1 | 0.990504147 | 99.87847443 |
| REACTOME_PATHWAY | R-HSA-372708:R-HSA-372708 | 3 | 0.190597205 | 0.351569214 | RAP1A, APBB1IP, FN1 | 745 | 15 | 9075 | 2.436241611 | 1 | 0.990504147 | 99.87847443 |
| REACTOME_PATHWAY | R-HSA-354192:R-HSA-354192 | 3 | 0.190597205 | 0.573300387 | RAP1A, APBB1IP, FN1 | 745 | 23 | 9075 | 1.588853224 | 1 | 0.998865833 | 99.99981446 |
| Annotation Cluster 132 | Enrichment Score: 0.4460029406474936 | | | | | | | | | | | |
| Category | Term | Count | % | PValue | Genes | List Total | Pop Hits | Pop Total | Fold Enrichment | Bonferroni | Benjamini | FDR |
| UP_KEYWORDS | Zymogen | 20 | 1.27064803 | 0.24591753 | CELA3A, CORIN, ADAMTS14, CFB, MMP8, MMP27, MMP16, PCSK2, GGT5, GZMK, CASP4, CASP9, BA | 1535 | 214 | 20581 | 1.253067064 | 1 | 0.677389287 | 98.26906801 |
| UP_KEYWORDS | Protease | 45 | 2.858958069 | 0.308723355 | CPO, METAP2, ADAMTS14, CORIN, MASP1, USP2, MMP8, MMP27, C1S, ENPEP, PCSK2, IMMP2L, API | 1535 | 544 | 20581 | 1.109102558 | 1 | 0.73259953 | 99.50398103 |
| GOTERM_BP_DIRECT | GO:0006508~proteolysis | 43 | 2.731893266 | 0.390220968 | CPO, ADAMTSL1, ADAMTS14, MASP1, UBE3A, MMP8, DPP10, MMP27, C1S, PCSK2, CASP4, CASP9, P | 1342 | 500 | 16792 | 1.076089419 | 1 | 0.999593805 | 99.98996145 |
| GOTERM_MF_DIRECT | GO:0004252~serine-type endopeptidase activity | 21 | 1.334180432 | 0.555032815 | CELA3A, MASP1, C4A, CORIN, C4B, CFB, MMP8, COLEC11, C1S, HGF, PCSK2, IMMP2L, GZMK, PARL, T | 1340 | 255 | 16881 | 1.037462687 | 1 | 0.998568936 | 99.99981564 |
| Annotation Cluster 133 | Enrichment Score: 0.44362994472418454 | | | | | | | | | | | |
| Category | Term | Count | % | PValue | Genes | List Total | Pop Hits | Pop Total | Fold Enrichment | Bonferroni | Benjamini | FDR |
| INTERPRO | IPR013806:Kringle-like fold | 5 | 0.317662008 | 0.187653892 | MRC2, ROR1, HGF, PRSS12, FN1 | 1452 | 29 | 18559 | 2.203738007 | 1 | 0.985407354 | 97.21320178 |
| UP_KEYWORDS | Kringle | 3 | 0.190597205 | 0.392480578 | ROR1, HGF, PRSS12 | 1535 | 18 | 20581 | 2.234636265 | 1 | 0.803507742 | 99.92249038 |
| INTERPRO | IPR000001:Kringle | 3 | 0.190597205 | 0.416411215 | ROR1, HGF, PRSS12 | 1452 | 18 | 18559 | 2.130280073 | 1 | 0.999358607 | 99.99065213 |
| INTERPRO | IPR018056:Kringle, conserved site | 3 | 0.190597205 | 0.416411215 | ROR1, HGF, PRSS12 | 1452 | 18 | 18559 | 2.130280073 | 1 | 0.999358607 | 99.99065213 |
| SMART | SM00130:KR | 3 | 0.190597205 | 0.473837792 | ROR1, HGF, PRSS12 | 878 | 18 | 10057 | 1.909073652 | 1 | 0.98013861 | 99.9877164 |
| Annotation Cluster 134 | Enrichment Score: 0.4353944338273869 | | | | | | | | | | | |
| Category | Term | Count | % | PValue | Genes | List Total | Pop Hits | Pop Total | Fold Enrichment | Bonferroni | Benjamini | FDR |
| UP_SEQ_FEATURE | domain:SAM | 9 | 0.571791614 | 0.278740594 | SAMD7, L3MBTL4, MAGI1, SAMD12, TP63, EPHB3, EPHB1, TP73, TNKS2 | 1499 | 83 | 20063 | 1.451304886 | 1 | 0.999583206 | 99.77211996 |
| INTERPRO | IPR013761:Sterile alpha motif/pointed domain | 12 | 0.762388818 | 0.306929653 | SAMD7, MAP3K5, L3MBTL4, FLI1, SAMD12, PPFIBP1, TP63, TFCP2, EPHB3, EPHB1, TP73, TNKS2 | 1452 | 117 | 18559 | 1.310941584 | 1 | 0.997049796 | 99.81925639 |
| INTERPRO | IPR001660:Sterile alpha motif domain | 9 | 0.571791614 | 0.455342884 | SAMD7, L3MBTL4, SAMD12, PPFIBP1, TP63, EPHB3, EPHB1, TP73, TNKS2 | 1452 | 94 | 18559 | 1.223777915 | 1 | 0.999557956 | 99.99715435 |
| SMART | SM00454:SAM | 9 | 0.571791614 | 0.465419571 | SAMD7, L3MBTL4, SAMD12, PPFIBP1, TP63, EPHB3, EPHB1, TP73, TNKS2 | 878 | 85 | 10057 | 1.212823261 | 1 | 0.980511471 | 99.98465417 |
| Annotation Cluster 135 | Enrichment Score: 0.43147775865383137 | | | | | | | | | | | |
| Category | Term | Count | % | PValue | Genes | List Total | Pop Hits | Pop Total | Fold Enrichment | Bonferroni | Benjamini | FDR |
| BIOCARTA | h_bard1Pathway:BRCA1-dependent Ub-ligase activity | 3 | 0.190597205 | 0.155150437 | FANCD2, FANCE, FANCC | 146 | 8 | 1625 | 4.17380137 | 1 | 0.995461208 | 88.37899547 |
| UP_KEYWORDS | Fanconi anemia | 4 | 0.254129606 | 0.164574729 | RAD51C, FANCD2, FANCE, FANCC | 1535 | 19 | 20581 | 2.82269844 | 1 | 0.559929057 | 92.45487267 |
| BIOCARTA | h_atrbrcaPathway:Role of BRCA1, BRCA2 and ATR in Cancer Susceptibility | 4 | 0.254129606 | 0.312403953 | FANCD2, FANCE, CHEK2, FANCC | 146 | 22 | 1625 | 2.02366127 | 1 | 0.997503486 | 99.16179727 |
| REACTOME_PATHWAY | R-HSA-6783310:R-HSA-6783310 | 4 | 0.254129606 | 0.648053597 | FANCD2, FANCE, FANCC, RPA3 | 745 | 40 | 9075 | 1.218120805 | 1 | 0.999364623 | 99.99999062 |
| KEGG_PATHWAY | hsa03460:Fanconi anemia pathway | 5 | 0.317662008 | 0.654743534 | RAD51C, FANCD2, FANCE, FANCC, RPA3 | 574 | 53 | 6879 | 1.130596279 | 1 | 0.908626745 | 99.9999212 |
| GOTERM_BP_DIRECT | GO:0036297~interstrand cross-link repair | 4 | 0.254129606 | 0.761410101 | FANCD2, FANCE, FANCC, RPA3 | 1342 | 49 | 16792 | 1.021442258 | 1 | 0.999999734 | 100 |
| Annotation Cluster 136 | Enrichment Score: 0.42528916970144476 | | | | | | | | | | | |
| Category | Term | Count | % | PValue | Genes | List Total | Pop Hits | Pop Total | Fold Enrichment | Bonferroni | Benjamini | FDR |
| GOTERM_MF_DIRECT | GO:0032266~phosphatidylinositol-3-phosphate binding | 5 | 0.317662008 | 0.246175994 | BBS5, SESTD1, SNX27, PHLDA3, HIP1 | 1340 | 32 | 16881 | 1.968400187 | 1 | 0.983149223 | 99.00281491 |

| | | | | | | | | | | | | |
|---|---|---|---|---|---|---|---|---|---|---|---|---|
| GOTERM_MF_DIRECT | GO:0080025~phosphatidylinositol-3,5-bisphosphate binding | 4 | 0.254129606 | 0.273703815 | SESTD1, CLVS2, PHLDA3, HIP1 | 1340 | 23 | 16881 | 2.19091499 | 1 | 0.987991546 | 99.45633131 |
| GOTERM_MF_DIRECT | GO:0005546~phosphatidylinositol-4,5-bisphosphate binding | 6 | 0.381194409 | 0.49169244 | SESTD1, ASAP1, SYTL2, KCNQ1, PHLDA3, HIP1 | 1340 | 58 | 16881 | 1.303216675 | 1 | 0.997091856 | 99.99838517 |
| GOTERM_MF_DIRECT | GO:0043325~phosphatidylinositol-3,4-bisphosphate binding | 3 | 0.190597205 | 0.600652431 | SESTD1, PHLDA3, HIP1 | 1340 | 25 | 16881 | 1.511731343 | 1 | 0.998995446 | 99.9999684 |

Annotation Cluster 137 Enrichment Score: 0.4185769503242083

| Category | Term | Count | % | PValue | Genes | List Total | Pop Hits | Pop Total | Fold Enrichment | Bonferroni | Benjamini | FDR |
|---|---|---|---|---|---|---|---|---|---|---|---|---|
| INTERPRO | IPR000242:Protein-tyrosine phosphatase, receptor/non-receptor type | 6 | 0.381194409 | 0.17248874 | PTPRE, PTPRM, PTPN18, PTPRT, PTPRO, PTPN11 | 1452 | 38 | 18559 | 2.01816007 | 1 | 0.981128987 | 96.16745643 |
| UP_KEYWORDS | Protein phosphatase | 14 | 0.889453621 | 0.1919481 | PDP1, PTPRM, PTPRE, PTPN18, PTPRT, PTPRO, PTPN11, PPM1D, PPM1E, DUSP14, PPM1H, CTDP1, M | 1535 | 133 | 20581 | 1.41134922 | 1 | 0.608648108 | 95.32556269 |
| SMART | SM00194:PTPc | 6 | 0.381194409 | 0.217221473 | PTPRE, PTPRM, PTPN18, PTPRT, PTPRO, PTPN11 | 878 | 37 | 10057 | 1.857477067 | 1 | 0.911069431 | 96.77504913 |
| UP_SEQ_FEATURE | domain:Tyrosine-protein phosphatase 1 | 3 | 0.190597205 | 0.224314344 | PTPRE, PTPRM, PTPRT | 1499 | 12 | 20063 | 3.346064043 | 1 | 0.998506684 | 99.11695651 |
| UP_SEQ_FEATURE | domain:Tyrosine-protein phosphatase 2 | 3 | 0.190597205 | 0.224314344 | PTPRE, PTPRM, PTPRT | 1499 | 12 | 20063 | 3.346064043 | 1 | 0.998506684 | 99.11695651 |
| GOTERM_BP_DIRECT | GO:0006470~protein dephosphorylation | 13 | 0.82592122 | 0.305249469 | CTTNBP2NL, PPM1D, PTPRE, PTPRM, PTPN18, CPPED1, PPM1H, CAMK2G, PTPRT, PTPRO, CTDP1, M | 1342 | 126 | 16792 | 1.29098952 | 1 | 0.998861269 | 99.88619818 |
| INTERPRO | IPR016130:Protein-tyrosine phosphatase, active site | 8 | 0.508259212 | 0.383844206 | PTPRE, PTPRM, PTPN18, DUSP14, PTPRT, PTPRO, MTMR4, PTPN11 | 1452 | 76 | 18559 | 1.345440046 | 1 | 0.999074931 | 99.97617744 |
| GOTERM_MF_DIRECT | GO:0005001~transmembrane receptor protein tyrosine phosphatase activity | 3 | 0.190597205 | 0.395270313 | PTPRE, PTPRM, PTPRO | 1340 | 17 | 16881 | 2.223134328 | 1 | 0.993861065 | 99.9725742 |
| INTERPRO | IPR000387:Protein-tyrosine/Dual specificity phosphatase | 8 | 0.508259212 | 0.46216047 | PTPRE, PTPRM, PTPN18, DUSP14, PTPRT, PTPRO, MTMR4, PTPN11 | 1452 | 82 | 18559 | 1.246993214 | 1 | 0.999584859 | 99.99770944 |
| UP_SEQ_FEATURE | active site:Phosphocysteine intermediate | 8 | 0.508259212 | 0.463711499 | PTPRE, PTPRM, PTPN18, DUSP14, PTPRT, PTPRO, MTMR4, PTPN11 | 1499 | 86 | 20063 | 1.245047086 | 1 | 0.999986047 | 99.99908487 |
| INTERPRO | IPR003595:Protein-tyrosine phosphatase, catalytic | 6 | 0.381194409 | 0.55334423 | PTPRE, PTPRM, PTPN18, PTPRT, PTPRO, PTPN11 | 1452 | 63 | 18559 | 1.217302899 | 1 | 0.999877493 | 99.99990666 |
| SMART | SM00404:PTPc_motif | 6 | 0.381194409 | 0.652910078 | PTPRE, PTPRM, PTPN18, PTPRT, PTPRO, PTPN11 | 878 | 63 | 10057 | 1.09089923 | 1 | 0.993195969 | 99.99996405 |
| GOTERM_BP_DIRECT | GO:0035335~peptidyl-tyrosine dephosphorylation | 8 | 0.508259212 | 0.685671688 | PTPRE, PTPRM, PTPN18, DUSP14, PTPRT, PTPRO, MTMR4, PTPN11 | 1342 | 99 | 16792 | 1.011124659 | 1 | 0.999998058 | 99.99999996 |
| GOTERM_MF_DIRECT | GO:0004725~protein tyrosine phosphatase activity | 8 | 0.508259212 | 0.688908056 | PTPRE, PTPRM, PTPN18, DUSP14, PTPRT, PTPRO, MTMR4, PTPN11 | 1340 | 100 | 16881 | 1.007820896 | 1 | 0.999674468 | 99.99999946 |
| UP_SEQ_FEATURE | domain:Tyrosine-protein phosphatase | 4 | 0.254129606 | 0.858842095 | PTPN18, DUSP14, PTPRO, PTPN11 | 1499 | 63 | 20063 | 0.849794043 | 1 | 1 | 100 |

Annotation Cluster 138 Enrichment Score: 0.4146799212955179

| Category | Term | Count | % | PValue | Genes | List Total | Pop Hits | Pop Total | Fold Enrichment | Bonferroni | Benjamini | FDR |
|---|---|---|---|---|---|---|---|---|---|---|---|---|
| GOTERM_BP_DIRECT | GO:0006298~mismatch repair | 5 | 0.317662008 | 0.304824482 | POLD3, MSH2, TDG, TP73, RPA3 | 1342 | 35 | 16792 | 1.787523951 | 1 | 0.998883596 | 99.88489553 |
| REACTOME_PATHWAY | R-HSA-5358565:R-HSA-5358565 | 3 | 0.190597205 | 0.351569214 | POLD3, MSH2, RPA3 | 745 | 15 | 9075 | 2.436241611 | 1 | 0.990504147 | 99.87847443 |
| REACTOME_PATHWAY | R-HSA-5358606:R-HSA-5358606 | 3 | 0.190597205 | 0.351569214 | POLD3, MSH2, RPA3 | 745 | 15 | 9075 | 2.436241611 | 1 | 0.990504147 | 99.87847443 |
| KEGG_PATHWAY | hsa03430:Mismatch repair | 3 | 0.190597205 | 0.582383141 | POLD3, MSH2, RPA3 | 574 | 23 | 6879 | 1.563172247 | 1 | 0.899571001 | 99.999026 |

Annotation Cluster 139 Enrichment Score: 0.4008607145149934

| Category | Term | Count | % | PValue | Genes | List Total | Pop Hits | Pop Total | Fold Enrichment | Bonferroni | Benjamini | FDR |
|---|---|---|---|---|---|---|---|---|---|---|---|---|
| GOTERM_BP_DIRECT | GO:0000086~G2/M transition of mitotic cell cycle | 15 | 0.952986023 | 0.204414223 | CEP72, BRSK2, ALMS1, CHEK2, PPM1D, CEP57, CLASP1, PAFAH1B1, CDK5RAP2, FBXL15, DYNC1I2, M | 1342 | 137 | 16792 | 1.370000109 | 1 | 0.994388351 | 98.58223106 |
| REACTOME_PATHWAY | R-HSA-380270:R-HSA-380270 | 9 | 0.571791614 | 0.307537019 | CEP72, TUBGCP3, TUBGCP5, CEP57, ALMS1, PAFAH1B1, CLASP1, CDK5RAP2, DYNC1I2 | 745 | 78 | 9075 | 1.405524006 | 1 | 0.987594177 | 99.6636359 |
| REACTOME_PATHWAY | R-HSA-5620912:R-HSA-5620912 | 10 | 0.635324015 | 0.412337549 | CEP72, SCLT1, CEP57, CEP89, AHI1, ALMS1, PAFAH1B1, CLASP1, CDK5RAP2, DYNC1I2 | 745 | 98 | 9075 | 1.242980414 | 1 | 0.994123734 | 99.973549 |
| REACTOME_PATHWAY | R-HSA-380259:R-HSA-380259 | 7 | 0.444726811 | 0.517245716 | CEP72, CEP57, ALMS1, PAFAH1B1, CLASP1, CDK5RAP2, DYNC1I2 | 745 | 70 | 9075 | 1.218120805 | 1 | 0.997972434 | 99.99874377 |
| REACTOME_PATHWAY | R-HSA-2565942:R-HSA-2565942 | 7 | 0.444726811 | 0.738478966 | CEP72, CEP57, ALMS1, PAFAH1B1, CLASP1, CDK5RAP2, DYNC1I2 | 745 | 88 | 9075 | 0.968959732 | 1 | 0.999751066 | 99.99999991 |

Annotation Cluster 140 Enrichment Score: 0.3954059612213853

| Category | Term | Count | % | PValue | Genes | List Total | Pop Hits | Pop Total | Fold Enrichment | Bonferroni | Benjamini | FDR |
|---|---|---|---|---|---|---|---|---|---|---|---|---|
| UP_KEYWORDS | Integrin | 7 | 0.444726811 | 0.115464906 | ITGB1BP1, ITGAE, ITGA8, ITGA11, ADAM33, ADAM12, ITGBL1 | 1535 | 45 | 20581 | 2.085660514 | 1 | 0.474542568 | 82.85268699 |
| INTERPRO | IPR000413:Integrin alpha chain | 3 | 0.190597205 | 0.416411215 | ITGAE, ITGA8, ITGA11 | 1452 | 18 | 18559 | 2.130280073 | 1 | 0.999358607 | 99.99065213 |
| INTERPRO | IPR013649:Integrin alpha-2 | 3 | 0.190597205 | 0.416411215 | ITGAE, ITGA8, ITGA11 | 1452 | 18 | 18559 | 2.130280073 | 1 | 0.999358607 | 99.99065213 |
| UP_SEQ_FEATURE | repeat:FG-GAP 7 | 3 | 0.190597205 | 0.420182925 | ITGAE, ITGA8, ITGA11 | 1499 | 19 | 20063 | 2.113303606 | 1 | 0.999963936 | 99.99608664 |
| UP_SEQ_FEATURE | repeat:FG-GAP 6 | 3 | 0.190597205 | 0.420182925 | ITGAE, ITGA8, ITGA11 | 1499 | 19 | 20063 | 2.113303606 | 1 | 0.999963936 | 99.99608664 |
| UP_SEQ_FEATURE | repeat:FG-GAP 5 | 3 | 0.190597205 | 0.420182925 | ITGAE, ITGA8, ITGA11 | 1499 | 19 | 20063 | 2.113303606 | 1 | 0.999963936 | 99.99608664 |
| UP_SEQ_FEATURE | repeat:FG-GAP 4 | 3 | 0.190597205 | 0.420182925 | ITGAE, ITGA8, ITGA11 | 1499 | 19 | 20063 | 2.113303606 | 1 | 0.999963936 | 99.99608664 |
| UP_SEQ_FEATURE | repeat:FG-GAP 3 | 3 | 0.190597205 | 0.420182925 | ITGAE, ITGA8, ITGA11 | 1499 | 19 | 20063 | 2.113303606 | 1 | 0.999963936 | 99.99608664 |
| UP_SEQ_FEATURE | repeat:FG-GAP 2 | 3 | 0.190597205 | 0.420182925 | ITGAE, ITGA8, ITGA11 | 1499 | 19 | 20063 | 2.113303606 | 1 | 0.999963936 | 99.99608664 |
| UP_SEQ_FEATURE | repeat:FG-GAP 1 | 3 | 0.190597205 | 0.420182925 | ITGAE, ITGA8, ITGA11 | 1499 | 19 | 20063 | 2.113303606 | 1 | 0.999963936 | 99.99608664 |
| INTERPRO | IPR013517:FG-GAP repeat | 3 | 0.190597205 | 0.443998406 | ITGAE, ITGA8, ITGA11 | 1452 | 19 | 18559 | 2.01816007 | 1 | 0.999477294 | 99.99594106 |
| INTERPRO | IPR013519:Integrin alpha beta-propellor | 3 | 0.190597205 | 0.443998406 | ITGAE, ITGA8, ITGA11 | 1452 | 19 | 18559 | 2.01816007 | 1 | 0.999477294 | 99.99594106 |
| SMART | SM00191:Int_alpha | 3 | 0.190597205 | 0.502888201 | ITGAE, ITGA8, ITGA11 | 878 | 19 | 10057 | 1.808596092 | 1 | 0.984136235 | 99.99446081 |
| GOTERM_CC_DIRECT | GO:0008305~integrin complex | 3 | 0.190597205 | 0.63462713 | ITGAE, ITGA8, ITGA11 | 1425 | 27 | 18224 | 1.420974659 | 1 | 0.982902201 | 99.99997356 |

Annotation Cluster 141 Enrichment Score: 0.3940018247061543

| Category | Term | Count | % | PValue | Genes | List Total | Pop Hits | Pop Total | Fold Enrichment | Bonferroni | Benjamini | FDR |
|---|---|---|---|---|---|---|---|---|---|---|---|---|
| REACTOME_PATHWAY | R-HSA-3299685:R-HSA-3299685 | 6 | 0.381194409 | 0.141723066 | NOX4, CYBA, GPX6, GPX5, CAT, SOD2 | 745 | 34 | 9075 | 2.149624951 | 1 | 0.956253548 | 90.63514575 |
| UP_KEYWORDS | Peroxidase | 3 | 0.190597205 | 0.565568794 | GPX6, GPX5, CAT | 1535 | 25 | 20581 | 1.608938111 | 1 | 0.892881507 | 99.99937447 |
| GOTERM_BP_DIRECT | GO:0098869~cellular oxidant detoxification | 5 | 0.317662008 | 0.820481035 | GPX6, GPX5, TXNRD3, CAT, TP53INP1 | 1342 | 70 | 16792 | 0.893761976 | 1 | 0.999999951 | 100 |

Annotation Cluster 142 Enrichment Score: 0.3912930254166313

| Category | Term | Count | % | PValue | Genes | List Total | Pop Hits | Pop Total | Fold Enrichment | Bonferroni | Benjamini | FDR |
|---|---|---|---|---|---|---|---|---|---|---|---|---|
| UP_SEQ_FEATURE | domain:CARD | 4 | 0.254129606 | 0.369613412 | BCL10, CASP4, CASP9, CARD10 | 1499 | 29 | 20063 | 1.846104299 | 1 | 0.999908924 | 99.98143299 |
| INTERPRO | IPR001315:Caspase Recruitment | 4 | 0.254129606 | 0.398138933 | BCL10, CASP4, CASP9, CARD10 | 1452 | 29 | 18559 | 1.762990406 | 1 | 0.999231959 | 99.98410096 |
| INTERPRO | IPR011029:Death-like domain | 9 | 0.571791614 | 0.455342884 | BCL10, NLRP5, CASP4, CASP9, ANK3, NLRP9, UNC5C, ANKDD1A, CARD10 | 1452 | 94 | 18559 | 1.223777915 | 1 | 0.999557956 | 99.99715435 |

Annotation Cluster 143 Enrichment Score: 0.3907997685543554

| Category | Term | Count | % | PValue | Genes | List Total | Pop Hits | Pop Total | Fold Enrichment | Bonferroni | Benjamini | FDR |
|---|---|---|---|---|---|---|---|---|---|---|---|---|
| UP_KEYWORDS | Helicase | 14 | 0.889453621 | 0.268407569 | XRCC5, BTAF1, CHD8, MCM7, DDX23, SKIV2L, DDX25, DDX24, DDX60L, EIF4A1, DDX10, DHX30, MCM | 1535 | 143 | 20581 | 1.31265347 | 1 | 0.694319978 | 98.87980117 |
| GOTERM_MF_DIRECT | GO:0003724~RNA helicase activity | 3 | 0.190597205 | 0.275975666 | SKIV2L, DDX24, DDX10 | 1340 | 13 | 16881 | 2.90717566 | 1 | 0.987941205 | 99.48340702 |
| UP_SEQ_FEATURE | short sequence motif:DEAD box | 5 | 0.317662008 | 0.280328105 | DDX23, DDX25, DDX24, EIF4A1, DDX10 | 1499 | 36 | 20063 | 1.858924468 | 1 | 0.999586083 | 99.78128014 |
| INTERPRO | IPR011545:DNA/RNA helicase, DEAD/DEAH box type, N-terminal | 8 | 0.508259212 | 0.293286734 | DDX23, SKIV2L, DDX25, DDX24, DDX60L, EIF4A1, DDX10, DHX30 | 1452 | 69 | 18559 | 1.481933964 | 1 | 0.996315698 | 99.74712501 |
| INTERPRO | IPR014014:RNA helicase, DEAD-box type, Q motif | 5 | 0.317662008 | 0.32759312 | DDX23, DDX25, DDX24, EIF4A1, DDX10 | 1452 | 37 | 18559 | 1.727254114 | 1 | 0.997792157 | 99.89269734 |
| UP_SEQ_FEATURE | short sequence motif:Q motif | 5 | 0.317662008 | 0.349511909 | DDX23, DDX25, DDX24, EIF4A1, DDX10 | 1499 | 40 | 20063 | 1.673032021 | 1 | 0.999868478 | 99.96669013 |
| INTERPRO | IPR000629:RNA helicase, ATP-dependent, DEAD-box, conserved site | 4 | 0.254129606 | 0.354431796 | DDX23, DDX24, EIF4A1, DDX10 | 1452 | 27 | 18559 | 1.893582288 | 1 | 0.998492474 | 99.94680587 |
| GOTERM_MF_DIRECT | GO:0004004~ATP-dependent RNA helicase activity | 7 | 0.444726811 | 0.412435231 | DDX23, SKIV2L, DDX25, DDX24, EIF4A1, DDX10, DHX30 | 1340 | 65 | 16881 | 1.356681975 | 1 | 0.994594102 | 99.98285075 |
| UP_SEQ_FEATURE | domain:Helicase C-terminal | 10 | 0.635324015 | 0.416635584 | BTAF1, CHD8, DDX23, SKIV2L, DDX25, DDX24, DDX60L, EIF4A1, DDX10, DHX30 | 1499 | 108 | 20063 | 1.239282979 | 1 | 0.999963194 | 99.99561598 |
| GOTERM_BP_DIRECT | GO:0010501~RNA secondary structure unwinding | 5 | 0.317662008 | 0.471057999 | DDX23, DDX25, DDX24, EIF4A1, DDX10 | 1342 | 44 | 16792 | 1.421894052 | 1 | 0.999891641 | 99.99928858 |
| INTERPRO | IPR001650:Helicase, C-terminal | 10 | 0.635324015 | 0.471773324 | BTAF1, CHD8, DDX23, SKIV2L, DDX25, DDX24, DDX60L, EIF4A1, DDX10, DHX30 | 1452 | 108 | 18559 | 1.18348893 | 1 | 0.999598474 | 99.99832115 |
| UP_SEQ_FEATURE | domain:Helicase ATP-binding | 10 | 0.635324015 | 0.481408147 | BTAF1, CHD8, DDX23, SKIV2L, DDX25, DDX24, DDX60L, EIF4A1, DDX10, DHX30 | 1499 | 114 | 20063 | 1.174057559 | 1 | 0.999990629 | 99.99951006 |
| INTERPRO | IPR014001:Helicase, superfamily 1/2, ATP-binding domain | 10 | 0.635324015 | 0.494174807 | BTAF1, CHD8, DDX23, SKIV2L, DDX25, DDX24, DDX60L, EIF4A1, DDX10, DHX30 | 1452 | 110 | 18559 | 1.161970949 | 1 | 0.99973921 | 99.99920421 |
| SMART | SM00490:HELICc | 10 | 0.635324015 | 0.595662636 | BTAF1, CHD8, DDX23, SKIV2L, DDX25, DDX24, DDX60L, EIF4A1, DDX10, DHX30 | 878 | 107 | 10057 | 1.07050859 | 1 | 0.990880231 | 99.99969418 |
| SMART | SM00487:DEXDc | 10 | 0.635324015 | 0.618314891 | BTAF1, CHD8, DDX23, SKIV2L, DDX25, DDX24, DDX60L, EIF4A1, DDX10, DHX30 | 878 | 109 | 10057 | 1.050866231 | 1 | 0.991899234 | 99.99986375 |
| GOTERM_MF_DIRECT | GO:0004386~helicase activity | 7 | 0.444726811 | 0.675656074 | BTAF1, DDX23, SKIV2L, DDX24, DDX60L, EIF4A1, DDX10 | 1340 | 85 | 16881 | 1.037462687 | 1 | 0.999617743 | 99.99999894 |

Annotation Cluster 144 Enrichment Score: 0.39076687996886567

| Category | Term | Count | % | PValue | Genes | List Total | Pop Hits | Pop Total | Fold Enrichment | Bonferroni | Benjamini | FDR |
|---|---|---|---|---|---|---|---|---|---|---|---|---|
| INTERPRO | IPR002073:3'5'-cyclic nucleotide phosphodiesterase, catalytic domain | 4 | 0.254129606 | 0.244762321 | PDE3B, PDE10A, PDE8A, RUNX1 | 1452 | 22 | 18559 | 2.323941898 | 1 | 0.99332032 | 99.20614539 |
| GOTERM_MF_DIRECT | GO:0004114~3',5'-cyclic-nucleotide phosphodiesterase activity | 4 | 0.254129606 | 0.273703815 | PDE3B, PDE10A, PDE8A, RUNX1 | 1340 | 23 | 16881 | 2.19091499 | 1 | 0.987991546 | 99.45633131 |
| GOTERM_MF_DIRECT | GO:0004115~3',5'-cyclic-AMP phosphodiesterase activity | 3 | 0.190597205 | 0.336423204 | PDE3B, PDE10A, PDE8A | 1340 | 15 | 16881 | 2.519552239 | 1 | 0.992081501 | 99.87532631 |
| GOTERM_BP_DIRECT | GO:0006198~cAMP catabolic process | 3 | 0.190597205 | 0.339492422 | PDE3B, PDE10A, PDE8A | 1342 | 15 | 16792 | 2.502533532 | 1 | 0.999174642 | 99.95557782 |
| UP_SEQ_FEATURE | metal ion-binding site:Divalent metal cation 2 | 4 | 0.254129606 | 0.348675573 | CPPED1, PDE3B, PDE10A, PDE8A | 1499 | 28 | 20063 | 1.912036596 | 1 | 0.999870952 | 99.96588361 |
| UP_SEQ_FEATURE | metal ion-binding site:Divalent metal cation 1 | 4 | 0.254129606 | 0.348675573 | CPPED1, PDE3B, PDE10A, PDE8A | 1499 | 28 | 20063 | 1.912036596 | 1 | 0.999870952 | 99.96588361 |
| INTERPRO | IPR023174:3'5'-cyclic nucleotide phosphodiesterase, conserved site | 3 | 0.190597205 | 0.496890596 | PDE3B, PDE10A, PDE8A | 1452 | 21 | 18559 | 1.825954349 | 1 | 0.99974334 | 99.99927469 |
| UP_KEYWORDS | cAMP | 4 | 0.254129606 | 0.508913961 | PDE3B, PDE10A, CNGB1, PDE8A | 1535 | 36 | 20581 | 1.48975751 | 1 | 0.865120233 | 99.99635785 |
| INTERPRO | IPR003607:HD/PDEase domain | 3 | 0.190597205 | 0.546452177 | PDE3B, PDE10A, PDE8A | 1452 | 23 | 18559 | 1.66717571 | 1 | 0.999868223 | 99.99987849 |
| SMART | SM00471:HDc | 3 | 0.190597205 | 0.608205302 | PDE3B, PDE10A, PDE8A | 878 | 23 | 10057 | 1.494057641 | 1 | 0.991280747 | 99.99980341 |
| KEGG_PATHWAY | hsa05032:Morphine addiction | 8 | 0.508259212 | 0.643387072 | GABRA2, GABRG3, ARRB1, PDE3B, PDE10A, GABBR2, PDE8A, KCNJ3 | 574 | 91 | 6879 | 1.053566642 | 1 | 0.90539238 | 99.99987915 |

Annotation Cluster 145 Enrichment Score: 0.3804835154339204

| Category | Term | Count | % | PValue | Genes | List Total | Pop Hits | Pop Total | Fold Enrichment | Bonferroni | Benjamini | FDR |
|---|---|---|---|---|---|---|---|---|---|---|---|---|
| REACTOME_PATHWAY | R-HSA-3000171:R-HSA-3000171 | 6 | 0.381194409 | 0.227483216 | COL3A1, NRXN1, FGF2, COL5A1, COL10A1, FN1 | 745 | 40 | 9075 | 1.827181208 | 1 | 0.973716361 | 98.1675235 |
| REACTOME_PATHWAY | R-HSA-1474244:R-HSA-1474244 | 3 | 0.190597205 | 0.351569214 | COL3A1, COL5A1, FN1 | 745 | 15 | 9075 | 2.436241611 | 1 | 0.990504147 | 99.87847443 |
| REACTOME_PATHWAY | R-HSA-3000170:R-HSA-3000170 | 4 | 0.254129606 | 0.383322497 | COL3A1, FGF2, COL5A1, FN1 | 745 | 27 | 9075 | 1.804623415 | 1 | 0.992910838 | 99.94418221 |
| KEGG_PATHWAY | hsa05146:Amoebiasis | 5 | 0.317662008 | 0.980711874 | C8B, PIK3CB, COL3A1, COL5A1, FN1 | 574 | 106 | 6879 | 0.56529814 | 1 | 0.996055796 | 100 |

Annotation Cluster 146 Enrichment Score: 0.3738148189948011

| Category | Term | Count | % | PValue | Genes | List Total | Pop Hits | Pop Total | Fold Enrichment | Bonferroni | Benjamini | FDR |
|---|---|---|---|---|---|---|---|---|---|---|---|---|
| BIOCARTA | h_fasPathway:FAS signaling pathway ( CD95 ) | 5 | 0.317662008 | 0.318139266 | PAK2, DFFA, FASLG, RB1, ARHGDIB | 146 | 32 | 1625 | 1.739083904 | 1 | 0.9967151 | 99.24680188 |
| BIOCARTA | h_hivnefPathway:HIV-I Nef: negative effector of Fas and TNF | 7 | 0.444726811 | 0.466049828 | MAP3K5, PAK2, CASP9, DFFA, FASLG, RB1, ARHGDIB | 146 | 61 | 1625 | 1.277228834 | 1 | 0.998319409 | 99.66679583 |
| BIOCARTA | h_tnfr1Pathway:TNFR1 Signaling Pathway | 4 | 0.254129606 | 0.509924543 | PAK2, DFFA, RB1, ARHGDIB | 146 | 30 | 1625 | 1.484018265 | 1 | 0.998714355 | 99.9888877 |

Annotation Cluster 147 Enrichment Score: 0.3719109851658836

| Category | Term | Count | % | PValue | Genes | List Total | Pop Hits | Pop Total | Fold Enrichment | Bonferroni | Benjamini | FDR |
|---|---|---|---|---|---|---|---|---|---|---|---|---|
| INTERPRO | IPR016160:Aldehyde dehydrogenase, conserved site | 3 | 0.190597205 | 0.359292003 | ALDH1L1, ALDH2, ALDH3B2 | 1452 | 16 | 18559 | 2.396565083 | 1 | 0.998586361 | 99.95329916 |
| GOTERM_MF_DIRECT | GO:0004029~aldehyde dehydrogenase (NAD) activity | 3 | 0.190597205 | 0.395270313 | ALDH1L1, ALDH2, ALDH3B2 | 1340 | 17 | 16881 | 2.223134328 | 1 | 0.993861065 | 99.9725742 |
| INTERPRO | IPR016162:Aldehyde dehydrogenase, N-terminal | 3 | 0.190597205 | 0.443998406 | ALDH1L1, ALDH2, ALDH3B2 | 1452 | 19 | 18559 | 2.01816007 | 1 | 0.999477294 | 99.99594106 |
| INTERPRO | IPR016161:Aldehyde/histidinol dehydrogenase | 3 | 0.190597205 | 0.443998406 | ALDH1L1, ALDH2, ALDH3B2 | 1452 | 19 | 18559 | 2.01816007 | 1 | 0.999477294 | 99.99594106 |
| INTERPRO | IPR015590:Aldehyde dehydrogenase domain | 3 | 0.190597205 | 0.443998406 | ALDH1L1, ALDH2, ALDH3B2 | 1452 | 19 | 18559 | 2.01816007 | 1 | 0.999477294 | 99.99594106 |
| INTERPRO | IPR016163:Aldehyde dehydrogenase, C-terminal | 3 | 0.190597205 | 0.443998406 | ALDH1L1, ALDH2, ALDH3B2 | 1452 | 19 | 18559 | 2.01816007 | 1 | 0.999477294 | 99.99594106 |
| GOTERM_MF_DIRECT | GO:0016620~oxidoreductase activity, acting on the aldehyde or oxo group of donors, NAD or NADP | 3 | 0.190597205 | 0.451601849 | ALDH1L1, ALDH2, ALDH3B2 | 1340 | 19 | 16881 | 1.989120189 | 1 | 0.995827395 | 99.99443168 |

Annotation Cluster 148 Enrichment Score: 0.3712115136900533

| Category | Term | Count | % | PValue | Genes | List Total | Pop Hits | Pop Total | Fold Enrichment | Bonferroni | Benjamini | FDR |
|---|---|---|---|---|---|---|---|---|---|---|---|---|
| INTERPRO | IPR008197:Whey acidic protein-type 4-disulphide core | 4 | 0.254129606 | 0.142873304 | WFDC10A, WFDC10B, EPPIN, EPPIN-WFDC6 | 1452 | 17 | 18559 | 3.007454221 | 1 | 0.975007669 | 92.97634335 |
| GOTERM_BP_DIRECT | GO:0010466~negative regulation of peptidase activity | 4 | 0.254129606 | 0.14961134 | WFDC10B, EPPIN, EPPIN-WFDC6, PI16 | 1342 | 17 | 16792 | 2.944157097 | 1 | 0.98677382 | 95.10160843 |
| INTERPRO | IPR020901:Proteinase inhibitor I2, Kunitz, conserved site | 3 | 0.190597205 | 0.416411215 | APP, EPPIN, EPPIN-WFDC6 | 1452 | 18 | 18559 | 2.130280073 | 1 | 0.999358607 | 99.99065213 |
| INTERPRO | IPR002223:Proteinase inhibitor I2, Kunitz metazoa | 3 | 0.190597205 | 0.443998406 | APP, EPPIN, EPPIN-WFDC6 | 1452 | 19 | 18559 | 2.01816007 | 1 | 0.999477294 | 99.99594106 |
| SMART | SM00131:KU | 3 | 0.190597205 | 0.473837792 | APP, EPPIN, EPPIN-WFDC6 | 878 | 18 | 10057 | 1.909073652 | 1 | 0.98013861 | 99.9877164 |
| UP_KEYWORDS | Serine protease inhibitor | 6 | 0.381194409 | 0.777418849 | WFDC10A, APP, CPAMD8, EPPIN, SERPINB12, EPPIN-WFDC6 | 1535 | 86 | 20581 | 0.935429134 | 1 | 0.963365367 | 99.99999996 |
| UP_KEYWORDS | Protease inhibitor | 8 | 0.508259212 | 0.825661959 | WFDC10A, APP, CPAMD8, EPPIN, SERPINB12, EPPIN-WFDC6, TIMP2, PI16 | 1535 | 124 | 20581 | 0.86502049 | 1 | 0.97595749 | 100 |
| GOTERM_MF_DIRECT | GO:0004867~serine-type endopeptidase inhibitor activity | 6 | 0.381194409 | 0.892075114 | WFDC10A, APP, CPAMD8, EPPIN, SERPINB12, EPPIN-WFDC6 | 1340 | 97 | 16881 | 0.77924296 | 1 | 0.99999667 | 100 |

Annotation Cluster 149 Enrichment Score: 0.37091118068039514

| Category | Term | Count | % | PValue | Genes | List Total | Pop Hits | Pop Total | Fold Enrichment | Bonferroni | Benjamini | FDR |
|---|---|---|---|---|---|---|---|---|---|---|---|---|
| UP_SEQ_FEATURE | domain:PI-PLC Y-box | 3 | 0.190597205 | 0.309852953 | PLCL1, PLCG1, PLCH1 | 1499 | 15 | 20063 | 2.676851234 | 1 | 0.999751312 | 99.89973676 |
| INTERPRO | IPR001711:Phospholipase C, phosphatidylinositol-specific, Y domain | 3 | 0.190597205 | 0.329946531 | PLCL1, PLCG1, PLCH1 | 1452 | 15 | 18559 | 2.556336088 | 1 | 0.997808954 | 99.89898659 |
| INTERPRO | IPR001192:Phosphoinositide phospholipase C | 3 | 0.190597205 | 0.329946531 | PLCL1, PLCG1, PLCH1 | 1452 | 15 | 18559 | 2.556336088 | 1 | 0.997808954 | 99.89898659 |
| SMART | SM00149:PLCYc | 3 | 0.190597205 | 0.380967216 | PLCL1, PLCG1, PLCH1 | 878 | 15 | 10057 | 2.290888383 | 1 | 0.961720177 | 99.87998128 |
| UP_SEQ_FEATURE | domain:PI-PLC X-box | 3 | 0.190597205 | 0.420182925 | PLCL1, PLCG1, PLCH1 | 1499 | 19 | 20063 | 2.113303606 | 1 | 0.999963936 | 99.99608664 |
| INTERPRO | IPR000909:Phospholipase C, phosphatidylinositol-specific , X domain | 3 | 0.190597205 | 0.47084235 | PLCL1, PLCG1, PLCH1 | 1452 | 20 | 18559 | 1.917252066 | 1 | 0.999608657 | 99.99826944 |
| SMART | SM00148:PLCXc | 3 | 0.190597205 | 0.473837792 | PLCL1, PLCG1, PLCH1 | 878 | 18 | 10057 | 1.909073652 | 1 | 0.98013861 | 99.9877164 |
| INTERPRO | IPR017946:PLC-like phosphodiesterase, TIM beta/alpha-barrel domain | 3 | 0.190597205 | 0.592489406 | PLCL1, PLCG1, PLCH1 | 1452 | 25 | 18559 | 1.533801653 | 1 | 0.999936269 | 99.99998078 |
| GOTERM_MF_DIRECT | GO:0004435~phosphatidylinositol phospholipase C activity | 3 | 0.190597205 | 0.643050074 | PLCL1, PLCG1, PLCH1 | 1340 | 27 | 16881 | 1.399751244 | 1 | 0.999414249 | 99.99999493 |

Annotation Cluster 150 Enrichment Score: 0.3690484914761442

| Category | Term | Count | % | PValue | Genes | List Total | Pop Hits | Pop Total | Fold Enrichment | Bonferroni | Benjamini | FDR |
|---|---|---|---|---|---|---|---|---|---|---|---|---|
| GOTERM_BP_DIRECT | GO:0035235~ionotropic glutamate receptor signaling pathway | 4 | 0.254129606 | 0.299616985 | APP, GRIN2B, GRIA2, GRID1 | 1342 | 24 | 16792 | 2.08544461 | 1 | 0.998830821 | 99.86774394 |
| INTERPRO | IPR019594:Glutamate receptor, L-glutamate/glycine-binding | 3 | 0.190597205 | 0.416411215 | GRIN2B, GRIA2, GRID1 | 1452 | 18 | 18559 | 2.130280073 | 1 | 0.999358607 | 99.99065213 |
| INTERPRO | IPR001320:Ionotropic glutamate receptor | 3 | 0.190597205 | 0.416411215 | GRIN2B, GRIA2, GRID1 | 1452 | 18 | 18559 | 2.130280073 | 1 | 0.999358607 | 99.99065213 |
| INTERPRO | IPR001508:NMDA receptor | 3 | 0.190597205 | 0.416411215 | GRIN2B, GRIA2, GRID1 | 1452 | 18 | 18559 | 2.130280073 | 1 | 0.999358607 | 99.99065213 |
| GOTERM_MF_DIRECT | GO:0005234~extracellular-glutamate-gated ion channel activity | 3 | 0.190597205 | 0.423794435 | GRIN2B, GRIA2, GRID1 | 1340 | 18 | 16881 | 2.099626866 | 1 | 0.994962 | 99.9875262 |
| SMART | SM00918:SM00918 | 3 | 0.190597205 | 0.473837792 | GRIN2B, GRIA2, GRID1 | 878 | 18 | 10057 | 1.909073652 | 1 | 0.98013861 | 99.9877164 |
| SMART | SM00079:PBPe | 3 | 0.190597205 | 0.473837792 | GRIN2B, GRIA2, GRID1 | 878 | 18 | 10057 | 1.909073652 | 1 | 0.98013861 | 99.9877164 |
| INTERPRO | IPR001828:Extracellular ligand-binding receptor | 4 | 0.254129606 | 0.542080452 | GRIN2B, GRIA2, GABBR2, GRID1 | 1452 | 36 | 18559 | 1.420186716 | 1 | 0.999860201 | 99.99985665 |

Annotation Cluster 151 Enrichment Score: 0.36003025561822616

| Category | Term | Count | % | PValue | Genes | List Total | Pop Hits | Pop Total | Fold Enrichment | Bonferroni | Benjamini | FDR |
|---|---|---|---|---|---|---|---|---|---|---|---|---|
| REACTOME_PATHWAY | R-HSA-983168:R-HSA-983168 | 29 | 1.842439644 | 0.299989283 | UBE3A, LMO7, UBE3D, ZNRF1, KLHL2, ZNRF2, FBXW9, RNF123, KLHL25, TPP2, PSMD1, KBTBD8, FBX[illegible] | 745 | 308 | 9075 | 1.146931927 | 1 | 0.986701481 | 99.60210722 |
| GOTERM_MF_DIRECT | GO:0016874~ligase activity | 25 | 1.588310038 | 0.309665143 | XRCC4, UBE3A, UBE3D, TTLL6, CBLL1, ZNRF1, TTLL1, TRIML1, ZNRF2, ZFP91, RNF123, RNF103, HECT[illegible] | 1340 | 270 | 16881 | 1.16645937 | 1 | 0.991071036 | 99.76246667 |
| GOTERM_MF_DIRECT | GO:0004842~ubiquitin-protein transferase activity | 29 | 1.842439644 | 0.389133044 | UBE3A, LMO7, CBLL1, ZNRF1, KLHL2, ZNRF2, ZFP91, RNF103, KBTBD8, KLHL24, HECTD4, FBXO24, KL[illegible] | 1340 | 329 | 16881 | 1.110440956 | 1 | 0.993722034 | 99.96766549 |
| GOTERM_BP_DIRECT | GO:0016567~protein ubiquitination | 30 | 1.905972046 | 0.504195505 | LMO7, UBE3D, CBLL1, KLHL2, RAB40B, TRIML1, RNF123, RNF103, KLHL25, KBTBD8, KLHL24, NEURL1[illegible] | 1342 | 359 | 16792 | 1.045626824 | 1 | 0.999937556 | 99.99978661 |
| UP_KEYWORDS | Ligase | 27 | 1.715374841 | 0.520656104 | UBE3A, FARS2, UBE3D, TTLL6, CBLL1, TTLL7, ZNRF1, TTLL1, TRIML1, ZNRF2, ZFP91, RNF123, RNF103[illegible] | 1535 | 347 | 20581 | 1.043259582 | 1 | 0.871025688 | 99.9974278 |
| UP_KEYWORDS | Ubl conjugation pathway | 48 | 3.049555273 | 0.728739583 | USP2, UBE3A, UBE3D, CBLL1, ZNRF1, RAB40B, KLHL2, TRIML1, ZNRF2, FBXW9, ZFP91, RNF123, RNF1[illegible] | 1535 | 680 | 20581 | 0.946434183 | 1 | 0.951876615 | 99.99999928 |

Annotation Cluster 152 Enrichment Score: 0.3558091086448179

| Category | Term | Count | % | PValue | Genes | List Total | Pop Hits | Pop Total | Fold Enrichment | Bonferroni | Benjamini | FDR |
|---|---|---|---|---|---|---|---|---|---|---|---|---|
| GOTERM_BP_DIRECT | GO:0007156~homophilic cell adhesion via plasma membrane adhesion molecules | 19 | 1.207115629 | 0.079979839 | ME3, PTPRM, PIK3CB, SDK1, PCDH9, DSCAML1, CDHR4, PCDH15, CDHR5, PTPRT, CDH3, CDH12, CDH[illegible] | 1342 | 158 | 16792 | 1.504687883 | 1 | 0.971558266 | 78.80648277 |
| INTERPRO | IPR014868:Cadherin prodomain | 3 | 0.190597205 | 0.124818096 | CDH13, DSC1, CDH3 | 1452 | 8 | 18559 | 4.793130165 | 1 | 0.969089745 | 89.94168933 |
| SMART | SM01055:SM01055 | 3 | 0.190597205 | 0.149671168 | CDH13, DSC1, CDH3 | 878 | 8 | 10057 | 4.295415718 | 1 | 0.909556504 | 89.70530327 |

| | | | | | | | | | | | | |
|---|---|---|---|---|---|---|---|---|---|---|---|---|
| INTERPRO | IPR027397:Catenin binding domain | 4 | 0.254129606 | 0.398138933 | CDH12, DSC1, CDH3, TCF7L2 | 1452 | 29 | 18559 | 1.762990406 | 1 | 0.999231959 | 99.98410096 |
| UP_SEQ_FEATURE | domain:Cadherin 7 | 3 | 0.190597205 | 0.420182925 | PCDH9, PCDH15, CDH23 | 1499 | 19 | 20063 | 2.113303606 | 1 | 0.999963936 | 99.99608664 |
| INTERPRO | IPR020894:Cadherin conserved site | 10 | 0.635324015 | 0.527268068 | CDH12, CDH13, ME3, PCDH9, CDHR4, PCDH15, CDHR5, DSC1, CDH3, CDH23 | 1452 | 113 | 18559 | 1.131122163 | 1 | 0.999826386 | 99.99975193 |
| INTERPRO | IPR000233:Cadherin, cytoplasmic domain | 3 | 0.190597205 | 0.546452177 | CDH12, DSC1, CDH3 | 1452 | 23 | 18559 | 1.66717571 | 1 | 0.999868223 | 99.99987849 |
| INTERPRO | IPR002126:Cadherin | 10 | 0.635324015 | 0.58060392 | CDH12, CDH13, ME3, PCDH9, CDHR4, PCDH15, CDHR5, DSC1, CDH3, CDH23 | 1452 | 118 | 18559 | 1.083193258 | 1 | 0.999921425 | 99.99996845 |
| INTERPRO | IPR015919:Cadherin-like | 10 | 0.635324015 | 0.601161144 | CDH12, CDH13, ME3, PCDH9, CDHR4, PCDH15, CDHR5, DSC1, CDH3, CDH23 | 1452 | 120 | 18559 | 1.065140037 | 1 | 0.99994392 | 99.99998673 |
| UP_SEQ_FEATURE | domain:Cadherin 3 | 9 | 0.571791614 | 0.603932035 | CDH12, CDH13, PCDH9, CDHR4, PCDH15, CDHR5, DSC1, CDH3, CDH23 | 1499 | 112 | 20063 | 1.075520585 | 1 | 0.999999624 | 99.99999676 |
| UP_SEQ_FEATURE | domain:Cadherin 4 | 9 | 0.571791614 | 0.603932035 | CDH12, CDH13, PCDH9, CDHR4, PCDH15, CDHR5, DSC1, CDH3, CDH23 | 1499 | 112 | 20063 | 1.075520585 | 1 | 0.999999624 | 99.99999676 |
| UP_SEQ_FEATURE | domain:Cadherin 1 | 9 | 0.571791614 | 0.633967762 | CDH12, CDH13, PCDH9, CDHR4, PCDH15, CDHR5, DSC1, CDH3, CDH23 | 1499 | 115 | 20063 | 1.047463526 | 1 | 0.99999978 | 99.99999925 |
| UP_SEQ_FEATURE | domain:Cadherin 2 | 9 | 0.571791614 | 0.633967762 | CDH12, CDH13, PCDH9, CDHR4, PCDH15, CDHR5, DSC1, CDH3, CDH23 | 1499 | 115 | 20063 | 1.047463526 | 1 | 0.99999978 | 99.99999925 |
| SMART | SM00112:CA | 10 | 0.635324015 | 0.692049033 | CDH12, CDH13, ME3, PCDH9, CDHR4, PCDH15, CDHR5, DSC1, CDH3, CDH23 | 878 | 116 | 10057 | 0.987451889 | 1 | 0.994483031 | 99.99999328 |
| UP_SEQ_FEATURE | domain:Cadherin 5 | 7 | 0.444726811 | 0.804999773 | CDH12, CDH13, PCDH9, PCDH15, DSC1, CDH3, CDH23 | 1499 | 105 | 20063 | 0.892283745 | 1 | 1 | 100 |
| UP_SEQ_FEATURE | domain:Cadherin 6 | 3 | 0.190597205 | 0.987841772 | PCDH9, PCDH15, CDH23 | 1499 | 83 | 20063 | 0.483768295 | 1 | 1 | 100 |

Annotation Cluster 153 Enrichment Score: 0.35088031068691933

| Category | Term | Count | % | PValue | Genes | List Total | Pop Hits | Pop Total | Fold Enrichment | Bonferroni | Benjamini | FDR |
|---|---|---|---|---|---|---|---|---|---|---|---|---|
| GOTERM_CC_DIRECT | GO:0005777~peroxisome | 12 | 0.762388818 | 0.188309972 | HAO1, FAR1, TRIM37, MYO5A, TMEM173, MLYCD, ABCD4, CAT, HSD17B4, URAD, ACAD11, AOC1 | 1425 | 104 | 18224 | 1.47562753 | 1 | 0.840802277 | 95.66511512 |
| UP_KEYWORDS | Peroxisome | 9 | 0.571791614 | 0.436827242 | HAO1, FAR1, TRIM37, MLYCD, ABCD4, CAT, HSD17B4, URAD, ACAD11 | 1535 | 97 | 20581 | 1.244024312 | 1 | 0.822006378 | 99.97392316 |
| GOTERM_CC_DIRECT | GO:0005782~peroxisomal matrix | 5 | 0.317662008 | 0.488868741 | HAO1, FAR1, MLYCD, CAT, HSD17B4 | 1425 | 46 | 18224 | 1.390083905 | 1 | 0.961106973 | 99.99587409 |
| KEGG_PATHWAY | hsa04146:Peroxisome | 8 | 0.508259212 | 0.544158085 | HAO1, FAR1, MLYCD, ABCD4, CAT, HSD17B4, ACSL5, SOD2 | 574 | 83 | 6879 | 1.155115234 | 1 | 0.893176925 | 99.99690108 |
| GOTERM_CC_DIRECT | GO:0005778~peroxisomal membrane | 4 | 0.254129606 | 0.804448151 | FAR1, ABCD4, CAT, HSD17B4 | 1425 | 54 | 18224 | 0.947316439 | 1 | 0.99717662 | 100 |

Annotation Cluster 154 Enrichment Score: 0.34817999441228853

| Category | Term | Count | % | PValue | Genes | List Total | Pop Hits | Pop Total | Fold Enrichment | Bonferroni | Benjamini | FDR |
|---|---|---|---|---|---|---|---|---|---|---|---|---|
| UP_KEYWORDS | Transcription | 188 | 11.94409149 | 0.25087894 | XRCC5, MMS19, FHIT, CDX1, TMEM18, ZNF821, CNOT2, ZNF782, MED23, PDX1, RORA, CNOT6, CSNK | 1535 | 2398 | 20581 | 1.051155007 | 1 | 0.682865816 | 98.42573376 |
| UP_KEYWORDS | Transcription regulation | 183 | 11.62642948 | 0.25986923 | XRCC5, MMS19, FHIT, CDX1, ZNF821, CNOT2, ZNF782, MED23, PDX1, RORA, CNOT6, CSNK2A2, BSX, | 1535 | 2332 | 20581 | 1.052157212 | 1 | 0.689064829 | 98.67651618 |
| UP_KEYWORDS | Nucleus | 403 | 25.60355781 | 0.271941351 | XRCC5, FHIT, XRCC4, HRAS, RUSC1, ZNF782, MED23, CLK1, RORA, CIAPIN1, BSX, INTS8, EPC2, FLI1, S | 1535 | 5244 | 20581 | 1.030387202 | 1 | 0.691622387 | 98.95510319 |
| GOTERM_BP_DIRECT | GO:0006351~transcription, DNA-templated | 155 | 9.847522236 | 0.588602364 | MMS19, XRCC5, FHIT, TMEM18, ZNF821, CNOT2, ZNF782, RORA, CNOT6, CSNK2A2, BSX, GATA2, ZN | 1342 | 1955 | 16792 | 0.992052935 | 1 | 0.999982488 | 99.99999338 |
| UP_KEYWORDS | DNA-binding | 144 | 9.14866582 | 0.818822601 | XRCC5, RAD51C, CDX1, RAD51B, TMEM18, ZNF821, ZNF782, PDX1, RORA, BSX, GATA2, ZNF184, TOP | 1535 | 2050 | 20581 | 0.941817431 | 1 | 0.974310119 | 100 |
| GOTERM_BP_DIRECT | GO:0006355~regulation of transcription, DNA-templated | 105 | 6.67090216 | 0.953265538 | FHIT, CDX1, CNOT2, ZNF782, MED23, RORA, CNOT6, CSNK2A2, ZNF184, SND1, SAP30L, NUP35, INSR | 1342 | 1504 | 16792 | 0.87355725 | 1 | 1 | 100 |

Annotation Cluster 155 Enrichment Score: 0.3345056130055581

| Category | Term | Count | % | PValue | Genes | List Total | Pop Hits | Pop Total | Fold Enrichment | Bonferroni | Benjamini | FDR |
|---|---|---|---|---|---|---|---|---|---|---|---|---|
| UP_SEQ_FEATURE | domain:Bromo | 4 | 0.254129606 | 0.369613412 | TRIM66, CECR2, SP110, BRD9 | 1499 | 29 | 20063 | 1.846104299 | 1 | 0.999908924 | 99.98143299 |
| INTERPRO | IPR001487:Bromodomain | 5 | 0.317662008 | 0.436552905 | TRIM66, SP100, CECR2, SP110, BRD9 | 1452 | 43 | 18559 | 1.486241912 | 1 | 0.999487941 | 99.99489572 |
| SMART | SM00297:BROMO | 5 | 0.317662008 | 0.504616846 | TRIM66, SP100, CECR2, SP110, BRD9 | 878 | 42 | 10057 | 1.363624037 | 1 | 0.98352179 | 99.99472488 |
| UP_KEYWORDS | Bromodomain | 4 | 0.254129606 | 0.563937791 | TRIM66, CECR2, SP110, BRD9 | 1535 | 39 | 20581 | 1.375160778 | 1 | 0.894412621 | 99.99933986 |

Annotation Cluster 156 Enrichment Score: 0.3318758874831108

| Category | Term | Count | % | PValue | Genes | List Total | Pop Hits | Pop Total | Fold Enrichment | Bonferroni | Benjamini | FDR |
|---|---|---|---|---|---|---|---|---|---|---|---|---|
| REACTOME_PATHWAY | R-HSA-1250347:R-HSA-1250347 | 3 | 0.190597205 | 0.320594134 | HRAS, ERBB4, HBEGF | 745 | 14 | 9075 | 2.610258869 | 1 | 0.987398843 | 99.74956138 |
| REACTOME_PATHWAY | R-HSA-1963640:R-HSA-1963640 | 3 | 0.190597205 | 0.382038787 | HRAS, ERBB4, HBEGF | 745 | 16 | 9075 | 2.28397651 | 1 | 0.993175733 | 99.94235427 |
| REACTOME_PATHWAY | R-HSA-1250196:R-HSA-1250196 | 3 | 0.190597205 | 0.469278262 | HRAS, ERBB4, HBEGF | 745 | 19 | 9075 | 1.92334864 | 1 | 0.996570224 | 99.9945477 |
| GOTERM_BP_DIRECT | GO:0038128~ERBB2 signaling pathway | 3 | 0.190597205 | 0.818470409 | HRAS, ERBB4, HBEGF | 1342 | 38 | 16792 | 0.987842184 | 1 | 0.99999995 | 100 |

Annotation Cluster 157 Enrichment Score: 0.32632183792624225

| Category | Term | Count | % | PValue | Genes | List Total | Pop Hits | Pop Total | Fold Enrichment | Bonferroni | Benjamini | FDR |
|---|---|---|---|---|---|---|---|---|---|---|---|---|
| REACTOME_PATHWAY | R-HSA-1810476:R-HSA-1810476 | 4 | 0.254129606 | 0.244976845 | HMGB1, APP, NFKBIB, ZBP1 | 745 | 21 | 9075 | 2.320230105 | 1 | 0.97971426 | 98.71503557 |
| REACTOME_PATHWAY | R-HSA-933542:R-HSA-933542 | 3 | 0.190597205 | 0.596876633 | HMGB1, APP, NFKBIB | 745 | 24 | 9075 | 1.522651007 | 1 | 0.999066042 | 99.9999231 |
| REACTOME_PATHWAY | R-HSA-445989:R-HSA-445989 | 3 | 0.190597205 | 0.7178354 | HMGB1, APP, NFKBIB | 745 | 30 | 9075 | 1.218120805 | 1 | 0.999664852 | 99.99999969 |

Annotation Cluster 158 Enrichment Score: 0.32594719981611664

| Category | Term | Count | % | PValue | Genes | List Total | Pop Hits | Pop Total | Fold Enrichment | Bonferroni | Benjamini | FDR |
|---|---|---|---|---|---|---|---|---|---|---|---|---|
| UP_SEQ_FEATURE | domain:Ricin B-type lectin | 4 | 0.254129606 | 0.327665536 | GALNT3, MRC2, GALNT15, GALNT12 | 1499 | 27 | 20063 | 1.982852766 | 1 | 0.999832316 | 99.93838427 |
| GOTERM_BP_DIRECT | GO:0016266~O-glycan processing | 7 | 0.444726811 | 0.345773522 | GALNT3, B3GNT8, GALNT15, MUC5AC, GALNT12, MUC6, MUC17 | 1342 | 60 | 16792 | 1.459811227 | 1 | 0.999185707 | 99.9628151 |
| REACTOME_PATHWAY | R-HSA-913709:R-HSA-913709 | 7 | 0.444726811 | 0.353964964 | GALNT3, B3GNT8, GALNT15, MUC5AC, GALNT12, MUC6, MUC17 | 745 | 59 | 9075 | 1.445228074 | 1 | 0.990329814 | 99.8852488 |
| UP_SEQ_FEATURE | region of interest:Catalytic subdomain B | 3 | 0.190597205 | 0.446401725 | GALNT3, GALNT15, GALNT12 | 1499 | 20 | 20063 | 2.007638426 | 1 | 0.999979603 | 99.99834666 |
| UP_SEQ_FEATURE | region of interest:Catalytic subdomain A | 3 | 0.190597205 | 0.446401725 | GALNT3, GALNT15, GALNT12 | 1499 | 20 | 20063 | 2.007638426 | 1 | 0.999979603 | 99.99834666 |
| SMART | SM00458:RICIN | 4 | 0.254129606 | 0.446461534 | GALNT3, MRC2, GALNT15, GALNT12 | 878 | 28 | 10057 | 1.636348845 | 1 | 0.977086178 | 99.97498379 |
| GOTERM_MF_DIRECT | GO:0004653~polypeptide N-acetylgalactosaminyltransferase activity | 3 | 0.190597205 | 0.478626757 | GALNT3, GALNT15, GALNT12 | 1340 | 20 | 16881 | 1.889664179 | 1 | 0.996646236 | 99.99755739 |
| INTERPRO | IPR000772:Ricin B lectin domain | 4 | 0.254129606 | 0.5028588 | GALNT3, MRC2, GALNT15, GALNT12 | 1452 | 34 | 18559 | 1.503727111 | 1 | 0.999766402 | 99.99940947 |
| INTERPRO | IPR001173:Glycosyl transferase, family 2 | 3 | 0.190597205 | 0.634941849 | GALNT3, GALNT15, GALNT12 | 1452 | 27 | 18559 | 1.420186716 | 1 | 0.999968065 | 99.99999711 |
| REACTOME_PATHWAY | R-HSA-977068:R-HSA-977068 | 3 | 0.190597205 | 0.641057799 | MUC5AC, MUC6, MUC17 | 745 | 26 | 9075 | 1.405524006 | 1 | 0.999312355 | 99.99998727 |
| KEGG_PATHWAY | hsa00512:Mucin type O-Glycan biosynthesis | 3 | 0.190597205 | 0.74316985 | GALNT3, GALNT15, GALNT12 | 574 | 31 | 6879 | 1.159772957 | 1 | 0.922891378 | 99.99999842 |

Annotation Cluster 159 Enrichment Score: 0.322832226168712

| Category | Term | Count | % | PValue | Genes | List Total | Pop Hits | Pop Total | Fold Enrichment | Bonferroni | Benjamini | FDR |
|---|---|---|---|---|---|---|---|---|---|---|---|---|
| GOTERM_MF_DIRECT | GO:0005160~transforming growth factor beta receptor binding | 6 | 0.381194409 | 0.237690041 | INHBB, BMP10, INHBC, SMAD3, TGFBR3, GDF10 | 1340 | 42 | 16881 | 1.799680171 | 1 | 0.982723485 | 98.80312431 |
| GOTERM_BP_DIRECT | GO:0010862~positive regulation of pathway-restricted SMAD protein phosphorylation | 6 | 0.381194409 | 0.337056282 | INHBB, BMP10, DAB2, RBPMS, INHBC, GDF10 | 1342 | 48 | 16792 | 1.564083458 | 1 | 0.999172051 | 99.95242734 |
| GOTERM_BP_DIRECT | GO:0032924~activin receptor signaling pathway | 3 | 0.190597205 | 0.369341045 | INHBB, BMP10, SMAD3 | 1342 | 16 | 16792 | 2.346125186 | 1 | 0.999452961 | 99.98121486 |
| GOTERM_BP_DIRECT | GO:0043408~regulation of MAPK cascade | 5 | 0.317662008 | 0.416479686 | INHBB, BMP10, INHBC, GDF10, SYNGAP1 | 1342 | 41 | 16792 | 1.525935081 | 1 | 0.999761527 | 99.99557536 |
| INTERPRO | IPR015615:Transforming growth factor-beta-related | 4 | 0.254129606 | 0.461977118 | INHBB, BMP10, INHBC, GDF10 | 1452 | 32 | 18559 | 1.597710055 | 1 | 0.999602753 | 99.99769595 |
| INTERPRO | IPR017948:Transforming growth factor beta, conserved site | 4 | 0.254129606 | 0.461977118 | INHBB, BMP10, INHBC, GDF10 | 1452 | 32 | 18559 | 1.597710055 | 1 | 0.999602753 | 99.99769595 |
| GOTERM_BP_DIRECT | GO:0060395~SMAD protein signal transduction | 6 | 0.381194409 | 0.558275182 | INHBB, BMP10, MAGI2, INHBC, SMAD3, GDF10 | 1342 | 62 | 16792 | 1.210903322 | 1 | 0.999970275 | 99.99997511 |
| INTERPRO | IPR001839:Transforming growth factor-beta, C-terminal | 4 | 0.254129606 | 0.561005805 | INHBB, BMP10, INHBC, GDF10 | 1452 | 37 | 18559 | 1.381803291 | 1 | 0.999888413 | 99.99993072 |
| SMART | SM00204:TGFB | 4 | 0.254129606 | 0.618822903 | INHBB, BMP10, INHBC, GDF10 | 878 | 36 | 10057 | 1.272715768 | 1 | 0.991477502 | 99.99986627 |
| GOTERM_BP_DIRECT | GO:0048468~cell development | 4 | 0.254129606 | 0.629622755 | DCT, INHBB, INHBC, GDF10 | 1342 | 40 | 16792 | 1.251266766 | 1 | 0.999992094 | 99.99999906 |
| GOTERM_MF_DIRECT | GO:0005179~hormone activity | 6 | 0.381194409 | 0.875833 | INHBB, BMP10, INHBC, CRH, EPO, THPO | 1340 | 94 | 16881 | 0.804112417 | 1 | 0.999994654 | 100 |

Annotation Cluster 160 Enrichment Score: 0.32246714098129947

| Category | Term | Count | % | PValue | Genes | List Total | Pop Hits | Pop Total | Fold Enrichment | Bonferroni | Benjamini | FDR |
|---|---|---|---|---|---|---|---|---|---|---|---|---|
| KEGG_PATHWAY | hsa04012:ErbB signaling pathway | 10 | 0.635324015 | 0.298278021 | HRAS, PAK2, ERBB4, PLCG1, PIK3CB, CAMK2G, HBEGF, MYC, AKT3, SHC4 | 574 | 87 | 6879 | 1.37750811 | 1 | 0.8126976 | 99.07303648 |
| KEGG_PATHWAY | hsa05221:Acute myeloid leukemia | 6 | 0.381194409 | 0.504669684 | HRAS, PIK3CB, RUNX1, MYC, TCF7L2, AKT3 | 574 | 56 | 6879 | 1.284034345 | 1 | 0.881222098 | 99.99070994 |
| KEGG_PATHWAY | hsa04917:Prolactin signaling pathway | 6 | 0.381194409 | 0.71609412 | HRAS, PRLR, PIK3CB, TH, AKT3, SHC4 | 574 | 71 | 6879 | 1.012759484 | 1 | 0.921573784 | 99.99999406 |

| Annotation Cluster 161 | Enrichment Score: 0.3193442923364121 | | | | | | | | | | | |
|---|---|---|---|---|---|---|---|---|---|---|---|---|
| Category | Term | Count | % | PValue | Genes | List Total | Pop Hits | Pop Total | Fold Enrichment | Bonferroni | Benjamini | FDR |
| REACTOME_PATHWAY | R-HSA-186763:R-HSA-186763 | 4 | 0.254129606 | 0.428718955 | HRAS, PLCG1, PIK3CB, PTPN11 | 745 | 29 | 9075 | 1.680166628 | 1 | 0.995251408 | 99.98293202 |
| REACTOME_PATHWAY | R-HSA-2424491:R-HSA-2424491 | 4 | 0.254129606 | 0.450974503 | HRAS, PLCG1, PIK3CB, VAV2 | 745 | 30 | 9075 | 1.624161074 | 1 | 0.995854771 | 99.99077926 |
| KEGG_PATHWAY | hsa04650:Natural killer cell mediated cytotoxicity | 11 | 0.698856417 | 0.569695055 | HRAS, PLCG1, PIK3CB, CD247, FASLG, VAV2, SH2D1B, SHC4, KIR2DL4, SH3BP2, PTPN11 | 574 | 122 | 6879 | 1.080553493 | 1 | 0.898141776 | 99.99855345 |

| Annotation Cluster 162 | Enrichment Score: 0.31748774431193869 | | | | | | | | | | | |
|---|---|---|---|---|---|---|---|---|---|---|---|---|
| Category | Term | Count | % | PValue | Genes | List Total | Pop Hits | Pop Total | Fold Enrichment | Bonferroni | Benjamini | FDR |
| UP_KEYWORDS | Glycolysis | 5 | 0.317662008 | 0.197013218 | TPI1, HKDC1, ENO2, PGAM1, OGDH | 1535 | 31 | 20581 | 2.162551224 | 1 | 0.612826513 | 95.72946981 |
| KEGG_PATHWAY | hsa01200:Carbon metabolism | 13 | 0.82592122 | 0.229792476 | ME3, SHMT2, HKDC1, ESD, PGAM1, OGDH, HAO1, MTHFR, TPI1, PHGDH, ENO2, HIBCH, CAT | 574 | 113 | 6879 | 1.378727144 | 1 | 0.816232404 | 96.82667205 |
| GOTERM_BP_DIRECT | GO:0006096~glycolytic process | 5 | 0.317662008 | 0.286367882 | TPI1, HKDC1, ENO2, PGAM1, OGDH | 1342 | 34 | 16792 | 1.840098185 | 1 | 0.998477068 | 99.81254671 |
| REACTOME_PATHWAY | R-HSA-70263:R-HSA-70263 | 4 | 0.254129606 | 0.494350406 | SLC25A12, TPI1, ENO2, PGAM1 | 745 | 32 | 9075 | 1.522651007 | 1 | 0.997324648 | 99.99742427 |
| KEGG_PATHWAY | hsa01230:Biosynthesis of amino acids | 7 | 0.444726811 | 0.561712195 | SHMT2, TPI1, MAT1A, PHGDH, ENO2, PGAM1, PAH | 574 | 72 | 6879 | 1.165142276 | 1 | 0.897575314 | 99.99815571 |
| GOTERM_BP_DIRECT | GO:0061621~canonical glycolysis | 3 | 0.190597205 | 0.626127408 | TPI1, ENO2, PGAM1 | 1342 | 26 | 16792 | 1.443769345 | 1 | 0.999991748 | 99.99999888 |
| KEGG_PATHWAY | hsa00010:Glycolysis / Gluconeogenesis | 6 | 0.381194409 | 0.666614256 | TPI1, HKDC1, ENO2, PGAM1, ALDH2, ALDH3B2 | 574 | 67 | 6879 | 1.073222736 | 1 | 0.912164837 | 99.99995037 |
| GOTERM_BP_DIRECT | GO:0006094~gluconeogenesis | 4 | 0.254129606 | 0.693627682 | SLC25A12, TPI1, ENO2, PGAM1 | 1342 | 44 | 16792 | 1.137515242 | 1 | 0.999998436 | 99.99999997 |
| REACTOME_PATHWAY | R-HSA-70171:R-HSA-70171 | 3 | 0.190597205 | 0.750755119 | TPI1, ENO2, PGAM1 | 745 | 32 | 9075 | 1.141988255 | 1 | 0.999791143 | 99.99999996 |
| KEGG_PATHWAY | hsa01130:Biosynthesis of antibiotics | 15 | 0.952986023 | 0.854390284 | BCKDHA, SHMT2, HKDC1, PGAM1, OGDH, CMBL, FNTB, HAO1, PGM3, TPI1, PHGDH, ALDH2, ENO2, A | 574 | 212 | 6879 | 0.847947209 | 1 | 0.956879976 | 100 |

| Annotation Cluster 163 | Enrichment Score: 0.30656865364970676 | | | | | | | | | | | |
|---|---|---|---|---|---|---|---|---|---|---|---|---|
| Category | Term | Count | % | PValue | Genes | List Total | Pop Hits | Pop Total | Fold Enrichment | Bonferroni | Benjamini | FDR |
| INTERPRO | IPR008979:Galactose-binding domain-like | 9 | 0.571791614 | 0.310416879 | PCSK2, NGLY1, CNTNAP5, NR2C2AP, CNTNAP2, EPHB3, PCSK5, NTN1, EPHB1 | 1452 | 82 | 18559 | 1.402867365 | 1 | 0.997018042 | 99.83429953 |
| INTERPRO | IPR011641:Tyrosine-protein kinase ephrin type A/B receptor-like | 3 | 0.190597205 | 0.592489406 | EPHB3, PCSK5, EPHB1 | 1452 | 25 | 18559 | 1.533801653 | 1 | 0.999936269 | 99.99998078 |
| SMART | SM01411:SM01411 | 3 | 0.190597205 | 0.654136106 | EPHB3, PCSK5, EPHB1 | 878 | 25 | 10057 | 1.37453303 | 1 | 0.992921335 | 99.99996579 |

| Annotation Cluster 164 | Enrichment Score: 0.30379657860282017 | | | | | | | | | | | |
|---|---|---|---|---|---|---|---|---|---|---|---|---|
| Category | Term | Count | % | PValue | Genes | List Total | Pop Hits | Pop Total | Fold Enrichment | Bonferroni | Benjamini | FDR |
| INTERPRO | IPR022272:Lipocalin conserved site | 3 | 0.190597205 | 0.240343159 | LCN6, PTGDS, LCN10 | 1452 | 12 | 18559 | 3.19542011 | 1 | 0.993507746 | 99.12220937 |
| INTERPRO | IPR012674:Calycin | 4 | 0.254129606 | 0.561005805 | LCN6, PTGDS, LCN10, LCNL1 | 1452 | 37 | 18559 | 1.381803291 | 1 | 0.999888413 | 99.99993072 |
| INTERPRO | IPR011038:Calycin-like | 4 | 0.254129606 | 0.597393863 | LCN6, PTGDS, LCN10, LCNL1 | 1452 | 39 | 18559 | 1.310941584 | 1 | 0.999941097 | 99.9999844 |
| INTERPRO | IPR000566:Lipocalin/cytosolic fatty-acid binding protein domain | 3 | 0.190597205 | 0.756403985 | LCN6, PTGDS, LCNL1 | 1452 | 34 | 18559 | 1.127795333 | 1 | 0.99999861 | 100 |

| Annotation Cluster 165 | Enrichment Score: 0.2998934890478816 | | | | | | | | | | | |
|---|---|---|---|---|---|---|---|---|---|---|---|---|
| Category | Term | Count | % | PValue | Genes | List Total | Pop Hits | Pop Total | Fold Enrichment | Bonferroni | Benjamini | FDR |
| REACTOME_PATHWAY | R-HSA-5218921:R-HSA-5218921 | 5 | 0.317662008 | 0.0877009 | HRAS, PLCG1, VEGFA, CALM2, ITPR2 | 745 | 21 | 9075 | 2.900287632 | 1 | 0.928010736 | 75.88468742 |
| KEGG_PATHWAY | hsa05214:Glioma | 9 | 0.571791614 | 0.17057648 | HRAS, PLCG1, PIK3CB, CAMK2G, RB1, CDK4, AKT3, CALM2, SHC4 | 574 | 65 | 6879 | 1.659367462 | 1 | 0.746692885 | 91.55451237 |
| BIOCARTA | h_biopeptidesPathway:Bioactive Peptide Induced Signaling Pathway | 5 | 0.317662008 | 0.444384447 | HRAS, PLCG1, CAMK2G, MYLK, CALM2 | 146 | 38 | 1625 | 1.464491709 | 1 | 0.999020455 | 99.44482867 |
| BIOCARTA | h_fcer1Pathway:Fc Epsilon Receptor I Signaling in Mast Cells | 5 | 0.317662008 | 0.464944323 | FCER1A, HRAS, PAK2, PLCG1, CALM2 | 146 | 39 | 1625 | 1.426940639 | 1 | 0.998732575 | 99.96590741 |
| BIOCARTA | h_pyk2Pathway:Links between Pyk2 and Map Kinases | 3 | 0.190597205 | 0.729267819 | HRAS, PLCG1, CALM2 | 146 | 28 | 1625 | 1.192514677 | 1 | 0.999548173 | 99.9999943 |
| BIOCARTA | h_tcrPathway:T Cell Receptor Signaling Pathway | 4 | 0.254129606 | 0.768379979 | HRAS, PLCG1, CD247, CALM2 | 146 | 44 | 1625 | 1.011830635 | 1 | 0.999661539 | 99.99999922 |
| BIOCARTA | h_bcrPathway:BCR Signaling Pathway | 3 | 0.190597205 | 0.822529059 | HRAS, PLCG1, CALM2 | 146 | 34 | 1625 | 0.982070911 | 1 | 0.999849556 | 99.99999997 |
| BIOCARTA | h_gpcrPathway:Signaling Pathway from G-Protein Families | 3 | 0.190597205 | 0.822529059 | HRAS, PLCG1, CALM2 | 146 | 34 | 1625 | 0.982070911 | 1 | 0.999849556 | 99.99999997 |
| REACTOME_PATHWAY | R-HSA-2871809:R-HSA-2871809 | 6 | 0.381194409 | 0.858205276 | FCER1A, PLCG1, TXK, VAV2, CALM2, ITPR2 | 745 | 88 | 9075 | 0.830536913 | 1 | 0.999966573 | 100 |
| REACTOME_PATHWAY | R-HSA-983695:R-HSA-983695 | 3 | 0.190597205 | 0.99683974 | PLCG1, CALM2, ITPR2 | 745 | 93 | 9075 | 0.392942195 | 1 | 1 | 100 |

| Annotation Cluster 166 | Enrichment Score: 0.29919687688718133 | | | | | | | | | | | |
|---|---|---|---|---|---|---|---|---|---|---|---|---|
| Category | Term | Count | % | PValue | Genes | List Total | Pop Hits | Pop Total | Fold Enrichment | Bonferroni | Benjamini | FDR |
| UP_KEYWORDS | Collagen degradation | 4 | 0.254129606 | 0.146228254 | ADAMTS14, MMP8, MMP27, MMP16 | 1535 | 18 | 20581 | 2.97951502 | 1 | 0.534251002 | 89.69012956 |
| INTERPRO | IPR021158:Peptidase M10A, cysteine switch, zinc binding site | 3 | 0.190597205 | 0.329946531 | MMP8, MMP27, MMP16 | 1452 | 15 | 18559 | 2.556336088 | 1 | 0.997808954 | 99.89898659 |
| INTERPRO | IPR018486:Hemopexin, conserved site | 3 | 0.190597205 | 0.38814783 | MMP8, MMP27, MMP16 | 1452 | 17 | 18559 | 2.255590666 | 1 | 0.999121106 | 99.97888706 |
| INTERPRO | IPR016293:Peptidase M10A, metazoans | 3 | 0.190597205 | 0.416411215 | MMP8, MMP27, MMP16 | 1452 | 18 | 18559 | 2.130280073 | 1 | 0.999358607 | 99.99065213 |
| UP_SEQ_FEATURE | short sequence motif:Cysteine switch | 5 | 0.317662008 | 0.435957577 | MMP8, MMP27, ADAM33, MMP16, ADAM12 | 1499 | 45 | 20063 | 1.487139575 | 1 | 0.999975119 | 99.99765847 |
| INTERPRO | IPR002477:Peptidoglycan binding-like | 3 | 0.190597205 | 0.47084235 | MMP8, MMP27, MMP16 | 1452 | 20 | 18559 | 1.917252066 | 1 | 0.999608657 | 99.99826944 |
| PIR_SUPERFAMILY | PIRSF001191:matrix metalloproteinase, stromelysin type | 3 | 0.190597205 | 0.494432569 | MMP8, MMP27, MMP16 | 154 | 18 | 1692 | 1.831168831 | 1 | 1 | 99.96736302 |
| UP_SEQ_FEATURE | domain:Hemopexin-like 3 | 3 | 0.190597205 | 0.496754612 | MMP8, MMP27, MMP16 | 1499 | 22 | 20063 | 1.825125841 | 1 | 0.999992869 | 99.99971995 |
| UP_SEQ_FEATURE | domain:Hemopexin-like 4 | 3 | 0.190597205 | 0.496754612 | MMP8, MMP27, MMP16 | 1499 | 22 | 20063 | 1.825125841 | 1 | 0.999992869 | 99.99971995 |
| UP_SEQ_FEATURE | domain:Hemopexin-like 1 | 3 | 0.190597205 | 0.52080984 | MMP8, MMP27, MMP16 | 1499 | 23 | 20063 | 1.745772544 | 1 | 0.999996216 | 99.9998875 |
| UP_SEQ_FEATURE | domain:Hemopexin-like 2 | 3 | 0.190597205 | 0.52080984 | MMP8, MMP27, MMP16 | 1499 | 23 | 20063 | 1.745772544 | 1 | 0.999996216 | 99.9998875 |
| INTERPRO | IPR001818:Peptidase M10, metallopeptidase | 3 | 0.190597205 | 0.546452177 | MMP8, MMP27, MMP16 | 1452 | 23 | 18559 | 1.66717571 | 1 | 0.999868223 | 99.99987849 |
| INTERPRO | IPR021190:Peptidase M10A | 3 | 0.190597205 | 0.546452177 | MMP8, MMP27, MMP16 | 1452 | 23 | 18559 | 1.66717571 | 1 | 0.999868223 | 99.99987849 |
| INTERPRO | IPR018487:Hemopexin-like repeats | 3 | 0.190597205 | 0.546452177 | MMP8, MMP27, MMP16 | 1452 | 23 | 18559 | 1.66717571 | 1 | 0.999868223 | 99.99987849 |
| INTERPRO | IPR000585:Hemopexin-like domain | 3 | 0.190597205 | 0.546452177 | MMP8, MMP27, MMP16 | 1452 | 23 | 18559 | 1.66717571 | 1 | 0.999868223 | 99.99987849 |
| SMART | SM00120:HX | 3 | 0.190597205 | 0.608205302 | MMP8, MMP27, MMP16 | 878 | 23 | 10057 | 1.494057641 | 1 | 0.991280747 | 99.99980341 |
| INTERPRO | IPR006026:Peptidase, metallopeptidase | 3 | 0.190597205 | 0.673846737 | MMP8, MMP27, MMP16 | 1452 | 29 | 18559 | 1.322242804 | 1 | 0.999985922 | 99.99999959 |
| SMART | SM00235:ZnMc | 3 | 0.190597205 | 0.732978514 | MMP8, MMP27, MMP16 | 878 | 29 | 10057 | 1.184942267 | 1 | 0.996054264 | 99.99999909 |
| INTERPRO | IPR024079:Metallopeptidase, catalytic domain | 6 | 0.381194409 | 0.768896092 | ADAMTS14, MMP8, MMP27, ADAM33, MMP16, ADAM12 | 1452 | 81 | 18559 | 0.946791144 | 1 | 0.999998979 | 100 |
| GOTERM_MF_DIRECT | GO:0004222~metalloendopeptidase activity | 6 | 0.381194409 | 0.951086802 | ADAMTS14, MMP8, MMP27, ADAM33, MMP16, ADAM12 | 1340 | 113 | 16881 | 0.668907674 | 1 | 0.999999905 | 100 |

| Annotation Cluster 167 | Enrichment Score: 0.2988350898658696 | | | | | | | | | | | |
|---|---|---|---|---|---|---|---|---|---|---|---|---|
| Category | Term | Count | % | PValue | Genes | List Total | Pop Hits | Pop Total | Fold Enrichment | Bonferroni | Benjamini | FDR |
| INTERPRO | IPR002049:EGF-like, laminin | 5 | 0.317662008 | 0.36410451 | CRELD1, ATRNL1, STAB1, MEGF11, NTN1 | 1452 | 39 | 18559 | 1.63867698 | 1 | 0.998672834 | 99.95898748 |
| UP_KEYWORDS | Laminin EGF-like domain | 4 | 0.254129606 | 0.4099417 | ATRNL1, STAB1, MEGF11, NTN1 | 1535 | 31 | 20581 | 1.730040979 | 1 | 0.811214248 | 99.94902791 |
| UP_SEQ_FEATURE | domain:Laminin EGF-like 1 | 3 | 0.190597205 | 0.588210376 | ATRNL1, STAB1, NTN1 | 1499 | 26 | 20063 | 1.54433725 | 1 | 0.999999408 | 99.99999331 |
| SMART | SM00180:EGF_Lam | 4 | 0.254129606 | 0.599360253 | ATRNL1, STAB1, MEGF11, NTN1 | 878 | 35 | 10057 | 1.309079076 | 1 | 0.990780074 | 99.99973115 |
| UP_SEQ_FEATURE | domain:Laminin EGF-like 2 | 3 | 0.190597205 | 0.609053963 | ATRNL1, STAB1, NTN1 | 1499 | 27 | 20063 | 1.487139575 | 1 | 0.999999615 | 99.99999746 |

| Annotation Cluster 168 | Enrichment Score: 0.28691251632189596 | | | | | | | | | | | |
|---|---|---|---|---|---|---|---|---|---|---|---|---|
| Category | Term | Count | % | PValue | Genes | List Total | Pop Hits | Pop Total | Fold Enrichment | Bonferroni | Benjamini | FDR |
| GOTERM_BP_DIRECT | GO:0034968~histone lysine methylation | 4 | 0.254129606 | 0.095362747 | PRDM6, SMYD3, MECOM, EHMT2 | 1342 | 14 | 16792 | 3.575047903 | 1 | 0.979190754 | 84.51500586 |
| GOTERM_MF_DIRECT | GO:0018024~histone-lysine N-methyltransferase activity | 5 | 0.317662008 | 0.374463485 | RBBP5, PRDM6, SMYD3, MECOM, EHMT2 | 1340 | 39 | 16881 | 1.615097589 | 1 | 0.992882906 | 99.9523881 |
| REACTOME_PATHWAY | R-HSA-3214841:R-HSA-3214841 | 6 | 0.381194409 | 0.611338646 | RBBP5, HIST4H4, SMYD3, MECOM, EHMT2, HIST1H3H | 745 | 64 | 9075 | 1.141988255 | 1 | 0.999147474 | 99.99995634 |
| SMART | SM00317:SET | 4 | 0.254129606 | 0.655783666 | PRDM6, SMYD3, MECOM, EHMT2 | 878 | 38 | 10057 | 1.205730728 | 1 | 0.99268392 | 99.999968 |
| INTERPRO | IPR001214:SET domain | 4 | 0.254129606 | 0.772276381 | PRDM6, SMYD3, MECOM, EHMT2 | 1452 | 51 | 18559 | 1.00248474 | 1 | 0.999999052 | 100 |
| UP_SEQ_FEATURE | domain:SET | 3 | 0.190597205 | 0.889736418 | PRDM6, SMYD3, EHMT2 | 1499 | 49 | 20063 | 0.819444255 | 1 | 1 | 100 |
| UP_KEYWORDS | S-adenosyl-L-methionine | 6 | 0.381194409 | 0.997120378 | PRDM6, SMYD3, LIAS, TRMT10B, EHMT2, METTL21C | 1535 | 174 | 20581 | 0.462338538 | 1 | 0.999958269 | 100 |

Annotation Cluster 169 — Enrichment Score: 0.28142274690714764

| Category | Term | Count | % | PValue | Genes | List Total | Pop Hits | Pop Total | Fold Enrichment | Bonferroni | Benjamini | FDR |
|---|---|---|---|---|---|---|---|---|---|---|---|---|
| INTERPRO | IPR003593:AAA+ ATPase domain | 14 | 0.889453621 | 0.26528852 | DNAH11, RAD51B, SPATA5L1, CLPX, ABCG2, MCM7, TAP2, DYNC2H1, ABCD4, FIGNL2, KATNAL2, ABC | 1452 | 136 | 18559 | 1.315761222 | 1 | 0.994852335 | 99.50615812 |
| UP_SEQ_FEATURE | domain:ABC transporter | 3 | 0.190597205 | 0.338081545 | TAP2, ABCD4, ABCG2 | 1499 | 16 | 20063 | 2.509548032 | 1 | 0.99984201 | 99.95392837 |
| SMART | SM00382:AAA | 14 | 0.889453621 | 0.406965387 | DNAH11, RAD51B, SPATA5L1, CLPX, ABCG2, MCM7, TAP2, DYNC2H1, ABCD4, FIGNL2, KATNAL2, ABC | 878 | 136 | 10057 | 1.179133726 | 1 | 0.967995592 | 99.93424114 |
| GOTERM_MF_DIRECT | GO:0042626~ATPase activity, coupled to transmembrane movement of substances | 5 | 0.317662008 | 0.465697468 | TAP2, ABCD4, ABCA13, ABCC5, ABCG2 | 1340 | 44 | 16881 | 1.431563772 | 1 | 0.996293289 | 99.99635813 |
| KEGG_PATHWAY | hsa02010:ABC transporters | 5 | 0.317662008 | 0.505019755 | TAP2, ABCD4, ABCA13, ABCC5, ABCG2 | 574 | 44 | 6879 | 1.361854609 | 1 | 0.878727146 | 99.99079633 |
| INTERPRO | IPR003439:ABC transporter-like | 5 | 0.317662008 | 0.556230375 | TAP2, ABCD4, ABCA13, ABCC5, ABCG2 | 1452 | 50 | 18559 | 1.278168044 | 1 | 0.999880259 | 99.99991652 |
| INTERPRO | IPR003959:ATPase, AAA-type, core | 5 | 0.317662008 | 0.572211454 | FIGNL2, SPATA5L1, KATNAL2, CLPX, TRIP13 | 1452 | 51 | 18559 | 1.253105926 | 1 | 0.999911791 | 99.99995562 |
| INTERPRO | IPR011527:ABC transporter, transmembrane domain, type 1 | 3 | 0.190597205 | 0.673846737 | TAP2, ABCD4, ABCC5 | 1452 | 29 | 18559 | 1.322242804 | 1 | 0.999985922 | 99.99999959 |
| INTERPRO | IPR017871:ABC transporter, conserved site | 3 | 0.190597205 | 0.868666533 | TAP2, ABCD4, ABCC5 | 1452 | 44 | 18559 | 0.871478212 | 1 | 0.999999983 | 100 |
| GOTERM_MF_DIRECT | GO:0016887~ATPase activity | 10 | 0.635324015 | 0.959041554 | DNAH11, MSH2, SWSAP1, DYNC2H1, ABCD4, ASNA1, KIF26A, CLPX, ABCA13, ABCG2 | 1340 | 183 | 16881 | 0.688402251 | 1 | 0.999999951 | 100 |

Annotation Cluster 170 — Enrichment Score: 0.27975479773726986

| Category | Term | Count | % | PValue | Genes | List Total | Pop Hits | Pop Total | Fold Enrichment | Bonferroni | Benjamini | FDR |
|---|---|---|---|---|---|---|---|---|---|---|---|---|
| UP_KEYWORDS | Ciliopathy | 12 | 0.762388818 | 0.338017025 | DNAH11, TRAF3IP1, BBS5, EVC, HYDIN, WDPCP, DYNC2H1, ARMC4, AHI1, ALMS1, CCDC39, IFT140 | 1535 | 126 | 20581 | 1.276935009 | 1 | 0.761213131 | 99.73378383 |
| GOTERM_CC_DIRECT | GO:0036064~ciliary basal body | 9 | 0.571791614 | 0.502421664 | TRAF3IP1, BBS5, EVC, EZR, AHI1, ALMS1, TTLL6, SSX2IP, IFT140 | 1425 | 98 | 18224 | 1.174479055 | 1 | 0.963377287 | 99.99724605 |
| GOTERM_BP_DIRECT | GO:0042384~cilium assembly | 11 | 0.698856417 | 0.534990656 | TRAF3IP1, SCLT1, BBS5, WDPCP, RFX4, CEP89, DYNC2H1, AHI1, ALMS1, SSX2IP, IFT140 | 1342 | 124 | 16792 | 1.109994712 | 1 | 0.999959681 | 99.99993531 |
| GOTERM_BP_DIRECT | GO:0060271~cilium morphogenesis | 11 | 0.698856417 | 0.653709976 | SCLT1, BBS5, WDPCP, CEP89, DYNC2H1, AHI1, ASAP1, SSX2IP, TBC1D32, PARVA, IFT140 | 1342 | 136 | 16792 | 1.012054002 | 1 | 0.999995573 | 99.99999973 |
| UP_KEYWORDS | Cilium biogenesis/degradation | 11 | 0.698856417 | 0.672183379 | TRAF3IP1, BBS5, WDPCP, CEP89, DYNC2H1, ARMC4, AHI1, ASAP1, SSX2IP, PARVA, IFT140 | 1535 | 148 | 20581 | 0.996526983 | 1 | 0.935140964 | 99.99998907 |

Annotation Cluster 171 — Enrichment Score: 0.27652170788171015

| Category | Term | Count | % | PValue | Genes | List Total | Pop Hits | Pop Total | Fold Enrichment | Bonferroni | Benjamini | FDR |
|---|---|---|---|---|---|---|---|---|---|---|---|---|
| KEGG_PATHWAY | hsa04660:T cell receptor signaling pathway | 11 | 0.698856417 | 0.320384961 | BCL10, HRAS, PAK2, PLCG1, PIK3CB, NFKBIB, CD247, MAP3K8, VAV2, CDK4, AKT3 | 574 | 100 | 6879 | 1.318275261 | 1 | 0.814317305 | 99.39278065 |
| KEGG_PATHWAY | hsa04664:Fc epsilon RI signaling pathway | 8 | 0.508259212 | 0.337105544 | FCER1A, JMJD7-PLA2G4B, HRAS, PLCG1, PIK3CB, VAV2, PLA2G4B, AKT3 | 574 | 68 | 6879 | 1.409920066 | 1 | 0.819506134 | 99.56310482 |
| REACTOME_PATHWAY | R-HSA-2424491:R-HSA-2424491 | 4 | 0.254129606 | 0.450974503 | HRAS, PLCG1, PIK3CB, VAV2 | 745 | 30 | 9075 | 1.624161074 | 1 | 0.995854771 | 99.09077926 |
| GOTERM_BP_DIRECT | GO:0038095~Fc-epsilon receptor signaling pathway | 11 | 0.698856417 | 0.910839818 | FCER1A, PSMA1, BCL10, HRAS, PAK2, PLCG1, PIK3CB, PSMD1, FER, VAV2, CALM2 | 1342 | 178 | 16792 | 0.773254743 | 1 | 0.999999999 | 100 |
| REACTOME_PATHWAY | R-HSA-2871796:R-HSA-2871796 | 5 | 0.317662008 | 0.934027036 | FCER1A, HRAS, PAK2, PLCG1, VAV2 | 745 | 87 | 9075 | 0.700069428 | 1 | 0.999996026 | 100 |

Annotation Cluster 172 — Enrichment Score: 0.2722004844605177

| Category | Term | Count | % | PValue | Genes | List Total | Pop Hits | Pop Total | Fold Enrichment | Bonferroni | Benjamini | FDR |
|---|---|---|---|---|---|---|---|---|---|---|---|---|
| UP_SEQ_FEATURE | domain:AGC-kinase C-terminal | 6 | 0.381194409 | 0.407853281 | MAST4, PKN2, GRK7, PRKCE, AKT3, DMPK | 1499 | 56 | 20063 | 1.434027447 | 1 | 0.99995953 | 99.99420968 |
| INTERPRO | IPR000961:AGC-kinase, C-terminal | 6 | 0.381194409 | 0.447651727 | MAST4, PKN2, GRK7, PRKCE, AKT3, DMPK | 1452 | 56 | 18559 | 1.369465762 | 1 | 0.999496184 | 99.99637681 |
| SMART | SM00133:S_TK_X | 5 | 0.317662008 | 0.645130587 | PKN2, GRK7, PRKCE, AKT3, DMPK | 878 | 50 | 10057 | 1.145444191 | 1 | 0.992858732 | 99.99995094 |
| INTERPRO | IPR017892:Protein kinase, C-terminal | 3 | 0.190597205 | 0.692000175 | PKN2, PRKCE, AKT3 | 1452 | 30 | 18559 | 1.278168044 | 1 | 0.999990571 | 99.99999985 |

Annotation Cluster 173 — Enrichment Score: 0.271156240744315

| Category | Term | Count | % | PValue | Genes | List Total | Pop Hits | Pop Total | Fold Enrichment | Bonferroni | Benjamini | FDR |
|---|---|---|---|---|---|---|---|---|---|---|---|---|
| KEGG_PATHWAY | hsa03440:Homologous recombination | 5 | 0.317662008 | 0.218634985 | POLD3, RAD51C, RAD51B, SYCP3, RPA3 | 574 | 29 | 6879 | 2.066262165 | 1 | 0.805802258 | 96.16240695 |
| GOTERM_BP_DIRECT | GO:0006310~DNA recombination | 9 | 0.571791614 | 0.343502482 | XRCC5, RAD51C, HMGB1, XRCC4, RAD51B, THAP9, TSN, RBPJ, RPA3 | 1342 | 83 | 16792 | 1.356795288 | 1 | 0.999186275 | 99.96033779 |
| UP_KEYWORDS | DNA recombination | 8 | 0.508259212 | 0.399835206 | XRCC5, RAD51C, HMGB1, XRCC4, RAD51B, THAP9, SWSAP1, RPA3 | 1535 | 81 | 20581 | 1.324228898 | 1 | 0.802867972 | 99.93493709 |
| GOTERM_BP_DIRECT | GO:0000722~telomere maintenance via recombination | 4 | 0.254129606 | 0.476310875 | POLD3, RAD51C, TEP1, RPA3 | 1342 | 32 | 16792 | 1.564083458 | 1 | 0.999899772 | 99.99940919 |
| REACTOME_PATHWAY | R-HSA-5685942:R-HSA-5685942 | 4 | 0.254129606 | 0.765450445 | POLD3, RAD51C, RAD51B, RPA3 | 745 | 48 | 9075 | 1.015100671 | 1 | 0.999836845 | 99.99999998 |
| GOTERM_BP_DIRECT | GO:0000731~DNA synthesis involved in DNA repair | 3 | 0.190597205 | 0.781017775 | POLD3, RAD51C, RAD51B | 1342 | 35 | 16792 | 1.072514371 | 1 | 0.999999851 | 100 |
| REACTOME_PATHWAY | R-HSA-5693616:R-HSA-5693616 | 3 | 0.190597205 | 0.841110498 | RAD51C, RAD51B, RPA3 | 745 | 39 | 9075 | 0.937016004 | 1 | 0.999949197 | 100 |
| GOTERM_BP_DIRECT | GO:0000724~double-strand break repair via homologous recombination | 4 | 0.254129606 | 0.941668313 | RAD51C, RAD51B, SWSAP1, RPA3 | 1342 | 74 | 16792 | 0.676360414 | 1 | 1 | 100 |

Annotation Cluster 174 — Enrichment Score: 0.2707428173341809

| Category | Term | Count | % | PValue | Genes | List Total | Pop Hits | Pop Total | Fold Enrichment | Bonferroni | Benjamini | FDR |
|---|---|---|---|---|---|---|---|---|---|---|---|---|
| INTERPRO | IPR009071:High mobility group (HMG) box domain | 6 | 0.381194409 | 0.463239733 | HMGB1, SP100, SOX15, SOX6, SOX8, TCF7L2 | 1452 | 57 | 18559 | 1.345440046 | 1 | 0.999575325 | 99.99778735 |
| SMART | SM00398:HMG | 6 | 0.381194409 | 0.51335212 | HMGB1, SP100, SOX15, SOX6, SOX8, TCF7L2 | 878 | 54 | 10057 | 1.272715768 | 1 | 0.984255058 | 99.99588958 |
| UP_SEQ_FEATURE | DNA-binding region:HMG box | 4 | 0.254129606 | 0.647963265 | SOX15, SOX6, SOX8, TCF7L2 | 1499 | 44 | 20063 | 1.216750561 | 1 | 0.999999871 | 99.99999964 |

Annotation Cluster 175 — Enrichment Score: 0.2705653814478511

| Category | Term | Count | % | PValue | Genes | List Total | Pop Hits | Pop Total | Fold Enrichment | Bonferroni | Benjamini | FDR |
|---|---|---|---|---|---|---|---|---|---|---|---|---|
| KEGG_PATHWAY | hsa04022:cGMP-PKG signaling pathway | 16 | 1.016518424 | 0.336316029 | SLC8A1, ATP1B2, PDE3B, CNGB1, PRKCE, CNGA1, ITPR2, ADRB2, ATP2B4, ATF6B, GUCY1A2, CACNA1 | 574 | 158 | 6879 | 1.213602082 | 1 | 0.827891135 | 99.55617816 |
| KEGG_PATHWAY | hsa04924:Renin secretion | 6 | 0.381194409 | 0.626008775 | ADRB2, GUCY1A2, PDE3B, CACNA1C, CALM2, ITPR2 | 574 | 64 | 6879 | 1.123530052 | 1 | 0.909918728 | 99.99977336 |
| KEGG_PATHWAY | hsa04970:Salivary secretion | 7 | 0.444726811 | 0.732783814 | ADRB2, ATP2B4, ATP1B2, RYR3, GUCY1A2, CALM2, ITPR2 | 574 | 86 | 6879 | 0.975467952 | 1 | 0.921076761 | 99.99999733 |

Annotation Cluster 176 — Enrichment Score: 0.26950008959601796

| Category | Term | Count | % | PValue | Genes | List Total | Pop Hits | Pop Total | Fold Enrichment | Bonferroni | Benjamini | FDR |
|---|---|---|---|---|---|---|---|---|---|---|---|---|
| KEGG_PATHWAY | hsa04750:Inflammatory mediator regulation of TRP channels | 11 | 0.698856417 | 0.298155374 | JMJD7-PLA2G4B, PLCG1, PIK3CB, CAMK2G, TRPV2, ASIC2, PRKCE, PLA2G4B, CALM2, HTR2A, ITPR2 | 574 | 98 | 6879 | 1.345178838 | 1 | 0.817876458 | 99.07089316 |
| KEGG_PATHWAY | hsa04724:Glutamatergic synapse | 11 | 0.698856417 | 0.480888806 | SLC17A7, PLD2, SLC1A2, DLGAP1, JMJD7-PLA2G4B, GRIN2B, GRIA2, CACNA1C, KCNJ3, PLA2G4B, ITP | 574 | 114 | 6879 | 1.156381808 | 1 | 0.874903251 | 99.98273567 |
| KEGG_PATHWAY | hsa04912:GnRH signaling pathway | 9 | 0.571791614 | 0.491219295 | PLD2, JMJD7-PLA2G4B, HRAS, CAMK2G, HBEGF, CACNA1C, PLA2G4B, CALM2, ITPR2 | 574 | 91 | 6879 | 1.185262473 | 1 | 0.876903711 | 99.98676313 |
| KEGG_PATHWAY | hsa04270:Vascular smooth muscle contraction | 9 | 0.571791614 | 0.769828205 | JMJD7-PLA2G4B, PLA2G12B, GUCY1A2, PRKCE, CACNA1C, PLA2G4B, MYLK, CALM2, ITPR2 | 574 | 117 | 6879 | 0.921870812 | 1 | 0.931343248 | 99.99999963 |
| KEGG_PATHWAY | hsa04726:Serotonergic synapse | 8 | 0.508259212 | 0.828596333 | APP, JMJD7-PLA2G4B, HRAS, CACNA1C, KCNJ3, PLA2G4B, HTR2A, ITPR2 | 574 | 111 | 6879 | 0.863734815 | 1 | 0.950237744 | 99.99999999 |

Annotation Cluster 177 — Enrichment Score: 0.2680805824834939

| Category | Term | Count | % | PValue | Genes | List Total | Pop Hits | Pop Total | Fold Enrichment | Bonferroni | Benjamini | FDR |
|---|---|---|---|---|---|---|---|---|---|---|---|---|
| UP_SEQ_FEATURE | domain:HECT | 4 | 0.254129606 | 0.348675573 | NEDD4, UBE3A, HECTD4, NEDD4L | 1499 | 28 | 20063 | 1.912036596 | 1 | 0.999870952 | 99.96588361 |
| INTERPRO | IPR000569:HECT | 4 | 0.254129606 | 0.376364618 | NEDD4, UBE3A, HECTD4, NEDD4L | 1452 | 28 | 18559 | 1.825954349 | 1 | 0.998951335 | 99.97067339 |
| SMART | SM00119:HECTc | 4 | 0.254129606 | 0.446461534 | NEDD4, UBE3A, HECTD4, NEDD4L | 878 | 28 | 10057 | 1.636348845 | 1 | 0.977086178 | 99.97498379 |
| UP_SEQ_FEATURE | active site:Glycyl thioester intermediate | 5 | 0.317662008 | 0.786219266 | NEDD4, UBE3A, HECTD4, NEDD4L, UBE2U | 1499 | 71 | 20063 | 0.942553251 | 1 | 1 | 100 |
| KEGG_PATHWAY | hsa04120:Ubiquitin mediated proteolysis | 6 | 0.381194409 | 0.991377474 | TRIM37, NEDD4, UBE3A, NEDD4L, PARK2, UBE2U | 574 | 137 | 6879 | 0.524860754 | 1 | 0.998636869 | 100 |

Annotation Cluster 178 — Enrichment Score: 0.26674162394194983

| Category | Term | Count | % | PValue | Genes | List Total | Pop Hits | Pop Total | Fold Enrichment | Bonferroni | Benjamini | FDR |
|---|---|---|---|---|---|---|---|---|---|---|---|---|
| UP_SEQ_FEATURE | domain:DAPIN | 3 | 0.190597205 | 0.496754612 | PYHIN1, NLRP5, NLRP9 | 1499 | 22 | 20063 | 1.825125841 | 1 | 0.999992869 | 99.99971995 |
| INTERPRO | IPR004020:DAPIN domain | 3 | 0.190597205 | 0.546452177 | PYHIN1, NLRP5, NLRP9 | 1452 | 23 | 18559 | 1.66717571 | 1 | 0.999868223 | 99.99987849 |
| SMART | SM01289:SM01289 | 3 | 0.190597205 | 0.583554014 | PYHIN1, NLRP5, NLRP9 | 878 | 22 | 10057 | 1.561969352 | 1 | 0.990540742 | 99.99953745 |

Annotation Cluster 179 — Enrichment Score: 0.26265625730224174

| Category | Term | Count | % | PValue | Genes | List Total | Pop Hits | Pop Total | Fold Enrichment | Bonferroni | Benjamini | FDR |
|---|---|---|---|---|---|---|---|---|---|---|---|---|
| GOTERM_BP_DIRECT | GO:0009083~branched-chain amino acid catabolic process | 3 | 0.190597205 | 0.455189596 | BCKDHA, HIBCH, AUH | 1342 | 19 | 16792 | 1.975684367 | 1 | 0.999853888 | 99.99876676 |

| | | | | | | | | | | | | |
|---|---|---|---|---|---|---|---|---|---|---|---|---|
| REACTOME_PATHWAY | R-HSA-70895:R-HSA-70895 | 3 | 0.190597205 | 0.469278262 | BCKDHA, HIBCH, AUH | 745 | 19 | 9075 | 1.92334864 | 1 | 0.996570224 | 99.9945477 |
| KEGG_PATHWAY | hsa00280:Valine, leucine and isoleucine degradation | 4 | 0.254129606 | 0.762796099 | BCKDHA, ALDH2, HIBCH, AUH | 574 | 47 | 6879 | 1.019942175 | 1 | 0.928711468 | 99.99999945 |
| Annotation Cluster 180 | Enrichment Score: 0.2429648255857971 | | | | | | | | | | | |
| Category | Term | Count | % | PValue | Genes | List Total | Pop Hits | Pop Total | Fold Enrichment | Bonferroni | Benjamini | FDR |
| REACTOME_PATHWAY | R-HSA-1169408:R-HSA-1169408 | 9 | 0.571791614 | 0.247127637 | IFIT1, EIF4E, PLCG1, NEDD4, NUP88, EIF4A1, JAK1, KPNA3, NUP35 | 745 | 73 | 9075 | 1.501792774 | 1 | 0.977948242 | 98.77060041 |
| REACTOME_PATHWAY | R-HSA-168276:R-HSA-168276 | 3 | 0.190597205 | 0.818882779 | NUP88, KPNA3, NUP35 | 745 | 37 | 9075 | 0.987665518 | 1 | 0.999928198 | 100 |
| GOTERM_BP_DIRECT | GO:0075733~intracellular transport of virus | 3 | 0.190597205 | 0.922491705 | NUP88, KPNA3, NUP35 | 1342 | 51 | 16792 | 0.736039274 | 1 | 1 | 100 |
| Annotation Cluster 181 | Enrichment Score: 0.23612144031204094 | | | | | | | | | | | |
| Category | Term | Count | % | PValue | Genes | List Total | Pop Hits | Pop Total | Fold Enrichment | Bonferroni | Benjamini | FDR |
| KEGG_PATHWAY | hsa00564:Glycerophospholipid metabolism | 11 | 0.698856417 | 0.265634372 | PLD2, CRLS1, JMJD7-PLA2G4B, PLB1, DGKG, PLA2G12B, DGKZ, CHPT1, PLA2G4B, AGPAT1, PTDSS2 | 574 | 95 | 6879 | 1.38765817 | 1 | 0.815302238 | 98.30946624 |
| GOTERM_BP_DIRECT | GO:0008654~phospholipid biosynthetic process | 4 | 0.254129606 | 0.629622755 | CRLS1, CHPT1, AGPAT1, PTDSS2 | 1342 | 40 | 16792 | 1.251266766 | 1 | 0.999992094 | 99.99999906 |
| UP_KEYWORDS | Phospholipid biosynthesis | 4 | 0.254129606 | 0.661911441 | CRLS1, CHPT1, AGPAT1, PTDSS2 | 1535 | 45 | 20581 | 1.191806008 | 1 | 0.931887589 | 99.99998297 |
| UP_KEYWORDS | Phospholipid metabolism | 4 | 0.254129606 | 0.717763335 | CRLS1, CHPT1, AGPAT1, PTDSS2 | 1535 | 49 | 20581 | 1.094515722 | 1 | 0.949408381 | 99.99999873 |
| UP_KEYWORDS | Lipid biosynthesis | 10 | 0.635324015 | 0.830322786 | FAR1, CRLS1, PTGDS, MLYCD, FAM213B, OSBPL10, CHPT1, OXSM, AGPAT1, PTDSS2 | 1535 | 156 | 20581 | 0.859475487 | 1 | 0.976935184 | 100 |
| Annotation Cluster 182 | Enrichment Score: 0.23306137284384842 | | | | | | | | | | | |
| Category | Term | Count | % | PValue | Genes | List Total | Pop Hits | Pop Total | Fold Enrichment | Bonferroni | Benjamini | FDR |
| GOTERM_BP_DIRECT | GO:0007223~Wnt signaling pathway, calcium modulating pathway | 5 | 0.317662008 | 0.379379518 | WNT5A, TNRC6C, TNRC6B, TCF7L2, CALM2 | 1342 | 39 | 16792 | 1.604188162 | 1 | 0.999537976 | 99.98606462 |
| REACTOME_PATHWAY | R-HSA-4086398:R-HSA-4086398 | 6 | 0.381194409 | 0.567792511 | WNT5A, TNRC6C, TNRC6B, TCF7L2, CALM2, ITPR2 | 745 | 61 | 9075 | 1.198151612 | 1 | 0.998892622 | 99.99977367 |
| KEGG_PATHWAY | hsa04916:Melanogenesis | 6 | 0.381194409 | 0.928009326 | DCT, WNT5A, HRAS, CAMK2G, TCF7L2, CALM2 | 574 | 100 | 6879 | 0.719059233 | 1 | 0.98109185 | 100 |
| Annotation Cluster 183 | Enrichment Score: 0.2304673188129147 | | | | | | | | | | | |
| Category | Term | Count | % | PValue | Genes | List Total | Pop Hits | Pop Total | Fold Enrichment | Bonferroni | Benjamini | FDR |
| UP_SEQ_FEATURE | repeat:I | 3 | 0.190597205 | 0.588210376 | CACNA1C, SCN4A, ITGBL1 | 1499 | 26 | 20063 | 1.54433725 | 1 | 0.999999408 | 99.99999331 |
| UP_SEQ_FEATURE | repeat:II | 3 | 0.190597205 | 0.588210376 | CACNA1C, SCN4A, ITGBL1 | 1499 | 26 | 20063 | 1.54433725 | 1 | 0.999999408 | 99.99999331 |
| UP_SEQ_FEATURE | repeat:III | 3 | 0.190597205 | 0.588210376 | CACNA1C, SCN4A, ITGBL1 | 1499 | 26 | 20063 | 1.54433725 | 1 | 0.999999408 | 99.99999331 |
| UP_SEQ_FEATURE | repeat:IV | 3 | 0.190597205 | 0.588210376 | CACNA1C, SCN4A, ITGBL1 | 1499 | 26 | 20063 | 1.54433725 | 1 | 0.999999408 | 99.99999331 |
| Annotation Cluster 184 | Enrichment Score: 0.2276194808181037 | | | | | | | | | | | |
| Category | Term | Count | % | PValue | Genes | List Total | Pop Hits | Pop Total | Fold Enrichment | Bonferroni | Benjamini | FDR |
| INTERPRO | IPR018253:DnaJ domain, conserved site | 4 | 0.254129606 | 0.354431796 | DNAJC16, DNAJC5, DNAJB1, DNAJB4 | 1452 | 27 | 18559 | 1.893582288 | 1 | 0.998492474 | 99.94680587 |
| INTERPRO | IPR001623:DnaJ domain | 5 | 0.317662008 | 0.572211454 | SLC8A1, DNAJC16, DNAJC5, DNAJB1, DNAJB4 | 1452 | 51 | 18559 | 1.253105926 | 1 | 0.999911791 | 99.99995562 |
| UP_SEQ_FEATURE | domain:J | 4 | 0.254129606 | 0.755833161 | DNAJC16, DNAJC5, DNAJB1, DNAJB4 | 1499 | 52 | 20063 | 1.029558167 | 1 | 0.999999998 | 100 |
| SMART | SM00271:DnaJ | 4 | 0.254129606 | 0.801690824 | DNAJC16, DNAJC5, DNAJB1, DNAJB4 | 878 | 48 | 10057 | 0.954536826 | 1 | 0.998615957 | 99.99999999 |
| Annotation Cluster 185 | Enrichment Score: 0.2239779710197233 | | | | | | | | | | | |
| Category | Term | Count | % | PValue | Genes | List Total | Pop Hits | Pop Total | Fold Enrichment | Bonferroni | Benjamini | FDR |
| INTERPRO | IPR017970:Homeobox, conserved site | 17 | 1.080050826 | 0.416240684 | CDX1, IRX6, SIX5, PDX1, BSX, BARX2, NKX1-1, POU5F1, LHX3, MNX1, MKX, CUX2, CUX1, LHX9, ALX3, | 1452 | 190 | 18559 | 1.143624039 | 1 | 0.999387234 | 99.99060496 |
| INTERPRO | IPR009057:Homeodomain-like | 28 | 1.778907243 | 0.463783064 | IRX6, CDX1, PAX5, PDX1, BSX, BARX2, CENPBD1, NKX1-1, POU5F1, LHX3, MKX, ALX3, LHX9, PITX1, SA | 1452 | 336 | 18559 | 1.065140037 | 1 | 0.999560294 | 99.99782562 |
| UP_KEYWORDS | Homeobox | 20 | 1.27064803 | 0.583976217 | SATB1, CDX1, IRX6, ZHX2, ZHX3, SIX5, PDX1, BSX, BARX2, NKX1-1, POU5F1, LHX3, MNX1, MKX, CUX2 | 1535 | 262 | 20581 | 1.023497526 | 1 | 0.898496079 | 99.99966425 |
| INTERPRO | IPR001356:Homeodomain | 20 | 1.27064803 | 0.627986923 | SATB1, CDX1, IRX6, ZHX2, ZHX3, SIX5, PDX1, BSX, BARX2, NKX1-1, POU5F1, LHX3, MNX1, MKX, CUX2 | 1452 | 256 | 18559 | 0.998568784 | 1 | 0.99996505 | 99.999996 |
| UP_SEQ_FEATURE | DNA-binding region:Homeobox | 14 | 0.889453621 | 0.676113026 | BSX, SATB1, CDX1, BARX2, NKX1-1, POU5F1, LHX3, MNX1, SIX5, PDX1, CUX1, ALX3, LHX9, PITX1 | 1499 | 191 | 20063 | 0.981044955 | 1 | 0.999999944 | 99.99999992 |
| INTERPRO | IPR020479:Homeodomain, metazoa | 7 | 0.444726811 | 0.734660814 | BSX, CDX1, BARX2, NKX1-1, MNX1, PDX1, PITX1 | 1452 | 92 | 18559 | 0.972519164 | 1 | 0.99999721 | 99.99999999 |
| SMART | SM00389:HOX | 20 | 1.27064803 | 0.769206334 | SATB1, CDX1, IRX6, ZHX2, ZHX3, SIX5, PDX1, BSX, BARX2, NKX1-1, POU5F1, LHX3, MNX1, MKX, CUX2 | 878 | 250 | 10057 | 0.916355353 | 1 | 0.997581513 | 99.99999988 |
| Annotation Cluster 186 | Enrichment Score: 0.2237599720525015 | | | | | | | | | | | |
| Category | Term | Count | % | PValue | Genes | List Total | Pop Hits | Pop Total | Fold Enrichment | Bonferroni | Benjamini | FDR |
| UP_SEQ_FEATURE | calcium-binding region:3 | 5 | 0.317662008 | 0.332112409 | SLC25A12, CAPS2, EFCAB1, CALM2, KCNIP4 | 1499 | 39 | 20063 | 1.715930278 | 1 | 0.999833156 | 99.94554602 |
| INTERPRO | IPR011992:EF-hand-like domain | 24 | 1.524777637 | 0.42699846 | MICU1, TESC, TBC1D8, UTRN, MYL12A, KCNIP4, SLC25A12, PLCL1, CAPS2, PLCG1, CAPN13, DGKG, RY | 1452 | 279 | 18559 | 1.099499393 | 1 | 0.99941958 | 99.99318073 |
| INTERPRO | IPR002048:EF-hand domain | 19 | 1.207115629 | 0.518608833 | MICU1, TESC, MYL12A, KCNIP4, SLC25A12, CAPS2, GNPTAB, PLCG1, DGKG, RYR3, PLCH1, NUCB2, MC | 1452 | 228 | 18559 | 1.065140037 | 1 | 0.999824064 | 99.99966086 |
| UP_SEQ_FEATURE | calcium-binding region:2 | 10 | 0.635324015 | 0.52366306 | SLC25A12, MICU1, CAPS2, DGKG, MCFD2, NUCB2, PLS1, EFCAB1, CALM2, KCNIP4 | 1499 | 118 | 20063 | 1.134258998 | 1 | 0.999996397 | 99.99989934 |
| INTERPRO | IPR018247:EF-Hand 1, calcium-binding site | 15 | 0.952986023 | 0.531132512 | MICU1, TESC, MYL12A, KCNIP4, SLC25A12, GNPTAB, PLCG1, DGKG, PLCH1, NUCB2, MCFD2, PLS1, EF | 1452 | 178 | 18559 | 1.077107902 | 1 | 0.999834093 | 99.99978464 |
| UP_SEQ_FEATURE | calcium-binding region:1 | 10 | 0.635324015 | 0.623197488 | SLC25A12, MICU1, CAPS2, DGKG, MCFD2, NUCB2, PLS1, EFCAB1, CALM2, KCNIP4 | 1499 | 128 | 20063 | 1.045645013 | 1 | 0.999999743 | 99.99999872 |
| UP_SEQ_FEATURE | domain:EF-hand 2 | 14 | 0.889453621 | 0.630895579 | MICU1, MYL12A, KCNIP4, SLC25A12, CAPS2, DGKG, CAPN13, PLCH1, NUCB2, MCFD2, PLS1, EFCAB1, | 1499 | 185 | 20063 | 1.012862629 | 1 | 0.999999778 | 99.99999913 |
| SMART | SM00054:EFh | 12 | 0.762388818 | 0.721183548 | SLC25A12, MICU1, TESC, CAPS2, DGKG, NUCB2, PLCH1, PLS1, EFCAB1, MYL12A, CALM2, KCNIP4 | 878 | 144 | 10057 | 0.954536826 | 1 | 0.995764763 | 99.99999833 |
| UP_SEQ_FEATURE | domain:EF-hand 1 | 13 | 0.82592122 | 0.734613261 | MICU1, MYL12A, SLC25A12, CAPS2, CAPN13, DGKG, PLCH1, NUCB2, MCFD2, EFCAB1, PLS1, RYR2, CA | 1499 | 185 | 20063 | 0.940515298 | 1 | 0.999999994 | 100 |
| UP_SEQ_FEATURE | domain:EF-hand 3 | 6 | 0.381194409 | 0.85268835 | SLC25A12, CAPS2, EFCAB1, MYL12A, CALM2, KCNIP4 | 1499 | 96 | 20063 | 0.836516011 | 1 | 1 | 100 |
| UP_SEQ_FEATURE | domain:EF-hand 4 | 3 | 0.190597205 | 0.951384514 | SLC25A12, CALM2, KCNIP4 | 1499 | 62 | 20063 | 0.647625299 | 1 | 1 | 100 |
| Annotation Cluster 187 | Enrichment Score: 0.21879103364903604 | | | | | | | | | | | |
| Category | Term | Count | % | PValue | Genes | List Total | Pop Hits | Pop Total | Fold Enrichment | Bonferroni | Benjamini | FDR |
| UP_SEQ_FEATURE | DNA-binding region:Fork-head | 5 | 0.317662008 | 0.51884834 | FOXJ3, FOXI1, FOXP1, FOXN3, FOXP2 | 1499 | 50 | 20063 | 1.338425617 | 1 | 0.99999615 | 99.99987861 |
| INTERPRO | IPR001766:Transcription factor, fork head | 5 | 0.317662008 | 0.539934151 | FOXJ3, FOXI1, FOXP1, FOXN3, FOXP2 | 1452 | 49 | 18559 | 1.304253106 | 1 | 0.999859701 | 99.99984462 |
| SMART | SM00339:FH | 5 | 0.317662008 | 0.629072684 | FOXJ3, FOXI1, FOXP1, FOXN3, FOXP2 | 878 | 49 | 10057 | 1.168820603 | 1 | 0.992109292 | 99.99990875 |
| INTERPRO | IPR018122:Transcription factor, fork head, conserved site | 3 | 0.190597205 | 0.756403985 | FOXJ3, FOXI1, FOXN3 | 1452 | 34 | 18559 | 1.127795333 | 1 | 0.99999861 | 100 |
| Annotation Cluster 188 | Enrichment Score: 0.21535245216239604 | | | | | | | | | | | |
| Category | Term | Count | % | PValue | Genes | List Total | Pop Hits | Pop Total | Fold Enrichment | Bonferroni | Benjamini | FDR |
| INTERPRO | IPR002209:Heparin-binding growth factor/Fibroblast growth factor | 3 | 0.190597205 | 0.522103424 | FGF14, FGF23, FGF2 | 1452 | 22 | 18559 | 1.742956424 | 1 | 0.999830232 | 99.99970086 |
| SMART | SM00442:FGF | 3 | 0.190597205 | 0.583554014 | FGF14, FGF23, FGF2 | 878 | 22 | 10057 | 1.561969352 | 1 | 0.990540742 | 99.99953745 |
| INTERPRO | IPR008996:Cytokine, IL-1-like | 3 | 0.190597205 | 0.741489563 | FGF14, FGF23, FGF2 | 1452 | 33 | 18559 | 1.161970949 | 1 | 0.999997693 | 99.99999999 |
| Annotation Cluster 189 | Enrichment Score: 0.2132758103335207 | | | | | | | | | | | |
| Category | Term | Count | % | PValue | Genes | List Total | Pop Hits | Pop Total | Fold Enrichment | Bonferroni | Benjamini | FDR |
| REACTOME_PATHWAY | R-HSA-418990:R-HSA-418990 | 4 | 0.254129606 | 0.494350406 | CDH12, CDH13, ANG, CDH3 | 745 | 32 | 9075 | 1.522651007 | 1 | 0.997324648 | 99.99742427 |
| GOTERM_BP_DIRECT | GO:0034332~adherens junction organization | 4 | 0.254129606 | 0.575893154 | CDH12, CDH13, ANG, CDH3 | 1342 | 37 | 16792 | 1.352720828 | 1 | 0.999977626 | 99.99998834 |
| UP_SEQ_FEATURE | domain:Cadherin 5 | 7 | 0.444726811 | 0.804999773 | CDH12, CDH13, PCDH9, PCDH15, DSC1, CDH3, CDH23 | 1499 | 105 | 20063 | 0.892283745 | 1 | 1 | 100 |
| Annotation Cluster 190 | Enrichment Score: 0.19968549712746375 | | | | | | | | | | | |
| Category | Term | Count | % | PValue | Genes | List Total | Pop Hits | Pop Total | Fold Enrichment | Bonferroni | Benjamini | FDR |
| INTERPRO | IPR000834:Peptidase M14, carboxypeptidase A | 3 | 0.190597205 | 0.546452177 | CPO, AGBL3, AGBL1 | 1452 | 23 | 18559 | 1.66717571 | 1 | 0.999868223 | 99.99987849 |
| GOTERM_MF_DIRECT | GO:0004181~metallocarboxypeptidase activity | 3 | 0.190597205 | 0.643050074 | CPO, AGBL3, AGBL1 | 1340 | 27 | 16881 | 1.399751244 | 1 | 0.999414249 | 99.99999493 |
| UP_KEYWORDS | Carboxypeptidase | 3 | 0.190597205 | 0.716385119 | CPO, AGBL3, AGBL1 | 1535 | 33 | 20581 | 1.218892508 | 1 | 0.949549843 | 99.99999864 |

Annotation Cluster 191 Enrichment Score: 0.19750119961293414

| Category | Term | Count | % | PValue | Genes | List Total | Pop Hits | Pop Total | Fold Enrichment | Bonferroni | Benjamini | FDR |
|---|---|---|---|---|---|---|---|---|---|---|---|---|
| UP_KEYWORDS | Flavoprotein | 11 | 0.698856417 | 0.425301849 | HAO1, FMO4, RNLS, SQRDL, MTHFR, TXNRD3, ACAD11, QSOX2, PPCDC, QSOX1, ACAD10 | 1535 | 122 | 20581 | 1.208901586 | 1 | 0.818542935 | 99.96511 |
| GOTERM_MF_DIRECT | GO:0050660~flavin adenine dinucleotide binding | 6 | 0.381194409 | 0.580880314 | FMO4, NOX4, MTHFR, TXNRD3, ACAD11, ACAD10 | 1340 | 64 | 16881 | 1.181040112 | 1 | 0.998764687 | 99.99993052 |
| UP_KEYWORDS | FAD | 9 | 0.571791614 | 0.581202228 | FMO4, RNLS, SQRDL, MTHFR, TXNRD3, ACAD11, QSOX2, QSOX1, ACAD10 | 1535 | 110 | 20581 | 1.097003257 | 1 | 0.89919872 | 99.9996306 |
| INTERPRO | IPR023753:Pyridine nucleotide-disulphide oxidoreductase, FAD/NAD(P)-binding domain | 4 | 0.254129606 | 0.841916866 | FMO4, RNLS, SQRDL, TXNRD3 | 1452 | 58 | 18559 | 0.881495203 | 1 | 0.999999934 | 100 |
| UP_SEQ_FEATURE | nucleotide phosphate-binding region:FAD | 4 | 0.254129606 | 0.851360687 | FMO4, TXNRD3, ACAD11, ACAD10 | 1499 | 62 | 20063 | 0.863500398 | 1 | 1 | 100 |

Annotation Cluster 192 Enrichment Score: 0.1971132627306219

| Category | Term | Count | % | PValue | Genes | List Total | Pop Hits | Pop Total | Fold Enrichment | Bonferroni | Benjamini | FDR |
|---|---|---|---|---|---|---|---|---|---|---|---|---|
| GOTERM_BP_DIRECT | GO:0002474~antigen processing and presentation of peptide antigen via MHC class I | 4 | 0.254129606 | 0.433527357 | HLA-DQB1, TAP2, HLA-C, TAPBP | 1342 | 30 | 16792 | 1.668355688 | 1 | 0.99982233 | 99.99745192 |
| REACTOME_PATHWAY | R-HSA-983170:R-HSA-983170 | 3 | 0.190597205 | 0.641057799 | TAP2, HLA-C, TAPBP | 745 | 26 | 9075 | 1.405524006 | 1 | 0.999312355 | 99.99998727 |
| REACTOME_PATHWAY | R-HSA-1236974:R-HSA-1236974 | 5 | 0.317662008 | 0.922033107 | PSMA1, TAP2, PSMD1, HLA-C, TAPBP | 745 | 84 | 9075 | 0.725071908 | 1 | 0.999994193 | 100 |

Annotation Cluster 193 Enrichment Score: 0.1949228366423763

| Category | Term | Count | % | PValue | Genes | List Total | Pop Hits | Pop Total | Fold Enrichment | Bonferroni | Benjamini | FDR |
|---|---|---|---|---|---|---|---|---|---|---|---|---|
| INTERPRO | IPR000299:FERM domain | 5 | 0.317662008 | 0.572211454 | EPB41L3, EZR, MYO7B, FRMD8, JAK1 | 1452 | 51 | 18559 | 1.253105926 | 1 | 0.999911791 | 99.99995562 |
| INTERPRO | IPR019748:FERM central domain | 5 | 0.317662008 | 0.572211454 | EPB41L3, EZR, MYO7B, FRMD8, JAK1 | 1452 | 51 | 18559 | 1.253105926 | 1 | 0.999911791 | 99.99995562 |
| INTERPRO | IPR019749:Band 4.1 domain | 5 | 0.317662008 | 0.572211454 | EPB41L3, EZR, MYO7B, FRMD8, JAK1 | 1452 | 51 | 18559 | 1.253105926 | 1 | 0.999911791 | 99.99995562 |
| SMART | SM00295:B41 | 5 | 0.317662008 | 0.66072317 | EPB41L3, EZR, MYO7B, FRMD8, JAK1 | 878 | 51 | 10057 | 1.122984501 | 1 | 0.992770291 | 99.99997387 |
| UP_SEQ_FEATURE | domain:FERM | 4 | 0.254129606 | 0.718902673 | EPB41L3, EZR, FRMD8, JAK1 | 1499 | 49 | 20063 | 1.09259234 | 1 | 0.999999988 | 99.99999999 |
| INTERPRO | IPR014352:FERM/acyl-CoA-binding protein, 3-helical bundle | 4 | 0.254129606 | 0.760507856 | EPB41L3, EZR, MYO7B, FRMD8 | 1452 | 50 | 18559 | 1.022534435 | 1 | 0.99999874 | 100 |

Annotation Cluster 194 Enrichment Score: 0.19452875374859024

| Category | Term | Count | % | PValue | Genes | List Total | Pop Hits | Pop Total | Fold Enrichment | Bonferroni | Benjamini | FDR |
|---|---|---|---|---|---|---|---|---|---|---|---|---|
| GOTERM_MF_DIRECT | GO:0005549~odorant binding | 16 | 1.016518424 | 0.005667502 | OR5I1, OR5M10, OR5M11, OR8H3, OR5M3, OR5M9, OR5A1, OR5B2, OR5M8, OR8K5, OR5AS1, OR8I | 1340 | 92 | 16881 | 2.19091499 | 0.999330607 | 0.839150274 | 8.851178045 |
| KEGG_PATHWAY | hsa04740:Olfactory transduction | 32 | 2.033036849 | 0.689121495 | OR10A3, CAMK2G, OR8H3, OR4D10, CNGB1, OR5A1, OR52D1, OR8K5, OR8I2, OR5AS1, OR5F1, OR5I | 574 | 399 | 6879 | 0.961148516 | 1 | 0.920093911 | 99.9999803 |
| UP_KEYWORDS | Sensory transduction | 43 | 2.731893266 | 0.724492095 | OR10A3, BBS5, KERA, OR8H3, CNGB1, OR4D10, OR5A1, CDH3, RIMS1, OR52D1, HMCN2, SLC24A4, O | 1535 | 609 | 20581 | 0.946693196 | 1 | 0.950809955 | 99.9999991 |
| UP_KEYWORDS | Olfaction | 31 | 1.969504447 | 0.735264787 | OR10A3, OR8H3, OR4D10, CNGB1, OR5A1, OR52D1, OR8K5, SLC24A4, OR8I2, OR5AS1, OR5F1, OR6J | 1535 | 443 | 20581 | 0.938244572 | 1 | 0.952521845 | 99.99999949 |
| INTERPRO | IPR000725:Olfactory receptor | 29 | 1.842439644 | 0.868508505 | OR10A3, OR8H3, OR4D10, OR5A1, OR52D1, OR8K5, OR8I2, OR5AS1, OR5F1, OR6J1, OR5I1, OR2AG2, | 1452 | 430 | 18559 | 0.862020309 | 1 | 0.999999984 | 100 |
| GOTERM_BP_DIRECT | GO:0050911~detection of chemical stimulus involved in sensory perception of smell | 29 | 1.842439644 | 0.880580603 | OR10A3, OR8H3, OR4D10, OR5A1, OR52D1, OR8K5, OR8I2, OR5AS1, OR5F1, OR6J1, OR5I1, OR2AG2, | 1342 | 425 | 16792 | 0.853805558 | 1 | 0.999999996 | 100 |
| GOTERM_MF_DIRECT | GO:0004984~olfactory receptor activity | 29 | 1.842439644 | 0.888870287 | OR10A3, OR8H3, OR4D10, OR5A1, OR52D1, OR8K5, OR8I2, OR5AS1, OR5F1, OR6J1, OR5I1, OR2AG2, | 1340 | 431 | 16881 | 0.847645185 | 1 | 0.99999628 | 100 |
| REACTOME_PATHWAY | R-HSA-381753:R-HSA-381753 | 29 | 1.842439644 | 0.913811883 | OR10A3, OR8H3, OR4D10, OR5A1, OR52D1, OR8K5, OR8I2, OR5AS1, OR5F1, OR6J1, OR5I1, OR2AG2, | 745 | 426 | 9075 | 0.829237168 | 1 | 0.999992811 | 100 |
| UP_KEYWORDS | Receptor | 107 | 6.797966963 | 0.957159471 | OSMR, RORA, OR4D10, VIPR1, OR52D1, IL31RA, GRIN2B, OR8K5, ROBO1, S1PR4, ROBO2, UNC5C, RC | 1535 | 1648 | 20581 | 0.870531846 | 1 | 0.997150729 | 100 |
| INTERPRO | IPR000276:G protein-coupled receptor, rhodopsin-like | 42 | 2.668360864 | 0.989582102 | OR10A3, OR8H3, DRD4, CXCR1, OR4D10, OR5A1, OR52D1, OR8K5, HRH3, S1PR4, MAS1, OR8I2, OR5 | 1452 | 719 | 18559 | 0.746635019 | 1 | 1 | 100 |
| INTERPRO | IPR017452:GPCR, rhodopsin-like, 7TM | 42 | 2.668360864 | 0.993571054 | OR10A3, OR8H3, DRD4, CXCR1, OR4D10, OR5A1, OR52D1, OR8K5, HRH3, S1PR4, MAS1, OR8I2, OR5 | 1452 | 736 | 18559 | 0.729389373 | 1 | 1 | 100 |
| UP_KEYWORDS | G-protein coupled receptor | 45 | 2.858958069 | 0.996494313 | OR10A3, OR8H3, DRD4, CXCR1, GABBR2, OR4D10, OR5A1, VIPR1, OR52D1, OR8K5, HRH3, S1PR4, M | 1535 | 842 | 20581 | 0.716569824 | 1 | 0.999943391 | 100 |
| UP_KEYWORDS | Transducer | 48 | 3.049555273 | 0.997307453 | OR10A3, OR8H3, DRD4, CXCR1, GABBR2, OR4D10, OR5A1, VIPR1, OR52D1, PLCL1, OR8K5, HRH3, S1 | 1535 | 899 | 20581 | 0.715879026 | 1 | 0.999961487 | 100 |
| GOTERM_MF_DIRECT | GO:0004930~G-protein coupled receptor activity | 36 | 2.287166455 | 0.999495222 | OR10A3, OR8H3, CXCR1, OR4D10, OR5A1, OR52D1, OR8K5, S1PR4, MAS1, OR8I2, OR5AS1, OR5F1, C | 1340 | 710 | 16881 | 0.638759723 | 1 | 1 | 100 |
| GOTERM_BP_DIRECT | GO:0007186~G-protein coupled receptor signaling pathway | 46 | 2.92249047 | 0.999875149 | OR10A3, C5, OR8H3, CXCR1, AKAP13, GABBR2, OR4D10, OR5A1, VIPR1, OR52D1, OR8K5, INPP5K, S1 | 1342 | 899 | 16792 | 0.640247733 | 1 | 1 | 100 |

Annotation Cluster 195 Enrichment Score: 0.19024902322629905

| Category | Term | Count | % | PValue | Genes | List Total | Pop Hits | Pop Total | Fold Enrichment | Bonferroni | Benjamini | FDR |
|---|---|---|---|---|---|---|---|---|---|---|---|---|
| UP_KEYWORDS | Host cell receptor for virus entry | 6 | 0.381194409 | 0.523982222 | NCAM1, LAMP1, HLA-DRB1, SCARB1, CLEC5A, HTR2A | 1535 | 64 | 20581 | 1.256982899 | 1 | 0.872025083 | 99.99767275 |
| GOTERM_MF_DIRECT | GO:0001618~virus receptor activity | 6 | 0.381194409 | 0.660959035 | NCAM1, LAMP1, HLA-DRB1, SCARB1, CLEC5A, HTR2A | 1340 | 70 | 16881 | 1.079808102 | 1 | 0.999559534 | 99.99999781 |
| GOTERM_BP_DIRECT | GO:0046718~viral entry into host cell | 6 | 0.381194409 | 0.775821543 | NCAM1, LAMP1, SCARB1, KPNA3, CLEC5A, HTR2A | 1342 | 80 | 16792 | 0.938450075 | 1 | 0.999999824 | 100 |

Annotation Cluster 196 Enrichment Score: 0.18166971690155279

| Category | Term | Count | % | PValue | Genes | List Total | Pop Hits | Pop Total | Fold Enrichment | Bonferroni | Benjamini | FDR |
|---|---|---|---|---|---|---|---|---|---|---|---|---|
| UP_SEQ_FEATURE | DNA-binding region:ETS | 3 | 0.190597205 | 0.609053963 | FLI1, ELK3, ETV5 | 1499 | 27 | 20063 | 1.487139575 | 1 | 0.999999615 | 99.99999746 |
| INTERPRO | IPR000418:Ets domain | 3 | 0.190597205 | 0.654832406 | FLI1, ELK3, ETV5 | 1452 | 28 | 18559 | 1.369465762 | 1 | 0.999978858 | 99.9999989 |
| SMART | SM00413:ETS | 3 | 0.190597205 | 0.714833525 | FLI1, ELK3, ETV5 | 878 | 28 | 10057 | 1.227261634 | 1 | 0.995590025 | 99.99999771 |

Annotation Cluster 197 Enrichment Score: 0.17373939613639452

| Category | Term | Count | % | PValue | Genes | List Total | Pop Hits | Pop Total | Fold Enrichment | Bonferroni | Benjamini | FDR |
|---|---|---|---|---|---|---|---|---|---|---|---|---|
| INTERPRO | IPR002165:Plexin | 3 | 0.190597205 | 0.614163327 | ATRNL1, PLXDC2, SEMA4A | 1452 | 26 | 18559 | 1.474809282 | 1 | 0.999956231 | 99.9999925 |
| INTERPRO | IPR016201:Plexin-like fold | 4 | 0.254129606 | 0.679356607 | SEMA5A, ATRNL1, PLXDC2, SEMA4A | 1452 | 44 | 18559 | 1.161970949 | 1 | 0.999987446 | 99.99999969 |
| SMART | SM00423:PSI | 4 | 0.254129606 | 0.721772163 | SEMA5A, ATRNL1, PLXDC2, SEMA4A | 878 | 42 | 10057 | 1.09089923 | 1 | 0.995561803 | 99.99999838 |

Annotation Cluster 198 Enrichment Score: 0.17330036332842266

| Category | Term | Count | % | PValue | Genes | List Total | Pop Hits | Pop Total | Fold Enrichment | Bonferroni | Benjamini | FDR |
|---|---|---|---|---|---|---|---|---|---|---|---|---|
| UP_KEYWORDS | Collagen degradation | 4 | 0.254129606 | 0.146228254 | ADAMTS14, MMP8, MMP27, MMP16 | 1535 | 18 | 20581 | 2.97951502 | 1 | 0.534251002 | 89.69012956 |
| UP_SEQ_FEATURE | short sequence motif:Cysteine switch | 5 | 0.317662008 | 0.435957577 | MMP8, MMP27, ADAM33, MMP16, ADAM12 | 1499 | 45 | 20063 | 1.487139575 | 1 | 0.999975119 | 99.99765847 |
| INTERPRO | IPR006586:ADAM, cysteine-rich | 3 | 0.190597205 | 0.654832406 | ADAMTS14, ADAM33, ADAM12 | 1452 | 28 | 18559 | 1.369465762 | 1 | 0.999978858 | 99.9999989 |
| UP_KEYWORDS | Metalloprotease | 11 | 0.698856417 | 0.704230427 | CPO, AGBL3, AGBL1, ADAMTS14, MMP8, MMP27, ADAM33, MMP16, ENPEP, ADAM12, C9ORF3 | 1535 | 152 | 20581 | 0.970302589 | 1 | 0.946579534 | 99.99999751 |
| SMART | SM00608:ACR | 3 | 0.190597205 | 0.714833525 | ADAMTS14, ADAM33, ADAM12 | 878 | 28 | 10057 | 1.227261634 | 1 | 0.995590025 | 99.99999771 |
| UP_SEQ_FEATURE | metal ion-binding site:Zinc; in inhibited form | 3 | 0.190597205 | 0.747333195 | ADAM33, MMP16, ADAM12 | 1499 | 35 | 20063 | 1.147221957 | 1 | 0.999999997 | 100 |
| UP_SEQ_FEATURE | metal ion-binding site:Zinc; catalytic | 9 | 0.571791614 | 0.755829495 | ADAMTS14, CA9, ADAM33, MMP16, ENPEP, CA2, ADAM12, CA1, C9ORF3 | 1499 | 129 | 20063 | 0.933785314 | 1 | 0.999999998 | 100 |
| INTERPRO | IPR024079:Metallopeptidase, catalytic domain | 6 | 0.381194409 | 0.768896092 | ADAMTS14, MMP8, MMP27, ADAM33, MMP16, ADAM12 | 1452 | 81 | 18559 | 0.946791144 | 1 | 0.999998979 | 100 |
| INTERPRO | IPR001590:Peptidase M12B, ADAM/reprolysin | 3 | 0.190597205 | 0.831064732 | ADAMTS14, ADAM33, ADAM12 | 1452 | 40 | 18559 | 0.958626033 | 1 | 0.999999896 | 100 |
| INTERPRO | IPR002870:Peptidase M12B, propeptide | 3 | 0.190597205 | 0.831064732 | ADAMTS14, ADAM33, ADAM12 | 1452 | 40 | 18559 | 0.958626033 | 1 | 0.999999896 | 100 |
| UP_SEQ_FEATURE | domain:Peptidase M12B | 3 | 0.190597205 | 0.831671232 | ADAMTS14, ADAM33, ADAM12 | 1499 | 42 | 20063 | 0.956018298 | 1 | 1 | 100 |
| UP_SEQ_FEATURE | domain:Disintegrin | 3 | 0.190597205 | 0.841394622 | ADAMTS14, ADAM33, ADAM12 | 1499 | 43 | 20063 | 0.933785314 | 1 | 1 | 100 |
| GOTERM_MF_DIRECT | GO:0008237~metallopeptidase activity | 5 | 0.317662008 | 0.893351752 | ADAMTSL1, ADAM33, ENPEP, ADAM12, C9ORF3 | 1340 | 81 | 16881 | 0.77763958 | 1 | 0.999996709 | 100 |
| GOTERM_MF_DIRECT | GO:0004222~metalloendopeptidase activity | 6 | 0.381194409 | 0.951086802 | ADAMTS14, MMP8, MMP27, ADAM33, MMP16, ADAM12 | 1340 | 113 | 16881 | 0.668907674 | 1 | 0.999999905 | 100 |

Annotation Cluster 199 Enrichment Score: 0.17098114732882475

| Category | Term | Count | % | PValue | Genes | List Total | Pop Hits | Pop Total | Fold Enrichment | Bonferroni | Benjamini | FDR |
|---|---|---|---|---|---|---|---|---|---|---|---|---|
| REACTOME_PATHWAY | R-HSA-1368108:R-HSA-1368108 | 5 | 0.317662008 | 0.473861631 | TBL1XR1, HDAC3, AVP, NR3C1, RORA | 745 | 43 | 9075 | 1.416419541 | 1 | 0.996632729 | 99.99523341 |
| REACTOME_PATHWAY | R-HSA-1368082:R-HSA-1368082 | 3 | 0.190597205 | 0.661676623 | TBL1XR1, HDAC3, RORA | 745 | 27 | 9075 | 1.353467562 | 1 | 0.999422165 | 99.99999491 |
| REACTOME_PATHWAY | R-HSA-1989781:R-HSA-1989781 | 9 | 0.571791614 | 0.718361738 | TBL1XR1, UGT1A9, HDAC3, MED30, FHL2, TEAD1, MED23, TEAD3, RORA | 745 | 113 | 9075 | 0.970184712 | 1 | 0.999648836 | 99.9999997 |
| REACTOME_PATHWAY | R-HSA-400253:R-HSA-400253 | 3 | 0.190597205 | 0.919252631 | TBL1XR1, HDAC3, RORA | 745 | 49 | 9075 | 0.745788248 | 1 | 0.999993995 | 100 |

Annotation Cluster 200 Enrichment Score: 0.17025377743414752

| Category | Term | Count | % | PValue | Genes | List Total | Pop Hits | Pop Total | Fold Enrichment | Bonferroni | Benjamini | FDR |
|---|---|---|---|---|---|---|---|---|---|---|---|---|
| UP_SEQ_FEATURE | binding site:NAD | 5 | 0.317662008 | 0.580782325 | ME3, CTBP2, PHGDH, DHPS, HSD17B4 | 1499 | 54 | 20063 | 1.239282979 | 1 | 0.999999294 | 99.99999067 |
| GOTERM_MF_DIRECT | GO:0051287~NAD binding | 4 | 0.254129606 | 0.589585575 | ME3, CTBP2, PHGDH, AHCYL2 | 1340 | 38 | 16881 | 1.326080126 | 1 | 0.998813841 | 99.99995065 |
| INTERPRO | IPR016040:NAD(P)-binding domain | 11 | 0.698856417 | 0.900903817 | FAR1, VAT1L, ME3, CTBP2, KCNT2, GMDS, PHGDH, HSD17B4, AHCYL2, DHRS7C, WWOX | 1452 | 179 | 18559 | 0.785466396 | 1 | 0.999999998 | 100 |

Annotation Cluster 201 Enrichment Score: 0.16578464617498867

| Category | Term | Count | % | PValue | Genes | List Total | Pop Hits | Pop Total | Fold Enrichment | Bonferroni | Benjamini | FDR |
|---|---|---|---|---|---|---|---|---|---|---|---|---|
| UP_SEQ_FEATURE | domain:Leucine-zipper | 11 | 0.698856417 | 0.335685534 | XRCC5, FOSL2, CEBPE, MAFB, ATF6B, LUZP2, TSN, TCF3, MYC, FOXP1, FOXP2 | 1499 | 113 | 20063 | 1.302892194 | 1 | 0.999836999 | 99.95072212 |
| INTERPRO | IPR004827:Basic-leucine zipper domain | 4 | 0.254129606 | 0.804758245 | FOSL2, CEBPE, MAFB, ATF6B | 1452 | 54 | 18559 | 0.946791144 | 1 | 0.999999704 | 100 |
| SMART | SM00338:BRLZ | 4 | 0.254129606 | 0.833843884 | FOSL2, CEBPE, MAFB, ATF6B | 878 | 51 | 10057 | 0.898387601 | 1 | 0.999224538 | 100 |
| UP_SEQ_FEATURE | DNA-binding region:Basic motif | 8 | 0.508259212 | 0.964225485 | FOSL2, CEBPE, MAFB, ATF6B, MGA, TCF3, MYC, TWIST1 | 1499 | 163 | 20063 | 0.656896008 | 1 | 1 | 100 |

Annotation Cluster 202 Enrichment Score: 0.16453286381868154

| Category | Term | Count | % | PValue | Genes | List Total | Pop Hits | Pop Total | Fold Enrichment | Bonferroni | Benjamini | FDR |
|---|---|---|---|---|---|---|---|---|---|---|---|---|
| INTERPRO | IPR015422:Pyridoxal phosphate-dependent transferase, major region, subdomain 2 | 4 | 0.254129606 | 0.579448528 | MOCOS, SHMT2, GADL1, PDXDC1 | 1452 | 38 | 18559 | 1.345440046 | 1 | 0.999923037 | 99.99996692 |
| INTERPRO | IPR015421:Pyridoxal phosphate-dependent transferase, major region, subdomain 1 | 4 | 0.254129606 | 0.664014634 | MOCOS, SHMT2, GADL1, PDXDC1 | 1452 | 43 | 18559 | 1.188993529 | 1 | 0.999983003 | 99.99999931 |
| INTERPRO | IPR015424:Pyridoxal phosphate-dependent transferase | 4 | 0.254129606 | 0.664014634 | MOCOS, SHMT2, GADL1, PDXDC1 | 1452 | 43 | 18559 | 1.188993529 | 1 | 0.999983003 | 99.99999931 |
| UP_KEYWORDS | Pyridoxal phosphate | 5 | 0.317662008 | 0.700394528 | MOCOS, SHMT2, GADL1, PHOSPHO2, PDXDC1 | 1535 | 63 | 20581 | 1.064112507 | 1 | 0.946449485 | 99.999997 |
| GOTERM_MF_DIRECT | GO:0030170~pyridoxal phosphate binding | 4 | 0.254129606 | 0.84066462 | MOCOS, SHMT2, GADL1, PDXDC1 | 1340 | 57 | 16881 | 0.884053417 | 1 | 0.999983068 | 100 |

Annotation Cluster 203 Enrichment Score: 0.1604405001092156

| Category | Term | Count | % | PValue | Genes | List Total | Pop Hits | Pop Total | Fold Enrichment | Bonferroni | Benjamini | FDR |
|---|---|---|---|---|---|---|---|---|---|---|---|---|
| UP_SEQ_FEATURE | repeat:30 | 3 | 0.190597205 | 0.393340669 | ALMS1, NELFE, MUC17 | 1499 | 18 | 20063 | 2.230709362 | 1 | 0.999938317 | 99.99091136 |
| UP_SEQ_FEATURE | repeat:15 | 7 | 0.444726811 | 0.479012988 | PRB4, KRTAP9-3, ALMS1, MAP4, NELFE, SCEL, MUC17 | 1499 | 74 | 20063 | 1.266078286 | 1 | 0.999990331 | 99.99946617 |
| UP_SEQ_FEATURE | repeat:16 | 6 | 0.381194409 | 0.482408606 | KRTAP9-3, ALMS1, MAP4, NELFE, SCEL, MUC17 | 1499 | 61 | 20063 | 1.316484214 | 1 | 0.999990478 | 99.99952737 |
| UP_SEQ_FEATURE | repeat:29 | 3 | 0.190597205 | 0.544081123 | ALMS1, NELFE, MUC17 | 1499 | 24 | 20063 | 1.673032021 | 1 | 0.999998084 | 99.99995548 |
| UP_SEQ_FEATURE | repeat:14 | 7 | 0.444726811 | 0.556422923 | PRB4, KRTAP9-3, ALMS1, MAP4, NELFE, SCEL, MUC17 | 1499 | 80 | 20063 | 1.171122415 | 1 | 0.999998721 | 99.99997329 |
| UP_SEQ_FEATURE | repeat:2 | 20 | 1.27064803 | 0.573393527 | SRRD, PLB1, PRB4, KRTAP9-3, DRD4, COL15A1, ALMS1, SPRR2G, CRIPAK, CDHR5, MUC6, HNRNPR, S | 1499 | 260 | 20063 | 1.029558167 | 1 | 0.999999112 | 99.99998708 |
| UP_SEQ_FEATURE | repeat:28 | 3 | 0.190597205 | 0.588210376 | ALMS1, NELFE, MUC17 | 1499 | 26 | 20063 | 1.54433725 | 1 | 0.999999408 | 99.99999331 |
| UP_SEQ_FEATURE | repeat:10 | 8 | 0.508259212 | 0.60412117 | PRB4, KRTAP9-3, ALMS1, MAP4, CRIPAK, NELFE, SCEL, MUC17 | 1499 | 98 | 20063 | 1.09259234 | 1 | 0.999999604 | 99.99999679 |
| UP_SEQ_FEATURE | repeat:13 | 7 | 0.444726811 | 0.604913545 | PRB4, KRTAP9-3, ALMS1, MAP4, NELFE, SCEL, MUC17 | 1499 | 84 | 20063 | 1.115354681 | 1 | 0.999999594 | 99.99999691 |
| UP_SEQ_FEATURE | repeat:3 | 17 | 1.080050826 | 0.617637647 | SRRD, PLB1, PRB4, KRTAP9-3, DRD4, COL15A1, SPRR2G, ALMS1, CRIPAK, CDHR5, SCEL, ZNF628, RYR | 1499 | 225 | 20063 | 1.011254911 | 1 | 0.999999695 | 99.99999832 |
| UP_SEQ_FEATURE | repeat:12 | 7 | 0.444726811 | 0.628039527 | PRB4, KRTAP9-3, ALMS1, MAP4, NELFE, SCEL, MUC17 | 1499 | 86 | 20063 | 1.0894162 | 1 | 0.999999777 | 99.99999899 |
| UP_SEQ_FEATURE | repeat:27 | 3 | 0.190597205 | 0.648303045 | ALMS1, NELFE, MUC17 | 1499 | 29 | 20063 | 1.384578225 | 1 | 0.999999866 | 99.99999965 |
| UP_SEQ_FEATURE | repeat:11 | 7 | 0.444726811 | 0.66120845 | PRB4, KRTAP9-3, ALMS1, MAP4, NELFE, SCEL, MUC17 | 1499 | 89 | 20063 | 1.052694306 | 1 | 0.999999914 | 99.99999982 |
| UP_SEQ_FEATURE | repeat:4 | 14 | 0.889453621 | 0.68333334 | PLB1, PRB4, DRD4, KRTAP9-3, COL15A1, ALMS1, CRIPAK, SCEL, ZNF628, RYR3, RYR2, MAP4, NELFE, [illegible] | 1499 | 192 | 20063 | 0.975935346 | 1 | 0.999999955 | 99.99999995 |
| UP_SEQ_FEATURE | repeat:26 | 3 | 0.190597205 | 0.684355102 | ALMS1, NELFE, MUC17 | 1499 | 31 | 20063 | 1.295250597 | 1 | 0.999999954 | 99.99999995 |
| UP_SEQ_FEATURE | repeat:9 | 8 | 0.508259212 | 0.695424403 | PRB4, KRTAP9-3, ALMS1, MAP4, CRIPAK, NELFE, SCEL, MUC17 | 1499 | 107 | 20063 | 1.00069205 | 1 | 0.999999971 | 99.99999998 |
| UP_SEQ_FEATURE | repeat:24 | 3 | 0.190597205 | 0.732683833 | ALMS1, NELFE, MUC17 | 1499 | 34 | 20063 | 1.18096378 | 1 | 0.999999994 | 100 |
| UP_SEQ_FEATURE | repeat:25 | 3 | 0.190597205 | 0.732683833 | ALMS1, NELFE, MUC17 | 1499 | 34 | 20063 | 1.18096378 | 1 | 0.999999994 | 100 |
| UP_SEQ_FEATURE | repeat:23 | 3 | 0.190597205 | 0.774567359 | ALMS1, NELFE, MUC17 | 1499 | 37 | 20063 | 1.08520996 | 1 | 0.999999999 | 100 |
| UP_SEQ_FEATURE | repeat:17 | 4 | 0.254129606 | 0.79886594 | ALMS1, MAP4, NELFE, MUC17 | 1499 | 56 | 20063 | 0.956018298 | 1 | 1 | 100 |
| UP_SEQ_FEATURE | repeat:1 | 17 | 1.080050826 | 0.809046345 | SRRD, PLB1, PRB4, KRTAP9-3, COL15A1, SPRR2G, ALMS1, CRIPAK, CDHR5, SCEL, ZNF628, AMBN, RY | 1499 | 257 | 20063 | 0.885339902 | 1 | 1 | 100 |
| UP_SEQ_FEATURE | repeat:22 | 3 | 0.190597205 | 0.821412827 | ALMS1, NELFE, MUC17 | 1499 | 41 | 20063 | 0.979335817 | 1 | 1 | 100 |
| UP_SEQ_FEATURE | repeat:21 | 3 | 0.190597205 | 0.841394622 | ALMS1, NELFE, MUC17 | 1499 | 43 | 20063 | 0.933785314 | 1 | 1 | 100 |
| UP_SEQ_FEATURE | repeat:6 | 9 | 0.571791614 | 0.866078465 | PRB4, KRTAP9-3, DRD4, ALMS1, MAP4, CRIPAK, NELFE, SCEL, MUC17 | 1499 | 147 | 20063 | 0.819444255 | 1 | 1 | 100 |
| UP_SEQ_FEATURE | repeat:20 | 3 | 0.190597205 | 0.867579071 | ALMS1, NELFE, MUC17 | 1499 | 46 | 20063 | 0.872886272 | 1 | 1 | 100 |
| UP_SEQ_FEATURE | repeat:7 | 8 | 0.508259212 | 0.875963868 | PRB4, KRTAP9-3, ALMS1, MAP4, CRIPAK, NELFE, SCEL, MUC17 | 1499 | 133 | 20063 | 0.80506804 | 1 | 1 | 100 |
| UP_SEQ_FEATURE | repeat:19 | 3 | 0.190597205 | 0.882763612 | ALMS1, NELFE, MUC17 | 1499 | 48 | 20063 | 0.836516011 | 1 | 1 | 100 |
| UP_SEQ_FEATURE | repeat:18 | 3 | 0.190597205 | 0.889736418 | ALMS1, NELFE, MUC17 | 1499 | 49 | 20063 | 0.819444255 | 1 | 1 | 100 |
| UP_SEQ_FEATURE | repeat:8 | 7 | 0.444726811 | 0.908527524 | PRB4, KRTAP9-3, ALMS1, CRIPAK, NELFE, SCEL, MUC17 | 1499 | 124 | 20063 | 0.755562848 | 1 | 1 | 100 |
| UP_SEQ_FEATURE | repeat:5 | 8 | 0.508259212 | 0.960912317 | PRB4, KRTAP9-3, DRD4, ALMS1, MAP4, CRIPAK, SCEL, MUC17 | 1499 | 161 | 20063 | 0.665056207 | 1 | 1 | 100 |

Annotation Cluster 204 Enrichment Score: 0.14769082184986942

| Category | Term | Count | % | PValue | Genes | List Total | Pop Hits | Pop Total | Fold Enrichment | Bonferroni | Benjamini | FDR |
|---|---|---|---|---|---|---|---|---|---|---|---|---|
| GOTERM_CC_DIRECT | GO:0071013~catalytic step 2 spliceosome | 8 | 0.508259212 | 0.585482445 | TFIP11, WDR83, DDX23, SNRPN, PPIL1, SNRPB, HNRNPR, SNURF | 1425 | 92 | 18224 | 1.112067124 | 1 | 0.977205769 | 99.99982348 |
| UP_KEYWORDS | Viral nucleoprotein | 3 | 0.190597205 | 0.628100962 | SNRPN, SNRPB, HNRNPR | 1535 | 28 | 20581 | 1.436551885 | 1 | 0.920073282 | 99.99993298 |
| GOTERM_CC_DIRECT | GO:0019013~viral nucleocapsid | 3 | 0.190597205 | 0.654520255 | SNRPN, SNRPB, HNRNPR | 1425 | 28 | 18224 | 1.370225564 | 1 | 0.98494901 | 99.99998861 |
| GOTERM_CC_DIRECT | GO:0005681~spliceosomal complex | 6 | 0.381194409 | 0.867364989 | TFIP11, WDR83, SNRPN, SNRPB, HNRNPR, SNURF | 1425 | 94 | 18224 | 0.816304591 | 1 | 0.998962067 | 100 |
| UP_KEYWORDS | Virion | 3 | 0.190597205 | 0.874737159 | SNRPN, SNRPB, HNRNPR | 1535 | 47 | 20581 | 0.855818144 | 1 | 0.985475792 | 100 |

Annotation Cluster 205 Enrichment Score: 0.13892523305551854

| Category | Term | Count | % | PValue | Genes | List Total | Pop Hits | Pop Total | Fold Enrichment | Bonferroni | Benjamini | FDR |
|---|---|---|---|---|---|---|---|---|---|---|---|---|
| GOTERM_BP_DIRECT | GO:0000413~protein peptidyl-prolyl isomerization | 4 | 0.254129606 | 0.678452476 | PPIG, FKBP4, PPIL1, FKBPL | 1342 | 43 | 16792 | 1.163969085 | 1 | 0.999997747 | 99.99999993 |
| GOTERM_MF_DIRECT | GO:0003755~peptidyl-prolyl cis-trans isomerase activity | 4 | 0.254129606 | 0.7179385 | PPIG, FKBP4, PPIL1, FKBPL | 1340 | 46 | 16881 | 1.095457495 | 1 | 0.999800853 | 99.99999989 |
| UP_KEYWORDS | Rotamase | 3 | 0.190597205 | 0.786353141 | PPIG, FKBP4, PPIL1 | 1535 | 38 | 20581 | 1.058511915 | 1 | 0.966014741 | 99.99999998 |

Annotation Cluster 206 Enrichment Score: 0.12606682812289685

| Category | Term | Count | % | PValue | Genes | List Total | Pop Hits | Pop Total | Fold Enrichment | Bonferroni | Benjamini | FDR |
|---|---|---|---|---|---|---|---|---|---|---|---|---|
| BIOCARTA | h_eifPathway:Eukaryotic protein translation | 4 | 0.254129606 | 0.165488103 | EIF4E, EIF4A1, EEF2, EIF1 | 146 | 16 | 1625 | 2.782534247 | 1 | 0.993688815 | 90.0692016 |
| GOTERM_BP_DIRECT | GO:0006446~regulation of translational initiation | 5 | 0.317662008 | 0.323400724 | EIF3D, KLHL25, EIF4A1, EIF1, EIF3I | 1342 | 36 | 16792 | 1.737870508 | 1 | 0.999071796 | 99.93047155 |
| UP_KEYWORDS | Protein biosynthesis | 13 | 0.82592122 | 0.461582068 | EEF1B2, AIMP1, FARS2, EEF2, LARS2, EIF3D, EIF4E, ELOF1, EIF4A1, LARS, EIF3I, EIF1, RMND1 | 1535 | 152 | 20581 | 1.146721241 | 1 | 0.839217399 | 99.98633261 |
| REACTOME_PATHWAY | R-HSA-72702:R-HSA-72702 | 6 | 0.381194409 | 0.567792511 | EIF3D, RPS18, EIF4E, EIF4A1, EIF3I, RPS7 | 745 | 61 | 9075 | 1.198151612 | 1 | 0.998892622 | 99.99977367 |
| REACTOME_PATHWAY | R-HSA-72649:R-HSA-72649 | 6 | 0.381194409 | 0.567792511 | EIF3D, RPS18, EIF4E, EIF4A1, EIF3I, RPS7 | 745 | 61 | 9075 | 1.198151612 | 1 | 0.998892622 | 99.99977367 |
| REACTOME_PATHWAY | R-HSA-156827:R-HSA-156827 | 10 | 0.635324015 | 0.597359206 | EIF3D, RPS18, EIF4E, RPL6, RPL31, EIF4A1, RPL27A, EIF3I, RPL10A, RPS7 | 745 | 114 | 9075 | 1.068527022 | 1 | 0.999015582 | 99.99992451 |
| REACTOME_PATHWAY | R-HSA-72706:R-HSA-72706 | 10 | 0.635324015 | 0.608047657 | EIF3D, RPS18, EIF4E, RPL6, RPL31, EIF4A1, RPL27A, EIF3I, RPL10A, RPS7 | 745 | 115 | 9075 | 1.059235483 | 1 | 0.999146096 | 99.99995025 |
| UP_KEYWORDS | Initiation factor | 5 | 0.317662008 | 0.636469615 | EIF3D, EIF4E, EIF4A1, EIF1, EIF3I | 1535 | 58 | 20581 | 1.155846344 | 1 | 0.923576001 | 99.99995168 |
| GOTERM_BP_DIRECT | GO:0006413~translational initiation | 11 | 0.698856417 | 0.662806564 | EIF3D, RPS18, EIF4E, RPL6, RPL31, EIF4A1, RPL27A, EIF1, EIF3I, RPL10A, RPS7 | 1342 | 137 | 16792 | 1.004666746 | 1 | 0.999996381 | 99.99999984 |
| GOTERM_MF_DIRECT | GO:0003743~translation initiation factor activity | 5 | 0.317662008 | 0.723403504 | EIF3D, EIF4E, EIF4A1, EIF1, EIF3I | 1340 | 61 | 16881 | 1.032603377 | 1 | 0.999817385 | 99.99999992 |
| REACTOME_PATHWAY | R-HSA-72689:R-HSA-72689 | 8 | 0.508259212 | 0.759551714 | EIF3D, RPS18, RPL6, RPL31, RPL27A, EIF3I, RPL10A, RPS7 | 745 | 104 | 9075 | 0.937016004 | 1 | 0.999821656 | 99.99999997 |
| REACTOME_PATHWAY | R-HSA-156902:R-HSA-156902 | 7 | 0.444726811 | 0.767203406 | RPS18, RPL6, RPL31, RPL27A, EEF2, RPL10A, RPS7 | 745 | 91 | 9075 | 0.937016004 | 1 | 0.999823998 | 99.99999998 |
| GOTERM_BP_DIRECT | GO:0019083~viral transcription | 8 | 0.508259212 | 0.800305874 | RPS18, NUP88, RPL6, RPL31, RPL27A, RPL10A, NUP35, RPS7 | 1342 | 112 | 16792 | 0.893761976 | 1 | 0.999999913 | 100 |
| REACTOME_PATHWAY | R-HSA-72695:R-HSA-72695 | 4 | 0.254129606 | 0.830866211 | EIF3D, RPS18, EIF3I, RPS7 | 745 | 54 | 9075 | 0.902311708 | 1 | 0.999940779 | 100 |
| REACTOME_PATHWAY | R-HSA-192823:R-HSA-192823 | 6 | 0.381194409 | 0.87706745 | RPS18, RPL6, RPL31, RPL27A, RPL10A, RPS7 | 745 | 91 | 9075 | 0.803156575 | 1 | 0.999979486 | 100 |
| GOTERM_BP_DIRECT | GO:0006614~SRP-dependent cotranslational protein targeting to membrane | 6 | 0.381194409 | 0.879529962 | RPS18, RPL6, RPL31, RPL27A, RPL10A, RPS7 | 1342 | 94 | 16792 | 0.798680914 | 1 | 0.999999996 | 100 |

| | | | | | | | | | | | | |
|---|---|---|---|---|---|---|---|---|---|---|---|---|
| REACTOME_PATHWAY | R-HSA-2408557:R-HSA-2408557 | 6 | 0.381194409 | 0.893753329 | RPS18, RPL6, RPL31, RPL27A, RPL10A, RPS7 | 745 | 94 | 9075 | 0.777523918 | 1 | 0.999988134 | 100 |
| REACTOME_PATHWAY | R-HSA-72764:R-HSA-72764 | 6 | 0.381194409 | 0.898863641 | RPS18, RPL6, RPL31, RPL27A, RPL10A, RPS7 | 745 | 95 | 9075 | 0.769339456 | 1 | 0.999989419 | 100 |
| REACTOME_PATHWAY | R-HSA-975956:R-HSA-975956 | 6 | 0.381194409 | 0.908447854 | RPS18, RPL6, RPL31, RPL27A, RPL10A, RPS7 | 745 | 97 | 9075 | 0.753476787 | 1 | 0.99999156 | 100 |
| GOTERM_CC_DIRECT | GO:0022625~cytosolic large ribosomal subunit | 4 | 0.254129606 | 0.908907694 | RPL6, RPL31, RPL27A, RPL10A | 1425 | 68 | 18224 | 0.752280702 | 1 | 0.999620045 | 100 |
| GOTERM_CC_DIRECT | GO:0005840~ribosome | 9 | 0.571791614 | 0.952526237 | RPS18, RPL6, RPL31, MRPL54, RPL27A, MRPL9, RPL10A, METTL17, RPS7 | 1425 | 166 | 18224 | 0.693367153 | 1 | 0.999935466 | 100 |
| GOTERM_BP_DIRECT | GO:0006364~rRNA processing | 12 | 0.762388818 | 0.960079528 | KRR1, SENP3, RPS18, NOLC1, RPL6, RPL31, EXOSC5, RPL27A, RPL10A, PWP2, WDR46, RPS7 | 1342 | 214 | 16792 | 0.701644916 | 1 | 1 | 100 |
| UP_KEYWORDS | Ribonucleoprotein | 16 | 1.016518424 | 0.960407603 | KRR1, SNRPN, RPL27A, MRPL9, HNRNPR, SNURF, RPS7, RPS18, PCBP3, RPL31, RPL6, MRPL54, SNRP | 1535 | 296 | 20581 | 0.724746897 | 1 | 0.997426219 | 100 |
| REACTOME_PATHWAY | R-HSA-1799339:R-HSA-1799339 | 6 | 0.381194409 | 0.962543618 | RPS18, RPL6, RPL31, RPL27A, RPL10A, RPS7 | 745 | 114 | 9075 | 0.641116213 | 1 | 0.999999212 | 100 |
| REACTOME_PATHWAY | R-HSA-975957:R-HSA-975957 | 6 | 0.381194409 | 0.964547066 | RPS18, RPL6, RPL31, RPL27A, RPL10A, RPS7 | 745 | 115 | 9075 | 0.63554129 | 1 | 0.999999283 | 100 |
| GOTERM_BP_DIRECT | GO:0000184~nuclear-transcribed mRNA catabolic process, nonsense-mediated decay | 6 | 0.381194409 | 0.965790579 | RPS18, RPL6, RPL31, RPL27A, RPL10A, RPS7 | 1342 | 119 | 16792 | 0.630890806 | 1 | 1 | 100 |
| UP_KEYWORDS | Ribosomal protein | 9 | 0.571791614 | 0.969878761 | RPS18, RPL6, RPL31, MRPL54, RPL27A, MRPL9, RPL10A, METTL17, RPS7 | 1535 | 185 | 20581 | 0.652272207 | 1 | 0.9983307 | 100 |
| GOTERM_BP_DIRECT | GO:0006412~translation | 14 | 0.889453621 | 0.970612215 | CPEB2, CPEB3, RPL27A, MRPL9, RPS7, SLC25A12, RPS18, SLC25A34, RPL31, RPL6, DHPS, RPL10A, RM | 1342 | 253 | 16792 | 0.622400582 | 1 | 1 | 100 |
| KEGG_PATHWAY | hsa03010:Ribosome | 7 | 0.444726811 | 0.975106101 | RPS18, RPL6, RPL31, RPL27A, MRPL9, RPL10A, RPS7 | 574 | 136 | 6879 | 0.616840029 | 1 | 0.994928002 | 100 |
| KEGG_PATHWAY | hsa03013:RNA transport | 9 | 0.571791614 | 0.979221509 | EIF3D, EIF4E, NUP88, EIF4A1, CYFIP1, THOC7, EIF1, EIF3I, NUP35 | 574 | 172 | 6879 | 0.627086541 | 1 | 0.995740436 | 100 |
| GOTERM_MF_DIRECT | GO:0003735~structural constituent of ribosome | 9 | 0.571791614 | 0.997336246 | SLC25A12, RPS18, SLC25A34, RPL6, RPL31, RPL27A, MRPL9, RPL10A, RPS7 | 1340 | 222 | 16881 | 0.510720048 | 1 | 1 | 100 |

Annotation Cluster 207 Enrichment Score: 0.12086381166412963

| Category | Term | Count | % | PValue | Genes | List Total | Pop Hits | Pop Total | Fold Enrichment | Bonferroni | Benjamini | FDR |
|---|---|---|---|---|---|---|---|---|---|---|---|---|
| GOTERM_MF_DIRECT | GO:0003730~mRNA 3'-UTR binding | 5 | 0.317662008 | 0.551706263 | CPEB2, CPEB3, RBMS3, HNRNPR, AUH | 1340 | 49 | 16881 | 1.285485836 | 1 | 0.998658392 | 99.99979183 |
| UP_SEQ_FEATURE | domain:RRM | 11 | 0.698856417 | 0.557673949 | RBFOX1, RALYL, RBPMS, TNRC6C, TRA2B, RBM20, PPRC1, SSB, NELFE, TNRC6B, LARP4B | 1499 | 135 | 20063 | 1.090569021 | 1 | 0.99999871 | 99.99997466 |
| INTERPRO | IPR012677:Nucleotide-binding, alpha-beta plait | 20 | 1.27064803 | 0.68075677 | RBFOX1, RALYL, CPEB2, CPEB3, RBFOX3, RBM20, TRA2B, SSB, LARP4B, HNRNPR, RBPMS, TNRC6C, C | 1452 | 264 | 18559 | 0.968309124 | 1 | 0.9999873 | 99.99999971 |
| INTERPRO | IPR000504:RNA recognition motif domain | 17 | 1.080050826 | 0.69707717 | RBFOX1, RALYL, CPEB2, CPEB3, RBM20, TRA2B, RBFOX3, SSB, HNRNPR, RBPMS, TNRC6C, CELF3, PP | 1452 | 226 | 18559 | 0.961453838 | 1 | 0.999991527 | 99.99999988 |
| SMART | SM00360:RRM | 16 | 1.016518424 | 0.836733231 | RBFOX1, RALYL, CPEB2, CPEB3, RBM20, TRA2B, RBFOX3, SSB, HNRNPR, RBPMS, CELF3, PPRC1, RBM | 878 | 212 | 10057 | 0.864486182 | 1 | 0.999171256 | 100 |
| GOTERM_MF_DIRECT | GO:0000166~nucleotide binding | 24 | 1.524777637 | 0.847411223 | NOX4, RBFOX1, FHIT, RALYL, CPEB2, CPEB3, RBFOX3, RBM20, TRA2B, SSB, LARP4B, HNRNPR, RBPM | 1340 | 348 | 16881 | 0.868811117 | 1 | 0.99998623 | 100 |
| UP_SEQ_FEATURE | domain:RRM 2 | 7 | 0.444726811 | 0.912328707 | CPEB2, CPEB3, CELF3, RBMS3, CELF2, MSI2, HNRNPR | 1499 | 125 | 20063 | 0.749518346 | 1 | 1 | 100 |
| UP_SEQ_FEATURE | domain:RRM 1 | 7 | 0.444726811 | 0.912328707 | CPEB2, CPEB3, CELF3, RBMS3, CELF2, MSI2, HNRNPR | 1499 | 125 | 20063 | 0.749518346 | 1 | 1 | 100 |
| UP_SEQ_FEATURE | domain:RRM 3 | 3 | 0.190597205 | 0.948157939 | CELF3, CELF2, HNRNPR | 1499 | 61 | 20063 | 0.658242107 | 1 | 1 | 100 |

Annotation Cluster 208 Enrichment Score: 0.11901676935910296

| Category | Term | Count | % | PValue | Genes | List Total | Pop Hits | Pop Total | Fold Enrichment | Bonferroni | Benjamini | FDR |
|---|---|---|---|---|---|---|---|---|---|---|---|---|
| UP_SEQ_FEATURE | domain:PX | 4 | 0.254129606 | 0.705665675 | PLD2, SNX29, SNX27, PIK3C2B | 1499 | 48 | 20063 | 1.115354681 | 1 | 0.99999998 | 99.99999999 |
| SMART | SM00312:PX | 4 | 0.254129606 | 0.764473413 | PLD2, SNX29, SNX27, PIK3C2B | 878 | 45 | 10057 | 1.018172615 | 1 | 0.997522773 | 99.99999984 |
| INTERPRO | IPR001683:Phox homologous domain | 4 | 0.254129606 | 0.81468236 | PLD2, SNX29, SNX27, PIK3C2B | 1452 | 55 | 18559 | 0.929576759 | 1 | 0.99999979 | 100 |

Annotation Cluster 209 Enrichment Score: 0.11564090002330855

| Category | Term | Count | % | PValue | Genes | List Total | Pop Hits | Pop Total | Fold Enrichment | Bonferroni | Benjamini | FDR |
|---|---|---|---|---|---|---|---|---|---|---|---|---|
| GOTERM_CC_DIRECT | GO:0031093~platelet alpha granule lumen | 5 | 0.317662008 | 0.632290672 | APP, VEGFA, HGF, QSOX1, FN1 | 1425 | 55 | 18224 | 1.16261563 | 1 | 0.982870771 | 99.99997089 |
| GOTERM_BP_DIRECT | GO:0002576~platelet degranulation | 7 | 0.444726811 | 0.84052583 | APP, VEGFA, WDR1, HGF, QSOX1, CALM2, FN1 | 1342 | 103 | 16792 | 0.850375472 | 1 | 0.999999981 | 100 |
| REACTOME_PATHWAY | R-HSA-114608:R-HSA-114608 | 9 | 0.571791614 | 0.846465887 | APP, CHID1, VEGFA, WDR1, HGF, RAB27B, QSOX1, CALM2, FN1 | 745 | 130 | 9075 | 0.843314404 | 1 | 0.99995502 | 100 |

Annotation Cluster 210 Enrichment Score: 0.1083762029372473

| Category | Term | Count | % | PValue | Genes | List Total | Pop Hits | Pop Total | Fold Enrichment | Bonferroni | Benjamini | FDR |
|---|---|---|---|---|---|---|---|---|---|---|---|---|
| UP_SEQ_FEATURE | domain:FHA | 3 | 0.190597205 | 0.747333195 | TIFA, TCF19, CHEK2 | 1499 | 35 | 20063 | 1.147221957 | 1 | 0.999999997 | 100 |
| INTERPRO | IPR008984:SMAD/FHA domain | 4 | 0.254129606 | 0.794390416 | TIFA, SMAD3, TCF19, CHEK2 | 1452 | 53 | 18559 | 0.964655128 | 1 | 0.999999583 | 100 |
| INTERPRO | IPR000253:Forkhead-associated (FHA) domain | 3 | 0.190597205 | 0.796751926 | TIFA, TCF19, CHEK2 | 1452 | 37 | 18559 | 1.036352468 | 1 | 0.9999996 | 100 |

Annotation Cluster 211 Enrichment Score: 0.10499567239551032

| Category | Term | Count | % | PValue | Genes | List Total | Pop Hits | Pop Total | Fold Enrichment | Bonferroni | Benjamini | FDR |
|---|---|---|---|---|---|---|---|---|---|---|---|---|
| GOTERM_BP_DIRECT | GO:0006418~tRNA aminoacylation for protein translation | 4 | 0.254129606 | 0.629622755 | AIMP1, FARS2, LARS, LARS2 | 1342 | 40 | 16792 | 1.251266766 | 1 | 0.999992094 | 99.99999906 |
| UP_KEYWORDS | Aminoacyl-tRNA synthetase | 3 | 0.190597205 | 0.786353141 | FARS2, LARS, LARS2 | 1535 | 38 | 20581 | 1.058511915 | 1 | 0.966014741 | 99.99999998 |
| KEGG_PATHWAY | hsa00970:Aminoacyl-tRNA biosynthesis | 3 | 0.190597205 | 0.977946151 | FARS2, LARS, LARS2 | 574 | 66 | 6879 | 0.544741844 | 1 | 0.995614686 | 100 |

Annotation Cluster 212 Enrichment Score: 0.09830099785596164

| Category | Term | Count | % | PValue | Genes | List Total | Pop Hits | Pop Total | Fold Enrichment | Bonferroni | Benjamini | FDR |
|---|---|---|---|---|---|---|---|---|---|---|---|---|
| UP_SEQ_FEATURE | domain:Ubiquitin-like | 4 | 0.254129606 | 0.663106529 | MIDN, PARK2, UBL3, HERPUD2 | 1499 | 45 | 20063 | 1.18971166 | 1 | 0.999999916 | 99.99999984 |
| INTERPRO | IPR000626:Ubiquitin | 4 | 0.254129606 | 0.8501917 | MIDN, PARK2, UBL3, HERPUD2 | 1452 | 59 | 18559 | 0.866554606 | 1 | 0.999999956 | 100 |
| SMART | SM00213:UBQ | 3 | 0.190597205 | 0.89949104 | MIDN, PARK2, HERPUD2 | 878 | 43 | 10057 | 0.79914711 | 1 | 0.999841125 | 100 |

Annotation Cluster 213 Enrichment Score: 0.09591736681932414

| Category | Term | Count | % | PValue | Genes | List Total | Pop Hits | Pop Total | Fold Enrichment | Bonferroni | Benjamini | FDR |
|---|---|---|---|---|---|---|---|---|---|---|---|---|
| UP_KEYWORDS | Nuclear pore complex | 4 | 0.254129606 | 0.730550123 | MYO1C, NUP88, XPO7, NUP35 | 1535 | 50 | 20581 | 1.072625407 | 1 | 0.951937035 | 99.99999935 |
| UP_KEYWORDS | Translocation | 6 | 0.381194409 | 0.759586568 | MYO1C, NUP88, XPO7, NUP35, SEC62, TIMM21 | 1535 | 84 | 20581 | 0.957701256 | 1 | 0.958999912 | 99.99999987 |
| GOTERM_CC_DIRECT | GO:0005643~nuclear pore | 5 | 0.317662008 | 0.824016194 | MYO1C, NUP88, KPNA6, KPNA3, XPO7 | 1425 | 72 | 18224 | 0.888109162 | 1 | 0.997803444 | 100 |
| UP_KEYWORDS | mRNA transport | 6 | 0.381194409 | 0.903998045 | MYO1C, NUP88, DDX25, THOC7, XPO7, NUP35 | 1535 | 106 | 20581 | 0.758933071 | 1 | 0.989975745 | 100 |

Annotation Cluster 214 Enrichment Score: 0.09280348917900104

| Category | Term | Count | % | PValue | Genes | List Total | Pop Hits | Pop Total | Fold Enrichment | Bonferroni | Benjamini | FDR |
|---|---|---|---|---|---|---|---|---|---|---|---|---|
| UP_SEQ_FEATURE | domain:Thioredoxin | 3 | 0.190597205 | 0.717317606 | DNAJC16, QSOX2, QSOX1 | 1499 | 33 | 20063 | 1.216750561 | 1 | 0.999999988 | 99.99999999 |
| INTERPRO | IPR013766:Thioredoxin domain | 3 | 0.190597205 | 0.841286034 | DNAJC16, QSOX2, QSOX1 | 1452 | 41 | 18559 | 0.93524491 | 1 | 0.999999936 | 100 |
| GOTERM_BP_DIRECT | GO:0045454~cell redox homeostasis | 5 | 0.317662008 | 0.872839442 | CYBA, DNAJC16, TXNRD3, QSOX2, QSOX1 | 1342 | 77 | 16792 | 0.812510887 | 1 | 0.999999996 | 100 |

Annotation Cluster 215 Enrichment Score: 0.0865554882750845

| Category | Term | Count | % | PValue | Genes | List Total | Pop Hits | Pop Total | Fold Enrichment | Bonferroni | Benjamini | FDR |
|---|---|---|---|---|---|---|---|---|---|---|---|---|
| GOTERM_MF_DIRECT | GO:0000149~SNARE binding | 4 | 0.254129606 | 0.7179385 | NBAS, SEC22A, VAMP5, TSNARE1 | 1340 | 46 | 16881 | 1.095457495 | 1 | 0.999800853 | 99.99999989 |
| GOTERM_MF_DIRECT | GO:0005484~SNAP receptor activity | 3 | 0.190597205 | 0.826659157 | SEC22A, VAMP5, TSNARE1 | 1340 | 39 | 16881 | 0.969058553 | 1 | 0.999975837 | 100 |
| GOTERM_CC_DIRECT | GO:0031201~SNARE complex | 3 | 0.190597205 | 0.926658707 | SEC22A, VAMP5, TSNARE1 | 1425 | 53 | 18224 | 0.723892751 | 1 | 0.99978958 | 100 |

Annotation Cluster 216 Enrichment Score: 0.08646345295075215

| Category | Term | Count | % | PValue | Genes | List Total | Pop Hits | Pop Total | Fold Enrichment | Bonferroni | Benjamini | FDR |
|---|---|---|---|---|---|---|---|---|---|---|---|---|
| GOTERM_BP_DIRECT | GO:0008380~RNA splicing | 14 | 0.889453621 | 0.571138894 | RBFOX1, TFIP11, SRPK2, SNRPN, RBM20, RBFOX3, SNURF, PPP1R9B, PPIG, CIR1, DDX23, CELF3, SNR | 1342 | 166 | 16792 | 1.055285224 | 1 | 0.999976697 | 99.99998565 |
| GOTERM_BP_DIRECT | GO:0000398~mRNA splicing, via spliceosome | 16 | 1.016518424 | 0.786114731 | POLR2H, TFIP11, CSTF3, SNRPN, PPIL1, TRA2B, POLR2C, HNRNPR, POLR2B, SNURF, WDR83, PAPOLA | 1342 | 222 | 16792 | 0.901813885 | 1 | 0.999999863 | 100 |
| GOTERM_BP_DIRECT | GO:0006397~mRNA processing | 11 | 0.698856417 | 0.914134214 | TFIP11, RBFOX1, PHRF1, CIR1, KHDRBS2, ADARB2, RBFOX3, RBM20, CELF2, THOC7, HNRNPR | 1342 | 179 | 16792 | 0.768934884 | 1 | 0.999999999 | 100 |
| UP_KEYWORDS | mRNA processing | 19 | 1.207115629 | 0.942119375 | RBFOX1, TFIP11, SRPK2, CSTF3, KHDRBS2, ADARB2, PPIL1, RBFOX3, RBM20, TRA2B, HNRNPR, WDR8 | 1535 | 332 | 20581 | 0.767314862 | 1 | 0.995589244 | 100 |
| UP_KEYWORDS | mRNA splicing | 14 | 0.889453621 | 0.955738685 | RBFOX1, TFIP11, SRPK2, RBM20, TRA2B, RBFOX3, PPIL1, HNRNPR, WDR83, CIR1, DDX23, CELF3, SN | 1535 | 260 | 20581 | 0.721959409 | 1 | 0.997038219 | 100 |

Annotation Cluster 217 Enrichment Score: 0.08528442476363163

| Category | Term | Count | % | PValue | Genes | List Total | Pop Hits | Pop Total | Fold Enrichment | Bonferroni | Benjamini | FDR |
|---|---|---|---|---|---|---|---|---|---|---|---|---|
| UP_KEYWORDS | Cell cycle | 46 | 2.92249047 | 0.721064362 | CLTA, CHMP3, PRC1, USP2, BRSK2, TP63, NR3C1, CHEK2, TTC28, CSNK2A2, MCM7, BRINP2, MAP3K8 | 1535 | 650 | 20581 | 0.948860937 | 1 | 0.950082292 | 99.99999893 |
| UP_KEYWORDS | Cell division | 26 | 1.65184244 | 0.804002947 | CLTA, CHMP3, PRC1, BRSK2, NR3C1, TTC28, CHEK2, CABLES2, SKA2, PAFAH1B1, CLASP1, CDK14, CDC | 1535 | 388 | 20581 | 0.898462003 | 1 | 0.970966246 | 99.99999999 |
| GOTERM_BP_DIRECT | GO:0051301~cell division | 25 | 1.588310038 | 0.811032201 | CLTA, BRSK2, NR3C1, TTC28, CHEK2, CABLES2, CLASP1, SKA2, CDK14, TUBA1C, CDCA3, PDS5B, CCNF, | 1342 | 350 | 16792 | 0.893761976 | 1 | 0.999999936 | 100 |
| UP_KEYWORDS | Mitosis | 16 | 1.016518424 | 0.88643031 | CLTA, PDS5B, CCNF, BRSK2, NR3C1, TTC28, CHEK2, SIRT2, TEX14, BUB1B, MAPRE2, SKA2, CLASP1, PA | 1535 | 262 | 20581 | 0.818798021 | 1 | 0.987664252 | 100 |
| GOTERM_BP_DIRECT | GO:0007067~mitotic nuclear division | 16 | 1.016518424 | 0.898800922 | CLTA, TADA2A, NUP88, CCNF, BRSK2, NR3C1, TTC28, HGF, SIRT2, NOLC1, BUB1B, MAPRE2, SKA2, PA | 1342 | 248 | 16792 | 0.807268881 | 1 | 0.999999998 | 100 |

Annotation Cluster 218 Enrichment Score: 0.07909866893147001

| Category | Term | Count | % | PValue | Genes | List Total | Pop Hits | Pop Total | Fold Enrichment | Bonferroni | Benjamini | FDR |
|---|---|---|---|---|---|---|---|---|---|---|---|---|
| UP_KEYWORDS | DNA repair | 20 | 1.27064803 | 0.768643878 | MMS19, XRCC5, RAD51C, XRCC4, HMGB1, RAD51B, MSH2, SWSAP1, DTX3L, MGMT, CHEK2, RPA3, C | 1535 | 293 | 20581 | 0.915209392 | 1 | 0.960683362 | 99.99999993 |
| UP_KEYWORDS | DNA damage | 23 | 1.461245235 | 0.833677098 | MMS19, XRCC5, RAD51C, HMGB1, XRCC4, RAD51B, MSH2, SWSAP1, DTX3L, MGMT, MACROD2, CHE | 1535 | 352 | 20581 | 0.87607899 | 1 | 0.977161242 | 100 |
| GOTERM_BP_DIRECT | GO:0006281~DNA repair | 15 | 0.952986023 | 0.903609835 | MMS19, RAD51C, RAD51B, PDS5B, MSH2, MGMT, EEPD1, RPA3, EPC2, FANCD2, TDG, USP45, UBE2L | 1342 | 235 | 16792 | 0.798680914 | 1 | 0.999999999 | 100 |

Annotation Cluster 219 Enrichment Score: 0.07633910021630046

| Category | Term | Count | % | PValue | Genes | List Total | Pop Hits | Pop Total | Fold Enrichment | Bonferroni | Benjamini | FDR |
|---|---|---|---|---|---|---|---|---|---|---|---|---|
| GOTERM_CC_DIRECT | GO:0071013~catalytic step 2 spliceosome | 8 | 0.508259212 | 0.585482445 | TFIP11, WDR83, DDX23, SNRPN, PPIL1, SNRPB, HNRNPR, SNURF | 1425 | 92 | 18224 | 1.112067124 | 1 | 0.977205769 | 99.99982348 |
| GOTERM_BP_DIRECT | GO:0000398~mRNA splicing, via spliceosome | 16 | 1.016518424 | 0.786114731 | POLR2H, TFIP11, CSTF3, SNRPN, PPIL1, TRA2B, POLR2C, HNRNPR, POLR2B, SNURF, WDR83, PAPOLA | 1342 | 222 | 16792 | 0.901813885 | 1 | 0.999999863 | 100 |
| REACTOME_PATHWAY | R-HSA-72163:R-HSA-72163 | 13 | 0.82592122 | 0.794526712 | POLR2H, TFIP11, CSTF3, SNRPN, TRA2B, PPIL1, POLR2C, HNRNPR, POLR2B, SNURF, PAPOLA, DDX23, | 745 | 177 | 9075 | 0.894664998 | 1 | 0.999892018 | 100 |
| GOTERM_CC_DIRECT | GO:0005681~spliceosomal complex | 6 | 0.381194409 | 0.867364989 | TFIP11, WDR83, SNRPN, SNRPB, HNRNPR, SNURF | 1425 | 94 | 18224 | 0.816304591 | 1 | 0.998962067 | 100 |
| UP_KEYWORDS | mRNA splicing | 14 | 0.889453621 | 0.955738685 | RBFOX1, TFIP11, SRPK2, RBM20, TRA2B, RBFOX3, PPIL1, HNRNPR, WDR83, CIR1, DDX23, CELF3, SNF | 1535 | 260 | 20581 | 0.721959409 | 1 | 0.997038219 | 100 |
| UP_KEYWORDS | Spliceosome | 6 | 0.381194409 | 0.964499139 | TFIP11, WDR83, DDX23, PPIL1, SNRPB, HNRNPR | 1535 | 127 | 20581 | 0.633440201 | 1 | 0.997846779 | 100 |
| KEGG_PATHWAY | hsa03040:Spliceosome | 4 | 0.254129606 | 0.999252139 | DDX23, PPIL1, TRA2B, SNRPB | 574 | 133 | 6879 | 0.360430693 | 1 | 0.999935922 | 100 |

Annotation Cluster 220 Enrichment Score: 0.07373510196998913

| Category | Term | Count | % | PValue | Genes | List Total | Pop Hits | Pop Total | Fold Enrichment | Bonferroni | Benjamini | FDR |
|---|---|---|---|---|---|---|---|---|---|---|---|---|
| UP_KEYWORDS | Fatty acid biosynthesis | 4 | 0.254129606 | 0.754749156 | PTGDS, MLYCD, FAM213B, OXSM | 1535 | 52 | 20581 | 1.031370584 | 1 | 0.957740257 | 99.99999983 |
| UP_KEYWORDS | Lipid biosynthesis | 10 | 0.635324015 | 0.830322786 | FAR1, CRLS1, PTGDS, MLYCD, FAM213B, OSBPL10, CHPT1, OXSM, AGPAT1, PTDSS2 | 1535 | 156 | 20581 | 0.859475487 | 1 | 0.976935184 | 100 |
| UP_KEYWORDS | Fatty acid metabolism | 6 | 0.381194409 | 0.958837701 | PTGDS, MLYCD, FAM213B, HSD17B4, OXSM, ACSL5 | 1535 | 124 | 20581 | 0.648765367 | 1 | 0.997295192 | 100 |

Annotation Cluster 221 Enrichment Score: 0.07360260143350028

| Category | Term | Count | % | PValue | Genes | List Total | Pop Hits | Pop Total | Fold Enrichment | Bonferroni | Benjamini | FDR |
|---|---|---|---|---|---|---|---|---|---|---|---|---|
| INTERPRO | IPR004087:K Homology domain | 3 | 0.190597205 | 0.808824337 | KRR1, KHDRBS2, PCBP3 | 1452 | 38 | 18559 | 1.009080035 | 1 | 0.999999739 | 100 |
| SMART | SM00322:KH | 3 | 0.190597205 | 0.85602194 | KRR1, KHDRBS2, PCBP3 | 878 | 38 | 10057 | 0.904298046 | 1 | 0.999456189 | 100 |
| INTERPRO | IPR004088:K Homology domain, type 1 | 3 | 0.190597205 | 0.868666533 | KRR1, KHDRBS2, PCBP3 | 1452 | 44 | 18559 | 0.871478212 | 1 | 0.999999983 | 100 |

Annotation Cluster 222 Enrichment Score: 0.05582167239788713

| Category | Term | Count | % | PValue | Genes | List Total | Pop Hits | Pop Total | Fold Enrichment | Bonferroni | Benjamini | FDR |
|---|---|---|---|---|---|---|---|---|---|---|---|---|
| REACTOME_PATHWAY | R-HSA-5617472:R-HSA-5617472 | 11 | 0.698856417 | 0.548524519 | POLR2H, HDAC3, RBBP5, HIST4H4, MAFB, H2AFJ, CNOT6, POLR2C, POLR2B, HIST1H3H, ZNF335 | 745 | 122 | 9075 | 1.098305644 | 1 | 0.998606554 | 99.9995551 |
| REACTOME_PATHWAY | R-HSA-3214858:R-HSA-3214858 | 7 | 0.444726811 | 0.612537021 | HIST4H4, SMARCC1, SMARCD1, H2AFJ, ARID1B, CDK4, HIST1H3H | 745 | 77 | 9075 | 1.10738255 | 1 | 0.999114428 | 99.99995838 |
| REACTOME_PATHWAY | R-HSA-5578749:R-HSA-5578749 | 9 | 0.571791614 | 0.648915037 | POLR2H, HIST4H4, NUP88, IPO8, H2AFJ, NUP35, POLR2C, POLR2B, HIST1H3H | 745 | 106 | 9075 | 1.034253514 | 1 | 0.999336896 | 99.99999097 |
| REACTOME_PATHWAY | R-HSA-2559580:R-HSA-2559580 | 9 | 0.571791614 | 0.800601902 | MAP3K5, HIST4H4, TNRC6C, MAPKAPK5, MINK1, H2AFJ, TNRC6B, CDK4, HIST1H3H | 745 | 123 | 9075 | 0.891307906 | 1 | 0.99989728 | 100 |
| REACTOME_PATHWAY | R-HSA-912446:R-HSA-912446 | 6 | 0.381194409 | 0.851402644 | RAD51C, HIST4H4, H2AFJ, CDK4, HIST1H3H, RPA3 | 745 | 87 | 9075 | 0.840083314 | 1 | 0.999959774 | 100 |
| REACTOME_PATHWAY | R-HSA-2299718:R-HSA-2299718 | 5 | 0.317662008 | 0.86681885 | HIST4H4, SET, RB1, H2AFJ, HIST1H3H | 745 | 74 | 9075 | 0.823054598 | 1 | 0.999972679 | 100 |
| GOTERM_BP_DIRECT | GO:0045815~positive regulation of gene expression, epigenetic | 4 | 0.254129606 | 0.8820553 | POLR2H, HIST4H4, MYO1C, HIST1H3H | 1342 | 62 | 16792 | 0.807268881 | 1 | 0.999999996 | 100 |
| REACTOME_PATHWAY | R-HSA-201722:R-HSA-201722 | 5 | 0.317662008 | 0.937637596 | RBBP5, HIST4H4, H2AFJ, TCF7L2, HIST1H3H | 745 | 88 | 9075 | 0.692114094 | 1 | 0.999996715 | 100 |
| REACTOME_PATHWAY | R-HSA-212300:R-HSA-212300 | 4 | 0.254129606 | 0.945391921 | HIST4H4, PHF1, H2AFJ, HIST1H3H | 745 | 73 | 9075 | 0.667463455 | 1 | 0.999997638 | 100 |
| REACTOME_PATHWAY | R-HSA-5250924:R-HSA-5250924 | 5 | 0.317662008 | 0.94741764 | POLR2H, HIST4H4, MYO1C, H2AFJ, HIST1H3H | 745 | 91 | 9075 | 0.669297146 | 1 | 0.99999756 | 100 |
| REACTOME_PATHWAY | R-HSA-2559582:R-HSA-2559582 | 6 | 0.381194409 | 0.953448442 | HIST4H4, IGFBP7, H2AFJ, EHMT2, CDK4, HIST1H3H | 745 | 110 | 9075 | 0.66442953 | 1 | 0.999998249 | 100 |
| REACTOME_PATHWAY | R-HSA-73728:R-HSA-73728 | 3 | 0.190597205 | 0.970131307 | HIST4H4, H2AFJ, HIST1H3H | 745 | 63 | 9075 | 0.580057526 | 1 | 0.999999569 | 100 |
| REACTOME_PATHWAY | R-HSA-5334118:R-HSA-5334118 | 3 | 0.190597205 | 0.974175946 | HIST4H4, H2AFJ, HIST1H3H | 745 | 65 | 9075 | 0.562209602 | 1 | 0.999999708 | 100 |
| REACTOME_PATHWAY | R-HSA-5625886:R-HSA-5625886 | 3 | 0.190597205 | 0.977689149 | HIST4H4, H2AFJ, HIST1H3H | 745 | 67 | 9075 | 0.545427226 | 1 | 0.999999828 | 100 |
| REACTOME_PATHWAY | R-HSA-427413:R-HSA-427413 | 5 | 0.317662008 | 0.978364203 | POLR2H, HIST4H4, SAP30L, H2AFJ, HIST1H3H | 745 | 106 | 9075 | 0.574585286 | 1 | 0.999999836 | 100 |
| REACTOME_PATHWAY | R-HSA-427359:R-HSA-427359 | 3 | 0.190597205 | 0.979267673 | HIST4H4, H2AFJ, HIST1H3H | 745 | 68 | 9075 | 0.537406238 | 1 | 0.999999852 | 100 |
| GOTERM_CC_DIRECT | GO:0000786~nucleosome | 4 | 0.254129606 | 0.980956474 | HIST1H1E, HIST4H4, H2AFJ, HIST1H3H | 1425 | 94 | 18224 | 0.544203061 | 1 | 0.999995252 | 100 |
| REACTOME_PATHWAY | R-HSA-73777:R-HSA-73777 | 4 | 0.254129606 | 0.981821221 | POLR2H, HIST4H4, H2AFJ, HIST1H3H | 745 | 90 | 9075 | 0.541387025 | 1 | 0.999999899 | 100 |
| UP_KEYWORDS | Nucleosome core | 3 | 0.190597205 | 0.991810683 | HIST4H4, H2AFJ, HIST1H3H | 1535 | 89 | 20581 | 0.451948908 | 1 | 0.999775155 | 100 |
| INTERPRO | IPR009072:Histone-fold | 4 | 0.254129606 | 0.99404799 | HIST4H4, ABTB2, H2AFJ, HIST1H3H | 1452 | 112 | 18559 | 0.456488587 | 1 | 1 | 100 |

Annotation Cluster 223 Enrichment Score: 0.05561086844389616

| Category | Term | Count | % | PValue | Genes | List Total | Pop Hits | Pop Total | Fold Enrichment | Bonferroni | Benjamini | FDR |
|---|---|---|---|---|---|---|---|---|---|---|---|---|
| KEGG_PATHWAY | hsa04923:Regulation of lipolysis in adipocytes | 5 | 0.317662008 | 0.697322905 | ADRB2, PIK3CB, PDE3B, INSR, AKT3 | 574 | 56 | 6879 | 1.070028621 | 1 | 0.920020346 | 99.99998616 |
| KEGG_PATHWAY | hsa04931:Insulin resistance | 5 | 0.317662008 | 0.982988904 | PIK3CB, PRKCE, INSR, AKT3, PTPN11 | 574 | 108 | 6879 | 0.554829655 | 1 | 0.996596925 | 100 |
| KEGG_PATHWAY | hsa04152:AMPK signaling pathway | 5 | 0.317662008 | 0.993538838 | MLYCD, PIK3CB, EEF2, INSR, AKT3 | 574 | 123 | 6879 | 0.487167502 | 1 | 0.999054575 | 100 |

Annotation Cluster 224 Enrichment Score: 0.05459185415706848

| Category | Term | Count | % | PValue | Genes | List Total | Pop Hits | Pop Total | Fold Enrichment | Bonferroni | Benjamini | FDR |
|---|---|---|---|---|---|---|---|---|---|---|---|---|
| REACTOME_PATHWAY | R-HSA-383280:R-HSA-383280 | 4 | 0.254129606 | 0.800393583 | ESRRG, NR2C2AP, NR3C1, RORA | 745 | 51 | 9075 | 0.955388867 | 1 | 0.9999028 | 100 |
| INTERPRO | IPR013088:Zinc finger, NHR/GATA-type | 4 | 0.254129606 | 0.841916866 | GATA2, ESRRG, NR3C1, RORA | 1452 | 58 | 18559 | 0.881495203 | 1 | 0.999999934 | 100 |
| UP_SEQ_FEATURE | zinc finger region:NR C4-type | 3 | 0.190597205 | 0.859326528 | ESRRG, NR3C1, RORA | 1499 | 45 | 20063 | 0.892283745 | 1 | 1 | 100 |
| UP_SEQ_FEATURE | DNA-binding region:Nuclear receptor | 3 | 0.190597205 | 0.859326528 | ESRRG, NR3C1, RORA | 1499 | 45 | 20063 | 0.892283745 | 1 | 1 | 100 |
| INTERPRO | IPR001628:Zinc finger, nuclear hormone receptor-type | 3 | 0.190597205 | 0.884433385 | ESRRG, NR3C1, RORA | 1452 | 46 | 18559 | 0.833587855 | 1 | 0.999999994 | 100 |
| INTERPRO | IPR001723:Steroid hormone receptor | 3 | 0.190597205 | 0.891642105 | ESRRG, NR3C1, RORA | 1452 | 47 | 18559 | 0.815851943 | 1 | 0.999999996 | 100 |
| INTERPRO | IPR000536:Nuclear hormone receptor, ligand-binding, core | 3 | 0.190597205 | 0.898431018 | ESRRG, NR3C1, RORA | 1452 | 48 | 18559 | 0.798855028 | 1 | 0.999999997 | 100 |
| SMART | SM00399:ZnF_C4 | 3 | 0.190597205 | 0.919345976 | ESRRG, NR3C1, RORA | 878 | 46 | 10057 | 0.74702882 | 1 | 0.999925011 | 100 |
| SMART | SM00430:HOLI | 3 | 0.190597205 | 0.930466101 | ESRRG, NR3C1, RORA | 878 | 48 | 10057 | 0.71590262 | 1 | 0.999953089 | 100 |
| GOTERM_MF_DIRECT | GO:0003707~steroid hormone receptor activity | 3 | 0.190597205 | 0.943348676 | ESRRG, NR3C1, RORA | 1340 | 56 | 16881 | 0.674880064 | 1 | 0.999999817 | 100 |

Annotation Cluster 225 Enrichment Score: 0.048870377725830826

| Category | Term | Count | % | PValue | Genes | List Total | Pop Hits | Pop Total | Fold Enrichment | Bonferroni | Benjamini | FDR |
|---|---|---|---|---|---|---|---|---|---|---|---|---|
| UP_SEQ_FEATURE | metal ion-binding site:Iron (heme axial ligand) | 7 | 0.444726811 | 0.75768065 | FRRS1, CYBA, DGCR8, CYP2B6, CYP21A2, HMOX1, CAT | 1499 | 99 | 20063 | 0.946361547 | 1 | 0.999999998 | 100 |
| GOTERM_MF_DIRECT | GO:0020037~heme binding | 8 | 0.508259212 | 0.925085304 | NOX4, CYBA, DGCR8, CYP2B6, CYP21A2, HMOX1, GUCY1A2, CAT | 1340 | 137 | 16881 | 0.73563569 | 1 | 0.999999306 | 100 |
| UP_KEYWORDS | Heme | 7 | 0.444726811 | 0.934599133 | FRRS1, CYBA, DGCR8, CYP2B6, CYP21A2, HMOX1, CAT | 1535 | 132 | 20581 | 0.71102063 | 1 | 0.994654995 | 100 |
| UP_KEYWORDS | Iron | 17 | 1.080050826 | 0.973251625 | CYP2B6, CYP21A2, TH, HGD, ACP5, PAH, CIAPIN1, BBOX1, FRRS1, CYBA, APP, DGCR8, KDM2A, HMOX | 1535 | 324 | 20581 | 0.703496602 | 1 | 0.998622924 | 100 |

Annotation Cluster 226 Enrichment Score: 0.048220165031033047

| Category | Term | Count | % | PValue | Genes | List Total | Pop Hits | Pop Total | Fold Enrichment | Bonferroni | Benjamini | FDR |
|---|---|---|---|---|---|---|---|---|---|---|---|---|
| INTERPRO | IPR018000:Neurotransmitter-gated ion-channel, conserved site | 3 | 0.190597205 | 0.884433385 | GABRA2, GABRG3, CHRNE | 1452 | 46 | 18559 | 0.833587855 | 1 | 0.999999994 | 100 |
| INTERPRO | IPR006202:Neurotransmitter-gated ion-channel ligand-binding | 3 | 0.190597205 | 0.898431018 | GABRA2, GABRG3, CHRNE | 1452 | 48 | 18559 | 0.798855028 | 1 | 0.999999997 | 100 |
| INTERPRO | IPR006201:Neurotransmitter-gated ion-channel | 3 | 0.190597205 | 0.898431018 | GABRA2, GABRG3, CHRNE | 1452 | 48 | 18559 | 0.798855028 | 1 | 0.999999997 | 100 |
| INTERPRO | IPR006029:Neurotransmitter-gated ion-channel transmembrane domain | 3 | 0.190597205 | 0.898431018 | GABRA2, GABRG3, CHRNE | 1452 | 48 | 18559 | 0.798855028 | 1 | 0.999999997 | 100 |

Annotation Cluster 227 Enrichment Score: 0.04316164380120197

| Category | Term | Count | % | PValue | Genes | List Total | Pop Hits | Pop Total | Fold Enrichment | Bonferroni | Benjamini | FDR |
|---|---|---|---|---|---|---|---|---|---|---|---|---|
| GOTERM_BP_DIRECT | GO:0007017~microtubule-based process | 4 | 0.254129606 | 0.556932795 | TUBAL3, PAFAH1B1, TUBA1C, MAP3K11 | 1342 | 36 | 16792 | 1.390296407 | 1 | 0.999971082 | 99.99997369 |
| GOTERM_CC_DIRECT | GO:0000776~kinetochore | 5 | 0.317662008 | 0.886377118 | TEX14, BUB1B, PAFAH1B1, CLASP1, WDR81 | 1425 | 81 | 18224 | 0.789430366 | 1 | 0.99934061 | 100 |
| GOTERM_BP_DIRECT | GO:0007062~sister chromatid cohesion | 6 | 0.381194409 | 0.921915379 | RAD51C, PDS5B, BUB1B, SKA2, PAFAH1B1, CLASP1 | 1342 | 103 | 16792 | 0.728893262 | 1 | 1 | 100 |
| REACTOME_PATHWAY | R-HSA-2500257:R-HSA-2500257 | 7 | 0.444726811 | 0.939030563 | PDS5B, TUBAL3, BUB1B, SKA2, PAFAH1B1, CLASP1, TUBA1C | 745 | 121 | 9075 | 0.704697987 | 1 | 0.999996613 | 100 |
| REACTOME_PATHWAY | R-HSA-68877:R-HSA-68877 | 6 | 0.381194409 | 0.948183865 | TUBAL3, BUB1B, SKA2, PAFAH1B1, CLASP1, TUBA1C | 745 | 108 | 9075 | 0.676733781 | 1 | 0.999997556 | 100 |
| REACTOME_PATHWAY | R-HSA-5663220:R-HSA-5663220 | 7 | 0.444726811 | 0.970159847 | DIAPH1, TUBAL3, BUB1B, SKA2, PAFAH1B1, CLASP1, TUBA1C | 745 | 135 | 9075 | 0.631618195 | 1 | 0.999999539 | 100 |
| UP_KEYWORDS | Centromere | 6 | 0.381194409 | 0.978586168 | TEX14, SYCP3, BUB1B, STAG3, SKA2, CLASP1 | 1535 | 137 | 20581 | 0.58720369 | 1 | 0.999032911 | 100 |
| UP_KEYWORDS | Kinetochore | 4 | 0.254129606 | 0.981299078 | TEX14, BUB1B, SKA2, CLASP1 | 1535 | 99 | 20581 | 0.541730004 | 1 | 0.999181259 | 100 |
| REACTOME_PATHWAY | R-HSA-2467813:R-HSA-2467813 | 9 | 0.571791614 | 0.987399195 | PSMA1, PDS5B, PSMD1, TUBAL3, BUB1B, SKA2, PAFAH1B1, CLASP1, TUBA1C | 745 | 185 | 9075 | 0.592599311 | 1 | 0.999999973 | 100 |
| GOTERM_CC_DIRECT | GO:0000777~condensed chromosome kinetochore | 3 | 0.190597205 | 0.993029973 | TEX14, BUB1B, CLASP1 | 1425 | 87 | 18224 | 0.440992136 | 1 | 0.99999974 | 100 |

Annotation Cluster 228 Enrichment Score: 0.03971089195620468

| Category | Term | Count | % | PValue | Genes | List Total | Pop Hits | Pop Total | Fold Enrichment | Bonferroni | Benjamini | FDR |
|---|---|---|---|---|---|---|---|---|---|---|---|---|
| UP_SEQ_FEATURE | zinc finger region:C2H2-type 1 | 39 | 2.477763659 | 0.7324119 | ZFP62, ZNF430, ZNF552, ZNF821, GLIS1, ZSCAN5A, GLI3, ZNF652, ZNF343, GLI1, ZNF184, ZFP91, BCL | 1499 | 554 | 20063 | 0.94221298 | 1 | 0.999999994 | 100 |
| UP_SEQ_FEATURE | zinc finger region:C2H2-type 2 | 42 | 2.668360864 | 0.7991525 | ZFP62, ZNF430, ZNF821, ZNF510, ZSCAN5A, GLI3, ZNF343, GLI1, ZNF184, ZFP91, BCL11B, BCL11A, ZN | 1499 | 615 | 20063 | 0.914046763 | 1 | 1 | 100 |
| UP_SEQ_FEATURE | zinc finger region:C2H2-type 9 | 25 | 1.588310038 | 0.848435046 | ZBTB48, ZFP62, ZNF430, ZNF552, ZNF354A, ZNF532, ZNF479, ZNF510, ZNF648, ZNF791, ZNF782, ZNI | 1499 | 385 | 20063 | 0.869107544 | 1 | 1 | 100 |
| UP_SEQ_FEATURE | zinc finger region:C2H2-type 10 | 22 | 1.397712834 | 0.849435716 | ZBTB48, ZFP62, ZNF430, ZNF354A, ZNF479, ZNF510, ZNF648, ZNF791, ZNF782, ZNF141, MECOM, ZN | 1499 | 341 | 20063 | 0.863500398 | 1 | 1 | 100 |
| UP_SEQ_FEATURE | zinc finger region:C2H2-type 5 | 36 | 2.287166455 | 0.860982116 | ZFP62, ZNF430, ZNF552, ZNF532, GLIS1, ZNF510, ZSCAN5A, ZNF782, GLI3, ZNF652, GLI1, ZNF343, ZFI | 1499 | 550 | 20063 | 0.876060404 | 1 | 1 | 100 |
| UP_SEQ_FEATURE | zinc finger region:C2H2-type 11 | 19 | 1.207115629 | 0.865977316 | ZBTB48, ZFP62, ZNF430, ZNF354A, ZNF532, ZNF479, ZNF791, ZNF782, ZNF141, ZNF627, ZNF335, ZNI | 1499 | 301 | 20063 | 0.84485338 | 1 | 1 | 100 |
| INTERPRO | IPR015880:Zinc finger, C2H2-like | 53 | 3.367217281 | 0.86818317 | ZNF532, ZNF821, ZNF782, GLI3, GLI1, ZNF343, ZFP91, ZNF184, ZNF681, ZNF490, ZNF391, ZBTB22, ZN | 1452 | 762 | 18559 | 0.889014519 | 1 | 0.999999985 | 100 |
| UP_SEQ_FEATURE | zinc finger region:C2H2-type 8 | 27 | 1.715374841 | 0.884975288 | ZFP62, ZNF430, ZNF552, ZNF532, ZNF510, ZNF782, ZNF652, ZNF343, ZNF184, ZNF681, CASZ1, ZNF49 | 1499 | 427 | 20063 | 0.84631128 | 1 | 1 | 100 |
| INTERPRO | IPR007087:Zinc finger, C2H2 | 54 | 3.430749682 | 0.91283633 | ZNF532, ZNF821, ZNF782, GLI3, GLI1, ZNF343, ZFP91, ZNF184, ZNF681, ZNF490, ZNF391, ZBTB22, ZN | 1452 | 799 | 18559 | 0.863843234 | 1 | 0.999999999 | 100 |
| UP_SEQ_FEATURE | zinc finger region:C2H2-type 3 | 40 | 2.541296061 | 0.919032362 | ZFP62, ZNF430, ZNF532, GLIS1, ZNF510, ZSCAN5A, GLI3, ZNF652, ZNF343, GLI1, ZFP91, ZNF184, BCL | 1499 | 636 | 20063 | 0.841777118 | 1 | 1 | 100 |
| UP_SEQ_FEATURE | zinc finger region:C2H2-type 7 | 28 | 1.778907243 | 0.929784186 | ZFP62, ZNF430, ZNF552, ZNF532, ZNF510, ZNF782, ZNF652, ZNF343, ZNF184, ZNF681, CASZ1, ZNF49 | 1499 | 463 | 20063 | 0.80941506 | 1 | 1 | 100 |
| GOTERM_MF_DIRECT | GO:0003676~nucleic acid binding | 67 | 4.256670902 | 0.940281514 | CPEB2, ZNF532, CPEB3, ZNF821, ZNF782, GLI3, ZNF343, ZNF184, ZFP91, DDX23, ANG, SND1, DDX25, | 1340 | 985 | 16881 | 0.856903553 | 1 | 0.999999771 | 100 |
| UP_SEQ_FEATURE | zinc finger region:C2H2-type 6 | 30 | 1.905972046 | 0.941743855 | ZFP62, ZNF430, ZNF552, ZNF532, ZNF510, ZNF782, ZNF652, ZNF343, ZNF184, BCL11B, BCL11A, CASZ | 1499 | 501 | 20063 | 0.801452465 | 1 | 1 | 100 |
| INTERPRO | IPR013087:Zinc finger C2H2-type/integrase DNA-binding domain | 46 | 2.92249047 | 0.947540061 | ZFP62, ZNF430, ZNF552, ZNF532, ZNF821, GLIS1, ZNF510, ZSCAN5A, ZNF782, CBLL1, GLI3, ZNF652, Z | 1452 | 712 | 18559 | 0.825782725 | 1 | 1 | 100 |
| UP_SEQ_FEATURE | zinc finger region:C2H2-type 12 | 14 | 0.889453621 | 0.949997154 | ZFP62, ZNF354A, ZNF532, ZNF479, ZNF791, ZNF782, ZNF628, ZNF335, ZNF343, ZNF184, ZNF714, ZNI | 1499 | 256 | 20063 | 0.731951509 | 1 | 1 | 100 |
| UP_SEQ_FEATURE | zinc finger region:C2H2-type 4 | 35 | 2.223634053 | 0.955367195 | ZFP62, ZNF430, ZNF532, GLIS1, ZNF510, ZSCAN5A, GLI3, ZNF652, GLI1, ZNF343, ZFP91, ZNF184, BCL | 1499 | 588 | 20063 | 0.796681915 | 1 | 1 | 100 |
| UP_SEQ_FEATURE | zinc finger region:C2H2-type 13 | 10 | 0.635324015 | 0.972050332 | ZNF184, ZFP62, ZNF354A, ZNF714, ZNF681, ZNF791, ZNF782, ZNF490, ZNF628, ZNF335 | 1499 | 204 | 20063 | 0.656090989 | 1 | 1 | 100 |
| SMART | SM00355:ZnF_C2H2 | 53 | 3.367217281 | 0.979779548 | ZNF532, ZNF821, ZNF782, GLI3, GLI1, ZNF343, ZFP91, ZNF184, ZNF681, ZNF490, ZNF391, ZBTB22, ZN | 878 | 762 | 10057 | 0.796700028 | 1 | 0.99999932 | 100 |
| UP_SEQ_FEATURE | domain:KRAB | 17 | 1.080050826 | 0.986626091 | ZNF430, ZNF552, ZNF354A, ZNF479, ZNF510, ZNF791, ZNF782, ZNF141, ZNF627, ZNF343, ZNF184, ZI | 1499 | 343 | 20063 | 0.663359635 | 1 | 1 | 100 |
| REACTOME_PATHWAY | R-HSA-212436:R-HSA-212436 | 19 | 1.207115629 | 0.993231026 | ZNF430, ZNF552, ZNF354A, ZNF479, ZNF510, ZNF791, ZNF782, ZNF141, MED23, ZNF627, ZNF343, ZN | 745 | 358 | 9075 | 0.646488696 | 1 | 0.999999997 | 100 |
| INTERPRO | IPR001909:Krueppel-associated box | 18 | 1.143583227 | 0.99877873 | ZNF430, ZNF552, ZNF354A, ZNF479, ZNF510, ZNF791, ZNF782, ZNF141, ZNF627, ZNF343, ZNF184, ZI | 1452 | 402 | 18559 | 0.57231405 | 1 | 1 | 100 |
| SMART | SM00349:KRAB | 18 | 1.143583227 | 0.999785867 | ZNF430, ZNF552, ZNF354A, ZNF479, ZNF510, ZNF791, ZNF782, ZNF141, ZNF627, ZNF343, ZNF184, ZI | 878 | 392 | 10057 | 0.525969272 | 1 | 1 | 100 |

Annotation Cluster 229 Enrichment Score: 0.03819103739684927

| Category | Term | Count | % | PValue | Genes | List Total | Pop Hits | Pop Total | Fold Enrichment | Bonferroni | Benjamini | FDR |
|---|---|---|---|---|---|---|---|---|---|---|---|---|
| GOTERM_BP_DIRECT | GO:0006369~termination of RNA polymerase II transcription | 4 | 0.254129606 | 0.894783958 | PAPOLA, CSTF3, SNRPB, THOC7 | 1342 | 64 | 16792 | 0.782041729 | 1 | 0.999999998 | 100 |
| REACTOME_PATHWAY | R-HSA-109688:R-HSA-109688 | 4 | 0.254129606 | 0.905240317 | PAPOLA, CSTF3, SNRPB, THOC7 | 745 | 64 | 9075 | 0.761325503 | 1 | 0.999990649 | 100 |
| GOTERM_BP_DIRECT | GO:0031124~mRNA 3'-end processing | 3 | 0.190597205 | 0.917095703 | PAPOLA, CSTF3, THOC7 | 1342 | 50 | 16792 | 0.75076006 | 1 | 0.999999999 | 100 |
| REACTOME_PATHWAY | R-HSA-72187:R-HSA-72187 | 3 | 0.190597205 | 0.946976309 | PAPOLA, CSTF3, THOC7 | 745 | 55 | 9075 | 0.66442953 | 1 | 0.999997784 | 100 |

Annotation Cluster 230 Enrichment Score: 0.03299871831404647

| Category | Term | Count | % | PValue | Genes | List Total | Pop Hits | Pop Total | Fold Enrichment | Bonferroni | Benjamini | FDR |
|---|---|---|---|---|---|---|---|---|---|---|---|---|
| INTERPRO | IPR020849:Small GTPase superfamily, Ras type | 4 | 0.254129606 | 0.597393863 | HRAS, RAP1A, RIT2, AGAP1 | 1452 | 39 | 18559 | 1.310941584 | 1 | 0.999941097 | 99.9999844 |
| INTERPRO | IPR005225:Small GTP-binding protein domain | 9 | 0.571791614 | 0.954719635 | GTPBP4, HRAS, MTG2, RAP1A, EEF2, RIT2, RAB27B, ARL4C, RAB40B | 1452 | 167 | 18559 | 0.688833078 | 1 | 1 | 100 |
| GOTERM_MF_DIRECT | GO:0003924~GTPase activity | 12 | 0.762388818 | 0.98152243 | GTPBP4, HRAS, MTG2, TUBAL3, TGM2, RAP1A, EEF2, RIT2, RAB27B, ARL4C, TUBA1C, ARHGDIB | 1340 | 234 | 16881 | 0.646039036 | 1 | 0.999999999 | 100 |
| INTERPRO | IPR001806:Small GTPase superfamily | 6 | 0.381194409 | 0.986615295 | HRAS, RAP1A, RIT2, AGAP1, RAB27B, RAB40B | 1452 | 139 | 18559 | 0.551727213 | 1 | 1 | 100 |
| GOTERM_MF_DIRECT | GO:0005525~GTP binding | 20 | 1.27064803 | 0.992211567 | GTPBP4, HRAS, FKBP4, SRL, EEF2, RAB40B, NOLC1, MTG2, TUBAL3, TGM2, GUCY1A2, RAP1A, SCG5, | 1340 | 384 | 16881 | 0.656133396 | 1 | 1 | 100 |
| UP_SEQ_FEATURE | short sequence motif:Effector region | 3 | 0.190597205 | 0.996444919 | HRAS, RAP1A, RAB27B | 1499 | 101 | 20063 | 0.397552163 | 1 | 1 | 100 |
| UP_KEYWORDS | GTP-binding | 14 | 0.889453621 | 0.998410651 | GTPBP4, HRAS, NOLC1, MTG2, TUBAL3, GUCY1A2, RAP1A, EEF2, RIT2, AGAP1, RAB27B, ARL4C, RAB | 1535 | 343 | 20581 | 0.547257861 | 1 | 0.999981869 | 100 |
| UP_SEQ_FEATURE | nucleotide phosphate-binding region:GTP | 12 | 0.762388818 | 0.998754574 | GTPBP4, HRAS, MTG2, TUBAL3, RAP1A, EEF2, RIT2, AGAP1, RAB27B, ARL4C, RAB40B, TUBA1C | 1499 | 310 | 20063 | 0.518100239 | 1 | 1 | 100 |

Annotation Cluster 231 Enrichment Score: 0.02777507371605624

| Category | Term | Count | % | PValue | Genes | List Total | Pop Hits | Pop Total | Fold Enrichment | Bonferroni | Benjamini | FDR |
|---|---|---|---|---|---|---|---|---|---|---|---|---|
| GOTERM_MF_DIRECT | GO:0070888~E-box binding | 3 | 0.190597205 | 0.763709108 | TCF3, MYC, TWIST1 | 1340 | 34 | 16881 | 1.111567164 | 1 | 0.999901982 | 99.99999999 |
| UP_SEQ_FEATURE | DNA-binding region:Basic motif | 8 | 0.508259212 | 0.964225485 | FOSL2, CEBPE, MAFB, ATF6B, MGA, TCF3, MYC, TWIST1 | 1499 | 163 | 20063 | 0.656896008 | 1 | 1 | 100 |
| UP_SEQ_FEATURE | domain:Helix-loop-helix motif | 4 | 0.254129606 | 0.993060054 | MGA, TCF3, MYC, TWIST1 | 1499 | 115 | 20063 | 0.465539345 | 1 | 1 | 100 |
| INTERPRO | IPR011598:Myc-type, basic helix-loop-helix (bHLH) domain | 4 | 0.254129606 | 0.99542905 | MGA, TCF3, MYC, TWIST1 | 1452 | 116 | 18559 | 0.440747601 | 1 | 1 | 100 |
| SMART | SM00353:HLH | 4 | 0.254129606 | 0.997774114 | MGA, TCF3, MYC, TWIST1 | 878 | 113 | 10057 | 0.40546697 | 1 | 1 | 100 |

Annotation Cluster 232 Enrichment Score: 0.022544825490825766

| Category | Term | Count | % | PValue | Genes | List Total | Pop Hits | Pop Total | Fold Enrichment | Bonferroni | Benjamini | FDR |
|---|---|---|---|---|---|---|---|---|---|---|---|---|
| UP_KEYWORDS | Monooxygenase | 5 | 0.317662008 | 0.90161097 | FMO4, CYP2B6, CYP21A2, TH, PAH | 1535 | 88 | 20581 | 0.761807818 | 1 | 0.989858194 | 100 |
| GOTERM_MF_DIRECT | GO:0004497~monooxygenase activity | 3 | 0.190597205 | 0.950581249 | CYP2B6, CYP21A2, PAH | 1340 | 58 | 16881 | 0.651608338 | 1 | 0.999999906 | 100 |
| GOTERM_MF_DIRECT | GO:0005506~iron ion binding | 5 | 0.317662008 | 0.998520748 | CYP2B6, CYP21A2, TH, PAH, BBOX1 | 1340 | 153 | 16881 | 0.411691542 | 1 | 1 | 100 |

Annotation Cluster 233 Enrichment Score: 0.02245872337066537

| Category | Term | Count | % | PValue | Genes | List Total | Pop Hits | Pop Total | Fold Enrichment | Bonferroni | Benjamini | FDR |
|---|---|---|---|---|---|---|---|---|---|---|---|---|
| UP_KEYWORDS | Thiol protease | 9 | 0.571791614 | 0.891201561 | SENP3, PGPEP1L, CASP4, CASP9, USP2, CAPN13, USP10, USP42, USP45 | 1535 | 153 | 20581 | 0.788695152 | 1 | 0.988269159 | 100 |
| INTERPRO | IPR018200:Peptidase C19, ubiquitin carboxyl-terminal hydrolase 2, conserved site | 4 | 0.254129606 | 0.942896561 | USP2, USP10, USP42, USP45 | 1452 | 76 | 18559 | 0.672720023 | 1 | 1 | 100 |
| INTERPRO | IPR001394:Peptidase C19, ubiquitin carboxyl-terminal hydrolase 2 | 4 | 0.254129606 | 0.955020285 | USP2, USP10, USP42, USP45 | 1452 | 80 | 18559 | 0.639084022 | 1 | 1 | 100 |
| GOTERM_MF_DIRECT | GO:0004843~thiol-dependent ubiquitin-specific protease activity | 4 | 0.254129606 | 0.9581778 | USP2, USP10, USP42, USP45 | 1340 | 80 | 16881 | 0.62988806 | 1 | 0.999999953 | 100 |
| GOTERM_BP_DIRECT | GO:0016579~protein deubiquitination | 5 | 0.317662008 | 0.961159363 | USP2, USP10, DMWD, USP42, USP45 | 1342 | 99 | 16792 | 0.631952912 | 1 | 1 | 100 |
| GOTERM_BP_DIRECT | GO:0006511~ubiquitin-dependent protein catabolic process | 8 | 0.508259212 | 0.992095702 | PSMA1, NPLOC4, UBE3A, USP2, UBR3, USP10, USP42, USP45 | 1342 | 182 | 16792 | 0.55000737 | 1 | 1 | 100 |

Annotation Cluster 234 Enrichment Score: 0.018985017271020370

| Category | Term | Count | % | PValue | Genes | List Total | Pop Hits | Pop Total | Fold Enrichment | Bonferroni | Benjamini | FDR |
|---|---|---|---|---|---|---|---|---|---|---|---|---|
| UP_KEYWORDS | mRNA transport | 6 | 0.381194409 | 0.903998045 | MYO1C, NUP88, DDX25, THOC7, XPO7, NUP35 | 1535 | 106 | 20581 | 0.758933071 | 1 | 0.989975745 | 100 |
| GOTERM_BP_DIRECT | GO:0006406~mRNA export from nucleus | 5 | 0.317662008 | 0.963310569 | EIF4E, NUP88, DDX25, THOC7, NUP35 | 1342 | 100 | 16792 | 0.625633383 | 1 | 1 | 100 |
| KEGG_PATHWAY | hsa03013:RNA transport | 9 | 0.571791614 | 0.979221509 | EIF3D, EIF4E, NUP88, EIF4A1, CYFIP1, THOC7, EIF1, EIF3I, NUP35 | 574 | 172 | 6879 | 0.627086541 | 1 | 0.995740436 | 100 |
| REACTOME_PATHWAY | R-HSA-159236:R-HSA-159236 | 3 | 0.190597205 | 0.984566761 | NUP88, THOC7, NUP35 | 745 | 72 | 9075 | 0.507550336 | 1 | 0.999999943 | 100 |

Annotation Cluster 235 Enrichment Score: 0.01894134715670335

| Category | Term | Count | % | PValue | Genes | List Total | Pop Hits | Pop Total | Fold Enrichment | Bonferroni | Benjamini | FDR |
|---|---|---|---|---|---|---|---|---|---|---|---|---|
| INTERPRO | IPR001841:Zinc finger, RING-type | 19 | 1.207115629 | 0.891414671 | RNF144B, PHRF1, DTX3L, RUFY1, TRIM52, CBLL1, NFXL1, ZNRF1, TRIML1, ZNRF2, TRIM37, RNF43, RN | 1452 | 295 | 18559 | 0.823226876 | 1 | 0.999999996 | 100 |
| INTERPRO | IPR017907:Zinc finger, RING-type, conserved site | 9 | 0.571791614 | 0.958511589 | RNF144B, PHRF1, DTX3L, RNFT2, SH3RF3, TRIM52, CBLL1, LONRF1, TRIML1 | 1452 | 169 | 18559 | 0.680681207 | 1 | 1 | 100 |
| UP_SEQ_FEATURE | zinc finger region:RING-type | 10 | 0.635324015 | 0.991029245 | RNF123, CHMP3, RNF103, DTX3L, RNFT2, SH3RF3, TRIM52, NEURL1B, CBLL1, TRIML1 | 1499 | 230 | 20063 | 0.581924181 | 1 | 1 | 100 |
| SMART | SM00184:RING | 13 | 0.82592122 | 0.991907091 | RNF144B, PHRF1, DTX3L, TRIM52, ZNRF1, TRIML1, ZNRF2, RNF43, RNF123, RNF103, RNFT2, SH3RF3 | 878 | 244 | 10057 | 0.610277643 | 1 | 0.999999962 | 100 |

Annotation Cluster 236 Enrichment Score: 0.01790198316430085

| Category | Term | Count | % | PValue | Genes | List Total | Pop Hits | Pop Total | Fold Enrichment | Bonferroni | Benjamini | FDR |
|---|---|---|---|---|---|---|---|---|---|---|---|---|
| REACTOME_PATHWAY | R-HSA-1169091:R-HSA-1169091 | 4 | 0.254129606 | 0.92092189 | PSMA1, BCL10, NFKBIB, PSMD1 | 745 | 67 | 9075 | 0.727236302 | 1 | 0.999994189 | 100 |
| GOTERM_BP_DIRECT | GO:0002223~stimulatory C-type lectin receptor signaling pathway | 6 | 0.381194409 | 0.929314831 | PSMA1, BCL10, HRAS, PAK2, PSMD1, CLEC4C | 1342 | 105 | 16792 | 0.715009581 | 1 | 1 | 100 |
| REACTOME_PATHWAY | R-HSA-5607764:R-HSA-5607764 | 3 | 0.190597205 | 0.99150931 | PSMA1, BCL10, PSMD1 | 745 | 80 | 9075 | 0.456795302 | 1 | 0.999999993 | 100 |
| REACTOME_PATHWAY | R-HSA-2871837:R-HSA-2871837 | 4 | 0.254129606 | 0.999331568 | FCER1A, PSMA1, BCL10, PSMD1 | 745 | 137 | 9075 | 0.35565571 | 1 | 1 | 100 |

Annotation Cluster 237 Enrichment Score: 0.016390988347218853

| Category | Term | Count | % | PValue | Genes | List Total | Pop Hits | Pop Total | Fold Enrichment | Bonferroni | Benjamini | FDR |
|---|---|---|---|---|---|---|---|---|---|---|---|---|
| REACTOME_PATHWAY | R-HSA-5362768:R-HSA-5362768 | 3 | 0.190597205 | 0.950609174 | PSMA1, PSMD1, ERLEC1 | 745 | 56 | 9075 | 0.652564717 | 1 | 0.999997882 | 100 |
| REACTOME_PATHWAY | R-HSA-5358346:R-HSA-5358346 | 3 | 0.190597205 | 0.962899961 | PSMA1, PSMD1, ERLEC1 | 745 | 60 | 9075 | 0.609060403 | 1 | 0.999999189 | 100 |
| REACTOME_PATHWAY | R-HSA-5678895:R-HSA-5678895 | 3 | 0.190597205 | 0.965479651 | PSMA1, PSMD1, ERLEC1 | 745 | 61 | 9075 | 0.599075806 | 1 | 0.999999264 | 100 |
| REACTOME_PATHWAY | R-HSA-382556:R-HSA-382556 | 4 | 0.254129606 | 0.97299308 | PSMA1, PSMD1, ERLEC1, ABCC5 | 745 | 84 | 9075 | 0.580057526 | 1 | 0.999999673 | 100 |

Annotation Cluster 238 Enrichment Score: 0.015689870781441540

| Category | Term | Count | % | PValue | Genes | List Total | Pop Hits | Pop Total | Fold Enrichment | Bonferroni | Benjamini | FDR |
|---|---|---|---|---|---|---|---|---|---|---|---|---|
| INTERPRO | IPR003877:SPla/RYanodine receptor SPRY | 5 | 0.317662008 | 0.94241077 | RNF123, RYR3, SPSB2, RYR2, TRIML1 | 1452 | 94 | 18559 | 0.679876619 | 1 | 1 | 100 |
| INTERPRO | IPR001870:B30.2/SPRY domain | 5 | 0.317662008 | 0.956146054 | RNF123, RYR3, SPSB2, RYR2, TRIML1 | 1452 | 99 | 18559 | 0.645539416 | 1 | 1 | 100 |
| SMART | SM00449:SPRY | 5 | 0.317662008 | 0.964973163 | RNF123, RYR3, SPSB2, RYR2, TRIML1 | 878 | 92 | 10057 | 0.622524017 | 1 | 0.99999549 | 100 |
| UP_SEQ_FEATURE | domain:B30.2/SPRY | 3 | 0.190597205 | 0.995315585 | RNF123, SPSB2, TRIML1 | 1499 | 97 | 20063 | 0.413946067 | 1 | 1 | 100 |

Annotation Cluster 239 Enrichment Score: 0.013146039861133692

| Category | Term | Count | % | PValue | Genes | List Total | Pop Hits | Pop Total | Fold Enrichment | Bonferroni | Benjamini | FDR |
|---|---|---|---|---|---|---|---|---|---|---|---|---|
| GOTERM_MF_DIRECT | GO:0004519~endonuclease activity | 3 | 0.190597205 | 0.925756918 | ANG, RNASE4, TSN | 1340 | 52 | 16881 | 0.726793915 | 1 | 0.999999298 | 100 |
| UP_KEYWORDS | Endonuclease | 3 | 0.190597205 | 0.988515521 | ANG, RNASE4, TSN | 1535 | 84 | 20581 | 0.478850628 | 1 | 0.999626605 | 100 |
| UP_KEYWORDS | Nuclease | 4 | 0.254129606 | 0.99788713 | ANG, RNASE4, TSN, CNOT6 | 1535 | 134 | 20581 | 0.400233361 | 1 | 0.999972676 | 100 |

Annotation Cluster 240 Enrichment Score: 0.012561614132713528

| Category | Term | Count | % | PValue | Genes | List Total | Pop Hits | Pop Total | Fold Enrichment | Bonferroni | Benjamini | FDR |
|---|---|---|---|---|---|---|---|---|---|---|---|---|
| UP_SEQ_FEATURE | zinc finger region:C2H2-type 12 | 14 | 0.889453621 | 0.949997154 | ZFP62, ZNF354A, ZNF532, ZNF479, ZNF791, ZNF782, ZNF628, ZNF335, ZNF343, ZNF184, ZNF714, ZNF | 1499 | 256 | 20063 | 0.731951509 | 1 | 1 | 100 |
| UP_SEQ_FEATURE | zinc finger region:C2H2-type 16 | 5 | 0.317662008 | 0.954291607 | ZNF184, ZFP62, ZNF681, ZNF791, ZNF628 | 1499 | 103 | 20063 | 0.649721173 | 1 | 1 | 100 |
| UP_SEQ_FEATURE | zinc finger region:C2H2-type 13 | 10 | 0.635324015 | 0.972050332 | ZNF184, ZFP62, ZNF354A, ZNF714, ZNF681, ZNF791, ZNF782, ZNF490, ZNF628, ZNF335 | 1499 | 204 | 20063 | 0.656090989 | 1 | 1 | 100 |
| UP_SEQ_FEATURE | zinc finger region:C2H2-type 15 | 6 | 0.381194409 | 0.974053096 | ZNF184, ZFP62, ZNF681, ZNF791, ZNF782, ZNF628 | 1499 | 133 | 20063 | 0.60380103 | 1 | 1 | 100 |
| UP_SEQ_FEATURE | zinc finger region:C2H2-type 17 | 3 | 0.190597205 | 0.986994853 | ZNF184, ZFP62, ZNF791 | 1499 | 82 | 20063 | 0.489667909 | 1 | 1 | 100 |
| UP_SEQ_FEATURE | zinc finger region:C2H2-type 14 | 6 | 0.381194409 | 0.992294615 | ZNF184, ZFP62, ZNF681, ZNF791, ZNF782, ZNF628 | 1499 | 156 | 20063 | 0.514779083 | 1 | 1 | 100 |

Annotation Cluster 241 Enrichment Score: 0.011350616047903906

| Category | Term | Count | % | PValue | Genes | List Total | Pop Hits | Pop Total | Fold Enrichment | Bonferroni | Benjamini | FDR |
|---|---|---|---|---|---|---|---|---|---|---|---|---|
| UP_SEQ_FEATURE | domain:Rho-GAP | 3 | 0.190597205 | 0.967060734 | SYDE2, ARHGAP10, ARHGAP9 | 1499 | 68 | 20063 | 0.59048189 | 1 | 1 | 100 |
| INTERPRO | IPR000198:Rho GTPase-activating protein domain | 3 | 0.190597205 | 0.971610119 | SYDE2, ARHGAP10, ARHGAP9 | 1452 | 67 | 18559 | 0.57231405 | 1 | 1 | 100 |
| INTERPRO | IPR008936:Rho GTPase activation protein | 4 | 0.254129606 | 0.977089214 | SYDE2, SYNGAP1, ARHGAP10, ARHGAP9 | 1452 | 91 | 18559 | 0.561832107 | 1 | 1 | 100 |
| SMART | SM00324:RhoGAP | 3 | 0.190597205 | 0.981109628 | SYDE2, ARHGAP10, ARHGAP9 | 878 | 65 | 10057 | 0.52866655 | 1 | 0.999999398 | 100 |

Annotation Cluster 242 Enrichment Score: 0.010544674195411378

| Category | Term | Count | % | PValue | Genes | List Total | Pop Hits | Pop Total | Fold Enrichment | Bonferroni | Benjamini | FDR |
|---|---|---|---|---|---|---|---|---|---|---|---|---|
| GOTERM_CC_DIRECT | GO:0005759~mitochondrial matrix | 19 | 1.207115629 | 0.958675472 | BCKDHA, PDP1, ME3, SHMT2, ERBB4, FARS2, NR3C1, LARS2, OGDH, CLPX, SOD2, AUH, MLYCD, MTG | 1425 | 327 | 18224 | 0.743078491 | 1 | 0.999956434 | 100 |
| UP_KEYWORDS | Mitochondrion | 68 | 4.320203304 | 0.978656365 | PDP1, FHIT, RAD51C, FASTKD1, OGDH, CLYBL, WDR81, CIAPIN1, AUH, DMPK, IMMP2L, TMEM173, A | 1535 | 1119 | 20581 | 0.814773544 | 1 | 0.999013942 | 100 |
| UP_SEQ_FEATURE | transit peptide:Mitochondrion | 26 | 1.65184244 | 0.982214564 | PDP1, ME3, FARS2, OGDH, CLYBL, AUH, PARL, MRPL54, LIAS, BCKDHA, SQRDL, SHMT2, CEP89, C21O | 1499 | 482 | 20063 | 0.721972325 | 1 | 1 | 100 |
| UP_KEYWORDS | Transit peptide | 29 | 1.842439644 | 0.984720758 | PDP1, FASTKD1, ME3, FARS2, OGDH, WDR81, CLYBL, AUH, PARL, MRPL54, LIAS, RMND1, BCKDHA, S | 1535 | 536 | 20581 | 0.725422967 | 1 | 0.999413734 | 100 |

Annotation Cluster 243 Enrichment Score: 0.007605647535700551

| Category | Term | Count | % | PValue | Genes | List Total | Pop Hits | Pop Total | Fold Enrichment | Bonferroni | Benjamini | FDR |
|---|---|---|---|---|---|---|---|---|---|---|---|---|
| UP_KEYWORDS | Citrullination | 4 | 0.254129606 | 0.978923638 | HIST1H1E, HIST4H4, TMPO, HIST1H3H | 1535 | 97 | 20581 | 0.552899694 | 1 | 0.999011395 | 100 |
| GOTERM_CC_DIRECT | GO:0000786~nucleosome | 4 | 0.254129606 | 0.980956474 | HIST1H1E, HIST4H4, H2AFJ, HIST1H3H | 1425 | 94 | 18224 | 0.544203061 | 1 | 0.999995252 | 100 |
| GOTERM_BP_DIRECT | GO:0006334~nucleosome assembly | 5 | 0.317662008 | 0.988062699 | HIST1H1E, HIST4H4, SET, SMYD3, HIST1H3H | 1342 | 119 | 16792 | 0.525742339 | 1 | 1 | 100 |

Annotation Cluster 244 Enrichment Score: 0.0062648315867155305

| Category | Term | Count | % | PValue | Genes | List Total | Pop Hits | Pop Total | Fold Enrichment | Bonferroni | Benjamini | FDR |
|---|---|---|---|---|---|---|---|---|---|---|---|---|
| GOTERM_BP_DIRECT | GO:0051436~negative regulation of ubiquitin-protein ligase activity involved in mitotic cell cycle | 3 | 0.190597205 | 0.980734041 | PSMA1, PSMD1, BUB1B | 1342 | 71 | 16792 | 0.528704267 | 1 | 1 | 100 |
| REACTOME_PATHWAY | R-HSA-174184:R-HSA-174184 | 3 | 0.190597205 | 0.984566761 | PSMA1, PSMD1, BUB1B | 745 | 72 | 9075 | 0.507550336 | 1 | 0.999999943 | 100 |
| GOTERM_BP_DIRECT | GO:0051437~positive regulation of ubiquitin-protein ligase activity involved in regulation of mitotic | 3 | 0.190597205 | 0.986543742 | PSMA1, PSMD1, BUB1B | 1342 | 76 | 16792 | 0.493921092 | 1 | 1 | 100 |
| REACTOME_PATHWAY | R-HSA-2467813:R-HSA-2467813 | 9 | 0.571791614 | 0.987399195 | PSMA1, PDS5B, PSMD1, TUBAL3, BUB1B, SKA2, PAFAH1B1, CLASP1, TUBA1C | 745 | 185 | 9075 | 0.592599311 | 1 | 0.999999973 | 100 |
| GOTERM_BP_DIRECT | GO:0031145~anaphase-promoting complex-dependent catabolic process | 3 | 0.190597205 | 0.989168567 | PSMA1, PSMD1, BUB1B | 1342 | 79 | 16792 | 0.475164595 | 1 | 1 | 100 |
